## Supplementary Data S3 for "Expanding the Landscape of BREX Diversity: Uncovering Multi-Layered Functional Frameworks and Identification of Novel BREX-Related Defense Systems"

*The PDF presents three distinct datasets organized as follows:*

**i. Representative MSAs of all analysed domains present in various protein components of BREX and their related systems.** Alignments are color-coded and annotated as in Supplementary Data S2. The corresponding topology denoting the boundaries of secondary structural elements are marked at the top of each alignment. Domain architecture and their boundaries are marked accordingly on top of each alignment. Consensus for each alignment at percent identity (70% to 100%) are marked below the alignments.

**ii. Shannon entropy plots for fast-evolving domains along with their corresponding alignment.** The bar plots represent positional entropy for each corresponding alignment: yellow-to-red indicates absolute entropy, while light-to-dark blue represents amino acid property-based entropy.

**iii. Synapomorphy table** with description of sequence-structure features of major domains associated with various protein components of BREX and related systems.


1. Type-1 BREX; BrxC-ATPase….4
2. Type-2 BREX; BrxC-ATPase….5
3. Type-3 BREX; BrxC-ATPase….6
4. Type-4 BREX; BrxC-ATPase (Version 1 and 2)….7
5. Type-1 BREX; BrxX/PglX-MTase….8
6. Type-2 BREX; BrxX/PglX-MTase….9
7. Type-3 BREX; PglXI-MTase….10
8. Type-1 BREX; PglZ (Representative 1, 2 and 3)….11
9. Type-2 BREX; PglZ (Representative 1 and 2)….12
10. Type-3 BREX; PglZ (Representative 1 and 2)….13
11. Type-4 BREX; PglZ (Representative 1, 2, 3 and 4)….14
12. BREX-PglZ and PorX PglZ core NPP phosphatase; comparison of catalytic residues….15
13. BREX-PglZ, PorX PglZ, and Type-3 BR-system standalone PglZ; comparison of catalytic residues….16
14. Type-1 BREX; BrxL (C-terminal inactive LonP version)….17
15. Type-1 BREX; BrxL (C-terminal active HKD-DNase version)….18
16. Type-4 BREX; BrxL….19
17. Type-1 BREX; BrxA (Representative 1)….20
18. Type-1 BREX; BrxA (Representative 2)….20
19. Type-1 BREX; BrxA (Representative 3)….20
20. Type-2 BREX; PglW BrxA-like Tripartite DNA-binding component….21
21. Type-3 BREX; BrxA….22
22. Type-4 BREX; DUF4007 (BrxA homolog; Representative 1)….23
23. Type-4 BREX; DUF4007 (BrxA homolog; Representative 2)….23
24. Type-1 BREX; BrxB-iSTAND….24
25. Type-2 BREX; PglW C-terminal iSTAND….25
26. Type-3 BREX; BrxF iSTAND (Representative 1 and 2)….26
27. Type-3 BREX; BrxF STAND (Representative 3, with Walker A & B intact)….26
28. Type-4 BREX; PglZ N-terminal iSTAND (Representative 1 and 2)….27

29. Type-1 BrxB-iSTAND and Type-2 BREX PglW C-terminal iSTAND (subset) alignment comparison….28
30. vWA-MoxR system; iSTAND….29
31. Type-2 BREX; PglW complete protein (Representative 1)….30
32. Type-2 BREX; PglW complete protein (Representative 2)….31
33. Type-2 BREX; BrxHI-Helicase….32
34. Type-2 BREX BrxHI-Helicase and Ski2-like Helicase alignment comparison….33
35. Type-2 BREX; BrxD AAA+ ATPase….34
36. Type-3 BREX; BrxHII-Helicase….35
37. Type-3 BREX BrxHII-Helicase and Swi2/Snf2-Helicase alignment comparison….36
38. **Representative alignments of core protein components associated with DUF499-centered Type-1 BREX-related systems**
(i) DUF499-ATPase (Representative 1)….37
(ii) DUF499-ATPase (Representative 2; HEPN fusion variant)….37
(iii) DUF499-ATPase (Representative 3)….37
(iv) Methyltransferase (PglXI-homolog)….38
(v) Helicase-nuclease fusion component (BrxHII homolog)….39
39. **Representative alignments of core protein components associated with DUF499-centered Type-2 BREX-related systems**
(i) DUF499-ATPase (Representative 1)….40
(ii) DUF499-ATPase (Representative 2)….40
(iii) DUF499-ATPase (Representative 3)….40
(iv) Methyltransferase (PglXI-homolog)….41
(v) Nuclease-Helicase fusion component (BrxHII homolog)….42
(vi) DUF3780….42
(vii) DUF3780+RAMA….42
40. **Representative alignments of core protein components associated with DUF499-centered Type-2 BREX-related systems**
(i) DUF499-ATPase (Representative 1)….44
(ii) Methyltransferase (PglXI-homolog; Representative 1 and 2)….45
(iii) Standalone PglZ….46
(iv) Inactive STAND (BrxB/BrxF-homolog; Representative 1 and 2)….47
(v) Helicase without nuclease fusion (BrxHII homolog)….48
41. Type-4 BREX; BrxP (DndC homolog; PAPS-reduct)….49
42. Dnd-system PAPS-reductase; DndC….49
43. SSP-systems PAPS-reductase; SspD….49
44. OLD-ABC+TOPRIM (Previously classified OLD-ABCs associated with ABC-ATPase centric conflict systems)….50
45. BREX-systems OLD-ABCs (TOPRIM associated)….50
46. **Representative alignments of core protein components associated with the novel HerA/FtsK-PglZ anchored Capture systems**
(i) HerA translocase….51
(ii) GNAT….52

(iii) tRNA-guanine transglycosylase (TGT)….53
(iv) DUF6079 (BrxC-counterpart)….54
(v) PglZ (BrxZ/PglZ counterpart)….55
(vi) DUF4007 (BrxA counterpart)….56
(vii) STAND-NTPase (BrxB counterpart)…57.

47. **Fast-evolving domain candidates with high entropy (Represented in the shannon entropy plot alongside their corresponding sequence alignment)**
(i) Type-1 BREX; BrxC (α+β-domain + wHTH)….59
(ii) Type-3 BREX; BrxC (α+β-domain + wHTH)….61
(iii) Type-4 BREX; BrxC (α+β-domain + wHTH)….63
(iv) BR-systems; DUF499-ATPase (α+β-domain + wHTH)….65
(v) Type-1 BREX; PglZ iSwi1/Snf2….67
(vi) Type-2 BREX; PglZ iSwi1/Snf2….69
(vii) Type-3 BREX; PglZ iSwi1/Snf2….71
(viii) Type-4 BREX; PglZ iSwi1/Snf2 (Representative 1)….73
(ix) Type-4 BREX; PglZ iSwi1/Snf2 (Representative 2)….75
(x) BRC-systems; PglZ iSwi1/Snf2….77
(xi) Type-1 BREX; PglZ C-terminal β-sandwich IG-like….79
(xii) Type-3 BREX; PglZ C-terminal β-sandwich IG-like (Representative 1)….81
(xiii) Type-3 BREX; PglZ C-terminal β-sandwich IG-like (Representative 2)….83
(xiv) Type-1 BREX; BrxB-iSTAND….85
(xv) Type-2 BREX; PglW C-terminal iSTAND….87
(xvi) Type-3 BREX; BrxF-iSTAND….89
(xvii) Type-4 BREX; PglZ N-terminal iSTAND….91
(xviii) Type-3 BR-systems; iSTAND….93
(xix) MoxR-vWA Ternary systems; iSTAND….95
(xx) Type-3 BREX; BrxHII-Helicase C-terminal iREase….97
(xxi) BR-systems; Helicase iREase (Representative 1)….99
(xxii) BR-systems; Helicase iREase (Representative 2)….101
(xxiii) BR-system; DUF499 C-terminal RRM/Ferredoxin (Representative 1)….103
(xxiii) BR-system; DUF499 C-terminal RRM/Ferredoxin (Representative 2)….105
(xxiv) BR-system; DUF499 C-terminal RRM/Ferredoxin (Representative 3)….107
(xxv) BR-system; DUF499 C-terminal RRM/Ferredoxin (Representative 4)….109
(xxvi) BR-system; DUF499 C-terminal RRM/Ferredoxin (Representative 5)….111
(xxvii) Type-2 BR-system; DUF499 C-terminal FnIII+FnIII….113

48. **Protein synapomorphy table**….114

```
# ; Type1_BrxC-ATPase; All-Merged Reference Alignment (ORC-CDC6-AAA+ATPase + wHTH + Alpha+Beta-Domain + wHTH + Coiled-Coil)

OrigSeq                                       [illegible]
MBR4615907.1_Kiritimatiellia_bacterium        [illegible]
MCI9358063.1_Lachnospiraceae_bacterium        [illegible]
NLJ79432.1_Bacillota_bacterium                [illegible]
MBQ2845004.1_Alphaproteobacteria_bacterium    [illegible]
NMA18734.1_Clostridiaceae_bacterium           [illegible]
WP_179400143.1_Macellibacteroides_fermentans  [illegible]
WP_118027524.1_Lactobacillus_amylovorus       [illegible]
KAF3303185.1_Carnobacterium_sp.               [illegible]
MSB67176.1_Leuconostoc_lactis                 [illegible]
WP_273096524.1_Dialister_succinatiphilus      [illegible]
MCD7882907.1_Lachnospiraceae_bacterium        [illegible]
WP_022770173.1_Butyrivibrio_sp._NC2007        [illegible]
MBQ8347028.1_Alphaproteobacteria_bacterium    [illegible]
WP_147675231.1_Numidum_massiliense            [illegible]
MBR4606065.1_Lachnospiraceae_bacterium        [illegible]
WP_219938189.1_Succinivibrio_faecicola        [illegible]
WP_091835632.1_Marininema_halotolerans        [illegible]
WP_019974718.1_Empedobacter_brevis            [illegible]
TVQ28583.1_Spirochaetaceae_bacterium          [illegible]
WP_091403827.1_Geotoga_petraea                [illegible]
WP_089525998.1_Paenibacillus_herberti         [illegible]
MBP6234856.1_Saprospiraceae_bacterium         [illegible]
PKK90022.1_Candidatus_Wallbacteria            [illegible]
MAE16011.1_Deltaproteobacteria_bacterium      [illegible]
WP_094389547.1_Prevotella_sp._P4-119          [illegible]
WP_257712004.1_Aeromonas_caviae               [illegible]
WP_098060820.1_Longimonas_halophila           [illegible]
MBZ5596030.1_Acidobacteriia_bacterium         [illegible]
MBN1458231.1_Armatimonadota_bacterium         [illegible]
MCR9098968.1_bacterium                        [illegible]
MBU6197215.1_Cyanobacteria_bacterium_REEB446  [illegible]
HJF93068.1_Mediterranea_massiliensis          [illegible]
HCY74134.1_Rikenellaceae_bacterium            [illegible]
WP_092090777.1_Pisciglobus_halotolerans       [illegible]
NLJ85301.1_Bacillota_bacterium                [illegible]
WP_051485152.1_Cloacibacillus_evryensis       [illegible]
MBR1685959.1_Clostridia_bacterium             [illegible]
WP_262567770.1_Endozoicomonas_gorgoniicola    [illegible]
WP_227411048.1_Bifidobacterium_pullorum       [illegible]
MCL6598118.1_Alicyclobacillus_macrosporangiidus [illegible]
WP_024332155.1_Winkia_neuii                   [illegible]
RHB16634.1_Collinsella_sp._AM40-7AC           [illegible]
WP_044073670.1_Prevotella_pectinovora         [illegible]
WP_270462427.1_Gemmiger_formicilis            [illegible]
OIO89050.1_Anaerolineae_bacterium_CG2_30_64_16 [illegible]
MCK6587209.1_Polyangiaceae_bacterium          [illegible]
MBF0258414.1_Desulfamplus_sp.                 [illegible]
MCK6575226.1_Myxococcota_bacterium            [illegible]
MCA6597699.1_Pseudanabaena_sp.                [illegible]
MBR6253006.1_Clostridia_bacterium             [illegible]
MCK6605711.1_Ignavibacteriaceae_bacterium     [illegible]
OQA19974.1_bacterium_ADurb.Bin363             [illegible]
MBP7846551.1_Burkholderiales_bacterium        [illegible]
MBR5630362.1_Bacteroidales_bacterium          [illegible]
NBJ07681.1_Alistipes_sp._Z76                  [illegible]
OJV92258.1_Bacteroidia_bacterium_44-10        [illegible]
MCH2449130.1_Gracilimonas_sp.                 [illegible]
MCH7965547.1_Bacteroidota_bacterium           [illegible]
WP_213195478.1_Cloacibacterium_caeni          [illegible]
ADB40822.1_Spirosoma_linguale_DSM_74          [illegible]
MCH8487209.1_Cyclonatronum_sp.                [illegible]
RGZ88026.1_Eubacterium_rectale                [illegible]
ETI91650.1_Clostridium_butyricum_DORA_1       [illegible]
APC47779.1_Virgibacillus_halodenitrificans    [illegible]
WP_048125616.1_Methanosarcina_lacustris       [illegible]
PTY02769.1_Verrucomicrobia_bacterium_LW23     [illegible]
TXT19373.1_Erysipelotrichaceae_bacterium      [illegible]
WP_146995455.1_Schleiferilactobacillus_harbinensis [illegible]
RKX87882.1_Spirochaetota_bacterium            [illegible]
MCH3907669.1_Sphaerochaeta_sp.                [illegible]
TWS24839.1_Tsukamurella_sputi                 [illegible]
MBP9037128.1_Candidatus_Cloacimonas_sp.       [illegible]
MBR4608207.1_Lachnospiraceae_bacterium        [illegible]
MBD1206697.1_Paracoccaceae_bacterium          [illegible]
MCI6014853.1_Coprobacillus_sp.                [illegible]
MBP8934558.1_Prevotella_sp.                   [illegible]
OYE05053.1_Nostoc_sp.                         [illegible]
MBQ6502300.1_Flexilinea_sp.                   [illegible]
WP_013460546.1_Sulfuricurvum_kujiense         [illegible]
WP_237187176.1_Rothia_nasimurium              [illegible]
MBQ6141563.1_Kiritimatiellia_bacterium        [illegible]
WP_190817045.1_Coleofasciculus_sp._FACHB-1120 [illegible]
WP_271768605.1_Candidatus_Pseudothioglobus_singularis [illegible]
SFW99921.1_Olsenella_sp._kh2p3                [illegible]
MCB0069888.1_Caldilineaceae_bacterium         [illegible]
OQC05381.1_Candidatus_Cloacimonetes_bacterium [illegible]
WP_193813001.1_Kaistella_flava                [illegible]
OQB98767.1_Spirochaetes_bacterium_ADurb.Bin110 [illegible]
MBV6422081.1_Ignavibacteriaceae_bacterium     [illegible]
MBQ0129310.1_bacterium                        [illegible]
consensus/100%                                [illegible]
consensus/95%                                 [illegible]
consensus/90%                                 [illegible]
consensus/85%                                 [illegible]
```

### ; Type2_BrxC-ATPase; ORC-Cdc6-AAA+ATPase + wHTH + Alpha+Beta-Domain + 3S-wHTH1 + 3SwHTH2 + Coiled-Coil

```
OrigSeq MS-----EYLRDVIDIPDRVTSNDYVLRLTDSVGATQRTLDNYVVTPALEQAFDQALGLVGDALQTHESRGAFLAGSFGSGKSHFMAVLHAMLTHQPAARAKAGLQGVIAKHEDALKDAKVLPLAFHLLGATTMEQALFDGYLRQIQEMHPGAPLPALYKSDALLDDADTMRRTLGDERFFAGLNGDMAAEDTVW [illegible]
--Walker-A--
S1 S2 <---HEH-Ins [illegible]

WP_038566393.1_Dermacoccus_nishinomiyaensis [illegible]
WP_015443641.1_Ilumatobacter_coccineus [illegible]
WP_276096852.1_Streptomyces_silvisoli [illegible]
EEP73860.1_Micromonospora_sp._ATCC_39149 [illegible]
HUP68595.1_Acidimicrobiales_bacterium [illegible]
HUY49759.1_Streptosporangiaceae_bacterium [illegible]
MBK6942983.1_Planctomycetota_bacterium [illegible]
MBK9578953.1_Fibrobacterota_bacterium [illegible]
MBL8678385.1_Myxococcales_bacterium [illegible]
MBL8842560.1_Planctomycetota_bacterium [illegible]
MBX3168148.1_Candidatus_Eremiobacteraeota_bacterium [illegible]
MBX3732320.1_Verrucomicrobiae_bacterium [illegible]
MCB9398403.1_Acidobacteriota_bacterium [illegible]
MCB9544268.1_Myxococcales_bacterium [illegible]
MCB9660515.1_Sandaracinaceae_bacterium [illegible]
MCE9632143.1_Planctomycetia_bacterium [illegible]
MCG5052490.1_Myxococcales_bacterium [illegible]
MCP5560212.1_Verrucomicrobiaceae_bacterium [illegible]
MDJ0841533.1_Acidobacteriota_bacterium [illegible]
MDX2267520.1_Bryobacter_sp. [illegible]
MSW49533.1_Actinomycetia_bacterium [illegible]
WP_030506727.1_Microbispora_rosea [illegible]
WP_052424042.1_Nonomuraea_candida [illegible]
WP_061229212.1_Dietzia_cinnamea [illegible]
WP_071086063.1_Parafrankia_colletiae [illegible]
WP_073883613.1_Corynebacterium_variabile [illegible]
WP_075083674.1_Mariniblastus_fucicola [illegible]
WP_123558026.1_Micromonospora_sp._HM5-17 [illegible]
WP_123661993.1_Actinocorallia_herbida [illegible]
WP_131153514.1_Egibacter_rhizosphaerae [illegible]
WP_144981417.1_Gimesia_aquarii [illegible]
WP_161103223.1_Actinomadura_rayongensis [illegible]
WP_165632998.1_unclassified_Actinopolyspora [illegible]
WP_184886687.1_Sphaerisporangium_siamense [illegible]
WP_191733017.1_Corynebacterium_gallinarum [illegible]
WP_199044191.1_Glycomyces_salinus [illegible]
WP_200346143.1_Halochromatium_glycolicum [illegible]
WP_222632854.1_unclassified_Rhodococcus [illegible]
WP_245209875.1_Ralstonia_solanacearum [illegible]
WP_247539401.1_Bradyrhizobium_sp._168 [illegible]
WP_256246191.1_Pseudarthrobacter_sp._R1 [illegible]
WP_267686051.1_Nannocystis_sp._SCPEA4 [illegible]
WP_270679261.1_Nocardiopsis_sp._LSu2-4 [illegible]
WP_296554869.1_Rhodococcus_sp. [illegible]
WP_322753649.1_Frankia_sp._Cas3 [illegible]
WUA45283.1_Streptomyces_sp._NBC_01367 [illegible]
consensus/100% [illegible]
consensus/95% [illegible]
consensus/90% [illegible]
consensus/85% [illegible]
consensus/80% [illegible]
consensus/75% [illegible]
consensus/70% [illegible]
```

### ; Type3_BrxC-ATPase; STAND/ORC-Cdc6-AAA+ATPase + wHTH + Alpha+Beta-Domain + 4S-wHTH + Coiled-Coil

```
                                      1---------11--------21--------31--------41--------51--------61--------71--------81--------91--------101-------111-------121-------131-------141-------151-------161-------171-------181-------191-
OrigSeq                               MKYRDLVSFEPIESVIQLLAANKADEGRRLVASYVISDAMADRIADVIFPQLDFDPAVDHKGILVVGNYGTGKSHLMSVLSTIAEDAAVLPEVRNEKVRSAAGPISGGFQVLRIEISSRMSLRDILVQELEGFLASKGLTFKFPPEDKVTQNKTALIEMMDVFEQKFSGQGLLVVVDEMLDYLRSRRDTELVHD
                                                                                                --Walker-A--                                                                                                  --Walker-B--
                                                                                        S1                                                                         S2         ---------------HEH-Insert-Region---------------       S3
                                      <<-------------------------------------------------------------------------------------------------------------------------------------------------------------------------------ORC-Cdc
                                      --HHHH----------HHHHH-HHHHHHHHHH----HHHHHHHHHHHHHHH---------EEEEE------HHHHHHHHHHHHH-HHHHHH---HHHHHHHHHH---EEEEEEE--|-HHHHHHHHHHHHHHHHH------------HHHHHHHHHHHHHHH--|---EEEEEEE-HHHHHHH--HHHHHHH
                                      --HHHH----------HHHHH-HHHHHHHHHH----HHHHHHHHHHHHHHH---------EEEEE------HHHHHHHHHHHHH-HHHHHH---HHHHHHHHHH---EEEEEEE--|-HHHHHHHHHHHHHHHHH------------HHHHHHHHHHHHHHH--|---EEEEEEE-HHHHHHH--HHHHHHH
                                      --HHHH----------HHHHH-HHHHHHHHHH----HHHHHHHHHHHHHHH---------EEEEE------HHHHHHHHHHHHH-HHHHHH---HHHHHHHHHH---EEEEEEE--|-HHHHHHHHHHHHHHHHH------------HHHHHHHHHHHHHHH--|---EEEEEEE-HHHHHHH--HHHHHHH
WP_164569833.1_Rhizobium_leguminosarum    MKYRDLVSFEPIESVIQLLAANKADEGRRLVASYVISDAMADRIADVifpqLDFDPAVDHKGILVVGNYGTGKSHLMSVLSTIAEDAAVLPEVRNEKVRSAAGPISGGFQVLRIEISSRMSLRDILVQELEGFLASKGLTFKFPPEDKVTQNKTALIEMMDVFEQKFSGQGLLVVVDEMLDYLRSRRDTELVHD
SDB48716.1_Belnapia_rosea                 MRYGDLIQFEPIESVIQLLDANRPDEAKKLVSTYVISDDMAERIAKLmipgLSFDEAVDHKGVLIVGNYGTGKSHLMSVLSLVAEDAAYVSMIRHPRVADAANAIAGKFKVHRIEISSQMSLRDIITQQLEVFLEKHGVSYSFPPADKIINNKAAFEEMMAAFAEVHPNHGVLLVVDELFLEYLRSRRDTELLD
MBA4157055.1_Gemmatimonadota_bacterium    MRYGDLIQYEPLRTVIELRKADRME--RELVSTYVISDEMADKMRDPiiqnLQFETPADNKGVLVVGNYGTGKSHLMSVVSAVAEHAELAELLSHPAVREAAAAIAGRFKVVRVEL-STAALRDVLCRQLERALAEWGVEYVFPPQDQVANNKEALAEMMEAFHQRFPDRGLLLVVDELLDFLRHRMDAGIILD
WP_190509459.1_Oscillatoria_sp.           MRYGDLIQFEPIETVVQLRDADRLQEAKRLVSTYVISDEMAEKLAVTlfpnLRLDG-HDSKGVLIVGNYGTGKSHLMSVISAIAEHSDLAKALPNSSIAEAATTIAGKFQVVRAEIGTKMSLRDILVSVLEEYLDKLGIPHNFPSIEEVVNTKPAFEEMMGKFNNEYRDKGLLLVVDELLDYLGSRTDQELNLD
RJQ83728.1_Desulfobacteraceae_bacterium   MRYGDLIQFEPIESVVQLRDADDAAVARQLVQTYVISGEMAERLVSLvipqLQFDRPVDNKGLMVVGNYGTGKSHLMSVLSAVAENGELSSHFNDGSVAGAAGKIGGRFQVARTEIGATMSLRDIIVAELEERLAALGVTYTFPPAAQISNNKRAFEEMMSAFHQEYPDHGLLLVVDELLDYLRSKRDQELILD
WP_013219148.1_Nitrosococcus_watsonii     LKYGDLIQFDPIESVVQLRDADKSSAAHTLVNTYVISEEMAERLTQLvipqMQFDQPVDNKGLLVVGNYGTGKSHLMSVVSSLAADASLLEGLNHAGVRDTASQIAGRFKVIRTEIGATMSLRDILVAELEEHLEKLGVEYVFPEAGTITSHKRAFEDMMAKFGEVFPEHGLLLVVDELLDYLRTRKDQELILD
GHU24474.1_Spirochaetia_bacterium         MKYGELIQFEALESVIQLRNADKSDNARRLVETFIISDEMAERITGIiipqMQYTVPADNKGILIVGNYGTGKSHLMSVLSSVAEDASLAHILYNDNWKSSLQKIAGQFKVIRAEISSQMSLRDIIVSLLEESFEKWNIDYHFPPAHSIVNNKSNFEDMMAAFEKVFPDQGLLIIIDELLDYLRSRNDQELILD
WP_209691006.1_Methanomicrobium_sp.       MKYKDLINFQPIESVIKLVDSEKKSTAIQLVRDYVISGEMSERLTSLvfpqVQFGSPCDNKGIFIVGNYGTGKSHLMSVISSIAEDETLLEELSNKDISESAKSIAGKFIAIRSEIGAEMSLRDIITGELEEKLRNYGINYSFPPANTIPNHISSFEEMMSLFHQKYPDKGLLVVVDELLDYLRTKEQALILD
NMB01453.1_Bacillota_bacterium            MKYHELFQFEPITSVIQLTKTGEKSFAKDLVRSFVVSDQMENLFVNrilpnLQFDSYVDNKALMIVGNYGTGKSHMMGVIASIAEDAGYLQELQYPRLREKAECIAGKFKVLRIEIGGATPLREHIILRELEPFLQNHGIAFDFPQSDTISNFKRGFEMMMEAFSQKYPNTGLLLVIDELLEFLQTKEERDLIRD
MCK9328444.1_Candidatus_Cloacimonadota_bacterium MKYKDLVSFEPITSVVQLRNADKPQMAQELVSSFVISKSMEENFTKLifphLQFEHPHDNKALMIIGNYGTGKSHMLSVISALAEDETSVHFVTNPSVKASALQIAGKFKVIRMEVGGQTPLETIILRTLEPNLATWGVPFKFSASNTISNFKDGFLSMMDAFAQKYPDQGILIVIDELLEFFKGKNQQEIQID
WP_047754172.1_Kosmotoga_pacifica         MKYGDLIQFDPVETVAQLKDANDKSKAANFVSTYVISNQMADNLHNVvfenLQFETPQDNKGLMIVGNYGTGKSHLMAVISSIAEDPSLIDDLLTNDRVKESASKIAGKFKVAIIAGRFKVIRTEIGTTMYLRDIILRELKENLAEMGIDFNYPPMDQVVNNKGILHEMMSLFNEKYPDKGLLMVVDELLDYLRHRNEQEIILD
MCB5228480.1_Candidatus_Cloacimonadota_bacterium MFYKDLIQYEPIESLVQLKAAVTQKAANLVSTYVISEEMSDRISNLivpqLQFDSPTDNKGLLIVGNYGTGKSHLLSVISAICGDAALAEHLTDEQVREKVKAIAGRFKVIRTELGSTMPLRDIVLRELENLVKEDGIDYTFPPMDKSFSIKKDLEDMMQAFHEVYPDDGILLVIDELLDYLRSRKGQDLVID
OBW63497.1_Dehalococcoides_mccartyi       MKFKELIQFEPIDTVIQLKESDNPTKAQSLVASYVISDEMAEKLSEIvaqqLQFDKPGDKMGIMVVGNYGTGKSHLMAVISSICENPELAKSVKNERVRNAFTPLAGRFKVVRAEIGTTMSLRDIVVATLQDSLSEMKVDFTFPNLNQTIGQKAPLEKMMNKFHEVYPEHGLLLVVDELLDYLRSRRDQELVLD
MBX3742594.1_Akkermansiaceae_bacterium    MRYADLIQFDPIESIVQLRESDKRDKALELVRSYVISKDMAQRLSSVffpnLALEGNPDTKGVMVVGNYGSGKSHLMSVVSALAEDASALDSLENTDVRKAAAVFAGRFKVIRMEIGSTMGLRELLTTSLTKGLAALGIEYVFPEAHVITENKTSLEDMMSAFHAKYPNLGLLLVVDELLDYLRSRKEQALVLD
MBT3747344.1_Bacteroidota_bacterium       MKYKDIVKFEAIESIIQLTDSAVETKAQELVSTYVISDRMADQLTEIvipqLQFNNPADNKGVLIIGNYGTGKSHLMSVISALAEYPSVGKKLNSDIVMESSSDISGKFKVIRSELGATMPLSQFVTGLIEEEMAKWGIDYEFPPSDQLRTFKEPIQEVMGLFNEKFPEKGLLFILDELLDYLRRRDEQAMILD
WP_106343732.1_Planifilum_fimeticola      MKYRDLVQFEPIESVVQLIEADAVDYAEKLVTTYVISDGMAERLTEVvfeqLQFERPADNKGLLVVGNYGTGKSHLMSVVSAVAEHGELAEKIRHPAVREKALEIGGKFRVIRTEIGSTMSLRNFVVGELEAGLAKLGVDFTFPPEEEVRNNKDAFHAMMGAFETRYPDQGLLLVVDELLDYLRTRKEHDLIRD
WP_254843978.1_Virgibacillus_dokdonensis  LRYDELIAFDPVESVVQLREADDQSRAISLLKTYVISDNMAEKLMNDifenIQFERMVDNKGMLIVGNYGSGKSHLMSVVSTIAELPGASKHLRNKQVAEKAKEIEGKFQVIRAEFGATMPLREIVCRQLEKGLANMGIDYTFPTADQVTNNKDMMYEMMELFHEEYPDKGLLLVIDELLDYLRGRKEQDLTLD
WP_235850150.1_Niallia_circulans          MLYQDLLQFEPIESIIKLTEANVEEYANNLLKTYVISERMAEQLNDLifeqLQFSYPIENKGLFIVGNYGTGKSHLMSVISTIAERENVSQFITNPFVTEKAKEIEGKFKVIRFEIGAKTPLRDIVLHALEDNLAKWGVDFSFPPVDQITNNKEALEEMMGLFNEKYPDQGLLLVIDELLDYLRGRREQEALLD
WP_146342557.1_Dellaglioa_algida          MKYRDLVNFEPIDSVIQLTDANNKDRALHLLDTFVISDRMADTINNTiinqLQFLRSADNKGLMIVGNYGTGKSHLMSVISTIAEHSGSSRHIQNKSVTESAQEIEGKFKVIRFEIGSEMSLREIITQQLELGLLELGINFEFPPDADQIVSNKNSLLEMMSIFNDQYSDKGLLLVVDELLDYLRGRKEQELTLD
WP_229736080.1_Kroppenstedtia_guangzhouensis MKYRELVHFEPIESIIRLTDANQQEYAYQLLDTYVISDRMAELLDELvieqLQMDRYADNKGLLIVGNYGTGKSHLMSVISTVAEMKGTCAHLSNQRVAQKAKEIEGKFKVIRVEIGSKMSLRDIICSELEDSLERIDVDFQFPPADQVHNNKNSLMEMMAAFHEVYPEQGLLLVVDELLDHLRTRKEQELALD
MCP4110753.1_Desulfobacteraceae_bacterium DAIGRLFFNFQQIKDVIDIDSDNESGKRADLVKSYVISELEMQIAEF----FDNLEKPSHKACTVIGNYGSGKSHLFGLISLTEEPALVRFIRSEKIRQAVGLSRKFSVQFELQAQVDLKEWFFGSVRRQLKKYQIDIPVFDPVKDFDDKENIKTVLDIVKTKDPSVGLLVIIDEISDFLATRQKEAMKALD
NUM38593.1_Armatimonadota_bacterium       MKYGDLFELNEITEVIKIAKADTASGAQELVKRFVLTKSLSDSIDGIaipqLNFESGVEGKGIFVVGNYGTGKSHVMSFLSILAENADMLRYVQDDEWRAKLAAFAGKYVVIRQEIAVSVTLFSIVCNSLEELASKNGFEFKFRR-EGVDNIKKEFARFMAAFEGVHPNKGALLVIDEVLDYLRFRDSSRVGLD
MCM1566961.1_Dehalobacter_sp.             IKYGDLFQLNPIETIIKLTHADEQSEAKRLVSSFVVTPSLAKSFIDVilpqLTFSKDHEKKGIFVVGNYGTGKSHVMSFVSILAENAGLLDHVMDPDIRNEFVAIAGKYIVVRRTEIGGQMNLYGIVTHELSMVAKQLGFSFEWQPLGQIVNVKEEFRRFMQAFEDYHPTKGLLLVIDELLHYLETRDDQALQLD
MBU0460852.1_Nanoarchaeota_archaeon       MKVNELINFEEIQDVIEIGAI---QNEEDLVEKFVISQTLQEEIIEL----IDVLSSQKHKSANIIGNYGTGKSHLLAFLSLVLSKPELINLIQNKKVKEKLENLKREFFIIKYELNAKTPLARIFFYRVRKQLKVYNIDIRDIDPEKEKNIKELIQEIMDVIKKKNPKKGLLVVFDEFSDFLRQKKREDMNYD
KFI36375.1_Peptococcaceae_bacterium       AIKIEFFNFEKIKEVVDIDAI---TDKKAMVENYIISPSLEEYLIHL----FRDINSDTHKAAQIIGGYGSGKSHLLAFIISLLTERDLYRYVQNDRVREIIEGPQRDFTVIHWELQPDVRLSEYFYDQLEIQLRKYGIEMALP-MSGVVDHKKTILQVLDKIKEDNPTRGLVVVVDEISDFLKQKTEKINHD
EMR73852.1_Thermoplasmatales_archaeon     VKVKELIDFVEIKDVIDIDSDDSTEGKKNIVKDYIISDNLKEHILDV----AENISKPKHKSVQIIGGYGSGKSHLLAWFVSILENKELIDNITNDGVKKRFEVLDREFAVVQFELQPVSALSDFFFDRIEQQLNKHDIEVPKRDSSKVEDFKKDIKDIIDAVKRKDPKMGLVVVIDEISDFLKQKTKMQQIHRD
WP_114474457.1_Halanaerobium_sp.          MKVKDLLHLEELDKVIDIDTYETEDKAKDTIEKYVITDDLKVHIQRI----AEDLKKPKHKSYQIIGNYGSGKSHFLAILAAVIDNHRLADSLQDEETKEIFKNLDREFAVVHFELQPQHTLSEFFYDRLTVKLKNYDIEIPEIDAGDIYDHKEKVREILDIIKSENPKMGLVVIIDEISDFLKSKPSKEMKAD
WP_098658108.1_Bacillus_toyonensis        NKIRDLIHFEDVEEVIKIRKQ---GQEDNIVSKYVISKKIKDHLIYM----FELLNSDTHKSFNIIGNYGTGKSHFLAFIAALLEKKDTRKLISEEEIEAALKFKREFVVVKFECPAEVPLRKIFYDNVQRQLEKYDFEIDEDINLEIHYDNKENIGWIIDQIKEENPERGLLVIVDEISDFLKQKSETKNKSD
MCL4298469.1_Anaerolineae_bacterium       FFIRDLLHFEEVEEVIKLRKD---EQAQEIVEKYVISDGLRRNLLYM----LELLSGMTHKSFNVVGNYGTGKSHFLAFVAALLEHPHLRSLVSEPEVRARAEALQRRYLVVKFELGAEVSLRHIFFDQVYQQLQKYDTDVKQVDISRDYDNKQNVLDILAIKNEDPEAGLVVIVDEISDFLKQKSEKEMAYD
MBP7946969.1_Verrucomicrobiota_bacterium  RELNKIISVPEPETTPSLSDIIQKEKAEKTVAEYLFTPALRGHGKRI----FDCVVNRKGQGFWVQAEYGAGKTHFLGTLVDLLVDLKVWDALRDQELKSEYGALKVKWFPVAFSLRGGDSLMRVFEEQIRESLKAFAPELE----GQIGKFKERFAYIYDQITKLGGYDGIVFVVDEFRSWQDRHPTAAYAED
MBT9159326.1_Dehalococcoidia_bacterium    TPIDKLIAVERPETTLSLAVA--CKNPIKTVSTVFTDSIRGYFSTI----LESVATGRGQGFWVQAEWGYGKTHFLATIAALLSSSQELWSSVQDREITRQVIHHRLNAKLFPVILSLRGGRSLLQIIEKQIGETLDKCHLQ------DRVVSTKERFSKVYDQLTD--GYNGLLFVIDEFAFWQDSHPSPEYAHD
NLF03150.1_Anaerolineales_bacterium       LPIGELITPARPETTLSLVPA-LQQQARDLVESYVLTDTIRGHFVKI----LENVARGDGQGFWVQAEYGAGKTHFLVTIAALLSDGELWDLVHDDADLAQPYRHRLARRLFPVVFSLRGGHTLLDVLLEGFERSLEHAGLE------DQVTSVKARLAHIYEQITAGLGFDGLLVVIDEYEGWQKGHAEGELSADA
GFP32117.1_Candidatus_Hakubanella_thermoalkaliphilus VPIGSLMKRDRPETTISLFQA--LNTSRTTVGTYFFTPSIQEFFEKI----LGAAGSKHGGGYWVQAEYGAGKTHFLATLSCLLMSEELWNELKNESVRNCRRRLGKKLFPVILSLKGADNLLNVVLDQIKEELGDRNLS------GKISSTKKRIAHIYNQLLD--GYDALLIVIDEYEFAAWQRRHATAESAQD
HIH77326.1_Halobacteria_archaeon          TRIGDLIQIEGLEATLSLANA--IKTPDITVGSYIFTNSISRYMKII----LDAVIKGHGAGFWIQAEYGAGKTHFLSTLASLLADEKVWNKVQDDEIRNYRQSLNYKLFPVLINLKGNDDLFEVIEKKIEDSLEECGIR------DKVTSIKETLKSVYDQLNG--GYNGMLFVIDEFAGWQRRHPSKAYATD
MBP6974014.1_Syntrophorhabdus_sp.         MKYSDLIQFDPIVTVIELKSADDNEKAKELVCSYVMSDNMADLISTKiisqLKLEDVVDNKGVLVVGNYGTGKSHLMSVISAIAANSAYLEEAQNEKFRASAKSIAGKFEVLRIEIGSEASLRAIITGNIEQDLARRGITYKFPDVGNITNNKDALMAMMAAFAEKYGEKGYLIIVIDELLDYLKGRKDNELMQD
HHW29753.1_Syntrophomonadaceae_bacterium  MKYSDLIQFEPVETVIQLISSEDTDYASQLVKTYVISERMAEVIVEViipqLQFHYPRDNKGILVVGNYGTGKSHLLSVLTSVAEDSALLPHLTNELVKENMHDIAGQFRVLRLEIGATNDLRSIICSNLERFLARIGVEYRFFPPASEITNNKDCLYEMMEAFHSVYPEQGLLIAIDELLDYLHSRKDFNLVLD
MBT6646600.1_Nitrososphaerota_archaeon    MKYSEIINFQPIDSVVQIKDSNDKAKATEYVQTFVISDKLAEDLSDLvipnLQFEEPLDNKGLVVIGNYGTGKSHLLSVISSLAEYPGMINNLQNQNIVETCKKIDGKFIVLRLELGARQSLRDIICHEIEKFLETKEVSYNFPTTG-IANNKTALQDMMISFQQKFPEQGFLLVIDELLDYLNSRQDQELITD
WP_053959405.1_Sulfobacillus_thermosulfidooxidans MKYHDLIHFQPIESVIKVQEANEKTKAMELVGNYVISERMEEEIRDHivpnLQFDAPFDNKGLLIVGNYGTGKSHLMAVISTIAENAEALTVLQNQRVAKSFEAIAGKFKVHVRMELGSKIDLRDAICLELEKALKEIGIDYPFPPVNRIINNKESFVDMMGIFHEFPEQGLLIVVIDELLDFLRARNEQEIVLD
WP_020614466.1_Sediminispirochaeta_bajacaliforniensis MKYKDLIQFDPIESVITLTEADNEQKAADLVKTFVVSQSMEINLNEIiipnLQFETPRDQKGIFVVGNYGTGKSHLMAFISALAENKDYLDLVGNLSVREQAAQIAGKFKVIRSELGGSTTLRNAICMELEKHLKSLGINFTFPSTDKIANNKDSIHEMMALFQEKYPDQGLLLILDELLDFLRGREDQKLIQD
WP_149544905.1_Calorimonas_adulescens     MKYRELMEFEPIETVISLSDASNMDKAAGMVESYVTSEKMTSNIDRLiisnLNLDG-AGRKGIFVVGNYGSGKSHLMSVVSSIAEHKGLLEHLKDEKLREYMDRIAGRFEVLRIELGALISLRDAVTKEIEKDLLKRGITYRFPPMDSIPNNKQCTLEMMSLFKKHYPEKGYLIVVIDELLDFIRGKKENEVMILD
NMA94940.1_Clostridiales_bacterium        MKYLELISFDPVESIIELKHADNADKAKELVQSYVMSDGMADSLDYGllrhLQLDEIVNNKGVFIVGNYGSGKSHLMSVVSAVAEDSALLNELRNERFAGYMEPIAGEFEVLRIELGSESSLSDSIVREIEKDLLERGIDYKFPDSSTITNHKDAFMMYAFETKYLGKGYLIVVIDELLDYLRARTEREMIALD
WP_209451817.1_Lactimicrobium_massiliense MKYSELISFHPIEDIIQLTTANDDNRAKEYVKSYVMSDQMAENLKAPvidqLQMDEVVDNHGVLIVGNYGTGKSHLMSVISAIASNADDLQYLQNKKFANEMQCVAGKFEILRMEIGGTMSLRDIIFGYIEDDFADRGIDFTVPDLDKVRDNKKLITDMMLAFNEKYPGKGYLIVVIDELLGYLSSRNEKQIVLD
EGD52432.1_Thermoanaerobacter_ethanolicus MKYGDLIQYEPIETVIKLKDADKIDEAKRLVKSFVVSEGMADSIINIcipqLQFDETIDNKGIFIVGNYGTGKSHLMMSVLSTVAEENKEMLDYVSHEELKKEMGKIAGKFKVLRFDIGANTPLRNIITKEIEKDLARYGIEFYFPPIDEVTNNKDAFMEMMSKFEDKFPDKGYLIVVIDELLDFLKARKEQELILD
NLV27395.1_Methanomicrobiales_archaeon    MKYKDIVQFDSIEGTIQLRDADELAGAERLVKTFVISQGMAEKLVGQiipqLSLTTTIDRKGLFIVGNYGTGKSHVMSFLTSVLSVLEYPNIYKYIANEVRNEIEPLSGKYKVIRTEIGAKRPIRDIFCSVLETELEKIGVNYKFPAADETANNKDDLIAMMGAFQVKYPDTGLIFAIDEMLEYLRQRPELDLIYD
consensus/100%                            ..h.phh.........pl............l.pah.s..h...................uh.l.u.aG.GKoHhhs.h..l..........h..........h......ph.s....l...h...h...h.................bo....hh..h.......uhlhhhDEh..ab..+.......D
consensus/95%                             ..h.clhph..ppp.lpl...........l.palho..h....h.........hp.....p..uh.l.upYGsGKoHhhu.ls.lh.........l.p..h.........h..ph.sh..ph.s..sL.phh....lp..h...............p.K..h.....hh..h.....s..GhlhllDEh..abp.+.p..b..D
consensus/90%                             .bh.-lhphp.lpp.lpl.........p.palho..h...h.........hp.....p.puh.l.upYGsGKoHhhu.lsslh.......p.l.p..h.......h..ca.slb.pl.s..sL.phh....lp..h...shp........s.K..h..hh..h......s..GhlhllDEh..abp.+.p..h..D
consensus/85%                             hbh.-Lhphp.lcsllpl.ps...p.s.phVpsaVhop.h.p.h.......hph....cppuhbllGsYGoGKSHhhuhluslhpp..h.p.lpp..l...h..l..cabslb.-l.s..sL.bphh...lp..L.p.slph......p.s.Kp.h..hh..hpp..s..GhllllDEh.calc.+.p..h..D
consensus/80%                             h+h.-Llpac.l-sllpl.ps...p.u.phVpsaVlSpph.ppl.......lph....cpKuhhllGNYGoGKSHhhuhluslhpp..hhp.lps.pl.p.h..l..+Fbll+.El.up.sLbphl..plcp.L.pbslph.b....spl.s.Kp.h.phh..hpp..sppGlLlllDEhb-aL+.+pppph..D
consensus/75%                             h+h.-Llpa-.l-sllpLppu...p.AbplVpoaVlSpph.cpl.ph.....lph....cpKGhhllGNYGoGKSHlhuhluslhcp..hhp.lpsppl.pp.h..lss+FbVl+hElsup.sLbcll..plcp.L.pbslph.h.s.spl.s.Kpsh.chhs.hpp..PppGlLlVlDElhDaL+.+pcpphhbD
consensus/70%                             M+a.-LlpF-sl-ollpLppusp.ppAbplVpoYVlSqph.-pl.plhh.pLph.ps.-pKGlhllGNYGoGKSHLhuhloulhEp..hhp.lpsppl.p.h..lsG+FbVlRhEluup.sLccllhpplEc.L.pbGlpaphss.spl.spKpshbchMshFpp.aPppGLLlVlDElLDaL+s+pcpchhbD
```

[illegible]

### ; Type4_BrxC-ATPase (Version 1); ORC-Cdc6-AAA+ATPase + wHTH + Alpha+Beta-Domain + wHTH + Coiled-Coil

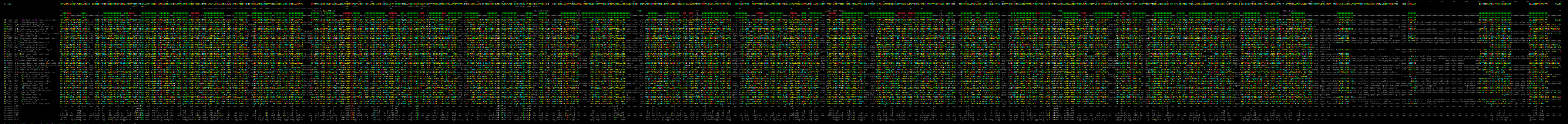


### ; Type4_BrxC-ATPase (Version 2); ORC-Cdc6-AAA+ATPase + wHTH + Alpha+Beta-Domain + wHTH + Extended-Coiled-Coil

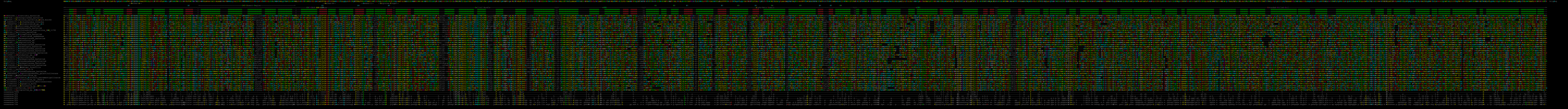

### ; Type 1 BREX MTase; NTD-Dimerisation-Domain + N6_METHYLASE + Target-Recognition-Domain(TRD) + Double_Helical_Spacer + wHTH + Coiled-Coil(CC)

OrigSeq [illegible]

MBZ5596028.1_Acidobacteriia_bacterium [illegible]
WP_251961449.1_Salinibacter_ruber [illegible]
CAG0947532.1_Anaerolineae_bacterium [illegible]
MCR5643212.1_Prevotella_sp. [illegible]
MCK6527599.1_Myxococcota_bacterium [illegible]
WP_191006192.1_Hymenobacter_montanus [illegible]
WP_263326061.1_Neobacillus_sp. [illegible]
MCI0601820.1_bacterium [illegible]
TXH34389.1_Burkholderiaceae_bacterium [illegible]
MBN2413377.1_candidate_division_KSB1_bacterium [illegible]
PIV21172.1_Deltaproteobacteria_bacterium [illegible]
WP_259215310.1_Bradyrhizobium_centrosematis [illegible]
MCC6132529.1_Acidobacteriota_bacterium [illegible]
MCL5952138.1_Chloroflexota_bacterium [illegible]
MCG3192297.1_Thermoanaerobaculia_bacterium [illegible]
MCH7912324.1_Deltaproteobacteria_bacterium [illegible]
WP_204660266.1_Dyadobacter_sandarakinus [illegible]
BAY35693.1_Nostoc_sp. [illegible]
WP_166386806.1_Polaribacter_sp._11A2H [illegible]
MBS1808656.1_Acidobacteriota_bacterium [illegible]
MCL4264722.1_Anaerolineae_bacterium [illegible]
MCG8407730.1_Phycisphaerales_bacterium [illegible]
MBP7797363.1_Thermoanaerobaculaceae_bacterium [illegible]
MCA9231626.1_Planctomycetales_bacterium [illegible]
MCC7280697.1_Chromatiaceae_bacterium [illegible]
WP_201065307.1_Thiorhodovibrio_winogradskyi [illegible]
WP_254579407.1_Marinobacterium_sedimentorum [illegible]
MCP5535435.1_Akkermansiaceae_bacterium [illegible]
GDY21841.1_Verrucomicrobiota_bacterium [illegible]
PZV04976.1_Cyanobium_sp. [illegible]
WP_219012196.1_Shewanella_algae [illegible]
MCC6190562.1_Anaerolineales_bacterium [illegible]
WP_259228203.1_Salinibacter_ruber [illegible]
WP_099325980.1_Candidatus_Kuenenia_stuttgartiensis [illegible]
WP_055669756.1_Desnuesiella_massiliensis [illegible]
WP_131821447.1_Desulfofundulus_thermosubterraneus [illegible]
MBV6422082.1_Ignavibacteriaceae_bacterium [illegible]
RJQ12088.1_Bacillota_bacterium [illegible]
MBS1828810.1_Acidobacteriota_bacterium [illegible]
NLW72145.1_Chloroflexota_bacterium [illegible]
MBN1450023.1_Anaerolineales_bacterium [illegible]
RLC10932.1_Deltaproteobacteria_bacterium [illegible]
WP_190817049.1_Coleofasciculus_sp._FACHB-1120 [illegible]
MBK9096487.1_Anaerolineae_bacterium [illegible]
MBA3548103.1_Nannocystis_sp. [illegible]
WP_012494684.1_Rhodopseudomonas [illegible]
MBI3732788.1_Chloroflexota_bacterium [illegible]
MBO6574435.1_Rhodothermales_bacterium [illegible]
WP_143101638.1_Stigmatella_aurantiaca [illegible]
MCD6541518.1_Candidatus_Bipolaricaulota_bacterium [illegible]
QRK08523.1_Archangium_violaceum [illegible]
MBK8030402.1_Chloroflexota_bacterium [illegible]
WP_129893876.1_Ktedonosporobacter_rubrisoli [illegible]
RRR74016.1_Candidatus_Viridilinea_halotolerans [illegible]
RWC69661.1_Mesorhizobium_sp. [illegible]
MCE8425367.1_Candidatus_Methanoperedens_sp. [illegible]
RGZ88025.1_Agathobacter_rectalis [illegible]
WP_250714473.1_Bacteroides_ovatus [illegible]
WP_110300685.1_Halanaerobium_congolense [illegible]
WP_087726353.1_Proteus_mirabilis [illegible]
MCB9043398.1_Chitinophagales_bacterium [illegible]
consensus/100% [illegible]
consensus/95% [illegible]
consensus/90% [illegible]
consensus/85% [illegible]
consensus/80% [illegible]
consensus/75% [illegible]
consensus/70% [illegible]

```
# ; Type 2 BREX MTase; NTD(Dimerisation-Domain) + N6_METHYLASE + Target-Recognition-Domain(TRD) + Coiled-Coil(CC)

                                                     1--------11--------21--------31--------41--------51--------61--------71--------81--------91--------101-------111-------121-------131-------141-------151-------161------- [illegible] -----921-------931-------941-------951----
OrigSeq                                              MIDIKALVPQLQKQVELLTEDLIGQAKAVEATSAFLREHYDKAIETGRLSQSFEEWSEDYLEQVAVSWVLSCVFVRFLEDNHLIDECWIASEGDRGKLADGHYEIYFRSYPHHSDREYLKYVFKEVGKIPAARELFDDEKTPLWAVDPSGDAARRLLMFFKEIDAERGGL [illegible] NKEIGLIERPEYKRRWSTELWADQVRRASENWLLDRLEAYS  OrigSeq

                                                     <<------------------------------------------------------------------------------------------------------NTD-Dimersiation-Domain--------------- [illegible] -------------------------------------->>
                                                     ---HHHHHHHHHHHHHHHHHHHHHHHHHH-HHHHHHHHHHHHHHHHH------HHHHHHHHHHHHHHHHHHHHHHHHHHHHHH------------HHHHHHHHHHHHHHHHH-----HHHHHHHHHHHHHH-HHHHHHH-------------HHHHHHHHHHH [illegible] -HHHHHHH------------HHHHHHHHHHHHHHHHHHH-
                                                     ---HHHHHHHHHHHHHHHHHHHHHHHHHH-HHHHHHHHHHHHHHHHH------HHHHHHHHHHHHHHHHHHHHHHHHHHHHHH------------HHHHHHHHHHHHHHHHH-----HHHHHHHHHHHHHH-HHHHHHH-------------HHHHHHHHHHH [illegible] -HHHHHHH------------HHHHHHHHHHHHHHHHHHH-
                                                     ---HHHHHHHHHHHHHHHHHHHHHHHHHH-HHHHHHHHHHHHHHHHH------HHHHHHHHHHHHHHHHHHHHHHHHHHHHHH------------HHHHHHHHHHHHHHHHH-----HHHHHHHHHHHHHH-HHHHHHH-------------HHHHHHHHHHH [illegible] -HHHHHHH------------HHHHHHHHHHHHHHHHHHH-
MBN9519353.1_bacterium                               MIDIKALVPQLQKQVELLTEDLIGQAKAVEATSAFLREHYDKAIETGRLSQSFEEWSEDYLEQVAVSWVLSCVFVRFLEDNHLIDECWIASEGDRGKLADGHYEIYFRSYPHHSDREYLKYVFKEVGKIPAARELFDDEKTPLWAVDPSGDAARRLLMFFKEIDAERGGL [illegible] NKEIGLIERPEYKRRWSTELWADQVRRASENWLLDRLEAYS
MCC6750663.1_Deltaproteobacteria_bacterium           ALDGAGLLRAAQPVLKVLNADLLARAKGSPAVRAALEARHAADRAARRTADSFLEWQGRFIAQVGAAWLLSCVFVRTLEDRGLLGRARLAGPG-----AADSQRLFFELAPSLTERDYLLTVFRELCGFPAARDLFDARHNPVWLLAPSAEAAKALLSFRTPTIEAPAL [illegible]
OZD41666.1_Rhodococcus_sp._06-1477-1B                TTAGPQLTAALKPQLEILVDDIRSRVRSNDLESGRWQTEYRRAVKERSTAVSWTEWSEDRFTQAAVGWLLASVFIRFCEDNLMLGEVWIAGPNRLRQRAEDAENDFYRQNQDASYREWLESAFAALANCRATRGLVD-DHAAFRITSPSSDAIQQLIAFWRKTDDA-QEL [illegible]
WP_172812478.1_Corallococcus_exiguus                 ANDRARLTTALRDHVAKIAADLRAKMRA----PGAARAAAEQLHKDERVAEDFEVWTDLLSRRAAVLWVLKSVYVRVLEDRRLLSPGRLLDP--------EAQQLFEKLAPNLGETSFLRWIYKDLASSGGVPELF--SRQPAEVALPSDELSRALIAFWRHDADTG-AV [illegible]
WP_188543682.1_Rhodococcus_trifolii                  AFKSEDLVSALRRVVLDVEYDLQVRVEGQDGVIEAWRTEYQTLLSAERTGVSWIEWRNDRITQAAVSWVLLTVFARYCEDNHLVTHRWISGPDDERGHALDTRTAYFRAHPEHTDRDWLTSIAHHFARYPATAGLVD-SYAPMWQVAPSGDAARALLEFWWDKDDT-GS- [illegible]
MBK9578957.1_Fibrobacterota_bacterium                MIATPDLTRQCQKLVRDFTPFLRQRLKALPAEDQALRARYRELRQHDQTSETEVDWLDAQLDQAAVAWVLACVFVRFLEDNRWLEHTWIAGVGNRLQWALDRQTQWFQENPSKSDLDYLEHVFKNMALYPGVSDLFGAGHNPLWSLKPTADGATFLLRFWRERDGSTGHL [illegible]
WP_104981127.1_Sorangium_cellulosum                  KVVERNLTRELSTLLKTLLPDLKERARP-EN [illegible]
GDX78542.1_Deltaproteobacteria_bacterium             AETYKTLLKDLQKLEKRIAEAHIAAVRADPV [illegible]
WP_193346318.1_Corallococcus_soli                    -MSIANLPKLLQPALKALEKDLLARAKT-ER [illegible]
WP_277955350.1_Saccharopolyspora_soli                ------MLADLQPVLKNLEDDLRERCERPGE [illegible]
WP_150112692.1_Dermacoccus_nishinomiyaensis          PVSSSALTPALKKLVLSTEADLRAWVEADGT [illegible]
WP_052413696.1_Streptomyces_glaucescens              SLDRAALLAGLRKQVRALEADLRERSDDVDR [illegible]
NMM16637.1_Cellulomonas_sp.                          MISSALLLADLKRRLALLESDLRAQSEDDVE [illegible]
GII83982.1_Sphaerisporangium_siamense                SVVSAEMLKDLQRQARLLEADLRAQAEEREE [illegible]
WP_020868705.1_Streptomyces_rapamycinicus            SLDHDALLKDLQKQVAALEADLRERAHGDSE [illegible]
ALO95599.1_Streptomyces_hygroscopicus_subsp          SLDRTRLLKDLQAQARILEDDLRARTEDEPE [illegible]
MCE9575764.1_Deltaproteobacteria_bacterium           PLDGGELLAAARPVLNLLREDLGTRAED-PA [illegible]
WP_145372506.1_Maioricimonas_rarisocia               MINRQELLSDLQTLVRRIEADLIARSDSVPE [illegible]
WP_238634083.1_unclassified_Halorhodospira           PVELETLRRDLQQQVRQLTEDLRERSDTVDA [illegible]
WP_164104215.1_Candidatus_Laterigemmans_baculatus    MINRQQLLKDLQALLRRLEADLRERHDEMPE [illegible]
WP_246179180.1_Actinomadura_decatromicini            ----PSLLAEARGLVRLLEDDLQAARDR--A [illegible]
WP_259859971.1_Dactylosporangium_fulvum              -MTQPTFLRDIKDLVRILGDDLYDQFAD-PA [illegible]
WP_215792560.1_Actinoplanes_bogorensis               MIE----LKALQGQVRLLVDDLRDQVAAIAE [illegible]
WP_131899849.1_Actinomadura_rubrisoli                MIERSALLADLRTQVAVLEGDLRKRAEDDEE [illegible]
WP_034267095.1_Actinospica_robiniae                  MIDRAKLLVDLKNQVKLAEKDLTAQISEVEG [illegible]
WP_120798744.1_Thiocapsa_rosea                       MIQPQRLLADLQSLLPRLEQDILAYTRTRPE [illegible]
WP_200236871.1_Thiohalocapsa_halophila               MIDPQRLLADLQGLLPRLEQDILDFTQADPA [illegible]
MBH2042924.1_Comamonadaceae_bacterium                MINASQLLADLKRRLTLLESDLRERIDTLPD [illegible]
RBJ11686.1_Micromonospora_chalcea                    MAVSTGLTAALKKVVLDLEDDLRDRVASQPD [illegible]
MCB9398404.1_Acidobacteriota_bacterium               ------MLNELTGLLRLLQDDLRMRVTEKPE [illegible]
WP_232328247.1_Kibdelosporangium_sp._MJ126-NF4       LPPDQALTKDLQRLVTRLEIDLRARLDAEPE [illegible]
MBZ5659871.1_Acidobacteriia_bacterium                MINEPLLRQGLQKLLKETESAIRDRLSDEPT [illegible]
WP_250297829.1_Streptomyces_sp._A_4/2                MIDRTALKTDLEDRLTELSADRADAVRGAEA [illegible]
TVR43182.1_Planctomycetota_bacterium                 MIDRTQLLKDCQALLKKLEKDIRAYAQADAD [illegible]
MBA4142468.1_Nitrosospira_sp.                        MINRTKLLADLKVQVRDLESDLRSRFATHAE [illegible]
MSW49531.1_Actinomycetia_bacterium                   MIDARTLTVDLKRLVTELEDDIRSHAAEDQA [illegible]
MCE9632144.1_Planctomycetia_bacterium                MIDRTALLEDLKRLVTRVEADLLARSEDVPE [illegible]
WP_073883614.1_Corynebacterium_variabile             GTAEASLTAALQPVIRDLVADMRTRLTENEV [illegible]
MCA6589822.1_Pseudanabaena_sp._M109S1SP1A06QC        PIARPVLLKDLQVLLKKLEDDLIDRSTSIPE [illegible]
WP_095741470.1_Rhodopirellula_sp._SM50               MIDRQQLLSDLQALLRSVEADLLARSEDLPE [illegible]
TVQ88635.1_Deltaproteobacteria_bacterium             TQARADLVTTLQGHVATIAEDLHTRLLERER [illegible]
consensus/100%                                       ..........hp................... [illegible]
consensus/95%                                        ......h...hp..h..h..sh......... [illegible]
consensus/90%                                        ......h...hp..l..l..Dlb.b...... [illegible]
consensus/85%                                        ......L..sLp..l..l..Dlb.b.p.... [illegible]
consensus/80%                                        ..s...L..sLp..l..lp.Dlb.b.p.... [illegible]
consensus/75%                                        .hs...L..sLp.blp.lp.DLbsb.ps... [illegible]
consensus/70%                                        .hs...L..sLp.blp.LpsDLbsbsps.s. [illegible]
```

```
# ; Type 3 BREX MTase (DUF499 system like MTase); N-terminal-Preceding-Region + N6_METHYLASE(TRD-Insert) + HTH1 + wHTH + HTH2

                                           1---------11--------21--------31--------41--------51--------61--------71--------81--------91--------101-------111-------121-------131-------141-------151-------161-------[illegible]
OrigSeq                                    MATRGEALNLMRAKSTKKLQGPVECLGMKFDNDEARRDYFLNLLREGLEEPEWKDQVGFPHGEIEDILKLSDPPYYTACPNPFIADFINYYGKPYDPEDNYRREPFAADVSEGKNDPIYNAHSYHTKVPQKAIMRYILHYTEPGDLVFDGFCGTGMTGVAAQLCGNRKAV[illegible]

                                                                                                                                                                              S1
                                           <<----------------------------------------Preceding-Region--------------------------------------------|-----------------Core-MTase-------------|--[illegible]
                                           ---HHHHHHHHHHHHH------EEE--EEE--HHHHHHHHHHHHHHHHH-HHHHH--------HHHHHHH----EEEE----HHHHHHHHH------------------------HHHHH---|----HHHHHHHHHHH-----EEEEE-----HHHHHHHHHH-|-HH[illegible]
                                           ---HHHHHHHHHHHHH------EEE--EEE--HHHHHHHHHHHHHHHHH-HHHHH--------HHHHHHH----EEEE----HHHHHHHHH------------------------HHHHH---|----HHHHHHHHHHH-----EEEEE-----HHHHHHHHHH-|-HH[illegible]
                                           ---HHHHHHHHHHHHH------EEE--EEE--HHHHHHHHHHHHHHHHH-HHHHH--------HHHHHHH----EEEE----HHHHHHHHH------------------------HHHHH---|----HHHHHHHHHHH-----EEEEE-----HHHHHHHHHH-|-HH[illegible]
MBW2068239.1_Deltaproteobacteria_bacterium MATRGEALNLMRAKSTKKLQGPVECLGMKFDNDEARRDYFLNLLREGLEEPEWKDQVGFPHGEIEDILKLSDPPYYTACPNPFIADFINYYGKPYDPEDNYRREPFAADVSEGKNDPIYNAHSYHTKVPQKAIMRYILHYTEPGDLVFDGFCGTGMTGVAAQLCGNRKAV[illegible]
WP_052507579.1_Desulfonatronovibrio_magnus MKQAEKKLEWGVEPEEKEEPKPVTCLGITFESDDERRKYFTKLLREKLKDPEFRKIEGFPIGEDEDILALSDPPYYTACPNPWIADFIAEWEEELDPPPPYHREPFAADVSEGKNDPIYNAHSYHTKVPHKAIMRYILHYTEPGDIVFDGFCGTGMTGVAAQMCGDRDTV[illegible]
HBT46728.1_Peptococcaceae_bacterium        MTT----L-FPGCEFRGEKQKQVECVGLTFESEEARRAYFLDRLREGLEEPHWKDEVGFPHGEIEDILRLSDPPYYTACPNPFIQDFIRYYGKPYDPDDAYRREPFAADVSEGKNDPIYNAHSYHTKVPHKAIMRYILHYTESGDLVFDGFCGTGMTGVAASLCGDKKAV[illegible]
WP_273888496.1_Rubrobacter_naiadicus       MNR-QDNL-FDNQEPKAGNQKPLECLGKTFDSGEARREHFLRLLREGLEDSSWKDVVGFPNGEIEDILNLSDPPYYTACPNPFIADFIKHYGKPYDP-VPYSKEPFAADVSEGKNDPIYNAHSYHTKVPHKAIMRYILHYTEPGDVVFDGFCGTGMTGVAAQLCGDRKTV[illegible]
MBC7752532.1_Candidatus_Saccharibacteria_bacterium MHNKGPELDLDNPNQTVINDGPVECLGKTFVNDDARREHYLAILAEKLKDPEFRKIEGFPIGEDEDILKLSDPPYYTACPNPFIADFIAVYGRPYDS-EKYKREPYAADVTEGKSDPIYNAHGYHTKVPHKAIMRYILHYTNPGDLVLDSFCGTGMTGVAAQMCGDRAEV[illegible]
MBT4512246.1_Chloroflexota_bacterium       MSKRQLDL-VDKSMEQTLEKIQVECLGMTFEDEDNRREYFLNKLREKLGDSEFRKIEGFPKADDKDILKLSDPPYHTACPNPFLDEILNFLQNSKKENLDYTKEPFSTDVSEGKNDLVYNAHSYHTKVPYKAVMRYVLHYTKPGDFVLDGFCGTGMTGVATGYCGEPRT[illegible]
WP_209876429.1_Paenibacillus_silagei       MTDKVEQLEFQSED-QSPQTGPVTC[illegible]
MBP8282509.1_Chromatiaceae_bacterium       MN----DL-FDSQ--SRAASGPVEC[illegible]
WP_186776371.1_Rubripirellula_reticaptiva  M------------------------[illegible]
MYB24178.1_Acidimicrobiia_bacterium        MNQPQDSLPLDAAD--------VHG[illegible]
MCA9693369.1_Myxococcales_bacterium        VE-----------------AGPVEC[illegible]
MBV6489841.1_Fimbriimonadaceae_bacterium   MT--DKRL-FEVEI-QPPTPEEIE-[illegible]
MBQ9466674.1_Muribaculaceae_bacterium      MSKKQTSLDFDAATGGQPADHSVTV[illegible]
WP_132761831.1_Vibrio_crassostreae         MK--QENL-FGTGN-EKKDTGPVEC[illegible]
WP_080996766.1_Moorella_thermoacetica      MDYETGSL-FEAK--NDPKLEP---[illegible]
WP_195923519.1_Clostridium_tepidum         MANK-----FEEG------------[illegible]
MBK7577376.1_Ignavibacteria_bacterium      MAKKSGEL-FDVR--HGDESRPVEC[illegible]
MBU0460855.1_Nanoarchaeota_archaeon        MEN-QNKL-------EKEKDEFVEE[illegible]
WP_273281158.1_Methylibium_petroleiphilum  MSD----L-LDDQ--LEREPEPTTC[illegible]
WP_210727635.1_Sphingopyxis_sp._PAMC25046  MNG-QLKM-----N-AKKSSASVEC[illegible]
KPQ15861.1_Algoriphagus_marincola_HL-49    MN-----------------------[illegible]
MBP8005394.1_Acinetobacter_sp.             MSNKVQSLDLDNPNHNQQLDGPVEC[illegible]
WP_073324031.1_Marisediminitalea_aggregata MK--QQEL-LDGSSAENQKNQQVIC[illegible]
MBO5155856.1_Prevotella_sp.                MSNNQPNL-FNDVVMTSHRDEHVVC[illegible]
TNF32507.1_Deltaproteobacteria_bacterium   MSN-QQDLGFAK---PQAYDGPVEC[illegible]
WP_076874183.1_Bacillus_cereus_group       MQ--QLSL-LEE---AQTNNSTIVC[illegible]
MCM1566962.1_Dehalobacter_sp.              MAKDQTKL-WIEEI-QQKKVTKVSV[illegible]
WP_173806728.1_Pseudodesulfovibrio_profundus MK--ETSL-FDSLFEPQKPSGPVTC[illegible]
WP_214805992.1_Exiguobacterium_sp._s46     MDE-QEQLFFSSE--AKTDNQKVVC[illegible]
WP_047715702.1_Enterobacteriaceae          MQ--QDSL-FSNNNSQTQTSCPVTC[illegible]
WP_075107049.1_Psychrobacter_sp._C_20.9    ME--QESL-LGIES-NIKESGPVEC[illegible]
MBF0121087.1_Desulfobacterales_bacterium   MK-----------------------[illegible]
WP_132941356.1_Vibrio_crassostreae         MK--QENL-FDTEG-ENKASGPVEC[illegible]
MBQ9636771.1_Prevotella_sp.                MA--SDKF-FSDEYLKEAEHGPVTV[illegible]
OGN72028.1_Chloroflexi_bacterium_GWB2_49_20 MNSSQAPL-FEEK--SKREKGPVEC[illegible]
MBR3317260.1_Atopobiaceae_bacterium        MEP----------------------[illegible]
MBI3175731.1_Chloroflexota_bacterium       MPRKQPEL-IAEEAA----------[illegible]
WP_201781196.1_Ornatilinea_apprima         MVIKQDTL-FPGEDA----------[illegible]
MBC7326961.1_bacterium                     MW-----------------------[illegible]
WP_270130053.1_Algoriphagus_sp._TR-M9      MD-----------------------[illegible]
WP_013278343.1_Acetohalobium_arabaticum    MD--EDKL-----------------[illegible]
WP_133532121.1_Flavobacterium_dankookense  MFQ-----------DTTQNQTITV[illegible]
consensus/100%                             h........................[illegible]
consensus/95%                              M........................[illegible]
consensus/90%                              M........................[illegible]
consensus/85%                              M........................[illegible]
consensus/80%                              M........................[illegible]
consensus/75%                              Mp....h..................[illegible]
consensus/70%                              Mp.....L.h...........l...[illegible]
```

### ; Type_1_BREX PglZ Representative_1; iSwi2/SNF2-Helicase + 3-HTH + Helical-Connector + Alkaline-Phosphatase + All-Beta-Sandwich-IG-like

[illegible]

### ; Type_1_BREX PglZ Representative_2; iSwi2/SNF2-Helicase + 3-HTH + Helical-Connector + Alkaline-Phosphatase + All-Beta-Sandwich-IG-Like

[illegible]

### ; Type_1_BREX PglZ Representative_3; iSwi2/SNF2-Helicase + HTH + Helical-Connector + Alkaline-Phosphatase + All-Beta-Sandwich-IG-Like

[illegible]

```
# ; Type_2_BREX PglZ Representative_1; iSwi2/SNF2-Helicase + 3-HTH + Helical-Connector + Alkaline-Phosphatase + 3s-wHTH

                                   1---------11--------21--------31--------41--------51--------61--------71--------81--------91--------101-------111-------1[illegible]
OrigSeq                            MGEAVAVSQRPVDESQVRAIAAQIGERDRRMVAVAGAPTWEGPERLETDAGTVRVLAAPSTLAVRAALVAHRAAGDHELLVVVTDRDERELGEEVVAELWDHRVVRPYGWEACKRLFRAEM[illegible]

                                   <<---------------------------------------------------------------------------iSwi/SNF2-Helicase----------->><<[illegible]
                                   -------------HHHHHHHHHHH------EEEEE---------eee----EEEEEE---HHHHHHHHHHH-------EEEEE----HHHHHHHHHHH---EEEE-||HHHHHHHHHH[illegible]
                                   -------------HHHHHHHHHHH------EEEEE---------eee----EEEEEE---HHHHHHHHHHH-------EEEEE----HHHHHHHHHHH---EEEE-||HHHHHHHHHH[illegible]
                                   -------------HHHHHHHHHHH------EEEEE---------eee----EEEEEE---HHHHHHHHHHH-------EEEEE----HHHHHHHHHHH---EEEE-||HHHHHHHHHH[illegible]
QBI18515.1_Egibacter_rhizosphaerae          MgeavavsqrpVDESQVRAIAAQIGERDRRMVAVAGAPTWEGPERLETDAGTVRVLAAPSTLAVRAALVAHRaagdhELLVVVTDRDERELGEEVVAELWDHRVVRPYGWEACKRLFRAEM[illegible]
WP_188941324.1_Nakamurella_endophytica      MspttdnlaatVDRSTVIAQIRLMRRKRRRVLAILATPVWSGTDVLEADGSTYRVWPCPSRLAVRDALRHRHete--PAPVILTDLEAAELGDGICDHLVDHRPLSPHPLSALLDLFSADR[illegible]
WP_285664605.1_Actinorhabdospora_filicis    Mtv--tadvpvVQPHLILGEARGVRKRTG-VMLLRASADWTGDELLKIDGETVRVRPARSVLAVLDAIDSRSeh---ELLIVLSPFREEELGPNVLARVVEQRVFSLGLWDVLLGETGADR[illegible]
WP_332348679.1_Accumulibacter_sp.           Mnv-----llnPSVRQIVAQVDAVLRHDR-VIAIRAQHDESWPSTLAVGEREFSVAWCPSPIAVREQLLALAdgamdPGLVVLTPLPDHVLGADVLARFSRGKVFSVEAWDMLRHAFQARA[illegible]
MBT3378822.1_Lentisphaerota_bacterium       M--------agIRTSQVKALIKSVRNKSRDFIGISSPFLWLEDESVEVDGETYDIVQCCSELHIREALAAATrkg--RSTVVITSLDDPDLGEDLRLRFHKRSLRKLNAWDNVMAIHEAYR[illegible]
WP_153758472.1_Actinomarinicola_tropica     M------trpaLTAQQLRATVERALSSTR-LVLVSGDPDAPPPL-DEVSGRPVTVVGSASPLEIRRLARQTGd---TPLVVVTNCESDELGDDLLARAASRRIITPDRWATVCSLFAAER[illegible]
MCB0954592.1_Ilumatobacter_sp.              M--------paADAQQLLALVNSVRRQED-AIAVRVDPVWAGPDTLD-GPVPIRIVPCPSPLAVREALAEHErtdndQVVMLLTGCGDSELGPDVVARLAKGRVVSLDPFTSVLGLFRAAV[illegible]
MDJ0841534.1_Acidobacteriota_bacterium      Mnt---------RARAVKVLVRDIAKKSP-PLAVAWDGVW--PEAERDDGEVLLFRRAVGPLAVRRVLAEPReag--EMSVILTPTAEKDLGADVITRLWGRRLFQVHKWTLTLQVFGVER[illegible]
WP_242476961.1_Halochromatium_glycolicum    Mde-------tTLPKPILA[illegible]
MDQ3647620.1_Actinomycetota_bacterium       M------tpakIAESALRE[illegible]
HEY0513780.1_Thermoanaerobaculia_bacterium  Mrt---aagtpPSLGLLQA[illegible]
WP_005004490.1_Nitrococcus_mobilis          M---------sLTPTQLAT[illegible]
MSW49534.1_Actinomycetia_bacterium          MadgpalhvtlVSTSHLKR[illegible]
MCO6436557.1_Phycisphaerae_bacterium        M------ptaaPSLGVIRA[illegible]
MBO9539708.1_bacterium                      M--------saLSGPQLSA[illegible]
MCX6543497.1_Acidobacteriota_bacterium      Mt-----ptltLSPAHVRA[illegible]
WP_239491483.1_Luteitalea_sp._TBR-22        Mqh----------VRAIRS[illegible]
MBL9044562.1_Myxococcales_bacterium         M------ssgtLSESLLRA[illegible]
MCG5052491.1_Myxococcales_bacterium         M------kaphLTRQDLAV[illegible]
MBF0261196.1_Magnetococcales_bacterium      M---------eLTESQIVA[illegible]
TAF23832.1_Verrucomicrobiota_bacterium      Mte----tmpsLIPEQLRA[illegible]
MCE9632142.1_Planctomycetia_bacterium       M-------aaaLSFNQIKA[illegible]
WP_330928032.1_Synoicihabitans_lomoniglobus M--------ttVTAGQIRA[illegible]
MCP5560213.1_Verrucomicrobiaceae_bacterium  M--------paLTPPILSA[illegible]
WP_287381673.1_Mesorhizobium_sp.            MeapglspllaPELADIER[illegible]
MCI0393077.1_Acidobacteriota_bacterium      M------sqslSLPPQIRA[illegible]
MBK6942982.1_Planctomycetota_bacterium      M------tsvvVGEAELKA[illegible]
MBI4662621.1_Verrucomicrobiota_bacterium    M----------LTLKQVRA[illegible]
MBL8842559.1_Planctomycetota_bacterium      M------selaLTTATLRI[illegible]
MBK6462401.1_Myxococcales_bacterium         MeaagmiegplLSQADVRL[illegible]
MCZ2341280.1_Bacteroidales_bacterium        M------sigtLSSQQLRS[illegible]
WP_075083673.1_Mariniblastus_fucicola       M--------sdLKRSHLIA[illegible]
MBZ5659873.1_Acidobacteriia_bacterium       Man---sektaMKESVLLS[illegible]
WP_238634086.1_unclassified_Halorhodospira  MtnptaahasgATPRTIEA[illegible]
HUP68596.1_Acidimicrobiales_bacterium       M------svraATAELIGQ[illegible]
MBL8678386.1_Myxococcales_bacterium         Mat--nfqaplVSRDALVE[illegible]
MCB9660516.1_Sandaracinaceae_bacterium      MsaaaglvvpvVSRQDLAE[illegible]
WP_256246190.1_Pseudarthrobacter_sp._R1     Mtq-----lsvVSKAEILE[illegible]
WP_154827459.1_Dietzia_kunjamensis          MtpvsssslgtATPADLIP[illegible]
WP_105324909.1_Corynebacterium_sp._J010B-136 Mts--lqssftVTPRDIEQ[illegible]
WP_312777491.1_Corynebacterium_variabile    M-----sarlrIRQRDIDD[illegible]
WP_138248771.1_Mycolicibacterium_phocaicum  M------ttlvATAPIIRA[illegible]
WP_322753648.1_Frankia_sp._Cas3             MtvpapapsaeATVAIVRA[illegible]
WP_144825789.1_Micrococcus_luteus           MtrtrrgtagrITGAAVRS[illegible]
WP_259849919.1_Brachybacterium_muris        Mtv--ttapapVPRAAVLA[illegible]
WP_157528564.1_Kibdelosporangium_sp.        MtsraesplplATEQYIRS[illegible]
WP_243731729.1_Nocardioides_ochotonae       MtwtlagatarAEPGYVRQ[illegible]
consensus/100%                              M..............l..[illegible]
consensus/95%                               M.............l...h..h......h.h................h.h..s.s.L.hb..h...........hhls......ls.sh..ch...p........h...h.h..[illegible]
consensus/90%                               M.............l...h..h..p....h.h................h.l..s.o.L.hb..h...........hhls......ls.sl..+h...ph.......l...a.hp.[illegible]
consensus/85%                               M..........p...l...l..h..p....hsl.s...h...........h.l..s.o.L.hR..l...........lhlss..p.pls.sl.s+h...ch...p.....l...F.hp.[illegible]
consensus/80%                               M.........hp...l...lp.h.p+...hhul.s...h.ss..h..ss..h.l..s.o.L.hR..L...........lllosb.p.pls.slbs+h...cl...p..p.l...F.hp.[illegible]
consensus/75%                               M.........hp...lbs.lp.h.p+...hlul.s.s.h.ss..h..ss..h.l..s.SsL.lRp.L...........VllTsbsp.clu.DlhsRhh..+l...p.bp.l...F.Ap.[illegible]
consensus/70%                               M.........hs...lbsblp.hbp+p..hlulps.s.Wsusp.hphss..hpV..ssSsL.lRp.Lsp.pp......VlLTshs-.cLG.DlhuRhh..Rlbp.s.ap.lbpbF.Ap.[illegible]

# ; Type_2_BREX PglZ Representative_2; iSwi2/SNF2-Helicase + 3-HTH + Helical-Connector + Alkaline-Phosphatase + 3s-wHTH

                                   1---------11--------21--------31--------41--------51--------61--------71--------81--------91--------101-------111-------12[illegible]
OrigSeq                            MSAPATDAVTTLPRLKTADVAQYLSAALADALEGRGERTVVLLRAEPAWAGSAELAIGERRARVVPAPSALAVHELVLRHLPPKVLVVLTDREHEELDPAILARTHRKRISLVDRWEIVRNA[illegible]

                                   <<-------------------------------------------------------------iSwi/SNF2-Helicase--------------------------->><<[illegible]
                                   ----------------HHHHHHHHHHH-----------EEEEEE---------eeee--EEEEEEE---HHHHHHHHHHHH---EEEEEE---HHH-HHHHHHH--EEE--||--HHHHHH[illegible]
                                   ----------------HHHHHHHHHHH-----------EEEEEE---------eeee--EEEEEEE---HHHHHHHHHHHH---EEEEEE---HHH-HHHHHHH--EEE--||--HHHHHH[illegible]
                                   ----------------HHHHHHHHHHH-----------EEEEEE---------eeee--EEEEEEE---HHHHHHHHHHHH---EEEEEE---HHH-HHHHHHH--EEE--||--HHHHHH[illegible]
WP_280369379.1_Nocardia_wallacei                MSAPATDAVTTLPRLKTADVAQYLSAAladalegrgERTVVLLRAEPAWAGSAELAIGERRARVVPAPSALAVHELVLRHLPPKVLVVLTDREHEELDPAILARTHRKRISLVDRWEIVRNA[illegible]
WP_285664605.1_Actinorhabdospora_filicis        MTV-----TADVPVVQPHLILGEARGLvrkrtgshnVPPVMLLRASADWTGDELLKIDAHTVRVRPARSVLAVLDAIDSRSEHELLIVLSPFREEELGPNVLARVVEQRVFSLGLWDVLLGE[illegible]
WP_148767669.1_Actinomadura_decatromicini       MTEAAAPGRRAVPVATSGILIAEVRRArrrs----pEGGTLFLRAAAEWRGGDVLDLDGATVQVAPCTTLLGVLDAVSAKR-TDFLVVLSAFERRELGESLLAHAIGHDIVAVNRWDLVRHA[illegible]
WP_213171150.1_Natronoglycomyces_albus          M-------SGAVPAASRGIV[illegible]
WP_158645688.1_Stackebrandtia_albiflava         MSV-----GTVLPEARQTMV[illegible]
WP_285505839.1_Actinokineospora_sp.             M-------MAAPPSVNRRMI[illegible]
WP_005461573.1_Saccharomonospora_glauca         M--------SAPPEVDRRMI[illegible]
WP_169440218.1_Streptomyces_sp.                 M-------GVTVPELDQRVV[illegible]
WP_326854225.1_Actinocrinis_sp.                 MAVAASVAAASAPRVSRGMI[illegible]
WP_039917244.1_Amycolatopsis_azurea             M--------AQVPPITRAGI[illegible]
WP_018383907.1_Wenjunlia_vitaminophila          MT------AAAAPPVGRRTI[illegible]
WP_051042993.1_Nocardiopsis_xinjiangensis       MNSPARPTSPALGQVNAEAV[illegible]
HEU5155676.1_Streptosporangiaceae_bacterium     MTE-----RMNAPEANEKVV[illegible]
WP_084008591.1_Planomonospora_sphaerica         MDT-----SLQLPQASRAVV[illegible]
MBV9853173.1_Streptosporangiaceae_bacterium     MSTPGPVTNGTVPAANRTVV[illegible]
WP_270679260.1_Nocardiopsis_sp._LSu2-4          MSP-----GAAVRQVTEDAV[illegible]
WP_169739992.1_Actinospica_robiniae             MPTAAEQTSQRAPRISRSVI[illegible]
WP_193587652.1_Thermobifida_fusca               MTT-----TTTVPPVTVDAV[illegible]
WP_189653453.1_Nonomuraea_spiralis              MSD---------APLLNDLVV[illegible]
NDU73742.1_Actinomadura_lepetitiana             MSASAGATGSGAMRLTVAAV[illegible]
WP_084958882.1_Thermoactinospora_rubra          M--------STALRPSAAAI[illegible]
WP_329144527.1_Streptomyces_niveus              MS---TATSTSAVRLNAATI[illegible]
WP_066069029.1_Frankia_sp._EI5c                 MSVAAGPVGSGAPLVTTAMI[illegible]
WP_264762815.1_Frankia_gtarii                   MSPPAAPAGAGAPRVTAAVV[illegible]
QGN48447.1_Micromonospora_sp.                   MRQTQAATPSQALRISPIAI[illegible]
WP_158624162.1_Micromonospora_sp.               M--SPAASALRALPVNPHTI[illegible]
WP_034267102.1_Actinospica_robiniae             MSITSAP-RAAARPLHLAAI[illegible]
WP_276475785.1_Streptomyces_sp.                 MNSAPAPAALGAGQASPRAA[illegible]
NED01600.1_Streptomyces_sp.                     M----AQALQGIARLTPTAV[illegible]
WP_077964593.1_Streptomyces_tsukubensis         M--TSTTAAAGLARLTPSLI[illegible]
consensus/100%                                  M.................h[illegible]
consensus/95%                                   M.............hp...l[illegible]
consensus/90%                                   M..........h..hp...l[illegible]
consensus/85%                                   M..........h..hs...l[illegible]
consensus/80%                                   M.........shs.hs..hl[illegible]
consensus/75%                                   M.........shs.hs..hl[illegible]
consensus/70%                                   M.......ssshs.ls..hl[illegible]
```

```
# ; Type_3_BREX PglZ Representative_1; iSwi2/SNF2-Helicase + 3-HTH + Helical-Connector + Alkaline-Phosphatase

                                    1---------11--------21--------31--------41--------51--------61--------71--------81--------91--------101-------111------- [illegible]
OrigSeq                             MASWRDHILKEFTPRVSRLTLVADPDGLLLEEGILEGIRERGFELIPFEDHIAFRYAYESRFRSRWDAALVVVLRSGLEALDVLPYDLLQAGRKLSFNLGDIFPHLSYPVVAALDRADLD [illegible]

                                    <<-----------------------------------iSwi/SNF2-Helicase---------------------------------------------->><<----------- [illegible]
                                    ---HHHHHHHH-------EEEEE----HHH-HHHHHHHHHH-EEEEE---HHHHHHHHHHHHHH----EEEEEE---HHHHHH--HHHHHH-EEEEE-HHHHH--||HHHHHH--HHHHH [illegible]
                                    ---HHHHHHHH-------EEEEE----HHH-HHHHHHHHHH-EEEEE---HHHHHHHHHHHHHH----EEEEEE---HHHHHH--HHHHHH-EEEEE-HHHHH--||HHHHHH--HHHHH [illegible]
                                    ---HHHHHHHH-------EEEEE----HHH-HHHHHHHHHH-EEEEE---HHHHHHHHHHHHHH----EEEEEE---HHHHHH--HHHHHH-EEEEE-HHHHH--||HHHHHH--HHHHH [illegible]
NPV81619.1_Bacillota_bacterium      MASWRDHILKEFTPRVSRLTLVADPDGLLLEEGILEGIRERGFELIPFEDHIAFRYAYESRFRSRWDAALVVVLRSGLEALDVLPYDLLQAGRKLSFNLGDIFPHLSYPVVAALDRADLD [illegible]
WP_013074456.1_Kyrpidia_tusciae     MDTWREQILRDFQPP-PPLTLACDPDDLLLDEELLAALRARQIEMVDAGDPVAFRYLYEKTYRERVAGRRVFLLIRTRGHVEELPYDVIKQGRRLTFRLSSLFPGLSLPVIRQLDGGARN [illegible]
MYH51650.1_Gemmatimonadota_bacterium  MKTWREEVIGRLDGGADHLTVAADPDELLVEERMLHRIDELGFDLITFDDHVAFRFAFESSNRAQADRPRRIVIRTRATDPNELPFDILARSRRVSLSLEKLFPKLSYPVVASLNRGDLD [illegible]
WP_190509464.1_Oscillatoria_sp.     KVNWRDRILEQFTPHAARLALVADPDNLLTEEDIAQKIQTRGFETLLYEDSISFRFIYESRYRSRWQDDLIVIVPSQPSELESLPFDLLKESRRLSFSLNELFPNLSYPVLSTLDRNLLD [illegible]
PZV08798.1_Leptolyngbya_sp.         M-TWRDLILNEFATEVSPITWV [illegible]
MCK9221199.1_Limnochordia_bacterium  MSNWQRQVLKKLVPSIARLFIV [illegible]
WP_264656686.1_Azospirillum_canadense  M-NWVDAVTKDLNPEIAPLVAV [illegible]
PKO06218.1_Chloroflexi_bacterium_HGW  MNNWRNTILNEFTPEVATITVV [illegible]
NLJ38120.1_Candidatus_Atribacteria_bacterium  MVSWRDHILNELNSADNNLIIA [illegible]
OBW63499.1_Dehalococcoides_mccartyi  MSTWQSEILKEILHTDSKLVIV [illegible]
WP_209690995.1_Methanomicrobium_sp.  M-GWRDSILSEFITDISKLTLV [illegible]
WP_178366973.1_Desulfobacter_latus  MSEWRDLILDRFAPDVNALSIV [illegible]
HIP19028.1_Sulfurovum_sp.           MRNWREEILAELVPEVNKLTFV [illegible]
OQB94797.1_Spirochaetes_bacterium   MASWRDEILKDFPSGICKLTIV [illegible]
MCP5207858.1_Hahellaceae_bacterium  M-SWRDSVLSEFSIAISRLYIA [illegible]
WP_069455074.1_Shewanella_xiamenensis  MSSWVKELISTLNLDGAFVYLI [illegible]
HEC58784.1_Methylophaga_sp.         M-QWQDRLLNKLALNESLCFVI [illegible]
WP_247116183.1_Marinobacter_sp.     YESWRKPILRALEPGLSDFFVV [illegible]
WP_255556632.1_Methylococcus_sp.    STAWVDRILKEFPVDLARLWIV [illegible]
WP_008603683.1_Pacificimonas_flava  MKDWVDHVAGGFQAEFAKLWAV [illegible]
WP_168134693.1_Sphingomonas_corticis  MTAWSEHVTAPFQREFSRLWAV [illegible]
MBO8173524.1_Bacillaceae_bacterium  MRSWREYVLHHFKEP-PSVSLV [illegible]
WP_188432816.1_Kroppenstedtia_guangzhouensis  MSSWRDRILHHFDEP-LRLYLV [illegible]
WP_181742020.1_Thermoactinomyces_mirandus  M-TWRDEVLKYFADP-PAVMII [illegible]
WP_113032578.1_Paenibacillus_contaminans  MKDWRQKVLESIQAP-SLITIA [illegible]
MCD8511937.1_Bacillus_sp.           MKAWRQLILDKLNVS-YPLILV [illegible]
WP_102026200.1_Salirhabdus_sp._Marseille  MNGWRQKLISQIPFQ-HKLLFV [illegible]
WP_173598486.1_Brevibacillus        MPTWRDAVLNHFNNH-SHLKIV [illegible]
WP_036836014.1_Pontibacillus_litoralis  MNDWREEVLKQFERP-SSVYIV [illegible]
WP_169099399.1_Bacillus_sp.         MANWRDQVLKQFIAN-NALYLV [illegible]
WP_275191612.1_Anoxybacillus_rupiensis  MRTWRDVIIQKLENQ-NRLLLV [illegible]
WP_190997467.1_Peribacillus_faecalis  MNNWRDIIINKMKYC-ESLIFE [illegible]
WP_106343516.1_Planifilum_fimeticola  MGDWQARILQPFHPG-ARLTLV [illegible]
MBT5530409.1_Cytophagia_bacterium   MTDWRKYILNHFDGQ-YRSIIV [illegible]
WP_257961949.1_Lysinibacillus_capsici  M-NWPEKIMSFFTNQ-GSIYYV [illegible]
MCM3358660.1_Psychrobacillus_sp.    MLNWQERIINHFTSK-GSLYIV [illegible]
WP_185807743.1_Lysinibacillus_telephonicus  M-NWVEKITSYFSIK-GALYVV [illegible]
WP_047391752.1_Exiguobacterium_sp.  MNSWLETL---FLKT-NPLYLV [illegible]
WP_214788919.1_Exiguobacterium_sp.  MMNDLHRIQSFLLQS-NPYYVV [illegible]
consensus/100%                      .......l...h........hh [illegible]
consensus/95%                       b..W.p.l...h......h.hh.D.p.lh....h...h....h.h....p.h.h+..aE...c..................hPa-h........hp...hhs.h...hl..h....b. [illegible]
consensus/90%                       M.sW.p.l...h......h.hh.D.c.lh.p..l...l....h.h..h.-.h.hRh.aE..hc........h.........hPaDhh..u....hph..hFs.hs..hl..hs...h. [illegible]
consensus/85%                       M.sW.p.lhp.h....s.l.hh.D.D.lh.c..l...lp...hplh.h.-.h.hRh.aEp.hR........hh......p.lPaDhhp.up.hphpl..lFPphs..llp.ls...hs [illegible]
consensus/80%                       M.sWbc.llp.h....s.l.hl.DsD.ll.-..l.p.lppb.hplh.apDsl.hRh.aEp.hR........ll......p.lPaDlhp.up.lphpls.lFPphs.sllp.ls...hs [illegible]
consensus/75%                       M.sWbc.llpph....s.l.lV.DsD.LL.-c.l.p.lppb.hplh.apDsl.FRabaEpphR.p.p.....ll....p..pplPaDllppup.lphsls.lFPpls.sllppls...hs [illegible]
consensus/70%                       M.sWb-pllpph....s.l.lV.DPD.LL.-cplbp.lppbshplh.apDsl.FRahYEpphR.p.p.....ll....pphpplPaDllppucplphslsplFPpls.sllppls.p.h- [illegible]

# ; Type_3_BREX PglZ Representative_2; iSwi2/SNF2-Helicase + HTH + Helical-Connector + Alkaline-Phosphatase

                                    1---------11--------21--------31--------41--------51--------61--------71--------81--------91--------101-------111--- [illegible]
OrigSeq                             MMINYIVSKLNRCDNNVILVFDEDNILSNSDIISMLKENGYDIYDFKDPDLFRYYFESKFKIAGVKLLIRYDGKITIPYDIKMSSSNIAITLTELFPKLNYTVLKEMILKKLYSIY [illegible]

                                    <<-----------------------------------iSwi/SNF2-Helicase--------------------------------------->><<---------------- [illegible]
                                    -HHHHHHHHHHH----EEEEE----HHHHHHHHHHHHHH--EEEE---HHHHHHHHHHH-------EEEEE-------HHHHHH-EEEEE-HHHHH||--HHHHHH--HHHHHHHH [illegible]
                                    -HHHHHHHHHHH----EEEEE----HHHHHHHHHHHHHH--EEEE---HHHHHHHHHHH-------EEEEE-------HHHHHH-EEEEE-HHHHH||--HHHHHH--HHHHHHHH [illegible]
                                    -HHHHHHHHHHH----EEEEE----HHHHHHHHHHHHHH--EEEE---HHHHHHHHHHH-------EEEEE-------HHHHHH-EEEEE-HHHHH||--HHHHHH--HHHHHHHH [illegible]
WP_014759117.1_Thermoanaerobacterium  MMINYIVSKLNRCDNNVI [illegible]
MBI9032248.1_bacterium              DFVTSILKKVLSLPTELI [illegible]
MBE6339837.1_Bacteroidales_bacterium  DLIKKVAQRITVDYCNLS [illegible]
WP_123477185.1_Muribaculum_intestinale  ELINEILDSISRGFHPLT [illegible]
MBR2506173.1_Bacilli_bacterium      RWVQTIIEKVTPSTVKPI [illegible]
WP_217803628.1_Parabacteroides_johnsonii  NLPQGIINRLFGSEKRLV [illegible]
MBQ9466673.1_Muribaculaceae_bacterium  ALTQNIVNRLFAGDKRLT [illegible]
MBO5155853.1_Prevotella_sp.         ELAKRIEEKVTDRHFHLT [illegible]
MCI6526368.1_Bacteroidales_bacterium  AFYRDIAGKITVPYKRLS [illegible]
TNF50598.1_Bacteroidota_bacterium   TLIEHIAQDLKDQPSKFI [illegible]
WP_076420664.1_Pontibacter_lucknowensis  AFVNHIAEEFLASPKRLI [illegible]
WP_133532125.1_Flavobacterium_dankookense  SFIEHIVSYFNSFDKKII [illegible]
WP_172278715.1_Lentimicrobium_sp._S6  TLIKKIVDDFVSLNRRVI [illegible]
MCC5935593.1_Balneolales_bacterium  ELLRNIAEKLKSYGARTM [illegible]
MBP7508728.1_Prolixibacteraceae_bacterium  NLLETLRKTLQ-AYQKHI [illegible]
WP_094391621.1_Prevotella_sp._P4-119  KFVETILQKITRQGCNLS [illegible]
WP_083331763.1_Prevotella_bryantii  SLLQCIVKEITRQYRRLS [illegible]
MBP6412566.1_Bacteroidia_bacterium  NWIDKFKKVLISSHKEHI [illegible]
WP_073205016.1_Prevotella_ruminicola  SLVSNIVHNIIQENVSIS [illegible]
CEA16535.1_Fermentimonas_caenicola  KLITDIINKLFGNRKRVV [illegible]
MCM1450899.1_Clostridium_sp.        RLLADIVNRIVGGDKRLE [illegible]
MBR3317262.1_Atopobiaceae_bacterium  MLAELVYRNSNAAFSTSV [illegible]
WP_195245438.1_Clostridium_celatum  MFDEYLEDRLALKYEKYI [illegible]
WP_071060846.1_Andreesenia_angusta  MFKDHMIKMLSIDYESRI [illegible]
WP_226978794.1_                     MFADLLYEKSSAAYEKHV [illegible]
WP_207652642.1_Anaerovirgula_multivorans  MFFNLMREILSLEFERLI [illegible]
WP_153014722.1_Syntrophomonas_wolfei  ILRDLLFEKAGLPWVERI [illegible]
MBP3452732.1_Clostridia_bacterium   -MKNRIFEVTGILSQNNC [illegible]
MBQ3643692.1_Candidatus_Riflebacteria_bacterium  SFSQYLYETNNAEQNKRI [illegible]
WP_252226004.1_Caldicoprobacter_algeriensis  LLCQYILEKMAVDYENQI [illegible]
WP_162523517.1_Calorimonas_adulescens  MFYEELLNGISSRKGQII [illegible]
MCR4431941.1_Tepidanaerobacteraceae_bacterium  MIEDVIIQRLPLNQNNFI [illegible]
WP_156946541.1_Desulfitibacter_alkalitolerans  MLME-LLSRLPLKDCDRI [illegible]
WP_045176026.1_Caldicellulosiruptor_danielii  MIEDAIWEKINSNLYDFI [illegible]
consensus/100%                      .h...h........... [illegible]
consensus/95%                       .h.p.h.p......p.. [illegible]
consensus/90%                       .h.p.l.p......p.. [illegible]
consensus/85%                       .h.p.lhp.h....p.. [illegible]
consensus/80%                       .h.p.lhpph....p.. [illegible]
consensus/75%                       .h.p.lhpph...pp.h [illegible]
consensus/70%                       .hhp.lhppls...ppbh [illegible]
```

```
# ; Type_4_BREX PglZ Representative_1; iSTAND-NTPase + iSwi/SNF2-Helicase + 3-HTH + Helical-Connector + Alkaline-Phosphatase + All-Beta-Sandwich-IG-Like

OrigSeq                                         [illegible]

RGH34270.1_Ruminococcus_sp._AM47-2BH             [illegible]
WP_073324030.1_Fibrobacter_sp._UWOV1             [illegible]
WP_216471320.1_Butyricicoccus_intestinisimiae    [illegible]
WP_044080180.1_Prevotella_pectinovora            [illegible]
MCI8555501.1_Clostridia_bacterium                [illegible]
MBD8953983.1_Dialister_sp.                       [illegible]
MZQ76087.1_Peptoclostridium_sp.                  [illegible]
HIS35238.1_Candidatus_Scatousia_excrementigallinarum [illegible]
WP_103226892.1_Clostridium_sp._chh4-2            [illegible]
NBJ16967.1_Dehalobacter_sp._4CP                  [illegible]
CDC05526.1_Clostridium_leptum_CAG:27             [illegible]
MCL2016626.1_Defluviitaleaceae_bacterium         [illegible]
WP_235396406.1_Pseudoflavonifractor_phocaeensis  [illegible]
MBR3817930.1_Clostridia_bacterium                [illegible]
MBP3656735.1_Clostridia_bacterium                [illegible]
ONI45221.1_Epulopiscium_sp._SCG-C07WGA-EpuloA2   [illegible]
MBO6272669.1_bacterium                           [illegible]
WP_037355505.1_Selenomonas_sp._FC4001            [illegible]
MBS6586269.1_Eubacterium_sp.                     [illegible]
NBK77202.1_bacterium_D16-76                      [illegible]
NSW92343.1_Bacillota_bacterium                   [illegible]
WP_048095116.1_Archaeoglobus_fulgidus            [illegible]
NPV08138.1_Anaerolineae_bacterium                [illegible]
MBQ7110119.1_Thermoguttaceae_bacterium           [illegible]
HED00230.1_Pseudomonadota_bacterium              [illegible]
MBE6101264.1_Selenomonas_ruminantium             [illegible]
ROT09401.1_Muribaculaceae_bacterium_Isolate-037  [illegible]
MBR2693834.1_Thermoguttaceae_bacterium           [illegible]
WP_216880780.1_                                  [illegible]
MCM1514006.1_Anaeroplasma_bactoclasticum         [illegible]
HHW17415.1_Bacillota_bacterium                   [illegible]
WP_157832981.1_Desulfofundulus_thermocisternus   [illegible]
WP_166670004.1_Aminivibrio_pyruvatiphilus        [illegible]
HCH97635.1_Lachnospiraceae_bacterium             [illegible]
WP_195464804.1_Clostridium_jeddahense            [illegible]
HBQ28533.1_Desulfotomaculum_sp.                  [illegible]
MBK5260961.1_Peptostreptococcaceae_bacterium     [illegible]
WP_006692206.1_Selenomonas_infelix               [illegible]
WP_028051765.1_Carboxydothermus_ferrireducens    [illegible]
consensus/100%                                   [illegible]
consensus/95%                                    [illegible]
consensus/90%                                    [illegible]
consensus/85%                                    [illegible]
consensus/80%                                    [illegible]
consensus/75%                                    [illegible]
consensus/70%                                    [illegible]

# ; Type_4_BREX PglZ Representative_2; iSTAND-NTPase + iSwi/SNF2-Helicase + 3-HTH + Helical-Connector + Alkaline-Phosphatase + All-Beta-Sandwich-IG-Like

OrigSeq                                         [illegible]

MBQ0159480.1_Bacteroidales_bacterium             [illegible]
PMQ01343.1_Dictyoglomus_sp.                      [illegible]
KUK66682.1_Parcubacteria_bacterium               [illegible]
WP_018963723.1_Coprothermobacter_platensis       [illegible]
MBR3117632.1_Oceanobacillus_sp.                  [illegible]
MBQ4585185.1_Clostridia_bacterium                [illegible]
WP_024292379.1_Lacrimispora_indolis              [illegible]
HAR58249.1_Acholeplasmatales_bacterium           [illegible]
WP_117621172.1_Hungatella                        [illegible]
WP_191216537.1_Methanothermobacter_sp            [illegible]
WP_011944160.1_Thermotoga_petrophila             [illegible]
OQC39307.1_Bacteroidetes_bacterium               [illegible]
MBQ4420212.1_Bacteroidales_bacterium             [illegible]
WP_183546997.1_Methanococcus_maripaludis         [illegible]
MBU7048061.1_Theionarchaea_archaeon              [illegible]
HHW21152.1_Thermodesulfovibrio_thiophilus        [illegible]
WP_207678199.1_Desulfonema_magnum                [illegible]
MBR5399199.1_Bacteroidales_bacterium             [illegible]
WP_022459608.1_Bacteroidales                     [illegible]
MCQ2340464.1_Paludibacteraceae_bacterium         [illegible]
MBE6838777.1_Ruminococcus_sp.                    [illegible]
MBR3710351.1_Bacteroidales_bacterium             [illegible]
WP_205109179.1_Marseilla_massiliensis            [illegible]
WP_172204608.1_Prevotella_sp._PTAC               [illegible]
MCH4183501.1_Prevotella_sp.                      [illegible]
WP_196042834.1_Parabacteroides_merdae            [illegible]
MCI6981218.1_Akkermansia_muciniphila             [illegible]
MCD7797870.1_Akkermansiaceae_bacterium           [illegible]
MBO6251938.1_Muribaculaceae_bacterium            [illegible]
MBR4838437.1_Bacteroidales_bacterium             [illegible]
RGX95620.1_Prevotella_copri                      [illegible]
WP_254009599.1_Bacteroides                       [illegible]
consensus/100%                                   [illegible]
consensus/95%                                    [illegible]
consensus/90%                                    [illegible]
consensus/85%                                    [illegible]
consensus/80%                                    [illegible]
consensus/75%                                    [illegible]
consensus/70%                                    [illegible]

# ; Type_4_BREX PglZ Representative_3; iSwi/SNF2-Helicase + 3-HTH + Helical-Connector + Alkaline-Phosphatase + All-Beta-Sandwich-IG-Like

OrigSeq                                         [illegible]

WP_192822482.1_Rufibacter_sp._LB8                [illegible]
WP_105482027.1_Abditibacterium_utsteinense       [illegible]
WP_243797665.1_Hymenobacter_tibetensis           [illegible]
MBX7251230.1_Candidatus_Promineofilum_sp.        [illegible]
WP_162787804.1_Anaerolinea_thermolimosa          [illegible]
WP_012642882.1_Thermomicrobium_roseum            [illegible]
WP_157205838.1_Calidithermus_timidus             [illegible]
MCL6576921.1_Kyrpidia_sp.                        [illegible]
NPV84914.1_Anaerolineae_bacterium                [illegible]
WP_139401683.1_Deinococcus_radiopugnans          [illegible]
MCE5327915.1_Planctomycetaceae_bacterium         [illegible]
OGX83061.1_Hymenobacter_glacialis                [illegible]
MBN2678362.1_Anaerolineaceae_bacterium           [illegible]
MBV5327602.1_Chlorobium_sp.                      [illegible]
WP_062417760.1_Levilinea_saccharolytica          [illegible]
MCL4273182.1_Anaerolineales_bacterium            [illegible]
WP_133161745.1_Deinococcus_koreensis             [illegible]
MCC6188628.1_Anaerolineales_bacterium            [illegible]
MCA9963066.1_Anaerolineales_bacterium            [illegible]
WP_215594968.1_Hymenobacter_piscis               [illegible]
WP_168570764.1_Oxynema_aestuarii                 [illegible]
WP_006100361.1_Coleofasciculus_chthonoplastes    [illegible]
WP_254031239.1_Planktothrix_agardhii             [illegible]
WP_190414231.1_Coleofasciculus_sp._FACHB-125     [illegible]
consensus/100%                                   [illegible]
consensus/95%                                    [illegible]
consensus/90%                                    [illegible]
consensus/85%                                    [illegible]
consensus/80%                                    [illegible]
consensus/75%                                    [illegible]
consensus/70%                                    [illegible]

# ; Type_4_BREX PglZ Representative_4; iSwi/SNF2-Helicase + 2-HTH + Helical-Connector + Alkaline-Phosphatase + All-Beta-Sandwich-IG-Like

OrigSeq                                         [illegible]

WP_273058112.1_Shewanella_vesiculosa             [illegible]
MCK9609000.1_Methylomonas_sp.                    [illegible]
WP_017840083.1_Methylotuvimicrobium_buryatense   [illegible]
MCK5897804.1_Methylococcales_bacterium           [illegible]
MBC7570526.1_Spirosoma_sp.                       [illegible]
WP_198076877.1_Hymenobacter_negativus            [illegible]
WP_235137408.1_Dyadobacter_fanqingshengii        [illegible]
WP_194112254.1_Mucilaginibacter_myungsuensis     [illegible]
HBJ34341.1_Planctomycetaceae_bacterium           [illegible]
MBX3399563.1_Gemmataceae_bacterium               [illegible]
WP_099078666.1_Vibrio_sp._PID17_43               [illegible]
SEH06866.1_Thiotrichales_bacterium_HS_08         [illegible]
QEP42412.1_Ectothiorhodospiraceae_bacterium      [illegible]
WP_194715675.1_Noviherbaspirillum_soli           [illegible]
WP_201382003.1_Ktedonobacter_sp._SOSP1-85        [illegible]
WP_221704724.1_Desmonostoc_muscorum              [illegible]
WP_007916556.1_Ktedonobacter_racemifer           [illegible]
NJL05928.1_Chloroflexaceae_bacterium             [illegible]
WP_129675140.1_Candidatus_Chloroploca_sp.        [illegible]
MBU6335166.1_Chloroflexota_bacterium             [illegible]
MBK9943589.1_Kouleothrix_sp.                     [illegible]
WP_097643873.1_Candidatus_Viridilinea            [illegible]
NJO81810.1_Blastochloris_sp.                     [illegible]
consensus/100%                                   [illegible]
consensus/95%                                    [illegible]
consensus/90%                                    [illegible]
consensus/85%                                    [illegible]
consensus/80%                                    [illegible]
consensus/75%                                    [illegible]
consensus/70%                                    [illegible]
```

### ; BREX PglZ Phosphatase and PorX Phosphatase Comparison Alignment

```
                                                      Alkaline-Phosphatase
WP_238754868.1_Leptospira_bandrabouensis              [illegible]
MBF0258418.1_Desulfamplus_sp.                         [illegible]
MBX3247846.1_Myxococcales_bacterium                   [illegible]
MBD2549130.1_Microcystis_elabens_FACHB-917            [illegible]
MBY0113210.1_Phycisphaerales_bacterium                [illegible]
MBX6362428.1_Gemmatimonadota_bacterium                [illegible]
WP_083610988.1_Desulfatibacillum_alkenivorans         [illegible]
MBN1165673.1_Methanospirillaceae_archaeon             [illegible]
WP_156746889.1_Mycobacterium_sp._E2733                [illegible]
MBV6422080.1_Ignavibacteriaceae_bacterium             [illegible]
WP_083423498.1_Stigmatella_aurantiaca                 [illegible]
NLN06912.1_Bacillota_bacterium                        [illegible]
WP_242609850.1_Comamonas_thiooxydans                  [illegible]
WP_243520949.1_Hymenobacter_monticola                 [illegible]
OQB98762.1_Spirochaetes_bacterium_ADurb.Bin110        [illegible]
WP_166386808.1_Polaribacter_sp._11A2H                 [illegible]
WP_235164441.1_Dyadobacter_chenhuakuii                [illegible]
MCP5535438.1_Akkermansiaceae_bacterium                [illegible]
MCC6145260.1_Candidatus_Hydrogenedentes_bacterium     [illegible]
MBN9589012.1_Alphaproteobacteria_bacterium            [illegible]
WP_100919682.1_Candidatus_Thiodictyon_syntrophicum    [illegible]
WP_011448105.1_Methanospirillum_hungatei              [illegible]
NBT25026.1_Actinomycetia_bacterium                    [illegible]
WP_246481782.1_Natronogracilivirga_saccharolytica     [illegible]
MBQ6007748.1_Kiritimatiellia_bacterium                [illegible]
WP_091825284.1_Butyrivibrio_sp._ob235                 [illegible]
WP_219938192.1_Succinivibrio_faecicola                [illegible]
MCK6605714.1_Ignavibacteriaceae_bacterium             [illegible]
MCH3907672.1_Sphaerochaeta_sp.                        [illegible]
WP_251178637.1_Adlercreutzia_agrestimuris             [illegible]
MBQ6502306.1_Flexilinea_sp.                           [illegible]
WP_237187173.1_Rothia_nasimurium                      [illegible]
OFQ22229.1_Actinomyces_sp._HMSC062G12                 [illegible]
QNI65417.1_Synechococcus_sp._A15-44                   [illegible]
MCF7790535.1_Victivallales_bacterium                  [illegible]
MBW8002950.1_Planctomycetota_bacterium                [illegible]
WP_262567766.1_Endozoicomonas_gorgoniicola            [illegible]
MBQ2845006.1_Alphaproteobacteria_bacterium            [illegible]
WP_216328025.1_Deinococcus_aestuarii                  [illegible]
WP_151792240.1_Acinetobacter_seifertii                [illegible]
MBP7846546.1_Burkholderiales_bacterium                [illegible]
7PVK_1|Chains                                         [illegible]
7PVA_1|Chains                                         [illegible]
8TED_1|Chains                                         [illegible]
8TFF_1|Chains                                         [illegible]
8THP_1|Chains                                         [illegible]
consensus/100%                                        [illegible]
consensus/95%                                         [illegible]
consensus/90%                                         [illegible]
consensus/85%                                         ...................s...[illegible]
consensus/80%                                         .......p.......h.s...[illegible]
consensus/75%                                         .......p..ha.p.l.s...[illegible]
consensus/70%                                         .......p..happ.l.s.h.[illegible]
Zn+ and Mg2+ Metal-Co-ordinating Residues: ^          [illegible]
```

### ; BREX_PglZ - PorX_PglZ - DUF499_PglZ Comparison Alignment

```
WP_238754868.1_Leptospira_bandrabouensis [illegible]
MBF0258418.1_Desulfamplus_sp. [illegible]
MBX3247846.1_Myxococcales_bacterium [illegible]
MBD2549130.1_Microcystis_elabens_FACHB-917 [illegible]
MBY0113210.1_Phycisphaerales_bacterium [illegible]
MBX6362428.1_Gemmatimonadota_bacterium [illegible]
WP_083610988.1_Desulfatibacillum_alkenivorans [illegible]
MBN1165673.1_Methanospirillaceae_archaeon [illegible]
WP_156746889.1_Mycobacterium_sp._E2733 [illegible]
MBV6422080.1_Ignavibacteriaceae_bacterium [illegible]
WP_083423498.1_Stigmatella_aurantiaca [illegible]
NLN06912.1_Bacillota_bacterium [illegible]
WP_242609850.1_Comamonas_thiooxydans [illegible]
WP_243520949.1_Hymenobacter_monticola [illegible]
OQB98762.1_Spirochaetes_bacterium_ADurb.Bin110 [illegible]
WP_166386808.1_Polaribacter_sp._11A2H [illegible]
WP_235164441.1_Dyadobacter_chenhuakuii [illegible]
MCP5535438.1_Akkermansiaceae_bacterium [illegible]
MCC6145260.1_Candidatus_Hydrogenedentes_bacterium [illegible]
MBN9589012.1_Alphaproteobacteria_bacterium [illegible]
WP_100919682.1_Candidatus_Thiodictyon_syntrophicum [illegible]
WP_011448105.1_Methanospirillum_hungatei [illegible]
NBT25026.1_Actinomycetia_bacterium [illegible]
WP_246481782.1_Natronogracilivirga_saccharolytica [illegible]
MBQ6007748.1_Kiritimatiellia_bacterium [illegible]
WP_091825284.1_Butyrivibrio_sp._ob235 [illegible]
WP_219938192.1_Succinivibrio_faecicola [illegible]
MCK6605714.1_Ignavibacteriaceae_bacterium [illegible]
MCH3907672.1_Sphaerochaeta_sp. [illegible]
WP_251178637.1_Adlercreutzia_agrestimuris [illegible]
MBQ6502306.1_Flexilinea_sp. [illegible]
WP_237187173.1_Rothia_nasimurium [illegible]
OFQ22229.1_Actinomyces_sp._HMSC062G12 [illegible]
QNI65417.1_Synechococcus_sp._A15-44 [illegible]
MCF7790535.1_Victivallales_bacterium [illegible]
MBW8002950.1_Planctomycetota_bacterium [illegible]
WP_262567766.1_Endozoicomonas_gorgoniicola [illegible]
MBQ2845006.1_Alphaproteobacteria_bacterium [illegible]
WP_216328025.1_Deinococcus_aestuarii [illegible]
WP_151792240.1_Acinetobacter_seifertii [illegible]
MBP7846546.1_Burkholderiales_bacterium [illegible]
7PVK_1|Chains [illegible]
7PVA_1|Chains [illegible]
8TED_1|Chains [illegible]
8TFF_1|Chains [illegible]
8THP_1|Chains [illegible]
HCF99179.1_Chloroflexota_bacterium [illegible]
CAD6492703.1_Candidatus_Argoarchaeum_ethanivorans [illegible]
HDN59917.1_Candidatus_Marinimicrobia_bacterium [illegible]
HDQ70819.1_Chloroflexota_bacterium [illegible]
HEU5078765.1_Opitutaceae_bacterium [illegible]
HIE29205.1_Candidatus_Poribacteria_bacterium [illegible]
HLE25735.1_Thermodesulfobacteriota_bacterium [illegible]
HNQ22500.1_Phycisphaerae_bacterium [illegible]
HZX48547.1_Nitrospirota_bacterium [illegible]
KXA95579.1_candidate_division_MSBL1_archaeon [illegible]
MBA3936147.1_Planctomycetota_bacterium [illegible]
MBC7332925.1_Actinomycetia_bacterium [illegible]
MBC7429624.1_Bacteriovorax_sp. [illegible]
MBE7508222.1_Planctomycetia_bacterium [illegible]
MBI4523255.1_Deltaproteobacteria_bacterium [illegible]
MBK6914963.1_Ignavibacteriales_bacterium [illegible]
MBK8636967.1_Chromatiaceae_bacterium [illegible]
MBM4466104.1_Chloroflexota_bacterium [illegible]
MBN2145087.1_Candidatus_Auribacterota_bacterium [illegible]
MBP6702581.1_Vicinamibacteria_bacterium [illegible]
MBQ8111586.1_Kiritimatiellia_bacterium [illegible]
MCD6507459.1_Candidatus_Poribacteria_bacterium [illegible]
MCK8825904.1_Fuchsiella_alkaliacetigena [illegible]
MCK9588325.1_Terrimicrobiaceae_bacterium [illegible]
MCL0058393.1_Dehalococcoidia_bacterium [illegible]
MCL0097130.1_Dehalococcoidia_bacterium [illegible]
MDB6140354.1_Verrucomicrobiaceae_bacterium [illegible]
MDD3269498.1_Syntrophomonadaceae_bacterium [illegible]
MDD4099138.1_Lentisphaeria_bacterium [illegible]
MDT3473331.1_Stenotrophomonas_maltophilia [illegible]
MDU2146555.1_Paeniclostridium_sordellii [illegible]
MEA1926980.1_Candidatus_Auribacterota_bacterium [illegible]
MEA3344887.1_Chloroflexota_bacterium [illegible]
MEA3401059.1_Armatimonadota_bacterium [illegible]
NLX99245.1_Rhodopirellula_sp. [illegible]
NMX21981.1_ANME-1_cluster_archaeon_GoMg4 [illegible]
RLF12261.1_Thermoprotei_archaeon [illegible]
RLF82337.1_Thermococci_archaeon [illegible]
TEU13000.1_Anaerolineales_bacterium [illegible]
WP_083335243.1_Planococcus_glaciei [illegible]
WP_084067307.1_Desulfocicer_vacuolatum [illegible]
WP_121919201.1_Haloplanus_aerogenes [illegible]
WP_125672296.1_Candidatus_Methanodesulfokores [illegible]
WP_149537233.1_Siccirubricoccus_phaeus [illegible]
WP_218932320.1_Adhaeretor_mobilis [illegible]
WP_257627367.1_Haloplanus_salinarum [illegible]
WP_259101347.1_Candidatus_Fervidibacter_sacchari [illegible]
WP_260594794.1_Salinirubellus_salinus [illegible]
WP_283022679.1_Bradyrhizobium_sp. [illegible]
consensus/85% [illegible]
consensus/80% [illegible]
consensus/75% [illegible]
consensus/70% [illegible]
```

```
# ; Type_1 BREX BrxL Representative_1; (SIGMA-HTH + OB-Fold + MCM-family-AAA+ATPase + Inactive-LonP)

                                                   1---------11--------21--------31--------41--------51--------61--------71--------[illegible]
OrigSeq                                            MIDETDRKLNEHFSGRVVRKDLTKLIKIGHNVPVYVLEYLLGINCATNDEALIQEGIQKVKKILSDNYVSPHEAEKIKSK[illegible]

                                                   -HHHHHHHHHH--[illegible]
                                                   -HHHHHHHHHH--[illegible]
                                                   -HHHHHHHHHH--[illegible]
WP_270096522.1_Methanococcoides_alaskense          MIDETDRKLNEHFS[illegible]
WP_199585971.1_Collinsella_sp._AF11-11             MMDVLDGKLLEHFG[illegible]
MBI2765421.1_Chloroflexota_bacterium               MPSSATQNVREVFA[illegible]
MBN1450020.1_Anaerolineales_bacterium              MMDDLDFKLTTHFP[illegible]
KYF76802.1_Sorangium_cellulosum                    -MDALREKIEGVFG[illegible]
MCD6506894.1_Candidatus_Poribacteria_bacterium     -MNRLDEKVLNLFP[illegible]
MXY51051.1_Gammaproteobacteria_bacterium           MLNDLDTLATESYP[illegible]
MCI0407274.1_Acidobacteriota_bacterium             MVDELDRKLLEAFA[illegible]
WP_005980888.1_Fusobacterium_ulcerans              -MDIINEIGNEAFE[illegible]
WP_244056943.1_Aeromonas_caviae                    MAQDLDARLNRYFK[illegible]
SFX61200.1_Ruminococcus_flavefaciens               MMDTLDRKIIDAFP[illegible]
HBO56052.1_Leuconostoc_mesenteroides               MVDDLLDKLEINFP[illegible]
MBK7688412.1_Elusimicrobiota_bacterium             MPDNLDRLASKALE[illegible]
MBI5489026.1_Deltaproteobacteria_bacterium         M--PADSKAHEVFG[illegible]
MBI4023516.1_Verrucomicrobiota_bacterium           M-----SKAREQFG[illegible]
MCD6317877.1_Candidatus_Aerophobetes_bacterium     MLDSLDQKILALFP[illegible]
MBU1143922.1_Bacillota_bacterium                   -MDSLDVKVYEQFS[illegible]
CCU85339.1_Mesotoga_infera                         MLDALDLKVLKIYP[illegible]
OQC41885.1_Acidobacteria_bacterium.                MRDPLDDKANLIFA[illegible]
MBR4647499.1_Bacteroidales_bacterium               MMNEIDKKLNEYYG[illegible]
WP_242979527.1_Subdoligranulum_sp.                 MYDSYEQRLKTVFQ[illegible]
AKB55566.1_Methanosarcina_barkeri_MS               MLEETDRKLIKYFP[illegible]
OGO91754.1_Clostridiales_bacterium_GWF2_38_85      -MEELNTLLSQHFA[illegible]
MBC8432678.1_Desulfobacterales_bacterium           MLDSLDRKLNDVFA[illegible]
MBX9637029.1_Nitrosomonas_sp.                      MLDELDEKLLEHFR[illegible]
WP_110300682.1_Halanaerobium_congolense            MTDELDLKLLDNFN[illegible]
WP_254547351.1_Halomarina_sp._BND7                 -RRDLDQKALEHFS[illegible]
WP_264798996.1_Brevibacterium_sp._JSBI002          MLDSIDETAASAFE[illegible]
consensus/100%                                     .............h.[illegible]
consensus/95%                                      ........h...a.[illegible]
consensus/90%                                      ..p..p.bh...a.[illegible]
consensus/85%                                      ..p.hs.bh.p.F.[illegible]
consensus/80%                                      ..p.hc.bh.p.F.[illegible]
consensus/75%                                      MhcplDp+h.p.Fs[illegible]
consensus/70%                                      Mh-plDpKl.p.Fs[illegible]
```

```
# ; Type_1 BREX BrxL Representative_2; (SIGMA-HTH + OB-Fold + MCM-family-AAA+ATPase + HKD-DNase)

                                               1---------11--------21--------31--------41--------51--------61--------71-------- [illegible]
OrigSeq                                        MDVLDIKLNEAFPGKVVRKDLLHEIKRAVNVPSFVLEFLLSRYCASEDPEEIEEGKKAVLQTIEKCYVRPDESNKAQAIV [illegible]

                                                <<-----------------------SIGMA-HTH---------------------------->>---------- [illegible]
                                                --HHHHHHHHH--------HHHHHHHHH----HHHHHHHHHHH-----HHHHHHHHHHHHHHHHHH-||HHHHHHHHHH [illegible]
                                                --HHHHHHHHH--------HHHHHHHHH----HHHHHHHHHHH-----HHHHHHHHHHHHHHHHHH-||HHHHHHHHHH [illegible]
                                                --HHHHHHHHH--------HHHHHHHHH----HHHHHHHHHHH-----HHHHHHHHHHHHHHHHHH-||HHHHHHHHHH [illegible]
HAM28421.1_Prevotella_sp.                       MDVLDIKLNEAFPGKVVRKDLLHEIKRAVNVPSFVLEFLLSRYCASEDPEEIEEGKKAVLQTIEKCYVRPDESNKAQAI [illegible]
WP_168040586.1_Neolewinella_antarctica          MTPLDQQINAAFPGKVVRKDLTSLVKGNAVVPTFVLEYLLGQYCATDEEETIRQGVETVKKILARHYVHREESELLKSR [illegible]
OFQ22228.1_Actinomyces_sp.                      LAGYEETAARAYDGFVVRKDLVGQVKGNAIVPSYVLEFLLAQYCATTSESAIVEGIESVRRILAQHYVNRGDSELIKSR [illegible]
WP_108716022.1_unclassified_Actinobaculum       QSELDRKINRLFPGVVVRKDLVKAVKGNAIVPTYVLEYLLGQYAASDDEATIQAGIDSVRRILADHYVHRNESELIKST [illegible]
NKY16940.1_Tsukamurella_spumae                  TSELDAKANVQFPGFVVRKDLVGRIKGNAVVPGYVLEFLLAQYCATDDPATIEAGIESVRDILAKHYVHRGEAELIKST [illegible]
TET34870.1_Planctomycetota_bacterium            MMSFERKAKDYYGEVVINKGLMGRAGGARAIPVYVGEWIISQFM--DGDELTDDGRTEIVNTISKYLPQKADKNMILNR [illegible]
WP_217054563.1_Bifidobacterium_longum           MSNLDRKITDDFSQYTVRKDLVSEVKGNALVPSYVLEYLLSKYATTTDQESINAGVKRVRDILADNYVHREEANLIQSK [illegible]
WP_084146426.1_Hyphomonas_oceanitis             --SLDEKINRHFAGRVVRKDLVKAVKGNAIVPSYVLEYLLGQYCATDDEASIQTGIATVKEILRKHYVHRNEAKLVQSN [illegible]
WP_014697721.1_Bifidobacterium_animalis         LSPLDRKITDVFPQHTVRKDLVGEIKGNAVVPTYVLEFLLSQFATTTDALSIESGVRRVQEILAQHYVHRDEAMLIQSK [illegible]
MBQ6502307.1_Flexilinea_sp.                     MNATDEKLNEVFAGLVVRKDLAKIVKGNVSVPNYVLEYLLGQYCATSEEEDIQAGVEQVKDILAQHFVQRSEAELVKST [illegible]
MBQ6007747.1_Kiritimatiellia_bacterium          MDHLDEVAVKALDGYLVRKDLVRKFSKQFPVPTYVVEFLLGRYCASTDEAEIAEGLELVQKSLQNKAVSNGNEELFKAH [illegible]
WP_259087830.1_Salinibacter_ruber               TTGLQRKIVDHFEGKVVRKDLTQLVKGNAVVPSYVLEYLLGQYCPTDDEDLIRRGVEKVRDIVNRHYVHRDESELVKSE [illegible]
MCH2449133.1_Gracilimonas_sp.                   ISEFENKVKEHFEGKVVRKDLTKFVKGNAVVPTYVLEYLLGQHCASSDEEIVQEGIERVKSIISNHYVHRDESELVKST [illegible]
WP_255087757.1_unclassified_Synechococcus       PSALDQKILEHFPGLVVRKDLTTELKQNAVVPTYVLEYLLGQHCATDDPAFIAEGLESVRRILSKHYVHRNQAELIKST [illegible]
NLK72538.1_Clostridiales_bacterium              MESFEIKARDYFGEIVINKSLIHKAGGSRAIPTYVAEWIIAHFVG-DDMELNEESRGKIARFVQKYIPSKGEKEAIKND [illegible]
MBE7502036.1_Verrucomicrobiales_bacterium       LDVLDEKLNHVFAGKVVRKDLLLQVKKGTNVPSFVLEFLLARFCASDDPREIQEGMAAVLETIQKNYVRPDESNRAQMV [illegible]
MBI1207572.1_Azospirillum_sp.                   PDALDLKLNQIFAGKVVRKDLLHRIKAGTNVPTFVLEFLLARYCASDDPTEIQEGMEAVLATLQENYVRQNESNAAQSR [illegible]
MCP5535439.1_Akkermansiaceae_bacterium          MDPIDLKLNETFEGRCVRKDLVQRIKKGTNIPTFVLEFLLAKYCATDDPDEINAGLQAVTETIQLNYVRPNEANKAQSM [illegible]
QOW09781.1_Kaistella_flava                      MDELDLKLNEFYPGKVVRKDLVHDIKKAANVPAFVLEFLLAKYCATDDPEEIIAGKMAVLEVVEKNYVRPDEANKAQSM [illegible]
AZB21333.1_Kaistella_haifensis                  MKDLDIKLNQYFGGKVVRKDLTKLVKGNAIVPMYVLEYLLGQYCATDDDQTIVEGVETVKSVISKHFVHRDEAQIVKST [illegible]
MBU6197219.1_Cyanobacteria_bacterium            VEDLNKKINEFFPGLAVRKDLVGLVKGNAIVPSYVLEYLLGQYCATSDEDSINAGIERVKKILAEHYVNRNEANLIKSK [illegible]
WP_028665039.1_Runella_zeae                     --DLQQKIIRYFEGKVVRKDLTSLVKGNAVVPTYVLEYLLGQYCAIDDEKIVQEGIEKVKNVIKNNYVHRSDAEVVKSN [illegible]
WP_252760244.1_Prevotella_cerevisiae            --TLQEKIMDAFLGKVVRKDLAFQVKSGLPVPTYVLEYLLGQYCASDDEEVIKDGLEKVKTVIQNNYVHRADAERVKGI [illegible]
MCH8487216.1_Cyclonatronum_sp.                  ---MKDKLYQYFEGRVVRKDLTSHVRGSAVVPTFVLEYLLGQNCAIDDEEIIQMGIEKVKEIISRNYVHPNEAELVKSK [illegible]
WP_237187172.1_Rothia_nasimurium                YTPLDEKIIAEFTGNVVRKDLSAHIKSNAPVPTYVLEYLLGQYATSHDETQIAEGIERVKLIMQNNFVQRNQAELIKST [illegible]
MCI1665138.1_Atopobiaceae_bacterium             MDDLSRRLIDAFPGYTVRKDLVKQVRGNAAVPSYVLEYLLGQYCSTDDEQEVAEGIEAVKNILAKHYVQRSEATLVRSH [illegible]
MCC6106581.1_Atopobium_sp.                      MDDLDRKLNDNFAGYVVRKDLVKLVRGNAAVPSYVLEYLLGQNCATDDEEEIQEGVERVRDILAAHYVQRAEAGAIRSD [illegible]
WP_084634624.1_Propionicicella_superfundia      KSALDDKINEQFAGAVVRKDLVKAVRGNAVVPSYVLEYLLGQYAASDDEATIQAGIDSVRQILATHYVNRNEHMLVKSE [illegible]
MBQ7190498.1_Kiritimatiellia_bacterium          DKKLNVKVLERFAKSVVRKDLAFQVKGNLPVPTYVIEYLLAQFCQSSDEQEIAKGMEEVKEIVRERYFNRADNEIIKGR [illegible]
MCB9093207.1_Halobacteriovoraceae_bacterium     MKQFEEKAKDAFGEIIINKSFIHNAGGSRAIPTYVSEWIISHFVK-EGEEISEHARERIAAFANKFIPQKSEKESIKNR [illegible]
consensus/100%                                  ....p..h...h....lpKsh...h.....lP.aV.Eallup.h..p.......u...l...h...h.p..p...hb.. [illegible]
consensus/95%                                   ...hp.ph...a....lpKsL...h..s..lPsaV.Eallup.h..p.......u...l...h...hsp..p...hbs. [illegible]
consensus/90%                                   ...hp.+h.p.a...sVRKDL...hp.s..lPsaVlEaLLupahs.s-...l..G...V.p.l.pphsp.s-.p.hbs. [illegible]
consensus/85%                                   ...hp.Kh.p.F...sVRKDL...l+.s..VPsaVlEaLLupahs.s-...l..G.p.Vbp.l.ppaVp.s-.p.hbu. [illegible]
consensus/80%                                   .p.hcbKl.c.FsGblVRKDLs..l+ushsVPoaVLEaLLupahsosD...I..Ghc.Vbphl.ppaVpps-uphlbup [illegible]
consensus/75%                                   .p.l-bKl.c.FsGbVVRKDLs.bl+ushsVPoYVLEaLLupahuosD.p.I..GhcpVbphl.ppYVp+s-uphlbup [illegible]
consensus/70%                                   .psL-bKl.c.FsGbVVRKDLspblKGshsVPoYVLEaLLuQaCAo-Dpp.IppGl-pVbpll.ppYVp+sEuphl+Sp [illegible]
```

```
# ; Type_4 BREX BrxL Representative Alignment; (SIGMA-HTH + OB-Fold + MCM-family-AAA+ATPase)

                                                      1---------11--------21--------31--------41--------51--------61--------71--------81--------91--------101-------111-------121-------131-------141-------151-------161------ [illegible]
OrigSeq                                               METNHTQADIEEQLKRYFGDMLVYKSPSNSKFFSALSLPSFMRDWLVMRFSDSEGHIDKEEVTSYVKRTIPKKEQWNDFQVELHNNQPVRFLTKVKIDFDTANRQALFSLPEFGVPEAVVEWHVIEKNRDYLLAPTEVWGIVELLCDLDDGKKTIIKLVDFTPFCPYT [illegible]
                                                                                                                              S1          S2          S3                      S4            S5   [illegible]
                                                      <<-----------------------------------SIGMA-HTH-------------------------->><<--------------------------------------------OB-Fold------------------------------------------->>< [illegible]
                                                      ------HHHHHHHHHHHH---------HHHHHHHHHH---HHHHHHHHHHH---------HHHHHHHHHHH||HHHHHHHH [illegible]
                                                      ------HHHHHHHHHHHH---------HHHHHHHHHH---HHHHHHHHHHH---------HHHHHHHHHHH||HHHHHHHH [illegible]
                                                      ------HHHHHHHHHHHH---------HHHHHHHHHH---HHHHHHHHHHH---------HHHHHHHHHHH||HHHHHHHH [illegible]

BDF48892.1_Lachnospiraceae_bacterium                  METNHTQADIEEQLKRYFGDMLVYKSPSNSKFFSALSLPSFMRDWLVMRFSDSEGHIDKEEVTSYVKRTIPKKEQWNDF [illegible]
AFZ48537.1_Cyanobacterium_stanieri_PCC_7202           MFSNNYSTYDNELVTRVFGSLSIDKKRLPSSSLTSVGVPSFVGEWILEHIVAGTGELNDGEVRNFVDKAFPRKDDKEEI [illegible]
HAR58248.1_Acholeplasmatales_bacterium                M------TKLDSKVLEIFPDESIYKSPNIYSCFTGFNLPSFIKDWLLKRYSNEHGKPDILKIKSFLIKHIPSKDYD--I [illegible]
HCF89836.1_Bacillota_bacterium                        M-----RCVKLDKLREAFGAMVVLKNPERTKFFSDLSLPSYMRDWLVMKFSDEGGNIDYKGVSRYIKQYIPSREDYEQF [illegible]
HED00231.1_Pseudomonadota_bacterium                   M---QTVGHVKDKVKTQFPDMVVYKDPSKNKFFSSLGIPSYLRDWLLKRFSDEEGRINIDEVQSYVRKVIPRKEEWEAI [illegible]
KUK07752.1_Archaeoglobus_fulgidus                     M-----QDPLGEKIKLVFPNESVYKDPQKSKFFSALNIPSYLRDWLIRKFSFNDGNIDLHSVKKFIEEKIPRKEHTQLV [illegible]
MAU00661.1_Anaerolineaceae_bacterium                  MINPPLASDLQNRAKNVFGEVCIDKGLFLDSGMISRSIPTFVAEWILDRFC-PNGELTDDIINLFIRKHLPRKDQKEEI [illegible]
MBE6363380.1_Lentisphaerota_bacterium                 M---------ENKLKEAFPNTTVFKDQNLIATFKAASIPAFLRDWILKKKAGVDGRIDKDELRQYMNAIIPRREDVLEI [illegible]
MBH8560404.1_Hymenobacter_negativus                   M-----NDALRQKVTDVFESDSIDKSLYSGSGLNEQSIPVFVGEWLIDRQL-KGGEWDEDVVIAFVRKFMPSKDDVEDL [illegible]
MBI2377776.1_Deltaproteobacteria_bacterium            MGAEVVNSSFPDKIRSVFGNAVIDKRRLRSSQVEKRGVPAYVAEWILDTIVPGVGPLTTEEVREWCSRYIPAPGDQNVL [illegible]
MBI9047159.1_Anaerolineaceae_bacterium                MNSESMKSEVDRLIEASFPDVSVHKGLANGIGLSGRVIPAFVADWLVSRYSSNDGSVDGDAIRQFIARYLPDKKQRESL [illegible]
MBN2258341.1_Anaerolineaceae_bacterium                MSQTSFLSNLDVKVQKVFGDVSIDKRRLPTTKLDKRGIPAYVAEWAIDMIAPGKGLLTNEQIRDWANRYIPGPDESKVI [illegible]
MBR5399200.1_Bacteroidales_bacterium                  M------SELGRKIQQVFGSAAVYKNPANYDVFTGRNLPSFIKDYLISTHVAVDGTFYREKLIRFLDSHIPADNNS--V [illegible]
MBU6335165.1_Chloroflexota_bacterium                  MMENQRMTSYEARASDVFGDYCINKALVEHAGFGARSIPAFVGEWIISRHA-PDGILDDAAIRTFLDRHLPTRDQKEQL [illegible]
MCD4818677.1_Candidatus_Cloacimonadota_bacterium       M------TTLDAKIKEHFPEESIYKSPDRYNVFSGKILPSFIKDWLIKRYTNSNGNLDSNALLNFMEEHIPLQKNN--L [illegible]
MCH4147879.1_Prevotella_sp.                           M-----KQEICDKIRQYFSAMSIYKDPASTSLFAGRNLPAFVKDFLLKRFLIQSGEVDKIALTDFLNKIIPSAQGS--V [illegible]
MCI7180755.1_Lachnospiraceae_bacterium                MNSCEIDADFINRLKDAFDTMVVFKDLQQNNFISSFKLPSFMRDYVVKNFQDDDGEVDVEGASEFIRTYIPKKEDWKNL [illegible]
NBI83184.1_Clostridiaceae_bacterium                   M---------VDKLREYFPTTTVYKDPSVMSMFKAAKIESFLRDWILKRKTSTDGIVDIGSLSDYVAAIIPSRSDKERL [illegible]
NLC55396.1_Armatimonadota_bacterium                   MTPTSSNGVLAQKIREVLAEVVIDKCRLPASKLRERGIPGYVAEWVLDYVVPGEGPLTPEEVQEWAVRYIPKADEQNLV [illegible]
NLG95480.1_Acetomicrobium_flavidum                    M---STVNDVQSKVRSVFSDAVIYKNPEYHNFFSSKSIPSFLRDWIIMRFADEEGNIDIDKVANFIKSSLPGAKDWESL [illegible]
NLI70094.1_Bacillota_bacterium                        M---------IEKLRNCFDE [illegible]
RKY16067.1_Planctomycetota_bacterium                  M------KSLGEKLRAVFP [illegible]
WP_007916555.1_Ktedonobacter_racemifer                M-----MLHFEEKALQYYG [illegible]
WP_119090030.1_Candidatus_Cryosericum_hinesii         M------DTLDEKLVKIFP [illegible]
WP_219156485.1_Hymenobacter_profundi                  M------ETLEDKIDRVFT [illegible]
WP_219722098.1_Deinococcus_planocerae                 MLDAEALKHLNAKVTEVFR [illegible]
WP_251958558.1_Nostoc_commune                         MNSWEATTYNNDLVREVFG [illegible]
WP_259125643.1_Salinibacter_ruber                     MESVDEKKSLDKLIKNNFE [illegible]
consensus/100%                                        M...........bh...h.. [illegible]
consensus/95%                                         M...........bh...a.. [illegible]
consensus/90%                                         M...........bh...F.p [illegible]
consensus/85%                                         M..........pblpphFsp [illegible]
consensus/80%                                         M..........p+lpphFsp [illegible]
consensus/75%                                         M........h.p+lpphFss [illegible]
consensus/70%                                         M......p.h.pKlpphFss [illegible]
```

### ; Type1 BrxA DNA-Binding Component Representative_1; (HTH1 + HTH2)

```
OrigSeq  [illegible]

MBZ4201979.1_Methylovulum_sp.  [illegible]
MCP9304659.1_Lacticaseibacillus_paracasei  [illegible]
MSB67178.1_Leuconostoc_lactis  [illegible]
RGO23299.1_Dorea_sp._OM02-2LB  [illegible]
WP_042694048.1_Methanobrevibacter_oralis  [illegible]
WP_085559183.1_Carnobacterium_iners  [illegible]
WP_107841262.1_Metasolibacillus_meyeri  [illegible]
WP_125758086.1_Lacticaseibacillus_hegangensis  [illegible]
WP_145463944.1_Weissella  [illegible]
WP_170068775.1_Alkalibacterium_olivapovliticus  [illegible]
WP_183128193.1_Clostridium_autoethanogenum  [illegible]
WP_220727332.1_Marinilactibacillus_psychrotolerans  [illegible]
WP_255464098.1_unclassified_Adlercreutzia  [illegible]
WP_257351444.1_Pseudalkalibacillus_decolorationis  [illegible]
WP_262953492.1_Turicibacter_sp._1E2  [illegible]
MBX9598964.1_Burkholderiales_bacterium  [illegible]
MRI34255.1_Endozoicomonas_sp._OPT23  [illegible]
WP_104852219.1_Acinetobacter_sp._SWBY1  [illegible]
WP_116688138.1_Pelagibaculum_spongiae  [illegible]
WP_184518068.1_Tardiphaga_robiniae  [illegible]
WP_224385468.1_Klebsiella  [illegible]
WP_231941296.1_Burkholderia_sp._PAMC_28687  [illegible]
WP_262567772.1_Endozoicomonas_gorgoniicola  [illegible]
WP_274377168.1_Ectothiorhodospira_magna  [illegible]
MBF0232704.1_Desulfamplus_sp.  [illegible]
MBK8989555.1_Chloroflexota_bacterium  [illegible]
MBK9103292.1_Saprospiraceae_bacterium  [illegible]
MBP6234854.1_Saprospiraceae_bacterium  [illegible]
MBP9037130.1_Candidatus_Cloacimonas_sp.  [illegible]
MBQ7190492.1_Kiritimatiellia_bacterium  [illegible]
MBQ8347030.1_Alphaproteobacteria_bacterium  [illegible]
MBQ9367248.1_Victivallales_bacterium  [illegible]
MBS5529917.1_bacterium  [illegible]
MBT3348503.1_Thiotrichales_bacterium  [illegible]
MBT62432.1_Puniceicoccaceae_bacterium  [illegible]
MCB0987054.1_Acidimicrobiales_bacterium  [illegible]
MCC6106576.1_Atopobium_sp.  [illegible]
MCE2892636.1_Hyphomonadaceae_bacterium  [illegible]
MCH8487207.1_Cyclonatronum_sp.  [illegible]
NEX15946.1_Halochromatium_sp.  [illegible]
NLK13332.1_Spirochaetales_bacterium  [illegible]
NUM59089.1_Pseudobdellovibrionaceae_bacterium  [illegible]
OJV92259.1_Bacteroidia_bacterium_44-10  [illegible]
WP_072727065.1_Bifidobacterium_lemurum  [illegible]
WP_073424903.1_Fibrobacter_sp._UWB7  [illegible]
WP_079511495.1_Maribacter_arcticus  [illegible]
WP_082099990.1_Bacteroides_ovatus  [illegible]
WP_129366657.1_Lutibacter_sp._HS1-25  [illegible]
WP_197728946.1_Rhizobium  [illegible]
WP_217901576.1_Pseudanabaena_sp._SR411  [illegible]
WP_229027683.1_Bacteroidales  [illegible]
WP_236888846.1_Desulfoluna_limicola  [illegible]
AXF55599.1_Salicibibacter_kimchii  [illegible]
EHI56638.1_Johnsonella_ignava_ATCC_51276  [illegible]
MBQ9252306.1_Clostridia_bacterium  [illegible]
MBR1685961.1_Clostridia_bacterium  [illegible]
MCI2147017.1_Clostridiales_bacterium  [illegible]
MCI6194162.1_Faecalibacterium_sp.  [illegible]
consensus/100%  [illegible]
consensus/95%  [illegible]
consensus/90%  [illegible]
consensus/85%  [illegible]
consensus/80%  [illegible]
consensus/75%  [illegible]
consensus/70%  [illegible]
```

### ; Type1 BrxA DNA-Binding Component Representative_2; (HTH1 + HTH2 + wHTH1)

```
OrigSeq  [illegible]

BAY35696.1_Nostoc_sp._NIES-2111  [illegible]
BBE18571.1_Aquipluma_nitroreducens  [illegible]
CAJ71911.1_Candidatus_Kuenenia_stuttgartiensis  [illegible]
ETI91652.1_Clostridium_butyricum_DORA_1  [illegible]
HCZ06895.1_Thermotogota_bacterium  [illegible]
KUK86069.1_candidate_division_TA06_bacterium_34_109  [illegible]
KUK92980.1_Thermotogales_bacterium_46_20  [illegible]
MBK8482435.1_Pseudomonadota_bacterium  [illegible]
MBK9368668.1_Deltaproteobacteria_bacterium  [illegible]
MBN1450030.1_Anaerolineales_bacterium  [illegible]
MBP7775369.1_Candidatus_Saccharimonas_sp.  [illegible]
MBS1808659.1_Acidobacteriota_bacterium  [illegible]
MBX6362417.1_Gemmatimonadota_bacterium  [illegible]
MBX9573700.1_Candidatus_Obscuribacterales_bacterium  [illegible]
MCA9087265.1_Planctomycetaceae_bacterium  [illegible]
MCB1001741.1_Acidimicrobiales_bacterium  [illegible]
MCD6506893.1_Candidatus_Poribacteria_bacterium  [illegible]
MCE8425370.1_Candidatus_Methanoperedens_sp.  [illegible]
MCI0393101.1_Acidobacteriota_bacterium  [illegible]
MCI0407269.1_Acidobacteriota_bacterium  [illegible]
MCI6342426.1_Prevotella_sp.  [illegible]
MCP5535432.1_Akkermansiaceae_bacterium  [illegible]
MCQ4334812.1_Natronomonas_aquatica  [illegible]
NIM13885.1_Candidatus_Aminicenantes_bacterium  [illegible]
NOQ74048.1_Crocinitomix_sp.  [illegible]
OFV66971.1_Candidatus_Syntrophoarchaeum_butanivorans  [illegible]
OQB98769.1_Spirochaetes_bacterium_ADurb.Bin110  [illegible]
OYX86685.1_Flavobacteriales_bacterium_32-34-25  [illegible]
PLW98640.1_Marinilabiliales_bacterium  [illegible]
RRJ96053.1_Opitutaceae_bacterium_TAV4  [illegible]
WP_015282555.1_Thioflavicoccus_mobilis  [illegible]
WP_072866938.1_Desulfofundulus_thermosubterraneus  [illegible]
WP_145167967.1_Rubripirellula_lacrimiformis  [illegible]
WP_151706157.1_Weizmannia_acidilactici  [illegible]
WP_173247267.1_Hymenobacter_sp._BRD67  [illegible]
WP_200392183.1_Roseibacillus_ishigakijimensis  [illegible]
WP_200858941.1_Geobacter_sulfurreducens  [illegible]
WP_201591402.1_Psychrobacter_pulmonis  [illegible]
WP_207689228.1_Desulfonema_limicola  [illegible]
WP_218608644.1_Methanospirillum_hungatei  [illegible]
consensus/100%  [illegible]
consensus/95%  [illegible]
consensus/90%  [illegible]
consensus/85%  [illegible]
consensus/80%  [illegible]
consensus/75%  [illegible]
consensus/70%  [illegible]
```

### ; Type1 BrxA DNA-Binding Component Representative_3; (HTH1 + HTH2 + wHTH1)

```
OrigSeq  [illegible]

MBL9184317.1_Verrucomicrobiaceae_bacterium  [illegible]
MBQ9739438.1_Kiritimatiellia_bacterium  [illegible]
MBK8265324.1_Nannocystis_sp.  [illegible]
MBU4371659.1_Pseudomonadota_bacterium  [illegible]
MCC6145268.1_Candidatus_Hydrogenedentes_bacterium  [illegible]
MCU0772113.1_Verrucomicrobiota_bacterium  [illegible]
TGJ96614.1_Actinotalea_fermentans  [illegible]
WP_027288598.1_Rhodovibrio_salinarum  [illegible]
WP_081756111.1_Candidatus_Contendobacter_odensis  [illegible]
WP_154296282.1_Spiribacter_sp._2438  [illegible]
WP_175475300.1_Butyrivibrio_sp._ob235  [illegible]
WP_210512156.1_Natronogracilivirga_saccharolytica  [illegible]
WP_211530101.1_Methanocalculus_chunghsingensis  [illegible]
WP_215427514.1_Akkermansia_muciniphila  [illegible]
WP_218164137.1_Chitinophaga_sancti  [illegible]
WP_222013553.1_Rhizobium_laguerreae  [illegible]
WP_249395099.1_Deinococcus_sp._QL22  [illegible]
WP_257742632.1_Methanoplanus_endosymbiosus  [illegible]
consensus/100%  [illegible]
consensus/95%  [illegible]
consensus/90%  [illegible]
consensus/85%  [illegible]
consensus/80%  [illegible]
consensus/75%  [illegible]
consensus/70%  [illegible]
```

```
# ; Type2 BREX PglW Tripartite-DNA-binding component; (HTH + wHTH1 + wHTH2)

                                           <<------------HTH------------>><<------------wHTH1------------>><<------------wHTH2------------>>
WP_185095299.1_Streptomyces_sp._CEV_2-1            [illegible]
WP_268746228.1_Dermacoccus_nishinomiyaensis        [illegible]
GII83978.1_Sphaerisporangium_siamense              [illegible]
WP_030506725.1_Microbispora_rosea                  [illegible]
RJQ78831.1_Pseudonocardiaceae_bacterium            [illegible]
MCB0954589.1_Ilumatobacter_sp.                     [illegible]
WP_208577501.1_Micromonospora_tulbaghiae           [illegible]
MCP9485019.1_Candidatus_Gaiellasilicea_maunaloa    [illegible]
WP_193120346.1_Nocardiopsis_coralli                [illegible]
RKN35100.1_Streptomyces_hoynatensis                [illegible]
WP_212908910.1_Streptomyces_sp._TS71-3             [illegible]
GES07632.1_Acrocarpospora_macrocephala             [illegible]
OAA24551.1_Frankia_sp._EI5c                        [illegible]
WP_213171153.1_Natronoglycomyces_albus             [illegible]
WP_144820010.1_Micrococcus_luteus                  [illegible]
MBL8842562.1_Planctomycetota_bacterium             [illegible]
MBX7222607.1_Blastocatellia_bacterium              [illegible]
MBO9539711.1_bacterium                             [illegible]
MBN9688999.1_Verrucomicrobiota_bacterium           [illegible]
KAA0252897.1_Acidobacteriota_bacterium             [illegible]
MBK9387522.1_Planctomycetota_bacterium             [illegible]
WP_144981411.1_Gimesia_aquarii                     [illegible]
WP_090664774.1_Belnapia_rosea                      [illegible]
WP_136411732.1_Luteimonas_yindakuii                [illegible]
KRE61575.1_Tetrasphaera_sp._Soil756                [illegible]
WP_052424041.1_Nonomuraea_candida                  [illegible]
WP_093484991.1_unclassified_Streptomyces           [illegible]
WP_109776035.1_Quadrisphaera_granulorum            [illegible]
WP_090069475.1_Lentzea_flaviverrucosa              [illegible]
WP_257478501.1_Acidipropionibacterium_jensenii     [illegible]
MBL9044566.1_Myxococcales_bacterium                [illegible]
MCP5560207.1_Verrucomicrobiaceae_bacterium         [illegible]
MBK6942988.1_Planctomycetota_bacterium             [illegible]
WP_217637676.1_Actinopolyspora_saharensis          [illegible]
TAF23825.1_Verrucomicrobiota_bacterium             [illegible]
MBZ5659870.1_Acidobacteriia_bacterium              [illegible]
MBO0801654.1_Nocardiopsaceae_bacterium             [illegible]
WP_200346137.1_Halochromatium_glycolicum           [illegible]
MBK9978482.1_Gemmatimonadota_bacterium             [illegible]
WP_130480151.1_Leptothrix_mobilis                  [illegible]
MBI4602629.1_Planctomycetota_bacterium             [illegible]
MCO6436553.1_Phycisphaerae_bacterium               [illegible]
RLS37136.1_Planctomycetota_bacterium               [illegible]
WP_085387166.1_Cellulosimicrobium_sp._TH-20        [illegible]
consensus/100%                                     [illegible]
consensus/95%                                      [illegible]
consensus/90%                                      [illegible]
consensus/85%                                      [illegible]
consensus/80%                                      [illegible]
consensus/75%                                      [illegible]
consensus/70%                                      [illegible]
```

```
# ; Type3 BrxA DNA-Binding Component; (HTH + wHTH1 + wHTH2)

                                            1---------11--------21--------31--------41--------51--------61--------71--------81--------91--------101-------111-------121-------131-------141-------151-------161-------171-------181-------191-------201-------211-------221-------231-----
OrigSeq                                     MRASQIGFSQRIRLEWLEQTANLVLAGNDKTEINDSLQELLKNKVSGEANREKVISILMKTWLTVPRGLEALRDEGLQIHQGLPRKDRIAVHWGMALAAYPFWGAVAAHTGRLLRLQGTAAAAHVQRRVKEQYGERETASRAARRVLRSFIDWNVLNETDKGVYAQGIRYSIQDPRLISWMVEASLHARANGSAATKDLLDGPSIFPFRLAITAEHLASSPRLDLLRHGLDEDLMMLR

                                            <<--------|-------------------------HTH-------------------------|---------------------------------|---------------------------wHTH1-----------------------------|--------------------------------wHTH2--------------------------->>
                                            -hhh------|--HHHHHHHHHHHHH---HHHHHHHHHHHHHHH---HHHHHHHHHHHHHHHH--|--hhhhhhhhhhhhh--hhhhhhhhhhhhhhh|HHHHHHHHHHHHHHHHHH----HHHHHHHHHHHH---HHHHHHHHHHHHHHHHHH-EEEE----EEEE---|----HHHHHHHHHHHHHHH----EEHHHHHH-----------HHHHHHHH---EEEEEE---EEEEEE-
                                            -hhh------|--HHHHHHHHHHHHH---HHHHHHHHHHHHHHH---HHHHHHHHHHHHHHHH--|--hhhhhhhhhhhhh--hhhhhhhhhhhhhhh|HHHHHHHHHHHHHHHHHH----HHHHHHHHHHHH---HHHHHHHHHHHHHHHHHH-EEEE----EEEE---|----HHHHHHHHHHHHHHH----EEHHHHHH-----------HHHHHHHH---EEEEEE---EEEEEE-
                                            -hhh------|--HHHHHHHHHHHHH---HHHHHHHHHHHHHHH---HHHHHHHHHHHHHHHH--|--hhhhhhhhhhhhh--hhhhhhhhhhhhhhh|HHHHHHHHHHHHHHHHHH----HHHHHHHHHHHH---HHHHHHHHHHHHHHHHHH-EEEE----EEEE---|----HHHHHHHHHHHHHHH----EEHHHHHH-----------HHHHHHHH---EEEEEE---EEEEEE-
MBW1792999.1_Deltaproteobacteria_bacterium  MRASQIGFSQRIRLEWLEQTANLVLAGNDKTEINDSLQELLKNKVSGEANREKVISILMKTWLTVPRGLEALRDEGLQIHQGLPRKDRIAVHWGMALAAYPFWGAVAAHTGRLLRLQGTAAAAHVQRRVKEQYGERETASRAARRVLRSFIDWNVLNETDKGVYAQGIRYSIQDPRLISWMVEASLHARANGSAATKDLLDGPSIFPFRLAITAEHLASSPRLDLLRHGLDEDLMMLR
WP_013627486.1_Rubinisphaera_brasiliensis   -MTAVVGFDRKLKLEWLDDLADQVAKGEDVSTLRTFLHERLEEDHPGETARGKSITVLMRIWVFVPDEHHEIREQAFELLKRINAKDRIWLHWGMCILAYPLFRDTATAIGRLLKLQDEFTLAQLQRRLIDSWGERSTVVRAFQRVVRSMVEWHTLEDATPGHFRAPTKVTTKSKSLQIWFLRACHAATGKEMVEVDQLLSLPSSFPFKITVAKSDLRNSKDFSIHRQGLDMDMVTTT
MBT3379348.1_Lentisphaerota_bacterium       SRHEVIGIKQVVRREWYDFALDRLISGDSTAAIRAGLDEFISERLQGDQTYTKAVTQIMRSWVTPGNDLVPFRDAALDHARSADRFERVVLHWAVTIAAYPFWHRVAVVVGRLLNLQEAVSQSQIRQRCYEAFGERTTIERSARRVIRSFVTWGLLDDSTKGMYRRSAPVVVTNHDLAALLIEGVLLAIPEGRAPQSQVLSNPGLYPFQLPMNGGMIAASERIDVGRFSLDDELLRLR
WP_271916069.1_Polyangium_sp._rjm3          TERRTIGFDRPVQLEWLDALAYRLAAGASAKEAREVIWNLLDGVVAGETARGKTLTVVSRIWLTVPPGVSPMRDAALLLIRHASVEERQAIHWALMSAAYTFFVDLASSIGKLIALNGEISLSQLTRRLVEVWGDRSTLRPAAQRVVRSMIRWGVLREGRPGQYLPPTKRIVVSAEVSELLLEGLLLAAGRGGMPLSQLLMHPAAFPFELRFDPTRLRKSARLRLHRQGDQADYVERE
WP_189000175.1_Paenibacillus_nasutitermitis GLKKAVGFDQKLQLHQLDHIAREFSRVENRQQLYDLVDQNLMADIAGAKARLNARTILFKIWVLVDNENVELRDRGLQLFANASREERLLLHWGMKLLAYPFFRDVADQVGRLFQLHGEFSSLQLSRKIFALYGERRRVSVSMGAVLGTFKHLEVISEQ-KKLYVNAEKIPIHQLEFKLWLIEIMLKATEKSAIELKQIATEPCLFPFNLDITESELRN-SRLQVTRQGIDMVMVVLR
WP_156173096.1_Kosmotoga_pacifica           ISSKMIGFDRQLKKEWLDLVIEMLHDGETRDKITEALNKYIGNDLKGKDAIRKTRLVINRMWFNPNFQLEFLRRSALELFNEARENATSIIHWGMCIACYPYFRDIARICGRLINLQGYFKPSQVRQRTMEIWGERPVVRRSVDRVIQTIMYWGLIEQEASDKYVVSKFVRINNPKLVNWLIRAYLVSRESKTASIDDLRNDPVLFPFSININIDNLLLNKEIELFNQGLKSEILILR
MBX7228944.1_Burkholderiaceae_bacterium     MKQGKIGFDRALKLRWLNATAEWTAHGFSPNEIRVRLDQLLEGEVAGEDAKGKTKTVLLHIWTLVPDHLKPLRNEGLGLIGELPPNNQLAVHWGMCLATYPFFRVVVEATGRLIKLQREFTKAQIQRRVEEVYGTRETVTRATRRVLQSLVEWDALVEDERGLYQANSLLNIANAKLAIWMTEAILASSDFSSATLTAVTQSPALFPFILTLNPRDIEANERLELLRYGLDQEFVKLR
NMB01449.1_Bacillota_bacterium              MRHDKIAFKQIVRLEWMNETLSLVLAGYPESEIRVKLDQYLSTQRQGNPATYGMSLGLLSCWFRKEPELEDFRKSLLTQATRMYQEDWLPLHWAMLAASYPFFAWVCEHVGRLFSLQESITSAQVYERMKQMFGDKEMIARNTRYVVRTLVSWNLLEDERKGQYSQPSVFQIQDTQLTQLLLEGLLLSMPDERCELNALLRHRALYGFSIGTSPQELVRNGRIVATQVGLTTGMLEIN
OQB94798.1_Spirochaetes_bacterium_ADurb.Bin110 QRTGIIGIKQTIRLEWMEKTANLILAGLPAEKVRQELTRYLANRRGSEQDCKFAVAILMNCWVNPLPELEDLRTQGLLLLREG-VEDHTAIHFGILCAAYPYWFNIAVQTGRLFNLQDAVTASQIVTRLKEIYGDRQTIMRTSRYVLRSFIAWDLIRDSRKGNYIKSEPIHIHDMRNVAFLLLSAFYAMGNPQADFNVILHSPAFFAFNMHIAAETLHQSDNIEIAQIGADVLVMKK-
MSP14283.1_Chloroflexota_bacterium          -MDKGIGFNRNIFLPWLDAVAALRAGTADAAQIRAQLDPIVQQDIASAENRRKAIDILINIWVKTGGVSPRLHQQALHFFQTTPPGDRLWLHYGMTLLYYPFFRETAAAIGQLSRYTEGITTALVKQRLIADRGELGSLNKAVERVVFSLRKWGIVQDARRNTYVPQRTLTANTQDLEAWLLACALWSHPAEELPFNDLVRLPELFPFCFSIGIDCIRQNSAFIVHRQGVSMDMVRLS
MCF7763175.1_Verrucomicrobiota_bacterium    MRFVKVGIQRELLVDWFDHAAGIHAMGADRIEARKKCYTYLDTAPGSPPTKAYIANAMVKTWINPDRELISIRNRALDLLTQD-RNDRTAIHWALLCAAYPFWFNLASITGRLLNLQDQATQSQIVLRLKEKYGDRPTVSRRARYVIRSFIAWKTVRDTNKGSYERANVIPIAQPEVTLLVEAALQATPNGTCELAQLINCPAWFPFHVHPLRSDLIACSGIEVIGFGPDQDLLKLK
WP_209690993.1_Methanomicrobium_sp._W14     -RHDIIGIKQSVRLEWMQKTSDLLLEGKNEKEIRNELKEYLSERMGSETACNFSVSMLMRIWVVPDKDLLPFRDAGLKFLKNSSEDEQVAIHFCMLSAAYPFWYNVSKQIGMLLRLQENVTIQQIIQRTKEYYGDRQTVDRNTQFVIRSLYYWNLLEETLRGCYKKGINIEILSETLNSWIIEAQLHSISDGKAVLSSLQNDMALFSFNTKLNSTVISKNKWIEVFQSGMSDEIMMIK
MBC7752535.1_Candidatus_Saccharibacteria_bacterium SKQPLVGFERFVAFDWLDQTAKWVIEGKSPKEVHDLIDDYLASQISGDTSKRKTKNVLSGVWVKSDGVDPEYKEQAKNLYLMANRTEKIAINYGLFIASYPFFYSLGRILGRLFKLQDDVTNAEFYRRCVEANGDRESIKRAAARYLQSVVEWGLLDDAESAKVKPTKKIVIDNPSIVTWLMTSVLFSSDRERLSVDELLSDPIWFPFEISTRDFQIGESQLLEVVHQNVGSVLLALK
OGK77266.1_Candidatus_Rokubacteria_bacterium -KPGEIGFDRRLRREWLDFVADCAAAHVAPDVIRAKLHDLLGPVVAESGARSKTITVLLRLWVVFDPRTDGLRLDALRRLPTCAPGERLTLHWGLAMAVYPFFADVVSSIGRLLRLQPTVSLAEVVRRTKEKWGDRERVARSARHVLQSIRDWAVLAETSAGVYQAAPPQAALRGDLALWMIEAVLQGSGTSIGSLRQLERAPALFPFTLTVRSAEIERGSRLELVRQAGDEDTVGLR
WP_152691916.1_Aeromonas_hydrophila         -MVKTPGFCRHVEKEWLEQSLRWVANGAEYTQLNDNLDNFLSAYIVSKENRRKARNLLTSVWSQPVDQYKNFFDKGIVLHKEAE-SFSLPIHWGMLIARKPFFAEVARFIGRQSRLNNTFTYAQVFRRMSMLYGETEATSRALSAVLRTIMELGILKRNSPASYSIVNQRVSINSHLANWLLDALLISSSKESIALDDALNDQALFPFHLDINLSNLDM-SAFECLQQGNSIVIFKKG
MBL9185845.1_Opitutaceae_bacterium          -TNQRIGFGKKVRWDWLLLAVRLRAKAVSFDDAREELERLIEETNAGKVAITKTMSVLRQLVFLPTPGCRLFSEHGLELFRRRGAAVAMPVIWGLSISAYSFFASCAETVGRLLRLHGEFTAAEMLRRLTEREGDRAFVIRIARFNLSSMLDWGLLEFDGSKRYRRGKLTAAHDPEILGWLAEGILLASDKTSMPQSQLFASNLFFPFDTKIAATQLIGNPRLQVIRQSLNEEIVALS
PKO06219.1_Chloroflexi_bacterium_HGW        SRDGQIGYNRQVELEWLDFTASLIASGETPQSIKSSLDEYLKDRLSSDASRGKTISILLSIWVTVPKHLTKLRDDALLLMRHLSASEHLALHWGMTMTAYPFFGVVADTAGRMLGLQETVSTLTLKNRMREIYGQREQVIRSTRFVLYSFTEWGVLIETERGVYRLSETKNLKDHQLQEWLLKAFLISEGKTLAPVNTAIRSPKLFPFKFE-EPFRSENYSDLEFFRQGLDEEMVMIK
WP_044665230.1_Syntrophaceticus_schinkii    --MQRIGFDRMIKLDWIEYYAKISRHEEDTGKLKEQMHETLFEECPQMASRKKNITILQRIWTRVEAEHRLLRRGALNILDDVEMPERLALHWGMSLLAYPFFREIAACCGRFLKLQKNFSRQQIKEQITATWGARPTLERALPRVLESMVDWGALSHARKSLYRAQPEIQVTDEQLRLWLLYVCLE-IEGGCLSLERAASLPCLFPFSFNISLAEIQQANCFSVTRERNNSIVVTCQ
WP_257961950.1_Lysinibacillus_capsici       -MTKAVGFDQKIYLDHLDKTASLLRHHIADEKIMYNLDGHLIGTIKGDKSRKNAITMLMKIWNFVDDSVVDLRNLALEEYPYLTQIEKKFIHYCLACLAYPFFSEQVCYIGKQFKQADRIYSRTVLAHMKDLYGERRRVEVATGAVFSSIKEWEIVSMPKAGQYEVKKEISAHNLLFSSFLIEVLLKYYGAGSLTLETLNNSAIFFPFDYHVGIGDLDK-QRFTTFSTVRDTVIERNT
WP_106343515.1_Planifilum_fimeticola        -AKMIIGFDYRVTLEDMEFAAFQDPNR-SRGDLITLFREHVGRNRPGRTTQESTATIMLNIWHGRSGETSHLRMRARSLLTETEARQRRALHWGMALLVYPFFRHFAKQVGLALQLDREVLAERINRKIKRIYGERRKVEVASKAVISSMREWGALKMVRHGVYVPTERLAIDHPDLRLWLAEVLIYASEAEMVPLDVINRSPELFPFSVNLSIPDLRASERLEVDRQGLDLIMVGCR
KUK47748.1_Actinobacteria_bacterium_66_15   -MSQMIGFDRSLRLDWLDAAAGVSRETEDIEQIRNYLHAYLAESYPHYTARRKTITVLTRVWSRIPDSHRLLRERAFVLLPTLNQTDRIWLHWGMCLLAYPFFRDVVRTVGHSIRYYGSFTKQEIIQRMSETWGERATLPRAVQRVIESLNHWGVIAGCQKGRYRCAVPLSTSNKDVEIWLLKVALCANPNDSFPVDQVHTLSEAFAFSCSLSIPEILGSREFEVSPMGNNRFALRCS
NLI08746.1_Thermotogaceae_bacterium         MKTISVGADRFLALQWVNYAYELFISSLDQDSSRKALREYLENQMLGQVSTRKTTDQLFRLWLYSGDPYQTLREMTLSIAKPIDLAQMAVFHLGMAINVFPIYRETVRVIGTLERLTDQISTKAISTRVIEKFPTTTSIPRTVNRVLQTLDDWHFIELT-QGLVVVKE-ILLLDPKLVNWFILTTLRAIQQNEITLQELDTCPFKLGINFSHPRNVVNDSEDLFFGRNQMGLEVIRIK
MBX3742602.1_Akkermansiaceae_bacterium      SSARPIAFNRRLRASWLEEGLRLRSDGLDGPVWVERMEELIKVDISGKDSIAKSMRYLRHVWADSGED-EAMRSEAIAIFRSSPSETAIILSWGMVIATYPFLNDVAATVGRMIRVQPEVKLEQILRKLSETYGERETVRRSGRYALGLITDFGFVKRASPGCYVLGNPIKI-DPTLSGWFVRSWFKALGSTVIDRVGMAGAPSLVFFDSALISDAIER-RILVPERLSFSQEVLRLN
MBN1459904.1_Armatimonadota_bacterium       -DRSRIGIDRALHLDWLDAAAGLAANGATCDTARTALLEVLGRELSSGGAADKTASVLLRTWFKIPAEAAPLRKDATLAMRHADGSGRLAAHWSMLLAAYPFFIDCTSVAGRLLRLSGGFQRSEYSRRVAELWGEKPTANRASGRVLETLVALGVLRQ--EGAHCTQTGRFTVDGPAAAMLAEGLLLGEDRMAVPVESVSSHAAMFPFEMPPIATAVVCSSRLGIEVRGGDTEMLVRT
MBI3175734.1_Chloroflexota_bacterium        MSEYLVGFDRFIALDWANYTLGLLSYPGDEAAKMSELKSWLSLYISGKEASQKTANVLARLWLRATPEISIFKKYAVSLAPELRKDELMYLHWGMSLLVFPLFHEVCTQVGRLLFLQSSVSRREIQIRMAEKYSNQSTIPRSVNRILQTLVAWGLLQEISPRELSAKR-QAPSNILIKAWLIECAVFSAPSKKVSLHGLYKNPILFPFEFHEDLSWIVNSDKLFLERDGNNLEYVVWL
WP_075724812.1_Tissierella_creatinophila    -MSKQVGMSRNISLDYLNKTIEFMEIEDDFEKVRENLREYIGLYIDDNTNIRKTANILSNIWLEDDKENRKIKDYAREEVAKSSSNLRLVAHWCMILITYKVFEDIASVIGKLEYMQIEITSKLIREKMTDMWGERPTLIHAIPKNIRTMRDTSVLETVKQGVYKVKK-HRVDDERAIILIVATLIHLKDKLYLSFDELIDDSLMFPFDYSIDVGVLEESNMFSFDRFGGELAISLKE
WP_020614472.1_Sediminispirochaeta_bajacaliforniensis MREKTIGYDRLLKNDLLNKVAYFVGDGNNPEDVKSKLRASFEDQTS---ALRKTINLLMGIWSTVPEEIVPLRDQAIKLFLKAYDSEKRIIHYFMSMANYPFFFDVCLLTGRALRLSNSFTSRQILTKIQEKWGDRSTTIRAVQRTLKTIEQWGWLNSTKQRDYSLNDMKTIIPKEWHGFIANCVLFGSLRSSMLIEEITQNPSLFPFHIEFNIHEMKQWNGIRLEQRGRGDLLVSLK
KAF0196958.1_Bacillota_bacterium            -MSKQIGFDRFIKLQWAEDLAWRMRETKDEVSLKRIIEDILSKELPTQETRRKTSIVLLKAFYRIPPQHERLRNEALELLPQVGVNEAKALYWGLMLLAFPIFTDTARHIGQLFSLQSDASAALVQRRVIETWGQRGSLHSTLNKVVKTMAEWGCIDLTHKKVLVPKA-PTPLGKPASLWLTRVALTVSETPLVPVDQALRPSICFPFTMDIHAQDFSD-GEFLLQRMDGRTEYVGLR
WP_270641086.1_Limosilactobacillus_mucosae  -MKKEVGLFRPLRFDQLDLAASLLSNY-ELNDAFEIMNEKLSNELKGNVGRRKVINNITRVWSGGKKQPSNFQKNVLSRYCNANHDSKIAYQYLMTAIAFPFFRDTVNIVGKYLRMTGKISNAFLLKEMENSYGSAESVYKGTNAVVRGLIDWNLVDGMEKGKLIEIDKLRISDEFLKQLIVKVLMEHYNTEYIQLDQLNEDAIFFPFNYHLSESYFDNLTNYEVIDDHSGKYVHMIG
WP_270691668.1_Bacillus_cereus_group_sp.    -MTKPVGFDQKVLLHHLDFAANHTKKY-SRKDMYEVLDGYLRNDITGAKSRKNAVTMLMKIWYLVDKDLIEIRDRVLEVLPVLTKEERVFIHWGMTLAAYPFFKDTIHELGKLFQLQDTVPSTVIGKRMKEEYGDRRRVEVATSAVLMSIKAWGIIIPTKNRSYQLPDKVFISNPLLQVFIVKIMLHVLEHTALQIDILKNHPMLFPFDHEFDLLELRQNEWLTLHHQGVDTLIVECK
WP_140397540.1_Flavonifractor_sp._An135     -MAKAVGLSRKITLKWLNKAHEFFSEGLTEEEFKTQINDYLSFEIDSPTVLRKTREILMHVWFYEDNYHTLIRRDADDLLKRFP-EYDIAIHWCMLLSVYPVFADICKLIGRISDFTDIVTLSQLKQKLYDEWGERSTLYHSTDKIIATMKELDAISSEKPGKYTIKK-HTIARSEVALFMAMIAMKVDGNSYYSFSELHDFELLFPFEYKTSKEELMTNERFTVSTFGGEVSVSLSV
MBT6646604.1_Nitrososphaerota_archaeon      -MLKNVGFLQYVTKDWLDATVAFYQPHQNIDEVKASLDTLIKFDVPKNEERRKTIDVLTRIWARIPPKFDSLRLDALSFYDDLDPNERIILHWGMALLAYPFFRDIVFSIGSLLKIQDDFTSTQLRNKLISNWGHRTTIIRAIDRNIQSLKNWNILIIDKS--LKQNSKMIIKNESLCLWFLECIFRAENLDSMLFEKLVSSQTMFPFELKIPVYKIKQAKNFEYQIHGMNIEMISLR
HEC58782.1_Methylophaga_sp.                 -MKVSIGFNRHLETEWLTQTASWVAKDIEGDELKSQIDLMLAPVFTSQVAKDKTRNLLFGIWNKLPAHIKAFQDDAAKLIMEHS-ELNLVFHWGMMLAKYPFFYLVVGQIGRLTKLNDAFLYAQLEQRVVENFGDTSTIKRSMQFVVRTLINLDLLANPKQGLYQTQKPIKVDSSSVKAWLVEASLYSESQSSRSLTTISDAPVWFPFELHFQASDLSQNGRLEVHHQSSDVIVFINN
WP_243191721.1_Vibrio_parahaemolyticus      -MSKRVNFSRHLEIQWLDQTARWVASGLDKATLNENVDRMLEPHIKCKVNKGKTRNQLINLWFSVSRDMDSFQNKAIEFVKSSD-ELPLTLHWGQLVAKNVFFSDVARFVGRKLKHGDSFTYAQAQKHLVELYGDTETVKRALRSALKTLVEFRLVSRESNRSYSVNKLEMYVDTRYKSWLLLALMYNQGFHNRSLSNLLDDLVWFPFDFSVSFNEIDA-SLFELHQQGNELVLFRKE
WP_149544908.1_Calorimonas_adulescens       -MIKKVGFSRALKLEWLNSTVEIYENTKDLETTRKLVKEYLTYKIKGRVSLSKCTGMLMDTWVDVPDEYRSFINKGIELYDGSDSEQKIILHWGALMVAFPIFTDISSTIGRFLYLHDSFNASSVKKRIYELWGERSTVERVVRYSLVTMSDFGVISRDKAGEYKRGKCIVVDNPEHKLFLINSYMLATNRKTLQYDTIEEQYPLFPFKMNLATHEINGSNVLTLNRFDNQTFVVMK-
WP_045174404.1_Caldicellulosiruptor_danielii --MKAIGFLKPVRVEWLNITVEELIKGKEKNEIRDVLKRIVSNYYKSEDTIRKILAILMSTWVNVSEKNMCLRDRGVDLFKIANPDEKIALHFGMLMVTYPIFNDVVFIIGKAFYLENKFKLEYLKKRIFEKWGERETLRFSIDRIIKTLKDWGLIVSKNSGVYEPGTKVTITSKEIKAFLIACYLKANERNYININEIENLYSFFPFNISVNLNELMNVDNLSVNKIDSIITVAVNR
WP_010252938.1_Treponema_primitia           -MLKSVGFLRAVSNAWLDKTVELYLTGLDPDKIREELTNHISFELKGKESIRKTIDVLMATWININTENQQARKTAITAYKNCA-ENHNIQHYCMLLLAYPVFNDFCNLIGKISKMQDTFTKSWLRDKLFEQWGDRVSIDNSLKYLIQTLLSLQILERIGVGKYQIKK-HEIMDNQNKILLLHALALLGGTKYCVLSSANFQPILFPYIVEFSPELILNNSILELCIFQGKAIISLKK
WP_140396494.1_Gordonibacter                -MARMVGFTRIIRLPWLNKTIELAGEGLEPTQMREELEEYLKFEISSDTNRRKTREILLLPWVAEDAALEALRPKALELALSHP-YELLAVHWGMLVVAFPMFGDLVRLIGKMCEYQDEFTAAQLKLKILDEWGERSTVVKGSEKMLASLNAIGVIQRVKQGRYVLAP-RVDADEELSLFLLHADMLAYQSGYRSYTELQALPELFPFVLHASKERLVEDARFSIGNFGREFTVSLNQ
WP_122014531.1_Biomaibacter_acetigenes      -MMQTVGFNRFIKREWLNFTAFLAGREKDEKIIRKNLNEYLEKETKAEITRQKTLVELLRIWVNVPASHEDLKEKALDLFEKVPEDEKMLFHWSMIMLAFPIFIDVASSLGKIFSLQDRVSLKILEKRIFEKWGERSTVKYALPKIVGSMVEWGVITRAKVGEYTKAQTIEIKNKDCKLFFINCYMQALEKDYLRFIEANNLSAAFPFKLNLGLDDFNNSNTLSLSRMGSDVVIIKR-
WP_035270489.1_Desulfitibacter_alkalitolerans -MLKRVGFDRQIKREWLDKTVELLINKEDSKDIKNILDGIVKEDIASNDNRRKAISILMHSWINVETDHIAIRSKALKLWETGNSTERLAIHWCMLSLAYPLFFNISEIIGKLINLQDEFSLNMVRRRVYDVWGERSTLDYAINRVVWSMKDWGTIGQNKVGEYKKIERIVIEDKEVQLLLLEAYFIASQRSNIPCAELEMINALFPFKTDITINDLQSNHRFKINNMGNNLVVSL--
WP_028274685.1_Atopococcus_tabaci           -NKKTIGLFRVLTVEDLDRTARLLKKY-SPQDAYDILNEKLMEKAPGDVGRRKLVNNLMMIWGNGKKEPANYQKKAMEVYNELPKEEQLVLHYLLALIAFPFLTFEARLAGKYLGLMDTVSSKVFMDEVMNKYGNSGTITRSVSNGFGILREFGFLKKEKAGRYSLGNRIIVKSTLLKNYIVLAALLNTDSDTLSLEAVQHDKMFFGLDFILYSSDLDP-NMFEVTSDRVHTYIKKR-
WP_202065228.1_Enterococcus                 -MKKMVGLFRVMTFEELDKAAEVMVKY-SPTDAYQILDETFAKTVKGDTFRRKALNNLMKVWGNGQKPLSDFQEKVLERYAISNYDEKKSLQVLMMLYSFSFFSDTINIIGKYLRMQDTFQSKTIMEEVRSTYGMTETVNKGVLTVLGSLVDWGFLFRSKPGVYELTKKILITDEFIKNVLIRSILAHTDAEYLPLEIINNQAGLFMFDYQVSRLEVHL-DDLNVLRERTDTFI----
MBP3452731.1_Clostridia_bacterium           -MAERVGLSRNISEEYLNVVADCKLADKPLLEARDILNALIGTHIHAETNIKNTRVILLNTWYRNE---LWFQNTCIEIAKEVTVPERLPLHWALLMVAYPVFFDLCTIIGTMLDYRDEFSATQVKEKERIFETWSARTTLIHALSKNLQTMKDMRILEPQKNGVYKPIS-HHVDNKKTVCVLAAAILKCSGLEYMTWEEIVHHPALFPFNIEVTQADLAACDRFLLERMGDNIVIRFRT
MBQ3643691.1_Candidatus_Riflebacteria_bacterium -MTKNIGLSQKLKFSWLEKTVEIVSSE-EDTFIKKKISEYLSCEIKSPTNIRKITKILMNIWVSNN---NDLQINALELIKKYP-EYSLVINWCLILTAYPVFTDICTIIGKIAKFQEVITNNQIKEKMFEEWGERSTLFYSVEKTLGTIIELGALKPIKKGSYSLCK-YAINNDEIIDFMLKSILLSNTKESKTINELQNLACFFPFKYKINENRIINNNSFKTVNLGGNLNISVA-
consensus/100%                              .....sh.b.h...h.................h..h...............h..h.........h.....................h........hh......G.......h....h..p....s............h..h.....l.......................hh..................h......h.h..............h...............
consensus/95%                               .....luh.b.l...hp..h.............h..l........s.......l...W.........hb..s...h..........hpa.h....asha...s...Gp...h...h....h..c....bu.p..h..s....l.sh..h..l.......h.........p.....hhh...h............h.....hasa........h.....h..........h....
consensus/90%                               .....lGh.b.l..pbhp..h............h...hp..l........s..p....l...W.........hb..s...h..........hpash.h.sasha...s...Gph..hp..h....h.p+h.p.aGpp.pl..s.....l.oh..h.hl.......h.........p..h..hhh.s.h..........hp.h.....hFsFp.......h.....h.h.........h....
consensus/85%                               .....lGh.b.l..pahp.sh.b.....s..phb..lp..l.....s..sb.ps...L.p.W.........hbp.uh.bh..........haash.hhsaPha.p.s..hGph..hp..hp...l.p+h.p.aGpp.pl..s.....l.oh..h.hl......a.........s..h..hhh.shh.s......h.hp.h.p...hFsFphp.....l.....h.h.p.s.p..h....
consensus/80%                               .....lGhpb.lpbcalp.sh.h.....s..phbp.lppbl..ph.u..sb.ps.shL.p.W...s.....hbppuhphh.p.....pbhlHash.hhsaPhF.p.s..hG+hhphp.phs..bl.p+h.p.aGpp.sl..u.p.sl.oh.phshl...p.s.Y...p...h.s.ph..hhh.shh.s......h.hp.l.p...hFPFphph.....l...p.hph.p.s.p..h....
consensus/75%                               .b.p.lGhp+.lphcWLs.shphh.p..s.pphbp.lpphl..ph.up.sbpKs.shLhphWh.ss.p...h+ppulphh.p.s.ppphhlHauhhhhsYPhF.shsp.lG+hhphpspho..blbp+h.-.aG-Rpol.+u.pbsl.oh.pashlpp.p.u.Y...p.b.h.s.ph..allpshhbs..p..hshppl.p.s.hFPFphphs..pl..sp.hpl.pbu.s..hh..p
consensus/70%                               .b.p.lGasR.lph-WLsbshphh.ps.s.pph+p.lcphl..ph.upssbpKs.shLhphWhpss.p...h+ppulplh.p.s.ppphhlHWuhhhhsYPhF.clsp.lG+hhplpspho..blbp+hb-.aG-Rpol.+uhpbllboh.pashlppsp.G.Yp..p.b.l.s.pl..allpshlbu.spp.hshppl.p.s.hFPFphphs.ppl.pspphpl.pbu.s..hh.bp
```

```
# ; Type4 DUF4007 Representative_1 DNA-Binding Component; (HTH + wHTH1 + wHTH2)

OrigSeq   [illegible]

CDE63278.1_Alistipes_putredinis_CAG:67   [illegible]
CVI65225.1_Alistipes_sp._CHKCI003   [illegible]
GFI28163.1_Lachnospiraceae_bacterium   [illegible]
GHU63481.1_Clostridia_bacterium   [illegible]
HAR58252.1_Acholeplasmatales_bacterium   [illegible]
HBZ34376.1_Bacteroidales_bacterium   [illegible]
HED00227.1_Pseudomonadota_bacterium   [illegible]
HHW21154.1_Thermodesulfovibrio_thiophilus   [illegible]
HHY36030.1_Bacillota_bacterium   [illegible]
HIQ39048.1_Methanothermococcus_okinawensis   [illegible]
KUK07756.1_Archaeoglobus_fulgidus   [illegible]
KUK66685.1_Parcubacteria_bacterium_34_609   [illegible]
MBB6067916.1_Methanococcus_maripaludis   [illegible]
MBD5142972.1_Ruminococcus_sp.   [illegible]
MBD8953985.1_Dialister_sp.   [illegible]
MBE5813906.1_Clostridiales_bacterium   [illegible]
MBE6363377.1_Lentisphaerota_bacterium   [illegible]
MBE6838774.1_Ruminococcus_sp.   [illegible]
MBF0554174.1_Nitrospirota_bacterium   [illegible]
MBK5260963.1_Peptostreptococcaceae_bacterium   [illegible]
MBO6251941.1_Muribaculaceae_bacterium   [illegible]
MBQ6210587.1_Prevotella_sp.   [illegible]
MBQ7110121.1_Thermoguttaceae_bacterium   [illegible]
MBR2732786.1_Selenomonadaceae_bacterium   [illegible]
MBR4291499.1_Oscillospiraceae_bacterium   [illegible]
MBR5214694.1_Bacteroidales_bacterium   [illegible]
MBR6984230.1_Ruminococcus_sp.   [illegible]
MBR7025509.1_Selenomonadaceae_bacterium   [illegible]
MCC8021079.1_Akkermansia_sp.   [illegible]
MCI9323608.1_Lachnospiraceae_bacterium   [illegible]
MCI9677951.1_Oscillospiraceae_bacterium   [illegible]
MCL2016624.1_Defluviitaleaceae_bacterium   [illegible]
MCM1232923.1_Ruminococcus_flavefaciens   [illegible]
MZQ76089.1_Peptoclostridium_sp.   [illegible]
NBJ16969.1_Dehalobacter_sp._4CP   [illegible]
NLT83667.1_Coprothermobacter_proteolyticus   [illegible]
NPV08141.1_Anaerolineae_bacterium   [illegible]
NSW92345.1_Bacillota_bacterium   [illegible]
ONI45223.1_Epulopiscium_sp._SCG-C07WGA-EpuloA2   [illegible]
OQC39309.1_Bacteroidetes_bacterium_ADurb.Bin037   [illegible]
WP_024333181.1_Desulfotignum_balticum   [illegible]
WP_062281146.1_Flexilinea_flocculi   [illegible]
WP_071148138.1_Bacteroides_ndongoniae   [illegible]
WP_126850816.1_Candidatus_Cryosericum_hinesii   [illegible]
WP_138286703.1_Phocaeicola_vulgatus   [illegible]
WP_197538712.1_Methanothermobacter_sp._CaT2   [illegible]
WP_207678197.1_Desulfonema_magnum   [illegible]
WP_268058970.1_Caproiciproducens_galactitolivorans   [illegible]
consensus/100%   [illegible]
consensus/95%   [illegible]
consensus/90%   [illegible]
consensus/85%   [illegible]
consensus/80%   [illegible]
consensus/75%   [illegible]
consensus/70%   [illegible]

# ; Type4 BrxA DUF4007 Representative_2 DNA-Binding Component; (HTH + wHTH1 + wHTH2)

OrigSeq   [illegible]

KEF34343.1_Deinococcus_sp._RL   [illegible]
EBT6907987.1_Salmonella_enterica   [illegible]
GIV81896.1_Anaerolineae_bacterium   [illegible]
MAU00652.1_Anaerolineaceae_bacterium   [illegible]
MBH8560400.1_Hymenobacter_negativus   [illegible]
MBI2377783.1_Deltaproteobacteria_bacterium   [illegible]
MBI2842135.1_Armatimonadota_bacterium   [illegible]
MBM4284315.1_Deltaproteobacteria_bacterium   [illegible]
MBU6335169.1_Chloroflexota_bacterium   [illegible]
MBU6335172.1_Chloroflexota_bacterium   [illegible]
MBV5327605.1_Chlorobium_sp.   [illegible]
MCK5872694.1_Methylococcales_bacterium   [illegible]
MCK9608994.1_Methylomonas_sp.   [illegible]
MCL4273185.1_Anaerolineales_bacterium   [illegible]
MCL4273188.1_Anaerolineales_bacterium   [illegible]
NJL05931.1_Chloroflexaceae_bacterium   [illegible]
NJL05934.1_Chloroflexaceae_bacterium   [illegible]
NPV84910.1_Anaerolineae_bacterium   [illegible]
OAB59991.1_Phormidium_willei_BDU_130791   [illegible]
PIQ01002.1_Shewanella_sp.   [illegible]
PNY80835.1_Deinococcus_koreensis   [illegible]
PSB31010.1_Stenomitos_frigidus_ULC18   [illegible]
QEP42420.1_Ectothiorhodospiraceae_bacterium_BW-2   [illegible]
SEH06869.1_Thiotrichales_bacterium_HS_08   [illegible]
SEH06873.1_Thiotrichales_bacterium_HS_08   [illegible]
TPG64628.1_Hymenobacter_nivis   [illegible]
WP_027878022.1_Meiothermus_cerbereus   [illegible]
WP_062417763.1_Levilinea_saccharolytica   [illegible]
WP_083924616.1_Lamprocystis_purpurea   [illegible]
WP_086759844.1_Nostoc_sp._106C   [illegible]
WP_129675134.1_Candidatus_Chloroploca_sp._Khr17   [illegible]
WP_165774646.1_Candidatus_Viridilinea_mediisalina   [illegible]
WP_169366112.1_Pseudanabaena_yagii   [illegible]
WP_192822488.1_Rufibacter_sp._LB8   [illegible]
WP_194112251.1_Mucilaginibacter_myungsuensis   [illegible]
WP_194715673.1_Noviherbaspirillum_soli   [illegible]
WP_210653449.1_Gemmata_palustris   [illegible]
WP_219156497.1_Hymenobacter_profundi   [illegible]
WP_235137402.1_Dyadobacter_fanqingshengii   [illegible]
WP_235137405.1_Dyadobacter_fanqingshengii   [illegible]
WP_254031242.1_Planktothrix_agardhii   [illegible]
consensus/100%   [illegible]
consensus/95%   [illegible]
consensus/90%   [illegible]
consensus/85%   [illegible]
consensus/80%   [illegible]
consensus/75%   [illegible]
consensus/70%   [illegible]
```

### ; Type1 BrxB iSTAND ATPase; Representative Alignment

```
                                            1---------11--------21--------31--------41--------51--------61--------71--------81--------91--------101-------111-------121-------131-------141-------151-------161-------171-------181------
OrigSeq                                     MLQQQFENHYRVITSAGFLQRQGLANEVPFFISTFSADQQVEAEGLVNSLFMRLQTQGVDVLKIDLFEFCLELLEQQGVLEDYLAMESQIKKADLKDALVGALSVQDKIAPAIARKLAEQNWKVLFLCGVGRAYPILRTHTVLSNLQSIVVSQPTVLFFPGRYTFVSLDLFGNLNEERYYRAFNLNNYQL

                                                                    S1                                    S2   ------------HEH-Extension-----------                                S3                                   S4                         S5
                                            -HHHHHHHHHHHHH-HHHHH---------EEEEEE----HHHHHHHHHHHHHHHHHHH---EEEEE|HHHHHHHHHHH-HHHHHHHHHHH--HHHHHHHHHH-|HHHHHHHHHHHHHHHHH----EEEEE-HHHH-----HHHHHHHHHHH-----EEEEE-|EEE---EEE---EEE--|--EEEE-----
                                            -HHHHHHHHHHHHH-HHHHH---------EEEEEE----HHHHHHHHHHHHHHHHHHH---EEEEE|HHHHHHHHHHH-HHHHHHHHHHH--HHHHHHHHHH-|HHHHHHHHHHHHHHHHH----EEEEE-HHHH-----HHHHHHHHHHH-----EEEEE-|EEE---EEE---EEE--|--EEEE-----
                                            -HHHHHHHHHHHHH-HHHHH---------EEEEEE----HHHHHHHHHHHHHHHHHHH---EEEEE|HHHHHHHHHHH-HHHHHHHHHHH--HHHHHHHHHH-|HHHHHHHHHHHHHHHHH----EEEEE-HHHH-----HHHHHHHHHHH-----EEEEE-|EEE---EEE---EEE--|--EEEE-----
WP_102077887.1_unclassified_Psychrobacter   MLNTDFNELMERVRAG---REFGHASFEPIFYLIFDPQKILKIKRQLPAWAAKLRNEGWDVHIFSMAKAVQEVFDEMPVFQDSAALENRDQWQKTNKSLAEALTKKNALQNKLEAKLFGRPNSILLVSDIEALHPYLRIGSMESQLQGKFH-VPTIFFYPGMRTGKQLKFLGFYPEDGNYRSVHVGG---
MAE16012.1_Deltaproteobacteria_bacterium    MLQQQFENHYRVITSAGFLQRQGLANEVPFFISTFSADQQVEAEGLVNSLFMRLQTQGVDVLKIDLFEFCLELLEQQGVLEDYLAMESQIKKADLKDALVGALSVQDKIAPAIARKLAEQNWKVLFLCGVGRAYPILRTHTVLSNLQSIVVSQPTVLFFPGRYTFVSLDLFGNLNEERYYRAFNLNNYQL
MCF2581773.1_Bacteroides_caecigallinarum    MMDTVFEKVYQKLSSPDF--GKNLGGELPLYIQPIPVSGQTELNTQVERLVSRLSKLGKNSIVVDLYRLALEIIDEEGILETLLEDEKNIDKDDLNATFESIFDTKEILIPRMRSMIEENKYDFVFITGVGRVYPFIRSHSIVNNMEGLADNANIVLFFPGEYNRLQISLFGKLPADNHYRAHNLNDIET
AUX12765.1_Latilactobacillus_sakei          MTNERLNELQARISDNQFLTKQGLGNEVSFYIFDYNPADELLVRSRLPEIQRYTNKLDTPIKEFNLFDIATRFFEERQYMEKNYQMEQTKDSFTLYDRMQRALKTATEVIKYIREH--CEDNTITFITGVGSAYPLVRAHTVLNNLQPIIVQKPLILFYPGVYEGNKLSLFNQFKDDNYYRAFRIVE---
ETR72220.1_Candidatus_Magnetoglobus_multicellularis MIESRIKILKDDLLNPL-GPSISTNQNYPFAIFHYPPHEEFKMREKLIELLDTLRNKAWVILNVDLFATLIEYLSIQEDIDALVQEEKLQPLNVLKNSLDIVLKNTDFVFDKILNKIE-ISKTVIFLSRIGGLYPFYRTSTLLRMLDNGIQ-VPTIVLYPGERTEQYLRFMGEMDADRDYRPRIY-----
HIU24259.1_Candidatus_Coprovicinus_avistercoris MLQERLDGIAKRISSQEFLENKGLGNEVGIHVFAYDPHYEWQVRAAVANLMHASGKLPCCIHERNLWEAVLEVCRQKRILDKIDALEHKRGSKALQKSLQPIVTPEA-LVASMNWQPHEHGRDVLFITGVGQAYPFVRAHQILENGQHVFDDIPVVLFYPGTYNDRELALFGRHDQSNYYRAFSLI----
KAB1640464.1_Ellagibacter_isourolithinifaciens MFSHCCEALRAHLLDVNFLENKGIGNEIGFFTFCYDPALEMQARTFFAALERESGDKPYRIVSRNLYDVFLNICEKRRILKAIPVQEAKHGTASQLKQLSKICTPAA-FAEAIDYEPH-ERGDVLVLTGVGEVYPFLRIHTLLDNLHVRFSDIPVVIAYPGSYTGHSFSLFNSLSDGNYYRAFDLV----
KAF3303186.1_Carnobacterium_sp._PL17RED31   MINELLDKFEETIMEEKFIESRSIGDDNNYYVFDYPHEKEMEVRNRIESMIIKSGAYPSELILFDIYDFTIDLLEKRGFLEMAYQFEEKNGMAFVADTVQSLLSKDGIFIDHIERN--SNTDSVILVGGIGKIHPIIKAKTILSSLQSTSNQSSTILLFPGTFTGLQLNLFEKISEENYYRALKLLD---
MBD1206698.1_Paracoccaceae_bacterium        MKPGLKDHLRSIITGPRVLGGDTPGNDVPFFLCPYDPEGEAAMQRLIADLAAEIGAAGTRVLTLDLYRIAVDLLQRTGRWDRLLSREARDDKVRFLALLQSTLNEEEVLVPEIRDLVDRTPHDLIFLTGFAPAFPMIRSYTILSRLQTVLKVAPTVIFFPGRYTQDSLKMLGRLRERNYYRAHNILHMRG
MBD5357713.1_Bacteroides_sp.                MSEIVVDKLYERLCQPDFQDTPHGDLFYNYFIVPYDPAKEYETRERIQWYHDNLPTNFVNALHIDIFEEFCNYLDNKPFLKYLIEKESKDSSAGVARSLTQQAQCDAYIRDRIVAHKEENKTPYVFISGFGNIYPYLRLSTFLNKYEEHTHAYKIFVFYPGEPDGNSYKLFDRFADQHVYRAALILDFRN
MBE7502026.1_Verrucomicrobiales_bacterium   MLKDNFETLRQRLGHG---REIDSTGTEPIYYLVFPVAEILEVKRQTKTWIARLANDGWNVVPLSLADTIAGIFQNHKLRKLWLASEQQLDPSEINKTLAKALSDNSELRVRLDAAVAAQPNGLLLLTDLEALHPYLRINTIEDYLRGRIR-HPVIVLYPGQREGKSLRFLEFYPPDPNYRSDHIG----
MBP6234855.1_Saprospiraceae_bacterium       M---QFDKILNVLARKDFQDPSTGNLFFPVYIYTYDTKDEALYKEQIPEMISHLPSNHLHTLQIDVWKEIMIFFEEKTFLELIKNKEN--TGEDVTQWVRHIFNSQNWLKNKIDAHYRLKFKPYIIVNGFTEAFPYMRPGEFLKNTESLPEHYKIILFYPGKVEGENFKLFGTLESESIYRANILNN---
MBP6733458.1_Chromatiaceae_bacterium        MLAARFEHLVNVIGSRRFLKMQGLNNDLPFYICEFRAAEAVEMQRLRRQLIHKLQGRGVRILDINLYDLSIALLQAREGWDDLLAIETEVDKDQLLELLQNVLGIEEYLIPAIGERLAQAEYDVIFLSGIGEVFPFIRSHNVLNNLQSRIKGKPLVMFFPGEYRHSSLELFGLLRDDKYYRAFNIYEIQA
MBQ6248037.1_Kiritimatiellia_bacterium      MTTEIFHQILPVLSSKPFLDGSLAAPGIPFYVCPYAEREQETVAMIGNLVRQLAAKGVAVTVVDAFRLMVERMKENGDLDYDLEDEPGYTRAEFLEHLQGILDVPKEIVPAICARVEEEKPRLIFVTGVGAAWPVLRAHSLLMNLESAVK-IPLVLFFPGAYRQISLYLFDKLPGDGYYRAFNLLDIHP
MBQ6343446.1_Anaerolineaceae_bacterium      MLSKRIEYLYKVFTSDRFLSMQGIGRETSYYICPFSPSEANEFSKQRELLVKRLTTNGIGVCEIDLYDLSVEILKEEDAWDGIMEQETAIDKSDLLETIQATIDLGGTFLRKIQNHVSRGDHKIIFISGIGEVYPWIRTHNLLENMQTVINGEPVVLFFPGSYKQTSLELFDTFKNERYYRAYNIFNYGV
MBQ8347029.1_Alphaproteobacteria_bacterium  MIEEKYNHLKKIISSPDFLSGKVLCNDLPIFIADYDISEEKDFLIMEKNLIKTLEAMEIKVLSVDLYDLIIEILKKNQGWDWVMSPDR--NKEELLEELKGILGIEDTVIPAIANKMANGKYDVLFLNGCGNLFPFLRTHTILNNLHSHIQ-KPVIVFFPGINVSKAFNLFGRIDEGNDYRATNIFQIRG
MBR3160179.1_Atopobiaceae_bacterium         MLSSKKNHILKVLASPQVIRGDRTMSRSTAIIVPYPPEDEVEWDDMAHTLVGDLGAQGIAASIANLYDIVLRILDDDGLYDEKVEIERTHGKARLIQELKGAADIELDIVPVIEDIIDDEGPQLLFLTGVGACHPFLRTSQLLEKLDDTID-VPVVVFYPGRYQRKPLEILNIPRGNAFYQGRNVFD---
MBR4606064.1_Lachnospiraceae_bacterium      MFYARLEQVPDRLLSKDLLHNIHIGNEIGFYIFDYPPDKELVVREWLPSLPEQVSEQKITLLSINIFQEVIRYIKNTNKYEKIIQVEKERGNDATLRGLGSLLNTTA-ITSIFKDVIN-TNADIILMHGVGAAWPLIRIHSFLNILQPIVERVPLVLFYPGTYNSSEIKLFNRLPSENYYRAFRLLP---
MBR6253007.1_Clostridia_bacterium           MIENRLQEVYDKLKTKKLSETQSLGNGLKFFIFDYDPEDEEKVYNYFETYLN--VKDDFHLRIVDVYDCIINILKEKGYLEKVYKMEKEKGIDYTNGVITSTIGVGSLLLNEITKD--LKKDEILVLKGIGKCFGVVRAHTILSNLEKVVDENPVLLLYPGGYDENGLRLFNKLKPNDYYKAIQLVY---
MBU6197214.1_Cyanobacteria_bacterium_REEB446 MRKQRFDLLLRFTSNKDNLDYRKY-NDIPFFIVPYEISEQNKIKEDIPSLESHLIKNDIRHLSINLYELSLELLDNNLDLKYFLETETERDKKRFIEDLQGVLTSQK-VAEKIGDECEKFKPELLLISGVGEVFPYLRLSKLLENLSCKVKEIVVLIFFPGIYCQSYLRLFSCLDYDHHYRAFNIEDLNL
MBX6362418.1_Gemmatimonadota_bacterium      MLKQRLEQLEQKILAG---RPLGPSTDIPFGIFVYDPAEELEMRRQASLFATRLENGGKRVRSVDLGALLWACLDESPL-PRALEVEEAFELEAILHQAHTMVAGQHPLERRIIEQLADRVRDFALLSRAGELFPIYRTSALLERLIGHVR-IPTLLLYPGTMTGPELRFMGVCDPSPNYRPTIYA----
MCB1062256.1_Verrucomicrobiae_bacterium     MLDATFERLFSVMQSERFLTSTGIGNEVSLFVQPYEIEVQDPVEKRIKALASRLSAKGLGVLCINLLELVAEFTSREDRLKRLLEKEPTFTRDKMLSTMARWTDAKEHLVPAMKAKFDAEKWDITLVYGCGSVFPFLRTHTILENLQSDMDKRPIVFFFPGEYQHRDLRLFGKLSHKGYYRAFNLDHYHL
MCC6107939.1_Atopobium_sp.                  MVNEAELFLVSKLSSDRFLKSAGAFSNSSMFVAPYPPEQQEDAAKMVDALALDLRSRGVTASVVDLFSVFTSVLDDQGLWEPYCQIEGTVPTKDFIIALKDAADLETQICPRVVAQVKEEHAQLLIVTGMGACYPVIRTHRLVEVLDPQ---VPVLVMFPGRQTHKSLDILATPAGEGQYRARNIFE---
MCF0260591.1_Erysipelotrichaceae_bacterium  MVTKRLENLMEKISTSAFLNNKGISNEVGLHIFTYKPSEEMIIKDFVRRLKKDH---PCRIVEFDLYKSLLKALENDGILDEVGEVEKEISHEELLEQLDTNLDQ---LMEAAELPLK-LGEDVLLITGIGDVFPFIRAHYVLNSIQQVVS-VPVVVFYPGEYTGRDLKLFNRIDDKNYYRAFNIC---
MCH3907668.1_Sphaerochaeta_sp.              MYNERYAFLLDLLDDDKFIRKKYVGAEIPNYICAFPPEDVSSYNDMLENLAKALTSKGRKVLKVNVYEIMIEMLKDSGDFEDYLE-EPSLTHKKIMEDFEGVLDNDTEFAPKVASIINESVPGIVLMNGIGEAYPFIRIHTLLEKLPTLLKRVPIVVFYPGNYDKVALQLFGKLEHKNYYRAFNIFT---
MCI6194161.1_Faecalibacterium_sp.           MTDQKLIKLDDFLHTFSTETGDGLCGEAGYYVFSYDPKDEMKVRRWTECETQILQQSNRPVVEFDLYKIMTGLLEEKHPRQKYEDLELRTSMTRVVNAVNSVLHIGQLMLNYIVDRIP-KKDAIILLTGVGKCYPLLRSHKILNDLHDRLADGIVLMLFPGRYEDGYLYLFGDVKDGNNYRAKPI-----
MCI9596747.1_Bacillota_bacterium            MLAVRLDKMEAAIRKPAFRQSSGKANEVNYWIFDYPPEAELRVRERIQHLKNKNGNDVFELVVFDLYDIIIDFLENRNFLDKCFDFEKRHGLDRITKAVNNSMKINDLIIQRIKEN--TPKNAIVFLTGIGKCYPILRSHKVLNNLHQAFVSVPVVLFFPGTYNEQELILFNEIKDDNYYRAFRLVK---
MCK6605710.1_Ignavibacteriaceae_bacterium   MISDKFDNLFNYILNSDLTAEAKTGGQLPFYISCYDIESEALIDQHINALYNRLTSSGKIITVIDLFEIVTGVLTMNNELQHLLHEESSYDYEERKEFFTEIHNPEKYVAPLIKNKADETNAQLILIKGISKIYPHLRTGHLLAVLERFISDMPVIVFFPGTYDGINLSLFGRFTD--SYRAFVLDALVN
MCK7578718.1_Chromatiales_bacterium         MTSDPTEHLFGILSHPDFLAMKGLANEVPIFIHTYEPRDEDATRRRVANLAARLKTMGIVLAVVDLFDLVLEQLEEEGRLHRIIEKEAAMGKQKLLELLNNLSDPKTRLVPRLIQRIGGNEIQLTLLTGVGRVFPFLRTHTLLESLQPAMMRHPVVLFFPGQYTQEQLRLFGSLSNRPYYRAFNLAHYRL
NLA16136.1_Bacteroidales_bacterium          MLSASINQAFEKMRNPKFLQKEGLGGEIPFFIFPYDIKYERQIDDTISGLKKRLNREGICVLELNLYTICIDILQSKGGIEKVLKLEERHDRPNFLKALQSALNIQEVFIPHIKNIVDTIEHHLIFITGVAPVFPYIRSHVILNNLHSVTDKVPTVMFYPGTYTGKTLNLFDRLKSNNYYRAYNLNEYQL
PZV04978.1_Cyanobium_sp.                    MLQQQLEPV---LTAADSAAGLSTYTDMPFGIFLYPPEDEWELRGELGDLVIRLENAGKTVQRLSMAVLLDQAIEAAGLMEEIAQGEADLGIAIVRDTIRDVLSGKAPLDQLVAEAVDPTSSHIVFLERVGALFGLHRPSALLDNLYRPVT-HPLVLFYPGQRDGPGLRFMDEAEADHNYRPKLFA----
RKX87881.1_Spirochaetota_bacterium          -TARQFENLFSIMSHEKFLNMEGLGNEVPFFIHAYDIREQNEIYEHIHLITRRLTTAGVETKLIGLYDMVLDILGETGSLEDVFDLEQESSKKELKETFSTLFNQDE-IKSYFTSKLREGDQQIVMIYQVGEVFPFLRTHDILNQLQSIVTDTPLVVFFPGEYVMSHLNLFGKFSGP-YYRAFRLDDYFA
RRF95307.1_Coriobacteriaceae_bacterium      MINDARGFIVDKLSDDALLGRNGYMMRQALYIIDYDPAQQQYAADLVRAICEKDPRRGVTPVVVNLYDLVLGYLDEQDLWEPLVEAEPDTPRLDLIQMLQDTVGVKDVVAPRVNEAIASPEADIAFVTGVGETFPYVRTHTLLEEISSP---IPVVLVFPGRYEQHALNILGLTQASTYYRATRVFD---
TVQ28582.1_Spirochaetaceae_bacterium        MTAKRFDRVYQIISHPRFLARQGLGNEVPYFIETYEPANEFAVTTEVRTIHERLLHNGIPAVLLPMYDIVIECLESDGRLAQVFEKEPAMGRQRFFAMLDAFTRPDAPVHDAIVRRLDAPDHKLVLMHQLGTVFPFLRTHTLLTNLHSTIYQVPLVAFFPGTYVSSYLSLFGTFKGD-YYRAFQLSDYIE
UCC65223.1_Anaerolineae_bacterium           MLEADFEKLRQRLGDP---DALNPAHSDPIFYFVYPPSQILTMRKLLPGWIARLRNEGLKVETLSLSEIMWELIDASGRWDDWLELEPEHDLDAVNEAIRDVLRAGNALVERVAERAASRENTVLFITDVELLHPYFRSRVIENYLNNKVL-IPTVFFYPGRRTGQGLHFLEFYPEDPGYRSTLIGGLE-
WP_004808450.1_Actinomycetaceae             MLQETLDIALRVMTSKRFLNREGLGNEVPYYVLRYRIEWDRSFDQSLRQMLSHL-NESVPTLHIDVYQLAIQIWRDCGYWDQILAQEAGMDRQDFADGLAQILDAERVLAPAIAERIQAPDSRVVILSGVHHLFPIMRAHRLLNCLQPLTGDVPVVLTFPGSYRQSALVLFDQVSEDNYYRAFDLLDQQY
WP_012411903.1_Nostoc_punctiforme           MINQRLDALLPKLQDARLLSNRGIGNEIGFYIFDYDAEYEPLVQRYVDRLKLQLSPIQLILLEIDLYKTILDILSERGVLKKAFELEATKGNAALAKTVAPLVRPDQ-VIAYIQKKLR-GDEQLVIMTGVGASWPLLRSHSILNNLHPVLDKIPLVMFFPGSYDGHELRLFDTFKDDNYYRAFPLIPHQQ
WP_013452129.1_Calditerrivibrio_nitroreducens MLTSKLNCLINILKDENYFRLEHIANEVPFYIFDYPDEAELDVRKTIKIISDS-DQLKYRIEVVNIYDFIISYLQEKDFLEKIFEMEQKLSPEKFLTRLKSIIDPEL-IINKIVQLNDEKNPDVFILTGIGNSYPIIRIHQIINRLQEKVCNKVVIVMYPGTFDVNKLILFNSLESKAFYRAFKICG---
WP_019974717.1_Empedobacter_brevis          MIKQDFDTIFKITSSEGFLKMEALNGEIPFWIAPYEVKLHNEYEKQIENLRKKLKTNSIQALVIDLFQLASEIIFEDFTVEQIEKLEKRKSKEQFKRALQATLNIHDIFIPKIKECLESNDYNLLILKNVGSLYPFMRSHTILNNLQSAIKTKPVLIFYPGAYTGQSLNLFGELIDDNYYRANNILKHKK
WP_029256830.1_Rhodococcus_erythropolis     MLTHEEKHVYDVLRSGRFLRMEGLSKEVPFFIYHFPPAWALDVDGMRDRVTKKLSDDGINVVEINLYNLAVELLNERGVWDRVLQAEPTMDKADFRDMLQGMLDPHDHLAPAIRARLDEPNTTVVFLTGIGEVFPFIRTHTVLENLQSVVVGRPMLAWFPGTYEFTQLRALNLGASDSYYRAKDILEQET
WP_048345182.1_Lactobacillaceae             MLEERFQHLKDKIQSPSFQHNQGLSNEVGYYIFDYPVTETLKMRTLLKRFLDSPMAAQLNIRCFNVFDIMIKEIKSFDYLNELEKVEERNGMDYLIEQVGNIMEITKAIVKYIQNRLP-KNESIIFITGLGEVYPLLRTHKILNAMTQVIDNYPVIFFYPGKYDSFSLQAFGEVKDQNYYRAFRID----
WP_094203898.1_Finegoldia_magna             MLNERLDNLKKIIQEPDFLEGKGLSNEVNINIFCYDPKEEMKIRYFVEKLKQDNSLKNCNILAFDLFELFIEVSNDRKLIHKYVEKEKKSGKEALLNSIKKVVSYDK-IIDKMDLSNKHMNNEVVIIYGVGRVHPFLKLHMILNEIQKV-TDLPIIALYPGKYDGLTVSLFGEFESSSYYRAFN------
WP_118234408.1_Fusobacterium_mortiferum     MINLRFRELVTKIKSDEFYNNRGLANEVPFYIFDYQPEDELSMRDNIKNLLLPADDVRLKAIEIDLFELLIESLNNDGIFESVFKMEERRGTKILYEKLKKSFNMEV-ICNYFKEK--SKDKNFVIITGVGKIFPLVRTHALLNNLQVIFNETKVLLFFPGEYTTTDLRVFG-FKDNHYYRAFRID----
WP_126029123.1_Bifidobacterium_callimiconis MIEREFESLFAIMRRPSFRTGSDTAGEPANYIYAYPPVKELEVERRTDQLANRLAETAPGVLIIDLYETAIQVLRDSRIFERVLRKERPRLIPDMFTQSLIDKLSPDKSIAEQYRQAREHKQGDIVFITGVGKVYPYIRTHILMERIQLVFEQRPVVLFFPGTYAKTTMRLFDRLESSNYYRAFSLNAMED
WP_135754989.1_Leptospira_bouyouniensis     MLLKRLNDLKKDILHPE-GIQVTQSQNYPFSIFIYPPADEFTVRAKFVEMIQDIKKENIEILEINLAHECLELLKQRDGIDEIIQKEKEFTFALVNDVFSPILEDENGISKSILNIMEEGRKGIVFITRAGFLYPFYRTSSLLKFLTNTRG-LSVVFLYPGTRTSESLSYMGVMSPDSNYRPRMY-----
WP_151792237.1_Acinetobacter_seifertii      MVDDRLSLIFERLNNPKFLENQSLSNDMPYWVFDYNPEDELIVRDYLAHKVPRFEKEGLNFHTIDLYDVMLSILDSRNLLERTYNLELSKGFDHLKKTMDSVLEQKK-VAKHIVDHLK-DNPSFIILHGVGNCFPYIKAPALLNALHGATGCTPVLLFFPGNYDGNMFETLNVVEPDPYYRAFRLVPEKT
WP_163588065.1_Ligilactobacillus_murinus    MIKDSLSILDETIKSEDFLKNRGLGNEVGYYIFDYDPKDELLVREYVRELCEQNTKDGYTIKVFNLYKIMIELLEEKKYLDIISKWEKQHDIMFVATRINRLLKMEEYFTKYIMDR--VEENSVVILTGIGEIYPLVRAHSVLNKMHLVYTKRPVIMMYPGEYDNKRLTILNTNRDANYYRAFRIH----
WP_168675170.1_Hymenobacter_artigasi        MIPEKAERLFQTISGSRFLKRELLGGDIHFFISTHAAEQQTEMRQAIAALIKRLDNTGIQVLEINLFKLALSVLDSEIGLPALFEFEEQESAAEFREALHSAMDMKQVLLPAIERHVQSATPQVYFLTGIGEVYPFIRSHSILNNLHHLVERAPLVAFFPGTYSGEQLKLFGLLADDNYYRAFNLDTFTE
WP_187771726.1_Phascolarctobacterium_faecium MLSERLGKIKNIITNSNFLNKTGNANEVSYFIFDYPPDKDIIVEDYIQRLATEIMEKDMQIKIFDIYGLIMDYLKDKKYIDKCILLEQKKGSAGLIKAIKDTLHLDSYIVNHIDENTI-QDKDIVFLVGVGRAFPIVRVRELLSVLNNLYTKCPIVLMLPGSYDGQHVTAFNKLKMENYYRAFKLVQ---
WP_200392182.1_Roseibacillus_ishigakijimensis MFDQKIDELIRRLGAPE--GTTNARQGDQVFNVHYPPHLWAELRTRLPKLMARLKDQSYAPRHCSLLALMEKIFGDNGKIALLRRSEQQSHQA-FTDTLHSILANKKPIVAAIEEEIQSKPNGLLLITDIEMAHPLLRVSAFEQVLQGRFA-VPTVFFYPGERGNINPSYLGVYPSDGNYRSTHIY----
WP_217359511.1_Anabaena_sp._UHCC_0204       NLDTAIAALKNDLLDPG-SPKISTMRNYRFAILQYDPREEYKLRDRIRKLTDELKSQGWNVLSVSLKSLFLNRLKKEDTLDSIIRTEKRLGINHLKDKISLYIEGADGIAQDVINLINKADHTLILLERAGSLYPFFRSSALLKHIDGKTNNLPVVLLYPGERRDLALSFMGELPADRDYRPRIYS----
WP_222635590.1_unclassified_Rhodococcus     MLTARFERAYRVMRSPSFLEMKGQIGEVPYFVLAYKPSDEPRVGAEHKRLVTRLRTDGLEVCDIDLWALANELWSATGQWELILEQETALPKAVFSEGVGNVISPKDYLSPRIARIVERPRPQVVLLSNVGRLYPLVRAHTVLNTLQPLLADLPIVLVFPGTYRQSSLVLFDRLTDDDYYRAFDLLELEN
WP_224137561.1_Candidatus_Kaistella_beijingensis MPNNINDKIFSLLSEEEFKDPDTGLLFFPVYIYTYNPEKEFQIREEIKELDRKLPSNNLNCLVLNIYNEFIDYLKHNIFLEQIFELEEIDALDFIAEKAQEKNDNDAHIGEKLKAYFDSPDRVYLLLYGFGSIFPYLRLSEFLKKTEEYVRNYKLIAFYPGELKNDHYSLFGIFGDENVYRANHLNQISQ
WP_235341611.1_Bifidobacterium_boum         MIDEDFQHLFDVMKQSSFRHSKGIAGEQPFYIYDYPPECELEVDRHIPWLMQKLDDYAPVVRRVDLYDVVLGILKRRKILDRVLKIEPRYSQDKFLGMLESILGADSQLPNAVVEAMHTVKPGIIFITGVGAVYPYVRAHTLLNTLQGMIEDIPVVLFYPGTFQSSTMTLFNCLEPDNYYRVMRIRDLNV
WP_243520940.1_Hymenobacter_monticola       MLNEAFTQLLDALNTRD--RTINPTGAAQFYYLVFPPEQALEVKRQLHKWKKRLTDQRYKTREYSFAKEMPGLAEKHPLVSRSLQLEAAKTAAEKFAASPKLLHQLTKLAADITARFG-SEKTVLLLTDLEALHPAVRIGTIENQLTQTLK-GPIIVFYPGQRTAKALKFLGFYPEDGNYRSTHFEATTA
WP_245747814.1_Methanolobus_profundi        MLEERLELLKEELQKEDFLKARGLGNEVPFWIFDYPPEKELLVRDTITKVIPALEKKSIQVLEIDLYDVCLEILEKKISAEKIGSFEGNKGSDELLKKLKLMLKPDI-LKNAIQQKMNEDGFQLIFLTGVGKAWPMVRSHSVLNNLQPLLGNIPLVTFYPGKYSGFDLSLFGKFRDANYYRAFRLINTDN
WP_251490388.1_Otoolea_muris                MIKERLDKVRALIQKPEFLEGRGLSNEVNIRIFCYEPENEMMVRHFVNQLETDP---DGHLKICNLYKTFLSVCGDMDITDAIPGMEEADGSAYLLEQLSFAIGNRE-FIDKIQYEPH-EPGDVLMLTGVGEVFPFMRIHTLLEALQPYFSDVPILVMYPGEFDGRHLKLFNRLKPNDYYRAFSVIE---
WP_258185035.1_unclassified_Bifidobacterium MIEQDFDRLFRYMSSKRFRNGEDTGGEPANFIYAYPPEKELEVRRRIPMLVNRLDDVAPAVLVIDLYEMAIGIMKDNDVFDSAIEQEPEMQEDDFYQALAGMLGPDNALRKCYQQAAASSEADMVFLTGMGQVYPYIRTHALLNCIQDVVPSTPVVVFFPGTYEKTVMRLFDRLDADNYYRAFSLNTMEG
consensus/100%                              .............................................................sh.................-....................h...h.........h.hh..h...as..+...h............h..hPG.........h........Yps........
consensus/95%                               M.......h...h................h..a...........h............hsh.......h...........E..........h..........h...h.........h.hh..h...aPhh+...hbp..........lhhaPG.......h.hh.......YRs..h.....
consensus/90%                               M....b..h...h..............sh.h..a.......h.......h...........hslh..h..hhp.p..hp.....E..........h......h.....h...h.........hhhl..h..haPhhR...hhp.hp.......s.lhhaPG.bp...h.hhs.......YRs..h.....
consensus/85%                               M....hp.l...h.p............p.shalh.a.s.....hp..h..h..p.......h..hslhp.h..hhppp..hp.h...E...s...h.p.h....h....h..h..........llhlpshu.haPhhR.p.llp.lp.......s.lhhaPGpbp...hphhs.h..p..YRu.l......
consensus/80%                               Mhppphp.lbp.l.p....p..s.ssp.shalhsY.sp.p..hppbh..h..p....s..h..hslap.hhphhppp..hc.hhp.E...s...h.p.l....hp.p..l...l.........pllhlpslG.haPhlR.p.lLp.lp..h...PllhhaPGpbp....lphhs.h..p..YRu.pl.....
consensus/75%                               Mhppphp.lbp.lpp.phbp.ps.ssphshalhsYssppp.bhcpbl..l..ph.p.sh.h..hslachhhphhppp..hc.hhp.E.p.s...h.p.lp.hhp.pp.l...l....h....p.pllhlsGlG.haPhlRspplLppLp..h...PllhhaPGphp.p.Lphhs.h.sps.YRAbpl.....
consensus/70%                               Mlppphpplbp.lps.phbp.ps.ss-hshaIhsYssppp.bhcpbl.pl.pcl.p.shpl..lsLachhlphlcpp..h-phhpbE.p.s...h.p.lp.hls.pp.lhs.l.p.h....p.pllhlsGlGphaPhlRs+slLppLps.hp..PlllhaPGpas.ppLplFsphpsssaYRAbpl.....
```

### ; Type 2 BREX systems PglW C-terminal iSTAND-AAA+ATPase

| Sequence | Alignment |
| --- | --- |
|  | ---1211------1221------1231------1241------1251------1261------1271------1281------1291------1301------1311------1321------1331------1341------1351------1361------1371------1381------1391------1401-- |
| | SAVTQRRITASTPEQKLRYRANAQLLESAARPGFRLLTVRYDDQRLALLTLTEEWRAEPVDVSILFLTSLRRLVEARPRPWDTILEADNAVPGSRDAMKFGEYTAAAWGEVEKELASNSRPLLLHDASVMARYGGTGILQRLADHARRGGRGLWLLSPLNDATAVPRLEDWTVALEDREEWIRLNHSWVVNDPEDAPAA |
|  | S1 S2 -----------HEH-Extension-------- S3 S4 S5 |
| | \|------HHHHH-HHHHHHHHHHHHHHHHHH---EEEEEEE-HHHHHHHHHHHHHH--EEEE\|HHHHHHHHHHHHHH----HHHHHHHHH---\|-HHHHHHHHHHHHHHHHHHHHHHHHH----EEEE-HHHHHHH-HHHHHHHHHHHHHH----EEEEEEE-----------HHHHHHHHH-EEEE-HHHHH--------- |
| | \|------HHHHH-HHHHHHHHHHHHHHHHHH---EEEEEEE-HHHHHHHHHHHHHH--EEEE\|HHHHHHHHHHHHHH----HHHHHHHHH---\|-HHHHHHHHHHHHHHHHHHHHHHHHH----EEEE-HHHHHHH-HHHHHHHHHHHHHH----EEEEEEE-----------HHHHHHHHH-EEEE-HHHHH--------- |
| | \|------HHHHH-HHHHHHHHHHHHHHHHHH---EEEEEEE-HHHHHHHHHHHHHH--EEEE\|HHHHHHHHHHHHHH----HHHHHHHHH---\|-HHHHHHHHHHHHHHHHHHHHHHHHH----EEEE-HHHHHHH-HHHHHHHHHHHHHH----EEEEEEE-----------HHHHHHHHH-EEEE-HHHHH--------- |
| WP_185095299.1_Streptomyces_sp. | SAVTQRRITASTPEQKLRYRANAQLLESAARPGFRLLTVRYDDQRLALLTLTEEWRAEPVDVSILFLTSLRRLVEARPRPWDTILEADNAVPGSRDAMKFGEYTAAAWGEVEKELASNSRPLLLHDASVMARYGGTGILQRLADHARRGGRGLWLLSPLNDATAVPRLEDWTVALEDREEWIRLNHSWVVNDPEDAPAA |
| WP_268746228.1_Dermacoccus_nishinomiyaensis | -----SRRPTHDVASSITGRFGVRLDDSVSSRSFLALGARSERLVEAVETLRRAYGAEVVDLSELLVDALKQQTVGGRLTWDHVRAADAAAPHTKAAQGLGQLVRRAWPQIENHLASAERPLLLTDAAPLARYGHMDMLARWSDLAR--SRPVWLLVPQVSGNHGAVLDGAPLPLAAPSQFVPLDSDWIDTHAAAAKEH |
| GII83978.1_Sphaerisporangium_siamense | -----GQTSTLGGVGRNLAAFEQRLTRSIAEHAFVTLAVDWRFHARAAAALAERFGVTPVDVTADLLAAMRREAEEHQVDWSLVLRSDRPDAAPQDAANLRTLVAMAAASLESRLEAETRPLLIVEAAPLARYDRLAVLERLADQAR--PAARWLLLPVE-QDGVPSIDGRPAP--GALGALRLSTTWITRPRDEAVS- |
| KEZ35702.1_Frankia_sp._CeD | VTSRGRWSTRVS---DEVLAADDRLTRSLVDGGWLVLSVPPRRLARAERCLAA---VTVVDAEQALLAGMREFCAQHRVQWSIVLAADAADRSSRDWANLSRVAQAGLAGVRASIEAAGPAVLITNAGVVARYDPAVVLDELRASVRMPVRTVWLLVPWADVDKQPLLDGAPVPQ-FGNQGLALSEEWIVRHESRGAA- |
| WP_030506725.1_Microbispora_rosea | -----SASRTAAPRQVVDSAAMQRLTGAAGRGGFLALNVDLRRAVLAVRVIPESLDVMPIDVGGEFLRALTDLASERDRPWSDVLRADERFSDGILPRGLAGYVREAWQRLATRLSQPSGTVLLHNAGVFARYPGGDLLVEIQQAARFGPRGVWLLCPTGTPKQRPILDGLVVEVIGENEWLELTAELLRSLDEKAA-- |
| RJQ78831.1_Pseudonocardiaceae_bacterium | ----SISLTTVGGVLTAREVVASKLTSAVQRGGFLALTLRGGELPGTAAMLAARFPVQPVDVDQEFLRTFRALADERGADWQKVRTLDLRLSTGEPTPGLASFARKAWDGVRSRVGAGETVLLLHHLSLLQRYYQWEFLTELQNDARYSPHGLWLLCPAENEKHDPHLDDRIVVECTDSEWAVLSKKFLNALRGQGAE- |
| WP_212516928.1_Actinospica_acidithermotolerans | -----ASGTGGAASGAVVDAAVKRLGEAVEDGGFRALTVSVRHLPGLSAAIERRFPLISVDGNAEFLACFRELAAEQGLEWERVLRVDARLQRGEAPRSYAEYVAEVASRLHAKWSSPRSTVLLHNAGLLARYWDIDLVVGLQQSARRAPRGLWLLCPAESARSSPNLDGRLVETVVAGEWIVLGSAAVEELKCQDGHE |
| WP_208577501.1_Micromonospora_tulbaghiae | ------TATTHTPLLTAADEASARLAAAIERRSFLAVLAPWRRLGPARRALLARLPLTEVDVTAILLDRLRAL----GFPWEAIVAADNGNPQDADFRSLMDLVHQVVPAIGEALAAVDGPALLTEAAPLARYGQLGLLQELADPTR--AAARLLLVPARRPE-PAVLDGVQLPLTSPGQSLWLSGDWIEAAGVASADH |
| MCP9485019.1_Candidatus_Gaiellasilicea_maunaloa | --RHVRRPTYEDPEVLDAAQFERRLETSTRRGGFLCLVGHDRSLDDIRDELARRFDAQHVSLDRLLIHHLRLAAGRRNVVWQTVVDADAATRDSRAWGNLQQLFVEAVERAEGELLAYEGTVLATDPGLLARYGQLEMIDRLRDRAS-SLSTLWLLVPDDNQSDRPAIDGQAIPIIGAGQWARVPSAWIENRHRGRAA- |
| WP_193120346.1_Nocardiopsis_coralli | ------HTGHREPGRAVSQETWERLGRAVERGGYRALKTWLPDTERSAEQIGNMPGVVPFDVSAEFVQAMDEVRQGKGLTWRHLTQADSSRPRPRP---FQTMLELALQRVGERVAAGHGVVLLHRATPLARYEQGALLGRLAEEARESPHGLWLLCPMRGPQSPALLDGEPAGIIADAEQVLLPRGFSAPE------- |
| WP_103561562.1_Actinomadura_rubteroloni | -ASGVRRGPHSDPALAAAARAEDQLVAAASRDGFRVLTVRMMLFSEALEELTGRFGGTEVSATRLFLDALHSLTAPGKRPWETILRADNAEPNEKSAHKFAEYATTALGRVEPHLASGNGPIILTDATVFAHYNAMDTLSRIADAAR-GKRGLWLVCPQSDPAREPRLGRRAVPYQPASEWVELNDHWARNRHRSERSA |
| WP_248824789.1_Frankia_sp._Ag45/Mut15 | ATPSVRQPTYDSPEQAAASLTEQRLAAAAEAGGFRALTVRRSRYLAARTELALRFAAAPVDVAAVFLDALHAQVDPRPRPWATILRADTAEPGSRAAGNLAEYVRTAWERAAPLLGGGPAPLLLHDAGIFGRYAGRDVLFALAERAR-GGRPVWVLCPTTEPTRPPRLDGALVQVATESEWVALPDAWTTNWHRSGKAS |
| WP_213171153.1_Natronoglycomyces_albus | -------TTGVGSGADEAARFTYRIDKAVESGGFRAIKTGVAETCAAAKWLAARPGVVALDVQAVFLRLLRQYVAERNKPWERVLAADTVPPPAP----LTNLLVEVWNRFETALTAEGRVLFLYNATPLGRYSGGKVLHELLSAARRRPHGVWLLCPTDALRHVAHLDGYNVGAIDSGEQIRFPAGVLKPRLDEDSGQ |
| WP_144820010.1_Micrococcus_luteus | -----TREATITLPSSELGAIGQRLLDSAAQRGYLVLGVPAHRMNRSRELLAARFGAEHVDLADLFLTRLRETVDRSPVSWEMALSADSAPPGTKAGRALQAWEAAALPQVVEHVSQAGRPVFLTGVAVLTRLGPLQPVARWADLGR--SRAVWVGVPQAPGNIGPVLDGVPVPTSSRSQFVRLEPEWIDTAARTSPA- |
| MBN9688999.1_Verrucomicrobiota_bacterium | --STATKGNGVTAEKLTANQFTDRLQRSIQTAGYLVLVSTTTNYLVVEQRLREGFPVAVCDIDTLLLNELKAQAESLKVKWEKILEADQAVETSTDWNKLNQLVGYVMPKLKAHLVAQDGPVLLTNIGLLARFNQIPILEQLREAIL-DPRSLWVLVPSDDQNRLPTLHGRPVPVATRGQWEQIPEVWLNLTTSATS-- |
| KAA0252897.1_Acidobacteriota_bacterium | -----APTTHVPTRDDPAAEFDRRLAQAAREGGYLVLLAPERELTRAREAIAARAGIEIRSLDRLFLDALKATAESKKVKWDAVLRADAAPADGPEGTKLRTLVGNALPRLEEALAASGDRLLLTEPGLLVRYRRLDVLDRLRDRVS--PGAAWVLVPADEQTDRPVLDGVAIPVLTPNQWARVPLAWIEAQATAGEVS |
| QQS02053.1_Austwickia_sp. | -----TATPTAGGAASLRDPVAARLAESVATRSFLALGVGALDLARVQDVLRERYDAHLLDVTGLLIDAMRAAAARHGVPWDVVLASDAATPGTRDRANLEQLVGMCLPEVWEAIHAKDAPVVLTDVAPLARYRHLGELSRWSDVAR--ERAVWVLAPQTPTSQGPVIDGQPLPLSSPGQYVRLDLRRLYASDDQNARA |
| MBK9387522.1_Planctomycetota_bacterium | SSLLPKAPLILSDEATDHRLFEKKLRHYAKDGGLLVLSTAPKSFVAAEQALVTFLALERCDLDALIVRAMHAAALANGIDWQTVLLADAPDASASDRANLHQLFHLALPEIEAALLAPGPAILVRKPGLLARYGALRIFALLKERAR--RPSLWFLLPSSQQSSVPRLFEEPLPI-SVDRHLRIPESWPRQQLSPRSTT |
| WP_090664774.1_Belnapia_rosea | -ATLIRHATEATPDVLAARAIEDKIVFAAEKGVFLALTVEPRRARDAETELLRRFPREVVSLERLMLHAMRTRAEERRIAWPKALTADAAARDSTDFKNLLRLAAEAAPRVAKEVLALRQPALLTRPGLIARYDLMTMLDEFSQASA--SPSLWLLIPQATPG-RPEIDGAILPVISAANWTRLTDPWLANAHRAGVPS |
| WP_136411732.1_Luteimonas_yindakuii | TSTLRRMTTADSPEAREARAFDATVRRALDAGRVVLVTADLRRVEDAVPELSQQYDLAAISLDRLLIDALKAQTSQIGGDWRVVRQADAAVQSSADWRRLTALVERAMPAVVAQLRDDPRPLLLQHLGLLVRYGHTGVIQTLRDAAS--APARLLLLPGDSHQ-PPMLDGVVLPVITPADWTHLPRSWLENRHRAAPTA |
| WP_150933330.1_Microbispora_cellulosiformans | ----ATRTTTGLPGLLGPTDAINRLVTVADRGGFLVVKARIKQAAALPELLSAELPVSCVDVTKAFIAALREVRDEWGVPWPIVLATDSPDAIPEDRVGLAEMTTEAFALVEDRVRAMEGVVLLHDATPLARYAGGELLARLADAARSSPHGLWLLAPMADPREPAQLDGQAIPVIPGNEELALSSDTVDALEDSSKAS |
| WP_090069475.1_Lentzea_flaviverrucosa | ---PSTSTGTMSPPLTAASDVDARLVEAARHGGVRIITVRRKRWERARTAVEIALGMPVIDVSVAFVAAVRQVARQLNIPFDRVLLADAAEPGSNDQLNLHRVVAQACELLEAEWSR--HPVPLDALTPLGRYPAGALLERLASRARYGPDALVLISPAGDETKVPRIGDHVIRVNTPEEWVIARSPWPSGVQPRSQAG |
| WP_183357094.1_Catenulispora_koreensis | -----KTHHTIPVGRKQPERLVRELRESVGARSFLAIGIPARQVERTTSDLLSTYGGVQVNVTDLLLGGLRSQAEEKSVPWDAVLAADAAESGTRAADGLAALVRQSISTIETAISEATQPVVVTDAAPLARYGHLRVIARLADVAR--SNAVWLIVPQG-PEPGAHLDGTPIPLSYSSQFLNLDNRVAVPAAAEETQ- |
| WP_257478501.1_Acidipropionibacterium_jensenii | -----ARDATFGVDLVEDSATTRRLEESVRYRSYLVLGAEPTRLHRLVRALGSRFHAPELDLSKCFLDALHRLSDDRRVPWPVLVAADAQPEASRAHIGLSKVIELAMPEVRSTLSDSPAPLVLTNVAPLVRCGYVKILRELSDLTR--GRAVWVVVPQYGLHVGPEVDDVRL-VTSPNQFVEIDRTWIDAETDAETE- |
| MBL9044566.1_Myxococcales_bacterium | TPGLARLTHPSGPEEEALALSAKLDRASREGGLLILSVLPQYADAAKQKLAQRFALSVYSIEEAILRHLKAFAHSKQIQWPVILLADAADRKSADFRKLQTAMSAALAQMVDEIRHAPSPLLLIRPGLLARYDRLDVIESIRVSVEPPPPGVWLLLPSDELDLGPRLDGQTVPVLTPAYWRAISPHFLRAALPAVRSS |
| MCP5560207.1_Verrucomicrobiaceae_bacterium | EKSSHRMRTVMQEDVATALLLEEKLSSCATRGSFLVLSVPPDRLARAESELLARFPVQRVDGDALFLRLLHAQAAEYEIPWDTILLADAAKEGTEDNQNLRRLVDMIKPQLRQALVSPAQTLLLVNPGLFARYGLMGMLAELRNDVR-APHGLWLLIPAVVQSALPKLNGQPVPVISEHQHISLPQAWLDNAHGKA--- |
| MBK6942988.1_Planctomycetota_bacterium | TRSTPRAATEISDEVAAARAFEERLARAQKDGVFLVLAITPSHALDAERELLRGFALERRSVDVLLLDAMRRHATANEVPWSVVLNADRARPDSVDGKNLRRLVRMAIADVEQQLATSTRTILLTQPGLLARFDCIDVLERLRDRVRVTLFGVWVLVPCDDQREPPVLNGVAVPVITRAQWARIPISWVKNRHRGSESR |
| WP_217637676.1_Actinopolyspora_saharensis | -----SRVGTVPAPSSRGDYLGATLAESISARSFLALGVAVDEADRAAHVLALRHGGTVLDVTGMLVDEMKTVAGKKGLPWDLVRGADAAQEGSRDAQGLHALLNKAFPAVGERLRAGAGPLILTELSPLARYGQLKMVARWSELAR--SRAVWVVLPQLSWMRGALVDSKPVQLGSNGQFVELDADWLATQEAETATR |
| TAF23825.1_Verrucomicrobiota_bacterium | -TIGSRRGTPVSEEVAEAQRLEDKLTRARAQGSYLVLTIDPKYLNDAANQL-ERIGAKRLNASELFLTAMKDKAADLGVPWEVVLEADAAEKGSADWTRLQELIRRSMPGVLEPLRNSNEDILLTDAGLFARYGRMNELDALRNDTT-QPHSLWLLVPGHGAATTPTLCGEAVPISNEAQFEQLTPQWARNHHRGGVA- |
| MBO0801654.1_Nocardiopsaceae_bacterium | -----RRKPSTATLLDETARASRRLEQASRRGGFVAVKAPLRDAARIGDVIAGMDGVTGVNVTREFVRLLRDEASKRPKLWETVLSADSADAPPGARRGFGELLRAVWGALEARIAGDGGIVLLHDATPLARYTGGELLARLAVAARDKPFGLWLLCPMENPKGSPLLDKSTVGVIPGMEQLDLPEGFGSGAESRAS-- |
| WP_238634082.1_unclassified_Halorhodospira | -----TRDTTDDEQGERAAEVERKVRRVASEGRFLALRIGPRHMVKAQRRLASGYGLQAYNLEGELLRAMRDVADERRVQWSKVLEADAAGPKGPDWKKLLQVVGEAVRRMRPHVIEPPQPLLLLNPGLLVRYGHIALLNQVRDACE--QPGVLLLLPGSSATTMPVIDDQALPIVHASEWLHADRHWLDAPANA---- |
| MBK9978482.1_Gemmatimonadota_bacterium | --TQTHTPNILDDAMDEARMLEERIAAARSDGGFLVLSTEPARFRRAELALAERHALTPMSLERLFVESMRKTAEANRVHWPLVLQADAAPHHDRAWRNLRLLAEKARPMMEATLAALVRPTLLLHPGLLARYDLLAVFEPLQDRSR--GPGVVLLLPSDGMSAMPVIDLVPLPMVYPSDWARLPREWFRLARSA---- |
| WP_130480151.1_Leptothrix_mobilis | STYLGRHATARLARLADASALEQRLQHHLQQGGLLSLTVEPRLARHAEAELLRRFGLQRLSLDALLRELRTQAYARRVDWATVLRADAAPAGSRDWTNLLRLVQLTLPALQQALLNSPAPLLVVHAGLLARYGLMRLVTELEDQSR--PPAAWLLLPGQRPG-VAEIDGTPVPLVHAQAVAALQPAWIGNEHRSAERP |
| MCO6436553.1_Phycisphaerae_bacterium | SGTPSRFPTQIPPEIAEARQFEERLRYAEKDGAFLALTAKPNLYERARQELLNRFDVRPLDLERVFLDSLRQAAEEVGADWNVVVKADAAGPNSTDWRNLNHLISKVVPKVEEAIVQDGKTVVIYHLNWLGRYDQVVMLSRVQQAVQ--DHGTWLLIPGSPQTEMPLLDGKAVPVITNNQWAAVPKSWCQNLHRTDEKA |
| WP_181762535.1_Pseudarthrobacter_sp. | -------TTHTKLSSTQILEIERQLAQSLTRKSAVTLAVRPRDFIRTSKALTQVFGLKLVDVSAEVVKAIRATAQSKNVDWDVVLRADAK-SSPSDHANLAQLVRMAVGPLWVELVDDPNPLLLTNIGPLLKFGFGAEVARLFDMSVPRPAARWVVAPKRGSANTPSSDGRVPPM-GPDGFVDISPGLASALSKRESTM |
| MBK9741405.1_Actinomycetia_bacterium | -------HVLTPPEVPPYDEIAARLQGSLRAGGYLTLAVDPRRVDQAAAVLARTYGLTRVDVTAELLAAAKEYAVEVGADWSTILDADKE-VGTRDRALLDSFVKAALDRRFPDILAATDPLLITDASILGRYGQQHWLADLADLAGNRPAARWLLVPHHASGGRPTLDAQPAPL-GADGYLVINAEFLAARQASEQSA |
| NMM16638.1_Cellulomonas_sp. | -------TNLTRVATEPADVVDGYLRASLTRSSALTLCVHPREHDRAVATLTTLYGVTPVDVSERLLTAVRALADRDGVDWSLVLRSDAE-PSGGDWENLRGLVQDALTEPWQQILATRRPLLLLNAAPLARYGLTAWLSDLLDQSQPRPAARWVLVPRRGHQAVPTLDGRPVPL-GADRWIDLPYDLSSLRPSSGVSA |
| WP_136709615.1_Agromyces_sp._H66 | -------LSRTLAKSLPASDLHARLLGSIGSNAALTLAVNPRRYTEARRYLTNRYDVTTIDLAAELVRRMHAVADERGAKWAVVTRADAS-ESSNDFKRVSQIARQAFEPYWRELSAEPNPLLLTNAAPIARYGLTEYLAEMFDLARPRPAARWLLVPHKAAKPTPDLDGRPIPL-GPDRWVSLPSDLAELFPSSSGVV |
| WP_210768105.1_Cellulomonas_humilata | -------GSSVHRGLVPASQVAAQLVSSLRTKSAVTLCAHPDAYVSTTRMLGRVFEIRTVDVGAELVRAARADAKVVGASWEFVLGADVE-PTSPLGSDLRVFLEPTVTAVWDRLMADTSPLLLTNAGPLLRYGFTSALAEMLDLATDRPAARWLLVPRPRAVAIPTLDGHAIPL-GADRPVDVPLDLESAVSGA---- |
| WP_067228577.1_Microterricola_viridarii | -------RSTTLQGSLPNDTLAPTLAESVQHRSALVLTTTWGRYDDSATALAARFGVEVVDLAALAVEQLHATAASVGADWLVVVDADKP-DSTIDKRNLQALAKQAMSSAWAEINDRDTALLFTNAAVLAHLGLVDLIARTIDLSTKRAAARWFLLPKHGAAAVPDLDGTPMPI-GAGKWIELPTTLAAVLPIPSSIA |
| WP_129336322.1_Cellulomonas_endophytica | -------LTSIATRTSTVPQVLAVLQRSVETRSGLLLAVTPERHEHASQQLAAAVGLVVLDVADALVREARRVADEDGADWSFVL--GADAE-PGSPLRSQLHEFLEAAVDAVWEQVVSTDAPLLLVHAGPLLRHGFEDRLDHLLDLATARRAARWVLAPRPSSSGVPLLDGHPVPV-GAGRWTDLPADPLPTLGAPS- |
| RKN35100.1_Streptomyces_hoynatensis | LTSASWGASDEDTAVAAAVKAEQRLAHSARQDGFRVLTVPQRRAEEAVRLLTEPYGAQPLSLTGLFLAELRGVVGEAARPWETVLKADAAETGSRAHQQLRIRTDKAWGRLEPELTADDGPLLLTDTAVLARYDAFDLLGRLAEEAR-GGRALWLLVVQSDPARAPRLAGQSVPYQAGDEWIVVPDAWVARRHLVSRAI |
| WP_262103914.1_Arthrobacter_sp._Marseille | -------STYTFIASGSVSELERRLRDSLDRKSALTLAAHPRDYLKAASTLAQRFGVKRIDIAAEVLRVIKSTARQADVSWELVLRSDSE-PSSGDFTNLTRLAQAAIRPMWGEMMQVPGPVLLTNIGPLIRYGFGPDVSKLLDLSHQRPAARWILAHRRGYDETPKVDGKPVPM-GPDGWIDMPRNLSDLLTTNDLNS |
| MBZ5659870.1_Acidobacteriia_bacterium | --SLHRFQTELPREMEEAVEFERRLHAAYRSPSYLVLATEPKYLEMARQNLAKHFPMTIFHCEREFLAALQKEAESKHIRWDVILRADTSKPGARDWDNLRKLAANAAHRVAEQVQSRTQSMLLIYPGLLGRYGQLSILDELADSLT---YSLWVLAGSEHQAASPMMDGQAIPA-RPTQWAWIPAKWLDNEFRKSTA- |
| MCB0954589.1_Ilumatobacter_sp. | GASGVRTGTAVAHEREEALAVHARLQRQEAEGGYLVMTVEPRWQRRAIE-LLRGYGPHTVDVDGWLVGAMRAYTDRRTIKWPAILGTDAAGPEGERWPRLLTVAKDAMAPLQDGLAQ--RHLLFVHPGLLGRYNLLGMLDQLRERTRPALRTVWVLVAADDPAATPTVAGRAVPVSTPAEHLVLPEPWLHNLHKTEGVA |
| MBL8842562.1_Planctomycetota_bacterium | DATSSRHTTEVDPEVVNAAEVERRLTLSHRRGGFLALSVLPRHYLAARAELRARFALDGLSFDAALLARMKRIAQEKGVDWQLVRATDAAGEGAAGWRNLVTLASLATGELLAELSAAGRPLLLTELGLLKRYEQIERLAALRERARRPLHGLWLLLAQSSPGPRPQIDGRLVPVLANGEWLALDSAWVKNRHRSVPRP |
| MBO9539711.1_bacterium | -STLHRLSTLPSDEILDARAFETRLERAVTDRRFLLLAIEPKYLLRAEAEITKRFEVHRVSFESLLVRKLRSAAQAVGAKWEVVVSADGTPRDSNAWRNLRGLVKRAMGEVEQELSTSSTPVLMVENALIARYDHMELLERLRDDCR-SAPGYLVLIASDDQSTAPKLERQAVPLITASEWVRIPHAWLENRHRSTPTP |
| WP_144981411.1_Gimesia_aquarii | SSIGSRRDTSTSPETVEARQTEDRLQHAYRDGGFLVLTVKPSYLRPCEHNLVHRFPLERVSFDDLLFEQLRAKSKEYEFPWSDLYEADSEGTNTDDWANLLHLMSEVGPNIETELMNSERPLLLVHPGLIARYQMMSVLQTLRDRVH--DPTVWVLIATDSQNEMPYLDGVQIPLISKGQRAAVSEYWIDNLHRGRSGS |
| WP_093484991.1_unclassified_Streptomyces | -LSGSGSGTPLDPE-STAARADAQLAGAARRDGLRVLTVRARNAPVVGRELRRQYGARAVSVTELFLKGLRGELTPGAMPWEVVVKADVL-PAGQRPTGFTNLVGAVWQAVQGVLAADPAPVLLTDAAVFARYDGMHVLDRLTALAR-GGRPLWVLVAQDDETAVPKLAGRPVPYLEAQEWIPIPKSWTRMHEPNAATT |
| WP_200346137.1_Halochromatium_glycolicum | -YDSSRFGSAALAAEQRQQAFDDRVRTALDEHRFLALSVRPADAQRAAQRLQARFGLVAISFDALLLRQLRRLCDARPPDWQVVLRADAAAPGSIDWSRLQGLVQRVLPELTAALGAAGAPVLLTDTGLIGRYGLVSWLAELRRRLE--GHALLLLIASDGTGPGARIEGVSVPDPGRREHAAIPSQWLQAAADG---- |
| WP_252180113.1_Endozoicomonas_sp._4G | ----SRTATVDTTTLEEVKRLEQEIATAIETSRFLALTVKPQLMVTARDKLCKDYPLKHISFDELLFHHLQELFNPKAPRWDVVLKADAAQKNSVDWGRLQMLVSKVLPAMANEIKSAGQPVLLTEPGLIARYDLVSWLNELRQHLNAEVHGMILLVAADAQRTAAVIDGTIIPRTGAAEFARIASVWLEASNDSQQA- |
| RLS37136.1_Planctomycetota_bacterium | SSSVGRWSTTLAPGLETALHFEDRLSRSRRQPNFLTLLVEPRGYGKARAALVQRFQAEIIDLEGFFLAALRHTATALGVEWELVLRADSH-RLPQEWMNLMALVRRAAPDFEERLMATDKALLLVQPGLLARYGLMDQLDKLVDQAR--RPAVWLLLAGD----RALIDGQAVPLIHAGLQVRVPLEWLENRHRAAELS |
| WP_241482799.1_Kocuria_polaris | -------VSTTQFAPTPAAEVARKLTESLERHSALTLTVAPKRYVAAARALASKFDIDVLDVAELVVSATREQIEKAGGIWSGLLGYDAL-DDSKQWKALEGVVQRSVEPVWGERLSAHRPLLLTNAGPLVRYGMTSLLATLLDTGTKRPAARWLLVAKAGDARAPLLEGRSVPL-SPSGWIDLPRDMAQLADDKTIQL |
| consensus/100% | ........................l............h............l.........p....hh...p.........a..h...D...........h.......................h...s.h.+......h............s.hh..........s.................h............... |
| consensus/95% | .......................pl.........h..l............l........hsh...hl..hp.........W..l..sD.....s.....h..h...s...h...h......hhh...s.h.+h.....l..h.........s.hllhs.......s.l....h..........h............... |
| consensus/90% | ........s...........h..pl..s....shb.l.........s...l.....h..hsh...hl..h+.........W..l..sD.....s.....h..hh..sh..h...l.....slll.p.s.h.Ra.....l..h.p.sp....s.hllhs.......s.l.s..l..........ls..h........... |
| consensus/85% | ........s...........h..pL..u..p.shbsl.s....h..s...L...hsh..hsh...hl..h+..s......W..lh.uDs....s.....L..hh..sh..h...l.....slll.p.u.lsRYs....l.ph.p.sp....u.hlLhs.......s.lss..lsh......h.ls..h........... |
| consensus/80% | ........s........s..h..cL..uhpp.uhlsLss..p.h..s...L...hsh..lshs..hl..h+..s......Wp.ll.ADs...ss.s...L..hh..sh..h...l.s...slLl.psu.lsRYs...hl.pl.p.sp....u.WlLhs.......s.l-u.slsh.ssspbh.ls..hh.......... |
| consensus/75% | .......ss..ss....s..h..+L.puhpp.uhlsLss..pph..u.p.L...ashp.lsls.bhlp.h+..s....h.Wp.ll.ADs...ss.s...L..hh..shs.lb..l.sss.slLL.psu.luRYs...hL.cL.p.sp..s.uhWlLhP..s....P.l-G.slPh.ssspah.ls..hhps........ |
| consensus/70% | .......ss..ss....s..hp.+L.puhpp.uaLsLss.spph..u.p.L..pashp.lsls.hhlp.h+..s....hsWp.lL.ADus..ss.s..pL.plhp.uhs.lb.pl.sss.slLLppsu.LuRYs...hLscL.-.sp..s.uhWlLhP.ps....P.lDG.slPl.ssspah.ls..ahps..p.s... |

```
# ; Type 3 BrxF iSTAND ATPase; Representative_1

                                         1---------11--------21--------31--------41--------51--------61--------71--------81--------91--------101-------111-------121-------131-------141-------151-
OrigSeq                                  MLSFDIDNIQKAIATAQSRYYKLVLAIPGTLLDGWIGMMAALGTTPVNLGLLLSQSLLEVEPRFRSLEFQDLFARIIRSAEGDLVAFDHAEILFQRDLQLDPLRMLLQHSRNKVLIVRWPGVFSGDKLTYGEPHHPEYRCYEQVDAVVCCMNQG

                                                            S1                S2  -----------HEH-Extension--------    S3                            S4                          S5
                                         -HHHHHHHHHHHHHHHHHH----EEEEE-------HHHHHHHH-EEEE|HHHHHHHHHH---HHHHHHHHHHHHHHHHHHH-|--EEEE--HHHHH-HHH---HHHHHHHH----EEEEE--|EEE--EEEE--------EEE--|--EEEE----
                                         -HHHHHHHHHHHHHHHHHH----EEEEE-------HHHHHHHH-EEEE|HHHHHHHHHH---HHHHHHHHHHHHHHHHHHH-|--EEEE--HHHHH-HHH---HHHHHHHH----EEEEE--|EEE--EEEE--------EEE--|--EEEE----
                                         -HHHHHHHHHHHHHHHHHH----EEEEE-------HHHHHHHH-EEEE|HHHHHHHHHH---HHHHHHHHHHHHHHHHHHH-|--EEEE--HHHHH-HHH---HHHHHHHH----EEEEE--|EEE--EEEE--------EEE--|--EEEE----
CEO89248.1_Syntrophaceticus_schinkii     MLSFDIDNIQKAIATAQSRYYKLVLAIPGTLLDGWIGMMAALGTTPVNLGLLLSQSLLEVEPRFRSLEFQDLFARIIRSAEGDLVAFDHAEILFQRDLQLDPLRMLLQHSRNKVLIVRWPGVFSGDKLTYGEPHHPEYRCYEQVDAVVCCMNQG
CEA16532.1_Fermentimonas_caenicola       -MSE----ISKQITHFAAPRYKFVVAPNERM---------LKDFSAFDLGYNLSLVLKENKNEELTFIVHEELTKLLNDNITDYICLSNIGILFEKELKINILQTFQRLSKNTLLILLWDGEVKGNTLFFLSENSKNKIDLREINHIIL-----
HHY32855.1_Bacillota_bacterium           -MR-----LHDLIKQMETNYYKMLVIVDNGHQPKIIGPLQAEGWTAVDVTDAVLKLIEGIPGEKVRVRIGGKLKEWFHSLP-DRLVLYNTSILYSPELRLNPVGAFKYKSREKEVVVLLEGRLACNRIQYSQYGRPDYAEMDVSEVIHADIDG-
KUO63975.1_Lutibacter_sp._BRH_c52        -MDKP---IEKHITTYSQSRNKVVLFKEN----------IFDFKPINVGHILSQALYNLDKDKLPMLVSKELNTILNASITRMLAISNLGILLEPDLKQDINSLFEKYSNNNVLFVQWEGELDHNNLYFLTKEKGIIINIKNLSHIAI-----
MBI3973658.1_Chloroflexota_bacterium     MMASEVHTISAAVDQLDPGYYACLVLVHYDVTHQGSAVLATYRWPHLSVGQELSEALLQEQPSRYPTAASKWLTDRVSSLSPGPVQVTAIDLLFGPALDLDPLILFRQASRTTRLVILWPGTYLNGVLSYAVPEHGHYRTWPKPDIQITTLQ--
MBI9032244.1_bacterium                   -M------LKDRLKEIQKLRNKLCLIPVNGVTKIIYTFNNTYNINSINIGFELSKVLKNELGKNLPIIAVESFRKIIEGSTSETIMVYNLGILLEDDLNLNLESILLEISKNIVVILLWEGEISKQGLHFLSIKHGKLLELTEKHYLQLEI---
CCX79045.1_Parabacteroides_johnsonii     -MDIT------AINKYSAPRYRLVVLKDEMP---------PDDIPKFDIGKSISAYIISEHSNDLKMLVNAEFMRLFREIASNAIALCNFGIVFEEKLGLDMRTILKDLSRNYLMFIQWEGVVKDDRLCFLTSDSSNFIDLSEISHLTI-----
MBK9455368.1_Bacteroidota_bacterium      -MENN-RHIDTYLALANSLRFKLWAIVGEDLIKAVIQYLQKHGYLLVDVGAELGSLYKELTNEEPSHDIGQKIKDWFLSKP-EKLILTNASILYHNAFKISPVGAFKYNSRNKSTVLFLDERIISNRISYGQAGSEEYYDKDINDILITDIEDN
MBL9119763.1_Phycisphaerae_bacterium     MLRGNLKDLCAAVLAAEERHTRLVLVVGTDEGALIAEFGLRLTTRVANLGLVVAERLINLPARHRPVTAADAAADFIRDAEPGPALVSNLQLLFLPELMLDPLKLLSDSARSRVVVASWPGERTGSCLRYAAPSHPEFREYHNPDCLVVEAPPP
MBO8173527.1_Bacillaceae_bacterium       -MYEEKQSVQFALEQLEGRRHRLLLLVSEDQRKVLNDVASKHQIPILNMNLSLSEELINVPVDRRPRKVGEIVEKILRHQKKDLVIISHIELLFDHSLHLNPIKLFEELSKTKTLIVGWNGKYENNVLSYADPGHSDYRSYYDCDAIIFSIK--
MBQ9466676.1_Muribaculaceae_bacterium    -MKPL---IDQRLTKFSAPRHKVVVLLNDSV---------GASLPAIDVGRELAAYLEAYTSHRAMDEAKQFLTKLIAQKTQPWVALRNLGILFERDLDISPANLIDNLSRNTLFIIQWPGLVNDGKLYFLDRGSRYFIDLSQISHITL-----
MBT3747343.1_Bacteroidota_bacterium      -MEIHISAISELIADMSDVREKYVIVCKNKHQPDFMHIAEQLNASYININLNLELSRRLLDIPKKKIPFKANRYLEDIINNSNKNIILLDRIEIILFDPLISIDPLYFIESNSKPVTMIVNWPGEFSSKILSYAKPVHPEYKSYPNISGRVISIK--
MCA6513726.1_Chitinophagaceae_bacterium  -MQ--IQTMSTPIKDITTERYKLYTLMCNSNELDKIAFVKSQNINVINIGKELANYIDSLDFRYLNIDVYDHTKKLLDKHRSDVVAIYNLGILMEPALELNAVQLFKEFSKSTSLIIIWDNQSENDRLDWQTQKQNFFLDFSDIQLKKLQYAI-
MCF1431077.1_Shewanella_sp.              -MEPG---LRQAIEQAQNYHSQLVVVAGESASLLLEHTASTDGLAVVNLGLELSERLIEVPRQDRAKSAASLFADLLHERNTRVLLLNHIEILFDKTLSIEPLKLLQSSAKNLTLVVAWPGEKTASSLTYAVPSHPEYRYYKASDFGETFVDAG
MCM1450902.1_Clostridium_sp.             -MAETSESVIADLLRFNGPRHRLVFVKDTNR---------TNSNIGVDVGWEVAKYLHDNSLGNIKAESQNFIKKLIYDNISKYIYLRNIGVLFEPELGLNVDMFLTNISRNTLLLLDWDGQMKYPYMYFLHTDSKHRIKIDTLNYMTL-----
NLI12135.1_Peptococcaceae_bacterium      -MSGNIYNIKKELAHIGYRRHQLLIICGVEHAKIIQDITRELAIPCINLSLKLSEELLEVPVVQRGRKVNQLVDKIVSESTGEILCFEHIELLFHPQLQQDPMRILENISRNKIIVVSWSGRYANGRLTYAEPEHPEYRVYNELEASVITID--
OJX40547.1_Chloroflexi_bacterium_44-23   -MP--------KLEIIFHELFDCLILVNPSIQIETANKIKEENISVINISQELGGFLLNISAMDRSRSTQNWMVDKLASHPHDPILCVCPDLLFDPTLQIDPFALFRQIARIKHIVVMWPGNFSSGILSYAVPEHHHFRTWKITDTIINQFPTS
PAD23514.1_Niallia_circulans             -MNTHLEQIENAISAVSEQYFKQIFVYDYINGNSVRTFAESHSLPYINVNVEISLLLQKVPINRRAFLITEVFQGLIGKYSDGVICLDYYELLFDNSLAIDPMILIKNNSRNKTLIISWRGKIIGDALIHAEPGHPEYKKFIVQDAVIIK----
PKO06211.1_Chloroflexi_bacterium         -MQAFQDELLVRLAIAPQQYHRLIIILNPDSRISKLDIAGKFGLRYINLGLILSQMLLDLTERQKALKMPQLVDQIIGCIEDKPAVLDHIEILFEPSLQQDPLRLLQGMSRNRTILAIWNGQVNNGYLTYAATEHPEYRKYSIRDLNILSLSDY
PZV08804.1_Leptolyngbya_sp.              -MGNYIAPLADLLAIAQGQYHQLVFATVEADSTYWRSVAVALDGHYLNLNLALSQRLLDIPQRQWPHKVMELIDAVVLSTDLRVTVLHCIEVLFDPQLRLDPLKSLQQIARRHNLLVVWPGTYDGRYLAYAEPWHPEYCRYPAKDLIVFPSA--
ROS80359.1_Muribaculaceae_bacterium      --------MRDKIAQYYDPRNKLIFAPSL-----------DQHLHSVDVGFELAKSIEGDSSPHLPMIAEDYLNAILRRSIKDYIALENWGILFEPALKLNLVSIFDSHSKSNTLILVNCGQADYENFHLVNKHFNTTFPIGNLHPYILQ----
TDP26837.1_Halanaerobium_congolense      -MT----HIDDLIKSNRGSHYKLLTLVGSEENHNIIDYLKNNGWNVYDIEEVILDLIEDIPENKIGLKIGDKIKEWLSDQE-NKIVLTNTSIIYSPELLIDPVETFRYAMRDKEAVIFIEGKMRDDKVIYSTPDKQDHKDIDISRIVSEEVEVN
TNF50602.1_Bacteroidota_bacterium        --------MKDKLLTYGSSRNKLVITSEK-----------IPGVHYLDVGETLSLSLANSPISWNSMKIKSEISKLFAESIESILSIENIGILLESEVQFDFKHFLEQYSRDMAIFLKWDGDHDESTLFFLSKDKGIKIPLTNLSYIKL-----
WP_061808754.1_Rossellomorea_vietnamensis -MNFS-EQLETLIEAIKGQRHQFLLLSFQANNKKIIDTLKEIRATTFNVNLLLANKLIELPISRRKRAVKDTLREALNEVENEVVYFERIEYLFDEELNQDPLKLFEYLSGNKVLLINWPGDVRCNGLSYATPDHPEYYQVDGYRDHLIVI---
WP_073205029.1_Prevotella_ruminicola     -M------MNSNLSSYKNPKYRLVFDLSS-----------SLEVESVDVGTYLADALQTKQKANLNMYASDAIRKMIRQHTHRYVALCNIGILFEEPALSFNLETIVEQTAKDELLIIQAKGGIANERFYFLSQQNNYSFSLKNLTYTIVE---
WP_075981269.1_Bacillus_massilionigeriensis -MS--LQQIKSEIETLDSRWYKLLIICGLKPYQ-DKDMANQLDIPFININLKLSEKLKDISKAKYALKVPEILQSILEQYEESYYWLGDIEILFDKQLQQNPVRLLENLGKRYSIIVTWPGQYANGQLTYATPEHPEFFSCHETDGKILEL---
WP_086717961.1_Bacillus_cereus_group     -MD--LYILEKEMALMHNWWHKLIFICASNELGIGEKIATNEDIYTVNLNLIMSETLVNISEEKYPLYVEEVTRETFDN-PFKIYLLQHIDVLFDPVIKINPIRLLENISKKSKLIVIWPGEYKNNQLVYAQEGHPEYFLSGSFEGKVILI---
WP_106450119.1_Trichococcus_alkaliphilus -LNYQLNQLDREVPYLKNKYHKLLYLYEYKKGDTLKYYSEANKVPYINVNLYLSENLADVKPNRRPYKVVDLM-NLLGDTEDDILCLDYFEILFEPSLMLQAFDIFKNLSKRKVVLVAWRDEIKDGYLIHAKPGHPEYTKELIADTTIIY----
WP_113032580.1_Paenibacillus_contaminans -MSGKLERLRAEMAGVRSRRYKLLFLLPETASAAIAGEMEREGIAAINVSLLLSDKLRHVAPDKRPYEVAKHVKQAINEYKADVVCLYHIEYLFDLELKQNPIKLLEALSGNIVLLVAWPGGAEQGVLYYADPEHPEYDSNGEYGSCILPY---
WP_118194774.1_Pedobacter_indicus        -MID----CKDKIVSFLIPRNRLIISTDFFD---------TEKVNWMNIGKLLAHELGNTDKSKIVFRINQLLDDVIVKNKIETIAIENIGLLLEPELKFNVEKFLSDYSQNNTLILKWEGEIDTNQLYFLTKNDNHKIDLNNISHIVI-----
WP_119865508.1_Clostridium_isatidis      -MEAEIYRMTDAVAAAKSKRNKLIIVNTKKVA--FEKAIAAIGLDCINLNLLLSEKLLEIPVSKRSRSVDSFIKDLIKSNNMNTLALSNYELLFLPELKQDPMRLFEDLSKERVILIVWEGRYENNSLYYAEPWHREYREYKDLDAIIVQGNI-
WP_155743288.1_Scytonema_sp._UIC_10036   -MGTSSSQISSYFDQVVERYYRLIVVVVPATRSNFCQVAESTNSRYVNINLELSGRLLELRQLHRATQVQQLLAEIIGNTNNGAVFLDRLEILFDESLKLKPLERLQKLARDRTIVAVWNGNIENNSLIYAKSGHPEYCRCPTDDCIVINL---
MBR1791000.1_Bacteroidaceae_bacterium    -MS-----LSEQVISFSSSHHHIVFCREM-----------MPELTFVNVGLVLSEKILEEDSPMLDFATDDFIQEIMSRHESNYLALTNIGILFEPCLKYNIPELFKSHSRNKILFIHTQAVIKDDVLYFNDPSDNITADLSGVHYIIV-----
WP_169099405.1_Bacillus_sp._DNRA2        -MSADVKKIEQELEGISTRYYKMILVCELSKGTAVKKAAESFGIPLLNISMLLSEKLKEYPSKRMAGRVHSLLSDILRNAGSSTLCLEHIEVLFDPTLKQDVIQLLQPLSRNYTLVVSWKGSYDGRRFVYAVPGHPEYYECKEFDGIIVTE---
WP_173978270.1_Magnetospirillum_sp._UT-4 -MQD-IGVVLQEITDVAGTYHRLLLLVGRVPAECIEAVAAHLDRPVLNLNAELSSALLEVPHRQRQLKAHKVVGDIIDGGPADIVLFDHTDILFSPSLRLDALALLKSLSRNKTVVATWSGTAKDGKLRHAEEWHPEYQSHAIEDIKVITVDQG
WP_181742022.1_Thermoactinomyces_mirandus MINTQVEAILTSLKEITRHRHQLIFLAGDCGKYDFRTLAEKIDAIYLNLSLELSHMLMEIPVHKRPRKVDICFREILTGYSQSLFLLDHIDVLFDSSLQLNPGELLENMSRKVVFVIPWYGQVTDSELIYADPGHLEHVRIPINEAMIIQI---
WP_214689525.1_Exiguobacterium_sp._s163  -MNLSVEKIQSSLPQVTNQYYKLMYVNEYLHGNTISYYAKNFETGYLNLGIELSLALLDIPLKRRPFKVTDIMENLIQKQHNKIICIDRIEVLFEPSLKTNPFQMLKQLSKNHTLIIAWRGIVSDQTLMYAKPNHPEYFKYTATEEWIL-----
consensus/100%                           ...........h.........h...................hsh...h...h.....................h.........h....hlh...h..p....h.........hh..........h......................
consensus/95%                            ...........h..h.....phh..................hsls..l...l....................h.........h.h....hlh...l.bp....h...u....hhh...s......h.a....p.........p........
consensus/90%                            .h......h...l..h.....phhhh...............hsls..lu..l.p.........h.p.h.phh.......hhh.p..lLhp..l.bs....hp..upp..hhh.h.u......h.a....p.........p........
consensus/85%                            .h......h...l..h...b.phlhh...............hsls..lu..l.p....pb...h.p.h.phh.p....p.hhl.ph.lLap..L.bs...hhp..u+p..hll.h.G.h....l.a....p..b.ph...p...h.....
consensus/80%                            .M......hp..l..h...b.+hlhh..............h..lslsb.Lu..Lbp....pbs..h.p.h.phl.p...p.lhlpph.lLac.pLpbs...hhpp.u+p..lll.W.Gphpss.L.ahp..p.pbbph...p...h.....
consensus/75%                            .M......lpp.l..hp..ba+llhl.s.............sh..lslsbbLup.Lbp.s.ppbsbbh.p.hpchl.p...p.lhlpph.lLF-.pLphsshphhpp.S+sp.lll.W.Gphpss.L.ahp..cscabph...p..hl.....
consensus/70%                            .Mp.....lpp.l..hps.ba+Llhl.s...........p.sh.hlNlshbLup.Lbphs.pcbsbbh.phhpcllpp...phlhlppl.lLF-spLphsshphhcphS+sp.lll.W.Gphpss.L.ahp..+scabph...c.bll.....

# ; Type 3 BrxF iSTAND+ATPase; Representative_2

                                              1---------11--------21--------31--------41--------51--------61--------71--------81--------91--------101-------111-------121-------131-------141-----
OrigSeq                                       MKDELKRARASRAAEYCSHNLVLVVLEPEVRDSQIEDVELINVSKSLSQRIVCLSANERIKRLDVEFEEVAQECNENIWLTKLDLLFEPSLKANPILLLKWLSKSRPVVAIWPGQIYEKTLVYAEPGSEDYRTYSLQELKDIQVVNAC

                                                                S1                S2  -----------HEH-Extension--------   S3                            S4                           S5
                                              -HHHHHHHHHHHHHHHHH---EEEEE--HHHH-------EEE|HHHHHHHHHH---HHHHHHHHHHHHHHHHHHH--|-EEEE-HHHHH-HHH---HHHHHHHHHHH--EEEEE-|EEE--EEEE--------EEEE|HHH--EEEE---
                                              -HHHHHHHHHHHHHHHHH---EEEEE--HHHH-------EEE|HHHHHHHHHH---HHHHHHHHHHHHHHHHHHH--|-EEEE-HHHHH-HHH---HHHHHHHHHHH--EEEEE-|EEE--EEEE--------EEEE|HHH--EEEE---
                                              -HHHHHHHHHHHHHHHHH---EEEEE--HHHH-------EEE|HHHHHHHHHH---HHHHHHHHHHHHHHHHHHH--|-EEEE-HHHHH-HHH---HHHHHHHHHHH--EEEEE-|EEE--EEEE--------EEEE|HHH--EEEE---
EGR2106511.1_Vibrio_cholerae                  MKDELKRARASRAAEYCSHNLVLVVLEPEVRDSQIEDVELINVSKSLSQRIVCLSANERIKRLDVEFEEVAQECNENIWLTKLDLLFEPSLKANPILLLKWLSKSRPVVAIWPGQIYEKTLVYAEPGSEDYRTYSLQELKDIQVVNAC
CCU85050.1_Mesotoga_infera                    MVKGDYSKNSLECLQAKEPGLVLLSGGNDPEVANEMNTEVLNLNFVLSKEMKTLSINERPLRASRFLSELVNSRE-SLLLDHIEILFEPQLKLLPIQILRRASRTTPIVANWPGSIENEFAIYAQRNHPEFMEVRLDSEIIFKIE---
HBR02122.1_Ruminiclostridium_sp.              MLGTLIRISEIIETIRKVTIPIIFCSLAPRLKKLINDYQELSLNKELAKALVEKDVIIRSRFVADEVMKIVSLIQGPILLTDYEMLFDPRYNIDVLKLFYELSRRAKMVIKWCGALDDNRLVYATPAYSDFHSYNIHDYDIICVI---
HCH5614386.1_Vibrio_parahaemolyticus          MTEQLLQQNACLEAEFRSFKLVLVVIPEDSREILNNSIGFVNLSKELARKLISLSISERVRQLETEVKIIANECNSSVWLTKLDVLFEPALANDPMRLLKGIAKSKAIVAIWPGEITETSVVYSKPGKPDYKTYPLVELNDIQVIDAC
HHV38844.1_Tepidimicrobium_sp.                -MGIDCNIKDLRIWNINNHYPVFSILQVDRIANEFKDFYYIALGEILSSSLLKYGKDDRKDYVIYEMDNIFHNMDSKLIIDHIDILFNPDYRIDILGYFIQLARNRRIIVIWPGEYVSGTLTYATPEYRNYKKYSIKDYNIICLI---
MBD9124683.1_Lachnospiraceae_bacterium        -MGRIIQYRLLSENQTGLMRPVVYCDEKYCES-----LHQVSLNEKIAALLIKIKPERRTMRIERCVQEVLTNIPEKSCIRDFDVLFNPAYKIDVLQLLTVANRSKSFSVLWPGTVIDGKLVYAEDGYADYKEYDVEQYDITCVV---
MBL0521045.1_Aeromonas_enteropelogenes        MAYELQANKALLRMSVQSSALVLVINSPTQIARQTEGLTYLELSEQLCERLLPLSCNDRSRHVSDIIAQMLNGIASVVWLDRIQVLFEPTLELDPLRQLQDLARLKPIVAIWPGQITERFLTFSVPGRDDYQSYSANDLANVPIITEQ
MBN2003265.1_Anaerolineae_bacterium           -DALALSQQRI-MAQIRSHALLVLVGMTDLVTALPDDSVYINVGLELAKHLRQ-NLPTGSLRAAQIMEELLESSGASLILDRLEILFALTLHLDVLRLLLDLNRRIPIAAIWPGEYRDARLTYAMPGHPEYREYEYPEAIFPITILAP
MBO2493598.1_Clostridia_bacterium             -GDNELSLRQVQGYTSNAGNIMVACVDKELVKAFFSNWKWINLGLVVSEQLCRLPKEQRKRMVADVVINAIKKTPSNIVVENIDLLFSPEYSLDVIRTLQLAGKNKRLLVIWAGDYQDGVLTYAQPDCPDFHRYEIKNYGVCCITR--
MBP3452736.1_Clostridia_bacterium             -MGAKIRRSDFRQYPAVYQGLFVFSDA----KRLPKDQPVINLGYVLSERLMGIPADRRCFEIDNCLRSIVETYT-AVSLSHIEILFTPSLKLDAIGVLLSLCRNRKICLSWPGRISGNTLTYAEPSSPEYYEADYTRFIDTYVITE-
MBP8682476.1_Trichococcus_sp.                 -MGSTIKQRDLAVIKQTRQGVFVLAPN----VTAPVNASVVNVGLLGERMFRVKTEGSSTSITRELDDILDEEHRDILLTNIDILFAPEYELDVIKLLLQLGRNQRLYLPWPGELVGERLTYSEPIRFDFKEYIVKNYVDTYVV---
MBR6410258.1_Clostridia_bacterium             --MYLLKSSELPENLKQSGVPLVFCSESPRICKRVESYIELKINIILSERLCRIPRDERAKLVEFKLNNLLNYYHAPVLISDFEMLFDPRYKIDILRFFCEKARIINVAIKWPGQYADGKLTYANPADPDYHEFDCNAYQIRIVQ---
MBX3742593.1_Akkermansiaceae_bacterium        MIAEKIAQVDIARASGEHFRLVWLAGGTPSELAEAEDGVVVDVGKILSSSFIEVPVSLRSASVEDCFTACLGESPSLTCLDHLEILFEPSLRVNPVALIKGASRHAVLVASWPGTTSTGHLFFGAADHPSHMDMSEHELESVLHL---
MCC5856368.1_Idiomarina_sp.                   MEEQLPSNEQIDKLALQSHQMLIVVGANTEQLTLLDNIPRLAISELLCEKLLPLASQRRVDEVEALTADVIASQATSVLLERIHLLFEPSLQLDVLRCLKALSKQRTLIIDWPGVYNGAALSFSMPGKPDYFNYSETELSSVPVLLTG
MCD8293624.1_Clostridia_bacterium             -MGTIIPAKIFPTDLTGLLKPIIYCCSYDGL-----DARALPINVPLARELLKYESAARTMLMKECFTNVVEEFPDDCTIKNFDVMFNPAYKIDVLGIMVSVCKIKPFNLIWSGTLDSNKLIYAENGFKDHTVYEIDDYDITCIAKK-
MCP5271842.1_Burkholderiaceae_bacterium       ----MLDLRLVNDAGALQSRLVLILGGDDRRAAAHFGTQSLPLGIDFARRISAVARRQRPLQASSLLRDSVDAAFPPALLDRTEILFDRTLQFHALVELKRLAHSRTVVAAWPGQLRDQRLIYAPANHPEHCDHPAAGFLVHHIS---
MCR4442916.1_Peptococcaceae_bacterium         -MGKEILLREASEYGTGKGPVFVACTNREVIKQLFPDWPWFNLGLCISQKVLSYKKEKRPNYIVDIIQEMVSASESRVIIDNIDLLFAPYYQLDVLKLFIQLGRNKKLIIMWDGAFRDGLLSSSVPELPDYHLYDIKNYDVYCLTK--
NMM63085.1_Clostridium_muellerianum           MMGATFKFSSMKVEIEKLMKPLILCVRQDKITSYISEYNVLSVNGLLSKNLLRYEFDKRSIFITDEFNKIIDNINDPILIKDFEILFDPAFQLDVLKLFIMANRKKRVTVLWCGKYKNGRLIFAEPEYEDYKSYNISDYDISCIV---
NPE22819.1_Fusibacter_sp._A1                  -MGQIIKYYDLSYFITSTNSLIVICNNKKLKKDIGLQYHHFSLNQELAKKMNEIEVEKKALFVIDILNEMFE-INENIVVSDFEMLFNPSYQLDPLKYFINLSNKRKIILEWCGDFDGENLTYATPNYLDYIAYKVDKHNITCVI---
QQS04458.1_Fibrobacterota_bacterium           ----MHSAQILSELKERHFKLVLLVGINPQEWANDCLTSVTQLGRALSPRLAELSVRSRPLEVSGALEELAKEVTTLLFLDGLEVLFDDSLKIDPFDTLKRLSLRRLVFAVWPGDYLRGRLTYAKVGHPEFRDCAPEGVLLYPRSMQ-
RKL62728.1_Thermoanaerobacteraceae_bacterium_SP2 --MLKALQNAIKILPHKYYKLILVVGPKERQMSLEFGYDYINLNLKLSEKLIRIPFDERCFYVEDYVNEIVESNHGLLILDNIEILFEKHLKIDPLLLLKNLSRYKACIAHWNGKVEDGHLIYAVPQHPDFVKYRMDGHDYIVIDTAQ
RRF92483.1_Coriobacteriaceae_bacterium        -MGALISRRQFSHKQLNYARPLVFCCDINGLT-----EDSVPLNVPLARRLAGIPAPSRSRRLEECLVDCLESLPNGVAIRDFDVLFNPSYEVDIVTLLVSAYRRHSFSVIWPGTLDGDELVYAEEGCADYRRYDINRYDITCIF---
WP_013778924.1_Tepidanaerobacter_acetatoxydans --MGLTDTNALMIFSHKFYQLLIAGDDSHKIAESGNYAYVNLNLALSERLMMIPSHEHSLFVNQCIDEIINGISATVIFDHIEILFEKPLNTDPMVLLKNISRHRKLISDWPGIIKDDILIYAKPGHREYVKYPVE-VDYTIIDIDQ
WP_013787752.1_Thermoanaerobacterium_xylanolyticum --MLLCERDALAILSNEYHKLIIVLTQLDDKISNKYAYDVLNVNLRLSEKLISVPISERWIYVDEYFKKLLNDYKTVIVLNNIEILFEPHLKVDPLRLLKSVSRYRKIIAIWTGNYTDGVLTYAEPSHPEYRRYKNDDLDFIMVCSNC
WP_028243927.1_Pseudobutyrivibrio_ruminis     -MANVMEWYKFLDNISGLMKPIIFCTSSEIEEL----SEAYSMNIPVAKELQKSGSSMRTMKMESILSSIISKLGDNPIIKDIDVMFNPDYQLDVLRTLINVSKRKSFMVVWPGTYKDGKLIYAQEGYSDYKSFDLNDYDITVVI---
WP_052235486.1_Alteromonas_marina             MMDDIFTQQSLSSGEMQSSCLIFLILEDSELSTLFSDALFLNTSKLLAERLITMSRRERERANVASALKSIVLAGDNMCYLQRIQLLFARELNINPVKLLQQLSRKKPVVAVWPGGLDCSNLYYAKPGHPEYVSYQLSEFQDVQVISTC
WP_071138484.1_Bacillota                      -MAEIYTKYNFLDLIKRNKLLGLLCMSADEFHIEVKDLSIINLGLDLSHKLKEYPMEYRNSKVLDELTNILAKAQTYIVVKNIDILFNPDYKLNILSYFINLSRSRLIFVEWPGRLKGRMLEYADINSPDYHKYNIDDYKIILIK---
WP_108775977.1_Lactimicrobium_massiliense     -MGVITRIDKLDDK---YMKPVIYTARNYASG-----IEQISMNRPLAERLIKLTTPQRGMRMGKVLLDCLKDQPDDVVITDIDVMFNPDYPIDVLQVLIDTCRVKAFQVLWPGSYESGALIYSVPDRPDYKKYLIENYDVTVII---
WP_164569834.1_Rhizobium_leguminosarum        MPAAAAAKQLLEATGVSGGKPVLLVGSEHDRLATLSNTSSINLNIHLVQALIDASDA-----RVDIISTISALEPSVLLLDRIQILMLPQLMVKATDVLCRVARRRSVCASWPGRLDRGRLLYAEPNHPEFLDEDASRALVIDLS---
WP_191750182.1_Clostridium_gallinarum         -MGDVIGIREVEMLIQQENKVIILCTNYKSIQSIFKEFKRISLGVILSERLLRYPIEIRNSILIQELDKLLLNTDDKLLIKDIDILFNPEYKIDILKYFYSLARGKKVAIEWNGSIKNNYLQYSEIGYLDYKRYSINKYDIICIK---
WP_243187470.1_Clostridium_celatum            -MEKVSKYKELSKINKLTNGLILITNSFDKAEIKIQDYKYISLGESLSKKLIKNDKVIRNRKVKLELDNIILNNDNSLVVDNIDILFNPEYKLNILKYFENVSKKKKIILVWPGNLDKNELVYSKLEFNDYKRYKIKDYNFILLK---
WP_271679096.1_Thermomonas_sp._2C3345         MWSVALEQSLMADTAQLQSKLILLVGGTPAELQRSQHATPLHLGVSMASALTELPQRERHLQAGNLLREIAQSHEPPLLIEGVELLFDASLKVNVLDLLKRQSQARPVVAIWPGEWRNGRLTYAVLGNPEYQDHAANGVIVFEI----
consensus/100%                                ......................h....................s..hs..h...........................hp..phhh...h.......h........h...WsG......h..u.....pa...............
consensus/95%                                 .........h.........hhh.................h.hs..hs..h...........h...h..hh........lp.hphhF...h..p.h...h...sp...h.h.WsG.h.....l.au.....pa...............
consensus/90%                                 .........h.........hlh.................h.ls..lup.h.......+...h...h.phh.......hhlpphclLF...hphs.h..h..hs+...h.h.WsG.h.....L.au.....-a.ph..p.h.....l....
consensus/85%                                 ........p.h.........hlhh...............lsls..luppl..h.....+...h...h.phh.......hhlpph-lLF...hpls.l..h..hs+.p.h.h.WsG.h.....L.au.....-a.ph.hp.h.....l....
consensus/80%                                 ........p.h.........llhhs.....b........lslsb.Luppl.phs...R..bh.p.hpphh.....s.lhlpph-lLFpsphplssl.hhb.hs+.+.hhh.WsGphps..L.Yu..s..-a.pa.hpph.....l....
consensus/75%                                 .....hpbp.h.........llhhs.....b........lslsb.LuccLhphs.pbRs.bl.p.lppllp.....s.lhlppl-lLFpPphplcslphhbphu+p+.lhh.WsGphpss.L.Yu.ss.s-a.paphpph.h..l....
consensus/70%                                 ....hhpbp.h.....p.....llhhs..p.b.....sh..lslsb.LuccLhphs.pbRsbbl.p.lppllpp...sslhlppl-lLFpPphplDslphhbplu+p+.lhh.WPGphpss.L.YApss.s-appYphpph.h..l....

# ; Type 3 BrxF iSTAND ATPase; Representative_3 (Walker A and Walker B intact)

                                                  1---------11--------21--------31--------41--------51--------61--------71--------81--------91--------101-------111-------121-------131-------141-------151-----
OrigSeq                                           MAASLSSEIHDVLLKTQDTYYKLILAVGPARTGKTAALTELAAKHKWPRINVNLCLSEKLLELTHRQRAVRVAGILNDTVRDETSNVVLLDNIELLFAPELAQDPLKLLQSLSRNRTIVAAWPGNFDGSSLTYSEPGHPEARRYLTPQAVIVKASETN

                                                                  S1                S2  -----------HEH-Extension--------   S3                            S4                          S5
                                                  ----HHHHHHHHHHHHHH-----EEEEEE-----HHHHHHHHHHHH---EEE|HHHHHHHHHH---HHHHHHHHHHHHHHHHHHHH-|--EEEEE--HHHH-HHH---HHHHHHHHHHH--EEEEEEE|EEE--EEEE--------EEEE-|-EEEEE-----
                                                  ----HHHHHHHHHHHHHH-----EEEEEE-----HHHHHHHHHHHH---EEE|HHHHHHHHHH---HHHHHHHHHHHHHHHHHHHH-|--EEEEE--HHHH-HHH---HHHHHHHHHHH--EEEEEEE|EEE--EEEE--------EEEE-|-EEEEE-----
                                                  ----HHHHHHHHHHHHHH-----EEEEEE-----HHHHHHHHHHHH---EEE|HHHHHHHHHH---HHHHHHHHHHHHHHHHHHHH-|--EEEEE--HHHH-HHH---HHHHHHHHHHH--EEEEEEE|EEE--EEEE--------EEEE-|-EEEEE-----
WP_063830324.1_Bradyrhizobium_murdochi            MAASLSSEIHDVLLKTQDTYYKLILAVGPARTGKTAALTELAAKHKWPRINVNLCLSEKLLELTHRQRAVRVAGILNDTVRDETSNVVLLDNIELLFAPELAQDPLKLLQSLSRNRTIVAAWPGNFDGSSLTYSEPGHPEARRYLTQAVIVKASETN
OQZ06365.1_Planctomycetes_bacterium_UTPLA1        MVQSLLNRLSQLVEDATSQYARLVLIVGQHGTGKSAVLREFAHRGGHPVFDLGADLPQGIFEVPARRRPVAASDRVGEQLR--NDDVAFVDNIEILFQPGLNLDPLKLLQDNARNRVIVATWPGTFDGRDLTFAEPGHPEFRLYPAAEVRILTL----
WP_211295643.1_Planifilum_fimeticola              MDRTAFQALMEAVRQMQTRRHRLILVVGPSGSGKTALLRAAAKILAAPLLNINLRVSARLHEVPRSRHPYEVGRIVRHMIADTASGTVFLDNLEMLFAPELQLDPLLLLENESRHRLLVASWAGRLEGEVLVYAEPDHPEYRRYTNFHAHLISVGGKV
WP_254950516.1_unclassified_Cyanobium             M-RDLARAIAEQAELAAKSFYRLVVVAGPPGSGKTIAFNELQVARGWSLLNLNKVLSERLLDLTSKQRRLRTADIVRDLIEEQNQTVTMLDNIGLLFHPALQQEPMLALQRASRNHTVIAAWQGEVVAGKLIYGSPDHPEYRRLDVKDVQIVTANGHC
NMD20860.1_Verrucomicrobiota_bacterium            -TDVSFPQLLDTIRATQGGYYKLTILAGSARAGKTDLLKQVAAQLDLPLINLSLHLSQRLLSQNRRQRVLNAEDVATEVIDENYKSGLCLDNTELLFDSTLQLNPLGFLQGVSRNRLIVATWNGILAGGKLRFGHTGHPDFFRQPVNGYPVVSVAEDK
MBL9185852.1_Opitutaceae_bacterium                -SSGFPEKIDETIRHARSGRHRLVLVIGDAGSGKSAFLQSIAAARAMQFLSLGEPLGIRLLETSPRTRPLVIEDTVRQFID--TRAGVCIDGTDILFNPALRCDPLRLACSISQNTFVAFTLTGRIEGKRFVHGYPDHPEFYSEEVPSVPIITLDRSE
MBA4157056.1_Gemmatimonadota_bacterium            -SPSAMDRLLPLLAAAPRQFYRLVLVVGASGAGKTATLRALGEHLGVGVVNLGLRLSERLLDVPVRYRPVEVLGLLDDILATSGDLPVLLDNTEILFELALEQQPLQVLQRLSRNRTVVATWSGVVQGGRLTYAPVGHPARREYAVDNLLLISLPGE-
ACD82385.1_Methylacidiphilum_infernorum_V4        MAEPLVDQVLRKIREARELYHRLILVVCPAGSGKTGALWEVSASTSAPLVNVNLELSRRMLDLAKSQRARKLQRILGEIMGEATGELVLLDNIEILFDVHLEHDPLRLLQRLSRNKTIVAAWNGSIVDGYVNYAVPDHREYRRYPIRDFMVACPEMTT
SCX35221.1_Agrobacterium_rosae                    MPEPIQDKIRDAIKQSEGLYHRLVLLVGENGSGKTTAIRDAAHALDADVINVNLALSTKLLELTAKQRALRLAELLGEVVENS-APVALLDNTEILFDHNLQQDPLRLLQSMSRNRCVVVSWNGTIRDNKLTYAEAGHPEYRRYDLTETLVVEIRQSI
GGE39031.1_Primorskyibacter_flagellatus           ----MIDRLDRVVEDISTVHSKLVLLIGEPGAGKSKLLCQLAERRAAPILNVGAALGRELLTVPRARRHLHVAEIFRRLADDLSQRILTVDNIELLFDRTLKLSPLDLLRRQAQSRRIIAVWPGGMRDNRLTYAETGHPEHQDYAIDGLVPFIVH---
MBI1186275.1_Alphaproteobacteria_bacterium        ----MLDDLVRLVEDVAGLNSKLVLLIGPPRSGKSRLLEQLSMRRKVPVVRLGEALGRLLAPLPASQRHLMAPDLLSGLAGGAGRDLLLLDNIEILFDRTLRLSPLDLMRRHSHGRRVVAAWPGELRDGRLVYAVPGHPEHQNYGTEGWVPCSVC---
MBK6654764.1_Zoogloea_sp.                         ----MISRLETLTSEISALHSRLILLIGPPRSGKTALLNSFGERMGMVPLNIGSELGRQLMPVPHKQRHLQAGNLLRELADIHTQDLLLIDNIELLFDPTLQLSPLDLLKKQAHTKKVVAVWPGELRGSRLTYAEMGHPEQRDCATNGVVLFEIQQ--
MBQ0714976.1_Paraperlucidibaca_sp.                M---TIERALHLLTEVGNTNTQLILIVGSRGAGKSAFLRECSLKLGIQHRNVSYELSRQLGFASCSERSFLVGDMLRDLAEKNRNIPLLLDNIELLFEPSLKISPLDLLKRLSHSRPILAVWPGLLQGENLTYADMTHPEYRNYSLDGITILKIKNS-
MBS3911805.1_Hydrogenophaga_sp.                   --MLMLTKLERYVDQLGGLHTKLIVLAGPRGSGKTKLLQELGAKLGVKPLNVNLELGRRLSATPHAARGFSAGQLLRDIADKQKEEVLLLDNIELLFERGLQINPLELIKRLAHSKRVVAVWPGELRGNRLYYAEMTHPEYRDYDADGVVALEI----
NMB01454.1_Bacillota_bacterium                    -TNLLIPALQRAVTSIRSNRVRLILLVASSGKGKTKILQKYAVENDFPLLSLGLLITKAIQEGILPEKLII---YLRQEFE--LGETVLLDNIEVLFQPSLRLQVLQLLKLLSRERTLIATFPGRMIDGSLVYAEASYPEYKQYEIKDFVILDMNGEL
QCB54648.1_Sphingopyxis_sp._PAMC25046             MERRMLEHLQQLVADAAQLNSKLVLVVANSGGGKTALLAEFAKTHDLQIINVGSELSNLLLPIPSSQRSFEVLGVLRGMLSHHDRGVVVLDNIEVMFDAELKINPFDTIKRLAHAMTVVVAWPGDFKDGRLLYAKVGHPEHCDVAVDGAIVLELN---
WP_198322149.1_Methylococcus                      ----MLSRLDSLVDETGALQSKLILLVGRPGCGKTRLLAMLANRRGAKVMNVGAALGKRLAALPLRQRALQANVVMRELAQSYSGDLLLLDNIELLFDQMLKLDPLDLLKRHAKALRVVAVWPGELRDGRLIYAEMGHPEYQDYGLDGLVPFEIDKFA
WP_202949433.1_Pseudomonas_sp._GM55               MRMEVVEELEKLLADVPALHNQLILLIGPPGSGKTAALLGLAKRLGVMLLNVGSALGTRLATVPGKARSLEATHLLREMADEHRNSVLLLDNIELLFDRHLRLDPLDLLKRQAQSRPVVAAWSGALHDGRLTYAHMGHPEYQDYAAIGLVHLQIDA--
WP_210246760.1_Bradyrhizobium_sp._UNPA324         MSEETLGRLERLVDEIESLNCKLVLVIGRSDAGKTELLAELAKRRQTSSLSVGAALGKALLSVPASRRHLQAPDLLREMTGNSANGLILLDDIEVLFDRTLRLNPLDLLRRNAQTRRVVASWPGELQNNRLSYAAMGHPEHQDYDAEGIVAFRM----
QVL49523.1_Thiocapsa_sp.                          ----MLTRMSALVDDVASLQNKLLILVGPPGCGKTALLDAYAEHRGVERRPLGSALGRRLAALPRTQRHLLAGSLLREIAETTTASPLLVDNIELLFDASLKLNPLALLKRLAQARVIVAVWPGELRHPRLTYAEPGHPEYRDDALDGVVTLDLQD--
WP_210879911.1_Roseovarius_autotrophicus          ----MIERLEKLVDDITSLNSKLVLLIGPPRSGKTALLVQLSQHRNVPVLNVGAALGQQLLMVPSTRRHLHAVDLMKDLTDETREGLLLLDNIELLFDRTLLLNPLDFLKRHAHARRVICVWPGEVRDGRLSYGRAGHPEYQDYGINGLVPFEISDGG
PSR33880.1_Sulfobacillus_benefaciens              MMRDSWETARHFVSQIQELPQPLVIIAGARKTGKTPLAKALATQGLGTYINLSLLLSRYLATDPDADDNLTASTMVRLLRDISPTGPMIVDNIEAVFQPELRLSPLRWFRQIARDMPLVVVWPGAVVEGEFRYSMPNRPDYFYSREPSLVVVNLETEF
HBM74931.1_Clostridiaceae_bacterium               --MPVVEDIERIVKLFPQQYYKLLLLIGPSGSGKTFVLKKCMQVLMCKYINLNLELSRKLKEVPVVERCYYVQDYIDEIVKNNDSDIIIFDDIEILFSKHLKIDPLSTLKNISKYKKFIAAWPGYIKGGYLVYAEEGHEDYIRYNLNELECSYLNLGR
KUJ90861.1_Thermoanaerobacter_thermocopriae       MDMSIYKDIQDAFILLPYKYYKLILLVGPIGSGKTQILKKISSDYNYDYINLNLKLSEKLINIPFEERCYYVDDFMEEILSQSKTDVTILDNIEILFAPHLKIDPLKMLKNMSKYKKIITSWGGSYINGILTYAEPGYPEYRRYKKDDLDIIVINVEG
PKO02178.1_Chloroflexi_bacterium_HGW-Chloroflexi-5 -KSAIKGDLITAIQTVQSDRHKLIILLGKFGSGKTAILKQTAVEVRGNYLNLNMELSEKLLGIPTSQYLTTVHAIIDQLCEDAHQEPLLVDNLEILFSPELKINPIDTFKRIARERPVVIALPAYRQGSSAIYSSSGHEDYVVMPLEEYIVIEIPEE-
WP_026486464.1_Caldanaerobius_polysaccharolyticus -ELIRKEDIDEALKKLKYRYYKLILVVGPMGSGKTDFIKSISLKYAFDYINLNLELSEKLINIPVDERCYYVEDFIDEILAGKRTEILAIDNIVILFGPHLKLEPLRLFKNMSKYRGMIVAWGGDLDDGYLTYARPGHLEFRKYSQNEIECMVIKLGG
WP_051534279.1_Desulfitibacter_alkalitolerans     --MATKDEILGCINKLDSHYYKLILLVGKTGSGKTRMLKALAAATDGAVADLNVELSKMLKNLPKMERCHYVQDFLDVVLRQSSGQPLFLDNTEILFSSHLKVDPIGMLKNASRYQPLVVTINGVIDNNKLTYAEPNHPDFIVYKVSELGCRYYSFRG
WP_053959406.1_Sulfobacillus_thermosulfidooxidans -MQESWDVISQQVKQLPTQLHPFYVIAGKPGTGKTAMARRIADEGLGVYVNLSFALAQYLLRQQDID-LVSVHDLLMCCEEIYSSGLFVFDNIELLMTLKLQVVPL--FQEFSRYHPAVVIWPGTVEDGIFQYSMPNLDDYYYFRDMSVLVTTLEHKS
WP_241751529.1_Bacillus_cereus_group              -DIVKEEKLLKAIKGASNDHFKLIVISGRFKTGKTSLMKRIANDFNYMYLNLNLIMSKRLLQIKEEAYVTQSQDIIKGIFDEIDDKILFVDNIELLFSKEVSLNPVEVFKNLARDKVIILSLPGRVKGDRIEYSTIDRNDYKLMDVFGLTVINMEE--
consensus/100%                                    ........h............h.lh.s....GKo..h...........ls..hs..h.....................hDs..hhh....l...sh..hp..up...hh..h.u......h.au..s.......................
consensus/95%                                     ........h...h.........Lhllhu...sGKo.hh..h.........sls..ls..l...........................hhDs..hlF...L..pPh..hp..up...hlhshsG.h.....h.au..s..-.......h.....h....
consensus/90%                                     ........h...h........bLlllhu...sGKo.hh..hu........hsls..lup.l...............hh...........hhlDs.-lLF...Lp.pPl..hpp.up...hlhshsG.h.s..h.au..s..-.b.......h.....h....
consensus/85%                                     ......ppl.p.l..h.....+LlllhG...sGKo.hl..hu.......hsls..LupbL...s..p+.h.h..hhp.hh.......hhlDN.ElLFp..LpbsPl.hhpp.up.b.llhsasG.h.ss.l.Yu..s+s-ab.h....h..h.h....
consensus/80%                                     ......ppl.phlp.h.....+LlllhG..ssGKT.hl..hu.......lsls..LupbL...s..p+.h.h..hlpphh......hhlDNlElLFp..LphsPL.hhpp.u+.+.llhsWsG.h.ss.l.Yu..sHs-ab.h....h..hph....
consensus/75%                                     .....hpcl.phlpph.....+LlLllG..ssGKT.hLpphu.p.sh.hlNlsh.LupbLh.ls.ppR.h.h.shlcphhp...p..lhlDNlElLFs.pLplsPLphlpp.u+.+.llhsWsG.lpssbL.Yup.sHP-abphs...h.hhplp...
consensus/70%                                     ....hhpcl.phlpph.s...+LlLllG.sGuGKT.hLpphu.p.sh.hlNlsh.LupbLh.ls.ppR.h.h.shlcphhpp.pp..lhlDNlElLFs.pLplsPLphlppbu+s+.llhsWsG.lpssbL.YupssHPEabphshpsh.hhplp...
```

### ; Type 4 BREX systems PglZ N-terminal iSTAND-AAA+ATPase; Representative_1

```
                                                   1---------11--------21--------31--------41--------51--------61--------71--------81--------91--------101-------111-------121------131------
OrigSeq                                            MRYDAESCFNEIIAYFNSEVGYPFIANIDDATILQDICSKMQADSKKIIRVSDYCNGDNLPNESELLTHVASADNGVVLGLSSYYMLRGEQALKKSVSTLLQMSVHGHVVIIIYGCSSILSNISADGRPDHRTVILEK

                                                                       S1                   S2 -----HEH-Extension----   S3                                S4                   S5
                                                   ----HHHHHHHHHHHHH------EEEEE--HHHHHHHHHHHHHH--EEEEHHHH--------HHHHHHHHHH--EEEE-HHHHHHHHH-HHHHHHHHHHHHH-----EEEEEE--HHHHHHHH---------EEEE--
                                                   ----HHHHHHHHHHHHH------EEEEE--HHHHHHHHHHHHHH--EEEEHHHH--------HHHHHHHHHH--EEEE-HHHHHHHHH-HHHHHHHHHHHHH-----EEEEEE--HHHHHHHH---------EEEE--
                                                   ----HHHHHHHHHHHHH------EEEEE--HHHHHHHHHHHHHH--EEEEHHHH--------HHHHHHHHHH--EEEE-HHHHHHHHH-HHHHHHHHHHHHH-----EEEEEE--HHHHHHHH---------EEEE--
RGH34270.1_Ruminococcus_sp._AM47-2BH               MRYDAESCFNEIIAYFNSEVGYPFIANIDDATILQDICSKMQADSKKIIRVSDYCNGDNLPNESELLTHVASADNGVVLGLSSYYMLRGEQALKKSVSTLLQMSVHGHVVIIIYGCSSILSNISADGRPDHRTVILEK
WP_073324030.1_Fibrobacter_sp._UWOV1               --MNLESCVLEAKSYRKSAQSYPLWIDFANAEDMRSFMENFLA--DKKISVEKYCNEDSMPRFENLYSDFRMEDFAFICNLSGFLKLLGSRKTAEVLSTLLSIG--SKKIILTVQCGVDFRI--TDPRLKCQILSVEG
WP_216471320.1_Butyricicoccus_intestinisimiae      MRFSIEKCIQAIDRYLVRSASTLRLVNVDNPAQMNQLIERYRVKGNEFLTVAQYSKPDEKAQTEEMLYAISKMGNLFVSGFTTFWKLEGEAEVKKQLDRLLHMSVKGHVVVLCYQCEHLLNF--ADARIERLIYFVDG
WP_044080180.1_Prevotella_pectinovora              --MKYEEVIRRIDKYKDSDKHQPIIVDLPNVQVYNQMVKHYDVGSNVFKDVSSFCNGEGLPVMDKLQYCLSIGGVVFLTGLSFYLKLQGEVSLRSCLRQMLDLSCSGKLVILTLNCSCVLNH---DSRLSGRISIVDG
MCI8555501.1_Clostridia_bacterium                  --MTIQE-VNYVKEYLFEGDNETCILNIESLDMYNNIVKAIKELNINIVRLSSFCNDNVSPNMENFYNALEINQPTAVIGISQFFKFIGKDRLKNEISVALSKTLKNKALFITFQCFDVLKELSQDIRLHNRITILGE
MBD8953983.1_Dialister_sp.                         MENTMDQCFDRIEAYCQKEIGHPLLVNTENAHDFQCLVKRMQLDGKTCVFVHDYC-GKGLPQIDALFRDVKKKGCFVMIGLSSYVMLLGKKRAEELMARLQELSVAGHVIVLLSHMKEIRLSMQQDQHLSYRVLLLEG
MZQ76087.1_Peptoclostridium_sp.                    MT-----AIDCIKKFLKQSNSKFLIVNINNVEDYMNIKSSLDIE-LSTIKLSNFCEYDDMPDIEKFVKCCRTDKSSLVLGVSQYLKLKNKESFEMWTQVLLSSMTLAKVIMIYFDCENKLKELDRDVRFQNQIILVKG
HIS35238.1_Candidatus_Scatousia_excrementigallinarum --MQLNSCIEKIEKHIHRGSSKPIFINLNNSQDVKELIDHFNVGGNKIVNVASNDQIDELPTLDAILGLIHRDETIFITGISSFARLMGQSQFASILNQIINSQSVSKAIVLLYQCEDILHDISKDPRIDSHIFFIDG
WP_103226892.1_Clostridium_sp._chh4-2              ----MNDCIERINNYLQDENTYPLFVNVENATQKMEILDYYRVGDNKIVHIADFAKKDSLPNVMGVIDDIKASGNMFVFELTTYLKLISGSELQKTMQTLCNMSIAKKAVIVCFHCGDILKKIQKDLRVAKRIVAVDS
NBJ16967.1_Dehalobacter_sp._4CP                    MYSGWDRTLNEVTKYLSNTSNAP-FFLVVGDGNYESIKTQLSEAGLKSVKVSELCNTDKRPNLDSLIGNVDPDKKIVILGLGEYLALCGEEAAFDRLSKIKDMKVNARVVILLRGVTSVVLRQNADPRFDGRRVLFTD
CDC05526.1_Clostridium_leptum_CAG:27               M------LCKEVQNYIATAKGLP-FFYVVGDEDYATVLDELRQANVSIVRMSDFCKDDKFPSVDELVDYFRRNNKFVVVGLGEYLALRGTAIVDKELRRLKNTTLNARAILLLRGVSTQASQIRDDNKMEQQRAYISA
MCL2016626.1_Defluviitaleaceae_bacterium           -------MNKKIKNYFSSELSKP-YFCFLNDEEYRQTMSELQADGVKIIRVSDMCGTDRPPNMDKLIDIIKNPAKMAIVGLGEYLAISGFDTARDNLNRLQATETRQKVVLLLRGVFSQIPKMRNDRRFDERVHYTSA
WP_235396406.1_Pseudoflavonifractor_phocaeensis    --MTIQEVIKKIDRYLKKDNVEPLIVDVQNKADLEAIMLHYQLPQNVFLCASDFCKPDEFPVIPNILERLAENTNFFVSEITSFLKLKGEKALVQELKELLSMSTEGHAVILTFQCVEYLHTIKNDRRLDSRICVLEG
MBR3817930.1_Clostridia_bacterium                  --MLLDACINRINNFITSSSSQPLIINVNNSSQFNELVSHFNVGDNRVISSKDYCKNDEMPQIDLVFDLISADGIVFITNFTTYLRMYGEDELRKTLKTIINLSTRGKVVFISYQCEQYLSF--TNPAQKNRVCLVDG
MBP3656735.1_Clostridia_bacterium                  --MKISDIYGRIERYLRSDDTRPRIVNFQDRASLIAFRDHFNVGTAAFKMPSDYSKVDEDILTENLYAEIQFQGNLFLIGFTSQLKLKGRDTLHQFIAQIAHMTRSGKLILVCFCCAEMLHF--DDTRIQGLVYPADD
ONI45221.1_Epulopiscium_sp._SCG-C07WGA-EpuloA2     M---VDEYIRSIDTYFKGEKNSPFIVNFENILDLQEAIAHFEVDGNTIVKTSTLANKDNLPSLDNVLSNCDITGQIFVTEYSSFLKLRGHNSLIETFRILLHHN--KKLVIFTYKCERQLRELNTDLRHRNNIYFHCG
MBO6272669.1_bacterium                             M---IRNYIEKIDRYFVRADTQPLLIDFNNKQMLNEFIQHYNVGAYNFIKAAECCREDEILNIVDLFHKLKSSNPTFLVNISAYLHLLGEKEAKNILKDLMSNN-HQKLVIVGYQICSYLGE--TDPRLKNRMVLVND
WP_037355505.1_Selenomonas_sp._FC4001              MRLDVMQCINDVASYCNNVSGFPLIIDTEDYDSFHDLIFRLRIDVKKFIYVSDFCKEGAIPNIEQAYSSAFADGVHVLVGLSQACMLQSQSTLQEYLDRAVNETCREHTIILLMHCREYLQKCRQDVRLERRIRFLGN
MBS6586269.1_Eubacterium_sp.                       MINDVSECFEQISAYFSGDCGFFLLVDTEDYDTFQKVLQRLQADGKKCVYVSEHCSRNGLPDVDSAVRAVCGDGDSVLVGVSQARMLQSGEALDRSLDDLLSRPVSGHCVALLDHCRQVLQKLHRDIRLKNRVVLVEE
NBK77202.1_bacterium_D16-76                        MKASIQNVKQEIDSYEEHFTGAPLVVGIDGGYNYTALLSLLKDDYKQVIRMSDSCSVDFPPDPAFQISLVSKAKPVIWIGAAQAKMLYGQQPTEQFLVNLLGTCFSGPVTVICPFCCTILEGARNYAKLGYNAIVMSD
NSW92343.1_Bacillota_bacterium                     MKYTIEELKQELLRDKAHRDRFPVFILIKGRDNWHKVVTLLEDMTDYTYKLSERCGEDTIPNLNDLTTKIESYSNMLVIPLAEYLRIFENRR--AILRELATIENKCRAYIPLFEVENIFFEMKGVSRFCAKYFSINT
WP_048095116.1_Archaeoglobus_fulgidus              MAGSVSRLITKLKDEANIRERFPVFILIEGSNAWKELLDGLKSFTDKVVLLSSFCDNDTYPDVNKAISMAKTGKRVIVLPFSEILRVDSDAE--RLIYELLDFQASGRIYIPLYGVTSQFNKWRTYDKEHSPPYFVED
NPV08138.1_Anaerolineae_bacterium                  MTHDVETLLTEVSYEAVSYTRFPVFILVTGVNAWRELIPALRAQMDACIDLSSLCSEDVLPYLPKVLALAREVPRAVVLPVAEALRLHPASS--SLLSQLAQLEKHQRIYVPLLEVEDLFGEVARMDRYLHPGWRLGT
MBQ7110119.1_Thermoguttaceae_bacterium             MN-----ANKALLAYLASKEATP-FFYVVGDGEYLEAKKSLLEMGMQVVKTSDFCDADRAPRFDALLDALRAGGTRVLLGLGEYLALKGEETARRQLDILRNLQIGGKAILLLRGVAGVVAARARDPRFDGRRYWASE
HED00230.1_Pseudomonadota_bacterium                MIPDLRDLFHELSKDAAKRARFPIFILVERLDHWKEVIEHVKAPANKLIKLSHLCDEDRFPKIDELKEILEGAGSVVILPLSEYVRLTGEK---EILTDLAQWELKGRVYIPLYGFASEFIEFVHVARYIHPGWHLSG
MBE6101264.1_Selenomonas_ruminantium               M------DMSAVEAYFVSPQHKP-FYLVVGDEDYKDALVTLSNRVTSVLRVSNYCTPDRIPNLDKMKDDLQEHNRVLVLGLGEYLALEGEEYAKSYLQSLKDFNLAGKAVLLLRGVAEQVKSIKGDRRLESGRIAIAS
ROT09401.1_Muribaculaceae_bacterium                MQMTLEQVKIKIEKYLNSTDIMPRIVNVSKADDLVELKNYFEIGNIKFIDMGDFASVDSMPLFDKLLNSISNKDITFLYGISTFLKLLGSDIMNLMLRNILEIVPQNKVIIILVACDQYIKL--QDPRIVDRGAYIMC
MBR2693834.1_Thermoguttaceae_bacterium             --MNKTDLIEAVKRYAVSKSKYP-FFAVSANRDYSSLRENIHEYGFKKVFVSDFCGPDTIPDWDELIDCVKAGEKILVIGLSEYFAFQGEEFALRKLKELKDIRLGGRAVLLFRSIGAWIKSLKSDPRFDGARYCDAA
WP_216880780.1                                     --MNLDDTKKRVQKYLNSNKRYPLIVDVQTREDLANIAEYFKVGSNKFPAIEEFCNQDEAIKLDELYASVSNSGNTFITGLSGFLMLFGENVAKQALKTLITTSISGHVVIITYQCEKYLRF--TDPRISGRLVIVDG
MCM1514006.1_Anaeroplasma_bactoclasticum           ---MHERLKNKINSFIDSNDNRPLIIDVSNVEERDNLLQDFFATPSIFELVKS---ECELPEISSLYEYMSKEKTVILTDLGTYFKLLGQDSLQQNIHALLEKSFATKFIILTLQCGQYLNE--KNPKNASKIVVSNQ
HHW17415.1_Bacillota_bacterium                     MLSSVDELVTYLRAESDKGERFATFILLEGTEAWDRLTARLQYEVDSTVHLSEFCEDDVFPDMSRLISYLKRCKKLLITPLSEWVRLNPERTD-VIIQILAEWPPVRRIYVPLLSIKDSFYQIERVVRYLPQEWYLSE
WP_157832981.1_Desulfofundulus_thermocisternus     MRKEIASLLDFIKNEAKVPTRYPAFILVRGLDAWQELLKSLRLEVDEVFTLSTLCNDDTFPYVEGIIPILKKTTKILVLPLAECLRFTGESR--LVLRELATWEDYKRVYIPLMELDDIFGYMNMIAQSLPEVLSFND
WP_166670004.1_Aminivibrio_pyruvatiphilus          --MTLQEILQRLADEKKVESRFPVLIFVDDWSQYKELETALENACDVTLELADFCAPDVLPNFSKIFKKIEREKHILLLSLGEYLRLTIAREVQPQKANIPSLWARTRVIIPLPCCRDLWDRIPYDERQRDCIWELSD
HCH97635.1_Lachnospiraceae_bacterium               --MTITDILQQEDLERKIESRFPVVIFCESMEEYRKLVTRLRGACDCCWNLAEFCEPDKYPKFRRLIQKIEQDKHILLLSVGEYLRMATKFEVYNNSAQFYDLWVKTRIFIPIFAAREYFNRVGTIERQLDFLWELDT
WP_195464804.1_Clostridium_jeddahense              MVNNFSELRTYLRKDRAESSLNPIFINVESMEMWVEVKKLLLSMSSGHLFLSEFCTEDTTPNIRRLIAKMKSTDSICVLPISEYLRINPEIAQ-PTILDIITKERLFRIYIPMYRMRSILYTSDNDPRRKGSILLLET
HBQ28533.1_Desulfotomaculum_sp.                    MIFTFKEFESFIAEDIKRKDLRIFFINVESLTLWVKVKNFLAQKCHNQIRLSTFCGDDLSPKINRLCAKLEIDDKTLLFPLSEHLRINNTKSD-EVLSQIVSTEISVHVYIPMYRMKDCLQAIAQNSRLNDNIIFLAE
MBK5260961.1_Peptostreptococcaceae_bacterium       MLKDYSSFTQYIDLDCGKADLNPVFINVETMKMWAQVKSYLTSKSQVLIKLSDFCNEDIAPNMNQLKKKIRVKDKTLVVPLSEYLRINRPLAT-KTLYDILHANTDLRIYIILYRMSEVLST----PREMNAVLIISE
WP_006692206.1_Selenomonas_infelix                 MV-----DLTAVERYLQDRERTP-FFLFAGGSVYAQLRHRLQGLGLHLLTLSSLSEEDRLPLPDDIVMTIKSGEKILLIGLAEYLALCGEAAARRLFSRLKSMPCDGAALLLLRGFFRYASELSEDPRIGERRLALDA
WP_028051765.1_Carboxydothermus_ferrireducens       MVSSREDLIAYLREERKKKERFATFILVTSWQQWRELIKALEFEVDVVVSLSHFCGDDVFPYLQGVLDTLEEGKTVLLFPLAEWLRLDWEDGS-KLLTKIVEWPEIKRLYVPLLAAREMLEQLRAFNRYLPPVWHLKG
consensus/100%                                     ........................hh.............h..........p................h......hh..hs....h............h..........h.....h...................
consensus/95%                                      ..........h............hh.......b..h...h..........p.s..p.......h...h......hh..hs....h............h.........hh....h....h.......p.........
consensus/90%                                      ..........l............hh.h.s...h.ph...h..........p.s..s...s....h...hp.....hl.shs.hh.h.........h..h.p.....+hhh....h.p.h.......+.........
consensus/85%                                      ......h.p.l.......p..s.hh.h.s...h.ph...hp..s...h.huphs..-..P.hp.hh..hp.....hl.shuphh.l.s.......h..l.p.....+hhl....h.p.h.......+b....h.h..
consensus/80%                                      ......phbp.lp.b..p.p..P.hh.h.s.p.h.ph.p.hp..s...l.huphs..D...P.hp.hh..hp.....hl.slupah.l.sp....p.l.pl.p.....+hhl..b.h.p.h.....s.Rb....h.hp.
consensus/75%                                      ....hpphbpblppbh.p.p.bP.hl.hps.p.h.plbp.hp..s..hlpluphCp.D.hPphpplh..lp...p.hlhslupalbl.spp...p.lpplhp.p...+hhl.hb.h.phh.....s.Rb....hhlps
consensus/70%                                      ....hpphhpblppbh.p.p.bP.hl.lps.p.hpplbp.hp..spphlpluphCp.D.hPphsplhp.lp.s.phhlhslupalbL.upp...p.lpplhpbp..s+hhlhhh.h.phhpp..pssRh.sp.hhlss
```

### ; Type 4 BREX systems PglZ N-terminal iSTAND-AAA+ATPase; Representative_2

```
                                                   1---------11--------21--------31--------41--------51--------61--------71--------81--------91--------101-------111-------121-------131-------141-------151
OrigSeq                                            MLFWSFSQLTQEIERDKDSRDMLAQRYAVRFIMLNNFNEFKELAKFMTNIGVDMLDLENLIDEDDTWITKDTLKDAIKACTKSTFVTPFSEVARFYNDDDFRGFFNEIMLLEDIHNPNKRIYIPLIGLQNRFTDFLNHFARIKESAPVWRYDA

                                                                       S1                       S2 -----HEH-Extension----     S3                                     S4                        S5
                                                   -----HHHHHHHHHHHHHH---HHHHH--EEEEE--HHHHHHHHHHHHHHHH--EEEEHHHH---------HHHHHHHHHH----EEEE--HHHHHH--HHHHHHHHHHHHHH---------EEEEEEE--HHHHHHHHHH---HHH----EEE--
                                                   -----HHHHHHHHHHHHHH---HHHHH--EEEEE--HHHHHHHHHHHHHHHH--EEEEHHHH---------HHHHHHHHHH----EEEE--HHHHHH--HHHHHHHHHHHHHH---------EEEEEEE--HHHHHHHHHH---HHH----EEE--
                                                   -----HHHHHHHHHHHHHH---HHHHH--EEEEE--HHHHHHHHHHHHHHHH--EEEEHHHH---------HHHHHHHHHH----EEEE--HHHHHH--HHHHHHHHHHHHHH---------EEEEEEE--HHHHHHHHHH---HHH----EEE--
MBQ0159480.1_Bacteroidales_bacterium               MLFWSFSQLTQEIERDKDSRDMLAQRYAVRFIMLNNFNEFKELAKFMTNIGVDMLDLENLIDEDDTWITKDTLKDAIKACTKSTFVTPFSEVARFYNDDDFRGFFNEIMLLEDIHNPNKRIYIPLIGLQNRFTDFLNHFARIKESAPVWRYDA
PMQ01343.1_Dictyoglomus_sp._NZ13-RE01              MIFGNLDELIEYLNKDRVSRNTYSIRFPTRLIMLSNYKLLHELINKLN--KANVISIGDILLNEDRWLTQSNIYNLIKNLQNDFIILGISEFLRFFSDNELYSFINFLLEIEGSPERNRRIYIPLIGMRNRIEAFLNKYHRRFEWDPLWIIND
KUK66682.1_Parcubacteria_bacterium_34_609          MIPLEENKLISLIKDDQEKFGISYGRYPVRFILIDNFEDMKKIIETMAKNGVETLDLSNLENDSNGWVSTYELIDVIKKLEKDYIVFPISEILRFLDNDDFYSLLTSLMEIENSTNSKRRIYIPILGLFSRFRSFLNRYHRRSEFSFVWKIGN
WP_018963723.1_Coprothermobacter_platensis         MNVNSMESLIELIDEDKSDNTPSATRYPVRFIFTKSVWAVNDITMLLKLRELDILDASELFPFSDGFPSLSTILQAIREKKDDVVVVSLGNLVRFWPDEDFYSLMMGLMEIENSPKPDRRIYVILEGLIERFDVFWSRRTRQIGSDPRWTLES
MBR3117632.1_Oceanobacillus_sp.                    M--NDVSSIVKQIIEDKQKSG-SYDRYPIRFLFMDLNNDTQQDIVKIVGISIRIKDISEELSFNDAWITKSKMLSIVSDLNTDYFITGFSELARFYSRQDLEALLLSFTNIENTSGGKQRIYFVCFSLFDKIAQELRTNSRNESINPIINGNK
MBQ4585185.1_Clostridia_bacterium                  M--TEQQEIIKLILDDKKKSN-TYDRYPVRFLFMKLSQNSEDELITLVELDIQFINLHTLLSFEDGWITKSHLLNFIASLNKDYIITGFSELIRFYSRNDLEALIISFTNIESVSKNKQRIYFVCYSLFEKIAIELRSNSRNESIDPILKPST
WP_024292379.1_Lacrimispora_indolis                M---DIDSMVDIIQSDLNRQG-SYDRYPIRFFSMKLSNGTSNYLMLKRKVCVEILDFQNFLLHDDGWITIDRFKKVYALDKSYIIVGFSEYARFLSNAEFITLLLGLLELENSGYVKRRIYIPCFAIYSQVRIVKANNRRMDVYNPFLNEMD
HAR58249.1_Acholeplasmatales_bacterium             M--SYTDELLNLMKEDKNLSN-TYDRYPIRFLFTRLSQETYNDISELIKMKLEVIRLSDYLNFNDGWLTKNSVYNIIKCLDKDYLIFGFSELARFYSKADLQSLIISFTDIESNSKSKQRIYFVCFSLFEKLSEVLKINGRNENINPIIYHMD
WP_117621172.1_Hungatella                          M---DVRGMVELIKKDIGKTN-SYDRYPIRFFSVHYEEGVADKLIQLQQINAEICDVKDFLAREDGWFSPDAFCKKICGLPKNYIVVGFSEYVRFINEPKFVSMLLTLLEIENTANRKRRIYIPCFALYNQISTIRKMHRRMDVYNPLMNETD
WP_191216537.1_Methanothermobacter_sp._THM-1       MEFNSFNKLFLEMENDLQTDN----RYPVRFIFLNSFEELKKFMEYIGEI-SKTVSLDGML-DGSEWLSFDNILKNIKEADSRICVVPLSEFLRFSPKKDFYALLKGLSEIET--NDKTKIYIPLVGLYKRFSEFMEEFYRKDEWAPIWCLNG
WP_011944160.1_Thermotoga_petrophila               MNFYSLEKLYDRIKQDKNSKRISDRRFPIRLIFINSFEELQEIIRFLKDCGAESKEITELLSSKDQWLTPNEIVNWIKGIYNNAVVTSLSEFLRFQNKENFYITLKKLTEIET--QNNIRIYIPLVGLCERFKEFWTNFYRKEEWAPIWKLES
OQC39307.1_Bacteroidetes_bacterium_ADurb.Bin037    MKFESIEALISEIKYDKATNDILSRRYPVRLIFLQRFETFRVLIEKLSSIGIENYHLEKDLPHHDGWITKDTLISIVKKNSKDTAVVPFSEIVRFYSKEDFKNFFNQLLLIEN--ELSRRIYLPLIGVEERFEDFFQDFTRKNESAPYWKISR
MBQ4420212.1_Bacteroidales_bacterium               IDF-SFDELKAEIMQDASSKDMLPRRYPARFIMLDNFETFRELRRFLHNLGAKLLQFDSLLDDVDGWLTSDQLIEIISNTTEPTLITPFSELVRFYEENKFSGFMHDISLLEDLSHPNKRLYIPLIGLENRVEGFLKGFTRIAESAPIWRCRT
WP_183546997.1_Methanococcus_maripaludis           MEYRDIECLYDAISKDLNSKEYLANRFSLRFIFVPDFDHMAKLCNYLTSMNLKEINVSKMLPHDDGQVTPQRIVETIKNEDKNLLITSLSEISRFYGEDDFLSLFLDLAYIEK----KNKIYVPVVGIYERFREVMPVFSRSNESAPIWALSL
MBU7048061.1_Theionarchaea_archaeon                MEFDSIDELVSHIQDDKKTPRLSFRRYPVRFIFLPNLWILKDLVHALVNIGIYRIELEDSLPGPDGWLTIDDIMNIFKKLKKDHVVISFSELTRFCSDDDFIGILNKLIEIENTENQERRLYIPMIGIFERFS---TTFSCRREWAPIWRISG
HHW21152.1_Thermodesulfovibrio_thiophilus          MVFENINYLIQKIKEDKKTKNVTEKRFPVRYIFLSNFDTLRDLIKETSKIGITTFEISQFLPKDDGWITKQEFIDKIKSLNKDYLLLPFSEIARFYNKQDFNNLFSQLTELENIANSNQRIYIPLMGIKVRFEEFYKNFNRKIEYSFVWEIYE
WP_207678199.1_Desulfonema_magnum                  MNFKFIEELYSEIDSEKKDKSPAVQRYPIRLIFLNSFNSLNLITDYFSKQAVDVLELNKLLPHNDGWLTSDDIINSIKLLEDDTVIVPFSEILRFKNTAEFNTILTGLFEIENSDKKIKRIYIPLVGMKERFEDFWKTFHRRGNWAPIWYLEE
MBR5399199.1_Bacteroidales_bacterium               MKYQSFEELVDAIKEDIKTRDFLSRRYGVRFIMLNSFDILRKLVLELQNLDVEMVTLDGLISEPDSWISTDTLKERVFSCTKPSLITPFSELVRFYPDEQFRGFFSEIILSER--SKTNRIYIPIIGLENRFIDFLSHFGRLSESAPIWQYNE
WP_022459608.1_Bacteroidales                       MVFYSLDDLKEAISVDIKSHDVLAQRYCVRFIMLNNFDAFRELTKFLVELGVEKFDLEKLALGSDKTIDIDTLSDAVRNLKVSSIVTPFSELARFFKEDEFKGFFNDIILSEDIAHPQKRIYIPIIGLHNRFTDFLKSFGRIEESAPIWQYYT
MCQ2340464.1_Paludibacteraceae_bacterium           M---------KEIQQDAASKETIAQRYPVRFIMLDNFDTFRDLSNELTSAGVETFPLESILEEDDAWITTDELKDAICQCTSSTMITPFSELARFYSDEEFVGFFNEIILHEDLAHPGKRIYIPLIGLKNRVEGFLSRFGRITESAPIWRLNS
MBE6838777.1_Ruminococcus_sp.                      M---NLEELKQKIADDKAKNTSSYRRYPVRFLFMELNQDTQNDIQDLVSSNGELLELSDFIMKDDGWLTKSRFLQVVKTAGKDIFIVGFSELIRFYSRKDIESTVLSLFDIENIMSSSRRIYFICFSMMDNVYVLQNSFARKDLLDPFINSDY
MBR3710351.1_Bacteroidales_bacterium               MRFDSYEALLDEIKRDMTDGDPLRCRYPVRFILLNNFDVFTKFAQDLAHHGVSPLNLEELIETPDRWITTDELCQAIKDRKVRTLVTPFSELVRFYSENDFRGFFNEIINTEDVDRPEKRIYIPLIGLQNRFSNFLNSFARIEESAPVWAYLS
WP_205109179.1_Marseilla_massiliensis              -SFNSLDALFAYIQEDKHWTGSHYNRYPVRFVLFDNFSDFNEFIVNRP--GIYKHSIDTMIDSPDDFLSYTELSREIRAFINDFVIYPFSEMARFYDNEEFDALITTIRASQAPEEEHVRIYIPIVGMQKKMGRFMN-----DNSTFVWEFKS
WP_172204608.1_Prevotella_sp._PTAC                 -DFEHIDSLIEEVKKDKAADGSILNRYPVRFVLFDNFADSKDFVSELILGVTKMQKIVDWMDHPDQILTHSCLANCIRQYDSDCIIVPFSELARFYDNHTFETLVSDIKGIQSATNNRQRVYIPMIGQYGKMSKFFS-----DSQSVIWHLVG
MCH4183501.1_Prevotella_sp.                        -EVTDFDGLKALVKKDKASAGSTRNRYPIRLLLLDNFRDCSDFVDFVRELKTMVYSVEKWLDYPDIMLTHTQLADNIRKQGKDCVIVLFSELARFYDNNEFEALLRTIKAIEATWVNHQRIYVLIVGLEGKMEAFSN-----DSQSFIWRLKR
WP_196042834.1_Parabacteroides_merdae              -IFTNIEALLWEVQADRHIEGSTANRFPVRFVLFDNFRDCCAFVDDCLLPNIAIQRIDDWMDYPDQFRSHKFIADKILNLTEYRIIMPFSELARFYNNEPFDALIATVKGFDTNRQYKQRIYIPIVGLEGKIQHFRD-----DSQSFIWYFHN
MCI6981218.1_Akkermansia_muciniphila               M-------------DKEAHYSGLNRYPVRFILFNSVEDYSKFIDVCAARGAYVQRISDWLLGNDKLLTYTELAEHIKEKGEDFVITPFSEIARFFGNSKFRSLMKTLGAWEASPRAHQRVYIPLMGISGMLEKLIT-----EPGTNVWDCCS
MCD7797870.1_Akkermansiaceae_bacterium             MPYQSLKDLEAAINDDKKRGG---NRFPCRFVLFDDYKNFDTFIQTRRKDGAFMCPLAEAL-SDDKQLTMKSLCDWIKGKKNDLVVFPISEIARFYNDNEFKCFITTIKGIES--DEQQRVYLPFIGQQSRMYNFVN-----DPAIVIWDNCN
MBO6251938.1_Muribaculaceae_bacterium              MKFSNIDELIQAVVVDKNAEGTVQNRYPIRFVLFDNFKDSYAFTIRLVELHITIEDIQQWIDYPDIIISHQKLATEIERCGESKIIAPFSELARFYDNHDFDTLIRTIKNIQTSAEKKQRIYIPIVGLEGKMSTFFG-----DTQATIWYMRS
MBR4838437.1_Bacteroidales_bacterium               -CFSHIDELMDEVRTDKWCNGTTLDRYPVRFVLFDNFRDSREFVATMQEMGCLVKNVNDWLDYDDRMMTHSKLAKQIEAFVKDYVIAPFSELARFYDNKDFDALVKTIKGKESRYQSRYRLYIPIVGLEGKMSQFHN-----DSQIFIWYLKS
RGX95620.1_Prevotella_copri                        MTIASFEELKALVEEDKNINGATANRYPIRFVLFDNFHDSFEFVMYMLECGVAVQSVDKWIDYPDLMITYQELSEHIADYGRDSVITPFSELARFYDNQSFDALIKTMKSIEASVENQQRIYIPIIGLEGKMSTFEN-----DTQINVWHLHD
WP_254009599.1_Bacteroides                         MYVENIAQLIELVKEDKLTQNYLRDRYPIRFVLFDNFNDSFEFVSQLMDCGCILKSVEDWVDYPDIMLTYNDLSRRIKNFSNDYVIAPFSELARFYDNNEFESLLRTIKGIESTSNNHQRIYIPIVGLEGKMQSILP-----DTQIIIWQLKS
consensus/100%                                     ..............-.........Ras.Rhh......................h.......s...s...h.p.h.......hl..huph.RF..p..h...h..h...p.......+lYh.h.u....h................h....
consensus/95%                                      ............h..D........RashRhh......................h...h...s.b.o.p.h.p.l.......hl.shSEh.RF..p.ph.s.h..h...p.......+lYh.hhub..bh...............hh....
consensus/90%                                      ......p.h...l..Db.......pRaslRhlhhp..p....b...h........plpphh...D.hho.pph.p.l....ps.hl.shSEhhRFhsp.ph.shh..h...-s....ppRlYhshhuhb.bh..h............hh....
consensus/85%                                      ....p.p.l...l..Db.......pRaPlRhlhhp..p....pb...h.....h....plpphh...D.hlo.pph.p.l....ps.hl.shSEhhRFhsppph.shh..l...Es..p.ppRlYlshhuhb.+h..hbp.......s.lh....
consensus/80%                                      M...php.L...lppDb..ps...pRYPlRFlhhpphp..pphh..h...sh..bplpphl..sD.hlo.ppl.p.lp....ps.ll.shSElhRFhsppcF.shh.pl..hEs..p.ppRIYlPhhGlb.+hp.hbp.....pp.s.lW...p
consensus/75%                                      M...shp.LbpblppDb.pps...pRYPlRFlhhsshps.pphh..h.p.sh..bplpphl..sD.hlo.splhp.Ipph.ps.llssFSElhRFaspp-F.uhh.pl..lEs..p.ppRIYlPhhGlb.+hp.hhp.....pp.sslWphps
consensus/70%                                      M.h.shcpLhpblppDbppps...pRYPlRFlhhssaps.pchh..h.p.shb.bplpphls.sD.hlT.splhp.Icphpps.llssFSElhRFaspp-F.uhhpslb.lEs..p.ppRIYIPllGlbp+hpphhp.....pp.sslWphps
```

```
# ; Type 1 BrxB-iSTAND and Type 2 BREX/Pgl systems PglW C-terminal iSTAND(subset) alignment comparison

WP_102077887.1_unclassified_Psychrobacter          MLNTDFNELMERVRAGREFGH------ASFEPIFYLIFDPQKILKIKRQLPAWAAKLRNEGWDVHIFSMAKAVQEVFDEMPVFQDSAALENRDQWQKTNKSLAEALTKKNALQNKLEAKLFGRPNSILLVSDIEALHPYLRIGSMESQLQGKFH-VPT---IFFYPGMRTGKQ---------------LKFLGFYPEDGNYRSVHVGG
MCF2581773.1_Bacteroides_caecigallinarum           MMDTVFEKVYQKLSSPDF--GKNL---GGELPLYIQPIPVSGQTELNTQVERLVSRLSKLGKNSIVVDLYRLALEIIDEEGILETLLEDEKNIDKDDLNATFESIFDTKEILIPRMRSMIEENKYDFVFITGVGRVYPFIRSHSIVNNMEGLADNANI---VLFFPGEYNRLQ---------------ISLFGKLPADNHYRAHNLND
RRF95307.1_Coriobacteriaceae_bacterium             MINDARGFIVDKLSDDALLGRNGY---MMRQALYIIDYDPAQQQYAADLVRAICEKDPRRGVTPVVVNLYDLVLGYLDEQDLWEPLVEAEPDTPRLDLIQMLQDTVGVKDVVAPRVNEAIASPEADIAFVTGVGETFPYVRTHTLLEEISSP---IPV---VLVFPGRYEQHA---------------LNILGLTQASTYYRATRVFD
TVQ28582.1_Spirochaetaceae_bacterium               MTAKRFDRVYQIISHPRFLARQGL---GNEVPYFIETYEPANEFAVTTEVRTIHERLLHNGIPAVLLPMYDIVIECLESDGRLAQVFEKEPAMGRQRFFAMLDAFTRPDAPVHDAIVRRLDAPDHKLVLMHQLGTVFPFLRTHTLLTNLHSTIYQVPL---VAFFPGTYVSSY---------------LSLFGTFKGD-YYRAFQLSD
UCC65223.1_Anaerolineae_bacterium                  MLEADFEKLRQRLGDPDALNP------AHSDPIFYFVYPPSQILTMRKLLPGWIARLRNEGLKVETLSLSEIMWELIDASGRWDDWLELEPEHDLDAVNEAIRDVLRAGNALVERVAERAASRENTVLFITDVELLHPYFRSRVIENYLNNKVL-IPT---VFFYPGRRTGQG---------------LHFLEFYPEDPGYRSTLIGG
WP_004808450.1_Actinomycetaceae                    MLQETLDIALRVMTSKRFLNREGL---GNEVPYYVLRYRIEWDRSFDQSLRQMLSHL-NESVPTLHIDVYQLAIQIWRDCGYWDQILAQEAGMDRQDFADGLAQILDAERVLAPAIAERIQAPDSRVVILSGVHHLFPIMRAHRLLNCLQPLTGDVPV---VLTFPGSYRQSA---------------LVLFDQVSEDNYYRAFDLLD
WP_126029123.1_Bifidobacterium_callimiconis        MIEREFESLFAIMRRPSFRTGSDT---AGEPANYIYAYPPVKELEVERRTDQLANRLAETAPGVLIIDLYETAIQVLRDSRIFERVLRKEPRLIPDMFTQSLIDKLSPDKSIAEQYRQAREHKQGDIVFITGVGKVYPYIRTHILMERIQLVFEQRPV---VLFFPGTYAKTT---------------MRLFDRLESSNYYRAFSLNA
WP_135754989.1_Leptospira_bouyouniensis           MLLKRLNDLKKDILHPEGIQVTQ----SQNYPFSIFIYPPADEFTVRAKFVEMIQDIKKENIEILEINLAHECLELLKQRDGIDEIIQKEKEFTFALVNDVFSPILEDENGISKSILNIMEEGRKGIVFITRAGFLYPFYRTSSLLKFLTNTRG-LSV---VFLYPGTRTSES---------------LSYMGVMSPDSNYRPRMY--
WP_168675170.1_Hymenobacter_artigasi               MIPEKAERLFQTISGSRFLKRELL---GGDIHFFISTHAAEQQTEMRQAIAALIKRLDNTGIQVLEINLFKLALSVLDSEIGLPALFEFEEQESAAEFREALHSAMDMKQVLLPAIERHVQSATPQVYFLTGIGEVYPFIRSHSILNNLHHLVERAPL---VAFFPGTYSGEQ---------------LKLFGLLADDNYYRAFNLDT
WP_187771726.1_Phascolarctobacterium_faecium       MLSERLGKIKNIITNSNFLNKTGN---ANEVSYFIFDYPPDKDIIVEDYIQRLATEIMEKDMQIKIFDIYGLIMDYLKDKKYIDKCILLEQKKGSAGLIKAIKDTLHLDSYIVNHIDENTIQ-DKDIVFLVGVGRAFPIVRVRELLSVLNNLYTKCPI---VLMLPGSYDGQH---------------VTAFNKLKMENYYRAFKLVQ
WP_200392182.1_Roseibacillus_ishigakijimensis      MFDQKIDELIRRLGAPEGTTNA-----RQGDQVFNVHYPPHLWAELRTRLPKLMARLKDQSYAPRHCSLLALMEKIFGDNGKI-ALLRRSEQQSHQAFTDTLHSILANKKPIVAAIEEEIQSKPNGLLLITDIEMAHPLLRVSAFEQVLQGRFA-VPT---VFFYPGERGNIN---------------PSYLGVYPSDGNYRSTHIY-
WP_222635590.1_unclassified_Rhodococcus            MLTARFERAYRVMRSPSFLEMKGQ---IGEVPYFVLAYKPSDEPRVGAEHKRLVTRLRTDGLEVCDIDLWALANELWSATGQWELILEQETALPKAVFSEGVGNVISPKDYLSPRIARIVERPRPQVVLLSNVGRLYPLVRAHTVLNTLQPLLADLPI---VLVFPGTYRQSS---------------LVLFDRLTDDDYYRAFDLLE
WP_224137561.1_Candidatus_Kaistella_beijingensis   MPNNINDKIFSLLSEEEFKDPDTG---LLFFPVYIYTYNPEKEFQIREEIKELDRKLPSNNLNCLVLNIYNEFIDYLKHNIFLEQIFELEEIDALDFIAEKAQEKNDNDAHIGEKLKAYFDSPDRVYLLLYGFGSIFPYLRLSEFLKKTEEYVRNYKL---IAFYPGELKNDH---------------YSLFGIFGDENVYRANHLNQ
WP_235341611.1_Bifidobacterium_boum                MIDEDFQHLFDVMKQSSFRHSKGI---AGEQPFYIYDYPPECELEVDRHIPWLMQKLDDYAPVVRRVDLYDVVLGILKRRKILDRVLKIEPRYSQDKFLGMLESILGADSQLPNAVVEAMHTVKPGIIFITGVGAVYPYVRAHTLLNTLQGMIEDIPV---VLFYPGTFQSST---------------MTLFNCLEPDNYYRVMRIRD
G8417454.1_                                        --RDPVAKALGAERSPHEAVRDMLLDAAGSRGFRILVTPPEHHADIGPSVAR--------ALGGTWVSFADAFFRDHGAN--------IAALERAERFVAQRDALTEAAEDTMFRLLD--EHGRPDHTAVVGDTGLLGVCDALDLPRRLYDETLSGSRGFWVLVLVPGVIYKSQPHFNEGPSMWQLAGATFSLLTALPD----------
MBK8541194.1_Ardenticatenia_bacterium              --PDPLPTFLPALTSPGDRAAAWLRSSRGMSSWRLVVTPPERHAELGRQVAA--------ALDARYVSFEAAFLKRIEDR--------FADFETAERMSALRGRLKKEAEGVLEALLA--EHGRPGTATVLGDTALLGLCEAKHLVTRVYERALGGALGFWVVVVPGVVHQQQPLLNEVDNLGSWPGMVLPLAEDVGV----------
MDO9017995.1_Myxococcales_bacterium                --RDTLPAEFIAHRSPDEIVVGLLRAARKSRGFRMIVAPPERHAEVARSVAQ--------ALDARFVSFEHGLLAGMEPE--------FAVYERAERFKAQRSRLTRAAEGLFEALLA--EHGRPGTCTVLGDMGILGLCDAAHMIRRLYDETQTGARGFWVVVVPGVIFQRQPLFNEKQPLFHVDGTVLPVDGELGR----------
WP_075011136.1_Stigmatella_aurantiaca              --QDPLPSDLSTQRSYEEVVADRLREASRGRGYRLVVAPPEKHVEVAHSVRR--------ALSAELVSFEARLLQRMEPD--------FPAFERAERFLAQKPKLKKEAVALLDALLE--QHGRPGNVVVLADTGILGVCGVKDVPRRLYDRVTERDLGFWVLVIPGVVSERQPLFNEQEPVLNLQ--VLPVPEPLDV----------
MCA9600002.1_Myxococcales_bacterium                --RDPVPDDLAPDRTPEQVLVARLRDAAERREWRLLVTPPEHHADYGCSVAA--------AINAEYISFEQRLLERMEPS--------FADFESALRMKARRRKLTRAAEALLEELLR--ERGRPDCRTVLGDTALLQMCDAAHLVRALYDETQGGQRGFWAFVIPGVIQERQPLLNEKDPVFHLPGATLPLSGPLPP----------
TVQ88636.1_Deltaproteobacteria_bacterium           --PDEIPELLEERRTPEEVARDVLRDAVKRSSYRLIVADPGQVGPIAKSVIR--------ALDATPIDLADAWFQRHAAT--------VKVDAHAERFPAMKAVLAQKAHKLFEELVA--EHGERGRAIVVYNTGLLDAIGGVVQVRNLYERVQGGQHGFWVLVVPGVISGRTPLFNERTQLWHQPGLTLPLRKPLPD----------
KYF91864.1_Sorangium_cellulosum                    --PDELPPEL--VKDPAEAARDILRSVAKTRGYRLVVAPPDDHVEIGRSVAR--------ALEAAFVSFEHEFFRRVDAQ--------IEVFDRAERFAAQRPRLRKEAEALLDALVQ--EHGKPGSTVVLGDTALFGTCDALHLVRRLYDLTATGGKGFWALVIPGVLHKRQPLFNEKATVFSIDGAVLPLAREIPE----------
MBI4699810.1_Deltaproteobacteria_bacterium         --SDPVPPELVAAKSPEEVAADLLRGAASGDGFRLVVSPAELAPEVGRSLAR--------ALEVRFVSFEQELLGRMDGA--------FDGFVRAERFKAQRGKLTREADGLLADLLA--REGRSGATVVLGDTAIFAVCEALHLVRKLYDETSAGGRGFWIMVVPGVVYQKQPLFLERAPIFHPE-STLPILRAIEG----------
consensus/100%                                     .............................................h..............s.....hsh...h..........................................h.............h.........h......p..............hh.hPG........................h................
consensus/95%                                      .........b......................hbh....s.....h..............s.....hsh...hb.......................h.............l...h.............l.....h.shh....h.p.h............hhhhPG......................h.h...h............
consensus/90%                                      ...p.h...b......................hbh....s.....h...h..........s..s..hsh...hb..h..p.................h...b.........l...l.......p.....l.....h.shhcs..h.p.l............lhhhPG....p.................h.h...h............
consensus/85%                                      ...p.h...h...ppsp..............shbhh..ss.....h.p.l..........uh.s..lsh...hh..hp.p........b........h...b....p..p.l...l.....p.p....hl.s...lhshhcs..h.p.l.......s....lhhhPG.b..p.................h.h...h............
consensus/80%                                      ...p.h...h...ppspb.............shblhs.sP.p..ph.ppl..........uh.s..lsh...hhp.hc.p........b........h...b.ph.p..p.l...l....pp.c.s.hhl.s...lhshhcs.ph.p.l..b....s....lhhhPG.h..pp................hshh..h............
consensus/75%                                      ...p.h.phh..hpsspb.............shblhshsPppp.pl.ppl..........uh.s..lshbp.hhp.hc.p........bp.......h...b.ph.p..p.l...l....ppsc.s.hhl.sss.lhshhcs.plhppL.sb....s....VhhhPG.h.ppp................lslh..h............
consensus/70%                                      ..pc.hsphh..hpsspbhs.p.....s.p.shblhshsPppp.pl.ppl..........ulps..lshbp.hhpbhcsp........bcsb.p.p.h...b.ph.p..pslh..l.p..ppscsshhhl.sss.lashhcs.pllppLbsb....s....VhhhPGshpppp................lslhs.hss..........
```

```
# ; vWA-MoxR subsystem 2: the iSTAND ternary systems

                                                    1---------11--------21--------31--------41--------51--------61--------71--------81--------91--------101-------111-------121-------131-------141-------151-------161-------171-------181-------191-------201-------211-------221-------231-------241-------251-------261-------271-----
OrigSeq                                             TFLEICQRLVKEIADFAVHDSDDLDIQVCVNRLDSNQLHKALLCLNYTKQENLFREFV----RHQIGACLIHGEPEHGQRWLLNRLLGL-VPNGRTESVIPLNL-LVATRSSSIDALWRELGKAVGAARL-ATCDEIAQKVGNLWKSQTVILIFHNLDQLPEDYFNEFICKFCSSQAN-VRNYRLLAFLMDLSGCVD-SWGITWAETIDLNWEPHTPLKLPKITKLSHNELSYWLEKSPHPLPAEIDKIALDILNSDDGKP----EAVLDYICELCD

                                                                                                                            S1                              S2    -------------HEH-Extension---------------        S3                                  S4                           S5     -------------Degenerated-Helical-Bundle-------------
                                                    ------------------------------------HHHHHHHH--HHHHHHHHHHH-------EEEEEEE------HHHHHHHHHHH------HHHEEEEE-|HHHHHH--HHHHHHHHHHHH--------HHHHHHHHHHHHH|--EEEEEE-HHH--HHHHHHHHHHHHHH-HHHHH--EEEEEEEE-------HHH-------------EEEEEE---|---HHHHHHHHHH-----HHHH--HHHHHHHHHHH-----HHHHHHHHHHHH
                                                    ------------------------------------HHHHHHHH--HHHHHHHHHHH-------EEEEEEE------HHHHHHHHHHH------HHHEEEEE-|HHHHHH--HHHHHHHHHHHH--------HHHHHHHHHHHHH|--EEEEEE-HHH--HHHHHHHHHHHHHH-HHHHH--EEEEEEEE-------HHH-------------EEEEEE---|---HHHHHHHHHH-----HHHH--HHHHHHHHHHH-----HHHHHHHHHHHH
                                                    ------------------------------------HHHHHHHH--HHHHHHHHHHH-------EEEEEEE------HHHHHHHHHHH------HHHEEEEE-|HHHHHH--HHHHHHHHHHHH--------HHHHHHHHHHHHH|--EEEEEE-HHH--HHHHHHHHHHHHHH-HHHHH--EEEEEEEE-------HHH-------------EEEEEE---|---HHHHHHHHHH-----HHHH--HHHHHHHHHHH-----HHHHHHHHHHHH
OCQ97673.1_Nostoc_sp._MBR_210                       TFLEICQRLVKEIADFAVHDSDDLDIQVCVNRLDSNQLHKALLCLNYTKQENLFREFV----RHQIGACLIHGEPEHGQRWLLNRLLGL-VPNGRTESVIPLNL-LVATRSSSIDALWRELGKAVGAARL-ATCDEIAQKVGNLWKSQTVILIFHNLDQLPEDYFNEFICKFCSSQAN-VRNYRLLAFLMDLSGCVD-SWGITWAETIDLNWEPHTPLKLPKITKLSHNELSYWLEKSPHPLPAEIDKIALDILNSDDGKP----EAVLDYICELCD
BAI89278.1_Arthrospira_platensis_NIES-39            MMETGFEKLETNINQNLQTIQESLNELIQTLNLSEQSIKDKLLHLDYREQVRSLEDFL----QDRVGAFWIYGSNQSEPHWLVTSLIENKIETSNTSSRVSLSF-VGNTQSHCLDNIIDSISKAFQIENPTEKLTYIIEQICIRWQTKSVVILAEIKKGVDPEIVEEFINKLRQKHYN-IYNYSLMLFLVDVHGCTN-NWEINFTEDILSEQIPEIPIRLTEIQPIPREQLRQWFRTHL---PDQLIEKMLDLIYHNN--Q----EKVFYQICKLFE
BAY43400.1_Scytonema_sp._HK-05                      WVSLIFVKL-----------------IADDSQIKNQQLYHVLLKLDYTDQEQLFSRFI----TNQIAAFLIHGSQDYGQHWLTNRLLKK-IRRVSDKPVL-VDL-DCLVHNRNSQTMWRELGRHFGGIK--DSPDLIAGKVLKSWTTKNIYIVVDNVQSMSEQLLQELIQKFQSIAS--QTNFKLLMFLIDNVSKVESSWNVSFAENFEPTWRCHQPVILPKNNVFSDIALRDWINHESVYLPENIDGVIQTILNSEAGIP----IPAFREICNLCE
KPA13794.1_Candidatus_Magnetomorum_sp._HK-1         FRPNGIQRL-----------------------PEKLRENFPYSVNRTEQVQGIKDIG----EKKPLVWLIHGDRNQSHERFLECVQWFHWENIKKGSKIEPRF---YYSNNFENDCMTEIWKNFSQDGQKDNWRDLSPLIDNNCNK--PVILYTMIRCSQQRLFYVLLDIQKKLAGKRKYP-LFLCILIIIYTHKHKR-----YQRKITRQLKKYNKYTLPLLSNIESEDVDEWQNMSF----KNERKIVKDIFDDIEQIPMTELAKIFKHCLYHQD
KPA12842.1_Candidatus_Magnetomorum_sp._HK-1         TKKAIFSDITKEINDTTKDTTDDKIV-------THHCGKFVHLMCDRANQVVEFRKTFSNENHKKPQFYVIHGKERNGHLSLVERLKIQVIQPFVNEQAVSLLV-EKGDLDMKMDILLDALFAEIFNQNDSIQYSLKALIQSDPIKADHAVIIAHRIPANHLDLANRYISDFADLSIVDTIPLFIVFFNIEYPRLVKRMKELE--YKIRSMVKELTNQLIKELQTINTHHVIDWFNSHQKNM-PEKFKAVDSIFDGQDEC-MFEIEDLLKKY---VP
ETR67363.1_Candidatus_Magnetoglobus_multicellularis LENLLMKQLDRVIPETIPDIPPTDQQ---SQAPIATIGDDINLYCNRITHVGEFLFTYRDDCGNMPQFYFVYGCKDECPESLFLRLRDIELRDHFKPQVPPIDERSNSSFEARKDVLLKSLFKTLTGDYQSNKFKLTDLYQSNAIKDNNVIVVPHFLSFDQQKLMEWYIKDY-------DAPHFMLFFIITCIPKKNFPKHVQ--KKLKKFSSQLSAYLFDPLHKIDKSEVAPLFKQFK----HEINKHIDDLFKDQELMNMSDIEKYLLRLKQTIK
OQX97257.1_Bacteroidetes_bacterium                  LQDTSCRDV-----------------NSNNKENISLDRNYYRYSCNCYKQNNYFENVFKKNKNEKIQFYVIHAKAEQCPEILPQRFFNEHINDNDNNSIF-----QVDNIDSLKIKTLYQIFSFFNEE---KEACLESLFNSKTLQKKKYITINFIVESRQHEFIEFFINDFCKNQIKKTKLRFIFFWSIIYTSEIKDRETKK--DKIQKCIKNLEFDVLPEIQIINLDDIRTWIKSIE----SKINISLFNIFTNKKEFTMEEVLEMFDFLSEKYN
WP_020569053.1_Neolewinella_persica                 EQANINGSLLYVLGESSADLKQDGAK--------EFIPPFHAFGVDRAPQEGQFFDEQFEAFDQKVFFYYLHGDDRQQASSLAERLGFKLVGLQLTADDLDIEALNGMREEHCRALLVHAITQQFMGGAVGLNLTILDLLNRPKLKRHPIVCINVTVGHSYPGMLRKFYETFCKVELPEDAPTFYFFFGLEHCDNKPT-----VRKEVAEAIAGRQNSALPELEPVLISDIVNWFTHHG---PYDAEQLVAENFTEGDQLDMLVVEADLKRLIDKFN
RMG26468.1_Bacteroidota_bacterium                   WEAQQEEELPWQLYGRHTDWRLALPG-------KKRSGRLLHLMCDRDPQAESFLDRFLSREAGDPQYYFIHGPRGERHKSLVRRFREIEIRHHLGRMPALVDF-TTGDLDMRKRNLKRSIARSRELPGIVGSKWESPALMALLAQRGNLVVFQHQLAAEKFRLLEWYIHDF-------ASFHFLLFVNIVYPEKWGSRQDIW--EDMQALKRKTPETILAELQPITPEDVAAWVDTYF---PDNMQHLPGTIFDKQKRLPMKVVETQLKLAIERLE
KPA17452.1_Candidatus_Magnetomorum_sp._HK-1         EMNTVCKNIALEAKNKSIIINSYDEK--------KFDGRIIPTLCNRDIHVEKFYKFFEKKHKKLPLFFIIHGFENQGHRSLIKRLRFTLIRAYLQKQKIELPF-DKGKLSDRKQILARQLFNAFQKMDDIKTYDLKRLNQLKAIETNNIVIISHNIEASLKDLLSWYIENYVDNFKYDDIPQFIIFLNIQYPVAIKKHSTIKVKKELTVFTQNFSKKLIDELNSITQQHVKDWICKYKSNM-YEIEEEIESIFENSEHMSMIEIEKKLKNILLK-S
WP_028158837.1_Bradyrhizobium_japonicum             NLDELVAELHRFVNDGAAP-------DQLGKARSKKFARLLPYLCDRRMQRRHFRDAVMERDVVRPQIFFVLGADSECADKFVEQLFFVDMPELLQRNGLEDRLADVSDIGDYLGELQYEFADKLG-----LKPRSSVTEIERRLQRHTALFLYFDVDIRKRRLLREWIGWWNSLVVMGRRYPIIVVGRFCSANQGIWSPTMRLVDDLTALSTTAGKTVLPQLGRVSFEDIEAWINENADFAPPDLRQRLRRFFIGERKVSMIQAATLLKTIMNEMK
RKZ89871.1_Candidatus_Parabeggiatoa_sp.             APSESAKQLIELIEQYPQGLTHLSEVVAPQKKFKLNVDDLRPHLPNRRPQESKLGRAITHHHKQRPLLCLIHSDECQCSDRFVDRLVQYYLPKVVPGDGIFVHC-HFSQGIDLHQEISEQLASRLGI----NPFAQPSEIVEAIMRERSPIIFHTDMCTKNMKAVHNFIEFWAKLDIPSTHNHLLICLYFNYKDIKPSKKSIN--DQIRAEFRQLENIVLPELMDIEKEEVYAWARDHLSSF-DSVQPKIDDLFSPDETIPMCDLARSLREILEEFP
OCR02031.1_Oscillatoriales_cyanobacterium_USR001    AVKQAIKKLAKTLASWNVPQVIVQNTNLPITQQSHDKVHQALLRLNYTHQVRLFRAFF----HHPIGAYLIHGEEESGQKWLLNRLMKF-ITDSRTAKVIPVKL-SRKSCGTYTEALWRQLGGHVGLMNQ-PSQTQIIKNVSELCKNQIVVLIFDELNAMPEECMSQFIGQFRDLT---QSQSRLLMFLVDNIGCVE-QWNVAFAETLDPNWEPHIPIKLPKINPIEQSDLVSWVRNAVDDLPNQMEEIVQNFFNGETILP----EAAMIKICDECD
RMG84230.1_Bacteroidota_bacterium                   KIAQVKKKLKATLAPEPQP---AGSE-------RVVFHEYHRYTCNRQPQNVQFHSSFVSKDKQKTHFFYMYGLNVQSPEGLFYRFSYDRLLDYLNPDLESVRAQLSPDETYNKQEVLKNLFAALSVPYAHKKRLEDVCLSSPTIRKEDYVCIFLGVSSFDPALAHWFITRFCACTLPEDAPTFLFFLAIIYEEDDPQ-----IRAEVKETVENSDLIGLPELGDVSLQDVKRWFFNYQTIFPKTRQQLIEERFGDAQTFEMEEVEIRLERIIDEYN
WP_028091549.1_Dolichospermum_circinale             EMNSLYEEL-EKIDQETKGLKSPIKK-------LSAQRSFLSYLANRSSQEFELTEAIKNRDKSRLLVCIIHGDELQSHYKFLERMRNDFLPDVLDLDQNKIKKSELKNLNELPKYLLMNLAKNLV-----NKGNTSLEEINEILNKSSPMIVHTDLISDDLHILNKLLEFWQDWHNQNTLNKVIICICVKYIRIEWTYQQIN--KKMRKHIQEFSLIILSELEAVGRADVENWACEHTKNF-QQLKDKIGDMFKSSSKIPMRYLADNLLEILEHFH
WP_018400346.1_filamentous_cyanobacterium_ESFC-1    SLSQLFEALDRELINIENIALEDQDHHQTTASLSQDLSELLFLLNCIQQENAFR---ARATANPAGTFLVRAKDLSLQKWLVNRLVRL-VPGY-ENAYCPFNV-YSPQIRLNFNLLWENLAANLNTT---ATIENVAEALIERYCHQTVILVLYGVHKLQDHQIAEFIQ-FQERYTR-ARRSRLILFLVEDTEEH--NYSFDCVESIDYQCL-EKPLRLSPLPLGQISKAEALTWFDEVYNQL-EQIDLLLQDI-NDPN--P-NEVSISLDEICMTCG
KPA18269.1_Candidatus_Magnetomorum_sp._HK-1         RIDFDIERLINAIEKNKERSDNKSTDNKTTGFKQPNNPKKLHLLCNRKKQALDFTLYFKNLKKNTIQLIFFYGPENECPYSFIERLAIKEIKDYWENQYDKINKEKKNDVNDQKNTLIQELYNSFLLIA--NDYKLSAFYQHPTFDKHNVIILYHEISIDKLKLLEWYINEHSQFFCHNNVKPLLLFFYIKVDGFKEKCQKML--DKLKKKIDKKFKLLLEPLQPVDKDDVREWFSNLE---PSIVEPEIEQIFGKENQMPMRKIITKLDIFITNID
WP_066426097.1_Anabaena_sp._4-3                     TLESFFDKLVTKCQEKTSP-------KQSVENNSSENIINALKKLNYNEQEKDFRDYIKIVENKYAATFLIHGQYGYGQRWLVNRLCEQ-VDDGLTTAFCPINL---TKKHRDIQRIWDDIAAKVGSSS--RNPQTIVDVIYQHWQTKPILLCFHNVDVVGCQELNKFIEPFKHPNT--VNTQPLLLFLIDNYGQQK-FSNLF--NQISKEYDIENNLEIKELDKFSDAEIKIWTKCYQKIF-SNLSEITQEIIYNNT--P----IQAIEYICQCFD
KPQ40445.1_Phormidium_sp._OSCR                      SYESDVSRHTEAVLRAMDELRGTGQESFPVNSSNKNNFIKILGRFNYVQQKRRFQVFL-EDANPRIGAFWLRGERDSAQKWLLSQLWLQNVPNSTKAIKKTLTM----TTRQNTERILEKLATWLQVE---ATPEALIKRILENCKTETVALVFYQVEYLDKGTLEELINTFQKTYLRHSLDNPLLLFMVDLGKKGDH-----CPINHSSDYNAEHPEQFVELTQRFQDRDFRWLTDNQTELPVSLEAIKKDMIDDNH--P----EEVFTELCDYFG
WP_071187406.1_Trichormus_sp._NMC-1                 KLSRKAERLLKEIQEKLSELDKNSSN-----------LEKSFQKIDFSKSREIAESLKKQYETEGAILLFLQKSKKQLGKYCIEEFIDVIISNQIETCPRTINF-ESNTYKSNKYGFINAISGYLGIEISSSIINEIHEKIVSSLSLDTIFFEIKSINRLEEEFIDWFIKIFKSLFN--QKRNRLIFALIAESPFPINSSNFC--QEVTLNHDK---IEIPLL-DWTVEDISNWLISHHKLFPDEIHKEAKNI-EESEGTP-----ESVCSKLQEYFY
SEH05005.1_Candidatus_Venteria_ishoeyi              VFNGVAQRLLQWVPALTDVLYHARKSINSTPNISKIIPNFAPYLVNYTKQQTCFEDIVQPTLGNHPLIFLLHGQESNTTHGLAARLKQNLEENCLHHGETTVRL-CPFDCDERPGFLLTTIEKNLKDK---FAVENLEILHESAFEA--TTLFYSEVEAKHKKLLQDFLNFWNDDGWPEQNHLFIVCLFIDYRPGPLRKMQLT--QRLKAKNIDICNMLLPELTPIPREDFTPWYNLMKTYYSDELEIRVTELFDENNKLSMEYLGPKLSKLLADLC
WP_017306429.1_Spirulina_subsalsa                   TLRDIYKRIKDKCNNREQQFQAWLKGTTLDQELLEYLLNDTLKKLNYDHQKRPVKDSF-PEINQRINAFIVQGQSHHAPRWLTNQLLYY-VPHHDNAKKHSIVL----KTHFRFNRLCEGIASGMDCG---AEPQNIVSALAQNLETQSVVLALRDIHQAKNGILSSFINEIQTNESLSTGQHYLILFLVDGDSSSSK-----WSTELNPFIQTDSPCSLPPLENFTQGTITLWRGRKEKILPDQYPQAIDEFIDTEE--P----KGFLDFIYDSCL
OCR02924.1_Oscillatoriales_cyanobacterium           DFETAYSNLTEVLLKGINP-------AVQNSATVENKLYNALLSLDYIRQESLFQEFV----NNQIGACLIHGKQEYGQRWLLHRLLRA-IPNNNTASLTKISF-LRQVQNGSLDALWGEIKRKLKLKCQ-PLPDELAKAVCNLWQTQTVVLIFDDADQIEEAHLEKFINDFNSKPPK-SRNSRLLVFLIDFDGCAP-KWSIPFADALDTTWKPHIPLKLPPIKPLREMELAYWLEKAFNDLPDSIEATVQSILNSDNGLDLHEFELAMAHICELCD
KIF28928.1_Hassallia_byssoidea_VB512170             EMDELAQQI-----------------DRLERASPVERLYRALLKLNYREQTKKFRRII----DNSVAAFLIHGLPDYGQGWLLNRLVQQHVPNSITAKVVQFNL-NRLARKSDVPSLWRELSRGVGLGGQ-GLPPDIADRVYQWWQTQNVLLIFRDVDFLSEIYLRELLENFTSANAK-VSQFQLIMFLIDEKGCVS-KWNVPFAEEPNSNWQPDSPVKLPEISGFSQEELAKWLDYESDDLPEQLEQTVQTILNSDNGVP----ELAMEQICYLSG
CUR12819.1_Planktothrix_paucivesiculata_PCC         PDDMLWTQL-----------------CQQQVPQSEHKLFKSLLRIDFDEQEEAVMEVL-SPSYNKTAAFLVHGEEKFGQETLVTRLSQL--SKLRNGRRIKIKV----GGMEDISNLWNEVGKHLLDKNQTLPETKIIDQIFELLQTQHLIFIFSDAHRTYISFLPELIQEF-------QRETYLAMFLVDNKGYIC-KSGLSLAWQVNHPDLPRIPLSLPPASRFPVRKLNEWVAVAAEVVP------VETLLESQGGVP----DLVYQRICYYCG
OQW38836.1_Proteobacteria_bacterium_SG              DEEEAETRLLSKIQDRPRPVFKKSES----PQYLSNQAEQKLGQINRKKHRDHLAIQIWQNENRKTHGFVISGEQQECPEDIRFKLSYLLQEDLINHNPKGLRTKSYRIGKSAEQFLWELLAKRLGSC---NPEKAAIEMRLNQVDR--SYLFFRELPSDEQQFLVDLLVAWSNLSLASTSHSHFLLLIHAAENSGNSKPSWR--DQIESLLKQHEVALLPEFHPPSKKEIRDWKGLHI----SVLRDGIEELFLKHESIPLGKFKAALPLLKKHYS
KPQ40050.1_Phormidium_sp._OSCR                      RLDSGFYKLESEITEIWKKLDEDTKEQLILEDSLPKRVHQCLLNFDYQKQEYEFDEFI----STQLGICIVNGLQKCGQPWFLYRMFQT--KYYFDDTSTPFNK-ISCFSRTQKLSLKEEICHFFRLPVN-ESIERIYQEIANHLEHKSLVFIVKNIHLTTQEYLDYFVKQFQQQPPN-NPENRLLFLLVDLEGVSH-HWEMSSEQELLSHYRSHTIYRLKQIKPLSWAERQDCLKQLAWDLP---DEISQKVILTEKFVP----ETVLEYICNCCN
ETR71372.1_Candidatus_Magnetoglobus                 VLTSFADEIFQLLQSQPLE------------KKSTSFGSASVLMCNRIPQLRSFLTNY--RVHKMPQFYFIHGREQDQHDSLVTRYNQTIIHKRIQEIYDPYMCEYGKKSDDQQNTLLYDIGHELTGESTHDITSFEKLVNATIFEQQNAIIISHKIEEDFKDLLTWYIQEFLKKYKHERIPQFILFFNIIYE-----KKKMI--RWINEITPKNSTVLIPELEPVIRPDVQSWFNTYTSDLPSKRNECIKQLFPKPKSRPMAVVEEKLVGF-----
WP_020531248.1_Flexithrix_dorotheae                 YHKNFVKKIELALKKKVNSIFNTS---------KNLIPEKFYLGCNRTIQFQDLQYIL-EISTKKPCMCILYGNREDCHRSLIERFYYYLLNK-------LINI-GKWDVRNRKERIIKSSFEAFNLTDIKAVPDNAQMIIDSYNQKCDVLLIEIEIDCNEKSTLKWYINEF-----------LKVLHLIKLVSKPSNERRIE--KDLRDLYSSEIRKVLPKLSPIKKYHVKELL----KGL-ASYVDKIDKLFLKKSEITMLNFEMFVT--KEFLN
GAK59821.1_Candidatus_Vecturithrix_granuli          LDNPDAQRLLWAITGQRELIADASAS-------QPFLGDIVYKMCNRGPQENDFLRFFLRENPRQPHFYFLHGSDDECHESLIERLARTRIKKYVEQTCGDVQS-IRGDLETRQEDLALNFMKTFDSDDLMPNELISGLCHLAVLRKHPCVILRHNLDAARERFIRWYIKAWLQPEWTADLPQFLIFLNIIYPRRHHRQKTIS--KQLCRIHTSSGEQLLPELKMVTPFDVSDWFSYYR---PWQRKEKIALIFGNQERCSMSDVEKHLKMIIDECT
KIJ79200.1_Tolypothrix_campylonemoides              GYCVLFEDLVRKVQERVSSWVTSVGGPLRTAASNEQPTEAEIIQLGHKSLSRQFEDSLLSSEQAAVVCFLIHGASGYGHKEMVVRLRQI-LEQQRSVQARKAAF-SPLWREKSLAKLLELIGDQIEQGWIPDSPEVLAQRLHKMLETSDVILEITEIQRFNGSLLS-FIKDF-------PTQHRLVVLLTLEQTILPEQSPLA--MDDDSEIDSMRPIKLAELKEFTEKELTSWLSKWR---PNDALVLAKTLMDETKGNP----QLLYNKLRDDST
RAM51670.1_Hapalosiphonaceae_cyanobacterium         SLDRLFSSLNLDLERESS--------IDQKPQKPTKEFMIALYDLDYLHQRTSFKTAI----DVKAATFLIHGKPHYGQRWLLNQLKSQ-VPYHTNAWQKSIYI---KPHRKDIEVIWQDLAQELKTS---ASPKDVAKELYQHWLKSTVILAIHDVDLIAGSPLKQFMDEFKKAGQL-QSSYRMLLFLVDNINSKSKEKSLSFLENIDIN-EPHIPFVLKELEPFGKNLIEAWVGVQEQSF-ASIKEVMCEIVRDSQ--P----ISVFTDICQCFD
WP_088278871.1_Ideonella_sp._A_288                  AWSSVSRDLFPLIETSTITAPSRSTQ-----QPLTTVDLPLPYLCNQEPLTRAFEDAAQRGGAKHAMLVLLAGGHNDAPEHFARRLQRHHLPRRLKTIGVAVHTPAAGTTLRHSMPLLRALFEALDDDLVPTEFDQAAQVLCEHLRARTAAVLMAWLPDRDLPALLRFIDAA-----------RLVLICVGEARGSVW-------RAVRKSSQA---AALEPMAEIDVAALTKWHKEHG----RVDLDRCETLFRAGR---AREFARHFQTFCEGLR
OCQ93508.1_Nostoc_sp._MBR_210                       TQNLRYHQLADELERYITSIFTAEKDDQPLSLLFNSQLFDCLLSIDFKDQEACVTQAIWQKKRQRTVAFLINGDDKYGQSAFIERLFRK-IPEFKHEKEIKIKL----ESGWETSNILQEIAKFFGLDYISDIQDKIINKACECLQFKNLIFVFYIPQRFFVGLLADLINDF-------HKDTYLTMFLVESNHKEKGLQELS--NKLAPREDDSSYLYLPLLSKFSRDHLEKWLENGVEDE-TQQKNFVDTLLETEWGIP----ERVYKKICNHFS
KPV52199.1_Kouleothrix_aurantiaca                   QIKQTLRAVDDEPNDDVRAGGAALGQETQRQNSGDGKLYQALQGLNYSAQKSAFVLSV----ARTVAAYLIDGQRNHGQRWLLNLLLKR-YEHALQA---SIDM-RRLGRGKSIADLCQEIGRKAGVAA--STPEQLAPVLCRWWETQHTFLIFHSVDLLGEAGLREILEHFGDPQHH-NAANKLLLFLVDNQGVAA-KWNLPLAQNPA---NAGGPERLPTIARFSEDELDRWMMASLLDLPERTD--VQTILNTDDGIP----ESVLEEICVICD
PKO80078.1_Betaproteobacteria_bacterium             ---------LQTVKEWAADLGKALSSLAGRRQASGDIPLAELGRIGRDTQRMALVDALRSVKGERAAAFLVHGPENHGQREFAEFLNVW-DNEWEDAQPI--------ADTDTADALIRWTCGQLGTPMIDAAIPELAQTLAARLRRGPVIIIQRSIGHRQ-ERLTRFIGGF-----------RLYWFLIDHCPLPQPEGVIA--PEAGADTADYDDLALPALGDISAGQVRKWLKELRKSAP-------DEILTTPDGNP----SDVYNRLWASAS
RKZ83713.1_Candidatus_Parabeggiatoa_sp.             ADNRLDNSITDSLVKAWPKLDP----DRTPKPKRKYKRTLLPYLPNRRDQELELGELIAHDDKHRPLLCVVHGDEYQCSDKFLDRIREESLPNIIPEQMHGVDL-FVCDTFQSVNELHRKMLANLGELFINNMYATCDEVAQAIAQRQCPVILHTDMSTTDINIIHGFIQFWADWPVLSAQNHLLVCLCFNYRDIEKMRPSVN--QAIRNTFKDLASRVLPELPSIERTHVEQWAKIYLSEF-YELIPYIRELFPPIEKIAMEPLAQQLKQILKECC
RMG30280.1_Bacteroidota_bacterium                   YKEEAAEVLNWRLPQQATQ--------------KGNLSMLDKYTCNRAEQNSRFKLNFKSRGNKKFQYYIIHGEEQESPYGLFQRFVLEHISTAYQPYFY----EEATSLEEAKINFLTALFQALELNPNRDQLTLANLLKAPAAQDMQCIALKLRVYSSERDMIQWVLQEFRESELSPNAPDLLMFISIIYDLQQASSKNPY--EKLIKTLAEFPEGLLPVLPPVKISDVNKWFDRIT---PEEKKQQIATYFDAQPEYHMVTVEKKLEQIIEVYN
PZN77012.1_Candidatus_Methylumidiphilus_alinenensis SARDINGPV-----------------YTTQNPRIPPVLSLLPLMFGRETQEEQLQDYLCFHKGQQIIAVVLPGPHRESPAHFVERYAKLSLRDAYQRHWREYVWEKHSPLQQRLDFLLRDLCRNIDGVSF-GEEGHVRDRLVSQFADQSHQILYYFLTADRANLIGEWIGFWQKLKLDTAKNMVSIIFCLHSEPSILPEWSSRCVKALRTLCPNAPDLMLEKLPPIKEQDSYNWAKEHESTFPAILYQRAKNAFRKRR--PLLNLMTKLEEAVAKAA
WP_072720293.1_Planktothrix_tepida                  TTLNIYKRIVGEINKDKQEFSEWLQTGSNQEDEQAEQVYQSLLRLDFRKSSGIFADWE----QNKAACLLVYGKKKHGQEWIMRRLMTK--PKYSKRMFYTCSF---SSYISDSKKLWKWLSEKIGIQFNRENQLRIIKALKEQLQYNNIVLIFTDLQANSLEYPQEFIDQFFSQQAQPNSCNKLLVFLLADEPEIK-NWCQNWVSNINENWSITEPIQLPEPPICPEDDLIPWLKQETDYLPVNPRKTAQEILKTNQGVP----EAVLMYICQYDQ
WP_020536784.1_Lewinella_cohaerens                  GILYLLEECKKSVGTSTTDAGG------------AALHEYHAYTCDRVAHSDTFRQLF-GEQKQPVQFYFLYGGDLQSHEGLFRRISYDRLQDYLNPELKSLQLEFSRQLEHYKQNILRSFFSMLGLPPNEEKDLTYALERSPRLQNTDYVCVFLHISQYDPAAARWFIKEFCQEGKVNTSVAILFFLAIEYDEEDAD-----LKSEVVAALEMAERQPLPELDMVAQRDIGRWLEHYKQVSPRTRKELMKTVFGNSREHYMEDVELELKKIIDQYN
ETR70369.1_Candidatus_Magnetoglobus_multicellularis RLKEIATELSQYMQNQPAALPDRTAS--------MLSRNLLPYMPDRSELIQEMSLALKKNQAGKPLVWIIHGNDNQCHYDLMDCIEGCYWERFLYPTSKLLRY-YQYETNQWRDLVAMELKHKFDIQK--EAPEDVFDAINRRCDKASPVMLYDVVDTDQQTVCQRFIDFWNQWPGMCCQNPLIVCLLVRYKSFPVSKKDPN--LEFKETILTLKNCVFPELPDISKKDTEDWYHNYKNYIPKNVSQCIEDIFDANQKFPMQSLAREIEQ------
PZN72517.1_Candidatus_Methylumidiphilus_alinenensis SEDGIEPNFESSPNANPTAIPSTRDNLSSQSNGLENTPAFKAGLVDKDEQATHVIDHLKEETHNNPLVYLLYGASTQWPDALLNILYYQQLPRFNKNSSPTLKYRILRGNIDPVEYLYELLDEELTCGLNR-------KAIERRLSKEKPLIFYRILTAAEPALVQGMLVAWQKLELSINSPRHILIFYYEYIETPKPEKTLI--NHLKASLPTEMSIFLPRLKSPNKKLVHDWISIHIQKL-EEAKKYVEDKYNAKKAYAKDDYHHDLKNILNEFC
consensus/90%                                       ........l...................................hsb..p...h.............hhhl.G........hh.ph..................h...h...h................hp....hhh...h.......h..hl..a...........hhhh.h................h.........l..h..h...ph..W...............phh...p..........h..h.....
consensus/85%                                       .......pl.................................b.hsb..p...h...h.....p...hhhl.G.p......hhpph....h.p..p.....h.............lh..l...h................hpp....hhh...l.......h..hlp.a...........hhhh.h................ph..........l..l..h...pl..Wh...............h.phh...p..s.......h..h.....
consensus/80%                                       ..p...ppl...h.............................b.hsb..p...h...h.....p...hhhl.G.ppps.p.hhpph....h.p..p.....h........p....lhp.l.p.h................hpppp.hhh...lp.....hhp.hlp.a..........blhhh.l...............pphp..........Ls.l..h...cl..Whp.......p....hpphh..pp..s.......hp.h...h.
consensus/75%                                       .bp..hppl...h.............................b.hsb..Q...h.p.h.....pp..hhhl.G.ppps.c.hhpcl.b..l.p..p.....hph.....pp....lhp.l.p.h................hpppshlhl...lp..p..hlpbhlp.a.p........bllhhhl...............pplp..........Lsbl..hp..cl..Whp.......p.p..hpplh.ppp..s.......hpblh..h.
consensus/70%                                       .bpp.hppl...l.p........................b..h.hsbppQpp.h.p.h....ppps.hallaG.ppps.c.hhpRl.b..l.p..p.....lph.....ppp.p.lhppl.p.h.........p.......p.hpppshlllbpplp..p.phlpbaIp.a.p.....p..bllhhhlp......p.ph...pplp...p....b.Lsbl..hsbpcl.pWhpp......pbp..hpplhpspp..s.....b..hpblhp.hs
```

### ; Type 2 BREX/Pgl system PglW Representative_1; NERD-REase + iPKinase + PKinase + HTH + wHTH1 + wHTH2 + wHTH3 + CTD-iSTAND-NTPase

```
OrigSeq [illegible]

WP_185095299.1_Streptomyces_sp. [illegible]
WP_268746228.1_Dermacoccus_nishinomiyaensis [illegible]
GII83978.1_Sphaerisporangium_siamense [illegible]
KEZ35702.1_Frankia_sp._CeD [illegible]
WP_030506725.1_Microbispora_rosea [illegible]
RJQ78831.1_Pseudonocardiaceae_bacterium [illegible]
MCB0954589.1_Ilumatobacter_sp. [illegible]
WP_212516928.1_Actinospica_acidithermotolerans [illegible]
WP_208577501.1_Micromonospora_tulbaghiae [illegible]
MCP9485019.1_Candidatus_Gaiellasilicea_maunaloa [illegible]
WP_193120346.1_Nocardiopsis_coralli [illegible]
RKN35100.1_Streptomyces_hoynatensis [illegible]
WP_188830920.1_Nocardia_camponoti [illegible]
WP_212908910.1_Streptomyces_sp._TS71-3 [illegible]
WP_103561562.1_Actinomadura_rubteroloni [illegible]
GES07632.1_Acrocarpospora_macrocephala [illegible]
WP_248824789.1_Frankia_sp._Ag45/Mut15 [illegible]
OAA24551.1_Frankia_sp._EI5c [illegible]
WP_213171153.1_Natronoglycomyces_albus [illegible]
WP_144820010.1_Micrococcus_luteus [illegible]
MBL8842562.1_Planctomycetota_bacterium [illegible]
MBX7222607.1_Blastocatellia_bacterium [illegible]
MBO9539711.1_bacterium [illegible]
MBN9688999.1_Verrucomicrobiota_bacterium [illegible]
KAA0252897.1_Acidobacteriota_bacterium [illegible]
QQS02053.1_Austwickia_sp. [illegible]
MBK9387522.1_Planctomycetota_bacterium [illegible]
WP_144981411.1_Gimesia_aquarii [illegible]
WP_090664774.1_Belnapia_rosea [illegible]
WP_136411732.1_Luteimonas_yindakuii [illegible]
WP_150933330.1_Microbispora_cellulosiformans [illegible]
KRE61575.1_Tetrasphaera_sp._Soil756 [illegible]
WP_052424041.1_Nonomuraea_candida [illegible]
WP_093484991.1_unclassified_Streptomyces [illegible]
WP_109776035.1_Quadrisphaera_granulorum [illegible]
WP_090069475.1_Lentzea_flaviverrucosa [illegible]
WP_183357094.1_Garicola_koreensis [illegible]
WP_257478501.1_Acidipropionibacterium_jensenii [illegible]
MBL9044566.1_Myxococcales_bacterium [illegible]
MCP5560207.1_Verrucomicrobiaceae_bacterium [illegible]
MBK6942988.1_Planctomycetota_bacterium [illegible]
MSW49530.1_Actinomycetia_bacterium [illegible]
WP_217637676.1_Actinopolyspora_saharensis [illegible]
TAF23825.1_Verrucomicrobiota_bacterium [illegible]
MBZ5659870.1_Acidobacteriia_bacterium [illegible]
MBO0801654.1_Nocardiopsaceae_bacterium [illegible]
WP_200346137.1_Halochromatium_glycolicum [illegible]
WP_238634082.1_unclassified_Halorhodospira [illegible]
MBK9978482.1_Gemmatimonadota_bacterium [illegible]
WP_130480151.1_Leptothrix_mobilis [illegible]
MBI4602629.1_Planctomycetota_bacterium [illegible]
MCO6436553.1_Phycisphaerae_bacterium [illegible]
WP_252180113.1_Endozoicomonas_sp._4G [illegible]
RLS37136.1_Planctomycetota_bacterium [illegible]
WP_085387166.1_Cellulosimicrobium_sp. [illegible]
WP_241482799.1_Kocuria_polaris [illegible]
WP_262103914.1_Arthrobacter_sp._Marseille [illegible]
WP_181762535.1_Pseudarthrobacter_sp._B4EP4b [illegible]
MBK9741405.1_Actinomycetia_bacterium [illegible]
MCH5644430.1_Gordonia_sp._ABSL49_1 [illegible]
NMM16638.1_Cellulomonas_sp. [illegible]
WP_136709615.1_Agromyces_sp._H66 [illegible]
MBU6244323.1_Actinomycetales_bacterium [illegible]
WP_210768105.1_Cellulomonas_humilata [illegible]
WP_104090838.1_Cryobacterium_sp._N19 [illegible]
WP_056862666.1_Nocardioides_sp._Leaf285 [illegible]
WP_067228577.1_Microterricola_viridarii [illegible]
WP_129336322.1_Cellulomonas_endophytica [illegible]
GGD24649.1_Nocardioides_daphniae [illegible]
WP_029289175.1_Cellulomonas_sp._HZM [illegible]
WP_155100111.1_Cellulosimicrobium_composti [illegible]
consensus/100% [illegible]
consensus/95% [illegible]
consensus/90% [illegible]
consensus/85% [illegible]
consensus/80% [illegible]
consensus/75% [illegible]
consensus/70% [illegible]
```

### ; Type 2 BREX/Pgl system PglW Representative_2; NERD-REase + iPKinase + PKinase + HTH1 + HTH2 + wHTH1 + wHTH2 + wHTH3 + CTD-iSTAND-NTPase(Type1-BREX-like-iSTAND)

```
                                           1---------11--------21--------31--------41--------51--------61--------71--------81--------91--------101-------111-------121-------131-------141-------151-------161-------171-------181-------191-------201- [illegible] ----1281------1291------1301------1311------1321------1331------1341------1351-
OrigSeq                                    MAAHFISMGEPAHDAERQAIRFLVEGLPTDYTVFGNAWLVERSGVVYELDAVVAAPHAIYVVEIKSYRGRIEGTDHTWYIPEPVPSPIKRNRKTAQVLKEQLKRRSYQAGQVWVQGLVFLSATTNVGVKGPASRDRIHTRKTILAALQDPGLVERLSGRQSLVPTESAQSDLLYLVLTGTRSGPRPLRRVREYDIVDTLDHDTY [illegible] DEHGRPDHTAVVGDTGLLGVCDALDLPRRLYDETLSGSRGFWVLVVPGVIYKSQPHFNEGPSMWQLAGATFSLLTALPD

                                           [illegible] S3 S4 -Cap-like-insert-region- S5
                                           <<------------------------------------------------------------------------------------------NERD-REase------------------------------------------------------------------------------------>><<- [illegible] ---------------------------------------------------------------------->>
                                           [illegible]
                                           [illegible]
                                           [illegible]
MBL0196963.1_Myxococcales_bacterium        MTAKFVPIGEPAHDAERQAIRFLVEGLPDTYTVYGNPWLVERSGVIYELDAVVVAPQAVFVVEIKSYRGIIQGTDSDWWTPEKRPSPLRLNRKTAQVLKSHLRDTTYQAGQVWTEGFVFLSATTDSRVRGLASNDRVHTRLTLLAALQSPELIERLSGGRARTPSSAAERELLELFTGAQQGPKPVRRVREYEIVETLDASDTF [illegible] EEHGAPGRVIVLGETALLGLCEALDLPRRLYDEALSGSHGFWVLVVPGVIHNRQPRFNEGAAMWHLEGATLGLATGLAA
MBK8541194.1_Ardenticatenia_bacterium      MTARYQAVGKPAHRQEIDALKFLIRGLPESHMLYANSWLAERDGALYELDAVVVAPHAVFIVEIKGYTGAIRGTDHDWYTPSPMRSPLRNNRLTAQMLKSALERENAALRGLWVEGYVFLSHTDDVAISGIGSEQRVHTRGSILKALTDPQFLTEHITRGPAPVGQEAAEAVHRILTGGDRRHPPPRRVREYELGDIVERGEDY [illegible] AEHGRPGTATVLGDTALLGLCEAKHLVTRVYERALGGALGFWVVVVPGVVHQQQPLLNEVDNLGSWPGMVLPLAEDVGV
MDO9017995.1_Myxococcales_bacterium        MAAKHIPLGDPAHDAERIAILHLVGNLPGSYTIWTNAWLTDRNGTIHELDAVVAAPHAVYIVEIKSYRGHVEGNENDWYLDDPIRSPLRLNRLTAQKLKDALRRDDFNAAQVWVEGLVFLSHATSFSPLGTAAIGRVFLRNGICAELQNPDSVQQRNGRRTGTVEPHVLAALTRVFQSERGRRPPQRKIREYRLETAIDRTDRY [illegible] AEHGRPGTCTVLGDMGILGLCDAAHMIRRLYDETQTGARGFWVVVVPGVIFQRQPLFNEKQPLFHVDGTVLPVDGELGR
MBL8718633.1_Myxococcales_bacterium        MGAKFIHIGDPAHDAERQALRFLVEGLPESFTVYGNAWLVERSGVLYELDAVVVAPHAVFVVEIKSYRGRIDGTDNDWWLPEKIRSPLKLNRKTAQVLKAHLKNTNYQAGQVLWTEGLVFLSATTDVGVRGPASNDRVHTRKTILAALQDEALLKRVSGGRAATPSSAAERELLEALFTGAQNGPRPVRRVREYEIVETLDHETF [illegible] EEHGRPGNVIVLGDTALFGLCEALDLPRRLYDETLSGSRGFWILVVPGVIHNRQPRFNEGPAMWHLEGATLPLLNPLPV
MCB9654435.1_Deltaproteobacteria_bacterium MAAKFIPIGDPAHDAERQALRFLVDGLPADYTVYGNPWLVERNGSVYELDAVVVAPHALFVVEIKSYRGRIDGTDNDWYIPHPIKSPIRLNRRTAQVLKSQLKRESYQAGQVWVQGFVFLSATTNVGIKGPASDDRIHTRKSILAAIQDQALIKRLSQRHSLTPTDDAESDLLRLLTGAKNGPRPTRRVREYEVIDRVDLQEHF [illegible] DEHGKAGRITVLGDTGLFGLCEALDFPRRLYDETLSGGRGFWVLVVPGVILNRQPHFNEGKAMWHLEGATLPLLQELGS
WP_075011136.1_Stigmatella_aurantiaca      MSARHHSLGEPAHEAERQAIRLIVDGLPEDYTVYSNAWLVERTGVVHELDVVVVAPHGFYVVELKAYRGVVRGTDHDWDVPQPIRSPLKKNRQTAQVLASELKRLSAEAARRWVEHFVCLSHASEHKVEGPASRGRVHTRHTILEALQDARGFHDRCPRGCPEVDEHTQRTVHTLLTQQDPALRPPERIGGWRLEYTVEATERW [illegible] EQHGRPGNVVVLADTGILGVCGVKDVPRRLYDRVTERDLGFWVLVIPGVVSERQPLFNEQEPVLNLQ--VLPVPEPLDV
MCA9600002.1_Myxococcales_bacterium        MPAKFIKVGEPAHDAERQALRFLVDGLDERCTVYGNPWIVERSGGIYELDAVVVAPHALFVVEIKAYRGRIRGTDFDWYVPAPVRSPLRLNRKTAQVLASELRRSFDAGRPWVDFLVFLSHAAGVELQGPLSRLHVHTRKSVLDAISDHSPVYQRARSGSPPVDDHCARTLHELLTGVDKTLAPPRRIREYEELENVLDTTDDY [illegible] RERGRPDCRTVLGDTALLQMCDAAHLVRALYDETQGGQRGFWAFVIPGVIQERQPLLNEKDPVFHLPGATLPLSGPLPP
MBK7192658.1_Myxococcales_bacterium        MAARFIKLGEPAHDSERQALRFLVDSLPATFTVYGNAWLVERSGTVHELDAVVVGPHAVFVVEIKGYRGTIEGTDYDWYLPEPVKSPIEKNRKTAQILHGLMKRVSYEAGQLWTQGLVFLSATTRCNVRGPASKDRIHTRATIIAALTDPAFVDRIANRAATPVTARTLADVDKLITSSAASPRPTRRIREYEIQAALESHDTCG [illegible] EEHGRPGHTVVLGETALLGLCGALDVPRWLYDEAMSGALGFWVLVVPGVILNRQPRFNENESFFHLEGVTFPLPAVMPV
TVQ88636.1_Deltaproteobacteria_bacterium   MAARIIKVGEPAHANEQAAIEHLASELPSSVTVFANPWLIEPNGAVYELDAVVVMPHAIYVVEIKGWRGTITGDHDWYLPAPQRSPLHLARKTAQILNNLLRRKDATAGAVWVQELVYLAAASRVELHGDAAKRRVATRQTIHAILHDEQRIRSLSRRMLTMVDADVESALVKVLDSKPP-RPPQRTILGFEVIDTVDTNERY [illegible] AEHGERGRAIVVYNTGLLDAIGGVVQVRNLYERVQGGQHGFWVLVVPGVISGRTPLFNERTQLWHQPGLTLPLRKPLPD
MCA9674838.1_Myxococcales_bacterium        MPAKFIPLGQPAHSAEVQGLKFLVDGLPSTYTVYGNPWLVERSGVVYETDAIVVAPHAVFVVEIKAYRGVIDGTDYDWYTPHPVKSPIGLNRKTAQVLNSLMKRDSYEAGQIWIQGLVFLSAASGCQVRGPASRDRIHTRATIIAALTQPELVTRLASRATAHNTPSTQSALHRILTSEDTRARPARRVREYEIERALETHDNCD [illegible] EEHGRPGQVIVLGDTALLALCDALDVPRRLYDETLSGSQGLWILVVPGVIHNRQPRFNEDQPLWHLEGVTFPLLVPMPA
WP_153825183.1_Polyangium_spumosum         MPAKHIQLGETAHEVEKAAIRYLVDGLPASFTVYSNAFIAERTGAVYELDAVVAAPHAVYVVEIKGYRGEIFGNDNDWYVPEPIRSPLRLNRKTAQVLQSCLRRASEDAGRVWVEGLVFLYQARSVQIIGPASESRVHARETILSELQNPAALQKRSSRGTPPLDKHAAEELDRLLRGADRQRPTKRIREYEIEATLEQTERF [illegible] DRHGKPGNVVVVGDAAILGTCDALHLVRRLYDETLSGSRGFWALVVPGIIQKRQVLFNEKAPVFNLDGATLPLLGEIPG
MBI2893937.1_Deltaproteobacteria_bacterium MSAKHIPLGEPAHDAERQAQRFLVEGLPETFTVYTNAWLVERSGVVHELDAVVAAPHAVFVVEIKSNRGAVEGTDYDWYVPEPIRSPVPENRKTAQALHSLLKRESHDVGQIWVQGLVFLSASASCGVRGPGSRDRIHTRKSILAALQDQSFIERIAQRPIAAVSQSGAEALHRVLTSAPKSARPTRRIREYEIILGTNETHETS [illegible] EHHGRPGHTVVVGDTALLGLCDALDVPRRLYDETLAGSRGFWVIVVPGVIHERQPFFNEGPPMWHLEGVTLPLLEALPR
NUN14841.1_Myxococcales_bacterium          M-ATFFQIGQPAHDAERQAIHFLVEGLPDSYIVVGNPWIVERTGVIYEIDALVIAPHAIYVIEIKSYRGRIEGTDTNWYLPQPIASPLRKIRVCAQVLNSRLRHENYEAGRVWVQELVYLSATTDVGIKGPASDGRIFNRSNILAALQDESLVLRLSRRSQLVPTGEAMHELNRIFAANASGPTPARQIREYDIIDTLSFSEQFS [illegible] NDLGKVGSTVVVGDTALFGVCDALDLPRRFYDETLSGAKGFWVLVIPGVIHKRQPLFNEGPAMWHLEGATLPLTENLPE
KYF91864.1_Sorangium_cellulosum            M-ATFIPIGEPASDRERQALRYLVDGLPEDYAVYGNPWLVERSGAVYEIDALVVAPHALYVVEIKSHSGVVSGNDNDWYLPEPIRSPLKLNRKTAQIFADQLKRHSEFAGRPYVEGFVFLSHADGCRVTGPASAERLHTRRTILQALLDRAAFVRRHGE-REPVDEHVRRVVDELLTGVDRAHPPPRRIREYEIEATLERTERY [illegible] QEHGKPGSTVVLGDTALFGTCDALHLVRRLYDLTATGGKGFWALVIPGVLHKRQPLFNEKATVFSIDGAVLPLAREIPE
MBX7096841.1_Myxococcaceae_bacterium       MPAKFIQIAESRHDHERRALRLLVDGLPSTFTVYGNPWLVERTGVIHELDAVVSAPHGIFVVEIKSYRGRIEGTDTNWYIPETITSPIKLNRITAQVLKNQLKDHDSAAGRSWVEGFIFLSETLNCGVRGPASNDRIYTKDKILLALQDTELLKRLSRGMSTPVNPATDRALLAIFQANQNPLRPVRSVREYDVVRKVSVEETF [illegible] EEHGRPGRVTVLGDTALLGLCESVDLARLPFSE----GQGFWVLVVPGVIHNRQPLFNEGDPVFHLPGQTLPLLKPLPT
MBI4699810.1_Deltaproteobacteria_bacterium MPAKHIPIGEPAHDAERQGIRFLVDGLDERFTVYSNALAERSGVIHELDAVVVSPHGFYVVELKSWRGDIKALDHDWYVPHAIRSPLKTTRVLAQKLKDEIKRRSADAARPWVEFFVLLSHTGQVSVFGPLNTGRVHTRKTIIEALGNAKTFYDRAGGQMPKVDAHAAKVLDDLLRGVDKRFAPPRRIREYELGDVLERSNRY [illegible] AREGRSGATVVLGDTAIFAVCEALHLVRKLYDETSAGGRGFWIMVVPGVVYQKQPLFLERAPIFHPE-STLPILRAIEG
consensus/100%                             M.Ap...hups.p..E..ub..lh..Ls.p..lhsNsal.-.sGslaE.DhlV..PpuhallElKu.pG.l.s.-.sW..s....SPl...R..AQ.h...hcp.s..h...a.p.hlhL..s......G..s..+l..+..l...l.p...h.p...........s...l..lh.......P.cpl..acl....p..pp. [illegible] .p.G..s...Vl.p.ulh..h.s....p..apb......GhWhhVlPGll..pps.h.E...h.p....shsl...h..
consensus/95%                              M.Ap...hups.p..E..ub..lh..Ls.p..lhsNsal.-.sGslaE.DhlV..PpuhallElKu.pG.l.s.-.sW..s....SPl...R..AQ.h...hcp.s..h...a.p.hlhL..s......G..s..+l..+..l...l.p...h.p...........s...l..lh.......P.cpl..acl....p..pp. [illegible] .p.G..s...Vl.p.ulh..h.s....p..apb......GhWhhVlPGll..pps.h.E...h.p....shsl...h..
consensus/90%                              M.Apa..lGpsAHp.Eb.ulc.LlpsLs.ph.lasNsWlhERsGslaElDAlVhuPHuhalVElKuapG.l.usD.sWa.P....SPl..sR.sAQ.L.s.h+c.s..hu..Wsp.hVhLu.s....l.G..u..Rlas+.slh..l.s...h.p........ss..s...l..lhpu.....P.+plb.acl....lp.p-ph [illegible] .ccGcssp..Vlu-sulhs.C.u.chsR.hY-cs.....GFWhhVlPGVl.ppQP.hNE..shhp....shsl...hs.
consensus/85%                              MsA+ah.lGcPAHc.Eb.Al+aLVcuLP.sasVauNsWlhERsGslaElDAlVhuPHulaVVElKuaRG.lpGsD.sWa.P.sh.SPlcbsRbTAQhL.s.l+c.s..Aup.Wsp.hVaLS.ss.h.l.Gsuup.RlasR.sIl.hLps...h.pbs......ss..s.p.l.plhpu......P.RplREa-l...l-.p-ph [illegible] .c+G+sGps.VlG-TulhuhC.uhchsRblY-cs.sGsbGFWhlVlPGVl.p+QPbFNE..shaph.G.shPl...ls.
```

### ; Type 2 BREX/Pgl systems Helicase (BrxHI-Helicase); SF1-SF2-Helicase(Ski2-like-RNA-Helicase) + wHTH + Small-B-Barrel(HAS) + Lhr-Helicase-CTD

```
OrigSeq [illegible]
MBK7581083.1_Myxococcales_bacterium [illegible]
WP_291996290.1_Candidatus_Accumulibacter [illegible]
MTV27845.1_Nitriliruptoraceae_bacterium_ZYF776 [illegible]
WP_223844567.1_Streptomyces [illegible]
WP_155450775.1_Allochromatium_palmeri [illegible]
WP_015767810.1_Accumulibacter_sp. [illegible]
WP_242667013.1_Frankia_casuarinae [illegible]
KAA0252885.1_Acidobacteriota_bacterium [illegible]
WP_271637792.1_Microbacterium_sp._nov._GSS16 [illegible]
WP_133290010.1_Dankookia_rubra [illegible]
MDX2267517.1_Bryobacter_sp. [illegible]
WP_251969640.1_Sphaerotilus_sp._FB-5 [illegible]
MDR3405892.1_Chthoniobacter_sp. [illegible]
MCP5560215.1_Verrucomicrobiaceae_bacterium [illegible]
WP_059037528.1_Gordonia_desulfuricans [illegible]
WP_205528529.1_Desertimonas_flava [illegible]
WP_129345931.1_Sorangium_cellulosum [illegible]
WP_250928728.1_Rhodopirellula_aestuarii [illegible]
WP_254377936.1_Streptomyces_sp._MAR25Y5 [illegible]
MCB9398398.1_Acidobacteriota_bacterium [illegible]
MBT3378820.1_Lentisphaerota_bacterium [illegible]
HAI12818.1_Phycisphaerales_bacterium [illegible]
MSW49536.1_Actinomycetia_bacterium [illegible]
WP_040409684.1_Arhodomonas_sp._KWT [illegible]
MCB9544265.1_Myxococcales_bacterium [illegible]
WP_241255648.1_Candidatus_Frankia_californiensis [illegible]
MBK9578950.1_Fibrobacterota_bacterium [illegible]
MDZ4800304.1_Bryobacteraceae_bacterium [illegible]
WP_015443645.1_Ilumatobacter_coccineus [illegible]
MDQ3647618.1_Actinomycetota_bacterium [illegible]
WP_005004495.1_Nitrococcus_mobilis [illegible]
WP_091695618.1_Micrococcus_terreus [illegible]
WP_312898983.1_Corynebacterium_variabile [illegible]
TAF23834.1_Verrucomicrobiota_bacterium [illegible]
WP_059289751.1_Corynebacterium_glutamicum [illegible]
MBI4662623.1_Verrucomicrobiota_bacterium [illegible]
consensus/100% [illegible]
consensus/95% [illegible]
consensus/90% [illegible]
consensus/85% [illegible]
consensus/80% [illegible]
consensus/75% [illegible]
consensus/70% [illegible]
```

### ; Type 2 BREX helicase and Ski2-Like-RNA-Helicase; Core Components Comparison Alignment;

```
MBK7581083.1_Myxococcales_bacterium              [illegible]
WP_291996290.1_Candidatus_Accumulibacter         [illegible]
MTV27845.1_Nitriliruptoraceae_bacterium          [illegible]
WP_223844567.1_Streptomyces                      [illegible]
WP_155450775.1_Allochromatium_palmeri            [illegible]
WP_015767810.1_Accumulibacter_sp.                [illegible]
WP_242667013.1_Frankia_casuarinae                [illegible]
WP_271637792.1_Microbacterium_sp._nov.           [illegible]
WP_133290010.1_Dankookia_rubra                   [illegible]
MDX2267517.1_Bryobacter_sp.                      [illegible]
MDR3405892.1_Chthoniobacter_sp.                  [illegible]
MCP5560215.1_Verrucomicrobiaceae_bacterium       [illegible]
WP_205528529.1_Desertimonas_flava                [illegible]
WP_129345931.1_Sorangium_cellulosum              [illegible]
WP_250928728.1_Rhodopirellula_aestuarii          [illegible]
MCB9398398.1_Acidobacteriota_bacterium           [illegible]
HAI12818.1_Phycisphaerales_bacterium             [illegible]
MSW49536.1_Actinomycetia_bacterium               [illegible]
MCB9544265.1_Myxococcales_bacterium              [illegible]
MBK9578950.1_Fibrobacterota_bacterium            [illegible]
WP_015443645.1_Ilumatobacter_coccineus           [illegible]
MDQ3647618.1_Actinomycetota_bacterium            [illegible]
WP_005004495.1_Nitrococcus_mobilis               [illegible]
WP_091695618.1_Micrococcus_terreus               [illegible]
WP_059289751.1_Corynebacterium_glutamicum        [illegible]
Q8IY21-DDX60_HUMAN                               [illegible]
Q8N3C0-HELC1_HUMAN                               [illegible]
A2PYH4-HFM1_HUMAN                                [illegible]
Q8TDG4-HL308_HUMAN                               [illegible]
P42285-SK2L2_HUMAN                               [illegible]
Q15477-SKIV2_HUMAN                               [illegible]
O75643-U5200_HUMAN                               [illegible]
P32639-BRR2_YEAST                                [illegible]
P51979-HFM1_YEAST                                [illegible]
P47047-MTR4_YEAST                                [illegible]
P35207-SKI2_YEAST                                [illegible]
P53327-SLH1_YEAST                                [illegible]
Q6XGE6-Q6XGE6_ECOLX                              [illegible]
consensus/90%                                    [illegible]
consensus/85%                                    [illegible]
consensus/80%                                    [illegible]
consensus/75%                                    [illegible]
consensus/70%                                    [illegible]
```

### ; Type 2 BREX/Pgl systems BrxD ORC/Cdc6-AAA+ATPase (Type 2 BREX systems)

| Sequence | Alignment |
| --- | --- |
| OrigSeq | MDIVISPQRRLDILDALR [illegible] |
| (secondary structure) | Preceding-Helices; S1; H1; … [illegible] |
| MBK8233499.1_Candidatus_Eisenbacteria_bacterium | [illegible] |
| WP_264762807.1_Frankia_gtarii | [illegible] |
| MCB9654431.1_Deltaproteobacteria_bacterium | [illegible] |
| WP_175348386.1_Cellulomonas_humilata | [illegible] |
| WP_242518426.1_Halochromatium_roseum | [illegible] |
| WP_274519969.1_unclassified_Halorhodospira | [illegible] |
| WP_170320004.1_Polyangium_spumosum | [illegible] |
| WP_091695616.1_Micrococcus_terreus | [illegible] |
| MBK7077332.1_Myxococcales_bacterium | [illegible] |
| TVQ88632.1_Deltaproteobacteria_bacterium | [illegible] |
| MBK7069431.1_Deltaproteobacteria_bacterium | [illegible] |
| HBQ10062.1_Myxococcales_bacterium | [illegible] |
| RZO66311.1_Sandaracinaceae_bacterium | [illegible] |
| WP_012240294.1_Sorangium_cellulosum | [illegible] |
| MBK8541202.1_Ardenticatenia_bacterium | [illegible] |
| WP_014010194.1_Corynebacterium_variabile | [illegible] |
| MBI4699820.1_Deltaproteobacteria_bacterium | [illegible] |
| MCE9575758.1_Deltaproteobacteria_bacterium | [illegible] |
| WP_095346964.1_Micrococcus_luteus | [illegible] |
| MCG5052492.1_Myxococcales_bacterium | [illegible] |
| MBO9539707.1_bacterium | [illegible] |
| WP_165633000.1_unclassified_Actinopolyspora | [illegible] |
| WP_179642186.1_Spinactinospora_alkalitolerans | [illegible] |
| WP_201724034.1_Rhodopirellula_sp._SM50 | [illegible] |
| MBL8678387.1_Myxococcales_bacterium | [illegible] |
| MSW90815.1_Actinomycetia_bacterium | [illegible] |
| WP_006368783.1_Gordonia_polyisoprenivorans | [illegible] |
| CCD90938.1_Bradyrhizobium_sp._ORS_375 | [illegible] |
| MCP9485024.1_Candidatus_Gaiellasilicea_maunaloa | [illegible] |
| WED64128.1_Opitutaceae_bacterium_LMO-M01 | [illegible] |
| WP_015443644.1_Ilumatobacter_coccineus | [illegible] |
| MCP5560214.1_Verrucomicrobiaceae_bacterium | [illegible] |
| TAF23833.1_Verrucomicrobiota_bacterium | [illegible] |
| MBX3732322.1_Verrucomicrobiae_bacterium | [illegible] |
| MCM2371083.1_Rhodopirellula_aestuarii | [illegible] |
| WP_134500748.1_Microvirga_pakistanensis | [illegible] |
| MCB1051860.1_Acidobacteriota_bacterium | [illegible] |
| WP_253862669.1_Prauserella_halophila | [illegible] |
| MBK9578951.1_Fibrobacterota_bacterium | [illegible] |
| MCB9544266.1_Myxococcales_bacterium | [illegible] |
| WP_007626462.1_Dietzia_cinnamea | [illegible] |
| WP_206824863.1_Microbacterium_esteraromaticum | [illegible] |
| WP_256246189.1_Pseudarthrobacter_sp._R1 | [illegible] |
| WP_188543685.1_Rhodococcus_trifolii | [illegible] |
| consensus/100% | [illegible] |
| consensus/95% | [illegible] |
| consensus/90% | [illegible] |
| consensus/85% | [illegible] |
| consensus/80% | [illegible] |
| consensus/75% | [illegible] |
| consensus/70% | [illegible] |

### ; Type 3 BREX BrxHII-Helicase Representative Alignment; TUDOR + Swi2/SNF2-Helicase + Inactive-REase + CTD(Alpha+Beta)

| Sequence | Alignment |
|---|---|
|  | 1--------11--------21--------31--------41--------51--------61--------71--------81--------91--------101-------111-- [illegible] |
| OrigSeq | MGENVHWVFDRWNNKPVKAVERYELWNTTNCLVYDPTSGRIYTLPADGVGDINEGYVYDEHRIRYILAGAKLKELIARQSILSPIGSNIIPLPHQIYALVRAISSNRIRFLLADE [illegible] |
|  | <<------------------TUDOR--------------------------->><<------ [illegible] |
|  | ----------------Preceding-Helices--------------- S1 [illegible] |
| | -----EEEEE---EEEEEEEEEEEE--EEEEEEEE---EEEEEEE-HHH-----\|\|---HHHHHHHHHHHHHHHHHHH----HHHH------HHHHHHHHHHHH----EEEEE-- [illegible] |
| | -----EEEEE---EEEEEEEEEEEE--EEEEEEEE---EEEEEEE-HHH-----\|\|---HHHHHHHHHHHHHHHHHHH----HHHH------HHHHHHHHHHHH----EEEEE-- [illegible] |
| | -----EEEEE---EEEEEEEEEEEE--EEEEEEEE---EEEEEEE-HHH-----\|\|---HHHHHHHHHHHHHHHHHHH----HHHH------HHHHHHHHHHHH----EEEEE-- [illegible] |
| WP_162523516.1_Calorimonas_adulescens | MGENVHWVFDRWNNKPVKAVERYELWNTTNCLVYDPTS [illegible] |
| MCL4298460.1_Anaerolineae_bacterium | MLKIGDFA----TGQTGRVIGLATLFGQTYADIFIQPN [illegible] |
| WP_207425053.1_Pedobacter_sp._SYSU_D00535 | ----------VFNGDDAEVIELSKVFGVEKAQIKLLRS [illegible] |
| WP_218808307.1_Methylococcus_sp._Mc7 | -WSVGAWCWHARQASPCRVIDRQDVWGEVAYRVWLPAK [illegible] |
| NPV81618.1_Bacillota_bacterium | M--GGVWAFSPAHSEPCRILDSQVLWGQTICRIWLPGR [illegible] |
| ELY62516.1_Natronococcus_jeotgali_DSM_18795 | MLQSGQQIHID--GQPAEIIRTYSVGDIESLRAYIEDA [illegible] |
| WP_053959402.1_Sulfobacillus_thermosulfidooxidans | M----NWRYLPSYRQMVRILDCHTVFGQKTCQIWLPNE [illegible] |
| MCD8293627.1_Clostridia_bacterium | -MNTGDYVYDKVAKTRVRIIDRIEAWGYVSYEVYDPAN [illegible] |
| WP_162251488.1_unclassified_Nocardioides | WLIPGILV--------VEIVEVSHVGAGAFIKVRDDLR [illegible] |
| NMB01451.1_Bacillota_bacterium | MYQIGQWLMYQPDFSECQIVESFDFWETPNYRIWIPLK [illegible] |
| WP_009146898.1_Thiorhodovibrio_sp._970 | MSVGQEWRYSTVHNSACLCLEEQTLWGQTVCRVWLPNQ [illegible] |
| HIP19029.1_Sulfurovum_sp. | M--NSNWQYSTTYNSPCKILDTQTLWGQTICHIWLPDS [illegible] |
| MBT5530410.1_Cytophagia_bacterium | -------------------MEEHTLWDQKVYDVWITSE [illegible] |
| WP_200891843.1_Caldicellulosiruptor_danielii | M--KDHFVYVKSLKKYAKVLDIKELWGYKIYVVLDTSS [illegible] |
| NLE00371.1_Fibrobacter_sp. | MMQPGDLVTCLLHEDKGKIISVSDLFEERYAEVNFS-T [illegible] |
| EMR73855.1_Thermoplasmatales_archaeon | M--IGQKVISDLYAEQGIVINITKIFNKEFCTVFYKKN [illegible] |
| WP_073324029.1_Marisediminitalea_aggregata | MWQAGCWLWSKSQECYVQLVEEHDLWQIPYCTVLGSNE [illegible] |
| MBN1459902.1_Armatimonadota_bacterium | MYSSGEWVSIELTGQAARVLESSELWGTREYQLWIPEL [illegible] |
| WP_068347600.1_Kosmotoga_arenicorallina | M----EWYWLEEKKQPCVLLEERKMFGYEMARVWLPTE [illegible] |
| WP_202636300.1_Rugosibacter_aromaticivorans | MLGVGDWCWFTRHASPCRVVDRQDVWGEVSLRVWLPAK [illegible] |
| WP_020614470.1_Sediminispirochaeta_bajacaliforniensis | MYSIGELVWSRQHKKEAKILSSSELWDKVSYEILIPEE [illegible] |
| MBT6646602.1_Nitrososphaerota_archaeon | MYSEEQWVWSKVHQEPLKIIEINKQWNVTSYKLWSQSQ [illegible] |
| MBV6489839.1_Fimbriimonadaceae_bacterium | MLVVGQRLWSPTLRAWVQVLETKKLFGRDVVKVLVEGS [illegible] |
| WP_073205019.1_Prevotella_ruminicola | M----------RGKSIEIIGKRQMFDKTFCWIHILEN [illegible] |
| NSW90672.1_Bacillota_bacterium | MFDIGQWAYDTVKGVRVQVIGKEDLWGYVSYQVYEPVS [illegible] |
| WP_148854793.1_Moorella_thermoacetica | M--PGILVRSTIYPEKGLVLGNEEFFDQVYVHVFFEKT [illegible] |
| MBU0460857.1_Nanoarchaeota_archaeon | M--IGDLVRSPDYKDLGRIINVTEIDGTKYYEIYFSKT [illegible] |
| KAF0196956.1_Bacillota_bacterium | MYNTGDWVCYR-GKNTAKVIDRRSLWGVITYCIYISET [illegible] |
| consensus/100% | ...................h..........h.h..... [illegible] |
| consensus/95% | ..................hl...ph.....h.l..... [illegible] |
| consensus/90% | .................bhl...ph.....h.lb...p [illegible] |
| consensus/85% | h.....b.......p.sbllp..phas...hplh...p [illegible] |
| consensus/80% | M.....b.......p.sbllp.pphas...hplh...p [illegible] |
| consensus/75% | M...sphh......p.sbllp.pplas...hplah..p [illegible] |
| consensus/70% | M...sphh......p.sbllcppplasb..hplahs.p [illegible] |

### ; Type3 BREX BrxHII-Helicase and Eukaryotic Swi2/SNF2-Helicase Alignment Comparison

```
WP_162523516.1_Calorimonas_adulescens            ydehririyilagaklkeliarqsilspigsNIIPLPHQIYALVRA--ISSNRIRFLLADEVGLGKTIEAGLIMTEM[illegible]
MCL4298460.1_Anaerolineae_bacterium              tiaslfvarlaalqlramltqsgvlsaanfRVTPLPHQVLAVDFV--LNQLRPRALLADEVGLGKTIEAAMVYEEL[illegible]
WP_207425053.1_Pedobacter_sp._SYSU_D00535        fsnvkirflaiaakikaeianqamiaplesNIIPLPHQILALEKV--MNGQSLRFLIADEVGMGKTIETGLILKEL[illegible]
NPV81618.1_Bacillota_bacterium                   vtanhiayvaaaarvaetltqdvllapigaSVIPLPHQISALSRA--ISGGRVRYLLADEVGLGKTIEAGLIMREL[illegible]
ELY62516.1_Natronococcus_jeotgali_DSM_18795      eaaewfdlrmdalnltlaheqgqllsisnsLVRLEPYQLDCVNEV--MQSLRQRYLIADDVGLGKTIEAGLILKEL[illegible]
WP_053959402.1_Sulfobacillus_thermosulfidooxidans lslaeicyralagriiqalnqdvllaplegQIEPMPHQFAVLERV--IYGGQGRLLLADEVGLGKTIEAGLVIREL[illegible]
MCD8293627.1_Clostridia_bacterium                idenylryvvllakirne-aaagllsslasGIIPLPHQIHVLKRV--LEKNAIRYMLADEVGLGKTIEAGMIIKEL[illegible]
WP_162251488.1_unclassified_Nocardioides         adidefrlaieairlthahslnpllavsssNVEPLPHQVQAVDHL--LKSHPQRFLLADDPGAGKTVMAGLFLKEA[illegible]
NLE00371.1_Fibrobacter_sp.                       ygpspffsrnailrldanlsemkvsaatnfKIMPLPHQILTVHFV--LNRFQPRCLIADEVGLGKTIEAILAYQEF[illegible]
EMR73855.1_Thermoplasmatales_archaeon            qspiefklnllshdievlssgeqalapinfKITPLPHQLLALNFV--IDQFRPRCLLADEVGLGKTIEAALIMEEL[illegible]
MBT6646602.1_Nitrososphaerota_archaeon           ynlefliytiatarinqayatesiisplegNLIPLPHQLTALKKI--TQNNQIRFLLADEVGLGKTIEAGLVLKEM[illegible]
MBV6489839.1_Fimbriimonadaceae_bacterium         fdlddalsvtagariwsalgsdlflaplvsKAIALPHQFRVLRKA--MGGFPVRMMLADEVGMGKTIEAGLVLKEL[illegible]
P46100-ATRX_HUMAN                                ------------------------------VIKLKPHQVDGVQFMWDCCSPGSGCILAHCMGLGKTLQVVSFLHTV[illegible]
O14981-BTAF1_HUMAN                               ------------------------------NAELRKYQQDGVNWLAFLNKYKLHGILCDDMGLGKTLQSICILAGD[illegible]
Q12873-CHD3_HUMAN                                ------------------------------GGTLHMYQLEGLNWLRFSWAQGTDTILADEMGLGKTIQTIVFLYSL[illegible]
Q96L91-EP400_HUMAN                               ------------------------------YGALRDYQKIGLDWLAKLYRKNLNGILADEAGLGKTVQIIAFFAHL[illegible]
Q03468-ERCC6_HUMAN                               ------------------------------FKKLFKYQQTGVRWLWELHCQQAGGILGDEMGLGKTIQIIAFLAGL[illegible]
Q9NRZ9-HELLS_HUMAN                               ------------------------------GGVMRWYQVEGMEWLRMLWENGINGILADEMGLGKTVQCIATIALM[illegible]
Q14527-HLTF_HUMAN                                ------------------------------SKELPPFQRNDLYYITNFSENVHGGILADDMGLGKTLTAIAVILTN[illegible]
Q9Y620-RAD54B_HUMAN                              ------------------------------VYHLRPHQKEGIIFLYECVNGRCGAILADEMGLGKTLQCISLIWTL[illegible]
Q149N8-SHPRH_HUMAN                               ---------------------------------------TSEGLKLYYNPYPQLLGGILADEMGLGKTVEVLALILTH[illegible]
P51531-SMCA2_HUMAN                               ------------------------------NGTLKHYQLQGLEWMVSLYNNNLNGILADEMGLGKTIQTIALITYL[illegible]
Q9H4L7-SMRCD_HUMAN                               ------------------------------SLSLKPYQKVGLNWLALVHKHGLNGILADEMGLGKTIQAIAFLAYL[illegible]
Q6ZRS2-SRCAP_HUMAN                               ------------------------------RGQLREYQHIGLDWLVTMYEKKLNGILADEMGLGKTIQTISLLAHL[illegible]
Q9UNY4-TTF2_HUMAN                                ------------------------------KVPLLLHQKQALAWLLWREQKPQGGILADDMGLGKTLTMIALILTQ[illegible]
Q5FWF4-ZRAB3_HUMAN                               ------------------------------RAKLLPFQKDGI-----IFKRNGRCMVADEMGLGKTIQAIGITYFY[illegible]
P53115-INO80_YEAST                               ------------------------------ACTLKEYQLKGLNWLANLYDQGINGILADEMGLGKTVQSISVLAHL[illegible]
P38144-ISW1_YEAST                                ------------------------------NGQLRPYQIQGVNWLVSLHKNKIAGILADEMGLGKTLQTISFLGYL[illegible]
P31244-RAD16_YEAST                               ------------------------------TIKLLPFQLEGLHWLISQESIYAGGVLADEMGMGKTIQTIALLMND[illegible]
P38086-RDH54_YEAST                               ------------------------------GKFLRPHQREGVKFMYDCLSDISGCLLADDMGLGKTLMSITLIWTL[illegible]
Q08562-RIS1_YEAST                                ------------------------------TVNLLKHQRLGLHWLLQVESAKKGGLLADDMGLGKTIQAIALMLAN[illegible]
P32849-RAD5_YEAST                                ------------------------------AQNLQQDVEDGIFFYANLHTMIKGGILSDEMGLGKTVAAYSLVLSC[illegible]
Q05471-SWR1_YEAST                                ------------------------------RGNLRTYQKQGLNWLASLYNNHTNGILADEMGLGKTIQTISLLAYL[illegible]
P31380-YAB9_YEAST                                ------------------------------EISLKDYQQTGINWLNLLYQNKMSCILADDMGLGKTCQVISFFAYL[illegible]
P43610-YFK8_YEAST                                ------------------------------NCILKPYQLEGLNWLITLYENGLNGILADEMGLGKTVQSIALLAFI[illegible]
Q06554-YL247_YEAST                               ------------------------------GPSLLWNKLTGYILLYNQYPVCAKGVLAEEMGLGKTIEILSLILLN[illegible]
P60240-RAPA_ECOLI                                ------------------------------RTSLIPHQLNIA-----HDRHAPRVLLADEVGLGKTIEAGMILHQQ[illegible]
consensus/90%                                    ..............................b.aQ..sl.bh.........llAD-hGhGKTlb...hh....[illegible]
consensus/85%                                    ..............................b.aQ..sl.bh.........lLAD-hGLGKTlbs..hh....[illegible]
consensus/80%                                    ..............................b.aQb.ul.bh..h.p.....lLAD-hGLGKTlbs..hh....[illegible]
consensus/75%                                    ..............................b.aQb.ulpbh..h.p....slLADEhGLGKTlbs..hl..hh[illegible]
consensus/70%                                    ............................hb.aQbpulphl..h.p...pslLADEhGLGKTlbshhll..hh[illegible]
```

Representative Alignments of Core Protien Components in DUF499 Associated Type-1 BREX-related System;
1. DUF499 ATPase
2. METHYLASE
3. HELICASE

### ; DUF499 AAA-ATPase Representative_1; ORC-CDC6-AAA+ATPase + wHTH + Alpha+Beta-Domain + 3S-wHTH + Degenerated-RRM(Ferredoxin-Fold)

OrigSeq

[illegible]

WP_148590534.1_Aquisphaera_giovannonii
MBQ9433643.1_Synergistaceae_bacterium
WP_054571357.1_Frankia_sp._R43
EEO54231.1_Bacteroides_sp._2_2_4
MBM4287682.1_Deltaproteobacteria_bacterium

[illegible]

NET07313.1_Symploca_sp._SIO2B6
MCX7997938.1_Leptospiraceae_bacterium
consensus/100%
consensus/95%
consensus/90%
consensus/85%
consensus/80%
consensus/75%
consensus/70%

### ; DUF499 AAA-ATPase Representative_2 (HEPN fusion Version); HEPN-EndoRNase + ORC-CDC6-AAA+ATPase + Alpha+Beta-Domain + 3S-wHTH + Degenerated-RRM(Ferredoxin-Fold)

OrigSeq

[illegible]

WP_244804062.1_Corynebacterium_sp._LD5P10
WP_005520661.1_Corynebacterium_matruchotii
MCL5447091.1_Actinobacteria_bacterium
WP_052687173.1_Streptomyces_lydicus

[illegible]

MDE0020779.1_Candidatus_Poribacteria_bacterium
MCA9831553.1_Dehalococcoidia_bacterium
consensus/100%
consensus/95%
consensus/90%
consensus/85%
consensus/80%
consensus/75%
consensus/70%

### ; DUF499 AAA-ATPase Representative_3; ORC-CDC6-AAA+ATPase + wHTH + Alpha+Beta-Domain + 3-wHTH + Degenerated-RRM(Ferredoxin-Fold)

OrigSeq

[illegible]

HGS37418.1_Deltaproteobacteria_bacterium
MXX34234.1_Gemmatimonadetes_bacterium
OLS15779.1_Candidatus_Lokiarchaeota_archaeon
BBH93037.1_Thermogemmatispora_argillosa

[illegible]

```
# ; METHYLASE; NTD(DUF1156) + N6_METHYLASE(TRD-Insert) + HTH + wHTH + HTH

OrigSeq [illegible]

MDE0324948.1_Candidatus_Poribacteria_bacterium [illegible]
MBU1103407.1_Nanoarchaeota_archaeon [illegible]
MCP3817894.1_Streptomyces_sp._A3M-1-3 [illegible]
GIV04313.1_Fimbriimonadales_bacterium [illegible]
GIV55346.1_Candidatus_Kapabacteria_bacterium [illegible]
HDP95528.1_Candidatus_Aminicenantes_bacterium [illegible]
MAU11781.1_Anaerolineaceae_bacterium [illegible]
MBE3095473.1_Actinomycetia_bacterium [illegible]
MBE7469121.1_Anaerolineales_bacterium [illegible]
MBF0121420.1_Desulfobacterales_bacterium [illegible]
WP_131964567.1_Actinomadura_sp._KC06 [illegible]
WP_237889221.1_Dissulfurimicrobium_hydrothermale [illegible]
MBZ0275668.1_Anaerolineae_bacterium [illegible]
WP_247337806.1_unclassified_Bradyrhizobium [illegible]
WP_069791012.1_Cyanobacterium_sp._IPPAS_B-1200 [illegible]
MYC94582.1_Caldilineaceae_bacterium [illegible]
MBN2002244.1_Anaerolineae_bacterium [illegible]
WP_012120128.1_Roseiflexus_castenholzii [illegible]
WP_013179177.1_Truepera_radiovictrix [illegible]
PWV37727.1_Desulfurococcaceae_archaeon [illegible]
MCF7707716.1_Verrucomicrobiota_bacterium [illegible]
WP_259099627.1_Candidatus_Fervidibacter_sacchari [illegible]
WP_221380247.1_Actinoplanes_polyasparticus [illegible]
MBI3467148.1_Planctomycetota_bacterium [illegible]
MBR1604260.1_Synergistaceae_bacterium [illegible]
MCC6334308.1_Myxococcales_bacterium [illegible]
MCY3951957.1_bacterium [illegible]
MCY4442679.1_Deltaproteobacteria_bacterium [illegible]
MDC0253130.1_Chloroflexi_bacterium [illegible]
MSW27208.1_Actinomycetia_bacterium [illegible]
TAK68534.1_Actinomycetota_bacterium [illegible]
WP_145451933.1_Gimesia_panareensis [illegible]
MBM3143887.1_Chloroflexota_bacterium [illegible]
NLH75304.1_Verrucomicrobiota_bacterium [illegible]
MBM3454154.1_Bacteroidota_bacterium [illegible]
NLB94468.1_Armatimonadota_bacterium [illegible]
WP_008352255.1_Caballeronia_zhejiangensis [illegible]
MCY4474612.1_Chloroflexi_bacterium [illegible]
WP_165360625.1_Candidatus_Chloroploca_sp._Khr17 [illegible]
MCA7120528.1_Acidibrevibacterium_sp. [illegible]
WP_066790911.1_Chloroflexus_islandicus [illegible]
MCL1800683.1_Promicromonosporaceae_bacterium [illegible]
MBK8015116.1_Deltaproteobacteria_bacterium [illegible]
WP_019991368.1_Rudanella_lutea [illegible]
OLS21119.1_Candidatus_Heimdallarchaeota_archaeon [illegible]
consensus/100% [illegible]
consensus/95% [illegible]
consensus/90% [illegible]
consensus/85% [illegible]
consensus/80% [illegible]
consensus/75% [illegible]
consensus/70% [illegible]
```

### ; HELICASE (BrxHII-Helicase Counterpart); TUDOR + Swi2/SNF2-Helicase + Inactive-REase + CTD(Alpha+Beta) + Active-PDDEXK-REase

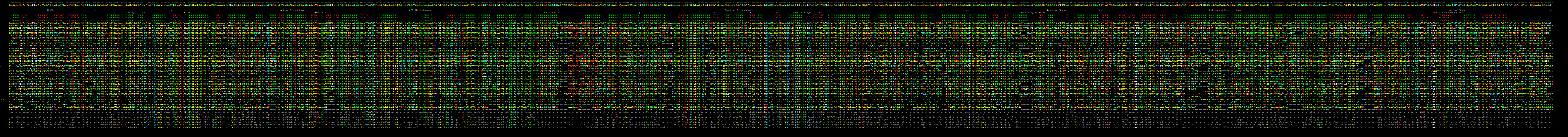

Representative Alignments of Core Protien Components in DUF499 Centered Type-2 Brex-Related System;
1. DUF499 ATPase
2. METHYLASE
3. HELICASE
4. DUF3780

### ; AAA-ATPase Representative_1; ORC-CDC6-AAA+ATPase + wHTH + Alpha+Beta-Domain + 4S-wHTH + RRM/Ferredoxin-Fold

OrigSeq MSLPTIFDKCTPRQDVLIGELAEDQFAASLADVAHSDAPEVYANPQLFFEKTFPTEGLQELLGRLATRFVSYHDQSNGVLRLDTSFGGGKTHNQIAAYHLAERPDAVPDLTDFLNDEQTALGLDVNTAVFVGTHVDGTSARCDDPNAPETKTMWGELAYQLFGQEGYEFLRENDEQQSPPGSQKLERLF[illegible]

[illegible]

WP_015408759.1_Natronomonas_moolapensis [illegible]
NHV98109.1_Thaumarchaeota_archaeon [illegible]
MBD5553419.1_Desulfovibrio_sp. [illegible]

[illegible]

WP_285255572.1_Halobacterium_salinarum [illegible]
consensus/100% [illegible]
consensus/95% [illegible]
consensus/90% [illegible]
consensus/85% [illegible]
consensus/80% [illegible]
consensus/75% [illegible]
consensus/70% [illegible]

### ; AAA-ATPase Representative_2; ORC-CDC6-AAA+ATPase + wHTH + Alpha+Beta-Domain + 2-wHTH + RRM/Ferredoxin-Fold

OrigSeq MTMSILDTCTPRKDLLAGTFNPEIFTANLMQVIGHyrgetgVVENIYTDPLAFFGEGTYPTEGMRLMLRNIFGRLKGDASFPAIQRSETAFGGGKTHTLIAATHLAYRGTEIADAARELIDADLLLAPGAVTVVGIAGDRVAIHETKGSRLIPYTLWGEIAYQIGGEDIYREIGSTATSFGSPGDEYFD[illegible]

[illegible]

ESQ16532.1_Thiohalocapsa_sp._PB-PSB1 [illegible]
KAE8440079.1_Halomonas_piezotolerans [illegible]
MBF6555079.1_Acidimicrobiales_bacterium [illegible]

[illegible]

WP_284505300.1_Streptomyces_cellostaticus [illegible]
consensus/100% [illegible]
consensus/95% [illegible]
consensus/90% [illegible]
consensus/85% [illegible]
consensus/80% [illegible]
consensus/75% [illegible]
consensus/70% [illegible]

### ; AAA-ATPase Representative_3; ORC-CDC6-AAA+ATPase + wHTH + Alpha+Beta-Domain + wHTH + Fn-III+Fn-III + RRM/Ferredoxin-Fold

[illegible]

### ; METHYLASE; NTD(DUF1156) + N6_METHYLASE(TRD-Insert) + HTH + wHTH + HTH

| Sequence | Alignment |
|---|---|
| Origseq | [illegible] |
| MCY4488570.1_Deltaproteobacteria_bacterium | [illegible] |
| WP_253156599.1_Stieleria_sp._TO1_6 | [illegible] |
| WP_171833829.1_Lichenicola_cladoniae | [illegible] |
| MCB1501368.1_Bauldia_sp. | [illegible] |
| MDA7889826.1_Akkermansiaceae_bacterium | [illegible] |
| MDD2765362.1_Opitutaceae_bacterium | [illegible] |
| WP_090142650.1_Limnohabitans_sp._DM1 | [illegible] |
| MBN8450448.1_Candidatus_Accumulibacter_sp. | [illegible] |
| AEW00272.1_Niastella_koreensis_GR20-10 | [illegible] |
| KOX56112.1_Streptomyces_purpurogeneiscleroticus | [illegible] |
| AGC68791.1_Thermoclostridium_stercorarium_subsp. | [illegible] |
| MBL0353837.1_Dechloromonas_sp. | [illegible] |
| WP_277247931.1_Phascolarctobacterium_succinatutens | [illegible] |
| MCP2383159.1_Akkermansia_muciniphila | [illegible] |
| WP_109646849.1_Rhodohalobacter_mucosus | [illegible] |
| WP_189055325.1_Deinococcus_daejeonensis | [illegible] |
| WP_224250596.1_Klebsiella_oxytoca | [illegible] |
| MCM8612466.1_Candidatus_Accumulibacter_sp. | [illegible] |
| MBC8243981.1_Verrucomicrobiota_bacterium | [illegible] |
| MAO03349.1_Citromicrobium_sp. | [illegible] |
| MBS0652529.1_Verrucomicrobiota_bacterium | [illegible] |
| WP_107325860.1_Phaeovulum_veldkampii | [illegible] |
| MCA3288089.1_Roseomonas_sp. | [illegible] |
| MCG2812995.1_Candidatus_Aminicenantes_bacterium | [illegible] |
| NLF73180.1_Candidatus_Anammoximicrobium_sp. | [illegible] |
| AEN07323.1_halophilic_archaeon_DL31 | [illegible] |
| CAG0963790.1_Methanosarcinales_archaeon | [illegible] |
| MBD6956364.1_Thermoplasmatales_archaeon | [illegible] |
| MBI3972514.1_Chloroflexota_bacterium | [illegible] |
| MBM3879662.1_Verrucomicrobiota_bacterium | [illegible] |
| MCC5947761.1_Nitriliruptoraceae_bacterium | [illegible] |
| MCL4868346.1_Anaerolineae_bacterium | [illegible] |
| MDE0525350.1_Thaumarchaeota_archaeon | [illegible] |
| MXZ98362.1_Acidimicrobiaceae_bacterium | [illegible] |
| MYC64341.1_Caldilineaceae_bacterium_SB0661. | [illegible] |
| OPY34215.1_Methanomassiliicoccales_archaeon | [illegible] |
| POG53811.1_Haloferax_marisrubri | [illegible] |
| QDV88972.1_Phycisphaerae_bacterium_RAS2 | [illegible] |
| RWX74209.1_Candidatus_Methanosuratincola_subterraneus | [illegible] |
| WAH99915.1_Candidatus_Methanoperedens_sp. | [illegible] |
| WP_013749026.1_Pyrococcus_sp._NA2 | [illegible] |
| WP_028962638.1_Sulfobacillus_thermosulfidooxidans | [illegible] |
| WP_082047906.1_Thauera_sp._SWB20 | [illegible] |
| WP_156785485.1_Synechococcus_sp._PCC_7002 | [illegible] |
| WP_233735706.1_Halonotius_aquaticus | [illegible] |
| WP_285255574.1_Halobacterium_salinarum | [illegible] |
| consensus/100% | [illegible] |
| consensus/95% | [illegible] |
| consensus/90% | [illegible] |
| consensus/85% | [illegible] |
| consensus/80% | [illegible] |
| consensus/75% | [illegible] |
| consensus/70% | [illegible] |

### ; HELICASE (BrxHII-Helicase Counterpart); HKD-EndoDNase + Swi2/SNF2-Helciase + Inactive-REase + CTD(Alpha+Beta-domain)

```
OrigSeq                                          [illegible]
                                                 -----HKD-Catalytic-Motif----- [illegible]
                                                 <<------HKD-EndoDNase------ [illegible]
MCK6447153.1_Planctomycetota_bacterium           [illegible]
AKC83094.1_Verrucomicrobia_bacterium             [illegible]
MCB9841664.1_Phycisphaeraceae_bacterium          [illegible]
WP_090215779.1_Pseudomonas_guangdongensis        [illegible]
WP_196479964.1_Burkholderia_pseudomultivorans    [illegible]
HCG7747318.1_Vibrio_parahaemolyticus             [illegible]
MCE9614803.1_Lentisphaerota_bacterium            [illegible]
MTE20051.1_Streptomyces_taklimakanensis          [illegible]
WP_206882508.1_Alicyclobacillus_fructus          [illegible]
WP_084460301.1_Aminiphilus_circumscriptus        [illegible]
MBI3972513.1_Chloroflexota_bacterium             [illegible]
KPC75468.1_Thermoactinomyces_vulgaris            [illegible]
MBX6754325.1_Thermorudis_peleae                  [illegible]
MCF8137375.1_Desulfotignum_sp.                   [illegible]
GIW50774.1_Gemmatimonadales_bacterium            [illegible]
MCI7613155.1_Selenomonadaceae_bacterium          [illegible]
MYD03943.1_Acidimicrobiia_bacterium              [illegible]
WP_073234049.1_Desulforamulus_putei              [illegible]
WP_036720040.1_Paracoccus_sphaerophysae          [illegible]
WP_161844016.1_Pseudoflavonifractor_sp.          [illegible]
WP_100917763.1_Candidatus_Thiodictyon_syntrophicum [illegible]
AXH34461.1_Humibacter_sp._BT305                  [illegible]
ELZ44607.1_Halorubrum_distributum_JCM_9100       [illegible]
MBD5553416.1_Desulfovibrio_sp.                   [illegible]
MBF6555076.1_Acidimicrobiales_bacterium          [illegible]
MBP7066719.1_Methanothrix_sp.                    [illegible]
MBV9397056.1_Bryobacterales_bacterium            [illegible]
MCD8487848.1_Desertifilum_sp.                    [illegible]
MCK6511424.1_Myxococcota_bacterium               [illegible]
MCY4068113.1_Acidimicrobiaceae_bacterium         [illegible]
MCY4434136.1_bacterium                           [illegible]
MDE0094453.1_Gammaproteobacteria_bacterium       [illegible]
MXW96682.1_Acidimicrobiaceae_bacterium           [illegible]
NLA99646.1_Methanomicrobiales_archaeon           [illegible]
NLJ36707.1_candidate_division_WS1_bacterium      [illegible]
OYY89820.1_Sphingomonas_sp.                      [illegible]
PZV04180.1_Cyanobium_sp.                         [illegible]
SNR42896.1_Halorubrum_vacuolatum                 [illegible]
WP_054466705.1_Planktothricoides_sp.             [illegible]
WP_103287095.1_Candidatus_Nitrosocaldus          [illegible]
WP_128478225.1_Halorussus_pelagicus              [illegible]
WP_164002139.1_Pyxidicoccus_caerfyrddinensis     [illegible]
WP_176737426.1_Micromonospora_citrea             [illegible]
WP_179052052.1_Nostoc_sp._TCL26-01               [illegible]
WP_285255575.1_Halobacterium_salinarum           [illegible]
consensus/100%                                   [illegible]
consensus/95%                                    [illegible]
consensus/90%                                    [illegible]
consensus/85%                                    [illegible]
consensus/80%                                    [illegible]
consensus/75%                                    [illegible]
consensus/70%                                    [illegible]
```

### ; DUF3780 Component

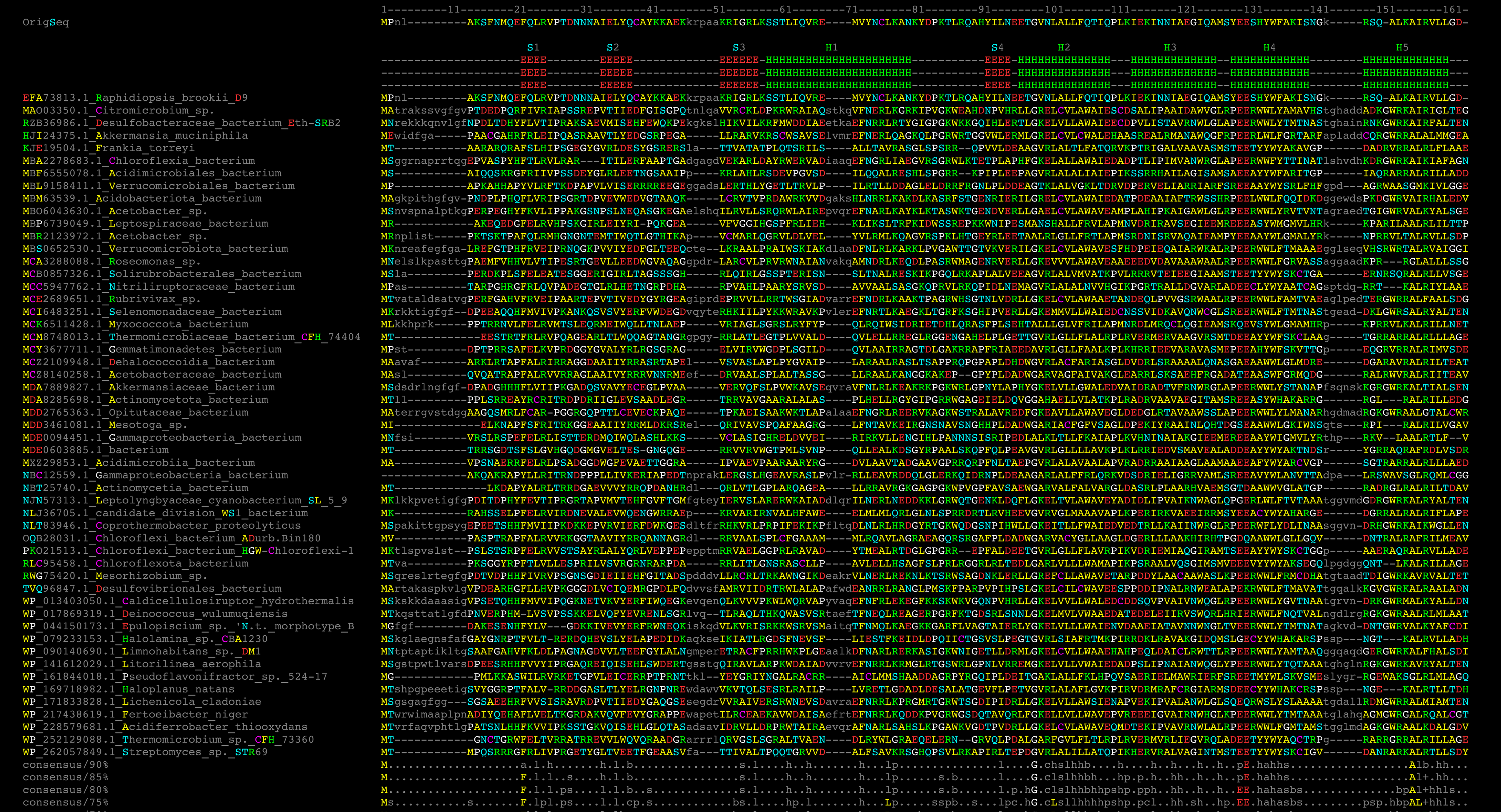

### ; DUF3780 + RAMA

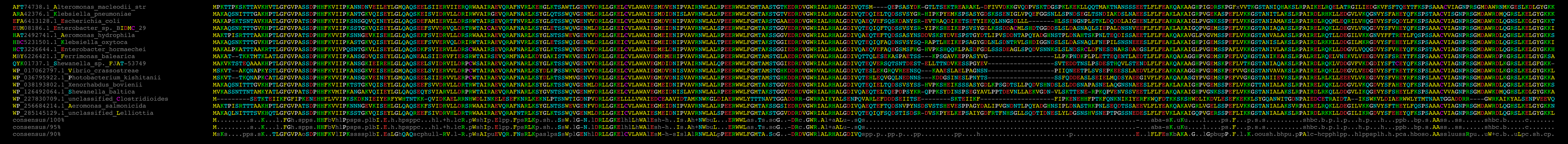

Representative Alignments of Core Protien Components in DUF499 Centered Type-3 Brex-Related System;
1. DUF499 ATPase
2. METHYLASE
3. PglZ-Phosphatase (Standalone-Domain)
4. Inactive STAND-NTPase
5. HELICASE

### ; DUF499 AAA-ATPase; ORC-CDC6-AAA+ATPase + wHTH + Alpha+Beta-Domain + wHTH + RRM/Ferredoxin-Fold

| | |
|---|---|
| OrigSeq | MPTIEDVLDLSEEVRNRHVKGNIELYKLISRGDEFEKNSEAIFNCTYPTGPIRSIVEFIARKIDPE [illegible] |
| CAD6492704.1_Candidatus_Argoarchaeum_ethanivorans | [illegible] |
| HCF99178.1_Chloroflexi_bacterium | [illegible] |
| HIE29204.1_Candidatus_Poribacteria_bacterium | [illegible] |
| HPN33675.1_bacterium | [illegible] |
| HUT59382.1_Phycisphaerae_bacterium | [illegible] |
| HVA45320.1_Pirellulales_bacterium | [illegible] |
| HXK27027.1_Candidatus_Binatia_bacterium | [illegible] |
| HZM03188.1_Candidatus_Saccharimonadales_bacterium | [illegible] |
| KOX91592.1_Haloarcula_rubripromontorii | [illegible] |
| KXA95606.1_candidate_division_MSBL1_archaeon | [illegible] |
| MBA3936148.1_Planctomycetota_bacterium | [illegible] |
| MBC7429623.1_Bacteriovorax_sp. | [illegible] |
| MBK6914962.1_Ignavibacteriales_bacterium | [illegible] |
| MBM4461756.1_Chloroflexi_bacterium | [illegible] |
| MBM4466103.1_Chloroflexi_bacterium | [illegible] |
| MBN2391962.1_Anaerolineae_bacterium | [illegible] |
| MBQ8111585.1_Kiritimatiellae_bacterium | [illegible] |
| MBU4312937.1_Candidatus_Omnitrophica_bacterium | [illegible] |
| MBX3129295.1_Polyangiaceae_bacterium | [illegible] |
| MBY8216331.1_Vibrio_fluvialis | [illegible] |
| MCE0498498.1_Methylacidiphilales_bacterium | [illegible] |
| MCG2864876.1_Vulcanisaeta_sp. | [illegible] |
| MCK8825903.1_Fuchsiella_alkaliacetigena | [illegible] |
| MCL0058394.1_Dehalococcoidia_bacterium | [illegible] |
| MDD3269499.1_Syntrophomonadaceae_bacterium | [illegible] |
| MDH4162607.1_Nitrospirota_bacterium | [illegible] |
| MDM7274691.1_Thermoprotei_archaeon | [illegible] |
| MDR0374944.1_Treponema_sp. | [illegible] |
| MDR0392109.1_Planctomycetaceae_bacterium | [illegible] |
| MDR2561287.1_Holophagales_bacterium | [illegible] |
| MDU2146556.1_Paeniclostridium_sordellii | [illegible] |
| MEA3344886.1_Chloroflexota_bacterium | [illegible] |
| MEA3401058.1_Armatimonadota_bacterium | [illegible] |
| MEA3428074.1_Thermodesulfobacteriota_bacterium | [illegible] |
| MEB3759607.1_Desulfurococcales_archaeon | [illegible] |
| NLX24424.1_Lentisphaerae_bacterium | [illegible] |
| NMX21982.1_ANME-1_cluster_archaeon_GoMg4 | [illegible] |
| OCQ93485.1_Nostoc_sp._MBR_210 | [illegible] |
| RLF82338.1_Thermococci_archaeon | [illegible] |
| SFC23913.1_Halobiforma_haloterrestris | [illegible] |
| TEU12999.1_Anaerolineales_bacterium | [illegible] |
| WP_011944180.1_Thermotoga_petrophila | [illegible] |
| WP_012608229.1_Desulfurococcus_amylolyticus | [illegible] |
| WP_125670094.1_Candidatus_Methanodesulfokores_washburnensis | [illegible] |
| WP_125672294.1_Candidatus_Methanodesulfokores_washburnensis | [illegible] |
| WP_259101344.1_Candidatus_Fervidibacter_sacchari | [illegible] |
| WP_260594793.1_Salinirubellus_salinus | [illegible] |
| WP_283022678.1_Bradyrhizobium_sp._CB1717 | [illegible] |
| WP_286167649.1_Ochrobactrum_sp._3-3 | [illegible] |
| consensus/100% | [illegible] |
| consensus/95% | [illegible] |
| consensus/90% | [illegible] |
| consensus/85% | [illegible] |
| consensus/80% | [illegible] |
| consensus/75% | [illegible] |
| consensus/70% | [illegible] |

### ; METHYLASE; NTD(DUF1156) + N6_METHYLASE(TRD-Insert) + HTH + wHTH + HTH

OrigSeq [illegible]

NTD(DUF1156)

MDB6140355.1_Verrucomicrobiaceae_bacterium [illegible]
MBC7429625.1_Bacteriovorax_sp. [illegible]
MSU35394.1_Pedosphaera_sp. [illegible]
WP_156915858.1_Desulfatirhabdium_butyrativorans [illegible]
HEX9782053.1_Opitutaceae_bacterium [illegible]
MBN2391964.1_Anaerolineae_bacterium [illegible]
HZM03184.1_Candidatus_Saccharimonadales_bacterium [illegible]
MDY0168857.1_Thermoguttaceae_bacterium [illegible]
MEA3344889.1_Chloroflexota_bacterium [illegible]
MBA3936146.1_Planctomycetota_bacterium [illegible]
GFP24747.1_Candidatus_Hakubanella_thermoalkaliphilus [illegible]
MDR6223668.1_Methanococcoides_alaskense [illegible]
MBI5901623.1_Rhodocyclales_bacterium [illegible]
MBX3394310.1_Phycisphaerae_bacterium [illegible]
MCC7203045.1_Nitrospirota_bacterium [illegible]
MCL5421040.1_Nitrospirota_bacterium [illegible]
WP_012508814.1_Pelodictyon_phaeoclathratiforme [illegible]
MBX3410399.1_Phycisphaeraceae_bacterium [illegible]
MBP7052833.1_Phycisphaerae_bacterium [illegible]
MBI4291278.1_Betaproteobacteria_bacterium [illegible]
MCL4198880.1_Phycisphaerales_bacterium [illegible]
WP_322248648.1_unclassified_Undibacterium [illegible]
WP_289404476.1_Xanthomonas_campestris [illegible]
MBN2145088.1_Candidatus_Auribacterota_bacterium [illegible]
MBR2124995.1_Acetobacter_sp. [illegible]
WP_002203285.1_Bacillus_cereus [illegible]
CAD6492700.1_Candidatus_Argoarchaeum_ethanivorans [illegible]
GAA0541230.1_Halorubrum_ejinorense [illegible]
HXK27028.1_Candidatus_Binatia_bacterium [illegible]
KXA95578.1_candidate_division_MSBL1_archaeon [illegible]
MBE0478556.1_Candidatus_Aerophobetes_bacterium [illegible]
MBM4461753.1_Chloroflexota_bacterium [illegible]
MBU7037417.1_Theionarchaea_archaeon [illegible]
MBX3129291.1_Polyangiaceae_bacterium [illegible]
MBX3129293.1_Polyangiaceae_bacterium [illegible]
MCK8825905.1_Fuchsiella_alkaliacetigena [illegible]
MCL0058392.1_Dehalococcoidia_bacterium [illegible]
MCL0075299.1_Dehalococcoidia_bacterium [illegible]
MEB3759608.1_Desulfurococcales_archaeon [illegible]
NMX21980.1_ANME-1_cluster_archaeon_GoMg4 [illegible]
RLF12262.1_Thermoprotei_archaeon [illegible]
WP_121816681.1_Halostella [illegible]
WP_125672293.1_Candidatus_Methanodesulfokores [illegible]
WP_147801106.1_Pelomicrobium_methylotrophicum [illegible]
WP_157687720.1_Haloplanus_rallus [illegible]
consensus/100% [illegible]
consensus/95% [illegible]
consensus/90% [illegible]
consensus/85% [illegible]
consensus/80% [illegible]
consensus/75% [illegible]

### ; METHYLASE; NTD(DUF1156) + N6_METHYLASE(TRD-Insert) + HTH + wHTH + HTH

OrigSeq [illegible]

NTD-(DUF1156)

ELY39842.1_Natronorubrum_tibetense_GA33 [illegible]
QCC56983.1_Natronorubrum_bangense [illegible]
WP_004516378.1_Haloarcula_vallismortis [illegible]
WP_017342168.1_Halorubrum_sp._T3 [illegible]
WP_049909043.1_Halorubrum_saccharovorum [illegible]
WP_049982437.1_Halorubrum_sp._BV1 [illegible]
WP_049996322.1_Halococcus_sediminicola [illegible]
WP_053969684.1_Haloarcula_rubripromontorii [illegible]
WP_089699567.1_Halogranum_gelatinilyticum [illegible]
WP_092663659.1_Halorientalis_persicus [illegible]
WP_120083766.1_Halonotius_pteroides [illegible]
WP_129069632.1_Halorientalis_pallida [illegible]
WP_135823129.1_Halostella_litorea [illegible]
WP_135823666.1_Halorussus_ruber [illegible]
WP_135823671.1_Halorussus_ruber [illegible]
WP_135853933.1_Halorussus_salinus [illegible]
WP_135853936.1_Halorussus_salinus [illegible]
WP_137692176.1_Halorubrum_sp._CGM5_25_10-8B [illegible]
WP_152120852.1_Halosegnis_rubeus [illegible]
WP_153952806.1_Halosegnis_longus [illegible]
WP_285254566.1_Halobacterium_salinarum [illegible]
consensus/100% [illegible]
consensus/95% [illegible]
consensus/90% [illegible]
consensus/85% [illegible]
consensus/80% [illegible]
consensus/75% [illegible]
consensus/70% [illegible]

```
# ; PglZ-Phosphatase; (Standalone-Domain)

                                              1---------11--------21--------31--------41--------51--------61--------71--------81--------91--------101-------111-------121-------131-------141-------151-------161-------171-------181-------191-------201-------211-------221-------231-------241-------251-------261-------271-------281-------291-------301-------311
OrigSeq                                       MYSPSDL-ELIFEGFDSVFGILRR-IWDPEN-KSISKYYRNGEIDTNRLEETLLELYPALYDELLERCMAENVAEITRKKKRYCLVIMDSLSLREAMLLENDLKDK-YSV-NLSYSFSSLPSETESFKQKVFGRT-------PDFKYLHDLD---GISVLPESDTLTIWTQAPDDKLHHTRSGHSEPWSMDDVYADTQQLVHEILRTCR-HDDVIITSDHGYVDLTAGCTFQISKEWKGVLGNQF-KNRWAKENNWELEQLYEKGFIRYAGEYYYVAGRYSWTSRGSVNARKHGGLSIMECMTPVLNVKVN

                                                                                                                  S1                  S2                  S3          S4                                          S5                  ----------------------Insert-Region---------------           S6
                                              -----HHHHHH---HHHHHHHHHHHHHHHH---------HHHHHHHHHHHHHHHHHH-HHHHHHHHHH---HHHHHHH-----EEEEEEE---HHHHHHHHHHHHHH-----EEEEEE------HHHHHHHH----HHHH------EEEE--------------EEEEEE--HHHHHHHH---HHHHHHHHHHHHH-HHHHHHHHHHHH---EEEEE-------HHH--------HHHHHHHHHHH---EEE----------EEEE-------EEEEE--------------------HHHH-EEEEEE---
                                              -----HHHHHH---HHHHHHHHHHHHHHHH---------HHHHHHHHHHHHHHHHHH-HHHHHHHHHH---HHHHHHH-----EEEEEEE---HHHHHHHHHHHHHH-----EEEEEE------HHHHHHHH----HHHH------EEEE--------------EEEEEE--HHHHHHHH---HHHHHHHHHHHHH-HHHHHHHHHHHH---EEEEE-------HHH--------HHHHHHHHHHH---EEE----------EEEE-------EEEEE--------------------HHHH-EEEEEE---
                                              -----HHHHHH---HHHHHHHHHHHHHHHH---------HHHHHHHHHHHHHHHHHH-HHHHHHHHHH---HHHHHHH-----EEEEEEE---HHHHHHHHHHHHHH-----EEEEEE------HHHHHHHH----HHHH------EEEE--------------EEEEEE--HHHHHHHH---HHHHHHHHHHHHH-HHHHHHHHHHHH---EEEEE-------HHH--------HHHHHHHHHHH---EEE----------EEEE-------EEEEE--------------------HHHH-EEEEEE---

HCF99179.1_Chloroflexota_bacterium            MTAVDWASLVLTGGTAERWQTLFDGVADLAMPG-------EFEQALDSGDRLIADAAFDLWTEFPSAAPSSVAELQSWGGSGRAVLVLDALSVREMHIILRVAAERGVTVTSSRVCGSAVPTDTQAFASALGLSGRAQLKASGRNFVFASSKPWTGFDHVPPAPNVFLWDPWPDKLLHDSANRSSGPKHFAATVQQRGDGFWKLVERLRQGRRVAITADHGYAVSHRFVDLPP--GLGEKMRLQFGASRSGAVPSALLH-PLYM----SVNGRAAVVGPWKWKSPGGYPHLNHGGLTLGEVCVPVITFEAI
CAD6492703.1_Candidatus_Argoarchaeum_ethanivorans MYSPSDL-ELIFEGFDSVFGILRR-IWDPEN-KSISKYYRNGEIDTNRLEETLLELYPALYDELLERCMAENVAEITRKKKRYCLVIMDSLSLREAMLLENDLKDK-YSV-NLSYSFSSLPSETESFKQKVFGRT-------PDFKYLHDLD---GISVLPESDTLTIWTQAPDDKLHHTRSGHSEPWSMDDVYADTQQLVHEILRTCR-HDDVIITSDHGYVDLTAGCTFQISKEWKGVLGNQF-KNRWAKENNWELEQLYEKGFIRYAGEYYYVAGRYSWTSRGSVNARKHGGLSIMECMTPVLNVKVN
HDN59917.1_Candidatus_Marinimicrobia_bacterium MVKINNINPLIRGDVDGIFISLNE-IWNYAW-SEDKKYYPLGEKLTSKFEEQVSKLYPEIYDSIMQNAERSNTISEIMSKDRFCLIIMDGMSLREVVPLLKDLKKYG-DV-RYRFSYSAIPSETEFFTKRYFDALRPNEK--YYYVHLQREDAIEDIP--LGKNKLLIWSLYPDDMFRYFRSGFE-TQDLQEVFIKTKNILFKLLEHLSSYNEIFITSDHGYFVD----TF-----SWKGLNDFPSGERY----------ESLKKYCKQFNDYWVLIGRYNTIKRGKYVHVRHGGLSFLETIVPLIEIKRN
HDQ70819.1_Chloroflexota_bacterium            M-DLSLIKQLIHIDLQTLYEVLNGEIWTPERHGDLHGYLREGEHAASRLEQAVYAIYRDVYALLEADDVRRDRVDVFRQREAYTLLLIDGLSLRELPLIQEVLNDRGLDC-EVDFALAPLPTETADFARRHYNASRNDHRYPFAFRHVANEDWRLDFS--PSERQRFIWYAFPDDYFGLKE-----TEYGRHVVQPIEAILAAVLGDPDLVRPLAITGDHSYLWQGNQCAWPVADRERAVLAEHFKLGRCTRTATEALA-RTGKAW--VAGSLGAARGRFAWGGRGPTSLFKHGGVSLMECLVPWVRI---
HEU5078765.1_Opitutaceae_bacterium            MLPHVTTQRLLTEDASAAWATIRSELQSQFFQGNVVEELIRREKAIEPLELLLGGAAWTLWEEFATAVPRGSARLLEWGHGGKAVLILDGLSAREMPVLMAEAAKRGFFVNLPELCGSELPAETNAYARALGFSHRGALDNNGGAHKVTG--AYTQLPSLSHESRLFVWHEWPDVRVHDLAGHGKGIRELLPEVVQRSDDFWTLVAKLATGREVLITSDHGYADAGSFRDAET--DEKNYLRDNFGANRYRQSELTT---PLALTLDCPSGRWSLALGRTKWAVAGGNKTLSHGGLTLLETFVPFLTLSKI
HIE29205.1_Candidatus_Poribacteria_bacterium  M-SVELLHRLIETEVHLIWEILTQTIWTYEQEHGADGYFAVCEQKTSEIERNINAVYFEIYERIFEQSLKEQKLKQLLNQDTYCLIIFDGLSLREAPVIANTLAEAGYRV-NQSYALAQIPSETEIFTQYVFGFDKSNLD--FQYYFIPDDSNT-EMS--RNTDKQVFWVRFPDVLLHGKNKGKTQILSYQEILQRTLTILKSLLK----AKEIVITSDHGYINLAPGCFWDLPGSEQKLIGKKFGWGRSTSTGLSELESLPNRV--KVIPGYASILGRYAWPTKGQASALNHGGLSFMEVIVPYLKARKT
HLE25735.1_Thermodesulfobacteriota_bacterium  MLILNNFASLIRPGIEWLEKWLLGEVWTTVRSLSPIEYLNDGESKVNELEEIIAQAAYRLYDEFLEEPRQGRNILGFINDESSALVIFDGLSLREIPVLLNLAQKSNFIVQEVGVSYSALPTETISYIEQRLKLGRREIKEKISAYYYDSQSQRHLLD--PDARKLFLWSAFPDNTYSDSGARF--AQHFEQIHTLLETAWLNTVQQIPRGRRILITSDHGYVYLGAGLTFARSNSELRPLSEYLGGERHRDLTEEGNQPEHPDLAVYPNHKVAVLRGRIQTHPPGATRLYKHGGLSIMEMLVPWIVLEAK
HNQ22500.1_Phycisphaerae_bacterium            MLTPTIFEKLIQPGLPWLADWLLGEVWSQHNQMPPGQYLQSGEQSVNEVEEMLCAAAARVYDEVLLPMPDARHLRQYLSGPPCAAVVFDGVSLREIPLLKRLAEQSGLRILEEGWSVAATPSETVDFIAARLGVGCAALRADITCHYLGQVNERCSLD--AAARAVLVWSFPDYRYTERDAKF--PELFENLHNMMMTAWQNSVQAVVQGRRILVTSDHGYVYFGGSHDNATVAPLTGYGDRFARLAEKPNPPDHRGVAVLVLADRGVAMIRGRVQTHPPRGARKLYKHGGLSLMEMLVPWLVLEAA
HZX48547.1_Nitrospirota_bacterium             MLTHELLDNLIRPGLPWIKKWLIEKVWTLPLSMSHIEYLKAGEDKVNRFEELASFTADRIYRELLSPPDPDKRLLNVL--EDTAVVVFDGLSLREIPILKLSERSGFNVKEVDCSIAAIPSETMDFVEREFQCGRSELKGRITAIYTNNITQGINAA--DGNSPLLVWSAFPDNTYTDSGSKF--ENHFENIHIQFETSWMHTVQQIKGRKTIIITSDHGYIFFGTGMDRTSSDREIRELNRYFGNNRNISFLDSPHPPNSDDIIIDESKGIAMIKGRIKTRSTGATKLYKHGGLSLMEMLTPWVVLEIG
KXA95579.1_candidate_division_MSBL1_archaeon  MIENKIVQELMESDFPILFEYLLEEIWTPK--PGIKEYYAQ-EQEMNKFERFLSNVYGEIYSELLPRYCMDKDKN---LDMEHSIILLDSLSLREAILLKKDLQEEGFDV-NLSFSYSSLPSDTKFYKEKISYEERK-----NKTKKIQDPD---SIS--LEGDEDIIWSDFPDAWLENISAGRTKLNKIEDMFNNCKNLLFNILDQMENTPAVTITSDHGYVRSEAAYSFKTNESDQKELRQVMGGSRYKPIENANGEECVKAGYLLNFNGYYLAKERHTWTIGGKYEIFQHGGVSLLECLIPRLEVEG-
MBA3936147.1_Planctomycetota_bacterium        MPSTAWRERIFGAGPSAIWSAIFGDV-----ACAKLGDGQRLETHCLDGDQLLADAGWDLWRSYADDADRTSDRLIAWPGPPPAVLVIDALSMRELPLLLQAGARRGVVPHTVDAAGAEVPTDTNAFAKALGLSQRAAIKNNPAAFRLQA--AFTRIPAIQHDPKIFVWHDLIDAHLSDIRS----QAQLDDLLRPDGDGFWRFVDHLRHGRRLVITGDHGYATKHHARPIDG--NLGAQVRDIFGGQRLESAAKALPNGPWVV---MTTGSHHVIVGNRHWRVPGSPKEAYHGGLSLLETIVPWIAFPAK
MBC7332925.1_Actinomycetia_bacterium          MLDTDFIVSLIKSEIYSVWESLLN-IWLNNT-SDLKEYYYHGEEDAFLFEDFLLKTYDEFFTSHLPKACLNEPSLFQKETHNVCFIVMDGMSIREGVLIFNALQKQGIAP-KMSYFLSGIPSDTQSFREKIKVDL------SDKFVEINNPK--KIR--ISEGECYIWSYFPDVLLDKIQVGHTVISDLENMYKTSEKIILEVLERIN-IKKIIILSDHGYIRSEPGFSFSVSDPAKKKLKEAFGSSRYIWPLIHGGISLMKSIRPDAEIRKL
MBC7429624.1_Bacteriovorax_sp.                MLSTEWLNKVLSKSPDEAWVDIFAHLKGLVPAQEAKTSYRKLEDDVAYIDHVLSASAWDLWRDFSS-APRASTYVQNFTVGPKAVLILDSLSIREINPICTEISKLGLKIKNIELAGSEIPSETDIYASAIGLSGRASLAKGPKGFLISTENTFVNTPKITSEKNIFIWHGWPDDSLHVYGKHDDAFSEFIEHVTDTSEGFGSCIQKLAQGRELLITSDHGYCNTSSFTSAVG--EHHQELKLL-GHTRAKKSADLVKGAPATMEMTSTNELYRIAVGRRR-PSDKGFPALTHGGLSMLECAVPLIHIASN
MBE7508222.1_Planctomycetia_bacterium         M--RPWEDIILGNLARDAWTEFASLCEIARPL-------ELHNKTIPPDRLLAEAGWDLWQAYPGAAPRSSEALREWIASGRAVLILDAFSLREMFVLLAAAEQRQIVPTQIRVTGSEVPSDTDNFAAAMGAQSRARLATGPAGFALASDDLAVNHPAIRPRRDLFVWHTWLDDILHVHERS---PDQVNSVARQQGPGFWRFIDALRQGRRLIITSDHGYALARSFATNETDEEVVTALRDTFSASRCKPASAAWN--PVVF----THGGHHVVMGQRKWKVQGGYPYLCHGGMSLLEVAVPFVELPAL
MBI4523255.1_Deltaproteobacteria_bacterium    MLDRALLDELTQPGLPRLINWLMTEVWTMDRGQKPAEYLLQGETLVHHRETLLTLAGESLYDELAAAPPPNRSIRAFLQYPGASVVILDGCSIRELPRLAELAAASGRPILEQSYGFAAVPSETEIFIGQRMGLGRRELREQIQAHWIRQPTDQIRIP--EGTETVLLWMRFPDMRFMDTLAAT--AALFDSIWDMLQTVWIRTVQKLPQARPVLVTSDHGYIFLGPGLSEPRLNGVDRPLE----GRRYREFSSAEELPKRPGLWIDPARRLAVIAGRVHNRPQASNSIYRHGGLSLMEMLTPWLVLETQ
MBK6914963.1_Ignavibacteriales_bacterium      MDFNTVFDLLTTPGIPLIHRWMKENVWSESASMDIKTYFESGEKQIQKLEELILTSSNGIYDELSALAEKNSSLLEEL-QSKTALIIFDGTSIRELPLLESLAVNTGFQIIESNYQASALPSDTVSFVEQRLVGKRKELSSRIKSYYYDTPIRHDFEIK--SSELNYLLVWSSFPDGTYTNFEARN--SLHFETFIKQFDLVWKNIVMAIPRGYRIIITSDHGYVYLNNGFTFSSVSGDMALRFLN----QERFRYYSEDEEIPNNSELKIISNRRLGILKGRIKNRPKGANKVKFRHGGLSMLEVLTPWLIIQRS
MBK8636967.1_Chromatiaceae_bacterium          M-DKALHADVFTLPADQAWPRIFDWAWSVCAAASVPSILGERERGLAPLDNLLAGSAWDLWDGFEAHAPRTSAALATFTPGGKAILILDALSLREVPWLLEQAQARGFTIHAARATASELPAETTPFARAIGFAQRSALAHNGKSPIFPG--ALTDLPLVPPHPGIVFWHTWPDERMHKLAEDSGGSRTLAKEAAQAGDDFWALIATLATGRRLVVTSDHGYANVGLFPDVEGADTQGQALKGVFKSGRSATGDDATAAGPLTRRLSTAHGDWLLVLGRKKWKSQGGYPTLAHGGLTLLEVAVPYIELSA-
MBM4466104.1_Chloroflexota_bacterium          MDESTLARDLITADLAVLFAALNQ-VWQADD-DDVNAFFTKRERETNQLERFVLETFFDVYDLLLAQVRQGRGRSRF-EGTTGPIIVLDGCSIREANLLMGQLQEAGYQVADYGYALSEVPSGTVDFNRNVFGTARQWHQ--YQVVSIESG---QLPLFPAETEVLVWISYPDELLHKVRGE---AVTPQEAFDKTAKVLLGTLDGLE-ADEFLVTSDHGYLYVENATLFKASGGDEKVLRQLFGGRRAVPAGDVGPQ-PKSRTYALFDDEGCYVRGRYHWSVQGPAPDIAHGGISLMECLVPVMRIRRG
MBN2145087.1_Candidatus_Auribacterota_bacterium MVPEELIKKLLQPGLGWLKEWLSREVWSSENELSPLDYLNRGEKIICDFEELLVSAAEQVYDELISSAEKQFSIPDAL-DKDTAVVIFDGLSVREMPMLKVLAEKSGMKIKTADYSLSSVPSETLDFIEQKIQVGRKEFKEKISAFYVSGHNETVTL---SGNNSMLLWSSFPDNTYKDSGARF--DSHFSNIRDVFETAWINCVQAIRNKKRIIITSDHGYIFLGSGLDFPRNTKDLRVLNDIFGNERCVSLDKLKGVETLEDVYLDKVRNLALLKGRVKTVGTGASRLYKHGGLSLMEMFTPWLELEIK
MBP6702581.1_Vicinamibacteria_bacterium       MVTAESLGRLLGGEPKEAWAEVFSLLWKGLSAPDVAAEMIRRDKSFESLDILLSGAAWDLWLSFEESVESTADALAAWASGGRAVLILDALSLRETPWILGEAAKRGYTVKSARPTASALPSDTTSFARALGLPQRSSLSAGVPSLRLAG--AATEQPTLPNAPDVLAWHDWPDSLMHDLSGAGPGLRQLAEDARKVGDDFWHFVDRMATGRRLVITSDHGYAASGLFADVPD--DQKTYFQQTFGAERHSATPGAAP--PVDLHLQTARGPHRFALVGRRWRVQSGRKHIPLTHGGLSVLETAVPFIELTKT
MBQ8111586.1_Kiritimatiellia_bacterium        MINAEWTDLLLAAHPAVTWHGVAQALAQLALPE-------ELHRMTVLPDRILAESAFDLWKAFPTAAPRTSARLAQWCTGSKAVLIMDALSLREMKVIVEEAKRRELAV-SAEVTFSEAPSDTTAFAKALGTPTRASLANNSGSFKLFGGRCSTSVPPVPPVPSVVIWHSFVDDLIHAGNKT---AAEVFKLTRQQDDGFWKLVDALRTGRQLVITSDHGYADAAKFSTEIKEPSAVDFLKANFHGKRFAENVSPQ---PLFL----VSNGFCMVTGQRKWPVQGGFPKICHGGLSLLEVASPWIELPAK
MCD6507459.1_Candidatus_Poribacteria_bacterium MAIDDFLKRLIEEDLIVIWQQMLE-LYQVDDPSDLIRFYNERETRLSQLEEALYHGYWEIYDLCFRRSQAQSGDYVFELREGYVVIIADSLSLREVGIMGKLLNDLGWYIEEKTFAVASFPTLTESLSRILLGTTKDVPA--FRYRYVAGPG--EVPNLPVGEPTLIWLRLPDKKLEQVTVAQ--VTTIADAFNATMETLIQLLEQIT-TRPVFITSDHGYLYATDISHYKMPGSIEKEVRKLFRESRAQPSDKEGMERQGNQRYFVRSDTHVGILGRYWWASAKGDRCTAHGGLSLIETLVPAIKVRKN
MCK8825904.1_Fuchsiella_alkaliacetigena       M-RLKLVRELLASEVEVVWKQLQE-IYITEKTDNILEYYSRTEKELLELEKFLFKSYFEFYQEDLKELANREDLTLVRDNPNLTYLLMDALSLRELGLFYDSLEQKGYQL-KIDYTFSALPSQTRFYKEKMRFDIRN-----IKFREIRDYQ---DIK--VRGNEQLIWSRYPDTLIEDIRSGQSLKSDLEETYKKTKQVLFTVLDNLD-KEKIIICSDHGYTRTEAGYNIKINNTVKKWLKELFGGSRFAAKEEANKELLLDRNLIVEVEDHYLPISHYSWPVSGKYSTFSHGGLSLLEVITPRIILTK-
MCK9588325.1_Terrimicrobiaceae_bacterium      MDIHFHHQGLTGEGLQAITGWLTETVWSRERGLSSEDYLRKGELAVCDLEALIAAGAPHVWDELESASAGATTLADWLLPSPHAAVVFDGLSLREMPMLLERAAASGFRVKSARAVATCLPTETRSFVEQRLLGKRREFAEKAQAFYFGQPNSRDNFP---VGRSLLVWSHYPDGLFFDDNARS--EQLFSTFASYIETIWKCTVQAIPAGVPIVVTSDHGYVFFGAGGETGALQADAGALL---EQSRWREFPPEDRFPWHPDLQLVPSRRLALLRGRIRTRKPGSNRLYQHGGFSLMETLVPWIELERK
MCL0058393.1_Dehalococcoidia_bacterium        MSTEVALRRLIGRELSVVWDTLVE-IWRVDSPKDLDGFYRDKEIESCRLEETLEQTYFEAYQELLNGTAVHKTSPWA-GEEERLAIIVDAMSVREACLLSPILEGQGYRVESLSFSLSALPSHTEAFARKHFNMGRQGI-------FVSPPGH--PVPQFPPYDLCFVWLRYPDRGLHAT------ILSPAEVFGKTSQYILEILRNSG-RNTFLLTSDHGYIYADTADHAPIAQSVQKEMRKVFGGSRYVQGPKEALATVYEKTL--SVGDTLLVRGRYLWRQ--RDKRYFHGGLSFVECLIPVIKAKTK
MCL0097130.1_Dehalococcoidia_bacterium        MNLSSLAAQLLTQAIGLLFQAINE-VWATSNANDIDAFYARQERETNHLEDFIKDTYFEIYELLVEWIKSHDRAAEF-ASYDSSIVILDGLSLREANWLMPRLRENGFEIRQYCYAFSRMPSTTQSFFNDVFGVARKWQG--FNFRSIKSG----DVPMFLDGPKSLIWLSFPDELMHPMRGK---GVTPSFALQRTEEFLLKILNLLE-TEDVVLTSDHGYMYAPSASSFHPAVSSRKVLQKLFGAKRGVLFDGKKVEEPKDQTYVMFDDKGCYVRGRYYWSAPGKQSDIAHGGVSLMECLVPVMHLRRK
MDB6140354.1_Verrucomicrobiaceae_bacterium    MLSSHTLQSLLTASEAEAWSSIRDLMAETWLAGAELSEVRRREAGLASLDLLLSHSIWPLWQNFPAGVPRAATQLVKFQLGGRAVLLLDSLSLRELPLLIMAAQSRGFSVHDSAVCASELPPETTTFAKALGFSSRGALDNNAASIAFPG--AFTNLPAVPPAPNIFCWHHWPDERLHQLDRADGGLEKLFPEMLRTGDDFWQFVQRLATGRRLVITSDHGYAATGAFPNVPA--EQKDYLRETFAAQRFRSGAIDA---PLTTTVTTASGLVTFVLGRQKWQVPGGFPALSHGGLTLMETFVPWIELSLS
MDD3269498.1_Syntrophomonadaceae_bacterium    MKQQALIIQLLDQRSKACWQSIIEYLGSMVYPA-------SLHRATVNLDRFLAESAWNLWQDYPASVTTTARTLQAWAPGGCAVLVLDALSLRELPLILQYGKEQGIEPLQAVVTGSEIPTETDAFAHALGASSRASLKNNGSSFILGGAAAYSSAPEIPPLKNLVVWHTWLDDRQIHEHRRE---PDELIRLVHKSSPGFWQLLQRLRQGRKLIVTSDHGYANCRLFASSETNGPAKEIMARIFGQTRSCPAQLDFP--PLAL----TMNNHHVVLGQRRWTVAGGYPHMTHGGLSLLEALVPFIEYAEV
MDD4099138.1_Lentisphaeria_bacterium          M-SQDWNSLILDSRPAEAWAGIFSALLSVARPV-------ALHEATVEPDRLLAEAAWELWHAFPGSAAHTAAELIRWAHHAQAILIIDALSLRELPLLSAASNRNILPLRVAVTGAECPSTTDHFAKAIGLPSRSAMANDPASFALFSEACYTSMPAVPSVPNVVLWHSWLDDLIHQQKL----PDQIASITATKGDGFWSFIDRLRKGRKLVITSDHGYAVSKLFSSEVADPDAIDILRSTFGASRNTAAATPWQ--PIVM----THNKQHVVMGQRKWKVQGGFPHVCHGGMSLLETVVPWIELPHR
MDT3473331.1_Stenotrophomonas_maltophilia     MLDAQILQRLTQDSPKEAWRIITDRVLEIASESNAPSEVVKRDREIAELDLFLASSGWDLWRVFGQGVERTSDRLARWHSVARAVLILDALSLRELPWLLQGAKERGFTLHTVGATASELPSETNPFARALGLTSRSQLQANGFAHKLQP--AQTDLPLVSGAKNWVFWHHWPDAKLHDYGA-GQGLELLTRDAAQSSDDFWSFIERLATGRRLVITSDHGYAATGYFPDADG--EVASHLKQTFSSGRSKAGNGDA---PVALQIDSPHGAHLLALGRRKWRSQGGYPTLTHGGLSLLEVLSPGFRRAAI
MDU2146555.1_Paeniclostridium_sordellii       MISLASLDNILSKRPEESWIKYISDCVWNIFSETCIENEASLRDSLTNDIESLLATSCSDLWDEFDRCISKTSDEVKSFSSDGKAVLILDGLSMRELPFILQGLESRNYTIDKACITGAELPSHTNSFAKAIGISSRSSLTNNSKKSIFNN--AYTGLSIISSQPNLFIWHEYPDKKVHELDSYGCTIQNLSKNMYSSDEKFWEFIKKISTGRDVIITSDHGYAHTGMFMDVTDN-DVKDYLKKQYKSARYRKTQNNE---AIEKSINSNHGDYTFVLNKIKWKVPGGYPTLSHGGLSLLEVMVPFIQFRQE
MEA1926980.1_Candidatus_Auribacterota_bacterium MLTTDILTKLVDPSLSWITDWLLDDVWSTERSTTPEKYLTAGENQVHALEELLAASADRFYNELAGAASQTKKLTERV--PDTAIVVFDGASIREIPLFLEKAHQSGYRVIERSYSFAALPSTTTDFVEQRLLGKRKELKERIKAFYFHDAISTNHIT--AGTEQLLLWSSFPDVTYQDSESRF--ARHFADMQQLYDTVWKNTVMQIPHDRRILITSDHGYIFFGTGVDSIRPNIACELLN----QNRFRIFDQDENLPEIKDLQIFPERRLAMLRGRLRNRPRGTNRVYRHGGLSLMEMLVPWIILEKE
MEA3344887.1_Chloroflexota_bacterium          MQLAELLSQLIQADMPVIWTYLQS-LWRPEE-ESGETYFKQGEKRTMYLEKVLGAAYPELYREILPERCARCNLDV--TRVEVPIIWMDALSLREALLLCSDLDADC----ELSFAYSRLPSETTFYKETIFYPW-------AKTVEIKDVN---WLS--LDGDERGVWCPIPDKELEDIKGRVK-VRTQTDIYEKSRQVLLNICEMLY-GNEFLVTSDHGYISTETHF-LTLPKRLKGAMQNLFDASRYVEVETKADA-LIEDGYIVEHDGYYLVSNRYAWPTRGKYKVMLHGGISLLECLVPVLRI---
MEA3401059.1_Armatimonadota_bacterium         M-KLDLLDRLLTEELVPPFEELRG-LWSVESAGDLQAYLSDNEHRASAFEREIWDVYADVYQVLEGEHPRKQFVSEVLQQDEFTLVVFDGLSLREVPCLLQVMEERDIEA-EVDFALAPVPTETSVFAQEHFGASQSGAE--LAFEHVKQTTWEPSFG--REQRRRLIWAPHPDNIFTLNSDT---VEYGGHVVEPVQAILAAILDSDP-VLPLYITSDHGYLWQGGACAWALDDDEEALLAREFKAGRSADDPTAELT-SGEKVWSDAERQVAAARGRFAWGGTGPPRNFKHGGVSLVECLVPALRLSGE
NLX99245.1_Rhodopirellula_sp.                 MPAEEWINLILNPAPTDTWTAIFARLRDFALSPIGKGWAESLHRETIVPDRLLAESAWDLWQDFARCAPTVVDELKKFTVSGTAVLILDGLSLRELPLIVQAGQSRGLPPVRVEVRGSEVPSETDRFAAALGLPGRSKLYNNPGTFAFAGPDVHTDAPSIPTSPRLFIWHKWPDELIHLHDDKKDGPAVVAAQTKAVSDGFWALVDRIRQGRRLVITGDHGYATSSDFSNEEKNEDTIKLLRNHFGASRCVQENPQSPWPPLVV----RIDGWLVVMGQRKWAVQGGFPHLCHRGLSLLEASVPVIEYPAK
NMX21981.1_ANME-1_cluster_archaeon_GoMg4      MLSNEFVEDLICEDLDRIWEYLVEDVWVHD--KSVSAYYNQGEKKTGSFERFFKDAYFEIYDLMAKESLGKRKKLRDIERSEYCLIVSDGLSIREANLLLRKLEENGYNIKKYTYEISNLPSDTIAFSRDFLNISRTSGEWRGKSVYIEDDKT-NYP-VPYSDQVIFSFPDAFDFDTVTAKI--GDSLEEIYLKASNIILKLQIENTN-HDQIVITSDHGYCTKHPGSNWSIPVSEKEIFQEVFGSARFKEESKIFGDMLDKTDRLKKVGDNVIVNGRYIWPVPGSKKQIYHGGISFMENFVPLIEVEK-
RLF12261.1_Thermoprotei_archaeon              M-LPSHVKLLVESKLCEVWRILTE-IWRPLD-DLKEHFSKDREKIASRFEYQLNRVFYEIYQHLHTSIHEST------KRFKSPLLIVDGLSVREGGLLARDLELKGYKVLRYWYSFALPSNTSSFRKVLK----------GRFREVKSG---RVP--YIDFSVPVWISYPDEILHHAGR----IISPYEAYERTRSVILKVLDRIVESCVVTITSDHGYVVRDVQP-WPISKAVRTFLRSIFGGSRFVKVEDIDKNKPEEDSYVFIDDEYCYVKGRYFWPIQGYRSNVDHGGLSTECIVPLIEVEV-
RLF82337.1_Thermococci_archaeon               MIPEALLEKIITTGTKEIGNFIVKDFSKA--ASPDFQYYSEFEQKVSLFEQWLYENWTEVYELLIPENKEEISIITAFSEINANVLIADGFSIRELIFLINEF---GKDRVTYEFSYAYPPTITSKAATIVFGSPKEIIITGWKTRLIETPM---DPPRIGHETGFAFYSYYPDAPLHNAQIHGTTLQNIGEVIKK----LINMVKELSRYTNLVITGDHGYIFLGNNPNL--------YLWKPIKSPRFGENLKKPP--------YLIVENEAVAIGRFHALK--RDSFITHGGISIMESLVPVVIIRRE
TEU13000.1_Anaerolineales_bacterium           MIDEQFITELIKDQLAAVWRQMER-VYTQTSCGDLRGLMDDRESALNDLERGVYRGFFEAYELLLERAGAQDENYVLSFDECPFVIITDSLSVREAVLLEKFWEGEAWSVERKGFAVAPFPPMTESLSNLLLGVPRDRPQ--FTYRYVAGPE---HIPAMPPSGSLLVWMRLPDTALEEVTVAQ--THTVADAFHSTVRTLEAVLETSG-RDAALATSDHGYLYARSPTQYPMPGGVEEIARSAFRASRVRPLTDERARGRVEQRFFAFGEAHAAMRGRHWWGSASNDRCTAHGGLSFVECLVPVLKIWRR
WP_083335243.1_Planococcus_glaciei            MIDKELLNSMLLGTDQRAWEAIEETLFNVFTEQSPVQEILKREGKISGLEALIAGAGWDLWHSFETNVSKTSELLIKFNHDGKAVFIIDGLSLREMPFFLTESQKRGYTIHQSKITASALPSDTTYFAKKLGFSQRSSLENNGLNHFFTG--AKTDMPLIGNDPNLIFWHTWAFDDRIHEYKVPGKGLRELSKSSETSGFWELIEKASLGRRVIITSDHGYASTGEFTNISG--EQGDYLKETYKSQRYTKNKERRP--PIEIQMENSQGLFSYVLGRRKWNSGGYPTLAHGGLSLMETMVPFIEITK-
WP_084067307.1_Desulfocicer_vacuolatum        MSSLEFVRKIQNESSKDAWHSILSYAWNFCSRTNPASEIIKRDRAVGNIDHYLATAGWDLWSTYEQSVPQTSNALINWHDTGRGVLILDAFSLREVPWLLQQAKERGFTIHKAGPVCAELPADTTPFAKALGFNQRSSLANNGSAHHLPG--AVTDMELIGSEPDWVFWHHFPDHRLHHHDAAGKGISSLVDEIKFTGDSFWSLIHRLTQGRRVIITSDHGYAASGLFPDANK--QQSDYLKKQFKSGRWHNNEMDTG--PIDLEIESRHGAYGYVNGRRKWKSAGGYPTLTHGGLTVLEIAVPYIEISRS
WP_121919201.1_Haloplanus_aerogenes           MLTSEWPKSLVEADVDVVWDALWD-IWWPPS-GPSDHYDHDREWASVELERTLVDIYWEFYSEVLPWRCVNEASIE--IPDDGAFLVMDAMSVREASLFAAALESDGYDV-KVSYSYATVPSETTPYKKRVGYADRE-----YPSATVRDL---DPS--LSGDERLVWSRYPDALLENIQEGKTELSSVEDAYDDSLRVFRSILDQLD-TDRIIVGADHGYVREESGYSFAISEHEKNRLRDTFGGQRFVGVDQADADDLVEDRLAVEADGYYMPVGRYTWPVRGKYSVYQHGGLSLMECLTPRLEVQR-
WP_125672296.1_Candidatus_Methanodesulfokores M-TENFVLSLIREKTSVIFSKMIE-LAESKLLSDLDKSYEDIEDELLRLEKWLFDAWVDLYRYHLAREKNKLAISKAFSEKGANVLISDGYSFREHVVVKNSIKAE----VDEDFGLSATPSITSEASKIFFDVPRRFIEGSWEMCIVDKIK---NPPKTGKKKGIAFLTYYPDAPLHEAKKHGIILQSITAVVGD----LVSLIDELSSNVPLVVTGDHGYIFTKGNPGL--------FLWKNWPKNRYGESYGEFQ---------LKIDETTVAIGRKHAPK--TESMITHGGVSLAESLVPITIIHKK
WP_149537233.1_Siccirubricoccus_phaeus        MLPLPLLRQLQEADPAAAWAALAQHALDVCDGKASDTAARLRDEALQAVDAWLAGAGWDLWAAFGDAMPRTADRLVDWHGAGAAVLILDGLSLRELPLLTQGALGHGFAVVRAEATAAELPGDTNSFAQALGVPARNTLANNPAGFRLLG--ATTDLPLVKAEPRWLFWHEWPDARFHA-----GSSFQALARNVADSDAAFWGFLRRLATGRRLLVTSDHGYAHIGGFHDTYE--DQKAWLRGVFAGGRTAADGET---PLALSLQGRQERVRLPLGRRKWPVPGGNANLGHGGLTLLEALSPFIELRLQ
WP_218932320.1_Adhaeretor_mobilis             MLSRGLLDTITSGSPQQAWDAIGQSLWETFSKQTATSEIVNRDANINELENVVSGSCWDLWQHFGDCVPRTSQSLIEFVGGGKAVLILDGLSLRESPWLLGALKERGYTIHKAGYRASELPCETTPFARSLGFQQRADLENNSKTHSLTG--AATGLPSVKSQEPFVYWHHWPDSRIHELAKPGEGLRKLAAEAQSSEDAFWEFVERLAHGRRLLVITADHGFAACGNFPDLGK--EQSNYMKALFKSGRTAVVPPSTEPTPIDLELETEHGKHRFALGRRKWKVSGKNRLLAHGGLSLLEVFVPFIELST-
WP_257627367.1_Haloplanus_salinarum           MYGSEQLSELVRGELDGAFECLRR-IWSRPD-ALDGQTLTDSEYRTRLLEHHLAAQEPKLYDELNASMGDHPIT----RLDSGCGVIMDALSLREGFQLLSDLDEHGWDVTLEWSGVERLPSETTFICREWFDAHR------DDYRFVGDL---DVPQLPASDPAFVWTRHPDRRLEESFKGNYSSEAMTDIYSDTKGLLEEIVAESV-HDEFLVTSDHGYVNVQDTNPYVLSDSNEDALSSKF-SGRHREVADDYALDLHDAGVIERAGGHYVVKGHYHPTKGPATKHIRHGGLSLPECMTPVLRITT-
WP_259101347.1_Candidatus_Fervidibacter_sacchari MLNESLVRELVEAELPVIWKAMVR-IWKPNGPETLKEFYDEREEQMFRLERFLSDTVSDLYNFLLQRAINQDGSYVLKLQGNNVVIIADSLSVREAALLTHWFPQINFAK-DQPFAVAPFPTLTESLAQKLLGVIRDTEK--FTYRYIAGPG---ALESFPEDRPLLIWLRLPDAELEQVTEAQ--TTKIADVLRVTVEALGKLFQRLC-GRKVVITSDHGYFYGASSKHF----S-FRGLEQLPRERRARGEASSLS-PLEQDAFIGHGEWIALKGRYWRSGSGNARHTSHGGLSLTEVLVPILEI---
WP_260594794.1_Salinirubellus_salinus         MYGIDEVRHLVNDDFDALFECLRR-IWDADD-AFEAKQVADREYRTRLLERESIRSQYPDLYDDLVDRVGDHPLM----ELTDGCGIIMDGMSLREGFRLAADLEERDWTVEYDWAACEGLPTETKFIAQPWFNANKH-----DHVAYIGEP----EVPPLPGTSPEYVWSRFPDKRLHNSQEGQYTLTELVEVYDEAKELVIDIIEESV-HDRFLVSSDHGYVNFSGNNPYRLAEEDEAILKEKF-SGRYREVEHSGLLRLERDGIIHRAGGHYLIRGHHSWTKRGATSRIDHGGLTLVETMTPLLTVDTT
WP_283022679.1_Bradyrhizobium_sp._CB1717      MLDRDVLATILTQTPSVAWSAIVERVHSICRDKDLIRSVREREREVGELDDLISTSAWDLWDTFQDGVPRTSALLQTFPAHGKAILILDGLSLRELPHLISGAEAHGLKLISETVTASEFPPETDPFAAALGFASRSSLRNAPTSKFFPG--AKTDMDAVMPDPNIILWHQMPDDRIHSLENDGDALGRLATELASQGVDFWGLVKKLTTGRSLIITSDHGYAAGGEFYELPD--GPTGFLRDNFKARRSLAGDGEL---PLVMRFDTPRGPYRLALGRRKWKVSGGFPKLTHGGLTLFEVLSPFVEITQ-
consensus/100%                                M.......h..........h...h...............c......-..h......ha......................hh.Du.ShRE...h.....................Ps.T..h..................................hh...D..h.........................h...h........h.h.uDHua.................h.......R.....................................H.Ghoh.c...P.......
consensus/95%                                 M.......l..........h...h...............c......-..l......ha...................shll.DuhShRE...h.....................s..Ps.T..h...................h.......................ha...D..h.........................h...l........hhlsuDHGYh.................h.......R..................................h..sp............HGGhoh.E.h.P.h.h...
consensus/90%                                 M.......l..........h...h...............c......-..l......phap.h.................shllhDuhSlRE...h.......s....p...h..u.hPs.T..h.......p.................h..............hhW..h.D..hp.........................h...l........hhlTuDHGYh.................h.......R..................................hh.up...........h.HGGlohhE.hsP.h.h...
consensus/85%                                 M.......lh.....hh..l..lhp...........p.-.....h-..l..s..clap.h...................shllhDuhSlREh.hl...h..psh...p..hs.u.hPspT..a.p.....+..........h.s......hs........hhW..hPD..hp..............h.............hb..lp.....p.lllTSDHGYh.................h...h......R...............................hh.G+b.....u.....h.HGGlolhEshsPhl.h...
consensus/80%                                 M.....hpplhp.....ha..l.p.lhp.........h.p.-p....h-p.l..sh.clap.h.......p..........s.slllhDuhSlREhshl...h..psh...p..hshu.hPopTp.a.p.b....+.........h.s.....phs.......hhW.aPD..hp..........ph.p.h.....hbphlp.h...pplllTSDHGYh.................lp..h....R..............................hh.G+b.......G....h.HGGlSlhEshsPhl.l...
consensus/75%                                 M...phhppllp.....hap.l.p.lhp.....s...bh.p.Ep....h-phl..sh.clap.h...s.ppp.....b....s.slllhDuhSlREhshlhp.hpppGh.l.p..hshu.lPocTp.asp.h..s.R..........l.s.....phs...s..phhlWppaPD.bhcp.........pph.p.h.....p.hhphlppl..scplllTSDHGYh......sh......p.Lpp.a.s.R.............................hhbGRb.h.s.Gs...h.HGGlSLhEshsPhlplp..
consensus/70%                                 M.s.phlppLlp.....hap.l.p.lap.....s...bh.p.Ep....h-phls.sh.-lap.h...s.ppp.....b....s.slllhDuLSlREhshlhp.hpppGhpl.p.phshu.lPo-TpsFsp.h..s.R.........bl.s.....phs...s..phhlWppaPD.bhcp.p......pph.p.hpp..p.hhphlppl..scplllTSDHGYh...s..sh......p.Lpp.F.upR.....................s...hhbGRbpW.s.Gs...hpHGGLSLhEshlPhlplp..
```

```
# ; Inactive-STAND-NTPase Representative_1;

                                            1---------11--------21--------31--------41--------51--------61--------71--------81--------91--------101-------111-------121-------131-------141-------151-------161----
OrigSeq                                     MSPINDKSFVRKGMSLIWDPELLAEIIAYHKAVTIRQFCQMKGKWPESLPSKDEHALVVVGLEGCMDVLKPSDAEKWLEQDIKPLILDFQDEYQGQCALIFWIPSGLKKIKHVSSTDNYVWQYSTPHSGKVEIGRILWAGAQEVRPIITGSADVADSNWKGLHHPRIS

                                                       S1            S2                 S3                                  S4     -------Insert-Region---------           S5            Terminal-Insert-Str
                                            -------HHHH--EEEE--HHHHHHHHHHH-EEE-HHHHHH--------------EEEEE-HHHHHHH--HHHHHHHHHHHHHHHHHHHHHHHHH----EEEEEE--|---EEEE---EEEEE--------EEE-HHH|-------EEE--------HHH-EEEEE----
                                            -------HHHH--EEEE--HHHHHHHHHHH-EEE-HHHHHH--------------EEEEE-HHHHHHH--HHHHHHHHHHHHHHHHHHHHHHHHH----EEEEEE--|---EEEE---EEEEE--------EEE-HHH|-------EEE--------HHH-EEEEE----
                                            -------HHHH--EEEE--HHHHHHHHHHH-EEE-HHHHHH--------------EEEEE-HHHHHHH--HHHHHHHHHHHHHHHHHHHHHHHHH----EEEEEE--|---EEEE---EEEEE--------EEE-HHH|-------EEE--------HHH-EEEEE----
MDO8527158.1_Deltaproteobacteria_bacterium  MSPINDKSFVRKGMSLIWDPELLAEIIAYHKAVTIRQFCQMKGKWPESLPSKDEHALVVVGLEGCMDVLKPSDAEKWLEQDIKPLILDFQDEYQGQCALIFWIPSGLKKIKHVSSTDNYVWqystphsgKVEIGRILWAGAQEVRPIITGSADvadsnwKGLHHPRIS
EFN9589256.1_Escherichia_coli               MSDVTNDKWKRRGVSVLWCGKTLAELNAASQVISLRQFIGYYEAWPDDMPLLNDDGLYVAGLDVAVDALSPEDALEWLESEIYEMIYDFQNH--ADAALIFWMPDQG-RWKEDLTTSTYHWclagkydaMFPLGQCIWNGAQDVRRIESTSGGkt-newLGLYLERIS
MBJ2281507.1_Pseudomonas_sp._MF6767         MRQVLDTVWQRRGTSWLWDEEARNQVCKAEEVWSLRQFLLSAQAWPDDLPSNNSRTLIVAGLEGSLDLLAPDQGEVWLGDTIKRAILSFQDEYAGEAALIFWLPQKHGRLKVQASTDAVSWlceapnrgQLDFGRLLWGEANYPQEILLREGAqt----AGLFHLRIT
WP_283022681.1_Bradyrhizobium_sp._CB1717    MRQIEDEVWRRRGFTVLWSPTALGLLCSTTEIISMREFLRVCTNWPDTLPTSNGNALVVTGLEGCLDRISPPDAEAWLDEVVQPLMLRFQRQYSDDAALIFWMPTGEKRVELVNLDDTFSWefappfkkPFSISRALFGGAHDAGKIVASEQDdrgqawIGLHLSRIA
MDU2146553.1_Paeniclostridium_sordellii     MRGRKSEIWKRSGISVLWELPPVIECIRNSEVITLNQLFRFSVEWPENLPSFSDSAIIVSGLEYILDLLSPSEAKEWLDQRLYPLLLEFQKEYEGYAALIFVIPQGDERIKYITVKDTFYWicncnd--EIPFSSSLWNGAEDAKYIDIDKYIk-----CGLYHPRIS
KYH32453.1_Moorella_mulderi_DSM_14980       MRE-FSSAWLRRGSSVVFDKYVLGHLIESASLVSLRQALAWLKAWPVEPP-AGRNTVLVSGLETLLEVLDPVEAEAFLRKRVKPLIMEFQYRW-DQCGLVFGFAAPERSFEVTVADEEVLFlr---rdgRVHLSFALWDGSTDVTRLVRDEPQtgr---VGYHVARIS
OPX92064.1_Pelotomaculum_sp._PtaB.Bin013    MRE-RGNGWLRSGSSLIWDAQLLSPLLENGVQIPLHKALNWIGGLPSTIPSGDGKTIFVVGLQTVLETLDSEAAFTFLRNRVQRLIVRVQDFY-NNVGIVFGCNCNWRQWRIDS-NEHAYLrl---rggELNVTFALWNGVAEAQIIMVEDKNvhreigGGFYVRRYS
MBE2220371.1_Anaerolineae_bacterium         MRE-SGTRWQRRGSSIVFDPETLSPLLIGQSLISLRAFLGWREDWPDDPP-GDGKTVLVGGLEPCLELYSPEEAESLLRYRIRPVVEEFQTRW-DQRGLVFGFGKPARAFQLDPTSDEMTYir---qdlVVRLAYSLWNGGADLTQIIRSTAKgse---IGYYVPRLS
MBP6702579.1_Vicinamibacteria_bacterium     MRAPEIDSWKRQGHTILWDAASLASLCHAEKAVTLRHFIRMHAAWPDDATLVKGRALVVAGLDAALDALDPDDALAWLEKTVYPAVISFEKNVASQAALVFWFADGR-RFKHSMADLVAWWdcslafgqQIPLTRGVWNGSSSMQEIRVKNSEgk-vvhVGYHVQKIS
HEU5078767.1_Opitutaceae_bacterium          MRADNDSVWTKRGVSVVWDAPELAKRCTLAKALSLRQWFRWQAEWPDYYTGKDGRTIIVAGLDAAIDSMTPETATEWMQAKLLAAIREFQAEVAQEAALIFWMVNRR-RFEHRRADDSVIWksapgyrgTIPISHCIWNGAQDIARIVPPGCAdl-ehgIGLFLQRVS
MBW4553277.1_Aphanocapsa_sp._GSE-SYN-MK-11-07L MQPVFGTGWTRKGFSLLWDVTALSKLVESDQIISMRRFFNLVGQWQDDLPAANGDALVVAGLDASLDALTADDASQWLEEKLRSALLSFQDYYQGDAALIFWLPGGRSRIEVQRATNTYLWlcgstsgrRLPLGQALWAGAEDARHIMAVNDPqtdkdwIGLHLPRIS
MBP7052832.1_Phycisphaerae_bacterium        MQE-TASKWKRTGSSIIWHPDLLADLIRDIEPTPLRTLIGWLKAFPEDPP-GGKQTVLVGGLQTAVETMTPDAVADWLRNHALAAVRAWKSHW-PDVGLVFVMDGPGSLFEFNEGDEIVYFgrgr-drsKVMLSLAIWNGAAAGAYQILVEGTgev---GGYFVRWLS
WP_149537235.1_Siccirubricoccus_phaeus      MP--LADAWARRGLTLLWGADAFRELASPDQVMPLRALFGERLNWPVDLVSNGGRALVATGLEASLDCLAPADAETWIATDLRDLVREFQDAYQGERALVLWLPSGAQRVTYDAPNDAWLWatadn---RLPLGRLLFSGAEDVERVSPAGWRgpapglIGLRQARLS
MCX6876430.1_Verrucomicrobia_bacterium      MNPPTDDVWKRKGISVLWDVAGLASLGSLQTAISLREFFLWNADWREDSTSGGARRIVVAGLEAALDCMEPEAAEEWMAQQLLPVINRCAEILFTGGALLFWMVKHD-RFRVKLENGEILWtcdgehrdKILFSHGLWNGAYDVQRIAPLRTEt--epgLGFYLQRIS
MDR4485162.1_Nitrospirales_bacterium        MNPDHDNTWLYCGTSLLWEAEALNHICSAEAVRSLREFLLLHQAWPDTLRLSNNRTLVVAGLEAAMDTLHPAKAIEWIEQKVYPAVLDFQESVAREAALILWLADHK-RIFHLPSENTYHWhcsaahrkSIPLGRCIWNGAESARRIVTKDTAkk-eiwAGLFQARIS
MCP3685978.1_bacterium                      MMVLRDDAWIRRGFTLLWGLKTLGKVVRPPQVVSMRKFFSMANAWPEELPASNGNALVVSGFEGCLDVLTGQDAERWIEKDIKDVILSFQDEYEGQAGLIFWVPSGHNRISMVGSSEEYHWkhrasksGLHIGRILWSGAEEVERLIDDQRStadkawVGLHHPRIS
MBI5511448.1_Deltaproteobacteria_bacterium  MLPPADRGWVRRGLSLVWNAEALGTVAGPDAAWTMRQLLTRSRAWPAALPSSNGEAVVVAGFDSCLDAFGPDAAEVWIQRELVPVMLSFQEEYDGQAALIFWLPEALRRLTHSSADGSYSWrwagrtsvPLALGRCLWAGAEDLRRIVVPAKGkvdpgwIGLHLARLS
MBQ8111589.1_Kiritimatiellia_bacterium      MKTTNYTLMLRGGSSVLFGKKELSELLTQIEPISLREFLLLGDDLPDDPPPEFEDGILVSGLETVLDAMPVGEAEEFLAGRLRGVIQQIQNHW-TRTGIVFGCSLSEKAFEKQGVSESVVFtt---kddELALSLALWGRSQDIMRLEHKEPGere---VGYHVPRIS
MBA3936145.1_Planctomycetota_bacterium      MKR-GGTSWMRQGSTVITHLETASMLLAQAGLEPLRTALRWAAEWPAE-LAN-VSTLVVNGLSAVLDRCTPEDSAEFLRTKAKLIIRRSQEQW-TRTGILFLFPTHPSTLSVDL-DDRVIYrl---pggSVALSHELWNGAADLREI-----Hadv---IGYHVLRLS
MBN8780446.1_Thiobacillus_sp.               MKILQGDVWKRRGVSLLWGGEALSVLAQPQQVVSIRQFFAMVGQWPDDLPCNDGNTLVVAGLEGCVDLMDPIEGEAWMKSDLLPAVLAFQDEYSLEAALVFWLPTGKNRVKMNRATEAYSWvcsaphsnHLDLGRILWAGAEDVGRIIDPDCTnsdpgwIGLHHPRIS
MBR2124994.1_Acetobacter_sp.                MKELRDTVWQRRGISWLWDNTALDHVCKAHEVGSLRQFLQLYGEYPKSLERSFGRAFVVGGLDTSLDLLTPEEAKTWLREDIKKAIKAFQQRWAEGAALIFWLPRGDSRFDI-SSTDVLTWrcrppyg-KIDFGRILWGQCD-PQHIMQGDKK------IGFFYPRLS
MBK8636965.1_Chromatiaceae_bacterium        MKASEDDVWTRRGINVLWDGAALARIGPVAAAISLRRFFTLDALWPDETPLVDDQALVVAGLDVALDALDPADIEDWIEQVLYPRIQEFQSVFAGHRALIFWMPDKG-RWLENSGEGAFEWrlgsgdqgRLPLARLLWDGAYDVQRIEEPGDKgv-nrwIGLYLARIS
OCQ93482.1_Nostoc_sp._MBR_210               MHK-SSSAWQRRGSSIVFHRDLLVPLIESGAMVSLREALSWANNWPLEPP-NGSNTVLIVGLETCLEVMVPSEAEIFLRRTIRPFLLESRNHW-DRCGLIFGFGCSEKRFCIDPQESILFVyl---dgtHIRLSEGLWNGAAELFYLTTPEQTknh---GGYHV-RLS
MBC7429626.1_Bacteriovorax_sp.              MEEIEAGVWFKRGCSIIWDNRCLTRLVKDNSITPISTALRMSKAWPDKLPASSGRAMVVVGLDGCLESLEDDQREKWLREHVKPLVLSFYRKFQNEASLIFFFERGTASF-IDHIDEAFVWqptn-kqgPLPLVKFLYGGRFEIKRLVSEHDPkke---WGLYVANPS
MBF0302144.1_Desulfamplus_sp.               MDT-SNTVWQRKGSSVIFDQDSLAPFVKDEAVVSLREALSWMKKLPTTPP-VKGKTIVVCGLETMIETLPEDEIDDFLTQRIRPLIIELQNRW-TGNGLVFGFSSHPNTFEESAMQEEVLFkr---rgnKVRLSEGLWDGSANMKRIITSEREsgq---IGYYVARIS
WP_145058250.1_Adhaeretor_mobilis           MDKPNDDLWKRRGISILWDADSLNSLCRSDEVISLRHFRQLFNDWSDDASMVDEETLVVAGLESCIDALPPEEATKWLQDVFYKAMVSYQGEVATQAALILWFVDSR-RLKYRTSDDGWSMhcagefkgQIPLGRCLFNGAADLKEIQDADGNs-----LGLYHPRIS
HCF99181.1_Chloroflexota_bacterium          MDE-AGDMWRRTGTSLVWDGDALHTAATEGVTVSLRTALMWAEVMPEDPP-DGKRVLVVTGLRTAIDVLTEAERDAVLEK-VARLVRAQSRYW-PEAAIVFALPETTRLL-VAPPTGTVVLdlgg---dSLDIGNRIWGGAGGANQLIESRLDskgrqaIGYWLRRVS
MBX3394309.1_Phycisphaerae_bacterium        MDEFDSKMWLTRGSSVIFDKVMLGPLLTEGALIPLRQALSWLSSWPSATP-TGGKTVLVVGGLETVLEMLAVSDAEDFVRRYIRPLIAEFQSCW-PEHGLVFGLNTEPGRLIEDPVSEMVNLkr---rddTICLSTELWNGSADMRRLVRTDRAsgq---GGYYAPR--
WP_270720981.1_unclassified_Bacillus_cereus_group MASSLGDTWTRRGISLLWDSNSLSLITEIKDVLSVREFLRYYQTWPNELHSANGYGMVVSGLETILDYTESQSLIDWMEKDFYPALLSFQSMYEGQCSLIFWLPDGQKRL-EQKTDGLIYWnctgayygQIPLSQCLWNGAADAKLIIKNTNVlsmkncLGIYHPRIS
consensus/100%                              M.......h...G.shl................shp.h......h...........hhhsGhp..h-.........hh....h....h..........ullh.hs......h.....p..h.h.........h.hs..las........................Gh......
consensus/95%                               M.....p.a.ppG.ohla....h..h.......shp.hh......h.........pshhlsGhps.h-.h.......hh...h..hh......h....ullh.hs......h.....p..h.h.........h.hs..las.s.....l.............Gha..p.s
consensus/90%                               M....sp.W.+pG.olla....h..h.......sh+phh.b....hs........pshlVsGLps.l-.h.s.....ah...hb.hh....p..h.....ullh.hs......h.....p..h.h.........l.hs..las.s.....l.............Gha..+ho
consensus/85%                               Mp...sp.W.RpG.ollap...Ls.lh....h.slRphh.h...aP.p.s....psllVsGL-shl-.h.s.p...alpp.lb.hlbphpp.h.spsuLlF.hsp...ph.....p..h.h.........l.lup.lWsGu.ph.bl.............Gha..RlS
consensus/80%                               Mp...ss.W.RpG.ollas...Ls.lh....hhslRphh.h...aPpp.s.ss.psllVsGL-shl-.hss.ph..alpplb.hlbphpp.h.spsuLlF.hsp...ph.....s..h.a.........l.lup.lWsGu.ph.bl....p........hGha..RlS
consensus/75%                               Mp...ss.WbRpG.SllWc.psLs.lhp...hholRphh.h...WPpp.s.ssspsllVsGL-shl-.hss.pup.alppplbshlbpaQppa.spsuLlFhhsp...php.p.ssp.h.a.......pl.lup.LWsGu.-hpbl...p........hGha..RlS
consensus/70%                               Mp..psssWbR+G.SllWc.psLs.lhps..hloLRphh.h...WPss.s.ssspsllVsGL-shlDhhsspcuppalcpplbshlbpaQppa.spsuLlFhhsp...php.p.ssp.h.a.......splslup.LWsGu.-hpcl..sp........hGha..RlS

# ; Inactive-STAND-NTPase Representative_2;

                                            1---------11--------21--------31--------41--------51--------61--------71--------81--------91--------101-------111-------121-------131
OrigSeq                                     ---MKGYEIFEQKAPYENKIGFLDFLRELKSGTQGIPSSFMVVGIDDVLYLAGRDERLALTIHKILQSSAKVLDQKIIEVQIVCKGRLYKGESFWSEYRGEKLPLDYIFGTPNKRRECPVYSTGFNLSS---

                                                   S1          S2                   S3                                    S4   ----Insert-Region-----      S5  Terminal-Insert-Str
                                            ------EEEE---------EEE-HHHHHHHHHH-------EEEEE-HHHHHHHH---HHHHHHHHHHHHHHHHHHHHHH--EEEEE-|-EEEE---EEEEE--EEEE-|HHH--EEE-----EEEE---------
                                            ------EEEE---------EEE-HHHHHHHHHH-------EEEEE-HHHHHHHH---HHHHHHHHHHHHHHHHHHHHHH--EEEEE-|-EEEE---EEEEE--EEEE-|HHH--EEE-----EEEE---------
                                            ------EEEE---------EEE-HHHHHHHHHH-------EEEEE-HHHHHHHH---HHHHHHHHHHHHHHHHHHHHHH--EEEEE-|-EEEE---EEEEE--EEEE-|HHH--EEE-----EEEE---------
CBX30016.1_uncultured_Desulfobacterium_sp.  ---MKGYEIFEQKAPYENKIGFLDFLRELKSGTQGIPSSFMVVGIDDVLYLAGRDERLALTIHKILQSSAKVLDQKIIEVQIVCKGRLYKGESFWSEYRGEKLPLDYIFGTPNKRRECPVYSTGFNLSS---
HIE51614.1_Armatimonadota_bacterium         --MRPGFYLITRRQRPAKTMSLIALLRALHRGE-ALPSAVCVVGLVGLLESASDLRQAARLLAQRFQRHRSALMRQSPVIVFPVEHLSPTPRGVQVGARGSNFIAEWIFP-RLEPKVPRAYYSPF-------
KXA95577.1_candidate_division_MSBL1_archaeon --MEEGFHIIEKSLPFKKMSIVNWIKNIRNEGFNQFESYTVVGLEDLLEKISNPQDFSEKLRKILIDQSNYLVSTGNIFQFLVNEKNPTVKTASGVIDLSAIFG-ELERKDVNWFYQQLNIQS---
MBM4461752.1_Chloroflexota_bacterium        EGLKAGYNLVERGEDDSEVMSIVQFLQRLQARD-TLPRRVKVAGLDRIMATMSDLKP----LHDILYEAGERLLRRSPVVLLPVD-KLTMDVEPKMYWRNQEIRLRPLFGNRLEPQGVGHFYSPFNIP---
MBN2391965.1_Anaerolineae_bacterium         MS-SSGYVLIEQRRSPRLQRGLLDFLRDLAEDELPYPSLVTVTGLDEVLYAAQPDEELAHEIHGRLNQAASRLESRLILIEMRLKGQIIRGDPFRLRHRGIDLPLHIIFDTPGKAADGNYYLVRCHLS----
MBP1716767.1_Deltaproteobacteria_bacterium  ---MGNYLLIEERLPNPKRMGFLDFLRELGQEVFPFSNRLCVVGLEEVLLQAGSGENVSLEIRKILTARANELEANLGQVQIVFRRPLKRSDDFWFDPGGNRMSLRPIFGSPIKQNGNEYYMTGFNLT----
MBU7037416.1_Theionarchaea_archaeon         MDLKRGFHVIEGQFTLIHKTSIINLLRLIETNSHQ--KLTIVEGLDRFLKYSTDE--AIEQARKVLNTGITDMLAIDASLIFVVECDIRIPDNPSIH----DRPLAMILPRPHDIMEPGYLYYPIM------
MBW1935302.1_Deltaproteobacteria_bacterium  ---MPGYQLLELREPYENKMGLLDFLREITQDPFPFPYRVQILGLEDVLFAARPQEELALKIQRILRSAAQNLEKKLVEIQIVFRGRLVRGESLWVEYRNAKLPIGHIFGSPIRQSRNQFYKKSFSLTNGVT
MCF8267322.1_Ignavibacteriales_bacterium    ---MPAYLLVEKKPANTQKIGFLDFLRDLEKDEFPYNSRLMIVGIEETLLAASPDIHAAIQIRNLLQKAANRLSNNCSDIQIVFRGTLQRGDQLVVRHVPQDIPIYLIFGTPIQEYGEKVYRPSFNRSSN--
MCK8825906.1_Fuchsiella_alkaliacetigena     --MQQGYNLMTQNDDFVERMSIMKFLKMVQEENKYIDYSLEIYGLEEFLCFLGKED-ANDLLTEILGKGVNYLTDNFATIKVITENKIIWNNKAVLKYSNEEFPLGNIFGSSLRQEDTEYWVANLNIDS---
NLY06148.1_Candidatus_Atribacteria_bacterium VEINIGFNIIHRSDIFRERISIIQFLGDLNRGS-KFPPDFAVFGLDSLLYYVEDREKMAKYIRHTLQDNANYLTSGNYIIQIVIEGDIESDEKPRVVYKDEEFPLYSVIG-RLKRVDLKHFYAPLNLQS---
NMX21979.1_ANME-1_cluster_archaeon_GoMg4    MNPTQGYNVLLKEPEGFAKMSIIKFLREIRRGQ-ELPKKVAIYGLESLLLTNKDARAGTKFIRNMFIENNRLFREKGYTFLFIPRKELYVNRDVYLKAEDKKANVSSLFGLRLQIKEVGIYYADFEF-----
WP_089768781.1_Halobellus_clavatus          MSLNAGYNLVTDREDTEEELPALAFFRRLANNG-TLPPRLTVTRLEDLLYETDEEERAVRELRDVLRESGSF--RGPKAIQFVFDGDLVDDDVFSVRIERGYLPVGNLFVEEPRVVEAGHAVA------R-K
WP_103424089.1_Salinigranum_rubrum          SDLDIGYHVIDEDGMFDEEMSIVEFLRRLKRRE-EIPLDMTVRGLDDYLLGVDDADVACDYIHRLLRDRVNYLSLRNPRVQFVVD-DVENWSGPVIPTGDEPIKLNRIFHGSMEQSGPGWYSSNLNVQS---
AKH97513.1_Halanaeroarchaeum_sulfurireducens MPLRPGYNIVSEPAETNSSLAALEFFRQVANNE-SIETPVTVTGLEDLLYNSAENERVLGRLRQALRDTGSI--GSMDAVQLHVDGKLVDDVEFRVRIEKAYLDIGQLFVEAPQLISPTHAVA------R-K
CAD6492699.1_Candidatus_Argoarchaeum_ethanivorans MRLKRGFNIVLNEDHFEDTMTLIEFLGDIRRNK-QIPSRVTVKGLDTLLLNSCDQEEMGLFIGELLRDGQSKLIRTSTVVQFIVDGKITKDIHTKIKVRNEYINLENLFYGKVSRLAPDWVHA------V-R
consensus/100%                              .....sa.lh.........shh.hh..l............l..l...h.............h...h....p........h.h..c..................h..lh....p......h..........
consensus/95%                               .....sa.lh.........shh.hh..l............l..l...h.............h...h....p........h.h..c..................h..lh....p......h..........
consensus/90%                               .....ua.lh........pbshl.hh..l.p....b...h.l.Gl-phL............l.p.h.p..p........hbh..c..l........h......h.l..lh....p......h..........
consensus/85%                               .....Ga.llppp....pphuhlpalp.l.p....h...h.l.GL-phLb.....p.h..blpp.L.p..p.h......lbh.hc.pl..s....h......h.l..lFs...p...s..h..........
consensus/80%                               .....Ga.llppp....pphuhlpFL+pl.p....hs..h.l.GL-phLh.s.p.c.hs.blpphL.p.sp.h..p...lbh.hc.pl..sp...l..s..hsl..lFs...p...s..h.s.h.......
consensus/75%                               ...p.Gapllppp....pphuhlpFL+plpppp..hs.ph..V.GL-clLh.sss.c.hs.bl+phLppsssbL.pp...lbhlhc.pl..spp..lc..s..hsl..lFs...cb..s.aa.s.h.......
consensus/70%                               ...p.Gapllppp....pphuhlpFL+plpppp..hs.ph..V.GL-clLh.sss.c.hs.bl+phLppsssbL.pp...lbhlhc.pl..spp..lc..s..hsl..lFs...cb..s.aa.s.h.......
```

```
# ; HELICASE (BrxHII-Helicase Counterpart); TUDOR + Swi2/SNF2-Helicase + Inactive-REase + CTD(Alpha+Beta);

OrigSeq                                              [illegible]

MBC7429627.1_Bacteriovorax_sp.                       [illegible]
MBE7508219.1_Planctomycetia_bacterium                [illegible]
MBI2192005.1_Planctomycetota_bacterium               [illegible]
MBI5511449.1_Deltaproteobacteria_bacterium           [illegible]
MBI5901621.1_Rhodocyclales_bacterium                 [illegible]
MBA3936144.1_Planctomycetota_bacterium               [illegible]
MBM4322262.1_Deltaproteobacteria_bacterium           [illegible]
MBN2145092.1_Candidatus_Auribacterota_bacterium      [illegible]
MBN2391966.1_Anaerolineae_bacterium                  [illegible]
MBP6702578.1_Vicinamibacteria_bacterium              [illegible]
MBQ8111590.1_Kiritimatiellia_bacterium               [illegible]
MBU4312941.1_Candidatus_Omnitrophota_bacterium       [illegible]
MBW1935303.1_Deltaproteobacteria_bacterium           [illegible]
MBW4553276.1_Aphanocapsa_sp._GSE-SYN-MK-11-07L       [illegible]
MBX3410397.1_Phycisphaeraceae_bacterium              [illegible]
MCC7337860.1_Pirellulaceae_bacterium                 [illegible]
MCX6876429.1_Verrucomicrobia_bacterium               [illegible]
MDR1607843.1_Deltaproteobacteria_bacterium           [illegible]
MDR2179037.1_Synergistaceae_bacterium                [illegible]
NLG01394.1_Lentisphaerota_bacterium                  [illegible]
NPV69259.1_Bacillota_bacterium                       [illegible]
UJS21265.1_Candidatus_Brocadia_sp.                   [illegible]
WP_026299088.1_Deinococcus_aquatilis                 [illegible]
WP_058554674.1_Thiohalocapsa_sp._ML1                 [illegible]
WP_277443845.1_Pelotomaculum_isophthalicicum         [illegible]
WP_282552852.1_Providencia_stuartii                  [illegible]
WP_283022682.1_Bradyrhizobium_sp._CB1717             [illegible]
WP_308999799.1_Paenibacillus_sp._LHD-38              [illegible]
WP_309896344.1_Archangium_sp.                        [illegible]
consensus/100%                                       [illegible]
consensus/95%                                        [illegible]
consensus/90%                                        [illegible]
consensus/85%                                        [illegible]
consensus/80%                                        [illegible]
consensus/75%                                        [illegible]
```

### ; Type-4 BREX PAPS-Reduct (DndC Homolog); (PAPS-reduct + A+B-Domain + Fer4_9 + DUF4007)

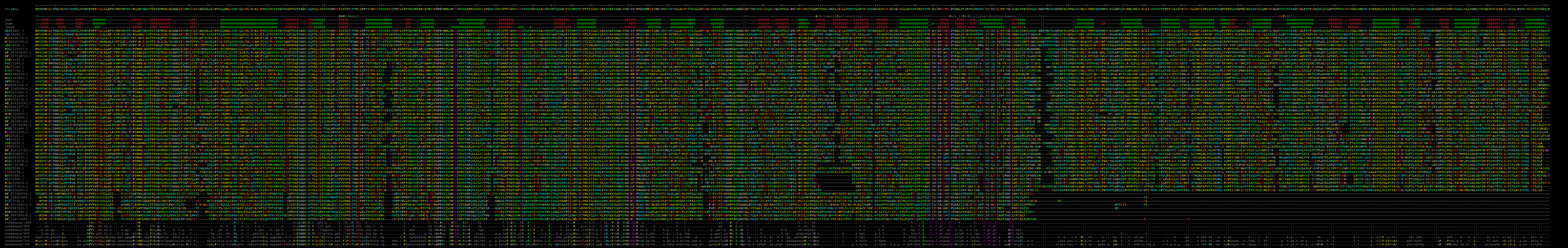

### ; Dnd-System PAPS-reduct DndC;

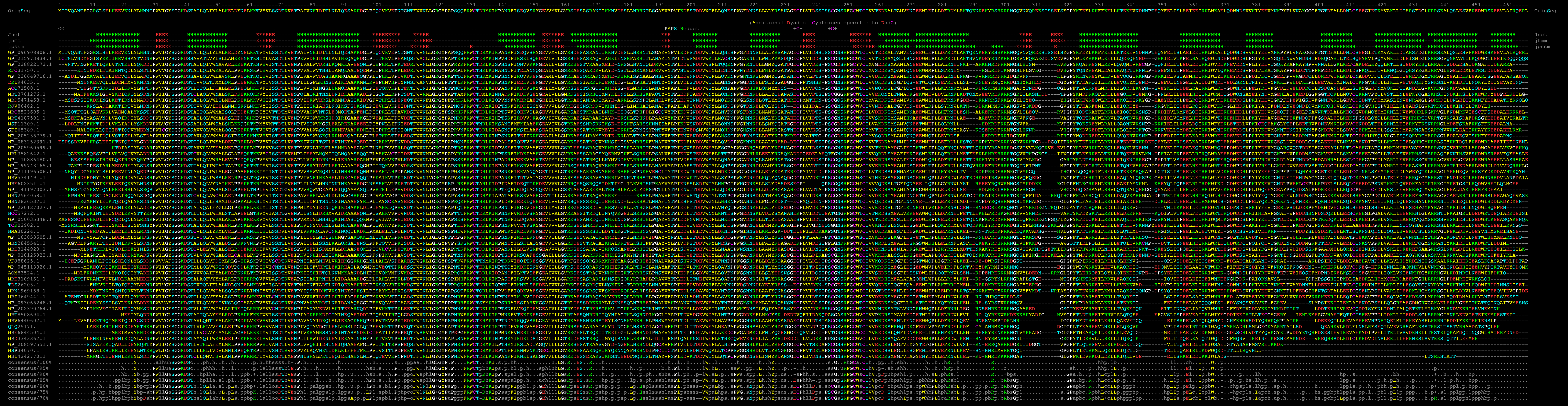

### ; SSP-systems PAPS-Reduct; (PAPS-reduct + A+B-Domain + Fer4_9)

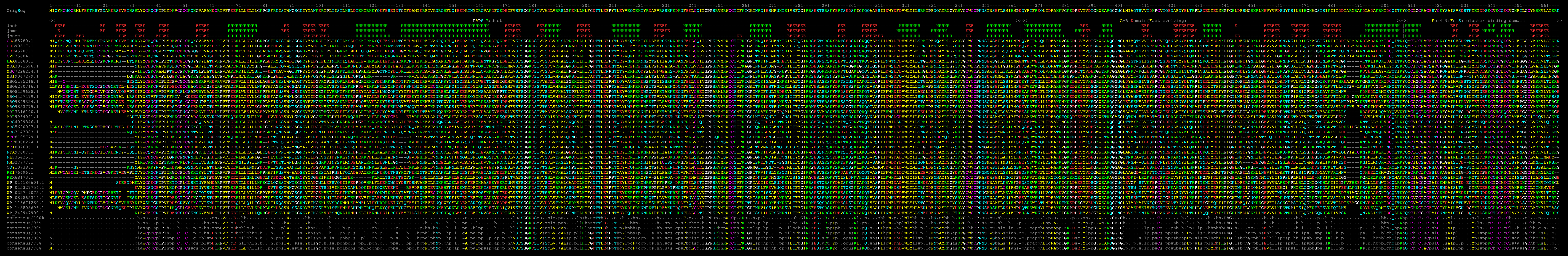

### ; OLD+TOPRIM (Previously Classified OLD-ABCs, associated with ABC-ATPase centric Conflict Systems)

[illegible]

### ; BREX-systems OLD-ABC (TOPRIM-linked); Representative_1;

[illegible]

### ; BREX-systems OLD-ABC (TOPRIM-linked); Representative_2;

[illegible]

### ; BREX-systems OLD-ABC (TOPRIM-linked); Representative_3;

[illegible]

### ; BREX-related (DUF499 centered) systems OLD-ABC (Associated with TOPRIM/REase/HNH)

[illegible]

Representative Alignments of Core Protien Components in Newly Identified PglZ-HerA/FtsK Anchored Conflict Systems;
1. HerA Translocase
2. GNAT
3. TGT; (Partially Conserved in System)
4. DUF6079 ATPase (BrxC Counterpart)
5. PglZ (BrxZ/PglZ Counterpart)
6. DUF4007 (BrxA Counterpart)
7. STAND-NTPase (BrxB Counterpart)

### ; HerA/FtsK Transloacase Representative Alignment;

```
                                                    1---------11--------21--------31--------41--------51--------61--------71--------81--------91--------101-------111-------121-------131-------141-------151-------161-------171-------181-------191-------201-------211-------221-------231-------241-------251-------261-------271-------281-------291-------301-------311-------321-------331-------341-------351-------361-------371-------381-------391-------
OrigSeq                                             MLLQVGRDIETREQVFLEGDRARTVFICGKRGSGKSYTLGVLVEELFEKSGGDVTVILVDPMGIFHTMVLPNKDQSRELLEWGLVEKGLPVRLLVPGDPVRRFGDSEIVSTLKSRGVDVLGLRLNPSDLTPENWCSLFDLSINDLMGIALYRAITGLQERKPYYLQDIERAIMRDTRAQDKTKEALTNRLSIAHRWDIFSDGPGANWGQLFRADSVNIVDVSVLDPGSNGLRSLVVDLITKQIFSVRISARRREEFGLQPQMPRVWLALDEAHQFVPSSGSSLCKESLTRWVKEGRQPGLGLIAATQQPSAMDSDLLSQCDLIIAHKITSLEDVQALNRLSATYMTGEMKSVLRNIGQRGEAVMVDDEKESVTTVAIRPRRSKHGGGETVEKKLRKLLF

                                                    |-Preceding-Hairpin-|                                                                      |-------------------------------------------------------Alpha-Helical-Insert(2-insert-Strands)---------------------------------------|                                                        -R-Finger- -Sensor-1-                |------------------------C-Terminal-Extension------------------------|
                                                                       S1             H1           S2                                                                                                                                                                                  H2                                    S3                                    H3             S4            H3          S5
                                                    -EEEEEEE---EEEEEEE----EEEEEE-------HHHHHHHHHHHHHHH---EEEEEEE---HHHHHH------HHHHHHH--------EEEEEE--HHHHH--HHHHHHHHHH--EEEEE---HHH--HHHHHHHH------HHHHHHHHHHHHHH------HHHHHHHHHH-----HHHHHHHHHHHHHHHHHHH------------------EEEEE----------HHHHHHHHHHHHHHHHHHHHHHHHHHH-------EEEEEE--------------HHHHHHHHHHHHHH--EEEEEEE--HHHHHHHHHHH--EEEEE----HHHHHHHHHH--------HHHHHHH-----HHHHHHHHHHHHHHHHH------------HHHHHHHH--
                                                    -EEEEEEE---EEEEEEE----EEEEEE-------HHHHHHHHHHHHHHH---EEEEEEE---HHHHHH------HHHHHHH--------EEEEEE--HHHHH--HHHHHHHHHH--EEEEE---HHH--HHHHHHHH------HHHHHHHHHHHHHH------HHHHHHHHHH-----HHHHHHHHHHHHHHHHHHH------------------EEEEE----------HHHHHHHHHHHHHHHHHHHHHHHHHHH-------EEEEEE--------------HHHHHHHHHHHHHH--EEEEEEE--HHHHHHHHHHH--EEEEE----HHHHHHHHHH--------HHHHHHH-----HHHHHHHHHHHHHHHHH------------HHHHHHHH--
                                                    -EEEEEEE---EEEEEEE----EEEEEE-------HHHHHHHHHHHHHHH---EEEEEEE---HHHHHH------HHHHHHH--------EEEEEE--HHHHH--HHHHHHHHHH--EEEEE---HHH--HHHHHHHH------HHHHHHHHHHHHHH------HHHHHHHHHH-----HHHHHHHHHHHHHHHHHHH------------------EEEEE----------HHHHHHHHHHHHHHHHHHHHHHHHHHH-------EEEEEE--------------HHHHHHHHHHHHHH--EEEEEEE--HHHHHHHHHHH--EEEEE----HHHHHHHHHH--------HHHHHHH-----HHHHHHHHHHHHHHHHH------------HHHHHHHH--
HHW27503.1_Candidatus_Fermentithermobacillaceae_bacterium  MLLQVGRDIETREQVFLEGDRARTVFICGKRGSGKSYTLGVLVEELFEKSGGDVTVILVDPMGIFHTMVLPNKDQSRELLEWGLVEKGLPVRLLVPGDPVRRFGDSEIVSTLKSRGVDVLGLRLNPSDLTPENWCSLFDLSINDLMGIALYRAITGLQERKPYYLQDIERAIMRDTRAQDKTKEALTNRLSIAHRWDIFSDGPGANWGQLFRADSVNIVDVSVLDPGSNGLRSLVVDLITKQIFSVRISARRREEFGLQPQMPRVWLALDEAHQFVPSSGSSLCKESLTRWVKEGRQPGLGLIAATQQPSAMDSDLLSQCDLIIAHKITSLEDVQALNRLSATYMTGEMKSVLRNIGQRGEAVMVDDEKESVTTVAIRPRRSKHGGGETVEKKLRKLLF
HQD80870.1_Bacillota_bacterium                      -FLNIGKDSETGKKLYIEADRSRAIFICGKRGSGKSYTLGVFLIEELFENAKGEALIVVCDPMGIYHTMVLPNREQEQVLRDWGLRAREAKVKLLVPGRDPVAQFGSTEIVERIKARGVELKRLRINAGSLTPDEWCYLLGLNLNDLMGIALYRAVDDLQEQRFFPFGELERTILKDRKSNDRTKQALANRLSGVKRWDVFQEGEEEPIQNMFTTDAINIIDLADLTPGPYSIRNLIMSVIGQQLFAMRIKERRKEEFGMHAHLPKVWLAIDEAHEFVPVSG-SLSKDLLVRWVKEGRQPGLSLIAATQQPAAVDSEVLSQCDVIISHKITSIEDINALNRLSATYMSGELRTLIRNLGRRGEAILVDDETEEVNHLMVRPRLSAHGGGETGEDEIRKSLF
GIK42764.1_Chloroflexota_bacterium                  -KLLLGKDAHNEEQVFLDARGSRAILICGKRGSGKSYTLGVLVIEELVSVGSTEVIPIVVDPMGVYHTMSQTNIIQQDELFKWGLSARSYPVRLLVPGDPLKLY-DRDILKIFADRDITVVPLRLNPSDLSPDGWCDLFDANINKPLGIALFRATHKLEGKQSYSIKDIIKEVEIDGRAIDTTKEALLNRLEATRSWQLFAENDYTPIHDLFVPGAVNVLDLSRLEPGPRGRRNLAVSVIARNLFKARSDARLREEFNLATPMPKVWILLDEAHQFAPNSGSTLSKDQLIRWAKEGRQPGLSLVVASQQPSAIDSEVLSQCDVILSHKMTTREDVNALNNLSQDYMGGELRTFIKNLKRTGEAVLVDDEQESVHTMRIRPRRSQHGGSSVAPVEEETFDI
RLC84906.1_Chloroflexota_bacterium                  -KLLLGRDAVHNGPVYLEAGGARAVLICGKRGSGKSYTLGVFVEELLTAAADAIIPIIVDPIGVYHTMSLPNELQRDDLFRWALNPRGYPVRMLVPGDPEALY-DPDVLRALRERQVEVVPLQLNAADLSPDGWCELFGANINQPLGIVLFRAVQSLIERRDFTVRDIARAVGTDERANETTQEALLNRLEAARTWRIFAENGYMPMDAIFRPGVVNIVDLSRLESGPRGRRNLTVSVIARNLFRARADARLREEFGLASPLPRIWLAIDEAHQFVPAGGTTLAKEQLVRWAKEGRQPGLSLIVASQRPSAIDREVLSQCDVVLSHKLTTRDDVSALNRLSQDYMGSELQALIKRLKRVGEAILVDDERETVHHLSIRPRRSRHGGSSVAPVEEERLDL
RKU21514.1_Candidatus_Poribacteria_bacterium        -NLRVGTDYETEDIAYIRATGSRAVFICGKRGSGKSYTLGVIVEELFAQERGKAVLIIADPMGIYHTMVSANESQSDEVRQALLTEGFPVRLIVPGDPLACFGDAQIVNRLQTNGVEVTSLWINAHDLSADAWCELFDLEINNAMGIAMWRAVDTLHETEHFNISDILASLRKDTRAKSTTVEALENRLTAAQRWGFFSE-ESTDLTDVFSLEHINVLDLSVVDAGTKGLRNLVLDIVARRLFTERTRMRRREEFGMETSLPRVWMAIDEAHQFVPQSRASLCKEMLIRWVKEGRQPGLSLIVATQQPSAIDSELLSQCDLIMAHKITTWDDINALDRLSATYMEGDLKGYIRQLNRRGEALLVDDETESVDTIRVRPRRSAHGGAEIQEKTASRSFF
MFZ1536133.1_Chromatiaceae_bacterium                -QLMLGRDLVSQQNVYLDASGSRAVLICGKRGSGKSYTLGVFIEELIAMGKKDVIPVIIDPMGIYHTMVMRNDRQSGELYRWGLPAQSFSVRLLIPGVARELY-DRDILDALKQRGVETVEMRLNPSDLSPDGWCDLFNANINQPLGIALFRAVQRLTKTADFTIEDIITAIEDDERAHDISKEALINRLEAADGWHLFNVGGYQPMDHIFRPGVVNIVDLSRLESGVRGRRNLILSVIGRNLFRARSDARLREEFGLAPELPRVWLAIDEAHQFIPSSGSSLAKPQMVRWAKEGRQPGLSLIVATQQPSAIDSEVLSQCDVILSHKLTIRDDVAALNALSQDYMGGELRGFITKLERTGQSVLVDDERESVSMLQIRPRQSLHGGGVAPSLAEEEYDI
MFN8469221.1_Caldilineaceae_bacterium               -LLNLGTKLEQSESCYLNVTGSRAIFICGKRGSGKSYSMGVIVEEMFERLKGEAVILIADPMGIYHTMVSPNTRQSGELRAIGRLAKGYPLRLIVPGDPVGRYGDASVVARFRENGAEVVALQINPSDMSPEAWCELFDVSINEAMGITLWRAVDTLRDKRPFSIHDIIKQVMNDARAKDASKEALENRLTAAGKWGIFAE-SPKALTDLFSVSHINVLDLSVLDAGTKGLRNMVLNVVAQRLFSERTRERRREEFGLGTQLPKVWLAIDEAHQFVPTGRTSLCKEMLIRWVKEGRQPGLGLIVATQQPAAMDSELLSQCDVILSHKITSWEDINALDRLSSTYLEGDLKTQIRALSRRGEALLVDDESESATFVKMRPRLSAHGGGETHEVQGKQTLF
MDQ2805419.1_Chloroflexota_bacterium                -KLFLGKDSKGNAPVYLEIGGSRAMLICGKRGSGKSYTMGVLIEELLN-GSEPVIPIVIDPMGIYHTMGGRNALQSNELYGWGLASRAYRVRLLIPGEPEHLY-DPEVLAAMQTRGVEVVPLRLNPADLSPDGWCDLFGLNINTPIGITLFRAVQHLAKQPGFRIADVMHNIELDGQAQDVSKQALLNRLEAAGAWNIFRTNQYEPIDTIFEPGIVNVVDVSRLESGPRGLRNLIVSVICRNLFRKRADARLREEFGLNAGLPRVWLTIDEAHQFVPAGSTTLAKEQIIRWAKEGRQPGLSLVVATQQPSAVDNEVLSQCDIILSHKLTTKADVAALNSLSQDYMGGELRTFIKQLNRSGESVLVDDERESVTLLKIRPRQSSHGGSSITPTEQEQIDL
MBW7959470.1_Candidatus_Promineofilum_sp.           -KLFLGQEPQSGTPTYLTIGGAHAVLICGKRGSGKSYTMGVLVEELLAGANTDIIPILVDPMGVYHTMAQANQAQQEDLFNWGLSAKGYTVRLLVPGPTEQLY-DPEVIDALQQRGVSIVPLQLNPSDLSPDGWCDLFDADINAPLGIVLFRAAQKLQRQRPFTIQELIVTIERDGRAKDTSKEALLNRLEAARGWQIFAEEGYTPIPELFTPSVVNILDLSRLEPGPRGLRNLTVSIIARNLFRARADARLREEFGLATPLPRVWMLIDEAHQFVPKGQSTLAKDQLIRWAKEGRQPGLSLAVASQQPSAIDPEVLSQCDIILSHKLTSREDVMALNSLSQDYMGGELRTYIRSLNRTGEAVIVNDEMETVQMLRIRPRRSQHGGSSLPPEEENAFDI
MFQ6042699.1_Candidatus_Poribacteria_bacterium      -MLTIGKEIDKDEYVYINAARSRAVLVCGKRGSGKSYTLGVFVEELI--ETEDVIVIIIDPMGIYHTMALPNREQEQVLRDWGLRAREAKVKLLVPGEAEELYGGAEIISAMQRRGVIFQALRINPSDMTPDGWCELFDLTLNEPMGIVLYRAVQELQRRQPFSIEQMISQVERDKRAVEKTREALLLRLEMAKDWGIFAD-EYQSILDFFEPKIVNVIDLSVIDPGRYGLRNLIVDVVCRDLFRKRTIDRRREELHLPTRVPKIWLAMDEAQQFIPSAKSSLSKETLIRWVKEGRQPGLSLIVASQQPAAIDGEVLSQCDILLAHKITALEDVQAINKLSQDYMSSELRNLIKRIKGSGFAVLVDDEEELAQTIKIRPRKSKHGGGESAVGRVATFKI
HHW40998.1_Syntrophomonadaceae_bacterium            -MLRIGKDIESNEEVVIEGDRSRVVFICGKRGSGKSYTMGVLIEELFEKAQGDAIIVLADPMGIFHTMALPNNDQTHDLYQWGLTERGFPVRLLVPGDPIRCYGDEEIVELLQERGVDVQSLRINPADLSPDAWCFLFNLNVNDMIGIALYRAVSDLQEKDFFSIADLERSVKRDPRAQEKTKEALLNRLNIASKWGIFEDSSSRAKEDMFTSSAINVLDLSIIDPGSHGLRNLVVDIVARNLFSIRTKARRKEEFGLSPELPRVWLAMDEAHQFVPQSGSSLCKEVLIRWVKEGRQPGLGLIAATQQPSAIDLELLSQCDLILAHKITSIDDIQSLNRLSATYMGGELKTFVRNLSRRGEAILVDDERESVSRITIRPRRSRHGGGETIEQKIKRTLF
WP_052723405.1_Methanosarcina_barkeri               -NLFIGKDKKTDEDAFIDASKSRAILICGKRGSGKSYTLGNLAEELQS-NCKDALCLIIDPMGIYWTMCNPNYDQENVLWEWGLSPRGLNVKILVPGDPKNRY-GDEVVNVMEEHGVQFQSISLNPSDISPDGWCDLFNLSINDAMGIVLYRAVQGLKKQNDFYIKDIINSIEYDPKANDKSREALLNRLEMANSWGLFGD-TYTNTWDSFDTNSVNIVDLSVLDPGRYGTRNLVVSIFCRDLFTKRTEARRREELGLGSTSTPKVWLMIDEAHQFAPSGKSTLSKELLVRWVKEGRQPGLSLVVATQQPSAIDHEVLSQCDIVISHKITTKEDVDALNKLSQDYMGGELKTYIRNIKGNGEAVMVDDENEKVSMIQIRPRKCKHGGGEHKEKNKY----
MFA0740147.1_Candidatus_Fervidibacter_sacchari      MMISLGRDRNSGEIVFVDDSRARAVLICGKRGSGKSYTMGVFLEELV--RRDDVLVIVLDPMRNFHTMVLPNVEQERLLWDWNMFAEGVPVRLLVPGDPYRRY-GQEVVEAFEKRGVEVVPLRLNPGDLPPDGWLELFGIDIATPMGLALYRAVSKLQRSRFFTVADIIAEIYSDPLIRETSIQALVLRLETANDWDLFST-RYYELLEVFQIGAVNVVDLGVLEVGRVSLRNLVAAVLVRDLFRKRKEARLQEELGIPTQVPKVWLAIDEAQEFVPAGKTALAKEPLIQWVKEGRQPGLSLIIASQQPGAIDKEVLSQCDLILCHKVTALEDVEAINKLSHDYMAHELRHYIRQLERTGEAVIVDDFRETVTVVQIRPRKSKHGGAEA----------
TEU15944.1_Anaerolineales_bacterium                 -SILLGKETEMGDMVRIEADRSQRILVCGKTGTGKSYSLGVCIEELA--IGSNIILVIDPQGIFWTMAEANTDEVDGLWDFDRSPRGFPLNLMVPGDPVERFGDPEIPAEMEKRGIQVQSLRLNPSDLTPEMWCDLFDLDINELRGIMLFKAVRQCKRRRDFLIDDISAEVDR-LKGLDKTKEAVQRRLDMAQDWQIFAEDQYREIWEILRPDQINVLDLSVIAQGRYGLRNLVIAVLANFIFRKRTIARRREALGLPSDTRKVWLVIDEAHNFCPSGKSTLSKEILIRWAKEGRQPGLSLIVASQQPAAIDREILTQCGIRIVHRITSKDDFQAVNALSQDYLTDGLHAQIKRLTGPGQTLMIDDARETVVRTQVRPRQSLHGGGSA----------
WP_089881367.1_Halogeometricum_limi                 -ELFIGKELDTDEEEYVDGSHSRAVFICGKRGSGKSYSMGDIIEEVHD-HTQGVVPLIVDPMGIYWTMVSENEEQQNSLWDWGLQSRAYRVNLLVPGNPKERY-GEEVLSEMKRRGLEVSAIKINPSDISPDGWCDLFDLNINKPMGITLYRAIRNLSRQNEFNLDDIIDTVEGDAQSNERTKEALLNRLEMSRDWGLFSD-TYTPIWDRLDETSLNVLDLSVVDPGKYGLRNLVVSVLCRDLFRKRIKARKREELELQVDVPRVWLFIDEAHNFVPSGESSLSKDILIRWVKEGRQPGVSLTVATQQPSALDHDMVSQCDVILCHKITTREDIQELNRLSQDYMSSELKTYIQKLDRVGQAVYVDDEDESISIVQMRPRRSRHGGGEN----------
RJP56335.1_Deltaproteobacteria_bacterium            -QLLIGKDVDDGNMFRISCEKSQRILVCGKTGTGKSYTMGVLVEELA--NIGDDIILVIDPQGIFWTMTQPNEKEVDNLWEYDRQAKGFRVNLLVPGDPVERYGGEDVVHEMERRGLNIQGLRLNPSDLSPEMWCELFNLDINELQGSMLFKAVRTCYQKHDFLLDDISSTVEK-LQGLDKTKEAIFRKIEMAHDWQLFEEFRYREIWEILQPNAVNILDLSVIAQGRYGLRNLVIAVLANFIFRMRTIARRREELGLPTDIKKVWIFIDEAHNFCPSGGSTLSKEILIRWAKEGRQPGLSLVVASQQPSALSSEVLTQCGIRIVHRITSKEDYRAVNALSQDFLPDDLQTYIKQLSGLGETLIINDESEKMAKVKIHPRQSSHGGGSA----------
NBT26026.1_Actinomycetota_bacterium                 -MLYVGRTTTTDEGFYLDASRSRAVFVCGKRGSGKSYTLGVILEELA--LAGNVLVIVADPMGIFHTMRLPNEPQRQALWDWNLGATGVPLRLLPPGDPEARY-GRDVLRVLRDRDVDVRSFRINPSDLSNEGWLELFSLDMAQPMGLILLRALQKSRARQHFFVPDIVSAVQNDPLSKEASAQALTLRLETANEWDLFTT-HYQDLSESFSTGMVNVLDLSVLDNSRFGLRNLVLSVLVRHLFARRRQARMQEELNLSPDVPRIWLAIDEAQEFVPSGKSTLSKEGLIQWVKEGRQPGLSLVVATQQPGAVDKELLSQCDLIISHKITAADDVDALNRLNQAYMGGEMRVMLRQLARTGEAVLIDDTAERATVLQVRPRKSRHGGGET----------
MCK4444003.1_Thermoplasmata_archaeon                -MIRIGKSLQDESDAYLDDDKSRVILICGKRGSGKSYTMGVMIEELL--RKKERIILVIDPMGIFHTMCLPNTDQEELLWNWGMHPEGSPIRVLVPGNPEERYGGQDIVAAMEKRGVRFVELRLNPSDISPEGWCETFGVGINDPMGIAIHKAVNKCRKKGSYFLDELVDLVEYDTRAASQTKDALTNRLEMAKEWGMFENEKYRLIEETLDPNSINILDLSTIESGRYGRRPLILDTLARSLFKRRSIARRKEELGLASEMKRLWMFIDEAQQFAPAGKSSISKETLVRWVKEGRQPGLSLVASSQQPSAITMEVLSQCDAVISHSLTTAVDKDYLNRLTKDYMRGELKTYINKIARTGEAVFVDDDKERVDTIRIQPRKSSHGGKEEAS--------
WP_423222230.1_Candidatus_Amarolinea_aalborgensis   -MLSIGRDIKTNDYVYLDVSRSRAALICGKRGSGKSYTLGVIVEELH--QAGNVLTVIIDPMGIYYPMAQPNHSQDRLLWDWGLSAQALPTLLLVPGDPEMLYGGREVLKAMEDMGVRFRQFRLNPADLSPDAWCELFDFSIADVMGIALFRAVQTLSRKEYFFIPDIIAEIEGDGLIADRTREALLNRLKMAQDWDLFSY-RYQEIWEIFDPKAINIVDLSTLDPGPRGRRNLVVDVLARDMFKRRTIARRKEELGLISELPRVWMLIDEAHQFAPSGRGTLAKEALIRWAKEGRQPGLSLTVASQQPSAIDREILTQCDVLLCQKITNSEDIQSINGLSQDYMGGELKTFIRKLERRGEAVLVDDEQEKVTMMQVRPRRSRHGGGEAAG--------
WP_338740222.1_Haloplanus_salilacus                 -RLFIGEELENGEDTYIDATSARSILACGKRGTGKSYALGDIVEEIHT-EGIDVVPLVIDPMGIYWTMAEENEEQRDLLWDWGLNEEGFPVNLLVPGEPEERF-DEEVLREFERRGIDINPLLLNPSDMSPDGWCDLFDLNINKPMGIALYRAVRKLNDEGDFYLDDIIKKIEMDGDANERTQEALLNRLEMAHDWGIFAD-SYQDIFRTVDEDRINILDVSVIEPGKYGLRNLVVEVLGKELFRRRQAARRREELQLGVDIPKVWLFIDEAHNFVPSGKSSLAKDTIIRWVKEGRQPGLSIVVATQQPSAIDSNVLSQCDVILCHKITTREDIKSLNKLSQDYMSGALKTYVQNIDNVGEAVYVDDEEETVKMVQIRPRKSKHGGGEA----------
WP_089309286.1_Halorubrum_ezzemoulense              -EIFIGEERDSGEETLIDTSKARALFVCGKRGTGKSYSLGNVIEEVNR-EMPDTVPLVVDPMGIYWTMAEPNEDQQDLLWDWGMQPTGFDTTLLVPGDPTERY-GPEVYDEMMRRGIEIESLRLNPSDISPDGWCDLFDLNINKPMGIALYRAVRSLNESDFFDLDELIEEVEYDEKANERTVDALVNRLEMAHEWGIFSD-SYTDIWNNLDPSRVNILDVSVLDPGKYGLRNLVVYVLTKKIFDERSKARRREELQLSSDIPKVWLFIDEAHNFAPSGSSSLAKDILIRWIKEGRQPGLSLGIATQQPSAVSSDVVSQMDLIFCHKITTRTDIQSLNELSQEYMGSQLKTYIKKIDKVGEAVCVDDEREVVKQVQMRPRMSQHGGSGA----------
MEI6180765.1_Chloroflexales_bacterium               -MINLGKDIKADTDAYIDDAKSRAVLICGKRGSGKSYTLGVIFEELI--EANNTLLILDPMGIYHPMSLANHEQERELWAWGLSKGMPVHLLVPGDPEQRYGGREVIAELERREISIRPLRINPSDLSPDAWCGLFDLSISDVMGIALYRAVQNLARKENFFLSDMMQAVELDGRVQDRTREALLNRLEMAVDWNLFSE-RYQEIWEVFDQRAVNIVDLSVLDPGQYGLRSLVVDVLARDIFRRRTIARRKEELGLVGDLPRVWVAIDEAHQFVPSGKATLAKEILIRWVKEGRQPGLSLVVASQQPSAIDSEVLSQCDVIIAQKLTNGEDIAALNKLSQDYMGGELKTYIRNLNRRGQAVVVDDEQESVRMVQVRPRRSRHGGGEAQG--------
WP_348254530.1_Leptolyngbya_sp._ST-U4               -MIFLGRDFETEEPIYVESDRSRAILICGKRGSGKSYTMGVFLEELY--GQEDSLVIVIDPLRNFYTMSRPNDDQEEALYRWNLSPRSVPVRLLIPGDPIKRY-GEKVIDALKSQGVQVESFRINPGDLTPDTWLDLFDLSLTEPMGLALYRSVSQLRHSVNFSLEDLIEKVKKSSRSKETTTEALLARLEVLDEWDLFSS-YYKEVTEIFDVRAVNVIDVGVLGSSRFGLRNLVVLILTQDLFRKRSQARLQEELGLPTGVPKVWLAIDEAQEFVPSGRASLSKESLIQWVKEGRQPGLSLVAATQQPAAIDRELLSQCDLVLAHKVTTSDDVDALNRLSHNYMASELKQYLRQIDRRGEALLVDDEEELVRVVQVRPRRSRHGGAELK---------
MBN1891047.1_Thermoflexales_bacterium               -MLFVGRNLDDNSDVHIDANKSRAVAICGKRGSGKSYTMGVLIEELC--REDSAVILVVDPMGIYHTMSLPNLEQERTLWDWGESPQGLPIKVLVPGDPVVRYGGREIVERLEERGVVFEQLRLNPSDISPEAWCDLFDFNINEPLGIALYKAIRTCHKKAEFYISQIMDIIELDARAAPKTVDALVNRLDTAQEWGFFEDSRYREVWEMLDPHVVNILDLSTIDSGRYGRRPLILSALTRNLFRKRSVARRREELGLEAGMRKIWMLIDEAHQFVPAGKSTLCKEALIQWVKEGRQPGLSLVVASQQPSALDMEALSQCDVIISHALTTTEDKSALNRLTKDYMGAELRVFINKIARTGQAAFVDDDVERVMMVQVKPRKSKPGGSEA----------
NPV62498.1_Methanotrichaceae_archaeon               -MINIGNEIKTKDHVYINSNKSHAVLICGKRGSGKSYTMGVIVEELL--DAEDVLVIIIDPMGIYYTMQQPNINQEQLLWDWGLSSKGVPVKVLVPGDPIELYGGREIVGAMERKGITFRTFKLNPSDLSPDAWCELFDVSISDVMGIGLFRAVQQLSRSNNFSVRDIIDSIGSDPLVADKTRQALVNRLEMAIDWGIFSE-WYQELWEIFDKHSVNIVDLSVLDPGPLGRRNLILNVLAKDIFRRRTIARRKEELKLSDDLPKVWMLIDEAHQFVPSGKSTLSKEILIRWAKEGRQPGLSLLVASQQPSAIDSEVVTQCDVVIAQKLTNKDDISALNALNQDYMGGEMKTYIRGLKRKGEAVVVNDEEEKVRIIQVRPRKSYHGGAEI----------
consensus/100%                                      ..h.lGpp........hp...up.hhhCGKpGoGKSYshGshhEEh..........llhDPb..aasM...N..b...lb.hsb..p.....hh.P..s...a.s.pl...h....h....h.lNs.shss-.Wh..hsh.hs..bG..h.+u.p....p..a.h.ph...l.........o.pAl..+lp....W.hF...........h....lNllDl..l..u..u.Rsh.h..hsp.hF..R...RbbE.h.h.s...+lWh.hDEAppFhP.u..slsK..h.bWhKEGRQPGlul.huoQbPuAh..........................................................................................
consensus/95%                                       ..l.lGpp........lp...up.hhlCGKpGoGKSYohGshhEEl........h.lhDPb..aasM...N..b...Lb.hsb..puh..plllPG.s...a.s.plh..hbp..l.h..hblNsuDhos-.Wh.LFshslsp.bG..la+Ahp.h..p...a.l.ph...l....b..p.o.pAl..+Lp.s..W.hF.........p.hp...lNllDlu.l..u..ubRsL.h.hlsp.lF..R...RbbE.h.h.s...+lWhhhDEAppFhP.u.soLsK-.l.bWhKEGRQPGLuL.hAoQQPuAhs.phloQhsh.hsp.hT...D...ls.Ls..ah...hp..lp.l...GbshhlsD..E.h....hpPRbs..GG............
consensus/90%                                       ..l.lGpp.p..p..blps..u+.hhlCGKpGoGKSYohGshlEEl........l.lhDPbGlaaTMs..N..Q.p.Lb.asb..puh..lplllPGps..ba.s.cll..hbppslph..hblNsuDhos-.Wh.LFshslsp.hGl.La+Alpph.pp..a.l.-l...l....bs.p.o.pAl..RLp.s..W.hF.......h.p.hp...lNllDlu.l..u..GbRsLhl.llsp.lFp.R...RbbEEh.l.s.h.+lWhhlDEAppFhP.u.soLsK-.llbWhKEGRQPGLuL.hAoQQPuAls.-hloQCsl.hspplTs..D...ls.Ls.sYh.s.hp.blp.l...GbslhlsD..E.h..h.h+PRbS.HGGu...........
consensus/85%                                       ..l.lGpp.pp.p..blpss.u+slhlCGKRGoGKSYohGshlEEl......psl.llhDPhGlaaTMs..N..Qpp.Lhpasb..puh.lplLlPGsP..ba.s.cll..hbppslph..hblNPuDloP-.WCpLFshslsp.hGlsLa+Alpph.pp..F.l.-l...lb.s.bs.-.o.pAL..RLp.u.pW.lF.p...p.h.p.hp...lNllDlu.lpsG.bGbRsLllsllspplFp.R..sRbbEEh.L.s.h.+lWhhlDEAppFhPsu.soLsK-.llbWhKEGRQPGLSLhhAoQQPuAls.-lloQCDlllsp+lTs.pDh.ulN.Lo.sYh.sph+shlppl...GbulhlsDp.E.h..hph+PRbS.HGGup.........
```

```
# ; GCN5-related N-acetyltransferases (GNAT) Representative Alignment;

                                                        1---------11--------21--------31--------41--------51--------61--------71--------81--------91--------101-------111-------121-------131-------141-------151
OrigSeq                                                 MVFGAETLVIRKATPADLDAIKAIADAHRHELGFVLRPALAESIGRGEVLVAENHQGLIGFAEYHHRRDAQTTLYHIAVIPQCRQQGVGRALVNALCAEASALGKLTVFLKCPADLSARGFYACLGFEVLGEEPGNGRSLIVWTLSLTENTQ


                                                        --------EEEE--HHHHHHHHHHHHHHHHHH----HHHHHHHHHH--EEEEE-----EEEEEEEE----EEEEEEEEE---------HHHHHHHHHHHHHHH---EEEEEEE---HHHHHHHHH--EEEEEEE----EEEEEEEEE-----
                                                        --------EEEE--HHHHHHHHHHHHHHHHHH----HHHHHHHHHH--EEEEE-----EEEEEEEE----EEEEEEEEE---------HHHHHHHHHHHHHHH---EEEEEEE---HHHHHHHHH--EEEEEEE----EEEEEEEEE-----
                                                        --------EEEE--HHHHHHHHHHHHHHHHHH----HHHHHHHHHH--EEEEE-----EEEEEEEE----EEEEEEEEE---------HHHHHHHHHHHHHHH---EEEEEEE---HHHHHHHHH--EEEEEEE----EEEEEEEEE-----
RLC84907.1_Chloroflexota_bacterium                      MVFGAETLVIRKATPADLDAIKAIADAHRHELGFVLRPALAESIGRGEVLVAENHQGLIGFAEYHHRRDAQTTLYHIAVIPQCRQQGVGRALVNALCAEASALGKLTVFLKCPADLSARGFYACLGFEVLGEEPGNGRSLIVWTLSLTENTQ
WP_048108637.1_Methanosarcina_barkeri                   MNNFNSDVEVTKARLEEVPQIKSIADKNKNLIGFILRGSLVESIEKENLVVLKYKKIIIGFINYHHRKDSQTTLYEICVDENNRNKGYGKILINYLLQEAKSQKKKTLLLNCPIDGIAHSFYLRCGFKIKSTRTHTKKKLVTWVYKLDDNNT
GAB4153040.1_Candidatus_Promineifilaceae_bacterium      MTSG-NEVLIRAGRSEDIDGIKQLADAHKRELGFLRRPALLEAIQRGELLVAQNGFTIVGFIEYRHRRDHQTTLYNVVVHPDYRRLGIGRQLVLALEKEAIQREKSQVVLKCPEDLPANDFYEQTGYERIHVEPGKLRRLNVWRKFLMNGCR
MBW7959469.1_Candidatus_Promineofilum_sp.               MPSG-SDLIIRKAVFTDLETVKRIADAHRHELGFVVLSALRERIERNEMLVAVRDNELIGFVDYHKRRDGQLTLYRIAVDEEDKGQRVGQALIDNLVAIARAQGCSQIILKCPVDLRANQFYEQHGFTPAGTVEGRRRSLNVWAMSLDHTTQ
HFD39725.1_Anaerolineae_bacterium                       M----SDPEIRQATMFDLNAIKELADAHRHELGFVRRPALARSIERGELFVAQNGQGVIGFVEYHHRRDGQTTLYHLAVQADYRRQGLGRRLVQALVHDAGQHSQEFIQLKCPVDLEANRFYQQIGFAQIDLQRGKRRDLAIWRLSLAPGCD
MBC8263372.1_Anaerolineales_bacterium                   M----TSLMIRKAIPADLDIVKGLADAHKNELGFVLRPALASSIDRGEVIVAENSRGIVGFVEYHHRQDEQTTLYHIAVAPQHRRQGIGQRLVNALVYDAGEHNQRFVLLKCPADLEANKFYERLGFSHVDTQPGKQRELSIWRLPLSSAIR
NLE46404.1_Chloroflexota_bacterium                      MTSGNQVSIVRKAVLDDLDTIKAIADAHRDELGFVIRRALQESIERGELSVAIDGDRLAGFVEYRHRRDQQTTLYHIAVVEQWRQRGIGRLLVEAVRRDALANGKNVIRLKCPESLPANEFYTRLGWQKIAVESGKRRRLVVWELPLVLEAK
MBV6438331.1_Anaerolineae_bacterium                     MISG-GNFVIRRARLDDIDNAKALVDRHKNELGFVIRSALVTSVSRGELLVALANDDIVGLVHFRHRRDKQSTLYSIVVSENVRSQSIGRALLAELVDECQRLGQERIVLKCPVELPANSIYPKFGFTLTGTENGKHRRLNIWQLELPGSN-
MBN2393031.1_Anaerolineae_bacterium                     MISGNAEITLRKATEEDLDNIKIMADAHRRELGFVRRPALLEAIHRKEIIVAQNSRHLAGFVHYRHRRDEQTTLYDIVVAAEYRLIGIGKALVQTLVAEIQALGKQTLILKCPAELPANTFYTHLGFERWKEEPGKHRPLIVWRLSIPSQQ-
MBI5668050.1_Chloroflexota_bacterium                    MISGNNNVTVRKATIEDLEAIKRLSDQHRAELGFVLKPALEKAILDGEVFVAV-APQIVGFVHYHHRRDSQTTLYHIVVNMEERRQGIGRALLSRLKVEATERGQSSILLKCPTELPANKFYEQCGYSHIDVEPGKHRRLNVWQISL-----
MFZ4828883.1_Phototrophicaceae_bacterium                MIFGSDDFVLRKASEIDLTSIKEIADENKKELGFINRGTIIESIKRYEVAIVEDGYGIVGFIHYRHRLDSQTTLYNLAVKHPFRNCGMGRRLIEFLKQDALASQKLTIRLKCPEELASNQFYERYGFKLMSVENGKHRRLNIWVLNL-----
GIK42763.1_Chloroflexota_bacterium                      MTSGVSEIIITKAEFADIGGIKQIADAHRNELGFVRRPALIEAIDRSEILIAKQNGNIVGFVEYRHRRDEQTTLYNIAVMPENRHTGIGRKLFQALVKEAEKRDKSHILLKCPEELAANKFYRALGLHLTELEPGKQRRLNIWQWEL-----
GAB4528674.1_Anaerolineae_bacterium                     MISGNSRFTLRKATLDDIDAIKALADAHRRELGFVRRPALIESINRQEIIVMQNSRDVIGFVEYHHRKDDQTTLYHICVEPAYRRRGVGRLLIEALQEEARRYGKHMIRIRCPANLPANQFYRRLGCFLRGKELGKKRCLAIWDIFV-----
MBP8947513.1_Promineifilum_sp.                          MTSG-NSLLVRPAQLADLTPIKALADAHRHELGFVLLPSLREQMERGQMLVAEQDGTLLGFVDYHRRRDGQITLYHIVVKPGVRGQGTGRALLAALEACGRDANCRRIVLKCPIDLASNHFYERYGFILDGTIEGRKRPLNVWMLNL-----
MDQ2805418.1_Chloroflexota_bacterium                    M----ESVIIRTATSADLDAVKALADRHKRDVGFVRRGALLESINRAHVLLALAGEDPVGFVQFHHRRDGQTTIHLIMVASEHRRRGIAALLLNQLRTACQGLGQQRIGLKCPVDLSANSFYQQAGFTLIAQEPGKIRALNLWQLPL-----
HET59441.1_Chloroflexota_bacterium                      M------IKIRKAKPVDLDAIKSLVDSHKHELGFVLRPALMKSIDRSEVFVANGGKTIVGVLDYHHRLDQQTTIYHFVVAPQLRRQGVGRLLVEALKEEAKEYGKSHILLKCPEDLESNDFYGRVGFQLTAIENGRQRRLNVWELVLIS---
NPV62500.1_Methanotrichaceae_archaeon                   -------MLIKKATIDDLEGIKLLADKNRNELGFVLNSALVRSINRGELIIAL-DETIVGFLEYHHRKDCQTTIYHFVIEENYRLRGISKILLNFLIRECESQGMRYVFLKCPADLNANNIYPKLGFKNVGLEGGKKRKLNLWNLNIYLS--
MFQ5758745.1_Candidatus_Bathyarchaeia_archaeon          M------VVVRKATLDDLDVIKALADSHRYELGFVRCAALQRAIEQNEVFVAQNRQYITGFVDYHHRRDGQTTLYHIGVSPALRGQGIGRSLVMALYEEAQQRGSRTIVLKCPADLPANDFYAALSFHLIGEERSKNRPLRIWSLSL-----
HHW27508.1_Candidatus_Fermentithermobacillaceae         MP------IVRLAEQADLRDVKSLADCHRKEIGFLPRAAFEESALHQCLIVACDDKGVVGFTRFRHRRDGVTVIYEIC--SQLSGSGAGGQMIDWLKADCREHGQRSIILKCPSDLPSNRFYESKGFRKVGVEAGKAQPLNIWSLEV-----
WP_310922608.1_Halogeometricum_sp._S1BR25-6             MTDQQSDFLVKTVESDRLGQLNDFFNSPKSELHFTHRDTLERAAVRNDLYYYR-FDEILGWCESRVLEDHQAQIRLVAVHPAYREYGIGRYLVKSAIKFAKVWEKSTMIADVAADASAVGFWKACGFHQVDQYQTKGREMYQMQRRI-----
WP_086215089.1_Halorubrum_sp._SD683                     MEKSGADFTVKAVNLSLVTAVNEFFNSTKSELHFTHRDTLERASERNDLLYIR-RETVIAWCKSRVLEDDQAQIRLVAVDPAYRNYGLGAYLVSAAVDFAQFFGKKEMIADVAAEAPAVTFWEHCDFAKKSEYSTEGRLMYRMWKPI-----
WP_232571137.1_Halobacterium_litoreum                   MSRGKKLMKVSSASPDDVERINAFFNSTKTDLHFTYRDTLDRAFEREDLLYVDDDGEIIGWCQSRVLDPGEAQIRLVAVSPDNRDEGLGRLLCECAEAFARRQSQDRMIADVVASAPAVEFWKSIGYDPLEEWETRGRSMLTVQKPL-----
WP_089673069.1_Halohasta_litchfieldiae                  MKSKLTVLPIQAASINEVEAINDFFNSSKNDLHFTYRDTLKRAFERGDLYYVETSGTLIGWCESRILDKHEAQIRLVAVSRVGRGAGIGRYLCEEAEEFARSHGKEQMIADVVKGSSAVEFWKSIGYEVQSGWETKKTEMLTVEKWL-----
WP_089881364.1_Halogeometricum_limi                     MTSN-ERSAIKTARPEDIDRIQRLYNHNKKELHYTYRDGLERAVSNEGILFTDETEEVIGWCESRVLKPGQAQVRLIAVHPDFRSTGLGADLIEASENYAKEAGKTVLIAETAASSDAKQFWLTMDFQVTSTRTTDGREMALMKKEI-----
WP_256533862.1_Halovivax_cerinus                        MKKP-EPITIKHAEEEDVTRINEFFNSKKQELHFTYRTTLSRAFSRDDLFYVE-NESLVGWCESRVLDDDEAQIRLVAVSPDRREEGIAARLCENAEEFATYHGQERVSADVACDSPAVNFWTSIEYDIEYEWQTDGREMYRMMKAL-----
WP_338740224.1_Haloplanus_salilacus                     MTQP-KSIAVEQVDEDDVGRVNDFFNLRKRDLHFTYRDTLDRAFCRDDLFYIE-DATIMAWCESRVLDDDEAQIRLVAVKPEHRGQGFGRALCERAEQFAFEFGEQKMSADVMAESQAVEFWQSLGYTIEEEWTTDNRSMYRVSKPI-----
WP_117592509.1_Haloprofundus_halophilus                 MS---DEFPVLSAEPETIEFLLNFFNSPKKELHFTHRATVERAFERDDLFYTVGDNTIIGWCESRVLEPHEAQIRLVAVDLDYRDCGIGKQLCDAAEGFAKECEKKAMIADVAEKSPAVSFWHACGYKTQKKWSTKNRPMLRVEKSL-----
consensus/100%                                          ........l..s....l..h..h.s..+..l.a....sh.p...p..h.h........uh.p.+.b......l..h......p....u..hh.............h.hps.....u..ha....h........p.p.h..h.b.l.....
consensus/95%                                           M........lp.s...pl..lp.hhs..+.-l.F...ssl.p.h.p.phhh......lhuhhp.+.b.s.b.plb.hhl....+..uhu..Lh..h...h....p..h.hcss.p..u..ha...sh.....b.sp.p.h..h.b.l.....
consensus/90%                                           M........lp.s...cl..lp.hhs.p+p-l.F..bssl.cuh.c.plhhh.....lhGahp.+.bps.bsplb.lhV....R..Ghu..Lh..h...h....pp.h.hcss.p..u.pFa...sa.....b.sc.+.h..h.b.l.....
consensus/85%                                           M......h.lp.Ap..-lp.lp.hhs.p+p-L.Fs.+ssL.cuhp+.clhhh.....llGahc.+.bcD.bsplb.lhV....R..GhG..Lhp.h...h....pp.h.hcss.p..AspFa.phGap....b.sc.R.h..h.b.l.....
```

```
# ; tRNA-guanine transglycosylase Representative Alignment; (TGT; Partially Conserved in System)

                                                   1---------11--------21--------31--------41--------51--------61--------71--------81--------91--------101-------111-------121-------131-------141-------151-------161-------171-------181-------191-------201-------211-------221-------231-------241-------251-------261-------271-------281-------291-------301-------311-------321-------331-----
OrigSeq                                            MILNFFTALTLCPSDIRKIRRLLDMHRIDMNPLRHILITPLFASPESLKLVRNLKEHNGSTVMFDSGGYYVQVGKLTYQELYYPLLEFYKANPWADIYTLPDHVPTSQDSEQVVWQKIRDTADYTALFYQEMPHHLQSRALAVVQGHTYEQVDLCLKTYLQLGVQRIGFGSFGTIGKNNGINIATNSAVDLARYVVDIAHKNSVKVHFFGLGAPALIAMIYGAGADSFDSSAWIKSAGFGQVYLPFTRGYNISHRSELQQGIQFKEFETLKILTGHSCPFCASIDELQAHKMHRALHNLIAIQQSVNMINSDAGDTIERIYRNGSPRYREEYQKWLNGI

                                                   ---EEEEE----HHHHHHHHHHHHHHH---------EEEE------HHHHHHHHHHHHHHH--EEEE---HHHHHH----HHHHHHHHHHHHHHH-----EEEE----------HHHHHHHHHHHHHHHHHHHHHH--HHHHHH-EEEE----HHHHHHHHHHHHHHH---EEEE---EEE----EEEE--HHHHHHHHHHHHHHHHHH--EEEE-----HHHHHHHHHH---EEEE-HHHHHHHH------------------------HHHHHHHHHHH---------HHHHHHHHHHHHHHHHHHHHHHHHHHHHH---HHHHHHHHH--HHHHHHHHHHH---
                                                   ---EEEEE----HHHHHHHHHHHHHHH---------EEEE------HHHHHHHHHHHHHHH--EEEE---HHHHHH----HHHHHHHHHHHHHHH-----EEEE----------HHHHHHHHHHHHHHHHHHHHHH--HHHHHH-EEEE----HHHHHHHHHHHHHHH---EEEE---EEE----EEEE--HHHHHHHHHHHHHHHHHH--EEEE-----HHHHHHHHHH---EEEE-HHHHHHHH------------------------HHHHHHHHHHH---------HHHHHHHHHHHHHHHHHHHHHHHHHHHHH---HHHHHHHHH--HHHHHHHHHHH---
                                                   ---EEEEE----HHHHHHHHHHHHHHH---------EEEE------HHHHHHHHHHHHHHH--EEEE---HHHHHH----HHHHHHHHHHHHHHH-----EEEE----------HHHHHHHHHHHHHHHHHHHHHH--HHHHHH-EEEE----HHHHHHHHHHHHHHH---EEEE---EEE----EEEE--HHHHHHHHHHHHHHHHHH--EEEE-----HHHHHHHHHH---EEEE-HHHHHHHH------------------------HHHHHHHHHHH---------HHHHHHHHHHHHHHHHHHHHHHHHHHHHH---HHHHHHHHH--HHHHHHHHHHH---
MBN2393039.1_Anaerolineae_bacterium                MILNFFTALTLCPSDIRKIRRLLDMHRIDMNPLRHILITPLFASPESLKLVRNLKEHNGSTVMFDSGGYYVQVGKLTYQELYYPLLEFYKANPWADIYTLPDHVPTSQDSEQVVWQKIRDTADYTALFYQEMPHHLQSRALAVVQGHTYEQVDLCLKTYLQLGVQRIGFGSFGTIGKNNGINIATNSAVDLARYVVDIAHKNSVKVHFFGLGAPALIAMIYGAGADSFDSSAWIKSAGFGQVYLPFTRGYNISHRSELQQGIQFKEFETLKILTGHSCPFCASIDELQAHKMHRALHNLIAIQQSVNMINSDAGDTIERIYRNGSPRYREEYQKWLNGI
MBV6438325.1_Anaerolineae_bacterium                M-LRFVASMTVCPSDIRKIRNHVGAAGTHLHPLQDILITPMFVQKQSLALARELAEEN-SNVTFDSGGYYVQLGRTKYEELYMPLLRAYRENRWASVYTLPDHVPTSQDTTEIVDRKVQDTIRFSTLFFQEMPDELKPRAMPVVQGHTYKQVEACLQAYIQLGVRQIGFGSFGTQGQNSEINVATKSAIEIARHVIAVAHQHDMRVHLFGLGTPSLVAMIKGIGADSFDSATWLKSAGFGQVFLPFMRAYNISYGSELQKGITFSQFQEWKALTGHRCRLCDDLDRLNSQKMYRAAHNLVVMAETVEMVNSGNPQMIKAIYKAGSPKYRDEFEKWLQRN
WP_086215083.1_Halorubrum_sp._SD683                M-IEFVASVTINKSQFNSLIDRLEEHGLVEDVLGNLMFCPLTAGESTSEIIERLATDYDIHVACDSGGYESQINKYSMRDIYQFDRKYYLETDWPTEYVMPDQVPLETDDEMTVEHKVQDTITLSRMLYRELPPEKQKRAVPVVQGHKREQIVNCLDAYRRLNVQKIGFGSFSTGGVNGGVNYLNEDNIELLQFVVKEAKKYGLDVHAFGIGGPTSIPILYECGVDTFDSTGWIRSAGYGNVFFPFKSRLNITHLDRSGPAVFREELSNLKTETNHSCPFCESFEDLQENREDRVLHNLVVMSEMADAIGEYSTDELLDIMNPRS-QYTKYLKKLTAI-
SDG37465.1_Halorientalis_regularis                 M-LTFYAALSVSPSSMRKTFARLEDGYGSLEPLENIMFTPISGGEKAHQTIRELNDHVGINIMFDSGGYEVQVGNKTFDELQAYLHDFYGENHWGHRYVLPDNVPVSQDSSEVVQRKVEQTADISALFYRQMPDPLKPRALAVVQGHTQDQLQHCLNKYASLGLQHVGFGSFGTSGVSNGVNMLTTEAYHNLEWAVNQAHEAGLSVHAFGVGGPTSIPLLYHAGVDSFDTTSWMRSSGYGNVFFPFKSRYNASHRKRGGNLLTKDELPHIKAETEHECPFCQEMSQLRSSRWNRIMHNLIVIHEMTNRVEDMTREEILKAMDEES-KYRRRLKKVTSSN
MBI5668044.1_Chloroflexota_bacterium               M-LQFVASVTICPSDIQKVRKHVEIAKTATDPLHSVLITPLFANPRSLMMVRDLAESG-KQVYFDSGGYYVQIGRLRYEELYMPLLQAYRANPWASIFTLPDHVPTTQDDPDTVECKVHDTITYSTLFFQELPDNLKPRAMPVVQGHTVRQIDACLQAYIRLGAKQIGFGSFGTGSKKSEINVATNGSVGLAKYVIQVAHSHGMKVHVFGLGAPALVAMLKGIKADSFDSSSWLKAAGFGQVFLPFMRAYNISHNSELQRGITFNQFDEWKTLTGHHCKLCEELDNLQKHKMYRAAHNLIVLAETIAMVNQGDFSRIKQIYVKGSGKYRQEFDRWLQPN
WP_114450123.1_Haloplanus_salinus                  M-LTFYAGMNISPSSMRKLFARLQNEHPLTAPLENIIFTPIAHGNGTHSEIESLSKELDLNLMFDSGGYEVQVGNKTFDELYEYLLEYYDENRWGQRYVLPDNVPMSDDDVQTVQEKVDETISATRMCFRRLPEDVREDSLAVIQGHTREQISRCIDEYKELGLTKVGFGSFATGGVNGGVNMITSEAFKNLRWATELAHEHGLSVHAFGVGGPTSIPLLYEAGVDSFDTTSWMRTSGYGNVYFPFKGRLNASHRHRSGNVLTRQELPHLKAETDHSCAFCEDISRLSNNRWDRILHNLIVTHEVVERIDDMTQQEVIDLMDQSS-RYRKRLENITPRK
MCZ7554452.1_Anaerolineales_bacterium              --MKFVASLTMDASLLKKLLALWDASGDSRNPFESVLVTPLFASASTLKMIRELKEKRGAQVYFDSGGYYVQQGKIGFEQLYQDVRAWYSENQWADWYVLPDNVPTSGDCGEVVEQKVNETVAAAKMFFAEAPAAVLERCMAVVHGHELWQVNRCIESYLGLGVKYLGFGSFGTGGGDNSINQLDKHSAEIVAHLASGLCSCDVLLHTFGISTPPAVYAFKKLGFSSFDSLGWQRSAGYGKVYMPFTRAYNVSHRSIHNSALTRDEFVRLCLLTGHECPFCVDFRQLSEDRLYRSLHNLVVLMETGNGTGESRQEKIARLIEDWSPRYYRLFQQVYNG-
GAB4528646.1_Anaerolineae_bacterium                M-LQFSTTLTLCPSDIRKICQLLDEHDIAATPLKHILVTPLFVQKESLQRIKAIKEQQGSTIMFDSGGYYVQIGKLTYEELYYPLLQFYRENPWADVYTLPDHVPTSQDTENDVWRKVRETVRYSILFHQELPADIQTRAMPVVQGHTLEQVDHCLEAYLSLSIRYIGFGSFGTIGKNSEVNVATNSAVNLAKYVVKVVQARGVKVHFFGLGAPALVAMIYGAGASSFDSSSWIKSAGFGQIYLPFMRGYNISHRSELQRRKVLTEFQDLRLLTGHHCPFCSSISELQRKKLYRALHNLLSIQETVDMINSGDHTIKTIYEQGSPRYKQEYEKWLAPA
MBC8263378.1_Anaerolineales_bacterium              --MQLVTSLTVDASLLRKLLRLWDESGDPRNPFATVLVTPLFASPSTLRLIRELKERRGATVYFDSGGYYVQQRKIAFEELCCTLRDYYQDNQWADWYVLPDNVPTSTDSPAIVEEKINDTVTAVRMFYAEMPSIVQERSMPVIQGHTFKQVNRCIEAYRELGLKYIGFGSFGTSGTNNSINVADSRSAATVAHIVRELDNDGIRLHTFGVSTPPVIYAFQRLGIYSFDSLAWQRSAGYGKVYMPFMRAYNVSHRSTRNSALSQVEFELIKERTGHHCPFCENFSHLSKRRLYRSLHNLVAVMDTVKVDQRQDSQSIASLIAWKSQRYYKMFKEIYLGQ
GAB4153075.1_Candidatus_Promineifilaceae_bacterium --MQFVASITRNASDIRKISRLWQGSGDTRNPLKNILITPLFTPPSTLKLIREMRERGEEQVYFDSGGYFVQMGRLSYEDMYWYLLQFYRANHWADWYVLPDYVPLSSDKEDEVWYKVRRTADGSKLFFLEMPSSLQDKALPVAQGSESKQIEYCLETYVRLGVQQIGFGSFGTSGKSSSVNAVTSNSLKMLGELSEILTQERLNLHAFGVGNPPVIYLLNQAGVDSFDSVGWMKTAGFGKIFMPLTRAYNITYRDPMARGLKEDAFYTLKELSNHDCYFCQSFQKLRESRDYRVMHNLTVVLDTLEMIDKGLSNLVERILQTYSPAYAKLGRRDRDA-
MBW7959475.1_Candidatus_Promineofilum_sp.          --MQFVASLTRGAAEIKRLIRHWNDSEDPRNPFANILVTPIFTPPSTLKLVRELKEEGVKQVFFDSGGFFVQMGRISYEDMYWQLLHFYRSNPWGDWYTLPDYVPLSSDDEETCWHKVRQTADGSRLFYQELPYEIKAKALAVIQGRTIAQVDYCLETYLRTGPERWGFGSFGTNGKQSNANIVTEEAFILLNYVSEIAVSNNRHLHAFGVSTPPAIYLLNESRVDSFDSIGWMKTAGYGKVLYLPFVRAYNITYRDKGARGMPPQVFEKLKIRSGHECSFCTDFRMLETSRTHRILHNLAVIQDMVDMIRERTGDDLLQIMEEYASTYARLRSIAES--
MBP7045033.1_Chloroflexota_bacterium               --MEFIASITLDVSYVHKIRRLWKMSGDPRNPLKNVLVSPLFTHPKTLAKFRELRENNEEQIYFDSGAYSVQKGRSSYEEMYWSLLNYYRENDWADWYVLPDYVPLSSDTPEDVWYKVQTTANMGSLFFAEMPEHLRLRALPVVQGHSIDQIEFCLESFLKMGIQRIGFGSFPTNGKDSSVNTLSPSALTFLSHLTKILDSKGIGLHAFGVGTPPVIFLLHQAGVKSFDSVGWMKTAGFGKIFMPFVRAYNITYRDRTARGLTKETFLELKRMTNHDCFFCQSFDTLAQSRDFRFMHNLIVVMDTIDQLGNLQSVDVGKILDIYSPYYAKMNIGANT--
HHW27507.1_Candidatus_Fermentithermobacillaceae    M-VRFVTALGIRPGDLAQIALAWGNVKDRNTPLANVLISPLFADPASIQMIRRLKEQGANTVVFDSGGYQVQQGKRIDYNTLYNRLLQYYEANDWADYYVMPDFPPVSGDSDSIVQEKVRITVDGTKRFFYDLPSTLRSRALAVVQGRRVEQVKFCLVNYHDLGISYVGFGSFPTNGAQSSINHLDAGAAEMLACIRRQYPS--VRIHAFGVGNPPVAYILNHMGIHSFDSSGWIKAAAYGNIYFPFTRAYNITYREPGRGALDAESFERIKTLTGHSCPYCSDFYELSRNRTSRFLHNLICMMETVEDISAPL---ASEVISHWAPAYKRFLTGRAL--
WP_117592495.1_Haloprofundus_halophilus            --MDFYAALSVNRSSIRKIAKRLEEEYSSFDPLQNIIFTPITQGETSHQEIRRVAEKYDVNLMFDSGGYEVQTGHFEFDDLYEYLIDYYSDNQWASRYVLPDNVPVSGDSPEVVEQKVTETLSASRSCFKRLPEFVQENAVGVIQGHTREQISRCVNEYSEFNLKHLGFGSFATSGGQNSVNMLTSESLQNLRWAVKLAKEHGFTIHAFGIGGPTSIPLLEEAGVDSFDTSSWMKSSGYGNVFFPFKSRYNASHRLRSGKVLMADNLEDLKEETNHDCPFCRDVAKLREDRLTRVMHNLVVTHELSNRVSGMARDQMAGQMDTNS-MYRRRLETM----
WP_089881382.1_Halogeometricum_limi                --MKLYASMTVSPSTVRKFKEHLLREYGTLEALDPLIYTPISTGDGLTEEMKQLSKE-GVNVMFDSGGYHVQTNNGDFDELYNYIMPFYDKNRWAHQYVLPDNVPLSDDDDETVERKVKETRTNAKLSYHFLPEELKSRAVPVIQGSKYPHITDCIDTYSELQIQTVGFGSFETFGKNNGINFISERVAETLRYAVNLAHERGLKIHAFGVGGPTAIPVLGKLGVDSFDCSSWLRAAGYGDIYFPFRTPRNVTHKDRTGPKVFRSDLEELKKETGHKCPFCASYDKLRDSRIIRILHNLIVLRDMAESINSYSTEEMLDLMNDKS-RYTTILRSIS---
MDW8029827.1_Armatimonadota_bacterium              L-PTFFVTLTHEATEIRKILRLWDESGDRRNPFAHLIFTPLFSKPATIATIRQLKDQRGSTVMFDSGGYQVQTGKATYEELFERLLRFYRENDWADFYVLPDHVPRSSDSDREVDFKVRETVDFARLFLKMMPSDFAAKAVGVVHGRTFEQVRYCVQAYVDMGISYIGFGSFGTSGPNGKVNMLSQKSLRLLKLVQTLACEFGLRLHIFGIGSPSHLIRLQQGAIDSFDSAGWWKAGGFGNIFFPAGRQLHITAINAEATLL---GIERQKRHSGHQCPFCDNSKLLRLSRVKRILHNLAAMLDTLKQVYGGQ-
WP_048108638.1_Methanosarcina_barkeri              --------MSAHPGLIKKMRDSLKDFEKIKWPLKNVLYTPVFSTKKTSEAIKDLKDNDGSKIMFDSGGFVQQGKITTQELFSKLTDFYSQNEWGDYYVLPDSPPVNGDSDLIVEYKVRDTITNAKIFHDQLKENIQKKCIPVVQGHNKEQIFRCVREYSDLNVSRIGFGSFGTCGSNNGINFLTSKSMENVEYMMNLLKDYSLEVHAFGVGGTSSLSKLCEAGFTSFDSAGWKRAAAYGDVLFPFHSTHNITHREKSCSILSRNNFEKVKTESNHACYFCEDFNKLQSQVYYRYLHNMIAIFDTVEFLKKYPISSHNDI------------------------------------------------
consensus/100%                                     ........hs...s.h.p....h.......sh..lhhsPh.........hc.h.pp...pl..hDSGua.sQ..p....pph......hY..s.Wsp.ashPD.sP.p.Dp...h..Klp.T....s......h......pshsVhpGpp..pl..Cl..a.....p.hGFGSF.T.G.ps.hN.hs..sh..h..h...h......lH.FGlusss.h..h....h.oFDs.sWb+suuaGplhhPh.....pho........h....h...p..o.H.C.hC.p...L..p...R.hHNh.s..p................................
consensus/95%                                      ........hs...s.h.p....h.......sh..lhhsPh.........hc.h.pp...pl..hDSGua.sQ..p....pph......hY..s.Wsp.ashPD.sP.p.Dp...h..Klp.T....s......h......pshsVhpGpp..pl..Cl..a.....p.hGFGSF.T.G.ps.hN.hs..sh..h..h...h......lH.FGlusss.h..h....h.oFDs.sWb+suuaGplhhPh.....pho........h....h...p..o.H.C.hC.p...L..p...R.hHNh.s..p................................
consensus/90%                                      ...ph.sshs.p.u.hpph...h........sh.plhhoPl.....s...hcpl.cp...plhFDSGGa.VQ.sp..hpphh..l..h.Y..N.Wup.YshPD.sP.s.Ds...V..KlppThp.s.bha.bhP..hb.cshsVlpGpp..Ql..Clp.Y.phshp.lGFGSFsT.G.ps.lN.hs..sh..h..h.p.h.p..h.lH.FGlusPs.l..h....hpSFDo.uWb+suuaGplahPh.p.bNhoa.p.....l....h.ph+.bosHpC.hC.ph..Lp.p+..R.hHNLhs.-.p..............................
consensus/85%                                      ..hphhssho.psu.hp+h...h.......pPh.pllhoPl.s...o...h+pl.cp...plhFDSGGYbVQ.sp.pap-ha..lbpaYp.N.WupbYsLPD.VP.o.Ds...Vb.KVppThs.s.hha.bhP..lb.+uhsVlQG+p..Qlp.Clp.Y.phslp.lGFGSFsT.G.ps.lN.hsppuh..h..h.p.hpp.shplHhFGlusPs.l.hl..huhpSFDo.uWb+suGaGplahPF.p.hNhoabp.....l....p.h.phK.bosHpC.aC.shppLppp+..RhhHNLhsh.-..p.l.......h..hh...u..Y.pb........
```

```
# ; DUF6079 ATPase Representative Alignment; ORC/CDC6-AAA-ATPase + wHTH + Alpha+Beta-Domain + 2-wHTH + Extended-Coiled-Coil

                                              1---------11--------21--------31--------41--------51--------61--------71--------81--------91--------101-------111-------121-------131-------141-------151-------161-------171-------181--- [illegible]
OrigSeq                                       MNTLWDQLDALRAIVDTAQADDIERQPEAIRTRYEPYAQAYFAIEAVKRVEERLIGELRAGRSVTGYLSADFGYGKTATAIYLWKECLDSGIVAVPPFLFRQLKGLMQATRGWLAYLLQHTQPALLPRLEETYRQHAERSVDELAKDIAARQGISEAKARAIVQEHIANRRDVTTTQSLLEFLRDA [illegible]

                                              |------------------------Preceding-Helices-----------------     S1                       S2       |-----------------------HEH-Insert-Region------------------------| [illegible]
                                              <<-----------------------------------------------------------------------------------------------------------------------------------------------ORC/CDC6-AAA-A [illegible]
                                              ---HHHHHHH---------HHHHHH-HHHHHHHHHHHHHHH----HHHHHHHHHHHHHHHHH---EEEEEE------HHHHHHHHHHHHHHH--EEEE------HHHHHHHHHHHHHHHHHHHHHHHHHH-HHHHHHHHHHHHHHHHHHHHHHHHHHHHHHHHHHH---HHHHHHHHHHHHHH------HHHHHHHHH [illegible]
                                              ---HHHHHHH---------HHHHHH-HHHHHHHHHHHHHHH----HHHHHHHHHHHHHHHHH---EEEEEE------HHHHHHHHHHHHHHH--EEEE------HHHHHHHHHHHHHHHHHHHHHHHHHH-HHHHHHHHHHHHHHHHHHHHHHHHHHHHHHHHHHH---HHHHHHHHHHHHHH------HHHHHHHHH [illegible]
                                              ---HHHHHHH---------HHHHHH-HHHHHHHHHHHHHHH----HHHHHHHHHHHHHHHHH---EEEEEE------HHHHHHHHHHHHHHH--EEEE------HHHHHHHHHHHHHHHHHHHHHHHHHH-HHHHHHHHHHHHHHHHHHHHHHHHHHHHHHHHHHH---HHHHHHHHHHHHHH------HHHHHHHHH [illegible]
BAL53915.1_uncultured_prokaryote              MNTLWDQLDALRAIVDTAQADDIERQPEAIRTRYEPYAQAYFAIEAVKRVEERLIGELRAGRSVTGYLSADFGYGKTATAIYLWKECLDSGIVAVPPFLFRQLKGLMQATRGWLAYLLQHTQPALLPRLEETYRQHAERSVDELAKDIAARQGISEAKARAIVQEHIANRRDVTTTQSLLEFLRDA [illegible]
MDT7893011.1_Armatimonadota_bacterium         LQRLMERLEQLQPIIATARAQELGEMPEQVRNNYEFTASVFYAMESVKDLRDPMLRRLTNNKPVLGYVSGEYGYGKTATMVWLWHELEREGFVAVPPFLFYSWDDLMLATTNWLSFRLAEKRPDLVDEAEKLYNSHRSKAVEELADEIAKRQRVSKEQARRIVEDLLQGKL-MTPSQMVDFLRDN [illegible]
MCS7186064.1_Armatimonadota_bacterium         MKRLLERLESLQPIVSTVARADDLYQQPEIVRNNYEFIASLFYPFESVKELREPLLRRLKDSKPVVGYVSGDYGYGKTATMVWLWQQCEHEGMISVPPFLFYDWDAILKATAFWLTFRLKERRPDIAKRVTELLDRFRSKAIDELAEEYVRRERVPFERAKRIIDDLLKRGML-SDPSQVVEFLQEA [illegible]
HEY9694182.1_Oculatellaceae_cyanobacterium    MKRLLKQLDSIEAIVDTVSPSQIEDDPQGVASCYELYASSYDPLERVKTLRDNLITGIHKGPVNGYLSADYGYGKTATLVYLWYECQQQQIVAVPPFKFKELGNLMVASYWWIKASLKLKQPELISEIEELYRKYGLSQSQQEQAAEIARKYRVSEVKALKIVQELKT---ETTNTDSILQFWQYS [illegible]
HIE11924.1_Desulfotomaculum_sp.               MPRLWEQIERLRAIVDTARAQDVEREPEAVRTRYEPYAESYFRIEAVKRLEQRLIGELHALQDGRSITGYLSADFGYGKTATAVYLWKRCLDSELVAVPPFLFRQLKDVMQATYGWLAYQLEQTYPSLIRRLEEAYQRHNVRSTEELVEAIARREGISEAKARRIVQEYIRRYKDVTTAEALLDFLREA [illegible]
MDX2239253.1_Leptolyngbyaceae_cyanobacterium  MKRLLEQLDAIKSILDTVTPAQLETDPEAVESGYEVYAQAYNPIERVKDLKDRLISEIRKGRPVNGYLSAGYGYGKTATLVYLWHECQKKQIAAVPPFKFREIGDLMIASYGWMKKCLSHSRPDLIAQVDKLYQTYGLKSQEAIAARIAKEDNISESKALKIVSRLVNYQ-QAINADSLLSFWRES [illegible]
HEY9694177.1_Oculatellaceae_cyanobacterium    MKRLLEQLDSIKAIVDTVSPSQIKDNPQGVANSYELYASAYEPIEQVKTLRDRLIAEIRKGKPVNGYLSADYGYGKTATLVYLWHQCQQKQIVAVPPFKFKELGDLMVASYGWIKACLR-SQPELIPEVETLYRKYGLKSQTRAAEMAGKYKVSEDKVLKIVQELKT---DTTNTDSILHFWHEI [illegible]
WP_310922611.1_Halogeometricum_sp._S1BR25-6   VGEVFDLVRKSEGIISTFSANEVAEEPQTVEAEYERFADTHISLGDTDGFEDRVYESLVNNKPTKGYLYGPFGYGKTSTSVSIWKEITDNDVIAVPPFTLTSFTAIMRATHGWMRYELSNKAPGYVDELEEIKDRYINQELEAYAEKKKDEYQFDKDQLVEMFHEMERNNDLSIQADTLIDFFSEC [illegible]
WP_338740219.1_Haloplanus_salilacus           VEELSDLVRKSEGIISTFEARSLKDEPETVEDDYRRYAQTHVSLGDTSEFENEIYETVIDESATKGYLYGPYGYGKTSTSVSIWHTLSDNDIIAVPPFTMDSFAAVMRATYGWLRYEFQNKAPGYVDDLDDIHERYLQQELQSVAKQKEGQHDLDFDKLVAMFEEMEQENDLSINADTLIDFFDEC [illegible]
WP_147270938.1_Haloplanus_salinus             TDELLDLIRKDEGIISTHTASDLETSAESVEEKYRTYAETHISLGDTSGFEQKVYDSLTENKPTKGYLYGPFGYGKTSTSVSIWHKLNQNEIIAVPPFTVSSFSSIMRATYGWMHYKLRTEAPPYVDELENIRESYLEKEIRSYAEEKKDAHDVGVDKLVTMFEEMERTGELSINADTLIDFFDEC [illegible]
MYC73120.1_Gemmatimonadota_bacterium          --MALDEIIQKNAVVDTPRASDLAAKPDEWADRFQLHVSTHVAMADMTDKWDAILKNLLRGRSATGLIHADTGYGKTSTAAALWHYAENRDVVTVPPFAWNSIADMLTATHGWVRHRLKVKRPDLISDLEQQYKTLVESSDEDRARRTSLEENIPLDHAKRVVQSLKAKGEDSISPNRLIDYLRQA [illegible]
MFN8469223.1_Caldilineaceae_bacterium         --MTLETIITKEPVIETPTARALAEEPKYAEFTFHVNTHVTVADLADKWDRMLHNLEQRKSVTGLIYADTGYGKTSTAAALWQYAEARGIVAVPPFLWASMTDMLIAAYGWSRFRLETKRPDLVEALDRRYGALMQSGLSDLSARLARTKGVSIADAQKIVEQLGSQEGQDVISASRLIDFLEE [illegible]
RKU21510.1_Candidatus_Poribacteria_bacterium  --MPLADIIEKDQTLYTPSAQEIADSPQEYADKFRFHVATYVTIKDIQSKWEATLGNLLKGRSATGLIYADTGYGKTSTGASLWHFAESKGIVTVPPFRWNSFADLLTATHGWVRYRLEDTRPDLIPNLEEKYREVIAIGEKVLAQRMSHEKHLSIAQAQEATEFFKSAGREELSPRDLLNYLRFT [illegible]
MYB94331.1_Candidatus_Poribacteria_bacterium  --MSLHEIHKKEPTVATPSVYNLVAAPEIWDRKFHFHVSTHVSIGNLGNRWDAIINNLLKKRSVTGLIYADKGYGKTSTGISLWQQAAEAKQIVTVPPFVWGSLADMLTATHGWVCYRLQDTRPELIPDIEQKHREVVEVDEEKLAQRLIQENGLSYDQARNTINHLREAGREVLSPHQLLDYLRFT [illegible]
HET59438.1_Chloroflexota_bacterium            MLTGLQAILDKEVVLSTVNADDLLHHTAKVEADYARHVATFIPMGDTSNFSRRIIKLTTNAKTPKGLVVADYGYGKTSTLAFLWHECEQQDIVAVPPFYCASLLDMLKATYGWVKFRLERRQPELVDDLDEVYQKYTAATVEEMAERYMREHGIAKVTAVGMLNDMLDGGSLELTPSNLLFFLDAA [illegible]
HFD39728.1_Anaerolineae_bacterium             MLEGLEAILDKQPVLSTVNADDLLQNPSKVEGDYARHVATFVPMGDVSGFAQRLVQRVIKAKTPKGMIVAPYGYGKTSTLAFLWHACEEQGLVAVPPFYCATWLDVLNATYGWVRFRLEHSQPELLADLDDVYQKYTASTVDEMAERYAEEHGLARITAKGMLQDMLEEGSLSLTPSNLLFFLDA [illegible]
MCZ7554455.1_Anaerolineales_bacterium         MIEGLQAILEKDIIVSTVNPRDLLEGAENVERVYQHHVSTFIPMGDMEGVAERLARKIQSEKTVKGLLTAPYGYGKTSSLAFLWRSCEDQGLLAVPPFYCASWLDILDATFGWVRFRLGNLQPGLVSEVERIYAKYTRTTLEDMARRYSEEHGISTNSAQSILDDLLEHGSLELSPASLLTFLDEI [illegible]
MBI5668047.1_Chloroflexota_bacterium          GLEGLDYILNKKEVVPTVSARDLLERAEQVDMDYKRHVRTYVPISNVEEFERKVIKQVKEARAPRGYITAEYGYGKTSTALYLWLRAEEDNLIAVPPFQMLELPDLVTATHGWTRYRLSHHRPDLVTSLDTLYETTMDRSIEKEAH----EKHVSVSVMREWVNQ----GRLDLQPTDYIHYFEDV [illegible]
GAB4528661.1_Anaerolineae_bacterium           SLDGLQAILHKEEIVATQRANDLLERPQFVDETYKRHVRTYIPLGSVIAFERRVIREVCQAGALRGYITAEFGHGKTSTALYLWDQARAENLLAVPPFQLNKLPDLIIACFGWIRYEVGRTRPNMIDEAQSLYKSLMERSAESLAK----RYDIDLTSAQRMAHD----RPLELTPADYIRFFEEM [illegible]
MBN2393034.1_Anaerolineae_bacterium           SLEGLQAILDKEEIVATQRATDLFERPEAVDEAYRRHVRTYVPLGNVASFERQVIQEVRQGGAMRGYITAEYGYGKTSTALYLWERARDANILAVPPFQLNRLTDLLVGCFGWARYEIGRTRPNLVSEAQAIYQAVMERNAETLAH----RYHIDVAAARRMAQD----KPLELGPADYIRFFEDM [illegible]
MBV6438328.1_Anaerolineae_bacterium           SIQGLDFITKKDEVVPTFTASDLLRDPEKVEESYKRHVRTYVPINSVDEFERKLIKAVKDARAPGGYITAEYGYGKTSTALYLWERAEESGVVVVPPFKLIHLPDLLTGIYAWVRYKLMSRAPHLVEKLEALYGAATGRSIEREAK----ANDVSEAVLRQWVEQ----GRLEPQASEYITFFENV [illegible]
MBP7045036.1_Chloroflexota_bacterium          NLEGLQAILDKEEIVATQRAEDLLTRPEIVDRVYQGHVRTYIPLSSVKDFERRVIREVKQAGAIRGYITAEYGHGKTSTALYLWQQARREENILAVPPFQLNKLTDFIRATYGWVRYEIKRTRPQSIQEADKLYDSLINRGAESIAR----QYNMSLADAKRMARE----KPLELTPADYIHFFEEM [illegible]
MBW7959472.1_Candidatus_Promineofilum_sp.     SIEGLQAILDKDEIVATHQAEDLIAHPDLVEGSYRHHVRTYVPFSSVPAFEKKVIRDIKDAKVVRGYITAEYGHGKTSTALYLWDRARQNNLLAVPPFKFNYLPDLARATYGWARYEIGRTRPGLLDKVKDAFTRFIDHSAEAYCN----KYDIPLATAQRLIRD----KPIELGANDYIAFFETI [illegible]
consensus/100%                                ......l...p..l.T.ps..l...s..h...a...s..a..h.......p..h.......s..Ghl.u..GaGKTuo.h.lW..h....hhssPPF.h..h.shh.u..W....h....P..h..hp..........p..s........hs...h....p..........s..hl.ahp.. [illegible]
consensus/95%                                 .....hp.l...p..l.T.ps.pl...sp.h...ap..s.sa.sh......p.hh..l.p..s..Ghl.u..GaGKTuT.h.lW..h..p.hlsVPPF.h.ph.shh.us..W....pl..p.P.hl.php..........p.bs........hs...h...hhp..........s.phl.ahp.. [illegible]
consensus/90%                                 ....hp.l.p.p.llsT.psppl...sp.h...ap.as.sa.sh.p...b.p.hh..l.p..s..Ghl.u..GaGKTuT.h.lWp.hp.p.llsVPPF.h.ph.shh.As.sWhphpl..p.P.hl.phpp.b........p.bu....p..ls...h..hhpp..........ps.phl.ahp.. [illegible]
consensus/85%                                 ..b.hp.l.pbp.llsT.psppl.ppsp.h...ap.aspsahsh.s...bbp.lhp.l.p.ps.pGhl.u.bGYGKTuT.h.LWp.hppp.llsVPPF.h.ph.shh.AsauWhpapl.pp.P.hl.phcp.app...ps.p.bA.....p..ls..ph..hhpp........ss.phlpahc.. [illegible]
```

```
# ; PglZ Representative Alignment; iSwi2/SNF2-Helicase + PglZ-Phosphatase

                                                        1---------11--------21--------31--------41--------51--------61--------71--------81--------91--------101-------111-------121-------131-------141-------151-------161-------171-------181-------191-------201-------211-------221-------231-------241-------251-------261-------271-------281-------291-------301-------311-------321-------331-------341-------351-------
OrigSeq                                                 MPTFTEIAEHAKVSPVVCSLPKLASFYLDDLAVYTSLDNFHHDLYDPDYRFEEDVVIDTDEHHPDYVRLSCDEEPVSPQLNEVVREHHDRLRRELLPQSGLKDYILRNVIGETVVVLLIVDGLSYESIRTTNLPVQPVIVDGITTTEPGFKRIIYGNNVSLYSELLHKKNFANKLGFTYWARGQEELSTDLHASMGDIRRIRDFSEAIEYLSNNSPTNRTYIQITRMGLDQESHNKKEAPNQEAVIENIVEDIRKLESELSKQERSYKIFATSDHGILWRTQLRPNVVIDDYKHQPRYVTGREKVPHSMTLTDSEGEIVSGLAYPYLARDFKHTEWGVHGGFSYYESVVPLIELASKHKD

                                                                                                                                              S1                                                         S2                    S3                         S4                                          S5                  --------------------Insert-Region--------------------         S6
                                                        <<-----------------------------------iSwi2/Snf2-Helicase------------------>>          <<------------------------------------------------------------------------PglZ-Phosphatase------------------------------------------------------------------------------------------------------------------>>
                                                        ---HHHHHHHHHH--EEE---HHHHHHHHH-EEE--HHHHHHHHH---------EEE---------EEE-----||HHHHHHH---HHHHHHH-||-HHHHHHHHHHHHH--EEEEEEEE---HHHHHHHH---EEEE------HHHHHHHHHH------HHHHHHHH---EEEEEE-------HHHHHHHH---EEE----HHHHHHHHHH------EEEEEEEE-HHHHH--------HHHHHHHHHHHHHHHHHHHHHH---EEEEEEE------HHH----------------EEE---------EEEEEE--EEEEEE----------------------HHHHEEEEE--------
                                                        ---HHHHHHHHHH--EEE---HHHHHHHHH-EEE--HHHHHHHHH---------EEE---------EEE-----||HHHHHHH---HHHHHHH-||-HHHHHHHHHHHHH--EEEEEEEE---HHHHHHHH---EEEE------HHHHHHHHHH------HHHHHHHH---EEEEEE-------HHHHHHHH---EEE----HHHHHHHHHH------EEEEEEEE-HHHHH--------HHHHHHHHHHHHHHHHHHHHHH---EEEEEEE------HHH----------------EEE---------EEEEEE--EEEEEE----------------------HHHHEEEEE--------
                                                        ---HHHHHHHHHH--EEE---HHHHHHHHH-EEE--HHHHHHHHH---------EEE---------EEE-----||HHHHHHH---HHHHHHH-||-HHHHHHHHHHHHH--EEEEEEEE---HHHHHHHH---EEEE------HHHHHHHHHH------HHHHHHHH---EEEEEE-------HHHHHHHH---EEE----HHHHHHHHHH------EEEEEEEE-HHHHH--------HHHHHHHHHHHHHHHHHHHHHH---EEEEEEE------HHH----------------EEE---------EEEEEE--EEEEEE----------------------HHHHEEEEE--------
MDS0297782.1_Halogeometricum_sp._S1BR25-6               MPTFTEIAEHAKVSPVVCSLPKLASFYLDDLAVYTSLDNFHHDLYDPDYRFEEDVVIDTDEHHPDYVRLSCDEEPVSPQLNEVVREHHDRLRRELLPQSGLKDYILRNVIGETVVVLLIVDGLSYESIRTTNLPVQPVIVDGITTTEPGFKRIIYGNNVSLYSELLHKKNFANKLGFTYWARGQEELSTDLHASMGDIRRIRDFSEAIEYLSNNSPTNRTYIQITRMGLDQESHNKKEAPNQEAVIENIVEDIRKLESELSKQERSYKIFATSDHGILWRTQLRPNVVIDDYKHQPRYVTGREKVPHSMTLTDSEGEIVSGLAYPYLARDFKHTEWGVHGGFSYYESVVPLIELASKHKD
WP_271955572.1_Halorubrum_ezzemoulense                  IESLSALAEATRDGPVVANIGGLESRVLQNIDTVESLEEFYRLYESPTNLIETDVIIRDQGPKTLYPSVEYTGTVYDRSLNRLISISWDDVESTTTCQSEVGDLIMEAASGVNVIALVIVDGLSYQDWTDSGYEAEPIYVDCPTRTDCGYPNVVRGANGKHLQYRLFNKGYKTRKAFTYWERKDNDLSDELHDGFSVIGNIQDFGDSISYLQTHDWRPKTYLQFTLTGPERVAHRMKEDPVIQSEVDAVHQKLSDLRETISKETDSFRIFAVADHGILWRMDTDFEALDEWKQGIRRDYEDPPASESVGETSFWAGQQYLRLYHPYIFNNLRSNEPGVHGGFSYQESIVPLIEYE-----
WP_139229723.1_Halogeometricum_limi                     MAKLEEVVDEAKSHPVVVDVHPLSYLYLQELELFESEKQFEDELDNPSKSFERNVAIKEPRVKTTYPSMSIDHDPLSPALNKIIGTKYEEVEEYLLTQRDIKRRILDYLTDESVVVLMIADGLGYKDVQGTELEGEPCLVNGPTITPIGYRNIVFGHDNKPIVDHLVKRGFMRRRGYSYWDKESNDLNKLVFENFSELTRVRTSEEIISDLENRRLDHRTFIQISRRALDGDAHDGKEVREINHEIKKLQQDIDAICEVLDEQVDSYLFILTADHGILWRKEFHMSILTTNGQSNMRYAKSKIGVQGDAKLLDAYGSTYTVFPYPFVSREFHHDEFGIHGGLSYEESLVPLLFRNSEDSE
ATW88247.1_Halohasta_litchfieldiae                      MAYFNEIKERAHQQPVIVPLPKLAQFYLEDFNIYSSIDDFQTDFYHPDFLFENDVIIDADEHHPDFDRLSCDEETVGPRLNTVVNTDHQRLKEDLLVQSQIKEHILDAVSTEDLVILIIVDGLSYESVRGTAIDARPVFVDGISTTEPGYRRVIYGKPTSVYADLLDAKRFYNDLSYTYWERGDEDLSTDLHSSMSSVTRISSFNEVVDALREEYPEEKTFVQITRMGLDQDSHNRKEEPNKSAVVQDILDDLGKMHETADKLSDRFRIFLTADHGILWRDQLEDSIVCEDYHPHARFVEGGMNIKEGRTIFETDGVKSIGLGYPHLTRKLANTEWGVHGGFSYYESIVPLIEVTEDSAL
WP_338740220.1_Haloplanus_salilacus                     MASFETLADRASEAPVISPLPALATFYLDGLSIYRSVDTFHRDLYDPDYQFEQDVVIDDDSRYHEYPYINCTTDPITPALNDVVEEHHDRIRQDLLQQSKLKNHILDAITGEDFVALIIIDGLSYDALRTVSNILQPVIVDGITTTEPGFRRIIYGGDPASIYASLTGEGFYTGYGFTYWERGQEDLSTDLHSAMGNIHRIRDFDEVISTLEADAPTEKTYVQVTRMGLDQDSHNRKENPNRNVVRDEILSDVKSLAETASTMSDNFRIFMTADHGILWRDQLDPPIVHDEWTNHARFLTGQHNLEHGMVVKDST-DPTTGLAYPYLTRELKNTEWGVHGGFSYQESIVPLIEVTSPGDL
WP_143414662.1_Halorubrum_sp._SD683                     MEDIKSIADRAHEEPIRADLPALAERILQQIDVISTTTEFYQRFEDPDHRFSADVLINDTTPKTTYPTIPYTGDPYTPELNSLIVLSWKDVEETTLSQSEVAEYVASESGSEDVIVVVIVDGLSYYDWIRAGYRPDPVYVDCPTITKCGYPNVVYGGSGESLANRLYNQGYKNRLAFSYWDKSDSDLTDQLHESFSVIGNVEDFDDIITYLNRHEWQKKTFIQVTLTGPERVAHKIKENPDTIHQAQLVQKKLTALNDVLSEQVSSHRIFATADHGILWRMDVDFETLTQWHHTKRRHIENPDVEESTGYTETYAGSKFLRLEYPYLFYDLKSNEPGVHGGYSFQESIVPLIEVQ-----
WP_048108644.1_Methanosarcina_barkeri                   -MKLSEIDNLLSKTPVYLSLDSLVSLCYLDITIFHNRRDFWRKIDQLEPLKLCRYVILDDPD-YRLPYVRLKHNPFTNNLNSILSKKYEEIQRLMLTQKDLSQEILFRSN--EIVVLILIIDGLSYIDCLKY--SPEPCIVNGTTTTESGFTNIVYGSENRPLSFKLYDKGYMRRFGFTYWERENNELTNKLFKGFNEVIKVSSFEDVIKHVRSMELKPHTFLQIIRNGLDRYAHNMHDKPLKEQEIEQIFKDVFELKETFESKGITANIFVTSDHGLLWKYEHEFKVIE-PYNESPRYLKGLHKIEGHSVISKLNEHYNTLLAYPYILRSFRNNEWGIHGGISFEESITPFIEYKVDK--
NPV62497.1_Methanotrichaceae_archaeon                   RVTLEELDALLNKAPIAADMNPVSRCYLPN--RFANLDDVVKELDSPRPVDPSRTLIIGPRKDEKFLNVRVVHEPFTSDLNEILFDHFDEIRSAYMLNQSEIAQKIYSDAD--DVIILYIVDGLSFHDVKKNPPVLEACLVDCPTVTQVAFPNII---GKPPLARLLFDKGYINRIGFSYWTRKDNKLTDELFSTIPTLHRAENFDKIISILSLDL-KRQTFVQIIRQGLDGLCHSQKLEINPSVILNQIYENLDQLIELCDQLKLSAKVYLTSDHGILWREEFEPKLIENCNNCHPRYIESNII-ENSGRNFLIGNQKFYCLNYPLTRRMLKIDERGVHGGISFQESIVPFLRME-----
MCK4444004.1_Thermoplasmata_archaeon                    -MRVESFLELLRRTPVVLKGNELPLLYVDRTRIIRDIEELNQQIDSPNPINDAEIVIVDDDKGWSFPVAEVSEDLMKEDLNSILNKDYDILQERMFSQTDIADRILSDL-DYDVIVLILVVDGLSYYDCRIVT-EAQPCLVNGPTLTRVGMRNII---NDPPIVSRLFDSGIRERRGFSHWNRNE-TLTNQVFEGFTQMHKIREFKDIMMRLEKEL-VKDTYIQIVLEGCDQISHKNRDRPLVSSRIEQLVEEIVALSNTIKSKNLSAVIHMTSDHGIWWKPELVPQLVHVPHARSRRYVKGYVD-GEHILHFSSYGMTYSVLKYPFIFSPYKSNEWGTHGGVSIYESIVPFCTAKVMAE-
MBI2842128.1_Armatimonadota_bacterium                   DALVSGLSATLNSYPPAFRGNPLNLLFLDDARPIREASAFLNLLDHPALCDERREYLLDDPAN-PFPAYNISRRPDPFLLSERLYDRFDSVKDVMTTQRDVGRILAEKAKEPDVVALMVVDGLSYYDLPDDP-ENHPCFVDGATTTEVGYREVI---GKPSISERLFSIGYRHQLGFTYFDTDSNTLASDLYGVFGQVCKIRRFDECVRMIQAEE-MSRGYIQIVAPGLDGLCHRHQDQPPVKHYLSELLARYDSLVEFLCRGNRSVLACLTADHGILWREQLAVADLHPDDSRHVRYIRGSRL-RHYACVRTCMGNAYSLLKLPFTTRELKSNEWGVHGGISAWESVVPLIIRTA----
HHW27504.1_Candidatus_Fermentithermobacillaceae          -MDVPRLIRMARETPVGIPNDLFLRLRLPDCNVFPNAQSFREALNGHRPMPNSPTLIVCESGDEQMPFEDTHYDFMDPAFNDVLCMRYEFLRATIPKQSQVPRIFVELVKQPRVAVIMVIDGLSYGEARDFI-PSTPCLVDGLSLTREGMLRIV---GSPTLAERLFSIGYRSRIGFSYWDRH-NDLGNEIFRVFAQSCVVKSLDDALMRLAEMKLEENTFVQIVRSGLDSYAHHHLDRPPVHELTAEICADIRRTAQVLASMADNYLLLVTSDHGILWEDTLTSSILPGGAGSPIRYYSANESVSMRKILVAVDEDGHKALPIRYLRRALGSREWGCHGGISVHESFVPIMVLAPDRGK
HIE11925.1_Desulfotomaculum_sp.                         KVDLEWLREMLRQTPVMAEIHPLLALLPKVVVFGDVWELIARLDSPELQPTRPMLCTQVKSEHRLPAIFCDHDPFLTTELNRVLLRRYEFIQRYVLCQSQVADEIVKQPT--DTVILMLVVDGLSYADVKRYAPKAVPVLVDGVSVTDQGMLRIV---GSPPIVQRLFGAGFRRMLGFTYWERAQEPLTDQIFKGFGCVRRVRSFGEILGCLEEID-LCGAYVQIVRAGLDGVAHRQREMPNVEAMVNDILNDFERLACLFEEKAISATLHLVSDHGVLWAWEHQLQLYEFSHAEHPRYYQHHKQ-GEHVLNVRFAEKEFAVLAYPYLRRELHANEWGVHGGLSFEESVVPWLVHRVSRHE
HHW40997.1_Syntrophomonadaceae_bacterium                 QLSLDEFIALLQKTPVITAWNPWLKVKLGKVVFPSLSEFRIAVNRFHPLADMPTVIFAGQNEHAFPKVQCTYDPKDPEFEFLLVQEFHELCPGVFKQSKIPDYFVESVEQPAVAVFMVVDGLSFADIPDY--SVLPCLVDGVSLTPEGMRRLV---GKPHIAERLFARGFSNRVGFSYWDRH-NELADQLFFTFTQLRKVEGFEQILEWLETERLEERTYVQIVRTGLDGYTHNHRDRPPKEHLLKAIRKDVEDLVDRLRKMRLTFLLFVTADHGILWDRDFEDAVFLEEATVPVRYYREKENINLRQLGVWHPRDNTFSLPVTHYRRRLRSTEWGCHGGVSVAESFVPLLRVF-----
HQD80871.1_Bacillota_bacterium                          MIRREHVPSIIDQYPVCVRFDPLLYMRLYGCEVLSSPEEFREHLDDPSYFCGPTLVMCDNLGESIYPLQTLDVV-LSTRSDEVILTYYRQFSPRLLQQSRIASSICHDAKLPDVLVLAIIDGLSYSDQLF--EDSSPCIVDGLTTTSYGMRRIVK--TGRPLAVELHMLGYTQRLGFSYWSPENERLTGELFEGFSQFRRLKDSRDVFSALDSL--SHRTFVQIVREGLDGYAHRHRDRPLREAVVVDLRNYLSELVATMRNTGLTANIYLTSDHGILWADDLNAVCLAGNQIGTVRYYDLRHDDEAYRVGVLVENHSFLALPKTHCRRKLSALEWGTHGGISFDECVTPFIVRRIR---
MBE2199248.1_Anaerolinea_sp.                            --MIELLLKQLNIAPVMGQFHPLWSLCFPPVVIAHSAKQVAEQLDGRERHEEKPILITEYNPADRFLSHHLPDCPPADLINRMLPLRMN-ARQPILTHKEIGKCIVSDTQGYETVILLLVVDGLCYDDTRHWSHPVTPCLIAGPSITDVGFPGIV---GQLPIARQLAQVGLPRSRGYSYWEREKNDVSEFLFSGMP-LQRVSGINTALTLLRDIN-LQGLYIQIVREGTDGLAHRRREVTAVEATVKAIWEDFQQLVELVAENGRRAAIYLVSDHGILWKYQHAFVRLPDENSAHARYTWERPFDPSCATLIPTGNGACYLYHYPFIGRQIPANDSGTHGGLSYWESIVPFVHVEVNV--
MBC8263374.1_Anaerolineales_bacterium                   MMSLDDLLTLLDCTPVIAEIPPVSICYLPEVTVLASAAEAAWQLDAPEPMLPGRLLITASDIQDRFPHVRCDHDPFTPALNTILFDRYDDIASRMLCQSQAVERIVVEAGQYDFVILFLVVDGLSYQDVRSWADSIEPCLVDVPTLTRVAFPNLI---GDPPVAVRLFDAGYHHRLGFTYWMREDNQLTDRLFHTIAEVKKIGHFSQILATLRRHLSQRKSYGQIMRTGLDGYAHAQKRRPPIAAIVEEVRREFEQLTELCSELGLRACLYLTADHGILWRDEFEPKVIGDPGKSSPRWCGWRDLYRDKGRRFLIDGEEYYCLDFPKLRRPLRIDEQGVHGGISFQESIVPFITVRIGD--
MCZ7554456.1_Anaerolineales_bacterium                   VLEWDDLLAALDASPMIVDMHPVLYAFLPDVRVVHDSAEASWLMDSPDPTLLSKMIIQREAALGRFPHVKCSRDPFSQELNRVMLQRFDLISESMLCQSVIADRILSRTSEIDVVVLMLVVDGLGYEDARQAVRELEPSLVDVPTMTRAAFPNIV---GSPPICVRLFRQGFYERLAFSYWTREDNALSDRIFQGFPHVQKTGQFDLITGSLREALTDAKKYVQILRTGLDGYVHAQKRFPPVQAIVDQVFAEFIELTELCAELGARALLFLTADHGILWRNQFTPVLVGAHGQSSPRYCGWREVHTDQYRQFKWETEDIYCLASPYIRRELRIDEQGVHGGISYEESIVPLVSMSV----
HID64303.1_Anaerolineae_bacterium                       SATMSQLDDTLDRTPVVYEGEPLLLAELDNLVVERDVWRYVQRVDNPAPVAVERTVLLGPVTSDRWPVVRMDARPDLAQFNRRLFQNYDQVRRFVFTQSDIAVRIVEDLD--DVIVLVLVVDGLSYTDWMDYP-GVQSCLVKGPTITPVGFRNIV---GKPSIARRLFKRGFRKRLGFSHWDRD-NQLTDVLFYGFDQMFKVSEFTEILLALDRLP-PEQTYVQILTNGLDAISHRHRGRPPVEAIARHLYEDLLALVERLQSMGATALVYATADHGILWKPEPELIVIRDERTDSHRYAKGTLI-IPHSRQFSCYGMNYTVLAYPYLFNRLTSLEWGMHGGISFQESVVPFVKLEVI---
MBN2393033.1_Anaerolineae_bacterium                     ETGFMRLLQALQRGPVMTEISPLALLCLPPIDIASDAETLVERLDILEPHSERPILIPCIDTRHRFQVESLTYTP--EAANEMLPRRLADARRLLLTHQRVAERIVSDTIGYCVVALLLVVDGLSYEDARHWPEAPEPCLIDGPSITTVGFPGIV---GAPTLARRLVDVGVPHSTGFSYWRRERNEVSAYLFEGVP-LTRVAGITTALETLEQQS-LEGTYVQLVREGLDGLAHSRREVTPIEAAVEAIHDNYRRLIEVVAATELPGVVYLTADHGILWKAQHDLRLVEELHSAHPRYQRGGGALSDYGVELSLTGQMYTLYRYPYLGATIHANDSGVHGGLSYWESLVPFVRVEVNV--
MBV6438329.1_Anaerolineae_bacterium                     ALSVDELEAMLSRTPVIAKVHPLVPTALSEVRIVSDANAFQIELDEHAPAPELPTLLSSNAAARFPRVTSLGLPPIERINAQMHQHLSYLRSHLVTTRTIADQIEADVVDWRIVVLLIIDGLSFEDVLDWDLVAQPCFVDGPSVTDVGFSAIV---GSPTLFWRLMRRGFRNAYGFTYWEPESNIISDHIFAQIP-TRSVANFEAILSELSNLSFEQPTYLQIMRQGLDGLAHSKRELARISGALSAIRNDIDRLAAVLNHRGNSACIYVVADHGILWKTEHQWVQLS-LKGSKPRFSESRPDQDEHYSRFECGSVPFYVLHHPYLGAKIPSNDSGVHGGLSYQESIVPLLRIGAKPS-
MBW7959471.1_Candidatus_Promineofilum_sp.               IEQIDTLRKQLSLAPVMGHFHPLLPLCLPAVMIVTEARRVALAMDARGPVPESQILITRIDPTDRFLSLSLTETPPAEIVNTGLITNPKTAR-HLLTNHEIGNRIVSDMLNYRCVTLLLVVDGLSYEDVRRWPEPSDPVFVDGPSITDVGFPAIT---GPAPLGLRLQHAGLTRSRGYTYWDRDQNDVSDLVFKGMH-QDRVGSISEALDRLAGQN-LDGTYIQIMRIGTDGLAHGRREVSNVMATVEAVHKDLQALVSLLRDAGTYGVVYLTSDHGVLWKSQHTLQKLDGYTTTHPRYTTEGLAHDPATVRITCGNTDFHLFKYPYIGTALRSNNGGVHGGLSYWESIIPLIKVEVKP--
MFN8469222.1_Caldilineaceae_bacterium                   ---MDW--SELNVIPHIVQLSCLWRAVLLQILVPITLHQWKGILDA-APRATAPTVFVDNPSNYGFPQVQMPLLPDEPDLSRTLVDALSSPIPDLLRQSQIAFEIARQAAGAHIIVLLVLDGLAYEDVLDWQYTRRPCFVDGITLTTTAMPRVI---GLPPLAHRLFQQGYKQHYGFTYWERDSNDLTDTLFAEFSQLTKVSEFSQILDRLADSNFSAPTYVQIVRNGLDQFVHSYRERPNVRNFLRELEHAVWDLLDLLATFQCSVRVYVTADHGILWYEKQPVTTIS-TNSPSARYVTTETHVDDRFTRICEASEDYTVAVGPYICRNRRVNEWGFHGGISGRESVIPFLTLDYRPQE
MBI5668048.1_Chloroflexota_bacterium                    ATTFNELAQLLAKIPVIGHVHPLIPLCLPPTQVVNDIQGFGYEIDSPELHAARPTLLIRHDKAARFPLVNVLNHPPIEAVNNNIAHNVRYARERHLLTHDKIAPHIERDVHPPDVVVLFLVVDGLSYADVLNWCWELQPCFVDGPSVTTVGFASII---NRPSVYERLYGLGYHQARGYTYWDRNENQIADFMFEAIP-VEGITNFEVALQWLKDEKSLRNCYIQIVREGLDGLAHGKRELRPVDIAVKAILQDIERLLNLLESKHLSVNLYLTADHGVLWKNEHSFRTLRGIRDSKPRFTSSCPAEREYTVRFENSYVPYYLYTYPYLGVPIRTNDSGVHGGLSYQESIVPFGKFKV----
NLE46402.1_Chloroflexota_bacterium                      VESINYLLDLLARAPVMGSFHPLLPLALPTVTIADSAQKVAEALDALEQHEPCPILINEYDVRDRFLVERFTELPPIESINAHLPERV-FTRVPILRQGEIAARITQDTLGYQVVIMILVVDGLSYDDVRQWHLEVEPCLVDGPSITDVGFPRIV---ANPPLARQLQAVGLRHSRGYTYWDREENDVTEYLFRGIP-VQRVQSISEAIDALYNEQ-LHGLYIQIIREGTDGLAHRRREVRQVESTTATIRDDLQALAKLLARKQLKGALYLTSDHGMLWKQEHEFHLLSSEGAISPRYAFECPAESDRASPITAADRTYYLYHYPYIGRRVRSNDSGIHGGLSYHESIVPFVRLEVN---
RLC84905.1_Chloroflexota_bacterium                      MTDPAELARLLMQAPVMGPLHPLQALCLPPTEVLADAAALAEELDDLAHHAVRPIWLTDVQPRDRFLQADASTPPPLDVVNRALPGRLDYVRRRWLTHRTVADRIVADAACYRTVALLLVVDGLGYADVADWPEPVQPCLVDGPSITEVGFPGII---GAPRLARRLIEVGLLHSRGFSYWSRETNAVSEYLFEGVP-LTRVARFSEAVAALRSAN-LECTYVQVVREGLDGLAHARREVTPVRQAVQAIRADALALLDVLREGPRPAALYLTADHGILWKAEHTFEMLETNDRVHPRYGTHLPAGTEHATRFTFGERTFYVYHWPYLGAAIRRNDSGVHGGLSVQESIVPFIRWEITE--
MDX2239252.1_Leptolyngbyaceae_cyanobacterium_bins.302    EVSLDKLPDYLEAGPVIAQMHPLSLAFLPILKVFDSAAELVAALDNPANECSDAMLALSYDANDRLLSLKINHQPLTPQLNQYLKERWELICSTLLCQSHLAELIAKEAS--ECIVLLVIVDGLSWADFSKYWQEAQPVVVDGASTTEQGMKRIV---GNPALAERLFAKGYERTYGFSYWTRDDNALTDHLFNRFSNVNKVRSFQDVLEQLKKFD-LMGTYIQIVRQGLDGLCHRHRDRPNIEHYVHDLAQDWAALLKLLKGKGISAQAFLTSDHGILWKEGNFRVQFLGKHTDPPRYFKQAYP-SPQTLEVSFEGSTYSLLTFPNLRRNFNVNEWGVHGGLSYEESVVPLLKARILHKD
MDE0397489.1_Candidatus_Poribacteria_bacterium          ------MAQQLCTIPCITEIRALWELCLTDVWCPVSLEEWRERIDSPREIDADSTLFVNLSPSFGFPAVKFPALPSIEQVNQALPSRFET--THLFKQSDVAAGITERIRLPQIVLLVFDGLSFYEITDWELRLEPCLVDGLSTTENAMQRLI---GSPVLTHRLFELEYVRRLGFSYWERSTNPLTDALFAEFSQLHRVTTFDEVLERIGASEWPTHTYVQIIRSGLDSACHSHRERPDVSSLLEGLRQDILSIFAVLQETRKSFRLFITADHGILWYNNQNILPLA-QEKFKTRYTEDNVVSSDNVLTLQSNERTYTVLTGDKIAKKRRSNEWGFHGGISAQESLIPFWDIRRSTDY
MYC73121.1_Gemmatimonadota_bacterium                    ------MIDILEKIPCLTTLQGLWDICLSDIWCPNSLAEWRDVVDAPQDDSTKATLFVNLPETFGFSTVDLPTFPPIEWVNQKLPEHFNALCSKLLKQSQLVQTIIDRVSLPDIMVFLVLDGLSLYDVADWAFVCEPCFVDGLSVTTSGMRRLI---GSPPLTHRLFALGYVQRFGFSYWERSQNQLTNVLFSEFPQLFRITTFEEVLDRLQNTDLPARTFIQIVRTGLDRVCHNHREKPDIEAILNQLELNIKQIVEAVGSSQKIVRIFVTSDHGILWFKKQKVMPFQ-SGTASARYVCAKSSSDMSGITIAESGETFTALTGDEIARNRRVTEWGFHGGISAQESFVPLFDITLNGR-
RJP56334.1_Deltaproteobacteria_bacterium                GADVTKLSTAIDAYPAAFRGNRLNLLFLDDAHPIDSSREFLRKVDSLQLQEERREYLISDKEG-HFPTISIESRPDPWKITEALFDNFEIAGKILPTQRDIGQILTEHALNPDIIILNVVDGLSYYDLPDNI-DAIPCLVRGVTTTDFGFRDVI---GKPNTSQRLFSLGYKSQLAFTFFDTDTRHLSAELHATFGQLNRINSIEEGIDIIRGNP-FWRGYIQIVAPGLDKLAHYHADRPMIKETLCRIFERLDKVIDSVSSKGRKVIACLTSDHGILWRDDAVVGDLFSEDTAHPRYVKGSLS-RNYARAVKLESKSYTLLKAPYMTRDWKHSEWGVHGGISAWESIVPIIIKEIG---
MFZ4828881.1_Phototrophicaceae_bacterium                GKNPEILGGVLQRSPAIVKAHPLFPLALPKVKIVTDGLDILEKIDNREHLVNHQTVILDQHPLDRFPTLEFPTHPSIDAINANMRYHLPYMREHLLTNHKVADYIQEDVSGYDVVVLLIIDGLSYGDVYTWGELVQPCFIDGVSVTSVGFASII---NSPSIQRRLYHLNYRNHLGFTYWKSSNNVVSRFLFDGIS-NITVESFKAIVHHLKSTTLLPSTYIQIMREGLDGLAHGKRELHPIESATHAINDDVKTLLNILEESGCHSVLYVVADHGILWKTEHDLRYLPSTINSHSRYTNNLPQDLEHTEHFKHDTLSYYLYVYPYIDAVIPKNHHGVHGGLSYQESLVPFAKFEVTSKW
consensus/100%                                          ......P......h...h......p...h...h..........hh........h..............p..h...........p..h.......h.h.lhDGLsh..........s.hl.s.o.T..uh..l...........L....h.p..uaoaa.....ls..ha..h.............l........a.Qh...u.-..sH.............l...h.....h.......h.hsuDHGhhW........h.......R.........................h...........p.G.HGGhS..Esh.Ph........
consensus/95%                                           ......h...h...P.......h....h.......p...h...h..........hh.........h.............hs..h............pp.l....h.......p.hhhhlhDGLua.p.........shhlps.o.T..uh..ll...s...h...L....h.p..uaoaa...p.ls..la..hs...h.....h..l..........alQl...G.-..sH.............l...h..h...h.......h.hsuDHGlLW..p....h.......Ra.........................h..s.....b...c.G.HGGhSh.ESh.Phh........
consensus/90%                                           .....h...hp..Phh.....l..h.L.......s...h...hp...........hh.........h.....ph...s....hs..l..p.p.h...h.ppppl...l.......phlhlhllDGLua.-h.......pPhhVss.o.Tp.uh..ll...s...l..bL...sh.p.buaoYWpp.pp.lsp.la..hs...pl.php.hl..l........salQl.b.G.D..sH..+c....p...p.l..ph..l.p.h.p......hhhsuDHGlLWb.p....h.......Ra.........................h..s.h...h..s-.GhHGGhSh.ESllPhh........
consensus/85%                                           .....h...hp..Phh.....l..hhL..h.hh.s..ph...hp...........lh.........a...ph...s....hsp.l..php.hp..hhpppppls..l.p....phlhlhllDGLua.-h.p.....pPhhVcs.o.Tp.uh..ll...s..sl..bL...Gh.ppbuaoYWppppp.lsp.la..hs.h.pl.phpphl..L........salQl.b.G.D.hsH..+-....p..hp.l..ph..l.p.h.p...p..lahsuDHGlLWb.p.....l.....p.Ra.p..........................h..sbh...h..s-.GhHGGhShbESllPhh..p.....
```

```
# ; DUF4007 DNA-Binding Component Representative Alignment; (HTH + HTH + wHTH)

                                                  1---------11--------21--------31--------41--------51--------61--------71--------81--------91--------101-------111-------121-------131-------141-------151-------161-------171-------181-------191-------201-------211-------221-------231-------241-------251-------261
OrigSeq                                           MSTKLHIEGTGRPDAIKSYLDLIERREYNSASAVMEDAYVEFNLTREVDTKGMLTTLTNLGLVEENNQQLTELGEEFIDVLIYNEDAFFDLFHLLYSTAYYRNSNTGISWSYFQISDAYRRRAPTNFRQEVIEEVMNRADPIFNDPGPLSKRSLNSYKRFVEKLVPPVLKDGTFDLRSFAKNELVIAAVDSLYRSDILRYGDRLELSNESREFLSTVLLVYEDDLPELLEHTASMDGRLSIESDYSIRIRLTEEVNIDDLA

                                                  <<--------------------------------HTH--------------------------------|---------------------------------|-----------------------------HTH------------------------------->><<----------------------------------------------wHTH-------------------------------------->>
                                                  ------------HHHHHHHHHHHHHH----HHHHHHHHHHHH-------HHHHHHHHHHHHH-------|-hhhhhhhhhhhh-hhhhhhhhhhhhhhhhhh-|-HHHHHHHHHHHHHHHHH-----HHHHHHHHHHHHHHH--------HHHHHHHHHHHHH-----||-----------HHHHHHHHHHHHH-------------HHHHHHHHHHH---HHHHHHHHHHHHHH---EEEE-----EEEEE---------
                                                  ------------HHHHHHHHHHHHHH----HHHHHHHHHHHH-------HHHHHHHHHHHHH-------|-hhhhhhhhhhhh-hhhhhhhhhhhhhhhhhh-|-HHHHHHHHHHHHHHHHH-----HHHHHHHHHHHHHHH--------HHHHHHHHHHHHH-----||-----------HHHHHHHHHHHHH-------------HHHHHHHHHHH---HHHHHHHHHHHHHH---EEEE-----EEEEE---------
                                                  ------------HHHHHHHHHHHHHH----HHHHHHHHHHHH-------HHHHHHHHHHHHH-------|-hhhhhhhhhhhh-hhhhhhhhhhhhhhhhhh-|-HHHHHHHHHHHHHHHHH-----HHHHHHHHHHHHHHH--------HHHHHHHHHHHHH-----||-----------HHHHHHHHHHHHH-------------HHHHHHHHHHH---HHHHHHHHHHHHHH---EEEE-----EEEEE---------
WP_162551688.1_Halohasta_litchfieldiae            MSTKLHIEGTGRPDAIKSYLDLIERREYNSASAVMEDAYVEFNLTREVDTKGMLTTLTNLGLVEENNQQLTELGEEFIDVLIYNEDAFFDLFHLLYSTAYYRNSNTGISWSYFQISDAYRRRAPTNFRQEVIEEVMNRADPIFNDPGPLSKRSLNSYKRFVEKLVPPVLKDGTFDLRSFAKNELVIAAVDSLYRSDILRYGDRLELSNESREFLSTVLLVYEDDLPELLEHTASMDGRLSIESDYSIRIRLTEEVNIDDLA
WP_310922613.1_Halogeometricum_sp._S1BR25-6       MSNKLHIVSAGRPEAIKRYLDLLDRRDYDTVESVMSDAYHEFGLSQHINTKDMTNTLARLGLVQEDNLALTEFGQELVEVLIHNEQRFYDLLHFVYASAYYRNPKRLISWSYYQITREYCERAPVDFKQEIVEAVLQTAETGFDNYGTLSKKSINNYRRFIDELEPPVLQEGQFVLREFVPSELVVASIDLLYRTDFQDYGGLMSLSGAPAEALQTLCLVQEDSLSDVVEQAVKMDSRLSLKADYTMHVRLSEPVNLNDLA
WP_117592497.1_Haloprofundus_halophilus           MSNKIHIEGDRRPVAMRDYLDLIEQYEYESEEDVITDAYQQ-GYEGEIPTKNMVSTLSSLGIVAESDQELTDFGRDLVDVLLYDEQLFYEMLHFRFSTVYGINPNNMMSWAYYEVSNILYNTAPIESKQDLADDIIYRSDPGFDEPGAFSRKSINGYQKFIEVLEPPVIEGGVIDLRSYAREELIISAIDYIYRSEVLSYGDLLELSNEIEEVIGTVCLLAPQSIDEMIEQTAAGYDSCSVESDHRVRVRLREEVNLSDLT
WP_338740217.1_Haloplanus_salilacus               M-AKLHIEAKHRPEAIKHYLDFIDQRDYDSVEGVMDDVYRETGLD-VVNTKNMQTTLAKLGLVDDSDQQLTDYGRELVEVLLYNDQLFYELLHFTYATAYYRDSDRAISWAYYQICDEFRRRAPVEFKQEIVESVDLTAEEGLSDSGPLSTTSLNNYARFIKELDPDVYDDSEVTLRSFAPTELVLATVDHLYRTDIRNYGDPVELSGDTTEIICNILLLQEEQLPDVIEHVASMDARLSIKSDYQLRVRLSDPVEINDLA
MDW8029828.1_Armatimonadota_bacterium             MRLPFHIQQNLTAERLWVIL-----CCGFDGTVDAAHIAEKCDLAVSVLEKVVLPFVRQLGLIRNTDFSLTDLGEQFYQLGKHSLILLPEAMHHLLYTLHHNDKGKHFSWAYAKVVDALWLSGERNLMAQLVGMVVDEAVQTFNEQIAFSRDSIRGVLNWLRALDPPVVKANKFRRRYFCPIPAFFWGLDFLYRINKIAYGVRMFLTSECIEQLCKVCVLDPSGLENVLMMAKRISDYYGTEGGFGRWLLLASPCPLPKLP
HIE11922.1_Desulfotomaculum_sp.                   MRVVLHIQHNIDPPKLAKIL------SGVATGGNILQIARELGIKEDSLPKRMFPFIRQLGMLDSRGCRLTNLGLRCHCLAQH--PLFPEAIHHLLYTGHLLDNTKAFSWAYATVADTLWATGSSVLKAQLVGHVVQEAQQTLGDRIAFSINSVRGALNWLGALSPQVVRANSFSRRFYCEPATFLWAVDFLYRFEGKPYGVRLFLTPERVERLCKLCILDPSGLHNVLLYTKNRSDYFGTEGGFGQWILLTRPCEVPSLP
WP_348254523.1_Leptolyngbya_sp._ST-U4             MKLTLHLQQNTTFERVVSIV-----EFCNQEETDLDELGVTCGLGTSVLQKVIFPFLRSISILSKKNPTLTNLGWVAARIHQENPAILSEFFHSLIYNLHYQEPDKRFSWVYSTVVQQLWQRKEVVLKNSLVSQVIEAATEKFDDEIAFSSNSITGVVNWLRALEPSVVKAEHFNLRYFCAAPAIVKAVDNLYQQGDRTYGVKIFLRESVQDTLCQMLLLDPSGLPGVLDNAKATYDYWGYEGGYGQWVMLNKPVDWLELI
MFA0782867.1_Candidatus_Fervidibacter_sacchari    MQLPFHIQHNTTLTVIVKTL-----RFPTNEQITPETLSGALGCAKGTVEKVIAPALRQLGVLDG--WRLSEFGERLKVLGQSHPDLLAEALHVHLYTLHWRQPDAYFSFAYATICDWLWERGTWVLKAELVGVVVNAAAEKFGNQIAFSQNSVHGAINWLKALNPSVIKSGRFSLRDACPVQTLLWSIDALYHHPKRDYGVRIILDDERLTQLCKICLVDLDGLEKMLDFAVRTCSRIGTEGGFGRWLSLTEPIPVGMLW
MBV6438326.1_Anaerolineae_bacterium               MKLSIHVPPDANPDNVSTTISILSENPSFSSSKEMLEYLTSGGVA---TRQELPYIAHAMGILERSKVTISENG---RALSDIREEARGDILHFLMYAGWDAEMTFPQSWGYRNCCDRYWAAHEVQLLDRQVEETIDEAYRTFEDEVAFSRKSLTGAHKWMLALDPPVLKDRVFRRRDFCQPELAVLALAWALRNESTVTGVDILLTTEKREAISKVCLLSPESLDQVLDWAIPTFPEAGTSAGYGRFVRLKRLPSLQDLA
MBI5668045.1_Chloroflexota_bacterium              MKLAIHIPPDANPVNITEMVVAIARDTVFESVRDLLEYTESQGIG---SRTEMQSTAVYLGLLEKSSLGLTRTG---FALAKIREDVRSDLLHYLMYTGWEESVNFLPSWSYRICCDRYWEGQEVTLLNRQVEETINQAREAFAAEVSFSRKSLEGAHKWLGALQPPVIENKVFTRRSFCPPELLLLAIGYVMKDETDATDIDILLTREKRDRICKVCLLDHNALDRALDWMLPIFPAPGTSAGYGRFVRLHKLPTLEDVV
GAB4528652.1_Anaerolineae_bacterium               MRLTFHVPQEGEPDKLIPLLEAMYAGMRFPTVKALLEFAREQGLG---GRTEMQILASACGLLDTDGDGWIIMSETAKVIVQLKPEVRPDLIHYLLYTGWQPESANTILWSYREITDSLWEKPSANVANVVAEEVRNRTQQVFIGGVSFSPKSIRGVRKWLEAVTPPVIEDDLFARRHFCPPELALLATGWVAQAMGGEIGIDFLLTPECREAICRVCLLELSALDRVLDWMLPLYVEPGTSAGYGRFLHFLKWPQVSDLA
HET59436.1_Chloroflexota_bacterium                MDFVFHVR-NAQPGDMIAILATLEDYPGLSTIAEIVDQADTLGFTRDRARLEALMTARELDFVEMDKNLLTDKGKALVRLEMNKPDLFPDIVHGYQYTLWDKKSIHCFSWTYREICKHLWQLGSYHMKKDLTSEIEGLARTHFESDIALSAKSVGGAFLWIEELVPGVIPSELFARRTFCPPELFILGVNFVYRGNEMDYGTNLLLGEEPRNTICQICLLEPDGFDRVLDYATAQFD-KGIGGGWGQYLTLYRAPVVEDFV
MCZ7554453.1_Anaerolineales_bacterium             MASSFHVR-NAQPDDLLAALTVLQAYPGLSSVADIAVRAETMGLSRDRLRLEALMTARELGLVHGDSNELTLDGEKALELEGQKPELFGDIVHGLQYMLWVPFWEHCFSWTYQAVCRSLWRQVRVEV---LASELEVEARTQFANDIALSGKSVRGVLMWLRELRPRVIEDDNYLSRHFCPPELFLLALDYVYRSEVADYGTNLLLNEERRDAICQMCLLDTEAFDRVLQYAVGQFP-RGIGGGWGHYMSLERRVTLGVLG
RKU21509.1_Candidatus_Poribacteria_bacterium      M-ARIHIL--RGFSDLDGALWVLKYHPELRTRSEIVATILENGLISDRTRFEILQLLQEWGIAESNGYQLTLKGKHFYSLWEMKPEIAIDVLHGLQYNLWTKHDQNQASWAYQRVCDYLWEYQILPEENDLTAYVYEMREENLGIANAFSSKSVKGAYDWLLPLDPPVLRRASFTRRTHCASALFLMGLSWVSRESEIGFGELVELTDERRIDVCRFCLIEESSFDIMLNQALTRFD-MQRIGR--LFVIIQREPQIADF-
MYD63886.1_Gemmatimonadota_bacterium              M-ARSHIL--KGFSDLESALWIHKYHPGLTIRRQMVKVALENGYSTDSARYEAVQLLSEWGLADGKSGSLTPEGQLFYSLWETKRSTAIDFLHGRQYGLWTQNDQNIASWAYKNICDYLWTRQTLPDKGDLVTYINDLRAEVVIMGNAFSNKSINDAYDWLRPLDPPVLNGATFAQRAFCSTLLFLMGLGYIVRETNTVFGDLVKIDDTRKQAVCCFCLISEIGFDLMLDEALRQVS-VQRGWD-GIYMELNQKPEVRDFI
MYB94330.1_Candidatus_Poribacteria_bacterium      M-SRIHIL--KGFSDVDSALWVLKYHPGTRSRADLVQIVRDNGYTANKD-YEIILLLTAWELVQRKPQGLTQKGMAFYHLWEVKRDVAIDVLHGLQYGLWTEHTQNTESWSYREVCNYLWDSDALPEKDDIISHIYSSLSLEIDEKAAFDGKSVNHAYDWLLPLSPPVLRNATFTRRSHCSTALFLMGLDWVARESGIIYGDLWMIGEEQRRQVCRFCLIDETRFDFMFNETLRRFSVVRRSGG--LYIVINNEPQISDF-
TEU15948.1_Anaerolineales_bacterium               MKEDVHFLDNAPPAEVEYVLACLELDIAYSTEDLRKRLLSEWGYSAQRNLTFSTRRLLDLGLADKANHVITQLGEKVRGILSIDRELYTEIMHFLHYTSYDGSSSRKLLWSYRTCCEIVWARKQVPPTPELVAEVQSRIAELFPVGGNFNAGGVSGWKPWLAQLQPSPFEGRALVPRHSHRLELAALALDHVYRSRGYRYGDPVILDEPLLDEIARVFFLDPICCRELIDLAARLTKASDTFTG--TSATLLAPYDVERL-
RJP56331.1_Deltaproteobacteria_bacterium          MFEDTHFYDNASPVEVTNVLRVLSPSHEFTPKEISDTLRDSYGFSMQKDYTLSPRRLFDLGLTGQNKYTLTELGSKVQNILGIDPALAMDLLHYLHYSGYTGKTQRKYLWSYRRCCEIVWGNMRALRSTELASTIQSEMREKFDTGARFNDKAAVQVYSWLRSLEPSPFSSSTVTPRKIDRYEIALLALDEAYQTSDYTYGDPVIMDNEFIKLVSGVFMLDPECCKGLLGIGARINQSSETLSG--ASINLMRPFKIDSI-
MBI2842131.1_Armatimonadota_bacterium             MEEDTHFYDSAPPDEVTRVLRVLAADVCLSPDEIADTLDSQYGEPMQRDHTYTPRRLFDLGLAEQMRYCLTSRGLRVQTVQASDAAFARDLLHYLHFSGYGGSTDRKYLWSYRRCCELMWANGKMIPAVELAALIQAEMVSAFPAGGRFNHKAASCVSSWLGCLTPPAFRGRHVQPRIVDRYELAALALDDVYRSRRYRYGDAVVMDDSLLNQVAGVFFLDRGCCANLLRLAASITPSSDTLAG--PSVNLTRGFTIEDL-
consensus/100%                                    M....Hh........h...l......................s.............h..h.h.......h...u.....l.........-hhH......a.........asY..hsp.h.......................h.....hs..uh..h..ah..l.P.sh....h..R.......hh.s.s.h.p......s..h.h.......l..hhhl....h..hh...................h.h.....h..h.
consensus/95%                                     M....Hh........h...l......................s.............h..h.h.......h...u.....l.........-hhH......a.........asY..hsp.h.......................h.....hs..uh..h..ah..l.P.sh....h..R.......hh.s.s.h.p......s..h.h.......l..hhhl....h..hh...................h.h.....h..h.
consensus/90%                                     M....Hh........h...l....................p.sh............h..hshh......lo..G.....l...p.....-hhH....s.a..p......WsY..hsp.hb......................h.....hs.puh..h..al..L.P.shp...h..R..s....hh.uls.hhp......G..h.hs....p.ls.hhhl....h..hl.............ss....h.l.p...l..h.
consensus/85%                                     M....Hh........l..hl.........p......p...p.Gh............h..hulhp.....lo..G..h..l...p..h..-hhH...hs.a..p......WsY.phsp.hh.........plh...l......h....shs.pulpsh..al..LpPsshps..h..R..s...hhhhuls.lhp.....hG..hbls.p..p.lsphhhlp..sh..hl..s....s.....us....h.l.p...l.ph.
```

### ; STAND-NTPase Representative Alignment; (Enzymatically Active)

```
                                          1---------11--------21--------31--------41--------51--------61--------71----- [illegible]
                                          MSRLVNFQNRIRRFFSTQWLTTNQKQTYDRLRQLLSFQDSVNLYGQSGVGKTFLIWVLSKEVSTDFYSNWEQLEEKE [illegible]
                                                                              S1                  S2 [illegible]
                                          --HHHHHHHHHHHH--HHH--HHHHHHHHHHHHHHHH--EEEEE------HHHHHHHHHHH--EEEE----------- [illegible]
                                          --HHHHHHHHHHHH--HHH--HHHHHHHHHHHHHHHH--EEEEE------HHHHHHHHHHH [illegible]
                                          --HHHHHHHHHHHH--HHH--HHHHHHHHHHHHHHHH--EEEEE------HHHHHHHHHHH [illegible]
                                          --HHHHHHHHHHHH--HHH--HHHHHHHHHHHHHHHH--EEEEE------HHHHHHHHHHH [illegible]
HEY9694178.1_Oculatellaceae_cyanobacterium        MSRLVNFQNRIRRFFSTQWLTTNQKQTYDRLRQLLSFQDSVNLYGQSGVGKTFLIWVLSKE [illegible]
HQD80873.1_Bacillota_bacterium                    VGKWFEIHNLIKARCSSSWLTPSQQKALRTIRELTSTFGRCNLHGNVGVGKTFVAWVLSKE [illegible]
MFA0782864.1_Candidatus_Fervidibacter_sacchari    MNRFVALLNRIRKEADWGWLTESQREASEQLRDFLGVSDVVNLFGFHGVGKTFLAWVWQKE [illegible]
RKU21511.1_Candidatus_Poribacteria_bacterium      MNWLTNHKRLSRLAETDETLTPSQLSARDVLLDTIHEETRINLWGGPGTGKTFLAHYLHHR [illegible]
WP_348254525.1_Leptolyngbya_sp._ST-U4             MSRLVEFRNRIYHRFRREWLSPSQEQAYTRLKNLLVFQDAINLCGSSGSGKTFMTWVLAKE [illegible]
MYD63885.1_Gemmatimonadota_bacterium              MHWQSYHRKLLALAETGETLTTSQQAVCEFLHQAIEYQQRINLWGSPGVGKTFLAHYLHYC [illegible]
MBP7045035.1_Chloroflexota_bacterium              MSQLLTLLNRLKAEARLERLTVSQQTAWHNIQQQLRFPERINLYGAAGSGKTFLAWALANE [illegible]
MBW7959473.1_Candidatus_Promineofilum_sp.         MSRLLAAHNHLKAQADIDRLASHQRQAYESIINQWRFPTWLNLHGSAGAGKTYLGWVVAES [illegible]
BAL53914.1_uncultured_prokaryote                  MSRFVDTLNRIRQSMKPEWMTPGQRAAYDLLRERLRFLDEVNLWGGPGVGKTFLGWVLHVQ [illegible]
HET59437.1_Chloroflexota_bacterium                MSEFIRLLNQIKTHRNRAWLTPSQVNAMRALQKTLRIPGTVNLFGSVGVGKTFLGWSLAEE [illegible]
MCZ7554454.1_Anaerolineales_bacterium             MSGFVRNLNRIKTLCNQDWLSPSQKAALDTLTQGILYPGTTNLWGPAGVGKTFLAWILQQE [illegible]
HFD39729.1_Anaerolineae_bacterium                 MSKFVRTLNSIKTHRCAEWLTPSQSRALAALRRELHAPGTVNLYGPAGVGKTFLAWILADE [illegible]
MBN2393037.1_Anaerolineae_bacterium               MSRLLVLLNRLKAEARHDWLTESQAATLSEIERLWRFPERINLYGPPGAGKTFLAWSVARA [illegible]
MBN1891044.1_Thermoflexales_bacterium             MISRTEVENQIKAQSDASWFFPSQVVAYQELLPFLGLHRVVNLYGAQGVGKTFLAHTLCKE [illegible]
GAB4528656.1_Anaerolineae_bacterium               MSRMLCLLNRIKAEARRDLLTGNQLIAFEEIKRLWQFPERVNLWGPPGSGKTMVGWVVGRS [illegible]
MBC8263376.1_Anaerolineales_bacterium             MSDLIRALNLIKTRRDPAWLTVSQQKALAALQEALRVPCTVNLFGATGVGKTFLAWTLADE [illegible]
WP_147270937.1_Haloplanus_salinus                 EADLLTRMNTLKQVDIKQHLTSTQKGALEEIRQHREGEPFINLHGPRNSGKTFLCWALRE- [illegible]
WP_338740218.1_Haloplanus_salilacus               EAPLLRQTNVLKEAADESLLTDAQKDARDAIRQHREGARFINLHGPRHAGKTFLCWVLHLS [illegible]
WP_232571133.1_Halobacterium_litoreum             NALLLQRMNALKDAASVERLSAAQQEAMDQIRKHRDDARFINLYGPEEAGKTFLCWALREA [illegible]
consensus/100%                                    ...b....p..........h...Q..s...l..........NL.G...sGKThh.a.h... [illegible]
consensus/95%                                     ...b....p..........h...Q..s...l..........NL.G...sGKThh.a.h... [illegible]
consensus/90%                                     ...b....p.lb...p...hs.sQ..s...l.p.b.....hNLaG...sGKTalsa.l... [illegible]
consensus/85%                                     bs.h....N.l+...p...LossQ..sbp.lbp.b.....lNLaGs.ssGKTFlsa.l... [illegible]
```

```
# ; Alpha+Beta-Domain + wHTH (BrxC-ATPase Type-1 BREX)

MBR4615907.1_Kiritimatiellia_bacterium [illegible]
MCI9358063.1_Lachnospiraceae_bacterium [illegible]
NLJ79432.1_Bacillota_bacterium [illegible]
MBQ2845004.1_Alphaproteobacteria_bacterium [illegible]
NMA18734.1_Clostridiaceae_bacterium [illegible]
WP_179400143.1_Macellibacteroides_fermentans [illegible]
WP_118027524.1_Lactobacillus_amylovorus [illegible]
KAF3303185.1_Carnobacterium_sp. [illegible]
MSB67176.1_Leuconostoc_lactis [illegible]
WP_273096524.1_Dialister_succinatiphilus [illegible]
MCD7882907.1_Lachnospiraceae_bacterium [illegible]
WP_022770173.1_Butyrivibrio_sp._NC2007 [illegible]
MBQ8347028.1_Alphaproteobacteria_bacterium [illegible]
WP_147675231.1_Numidum_massiliense [illegible]
MBR4606065.1_Lachnospiraceae_bacterium [illegible]
WP_219938189.1_Succinivibrio_faecicola [illegible]
WP_091835632.1_Marininema_halotolerans [illegible]
WP_019974718.1_Empedobacter_brevis [illegible]
TVQ28583.1_Spirochaetaceae_bacterium [illegible]
WP_091403827.1_Geotoga_petraea [illegible]
WP_089525998.1_Paenibacillus_herberti [illegible]
MBP6234856.1_Saprospiraceae_bacterium [illegible]
PKK90022.1_Candidatus_Wallbacteria [illegible]
MAE16011.1_Deltaproteobacteria_bacterium [illegible]
WP_094389547.1_Prevotella_sp._P4-119 [illegible]
WP_257712004.1_Aeromonas_caviae [illegible]
WP_098060820.1_Longimonas_halophila [illegible]
MBZ5596030.1_Acidobacteriia_bacterium [illegible]
MBN1458231.1_Armatimonadota_bacterium [illegible]
MCR9098968.1_bacterium [illegible]
MBU6197215.1_Cyanobacteria_bacterium_REEB446 [illegible]
HJF93068.1_Mediterranea_massiliensis [illegible]
HCY74134.1_Rikenellaceae_bacterium [illegible]
WP_092090777.1_Pisciglobus_halotolerans [illegible]
NLJ85301.1_Bacillota_bacterium [illegible]
WP_051485152.1_Cloacibacillus_evryensis [illegible]
MBR1685959.1_Clostridia_bacterium [illegible]
WP_262567770.1_Endozoicomonas_gorgoniicola [illegible]
WP_227411048.1_Bifidobacterium_pullorum [illegible]
MCL6598118.1_Alicyclobacillus_macrosporangiidus [illegible]
WP_024332155.1_Winkia_neuii [illegible]
RHB16634.1_Collinsella_sp._AM40-7AC [illegible]
WP_044073670.1_Prevotella_pectinovora [illegible]
WP_270462427.1_Gemmiger_formicilis [illegible]
OIO89050.1_Anaerolineae_bacterium_CG2_30_64_16 [illegible]
MCK6587209.1_Polyangiaceae_bacterium [illegible]
MBF0258414.1_Desulfamplus_sp. [illegible]
MCK6575226.1_Myxococcota_bacterium [illegible]
MCA6597699.1_Pseudanabaena_sp. [illegible]
MBR6253006.1_Clostridia_bacterium [illegible]
MCK6605711.1_Ignavibacteriaceae_bacterium [illegible]
OQA19974.1_bacterium_ADurb.Bin363 [illegible]
MBP7846551.1_Burkholderiales_bacterium [illegible]
MBR5630362.1_Bacteroidales_bacterium [illegible]
NBJ07681.1_Alistipes_sp._Z76 [illegible]
OJV92258.1_Bacteroidia_bacterium_44-10 [illegible]
MCH2449130.1_Gracilimonas_sp. [illegible]
MCH7965547.1_Bacteroidota_bacterium [illegible]
WP_213195478.1_Cloacibacterium_caeni [illegible]
ADB40822.1_Spirosoma_linguale_DSM_74 [illegible]
MCH8487209.1_Cyclonatronum_sp. [illegible]
RGZ88026.1_Eubacterium_rectale [illegible]
ETI91650.1_Clostridium_butyricum_DORA_1 [illegible]
APC47779.1_Virgibacillus_halodenitrificans [illegible]
WP_048125616.1_Methanosarcina_lacustris [illegible]
PTY02769.1_Verrucomicrobia_bacterium_LW23 [illegible]
TXT19373.1_Erysipelotrichaceae_bacterium [illegible]
WP_146995455.1_Schleiferilactobacillus_harbinensis [illegible]
RKX87882.1_Spirochaetota_bacterium [illegible]
MCH3907669.1_Sphaerochaeta_sp. [illegible]
TWS24839.1_Tsukamurella_sputi [illegible]
MBP9037128.1_Candidatus_Cloacimonas_sp. [illegible]
MBR4608207.1_Lachnospiraceae_bacterium [illegible]
MBD1206697.1_Paracoccaceae_bacterium [illegible]
MCI6014853.1_Coprobacillus_sp. [illegible]
MBP8934558.1_Prevotella_sp. [illegible]
OYE05053.1_Nostoc_sp. [illegible]
MBQ6502300.1_Flexilinea_sp. [illegible]
WP_013460546.1_Sulfuricurvum_kujiense [illegible]
WP_237187176.1_Rothia_nasimurium [illegible]
MBQ6141563.1_Kiritimatiellia_bacterium [illegible]
WP_190817045.1_Coleofasciculus_sp._FACHB-1120 [illegible]
WP_271768605.1_Candidatus_Pseudothioglobus_singularis [illegible]
SFW99921.1_Olsenella_sp._kh2p3 [illegible]
MCB0069888.1_Caldilineaceae_bacterium [illegible]
OQC05381.1_Candidatus_Cloacimonetes_bacterium [illegible]
WP_193813001.1_Kaistella_flava [illegible]
OQB98767.1_Spirochaetes_bacterium_ADurb.Bin110 [illegible]
MBV6422081.1_Ignavibacteriaceae_bacterium [illegible]
MBQ0129310.1_bacterium [illegible]
```

### Type-1 BREX BrxC Alpha+Beta-Domain + wHTH Entropy Plot

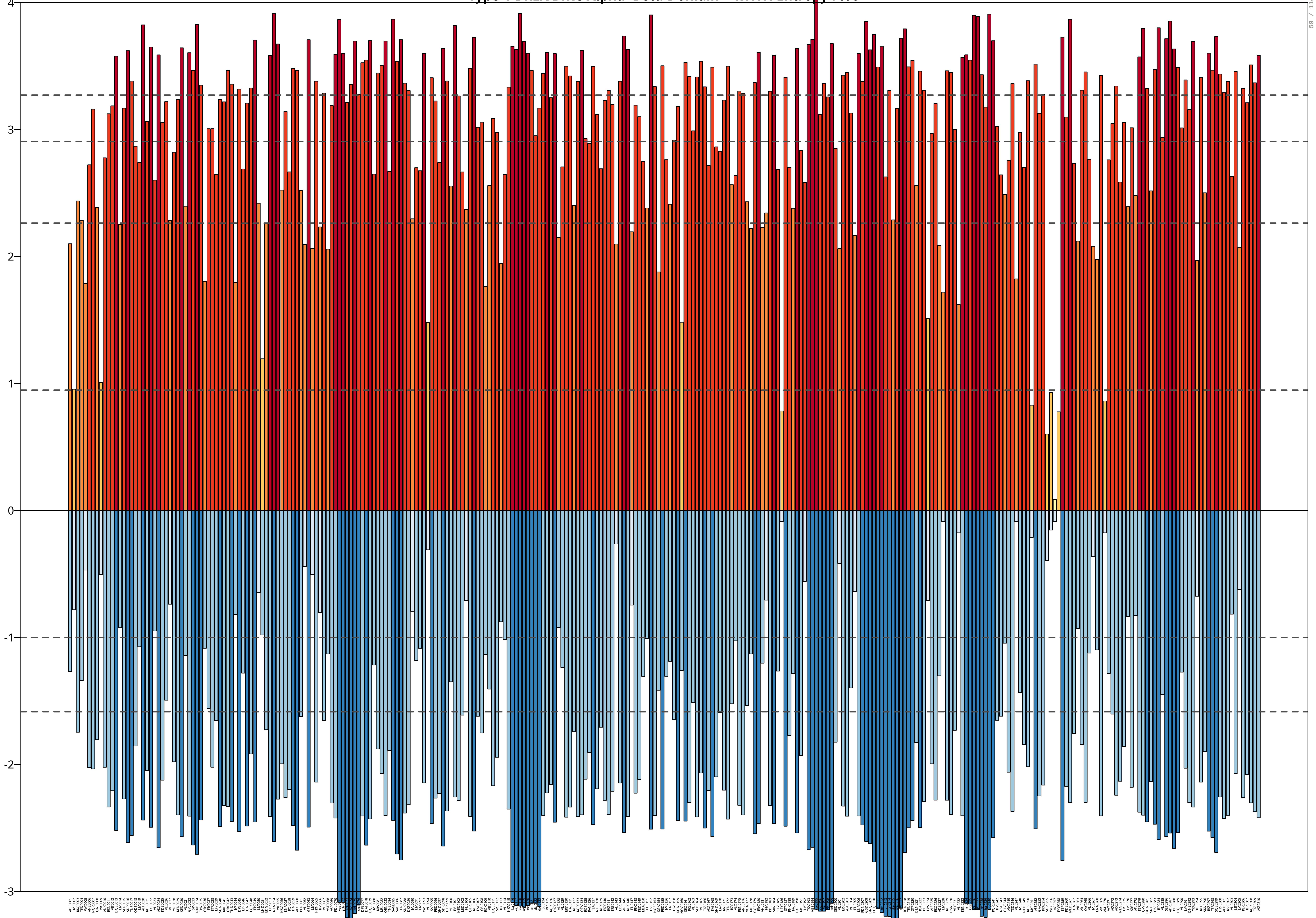

```
# ; Alpha+Beta-Domain + wHTH (BrxC-ATPase Type-3 BREX)

WP_164569833.1_Rhizobium_leguminosarum                 DVDYDAEIDKRGESLDDEDLDNAYYQAVAHLMERtdeathlpgHKIWQYPLEWFERRVHRDGYLFFGSPNDRPTAYPARDFYLYFVQPYDPPKFKDEEKPDEVFLRLASKDEALTAILRRFSSAAALAGLNSGNAKNVYTNKAEDALKSMRKWLRDQQATAFNVTYRGKTRTLGEWAKGFSVETINFREYINASAGFLLAPHFAEQYPQYPTFGTIVTDANRRMLSTTAVKALSSPARPKDAVMILDALKLLDGDKLDPRQSPYAQEITARLAAKGIGQVLNRGEIFeatsmaesfapGRFGLEPDLVAVVLAALVHSGDVVVSVAGRKVDSSNLGDGSTSLVEELAAFKHVEAP
SDB48716.1_Belnapia_rosea                              DVDYDAQIEKRAEALSDDSLDRAYFSAIRQLMERtdeaayvtgHQIWQYQIEWQDRRVERIGYLFFGAPNDRPTAQPERDFYIYFIQPFEPPRFRDDHKADEVFFRLKGRDDAIKRHLAFYAAAQDLESTASGGAKSIYLDKGKDALRNISKWLQEKQMAAYEVTYQGKTRNLQEWTKGVSLERINFRDVVNIISGVALNNRFAEVAPEYPTFSALVTESNRRQLVGNALRALAGGNRTKDAVVILDGLEVLDGDRIDAVRSRYAQDVLSRLKSKGHGQVLNRGELLsgdgdveyfapLKSRLEPDLLVAVLGGLVYAGDIVLAIAGDKIDSGKITLLAERSLDELKQFKHVEAP
MBA4157055.1_Gemmatimonadota_bacterium                 TVDYDAQIDRRASTLSEDVFDRRYFEALTIALERkdqptvatgYKIWEYELTWRERNASRLGYLVLGAPNDRNTAHPPRDFYLYFLQLYRPPRYQDEKRLDEVFFELTGADEAFRGSLTRYAAAMDLASTSAGNEKATYQRKADQYLRELVRWLQDNMLAAVEVIHQGRTKKLLDWVQGR-VETANVRDVINLVASTCLAPAFEEQAPDYPQFSILITARNREQAAQEAIRWFRGGTQTKQATAVLDALELLDGDQVVPERSRYASHIRQVLQQKQQGQVANRAELIgnvrgveymspDTFRLEPEWVAVLLAALVYNGDVVLAIPGRKFNAGDLDALASTPVADLVAFKHLEPP
WP_190509459.1_Oscillatoria_sp.                        TTDYDAQINKRAESLDGSKLDQAYFRALAQILYLsdry-ypgtHLAWEYELEWRSHHASRIGYMFFGTPSQRSTAQPPRDFYIFFLQPFSELPKLSKPEASDVFFELGRRDETFDTALNWYAAALDLEATSSGKAKETYASKAAGYLRNLVRWLETNITTAFKVTYRGQTKPLPEWIKGKLTTRAGVRDIVNTAASVCLEDHFRQLAPDYPGFSVLITNENRPQAAQDALRWMRGATQTKQATAVLDALELLDGDRLTPSRSKYTNYILELLRNKGQGQVLNRSELIqdefgiefmi--RYRLEPEWIVVLLAVLVYNGDVVLSVPGKKYDAGNLDSLITAPISELVNFKHIEQP
RJQ83728.1_Desulfobacteraceae_bacterium                TDDFDALIEKRADSLDGSQLDRYYYEALKRVMECtdht-yvtgFKIWQHEITWLERKAARQGYLFFGAPNERSTAVPPRDFYLYFIQPYDAPRFKDEKKPDEVFLRLKQKDEAFHKALRNYAAALDLASTASGYKKSTYESKASVCLQELTQWLKENIFTAFEVTCQGRARPLVDWAKDKSIERLNFRDLINTIAGVCLGSHFQDQAPLYPCFPVLITGANLAQAAQDALRAVAGQKPTKQATAVLDALEILDGDRLDPYQSKYARHILDIVKKKGHGQVVNRSELIqeilgveylapQTLRLESEWAVVVLAALVYSGELVLAVPGKKFDATGLPQLAGTSIEELTRFKHIERP
WP_013219148.1_Nitrosococcus_watsonii                  TDDFDALIDKRAESLGQAQLDRFYYEALKRVMECqdat-yvtgYKIWQHELVWQEHKAARSGYLFFGAPNERSTAVPQRDFYLYFIQPNDPPRFKDDKVNDEIFFRLKGTDEEFQTALKSYAAALDLAATSSGHAKATYESKANGFLKKLVQWLQKHMSDAFEVTYQGRAKSMTEWAKGKSIETINFRDLVNTIAGVCLAPNFENQAPDYPFFSVLITGNNRAQAAQDALRAIAGQNRTKQATAVLDALELLNGDKIDPYKSKYTKFILDTVKAKGHGQVVNRSEIIqddhgleymnpGGSRLEPEWVSVLVAALVYSGDIVLAIPGKKFDATGLQQLAATGMDELVRFKHLEQP
GHU24474.1_Spirochaetia_bacterium                      TEDFDTLIENRAASLSANQLDRYYFEALKRAMECqdvs-yvtgYHIWQHELMWREHKAARNGYLFFGTPNERSTAVPRQSFYLYFIQLNSPPHFDDTKSDDEIFFRLKHIDNDFQTVLKNYAAAVDLSLTSSANTKKIYLDKSNFYLKKLNEWIQNH-YDAFDVCYQGQSKSFYDWLQNKSLQTLNFRDKMNEIAGICLAPAFENQSPGYPVFSIVITGANRKQAAQDALRGIANQSRTKQAVAVLDALQLLDGDKFSPRKSKYAQFILELMKVKGAGRVVNRSEIIeniygveymdsTGCRLEPEFVIVILAALVYSGDIELSIPGKKFNATELSNIAAANFDDLLNFKHIEQP
WP_209691006.1_Methanomicrobium_sp.                    TEDYDAIIEKKAETLDDNILDRYYASALSQLMECstqt-yrtgFNIWEYELEWHEKRASRMGYLFFGVPNERSTTVPERDFYLYFIQPYDPPKFKDEKKSDEVFFRLKNPDTEFLLALKKYAGAGELAGTSSGSAKSTYHSKAEENLRKLVSWLSKNLYSAYEITYQGETKSLVEWKSNRSLKIVNFRDLINTVAGTCLEPHFSDIAPDYPVFSVMVRNDSRKQAAKDALRLIANQSRTKQAIAVSDALELLDGDERLAPERSRYVQFILNLMREKGAGQVLNRNEILnddlgleymdpEGARLEPEWVVVLLSAIVYSGDGVLAIPGQKFDSTSVDKLASVSVEDLINFKHIEPP
NMB01453.1_Bacillota_bacterium                         TVDFDQIIETKAQMISNDLLDQYYFELLKQMLECqdelsfqsnSKIWDYELQWLSHKMFRKGWLFFGLPNERPTAIPEKAFYLFFIPPMTSPRYLDEKRNDELILKLEQKEDEFVEKLRLYAATSELASMSSAQQKSMYQDKASEHFRFLAKWFLAHLQNSFTLTYRGSTRLLKNWLVGTDVATLNFKDAFDLIASYCLRDEFHQQASEYPTFSVVLTSQNYTSTLNETCRYLVNRGTSKLPMAILSGLGLVDNSYITPKNSKYAQWILEKLQEKPSGVVLRRDELFttihgeeylanPRYRLEKEFVSLLLVTLVSSGDIILKLRDKNYDASNLKDFINESVEAIADFDSINRP
MCK9328444.1_Candidatus_Cloacimonadota_bacterium       TFDFEQTIATRIEQMNADDVNNYYFAVMKLILECpkea-afmtSQIWDYELQFQGHKMFRKGWLFFGTPNERPTAVPEKDFYLFFIPPKGTHKYKDESKPDELIIKYVGQDEEFEAKLKEFAAASLLADSSSKTYKGTYQNKADTNLAYLTSWMRKNILNAFTFTYCGKTKTLTEWTAKTSLQVLSAKEAIDAIAGYCLKQNFEDLAPCYPSFTTRISYTNFIQTVNDAYKAVVFNNYSQTALSVLNGLQLLENGNLHPRSSIYAKWILAKLDNKPNGVVLNNDELLelihdd-yyclKDYRLEKEFVSVILVSLIASGDIVLTLKGKKYDASNIRDLISGGATELASFNSIDKP
WP_047754172.1_Kosmotoga_pacifica                      TIDFVALIEKKAEILNKDTLNRYYYDILAKLMECsdkt-rvprFKIWEYELRWYEKNTTRLGYLFFGQPNERSNAQPPRDFYIYFLPMYKIHDFKDDGRDDEVFFSLKTINEEFEKDLTNYAAAMELAKT-AGMYKSSYEDKARSFQRALTRWLNEHFLDSFEVTYQKRKKDLREALKEGSFKSDNFKETVDYIASTLFNNYFNDTAIGYPKFPSLITSQNRIQNIESTLKAIAMENFTKQARDILSALGLLDESNLNTSKSPYIKYIRSKLQENGINKVLNRNELIdevkdvey--fGPYRLEPEWVVILLAAMVYMGEIEVSLAGETIDASNVEKFAKIPLEQLLNFKNIKHP
MCK5228480.1_Candidatus_Cloacimonadota_bacterium       VDDFDAIIAKKRDTLGDASLNTYYNEALKRVLERmdvtphvtnYNIWQYEINWRERNAPRSGYLFFGSPNERSTAVPQRDYYIYFLRPYDNTPVKDQKLDDEVFIYLNGRDAEFDEALGLFAASHELSLTASGNAKNIYLQKSEQYLRKVVTWLQKSIWEAFDVVYQGKRQSLENWAKGKNIETANFRDIVNAIASICLAQHFENQAPKYPKFSVLITGYNRKQNITEALRSISGGSKTKQATAILDALQLLDGDRLTTAQSPIAQYVIDKLTAKGKGQVLNRREELIekvngieyldpNGFRLEPEWVVVVLAAMVYAGDIVLTVVGDKFDATKLQYLGATDLPTLVDFKHIEFP
OBW63497.1_Dehalococcoides_mccartyi                    VEDYEARIEQRAESLDKQQLDRHYYEALKSIMGCaedy-kftgYKIWQHELTWTEHSAPRPGWLFFGAPNERSTAVPPLDFYVYFLQPHDPPPFKDEKKSDEVFFRFVGGDETFCRSLKHYAAAKLLASTSSGNAKNNYESNAEISRRELVNWLQQHVTTAYEITYQGKTKSFAEWIKGK--KSVNVRDIVNLACSTILDNYFNETAPEYPKFSILLTSDNLSIAAQDAIRNIAAGTRSKQASAVLDALELLDGDQIKIANSRYAGYILDLISQKGIGQVINRNEIIqevysveymapVKYRLQPELAVVVLATLVYSGDIVLSIPGKNYDAANFTALIATPLADLVNFKHFEKP
MBX3742594.1_Akkermansiaceae_bacterium                 TDDYDALVEKKAETLSKDSLDRAYYQALVEILGCsdps-syqgFRIWERSIQWTERKVSKLGWLFFGTPSERSTAQPPRDFYLFFPQPFDAPNYIDEKKADEVFFKLDSRDVAFTEPLKLYAAALDLANTSTGQKKQAYGDKANIHFKTLTKWLREKFLTALEVIHEGKKKSLGQALAGANAGGMTPAEQVFTAASRQLSGHFASVCGDYPTFSRLITDGNAIAATQDALRGLSA-NATQTGIAVLDGLGLMNGEKIDPYVSPYARYVLDLLNQKGHGQVLNRSELVqdiygvpyfspGKFRLEPELLVVVLGALVHSGDIMFCLPGKEFAATDLKEMATRSLDDLVEFKHIKKP
MBT3747344.1_Bacteroidota_bacterium                    DIDYDEKIKQKASTLSKDQFDRYYFELLKIKAECydtntyvrgFKIWQHEVPWEDHKINRLGYLFFGSPNDRSTAHPPRDFFLYLLQPFNEPDFKDNQRPDEIYFRLKKKDDEFVKFIQLYAGAKELAQSATRAVRDTYEKKASDYLRDLSEWLDNNFTTAIEVTYKGVTKPLIDIAKGKTPPHATIRELINVVGSVSLSPYFKEISPDYPKFKGLVTAENISVYAQDSLKWISNMAKTKNARAILDAFGLLDGESLRPVHSKYSKWILGIKDKKGQGQVINRSELIeffggvey--tKQFRLEPEWLIVILTALVHHGDIILAIPGEKFNSADLVKLANTPIKSLIDFKHLESP
WP_106343732.1_Planifilum_fimeticola                   GVDVERLIEQKAETLSDAQLDRHYFDALEKALECsdtp-yrsgFRLWPHGLLWADRKVTRPGYLFLGTPNERSTAQPPRDFYLYMVQPFDPPRFRDEQRGDEVFFRLVHRDDTFMHHMTRYAAARELEINATTATKKRYVDHISRHLKSLNAWLRENFGTAFRITHKGVSKKASEYLSGLS-ANATLREMVDHVGAACLSAWFEERYPDYPSFRAVITVSNRDSYADSALKWLAG-HKTAGGEGVLKGLVLLHDGKVEPQRSAYAKWILDLLEKKGEGQVLNRSELIeelyggevvaaGPFRLEPEWVAVILAALVYAGLAEIQLPGEKIDATGMERLASLPVKKLVEFKYVKKP
WP_254843978.1_Virgibacillus_dokdonensis               SIPVDELIRQRGEGLKDSQLDSYYFDVLKQATEVsettayvfgYKIWLHEIPWVDRRVKREGYLFFGAPNERSTAQPERDFYIYMLQAFEEPNFKDEQKEDEVFFRLKKKNDTFVDLLRLYGGATEMYND-TTTNKKLYKPKMEEYKKKLVKWIKENFVDTYEIVYRGKSAGVLEHGMFLPSNPDTLVELVDSVSQDLLSQWFEVKYEEYPSFRKLLTKKNIATYVKDALDYLNG-RKTNQGEAILDGLLLLDQRNPTTQKSGYAKWVTDLLDSKGNGQVLNQSELIeviqgtpdqrtRKFTMEPELLVVILGSLIQDGQIVVTVKGTQYEAMNFSEFIRLPIQDITYFDHIKKP
WP_235850110.1_Niallia_circulans                       DIDVDSLIQAQAETIEDDKLDHYYFSLLQKAMELddnt-hvsyVKVWKYDLAWEAHHVMRPGYIFFGAPNERGTAQPERDFYIYMLRPYDKVTFKNEHKPDEIFFEYNRSNEDFDKYLKLYAAAQDLATDASPATKKLYTAKVEEYQKKLFKWLIDNFDSTFEITYKGKKSTVADLRMYIL-NNPTVKQIVDTAAEFYLTDWFEQKYEDYPSFGKILTKDNMETYVQKALQYING-AESKMGADILDGFVLLDKSRLTINNSGYAKWIINLLEAKGHGQVINHSDLIetikgipdlqtKQFSLEPELFVVLLGAMISAGAIEVTIGGKTYNALNLHEYVQRPIKELTQFSHIKKP
WP_146342557.1_Dellaglioa_algida                       DIDIESLISERTEIIENTTLDSYYYKILQNAVALddnt-yvsgYRIWQHELPWEQRRVMRRGYLFFGSPNERSTAQPERDFYVYMLRPFEKVQYKDEKLSDEVFFELSLEDKNFKKNLLSYAAANELYNETTSAQKRLYKDKIDSYFKKLNKWLTEHFVNTFKITYKGKTGYLDDFGMFLP-QEDDVKSLVENFSAQEFLAEHFSEKYPEYPSFRNYISSNNIKVVASDALQRING-RNNSQGESILEGLVLLSNQSDSVTNSAYARWIVETLELKGAGKVLNNDELFnivrgvedrrsKSFDLEPELVVVLLGALLSTGKIEVNIDGTTYTAMNFSEFAVLSIEKLTRFSYIKRP
WP_229736080.1_Kroppenstedtia_guangzhouensis           DIAVDDLIEQKSETLSDQLDRYYFKLLAEVTENskts-yrsgYRIWLHELPWHDRKVTRQGYLFFGAPNERSTAQPERDFYVYFLQPFDPPKFKDEEKPDEVFFKFVDRDEHFMRWLRLYAGAEEMKETSASSTRRLYEEKADGYRKQLVEWFRQHMPTAFKITYRGVTKKLAEWSIFAH-AHATVREIIDEAAGDCLKSWFEKYPEYPSFSKLITKASLPGYAVDAIKDISS-PRTQNGTAILSGLVLLGDKLNVRGSGFADWVLEKLEAKGPGQVLNRSELIetvqgtediqtVERNLEPEFFAVVLAALVYSGDIVITIDGTTYDAMKWDELIRLPLNQLTQFSHISKS
MBP4110753.1_Desulfobacteraceae_bacterium              GVDPEQKIEQKAAGVSDSEIEYELFYQLAEILELdrvn---ghSDVFHDECEWKSVKSFRKGYIIFVKENSEFSALPDRDYAIVFLSPFVTQFSRK-FACDQLCIKLKISVAENIERIKQIVAIKALIN--SNFNKSQMLRKLDGRINGFKHSFLKLLMNKAECDFNGRPESIKFHIGRER---NNVPEIIDELKASVFDGQFNKAYPLHPKYSIQLSSVNIVPSLNPIASELTKGNLSPSVRSFLRSVLLDQQYPDISQSKIVACLLDKIKANK-KKVTDIQELVep------lcaSQYGSEPEIVRFVLVMTVMGRIFLQAGGDKIDINNIKDKFK-SLAAFETIAYAKIQ
NUM38593.1_Armatimonadota_bacterium                    DRDYEQEVNAFNPG--KNDVQRHLNEILGIALELdnaq-kapqGTIWNYAITWEDKKVERPGWLFFGFPSERSTAKPPLDFYIFIMPSKRITEKEDANVPDESYVTLQDDPDTFLDLLRKYTSASLRADVVGSNDRVAFEGIAKKLRGMLAEEFRENAGEWLSITHGGNKKRFREWIQTIDPANFPLKSQFDSVAQGMFKARFETEYPDYPSFDVKVSESSRPGACQSAIEIVCQRQGSKEGWAVLEALGLAHGNALTLEKSDWLKKARNLLKGKGDGQVLNHSDLFekrkddkww-fKGECIEAEFLNVVIAAGVYAGDFIITTKNEQVDAANLQAKYE-EIRNFQNVKRLSRP
WP_111566961.1_Dehalobacter_sp.                        DKDYEQDIRNYSKTLSREVLQRYLNEIVLRTLLLdava-pvqeGRLWDYALLWPDNNVERPGWLFFGFPNQRSTAYPPRDFYIFFIPSKRITGREDNDNVDEVYCFLDDAQDTYLDKLAYYAAARELQSNSAGLDRQGYEQEAKRKLERIVEEFHSHVGQWISLRFNGQKRTLEQWITEQAPLNAPLRSQIDALTQGLLRAHFENKYPDYPIFTERIYENTRSAAAISAIEIICGVQKTAQGKSVLRALALMDGDLLTAEKSPWLKLIMERLEGLPPGQVLNQSDLFekreeriw--lVGTCLEAEWVHVTIAAGIEAGFLVVIGNNQKYDATVLKMFYQ-NIRTWQDIIRIARP
MBU0460852.1_Nanoarchaeota_archaeon                    TQDYDVIIQNKVANMNLKYVNEKFVEFLLKELDLdieyfetskKYVLNDSAYWEDRKSFKKGYLIIDVGYDLKID-SGEDYSITILGYGQKENKI--NSKNNILIKI-PYSDTFVESIKKLAAIEEFIR--TKAYVSIMGNKKRSVIDNLRPALTKALFEA-KISYQGKDYKAEEDLGIGT---EVTAEIFRQIKQNLLNEEFTKTYPKYPKLKSRISAENIKGTVESIFRDISSKNLLNQSVNILLPLGLYKDNRLDLSESEYAKVILDKLEDS--SKNVSIQDVLkd------likIPYGLQKELVYLVLAVLLRNGDIILSSRGQVYSASDFNTLFSNGLKAFDELSYLKE
KFI36375.1_Peptococcaceae_bacterium                    TVDYDQVIRRRAENLPENILDEEILALLKDQLLLtdgp----aPAVFWDTCRWPGRRAFREGLFIYETGRGETVG-ESGDYQVIFVSPFYTGNRYA-TAENRAQVSG-SLSREAVEQLKLAAAARTLAR--ENYHRSIMEKKYAGFKKEFTAMFMQAWLETGQVDTGKAKKGIKTLIAREF---ANFDELFSEIKPGLFEDYFNHKYPKHPRFTQLITRDNIKGEFGATTREIISKSLFSNARSILNALNLLDKKNLSTAGSEVAAKIIETARNAE-GKVVGVNELIke------fqePPFGYDRWMTAFVLIILTYNGEIVLKAGGKVISSSEVGETFGTGLDAFDENIKYLAME
EMR73852.1_Thermoplasmatales_archaeon                  SIDYDEVVKRKMKTLPEGMENEELLKILKYTDLIeadl-aesyVRSFKDSCPWSDKKSFRLGNFIFDDGSDQVKS-DDLDFNLVVQSPYVSSSKLD-SSKETAILSI-KYSDELDDALKYLAATKLLIN--ENYAKSVMIKKQNTCSDDARNLILQKMIDS-EIEIDGSKKKVKSIISKEP---DNIDEFFHHVKENVFNDYFSSTYSKYPKFLNQISYENVKGEVESTINELLSKNLFSNAKNILSSLDLMDIDNIDTSNSLYVKVIVEELEKNK-GKNVKIDDILek------lqkPPFGLDVEVIYLVFVVLTYNGEINLRKGGGTITSSDLKDVFNIGISAFRDIPYATLE
WP_114474457.1_Halanaerobium_sp.                       SVDYDVVIQRKAENLYVGAEDSVLLNILKDNFYLdsds---nhDRVFDDDSYWKEKKSFREGSFIYDDNSVVLDK-GDGDFNFVIVSPYQKSATIK-SDFDTAILNL-EYNEELDNLLSKIAAVNSLKS-GSNYPKSIMDKKKRNFLEEAKEKLLKLLLLT-NIKNGGDYDSTKKLLAQEP---DTLSAYYEAVKPAVFGHVFNDKYPEYPDLISRLSRENIQGEVERTIDSILTGISLSNSQNLLSSLNLLDDNYLDTRNSKYANIILDKLKDNK-GKNISVNGLVem------laePPFGLQKELAYLILLVITYNGEANLKHNRGTLTASDLKQFFSSGLNKFDEVSYVSLE
WP_098658108.1_Bacillus_toyonensis                     VQDFDVLIEKKIEKQDIDDWKETFKNLLISHLNLneqnydkafPVVFEDTAFWSSRRSYRQGYIIIGQRNSGPSVFNRRDYKLILHSPFANHDTKQ-TVNDELIIGV-EFNEEVLLKLKRISATESLIR--DGVFRQELQGIYNKEMKDFYNKYLQLLTTEGYAIHGGRKEFLCELPVNRF---DHVSDVLEHFKSHFFEGYFNDLYPKYPIFKRDITLQNIKNEIVRSITNLDKIGMDENAQSYLTSFHAQKDGKIMSDGSEYMQYILNLISQNTEGEMTQLEVVVdn------lskKPFGLQKELVYFLLVALLYNGEMNFVKGGVRLFAHDFSNHFKSNLQIFESIKYIEEE
MCI4298469.1_Anaerolineae_bacterium                    IDDYDAIIEKKALASVKQEINLAFRQIAETELGYkdpkplvagLSLFDDTAPWPSRRAFRPGVLVIGQTSDGANL-QRGDYRFVVQGPLAGSSSQ--PRQEEVILAF-SFTDELNSLLTRARAAELLAR--EGVHKKVMTDLQKKATDEFKARYLGHLLSQGYVLHGGHRTPLSDLPTTRPM--TTLADVVDHVKGALLDQTFSDKYPNYPSFRTIVTAANINSEISRALQALSKAALDLNSRGYLESFGAMKDGHFSASHSPACRLILERVEANDQNKLTDLDDLLre------ferPPWGLPRELVYLLLGALVFNGYLLLVRGGARLHAGDVGPLLKQGTDLFEEIRYLERD
MBP7946969.1_Verrucomicrobiota_bacterium               QVNPVKFFAPLKKKFLKKRQDEKWVESL--FWQLpditelgvnGGLFAAASTRKLYRVAYGGEIVVSDRWRDEFGDTDQHFRIVYLV-NKPADTDAKAALKETRVAVEDLSEGTRDALADLLAAEEMKRT------------AAPNQVSLRDYADGKRRDAVKALLKCQQDEFRRGQILTQ---ATPKEREESLAGRLLEKSYDTPLFSPKELKKDFTETDARKVFAGLFTKEPATAEKDAVVNFGVGLELTLKSELKPDASQALAEVRKELSGIPD---KPIADIHka------lcrPPYGLTQEMANLYLFALVKSGGYELALPATKLTAHTLGLCWNSQMDKALLGARIVAS
MBT9159326.1_Dehalococcoidia_bacterium                 GLSAPQRFNEIKQRITQHHIQKKWRESL----------emlgeSTLFADSIRIENRRLEYSGEVILSDTWRPELGQEDHHFKIVILM--ENTDAGG-IAIEDERIAVNGLSELAHDAVRDYMAICDMEE--------IYRHERGSDAEQIREWVRGKKPEVIRALISSQPKQYQSGKICTR-SQPSNERRFALIANALLDNAYTARTLDSRAFKKTLSSTDVGKVFNGFFRGAASAQDTSALDNFALGLHLAATDKFAPAANAVFDYLKGKLGENE-G-HLPTWQVYrd------lsnIPYGLPFPLISLYILCFTRYGAIDLYLPNHKLTSATVPELWKTGLDKY--FDAISLS
NLF03150.1_Anaerolineales_bacterium                    PLAVLDLFNQEKRRVARYRLQQVWTGSL----------dtggaPGLFGETRRFECRRIDYSGEVVVATSWRVDQGMGDTHFRVVILTPVAAQAVRA-TDLQDPRIAVGELNEEARDAAAGFLAWTAMRE--------AYKDRAGSDAEEIRSWIGSQRRAIVDALLATHLKLYQSGRVITR-RGGGNDARLAHVVNQVLLSAYQQVPLEHEQLRSTLSSTQVGNVFAGYFAGSSGAAVMAALRNFGVPLGLSHADRFAPQSPRAFELLEAMLAEKG-GAELPVWQVYdr------lsgAPFGLPYAVIQLYLLAFVRRGRVDLLLPKHRLTAATVVDLWKPGLQDS--FDALVPA
GFP32117.1_Candidatus_Hakubanella_thermoalkaliphilus   EAKPAQIFMSFQKKITISQIQEAWDKGL----------etggeESFFSGRATVEFRKLEYPGEVIVTRRWRAEYGENDLHFRIVILT--GPESIDV-KEIKDFRIGVSSLTEAARDAARDYLALSEMEE--------AYRDKTGSEAEQIRSWLQSKRPEVVRDLLRKQLGIYRNGTIITA-LLNKIDKVLDQLVPTILADAYAKPVVDSQLFKKNFSGREARNVFEGLFKKANVPAATSACINYAPALKLSTTDKFNPSPNKVFELIENRVK----AGEVQIWKLFee------lhnQ--GLTNELINLYLLAFVRNGNVEISLTGHKITSFNIPQVWRARLEII--LDTLYPS
HIW77326.1_Halobacteria_archaeon                       NSEVIKTFNRYKKKVSDAEIRDYWLRLL----------yidgfSSLFMSKRKVFFKNIEYPGEVILSRTLRDEYLGGDSHFRIVFLS--EPADIDP-AKLADVRIAVSDLPDTLKDRIREYCAISKTKD--------EYF-RSDQRSDDDKEWIKNKEKETIINIIREQRSVYRDGKVFTN-TSTDLDQILSSIVGTLLSSSYTKQPFD--TFKRKFSENEARKLFEGFFKANPKRNALDACRNYGPNLALSLDDKFHPMNDNIFQFLRAKLDEKPEG--VPVWRIYke------ledPPYGLLKNLITLYILAFVRANNTEIRLSGSRINSFNIEKIWN-NFEKD--LDILVKS
MBP6974014.1_Syntrophorhabdus_sp.                      DVDYDKKIQEKADFLGEDRLNQYFYTVMYNAADLqlte-hvtgRKIYEYRINWTEKNIFREGYLFMGLPSGRPTAQPPQDFYVYMLPLFG-SKIIDEPKEDEVYFTF-QNSEKFTSLLKEYAGAKEMEVMAQGETRSTYAKRATSTLRDIRRYLEENKTVCFQISYLGAEKPVLEYLRGARMNEITVRDVISIVASKALSSYFNERYPYYPVFKKPVTKDNQATMRAEAINAIAG-KATQLGQSLLESLGLLLDGKVSVENSKYATYYLGKLAKLPEGGVLNASDIMtslngqdmv-dGEFKLSVIWTSVIWTALVYSGNCVLVGDNKRYDASNVEEFIKNP-SVTYDFKRLEKP
HHW29753.1_Syntrophomonadaceae_bacterium               GEDPDVLIAERAEDLSDYMLDQYYFALLRRVMQCsdet-yvsgVSIWEYDLVWQEKKVTRTGYLFFGAPNERSTAQPPQDYYIYFLQPFDPPPFTDEERPDEVFFYLKERSPEFIEKLKLFGGAMEMLKIAPTPDRPIYREKGEEYLSDLSSYLLENISNAFMVKYQGSKKPLAERLQGK--RGGSFKEIIDQVASFCLSGYFAGEYPDYPSFPVLLTRQNLKPSIEEAIKYLAT-GQSRLGAQVLEGLQLLDGEEIVPEKSRYSKHFLGILKEKGGNQVVNREEIIkgpdraer--dRFFKLEPEFVVVVLAALVYRGDIVLNLAGREINASNLVDITRISLDELVNFRYYKTP
MBT6646600.1_Nitrososphaerota_archaeon                 NIDYNAKIEKKITSLDDDKKDRYFFEILQNAMETpsvs-avadRKIWQWEMEWEEKKITRKGYLFFGAPNERSTAQPPRDFYLYFAKFFNLSSFTDEKKSDEVFFKFKDIDSQIKEDILFYCASRDLEAVASGSTKQIYSNKSEQRLKNIVMWIKEHGVSNIQVTFQGNEKSLLEIPITKKGSNYSFREIINSIGSFYLKEHFNQKYPNYPVFEKLITNENKDQYVKETLKFITNNVPSKISTSVLSSLELLNSNDIDTTSSKYANVIKNKLQSKATNEVLNRDEFMifeqneyf--dSELHLENDWVLVVLCAMVYSGDLVIGYSGKKISAENLTEINSLSFEEELDFKHVQRP
WP_053959405.1_Sulfobacillus_thermosulfidooxidans      MVDYDGMLKDKASVLSPDTLNRHYFEALTRILQRtdlp-ykpnTRIWRYDVPWLSHNVERPGYLFLGTPNERSTAQPPREFYVYFLPPFKTSRFEDEGRDDDVFFRLAEMNDELTDMLRYYAAARELANDSNSPARREYTIRSDRALKNFNDWLGQHFITHFEVVYERQPRRLAGVVKNLNLSDTLIHEMVPKIAGYFLEKAFERQCAGYPSFTPFVTQSSLESYAKEAIDRIAGKSTSAATGMLLESLGLLEDNKIHPRRSPYAKWVLELMEQQPAAHLVNRGLLIeekngvedvatPQFHLEPVLLLVVLTALVYNGDLVIAAGKVKHDASRLSDWAKLPLVDKIHFSHIERP
WP_020614466.1_Sediminispirochaeta_bajacaliforniensis  GKDPNQEIRNKASKLSDKELTRYYFEILSLIMEQtdvt-yrtgFKIWEYPLMWYQKKSEIDGYLFFGNPNERSTTQPPREFYLYFIDPFRQNTTVSSGKDDEVFIQLKDVDDDFEKNLRLYAAAKELSLTAVQEYKTVYNQLAQKYQRELLSWLKNNISDRFQISYKNSTKLFKTWSASINLETLSFKQQIDTLASGLLSDYIADQKPDYPEFSIKITNQNRRQTVQEALRAVVRGQSSGQANAVLSALDLLSGEGISVEHASICSEILAIVNSKESSQVINRNELVserndkeyil--NGSLEIDFLPVYLAALISEGYINLSVGSDRYTASNLDSLLSIGFNQILDFNHIKKA
WP_149544905.1_Calorimonas_adulescens                  DIDYDAKIDEAVDVISNDALNRYFYEILLNKLELaesskiadaHLLWSYSLIWKEKNIERGVYIKFGLPEENS---PLGDMLCLFVGPYNDIE-PDQIRDNYLYFIMRNYDEKFENDLKHYSAARYLYQLATDDIKQIYDEKCRKYARGLNEWLDIHMRNAFDVRYGGKKIKIDR-------DKMSFKEIIDECINMGLYDYMATILPDYPRFNTVITNKNRNEVFRAAINFLSG-KRNELGKKILESLELWDGTEVNIHDSKYAGYFLGLLNDKPDG--IAREDIIdg------fddRRFRLETEWVAVVLLSLVYCGEAVLKLQKDEINLLNISGLSRYNVSEITLFSGYRKM
NMA94940.1_Clostridiales_bacterium                     DIDYDVEIESMADLLSDEGLNEYYYAIAYELLDWkgnr-avpgYEIYECSLNWDSHNIFRRGYLFFGIPTERSTAQPPRDYYIYILPPYGAMDYQDEKRKDEVFFSL-RPDDKFYRNLKLYAAARILQDRSEQGKKDIYRKEARKYESKIKKWLNNNKETRYDVVYRGVQQQLIQATKGRT-RGFNFKDTIDTTASLCLDEYFTERYPGFPVFVKTRVTQENNLELIDRVIKHFAG-IETEDTRAFLKSFGLIENGRIDVESSKYAMELVRQMEKLPRDGVLNRYDILeeergygelrdKSFKIGNEYFIAVILALVHTGYANLALKNITITASNMDRLADISGWDVKNFKHLIKP
WP_209451817.1_Lactimicrobium_massiliense              VVDYDEKIKQKASIMAPSEWNRYFYEVVYGCMEWsekq-yvpgFEIYQYDLNWESHNIFREGYLFMGIPEERSTAQPERDFYIHFMPPYDAKEDHVRNLDDEVYFYF-KPSAEFNDNMQLYAAASAMANESEGKDKDAYLTKKSTLKKKLIRYLSENRNTCFDVSYHQHRRQLIEILRGKHIVDSTFKGSIDLAASLCFDEYFDNKYPEFPVFRTKITRKNMAENVRAAFDYFAGRKNSQVAVALLKTFGVLDDHRIKPEGSKYAGYYIDLLKKLPQQGVLNYSDLFeyqdsvefv-dKHFKISMLFTPIIFLSMVYGGYAVITLDGKQVTASSLDQIPKTGALDLYEFKYLSKP
EGD52432.1_Thermoanaerobacter_ethanolicus              GINYDAKIQEKAEILSEDIFNRHYFELLIKVLDWnape-kvpnMRIWEYSLTWNEKNVKRYGYLFMGYPDERSTAQPPRDFYIYFLPPFVGVNYNKKGKEDEVFFEFVNTDEEIIEKIKKYAATMELSELSSSDQKKIYEDKANGFQKDITKWIRDNVSKCFNVSYKGETQSIIYWLKGKTANYDTIKSYIDEAASSCLSLYFSEKYSEYPTFTYTKITEANLNEVFKSAMNYLAG-KKTETGKKVLQSFDLILGDDIVPGNSKYANYFLNELYKLPPGKVLNREDIIenvngedi--dKRFKLEVQWVVLLLSALVYSGDIILRIKNKVYDATTIEALANMNVGELIDFDSFERP
NLV27395.1_Methanomicrobiales_archaeon                 SVDIDLLIEEKGAGLNDDQLDKYYFESLTRLMDRpdtt-yvsgYRIWEHELIWKDRNVGRMGYLFFGAPNERSTAQPPRDFYLYFIQPFNPPVFNDEHKPDEIFFRLIKKDDAFIDALRKYGGAREMSAITNVKDRQDYDRKAQGYLKDIVQWLSANMGHSLEVTYQSQTKKLVEWSKESTILGPEVHRAINEIASYCVSPHFNDISPDYPKFSVTITSTNRHDMVIDALSWIKG-RTTKSGSQVLAGLELMTEGVLTPEKSRYSQYILTTLSEKQ-GQVINRQDLIgllngvey--dKKFRIEPELFMVVLSALVWRGSIVISYVHGKIDASNVDTMAKMAGDDLIAFKHLEPP
```

**Type-3 BREX BrxC Alpha+Beta-Domain + wHTH Entropy Plot**

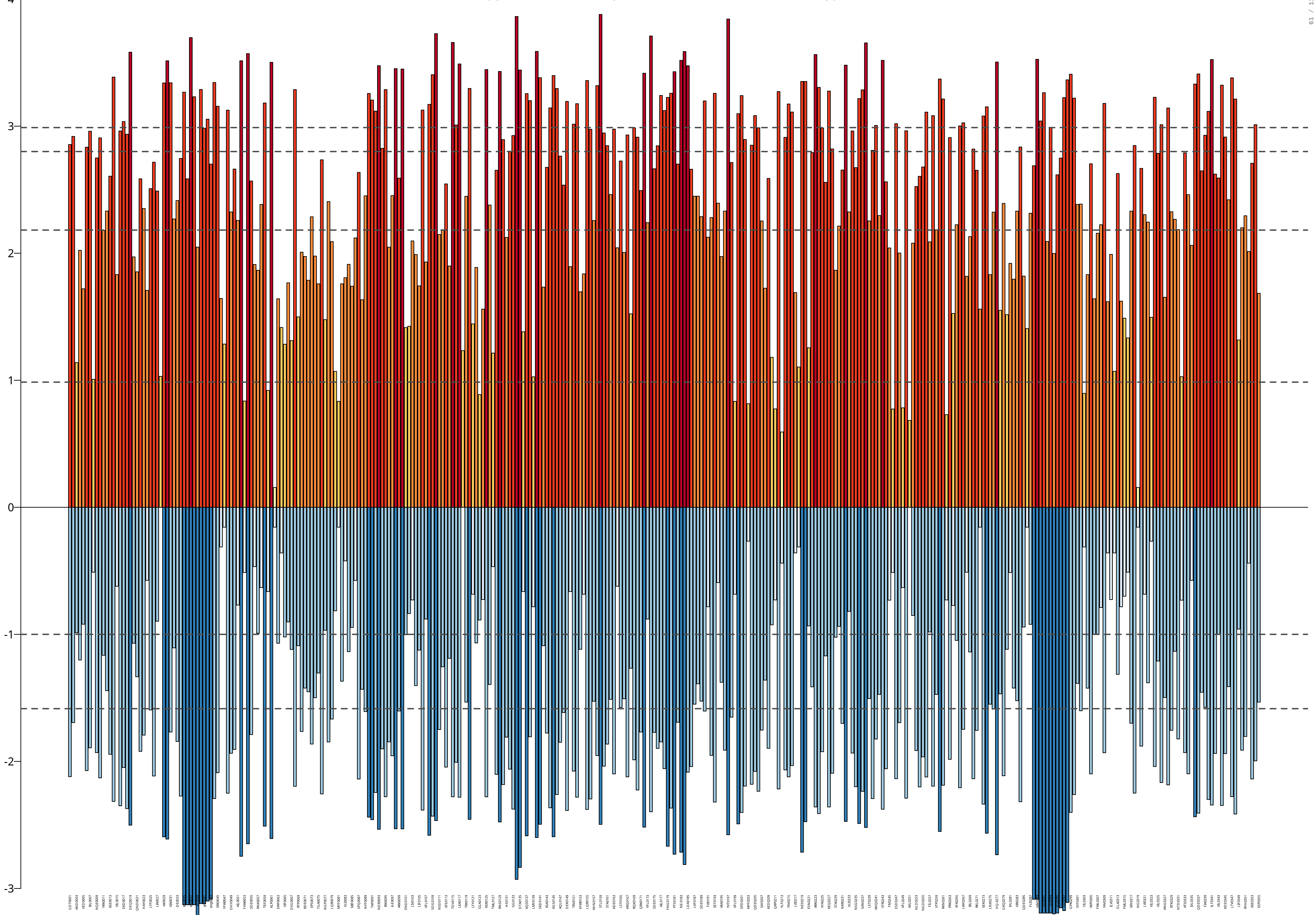

```
# ; Alpha+Beta-Domain + wHTH (BrxC-ATPase Type-4 BREX)

WP_165846692.1_Candidatus_Cryosericum_hinesii    [illegible]
WP_169366114.1_Pseudanabaena_yagii    [illegible]
HAR58250.1_Acholeplasmatales_bacterium    [illegible]
HHW21153.1_Thermodesulfovibrio_thiophilus    [illegible]
KUK66683.1_Parcubacteria_bacterium_34_609    [illegible]
MAU00653.1_Anaerolineaceae_bacterium    [illegible]
MBD3337553.1_Candidatus_Lokiarchaeota_archaeon    [illegible]
MBF0303406.1_Desulfamplus_sp.    [illegible]
MBF2075132.1_Synechococcales_cyanobacterium    [illegible]
MBK9943588.1_Kouleothrix_sp.    [illegible]
MBN2678363.1_Anaerolineaceae_bacterium    [illegible]
MBR3710350.1_Bacteroidales_bacterium    [illegible]
MBU7048060.1_Theionarchaea_archaeon    [illegible]
MBX7251231.1_Candidatus_Promineofilum_sp.    [illegible]
MCI9127450.1_Eubacterium_sp.    [illegible]
MCK4843012.1_Methylococcales_bacterium    [illegible]
MCL6576920.1_Kyrpidia_sp.    [illegible]
NJL05929.1_Chloroflexaceae_bacterium    [illegible]
NLT83665.1_Coprothermobacter_proteolyticus    [illegible]
OQC38485.1_Bacteroidetes_bacterium_ADurb.Bin041    [illegible]
OQC39308.1_Bacteroidetes_bacterium_ADurb.Bin037    [illegible]
PMQ01344.1_Dictyoglomus_sp._NZ13-RE01    [illegible]
WP_048175326.1_Methanothermobacter_sp.    [illegible]
WP_055301605.1_Roseburia_inulinivorans    [illegible]
WP_062196491.1_Anaerolinea_thermolimosa    [illegible]
WP_139515175.1_Hymenobacter_jejuensis    [illegible]
WP_140468254.1_Hymenobacter_nivis    [illegible]
WP_143714756.1_Thermomicrobium_roseum    [illegible]
WP_189070975.1_Deinococcus_radiotolerans    [illegible]
WP_195558544.1_Bacteroides    [illegible]
WP_201382002.1_Ktedonobacter_sp.    [illegible]
WP_214462063.1_Bacteroidaceae    [illegible]
WP_235137407.1_Dyadobacter_fanqingshengii    [illegible]
MCD8003208.1_Clostridia_bacterium    [illegible]
HCH97636.1_Lachnospiraceae_bacterium    [illegible]
OQB25825.1_Firmicutes_bacterium_ADurb.Bin182    [illegible]
WP_198469750.1_Acetomicrobium_sp._S15    [illegible]
NPV08139.1_Anaerolineae_bacterium    [illegible]
HED00229.1_Pseudomonadota_bacterium    [illegible]
SHE91973.1_Desulfofundulus_australicus_DSM_11792    [illegible]
NSW92344.1_Bacillota_bacterium    [illegible]
MBF0318625.1_Nitrospirota_bacterium    [illegible]
MBR4835363.1_Thermoguttaceae_bacterium    [illegible]
NLY01889.1_Rhodopirellula_sp.    [illegible]
MBR0237621.1_Thermoguttaceae_bacterium    [illegible]
MBP6982276.1_Thermotogota_bacterium    [illegible]
MBR2693833.1_Thermoguttaceae_bacterium    [illegible]
MBQ7111119.1_Thermoguttaceae_bacterium    [illegible]
MCL2016625.1_Defluviitaleaceae_bacterium    [illegible]
NBJ16968.1_Dehalobacter_sp._4CP    [illegible]
MBD5156842.1_Butyrivibrio_sp.    [illegible]
MBR2520311.1_Selenomonadaceae_bacterium    [illegible]
MBQ9371615.1_Thermoguttaceae_bacterium    [illegible]
MBQ7256248.1_Abditibacteriota_bacterium    [illegible]
WP_215692602.1_Gemmiger_formicilis    [illegible]
NBK77201.1_bacterium_D16-76    [illegible]
MBR2215718.1_Selenomonadaceae_bacterium    [illegible]
WP_268058969.1_Caproiciproducens_galactitolivorans    [illegible]
MZQ76088.1_Peptoclostridium_sp.    [illegible]
HIS35237.1_Candidatus_Scatousia_excrementigallinarum    [illegible]
WP_031492453.1_Succinivibrio_dextrinosolvens    [illegible]
WP_123481433.1_Lepagella_muris    [illegible]
WP_118016040.1_Mitsuokella_sp._AF21-1AC    [illegible]
ONI45222.1_Epulopiscium_sp._SCG-C07WGA    [illegible]
```

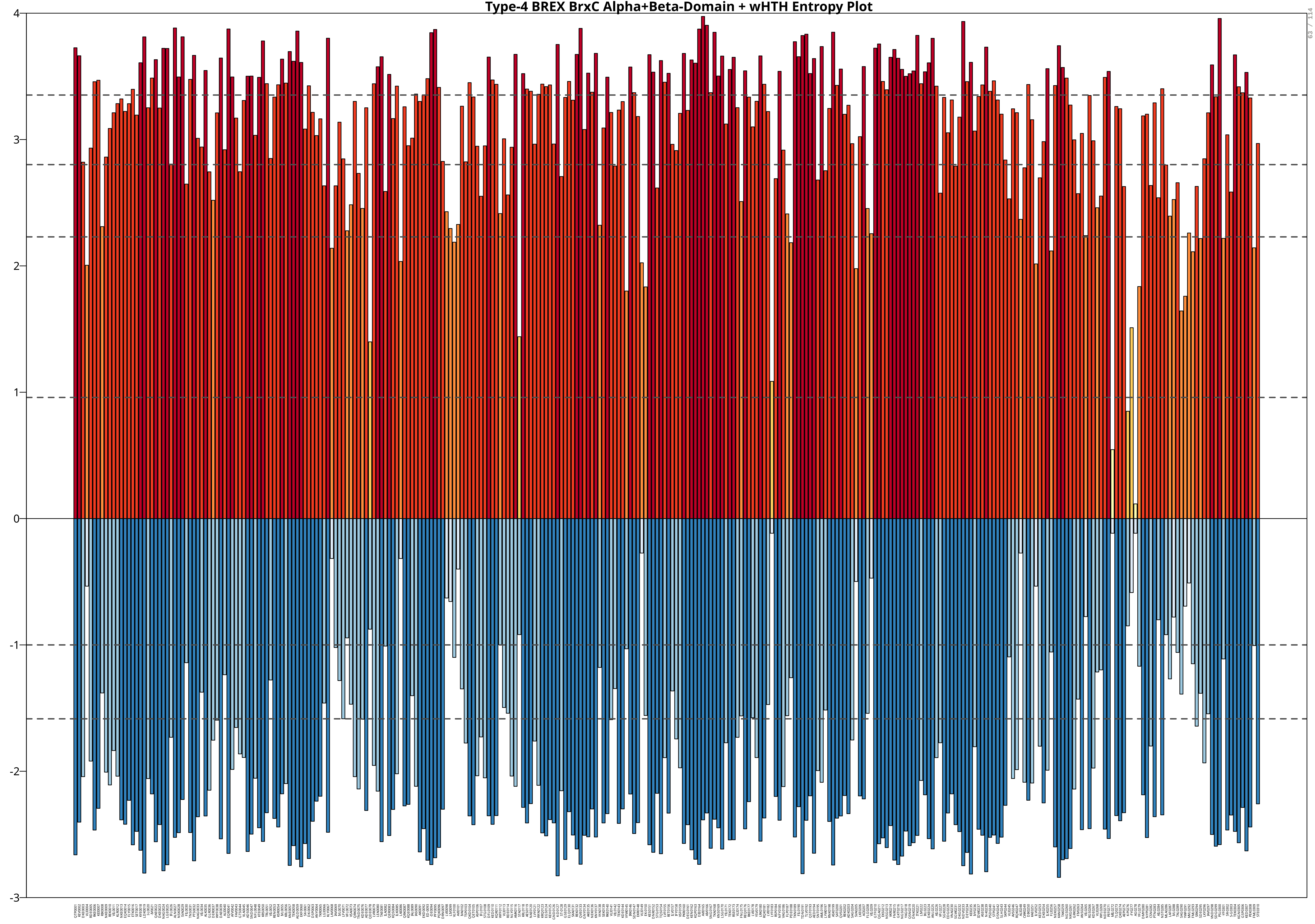
Type-4 BREX BrxC Alpha+Beta-Domain + wHTH Entropy Plot
4
3
2
1
0
-1
-2
-3

\# ; Alpha+Beta-Domain + wHTH (DUF499-ATPase BR-systems)

```
WP_148590534.1_Aquisphaera_giovannonii                 NLNQILVSRRGAVKPKDVEDRVRKTVEDLFGPKFLDRRFFPERSNDVPDRPQLTLVVLAPSGTRSLVESIVKECGTSGRTFKSALIFAAPSEATAVTDAARDLLAWEDVRDDEATQLDEPQKRSLNQSLGRAKADLREALWRSYRHILMLNKGNAIKDVDLGQINSSMASTLPDLIVGTLIKDDEITDKVGGSRLVRALTEWPTKAVRDAFYASPALPRLIDPNIIKRTISDAVSSKFVGYARKVGARTVLERFGEALGISEDYVLLRA
MBQ9433643.1_Synergistaceae_bacterium                  TLKKIAQELADEVKAYDVREEVTRRLKKLMQGAFVGVHVH-DKEIEVPDKQEVRLVVLGL-DSRSTALEVLRSKGQPREYMNMLVFLEADGVEGVRRNARSYLAWKKLESERESNLDARQQREVQRSVADFDGELNGAIVRAWNCVLSPHSEKKVDDIALEGLKLSDGDAVAE-ALRTLRSSDRLSDKWGAVHMST-REDIPVKTLWEYVRCYPMPRIERIEGVYAGISEGVAKGYFALADG--GKYTHLRMGESVLVNDSDFVVKK
WP_054571357.1_Frankia_sp._R43                         NLTKLLRDTSNSFGGEDVLAEIRTEFEDQFR----GVSIWPADPGAIVDEPVFTLAYLHPSGTRLPLSQFVEYVSRRLRSYRNGLALVLPESSDRVRQATRSRMAADHLLGSSKHRFNAEQLEELALRRDSARRDGAAALTLAYRTVVLPVRGERMEEIDLRSL-VSAGRSLQERVFDALSS--RVFSTITPSKLVSGKELIPCAELVDCFFSYYEFVKIRDKGVIADAIARAVSNGDAGYVVGSNEDRRAVRFELTLRLSEDAAIMSA
EEO54231.1_Bacteroides_sp._2_2_4                       NINILVNQAKAEISQADISGEIINRLNAQTVSKI-KVLVNP--ANDIPEQKSLTLVILGPTKTKKQVEQIAQNKGYSSRIYRNTILYLACSEIGMLHSKLLEYLACAKIQAEYS-QIEPEQKKDLIERKAEYDKQANALLIAAYNIVCKYSVSEGIEKIEIKDFAQDFNTQLSSNLFNNIKEEEWLLEKLGIGTLRS-EQPIQVNDLYEAFLRFDDKPMICGVETVSKSIQRYCENGDFNVACGEQGNYNHIYHHESPFVTDPQWLVDK
MBM4287682.1_Deltaproteobacteria_bacterium             NVIKLIEERMPNIPLEDAKSRVQAEVQGVFPGLM-RVVAWPSSPQQVLDTAQLQVALCD-ETVAQAVCRYADAEAPMPRRFRNGVVAVTAKADQEAVTRARKVLAAEDIESEHKPKLLREQLKTLMPTLKRG---LRLRSCQAFNTVVLAD--KGPYSLEYQVREETEKGQGLNQLLSFFQDKTLIYDTLDVDLFVSKPEVVTAKAVKERFWAAPGLKLLRDDAIVRQSILKGLQDGKVLVRLGDG-VYDAQYIQRRGSLDEATLLTPD
MBI3467145.1_Planctomycetota_bacterium                 NLNRILVDREEMVRSEDVRDFAKQTLNDMIDVAF-RVYRYPEEDRDVADEPRLSLVVLDLADTEQFVTQVLKQHGKGFRKHANVLIFLAPDQASEVIDAARRLLALRNIEEDKGKQLTEEQLKDLAGRLKEAEARLPAALATVCRFVLVPAKNKALRCFDLGITGYSGKTTISNRIAEKLIDEQQILEKLDPGILIGLQEVINVRTLADYFTQLTHLPRLNNAAVLPDCLSKGVHRGLFAYALGEKKEFDTIYFNDKVSITETAWLVRP
WP_259054315.1_Salinibacter_ruber                      NLNRMVVDKKGQIQPSDVRSKLESLIRDELRM---RTYLWPSSGDNLPDNEDLKLAVLDP---EQELRRWVNRSGQSYRSYKNTIFFAQADEDGRIQDDIKDFLALQDILNGVRKSGLEEKENEVQRRLENVEGEFSQKIRDLYHVALVPVMNGSLEEIDFGQSAT-GRENLDSWFRKKLSSHSKILERPSANLINKGSDTIELQAILDQFYKDPNLPALDQADLIATVANGVQDGSFGLARREDGRPTSVRTNEPLPFREEGVLVAD
MBQ9726423.1_Kiritimatiellae_bacterium                 NLREMESRKARFPAADVDALVRRVFTERFSPAFAGFHPF-VRSGDAPDDPGIRVVALPGGNAFDAARDILLHHGTGARVHRNRLVFLAADLNGRVEDIARTVLAWRQISDDVSRETTVTEKAQVDREKDGMLRTLSQTVAECWRLALVPRQV--PEPTFDVRKL-ATGTIAQALETALRQNEDVVERWSPAFLKA-ADEVSVGRVWNDLCNLCEFRRLSVDVFETTVREGLGDGLFGYAQAD-GYEGFVFRQSFSVNESALFVRG
CAI1753142.1_Serratia_quinivorans                      NLRREMETRKSKVGERELTPLLKKSVTACFQHCFAGIHVF-TPSSDVLDENGPRLVVLEPNQAFKAAEVILFKRGEQPRVKQNRLIFLAADYDNRLKDQGRTYLAWHSIVIDIEVNQDLAHLTQAKGNRDTAKKQFDSLVRDTYKWLLAPLQD---PQLEWEVSQI-SSPKLVQEIESKLAEEEWLIKEWSPVHLRN-AESVSAVKVWQDACHLYAPPRLVNDQVLRRSVEEAVKTDFFAFASGDD-RFEGFAFGKSTVLDESNVSLISR
MCL0064885.1_Dehalococcoidia_bacterium                 CLTKLLADRRALVSPSVEERIREVIRSTFPKTF-QRCYFPEELSDIPDLTFLILVIMRPERTKEIETFIKEYGSKPRIFKNSLFFLLPASHREIEKARDLLAVEALEHEVDVELPEEQKRILKDIKSKAQKELSEKVRQAYRWLFLEE--GLRWMDLGIFSPGVADTPVGFIEHRLKLEGILEEKVTYKFLKALIEWPTKKVKEDFYASPKFPRLTDWQNLKRTIAEGVRGGSFGYAIKPEEGYIEVSIEPFFTSPDTESKLE
MCD6101177.1_Candidatus_Marinimicrobia_bacterium       NVNKLLEDEAEGIRNDEIKNEIKNMLEKELK----SAVIWPQNSKDIPDKPAFQIAYLPIKNKEKIGIEFLMQYGDKPRVYKNGLALAIPDKNEPLRRAVRYLIAVQRVKGKNKSNLTEEQQEQLKERETEKASRDSSFRNLYNTVWLLNLEDVLETREVGGRAL-QATNIHERLMELLMRPPKVFDSLTAKRLSDIEEGIKTKDIIDTFFSSLDFPRIIDENVIKKAILKGVKDSFGLTNKVEGAKEHVIIGKEIPLPSGYIVNSK
WP_116051372.1_Pseudonocardiaceae                      THRMLVNNIRRTVTEAERDEVLFEFAQGLSGGPFRE--LMP--AADVLDAHTTRLIVLDPAPTMEALTVAMGGEGTKLVQWGSSAVYALVNTQGLARGVAVEYLARQKALAAPEQ--DDDLKATGTKDLASAKQQLEKAVKRAYQHVVYLTQ-PDDGERYLDQLAFDDTALDGTLVWKSLAARDKVFGQFDPKALLHDQDYRTLSDLRAAFYSAPRLPLLFAGRDLQQAIYDAAETGLVDIVDGSG----------RTALTSAGLRLKP
RME04985.1_Planctomycetota_bacterium                   NLNRILILREENVTDGEIYEERREVIRRHIESRF-RVYLHPFPKDIADNEELKLVVIE---DSFPSKEFLERYGEGPRVYRNTIMFLYGDEGFVSYLRKLIALRNVERDR-GRLTEGQRKELRSKIARQNQLEYEELRRFYKFLFVPSGGGPFQEIDMGLPTFG-ERELDREIEFTILKNLGEVLERLSPRVIRD-GDCVEVRVLYESLLKAPGLEKVASWEGFLEGIREGVEAGLFRVGEFSEDGGRVRWKGVSGEGGEGEVIRL
MDE0129995.1_Gammaproteobacteria_bacterium             NITLLVEQEAEAVGRDEVRERVRQMIDQRLQ----RAIVWPEKPGDVPDRPEFLIAYLPLGDQNQALAICEQYGNRQREYRNGLGLAVPSGDEILRRAVRYLLAVERVKGKWRENLTDAQRSQLKERDATEAAAAESAMLKLYGEVWLPSSGDEFDRVQLGGRPLQTRALIHQRLMELLTTHRRVFGTLAPGRIVEQGMGIATDTVLAGFYEFIDFPRLLTAEAVRKAIARGVETGLFGYTTGGEDDRSRVAFQRNVALDSGFLIVPP
PJF35651.1_Candidatus_Thermofonsia                     TVNRVAEERAQMLRAEQVCEEARRELRELRRPQGIRVHVAPRESGDVPDEQEACLIVLDPSKVAEVRKYYETRGDTARVHRNMLLFIAPEARANWESALRSYLAWQSIKNDREQNLDAQQERQVEARLKEANETVEARMNETYSLLIVPMQR---DTVEFVAYTLKG-DDFYRRAMSVLNQDELLIRSLSSQALLNHNQWHNLKQLWGHFTQYVYFPRLLNGEALLHAVRSVKEGLLYARRENGKYEGLVLGSDVQLSEEAVIVRP
MBQ3918676.1_Oscillospiraceae_bacterium                TLRKTVEDRISKFSLDDVYYEIEKRLNKIHEPPFSGIHICPRSSLDVPDDQSVRLVLLPPSDAMNAAREILDNRGTSPRLYRNMIAFVAADQNNELISEVRRYLAWDSVKQDTEINLDTTQIRETDRSIKRCSDTINLSSNEVYCHLLTPYIDRDIKEIVWEENRING-DSIIKKAAQKMVQNEALITNWAPALLKM-SNDITIKKLWDYLCTCYLPRLANVEVLYDAIRKGMPGEYFALAGTDD-RYIDLILAK---LSNSDLLVKV
WP_157574125.1_Nocardia_jejuensis                      RLTMMVRAMRAAVTDADRDQIIVERIRAKAQQFQDL-----RPRSLLDARLTRLVILDPSATVSATRALFG--GARIVGWSSSLVFLLATARKHARQFAADYLAREVRTGPEA--DLDLYSRVNEECEDFCARLDESIARAFQHVQYLSQ-PDGVERTVETIKIDSGTLDGDEVWEALAKAQKAFKAFTATALLHDGYRPLSEIRDAFWNTPRLGLLPGGSDLKQAILDAVQQGLLHIVRGDG----------ASDNAATYRLRP
MYJ80998.1_Acidimicrobiaceae_bacterium                 TLAMLTRAEKKTVSDRDRDRAITERAFDLASGPFDK--IVPDEGGAVLDKHQTRLVILDSTATREATTAAMG--GPPMVQWASSAVFACANTAAQARGLASEWIARTRVAEHPAK--DKDMHKQAQEEAREAKKRLDSMVKQCYKHIIYLA--PKDYERSVEFLRLRKSALNGSDVWEELREYSKVFGEFTRKALLHDNDYCPISEIRDGFWSNPHKPLLPTGAELRDAIFDAIAMGDIELVNPEG----------ARNLAADNIRLRP
TME11450.1_Chloroflexi_bacterium                       GVTRLAQDRAAQLDEEKVLEEVEKRLRVEQRGDFARVHVCPTSGADIADDESVRLVILKPSKAEAANEMMNLRGNTRRGYRNTVLFLAADRNEDLKDQGVRQFLAWDSINQESETNLDAFQRNQARTKRDEANKIVDARIPETYTWLIVPEQPDPRQPDELQEFKLQPLNSLAINASRRLKTDDLLITQYSGTLRR-GNHIGIKLAEYFARYVYLPRFKNIDVLIDSIGQGIQRETFAYADG--SRYFGLQRKQRQVINAETLLVKP
WP_247297741.1_Bradyrhizobium_sp._179                  NILKQIEERMSQVPRADALDRLKTDVQKSFQGALAKLLPWPPNAKAVPERPELQLALCE-EDIAKSVVTYSDAGGQVMRTYRNAIVAVAPDATEQAIQRVQRLKAAEEIEDETPAKLAREQLKKQKPELIKA---ARLQAARTFNRLVLAD--GSLLTIEFIVPPEGSLPSGQDAVRAFIEDRKLIYDSLAPDYFVDGTEVRSTASLQRRFLGAQGLKLVSDPSVIRASILRAVREGLVVRQEDG-AFDKLAVHQRRKMDETTHVAAT
HEC67877.1_Candidatus_Desulfofervidus_auxilii          NLNRVIIDREETLSEEEVNECLKNQLKKLI--EF-EIYLWPQETRDIPDTKHFKLALLPPRETEIFVKEIYEKAGIGFRIYKNTLFVIALDTSSVLKKSLKRYLALNQIQKIEMTLTPQRIAELKEKIKDTEKSLPFHILSAYRHLAFMSH-QGVLWYDMGLPVIGRMELIKRIKTFLQEEQKLLRHITPKYILQGEAEKGLVEVFEIFLKTPGLPLLENINVLKETIVNGAKTGIIGIRLE-----IYFKQTVEIPEETIILRP
MBR0288712.1_Selenomonadaceae_bacterium                TLRKLVDDKLEQFSDDDIDYEISQRLRTR-TPSWSAVHFCPKTSADVVDEQNARLVILAPNPALAFAKTILNTRGTVPRKYKNTLLFLAPDSDESLKKTVREFKAWSEVIDEADQNLDRVQLVDAKSNRKSTEKNFAMKISQAYCQLFAPHTFGDMTVLKWFFAEIDCEDNIS-VTAEKFSSNEMLLKSLGHEKLKN-RDTVNLNQLWEYFTTYYYMPRLTDKSVLLETVRKGVAEKTFALSDDLT----ELKFGD---ISTESFLVKS
MBM4285998.1_Deltaproteobacteria_bacterium             NLRRLHMERKAAINPEQVEKQIRAVVQQIFKAEI-NLSFFPQESTDIQDRPALNLVVLDPDELLTKLTAWTQYSGQSHRQFPGGIFWLVPASVRSVQEAVKDWLVWDSIKKDIDKLKEDEAMVPTELRKAQNMVEDQVWGLYTNLYLWNKDEMLKPVELRHMDKSEARSITGAVMARLRHAGELNQEVQPTYVERLTGAWPLASLAAFFQGRTLTLREKAHSLKQMIVKAVRQGSFGLIGSEEKFDRVWFKEELDFDHETFLLLP
NJK30457.1_Acaryochloris_sp._SU_5_25                   SVTRLAQDRAQQFEPNTVHAKIIEYLQSEARAEFAGLHVYTHEGDEIADEKEARLIVLGPSPARTIINTILEKRGNQPRLFKNMLVFLAPEQSSALEEATRLAIAWKSISEDT-ADLTRNQTTQANSKRKEAEAAVNRLIQNVWTWCLYPSQEPGTKDVEIEALKLTGTENLAAQVSKKLVDREDLLPEYGCDRLKL-RNHIQIQQLWEFFATHPYLPRLKDQYCLFQAIKDEFEIDGFAYAASATGKYLNLVTNRGSFVSLTGFIVKP
WP_048201423.1_Methanocaldococcus_bathoardescens       NLNKTVITKMTEIEMKQLEEEIKTLLKKYL--KL-LTYIYPKNSRDIPDTEEFKLVILP-TNDKDFVKEIIQNYGNNPRVNKNTLFFLCPMESHNFERFLEKLSAWESIAKDK-HVLTDQQKKEVKEKIEKLKKDEKDKLRYFYRIIYLPGKD--IKEIDLGIPTYGLKKTITEEIFDRLREEKEIVEKLAPILILNAKDYLPIQKYLALLTTPGEPRISKTNFI-KSIEEGVKNGYFGFGILKDGEVECIKIKATPKFEDYEVIIKK
MBF0121417.1_Desulfobacterales_bacterium               NLIKRFSDRKATIAENDIKELMNQEVNSVFTCSL-DCILFPTMTNQVPDRPTLSFFILSPAQTFLLIKDIFTKYGSNPRVYRNGIIGVCAENVQGIKEEARKQLAWISIEKDAIEQFEKEVVKNLKENIHRSKRELKEQVWKAYRHIFLIKKNGDVNHIDLGLLHSQSGNLIDFYLHRLIKQGEITEEVTPNFLVRALVQWTKQIRDAFFSSPSFPKLKNSSSIAFTISKGVANSLLGYSQLQSGKIIKIIGKMTYEEKVAILQA
MCS6856320.1_Sandaracinaceae_bacterium                 NLNRIIAERRERGVSSDAVEEEIRKCLEEESEKEF-GVILWPTDPQDVTDGPEPKLVLLSLTNSERFANSILEKSGSAFRIRKNAVLLLVAEENARVQGAKRYLAYRSIEEDKG-RLSEEQRKAVSRKLSDTKNEITHQLAAAYARLGKFKG----GVIEMKLEPMSTSTSIVDRVRNYLEKRDLMVRKLSPSYLLEQEQEKTVGSIYEIFASYTELPILKSKDVLLDAIAEGVQKGVFGVRLGER-----ILFKEVVNLDDEAVVIQK
MBT7692838.1_Gemmatimonadales_bacterium                NVTRVAADRAANYTDRDADDEVKARIATQTRGNFSAIQIFAEGPGDVPDDGVRLVILAPSPAIHLATELLSQRDGGPRLNRNLLVFTAAAANTELRAAARSLLAWKSIVDDNVADLTAHQKKQADSKVTETAQQVDSLIAETFTQVLTPTQEPGTADIKDWQTRATA-GDIGDRVSKKLATEEKLIAVYGGVRVKM-RHDIAVGELWATYARFLHMPVASSRGLSDAISDGTSQEFTYAEAGT-TWVGIHTSE---TSPGGFIVRP
MBU2110960.1_Actinomycetia_bacterium                   SLNRTVNERAANLHEDDVWAEVVTRLKQAGGADFATAIVAPTDASDVPEAEAVRLIYVHPSRAIRFARDVVANRGSASRERRNTLVFLAADEQAELESIVRQHLAWRSVVKDKDHDLTQQRVALASAKVDETNKVVAQRIGTSWIWALYPRDRGENEPFSISVVRADG-DRLGVRTGKKLVKEDVLREVIAPATIRHKQGRIRVGDLWDYYTKYPYLPRLRDKTVLIRAIEDVAMQTGFALAL---GDFKGLTVPEDPIVTDDTYLVAP
UGS34698.1_Solirubrobacterales_bacterium_0166_1        NLNRLIQQEIAEDTANVSRESVQIVRARLEDVLRGASRHVVLWPGDVPGGQVPDEEAFQVVYLAASPAQAALSRYVLESASGPRVARNALAFVPASGDDARAAARRALAVEALLSQGRSGLSGEQKEELRERLRVAEQELRAAALGQAYTRVHVPLGDDRFATRDLGTI-LSGGRSLHERVREALQT--YVADELFPTKIAASRELRWRVSEVAHALPRFFEAPKVWNQDALTTGIVKAVAQGALGYVARDGTGSVRLEDLDLEGGCALRV
RIK37466.1_Chloroflexi_bacterium                       NLTRLLARRKEEVTQADAGELVREVVQQEFQSSH-KPIYFPERSNDIPNAPALTFVVVAPPATLAMIDRYTREYGASARTFKNALVWAVADDSTPLYEAARKVRAWEEIDRDRDEQLSPDQKGQIEGHLLRAQGELRETVWRVYHLVVLLNKDNTLRQLDLGRHNSSAATSLTELVERELRKYDIVDGISPTFLVRAFKEWSVKAIRDAFFAAPQFPRITDDQVVREAVARGVTNGLIAVVSKTDGAVYSFYFQTPFPIDSDDVFVITA
MXV84967.1_Candidatus_Poribacteria_bacterium           NLRRIVQTMRNIDDKVVAGRNSTQLVRSFSYGFL-KTYIAPKGDTDIRDTDDLKLVILS-KRDDAFCQNLVENRGETPRIYRNTLFFIIPSAGNQLTKEVSVIAHEEIKKDN-SLSTTQKKEIDTALKQSDSALENAVRQDYRILVIPTK-EGFREEDLGLPAAGMSFTDEIVYEMLRVKSEILTSIGPRNISIKNDTVSTSQLFHSSLRSPGETRVLEVRAWI-NGIRQGVQDGLFAGEKAGGKLVPYFKEPLELDNEVIIQP
MBC7327984.1_bacterium                                 NLNRIFTMEKERVEKRELEEAESEILKRSLNEPL-RSYLYPENSSDIPDDRTLKLAVLK---DKSKAKEMWENCGARPRVYRNTLFFLCPYNSAGFEDYLRGLIAWRRIKSAE-NQLTPGQRGEVERRLREMEGQRYEQIRRLYRILLIPRQ-GGLEEKDLGVPIVGGRENLTGEVYETLRADERILERITPQFLFEKKEYLSTKDLLESFYRTPGEIILKNEEVLREAIREGVRKLFGLGKLEDSKPDCKFFGNDCSLGEGELIVSE
WP_013295752.1_Methanothermobacter_marburgensis         NLNAIVLRKKENIRDEDVRDEERRLLGEITKKIF-SVYIWPSSASDVPDDMKLKLVILE---DADDCTGFLEFKGEKPRVYKNTLIFLKADDTYQLVEHIKTKLAWMAVESDR-QLSTDEQRKEVREKIKSVSAEDRESIRNLYRMVFVPSR-SGLERIDLGMASYGADVKDVYEIKSRLIEEDKLLESLDPVIIERQEDYIETERILRSFHTTPGAMRIMSPSVLEEAIRKGVKRRLFGLVGLEDDPKCRHIGEDCSLPEIIVNP
NPV88406.1_Coprothermobacteraceae_bacterium            NLNKVMLDKMDGITEEEVKEFVRQLVESL--KF-ANYLWPERPADVPDDESLKLIILR-RKDEELMQLIRDNVAQVPRRFRNTLIFLCGKDDVVFTEKVRALAWKMVEADR-STLTQEQRKDVGEKVRSARDDARHYLRNVYRLVLLPGG-SGFEEVDLGIATYGDTSKIDEQVYNALRSQSKIVETLSPKLLFDGQNFVETAAIYRALLSAPGLRITSKRVLEEAIRKGVKEEIFGLGRKVGPSAECKHFGEDCEFDVEVIIQR
PWV37728.1_Desulfurococcaceae_archaeon                 NLNKILLVMMENVTEDEMRREERKILESIFGRF-KVYIWPRRSSDIPDDKKLKLVVMD-SDNEEAMGILKNHGKSLRTNPNTLIFLCPDSSNSFNQFLRSKIALEKILKDE-TKLTEQQRKEVEKRLSKLANDLYRAVRLVYVPSR-EGFKKIDLGVHTYGMPIEPDREIWDELKRQGKIEEEMPPEFLREGKDYVAVKDILESLYTTPGEVRIASEEVLVNCVRKGVAQGIFGYGVLKDNKPECLQIRDLELAENVILTKP
KEQ56705.1_Marine_Group_I_thaumarchaeote               NILKMKVDTMENIKEREIDEAQKSLIKNNVNQNF-RVFLWPNAPKEVENSNSLKLVIMK-ENDIRLIQSIYEYTIGESPRVFRNNILFLTPSDGGKFLNSLKSKIAWEKIKSDPESTLKEDQITSLNNELKKENERLNDLIKEYYSTLYIPEK-NGPSKVHLNIPLI--SKGIDDIIYDDLVDQDQIHPQLGALFLKSKQQYVETSNLFESMLSTPGERRPLNKNVVEQAIIDGVFKGEFGIGTLENGNPVPKLIKKQTRFDPGEIIIQA
MBC8384565.1_Candidatus_Cloacimonetes_bacterium        NLNHILNTKIENVPDNKVKEMEKDEIRKSLVGKF-KTYLYPRNSNDIVDDNQFKLVILS-QNDQEFIKHIIGNKGNNPRVYRNTLFFLVPMESHSFTNSVKKLIAYEKIKNDI-TNLSKQQIKEVDEKIKNLGNELKDLIRSYYRLVYIIQK-DGLKEIDLGIPTFGNVQKINDEIYDKLISNNEIVEKAVAPIYIFNTQDYVNIEKIYQDATLKTPGELRLQSKDVLVSSVTKGVENGIFGFGELEDEIKCLYFKKQFLSENIIILP
NET07313.1_Symploca_sp._SIO2B6                         NLNKLTADEESKITTDEVLDKIRDDLSKSLRG---KVVLWPKDSGAIIDRSQFSIVYLNPEVVESEAKVRLEQRGNDKRDYKNALAFVVPSKVDKARRAARTALAVASLIKQQAKKFTPENTEELKGKLKEATTGINAALGRLYRTLYIPLLPGTRELDLLHSQ-INTSHHLQQRVLEALKKH--VFESIRPAKLLQLTGYIKAEEKLVSYFFRFPTLPKMLDPSGIQTAIKAIEQGLCGYVPLSNNNPALIRFERIIKLENSGILSP
MCX7997938.1_Leptospiraceae_bacterium                  NLNKVILDKKENVSKKEILELEKKLLQKNLGKYF-KIYLWEKNSENIADTEDLKLIILQ-ESDLDVMQQIVSKKGNFTRIHCNTLFFLYPLESTNLQDTIKTYLAYKSLEEDT-TNLDDSQKKYIQAQQKTLKENLNQLIQNTYQMLVVPKE-NGLEKDNLGTPTFGDISKLDERVYQKLKGKN-IAENLSPVVIQDKKNQESIYQILQSFYRTPGEIRVLNREVFRKALQEGVKEIFGVARKEGETLKPLRANE---LLESEVLIPI
WP_244804062.1_Corynebacterium_sp._LD5P10              NITSEVRERAEALSPDDINEEISDRLRTLTHRDSVKVVCTPSNGDILDEEQLRLVVVPPGAGEWARSATNNVGLKPRTFKNTVVYLAADARDSLNDGVRQYLAWSSVQRDREKDLTQTSASQASTQTQKTQDTVDDRLMEAYQWVLEPVESSTHNYVDLDHQAIHGGNEIIRRAIDKLQTMSAFATQQAPVLIATKNGHYRFGDLWDMYPRYLYMQRVLKRSVVEDGLRHAGGGRPFAFADDVSRRYTRLRWDDGSELPDTTLLVVTP
WP_005520661.1_Corynebacterium_matruchotii             SVAKAARDHAEQLDPDTAWNEIVRRLKAAERGFFSHIHIAPDTAADIPDMDTVRLVIVHPSEVVKWIRAAVESKGAAQRTHRNTLVFLVADSDERLENTTRNYLGWKMVQDSAEQNLSKQQSNQADSWVKRLNDTINSNIRSTYMWMLYPEQIDPTRPFELAAEKTSDSDTLTERVFTKIKRDGQLITELAPTMLGMRVDDMTVGDLWGYFTQYAYMPRLASRTVLDDALRSVIDVEQFALATGKDAHYQGLILPSSAAITDNTLVVXW
MCL5447091.1_Actinobacteria_bacterium                  SVARTARERTERYVDGDVTEEILKRLRKDSRAEFTAVHLSPTSSGDVPDEDGVRLVILGPSAALRAALELSRGSSPRLYRNMIVFLAPSRQRDDLFQATAEYLAWSSIREDRTLNLNHDQSHVETKINQADEVVQSRLANTFQWLLVPSQPDPAGEVTIDAARVNGKGSLAARASAALVHGHYLNVEYNPYLLRL-GNHISVELWEMARYPYLPRLSDRKVLYSAVENTSTQEGFALADRFDQRYTGLTYGRSENVTGDTLVVQP
WP_052687173.1_Streptomyces_lydicus                    NITRLAKDEANRLRQEDVWAEIVRRLRSPLARADFKAVHAAPRGTRDIPDRDEARLVLLHPTLAMEFVAEALLTHGPSQRSHRNMLVFLAPDKRQELEEAVREFLGWRKVLADHLQDLSESQRRQAEARTAQAAEKVELRIGQSYHWALVPQQGEPGARVTWEDIKIDTGSDLAARTAERLKRKAQLHLQQAPNLIQQAEGHLSVGRLWELHTTYPYLNRLIGRSVLDAGVDDVLSLEGFALATGYEDRYQGLVLGGGARITDSTLLVRP
WP_053587850.1_Actinomyces_sp.                         SINRIVVDRAAALDAADVAEAGVEVLRAVAAPEFSAVDIAPTSTGDVADSRSVRLVILHPGPGMEFADELLRRRASAARLNANALIVLAPDAGEDADHALRLHLAWSQMARPSDALTQSQAAQARIADAERAAAERAVSAAWIWALHPDQPDPGGRPFVVGAMRVDGSERIAVRARGRKLQEDIVFTSAASATIALNEGRITAGELWGIFTRYPYMPRLRDERVFRAALASTMNDGGFALAAGYDGRFIDLRLPHDDAITDETVLVAP
WP_270528574.1_Mitsuokella_jalaludinii                 TLRKLVADQEQRIHEDAVDFEIQRRLHKWRSCHGLGLNILPTSSSDVPDDTKVALVLLPLNEAKKTAQEYLEQRGEAGRIHRNMLLFLAPDMQRLRQMTRHFLAWQMIRDEAEENLDQSQLREADNISTKQVEKDLQMKLSQCYCWLLAPYSDDDIRTIQWDVYEISCTDSNVTKAYERMKEEEIILEIKWAPRLLLMDKDHSLSGGLWEYLCNYCYLPRLKNEEVLYDAIRTDVSEGFFGLADTCIGRYEGLIFPEHGYIAPQQLLVKA
MCV7506294.1_Micrococcus_luteus                        SLNRTAADRARQLDLGLVWDEVVSRLRKSNTPEFGAVVVAPRSTADVPEDDSVRLVILHPSPATEFARELLTTGSSARAMPNTLIMLASDAGKDLESTVRSHLAWESIHEETADLTPTNEAQARQVETTNRTIDDQITSTWTWGLHAVQPDPRLPFDVGHVKCDGQMSLAERTGRRFVKEDLLRVESAAALVRMNSGHIRFGDLWDVHTRYPYMSRLRDRKVLAAAASLVTRDRGFALATGYDGLYSGLAVLEDLIDDTLLVRP
MCY3894210.1_Acidimicrobiaceae_bacterium               NVTRTAQDEIQSLERHEIQAEIVRRLREGASSPFVGLHIAPAGPADVADEPRARLVILEPSPALATAREILEQRGAGPRRYRNMLVFAAAEHREGLEQAAEYLAWDGICSRHEDNLDADQQKNLAGTLRERADQAAGLRLVEAYRYLLVPQQREPRGMVELEAVRIDPERSIAESSAGRRLLNDGTLAVQFPAVILRQADGHVPASVLWEDLARYVYLPRLRDQDVLLATVGAGPEGEGFGVASYDDRYVGLSTGQPAAIAPSTLVVRA
WP_238518868.1_Azospirillum_doebereinerae              TVNKLAEDRARTLPPEEVDAEIVRVLRDARRVFGRVHAAPDDPTDVDDTPELGLVILGPSAATRAAVEVVRLKGQTQRRKRNMLVFLAADVGEVARVATRKALAWRSVVKDVSTVLTGTQERDAEKKSSGGAAKAIRSAWIHAFYPTKGDPPFALEHAAVRPADPLPHVAVYDKLCMDNAVWKTIGGKNVRDDAPHLAVAEVWDYIYLPRLAGAIDLHDAINSALALPFAVARRTASGYCDIAPGVLYQEDSVLLRR
HBM97560.1_bacterium_UBP9_UBA11836                     TLQKLARERGSKIDSVDIQAEITSRLRKSQEEPLGGLHICPESSADVNDSSDVRLVILGLSELFEILDSSLLKTRGSGNRTNQNMLVFLAPDKGPTLKSAVVTYLAWKSIQEDKE-NLDKNQQKEVKQRIETYSDNIDSCLKETYRWLLVPSTKGEKQEITWHPYNL-GSGKLIEKVVRKLKDDELVITRWNPIFFREDSSVITVKQAWSQLCTYCYMQRLANYKVLEDCLKEGAASGCFIASSQDGYFKDLKFAEKIDISDNLLVPA
MBC6439887.1_Rhodospirillales_bacterium                SLNRIAADRAGQQEDAFILEEIDRALTTHVRGHFDAVQVAPSSSADVPDEGGVRAVLGVSQALIEARDILTKRGATPRVWRNTLVFLCADARDDLQRVARRVAWRNRAVDSIVKDVQRDLKQSDEATAKEKLLKEAGETITRRLRECWCYLLYPRQDDARAEVEWVARIKPGQDGLLSTASKLEKDELLITELGPTRLDRGKPLPKLKDWDHLNRYTWLPRLQHRSVLVKAVQAAVSQGFAYAGRFDGRYPGLIAAANAIDSSVVIRP
OZG52261.1_Pseudoscardovia_radai                       SINKTAQDIIRQLDPTSVYNTITEFMKANGKRLFERVCLAPKDTSEIPDVDEATLVILYPSETVRFMEDATKHRGNSRTNRNMLAFLAPDNTELAFTDTACSLAWKDICNRAKQNLQPTQVAQADEKLTMYRTQLGKRLAESYIWSFYLATDPKKEPVLQHSKVTAGDELAERTGARIQDRGQLDLGGIAPGILCAEKGSLSVGALWEVFTQYAYLPRLRNRGVLDFALQQMPQNHPYALATGYDGFTGLTMGEDQLITDSTLIVTG
MCF6470725.1_Nonomuraea_sp._MG754425                   SVNRLARDRAEEYHPEDVYVEIVRRLKERDKGLFGKV--FPLERGDIPETNEVALSIIHPSPARDFATQALEARGDTPRVNQNMLVFAAADREDYLKDATRDYLAWDSIVNDRD-PLVPDQAAMAKKRRKTADDTVWLRLTQAYNTVIYPIWDDKAGRIGLRSVTLDDEGVIPARVETKLRNLDQLRAVQGARLIRHEKGHISLGDLWQQYCRFPYLSRLRNRKVLLEEGIFRVLEEEGFAVAARYDDRYIELVIGTGEYLTDRSVTAA
MXX78159.1_Gemmatimonadales_bacterium                  TLNKLAADRARDVGDRADARIVEVLRRERQAGFPRVHAAPDNLTDIEDRRATALVVLPPSSAVVAAREALEERGSGQRSYRNALVFVAPDSDAARQNAKRERAWHSILKDPDRNLTQADVSLAEAKRSRALLARSIQRAWVHVLHPGSPDGAGYVVRSTLINRGGGIPGVVDWNKVSGDGTVIAEMGPQNVIPHPVIETIIIPPSVYQSEWATHDSAFVSYYMPRIRDDATLYRALKELEEDEPYVVPRAFDGYYDIDEEIEGALLVAE
MCI1646461.1_Olsenella_sp.                             TLEKLARDRASRQDAASIEARVVEILRQGERSGFANVNVCPQDGSEVPDEPQLTLVLVAPPSAAVAWAADCLTMRGGSMRARNMNLLFFLAAADSTDDARDAVRREAAWSSIRADRADSILTRHARDAHGKLQTATQRAQRLVGLAYKLHLVFPQQDGDASTIEWDAAPIPGNDAVSKRAYQRAKDDGLVVEVYTPFLLQRDRDCVAAPELWDDYCRYCYLPRLRDSSVLAESIANGATTDFFGYAQGDGRFGLHMGERPSDMRLGLVVAR
WP_094451589.1_Parenemella_sanctibonifatiensis         TVSQIVQERADAYHADQVFDELATVVRRETKGLFQRVSDLPVDEDDDVPGIGLVIFGPSNAEAAAREFLTKRGGANARIHKNSLVFLAPEVDDLLRSVRQAWAAEYILDKNDSNLDQHNIKATSRVAQAEQTVTDAIRLEAYKWLLIPHQEPGPDPKDIDLEAVLMRGDEKLDRLSLEVLRSAGQLQRTWSDVPLGLEQVSSQTLLPTAVPLEAVKVLADGIEVYSEPRESAEDGHVEHEVFRLMPALRVTPRHVAPLERPYQAI
MCG3151370.1_bacterium                                 TINRLAVERARYTDEEAISQELSNRVKLLFRGKWPKISVCPRDPSEVGDDAEARLVVLSPSPALASAKKILEERAGGPRRHKNVLFFQVPLKAEKLNEKAATYLAWHSIGDIANEVDGQQKKQAKDKLHEADATLLASLPLYLAHVLYPSMKEPGSPVVWGQKLSLSRDPAVESFRKLEQVSVLIDNLGATVLRMNLDHVEIRKLGDLFPVYLYLPKLSTPGVLERCIATGLEGETFAYAESYDGRYVGLVEGRLPYLDGAGLLVKA
WP_092673506.1_Siccationidurans_arizonensis            NVNRTAEERAGHIEKGDVEDYLKSLILDLLQGKLRRFYMMPASGADIPDQEQMKLIVLASSSQARKAEEVLQYRGSGSRMNRNTLVFMAPDKAQELHGAVKTYLGWESVKRDQLLNLDHFQKTQVEQVRVAAAKDAVKSQIPTVFACLLSPTQVRGSRDVEWIESRLNNGSLHKPLGLVNIEQGHVERLHLVVALERPRDNLPYLKRKCERRHSSGASDVRLLHLQDHQNLPYNRLLSRAVSSKHQNFSAEYGSKLRKEKERLIIKR
WP_201387653.1_Ktedonobacter_sp._SOSP1-52              SVLRLAQERAGQLEEYLVEEELEKRLRRVQRGLFARVHPTPASSADIADEAAVRLVILAKSPAIHAAQEILDKRGNASRHARNALIFLAAADRDIDALQQIRAYLAWNSIYQEQEVNLDAFQRQAVKSKLADAEKTVELRIPETYVWLLVPGKEGPHGAEFWEGIRLQPDDTLALAASTRSLKDGQLISAYAATSLRRGANHVSIETLKQDFANYLYLPRFKNTEVLLEALRSGLQSETFAYADRWDQRYTGVGLVKAGPGSNLLESGLLVKS
MBD2550060.1_Microcystis_elabens_FACHB-917             NLNRTAEDYRESYQREELLVEANRLLVQQARGAFAGVHSAQLDSSGIPDEPSTRLVILAASPALAEWAAQCLQSKGNSPRQYANTLVFLAPDQDEKLFQAIADVQARAWQKVIDEKDLQIYNQENQAAKIQEAITAIAEIPETWCHLLVPHQSPGNQGPSWDEFRLSAGSGLVKRASSKCQQEEALVQALGASRIRRNRDSVTEVQLWVAWCRKYLYPRVSTDDVILEGLTNAQAASTFYLAEGFDGSFLGLKPGQFSAATTAMILVKD
WP_188816795.1_Calditerricola_satsumensis              TVNRVAQERAGQWTDQDVEAEIIRRLREHVKGPFHAVHVAPYDQGEVADEPAVRLVVLAPEPNTPPTRALQLAARILEERGNMPRLYRNMLVFLATDAGDSLAQAVRLYLAWSVKEEDSEQNLDMHQQMHVKSIRHWDQTVNVRLEQAEYDWLLVPTQ-DGIGPVQWEALRLAGTDALARVRAGNKMEESGHLVLSFHPQLLAGDHRHLPIQEVWTCYARYLYLSRLRDASVLKAVEEGLASEHFAYAEKDGRFDGLVLGASRPSGERV
MCL1790896.1_Peptococcaceae_bacterium                  TLKKTATERARLQNADDVLYEIEKFLKSSCKEFFDSVHIAPSDSTDVPDQPQSRIVLLSPTPALDAAREYLEYRGSTPRMYRNMLVFLAPDRDEGSQLKQDMRALMAWCSIDRDADLDNSQKREKEARIQLETIVTDRVQETWQLLLVPIQ-HGTQPVVWDTYIVQGANPIA-KAAQKRLAADDLLTDVFSPKILAQDAESIRELWEDYTRYLYLYRLTGVSVITEALAEAGVRSDFYAYADGQGGRYLGLCFGDTGYTTLDGKVVS
MCY4445744.1_Rhodobacteraceae_bacterium                TVSRLAEDRAEEFKRENITYNRLNEQFRGIFRGLHIFPDSTGEVFDTEDLSLIILPTDPAIKSTWEFLQNKGNTPRTYLNTMVFLAPDKEDYLENDVRLYLAWKSISDEKEINLPPFQQKQVNKRMEDSLSSIKSKILDTFNILLVPTSVQEILLPTDNVEISSKKISVYGKITVEVIRRALNKNDQLVSEFLSPMLRKDGSHLRVTDLRQYEEYFLKYLYLPRLEKDEVLFQSIRYGSEDETFAYADSFDEKYSGLRTKN---YGKTGLLVVK
WP_127367955.1_unclassified_Mesorhizobium              TLNRLADDRAKALPDHEVDAAITEVLREDAKGGFHRVFAAPDDPVTIDEAHSLSLLVILNPSPASDAVADALMRCRASQRRLRNTLIFVAPDEADTAREVMRKAMAWASIDHDVERQTTRTQADDADKAKTNRESLAKIARAAWSHILYPVKSTPGKPFDLEHSLISSRDAIPHVVVDKTKADGIVLKELGTEKLWHDRLPHLSITEIAKWFAAYVYLPRLRDRLVLEQSVRIAAIKRPGFYAQFDGRYGHLRHLIWAKNPPASLTAMIVLE
MCZ0940408.1_Caldilineaceae_bacterium                  SLTKMAADRALSNSDDADIKLGERIQRTRKELGAVHVFPDGPGDVPDEDRVRLVLLSPSPAEKTAQDILEQRRGGPRINRNLLVFLAAEKAPELRQAIRSRMAWQSIDLDDQGANLTPADVQSAETRKREADRTVDLRIEETFSQVLYPVAPQGQSEIRWAAVRASSSNLAERVARKLESSEHLIPRYHGTRVRRGGTHIRIQLWSEYYCRYLYMPRLAGFGVLAAAISGGVASETFAYAESYDDRYDLGLQTGQHVDRSQEAVLVHP
WP_132287513.1_Paracraurococcus_sp._NE82               NVNRLAEERAAGYEDDDVNEKLVALLRDHVKGGFSEVYTCPEGTGGVGDETDVRLVIIGPRSASEAAIEAILTQRGTAGRVHRNNLVFLAADAASDLKTEVRSYLAWRSITCDEREANLDPFQSRTAAGRLKEAERTVDAQLPDCWQWLLAPEQPPTDPALVWAPLKLNGADPLPLALKKLGAEEKVITNYGRGKLRGSDFHVLTQTLWGYFARYPYLPRLRDASVLAEAMTKGLDLDWDYAAQVTGRYDVLGCGGRAEVNARSALVKP
CAG0999165.1_Planctomycetaceae_bacterium               TVARLAQDRAAQQKPDDIKAHINTCVDEAARGPFAAVQAVPSGHGEVPDERRARLVVLGASPAMAQAIEAILAHRGNSPRYEKNMIVFLAADTKPEMETAVRAHLAWKSIMRDENKLLTSQKNQADEKKTDTGKTIGLRLAQTWVWPLYPTQPAATGPVEIEQRLKGASSFVEDLGSKLEDADEMLFTTLGPKRLKMDADHLGIRKLWEELTKYLYLPRLRDQNVLMEAIRSGLAGDFFGYAGGFGYDGRYLGLKISGGAMWHGDDAQ
WP_151650193.1_Bradyrhizobium_betae                    SLNQEAEDRAKGLSGSDVEAELVSLIRLEEKGGFLRVHGAPDDPLSIDDAYEAALVLLPPTPAMTLAADIVEHKGPGQRRNRNRLAFLAADQSEDIQNVVRKKLAWGSIVRDARIQLPPAQEDDAKKKLAEQEAALNAVRRGWKHLLLPQEPPGSAGFDLEPVLTNGSDPLAQLAWKKCEDDGLIVARLG--VLDNNQQHVAVRQLRDWFAQFPYLSKLREPPVLAKAISRALARAKYGLADRFDGEYGLYGRLKLAQLVLSNSDVVLIRR
MDE2480824.1_bacterium                                 NLNRRAQDEASAYERYIHEEIVRRLQRFQRGDFVRVHTAPANSEPEERNARLVVLGPTEALKFSLECTLKIGEGNRVFKNALVFLAPDESEQLESINRQIALRQWAKIADQQQELTSPQQEMARQRAKKLGDGDAVRALNQCWCHALVPTQNTPGSPIELDEHKIGDSGPALATSEKLRDGLLNPTLGGTNLRSENSGIQIGRLFDMFAQYPYMPRLKNEGALRDGIQGGLVRDTFAYAENYDKRFVGLGVAGEDAADMTNGILVKSP
HDQ14361.1_Sediminispirochaeta_sp.                     TLRKLAEDRKAAFDASNIYEVLEKQLLKSPKSDFSRIHIAPHSASEVADQQSVGLVILSPSPAMLKAAALLDSRKSGPGRLNKNMIVFAAPDVDEAQDLEAAVRSWMAWGDILDHAEKEVTTGQLSSARSEKDEAYSQIQARIPETYRWLLIPEQADPKSPVTIEEYPIRGDKAIAQKAAEYMHRIEQLIPVMDGPRLKLKNNTVSLDALAEYFSRYIYLPRIASRETLTVHAVEDGVSSHTFALASGYDERFTDLHAGESIRSMEKVLILQP
MYD10935.1_Chloroflexi_bacterium                       TVNRIADRQDVDSYQVLDEAGARLRAQPRNFLGSVHFLPENSAEIPDEQRARVVVLGPSDALQFIGEIMTNRGNSPRHHRNTLVFIAPDETGEWASIRRYLAWRSIEADKESNLDVQQSKQVEQALKREAEEEAGGLLSHLVKRFEPDLAEVLEVFLDRLDRKPLVLLGVLSERLLGKQEYPRADRLLEDLYTLRKRNSGGTLIEQFGKALPAAPYLLEDSNQLRPDTEIPLASSPSKASRSMQSLLGKGHAD
WP_075830034.1_Deinococcus_marmoris                    SLNRRAADLAAHLPEDEVLHDIERRLQVHTRGDFAGVHVT-ADSGSVPDEPAVRLVILPPSPATTRAHEIVTRRGNADRLGRNNVVVLAVDKADELFGRIKQYLAWKSIHQDADANLTPSNKRQAESRMKEHDKGSDAQIGAAYTFLLTPYQTAGQGAILMDEERLAGDGLEIERVTKKLVNSGKLTRNFGALLLNMNEPHLSVKLLAYFAQYLYLPRLTSESVLLAALRSGVAAPYFGYADGFVGKYSNLKFGQDATADDLSVVSR
MYA05496.1_Caldilineaceae_bacterium                    SVAKIAQDRASGYTEEVIATRMLEHLEHQVRGDFDGVHVSPQHSADVPDSDSIRLVVLGSSEAHRVCRDYLLNAGSQRRFRNMLVFVAPDEAASLNDAVCNLLAWESSVVTEKETNTLDLQRNSAALQQRRAKAADSVETRTLIPTEVWRRVLPEKAVDADGVTGSHTKFSLKDAGGLLSKTLADRVSQRLIQEQELLTTRFGPKRMLMDQKHVSAKQVWEDLCSFLYLPRLRDRKVFLDTIRAGAAAVRNRETFAFAKGLDGGYTGLSVDGVGAQDDQSLVLQ
WP_106140933.1_Spirosoma_oryzae                        NLNILVNHARQEVSQEQVKQEVLKRLNE--RRQF-HPLVDP--ADDVPEQQRPTLIILGPRKAQERIERLATKRGNNERLYRNTLLFLLPLPAQGQLNDSVQTYLAGERVKGDFGSQLDADQRTDIKKRIDEASSQIDKALAATYSVLAKYSAKSGITKIEATQYRDRVDAQINDVLIPMLKDGSCLLDKVSYNTLAKPGHPVQASAIYEAFLRYDDKDMVSGVGALQESLQRYCTNGEYAIGAGDGKFRRIFYQETVPQDPTYWLVDK
WP_118201547.1_Prevotella_copri                        NVNILLQQAKAEIKDAEVAEIKRLQIALSAQKL-NVLVAP--TGDVPEQTKSLLVIMSPSSLEHAINQIARKKGNSDRIYRNTIFYLVCSEAAELNNQYCDLLACNKILEEYAGTLEADQKQDIQSRKKEYDRAVSQSDLAIRAYNIVASKSVKNGVEYVKMAYAMDFSSQVRQNMMGELEQEEWIKLSIGAKYLGIERPVQLYETFLRFDDKPMIASRDAINTVNKYCQEGLFNVGVGIKGKTIKYNHEEVPSWEDLWLLDT
WP_051018640.1_Saprospira_grandis                      NINILINQFQAAIKKEAIEQELMERLRLLSRGLF-RLIVDP--QEALPEQNKFTLVIMPPKAAEEKLRQIVGKKGQADRIYRNTLLFLFANPSGTVESAIALYLASKQTEREYK-NLDKGQEDDLRRRKQEANDKIERNLPGLYNYLVVKYSMKDGAKFLLLQEDAFSLKQLLEHKIIDRLKEEQWLLDSLGRSLLSKEGNAISVQMIYEAFLRYDDKPLITGPEAISQAITQFLHRSYCALAEGASPKVLEQFYEDSLPEEEQFYLIDL
RKU15614.1_Candidatus_Poribacteria_bacterium           NLNKVAIDRADEFSDREINEHILEIGEAVRTHRSKVITCPEDLEAVPDSDELRLVILPPDEATPAALHILTHCGTAGRTHKNTLLFLTAKTDRDMKEYLRMKTRDFLAWKSITQEGERRNLTGERSNQAQANLRKADESVNRMIPKAYRWALSTAPHVSMSISEQTRVLEDIVESAFDTFKSREVIEQYEQESLEADKDHISVGDLWNLMTQYVYMPRLINKEVLTAAVKEGVEDGTFGYAEVYDQAYRGLLFHAFQIVPQLNNLGGLIVKS
MCY4474618.1_Chloroflexi_bacterium                     NLNKVHADRVDATSNQVADRFIINELNQAIRGNSSVIVCPQSSADIAETDSVRFIITEPDTATETFIKDVIVNRGDTSRIKRNTVVFIITAKADRALRNTTRSHLAWDSMLNGEDKNSLSGRSMESVSANIRNTRSQLHNDLTRAYRWVVAPSRQPQAEYTLNPSMTQATEEIVRSADEKLVADEKLVDTIAPSQLARGNYHITIDSLWDLFTSHVYLPRLRNASVLEDAIKRGTANGDFGYADQYADNYENLRFKEPIDRDRARIIVEP
MDE0020779.1_Candidatus_Poribacteria_bacterium         NLVRVAQDRIDKLTREQLDAHIVQKLNEAT---HPGVVVCPDETFMTDDQNRVTLAVLSPDEATEAALKILTTRGGSPRSFRNNMIFVLAAKNDRLRRWAGQYLAWSSIVNGNEKGLKDDRLTEAKARLDKNKKTEDALIRAYQHALYPRQPNTNAKYEFKQVQTDAGDIVEAAFNKLIEEDAVADNMSAASLERTKDHIDIDKLWEFLAQFVYMPRLESKSVLANAVAASVEGLLGSASKYDGRYENLQYESKMSNDLRGLIVRK
MCA9831553.1_Dehalococcoidia_bacterium                 NLYRRIAETAQTQSPHQVTEKLHQVMTAASGEF-RVLQWAGEDGAIPDAPEPAVAILAPGQERAAIEGLWNKSGGGPRIYRNALIQVAPDREDRAEQATREVMAYETVLSGHSELSEGEKKDLQARKRDKEQSLSRTSLVTAYRWVFYPGDEGKLESIAL--PTATANETIARRVVDRLESHDKILRAISAMFFQAAGTELEVGELVRRFSEWTFLPILPDRGTIRACISDGIDKGAWALAYGIPGQARRLI-EKPGELDGASFLVEP
```

#### BR-system DUF499-ATPase Alpha+Beta-Domain + wHTH Entropy Plot

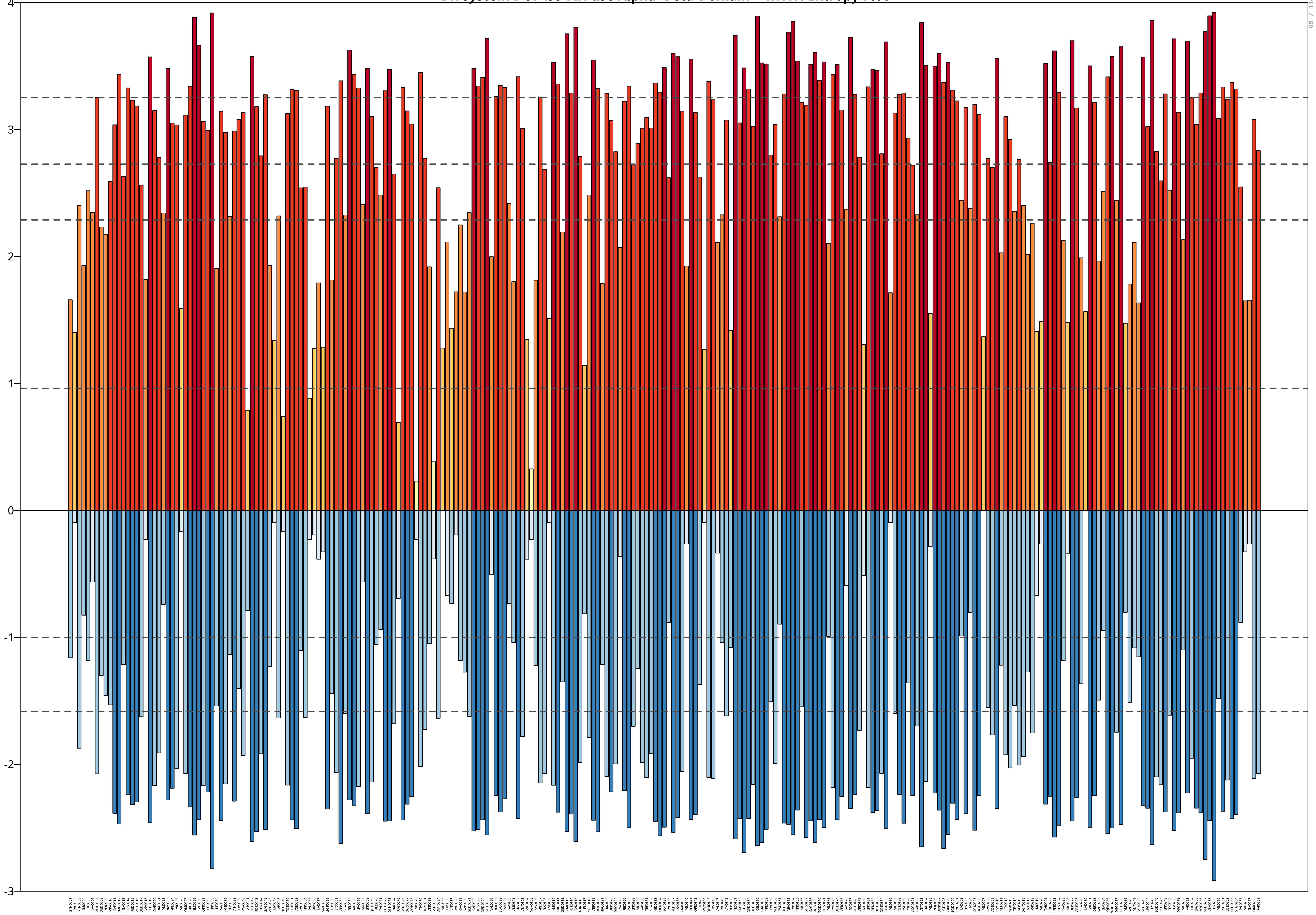

```
# ; Type_1 BREX PglZ NTD iSwi2/SNF2-Helicase

WP_238754868.1_Leptospira_bandrabouensis          MLYNAITSYIDLKINRECLVSWIDPTGEFSGYIDQLEKKKYHLVRFKGSFLESILALHQDVKKPDCLVYIPFISEKTPFLEVLKSSEQVPS
MBF0258418.1_Desulfamplus_sp.                     MIAEHIKRKLEQDLRTKGLLVWLDKENQFTPLVDTWILFHYDIFAFRGSFLELMVKSQSGRDIPKCVIHMPGFNKETPVLEAYKAGQRWRV
WP_009097832.1_Rhodopirellula_sp._SWK7            MVSKALESELRALIGQHDIVIWLDAGSIYTDFVDSLAKLPYAVHAFRGSHLQLLFDLRSSPSRHPMVIHMPGFNKQTPILELYLAGRSFEK
MCC6649976.1_Candidatus_Eisenbacteria_bacterium   MASRALAAEVAKKVRERGLVVWVDAERKFDAFVDAIRGFSYPVVAYRGSYLELMLALKNGLYPEHVLIHLPGINRETPVFELFKAGTVFEK
MBD2549130.1_Microcystis_elabens_FACHB-917        MLQARFCQELSRQLSLHRVVLLFDPAEQLRSLLDALAELQAQWVVAGTSLFALRHQLECADLPDPLLVYLPQRQGRATLLELIRAGTVFEA
WP_096450969.1_Thauera_sp._K11                    MLHQQIASTLDRLLRERRIVVFYDLREEFQPFLDELEDTLACLARFDGSFFALKAAVEAAPLPEPLLVYVPGYAVWNVLFELDCAGKSYEP
MBY3465673.1_Rhizobium_laguerreae                 MLHDYIAARIGDRVKDRRVVVIYDERKEYTAFFDEILSRKASVVRFEGSFLELRYAVEGGDNIEDVVIYLPGVREGSLLLEIEKLGTHYAP
MBY0113210.1_Phycisphaerales_bacterium            MFHDYLASRLAEKVAKRQVAVWYDPREEFRGFIAELRGRDVRLICKNGSLFETKLVAEDGETIEPLLIYLPGERTLNALLELEKAGTPVEW
WP_273378697.1_Symbiobacterium_thermophilum       MVSQYVAQLVRRKLDQHLVLVWYDPTRVFAELVDHLDPPGTRLARFAGSYFELRAEAEARMTAAPLLVYVPAAPQIDPLLGLAAAGGVLKE
WP_209466794.1_Symbiobacterium_terraclitae        MVSQWVVDHVSRVLKEESLVVWFDPERYYETLLPYLEEPNRVVIPYRGSYYEQRRAAENNLSGKQLLLYVPAKQDASPLLEFVLAGRALEV
MBW2308572.1_Deltaproteobacteria_bacterium        MILDELRRILEKQLDNAGIVVWYDPEGIYETVAHRLPIEGATFITFDGSYFKVRREIEISVNRARVLVYVKSPRKLPPLIELEAAGVVCET
WP_083610988.1_Desulfatibacillum_alkenivorans     MLSDRIRSMIARQVKDRGIVVWYDPEKAYAGLIQNLDLPETAVLQYADSFFRLRHELEEFVNAPNVVVYVPMERTSFALVEAETAGVVVES
MBN1165673.1_Methanospirillaceae_archaeon         MATEYLKTQITSSLKNHHLVVLFDPEKHYSDIIKNLDLNGIPIHTYTGSFIELRKEIATGEKPPDLLLYIPMAESNNALIGFTKPGVTLQT
MCI0601819.1_bacterium                            MIADHLQEMLARQVAEQGIVLWFDPDQTYSSLIETLKIPSCKIARYNTSYYDLRAQIERGPHRPHMLVYVPLHRTKQALAELAALGAVAQT
WP_062190142.1_Anaerolinea_thermolimosa           MVSQYLIQTLRQQIQEHHIVVWFDPEKQYQALVRAENFPNVQLFSYQKGFMALRRELEGTFEPPQLLIYVPTSESNHALVEYTTAGVTVQS
WP_156746889.1_Mycobacterium_sp._E2733            MLRDQLSAELAKAVRQAGLVVWQDEHREYASSARSMCPESVQVVAYDGSWYGLRWEVEAGDRAPELVVYVPSAAPEDPLAEIRSAGKEFKR
MBN2413376.1_candidate_division_KSB1_bacterium    MVNEFISAYLLDHIREYGIVLWYDPEQLYTEMLNDLTLPNVPVFIYEGSFLKLRYNIEAGEKKPNMLIYINLARTEYAMIEAESAGCYLMT
MBV6422080.1_Ignavibacteriaceae_bacterium         MLKTYLQDLISTQLDEKRIVLWYDEKGFFKNYIGDFVDSD--IIHFNESYFELRDLLKENPEESKILIYLITKRKNLPLNEIESIGFIFEL
WP_083423498.1_Stigmatella_aurantiaca             MVRDYVVQQVMERLRDRRILVWYDPEGDFYDLFREFPMPRLIRADARHSVLEARRAADRKTFPDPVLIYVPKVRRTEPFEAFALAGAAFGE
WP_218933729.1_Rubripirellula_lacrimiformis       MVSTFIHRRLVDLLEEQRVVVWYDAEQSFTELVAKLTVPNCIVVDTSLSTLQSRRESDAGINHKHLLVYCPRGRYEDPFEVFAVIGAAFGD
WP_055669757.1_Desnuesiella_massiliensis          MITEFLNNSLLEKLDKYRVVVWYDGKHEFKDFIDKFTAPNCVVINATKSIIIARREAEKKMNQKNMIIYLPFFPLDNQFELFSRVGTVFGE
WP_242609850.1_Comamonas_thiooxydans              MIAEFIRESLRPRLKQSGCLVVYDADRRYREQCFDLAADKVRVVDASESSIESREAALREVGLEGVLIYVPAKRQTDPFALYAECGAVFPD
WP_243520949.1_Hymenobacter_monticola             MIRAFIQNQIQQKLTKRRCVVVYDPQHLYAELVAELAAPQAVVVAALPGPTQALLSAWASLGGPPLVVYVATAARADPFWALAVAGCAFGE
WP_235164441.1_Dyadobacter_chenhuakuii            MINQYIADYLRQKLVDSPTLLVYDPNGRYHDLVQTLASGLVHVVDASKSFIKAREEAQTELPQDRLVLYLPLKSIQDPYFMFGLGGAIFPD
MCP5535438.1_Akkermansiaceae_bacterium            MFLKFIANHWQQRLSSNRSIVIYDGSGALRGALALLDLKELHVVDTSESSMRSRLEAWLAVGQTRLLVYVPQNLLADPYQPFRLAGCHFPD
MCC6145260.1_Candidatus_Hydrogenedentes_bacterium MVLDALVSRLRALEAAPVAILWMDEKREWEGVLPRLKEAMPELFTLTGPGVWLRMAADGFAPGETAILYLPGVGRTDPIAELQFRGCFWRQ
MBN9589012.1_Alphaproteobacteria_bacterium        MLIHGLARALIAGEGWPAALLWPDPERQWHFAFADLRRRLASLYALIGPAIWLRCLIEAPDDAKHSVILLPGISWRDMLVDLQYRGDVFRQ
MBW6486418.1_Syntrophobacterales_bacterium        MILESILKYLRHDLAPPRVILWPDEEGLWKRCIEPLRANYPALWSLTGPAVWLRYQLEVHEGEDIPVIYLPGIGRYAHLFALQFQGQFWTQ
WP_004078483.1_Methanoplanus_limicola             MFIEEVKKAILERLSAPVALIWTDHNREWEPVVEKLVKDM-PIATFKGPAIYIRCLISHNLKEGTPVIYLPGYSKDVQLFELQYRGIVWSQ
WP_011448105.1_Methanospirillum_hungatei          MLDEVICASLQKDVVAPLSVFWFDKTGEWQPVIDRLKKRF-TIYTYTGPAIYIRTSVMHHESRNIPILYLPGYSREEPLMGLYHRSVSWTS
WP_030150652.1_Oerskovia_turbata                  MVRAVVAESLRDAQERPVAVLWPDPDRTWEPAIRSLQEAV-PILVLQGPALWIRAVLAAPETANPWVLHLPGIHRHEPLADIALRSTWWGP
WP_248598775.1_Agromyces_sp._C10                  MFAESVHQSILADEIPPAAILWPDPIQAWKPVVSHLRRAGAAILTLRGPAIWIRSQLAAASDDMPRIVYLPGKDRATPIAEVQFRSNWWAE
MBA4013264.1_Phenylobacterium_sp.                 MPLDALIEALEDVQARPEAVVWCDPDASFGPVMPMLRARLLRLHSLTAPAFALRLVTERAEGEPPPILWLPNVGREAPLAWLAVAGAFFGH
NBT25026.1_Actinomycetia_bacterium                MLIQAMVAAWEDGVVPPVAVLWTDHAGEWNKLWPTVLDAFPAL-RLTGPAIWIRAVLAGALPGQCPVVYLPGIERNQPLVELLYRGTGWWH
WP_246481782.1_Natronogracilivirga_saccharolytica MLKQTLINSIRGNLTAPVVVLWPDPEKQWSSVIAILQEDMPELMVLTGPAIWLKCMVASTLASLVPVIYLPGISKNDPLMEYQYTGTLWTH
WP_091825284.1_Butyrivibrio_sp._ob235             MIIDILIRKIRSAEVKPAVILWPDAERRFEYAIKVLQTEMNELFVLTGPAIWLRCAMTGTVKDRVPVIYMPGVGREKPLIPLQYLGRFFSQ
WP_219938192.1_Succinivibrio_faecicola            MASSRITEALLNAFNKYRIVFWDDDKEKLIDEFNSLELPDVTKIKLDNNEFSVKYRILVEETDKKFLIYRPAGEPDVWLLDVQLASYKFVA
MCK6605714.1_Ignavibacteriaceae_bacterium         M---DLRPVLAEYFQKTDVVFWYDKEGEYNIEINTLLPEGVTLLRLDNNHFGIKYRIYNSAAGSRFLIYSCNEKPLPPLLGLLLTGREFLP
WP_251178637.1_Adlercreutzia_agrestimuris         MTAIEIAGRLAAYFGKHRLVFWEDEDGGYSEILDNLSVDGATIVDVTGRELSSKRRILRDERDGKFVIYRQGALPAPFLYDLKLAALPFNV
WP_149431761.1_Rhodococcus_cavernicola            MSAERIAELLADRFATTPILTWRDVEGEHADALDAVAAAELTTHRVSGNEFGIKHQLYAQHEWARHLLYRTGEPPQDWLLDIEAGYGLFTA
MBX3116434.1_Cryobacterium_sp.                    MGLGVLKKSLTDRFGDRHIAFWRDQEGEYEEDLDQLELGDVKVIRVDGDEFAVKHRLLKEEPEQAFVVYRKGDVPEGWLWDLELAYGLFKA
HJG42481.1_Corynebacterium_phoceense              MSLESVLPHLEEKFAAQRIVFWHDDEAEFSAEIDSLAIPGVTVVKVANNELGIKDKLITSADDEKFLVYRAGTVPKGWLLDQELAYGNFSA
MBU6197218.1_Cyanobacteria_bacterium_REEB446      MILRQIKQSLGKLLDLYDVFFWYDTKAEFRREYEKLNLENLIKYELNNNEFNLRYEIHAEKTEAKFLIYSNSARLADCLLDLLLSNHEFIA
QNI65417.1_Synechococcus_sp._A15-44               MPTSRIHESLDAALARKRVVLWYDPNGEWASEFDDYQSAAAEKLRVEGNEFSVKVAISRAPLEQRFLLYLPCAKPPEWLLDLLLAGHEFTA
WP_147932466.1_Neolewinella_aurantiaca            M--STLNQRLHKLFDDHRIVLWYNDSSEMTAEFEAMELAGVTKLNVQNQALKTKFSVQVTDDKDKFLLFSPGKVPSDWLLDLQLAYPLFST
WP_153112503.1_Prevotella_copri                   MTNEKIKQVLTSMVEQNRYVFWYDEGGQMKDIATSLDVPGISVLEMEDNAFSIKFQMQKKQPETGFVIYSAKPMPKDWLLDYQEEGALFSA
MBK9103302.1_Saprospiraceae_bacterium             MSSDKVKLTLEKIFQQHRLIFWYDEQAEMKDLFDDLSFENIGKLVIDKNEFGLKYHVLIEEPTKSFLIYKAGEKPEDWLLDLELSNYEFHT
MA064830.1_Balneola_sp.                           MTQNKLEKNLKGLFEKHRIIFWYDSEQEFSEAFSSLNLDAVEKVNCDGNPFFVKHLVSKKQPEQKFLLYFPYEQPKDWLLDLQLAHKVYHS
WP_262567766.1_Endozoicomonas_gorgoniicola        MDTGKVLEDIQRRFSSHRLVFWHDTDRVYADELDELMLPDVALIRLDNEMLGVKTRIEIDEPDQPFLLFSEKAKPEPWLLDVRLYSDEVAV
MBQ2845006.1_Alphaproteobacteria_bacterium        MNETQIKQALQKYFEDNRIVLWYDDNAEFSELVAEI---DVKVINLKEIALQARVMFDVDNPTEKFLLYAPFPKPEYWFLDVSKYAPIFKT
WP_156790427.1_Alcanivorax_sp._DG881              MSIERIEKGLAKLLAGNRIVWWSDPDGEFADAVADLSLEEADVVSLGDTALAVKRQIELESPASPFVLYEGQEAPDPWLLDIRLYAEPFAA
TXI44184.1_Methylophilus_sp.                      MLNQKIIDSIQSQLTKERFLFWYDTEKEYLECISSLALEGVRLVMTDNSALAIKLEIAQSDASTRFLFYSNNPEPERWLLSYRLKGKSFSA
WP_006086471.1_Shewanella_baltica                 MQISELAQGVAAKLEQSRIVFWYDPEQSFTEELEHLAPNDLTILNMDNEVLAVKKRIEVDEREAKFLIYFPSAEPEPWLLDVRLYSEQFFA
WP_151792240.1_Acinetobacter_seifertii           LDIDQLLNNILLLSQDSRILFWYDAGKQFEQALEQLSLESIELINLSGRTFEVKKILERDKPQQNFLIYAPFAEPEYPLLDIQKYSKQFHA
MBP7846546.1_Burkholderiales_bacterium            MDIDKIIKQLQELCKNNRIIFWQD-DGYFEKILAEIELSDVTIINTSLVDLEIKVRVELEQINDKFIIYRTTE-PLVWLLNIKCYAKAFSA
consensus/100%                                    h....h..h.........h.h.s.......h..................b...........lh.........h........h..
consensus/95%                                     M....l...l.........hhh.D....h...h..h.............h..b............lla.........h..h......h..
consensus/90%                                     M....l...l........hlhh.D....h...h..h.......h.h....h..+..h..........lla.s.......hh.h....s..h..
consensus/85%                                     M..p.l...l...h....hlhW.D.p..a...h..l......hh.hp.s.h.h+..h..........llY.s.......hh.h...u..h..
consensus/80%                                     M..p.l.p.l...h....hlhW.D.p..a..hh.pl......lh.hpss.h.h+..h.........hllY.s.......lh.h...u..a..
consensus/75%                                     M..p.l.p.l.p.h.p..hlhW.D.p.bap.hh.pl....h.lh.hpss.h.l+..h.........hllY.s.......lh.h...u..a..
consensus/70%                                     M..p.l.p.l.pbh.p..llhWbDsp.bap.hlppL....h.lhphpss.h.l+..lb.......hllYhP....ps.Lh-lbb.u..a..
```

### Type 1 BREX PglZ NTD iSwi2/SNF2-Helicase

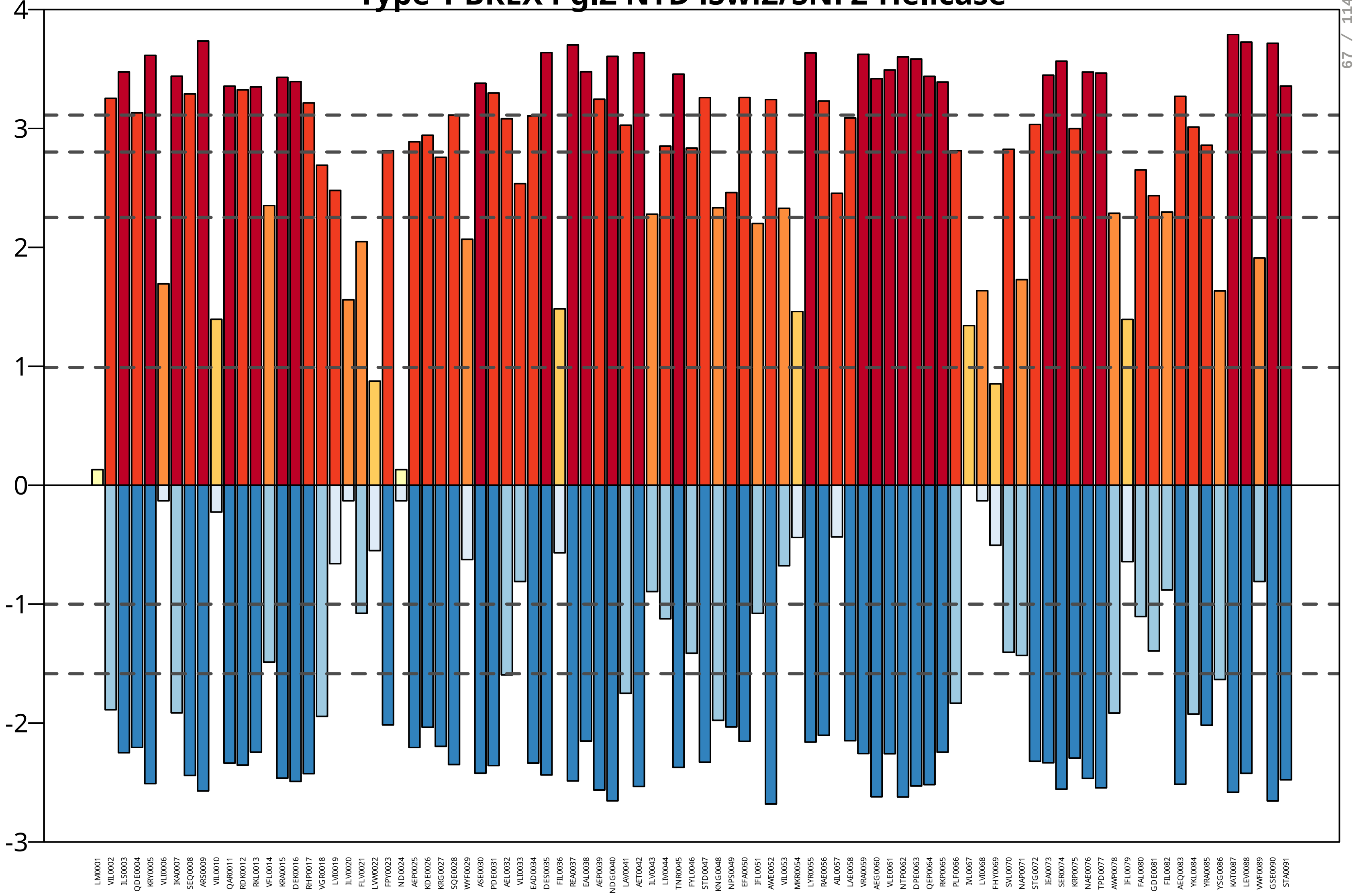

```
# ; Type_2 BREX PglZ NTD iSwi2/SNF2-Helicase

QBI18515.1_Egibacter_rhizosphaerae                MGEAVVDESQVRAIAAQIGERRRMVAVAGAPTWEGPERLETDAGTVRVLAAPSTLAVRAALVAHRAAELLVVVTDRDERELGEEVVAELWDHRVVRP
WP_188941324.1_Nakamurella_endophytica            MSPTTVDRSTVIAQIRLMRRKRRVLAILATPVWSGTDVLEADGSTYRVWPCPSRLAVRDALRHRHETPAPVILTDLEAAELGDGICDHLVDHRPLSP
WP_332348679.1_Accumulibacter_sp.                 MNVLLPSVRQIVAQVDAVLRHDRVIAIRAQHDESWPSTLAVGEREFSVAWCPSPIAVREQLLALADGPGLVVLTPLPDHVLGADVLARFSRGKVFSV
MBT3378822.1_Lentisphaerota_bacterium             MAG--IRTSQVKALIKSVRNKRDFIGISSPFLWLEDESVEVDGETYDIVQCCSELHIREALAAATRKRSTVVITSLDDPDLGEDLRLRFHKRSLRKL
MCB0954592.1_Ilumatobacter_sp.                    MPA--ADAQQLLALVNSVRRQEDAIAVRVDPVWAGPDTLD-GPVPIRIVPCPSPLAVREALAEHERTQVVMLLTGCGDSELGPDVVARLAKGRVVSL
MDJ0841534.1_Acidobacteriota_bacterium            MN---TRARAVKVLVRDIAKKSPPLAVAWDGVW--PEAERDDGEVLLFRRAVGPLAVRRVLAEPREAEMSVILTPTAEKDLGADVITRLWGRRLFQV
WP_005004490.1_Nitrococcus_mobilis                MS---LTPTQLATHAAKALQQATTLAYHWPQPWNGGDSIVVDGQTLPLRHCVSPLEIREALAEASD-VSRVLLVSQQETGLGQDVLARVFRHRLLHV
MSW49534.1_Actinomycetia_bacterium                MADGPVSTSHLKRHLERWLDGPPRFLIHGEWPAERPTELVAFGHTVDVVSCPSELSVREELVRDRPDRPLVMVTPID--RVGSDVIARLTKRRMISL
MBO9539708.1_bacterium                            MSA--LSGPQLSALIASLRAKRAVIGVHAPDGWQGASELRVGNETWTVAVCETPLAVSVALSERPEASPMAVLTRLTEADLTQDILVRFPRRELVRL
MCX6543497.1_Acidobacteriota_bacterium            MTPTLLSPAHVRAEALDLRKRVTVVGIRYPGQWSGPSAFETEHGTVSVLCSQSPLEIRERLVDHERTVPLVILTSLSEQDLGDDVTARLAGRRLRQP
MCG5052491.1_Myxococcales_bacterium               MKA--LTRQDLAVELQRFHKPPENTAFYGFGERDRVSVIGADSHDFEIVPVKCELELRKELGRIEREAQAAFLVDFTE-RLPLDLGGRLASGDLQWV
MBF0261196.1_Magnetococcales_bacterium            ME---LTESQIVAQVEAILKKDPAIGIRSMGPWRGGETIVAGDRQVRVRHCATPLAFRERLVAVKESEVSVLLTPLDTNSLGQDVLARIWRQQLIAI
TAF23832.1_Verrucomicrobiota_bacterium            MTETMLIPEQLRAQIDRWRDRRGVFAVRTDFDYAGPAQITCDGLAVTVADCRSDLEARDLLLGLNSQHGLLMLMRTDEDRIGEDVRARFAGKTVHTL
MCE9632142.1_Planctomycetia_bacterium             MAAA-LSFNQIKAEVTALGRKKQPIGIRAPSRWIGEAVHTDGEAGYLIRQCDSPLAMRLALREAPPPVVKVIVTNLPDAEISRDIFARLHRQRLFSI
WP_330928032.1_Synoicihabitans_lomoniglobus       MTT--VTAGQIRAQVAGLRKKRNAFALRVNGTWTGPDHMEIDGTAHLIRSCASDLQAREALHDATEKSPIILLCSIASDRLGEDVIARFARNRVLAP
MCP5560213.1_Verrucomicrobiaceae_bacterium        M----LTPPILSAQVKAARAKTRPLAFRSYDTWSGDETLAFEDKHLTVKECRSDLETREAMGLGAP-DELVLLVRASDGLLAEDTLSRCAKGRLLDL
WP_287381673.1_Mesorhizobium_sp.                  MEAPGPELADIERALDSLLLKDEILAMRSPIRRAWPEAVERHGRRFCIAWCPSELEVRERLDEVERNEGLVVVTPLDAATLGDDIVARFPRARLDQT
MCI0393077.1_Acidobacteriota_bacterium            MSQSLSLPPQIRAHVELLHKKHGVFGIRMSVPAPVSERSEVNGRECFFNWCDSVLQIRELLFEREKDALQVVLTNLSEADLGQDLLARFAKKRFFTD
MBK6942982.1_Planctomycetota_bacterium            MTSVVVGEAELKAHLQQFLRKTEVIALRTASTWNGPSSIEVDGVTFDVVACSSTLAVRESMAKFEDTHRLVVLTDVADRELGDDVLAHVAKGRGVPP
MBI4662621.1_Verrucomicrobiota_bacterium          M----LTLKQVRAQVEAIRRRSPAIGLVSAVRWTGENELKVDGERYDLRAAETELEMRTVLVEAKRKSRVVVLTNLDPSAIGKDLLARFARRRFFSI
MBL8842559.1_Planctomycetota_bacterium            MSELALTTATLRIAVNDVLDRRRVVAVQSSTPWRGGSEIDLGESRARVICCASPLEVREALTESEAALPLVVLTPADERQLGQDVLARLARAQLHPL
MBK6462401.1_Myxococcales_bacterium               MEAAGLSQADVRLEVERLFRRDHLLALFGRGQ---PGRFELDGLKWDIIPTACELSLRSRLPAPGEPTGRVYLVDWAEDALPLDVSCRLAGGRIYHV
MCZ2341280.1_Bacteroidales_bacterium              MSIGTLSSQQLRSLLEDKWQRDAVIGLHVAQPWKTPSVVDFEAGKAQVVRADTVFQVREVLGDAERNERIILLTKLRQGDLGNDVVGRLARCRLFAI
WP_075083673.1_Mariniblastus_fucicola             MSD--LKRSHLIAGVSSL---KKVVGVRSSISWSGDIESEVNGIRYCIRQCDSPLAVREFIRSAVDGAINIAVTNMDEETIGMDAMLRMASTRLLTI
MBZ5659873.1_Acidobacteriia_bacterium             MANSEMKESVLLSQIESIRSKREAIGVHSQSRWQGRDKILVAQETWRVAQCDSVLELRQRLSEQID-TPLVLITPLPTTEIGDDVRARLFKQQLLPV
WP_238634086.1_unclassified_Halorhodospira        MTNPTATPRTIEARVTGLVSRGEVVAIGMSGHWSAGETVEINSTTWPVRQCDSVLSMGLALTELETQGRGVVLTSLSESDLPEDMLARLQGRRLHRL
MBL8678386.1_Myxococcales_bacterium               MATNFVSRDALVEELKALHKRWHDTLLYARLERELPSIVDTNEGPFVVRTAECELELRAALVDGVREDPVVVLVDFDLPRVPADLLGRVATGKVLRI
MCB9660516.1_Sandaracinaceae_bacterium            MSAAAVSRQDLAEDLKRLYESPRLFAYYGTGS---EDAVDLNAARVTVVPVRSELELRMKLPLLDDDARVAFLVPFK-GEVPLDLQARFALGRVRRI
WP_154827459.1_Dietzia_kunjamensis                MTPVSATPADLIPLLTSYRDKRHLLVVHARPEWDG-TVLRVDDLPVPVHCCRTVLEIRDVLRHRNDPPWRVVLTDLHDDQIPQGVKEHFRPGRSLEL
WP_105324909.1_Corynebacterium_sp._J010B-136      MTSLQVTPRDIEQHVEKLRNKNYTVAIRTSADFVGRDQWVSNGQNIEVQRVNTPLLLRAALRRSDEAPWRFFLTTLPDSEMPLDLQEKLYPGRVFTP
WP_312777491.1_Corynebacterium_variabile          MSARLIRQRDIDDQIDYLTGKNYVLVLHTDQGWDGPASYTRSGDLVAVTTVRTPLALRHALVQAEQQPWNVFLTPLGDADLPADVRDRLRPHDVQVV
WP_322753648.1_Frankia_sp._Cas3                   MTVPAATVAIVRAMIDKALEKRYVIGVRARPVWSGPETFTHRDRPARVVVCRSALAVREALRERSAGRWLVVITDRDDADLGAGILAHLVFQRLRTP
WP_144825789.1_Micrococcus_luteus                 MTRTRITGAAVRSVIADAQRQYAVVGIYADPAGVAELGVVVDGAAVTVLPAASTLAALDQVHHAPRGSWSALVTNRPEEDLGVGVLSALVGGVLRTP
WP_259849919.1_Brachybacterium_muris              MTVTTVPRAAVLATLDAVRRRESVVGIRASRGLDDTTPSDAHHEPITVRWCPSVLSLWDALTGWNEQGWLVLLTDRTEEEIGSGLLARMVQQRLHRP
WP_243731729.1_Nocardioides_ochotonae             MTWTLAEPGYVRQRAAAVRNHDPVLILHAVPQWSGEPIFTVGDAVVHVRACESQLAVLDAYAQLPEGDYLVVLTDRTAQDLGDAVLLRAWRRRVEMP
WP_280369379.1_Nocardia_wallacei                  MSAPALKTADVAQYLSAARGERTVVLLRAEPAWAGSAELAIGERRARVVPAPSALAVHELVLRHLPPKVLVVLTDREHEELDPAILARTHRKRISL-
WP_285664605.1_Actinorhabdospora_filicis          MTVTAVQPHLILGEARGLRKRPPVMLLRASADWTGDELLKIDAHTVRVRPARSVLAVLDAIDSRSEHELLIVLSPFREEELGPNVLARVVEQRVFS-
WP_148767669.1_Actinomadura_decatromicini         MTEAAATSGILIAEVRRARRRGGTLFLRAAAEWRGGDVLDLDGATVQVAPCTTLLGVLDAVSAKRT-DFLVVLSAFERRELGESLLAHAIGHDIVA-
WP_158645688.1_Stackebrandtia_albiflava           MSVGTARQTMVHAKVVELPGESAVLLLQAAPVWNGPETFPVPGAEVRVVPADSPLAVMEALSGASKDEVTVIVSPYGD-ELGANVLARVYKRRAQS-
WP_169440218.1_Streptomyces_sp.                   MGVTVLDQRVVAGLLTAHLAKRRLVLVQGKYAAGSPEEFTLSRRSVRVVHSTSVLGITDAWQAHLRRDVLVVTTDVPDDRLGLDLVGEMMRSHVIK-
WP_326854225.1_Actinocrinis_sp.                   MAVAAVSRGMLESLVRRERKASRLLLVYAQYAAQGPARFRVQRYVVTVTACSSVLAVADAWESFQASRVLVITSTLAAQEIGLSLRAFAARMGTIA-
WP_018383907.1_Wenjunlia_vitaminophila            MTAAAVGRRTIEALLDTHAGRRRLMLVHGHYPPTAAPEFTVRTRRVHVVDRSSILGILQAWHDHQSRDLLVVTTAVDDAQLGWDIRGHAVRRRTLT-
WP_051042993.1_Nocardiopsis_xinjiangensis         MNSPAVNAEAVAGALERARSGATALALSANPRWDGSARLTVHPRSVRVVGADSVISLVYALSGHDDTDHVVVLTPLESADLGEALLGRFLGEEVVR-
MBV9853173.1_Streptosporangiaceae_bacterium       MSTPGANRTVVTAELRRAAGPSRVLLLRAAPEWLDEPRLTVSTVTANVAGCPTVLAVLDALATRPEDGYLVILTPCDDDELGDSVLARVIANQVHT-
WP_189653453.1_Nonomuraea_spiralis                MSD--LNDLVVREFVRTLMRRQDIILLYGRYEPGAPTELKVEERQVRVADCPSVLSVLEAEVGHPG-GLLVIVTPRSDDELGNTLRSRAVKQRALR-
WP_066069029.1_Frankia_sp._EI5c                   MSVAAVTTAMLDGRVARLGERARILLVRAQPEWPGPGELLVSGRTVRVVACVSPLGVLEEVARWGPDGWLVLLTDADEETLGDTVLAQVHRQRVDV-
QGN48447.1_Micromonospora_sp.                     MRQTQISPIALRQKVTQQRHRYPVLVVRAEPAWPHDPAMQLPGRRVRVVPCVSPLAIWEHLVAERGDEVLVLLTDLAESSLGAGVRGRIFRQQVIM-
WP_034267102.1_Actinospica_robiniae               MSITSAARPLHLAALRQLDDEPRVYLVRGAPSWNGPATLPGPGHDVRIATAVSPLEVWELIVRHADESLLAIITDAEEHDLGPNVLAHVFRYRVLI-
consensus/100%                                    M.............h...........h....................h..s...h...............hh........hs...............
consensus/95%                                     M.........l...h.........h.h..................h.l..s.s.l.h...h.........hh.s......ls.sh..ph...p....
consensus/90%                                     M....h....l...h.........h.h..................h.l..s.o.L.h...h.........hhlss.....ls.sl..ch...p....
consensus/85%                                     Ms...hp...l...l..h..p...h.l.s.........h......h.l..s.o.L.hb..l.........lhlss....pls.sl.s+h...ph...
consensus/80%                                     Ms...hp...l...lp.h..p..hh.l.s...h..s..h......hpl..s.o.L.lbp.l.........lllos..p.pls.slbs+h...ch...
consensus/75%                                     Ms...hs...l.s.lp.h.pc..hl.lpu...h.ss..h.hss..hpl..s.osL.lbc.l....p....VllTshsp.pLG.slbu+hh..cl...
consensus/70%                                     Ms...ls...l.sblp.hbp+p.hlhlpu...a.uss.hphss.phpV..ssSsL.lbchL....p....VllTshsc.cLG.slhuRhhp.+lb..
```

### Type 2 BREX PglZ NTD iSwi2/SNF2-Helicase

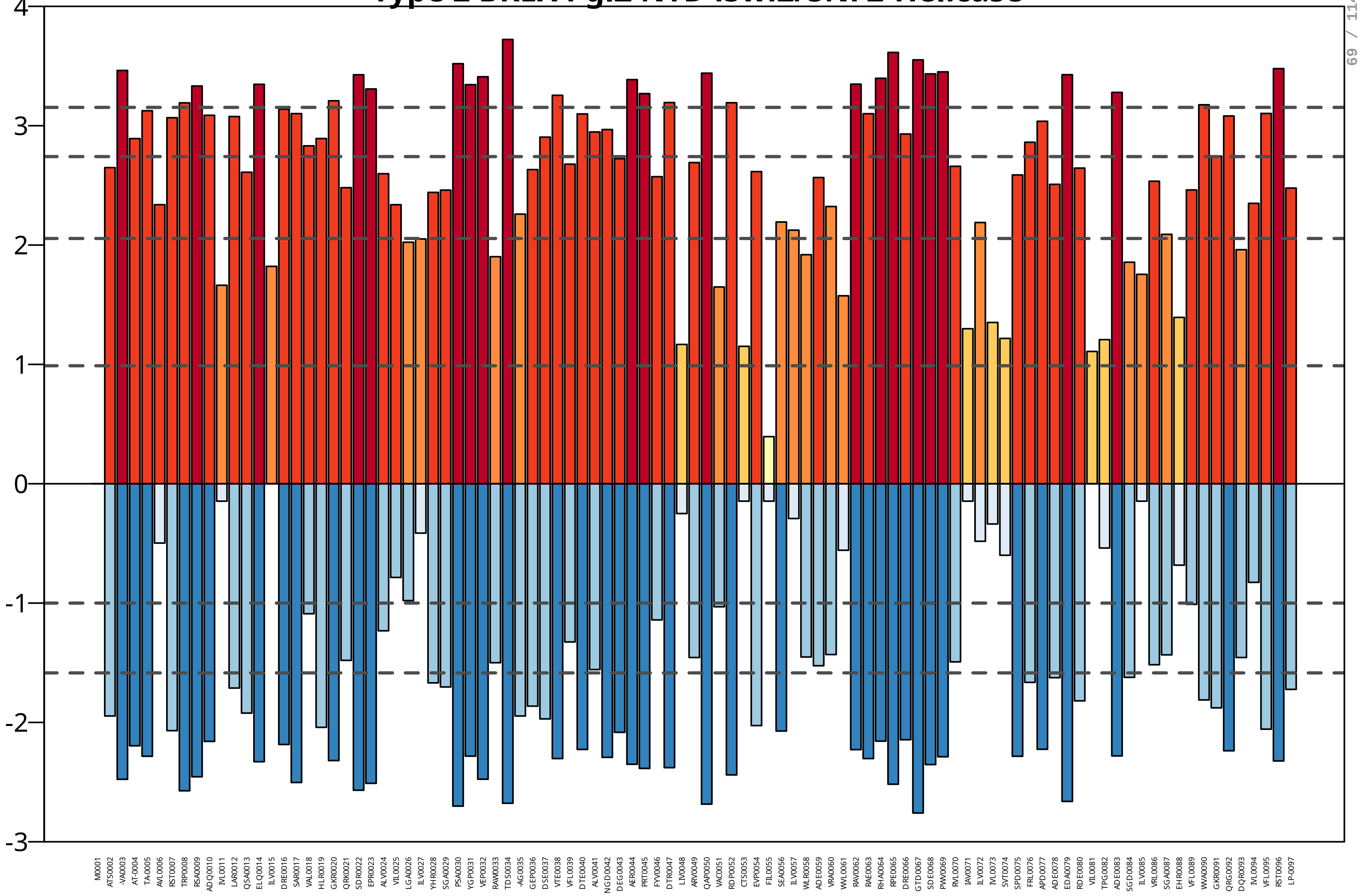

```
# ; Type_3 BREX PglZ NTD iSwi2/SNF2-Helicase

NPV81619.1_Bacillota_bacterium                      MWRDHILKEFTPRVSRLTLVADPDGLLLEEGILEGIRERGFELIPFEDHIAFRYAYESRFRSRDAALVVVLRSGDVLPYDLLQAGRKLSFNLGDIFP
WP_264656686.1_Azospirillum_canadense               MWVDAVTKDLNPEIAPLVAVIDPDGIFREEAVLAAVQARGYAVVQFEDEIQFRVEFEAKFRGRDRHLLIVHHGGEDLPYDLVSAATVVSASLPSLFP
WP_069455074.1_Shewanella_xiamenensis               MWVKELISTLNLDGAFVYLLDDPDGLCYEPQIASFFQDKRAVLIDEPDPLAIRLTYEEWLRENNQVSLIIRISDHSIPFDIALHAKRRSFNLAEICP
HEC58784.1_Methylophaga_sp.                         MWQDRLLNKLALNESLCFVINDPDRICYEPHISNAIKQSKAIIFDDVDPIALHLIYEEWLEKESGGAFLFRYSDINIPYELLRKSKIVDFHISEVIE
WP_247116183.1_Marinobacter_sp.                     YWRKPILRALEPGLSDFFVVEDTDGLLAEPGIQKMLQARDVQFYAYDDPIAFRHFYETRIRGDEVTLVISLGEDRSLPFDVINRARVESINISDCFP
WP_008603683.1_Pacificimonas_flava                  MWVDHVAGGFQAEFAKLWAVSDPDDVLFGDTVSRTLVERGIDVVEYAEPIAFRLLYERDLRENEAPTIIHVRGDQELPWDVLQSSRHVDLRLAEIFD
WP_181742020.1_Thermoactinomyces_mirandus           MWRDEVLKYFADP-PAVMIISDPDRLLQEKKILDQLLEKNLEIVAFEDSIRFRYLFEATYRDRGNFCLVVHTEEHHVPYFLYQHGEIVELRLADLFP
WP_113032578.1_Paenibacillus_contaminans            MWRQKVLESIQAP-SLITIAHDPDLLLQDEGIVCELQERGIEIADFQDRAVFRYWYESRNRLSGGRHVLIRQAGEDIPYDVWVQGHHVSLSKSALFP
MCD8511937.1_Bacillus_sp.                           MWRQLILDKLNVS-YPLILVADRDDLLSDEELILYLQENSIHVHEAMDRAEMRFIFEQFKEGKVGEKLLFKLGDEFFPYDFLTEGHFVSLSVSEIFP
WP_102026200.1_Salirhabdus_sp._Marseille            MWRQKLISQIPFQ-HKLLFVYDRDDILSDEELLSYLQQNEVNVFSFKDRAELRIHYEGQLEYSSGKWIYKVGEGFTFPFDMLNNQEPIELTLHIAFP
WP_036836014.1_Pontibacillus_litoralis              MWREEVLKQFERP-SSVYIVADPDHLISDEKILSYLQKEKLHVVDIKDTIDFRYIFESKFREKSKEYLVIRFSRTSIPYDFLQIGHQINVSLADVFP
WP_169099399.1_Bacillus_sp.                         MWRDQVLKQFIAN-NALYLVLDNDQLLLDELILGKLKEDNIQVVNFNESIIFRYEYERDYRTKEEICLLVREGTNVFPYDLIQSGQRTRLDVSIIFP
WP_190997467.1_Peribacillus_faecalis                MWRDIIINKMKYC-ESLIFFIDPDYLLSDEILLRKIEDLEFTITRYENSIQFRFFYEKYCRYSNKKFL-VYNNETELPFDIKQQSAILQIDISTIFP
MBT5530409.1_Cytophagia_bacterium                   MWRKYILNHFDGQ-YRSIIVEDPDDLFFDETIQDNIRRNSFKVYSYSTSLAFYYFYESVWRNNNGDVLIIVQSEEFAPYELHRDIQKIEVSLPILFP
MCM3358660.1_Psychrobacillus_sp.                    MWQERIINHFTSK-GSLYIVTDEHELLREKTVQEMLRIQNCTLHFYDDPVAFRYEYETTFRHIHRATLVIVLDHNHVPYDVYINANRSTLSFETLFP
WP_214788919.1_Exiguobacterium_sp.                  MDLHRI-QSFLLQSNPYYVVRDMNQLFNSPSFRNVVTNLGYTIHVADDPIAIRLAYETFRLSNKGTFLLISQDFSSVPFDILKSSLQVTITWGLLFP
WP_014759117.1_Thermoanaerobacterium                MMINYIVSKLNRCDNNVILVFDEDNILSNSDIISMLKENGYDIYDFKDPDLFRYYFESKFKIAGVKLLIRYDGKITIPYDIKMSSSNIAITLTELFP
MBI9032248.1_bacterium                              DFVTSILKKVLSLPTELIFINDIDNIFTYSEVLNRLNELSYKVSNYICPIAFRIEYELEIKPFGSKYIVRCSSSPYLLEDIAEHSDIMKISFNDLYP
MBE6339837.1_Bacteroidales_bacterium                DLIKKVAQRITVDYCNLSIVENPDGFLSQDVVRALLQDCGIEVV-VGSNLQLRIHYELHYKVSGRRYVYICKSKDTILADMKRSAHVVDFSISDVFP
WP_123477185.1_Muribaculum_intestinale              ELINEILDSISRGFHPLTIVSNPDGFLAHEDVIQELRESPTDVV-VGSDIQLRVHYELNLRRNEKSAIYILSTG-DMLPDISTKANNVRFKIAELFP
MBR2506173.1_Bacilli_bacterium                      RWVQTIIEKVTPSTVKPIIVIDHQRYLEIGEIQELLRKNGYTLVFVDCGILVRIKYELEVKGK-ENTILVIKGNYSLVDDMSASAFVIDLKSKEVFN
WP_217803628.1_Parabacteroides_johnsonii            NLPQGIINRLFGSEKRLVVVQTYDEFLAREDVRMMLTYASLRIS-EGDSLTLRLWWETTVKEENIRFVFVQREQFDVPDDIAIYADTTQFVARHIFP
MBQ9466673.1_Muribaculaceae_bacterium               ALTQNIVNRLFAGDKRLTIVINRDGFLSRTDVQNALRHAAVTVR-HGKAIDLRVLVELHIDDDTGKTLFICDDEFDIMPDILPRATVMTFQARNLFP
MBO5155853.1_Prevotella_sp.                         ELAKRIEEKVTDRHFHLTMVRNSDGFLSRSDVQQAVQECGVLLLPVCSALELRVRYELNDRLSEQKVCYIVNGNLTLLPDMKPYIKDKMINLSDLMP
TNF50598.1_Bacteroidota_bacterium                   TLIEHIAQDLKDQPSKFLLIENPDRFFQRKDIQVLLESQGIIFL-SGSSLYLRVSLETQDEKNDATVCYFISNENHVLEDIREKVFKKKLSINDYLK
WP_076420664.1_Pontibacter_lucknowensis             AFVNHIAEEFLASPKRLILVRNPDGFLRMPEVQKALKAYSIRLC-DKQGIDLRIDFELRYKATADKTVYLISSLEAILEDIASEGEYRVFHLADYFP
WP_133532125.1_Flavobacterium_dankookense           SFIEHIVSYFNSFDKKIILVENPDKFLLREDIIKIFEDNNISIV-NGTGIEQRIKFEL--KSSVTTLIFTSDDNENYLEDIKRNAAKTTFHLSNFIK
WP_172278715.1_Lentimicrobium_sp._S6                TLIKKIVDDFVSLNRRVIAVENNDGFLFREDVIEALFANGIELV-SGTSLDHRIAYELRDK--SKTLILLSKQKQDYVPDMGAQLYYYEFEMNQYLN
MCC5935593.1_Balneolales_bacterium                  ELLRNIAEKLKSYGARTMLIENPDGFLNRPDVAKELLKYGLTLH-RGTQLEQRIQYEL--RDPEERLVLLNKDNSSFLQDIKQNAFPVKFRLDDYLG
MBP7508728.1_Prolixibacteraceae_bacterium           NLLETLRKTLQAY-QKHILVFDKDGIFFHSEVQTEAQEVGFKLIFIEDAYKFRISFELNHNLN-GRILYITNCTFPVLPDIIREVKTITVSFRDFFP
WP_083331763.1_Prevotella_bryantii                  SLLQCIVKEITRQYRRLSVISNPDGFLRRKDTIRAMEQADITVL-SFSQLELRVWFETVFKEDANRFVVLMDDTTKLVADIYNQAFVTVFKTRDLLL
MBP6412566.1_Bacteroidia_bacterium                  NWIDKFKKVLISSHKEHIVVADIDDLFEYSELKQAFETDGYKIETCKSSLAVRLFFELNVRNSKDKFLIIAPAYYFPLPDIEMQVQFQAIGLLNLFP
WP_073205016.1_Prevotella_ruminicola                SLVSNIVHNIIQENVSISIIRYADSLLMREDIYSRLEESGVKFV-TGSNIELRFHYELHKNDDGQHYCYLLGTGQMLMPDIMRKCYVTDFKLTDLFS
MCM1450899.1_Clostridium_sp.                        RLLADIVNRIVGGDKRLFVVDNRDGFIMKYRQALETYGNRCRIF-GGTSLDLRIVRDYEMMLDESRFIFVPSEDFTVLDDVAADSERVTINVQRFFA
MBR3317262.1_Atopobiaceae_bacterium                 MLAELVYRNSNAAFSTSVLLVDH-GIEARCGYAEVFERNGFVVIHWVDDLRFRIEWEDQLKRG-ERLAVIAPDGAYVPYDLAQLMQRYDVSLRNLFP
WP_195245438.1_Clostridium_celatum                  MFDEYLEDRLALKYEKYIMVLDEDDILSRYNIIEKIKENKFNVEQYNDVEKFRYLFETKIKKNTEKYFVIVNSDIYIPYDIRNFFYQVNISFQNLFP
WP_071060846.1_Andreesenia_angusta                  MFKDHMIKMLSIDYESRILVFDYDDMESKYGYNEILVENGFRIIKYEDPTEFRFIYESEIKHSDEKFAVIVKTEGYVPYDIIKSFYSVELRLEDIFP
WP_226978794.1_                                     MFADLLYEKSSAAYEKHVLLIEEDALEDKTEYCAYFAGKGFRIIRYENDLVYRVKHEEAVRAGDSKILMLASSGVYIPYDVQKCFRVFKVSLAKLFP
WP_153014722.1_Syntrophomonas_wolfei                ILRDLLFEKAGLPWVERLIIYDTDNLVSRTAGTNGLYWDEYSVFEVKCSEELRFIYERDCRQTESTIFIVPSLDIKIPYDIYKQFTIVTLGLDTVFP
MBP3452732.1_Clostridia_bacterium                   M-KNRIFEVTGILSQNNCILFDIDGIWQEYDCSSNASEAGFQVINIHNEMELRYEHEKKLMHNEDRILLMHQLDMYIPVDIRRLFTITMLTYDRLFQ
MBQ3643692.1_Candidatus_Riflebacteria_bacterium     SFSQYLYETNNAEQNKRIIFIDEENLEEKFSISDYFKKQGYDVY-YSSKTPFVFVFVGELQESDKQILLIVRSDGDIPNEIRDICVVKRINIEEIFP
WP_252226004.1_Caldicoprobacter_algeriensis         LLCQYILEKMAVDYENQILLFDYDDLCERSGALRFAHNKGFQIIEYADPEEFRYFYELKLRDNDVKVIIVMNEKIYVPYDIRKRYYNVFVDYASVFP
WP_162523517.1_Calorimonas_adulescens               MFYEELLNGISSRKGQIIIIHDPDGFFSEVTV-SELFDGSIDVHYFNDPDLFRYEYERDYRQRDKVFMVICSNNRGIPYDVLSKGSLVDMELSRIFP
MCR4431941.1_Tepidanaerobacteraceae_bacterium       MIEDVIIQRLPLNQNNFILLYNPDDMVAGEKIVSRISSCGYTLINYEDPDMFRFYFEQILNKGGDKIILRLTRNTYVPYDIQSIFQEIRITIKDIFP
WP_045176026.1_Caldicellulosiruptor_danielii        MIEDAIWEKINSNLYDFLLVYDYDNLLCEFNVITKIFDEGYEVFYYKDIEEYRIFYETEIRYTDKKVILIVTENVYIPWDVKCDYHEVLIRVKDLFF
consensus/100%                                      .....h...........hh....h...............h...p..................h...........h.......h...hh.
consensus/95%                                       .h.p.l.p.........hh.s.p.h........h...h.h...p...+h..E.........hh......h..-h.......h...hh.
consensus/90%                                       .h.p.l.p.h.......hl.s.-.hh...h..h...h.l...s...hRh.aE...p.....hh.......h..Dh.......hph..hh.
consensus/85%                                       .h.p.lhp.h.......hl.s.D.hh.p..h..h...hpl...s...hRh.aE...p.....hh.......h..Dh.......hph.phhs
consensus/80%                                       .h.p.lhpph....p.hll.s.D.hh.c..l.p.hbp.shpl...s.h.hRh.aE...c.p..phlhh......l..Dl......phph.plFs
consensus/75%                                       .h.p.lhpph....p.hll.s.D.hh.c.pl.p.lbppshplh...ssh.hRh.aE.phc.p..phlhl.psp..l..Dlbpp....phph.plFs
consensus/70%                                       bh.p.lhpph....phhll.s.Dshh.c.pl.p.lbppshplh.h.ssh.hRhbaE.ph+.s..phlhl.psp..ls.Dlbppsp..phplpplFP
```

### Type 3 BREX PglZ NTD iSwi2/SNF2-Helicase

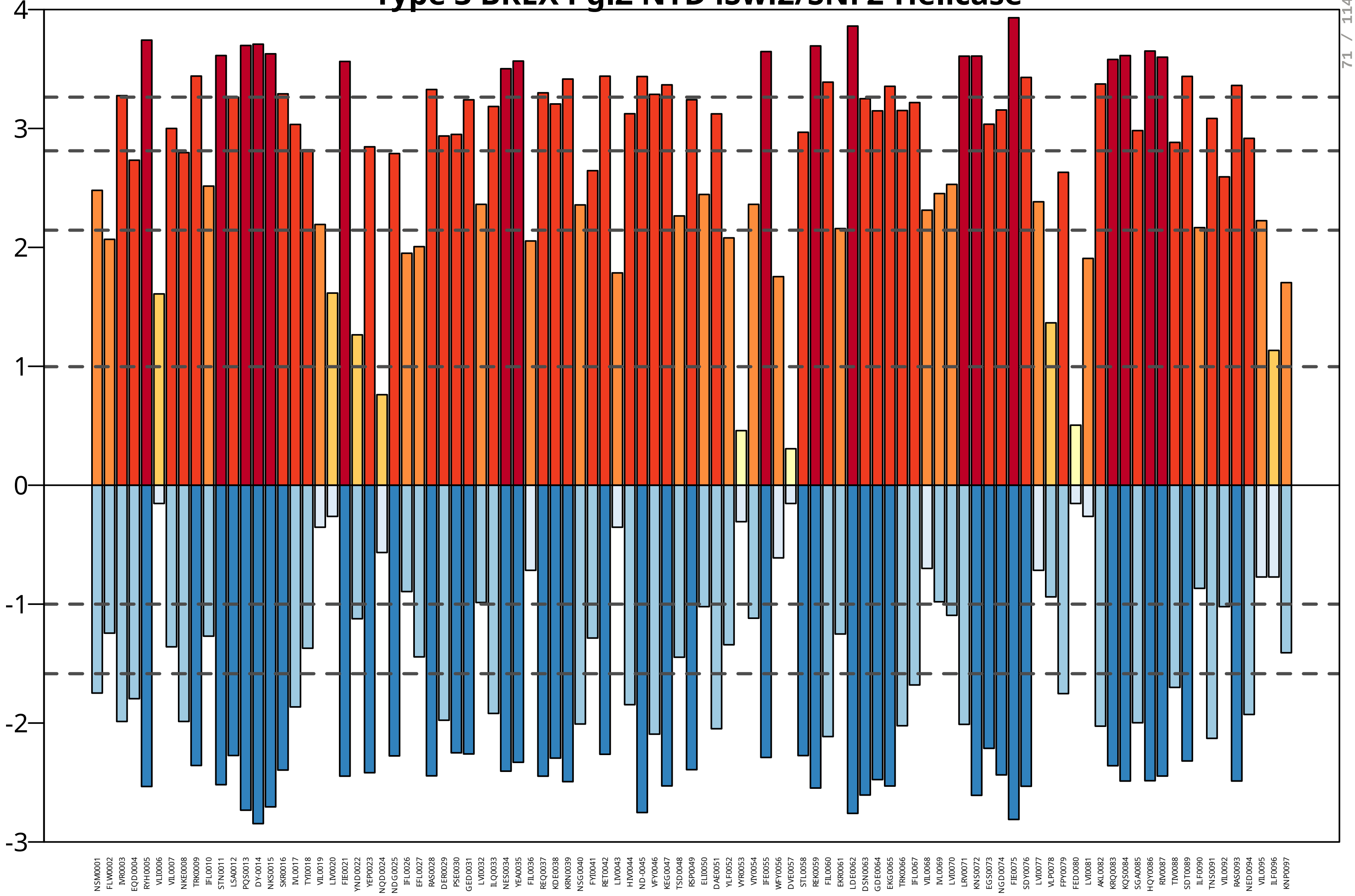

```
# ; Type_4 BREX PglZ NTD iSwi2/SNF2-Helicase Rep_1

RGH34270.1_Ruminococcus_sp._AM47-2BH                   PQICLPKITLTQSSELEKDSSVIKGISKLFSMLENMSKSELIVHTHYSSTLFKNAMMPVDTING
WP_073324030.1_Fibrobacter_sp._UWOV1                   MPQKYLDVCFVPAPEYLQGIRPYGGVHLFGSMFEQQDDAVAYILTSKKKRDFPDAKINIRELNN
WP_216471320.1_Butyricicoccus_intestinisimiae          EIIRRPEICFYREVDFVIETEFVVGIQAIPEAIETRVMMSLAVVTTKPKTAFPYAQFVIQQEES
WP_044080180.1_Prevotella_pectinovora                  EKLQPTTLFFVKPCLADSAQACIKGVNNLLKLLECGNGGVISIITSKTKRDYPYSMYDIEEYSS
MCI8555501.1_Clostridia_bacterium                      KSIDNFCFDFIKVDN---QSNEIRGFKNLLAKMEDNLSKDYRCFTTIDDRIFKEHCYAINIINN
MBD8953983.1_Dialister_sp.                             DPTPLPSIYLASPEENSFGQLVLPNIQALLEKLEQYDETSFAVRTSLHKEVFSESMYSIRSSKG
MZQ76087.1_Peptoclostridium_sp.                        DRLENATLNLVNSQITNKNGNVINGIKNYLDQCEEECIVKLPVSTRHYNYDYSEAGVNIRKISD
HIS35238.1_Candidatus_Scatousia_excrementigallinarum   DKQTLPKIIFVKENVAKLAQPIIIGMAALIKEVEYTNKKVLYVKSHFKKSNFTESIYYIEELNS
WP_103226892.1_Clostridium_sp._chh4-2                  TVDSVPHITFVAPEFSFDSANVVEGVEFLPVAIESAVQSEIYIRTAKKAAVFANGQYAIATMDT
CDC05526.1_Clostridium_leptum_CAG:27                   SPLTNISIINVPNDM---GVVTKSGVKYLLHNLEDGVFGNVYASTS---LILDNTLFPSSTLSG
MCL2016626.1_Defluviitaleaceae_bacterium               IQQNALNVTVLASDLQDLQGTILPNFKALLAELEKAAEDKILAITNL--SNVANSTFEINKINN
WP_235396406.1_Pseudoflavonifractor_phocaeensis        IPSARPKLIFTAEGIQPDASALVKGLRGMAKAVESAAAEILYIETAKSKDQYPFSLYTISDLKN
MBR3817930.1_Clostridia_bacterium                      EYQAIPQLIFISNSKFANASFMNHGLPSIGNSIEALSRDKIFIVTNHLTTNFTNSLIPISAIKS
MBP3656735.1_Clostridia_bacterium                      ATSNLPQLRFVGDDISIKDQKVTKGIDKVGLLVETVYADTLLIRTKKLAQDYPNSMLHIEELVD
ONI45221.1_Epulopiscium_sp._SCG-C07WGA-EpuloA2         KYELEKTIVFVTNNLRIDDVKILPGINYIAPEIEKSTEEKIYIYTAKSKDIYRNSLLEIVELNN
MBO6272669.1_bacterium                                 RTDKLPEIYFCEDSDYFENKNVINGINLLGAELENSSLLTVYVKTKKNKSDFPYSCLKIQNLKN
WP_037355505.1_Selenomonas_sp._FC4001                  KNQPLPQIIIERNPA---RGKTYNSIGELLSALEQMTSLMISLVTKYSPTLFNKAVYSVSQGPE
MBS6586269.1_Eubacterium_sp.                           NSSPLPKIRLVKSAELCVGAEPLNGIPGLLGYLEKMSQPVLNVLCGLNPGLFSRAAYYVSAADG
NBK77202.1_bacterium_D16-76                            GERNIPAIHVVPENVAYPGVDSVQGIKELLRILENGKGTDIYLTTSCKVVYLERSMYPVSEGMN
NSW92343.1_Bacillota_bacterium                         GEDEKVSLQIMPRNIETLCEDVIRGIKQYMQIWEKTSRQNSFLITGFAQAKETAGDYCIRICNS
WP_048095116.1_Archaeoglobus_fulgidus                  LEKTTVDVWIITDRLTKRNVVAVDGFKEYLKLWENQVIHDILVYSILLHSTPIAGNVKISPITT
MBQ7110119.1_Thermoguttaceae_bacterium                 RAECDIRVDLYDETTLGGLKHCIDGIKNALETLEAGRARTVAVRTR---EDFSSSLCSVRRVAS
HED00230.1_Pseudomonadota_bacterium                    -RPEEINISVFQRQIEGSRFFWINGIKNYMGLWENAKVSSICLSTRFADRGAQVSVIRIDVFKS
ROT09401.1_Muribaculaceae_bacterium_Isolate-037        DDCKPIKLTLINKSLAKFYETNTYGINQISKLIELIGLNEILIETKHKRHEFPDSIFEIHERNS
MBR2693834.1_Thermoguttaceae_bacterium                 DPKDSFSVIRVPLGS---NASFCDGFKAFLHEAEKGEPREYWVSSA---LTFEEAKFSVKTVSS
MCM1514006.1_Anaeroplasma_bactoclasticum               NYSSTPAIVFVNNEYQ--YVRAENGLNVALRNVERNTGVKIYVLTSYKKKDFKSPLFNIEECKD
WP_166670004.1_Aminivibrio_pyruvatiphilus              HSLEPFSLNVYSQEFGECGTDIMQGVQTWFYKWSGMIRKSCRLVTNLWRNQISSSLINIKIISD
HCH97635.1_Lachnospiraceae_bacterium                   DDEKTYSLTVYSEKFKNPLENIAHNLKEWLEKWQDCFFSQAMIMTGQIENEKTYGKVTIDIVEY
WP_195464804.1_Clostridium_jeddahense                  GEETDYSLTIIQKGLDVLSGNEIYGFKKYLQYWEQNPDKPLILHTSNAVYENNVFFDDVKVIVT
WP_006692206.1_Selenomonas_infelix                     AEDSVVSLSVLPPEP---MEVGIHGARSLLAALEQGRDGTHIVQS---RQSFPHALLRVHHVRD
WP_028051765.1_Carboxydothermus_ferrireducens          GEEEKCELIVAPFARKGKEFVSRNGVKDYFSFWEKDGVPSLWLVTSWASHPSQAINFKVKVYTD
MBQ0159480.1_Bacteroidales_bacterium                   -ETQSVEVFFSKYKNES-VQCQLNSLREWLMFWKKQAPQERIVCTSAPKYSKPDNIFNFTRIAN
KUK66682.1_Parcubacteria_bacterium                     -GSIKYNLVFCKFASIE-GFTIVHTTKDFLELWKKDDISSSILISSELNRAIDDELFTLTKIKD
WP_018963723.1_Coprothermobacter_platensis             -PRDKVQVYLIKFDYID-DSNALPDVKSLMKLWEKSKSDE-FICFSKTSCFCSDDVVSVNNVLT
MBR3117632.1_Oceanobacillus_sp.                        -YKDKICVYYTNSSLEV-LGNKISSSKEWLNIYKNGNYDKGIVCVSDAEQAKPDNFVDIEKLDN
WP_024292379.1_Lacrimispora_indolis                    -FEDLPRIFFIDDNLSA-YENEVTNSSEWFGMWRKSDVTKPIICTSKVEKASPDNVYNIKRIST
WP_191216537.1_Methanothermobacter_sp                  -TSDQLTIYHADFDFLE-EFEIIRNTHDWFNLWKKTYYNDRFLSFLPVEEWLPDEIFNLRIIKN
OQC39307.1_Bacteroidetes_bacterium                     -ETNSIKVYLTSQTLID-NYETIANTEEWLKFWKRKSPCD-VICYSKPKNTLPDTIFTIEQVDN
MBQ4420212.1_Bacteroidales_bacterium                   -EKQATRVYLMSETPNS-HITVLKNAKEWLQFWKTQAPQDVVICTSRPNYSQPDNIFSFLPIND
WP_183546997.1_Methanococcus_maripaludis               -QDTPNQIFIPKFEYND-QYHVVKNVRDWINLWNNQSITKNIVCTSRFENVFPDYFFKFNILDN
MBU7048061.1_Theionarchaea_archaeon                    -KNDKIEVFLTDFLVSV-HFSVIENTRDWLSLWKGREKRK-IICHSRSKNAIFDKAFEIEKVED
HHW21152.1_Thermodesulfovibrio_thiophilus              -SIQRTNIFLYKDLFIN-TIQKVKGSKEWLTLWKKSLSTP-ILVLSKTDNAKPDEFFDLKKINN
WP_207678199.1_Desulfonema_magnum                      -PFKKIKIYQISFEALC-NFKTINNTLGWLDLWKQNDLEV-VICKSGTKDFLPDMMFFPEEIST
MBE6838777.1_Ruminococcus_sp.                          -ELEYRQICFVSDEYNI-KTNKITTSVEWIGLWRSADFTKPIWCCSASQEASPDNAFQIDSISN
WP_205109179.1_Marseilla_massiliensis                  -EIGTYNLVVTNGTISD-KYTYVKTLYEWLKLWEKGKIKQTIISSSPNHFAQPDNAFTYTECKN
MCH4183501.1_Prevotella_sp.                            -PNFSYRLILTDGETET-HYTIVNNMHEWLSLWRDEQENPNIVCTSHSGYAQPDHAFSYIICPD
MCI6981218.1_Akkermansia_muciniphila                   -PADKYRIILTNGTTKD-SCVICHNTREWVKSWEDLDGRRTFICSSKTQNISPNDCFEYVRCET
MCD7797870.1_Akkermansiaceae_bacterium                 -QANCYSLILTDDVKSN-KYSICSSLKDWVNLWQPNKIKHKIILSSKTSLPTSDNAIDVKFYPD
consensus/100%                                         .......h...............s...h...hp.......h.....................
consensus/95%                                          ......pl...............s...h...hp.......h............h.h....s
consensus/90%                                          ......pl.h.............s...h...hc.......l............h.h....s
consensus/85%                                          ......pl.h...........h.s.p.hh..hcp.....hl.s..........h.hp.h.s
consensus/80%                                          ...p..pl.h...........h.s.p.hh..hcp.....hl.o..........h.lp.hps
consensus/75%                                          ...p..pl.hhp.p......p.hpshp.hh.bhcp...pp.hl.os..p..bspshhplp.hps
consensus/70%                                          ..pp..pl.hhp.p......p.lpshpphhpbhcp...pp.hl.os..pp.bssshhplp.hps
```

### Type 4 BREX PglZ NTD iSwi2/SNF2-Helicase Representative_1

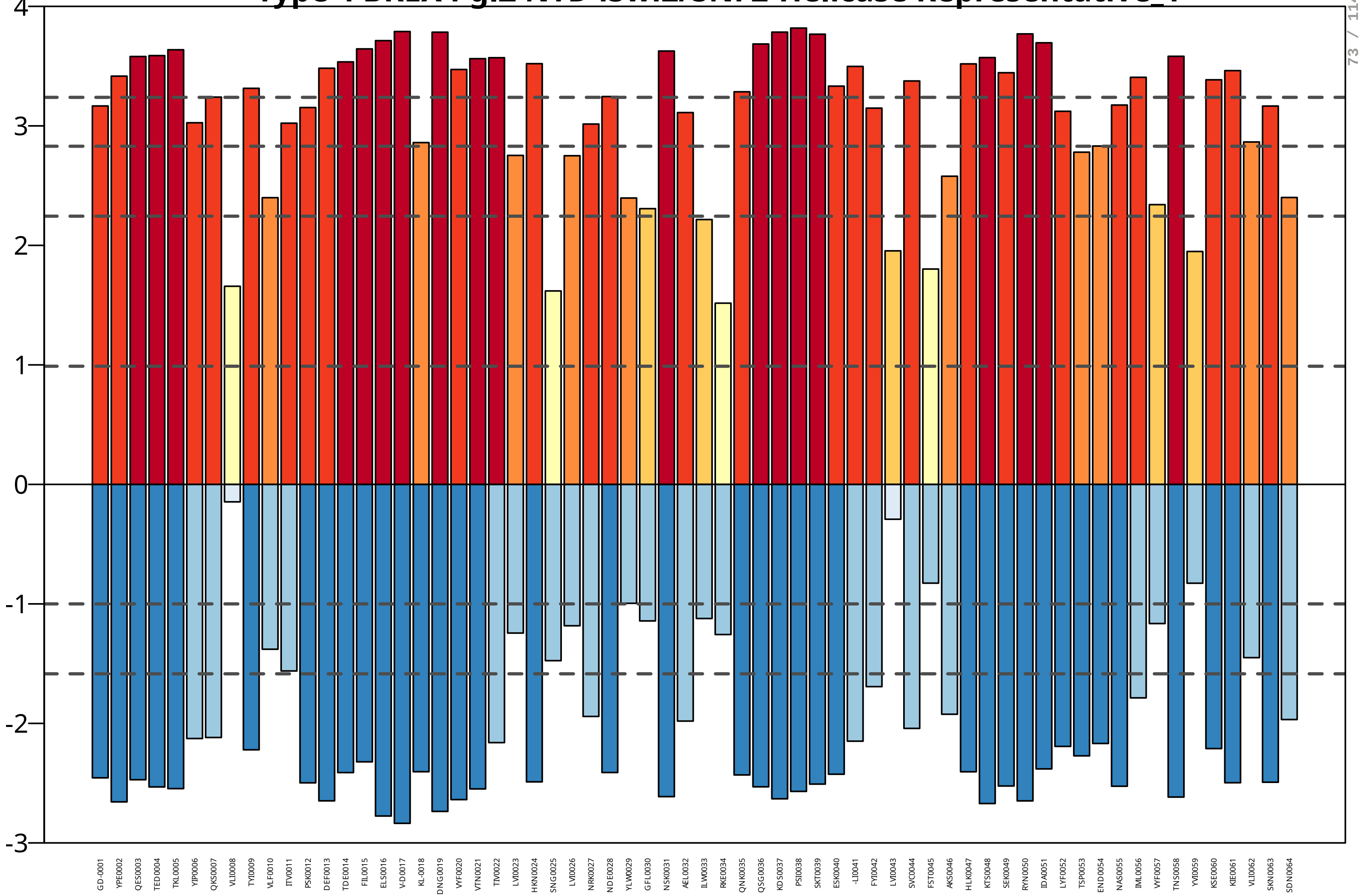

```
# ; Type_4 BREX PglZ NTD iSwi2/SNF2-Helicase Rep_2

WP_192822482.1_Rufibacter_sp._LB8                   MKALLLDPSGDAPAPEGYSVIDTDVAWLKAADNVPKVVVRGVERCNWAYIWWQALGGQTATLIS
WP_105482027.1_Abditibacterium_utsteinense          MSSLRLDWKGDAEIPEGARLVTNEVEWLRVAPDAQDVWVRGATVCKWASQWWQAAGGKCEEVRG
WP_243797665.1_Hymenobacter_tibetensis              MSFLLLDPHGDARVPVGYAVVATELEWLRETLSAPPIVVRGLELTRWVRKWWEARGGECQELVS
MBX7251230.1_Candidatus_Promineofilum_sp.           MTDVWLDRLGDADTPEGAAVVECEADFLRCALDEPRLLVRGERLCEWAGAFYRGRGLPVMETHS
WP_162787804.1_Anaerolinea_thermolimosa             MMEILLDHEQDCPVPDGYILVENEVDFLRLATSGQSLLIRGRELCAWAEGFYSLRNLPVRVVES
WP_012642882.1_Thermomicrobium_roseum               MREVWFDQDGLCPPRPGTVSISDDVSFLEEIDTDRPLLIRGRQRCEWARALADARNWRVHEVQP
MCL6576921.1_Kyrpidia_sp.                           MRKIYLNPNADAYKPEGTRVVSSETEFWIYALRPEPLVVQGLRLCEWAREFAEGRGLEVEELPS
WP_139401683.1_Deinococcus_radiopugnans             MARFYLDAAGGAEV-AGAHSVNSEAEFLRLALGDGPLLVHGEALCRWARRFGAGRGIPTQEVVA
MCE5327915.1_Planctomycetaceae_bacterium            MRELLLDIHGEANPVPGCRRIETELDFLQALSEGKPVLVCGEHLCEWAGAIWQARGWRVQRLTS
MBV5327602.1_Chlorobium_sp.                         MICFVFDPSGTESFESEYRLVEDDVSLLQGVLYQQSLFVRGERLCRWASLIADARGYDQVWRKA
MCC6188628.1_Anaerolineales_bacterium               MSEILLDREGTARIPPGFQVAETEVEFLRVAPEAEALVVRGARLCDWAATFFRGRRATVRTLVP
WP_168570764.1_Oxynema_aestuarii                    MSDVVLDPTELYDIPAEYTPISTEVEWLEHVITSNPLWIRGEFLYKWTREWLRVWNRTVEIKQP
WP_006100361.1_Coleofasciculus_chthonoplastes       MIEIVLDPSKTYAIPPGYHLIASELDWMRDFINGSPYWVQGESLCNWTQTWLRCWNRSDQIKRP
WP_273058112.1_Shewanella_vesiculosa                MIKFYLDIYQVFSGDSDVDRFRQLIYSNLSRTTDLCIYFHSKTLYSRFRDFEGLDGVAHTNIID
MCK9609000.1_Methylomonas_sp.                       MVMLVIDTYQIDTEQPAIEQLRQKISQHLKDEHLLRVRFKSKALYERFKDFEGLDNVLPTQQLL
WP_017840083.1_Methylotuvimicrobium_buryatense      MVQLILDYFGVDADDPSIAAVTRQIHQYLREPVLHRVRLTSQALYHRFEHLEGLDGVLKTQTLI
MCK5897804.1_Methylococcales_bacterium              MIRLIIDYFNIDKQDKAVNINRKKIDTHLEQDTLLRVRLNTPALFRHFSDYEGLNRVLLTQKLL
MBC7570526.1_Spirosoma_sp.                          M--LTLDPFRLSPPDTAYVEARRQILAYRLGSKAPTFVVYETPYLFWFNDLPDEERTD----VT
WP_198076877.1_Hymenobacter_negativus               M-KIYLDPFSLQPASASYVVARCLLLDFMRSQSNRDVIVRQKPWIEWFGDLPQTLIAR----VS
WP_235137408.1_Dyadobacter_fanqingshengii           M-KVIIDPYKIYKSASTYVISRTSLLDYFVSEDFENIHVTEHPFTKWYDDLGEHLQV-----IK
WP_194112254.1_Mucilaginibacter_myungsuensis        M-KVIIDRHHLYKTTQTYQKARGLLLEFLLSAGYDDIHVFKEPFVDWFDDIPELVAP-----VT
HBJ34341.1_Planctomycetaceae_bacterium              MITLIFDPFSIAEKQKAELLAFEEINDSLRRTVVPKVYIIPSSSIEHFRTYREVKGVNIQT-VT
MBX3399563.1_Gemmataceae_bacterium                  MIELVFDPYRIYTGESQEVAAFHTVGEMLRQTRRPKIYVLDGIAWEHFRSLRSVRGVRVRE-FT
WP_099078666.1_Vibrio_sp._PID17_43                  MVEFLVDPYNIDSDSASVREARLRINENLASIRNLRIRIVHESLLRRFSDYLGINGVKVTFTPS
SEH06866.1_Thiotrichales_bacterium_HS_08            MIQLLVDDFDLEDGEDDIPHIRQKIYAHLAYQGALSIRM-SSVFFQRFKSFEGGQGITVKR-IS
QEP42412.1_Ectothiorhodospiraceae_bacterium         MVILEADSWGISANADQISGARSKIRHYLLQAEELTIPV-KPAYRQRFADFIGFDGVE-WREVR
WP_194715675.1_Noviherbaspirillum_soli              MVEILVDSLKLSRKPSDLLHVRERIRDHLRRSTTLRVRL-DEARYRKVADLDGINDLLVVH-VT
WP_201382003.1_Ktedonobacter_sp._SOSP1-85           MITIYCDFYSILQVEQDYIPLRQRIRSALDKNTAYTVYVQNPILAQWLKDLRGYDPHVRWEDID
NJL05928.1_Chloroflexaceae_bacterium                MITIIADVHSCEAAPTEYVPLRQQIQRALTTQQALVVVIRHATLVHWLADLRQYPSSIVWRTVD
WP_129675140.1_Candidatus_Chloroploca_sp.           MITIIADPHNYKGDVRDYVAARQAIRRALDLDNSLTLHVTSHLLLHWLEDFNGYTGVD-WKQIA
MBU6335166.1_Chloroflexota_bacterium                MITILHDRLGVLPPPEDYVAVRRAVRVALDLGQALTVVVHDPQVSRALADLRTYPAAVVWRDAD
WP_097643873.1_Candidatus_Viridilinea               MITIIADRFAAAAAPADYVEVRCQIKAALVNFTSLTVYVTDPVVLCWLSDLQRYPASLCWHEVD
consensus/100%                                      M..h..s....................b......h.h..........h............
consensus/95%                                       M..h.hD................h...b......h.h..........h............
consensus/90%                                       M..hhhD............h.p.l...b......l.l......p.h..h............
consensus/85%                                       M..lhhD.........h..hpp.l...b......l.l....hhpbh..h............
consensus/80%                                       M.plhhD........sh..hpppl...b...p...l.l....hhpbh.ph....s.......s
consensus/75%                                       M.plhhD.......s.sh..hppplp.hb...p...l.lps..hhpah.sh..h.s.......s
consensus/70%                                       M.plhhD.......s.sh..hppplp.hb...ps.sl.lps..hhcahpsh..h.s....p..s
```

### Type 4 BREX PglZ NTD iSwi2/SNF2-Helicase Representative_2

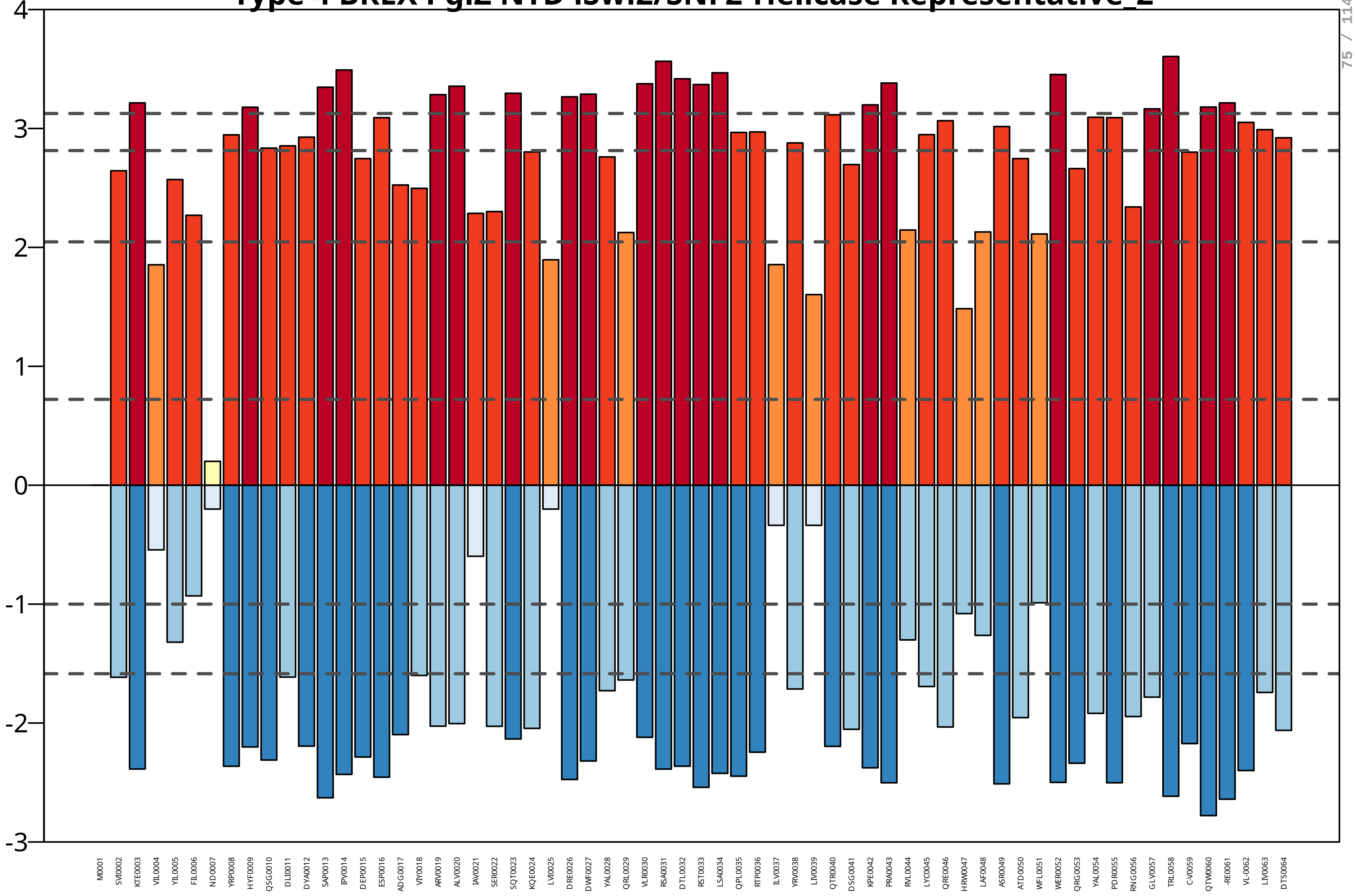

```
# ;BRC-system PglZ NTD iSwi2/SNF2-Helicase

MDS0297782.1_Halogeometricum_sp._S1BR25-6                MPTFTEIAEHAKVSPVVCSLPKLASFYLDDLAVYTSLDNFHHDLYDPDYRFEEDVVIDTDEHHPDYVRLSCDEEPV
WP_271955572.1_Halorubrum_ezzemoulense                   IESLSALAEATRDGPVVANIGGLESRVLQNIDTVESLEEFYRLYESPTNLIETDVIIRDQGPKTLYPSVEYTGTVY
WP_139229723.1_Halogeometricum_limi                      MAKLEEVVDEAKSHPVVVDVHPLSYLYLQELELFESEKQFEDELDNPSKSFERNVAIKEPRVKTTYPSMSIDHDPL
ATW88247.1_Halohasta_litchfieldiae                       MAYFNEIKERAHQQPVIVPLPKLAQFYLEDFNIYSSIDDFQTDFYHPDFLFENDVIIDADEHHPDFDRLSCDEETV
WP_338740220.1_Haloplanus_salilacus                      MASFETLADRASEAPVISPLPALATFYLDGLSIYRSVDTFHRDLYDPDYQFEQDVVIDDDSRYHEYPYINCTTDPI
WP_143414662.1_Halorubrum_sp._SD683                      MEDIKSIADRAHEEPIRADLPALAERILQQIDVISTTTEFYQRFEDPDHRFSADVLINDTTPKTTYPTIPYTGDPY
WP_048108644.1_Methanosarcina_barkeri                    -MKLSELDNLLSKTPVYLSLDSLVSLCYLDITIFHNRRDFWRKIDQLEPLKLCRYVILDDPD-YRLPYVRLKHNPF
NPV62497.1_Methanotrichaceae_archaeon                    RVTLEELDALLNKAPIAADMNPVSRCYLPN--RFANLDDVVKELDSPRPVDPSRTLIIGPRKDEKFLNVRVVHEPF
MCK4444004.1_Thermoplasmata_archaeon                     -MRVESFLELLRRTPVVLKGNELPLLYVDRTRIIRDIEELNQQIDSPNPINDAEIVIVDDDKGWSFPVAEVSEDLM
MBI2842128.1_Armatimonadota_bacterium                    DALVSGLSATLNSYPPAFRGNPLNLLFLDDARPIREASAFLNLLDHPALCDERREYLLDDPAN-PFPAYNISRRPD
HHW27504.1_Candidatus_Fermentithermobacillaceae           -MDVPRLIRMARETPVGIPNDLFLRLRLPDCNVFPNAQSFREALNGHRPMPNSPTLIVCESGDEQMPFEDTHYDFM
HIE11925.1_Desulfotomaculum_sp.                          KVDLEWLREMLRQTPVMAEIHPLLLALLPKVVVFGDVWELIARLDSPELQPTRPMLCTQVKSEHRLPAIFCDHDPL
HHW40997.1_Syntrophomonadaceae_bacterium                 QLSLDEFIALLQKTPVITAWNPWLKVKLGKVVVFPSLSEFRIAVNRFHPLADMPTVIFAGQNEHAFPKVQCTYDPK
HQD80871.1_Bacillota_bacterium                           MIRREHVPSIIDQYPVCVRFDPLLYMRLYGCEVLSSPEEFREHLDDPSYFCGPTLVMCDNLGESIYPLQTLDVV-L
MBE2199248.1_Anaerolinea_sp.                             --MIELLLKQLNIAPVMGQFHPLWSLCFPPVVIAHSAKQVAEQLDGRERHEEKPILITEYNPADRFLSHHLPDCPP
MBC8263374.1_Anaerolineales_bacterium                    MMSLDDLLTLLDCTPVIAEIPPVSICYLPEVTVLASAAEAAWQLDAPEPMLPGRLLITASDIQDRFPHVRCDHDPF
MCZ7554456.1_Anaerolineales_bacterium                    VLEWDDLLAALDASPMIVDMHPVLYAFLPDVRVVHDSAEASWLMDSPDPTLLSKMIIQREAALGRFPHVKCSRDPF
HID64303.1_Anaerolineae_bacterium                        SATMSQLDDTLDRTPVVYEGEPLLLAELDNLVVERDVWRYVQRVDNPAPVAVERTVLLGPVTSDRWPVVRMDARPD
MBN2393033.1_Anaerolineae_bacterium                      ETGFMRLLQALQRGPVMTEISPLALLCLPPIDIASDAETLVERLDILEPHSERPILIPCIDTRHRFQVESLTYTP-
MBV6438329.1_Anaerolineae_bacterium                      ALSVDELEAMLSRTPVIAKVHPLVPTALSEVRIVSDANAFQIELDEHAPAPELPTLLLSSNAAARFPRVTSLGLPP
MBW7959471.1_Candidatus_Promineofilum_sp.                IEQIDTLRKQLSLAPVMGHFHPLLPLCLPAVMIVTEARRVALAMDARGPVPESQILITRIDPTDRFLSLSLTETPP
MFN8469222.1_Caldilineaceae_bacterium                    ---MDW--SELNVIPHIVQLSCLWRAVLLQILVPITLHQWKGILDA-APRATAPTVFVDLSPNYGFPQVQMPLLPD
MBI5668048.1_Chloroflexota_bacterium                     ATTFNELAQLLAKIPVIGHVHPLIPLCLPPTQVVNDIQGFGYEIDSPELHAARPTLLIRHDKAARFPLVNVLNHPP
NLE46402.1_Chloroflexota_bacterium                       VESINYLLDLLARAPVMGSFHPLLPLALPTVTIADSAQKVAEALDALEQHEPCPILINEYDVRDRFLVERFTELPP
RLC84905.1_Chloroflexota_bacterium                       MTDPAELARLLMQAPVMGPLHPLQALCLPPTEVLADAAALAEELDDLAHHAVRPIWLTDVQPRDRFLQADASTPPP
MDX2239252.1_Leptolyngbyaceae_cyanobacterium_bins.302    EVSLDKLPDYLEAGPIMAQMHPLSLAFLPILKVFDSAAELVAALDNPANECSDAMLALSYDANDRLLSLKINHQPL
MDE0397489.1_Candidatus_Poribacteria_bacterium           ------MAQQLCTIPCITEIRALWELCLTDVWCPVSLEEWRERIDSPREIDADSTLFVNLSPSFGFPAVKFPALPS
MYC73121.1_Gemmatimonadota_bacterium                     ------MIDILEKIPCLTTLQGLWDICLSDIWCPNSLAEWRDVVDAPQDDSTKATLFVNLPETFGFSTVDLPTFPP
RJP56334.1_Deltaproteobacteria_bacterium                 GADVTKLSTAIDAYPAAFRGNRLNLLFLDDAHPIDSSREFLRKVDSLQLQEERREYLISDKEG-HFPTISIESRPD
MFZ4828881.1_Phototrophicaceae_bacterium                 GKNPEILGGVLQRSPAIVKAHPLFPLALPKVKIVTDGLDILEKIDNREHLVNHQTVILDQHPLDRFPTLEFPTHPS
consensus/100%                                           ..............P.......h....h.......p...h...h...........hh........h..........
consensus/95%                                            ......h...h...P.......h....h.......p...h...h...........hh........h..........
consensus/90%                                            ......h...hp..Phh.....l..h.L.......s...h...hp..........hh........h...ph...s.
consensus/85%                                            ......h...hp..Phh.....l..hhL..h.hh.s..ph...hp..........lh........a...ph...s.
```

### BRC-systems PglZ iSwi2/SNF2-Helicase

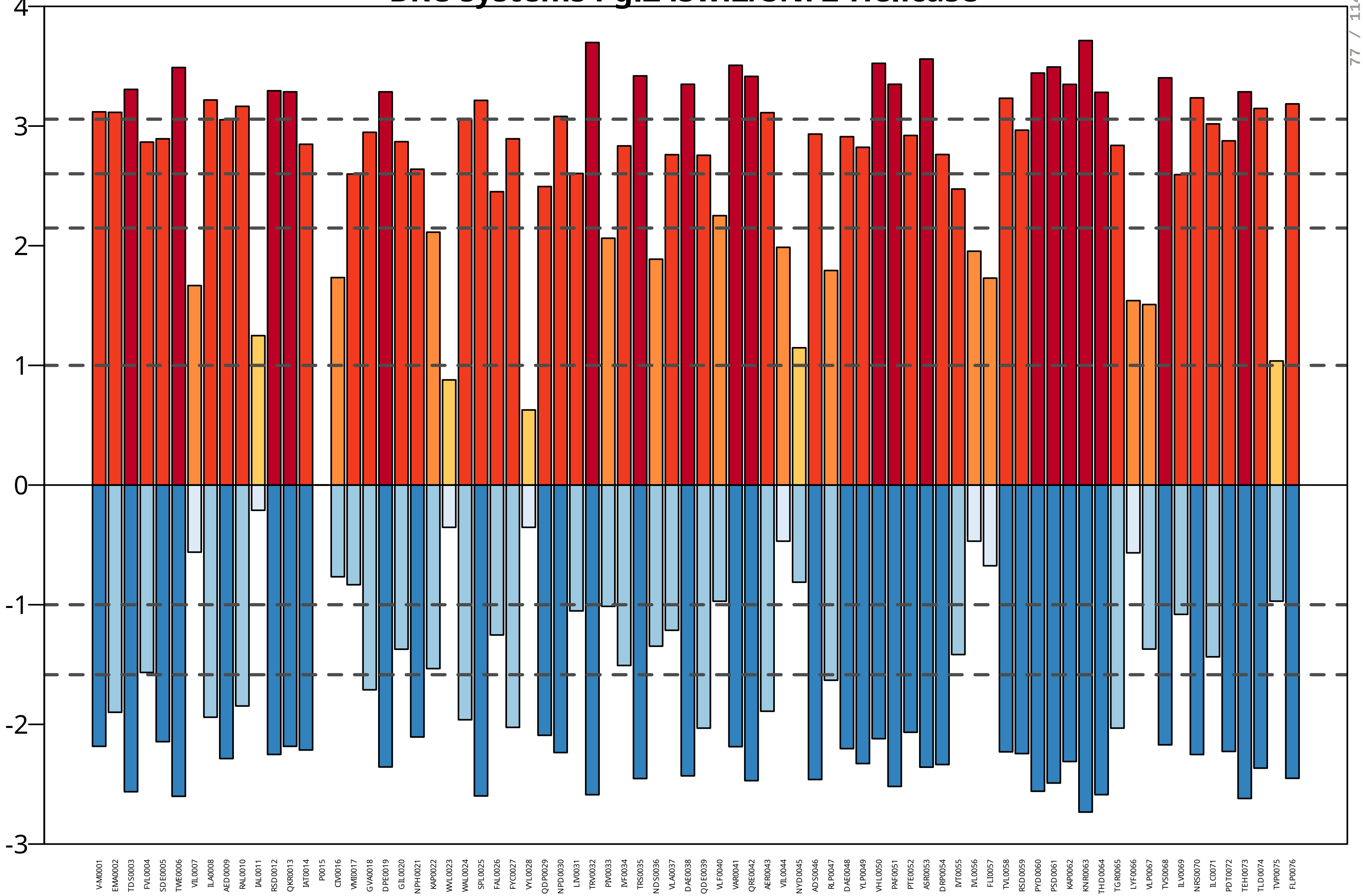

```
# ; Type_1 BREX PglZ CTD IG-like

WP_238754868.1_Leptospira_bandrabouensis       NDISYEILAELFIGKGKHRIELSLENVLFKTLIVELDSVDGARLKIISSNSYDNNGIVVEPNTKVNLEFEITTDYQEKASVKIWSPTLNSSFVKVPL
MBF0258418.1_Desulfamplus_sp.                  GTFELTIQKPDALGFHRIGVLPLSTPGFFSIPHVEVQMFSDQEIGDIVGAQRKGNILTLPLDRESEIYFKLSKGACSKAQIYFKATQQRVLLTNSEY
MBX3247846.1_Myxococcales_bacterium            LRYVVRGSREPLAGLQCVRAVVEVAQRSLEFLELALRATDDAELVDVRGGRTDAGTVLARVGQELEIFFRLATSAVERAQIELSHPSKMALYFEVMP
MBT9554533.1_Myxococcales_bacterium            AKYRTEVLPADVLGLRRVRVRVLPDLSALEFVDVELVATGRPTLKDATGGVVKGGRLAVRVGEWTEVFFGLEGPREEMVALEVVGPGPVAAFFSVSG
MCC6649976.1_Candidatus_Eisenbacteria          SKYEVAARAPAHLGRQRLRVAVRLETASLGFISLALRVPGRPSLLDAGPPTLSDGRVLVPPNREALIEFELEALADEKVRVEIYHPDAREVFFDAGR
KPQ33026.1_Phormidesmis_priestleyi_Ana         VPYVIEAEASAIAGFDRVQVKVKPAQGVLTFLNLSLRVPNRPNIAEATGAQLKNQQVQLAIDQWVEILFKLKGPKDERIKLEVYHSDGAEIYFEVAG
MBD2549130.1_Microcystis_elabens_FACHB-917     TDPAWPGALPEKITNRILMLPIELPVTLLNQRQVVLGAYDSKTPIQAIGADTGTNRLTLTPGQAATIGLMLPAESARKLYLELRDAATDLVLHKSPE
WP_096450969.1_Thauera_sp._K11                 ASSKVALEGPSVLTNRTFGMRVLVAAELFSQVPVRLILLADGRCGMALDADRASGVLMLQPGKPANVAMMLTRDEVNKIRIVAQDPATDAVLAQSAD
MBY3465673.1_Rhizobium_laguerreae              EKNAVSVKLFDAVTNRIFSVEISLSGGMFAERRVRPVAICDGSAKMTSNGAVEDGVVVVEPGKAINVAFLLSDDQIKTLKIQMLDADTDAVLYASPD
MBY0113210.1_Phycisphaerales_bacterium         ATLTATVGDPAKITNRTFRVTLTLADGLFTDASVRVSLVHKGSVGHAIDADAVSKTVKLEIGKAATILMLLQRDDAKQARLVIEDATSRAVLYQSDT
WP_273378697.1_Symbiobacterium_thermophilum    AGDVFKIRLPPRVTNRFFMVTVLYVRSLLSPRRVRLELVANGHAKASQYGDPSSEEITLESGKDNTVTLFFKADASGDLVIQMKDAESGAVLAEEGP
WP_209466794.1_Symbiobacterium_terraclitae     RGSDVRLAPSDPVTNRAFKVDLLYA-SVIEPRTVRLTVLLGDEVMVAEG-NDQTKEVTLVSQKPNTVFLKLSSDGQGEFTVQVVDSRTGLVLAQRN-
MBX6362428.1_Gemmatimonadota_bacterium         KKPAVTLELGRKVTNAIFQVTLSFALDLFDERRFRVTVVDPSKTALATDGREAGAEVELDTGRTSTLVLQLTDPGSGDLELHVADVDAGETILRKK-
MBW2308572.1_Deltaproteobacteria_bacterium     VDIGYELEITPRVTTRMFTVRVVYPAGLFVPVKVAFRVLKGRTAATSMYGQEGTREVILEKERENYVTVLLEGELTGKVSVRMLDADTDAELQRIDD
WP_083610988.1_Desulfatibacillum_alkenivorans  TSVTLKLEFKSAITNRFFSMTATLTHGLFESVRVRAAVLSNKFCAMAAYGEEGSREITMRVNEPNALTFMLSEDGIEKVTIQIVDCHTQLELATMAD
NIM12894.1_Candidatus_Aminicenantes_bacterium  ESIEIKLSISKKITSRHFSVTVIYLKSLFGPIRIKAQVKSGNNAVMSAYGEGGTKEILLKKDKENHITFMLTEEDIQHVFVQILDTKSQIELKRSDK
MBN1165673.1_Methanospirillaceae_archaeon      SSISWSLAIGDKISSRMVSVKINGITSITCDPKVRVEIRSGDNPLTAHYDREATGDISMRINEENTAIMQVHTIPKGQATIHLLDSETDVELTKIGP
MCI0601819.1_bacterium                         VKISWKLALTPKITSRFFSVTVTGESRVLEMPRIKVEARVGNIPVSGSYGNQTTADFALRESEPNTIALMLTEKASGSVTIHLIDSLTGMELVKPNP
WP_062190142.1_Anaerolinea_thermolimosa        KEFHWEVILSKQISTRFFSVTINGATELLATPKIRVELKSGNMPVAATYGDETTKDVKMKIEIPNTVTLFIDDIGTVTVDLDLINSETGLALTQVQN
WP_156746889.1_Mycobacterium_sp._E2733         QKLDIKLTIGGRITTGVFAATFEFLGDLFTDVTFRVVVRGETRIVSGDGYDSESGSVTIAQGKPSVLTFQVISNKDAHIGVQVLDARTGRRLAESVA
MBN2413376.1_candidate_division_KSB1           GDAMFRVSMKKEITNRFFTLQIRYTADLFLPRRVHLDIEYDKQIVTASYGDPRSGDILLEKNKDNSIAVMLNEPIPPTLDLILSDTETLAVLSKTTK
WP_164995432.1_Candidatus_Kuenenia             TAPEIKIILKAKITNRLFSVQLVWQRKLLGTKRVRVAVKSGKEAVHAVYGEDGTKDIILEQNKPNLVTIMLTGEAQNVLYLVLVDAETLVELARLEN
MBV6422080.1_Ignavibacteriaceae_bacterium      PAEDFSIKLKDQITNRIFMIDVLFKLDFVAEEKVKLTIHSQESVVAAKSGNESNRSLELKNNELNTITIMLSDVGISKFDITMYEYESDVLLASKKD
WP_083423498.1_Stigmatella_aurantiaca          QSVRVDLTFSGQFTNRILLITATAFGDMFTDVPVRITVQHASILGDLDERDDATGLVSLKPRQPVKLPIRIADDFRGDIVIRALHATPDVEYGRLV-
NLN06912.1_Bacillota_bacterium                 GKPVIEIRYSDRFITQVIGLKIYLQENLFKQEQVLVEAYDGKEAADCDARDERTRLVTLQAGQEIFVPVLIYPDFNGPVQIRVIDPQTRVVWAKKD-
WP_055669757.1_Desnuesiella_massiliensis       KNENVTISYRNFFTNRVFSINICF-DSLIPSLRTKVDVYDGTTVADCPSRDETTHEVILNSGCPVHIPILIDSDFNGDIEIRVIDSTSKVIWTRLK-
WP_242609850.1_Comamonas_thiooxydans           RKVVVELSYAKRITTRLPVIEVILADDMFSQVEILLEAQDSKEPRPGGDVNPATRTVTLMPGQRKQIALRMDDEFRGKFSVKALNPTTLAAYSNLA-
WP_243520949.1_Hymenobacter_monticola          KRTSLSLTYKPVVTTRQPILTAEAGGSLLDFVEFQLVVTAADKLGLHTAVDVATQLVRLSVQEKARLPVRLDEEFEGAFTVRALHPDTQVELASIT-
OQB98762.1_Spirochaetes_bacterium              HEVGVSLSYTTKITTMRPSFDLSVHTDPFKGIAIAIAARAGGRAQASAGYDPGSGCFRLKPGKSIKITLAMSEEYRGSFTVSAFDPVTHELYAKLD-
WP_166386808.1_Polaribacter_sp._11A2H          QDYQINLTYSGYVTTRRPSIEISCGDSFFDEIDIMLEAIGDNGPAASENVNSTSGFAEIKPGQSLKITMSLHDDFEGKFTIYAKAPATGVILSEIN-
WP_235164441.1_Dyadobacter_chenhuakuii         SKADVTLSHKGFITGLRPSIQVAASESLFAELVVRIEAVAGGTLADQTGVNLLAGTVELTPGTTSSLTLEMEEDFEGNFDVVAINPATGKQYHFLS-
MCP5535438.1_Akkermansiaceae_bacterium         FSFEVNLNYSKKIMTRRPSIEIGIQEGLFAGVRFQLEAVHKKRPAPCQYVDPSTNYVQMPPQSFAKLTLIMDEEFNGEFELRAIDPDTKMPLGKQAK
WP_219938192.1_Succinivibrio_faecicola         DDVSEVISRERIITSGQILITFYQEQPVVKGISIKAGFFRDGVPVSNIEDLEFNKETTNVEEREQKRQFKLSTGIDSGVILKLTKRVENTPYDEITF
MCK6605714.1_Ignavibacteriaceae_bacterium      ENISKVISRSTTLSSNRPVFELVQKSPITLPRELSVGIYSDGLLISDTVRVMFDSKSEDLRERTKKITLRMQSGIENKVILKLSEISEKVPYHDFTL
MCH3907672.1_Sphaerochaeta_sp.                 DDITDVLTNLKTITTGSVAVTFFQEDYVVQGYEATFGFYQDGTAISDIEKRTIASEAIDPKKREFQIVFYLNKESERRVFLRIQKILPSGMIAEKEC
WP_251178637.1_Adlercreutzia_agrestimuris      KKTYETFATGLSITGSTVAVDVYQVQPCILPVTVKVGLYPDGKLLSATKTLVLESENKASDARKSRVTLDVTNDIDRTAMVRISKQVGQTSVWEQEY
MCC6107936.1_Atopobium_sp.                     RSAAPVLTGGKVITGSFVRFEIFQTEPVVLPVSLQVGLYADGQLRSETKPIEAASSSRSAEERRIPVTLAVAQDVAGALTLRVNQRYGTTTVAQAEY
MCI1665139.1_Atopobiaceae_bacterium            RKAETVLTGGTLITGNAVALDLYQTDPVASKIHVRVFASAGEAAISPVEELDLASESEDTAARISHIRLPLASTVRGQVTVKVQRRVGETDLASQDY
MBQ6502306.1_Flexilinea_sp.                    HINQKVANNTRIITTSVKTVVFFQQDPVIQPLTVTAGFYPEGEAVSDEPVITFNLTEPEDRLRQVQQNFTFGNNAVGKVELRLTEQLSATTQTLGVY
WP_237187173.1_Rothia_nasimurium               NTAQTVAEVPQRMTTNLLTIKLLQQEPILKPLKLRAGIYAD-KLISDQPEATFHFTSEMPSDRQIKLTLELTRAADAGADIRLESYNTTTHYDKTSI
WP_149431761.1_Rhodococcus_cavernicola         KDTSKVNTSSPTITSSAVLVTLTQREPVRRGRSLSIGVFDDGTLLSNQRAVVVDSRADDIRERHIAVELVLGDDAMGQVQIRADEVSHDTGYKSTSV
OFQ22229.1_Actinomyces_sp._HMSC062G12          TDVRDVHYDDTSLKVNRFSPRVIQQEPVRRPLTVTIGLEATGERLSSTRVITLDADQRSEVSLKQSSELILNDGVLGQARIVAYKTVNGQAVAVHEL
MBX3116434.1_Cryobacterium_sp.                 SEVKPVKPKSDVLTTAIVRIGLLQAEPIAQARELRASLYGD-KLISDQVVVVFNATGDE-RDRIQEVKLQIRPDAGGAVEFRLEQQSTISTPYKHLF
HJG42481.1_Corynebacterium_phoceense           STNSVVFQNSTVITTQQLIIKMTQEKRVIQPAQVRLSLYKRQVLISDQPKITLGSTSHESRDWEVEVKLSLNSDADGTVTAVIEIPLPNTTLTTQTY
WP_219100161.1_Bifidobacterium_pluvialisilvae  ADVRPVLQRTDRITTGQLTVEFQQTEPVVAQRTVFAGLYVGGEPISNEVPLTFRSESPEARDRHVAATFLLTSEADDTVELRLSERIAGSLEQKADY
MBI9030469.1_bacterium                         DDVKEVISQAKNITSGQISVTLYQSEAVIKACYISVGLYQDDILLSDKHEVSFDSESELSRDREVKVTLRLNEKAEEQILLKLKKKMSGTDYKSDNY
MBD1206695.1_Paracoccaceae_bacterium           DDIEEVIGAGRIITTGQLTIRLYQAKPSRKSVRLRIGLEPDRTKISNEEDRIFDFTSEDPRQRETSVRLLLSKAAEGGVRLVMRRLDHKGVYRSETY
QNI65417.1_Synechococcus_sp._A15-44            DESREVLRVPEKITTARLKFSLFQDEPCFLPITLRISVVNDKELLCEPATLRLDSTAVEPREREQLIDLKLSNAAGNQLELRIDQMLEGVRLETKEV
WP_147932466.1_Neolewinella_aurantiaca         ADALPVIAGNGHITNNEYSVRFFQKQAVRGERRLRIGFYADEKLISDEKNIVFNSEAVEERNRETRLRFRFGADAGQRVKLRLRD-----IVFEHPF
WP_013460554.1_Sulfuricurvum_kujiense          DDVENVVKSFSRISSNQITLTLWQKTPAMLPRTLYAGFYGD-ELVSNEEQLVFGSTDEDSRNREVKVKFHFVQTISNQISLRLSELEPGTVYKQEQF
WP_260916970.1_Aliarcobacter_butzleri          NMINEVMPIS-TITTNNVNVSVYQEEPIIKPITISCGFYEDGIELSDKHKITLNSSDDENRNRENKLKFTFKEISNGKIKFVMKKVIENSIYKTYDV
MCF7790535.1_Victivallales_bacterium           DKVNQVLGKPERITSGQLSLKFYQLEPIRLSRDLRIGFYPDSKLISNREDVSFESVDPVGDNRTKRLTFIFSKDIDNKIELRLEEQIKNSTYKKYKF
WP_153112503.1_Prevotella_copri                QGVSEVLNKKSKITTNTQTISFYQNEPVVKEVTLRFGFYQADNLLSDSITLTFGSMSEDSTQREQKHRFMFKRQLSGQIYLRKEQQVAGSKLEDIAY
MBK9103302.1_Saprospiraceae_bacterium          SDIEQVISRSPNITSNTFGVIFYQTRPVLLPRKLRTGFFSSGQKISDLVTILFNSTDSDSAAREKRQTYIFSNEANAQVYLRMEELIEGTEYKKITY
WP_006745114.1_Bacteroides_coprosuis           DTLREVINQANRITTNQHRVQFYQKEPIVVARDIKAYFASEGKVISDIFTYTLDKKESRSQDREIDYLFTISSHSSE-VFLYIECRIRNSSLKKFRY
WP_053405041.1_Persicobacter_sp._CCB-QB2       NTVKAVIRSVERITTGALAVSFMQSEPVCLAQKIRAGIYQDGVLLSDVFTYNFDSEAAMERQREQTHHFKLNYEASNQVALRLETPVEGTKYKEFFY
MBW8002950.1_Planctomycetota_bacterium         DDVREVMSKSEAITTNQIAVSLFQEEPVIQSRKITLRFETEGQVLSNQRDIVFDSNDPNAQNRTLLASFVFSKEIETQIYLCFYQDRYGQIIDPVPY
WP_262567766.1_Endozoicomonas_gorgoniicola     QRVSDVLGDNHRITTPTHYFTMLQASVVVKAVSLRAEIRDSSKAVSDQQILTFDSASASMDERKKRVRFTLKALQFDRAQYYLVLEKADTEVMSQPV
MBQ2845006.1_Alphaproteobacteria_bacterium     ARVQDVMAAPARITNKIQSFRLIQTDAVNQPAEIKVAVYEDENPITNVVNMTFDSESANMDERIQVATLTLISKPYDKKDYFLRITNTNDILSEYNV
WP_156790427.1_Alcanivorax_sp._DG881           ARVSKVLGQQHRITTNFHVFTLLQTEPVVLATRITIELVNQHEGISSREILNLDSDATELGERQSKVSLSLAAQAVKPNDHFLVLSDAESELQRIPV
OYU71397.1_Alphaproteobacteria_bacterium_PA2   TSIRKVLFHSTRVTTARHRFTLTQTEAVVQPATARLAIYDGDQQISNTAVVTFDSAVADMNLWRKEVWLTLANLPFDAKQYRLIIRDNDDDLPPFPI
MBN1905583.1_Deltaproteobacteria_bacterium     TKVKPVLGTIHKITTSRHKFDIIQMESVVKPVTLKIAIYDAEEPVTNIESVKFESVSDKIDDRKKTVMLVLQNRKYDKRQYRLILRDAETEQESRNI
MCI7405010.1_Desulfovibrio_piger               DILKQVLGQPSRIVNNIQKFEFIQTEKVCLPRKLLVSLSDGKESISDEKKLIFDSVSEDMENRKKSVKLALKNRTFDSATYWLTLRDEETEYLRVGM
TXI44184.1_Methylophilus_sp.                   AKTKSVLGQVLKVVTNMQRFDLMQTESVVLPRTLKLSIQDGHQPISDEQTITFDKSGSMLDDLKVSTLITLKAGDYDRKMYYFVAKDIVSEVIRQPI
WP_006086471.1_Shewanella_baltica              HEKQPVATQPIKLVNNIDKVRFIQTDPVFVARLLDIYIVDVKNEVSSREMINFDSVSKVMDERTREARLKLIGSQFDRAQYTLVLENAETRYSQYAV
WP_216328025.1_Deinococcus_aestuarii           QASRKVVARSRRVTNTLFSVPFVQAEAVVRPRTVTVRLVDAAREVTDVKQLTFESASPHPSEREQVARLSVTLQGPDVATYFLSVTDTEDELLREPW
WP_151792240.1_Acinetobacter_seifertii         KEVKPVLDRNIRFMNPVSFVRTMQTSIVTKSSRVEFYIKDDNQIRSSRIIENYDATKAKKFEKHVRINLVGRDFDSGQYFLVATNLETGQ-VIEYPV
WP_186816219.1_Deinococcus_cellulosilyticus    GVSRKVTSTDRTVRNNTFTVMVVQEEPVIRPRRVRIGLYEKEIAVTNEVLLDLASESSYATEREFPVKLIIGSRKTSSTPYLLEVRDAEDVVTSEEW
MBP7846546.1_Burkholderiales_bacterium         NKISLVLSANFKVTSYNFNVDVFQCEAVILADEVSIEVFDEHISVSEVKRISLASISHEISQRTFKVSLALKQSIYARNKYRLTMRGKAQTKDNQFI
consensus/100%                                 ...........h..................................................h...........................
consensus/95%                                  ...........h.....h.h..........h.h.h...........................h.h........h.h............
consensus/90%                                  .....h....hss...h.h....h....h.h.h..............s.......b....h.h........h.h........h.p..
consensus/85%                                  .....h....lss...hph...shh...pl.h.h........s.....s.......b..h.h.h......h.h.h.......h.p..
consensus/80%                                  .p...h....lss...hph.b.pshh...plph.h.....hss.....s.......b..h.h.h.....ph.h.h.......h.p..
consensus/75%                                  .p.p.sl..s..los....lph.b.pslh...plph.h.ss.p.hss.....s........c..pl.h.lpp...sph.lbh.p......h.p..
consensus/70%                                  pphp.sl..s..lTsp.h.lph.b.pslh.s.plpl.hbss.p.hss.....ssps....pcp.pl.h.lppp..sphplbh.p..s.s.h.p.s.
```

### Type 1 BREX PglZ all-Beta IG-like

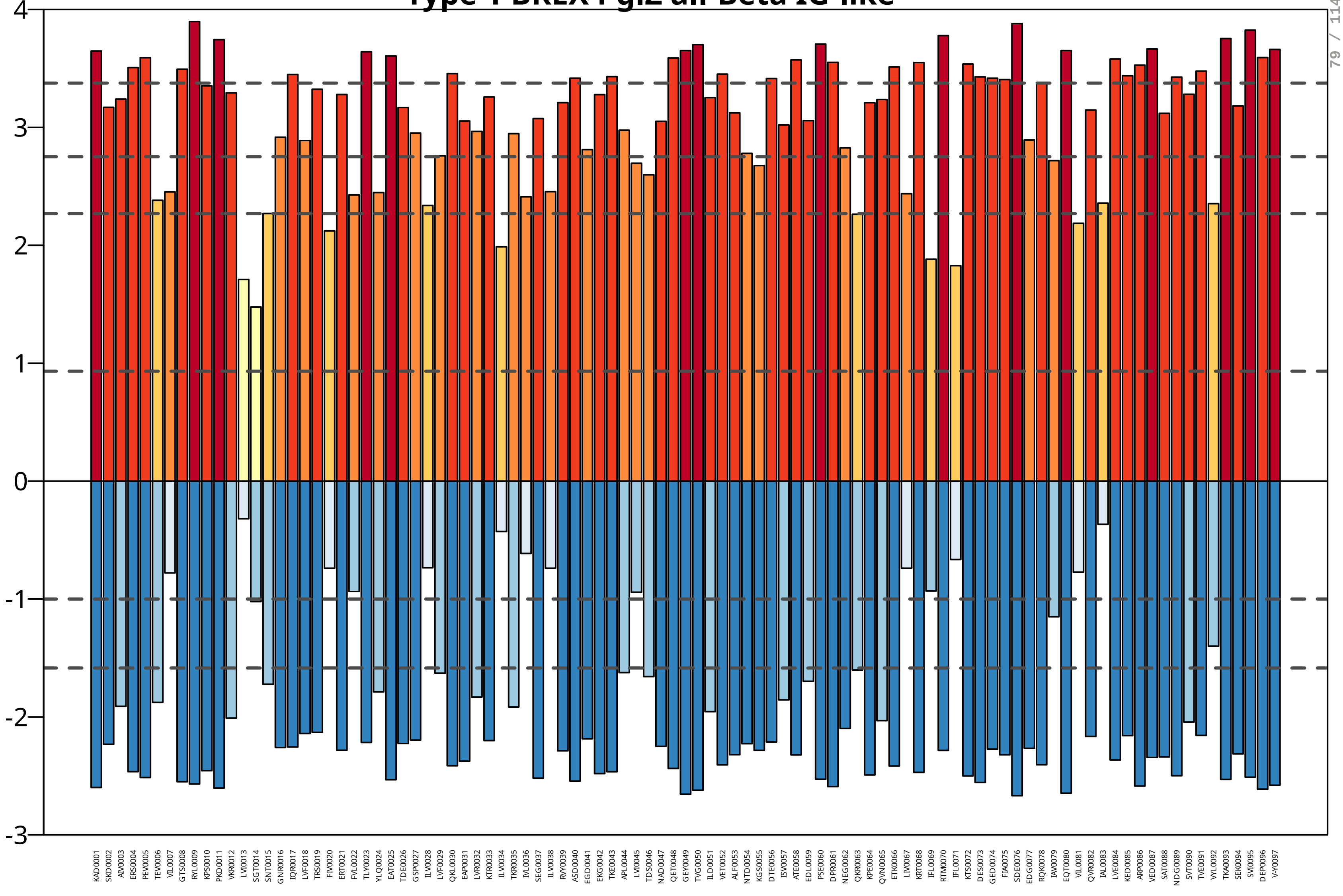

```
# ; Type_3 BREX PglZ CTD IG-Like Rep_1

RGH34270.1_Ruminococcus_sp._AM47-2BH          KTDIYFVSKYIEFHIVSVVVHSNLMIAAPKLIIKGPAFSCECDTKHYKFDIPEKRSGKFTADLYDGDSLIQMTFETKKAVGTKDFF
WP_073324030.1_Fibrobacter_sp._UWOV1          NVEIWIETPEVTVGNAKIKIASKTVLKNVFMEINGKKIPAIESGQKFSFDVPKIKAGMYDVRVFENGNLLKLKCEIKKKMAQRNLL
WP_216471320.1_Butyricicoccus_intestinisimiae EIEIIIENPVVESSPPVIYFYTKTKLENIAVDIQGTKYSVEWDEHSFCIVMDKTRKGMYKASVISDEHILVLEFKIEKAGMKKDIL
WP_044080180.1_Prevotella_pectinovora         NITCRLTDKPIMVSKARLKLFVDIDSDNLAVSVNGKFYALAKDISYYIAEMPEKTVGIYKFNVYEDESLIAMEFEVKKEGASRKFF
MCI8555501.1_Clostridia_bacterium             QINVKLINTLIKVSKAILGIETSIKLQNPYVIINGKKYNAENNEKLAEFILDDKKPGDYIAKLYDGGNQIGLNFIIESKVSSNTLF
MBD8953983.1_Dialister_sp.                    DPMVEFQPALVQLRVGELLLVTNFDMADPYIIVNEKIYRGKLHRRRAIIPLPDKRTHDYMADIYDGDTLLKVAFHVKKEVAKNLGF
MZQ76087.1_Peptoclostridium_sp.               IIRANLVEDVITVSKAVLKIFVTKKFEKIKVKLNNNEYESTEDGRYYTVPIYDIKLGEYEGEVYANEQYIDIKFNVIKPTSNKDYG
HIS35238.1_Candidatus_Scatousia               DINIEVLTKIIQTGKPILKFVSNHLLNNVNVLIGDKRYTAVTDGKHFEVELKGNREKEYSFIVLSNNNHIVLKFKIKTAGMSNDLL
WP_103226892.1_Clostridium_sp._chh4-2         VWEFKVMNRKVIFSQPVLVIWSKNELNNLSVKVNGKMYAGKQDKKTFRFELDKEKACDCSADIYVSSNCVKVKFKFEREGVKSQKF
NBJ16967.1_Dehalobacter_sp._4CP               DTSVEVVDDRLIASPLEFMLFSKTELQRVSVVIKGTPFVAKKDKNHYRIRTEIKRPGKYLADVFDSDNLIGINLDVLSETQKSGDF
CDC05526.1_Clostridium_leptum_CAG:27          TGVIVVIHNDITADGVTLNLYISDTSENIRLVIEDKIYQGISDESHYTFELKDKRAKTYTADVFDGADLIGVSFRVKGKTATKADF
MCL2016626.1_Defluviitaleaceae_bacterium      QLIVQITTEIIKADGIEFEVSLNKPPNPVILVLNNEKYTATPINNKYNVKMPNKKAGTYQAEVHIGNDLINISIKVQGAAAMDDFF
WP_235396406.1_Pseudoflavonifractor_phocaeensis AVEVKIMPPEIFVSKAAIKVFSTEKLVDVSVVIDGHTYEAKPADNFVVEEMTERRAKTYSVDVFAGGMPVALPLRVKKESGSKNIL
MBR3817930.1_Clostridia_bacterium             DNEFIVTTPIVTASEPVLKLFIKYPMDKISIVAEGTSYEGESGKGNYEFVLSDKKVGDYSFDVYYCNGIIALNFKLKKEGQNRKFF
MBP3656735.1_Clostridia_bacterium             EVEVQVETKIVEYSQPVLKLFSKTILTNASVLINGKWYEADADGQNFSFMLAGLKKGQYTAALYANRSEVALTFELRSKAGSTGKL
WP_037355505.1_Selenomonas_sp._FC4001         RLTYAFENDSIERKPTKLVLFCNKEMTNPQLRVKNKFYSGKLDNKHAEFILDDKRKSIYLAEIYDDNKATGLQFKIVKKTHEDLGF
MBS6586269.1_Eubacterium_sp.                  NVEFCFTEQVITLVVPQLTLYANVPMARPRLLIDGEFIDGELDSRHAKFLLPKKRRGEYLAEVYDGDVSRGLAFTAQKNTKEDLFF
NBK77202.1_bacterium_D16-76                   EQQVFFVETSVLCSGSRIRMFANPSLKKPRLVVNGQSYDGKFDKHNVIFDMPDRRKGHYEAEIYNGGIKVALTFETKRQTGTKQLL
NSW92343.1_Bacillota_bacterium                ENRFSLHSSTIRLNEAVLNLTLSDRVEKLSLVIRSVRFEARYKDNKWEITLRNLKPGQYRGTLWADNKNIGVTFELTKGITINDMG
WP_048095116.1_Archaeoglobus_fulgidus         IPLIKVLESKLPAYKPVLHVVVSPPVENVGLRILSELVSGKRDETLWEFDLKSLKPGIYTATVEVGSFKKEIQLEIESGIEEEELL
NPV08138.1_Anaerolineae_bacterium             ELVFKVLSPTVRVSEGHLETVTSLPLEELRLVVAGHWLKGSRGRDRWAFDVSRLRAGHYKGRLESPGQSPGVEFEVRRGIAEDDMG
MBQ7110119.1_Thermoguttaceae_bacterium        EGLVELKTTEVAVKAPVVELFALQRLNGVEIEVDGKRYTASSDGQHFSAELSDKKKGDYSICFGIGAEVLEFDIRVVQGGVKNAMF
HED00230.1_Pseudomonadota_bacterium           KKEFRLWPEKISLDRAVIEIRFDPDLDELHVLLSSPLIKAEKGVGVFRCEIQGLRSGRQWVDLISHGKNIGASFEVVVGMEEEDLG
MBE6101264.1_Selenomonas_ruminantium          SFSVKITNDNITADGIKFNLSLSKSNENLALGYDNQRYLAQRDDYNYEVEISAIKAGTYSLDAYLGDNLVELSIVVSGKSAMNDDF
ROT09401.1_Muribaculaceae_bacterium_Isolate-037 KIECNILDETIFISKAEIKLFISTLSEQYRILINGKFYSAKLDVKYYDFVIPDKKSGNYLFDLYDGDNIVVLSFIVKKEGSNRNLF
MBR2693834.1_Thermoguttaceae_bacterium        KDKIKLVNPEILLRTPELLLSASWSYKEITVEINGRRFTVSADGRQYRVHLGGGKKGVYTANVFSGEEFLAISFCIASKGMQKKDF
MCM1514006.1_Anaeroplasma_bactoclasticum      NVEIRVITEVIKVSIAVLKFYSSSKLSNVTVHVNGASYSAESDGFNFLSELIDKRSGNYSFEVWANNKLVSNNFKIEKESAKNNLW
HHW17415.1_Bacillota_bacterium                ELQFELVDNVVRLNEGVLTMRCNRVVHDLRLLLRGQIISGKSNGREWVFLLRMLKSSQHTAEVYCGTKPVGINFRTVKGITQSDLG
WP_157832981.1_Desulfofundulus_thermocisternus DGAIAIFTSVVKLNEGCLEVELTAPVEKVILQVAGQVYPGILGVQKWVIGIRNLKAGRYRGRLEYEGGFLGIEFEVTRGLMEEDLG
WP_166670004.1_Aminivibrio_pyruvatiphilus     LAPPTLLATFVYRESVSLRVKFSKKISVLSAFSGEKRADCSPSGREWQIHFSEMKQGKHLLRFEADGKMLECSFEVGTRGIADDVL
HCH97635.1_Lachnospiraceae_bacterium          PGRYKINNKKI-LSKATLRVEFNSPVQSLKIITNNGECECIQNVEEWTAHFSNLHEGEAELEIIADNKMIKKIMQVGSRGIQNSML
WP_195464804.1_Clostridium_jeddahense         KVKKKIVNKEFKISVVTIEFRLEVVMEAVSVFLRGKRIVCEKCDSDYTFKYKPDEETEATATIFAGSDEICTKFSIRRPLTTNKKF
HBQ28533.1_Desulfotomaculum_sp.               EKKITLFEAVIRLKLPKVKFKIDKPFTELYATVDAKKYFCHR-EDYWYFEPEVGKKERYVAKISSKGNIGEFEYRIIKGRTDSDKF
MBK5260961.1_Peptostreptococcaceae_bacterium  LSEITVISQLIKIPDIEVQFSVDMSLNKIGVIVDNQKFDCEYNNGIYSFIQHLSKKTDYVAKVVSDGHIIGFKYRVVRPMEKRQEF
WP_006692206.1_Selenomonas_infelix            QTVVSLADKNVFVDGLCLALFADAPINALGIVWEGKFYAGQMDAQHFVVQIPETHAGKYSLAVYEGDTPLNLVVEAQSKGMKNDGF
WP_028051765.1_Carboxydothermus_ferrireducens AVRINVLNERVKLDEGWLEIEVDKPLASLELRIGGKVFSGEESPQRWKFSLQKLGEKEYTAELWELAWKIGVSFTVERKGIKDDLG
MBQ0159480.1_Bacteroidales_bacterium          NHQVKITSEDIMISNPVVSLTIIPEPKGAILSCEGRSYRMNRCGTTWTVVLEGITEGRHEVEVKPGANSINKPINVVGGVNSNDMF
PMQ01343.1_Dictyoglomus_sp.                   KFEVHILKTDISIRNPILEVKIKPKPKVKKIEYSGRLYEMNKESDIYSIKLENIKVGNLSIKIKIGEFEKPFEINIIGGMKERDIL
KUK66682.1_Parcubacteria_bacterium            EYKIELLTKEINIRTPILELKIKPDPEKVSFLVKNRDLNADRDKEIYKINLSGYKAGEYTLTVKIGLWTKDLKFKIRGGLREKDIL
WP_018963723.1_Coprothermobacter_platensis    KYEVKLESATISVRRRVFRFRIKPEPKSVHVTVGNTRYEVSKDGDWFEIKLSNIKSGNYTAKITLPGFEQEVEFAVKGGIEER---
MBR3117632.1_Oceanobacillus_sp.               KHSVELKSGLLSGLNRSVTIEILPPTEKAVIVEDGSKHIMKVSNNIWVSPINS--IKSQVLTVKVGNSDYKVSVSSSLGASQDDGF
WP_024292379.1_Lacrimispora_indolis           TYKVNPKDLKVSGMKKELEFKISPKPDSALMAKDGTNINMDQEKKIWIGKLKR--GIEQTIEVCIENQYFCFTIPPTKMG--DDLF
HAR58249.1_Acholeplasmatales_bacterium        NYNINPIKTTVNGLDKIIKIEITPKPKNQLYIDFGNQGVFNEGNNVWSVKLQV--ITNQNIKIAIGSFSDKLNVCSSSGITKDDGF
WP_191216537.1_Methanothermobacter_sp         IYRLKESEIEVEISEPKVPFEIYPTPTETKVLLNERELNITIGKGCYEIDVSDLKAGEYNVKINLKKQFLDLKINLKAGFRETDLF
OQC39307.1_Bacteroidetes_bacterium            DFSITIEKTEIPKKESLVSINITPKPSIANIDIKGKITKLNEMTQKWEVNLDKSLSGKIPIKVKIGKCEKEFTINIISGIIEEDLF
MBQ4420212.1_Bacteroidales_bacterium          NYTIRIKKNKIPLSNPLVEIFVSPKPTEVKMKIGSVYIDMHETGDMWVASLPDAKEGNIRIAIQPGGAAQNFDIEVFGGFGNGNMF
WP_183546997.1_Methanococcus_maripaludis      EYEIRLQNDIINTK---ISLEITPWPKSVLKDQDDNEYILYSDESFWNLQNPEKKASVVTYAVVIGNIIKNIEINTKGFIE-KDLG
MBU7048061.1_Theionarchaea_archaeon           SYKVAPQEQDISIRNPTIHLSIEPFPDIEPLAYKDRTFDFDSTEDKWSAILEGLKTGKYELEIRIGDWTGKISIEIKGGMKERDLL
HHW21152.1_Thermodesulfovibrio_thiophilus     TYNIKLITDEISIKNPVVIFEISPVPNKKFLAYKKKEIELKNEGGQYTNKININKAGQYKIKVKIGDFQQEFNINIKSGFKEEDLF
WP_207678199.1_Desulfonema_magnum             IYEIKLLTKELSTKNPIMDILIKPQPSTPVLTFKGSFIKLKQKDEKWRASLKGFKPGNYQFKLNVQEQKFEMEVSIEGGMKEKELF
MBR5399199.1_Bacteroidales_bacterium          DFHIIPVNDKVAVSEALVEVTIIPQPTAAFISFNGVEYEMKRKNTQWTAKIDGVSEGINNFIIIPAGEPQNISIEFFGGAGSDDEF
WP_022459608.1_Bacteroidales                  KYTIVPVEAKVPISDRKVAFKIIPEPLSSKAIFNGQEIALIRHGMQWEGIIPNATEGKHHITVIPRGKPVQLEIEFFGGFNSLDLD
MCQ2340464.1_Paludibacteraceae_bacterium      PYVIQVIQNRLPLSKPVVSMQIIPKPDSCKLMVDGVEYDMNTTKDIWETTINNPEERTYKCIVKPKGTNHPVDVEFYGGFGSNDIN
MBE6838777.1_Ruminococcus_sp.                 TYKVFDENLQVNGIDRVVRFSIIPDVEAAVVESSGEKHSLVKVKGMYQAELSS--GKEQNISVLIEDKVFGFTVNTAKKNMEDDGF
MBR3710351.1_Bacteroidales_bacterium          RFKYELVSKELEVSEPVVSVVIMPQPSKVKLTIDNQEYNMTRSAGKWSVRVEGLEEGEYKVDISIGGTSHSETIKVIGGFGGNDFL
WP_205109179.1_Marseilla_massiliensis         TYSVTIDNDEIDGRQPNLKFTIKGLASVDTLEYNGVIYHLNLGGDKFESERLNLVDTATRAVVRINDIFGTFNIKISTGAVEDDLF
WP_172204608.1_Prevotella_sp._PTAC            EYSISLIDDAVNGNNPILMFRIKGVTNLEKLIYNNTGYNLNQGHFRFESDRLELTQEVDEVEIRIGSYSQRLYALVP-----NHVF
MCH4183501.1_Prevotella_sp.                   NWTAELLTTEISGANPTFQFVINELPSTEYIVYDGTRYELNTEDDIYESDPVIINEKACEVQLVIGSKLYIKNVTVLTGAKEEDLF
WP_196042834.1_Parabacteroides_merdae         TWRAVFKSFEVEEANPVVVYSIAGLDNNQMVEYDGKFYSMIT-GNTYTSERLPLNKDENKVILHIGTWQKEPIFTIKMAVIEDDPF
MCI6981218.1_Akkermansia_muciniphila          ACSVKLITNAPYARLSRLEFEIRGLKKSDCVVYNSKKYLLRDKGVSFMTENIPICTQCKGIRVMIGVQSWDFSLEIRGGAVEDDFF
MCD7797870.1_Akkermansiaceae_bacterium        RINAKILTKKISPIRSNIQYQIKGITSSAEIEYNCKKYTLKVDESTFECPKVNIQQTETKVILRIGASFEQVTLIVKTGAEETDLF
MBO6251938.1_Muribaculaceae_bacterium         TWEARLVEVEVSATHPVVQIHITGLSAMDLLIYKGNSYKLNKGNNIYESERMPLEENECTLDLHIGDYSEILKITVNTGGHMDDMF
MBR4838437.1_Bacteroidales_bacterium          DWTATLQTKTLSESDPIVRYVFKGDTTNLRIVYNGKQYALRINDTLFESDRLTLDKDATTIKLVVGDIEQKDSIVVNLGATEDDLF
RGX95620.1_Prevotella_copri                   HWSVVVLSQELTTVNPQVRLRIKGLSASDYVVYNNKSYPLKLGMDEFMTAPLEANENDSELSVLVGDKVIDRSILWKKAAQEEDLF
WP_254009599.1_Bacteroides                    NWKANIISSNVSANKPVVSIEIKNAPIDAYLIYDGKRYEMEKQNEIFESEELILNADVKRIALYVGPDKKDFHIDFSKAADVDNMF
consensus/100%                                ................h.........................h.............h.........h...........
consensus/95%                                 ...h......l.....h.h.........h.............h...............l.........h.......p...
consensus/90%                                 ...h...p..l.....l.h.........h...s..h.......a............h.l.........h.......p...
consensus/85%                                 ...h.h.p..l.....l.h.....p...l.hpsp.h.......a......p.....h.l.......hphph......pp..
consensus/80%                                 p.ph.l.p..l..p.s.l.h.hp...p...l.hpsp.h..p...p.a...b...p.sp.phpl..ss....hphph.....ppp.h
consensus/75%                                 p.phpl.sp.l..p.s.lph.hp...pp..l.hpspbhphp.s.p.aph.h...p.sp.phplb.ss....hphpl.....ppsbh
consensus/70%                                 phphpl.sp.lphpps.lph.hp...pp..l.hsspbaphp.s.p.aphph..bc.sphphplb.ss..b.hphpl.....ppchh
```

### Type 3 BREX PglZ all-Beta IG-like Representative_1

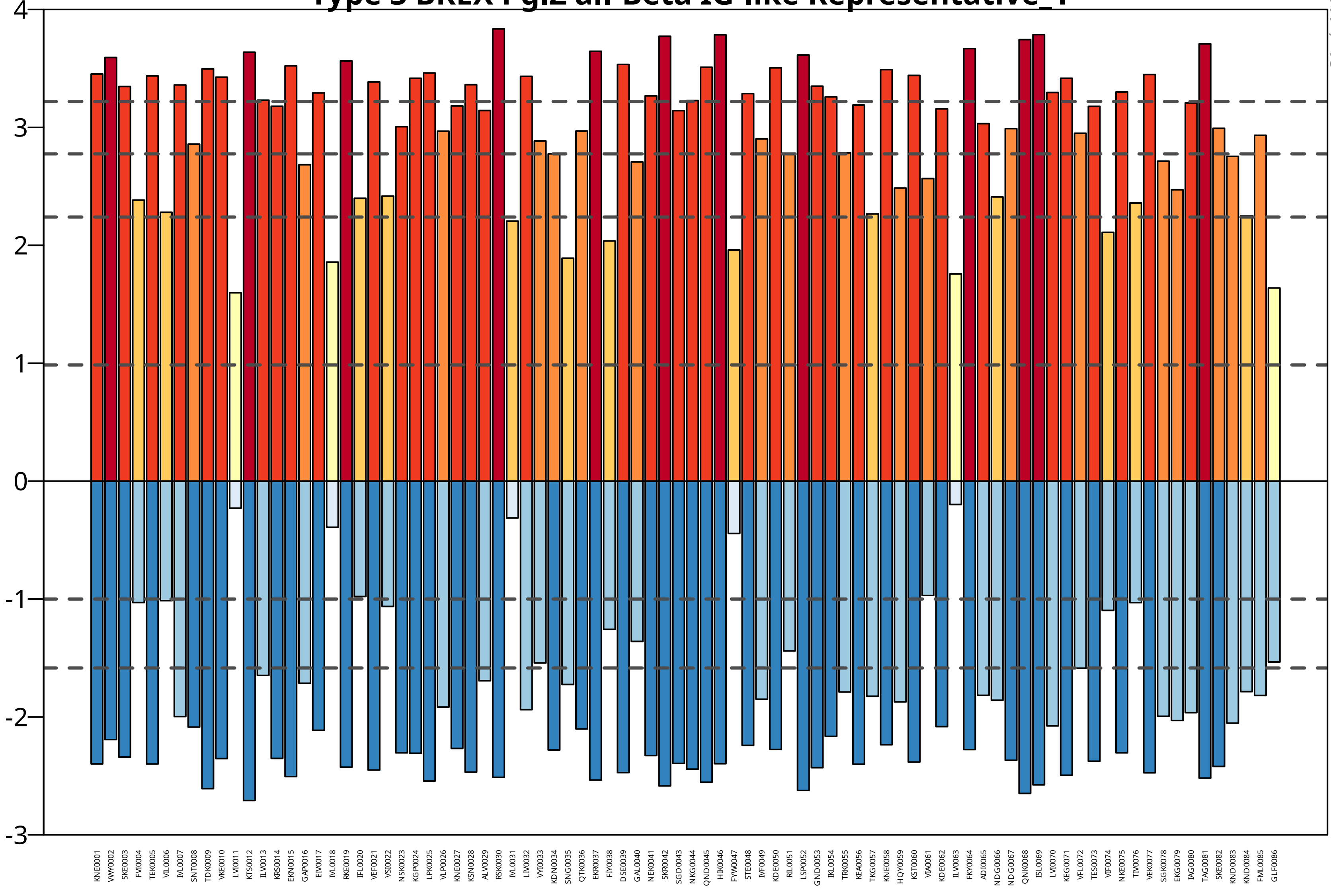

```
# ; Type_3 BREX PglZ CTD IG-Like Rep_2

WP_192822482.1_Rufibacter_sp._LB8         LMIGGFLSGESRSYRQGKAELHLNNSRPARLRKVKVQIPRVELLPGTIGISKSIEIPRWPDTAQAEKTVAQVTLETDGELHTFNLNTNLRNTDLRQSSTPLDDLF
WP_105482027.1_Abditibacterium            IRVEAVLSGKGRAGRLGKAALRLINFRALLMENIELRYGGKVTVAAFGHAEIEVQLDAWPETAQARDLTMLVLLRADGRLVQSVAEIHLQVEALQTRENILGDLI
WP_243797665.1_Hymenobacter_tibetensis    IKITGSATGRARAGSTGTITLQLFNERPVRLRRLQISWTGTTTLSGQQSTSVSVPLSNWPDGLRIGTATLFITVELDGREQQFSLTNQLSTDEFQTRQVLLGDL-
MBX7251230.1_Candidatus_Promineofilum_sp. PEIGVTIRGRSVAGRPGQATVQVLNPVSVTLTWIDLRRPKGLTIAPRSQAQAMLDIDPWPTPAEVDELTAEAVFHLSGKRFDVSVSLELDSEEMYGRDNILEDLV
WP_162787804.1_Anaerolinea_thermolimosa   PELDISISGEGEANMSSKVFISILNSMSLECQEIEISGENYKEIPPLDKVIFQEILAPWPSKIIERNLTAKVTFTLNGKIYTQNVTPDLKVQILYQQSDLLKDLL
WP_012642882.1_Thermomicrobium_roseum     PSPVVAIKGDGRAYGSGTLVVMVTNEIPLQLLKLELEFRQRETIGPLSQVTITRQIESWPTPQDCREARVRLTYRLSGAHPTIQEKIELTSRTLYDQTNILEDLQ
WP_157205838.1_Calidithermus_timidus      KDIEAELLGKGVVERAGQIVLQVRNPISVVVLELSFGAKQQREVSAMGGLEHALEIPSWPSKRDVQSLRAQVSFLRDQRPAYVEAKVRLEVEEMYQQGDPLEDLL
MCL6576921.1_Kyrpidia_sp.                 LPTKAYLVGTGVAGRINNAVLVVENPAPVELVQLELILGGTSMANPMSRTEWSIDLNPWPEKKVLKNGQARLIYRLGGKQDAAPVELRLTSLEMYEQNDILGDLL
NPV84914.1_Anaerolineae_bacterium         PKISIEITGHGEANKTGTLLVKITNQVTLECLEISLSAVINCQISPLQETTFKFNTISWPTKSALPSLKAKLLFRMNGKTFEQEIMPKLEADELYSQPT-LKDLG
WP_139401683.1_Deinococcus_radiopugnans   RLPTLTAEGDGQAEEPGTFILTLTNPVPLVVERLGGPLGAQWTVAPQSQQRLTVALGSWPTSAALSALTLRATVRARSAAQDVAVTLKLKSVELYTSTAILGDLL
MCE5327915.1_Planctomycetaceae_bacterium  PNAVAKITGKAREGAEGRLTVAVTNSISLQATWLEIAFGTAHEIAPLTKREYVISVADWPAKPNAISARAWIGIRPAGPEIRIETEVVLDSEGFYVQDAILGDLT
OGX83061.1_Hymenobacter_glacialis         VKVAASLTGAGRAGFPGVAQLRLRNDHHLTVVHVSIRFGAAQPSVPLKTIDVSIDLPSWPHSSEVQSGQLVVVVETDGKQASFPVDANLTNTGLQDRGVIGAEFG
MBN2678362.1_Anaerolineaceae_bacterium    PNIEFAFSGAGRATLPGKIIVSTINPIDLDLTKAEINFGGDKNVPGLKKTEFEIDITTWPSSEQVSLGKANFTLRLSGEEFGVSSLVGISVTELYTRDKILEGLD
MBV5327602.1_Chlorobium_sp.               IILEATLTGKGVAKSTGVYLLEILNTVRIQVLEIELIQIGYFEVAAMDKKLINIPVTYWPTTLESNSLEASILYSLTGERKVLVIKPSLETEEMYQQEDILSDL-
WP_062417760.1_Levilinea_saccharolytica   PDATITISGNGRARRDGTFAVTVLNKIDLFLETVTLTLRTGQKIPARTQIPAEIPYSPWPSAGELETVHATVRLRLNKMVYEYSAETSVQSEDIYTRPEILEDLD
MCC6188628.1_Anaerolineales_bacterium     PQVEVTLTGTGRARRTGILRLTVMNMVPVELRSLTLDVGRSLKIGQRAKAEWEGDLSPWPSPSDCRAASAIAHCVPNGVSFDVTVSASLQSEDLYIAPDILEDLT
MCA9963066.1_Anaerolineales_bacterium     PAVTVTLRGSGQAGQSGQFILQLRNPISVTAVALSLLIAGQQAITPYSEQTFTLSWSSWLKRAEVSQTTVTLSLQLNGLSFTVPANPEIESIEMYRRDNNLGDLD
WP_215594968.1_Hymenobacter_piscis        PLITATLSGAAPAGRSGIATLELLNPRTLTLRELHVRLGNKGSCPGLQTLSVSVQLPDWPDSAQLQTAQAKVLLQTDLQEHTCSVGLNLRNTGLQDRTDILADLF
WP_006100361.1_Coleofasciculus            APIIIKCSGEGRPGENSPLRVDIYNPVAIEDLRLRVNQLVSVTVQPHQRQLIEISISTWPELPPTHELTGKLEFQYNAERGWADPESSIEVHQIFSSGEGLDDFL
WP_254031239.1_Planktothrix_agardhii      YPVIVTCRGEGKAKQPGELTITIHNHVSLTNLCLYIDQLNDITLKPNDKVTHKISISEYPELAPDKNLSGQLTFSFNTDSSQSNSQSYIIIKQIFSSGQGLDDFL
WP_273058112.1_Shewanella_vesiculosa      SYNNFTLNAISSVSGKWCLRLKISTPVNIDNFRIKLGKHAIERINEGQTIEFDFAFECTVSQEGLTDIKFDLQFEVNGNKKIQTQSISVCFPPKSLASKKFEGMF
MCK9609000.1_Methylomonas_sp.             VKMPLELATLSDSDKSWQISLELSAGINVKNINLKLGEESISAISAGTSQKLILAFSALTEQSGLTEITIHLSYDREGAHEINTKQISCIFPAPDAATQGFEDMF
WP_017840083.1_Methylotuvimicrobium       LETHLEVSLENNDDKHWQIDLVLTSSVTASDICIKIGAETVDSLRVNKSQKVALRFNCSNGQEGLIRIELVISYHHSGANEQLIKFIDVNFPASDGGVQSFEGMF
MBC7570526.1_Spirosoma_sp.                VPLVLTYYGSSDGKTIKDFGIQVPNPFAIELLTLSIVEDLPMRIEPTASATLTGSIAQKKPQQGQLSLAVSITFRANAEESAQSATLTVPIQTSELDD--FDFDF
WP_198076877.1_Hymenobacter_negativus     VELAISYFGKKEGKTLKDVQFRIPNDFIVELRGLSLAEDLPAQLAPNSSVALKTKLPSSRPKEKSVSLTASLEYVVQGDPYTTQASVSVPIQQANELD-DFDFDI
WP_235137408.1_Dyadobacter_fanqingshengii VKLSIRPIGFNEGKSERNFKFRISNINSIQISKVTVLQELPLDIPANAEADLTGTLHPRHSKDGKLNLELAVKYTLLGHHYEDVINLDILTERD-----EFENDF
WP_194112254.1_Mucilaginibacter           QQLQIRYTGIRDGKTEREFKLKVVGTTAVELQEFTILEDMPVTIGPGADKPLHSALPQKNAKGGKLDLNLEVTCLILGVRVVQSLKCTVTTEKD-----DFEDDF
HBJ34341.1_Planctomycetaceae_bacterium    TWPSVSFPEAGYDRGVFRFSVTLSRSVPTTSCTLRIGEVKLPAMSCGDSTKLDIVLPLRTSDCDTVAIVVTMRAKETGELIDKCNDFIVPRKKLTNDQDDFENMF
MBX3399563.1_Gemmataceae_bacterium        AFPDVTVPNGATDRGQWILTLQLSHPIQTNGGRIRVGEESYPALRPNATHMLPLEVPGQLPNDQELVLEVTLSPRGDTPAPDLTRHVPVARVRQTKDQAAFEDMF
WP_099078666.1_Vibrio_sp._PID17_43        HEQPFILTSDEVGEKRWNITLSAIKNASEEHVTFKMGNGNW----DSANEQLHLSLHSSVNHEGLVEVAFTVGFKG---KHVTTVTLEIDFPKSDKASSAFDDML
SEH06866.1_Thiotrichales_bacterium_HS_08  AKIPIKVNVLKPLEKGWHLELSILPRIRLNKLQLQFANTNYSDLQAHSETKIPLQLVSEQAQSGLTQVDLTLRYLPQTTWTSLTVNLSLDLVAPTQEADNFNKMF
QEP42412.1_Ectothiorhodospiraceae         PLPPLQLQPKRYAEQQWLVQLVIQLHETVERVKIRAGQVRFNQID--EALNLDLLVVANQAQRGVISLPLTLETERNHEVMVLEESVEVEFFASDESNQLFEGMF
WP_194715675.1_Noviherbaspirillum_soli    KTPDVQISSEQDSGEGWTVTFKVASATHLRRVQIVCGHCTVENILEQQSKLVTFNIRSTVHQEGAVNLMLNVAYVLGVGFQHVQLPVSLKLRPHEQGSADFDNMF
WP_201382003.1_Ktedonobacter_sp._SOSP1-85 FEPVINISGFLFSSKVNALQVTLMNPFPITVLQLLVADSIQSYVEPNAMITGEVILPAVVYRESARLVTWSLTCEGGGIRQVFRGDVSIPIRRLQTSD--VDEMF
WP_221704724.1_Desmonostoc_muscorum       ISPVVKIEGNFKPNQVSTLRVLVLNPVPLRKVRLEISNAIWSDVPAFSKSSIEEILPA-SHNGLKQTVEWFINCEAVGKTWQFSGQIEVAIRRFQVNS--VDELF
WP_007916556.1_Ktedonobacter_racemifer    LAPILQIEGSLRTGTISKLETIIPNNFPLKIIQFSIDNVKPTRIGANSTILIPIQVQT-TSNQSDQYIKWFLSYERGGQHTSIEGSVNIQVRRLYANT--LDDMF
NJL05928.1_Chloroflexaceae_bacterium      WALDVALHGQLRANQAGTLELEIPNPFPVQVQQLTLDTDAPFAVAALAKDTFPVAVAPPTTRADQLHVQWHITTTSPAGQQTQHGAADVPIRRLQTTDPDFDAMF
WP_129675140.1_Candidatus_Chloroploca_sp. VNLQLTLDGQLQPQREGVLSLLLPNPAPIDGLVVRVADLTIERIAAQDRYETTIRFPARPINGTKLSLQWTLDGQVLGVAYHQQGEASIKVRRLQSGN-SIDDLF
MBU6335166.1_Chloroflexota_bacterium      HPPVLRLSGVPSAIDDATLVVELFNAGALEAVTLVLDGVPLPSLPGGGHARHAIAVPRCHSAAA--TIDWWLEYRCLGDAYRTPGPAPLAVRRLHHDDGGFDAMF
MBK9943589.1_Kouleothrix_sp.              LPLMIELSGTLRVGQPSQITVGLPNRFPIDTVTLDLGAGHIDRVESSARCDLVIEFPAIASALSELPIKWRMRYDTVAGTQTQDGRLLLPIRRLQAEDMGFDDFF
WP_097643873.1_Candidatus_Viridilinea     MPLELVVQGELRARRAGTLTMLIHNPFPIENLHWQISGGQIAILGPNASQTISISFPPINPAGDTFTINYTLDFSHFGKPEQNSGMCKIRVQHLYIQDEGLEDMF
NJO81810.1_Blastochloris_sp.              IPLDILLQGTLRAGQSGILTLIIINTFPVEQIVVTVTASEVARIEGLAKATAEIHIPMVTVSGTELPVTYEVQYYAFGNSYQTMGKTTIAVRRLQVEDTSFDDMF
consensus/100%                            ..................h.............h.....................................h...............h................
consensus/95%                             ....h.h.s...s.....h.h.l.........h.h......h..........h............h.h...............l..........h....
consensus/90%                             ....h.h.s...s.....h.h.l.s...hp...h.h......h.s..p.....h............h.h.h..............l..........h..h.
consensus/85%                             ....h.h.u...s.....h.l.l.s..slp..plpl.......l.s..p..h...h...........hph.l.h...s..........l..........h.sh.
consensus/80%                             ....h.hpu...s.p...h.l.l.s..slp..plpl.......l.s..p..h.h.hss.s.......hph.l.hp..s.......p.pl........s..hpsh.
consensus/75%                             ....hplpG..ps.p..ph.l.l.N.hslp..plpl.......l.s..p.ph.h.lss.s.p....plphplphp..u.....p.phpl..p.b..ps..hcshh
consensus/70%                             ....lplpGp.pu.p..ph.lpl.Nshslp..plpls.....plss.sp.phpl.lssbssp....plphplphp.su...p.p.phpl..pphb.ps..h--hh
```

### Type 3 BREX PglZ all-Beta IG-like Representative_2

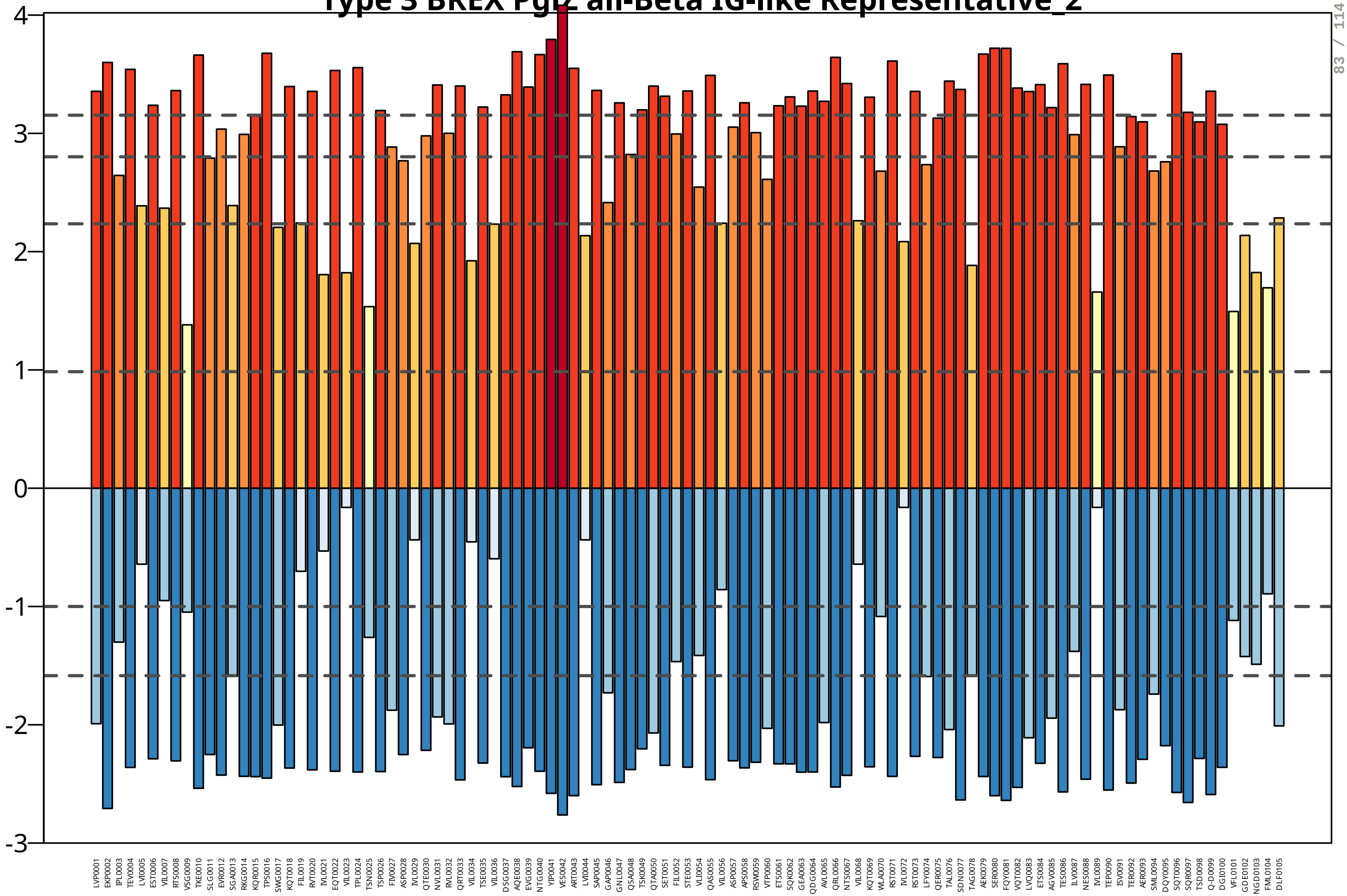

### ; Type 1 BREX BrxB-iSTAND NTPase; (For Supplementary ENTROPY)

```
WP_258176317.1_Halomonas_ventosae [illegible]
WP_262567771.1_Endozoicomonas_gorgoniicola [illegible]
WP_094203898.1_Finegoldia_magna [illegible]
MCK7578718.1_Chromatiales_bacterium [illegible]
MAE16012.1_Deltaproteobacteria_bacterium [illegible]
WP_165775784.1_Bifidobacterium [illegible]
HIU24259.1_Candidatus_Coprovicinus [illegible]
WP_168040576.1_Neolewinella_antarctica [illegible]
WP_151792237.1_Acinetobacter_seifertii [illegible]
MA064837.1_Balneola_sp. [illegible]
WP_245747814.1_Methanolobus_profundi [illegible]
WP_167658113.1_Pseudoalteromonas_sp._HF66 [illegible]
KAB1640464.1_Ellagibacter [illegible]
MBU6197214.1_Cyanobacteria_bacterium_REEB446 [illegible]
WP_235341611.1_Bifidobacterium_boum [illegible]
WP_012411903.1_Nostoc_punctiforme [illegible]
MBR4606064.1_Lachnospiraceae_bacterium [illegible]
MCB1062256.1_Verrucomicrobiae_bacterium [illegible]
MBD1206698.1_Paracoccaceae_bacterium [illegible]
WP_159471216.1_Dyadobacter_sp._3J3 [illegible]
WP_168675170.1_Hymenobacter_artigasi [illegible]
MBP6733458.1_Chromatiaceae_bacterium [illegible]
WP_231782752.1_Rhodococcus_qingshengii [illegible]
MCH3907668.1_Sphaerochaeta_sp. [illegible]
WP_013460545.1_Sulfuricurvum_kujiense [illegible]
MBQ6141564.1_Kiritimatiellia_bacterium [illegible]
WP_048345182.1_Lactobacillaceae [illegible]
MBQ6502299.1_Flexilinea_sp. [illegible]
MBW7869162.1_Brumimicrobium_sp. [illegible]
WP_019974717.1_Empedobacter_brevis [illegible]
WP_222635590.1_unclassified_Rhodococcus [illegible]
TVQ28582.1_Spirochaetaceae_bacterium [illegible]
RKX87881.1_Spirochaetota_bacterium [illegible]
WP_217359511.1_Anabaena_sp._UHCC_0204 [illegible]
WP_238754389.1_Leptospira_bandrabouensis [illegible]
MBY0113215.1_Phycisphaerales_bacterium [illegible]
PZV04978.1_Cyanobium_sp. [illegible]
WP_024332154.1_Winkia_neuii [illegible]
WP_251490388.1_Otoolea_muris [illegible]
WP_153861779.1_Fundicoccus_ignavus [illegible]
WP_196032769.1_Collinsella_aerofaciens [illegible]
KAF3303186.1_Carnobacterium_sp._PL17RED31 [illegible]
HDN19843.1_Candidatus_Acetothermia [illegible]
NBI63882.1_Clostridiales_bacterium [illegible]
MBQ7190493.1_Kiritimatiellia_bacterium [illegible]
ETR72220.1_Candidatus_Magnetoglobus [illegible]
WP_224137561.1_Candidatus_Kaistella [illegible]
WP_098060819.1_Longimonas_halophila [illegible]
MCF0260591.1_Erysipelotrichaceae_bacterium [illegible]
MBX6362418.1_Gemmatimonadota_bacterium [illegible]
WP_270462425.1_Gemmiger_formicilis [illegible]
WP_215020743.1_Bacillus_sp._ISL-40 [illegible]
MBR6253007.1_Clostridia_bacterium [illegible]
WP_273738558.1_Natrinema_soli [illegible]
MBK7566197.1_Bacteroidota_bacterium [illegible]
WP_200392182.1_Roseibacillus_ishigakijimensis [illegible]
WP_242820840.1_Deferrivibrio_essentukiensis [illegible]
WP_235327429.1_Bacteroides_caecigallinarum [illegible]
NMA18736.1_Clostridiaceae_bacterium [illegible]
WP_163588065.1_Ligilactobacillus_murinus [illegible]
AUX12765.1_Latilactobacillus_sakei [illegible]
WP_162921703.1_Listeria_costaricensis [illegible]
MBD5357713.1_Bacteroides_sp. [illegible]
MBP6234855.1_Saprospiraceae_bacterium [illegible]
WP_215708905.1_Bacteroides_uniformis [illegible]
MBQ0129311.1_bacterium [illegible]
WP_028665034.1_Runella_zeae [illegible]
MCI5856702.1_Agathobacter_sp. [illegible]
WP_061570111.1_Caldibacillus_debilis [illegible]
WP_271398743.1_Salinicoccus_roseus [illegible]
WP_091403824.1_Geotoga_petraea [illegible]
MCK6587210.1_Polyangiaceae_bacterium [illegible]
MBN1450029.1_Anaerolineales_bacterium [illegible]
MBE7502026.1_Verrucomicrobiales_bacterium [illegible]
WP_261427665.1_Serratia_proteamaculans [illegible]
MBQ8347029.1_Alphaproteobacteria_bacterium [illegible]
MBQ6248037.1_Kiritimatiellia_bacterium [illegible]
OI089049.1_Anaerolineae_bacterium_CG2_30_64_16 [illegible]
MBX9598965.1_Burkholderiales_bacterium [illegible]
WP_191006190.1_Hymenobacter_montanus [illegible]
MBR3160179.1_Atopobiaceae_bacterium [illegible]
MCH8487208.1_Cyclonatronum_sp. [illegible]
MBP2454131.1_Mycolicibacterium_lutetiense [illegible]
MCK6605710.1_Ignavibacteriaceae_bacterium [illegible]
GDY21839.1_Verrucomicrobiota_bacterium [illegible]
WP_025641434.1_Schnuerera_ultunensis [illegible]
WP_201561161.1_Psychrobacter_glacincola [illegible]
RKY59329.1_Candidatus_Latescibacteria_bacterium [illegible]
UCC65223.1_Anaerolineae_bacterium [illegible]
WP_154435942.1_Olsenella_porci [illegible]
WP_075010320.1_Stigmatella_aurantiaca [illegible]
WP_187771726.1_Phascolarctobacterium_faecium [illegible]
WP_252697946.1_Natribaculum_sp._CGA3 [illegible]
MCC6107939.1_Atopobium_sp. [illegible]
WP_255679049.1_Methanosarcina_sp._DH1 [illegible]
MCL4835317.1_Caldilineaceae_bacterium [illegible]
WP_012508723.1_Pelodictyon_phaeoclathratiforme [illegible]
MBL9184318.1_Verrucomicrobiaceae_bacterium [illegible]
WP_157203231.1_Methanofollis_liminatans [illegible]
MCC6785238.1_Planctomycetota_bacterium [illegible]
WP_050592628.1_Mesorhizobium_sp._L2C084A000 [illegible]
WP_100919676.1_Candidatus_Thiodictyon [illegible]
EMR73246.1_Thaumarchaeota_archaeon_SCGC [illegible]
MCC6224162.1_Thermoleophilia_bacterium [illegible]
WP_247863886.1_Gordonia_sp._PP30 [illegible]
WP_131908314.1_Flavobacterium_caseinilyticum [illegible]
WP_262301579.1_Prevotella_hominis [illegible]
WP_011448111.1_Methanospirillum_hungatei [illegible]
RDE50959.1_Candidatus_Accumulibacter_meliphilus [illegible]
MCC5847926.1_Verrucomicrobiota_bacterium [illegible]
MBA2375852.1_Rubrobacter_sp. [illegible]
WP_019935797.1_Oceanimonas_smirnovii [illegible]
WP_229574328.1_Rhizobium_sp. [illegible]
WP_210512158.1_Natronogracilivirga [illegible]
consensus/100% [illegible]
consensus/95% [illegible]
consensus/90% [illegible]
consensus/85% [illegible]
consensus/80% [illegible]
consensus/75% [illegible]
consensus/70% [illegible]
```

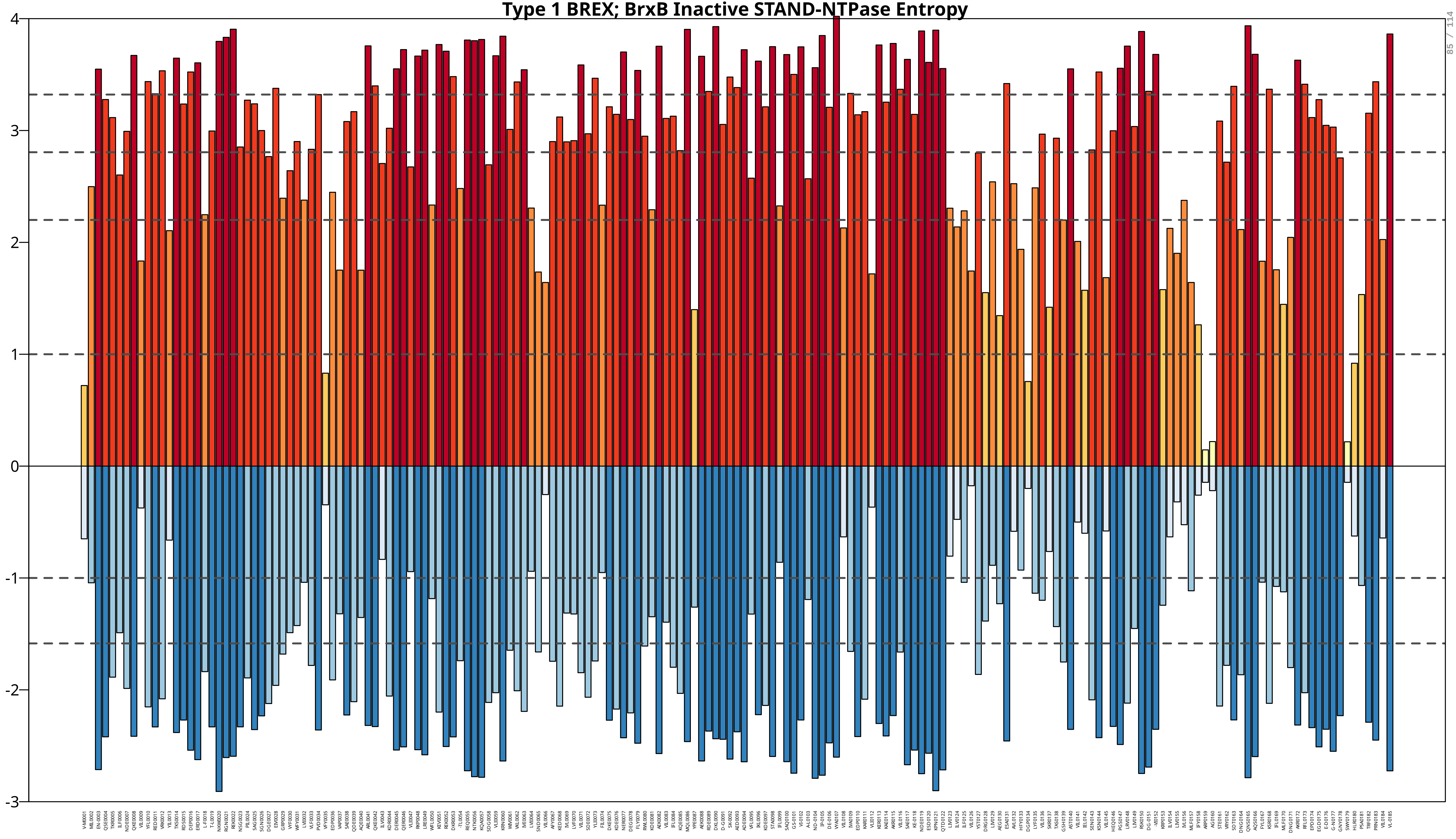
Type 1 BREX; BrxB Inactive STAND-NTPase Entropy
4
3
2
1
0
-1
-2
-3

```
# ; Type 2 BREX PglW C-terminal iSTAND NTPase; (For Supplementary ENTROPY)

WP_185095299.1_Streptomyces_sp.                   ITASTPEQKLRYRANAQLLESAARPGFRLLTVRYDDQRLALLTLTEEWRAEPVDVSILFLTSLRRLVEARPRPWDTILEADNAVPGSRDAMKFGEYTAAAWGEVEKELASNSRPLLLHDASVMARYGGTGILQRLADHARGRGLWLLSPLNDATAVPRLEDWTVALEDREEWIRLNHSWVVNDPEDAPA
WP_268746228.1_Dermacoccus_nishinomiyaensis       RPTHDVASSITGRFGVRLDDSVSSRSFLALGARSERLVEAVETLRRAYGAEVVDLSELLVDALKQQTVGGRLTWDHVRAADAAAPHTKAAQGLGQLVRRAWPQIENHLASAERPLLLTDAAPLARYGHMDMLARWSDLARSRPVWLLVPQVSGNHGAVLDGAPLPLAAPSQFVPLDSDWIDTHAAAAKE
GII83978.1_Sphaerisporangium_siamense             TSTLGGVGRNLAAFEQRLTRSIAEHAFVTLAVDWRFHARAAAALAERFGVTPVDVTADLLAAMRREAEEHQVDWSLVLRSDRPDAAPQDAANLRTLVAMAAASLESRLEAETRPLLIVEAAPLARYDRLAVLERLADQARPAARWLLLPVE-QDGVPSIDGRPAP--GALGALRLSTTWITRPRDEAVS
KEZ35702.1_Frankia_sp._CeD                        STRVS---DEVLAADDRLTRSLVDGGWLVLSVPPRRLARAERCLAA---VTVVDAEQALLAGMREFCAQHRVQWSIVLAADAADRSSRDWANLSRVAQAGLAGVRASIEAAGPAVLITNAGVVARYDPAVVLDELRASVRVRTVWLLVPWADVDKQPLLDGAPVPQ-FGNQGLALSEEWIVRHESRGAA
WP_030506725.1_Microbispora_rosea                 SRTAAPRQVVDSAAMQRLTGAAGRGGFLALNVDLRRAVLAVRVIPESLDVMPIDVGGEFLRALTDLASERDRPWSDVLRADERFSDGILPRGLAGYVREAWQRLATRLSQPSGTVLLHNAGVFARYPGGDLLVEIQQAARPRGVWLLCPTGTPKQRPILDGLVVEVIGENEWLELTAELLRSLDEKAA-
RJQ78831.1_Pseudonocardiaceae_bacterium           LTTVGGVLTAREVVASKLTSAVQRGGFLALTLRGGELPGTAAMLAARFPVQPVDVDQEFLRTFRALADERGADWQKVRTLDLRLSTGEPTPGLASFARKAWDGVRSRVGAGETVLLLHHLSLLQRYYQWEFLTELQNDARPHGLWLLCPAENEKHDPHLDDRIVVECTDSEWAVLSKKFLNALRGQGAE
WP_212516928.1_Actinospica_acidithermotolerans    GTGGAASGAVVDAAVKRLGEAVEDGGFRALTVSVRHLPGLSAAIERRFPLISVDGNAEFLACFRELAAEQGLEWERVLRVDARLQRGEAPRSYAEYVAEVASRLHAKWSSPRSTVLLHNAGLLARYWDIDLVVGLQQSARPRGLWLLCPAESARSSPNLDGRLVETVVAGEWIVLGSAAVEELKCQDGH
WP_208577501.1_Micromonospora_tulbaghiae          ATTHTPLLTAADEASARLAAAIERRSFLAVLAPWRRLGPARRALLARLPLTEVDVTAILLDRLRAL----GFPWEAIVAADNGNPQDADFRSLMDLVHQVVPAIGEALAAVDGPALLTEAAPLARYGQLGLLQELADPTRAAARLLLVPARRPE-PAVLDGVQLPLTSPGQSLWLSGDWIEAAGVASAD
MCP9485019.1_Candidatus_Gaiellasilicea_maunaloa   PTYEDPEVLDAAQFERRLETSTRRGGFLCLVGHDRSLDDIRDELARRFDAQHVSLDRLLIHHLRLAAGRRNVVWQTVVDADAATRDSRAWGNLQQLFVEAVERAEGELLAYEGTVLATDPGLLARYGQLEMIDRLRDRASLSTLWLLVPDDNQSDRPAIDGQAIPIIGAGQWARVPSAWIENRHRGRAA
WP_193120346.1_Nocardiopsis_coralli               TGHREPGRAVSQETWERLGRAVERGGYRALKTWLPDTERSAEQIGNMPGVVPFDVSAEFVQAMDEVRQGKGLTWRHLTQADSSRPRPRP---FQTMLELALQRVGERVAAGHGVVLLHRATPLARYEQGALLGRLAEEARPHGLWLLCPMRGPQSPALLDGEPAGIIADAEQVLLPRGFSAPE------
WP_103561562.1_Actinomadura_rubteroloni           GPHSDPALAAAARAEDQLVAAASRDGFRVLTVRMMLFSEALEELTGRFGGTEVSATRLFLDALHSLTAPGKRPWETILRADNAEPNEKSAHKFAEYATTALGRVEPHLASGNGPIILTDATVFAHYNAMDTLSRIADAARKRGLWLVCPQSDPAREPRLGRRAVPYQPASEWVELNDHWARNRHRSERS
WP_248824789.1_Frankia_sp._Ag45/Mut15             PTYDSPEQAAASLTEQRLAAAAEAGGFRALTVRRSRYLAARTELALRFAAAPVDVAAVFLDALHAQVDPRPRPWATILRADTAEPGSRAAGNLAEYVRTAWERAAPLLGGGPAPLLLHDAGIFGRYAGRDVLFALAERARGRPVWVLCPTTEPTRPPRLDGALVQVATESEWVALPDAWTTNWHRSGKA
WP_213171153.1_Natronoglycomyces_albus            TTGVGSGADEAARFTYRIDKAVESGGFRAIKTGVAETCAAAKWLAARPGVVALDVQAVFLRLLRQYVAERNKPWERVLAADTVPPPAP----LTNLLVEVWNRFETALTAEGRVLFLYNATPLGRYSGGKVLHELLSAARPHGVWLLCPTDALRHVAHLDGYNVGAIDSGEQIRFPAGVLKPRLDEDSG
WP_144820010.1_Micrococcus_luteus                 EATITLPSSELGAIGQRLLDSAAQRGYLVLGVPAHRMNRSRELLAARFGAEHVDLADLFLTRLRETVDRSPVSWEMALSADSAPPGTKAGRALQAWEAAALPQVVEHVSQAGRPVFLTGVAVLTRLGPLQPVARWADLGRSRAVWVGVPQAPGNIGPVLDGVPVPTSSRSQFVRLEPEWIDTAARTSPA
MBN9688999.1_Verrucomicrobiota_bacterium          GNGVTAEKLTANQFTDRLQRSIQTAGYLVLVSTTTNYLVVEQRLREGFPVAVCDIDTLLLNELKAQAESLKVKWEKILEADQAVETSTDWNKLNQLVGYVMPKLKAHLVAQDGPVLLTNIGLLARFNQIPILEQLREAILPRSLWVLVPSDDQNRLPTLHGRPVPVATRGQWEQIPEVWLNLTTSATS-
KAA0252897.1_Acidobacteriota_bacterium            TTHVPTRDDPAAEFDRRLAQAAREGGYLVLLAPERELTRAREAIAARAGIEIRSLDRLFLDALKATAESKKVKWDAVLRADAAPADGPEGTKLRTLVGNALPRLEEALAASGDRLLLTEPGLLVRYRRLDVLDRLRDRVSPGAAWVLVPADEQTDRPVLDGVAIPVLTPNQWARVPLAWIEAQATAGEV
QQS02053.1_Austwickia_sp.                         TPTAGGAASLRDPVAARLAESVATRSFLALGVGALDLARVQDVLRERYDAHLLDVTGLLIDAMRAAAARHGVPWDVVLASDAATPGTRDRANLEQLVGMCLPEVWEAIHAKDAPVVLTDVAPLARYRHLGELSRWSDVARERAVWVLAPQTPTSQGPVIDGQPLPLSSPGQYVRLDLRRLYASDDQNAR
MBK9387522.1_Planctomycetota_bacterium            PLILSDEATDHRLFEKKLRHYAKDGGLLVLSTAPKSFVAAEQALVTFLALERCDLDALIVRAMHAAALANGIDWQTVLLADAPDASASDRANLHQLFHLALPEIEAALLAPGPAILVRKPGLLARYGALRIFALLKERARRPSLWFLLPSSQQSSVPRLFEEPLPI-SVDRHLRIPESWPRQQLSPRST
WP_090664774.1_Belnapia_rosea                     ATEATPDVLAARAIEDKIVFAAEKGVFLALTVEPRRARDAETELLRRFPREVVSLERLMLHAMRTRAEERRIAWPKALTADAAARDSTDFKNLLRLAAEAAPRVAKEVLALRQPALLTRPGLIARYDLMTMLDEFSQASASPSLWLLIPQATPG-RPEIDGAILPVISAANWTRLTDPWLANAHRAGVP
WP_136411732.1_Luteimonas_yindakuii               TTADSPEAREARAFDATVRRALDAGRVVLVTADLRRVEDAVPELSQQYDLAAISLDRLLIDALKAQTSQIGGDWRVVRQADAAVQSSADWRRLTALVERAMPAVVAQLRDDPRPLLLQHLGLLVRYGHTGVIQTLRDAASAPARLLLLPGDSHQ-PPMLDGVVLPVITPADWTHLPRSWLENRHRAAPT
WP_150933330.1_Microbispora_cellulosiformans      TTTGLPGLLGPTDAINRLVTVADRGGFLVVKARIKQAAALPELLSAELPVSCVDVTKAFIAALREVRDEWGVPWPIVLATDSPDAIPEDRVGLAEMTTEAFALVEDRVRAMEGVVLLHDATPLARYAGGELLARLADAARPHGLWLLAPMADPREPAQLDGQAIPVIPGNEELALSSDTVDALEDSSKA
WP_090069475.1_Lentzea_flaviverrucosa             TGTMSPPLTAASDVDARLVEAARHGGVRIITVRRKRWERARTAVEIALGMPVIDVSVAFVAAVRQVARQLNIPFDRVLLADAAEPGSNDQLNLHRVVAQACELLEAEWSR--HPVPLDALTPLGRYPAGALLERLASRARPDALVLISPAGDETKVPRIGDHVIRVNTPEEWVIARSPWPSGVQPRSQA
WP_183357094.1_Garicola_koreensis                 HHTIPVGRKQPERLVRELRESVGARSFLAIGIPARQVERTTSDLLSTYGGVQVNVTDLLLGGLRSQAEEKSVPWDAVLAADAAESGTRAADGLAALVRQSISTIETAISEATQPVVVTDAAPLARYGHLRVIARLADVARSNAVWLIVPQG-PEPGAHLDGTPIPLSYSSQFLNLDNRVAVPAAAEETQ
WP_257478501.1_Acidipropionibacterium_jensenii    DATFGVDLVEDSATTRRLEESVRYRSYLVLGAEPTRLHRLVRALGSRFHAPELDLSKCFLDALHRLSDDRRVPWPVLVAADAQPEASRAHIGLSKVIELAMPEVRSTLSDSPAPLVLTNVAPLVRCGYVKILRELSDLTRGRAVWVVVPQYGLHVGPEVDDVRL-VTSPNQFVEIDRTWIDAETDAETE
MBL9044566.1_Myxococcales_bacterium               THPSGPEEEEALALSAKLDRASREGGLLILSVLPQYADAAKQKLAQRFALSVYSIEEAILRHLKAFAHSKQIQWPVILLADAADRKSADFRKLQTAMSAALAQMVDEIRHAPSPLLLIRPGLLARYDRLDVIESIRVSVEPPGVWLLLPSDELDLGPRLDGQTVPVLTPAYWRAISPHFLRAALPAVRS
MCP5560207.1_Verrucomicrobiaceae_bacterium        RTVMQEDVATALLLEEKLSSCATRGSFLVLSVPPDRLARAESELLARFPVQRVDGDALFLRLLHAQAAEYEIPWDTILLADAAKEGTEDNQNLRRLVDMIKPQLRQALVSPAQTLLLVNPGLFARYGLMGMLAELRNDVRPHGLWLLIPAVVQSALPKLNGQPVPVISEHQHISLPQAWLDNAHGKA--
MBK6942988.1_Planctomycetota_bacterium            ATEISDEVAAARAFEERLARAQKDGVFLVLAITPSHALDAERELLRGFALERRSVDVLLLDAMRRHATANEVPWSVVLNADRARPDSVDGKNLRRLVRMAIADVEQQLATSTRTILLTQPGLLARFDCIDVLERLRDRVRLFGVWVLVPCDDQREPPVLNGVAVPVITRAQWARIPISWVKNRHRGSES
WP_217637676.1_Actinopolyspora_saharensis         VGTVPAPSSRGDYLGATLAESISARSFLALGVAVDEADRAAHVLALRHGGTVLDVTGMLVDEMKTVAGKKGLPWDLVRGADAAQEGSRDAQGLHALLNKAFPAVGERLRAGAGPLILTELSPLARYGQLKMVARWSELARSRAVWVVLPQLSWMRGALVDSKPVQLGSNGQFVELDADWLATQEAETAT
TAF23825.1_Verrucomicrobiota_bacterium            GTPVSEEVAEAQRLEDKLTRARAQGSYLVLTIDPKYLNDAANQL-ERIGAKRLNASELFLTAMKDKAADLGVPWEVVLEADAAEKGSADWTRLQELIRRSMPGVLEPLRNSNEDILLTDAGLFARYGRMNELDALRNDTTPHSLWLLVPGHGAATTPTLCGEAVPISNEAQFEQLTPQWARNHHRGGVA
MBO0801654.1_Nocardiopsaceae_bacterium            KPSTATLLDETARASRRLEQASRRGGFVAVKAPLRDAARIGDVIAGMDGVTGVNVTREFVRLLRDEASKRPKLWETVLSADSADAPPGARRGFGELLRAVWGALEARIAGDGGIVLLHDATPLARYTGGELLARLAVAARPFGLWLLCPMENPKGSPLLDKSTVGVIPGMEQLDLPEGFGSGAESRAS-
WP_239634082.1_unclassified_Halorhodospira        DTTDDEQGERAAEVERKVRRVASEGRFLALRIGPRHMVKAQRRLASGYGLQAYNLEGELLRAMRDVADERRVQWSKVLEADAAGPKGPDWKKLLQVVGEAVRRMRPHVIEPPQPLLLLNPGLLVRYGHIALLNQVRDACEQPGVLLLLPGSSATTMPVIDDQALPIVHASEWLHADRHWLDAPANA---
MBK9978482.1_Gemmatimonadota_bacterium            PNILDDAMDEARMLEERIAAARSDGGFLVLSTEPARFRRAELALAERHALTPMSLERLFVESMRKTAEANRVHWPLVLQADAAPHHDRAWRNLRLLAEKARPMMEATLAALVRPTLLLHPGLLARYDLLAVFEPLQDRSRGPGVVLLLPSDGMSAMPVIDLVPLPMVYPSDWARLPREWFRLARSA---
WP_130480151.1_Leptothrix_mobilis                 ATARLARLADASALEQRLQHHLQQGGLLSLTVEPRLARHAEAELLRRFGLQRLSLDALLLRELRTQAYARRVDWATVLRADAAPAGSRDWTNLLRLVQLTLPALQQALLNSPAPLLVVHAGLLARYGLMRLVTELEDQSRPPAAWLLLPGQRPG-VAEIDGTPVPLVHAQAVAALQPAWIGNEHRSAER
MCO6436553.1_Phycisphaerae_bacterium              PTQIPPEIAEARQFEERLRYAEKDGAFLALTAKPNLYERARQELLNRFDVRPLDLERVFLDSLRQAAEEVGADWNVVVKADAAGPNSTDWRNLNHLISKVVPKVEEAIVQDGKTVVIYHLNWLGRYDQVVMLSRVQQAVQDHGTWLLIPGSPQTEMPLLDGKAVPVITNNQWAAVPKSWCQNLHRTDEK
WP_181762535.1_Pseudarthrobacter_sp.              TTHTKLSSTQILEIERQLAQSLTRKSAVTLAVRPRDFIRTSKALTQVFGLKLVDVSAEVVKAIRATAQSKNVDWDVVLRADAK-SSPSDHANLAQLVRMAVGPLWVELVDDPNPLLLTNIGPLLKFGFGAEVARLFDMSVPAARWVVAPKRGSANTPSSDGRVPPM-GPDGFVDISPGLASALSKREST
MBK9741405.1_Actinomycetia_bacterium              HVLTPPEVPPYDEIAARLQGSLRAGGYLTLAVDPRRVDQAAAVLARTYGLTRVDVTAELLAAAKEYAVEVGADWSTILDADKE-VGTRDRALLDSFVKAALDRRFPDILAATDPLLITDASILGRYGQQHWLADLADLAGPAARWLLVPHHASGGRPTLDAQPAPL-GADGYLVINAEFLAARQASEQS
NMM16638.1_Cellulomonas_sp.                       TNLTRVATEPADVVDGYLRASLTRSSALTLCVHPREHDRAVATLTTLYGVTPVDVSERLLTAVRALADRDGVDWSLVLRSDAE-PSGGDWENLRGLVQDALTEPWQQILATRRPLLLLNAAPLARYGLTAWLSDLLDQSQPAARWVLVPRRGHQAVPTLDGRPVPL-GADRWIDLPYDLSSLRPSSGVS
WP_136709615.1_Agromyces_sp._H66                  LSRTLAKSLPASDLHARLLGSIGSNAALTLAVNPRRYTEARRYLTNRYDVTTIDLAAELVRRMHAVADERGAKWAVVTRADAS-ESSNDFKRVSQIARQAFEPYWRELSAEPNPLLLTNAAPIARYGLTEYLAEMFDLARPAARWLLVPHKAAKPTPDLDGRPIPL-GPDRWVSLPSDLAELFPSSSGV
WP_210768105.1_Cellulomonas_humilata              GSSVHRGLVPASQVAAQLVSSLRTKSAVTLCAHPDAYVSTTRMLGRVFEIRTVDVGAELVRAARADAKVVGASWEFVLGADVE-PTSPLGSDLRVFLEPTVTAVWDRLMADTSPLLLTNAGPLLRYGFTSALAEMLDLATPAARWLLVPRPRAVAIPTLDGHAIPL-GADRPVDVPLDLESAVSGA---
WP_067228577.1_Microterricola_viridarii           RSTTLQGSLPNDTLAPTLAESVQHRSALVLTTTWGRYDDSATALAARFGVEVVDLAALAVEQLHATAASVGADWLVVVDADKP-DSTIDKRNLQALAKQAMSSAWAEINDRDTALLFTNAAVLAHLGLVDLIARTIDLSTAAARWFLLPKHGAAAVPDLDGTPMPI-GAGKWIELPTTLAAVLPIPSSI
WP_129336322.1_Cellulomonas_endophytica           LTSIATRTSTVPQVLAVLQRSVETRSGLLLAVTPERHEHASQQLAAAVGLVVLDVADALVREARRVADEDGADWSFVLGADAE-PGSPLRSQLHEFLEAAVDAVWEQVVSTDAPLLLVHAGPLLRHGFEDRLDHLLDLATRAARWVLAPRPSSSGVPLLDGHPVPV-GAGRWTDLPADPLPTLGAPS--
RKN35100.1_Streptomyces_hoynatensis               ASDEDTAVAAAVKAEQRLAHSARQDGFRVLTVPQRRAEEAVRLLTEPYGAQPLSLTGLFLAELRGVVGEAARPWETVLKADAAETGSRAHQQLRIRTDKAWGRLEPELTADDGPLLLTDTAVLARYDAFDLLGRLAEEARGRALWLLVVQSDPARAPRLAGQSVPYQAGDEWIVVPDAWVARRHLVSRA
WP_262103914.1_Arthrobacter_sp._Marseille         STYTFIASGSVSELERRLRDSLDRKSALTLAAHPRDYLKAASTLAQRFGVKRIDIAAEVLRVIKSTARQADVSWELVLRSDSE-PSSGDFTNLTRLAQAAIRPMWGEMMQVPGPVLLTNIGPLIRYGFGPDVSKLLDLSHPAARWILAHRRGYDETPKVDGKPVPM-GPDGWIDMPRNLSDLLTTNDLN
MBZ5659870.1_Acidobacteriia_bacterium             QTELPREMEEAVEFERRLHAAYRSPSYLVLATEPKYLEMARQNLAKHFPMTIFHCEREFLAALQKEAESKHIRWDVILRADTSKPGARDWDNLRKLAANAAHRVAEQVQSRTQSMLLIYPGLLGRYGQLSILDELADSLT-YSLWVLAGSEHQAASPMMDGQAIPA-RPTQWAWIPAKWLDNEFRKSTA
MCB0954589.1_Ilumatobacter_sp.                    GTAVAHEREEALAVHARLQRQEAEGGYLVMTVEPRWQRRAIE-LLRGYGPHTVDVDGWLVGAMRAYTDRRTIKWPAILGTDAAGPEGERWPRLLTVAKDAMAPLQDGLAQ--RHLLFVHPGLLGRYNLLGMLDQLRERTRLRTVWVLVAADDPAATPTVAGRAVPVSTPAEHLVLPEPWLHNLHKTEGV
MBL8842562.1_Planctomycetota_bacterium            TTEVDPEVVNAAEVERRLTLSHRRGGFLALSVLPRHYLAARAELRARFALDGLSFDAALLARMKRIAQEKGVDWQLVRATDAAGEGAAGWRNLVTLASLATGELLAELSAAGRPLLLTELGLLKRYEQIERLAALRERARLHGLWLLLAQSSPGPRPQIDGRLVPVLANGEWLALDSAWVKNRHRSVPR
MBO9539711.1_bacterium                            STLPSDEILDARAFETRLERAVTDRRFLLLAIEPKYLLRAEAEITKRFEVHRVSFESLLVRKLRSAAQAVGAKWEVVVSADGTPRDSNAWRNLRGLVKRAMGEVEQELSTSSTPVLMVENALIARYDHMELLERLRDDCRAPGYLVLIASDDQSTAPKLERQAVPLITASEWVRIPHAWLENRHRSTPT
WP_144981411.1_Gimesia_aquarii                    DTSTSPETVEARQTEDRLQHAYRDGGFLVLTVKPSYLRPCEHNLVHRFPLERVSFDDLLFEQLRAKSKEYEFPWSDLYEADSEGTNTDDWANLLHLMSEVGPNIETELMNSERPLLLVHPGLIARYQMMSVLQTLRDRVHDPTVWVLIATDSQNEMPYLDGVQIPLISKGQRAAVSEYWIDNLHRGRSG
WP_093484991.1_unclassified_Streptomyces          GTPLDPE-STAARADAQLAGAARRDGLRVLTVRARNAPVVGRELRRQYGARAVSVTELFLKGLRGELTPGAMPWEVVVKADVL-PAGQRPTGFTNLVGAVWQAVQGVLAADPAPVLLTDAAVFARYDGMHVLDRLTALARGRPLWVLVAQDDETAVPKLAGRPVPYLEAQEWIPIPKSWTRMHEPNAAT
WP_200346137.1_Halochromatium_glycolicum          GSAALAAEQRQQAFDDRVRTALDEHRFLALSVRPADAQRAAQRLQARFGLVAISFDALLLRQLRRLCDARPPDWQVVLRADAAAPGSIDWSRLQGLVQRVLPELTAALGAAGAPVLLTDTGLIGRYGLVSWLAELRRRLEGHALLLLIASDGTGPGARIEGVSVPDPGRREHAAIPSQWLQAAADG---
WP_252180113.1_Endozoicomonas_sp._4G              ATVDTTTLEEVKRLEQEIATAIETSRFLALTVKPQLMVTARDKLCKDYPLKHISFDELLFHHLQELFNPKAPRWDVVLKADAAQKNSVDWGRLQMLVSKVLPAMANEIKSAGQPVLLTEPGLIARYDLVSWLNELRQHLNVHGMILLVAADAQRTAAVIDGTIIPRTGAAEFARIASVWLEASNDSQQA
RLS37136.1_Planctomycetota_bacterium              STTLAPGLETALHFEDRLSRSRRQPNFLTLLVEPRGYGKARAALVQRFQAEIIDLEGFFLAALRHTATALGVEWELVLRADSH-RLPQEWMNLMALVRRAAPDFEERLMATDKALLLVQPGLLARYGLMDQLDKLVDQARRPAVWLLLAGD----RALIDGQAVPLIHAGLQVRVPLEWLENRHRAAEL
WP_241482799.1_Kocuria_polaris                    VSTTQFAPTPAAEVARKLTESLERHSALTLTVAPKRYVAAARALASKFDIDVLDVAELVVSATREQIEKAGGIWSGLLGYDAL-DDSKQWKALEGVVQRSVEPVWGERLSAHRPLLLTNAGPLVRYGMTSLLATLLDTGTPAARWLLVAKAGDARAPLLEGRSVPL-SPSGWIDLPRDMAQLADDKTIQ
G8417454.1_                                       AKALGAERSPHEAVRDMLLDAAGSRGFRILVTPPEHHADIGPSVARALGGTWVSFADAFFRDHGANIAAL----------ERAERFVAQRDALTEAAEDTMFRLLDEHGRPDHTAVVGDTGLLGVCDALDLPRRLYDETLSRGFWVLVPGVIYKSQPHFNGATFSLLTALPD-----------------
MBK8541194.1_Ardenticatenia_bacterium             PTFLPALTSPGDRAAAWLRSSRGMSSWRLVVTPPERHAELGRQVAAALDARYVSFEAAFLKRIEDRFADF----------ETAERMSALRGRLKKEAEGVLEALLAEHGRPGTATVLGDTALLGLCEAKHLVTRVYERALALGFWVVVPGVVHQQQPLLNGMVLPLAEDVGV-----------------
MDO9017995.1_Myxococcales_bacterium               PAEFIAHRSPDEIVVGLLRAARKSRGFRMIVAPPERHAEVARSVAQALDARFVSFEHGLLAGMEPEFAVY----------ERAERFKAQRSRLTRAAEGLFEALLAEHGRPGTCTVLGDMGILGLCDAAHMIRRLYDETQARGFWVVVPGVIFQRQPLFNGTVLPVDGELGR-----------------
WP_075011136.1_Stigmatella_aurantiaca             PSDLSTQRSYEEVVADRLREASRGRGYRLVVAPPEKHVEVAHSVRRALSAELVSFEARLLQRMEPDFPAF----------ERAERFLAQKPKLKKEAVALLDALLEQHGRPGNVVVLADTGILGVCGVKDVPRRLYDRVTDLGFWVLVPGVVSERQPLFN-QVLPVPEPLDV-----------------
MCA9600002.1_Myxococcales_bacterium               PDDLAPDRTPEQVLVARLRDAAERREWRLLVTPPEHHADYGCSVAAAINAEYISFEQRLLERMEPSFADF----------ESALRMKARRRKLTRAAEALLEELLRERGRPDCRTVLGDTALLQMCDAAHLVRALYDETQQRGFWAFVPGVIQERQPLLNGATLPLSGPLPP-----------------
TVQ88636.1_Deltaproteobacteria_bacterium          PELLEERRTPEEVARDVLRDAVKRSSYRLIVADPGQVGPIAKSVIRALDATPIDLADAWFQRHAATVKVD----------AHAERFPAMKAVLAQKAHKLFEELVAEHGERGRAIVVYNTGLLDAIGGVVQVRNLYERVQQHGFWVLVPGVISGRTPLFNGLTLPLRKPLPD-----------------
KYF91864.1_Sorangium_cellulosum                   PPELV--KDPAEAARDILRSVAKTRGYRLVVAPPDDHVEIGRSVARALEAAFVSFEHEFFRRVDAQIEVF----------DRAERFAAQRPRLRKEAEALLDALVQEHGKPGSTVVLGDTALFGTCDALHLVRRLYDLTAGKGFWALVPGVLHKRQPLFNGAVLPLAREIPE-----------------
MBI4699810.1_Deltaproteobacteria_bacterium        PPELVAAKSPEEVAADLLRGAASGDGFRLVVSPAELAPEVGRSLARALEVRFVSFEQELLGRMDGAFDGF----------VRAERFKAQRGKLTREADGLLADLLAREGRSGATVVLGDTAIFAVCEALHLVRKLYDETSGRGFWIMVPGVVYQKQPLFLESTLPILRAIEG-----------------
consensus/100%                                    .................l...........h.............l.........p....hh................................h.......................h...s.h...................s.hh..........s................................
consensus/95%                                     .................l..s.....h..l.............l........hsh...hh..hp................-...........h..........h...........lh.p.s.h..h........h..h.......s.hhlhs.......s.h....h.........................
consensus/90%                                     .s...........h..bl..s....shb.l.s...........l...h.h..hsh...hl..hc................-.....s.....l..hh...h..h.........shll.p.s.h..h.....l..h.p.s...s.hllhs.......s.h....ls........................
consensus/85%                                     .s...........h..bL..u..p.uhb.lss..p....h...l...hsh..lsh...hl..hc..h......W..h...D.....s.....L..hh..sh..h...h.....slll.p.u.ls+hs....l.pl.p.sp..u.Wllhs.......s.lss.slsh........h..............
consensus/80%                                     ss.............h..bL..u.pp.uhbhlss..p....h...l...hsh..lshp..hl..h+..h......W..l..sD.....s.....L..hh..sh..h...l...s.slll.psu.lsRas...hl.pl.p.sp..u.WlLhP.......P.lsu.slPh..s.....ls..h..........
consensus/75%                                     ss..ss....s..h..bL.puhpp.uabhLss.spph..h.p.L...hshp.lshp.bhlp.h+..h......W..lh.uDss...s.....L.phhp.sh..lb.pl..ss.sllL.psu.luRYs...hl.cL.-.sps.uhWlLhP.......P.lsG.slPl.ssspbh.ls..h..........
consensus/70%                                     ss..ss....s..h..+L.puhpp.ualhLss.scph.ph.p.L...hshp.lsls.bhlp.h+..h......Wp.ll.ADss..ss.s..pL.phhp.shs.lb.pl.sss.slLLspsu.LuRYs..phl.cL.-.sps.uhWlLlP.ss.p..P.l-G.slPl.ssspbh.ls..hh.........
```

#### Type 2 BREX; PglW C-terminal Inactive STAND-NTPase Entropy

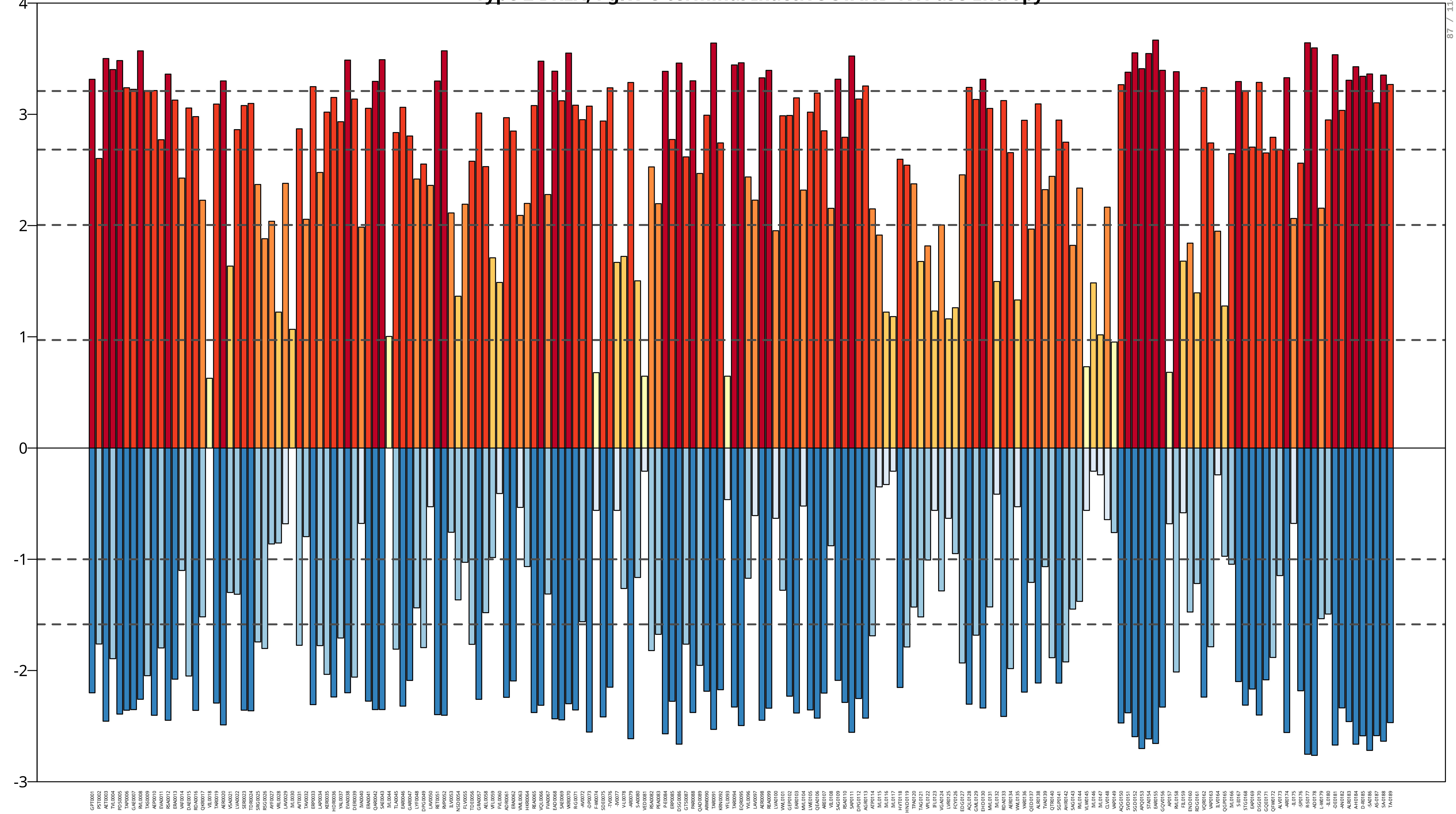

```
# ; Type 3 BREX BrxF-iSTAND NTPase; (For Supplementary ENTROPY)

MBI3973658.1_Chloroflexota_bacterium                 MAAAVDQLDPGYYACLVLVHYLATYRWPHLSVGQELSEALLQEQPSRYPTAASKWLTDRVSSLSPGPVQVTAIDLLFGPALDLDPLILFRQASRTTRLVILWPGTYLNGVLSYAVPEHGHYRTWPKPDIQITT
MBO8173527.1_Bacillaceae_bacterium                   MYFALEQLEGRRHRLLLLVSEASKHQIPILNMNLSLSEELINVPVDRRPRKVGEIVEKILRHQKKDLVIISHIELLFDHSLHLNPIKLFEELSKTKTLIVGWNGKYENNVLSYADPGHSDYRSYYDCDAIIFS
CEO89248.1_Syntrophaceticus_schinkii                 LSKAIATAQSRYYKLVLAIPGMAALGTTPVNLGLLLSQSLLEVEPRFRSLEFQDLFARIIRSAEGDLVAFDHAEILFQRDLQLDPLRMLLQHSRNKVLIVRWPGVFSGDKLTYGEPHHPEYRCYEQVDAVVCC
CEA16532.1_Fermentimonas_caenicola                   MSKQITHFAAPRYKFVVAPNERMLKDFSAFDLGYNLSLVLKENKNEELTFIVHEELTKLLNDNITDYICLSNIGILFEKELKINILQTFQRLSKNTLLILLWDGEVKGNTLFFLSENSKNKIDLREINHIIL-
HHY32855.1_Bacillota_bacterium                       MRDLIKQMETNYYKMLVIVDNLQAEGWTAVDVTDAVLKLIEGIPGEKVRVRIGGKLKEWFHS-LPDRLVLYNTSILYSPELRLNPVGAFKYKSREKEVVVLLEGRLACNRIQYSQYGRPDYAEMDVSEVIHAD
KUO63975.1_Lutibacter_sp._BRH_c52                    MDKHITTYSQSRNKVVLFK--ENIFDFKPINVGHILSQALYNLDKDKLPMLVSKELNTILNASITRMLAISNLGILLEPDLKQDINSLFEKYSNNNVLFVQWEGELDHNNLYFLTKEKGIIINIKNLSHIAI-
MBI9032244.1_bacterium                               M-DRLKEIQKLRNKLCLIPVNNNTYNINSINIGFELSKVLKNELGKNLPIIAVESFRKIIEGSTSETIMVYNLGILLEDDLNLNLESILLEISKNIVVILLWEGEISKQGLHFLSIKHGKLLELTEKHYLQLE
CCX79045.1_Parabacteroides_johnsonii                 MDTAINKYSAPRYRLVVLKDEMPPDDIPKFDIGKSISAYIISEHSNDLKMLVNAEFMRLFREIASNAIALCNFGIVFEEKLGLDMRTILKDLSRNYLMFIQWEGVVKDDRLCFLTSDSSNFIDLSEISHLTI-
MBK9455368.1_Bacteroidota_bacterium                  METYLALANSLRFKLWAIVGELQKHGYLLVDVGAELGSLYKELTNEEPSHDIGQKIKDWFLS-KPEKLILTNASILYHNAFKISPVGAFKYNSRNKSTVLFLDERIISNRISYGQAGSEEYYDKDINDILITD
MBL9119763.1_Phycisphaerae_bacterium                 LRAAVLAAEERHTRLVLVVGTGLRLTTRVANLGLVVAERLINLPARHRPVTAADAAADFIRDAEPGPALVSNLQLLFLPELMLDPLKLLSDSARSRVVVASWPGERTGSCLRYAAPSHPEFREYHNPDCLVVE
MBQ9466676.1_Muribaculaceae_bacterium                MKQRLTKFSAPRHKVVVLLNDSVGASLPAIDVGRELAAYLEAYTSHRAMDEAKQFLTKLIAQKTQPWVALRNLGILFERDLDISPANLIDNLSRNTLFIIQWPGLVNDGKLYFLDRGSRYFIDLSQISHITL-
MBT3747343.1_Bacteroidota_bacterium                  MEELIADMSDVREKYVIVCKNAEQLNASYININLELSRRLLDIPKKKIPFKANRYLEDIINNSNKNIILLDRIEILFDPLISIDPLYFIESNSKPVTMIVNWPGEFSSKILSYAKPVHPEYKSYPNISGRVIS
MCA6513726.1_Chitinophagaceae_bacterium              MQTPIKDITTERYKLYTLMCNVKSQNINVINIGKELANYIDSLDFRYLNIDVYDHTKKLLDKHRSDVVAIYNLGILMEPALELNAVQLFKEFSKSTSLIIIWDNQSENDRLDWQTQKQNFFLDFSDIQLKKLQ
MCF1431077.1_Shewanella_sp.                          MEQAIEQAQNYHSQLVVVAGEASTDGLAVVNLGLELSERLIEVPRQDRAKSAASLFADLLHERNTRVLLLNHIEILFDKTLSIEPLKLLQSSAKNLTLVVAWPGEKTASSLTYAVPSHPEYRYYKASDFGETF
MCM1450902.1_Clostridium_sp.                         MAADLLRFNGPRHRLVFVKDTNRTNSNIGVDVGWEVAKYLHDNSLGNIKAESQNFIKKLIYDNISKYIYLRNIGVLFEPELGLNVDMFLTNISRNTLLLLDWDGQMKYPYMYFLHTDSKHRIKIDTLNYMTL-
NLI12135.1_Peptococcaceae_bacterium                  MSKELAHIGYRRHQLLIICGVTRELAIPCINLSLKLSEELLEVPVVQRGRKVNQLVDKIVSESTGEILCFEHIELLFHPQLQQDPMRILENISRNKIIVVSWSGRYANGRLTYAEPEHPEYRVYNELEASVIT
OJX40547.1_Chloroflexi_bacterium_44-23               M-PKLEIIFHELFDCLILVNPIKEENISVINISQELGGFLLNISAMDRSRSTQNWMVDKLASHPHDPILCVCPDLLFDPTLQIDPFALFRQIARIKHIVVMWPGNFSSGILSYAVPEHHHFRTWKITDTIINQ
PAD23514.1_Niallia_circulans                         MNNAISAVSEQYFKQIFVYDYAESHSLPYINVNVEISLLLQKVPINRRAFLITEVFQGLIGKYSDGVICLDYYELLFDNSLAIDPMILIKNNSRNKTLIISWRGKIIGDALIHAEPGHPEYKKFIVQDAVIIK
PKO06211.1_Chloroflexi_bacterium                     MQVRLAIAPQQYHRLIIILNPAGKFGLRYINLGLILSQMLLDLTERQKALKMPQLVDQIIGCIEDKPAVLDHIEILFEPSLQQDPLRLLQGMSRNRTILAIWNGQVNNGYLTYAATEHPEYRKYSIRDLNILS
PZV08804.1_Leptolyngbya_sp.                          MGDLLAIAQGQYHQLVFATVEAVALDGHYLNLNLALSQRLLDIPQRQWPHKVMELIDAVVLSTDLRVTVLHCIEVLFDPQLRLDPLKSLQQIARRHNLLVVWPGTYDGRYLAYAEPWHPEYCRYPAKDLIVFP
ROS80359.1_Muribaculaceae_bacterium                  --DKIAQYYDPRNKLIFAP--SLDQHLHSVDVGFELAKSIEGDSSPHLPMIAEDYLNAILRRSIKDYIALENWGILFEPALKLNLVSIFDSHSKSNTLILVNCGQADYENFHLVNKHFNTTFPIGNLHPYILQ
TDP26837.1_Halanaerobium_congolense                  MTDLIKSNRGSHYKLLTLVGSLKNNGWNVYDIEEVILDLIEDIPENKIGLKIGDKIKEWLSD-QENKIVLTNTSIIYSPELLIDPVETFRYAMRDKEAVIFIEGKMRDDKVIYSTPDKQDHKDIDISRIVSEE
TNF50602.1_Bacteroidota_bacterium                    MKDKLLTYGSSRNKLVITS--EKIPGVHYLDVGETLSLSLANSPISWNSMKIKSEISKLFAESIESILSIENIGILLESEVQFDFKHFLEQYSRDMAIFLKWDGDHDESTLFFLSKDKGIKIPLTNLSYIKL-
WP_061808754.1_Rossellomorea_vietnamensis            MNTLIEAIKGQRHQFLLLSFQLKEIRATTFNVNLLLANKLIELPISRRKRAVKDTLREALNEVENEVVYFERIEYLFDEELNQDPLKLFEYLSGNKVLLINWPGDVRCNGLSYATPDHPEYYQVDGYRDHLIV
WP_073205029.1_Prevotella_ruminicola                 M-SNLSSYKNPKYRLVFDL--SSSLEVESVDVGTYLADALQTKQKANLNMYASDAIRKMIRQHTHRYVALCNIGILFEPALSFNLETIVEQTAKDELLIIQAKGGIANERFYFLSQQNNYSFSLKNLTYTIVE
WP_075981269.1_Bacillus_massilionigeriensis          MSSEIETLDSRWYKLLIICGLANQLDIPFININLKLSEKLKDISKAKYALKVPEILQSILEQYEESYYWLGDIEILFDKQLQQNPVRLLENLGKRYSIIVTWPGQYANGQLTYATPEHPEFFSCHETDGKILE
WP_086717961.1_Bacillus_cereus_group                 MDKEMALMHNWWHKLIFICASATNEDIYTVNLNLIMSETLVNISEEKYPLYVEEVTRETFDN-PFKIYLLQHIDVLFDPVIKINPIRLLENISKKSKLIVIWPGEYKNNQLVYAQEGHPEYFLSGSFEGKVIL
WP_106450119.1_Trichococcus_alkaliphilus             LNREVPYLKNKYHKLLYLYEYSEANKVPYINVNLYLSENLADVKPNRRPYKVVDLM-NLLGDTEDDILCLDYFEILFEPSLMLQAFDIFKNLSKRKVVLVAWRDEIKDGYLIHAKPGHPEYTKELIADTTIIY
WP_113032580.1_Paenibacillus_contaminans             MSAEMAGVRSRRYKLLFLLPEMEREGIAAINVSLLLSDKLRHVAPDKRPYEVAKHVKQAINEYKADVVCLYHIEYLFDLELKQNPIKLLEALSGNIVLLVAWPGGAEQGVLYYADPEHPEYDSNGEYGSCILP
WP_118194774.1_Pedobacter_indicus                    MIDKIVSFLIPRNRLIISTDFFDTEKVNWMNIGKLLAHELGNTDKSKIVFRINQLLDDVIVKNKIETIAIENIGLLLEPELKFNVEKFLSDYSQNNTLILKWEGEIDTNQLYFLTKNDNHKIDLNNISHIVI-
WP_119865508.1_Clostridium_isatidis                  MEDAVAAAKSKRNKLIIV--NIAAIGLDCINLNLLLSEKLLEIPVSKRSRSVDSFIKDLIKSNNMNTLALSNYELLFLPELKQDPMRLFEDLSKERVILIVWEGRYENNSLYYAEPWHREYREYKDLDAIIVQ
WP_155743288.1_Scytonema_sp._UIC_10036               MGSYFDQVVERYYRLIVVVVPAESTNSRYVNINLELSGRLLELRQLHRATQVQQLLAEIIGNTNNGAVFLDRLEILFDESLKLKPLERLQKLARDRTIVAVWNGNIENNSLIYAKSGHPEYCRCPTDDCIVIN
MBR1791000.1_Bacteroidaceae_bacterium                MSEQVISFSSSHHHIVFCR--EMMPELTFVNVGLVLSEKILEEDSPMLDFATDDFIQEIMSRHESNYLALTNIGILFEPCLKYNIPELFKSHSRNKILFIHTQAVIKDDVLYFNDPSDNITADLSGVHYIIV-
WP_169099405.1_Bacillus_sp._DNRA2                    MSQELEGISTRYYKMILVCELAESFGIPLLNISMLLSEKLKEYPSKRMAGRVHSLLSDILRNAGSSTLCLEHIEVLFDPTLKQDVIQLLQPLSRNYTLVVSWKGSYDGRRFVYAVPGHPEYYECKEFDGIIVT
WP_173978270.1_Magnetospirillum_sp._UT-4             MQQEITDVAGTYHRLLLLVGRAAHLDRPVLNLNAELSSALLEVPHRQRQLKAHKVVGDIIDGGPADIVLFDHTDILFSPSLRLDALALLKSLSRNKTVVATWSGTAKDGKLRHAEEWHPEYQSHAIEDIKVIT
WP_181742022.1_Thermoactinomyces_mirandus            INTSLKEITRHRHQLIFLAGDAEKIDAIYLNLSLELSHMLMEIPVHKRPRKVDICFREILTGYSQSLFLLDHIDVLFDSSLQLNPGELLENMSRKVVFVIPWYGQVTDSELIYADPGHLEHVRIPINEAMIIQ
WP_214689525.1_Exiguobacterium_sp._s163              MNSSLPQVTNQYYKLMYVNEYAKNFETGYLNLGIELSLALLDIPLKRRPFKVTDIMENLIQKQHNKIICIDRIEVLFEPSLKTNPFQMLKQLSKNHTLIIAWRGIVSDQTLMYAKPNHPEYFKYTATEEWIL-
EGR2106511.1_Vibrio_cholerae                         MKRASRAAEYCSHNLVLVVLESQIEDVELINVSKSLSQRIVCLSANERIKRLDVEFEEVAQE-CNENIWLTKLDLLFEPSLKANPILLLKWLSKSRPVVAIWPGQIYEKTLVYAEPGSEDYRTYSLQELKDIQ
CCU85050.1_Mesotoga_infera                           VKNSLECLQAKEPGLVLLSGGANEMNTEVLNLNFVLSKEMKTLSINERPLRASRFLSELVNS--RESLLLDHIEILFEPQLKLLPIQILRRASRTTPIVANWPGSIENEFAIYAQRNHPEFMEVRLDSEIIFK
HBR02122.1_Ruminiclostridium_sp.                     LGSEIIETIRKVTIPIIFCSLKLINDYQELSLNKELAKALVEKDVIIRSRFVADEVMKIVSL-IQGPILLTDYEMLFDPRYNIDVLKLFYELSRRAKMVIKWCGALDDNRLVYATPAYSDFHSYNIHDYDIIC
HHV38844.1_Tepidimicrobium_sp.                       MGKDLRIWNINNHYPVFSILQNEFKDFYYIALGEILSSSLLKYGKDDRKDYVIYEMDNIFHN-MDSKLIIDHIDILFNPDYRIDILGYFIQLARNRRIIVIWPGEYVSGTLTYATPEYRNYKKYSIKDYNIIC
MBD9124683.1_Lachnospiraceae_bacterium               MGRLLSENQTGLMRPVVYCDE--CESLHQVSLNEKIAALLIKIKPERRTMRIERCVQEVLTN-IPEKSCIRDFDVLFNPAYKIDVLQLLTVANRSKSFSVLWPGTVIDGKLVYAEDGYADYKEYDVEQYDITC
MBL0521045.1_Aeromonas_enteropelogenes               MAKALLRMSVQSSALVLVINSRQTEGLTYLELSEQLCERLLPLSCNDRSRHVSDIIAQMLNG-IASVVWLDRIQVLFEPTLELDPLRQLQDLARLKPIVAIWPGQITERFLTFSVPGRDDYQSYSANDLANVP
MBN2003265.1_Anaerolineae_bacterium                  --QQRIMAQIRSHALLVLVGMALPDDSVYINVGLELAKHLRQ-NLPTGSLRAAQIMEELLES-SGASLILDRLEILFALTLHLDVLRLLLDLNRRIPIAAIWPGEYRDARLTYAMPGHPEYREYEYPEAFPIT
MBO2493598.1_Clostridia_bacterium                    GDRQVQGYTSNAGNIMVACVDAFFSNWKWINLGLVVSEQLCRLPKEQRKRMVADVVINAIKK-TPSNIVVENIDLLFSPEYSLDVIRTLQLAGKNKRLLVIWAGDYQDGVLTYAQPDCPDFHRYEIKNYGVCC
MBP3452736.1_Clostridia_bacterium                    MGSDFRQYPAVYQGLFVFS--RLPKDQPVINLGYVLSERLMGIPADRRCFEIDNCLRSIVET--YTAVSLSHIEILFTPSLKLDAIGVLLSLCRNRKICLSWPGRISGNTLTYAEPSSPEYYEADYTRFIDTY
MBP8682476.1_Trichococcus_sp.                        MGRDLAVIKQTRQGVFVLA--TAPVNASVVNVGLLLGERMFRVKTEGSSTSITRELDDILDE-EHRDILLTNIDILFAPEYELDVIKLLLQLGRNQRLYLPWPGELVGERLTYSEPIRFDFKEYIVKNYVDTY
MBR6410258.1_Clostridia_bacterium                    MYSELPENLKQSGVPLVFCSEKRVESYIELKINIILSERLCRIPRDERAKLVEFKLNNLLNY-YHAPVLISDFEMLFDPRYKIDILRFFCEKARIINVAIKWPGQYADGKLTYANPADPDYHEFDCNAYQIRI
MBX3742593.1_Akkermansiaceae_bacterium               IAVDIARASGEHFRLVWLAGGAEAEDGVVVDVGKILSSSFIEVPVSLRSASVEDCFTACLGE-SPSLTCLDHLEILFEPSLRVNPVALIKGASRHAVLVASWPGTTSTGHLFFGAADHPSHMDMSEHELSVLH
MCC5856368.1_Idiomarina_sp.                          MEEQIDKLALQSHQMLIVVGATLLDNIPRLAISELLCEKLLPLASQRRVDEVEALTADVIAS-QATSVLLERIHLLFEPSLQLDVLRCLKALSKQRTLIIDWPGVYNGAALSFSMPGKPDYFNYSETELSSVP
MCD8293624.1_Clostridia_bacterium                    MGKIFPTDLTGLLKPIIYCCS--GLDARALPINVPLARELLKYESAARTMLMKECFTNVVEE-FPDDCTIKNFDVMFNPAYKIDVLGIMVSVCKIKPFNLIWSGTLDSNKLIYAENGFKDHTVYEIDDYDITC
MCP5271842.1_Burkholderiaceae_bacterium              MLRLVNDAGALQSRLVLILGGAAHFGTQSLPLGIDFARRISAVARRQRPLQASSLLRDSVDA-AFPPALLDRTEILFDRTLQFHALVELKRLAHSRTVVAAWPGQLRDQRLIYAPANHPEHCDHPAAGFLVHH
MCR4442916.1_Peptococcaceae_bacterium                MGREASEYGTGKGPVFVACTNQLFPDWPWFNLGLCISQKVLSYKKEKRPNYIVDIIQEMVSA-SESRVIIDNIDLLFAPYYQLDVLKLFIQLGRNKKLIIMWDGAFRDGLLSSSVPELPDYHLYDIKNYDVYC
NPE22819.1_Fusibacter_sp._A1                         MGYDLSYFITSTNSLIVICNNDIGLQYHHFSLNQELAKKMNEIEVEKKALFVIDILNEMFE--INENIVVSDFEMLFNPSYQLDPLKYFINLSNKRKIILEWCGDFDGENLTYATPNYLDYIAYKVDKHNITC
QQS04458.1_Fibrobacterota_bacterium                  MHQILSELKERHFKLVLLVGIANDCLTSVTQLGRALSPRLAELSVRSRPLEVSGALEELAKE-VTTLLFLDGLEVLFDDSLKIDPFDTLKRLSLRRLVFAVWPGDYLRGRLTYAKVGHPEFRDCAPEGVLLYP
RKL62728.1_Thermoanaerobacteraceae_bacterium_SP2     M-NAIKILPHKYYKLILVVGPSLEFGYDYINLNLKLSEKLIRIPFDERCFYVEDYVNEIVES-NHGLLILDNIEILFEKHLKIDPLLLLKNLSRYKACIAHWNGKVEDGHLIYAVPQHPDFVKYRMDGHDYIV
WP_028243927.1_Pseudobutyrivibrio_ruminis            MAYKFLDNISGLMKPIIFCTS-IEELSEAYSMNIPVAKELQKSGSSMRTMKMESILSSIISK-LGDNPIIKDIDVMFNPDYQLDVLRTLINVSKRKSFMVVWPGTYKDGKLIYAQEGYSDYKSFDLNDYDITV
WP_052235486.1_Alteromonas_marina                    MMQSLSSGEMQSSCLIFLILETLFSDALFLNTSKLLAERLITMSRREREANVASALKSIVLA-GDNMCYLQRIQLLFARELNINPVKLLQQLSRKKPVVAVWPGGLDCSNLYYAKPGHPEYVSYQLSEFQDVQ
WP_071138484.1_Bacillota                             MAYNFLDLIKRNKLLGLLCMSIEVKDLSIINLGLDLSHKLKEYPMEYRNSKVLDELTNILAK-AQTYIVVKNIDILFNPDYKLNILSYFINLSRSRLIFVEWPGRLKGRMLEYADINSPDYHKYNIDDYKIIL
WP_164569834.1_Rhizobium_leguminosarum               MPQLLEATGVSGGKPVLLVGSATLSNTSSINLNIHLVQALIDASDAR-----VDIISTISAL-EPSVLLLDRIQILMLPQLMVKATDVLCRVARRRSVCASWPGRLDRGRLLYAEPNHPEFLDEDASRALVID
WP_191750182.1_Clostridium_gallinarum                MGREVEMLIQQENKVIILCTNSIFKEFKRISLGVILSERLLRYPIEIRNSILIQELDKLLLN-TDDKLLIKDIDILFNPEYKIDILKYFYSLARGKKVAIEWNGSIKNNYLQYSEIGYLDYKRYSINKYDIIC
WP_271679096.1_Thermomonas_sp._2C3345                MWSLMADTAQLQSKLILLVGGQRSQHATPLHLGVSMASALTELPQRERHLQAGNLLREIAQS-HEPPLLIEGVELLFDASLKVNVLDLLKRQSQARPVVAIWPGEWRNGRLTYAVLGNPEYQDHAANGVIVFE
consensus/100%                                       ..............................h...h..........................h....hhh...h.......h..........h..........h......................
consensus/95%                                        h...h.........hh............h.ls..ls..h..............h...h.........h....hlh...h..p....h...sp...hhh.h.u.......h.a...................
consensus/90%                                        h...h.........hhh...........hsls..lu..l..........h...h.phh.......hhlpph.lLa...h.bs....h...up...hhh.W.G.h.....l.a.......p.......p.....
consensus/85%                                        h...h..h......hlhh..........hslsb.lu..l.p.....b...h.p.h.phh.......hhlpph.lLFp..hphs...hhb..u+p..hhh.W.G.h....L.a....p.pa.ph...p.....
consensus/80%                                        M.p.l..h.....phlhh..........lslsb.lup.l.p...ppbs..h.p.hpphl.p...p.lhlpphplLFp.phphssh.hhbphu+p..lhh.W.Gph.s..L.au...p.ca.ph...p.....
consensus/75%                                        M.p.l..h.....phlhhh......sh..lslsb.Lup.Lbphs.ppbs.bh.p.hpphl.p...p.lhlpphplLFpsplplsshphhbphu+pp.lll.WsGphpss.L.aup..psca.ph...p...h.
consensus/70%                                        M.p.l..h...b.pllllhs.....sh..lslsb.LupbLbphs.ppbsbbh.phlppllpp...s.lhlpplplLFpspLplsslphhbphu+pp.lll.WsGphpss.L.aups.ps-abphp..ph.lh.
```

### Type 3 BREX; BrxF Inactive STAND-NTPase Entropy

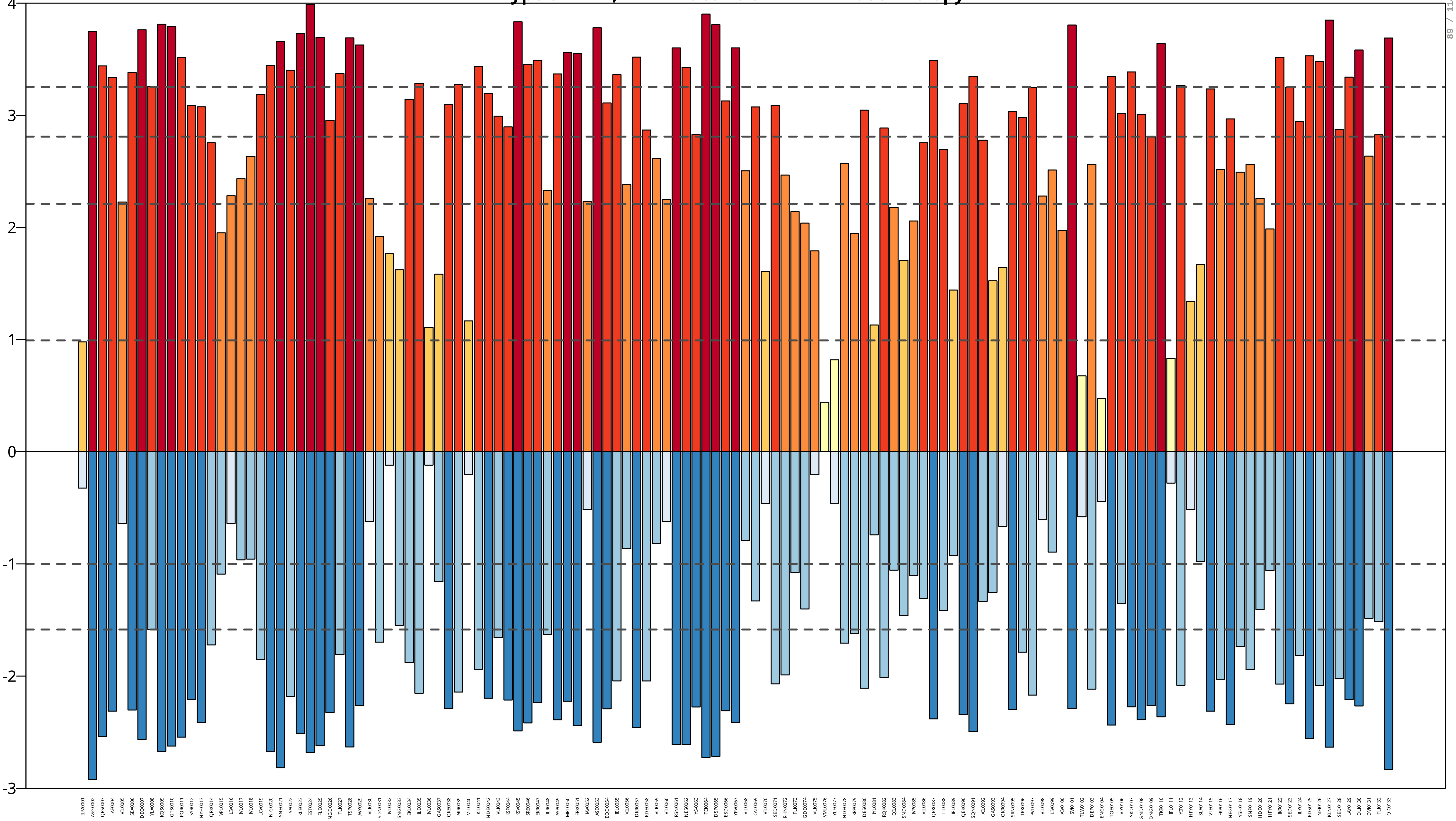

```
# ; Type 4 BREX PglZ N-terminal iSTAND NTPase; (For Supplementary ENTROPY)

RGH34270.1_Ruminococcus_sp._AM47-2BH                     MDAESCFNEIIAYFNSEVGYPFIANIDDATILQDICSKMQADSKKIIRVSDYCNGDNLPNESELLTHVASADNGVVLGLSSYYMLRGEQALKKSVSTLLQMSVHGHVVIIIYGCSSILSNISADGRPDHRTVILEK
MCI8555501.1_Clostridia_bacterium                        MTIQE-VNYVKEYLFEGDNETCILNIESLDMYNNIVKAIKELNINIVRLSSFCNDNVSPNMENFYNALEINQPTAVIGISQFFKFIGKDRLKNEISVALSKTLKNKALFITFQCFDVLKELSQDIRLHNRITILGE
MBD8953983.1_Dialister_sp.                               MTMDQCFDRIEAYCQKEIGHPLLVNTENAHDFQCLVKRMQLDGKTCVFVHDYCGK-GLPQIDALFRDVKKKGCFVMIGLSSYVMLLGKKRAEELMARLQELSVAGHVIVLLSHMKEIRLSMQQDQHLSYRVLLLEG
WP_073324030.1_Fibrobacter_sp._UWOV1                     MNLESCVLEAKSYRKSAQSYPLWIDFANAEDMRSFMENFLA--DKKISVEKYCNEDSMPRFENLYSDFRMEDFAFICNLSGFLKLLGSRKTAEVLSTLLSI--GSKKIILTVQCGVDFR--ITDPRLKCQILSVEG
WP_216471320.1_Butyricicoccus_intestinisimiae            MSIEKCIQAIDRYLVRSASTLRLVNVDNPAQMNQLIERYRVKGNEFLTVAQYSKPDEKAQTEEMLYAISKMGNLFVSGFTTFWKLEGEAEVKKQLDRLLHMSVKGHVVVLCYQCEHLLN--FADARIERLIYFVDG
WP_044080180.1_Prevotella_pectinovora                    MKYEEVIRRIDKYKDSDKHQPIIVDLPNVQVYNQMVKHYDVGSNVFKDVSSFCNGEGLPVMDKLQYCLSIGGVVFLTGLSFYLKLQGEVSLRSCLRQMLDLSCSGKLVILTLNCSCVLN---HDSRLSGRISIVDG
MZQ76087.1_Peptoclostridium_sp.                          M---TAIDCIKKFLKQSNSKFLIVNINNVEDYMNIKSSLDIE-LSTIKLSNFCEYDDMPDIEKFVKCCRTDKSSLVLGVSQYLKLKNKESFEMWTQVLLSSMTLAKVIMIYFDCENKLKELDRDVRFQNQIILVKG
HIS35238.1_Candidatus_Scatousia_excrementigallinarum     MQLNSCIEKIEKHIHRGSSKPIFINLNNSQDVKELIDHFNVGGNKIVNVASNDQIDELPTLDAILGLIHRDETIFITGISSFARLMGQSQFASILNQIINSQSVSKAIVLLYQCEDILHDISKDPRIDSHIFFIDG
WP_103226892.1_Clostridium_sp._chh4-2                    M--NDCIERINNYLQDENTYPLFVNVENATQKMEILDYYRVGDNKIVHIADFAKKDSLPNVMGVIDDIKASGNMFVFELTTYLKLISGSELQKTMQTLCNMSIAKKAVIVCFHCGDILKKIQKDLRVAKRIVAVDS
NBJ16967.1_Dehalobacter_sp._4CP                          MGWDRTLNEVTKYLSNTSNAPFFLVVGD-GNYESIKTQLSEAGLKSVKVSELCNTDKRPNLDSLIGNVDPDKKIVILGLGEYLALCGEEAAFDRLSKIKDMKVNARVVILLRGVTSVVLRQNADPRFDGRRVLFTD
CDC05526.1_Clostridium_leptum_CAG:27                     M----LCKEVQNYIATAKGLPFFYVVGD-EDYATVLDELRQANVSIVRMSDFCKDDKFPSVDELVDYFRRNNKFVVVGLGEYLALRGTAIVDKELRRLKNTTLNARAILLLRGVSTQASQIRDDNKMEQQRAYISA
MCL2016626.1_Defluviitaleaceae_bacterium                 M------KKIKNYFSSELSKPYFCFLND-EEYRQTMSELQADGVKIIRVSDMCGTDRPPNMDKLIDIIKNPAKMAIVGLGEYLAISGFDTARDNLNRLQATETRQKVVLLLRGVFSQIPKMRNDRRFDERVHYTSA
WP_235396406.1_Pseudoflavonifractor_phocaeensis          MTIQEVIKKIDRYLKKDNVEPLIVDVQNKADLEAIMLHYQLPQNVFLCASDFCKPDEFPVIPNILERLAENTNFFVSEITSFLKLKGEKALVQELKELLSMSTEGHAVILTFQCVEYLHTIKNDRRLDSRICVLEG
MBR3817930.1_Clostridia_bacterium                        MLLDACINRINNFITSSSSQPLIINVNNSSQFNELVSHFNVGDNRVISSKDYCKNDEMPQIDLVFDLISADGIVFITNFTTYLRMYGEDELRKTLKTIINLSTRGKVVFISYQCEQYLS--FTNPAQKNRVCLVDG
MBP3656735.1_Clostridia_bacterium                        MKISDIYGRIERYLRSDDTRPRIVNFQDRASLIAFRDHFNVGTAAFKMPSDYSKVDEDILTENLYAEIQFQGNLFLIGFTSQLKLKGRDTLHQFIAQIAHMTRSGKLILVCFCCAEMLH--FDDTRIQGLVYPADD
ONI45221.1_Epulopiscium_sp._SCG-C07WGA-EpuloA2           M-VDEYIRSIDTYFKGEKNSPFIVNFENILDLQEAIAHFEVDGNTIVKTSTLANKDNLPSLDNVLSNCDITGQIFVTEYSSFLKLRGHNSLIETFRILLHH--NKKLVIFTYKCERQLRELNTDLRHRNNIYFHCG
MBO6272669.1_bacterium                                   M-IRNYIEKIDRYFVRADTQPLLIDFNNKQMLNEFIQHYNVGAYNFIKAAECCREDEILNIVDLFHKLKSSNPTFLVNISAYLHLLGEKEAKNILKDLMSNN-HQKLVIVGYQICSYLG--ETDPRLKNRMVLVND
WP_037355505.1_Selenomonas_sp._FC4001                    MDVMQCINDVASYCNNVSGFPLIIDTEDYDSFHDLIFRLRIDVKKFIYVSDFCKEGAIPNIEQAYSSAFADGVHVLVGLSQACMLQSQSTLQEYLDRAVNETCREHTIILLMHCREYLQKCRQDVRLERRIRFLGN
MBS6586269.1_Eubacterium_sp.                             MDVSECFEQISAYFSGDCGFFLLVDTEDYDTFQKVLQRLQADGKKCVYVSEHCSRNGLPDVDSAVRAVCGDGDSVLVGVSQARMLQSGEALDRSLDDLLSRPVSGHCVALLDHCRQVLQKLHRDIRLKNRVVLVEE
NBK77202.1_bacterium_D16-76                              MSIQNVKQEIDSYEEHFTGAPLVVGIDGGYNYTALLSLLKDDYKQVIRMSDSCSVDFPPDPAFQISLVSKAKPVIWIGAAQAKMLYGQQPTEQFLVNLLGTCFSGPVTVICPFCCTILEGARNYAKLGYNAIVMSD
NSW92343.1_Bacillota_bacterium                           MTIEELKQELLRDKAHRDRFPVFILIKGRDNWHKVVTLLEDMTDYTYKLSERCGEDTIPNLNDLTTKIESYSNMLVIPLAEYLRIFEN--RRAILRELATIENKCRAYIPLFEVENIFFEMKGVSRFCAKYFSINT
WP_048095116.1_Archaeoglobus_fulgidus                    MSVSRLITKLKDEANIRERFPVFILIEGSNAWKELLDGLKSFTDKVVLLSSFCDNDTYPDVNKAISMAKTGKRVIVLPFSEILRVDSD--AERLIYELLDFQASGRIYIPLYGVTSQFNKWRTYDKEHSPPYFVED
NPV08138.1_Anaerolineae_bacterium                        MDVETLLTEVSYEAVSYTRFPVFILVTGVNAWRELIPALRAQMDACIDLSSLCSEDVLPYLPKVLALAREVPRAVVLPVAEALRLHPA--SSSLLSQLAQLEKHQRIYVPLLEVEDLFGEVARMDRYLHPGWRLGT
HED00230.1_Pseudomonadota_bacterium                      MDLRDLFHELSKDAAKRARFPIFILVERLDHWKEVIEHVKAPANKLIKLSHLCDEDRFPKIDELKEILEGAGSVVILPLSEYVRLTGE---KEILTDLAQWELKGRVYIPLYGFASEFIEFVHVARYIHPGWHLSG
MBE6101264.1_Selenomonas_ruminantium                     M----DMSAVEAYFVSPQHKPFYLVVGD-EDYKDALVTLSNRVTSVLRVSNYCTPDRIPNLDKMKDDLQEHNRVLVLGLGEYLALEGEEYAKSYLQSLKDFNLAGKAVLLLRGVAEQVKSIKGDRRLESGRIAIAS
ROT09401.1_Muribaculaceae_bacterium                      MTLEQVKIKIEKYLNSTDIMPRIVNVSKADDLVELKNYFEIGNIKFIDMGDFASVDSMPLFDKLLNSISNKDITFLYGISTFLKLLGSDIMNLMLRNILEIVPQNKVIIILVACDQYIK--LQDPRIVDRGAYIMC
MBR2693834.1_Thermoguttaceae_bacterium                   MNKTDLIEAVKRYAVSKSKYPFFAVSAN-RDYSSLRENIHEYGFKKVFVSDFCGPDTIPDWDELIDCVKAGEKILVIGLSEYFAFQGEEFALRKLKELKDIRLGGRAVLLFRSIGAWIKSLKSDPRFDGARYCDAA
WP_216880780.1                                           MNLDDTKKRVQKYLNSNKRYPLIVDVQTREDLANIAEYFKVGSNKFPAIEEFCNQDEAIKLDELYASVSNSGNTFITGLSGFLMLFGENVAKQALKTLITTSISGHVVIITYQCEKYLR--FTDPRISGRLVIVDG
MCM1514006.1_Anaeroplasma_bactoclasticum                 M-HERLKNKINSFIDSNDNRPLIIDVSNVEERDNLLQDFFATPSIFELVKSEC---ELPEISSLYEYMSKEKTVILTDLGTYFKLLGQDSLQQNIHALLEKSFATKFIILTLQCGQYLN--EKNPKNASKIVVSNQ
HHW17415.1_Bacillota_bacterium                           MSVDELVTYLRAESDKGERFATFILLEGTEAWDRLTARLQYEVDSTVHLSEFCEDDVFPDMSRLISYLKRCKKLLITPLSEWVRLNPE-RTDVIIQILAEWPPVRRIYVPLLSIKDSFYQIERVVRYLPQEWYLSE
WP_157832981.1_Desulfofundulus_thermocisternus           MEIASLLDFIKNEAKVPTRYPAFILVRGLDAWQELLKSLRLEVDEVFTLSTLCNDDTFPYVEGIIPILKKTTKILVLPLAECLRFTGE--SRLVLRELATWEDYKRVYIPLMELDDIFGYMNMIAQSLPEVLSFND
WP_166670004.1_Aminivibrio_pyruvatiphilus                MTLQEILQRLADEKKVESRFPVLIFVDDWSQYKELETALENACDVTLELADFCAPDVLPNFSKIFKKIEREKHILLLSLGEYLRLTIAREVQPQKANIPSLWARTRVIIPLPCCRDLWDRIPYDERQRDCIWELSD
WP_195464804.1_Clostridium_jeddahense                    MNFSELRTYLRKDRAESSLNPIFINVESMEMWVEVKKLLLSMSSGHLFLSEFCTEDTTPNIRRLIAKMKSTDSICVLPISEYLRINPE-IAQPTILDIITKERLFRIYIPMYRMRSILYTSDNDPRRKGSILLLET
HBQ28533.1_Desulfotomaculum_sp.                          MTFKEFESFIAEDIKRKDLRIFFINVESLTLWVKVKNFLAQKCHNQIRLSTFCGDDLSPKINRLCAKLEIDDKTLLFPLSEHLRINNT-KSDEVLSQIVSTEISVHVYIPMYRMKDCLQAIAQNSRLNDNIIFLAE
WP_006692206.1_Selenomonas_infelix                       M----DLTAVERYLQDRERTPFFLFAGG-SVYAQLRHRLQGLGLHLLTLSSLSEEDRLPLPDDIVMTIKSGEKILLIGLAEYLALCGEAAARRLFSRLKSMPCDGAALLLLRGFFRYASELSEDPRIGERRLALDA
MBQ0159480.1_Bacteroidales_bacterium                     MSFSQLTQEIERDKDSAQRYAVFIMLNNFNEFKELAKFMTNIGVDMLDLENIDEDDTWITKDTLKDAIKATKSTFVTPFSEVARFYNDDDFRGFFNEIMLLEDNKRIYIPLIGLQNRFTDFLNHFARIKEVWRYDA
PMQ01343.1_Dictyoglomus_sp._NZ13-RE01                    MNLDELIEYLNKDRVSSIRFPTLIMLSNYKLLHELINKLNK--ANVISIGDLLNEDRWLTQSNIYNLIKNQNDFIILGISEFLRFFSDNELYSFINFLLEIEGNRRIYIPLIGMRNRIEAFLNKYHRRFELWIIND
KUK66682.1_Parcubacteria_bacterium_34_609                MEENKLISLIKDDQEKYGRYPVFILIDNFEDMKKIIETMAKNGVETLDLSNENDSNGWVSTYELIDVIKKEKDYIVFPISEILRFLDNDDFYSLLTSLMEIENKRRIYIPILGLFSRFRSFLNRYHRRSEVWKIGN
WP_018963723.1_Coprothermobacter_platensis               MSMESLIELIDEDKSDATRYPVFIFTKSVWAVNDITMLLKLRELDILDASEFPFSDGFPSLSTILQAIREKDDVVVVSLGNLVRFWPDEDFYSLMMGLMEIENDRRIYVILEGLIERFDVFWSRRTRQIGRWTLES
MBR3117632.1_Oceanobacillus_sp.                          MDVSSIVKQIIEDKQKYDRYPIFLFMDLNNDTQQDIVKIVGISIRIKDISELSFNDAWITKSKMLSIVSDNTDYFITGFSELARFYSRQDLEALLLSFTNIENKQRIYFVCFSLFDKIAQELRTNSRNESIINGNK
WP_024292379.1_Lacrimispora_indolis                      MDIDSMVDIIQSDLNRYDRYPIFFSMKLSNGTSNYLMKLRKVCVEILDFQNLLHDDGWITIDRFRKKVYADKSYIIVGFSEYARFLSNAEFITLLLGLLELENKRRIYIPCFAIYSQVRIVKANNRRMDVFLNEMD
WP_183546997.1_Methanococcus_maripaludis                 MDIECLYDAISKDLNSANRFSLFIFVPDFDHMAKLCNYLTSMNLKEINVSKLPHDDGQVTPQRIVETIKNDKNLLITSLSEISRFYGEDDFLSLFLDLAYIEKKNKIYVPVVGIYERFREVMPVFSRSNEIWALSL
HHW21152.1_Thermodesulfovibrio_thiophilus                MNINYLIQKIKEDKKTEKRFPVYIFLSNFDTLRDLIKETSKIGITTFEISQLPKDDGWITKQEFIDKIKSNKDYLLLPFSEIARFYNKQDFNNLFSQLTELENNQRIYIPLMGIKVRFEEFYKNFNRKIEVWEIYE
WP_207678199.1_Desulfonema_magnum                        MFIEELYSEIDSEKKDVQRYPILIFLNSFNSLNLITDYFSKQAVDVLELNKLPHNDGWLTSDDIINSIKLEDDTVIVPFSEILRFKNTAEFNTILTGLFEIENIKRIYIPLVGMKERFEDFWKTFHRRGNIWYLEE
MCI6981218.1_Akkermansia_muciniphila                     M-----------DKEALNRYPVFILFNSVEDYSKFIDVCAARGAYVQRISDLLGNDKLLTYTELAEHIKEGEDFVITPFSEIARFFGNSKFRSLMKTLGAWEAHQRVYIPLMGISGMLEKLITEPGTN--VWDCCS
MCD7797870.1_Akkermansiaceae_bacterium                   MSLKDLEAAINDDKKRGNRFPCFVLFDDYKNFDTFIQTRRKDGAFMCPLAEALSDDKQLTMKSLCDWIKGKNDLVVFPISEIARFYNDNEFKCFITTIKGIESQQRVYLPFIGQQSRMYNFVNDPAIV--IWDNCN
MBO6251938.1_Muribaculaceae_bacterium                    MNIDELIQAVVVDKNAQNRYPIFVLFDNFKDSYAFTIRLVELHITIEDIQQIDYPDIIISHQKLATEIERGESKIIAPFSEIARFYDNHDFDTLIRTIKNIQTKQRIYIPIVGLEGKMSTFFGDTQAT--IWYMRS
consensus/100%                                           M.....................hh..........................p................h......hh..hs....h............h..........h...........................
consensus/95%                                            M........l............hh..........h...h...........p............h...h......hl..hs.h..h.s.......h..h.......+hhh....h...h..................
consensus/90%                                            M........l..........s.hh...s......h...h.........h.p.s..s.......h...hp.....hl.shsphh.h.s.......h..l.......+hhh....h...h.......p.....h....
consensus/85%                                            M...ph.p.l.....p.p.bshhl.h.s.p.h.ph...h.......h.h.phs..D.b.p.pphh..hp.....hl.shuphhbh.sp......h..l.p.....+hhl....h.pbh.......pb....h.h..
consensus/80%                                            M..pphbp.lpp.b.p.p.bPhhl.hps.p.h.ph...hp..s...h.huphs..D.b.p.pphh..lp...p.hl.shuphhbh.sp..h.p.h.pl.pbp...+hhl.hb.h.pbh.....s.pb....h.h..
consensus/75%                                            M.hpphhpblpp.bpp.ppbPhhl.hps.p.hpph.p.hp..s.p.lpluphsp.D.h.p.pplhp.lp...p.hl.shuphhbhbsp..hpp.lppl.pbp...+hhl.hh.h.pbh....ps.+b....h.hps
consensus/70%                                            Mphpphhpblpp.bpp.ppbPhhl.hps.pphpplhp.hp..s.phlpluphsppD.hsshsplhp.lp.spphhlhslophh+hbupp.hppblpplhphp.p.+lhl.hh.h.pbhp.h.ps.+bp.phh.lss
```

### Type 4 BREX; PglZ N-terminal Inactive STAND-NTPase Entropy

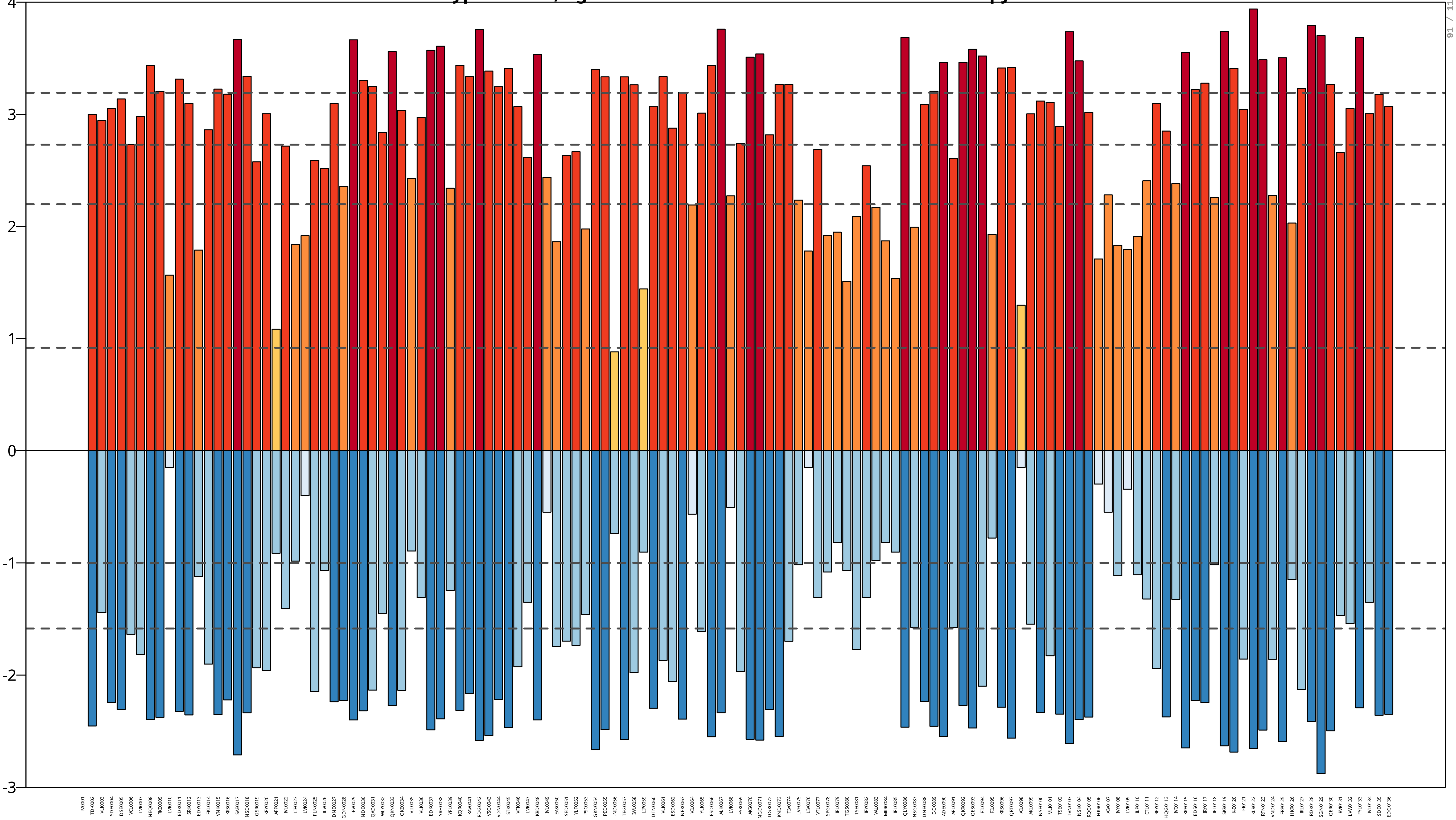

```
# ; Type-3 BRSystems iSTAND NTPase

MBP8199113.1_Nitrospira_sp.                     RPPLEIDTWKRRGHSILWDAASLASLCSSDQAVTLRQFLLHAPAAQTDVRGRALVVAGLDAALDDPDESLIWLEKFLYPAVISFENVADGALIFWFADVMSEICAHWHCSLAFIPLSRGLWNGSSVSMQEILETRTDGSMHVGYYVQKIS
MCX6876430.1_Verrucomicrobia_bacterium          NPPLTDDVWKRKGISVLWDVAGLASLGSLQTAISLREFFWNAREDSTHAGARRIVVAGLEAALDEPEAAEEWMAQQLLPVINRCAILFEGALLFWMVKHLENGEILWTCDGEHILFSHGLWNGAYRDVQRIAPLRTETEPGLGFYLQRIS
MCP3685978.1_bacterium                          VLTRRDDAWIRRGFTLLWGLKTLGKVVRPPQVVSMRKFFSMAPEELPASNGNALVVSGFEGCLDTGQDAERWIEKDIKDVILSFQDEYEGGLIFWVPSGGSSEEYHWKHRASKLHIGRLLWSGAEKEVERLIDDQRSTKAWVGLHHPRIS
WP_270720981.1_unclassified_Bacillus            SSVLLGDTWTRRGISLLWDSNSLSLITEIKDVLSVREFLYYQPNELHSANGYGMVVSGLETILDESQSLIDWMEKDFYPALLSFQSMYEGSLIFWLPDG-TDGLIYWNCTGAYIPLSQCLWNGAARDAKLIIKNTNVLKNCLGIYHPRIS
MSU35393.1_Pedosphaera_sp.                      NHDLLHNTWMRCGTSLLWDAQALNQVCSPEAVRSLREFLLHQPEGTLKIQNRTFVVAGLETAMDHPDEAVEWLEKTVYPVVLDFQEIVAGALIFWLADRASENTYHWHCTGEHIPLGRCIWNGAESDVRRIIAPDTQKTHHIGLFLQRIS
MBI5511448.1_Deltaproteobacteria_bacterium      LPPMADRGWVRRGLSLVWNAEALGTVAGPDAAWTMRQLLTRSPAALPSSNGEAVVVAGFDSCLDGPDAAEVWIQRELVPVMLSFQEEYDGALIFWLPEASADGSYSWRWAGRTLALGRCLWAGAEADLRRIVVPAKGKPGWIGLHLARLS
WP_263354237.1_Acidicapsa_acidisoli             MKVLHGDVWTRRGMNLLWDGKALSALAEPQSVVSIRQFFAMIPADLPCNEGKALVVAGLEGCLDKPGDAERWLDQHLGPAIDAFGRKYELALVFWMPTGAASEAYSWLCSAPHIEIGRILWAGAEADVRRIIDPKCANPAWIGLNLTRLS
WP_190375081.1_Leptolyngbya_sp._FACHB-17        MQPIFGTGWTRKGFSLLWDVTALSKIVESNQVISMRRFFGLVQDDLPAANGDALVVAGLDASLDTSDDASQWLEEKLRSAMLSFQDYYQGALIFWLPGGRATNTYLWLCGPTSLPLGQALWAGAEADAGHIIASSDFQKDWIGLHLPRIS
WP_125861692.1_Stutzerimonas_xanthomarina       MRQVLDTVWQRRGTCWLWDEEARNQVCVAGEVWSLRQFLAAQPDDLPSNNGRTLVVAGLEGSLDAPDQGEIWLGDTIKRAILSFQDEYAGALIFWLPQGASTDAVSWLCEAPHLDFGRLLWGEANEYPQEILLREGAK---AGLFHLRIT
OPX92064.1_Pelotomaculum_sp._PtaB.Bin013        MREIRGNGWLRSGSSLIWDAQLLSPLLENGVQIPLHKALWIGPSTIPSGDGKTIFVVGLQTVLEDSEAAFTFLRNRVQRLIVRVQDFYGNGIVFGCNCN-SNEHAYLRLRGGHLNVTFALWNGVAREAQIIMVEDKNVEIGGGFYVRRYS
WP_145058250.1_Adhaeretor_mobilis               DKPLNDDLWKRRGISILWDADSLNSLCRSDEVISLRHFRLFNSDVDASVDEETLVVAGLESCIDPPEEATKWLQDVFYKAMVSYQEVADGALILWFVDSTSDDGWSMHCAGEFIPLGRCLFNGAAKDLKEIQDADGNS---LGLYHPRIS
MBV5311590.1_Chromatium_okenii                  TAPLTDNVWTRRGINVLWDGAALARIGPVSGVISLRRFFLDQPEEATPTGGEALVVAGLDAAIDDAETVTHWIEQELYPRIQEFQSVFDGALIFWMADKTGKGCFEWRLGCGNLPLASLLWNGAHKDVRRIEASAEKGNRWIGLYLARIS
MBP7052832.1_Phycisphaerae_bacterium            MQEMTASKWKRTGSSIIWHPDLLADLIRDIEPTPLRTLIWLKPEDPPG-GKQTVLVGGLQTAVETPDAVADWLRNHALAAVRAWKSHWPDGLVFVMDGP-EGDEIVYFGRGRDVMLSLAIWNGAASGAGAYQILVEGTGEVGGYFVRWLS
MBQ8111589.1_Kiritimatiellia_bacterium          KTTTNYTLMLRGGSSVLFGKKELSELLTQIEPISLREFLLGDPDDPPPEFEDGILVSGLETVLDPVGEAEEFLAGRLRGVIQQIQNHWTRGIVFGCSLSGVSESVVFTTKDDTLALSDALWGRSQSGDIMRLEHKEPGSRRVGYHVPRIS
HCF99181.1_Chloroflexota_bacterium              MDEYAGDMWRRTGTSLVWDGDALHTAATEGVTVSLRTALWAEPEDPPD-GKRVLVVTGLRTAIDTEAERDAVL-EKVARLVRAQSRYWPEAIVFALPETPPTGTVVLDLGG--LDIGNRIWGGAGTGANQLIESRLDSRQAIGYWLRRVS
WP_149537235.1_Siccirubricoccus_phaeus          ----LADAWARRGLTLLWGADAFRELASPDQVMPLRALFGERPVDLVSNGGRALVATGLEASLDAPADAETWIATDLRDLVREFQDAYQGALVLWLPSGAPNDAWLWATAD--LPLGRLLFSGAERDVERVSPAGWRGPGLIGLRQARLS
MBF0302144.1_Desulfamplus_sp.                   MDTISNTVWQRKGSSVIFDQDSLAPFVKDEAVVSLREALWMKPTTPPV-KGKTIVVCGLETMIEPEDEIDDFLTQRIRPLIIELQNRWTGGLVFGFSSHAMQEEVLFKRRGNRVRLSEGLWDGSAAVNMKRIITSEREETIIGYYVARIS
MBX3394309.1_Phycisphaerae_bacterium            DEFVDSKMWLTRGSSVIFDKVMLGPLLTEGALIPLRQALWLSPSATPT-GGKTVLVGGLETVLEAVSDAEDFVRRYIRPLIAEFQSCWPEGLVFGLNTEPVSEMVNLKRRDDTICLSTELWNGSAVTDMRRLVRTDRAKVRGGYYAPRS-
MBN2415250.1_bacterium                          NPVLLDDVWLRKGISVIWDNSMLTKLVRNNRAVSLREFFYYKPEENMHINEEMLLVAGLDAALDDAQSAEEWVTSEVYERIYDFQNWAEGALVFWMSQKIESSRYTWLCDGKDIELGRGIWNGAQLSVSRIESES----RWIGLFLARIS
MBA3936145.1_Planctomycetota_bacterium          MKRIGGTSWMRQGSTVITHLETASMLLAQAGLEPLRTALWAAPAE---ANVSTLVVNGLSAVLDTPEDSAEFLRTKAKLIIRRSQEQWTRGILFLFPTH-LDDRVIYRLPG--VALSHELWNGAAKDLREIHADV---DTIIGYHVLRLS
OCQ93482.1_Nostoc_sp._MBR_210                   MHKMSSSAWQRRGSSIVFHRDLLVPLIESGAMVSLREALWANPLEPPN-GSNTVLIVGLETCLEVPSEAEIFLRRTIRPFLLESRNHWDRGLIFGFGCSPQESILFVYLDGTTIRLSEGLWNGAARNELFYLTTPEQTDVVGGYHV-RLS
MBE2220371.1_Anaerolineae_bacterium             MREESGTRWQRRGSSIVFDPETLSPLLIGQSLISLRAFLWREPDDPPG-DGKTVLVGGLEPCLESPEEAESLLRYRIRPVVEEFQTRWDQGLVFGFGKPPTSDEMTYIRQDLKVRLAYSLWNGGAAMDLTQIIRSTAK--VIGYYVPRLS
MBE7511465.1_Anaerolineales_bacterium           MRDMSASSWLRQGSSIVFDKTKLGPLVTTGCLISLREALWVDPAAPKN-GADTVLVGGLETCLDPAPDAETWIRTHVRRFIVKFQNEWDRGLVFGFSTP-SDDEVLFQSNN--IRLSHALWNGTATVDVARLVTDDRE---VGYHVRRIS
RKY62549.1_Candidatus_Latescibacteria           -------MILTYGLYAFEKQI-LTKLAQ---HRSILAFLLRH-RE---PLADEVLVTGLDRMLYAEDKAKD-AVKIFGSELYRHRADLAR-VLFPLEYVYWKAGIRYRPTGEPFRLEWFF-PRCEVTEIAG----------EPACYSMF-
OIO18554.1_Ignavibacteria_bacterium_CG1         ----------MAGYMLVEQRSFAVPQTPNRGKIGIADFLLEQ-EEFPFDENSSLMVTGIEEYLLSRPDMEV-TAREIRMKLQKAAGFFNDNVQIVFRQPKRGEHLIVDHVTQSIPIYLIFNTPIQTDIGG-----------QTVFISQFN
MBW1935302.1_Deltaproteobacteria_bacterium      ----------MPGYQLLELRH-LRPNMV-EGKMGLLDFLITQ-DPFPFPKFYRVQILGLEDVLFARPQEEE-LALKIQRILRSAAQNLEK-IQIVFRGRVRGESLWVEYRNAKLPIGHIFGSPIRQTDASR----------NQFYKKSFS
MBU1206361.1_Pseudomonadota_bacterium           ----------MGNYLLIEERM-LQPYRP-KGRMGFLDFLLRQ-EVFPFSRGNKLCVVGLEEVLLGSGENFMKVSLGIRNILTSRANELEA-VQIVFRRSKRADDFWFDPGGRRVSLRPIFGSPIKQSDQNG----------NEYFMTGFN
CBX30016.1_uncultured_Desulfobacterium_sp.      ----------MKGYEIFEQKY-LKDYFGEEGKIGFLDFLLKS-GTQGIPSSSSFMVVGIDDVLYGRDERKM-LALTIHKILQSSAKVLDQ-VQIVCKGRYKGESFWSEYRGEKLPLDYIFGTPNKRDIRE-----------CPVYSTGFN
MBN2391965.1_Anaerolineae_bacterium             --------MSSSGYVLIEQRE-PTPTILRDDQRGLLDFLLAE-DELPYPKWSLVTVTGLDEVLYAQPDEEA-LAHEIHGRLNQAASRLES-IEMRLKGQIRGDPFRLRHRGIDLPLHIIFDTPGKAADGN-------------YYLVRCH
WP_248664448.1_Fuchsiella_alkaliacetigena       ---------MQQGYNLMTQNY-NQQYYQ---RMSIMKFLVQE-EN---KIDYSLEIYGLEEFL-GKEDA----NDLLTEILGKGVNYLTD-IKVITENKIWNNKAVLKYSNEEFPLGNIFGSSLRQEDTE-------------YWVANLN
WP_103424089.1_Salinigranum_rubrum              -----MASDLDIGYHVIDEDA-LDD------EMSIVEFLLKR-RE---EIPLDMTVRGLDDYL-GVDDADV-ACDYIHRLLRDRVNYLSL-VQFVVDD-ENWSGPVIPTGDEPIKLNRIFHGSMEQSGPG-------------WYSSNLN
KXA95577.1_candidate_division_MSBL1_archaeon    ---------MEEGFHIIEKSS-VKERDL---KMSIVNWIIRN-EG---NQFESYTVVGLEDLL-KISNPQD-FSEKLRKILIDQSNYLVS-FQFLIDNSVWEKNPTVKTASEVIDLSAIFG-ELERKDVN-------------WFYQQLN
NLY06148.1_Candidatus_Atribacteria_bacterium    ---M---VEINIGFNIIHRSN----------RISIIQFLLNR-GS---KFPPDFAVFGLDSLL-YVEDREK-MAKYIRHTLQDNANYLTS-IQIVIEGDESDEKPRVVYKDEEFPLYSVIG-RLKRVDLK-------------HFYAPLN
HIE51614.1_Armatimonadota_bacterium             ---------MRPGFYLITRRQ-LGRAIQ---TMSLIALLLHR-GE---ALPSAVCVVGLVGLL-SASDLRQ-AARLLAQRFQRHRSALMR-IVFPVEHLPTPRGVQVGARGSQNFIAEWIFPRLEPLEKV-----------PRAYYSPF-
WP_285254567.1_Halobacterium_salinarum          ---MSDIDAVAGMYNIIVSSE----------RVPIEEFVLET-RD----LPDEICVAGLEEVL--TEDEDL--RDQLVSALRQEMDYLNSAIQFVVDGDGAGDSFDVEIDDEFHSLEPIFGRQIKKRDS-------------GWLVAPFR
MCR8454635.1_Thermoproteota_archaeon            ---MKSLKTMEVGYNIVEKSL-VGKEFE---KISIIEFLLYR-KQ---PVNRKIAVIGLEEAL-AGEET----ARYIRRILVDSTYMLRTIIQFQINGEILDKEPKIRYRSKEINLIPIFGNRIKPKTVG-------------YFHSPPN
WP_135823665.1_Halorussus_ruber                 ---MPEAGRLRGVYNIVTAGD----------QIPVDEFLLED-DD----VPEEVTVVGLEDAY---DDESF--IDELSRAMDRKADDLENMVQFEIDADRGQVAFDIFHEGEIYPLHEVFGRQIKPKDQT-----------GSWYTTPF-
WP_153952804.1_Halosegnis_longus                ---MPTDGQLRGVYNIVEHGT----------QLPIEAFELAE-GD----VPNQVTVVNLGEAF---TDQEF--INSLAKTMADHGNALATIIQFVVQGEHRGEPFDLFIDGEIHSLRPLFGRQMQALTDD-----------HSWLKAPF-
NMX21979.1_ANME-1_cluster_archaeon_GoMg4        -------MNPTQGYNVLLKE----DEPP---KMSIIKFLIRR-GQ---ELPKKVAIYGLESLL-TNKDARA-GTKFIRNMFIENNRLFRE-FLFIPRKEYVNRDVYLKAEDKKANVSSLFGLRLQIKEVG-------------IYYADFE
MCL4459198.1_Chloroflexota_bacterium            ---M---EGLKAGYNLVERAL-IGQRYE---VMSIVQFLLES-GE---NLPRRFKVTGLDRLM-AIDD-----SGTIRSILYEAGEKLLR-VLFPVDK-TMDAEPKMYWRNQEIRLRPLFGNRLEPQGVG-------------YFYSPFN
consensus/100%                                  ..............hh................sh..h...............h.h.sh...h.............h...h......h...h.h....................h...h............................
consensus/95%                                   ............s..lh...............sh..hh..............h.l.Glp..h.............h...h......h...l.h.h..........h...s...l...h.......................h....h.
consensus/90%                                   ...........G..ll................sh..hhh.............h.l.GLp.hl....p........h...h.p..p.h...l.h.h..........h...s..h.l...h.......................h....h.
consensus/85%                                   .........p.G.pll.p.............hslbphhh..........p.h.VsGL-.hl....p.....p.l...h.p..p.h...l.h.h........h.h...s..h.l...h.s......................ha...hp
consensus/80%                                   .........p.G.sllbp...h.........hslbphhh.p.........sp.lhVsGL-.hl....p....h.p.l..hl.p..p.h...l.h.h........h.h...s..l.l...h.s.h..................ha...hs
consensus/75%                                   .........p.G.sllbp...h.........hulbphlh.p..p.....sp.lhVsGL-shL.s.pc.p..h.p.l..hl.p..p.h...l.F.h.....sp.h.h...s..l.Ls.hh.s.h...s.................ha...hs
consensus/70%                                   .......b.c.G.sllbc.p.ls.hh.....hulbpFLh.p..p.....spslhVsGL-shL.ssp-.p..h.pplb.hlbp..s.hp..lbF.hsp...sp.h.h..ps..l.Ls.hh.sshp.ps.............shahsphs
```

### Type-3 BR-systems; Inactive STAND-NTPase entropy

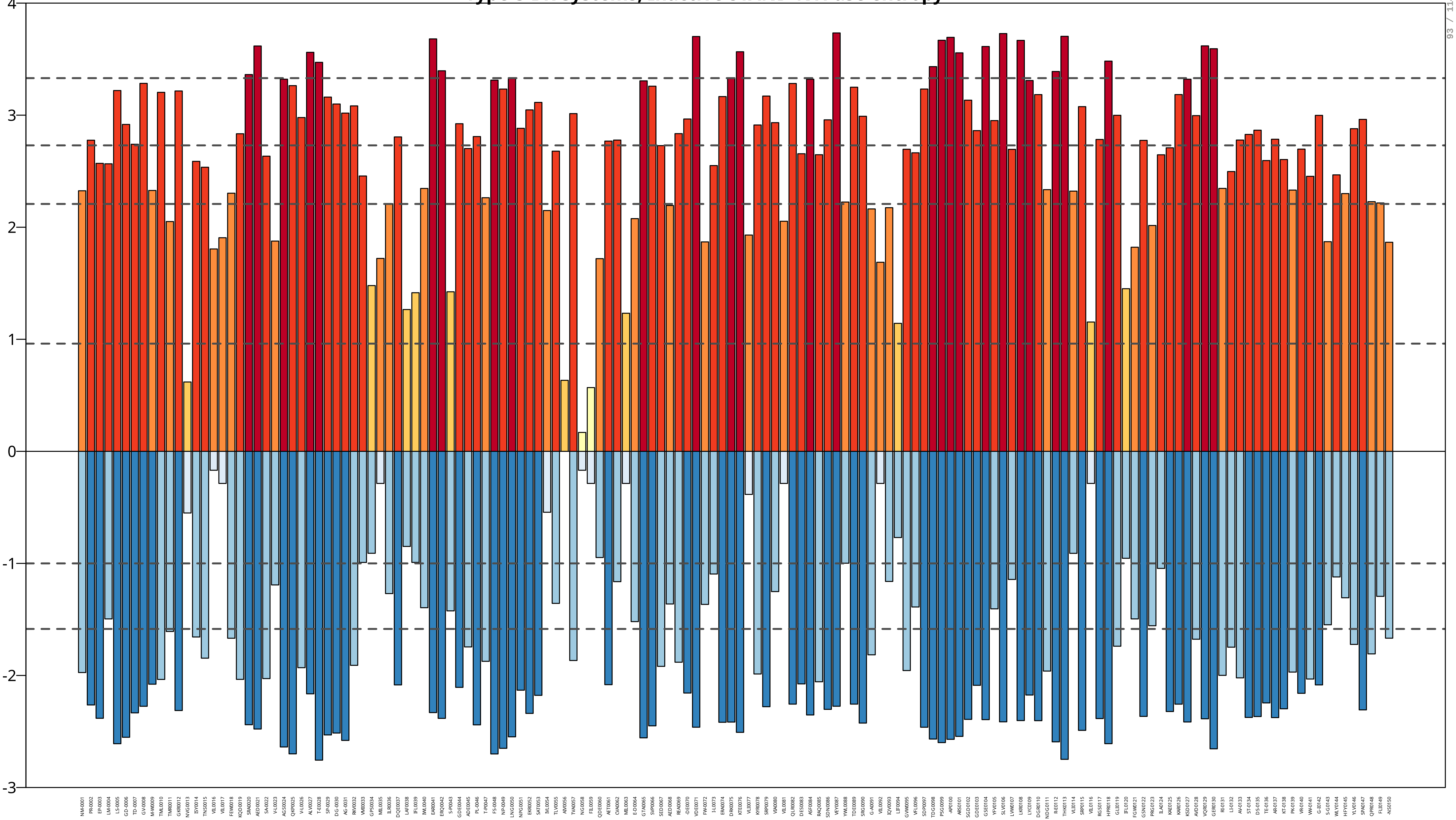

```
# ; vWA-MoxR subsystem 2: the iSTAND ternary systems; (For Supplementary ENTROPY)

OCQ97673.1_Nostoc_sp._MBR_210                         LHKALLCLNYTKQENLFREFVRHQIGACLIHGEPEHGQRWLLNRLLGLVPNGRTESVIPLNLTRSSSIDALWRELGKAVGAAATCDEIAQKVGNLWKSQTVILIFHNLDQLPEDYFNEFICKFCVRNYRLLAFLMDLSGCVDETIDLNWEPHTPLKLPKITKLSHNELSYWLEKSPAEIDKIALDILNSDDEAVLDYICELCD
BAI89278.1_Arthrospira_platensis_NIES-39              IKDKLLHLDYREQVRSLEDFLQDRVGAFWIYGSNQSEPHWLVTSLIENIETSNTSSRVSLSFTQSHCLDNIIDSISKAFQIEEKLTYIIEQICIRWQTKSVVILAEIKKGVDPEIVEEFINKLRIYNYSLMLFLVDVHGCTNEDILSEQIPEIPIRLTEIQPIPREQLRQWFRTHLDQLIEKMLDLIYHNNEKVFYQICKLFE
BAY43400.1_Scytonema_sp._HK-05                        LYHVLLKLDYTDQEQLFSRFITNQIAAFLIHGSQDYGQHWLTNRLLKKIRRVSDKPVL-VDLVHNRNSQTMWRELGRHFGGIDSPDLIAGKVLKSWTTKNIYIVVDNVQSMSEQLLQELIQKFQQTNFKLLMFLIDNVSKVEENFEPTWRCHQPVILPKNNVFSDIALRDWINHESENIDGVIQTILNSEAIPAFREICNLCE
KPA13794.1_Candidatus_Magnetomorum_sp._HK-1           RENFPYSVNRTEQVQGIKDIGEKKPLVWLIHGDRNQSHERFLECVQWFWENIKKGSKIEPRFYSNNFENDCMTEIWKNFSQDDNWRDLSPLIDNNCNK--PVILYTMIRCSQQRLFYVLLDIQKKYP-LFLCILIIYTHKHKRKITRQLKKYNKYTLPLLSNIESEDVDEWQNMSFKNERKIVKDIFDDIEAKIFKHCLYHQD
KPA12842.1_Candidatus_Magnetomorum_sp._HK-1           GKFVHLMCDRANQVVEFRKTFHKKPQFYVIHGKERNGHLSLVERLKIQIQPFVNEQAVSLLVDLDMKMDILLDALFAEIFNQIQYSLKALIQSDPIKADHAVIIAHRIPANHLDLANRYISDFATIPLFIVFFNIEYPRLVKYKIRSMVKELTNQLIKELQTINTHHVIDWFNSHQPEKFKAVDSIFDGQDEDLLKKY---VP
ETR67363.1_Candidatus_Magnetoglobus_multicellularis   GDDINLYCNRITHVGEFLFTYGNMPQFYFVYGCKDECPESLFLRLRDILRDHFKPQVPPIDESFEARKDVLLKSLFKTLTGDNKFKLTDLYQSNAIKDNNVIVVPHFLSFDQQKLMEWYIKDY-DAPHFMLFFIITCIPKKNKKLKKFSSQLSAYLFDPLHKIDKSEVAPLFKQFKHEINKHIDDLFKDQEEKYLLRLKQTIK
OQX97257.1_Bacteroidetes_bacterium                    RNYYRYSCNCYKQNNYFENVFNEKIQFYVIHAKAEQCPEILPQRFFNEINDNDNNSIF----NIDSLKIKTLYQIFSFFNEEKEACLESLFNSKTLQKKKYITINFIVESRQHEFIEFFINDFCTKLRFIFFWSIIYTSEIKDKIQKCIKNLEFDVLPEIQIINLDDIRTWIKSIESKINISLFNIFTNKKLEMFDFLSEKYN
WP_020569053.1_Neolewinella_persica                   PPFHAFGVDRAPQEGQFFDEQDQKVFFYYLHGDDRQQASSLAERLGFKVGLQLTADDLDIEAREEHCRALLVHAITQQFMGGLNLTILDLLNRPKLKRHPIVCINVTVGHSYPGMLRKFYETFCDAPTFYFFFGLEHCDNKPKEVAEAIAGRQNSALPELEPVLISDIVNWFTHHGYDAEQLVAENFTEGDEADLKRLIDKFN
RMG26468.1_Bacteroidota_bacterium                     GRLLHLMCDRDPQAESFLDRFAGDPQYYFIHGPRGERHKSLVRRFREIIRHHLGRMPALVDFDLDMRKRNLKRSIARSRELPGSKWESPALMALLAQRGNLVVFQHQLAAEKFRLLEWYIHDF-ASFHFLLFVNIVYPEKWGEDMQALKRKTPETILAELQPITPEDVAAWVDTYFDNMQHLPGTIFDKQKETQLKLAIERLE
KPA17452.1_Candidatus_Magnetomorum_sp._HK-1           GRIIPTLCNRDIHVEKFYKFFKKLPLFFIIHGFENQGHRSLIKRLRFTIRAYLQKQKIELPFKLSDRKQILARQLFNAFQKMKTYDLKRLNQLKAIETNNIVIISHNIEASLKDLLSWYIENYVDIPQFIIFLNIQYPVAIKKELTVFTQNFSKKLIDELNSITQQHVKDWICKYKYEIEEEIESIFENSEEKKLKNILLK-S
WP_028158837.1_Bradyrhizobium_japonicum               ARLLPYLCDRRMQRRHFRDAVVVRPQIFFVLGADSECADKFVEQLFFVMPELLQRNGLEDRLDIGDYLGELQYEFADKLG--LKPRSSVTEIERRLQRHTALFLYFDVDIRKRRLLREWIGWWNRRYPIIVVGRFCSANQGIDDLTALSTTAGKTVLPQLGRVSFEDIEAWINENAPDLRQRLRRFFIGERATLLKTIMNEMK
RKZ89871.1_Candidatus_Parabeggiatoa_sp.               DDLRPHLPNRRPQESKLGRAIKQRPLLCLIHSDECQCSDRFVDRLVQYLPKVVPGDGIFVHCQGIDLHQEISEQLASRLGI-NPFAQPSEIVEAIMRERSPIIFHTDMCTKNMKAVHNFIEFWATHNHLLICLYFNYKDIKPDQIRAEFRQLENIVLPELMDIEKEEVYAWARDHLDSVQPKIDDLFSPDEARSLREILEEFP
OCR02031.1_Oscillatoriales_cyanobacterium_USR001      VHQALLRLNYTHQVRLFRAFFHHPIGAYLIHGEEESGQKWLLNRLMKFITDSRTAKVIPVKLSCGTYTEALWRQLGGHVGLMPSQTQIIKNVSELCKNQIVVLIFDELNAMPEECMSQFIGQFRQSQSRLLMFLVDNIGCVEETLDPNWEPHIPIKLPKINPIEQSDLVSWVRNAVNQMEEIVQNFFNGETEAAMIKICDECD
RMG84230.1_Bacteroidota_bacterium                     HEYHRYTCNRQPQNVQFHSSFKQKTHFFYMYGLNVQSPEGLFYRFSYDLLDYLNPDLESVRADETYNKQEVLKNLFAALSVPKKRLEDVCLSSPTIRKEDYVCIFLGVSSFDPALAHWFITRFCDAPTFLFFLAIIYEEDDPAEVKETVENSDLIGLPELGDVSLQDVKRWFFNYQKTRQQLIEERFGDAQEIRLERIIDEYN
WP_028091549.1_Dolichospermum_circinale               RSFLSYLANRSSQEFELTEAIKSRLLVCIIHGDELQSHYKFLERMRNDLPDVLDLDQNKIKKNLNELPKYLLMNLAKNLV--NKGNTSLEEINEILNKSSPMIVHTDLISDDLHILNKLLEFWQTLNKVIICICVKYIRIEWKKMRKHIQEFSLIILSELEAVGRADVENWACEHTQQLKDKIGDMFKSSSADNLLEILEHFH
WP_018400346.1_filamentous_cyanobacterium_ESFC-1      LSELLFLLNCIQQENAFR---ANPAGTFLVRAKDLSLQKWLVNRLVRLVPGY-ENAYCPFNVQIRLNFNLLWENLAANLNTTATIENVAEALIERYCHQTVILVLYGVHKLQDHQIAEFIQ-FQARRSRLILFLVEDTEEH-ESIDYQCL-EKPLRLSPLGQISKAEALTWFDEVYEQIDLLLQDI-NDPNSISLDEICMTCG
KPA18269.1_Candidatus_Magnetomorum_sp._HK-1           PKKLHLLCNRKKQALDFTLYFKNTIQLIFFYGPENECPYSFIERLAIKIKDYWENQYDKINKDVNDQKNTLIQELYNSFLLINDYKLSAFYQHPTFDKHNVIILYHEISIDKLKLLEWYINEHSNVKPLLLFFYIKVDGFKEDKLKKKIDKKFKLLLEPLQPVDKDDVREWFSNLESIVEPEIEQIFGKENITKLDIFITNID
WP_066426097.1_Anabaena_sp._4-3                       IINALKKLNYNEQEKDFRDYINKYAATFLIHGQYGYGQRWLVNRLCEQVDDGLTTAFCPINLKKHRDIQRIWDDIAAKVGSSRNPQTIVDVIYQHWQTKPILLCFHNVDVVGCQELNKFIEPFKVNTQPLLLFLIDNYGQQKNQISKEYDIENNLEIKELDKFSDAEIKIWTKCYQSNLSEITQEIIYNNTIQAIEYICQCFD
KPQ40445.1_Phormidium_sp._OSCR                        FIKILGRFNYVQQKRRFQVFLNPRIGAFWLRGERDSAQKWLLSQLWLQVPNSTKAIKKTLTMTTRQNTERILEKLATWLQVEATPEALIKRILENCKTETVALVFYQVEYLDKGTLEELINTFQSLDNPLLLFMVDLGKKGDINHSSDYNAEHPEQFVELTQRFQDRDFRWLTDNQVSLEAIKKDMIDDNHEEVFTELCDYFG
WP_071187406.1_Trichormus_sp._NMC-1                   LEKSFQKIDFSKSREIAESLKTEGAILLFLQKSKKQLGKYCIEEFIDVISNQIETCPRTINFTYKSNKYGFINAISGYLGIESIINEIHEKIVSSLSLDTIFFEIKSINRLEEEFIDWFIKIFKQKRNRLIFALIAESPFPIQEVTLNHDK---IEIPLL-DWTVEDISNWLISHHDEIHKEAKNI-EESEESVCSKLQEYFY
SEH05005.1_Candidatus_Venteria_ishoeyi                PNFAPYLVNYTKQQTCFEDIVGNHPLIFLLHGQESNTTHGLAARLKQNEENCLHHGETTVRLDCDERPGFLLTTIEKNLKDKFAVENLEILHESAFEA--TTLFYSEVEAKHKKLLQDFLNFWNQNHLFIVCLFIDYRPGPLQRLKAKNIDICNMLLPELTPIPREDFTPWYNLMKDELEIRVTELFDENNGPKLSKLLADLC
WP_017306429.1_Spirulina_subsalsa                     LNDTLKKLNYDHQKRPVKDSFNQRINAFIVQGQSHHAPRWLTNQLLYYVPHHDNAKKHSIVLKTHFRFNRLCEGIASGMDCGAEPQNIVSALAQNLETQSVVLALRDIHQAKNGILSSFINEIQTGQHYLILFLVDGDSSSSTELNPFIQTDSPCSLPPLENFTQGTITLWRGRKEDQYPQAIDEFIDTEEKGFLDFIYDSCL
OCR02924.1_Oscillatoriales_cyanobacterium             LYNALLSLDYIRQESLFQEFVNNQIGACLIHGKQEYGQRWLLHRLLRAIPNNNTASLTKISFVQNGSLDALWGEIKRKLKLKPLPDELAKAVCNLWQTQTVVLIFDDADQIEEAHLEKFINDFNSRNSRLLVFLIDFDGCAPDALDTTWKPHIPLKLPPIKPLREMELAYWLEKAFDSIEATVQSILNSDNELAMAHICELCD
KIF28928.1_Hassallia_byssoidea_VB512170               LYRALLKLNYREQTKKFRRIIDNSVAAFLIHGLPDYGQGWLLNRLVQQVPNSITAKVVQFNLARKSDVPSLWRELSRGVGLGGLPPDIADRVYQWWQTQNVLLIFRDVDFLSEIYLRELLENFTVSQFQLIMFLIDEKGCVSEEPNSNWQPDSPVKLPEISGFSQEELAKWLDYESEQLEQTVQTILNSDNELAMEQICYLSG
CUR12819.1_Planktothrix_paucivesiculata_PCC           LFKSLLRIDFDEQEEAVMEVLYNKTAAFLVHGEEKFGQETLVTRLSQL-SKLRNGRRIKIKVGGMEDISNLWNEVGKHLLDKLPETKIIDQIFELLQTQHLIFIFSDAHRTYISFLPELIQEF-QRETYLAMFLVDNKGYICWQVNHPDLPRIPLSLPPASRFPVRKLNEWVAVAA------VETLLESQGDLVYQRICYYCG
OQW38836.1_Proteobacteria_bacterium_SG                AEQKLGQINRKKHRDHLAIQINRKTHGFVISGEQQECPEDIRFKLSYLQEDLINHNPKGLRTIGKSAEQFLWELLAKRLGSCNPEKAAIEMRLNQVDR--SYLFFRELPSDEQQFLVDLLVAWSTSHSHFLLLIHAAENSGNDQIESLLKQHEVALLPEFHPPSKKEIRDWKGLHISVLRDGIEELFLKHEKAALPLLKKHYS
KPQ40050.1_Phormidium_sp._OSCR                        VHQCLLNFDYQKQEYEFDEFISTQLGICIVNGLQKCGQPWFLYRMFQT-KYYFDDTSTPFNKFSRTQKLSLKEEICHFFRLPESIERIYQEIANHLEHKSLVFIVKNIHLTTQEYLDYFVKQFQNPENRLLFLLVDLEGVSHQELLSHYRSHTIYRLKQIKPLSWAERQDCLKQLA---DEISQKVILTEKETVLEYICNCCN
ETR71372.1_Candidatus_Magnetoglobus                   GSASVLMCNRIPQLRSFLTNYHKMPQFYFIHGREQDQHDSLVTRYNQTIHKRIQEIYDPYMCKSDDQQNTLLYDIGHELTGEDITSFEKLVNATIFEQQNAIIISHKIEEDFKDLLTWYIQEFLRIPQFILFFNIIYE----RWINEITPKNSTVLIPELEPVIRPDVQSWFNTYTSKRNECIKQLFPKPKEEKLVGF-----
WP_020531248.1_Flexithrix_dorotheae                   PEKFYLGCNRTIQFQDLQYILTKKPCMCILYGNREDCHRSLIERFYYYLNK-------LINIDVRNRKERIIKSSFEAFNLTAVPDNAQMIIDSYNQKCDVLLIEIEIDCNEKSTLKWYINEF-----LKVLHLIKLVSKPSKDLRDLYSSEIRKVLPKLSPIKKYHVKELL----ASYVDKIDKLFLKKSEMFVT--KEFLN
GAK59821.1_Candidatus_Vecturithrix_granuli            GDIVYKMCNRGPQENDFLRFFPRQPHFYFLHGSDDECHESLIERLARTIKKYVEQTCGDVQSDLETRQEDLALNFMKTFDSDPNELISGLCHLAVLRKHPCVILRHNLDAARERFIRWYIKAWLDLPQFLIFLNIIYPRRHHKQLCRIHTSSGEQLLPELKMVTPFDVSDWFSYYRWQRKEKIALIFGNQEEKHLKMIIDECT
KIJ79200.1_Tolypothrix_campylonemoides                TEAEIIQLGHKSLSRQFEDSLQAAVVCFLIHGASGYGHKEMVVRLRQILEQQRSVQARKAAFWREKSLAKLLELIGDQIEQGDSPEVLAQRLHKMLETSDVILEITEIQRFNGSLLS-FIKDF-PTQHRLVVLLTLEQTILPMDDDSEIDSMRPIKLAELKEFTEKELTSWLSKWRNDALVLAKTLMDETKQLLYNKLRDDST
RAM51670.1_Hapalosiphonaceae_cyanobacterium           FMIALYDLDYLHQRTSFKTAIDVKAATFLIHGKPHYGQRWLLNQLKSQVPYHTNAWQKSIYIPHRKDIEVIWQDLAQELKTSASPKDVAKELYQHWLKSTVILAIHDVDLIAGSPLKQFMDEFKQSSYRMLLFLVDNINSKSENIDIN-EPHIPFVLKELEPFGKNLIEAWVGVQEASIKEVMCEIVRDSQISVFTDICQCFD
WP_088278871.1_Ideonella_sp._A_288                    DLPLPYLCNQEPLTRAFEDAAAKHAMLVLLAGGHNDAPEHFARRLQRHLPRRLKTIGVAVHTTTLRHSMPLLRALFEALDDDTEFDQAAQVLCEHLRARTAAVLMAWLPDRDLPALLRFIDAA-----RLVLICVGEARGSVRAVRKSSQA---AALEPMAEIDVAALTKWHKEHGRVDLDRCETLFRAGRARHFQTFCEGLR
OCQ93508.1_Nostoc_sp._MBR_210                         LFDCLLSIDFKDQEACVTQAIRQRTVAFLINGDDKYGQSAFIERLFRKIPEFKHEKEIKIKLESGWETSNILQEIAKFFGLDDIQDKIINKACECLQFKNLIFVFYIPQRFFVGLLADLINDF-HKDTYLTMFLVESNHKEKNKLAPREDDSSYLYLPLLSKFSRDHLEKWLENGVTQQKNFVDTLLETEWERVYKKICNHFS
KPV52199.1_Kouleothrix_aurantiaca                     LYQALQGLNYSAQKSAFVLSVARTVAAYLIDGQRNHGQRWLLNLLLKRYEHALQA---SIDMGRGKSIADLCQEIGRKAGVASTPEQLAPVLCRWWETQHTFLIFHSVDLLGEAGLREILEHFGNAANKLLLFLVDNQGVAAQNPA---NAGGPERLPTIARFSEDELDRWMMASLERTD--VQTILNTDDESVLEEICVICD
PKO80078.1_Betaproteobacteria_bacterium               PLAELGRIGRDTQRMALVDALGERAAAFLVHGPENHGQREFAEFLNVWDNEWEDAQPI----ADTDTADALIRWTCGQLGTPAAIPELAQTLAARLRRGPVIIIQRSIGHRQ-ERLTRFIGGF-----RLYWFLIDHCPLPQPEAGADTADYDDLALPALGDISAGQVRKWLKELR-------DEILTTPDSDVYNRLWASAS
RKZ83713.1_Candidatus_Parabeggiatoa_sp.               RTLLPYLPNRRDQELELGELIKHRPLLCVVHGDEYQCSDKFLDRIREELPNIIPEQMHGVDLDTFQSVNELHRKMLANLGELNMYATCDEVAQAIAQRQCPVILHTDMSTTDINIIHGFIQFWAAQNHLLVCLCFNYRDIEKQAIRNTFKDLASRVLPELPSIERTHVEQWAKIYLYELIPYIRELFPPIEAQQLKQILKECC
RMG30280.1_Bacteroidota_bacterium                     SMLDKYTCNRAEQNSRFKLNFNKKFQYYIIHGEEQESPYGLFQRFVLEISTAYQPYFY----SLEEAKINFLTALFQALELNDQLTLANLLKAPAAQDMQCIALKLRVYSSERDMIQWVLQEFRNAPDLLMFISIIYDLQQAEKLIKTLAEFPEGLLPVLPPVKISDVNKWFDRITEEKKQQIATYFDAQPEKKLEQIIEVYN
PZN77012.1_Candidatus_Methylumidiphilus_alinenensis   LSLLPLMFGRETQEEQLQDYLGQQIIAVVLPGPHRESPAHFVERYAKLLRDAYQRHWREYVWPLQQRLDFLLRDLCRNIDGVGEEGHVRDRLVSQFADQSHQILYYFLTADRANLIGEWIGFWQAKNMVSIIFCLHSEPSILKALRTLCPNAPDLMLEKLPPIKEQDSYNWAKEHEAILYQRAKNAFRKRRMTKLEEAVAKAA
WP_072720293.1_Planktothrix_tepida                    VYQSLLRLDFRKSSGIFADWEQNKAACLLVYGKKKHGQEWIMRRLMTK-PKYSKRMFYTCSFSYISDSKKLWKWLSEKIGIQENQLRIIKALKEQLQYNNIVLIFTDLQANSLEYPQEFIDQFFNSCNKLLVFLLADEPEIKSNINENWSITEPIQLPEPPICPEDDLIPWLKQETVNPRKTAQEILKTNQEAVLMYICQYDQ
WP_020536784.1_Lewinella_cohaerens                    HEYHAYTCDRVAHSDTFRQLFKQPVQFYFLYGGDLQSHEGLFRRISYDLQDYLNPELKSLQLQLEHYKQNILRSFFSMLGLPEKDLTYALERSPRLQNTDYVCVFLHISQYDPAAARWFIKEFCTSVAILFFLAIEYDEEDASEVVAALEMAERQPLPELDMVAQRDIGRWLEHYKRTRKELMKTVFGNSRELELKKIIDQYN
ETR70369.1_Candidatus_Magnetoglobus_multicellularis   RNLLPYMPDRSELIQEMSLALAGKPLVWIIHGNDNQCHYDLMDCIEGCWERFLYPTSKLLRYETNQWRDLVAMELKHKFDIQEAPEDVFDAINRRCDKASPVMLYDVVDTDQQTVCQRFIDFWNCQNPLIVCLLVRYKSFPVLEFKETILTLKNCVFPELPDISKKDTEDWYHNYKKNVSQCIEDIFDANQAREIEQ------
PZN72517.1_Candidatus_Methylumidiphilus_alinenensis   PAFKAGLVDKDEQATHVIDHLHNNPLVYLLYGASTQWPDALLNILYYQLPRFNKNSSPTLKYGNIDPVEYLYELLDEELTCG-------KAIERRLSKEKPLIFYRILTAAEPALVQGMLVAWQNSPRHILIFYYEYIETPKNHLKASLPTEMSIFLPRLKSPNKKLVHDWISIHIEEAKKYVEDKYNAKKHHDLKNILNEFC
consensus/90%                                         ......hsb..p...h.........hhhl.G........hh.ph.........................h...h...h...............hp....hhh...h.......h..hl..a......hhhh.h.........h...........l..h..h...ph..W............phh...p...h..h.....
consensus/85%                                         .....b.hsb..p...h...h.p...hhhl.G.p......hhpph...h.p..p.....h..........lh..l...h...............hpp...hhh...l.......h..hlp.a......hhhh.h........ph...........l..l..h...pl..Wh..........h.phh...p...h..h.....
consensus/80%                                         .....b.hsb..p...h...h.p...hhhl.G.ppps.p.hhpph...h.p..p.....h.....p....lhp.l.p.h...............hpppp.hhh...lp.....hhp.hlp.a.....blhhh.l.......pphp..........Ls.l..h...cl..Whp....p....hpphh..pp...hp.h...h.
consensus/75%                                         .....b.hsb..Q...h.p.h.pp..hhhl.G.ppps.c.hhpcl.b.l.p..p.....hph..pp....lhp.l.p.h...............hpppshlhl...lp..p..hlpbhlp.a.....bllhhhl.......pplp..........Lsbl..hp..cl..Whp....p.p..hpplh.ppp...hpblh..h.
consensus/70%                                         ..b..h.hsbppQpp.h.p.hppps.hallaG.ppps.c.hhpRl.b.l.p..p.....lph..ppp.p.lhppl.p.h......p.......p.hpppshlllbpplp..p.phlpbaIp.a.p...bllhhhlp......pplp...p....b.Lsbl..hsbpcl.pWhpp...pbp..hpplhpsppb..hpblhp.hs
```

MoxR vWA Ternery systems; Inactive STAND-NTPase Entropy

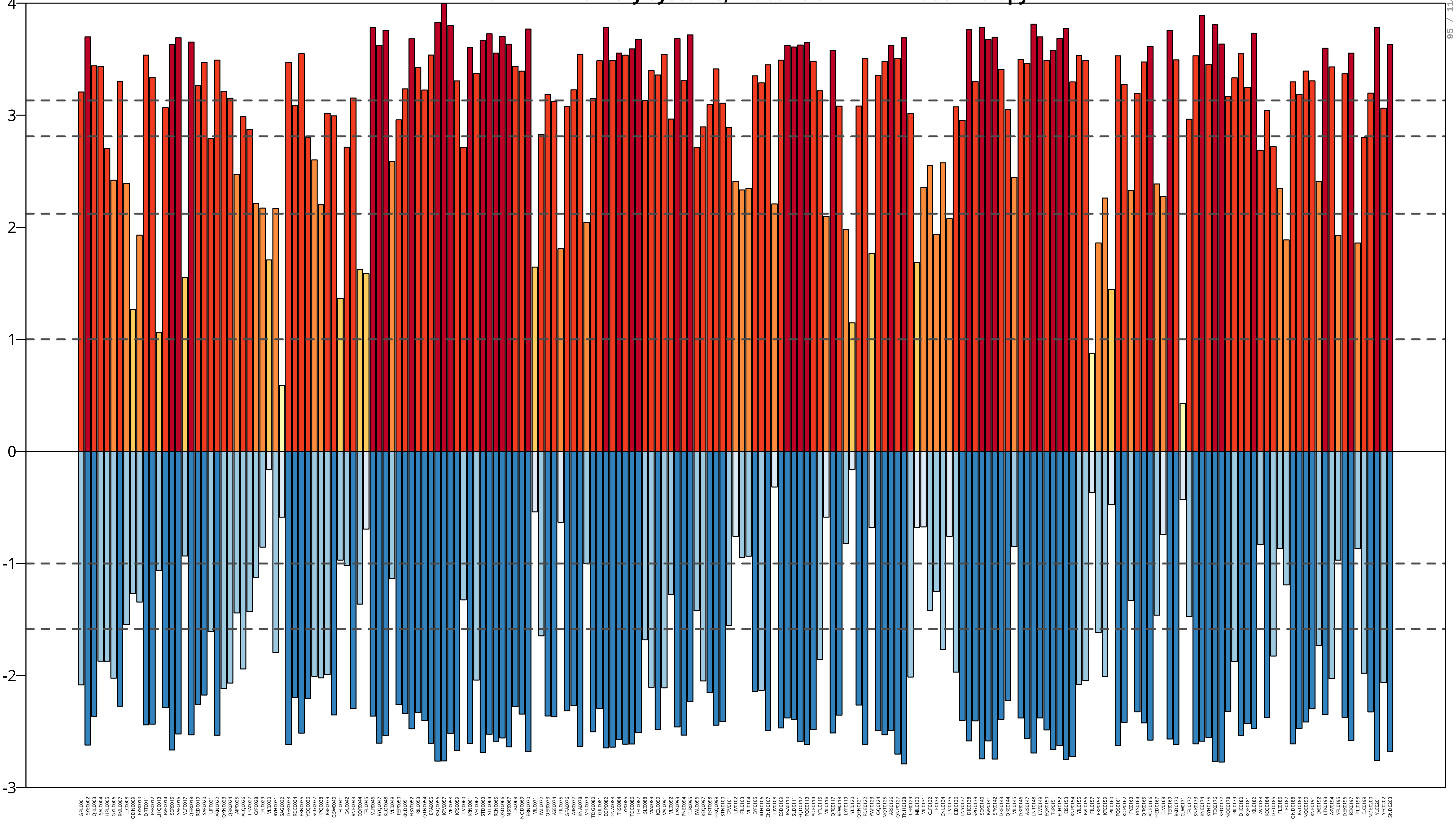

### ; Type_3 BREX Helicase CTD Inactive REase

| | |
|---|---|
| WP_162523516.1_Calorimonas_adulescens | DEKELSIDILKNIKYGEIYELLRTMFINYVLYNGGTVKEGKNGFDITIDGVRRYKGVTFDPSGNGANAITLNHEYVKKMIQELPVYPE |
| MCL4298460.1_Anaerolineae_bacterium | PQTQAEVEQYRQRLIEISQAQVKAMLGGYLAAHGESLNEYSRRPGVYYFDKTRYSNVVFDRADDGLTYLHLNHPAVQTILTHLADD-R |
| WP_207425053.1_Pedobacter_sp._SYSU_D00535 | TLAEIKADKVEAIKHSPLPKWLEGLTKSYLQIKGIGYKYLNDGINFKFP-GHQESIYTFNINNPIPEPISLQHEIIQTILKDAIHYTS |
| WP_218808307.1_Methylococcus_sp._Mc7 | EDHALEADDARKWRDHPAQYWLERAITAGLPARGGMAVREGNCWRIRWPDATESPRVCFDAQNPEMEWITLEDPRARAVISELPRCVA |
| NPV81618.1_Bacillota_bacterium | IAQELDPGETQRLLSYPIPYWVERMTVSYLRAHGGKAEQNGHLWDLTWPDGKTQKGIVFSGEFPSGRHLTLEDPKIRGLIMRLPCFVP |
| ELY62516.1_Natronococcus_jeotgali_DSM_18795 | STERIHEVMDESEDVFGSEAAIRSFVERSLDIFGGNLDRAGNLYRIEAPNSETRGPVTFSRNNEGIEHLSPDDPLIQRLRERILEDDR |
| WP_053959402.1_Sulfobacillus_thermosulfidooxidans | AL-PPDLEKVSKALGHPFPEWVEAMTEQYVMARGGMVERRLFGIDIHFPDGEVISQAVFRKRAAQSHYVGIHHPRIASLIDKVPRVTP |
| MCD8293627.1_Clostridia_bacterium | ENKDLTQLVGTK-PNFDIDSALRSMLSHYESWQGR---------STQWID-----------------KISIADNEIVQHLRSKILQDE |
| WP_162251488.1_unclassified_Nocardioides | EEAVLRQEMELARAASFQPDVVKDFTLTALARFRGDIAQAGSTWQVRHVPLPRYDRLIFGRSFVTPEMMSPGHPLLAALVAAVDT--- |
| NMB01451.1_Bacillota_bacterium | SIMKPTYDEIRKLQESLVPFWTENAVLGYLRYMGGSAVKSSKGWSLNWPDGTNQKFVGFQG---PKQLLSTASPKVLQLIQQAPDFHI |
| WP_009146898.1_Thiorhodovibrio_sp._970 | LSEEPDVQAAERLRSHPLPHWVERMTVAYINSHGGAATRKRSWWDLNWPDGEEHRKAVFSARLTDATLLNPENSRVRGLAMNLPQVAP |
| HIP19029.1_Sulfurovum_sp. | ISDEHDPSAVKKLRTHPLPHWVERMTTSYIEAYGGQVTRKQHWWELKWPDDTLHKKVVFNIHYNNTNFLNLENDKLRGLTKNLPQVTD |
| MBT5530410.1_Cytophagia_bacterium | DSDENMLELSKSYLDNQLPFWVEQMTTSFLDWSGGKVKKDLFGYSLLWPDGEHFKKIVFGWQSTGYRHLTLESPKVQELLGKDNRFIP |
| WP_200891843.1_Caldicellulosiruptor_danielii | DEKVLSVNLLAELPSDKVDEYIRKMYLYKEISLGN--------------------------LNISEETLSLENDKIKELLN-IPEYPI |
| NLE00371.1_Fibrobacter_sp. | PF----STDAKELINTGINDIVACLVTSFLRYREIRINRYKENADMCYFSDPIIRNATFYNVSEKAELINADHHIVKNIRAFFASNPS |
| EMR73855.1_Thermoplasmatales_archaeon | PFDQENISNIQISEIQNISKKIQKFTELFLQNRNLIITEYKDKTGVYYFKEKLYSKIIFDQDDEDAELLSFTHPYIKESINHSKQNGR |
| WP_073324029.1_Marisediminitalea_aggregata | GDIPLQLDEYKNALNHPMPYWVEQMVSSFLKALGKSINRNAKFLDITWPDGSTSNNVTFSGQTTNVNLLTLEEPKVSELLQKLAPWVP |
| MBN1459902.1_Armatimonadota_bacterium | SA-DVQADTAARLLEGQIPIAVRTMTLAHLAERGREVGDSEQVYRLEWPDGSIVERAVFDNQFDDVTLVGLDDARVRGLLDRVVPVVD |
| WP_068347600.1_Kosmotoga_arenicorallina | AYEITNPKEAQKLVQQPVNYWLERAVISYIKAYGGKATQIGDNWKIRYLETGREELITFDRKNPDVKKIAFNDQLVNDAIKHLPELSE |
| WP_202636300.1_Rugosibacter_aromaticivorans | DGHALEADDARKWRDHPAQFWLERAITSGLPARGGSAVKVDQAWRVKWVDGSESAQVCFDAQNPELEWITLEDPRARALISELPRCVA |
| WP_020614470.1_Sediminispirochaeta_bajacaliforniensis | STGSMDPGIAKEIEQHPIGFWIQQMVINYLNQNNARIEEGKTGYNIYWPDGDYSEEVTFDRDTYSLDLLSLESSIIQELLEKSTPLPE |
| MBT6646602.1_Nitrososphaerota_archaeon | QDKNLDVEIAKTVNEHPLPLWIESMVLNYLKSNNGSIIEKNRGFDIVWPDGETSLSISFEKENKNLKFLSLEDEHVRKISKYISTFVK |
| MBV6489839.1_Fimbriimonadaceae_bacterium | GQ----LADGDKIPTVPLRAWLDTLRGAHQ-------------------------RTLMDEE--------LSELVVERINSLRPYFCD |
| WP_073205019.1_Prevotella_ruminicola | SIKEITKTAASEVKYSPLPIWLEDLMDLHCINVKGFYDKMLNLTLYHLPDGNL--GAIFDSDNPAAIHITLQHPIIQHIIRDIDGGI- |
| NSW90672.1_Bacillota_bacterium | DDKTFDVALISNIKNNNVIDLLRRMYVNYKEW----------------------------KENIDQLMIDLNHEEIKTLVNQVLHWDK |
| WP_148854793.1_Moorella_thermoacetica | PILAPEYVLASQRDLEKRAKRVQRLVEQYLQVYGASLHPYKLREGVYYFQGRRLHNVIFQQANEGAELLSFQHPYMVELLAHLEDALR |
| MBU0460857.1_Nanoarchaeota_archaeon | PFDKKDIKEIEKSTVENESNIIKNLVETYAKFNNFDFKEYSKKKNIFYTDETKIKNAVFNKEDESLDLINITHPLVKEITNELIK--R |
| KAF0196956.1_Bacillota_bacterium | ASEPLSPALAQQVAESPIPYWVEQMTKSFLTAEGAHIRYVGESYEVIWPGETTVEKVSFRETTEASGVASLEHPRIRALLDRLSPVVP |
| consensus/100% | ........................h...............................................p..h............ |
| consensus/95% | .....................lp.h...........................................h...p..h............ |
| consensus/90% | .....................lp.h...........................................hs..p..h..h......... |
| consensus/85% | .....p....pp........hlcph....h......h........h.h........h.F.........lshpp.bl..l......... |
| consensus/80% | .....p....pp.....h..hlcph...ah...s..h........h.h........h.F.......p.lshpp.blp.l..ph..... |
| consensus/75% | s....p...hpph..p.hs.hlcph..sal...s..h.b..p..pl.a.s.p....h.Fp.p..s.p.lshpcsblp.llpphs..s. |
| consensus/70% | s....p.p.hpph..pshs.hlcph..sal...s..h.b..p.bpl.a.s.p....hsFp.ps.s.p.lslpcsblppllpplsphs. |

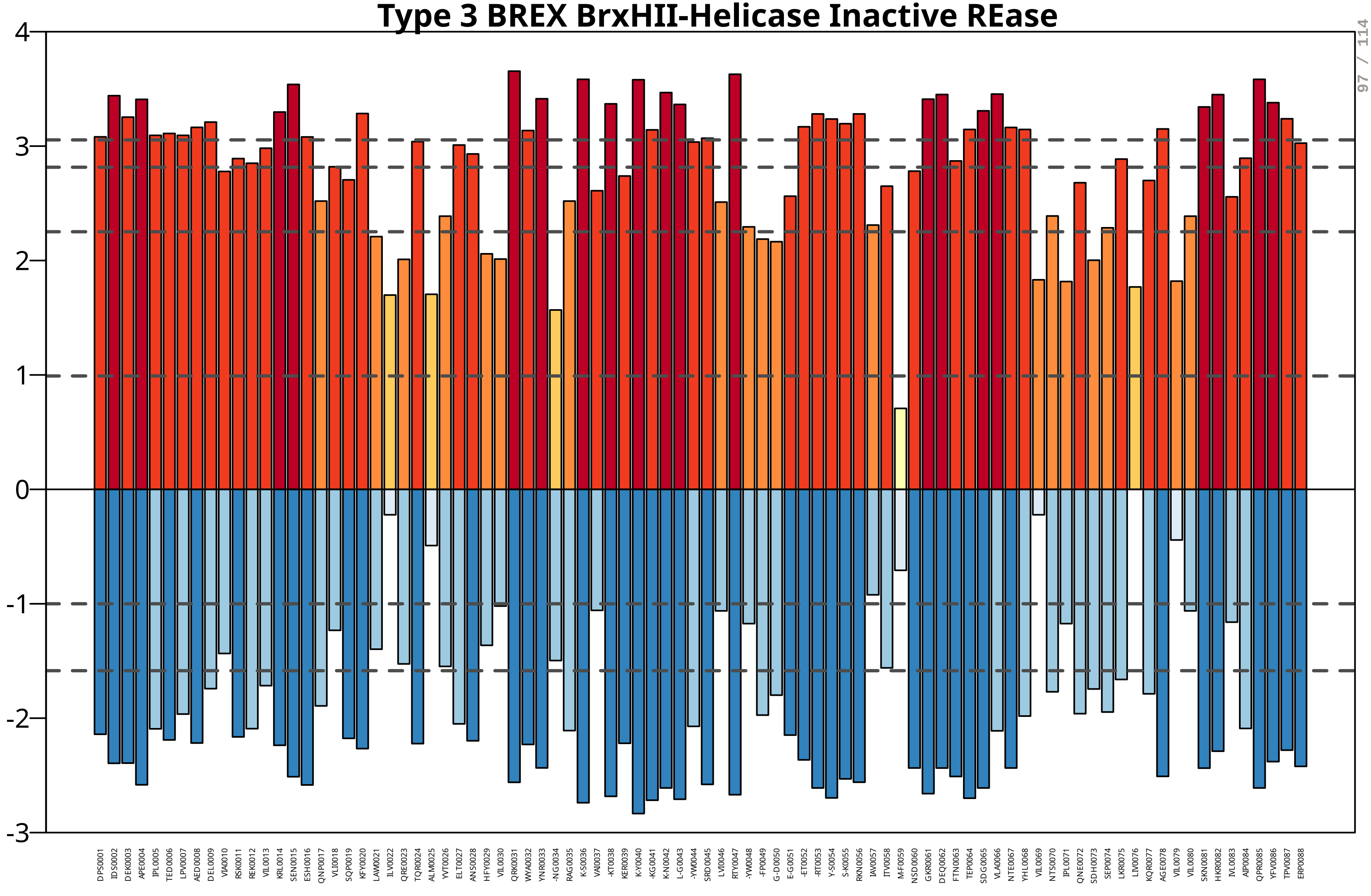
Type 3 BREX BrxHII-Helicase Inactive REase
4
3
2
1
0
-1
-2
-3
DPS0001
IDS0002
DEK0003
APE0004
IPL0005
TED0006
LPV0007
AED0008
DEL0009
VIA0010
RSK0011
REK0012
VIL0013
KRL0014
SEN0015
ESH0016
QNP0017
VLI0018
SQP0019
KFY0020
LAW0021
ILV0022
QRE0023
TQR0024
ALM0025
YVT0026
ELT0027
ANS0028
HFY0029
VIL0030
QRK0031
WYA0032
YNR0033
-NG0034
RAG0035
K-S0036
VAI0037
-KT0038
KER0039
K-Y0040
-KG0041
K-N0042
L-G0043
-YW0044
SRD0045
LVI0046
RTY0047
-YW0048
-FP0049
G-D0050
E-G0051
-ET0052
-RT0053
Y-S0054
S-K0055
RKN0056
IAV0057
ITV0058
M-F0059
NSD0060
GKR0061
DEQ0062
FTN0063
TEP0064
SDG0065
VLA0066
NTE0067
YHL0068
VIL0069
NTS0070
IPL0071
QNE0072
SDH0073
SEP0074
LKR0075
LIV0076
KQR0077
AGE0078
VIL0079
VIL0080
SKN0081
HKR0082
IVL0083
AIP0084
QPR0085
YFV0086
TPV0087
ERP0088

```
BR-system  Helicase CTD Inactive REase Rep_1

WP_169116183.1_Tepidiphilus_baoligensis          QPFFIRSFFTQAFRRFQGAMHPREPGRYEITHPVLRRYERICFEKRHVRGAPMASLIHPGHPLMQALTDLVLEEHRPK
MCY3585835.1_Acidimicrobiaceae_bacterium         NPVIVEHFLERVASASLIGRAPLGDGFYYLSALVVTRHERSSAIAQGVDRAADAHMLGPSDTAFRALVEALRTRLETD
WP_094451591.1_Parenemella_sanctibonifatiensis   QPHHVEAFFHKAFKDLGGNLRMRESHRFEIKSPIVNTYHRVAFDRANVKGNHVATLLHPGHPLMAATIGCITDRHHDA
WP_042699659.1_Methanocorpusculum_bavaricum      QPYFVRSYFLNAFSKVGGVLNQRETGRYEISHPVVHKYDRVCFDKEDIEKNKQAVMLHPGHPLMRAVTSIILEKYGGL
SCM57150.1_Methanothermobacter_wolfeii           APEYVEEFFRTVFRRMGWKYTER-KGLYTLRTALRREYRNLTFNP----GTDEHEFISFGHPLLEATIRYTGKRFGDA
AXV39096.1_Methanobacterium_sp._BAmetb5          VPEYLEEFFKKAFKKVGGKFTTRKNGTLNIASRLNKSYPRLTFDKNFASENSDYDFLSFGHPLLEAVLKWVELNYSNT
WP_106140929.1_Spirosoma_oryzae                  QPVYIRQFFEKAFTYMGGKLQELQPQLFRITTLINPAELYLCFDKRLFLNLTKTIYLNPGNALFDALLSLIQQQCRPD
GIV15258.1_Armatimonadota_bacterium              VPWDVERFTRLAVQTVGGQFSERRPNVYRLSVDAFARGLRVAFQRAVAR--ENAEFFAPGHPLLEALIDHFLQPNLPV
NNN07583.1_Acidimicrobiaceae_bacterium           QPGFVEAFFTAALTDLGGRITPRERGRFEITRPLQSAYERVTFDKDRITDRVRAELLAPGNPLVKAVIETVLERYGAT
MDE0136609.1_Acidimicrobiaceae_bacterium         QPHYVQAFFEAALSDLGGTMREVEPGRFQIARAVLERYTRITFDKDLTEGAPDADLLAPGHRLLDVVVDLTLERHRDQ
OZG52264.1_Pseudoscardovia_radai                 QPGYIEAFFVEAILDLGGSISRRETGRWEILRPLANRYERVTFDPAHVDNQTKALLVTPGTPILNAVIAVVLRKYAGE
WP_259102394.1_Candidatus_Fervidibacter_sacchari IPEYIERFFLRACESLKVTVKPREKGVFQVESEVKMAYRSLTFRKELAK--DTVEFVAPGHPLLEAIVDAVMAEGKEE
PWV37726.1_Desulfurococcaceae_archaeon           NPSHVEEFFRRAFEKAGGRIKILGDGFISIGSPVKDKYPKITFYKDLALKDPEAELVTLGHPLFEATLEWVRRKFFLS
RLC12746.1_Deltaproteobacteria_bacterium         MPRYVEAQFVAAARQVGLRVEPRADGLWRIEHKADSSYRKITFHKHHLEAHLDAVLMGPGHPLYAAVDEELNDRLALL
PZR94482.1_Chloroflexota_bacterium               VPYFIASFFLEAFRNLGGEAVEREAGRYEIKFHIVTKYARICFDKADIAGHLKAEFIAPGHSLLDTVVELLLEREGSL
NJL06253.1_Chloroflexaceae_bacterium             VPEYIQDFVVQAFNTLAADLQRRNDAVWRIESTLDSSYPRLVFKRDLLEQYPNAEFVAPGHPLFVALLNLVRDRYGSL
MCY4618413.1_Chloroflexi_bacterium               TPEFVERFFVDGLRYLGGRIARGDDRDWRLDHEFGEDGRIITFNKDRLRREPPAEFAAPGHPLFDTVLNRILDQGRPA
MCS6856322.1_Sandaracinaceae_bacterium           RQEEIHRFLEEAFAHLGVATEVIKKG--SILRTGKKEWVEVALDRASAI--EGVELVGPGHPLLVEVTKRVLREAKPH
GAK56805.1_Candidatus_Vecturithrix_granuli       MPAFIRYYIEELAPYLGLTLAGDIRGLFRFEDPVLLTYPRLTFEKTLALSILQAIYLHPGEIVFDRISSRFLAKYSRD
MCP9446721.1_Nitrospira_sp.                      VPETIARFMQEAAEFVPLTLKPIPPHTFEPGRALATKYPRCSTDRETAE--HNLEWVTPGHPLFEAIRRHTYDKARDA
MBN1909331.1_Pirellulales_bacterium              MPEYIERFFVEAYRSFGGTITPTKKGVWSIGRKIGNSYPQITFDKELLVGYSDMEFIGPGHPLFEGIVERVLTQYGPS
WP_059053655.1_Tractidigestivibacter_scatoligenes QPHYIQSFFAWAVEDLGGHLASNRDGTWRIRKPLADSYSSLAFDKEDVADPGRTDLVAVGHPLLDAAVEETLAKHGGA
MCL5028417.1_Bacteroidota_bacterium              QPLYIERFFTKAFSYAGGFIKQHEQPVIYLGQPIRDYYNPFVFNKSLVSSPDGTKLLGPGHPLFDCLLNYLKKQAAEV
WP_090584037.1_Arthrobacter_sp._ov407            NPAIIEQYLHRLNLAGVLSVTAAGDGILRIQGLIATSGKALSDAQRGGALLSNVFTLGPGEPSFNELVDYAGAELEPD
WP_159000922.1_Streptomyces_sp._SBT349           NPVIVDAFLDQLARAQGWALAP---GLRLLTSLVAADGARLAINDGATD--DDVIVLGPTEEPFGELVELALRSGEMD
MBC7429627.1_Bacteriovorax_sp.                   DLKRLFSAFQAGLSLFDGSYSNLDNKTFEVSYPKGSNKTLVTFDRDLANGDSKLTLLGFDHPIVANMINTIESRKNL-
MBE7508219.1_Planctomycetia_bacterium            GMRRLVAFVQRAAALFGGQFVSKSGTRYVLIDP-DKAETPFTTDRDEALREENVELLGLEHPAVSRLIERCRQLEAE-
MBI2192005.1_Planctomycetota_bacterium           GLQRLAAFIKRAARQQDATLREREDGLWELLLR-GREALLCTLDRERALQSEGVDLLGLDHPIVKQWLEQCRALPAR-
MBI5511449.1_Deltaproteobacteria_bacterium       SVERMLRYFGAATADTGKTLATTGPGRFEVRDGSGAALLRFTTDRDLAREDDALDLFGLDHPLMQEALRRWREAAPE-
MBI5901621.1_Rhodocyclales_bacterium             TLAELGAWVRGMILQLGGAALP-DGQFWSFLLPEAPRYEQVCFDRDLALRARQSELGGIGHPLVDALLHEAKSPGFP-
MBA3936144.1_Planctomycetota_bacterium           SMMALLDFARRAAVAEGGRFDSAADGTHDFT-PVGQPPLRFSTDRAVALANGGIQLLGIDHPEVQKWLEKFASLPPE-
MBN2145092.1_Candidatus_Auribacterota_bacterium  TLSDLRLFVEKAVLRLGGGFIP-SGEIVEILVPESAVYKNSTFNRKMAVRKKGVELLGIGHPLINALLDYFKSDLGA-
MBP6702578.1_Vicinamibacteria_bacterium          GLGALDRFVAEAAREDGGRLEPMGCGLSRLVDPDGSTAAVFCHDRHRAMADEGLALMGLDHPAVDAWLRSSRDTPPE-
MBQ8111590.1_Kiritimatiellia_bacterium           NMARLVKFLREASSFAGYAYREEEGGDFTLTS--ANDTWSFTCEREKALKDDSLQLAGLEHPVVTAIIDACRRALAE-
MBW4553276.1_Aphanocapsa_sp._GSE-SYN-MK-11-07L   SMDRLIRFLSAALEEKKQTIVRVSNGIYELHNELGEKVLRFTTNRELAREQDGLDLMGLDHPIMQKAIEKWQNVVPE-
MBX3410397.1_Phycisphaeraceae_bacterium          GMQRLLAFTRECARAAGGAMRDAGGSIHEVSLP-GGRIERFTTDRAKAKDDDALALLGLEHPLVREFMDEYRALPAE-
MCC7337860.1_Pirellulaceae_bacterium             GMSRIVRFAQLATEETGGTWNTVGDGTFEFRGKDGRWPAVFTDSRERSLEDEHLQLLGLDHALIQGYLEDYRNVPAN-
MCX6876429.1_Verrucomicrobia_bacterium           DRKDIIGFAAQATTASGGNWSTISPDRFSCAVP-GETELVFTTDREQANASVSLELLGLDHPFVANQLRKWQSLPPE-
MDR1607843.1_Deltaproteobacteria_bacterium       GMRRLLFFAQKAAHLNGEEFLPIGGDIFELSRP-NSETVRFTTDRDQALEAEYLSLLGLEHPIVKQWLDSSASLQPR-
MDR2179037.1_Synergistaceae_bacterium            AMDRLVAFFSSALAAHGQKLLKTADGLYQLSSPDGFEKIIFTLDRDEATSREDIELLGLDHPLVKKELEAWRDLPPE-
NPV69259.1_Bacillota_bacterium                   GMERLVSFMSRVERLTERQMKQRDATLWELWQ--SDTPLLFTSDRERALQNEDLHLMGLEHPVVRERFKTCLGLPEA-
WP_026299088.1_Deinococcus_aquatilis             SFERMVQFLCLAIAEDDRHIVKIDQHIYAITASDGQQELMFTTDRDLAIRETQLELIGIDHPIMQQEIKRWQSTPPE-
WP_058554674.1_Thiohalocapsa_sp._ML1             SKHRLLAFLATAAEAAGGGLRRLDGQRFELHAGAGAEPMTCTLDRELAQAEEGLTLIGIDHPLVARFCAALQGMDPA-
WP_277443845.1_Pelotomaculum_isophthalicicum     GMLRLIHFATIALQYKNGKLIEVEKGKTYIADIGNGQSVRFTTQRDIAIEDENLSLLGLEHPLIQQLLEAYTNLPPE-
WP_282552852.1_Providencia_stuartii              SIDDLIRFMQKALPYEGEAIEIRGQKQWALVKD-GLEQMLFTTVREDATNSEKLHLIGLDHPVIAKQLDKYRNLDPE-
WP_283022682.1_Bradyrhizobium_sp._CB1717         GFRRLIDFLAASLPDQGRRLVRKEEDLYEVVDNAGKRTSFLTTDRAKAADREDLELVGLDHPLVEAELAKWRDLNPA-
WP_308999799.1_Paenibacillus_sp._LHD-38          GLEQIEMFIREAVTIEGGEYRKKSDDLIEISTNSGLSSIIFTTDRESYIEDDKVELLGIDHPLIANYIAQYQGLDPD-
WP_309896344.1_Archangium_sp.                    GLERVLEFLRTSLAADGRRVEAQDDGTYVVKDAEQQTVCRFTLDREAARARADLDLLGLDHPLVQEAIKRWQTTSPE-
consensus/100%                                   ....h....................................s..................p..h..............
consensus/95%                                    ....l..ah..........h.........h..........hs.............hh...p..h..............
consensus/90%                                    ....l..ah..s.......h.........h..........hs.pc..........hhs..cshh..............
consensus/85%                                    ....l..Fh..sh...s..h.......h.l..........hs.p+..h.....h.hhs.sHPlh...hp.hb......
consensus/80%                                    ...bl..Fh..uh...s..h.......hpl......p...hs.p+..h.....hphlu.sHPlh..hlp.hb......
consensus/75%                                    ...bl..Fh..Ah...G.ph....ss.apl....s.p..bhs.s+p.h.....hphlu.sHPlhp.hlc.hbpb....
consensus/70%                                    s..blb.FhppAhp..G.ph....ss.apl.p..spp..bho.c+c.h...pphpllu.sHPlhp.hlc.hbpb....
```

### BR-systems Helicase inactive REase Representative_1



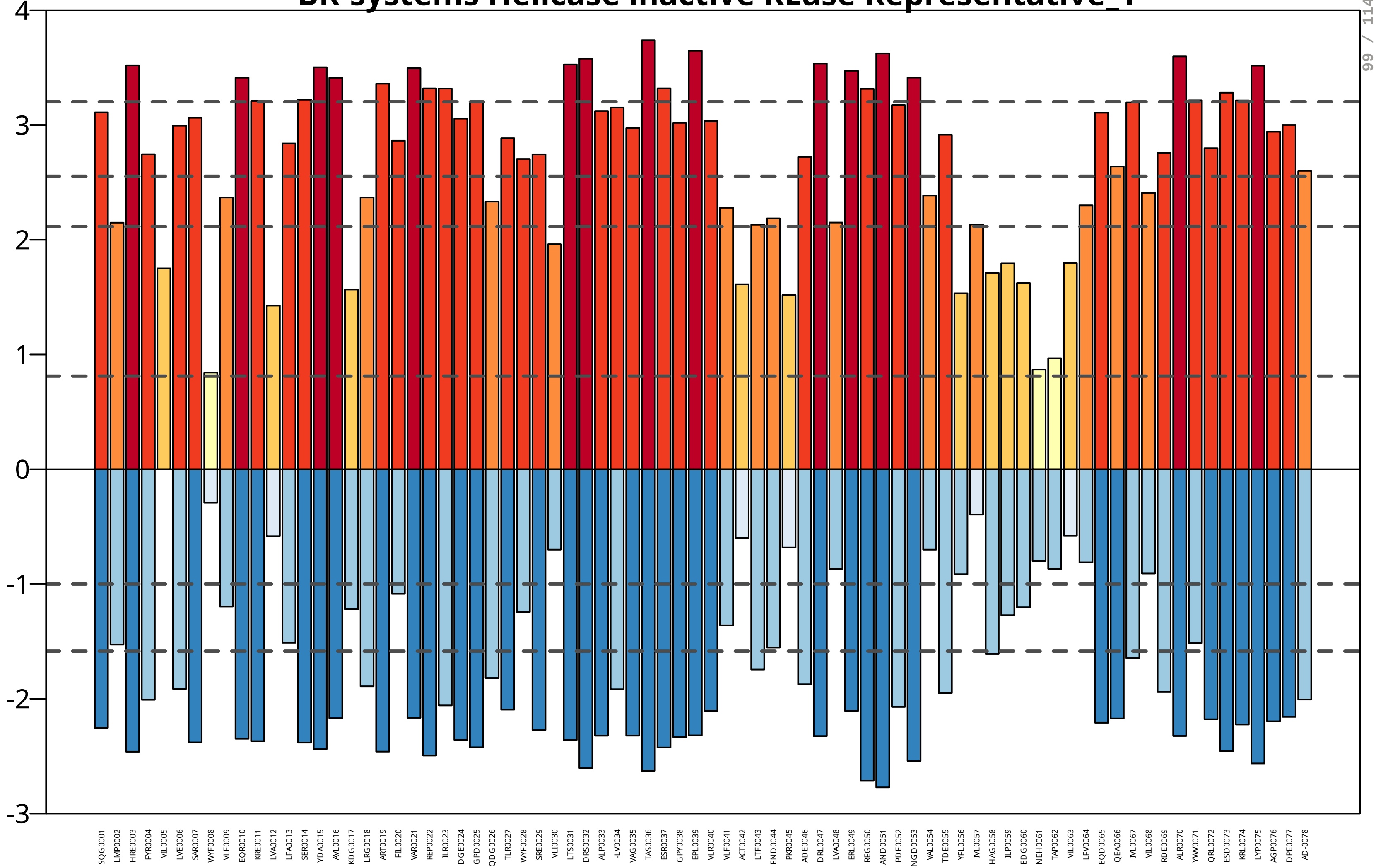

```
BR-systems Helicase CTD Inactive REase Rep_2

MCK6447153.1_Planctomycetota_bacterium                  PPVTLQQLERAVVGSKTLGSKFTPHNQIAGAHALAWG--ASETVTFSPEVFDVHPNVKLLS
AKC83094.1_Verrucomicrobia_bacterium                    PLATRDSVRCALYRLA----GTHATVSDSNRVGVS-GLGDAFSVGLTQEALEGDSDALPPT
MCB9841664.1_Phycisphaeraceae_bacterium                 RVITLDDIVHALSESEYLRAGSKVRETGQPWVEIRGGLYTRFALTAKQDLYERGLALRFAS
WP_090215779.1_Pseudomonas_guangdongensis               LPVTLAQIWRSLIESEYLKSGCQFSPC-ARYITLNGPLIDGSNLTVDRELMDVGLPLHFAS
WP_196479964.1_Burkholderia_pseudomultivorans           LPITLEQIYSTLAASMTLKSGSEVIPSYGEALLVRGGS-ESFMLTVSRALYERGIALHFAT
HCG7747318.1_Vibrio_parahaemolyticus                    IPVRLNDIWDAIVNSAFLKSGAKLESFNGEAVEINGGIPGRTLFTVSRELFEEGIDLVFAT
MCE9614803.1_Lentisphaerota_bacterium                   PAITRADVIAALVTLQP---GLQTVGHAKGTVTIK-GLTKPIRLAFTDPALDADTAARPLT
MTE20051.1_Streptomyces_taklimakanensis                 SPVTLEDLREVLVERFAAVL-------------------EGRPVTFDPSRASRDGETALAT
WP_206882508.1_Alicyclobacillus_fructus                 GPVQLDDIWRALVESPFLRHGCHTEETHGPVFVVRGGVEDGTRLTVSPELYERGLPVHFAS
WP_084460301.1_Aminiphilus_circumscriptus               MPVTLESIWGLLANCRYLCDGCELVPGNEKVLRLAGGIPRGCCLTGDRAVLETGIPLHLAT
MBI3972513.1_Chloroflexota_bacterium                    PPVEKEQLKAALARLVEADVGVSAEALDDGAWQIRT---NATTVGLDSAVLEAREDLRELP
KPC75468.1_Thermoactinomyces_vulgaris                   APIDLQSIWSTLTESRYLQEGCKIFDENNLILEINHGVPDGSVLTVSRELYESGLPVHFAS
MBX6754325.1_Thermorudis_peleae                         SPVSLEDLARVVPSLPGLAERFSPHPTIPNAWLVSW---QLHTVTFDREVFDMHSQVRLLT
MCF8137375.1_Desulfotignum_sp.                          SPVGLDAFWQAVAGSRYLKDGCQTLDAADRMVCLSGKIIDGTFMTFSRKVYEEHPRLAFGT
GIW50774.1_Gemmatimonadales_bacterium                   PPVSLSDLERVLLTSS-LRERFVAHPELAGAYRLDLG--RLWEVTFDPAVADAHPGVRLLT
MCI7613155.1_Selenomonadaceae_bacterium                 LPATVDDIWQAFADSVALQAGLQVDGD---------GKMLEVAFTKDRGEVD----VHYLT
MYD03943.1_Acidimicrobiia_bacterium                     PPATPSELEAFYTRSHSFGRLFRPDADLDGAYLL-----EGRSVTFRPELADQHPDLHFLT
WP_073234049.1_Desulforamulus_putei                     PPVTPLELERIFTAT-LLPEGVSLQPGEKGVYLLSF---QSYRVTFNPRVYDRRPDVKLLA
WP_036720040.1_Paracoccus_sphaerophysae                 APVSLADIDVALTSLAKVAPLLSVT--HPGIFALRYG--RVSHITASRELLDRPPPLEFAS
WP_161844016.1_Pseudoflavonifractor_sp.                 PSACLEDLWAALTQSPYLQSGAFCSGP---VWRLPE---PALEGTIQRDAIT----VSFLT
WP_100917763.1_Candidatus_Thiodictyon_syntrophicum      LPATLDGLWGVLTGSAYLKAGCACAVNGREPLLLRGGVLDGTALTTDRRLYEEGIELRFAT
AXH34461.1_Humibacter_sp._BT305                         GATTPEVLERLLTTNGLSRAAFEVDNQRPKVYRLKPGVRETLPVTFDRATWDESDDLVFLT
ELZ44607.1_Halorubrum_distributum_JCM_9100              APYQKSSLEAVLVENETLADGVTFTPVEETTYRLNWGMEDSVAVTFSSDCADEFPSIHHFA
MBD5553416.1_Desulfovibrio_sp.                          PAWDLSLLRRVLAEPRLMPPGYAASQLGSHDWQLIAGAAQPWRVSVDAEFYANHVDVAFWT
MBF6555076.1_Acidimicrobiales_bacterium                 PVVTLEDVQRILLDSPATREQLSEHPMIDGAWLVEDG--NKIAVTFDRIVLDENSPVRLLS
MBP7066719.1_Methanothrix_sp.                           DDVLVERLRGFVVSAA----GLSTHELNDKLFV---GVPNGAALTSEK--------ANALT
MBV9397056.1_Bryobacterales_bacterium                   PAYDLPVLDQAMILDASRPMDLEWRPLDARTYAARLGLPEPIRVTTSADVFEESSDHEFFS
MCD8487848.1_Desertifilum_sp.                           APMTPETIEQLFTTSTILRSGALFECQAHRTWQLTYG--QNYNVTFYPEVFDERPSLRLMT
MCK6511424.1_Myxococcota_bacterium                      PPYDLRDLQRILAQPDLMPNGVTIRTLGPEEVGYRSQHVEEVRVTTSPTYFDQHTEVELWS
MCY4068113.1_Acidimicrobiaceae_bacterium                PAMTLEGLRERLLAVPAAARLLKPDPQRCGVWLLDLGADAPVPVAFDRSVCEAHDDVTLLT
MCY4434136.1_bacterium                                  PAMTLNDLQDALLSIPWCRNRFTPHPDFSDAWLLSLGT-STQPVTFNPTRYQDTPHITLLT
MDE0094453.1_Gammaproteobacteria_bacterium              PAIDFNYLERALEDETLLPSGVNAIRKGVKEFHIKVGTHDTVRVTTNPSYYEAHSEVELWS
MXW96682.1_Acidimicrobiaceae_bacterium                  PAMTLDGLCDRLLSLPAVATRLVADPQRVGVWLLSLGAEAGVPVTFDRSICAGYDDVALLT
NLA99646.1_Methanomicrobiales_archaeon                  DDAVILRTAAFIWTAA----GFTVEERDARLYIVGSGIPNGASLAAGR--------ANALT
NLJ36707.1_candidate_division_WS1_bacterium             PLCQPEDIRDLILTSQAITANWTLEEHVAGVYTLTEKTDKTTTVTFRQDILEARPEHHCLA
OYY89820.1_Sphingomonas_sp.                             PALSLTDLARIVADHRLLPPGYAMNSLGEQDFAVEQESRRRVRATISREFYASHFDTDFWT
PZV04180.1_Cyanobium_sp.                                PPLTLADLEAILKQPALLPPGCSARSIGAYDFAWSQGLDKEMRVTCNASYYEDNSDCELWV
SNR42896.1_Halorubrum_vacuolatum                        PLLNPEVVKHLFTQSQILRNGWTFEMLYQKLYRLEIASPDEVLVSFDPEVLEWYPSVIIPL
WP_054466705.1_Planktothricoides_sp.                    SPITPAEIEQLFTQSEIMAEGIVFTPIGDKIWQLTYN--QEISVTFYPEVFDEHPSLRLMT
WP_103287095.1_Candidatus_Nitrosocaldus                 PTFTEVEVSRVLERFVRLCCGMRLVKRGDLLFVEPL---ERLPVTV-RSFVENGLSPDALV
WP_128478225.1_Halorussus_pelagicus                     PVFTVKAIKGVFIESETLNEGVHFDSLAESIYCLQHGDAEGIGVTFDSETADEYPSLRFLA
WP_164002139.1_Pyxidicoccus_caerfyrddinensis            PPATQAELAYWCLQHPA---GMQIRTGEPGTDRLATGTNEEVLATFSGEVADRHPPVRLLT
WP_176737426.1_Micromonospora_citrea                    PSDTLADLAAAIVGNKAVAPWLTADPTEEGAWELALGA-PPQLVSFDRAVVDASKRVALFT
WP_179052052.1_Nostoc_sp._TCL26-01                      PLYDLKTLDQILQSPSLLPPGIDVQLMQDGEYKFSMGMKETLRVTTKPDYFDEHPDTELWS
WP_285255575.1_Halobacterium_salinarum                  PPYDPPIAASLLLDSQFLAIDVECEPLGDSEYRLQW---QVAVVTTNPE--QTRMETRLLT
consensus/100%                                          ...........h................................s................
consensus/95%                                           ..h....h...h...............................hs................
consensus/90%                                           ..hp...l...h...................h.h.........hs.p...h.......hhs
consensus/85%                                           .shs...l..hh.........h.........h.h.....p...lo.p...hp....h.hhs
consensus/80%                                           sshs...l..hh.p...h...hp.p......h.l.....p...lT.s..hhp....h.hhs
consensus/75%                                           sshsbp.l..hl.ps..h..shphps.....h.l..s..p...lThs.phh-....lphho
consensus/70%                                           sshsbpplb.hl.ps..h..shphps.....a.lp.u..ps..lThs.phh-...slphho
```

### BR-systems Helicase inactive REase Representative_2

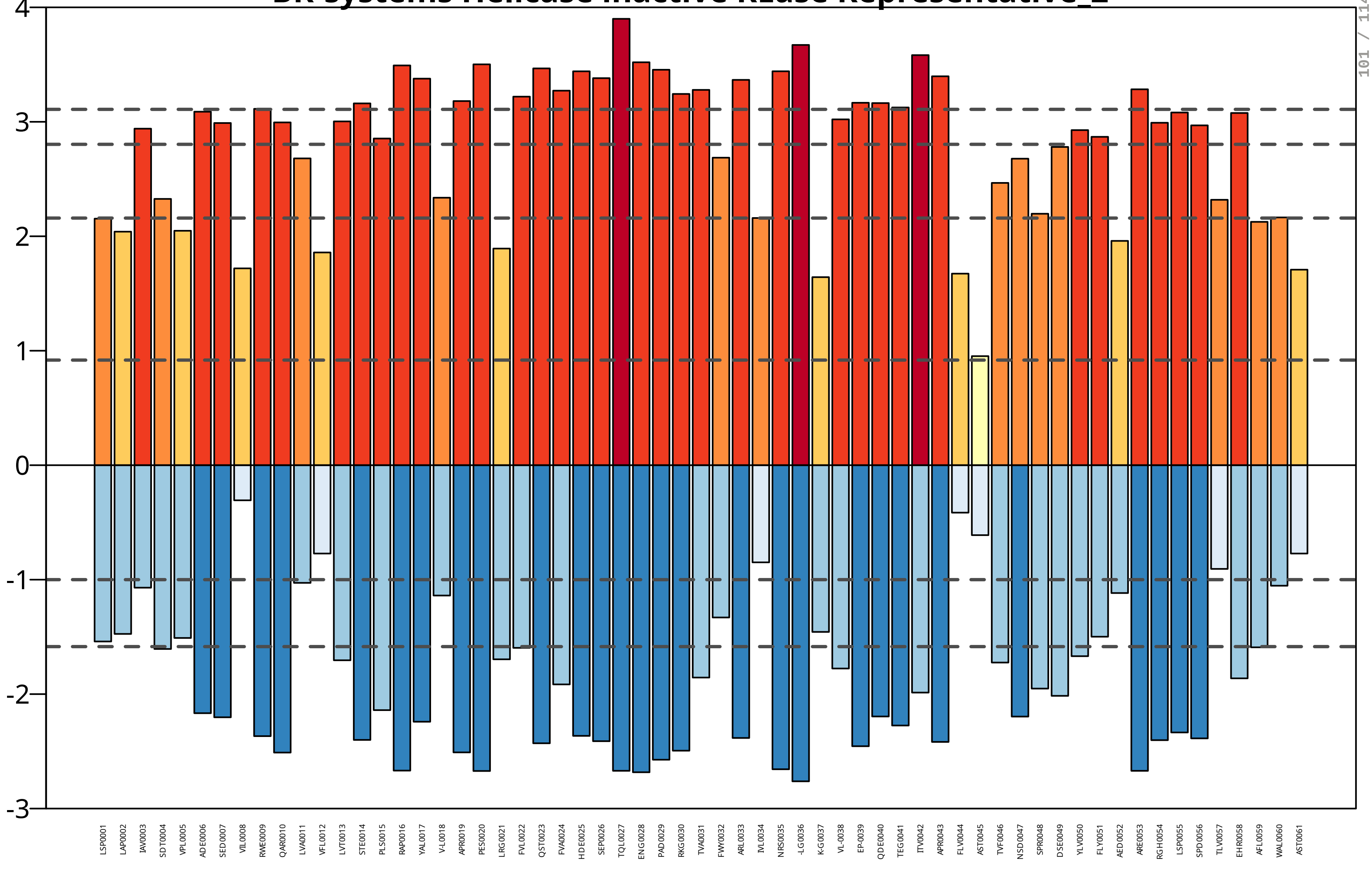

Type-2 BR-systems

### ; DUF499_2 C-Terminal RRM Rep_1 (MAIN_FIG)

```
WP_015408759.1_Natronomonas_moolapensis            ASTSSPYHVPRALKEMRADHPDEIITAVEGVWIRIEGADGWKGAWFTANKLGDSHTLVNFD--YTASESMVDITYRGDPSAFADFNMEPEDFSTPDGERTAEADFSIEFEGDPMLYGETFEALDDLLAVDNAFTVTMETEVEIRARD
NHV98109.1_Thaumarchaeota_archaeon                 LIVLAESSV-ETLNDIEKRAIRDRFTKVNSVILKISSNLEIKNLLTRLG--PERNCNIRLSGSITRRSPSYTISFEINKEDLQKGKSMLDLLWRLKGIDSLQVRLELKWGEPVKPEDAA-----GVLRSLGDIVAGLEANVGA----
MBD5553419.1_Desulfovibrio_sp.                     LEAGANGAVAEAFAELEEKMKAAGMREIASLTAEAFLPRDGFGLTRLAAGVP-GRKQVALSASWATAGANFSLELEGSCQEAAEAGQFVQGQMRGAAEQSFSLALTVSFEPPLS----DLAPVRTQFCKLADAALFLVCRGRQ----
MCK6511429.1_Myxococcota_bacterium                 FTLDHENNLPEILRFLEQSLQQRKARSIRSFSCQLFHLPDFTQLTQSSNVFKDASPHLEFRATFAKNNSEGILSFEGHPDDIRSVLPFLQSQSQRRAEVEINATFSFDFSTPLPLPDPQ--AFSARIRPLEPNKCKISLTVSTKDT-
MBY0489937.1_Gemmatimonadaceae_bacterium           -------------SDLAAAIAAAAVRPLERLVLTARTTADAKLLTQFAQPFGATSLGLTVSVSGDCGGGTMALMVDGVKPTHPTPLDMASAIANSTSATSFEASLTMDFEPGLLLAGDQLQQL-------AASSVTVRCRFGPANGS
OYR54791.1_Halorubrum_halodurans                   DTATSQMAASRAFNEVRTHAPSDGTPGVSQVTVEVMGEDRLSKGSFIAQ---RDAFAVSVRMSYDVTGTSYSASYQGGLDDFSRVTNQPDPF--------GSINVTLKFRVDLDETDDEDDVLAELQAGLGQTNIDVKVQARGPVEV
GBD14658.1_bacterium_HR25                          ALLEATGDPGLALADLAARAKDARWTAIGQARVFWQGDPRLSALRTVLGHLAHARARVEVDLAVRYDGDEWESHFRGAAARYQSLAPTLEAQAGQAQDAHVNIALALEFPSGLPPEGPDYGDIRDVLDLAGLGHIRIVAWPLQGGDS
MBQ7457436.1_Desulfovibrio_sp.                     TEFSKEGILKEAFVRLWEEIRHAKVAVISEFHIEAVEMADALRLMPLVQAIRNADKTLTITGILETHGAKLTLEYSGSIEEAQALRDYLSIQLRQSTSHSLEVAFGCLFPQGLDMNSNMPEAILDRLSKAAASTAHIRAKAKV----
MBA3046419.1_Euryarchaeota_archaeon                IPPITEKTLQGIIGELLPILEDKNIEVIQSIGLKAKGRDDLARLLSAIPQFLGAKAYYIIEATLNEKGGFMQLDYRGDDAGFKVVKPVLLNYQPKEQFSSCDLVLTFSWDGGIS-RDEFVGILRDKVANFTGDSIYTAKVTPKEAEK
MYC64339.1_Caldilineaceae_bacterium_SB0661_bin_34  SEFQSTRAPAEAIEDVVAQAQDASATRMEELILEWDAEGCVAKALHTLAQHPNVDPSLSVD--LDTEGKPFTLKMTCSAPQAQMMGDSLKAILRQGEGTACTTRIVLAFPASTSVEGNEKRTIQDAVRGTMSRSLTILKTA------
MCA3396953.1_Roseomonas_sp.                        ELLHAEGVLKEALNRIFEQAAARKVSAISRLTVRVFEPADGFRLLGVVSAVRTAEVRASIKGEFETAGSTVMIGFEGVPQDALPLRDFLDPQLKLAAEKDVSFVLSLGYKDGLPTSGEAPTKLIEQLTRFVTGAAYVEATAEPKP--
MBR2123973.1_Acetobacter_sp.                       NPISGEGVFKQILRSLLEETKRRSIEEITQIDIKITEPTDGFRFLGTIGNISLVQKELQMAVDYETTKGCFKVGYKGQAAEAIIFKEFLEPQLRTAEETNFTMVFSLKFEPSFSLRDEENRKLSDQLTRFIAAQVFVTIFPSS----
QES21410.1_Streptomyces_venezuelae                 TVFTAAGAAGSAFAQARSAAADTQRESLVELKIEVDHLPELLRLHSIIPPTLGADLTYDVDVVVTLDGQTAKISYVGTPRDYAPLREALKQLLPR--ESVLKASVTAAFADPLPLSGDVVERFAQAARDTGPTKCRISMRTESDG--
NLV45647.1_Candidatus_Hydrogenedentes_bacterium    TLLSAEGPLRQALVELFDKARSQKVEYLKSISMRFYEAKGAWNAHQAVATYRDSETICQLSVGIEGGIESFEVQFSGTLNKANSVKSFLEPQLRSATNATFEANYTLTFATPFATVKEKTNDFIAAMTKYGASEAYVEAEADRAEGR
MCY4162822.1_bacterium                             IIVRRSGTVGKAFRKIEDALGDRGGGPVAGLTVIAAADRDLRALGYCVSQLPRFDCRVTANIVVEYLDGSLSADLAGAATGWRQVEDALLGLADRGSAVGGTLTLEFTPAVPIPYGGPDWDQFRSVVTNNDPGAIEITVTLADGDSR
MYE76343.1_Acidimicrobiaceae_bacterium             RVLEAHGSAAAALQKLDDAMRDAGAQHLSQVAVRVSAEQPFRVLGMCVPQLARFDVALKLDATAEFPSGAASIRFEGSAAACTPVFEAFWGALSGASDVAGSLTLTASQADAIAYDGPDFRQFAEVLRRAAPGHVTLSAKVAETAAA
MCY3677712.1_Gemmatimonadetes_bacterium            LVVTVDAAPGRAFQRIADGFQDAKREFIYTLKITCEDLSDMRALGMAVPQFPKGEYRVDLQLVAEFADGSVEVRYRGLWDRYKRLRSTLEAQAGEATKVDVKIVLTASYPEGLAPGSRAFQTLRDVMVQLDMGRIEVRAEERPASDP
GBD21619.1_bacterium_HR28                          ELISLDGTLGQVSKGLLDVVRDKQLERLRDVTLRLETASEVRAFGLWVAQLGQLQPELEASLTATLQGGDLRLTFTGPPEQYVRYRSLFETPLRDARDLDARFLLRVT--NLVAKDVQQLVLLGEDELAAELGKVRLTAQGYREHA-
HHV54946.1_Firmicutes_bacterium                    EALEASGTAQQALQGILDRMHDRNVAALAVLQLAVDGDRELRSLGLAVPQLGPGQFRVEVSATGELEGGSIRLEFRGPWERYKDLKDAVVSVLERASAAQVRARLDWRARDPLTPEG--INQVRDVLAHLIPGRVRITAEPATGEVD
OQA44952.1_Chloroflexi_bacterium_ADurb.Bin325      TRFQGQGAAAQAFGQLVDLCAEHKVAHLRRLTITVDALADVRGLGLAIPQFGKGRYVVEQDVKAEFAAESFTLNFRGGWDRYKRFRSLVEAFGSEADELKATMRVAGEFEDGLEVAGAQFAAIRDVLTTLELGRISVEGIPLP----
WP_014555848.1_Haloquadratum_walsbyi               GKTDSPAVVTEAVDTITAESTHEIKQSLDGLTLKLQGDDAWKKGEWIAKQMPEQNVAVELE--YEADDSHVETHFDGTGKMFADLKHTNKP--------SGDQYIEILIKADTSLELNRLDILEDIIPGATTIEMSVRTALELTEDE
WP_073234041.1_Desulforamulus_putei                KPLVAAGVPNKAFADLRDLCEKEKVEALASLTIKLKDIKSVQAVALMLPQYQKQNPVIEQDFTVELPDGSLELKFSGGWQNYREVKNFTERLAKSARETYVETRLTFEYPEPVSPHGEELAKMNESLNTFKVEKIEIEAKETKDE--
SBW09990.1_uncultured_Desulfovibrio_sp.            AAVSQEGVLKEALAILWEKARQKHFLAIATLDLKIYDAQDAFTLLAAVASLSTYDKRVQLEVNFETDHSTISISLNGSTSSAQQLKDFLQPQLRAALDKEFSAIFTIYFANGLSLAETAPEDLAEKLTRVGGVTAYVTATAQRQGA-
NLJ36704.1_candidate_division_WS1_bacterium        QQFSSDISPQRAFAEMQDWVTDHKIERLEWLQIEVRDRQAWTGMTYFV----ELSIDVTFEGSIELQGGRIALDFAGSGQNYQRVYDFFHALAREQEGVQFVAIFRANLPAPLAPAAGEISDLRRRFADGDVQQIKLRAGAPREGAS
MDA8372907.1_Actinomycetota_bacterium              RFSNASMSPSAAADQVAASL---DASARKRLRVKLQENPGSTAWTVLISQSAKVNVLVVVE--LQGSGARLRVNLQGTAEEFKRASALIVEFAKRDVKTEGRVEV------GFVVTGAELVGFMESVKLLELQTIGLDLEEIQ----
MBP6739048.1_Leptospiraceae_bacterium              KIFISEGPLVLALESLWTKLREAKIKSISKVRIKTYDSNSVSKIQQGAINYNLATKTISYNGKISTSIPDFSVNYSGDVGKFSSIRNFFESQIRSGGESSVEGEFQFEFNEPMKLSDSNPENFAKELSSYGGAEVYIEAEVKE----
PSB37932.1_Aphanothece_cf._minutissima             GVIERTDNVKTALTQLAQQVSSSG-KAIKNLTWRMNSGDGFLPM-ALLKAEPGAMVRVEMEGGWSSDGSEWSFDFNGTPEEAAHLRGFIEGQWRRGGEKSLDITYQLRYEPPLRWEDGG-EGWITRLTRAGTATIRLELTAAAEGQA
MBY0527363.1_Gemmataceae_bacterium                 VRLHAEGTPSQTFQTVTDQCHDQKVKALKRMFIRVEGLRDARSLGLAIPQIGKVTIALDQKMVLEFDGEKFTVEFAGSWERYKRIKALTDALGQEASNASVKLVLRADFDGGLDVSGDQYQTIRDVLGNLNIGRIAVDGEPMAEGQG
MCW5609142.1_Rubrivivax_sp.                        GVVEAEDVLKAALNRVFELARHNGALTFGTMAIRPFDKGDAMKLLALVKSVPNATKRIELEAQFETSQSEATITFKGDLEDATPLKDYLEPQFRAASDSDASVTYRLHFEPPLSVNEKAQEALVQRLTRLVSPVAHVLAVVAEKRS-
SPE38928.1_Candidatus_Sulfopaludibacter_sp.        TELHAEGPLLEALREIWEKARKAKHARIQKVIIRLYDALAIFKVHGAMATLKDATVNCELEAELQFGVKEFLVRFHGPIDKANAVKTFLEPQARSASDRRVEAVYTLEFAVGLPLAGDGPESLTTSLTRYGGGEAYVEAHAAAPEPK
WP_091105663.1_Micromonospora_citrea               RIEAKTGQAGVALSNFRDELMDKG-KPLVEMRLTVRAEQDIDLLGMAVAQMPKQPAKVTIDVAASMLEGVIEIRLEGDKAKYQQASSKLGPLLKAAEEIDGTLTLTLDFGEGVTVTDNAFSTVDSVIRGLGPSELTVEGTLA-----
WP_142649215.1_Microbispora_sp._SCL1-1             KTFTATGPGGVAFQHIVDQVTDFTIQTISRLTLKVTADHDIDLAVAALGMLQKQSITVRSTIRAEYLNGGLQFMGTADRMDFQSTYNHVKKALVSATKVAGDLTLNITFQPAVAVNDAVISQIHTVIKNLQMQQTSMTAEVTK----
WP_164002142.1_Pyxidicoccus_caerfyrddinensis       QPVVQVGTPREALTAAWDALATARNVRIAELGLSVEDRESFDNLRVAWADRPKVAVHATLN------GQAVNVTYEG---RFETLKDLLAPLWRSQGELQVTITVALKFAEPPAVDDAALGTFRTALMNAGQGRVEVRLVPVRPRKT
WP_168134876.1_Sphingomonas_corticis               -VFVHEGVLKEALTVVFQKARSAKAVSLASVTISPFDYADAFKLLPVAEGVPDATKHVEMEGDLATAGGELSIRAEGTLTEMKGLREFVEPQLRAASEKRLEVVIRLDFANGLALDGDAPERIVDRLTRFGAAAAFVEASAKVNS--
WP_199338599.1_Nostoc_sp._FACHB-145                KIFKAEAPLREALSKIWEDARKAQIKKISRLSLRVFDVSDGFKLLGVINTVSGADKQVKLSAEYETNSSSFSIEFEGQVSDAQPIKDFLQPQLRAASETHLEITFTLTYANGLELNTDEPEKITEKLARFATGAAFVEAYAEA----
WP_233194430.1_Haloferax_marisrubri                SHTSNTASAKRAFSEVREEAGNNATPTVESVSIEVSGDEVLKHAFFLA----QTTLHTTVSMIYRAGDDEFKANFNGSLASFNTLNQRLEGFADKSGQRNVDLTFTLSLPSSTEDETDVLAVLRDKLSG---TDVTLQVEGSGDIDT
WP_239638853.1_Halorubrum_californiense            VSTDSPSAITGAFSDVNSASSSDIVVEVDTIEIELEGDQVWERAWFISNKLKDLATSIVFEYVAQGSSSVFEAGFEGSAAEFADTNHIPESFISAGGDSEGEAVLIIDTESQDNTREATLDVLQEALEGLGNVELVVSGVVKEASEQ
WP_242016564.1_Pseudanabaena_sp._FACHB-1277        EIIDLSGTVNATFTGFSDRCADYKVKAIHSIEIMIDQVTDYRKLMTAFPLLSKFVFEIDQKATIQMDKQFVRFDYQGGLRGFQSFSNPINALLAEGVRANVSLKLTFTFDPAIAPNGNELQTLQQALMRNPVDRLSLTTRVSY----
WP_257121428.1_Dolichospermum_heterosporum         ETFNAEGTHNSVFTQLADFIKDNQAEGIELLEISVAEVMDYRKLLTAFPLLMKLPLQIDQTASITLGEQFIRLEYQGPYKGFNSFQNTLNTLLSKDVKADVFLKLEFKFLSPILPEGTEISNIKSTLNRNPVDRLNLMAKLTY----
WP_285255572.1_Halobacterium_salinarum             RANDKPMGVLRALTELRDQTFENGAPVLDDLDITFTGNNAWDDAYDFV----DYNPFADVEGTDEFRGDEVECTASGDVEDMPSLS-------QNGGEQQISLSTTVE-PDAIVGDTSLLDTLIGSLEDGSRGTVIVEAEVHNVPEG
consensus/100%                                     ...............h..............h.h..................................................................................................................
consensus/95%                                      ............h..h...........h..h.h..........h...h.........h..p.........h.h.h.u....h...................p..h.h....s..........h...........h.h..........
consensus/90%                                      .......s...hh..h.....p.....l..h.lph........h...h.........hp.p..hp.....h.h.hpG....h..h................ph.h.h....s..........h...h.......h.l..........
consensus/85%                                      .......s...hh..l...h.p.....l.pl.lph....sh..h..hh.........hphp..hp.....hphphpGs...h..h......b.........ph.h.h.h..sh...s.....h.p.h.......h.lp.........
consensus/80%                                      .....pss...hh.pl.p.h.p.....l.pl.lph....sh..h..hh......p..lphp..hp.....hphphpGs..ph..hpp.hpsbhp.s...phph.hphph.ssh...s.....h.p.l.....sph.lps.s......
consensus/75%                                      ..hp.pss...Ahppl.p.h.p..h..l.plplph.s..sh..h..hh..h.p.p..lphp..hp..ssphplpapGs..ph..hps.hpsbhp.s.p.phph.hphpa.ssls..s.....h.p.l..hs.sph.lpsps......
consensus/70%                                      ..hpspush.pAhppl.p.h.p..h..lpplplph.s..shb.h..hhs.h.p.s.plphp..hp.sssphplpapGs.pphpphpshlpsbhp.usp.phph.hphpa.ssls.pss..p.l.p.l.phsssph.lpsps......
```

### Type-2 BR-system DUF499-ATPase C-terminal RRM/Ferredoxin Fold Representative_1

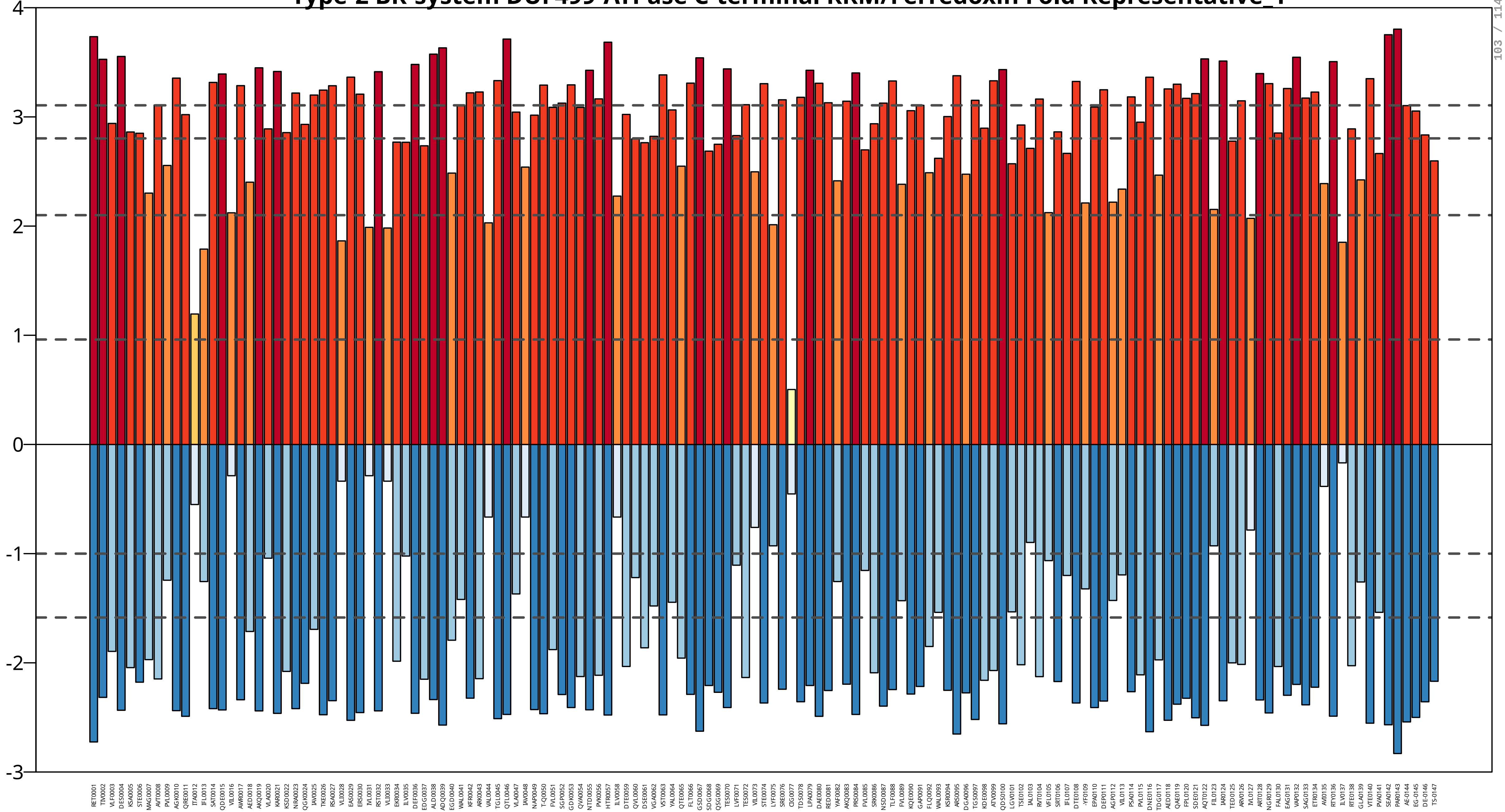

### Type-2 BR-systems

```
# ; DUF499_2 C-Terminal RRM Rep_2 (MAIN_FIG)

AFM14204.1_Turneriella_parva_DSM_21527                ESKAAIVIRRHKFDGTDKCYQLIESMRRYQISVSQVEVAIEAADTFVSYSTGGQIIRDATKLSELIDNLRKPFIEEYRIRLMIGEMHYRQGGDIQHFAGEMNLPGEPHEVRQ
MBX3108928.1_Fimbriimonadaceae_bacterium              PTKAYDWDRKFDKDSTMEAYEFLSLCRKHDVGMRGVQANADISHRFAEFLFDAKFPYMSDNVQSVLDLLQS-LLPESKLSLRVSGLNFANGQAFLDMLAELKVQLPPHEVKE
AGC68793.1_Thermoclostridium_stercorarium             PAKPVKYQPRVRFSDKVAYYYEIQKLKSSNAKPVKIYLDIRGSKREMTVILSEGEGIDIDLLQNLVENLSTEIIRDAQVSGRFETMRFETGKDFLEWLKLQNKKIESGEVEQ
EAZ58601.1_Pseudomonas_aeruginosa_2192                DAKPAKLNKKVSLDSTEKVFGVLNSFKSSPATFKGVRIQIGEGENTVSINFVER-EVTASVIEAAINGLRKALGNEETVTVQIRSAAFENGFAAKEFAKLSGVELRPGDVIQ
MBK7550235.1_Syntrophaceae_bacterium                  RSKAAVWRCNLKKDSTGETYRLLETAKKHGAELGGVRLIIAREGRWIELNSPDDAYHAVQHFEHGADLLKE-FIPEGAISIDVGTLKFDSGQLLLEMVADLKTELKEGDVRQ
MBK8067117.1_Rhodanobacteraceae_bacterium             LNKPVAFQRNAQITSREKVFTALAKATERNIEFTDVVLRIQEGSAFGQFGLQGQ-SAKAERVQQALTMLAVGFSPSTPVTMQFRA-QFPSGQDLIDFAGTLELSY-AGDWKE
MBR2257284.1_Blautia_sp.                              DEKGLEYKSMKKCPDTASAYSEFGRLQKLAGTYIRHTVVITEADNYMEITTARV-PYDASNLQATVDIIRESAFANVVVEFEYKSILFTKGAIFKQWVDMNKLDMNTNEIIQ
MBS0652531.1_Verrucomicrobia_bacterium                PLRPAWLNQNNGTFADAQAWKLLAEARTLGATLYVRSVDLKDGSKTLALRALDAWETDPAHWEEAIQLHARHFGDKPILTLQISEIRFQTPQDLELLAERLQLSYKNAEVKQ
MBX3024939.1_bacterium                                PAAPARLAKLVSAPSTAEAFRVLAQARRDGIRLSEVSLEVGSGTRTASLRFGRDGVLTAEQLEQVIAALRAALGEDAEVRLSARRLEFTRGHDLSEFVKAWGLSVDANEVEQ
MCA3288087.1_Roseomonas_sp.                           AGKASSLLKRVERTDTQGAFELINQIKSARAQAFGAILTVGLGDASAMLRIGRDAGAEGAAIERAATLLRELVSDEAPVTLRLDRLDFPTGADLLALVDALHEPVEPAMVKQ
MCF7997122.1_Chromatiaceae_bacterium                  TTQPATWASGFSFTTTRTAYGFVERLRRRNGKAAVTRIAVL-NNCWVDLNLADDVLLDPDQIQTTIEHLRS-LLEEGEVSIEAPSLWFPTGQDLLDYWAQR----EAAAVEN
MCK9217630.1_Firmicutes_bacterium                     KDKALKITEPVNYSNATDLYRELELLEKFNVSLSGLTLVVSDNKGYVQLMIGGGFVVSSQKIINELNHILNNLFEGFKANLTINEIEFKKGEDFERWISAKKEKIEKGKIRQ
MCL6686250.1_Akkermansia_muciniphila                  PDAPARLDAAGYKSPGVSAAKALDVAEECGVTFREAQLLFGSGEQSVQLKLGRL-EIPAGRLKGLAAAILAPGDGGAGMELRFKTCLAPAARELERFLKNAAIAYQPEELRQ
MCP4350268.1_Desulfobacterales_bacterium              KDKALTLARRIKTNDTSETYGELNQVKKHSALLSQVFVTFYKGKGWIELSFDGTTKVDPKKLEIEMDNIRSNFMNEIDISLEYTGICFRTGQNFLDWVAEKKLALNEEEIIQ
MCX6251131.1_Bacteroidetes_bacterium                  KMIPLTWKVRLKRDSTSESYKLVDALKKTGAKVSGIDINSH-AEDWVSLQTSPEMTFEASLVEEYLKFIQDHIYGKGMLGLQIDACHFETGQKFLDLIRELKTDYKENEIEQ
MCZ8170923.1_Brevundimonas_sp.                        NERPAKAKRDFQFASKGEVLRVVRAAKGKSVRFQGVSVTVGEGERSLRVRSGGDVSLSGADIETMIEGLRAALGKPAEVQLRFREVDFPDGYTLKDFATQVGIDIPVEDVEQ
MDA7889828.1_Akkermansiaceae_bacterium                PNIPATWKHTLTAATTAEVYNLLDHLQSCNAKISDIQITAMKGADCAELMITGPTKRAILEAKDTVKFLQD-LIPQAEVNIDIKASYFELGSELKKVLELLQVAAKAEEIKQ
MDD2765364.1_Opitutaceae_bacterium                    PDKPICWKPKEKFDASTG--RLVALLGQRNARLKKVGMGLAVGRSFGTIEFGDA-EVSAGFVDRLASMVRE-ETGATVTSLRFGEARFEKGEDLVALVKEMNLDLPPEEWLQ
MDD5268641.1_Methylococcales_bacterium                PSKPLTWNTHQGTLSAKAAFDFIERLIKFEGTVQGVIIDVFAESASLSYSAEGSTQYTGEEIKTLLEKLQG-VMKGSQVSISVEDVKFQRGQLLIDWLAEVGRQLQPGEVRQ
NCB71009.1_Clostridia_bacterium                       KDKPLVLSSVIQTGSTKQTYELMSNLIKVEASINGLSTNFSIGSEWFEFNSGGR-DINASEARNVIDSIVSLFGREVSITTQIESLKFPSGERFESWIAAEKKEITTMNIVQ
ORC33988.1_Marispirochaeta_aestuarii                  PDKPLDWKTEYYNIPQTDAYSVFEKCTKYNARLANVFMTIIHSGETLSYTLPEGIGKTAHELSSLMAALQE-LIGEASVQVAIGKISFETGQAFLDWAQEEKFTPDPQEVEQ
PYG00798.1_Thioalkalivibrio_sp._ALE21                 PHQPVRLQNSVRLDTTQSVFQVLVDFKDRNVSFTQVRLQVGEGEDAIEVNFNSR-AVTPAIIESTIESLRKATNDSAEVQMFIRNGDFATGEDLKRFAEIAGLQLKHDNVSQ
RDE52523.1_Candidatus_Accumulibacter_meliphilus       PDIPASVNGTAHLVTRSATYHLLGSLPADRLQMVQVKSTVAATDTTVTMTWDSKTQLAPQAVMEAFEFLDK-QVEGGEWSLRFKHLHFATGKSLLQWQVDTSSKIELSQVSQ
RQH12683.1_Bradyrhizobium_sp._RP6                     PDRPVVIKKRLKIASTKDVFVVINALKAAHGKVRGSSATVGQGDVNATTRFGPKTELSAEILEGFLSAGRAALANEAEVEVGFSEVQFSTGREMEDFIAAVGWEVQPNEVEQ
TPG55477.1_Roseomonas_nepalensis                      EAKPATLVLDIRQSSTEDFFSLVTALKAASAEAVVPQVTVGESTAALSMRLGGDLRLKGEGLSAVVEAVRASMGRPATVTSAVKSVSFPTGRDLIAFVDALKVDVDPRCVTQ
WP_005488051.1_Halanaerobium_saccharolyticum          RDKPVKLTEPQTPKGTKQTYEELEFLNETNATFKQAQFIITGGDFSITFMIDKVEIDDMNMLEEQLKNIKDNFVGEHDLTANITGIKFKSGRDFLRWLEKNEFNLEKNRFSQ
WP_012692622.1_Deinococcus_deserti                    PVTPARWARTHKTTATADSVKLMDRLEKHGAKAAGVKVMVDGGDQYGTATFSEGAAFTADALRELSEYVRKLVGPETNLHLDVKEIRFERGQNLSDWTAETGDKLNMSEVAQ
WP_018962942.1_Coprothermobacter_platensis            QAKPLLFSPKITAPSLRDLRTLVDTLARYNASIAEAYITADTQNATIYLRTKEGIYQKPDQLFDIVEKMSDLLKTDVNISAQIRTIYFEHGRDFSSWVNGQNFKFEDSEWEQ
WP_045169275.1_Caldicellulosiruptor_morganii          YEKPVRLKIAIQLYNMEEVNREIDLLRSFGGKPVKYSIDIYKDDNYIVITNNIPKGIEADEFLRLIEKVKSEFYNNQNVAAVITELYFEDARNFEEWLKQKGKTLQKEAITQ
WP_085411740.1_Fibrella_sp._ES10-3-2-2                KQKPARWNRTIQFDTTQKTYDFLTKMDKAQVNVVLAGAAIT-GIEYVELMLSEKLSFKPAQIQAAIDALRTNLYADGQVTLDVRALLFPDGQALEEYAQELRIDLKSHEIEQ
WP_090663037.1_Belnapia_rosea                         PARPAQWSSRFMFSERRKAFTALECLKRLGGQLGGARLNVQVSTRFMELTHGPAVFRKPDEIEALATDLETKLGAPPNLQLFAARLSFPTGRALTEAAKELGIALDTGEITQ
WP_119720856.1_Methanococcus_maripaludis              RDYPLTITKNEKLNSSQETYRCIDLFEKYCVKTSGNTIYIKGNEKWGQIELGTDVELNMQDLKKLIDVIRDQISNEINVSLRYTKAKFEKGSDFIRWTKDAEKDIEEGHTKQ
WP_171833827.1_Lichenicola_cladoniae                  PNRPAKWKARLKFRTTEDTYNAVKAAKDAQATIMGLEINVTSSEAFASLVMGEGLGLPASDMEGVLADLQA-RVPGGEVTLAAPVVSFETGTDLNGFSNAIGQAYDVKDVVQ
WP_190610329.1_Synechococcus_sp._FACHB-909            PTRPARWEKGGKLDDSGAVWDLITRLEQNPAVLAYDHLTAQSGEQIVEYTGSLANGYCGAELKTVAGKLQN-IVQDGTLRMEIGELGFPTGQALIDWLNVVKEPFNLSYVSQ
WP_231170937.1_Bradyrhizobium                         DGLPVFVNYVVESKSTPQAYEIIEAIKAATGSLVKGRIEVMGEHDFLAATFGDTIALDSETIETLVRTLAGKAGTLPTVTLQINSIRFGTGRDFKNFADKFAIDFEKANWSQ
WP_274435612.1_Alicyclobacillus_sp._ALC3              RESPAIWHSPKRLDNSSKTYEGLKMAKEKSIEFEQVMLMVGSAPRAIHLSLGEI-KIDAEFIETELGHLQSLVGSEAPVIMTFKKAYMPTGYDLEQFAKQLGIEIGNGEVEQ
WP_277247933.1_Phascolarctobacterium_succinatutens    NAKSLKLKINKRYGNNADVYQYLSDMKKIGFTATLDSINVADDDCYFDITCSGQ-NLTAEQIENMLDTIRQVVMKDTKVELGIAEMHFTSGQSFRDWIAASKDEIKKGCIEQ
consensus/100%                                        ...s..h..............h..h..............................h....................h....h..s..h..h..............p
consensus/95%                                         ...sh.h.......s.......h..h...........h.......h...........h...h...........h...h....h..u..h..h............h.p
consensus/90%                                         ...sh.h.......s....h..h..h.....ph....h.h.........h...........h...h..l.........hph.h..h.F.pG..h..h.......h....h.Q
consensus/85%                                         ..bsh.h....p..sp..shp.l..hpp...ph....h.h..sp..hph..s......s..hb..h..l...h.....lph.h..h.F.pG..h.phh......hp..ph.Q
consensus/80%                                         ..bsh.hp...p.ssp.psaphlp.hpp..sph....h.h..sp..hph..s.....ps..lb..hp.lp..h...s.lph.h..h.F.pGp.hbphh......hp..plpQ
consensus/75%                                         .sbPhphp...p.sss.psaphlp.hpp.ssph..h.l.l..sp..hphp.s....hssp.lbphhp.lpp.h...splphphpph.F.pGpshbpah.....php..plpQ
consensus/70%                                         .s+Phphppp.p.sss.psaphlp.hcp.ssph..hplsl..upp.hphphs....hssp.lbphlp.lpp.h..psplplphpphpF.sGpshbcah....bphp..clpQ
```

### Type-2 BR-system DUF499-ATPase C-terminal RRM/FerredoxinFold Representative_2

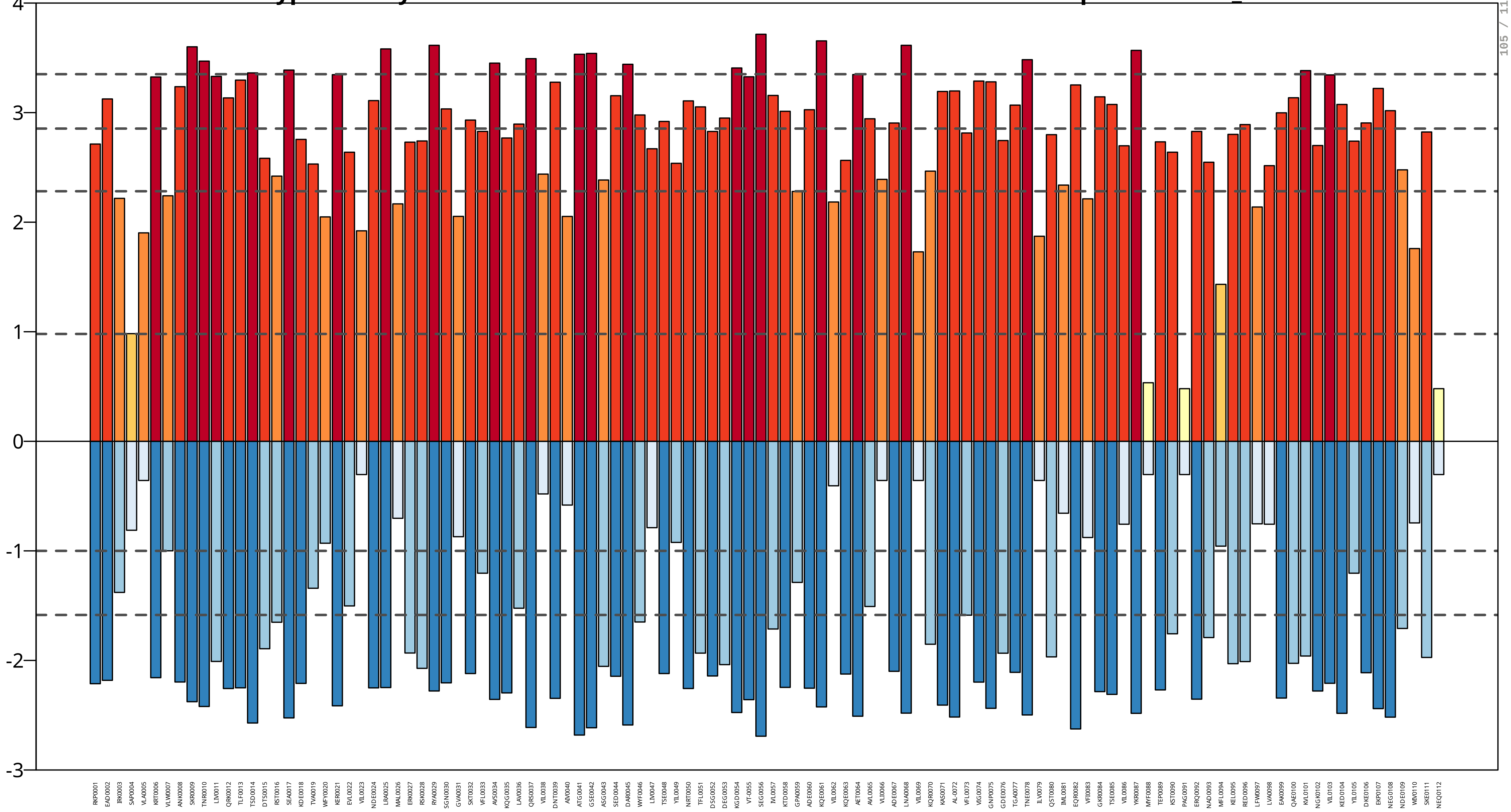

Type-1 BR-systems

```
# ; DUF499_1 C-Terminal Degenerated RRM Rep_3

WP_148590534.1_Aquisphaera_giovannonii              AGGGTATKP---KMIRWRGTIPPQKWTNFYAKVVSPFALAQSLTAEIVIPADAGLSSAQVEKIRAALKDLKLDEQDLS
MBQ9433643.1_Synergistaceae_bacterium               ------TPAKLNRKCKVDAGKFQRDTRQIHDNIILCLQESPSIRVDIYMETPESISAAVESAIRDNIKSMDIDYC---
WP_054571357.1_Frankia_sp._R43                      VDQGPALPHDTVRTAQISAHFDRSGLFRLVRALSK-LRDVDRVEVDVKLRVEGELDRAEFRSVIEPIDEDGVDIQQLT
EEO54231.1_Bacteroides_sp._2_2_4                    -KSEHTAPSRKFKSIKVSGKVPVERWTDLFSSFVVPLKNNGEISFKAKTTSLNPLDESAYKVVKESAMQLGLNLEEE-
MBM4287682.1_Deltaproteobacteria_bacterium          --GGGAPPPAAARPLQLQLRVPNPVTANKLLALAQPLT----LDVTVTGPLKDGGTMNFAGEVKPTLQAIELSLNEAR
MBI3467145.1_Planctomycetota_bacterium              GTGGGSTVARRVSKVRVEMSLPWENWHDVYNEVIQPLANEGFCQVVVIAKGDAAIRENTVEGIRESLSQRDIEAD---
MBI4291280.1_Betaproteobacteria_bacterium           PLPQPEPPSAQPRSIG-------AKGVNLLGELEKWLP---HASLTFNGVSKE---------LRDLCMRL-LQLTEED
WP_259054315.1_Salinibacter_ruber                   GQEGEEEPLV--HRFSLRADIPTRKLTDLNRGVLMPITRDATFTIEFEVASTDGISERKIETVMETLRQLGATVEKDT
MCX5685413.1_Planctomycetota_bacterium              GVGGGTTPVVKTQRAD-----EPQAINTIVTAGDTWVNTILEAWTAITATSAGGAGRAVIDSLRELLSEVGLDYKEEK
MBA3936148.1_Planctomycetota_bacterium              PPGGPQTPPTESRR-------SPPAKTNLLSWIEQQIE----ASLRLTGVSIE---------LRKLLQSL-VEVQQGG
WP_286167649.1_Ochrobactrum_sp._3-3                 PGDSGETPSQQLTTLS-----NPATPLNLIGKLEGWIG---EVMLKVPAATAQ---------LKELLKKL-LSLD---
MBQ9726423.1_Kiritimatiellae_bacterium              ------QPPQAFRHYALDPQLAVARVAQIQDELLRLFNAKPSVKLDIEASSATPFDPNLVRAVRENASPSNLNLSTE-
CAI1753142.1_Serratia_quinivorans                   SGTGSVSPQSQKTRFRLDAIRAKMDFANIMDEVVQNFASQYEISIEINARSEKGFDANLQRAIKENCSMLKFGNAEE-
MCC7203047.1_Nitrospirae_bacterium                  TPRCGQRPCSLAFRLEYDG-------AEITKATYKIFQKDGRGNLEISITKNGRFVKN---QIEQQVESLGLIVENK-
MCL0064885.1_Dehalococcoidia_bacterium              KEEKQIEPKIEIVGIRWKGEVPYKQLMNFSTRVLSPFA--KSLQIDFEVYPPEGVSPEKVQEVKQALRELGLDETEMV
MCD6101177.1_Candidatus_Marinimicrobia_bacterium    PEEETEKPEGKITQITYTLKVNRQQLYKCFNALGNLVE--KTLSLKIEAQTKEGIDTNWLRAVEEPIEEAGVEIKEEK
WP_116051372.1_Pseudonocardiaceae                   ATSGAATGSKQAASFLANSG-NSDAFAALFKTLYMALDERQQGTLQLVVDADD---------LQQNMDELGISGTDI-
RME04985.1_Planctomycetota_bacterium                ELEGERRVERVRRDLELVLGVEVGQMSTVSRVVRY-LKERF--------------CEAEYEKVLEALYQEGIRVEG--
MDE0129995.1_Gammaproteobacteria_bacterium          DSGAGGRPGSGSREITFSFVADQRSLYQAWNAMANLAD--LAVAINAKATAPEGYDESKVEGVLEPLRELGLLDEES-
PJF35651.1_Candidatus_Thermofonsia                  SEVHTAQPLIVKRRYKLNPLHANAEMSSVVEEILKHFLNFSELELEISARRAEGFDESLMRDINENSRSLKFTSYEED
MBQ3918676.1_Oscillospiraceae_bacterium             DDHGGETPPQKKTRFTLDNTRINRDVQNLVDEVISHISA-QKITLDVEAECSEGYTQPTIRAVNENCKTLHVKTFED-
WP_157574125.1_Nocardia_jejuensis                   VASNSGKPAQILATLANDPS-STDLLAQVFRRLFDLIDREAQGTMQLHLPSDD---------LATLIRELGIAVNDK-
MYJ80998.1_Acidimicrobiaceae_bacterium              VGDSTGQPDRHWQTLAVNPDDPDADLVHLLRELSNRLEEGNNQTTQITVTGQE--------AIEDLAKEAGTAVNQL-
TME11450.1_Chloroflexi_bacterium                    IADIQELPVPKYHRFIIDERRMGTQAGNIMEEVVKHLTSLYKVTLEIQAELQNGISEATVQTVMENCRTLKFESFEEE
WP_247297741.1_Bradyrhizobium_sp._179               --DLPLPPPADKRRLQLRISCVSAGDAQKVLGATQPLQ----IEAELTGDLKDGGRLSFVAEAKISLQTLGVVLEAR-
HEC67877.1_Candidatus_Desulfofervidus_auxilii       KTEDGKTPTTLPTTLQFKAGIPWDKLHTIIAGIITPLKDKGEITLEIKAHTSTGLDRTTLDKIKETLLQIGAKIEEE-
MBR0288712.1_Selenomonadaceae_bacterium             -GDAGTDPSLLPKHFELDKTRLNKSFNACIDEVASYLMHLPSIRLAINISAPEGIPEDLKEVISENCHTLKVRNFEN-
MDD3269499.1_Syntrophomonadaceae_bacterium          PMPGADRPEPVVKR-------SEEKILNLGGEFDVWME---QARLEFNGLTQD---------IKQLLRRL-LEIKE--
MBM4285998.1_Deltaproteobacteria_bacterium          PVGTGPTPPKQPVDIEWRGTMPKEKWNLFSHRVLARLSTAEQIEVTVKAKVPD---PTVKQQLNMALQDLDLEGE---
NJK30457.1_Acaryochloris_sp._SU_5_25                ETSSTSYPQRQINRFSLDAYRVGKDAGRIAEEIIQHLSEVAEVKLDIQVQFSNQPTDQVIRTITENCKTLKFDDLESD
WP_048201423.1_Methanocaldococcus_bathoardescens    TIQKEDKIKPTINELTLEVKIPRGKLSEFYRYVMRNLENVFEITIKIKATNGN-LKKSDYEKIKETLNQINAEVLEDI
MBF0121417.1_Desulfobacterales_bacterium            FVRKELKVQIRKKSIRWSGEVPNSKWMVFYNKVLAKQVSKSKVRVSFEFETEESEQEASIESIKSELNELGFNTEEI-
MBN2391962.1_Anaerolineae_bacterium                 PPEASVSPSQLAFQFDAGD-------VRVLRVRFSIVEQSRRGSLEIVLTFDGEYAKS---AVEQMAEQLGLTLQQNS
MCS6856320.1_Sandaracinaceae_bacterium              EVSQESKSQGRLKKYELSAVIPAGMLSHFAKGVLMPLQMAEELEIHLRASSEVGIPLYVIKQVEETIRQIGGKIHEGQ
MBT7692838.1_Gemmatimonadales_bacterium             EGDGGGKPGTAKHQFDLDPVRSIKQLG----EILEHVGAKLEISLEVRAKNAEGFDDRTRRTVSENANNLGANEAE--
MBU2110960.1_Actinomycetia_bacterium                -SNGPAAPAVERARYEVDASDADARLREVIDEVVRHIVSAPSIVLEVSAENSDGFSESVGRTVRENSRVLGFDKSEDV
UGS34698.1_Solirubrobacterales_bacterium_0166_1     PQNDPPAPQGDATGLALSITATEDDLFTLSQALGKLLLGEGGLQIEVRSATGEAIDRLKARTVVEPLEENGVTLSQTR
RIK37466.1_Chloroflexi_bacterium                    ASSSRTTPR---QSLEWTGELPHQKWMNFYMKVLTRFAGQHHIRLNVEISAPSGVSTQKIAEMQAALRELGLDDKE--
MXV84967.1_Candidatus_Poribacteria_bacterium        RQSGPETPPLPRKSIGIRFTLPQGKVSNVAQHFNQ-LQTNF-MQIELKASDGE-ISQDAYEELKENLRALGIEIEEI-
MBC7327984.1_bacterium                              KMKEEISAPPPLKEIYLKLQVPFGKLSDVIKMVNL-IKSKYEMFMEVKAKQGEGMTNTEYENLKEGLRQAGVDIEEK-
WP_013295752.1_Methanothermobacter_marburgensis     RVDGKDMPTGHIKWIELEFTTALENMSNIIRIKRT-ISEDFVAEIRIVARDGE-MDDTKYERILETFEQLGIDLKKVR
NPV88406.1_Coprothermobacteraceae_bacterium         DGGGEGEGEAVVKELELDLDVPVGSFSAIAQIVPF-MKLKFEIHVRIVAHGGS-LTNKDYEKVIEALKLANIKIVEEK
PWV37728.1_Desulfurococcaceae_archaeon              PSERRDKPELTYKELHLELKAKPGMISEIMRLVSSILKPKFTITVKIQAGEGE-ITREDYEEVREMLRQLGVEIKEEP
KEQ56705.1_Marine_Group_I_thaumarchaeote            -GEGGSMPDDSIENLDFGFTIPEGQINNVSTMLLK-IASHY-LKLKVEASDGK-MTKHDLDLIKETLSQIGSDL----
MBC8384565.1_Candidatus_Cloacimonetes_bacterium     KLEKPKQEETKKDHLNLSFEIPKGKVAGILGILNF-LQLKF-VKVDITSTNGE-ITESDYEKIKEALMQLGINFK---
NET07313.1_Symploca_sp._SIO2B6                      SEYKSETSSKKTARFKLNSTVNKAQVFQLFEVLQT-LSDKATIQIEIRAYTNQEFDPNWIRAIEEPLDELDIKASE--
MCX7997938.1_Leptospiraceae_bacterium               PTPQPFTPTSTKKEVKLEFSVPKGKVYEIAKMLHS-LHEKF-ITLTIHSIDGS-ITTQDYEQILETLKQLGIQYREE-
consensus/100%                                      ..............................................................h...............
consensus/95%                                       ..............h.....................h.........h...............l...h...........
consensus/90%                                       .......s......h.............h...h...h.......hph...............l...h....h......
consensus/85%                                       .......s.....ph.h..........ph...h...h.....hphph...............l.p.hpph.hp.....
consensus/80%                                       .......s....pph.hp.......h.ph...l...h.....hplplp....p.........lbp.hpph.hp..p..
consensus/75%                                       ......pP....pphphp...s..ph.phhp.l...l.....hplplph.s.p.........lbpshppl.hp.pp..
consensus/70%                                       ......pP....pphphp...s..ph.plhp.l.p.lp...phplclphpssp.hs....p.lb-shpplslp.pp..
```

#### Type-1 BR-system DUF499-ATPase C-terminal RRM/Ferredoxin Fold Representative_3

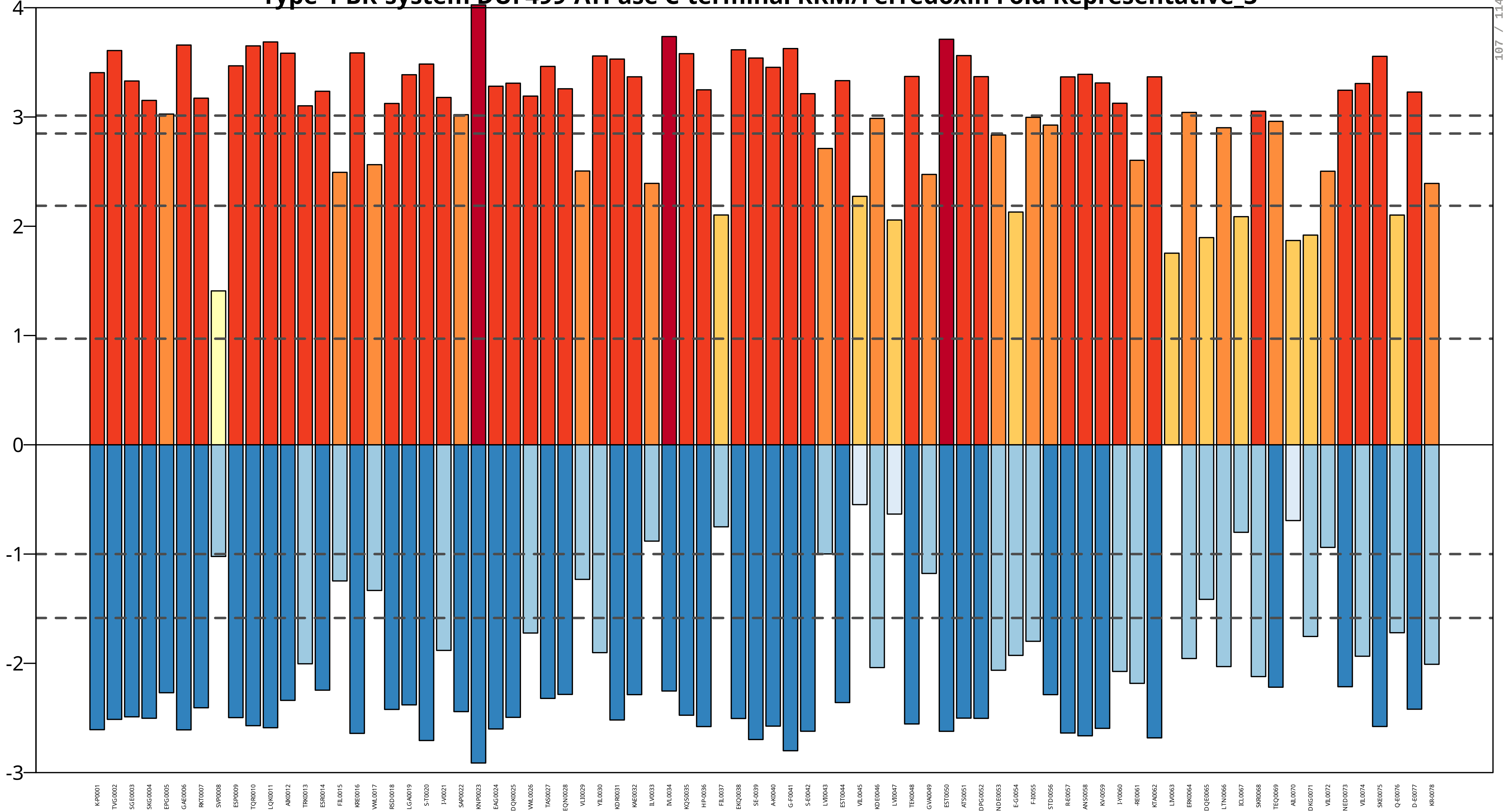

#### Type-1 BR-systems

```
# ; DUF499_1 C-Terminal Degenerated RRM Rep_4

WP_244804062.1_Corynebacterium_sp._LD5P10            TTPPRPSGEPGATRFTATFDLPLTGGVGRYAIIDDEVFTHLKAGSGGSVSISLDVEATNPEGFSPAVQKMINGSLAEMSDADGAFE
WP_005520661.1_Corynebacterium_matruchotii           LSGGPTVVELPDTHYTGSVVINSDRYVRMVNNIIEEVIDRLA-GSGADLEITMNIHATKPKGFTETEKRIISENSQTLKFGYYGFE
MCL5447091.1_Actinobacteria_bacterium                PIGTPGSGSRRPRRFHGSIVLHSDRLNRDFGQVVQEIVQQLQNANSADVEVTVEISATSNDGFDDAVMRTVTENARTLRFRDQGFE
WP_052687173.1_Streptomyces_lydicus                  GISVQPILPRTLNRFYATARLTPEQYSKNATDYVFEILQHLA-LSGTDVEVTVEIQASRPEGFPEDKVRILRENSNTLNLQAE-FE
WP_053587850.1_Actinomyces_sp.                       GGADPEAARPKNTRFTGSVELDPGDIAAQLASLAEEIIVHLRRGGADGLEINVNIDATRYAGFDQATVRTVGENARVLGLRPGRFE
WP_270528574.1_Mitsuokella_jalaludinii               NGAASAHPGPQYSHFSMTQDLDSLRATKQVNDCLQEVVNHLSDLKGSQVTIKLEVQADLPANLPKDVVRTITENCQALGIKFT---
MCV7506294.1_Micrococcus_luteus                      PETEKSAARVPNARYSGRVEVPGADLPSTFQTLIDEVLSHLQNADPDTLEILVEVHAEKSSGFSTATSRTVGENARQLGFIKTHFL
MCY3894210.1_Acidimicrobiaceae_bacterium             PRDDGDAVQRGLRVFRGSVKLDVSRPTQAFTRLSDGVLTRVLGHPDAVVDIRVEIEVHNPAGFDSDVIRNVNENTRELDFEEGGFT
WP_238518868.1_Azospirillum_doebereinerae            SPQPIAAAEKPKTRFFATLDLDPVRMMAKLPQVAQGVVSELARPEGARIRISLEIHVDAPDGFPNDVMDIVRDNLKDLGFPAGRFE
HBM97560.1_bacterium_UBP9_UBA11836                   KPLETEKSESLKTSLYASVKIDPNCCNQVVKKILENIVYDLVNLEDCELNLTFSVEANAKQGIPAEIVSAIEENSKSLNLDDFDLH
MBC6439887.1_Rhodospirillales_bacterium              -NTDSPDPELPPTRFTGTVSISAERPARDMNRIVDEIVDHLKTQPSARISLRLEIDADLPDGMDPAKVRTLRENATTLGFIDQTIE
OZG52261.1_Pseudoscardovia_radai                     PSVPGPAPEPKQTRFYGSVVLDPDQYNRQIAKINEMIIDQLR-FSHAKMELRLDLQATQPDGFDPSIVQRISSSAETLGFTASGFD
MCF6470725.1_Nonomuraea_sp._MG754425                 PGPNPPAPPVRNTRFWGSLKVNPERYSRDLNSLAKDLLALLTAPDDVELEVEVQITARRTEGFSDDTVMKVLENLGHLKVEGK-FE
MXX78159.1_Gemmatimonadales_bacterium                GETEEEAVEKPPARFFASIPVEPDRAGFVVARIMDALLVELTRSPGSNVRVTLDLEGNAGNGYPKDVVDIVKANARDLKIDKNGFE
MCI1646461.1_Olsenella_sp.                           GGGDRPAPEPRKHDVELEATLDQWSAGSDVGAIVEEILQRLYDLRGSSAHLTLSAEVSVPGGIDDTTLRIVTENAKALGVTLR---
WP_094451589.1_Parenemella_sanctibonifatiensis       GTGASSAAPAVPTRFHATKDLNTTRVVRDASQIYEEVISHFV-ASGINVTVTLDVESDQLDKLTGDQRTAIRENLKTLGFADHDWS
MCG3151370.1_bacterium                               PTMNEPVAPALPTRFHGSVLLDPTRVGREAGKIADEVITHLEKLIGAKVEVRLEIQATLPEGYPEDVQRTVSENSRTLKFTGFGFE
WP_092673506.1_Siccationidurans_arizonensis          DDNTAKDSGTKRTRFYADANLDVTRFFRDAEALEKEIIKHLP----KGATVTMHIEARNVDGFTLDLQRVLQENMRTLGLGSAEFE
WP_201387653.1_Ktedonobacter_sp._SOSP1-52            AVPVTVKNETIKRRFYGSVKLNPRMMAGDAGKIMDEIVTHLTTLSKADVSITLEIQANIPGGVPAETIRTVMENSNTLKFESAEFE
MBD2550060.1_Microcystis_elabens_FACHB-917           PAPLPPPPAPQRTTFTGSLKLDPVRAGLQMGQFLEEVMSHLQALPGAEVNLSVEVHVKAPNGIDDQTARIVLQNAADLKLDNP---
WP_188816795.1_Calditerricola_satsumensis            VDPPIVPPPPKVRRLNLFANVPPERLIGKSSEIFQEVLTHLLSLDGAEVEVNLDITVRIPNGLPGHVRRVINENARVLGVQVVEYE
MCL1790896.1_Peptococcaceae_bacterium                SVQPDIPPAGKNTHFYGSVRIDASKLGSTAGTINTEVLQHLNRLTGARTRITLDIQVSVPDGVPDHVAHTIRENCKFLKFDNAEFG
MCY4445744.1_Rhodobacteraceae_bacterium              DNGIKPIKEKKYRKYKATTTLNPESPNPKFNDIANYILDPLL-RDVANVQVKVEIVAIMPNGFSDETIRTLNENSPLIDLDGEFLK
WP_127367955.1_unclassified_Mesorhizobium            PGGTRPSAPGKPRRFYGSVELDMVRPVKSFDAILNAVVMELQRTHSAKVKLTLEIEAEASDGFADSEVSVVRDNARQLKFKSEGFE
MCZ0940408.1_Caldilineaceae_bacterium                VDVGSGREEPSLTRFYGQVELDSVRAIRDLESILKEVVSHLR-RAGKKVTISVEVNAEA-DGFDTHTVRVVSENATQLGFTSHEFE
WP_132287513.1_Paracraurococcus_sp._NE82             SPGPAPNSPKPLTRYFGTAEIGADRAMRDFGTISDAVLAHLVTQRGAKVKLRIEIEAELPSGFTTDLVRVVTENGNTLKFTQQGFE
CAG0999165.1_Planctomycetaceae_bacterium             TQSGKPVAPKLPTQFFGTVELSPDRVGSQAGKVAMEVLLHLMNQPDSKVKVTLDIEATLPKGADEAIQRTVLENCAALKFVNREFT
WP_151650193.1_Bradyrhizobium_betae                  PLPTGGTRAARPRRFYAKIVLDPNRPTPQVSNIAQSILSELDRVRGIRMTLTLDIDAEAADGFPEDVESVVRDNAASLRITDFGFE
MDE2480824.1_bacterium                               VAVEQPQASGPPKHFFLRVKADDTKLPRVATQLANDIVAHLASIGGASVEVNIEVKADISNGINAQLEDRLRKNAAAHQFPDPEFN
HDQ14361.1_Sediminispirochaeta_sp.                   QRPVKPAEAAAKKRFYGTVDLNPHAPSGKMGTIAEEVLAHLA-ADGARVKVHLEIEAESAEGFSEETVNTVLENTRTLGFEQKGFE
MYD10935.1_Chloroflexi_bacterium                     PDALLPDPPKPKTRYFGSVRVDPQRAMRDFNQIADEIILRLASLPGADVSVTVEIKCERSDGFDDVTVRAVSENSNTLNFSAHGFE
WP_075830034.1_Deinococcus_marmoris                  PEPKPVVPPPLPQQVFAEGKLDDARMVRKFQTIYEEIIQQVI-DAGGDVSVELVIRGDVRGGLNTTQQRNLSENARNLGLKLQ---
MYA05496.1_Caldilineaceae_bacterium                  IIDPPPPPPPSSQRFFGAVKVDPERLGLEAARIADEVLTHLTTQPGVQVEVSLEINAVLPQGLTEDELRDIRENCSTLRFASFDFE
WP_106140933.1_Spirosoma_oryzae                      GSTTSPNSVPTGPRPIHTVTVHGKVDIANYSQVFQTFIMPLA-QQEVEIEIRIKGKSTTAKPLTETSPEVVIEGARQLGLRVE--E
WP_118201547.1_Prevotella_copri                      TNPLNSDSSENEVKTYSKVTISGSVPMENWVQLFPSFITTLK-KNNLEIEVKFTAKSTEQNPLTENSPTSIKESASQLGLDFE-ME
WP_051018640.1_Saprospira_grandis                    LSVQKVDEELKYTAFQTTG----QVGPKEYTKLFDYFIRPFL-REGDEISIDLSFNIKTGTSFDEKDERRAKEAAKQLGFDIRP--
RKU15614.1_Candidatus_Poribacteria_bacterium         ASVKKDKAKLGSKRIVARKIVEGELSLDDIGDLCQEIIGPLS-TDGGDVTIEIIITAYKEDGFSQNIERSIKENSIQLNIEVQSTN
MCY4474618.1_Chloroflexi_bacterium                   GEGMQPPPQTGPSRITATKQLQGNISFDEVSGIREEIIRNLT-DDGGDVTVTITVSASKEGGFSQNTIRAVRENGDALQLVLQQTE
MDE0020779.1_Candidatus_Poribacteria_bacterium       RRRKELDPPKLPKLFTMTAEAKPDTLKQAVEQLENEIMPSLR---NGEVSMRIIVTAKHARGFDENDARAVRENCKELGIELDEES
MCA9831553.1_Dehalococcoidia_bacterium               VQTPVPQPATRHSRV----RIRVDMPVAKSTNLQPYLWKVLQ-GVDIDTKLSITVEVESAAGVPEDVLNRIVEGLEQLGIQVE---
consensus/100%                                       ................................h...hh..h........ph.h.........hs......h..s............
consensus/95%                                        ..............h.....h........h..h...hh..l.......hplph.hps....shs....p.l.ps...l.h......
consensus/90%                                        ............pph..p..l.s......h..l.p.ll..L....s.phplplplpsp...uhs....p.l.cs...L.h......
consensus/85%                                        ..s..s.s....pph..p..lps.....phspl.p.ll.pL....ssplplplplpsp...Ghs....p.l.-Ns..L.h......
consensus/80%                                        s.s..s.s....p+h.uphplss.....phspl.p.ll.pL....ssplplplplpsp.spGhs.p..p.l.ENsp.L.h.....p
consensus/75%                                        s.s..sss....p+a.ushplsspp...phsplhppll.pL....ssplplplplpup.spGhsps..c.lpENsppL.h....hp
consensus/70%                                        s.s..sss....p+a.uohplsspp.s.phsplhppll.cL...sssclplplclpup.ssGhsps..cslpENsppL.hp...hp
```

### Type-1 BR-system DUF499-ATPase C-terminal RRM/Ferredoxin Fold Representative_4

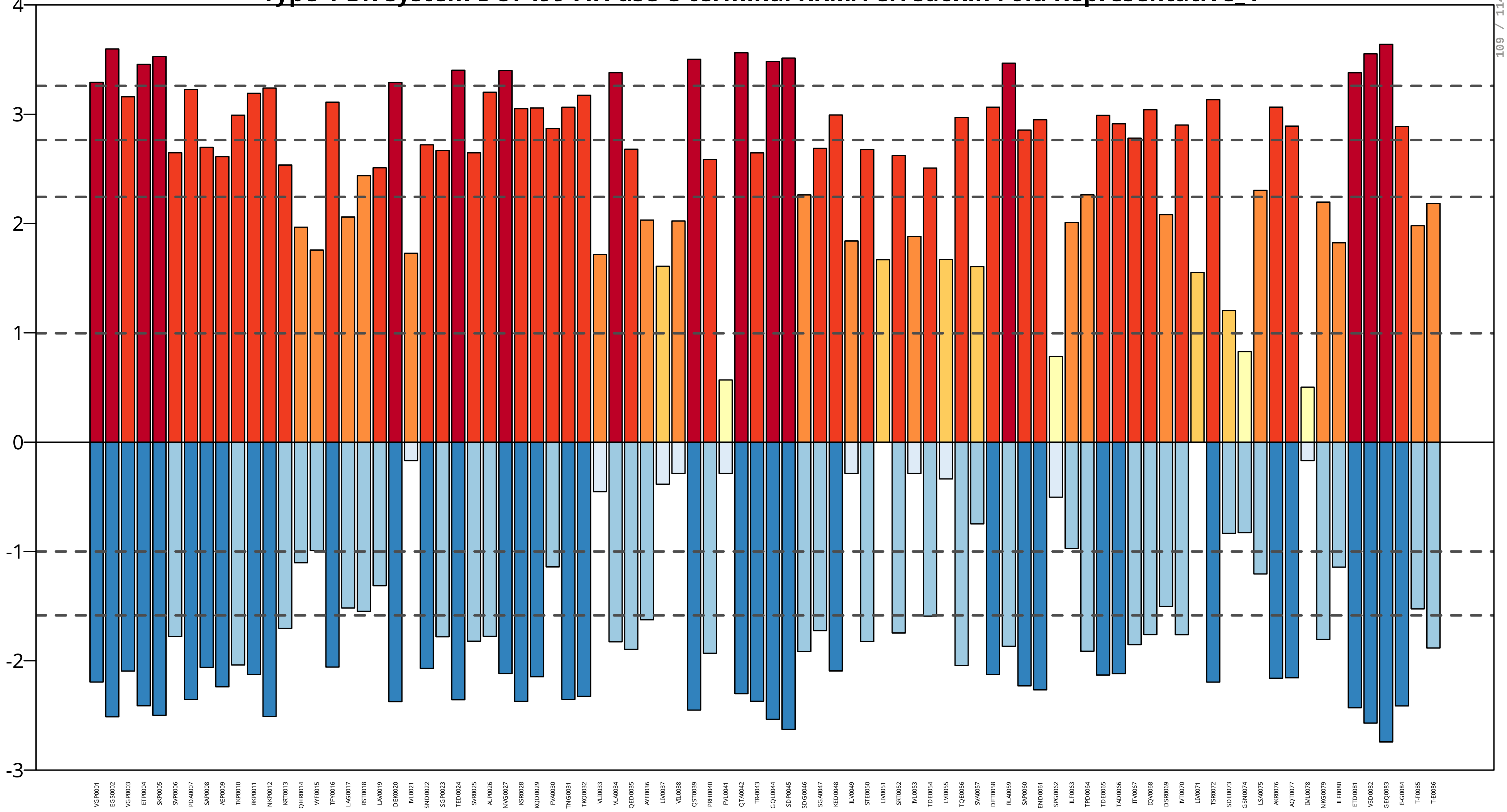

### Type-3 BR-systems

```
# ; DUF499_3 C-Terminal RRM Rep_5 (For MAIN_FIG)

CAD6492704.1_Candidatus_Argoarchaeum_ethanivorans            IGDEVAKTPPSKTVTVPPITAKPELIDKIDREINSGDILKEIDLSISGVDKNGFISFLRELKTPKIATFEVQMKVVRG
HCF99178.1_Chloroflexi_bacterium                             TPPTGGTSPAAKTLKSDDARIGLNLLGDVAKWGVQDREILSARLALEGLSYQELKELLQRIPPKLKASLEIDVEDQTP
HIE29204.1_Candidatus_Poribacteria_bacterium                 VKEGGSDEPTTETVQTAIFSNIGGLIDQLEKTVATTGTIRGVTIKMNTYTKNELLQNLNKLPNIPSGEIQAIVKVERN
HPN33675.1_bacterium                                         -LKTTSEVPQQSKIQTPFTNSLGQLRQEVAGRLLDADQITDLEFRIFAFSKAQVEQLVEALPVYQGATYKAEMSVLDQ
HUT59382.1_Phycisphaerae_bacterium                           SPSVATVSPAGETISIPFKSSKGLLRQEVASRLAQEGQVTSVRFQIFQLSKAQVEQFCEQLPDFPNAQYMARLEITGA
HVA45320.1_Pirellulales_bacterium                            APLAGGVQPAGVIHRSLGAKSGINLLGDLERWALPGQRVIQASLTISGATVKELRELCSRIPAKIVAELQITLPPEGD
HZM03188.1_Candidatus_Saccharimonadales_bacterium            TPAPGGGQTSGVTRRTDSESNTISLIGQLELWGVTDKMLTSAKVEFSNLTAQQLRRILQQIPSAHRANLEVSYQQQQT
KOX91592.1_Haloarcula_rubripromontorii                       GGSSEEDGGSSKQVT----PSARVLRSHAESRVNETDTVTHVDLSYDVLSKEELIELVEQLPGANHIEATVVIEREVQ
MBA3936148.1_Planctomycetota_bacterium                       QPAGGPQTPPSTESRRSPPASKTNLLSWIEQQRDIAGAPLEASLRLTGVSVIELRKLLQSLPVSVQPSVEVQWTADGQ
MBC7429623.1_Bacteriovorax_sp.                               TPIGAEASPSQKLMRLGYGRAPLQILDQLERWGVSETKLHDVSLVFTALSGEQLRKIIQTLDSDIELNLKLIQEEDK-
MBK6914962.1_Ignavibacteriales_bacterium                     TDNGISGIPGAINISTANFKSIGELRQEIASRLNEENVIENIRFTIFLFSKSQIEQLSEQLPSYQDAVYKATLKGYVK
MBM4461756.1_Chloroflexi_bacterium                           -TTEYERTPEGVPLASQEHTTPFKMVTELEGRLSSGDKVASVTISLRGMAKDELMKLLGKLPVPSEGIIKATLEVEKH
MBM4466103.1_Chloroflexi_bacterium                           KPPEEAEEPKAVVVRTEAHATPFSLQTDVEGKLRSGDRVTGLEVTIEGQQKQEILKLLDRLPIPVDGQVRATLEVESS
MBN2391962.1_Anaerolineae_bacterium                          EPPEASVSPSVRELAIRYQPTPGQLRQEIAFQFQDGDRVLRVRFSILVYAKSAVEQMAEQLPAIPGGEYTATLTLQAS
MBQ8111585.1_Kiritimatiellae_bacterium                       NPPTVLVAPKK----TAAPANGSTLQGKFEEWGLAGKKLKSATLSVEGISVSDLKTFLMRLPSKYKASMEVEEEA---
MBU4312937.1_Candidatus_Omnitrophica_bacterium               EPQIGTVTPAQQTISVPSQPGIGALRQAVAVRLQESEKVIKAVFTAYLFTKGQIESHIESLPNVKNAEYSARLDILIS
MBY8216331.1_Vibrio_fluvialis                                EPTGNGSAASGKTVASAQNTSTLSLLGKLEQWGINGSKVKDVKLSTSQLTGAQLEHLLKNLPDGVTYSLDLDKEQ---
MCE0498498.1_Methylacidiphilales_bacterium                   SPNPGNYQPLPETDSTGFLKSRQEVRQEVARRLDAENKIVHVRIGLTYYTKAQVEQLMERLPDFTPGACRVTIQLQLR
MCK8825903.1_Fuchsiella_alkaliacetigena                      --ETAEQSNASVEIH----NSNRLLISQVETQLRAADQINKVVITINFFSKEDLKELLLKLPDKEG-SYSLEVRGERE
MCL0058394.1_Dehalococcoidia_bacterium                       EPPPTPLTPVR----TESRATPPGLRAEVYNKLEEKAKVIQFSIDITALSKANIIDLINKLPMPVADTITLNLEVEIE
MDD3269499.1_Syntrophomonadaceae_bacterium                   ENQVGIVMPGDTVVKRSEEKTILNLGGEFDVWGMETRKIAQARLEFNGLTVQDIKQLLRRLPPTTRAVLEIKLPGGAE
MDH4162607.1_Nitrospirota_bacterium                          QPAESDVEPGEEERSTPSCRSLGELRQQTAARLTDDDIVRTVRFTIFAMTKAGLEQHCEKLPNLSGASYMARFSVELQ
MDM7274691.1_Thermoprotei_archaeon                           STIAPPPPPPS--------RTLNEFRQRLDLLYHTGQLVNYMAVKADGHEVSHILNFLKHLSEKYKATLQLEIGFAEP
MDR0374944.1_Treponema_sp.                                   VPVLGTMSPPPVRTKSSAPRSSINLIGQFESWGLSDKTINKATLEFDGLSVFQIKQILQRIPIAYQAGLSISYDDIEN
MDR0392109.1_Planctomycetaceae_bacterium                     PP-----PPPTKHYATNDSTSGLNLLGRIEQWGISTTSVSNVKLTITQMSGKNLDLLLHVLPSGVKYELELDTEE---
MDR2561287.1_Holophagales_bacterium                          ASAVTTTGPATTVHKECPATSPLNLLGNMETWGVMNSITEKITLSISCATGAQLKEIIKKLPEDSSCELKLYIRAKD-
MEA3344886.1_Chloroflexota_bacterium                         EGEAVEGVPVSFTIS----LTPLGL-----QELGPQDTVREVAIVVDELDKATLEKLIAGLPEGTAIRLKLKVVRRG-
MEA3401058.1_Armatimonadota_bacterium                        ARAEQQQLPQPKTERNWVHLPKGAMVTAVVQEINKTDLLRNVNITLQKSDRDGLVALLNALPDDESMQVSLSLERMER
MEA3428074.1_Thermodesulfobacteriota_bacterium               TPPICGKDPCELTIRIPPQNSIGSLRQETAFKLQDEDIITRVTCKIFLFSKSQIEQHIESLPVISGADFSVDMEVEVI
MEB3759607.1_Desulfurococcales_archaeon                      LPPQKPFTPATISLE----PSGDDVQQKLTVFIEHKATVNELVFQLETVDVDTLRNLISRLSSKETGTVKVSITLGEP
NLX24424.1_Lentisphaerae_bacterium                           QTGETEKSPVPVTVETPFVHGMGALRQEVAIKLSDADIVKEMTFRIFYLGKSDVERVVESLPQFPGGEYKAELKILRP
NMX21982.1_ANME-1_cluster_archaeon_GoMg4                     -EEDEVVTPPSFKIE----STPWSLQGRLGNTLDESVDVGEVSVTMKRLKGKDVKGIVDNILEEHEDNLRMKLDSKVR
OCQ93485.1_Nostoc_sp._MBR_210                                TPVGNGSIPPVKSQRTHEQSTGINLSGQFEKWGLAATPLSTAKLEFTDMTVQQLKQVLQKLPPVLKAHLEVSYLEDEQ
RLF82338.1_Thermococci_archaeon                              EVKKKEKKPKIKTVD----DGNINAPIIVESEEIGESGFGHLEISISEIERWKLIKIIRKLSLKELGFEKITLRIERE
SFC23913.1_Halobiforma_haloterrestris                        GAGQIFTCPQDTTTRFPVMDDKPSLIDTLERELGIEWTIHTVSLSYTGLSRDALLDVLLRLDVPDDASFEVWMEVEKG
TEU12999.1_Anaerolineales_bacterium                          PSGQRFPAPPDPSLIHDVSDTVDQFLRDLRARLREKARIQSARFQVSPVSKTAFLNLIGQLEQIEGVLVSVEVSGDVP
WP_011944180.1_Thermotoga_petrophila                         ---TGDDKNENDELS----NSPRVILSQIEARTSEKDEFEEILVEYRFLSKQEIMKFVKQLPFQEDCEIKAKIILWRG
WP_012608229.1_Desulfurococcus_amylolyticus                  -----------VEIQ----DAKSGLLEVLIKYLNRRNSITSCRLVVQLYEKDEALQMVKSMGLSPGGVRLIESREQ--
WP_125670094.1_Candidatus_Methanodesulfokores_washburnensis  KVPEIRRTPAQWTIETSEYQTPFRLSEEMERRIPESVKIQQIKIGFSKMDKKEAIKLIEKLPASLGGIIKASIEVKKV
WP_125672294.1_Candidatus_Methanodesulfokores_washburnensis  ELGKKEEIPPEIDIK----INGEKYRTPVQLKFERGEKVIVDSLEFDGRSVSVVANKIKNIPPEEISISTFKLEMSAR
WP_259101344.1_Candidatus_Fervidibacter_sacchari             TNGERYPEPPDQVLEISVTNAKDLWLGELENKLDEDAQVERVSCEIQCAGKEELLRWIESLPEGMSFVVSTKIEGKRP
WP_260594733.1_Salinirubellus_salinus                        TGTEVTDGPADSTVTFPAMDNKFSLVDKLERRMGADWEVHHLTLKIEALSKRRVLDIVEDVSVPRVATLSIRLEVMKH
WP_283022678.1_Bradyrhizobium_sp._CB1717                     TP-----APASRPRAVESPTSSLNLVGQLETWQVKATRVKNVTVTISDATGAQVEALLKKLPDGMVYGLQLDAEDDN-
WP_286167649.1_Ochrobactrum_sp._3-3                          TPMDSGETPSGRTTLSNPATSPLNLIGKLEGWGIGATPVKEVMLKVPAATGAQLKELLKKLPDGMTFELSLDKEDG--
consensus/100%                                               ......................h...................h.........h......h.................
consensus/95%                                                ........s............s...h......b......h..h.h.h.......h...h..l.........h.......
consensus/90%                                                ........s............s...h...h..b......l.ph.h.h..hp...lbp.h.pls......hph.......
consensus/85%                                                ........P............s...h...h..b......l.ph.hph..hs...lbphlppls......hphphp....
consensus/80%                                                .s......P............s...l..ph..b......l.phphph..hs..plbphlppLs......hphphp....
consensus/75%                                                .s......Pss.p.p....ps...Lb.pl..b....s.l.phplphp.hs..plbphlppLP....s.hphphp....
consensus/70%                                                .s......Pss.p.p....ps..pLb.pl..b..pssplpphplphp.hs..plbpllppLPs...sphphphp....
```

### Type-3 BR-system DUF499-ATPase C-terminal RRM/Ferredoxin Fold Representative_5

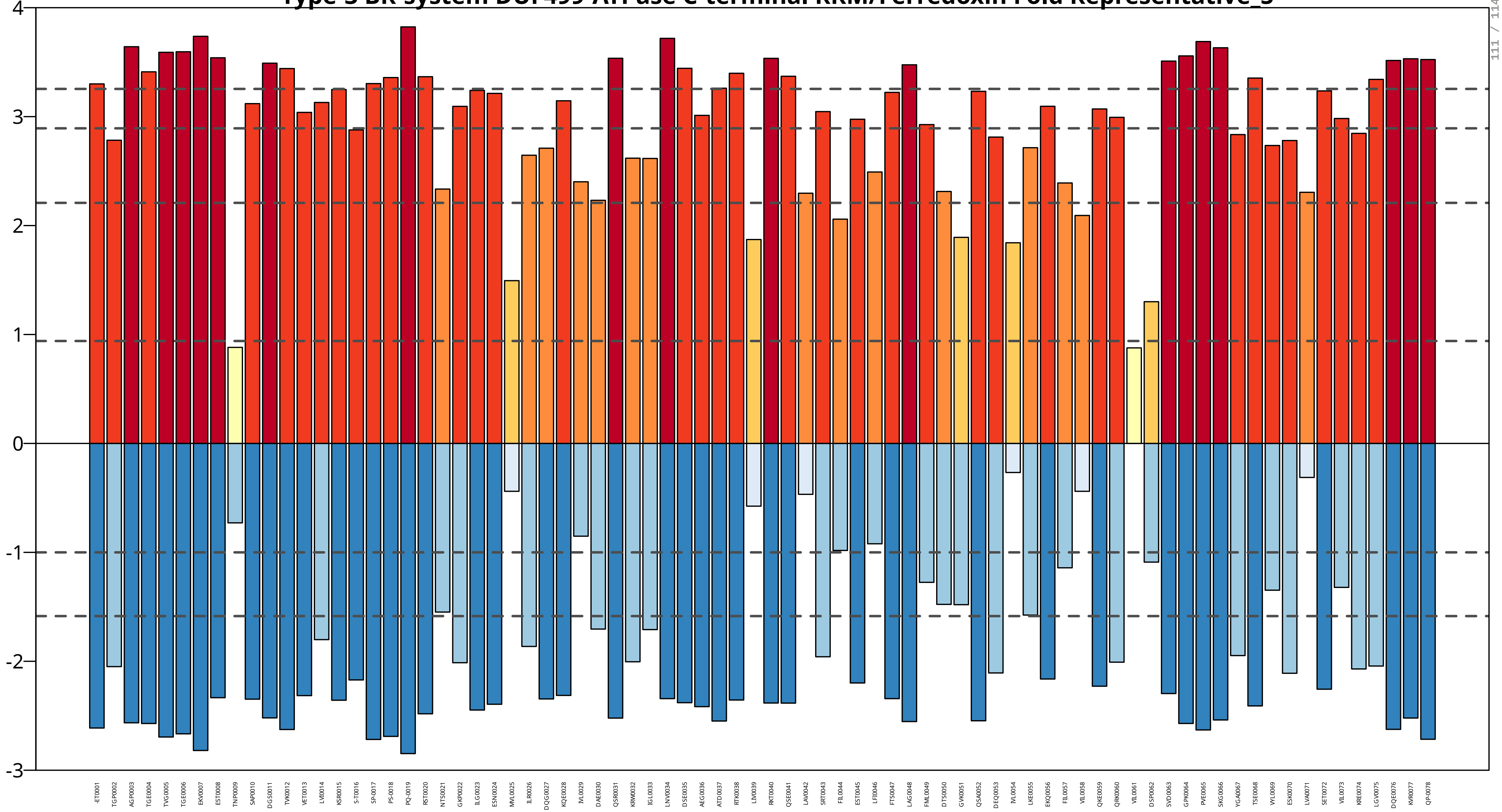

#### Type-2 BR-systems

### ; DUF499_2 C-Terminal Fn-III + Fn-III

| | |
|---|---|
| AFM14204.1_Turneriella_parva_DSM_21527 | AAKTDVIINQVLRDEETGAVKLRVEPQHGDKVYVDYGADPTTASELIHHGNFETARMEVRFLCVDSTGKHPQGPVRIWKNTIALKYDTTPRDGKVFVTLRAIPDADIRYTTDGSAPRELGGEYTGVFEVKKKCT-VMAIATKEGIESQILSFEADPTASKGARI |
| MBX3108928.1_Fimbriimonadaceae_bacterium | QPQTEVRIQQVYRDSETGEATLKVTALHGDQVFVEVGGPATPGSRKLDSNDYKTKEMEVSFLAVDTTKVHEAGPAQTWKNTVTLKHRFFQDGEVMRCEVKAAPGGQIKFTTDGSNPTQNGGTYTEPFIVPPGSKFIQAVGTNGSVQSEVLRVSVPA-EPEKLVI |
| AGC68793.1_Thermoclostridium_stercorarium | KAAVTVD-SDVEYDFENDEIVLKLNVKNADTVYYNPGTTIDVTAKNVDWDEFRLKPGKFTFVAMDSSGANIDGDPYVFTCPIKLRKAEEADQGLKHIKVKSLPEADIYYTTDGSSPVTSDTLLEGEIIVPAQVQLLLIVAQNDKGYSNVLEINTKM----GLVI |
| EAZ58601.1_Pseudomonas_aeruginosa_2192 | AEKTAVNVTVHAANEATGETTLALIPRNAPVVHYAPTAKVSTSDPKVELDNFTTDAATLYFLAVDSEGKHETGEPQRWVAELRVRHQVQAIADKRMVTLSCMPAADMRYTLDGSNPKD-GELYGDPFEVGPQAARLMVYACSGEATRTA-DFQIPAANDKQVQI |
| MBK7550235.1_Syntrophaceae_bacterium | KPATSVRVDVLSRDDQTGQATLRIRPLNGDTVYMETKGVATVSSKKIEEYDIKTTELKLSFLCVDSKGEHATGEAVMWTNSLSLKHRFFQKGAKRMCELKAVPAGRIRYTTDGSGVETKGIAYEKPFEIPADCRVILVVAEETGIPPQKASIPAPQGVDVAPTI |
| MBK8067117.1_Rhodanobacteraceae_bacterium | -KQAGVQVTPVGKMQDDGSTVLQIGIINATSVHYAEDGEVTEASPKLNDDKLTTKALRVAFLAVDHSGRTQKGAPYIWPNKLMVQWEIAPGQGKRVVTLKVLPRDVLRYTLNGAEPRN-GIGYTAPFTVGPEAAILLVFAEASGLETKA-DFRIPQAGGAEPEP |
| MBR2257284.1_Blautia_sp. | KEPTEVSVEQLSYDEGTGEFTLKVRGVYGNTVYYDVGAEPTEASAIVP-TPFITRETDLWFLCVDQSDPHPTGDAKHFMCKVPLRAEQREAAVGTVMQLATHPGFEVRYTTDGSNPKESGGKYSGEFVIPKGCKYVRTAVYNHGELVEEKDIPITEQSSKKAQI |
| MBS0652531.1_Verrucomicrobia_bacterium | -KKPTVRITQVAPLRDDGTYELEVVAVHAPEIYYSEHPNVPKQRTQLNGQRFSTQKLRLYFLAVDPTGEAGEGAPVEHKGSIKITAMPPKVQGKRLVELKALPYGLLKYTLDGSDPKLRGSSVDGALDIGSEAVTLLVYAEADGLESHWKEM-YPSLGNGETVL |
| MBX3024939.1_bacterium | KPKTSVILDERAYDEATGEATLEVTVRDAPRIYWDTTPNVTARSHRLTDSKLVTDATCVWVVAEDPAGGHERGDPVLWRNRLTITHQPREGVGSRSVELHVVPRGTIRYTLNGANPAE-GTVYDGPIEIGDGETTIWCHAEHEGVTERR-TFTIPPAGNRGIRI |
| MCA3288087.1_Roseomonas_sp. | KARTTVSVSRIAYDSNSGEARIEVTARDAPVIRWSDTGPALPSSPELKELSFATSKLRLWFLAEDSTGDHPTGDAVLWRNEITILHEPRETSGGRVVELKAVPAGSLRWTLGGTSPRDMGAAYVGAVPIPLSGGLLRVLAEDGDISAEN-SFDFAPKGVEEKYV |
| MCF7997122.1_Chromatiaceae_bacterium | KPATQVKVQELSRDNDSGRVKLRATPINGDTIYVEVGAAATPASHKLSGRDYETDELEVSFLAVDSTSEHDTGDPLAWKNRITLQSRVYDAGGDKMIELRAAPPVPIRYTTDGSDPKVSGGTYDDAFVVPEGTRLVLMVAEDKGVASEVLQVPVNWDPNVEEPI |
| MCK9217630.1_Firmicutes_bacterium | APETSLTVITVYEDEDIGEVTLKLIPENGDTIYYEIGQTATKASLKVETNNFKTKELKLSFLCVDSRGEHEEGPAVEWSKNVKVKYKPYDKEGKQYIELKATADVKILYSTDGSNPRESGGIYEDDFIVPVGAKFVQAIAEKLEVFSEPLQIEI---KEREFEI |
| MCL6686250.1_Akkermansia_muciniphila | -RRASLRWTREYGPDDSGLVRLAVDPVDGPEIHYEEDGAVTEESPCLTSDRLETRALRVEFLVKDPTGKFETGDPQCWRNELRLRVNRRPGTGAELLLAPARPDIPIRYTLDGSNPRD-GLLYEGPLEVGAGEVRIEAFAEHEGLETRE-SFTLPAESDPMAVL |
| MCP4350268.1_Desulfobacterales_bacterium | KEKTDVLIRELHRNTETGEVILMITPRYGDRVLYETGSAATPDSPPAELNKFRTSELKLSFLCIDTKGEHPAGEPKEWCNKIAIKYRVYDKEDDKVLELKSVPDIPIRYTVDGSNPKDYGVPYDGELKVPSDVTHVQAVAEEQDVYSEVLTVRIDRTKGATVGI |
| MCX6251131.1_Bacteroidetes_bacterium | KPPTKVTVQQIERNPDTGKVKLRIKPVHGDKILYQYGKGVTEQCPEWDQNIMETDAMDVEFLCIDSKNEHEKGKPVLWTNEITLKYGWIPQGDETMMEINVAPKAQVKYTTDGSEPLLNGGIYLSPFLVP--MTLVSIIGVTDSFHSEIKYVKVPA-KPTKFEI |
| MCZ8170923.1_Brevundimonas_sp. | -PQPLIRVTRIGGEEATGESELELAVSNAPEVLVAPTRDGLDAGEVITVGTYKTTEVELWFQARNPE-TKEVSDPYRWSGSITITHERRDNAGMWQVTLAARPEAELRWNTTGINPKD-GTVYEGPIEIGRQKTTLYVYAVKGGVSAQR-SFTFDA-VGARRTI |
| MDA7889828.1_Akkermansiaceae_bacterium | KDKTSIRWTE-NRDPETQEVILNLVPEFGDTIHYEEGGVATESSPKIDKRTHKTTAMEISFLVVDSKGEHKMGDPMPWTNKLVIKHRVLENGAKRKCELKAYPDAVISYTTDNMDPAKSGSTYSAPFELPDTCKIVQAIATLGSHKTATPEINIAE-KGKKFEI |
| MDD2765364.1_Opitutaceae_bacterium | TPQAQVRNAIVSADEKRGLCRVSVQPVNGDSVVWEREALPGDSSPVWKGDAAAHDWLWAGFLGRDAANPAAPKGEVSWNPNVSIKHDFKKQPNTVLVLEAAPPRIRIFYTFDGSAPTNAQAAYDGPAEVPADCRAILVVGEFDGPEGVL-RTEVQKFPGNPLPP |
| MDD5268641.1_Methylococcales_bacterium | KPETQVNVQLLSRNPDTGEVNLKITPVNGNKVLYELGHSPSSASMDVAYSRFSTSDMRVTFACIDSRGEHDQGSVYVWENTLQLKHEVIQQGDKWRVELKAVPGAEIRYTTDGSDPSSLGASYDSVFEVPASSRFVQAIATKDGIQSRLEKIDMDQYRDKKVVI |
| NCB71009.1_Clostridia_bacterium | QPQTDVGVHEVARTDD--TVILRISPLNGDTVYYEVGQPATTASMKINLNGFRTNELELHFLCIDSIGEHETGTEKIWRNITNCRYNTYDNHGDKMLELKSEAPLTIRYTLDGSDPRSSAATYVAAVPLPKGKVFVQAITSKYDFWGNALGVWIVNKPDTEIVI |
| ORC33988.1_Marispirochaeta_aestuarii | KDPTGVKVIVKNRDEHTGETILQITPEGGRVVYYEIGKKPSSASLQVTYQNFRTRELSLTFLCVDDSDPRPTGEPLYWKNQVTIKGQFYQQGDEIMFKALSIPPIPLKYTTDGSNPLSHGVPYNGDFLVPNGAQLIQVLAQKDDIQAKE-TFTVK--SATGPVV |
| PYG00798.1_Thioalkalivibrio_sp._ALE21 | KEQTSVSVTPKSTDEATGELILQLVPKHSPRVHYCTDEGVSEQDPVADLEEFRTTEPTLYFLAVDPDGNHETGDATLWKAPIKIRHEVHTKPDHRVVELQSTPKAALYYTTDGSGPRE-GFEYTGPFPIEDGRVLVQVYAKAGASEQTE-KFTIQPSNDGRPEI |
| RDE52523.1_Candidatus_Accumulibacter | -EITGVSI-EYARDKDSGVITTTINLVHADKLFVREDAAEYRVHP---DQALVSGAMLIEFKAVDSSGKNKEGKPYRIENSIDLAHDFVPPAGHQVVKVKVVPPSTLLYSLDGSNPANNGKPYAKPGIEAPEGSTVRLHAAKGGVVRDI-SITVPK-GGTGPAI |
| RQH12683.1_Bradyrhizobium_sp._RP6 | AAKTSVKVLTRNYDDQTGVATIELTATDAGKIHFAATSDVSAKSPIVPDLITDRDETVLWFVAVDPDGKHETGEPVKWTNTLTLTYEPKEVMGKRSVALTVKPRGNIRWNTDGTNARE-GKPYTGPIPIGNDEVKIYAYAEDAGVETQK-IFTIRPVKGGEVQI |
| TPG55477.1_Roseomonas_nepalensis | VDKTRVSVTPESRSLD-GEVRLSVVALDAPRIHFAKGTTVSTSDPEMTELSFSTTDLRIAFLAVDPKGVHPTGEPYVWRNEITIQHTVSPSADGKRVELFARPSATIRYTLNGTLARE-GTVYDGSFLVTGGSVLLRAIAIDGDLEGEE-TFHIPMTEGTDMFV |
| WP_005488051.1_Halanaerobium_saccharolyticum | KKETNVNITKMSTVSENGISTLRITPVNADEVYYEIGQEPTKASKKVEHNSFEATDLVYYFLAVDPDGVNETGDPVRWENDINLQYKELTIKGKDALKLQATPSCEIRYTTDGASPKENGGVYQEPIIIPKNAKYIQAVAEEHDITSDVLQYEV---KGKEVTV |
| WP_012692622.1_Deinococcus_deserti | LEPTSVTVRHDILDAAADRYRLTVTPKYGTQVHYAEGSTVDIAAPVMTDASMEVTGQTFTFLAVDPSKGHPTGQSFTWQAPTFLKGVWESG----EVRFEASPDVEIRVTFDGSAPE-KGQIYEPPITPPPGTHVVQAVATRGGVNSTPITL-----KPDVIPL |
| WP_018962942.1_Coprothermobacter_platensis | KIPEILSYDIAEYDFENETVTIRLYTRDADAIYWTFGNRLEISDANRELTELTINPENNTQITIIPAFGETLGDPTTINIPIKVKQTEKTDNGNIHVQLKVLPKDKLVYEYGSSTETFTASEATSELVVPTEASQVQVKAPQGEEYLPLTTVAVDK------HV |
| WP_045169275.1_Caldicellulosiruptor_morganii | KDKTTVRVQFVSADFESGEVVLKLIPMFGDVIHYEEGDEVTENSPVVQINEFRTKSAKLKFLCIDSTGYHPKGEIVKYECPIMLDDNIKYDEGKVHIKVRTAPQATVKYTTDGTNPRYNGKAENGDIVIPDDAKVIIVVAEKDNVFSELKHIPVKKDVVKTVEL |
| WP_085411740.1_Fibrella_sp._ES10-3-2-2 | LPQTDLRFQQVARDTDTGEVILKLQPVHGDEVYYEIGAVATTASARVALKNFVTTELEISFLCHDSTGEHETGPAVTWNNQITLKYDLYDKGGQKMLELKAAPNALIRYTTDGSNPLTTGGTYTAPLAVPPATRLVLAIAEKKGIQSNALNIDIDWKAPNTVTL |
| WP_090663037.1_Belnapia_rosea | PEPTEVLVTERAPQDDRGRVVLSVTAKRGDSVHWEVGSPPTAASAKVENGELATDEMRVTFIAVDSTGQVPTGNHFTWENRLRVKSETPYRDGTFRLRLKAVPESKLRYTFDGSDPKLGGGAYDGETPVPEGTRVILAVAENSGIYSRVERIDLPDAGDDGPAV |
| WP_119720856.1_Methanococcus_maripaludis | EPKTAVSLN-VKSELNNGEVILRVVPQYGEKVYYDIDSDPTTASSEVEINSFKTDELEVRFLCVDPTGKHETGDINIWRNIPKIKYDLISKGNKKQLELFALPKLEIKYTTDGSNPKEHGAVYVEPVLLPENCDTVCAIAKKYNLTSEKLEIKIPKNSGPKVKI |
| WP_171833827.1_Lichenicola_cladoniae | PDRTGVICRTRTRDETTGKTSLEVKAVHGDRVHWAPTSSVSAASPIVTGEVLDSDKVSLWFLCVDTKGHHETGDPVHWTGKIEVKIDPQSGATTRRMALSSIPPAAIRFTMDDSEPASSGTPYTGPIEIGDGGCTFAAIAEREGVRSEKVTYRFPPRGAEAPEI |
| WP_190610329.1_Synechococcus_sp._FACHB-909 | LPTPSIDIRELSRDDDDGRTYLQLVPLHCDSIVFESGEEPTTASSPVPTAKFEAKGLQYKFLAFDSDNPERQSEVKTWTATLRLKKQLHDRGDHYEVELLAFPRVSIHYTTNGSSPITNAAVYDGPIRVPDSCRKVCAVAPAYSLTSQGIVQDIPKRGEEARTI |
| WP_231170937.1_Bradyrhizobium | VTSVTARVEPAGDAHVRGTYILSLSSEDADTIYVSETGVPDARSEKVSGRQYETAAATVWFMAVDSTGKAATGPAYPWYAPIDVKPTLNSSSSGMKLSVSVMPRAVVLATFDGS-PAREGAEITTETKVPPGTEMVRVIGNVDGRYGQEISIVVQRPVPAKPEL |
| WP_274435612.1_Alicyclobacillus_sp._ALC3 | -KTTEVIIGEDSSPDDSGTVRLKIDVANAPRIHYAEDSEVSESSPVLSDNTLVTKALRVQFLAVDPTGKNLTGAPKTWKNRLTLRNRFDEV--SRTVELFVAPKGSIKYTLDGSEARN-GTDYAGPIQLGDDETTVYVFAECDGLEEKR-NFTFAKSGSKEVPI |
| WP_277247933.1_Phascolarctobacterium | VPKTGLRIVETSRNGE--DVTLKLVPENGDVIRWEIEQDPTSASPIVKLGAFRTNEMSLSFICEDSTGKHEVGEVKTWTNKVEVQYNFFDDNGIKRCELEATNSVKIMYSTNGSNPR-NGAVYNGSFVVPKDASILLAVSEPLDLYGAELRAAIPAFPEPEIVI |
| consensus/100% | .....h.................h....s..l.h............................s....................h............h...s.s...h.hp.ss......................h...s........................ |
| consensus/95% | ...s.l...............l.l....u..l.h........................h.h.s.s.......u....h.....l............h.h.s.s...l.hs.ss..s...s..h............l.h.s........................ |
| consensus/90% | ...s.l.h.............lpl.s..us.l.h.....s...s..h....h..p...h.Fhs.Ds......G....a.s.l.lp.......s...h.h.s.P...lbao.sGs.s...u..h...h.ls.....l.hhu...........h.h.........l |
| consensus/85% | ...s.l.h.....s..ps...Lpl.s..us.l.h..s..ss..s..h....h.sp..ph.Fls.Dsps.p..G.s..aps.l.lp.p.....s...hpl.s.P...lbao.sGo.P...G..Y.s.h.ls.....lbhhu...s...p..ph.h....s....l |
| consensus/80% | ...s.lplp....s.psGph.Lpl.s.puspl.a..s..ss.su..hp...h.op.hpl.FlshDsps.p.pG.s..Wps.lplp.p.....s.b.hpl.s.P.s.lbYT.sGosP...G..Ypssh.ls.....lbhhAp..sh.sp..ph.h....s....l |
| consensus/75% | ...TpVplp....s.psGph.Lpl.s.pusplaap.s..so.su..lp..phpTp.hpl.FlslDspu.p.pGps..Wpsplplp.p.....s.b.lplps.P.s.l+YT.DGSsP.p.Gs.Ysssh.ls.ssp.lbhhApp.sh.sp..ph.h....s....l |
| consensus/70% | ..pTsVplp..s.sppsGph.Lplss.pusplaap.sssso.uS..lp.pphpTs.hpl.FLslDssG.c.pGps..Wpsplplc.p.bp.sspb.lcLpu.P.spl+YT.DGSsPbp.GssYsush.lssssp.lbhhApp.sl.sp..phpls...s....l |

### Type-2 BR-system Type 2 DUF499 systems; DUF499-ATPase C-terminal FnIII+FnIII

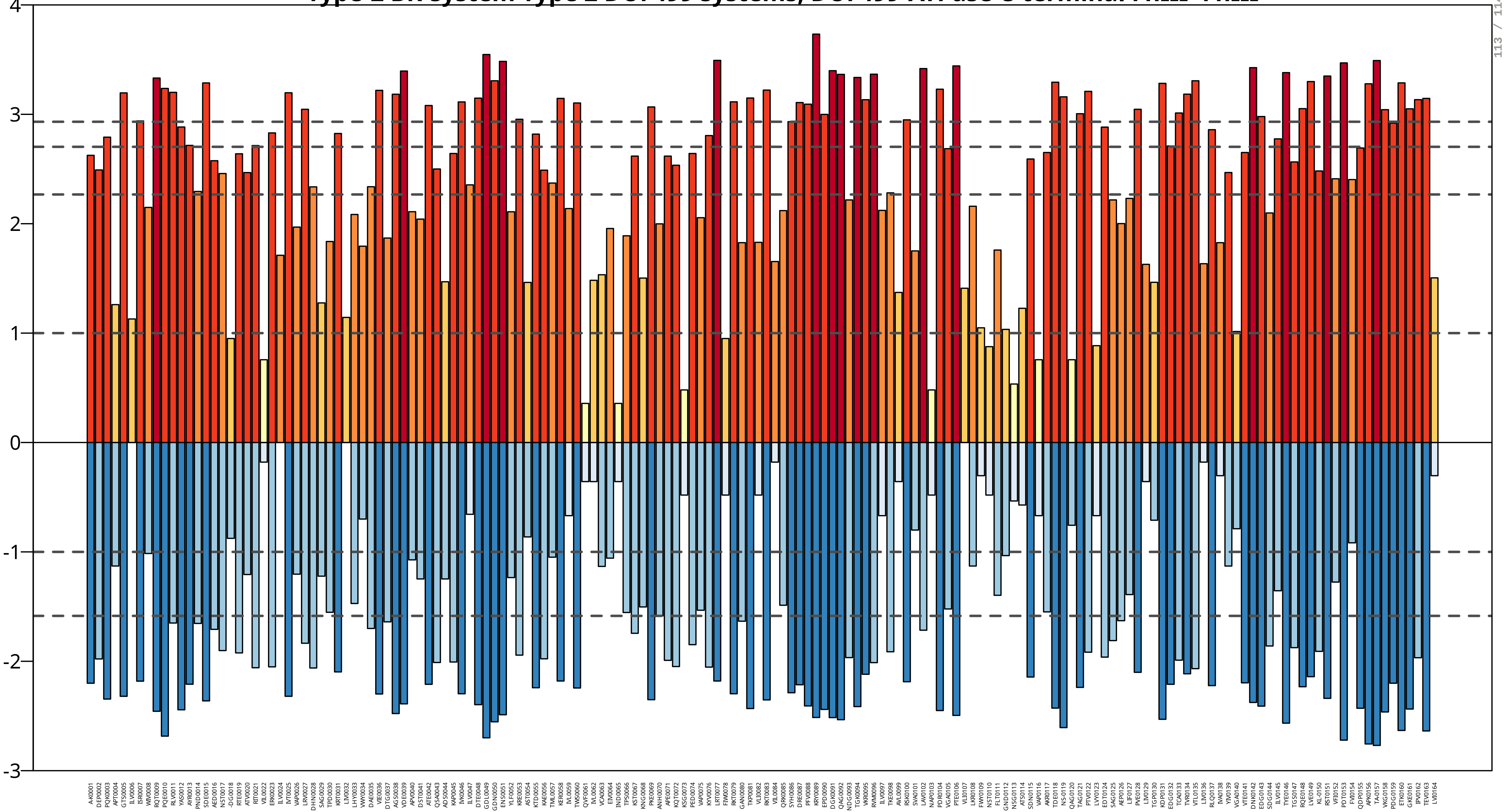

| SYSTEMS-->> | Type 1 BREX | Type 2 BREX/Pgl-systems | Type 3 BREX | Type 4 BREX | Type-1 BR-system | Type-2 BR-system | Type-3 BR-system |
|---|---|---|---|---|---|---|---|
| **Protein Components** | | | | | | | |
| **Core-ATPase** | ✔ | ✔ | ✔ | ✔ | ✔ | ✔ | ✔ |
| Designated-name | **BrxC-ATPase (DUF6079)** | **BrxC/PglY-ATPase (DUF6079)** | **BrxC-ATPase (DUF6079)** | **BrxC-ATPase (DUF6079)** | **DUF499 ATPase (DUF499)** | --- | --- |
| Domain architecture | STAND/ORC-Cdc6-AAA+ATPase + wHTH + α+β-Domain + wHTH + Coiled-Coil | STAND/ORC-Cdc6-AAA+ATPase + wHTH + α+β-Domain + wHTH + Coiled-Coil | STAND/ORC-Cdc6-AAA+ATPase + wHTH + α+β-Domain + wHTH + Coiled-Coil | STAND/ORC-Cdc6-AAA+ATPase + wHTH + α+β-Domain + wHTH + Coiled-Coil | STAND/ORC-Cdc6-AAA+ATPase + wHTH + α+β-Domain + wHTH + RRM<br>or<br>STAND/ORC-Cdc6-AAA+ATPase + wHTH + α+β-Domain + 3wHTH + RRM<br>or<br>HEPN + STAND/ORC-Cdc6-AAA+ATPase + wHTH + α+β-Domain + wHTH + RRM | STAND/ORC-Cdc6-AAA+ATPase + wHTH + α+β-Domain + wHTH + RRM<br>or<br>STAND/ORC-Cdc6-AAA+ATPase + wHTH + α+β-Domain + 2wHTH + RRM<br>or<br>STAND/ORC-Cdc6-AAA+ATPase + wHTH + α+β-Domain + wHTH + Fn-III+Fn-III + RRM | STAND/ORC-Cdc6-AAA+ATPase + wHTH + α+β-Domain + wHTH + RRM |
| Type-specific Synapomorphy | **AAA+ ATPase:**<br>Canonical features - Walker A and Walker B residues, Sensor-1 atop strand-4, Arginine-Finger at the base of strand-5, Sensor-2-Arginine at distal end of helical-bundle, elaborated HEH-insert region, C-terminal helical bundle.<br>**Conserved canonical wHTH:**<br>**α+β-Domain:**<br>Average length 185 residues;<br>Topology - (αH1-αH2 bi-helical connector)-βS1-βS2-βS3-αH3-βS4-(αH4-αH5 long helices forming bundled extension with bi-helical connector)-βS5-βS6-αH6.<br>Structurally distinguished into an α-helical region (formed by the bi-helical connector and αH4-αH5 helix extension), and an α/β/α globular region comprising of a six stranded central β-sheet;<br>Central β-sheet is flanked by two β-hairpins on either end, βS1-βS2 (N-terminal hairpin) and βS5-βS6 (C-terminal hairpin).<br>**wHTH:**<br>Average length 118 residues; Elaborated helical bundle and three stranded wHTH.<br>**Coiled-coil**:<br>Extended coiled-coil helices with an average length of 297 residues. | **AAA+ ATPase:**<br>Canonical features same as Type-1 BREX.<br>**Conserved canonical wHTH:**<br>**α+β-Domain:**<br>Average length 237 residues; Structural and topological features are same as Type-1 BREX;<br>Sequence profile is markedly distinct and relatively conserved within this subtype (primarily due to lesser phyletic distribution of Type-2 BREX);<br>However, like the α+β domain for every other BREX subtype, the sequence profile of α+β domain in Type-2 BREX systems is also unique, with no similarities identified across other variants of α+β domain;<br>**wHTH:**<br>*Elaborated into two closely packed three-stranded wHTH;*<br>wHTH1 average length is 57 residues, wHTH2 average length is 79 residues.<br>**Coiled-coil:**<br>Extended coiled-coil helices with an average length of 241 residues. | **AAA+ ATPase:**<br>Canonical features same as Type-1 BREX, HEH-insert region consists of usual 2-helix "helix-extension-helix" element.<br>**Conserved canonical wHTH:**<br>**α+β-Domain:**<br>Average length 209 residues; Structural and topological features are same as Type-1 BREX;<br>H3 after core S4 is either degenerate or not predicted in AF3 models;<br>**wHTH:**<br>Single wHTH domain, elaborated into a four-stranded wHTH domain; Average length is 137 residues.<br>**Coiled-coil:**<br>Extended coiled-coil helices with an average length of 395 residues.<br>*Features a Type-3 specific 3C-H type of Zn-finger motif highlighted in the reference alignments and AF3 models.* | **AAA+ ATPase:**<br>Canonical features same as Type-1 BREX.<br>**Conserved canonical wHTH:**<br>**α+β-Domain:**<br>Average length 196 residues: Structural and topological features are same as Type-1 BREX; Absence of first βS1 hairpin;<br>**wHTH:**<br>Elaborated solitary three-stranded wHTH with an extended helical section, averaging over 150 residues in length.<br>**Coiled-coil:**<br>Extended coiled-coil helices.<br>Two distinct versions of coiled-coil observed within Type-4 BREX systems;<br>One is averaging 279 residues, other is markedly extended, averaging 497 residues in length. | **AAA+ ATPase:**<br>Canonical features same as Type-1 BREX. HEH-insert region consists of usual 2-helix "helix-insert-helix" element. N-terminal;<br>HEPN fusion occuring in a subset.<br>**Conserved canonical wHTH:**<br>**α+β-Domain:**<br>Average length 184 residues; Structural and topological features are same as Type-1 BREX;<br>Absence of first βS1 hairpin;<br>A conserved motif consisting of a glycine, followed by a highly conserved stretch of [Rxx+N] (+ denotes positively charged residues).<br>*The Rxx+N motif (particularly the conserved N) is a synapomorphy specifically found in DUF499 associated α+β-Domains;* Conserved alanine in H4.<br>**wHTH:**<br>Single three-stranded canonical wHTH, typified by a usual three-helical triangular core, with an average length of 103 residues;<br>Expanded into three distinct three-stranded wHTH, with a total average length of 268 residues, in a subset of Type-3 DUF499-ATPase.<br>**RRM:**<br>Degenerated RRM domain with hypervariable sequence profile; Average length of 103 residues;<br>*AF3 structure prediction is inconsistent due to divergent and degenerate sequence features.* | **AAA+ ATPase:**<br>Canonical features same as Type-1 BREX. HEH-insert region consists of usual 2-helix "helix-insert-helix" element.<br>**Conserved canonical wHTH:**<br>**α+β-Domain:** Average length of 197 residues;<br>Structural and topological features are same as Type-1 BREX;<br>Absence of first βS1 hairpin;<br>Similar sequence profile to the Type-1 DUF499 α+β-Domain.<br>**wHTH:**<br>Four-stranded wHTH with an average length of 92 residues;<br>In a subset, the 4S-wHTH is followed by another 3S-wHTH, separated by a short loop region;<br>*Features an additional downstream β-sandwich (Fn-III associated repeat) in a subset of Type-2 DUF499-ATPase.*<br>**RRM:**<br>*Structurally complete RRM domain with hypervariable sequence profile, maintaning the characteristic β-α-β-β-α-β core topology;* Average length of 143 residues. | **AAA+ ATPase:**<br>Canonical features same as Type-1 BREX;<br>HEH-insert region consists of usual 2-helix "helix-insert-helix" element.<br>**Conserved canonical wHTH:**<br>**α+β-Domain:**<br>Average length 170 residues;<br>Structural and topological features are same as Type-1 BREX;<br>Absence of first βS1 hairpin;<br>Sequence profile is significantly more divergent relative to the Type-1 and Type-2 BR systems;<br>The Rxxx+N motif is divergent, characterized by the loss of first Arginine, and the terminal "+N" is usually represented by a conserved [RN];<br>**wHTH:**<br>Single three-stranded canonical wHTH with an average length of 107 residues, typified by a usual three-helical triangular core.<br>**RRM:**<br>Structurally complete RRM domain with hypervariable sequence profile;<br>Relatively smaller with average length of 96 residues;<br>Maintains the characteristic β-α-β-β-α-β core topology. |
| **Methylase** | ✔ | ✔ | ✔ | --- | ✔ | ✔ | ✔ |
| Designated-name | **BrxX/PglX-MTase** | **BrxX/PglX-MTase** | **PglXI-MTase** | --- | **MTase** | --- | --- |
| Domain architecture | NTD(Dimerisation) + N6-MTase + TRD + wHTH + CC-like helical ext | NTD(Dimerisation) + N6-MTase + TRD + CC-like helical ext | N-terminal ext + N6-MTase(TRD-Insert) + HTH1 + wHTH + HTH2 | --- | DUF1156 + N6-MTase(TRD-Insert) + HTH1 + wHTH + HTH2 | DUF1156 + N6-MTase(TRD-Insert) + HTH1 + wHTH + HTH2 | DUF1156 + N6-MTase(TRD-Insert) + HTH1 + wHTH + HTH2 |
| Type-specific Synapomorphy | **NTD (Dimerisation)**:<br>Average length 206 residues;<br>Conserved stretch in αH3 represented by [ExhhT] and [NRhh] in the middle of the helix, followed by a conserved [EsR] in the terminal end of the helix;<br>A conserved proline situated in the interior region of the core structure;<br>Terminal helices consists a stretch of conserved motifs represented by a dyad of negatively charged residues [DE], followed by a distinct motif [GWIYQ], which is shared with the Type-2 BREX-MTase NTD (dimerisation) domains.<br>**N6-MTase:**<br>Central β-sheet comprise of nine β-strands;<br>Two additional anti-parallel β-strands are situated C-terminally to the βS6-βS7 MTase hairpin;<br>Large insert region after core βS3, forming 5 packed helices protruding outside from the core;<br>Core catalytic motifs for SAM and substrate binding includes:<br>(i) glycine rich loop after βS1, represented by conserved [DP] on top of βS1, followed by [CGxGHF] in the loop region;<br>(ii) Conserved [DID] on top of βS2 and<br>(iii) [NPP(Y/F)] motif on top of βS4.<br>**TRD:**<br>*Separate domain following the N6-MTase*, with an average length of 273 amino acid residues;<br>Conserved residues includes a glycine on top of βS1, followed by a threonine, following which the βS2 includes a conserved stretch of residues represented by [RFIRhhWE], extending throughout the β-strand. Top of βS3 includes a glycine rich loop represented by [KGG];<br>**C-terminal end:**<br>Three distinct region at the C-terminal end, following the TRD-domain;<br>(i) Two extended helical region with an average length of 91 residues, termed as double-helical spacer, suggested to mimic the coiled-coil spacers found in the specificity subunit (HsdS) of M.EcoKI;<br>(ii) *A three-stranded wHTH with an average length of 155 residues, restricted to only Type-1 BREX MTases;*<br>(iii) A C-terminal CC-like helical extension with an average length of 134 residues, and a sequence profile with multiple conserved residues specific to Type-1 BREX MTases. | **NTD (Dimerisation):**<br>Average length 197 residues; Conserved stretch of amino acid residues in αH3, containing a glutamine, followed by a stretch of hydrophobic residues, ends with a conserved motif of [RhhEDN];<br>Terminal helices consists of a conserved stretch containing the motif [TRFL], immediatedly followed by a conserved [GDLYQ] comparable to that of Type-1 BREX-MTase NTD.<br>**N6-MTase:**<br>Sequence and structural profile is similar to the Type-1 BREX core N6-MTase;<br>Central β-sheet comprising nine β-strands with two additional anti-parallel β-strands, situated C-terminally to the βS6-βS7 MTase hairpin;<br>Relatively smaller insert region after core βS3, consisting of a β-hairpin and two small α-helices, leading into the core βS4; Core catalytic motifs for SAM and substrate binding are similar to those found in Type-1 BREX N6-MTases.<br>**TRD:**<br>*Separate domain following N6-MTase*, with an average length of 263 residues;<br>Structurally similar to the TRD of Type-1 BREX MTase with markedly distinct sequence profile;<br>Begins N-terminally with a glycine rich loop; Conserved glycine at the top of βS1;<br>Conserved dyad of negatively charged residues [D/E] at the base of βS2;<br>Conserved glycine followed by a downstream conserved motif [RDW] in the loop connecting βS3 and βS4;<br>Conserved motif [HNH] at the base of βS6;<br>Conserved stretch of polar residues [NSS] buried in the structural core, followed by a motif [FWhKQ] and [KG] in the downstream α-helix.<br>**C-terminal end:**<br>*Absence of the double helical spacer and the downstream three-stranded wHTH found in Type-1 BREX MTases;*<br>Features a C-terminal CC-like helical extension with an average length of 150 residues. | **N-terminal preceding extension:**<br>Average length 111 resdiues;<br>Structurally consists of a small incosistently occuring β-hairpin in its N-termini, followed by two fairly conserved helices and a β-strand lodged in the rightmost end of the central β-sheet of MTase rossmann fold;<br>Overall sequence profile is fairly conserved with multiple conserved stretches of amino acid residues, which are not shared with any other MTase NTD across all BREX subtypes.<br>**N6-MTase:**<br>Distinct from Type-1 and Type-2 BREX MTases;<br>Central β-sheet comprises eight β-strands, with one additional β-strands occuring from the N-terminal preceding extension, lodged in the rightmost end of the central β-sheet;<br>*Two distinct insert region, one small inconstently retained insert region occuring immediately downstream of the glycine rich loop, whereas the second insert region occurs after core βS2, forming a large TRD insert;*<br>Core catalytic motifs for SAM binding and substrate binding are retained in their usual structural positions, including the<br>(i) Conserved [DG] on top of βS1, followed by glycine rich loop [CGxGMTG] after βS1;<br>(ii) Conserved D on top of βS2 and<br>(iii) [DPPF] motif on top of βS4.<br>**Insert TRD:**<br>Large insert region after core βS2 with an average length of 254 residues;<br>Structurally composed of α-helices at the N-terminal and C-terminal end, whereas the β-sheet elements in the middle of the insert-TRD is protruding outside, potentially forming the specific DNA-binding interfaces;<br>4C Zinc finger at the apex of the insert-TRD;<br>A conserved motif R[N/T/Q] positioned atop the 4th α-helix; and a conserved negatively charged aspartate in the terminal β-sheet of the insert TRD.<br>**C-terminal end:**<br>*Consist of three distinct region, each encoding a DNA binding HTH-like domain,* sequentially occuring as HTH1, a canonical wHTH and the C-terminal most HTH2;<br>Structurally homologous domains also occur in the C-terminal end of DUF499 associated MTases;<br>However, the seqeunce profile in Type-3 BREX MTase, and in each subtype of DUF499 associated MTases are unique with no significant conservation of residues retained across these multiple subtypes. | --- | **N-terminal DUF1156:**<br>Average length 101 residues; *Distinct from the N-terminal extension of Type-3 BREX MTase, the N-terminal all α-helical bundle in DUF499 associated MTases, is positioned in the anterior end of the core MTase, organised in four tightly packed α-helices;*<br>Conserved residues include two aspartate in the N-termini, followed by a motif [HxWWARR], where H, W, and R are positioned internally towards the interface of the core MTase and insert-TRD domain;<br>These conserved residues are retained across all DUF499 associated MTases in structurally superimposable regions.<br>**N6-MTase:**<br>Similar to the Type-3 BREX MTase;<br>Central β-sheet comprises seven β-strands;<br>The small insert region found in Type-3 BREX MTases, are absent in DUF499 associated MTases;<br>The core βS2 and the downstream helix leads to the large insert-TRD;<br>Core catalytic motifs for SAM binding and substrate binding are retained in their usual structural positions, including the<br>(i) Conserved [DP] on top of βS1, followed by glycine rich loop [sGGG] after βS1;<br>(ii) Conserved D on top of βS2 and<br>(iii) [DPPY] motif on top of βS4.<br>**Insert TRD:**<br>*Large insert region after core βS2 with an average length of 318 residues;*<br>*Structurally homologous to the insert-TRD of Type-3 BREX MTase;*<br>Also retains the 4C Zinc finger at the apex of the TRD, the conserved motif R[N/T/Q] atop the 4th α-helix, and a conserved negatively charged aspartate in the terminal β-sheet of the insert TRD.<br>**C-terminal end:**<br>Consist of three distinct globular region, each encoding a DNA binding HTH-like domain, sequentially occuring as HTH1, a canonical wHTH and the C-terminal most HTH2. | **N-terminal DUF1156:**<br>Average length 150 residues; extended α-helical bundle in a subset of representatives;<br>Sequence and structural profile same as Type-1 DUF499 associated MTases.<br>**N6-MTase:**<br>Similar to Type-1 DUF499 associated MTase, sharing all major sequence and structural synapomorphies.<br>**Insert TRD:**<br>Large insert region after core βS2; Structurally similar to the insert-TRD found in other DUF499 associated MTases and Type-3 BREX MTase;<br>Average length of 324 residues;<br>The 4C Zinc finger is inconsistely retained;<br>The conserved motif R[N/T/Q] atop the 4th α-helix is retained, however the aspartic acid in the terminal β-sheet is inconsistently retained.<br>**C-terminal end:**<br>Structurally and topologically similar to Type-3 BREX and Type-1 DUF499 associated MTases, with a distinctive sequence profile. | **N-terminal DUF1156:**<br>Average length 85 residues; Sequence and structural profile same as Type-1 DUF499 associated MTases.<br>**N6-MTase:**<br>Similar to Type-1 DUF499 associated MTase, sharing all major sequence and structural synapomorphies.<br>**Insert TRD:**<br>Large insert region after core βS2; Structurally similar to the insert-TRD found in other DUF499 associated MTases and Type-3 BREX MTase;<br>*Relatively more extended β-sheet elements, increasing the average length of the insert-TRD to 433 residues;*<br>*Often features two or three 4C Zinc finger compared to the single 4C zinc finger found in other subtypes;*<br>The conserved motif R[N/T/Q] atop the 4th α-helix; and a conserved negatively charged aspartate in the terminal β-sheet of the insert TRD, are both consistently retained.<br>**C-terminal end:**<br>Structurally and topologically similar to Type-3 BREX, Type-1 DUF499, and Type-2 DUF499 associated MTases, with a distinctive sequence profile. |
| **PglZ component** | ✔ | ✔ | ✔ | ✔ | --- | --- | ✔ |
| Designated-name | **BrxZ/PglZ-Phosphatase** | **PglZ-Phosphatase** | **PglZ-Phosphatase** | **PglZ-Phosphatase** | --- | --- | --- |
| Domain architecture | iSwi2/Snf2-Helicase + 3HTH + helical-connector + PglZ-Phosphatase + IG-like | iSwi2/Snf2-Helicase + 3HTH + helical-connector + PglZ-Phosphatase + 3S-wHTH | iSwi2/Snf2-Helicase + 3HTH + helical-connector + PglZ-Phosphatase<br>or<br>iSwi2/Snf2-Helicase + HTH + helical-connector + PglZ-Phosphatase | iSTAND + iSwi2/Snf2-Helicase + 3HTH + helical-connector + PglZ-Phosphatase + IG-like<br>or<br>iSwi2/Snf2-Helicase + 3HTH + helical-connector + PglZ-Phosphatase + IG-like | --- | --- | PglZ-Phosphatase |
| Type-specific Synapomorphy | **iSwi2/Snf2-Helicase:**<br>*Small truncated inactive P-loop domain of the Swi2/Snf2-Helicases;*<br>Average length of 101 residues, typically represented by 4 short core-strands;<br>Highly variable at the sequence level with very few conserved residues, including a conserved stretch of hydrophobic residues in βS1, ending with the conserved D on top of βS1;<br>Conserved hydrophobic stretch in βS3, across all variants of Type-1 BREX PglZ.<br>**3HTH-like:**<br>Three structurally conserved HTH-like units arranged tandemly;<br>Highly divergent sequence profile with no conservation of residues;<br>Combined average length of 167 residues.<br>**Helical-connector:**<br>Five to six extended α-helix forming the connector region; Average length of 159 residues;<br>Conserved residues include an ariginine and tryptophan in αH2; tyrosine, aspartate and arginine in αH4;<br>Most conserved residues are pointing inwards towards the structural core, potentially forming stabilizing interactions.<br>**Phosphatase:**<br>Core topology in comparison with other nucleotide pyrophosphatase superfamily of enzymes - [Preceding(αH)-βS1-αH1-βS2-αH2-(β-hairpin *insert-1*)-αH3-βS3-αH4-βS4-αH5-αH6-βS5-(*Insert-Region-2*)-βS6].<br>Structurally includes a six-stranded central β-sheet of the core-domain and two distinct insert regions, insert region-1 forming a β-hairpin and insert region-2 with an elaborated cap-like subdomain;<br>Insert-region-2 forms a large elaborate cap-like subdomain consisting of six β-strands, βS1 to βS6, where βS2-βS3-βS4-βS5 forms a four stranded anti-parallel β-sheet, and βS1 and βS6 are arranged anti-parallely and often predicted as a loop region in AF3 predicted models;<br>Conservation of residues involved in coordination of $Mg^{2+}$ and $Zn^{+}$ ions includes;<br>(i) Aspartate on top of βS1, (ii) threonine at the base of αH2, (iii) aspartate in αH5,<br>(iv) Conserved motif [DHG] on top of βS5, and (v) conserved histidine in the loop preceding βS6. | **iSwi2/Snf2-Helicase:**<br>Average length of 102 residues;<br>Structurally and topologically similar to Type-1 BREX PglZ iSwi2/Snf2-helicases, with an additional small insert-strand placed in-between βS1 and βS2;<br>Highly divergent sequence profile distinguished from those of Type-1 BREX PglZ iSwi2/Snf2-helicases;<br>Conserved motifs unique to Type-2 occurs in the beginning of αH1, represented by [SxLhhR] and conserved strech in βS3 represented by [VIIT] (I-aliphatic amino acid).<br>**3HTH-like:**<br>Three structurally conserved HTH-like units arranged tandemly;<br>Highly divergent sequence profile with no significant conservation of sequences other than few sporadically conserved hydrophobic residues;<br>Combined average length of 222 residues.<br>**Helical-connector:**<br>Five extended α-helix forming the connector region; Average length of 135 residues;<br>Conserved residues include an arginine in αH1, histidine in αH2, alanine and arginine in [AhRL] motif in αH3, aspartate and alanine in [DbA] motif in αH4, arginine followed by [FA] in αH5;<br>Like Type-1 BREX PglZ, most conserved residues are pointing inwards towards the structural core.<br>**Phosphatase:**<br>Similar to Type-1 BREX PglZ phosphatase domains with minimal variation in secondary structural elements; Core topology - [Preceding region(βS-αH)-βS1-αH1-βS2-(insert βS)-αH2-αH3-βS3-αH4-βS4-αH5-αH6-βS5-(Insert-Region-2)-βS6-terminal strand];<br>Structurally retains the the six-stranded central β-sheet with a small additional terminal β-strand lodged beside the left-most core βS2;<br>Insert β-hairpin found after αH2 in Type-1 variants are absent;<br>Cap-like sub-domain is retained;<br>Catalytic residues involved in metal ion coordination are conserved.<br>**3S-wHTH:** Canonical wHTH, with an average length of 105 residues;<br>*Only in Type-2 BREX system's PglZ component, the C-terminal regions harbors a wHTH domain, instead of the IG-like domains found in other BREX subtypes;* | **iSwi2/Snf2-Helicase:** Average length of 100 residues;<br>Structurally and topologically similar to Type-1 BREX PglZ iSwi2/Snf2-helicases;<br>Divergent sequence profile distinguished from those of Type-1 and Type-2 PglZ iSwi2/Snf2-helicases;<br>Conserved motifs includes (i) Aspartate on top of βS1, similar to Type-1 PglZ;<br>(ii) Conserved motf [RhxaE] (a-aromatic amino acid) in αH2 and<br>(iii) conserved aspartate in αH3.<br>**3HTH-like:**<br>Three structurally conserved HTH-like units arranged tandemly;<br>Highly divergent sequence profile with no significant conservation of sequences apart from few sporadically conserved hydrophobic residues;<br>Combined average length of 154 residues;<br>*In a subset of Type-3 PglZ, second and third HTH-like elements are lost, retaining only the N-terminal HTH1-like unit, before leading to the helical connector.*<br>**Helical-connector:**<br>*Relatively shorter version of the helical connector made up of three helices.*<br>Average length of 78 residues;<br>Conserved residues include a tryptophan in αH2, and [FppW] motif in αH3, all of which are pointing inwards towards the structural core, potentially forming stabilizing interactions.<br>**Phosphatase:**<br>Similar to Type-1 BREX PglZ phosphatase domains;<br>Core topology: Same as Type-1 and Type-2 BREX PglZ Phosphatase;<br>Structurally retains the six-stranded central β-sheet βS2;<br>Insert β-hairpin found after αH2 in Type-1 variants are absent;<br>Cap-like sub-domain is retained;<br>Catalytic residues involved in metal ion coordination are conserved;<br><br>*Type-3 BREX PglZ is the only version across all BREX subtypes which does not feature any additional domain fusion towards the C-terminal end of the core phosphatase domain, making it the smallest version among all BREX subtypes.* | **iSwi2/Snf2-Helicase:**<br>Three distinct sub-variants of iSwi2/SNF2-Helicase are observed within Type-4 BREX PglZ N-termini, with extreme diversity in their sequence profile and no conservation of residues across the sub-variants;<br>Average length varies from 78 residues to 116 residues across these variants;<br>Most variants exhibit further reduction in its core structural elements typified by the loss of αH1 and αH2 usually found in other subtypes;<br>**iSTAND:**<br>*A subset of Type-4 PglZ, contains an iSTAND domain fused N-terminally to the iSwi2/Snf2-helicase;*<br>Features the typical iSTAND topology with the loss of walker A, walker B and C-terminal helical bundle.<br>**3HTH-like:**<br>Three HTH-like units arranged tandemly;<br>Divergent sequence profile with no significant conservation of sequences other than few sporadically conserved hydrophobic residues;<br>Combined average length of 164 residues;<br>*Occasionally, third HTH-like element is lost.*<br>**Helical-connector:**<br>Highly divergent helical connector, ranging from three to four helix extension;<br>No significant conservation of residues, except few stretches of tyrosine;<br>Average length of 61 residues;<br>In a subset of these, the region is highly elaborated to form eight helices.<br>**Phosphatase:**<br>Two distinct version of phosphatase domains are observed in Type-4 BREX PglZ;<br>*In the canonical versions of domain architecture with no N-terminal iSTAND, the structural and topological features are similar to other variants* found in Type-1, Type-2 and Type-3 BREX systems;<br>*The other version with an iSTAND fusion in its N-termini, is typified by the loss of βS3-αH4, occuring in the rightmost end of the core structure, thereby reducing the central β-sheet to a five-stranded sheet instead of the usual six-stranded β-sheet;*<br>Cap-like subdomain is retained;<br>Catalytic residues involved in metal ion coordination are conserved. | --- | --- | **Phosphatase:**<br>*Standalone PglZ Phosphatase domain with no additional domain fusions in its N-termini or C-termini;*<br>Core topology is similar to those of BREX systems;<br>Structurally retains the the six-stranded central β-sheet with an additional four-helix bundle in the N-terminal of the core domain;<br>Cap-like sub-domain is retained;<br>Catalytic residues involved in metal ion coordination are conserved. |
| **Inactive STAND** | ✔ | --- | ✔ | --- | --- | --- | ✔ |
| Designated-name | **BrxB** | --- | **BrxF** | --- | --- | --- | --- |
| Domain architecture | Inactive STAND-NTPase | --- | Inactive STAND-NTPase | --- | --- | --- | Inactive STAND-NTPase |
| Type-specific Synapomorphy | Average length 186 residues; Absence of walker A, walker B and C-terminal helical bundle;<br>Central β-sheet consists of the usual five core-strands found in AAA+ ATPases;<br>Retention of STAND/ORC-Cdc6 specific HEH (Helix-extension-helix) insert region after core βS2;<br>The top of strand-4 consistently extends into a three-stranded antiparallel β-sheet, protruding outward to form a cap-like subdomain;<br>Conserved sequence elements includes a hydrophobic stretch ending with a tyrosine in βS1;<br>Conserved aspartate in the HEH insert region, conserved motif [GxG] on top of βS3;<br>Conserved motif [PhhR] in loop connecting βS3 and αH4;<br>*A stretch of conserved hydrophobic residues in βS4 which leads to a highly conserved proline-glycine dyad [PG] on top of βS4, leading into the cap-like subdomain;*<br>The conserved motifs of βS4 are also retained across Type-2 and Type-3 iSTAND components; | --- | Average length 147 residues; Absence of Walker A, Walker B and C-terminal helical bundle;<br>Structurally similar to Type-1 BREX iSTAND(BrxB);<br>HEH insert region is smaller consisting of 'one-helix insert';<br>Strand-4 leads to the three-stranded antiparallel β-sheet insert region, protruding outward to form a cap-like subdomain;<br>Sequence profile is highly divergent, with multiple versions observed (see supplementary alignments);<br>Conserved motifs include a stretch of polar residues (represented by [NhN] motif) in loop connecting βS2 and HEH insert region;<br>Hydrophobic residues in αH3;<br>*Conserved tryptophan followed by proline-glycine dyad [WPG] on top of βS4 is somewhat retained, similar to the Type-1 BrxB component;*<br>*In a very limited subset of BrxF(iSTAND) specific to Type-3 BREX systems, the walker A and walker B residues are intact* (represented in supplementary alignment). | --- | --- | --- | Average length 163 residues; Absence of Walker A, Walker B and C-terminal helical bundle;<br>Structurally similar to Type-1 BREX iSTAND(BrxB); Includes an additional terminal β-strand, inserted antiparallelly into the central five-stranded β-sheet, making the central sheet a six-stranded β-sheet;<br>Strand-4 leads to the three-stranded antiparallel β-sheet insert region, protruding outward to form a cap-like subdomain;<br>Overall sequence profile is markedly distinct from those of Type-2 and Type-3 BREX MTase, however, similar to Type-3 BREX iSTAND, the [WxG] motif on top of βS4 is retained; |
| **DNA-binding HTH** | ✔ | --- | ✔ | ✔ | --- | --- | --- |
| Designated-name | **BrxA** | --- | **BrxA** | **DUF4007** | --- | --- | --- |
| Domain architecture | HTH + wHTH1 (Version 1)<br>or<br>HTH + wHTH1 + wHTH2 (Version 2) | --- | HTH + wHTH1 + wHTH2 | PAPS_Reduct + HTH + wHTH1 + wHTH2 (Version 1; DUF4007)<br>or<br>HTH + wHTH1 + wHTH2 (Version 2; DUF4007) | --- | --- | --- |
| Type-specific Synapomorphy | Two distinct versions of domain architecture found in BrxA component within Type-1 BREX systems;<br>One version has a single wHTH domain (wHTH1), whereas the second version has two wHTH domains towards the C-termini (wHTH1 and wHTH2);<br>***Version 1:*** Average length of 199 residues; Topologically contains a preceding helix region, followed by HTH, a small helical spacer, a wHTH and a C-terminal helix region;<br>***Version 2:*** Average sequence length is 254 residues;<br>Topologically retains the preceding helix region, the small helical connector between HTH and wHTH1, and the C-terminal most helical extension; | --- | Average length 235 residues; Topologically contains a small preceding helix, followed by a three-helical HTH domain, a two-helical spacer region, wHTH1, and C-terminal wHTH2;<br>Absence of the terminal helical extension observed in Type-1 and Type-4 BrxA components;<br>The N-terminal HTH domain is clustered with HTH domain of Type-1 BrxA; | The BrxA component (denoted by the architechture of HTH+wHTH1+wHTH2, is often described as DUF4007 in the literature, because of the DUF4007 (UPF0758 entry) top hit, recovered from HHpred searches;<br>BrxA component found in Type-1 and Type-3 BREX systems, and DUF4007 component found in Type-4 BREX systems, both recover the BrxA PDB structure (7RZB) and DUF4007 profile as their top hits, suggesting the occurence of shared domains among these components;<br>Our analysis of domain architecture and sequence-structure features suggests that BrxA and DUF4007 represents homologous counterparts in various BREX subtypes;<br>Two distinct versions of domain architecture found in DUF4007 component within Type-4 BREX;<br>Version 1 is fused to an N-terminal PAPS reductase; Average length 248 residues;<br>Topology same as Type-1 BrxA;<br>Version 2 is standalone (HTH + wHTH1 + wHTH2) BrxA; | --- | --- | --- |
| **AAA+ Helicase** | ✔ | --- | --- | ✔ | --- | --- | --- |
| Designated-name | **BrxL** | --- | --- | **BrxL** | --- | --- | --- |
| Domain architecture | SIGMA-HTH + OB-Fold + MCM-family-AAA+ATPase + iLon-Protease<br>or<br>SIGMA-HTH + OB-Fold + MCM-family-AAA+ATPase + HKD-DNase | --- | --- | SIGMA-HTH + OB-Fold + MCM-family-AAA+ATPase | --- | --- | --- |
| Type-specific Synapomorphy | **SIGMA-HTH:**<br>Average length 67 residues;<br>Begins N-terminally with a small α-helix, followed by a tri-helical SIGMA-HTH like domain;<br>Topologically the SIGMA-HTH includes [Preceding αH-Insert β-strand-αH1-αH2-αH3]; The small insert β-strand together with a small β-strand present in the C-terminal helical bundle of the downstream MCM-AAA+ATPase, forms a small β-bridge element;<br>The sequence profile of SIGMA-HTH is highly conserved, including (i) a stretch of highly conserved positively charged residues [lysine and arginine], in the insert β-strand and following loop region;<br>(ii) Hydrophobic stretch of residues and a highly conserved negatively charged residues [D/E] in the middle of αH1;<br>**OB-Fold:**<br>[αH1-βS1-βS2-βS3-αH2-βS4-βS5]; Average length 93 residues;<br>Typical topology associated with OB-fold from small β-barrel assemblage of proteins;<br>αH1 is followed by βS1-βS2-βS3, together forming the β-meander sheet-A;<br>βS3 is followed by a helix and βS4-βS5, which forms the three stranded β-sheet B, along with the N-terminal half of βS1.<br>Overall sequence profile within Type-1 BrxL is highly conserved with multiple conserved, residues but is relatively distinctive from the OB-fold of Type-4 BrxL component.<br>**MCM-family-AAA+ATPase:**<br>Average length 329 residues; Structural and sequence features are typical to those of MCM-clade of AAA+ ATPases, including the clade-specific H2-insert in core αH2, and the PSI (Pre-sensor 1) insert region after core αH3;<br>C-terminal helical bundle terminates with a three-stranded wHTH-like configuration; The small β-strand occuring after core βS5 towardds the C-terminal end, pairs along with the insert βS from the N-terminal SIGMA-HTH to form a β-bridge.<br>**iLon-Protease:**<br>Average length 191 residues;<br>Structurally and topologically similar to canonical Lon-Proteases from E.coli;<br>*Absence of catalytic serine-lysine dyad residues, suggesting that it is enzymatically inactive;*<br>**HKD-DNase:**<br>In a subset of Type-1 BrxL, the C-terminal iLonP is replaced with an HKD-DNase component; Average length of BrxL HKD-DNase is 171 residues;<br>Structurally and topologically similar to the canonical versions of HKD-DNases belonging to PLD superfamily of endonucleases in RM systems;<br>*Retains a catalytic triad represented by [HRD], instead of the usual [HKD], found inType-I/Type-III-like RM system's endonucleases;* | --- | --- | The sequence and structural profile of all individual domains including the N-terminal SIGMA-HTH, the following OB-fold and the MCM-family AAA+ATPases, are similar to those found in Type-1 BrxL component;<br>*The key distinction lies in the C-terminal end, where unlike the Type-1 BrxL component, no additional domain fusions (inactive LonP or HKD-endoDNase) are retained, thus reducing the overall domain architecture to a three-domain containing BrxL component.* | --- | --- | --- |
| **PglW (NERD + Kinase)** | --- | ✔ | --- | --- | --- | --- | --- |
| Designated-name | --- | **PglW** | --- | --- | --- | --- | --- |
| Domain architecture | --- | NERD + iPkinase + Pkinase + HTH + wHTH1 + wHTH2 + wHTH3 + iSTAND(BrxB) | --- | --- | --- | --- | --- |
| Type-specific Synapomorphy | --- | **NERD:**<br>Average length 198 residues; Structurally and topologically similar to canonical PDDEXK superfamily of nuclease and related domain (NERD);<br>Retention of PDDEXK catalytic residues.<br>**iPkinase:**<br>Average length 289 residues; Belongs to Serine/Threonine/Tyrosine (S/T/Y) class of protein kinases;<br>Retains the typical structural and topological features of S/T/Y kinases;<br>Absence of the glycine rich loop between S1-S2, used for ATP localization;<br>Degeneration of the HRD motif of the catalytic loop and DFG motif of the Mg+ binding loop in βS6 and βS9 respectively;<br>No conserved D in Mg+ binding loop;<br>Lack of all catalytic motifs and loop features makes this S/T/Y kinase enzymatically inactive.<br>**Pkinase:**<br>Average length 278 residues; Belongs to the same S/T/Y class of protein kinases as the previous domain;<br>Retains the typical structural, topological and sequence features of S/T/Y kinases;<br>Glycine rich loop is conserved;<br>Catalytic loop with HRD motif and Mg+ binding loop with DFx motif are both conserved, followed by the activation loop;<br>This is a functional copy of S/T/Y kinase domain which is preceded by a enzymatically inactive copy of S/T/Y kinase.<br>**HTH + wHTH1 + wHTH2 + wHTH3:**<br>*The first HTH domain resembles the CTD of the RNA polymerase α-subunit, which is followed by three set of wHTH domains, together forming a large DNA-binding module similar to BrxA architecture, found in Type-1 and Type-4 BREX systems;*<br>**iSTAND:**<br>The C-terminal most region encodes an iSTAND, characterized by the absence of Walker A, Walker B, and the C-terminal helical-bundle; Average length of 192 residues;<br>Central β-sheet consists of the usual five core-strands found in AAA+ ATPases;<br>Retention of STAND/ORC-Cdc6 specific HEH (Helix-extension-helix) insert region after core βS2;<br>Conserved residues includes a Tryptophan and Aspartate in HEH insert region, a conserved arginine (not arginine finger) and tyrosine in αH3, a conserved stretch of hydrophobic residues in βS4.<br>*Although the sequence profile of majority of PglW C-terminal iSTAND is distinct from the iSTAND BrxB or BrxF components, found in other BREX systems, one subset of iSTAND from Type-2 PglW shares its sequence and structural features with those of BrxB/BrxF;*<br>*This includes stretch of hydrophobic residues and a highly conserved [PG] residues in βS4, which is followed by the three-stranded cap-like subdomain insert region.* | --- | --- | --- | --- | --- |
| **Helicase-I** | --- | ✔ | --- | --- | --- | --- | --- |
| Designated-name | --- | **BrxHI-Helicase** | --- | --- | --- | --- | --- |
| Domain architecture | --- | Ski2-like-RNA-Helicase + wHTH + Small-β-Barrel (HAS-barrel) + Lhr-Helicase-CTD | --- | --- | --- | --- | --- |
| Type-specific Synapomorphy | --- | **Ski2-like-RNA-Helicase:**<br>SF2-helicase with all typical sequence and structural features, including two tandem P-loop domains (an active P-loop, followed by an inactive P-loop);<br>*Belongs to the Ski2-like helicase family, within SF2-Helicase superfamily;*<br>**wHTH:**<br>Canonical wHTH domain with an average length of 89 residues;<br>**HAS-barrel:**<br>Small All-β protein domain, belonging to the small β-barrel assemblage of proteins, with structural and topological features identifying it as a homolog of HAS-barrel domains;<br>**Lhr-helicase-CTD:**<br>A C-terminal α+β domain with an average length of 169 residues;<br>No structural homologs detected, apart from the C-terminal domain found in Lhr-helicases; five-stranded central β-sheet flanked by α-helices<br>and a distinct three-helical bundle extending outward;<br>Conserved residues includes:<br>(i) an arginine at the base of αH2;<br>(ii) a conserved motif [WWTaAG] (a-aromatic) in the loop connecting βS2-αH3;<br>(iii) a conserved asparagine in the beginning of αH3;<br>(iv) Conserved dyad [KF] in the beginning of αH6;<br>(v) A conserved arginine or lysing at the end of αH6.<br>Only the conserved arginine in αH2(i), conserved dyad [KF](iv) and the conserved lysine or arginine(v) are pointing outwards from the structure, rest all are buried within the central β-sheet and surrounding helices of the core structure;<br>A conserved dyad of lysine/arginine residues [KR] in the CTD of Lhr-helicase has been shown to be involved in DNA-binding. | --- | --- | --- | --- | --- |
| **Standalone-ATPase** | --- | ✔ | --- | --- | --- | --- | --- |
| Designated-name | --- | **BrxD-ATPase** | --- | --- | --- | --- | --- |
| Domain architecture | --- | STAND/ORC-Cdc6-AAA+ATPase | --- | --- | --- | --- | --- |
| Type-specific Synapomorphy | --- | **STAND/ORC-Cdc6-AAA+ATPase:**<br>Standalone canonical STAND NTPase with an average length of 435 residues;<br>Canonical features - Walker A and Walker B residues, Sensor-1 atop strand-4, Arginine-Finger at the base of strand-5, Sensor-2-Arginine at distal end of helical-bundle, C-terminal helical bundle, are all retained;<br>Similar to the BrxC-STAND/ORC-Cdc6**,** the HEH insert region after core βS2 is elaborated forming four closely pacled α-helices. | --- | --- | --- | --- | --- |
| **Helicase-II** | --- | --- | ✔ | --- | ✔ | ✔ | ✔ |
| Designated-name | --- | --- | **BrxHII-Helicase** | --- | --- | --- | --- |
| Domain architecture | --- | --- | TUDOR + Swi2/Snf2-Helicase + iREase + CTD | --- | TUDOR + Swi2/Snf2-Helicase + iREase + CTD + REase(PDDEXK) | HKD-DNase + Swi2/Snf2-Helicase + iREase + CTD | TUDOR + Swi2/Snf2-Helicase + iREase + CTD |
| Type-specific Synapomorphy | --- | --- | **TUDOR:**<br>A Small-β-Barrel domain with homology to TUDOR domains;<br>**Swi2/SNF2-Helicase:**<br>Average length of 601 residues;<br>SF2-Helicase belonging to the Swi2/SNF2-helicase superfamily, with retention of all canonical features associated with these helicases, including two tandem P-loop domains (an active P-loop, followed by an inactive P-loop);<br>A large all-α helical insert region (average length 115 residues) occuring after the core βS2 of the 2nd inactive P-loop domain;<br>**iREase:**<br>Small degenerated α+β domain with an average length of 82 residues;<br>Topologically represented by [αH1-βS1-βS2-βS3-(small-degenerate βS)-αH2];<br>The central β-sheet consists of three-stranded β-sheet, and both αH1 and αH2 are located behind the central β-sheet;<br>The domain is structurally and sequentially homologous to the first small α+β domain of RapA;<br>Our comparative structural analysis reveals that both the first small α+β domain of RapA and BrxHII-Helicase adopts a small REase like fold;<br>*Structurally and topologically it is similar to various endonucleases related to the virus-type replication-repair (VRR) nuclease and holliday junction resolvases (HJC), which belongs to the PD-(D/E)-XK superfamily of nucleases;*<br>*Despite the retention of the PDDEXK-fold, the core β-sheet (particularly βS4) exhibits partial deterioration;*<br>βS4 in RapA is represented as a small degenerated β-strand, but in BrxHII-iREase, it takes the form of a loop section which leads directly to αH4 behind the central β-sheet;<br>The catalytic PD-(D/E)-XK motif is entirely degenerate, with no conservation of residues across all known representatives, and a fast-evolving sequence profile, suggesting that this REase module is potentially inactive.<br>**CTD α+β:**<br>An α+β domain with an average length of 166 residues;<br>Topologically represented by [βS1-βS2-βS3-αH1-βS4-αH2-αH3-βS4];<br>The five β-strands are arranged anti-parallely to form a five-stranded β-sheet;<br>Structural searches did not identify any true homologs beyond the C-terminal most α+β domain of RapA Swi2/Snf2-helicase, suggesting that this is unique to Type-2 helicases. | --- | Overall features of all individual domains are same as those found in Type-3 BrxHII-Helicase;<br>The sequence profile of the C-terminal α+β domain is distinct from other variants found in Type-3 BREX and Type-2, Type-3 BR system's helicase components;<br>*The key distinction lies in the occurence of a canonical REase domain in the C-terminal end of the helicase component, belonging to the PDDEXK superfamily;*<br>The REase features the typical sequence and structural features associated with PDDEXK REases, and also features the catalytic residues, suggesting that unlike the degenerated version of an iREase found in the preceding region, this REase is enzymatically active and can potentially faciliate DNA degradation restriction activity. | **HKD-DNase:**<br>N-terminal TUDOR domain is absent;<br>*Instead a enzymatically active restriction endonuclease from the HKD-superfamily (also known as Phospholipase-D), is found in the N-terminal region;*<br>Catalytic residues [HKD] intact, resembling the canonical versions found in Type-I/Type-III RM-like systems;<br>**Swi2/SNF2-Helicase:**<br>Overall sequence and structural features are similar to those found in Type-3 BREX BrxHII-Helicase and the helicase component of Type-1 DUF499 associated systems;<br>**CTD α+β:**<br>The C-terminal α+β domain is slightly shortened;<br>Sequence profile is highly divergent and distinctive from other versions of C-terminal α+β domains, found across all variants of this helicase component. | *Structurally and architecturally similar to the Type-3 BREX system's helicase BrxHII;*<br>Begins with a standalone N-terminal TUDOR domain, followed by a Swi2/SNF2-Helicase, a degenerated inactive REase, and a C-terminal α+β domain;<br>*Compared to the helicase component of Type-1 and Type-2 BR systems, this lacks an active endonuclease domain;*<br>The C-terminal α+β domain of Type-3 associated DUF499 systems are relatively smaller with an average length of 108 residues, primarily due to presence of a four antiparallel strands in the central b-sheet, and reduction in size of the coiled-coi |
