## Supplementary Data S4 for "Expanding the Landscape of BREX Diversity: Uncovering Multi-Layered Functional Frameworks and Identification of Novel BREX-Related Defense Systems"

*The PDF presents three distinct datasets organized as follows:*

**(i) Gene neighbourhoods of BREX and related systems containing auxiliary effectors from diverse superfamilies.** The file follows the format of Supplementary Data S1, with column-specific information marked at the top of each page. Within each neighbourhood, core components of a given BREX subtype are highlighted in light blue, while nearby auxiliary nuclease effectors are shown in distinct colors for clarity. Nucleases occurring as fusion components or in consistent genomic association with specific core proteins (as discussed in the manuscript) are also color-coded for visualization.

**(ii) Domain architectures and corresponding NCBI node information** for all identified nuclease superfamilies associated with BREX and related systems.

**(iii) Representative MSAs and domain boundary characterizations** for all auxiliary nuclease superfamilies linked to BREX and related systems. Alignments are color-coded and annotated as in Supplementary Data S2 and S3.


1. **Backup effector nucleases associated with Type-1/5/6 BREX**
   - (i) HEPN-endoRNase……4-8
   - (ii) HKD-DNase(BrxL-Fusion)……9-24
   - (iii) HKD-DNase+OGG-like……24-28
   - (iv) ParB+HNH(GmrSD)……29-39
   - (v) OLD-ABC associated HNH……39-40
   - (vi) wHTH(SBD)+HNH……40-41
   - (vii) HNH (Standalone)……41-42
   - (viii) PIN-EndoRNase……43-44
   - (ix) BrxHII-Helicase+REase (Type-5 BREX)……45
   - (x) CR-ATPase+REase……45-46
   - (xi) (w)HTH+REase/HARE-HTH+REase……46-49
   - (xii) SF2-Helicase+REase……49-51
   - (xiii) Standalone REase; OLD-ABC associated REase……51-57
   - (xiv) Schlafen+HATPase+wHTH……58-67
   - (xv) OLD-ABC associated TOPRIM……68-79
2. **Backup effector nucleases associated with Type-2 BREX**
   - (i) HEPN-endoRNase……80-81
   - (ii) SF2-Helicase+HKD-DNase……82
   - (iii) ParB+HNH (GmrSD)……83-84
   - (iv) OLD-ABC associated HNH and SAD(SRA)+HNH……84
   - (v) PglW component……85-97
   - (vi) PIN-EndoRNase……98
   - (vii) REase; SF2-Helicase+REase; REase+SF2-Helicase……99-100
   - (viii) Schlafen+HATPase+wHTH……101
   - (ix) OLD-ABC associated TOPRIM……102
3. **Backup effector nucleases associated with Type-3 BREX**
   - (i) HEPN-endoRNase……103-104
   - (ii) HKD-DNase……105
   - (iii) ParB+HNH(GmrSD)……106
   - (iv) HNH(standalone) and wHTH(SBD)+HNH……106-107
   - (v) PIN-EndoRNase……108
   - (vi) BrxHII-Helicase+REase……109
   - (vii) (w)HTH+REase……109-110
   - (viii) Standalone REase; OLD-ABC associated REase; SF1-SF2-Helicase+REase……110-111
   - (ix) Schlafen+HATPase+wHTH……112-115
   - (x) OLD-ABC associated TOPRIM……116-117
4. **Backup effector nucleases associated with Type-4 BREX**

(i) HEPN-endoRNase……118
(ii) ParB+HNH(GmrSD)……119
(iii) wHTH(SBD)+HNH……119
(iv) HNH (standalone)……119-120
(v) PIN-EndoRNase……121
(vi) Standalone REase; HTH+REase……122
(vii) CR-ATPase+REase……122
(viii) Schlafen+HATPase+wHTH……123
(ix) OLD-ABC associated TOPRIM……124
5. **Backup effector nucleases associated with Type-1 BREX-Related systems**
(i) HEPN+DUF499……125-146
(ii) Standalone HEPN; DNA-PolB associated HEPN……146-148
(iii) HKD-DNase……149
(iv) ParB+HNH(GmrSD)……150-152
(v) wHTH(SBD)+HNH……153
(vi) HNH standalone; OLD-ABC associated HNH……153-155
(vii) PIN-EndoRNase; OLD-ABC associated PIN……156-161
(viii) Additional PDDEXK-REase alongside primary REase effector; OLD-ABC associated REase……162-167
(ix) REase, SF2-Helicase+REase……167-170
(x) Schlafen+HATPase+wHTH……171-174
(xi) OLD-ABC associated TOPRIM……175-178
6. **Backup effector nucleases associated with Type-2 BREX-Related systems**
(i) HEPN-EndoRNase……179
(ii) ParB+HNH(GmrSD)……180
(iii) HNH; Standalone versions, OLD-ABC associated and wHTH(SBD)+HNH……181
(iv) PIN-EndoRNase……182-183
(v) (w)HTH+REase……184-186
(vi) REase……186-187
(vii) Schlafen+HATPase+wHTH……188-189
(viii) OLD-ABC associated TOPRIM……190
7. **Backup effector nucleases associated with Type-2 BREX-Related systems**
(i) HEPN-EndoRNase……191
(ii) HKD-DNase……192
(iii) ParB+HNH(GmrSD)……193
(iv) HNH……193
(v) PIN-EndoRNase……194
(vi) REase……195
(vii) Schlafen+HATPase+wHTH……196
(viii) OLD-ABC associated TOPRIM……197
8. **OLD-ABC associated nucleases other than TOPRIM (OLD ABC associated with HNH/REase/PIN)....198-207**
9. **Representative domain architectures and corresponding NCBI node information**
(i) HKD-DNase+OGG-like……208-210
(ii) HNH-standalone……211-216
(iii) ParB+HNH……216-231
(iv) wHTH(SBD)+HNH……232
(v) SAD(SRA)+wHTH(SBD)+HNH……233

(vi) PIN-EndoRNase.........234-242
(vii) CR-ATPase+REase.........243
(viii) (w)HTH+REase.........244-248
(ix) McrC-NTD+REase.........248
(x) REase+SF2-Helicase.........249-250
(xi) REase-standalone.........250-260
(xii) SF2-Helicase+REase.........261-263
(xiii) TUDOR+SF1-SF2-Helicase+iREase+CTD+REase.........263-264
(xiv) Schlafen+HATPase+wHTH.........265-278
(xv) OLD-ABC.........279-291
(xvi) OLD-ABC+TOPRIM.........291-294
(xv) TOPRIM.........294-306
10. Representative alignments with topology and domain architectures of backup nuclease effectors
(i) HEPN-EndoRNase.........307-309
(ii) CBS+HEPN.........309
(iii) HKD-DNase+OGG-like.........310
(iv) SIGMA-HTH+OB-Fold+MCM-AAA+HKD-DNase.........310
(v) ParB+HNH.........311-312
(vi) wHTH(SBD)+HNH.........312
(vii) SAD(SRA)+wHTH(SBD)+HNH.........312-313
(viii) Standalone HNH EndoDNase.........313
(ix) PIN-EndoRNase.........314-316
(x) Type-5 BREX and Type-3 BREX BrxHII-Helicase with C-terminal Active PDDEXK REase; (TUDOR+SF1-SF2-Helicase+iREase+CTD+REase).........317
(xi) REase+SF2-Helicase.........317
(xii) SF2-Helicase+REase.........318
(xiii) CR-ATPase+REase.........318-319
(xiv) (w)HTH+REase.........319-320
(xv) McrC-NTD+REase.........320
(xvi) Standalone PDDEXK-REase.........321-323
(xvii) Schlafen+HisKin+wHTH.........324
(xviii) OLD-ABC+TOPRIM (Previously classified OLD-ABCs in association with ABC-ATPase centric conflict systems).........325
(xix) OLD-ABC (TOPRIM-linked).........325-327
(xx) BR-systems OLD-ABC.........327
(xxi) OLD-ABC+TOPRIM.........327
(xxii) TOPRIM-nuclease.........328

*#Titles are internally hyperlinked.*
*Click on the titles to access the material.*

| Protein-ID | System | Gene-Neighbourhoods | Length | Taxonomy | Assembly-ID | Strain | Locus-Tag | Genomic-coordinates | TaxID | Fasta-defline |
|---|---|---|---|---|---|---|---|---|---|---|
| -- | | | | | | | | | | |
| --HEPN-EndoRNase | | | | | | | | | | |
| -- | | | | | | | | | | |
| WP_015789649.1 | Type1.BREX | <-REase<-?<-?\|\|DOC+zhth-><-BirA-HTH<-?\|\|?->BrxA-DNA-binding->DUF6079*->BrxB-BrxF-iSTAND->DUF6079-><-?<-?\|\|?-><-OLD-ABC+TOPRIM\|\|N6-METHYLASE->TBP+TBP->PglZ-> | 1236 | Archaea>Euryarchaeota>Stenosarchaea_group>Halobacteria>Halobacteriales>Haloarculaceae | GCF_000023945.1 | Halorhabdus_utahensis | HUTA_RS09330 | 1919731..1923441 | 146826 | hypothetical-protein-[Halorhabdus-utahensis] |
| WP_015789651.1 | Type1.BREX | DOC+zhth-><-BirA-HTH<-?\|\|?->BrxA-DNA-binding->DUF6079->BrxB-BrxF-iSTAND->DUF6079*-><-?<-?\|\|?-><-OLD-ABC+TOPRIM\|\|N6-METHYLASE->TBP+TBP->PglZ->HEPN->TUDOR+SF1-SF2-Helicase+iREase+CTD-> | 1280 | Archaea>Euryarchaeota>Stenosarchaea_group>Halobacteria>Halobacteriales>Haloarculaceae | GCF_000023945.1 | Halorhabdus_utahensis | HUTA_RS09340 | 1924016..1927858 | 146826 | hypothetical-protein-[Halorhabdus-utahensis] |
| WP_205252856.1 | Type1.BREX | <-SF1-SF2-Helicase+REase<-TUDOR+SF1-SF2-Helicase+iREase+CTD<-HEPN<-PglZ<-?<-N6-METHYLASE\|\|?->?-><-REase<-DUF6079*<-BrxB-BrxF-iSTAND<-DUF6079<-BrxA-DNA-binding\|\|ORC-CDC6->ORC-CDC6+HTH-><-?\|\|MarR-HTH-> | 1280 | Archaea>Euryarchaeota>Stenosarchaea_group>Halobacteria>Halobacteriales>Halobacteriaceae | GCF_016917835.2 | Halobacterium_sp._BOL4-2 | JRZ79_RS24035 | 52137..55979 | 2810537 | hypothetical-protein-[Halobacterium-sp.-BOL4-2] |
| WP_205252782.1 | Type1.BREX | <-HEPN<-PglZ<-?<-N6-METHYLASE\|\|?->?-><-REase<-DUF6079<-BrxB-BrxF-iSTAND<-DUF6079*<-BrxA-DNA-binding\|\|ORC-CDC6->ORC-CDC6+HTH-><-?\|\|MarR-HTH-><-?<-?\|\|HTH_28->NTP_transf_5-> | 1227 | Archaea>Euryarchaeota>Stenosarchaea_group>Halobacteria>Halobacteriales>Halobacteriaceae | GCF_016917835.2 | Halobacterium_sp._BOL4-2 | JRZ79_RS24045 | 56554..60237 | 2810537 | hypothetical-protein-[Halobacterium-sp.-BOL4-2] |
| WP_004047662.1 | Type5.BREX | Peptidase_M29-><-?<-BirA-HTH\|\|?-><-?<-ParA-Soj-PloopNTPase<-?\|\|?->BrxA-DNA-binding->DUF6079*->BrxB-BrxF-iSTAND->DUF6079->?->HEPN->N6-METHYLASE->PglZ->N6-METHYLASE->TUDOR+SF1-SF2-Helicase+iREase+CTD-> | 1249 | Archaea>Euryarchaeota>Stenosarchaea_group>Halobacteria>Haloferacales>Halorubraceae | GCF_000337915.1 | Halorubrum_saccharovorum | C471_RS07440 | 145399..149148 | 2248 | hypothetical-protein-[Halorubrum-saccharovorum] |
| WP_004047666.1 | Type5.BREX | <-BirA-HTH\|\|?-><-?<-ParA-Soj-PloopNTPase<-?\|\|?->BrxA-DNA-binding->DUF6079->BrxB-BrxF-iSTAND->DUF6079*->?->HEPN->N6-METHYLASE->PglZ->N6-METHYLASE->TUDOR+SF1-SF2-Helicase+iREase+CTD->?->?-><-DDE_Tnp_IS240 | 1276 | Archaea>Euryarchaeota>Stenosarchaea_group>Halobacteria>Haloferacales>Halorubraceae | GCF_000337915.1 | Halorubrum_saccharovorum | C471_RS07450 | 149762..153592 | 2248 | hypothetical-protein-[Halorubrum-saccharovorum] |
| WP_137705565.1 | Type1.BREX | <-ParA-Soj-PloopNTPase\|\|?-><-PADR-HTH<-?<-?<-?<-?\|\|BrxA-DNA-binding->DUF6079*->BrxB-BrxF-iSTAND->DUF6079->REase-><-?<-?\|\|N6-METHYLASE->?->PglZ->HEPN-> | 1227 | Archaea>Euryarchaeota>Stenosarchaea_group>Halobacteria>Haloferacales>Halorubraceae | GCF_005435115.1 | Halorubrum_sp._ARQ200 | EXE50_RS16210 | 7684..11367 | 1855872 | hypothetical-protein-[Halorubrum-sp.-ARQ200] |
| WP_137705567.1 | Type1.BREX | <-PADR-HTH<-?<-?<-?<-?\|\|BrxA-DNA-binding->DUF6079->BrxB-BrxF-iSTAND->DUF6079*->REase-><-?<-?\|\|N6-METHYLASE->?->PglZ->HEPN->TUDOR+SF1-SF2-Helicase+iREase+CTD-><-BirA-HTH | 1280 | Archaea>Euryarchaeota>Stenosarchaea_group>Halobacteria>Haloferacales>Halorubraceae | GCF_005435115.1 | Halorubrum_sp._ARQ200 | EXE50_RS16220 | 11942..15784 | 1855872 | hypothetical-protein-[Halorubrum-sp.-ARQ200] |
| WP_138006592.1 | Type5.BREX | <-TUDOR+SF1-SF2-Helicase+iREase+CTD<-PglZ<-N6-METHYLASE<-?<-HEPN<-Peptidase_M1<-DUF6079*<-BrxB-BrxF-iSTAND<-DUF6079<-BrxA-DNA-binding | 1272 | Archaea>Euryarchaeota>Stenosarchaea_group>Halobacteria>Haloferacales>Halorubraceae | GCF_005543295.1 | Halalkalirubrum_salinum | E5K03_RS09380 | 11807..15625 | 2563889 | hypothetical-protein-[Halalkalirubrum-salinum] |
| WP_138006594.1 | Type5.BREX | <-TUDOR+SF1-SF2-Helicase+iREase+CTD<-PglZ<-N6-METHYLASE<-?<-HEPN<-Peptidase_M1<-DUF6079<-BrxB-BrxF-iSTAND<-DUF6079*<-BrxA-DNA-binding | 1258 | Archaea>Euryarchaeota>Stenosarchaea_group>Halobacteria>Haloferacales>Halorubraceae | GCF_005543295.1 | Halalkalirubrum_salinum | E5K03_RS09390 | 16209..19985 | 2563889 | hypothetical-protein-[Halalkalirubrum-salinum] |
| WP_121744813.1 | Type6.BREX | Thermonuclease->?->?->wHTH+REase->SF1-SF2-Helicase->BrxE->BrxA-DNA-binding->BrxB-BrxF-iSTAND->DUF6079*-><-?\|\|N6-METHYLASE-><-HAD<-HTH+PADR-HTH\|\|?-><-?\|\|PglZ->SIGMA+OB-Fold+AAA-ATPase+Lon-Protease-BrxL->HEPN-> | 1206 | Archaea>Euryarchaeota>Stenosarchaea_group>Halobacteria>Natrialbales>Natrialbaceae | GCF_003670115.1 | Natronorubrum_halophilum | DWB23_RS21360 | 42652..46272 | 1702106 | BREX-system-P-loop-protein-BrxC-[Natronorubrum-halophilum] |
| MBU4220824.1 | Type1.BREX | PAIREDC-HTH->?->?->?->?-><-?\|\|BrxB-BrxF-iSTAND->DUF6079*->N6-METHYLASE->TIR->Schlafen-p->HEPN->Schlafen+HATPase+wHTH->?->?->PglZ-p-> | 1180 | Archaea>Euryarchaeota>unclassified_Euryarchaeota | GCA_018898145.1 | Euryarchaeota_archaeon | KKA10_04250 | 6252..9794 | 2026739 | BREX-system-P-loop-protein-BrxC-[Euryarchaeota-archaeon] |
| MBD0371455.1 | Type1.BREX | VirB11->PSBP->?-><-SIGMA+OB-Fold+AAA-ATPase+Lon-Protease-BrxL<-PglZ<-ParB+HNH<-NERD<-N6-METHYLASE<-HEPN<-DUF6079*<-BrxB-BrxF-iSTAND<-BrxA-DNA-binding | 1218 | Bacteria>Acidobacteriota>Blastocatellia>Blastocatellales>Pyrinomonadaceae>unclassified_Pyrinomonadaceae | GCA_014534425.1 | Pyrinomonadaceae_bacterium | ICV60_11505 | 29495..33151 | 2283092 | BREX-system-P-loop-protein-BrxC-[Pyrinomonadaceae-bacterium] |
| MCI0490100.1 | Type1.BREX | BrxA-DNA-binding->BrxB-BrxF-iSTAND->DUF6079*->HEPN->Four-helical-protein-><-?<-?\|\|N6-METHYLASE->Schlafen+HATPase+wHTH-p-> | 1218 | Bacteria>Acidobacteriota>Blastocatellia>unclassified_Blastocatellia | GCA_022615215.1 | Blastocatellia_bacterium | L0229_26200 | 2085..5741 | 2052146 | BREX-system-P-loop-protein-BrxC-[Blastocatellia-bacterium] |
| MBK9165938.1 | Type1.BREX | XIS-HTH-><-?<-SIGMA+OB-Fold+AAA-ATPase+Lon-Protease-BrxL<-PglZ<-PglZ<-PglZ<-?<-N6-METHYLASE<-HEPN<-DUF6079*<-BrxB-BrxF-iSTAND<-BrxA-DNA-binding<-CopG/RHH\|\|?-><-ParD_antitoxin<-HTH_37\|\|?->?->Tox-RelE-> | 1234 | Bacteria>Acidobacteriota>Terriglobia>Bryobacterales>unclassified_Bryobacterales | GCA_016716385.1 | Bryobacterales_bacterium | IPM24_00570 | 108049..111753 | 2026791 | BREX-system-P-loop-protein-BrxC-[Bryobacterales-bacterium] |
| OFW17305.1 | Type1.BREX | <-SF1-SF2-Helicase<-SIGMA+OB-Fold+AAA-ATPase+Lon-Protease-BrxL<-PglZ<-HATPase+wHTH<-HATPase_c_4<-Schlafen-p<-N6-METHYLASE<-HEPN<-PolB<-DUF6079*<-BrxB-BrxF-iSTAND<-BrxA-DNA-binding\|\|wHTH+AbiEi_2->AbiEii-><-?<-?<-PIN | 1156 | Bacteria>Acidobacteriota>unclassified_Acidobacteriota | GCA_001767195.1 | Acidobacteria_bacterium_RIFCSPLOWO2_02_FULL_67_21 | A3H29_10520 | 11544..15014 | 1797183 | hypothetical-protein-A3H29_10520-[Acidobacteria-bacterium-RIFCSPLOWO2_02_FULL_67_21] |
| OFW39743.1 | Type1.BREX | PIN->?->?-><-AbiEii<-wHTH+AbiEi_2\|\|BrxA-DNA-binding->BrxB-BrxF-iSTAND->DUF6079*->PolB->HEPN->N6-METHYLASE->Schlafen+HATPase+wHTH->Schlafen+HATPase+wHTH->PglZ->SIGMA+OB-Fold+AAA-ATPase+Lon-Protease-BrxL->SF1-SF2-Helicase->DDE_Tnp_1_4-> | 1156 | Bacteria>Acidobacteriota>unclassified_Acidobacteriota | GCA_001767515.1 | Acidobacteria_bacterium_RIFCSPLOWO2_12_FULL_67_14 | A3F70_01760 | 7438..10908 | 1797192 | hypothetical-protein-A3F70_01760-[Acidobacteria-bacterium-RIFCSPLOWO2_12_FULL_67_14] |
| MBS1828808.1 | Type1.BREX | D5N+D5ATPase-><-?\|\|ParB-><-?<-Phage_integrase+Phage_integrase<-?\|\|CopG/RHH->BrxA-DNA-binding->BrxB-BrxF-iSTAND->DUF6079*->HEPN->N6-METHYLASE->PglZ->SIGMA+OB-Fold+AAA-ATPase+Lon-Protease-BrxL->REase->HEPN->?->Phage_integrase-> | 1226 | Bacteria>Acidobacteriota>unclassified_Acidobacteriota | GCA_018268935.1 | Acidobacteriota_bacterium | JST93_26130 | 24024..27704 | 1978231 | BREX-system-P-loop-protein-BrxC-[Acidobacteriota-bacterium] |
| MCI0393103.1 | Type1.BREX | BrxA-DNA-binding->BrxB-BrxF-iSTAND->DUF6079*-><-?\|\|HEPN->N6-METHYLASE->Schlafen+HATPase+wHTH+HTH->PglZ->SIGMA+OB-Fold+AAA-ATPase+Lon-Protease-BrxL-><-MPTase<-HTH_3 | 1220 | Bacteria>Acidobacteriota>unclassified_Acidobacteriota | GCA_022614935.1 | Acidobacteriota_bacterium | MOB07_30595 | 1910..5572 | 1978231 | BREX-system-P-loop-protein-BrxC-[Acidobacteriota-bacterium] |
| MCI0698477.1 | Type1.BREX | <-Schlafen+HATPase+wHTH<-HEPN<-DUF6079*<-BrxB-BrxF-iSTAND<-BrxA-DNA-binding<-AbiEii+ZnR<-AbiEi_4+AbiEi_1 | 1215 | Bacteria>Bacteria_incertae_sedis>Bacteria_candidate_phyla>candidate_division_KSB1>unclassified_candidate_division_KSB1 | GCA_022616415.1 | candidate_division_KSB1_bacterium | L0337_41575 | 2047..5694 | 2172550 | BREX-system-P-loop-protein-BrxC-[candidate-division-KSB1-bacterium] |
| MBI2566519.1 | Type1.BREX | NtrC-AAA+FIS-HTH->?->?-><-?<-?\|\|BrxA-DNA-binding->BrxB-BrxF-iSTAND->DUF6079*->?->N6-METHYLASE->HEPN->ParA-Soj-PloopNTPase->?->PglZ->HNH-><-?<-NtrC-AAA+FIS-HTH | 1109 | Bacteria>Bacteria_incertae_sedis>Bacteria_candidate_phyla>Candidatus_Schekmanbacteria | GCA_016188575.1 | Candidatus_Schekmanbacteria_bacterium | HYV63_05760 | 102186..105515 | 2419984 | BREX-system-P-loop-protein-BrxC-[Candidatus-Schekmanbacteria-bacterium] |
| PKK90022.1 | Type1.BREX | <-Amidohydro_1<-?<-?\|\|BrxA-DNA-binding->BrxB-BrxF-iSTAND->DUF6079*->HEPN->N6-METHYLASE-p->?->HEPN-> | 1176 | Bacteria>Bacteria_incertae_sedis>Bacteria_candidate_phyla>Candidatus_Wallbacteria | GCA_002838975.1 | Candidatus_Wallbacteria_bacterium_HGW-Wallbacteria-1 | CVV64_11925 | 128140..131670 | 2013854 | BREX-system-P-loop-protein-BrxC-[Candidatus-Wallbacteria-bacterium-HGW-Wallbacteria-1] |
| WP_046997142.1 | Type1.BREX | wHTH+WYL+WCX->BrxA-DNA-binding->BrxB-BrxF-iSTAND->DUF6079*->N6-METHYLASE->?->HEPN->Schlafen+HATPase+wHTH->PglZ->SIGMA+OB-Fold+AAA-ATPase+Lon-Protease-BrxL-> | 1202 | Bacteria>Campylobacterota>Epsilonproteobacteria>Campylobacterales>Arcobacteraceae>Aliarcobacter | GCF_001010585.1 | Aliarcobacter_butzleri | AA20_RS09495 | 10373..13981 | 28197 | BREX-system-P-loop-protein-BrxC-[Aliarcobacter-butzleri] |
| WP_260968564.1 | Type1.BREX | ABC-ATPase+SbcC->?->LuxR-HTH-><-?\|\|Phage_integrase->?->wHTH+WYL+WCX->BrxA-DNA-binding->BrxB-BrxF-iSTAND->DUF6079*->N6-METHYLASE-p->HEPN+RES->N6-METHYLASE-p->Schlafen+HATPase+wHTH->Schlafen+HATPase+wHTH->PglZ->SIGMA+OB-Fold+AAA-ATPase+Lon-Protease-BrxL->Calcineurin-><-SIGMA-HTH | 1162 | Bacteria>Campylobacterota>Epsilonproteobacteria>Campylobacterales>Arcobacteraceae>Aliarcobacter | GCF_025335945.1 | Aliarcobacter_cryaerophilus | N5S77_RS10385 | 37284..40772 | 28198 | BREX-system-P-loop-protein-BrxC-[Aliarcobacter-cryaerophilus] |
| OZA03558.1 | Type1.BREX | <-SAM-METHYLASE\|\|?-><-TPR\|\|?->BrxB-BrxF-iSTAND->ACET->XIS-HTH->BrxA-DNA-binding->BrxB-BrxF-iSTAND->DUF6079*->HEPN->N6-METHYLASE->?->ZnR+HEPN->?->PglZ->SIGMA+OB-Fold+AAA-ATPase+Lon-Protease-BrxL-> | 1158 | Bacteria>Campylobacterota>Epsilonproteobacteria>Campylobacterales>Thiovulaceae>Sulfuricurvum | GCA_002282015.1 | Sulfuricurvum_sp._17-40-25 | B7X89_02520 | 35980..39456 | 1970439 | MAG:-hypothetical-protein-B7X89_02520-[Sulfuricurvum-sp.-17-40-25] |
| WP_013460546.1 | Type1.BREX | Phage_integrase-><-XIS-HTH<-HTH_36<-?\|\|HTH_3-><-XIS-HTH\|\|BrxA-DNA-binding->BrxB-BrxF-iSTAND->DUF6079*-><-?\|\|N6-METHYLASE-p->?->?->HEPN->BrxD->N6-METHYLASE-p->CR-ATPase5+REase->PglZ-> | 1163 | Bacteria>Campylobacterota>Epsilonproteobacteria>Campylobacterales>Thiovulaceae>Sulfuricurvum | GCF_000183725.1 | Sulfuricurvum_kujiense | SULKU_RS08475 | 1692937..1696428 | 148813 | BREX-system-P-loop-protein-BrxC-[Sulfuricurvum-kujiense] |
| QFR43750.1 | Type1.BREX | Glycos_transf_4->?->?->tRNA->tRNA->XIS-HTH->BrxA-DNA-binding->BrxB-BrxF-iSTAND->DUF6079*->HEPN->N6-METHYLASE-p->?->DUF3644->?->N6-METHYLASE-p->?->PglZ->SIGMA+OB-Fold+AAA-ATPase+Lon-Protease-BrxL-> | 1163 | Bacteria>Campylobacterota>Epsilonproteobacteria>Campylobacterales>Thiovulaceae>Sulfurimonas | GCF_009258045.1 | Sulfurimonas_xiamenensis | FJR47_07440 | 1472966..1476457 | 2590021 | BREX-system-P-loop-protein-BrxC-[Sulfurimonas-xiamenensis] |
| WP_188093712.1 | Type1.BREX | Glycos_transf_4->?->?->tRNA->tRNA->XIS-HTH->BrxA-DNA-binding->BrxB-BrxF-iSTAND->DUF6079*->HEPN->N6-METHYLASE-p->?->DUF3644->?->N6-METHYLASE-p->?->PglZ->SIGMA+OB-Fold+AAA-ATPase+Lon-Protease-BrxL-> | 1160 | Bacteria>Campylobacterota>Epsilonproteobacteria>Campylobacterales>Thiovulaceae>Sulfurimonas | GCF_009258045.1 | Sulfurimonas_xiamenensis | FJR47_RS07440 | 1472975..1476457 | 2590021 | BREX-system-P-loop-protein-BrxC-[Sulfurimonas-xiamenensis] |
| WP_193150356.1 | Type1.BREX | <-Phage_integrase<-SIGMA+OB-Fold+AAA-ATPase+Lon-Protease-BrxL<-PglZ<-?<-SKI<-N6-METHYLASE-p<-RVT<-N6-METHYLASE-p<-HEPN<-DUF6079*<-BrxB-BrxF-iSTAND<-BrxA-DNA-binding<-DDE_Tnp<-?<-DDE_Tnp | 1160 | Bacteria>Campylobacterota>Epsilonproteobacteria>Campylobacterales>Thiovulaceae>Sulfurimonas | GCF_014905115.1 | Sulfurimonas_sediminis | FJR45_RS09735 | 1812721..1816203 | 2590020 | BREX-system-P-loop-protein-BrxC-[Sulfurimonas-sediminis] |
| OYZ34277.1 | Type1.BREX | <-SIGMA+OB-Fold+AAA-ATPase+Lon-Protease-BrxL<-PglZ<-?<-ZnR+HEPN<-?<-N6-METHYLASE<-HEPN<-DUF6079*<-DUF6079-p<-BrxB-BrxF-iSTAND<-BrxA-DNA-binding<-XIS-HTH<-ACET<-BrxB-BrxF-iSTAND<-?\|\|TPR-> | 1013 | Bacteria>Campylobacterota>Epsilonproteobacteria>Campylobacterales>unclassified_Campylobacterales | GCA_002281685.1 | Campylobacterales_bacterium_16-40-21 | B7Y30_07055 | 21520..24561 | 1970501 | hypothetical-protein-B7Y30_07055-[Campylobacterales-bacterium-16-40-21] |
| WP_013452128.1 | Type1.BREX | CR-ATPase8+REase-><-?<-SIGMA+OB-Fold+AAA-ATPase+Lon-Protease-BrxL<-PglZ<-N6-METHYLASE<-HEPN<-N6-METHYLASE-p<-DUF6079*<-BrxB-BrxF-iSTAND<-BrxA-DNA-binding\|\|CR-ATPase8+REase-> | 1161 | Bacteria>Deferribacterota>Deferribacteres>Deferribacterales>Calditerrivibrionaceae>Calditerrivibrio | GCF_000183405.1 | Calditerrivibrio_nitroreducens | CALNI_RS10260 | 2104217..2107702 | 477976 | BREX-system-P-loop-protein-BrxC-[Calditerrivibrio-nitroreducens] |
| MBZ4644075.1 | Type1.BREX | <-TPR<-ARG<-?\|\|McrB-AAA->wHTH+REase->BrxA-DNA-binding->BrxB-BrxF-iSTAND->DUF6079*->N6-METHYLASE-p->HEPN->N6-METHYLASE->PglZ->SIGMA+OB-Fold+AAA-ATPase+Lon-Protease-BrxL->?->SF1-SF2-Helicase-> | 1168 | Bacteria>Deferribacterota>Deferribacteres>Deferribacterales>Deferribacteraceae>unclassified_Deferribacteraceae | GCA_020054865.1 | Deferribacteraceae_bacterium | JG767_1684 | 7812..11318 | 2053543 | hypothetical-protein-[Deferribacteraceae-bacterium] |
| WP_242820841.1 | Type1.BREX | NUDIX-><-TPR+S1<-ARG<-?<-?\|\|McrB-AAA->wHTH+REase->BrxA-DNA-binding->BrxB-BrxF-iSTAND->DUF6079*->N6-METHYLASE-p->HEPN->N6-METHYLASE->PglZ->SIGMA+OB-Fold+AAA-ATPase+Lon-Protease-BrxL->?->SF1-SF2-Helicase-> | 1168 | Bacteria>Deferribacterota>Deferribacteres>Deferribacterales>Deferrivibrionaceae>Deferrivibrio | GCF_020480685.1 | Deferrivibrio_essentukiensis | LF845_RS09815 | 29861..33367 | 2880922 | BREX-system-P-loop-protein-BrxC-[Deferrivibrio-essentukiensis] |
| EFK06105.1 | Type1.BREX | <-Acyltransferase<-?<-?<-SIGMA+OB-Fold+AAA-ATPase+Lon-Protease-BrxL<-PglZ<-TOPRIM<-OLD-ABC<-N6-METHYLASE<-HEPN<-DUF6079*<-BrxB-BrxF-iSTAND-p\|\|?-><-BrxA-DNA-binding\|\|?->?->?-><-DnaJ | 1215 | Bacteria>delta/epsilon_subdivisions>Deltaproteobacteria>unclassified_Deltaproteobacteria | GCA_000179315.1 | delta_proteobacterium_NaphS2 | NPH_2628 | 46790..50437 | 88274 | conserved-hypothetical-protein-[delta-proteobacterium-NaphS2] |
| MBI5543786.1 | Type6.BREX | XIS-HTH->Classical-AAA->Peptidase_S8-><-SF1-SF2-Helicase+wHTH+Small-B-barrel+DUF1998<-BrxD-AAA<-PglZ<-HEPN+RES<-N6-METHYLASE<-DUF6079*<-BrxB-BrxF-iSTAND<-BrxA-DNA-binding<-BrxE | 1206 | Bacteria>delta/epsilon_subdivisions>Deltaproteobacteria>unclassified_Deltaproteobacteria | GCA_016218885.1 | Deltaproteobacteria_bacterium | HY901_07870 | 25804..29424 | 2026735 | BREX-system-P-loop-protein-BrxC-[Deltaproteobacteria-bacterium] |
| MCS4056789.1 | Type1.BREX | <-Calcineurin-SbcD<-SIGMA+OB-Fold+AAA-ATPase+HKD-DNase-BrxL<-PglZ<-HEPN<-PolB\|\|?-><-?<-N6-METHYLASE<-DOC+HTH<-DUF6079*<-BrxB-BrxF-iSTAND<-BrxA-DNA-binding<-?<-viral-protein<-?<-?\|\|?-><-RHH | 1165 | Bacteria>FCB_group>Bacteroidetes/Chlorobi_group>Bacteroidota>Bacteroidetes_incertae_sedis>Rhodothermaceae | GCA_024809315.1 | Salinibacter_ruber | GGP75_002131 | 31231..34728 | 146919 | hypothetical-protein-[Salinibacter-ruber] |
| WP_259115052.1 | Type1.BREX | <-SbcC+SbcC<-Calcineurin-SbcD<-SIGMA+OB-Fold+AAA-ATPase+HKD-DNase-BrxL<-PglZ<-HEPN<-PolB<-?<-N6-METHYLASE<-DOC+HTH<-DUF6079*<-BrxB-BrxF-iSTAND<-BrxA-DNA-binding<-?<-viral-protein<-?<-?\|\|?-><-RHH\|\|HTH_3+Peptidase_S24-> | 1167 | Bacteria>FCB_group>Bacteroidetes/Chlorobi_group>Bacteroidota>Bacteroidetes_incertae_sedis>Rhodothermaceae | GCF_024809315.1 | Salinibacter_ruber | GGP75_RS10390 | 31231..34734 | 146919 | BREX-system-P-loop-protein-BrxC-[Salinibacter-ruber] |
| CDE09669.1 | Type1.BREX | ArdA->HTH_3->BrxA-DNA-binding->BrxB-BrxF-iSTAND->DUF6079*->Pkinase->N6-METHYLASE->HEPN->?->?->PglZ->SIGMA+OB-Fold+AAA-ATPase+Lon-Protease-BrxL-> | 1221 | Bacteria>FCB_group>Bacteroidetes/Chlorobi_group>Bacteroidota>Bacteroidia>Bacteroidales | GCA_000437495.1 | Prevotella_sp._CAG:485 | BN677_00784 | 6933..10598 | 1262927 | putative-uncharacterized-protein-[Prevotella-sp.-CAG:485] |
| WP_191006191.1 | Type6.BREX | HEPN->Phage_integrase+Phage_integrase-><-?<-HEPN<-TIR\|\|?->BrxE->BrxA-DNA-binding->BrxB-BrxF-iSTAND->DUF6079*->N6-METHYLASE->HEPN->PglZ->SIGMA+OB-Fold+AAA-ATPase+HKD-DNase-BrxL->N6_N4_METHYLASE->MORC+HISKIN->?-><-DOC+HTH | 1204 | Bacteria>FCB_group>Bacteroidetes/Chlorobi_group>Bacteroidota>Cytophagia>Cytophagales | GCF_014699115.1 | Hymenobacter_montanus | IC235_RS15945 | 31815..35429 | 2771359 | BREX-system-P-loop-protein-BrxC-[Hymenobacter-montanus] |
| WP_243520943.1 | Type6.BREX | Glyoxalase-><-?<-HEPN<-?\|\|?->?->BrxE->BrxA-DNA-binding->BrxB-BrxF-iSTAND->DUF6079*->N6-METHYLASE->HEPN->PglZ->SIGMA+OB-Fold+AAA-ATPase+HKD-DNase-BrxL->?->HKD-DNase->HTH+REase-><-CopG_antitoxin | 1204 | Bacteria>FCB_group>Bacteroidetes/Chlorobi_group>Bacteroidota>Cytophagia>Cytophagales | GCF_022811645.1 | Hymenobacter_monticola | MTP16_RS25545 | 68480..72094 | 1705399 | BREX-system-P-loop-protein-BrxC-[Hymenobacter-monticola] |
| WP_059014619.1 | Type1.BREX | <-HEPN<-ParB+HNH<-N6-METHYLASE<-DUF6079*<-BrxB-BrxF-iSTAND<-BrxA-DNA-binding<-HTH_3<-Classical-AAA<-HTH_36<-HTH_36<-XIS-HTH<-Arm-DNA-bind_5+Phage_int_SAM_5+Phage_integrase | 1169 | Bacteria>FCB_group>Bacteroidetes/Chlorobi_group>Bacteroidota>Flavobacteriia>Flavobacteriales | GCF_001485515.1 | Flavobacterium_psychrophilum | FPK15_RS07995 | 6803..10312 | 96345 | BREX-system-P-loop-protein-BrxC-[Flavobacterium-psychrophilum] |
| WP_133546801.1 | Type1.BREX | <-XIS-HTH<-?<-?<-?<-AraC-HTH<-Arm-DNA-bind_5+Phage_int_SAM_5+Phage_integrase\|\|?->BrxA-DNA-binding->BrxB-BrxF-iSTAND->DUF6079*->N6-METHYLASE->REase+DUF5655->NERD+SF1-SF2-Helicase->TPR+S1->PglZ->SIGMA+OB-Fold+AAA-ATPase+HKD-DNase-BrxL->HEPN->?-><-HTH_36 | 1167 | Bacteria>FCB_group>Bacteroidetes/Chlorobi_group>Bacteroidota>Flavobacteriia>Flavobacteriales | GCF_004362395.1 | Salegentibacter_sp._24 | DET49_RS03950 | 90042..93545 | 2183986 | BREX-system-P-loop-protein-BrxC-[Salegentibacter-sp.-24] |
| WP_165150638.1 | Type1.BREX | <-ABhydrolase<-REase+RNase<-?<-wHTH+WYL+WCX\|\|?->?->?->BrxA-DNA-binding->BrxB-BrxF-iSTAND->DUF6079*->N6-METHYLASE->ParB+HNH->PglZ->SIGMA+OB-Fold+AAA-ATPase+Lon-Protease-BrxL->HEPN->?->?-><-FGase<-tRNA\|\|SKI-> | 1198 | Bacteria>FCB_group>Bacteroidetes/Chlorobi_group>Bacteroidota>Flavobacteriia>Flavobacteriales | GCF_011065475.1 | Aequorivita_sp._KMM_9714 | G3A46_RS13570 | 21223..24819 | 2707173 | BREX-system-P-loop-protein-BrxC-[Aequorivita-sp.-KMM-9714] |
| WP_163432971.1 | Type1.BREX | <-LexA-protease<-?\|\|?->XIS-HTH->PIN->GDResolvase+HIN-HTH->HTH_3->BrxA-DNA-binding->BrxB-BrxF-iSTAND->DUF6079*->N6-METHYLASE->PglZ->HEPN-><-?\|\|?->Relaxase->VirD4-TraG+VirD4-FtsK-> | 1189 | Bacteria>FCB_group>Bacteroidetes/Chlorobi_group>Bacteroidota>Flavobacteriia>Flavobacteriales | GCF_014050215.1 | unclassified_Flavobacterium | G7B00_RS14255 | 32052..35621 | 196869 | MULTISPECIES:-BREX-system-P-loop-protein-BrxC-[unclassified-Flavobacterium] |
| WP_095606906.1 | Type1.BREX | <-REase\|\|?->?-><-SIGMA+OB-Fold+AAA-ATPase+Lon-Protease-BrxL<-PglZ<-N6-METHYLASE<-DUF6079*<-BrxB-BrxF-iSTAND<-BrxA-DNA-binding<-HEPN\|\|?-><-XIS-HTH<-?<-?<-?<-Phage_int_SAM_5+Phage_integrase | 1213 | Bacteria>FCB_group>Bacteroidetes/Chlorobi_group>Balneolota>Balneolia>Balneolales | GCF_002287075.1 | Aliifodinibius_sp._WN023 | CK503_RS11215 | 230173..233814 | 2032627 | BREX-system-P-loop-protein-BrxC-[Aliifodinibius-sp.-WN023] |
| WP_265763895.1 | Type1.BREX | <-HEPN<-SIGMA+OB-Fold+AAA-ATPase+HKD-DNase-BrxL<-PglZ<-?<-Schlafen+HATPase+wHTH<-REase<-?<-N6-METHYLASE<-DUF6079*<-BrxB-BrxF-iSTAND<-BrxA-DNA-binding<-tRNA\|\|?->?->?-><-?<-?<-ABC-ATPase+SbcC | 1165 | Bacteria>FCB_group>Bacteroidetes/Chlorobi_group>Balneolota>Balneolia>Balneolales | GCF_026229185.1 | Aliifodinibius_sp._1BSP15-2V2 | J6I44_RS00130 | 34405..37902 | 2820877 | BREX-system-P-loop-protein-BrxC-[Aliifodinibius-sp.-1BSP15-2V2] |
| OQC05381.1 | Type1.BREX | BrxA-DNA-binding->BrxB-BrxF-iSTAND->DUF6079*->OLD-ABC->REase->N6-METHYLASE->PglZ->SIGMA+OB-Fold+AAA-ATPase+HKD-DNase-BrxL->HEPN-> | 1162 | Bacteria>FCB_group>Candidatus_Cloacimonetes>unclassified_Candidatus_Cloacimonetes | GCA_002070875.1 | Candidatus_Cloacimonetes_bacterium_ADurb.Bin089 | BWX76_00750 | 1812..5300 | 1852836 | MAG:-hypothetical-protein-BWX76_00750-[Candidatus-Cloacimonetes-bacterium-ADurb.Bin089] |
| NLV46851.1 | Type1.BREX | Mrr_N->Mrr_N->Mrr_N->DUF4007->LON-AAA->N6-METHYLASE->DUF6079*->PglZ->SIGMA+OB-Fold+AAA-ATPase-BrxL->HEPN->?->Radical-SAM->MPTase->?->?->REase-> | 1339 | Bacteria>FCB_group>Candidatus_Hydrogenedentes>unclassified_Candidatus_Hydrogenedentes | GCA_012730105.1 | Candidatus_Hydrogenedentes_bacterium | GXY07_20415 | 9749..13768 | 2030809 | hypothetical-protein-[Candidatus-Hydrogenedentes-bacterium] |
| NUM53512.1 | Type1.BREX | <-SF1-SF2-Helicase<-OLD-ABC+TOPRIM<-?<-SIGMA+OB-Fold+AAA-ATPase+Lon-Protease-BrxL<-PglZ<-ArdCN+MPTase<-?<-N6-METHYLASE<-HEPN<-DUF6079*<-BrxB-BrxF-iSTAND<-BrxA-DNA-binding\|\|?->HTH_3->MPTase-> | 1149 | Bacteria>FCB_group>Candidatus_Hydrogenedentes>unclassified_Candidatus_Hydrogenedentes | GCA_013359265.1 | Candidatus_Hydrogenedentes_bacterium | HUU46_07705 | 154693..158142 | 2030809 | BREX-system-P-loop-protein-BrxC-[Candidatus-Hydrogenedentes-bacterium] |
| MCC6156024.1 | Type1.BREX | <-SIGMA+OB-Fold+AAA-ATPase+Lon-Protease-BrxL<-PglZ<-ArdCN+MPTase<-N6-METHYLASE<-HEPN<-DUF6079* | 1142 | Bacteria>FCB_group>Candidatus_Hydrogenedentes>unclassified_Candidatus_Hydrogenedentes | GCA_020846215.1 | Candidatus_Hydrogenedentes_bacterium | IT367_19820 | 10018..13447 | 2030809 | BREX-system-P-loop-protein-BrxC-[Candidatus-Hydrogenedentes-bacterium] |
| ODT04869.1 | Type6.BREX | <-SIGMA+OB-Fold+AAA-ATPase+Lon-Protease-BrxL<-PglZ<-N6-METHYLASE<-N6-METHYLASE-p<-ORC-CDC6<-ParA-Soj-PloopNTPase<-HEPN<-N6-METHYLASE<-DUF6079*<-BrxB-BrxF-iSTAND<-BrxA-DNA-binding<-BrxE\|\|McrB-AAA->wHTH+REase->REase->?->EVE+HNH-> | 1181 | Bacteria>FCB_group>Gemmatimonadota>unclassified_Gemmatimonadota | GCA_001724275.1 | Gemmatimonadetes_bacterium_SCN_70-22 | ABS52_01570 | 17993..21538 | 1660163 | hypothetical-protein-ABS52_01570-[Gemmatimonadetes-bacterium-SCN-70-22] |
| WP_098702328.1 | Type1.BREX | SKI->JAB+sGTP->wHTH+WYL+WCX->?->?->ABC-ATPase+SbcC->?-><-?\|\|DUF6079*->OLD-ABC->TOPRIM->N6-METHYLASE->HEPN->CR-ATPase8+REase->Classical-AAA->Peptidase_S8->PglZ->SIGMA+OB-Fold+AAA-ATPase+Lon-Protease-BrxL-> | 1250 | Bacteria>Fusobacteriota>Fusobacteriia>Fusobacteriales>Fusobacteriaceae>Fusobacterium | GCF_002573625.1 | Fusobacterium_nucleatum | RN96_RS03470 | 669523..673275 | 851 | BREX-system-P-loop-protein-BrxC-[Fusobacterium-polymorphum] |
| WP_099959237.1 | Type1.BREX | <-SIGMA+OB-Fold+AAA-ATPase+Lon-Protease-BrxL<-PglZ<-Peptidase_S8<-Classical-AAA<-CR-ATPase8+REase<-HEPN<-N6-METHYLASE<-TOPRIM<-OLD-ABC<-DUF6079*\|\|?-><-?<-?<-Peptidase_M3_N+Peptidase_M3\|\|RNaseH->?->?->Peptidase_M20-> | 1250 | Bacteria>Fusobacteriota>Fusobacteriia>Fusobacteriales>Fusobacteriaceae>Fusobacterium | GCF_002761955.1 | Fusobacterium_pseudoperiodonticum | CTM71_RS09955 | 1967192..1970944 | 2663009 | BREX-system-P-loop-protein-BrxC-[Fusobacterium-pseudoperiodonticum] |
| WP_100024172.1 | Type1.BREX | <-Peptidase_M20<-?<-?<-RNaseH\|\|Peptidase_M3_N+Peptidase_M3->?->?-><-?\|\|DUF6079*->OLD-ABC->TOPRIM->N6-METHYLASE->HEPN->CR-ATPase8+REase->PglZ->SIGMA+OB-Fold+AAA-ATPase+Lon-Protease-BrxL->BrxA-DNA-binding->BrxB-BrxF-iSTAND-> | 1250 | Bacteria>Fusobacteriota>Fusobacteriia>Fusobacteriales>Fusobacteriaceae>Fusobacterium | GCF_002763625.1 | Fusobacterium_pseudoperiodonticum | CTM72_RS01085 | 199877..203629 | 2663009 | BREX-system-P-loop-protein-BrxC-[Fusobacterium-pseudoperiodonticum] |
| WP_238959505.1 | Type1.BREX | <-BrxB-BrxF-iSTAND<-BrxA-DNA-binding<-SIGMA+OB-Fold+AAA-ATPase+Lon-Protease-BrxL<-PglZ<-CR-ATPase8+REase<-HEPN<-MORC<-N6-METHYLASE<-Schlafen+HATPase+wHTH<-DUF6079*\|\|?->?-><-?<-ABC-ATPase+SbcC<-?<-?<-wHTH+WYL+WCX<-JAB+sGTP<-SKI | 1250 | Bacteria>Fusobacteriota>Fusobacteriia>Fusobacteriales>Fusobacteriaceae>Fusobacterium | GCF_022340045.1 | Fusobacterium_nucleatum | K5I20_00280 | 65514..69266 | 851 | BREX-system-P-loop-protein-BrxC-[Fusobacterium-nucleatum] |
| WP_238972693.1 | Type1.BREX | SKI->JAB+sGTP->wHTH+WYL+WCX->?->ABC-ATPase+SbcC->?-><-?<-?\|\|DUF6079*->OLD-ABC->TOPRIM->Schlafen+HATPase+wHTH->N6-METHYLASE->MORC->HEPN->CR-ATPase8+REase->PglZ->SIGMA+OB-Fold+AAA-ATPase+Lon-Protease-BrxL-> | 1250 | Bacteria>Fusobacteriota>Fusobacteriia>Fusobacteriales>Fusobacteriaceae>Fusobacterium | GCF_022340165.1 | Fusobacterium_nucleatum | K5I19_RS02505 | 127932..131684 | 851 | BREX-system-P-loop-protein-BrxC-[Fusobacterium-nucleatum] |
| WP_060798221.1 | Type1.BREX | <-BrxA-DNA-binding<-SIGMA+OB-Fold+AAA-ATPase+Lon-Protease-BrxL<-PglZ<-CR-ATPase8+REase<-HEPN<-MORC<-N6-METHYLASE<-TOPRIM<-OLD-ABC<-DUF6079*<-?<-SWIM+DUF3670+SF1-SF2-Helicase<-?<-?<-?<-SF1-SF2-Helicase | 1250 | Bacteria>Fusobacteriota>Fusobacteriia>Fusobacteriales>Fusobacteriaceae>Fusobacterium | GCF_023383525.1 | Fusobacterium_nucleatum | YWH7054_RS02310 | 21716..25468 | 851 | BREX-system-P-loop-protein-BrxC-[Fusobacterium-nucleatum] |
| WP_249524183.1 | Type1.BREX | <-BrxA-DNA-binding<-SIGMA+OB-Fold+AAA-ATPase+Lon-Protease-BrxL<-PglZ<-CR-ATPase8+REase<-HEPN<-MORC<-N6-METHYLASE<-TOPRIM<-OLD-ABC<-DUF6079*<-?<-SWIM+DUF3670+SF1-SF2-Helicase<-?<-?<-?<-SF1-SF2-Helicase | 1250 | Bacteria>Fusobacteriota>Fusobacteriia>Fusobacteriales>Fusobacteriaceae>Fusobacterium | GCF_023383605.1 | Fusobacterium_nucleatum | YWH7056_RS07045 | 28639..32391 | 851 | BREX-system-P-loop-protein-BrxC-[Fusobacterium-nucleatum] |
| WP_249526950.1 | Type1.BREX | <-BrxB-BrxF-iSTAND<-BrxA-DNA-binding<-SIGMA+OB-Fold+AAA-ATPase+Lon-Protease-BrxL<-PglZ<-HEPN<-MORC<-N6-METHYLASE<-TOPRIM<-OLD-ABC<-DUF6079*<-?<-SWIM+DUF3670+SF1-SF2-Helicase<-?<-?<-?<-SF1-SF2-Helicase | 1250 | Bacteria>Fusobacteriota>Fusobacteriia>Fusobacteriales>Fusobacteriaceae>Fusobacterium | GCF_023383645.1 | Fusobacterium_nucleatum | YWH7199_RS06535 | 22791..26543 | 851 | BREX-system-P-loop-protein-BrxC-[Fusobacterium-nucleatum] |
| WP_008793539.1 | Type1.BREX | <-RNaseH\|\|Peptidase_M3_N+Peptidase_M3->?->?-><-?<-HTH_OrfB_IS605+OrfB_IS605+OrfB_Zn_ribbon\|\|DUF6079*->OLD-ABC->RNase+PHP->Schlafen+HATPase+wHTH->N6-METHYLASE->HEPN->CR-ATPase8+REase->PglZ->SIGMA+OB-Fold+AAA-ATPase+Lon-Protease-BrxL->BrxA-DNA-binding-> | 1250 | Bacteria>Fusobacteriota>Fusobacteriia>Fusobacteriales>Fusobacteriaceae>Fusobacterium | GCF_902373915.1 | Fusobacterium_periodonticum | FY302_RS02975 | 615184..618936 | 860 | BREX-system-P-loop-protein-BrxC-[Fusobacterium-periodonticum] |
| WP_026745399.1 | Type1.BREX | Peptidase_M24->?->?->Peptidase_M18+Peptidase_M42->?->BrxA-DNA-binding->BrxB-BrxF-iSTAND->DUF6079*->REase->N6-METHYLASE->HEPN->?->HicA_toxin->HicB_lk_antitox->?->PglZ->SIGMA+OB-Fold+AAA-ATPase+Lon-Protease-BrxL-> | 1193 | Bacteria>Fusobacteriota>Fusobacteriia>Fusobacteriales>Leptotrichiaceae>Leptotrichia | GCF_007990525.1 | Leptotrichia_hofstadii | FVE77_RS00755 | 171012..174593 | 157688 | BREX-system-P-loop-protein-BrxC-[Leptotrichia-hofstadii] |
| WP_151554984.1 | Type1.BREX | <-REC+NtrC-AAA+FIS-HTH<-HAMP+PAS+HISKIN<-ACET<-?<-SIGMA+OB-Fold+AAA-ATPase+Lon-Protease-BrxL<-PglZ<-HEPN<-N6-METHYLASE<-DUF6079*<-BrxB-BrxF-iSTAND<-BrxA-DNA-binding\|\|TRPR-HTH->Couple_hipA+HipA_C->SF1-SF2-Helicase->?-><-wHTH+REase<-McrB-AAA<-TraI-VirD2-RCRE | 1205 | Bacteria>Pseudomonadota>Alphaproteobacteria>Hyphomicrobiales>Brucellaceae>Brucella/Ochrobactrum_group | GCF_008932275.1 | Brucella_tritici | F9K85_RS05045 | 164979..168596 | 94626 | BREX-system-P-loop-protein-BrxC-[Brucella-tritici] |
| WP_080921652.1 | Type1.BREX | HEPN->HEPN->?-><-DEDD_Tnp_IS110+Transposase_20\|\|<-?<-HEPN<-SIGMA+OB-Fold+AAA-ATPase+HKD-DNase-BrxL<-PglZ<-N6-METHYLASE<-DUF6079*<-BrxB-BrxF-iSTAND<-BrxA-DNA-binding<-wHTH+WYL+WCX\|\|Calcineurin-SbcD->SbcC+CCDC158->?->HTH_3->Peptidase+SF1-SF2-Helicase+REase-> | 1179 | Bacteria>Pseudomonadota>Alphaproteobacteria>Hyphomicrobiales>Phyllobacteriaceae>Pseudaminobacter | GCF_002075885.1 | Pseudaminobacter_manganicus | BFN67_RS22220 | 17195..20734 | 1873176 | BREX-system-P-loop-protein-BrxC-[Pseudaminobacter-manganicus] |
| WP_088677288.1 | Type1.BREX | <-SPOUT\|\|tRNA->tRNA->?->?->?->tRNA->?->Classical-AAA+S2P->HEPN->BrxA-DNA-binding->BrxB-BrxF-iSTAND->DUF6079*->N6-METHYLASE->PglZ->SIGMA+OB-Fold+AAA-ATPase+HKD-DNase-BrxL-><-?\|\|?->?->MORC+REase->RNase-> | 1194 | Bacteria>Pseudomonadota>Alphaproteobacteria>Hyphomicrobiales>Rhizobiaceae>Rhizobium/Agrobacterium_group | GCF_002211395.1 | Rhizobium_sp._R339 | ATY76_RS24585 | 353019..356603 | 1764273 | BREX-system-P-loop-protein-BrxC-[Rhizobium-sp.-R339] |
| WP_222013551.1 | Type1.BREX | <-SIGMA+OB-Fold+AAA-ATPase+Lon-Protease-BrxL<-PglZ<-N6-METHYLASE<-DUF6079*<-BrxB-BrxF-iSTAND<-BrxA-DNA-binding<-HEPN<-?<-PIWI<-SIR2<-HARE-HTH\|\|Phage_integrase->Phage_Integr_2+Phage_integrase-> | 1137 | Bacteria>Pseudomonadota>Alphaproteobacteria>Hyphomicrobiales>Rhizobiaceae>Rhizobium/Agrobacterium_group | GCF_019779115.1 | Rhizobium_laguerreae | HFO05_RS34495 | 7944..11357 | 1076926 | BREX-system-P-loop-protein-BrxC-[Rhizobium-laguerreae] |
| PSL17302.1 | Type6.BREX | <-HAMP+HISKIN\|\|?->tRNA-><-SIR2<-SIGMA+OB-Fold+AAA-ATPase+HKD-DNase-BrxL<-PglZ<-N6-METHYLASE<-N6-METHYLASE<-DUF6079*<-BrxB-BrxF-iSTAND<-BrxA-DNA-binding<-BrxE<-XIS-HTH<-HEPN<-Polbeta<-wHTH+AbiEi_2\|\|REase-> | 1216 | Bacteria>Pseudomonadota>Alphaproteobacteria>Rhodobacterales>Roseobacteraceae>Shimia | GCA_003014475.1 | Shimia_abyssi | CLV88_11975 | 80664..84314 | 1662395 | hypothetical-protein-CLV88_11975-[Shimia-abyssi] |
| WP_207797113.1 | Type6.BREX | <-HAMP+HISKIN\|\|?->tRNA-><-SIR2<-SIGMA+OB-Fold+AAA-ATPase+HKD-DNase-BrxL<-PglZ<-N6-METHYLASE<-N6-METHYLASE<-DUF6079*<-BrxB-BrxF-iSTAND<-BrxA-DNA-binding<-BrxE<-XIS-HTH<-HEPN<-Polbeta<-wHTH+AbiEi_2\|\|REase-> | 1207 | Bacteria>Pseudomonadota>Alphaproteobacteria>Rhodobacterales>Roseobacteraceae>Shimia | GCF_003014475.1 | Shimia_abyssi | CLV88_RS20200 | 80664..84287 | 1662395 | BREX-system-P-loop-protein-BrxC-[Shimia-abyssi] |
| QKS29579.1 | Type6.BREX | <-SIGMA+OB-Fold+AAA-ATPase+Lon-Protease-BrxL<-PglZ<-HNH<-OLD-ABC<-N6-METHYLASE<-ORC-CDC6<-ParA-Soj-PloopNTPase<-HEPN<-N6-METHYLASE<-DUF6079*<-BrxB-BrxF-iSTAND<-BrxA-DNA-binding<-BrxE<-XIS-HTH | 1180 | Bacteria>Pseudomonadota>Betaproteobacteria>Betaproteobacteria_incertae_sedis>Candidatus_Accumulibacter | GCA_013347225.1 | Candidatus_Accumulibacter_similis | HT579_12075 | 2666587..2670129 | 2954388 | MAG:-BREX-system-P-loop-protein-BrxC-[Candidatus-Accumulibacter-similis] |
| HCN68765.1 | Type1.BREX | HTH_3-><-?<-?\|\|?->wHTH+AbiEi_2->Polbeta->REase->BrxA-DNA-binding->BrxB-BrxF-iSTAND->DUF6079*->HEPN->N6-METHYLASE->OLD-ABC+TOPRIM->SF1-SF2-Helicase->N6-METHYLASE->PglZ->SIGMA+OB-Fold+AAA-ATPase+Lon-Protease-BrxL-> | 1154 | Bacteria>Pseudomonadota>Betaproteobacteria>Betaproteobacteria_incertae_sedis>Candidatus_Accumulibacter>unclassified_Candidatus_Accumulibacter | GCA_003535635.1 | Candidatus_Accumulibacter_sp. | DIT03_10970 | 31798..35262 | 2053492 | BREX-system-P-loop-protein-BrxC-[Candidatus-Accumulibacter-sp.] |
| WP_043683814.1 | Type1.BREX | DDE_Tnp_1_4->?->VirB5-><-DUF772+DDE_Tnp_1+DDE_Tnp_1_6<-?\|\|wHTH+WYL+WCX->BrxA-DNA-binding->BrxB-BrxF-iSTAND->DUF6079*->HEPN->N6-METHYLASE->OLD-ABC->HNH->DUF772+DDE_Tnp_1+DDE_Tnp_1_6->HNH->PglZ->SIGMA+OB-Fold+AAA-ATPase+Lon-Protease-BrxL-><-YdaS_antitoxin | 1153 | Bacteria>Pseudomonadota>Betaproteobacteria>Burkholderiales>Alcaligenaceae>Castellaniella | GCF_000612685.1 | Castellaniella_defragrans | BN940_RS13240 | 2967883..2971344 | 75697 | BREX-system-P-loop-protein-BrxC-[Castellaniella-defragrans] |
| WP_183964658.1 | Type6.BREX | <-SWIM<-?\|\|NUDIX+NAAR1+NAAR2->?->?->?->HEPN->BrxE->BrxB-BrxF-iSTAND->DUF6079*->N6-METHYLASE->N6-METHYLASE-p->PglZ->SIGMA+OB-Fold+AAA-ATPase+HKD-DNase-BrxL-><-RVT<-?<-SAM-METHYLASE+wHTH+HNH<-BirA-HTH | 1210 | Bacteria>Pseudomonadota>Betaproteobacteria>Burkholderiales>Burkholderiaceae>Quisquiliibacterium | GCF_014202215.1 | Quisquiliibacterium_transsilvanicum | HNQ70_RS04390 | 209103..212735 | 1549638 | BREX-system-P-loop-protein-BrxC-[Quisquiliibacterium-transsilvanicum] |
| WP_086927320.1 | Type1.BREX | <-SIGMA+OB-Fold+AAA-ATPase+Lon-Protease-BrxL<-PglZ<-ParB+HNH<-SF1-SF2-Helicase<-OLD-ABC+TOPRIM<-N6-METHYLASE<-HEPN<-DUF6079*<-BrxB-BrxF-iSTAND<-BrxA-DNA-binding<-RHH_5<-?<-XIS-HTH<-tRNA<-tRNA<-35exo<-?<-?<-DnaJ | 1153 | Bacteria>Pseudomonadota>Betaproteobacteria>Burkholderiales>Comamonadaceae>Acidovorax | GCF_002157125.1 | Acidovorax_carolinensis | CBP36_RS10055 | 2162910..2166371 | 553814 | BREX-system-P-loop-protein-BrxC-[Acidovorax-carolinensis] |
| WP_094097616.1 | Type1.BREX | <-Peptidase_C39_2<-?\|\|?-><-?\|\|XIS-HTH->Phage_X->BrxA-DNA-binding->BrxB-BrxF-iSTAND->DUF6079*->HNH->HEPN->ParD_antitoxin->RelE-ParE->N6-METHYLASE->OLD-ABC->TOPRIM->ZnR+TOPRIM->PglZ-> | 1214 | Bacteria>Pseudomonadota>Betaproteobacteria>Burkholderiales>Comamonadaceae>Acidovorax | GCF_002157145.1 | Acidovorax_carolinensis | CBP34_RS06480 | 1379414..1383058 | 553814 | BREX-system-P-loop-protein-BrxC-[Acidovorax-carolinensis] |
| WP_204247869.1 | Type1.BREX | DnaJ->?->?->tRNA->tRNA->XIS-HTH->?->RHH_5->BrxA-DNA-binding->BrxB-BrxF-iSTAND->DUF6079-p*->DUF6079-p->HEPN->N6-METHYLASE->OLD-ABC+TOPRIM->SF1-SF2-Helicase->ParB+HNH->PglZ->SIGMA+OB-Fold+AAA-ATPase+Lon-Protease-BrxL-> | 590 | Bacteria>Pseudomonadota>Betaproteobacteria>Burkholderiales>Comamonadaceae>Acidovorax | GCF_002157185.1 | Acidovorax_carolinensis | CBP35_RS21920 | 1914645..1916417 | 553814 | BREX-system-P-loop-protein-BrxC-[Acidovorax-carolinensis] |
| GAO25248.1 | Type1.BREX | HTH_3-><-SIGMA+OB-Fold+AAA-ATPase+Lon-Protease-BrxL<-PglZ<-ZnR+TOPRIM<-TOPRIM<-OLD-ABC<-N6-METHYLASE<-HEPN<-HNH<-DUF6079*<-BrxB-BrxF-iSTAND<-BrxA-DNA-binding<-HAD+CHTH<-CHTH\|\|?->AAA-ATPase-p->?-><-AAA-ATPase | 1205 | Bacteria>Pseudomonadota>Betaproteobacteria>Burkholderiales>Comamonadaceae>Alicycliphilus | GCA_000974645.1 | Alicycliphilus_sp._B1 | ALISP_5068 | 1538304..1541921 | 1603291 | ATPase-like-protein-[Alicycliphilus-sp.-B1] |
| WP_103018746.1 | Type1.BREX | APATPase->?-><-?\|\|?->?->?->RHH_5->BrxA-DNA-binding->BrxB-BrxF-iSTAND->DUF6079*->HEPN-><-?\|\|N6-METHYLASE->SF1-SF2-Helicase+PRTase->SLOG+BirA-HTH->?->TPR+REase->?->ParB+HNH-> | 1153 | Bacteria>Pseudomonadota>Betaproteobacteria>Burkholderiales>Comamonadaceae>Alicycliphilus | GCF_002894355.1 | Alicycliphilus_denitrificans | CUB93_RS20845 | 10909..14370 | 179636 | BREX-system-P-loop-protein-BrxC-[Alicycliphilus-denitrificans] |
| WP_169107995.1 | Type1.BREX | SIGMA+OB-Fold+AAA-ATPase+Lon-Protease-BrxL->tRNA->tRNA->tRNA->tRNA->Phage_integrase->?->?-><-XIS-HTH\|\|?->RHH_5->BrxA-DNA-binding->BrxB-BrxF-iSTAND->DUF6079*->HEPN->N6-METHYLASE->PHP+SbcC->MarR-HTH->Polbeta->PglZ->SIGMA+OB-Fold+AAA-ATPase+Lon-Protease-BrxL->SF1-SF2-Helicase->PIN-> | 1153 | Bacteria>Pseudomonadota>Betaproteobacteria>Burkholderiales>Comamonadaceae>Comamonas | GCF_012844455.2 | Comamonas_suwonensis | HF327_RS09545 | 49446..52907 | 2606214 | BREX-system-P-loop-protein-BrxC-[Comamonas-suwonensis] |
| WP_172207178.1 | Type1.BREX | <-OLD-ABC\|\|<-N6-METHYLASE-p<-RelE-ParE<-ParD_antitoxin<-PIN<-MazE_antitoxin\|\|?-><-HEPN<-HNH<-DUF6079*<-BrxB-BrxF-iSTAND<-BrxA-DNA-binding<-Phage_X<-XIS-HTH\|\|?-><-?\|\|?->Peptidase_C39_2-> | 1213 | Bacteria>Pseudomonadota>Betaproteobacteria>Burkholderiales>Comamonadaceae>Diaphorobacter | GCF_013154915.1 | Diaphorobacter_sp._JS3050 | HND92_RS11875 | 2478331..2481972 | 2735554 | BREX-system-P-loop-protein-BrxC-[Diaphorobacter-sp.-JS3050] |
| WP_226462338.1 | Type1.BREX | <-SIGMA+OB-Fold+AAA-ATPase+Lon-Protease-BrxL<-PglZ<-ZnR+TOPRIM<-?<-HEPN<-N6-METHYLASE<-N6-METHYLASE<-Couple_hipA+HipA_C<-HTH_3<-DUF6079*<-BrxB-BrxF-iSTAND<-BrxA-DNA-binding<-SbcC+CCDC158<-Calcineurin-SbcD<-?<-MoxR-AAA<-HEPN | 1204 | Bacteria>Pseudomonadota>Betaproteobacteria>Burkholderiales>Comamonadaceae>Hydrogenophaga | GCF_020510545.1 | Hydrogenophaga_taeniospiralis | KI616_RS15915 | 3311609..3315223 | 65656 | BREX-system-P-loop-protein-BrxC-[Hydrogenophaga-taeniospiralis] |
| WP_101103324.1 | Type1.BREX | HTH_Tnp_1+HTH_21+rve+rve_3->RHH_5->BrxA-DNA-binding->BrxB-BrxF-iSTAND->DUF6079*->HEPN->N6-METHYLASE->OLD-ABC+TOPRIM->SF1-SF2-Helicase->OLD-ABC->HNH->ParB+HNH->PglZ->SIGMA+OB-Fold+AAA-ATPase+Lon-Protease-BrxL-> | 1153 | Bacteria>Pseudomonadota>Betaproteobacteria>Burkholderiales>Comamonadaceae>Macromonas | GCF_002837135.1 | Macromonas_bipunctata | CON76_RS10585 | 3906..7367 | 183670 | BREX-system-P-loop-protein-BrxC-[Macromonas-bipunctata] |
| OOG58024.1 | Type1.BREX | <-HTH_3<-?<-?\|\|?-><-SIGMA+OB-Fold+AAA-ATPase+Lon-Protease-BrxL<-PglZ<-HEPN<-N6-METHYLASE<-DUF6079*<-BrxB-BrxF-iSTAND<-BrxA-DNA-binding\|\|?->HD+PspB-> | 1207 | Bacteria>Pseudomonadota>Betaproteobacteria>Burkholderiales>Comamonadaceae>Polaromonas | GCA_002002705.1 | Polaromonas_sp._CO4 | BOE49_04110 | 64714..68337 | 1945857 | ATPase-[Polaromonas-sp.-CO4] |
| WP_011485879.1 | Type1.BREX | N6-METHYLASE-><-?<-?<-?<-REC<-HISKIN\|\|HTH_3->BrxA-DNA-binding->BrxB-BrxF-iSTAND->DUF6079*->CHTH->Tox-RES->N6-METHYLASE->HEPN->NERD+Pkinase+Pkinase+SF1-SF2-Helicase->?->PglZ->SIGMA+OB-Fold+AAA-ATPase+Lon-Protease-BrxL-><-PHP+SbcC | 1202 | Bacteria>Pseudomonadota>Betaproteobacteria>Burkholderiales>Comamonadaceae>Polaromonas | GCF_000013865.1 | Polaromonas_sp._JS666 | BPRO_RS25145 | 121747..125355 | 296591 | BREX-system-P-loop-protein-BrxC-[Polaromonas-sp.-JS666] |
| APW46516.1 | Type1.BREX | Glyco_transf_9-><-?\|\|?->?->TPR+TPR+TPR->?->tRNA-><-?\|\|DUF6079-p*->MarR-HTH+Polbeta->HEPN->HEPN->N6-METHYLASE-p->N6-METHYLASE->?->?-><-DDE_Tnp_1+DDE_Tnp_1 | 269 | Bacteria>Pseudomonadota>Betaproteobacteria>Burkholderiales>Comamonadaceae>Rhodoferax | GCA_001955735.1 | Rhodoferax_antarcticus | RA876_09190 | 1894413..1895222 | 81479 | hypothetical-protein-RA876_09190-[Rhodoferax-antarcticus] |
| MBP9058688.1 | Type1.BREX | <-PglZ<-SLOG+BirA-HTH<-SF1-SF2-Helicase+PRTase<-?<-TPR+REase<-?<-N6-METHYLASE<-HEPN<-MarR-HTH+Polbeta<-DUF6079*<-BrxB-BrxF-iSTAND<-BrxA-DNA-binding<-RHH_5\|\|N6-METHYLASE->ORC-CDC6+ZnR-> | 1153 | Bacteria>Pseudomonadota>Betaproteobacteria>Burkholderiales>Comamonadaceae>Rhodoferax | GCA_018057375.1 | Rhodoferax_sp. | KBF98_00065 | 15882..19343 | 50421 | BREX-system-P-loop-protein-BrxC-[Rhodoferax-sp.] |
| MCF8211220.1 | Type1.BREX | BrxA-DNA-binding->BrxB-BrxF-iSTAND->DUF6079*->MarR-HTH+Polbeta->OLD-ABC->REase->HEPN->N6-METHYLASE-p-> | 1153 | Bacteria>Pseudomonadota>Betaproteobacteria>Burkholderiales>Comamonadaceae>Rhodoferax | GCA_021738345.1 | Rhodoferax_sp. | K9K38_17735 | 1925..5386 | 50421 | BREX-system-P-loop-protein-BrxC-[Rhodoferax-sp.] |
| WP_139313405.1 | Type1.BREX | ABC-ATPase+SbcC->?->Glyco_transf_9-><-?\|\|?->?->TPR+TPR+TPR->?->tRNA->\|\|DUF6079-p*->MarR-HTH+Polbeta->HEPN->?->HEPN->N6-METHYLASE-p->N6-METHYLASE->?->?->RAMA-> | 323 | Bacteria>Pseudomonadota>Betaproteobacteria>Burkholderiales>Comamonadaceae>Rhodoferax | GCF_001955735.1 | Rhodoferax_antarcticus | RA876_RS09190 | 1894251..1895222 | 81479 | hypothetical-protein-[Rhodoferax-antarcticus] |
| WP_264569449.1 | Type1.BREX | <-Glyco_transf_9\|\|?->?->TPR+TPR+TPR->?->tRNA->\|\|DUF6079-p*->MarR-HTH+Polbeta->HEPN->?->HEPN->N6-METHYLASE-p->N6-METHYLASE-p-> | 323 | Bacteria>Pseudomonadota>Betaproteobacteria>Burkholderiales>Comamonadaceae>Rhodoferax | GCF_025961535.1 | Rhodoferax_antarcticus | M2244_RS19480 | 9793..10764 | 81479 | hypothetical-protein-[Rhodoferax-antarcticus] |
| MBH2043138.1 | Type1.BREX | RHH_5->BrxA-DNA-binding->BrxB-BrxF-iSTAND->DUF6079*->MarR-HTH+Polbeta->HEPN->OLD-ABC->HNH->PglZ->SIGMA+OB-Fold+AAA-ATPase+Lon-Protease-BrxL-> | 1153 | Bacteria>Pseudomonadota>Betaproteobacteria>Burkholderiales>Comamonadaceae>unclassified_Comamonadaceae | GCA_016000285.1 | Comamonadaceae_bacterium | I8H87_06315 | 1878..5339 | 1871071 | BREX-system-P-loop-protein-BrxC-[Comamonadaceae-bacterium] |
| WP_096234088.1 | Type1.BREX | HTH_3->?->?->?->BrxA-DNA-binding->BrxB-BrxF-iSTAND->DUF6079*->N6-METHYLASE->OLD-ABC->TOPRIM->PglZ->SIGMA+OB-Fold+AAA-ATPase+Lon-Protease-BrxL->HEPN-><-TPR | 1204 | Bacteria>Pseudomonadota>Betaproteobacteria>Burkholderiales>Oxalobacteraceae>Janthinobacterium | GCF_002327145.1 | Janthinobacterium_svalbardensis | CNX70_RS06880 | 1564323..1567937 | 368607 | BREX-system-P-loop-protein-BrxC-[Janthinobacterium-svalbardensis] |
| WP_203569835.1 | Type1.BREX | <-HAD+HAD\|\|?->HTH_Tnp_1+HTH_21+rve+rve_3->wHTH+WYL+WCX->BrxA-DNA-binding->BrxB-BrxF-iSTAND->DUF6079*->HEPN->N6-METHYLASE->?->?->?->?->PglZ->SIGMA+OB-Fold+AAA-ATPase+Lon-Protease-BrxL-> | 1156 | Bacteria>Pseudomonadota>Betaproteobacteria>Neisseriales>Chromobacteriaceae | GCF_014042245.2 | Deefgea | H1229_RS02985 | 646302..649772 | 400947 | MULTISPECIES:-BREX-system-P-loop-protein-BrxC-[Deefgea] |
| MBK7491769.1 | Type1.BREX | HIN-HTH->?-><-SF1-SF2-Helicase<-?\|\|?->DUF772+DDE_Tnp_1->PIN+OST-HTH->BrxA-DNA-binding->BrxB-BrxF-iSTAND->DUF6079*->N6-METHYLASE->HEPN->HTH+REase->PglZ->SIGMA+OB-Fold+AAA-ATPase+Lon-Protease-BrxL->HIN-HTH-> | 1203 | Bacteria>Pseudomonadota>Betaproteobacteria>Nitrosomonadales>Nitrosomonadaceae>Nitrosomonas | GCA_016707165.1 | Nitrosomonas_sp. | IPI17_07005 | 52164..55775 | 42353 | BREX-system-P-loop-protein-BrxC-[Nitrosomonas-sp.] |
| MBK7235806.1 | Type6.BREX | <-PhdYeFM_antitox\|\|?-><-tRNA<-SIGMA+OB-Fold+AAA-ATPase+Lon-Protease-BrxL<-PglZ<-N6-METHYLASE<-ORC-CDC6<-ParA-Soj-PloopNTPase<-HEPN<-N6-METHYLASE<-DUF6079*<-BrxB-BrxF-iSTAND<-BrxA-DNA-binding<-BrxE<-XIS-HTH<-NUDIX\|\|?->?->Trypsin+PDZ-> | 1180 | Bacteria>Pseudomonadota>Betaproteobacteria>Nitrosomonadales>Sterolibacteriaceae>unclassified_Sterolibacteriaceae | GCA_016707455.1 | Sterolibacteriaceae_bacterium | IPI02_09355 | 160184..163726 | 2682146 | BREX-system-P-loop-protein-BrxC-[Sterolibacteriaceae-bacterium] |
| MBK6972675.1 | Type6.BREX | <-Trypsin+PDZ<-?<-?\|\|NUDIX->XIS-HTH->BrxE->BrxA-DNA-binding->BrxB-BrxF-iSTAND->DUF6079*->N6-METHYLASE->HEPN->ParA-Soj-PloopNTPase->ORC-CDC6->N6-METHYLASE->PglZ->SIGMA+OB-Fold+AAA-ATPase+Lon-Protease-BrxL->tRNA-><-?\|\|PhdYeFM_antitox-> | 1180 | Bacteria>Pseudomonadota>Betaproteobacteria>Nitrosomonadales>Sterolibacteriaceae>unclassified_Sterolibacteriaceae | GCA_016708905.1 | Sterolibacteriaceae_bacterium | IPH26_06870 | 229623..233165 | 2682146 | BREX-system-P-loop-protein-BrxC-[Sterolibacteriaceae-bacterium] |
| MBL8465274.1 | Type6.BREX | ABC-ATPase+SbcC->BrxE->BrxA-DNA-binding->BrxB-BrxF-iSTAND->DUF6079*->MNS+MarR-HTH->N6-METHYLASE->N6-METHYLASE->HEPN->ParA-Soj-PloopNTPase->ORC-CDC6->PglZ->SIGMA+OB-Fold+AAA-ATPase+HKD-DNase-BrxL-> | 1209 | Bacteria>Pseudomonadota>Betaproteobacteria>Rhodocyclales>Zoogloeaceae>Thauera | GCA_016790365.1 | Thauera_sp. | JNL37_10520 | 3981..7610 | 1905334 | BREX-system-P-loop-protein-BrxC-[Thauera-sp.] |
| WP_096450985.1 | Type1.BREX | <-SIGMA+OB-Fold+AAA-ATPase+Lon-Protease-BrxL<-PglZ<-HNH<-OLD-ABC<-N6-METHYLASE<-ORC-CDC6<-ParA-Soj-PloopNTPase<-HEPN<-N6-METHYLASE<-DUF6079*<-BrxB-BrxF-iSTAND<-BrxA-DNA-binding<-BrxE<-XIS-HTH<-wHTH_3\|\|?-><-ParB<-?\|\|HTH_3-> | 1180 | Bacteria>Pseudomonadota>Betaproteobacteria>Rhodocyclales>Zoogloeaceae>Thauera | GCF_002354895.1 | Thauera_sp._K11 | CCZ27_RS19540 | 4450815..4454357 | 2005884 | BREX-system-P-loop-protein-BrxC-[Thauera-sp.-K11] |
| WP_247725420.1 | Type6.BREX | <-wHTH<-ORC-CDC6<-ParA-Soj-PloopNTPase<-HEPN<-N6-METHYLASE-p<-PIN<-?<-N6-METHYLASE<-MNS+MarR-HTH<-DUF6079*<-BrxB-BrxF-iSTAND<-BrxA-DNA-binding<-BrxE<-?\|\|ABC-ATPase+SbcC+ABC-ATPase+SbcC->HAD-> | 1209 | Bacteria>Pseudomonadota>Betaproteobacteria>Rhodocyclales>Zoogloeaceae>Thauera | GCF_023093595.1 | Thauera_aromatica | MXC99_RS10610 | 105350..108979 | 59405 | BREX-system-P-loop-protein-BrxC-[Thauera-aromatica] |
| BDD08311.1 | Type1.BREX | ARG->RHS->HTH_23+HTH_32+HTH_33->DDE_3->RHS->BrxA-DNA-binding->BrxB-BrxF-iSTAND->DUF6079-p-><-DDE_Tnp_1\|\|DUF6079*->ParB+HNH->N6-METHYLASE->HEPN->PAIREDC-HTH+HIN-HTH->N6-METHYLASE-p->DUF3944->?->PglZ->SIGMA+OB-Fold+AAA-ATPase+Lon-Protease-BrxL-> | 1089 | Bacteria>Pseudomonadota>Gammaproteobacteria>Aeromonadales>Aeromonadaceae>Aeromonas | GCA_024347775.1 | Aeromonas_caviae | KAM643c_18840 | 2028863..2032132 | 648 | hypothetical-protein-KAM643c_18840-[Aeromonas-caviae] |
| WP_059169685.1 | Type1.BREX | <-PglZ<-N6-METHYLASE-p<-HEPN<-RVT<-N6-METHYLASE<-?<-HNH<-TOPRIM<-OLD-ABC<-DUF6079*<-BrxB-BrxF-iSTAND<-BrxA-DNA-binding<-RHS<-ARG<-RHS\|\|LexA-protease->?-><-HDRnase<-Arm-DNA-bind_2+Phage_int_SAM_3+Phage_integrase | 1240 | Bacteria>Pseudomonadota>Gammaproteobacteria>Aeromonadales>Aeromonadaceae>Aeromonas | GCF_001481535.2 | Aeromonas_salmonicida | VO70_RS19205 | 18975..22697 | 645 | BREX-system-P-loop-protein-BrxC-[Aeromonas-salmonicida] |
| WP_101617899.1 | Type1.BREX | <-TetR-HTH<-Phage_integrase+Phage_integrase<-Phage_integrase+Phage_integrase<-Phage_integrase<-LexA-protease\|\|?->RHS->BrxA-DNA-binding->BrxB-BrxF-iSTAND->DUF6079*->OLD-ABC->TOPRIM->N6-METHYLASE->N6-METHYLASE-p->HEPN->N6-METHYLASE-p-><-AAA-ATPase<-HTH+rve\|\|DUF3944-> | 1231 | Bacteria>Pseudomonadota>Gammaproteobacteria>Aeromonadales>Aeromonadaceae>Aeromonas | GCF_002850695.3 | Aeromonas_hydrophila | C1A23_RS07645 | 1215070..1218765 | 644 | BREX-system-P-loop-protein-BrxC-[Aeromonas-hydrophila] |
| WP_160837717.1 | Type1.BREX | <-N6-METHYLASE-p<-TIR<-HEPN<-N6-METHYLASE<-TOPRIM<-OLD-ABC<-DUF6079*<-BrxB-BrxF-iSTAND<-BrxA-DNA-binding<-RHS<-ZnR<-ARG<-RHS\|\|LexA-protease->?-><-HDRnase | 1231 | Bacteria>Pseudomonadota>Gammaproteobacteria>Aeromonadales>Aeromonadaceae>Aeromonas | GCF_009831085.1 | Aeromonas_caviae | GR302_RS08560 | 7464..11159 | 648 | BREX-system-P-loop-protein-BrxC-[Aeromonas-caviae] |
| WP_171280306.1 | Type1.BREX | AAA-ToxA+Classical-AAA->Classical-AAA->ARG->?->ZnR->RHS->BrxA-DNA-binding->BrxB-BrxF-iSTAND->DUF6079*->N6-METHYLASE->ParB+HNH->PglZ->HEPN+Sel1+SEC-C-><-SF1-SF2-Helicase+REase<-SF1-SF2-Helicase+REase\|\|HIN-HTH-> | 1157 | Bacteria>Pseudomonadota>Gammaproteobacteria>Aeromonadales>Aeromonadaceae>Aeromonas | GCF_013085505.1 | Aeromonas_sp._2692-1 | E5E97_RS12155 | 2531425..2534898 | 2560029 | BREX-system-P-loop-protein-BrxC-[Aeromonas-sp.-2692-1] |
| WP_210557479.1 | Type1.BREX | DEDD_Tnp_IS110+Transposase_20->\|\|BrxA-DNA-binding->BrxB-BrxF-iSTAND->DUF6079*->HNH->?->N6-METHYLASE->N6-METHYLASE-p->HEPN->N6-METHYLASE-p-> | 1218 | Bacteria>Pseudomonadota>Gammaproteobacteria>Aeromonadales>Aeromonadaceae>Aeromonas | GCF_017939245.1 | Aeromonas_sp._PrichA-15 | J9889_RS17255 | 4362..8018 | 2823360 | BREX-system-P-loop-protein-BrxC-[Aeromonas-sp.-PrichA-15] |
| WP_213387487.1 | Type1.BREX | <-LexA-protease\|\|?-><-wHTH+WYL+WCX\|\|ARG->?->ZnR->RHS->BrxA-DNA-binding->BrxB-BrxF-iSTAND->DUF6079*->N6-METHYLASE->OLD-ABC->HNH->PglZ->HEPN+Sel1+SEC-C->HIN-HTH-><-SF1-SF2-Helicase+REase<-SF1-SF2-Helicase+REase | 1157 | Bacteria>Pseudomonadota>Gammaproteobacteria>Aeromonadales>Aeromonadaceae>Aeromonas | GCF_018359765.1 | Aeromonas_veronii | J4G64_RS11995 | 290568..294041 | 654 | BREX-system-P-loop-protein-BrxC-[Aeromonas-veronii] |
| WP_234578020.1 | Type1.BREX | <-SIGMA+OB-Fold+AAA-ATPase+Lon-Protease-BrxL<-PglZ<-?<-DUF3944<-N6-METHYLASE-p<-HEPN<-N6-METHYLASE<-TOPRIM<-OLD-ABC<-DUF6079*<-BrxB-BrxF-iSTAND<-BrxA-DNA-binding<-RHS<-ZnR+ZnR\|\|wHTH+WYL+WCX-><-?\|\|LexA-protease->?-><-SF1-SF2-Helicase | 1231 | Bacteria>Pseudomonadota>Gammaproteobacteria>Aeromonadales>Aeromonadaceae>Aeromonas | GCF_021440865.1 | Aeromonas_salmonicida | LZ632_RS15270 | 23400..27095 | 645 | BREX-system-P-loop-protein-BrxC-[Aeromonas-salmonicida] |
| WP_139392567.1 | Type1.BREX | <-Phage_integrase+Phage_integrase<-Phage_integrase<-LexA-protease\|\|?-><-wHTH+WYL+WCX\|\|?->RHS->BrxA-DNA-binding->BrxB-BrxF-iSTAND->DUF6079*->OLD-ABC->TOPRIM->N6-METHYLASE->RVT->HEPN->N6-METHYLASE-p->DUF3944->?->PglZ-> | 1225 | Bacteria>Pseudomonadota>Gammaproteobacteria>Aeromonadales>Aeromonadaceae>Aeromonas | GCF_021654355.1 | Aeromonas_hydrophila | L6424_RS03515 | 743150..746827 | 644 | BREX-system-P-loop-protein-BrxC-[Aeromonas-hydrophila] |
| WP_261925329.1 | Type1.BREX | <-LexA-protease\|\|ARG->ZnR->RHS->RHS->BrxA-DNA-binding->BrxB-BrxF-iSTAND->DUF6079-p-><-DDE_Tnp_1\|\|DUF6079*->ParB+HNH->N6-METHYLASE->HEPN->N6-METHYLASE-p->PAIREDC-HTH+HIN-HTH->N6-METHYLASE-p->DUF3944->?->PglZ-> | 1089 | Bacteria>Pseudomonadota>Gammaproteobacteria>Aeromonadales>Aeromonadaceae>Aeromonas | GCF_024347775.1 | Aeromonas_caviae | OCX50_RS09760 | 2028863..2032132 | 648 | BREX-system-P-loop-protein-BrxC-[Aeromonas-caviae] |
| WP_257712004.1 | Type1.BREX | <-PglZ<-TOPRIM<-OLD-ABC<-N6-METHYLASE-p<-HEPN<-RVT<-N6-METHYLASE<-TOPRIM<-OLD-ABC<-DUF6079*<-BrxB-BrxF-iSTAND<-BrxA-DNA-binding<-RHS<-?\|\|wHTH+WYL+WCX-> | 1281 | Bacteria>Pseudomonadota>Gammaproteobacteria>Aeromonadales>Aeromonadaceae>Aeromonas | GCF_024649525.1 | Aeromonas_caviae | NUK31_RS14230 | 17564..21409 | 648 | BREX-system-P-loop-protein-BrxC-[Aeromonas-caviae] |
| WP_265783524.1 | Type1.BREX | <-TOPRIM<-OLD-ABC<-N6-METHYLASE-p<-TIR<-HEPN<-?<-HEPN<-TOPRIM<-OLD-ABC<-N6-METHYLASE<-DUF6079*<-BrxB-BrxF-iSTAND<-BrxA-DNA-binding<-RHS<-?\|\|LexA-protease->?->PBPB+PBPB+PAS+PAS+HISKIN+REC+HPT->REC+EAL-> | 1235 | Bacteria>Pseudomonadota>Gammaproteobacteria>Aeromonadales>Aeromonadaceae>Aeromonas | GCF_026230585.1 | Aeromonas_veronii | OD810_RS03320 | 107717..111424 | 654 | BREX-system-P-loop-protein-BrxC-[Aeromonas-veronii] |
| WP_270698108.1 | Type1.BREX | <-LexA-protease\|\|?-><-wHTH+WYL+WCX\|\|?->RHS->BrxA-DNA-binding->BrxB-BrxF-iSTAND->DUF6079*->OLD-ABC->TOPRIM->N6-METHYLASE->RVT->HEPN->N6-METHYLASE->OLD-ABC->TOPRIM->PglZ-> | 1280 | Bacteria>Pseudomonadota>Gammaproteobacteria>Aeromonadales>Aeromonadaceae>Aeromonas | GCF_027705185.1 | Aeromonas_sp._QDB05 | OKC32_RS09690 | 4766..8608 | 2990478 | BREX-system-P-loop-protein-BrxC-[Aeromonas-sp.-QDB05] |
| WP_201934227.1 | Type1.BREX | RHS->BrxA-DNA-binding->BrxB-BrxF-iSTAND->DUF6079*->ParB+HNH->N6-METHYLASE->HEPN->N6-METHYLASE-p->DUF3944->?->PglZ->SIGMA+OB-Fold+AAA-ATPase+Lon-Protease-BrxL-> | 1231 | Bacteria>Pseudomonadota>Gammaproteobacteria>Aeromonadales>Aeromonadaceae | GCF_019972675.1 | Aeromonas | MWG22_RS17445 | 1818..5513 | 642 | MULTISPECIES:-BREX-system-P-loop-protein-BrxC-[Aeromonas] |
| WP_171273737.1 | Type1.BREX | Arm-DNA-bind_2+Phage_int_SAM_3+Phage_integrase->HDRnase-><-?\|\|RHS->ARG->ZnR->RHS->BrxA-DNA-binding->BrxB-BrxF-iSTAND->DUF6079*->OLD-ABC->TOPRIM->N6-METHYLASE->HEPN->TIR->N6-METHYLASE-p->DUF3944->?->?->\|\|PglZ-> | 1231 | Bacteria>Pseudomonadota>Gammaproteobacteria>Aeromonadales>Aeromonadaceae | GCF_021654375.1 | Aeromonas | E4185_RS15845 | 3355159..3358854 | 642 | MULTISPECIES:-BREX-system-P-loop-protein-BrxC-[Aeromonas] |
| GGW57088.1 | Type1.BREX | <-wHTH+WYL+WCX-p\|\|SIR2->?-><-?\|\|HEPN-p->JAB->BrxA-DNA-binding->BrxB-BrxF-iSTAND->DUF6079*->N6-METHYLASE->OLD-ABC->HNH->PglZ->HEPN+Sel1+SEC-C->Immank+ANK+ANK->REase-><-?<-SF1-SF2-Helicase+REase | 1114 | Bacteria>Pseudomonadota>Gammaproteobacteria>Alteromonadales>Alteromonadaceae>Alishewanella | GCA_014652375.1 | Alishewanella_tabrizica | GCM10008111_11430 | 80462..83806 | 671278 | hypothetical-protein-GCM10008111_11430-[Alishewanella-tabrizica] |
| WP_229796961.1 | Type1.BREX | <-wHTH+WYL+WCX-p\|\|SIR2->?-><-?\|\|HEPN-p->JAB->BrxA-DNA-binding->BrxB-BrxF-iSTAND->DUF6079*->N6-METHYLASE->OLD-ABC->HNH->PglZ->HEPN+Sel1+SEC-C->Immank+ANK+ANK->REase-><-?<-SF1-SF2-Helicase+REase | 1125 | Bacteria>Pseudomonadota>Gammaproteobacteria>Alteromonadales>Alteromonadaceae>Alishewanella | GCF_014652375.1 | Alishewanella_tabrizica | IEO70_RS05825 | 80429..83806 | 671278 | BREX-system-P-loop-protein-BrxC-[Alishewanella-tabrizica] |
| WP_240919947.1 | Type1.BREX | BrxA-DNA-binding->BrxB-BrxF-iSTAND->DUF6079*->N6-METHYLASE->ParB+HNH->OLD-ABC->Tox-RelE->PglZ->SIGMA+OB-Fold+AAA-ATPase+Lon-Protease-BrxL->HEPN-> | 1180 | Bacteria>Pseudomonadota>Gammaproteobacteria>Alteromonadales>Alteromonadaceae>Paraglaciecola | GCF_011378905.1 | Paraglaciecola_sp._20A4 | GQS54_RS13935 | 91436..94978 | 2687288 | BREX-system-P-loop-protein-BrxC-[Paraglaciecola-sp.-20A4] |
| WP_058373560.1 | Type1.BREX | <-SIGMA+OB-Fold+AAA-ATPase+Lon-Protease-BrxL<-PglZ<-SF1-SF2-Helicase<-?<-N6-METHYLASE<-HEPN<-N6-METHYLASE<-DUF6079*<-BrxB-BrxF-iSTAND<-BrxA-DNA-binding<-wHTH+WYL+WCX<-HTH_26\|\|Phage_AlpA->Arm-DNA-bind_2+Phage_int_SAM_3+Phage_integrase->HDRnase-><-Phage_integrase<-Phage_integrase+Phage_integrase | 1229 | Bacteria>Pseudomonadota>Gammaproteobacteria>Alteromonadales>Pseudoalteromonadaceae>Pseudoalteromonas | GCF_001465295.1 | Pseudoalteromonas_translucida | PTRA_a2157 | 1987847..1991536 | 166935 | BREX-system-P-loop-protein-BrxC-[Pseudoalteromonas-translucida] |
| WP_233998422.1 | Type1.BREX | <-PglZ<-?<-?<-SF1-SF2-Helicase<-HEPN<-N6-METHYLASE<-N6-METHYLASE-p<-HATPase+wHTH<-Schlafen+HATPase+wHTH<-DUF6079*<-BrxB-BrxF-iSTAND<-BrxA-DNA-binding<-wHTH+WYL+WCX<-?<-?\|\|?-><-?\|\|?-><-SIR2 | 1252 | Bacteria>Pseudomonadota>Gammaproteobacteria>Alteromonadales>Pseudoalteromonadaceae>Pseudoalteromonas | GCF_002964785.1 | Pseudoalteromonas_sp._T11g48 | C2M29_RS12220 | 655771..659529 | 2077100 | BREX-system-P-loop-protein-BrxC-[Pseudoalteromonas-sp.-T11g48] |

<table>
<tr><th>Protein-ID</th><th>System</th><th>Gene-Neighbourhoods</th><th>Length</th><th>Taxonomy</th><th>Assembly-ID</th><th>Strain</th><th>Locus-Tag</th><th>Genomic-coordinates</th><th>TaxID</th><th>Fasta-defline</th></tr>
<tr><td>WP_182739649.1</td><td>Type1.BREX</td><td><-Phage_integrase<-DUF2787<-?||XIS-HTH->HEPN->?->wHTH+WYL+WCX->BrxA-DNA-binding->BrxB-BrxF-iSTAND->DUF6079*->N6-METHYLASE-p-></td><td>1235</td><td>Bacteria>Pseudomonadota>Gammaproteobacteria>Alteromonadales>Pseudoalteromonadaceae>Pseudoalteromonas</td><td>GCF_014163545.1</td><td>Pseudoalteromonas_sp._SR45-6</td><td>H5158_RS05620</td><td>135410..139117</td><td>2760927</td><td>BREX-system-P-loop-protein-BrxC-[Pseudoalteromonas-sp.-SR45-6]</td></tr>
<tr><td>WP_193038371.1</td><td>Type1.BREX</td><td><-ZetaToxin<-?<-PglZ<-ParB+HNH<-N6-METHYLASE<-DUF6079*<-BrxB-BrxF-iSTAND<-BrxA-DNA-binding<-wHTH+WYL+WCX<-?<-HEPN<-XIS-HTH||?->DUF2787->Phage_integrase-></td><td>1159</td><td>Bacteria>Pseudomonadota>Gammaproteobacteria>Alteromonadales>Pseudoalteromonadaceae>Pseudoalteromonas</td><td>GCF_014897925.1</td><td>Pseudoalteromonas_nigrifaciens</td><td>INT11_RS00030</td><td>7982..11461</td><td>28109</td><td>BREX-system-P-loop-protein-BrxC-[Pseudoalteromonas-nigrifaciens]</td></tr>
<tr><td>WP_248533819.1</td><td>Type1.BREX</td><td><-MarR-HTH||?-><-?<-?||BrxA-DNA-binding->BrxB-BrxF-iSTAND->DUF6079*->SSB->GP+REase->SF1-SF2-Helicase+PRTase->SLOG+BirA-HTH->N6-METHYLASE->HEPN->N6-METHYLASE->DnaB+McrB-AAA->wHTH+REase+ZnR-></td><td>1229</td><td>Bacteria>Pseudomonadota>Gammaproteobacteria>Alteromonadales>Pseudoalteromonadaceae>Pseudoalteromonas</td><td>GCF_023161935.1</td><td>Pseudoalteromonas_sp._2CM28B</td><td>MTN96_RS29910</td><td>10677..14366</td><td>2929851</td><td>BREX-system-P-loop-protein-BrxC-[Pseudoalteromonas-sp.-2CM28B]</td></tr>
<tr><td>WP_037429580.1</td><td>Type1.BREX</td><td><-PglZ<-ParB+HNH<-N6-METHYLASE<-DUF6079*<-BrxB-BrxF-iSTAND<-BrxA-DNA-binding<-?<-HEPN<-?<-MoxR-AAA<-NtrC-AAA+FIS-HTH<-XIS-HTH</td><td>1157</td><td>Bacteria>Pseudomonadota>Gammaproteobacteria>Alteromonadales>Shewanellaceae</td><td>GCF_000282755.1</td><td>Shewanella</td><td>SHEWPOL2_RS15310</td><td>7136..10609</td><td>22</td><td>MULTISPECIES:-BREX-system-P-loop-protein-BrxC-[Shewanella]</td></tr>
<tr><td>HAY94623.1</td><td>Type1.BREX</td><td><-DUF2787<-CbeA_antitoxin<-?<-?||XIS-HTH->wHTH+WYL+WCX-p->wHTH+WYL+WCX->BrxA-DNA-binding->BrxB-BrxF-iSTAND->DUF6079*->HNH->?->?->ParB+HNH->N6-METHYLASE->HEPN->N6-METHYLASE-p-></td><td>1224</td><td>Bacteria>Pseudomonadota>Gammaproteobacteria>Alteromonadales>Shewanellaceae>Shewanella</td><td>GCA_003486485.1</td><td>Shewanella_sp.</td><td>DCY70_12125</td><td>4933..8607</td><td>50422</td><td>BREX-system-P-loop-protein-BrxC-[Shewanella-sp.]</td></tr>
<tr><td>MCD8561045.1</td><td>Type1.BREX</td><td><-N6-METHYLASE-p<-HEPN<-N6-METHYLASE-p<-N6-METHYLASE-p<-N6-METHYLASE<-?<-HNH<-DUF6079*<-BrxB-BrxF-iSTAND<-BrxA-DNA-binding<-wHTH+WYL+WCX<-XIS-HTH</td><td>1231</td><td>Bacteria>Pseudomonadota>Gammaproteobacteria>Alteromonadales>Shewanellaceae>Shewanella</td><td>GCA_021209305.1</td><td>Shewanella_xiamenensis</td><td>LRY75_19995</td><td>6155..9850</td><td>332186</td><td>BREX-system-P-loop-protein-BrxC-[Shewanella-xiamenensis]</td></tr>
<tr><td>WP_011622483.1</td><td>Type1.BREX</td><td>XIS-HTH->NtrC-AAA+FIS-HTH->MoxR-AAA->?->HEPN->?->BrxA-DNA-binding->BrxB-BrxF-iSTAND->DUF6079*->N6-METHYLASE->ParB+HNH->PglZ->ZetaToxin->?-><-SF1-SF2-Helicase+REase<-SF1-SF2-Helicase+REase||HerA/FtsK-></td><td>1157</td><td>Bacteria>Pseudomonadota>Gammaproteobacteria>Alteromonadales>Shewanellaceae>Shewanella</td><td>GCF_000014685.1</td><td>Shewanella_sp._MR-4</td><td>SHEWMR4_RS09030</td><td>2019364..2022837</td><td>60480</td><td>BREX-system-P-loop-protein-BrxC-[Shewanella-sp.-MR-4]</td></tr>
<tr><td>WP_011716828.1</td><td>Type1.BREX</td><td>XIS-HTH->NtrC-AAA+FIS-HTH->MoxR-AAA->?->HEPN->?->BrxA-DNA-binding->BrxB-BrxF-iSTAND->DUF6079*->N6-METHYLASE->ParB+HNH->PglZ->ZetaToxin->?-><-SF1-SF2-Helicase+REase<-SF1-SF2-Helicase+REase||HerA/FtsK-></td><td>1157</td><td>Bacteria>Pseudomonadota>Gammaproteobacteria>Alteromonadales>Shewanellaceae>Shewanella</td><td>GCF_000203935.1</td><td>Shewanella_sp._ANA-3</td><td>SHEWANA3_RS09590</td><td>2144277..2147750</td><td>94122</td><td>BREX-system-P-loop-protein-BrxC-[Shewanella-sp.-ANA-3]</td></tr>
<tr><td>WP_028764576.1</td><td>Type1.BREX</td><td>BrxA-DNA-binding->BrxB-BrxF-iSTAND->DUF6079*->N6-METHYLASE->ParB+HNH->PglZ->HEPN+Sel1+SEC-C-><-SF1-SF2-Helicase+REase<-SF1-SF2-Helicase+REase<-?<-MerR-HTH</td><td>1157</td><td>Bacteria>Pseudomonadota>Gammaproteobacteria>Alteromonadales>Shewanellaceae>Shewanella</td><td>GCF_000518705.1</td><td>Shewanella_colwelliana</td><td>L876_RS0115155</td><td>4861..8334</td><td>23</td><td>BREX-system-P-loop-protein-BrxC-[Shewanella-colwelliana]</td></tr>
<tr><td>WP_055648628.1</td><td>Type1.BREX</td><td><-PVC-AAA<-?||SF1-SF2-Helicase+REase->SF1-SF2-Helicase+REase-><-?<-ZetaToxin<-PglZ<-ParB+HNH<-N6-METHYLASE<-DUF6079*<-BrxB-BrxF-iSTAND<-BrxA-DNA-binding<-?<-HEPN</td><td>1157</td><td>Bacteria>Pseudomonadota>Gammaproteobacteria>Alteromonadales>Shewanellaceae>Shewanella</td><td>GCF_001308045.1</td><td>Shewanella_sp._Sh95</td><td>AEA42_RS19845</td><td>36695..40168</td><td>1689868</td><td>BREX-system-P-loop-protein-BrxC-[Shewanella-sp.-Sh95]</td></tr>
<tr><td>WP_244159705.1</td><td>Type1.BREX</td><td><-HDRnase<-DDE_Tnp_1<-Arm-DNA-bind_2+Phage_int_SAM_3+Phage_integrase<-Phage_AlpA||HTH_26->wHTH+WYL+WCX->BrxA-DNA-binding->BrxB-BrxF-iSTAND->DUF6079*->HNH->?->N6-METHYLASE->N6-METHYLASE-p->HEPN->N6-METHYLASE-p->MULE-TRANSPOSASE->N6-METHYLASE-p-></td><td>1254</td><td>Bacteria>Pseudomonadota>Gammaproteobacteria>Alteromonadales>Shewanellaceae>Shewanella</td><td>GCF_001723195.1</td><td>Shewanella_xiamenensis</td><td>ABT47_RS18205</td><td>4085889..4089653</td><td>332186</td><td>BREX-system-P-loop-protein-BrxC-[Shewanella-xiamenensis]</td></tr>
<tr><td>WP_069454895.1</td><td>Type1.BREX</td><td><-HDRnase<-Arm-DNA-bind_2+Phage_int_SAM_3+Phage_integrase<-Phage_AlpA||HTH_26->wHTH+WYL+WCX->BrxA-DNA-binding->BrxB-BrxF-iSTAND->DUF6079*->HNH->?->N6-METHYLASE->N6-METHYLASE-p->HEPN->N6-METHYLASE-p->MULE-TRANSPOSASE->N6-METHYLASE-p-></td><td>1240</td><td>Bacteria>Pseudomonadota>Gammaproteobacteria>Alteromonadales>Shewanellaceae>Shewanella</td><td>GCF_002074855.1</td><td>Shewanella_xiamenensis</td><td>V000_RS12940</td><td>4702..8424</td><td>332186</td><td>BREX-system-P-loop-protein-BrxC-[Shewanella-xiamenensis]</td></tr>
<tr><td>WP_080916542.1</td><td>Type1.BREX</td><td><-Pkinase+STYKIN||?->?->?->wHTH+WYL+WCX->BrxA-DNA-binding->BrxB-BrxF-iSTAND->DUF6079-p->DUF6079*->ParB+HNH->ParB+HNH-p->HEPN->PglZ->SIGMA+OB-Fold+AAA-ATPase+Lon-Protease-BrxL-><-MarR-HTH||?->?->Phage_int_SAM_4-></td><td>1234</td><td>Bacteria>Pseudomonadota>Gammaproteobacteria>Alteromonadales>Shewanellaceae>Shewanella</td><td>GCF_002075795.1</td><td>Shewanella_japonica</td><td>SJ2017_RS16905</td><td>3936240..3939944</td><td>93973</td><td>BREX-system-P-loop-protein-BrxC-[Shewanella-japonica]</td></tr>
<tr><td>WP_088585687.1</td><td>Type1.BREX</td><td>ZnR-><-HEPN-p<-HEPN-p<-SIGMA+OB-Fold+AAA-ATPase+Lon-Protease-BrxL<-PglZ<-N6-METHYLASE-p||DDE_Tnp_1-><-N6-METHYLASE<-DUF6079*<-BrxB-BrxF-iSTAND<-BrxA-DNA-binding<-DDE_Tnp_1<-Prok-JAB<-wHTH+WYL+WCX<-?<-XIS-HTH</td><td>1229</td><td>Bacteria>Pseudomonadota>Gammaproteobacteria>Alteromonadales>Shewanellaceae>Shewanella</td><td>GCF_002209245.2</td><td>Shewanella_sp._FDAARGOS_354</td><td>CEQ32_RS05910</td><td>1298859..1302548</td><td>1930557</td><td>BREX-system-P-loop-protein-BrxC-[Shewanella-sp.-FDAARGOS_354]</td></tr>
<tr><td>WP_208124435.1</td><td>Type1.BREX</td><td>XIS-HTH->NtrC-AAA+FIS-HTH->MoxR-AAA->?->HEPN->BrxA-DNA-binding->BrxB-BrxF-iSTAND->DUF6079*->N6-METHYLASE->ParB+HNH->PglZ->ZetaToxin-><-SF1-SF2-Helicase+REase<-SF1-SF2-Helicase+REase||REase->HIN-HTH-><-GDResolvase+HIN-HTH</td><td>1157</td><td>Bacteria>Pseudomonadota>Gammaproteobacteria>Alteromonadales>Shewanellaceae>Shewanella</td><td>GCF_017570045.1</td><td>Shewanella_algae</td><td>JKK45_RS07515</td><td>1709365..1712838</td><td>38313</td><td>BREX-system-P-loop-protein-BrxC-[Shewanella-algae]</td></tr>
<tr><td>WP_144191359.1</td><td>Type1.BREX</td><td>XIS-HTH->NtrC-AAA+FIS-HTH->MoxR-AAA->?->HEPN->BrxA-DNA-binding->BrxB-BrxF-iSTAND->DUF6079*->N6-METHYLASE->ParB+HNH->PglZ->ZetaToxin->?-><-SF1-SF2-Helicase+REase<-SF1-SF2-Helicase+REase</td><td>1157</td><td>Bacteria>Pseudomonadota>Gammaproteobacteria>Alteromonadales>Shewanellaceae>Shewanella</td><td>GCF_017580415.1</td><td>Shewanella_algae</td><td>I6M45_RS07395</td><td>495996..499469</td><td>38313</td><td>BREX-system-P-loop-protein-BrxC-[Shewanella-algae]</td></tr>
<tr><td>WP_224020606.1</td><td>Type1.BREX</td><td><-HerA/FtsK||SF1-SF2-Helicase+REase->SF1-SF2-Helicase+REase-><-?<-ZetaToxin<-PglZ<-ParB+HNH<-N6-METHYLASE<-DUF6079*<-BrxB-BrxF-iSTAND<-BrxA-DNA-binding<-?<-HEPN<-?<-MoxR-AAA<-NtrC-AAA+FIS-HTH<-XIS-HTH</td><td>1157</td><td>Bacteria>Pseudomonadota>Gammaproteobacteria>Alteromonadales>Shewanellaceae>Shewanella</td><td>GCF_019973655.1</td><td>Shewanella_xiamenensis</td><td>LDO18_RS12250</td><td>2746465..2749938</td><td>332186</td><td>BREX-system-P-loop-protein-BrxC-[Shewanella-xiamenensis]</td></tr>
<tr><td>WP_248084385.1</td><td>Type1.BREX</td><td>SF1-SF2-Helicase-><-DUF2787<-CbeA_antitoxin<-?<-?||XIS-HTH->wHTH+WYL+WCX->BrxA-DNA-binding->BrxB-BrxF-iSTAND->DUF6079*->HNH->?->N6-METHYLASE->N6-METHYLASE-p->HEPN->N6-METHYLASE-p-></td><td>1231</td><td>Bacteria>Pseudomonadota>Gammaproteobacteria>Alteromonadales>Shewanellaceae>Shewanella</td><td>GCF_023159095.1</td><td>unclassified_Shewanella</td><td>MOD77_RS02670</td><td>216752..220447</td><td>196818</td><td>MULTISPECIES:-BREX-system-P-loop-protein-BrxC-[unclassified-Shewanella]</td></tr>
<tr><td>WP_249553736.1</td><td>Type1.BREX</td><td><-SIGMA+OB-Fold+AAA-ATPase+Lon-Protease-BrxL<-PglZ<-?<-N6-METHYLASE<-HEPN<-RVT<-N6-METHYLASE-p<-N6-METHYLASE<-DUF6079*<-BrxB-BrxF-iSTAND<-BrxA-DNA-binding<-wHTH+WYL+WCX||?->?->CbeA_antitoxin->DUF2787->DUF2787->Phage_integrase-></td><td>1268</td><td>Bacteria>Pseudomonadota>Gammaproteobacteria>Alteromonadales>Shewanellaceae>Shewanella</td><td>GCF_023475945.1</td><td>Shewanella_sp._8A</td><td>M2897_RS12020</td><td>2756867..2760673</td><td>2943323</td><td>BREX-system-P-loop-protein-BrxC-[Shewanella-sp.-8A]</td></tr>
<tr><td>WP_259573703.1</td><td>Type1.BREX</td><td><-CHTH<-REase<-SIGMA+OB-Fold+AAA-ATPase+Lon-Protease-BrxL<-PglZ<-N6-METHYLASE-p<-HEPN<-RVT<-N6-METHYLASE-p<-N6-METHYLASE<-DUF6079*<-BrxB-BrxF-iSTAND<-BrxA-DNA-binding<-wHTH+WYL+WCX<-HTH_26||Phage_AlpA->Arm-DNA-bind_2+Phage_int_SAM_3+Phage_integrase->HDRnase->?-><-SF1-SF2-Helicase</td><td>1240</td><td>Bacteria>Pseudomonadota>Gammaproteobacteria>Alteromonadales>Shewanellaceae>Shewanella</td><td>GCF_024999165.1</td><td>Shewanella_baltica</td><td>G7A68_RS06675</td><td>25637..29359</td><td>62322</td><td>BREX-system-P-loop-protein-BrxC-[Shewanella-baltica]</td></tr>
<tr><td>WP_263197229.1</td><td>Type1.BREX</td><td><-SIGMA+OB-Fold+AAA-ATPase+Lon-Protease-BrxL<-PglZ<-N6-METHYLASE-p<-HEPN<-N6-METHYLASE-p<-?<-HNH<-DUF6079*<-BrxB-BrxF-iSTAND<-BrxA-DNA-binding<-wHTH+WYL+WCX<-HTH_26||Phage_AlpA->Arm-DNA-bind_2+Phage_int_SAM_3+Phage_integrase->HDRnase-></td><td>1246</td><td>Bacteria>Pseudomonadota>Gammaproteobacteria>Alteromonadales>Shewanellaceae>Shewanella</td><td>GCF_025654545.1</td><td>Shewanella_sp._SM20</td><td>L5M18_RS13095</td><td>42119..45859</td><td>2912792</td><td>BREX-system-P-loop-protein-BrxC-[Shewanella-sp.-SM20]</td></tr>
<tr><td>WP_101520955.1</td><td>Type1.BREX</td><td>HEPN->HNH+Polbeta->?->?->Peptidase_M29+DUF2971->XIS-HTH->BrxA-DNA-binding->BrxB-BrxF-iSTAND->DUF6079*->N6-METHYLASE->MarR-HTH->Polbeta->PglZ->SIGMA+OB-Fold+AAA-ATPase+Lon-Protease-BrxL->HEPN+SF1-SF2-Helicase+RNase+RNase->TPR-><-?<-Arm-DNA-bind_3+Phage_integrase</td><td>1151</td><td>Bacteria>Pseudomonadota>Gammaproteobacteria>Cellvibrionales>Halieaceae>Kineobactrum</td><td>GCF_002868715.1</td><td>Kineobactrum_sediminis</td><td>CWI75_RS08140</td><td>627029..630484</td><td>1905677</td><td>BREX-system-P-loop-protein-BrxC-[Kineobactrum-sediminis]</td></tr>
<tr><td>MBN49711.1</td><td>Type1.BREX</td><td><-SIGMA+OB-Fold+AAA-ATPase+Lon-Protease-BrxL<-PglZ<-DOC+HTH<-?<-ZNR+RmuC<-HEPN<-?<-N6-METHYLASE<-?<-DUF6079*<-BrxB-BrxF-iSTAND<-BrxA-DNA-binding||wHTH+WYL+WCX->?-><-?<-SF1-SF2-Helicase||?->?->Phage_integrase-></td><td>1193</td><td>Bacteria>Pseudomonadota>Gammaproteobacteria>Cellvibrionales>Spongiibacteraceae>unclassified_Spongiibacteraceae</td><td>GCA_002723235.1</td><td>Spongiibacteraceae_bacterium</td><td>CMN85_09225</td><td>177592..181173</td><td>2026793</td><td>hypothetical-protein-[Spongiibacteraceae-bacterium]</td></tr>
<tr><td>QVL47244.1</td><td>Type6.BREX</td><td><-PIN<-RHH<-N6-METHYLASE<-TOPRIM<-OLD-ABC<-N6-METHYLASE-p<-ORC-CDC6<-ParA-Soj-PloopNTPase<-HEPN<-DUF6079*<-?<-BrxB-BrxF-iSTAND<-BrxA-DNA-binding<-BrxE<-?<-NTP_transf_3+Hexapep+Hexapep</td><td>1179</td><td>Bacteria>Pseudomonadota>Gammaproteobacteria>Chromatiales>Chromatiaceae>Thiocapsa</td><td>GCA_018399035.1</td><td>Thiocapsa_sp.</td><td>KFB96_16145</td><td>3553103..3556642</td><td>2024551</td><td>MAG:-BREX-system-P-loop-protein-BrxC-[Thiocapsa-sp.]</td></tr>
<tr><td>MBK8639641.1</td><td>Type6.BREX</td><td>REC-><-?||BrxE->BrxA-DNA-binding->BrxB-BrxF-iSTAND->DUF6079*->?->N6-METHYLASE->RHH->PIN->PglZ->PolB->HEPN->SIGMA+OB-Fold+AAA-ATPase+Lon-Protease-BrxL-></td><td>1178</td><td>Bacteria>Pseudomonadota>Gammaproteobacteria>Chromatiales>Chromatiaceae>unclassified_Chromatiaceae</td><td>GCA_016714645.1</td><td>Chromatiaceae_bacterium</td><td>IPN92_15725</td><td>846355..849891</td><td>2049432</td><td>BREX-system-P-loop-protein-BrxC-[Chromatiaceae-bacterium]</td></tr>
<tr><td>MCF7977306.1</td><td>Type6.BREX</td><td><-ParA-Soj-PloopNTPase<-HEPN<-DUF6079*<-BrxB-BrxF-iSTAND<-BrxA-DNA-binding<-BrxE||?->MlaE->ABC-ATPase+SbcC-></td><td>1182</td><td>Bacteria>Pseudomonadota>Gammaproteobacteria>Chromatiales>Chromatiaceae>unclassified_Chromatiaceae</td><td>GCA_021736685.1</td><td>Chromatiaceae_bacterium</td><td>K9L82_04715</td><td>1611..5159</td><td>2049432</td><td>BREX-system-P-loop-protein-BrxC-[Chromatiaceae-bacterium]</td></tr>
<tr><td>RJT19525.1</td><td>Type1.BREX</td><td><-PglZ<-RVT<-HEPN<-N6-METHYLASE<-ParB+HNH<-DUF6079*<-BrxB-BrxF-iSTAND<-BrxA-DNA-binding<-?<-XIS-HTH||?->?->DUF932->?->wHTH+WYL-></td><td>1233</td><td>Bacteria>Pseudomonadota>Gammaproteobacteria>Enterobacterales>Enterobacteriaceae>Buttiauxella</td><td>GCA_003601925.1</td><td>Buttiauxella_izardii</td><td>D6029_18525</td><td>11062..14763</td><td>82991</td><td>BREX-system-P-loop-protein-BrxC-[Buttiauxella-izardii]</td></tr>
<tr><td>WP_244212427.1</td><td>Type1.BREX</td><td><-PglZ<-N6-METHYLASE-p<-N6-METHYLASE-p<-RVT<-HEPN<-N6-METHYLASE<-ParB+HNH<-DUF6079*<-BrxB-BrxF-iSTAND<-BrxA-DNA-binding<-?<-XIS-HTH||?->?->DUF932->?->wHTH+WYL-></td><td>1237</td><td>Bacteria>Pseudomonadota>Gammaproteobacteria>Enterobacterales>Enterobacteriaceae>Buttiauxella</td><td>GCF_003601925.1</td><td>Buttiauxella_izardii</td><td>D6029_RS18475</td><td>11062..14775</td><td>82991</td><td>BREX-system-P-loop-protein-BrxC-[Buttiauxella-izardii]</td></tr>
<tr><td>WP_181630678.1</td><td>Type1.BREX</td><td><-Sulfatase<-?<-?<-?<-Schlafen+HATPase+wHTH<-PHP+SbcC<-?||BrxA-DNA-binding->BrxB-BrxF-iSTAND->DUF6079*->ParB+HNH->N6-METHYLASE->HEPN-><-HIN-HTH||N6-METHYLASE-p->N6-METHYLASE-p->PglZ->SIGMA+OB-Fold+AAA-ATPase+Lon-Protease-BrxL-><-HTH_23+HTH_32+HTH_33+DDE_3</td><td>1233</td><td>Bacteria>Pseudomonadota>Gammaproteobacteria>Enterobacterales>Enterobacteriaceae>Citrobacter</td><td>-</td><td>Citrobacter_sp._RHBSTW-01013</td><td>HV350_21470</td><td>4470372..4474073</td><td>2742677</td><td>BREX-system-P-loop-protein-BrxC-[Citrobacter-sp.-RHBSTW-01013]</td></tr>
<tr><td>QLR26611.1</td><td>Type1.BREX</td><td><-Sulfatase<-?<-?<-?<-Schlafen+HATPase+wHTH<-PHP+SbcC<-?||BrxA-DNA-binding->BrxB-BrxF-iSTAND->DUF6079*->ParB+HNH->N6-METHYLASE->HEPN-><-HIN-HTH||N6-METHYLASE-p->N6-METHYLASE-p->PglZ->SIGMA+OB-Fold+AAA-ATPase+Lon-Protease-BrxL-><-HTH_23+HTH_32+HTH_33+DDE_3</td><td>1233</td><td>Bacteria>Pseudomonadota>Gammaproteobacteria>Enterobacterales>Enterobacteriaceae>Citrobacter</td><td>GCA_013781985.1</td><td>Citrobacter_sp._RHBSTW-01013</td><td>HV350_21470</td><td>4470372..4474073</td><td>2742677</td><td>BREX-system-P-loop-protein-BrxC-[Citrobacter-sp.-RHBSTW-01013]</td></tr>
<tr><td>MBM9568541.1</td><td>Type1.BREX</td><td><-wHTH+WYL<-DUF932<-nSTAND1+GTPase-AIG+sGTP<-CopG/RHH||XIS-HTH->?->?->BrxA-DNA-binding->BrxB-BrxF-iSTAND->DUF6079*->ParB+HNH->N6-METHYLASE->HEPN->RVT->N6-METHYLASE-p->PglZ->SIGMA+OB-Fold+AAA-ATPase+Lon-Protease-BrxL-><-Arm-DNA-bind_3+Phage_integrase</td><td>1233</td><td>Bacteria>Pseudomonadota>Gammaproteobacteria>Enterobacterales>Enterobacteriaceae>Citrobacter</td><td>GCA_014931625.2</td><td>Citrobacter_sedlakii</td><td>INP82_14060</td><td>158441..162142</td><td>67826</td><td>BREX-system-P-loop-protein-BrxC-[Citrobacter-sedlakii]</td></tr>
<tr><td>WP_249418019.1</td><td>Type1.BREX</td><td><-Sulfatase<-?<-?<-?<-Schlafen+HATPase+wHTH<-PHP+SbcC<-?||BrxA-DNA-binding->BrxB-BrxF-iSTAND->DUF6079*->ParB+HNH->N6-METHYLASE->HEPN->RVT-><-HIN-HTH||RVT->N6-METHYLASE-p->N6-METHYLASE-p->PglZ-></td><td>1240</td><td>Bacteria>Pseudomonadota>Gammaproteobacteria>Enterobacterales>Enterobacteriaceae>Citrobacter</td><td>GCF_013781985.1</td><td>Citrobacter_sp._RHBSTW-01013</td><td>HV350_RS21465</td><td>4470351..4474073</td><td>2742677</td><td>BREX-system-P-loop-protein-BrxC-[Citrobacter-sp.-RHBSTW-01013]</td></tr>
<tr><td>WP_275379687.1</td><td>Type1.BREX</td><td>Arm-DNA-bind_3+Phage_integrase-><-SIGMA+OB-Fold+AAA-ATPase+Lon-Protease-BrxL<-PglZ<-N6-METHYLASE-p<-RVT<-HEPN<-N6-METHYLASE<-ParB+HNH<-DUF6079*<-BrxB-BrxF-iSTAND<-BrxA-DNA-binding<-?<-?<-XIS-HTH<-?||?->nSTAND1+GTPase-AIG+sGTP->DUF932-></td><td>1240</td><td>Bacteria>Pseudomonadota>Gammaproteobacteria>Enterobacterales>Enterobacteriaceae>Citrobacter</td><td>GCF_029104295.1</td><td>Citrobacter_freundii</td><td>L2W90_RS09965</td><td>166217..169939</td><td>546</td><td>BREX-system-P-loop-protein-BrxC-[Citrobacter-freundii]</td></tr>
<tr><td>CAF2998376.1</td><td>Type1.BREX</td><td>Arm-DNA-bind_3+Phage_integrase-><-SIGMA+OB-Fold+AAA-ATPase+Lon-Protease-BrxL<-PglZ<-N6-METHYLASE-p<-RVT<-HEPN<-N6-METHYLASE<-ParB+HNH<-DUF6079*<-BrxB-BrxF-iSTAND<-BrxA-DNA-binding<-Glyoxalase<-ZnR<-SAM-METHYLASE<-YjhX_toxin<-Glyoxalase<-?<-Glycoside_Hydrolase</td><td>1240</td><td>Bacteria>Pseudomonadota>Gammaproteobacteria>Enterobacterales>Enterobacteriaceae>Enterobacter</td><td>-</td><td>Enterobacter_cloacae</td><td>AI2972V1_0508</td><td>557695..561417</td><td>550</td><td>hypothetical-protein-AI2972V1_0508-[Enterobacter-cloacae]</td></tr>
<tr><td>WP_039268698.1</td><td>Type1.BREX</td><td>Arm-DNA-bind_3+Phage_integrase-><-SIGMA+OB-Fold+AAA-ATPase+Lon-Protease-BrxL<-PglZ<-N6-METHYLASE-p<-RVT<-HEPN<-ParB+HNH<-DUF6079*<-BrxB-BrxF-iSTAND<-BrxA-DNA-binding<-Glyoxalase<-ZnR<-SAM-METHYLASE<-YjhX_toxin<-Glyoxalase<-?<-Glycoside_Hydrolase</td><td>1233</td><td>Bacteria>Pseudomonadota>Gammaproteobacteria>Enterobacterales>Enterobacteriaceae>Enterobacter</td><td>-</td><td>Enterobacter_hormaechei</td><td>LI66_02640</td><td>557969..561670</td><td>158836</td><td>MULTISPECIES:-BREX-system-P-loop-protein-BrxC-[Enterobacter]</td></tr>
<tr><td>AFP68425.1</td><td>Type1.BREX</td><td>Arm-DNA-bind_3+Phage_integrase-><-SIGMA+OB-Fold+AAA-ATPase+Lon-Protease-BrxL<-PglZ<-N6-METHYLASE-p<-RVT<-HEPN<-N6-METHYLASE<-ParB+HNH<-DUF6079*<-BrxB-BrxF-iSTAND<-BrxA-DNA-binding||?->?-><-?<-?<-Glyoxalase<-ZnR<-METHYLASE</td><td>1233</td><td>Bacteria>Pseudomonadota>Gammaproteobacteria>Enterobacterales>Enterobacteriaceae>Enterobacter</td><td>GCA_000286275.1</td><td>Enterobacter_kobei</td><td>ECENHK_02660</td><td>581253..584954</td><td>208224</td><td>hypothetical-protein-ECENHK_02660-[Enterobacter-kobei]</td></tr>
<tr><td>AJB80307.1</td><td>Type1.BREX</td><td>Arm-DNA-bind_3+Phage_integrase-><-SIGMA+OB-Fold+AAA-ATPase+Lon-Protease-BrxL<-PglZ<-N6-METHYLASE-p<-RVT<-HEPN<-ParB+HNH<-DUF6079*<-BrxB-BrxF-iSTAND<-BrxA-DNA-binding<-Glyoxalase<-ZnR<-SAM-METHYLASE<-YjhX_toxin<-Glyoxalase<-?<-Glycoside_Hydrolase</td><td>1233</td><td>Bacteria>Pseudomonadota>Gammaproteobacteria>Enterobacterales>Enterobacteriaceae>Enterobacter</td><td>GCA_000814225.1</td><td>Enterobacter_hormaechei_subsp._xiangfangensis</td><td>LI66_02640</td><td>557969..561670</td><td>1296536</td><td>DNA-repair-protein-[Enterobacter-hormaechei-subsp.-xiangfangensis]</td></tr>
<tr><td>MBG0523800.1</td><td>Type1.BREX</td><td>Arm-DNA-bind_3+Phage_integrase-><-SIGMA+OB-Fold+AAA-ATPase+Lon-Protease-BrxL<-PglZ<-N6-METHYLASE-p<-RVT<-HEPN<-N6-METHYLASE<-ParB+HNH<-DUF6079*<-BrxB-BrxF-iSTAND<-BrxA-DNA-binding<-HIN-HTH<-REase<-wHTH+SAD+HNH||SymE_toxin-></td><td>1233</td><td>Bacteria>Pseudomonadota>Gammaproteobacteria>Enterobacterales>Enterobacteriaceae>Enterobacter</td><td>GCA_015704905.1</td><td>Enterobacter_cloacae</td><td>IOR24_17915</td><td>3774454..3778155</td><td>550</td><td>BREX-system-P-loop-protein-BrxC-[Enterobacter-cloacae]</td></tr>
<tr><td>MBK3145473.1</td><td>Type1.BREX</td><td>Glyoxalase->ACET->YjhX_toxin->SAM-METHYLASE->ZnR->Glyoxalase->BrxA-DNA-binding->BrxB-BrxF-iSTAND->DUF6079*->ParB+HNH->N6-METHYLASE->HEPN->RVT->N6-METHYLASE-p->PglZ->SIGMA+OB-Fold+AAA-ATPase+Lon-Protease-BrxL-><-Arm-DNA-bind_3+Phage_integrase</td><td>1233</td><td>Bacteria>Pseudomonadota>Gammaproteobacteria>Enterobacterales>Enterobacteriaceae>Enterobacter</td><td>GCA_016616795.1</td><td>Enterobacter_hormaechei</td><td>FKK11_12155</td><td>273388..277089</td><td>158836</td><td>BREX-system-P-loop-protein-BrxC-[Enterobacter-hormaechei]</td></tr>
<tr><td>WP_228371847.1</td><td>Type1.BREX</td><td>Arm-DNA-bind_3+Phage_integrase-><-SIGMA+OB-Fold+AAA-ATPase+Lon-Protease-BrxL<-PglZ<-N6-METHYLASE-p<-RVT<-HEPN<-N6-METHYLASE<-ParB+HNH<-DUF6079*<-BrxB-BrxF-iSTAND<-BrxA-DNA-binding||?-><-?<-Glyoxalase<-ZnR<-METHYLASE<-YjhX_toxin</td><td>1240</td><td>Bacteria>Pseudomonadota>Gammaproteobacteria>Enterobacterales>Enterobacteriaceae>Enterobacter</td><td>GCF_000286275.1</td><td>Enterobacter_kobei</td><td>ECENHK_RS02760</td><td>581253..584975</td><td>208224</td><td>BREX-system-P-loop-protein-BrxC-[Enterobacter-kobei]</td></tr>
<tr><td>WP_234704204.1</td><td>Type1.BREX</td><td>Arm-DNA-bind_3+Phage_integrase-><-SIGMA+OB-Fold+AAA-ATPase+Lon-Protease-BrxL<-PglZ<-N6-METHYLASE-p<-RVT<-HEPN<-N6-METHYLASE<-ParB+HNH<-DUF6079*<-BrxB-BrxF-iSTAND<-BrxA-DNA-binding<-Glyoxalase<-ZnR<-SAM-METHYLASE<-YjhX_toxin<-ACET<-Glyoxalase</td><td>1240</td><td>Bacteria>Pseudomonadota>Gammaproteobacteria>Enterobacterales>Enterobacteriaceae>Enterobacter</td><td>GCF_000814225.1</td><td>Enterobacter_cloacae_complex</td><td>LI66_RS02640</td><td>557969..561691</td><td>354276</td><td>MULTISPECIES:-BREX-system-P-loop-protein-BrxC-[Enterobacter-cloacae-complex]</td></tr>
<tr><td>WP_230863208.1</td><td>Type1.BREX</td><td>Arm-DNA-bind_3+Phage_integrase-><-SIGMA+OB-Fold+AAA-ATPase+Lon-Protease-BrxL<-PglZ<-N6-METHYLASE-p<-RVT<-HEPN<-N6-METHYLASE<-ParB+HNH<-DUF6079*<-BrxB-BrxF-iSTAND<-BrxA-DNA-binding<-REase<-wHTH+SAD+HNH||SymE_toxin-></td><td>1240</td><td>Bacteria>Pseudomonadota>Gammaproteobacteria>Enterobacterales>Enterobacteriaceae>Enterobacter</td><td>GCF_015704905.1</td><td>Enterobacter_cloacae</td><td>IOR24_RS17915</td><td>3774454..3778176</td><td>550</td><td>BREX-system-P-loop-protein-BrxC-[Enterobacter-cloacae]</td></tr>
<tr><td>WP_234450174.1</td><td>Type1.BREX</td><td>Glycoside_Hydrolase->?->Glyoxalase->YjhX_toxin->SAM-METHYLASE->ZnR->Glyoxalase->BrxA-DNA-binding->BrxB-BrxF-iSTAND->DUF6079*->ParB+HNH->N6-METHYLASE->HEPN->RVT->N6-METHYLASE-p->PglZ->SIGMA+OB-Fold+AAA-ATPase+Lon-Protease-BrxL-><-Arm-DNA-bind_3+Phage_integrase</td><td>1240</td><td>Bacteria>Pseudomonadota>Gammaproteobacteria>Enterobacterales>Enterobacteriaceae>Enterobacter</td><td>GCF_016616795.1</td><td>Enterobacter_hormaechei</td><td>FKK11_RS12010</td><td>273367..277089</td><td>158836</td><td>BREX-system-P-loop-protein-BrxC-[Enterobacter-hormaechei]</td></tr>
<tr><td>WP_174580485.1</td><td>Type1.BREX</td><td>DDE_Tnp_IS1->?->ZnR->BrxA-DNA-binding->BrxB-BrxF-iSTAND->DUF6079*->ParB+HNH->N6-METHYLASE->HEPN->RVT->N6-METHYLASE-p->PglZ->SIGMA+OB-Fold+AAA-ATPase+Lon-Protease-BrxL-><-Arm-DNA-bind_3+Phage_integrase||Phage_integrase-></td><td>1233</td><td>Bacteria>Pseudomonadota>Gammaproteobacteria>Enterobacterales>Enterobacteriaceae>Escherichia</td><td>-</td><td>Escherichia_coli</td><td>HUC40_15465</td><td>2945..6646</td><td>562</td><td>BREX-system-P-loop-protein-BrxC-[Escherichia-coli]</td></tr>
<tr><td>EEZ5714529.1</td><td>Type1.BREX</td><td><-wHTH+WYL<-nSTAND1+sGTP<-?||XIS-HTH->?->?->ZnR->BrxA-DNA-binding->BrxB-BrxF-iSTAND->DUF6079*->ParB+HNH->N6-METHYLASE->HEPN->RVT->N6-METHYLASE-p->PglZ->SIGMA+OB-Fold+AAA-ATPase+Lon-Protease-BrxL-><-Arm-DNA-bind_3+Phage_integrase</td><td>1233</td><td>Bacteria>Pseudomonadota>Gammaproteobacteria>Enterobacterales>Enterobacteriaceae>Escherichia</td><td>GCA_012215525.1</td><td>Escherichia_coli_O25</td><td>DBU27_000026</td><td>20065..23766</td><td>1095709</td><td>BREX-system-P-loop-protein-BrxC-[Escherichia-coli-O25]</td></tr>
<tr><td>EFD0312067.1</td><td>Type1.BREX</td><td>GDResolvase+HIN-HTH-><-SIGMA+OB-Fold+AAA-ATPase+Lon-Protease-BrxL<-PglZ<-N6-METHYLASE-p<-RVT<-HEPN<-N6-METHYLASE<-ParB+HNH<-DUF6079*<-BrxB-BrxF-iSTAND<-BrxA-DNA-binding<-?<-?<-XIS-HTH<-?||?->nSTAND1+GTPase-AIG+sGTP->DUF932-></td><td>1233</td><td>Bacteria>Pseudomonadota>Gammaproteobacteria>Enterobacterales>Enterobacteriaceae>Escherichia</td><td>GCA_012380475.1</td><td>Escherichia_coli</td><td>E6D44_20535</td><td>19474..23175</td><td>562</td><td>BREX-system-P-loop-protein-BrxC-[Escherichia-coli]</td></tr>
<tr><td>EFN5198232.1</td><td>Type1.BREX</td><td><-TIR||BrxA-DNA-binding->BrxB-BrxF-iSTAND->DUF6079*->ParB+HNH->N6-METHYLASE->HEPN->RVT->N6-METHYLASE-p->PglZ->SIGMA+OB-Fold+AAA-ATPase+Lon-Protease-BrxL->wHTH+WYL+WCX-></td><td>1233</td><td>Bacteria>Pseudomonadota>Gammaproteobacteria>Enterobacterales>Enterobacteriaceae>Escherichia</td><td>GCA_012993025.1</td><td>Escherichia_coli</td><td>FQG10_00765</td><td>2767..6468</td><td>562</td><td>BREX-system-P-loop-protein-BrxC-[Escherichia-coli]</td></tr>
<tr><td>NUC65042.1</td><td>Type1.BREX</td><td>DDE_Tnp_IS1->?->ZnR->BrxA-DNA-binding->BrxB-BrxF-iSTAND->DUF6079*->ParB+HNH->N6-METHYLASE->HEPN->RVT->N6-METHYLASE-p->PglZ->SIGMA+OB-Fold+AAA-ATPase+Lon-Protease-BrxL-><-Arm-DNA-bind_3+Phage_integrase||Phage_integrase-></td><td>1233</td><td>Bacteria>Pseudomonadota>Gammaproteobacteria>Enterobacterales>Enterobacteriaceae>Escherichia</td><td>GCA_013342565.1</td><td>Escherichia_coli</td><td>HUC40_15465</td><td>2945..6646</td><td>562</td><td>BREX-system-P-loop-protein-BrxC-[Escherichia-coli]</td></tr>
<tr><td>WP_250907975.1</td><td>Type1.BREX</td><td>DDE_Tnp_IS1->?->ZnR->BrxA-DNA-binding->BrxB-BrxF-iSTAND->DUF6079*->ParB+HNH->N6-METHYLASE->HEPN->RVT->N6-METHYLASE-p->PglZ->SIGMA+OB-Fold+AAA-ATPase+Lon-Protease-BrxL-><-Arm-DNA-bind_3+Phage_integrase||Phage_integrase-></td><td>1240</td><td>Bacteria>Pseudomonadota>Gammaproteobacteria>Enterobacterales>Enterobacteriaceae>Escherichia</td><td>GCA_030820635.1</td><td>Escherichia_coli</td><td>HUC40_RS15465</td><td>2924..6646</td><td>562</td><td>BREX-system-P-loop-protein-BrxC-[Escherichia-coli]</td></tr>
<tr><td>WP_001019654.1</td><td>Type1.BREX</td><td>LexA-protease->||Arm-DNA-bind_3+Phage_integrase->N6-METHYLASE->ORC-CDC6+ZnR->HEPN-><-Zn_Tnp_IS1+HTH_Tnp_IS1+DDE_Tnp_IS1<-REase||BrxA-DNA-binding->BrxB-BrxF-iSTAND->DUF6079*->N6-METHYLASE->PglZ->SIGMA+OB-Fold+AAA-ATPase+Lon-Protease-BrxL-><-ZnR<-ZnR<-?||HTH_Tnp_1+HTH_21+rve+rve_3-><-LuxR-HTH</td><td>1213</td><td>Bacteria>Pseudomonadota>Gammaproteobacteria>Enterobacterales>Enterobacteriaceae>Escherichia</td><td>GCF_000017765.1</td><td>Escherichia_coli</td><td>ECHS_RS01710</td><td>342111..345752</td><td>562</td><td>BREX-system-P-loop-protein-BrxC-[Escherichia-coli]</td></tr>
<tr><td>WP_233470897.1</td><td>Type1.BREX</td><td><-DUF932<-nSTAND1+GTPase-AIG+sGTP<-CopG/RHH||?->XIS-HTH->?->?->BrxA-DNA-binding->BrxB-BrxF-iSTAND->DUF6079*->ParB+HNH->N6-METHYLASE->HEPN->RVT->N6-METHYLASE->PglZ->SIGMA+OB-Fold+AAA-ATPase+Lon-Protease-BrxL-><-Arm-DNA-bind_3+Phage_integrase</td><td>1240</td><td>Bacteria>Pseudomonadota>Gammaproteobacteria>Enterobacterales>Enterobacteriaceae</td><td>GCF_014931625.2</td><td>Citrobacter</td><td>INP82_RS14360</td><td>158420..162142</td><td>544</td><td>MULTISPECIES:-BREX-system-P-loop-protein-BrxC-[Citrobacter]</td></tr>
<tr><td>WP_225990544.1</td><td>Type1.BREX</td><td>Glyoxalase->ACET->YjhX_toxin->SAM-METHYLASE->ZnR->Glyoxalase->BrxA-DNA-binding->BrxB-BrxF-iSTAND->DUF6079*->ParB+HNH->N6-METHYLASE->HEPN->RVT->N6-METHYLASE-p->PglZ->SIGMA+OB-Fold+AAA-ATPase+Lon-Protease-BrxL-><-Arm-DNA-bind_3+Phage_integrase</td><td>1240</td><td>Bacteria>Pseudomonadota>Gammaproteobacteria>Enterobacterales>Enterobacteriaceae</td><td>GCF_954736865.1</td><td>Enterobacter</td><td>CU081_RS00925</td><td>173954..177676</td><td>547</td><td>MULTISPECIES:-BREX-system-P-loop-protein-BrxC-[Enterobacter]</td></tr>
<tr><td>AUN75366.1</td><td>Type1.BREX</td><td>Arm-DNA-bind_3+Phage_integrase-><-SIGMA+OB-Fold+AAA-ATPase+Lon-Protease-BrxL<-PglZ<-N6-METHYLASE-p<-RVT<-HEPN<-N6-METHYLASE<-ParB+HNH<-DUF6079*<-BrxB-BrxF-iSTAND<-BrxA-DNA-binding<-?<-?<-XIS-HTH||?->nSTAND1+sGTP->wHTH+WYL-></td><td>1233</td><td>Bacteria>Pseudomonadota>Gammaproteobacteria>Enterobacterales>Enterobacteriaceae>Klebsiella/Raoultella_group</td><td>GCA_002870985.1</td><td>Klebsiella_pneumoniae</td><td>C0077_01580</td><td>326913..330614</td><td>573</td><td>BREX-system-P-loop-protein-BrxC-[Klebsiella-pneumoniae]</td></tr>
<tr><td>QHX00137.1</td><td>Type1.BREX</td><td>YjhX_toxin->ACET->METHYLASE->ZnR->Glyoxalase->BrxA-DNA-binding->BrxB-BrxF-iSTAND->DUF6079*->ParB+HNH->N6-METHYLASE->HEPN->RVT->N6-METHYLASE-p->PglZ->SIGMA+OB-Fold+AAA-ATPase+Lon-Protease-BrxL-><-Arm-DNA-bind_3+Phage_integrase</td><td>1233</td><td>Bacteria>Pseudomonadota>Gammaproteobacteria>Enterobacterales>Enterobacteriaceae>Klebsiella/Raoultella_group</td><td>GCA_010365565.1</td><td>Klebsiella_variicola</td><td>GZS05_24295</td><td>4949707..4953408</td><td>244366</td><td>BREX-system-P-loop-protein-BrxC-[Klebsiella-variicola]</td></tr>
<tr><td>QJM45197.1</td><td>Type1.BREX</td><td><-wHTH+WYL<-sGTP<-?||XIS-HTH->?->?->BrxA-DNA-binding->BrxB-BrxF-iSTAND->DUF6079*->ParB+HNH->N6-METHYLASE->HEPN->RVT->N6-METHYLASE-p->PglZ->SIGMA+OB-Fold+AAA-ATPase+Lon-Protease-BrxL-><-Arm-DNA-bind_3+Phage_integrase</td><td>1233</td><td>Bacteria>Pseudomonadota>Gammaproteobacteria>Enterobacterales>Enterobacteriaceae>Klebsiella/Raoultella_group</td><td>GCA_012970665.1</td><td>Klebsiella_pneumoniae</td><td>HJW71_23035</td><td>4620464..4624165</td><td>573</td><td>BREX-system-P-loop-protein-BrxC-[Klebsiella-pneumoniae]</td></tr>
<tr><td>EJG2380286.1</td><td>Type1.BREX</td><td><-DUF905<-DUF932<-CopG/RHH||?->XIS-HTH-><-TIR<-TIR||BrxA-DNA-binding->BrxB-BrxF-iSTAND->DUF6079*->ParB+HNH->N6-METHYLASE->HEPN->RVT->N6-METHYLASE-p->PglZ->SIGMA+OB-Fold+AAA-ATPase+Lon-Protease-BrxL->wHTH+WYL+WCX-></td><td>1233</td><td>Bacteria>Pseudomonadota>Gammaproteobacteria>Enterobacterales>Enterobacteriaceae>Klebsiella/Raoultella_group</td><td>GCA_016088515.2</td><td>Raoultella_ornithinolytica</td><td>AM381_RS08835</td><td>19475..23176</td><td>54291</td><td>BREX-system-P-loop-protein-BrxC-[Raoultella-ornithinolytica]</td></tr>
<tr><td>MBN7733855.1</td><td>Type1.BREX</td><td><-wHTH+WYL<-sGTP<-?||XIS-HTH->?->?->BrxA-DNA-binding->BrxB-BrxF-iSTAND->DUF6079*->ParB+HNH->N6-METHYLASE->HEPN->RVT->N6-METHYLASE-p->PglZ->SIGMA+OB-Fold+AAA-ATPase+Lon-Protease-BrxL-><-Arm-DNA-bind_3+Phage_integrase</td><td>1233</td><td>Bacteria>Pseudomonadota>Gammaproteobacteria>Enterobacterales>Enterobacteriaceae>Klebsiella/Raoultella_group</td><td>GCA_017255495.1</td><td>Klebsiella_pneumoniae</td><td>JX616_18220</td><td>31914..35615</td><td>573</td><td>BREX-system-P-loop-protein-BrxC-[Klebsiella-pneumoniae]</td></tr>
<tr><td>EIV3098057.1</td><td>Type1.BREX</td><td><-wHTH+WYL<-nSTAND1+sGTP<-?||XIS-HTH->?->?->BrxA-DNA-binding->BrxB-BrxF-iSTAND->DUF6079*->ParB+HNH->N6-METHYLASE->HEPN->RVT->N6-METHYLASE-p->PglZ->SIGMA+OB-Fold+AAA-ATPase+Lon-Protease-BrxL-><-Arm-DNA-bind_3+Phage_integrase</td><td>1233</td><td>Bacteria>Pseudomonadota>Gammaproteobacteria>Enterobacterales>Enterobacteriaceae>Klebsiella/Raoultella_group</td><td>GCA_021854435.1</td><td>Klebsiella_pneumoniae</td><td>L7W77_004838</td><td>45573..49274</td><td>573</td><td>BREX-system-P-loop-protein-BrxC-[Klebsiella-pneumoniae]</td></tr>
<tr><td>HBR1784209.1</td><td>Type1.BREX</td><td><-wHTH+WYL<-sGTP<-?||XIS-HTH->?->?->BrxA-DNA-binding->BrxB-BrxF-iSTAND->DUF6079*->ParB+HNH->N6-METHYLASE->HEPN->RVT->N6-METHYLASE-p->PglZ->SIGMA+OB-Fold+AAA-ATPase+Lon-Protease-BrxL-><-Arm-DNA-bind_3+Phage_integrase</td><td>1233</td><td>Bacteria>Pseudomonadota>Gammaproteobacteria>Enterobacterales>Enterobacteriaceae>Klebsiella/Raoultella_group</td><td>GCA_021911385.1</td><td>Klebsiella_quasipneumoniae_subsp._quasipneumoniae</td><td>L9M28_003408</td><td>20306..24007</td><td>1667327</td><td>BREX-system-P-loop-protein-BrxC-[Klebsiella-quasipneumoniae-subsp.-quasipneumoniae]</td></tr>
<tr><td>HCD1392381.1</td><td>Type1.BREX</td><td><-wHTH+WYL+WCX<-SIGMA+OB-Fold+AAA-ATPase+Lon-Protease-BrxL<-PglZ<-N6-METHYLASE-p<-RVT<-HEPN<-N6-METHYLASE<-ParB+HNH<-DUF6079*<-BrxB-BrxF-iSTAND<-BrxA-DNA-binding||TIR->TIR-><-XIS-HTH<-?||CopG/RHH->DUF932->DUF905-></td><td>1233</td><td>Bacteria>Pseudomonadota>Gammaproteobacteria>Enterobacterales>Enterobacteriaceae>Klebsiella/Raoultella_group</td><td>GCA_023530865.1</td><td>Klebsiella_pneumoniae_subsp._pneumoniae</td><td>M9022_004645</td><td>23383..27084</td><td>72407</td><td>BREX-system-P-loop-protein-BrxC-[Klebsiella-pneumoniae-subsp.-pneumoniae]</td></tr>
<tr><td>MCS5967892.1</td><td>Type1.BREX</td><td><-nSTAND1+sGTP<-?<-?||?->BrxA-DNA-binding->BrxB-BrxF-iSTAND->DUF6079-p*->DUF6079-p->?->ParB+HNH->ParB+HNH->N6-METHYLASE->HEPN->?->RVT-></td><td>389</td><td>Bacteria>Pseudomonadota>Gammaproteobacteria>Enterobacterales>Enterobacteriaceae>Klebsiella/Raoultella_group</td><td>GCA_024990005.1</td><td>Klebsiella_variicola_subsp._variicola</td><td>LNP26_07865</td><td>1516930..1518099</td><td>2590157</td><td>BREX-system-P-loop-protein-BrxC-[Klebsiella-variicola-subsp.-variicola]</td></tr>
<tr><td>WEJ89011.1</td><td>Type1.BREX</td><td><-wHTH+HNH||tRNA->Arm-DNA-bind_3+Phage_integrase-><-SIGMA+OB-Fold+AAA-ATPase+Lon-Protease-BrxL<-PglZ<-N6-METHYLASE-p<-RVT<-HEPN<-N6-METHYLASE<-ParB+HNH<-DUF6079*<-BrxB-BrxF-iSTAND<-BrxA-DNA-binding<-ZnR<-XIS-HTH<-?||HIN-HTH->?->LZ_Tnp_IS66+zf-IS66+DDE_Tnp_IS66+DDE_Tnp_IS66_C-></td><td>1240</td><td>Bacteria>Pseudomonadota>Gammaproteobacteria>Enterobacterales>Enterobacteriaceae>Klebsiella/Raoultella_group</td><td>GCA_029202365.1</td><td>Klebsiella_huaxiensis</td><td>POY63_27795</td><td>5833350..5837072</td><td>2153354</td><td>MAG:-BREX-system-P-loop-protein-BrxC-[Klebsiella-huaxiensis]</td></tr>
<tr><td>WP_256991583.1</td><td>Type1.BREX</td><td><-DUF905<-DUF932<-CopG/RHH||?->XIS-HTH->ZnR-><-TIR||BrxA-DNA-binding->BrxB-BrxF-iSTAND->DUF6079*->ParB+HNH->N6-METHYLASE->HEPN->RVT->N6-METHYLASE-p->PglZ->SIGMA+OB-Fold+AAA-ATPase+Lon-Protease-BrxL->wHTH+WYL+WCX-></td><td>1240</td><td>Bacteria>Pseudomonadota>Gammaproteobacteria>Enterobacterales>Enterobacteriaceae>Klebsiella/Raoultella_group</td><td>GCF_002264145.1</td><td>Raoultella_planticola</td><td>CIG23_RS11860</td><td>59081..62803</td><td>575</td><td>BREX-system-P-loop-protein-BrxC-[Raoultella-planticola]</td></tr>
<tr><td>WP_264757344.1</td><td>Type1.BREX</td><td><-wHTH+WYL<-nSTAND1+sGTP<-?||XIS-HTH->?->?->BrxA-DNA-binding->BrxB-BrxF-iSTAND-><-DDE_Tnp_1_5+DDE_Tnp_1||DUF6079-p*->ParB+HNH->N6-METHYLASE->HEPN->RVT->N6-METHYLASE-p->PglZ->SIGMA+OB-Fold+AAA-ATPase+Lon-Protease-BrxL-><-Arm-DNA-bind_3+Phage_integrase</td><td>995</td><td>Bacteria>Pseudomonadota>Gammaproteobacteria>Enterobacterales>Enterobacteriaceae>Klebsiella/Raoultella_group</td><td>GCF_003660185.1</td><td>Klebsiella_pneumoniae</td><td>D9K63_RS23205</td><td>4667642..4670629</td><td>573</td><td>BREX-system-P-loop-protein-BrxC-[Klebsiella-pneumoniae]</td></tr>
<tr><td>WP_223814682.1</td><td>Type1.BREX</td><td><-wHTH+WYL<-nSTAND1+sGTP<-?||XIS-HTH->?->?->BrxA-DNA-binding->BrxB-BrxF-iSTAND->DUF6079*->ParB+HNH->N6-METHYLASE->HEPN->RVT->N6-METHYLASE-p->PglZ->SIGMA+OB-Fold+AAA-ATPase+Lon-Protease-BrxL-><-Arm-DNA-bind_3+Phage_integrase</td><td>1240</td><td>Bacteria>Pseudomonadota>Gammaproteobacteria>Enterobacterales>Enterobacteriaceae>Klebsiella/Raoultella_group</td><td>GCF_004313365.1</td><td>Klebsiella</td><td>EO018_RS24630</td><td>24494..28216</td><td>570</td><td>MULTISPECIES:-BREX-system-P-loop-protein-BrxC-[Klebsiella]</td></tr>
<tr><td>WP_235410418.1</td><td>Type1.BREX</td><td><-wHTH+WYL<-sGTP<-?||XIS-HTH->?->?->BrxA-DNA-binding->BrxB-BrxF-iSTAND->DUF6079*->ParB+HNH->N6-METHYLASE->HEPN->RVT->N6-METHYLASE-p->PglZ->SIGMA+OB-Fold+AAA-ATPase+Lon-Protease-BrxL-><-Arm-DNA-bind_3+Phage_integrase</td><td>1240</td><td>Bacteria>Pseudomonadota>Gammaproteobacteria>Enterobacterales>Enterobacteriaceae>Klebsiella/Raoultella_group</td><td>GCF_021532695.1</td><td>Klebsiella_pneumoniae</td><td>LZ602_RS12600</td><td>31915..35637</td><td>573</td><td>BREX-system-P-loop-protein-BrxC-[Klebsiella-pneumoniae]</td></tr>
<tr><td>WP_223339928.1</td><td>Type1.BREX</td><td><-wHTH+WYL<-sGTP<-?||XIS-HTH->?->?->BrxA-DNA-binding->BrxB-BrxF-iSTAND->DUF6079*->ParB+HNH->N6-METHYLASE->HEPN->RVT->N6-METHYLASE-p->PglZ->SIGMA+OB-Fold+AAA-ATPase+Lon-Protease-BrxL-><-Arm-DNA-bind_3+Phage_integrase</td><td>1240</td><td>Bacteria>Pseudomonadota>Gammaproteobacteria>Enterobacterales>Enterobacteriaceae>Klebsiella/Raoultella_group</td><td>GCF_022353805.1</td><td>Klebsiella_pneumoniae</td><td>JX616_RS18220</td><td>31893..35615</td><td>573</td><td>BREX-system-P-loop-protein-BrxC-[Klebsiella-pneumoniae]</td></tr>
<tr><td>WP_227643422.1</td><td>Type1.BREX</td><td>Arm-DNA-bind_3+Phage_integrase-><-SIGMA+OB-Fold+AAA-ATPase+Lon-Protease-BrxL<-PglZ<-N6-METHYLASE-p<-RVT<-HEPN<-N6-METHYLASE<-ParB+HNH<-DUF6079*<-BrxB-BrxF-iSTAND<-BrxA-DNA-binding<-?<-?<-XIS-HTH<-?||CopG/RHH->nSTAND1+GTPase-AIG+sGTP->DUF932-></td><td>1240</td><td>Bacteria>Pseudomonadota>Gammaproteobacteria>Enterobacterales>Enterobacteriaceae>Klebsiella/Raoultella_group</td><td>GCF_900173735.1</td><td>Klebsiella_pneumoniae</td><td>B7E22_RS15515</td><td>109572..113294</td><td>573</td><td>BREX-system-P-loop-protein-BrxC-[Klebsiella-pneumoniae]</td></tr>
<tr><td>WP_227635281.1</td><td>Type1.BREX</td><td>XIS-HTH-><-TIR<-TIR||BrxA-DNA-binding->BrxB-BrxF-iSTAND->DUF6079*->ParB+HNH->N6-METHYLASE->HEPN->RVT->N6-METHYLASE-p->PglZ->SIGMA+OB-Fold+AAA-ATPase+Lon-Protease-BrxL->wHTH+WYL+WCX-></td><td>1240</td><td>Bacteria>Pseudomonadota>Gammaproteobacteria>Enterobacterales>Enterobacteriaceae>Klebsiella/Raoultella_group</td><td>GCF_902159715.1</td><td>Klebsiella</td><td>PIK69_RS17535</td><td>2737..6459</td><td>570</td><td>MULTISPECIES:-BREX-system-P-loop-protein-BrxC-[Klebsiella]</td></tr>
<tr><td>WP_227627429.1</td><td>Type1.BREX</td><td><-wHTH+WYL+WCX<-?<-SIGMA+OB-Fold+AAA-ATPase+Lon-Protease-BrxL<-PglZ<-N6-METHYLASE-p<-RVT<-HEPN<-N6-METHYLASE<-ParB+HNH<-DUF6079*<-BrxB-BrxF-iSTAND<-BrxA-DNA-binding||TIR-><-ZnR<-XIS-HTH<-?||?->DUF932->DUF905-></td><td>1240</td><td>Bacteria>Pseudomonadota>Gammaproteobacteria>Enterobacterales>Enterobacteriaceae>Klebsiella/Raoultella_group</td><td>-</td><td>Klebsiella_michiganensis</td><td>G4403_RS10440</td><td>98242..101964</td><td>1134687</td><td>BREX-system-P-loop-protein-BrxC-[Klebsiella-michiganensis]</td></tr>
<tr><td>WP_117042583.1</td><td>Type1.BREX</td><td><-DUF905<-DUF932<-CopG/RHH||?->XIS-HTH-><-TIR<-TIR||BrxA-DNA-binding->BrxB-BrxF-iSTAND->DUF6079*->ParB+HNH->N6-METHYLASE->HEPN->RVT->N6-METHYLASE-p->PglZ->SIGMA+OB-Fold+AAA-ATPase+Lon-Protease-BrxL->wHTH+WYL+WCX-></td><td>1233</td><td>Bacteria>Pseudomonadota>Gammaproteobacteria>Enterobacterales>Enterobacteriaceae>Klebsiella/Raoultella_group</td><td>-</td><td>Klebsiella_pneumoniae</td><td>AM381_RS08835</td><td>19475..23176</td><td>573</td><td>MULTISPECIES:-BREX-system-P-loop-protein-BrxC-[Klebsiella]</td></tr>
<tr><td>WP_114263049.1</td><td>Type1.BREX</td><td><-DUF932<-GspA-ATPase+GTPase<-CopG/RHH||?->XIS-HTH->?->?->BrxA-DNA-binding->BrxB-BrxF-iSTAND->DUF6079*->ParB+HNH->N6-METHYLASE->HEPN->RVT->N6-METHYLASE-p->PglZ->SIGMA+OB-Fold+AAA-ATPase+Lon-Protease-BrxL-><-Arm-DNA-bind_3+Phage_integrase</td><td>1233</td><td>Bacteria>Pseudomonadota>Gammaproteobacteria>Enterobacterales>Enterobacteriaceae>Klebsiella/Raoultella_group</td><td>-</td><td>Klebsiella_pneumoniae</td><td>AN2353V1_3685</td><td>3967252..3970953</td><td>573</td><td>MULTISPECIES:-BREX-system-P-loop-protein-BrxC-[Enterobacteriaceae]</td></tr>
<tr><td>WP_062729754.1</td><td>Type1.BREX</td><td>Arm-DNA-bind_3+Phage_integrase-><-SIGMA+OB-Fold+AAA-ATPase+Lon-Protease-BrxL<-PglZ<-N6-METHYLASE-p<-RVT<-HEPN<-N6-METHYLASE<-ParB+HNH<-DUF6079*<-BrxB-BrxF-iSTAND<-BrxA-DNA-binding<-?<-?<-XIS-HTH||?->nSTAND1+sGTP->wHTH+WYL-></td><td>1233</td><td>Bacteria>Pseudomonadota>Gammaproteobacteria>Enterobacterales>Enterobacteriaceae>Klebsiella/Raoultella_group</td><td>-</td><td>Klebsiella_pneumoniae</td><td>C0077_01580</td><td>326913..330614</td><td>573</td><td>MULTISPECIES:-BREX-system-P-loop-protein-BrxC-[Enterobacteriaceae]</td></tr>
<tr><td>WP_073511310.1</td><td>Type1.BREX</td><td><-wHTH+WYL<-sGTP<-?||XIS-HTH->?->?->BrxA-DNA-binding->BrxB-BrxF-iSTAND->DUF6079*->ParB+HNH->N6-METHYLASE->HEPN->RVT->N6-METHYLASE-p->PglZ->SIGMA+OB-Fold+AAA-ATPase+Lon-Protease-BrxL-><-Arm-DNA-bind_3+Phage_integrase</td><td>1233</td><td>Bacteria>Pseudomonadota>Gammaproteobacteria>Enterobacterales>Enterobacteriaceae>Klebsiella/Raoultella_group</td><td>-</td><td>Klebsiella_pneumoniae</td><td>L7V03_004584</td><td>32186..35887</td><td>573</td><td>BREX-system-P-loop-protein-BrxC-[Klebsiella-pneumoniae]</td></tr>
<tr><td>WP_117247358.1</td><td>Type1.BREX</td><td><-wHTH+WYL<-nSTAND1+sGTP<-?||XIS-HTH->?->?->BrxA-DNA-binding->BrxB-BrxF-iSTAND->DUF6079*->ParB+HNH->N6-METHYLASE->HEPN->RVT->N6-METHYLASE-p->PglZ->SIGMA+OB-Fold+AAA-ATPase+Lon-Protease-BrxL-><-Arm-DNA-bind_3+Phage_integrase</td><td>1233</td><td>Bacteria>Pseudomonadota>Gammaproteobacteria>Enterobacterales>Enterobacteriaceae>Klebsiella/Raoultella_group</td><td>-</td><td>Klebsiella_pneumoniae</td><td>L7W77_004838</td><td>45573..49274</td><td>573</td><td>MULTISPECIES:-BREX-system-P-loop-protein-BrxC-[Klebsiella]</td></tr>
<tr><td>WP_094898295.1</td><td>Type1.BREX</td><td><-TIR||BrxA-DNA-binding->BrxB-BrxF-iSTAND->DUF6079*->ParB+HNH->N6-METHYLASE->HEPN->RVT->N6-METHYLASE-p->PglZ->SIGMA+OB-Fold+AAA-ATPase+Lon-Protease-BrxL->wHTH+WYL+WCX-></td><td>1233</td><td>Bacteria>Pseudomonadota>Gammaproteobacteria>Enterobacterales>Enterobacteriaceae>Klebsiella/Raoultella_group</td><td>-</td><td>Raoultella_planticola</td><td>FQG10_00765</td><td>2767..6468</td><td>575</td><td>BREX-system-P-loop-protein-BrxC-[Raoultella-planticola]</td></tr>
<tr><td>QXB23949.1</td><td>Type1.BREX</td><td>XIS-HTH->?->ZnR->ZnR->?->||BrxA-DNA-binding->BrxB-BrxF-iSTAND->DUF6079*->ParB+HNH->N6-METHYLASE->HEPN->RVT->N6-METHYLASE-><-HIN-HTH||PglZ->SIGMA+OB-Fold+AAA-ATPase+Lon-Protease-BrxL-></td><td>1233</td><td>Bacteria>Pseudomonadota>Gammaproteobacteria>Enterobacterales>Enterobacteriaceae>Lelliottia</td><td>GCA_019048185.1</td><td>Lelliottia_amnigena</td><td>I6L76_15755</td><td>3242307..3246008</td><td>61646</td><td>BREX-system-P-loop-protein-BrxC-[Lelliottia-amnigena]</td></tr>
<tr><td>WP_254208058.1</td><td>Type1.BREX</td><td>XIS-HTH->?->ZnR->ZnR->?->||BrxA-DNA-binding->BrxB-BrxF-iSTAND->DUF6079*->ParB+HNH->N6-METHYLASE->HEPN->RVT->N6-METHYLASE-><-HIN-HTH||PglZ->SIGMA+OB-Fold+AAA-ATPase+Lon-Protease-BrxL-></td><td>1237</td><td>Bacteria>Pseudomonadota>Gammaproteobacteria>Enterobacterales>Enterobacteriaceae>Lelliottia</td><td>GCF_019048185.1</td><td>Lelliottia_amnigena</td><td>I6L76_RS15755</td><td>3242295..3246008</td><td>61646</td><td>BREX-system-P-loop-protein-BrxC-[Lelliottia-amnigena]</td></tr>
<tr><td>WP_216981896.1</td><td>Type1.BREX</td><td>XIS-HTH->?->ZnR->ZnR->?->||BrxA-DNA-binding->BrxB-BrxF-iSTAND->DUF6079*->ParB+HNH->N6-METHYLASE->HEPN->RVT->N6-METHYLASE-><-HIN-HTH||PglZ->SIGMA+OB-Fold+AAA-ATPase+Lon-Protease-BrxL-></td><td>1233</td><td>Bacteria>Pseudomonadota>Gammaproteobacteria>Enterobacterales>Enterobacteriaceae>Lelliottia</td><td>-</td><td>Lelliottia_amnigena</td><td>I6L76_15755</td><td>3242307..3246008</td><td>61646</td><td>BREX-system-P-loop-protein-BrxC-[Lelliottia-amnigena]</td></tr>
<tr><td>EAA4435938.1</td><td>Type1.BREX</td><td>XIS-HTH->?->?-><-?<-ZnR||BrxA-DNA-binding->BrxB-BrxF-iSTAND->DUF6079*->ParB+HNH->N6-METHYLASE->HEPN->RVT->N6-METHYLASE-p->PglZ->SIGMA+OB-Fold+AAA-ATPase+Lon-Protease-BrxL-><-tRNA||?-><-SKI</td><td>1233</td><td>Bacteria>Pseudomonadota>Gammaproteobacteria>Enterobacterales>Enterobacteriaceae>Salmonella</td><td>GCA_004179155.1</td><td>Salmonella_enterica_subsp._salamae</td><td>DPE44_05505</td><td>71371..75072</td><td>59202</td><td>BREX-system-P-loop-protein-BrxC-[Salmonella-enterica-subsp.-salamae]</td></tr>
<tr><td>WP_079805580.1</td><td>Type1.BREX</td><td>SLOG->HAD-><-SIGMA+OB-Fold+AAA-ATPase+Lon-Protease-BrxL<-PglZ<-N6-METHYLASE-p<-RVT<-HEPN<-N6-METHYLASE<-ParB+HNH<-DUF6079*<-BrxB-BrxF-iSTAND<-BrxA-DNA-binding||Schlafen+HATPase+wHTH-><-?<-?<-XIS-HTH</td><td>1240</td><td>Bacteria>Pseudomonadota>Gammaproteobacteria>Enterobacterales>Enterobacteriaceae>Salmonella</td><td>GCF_016903075.1</td><td>Salmonella_enterica</td><td>AJH70_RS01330</td><td>291078..294800</td><td>28901</td><td>BREX-system-P-loop-protein-BrxC-[Salmonella-enterica]</td></tr>
<tr><td>UCB31815.1</td><td>Type1.BREX</td><td><-wHTH+WYL<-?||XIS-HTH->?->BrxA-DNA-binding->BrxB-BrxF-iSTAND->DUF6079*->ParB+HNH->N6-METHYLASE->HEPN->RVT->N6-METHYLASE-p->PglZ->SIGMA+OB-Fold+AAA-ATPase+Lon-Protease-BrxL-><-Arm-DNA-bind_3+Phage_integrase</td><td>1240</td><td>Bacteria>Pseudomonadota>Gammaproteobacteria>Enterobacterales>Erwiniaceae>Duffyella</td><td>GCA_020342335.1</td><td>Duffyella_gerundensis</td><td>J9874_02366</td><td>2569295..2573017</td><td>1619313</td><td>hypothetical-protein-J9874_02366-[Duffyella-gerundensis]</td></tr>
<tr><td>WP_225700910.1</td><td>Type1.BREX</td><td><-wHTH+WYL<-?||XIS-HTH->?->BrxA-DNA-binding->BrxB-BrxF-iSTAND->DUF6079*->ParB+HNH->N6-METHYLASE->HEPN->RVT->N6-METHYLASE-p->PglZ->SIGMA+OB-Fold+AAA-ATPase+Lon-Protease-BrxL-><-Arm-DNA-bind_3+Phage_integrase</td><td>1237</td><td>Bacteria>Pseudomonadota>Gammaproteobacteria>Enterobacterales>Erwiniaceae>Duffyella</td><td>GCF_020342335.1</td><td>Duffyella_gerundensis</td><td>J9874_RS11820</td><td>2569304..2573017</td><td>1619313</td><td>BREX-system-P-loop-protein-BrxC-[Duffyella-gerundensis]</td></tr>
<tr><td>WP_223177211.1</td><td>Type1.BREX</td><td><-wHTH+WYL<-nSTAND1+sGTP<-?||XIS-HTH->?->?->BrxA-DNA-binding->BrxB-BrxF-iSTAND->DUF6079*->ParB+HNH->N6-METHYLASE->HEPN->RVT->N6-METHYLASE-p->PglZ->SIGMA+OB-Fold+AAA-ATPase+Lon-Protease-BrxL-><-Arm-DNA-bind_3+Phage_integrase</td><td>1240</td><td>Bacteria>Pseudomonadota>Gammaproteobacteria>Enterobacterales</td><td>GCF_004312085.1</td><td>Enterobacteriaceae</td><td>EOM53_RS14900</td><td>53286..57008</td><td>543</td><td>MULTISPECIES:-BREX-system-P-loop-protein-BrxC-[Enterobacteriaceae]</td></tr>
<tr><td>WP_175320436.1</td><td>Type1.BREX</td><td><-DUF932<-nSTAND1+GTPase-AIG+sGTP<-CopG/RHH||?->XIS-HTH->?->?->BrxA-DNA-binding->BrxB-BrxF-iSTAND->DUF6079*->ParB+HNH->N6-METHYLASE->HEPN->RVT->N6-METHYLASE-p->PglZ->SIGMA+OB-Fold+AAA-ATPase+Lon-Protease-BrxL-><-Arm-DNA-bind_3+Phage_integrase</td><td>1233</td><td>Bacteria>Pseudomonadota>Gammaproteobacteria>Enterobacterales>Hafniaceae>Hafnia</td><td>GCF_013360205.1</td><td>Hafnia_paralvei</td><td>G4F99_RS14350</td><td>55379..59080</td><td>546367</td><td>BREX-system-P-loop-protein-BrxC-[Hafnia-paralvei]</td></tr>
<tr><td>WP_234619586.1</td><td>Type1.BREX</td><td><-nSTAND1+GTPase-AIG+sGTP<-CopG/RHH||XIS-HTH->?->?->ZnR->?->BrxA-DNA-binding->BrxB-BrxF-iSTAND->DUF6079*->ParB+HNH->N6-METHYLASE->HEPN->RVT->N6-METHYLASE-p->PglZ->SIGMA+OB-Fold+AAA-ATPase+Lon-Protease-BrxL-><-HTH+MPTase</td><td>1237</td><td>Bacteria>Pseudomonadota>Gammaproteobacteria>Enterobacterales>Hafniaceae>Hafnia</td><td>GCF_021440285.1</td><td>Hafnia_alvei</td><td>LZ667_RS17495</td><td>58169..61882</td><td>569</td><td>BREX-system-P-loop-protein-BrxC-[Hafnia-alvei]</td></tr>
<tr><td>AUR32853.1</td><td>Type1.BREX</td><td>HTH->Pertactin+Autotransporter->?->wHTH+SAD+HNH->?->ZnR->BrxA-DNA-binding->BrxB-BrxF-iSTAND->DUF6079*->ParB+HNH->N6-METHYLASE->HEPN->RVT->N6-METHYLASE-p->PglZ->SIGMA+OB-Fold+AAA-ATPase+Lon-Protease-BrxL-><-Arm-DNA-bind_3+Phage_integrase</td><td>1233</td><td>Bacteria>Pseudomonadota>Gammaproteobacteria>Enterobacterales>Morganellaceae>Morganella</td><td>GCA_002891475.1</td><td>Morganella_morganii</td><td>C1070_15730</td><td>3342273..3345974</td><td>582</td><td>BREX-system-P-loop-protein-BrxC-[Morganella-morganii]</td></tr>
<tr><td>WP_225973345.1</td><td>Type1.BREX</td><td>HTH->Pertactin+Autotransporter->wHTH+SAD+HNH->?->ZnR->BrxA-DNA-binding->BrxB-BrxF-iSTAND->DUF6079*->ParB+HNH->N6-METHYLASE->HEPN->RVT->N6-METHYLASE-p->PglZ->SIGMA+OB-Fold+AAA-ATPase+Lon-Protease-BrxL-><-Arm-DNA-bind_3+Phage_integrase</td><td>1240</td><td>Bacteria>Pseudomonadota>Gammaproteobacteria>Enterobacterales>Morganellaceae>Morganella</td><td>GCF_002891475.1</td><td>Morganella_morganii</td><td>C1070_RS15730</td><td>3342252..3345974</td><td>582</td><td>BREX-system-P-loop-protein-BrxC-[Morganella-morganii]</td></tr>
<tr><td>EGS9996488.1</td><td>Type1.BREX</td><td><-nSTAND1+GTPase-AIG+sGTP<-CopG/RHH||XIS-HTH->?->?->BrxA-DNA-binding->BrxB-BrxF-iSTAND->DUF6079*->ParB+HNH->N6-METHYLASE->HEPN->RVT->N6-METHYLASE-p->PglZ->SIGMA+OB-Fold+AAA-ATPase+Lon-Protease-BrxL-><-Arm-DNA-bind_3+Phage_integrase</td><td>1233</td><td>Bacteria>Pseudomonadota>Gammaproteobacteria>Enterobacterales>Yersiniaceae>Serratia</td><td>GCA_015929585.1</td><td>Serratia_marcescens</td><td>I9Y15_003567</td><td>178923..182624</td><td>615</td><td>BREX-system-P-loop-protein-BrxC-[Serratia-marcescens]</td></tr>
<tr><td>WP_260965787.1</td><td>Type1.BREX</td><td>Arm-DNA-bind_3+Phage_integrase-><-SIGMA+OB-Fold+AAA-ATPase+Lon-Protease-BrxL<-PglZ<-N6-METHYLASE-p<-RVT<-HEPN<-N6-METHYLASE<-ParB+HNH<-DUF6079*<-BrxB-BrxF-iSTAND<-BrxA-DNA-binding<-?<-XIS-HTH<-?||?->GspA-ATPase+GTPase-></td><td>1240</td><td>Bacteria>Pseudomonadota>Gammaproteobacteria>Enterobacterales>Yersiniaceae>Serratia</td><td>GCF_025336405.1</td><td>Serratia_marcescens</td><td>N7751_RS05105</td><td>1087639..1091361</td><td>615</td><td>BREX-system-P-loop-protein-BrxC-[Serratia-marcescens]</td></tr>
<tr><td>WP_262254721.1</td><td>Type1.BREX</td><td><-GTPase-AIG<-CopG/RHH||?->XIS-HTH-><-?<-?||BrxA-DNA-binding->BrxB-BrxF-iSTAND->DUF6079*->ParB+HNH->N6-METHYLASE->HEPN->RVT->N6-METHYLASE-p->PglZ->SIGMA+OB-Fold+AAA-ATPase+Lon-Protease-BrxL-><-Arm-DNA-bind_3+Phage_integrase</td><td>1240</td><td>Bacteria>Pseudomonadota>Gammaproteobacteria>Enterobacterales>Yersiniaceae>Serratia</td><td>GCF_946228755.1</td><td>Serratia_proteamaculans</td><td>OC575_RS10145</td><td>2133622..2137344</td><td>28151</td><td>BREX-system-P-loop-protein-BrxC-[Serratia-proteamaculans]</td></tr>
<tr><td>WP_132542053.1</td><td>Type1.BREX</td><td><-SIGMA+OB-Fold+AAA-ATPase+Lon-Protease-BrxL<-PglZ<-HNH<-OLD-ABC<-N6-METHYLASE<-HEPN<-PolB<-DUF6079*<-BrxB-BrxF-iSTAND<-BrxA-DNA-binding||ABhydrolase-><-Acyl_transf_3<-?<-?||Peptidase_M3-><-ABhydrolase</td><td>1153</td><td>Bacteria>Pseudomonadota>Gammaproteobacteria>Gammaproteobacteria_incertae_sedis>Candidatus_Competibacteraceae>Plasticicumulans</td><td>GCF_004341245.1</td><td>Plasticicumulans_lactativorans</td><td>EV699_RS09620</td><td>184945..188406</td><td>1133106</td><td>BREX-system-P-loop-protein-BrxC-[Plasticicumulans-lactativorans]</td></tr>
<tr><td>WP_223374968.1</td><td>Type1.BREX</td><td>Arm-DNA-bind_3+Phage_integrase-><-SIGMA+OB-Fold+AAA-ATPase+Lon-Protease-BrxL<-PglZ<-N6-METHYLASE-p<-RVT<-HEPN<-N6-METHYLASE<-ParB+HNH<-DUF6079*<-BrxB-BrxF-iSTAND<-BrxA-DNA-binding||?->PHP+SbcC->Schlafen+HATPase+wHTH->?->?->?->Sulfatase-></td><td>1240</td><td>Bacteria>Pseudomonadota>Gammaproteobacteria</td><td>GCF_003051945.2</td><td>Enterobacterales</td><td>N5D94_RS08195</td><td>82368..86090</td><td>91347</td><td>MULTISPECIES:-BREX-system-P-loop-protein-BrxC-[Enterobacterales]</td></tr>
<tr><td>WP_099433763.1</td><td>Type1.BREX</td><td><-Schlafen+HATPase+wHTH<-PHP+SbcC<-?||BrxA-DNA-binding->BrxB-BrxF-iSTAND->DUF772+DDE_Tnp_1->DUF6079*->ParB+HNH->N6-METHYLASE->HEPN->RVT->N6-METHYLASE-p->PglZ->SIGMA+OB-Fold+AAA-ATPase+Lon-Protease-BrxL-><-Arm-DNA-bind_3+Phage_integrase</td><td>1233</td><td>Bacteria>Pseudomonadota>Gammaproteobacteria</td><td>GCF_014168815.1</td><td>Enterobacterales</td><td>H7R56_RS21645</td><td>4484852..4488553</td><td>91347</td><td>MULTISPECIES:-BREX-system-P-loop-protein-BrxC-[Enterobacterales]</td></tr>
<tr><td>WP_253145790.1</td><td>Type1.BREX</td><td>DDE_Tnp_IS66+DDE_Tnp_IS66_C-><-HEPN<-?<-SIGMA+OB-Fold+AAA-ATPase+Lon-Protease-BrxL<-PglZ<-?<-N6-METHYLASE<-DUF6079*<-BrxB-BrxF-iSTAND<-BrxA-DNA-binding<-wHTH+WYL+WCX||DUF4236-><-?<-ParB+HNH-p<-CR-ATPase5+REase||HicA->HicB-></td><td>1222</td><td>Bacteria>Pseudomonadota>Gammaproteobacteria>Moraxellales>Moraxellaceae>Acinetobacter</td><td>GCF_024129855.1</td><td>Acinetobacter_lwoffii</td><td>NI465_RS14465</td><td>11210..14878</td><td>28090</td><td>BREX-system-P-loop-protein-BrxC-[Acinetobacter-lwoffii]</td></tr>
<tr><td>WP_161384457.1</td><td>Type1.BREX</td><td><-wHTH+WYL+WCX||?-><-RelE-ParE<-RHH_1||?->HEPN->?->BrxA-DNA-binding->BrxB-BrxF-iSTAND->DUF6079*->ParB+HNH->N6-METHYLASE->?->HEPN->ZNR+RmuC->PglZ->SIGMA+OB-Fold+AAA-ATPase+Lon-Protease-BrxL-><-?<-HATPase_c</td><td>1192</td><td>Bacteria>Pseudomonadota>Gammaproteobacteria>Oceanospirillales>Alcanivoracaceae>Alcanivorax</td><td>-</td><td>Alcanivorax_sp._ZXX171</td><td>GN155_RS10555</td><td>118232..121810</td><td>2695858</td><td>BREX-system-P-loop-protein-BrxC-[Alcanivorax-sp.-ZXX171]</td></tr>
<tr><td>WP_153499070.1</td><td>Type6.BREX</td><td><-PHP<-wHTH+WYL+WCX||?->?->HEPN-p->BrxE->BrxA-DNA-binding->BrxB-BrxF-iSTAND->DUF6079*->N6-METHYLASE->N6-METHYLASE->?->ParB+HNH->PglZ->SIGMA+OB-Fold+AAA-ATPase+HKD-DNase-BrxL-><-?<-HTH_3</td><td>1214</td><td>Bacteria>Pseudomonadota>Gammaproteobacteria>Oceanospirillales>Alcanivoracaceae>Alcanivorax</td><td>GCF_009601165.1</td><td>Alcanivorax_sediminis</td><td>GFN93_RS03795</td><td>861896..865540</td><td>2663008</td><td>BREX-system-P-loop-protein-BrxC-[Alcanivorax-sediminis]</td></tr>
<tr><td>WP_145953601.1</td><td>Type1.BREX</td><td><-RelE-ParE<-RelE-ParE<-RelB<-SIGMA+OB-Fold+AAA-ATPase+Lon-Protease-BrxL<-PglZ<-ZNR+RmuC<-HEPN<-?<-N6-METHYLASE<-DUF6079*<-BrxB-BrxF-iSTAND<-BrxA-DNA-binding<-?<-HEPN||?->wHTH+WYL+WCX-></td><td>1192</td><td>Bacteria>Pseudomonadota>Gammaproteobacteria>Oceanospirillales>Alcanivoracaceae>Alloalcanivorax</td><td>GCF_002072815.1</td><td>Alloalcanivorax_xenomutans</td><td>P40_RS14805</td><td>3223957..3227535</td><td>1094342</td><td>MULTISPECIES:-BREX-system-P-loop-protein-BrxC-[Gammaproteobacteria]</td></tr>
<tr><td>MBR2514422.1</td><td>Type6.BREX</td><td>Arm-DNA-bind_3+Phage_integrase-><-?<-SIGMA+OB-Fold+AAA-ATPase+HKD-DNase-BrxL<-PglZ<-ORC-CDC6<-ParA-Soj-PloopNTPase<-HEPN<-N6-METHYLASE<-DUF6079*<-BrxB-BrxF-iSTAND<-BrxA-DNA-binding<-BrxE<-SIGMA-HTH<-?<-?<-?<-?<-XIS-HTH</td><td>1224</td><td>Bacteria>Pseudomonadota>Gammaproteobacteria>Oceanospirillales>Halomonadaceae>Halomonas</td><td>GCA_017515565.1</td><td>Halomonas_sp.</td><td>IKE45_10465</td><td>23089..26763</td><td>1486246</td><td>BREX-system-P-loop-protein-BrxC-[Halomonas-sp.]</td></tr>
<tr><td>WP_062374360.1</td><td>Type1.BREX</td><td><-ABC-ATPase+SbcC<-ABC-ATPase+SbcC<-?<-?<-?<-TetR-HTH||?->tRNA->ncRNA->BrxA-DNA-binding->BrxB-BrxF-iSTAND->DUF6079*->Phd-YefM->PIN->Schlafen+HATPase+wHTH->N6-METHYLASE->N6-METHYLASE-p->HEPN->OLD-ABC->TOPRIM-></td><td>1227</td><td>Bacteria>Pseudomonadota>Gammaproteobacteria>Oceanospirillales>Halomonadaceae>Halomonas</td><td>GCF_001596805.1</td><td>Halomonas_sp._KX33721</td><td>A3220_RS08340</td><td>63501..67184</td><td>1819251</td><td>BREX-system-P-loop-protein-BrxC-[Halomonas-sp.-KX33721]</td></tr>
<tr><td>WP_101147206.1</td><td>Type1.BREX</td><td><-DiS_P_DiS+Peptidase_A24<-?<-VirB11||?->?->?->tRNA->BrxA-DNA-binding->BrxB-BrxF-iSTAND->DUF6079*->Phd-YefM->PIN->Schlafen+HATPase+wHTH->N6-METHYLASE->HEPN-><-?||OLD-ABC->TOPRIM-></td><td>1230</td><td>Bacteria>Pseudomonadota>Gammaproteobacteria>Oceanospirillales>Halomonadaceae>Halomonas</td><td>GCF_002836035.1</td><td>Halomonas_sp._MES3-P3E</td><td>CXF87_RS15340</td><td>100201..103893</td><td>2058321</td><td>BREX-system-P-loop-protein-BrxC-[Halomonas-sp.-MES3-P3E]</td></tr>
<tr><td>WBU42642.1</td><td>Type1.BREX</td><td><-Mrr-N||?->?->?->SWIM+DUF3670+SF1-SF2-Helicase->HTH_3->BrxA-DNA-binding->BrxB-BrxF-iSTAND->DUF6079*->DOC+HTH->N6-METHYLASE->?->N6-METHYLASE-p->HEPN->?->TIR->?->TIR-></td><td>1221</td><td>Bacteria>Pseudomonadota>Gammaproteobacteria>Pseudomonadales>Marinobacteraceae>Marinobacter</td><td>GCA_027912435.1</td><td>Marinobacter_alkaliphilus</td><td>PBN92_07055</td><td>1547680..1551345</td><td>254719</td><td>BREX-system-P-loop-protein-BrxC-[Marinobacter-alkaliphilus]</td></tr>
<tr><td>WP_058091394.1</td><td>Type1.BREX</td><td>wHTH+WYL+WCX->?-><-Mrr-N||HTH_3->BrxA-DNA-binding->BrxB-BrxF-iSTAND->DUF6079*->DOC+HTH->SF1-SF2-Helicase->N6-METHYLASE->HEPN->?->TIR->OLD-ABC->TOPRIM-></td><td>1221</td><td>Bacteria>Pseudomonadota>Gammaproteobacteria>Pseudomonadales>Marinobacteraceae>Marinobacter</td><td>GCF_001447155.2</td><td>Marinobacter_sp._LQ44</td><td>ASQ50_RS18570</td><td>4052781..4056446</td><td>1749259</td><td>BREX-system-P-loop-protein-BrxC-[Marinobacter-sp.-LQ44]</td></tr>
<tr><td>WP_273917578.1</td><td>Type1.BREX</td><td><-SIGMA+OB-Fold+AAA-ATPase+Lon-Protease-BrxL<-PglZ<-HEPN<-TPR<-NERD+GspA-ATPase<-N6-METHYLASE<-N6-METHYLASE<-TOPRIM<-OLD-ABC<-DUF6079*<-BrxB-BrxF-iSTAND<-BrxA-DNA-binding<-wHTH+WYL+WCX<-wHTH+WYL+WCX||?->ARG-></td><td>1223</td><td>Bacteria>Pseudomonadota>Gammaproteobacteria>Pseudomonadales>Pseudomonadaceae>Pseudomonas</td><td>GCF_028656895.1</td><td>Pseudomonas_sp._TNT2022_ID1044</td><td>M5G20_RS01570</td><td>347595..351266</td><td>2942636</td><td>BREX-system-P-loop-protein-BrxC-[Pseudomonas-sp.-TNT2022-ID1044]</td></tr>
<tr><td>WP_053105412.1</td><td>Type1.BREX</td><td>METHYLASE->?->?-><-?<-?||wHTH+WYL+WCX->BrxA-DNA-binding->BrxB-BrxF-iSTAND->DUF6079*->N6-METHYLASE->ParB+HNH->OLD-ABC->TOPRIM->HEPN->PglZ->SIGMA+OB-Fold+AAA-ATPase+Lon-Protease-BrxL->Arm-DNA-bind_3+Phage_integrase-><-SIGMA+OB-Fold+AAA-ATPase+Lon-Protease-BrxL</td><td>1155</td><td>Bacteria>Pseudomonadota>Gammaproteobacteria>Pseudomonadales>Pseudomonadaceae>Thiopseudomonas</td><td>GCF_001267275.1</td><td>Thiopseudomonas_alkaliphila</td><td>AKN90_RS00615</td><td>123321..126788</td><td>1697053</td><td>BREX-system-P-loop-protein-BrxC-[Thiopseudomonas-alkaliphila]</td></tr>
<tr><td>WP_101840169.1</td><td>Type1.BREX</td><td>ZnR+TOPRIM+TOPRIM->SIGMA-HTH+SIGMA-HTH+SIGMA-HTH-><-?<-wHTH+WYL+WCX||?->HEPN->BrxA-DNA-binding->BrxB-BrxF-iSTAND->DUF6079*->N6-METHYLASE-></td><td>1210</td><td>Bacteria>Pseudomonadota>Gammaproteobacteria>Thiotrichales>Piscirickettsiaceae>Hydrogenovibrio</td><td>GCF_002848135.1</td><td>Hydrogenovibrio_sp._SC-1</td><td>CYQ88_RS04565</td><td>151062..154694</td><td>2065820</td><td>BREX-system-P-loop-protein-BrxC-[Hydrogenovibrio-sp.-SC-1]</td></tr>
<tr><td>MBK85426.1</td><td>Type1.BREX</td><td>HHH_5+SF1-SF2-Helicase->BrxA-DNA-binding->BrxB-BrxF-iSTAND->DUF6079*->N6-METHYLASE->HEPN->Schlafen+HATPase+wHTH->TIR->PglZ->SIGMA+OB-Fold+AAA-ATPase+Lon-Protease-BrxL->?->RNase-><-SWIM+DUF3670+SF1-SF2-Helicase</td><td>1217</td><td>Bacteria>Pseudomonadota>Gammaproteobacteria>unclassified_Gammaproteobacteria</td><td>GCA_002721025.1</td><td>Gammaproteobacteria_bacterium</td><td>CMD81_16515</td><td>7659..11312</td><td>1913989</td><td>ATPase-[Gammaproteobacteria-bacterium]</td></tr>
<tr><td>NLY14765.1</td><td>Type1.BREX</td><td><-SIGMA+OB-Fold+AAA-ATPase+Lon-Protease-BrxL<-PglZ<-HEPN<-TOPRIM<-OLD-ABC<-ParB+HNH<-N6-METHYLASE<-DUF6079-p*</td><td>438</td><td>Bacteria>Pseudomonadota>Gammaproteobacteria>unclassified_Gammaproteobacteria</td><td>GCA_012728415.1</td><td>Gammaproteobacteria_bacterium</td><td>GXZ05_00075</td><td>12795..14111</td><td>1913989</td><td>hypothetical-protein-[Gammaproteobacteria-bacterium]</td></tr>
<tr><td>MBK7728002.1</td><td>Type1.BREX</td><td>Peptidase-><-DDE_Tnp<-SIGMA+OB-Fold+AAA-ATPase+Lon-Protease-BrxL<-PglZ<-HEPN<-?||?-><-DUF6079*<-BrxB-BrxF-iSTAND<-BrxA-DNA-binding||URI-DNase->?-><-tRNA<-MerR-HTH</td><td>1048</td><td>Bacteria>Pseudomonadota>Gammaproteobacteria>unclassified_Gammaproteobacteria</td><td>GCA_016709565.1</td><td>Gammaproteobacteria_bacterium</td><td>IPJ33_05745</td><td>303360..306506</td><td>1913989</td><td>BREX-system-P-loop-protein-BrxC-[Gammaproteobacteria-bacterium]</td></tr>
<tr><td>MBT5223355.1</td><td>Type1.BREX</td><td>BrxA-DNA-binding->BrxB-BrxF-iSTAND->DUF6079*->MazE_antitoxin->PIN->N6-METHYLASE->N6-METHYLASE->HEPN->GrpE->?->?->RelE-ParE-></td><td>1192</td><td>Bacteria>Pseudomonadota>Gammaproteobacteria>unclassified_Gammaproteobacteria</td><td>GCA_018661125.1</td><td>Gammaproteobacteria_bacterium</td><td>HOM15_11165</td><td>1377..4955</td><td>1913989</td><td>BREX-system-P-loop-protein-BrxC-[Gammaproteobacteria-bacterium]</td></tr>
<tr><td>MCG3145056.1</td><td>Type1.BREX</td><td>HTH_23+HTH_32+HTH_33->DDE_3-><-SIGMA+OB-Fold+AAA-ATPase+Lon-Protease-BrxL<-PglZ<-Schlafen+HATPase+wHTH+HTH<-N6-METHYLASE<-HEPN<-DUF6079*<-BrxB-BrxF-iSTAND<-BrxA-DNA-binding<-?||?-><-ParB<-ParA-Soj-PloopNTPase||Glyoxalase-><-?||AbiEi_4+AbiEi_1-></td><td>1220</td><td>Bacteria>Pseudomonadota>Gammaproteobacteria>unclassified_Gammaproteobacteria</td><td>GCA_022072255.1</td><td>Gammaproteobacteria_bacterium</td><td>HONDAALG_02597</td><td>52539..56201</td><td>1913989</td><td>hypothetical-protein-[Gammaproteobacteria-bacterium]</td></tr>
<tr><td>WP_190395653.1</td><td>Type1.BREX</td><td><-Couple_hipA+HipA_C<-HTH_3||?-><-RNase<-PALP||wHTH+WYL+WCX->BrxA-DNA-binding->BrxB-BrxF-iSTAND->DUF6079*->N6-METHYLASE->OLD-ABC->TOPRIM->PglZ->HEPN+SF1-SF2-Helicase+RNase+RNase->TPR-><-SF1-SF2-Helicase+REase<-Peptidase_M32</td><td>1157</td><td>Bacteria>Pseudomonadota>Gammaproteobacteria>Vibrionales>Vibrionaceae>Aliivibrio</td><td>GCF_014694495.1</td><td>Aliivibrio_sp._S10_S31</td><td>HC728_RS13680</td><td>57939..61412</td><td>2720224</td><td>BREX-system-P-loop-protein-BrxC-[Aliivibrio-sp.-S10_S31]</td></tr>
<tr><td>WP_159406353.1</td><td>Type1.BREX</td><td><-GspA-ATPase<-?||CR-ATPase4->?-><-?<-LexA-protease||wHTH+WYL+WCX->BrxA-DNA-binding->BrxB-BrxF-iSTAND->DUF6079*->N6-METHYLASE->ParB+HNH->PglZ->HEPN+SF1-SF2-Helicase+RNase+RNase->TPR-><-SF1-SF2-Helicase+REase<-SF1-SF2-Helicase+REase<-viral-protein<-AbiEi</td><td>1157</td><td>Bacteria>Pseudomonadota>Gammaproteobacteria>Vibrionales>Vibrionaceae</td><td>GCF_009883895.1</td><td>Vibrio</td><td>O8X55_RS08175</td><td>512098..515571</td><td>662</td><td>MULTISPECIES:-BREX-system-P-loop-protein-BrxC-[Vibrio]</td></tr>
<tr><td>WP_086713494.1</td><td>Type1.BREX</td><td><-GrpE<-ANK+ANK+ANK||?->SF1-SF2-Helicase->?->?->wHTH+WYL+WCX->BrxA-DNA-binding->BrxB-BrxF-iSTAND->DUF6079*->N6-METHYLASE->ParB+HNH->PglZ->HEPN+SF1-SF2-Helicase+RNase+RNase->TPR-><-SF1-SF2-Helicase+REase<-SF1-SF2-Helicase+REase<-?||AbiEi_4-></td><td>1157</td><td>Bacteria>Pseudomonadota>Gammaproteobacteria>Vibrionales>Vibrionaceae</td><td>GCF_024347515.1</td><td>Vibrio</td><td>EDB41_RS15380</td><td>67742..71215</td><td>662</td><td>MULTISPECIES:-BREX-system-P-loop-protein-BrxC-[Vibrio]</td></tr>
<tr><td>WP_232606822.1</td><td>Type1.BREX</td><td>ROK-HTH-><-?<-sGTP+sGTP||?->ANK+ANK->wHTH+WYL+WCX-p->BrxA-DNA-binding->BrxB-BrxF-iSTAND->DUF6079*->N6-METHYLASE->N6-METHYLASE->PglZ->HEPN+SF1-SF2-Helicase+RNase+RNase->TPR-><-SF1-SF2-Helicase+REase<-SF1-SF2-Helicase+REase||McrB-AAA->wHTH+REase-></td><td>1157</td><td>Bacteria>Pseudomonadota>Gammaproteobacteria>Vibrionales>Vibrionaceae>Photobacterium</td><td>GCF_021237045.1</td><td>Photobacterium_phosphoreum</td><td>CJF26_RS18915</td><td>30954..34427</td><td>659</td><td>BREX-system-P-loop-protein-BrxC-[Photobacterium-phosphoreum]</td></tr>
<tr><td>MBE4310404.1</td><td>Type1.BREX</td><td><-PIN+OST-HTH||CR-ATPase4->?-><-?<-DUF4113||wHTH-><-HTH_Tnp_1+HTH_21+rve+rve_3||wHTH+WYL+WCX->BrxA-DNA-binding->BrxB-BrxF-iSTAND->DUF6079*->ParB+HNH->PglZ->HEPN+SF1-SF2-Helicase+RNase+RNase->TPR-><-SF1-SF2-Helicase+REase<-SF1-SF2-Helicase+REase<-viral-protein<-AbiEi</td><td>1157</td><td>Bacteria>Pseudomonadota>Gammaproteobacteria>Vibrionales>Vibrionaceae>Vibrio</td><td>GCA_014921165.1</td><td>Vibrio_parahaemolyticus</td><td>HJ064_04290</td><td>963533..967006</td><td>670</td><td>BREX-system-P-loop-protein-BrxC-[Vibrio-parahaemolyticus]</td></tr>
<tr><td>EGR0719595.1</td><td>Type1.BREX</td><td>HTH_20-><-HEPN<-SbcC||CR-ATPase4->?-><-DUF4113||wHTH+WYL+WCX->BrxA-DNA-binding->BrxB-BrxF-iSTAND->DUF6079*->N6-METHYLASE->ParB+HNH->PglZ->HEPN+SF1-SF2-Helicase+RNase+RNase->TPR-><-SF1-SF2-Helicase+REase<-SF1-SF2-Helicase+REase<-viral-protein<-AbiEi</td><td>1157</td><td>Bacteria>Pseudomonadota>Gammaproteobacteria>Vibrionales>Vibrionaceae>Vibrio</td><td>GCA_015789455.1</td><td>Vibrio_alginolyticus</td><td>EX464_05795</td><td>8586..12059</td><td>663</td><td>BREX-system-P-loop-protein-BrxC-[Vibrio-alginolyticus]</td></tr>
<tr><td>EGR4188113.1</td><td>Type1.BREX</td><td>AbiEi->viral-protein->SF1-SF2-Helicase+REase->SF1-SF2-Helicase+REase-><-TPR<-HEPN+SF1-SF2-Helicase+RNase+RNase<-PglZ<-ParB+HNH<-N6-METHYLASE<-DUF6079*<-BrxB-BrxF-iSTAND<-BrxA-DNA-binding<-wHTH+WYL+WCX||DDE_Tnp_1_5+DDE_Tnp_1->PIN+OST-HTH-><-?<-HTH_20</td><td>1157</td><td>Bacteria>Pseudomonadota>Gammaproteobacteria>Vibrionales>Vibrionaceae>Vibrio</td><td>GCA_015809855.1</td><td>Vibrio_cholerae</td><td>DDM99_15775</td><td>19819..23292</td><td>666</td><td>BREX-system-P-loop-protein-BrxC-[Vibrio-cholerae]</td></tr>
<tr><td>EGR4671255.1</td><td>Type1.BREX</td><td>AbiEi->viral-protein->SF1-SF2-Helicase+REase->SF1-SF2-Helicase+REase-><-TPR<-HEPN+SF1-SF2-Helicase+RNase+RNase<-PglZ<-ParB+HNH<-N6-METHYLASE<-DUF6079*<-BrxB-BrxF-iSTAND<-BrxA-DNA-binding<-wHTH+WYL+WCX||DUF4113-><-?<-CR-ATPase4||SbcC-></td><td>1157</td><td>Bacteria>Pseudomonadota>Gammaproteobacteria>Vibrionales>Vibrionaceae>Vibrio</td><td>GCA_015814545.1</td><td>Vibrio_parahaemolyticus</td><td>C4G45_02530</td><td>555237..558710</td><td>670</td><td>BREX-system-P-loop-protein-BrxC-[Vibrio-parahaemolyticus]</td></tr>
</table>

| Protein-ID | System | Gene-Neighbourhoods | Length | Taxonomy | Assembly-ID | Strain | Locus-Tag | Genomic-coordinates | TaxID | Fasta-defline |
|---|---|---|---|---|---|---|---|---|---|---|
| MBM4838763.1 | Type1.BREX | AbiEi->viral-protein->SF1-SF2-Helicase+REase->SF1-SF2-Helicase+REase-><-TPR<-HEPN+SF1-SF2-Helicase+RNase+RNase<-PglZ<-ParB+HNH<-N6-METHYLASE<-DUF6079*<-BrxB-BrxF-iSTAND<-BrxA-DNA-binding<-wHTH+WYL+WCX\|\|HTH_Tnp_1+HTH_21+rve+rve_3-><-wHTH\|\|DUF4113-><-?<-CR-ATPase4\|\|PIN+OST-HTH-> | 1157 | Bacteria>Pseudomonadota>Gammaproteobacteria>Vibrionales>Vibrionaceae>Vibrio | GCA_016876235.1 | Vibrio_parahaemolyticus | HY055_06530 | 138658..142131 | 670 | BREX-system-P-loop-protein-BrxC-[Vibrio-parahaemolyticus] |
| MBM4875012.1 | Type1.BREX | <-AAA-ATPase-p<-HEPN\|\|CR-ATPase4->?->\|\|wHTH+WYL+WCX->BrxA-DNA-binding->BrxB-BrxF-iSTAND->DUF6079*->N6-METHYLASE->ParB+HNH->PglZ->HEPN+SF1-SF2-Helicase+RNase+RNase->TPR-><-SF1-SF2-Helicase+REase<-SF1-SF2-Helicase+REase<-viral-protein<-AbiEi | 1157 | Bacteria>Pseudomonadota>Gammaproteobacteria>Vibrionales>Vibrionaceae>Vibrio | GCA_016876415.1 | Vibrio_parahaemolyticus | HY045_05440 | 517620..521093 | 670 | BREX-system-P-loop-protein-BrxC-[Vibrio-parahaemolyticus] |
| HBC3445072.1 | Type1.BREX | <-ZnR<-?\|\|AbiEi->viral-protein->wHTH+WYL+WCX->BrxA-DNA-binding->BrxB-BrxF-iSTAND->DUF6079*->N6-METHYLASE->?->?->PglZ->SIGMA+OB-Fold+AAA-ATPase+Lon-Protease-BrxL-><-TPR<-HEPN+SF1-SF2-Helicase+RNase+RNase\|\|<-?<-CR-ATPase4 | 1225 | Bacteria>Pseudomonadota>Gammaproteobacteria>Vibrionales>Vibrionaceae>Vibrio | GCA_018119025.1 | Vibrio_parahaemolyticus | KDY48_003788 | 6147..9824 | 670 | BREX-system-P-loop-protein-BrxC-[Vibrio-parahaemolyticus] |
| WP_021709035.1 | Type1.BREX | <-Thioredoxin<-?<-Phage_integrase\|\|?->?-><-?\|\|wHTH+WYL+WCX->BrxA-DNA-binding->BrxB-BrxF-iSTAND->DUF6079*->N6-METHYLASE->HEPN->PglZ->SIGMA+OB-Fold+AAA-ATPase+Lon-Protease-BrxL->GNAT-><-Peptidase | 1229 | Bacteria>Pseudomonadota>Gammaproteobacteria>Vibrionales>Vibrionaceae>Vibrio | GCF_000400265.1 | Vibrio_azureus | VAN_RS15510 | 14465..18154 | 512649 | BREX-system-P-loop-protein-BrxC-[Vibrio-azureus] |
| WP_053027640.1 | Type1.BREX | DUF6079-p*->N6-METHYLASE->PglZ->SIGMA+OB-Fold+AAA-ATPase+Lon-Protease-BrxL->HEPN->HDRnase->VirD4-TraG+VirB4-FtsK->?->?-><-Glyoxalase_8 | 848 | Bacteria>Pseudomonadota>Gammaproteobacteria>Vibrionales>Vibrionaceae>Vibrio | GCF_001259875.1 | Vibrio_cholerae | ALV71_RS07765 | 696..3240 | 666 | BREX-system-P-loop-protein-BrxC,-partial-[Vibrio-cholerae] |
| WP_096070868.1 | Type1.BREX | HTH_20->CR-ATPase4->?-><-?<-LexA-protease\|\|wHTH+WYL+WCX->BrxA-DNA-binding->BrxB-BrxF-iSTAND->DUF6079*->N6-METHYLASE->ParB+HNH->PglZ->HEPN+SF1-SF2-Helicase+RNase+RNase->TPR-><-LZ_Tnp_IS66+zf-IS66+DDE_Tnp_IS66+DDE_Tnp_IS66_C<-?<-HTH_Tnp_1<-AAA-ATPase | 1157 | Bacteria>Pseudomonadota>Gammaproteobacteria>Vibrionales>Vibrionaceae>Vibrio | GCF_002313025.1 | Vibrio_cholerae | FORC55_RS15555 | 276972..280445 | 666 | BREX-system-P-loop-protein-BrxC-[Vibrio-cholerae] |
| WP_148513618.1 | Type1.BREX | AbiEi->viral-protein->SF1-SF2-Helicase+REase->SF1-SF2-Helicase+REase-><-TPR<-HEPN+SF1-SF2-Helicase+RNase+RNase<-PglZ<-ParB+HNH<-N6-METHYLASE<-DUF6079*<-BrxB-BrxF-iSTAND<-BrxA-DNA-binding<-wHTH+WYL+WCX\|\|DDE_Tnp_1_5+DDE_Tnp_1-> | 1157 | Bacteria>Pseudomonadota>Gammaproteobacteria>Vibrionales>Vibrionaceae>Vibrio | GCF_008083315.1 | Vibrio_cholerae | FXF10_RS00340 | 29414..32887 | 666 | BREX-system-P-loop-protein-BrxC-[Vibrio-cholerae] |
| WP_148490856.1 | Type1.BREX | <-DDE_Tnp_1_5+DDE_Tnp_1\|\|HTH_Tnp_1+HTH_21+rve+rve_3-><-SbcC\|\|CR-ATPase4->?-><-DUF4113\|\|wHTH+WYL+WCX->BrxA-DNA-binding->BrxB-BrxF-iSTAND->DUF6079*->N6-METHYLASE->ParB+HNH->PglZ->HEPN+SF1-SF2-Helicase+RNase+RNase->TPR-><-SF1-SF2-Helicase+REase<-SF1-SF2-Helicase+REase<-viral-protein<-AbiEi | 1157 | Bacteria>Pseudomonadota>Gammaproteobacteria>Vibrionales>Vibrionaceae>Vibrio | GCF_008084645.1 | Vibrio_cholerae | FXE54_RS16715 | 26974..30447 | 666 | BREX-system-P-loop-protein-BrxC-[Vibrio-cholerae] |
| WP_213899843.1 | Type1.BREX | AbiEi->viral-protein->SF1-SF2-Helicase+REase->SF1-SF2-Helicase+REase-><-TPR<-HEPN+SF1-SF2-Helicase+RNase+RNase<-PglZ<-ParB+HNH<-N6-METHYLASE<-DUF6079*<-BrxB-BrxF-iSTAND<-BrxA-DNA-binding<-wHTH+WYL+WCX\|\|<-?<-CR-ATPase4\|\|SbcC->HEPN-><-HTH_20 | 1157 | Bacteria>Pseudomonadota>Gammaproteobacteria>Vibrionales>Vibrionaceae>Vibrio | GCF_018458525.1 | Vibrio_alginolyticus | J4H25_RS19085 | 22843..26316 | 663 | BREX-system-P-loop-protein-BrxC-[Vibrio-alginolyticus] |
| WP_248354663.1 | Type1.BREX | AbiEi->viral-protein->SF1-SF2-Helicase+REase->SF1-SF2-Helicase+REase-><-TPR<-HEPN+SF1-SF2-Helicase+RNase+RNase<-PglZ<-ParB+HNH<-N6-METHYLASE<-DUF6079*<-BrxB-BrxF-iSTAND<-BrxA-DNA-binding<-wHTH+WYL+WCX\|\|DDE_Tnp_1_5+DDE_Tnp_1->PIN+OST-HTH-><-?<-HTH_20 | 1157 | Bacteria>Pseudomonadota>Gammaproteobacteria>Vibrionales>Vibrionaceae>Vibrio | GCF_024745175.1 | Vibrio_parahaemolyticus | DDM99_15775 | 19819..23292 | 670 | BREX-system-P-loop-protein-BrxC-[Vibrio-parahaemolyticus] |
| WP_272236739.1 | Type1.BREX | TraJ-RHH->TraI-VirD2-RCRE->REase-><-AbiEi_4<-?\|\|?->wHTH+WYL+WCX->BrxA-DNA-binding->BrxB-BrxF-iSTAND->DUF6079*->N6-METHYLASE->HEPN->PglZ->SIGMA+OB-Fold+AAA-ATPase+Lon-Protease-BrxL-><-?<-?<-SF1-SF2-Helicase<-?<-Arm-DNA-bind_3 | 1229 | Bacteria>Pseudomonadota>Gammaproteobacteria>Vibrionales>Vibrionaceae>Vibrio | GCF_028446685.1 | Vibrio_europaeus | OPW18_RS21520 | 351695..355384 | 300876 | BREX-system-P-loop-protein-BrxC-[Vibrio-europaeus] |
| WP_229611425.1 | Type1.BREX | REase+SPF-cons-><-PIN+OST-HTH\|\|CR-ATPase4->?->\|\|wHTH+WYL+WCX->BrxA-DNA-binding->BrxB-BrxF-iSTAND->DUF6079*->N6-METHYLASE->ParB+HNH->PglZ->HEPN+SF1-SF2-Helicase+RNase+RNase->TPR-><-SF1-SF2-Helicase+REase<-SF1-SF2-Helicase+REase<-viral-protein<-AbiEi | 1157 | Bacteria>Pseudomonadota>Gammaproteobacteria>Vibrionales>Vibrionaceae>Vibrio | GCF_905331955.1 | Vibrio_parahaemolyticus | LPR99_RS16740 | 18511..21984 | 670 | BREX-system-P-loop-protein-BrxC-[Vibrio-parahaemolyticus] |
| WP_107394298.1 | Type1.BREX | <-HisKin<-SF1-SF2-Helicase+REase<-35Exo+REase\|\|wHTH+WYL+WCX->BrxA-DNA-binding->BrxB-BrxF-iSTAND->DUF6079*->N6-METHYLASE->HEPN->PglZ->SIGMA+OB-Fold+AAA-ATPase+Lon-Protease-BrxL-><-AbiEii | 1140 | Bacteria>Pseudomonadota | GCF_003034225.1 | Alphaproteobacteria | CV103_RS05945 | 27177..30599 | 28211 | MULTISPECIES:-BREX-system-P-loop-protein-BrxC-[Alphaproteobacteria] |
| WP_053322564.1 | Type1.BREX | HNH->?->?->LON-AAA->Calcineurin->?->BrxA-DNA-binding->BrxB-BrxF-iSTAND->DUF6079*->N6-METHYLASE-p->Schlafen+HATPase+wHTH->HEPN->REase->N6-METHYLASE-p->HerA/FtsK->PglZ->SIGMA+OB-Fold+AAA-ATPase+HKD-DNase-BrxL-> | 1181 | Bacteria>Pseudomonadota>Zetaproteobacteria>Mariprofundales>Mariprofundaceae>Mariprofundus | GCF_001273615.1 | Mariprofundus_ferrooxydans | AL013_RS02410 | 88551..92096 | 314344 | BREX-system-P-loop-protein-BrxC-[Mariprofundus-ferrooxydans] |
| MCF7790541.1 | Type1.BREX | <-HEPN<-SIGMA+OB-Fold+AAA-ATPase+HKD-DNase-BrxL<-PglZ<-TOPRIM<-OLD-ABC<-?<-?<-?<-DUF6079*<-BrxB-BrxF-iSTAND<-BrxA-DNA-binding<-wHTH+WYL+WCX<-?\|\|?-><-?<-?\|\|S2P+PDZ-> | 1176 | Bacteria>PVC_group>Lentisphaerota>Lentisphaeria>Victivallales>unclassified_Victivallales | GCA_021733985.1 | Victivallales_bacterium | K9M56_00960 | 14578..18108 | 2212475 | BREX-system-P-loop-protein-BrxC-[Victivallales-bacterium] |
| CAJ71903.1 | Type1.BREX | Unstab_antitox->RelE-ParE->RelE-ParE->HEPN->ABhydrolase-><-HEPN<-Polbeta<-SIGMA+OB-Fold+AAA-ATPase+Lon-Protease-BrxL<-PglZ<-DUF6079*<-?<-?<-N6-METHYLASE-p\|\|?-><-DDE_Tnp_1_4<-N6-METHYLASE<-BrxB-BrxF-iSTAND<-BrxA-DNA-binding | 1220 | Bacteria>PVC_group>Planctomycetota>Candidatus_Brocadiia>Candidatus_Brocadiales>Candidatus_Brocadiaceae | - | Candidatus_Kuenenia_stuttgartiensis | kustc1158 | 789101..792763 | 174633 | hypothetical-protein-kustc1158-[Candidatus-Kuenenia-stuttgartiensis] |
| MBQ9812620.1 | Type1.BREX | OLD-ABC->HEPN->BrxA-DNA-binding->BrxB-BrxF-iSTAND->DUF6079-p*->DUF6079-p->N6-METHYLASE->?->PglZ->SIGMA+OB-Fold+AAA-ATPase+Lon-Protease-BrxL-><-HNH<-OLD-ABC | 596 | Bacteria>PVC_group>Planctomycetota>Planctomycetia>Pirellulales>Thermoguttaceae | GCA_017441335.1 | Thermoguttaceae_bacterium | IJM54_04830 | 4203..5993 | 2691551 | BREX-system-P-loop-protein-BrxC-[Thermoguttaceae-bacterium] |
| WP_145192991.1 | Type1.BREX | <-SIGMA+OB-Fold+AAA-ATPase+Lon-Protease-BrxL<-PglZ<-?<-NTP_transf<-N6-METHYLASE<-HEPN<-DUF6079*<-BrxB-BrxF-iSTAND<-BrxA-DNA-binding<-XIS-HTH\|\|ABhydrolase-><-?<-?<-SIGMA-HTH\|\|GDResolvase+HIN-HTH-><-SIR2 | 1220 | Bacteria>PVC_group>Planctomycetota>Planctomycetia>Planctomycetales>Planctomycetaceae | GCF_007744075.1 | Gimesia_chilikensis | HG66A1_RS30060 | 7903021..7906683 | 2605989 | BREX-system-P-loop-protein-BrxC-[Gimesia-chilikensis] |
| TET34868.1 | Type1.BREX | <-HNH\|\|?-><-?\|\|XIS-HTH->DUF4007->LON-AAA->N6-METHYLASE->ParB+HNH->HEPN->DUF6079*->PglZ->SIGMA+OB-Fold+AAA-ATPase-BrxL->SbcC->HISKIN->wHTH+WYL->RADICAL-SAM->DEOR-HTH-><-GDResolvase+Recombinase+Zn_ribbon_recom | 1347 | Bacteria>PVC_group>Planctomycetota>unclassified_Planctomycetota | GCA_004376375.1 | Planctomycetota_bacterium | E3J72_12085 | 41890..45933 | 2026780 | MAG:-hypothetical-protein-E3J72_12085-[Planctomycetota-bacterium] |
| MBI3468946.1 | Type1.BREX | <-N6-METHYLASE-p<-HEPN<-DUF6079*<-BrxB-BrxF-iSTAND<-BrxA-DNA-binding\|\|?->Amidohydro_2->?-><-Phage_integrase<-wHTH+WYL+WCX\|\|SIGMA-HTH->Pkinase+CR-Kinase-> | 1222 | Bacteria>PVC_group>Planctomycetota>unclassified_Planctomycetota | GCA_016196255.1 | Planctomycetota_bacterium | HY000_38570 | 1768..5436 | 2026780 | BREX-system-P-loop-protein-BrxC-[Planctomycetota-bacterium] |
| MBL1217602.1 | Type1.BREX | BrxA-DNA-binding->BrxB-BrxF-iSTAND->DUF6079*->HEPN->N6-METHYLASE->RNase->PglZ->SIGMA+OB-Fold+AAA-ATPase+Lon-Protease-BrxL-><-?<-?<-Phage_integrase | 1165 | Bacteria>PVC_group>Planctomycetota>unclassified_Planctomycetota | GCA_016743455.1 | Planctomycetota_bacterium | D8M59_08910 | 1591..5088 | 2026780 | BREX-system-P-loop-protein-BrxC-[Planctomycetota-bacterium] |
| MBW8002946.1 | Type1.BREX | BrxA-DNA-binding->BrxB-BrxF-iSTAND->DUF6079*->N6-METHYLASE->Ntox45->HEPN->PglZ->SIGMA+OB-Fold+AAA-ATPase+HKD-DNase-BrxL-><-HEPN<-DDE_Tnp_ISAZ013<-DDE_Tnp_ISAZ013 | 1164 | Bacteria>PVC_group>Planctomycetota>unclassified_Planctomycetota | GCA_019456255.1 | Planctomycetota_bacterium | FVQ80_13155 | 4626..8120 | 2026780 | BREX-system-P-loop-protein-BrxC-[Planctomycetota-bacterium] |
| MCC6785237.1 | Type1.BREX | <-RAMPs<-?<-?<-RAMPs<-HEPN<-Schlafen+HATPase+wHTH<-N6-METHYLASE<-SbcC<-DUF6079*<-BrxB-BrxF-iSTAND<-BrxA-DNA-binding | 1150 | Bacteria>PVC_group>Planctomycetota>unclassified_Planctomycetota | GCA_020848735.1 | Planctomycetota_bacterium | IT457_20475 | 22318..25770 | 2026780 | BREX-system-P-loop-protein-BrxC-[Planctomycetota-bacterium] |
| MBI4023965.1 | Type1.BREX | CBS->BrxA-DNA-binding->BrxB-BrxF-iSTAND->DUF6079*->HEPN->N6-METHYLASE->REase+AIPR->PglZ->SIGMA+OB-Fold+AAA-ATPase+Lon-Protease-BrxL-> | 1148 | Bacteria>PVC_group>Verrucomicrobiota>unclassified_Verrucomicrobiota | GCA_016198745.1 | Verrucomicrobiota_bacterium | HY360_03230 | 10391..13837 | 2026799 | BREX-system-P-loop-protein-BrxC-[Verrucomicrobiota-bacterium] |
| NLV85368.1 | Type1.BREX | DUF6079*->N6-METHYLASE->HEPN->N6-METHYLASE-p->PglZ->SIGMA+OB-Fold+AAA-ATPase+Lon-Protease-BrxL-> | 1160 | Bacteria>Spirochaetota>Spirochaetia>Spirochaetales>unclassified_Spirochaetales | GCA_012729295.1 | Spirochaetales_bacterium | GXY60_12460 | 48..3530 | 2026792 | BREX-system-P-loop-protein-BrxC-[Spirochaetales-bacterium] |
| RKX87882.1 | Type1.BREX | <-MarR-HTH\|\|?->tRNA->BrxA-DNA-binding->BrxB-BrxF-iSTAND->DUF6079*->PolB->HEPN->Polbeta-><-?\|\|N6-METHYLASE-p-> | 1168 | Bacteria>Spirochaetota>unclassified_Spirochaetota | GCA_003644105.1 | Spirochaetota_bacterium | DRP70_07570 | 3107..6613 | 2202144 | BREX-system-P-loop-protein-BrxC-[Spirochaetota-bacterium] |
| WP_009648402.1 | Type1.BREX | REase-><-?<-DDE_Tnp<-DDE_Tnp<-?\|\|?->BrxA-DNA-binding->BrxB-BrxF-iSTAND->DUF6079*->N6-METHYLASE->RVT->PglZ->SIGMA+OB-Fold+AAA-ATPase+HKD-DNase-BrxL->HEPN-> | 1169 | Bacteria>Terrabacteria_group>Actinomycetota>Actinomycetes>Actinomycetales>Actinomycetaceae | GCF_000278725.1 | Actinomyces_sp._ICM47 | HMPREF1136_RS05845 | 15295..18804 | 936548 | BREX-system-P-loop-protein-BrxC-[Actinomyces-sp.-ICM47] |
| KFI62072.1 | Type1.BREX | <-XIS-HTH+GDResolvase\|\|REase->?->BrxA-DNA-binding->Schlafen+HATPase+wHTH+HTH-><-XIS-HTH\|\|BrxA-DNA-binding->BrxB-BrxF-iSTAND->DUF6079*->HEPN->N6-METHYLASE->PglZ->SIGMA+OB-Fold+AAA-ATPase+HKD-DNase-BrxL-><-?<-ZnR+GP<-?<-MPTase | 1219 | Bacteria>Terrabacteria_group>Actinomycetota>Actinomycetes>Bifidobacteriales>Bifidobacteriaceae | GCA_000741575.1 | Bifidobacterium_cuniculi | BCUN_1388 | 51093..54752 | 1688 | putative-ATP-binding-protein-[Bifidobacterium-cuniculi] |
| GDY95791.1 | Type1.BREX | <-HTH_3<-tRNA<-ACET\|\|Acyl_transf_3->Acyl_transf_3->?-><-?\|\|?->BrxA-DNA-binding->BrxB-BrxF-iSTAND->DUF6079*->N6-METHYLASE->HEPN->Schlafen+HATPase+wHTH->PglZ->SIGMA+OB-Fold+AAA-ATPase+HKD-DNase-BrxL->ZnR+GP->ZnR->ParB+HNH+RAMA-> | 1223 | Bacteria>Terrabacteria_group>Actinomycetota>Actinomycetes>Bifidobacteriales>Bifidobacteriaceae | GCA_018326565.1 | Bifidobacteriaceae_bacterium_MCC01943 | MCC01943_10780 | 5409..9080 | 2569488 | hypothetical-protein-MCC01943_10780-[Bifidobacteriaceae-bacterium-MCC01943] |
| WP_032738770.1 | Type1.BREX | HTH_28->BrxA-DNA-binding->BrxB-BrxF-iSTAND->DUF6079*->N6-METHYLASE->HEPN->?->Schlafen+HATPase+wHTH->PglZ->SIGMA+OB-Fold+AAA-ATPase+HKD-DNase-BrxL-> | 1218 | Bacteria>Terrabacteria_group>Actinomycetota>Actinomycetes>Bifidobacteriales>Bifidobacteriaceae | GCF_000730055.1 | Bifidobacterium_longum | BL71B_RS01495 | 365977..369633 | 216816 | BREX-system-P-loop-protein-BrxC-[Bifidobacterium-longum] |
| WP_051922710.1 | Type1.BREX | <-His_Phos_1<-?\|\|ZnR->?->ABC-ATPase+SbcC-><-?\|\|BrxA-DNA-binding->BrxB-BrxF-iSTAND->DUF6079*->N6-METHYLASE->?->HEPN+RES->Schlafen+HATPase+wHTH->PglZ->SIGMA+OB-Fold+AAA-ATPase+HKD-DNase-BrxL-> | 1249 | Bacteria>Terrabacteria_group>Actinomycetota>Actinomycetes>Bifidobacteriales>Bifidobacteriaceae | GCF_000741785.1 | Bifidobacterium_stellenboschense | BSTEL_1465 | 61247..64996 | 762211 | BREX-system-P-loop-protein-BrxC-[Bifidobacterium-stellenboschense] |
| WP_160257632.1 | Type1.BREX | <-XIS-HTH+GDResolvase\|\|REase->?->BrxA-DNA-binding->Schlafen+HATPase+wHTH+HTH-><-XIS-HTH\|\|BrxA-DNA-binding->BrxB-BrxF-iSTAND->DUF6079*->HEPN->N6-METHYLASE->PglZ->SIGMA+OB-Fold+AAA-ATPase+HKD-DNase-BrxL-><-?<-ZnR+GP<-MPTase | 1215 | Bacteria>Terrabacteria_group>Actinomycetota>Actinomycetes>Bifidobacteriales>Bifidobacteriaceae | GCF_000771465.1 | Bifidobacterium_cuniculi | BCUN_RS07325 | 51105..54752 | 1688 | BREX-system-P-loop-protein-BrxC-[Bifidobacterium-cuniculi] |
| WP_100495666.1 | Type1.BREX | <-ABhydrolase\|\|TPR+TPR+TPR+TPR->BrxA-DNA-binding->BrxB-BrxF-iSTAND->DUF6079*-><-?<-?<-?\|\|HNH->?->SbcC->N6-METHYLASE->HEPN->Schlafen+HATPase+wHTH-> | 1240 | Bacteria>Terrabacteria_group>Actinomycetota>Actinomycetes>Bifidobacteriales>Bifidobacteriaceae | GCF_002802865.1 | Bifidobacterium_scaligerum | CUU80_RS01790 | 434556..438278 | 2052656 | BREX-system-P-loop-protein-BrxC-[Bifidobacterium-scaligerum] |
| WP_180326342.1 | Type1.BREX | DUF6079-p*->N6-METHYLASE->PglZ->SIGMA+OB-Fold+AAA-ATPase+HKD-DNase-BrxL-><-HEPN<-PRTase | 865 | Bacteria>Terrabacteria_group>Actinomycetota>Actinomycetes>Bifidobacteriales>Bifidobacteriaceae | GCF_002846775.1 | Bifidobacterium_pseudolongum | CQR50_RS05955 | 514..3111 | 1694 | BREX-system-P-loop-protein-BrxC-[Bifidobacterium-pseudolongum] |
| WP_109056521.1 | Type1.BREX | <-ABhydrolase\|\|BrxA-DNA-binding->BrxB-BrxF-iSTAND->DUF6079*->N6-METHYLASE->HEPN->Schlafen+wHTH->CR-ATPase5+REase->PglZ->SIGMA+OB-Fold+AAA-ATPase+HKD-DNase-BrxL->METHYLASE-> | 1246 | Bacteria>Terrabacteria_group>Actinomycetota>Actinomycetes>Bifidobacteriales>Bifidobacteriaceae | GCF_003129905.1 | Bifidobacterium_callitrichidarum | DF196_RS03600 | 108404..112144 | 2052941 | BREX-system-P-loop-protein-BrxC-[Bifidobacterium-callitrichidarum] |
| WP_115955260.1 | Type1.BREX | <-OLD-ABC+TOPRIM<-ZnR<-ZnR+GP<-SIGMA+OB-Fold+AAA-ATPase+HKD-DNase-BrxL<-PglZ<-Schlafen+HATPase+wHTH<-?<-HEPN<-N6-METHYLASE<-DUF6079*<-BrxB-BrxF-iSTAND<-BrxA-DNA-binding<-?<-?<-?<-ABC-ATPase+SbcC<-?<-ZnR | 1217 | Bacteria>Terrabacteria_group>Actinomycetota>Actinomycetes>Bifidobacteriales>Bifidobacteriaceae | GCF_003390735.1 | Bifidobacterium_bifidum | CJD48_RS06210 | 1417204..1420857 | 1681 | BREX-system-P-loop-protein-BrxC-[Bifidobacterium-bifidum] |
| WP_117759621.1 | Type1.BREX | <-ParB+HNH+RAMA<-ZnR<-ZnR+GP<-SIGMA+OB-Fold+AAA-ATPase+HKD-DNase-BrxL<-PglZ<-Schlafen+HATPase+wHTH<-HEPN<-N6-METHYLASE<-DUF6079*<-BrxB-BrxF-iSTAND<-BrxA-DNA-binding<-?\|\|?-><-Acyl_transf_3\|\|ACET->tRNA-><-PIN<-XIS-HTH | 1223 | Bacteria>Terrabacteria_group>Actinomycetota>Actinomycetes>Bifidobacteriales>Bifidobacteriaceae | GCF_003458805.1 | Bifidobacterium_adolescentis | DWX79_RS04615 | 1022537..1026208 | 1680 | BREX-system-P-loop-protein-BrxC-[Bifidobacterium-adolescentis] |
| WP_126029122.1 | Type1.BREX | <-SIGMA+OB-Fold+AAA-ATPase+HKD-DNase-BrxL<-PglZ<-HEPN<-N6-METHYLASE<-DUF6079*<-BrxB-BrxF-iSTAND<-BrxA-DNA-binding<-?\|\|SIGMA-HTH-><-?<-?<-?<-?<-METHYLASE | 1235 | Bacteria>Terrabacteria_group>Actinomycetota>Actinomycetes>Bifidobacteriales>Bifidobacteriaceae | GCF_003952005.1 | Bifidobacterium_callimiconis | D2E23_RS00850 | 182499..186206 | 2306973 | BREX-system-P-loop-protein-BrxC-[Bifidobacterium-callimiconis] |
| WP_150379196.1 | Type1.BREX | <-SbcD\|\|?-><-ABhydrolase\|\|BrxA-DNA-binding->BrxB-BrxF-iSTAND->DUF6079*->CR-ATPase5+REase->TPR+TPR->N6-METHYLASE->HEPN->MazE_antitoxin->PglZ->SIGMA+OB-Fold+AAA-ATPase+HKD-DNase-BrxL->METHYLASE-> | 1267 | Bacteria>Terrabacteria_group>Actinomycetota>Actinomycetes>Bifidobacteriales>Bifidobacteriaceae | GCF_008698155.1 | Bifidobacterium_myosotis | EM091_RS06000 | 83624..87427 | 1630166 | BREX-system-P-loop-protein-BrxC-[Bifidobacterium-myosotis] |
| WP_152233646.1 | Type1.BREX | REase+Bacteriocin_IIc->?->?->?->?-><-?\|\|BrxA-DNA-binding->BrxB-BrxF-iSTAND->DUF6079*-><-?\|\|CR-ATPase5+REase->N6-METHYLASE->?->?->HEPN+RES->Schlafen+HATPase+wHTH->PglZ->SIGMA+OB-Fold+AAA-ATPase+HKD-DNase-BrxL-> | 1235 | Bacteria>Terrabacteria_group>Actinomycetota>Actinomycetes>Bifidobacteriales>Bifidobacteriaceae | GCF_009193355.1 | Bifidobacterium_leontopitheci | F7D09_RS01345 | 123122..126829 | 2650774 | BREX-system-P-loop-protein-BrxC-[Bifidobacterium-leontopitheci] |
| WP_117762529.1 | Type1.BREX | <-ParB+HNH+RAMA<-ZnR<-ZnR+GP<-SIGMA+OB-Fold+AAA-ATPase+HKD-DNase-BrxL<-PglZ<-Schlafen+HATPase+wHTH<-HEPN<-N6-METHYLASE<-DUF6079*<-BrxB-BrxF-iSTAND<-BrxA-DNA-binding<-?\|\|?-><-Acyl_transf_3\|\|ACET-> | 1223 | Bacteria>Terrabacteria_group>Actinomycetota>Actinomycetes>Bifidobacteriales>Bifidobacteriaceae | GCF_009832825.1 | Bifidobacterium_adolescentis | DWZ00_RS05085 | 1174565..1178236 | 1680 | BREX-system-P-loop-protein-BrxC-[Bifidobacterium-adolescentis] |
| WP_196336889.1 | Type1.BREX | 35exo+HRDC->35exo+BRCT-><-HTH\|\|?-><-ACET\|\|?->BrxA-DNA-binding->BrxB-BrxF-iSTAND->DUF6079*->N6-METHYLASE->HEPN->?->Schlafen+HATPase+wHTH->PglZ->SIGMA+OB-Fold+AAA-ATPase+HKD-DNase-BrxL->ZnR-><-DSBH+AraC-HTH+AraC-HTH\|\|Glyoxalase-> | 1217 | Bacteria>Terrabacteria_group>Actinomycetota>Actinomycetes>Bifidobacteriales>Bifidobacteriaceae | GCF_015703835.1 | Bifidobacterium_dentium | IHQ82_RS06050 | 179848..183501 | 1689 | BREX-system-P-loop-protein-BrxC-[Bifidobacterium-dentium] |
| WP_214374600.1 | Type1.BREX | ZnR->?->ABC-ATPase+SbcC-><-?\|\|?-><-ABhydrolase\|\|BrxA-DNA-binding->BrxB-BrxF-iSTAND->DUF6079*->CR-ATPase5+REase->N6-METHYLASE->?->?->HEPN+RES->Schlafen+HATPase+wHTH->PglZ->MIT_C->METHYLASE-> | 1249 | Bacteria>Terrabacteria_group>Actinomycetota>Actinomycetes>Bifidobacteriales>Bifidobacteriaceae | GCF_018555335.1 | Bifidobacterium | JS541_RS11260 | 52363..56112 | 1678 | MULTISPECIES:-BREX-system-P-loop-protein-BrxC-[Bifidobacterium] |
| WP_217054570.1 | Type1.BREX | <-SIGMA+OB-Fold+AAA-ATPase+HKD-DNase-BrxL<-PglZ<-HEPN+RES<-?<-?<-REase<-N6-METHYLASE<-DUF6079*<-BrxB-BrxF-iSTAND<-BrxA-DNA-binding\|\|?->CR-Kinase-><-CR-ATPase5+REase-p<-REase<-HTH_3 | 1229 | Bacteria>Terrabacteria_group>Actinomycetota>Actinomycetes>Bifidobacteriales>Bifidobacteriaceae | GCF_019041665.1 | Bifidobacterium_longum | KTF47_RS04030 | 15252..18941 | 216816 | BREX-system-P-loop-protein-BrxC-[Bifidobacterium-longum] |
| WP_270561184.1 | Type1.BREX | 35exo+HRDC->?->35exo+BRCT-><-HTH\|\|?-><-ACET\|\|?->BrxA-DNA-binding->BrxB-BrxF-iSTAND->DUF6079*->SIR2->HerA/FtsK->N6-METHYLASE-p->HEPN->?->Schlafen+HATPase+wHTH->PglZ->SIGMA+OB-Fold+AAA-ATPase+HKD-DNase-BrxL->ZnR-> | 1217 | Bacteria>Terrabacteria_group>Actinomycetota>Actinomycetes>Bifidobacteriales>Bifidobacteriaceae | GCF_027698005.1 | Bifidobacterium_dentium | PG823_RS09640 | 38589..42242 | 1689 | BREX-system-P-loop-protein-BrxC-[Bifidobacterium-dentium] |
| WP_029253807.1 | Type1.BREX | ABC-ATPase+SbcC-><-?<-HEPN<-ParB+HNH<-SIGMA+OB-Fold+AAA-ATPase+HKD-DNase-BrxL<-PglZ<-N6-METHYLASE<-DUF6079*<-BrxB-BrxF-iSTAND<-BrxA-DNA-binding<-AbiEi<-AbiEi<-REase | 1160 | Bacteria>Terrabacteria_group>Actinomycetota>Actinomycetes>Micrococcales>Cellulomonadaceae | GCF_000701465.1 | Paraoerskovia_marina | Q343_RS0111945 | 114651..118133 | 545619 | BREX-system-P-loop-protein-BrxC-[Paraoerskovia-marina] |
| WP_156999272.1 | Type1.BREX | <-HEPN<-PolB<-HEPN+SF1-SF2-Helicase+REase\|\|?->CBS+HEPN+HEPN->BrxA-DNA-binding->BrxB-BrxF-iSTAND->DUF6079*->N6-METHYLASE->PglZ->SIGMA+OB-Fold+AAA-ATPase+HKD-DNase-BrxL->sGTP+sGTP+GTPase-AIG->?-><-ParB+HNH\|\|SF1-SF2-Helicase+REase->REase->REase-> | 1173 | Bacteria>Terrabacteria_group>Actinomycetota>Actinomycetes>Micrococcales>Microbacteriaceae | GCF_024172065.1 | Agromyces_terreus | BJ978_RS07845 | 1788329..1791850 | 424795 | BREX-system-P-loop-protein-BrxC-[Agromyces-terreus] |
| WP_085152401.1 | Type1.BREX | <-ABhydrolase+ABhydrolase<-Preader7<-HEPN+SF1-SF2-Helicase+REase<-?<-SIGMA+OB-Fold+AAA-ATPase+HKD-DNase-BrxL<-PglZ<-N6-METHYLASE<-DUF6079*<-BrxB-BrxF-iSTAND<-BrxA-DNA-binding<-XIS-HTH<-HKD-DNase+SF1-SF2-Helicase+HTH+HTH+SRA\|\|35exo+SF1-SF2-Helicase->SF1-SF2-Helicase-> | 1156 | Bacteria>Terrabacteria_group>Actinomycetota>Actinomycetes>Mycobacteriales>Mycobacteriaceae | GCF_010729895.1 | Mycolicibacterium_confluentis | H7K00_RS24370 | 129281..132751 | 28047 | BREX-system-P-loop-protein-BrxC-[Mycolicibacterium-confluentis] |
| WP_094721084.1 | Type1.BREX | SF1-SF2-Helicase+REase->SF1-SF2-Helicase+HTH-><-N6-METHYLASE\|\|MORC+HISKIN->?-><-SIGMA+OB-Fold+AAA-ATPase+HKD-DNase-BrxL<-PglZ<-N6-METHYLASE<-DUF6079*<-BrxB-BrxF-iSTAND<-BrxA-DNA-binding<-HEPN+SF1-SF2-Helicase+REase\|\|HIRAN->?-><-TIR<-HD<-?<-MarR-HTH | 1174 | Bacteria>Terrabacteria_group>Actinomycetota>Actinomycetes>Mycobacteriales>Nocardiaceae | GCF_002259415.1 | Rhodococcus_sp._14-2470-1b | CH293_RS18620 | 24333..27857 | 2023149 | BREX-system-P-loop-protein-BrxC-[Rhodococcus-sp.-14-2470-1b] |
| MBW3578771.1 | Type1.BREX | <-HTH_3+HTH_3\|\|?->?->BrxA-DNA-binding->BrxB-BrxF-iSTAND->DUF6079*->N6-METHYLASE->PglZ->SIGMA+OB-Fold+AAA-ATPase+Lon-Protease-BrxL-><-HEPN\|\|?-><-Phage_int_SAM_1+Phage_integrase<-?<-?<-Pentapeptide+Pentapeptide+Pentapeptide+Pentapeptide | 1145 | Bacteria>Terrabacteria_group>Actinomycetota>Actinomycetes>unclassified_Actinomycetes | GCA_019347675.1 | Actinomycetia_bacterium | KY462_13720 | 21398..24835 | 1883427 | BREX-system-P-loop-protein-BrxC-[Actinomycetia-bacterium] |
| MBR3314870.1 | Type1.BREX | <-HEPN\|\|?-><-?<-?<-?<-SIGMA+OB-Fold+AAA-ATPase+Lon-Protease-BrxL<-PglZ<-DOC+HTH<-N6-METHYLASE<-DUF6079*<-BrxB-BrxF-iSTAND<-BrxA-DNA-binding\|\|MACRODOMAIN->ACET-><-?<-Schlafen+HATPase+wHTH | 1189 | Bacteria>Terrabacteria_group>Actinomycetota>Coriobacteriia>Coriobacteriales>Atopobiaceae | GCA_017510745.1 | Atopobiaceae_bacterium | IKG18_12110 | 33041..36610 | 2485926 | BREX-system-P-loop-protein-BrxC-[Atopobiaceae-bacterium] |
| HIZ18751.1 | Type1.BREX | <-Thr_synth_N+PALP\|\|NAD_binding_3+Homoserine_dh->AA_kinase+ACT_7->?->ABhydrolase->?->BrxA-DNA-binding->BrxB-BrxF-iSTAND->DUF6079*->N6-METHYLASE->HEPN-> | 1177 | Bacteria>Terrabacteria_group>Actinomycetota>Coriobacteriia>Coriobacteriales>Atopobiaceae | GCA_019116265.1 | Candidatus_Olsenella_stercoravium | IAA22_06560 | 20591..24124 | 2838713 | BREX-system-P-loop-protein-BrxC-[Candidatus-Olsenella-stercoravium] |
| WP_204537521.1 | Type1.BREX | Cysteine-Desulfurase->?-><-SIGMA+OB-Fold+AAA-ATPase+HKD-DNase-BrxL<-PglZ<-Schlafen+HATPase+wHTH<-N6-METHYLASE<-HEPN<-DUF6079*<-BrxB-BrxF-iSTAND<-BrxA-DNA-binding<-TraJ-RHH | 1171 | Bacteria>Terrabacteria_group>Actinomycetota>Coriobacteriia>Coriobacteriales>Coriobacteriaceae | - | Collinsella_tanakaei | H6A33_RS00145 | 33492..37007 | 626935 | BREX-system-P-loop-protein-BrxC-[Collinsella-tanakaei] |
| MBD9244247.1 | Type1.BREX | <-LRR_5<-PglZ<-SF1-SF2-Helicase<-HEPN+ParB<-?<-N6-METHYLASE-p<-HEPN<-N6-METHYLASE<-DUF6079*<-BrxB-BrxF-iSTAND<-BrxA-DNA-binding | 1191 | Bacteria>Terrabacteria_group>Actinomycetota>Coriobacteriia>Coriobacteriales>Coriobacteriaceae | GCA_014847015.1 | Coriobacteriaceae_bacterium | EGQ66_00120 | 16164..19739 | 2011094 | BREX-system-P-loop-protein-BrxC-[Coriobacteriaceae-bacterium] |
| MBN2945151.1 | Type1.BREX | <-Cysteine-Desulfurase\|\|BrxA-DNA-binding->BrxB-BrxF-iSTAND->DUF6079*->N6-METHYLASE->HEPN->N6-METHYLASE-p->PglZ->SIGMA+OB-Fold+AAA-ATPase+Lon-Protease-BrxL->?->Nif11-> | 1220 | Bacteria>Terrabacteria_group>Actinomycetota>Coriobacteriia>Coriobacteriales>Coriobacteriaceae | GCA_016939135.1 | Collinsella_sp. | JTJ24_07700 | 2990..6652 | 1965294 | BREX-system-P-loop-protein-BrxC-[Collinsella-sp.] |
| MBN2939679.1 | Type1.BREX | <-Nif11<-PglZ<-SF1-SF2-Helicase<-HEPN+ParB<-?<-N6-METHYLASE-p<-HEPN<-N6-METHYLASE<-DUF6079*<-BrxB-BrxF-iSTAND<-BrxA-DNA-binding\|\|PhdYeFM_antitox-> | 1196 | Bacteria>Terrabacteria_group>Actinomycetota>Coriobacteriia>Coriobacteriales>Coriobacteriaceae | GCA_016939175.1 | Collinsella_sp. | JTJ11_06140 | 12590..16180 | 1965294 | BREX-system-P-loop-protein-BrxC-[Collinsella-sp.] |
| MBS5295771.1 | Type1.BREX | His_Phos_1->His_Phos_1-><-?<-Peptidase_M20<-?<-Cysteine-Desulfurase\|\|BrxA-DNA-binding->BrxB-BrxF-iSTAND->DUF6079*->N6-METHYLASE->HEPN->N6-METHYLASE-p->PglZ->SIGMA+OB-Fold+AAA-ATPase+Lon-Protease-BrxL-> | 1237 | Bacteria>Terrabacteria_group>Actinomycetota>Coriobacteriia>Coriobacteriales>Coriobacteriaceae | GCA_018367145.1 | Collinsella_sp. | KHY16_04805 | 17070..20783 | 1965294 | BREX-system-P-loop-protein-BrxC-[Collinsella-sp.] |
| MBS6611921.1 | Type1.BREX | DNA_METHYLASE-><-PglZ<-?<-N6-METHYLASE-p\|\|RNase-><-35exo+wHTH+SF1-SF2-Helicase+HTH<-?<-HEPN<-N6-METHYLASE<-DUF6079*<-BrxB-BrxF-iSTAND<-BrxA-DNA-binding\|\|SIGMA+OB-Fold+AAA-ATPase+Lon-Protease-BrxL->?-><-?<-?<-ZnR\|\|?-><-STYKIN | 1187 | Bacteria>Terrabacteria_group>Actinomycetota>Coriobacteriia>Coriobacteriales>Coriobacteriaceae | GCA_018377935.1 | Collinsella_intestinalis | KH279_01485 | 57380..60943 | 147207 | BREX-system-P-loop-protein-BrxC-[Collinsella-intestinalis] |
| MBS6608517.1 | Type1.BREX | <-RNase<-Nif11<-?<-SIGMA+OB-Fold+AAA-ATPase+Lon-Protease-BrxL<-PglZ<-N6-METHYLASE-p<-HEPN<-?<-N6-METHYLASE<-DUF6079*<-BrxB-BrxF-iSTAND<-BrxA-DNA-binding\|\|PhdYeFM_antitox->Cysteine-Desulfurase->?-><-AbiEi+REase\|\|?-><-His_Phos_1<-His_Phos_1 | 1220 | Bacteria>Terrabacteria_group>Actinomycetota>Coriobacteriia>Coriobacteriales>Coriobacteriaceae | GCA_018377985.1 | Collinsella_sp. | KH291_01665 | 76425..80087 | 1965294 | BREX-system-P-loop-protein-BrxC-[Collinsella-sp.] |
| MBS6695654.1 | Type1.BREX | RNase-><-Nif11<-?<-SIGMA+OB-Fold+AAA-ATPase+Lon-Protease-BrxL<-PglZ<-N6-METHYLASE-p<-HEPN<-N6-METHYLASE<-DUF6079*<-BrxB-BrxF-iSTAND<-BrxA-DNA-binding\|\|PhdYeFM_antitox-> | 1220 | Bacteria>Terrabacteria_group>Actinomycetota>Coriobacteriia>Coriobacteriales>Coriobacteriaceae | GCA_018378775.1 | Collinsella_sp. | KH246_00985 | 77204..80866 | 1965294 | BREX-system-P-loop-protein-BrxC-[Collinsella-sp.] |
| MBS6858450.1 | Type1.BREX | His_Phos_1->His_Phos_1-><-?<-Peptidase_M20<-?<-Cysteine-Desulfurase\|\|BrxA-DNA-binding->BrxB-BrxF-iSTAND->DUF6079*->N6-METHYLASE->HEPN->N6-METHYLASE-p->PglZ->SIGMA+OB-Fold+AAA-ATPase+Lon-Protease-BrxL->?->Nif11->RNase-> | 1220 | Bacteria>Terrabacteria_group>Actinomycetota>Coriobacteriia>Coriobacteriales>Coriobacteriaceae | GCA_018380015.1 | Collinsella_sp. | KH187_04800 | 313170..316832 | 1965294 | BREX-system-P-loop-protein-BrxC-[Collinsella-sp.] |
| MBS7081308.1 | Type1.BREX | His_Phos_1->His_Phos_1-><-?<-Peptidase_M20<-?<-Cysteine-Desulfurase<-PhdYeFM_antitox\|\|BrxA-DNA-binding->BrxB-BrxF-iSTAND->DUF6079*->N6-METHYLASE->HEPN->N6-METHYLASE-p->PglZ->SIGMA+OB-Fold+AAA-ATPase+Lon-Protease-BrxL->?->Nif11-><-RNase | 1220 | Bacteria>Terrabacteria_group>Actinomycetota>Coriobacteriia>Coriobacteriales>Coriobacteriaceae | GCA_018382125.1 | Collinsella_sp. | KH077_07095 | 14975..18637 | 1965294 | BREX-system-P-loop-protein-BrxC-[Collinsella-sp.] |
| WP_006235452.1 | Type1.BREX | His_Phos_1->His_Phos_1-><-?<-Peptidase_M20<-?<-Cysteine-Desulfurase<-PhdYeFM_antitox\|\|BrxA-DNA-binding->BrxB-BrxF-iSTAND->DUF6079*->N6-METHYLASE->HEPN->N6-METHYLASE-p->PglZ->SIGMA+OB-Fold+AAA-ATPase+Lon-Protease-BrxL->?->Nif11-><-RNase | 1220 | Bacteria>Terrabacteria_group>Actinomycetota>Coriobacteriia>Coriobacteriales>Coriobacteriaceae | GCF_000169035.1 | Collinsella_aerofaciens | COLAER_RS06810 | 43874..47536 | 74426 | BREX-system-P-loop-protein-BrxC-[Collinsella-aerofaciens] |
| WP_087204139.1 | Type1.BREX | <-Glycos_transf<-?<-?<-?\|\|Peptidase-C39-><-?<-?\|\|BrxA-DNA-binding->BrxB-BrxF-iSTAND->DUF6079*->N6-METHYLASE->HEPN->?->SF1-SF2-Helicase->N6-METHYLASE-p->PglZ->SIGMA+OB-Fold+AAA-ATPase+Lon-Protease-BrxL-><-?<-Pkinase | 1199 | Bacteria>Terrabacteria_group>Actinomycetota>Coriobacteriia>Coriobacteriales>Coriobacteriaceae | GCF_002159765.1 | Collinsella_sp._An268 | B5F70_RS03250 | 39703..43302 | 1965612 | BREX-system-P-loop-protein-BrxC-[Collinsella-sp.-An268] |
| WP_099431905.1 | Type1.BREX | <-Nif11<-?<-SIGMA+OB-Fold+AAA-ATPase+Lon-Protease-BrxL<-PglZ<-N6-METHYLASE-p<-HEPN<-N6-METHYLASE<-DUF6079*<-BrxB-BrxF-iSTAND<-BrxA-DNA-binding\|\|PhdYeFM_antitox->Cysteine-Desulfurase->?-><-AbiEi+REase\|\|Peptidase_M20->?-><-His_Phos_1 | 1219 | Bacteria>Terrabacteria_group>Actinomycetota>Coriobacteriia>Coriobacteriales>Coriobacteriaceae | GCF_002736145.1 | Collinsella_aerofaciens | FX111_RS04100 | 928827..932486 | 74426 | BREX-system-P-loop-protein-BrxC-[Collinsella-aerofaciens] |
| WP_117639687.1 | Type1.BREX | <-Nif11<-?<-SIGMA+OB-Fold+AAA-ATPase+Lon-Protease-BrxL<-PglZ<-N6-METHYLASE-p<-HEPN<-N6-METHYLASE<-DUF6079*<-BrxB-BrxF-iSTAND<-BrxA-DNA-binding\|\|PhdYeFM_antitox->Cysteine-Desulfurase->?->Peptidase_M20->?-><-His_Phos_1<-His_Phos_1 | 1220 | Bacteria>Terrabacteria_group>Actinomycetota>Coriobacteriia>Coriobacteriales>Coriobacteriaceae | GCF_003436395.1 | Collinsella_sp._TM06-3 | DXD56_RS01895 | 14278..17940 | 2292342 | BREX-system-P-loop-protein-BrxC-[Collinsella-sp.-TM06-3] |
| WP_117735584.1 | Type1.BREX | His_Phos_1->His_Phos_1-><-?<-Peptidase_M20\|\|AbiEi+REase-><-?<-Cysteine-Desulfurase\|\|BrxA-DNA-binding->BrxB-BrxF-iSTAND->DUF6079*->N6-METHYLASE->HEPN->N6-METHYLASE-p->?->HEPN+ParB->SF1-SF2-Helicase->PglZ->Nif11-> | 1196 | Bacteria>Terrabacteria_group>Actinomycetota>Coriobacteriia>Coriobacteriales>Coriobacteriaceae | GCF_003436895.1 | Collinsella_sp._TF09-1AT | DXC90_RS08415 | 29935..33525 | 2292334 | BREX-system-P-loop-protein-BrxC-[Collinsella-sp.-TF09-1AT] |
| WP_117716812.1 | Type1.BREX | His_Phos_1-><-?<-Peptidase_M20\|\|AbiEi+REase-><-?<-Cysteine-Desulfurase<-PhdYeFM_antitox\|\|BrxA-DNA-binding->BrxB-BrxF-iSTAND->DUF6079*->N6-METHYLASE->?->HEPN->N6-METHYLASE-p->PglZ->SIGMA+OB-Fold+AAA-ATPase+Lon-Protease-BrxL->?->Nif11-> | 1220 | Bacteria>Terrabacteria_group>Actinomycetota>Coriobacteriia>Coriobacteriales>Coriobacteriaceae | GCF_003438115.1 | Collinsella_sp._OM08-14AT | DXC18_RS07380 | 14092..17754 | 2292329 | BREX-system-P-loop-protein-BrxC-[Collinsella-sp.-OM08-14AT] |
| WP_117746709.1 | Type1.BREX | RNase-><-Nif11<-?<-SIGMA+OB-Fold+AAA-ATPase+Lon-Protease-BrxL<-PglZ<-N6-METHYLASE-p<-HEPN<-N6-METHYLASE<-DUF6079*<-BrxB-BrxF-iSTAND<-BrxA-DNA-binding\|\|PhdYeFM_antitox->Cysteine-Desulfurase->?->Peptidase_M20->?-><-His_Phos_1<-His_Phos_1 | 1220 | Bacteria>Terrabacteria_group>Actinomycetota>Coriobacteriia>Coriobacteriales>Coriobacteriaceae | GCF_003438495.1 | Collinsella_sp._TF05-9AC | DXC52_RS07155 | 15430..19092 | 2292330 | BREX-system-P-loop-protein-BrxC-[Collinsella-sp.-TF05-9AC] |
| WP_117763119.1 | Type1.BREX | <-RNase<-Nif11<-?<-SIGMA+OB-Fold+AAA-ATPase+Lon-Protease-BrxL<-PglZ<-N6-METHYLASE-p<-HEPN<-N6-METHYLASE<-DUF6079*<-BrxB-BrxF-iSTAND<-BrxA-DNA-binding\|\|PhdYeFM_antitox->Cysteine-Desulfurase->?-><-AbiEi+REase\|\|Peptidase_M20->?-><-His_Phos_1 | 1220 | Bacteria>Terrabacteria_group>Actinomycetota>Coriobacteriia>Coriobacteriales>Coriobacteriaceae | GCF_003466125.1 | Collinsella_sp._TM10-22 | DXD94_RS02350 | 80369..84031 | 2292344 | BREX-system-P-loop-protein-BrxC-[Collinsella-sp.-TM10-22] |
| WP_117786732.1 | Type1.BREX | <-RNase<-Nif11<-?<-SIGMA+OB-Fold+AAA-ATPase+Lon-Protease-BrxL<-PglZ<-N6-METHYLASE-p<-HEPN<-N6-METHYLASE<-DUF6079*<-BrxB-BrxF-iSTAND<-BrxA-DNA-binding\|\|PhdYeFM_antitox->Cysteine-Desulfurase-> | 1224 | Bacteria>Terrabacteria_group>Actinomycetota>Coriobacteriia>Coriobacteriales>Coriobacteriaceae | GCF_003467795.1 | Collinsella_sp._AM36-4AA | DW850_RS07585 | 35525..39199 | 2292317 | BREX-system-P-loop-protein-BrxC-[Collinsella-sp.-AM36-4AA] |
| WP_118226697.1 | Type1.BREX | His_Phos_1-><-?<-Peptidase_M20\|\|AbiEi+REase-><-?<-Cysteine-Desulfurase<-PhdYeFM_antitox\|\|BrxA-DNA-binding->BrxB-BrxF-iSTAND->DUF6079*->N6-METHYLASE->HEPN->N6-METHYLASE-p->PglZ->SIGMA+OB-Fold+AAA-ATPase+Lon-Protease-BrxL->?->Nif11->RNase-> | 1224 | Bacteria>Terrabacteria_group>Actinomycetota>Coriobacteriia>Coriobacteriales>Coriobacteriaceae | GCF_003470945.1 | Collinsella_sp._AM23-17 | DW645_RS05360 | 15377..19051 | 2292030 | BREX-system-P-loop-protein-BrxC-[Collinsella-sp.-AM23-17] |
| WP_117583764.1 | Type1.BREX | His_Phos_1-><-?<-Peptidase_M20\|\|AbiEi+REase-><-?<-Cysteine-Desulfurase<-PhdYeFM_antitox\|\|BrxA-DNA-binding->BrxB-BrxF-iSTAND->DUF6079*->N6-METHYLASE->?->HEPN->N6-METHYLASE-p->PglZ->SIGMA+OB-Fold+AAA-ATPase+Lon-Protease-BrxL->?->Nif11-> | 1220 | Bacteria>Terrabacteria_group>Actinomycetota>Coriobacteriia>Coriobacteriales>Coriobacteriaceae | GCF_003472325.1 | unclassified_Collinsella | DW136_RS06205 | 18574..22236 | 2637548 | MULTISPECIES:-BREX-system-P-loop-protein-BrxC-[unclassified-Collinsella] |
| WP_118273143.1 | Type1.BREX | RNase-><-Nif11<-?<-SIGMA+OB-Fold+AAA-ATPase+Lon-Protease-BrxL<-PglZ<-N6-METHYLASE-p<-HEPN<-N6-METHYLASE<-DUF6079*<-BrxB-BrxF-iSTAND<-BrxA-DNA-binding\|\|PhdYeFM_antitox->Cysteine-Desulfurase->?-><-AbiEi+REase<-?\|\|Peptidase_M20-> | 1220 | Bacteria>Terrabacteria_group>Actinomycetota>Coriobacteriia>Coriobacteriales>Coriobacteriaceae | GCF_003472465.1 | unclassified_Collinsella | DW129_RS04200 | 27076..30738 | 2637548 | MULTISPECIES:-BREX-system-P-loop-protein-BrxC-[unclassified-Collinsella] |
| WP_117629077.1 | Type1.BREX | His_Phos_1-><-?<-Peptidase_M20\|\|AbiEi+REase-><-?<-Cysteine-Desulfurase<-PhdYeFM_antitox\|\|BrxA-DNA-binding->BrxB-BrxF-iSTAND->DUF6079*->N6-METHYLASE->?->HEPN->N6-METHYLASE-p->PglZ->SIGMA+OB-Fold+AAA-ATPase+Lon-Protease-BrxL->?->Nif11-> | 1220 | Bacteria>Terrabacteria_group>Actinomycetota>Coriobacteriia>Coriobacteriales>Coriobacteriaceae | GCF_003474585.1 | unclassified_Collinsella | DW037_RS01545 | 67055..70717 | 2637548 | MULTISPECIES:-BREX-system-P-loop-protein-BrxC-[unclassified-Collinsella] |
| WP_118384652.1 | Type1.BREX | His_Phos_1->His_Phos_1-><-?<-Peptidase_M20<-?<-Cysteine-Desulfurase\|\|BrxA-DNA-binding->BrxB-BrxF-iSTAND->DUF6079*->N6-METHYLASE->HEPN->N6-METHYLASE-p->PglZ->SIGMA+OB-Fold+AAA-ATPase+Lon-Protease-BrxL->?->Nif11-> | 1220 | Bacteria>Terrabacteria_group>Actinomycetota>Coriobacteriia>Coriobacteriales>Coriobacteriaceae | GCF_003475325.1 | Collinsella_sp._AF33-16 | DWZ52_RS00745 | 185923..189585 | 2292012 | BREX-system-P-loop-protein-BrxC-[Collinsella-sp.-AF33-16] |
| WP_124786083.1 | Type1.BREX | His_Phos_1->His_Phos_1-><-?<-Peptidase_M20\|\|AbiEi+REase-><-?<-Cysteine-Desulfurase\|\|BrxA-DNA-binding->BrxB-BrxF-iSTAND->DUF6079*->N6-METHYLASE->HEPN->N6-METHYLASE-p->N6-METHYLASE-p->PglZ->SIGMA+OB-Fold+AAA-ATPase+Lon-Protease-BrxL->?->Nif11-> | 1220 | Bacteria>Terrabacteria_group>Actinomycetota>Coriobacteriia>Coriobacteriales>Coriobacteriaceae | GCF_003856815.1 | Collinsella_aerofaciens | CV096_RS03705 | 854606..858268 | 74426 | BREX-system-P-loop-protein-BrxC-[Collinsella-aerofaciens] |
| WP_129423162.1 | Type1.BREX | <-SIGMA+OB-Fold+AAA-ATPase+Lon-Protease-BrxL<-PglZ<-N6-METHYLASE-p\|\|NTP_transf-><-HEPN<-N6-METHYLASE\|\|rve->AAA-ATPase-><-ACET<-DUF6079*<-BrxB-BrxF-iSTAND<-BrxA-DNA-binding\|\|?->PNPase->?-><-MODE-HTH+PBPII<-?\|\|?->Phos_pyr_kin-> | 1205 | Bacteria>Terrabacteria_group>Actinomycetota>Coriobacteriia>Coriobacteriales>Coriobacteriaceae | GCF_004135645.1 | Senegalimassilia_faecalis | ET524_RS02380 | 534170..537787 | 2509433 | BREX-system-P-loop-protein-BrxC-[Senegalimassilia-faecalis] |
| WP_154515691.1 | Type1.BREX | <-Nif11<-?<-SIGMA+OB-Fold+AAA-ATPase+Lon-Protease-BrxL<-PglZ<-N6-METHYLASE-p<-PSBP<-HEPN<-N6-METHYLASE<-DUF6079*<-BrxB-BrxF-iSTAND<-BrxA-DNA-binding<-?<-SF1-SF2-Helicase<-HEPN+ParB<-SIGMA-HTH+SIGMA-HTH\|\|SF1-SF2-Helicase->Cysteine-Desulfurase-> | 1219 | Bacteria>Terrabacteria_group>Actinomycetota>Coriobacteriia>Coriobacteriales>Coriobacteriaceae | GCF_009696325.1 | unclassified_Collinsella | FYJ40_RS02790 | 33433..37092 | 2637548 | MULTISPECIES:-BREX-system-P-loop-protein-BrxC-[unclassified-Collinsella] |
| WP_204559883.1 | Type1.BREX | Cysteine-Desulfurase->?-><-SIGMA+OB-Fold+AAA-ATPase+HKD-DNase-BrxL<-PglZ<-Schlafen+HATPase+wHTH<-REase<-N6-METHYLASE<-HEPN<-DUF6079*<-BrxB-BrxF-iSTAND<-BrxA-DNA-binding<-TraJ-RHH | 1171 | Bacteria>Terrabacteria_group>Actinomycetota>Coriobacteriia>Coriobacteriales>Coriobacteriaceae | GCF_016902615.1 | Collinsella_tanakaei | H6A21_RS01790 | 55502..59017 | 626935 | BREX-system-P-loop-protein-BrxC-[Collinsella-tanakaei] |
| WP_215694813.1 | Type1.BREX | His_Phos_1->His_Phos_1-><-?<-Peptidase_M20<-?<-Cysteine-Desulfurase<-PhdYeFM_antitox\|\|BrxA-DNA-binding->BrxB-BrxF-iSTAND->DUF6079*->N6-METHYLASE->HEPN->N6-METHYLASE-p->PglZ->SIGMA+OB-Fold+AAA-ATPase+Lon-Protease-BrxL->?->Nif11-> | 1220 | Bacteria>Terrabacteria_group>Actinomycetota>Coriobacteriia>Coriobacteriales>Coriobacteriaceae | GCF_018785265.1 | Collinsella_aerofaciens | GPK42_RS08810 | 321305..324967 | 74426 | BREX-system-P-loop-protein-BrxC-[Collinsella-aerofaciens] |
| WP_270471419.1 | Type1.BREX | RuvB-AAA+RuvB-HTH-><-SIGMA+OB-Fold+AAA-ATPase+Lon-Protease-BrxL<-PglZ<-N6-METHYLASE-p\|\|RNase->?-><-HEPN<-N6-METHYLASE<-DUF6079*<-BrxB-BrxF-iSTAND<-BrxA-DNA-binding\|\|ParA-Soj-PloopNTPase->?->Peptidase_C69->?->Pkinase+STYKIN-><-METHYLASE<-ACET | 1184 | Bacteria>Terrabacteria_group>Actinomycetota>Coriobacteriia>Coriobacteriales>Coriobacteriaceae | GCF_027683425.1 | Collinsella_intestinalis | PGX73_RS02685 | 38070..41624 | 147207 | BREX-system-P-loop-protein-BrxC-[Collinsella-intestinalis] |
| WP_271743294.1 | Type1.BREX | <-Peptidase_M20\|\|AbiEi+REase-><-?<-Cysteine-Desulfurase\|\|BrxA-DNA-binding->BrxB-BrxF-iSTAND->DUF6079*->N6-METHYLASE->HEPN->N6-METHYLASE-p->PglZ->SIGMA+OB-Fold+AAA-ATPase+Lon-Protease-BrxL->?->Nif11-> | 1220 | Bacteria>Terrabacteria_group>Actinomycetota>Coriobacteriia>Coriobacteriales>Coriobacteriaceae | GCF_028206795.1 | Collinsella_aerofaciens | PMX70_RS07965 | 7863..11525 | 74426 | BREX-system-P-loop-protein-BrxC-[Collinsella-aerofaciens] |
| WP_271729196.1 | Type1.BREX | <-RNase<-Nif11<-?<-SIGMA+OB-Fold+AAA-ATPase+Lon-Protease-BrxL<-PglZ<-N6-METHYLASE-p<-HEPN<-N6-METHYLASE<-DUF6079*<-BrxB-BrxF-iSTAND<-BrxA-DNA-binding\|\|PhdYeFM_antitox->Cysteine-Desulfurase->?-><-AbiEi+REase\|\|Peptidase_M20->?-><-His_Phos_1 | 1220 | Bacteria>Terrabacteria_group>Actinomycetota>Coriobacteriia>Coriobacteriales>Coriobacteriaceae | GCF_028206955.1 | Collinsella_aerofaciens | PMX73_RS03570 | 36978..40640 | 74426 | BREX-system-P-loop-protein-BrxC-[Collinsella-aerofaciens] |
| WP_196032768.1 | Type1.BREX | <-Nif11<-PglZ<-SF1-SF2-Helicase<-HEPN+ParB<-N6-METHYLASE-p<-HEPN<-N6-METHYLASE<-DUF6079*<-BrxB-BrxF-iSTAND<-BrxA-DNA-binding\|\|PhdYeFM_antitox->RelE-ParE->Cysteine-Desulfurase->?->Peptidase_M20->?-><-His_Phos_1 | 1196 | Bacteria>Terrabacteria_group>Actinomycetota>Coriobacteriia>Coriobacteriales>Coriobacteriaceae | GCF_028209425.1 | Collinsella_aerofaciens | PMX18_RS06205 | 35659..39249 | 74426 | BREX-system-P-loop-protein-BrxC-[Collinsella-aerofaciens] |
| WP_261981442.1 | Type1.BREX | His_Phos_1->His_Phos_1-><-?<-Peptidase_M20\|\|AbiEi+REase-><-?<-Cysteine-Desulfurase\|\|BrxA-DNA-binding->BrxB-BrxF-iSTAND->DUF6079*->N6-METHYLASE->HEPN->N6-METHYLASE-p->HEPN+ParB->SF1-SF2-Helicase->PglZ->Nif11-> | 1191 | Bacteria>Terrabacteria_group>Actinomycetota>Coriobacteriia>Coriobacteriales>Coriobacteriaceae | GCF_901212495.1 | Collinsella_aerofaciens | SFLMC1_RS10875 | 26184..29759 | 74426 | BREX-system-P-loop-protein-BrxC-[Collinsella-aerofaciens] |
| WP_118094641.1 | Type1.BREX | RNase-><-Nif11<-?<-SIGMA+OB-Fold+AAA-ATPase+Lon-Protease-BrxL<-PglZ<-N6-METHYLASE-p<-HEPN<-N6-METHYLASE<-DUF6079*<-BrxB-BrxF-iSTAND<-BrxA-DNA-binding\|\|PhdYeFM_antitox->Cysteine-Desulfurase->?-><-AbiEi+REase\|\|Peptidase_M20->?-><-His_Phos_1 | 1220 | Bacteria>Terrabacteria_group>Actinomycetota>Coriobacteriia>Coriobacteriales>Coriobacteriaceae | GCF_902364885.1 | Collinsella | FY305_RS06610 | 68794..72456 | 102106 | MULTISPECIES:-BREX-system-P-loop-protein-BrxC-[Collinsella] |
| WP_226814226.1 | Type1.BREX | His_Phos_1->His_Phos_1-><-?<-Peptidase_M20\|\|AbiEi+REase-><-?<-Cysteine-Desulfurase\|\|BrxA-DNA-binding->BrxB-BrxF-iSTAND->DUF6079*->N6-METHYLASE->HEPN->N6-METHYLASE-p->HEPN+ParB->SF1-SF2-Helicase->PglZ->Nif11-> | 1191 | Bacteria>Terrabacteria_group>Actinomycetota>Coriobacteriia>Coriobacteriales>Coriobacteriaceae | GCF_902501535.1 | Collinsella_aerofaciens | GDK10_RS03455 | 524536..528111 | 74426 | BREX-system-P-loop-protein-BrxC-[Collinsella-aerofaciens] |
| WP_136846141.1 | Type1.BREX | Nif11->ParD_antitoxin-><-?\|\|ParD_antitoxin->35exo+wHTH+SF1-SF2-Helicase+HTH->REase+SF1-SF2-Helicase->BrxA-DNA-binding->BrxB-BrxF-iSTAND->DUF6079*->N6-METHYLASE->?->HEPN-><-?<-?\|\|ParD_antitoxin->N6-METHYLASE-p->PglZ->SIGMA+OB-Fold+AAA-ATPase+Lon-Protease-BrxL-> | 1192 | Bacteria>Terrabacteria_group>Actinomycetota>Coriobacteriia>Coriobacteriales>Coriobacteriaceae | GCF_948466405.1 | Parvibacter_caecicola | E5982_RS08750 | 41748..45326 | 747645 | BREX-system-P-loop-protein-BrxC-[Parvibacter-caecicola] |
| HIT45392.1 | Type1.BREX | <-N6-METHYLASE-p<-ParD_antitoxin<-HEPN<-N6-METHYLASE<-DUF6079*<-BrxB-BrxF-iSTAND<-BrxA-DNA-binding\|\|HTH_3-> | 1225 | Bacteria>Terrabacteria_group>Actinomycetota>Coriobacteriia>Coriobacteriia_incertae_sedis>Candidatus_Aphodovivens | GCA_018715935.1 | Candidatus_Aphodovivens_excrementavium | IAC28_04775 | 5578..9255 | 2840682 | BREX-system-P-loop-protein-BrxC-[Candidatus-Aphodovivens-excrementavium] |
| CDB32945.1 | Type1.BREX | <-PglZ-p<-LRR_5<-?<-HEPN<-HEPN<-N6-METHYLASE-p<-N6-METHYLASE-p<-ACET<-DUF6079-p<-DUF6079-p*<-BrxB-BrxF-iSTAND<-BrxA-DNA-binding\|\|?-><-SAM-METHYLASE<-DADA-Ligase | 819 | Bacteria>Terrabacteria_group>Actinomycetota>Coriobacteriia>Eggerthellales>Eggerthellaceae | GCA_000436075.1 | Eggerthella_sp._CAG:209 | BN534_00740 | 18377..20836 | 1262875 | uncharacterized-protein-BN534_00740-[Eggerthella-sp.-CAG:209] |
| MCI9261625.1 | Type1.BREX | BrxA-DNA-binding->BrxB-BrxF-iSTAND->DUF6079*->N6-METHYLASE->HEPN-><-RNase\|\|N6-METHYLASE-p->?->PglZ->SIGMA+OB-Fold+AAA-ATPase+Lon-Protease-BrxL->LRR_5-> | 1223 | Bacteria>Terrabacteria_group>Actinomycetota>Coriobacteriia>Eggerthellales>Eggerthellaceae | GCA_022798975.1 | Eggerthellaceae_bacterium | HFJ71_04015 | 1625..5296 | 1972561 | BREX-system-P-loop-protein-BrxC-[Eggerthellaceae-bacterium] |
| WP_160709085.1 | Type1.BREX | BrxA-DNA-binding->BrxB-BrxF-iSTAND->DUF6079*->HEPN-><-?\|\|SIGMA-HTH+SIGMA-HTH->N6-METHYLASE-p->?->?->PglZ->SIGMA+OB-Fold+AAA-ATPase+Lon-Protease-BrxL-> | 1222 | Bacteria>Terrabacteria_group>Actinomycetota>Coriobacteriia>Eggerthellales>Eggerthellaceae | GCF_009911695.1 | Enterorhabdus_sp._P55 | D1643_RS04550 | 2737..6405 | 2304571 | BREX-system-P-loop-protein-BrxC-[Enterorhabdus-sp.-P55] |
| WP_165048948.1 | Type1.BREX | <-URI-DNase+UVR+UvrC_HhH_N+HHH<-Nif11<-SIGMA+OB-Fold+AAA-ATPase+Lon-Protease-BrxL<-PglZ<-N6-METHYLASE-p\|\|?->RNase-><-HEPN<-N6-METHYLASE<-DUF6079*<-BrxB-BrxF-iSTAND<-BrxA-DNA-binding<-?<-?<-?\|\|MODE-HTH+PBPII-><-HD\|\|BrnT_toxin->CopG_antitoxin-> | 1188 | Bacteria>Terrabacteria_group>Actinomycetota>Coriobacteriia>Eggerthellales>Eggerthellaceae | GCF_011038975.1 | unclassified_Adlercreutzia | G7Y50_RS02135 | 57415..60981 | 2636013 | MULTISPECIES:-BREX-system-P-loop-protein-BrxC-[unclassified-Adlercreutzia] |
| WP_195347157.1 | Type1.BREX | <-HTH_3\|\|HTH->Arm-DNA-bind_4+Phage_int_SAM_3+Phage_integrase->TraJ-RHH-><-HEPN<-SIGMA+OB-Fold+AAA-ATPase+Lon-Protease-BrxL<-PglZ<-Schlafen+HATPase+wHTH<-N6-METHYLASE<-DUF6079*<-BrxB-BrxF-iSTAND<-BrxA-DNA-binding<-TraJ-RHH\|\|?->?->ParA-Soj-PloopNTPase->ParB->TraJ-RHH->TraI-VirD2-RCRE-> | 1201 | Bacteria>Terrabacteria_group>Actinomycetota>Coriobacteriia>Eggerthellales>Eggerthellaceae | GCF_028209285.1 | Eggerthella_lenta | PMV52_RS07270 | 251002..254607 | 84112 | unnamed-protein-product-[Eggerthella-lenta] |
| WP_251231669.1 | Type1.BREX | <-SIGMA+OB-Fold+AAA-ATPase+Lon-Protease-BrxL<-PglZ<-?<-?<-N6-METHYLASE-p<-SIGMA-HTH+SIGMA-HTH<-HEPN<-REase<-N6-METHYLASE<-DUF6079*<-BrxB-BrxF-iSTAND<-BrxA-DNA-binding<-Nif11<-CR-ATPase5+REase<-?<-Pkinase<-?<-PspA | 1213 | Bacteria>Terrabacteria_group>Actinomycetota>Coriobacteriia>Eggerthellales>Eggerthellaceae | GCF_943193145.1 | Adlercreutzia_aquisgranensis | NFR41_RS09570 | 19774..23415 | 2941323 | BREX-system-P-loop-protein-BrxC-[Adlercreutzia-aquisgranensis] |
| WP_227070741.1 | Type1.BREX | <-TraI-VirD2-RCRE<-TraJ-RHH<-ParB<-ParA-Soj-PloopNTPase<-?<-?\|\|TraJ-RHH->BrxA-DNA-binding->BrxB-BrxF-iSTAND->DUF6079*->N6-METHYLASE->Schlafen+HATPase+wHTH->SIGMA+OB-Fold+AAA-ATPase+Lon-Protease-BrxL->HEPN-><-TraJ-RHH<-Arm-DNA-bind_4+Phage_int_SAM_3+Phage_integrase<-HTH\|\|HTH_3-> | 1199 | Bacteria>Terrabacteria_group>Actinomycetota>Coriobacteriia>Eggerthellales | GCF_015551405.1 | Eggerthellaceae | I2D50_RS06370 | 44195..47794 | 1643826 | MULTISPECIES:-BREX-system-P-loop-protein-BrxC-[Eggerthellaceae] |
| MBA3474041.1 | Type1.BREX | <-HTH+REase<-PemK_toxin<-?<-METHYLASE<-?<-HEPN<-HEPN<-DUF6079*<-REase<-?<-?<-N6-METHYLASE-p | 1219 | Bacteria>Terrabacteria_group>Actinomycetota>Rubrobacteria>Rubrobacterales>Rubrobacteraceae | GCA_013813465.1 | Rubrobacter_sp. | HOT57_12585 | 3865..7524 | 2036010 | BREX-system-P-loop-protein-BrxC-[Rubrobacter-sp.] |
| MCL6447596.1 | Type1.BREX | <-AAA+<-DEOR-HTH<-?<-HTH_3\|\|?->BrxA-DNA-binding->BrxB-BrxF-iSTAND->DUF6079*->HEPN->N6-METHYLASE->PglZ->SIGMA+OB-Fold+AAA-ATPase+Lon-Protease-BrxL-><-ARG<-?<-?<-?<-35exo+T7DNApol | 1222 | Bacteria>Terrabacteria_group>Armatimonadota>unclassified_Armatimonadota | GCA_023511535.1 | Armatimonadota_bacterium | K6U04_05490 | 13164..16832 | 2033014 | BREX-system-P-loop-protein-BrxC-[Armatimonadota-bacterium] |
| MCI1589808.1 | Type1.BREX | ParD_antitoxin-><-REase<-SIGMA+OB-Fold+AAA-ATPase+Lon-Protease-BrxL<-PglZ<-?<-?<-HEPN<-N6-METHYLASE<-DUF6079*<-BrxB-BrxF-iSTAND<-BrxA-DNA-binding<-?<-?<-HTH_31<-?<-TRAM+METHYLASE\|\|MerR-HTH-> | 1190 | Bacteria>Terrabacteria_group>Bacillota>Bacilli>Bacillales>Bacillaceae | GCA_022646685.1 | Heyndrickxia_oleronia | LKH79_04555 | 33937..37509 | 38875 | BREX-system-P-loop-protein-BrxC-[Heyndrickxia-oleronia] |
| WP_031537547.1 | Type1.BREX | REase+DUF4268-><-?\|\|ZnR+RHH_7->XIS-HTH->MerR-HTH->?->?->BrxA-DNA-binding->BrxB-BrxF-iSTAND->DUF6079*->N6-METHYLASE->PglZ->SIGMA+OB-Fold+AAA-ATPase+Lon-Protease-BrxL->HEPN->?->MarR-HTH-> | 1187 | Bacteria>Terrabacteria_group>Bacillota>Bacilli>Bacillales>Bacillaceae | GCF_000701605.1 | Bacillus_sp._MB2021 | T349_RS0112475 | 16440..20003 | 1408303 | BREX-system-P-loop-protein-BrxC-[Bacillus-sp.-MB2021] |
| WP_048824801.1 | Type1.BREX | Preader17->?->HKD-DNase+SF1-SF2-Helicase+HTH+HTH+SRA->SIGMA-HTH+SIGMA-HTH->35exo+wHTH+SF1-SF2-Helicase+HTH->?->BrxA-DNA-binding->BrxB-BrxF-iSTAND->DUF6079*->N6-METHYLASE->HEPN->PglZ->SIGMA+OB-Fold+AAA-ATPase+Lon-Protease-BrxL->AbiGi->JetA-> | 1190 | Bacteria>Terrabacteria_group>Bacillota>Bacilli>Bacillales>Bacillaceae | GCF_000821085.1 | Bacillus_sp._B-jedd | BN1002_01993 | 1962591..1966163 | 1476857 | BREX-system-P-loop-protein-BrxC-[Bacillus-sp.-B-jedd] |
| WP_062353349.1 | Type1.BREX | HTH+HTH+SF1-SF2-Helicase->HTH_31->?->?->BrxA-DNA-binding->BrxB-BrxF-iSTAND->DUF6079*->N6-METHYLASE->HEPN->PglZ->SIGMA+OB-Fold+AAA-ATPase+Lon-Protease-BrxL->REase->REase-><-HTH_Tnp_1+HTH_21+rve+rve_3\|\|?->Schlafen+HATPase+wHTH-> | 1190 | Bacteria>Terrabacteria_group>Bacillota>Bacilli>Bacillales>Bacillaceae | GCF_001375515.1 | Bacillus_kwashiorkori | BN1242_RS07000 | 23447..27019 | 1522318 | BREX-system-P-loop-protein-BrxC-[Bacillus-kwashiorkori] |
| WP_108713788.1 | Type1.BREX | TPR-><-?<-Phage_CRI+Phage_X<-ParB+HNH<-SIGMA+OB-Fold+AAA-ATPase+Lon-Protease-BrxL<-PglZ<-HEPN<-AbiGi+seqA+RAMA<-N6-METHYLASE<-DUF6079*<-BrxB-BrxF-iSTAND<-BrxA-DNA-binding<-?\|\|XIS-HTH->Arm-DNA-bind_4+Phage_int_SAM_3+Phage_integrase-><-GDResolvase+HIN-HTH | 1191 | Bacteria>Terrabacteria_group>Bacillota>Bacilli>Bacillales>Bacillaceae | GCF_003073375.1 | unclassified_Lysinibacillus | DCE79_RS14920 | 3050129..3053704 | 2636778 | MULTISPECIES:-BREX-system-P-loop-protein-BrxC-[unclassified-Lysinibacillus] |
| WP_114371647.1 | Type1.BREX | <-Polbeta<-?<-RAMA<-?<-?<-HEPN<-PglZ<-N6-METHYLASE<-DUF6079*<-BrxB-BrxF-iSTAND<-BrxA-DNA-binding<-SF1-SF2-Helicase+wHTH+Small-B-barrel+DUF1998<-BrxD-AAA<-?<-HTH_36+HTH_Crp_2 | 1205 | Bacteria>Terrabacteria_group>Bacillota>Bacilli>Bacillales>Bacillaceae | GCF_003336365.1 | Salicibibacter_kimchii | DT065_05890 | 1163447..1167064 | 2099786 | BREX-system-P-loop-protein-BrxC-[Salicibibacter-kimchii] |
| WP_126293910.1 | Type1.BREX | TRAM+METHYLASE-><-?\|\|HTH_31->?->?->BrxA-DNA-binding->BrxB-BrxF-iSTAND->DUF6079*->N6-METHYLASE->HEPN->PglZ->SIGMA+OB-Fold+AAA-ATPase+Lon-Protease-BrxL->REase-><-ParD_antitoxin<-?\|\|?->HTH_3-> | 1190 | Bacteria>Terrabacteria_group>Bacillota>Bacilli>Bacillales>Bacillaceae | GCF_003966145.1 | Lysinibacillus_telephonicus | EKG35_RS07955 | 47066..50638 | 1714840 | BREX-system-P-loop-protein-BrxC-[Lysinibacillus-telephonicus] |
| WP_144538762.1 | Type1.BREX | <-Phage_CRI+Phage_X<-ParB+HNH<-SIGMA+OB-Fold+AAA-ATPase+Lon-Protease-BrxL<-PglZ<-HEPN<-HEPN<-?<-?<-N6-METHYLASE<-DUF6079*<-BrxB-BrxF-iSTAND<-BrxA-DNA-binding<-?<-?<-HEPN<-?<-?<-RhoN+RhoN+RNase+S1<-ABhydrolase | 1191 | Bacteria>Terrabacteria_group>Bacillota>Bacilli>Bacillales>Bacillaceae | GCF_007677425.1 | Bacillus_sp._FJAT-22090 | FS503_RS11600 | 136183..139758 | 1581038 | BREX-system-P-loop-protein-BrxC-[Bacillus-sp.-FJAT-22090] |
| WP_216486703.1 | Type1.BREX | <-SIGMA+OB-Fold+AAA-ATPase+Lon-Protease-BrxL<-PglZ<-?<-SIR2<-?<-SbcC<-?<-HEPN<-N6-METHYLASE<-DUF6079*<-BrxB-BrxF-iSTAND<-BrxA-DNA-binding<-ParB+HNH<-?<-Peptidase_S8\|\|?->PhdYeFM_antitox->RelE-ParE-> | 1194 | Bacteria>Terrabacteria_group>Bacillota>Bacilli>Bacillales>Bacillaceae | GCF_018918075.1 | Virgibacillus_sp._MSJ-26 | KQI49_RS03110 | 240564..244148 | 2841522 | BREX-system-P-loop-protein-BrxC-[Virgibacillus-sp.-MSJ-26] |
| WP_220158113.1 | Type1.BREX | <-SIGMA+OB-Fold+AAA-ATPase+Lon-Protease-BrxL<-PglZ<-RVT<-HEPN<-N6-METHYLASE<-DUF6079*<-BrxB-BrxF-iSTAND<-BrxA-DNA-binding | 1190 | Bacteria>Terrabacteria_group>Bacillota>Bacilli>Bacillales>Bacillaceae | GCF_019459225.1 | Bacillus_sp._IITD106 | KOH71_20660 | 11529..15101 | 2864214 | BREX-system-P-loop-protein-BrxC-[Bacillus-sp.-IITD106] |
| WP_259548277.1 | Type1.BREX | <-MarR-HTH<-?<-?<-HEPN<-HEPN<-PIN<-SIGMA+OB-Fold+AAA-ATPase+Lon-Protease-BrxL<-PglZ<-N6-METHYLASE<-DUF6079*<-BrxB-BrxF-iSTAND<-BrxA-DNA-binding<-?<-?<-MerR-HTH<-XIS-HTH<-ZnR+RHH_7\|\|?->UDG-> | 1187 | Bacteria>Terrabacteria_group>Bacillota>Bacilli>Bacillales>Bacillaceae | GCF_024971915.1 | Heyndrickxia_oleronia | KXF32_RS03880 | 741080..744643 | 38875 | BREX-system-P-loop-protein-BrxC-[Heyndrickxia-oleronia] |
| WP_131470342.1 | Type1.BREX | <-REase+DUF5655<-SF1-SF2-Helicase\|\|SF1-SF2-Helicase+REase-><-?\|\|HTH_3-><-SIGMA+OB-Fold+AAA-ATPase+Lon-Protease-BrxL<-PglZ<-HEPN<-N6-METHYLASE<-DUF6079*<-BrxB-BrxF-iSTAND<-BrxA-DNA-binding<-?<-AAA-ATPase<-rve | 1188 | Bacteria>Terrabacteria_group>Bacillota>Bacilli>Bacillales>Bacillales_incertae_sedis | GCF_004337105.1 | Exiguobacterium_sp._SH3S1 | EVJ26_RS09655 | 119198..122764 | 2510955 | BREX-system-P-loop-protein-BrxC-[Exiguobacterium-sp.-SH3S1] |
| WP_214711985.1 | Type1.BREX | wHTH+HNH->?->?->BrxA-DNA-binding->BrxB-BrxF-iSTAND->DUF6079*->N6-METHYLASE->HEPN->SIGMA+OB-Fold+AAA-ATPase+Lon-Protease-BrxL-><-HTH_3\|\|?-><-SF1-SF2-Helicase+REase\|\|SF1-SF2-Helicase->REase+DUF5655->PIWI-> | 1188 | Bacteria>Terrabacteria_group>Bacillota>Bacilli>Bacillales>Bacillales_incertae_sedis | GCF_018617985.1 | Exiguobacterium_sp._s55 | KNP09_RS10355 | 38338..41904 | 2751245 | BREX-system-P-loop-protein-BrxC-[Exiguobacterium-sp.-s55] |
| WP_148456431.1 | Type1.BREX | Resolvase+Recombinase-><-?\|\|REase->wHTH+WYL+WCX-><-35exo\|\|HEPN+HTH->BrxA-DNA-binding->BrxB-BrxF-iSTAND->DUF6079*->N6-METHYLASE->SIR2->PglZ->SIGMA+OB-Fold+AAA-ATPase+Lon-Protease-BrxL->SLATT->UDG-conflict->JAB->?->REase-> | 1206 | Bacteria>Terrabacteria_group>Bacillota>Bacilli>Bacillales>Paenibacillaceae | GCF_008084145.1 | Paenibacillus_faecis | FRY98_RS22745 | 231357..234977 | 862114 | BREX-system-P-loop-protein-BrxC-[Paenibacillus-faecis] |
| WP_212938911.1 | Type1.BREX | <-pPIWI<-pPIWI+pPIWI<-?<-RVT<-?<-SIGMA+OB-Fold+AAA-ATPase+Lon-Protease-BrxL<-PglZ<-SIR2<-N6-METHYLASE<-DUF6079*<-BrxB-BrxF-iSTAND<-BrxA-DNA-binding<-HEPN+HTH\|\|35exo-><-SF1-SF2-Helicase+REase<-SF1-SF2-Helicase+REase<-wHTH+WYL+WCX<-?<-REase | 1206 | Bacteria>Terrabacteria_group>Bacillota>Bacilli>Bacillales>Paenibacillaceae | - | Paenibacillus_antibioticophila | KI368_RS07085 | 201985..205605 | 1274374 | BREX-system-P-loop-protein-BrxC-[Paenibacillus-antibioticophila] |
| PIC94624.1 | Type1.BREX | Phage_CRI+Phage_X->?->BrxA-DNA-binding->BrxB-BrxF-iSTAND->DUF6079*->N6-METHYLASE->HEPN-> | 1186 | Bacteria>Terrabacteria_group>Bacillota>Bacilli>Bacillales>Planococcaceae | GCA_002743245.1 | Sporosarcina_sp._P26b | CSV69_15785 | 3496..7056 | 2048253 | hypothetical-protein-CSV69_15785-[Sporosarcina-sp.-P26b] |
| WP_233192413.1 | Type1.BREX | Phage_CRI+Phage_X->?->BrxA-DNA-binding->BrxB-BrxF-iSTAND->DUF6079*->N6-METHYLASE->HEPN-> | 1188 | Bacteria>Terrabacteria_group>Bacillota>Bacilli>Bacillales>Planococcaceae | GCA_002743245.1 | Sporosarcina_sp._P26b | CSV69_RS15805 | 3490..7056 | 2048253 | BREX-system-P-loop-protein-BrxC-[Sporosarcina-sp.-P26b] |
| WP_010024517.1 | Type1.BREX | <-HEPN<-?<-N6-METHYLASE<-DUF6079*<-BrxB-BrxF-iSTAND<-BrxA-DNA-binding<-HEPN+PAS+DUF4011+SF1-SF2-Helicase+REase<-nSTAND1+HNH | 1189 | Bacteria>Terrabacteria_group>Bacillota>Bacilli>Bacillales>Sporolactobacillaceae | GCF_000222445.2 | Sporolactobacillus_inulinus | SINU_RS07430 | 4602..8171 | 2078 | BREX-system-P-loop-protein-BrxC-[Sporolactobacillus-inulinus] |
| HJF20832.1 | Type1.BREX | <-TetR-HTH<-PglZ<-HEPN-p<-nSTAND3<-N6-METHYLASE<-DUF6079*<-BrxB-BrxF-iSTAND<-BrxA-DNA-binding<-?\|\|<-?<-RFC-AAA | 1239 | Bacteria>Terrabacteria_group>Bacillota>Bacilli>Bacillales>Staphylococcaceae | GCA_020742605.1 | Mammaliicoccus_lentus | K8U93_01665 | 12906..16625 | 42858 | BREX-system-P-loop-protein-BrxC-[Mammaliicoccus-lentus] |

| Protein-ID | System | Gene-Neighbourhoods | Length | Taxonomy | Assembly-ID | Strain | Locus-Tag | Genomic-coordinates | TaxID | Fasta-defline |
|---|---|---|---|---|---|---|---|---|---|---|
| MBK0347250.1 | Type1.BREX | HTH_36->?->HTH_26-> BrxA-DNA-binding -> BrxB-BrxF-iSTAND -> DUF6079 *->Schlafen+HATPase+wHTH-> N6-METHYLASE -> HEPN -> PglZ -> wHTH+WYL+WCX ->?->ParA-Soj-PloopNTPase-><-DEOR-HTH | 1198 | Bacteria>Terrabacteria_group>Bacillota>Bacilli>Lactobacillales>Aerococcaceae | GCA_016522115.1 | Aerococcaceae_bacterium_zg-1578 | JDW15_01195 | 55878..59474 | 2796350 | BREX-system-P-loop-protein-BrxC-[Aerococcaceae-bacterium-zg-1578] |
| WP_274731953.1 | Type1.BREX | LACI-HTH+PBPI->?-> HEPN ->HTH_26-> BrxA-DNA-binding -> BrxB-BrxF-iSTAND -> DUF6079 *-> N6-METHYLASE -p-> | 1193 | Bacteria>Terrabacteria_group>Bacillota>Bacilli>Lactobacillales>Carnobacteriaceae | GCF_028863945.1 | Jeotgalibaca_sp._YN-L-12 | PVE40_RS10850 | 8874..12455 | 3028623 | BREX-system-P-loop-protein-BrxC-[Jeotgalibaca-sp.-YN-L-12] |
| TXJ87453.1 | Type1.BREX | HDRnase->CHC2+TOPRIM-DNAG->SIGMA-HTH+SIGMA-HTH+SIGMA-HTH->SIGMA-HTH+SIGMA-HTH->\|\|HAD+HAD->\|\|?-> BrxA-DNA-binding -> BrxB-BrxF-iSTAND ->\|\| DUF6079 -p*-> N6-METHYLASE -> SIGMA+OB-Fold+AAA-ATPase+Lon-Protease-BrxL -> HEPN -> | 917 | Bacteria>Terrabacteria_group>Bacillota>Bacilli>Lactobacillales>Lactobacillaceae | GCA_008016355.1 | Lactobacillus_delbrueckii | FU655_07470 | 24596..27349 | 1584 | BREX-system-P-loop-protein-BrxC-[Lactobacillus-delbrueckii] |
| MBT8803912.1 | Type1.BREX | ParB+HNH-> BrxA-DNA-binding -> BrxB-BrxF-iSTAND -> DUF6079 *-> N6-METHYLASE -> PglZ -> SIGMA+OB-Fold+AAA-ATPase+Lon-Protease-BrxL -> HEPN ->?->O6MGT-HTH-><-?<-NUDIX\|\|METHYLASE-> | 1178 | Bacteria>Terrabacteria_group>Bacillota>Bacilli>Lactobacillales>Lactobacillaceae | GCA_018719835.1 | Lactobacillus_delbrueckii_subsp._bulgaricus | BTJ20_08000 | 3864..7400 | 1585 | ATPase-[Lactobacillus-delbrueckii-subsp.-bulgaricus] |
| MBT8901771.1 | Type1.BREX | <-METHYLASE\|\|NUDIX->?->\|\|<-AbiGi<- HEPN <- SIGMA+OB-Fold+AAA-ATPase+Lon-Protease-BrxL <- PglZ <- N6-METHYLASE <- DUF6079 *<- BrxB-BrxF-iSTAND <- BrxA-DNA-binding <-ParB+HNH<-?\|\|<-HAD+HAD<-SIGMA-HTH+SIGMA-HTH<-SIGMA-HTH+SIGMA-HTH | 1178 | Bacteria>Terrabacteria_group>Bacillota>Bacilli>Lactobacillales>Lactobacillaceae | GCA_018720695.1 | Lactobacillus_delbrueckii_subsp._bulgaricus | BTI52_06840 | 21471..25007 | 1585 | ATPase-[Lactobacillus-delbrueckii-subsp.-bulgaricus] |
| MBT8914213.1 | Type1.BREX | ParB+HNH-> BrxA-DNA-binding -> BrxB-BrxF-iSTAND -> DUF6079 *-> N6-METHYLASE -> PglZ -> SIGMA+OB-Fold+AAA-ATPase+Lon-Protease-BrxL -> HEPN -> | 1178 | Bacteria>Terrabacteria_group>Bacillota>Bacilli>Lactobacillales>Lactobacillaceae | GCA_018720815.1 | Lactobacillus_delbrueckii_subsp._bulgaricus | BTI61_07605 | 4087..7623 | 1585 | ATPase-[Lactobacillus-delbrueckii-subsp.-bulgaricus] |
| MCT7861392.1 | Type1.BREX | <- PglZ <- HEPN <-?<-?<- N6-METHYLASE <-?<- DUF6079 *<- BrxB-BrxF-iSTAND <- BrxA-DNA-binding | 1165 | Bacteria>Terrabacteria_group>Bacillota>Bacilli>Lactobacillales>Lactobacillaceae | GCA_025347225.1 | Lactobacillus_crispatus | N4Q24_00945 | 8507..12000 | 47770 | BREX-system-P-loop-protein-BrxC-[Lactobacillus-crispatus] |
| WP_025283578.1 | Type1.BREX | ASCH->Glyoxalase->Glyoxalase->ParB+HNH-> BrxA-DNA-binding -> BrxB-BrxF-iSTAND -> DUF6079 *-> N6-METHYLASE -> HEPN -> HEPN -> PglZ -><-?\|\|DDE_Tnp_1-><-Calcineurin<-Peptidase_M13_N+Peptidase_M13 | 1243 | Bacteria>Terrabacteria_group>Bacillota>Bacilli>Lactobacillales>Lactobacillaceae | GCF_000525715.1 | Lactobacillus_helveticus | LBH_0939 | 971853..975584 | 1587 | BREX-system-P-loop-protein-BrxC-[Lactobacillus-helveticus] |
| WP_060598044.1 | Type1.BREX | <-METHYLASE\|\|NUDIX->?-><-AraC-HTH+O6MGT-HTH<-?<- HEPN <- SIGMA+OB-Fold+AAA-ATPase+Lon-Protease-BrxL <- PglZ <- N6-METHYLASE <- DUF6079 *<- BrxB-BrxF-iSTAND <- BrxA-DNA-binding <-ParB+HNH<-?<-?\|\|<-HAD+HAD\|\|<-SIGMA-HTH+SIGMA-HTH<-SIGMA-HTH+SIGMA-HTH+SIGMA-HTH<-SIGMA-HTH+SIGMA-HTH+SIGMA-HTH | 1178 | Bacteria>Terrabacteria_group>Bacillota>Bacilli>Lactobacillales>Lactobacillaceae | GCF_001469775.1 | Lactobacillus_delbrueckii | AT236_RS05345 | 1068532..1072068 | 1584 | BREX-system-P-loop-protein-BrxC-[Lactobacillus-delbrueckii] |
| WP_086355469.1 | Type1.BREX | LZ_Tnp_IS66+zf-IS66+DDE_Tnp_IS66+DDE_Tnp_IS66_C-><-AbiGi<-?<- HEPN <- SIGMA+OB-Fold+AAA-ATPase+Lon-Protease-BrxL <- PglZ <- N6-METHYLASE <- DUF6079 *<- BrxB-BrxF-iSTAND <- BrxA-DNA-binding <-ParB+HNH\|\|<-ParB+HNH<-?<-?<-REase+N6_METHYLASE<-ABhydrolase+ABhydrolase<-ABhydrolase | 1183 | Bacteria>Terrabacteria_group>Bacillota>Bacilli>Lactobacillales>Lactobacillaceae | GCF_002142575.1 | Lactobacillus_delbrueckii | B9N98_RS00590 | 112113..115664 | 1584 | BREX-system-P-loop-protein-BrxC-[Lactobacillus-delbrueckii] |
| WP_087363710.1 | Type1.BREX | Peptidase_M13_N+Peptidase_M13->Acetate_kinase-><-TerBN+TPR<- PglZ <- HEPN <- HEPN <- N6-METHYLASE <- DUF6079 *<- BrxB-BrxF-iSTAND <- BrxA-DNA-binding <-ParB+HNH<-?<-OrfB_Zn_ribbon | 1244 | Bacteria>Terrabacteria_group>Bacillota>Bacilli>Lactobacillales>Lactobacillaceae | GCF_002160975.1 | Lactobacillus_gallinarum | B5E63_RS08425 | 13791..17525 | 52242 | BREX-system-P-loop-protein-BrxC-[Lactobacillus-gallinarum] |
| WP_113532397.1 | Type1.BREX | YTCD-HTH-><-?<-?<-TetR-HTH<-TetR-HTH<- PglZ <- HEPN +REase<-nSTAND3<- N6-METHYLASE <- DUF6079 *<- BrxB-BrxF-iSTAND <- BrxA-DNA-binding <-?<-?<-?<-NUDIX<-HAD+HAD | 1239 | Bacteria>Terrabacteria_group>Bacillota>Bacilli>Lactobacillales>Lactobacillaceae | GCF_003584685.1 | Lactobacillus_paragasseri | LpgJCM5343_RS05125 | 1058950..1062669 | 2107999 | BREX-system-P-loop-protein-BrxC-[Lactobacillus-paragasseri] |
| WP_172987666.1 | Type1.BREX | ParB+HNH-> BrxA-DNA-binding -> BrxB-BrxF-iSTAND -> DUF6079 *-> N6-METHYLASE -> HEPN -> HEPN -> PglZ ->TerBN+TPR->ParB->ParB+HNH-> | 1243 | Bacteria>Terrabacteria_group>Bacillota>Bacilli>Lactobacillales>Lactobacillaceae | GCF_013280955.1 | Lactobacillus_helveticus | IMAU50022_RS01295 | 3445..7176 | 1587 | BREX-system-P-loop-protein-BrxC-[Lactobacillus-helveticus] |
| WP_173006741.1 | Type1.BREX | Peptidase_M13_N+Peptidase_M13->?-><-TerBN+TPR<- PglZ <- HEPN <- HEPN <- N6-METHYLASE <- DUF6079 *<- BrxB-BrxF-iSTAND <- BrxA-DNA-binding <-ParB+HNH\|\|MODE-HTH+PBPII-><-?<-?<-wHTH+WYL | 1241 | Bacteria>Terrabacteria_group>Bacillota>Bacilli>Lactobacillales>Lactobacillaceae | GCF_013281475.1 | Lactobacillus_helveticus | IMAU30014_RS01855 | 25052..28777 | 1587 | BREX-system-P-loop-protein-BrxC-[Lactobacillus-helveticus] |
| WP_220735234.1 | Type1.BREX | Calcineurin-><-?\|\|?->?->?-><-?\|\| BrxA-DNA-binding -> BrxB-BrxF-iSTAND -> DUF6079 *-> N6-METHYLASE -> HEPN -> HEPN -><-Phage_int_SAM_5+Phage_integrase<- N6-METHYLASE \|\| PglZ -> SIGMA+OB-Fold+AAA-ATPase+Lon-Protease-BrxL -> | 1212 | Bacteria>Terrabacteria_group>Bacillota>Bacilli>Lactobacillales>Lactobacillaceae | GCF_019656015.1 | Leuconostoc_inhae | K5Q14_RS07795 | 32108..35746 | 178001 | BREX-system-P-loop-protein-BrxC-[Leuconostoc-inhae] |
| WP_223660616.1 | Type1.BREX | BrxA-DNA-binding -> BrxB-BrxF-iSTAND -> DUF6079 *-> N6-METHYLASE ->nSTAND3-> HEPN -p-> PglZ ->TetR-HTH->?-><-?\|\|MORC+DNA_gyrase+TOPRIM-> | 1239 | Bacteria>Terrabacteria_group>Bacillota>Bacilli>Lactobacillales>Lactobacillaceae | GCF_020026535.1 | Lactobacillus_johnsonii | K7D69_RS01700 | 3964..7683 | 33959 | BREX-system-P-loop-protein-BrxC-[Lactobacillus-johnsonii] |
| WP_231523637.1 | Type1.BREX | GGDEF+PAS-><-ABhydrolase\|\|ABhydrolase->ABhydrolase+ABhydrolase->ParB+HNH->\|\|ParB+HNH->ParB+HNH-> BrxA-DNA-binding -> BrxB-BrxF-iSTAND -> DUF6079 *-> N6-METHYLASE -> PglZ -> SIGMA+OB-Fold+AAA-ATPase+Lon-Protease-BrxL -> HEPN ->AbiGi-><-AraC-HTH+AraC-HTH\|\|?->AraC-HTH+O6MGT-HTH-> | 1183 | Bacteria>Terrabacteria_group>Bacillota>Bacilli>Lactobacillales>Lactobacillaceae | GCF_021134645.1 | Lactobacillus_delbrueckii | L0B39_RS07020 | 33020..36571 | 1584 | BREX-system-P-loop-protein-BrxC-[Lactobacillus-delbrueckii] |
| WP_236153484.1 | Type1.BREX | ABhydrolase->ABhydrolase+ABhydrolase->?->OrfB_Zn_ribbon->HNH->ParB+HNH->ParB+HNH-> BrxA-DNA-binding -> BrxB-BrxF-iSTAND -> DUF6079 *-> N6-METHYLASE -> PglZ -> SIGMA+OB-Fold+AAA-ATPase+Lon-Protease-BrxL -> HEPN ->AbiGi-><-AraC-HTH+AraC-HTH | 1183 | Bacteria>Terrabacteria_group>Bacillota>Bacilli>Lactobacillales>Lactobacillaceae | GCF_021600265.1 | Lactobacillus_delbrueckii | ME802_14840 | 21460..25011 | 1584 | BREX-system-P-loop-protein-BrxC-[Lactobacillus-delbrueckii] |
| WP_236149793.1 | Type1.BREX | ABhydrolase->ABhydrolase+ABhydrolase->REase+N6_METHYLASE->?->?->ParB+HNH-p->ParB+HNH-> BrxA-DNA-binding -> BrxB-BrxF-iSTAND -> DUF6079 *-> N6-METHYLASE -> PglZ -> SIGMA+OB-Fold+AAA-ATPase+Lon-Protease-BrxL -> HEPN ->AbiGi-><-AraC-HTH+AraC-HTH | 1188 | Bacteria>Terrabacteria_group>Bacillota>Bacilli>Lactobacillales>Lactobacillaceae | GCF_021600925.1 | Lactobacillus_delbrueckii | L5981_RS02870 | 31328..34894 | 1584 | BREX-system-P-loop-protein-BrxC-[Lactobacillus-delbrueckii] |
| WP_050952512.1 | Type1.BREX | <-AraC-HTH+O6MGT-HTH<-?\|\|AraC-HTH+AraC-HTH-><-AbiGi<- HEPN <- SIGMA+OB-Fold+AAA-ATPase+Lon-Protease-BrxL <- PglZ <- N6-METHYLASE <- DUF6079 *<- BrxB-BrxF-iSTAND <- BrxA-DNA-binding <-ParB+HNH<-ParB+HNH-p<-?<-?<-REase+N6_METHYLASE<-ABhydrolase+ABhydrolase<-ABhydrolase | 1188 | Bacteria>Terrabacteria_group>Bacillota>Bacilli>Lactobacillales>Lactobacillaceae | GCF_021601025.1 | Lactobacillus_delbrueckii | L5988_RS00790 | 158168..161734 | 1584 | BREX-system-P-loop-protein-BrxC-[Lactobacillus-delbrueckii] |
| WP_268886389.1 | Type1.BREX | <-AraC-HTH+O6MGT-HTH<-?\|\|AraC-HTH+AraC-HTH-><-AbiGi<- HEPN <- SIGMA+OB-Fold+AAA-ATPase+Lon-Protease-BrxL <- PglZ <- N6-METHYLASE <- DUF6079 *<- BrxB-BrxF-iSTAND <- BrxA-DNA-binding <-ParB+HNH<-ParB+HNH\|\|<-ParB+HNH<-ABhydrolase+ABhydrolase<-ABhydrolase\|\|ABhydrolase-><-GGDEF+PAS | 1183 | Bacteria>Terrabacteria_group>Bacillota>Bacilli>Lactobacillales>Lactobacillaceae | GCF_026891475.1 | Lactobacillus_delbrueckii | O0549_RS05810 | 14866..18417 | 1584 | BREX-system-P-loop-protein-BrxC-[Lactobacillus-delbrueckii] |
| WP_270447891.1 | Type1.BREX | ABhydrolase+ABhydrolase-><-?\|\|?->?->?->HTH+REase-> BrxA-DNA-binding -> BrxB-BrxF-iSTAND -> DUF6079 *-> N6-METHYLASE -> PglZ -> SIGMA+OB-Fold+AAA-ATPase+Lon-Protease-BrxL -> HEPN -> HEPN ->SF1-SF2-Helicase->?-><-AraC-HTH | 1188 | Bacteria>Terrabacteria_group>Bacillota>Bacilli>Lactobacillales>Lactobacillaceae | GCF_027681365.1 | Lactobacillus_delbrueckii | PGX26_RS03415 | 69740..73306 | 1584 | BREX-system-P-loop-protein-BrxC-[Lactobacillus-delbrueckii] |
| WP_271018243.1 | Type1.BREX | BrxA-DNA-binding -> BrxB-BrxF-iSTAND -> DUF6079 *-> N6-METHYLASE -> PglZ -> SIGMA+OB-Fold+AAA-ATPase+Lon-Protease-BrxL -> HEPN -> HEPN ->AbiGi-><-AraC-HTH+AraC-HTH\|\|?->AraC-HTH+O6MGT-HTH-> | 1183 | Bacteria>Terrabacteria_group>Bacillota>Bacilli>Lactobacillales>Lactobacillaceae | GCF_027858035.1 | Lactobacillus_delbrueckii | PF599_RS07175 | 1642..5193 | 1584 | BREX-system-P-loop-protein-BrxC-[Lactobacillus-delbrueckii] |
| WP_271873858.1 | Type1.BREX | <-TerBN+TPR<- PglZ <- HEPN <- HEPN <- N6-METHYLASE <- DUF6079 *<- BrxB-BrxF-iSTAND <- BrxA-DNA-binding <-ParB+HNH<-?<-?<-AAA_33 | 1244 | Bacteria>Terrabacteria_group>Bacillota>Bacilli>Lactobacillales>Lactobacillaceae | GCF_028308865.1 | Lactobacillus_amylovorus | ODV08_RS03455 | 54390..58124 | 1604 | BREX-system-P-loop-protein-BrxC-[Lactobacillus-amylovorus] |
| WP_271897403.1 | Type1.BREX | <-HIN-HTH+PAIREDC-HTH\|\|?->CR-ATPase5+REase-><-TerBN+TPR<- PglZ <- HEPN <- HEPN <- N6-METHYLASE <- DUF6079 *<- BrxB-BrxF-iSTAND <- BrxA-DNA-binding <-ParB+HNH<-?<-?<-AAA_33 | 1233 | Bacteria>Terrabacteria_group>Bacillota>Bacilli>Lactobacillales>Lactobacillaceae | GCF_028308945.1 | Lactobacillus_amylovorus | ODV12_RS07245 | 16097..19798 | 1604 | BREX-system-P-loop-protein-BrxC-[Lactobacillus-amylovorus] |
| WP_064081740.1 | Type1.BREX | ParB+HNH-> BrxA-DNA-binding -> BrxB-BrxF-iSTAND -> DUF6079 *-> N6-METHYLASE -> PglZ -> SIGMA+OB-Fold+AAA-ATPase+Lon-Protease-BrxL -> HEPN ->?->?->O6MGT-HTH-><-?<-NUDIX | 1178 | Bacteria>Terrabacteria_group>Bacillota>Bacilli>Lactobacillales>Lactobacillaceae | GCF_925300675.1 | Lactobacillus_delbrueckii | NGL01_RS07850 | 3981..7517 | 1584 | BREX-system-P-loop-protein-BrxC-[Lactobacillus-delbrueckii] |
| WP_118669593.1 | Type1.BREX | <- SIGMA+OB-Fold+AAA-ATPase+Lon-Protease-BrxL <-OLD-ABC+TOPRIM<- PglZ <-Schlafen+HATPase+wHTH<- HEPN <-TIR<- N6-METHYLASE <- DUF6079 *<- BrxB-BrxF-iSTAND <- BrxA-DNA-binding <-HTH_26 | 1196 | Bacteria>Terrabacteria_group>Bacillota>Clostridia>Eubacteriales>Clostridiaceae | - | Clostridium_sp._TF08-15 | DXC82_15795 | 32814..36404 | 2293052 | BREX-system-P-loop-protein-BrxC-[Clostridium-sp.-TF08-15] |
| CCZ63532.1 | Type1.BREX | HTH_26-> BrxA-DNA-binding -> BrxB-BrxF-iSTAND -> DUF6079 *-> N6-METHYLASE -> PglZ -> SIGMA+OB-Fold+AAA-ATPase+Lon-Protease-BrxL -> HEPN ->?->?->NUDIX->?->HTH+HisKin-> | 1215 | Bacteria>Terrabacteria_group>Bacillota>Clostridia>Eubacteriales>Clostridiaceae | GCA_000433395.1 | Clostridium_hathewayi_CAG:224 | BN544_01435 | 55563..59210 | 1263067 | putative-uncharacterized-protein-[Clostridium-hathewayi-CAG:224] |
| CDD75188.1 | Type1.BREX | MACRODOMAIN->?->PIN->ZnR+ZnR+ZnR->HTH_26-> BrxA-DNA-binding -> BrxB-BrxF-iSTAND -> DUF6079 *->OLD-ABC->PIN-> N6-METHYLASE ->OLD-ABC->NERD-> HEPN ->Schlafen+HATPase+wHTH-> PglZ ->DUF4365+REase-> | 1200 | Bacteria>Terrabacteria_group>Bacillota>Clostridia>Eubacteriales>Clostridiaceae | GCA_000436175.1 | Clostridium_sp._CAG:62 | BN737_01375 | 69547..73149 | 1262828 | uncharacterized-protein-BN737_01375-[Clostridium-sp.-CAG:62] |
| RGF28780.1 | Type1.BREX | <-OLD-ABC+TOPRIM<- PglZ <-Schlafen+HATPase+wHTH<- HEPN <- N6-METHYLASE <- DUF6079 *<- BrxB-BrxF-iSTAND <- BrxA-DNA-binding <-HTH_26 | 1196 | Bacteria>Terrabacteria_group>Bacillota>Clostridia>Eubacteriales>Clostridiaceae | GCA_003435925.1 | Clostridium_sp._AF46-9NS | DW081_14990 | 34449..38039 | 2293020 | BREX-system-P-loop-protein-BrxC-[Clostridium-sp.-AF46-9NS] |
| RHU61449.1 | Type1.BREX | <- SIGMA+OB-Fold+AAA-ATPase+Lon-Protease-BrxL <-OLD-ABC+TOPRIM<- PglZ <-Schlafen+HATPase+wHTH<- HEPN <-TIR<- N6-METHYLASE <- DUF6079 *<- BrxB-BrxF-iSTAND <- BrxA-DNA-binding <-HTH_26 | 1196 | Bacteria>Terrabacteria_group>Bacillota>Clostridia>Eubacteriales>Clostridiaceae | GCA_003481275.1 | Clostridium_sp._TF08-15 | DXC82_15795 | 32814..36404 | 2293052 | BREX-system-P-loop-protein-BrxC-[Clostridium-sp.-TF08-15] |
| MBS5781623.1 | Type1.BREX | <-SMODS<-?<-SAVED<- SIGMA+OB-Fold+AAA-ATPase+Lon-Protease-BrxL <- PglZ <- HEPN <- N6-METHYLASE -p<- N6-METHYLASE -p<- N6-METHYLASE <- DUF6079 *<- BrxB-BrxF-iSTAND <- BrxA-DNA-binding \|\|?->?->?-><-?\|\|XIS-HTH->ZnR+TOPRIM+AAA-helicase+wHTH-> | 1171 | Bacteria>Terrabacteria_group>Bacillota>Clostridia>Eubacteriales>Clostridiaceae | GCA_018371215.1 | Clostridium_sp. | KID04_01935 | 432250..435765 | 1506 | BREX-system-P-loop-protein-BrxC-[Clostridium-sp.] |
| MCI9636406.1 | Type1.BREX | <-SF1-SF2-Helicase+REase<-ParB<-?<- wHTH+WYL+WCX <- SIGMA+OB-Fold+AAA-ATPase+Lon-Protease-BrxL <- PglZ <- N6-METHYLASE -p<- HEPN <- N6-METHYLASE <- DUF6079 *<- BrxB-BrxF-iSTAND <- BrxA-DNA-binding <-REase<-Sulfatase<-REC+AraC-HTH+AraC-HTH<-dCache_1+HAMP+HisKin+HATPase_c<-Sulfatase+DUF4976<-ABC-ATPase+SbcC | 1182 | Bacteria>Terrabacteria_group>Bacillota>Clostridia>Eubacteriales>Clostridiaceae | GCA_022802035.1 | Hungatella_sp. | HFG77_08430 | 22407..25955 | 2613924 | BREX-system-P-loop-protein-BrxC-[Hungatella-sp.] |
| WP_003411890.1 | Type1.BREX | <- HEPN <- N6-METHYLASE <- DUF6079 *<- BrxB-BrxF-iSTAND <- BrxA-DNA-binding <-?<-MORC\|\|HTH_3-><-?<-Phage_integrase<-?<-Phage_integrase | 1215 | Bacteria>Terrabacteria_group>Bacillota>Clostridia>Eubacteriales>Clostridiaceae | GCF_000171115.1 | Clostridium_butyricum | CBY_RS17320 | 4352..7999 | 1492 | BREX-system-P-loop-protein-BrxC-[Clostridium-butyricum] |
| WP_002598576.1 | Type1.BREX | <-CRISPR_Cas2<-Cas_Cas1<-?<-Cas9+HNH+RuvC<- N6-METHYLASE <-Ntox45<- HEPN <- DUF6079 *<- BrxB-BrxF-iSTAND <- BrxA-DNA-binding <-?\|\|TetR-HTH->?-><-DnaB+TnsC-ATPase<-?\|\|HAD-> | 1208 | Bacteria>Terrabacteria_group>Bacillota>Clostridia>Eubacteriales>Clostridiaceae | GCF_000371465.1 | Clostridium_thermobutyricum | HMPREF1092_RS10115 | 2173864..2177490 | 29372 | BREX-system-P-loop-protein-BrxC-[Clostridium-thermobutyricum] |
| WP_065077880.1 | Type1.BREX | <-GNTR-HTH+UTRA\|\|?->?->tRNA-> HEPN -> BrxA-DNA-binding -> BrxB-BrxF-iSTAND -> DUF6079 *-> N6-METHYLASE -> | 1188 | Bacteria>Terrabacteria_group>Bacillota>Clostridia>Eubacteriales>Clostridiaceae | GCF_001675165.1 | Clostridium_ragsdalei | CLRAG_RS07695 | 80363..83929 | 217158 | BREX-system-P-loop-protein-BrxC-[Clostridium-ragsdalei] |
| WP_118731364.1 | Type1.BREX | PALP->?->RVT->HTH_26-> BrxA-DNA-binding -> BrxB-BrxF-iSTAND -> DUF6079 *-> N6-METHYLASE -> HEPN ->Schlafen+HATPase+wHTH-> PglZ ->OLD-ABC+TOPRIM-> SIGMA+OB-Fold+AAA-ATPase+Lon-Protease-BrxL -> | 1196 | Bacteria>Terrabacteria_group>Bacillota>Clostridia>Eubacteriales>Clostridiaceae | GCF_003480225.1 | Clostridium_sp._AM34-9AC | DW835_RS14805 | 20266..23856 | 2293030 | BREX-system-P-loop-protein-BrxC-[Clostridium-sp.-AM34-9AC] |
| WP_118742502.1 | Type1.BREX | 35exo+T7DNApol->?->?->?->HTH_26-> BrxA-DNA-binding -> BrxB-BrxF-iSTAND -> DUF6079 *-> N6-METHYLASE -> HEPN ->Schlafen+HATPase+wHTH-> PglZ -> SIGMA+OB-Fold+AAA-ATPase+Lon-Protease-BrxL -><-MODE-HTH+PBPII | 1182 | Bacteria>Terrabacteria_group>Bacillota>Clostridia>Eubacteriales>Clostridiaceae | GCF_003480665.1 | Clostridium_sp._AM28-20LB | DW739_RS01385 | 291245..294793 | 2293027 | BREX-system-P-loop-protein-BrxC-[Clostridium-sp.-AM28-20LB] |
| WP_118744885.1 | Type1.BREX | ACET->REase->?->?->?->?->HTH_26-> BrxA-DNA-binding -> BrxB-BrxF-iSTAND -> DUF6079 *->OLD-ABC->PIN-> N6-METHYLASE -> HEPN ->?->OLD-ABC->HNH->?->TIR-> | 1203 | Bacteria>Terrabacteria_group>Bacillota>Clostridia>Eubacteriales>Clostridiaceae | GCF_003480745.1 | Clostridium_sp._AM27-31LB | DW721_RS03885 | 61324..64935 | 2293026 | BREX-system-P-loop-protein-BrxC-[Clostridium-sp.-AM27-31LB] |
| WP_164950923.1 | Type5.BREX | <-GDResolvase<-?<-?<- TUDOR+SF1-SF2-Helicase+iREase+CTD +REase<- PglZ <-TOPRIM<-OLD-ABC<- HEPN <- N6-METHYLASE <- DUF6079 *<- BrxB-BrxF-iSTAND <- BrxA-DNA-binding <-?<-Calcineurin<-PARG+MACRODOMAIN<-Calcineurin<-HAD<-LRR+LRR | 1191 | Bacteria>Terrabacteria_group>Bacillota>Clostridia>Eubacteriales>Clostridiaceae | GCF_011065955.2 | Clostridium_sp._FP2 | G9F71_RS16320 | 3398773..3402348 | 2724481 | BREX-system-P-loop-protein-BrxC-[Clostridium-sp.-FP2] |
| WP_164949146.1 | Type1.BREX | <- HEPN <- SIGMA+OB-Fold+AAA-ATPase+Lon-Protease-BrxL <- PglZ <-HNH<-OLD-ABC<- N6-METHYLASE <- DUF6079 *<- BrxB-BrxF-iSTAND <- BrxA-DNA-binding <-?<-CR-ATPase8+REase\|\|?->?-><-?<-?\|\|METHYLASE-> | 1215 | Bacteria>Terrabacteria_group>Bacillota>Clostridia>Eubacteriales>Clostridiaceae | GCF_011065955.2 | Clostridium_sp._FP2 | G9F71_RS17965 | 3736090..3739737 | 2724481 | BREX-system-P-loop-protein-BrxC-[Clostridium-sp.-FP2] |
| WP_077854163.1 | Type1.BREX | <-HAD+HAD<-?<-Peptidase_M42<- HEPN <- SIGMA+OB-Fold+AAA-ATPase+Lon-Protease-BrxL <-?<- PglZ <- N6-METHYLASE <- DUF6079 *<- BrxB-BrxF-iSTAND <- BrxA-DNA-binding <-MORC\|\|HTH_3-><-?<-Phage_integrase<-?<-Phage_integrase<-Aminotran_1_2 | 1216 | Bacteria>Terrabacteria_group>Bacillota>Clostridia>Eubacteriales>Clostridiaceae | GCF_013297045.1 | Clostridium_beijerinckii | BCD95_RS19475 | 4461966..4465616 | 1520 | BREX-system-P-loop-protein-BrxC-[Clostridium-beijerinckii] |
| WP_195540766.1 | Type1.BREX | <-HNH<-ClpABC-AAA<-?<- SIGMA+OB-Fold+AAA-ATPase+Lon-Protease-BrxL <- PglZ <-Schlafen+HATPase+wHTH<- HEPN <- N6-METHYLASE <-?<- DUF6079 *<- BrxB-BrxF-iSTAND <- BrxA-DNA-binding <-HTH_26<-?<-?<-?<- wHTH+WYL+WCX \|\|?-><-NUDIX | 1182 | Bacteria>Terrabacteria_group>Bacillota>Clostridia>Eubacteriales>Clostridiaceae | GCF_015558125.1 | Clostridium_sp._D43t1_170807_D5 | I2G39_RS02195 | 465793..469341 | 2787150 | BREX-system-P-loop-protein-BrxC-[Clostridium-sp.-D43t1_170807_D5] |
| WP_216574650.1 | Type1.BREX | <-ABhydrolase\|\|IG+IG->?->MazE_antitoxin+SpoVT_C->HTH_26-> BrxA-DNA-binding -> BrxB-BrxF-iSTAND -> DUF6079 *-> N6-METHYLASE -> HEPN ->Schlafen+HATPase+wHTH-> PglZ -> SIGMA+OB-Fold+AAA-ATPase+Lon-Protease-BrxL ->TIR->?->MazE_antitoxin+SpoVT_C-> | 1182 | Bacteria>Terrabacteria_group>Bacillota>Clostridia>Eubacteriales>Clostridiaceae | GCF_018918265.1 | Falcatimonas_sp._MSJ-15 | KQI85_RS03175 | 188313..191861 | 2841515 | BREX-system-P-loop-protein-BrxC-[Falcatimonas-sp.-MSJ-15] |
| WP_270807575.1 | Type1.BREX | <-LRR_5+LRR_8<- SIGMA+OB-Fold+AAA-ATPase+Lon-Protease-BrxL <-?<-PIN<-SbcC<- PglZ <- HEPN <- N6-METHYLASE <- DUF6079 *<- BrxB-BrxF-iSTAND <- BrxA-DNA-binding <-HTH_26<-PIN<-?<-MACRODOMAIN | 1201 | Bacteria>Terrabacteria_group>Bacillota>Clostridia>Eubacteriales>Clostridiaceae | GCF_027723425.1 | Clostridium_fessum | PG965_RS01510 | 334129..337734 | 2126740 | BREX-system-P-loop-protein-BrxC-[Clostridium-fessum] |
| WP_072525015.1 | Type1.BREX | TRAM+METHYLASE->SymE_toxin->ParA-Soj-PloopNTPase->ParB->?->HTH_26-> BrxA-DNA-binding -> BrxB-BrxF-iSTAND -> DUF6079 *-> N6-METHYLASE -> PglZ -> SIGMA+OB-Fold+AAA-ATPase+Lon-Protease-BrxL -> HEPN -> HEPN -><-XIS-HTH<-PemK_toxin | 1199 | Bacteria>Terrabacteria_group>Bacillota>Clostridia>Eubacteriales>Clostridiaceae | GCF_900120155.1 | Clostridium_sp._Marseille-P3244 | BQ7367_RS08330 | 1368431..1372030 | 1871020 | BREX-system-P-loop-protein-BrxC-[Clostridium-sp.-Marseille-P3244] |
| WP_073007111.1 | Type1.BREX | PAS+GGDEF->?->?->?->?-> BrxA-DNA-binding -> BrxB-BrxF-iSTAND -> DUF6079 *-> N6-METHYLASE -> HEPN ->HipA_C+STYKinase->Couple_hipA+HIRAN->Cas9+HNH+RuvC->Cas_Cas1->CRISPR_Cas2-> PglZ -> SIGMA+OB-Fold+AAA-ATPase+Lon-Protease-BrxL -> | 1214 | Bacteria>Terrabacteria_group>Bacillota>Clostridia>Eubacteriales>Clostridiaceae | GCF_900142075.1 | Clostridium_amylolyticum | BUA82_RS09870 | 579511..583155 | 1121298 | BREX-system-P-loop-protein-BrxC-[Clostridium-amylolyticum] |
| WP_117466987.1 | Type1.BREX | <-OLD-ABC+TOPRIM<- PglZ <-Schlafen+HATPase+wHTH<- HEPN <- N6-METHYLASE <- DUF6079 *<- BrxB-BrxF-iSTAND <- BrxA-DNA-binding <-HTH_26 | 1196 | Bacteria>Terrabacteria_group>Bacillota>Clostridia>Eubacteriales>Clostridiaceae | - | unclassified_Clostridium | DW081_14990 | 34449..38039 | 2614128 | MULTISPECIES:-BREX-system-P-loop-protein-BrxC-[unclassified-Clostridium] |
| WP_014901355.1 | Type1.BREX | ANK+ANK->?->?->SMODS->SLATT->?-> BrxA-DNA-binding -> BrxB-BrxF-iSTAND -> DUF6079 *-> N6-METHYLASE -> HEPN -> PglZ -> SIGMA+OB-Fold+AAA-ATPase+Lon-Protease-BrxL ->?-> HEPN +PAS+DUF4011+SF1-SF2-Helicase+REase->?->TniQ->TniQ-> | 1203 | Bacteria>Terrabacteria_group>Bacillota>Clostridia>Eubacteriales>Desulfitobacteriaceae | GCF_000231385.2 | Desulfosporosinus_meridiei | Desmer_0377 | 362087..365698 | 79209 | BREX-system-P-loop-protein-BrxC-[Desulfosporosinus-meridiei] |
| MCH3955225.1 | Type1.BREX | <-RVT+ HEPN \|\|HIN-HTH+PAIREDC-HTH-><-RVT+ HEPN <- SIGMA+OB-Fold+AAA-ATPase+Lon-Protease-BrxL <- PglZ <- N6-METHYLASE <- DUF6079 *<- BrxB-BrxF-iSTAND <- BrxA-DNA-binding <-?<-?<-HNH+3C-ABC\|\|SIGMA-HTH-><-Recombinase<-Resolvase+Recombinase+ZnR | 1171 | Bacteria>Terrabacteria_group>Bacillota>Clostridia>Eubacteriales>Eubacteriaceae | GCA_022482865.1 | Eubacterium_sp. | LKE44_08530 | 1944805..1948320 | 142586 | BREX-system-P-loop-protein-BrxC-[Eubacterium-sp.] |
| MCH4073094.1 | Type1.BREX | <-ClpABC-AAA<-?<- HEPN <- HEPN <- SIGMA+OB-Fold+AAA-ATPase+Lon-Protease-BrxL <- PglZ <-Schlafen+HATPase+wHTH<-Schlafen+HATPase+wHTH<- N6-METHYLASE <- DUF6079 *<- BrxB-BrxF-iSTAND <- BrxA-DNA-binding <-?<-TRAM+METHYLASE<-DTXR-HTH<-RADICAL-SAM<-?<-?<-HAD+HAD | 1171 | Bacteria>Terrabacteria_group>Bacillota>Clostridia>Eubacteriales>Eubacteriaceae | GCA_022483875.1 | Pseudoramibacter_sp. | LKF02_09135 | 1854333..1857848 | 2034862 | BREX-system-P-loop-protein-BrxC-[Pseudoramibacter-sp.] |
| MCI5510134.1 | Type1.BREX | CopG/RHH->?->TraJ-RHH->TraI-VirD2-RCRE->?->HTH_26-> BrxA-DNA-binding -> BrxB-BrxF-iSTAND -> DUF6079 *->?-> N6-METHYLASE -> HEPN ->Schlafen+HATPase+wHTH-> PglZ -> SIGMA+OB-Fold+AAA-ATPase+Lon-Protease-BrxL ->OLD-ABC+TOPRIM->SF1-SF2-Helicase-><-HTH | 1182 | Bacteria>Terrabacteria_group>Bacillota>Clostridia>Eubacteriales>Eubacteriaceae | GCA_022764165.1 | Eubacterium_sp. | MR436_06790 | 9461..13009 | 142586 | BREX-system-P-loop-protein-BrxC-[Eubacterium-sp.] |
| CVI69463.1 | Type1.BREX | <-Schlafen+wHTH<- HEPN <- N6-METHYLASE <-TIR<- DUF6079 *<- BrxB-BrxF-iSTAND <- BrxA-DNA-binding <-HTH_26<-ParB+HNH | 1179 | Bacteria>Terrabacteria_group>Bacillota>Clostridia>Eubacteriales>Eubacteriaceae | GCA_900016875.1 | Eubacteriaceae_bacterium_CHKCI004 | BN3660_01565 | 6017..9556 | 1780380 | hypothetical-protein-BN3660_01565-[Eubacteriaceae-bacterium-CHKCI004] |
| SEG33478.1 | Type1.BREX | <-SbcC<-Calcineurin<-?<-?<- SIGMA+OB-Fold+AAA-ATPase+Lon-Protease-BrxL <- PglZ <- N6-METHYLASE <- HEPN <- DUF6079 *<- BrxB-BrxF-iSTAND <- BrxA-DNA-binding <-?<- wHTH+WYL+WCX <-35exo<-HTH_3<-Tox-RelA<-PTSEIIA1 | 1157 | Bacteria>Terrabacteria_group>Bacillota>Clostridia>Eubacteriales>Eubacteriaceae | GCA_900108205.1 | Eubacterium_ruminantium | SAMN04487934_11616 | 23559..27032 | 42322 | hypothetical-protein-SAMN04487934_11616-[Eubacterium-ruminantium] |
| WP_055289382.1 | Type1.BREX | HTH-><-?<-PHP+SbcC<- SIGMA+OB-Fold+AAA-ATPase+Lon-Protease-BrxL <- PglZ <-Schlafen+HATPase+wHTH<- HEPN <- N6-METHYLASE <-?<- DUF6079 *<- BrxB-BrxF-iSTAND <- BrxA-DNA-binding <-HTH_26<-TraI-VirD2-RCRE<-TraJ-RHH<-?<-CopG/RHH<-?<-HTH_36 | 1183 | Bacteria>Terrabacteria_group>Bacillota>Clostridia>Eubacteriales>Eubacteriaceae | GCF_001406295.1 | Eubacterium_ramulus | ARA08_RS03325 | 204677..208228 | 39490 | BREX-system-P-loop-protein-BrxC-[Eubacterium-ramulus] |
| WP_078766837.1 | Type1.BREX | AbiEi_4->AbiEii->HTH_3->?->REase->wHTH+HNH->HTH_26-> BrxA-DNA-binding -> BrxB-BrxF-iSTAND -> DUF6079 *-> N6-METHYLASE -> HEPN -> PglZ ->SLATT->?-> SIGMA+OB-Fold+AAA-ATPase+Lon-Protease-BrxL ->RVT->SbcC-> | 1203 | Bacteria>Terrabacteria_group>Bacillota>Clostridia>Eubacteriales>Eubacteriaceae | GCF_900167115.1 | Eubacterium_uniforme | B5D16_RS09945 | 14532..18143 | 39495 | BREX-system-P-loop-protein-BrxC-[Eubacterium-uniforme] |
| MCI6150868.1 | Type1.BREX | <-wHTH+WYL<-?<-HTH+MPTase<-ZnR<- SIGMA+OB-Fold+AAA-ATPase+Lon-Protease-BrxL <- PglZ <- HEPN <-HKD-DNase<- N6-METHYLASE <- DUF6079 *<- BrxB-BrxF-iSTAND <- BrxA-DNA-binding <-HMG+NLPC+NLPC<-VirB4-FtsK<-?\|\|?-><-TrwI<-TraM<-VirD4-TraG+VirD4-FtsK | 1179 | Bacteria>Terrabacteria_group>Bacillota>Clostridia>Eubacteriales>Eubacteriales_incertae_sedis | GCA_022772165.1 | Flintibacter_sp. | MR705_10640 | 43131..46670 | 1918624 | BREX-system-P-loop-protein-BrxC-[Flintibacter-sp.] |
| WP_118637679.1 | Type1.BREX | <-LexA-protease<-MNS+REase<- wHTH+WYL+WCX <- N6-METHYLASE <- HEPN <-ParB+HNH<- SIGMA+OB-Fold+AAA-ATPase+Lon-Protease-BrxL <- PglZ <- N6-METHYLASE <- DUF6079 *<- BrxB-BrxF-iSTAND <- BrxA-DNA-binding <-HTH_26\|\|TraJ-RHH->TraI-VirD2-RCRE-> | 1215 | Bacteria>Terrabacteria_group>Bacillota>Clostridia>Eubacteriales | GCF_027924705.1 | Oscillospiraceae | H6A32_RS13625 | 30472..34119 | 216572 | MULTISPECIES:-BREX-system-P-loop-protein-BrxC-[Oscillospiraceae] |
| WP_256171760.1 | Type1.BREX | HTH_36->?->CopG/RHH->?->TraJ-RHH->TraI-VirD2-RCRE->HTH_26-> BrxA-DNA-binding -> BrxB-BrxF-iSTAND -> DUF6079 *->?-> N6-METHYLASE -> HEPN ->Schlafen+HATPase+wHTH-> PglZ -> SIGMA+OB-Fold+AAA-ATPase+Lon-Protease-BrxL ->OLD-ABC+TOPRIM->SF1-SF2-Helicase-><-HTH | 1182 | Bacteria>Terrabacteria_group>Bacillota>Clostridia>Eubacteriales>Lachnospiraceae | - | Dorea_longicatena | NE607_07100 | 21358..24906 | 88431 | BREX-system-P-loop-protein-BrxC-[Dorea-longicatena] |
| OOOO0080.1 | Type1.BREX | OLD-ABC->?->RADICAL-SAM->?->?->?-> BrxA-DNA-binding -> BrxB-BrxF-iSTAND -> DUF6079 *-><- HEPN | 1209 | Bacteria>Terrabacteria_group>Bacillota>Clostridia>Eubacteriales>Lachnospiraceae | GCA_002007315.1 | Epulopiscium_sp._Nele67-Bin004 | ATN35_09340 | 17804..21433 | 1764962 | MAG:-ATPase-[Epulopiscium-sp.-Nele67-Bin004] |
| RGI65884.1 | Type1.BREX | HTH_26-> BrxA-DNA-binding -> BrxB-BrxF-iSTAND -> DUF6079 *->OLD-ABC->PIN-> N6-METHYLASE ->TIR-> HEPN ->Schlafen+HATPase+wHTH-> PglZ ->Tox-ART-HYD3-> SIGMA+OB-Fold+AAA-ATPase+Lon-Protease-BrxL -> | 1196 | Bacteria>Terrabacteria_group>Bacillota>Clostridia>Eubacteriales>Lachnospiraceae | GCA_003436035.1 | [Eubacterium]_rectale | DXD95_13285 | 1619..5209 | 39491 | BREX-system-P-loop-protein-BrxC-[[Eubacterium] |
| HBA06127.1 | Type1.BREX | MNS+REase->TOPRIM+Topoisom_bac+TOPRIM_Crpt+TOPRIM_Crpt+TOPRIM_Crpt+TOPRIM_Crpt->?->?->?->HTH_26-> BrxA-DNA-binding -> BrxB-BrxF-iSTAND -> DUF6079 *->?-> N6-METHYLASE -> HEPN ->Schlafen+HATPase+wHTH-> PglZ -> SIGMA+OB-Fold+AAA-ATPase+Lon-Protease-BrxL ->?->Tox-ART-HYD3->AT-HTH-HYD2-> | 1182 | Bacteria>Terrabacteria_group>Bacillota>Clostridia>Eubacteriales>Lachnospiraceae | GCA_003491515.1 | Roseburia_sp. | DC452_03320 | 12130..15678 | 2049040 | BREX-system-P-loop-protein-BrxC-[Roseburia-sp.] |
| TGX97927.1 | Type1.BREX | <-OLD-ABC+TOPRIM-p<-OLD-ABC+TOPRIM-p<- PglZ <-HNH<-OLD-ABC<- N6-METHYLASE <- HEPN <-ParB+HNH<- N6-METHYLASE <- DUF6079 *<- BrxB-BrxF-iSTAND <- BrxA-DNA-binding <-HTH_26<-ZnR+ZnR+ZnR<-35exo+ADA-DBD<-?<-MACRODOMAIN | 1201 | Bacteria>Terrabacteria_group>Bacillota>Clostridia>Eubacteriales>Lachnospiraceae | GCA_004793585.1 | Lachnospiraceae_bacterium | E5357_10825 | 35909..39514 | 1898203 | BREX-system-P-loop-protein-BrxC-[Lachnospiraceae-bacterium] |
| TYL56933.1 | Type1.BREX | HTH_26-> BrxA-DNA-binding -> BrxB-BrxF-iSTAND -> DUF6079 *-> N6-METHYLASE ->REase->Schlafen+HATPase+wHTH-> PglZ ->?->?-> HEPN -> SIGMA+OB-Fold+AAA-ATPase+Lon-Protease-BrxL ->?->LRR_5+LRR_8-> | 1196 | Bacteria>Terrabacteria_group>Bacillota>Clostridia>Eubacteriales>Lachnospiraceae | GCA_008123415.1 | [Eubacterium]_rectale | FYL31_13935 | 1744..5334 | 39491 | BREX-system-P-loop-protein-BrxC-[[Eubacterium] |
| QNM03383.1 | Type1.BREX | GNTR-HTH+Aminotran_1_2->?->HTH_26-> BrxA-DNA-binding -> BrxB-BrxF-iSTAND -> DUF6079 *->Tox-ART-HYD2->AT-HTH-HYD2-1-> N6-METHYLASE -> HEPN ->Schlafen+HATPase+wHTH-> PglZ -> SIGMA+OB-Fold+AAA-ATPase+Lon-Protease-BrxL ->?->RNase-> | 1215 | Bacteria>Terrabacteria_group>Bacillota>Clostridia>Eubacteriales>Lachnospiraceae | GCA_014337175.1 | Simiaoa_sunii | H9Q77_04470 | 956158..959805 | 2763672 | BREX-system-P-loop-protein-BrxC-[Simiaoa-sunii] |
| MBR3244379.1 | Type1.BREX | TraJ-RHH->TraI-VirD2-RCRE->XIS-HTH-> BrxA-DNA-binding -> BrxB-BrxF-iSTAND -> DUF6079 *-> N6-METHYLASE ->?-> PglZ -> HEPN +SH3-like+SF1-SF2-Helicase-> | 1207 | Bacteria>Terrabacteria_group>Bacillota>Clostridia>Eubacteriales>Lachnospiraceae | GCA_017511245.1 | Parasporobacterium_sp. | IKF90_17090 | 4939..8562 | 2740533 | BREX-system-P-loop-protein-BrxC-[Parasporobacterium-sp.] |
| MBR4345194.1 | Type1.BREX | RelB->PIN-> BrxA-DNA-binding -> BrxB-BrxF-iSTAND -> DUF6079 *->VbhA+DOC+HTH-> N6-METHYLASE -> N6-METHYLASE -p-> HEPN ->SbcC+SbcC->Schlafen+HATPase+wHTH-> PglZ -> SIGMA+OB-Fold+AAA-ATPase+Lon-Protease-BrxL ->TIR-> | 1173 | Bacteria>Terrabacteria_group>Bacillota>Clostridia>Eubacteriales>Lachnospiraceae | GCA_017542245.1 | Lachnospiraceae_bacterium | IKP88_21230 | 56947..60468 | 1898203 | BREX-system-P-loop-protein-BrxC-[Lachnospiraceae-bacterium] |
| MBP3802955.1 | Type1.BREX | RelB-><-?\|\|HTH_26-> BrxA-DNA-binding -> BrxB-BrxF-iSTAND -> DUF6079 *-> N6-METHYLASE ->?->Schlafen+BirA-HTH-> PglZ -> SIGMA+OB-Fold+AAA-ATPase+Lon-Protease-BrxL -> HEPN -> | 1204 | Bacteria>Terrabacteria_group>Bacillota>Clostridia>Eubacteriales>Lachnospiraceae | GCA_017940955.1 | Oribacterium_sp. | J6I76_03520 | 9410..13024 | 1969407 | BREX-system-P-loop-protein-BrxC-[Oribacterium-sp.] |
| MCB7506159.1 | Type1.BREX | <-ClpABN-AAA+ClpABC-AAA\|\|HTH-><-MULE-TRANSPOSASE<-PHP+SbcC<- SIGMA+OB-Fold+AAA-ATPase+Lon-Protease-BrxL <- PglZ <-Schlafen+HATPase+wHTH<- HEPN <- N6-METHYLASE <- DUF6079 *<- BrxB-BrxF-iSTAND <- BrxA-DNA-binding <-HTH_26<-HTH_36 | 1182 | Bacteria>Terrabacteria_group>Bacillota>Clostridia>Eubacteriales>Lachnospiraceae | GCA_020553125.1 | Blautia_sp._MSK20_18 | LI091_01875 | 117354..120902 | 2883186 | BREX-system-P-loop-protein-BrxC-[Blautia-sp.-MSK20_18] |
| MCC2229908.1 | Type1.BREX | HTH_36->?->CopG/RHH->TraJ-RHH->TraI-VirD2-RCRE->?->HTH_26-> BrxA-DNA-binding -> BrxB-BrxF-iSTAND -> DUF6079 *->?-> N6-METHYLASE -> HEPN ->Schlafen+HATPase+wHTH-> PglZ -> SIGMA+OB-Fold+AAA-ATPase+Lon-Protease-BrxL ->OLD-ABC+TOPRIM->SF1-SF2-Helicase-><-HTH | 1182 | Bacteria>Terrabacteria_group>Bacillota>Clostridia>Eubacteriales>Lachnospiraceae | GCA_020687485.1 | Lachnospiraceae_bacterium_CLA-AA-H215 | LKD81_02670 | 26793..30341 | 2885348 | BREX-system-P-loop-protein-BrxC-[Lachnospiraceae-bacterium-CLA-AA-H215] |
| MCI8732218.1 | Type1.BREX | <-STYKinase-like\|\|OLD-ABC->MerR-HTH-><-?<-?\|\| BrxA-DNA-binding -> BrxB-BrxF-iSTAND -> DUF6079 *-> N6-METHYLASE -> HEPN -> | 1210 | Bacteria>Terrabacteria_group>Bacillota>Clostridia>Eubacteriales>Lachnospiraceae | GCA_022794815.1 | Lachnospiraceae_bacterium | HFH57_14060 | 7970..11602 | 1898203 | BREX-system-P-loop-protein-BrxC-[Lachnospiraceae-bacterium] |
| MCI9358063.1 | Type1.BREX | <-STYKinase-like\|\|Schlafen+HATPase+wHTH-p->BirA-HTH-><-Peptidase_M50<-?\|\| BrxA-DNA-binding -> BrxB-BrxF-iSTAND -> DUF6079 *-> N6-METHYLASE -> PolB -> HEPN ->OLD-ABC-> PglZ ->?->nSTAND3->?->Calcineurin-> | 1213 | Bacteria>Terrabacteria_group>Bacillota>Clostridia>Eubacteriales>Lachnospiraceae | GCA_022799775.1 | Lachnospiraceae_bacterium | HFH59_11085 | 6660..10301 | 1898203 | BREX-system-P-loop-protein-BrxC-[Lachnospiraceae-bacterium] |
| MCM1125306.1 | Type1.BREX | MoxR-AAA->LRR_5->Glyco_hydro_3_C+Fn3-like+Glyco_hydro_3->Peptidase_C26->tRNA->tRNA-><-?<-MerR-HTH\|\|HTH_26-> BrxA-DNA-binding -> BrxB-BrxF-iSTAND -> DUF6079 *->ParB+HNH-> HEPN -> N6-METHYLASE -p->?-><-tRNA\|\|?->HAD+HAD-> | 1205 | Bacteria>Terrabacteria_group>Bacillota>Clostridia>Eubacteriales>Lachnospiraceae | GCA_023663945.1 | Lachnospiraceae_bacterium | NC429_02425 | 28301..31918 | 1898203 | BREX-system-P-loop-protein-BrxC-[Lachnospiraceae-bacterium] |
| MCQ4892848.1 | Type1.BREX | HTH_36->?->CopG/RHH->?->TraJ-RHH->TraI-VirD2-RCRE->HTH_26-> BrxA-DNA-binding -> BrxB-BrxF-iSTAND -> DUF6079 *->?-> N6-METHYLASE -> HEPN ->Schlafen+HATPase+wHTH-> PglZ -> SIGMA+OB-Fold+AAA-ATPase+Lon-Protease-BrxL ->OLD-ABC+TOPRIM->SF1-SF2-Helicase-><-HTH | 1182 | Bacteria>Terrabacteria_group>Bacillota>Clostridia>Eubacteriales>Lachnospiraceae | GCA_024461355.1 | Dorea_longicatena | NE607_07100 | 21358..24906 | 88431 | BREX-system-P-loop-protein-BrxC-[Dorea-longicatena] |
| MCR5727513.1 | Type1.BREX | BrxA-DNA-binding -> BrxB-BrxF-iSTAND -> DUF6079 *->VbhA+DOC+HTH-> N6-METHYLASE -> HEPN ->SbcC+SbcC->?-> PglZ -> SIGMA+OB-Fold+AAA-ATPase+Lon-Protease-BrxL ->?->SF1-SF2-Helicase+REase-> | 1173 | Bacteria>Terrabacteria_group>Bacillota>Clostridia>Eubacteriales>Lachnospiraceae | GCA_024699435.1 | Lachnospiraceae_bacterium | K6G24_08610 | 2669..6190 | 1898203 | BREX-system-P-loop-protein-BrxC-[Lachnospiraceae-bacterium] |
| SCW50280.1 | Type1.BREX | Arm-DNA-bind_4+Phage_int_SAM_3+Phage_integrase->TraJ-RHH->ParA-Soj-PloopNTPase->ParB->?->HTH_26-> BrxA-DNA-binding -> BrxB-BrxF-iSTAND -> DUF6079 *-> N6-METHYLASE ->?-> PglZ -> HEPN -> SIGMA+OB-Fold+AAA-ATPase+Lon-Protease-BrxL ->Peptidase_C14-> | 1181 | Bacteria>Terrabacteria_group>Bacillota>Clostridia>Eubacteriales>Lachnospiraceae | GCA_900100095.1 | Lachnospiraceae_bacterium_C10 | SAMN02910400_01008 | 22627..26172 | 1520822 | hypothetical-protein-SAMN02910400_01008-[Lachnospiraceae-bacterium-C10] |
| SNU04610.1 | Type1.BREX | HTH_3->?->?->35exo->SbcC->?-> wHTH+WYL+WCX -> BrxA-DNA-binding -> BrxB-BrxF-iSTAND -> DUF6079 *->KAP-NTPase-> N6-METHYLASE -> HEPN -> PglZ -> HEPN +SH3-like+SF1-SF2-Helicase+wHTH->MORC+DUF3684+HNH->MORC+DUF3684+HNH->SF1-SF2-Helicase->SF1-SF2-Helicase+REase-> | 1206 | Bacteria>Terrabacteria_group>Bacillota>Clostridia>Eubacteriales>Lachnospiraceae | GCA_900199595.1 | Lachnospiraceae_bacterium | SAMN06297422_101152 | 246659..250279 | 1898203 | hypothetical-protein-SAMN06297422_101152-[Lachnospiraceae-bacterium] |
| WP_009536434.1 | Type1.BREX | <-SF1-SF2-Helicase<-ParB<-SbcC<- SIGMA+OB-Fold+AAA-ATPase+Lon-Protease-BrxL <- PglZ <-Schlafen+HATPase+wHTH<- N6-METHYLASE -p<- HEPN <- N6-METHYLASE <- DUF6079 *<- BrxB-BrxF-iSTAND <- BrxA-DNA-binding <-HTH_26<-HTH+rve<-Sigma-HTH<-TIR<-Peptidase_C14<-TIR\|\|DEDD_Tnp_IS110-> | 1217 | Bacteria>Terrabacteria_group>Bacillota>Clostridia>Eubacteriales>Lachnospiraceae | GCF_000238075.1 | Oribacterium_asaccharolyticum | HMPREF9624_00544 | 316049..319702 | 1501332 | BREX-system-P-loop-protein-BrxC-[Oribacterium-asaccharolyticum] |
| WP_002578404.1 | Type1.BREX | HTH-><-SF1-SF2-Helicase<-OLD-ABC+TOPRIM<- SIGMA+OB-Fold+AAA-ATPase+Lon-Protease-BrxL <- PglZ <-Schlafen+HATPase+wHTH<- HEPN <- N6-METHYLASE <-?<- DUF6079 *<- BrxB-BrxF-iSTAND <- BrxA-DNA-binding <-HTH_26<-HTH_36<-TraI-VirD2-RCRE<-TraJ-RHH<-CopG/RHH<-?<-HTH_36 | 1182 | Bacteria>Terrabacteria_group>Bacillota>Clostridia>Eubacteriales>Lachnospiraceae | GCF_000371725.1 | Enterocloster_bolteae | HMPREF1095_RS19895 | 1377442..1380990 | 208479 | BREX-system-P-loop-protein-BrxC-[Enterocloster-bolteae] |
| WP_016216871.1 | Type1.BREX | <-HTH\|\|\|\|HTH->\|\|HTH_26-> BrxA-DNA-binding -> BrxB-BrxF-iSTAND -> DUF6079 *-><-DEDD_Tnp_IS110+Transposase_20\|\| HEPN -> N6-METHYLASE ->HTH+REase->Schlafen+HATPase+wHTH-> PglZ -> | 1182 | Bacteria>Terrabacteria_group>Bacillota>Clostridia>Eubacteriales>Lachnospiraceae | GCF_000403455.2 | Dorea_sp._5-2 | C817_RS00440 | 102069..105617 | 1235798 | BREX-system-P-loop-protein-BrxC-[Dorea-sp.-5-2] |
| WP_026519395.1 | Type1.BREX | <-LRR_5<-BrxL<- SIGMA+OB-Fold+AAA-ATPase+Lon-Protease-BrxL <-PHP+SbcC<- PglZ <- HEPN <- N6-METHYLASE <- DUF6079 *<- BrxB-BrxF-iSTAND <- BrxA-DNA-binding <-HTH_26<-?<-?<-?<-TraJ-RHH<-LPD2<-ZnR+TOPRIM | 1194 | Bacteria>Terrabacteria_group>Bacillota>Clostridia>Eubacteriales>Lachnospiraceae | GCF_000424525.1 | Butyrivibrio_sp._FCS006 | G633_RS0106250 | 43812..47396 | 1280684 | BREX-system-P-loop-protein-BrxC-[Butyrivibrio-sp.-FCS006] |
| WP_029067206.1 | Type1.BREX | HTH_26-> BrxA-DNA-binding -> BrxB-BrxF-iSTAND -> DUF6079 *-> N6-METHYLASE -> PglZ ->ParB+HNH-> SIGMA+OB-Fold+AAA-ATPase+Lon-Protease-BrxL ->RVT-> HEPN ->HTH_38->HTH_26-> BrxA-DNA-binding -> BrxB-BrxF-iSTAND -> | 1182 | Bacteria>Terrabacteria_group>Bacillota>Clostridia>Eubacteriales>Lachnospiraceae | GCF_000687515.1 | Lachnobacterium_bovis | CC91_RS0108435 | 378442..381990 | 140626 | BREX-system-P-loop-protein-BrxC-[Lachnobacterium-bovis] |
| WP_055216182.1 | Type1.BREX | <-PIN<-ClpABC-AAA<-?<- SIGMA+OB-Fold+AAA-ATPase+Lon-Protease-BrxL <- PglZ <- HEPN <- N6-METHYLASE <- DUF6079 *<- BrxB-BrxF-iSTAND <- BrxA-DNA-binding <-HTH_26<-?<-?<-?<-ArdCN+LPD32+LPD36+N6_Mtase+SF1-SF2-Helicase | 1186 | Bacteria>Terrabacteria_group>Bacillota>Clostridia>Eubacteriales>Lachnospiraceae | GCF_001404435.1 | Lachnospira_eligens | ERS852490_02366 | 200381..203941 | 39485 | BREX-system-P-loop-protein-BrxC-[Lachnospira-eligens] |
| WP_118348289.1 | Type1.BREX | <- SIGMA+OB-Fold+AAA-ATPase+Lon-Protease-BrxL <- PglZ <-Schlafen+HATPase+wHTH<- HEPN <- N6-METHYLASE <-?<- DUF6079 *<- BrxB-BrxF-iSTAND <- BrxA-DNA-binding <-HTH_26<-?<-TraI-VirD2-RCRE<-TraJ-RHH<-?<-CopG/RHH<-HTH_36 | 1182 | Bacteria>Terrabacteria_group>Bacillota>Clostridia>Eubacteriales>Lachnospiraceae | GCF_003458885.1 | Coprococcus_sp._AF21-14LB | DWX73_RS07815 | 11153..14701 | 2292231 | BREX-system-P-loop-protein-BrxC-[Coprococcus-sp.-AF21-14LB] |
| WP_118051545.1 | Type1.BREX | <-Phage_int_SAM_5+Phage_integrase\|\|TraJ-RHH->?-><-?\|\|HTH_26-> BrxA-DNA-binding -> BrxB-BrxF-iSTAND -> DUF6079 *->?-> N6-METHYLASE -> HEPN ->Schlafen+HATPase+wHTH-> PglZ ->PHP+SbcC-> | 1183 | Bacteria>Terrabacteria_group>Bacillota>Clostridia>Eubacteriales>Lachnospiraceae | GCF_003464645.1 | Blautia_obeum | DWW07_RS08215 | 33645..37196 | 40520 | BREX-system-P-loop-protein-BrxC-[Blautia-obeum] |
| WP_118592376.1 | Type1.BREX | <-Schlafen+HATPase+wHTH-p<-ClpABC-AAA<-?<- SIGMA+OB-Fold+AAA-ATPase+Lon-Protease-BrxL <- PglZ <- HEPN <- N6-METHYLASE <- DUF6079 *<- BrxB-BrxF-iSTAND <- BrxA-DNA-binding <-HTH_26<-?<-?<-?<-?<-SF1-SF2-Helicase<-N6_N4_METHYLASE | 1186 | Bacteria>Terrabacteria_group>Bacillota>Clostridia>Eubacteriales>Lachnospiraceae | GCF_003467035.1 | Roseburia_intestinalis | DW927_RS18160 | 14462..18022 | 166486 | BREX-system-P-loop-protein-BrxC-[Roseburia-intestinalis] |
| WP_118198114.1 | Type1.BREX | <- SIGMA+OB-Fold+AAA-ATPase+Lon-Protease-BrxL <-?<-PHP<- PglZ <-Schlafen+HATPase+wHTH<- HEPN <- N6-METHYLASE <-?<- DUF6079 *<- BrxB-BrxF-iSTAND <- BrxA-DNA-binding <-HTH_26\|\|?-><-?<-TraJ-RHH\|\|Phage_int_SAM_5+Phage_integrase-> | 1183 | Bacteria>Terrabacteria_group>Bacillota>Clostridia>Eubacteriales>Lachnospiraceae | GCF_003471125.1 | Blautia_obeum | DW272_RS15845 | 24197..27748 | 40520 | BREX-system-P-loop-protein-BrxC-[Blautia-obeum] |
| WP_118742111.1 | Type1.BREX | ZnR->?->HTH_26-> BrxA-DNA-binding -> BrxB-BrxF-iSTAND -> DUF6079 *-> N6-METHYLASE ->RAMA-> HEPN ->Schlafen+HATPase+wHTH-> PglZ -> SIGMA+OB-Fold+AAA-ATPase+Lon-Protease-BrxL ->OLD-ABC+TOPRIM->SF1-SF2-Helicase-> | 1182 | Bacteria>Terrabacteria_group>Bacillota>Clostridia>Eubacteriales>Lachnospiraceae | GCF_003480245.1 | Blautia_sp._AM42-2 | DW915_10705 | 4262..7810 | 2292976 | BREX-system-P-loop-protein-BrxC-[Blautia-sp.-AM42-2] |
| WP_154780312.1 | Type1.BREX | Peptidase->?->?->MACRODOMAIN->?->ZnR+ZnR+ZnR->HTH_26-> BrxA-DNA-binding -> BrxB-BrxF-iSTAND -> DUF6079 *-> N6-METHYLASE ->RVT->?->OLD-ABC->HNH->?-> PglZ -> HEPN -> SIGMA+OB-Fold+AAA-ATPase+Lon-Protease-BrxL -> | 1201 | Bacteria>Terrabacteria_group>Bacillota>Clostridia>Eubacteriales>Lachnospiraceae | GCF_009707925.1 | Blautia_luti | GKZ57_RS08760 | 74022..77627 | 89014 | BREX-system-P-loop-protein-BrxC-[Blautia-luti] |
| WP_249326641.1 | Type1.BREX | GNTR-HTH+Aminotran_1_2->?->HTH_26-> BrxA-DNA-binding -> BrxB-BrxF-iSTAND -> DUF6079 *->Tox-ART-HYD2->AT-HTH-HYD2-1-> N6-METHYLASE -> HEPN ->Schlafen+HATPase+wHTH-> PglZ -> SIGMA+OB-Fold+AAA-ATPase+Lon-Protease-BrxL ->?->RNase-> | 1215 | Bacteria>Terrabacteria_group>Bacillota>Clostridia>Eubacteriales>Lachnospiraceae | GCF_014337175.1 | Simiaoa_sunii | H9Q77_RS04465 | 956158..959805 | 2763672 | BREX-system-P-loop-protein-BrxC-[Simiaoa-sunii] |
| WP_128132381.1 | Type1.BREX | MerR-HTH-><-?<-REase<-?<- SIGMA+OB-Fold+AAA-ATPase+Lon-Protease-BrxL <- PglZ <- HEPN <- N6-METHYLASE <- DUF6079 *<- BrxB-BrxF-iSTAND <- BrxA-DNA-binding <-HTH_26<-HTH\|\|?-><-?<-TraJ-RHH<-SF1-SF2-Helicase | 1182 | Bacteria>Terrabacteria_group>Bacillota>Clostridia>Eubacteriales>Lachnospiraceae | GCF_015547745.1 | Enterocloster_bolteae | I2D45_RS23920 | 73790..77338 | 208479 | BREX-system-P-loop-protein-BrxC-[Enterocloster-bolteae] |
| WP_195251421.1 | Type1.BREX | MACRODOMAIN->?->35exo+ADA-DBD->HTH_26-> BrxA-DNA-binding -> BrxB-BrxF-iSTAND -> DUF6079 *-> N6-METHYLASE ->ParB+HNH-> HEPN -> | 1201 | Bacteria>Terrabacteria_group>Bacillota>Clostridia>Eubacteriales>Lachnospiraceae | GCF_015550485.1 | Coprococcus_comes | I2A60_RS13430 | 4839..8444 | 410072 | BREX-system-P-loop-protein-BrxC-[Coprococcus-comes] |
| WP_215726767.1 | Type1.BREX | ParB->?-><-?\|\|HTH_3->HTH_26-> BrxA-DNA-binding -> BrxB-BrxF-iSTAND -> DUF6079 *->?-> N6-METHYLASE -> HEPN +ZNR-> HEPN -> PglZ ->?->?-> SIGMA+OB-Fold+AAA-ATPase+Lon-Protease-BrxL -> wHTH+WYL+WCX ->HTH+REase-> | 1205 | Bacteria>Terrabacteria_group>Bacillota>Clostridia>Eubacteriales>Lachnospiraceae | GCF_018784865.1 | Dorea_longicatena | GPK48_RS02600 | 18956..22573 | 88431 | BREX-system-P-loop-protein-BrxC-[Dorea-longicatena] |
| WP_252210442.1 | Type1.BREX | <-TIR<- SIGMA+OB-Fold+AAA-ATPase+Lon-Protease-BrxL <- PglZ <-Schlafen+HATPase+wHTH<- HEPN <-AT-HTH-HYD2<-Tox-ART-HYD2<-AT-HTH-HYD2<-SIR2<- DUF6079 *<- BrxB-BrxF-iSTAND <- BrxA-DNA-binding <-HTH_26<-HTH_36<-ABhydrolase<-?<-SAM-METHYLASE | 1099 | Bacteria>Terrabacteria_group>Bacillota>Clostridia>Eubacteriales>Lachnospiraceae | GCF_018785625.1 | Blautia_faecis | GPK86_RS03045 | 13256..16555 | 871665 | BREX-system-P-loop-protein-BrxC-[Blautia-faecis] |
| WP_216429335.1 | Type1.BREX | HTH_26-> BrxA-DNA-binding -> BrxB-BrxF-iSTAND -> DUF6079 *-> N6-METHYLASE ->RAMA-> HEPN -> PglZ -> SIGMA+OB-Fold+AAA-ATPase+Lon-Protease-BrxL ->?->REase->?-><-HTH_3 | 1182 | Bacteria>Terrabacteria_group>Bacillota>Clostridia>Eubacteriales>Lachnospiraceae | GCF_018918145.1 | Anaerotignum_sp._MSJ-24 | KQI66_RS10650 | 65878..69426 | 2841521 | BREX-system-P-loop-protein-BrxC-[Anaerotignum-sp.-MSJ-24] |
| WP_265190723.1 | Type1.BREX | <-ClpABN-AAA+ClpABC-AAA\|\|HTH-><-MULE-TRANSPOSASE<-PHP+SbcC<- SIGMA+OB-Fold+AAA-ATPase+Lon-Protease-BrxL <- PglZ <-Schlafen+HATPase+wHTH<- HEPN <- N6-METHYLASE <- DUF6079 *<- BrxB-BrxF-iSTAND <- BrxA-DNA-binding <-HTH_26<-HTH_36 | 1182 | Bacteria>Terrabacteria_group>Bacillota>Clostridia>Eubacteriales>Lachnospiraceae | GCF_020553125.1 | Blautia_sp._MSK20_18 | LI091_RS01865 | 117354..120902 | 2883186 | BREX-system-P-loop-protein-BrxC-[Blautia-sp.-MSK20_18] |
| WP_227165162.1 | Type1.BREX | MACRODOMAIN->?->35exo+ADA-DBD->ZnR+ZnR+ZnR->HTH_26-> BrxA-DNA-binding -> BrxB-BrxF-iSTAND -> DUF6079 *-> N6-METHYLASE -> HEPN -> PglZ ->OLD-ABC+TOPRIM-> SIGMA+OB-Fold+AAA-ATPase+Lon-Protease-BrxL ->LRR_5+LRR_8->?->STYKIN-> | 1201 | Bacteria>Terrabacteria_group>Bacillota>Clostridia>Eubacteriales>Lachnospiraceae | GCF_020560115.1 | Mediterraneibacter_sp._210702-DFI.5.30 | LI126_RS08100 | 75769..79374 | 2883232 | BREX-system-P-loop-protein-BrxC-[Mediterraneibacter-sp.-210702-DFI.5.30] |
| WP_158358547.1 | Type1.BREX | <- SIGMA+OB-Fold+AAA-ATPase+Lon-Protease-BrxL <- PglZ <-Schlafen+HATPase+wHTH<- HEPN + HEPN <-ParB+HNH<-AT-HTH-HYD2<-Tox-ART-HYD2<-AT-HTH-HYD2<- N6-METHYLASE <- DUF6079 *<- BrxB-BrxF-iSTAND <- BrxA-DNA-binding <-HTH_26<-?<-?<-?<-DOC+HTH<-HTH_3<-PDZ_6+Peptidase_S55 | 1182 | Bacteria>Terrabacteria_group>Bacillota>Clostridia>Eubacteriales>Lachnospiraceae | GCF_025567195.1 | Alitiscatomonas_aceti | OCV69_RS08300 | 70137..73685 | 2981724 | BREX-system-P-loop-protein-BrxC-[Alitiscatomonas-aceti] |
| WP_270422341.1 | Type1.BREX | CopG/RHH->?->TraJ-RHH->TraI-VirD2-RCRE->?->HTH_26-> BrxA-DNA-binding -> BrxB-BrxF-iSTAND -> DUF6079 *->?-> N6-METHYLASE -> HEPN ->Schlafen+HATPase+wHTH-> PglZ -> SIGMA+OB-Fold+AAA-ATPase+Lon-Protease-BrxL ->OLD-ABC+TOPRIM->SF1-SF2-Helicase-><-HTH | 1182 | Bacteria>Terrabacteria_group>Bacillota>Clostridia>Eubacteriales>Lachnospiraceae | GCF_027661365.1 | Fusicatenibacter_saccharivorans | PGQ16_RS03035 | 254369..257917 | 1150298 | BREX-system-P-loop-protein-BrxC-[Fusicatenibacter-saccharivorans] |
| WP_119247667.1 | Type1.BREX | HTH_36->?->CopG/RHH->TraJ-RHH->TraI-VirD2-RCRE->HTH_26-> BrxA-DNA-binding -> BrxB-BrxF-iSTAND -> DUF6079 *->?-> N6-METHYLASE -> HEPN ->Schlafen+HATPase+wHTH-> PglZ -> SIGMA+OB-Fold+AAA-ATPase+Lon-Protease-BrxL ->OLD-ABC+TOPRIM->SF1-SF2-Helicase-><-HTH | 1182 | Bacteria>Terrabacteria_group>Bacillota>Clostridia>Eubacteriales>Lachnospiraceae | GCF_027674485.1 | Dorea_longicatena | PGV47_RS07575 | 41898..45446 | 88431 | BREX-system-P-loop-protein-BrxC-[Dorea-longicatena] |
| WP_099385222.1 | Type1.BREX | <-Tox-RES<-?<-URI-DNase<- SIGMA+OB-Fold+AAA-ATPase+Lon-Protease-BrxL <-?<- PglZ <- HEPN <- N6-METHYLASE <- N6-METHYLASE <- DUF6079 *<- BrxB-BrxF-iSTAND <- BrxA-DNA-binding <-HTH_26<-ACET | 1207 | Bacteria>Terrabacteria_group>Bacillota>Clostridia>Eubacteriales>Lachnospiraceae | GCF_028539255.1 | Agathobacter_ruminis | CSX02_RS00500 | 90797..94420 | 1712665 | BREX-system-P-loop-protein-BrxC-[Agathobacter-ruminis] |
| WP_090167800.1 | Type1.BREX | <-HKD-DNase<-?<- SIGMA+OB-Fold+AAA-ATPase+Lon-Protease-BrxL <- BrxD-AAA <-?<- PglZ <- HEPN <- N6-METHYLASE <- N6-METHYLASE <- DUF6079 *<- BrxB-BrxF-iSTAND <- BrxA-DNA-binding <-HTH_26<-?<-?<- wHTH+WYL+WCX <-HNH<-?<-ARG | 1201 | Bacteria>Terrabacteria_group>Bacillota>Clostridia>Eubacteriales>Lachnospiraceae | GCF_900100545.1 | Pseudobutyrivibrio_sp._49 | BLP54_RS06100 | 185622..189227 | 1855344 | BREX-system-P-loop-protein-BrxC-[Pseudobutyrivibrio-sp.-49] |
| WP_090393802.1 | Type1.BREX | wHTH+WYL+WCX ->?->?->HKD-DNase->HTH_26-> BrxA-DNA-binding -> BrxB-BrxF-iSTAND -> DUF6079 *-> N6-METHYLASE ->?-> PglZ -> SIGMA+OB-Fold+AAA-ATPase+Lon-Protease-BrxL -> HEPN ->?->HKD-DNase->SF1-SF2-Helicase->REase-> | 1182 | Bacteria>Terrabacteria_group>Bacillota>Clostridia>Eubacteriales>Lachnospiraceae | GCF_900111635.1 | Pseudobutyrivibrio_sp._C4 | BMY66_RS00580 | 126012..129560 | 1520803 | BREX-system-P-loop-protein-BrxC-[Pseudobutyrivibrio-sp.-C4] |
| WP_090545210.1 | Type1.BREX | <-HKD-DNase<-?<- SIGMA+OB-Fold+AAA-ATPase+Lon-Protease-BrxL <- BrxD-AAA <-?<- PglZ <- HEPN <- N6-METHYLASE <- N6-METHYLASE <- DUF6079 *<- BrxB-BrxF-iSTAND <- BrxA-DNA-binding <-HTH_26<-?<-?<- wHTH+WYL+WCX <-HNH<-?<-ARG | 1201 | Bacteria>Terrabacteria_group>Bacillota>Clostridia>Eubacteriales>Lachnospiraceae | GCF_900113655.1 | Pseudobutyrivibrio_sp._OR37 | BM097_RS01535 | 134498..138103 | 1798186 | BREX-system-P-loop-protein-BrxC-[Pseudobutyrivibrio-sp.-OR37] |
| WP_072699981.1 | Type1.BREX | <- HEPN <-?<- SIGMA+OB-Fold+AAA-ATPase+Lon-Protease-BrxL <-TIR<- PglZ <-RAMA<- N6-METHYLASE <- DUF6079 *<- BrxB-BrxF-iSTAND <- BrxA-DNA-binding <-HTH_26<-?<-TraJ-RHH<-LPD2<-TOPRIM+Topoisom_bac<-?<-ArdCN+MPTase-inact+LPD19 | 1185 | Bacteria>Terrabacteria_group>Bacillota>Clostridia>Eubacteriales>Lachnospiraceae | GCF_900143205.1 | Butyrivibrio_hungatei | BUA02_RS00135 | 23142..26699 | 185008 | BREX-system-P-loop-protein-BrxC-[Butyrivibrio-hungatei] |
| WP_102289727.1 | Type1.BREX | <-TnpB_IS66\|\|?-><- SIGMA+OB-Fold+AAA-ATPase+Lon-Protease-BrxL <-Schlafen+HATPase+wHTH<- PglZ <- HEPN <- N6-METHYLASE <- DUF6079 *<- BrxB-BrxF-iSTAND <- BrxA-DNA-binding <-HTH_26<-?<-?<-?<-?<-?\|\|HAMP+HisKin+HATPase_c-> | 1215 | Bacteria>Terrabacteria_group>Bacillota>Clostridia>Eubacteriales>Lachnospiraceae | GCF_900243045.1 | Eisenbergiella_massiliensis | C1733_RS26410 | 2142071..2145718 | 1720294 | BREX-system-P-loop-protein-BrxC-[Eisenbergiella-massiliensis] |
| WP_055192172.1 | Type1.BREX | <-MORC+DUF3684+HNH<- HEPN +SH3-like+SF1-SF2-Helicase+wHTH<- PglZ <-Schlafen+HATPase+wHTH<- HEPN <-?<-?<- N6-METHYLASE <- DUF6079 *<- BrxB-BrxF-iSTAND <- BrxA-DNA-binding <-?<-35exo+wHTH+SF1-SF2-Helicase+HTH<-?<-35exo | 1204 | Bacteria>Terrabacteria_group>Bacillota>Clostridia>Eubacteriales>Oscillospiraceae | - | Faecalibacterium_prausnitzii | FXW87_RS13010 | 15894..19508 | 853 | BREX-system-P-loop-protein-BrxC-[Faecalibacterium-prausnitzii] |
| MCL2124490.1 | Type1.BREX | <-sGTP<-?\|\|?->?->?-> BrxA-DNA-binding -> BrxB-BrxF-iSTAND -> DUF6079 *-> N6-METHYLASE ->NERD+SF1-SF2-Helicase->OLD-ABC->HNH-> PolB -> HEPN -> PglZ -> SIGMA+OB-Fold+AAA-ATPase+Lon-Protease-BrxL -> | 1207 | Bacteria>Terrabacteria_group>Bacillota>Clostridia>Eubacteriales>Oscillospiraceae | GCA_009776905.1 | Oscillospiraceae_bacterium | FWH33_00685 | 36555..40178 | 2485925 | BREX-system-P-loop-protein-BrxC-[Oscillospiraceae-bacterium] |
| MBS7028836.1 | Type1.BREX | <-RVT<- SIGMA+OB-Fold+AAA-ATPase+Lon-Protease-BrxL <- PglZ <- HEPN <- N6-METHYLASE <-?<-?<- DUF6079 *<- BrxB-BrxF-iSTAND <- BrxA-DNA-binding <-DOC+HTH\|\|DUF932->?->?-><-CR-ATPase5+REase->?->CR-ATPase8+REase-> | 1182 | Bacteria>Terrabacteria_group>Bacillota>Clostridia>Eubacteriales>Oscillospiraceae | GCA_018381555.1 | Faecalibacterium_prausnitzii | KH132_10095 | 50879..54427 | 853 | BREX-system-P-loop-protein-BrxC-[Faecalibacterium-prausnitzii] |
| MCI6921905.1 | Type1.BREX | ABC-ATPase+SbcC+ABC-ATPase+SbcC->?->?->?->?->LRR_5-> HEPN -> BrxA-DNA-binding -> BrxB-BrxF-iSTAND -> DUF6079 *-> N6-METHYLASE -p-> | 1161 | Bacteria>Terrabacteria_group>Bacillota>Clostridia>Eubacteriales>Oscillospiraceae | GCA_022778185.1 | Oscillospiraceae_bacterium | MR756_00805 | 25286..28771 | 2485925 | BREX-system-P-loop-protein-BrxC-[Oscillospiraceae-bacterium] |
| UYJ36126.1 | Type1.BREX | TraJ-RHH-><-CR-ATPase5+REase\|\|TraI-VirD2-RCRE->?->?->?-> BrxA-DNA-binding -> BrxB-BrxF-iSTAND -> DUF6079 *-> N6-METHYLASE -> PglZ -> HEPN +SH3-like+SF1-SF2-Helicase+wHTH->MORC+DUF3684+HNH->?->Tox-RelA->HKD-DNase+HKD-DNase->HKD-DNase+HKD-DNase->REC+OmpR-HTH-> | 1205 | Bacteria>Terrabacteria_group>Bacillota>Clostridia>Eubacteriales>Oscillospiraceae | GCA_025757985.1 | Oscillospiraceae_bacterium | OGM67_07425 | 1479517..1483134 | 2485925 | MAG:-BREX-system-P-loop-protein-BrxC-[Oscillospiraceae-bacterium] |
| UYJ36609.1 | Type1.BREX | HTH-><-TIR<- SIGMA+OB-Fold+AAA-ATPase+Lon-Protease-BrxL <- PglZ <-Schlafen+HATPase+wHTH<- HEPN <- N6-METHYLASE <-?<- DUF6079 *<- BrxB-BrxF-iSTAND <- BrxA-DNA-binding <-HTH_26<-TraI-VirD2-RCRE<-TraJ-RHH<-?<-CopG/RHH<-?<-HTH_36 | 1182 | Bacteria>Terrabacteria_group>Bacillota>Clostridia>Eubacteriales>Oscillospiraceae | GCA_025758005.1 | Oscillospiraceae_bacterium | OGM10_10415 | 2214157..2217705 | 2485925 | MAG:-BREX-system-P-loop-protein-BrxC-[Oscillospiraceae-bacterium] |
| WP_044960386.1 | Type1.BREX | <-CR-ATPase8+REase<-?<-?<-DUF932\|\|DOC+HTH-> BrxA-DNA-binding -> BrxB-BrxF-iSTAND -> DUF6079 *->?->?-> N6-METHYLASE -> HEPN -> PglZ -> SIGMA+OB-Fold+AAA-ATPase+Lon-Protease-BrxL ->RVT->RVT-> | 1182 | Bacteria>Terrabacteria_group>Bacillota>Clostridia>Eubacteriales>Oscillospiraceae | GCF_000209855.1 | Faecalibacterium_prausnitzii | FPR_RS00845 | 148257..151805 | 853 | BREX-system-P-loop-protein-BrxC-[Faecalibacterium-prausnitzii] |
| WP_087369154.1 | Type1.BREX | <-GspA-ATPase<-SIR2<-?<-?<-?\|\| BrxA-DNA-binding -> BrxB-BrxF-iSTAND -> DUF6079 *-> N6-METHYLASE ->?-> HEPN -> PglZ -> SIGMA+OB-Fold+AAA-ATPase+Lon-Protease-BrxL -><-Lysozyme+NLPC<-VirB4-FtsK | 1178 | Bacteria>Terrabacteria_group>Bacillota>Clostridia>Eubacteriales>Oscillospiraceae | GCF_002161675.1 | Pseudoflavonifractor_sp._An44 | B5F98_RS00420 | 121430..124966 | 1965635 | BREX-system-P-loop-protein-BrxC-[Pseudoflavonifractor-sp.-An44] |
| WP_113992229.1 | Type1.BREX | <-CR-ATPase5+REase<-?<-?<-DUF932<-NUDIX<-HKD-DNase+SF1-SF2-Helicase+HTH+HTH+SRA\|\|XIS-HTH-> BrxA-DNA-binding -> BrxB-BrxF-iSTAND -> DUF6079 *->wHTH+ZnR->Four-helical-protein-> N6-METHYLASE -> HEPN -> PglZ -> SIGMA+OB-Fold+AAA-ATPase+Lon-Protease-BrxL ->ParB+HNH->HKD-DNase+SF1-SF2-Helicase+HTH+HTH+SRA->nSTAND-> | 1218 | Bacteria>Terrabacteria_group>Bacillota>Clostridia>Eubacteriales>Oscillospiraceae | GCF_003324185.1 | Faecalibacterium_prausnitzii | C7J97_RS04695 | 32898..36554 | 853 | BREX-system-P-loop-protein-BrxC-[Faecalibacterium-prausnitzii] |
| WP_158398728.1 | Type1.BREX | <-ParB+HNH<- SIGMA+OB-Fold+AAA-ATPase+Lon-Protease-BrxL <- PglZ <- HEPN <- N6-METHYLASE <-TIR<-?<- DUF6079 *<- BrxB-BrxF-iSTAND <- BrxA-DNA-binding <-HTH_3<-?<-RepA_N+HTH\|\|<-Peptidase_S51\|\|SIGMA-HTH+SIGMA-HTH->ncRNA->RFC-AAA+RuvB-AAA-> | 1182 | Bacteria>Terrabacteria_group>Bacillota>Clostridia>Eubacteriales>Oscillospiraceae | GCF_902167845.1 | Faecalibacterium_prausnitzii | FQ260_RS07280 | 12413..15961 | 853 | BREX-system-P-loop-protein-BrxC-[Faecalibacterium-prausnitzii] |
| WP_118142807.1 | Type1.BREX | wHTH+WYL+WCX ->?->?->?->HTH_26-> BrxA-DNA-binding -> BrxB-BrxF-iSTAND -> DUF6079 *->?-> N6-METHYLASE ->DUF4365+REase-> PglZ -> HEPN +SH3-like+SF1-SF2-Helicase+wHTH->MORC+DUF3684+HNH->SF1-SF2-Helicase->PIN->?->CHTH-> | 1203 | Bacteria>Terrabacteria_group>Bacillota>Clostridia>Eubacteriales>Oscillospiraceae | - | Ruminococcus_sp._OM07-7 | DXC06_07755 | 161101..164712 | 2293234 | BREX-system-P-loop-protein-BrxC-[Ruminococcus-sp.-OM07-7] |
| WP_066638545.1 | Type1.BREX | <-Glyoxalase<-?<- SIGMA+OB-Fold+AAA-ATPase+Lon-Protease-BrxL <- PglZ <- HEPN <- PolB <- N6-METHYLASE <- DUF6079 *<- BrxB-BrxF-iSTAND <- BrxA-DNA-binding <-?<-SLATT<-MazG<-NUDIX<-35exo<-35exo | 1184 | Bacteria>Terrabacteria_group>Bacillota>Clostridia>Eubacteriales>Peptococcaceae | - | Desulfolucianica_intricata | DIN01_RS10945 | 45737..49291 | 1285191 | BREX-system-P-loop-protein-BrxC-[Desulfolucianica-intricata] |
| WP_057552934.1 | Type1.BREX | TetR-HTH->?-><-?\|\| BrxA-DNA-binding -> BrxB-BrxF-iSTAND -> DUF6079 *-> N6-METHYLASE ->Preader17-> HEPN ->?->?->OLD-ABC-> HEPN -> PglZ -> SIGMA+OB-Fold+AAA-ATPase+Lon-Protease-BrxL -> | 1185 | Bacteria>Terrabacteria_group>Bacillota>Clostridia>Eubacteriales>Peptostreptococcaceae | GCF_001372915.1 | Paeniclostridium_sordellii | SSCC32135_RS03600 | 682084..685641 | 1505 | BREX-system-P-loop-protein-BrxC-[Paeniclostridium-sordellii] |

| Protein-ID | System | Gene-Neighbourhoods | Length | Taxonomy | Assembly-ID | Strain | Locus-Tag | Genomic-coordinates | TaxID | Fasta-define |
|---|---|---|---|---|---|---|---|---|---|---|
| -- | | | | | | | | | | |
| --HKD-DNase (BrxL-Fusion) | | | | | | | | | | |
| -- | | | | | | | | | | |
| BDC48932.1 | Type6.BREX | <-HKD-DNase+HKD-Rease<-?<-SIGMA+OB-Fold+AAA-ATPase+HKD-DNase-BrxL<-PglZ<-N6-METHYLASE<-DUF6079*<-BrxB-BrxF-iSTAND<-BrxA-DNA-binding<-BrxE<-NACHT+GTPase-AIG+sGTP<-NTP_transf_8<-REase+HTH | 1202 | Bacteria>Acidobacteriota>Terriglobia>Bryobacterales>unclassified_Bryobacterales | GCA_016865505.2 | Bryobacterales_bacterium_F-183 | F183_A12480 | 1423409..1427017 | 2802972 | hypothetical-protein-F183_A12480-[Bryobacterales-bacterium-F-183] |
| NLV30311.1 | Type1.BREX | DNA-glycosylase-><-?<-SH3\|\|wHTH+WYL+WCX->SF1-SF2-Helicase->BrxA-DNA-binding->BrxB-BrxF-iSTAND->DUF6079*->Schlafen+HATPase+wHTH+HTH->N6-METHYLASE->PglZ->SIGMA+OB-Fold+AAA-ATPase+HKD-DNase-BrxL->HigB_toxin->HTH+MPTase->SF1-SF2-Helicase->?-><-DSBA+Thioredoxin | 1169 | Bacteria>Acidobacteriota>unclassified_Acidobacteriota | GCA_012729555.1 | Acidobacteriota_bacterium | GXY47_04070 | 27594..31103 | 1978231 | BREX-system-P-loop-protein-BrxC-[Acidobacteriota-bacterium] |
| MBS1808657.1 | Type6.BREX | <-SF1-SF2-Helicase+DUF1998<-?<-35exo\|\|NtrC-AAA+FIS-HTH-><-SIGMA+OB-Fold+AAA-ATPase+HKD-DNase-BrxL<-PglZ<-N6-METHYLASE<-DUF6079*<-BrxB-BrxF-iSTAND<-BrxA-DNA-binding<-BrxE\|\|?->Cysteine-Desulfurase-><-Peptidase | 1212 | Bacteria>Acidobacteriota>unclassified_Acidobacteriota | GCA_018268995.1 | Acidobacteriota_bacterium | JST84_10740 | 129628..133266 | 1978231 | BREX-system-P-loop-protein-BrxC-[Acidobacteriota-bacterium] |
| MCK9474444.1 | Type1.BREX | BrxA-DNA-binding->BrxB-BrxF-iSTAND->DUF6079*->OLD-ABC->NERD->N6-METHYLASE->PglZ->SIGMA+OB-Fold+AAA-ATPase+HKD-DNase-BrxL->?->ZnR->?->PadR-HTH-> | 1170 | Bacteria>Bacteria_incertae_sedis>Bacteria_candidate_phyla>Candidatus_Muirbacteria>Candidatus_Muirbacterium | GCA_023228995.1 | Candidatus_Muirbacterium_halophilum | MOR46_00860 | 28955..32467 | 2053465 | BREX-system-P-loop-protein-BrxC-[Candidatus-Muirbacterium-halophilum] |
| PKL50472.1 | Type1.BREX | <-MPTase<-HTH_19<-?\|\|BrxA-DNA-binding->BrxB-BrxF-iSTAND->DUF6079*->Schlafen+HATPase+wHTH+HTH->OLD-ABC->REase->N6-METHYLASE->PglZ->SIGMA+OB-Fold+AAA-ATPase+HKD-DNase-BrxL-> | 1185 | Bacteria>Bacteria_incertae_sedis>Bacteria_candidate_phyla>Candidatus_Riflebacteria>unclassified_Candidatus_Riflebacteria | GCA_002839435.1 | Candidatus_Riflebacteria_bacterium_HGW-Riflebacteria-2 | CVV42_02960 | 62937..66494 | 2013830 | MAG:-BREX-system-P-loop-protein-BrxC-[Candidatus-Riflebacteria-bacterium-HGW-Riflebacteria-2] |
| NUM59087.1 | Type1.BREX | Phage_integrase->Phage_integrase->Phage_int_SAM_1+Phage_integrase->?-><-?<-SbcC+SbcC<-SIGMA+OB-Fold+AAA-ATPase+HKD-DNase-BrxL<-PglZ<-N6-METHYLASE<-DUF6079*<-BrxB-BrxF-iSTAND<-BrxA-DNA-binding<-XIS-HTH<-?<-?<-ABhydrolase<-REC+NtrC-AAA+FIS-HTH<-HISKIN+MORC | 1157 | Bacteria>Bdellovibrionota>Bdellovibrionia>Bdellovibrionales>Pseudobdellovibrionaceae>unclassified_Pseudobdellovibrionaceae | GCA_013359135.1 | Pseudobdellovibrionaceae_bacterium | HUU56_10670 | 15517..18990 | 2026715 | BREX-system-P-loop-protein-BrxC-[Pseudobdellovibrionaceae-bacterium] |
| PIT99446.1 | Type1.BREX | <-ABhydrolase<-REC+NtrC-AAA+FIS-HTH<-HISKIN+MORC<-?<-?\|\|XIS-HTH->BrxA-DNA-binding->BrxB-BrxF-iSTAND->DUF6079*->N6-METHYLASE->PglZ->SIGMA+OB-Fold+AAA-ATPase+HKD-DNase-BrxL->SIR2-><-MuB-ATPase<-HTH_28+HTH_32+rve+rve_3 | 1164 | Bacteria>Bdellovibrionota>Bdellovibrionia>Bdellovibrionales>unclassified_Bdellovibrionales | GCA_002778785.1 | Bdellovibrionales_bacterium_CG10_big_fil_rev_8_21_14_0_10_45_34 | COT74_10625 | 394529..398023 | 1973927 | BREX-system-P-loop-protein-BrxC-[Bdellovibrionales-bacterium-CG10_big_fil_rev_8_21_14_0_10_45_34] |
| KAA3613611.1 | Type1.BREX | <-PIN\|\|?->?->BrxA-DNA-binding->BrxB-BrxF-iSTAND->DUF6079*->N6-METHYLASE->?->PglZ->SIGMA+OB-Fold+AAA-ATPase+HKD-DNase-BrxL->REase->CARF+NtrC-AAA+FIS-HTH->REase-> | 1159 | Bacteria>Calditrichota>unclassified_Calditrichota | GCA_008501795.1 | Calditrichota_bacterium | DWQ03_12295 | 106728..110207 | 2212469 | BREX-system-P-loop-protein-BrxC-[Calditrichota-bacterium] |
| WP_187475233.1 | Type1.BREX | METHYLASE->Peptidase_M3->?->?-><-DDE_Tnp_1_4\|\|?->XIS-HTH->BrxA-DNA-binding->BrxB-BrxF-iSTAND->DUF6079*->OLD-ABC->REase->N6-METHYLASE->DrHyd->Schlafen+HATPase+wHTH->Trypsin->?->PglZ->SIGMA+OB-Fold+AAA-ATPase+HKD-DNase-BrxL-> | 1166 | Bacteria>Campylobacterota>Epsilonproteobacteria>Campylobacterales>Arcobacteraceae>Aliarcobacter | GCF_014352915.1 | Aliarcobacter_cryaerophilus | HOD34_RS05640 | 1060903..1064403 | 28198 | BREX-system-P-loop-protein-BrxC-[Aliarcobacter-cryaerophilus] |
| WP_260896239.1 | Type1.BREX | Peptidase->Phage_integrase->?->XIS-HTH->BrxA-DNA-binding->BrxB-BrxF-iSTAND->DUF6079*->?->N6-METHYLASE->DrHyd->Schlafen+HATPase+wHTH->REase->ACET->PglZ->SIGMA+OB-Fold+AAA-ATPase+HKD-DNase-BrxL-> | 1166 | Bacteria>Campylobacterota>Epsilonproteobacteria>Campylobacterales>Arcobacteraceae>Aliarcobacter | GCF_025335025.1 | Aliarcobacter_cryaerophilus | N5T57_RS09580 | 13633..17133 | 28198 | BREX-system-P-loop-protein-BrxC-[Aliarcobacter-cryaerophilus] |
| TDB40153.1 | Type1.BREX | Phage_integrase->SIR2-><-Thermonuclease<-?<-LexA-protease\|\|?->BrxA-DNA-binding->BrxB-BrxF-iSTAND->DUF6079*->Schlafen+HATPase+wHTH+HTH->N6-METHYLASE->REase->PglZ->SIGMA+OB-Fold+AAA-ATPase+HKD-DNase-BrxL-><-?<-?<-35exo<-Calcineurin | 1180 | Bacteria>delta/epsilon_subdivisions>Deltaproteobacteria>unclassified_Deltaproteobacteria | GCA_004348915.1 | Deltaproteobacteria_bacterium | D9V46_00535 | 112112..115654 | 2026735 | BREX-system-P-loop-protein-BrxC-[Deltaproteobacteria-bacterium] |
| MBC8466717.1 | Type1.BREX | <-RAMA<-SIGMA+OB-Fold+AAA-ATPase+HKD-DNase-BrxL<-PglZ<-N6-METHYLASE<-REase<-OLD-ABC<-DUF6079*<-BrxB-BrxF-iSTAND<-BrxA-DNA-binding | 1172 | Bacteria>delta/epsilon_subdivisions>Deltaproteobacteria>unclassified_Deltaproteobacteria | GCA_014384065.1 | Deltaproteobacteria_bacterium | H8D55_02650 | 10326..13844 | 2026735 | BREX-system-P-loop-protein-BrxC-[Deltaproteobacteria-bacterium] |
| MCC6503799.1 | Type1.BREX | <-SbcC<-Calcineurin+Calcineurin-SbcD<-SbcC+APATPase+SbcC<-SIGMA+OB-Fold+AAA-ATPase+HKD-DNase-BrxL<-PglZ<-N6-METHYLASE<-REase<-OLD-ABC<-DUF6079*<-BrxB-BrxF-iSTAND<-BrxA-DNA-binding<-XIS-HTH | 1181 | Bacteria>delta/epsilon_subdivisions>Deltaproteobacteria>unclassified_Deltaproteobacteria | GCA_020846435.1 | Deltaproteobacteria_bacterium | IT362_11790 | 16699..20244 | 2026735 | BREX-system-P-loop-protein-BrxC-[Deltaproteobacteria-bacterium] |
| MCP4158880.1 | Type1.BREX | <-CARF<-HKD-DNase+HKD-DNase<-?<-S1+SF1-SF2-Helicase+S1<-SIGMA+OB-Fold+AAA-ATPase+HKD-DNase-BrxL<-PglZ<-Couple_hipA+HipA_C<-HTH_3<-N6-METHYLASE<-DUF6079* | 1006 | Bacteria>delta/epsilon_subdivisions>Deltaproteobacteria>unclassified_Deltaproteobacteria | GCA_024224625.1 | Deltaproteobacteria_bacterium | GY760_02295 | 25697..28718 | 2026735 | BREX-system-P-loop-protein-BrxC-[Deltaproteobacteria-bacterium] |
| BAH89618.1 | Type6.BREX | CHTH-><-?\|\|?->ZnR+TOPRIM+AAA-helicase+wHTH-><-?\|\|BrxE->BrxA-DNA-binding->BrxB-BrxF-iSTAND->DUF6079*->GreA_GreB+SF1-SF2-Helicase->N6-METHYLASE->N6-METHYLASE->DOC+HTH->PglZ->SIGMA+OB-Fold+AAA-ATPase+HKD-DNase-BrxL->?->?->RADICAL-SAM-> | 1211 | Bacteria>environmental_samples | - | uncultured_bacterium | . | 12703..16338 | 77133 | ATPase-[uncultured-bacterium] |
| MCS4056789.1 | Type1.BREX | <-Calcineurin-SbcD<-SIGMA+OB-Fold+AAA-ATPase+HKD-DNase-BrxL<-PglZ<-HEPN<-PolB\|\|?-><-?<-N6-METHYLASE<-DOC+HTH<-DUF6079*<-BrxB-BrxF-iSTAND<-BrxA-DNA-binding<-?<-viral-protein<-?<-?\|\|?-><-RHH | 1165 | Bacteria>FCB_group>Bacteroidetes/Chlorobi_group>Bacteroidota>Bacteroidetes_incertae_sedis>Rhodothermaceae | GCA_024809315.1 | Salinibacter_ruber | GGP75_002131 | 31231..34728 | 146919 | hypothetical-protein-[Salinibacter-ruber] |
| WP_119843039.1 | Type1.BREX | <-HD<-SbcC+SbcC<-Calcineurin-SbcD<-HORMA<-SIGMA+OB-Fold+AAA-ATPase+HKD-DNase-BrxL<-PglZ<-N6-METHYLASE<-DUF6079*<-BrxB-BrxF-iSTAND<-BrxA-DNA-binding<-REase+RNase<-viral-protein\|\|?-><-Phage_holin_2_4<-?<-PAS+PAS+PAS+GAF+HISKIN | 1201 | Bacteria>FCB_group>Bacteroidetes/Chlorobi_group>Bacteroidota>Bacteroidetes_incertae_sedis>Rhodothermaceae | GCF_002290345.1 | Salinibacter_ruber | B7W84_RS15115 | 76596..80201 | 146919 | BREX-system-P-loop-protein-BrxC-[Salinibacter-ruber] |
| WP_259091014.1 | Type6.BREX | SF1-SF2-Helicase+REase-><-HORMA\|\|BrxE->BrxA-DNA-binding->BrxB-BrxF-iSTAND->DUF6079*->N6-METHYLASE->PglZ->SIGMA+OB-Fold+AAA-ATPase+HKD-DNase-BrxL->MULE-TRANSPOSASE->HORMA->Calcineurin-SbcD->SbcC+SbcC->HD->HNH-> | 1192 | Bacteria>FCB_group>Bacteroidetes/Chlorobi_group>Bacteroidota>Bacteroidetes_incertae_sedis>Rhodothermaceae | GCF_024807395.1 | Salinibacter_ruber | GGP54_RS13575 | 8828..12406 | 146919 | BREX-system-P-loop-protein-BrxC-[Salinibacter-ruber] |
| WP_259130005.1 | Type6.BREX | <-SbcC+SbcC<-Calcineurin-SbcD<-HORMA<-SIGMA+OB-Fold+AAA-ATPase+HKD-DNase-BrxL<-PglZ<-N6-METHYLASE<-DUF6079*<-BrxB-BrxF-iSTAND<-BrxA-DNA-binding<-BrxE<-REase+RNase<-viral-protein<-?<-?\|\|?-><-HKD-DNase+MED12 | 1199 | Bacteria>FCB_group>Bacteroidetes/Chlorobi_group>Bacteroidota>Bacteroidetes_incertae_sedis>Rhodothermaceae | GCF_024807455.1 | Salinibacter_ruber | GGQ04_003012 | 22826..26425 | 146919 | BREX-system-P-loop-protein-BrxC-[Salinibacter-ruber] |
| WP_259115052.1 | Type1.BREX | <-SbcC+SbcC<-Calcineurin-SbcD<-SIGMA+OB-Fold+AAA-ATPase+HKD-DNase-BrxL<-PglZ<-HEPN<-PolB<-?<-N6-METHYLASE<-DOC+HTH<-DUF6079*<-BrxB-BrxF-iSTAND<-BrxA-DNA-binding<-?<-viral-protein<-?<-?\|\|?-><-RHH\|\|HTH_3+Peptidase_S24-> | 1167 | Bacteria>FCB_group>Bacteroidetes/Chlorobi_group>Bacteroidota>Bacteroidetes_incertae_sedis>Rhodothermaceae | GCF_024809315.1 | Salinibacter_ruber | GGP75_RS10390 | 31231..34734 | 146919 | BREX-system-P-loop-protein-BrxC-[Salinibacter-ruber] |
| WP_259076695.1 | Type1.BREX | TraJ-RHH->RHH->Phage_holin_2_4->\|\|viral-protein->?->BrxA-DNA-binding->BrxB-BrxF-iSTAND->DUF6079*->DOC+HTH->N6-METHYLASE->?->?->PglZ->SIGMA+OB-Fold+AAA-ATPase+HKD-DNase-BrxL->Calcineurin-SbcD->SbcC+SbcC-> | 1167 | Bacteria>FCB_group>Bacteroidetes/Chlorobi_group>Bacteroidota>Bacteroidetes_incertae_sedis>Rhodothermaceae | GCF_024809595.1 | Salinibacter_ruber | GGP73_RS15155 | 19341..22844 | 146919 | BREX-system-P-loop-protein-BrxC-[Salinibacter-ruber] |
| WP_259087839.1 | Type1.BREX | URI-DNase-><-SIGMA+OB-Fold+AAA-ATPase+HKD-DNase-BrxL<-PglZ<-PIN<-XIS-HTH<-?<-?<-N6-METHYLASE<-DUF6079*<-BrxB-BrxF-iSTAND<-BrxA-DNA-binding<-?\|\|Phage_integrase-><-TOPRIM<-?\|\|XIS-HTH->Phage_int_SAM_4+Phage_integrase-><-tRNA\|\|BetaPropeller+ABhydrolase-> | 1168 | Bacteria>FCB_group>Bacteroidetes/Chlorobi_group>Bacteroidota>Bacteroidetes_incertae_sedis>Rhodothermaceae | GCF_024809725.1 | Salinibacter_ruber | GGQ07_RS07865 | 19343..22849 | 146919 | BREX-system-P-loop-protein-BrxC-[Salinibacter-ruber] |
| WP_259228204.1 | Type6.BREX | <-REC+OmpR-HTH<-PAS+PAS+PAS+GAF+HISKIN<-Phage_integrase<-?<-?<-?<-SIGMA+OB-Fold+AAA-ATPase+HKD-DNase-BrxL<-PglZ<-N6-METHYLASE<-DUF6079*<-BrxB-BrxF-iSTAND<-BrxA-DNA-binding<-BrxE\|\|?->?->?-><-viral-protein | 1201 | Bacteria>FCB_group>Bacteroidetes/Chlorobi_group>Bacteroidota>Bacteroidetes_incertae_sedis>Rhodothermaceae | GCF_024809755.1 | Salinibacter_ruber | GGP93_RS14280 | 71185..74790 | 146919 | BREX-system-P-loop-protein-BrxC-[Salinibacter-ruber] |
| WP_263820582.1 | Type1.BREX | <-SbcC+SbcC<-Calcineurin-SbcD<-SIGMA+OB-Fold+AAA-ATPase+HKD-DNase-BrxL<-PglZ<-?<-N6-METHYLASE<-DOC+HTH<-DUF6079*<-BrxB-BrxF-iSTAND<-BrxA-DNA-binding<-DUF4258<-REase+RNase<-viral-protein<-?\|\|CHTH-><-Phage_holin_2_4<-RHH | 1167 | Bacteria>FCB_group>Bacteroidetes/Chlorobi_group>Bacteroidota>Bacteroidetes_incertae_sedis>Rhodothermaceae | GCF_947077735.1 | Salinibacter_sp. | OJA43_RS08805 | 45420..48923 | 2065818 | BREX-system-P-loop-protein-BrxC-[Salinibacter-sp.] |
| WP_259248797.1 | Type1.BREX | <-SbcC+SbcC<-Calcineurin-SbcD<-SIGMA+OB-Fold+AAA-ATPase+HKD-DNase-BrxL<-PglZ<-CHTH<-RVT<-N6-METHYLASE<-DOC+HTH<-DUF6079*<-BrxB-BrxF-iSTAND<-BrxA-DNA-binding<-DUF4258<-REase+RNase<-HKD-DNase+SF1-SF2-Helicase<-RHH\|\|HTH_23+HTH_32+HTH_33->DDE_3-><-TraJ-RHH | 1167 | Bacteria>FCB_group>Bacteroidetes/Chlorobi_group>Bacteroidota>Bacteroidetes_incertae_sedis>Rhodothermaceae | - | Salinibacter_ruber | GGP55_RS14300 | 49306..52809 | 146919 | BREX-system-P-loop-protein-BrxC-[Salinibacter-ruber] |
| WP_117964426.1 | Type6.BREX | <-TRAM+METHYLASE\|\|?-><-UDG<-?<-ARG\|\|RHH+MobC->?-><-SIGMA+OB-Fold+AAA-ATPase+HKD-DNase-BrxL<-PglZ<-DUF6079-p*<-DUF6079-p<-BrxB-BrxF-iSTAND<-BrxA-DNA-binding<-BrxE<-HTH_3<-?\|\|<-XIS-HTH<-AbiEii\|\|HTH_3-> | 938 | Bacteria>FCB_group>Bacteroidetes/Chlorobi_group>Bacteroidota>Bacteroidia>Bacteroidales | - | Bacteroides_uniformis | IYV59_08305 | 23505..26321 | 820 | MULTISPECIES:-BREX-system-P-loop-protein-BrxC-[Bacteroides] |
| WP_118696414.1 | Type1.BREX | <-Arm-DNA-bind_5+Phage_int_SAM_5+Phage_integrase\|\|DUF4007->BrxB-BrxF-iSTAND->DUF6079-p->DUF6079-p*->N6-METHYLASE->PglZ->SIGMA+OB-Fold+AAA-ATPase+HKD-DNase-BrxL-><-?<-RHH+MobC\|\|ARG->?->UDG-> | 938 | Bacteria>FCB_group>Bacteroidetes/Chlorobi_group>Bacteroidota>Bacteroidia>Bacteroidales | - | Bacteroides_uniformis | KSX92_08935 | 3061..5877 | 820 | BREX-system-P-loop-protein-BrxC-[Bacteroides-uniformis] |
| EFG14417.1 | Type6.BREX | PRTase->?->?->tRNA->tRNA-><-?<-SIGMA+OB-Fold+AAA-ATPase+HKD-DNase-BrxL<-PglZ<-HNH<-OLD-ABC<-N6-METHYLASE<-DUF6079-p*<-DUF6079-p<-BrxB-BrxF-iSTAND<-BrxA-DNA-binding<-BrxE<-HTH_3<-?<-TraJ-RHH<-ZnR+TOPRIM<-CR-ATPase4 | 762 | Bacteria>FCB_group>Bacteroidetes/Chlorobi_group>Bacteroidota>Bacteroidia>Bacteroidales | GCA_000178215.1 | Bacteroides_xylanisolvens_SD_CC_1b | CW3_0497 | 100900..103188 | 702447 | hypothetical-protein-CW3_0497-[Bacteroides-xylanisolvens-SD-CC-1b] |
| EFF57551.1 | Type6.BREX | CR-ATPase4->ZnR+TOPRIM->TraJ-RHH->?->HTH_3->BrxE->BrxA-DNA-binding->BrxB-BrxF-iSTAND->DUF6079-p->DUF6079-p*->N6-METHYLASE->OLD-ABC->HNH->PglZ->SIGMA+OB-Fold+AAA-ATPase+HKD-DNase-BrxL-> | 706 | Bacteria>FCB_group>Bacteroidetes/Chlorobi_group>Bacteroidota>Bacteroidia>Bacteroidales | GCA_000178295.1 | Bacteroides_xylanisolvens_SD_CC_2a | CW1_2295 | 25339..27459 | 702444 | hypothetical-protein-CW1_2295-[Bacteroides-xylanisolvens-SD-CC-2a] |
| CDD81792.1 | Type1.BREX | <-HIRAN<-RADICAL-SAM<-NADAR<-HTH_3<-SIGMA+OB-Fold+AAA-ATPase+HKD-DNase-BrxL<-PglZ<-Schlafen+HATPase+wHTH<-N6-METHYLASE<-Four-helical-protein<-DUF6079*<-BrxB-BrxF-iSTAND<-BrxA-DNA-binding<-HTH_3\|\|?->?->?->?->?-><-XIS-HTH | 1217 | Bacteria>FCB_group>Bacteroidetes/Chlorobi_group>Bacteroidota>Bacteroidia>Bacteroidales | GCA_000435095.1 | Bacteroides_sp._CAG:462 | BN666_01223 | 46564..50217 | 1262740 | putative-uncharacterized-protein-[Bacteroides-sp.-CAG:462] |
| CDM00531.1 | Type6.BREX | PRTase->?->?->tRNA->tRNA-><-?<-SIGMA+OB-Fold+AAA-ATPase+HKD-DNase-BrxL<-PglZ<-HNH<-OLD-ABC<-N6-METHYLASE<-DUF6079-p*<-DUF6079-p<-BrxB-BrxF-iSTAND<-BrxA-DNA-binding<-BrxE<-HTH_3<-?<-TraJ-RHH<-ZnR+TOPRIM | 689 | Bacteria>FCB_group>Bacteroidetes/Chlorobi_group>Bacteroidota>Bacteroidia>Bacteroidales | GCA_000577955.1 | Bacteroides_xylanisolvens_SD_CC_2a | BN891_34560 | 156968..159037 | 702444 | hypothetical-protein-BN891_34560-[Bacteroides-xylanisolvens-SD-CC-2a] |
| OCW94638.1 | Type1.BREX | MarR-HTH->Polbeta->Calcineurin-><-DOC+HTH\|\|?->HTH_3->BrxA-DNA-binding->BrxB-BrxF-iSTAND->DUF6079*->N6-METHYLASE->PglZ->SIGMA+OB-Fold+AAA-ATPase+HKD-DNase-BrxL->?->?->XIS-HTH->TOPRIM->XIS-HTH-><-AbiEii | 1202 | Bacteria>FCB_group>Bacteroidetes/Chlorobi_group>Bacteroidota>Bacteroidia>Bacteroidales | GCA_001704385.2 | Macellibacteroides_sp._HH-ZS | A9168_06455 | 108865..112473 | 1857568 | hypothetical-protein-A9168_06455-[Macellibacteroides-sp.-HH-ZS] |
| OKZ04598.1 | Type1.BREX | <-SIGMA-HTH<-?\|\|35exo+SF1-SF2-Helicase-><-SIGMA+OB-Fold+AAA-ATPase+HKD-DNase-BrxL<-PglZ<-Schlafen+HATPase+wHTH<-SIR2+SIR2<-N6-METHYLASE<-DUF6079*<-BrxB-BrxF-iSTAND<-BrxA-DNA-binding<-?\|\|<-REase\|\|HTH_3->DNA_METHYLASE->McrC-NTD+REase->SbcC->REase-> | 1202 | Bacteria>FCB_group>Bacteroidetes/Chlorobi_group>Bacteroidota>Bacteroidia>Bacteroidales | GCA_001915435.1 | Bacteroides_sp._41_26 | BHV71_10470 | 85707..89315 | 1896973 | MAG:-hypothetical-protein-BHV71_10470-[Bacteroides-sp.-41_26] |
| OKZ20119.1 | Type1.BREX | <-REase<-?<-?<-?<-?<-SIGMA+OB-Fold+AAA-ATPase+HKD-DNase-BrxL<-PglZ<-Schlafen+HATPase+wHTH<-N6-METHYLASE<-DUF6079*<-BrxB-BrxF-iSTAND<-BrxA-DNA-binding<-HTH_3<-?<-?<-?\|\|?->?->SIGMA-HTH-> | 1185 | Bacteria>FCB_group>Bacteroidetes/Chlorobi_group>Bacteroidota>Bacteroidia>Bacteroidales | GCA_001915545.1 | Bacteroides_sp._43_108 | BHV77_10980 | 38384..41941 | 1896974 | MAG:-hypothetical-protein-BHV77_10980-[Bacteroides-sp.-43_108] |
| HAH91106.1 | Type1.BREX | <-HIRAN<-RADICAL-SAM<-ARG<-?<-?\|\|HTH_3->wHTH+WYL+WCX->BrxA-DNA-binding->BrxB-BrxF-iSTAND->DUF6079*->OLD-ABC->TOPRIM->?->SF1-SF2-Helicase->?->PglZ->SIGMA+OB-Fold+AAA-ATPase+HKD-DNase-BrxL->McrB-AAA->McrC-NTD+REase-> | 1202 | Bacteria>FCB_group>Bacteroidetes/Chlorobi_group>Bacteroidota>Bacteroidia>Bacteroidales | GCA_003450425.1 | Prevotella_sp. | DCL96_05230 | 7787..11395 | 59823 | BREX-system-P-loop-protein-BrxC-[Prevotella-sp.] |
| HAM28416.1 | Type6.BREX | ChlI+Lon_C+AAA-><-RNase<-AraC-HTH\|\|?-><-?\|\|HTH_3->BrxE->BrxA-DNA-binding->BrxB-BrxF-iSTAND->DUF6079*->N6-METHYLASE->OLD-ABC->HNH->PglZ->SIGMA+OB-Fold+AAA-ATPase+HKD-DNase-BrxL->?->?->35exo-> | 1218 | Bacteria>FCB_group>Bacteroidetes/Chlorobi_group>Bacteroidota>Bacteroidia>Bacteroidales | GCA_003453655.1 | Prevotella_sp. | DCP79_00675 | 66224..69880 | 59823 | BREX-system-P-loop-protein-BrxC-[Prevotella-sp.] |
| RHC75801.1 | Type1.BREX | sGTP->Phage_int_SAM_5+Phage_integrase->?->XIS-HTH->SMS-KaiC->HTH_3->BrxA-DNA-binding->BrxB-BrxF-iSTAND->DUF6079-p*->DUF6079-p->OLD-ABC->REase->N6-METHYLASE->PglZ->SIGMA+OB-Fold+AAA-ATPase+HKD-DNase-BrxL-><-REC+AraC-HTH+AraC-HTH | 691 | Bacteria>FCB_group>Bacteroidetes/Chlorobi_group>Bacteroidota>Bacteroidia>Bacteroidales | GCA_003468465.1 | Bacteroides_uniformis | DW831_02855 | 19890..21965 | 820 | BREX-system-P-loop-protein-BrxC-[Bacteroides-uniformis] |
| RHK71295.1 | Type6.BREX | <-ARG\|\|RHH+MobC->?-><-?<-SIGMA+OB-Fold+AAA-ATPase+HKD-DNase-BrxL<-PglZ<-N6-METHYLASE<-DUF6079-p<-?<-DUF6079-p*<-BrxB-BrxF-iSTAND<-BrxA-DNA-binding<-BrxE<-HTH_3\|\|<-XIS-HTH<-AbiEii<-?\|\|HTH_3->HISKIN-> | 145 | Bacteria>FCB_group>Bacteroidetes/Chlorobi_group>Bacteroidota>Bacteroidia>Bacteroidales | GCA_003474125.1 | Bacteroides_uniformis | DW051_06915 | 97313..97750 | 820 | hypothetical-protein-DW051_06915-[Bacteroides-uniformis] |
| RHR54760.1 | Type1.BREX | ACET-><-?<-Cysteine-Desulfurase\|\|tRNA-><-?<-?<-?<-?\|\|BrxA-DNA-binding->BrxB-BrxF-iSTAND->DUF6079*->N6-METHYLASE->Schlafen+HATPase+wHTH->PglZ->SIGMA+OB-Fold+AAA-ATPase+HKD-DNase-BrxL-><-?\|\|?->PolB->HEPN-> | 1216 | Bacteria>FCB_group>Bacteroidetes/Chlorobi_group>Bacteroidota>Bacteroidia>Bacteroidales | GCA_003479205.1 | Parabacteroides_sp._AF17-28 | DWW90_15085 | 22918..26568 | 2292241 | BREX-system-P-loop-protein-BrxC-[Parabacteroides-sp.-AF17-28] |
| HBQ57053.1 | Type1.BREX | <-SF1-SF2-Helicase\|\|tRNA-><-?<-?<-SIGMA+OB-Fold+AAA-ATPase+HKD-DNase-BrxL<-PglZ<-N6-METHYLASE<-DUF6079*<-BrxB-BrxF-iSTAND<-BrxA-DNA-binding<-?<-REase | 1176 | Bacteria>FCB_group>Bacteroidetes/Chlorobi_group>Bacteroidota>Bacteroidia>Bacteroidales | GCA_003511405.1 | Porphyromonadaceae_bacterium | DD786_07890 | 28635..32165 | 2049046 | BREX-system-P-loop-protein-BrxC-[Porphyromonadaceae-bacterium] |
| ROS89081.1 | Type1.BREX | <-SF1-SF2-Helicase<-SIGMA+OB-Fold+AAA-ATPase+HKD-DNase-BrxL<-PglZ\|\|?-><-Schlafen+HATPase+wHTH<-N6-METHYLASE<-DUF6079*<-BrxB-BrxF-iSTAND<-BrxA-DNA-binding<-HTH_3 | 1215 | Bacteria>FCB_group>Bacteroidetes/Chlorobi_group>Bacteroidota>Bacteroidia>Bacteroidales | GCA_003762635.1 | Muribaculaceae_bacterium_Isolate-043_(Harlan) | EEL36_13385 | 16553..20200 | 2486463 | BREX-system-P-loop-protein-BrxC-[Muribaculaceae-bacterium-Isolate-043-(Harlan)] |
| ROT16603.1 | Type1.BREX | <-SIGMA+OB-Fold+AAA-ATPase+HKD-DNase-BrxL<-PglZ<-MORC+DUF3684+SF1-SF2-Helicase<-ACET+TPR<-AT-HTH-HYD2<-Tox-ART-HYD2<-AT-HTH-HYD2<-N6-METHYLASE<-DUF6079*<-BrxB-BrxF-iSTAND<-BrxA-DNA-binding<-HTH_3\|\|SAVED->SMODS->?->?-><-Phage_lysozyme | 1195 | Bacteria>FCB_group>Bacteroidetes/Chlorobi_group>Bacteroidota>Bacteroidia>Bacteroidales | GCA_003762745.1 | Muribaculaceae_bacterium_Isolate-105_(HZI) | EEL50_01930 | 18926..22513 | 2486472 | BREX-system-P-loop-protein-BrxC-[Muribaculaceae-bacterium-Isolate-105-(HZI)] |
| ROT18783.1 | Type1.BREX | <-SIGMA+OB-Fold+AAA-ATPase+HKD-DNase-BrxL<-PglZ<-Schlafen+HATPase+wHTH<-REase<-N6-METHYLASE<-?<-Tox-ART-HYD2<-?<-DUF6079*<-Four-helical-protein<-BrxB-BrxF-iSTAND<-BrxA-DNA-binding<-HTH_3<-?<-AbiEii<-AbiEii | 1223 | Bacteria>FCB_group>Bacteroidetes/Chlorobi_group>Bacteroidota>Bacteroidia>Bacteroidales | GCA_003833095.1 | Muribaculaceae_bacterium_Isolate-114_(HZI) | EEL53_11930 | 25441..29112 | 2486475 | BREX-system-P-loop-protein-BrxC-[Muribaculaceae-bacterium-Isolate-114-(HZI)] |
| KAA3824840.1 | Type1.BREX | DUF6079*->N6-METHYLASE->SIR2+SIR2->Schlafen+HATPase+wHTH->PglZ->SIGMA+OB-Fold+AAA-ATPase+HKD-DNase-BrxL->\|\|BACON+Peptidase_M64-> | 1065 | Bacteria>FCB_group>Bacteroidetes/Chlorobi_group>Bacteroidota>Bacteroidia>Bacteroidales | GCA_008565655.1 | Bacteroides_ovatus | F3F58_10380 | 1..3200 | 28116 | BREX-system-P-loop-protein-BrxC,-partial-[Bacteroides-ovatus] |
| KAA4088264.1 | Type1.BREX | <-SIGMA+OB-Fold+AAA-ATPase+HKD-DNase-BrxL<-PglZ<-SWIM+DUF3670+SF1-SF2-Helicase<-?<-?<-?<-Peptidase_C39_2<-ACET+TPR<-N6-METHYLASE<-DUF6079* | 1069 | Bacteria>FCB_group>Bacteroidetes/Chlorobi_group>Bacteroidota>Bacteroidia>Bacteroidales | GCA_008566385.1 | Bacteroides_ovatus | F3F49_00290 | 70210..73420 | 28116 | BREX-system-P-loop-protein-BrxC,-partial-[Bacteroides-ovatus] |
| KAA4007861.1 | Type1.BREX | <-NUDIX+NUDIX<-?<-?<-?\|\|HTH_3->wHTH+WYL+WCX->BrxA-DNA-binding->BrxB-BrxF-iSTAND->DUF6079*->N6-METHYLASE->ACET+TPR->Peptidase_C39_2->?->?->?->SWIM+DUF3670+SF1-SF2-Helicase->PglZ->SIGMA+OB-Fold+AAA-ATPase+HKD-DNase-BrxL-> | 1208 | Bacteria>FCB_group>Bacteroidetes/Chlorobi_group>Bacteroidota>Bacteroidia>Bacteroidales | GCA_008566415.1 | Bacteroides_ovatus | F3F37_14290 | 84100..87726 | 28116 | BREX-system-P-loop-protein-BrxC-[Bacteroides-ovatus] |
| KAA4743085.1 | Type1.BREX | AbiEi_4-><-Phage_int_SAM_5+Phage_integrase\|\|STYKinase-like->?->?->\|\|HTH_3->BrxA-DNA-binding->BrxB-BrxF-iSTAND->DUF6079*->DUF6079-p->N6-METHYLASE->Schlafen+HATPase+wHTH->PglZ->SIGMA+OB-Fold+AAA-ATPase+HKD-DNase-BrxL->?->URI-DNase+RAMA->DOC+HTH-> | 1056 | Bacteria>FCB_group>Bacteroidetes/Chlorobi_group>Bacteroidota>Bacteroidia>Bacteroidales | GCA_008569225.1 | Bacteroides_fragilis | F3B36_09960 | 278156..281326 | 817 | BREX-system-P-loop-protein-BrxC-[Bacteroides-fragilis] |
| KAB5317657.1 | Type1.BREX | <-Phage_int_SAM_5<-Phage_integrase<-Phage_int_SAM_5+Phage_integrase<-SIGMA+OB-Fold+AAA-ATPase+HKD-DNase-BrxL<-PglZ<-N6-METHYLASE<-REase<-OLD-ABC<-DUF6079*<-BrxB-BrxF-iSTAND<-BrxA-DNA-binding<-HTH_3<-SMS-KaiC<-XIS-HTH | 1196 | Bacteria>FCB_group>Bacteroidetes/Chlorobi_group>Bacteroidota>Bacteroidia>Bacteroidales | GCA_009025145.1 | Bacteroides_stercoris | F9949_11590 | 61674..65264 | 46506 | BREX-system-P-loop-protein-BrxC-[Bacteroides-stercoris] |
| NBJ07681.1 | Type1.BREX | LRR_5->HTH_3->PSBP->?->?->?->HTH_3->BrxA-DNA-binding->BrxB-BrxF-iSTAND->DUF6079*->N6-METHYLASE->REase->Schlafen+HATPase+wHTH->PglZ->SIGMA+OB-Fold+AAA-ATPase+HKD-DNase-BrxL->SF1-SF2-Helicase->?->?->MPTase-> | 1223 | Bacteria>FCB_group>Bacteroidetes/Chlorobi_group>Bacteroidota>Bacteroidia>Bacteroidales | GCA_009911575.1 | Alistipes_sp._Z76 | D1647_16085 | 15947..19618 | 2304565 | BREX-system-P-loop-protein-BrxC-[Alistipes-sp.-Z76] |
| NLA16135.1 | Type1.BREX | <-HTH_3\|\|HTH+REase-><-SIGMA+OB-Fold+AAA-ATPase+HKD-DNase-BrxL<-PglZ<-N6-METHYLASE<-DUF6079*<-BrxB-BrxF-iSTAND<-BrxA-DNA-binding<-HTH_3<-?\|\|?->tRNA-><-?<-?\|\|?->LuxR-HTH-> | 1174 | Bacteria>FCB_group>Bacteroidetes/Chlorobi_group>Bacteroidota>Bacteroidia>Bacteroidales | GCA_012510445.1 | Bacteroidales_bacterium | GX877_06335 | 9723..13247 | 2030927 | BREX-system-P-loop-protein-BrxC-[Bacteroidales-bacterium] |
| NLN95595.1 | Type1.BREX | PolB->HEPN->?-><-?<-SIGMA+OB-Fold+AAA-ATPase+HKD-DNase-BrxL<-PglZ<-?<-N6-METHYLASE<-?<-DUF6079*<-BrxB-BrxF-iSTAND<-BrxA-DNA-binding<-HTH_3\|\|?->TRD+TRD-> | 1176 | Bacteria>FCB_group>Bacteroidetes/Chlorobi_group>Bacteroidota>Bacteroidia>Bacteroidales | GCA_012522795.1 | Bacteroidales_bacterium | GX128_05430 | 111716..115246 | 2030927 | BREX-system-P-loop-protein-BrxC-[Bacteroidales-bacterium] |
| MBD5181931.1 | Type1.BREX | <-Acyltransferase<-?<-?\|\|HISKIN+MORC->REC+OmpR-HTH->BrxA-DNA-binding->BrxB-BrxF-iSTAND->DUF6079*->N6-METHYLASE->OLD-ABC->TOPRIM->PglZ->SIGMA+OB-Fold+AAA-ATPase+HKD-DNase-BrxL-> | 1193 | Bacteria>FCB_group>Bacteroidetes/Chlorobi_group>Bacteroidota>Bacteroidia>Bacteroidales | GCA_014800235.1 | Bacteroidales_bacterium | HDS98_05210 | 20716..24297 | 2030927 | BREX-system-P-loop-protein-BrxC-[Bacteroidales-bacterium] |
| MBD5263514.1 | Type1.BREX | <-Calcineurin<-?<-?<-?<-Aminotran_1_2\|\|?-><-?\|\|BrxA-DNA-binding->BrxB-BrxF-iSTAND->DUF6079*->N6-METHYLASE->Schlafen+HATPase+wHTH->SF1-SF2-Helicase->BirA-HTH->PglZ->SIGMA+OB-Fold+AAA-ATPase+HKD-DNase-BrxL-> | 1221 | Bacteria>FCB_group>Bacteroidetes/Chlorobi_group>Bacteroidota>Bacteroidia>Bacteroidales | GCA_014801365.1 | Bacteroides_sp. | HDS39_05520 | 20086..23751 | 29523 | BREX-system-P-loop-protein-BrxC-[Bacteroides-sp.] |
| MBD5349669.1 | Type1.BREX | <-ABhydrolase<-JAB<-?<-?<-SIGMA+OB-Fold+AAA-ATPase+HKD-DNase-BrxL<-Schlafen+HATPase+wHTH<-?<-N6-METHYLASE<-Four-helical-protein<-DUF6079*<-BrxB-BrxF-iSTAND<-BrxA-DNA-binding<-HTH_3<-TraJ-RHH\|\|Zn_Tnp_IS1595+DDE_Tnp_IS1595->?-><-?<-HTH_58<-METHYLASE | 1219 | Bacteria>FCB_group>Bacteroidetes/Chlorobi_group>Bacteroidota>Bacteroidia>Bacteroidales | GCA_014802205.1 | Bacteroides_sp. | HDR89_02130 | 36441..40100 | 29523 | BREX-system-P-loop-protein-BrxC-[Bacteroides-sp.] |
| MBD5357714.1 | Type1.BREX | AAA+->?->XIS-HTH-><-?<-?<-?\|\|BrxA-DNA-binding->BrxB-BrxF-iSTAND->DUF6079*->N6-METHYLASE->HU-HIG+REase->PglZ->SIGMA+OB-Fold+AAA-ATPase+HKD-DNase-BrxL->REase->?->CASPASE-> | 1203 | Bacteria>FCB_group>Bacteroidetes/Chlorobi_group>Bacteroidota>Bacteroidia>Bacteroidales | GCA_014802265.1 | Bacteroides_sp. | HDR88_12035 | 22916..26527 | 29523 | BREX-system-P-loop-protein-BrxC-[Bacteroides-sp.] |
| MBD5419378.1 | Type1.BREX | BrxB-BrxF-iSTAND->DUF6079*->N6-METHYLASE->Schlafen+HATPase+wHTH->PglZ->SIGMA+OB-Fold+AAA-ATPase+HKD-DNase-BrxL->?->Calcineurin->Glyco_hydro_3-> | 1201 | Bacteria>FCB_group>Bacteroidetes/Chlorobi_group>Bacteroidota>Bacteroidia>Bacteroidales | GCA_014803765.1 | Bacteroides_sp. | HDR48_05020 | 1391..4996 | 29523 | BREX-system-P-loop-protein-BrxC-[Bacteroides-sp.] |
| MBE6254618.1 | Type1.BREX | <-SF1-SF2-Helicase<-?<-SIGMA+OB-Fold+AAA-ATPase+HKD-DNase-BrxL<-PglZ<-SIR2<-N6-METHYLASE<-DUF6079*<-BrxB-BrxF-iSTAND<-BrxA-DNA-binding<-wHTH+WYL+WCX | 1211 | Bacteria>FCB_group>Bacteroidetes/Chlorobi_group>Bacteroidota>Bacteroidia>Bacteroidales | GCA_015060555.1 | Prevotella_sp. | E7105_03785 | 25107..28742 | 59823 | BREX-system-P-loop-protein-BrxC-[Prevotella-sp.] |
| MBF7062545.1 | Type6.BREX | <-TRAM+METHYLASE\|\|?-><-UDG<-?<-ARG\|\|RHH+MobC->?-><-SIGMA+OB-Fold+AAA-ATPase+HKD-DNase-BrxL<-PglZ<-DUF6079-p*<-DUF6079-p<-BrxB-BrxF-iSTAND<-BrxA-DNA-binding<-BrxE<-HTH_3<-?\|\|<-XIS-HTH<-AbiEii\|\|HTH_3-> | 938 | Bacteria>FCB_group>Bacteroidetes/Chlorobi_group>Bacteroidota>Bacteroidia>Bacteroidales | GCA_015594615.1 | Bacteroides_sp._HF-5613 | ITJ87_11475 | 2819894..2822710 | 2785794 | BREX-system-P-loop-protein-BrxC-[Bacteroides-sp.-HF-5613] |
| MBM6671973.1 | Type6.BREX | <-SIGMA+OB-Fold+AAA-ATPase+HKD-DNase-BrxL<-PglZ<-N6-METHYLASE<-DUF6079*<-BrxB-BrxF-iSTAND<-BrxA-DNA-binding<-BrxE<-GNAT | 1224 | Bacteria>FCB_group>Bacteroidetes/Chlorobi_group>Bacteroidota>Bacteroidia>Bacteroidales | GCA_016900015.1 | Phocaeicola_coprophilus | H6B14_14560 | 8845..12519 | 387090 | BREX-system-P-loop-protein-BrxC-[Phocaeicola-coprophilus] |
| MBN2915288.1 | Type1.BREX | <-McrC-NTD+REase<-McrB-AAA<-SIGMA+OB-Fold+AAA-ATPase+HKD-DNase-BrxL<-PglZ<-SIR2+SIR2<-N6-METHYLASE<-TOPRIM<-OLD-ABC<-DUF6079-p* | 971 | Bacteria>FCB_group>Bacteroidetes/Chlorobi_group>Bacteroidota>Bacteroidia>Bacteroidales | GCA_016939035.1 | Prevotella_sp. | JTJ26_04215 | 25012..27927 | 59823 | BREX-system-P-loop-protein-BrxC-[Prevotella-sp.] |
| MBQ6178640.1 | Type1.BREX | ArdCN+MPTase->?->?->?-><-wHTH+WYL+WCX<-?\|\|HD->BrxA-DNA-binding->BrxB-BrxF-iSTAND->DUF6079*->N6-METHYLASE->REase->PglZ->SIGMA+OB-Fold+AAA-ATPase+HKD-DNase-BrxL-> | 1202 | Bacteria>FCB_group>Bacteroidetes/Chlorobi_group>Bacteroidota>Bacteroidia>Bacteroidales | GCA_017429185.1 | Bacteroidales_bacterium | IJK32_03880 | 18867..22475 | 2030927 | BREX-system-P-loop-protein-BrxC-[Bacteroidales-bacterium] |
| MBQ6872144.1 | Type1.BREX | BrxA-DNA-binding->BrxB-BrxF-iSTAND->DUF6079*->N6-METHYLASE->PglZ->SIGMA+OB-Fold+AAA-ATPase+HKD-DNase-BrxL-> | 1190 | Bacteria>FCB_group>Bacteroidetes/Chlorobi_group>Bacteroidota>Bacteroidia>Bacteroidales | GCA_017435685.1 | Bacteroidales_bacterium | IJD23_07295 | 3964..7536 | 2030927 | BREX-system-P-loop-protein-BrxC-[Bacteroidales-bacterium] |
| MBQ9588226.1 | Type1.BREX | Peptidase->?-><-HD\|\|BrxA-DNA-binding->BrxB-BrxF-iSTAND->DUF6079*->N6-METHYLASE->HTH+REase->N6-METHYLASE-p->PglZ->SIGMA+OB-Fold+AAA-ATPase+HKD-DNase-BrxL-> | 1183 | Bacteria>FCB_group>Bacteroidetes/Chlorobi_group>Bacteroidota>Bacteroidia>Bacteroidales | GCA_017446745.1 | Bacteroidales_bacterium | IJR26_10275 | 19779..23330 | 2030927 | BREX-system-P-loop-protein-BrxC-[Bacteroidales-bacterium] |
| MBQ7672328.1 | Type1.BREX | REase+35exo->?->?->LRR_5->LRR_5->BrxA-DNA-binding->BrxB-BrxF-iSTAND->DUF6079*->N6-METHYLASE->PglZ->SIGMA+OB-Fold+AAA-ATPase+HKD-DNase-BrxL->?->DOC+HTH-> | 1193 | Bacteria>FCB_group>Bacteroidetes/Chlorobi_group>Bacteroidota>Bacteroidia>Bacteroidales | GCA_017450385.1 | Paludibacteraceae_bacterium | IJS49_04535 | 76772..80353 | 2748031 | BREX-system-P-loop-protein-BrxC-[Paludibacteraceae-bacterium] |
| MBR6863890.1 | Type1.BREX | <-ParB+HNH<-wHTH+WYL+WCX<-?\|\|?->Couple_hipA+HIRAN->EVE->HD->BrxA-DNA-binding->BrxB-BrxF-iSTAND->DUF6079*->N6-METHYLASE->PglZ->SIGMA+OB-Fold+AAA-ATPase+HKD-DNase-BrxL->?->SF1-SF2-Helicase-> | 1202 | Bacteria>FCB_group>Bacteroidetes/Chlorobi_group>Bacteroidota>Bacteroidia>Bacteroidales | GCA_017531205.1 | Bacteroidales_bacterium | IKM75_03420 | 106682..110290 | 2030927 | BREX-system-P-loop-protein-BrxC-[Bacteroidales-bacterium] |
| MBR5893775.1 | Type1.BREX | <-SIGMA+OB-Fold+AAA-ATPase+HKD-DNase-BrxL<-PglZ<-DOC+HTH<-N6-METHYLASE<-?<-DUF6079*<-BrxB-BrxF-iSTAND<-BrxA-DNA-binding<-?<-HAD+HAD | 1185 | Bacteria>FCB_group>Bacteroidetes/Chlorobi_group>Bacteroidota>Bacteroidia>Bacteroidales | GCA_017554605.1 | Bacteroidaceae_bacterium | IKZ37_09095 | 11422..14979 | 2212467 | BREX-system-P-loop-protein-BrxC-[Bacteroidaceae-bacterium] |
| MBR5630362.1 | Type1.BREX | <-CopG/RHH\|\|BrxA-DNA-binding->BrxB-BrxF-iSTAND->DUF6079*->N6-METHYLASE->PglZ->SIGMA+OB-Fold+AAA-ATPase+HKD-DNase-BrxL-> | 1171 | Bacteria>FCB_group>Bacteroidetes/Chlorobi_group>Bacteroidota>Bacteroidia>Bacteroidales | GCA_017557685.1 | Bacteroidales_bacterium | IKW82_01820 | 70807..74322 | 2030927 | BREX-system-P-loop-protein-BrxC-[Bacteroidales-bacterium] |
| MBR5576788.1 | Type1.BREX | <-LRR_5+LRR_5+LRR_5+LRR_5<-?<-?<-?<-?<-?<-?\|\|BrxA-DNA-binding->BrxB-BrxF-iSTAND->DUF6079*->N6-METHYLASE->SbcC+SbcC->?->ACET+TPR->?->SF1-SF2-Helicase->PglZ->SIGMA+OB-Fold+AAA-ATPase+HKD-DNase-BrxL-> | 1209 | Bacteria>FCB_group>Bacteroidetes/Chlorobi_group>Bacteroidota>Bacteroidia>Bacteroidales | GCA_017558255.1 | Bacteroidaceae_bacterium | IKW22_05890 | 14217..17846 | 2212467 | BREX-system-P-loop-protein-BrxC-[Bacteroidaceae-bacterium] |
| MBO7470104.1 | Type1.BREX | BrxA-DNA-binding->BrxB-BrxF-iSTAND->DUF6079*->N6-METHYLASE->?->?->PglZ->SIGMA+OB-Fold+AAA-ATPase+HKD-DNase-BrxL-> | 1169 | Bacteria>FCB_group>Bacteroidetes/Chlorobi_group>Bacteroidota>Bacteroidia>Bacteroidales | GCA_017649945.1 | Bacteroidales_bacterium | J6T81_08275 | 1593..5102 | 2030927 | BREX-system-P-loop-protein-BrxC-[Bacteroidales-bacterium] |
| MBS6238336.1 | Type1.BREX | <-PAS+HISKIN\|\|?-><-AraC-HTH+AraC-HTH<-SIGMA+OB-Fold+AAA-ATPase+HKD-DNase-BrxL<-PglZ<-Schlafen+HATPase+wHTH<-ParB+HNH<-ParB+HNH<-N6-METHYLASE<-DUF6079*<-BrxB-BrxF-iSTAND<-BrxA-DNA-binding<-HTH_3<-?<-?<-?<-?\|\|SIGMA-HTH-> | 1205 | Bacteria>FCB_group>Bacteroidetes/Chlorobi_group>Bacteroidota>Bacteroidia>Bacteroidales | GCA_018374845.1 | Bacteroides_sp. | KH897_08200 | 204452..208069 | 29523 | BREX-system-P-loop-protein-BrxC-[Bacteroides-sp.] |
| MBV3827782.1 | Type6.BREX | <-UDG<-?<-ARG\|\|RHH+MobC->?-><-?<-SIGMA+OB-Fold+AAA-ATPase+HKD-DNase-BrxL<-PglZ<-N6-METHYLASE<-DUF6079-p*<-?<-DUF6079-p<-BrxB-BrxF-iSTAND<-BrxA-DNA-binding<-BrxE<-HTH_3\|\|<-XIS-HTH<-AbiEii | 938 | Bacteria>FCB_group>Bacteroidetes/Chlorobi_group>Bacteroidota>Bacteroidia>Bacteroidales | GCA_019129685.1 | Bacteroides_uniformis | DW051_06905 | 94105..96921 | 820 | BREX-system-P-loop-protein-BrxC-[Bacteroides-uniformis] |
| MCB9017748.1 | Type1.BREX | <-ACET\|\|?-><-SIGMA+OB-Fold+AAA-ATPase+HKD-DNase-BrxL<-PglZ<-N6-METHYLASE-p<-HTH+REase<-?<-Schlafen+HATPase+wHTH<-DUF6079*<-BrxB-BrxF-iSTAND<-BrxA-DNA-binding<-?<-PALP | 1207 | Bacteria>FCB_group>Bacteroidetes/Chlorobi_group>Bacteroidota>Bacteroidia>Bacteroidales | GCA_020636675.1 | Prevotellaceae_bacterium | K6544_03995 | 62697..66320 | 2049047 | BREX-system-P-loop-protein-BrxC-[Prevotellaceae-bacterium] |
| HJF82219.1 | Type1.BREX | <-SF1-SF2-Helicase-p<-HTH_3<-?<-?\|\|BrxA-DNA-binding->BrxB-BrxF-iSTAND->DUF6079*->OLD-ABC->TOPRIM->N6-METHYLASE->SIR2+SIR2->PglZ->SIGMA+OB-Fold+AAA-ATPase+HKD-DNase-BrxL-> | 1195 | Bacteria>FCB_group>Bacteroidetes/Chlorobi_group>Bacteroidota>Bacteroidia>Bacteroidales | GCA_020742805.1 | Phocaeicola_plebeius | K8V40_11345 | 4107..7694 | 310297 | BREX-system-P-loop-protein-BrxC-[Phocaeicola-plebeius] |
| MCC8037087.1 | Type1.BREX | <-HAD+PRTase+PRTase+REase+SF1-SF2-Helicase<-?<-?<-?<-STYKIN<-?<-SIGMA+OB-Fold+AAA-ATPase+HKD-DNase-BrxL<-PglZ<-DUF6079*<-BrxB-BrxF-iSTAND<-BrxA-DNA-binding<-?<-?<-?<-?<-Acetate_kinase | 1219 | Bacteria>FCB_group>Bacteroidetes/Chlorobi_group>Bacteroidota>Bacteroidia>Bacteroidales | GCA_020859755.1 | Bacteroidales_bacterium | LIP02_02980 | 18317..21976 | 2030927 | BREX-system-P-loop-protein-BrxC-[Bacteroidales-bacterium] |
| MCC8115076.1 | Type1.BREX | PglZ->SIGMA+OB-Fold+AAA-ATPase+HKD-DNase-BrxL->BrxA-DNA-binding->BrxB-BrxF-iSTAND->DUF6079-p*->DUF6079->N6-METHYLASE-> | 159 | Bacteria>FCB_group>Bacteroidetes/Chlorobi_group>Bacteroidota>Bacteroidia>Bacteroidales | GCA_020860445.1 | Bacteroidales_bacterium | LIP03_14010 | 6329..6808 | 2030927 | hypothetical-protein-[Bacteroidales-bacterium] |
| MCC8115077.1 | Type1.BREX | PglZ->SIGMA+OB-Fold+AAA-ATPase+HKD-DNase-BrxL->BrxA-DNA-binding->BrxB-BrxF-iSTAND->DUF6079-p->DUF6079*->N6-METHYLASE-> | 1040 | Bacteria>FCB_group>Bacteroidetes/Chlorobi_group>Bacteroidota>Bacteroidia>Bacteroidales | GCA_020860445.1 | Bacteroidales_bacterium | LIP03_14015 | 6789..9911 | 2030927 | BREX-system-P-loop-protein-BrxC-[Bacteroidales-bacterium] |
| MCH4018978.1 | Type1.BREX | XIS-HTH->?->AAA+-><-?\|\|wHTH+WYL+WCX->HTH_3->BrxA-DNA-binding->BrxB-BrxF-iSTAND->DUF6079*->N6-METHYLASE->Schlafen+HATPase+wHTH->SIR2->PglZ->SIGMA+OB-Fold+AAA-ATPase+HKD-DNase-BrxL->SIR2+SIR2->ZnR->DDE_Tnp_ISL3->Tox-RelE-> | 1198 | Bacteria>FCB_group>Bacteroidetes/Chlorobi_group>Bacteroidota>Bacteroidia>Bacteroidales | GCA_022483425.1 | Prevotella_sp. | LKE74_11130 | 2793807..2797403 | 59823 | BREX-system-P-loop-protein-BrxC-[Prevotella-sp.] |
| MCI1742488.1 | Type1.BREX | <-HTH_31\|\|tRNA-><-?<-SIR2+SIR2<-SIGMA+OB-Fold+AAA-ATPase+HKD-DNase-BrxL<-PglZ<-HTH+REase<-Schlafen+HATPase+wHTH<-N6-METHYLASE<-DUF6079*<-BrxB-BrxF-iSTAND<-BrxA-DNA-binding<-HTH_3<-wHTH+WYL+WCX\|\|?-><-AAA+<-?<-XIS-HTH | 1206 | Bacteria>FCB_group>Bacteroidetes/Chlorobi_group>Bacteroidota>Bacteroidia>Bacteroidales | GCA_022647785.1 | Prevotella_sp. | LKI18_09155 | 39395..43015 | 59823 | BREX-system-P-loop-protein-BrxC-[Prevotella-sp.] |
| MCI2087046.1 | Type1.BREX | <-SIGMA+OB-Fold+AAA-ATPase+HKD-DNase-BrxL<-PglZ<-HTH+REase<-Schlafen+HATPase+wHTH<-N6-METHYLASE<-DUF6079*<-BrxB-BrxF-iSTAND<-BrxA-DNA-binding<-HTH_3<-wHTH+WYL+WCX\|\|?-><-AAA+<-?<-XIS-HTH | 1198 | Bacteria>FCB_group>Bacteroidetes/Chlorobi_group>Bacteroidota>Bacteroidia>Bacteroidales | GCA_022653375.1 | Prevotella_sp. | LKK10_02020 | 11419..15015 | 59823 | BREX-system-P-loop-protein-BrxC-[Prevotella-sp.] |
| MCI7017346.1 | Type1.BREX | <-SIGMA+OB-Fold+AAA-ATPase+HKD-DNase-BrxL<-PglZ<-?<-Schlafen+HATPase+wHTH<-N6-METHYLASE<-AT-HTH-HYD2<-Tox-ART-HYD2<-AT-HTH-HYD2<-DUF6079*<-BrxB-BrxF-iSTAND<-BrxA-DNA-binding<-wHTH+WYL+WCX | 1189 | Bacteria>FCB_group>Bacteroidetes/Chlorobi_group>Bacteroidota>Bacteroidia>Bacteroidales | GCA_022775925.1 | Prevotella_sp. | MR923_09655 | 21723..25292 | 59823 | BREX-system-P-loop-protein-BrxC-[Prevotella-sp.] |
| MCI6342424.1 | Type6.BREX | DnaB+TnsC-ATPase->?-><-SIGMA+OB-Fold+AAA-ATPase+HKD-DNase-BrxL<-PglZ<-HNH<-OLD-ABC<-N6-METHYLASE<-DUF6079*<-BrxB-BrxF-iSTAND<-BrxA-DNA-binding<-BrxE<-HTH_3 | 1217 | Bacteria>FCB_group>Bacteroidetes/Chlorobi_group>Bacteroidota>Bacteroidia>Bacteroidales | GCA_022777695.1 | Prevotella_sp. | MR788_07720 | 61686..65339 | 59823 | BREX-system-P-loop-protein-BrxC-[Prevotella-sp.] |
| MCI7387531.1 | Type6.BREX | Arm-DNA-bind_5-><-UDG<-?<-ARG\|\|?-><-SIGMA+OB-Fold+AAA-ATPase+HKD-DNase-BrxL<-PglZ<-N6-METHYLASE<-DUF6079*<-BrxB-BrxF-iSTAND<-BrxA-DNA-binding<-BrxE<-HTH_3<-?\|\|?-><-XIS-HTH<-AbiEii | 1215 | Bacteria>FCB_group>Bacteroidetes/Chlorobi_group>Bacteroidota>Bacteroidia>Bacteroidales | GCA_022784345.1 | Bacteroides_uniformis | MSH64_12855 | 232214..235861 | 820 | BREX-system-P-loop-protein-BrxC-[Bacteroides-uniformis] |
| MCK9398992.1 | Type1.BREX | PIN-><-?\|\|DDE_Tnp_1-><-?<-?\|\|HTH_3->BrxA-DNA-binding->BrxB-BrxF-iSTAND->DUF6079*->N6-METHYLASE->PglZ->SIGMA+OB-Fold+AAA-ATPase+HKD-DNase-BrxL->?->?->?-><-tRNA\|\|RADICAL-SAM-> | 1163 | Bacteria>FCB_group>Bacteroidetes/Chlorobi_group>Bacteroidota>Bacteroidia>Bacteroidales | GCA_023229505.1 | Bacteroidales_bacterium | MOQ51_03210 | 70739..74230 | 2030927 | BREX-system-P-loop-protein-BrxC-[Bacteroidales-bacterium] |
| MCQ2071598.1 | Type1.BREX | <-TOPRIM<-?<-?\|\|Peptidase_M23->BrxA-DNA-binding->BrxB-BrxF-iSTAND->DUF6079*->N6-METHYLASE->Schlafen+HATPase+wHTH->PglZ->SIGMA+OB-Fold+AAA-ATPase+HKD-DNase-BrxL->XIS-HTH->wHTH+WYL+WCX->?->HHH_5+SF1-SF2-Helicase-> | 1190 | Bacteria>FCB_group>Bacteroidetes/Chlorobi_group>Bacteroidota>Bacteroidia>Bacteroidales | GCA_024398655.1 | Bacteroidaceae_bacterium | MJY96_00535 | 107887..111459 | 2212467 | BREX-system-P-loop-protein-BrxC-[Bacteroidaceae-bacterium] |
| WP_156315612.1 | Type1.BREX | XIS-HTH-><-?<-?\|\|?->?->?->HTH_3->BrxA-DNA-binding->BrxB-BrxF-iSTAND->DUF6079-p*->DUF6079-p->N6-METHYLASE-p->N6-METHYLASE->PglZ-p->PglZ->SIGMA+OB-Fold+AAA-ATPase+HKD-DNase-BrxL->KAP-NTPase-> | 925 | Bacteria>FCB_group>Bacteroidetes/Chlorobi_group>Bacteroidota>Bacteroidia>Bacteroidales | GCA_030820635.1 | Phocaeicola_coprophilus | JCM13816_RS17810 | 17247..20024 | 387090 | BREX-system-P-loop-protein-BrxC-[Phocaeicola-coprophilus] |
| SHK68061.1 | Type1.BREX | <-Classical-AAA<-?\|\|tRNA-><-?<-HTH+DOC\|\|?->HTH_3->BrxA-DNA-binding->BrxB-BrxF-iSTAND->DUF6079*->N6-METHYLASE->SbcC+SbcC->?->SIR2+SIR2->PglZ->SIGMA+OB-Fold+AAA-ATPase+HKD-DNase-BrxL->?->ParB+HNH->ParB+HNH-> | 1206 | Bacteria>FCB_group>Bacteroidetes/Chlorobi_group>Bacteroidota>Bacteroidia>Bacteroidales | GCA_900142525.1 | Prevotella_ruminicola | SAMN05216463_10942 | 46265..49885 | 839 | hypothetical-protein-SAMN05216463_10942-[Prevotella-ruminicola] |
| WP_008760280.1 | Type1.BREX | D5ATPase+DUF3874-><-?\|\|ADH_N+ADH_zinc_N+METHYLASE->SPOUT-><-SIGMA+OB-Fold+AAA-ATPase+HKD-DNase-BrxL<-PglZ<-HTH+REase<-N6-METHYLASE-p<-N6-METHYLASE<-DUF6079*<-BrxB-BrxF-iSTAND<-BrxA-DNA-binding<-HTH_3<-?<-?<-?<-TIR<-?<-TIR | 1193 | Bacteria>FCB_group>Bacteroidetes/Chlorobi_group>Bacteroidota>Bacteroidia>Bacteroidales | GCF_000162515.1 | Bacteroides_thetaiotaomicron | HMPREF9007_RS20215 | 181866..185447 | 818 | MULTISPECIES:-BREX-system-P-loop-protein-BrxC-[Bacteroides] |
| WP_232506819.1 | Type6.BREX | PRTase->?->?->tRNA->tRNA-><-?<-SIGMA+OB-Fold+AAA-ATPase+HKD-DNase-BrxL<-PglZ<-HNH<-OLD-ABC<-N6-METHYLASE<-DUF6079-p*<-DUF6079-p<-BrxB-BrxF-iSTAND<-BrxA-DNA-binding<-BrxE<-HTH_3<-?<-TraJ-RHH<-ZnR+TOPRIM | 794 | Bacteria>FCB_group>Bacteroidetes/Chlorobi_group>Bacteroidota>Bacteroidia>Bacteroidales | GCF_000178215.1 | Bacteroides_xylanisolvens | CW3_RS0123000 | 100900..103284 | 371601 | hypothetical-protein-[Bacteroides-xylanisolvens] |
| WP_008016782.1 | Type6.BREX | <-SIGMA+OB-Fold+AAA-ATPase+HKD-DNase-BrxL<-PglZ<-HNH<-OLD-ABC<-N6-METHYLASE<-DUF6079-p<-DUF6079-p*<-BrxB-BrxF-iSTAND<-BrxA-DNA-binding<-BrxE<-HTH_3<-?<-TraJ-RHH<-ZnR+TOPRIM<-CR-ATPase4 | 425 | Bacteria>FCB_group>Bacteroidetes/Chlorobi_group>Bacteroidota>Bacteroidia>Bacteroidales | GCF_000178215.1 | Bacteroides_xylanisolvens | CW3_RS26225 | 103281..104558 | 371601 | DUF6079-family-protein-[Bacteroides-xylanisolvens] |
| WP_004312187.1 | Type1.BREX | <-SIGMA+OB-Fold+AAA-ATPase+HKD-DNase-BrxL<-PglZ<-SWIM+DUF3670+SF1-SF2-Helicase<-?<-?<-?<-Peptidase_C39_2<-ACET+TPR<-N6-METHYLASE<-DUF6079*<-BrxB-BrxF-iSTAND<-BrxA-DNA-binding<-wHTH+WYL+WCX<-HTH_3\|\|?->?->?->NUDIX+NUDIX-> | 1208 | Bacteria>FCB_group>Bacteroidetes/Chlorobi_group>Bacteroidota>Bacteroidia>Bacteroidales | GCF_000178275.1 | Bacteroidaceae | K0G46_RS13725 | 52737..56363 | 815 | MULTISPECIES:-BREX-system-P-loop-protein-BrxC-[Bacteroidaceae] |
| WP_004314649.1 | Type6.BREX | XIS-HTH->CR-ATPase4->ZnR+TOPRIM->TraJ-RHH->?->?->BrxE->BrxA-DNA-binding->BrxB-BrxF-iSTAND->DUF6079-p*->DUF6079-p->N6-METHYLASE->OLD-ABC->HNH->PglZ->SIGMA+OB-Fold+AAA-ATPase+HKD-DNase-BrxL-> | 515 | Bacteria>FCB_group>Bacteroidetes/Chlorobi_group>Bacteroidota>Bacteroidia>Bacteroidales | GCF_000178295.1 | Bacteroides_xylanisolvens | CW1_RS26610 | 23799..25346 | 371601 | BREX-system-P-loop-protein-BrxC-[Bacteroides-xylanisolvens] |
| WP_232505074.1 | Type6.BREX | CR-ATPase4->ZnR+TOPRIM->TraJ-RHH->?->?->BrxE->BrxA-DNA-binding->BrxB-BrxF-iSTAND->DUF6079-p->DUF6079-p*->N6-METHYLASE->OLD-ABC->HNH->PglZ->SIGMA+OB-Fold+AAA-ATPase+HKD-DNase-BrxL-> | 711 | Bacteria>FCB_group>Bacteroidetes/Chlorobi_group>Bacteroidota>Bacteroidia>Bacteroidales | GCF_000178295.1 | Bacteroides_xylanisolvens | CW1_RS26615 | 25324..27459 | 371601 | hypothetical-protein-[Bacteroides-xylanisolvens] |
| WP_006745116.1 | Type1.BREX | <-MODE-HTH+PBPII\|\|?-><-?<-NUDIX\|\|ABhydrolase->DNA-glycosylase-><-SIGMA+OB-Fold+AAA-ATPase+HKD-DNase-BrxL<-PglZ<-N6-METHYLASE<-DUF6079*<-BrxB-BrxF-iSTAND<-BrxA-DNA-binding<-?\|\|?->REase-><-?<-?<-?<-SIGMA-HTH | 1192 | Bacteria>FCB_group>Bacteroidetes/Chlorobi_group>Bacteroidota>Bacteroidia>Bacteroidales | GCF_000212915.1 | Bacteroides_coprosuis | BCOP_RS09060 | 2132867..2136445 | 151276 | BREX-system-P-loop-protein-BrxC-[Bacteroides-coprosuis] |
| WP_018109295.1 | Type1.BREX | <-FURR-HTH<-?<-Peptidase_M49<-AraC-HTH\|\|SF1-SF2-Helicase+PRTase->tRNA->HTH_3->BrxA-DNA-binding->BrxB-BrxF-iSTAND->DUF6079*->N6-METHYLASE->PglZ->SIGMA+OB-Fold+AAA-ATPase+HKD-DNase-BrxL->?->?->XIS-HTH->HTH_3->MerR-HTH->TOPRIM-> | 1202 | Bacteria>FCB_group>Bacteroidetes/Chlorobi_group>Bacteroidota>Bacteroidia>Bacteroidales | GCF_000375405.1 | Bacteroides_propionicifaciens | B091_RS0106980 | 23999..27607 | 392838 | BREX-system-P-loop-protein-BrxC-[Bacteroides-propionicifaciens] |
| WP_002561973.1 | Type1.BREX | 35exo+SF1-SF2-Helicase-><-DDE_Tnp\|\|<-SIGMA+OB-Fold+AAA-ATPase+HKD-DNase-BrxL<-PglZ<-Schlafen+HATPase+wHTH<-SF1-SF2-Helicase<-OLD-ABC+TOPRIM<-N6-METHYLASE<-DUF6079*<-BrxB-BrxF-iSTAND<-BrxA-DNA-binding | 1202 | Bacteria>FCB_group>Bacteroidetes/Chlorobi_group>Bacteroidota>Bacteroidia>Bacteroidales | GCF_000382465.1 | Bacteroides_sp._HPS0048 | HMPREF1214_RS19625 | 483033..486641 | 1078089 | BREX-system-P-loop-protein-BrxC-[Bacteroides-sp.-HPS0048] |
| WP_155941928.1 | Type6.BREX | Glyoxalase-><-?<-?\|\|HD->wHTH+WYL+WCX->HTH_3->BrxE->BrxA-DNA-binding->BrxB-BrxF-iSTAND->DUF6079*->N6-METHYLASE->PglZ->SIGMA+OB-Fold+AAA-ATPase+HKD-DNase-BrxL->KAP-NTPase->?->?->?->SWIM+DUF3670+SF1-SF2-Helicase->ACET-> | 1198 | Bacteria>FCB_group>Bacteroidetes/Chlorobi_group>Bacteroidota>Bacteroidia>Bacteroidales | GCF_000424185.1 | Prevotella_sp._AGR2160 | G604_RS15270 | 9168..12764 | 1280674 | BREX-system-P-loop-protein-BrxC-[Prevotella-sp.-AGR2160] |
| WP_024993540.1 | Type1.BREX | <-wHTH+WYL+WCX<-wHTH+WYL+WCX<-SIGMA+OB-Fold+AAA-ATPase+HKD-DNase-BrxL<-PglZ<-Schlafen+HATPase+wHTH<-N6-METHYLASE<-DUF6079*<-BrxB-BrxF-iSTAND<-BrxA-DNA-binding<-HTH_3 | 1210 | Bacteria>FCB_group>Bacteroidetes/Chlorobi_group>Bacteroidota>Bacteroidia>Bacteroidales | GCF_000613805.1 | Phocaeicola_paurosaccharolyticus | JCM15092_RS04015 | 53285..56917 | 732242 | BREX-system-P-loop-protein-BrxC-[Phocaeicola-paurosaccharolyticus] |
| WP_042520323.1 | Type1.BREX | DDE_Tnp_ISL3-><-PARG+MACRODOMAIN<-REase<-SIGMA+OB-Fold+AAA-ATPase+HKD-DNase-BrxL<-PglZ<-N6-METHYLASE<-DUF6079*<-BrxB-BrxF-iSTAND<-BrxA-DNA-binding<-HTH_3 | 1204 | Bacteria>FCB_group>Bacteroidetes/Chlorobi_group>Bacteroidota>Bacteroidia>Bacteroidales | GCF_000833975.1 | Prevotella_pectinovora | ST44_RS13175 | 10747..14361 | 1602169 | BREX-system-P-loop-protein-BrxC-[Prevotella-pectinovora] |
| WP_044064719.1 | Type6.BREX | <-REase<-?<-SIGMA+OB-Fold+AAA-ATPase+HKD-DNase-BrxL<-PglZ<-HNH<-OLD-ABC<-N6-METHYLASE<-DUF6079*<-BrxB-BrxF-iSTAND<-BrxA-DNA-binding<-BrxE<-HTH_3<-AAA+<-?<-XIS-HTH<-?<-Phage_int_SAM_5+Phage_integrase | 1217 | Bacteria>FCB_group>Bacteroidetes/Chlorobi_group>Bacteroidota>Bacteroidia>Bacteroidales | GCF_000833995.1 | Prevotella_pectinovora | ST45_RS02505 | 69518..73171 | 1602169 | BREX-system-P-loop-protein-BrxC-[Prevotella-pectinovora] |
| WP_044073670.1 | Type1.BREX | HTH_3->BrxA-DNA-binding->BrxB-BrxF-iSTAND->DUF6079*->N6-METHYLASE->PglZ->SIGMA+OB-Fold+AAA-ATPase+HKD-DNase-BrxL->REase->PARG+MACRODOMAIN-><-DDE_Tnp_ISL3 | 1206 | Bacteria>FCB_group>Bacteroidetes/Chlorobi_group>Bacteroidota>Bacteroidia>Bacteroidales | GCF_000834015.1 | Prevotella_pectinovora | ST42_RS12125 | 1627..5247 | 1602169 | BREX-system-P-loop-protein-BrxC-[Prevotella-pectinovora] |
| WP_008139697.1 | Type6.BREX | HTH_3->BrxE->BrxA-DNA-binding-p->BrxB-BrxF-iSTAND->DUF6079*->N6-METHYLASE-p->N6-METHYLASE-p->N6-METHYLASE-p->PglZ->PglZ->SIGMA+OB-Fold+AAA-ATPase+HKD-DNase-BrxL->KAP-NTPase-> | 1215 | Bacteria>FCB_group>Bacteroidetes/Chlorobi_group>Bacteroidota>Bacteroidia>Bacteroidales | GCF_001315785.1 | Phocaeicola_coprophilus | JCM13818_RS17975 | 17242..20889 | 387090 | BREX-system-P-loop-protein-BrxC-[Phocaeicola-coprophilus] |
| WP_140392864.1 | Type6.BREX | <-TatD_DNase<-?<-?<-KAP-NTPase<-SIGMA+OB-Fold+AAA-ATPase+HKD-DNase-BrxL<-PglZ<-N6-METHYLASE<-DUF6079*<-BrxB-BrxF-iSTAND<-BrxA-DNA-binding<-BrxE<-HTH_3 | 1211 | Bacteria>FCB_group>Bacteroidetes/Chlorobi_group>Bacteroidota>Bacteroidia>Bacteroidales | GCF_002159645.1 | Parabacteroides_sp._An277 | B5F77_RS15595 | 54457..58092 | 1965619 | BREX-system-P-loop-protein-BrxC-[Parabacteroides-sp.-An277] |
| WP_094389547.1 | Type1.BREX | LRR_5->?->?->?-><-?\|\|?->HTH_3->BrxA-DNA-binding->BrxB-BrxF-iSTAND->DUF6079*->N6-METHYLASE->PglZ->SIGMA+OB-Fold+AAA-ATPase+HKD-DNase-BrxL->REase-> | 1228 | Bacteria>FCB_group>Bacteroidetes/Chlorobi_group>Bacteroidota>Bacteroidia>Bacteroidales | GCF_002251345.1 | Prevotella_sp._P4-119 | CIK89_RS00860 | 20388..24074 | 2024218 | BREX-system-P-loop-protein-BrxC-[Prevotella-sp.-P4-119] |
| WP_118441918.1 | Type1.BREX | HTH_Tnp_1+HTH_21+rve+rve_3-><-?<-?\|\|BrxA-DNA-binding->BrxB-BrxF-iSTAND->DUF6079*->OLD-ABC->TOPRIM->N6-METHYLASE->?->PglZ->SIGMA+OB-Fold+AAA-ATPase+HKD-DNase-BrxL->?->?->Peptidase_M28-> | 1195 | Bacteria>FCB_group>Bacteroidetes/Chlorobi_group>Bacteroidota>Bacteroidia>Bacteroidales | GCF_003474535.1 | Phocaeicola_plebeius | DW041_RS13375 | 4564..8151 | 310297 | BREX-system-P-loop-protein-BrxC-[Phocaeicola-plebeius] |

| Protein-ID | System | Gene-Neighbourhoods | Length | Taxonomy | Assembly-ID | Strain | Locus-Tag | Genomic-coordinates | TaxID | Fasta-defline |
|---|---|---|---|---|---|---|---|---|---|---|
| WP_135993587.1 | Type1.BREX | <-SF1-SF2-Helicase<- SIGMA+OB-Fold+AAA-ATPase+ HKD-DNase -BrxL <- PglZ <-Schlafen+HATPase+wHTH<-SIR2<- N6-METHYLASE <- DUF6079 *<- BrxB-BrxF-iSTAND <- BrxA-DNA-binding <-HTH_3<-?\|\|Phage_int [illegible] | 1221 | Bacteria>FCB_group>Bacteroidetes/Chlorobi_group>Bacteroidota>Bacteroidia>Bacteroidales | GCF_004793655.1 | Muribaculum_intestinale | E5330_RS10970 | 18051..21716 | 1796646 | BREX-system-P-loop-protein-BrxC-[Muribaculum-intestinale] |
| WP_172774764.1 | Type1.BREX | OLD-ABC->TOPRIM->XIS-HTH-><-?<-?<-?\|\|HTH_3-> BrxA-DNA-binding -> BrxB-BrxF-iSTAND -> DUF6079 *-> N6-METHYLASE ->OLD-ABC+TOPRIM->SF1-SF2-Helicase->Schlafen+HATPase+wHTH-> PglZ -> [illegible] | 1217 | Bacteria>FCB_group>Bacteroidetes/Chlorobi_group>Bacteroidota>Bacteroidia>Bacteroidales | GCF_012976275.1 | Phocaeicola_vulgatus | HKQ53_RS16515 | 79396..83049 | 821 | BREX-system-P-loop-protein-BrxC-[Phocaeicola-vulgatus] |
| WP_175324259.1 | Type1.BREX | <- SIGMA+OB-Fold+AAA-ATPase+ HKD-DNase -BrxL <- PglZ <-SWIM+DUF3670+SF1-SF2-Helicase<-?<-?<-?<-Peptidase_C39_2<-ACET+TPR<- N6-METHYLASE <- DUF6079 *<- BrxB-BrxF-iSTAND <- BrxA-DNA-binding [illegible] | 1208 | Bacteria>FCB_group>Bacteroidetes/Chlorobi_group>Bacteroidota>Bacteroidia>Bacteroidales | GCF_013361175.1 | Phocaeicola_vulgatus | HUV02_RS04070 | 75655..79281 | 821 | BREX-system-P-loop-protein-BrxC-[Phocaeicola-vulgatus] |
| WP_181757796.1 | Type6.BREX | <-DUF772+DDE_Tnp_1+DDE_Tnp_1_6\|\|Arm-DNA-bind_5-><-UDG<-?<-ARG\|\|?-><- SIGMA+OB-Fold+AAA-ATPase+ HKD-DNase -BrxL <- PglZ <- N6-METHYLASE <- DUF6079 -p*<- DUF6079 -p<- BrxB-BrxF-iSTAND [illegible] | 722 | Bacteria>FCB_group>Bacteroidetes/Chlorobi_group>Bacteroidota>Bacteroidia>Bacteroidales | GCF_013874375.1 | Bacteroides_sp._CACC_737 | H1A11_RS04980 | 1170653..1172821 | 2755405 | hypothetical-protein-[Bacteroides-sp.-CACC-737] |
| WP_181757797.1 | Type6.BREX | Arm-DNA-bind_5-><-UDG<-?<-ARG\|\|?-><- SIGMA+OB-Fold+AAA-ATPase+ HKD-DNase -BrxL <- PglZ <- N6-METHYLASE <- DUF6079 -p<- DUF6079 -p*<- BrxB-BrxF-iSTAND <- BrxA-DNA-binding <- BrxE <-HTH_3<-?\|\| [illegible] | 496 | Bacteria>FCB_group>Bacteroidetes/Chlorobi_group>Bacteroidota>Bacteroidia>Bacteroidales | GCF_013874375.1 | Bacteroides_sp._CACC_737 | H1A11_RS04985 | 1172743..1174233 | 2755405 | DUF6079-family-protein-[Bacteroides-sp.-CACC-737] |
| WP_186905779.1 | Type6.BREX | <-TRAM+METHYLASE\|\|?->Arm-DNA-bind_5-><-UDG<-?<-ARG\|\|<- SIGMA+OB-Fold+AAA-ATPase+ HKD-DNase -BrxL <- PglZ <- N6-METHYLASE <- DUF6079 *<- BrxB-BrxF-iSTAND <- BrxA-DNA-binding <- BrxE <-HTH_3 [illegible] | 1213 | Bacteria>FCB_group>Bacteroidetes/Chlorobi_group>Bacteroidota>Bacteroidia>Bacteroidales | GCF_014287575.1 | Bacteroides_sp._NSJ-21 | H8S53_RS11040 | 2778988..2782629 | 2763025 | BREX-system-P-loop-protein-BrxC-[Bacteroides-sp.-NSJ-21] |
| WP_115498725.1 | Type1.BREX | <- SIGMA+OB-Fold+AAA-ATPase+ HKD-DNase -BrxL <- PglZ <-Schlafen+HATPase+wHTH<- N6-METHYLASE <- DUF6079 *<- BrxB-BrxF-iSTAND <- BrxA-DNA-binding \|\|?->?->?->?-><-tRNA\|\|Cysteine-Desulfurase->? [illegible] | 1211 | Bacteria>FCB_group>Bacteroidetes/Chlorobi_group>Bacteroidota>Bacteroidia>Bacteroidales | GCF_014385205.1 | Parabacteroides_acidifaciens | H8784_RS05825 | 88295..91930 | 2290935 | BREX-system-P-loop-protein-BrxC-[Parabacteroides-acidifaciens] |
| WP_004319896.1 | Type1.BREX | BrxA-DNA-binding -> BrxB-BrxF-iSTAND -> DUF6079 *-> N6-METHYLASE ->SIR2+SIR2->Schlafen+HATPase+wHTH-> PglZ -> SIGMA+OB-Fold+AAA-ATPase+ HKD-DNase -BrxL -><-35exo+SF1-SF2-Helicase\|\|BACON+Pep [illegible] | 1202 | Bacteria>FCB_group>Bacteroidetes/Chlorobi_group>Bacteroidota>Bacteroidia>Bacteroidales | GCF_015548155.1 | Bacteroides | P0193_RS10880 | 44218..47826 | 816 | MULTISPECIES:-BREX-system-P-loop-protein-BrxC-[Bacteroides] |
| WP_195290998.1 | Type1.BREX | Calcineurin-><-?<- SIGMA+OB-Fold+AAA-ATPase+ HKD-DNase -BrxL <- PglZ <-Cas4+SF1-SF2-Helicase<-ACET+TPR<- N6-METHYLASE <- DUF6079 *<- BrxB-BrxF-iSTAND <- BrxA-DNA-binding <-HTH_3<-Arm-DNA-bi [illegible] | 1188 | Bacteria>FCB_group>Bacteroidetes/Chlorobi_group>Bacteroidota>Bacteroidia>Bacteroidales | GCF_015548275.1 | Alistipes_finegoldii | I2C47_RS09460 | 177258..180824 | 214856 | BREX-system-P-loop-protein-BrxC-[Alistipes-finegoldii] |
| WP_195366148.1 | Type6.BREX | <-TatD_DNase<-?<-KAP-glycosylase<-KAP-NTPase<- SIGMA+OB-Fold+AAA-ATPase+ HKD-DNase -BrxL <- PglZ <- N6-METHYLASE <- DUF6079 -p<- DUF6079 -p*<- BrxB-BrxF-iSTAND <- BrxA-DNA-binding <- BrxE [illegible] | 921 | Bacteria>FCB_group>Bacteroidetes/Chlorobi_group>Bacteroidota>Bacteroidia>Bacteroidales | GCF_015550085.1 | Bacteroides_ovatus | P0368_RS20360 | 46610..49375 | 28116 | BREX-system-P-loop-protein-BrxC-[Bacteroides-ovatus] |
| WP_118403962.1 | Type1.BREX | <-Phage_int_SAM_5+Phage_integrase\|\|STYKinase-like->?->?-><-HEPN\|\|HTH_3-> BrxA-DNA-binding -> BrxB-BrxF-iSTAND -> DUF6079 *-> N6-METHYLASE ->Schlafen+HATPase+wHTH-> PglZ -> [illegible] | 1211 | Bacteria>FCB_group>Bacteroidetes/Chlorobi_group>Bacteroidota>Bacteroidia>Bacteroidales | GCF_015553065.1 | Bacteroides_cellulosilyticus | I2F01_RS18130 | 73980..77615 | 246787 | BREX-system-P-loop-protein-BrxC-[Bacteroides-cellulosilyticus] |
| WP_195571580.1 | Type1.BREX | BrxA-DNA-binding -> BrxB-BrxF-iSTAND -> DUF6079 *-> N6-METHYLASE ->SIR2+SIR2->Schlafen+HATPase+wHTH-> PglZ -> SIGMA+OB-Fold+AAA-ATPase+ HKD-DNase -BrxL -><-35exo+SF1-SF2-Helicase\|\|?->SIGMA- [illegible] | 1202 | Bacteria>FCB_group>Bacteroidetes/Chlorobi_group>Bacteroidota>Bacteroidia>Bacteroidales | GCF_015557365.1 | Bacteroides_nordii | I2G99_RS18035 | 12742..16350 | 291645 | BREX-system-P-loop-protein-BrxC-[Bacteroides-nordii] |
| WP_195628684.1 | Type1.BREX | [illegible] | 769 | Bacteria>FCB_group>Bacteroidetes/Chlorobi_group>Bacteroidota>Bacteroidia>Bacteroidales | GCF_015558155.1 | Bacteroides_finegoldii | I2H47_RS02655 | 98961..101270 | 338188 | hypothetical-protein-[Bacteroides-finegoldii] |
| WP_195663259.1 | Type1.BREX | [illegible] | 1206 | Bacteria>FCB_group>Bacteroidetes/Chlorobi_group>Bacteroidota>Bacteroidia>Bacteroidales | GCF_015559525.1 | Bacteroides_congonensis | I2I22_RS15360 | 96982..100602 | 1871006 | BREX-system-P-loop-protein-BrxC-[Bacteroides-congonensis] |
| WP_117964425.1 | Type6.BREX | [illegible] | 265 | Bacteria>FCB_group>Bacteroidetes/Chlorobi_group>Bacteroidota>Bacteroidia>Bacteroidales | GCF_015679285.1 | Bacteroides | ITJ87_RS11480 | 2822744..2823541 | 816 | MULTISPECIES:-DUF6079-family-protein-[Bacteroides] |
| WP_203041477.1 | Type1.BREX | [illegible] | 1207 | Bacteria>FCB_group>Bacteroidetes/Chlorobi_group>Bacteroidota>Bacteroidia>Bacteroidales | GCF_016803185.1 | Prevotella_copri | F8Q96_RS13415 | 22671..26294 | 165179 | BREX-system-P-loop-protein-BrxC-[Prevotella-copri] |
| WP_203049164.1 | Type6.BREX | [illegible] | 1216 | Bacteria>FCB_group>Bacteroidetes/Chlorobi_group>Bacteroidota>Bacteroidia>Bacteroidales | GCF_016803255.1 | Prevotella_copri | F8Q89_RS01960 | 71962..75612 | 165179 | BREX-system-P-loop-protein-BrxC-[Prevotella-copri] |
| WP_204430319.1 | Type1.BREX | [illegible] | 1217 | Bacteria>FCB_group>Bacteroidetes/Chlorobi_group>Bacteroidota>Bacteroidia>Bacteroidales | GCF_016899805.1 | Bacteroides_gallinaceum | H6B13_RS12495 | 13542..17195 | 1462571 | BREX-system-P-loop-protein-BrxC-[Bacteroides-gallinaceum] |
| WP_204392825.1 | Type1.BREX | [illegible] | 1185 | Bacteria>FCB_group>Bacteroidetes/Chlorobi_group>Bacteroidota>Bacteroidia>Bacteroidales | GCF_016901995.1 | Bacteroides_caecigallinarum | H6A61_RS08410 | 10706..14263 | 1411144 | BREX-system-P-loop-protein-BrxC-[Bacteroides-caecigallinarum] |
| WP_204506195.1 | Type1.BREX | [illegible] | 1187 | Bacteria>FCB_group>Bacteroidetes/Chlorobi_group>Bacteroidota>Bacteroidia>Bacteroidales | GCF_016902095.1 | Phocaeicola_plebeius | H6A37_RS09755 | 27157..30720 | 310297 | BREX-system-P-loop-protein-BrxC-[Phocaeicola-plebeius] |
| WP_211451926.1 | Type1.BREX | [illegible] | 1208 | Bacteria>FCB_group>Bacteroidetes/Chlorobi_group>Bacteroidota>Bacteroidia>Bacteroidales | GCF_018289355.1 | Phocaeicola_vulgatus | INE85_RS07560 | 1903040..1906666 | 821 | BREX-system-P-loop-protein-BrxC-[Phocaeicola-vulgatus] |
| WP_215652320.1 | Type1.BREX | [illegible] | 1206 | Bacteria>FCB_group>Bacteroidetes/Chlorobi_group>Bacteroidota>Bacteroidia>Bacteroidales | GCF_018784085.1 | Prevotella_copri | GPK65_RS08775 | 33741..37361 | 165179 | BREX-system-P-loop-protein-BrxC-[Prevotella-copri] |
| WP_215708904.1 | Type1.BREX | [illegible] | 1215 | Bacteria>FCB_group>Bacteroidetes/Chlorobi_group>Bacteroidota>Bacteroidia>Bacteroidales | GCF_018785865.1 | Bacteroides_uniformis | GRD73_RS11230 | 15032..18679 | 820 | BREX-system-P-loop-protein-BrxC-[Bacteroides-uniformis] |
| WP_215758559.1 | Type6.BREX | [illegible] | 1217 | Bacteria>FCB_group>Bacteroidetes/Chlorobi_group>Bacteroidota>Bacteroidia>Bacteroidales | GCF_018789675.1 | Hallella_faecis | KM934_00340 | 78197..81850 | 2841596 | BREX-system-P-loop-protein-BrxC-[Hallella-faecis] |
| WP_216951216.1 | Type1.BREX | [illegible] | 1208 | Bacteria>FCB_group>Bacteroidetes/Chlorobi_group>Bacteroidota>Bacteroidia>Bacteroidales | GCF_019042015.1 | Phocaeicola_vulgatus | KTG13_RS11225 | 6191..9817 | 821 | BREX-system-P-loop-protein-BrxC-[Phocaeicola-vulgatus] |
| WP_217745553.1 | Type1.BREX | [illegible] | 1206 | Bacteria>FCB_group>Bacteroidetes/Chlorobi_group>Bacteroidota>Bacteroidia>Bacteroidales | GCF_019127365.1 | Prevotella_copri | KSW85_RS06175 | 41108..44728 | 165179 | BREX-system-P-loop-protein-BrxC-[Prevotella-copri] |
| WP_217718390.1 | Type1.BREX | [illegible] | 1217 | Bacteria>FCB_group>Bacteroidetes/Chlorobi_group>Bacteroidota>Bacteroidia>Bacteroidales | GCF_019127825.1 | Bacteroides_uniformis | KSX11_RS12350 | 118043..121696 | 820 | BREX-system-P-loop-protein-BrxC-[Bacteroides-uniformis] |
| WP_217725669.1 | Type1.BREX | [illegible] | 1202 | Bacteria>FCB_group>Bacteroidetes/Chlorobi_group>Bacteroidota>Bacteroidia>Bacteroidales | GCF_019128915.1 | unclassified_Bacteroides | KSX87_RS10005 | 229853..233461 | 2646097 | MULTISPECIES:-BREX-system-P-loop-protein-BrxC-[unclassified-Bacteroides] |
| WP_217719995.1 | Type1.BREX | [illegible] | 265 | Bacteria>FCB_group>Bacteroidetes/Chlorobi_group>Bacteroidota>Bacteroidia>Bacteroidales | GCF_019412375.1 | Bacteroides_uniformis | KSX92_RS08930 | 2230..3027 | 820 | DUF6079-family-protein-[Bacteroides-uniformis] |
| WP_138333738.1 | Type6.BREX | [illegible] | 1224 | Bacteria>FCB_group>Bacteroidetes/Chlorobi_group>Bacteroidota>Bacteroidia>Bacteroidales | GCF_019412725.1 | Bacteroides | GPL18_RS15350 | 126249..129923 | 816 | MULTISPECIES:-BREX-system-P-loop-protein-BrxC-[Bacteroides] |
| WP_115473019.1 | Type1.BREX | [illegible] | 1211 | Bacteria>FCB_group>Bacteroidetes/Chlorobi_group>Bacteroidota>Bacteroidia>Bacteroidales | GCF_019733435.1 | Bacteroides_fragilis | G7060_RS07670 | 242068..245703 | 817 | BREX-system-P-loop-protein-BrxC-[Bacteroides-fragilis] |
| WP_120489887.1 | Type1.BREX | [illegible] | 1202 | Bacteria>FCB_group>Bacteroidetes/Chlorobi_group>Bacteroidota>Bacteroidia>Bacteroidales | GCF_019733535.1 | Bacteroides_ovatus | G7086_RS03160 | 85646..89254 | 28116 | BREX-system-P-loop-protein-BrxC-[Bacteroides-ovatus] |
| WP_130041244.1 | Type1.BREX | [illegible] | 1175 | Bacteria>FCB_group>Bacteroidetes/Chlorobi_group>Bacteroidota>Bacteroidia>Bacteroidales | GCF_019733625.1 | Bacteroides_thetaiotaomicron | EAJ10_RS04945 | 124743..128270 | 818 | BREX-system-P-loop-protein-BrxC-[Bacteroides-thetaiotaomicron] |
| WP_221729077.1 | Type1.BREX | [illegible] | 1206 | Bacteria>FCB_group>Bacteroidetes/Chlorobi_group>Bacteroidota>Bacteroidia>Bacteroidales | GCF_019733655.1 | Parabacteroides_merdae | HCH02_RS20805 | 4560..8180 | 46503 | BREX-system-P-loop-protein-BrxC-[Parabacteroides-merdae] |
| WP_225659484.1 | Type1.BREX | [illegible] | 1175 | Bacteria>FCB_group>Bacteroidetes/Chlorobi_group>Bacteroidota>Bacteroidia>Bacteroidales | GCF_020313445.1 | Bacteroides_thetaiotaomicron | KFX61_RS09580 | 34477..38004 | 818 | BREX-system-P-loop-protein-BrxC-[Bacteroides-thetaiotaomicron] |
| WP_217720628.1 | Type1.BREX | [illegible] | 1208 | Bacteria>FCB_group>Bacteroidetes/Chlorobi_group>Bacteroidota>Bacteroidia>Bacteroidales | GCF_020827675.1 | Bacteroidaceae | HZY58_RS04175 | 119839..123465 | 815 | MULTISPECIES:-BREX-system-P-loop-protein-BrxC-[Bacteroidaceae] |
| WP_234117060.1 | Type1.BREX | [illegible] | 1202 | Bacteria>FCB_group>Bacteroidetes/Chlorobi_group>Bacteroidota>Bacteroidia>Bacteroidales | GCF_021410745.1 | Bacteroides_ovatus | K0G75_RS22190 | 19952..23560 | 28116 | BREX-system-P-loop-protein-BrxC-[Bacteroides-ovatus] |
| WP_237798701.1 | Type6.BREX | [illegible] | 1216 | Bacteria>FCB_group>Bacteroidetes/Chlorobi_group>Bacteroidota>Bacteroidia>Bacteroidales | GCF_022024195.1 | Prevotella_bryantii | L6467_12860 | 2115705..2119355 | 77095 | BREX-system-P-loop-protein-BrxC-[Prevotella-bryantii] |
| WP_249093422.1 | Type1.BREX | [illegible] | 1217 | Bacteria>FCB_group>Bacteroidetes/Chlorobi_group>Bacteroidota>Bacteroidia>Bacteroidales | GCF_023337025.1 | Mediterranea_sp._ET5 | M3087_RS02205 | 173446..177099 | 2939419 | BREX-system-P-loop-protein-BrxC-[Mediterranea-sp.-ET5] |
| WP_250803006.1 | Type1.BREX | [illegible] | 1208 | Bacteria>FCB_group>Bacteroidetes/Chlorobi_group>Bacteroidota>Bacteroidia>Bacteroidales | GCF_023668475.1 | Phocaeicola_massiliensis | NCY86_RS00900 | 193015..196641 | 204516 | BREX-system-P-loop-protein-BrxC-[Phocaeicola-massiliensis] |
| WP_252760249.1 | Type1.BREX | [illegible] | 1205 | Bacteria>FCB_group>Bacteroidetes/Chlorobi_group>Bacteroidota>Bacteroidia>Bacteroidales | GCF_024054555.1 | Prevotella_cerevisiae | NG821_RS03330 | 12038..15655 | 2053716 | BREX-system-P-loop-protein-BrxC-[Prevotella-cerevisiae] |
| WP_252760927.1 | Type1.BREX | [illegible] | 1206 | Bacteria>FCB_group>Bacteroidetes/Chlorobi_group>Bacteroidota>Bacteroidia>Bacteroidales | GCF_024054555.1 | Prevotella_cerevisiae | NG821_RS06905 | 6531..10151 | 2053716 | BREX-system-P-loop-protein-BrxC-[Prevotella-cerevisiae] |
| WP_258327305.1 | Type1.BREX | [illegible] | 1217 | Bacteria>FCB_group>Bacteroidetes/Chlorobi_group>Bacteroidota>Bacteroidia>Bacteroidales | GCF_024733785.1 | Barnesiella_sp._ET7 | NW211_RS06370 | 166743..170396 | 2972460 | BREX-system-P-loop-protein-BrxC-[Barnesiella-sp.-ET7] |
| WP_258898999.1 | Type1.BREX | [illegible] | 1208 | Bacteria>FCB_group>Bacteroidetes/Chlorobi_group>Bacteroidota>Bacteroidia>Bacteroidales | GCF_024759365.1 | Bacteroides_ovatus | NXX44_RS13110 | 3467956..3471582 | 28116 | BREX-system-P-loop-protein-BrxC-[Bacteroides-ovatus] |
| WP_258991265.1 | Type1.BREX | [illegible] | 1208 | Bacteria>FCB_group>Bacteroidetes/Chlorobi_group>Bacteroidota>Bacteroidia>Bacteroidales | GCF_024790585.1 | Phocaeicola_vulgatus | NXX11_RS15345 | 3784105..3787731 | 821 | BREX-system-P-loop-protein-BrxC-[Phocaeicola-vulgatus] |
| WP_259011403.1 | Type1.BREX | [illegible] | 1202 | Bacteria>FCB_group>Bacteroidetes/Chlorobi_group>Bacteroidota>Bacteroidia>Bacteroidales | GCF_024791585.1 | Bacteroides_sp._BFG-638 | NXX54_RS14145 | 3701947..3705555 | 2972765 | BREX-system-P-loop-protein-BrxC-[Bacteroides-sp.-BFG-638] |
| WP_258973412.1 | Type1.BREX | [illegible] | 1133 | Bacteria>FCB_group>Bacteroidetes/Chlorobi_group>Bacteroidota>Bacteroidia>Bacteroidales | GCF_024793205.1 | Bacteroides_thetaiotaomicron | NXX32_RS11560 | 3197619..3201020 | 818 | BREX-system-P-loop-protein-BrxC-[Bacteroides-thetaiotaomicron] |
| WP_258979859.1 | Type1.BREX | [illegible] | 780 | Bacteria>FCB_group>Bacteroidetes/Chlorobi_group>Bacteroidota>Bacteroidia>Bacteroidales | GCF_024793605.1 | Bacteroides_sp._BFG-637 | NXX38_RS14295 | 3701972..3704314 | 2972764 | BREX-system-P-loop-protein-BrxC-[Bacteroides-sp.-BFG-637] |
| WP_009000708.1 | Type1.BREX | [illegible] | 1202 | Bacteria>FCB_group>Bacteroidetes/Chlorobi_group>Bacteroidota>Bacteroidia>Bacteroidales | GCF_025147085.1 | Bacteroides | NQ505_RS12870 | 3648371..3651979 | 816 | MULTISPECIES:-BREX-system-P-loop-protein-BrxC-[Bacteroides] |
| WP_195661285.1 | Type1.BREX | [illegible] | 1202 | Bacteria>FCB_group>Bacteroidetes/Chlorobi_group>Bacteroidota>Bacteroidia>Bacteroidales | GCF_025914175.1 | Bacteroides_nordii | KQP55_RS20540 | 5436855..5440463 | 291645 | BREX-system-P-loop-protein-BrxC-[Bacteroides-nordii] |
| WP_264912090.1 | Type1.BREX | [illegible] | 1206 | Bacteria>FCB_group>Bacteroidetes/Chlorobi_group>Bacteroidota>Bacteroidia>Bacteroidales | GCF_026015575.1 | Prevotella_copri | ONS98_RS12175 | 17740..21360 | 165179 | BREX-system-P-loop-protein-BrxC-[Prevotella-copri] |
| WP_264965401.1 | Type1.BREX | [illegible] | 1194 | Bacteria>FCB_group>Bacteroidetes/Chlorobi_group>Bacteroidota>Bacteroidia>Bacteroidales | GCF_026015685.1 | Prevotella_copri | ONT16_RS03155 | 729578..733162 | 165179 | BREX-system-P-loop-protein-BrxC-[Prevotella-copri] |
| WP_270558549.1 | Type1.BREX | [illegible] | 1216 | Bacteria>FCB_group>Bacteroidetes/Chlorobi_group>Bacteroidota>Bacteroidia>Bacteroidales | GCF_027686315.1 | Bacteroides_uniformis | PG022_RS08930 | 563554..567204 | 820 | BREX-system-P-loop-protein-BrxC-[Bacteroides-uniformis] |
| WP_271687043.1 | Type1.BREX | [illegible] | 1208 | Bacteria>FCB_group>Bacteroidetes/Chlorobi_group>Bacteroidota>Bacteroidia>Bacteroidales | GCF_028199265.1 | Bacteroides_xylanisolvens | PL435_RS07875 | 93897..97523 | 371601 | BREX-system-P-loop-protein-BrxC-[Bacteroides-xylanisolvens] |
| WP_271704578.1 | Type1.BREX | [illegible] | 1209 | Bacteria>FCB_group>Bacteroidetes/Chlorobi_group>Bacteroidota>Bacteroidia>Bacteroidales | GCF_028200085.1 | Bacteroides_xylanisolvens | PL493_RS08910 | 20513..24142 | 371601 | BREX-system-P-loop-protein-BrxC-[Bacteroides-xylanisolvens] |
| WP_272709339.1 | Type1.BREX | [illegible] | 1208 | Bacteria>FCB_group>Bacteroidetes/Chlorobi_group>Bacteroidota>Bacteroidia>Bacteroidales | GCF_028528025.1 | Bacteroides | PQG94_RS11850 | 55098..58724 | 816 | MULTISPECIES:-BREX-system-P-loop-protein-BrxC-[Bacteroides] |
| WP_272706625.1 | Type6.BREX | [illegible] | 191 | Bacteria>FCB_group>Bacteroidetes/Chlorobi_group>Bacteroidota>Bacteroidia>Bacteroidales | GCF_028528065.1 | Bacteroides_faecis | PQH00_RS23500 | 20443..21015 | 674529 | DUF6079-family-protein,-partial-[Bacteroides-faecis] |
| WP_272706631.1 | Type6.BREX | [illegible] | 862 | Bacteria>FCB_group>Bacteroidetes/Chlorobi_group>Bacteroidota>Bacteroidia>Bacteroidales | GCF_028528065.1 | Bacteroides_faecis | PQH00_RS23505 | 21026..23611 | 674529 | BREX-system-P-loop-protein-BrxC,-partial-[Bacteroides-faecis] |
| WP_274990748.1 | Type6.BREX | [illegible] | 772 | Bacteria>FCB_group>Bacteroidetes/Chlorobi_group>Bacteroidota>Bacteroidia>Bacteroidales | GCF_028993715.1 | Bacteroides_uniformis | PX688_RS13675 | 79793..82111 | 820 | hypothetical-protein-[Bacteroides-uniformis] |
| WP_274990749.1 | Type6.BREX | [illegible] | 411 | Bacteria>FCB_group>Bacteroidetes/Chlorobi_group>Bacteroidota>Bacteroidia>Bacteroidales | GCF_028993715.1 | Bacteroides_uniformis | PX688_RS13680 | 82199..83434 | 820 | DUF6079-family-protein-[Bacteroides-uniformis] |
| WP_032560932.1 | Type1.BREX | [illegible] | 1211 | Bacteria>FCB_group>Bacteroidetes/Chlorobi_group>Bacteroidota>Bacteroidia>Bacteroidales | GCF_029872615.1 | Bacteroides_fragilis | M068_RS14675 | 124538..128173 | 817 | BREX-system-P-loop-protein-BrxC-[Bacteroides-fragilis] |
| WP_074557116.1 | Type1.BREX | [illegible] | 1202 | Bacteria>FCB_group>Bacteroidetes/Chlorobi_group>Bacteroidota>Bacteroidia>Bacteroidales | GCF_900102645.1 | Bacteroides_ovatus | BLR14_RS05840 | 106864..110472 | 28116 | BREX-system-P-loop-protein-BrxC-[Bacteroides-ovatus] |
| WP_072542576.1 | Type1.BREX | [illegible] | 1195 | Bacteria>FCB_group>Bacteroidetes/Chlorobi_group>Bacteroidota>Bacteroidia>Bacteroidales | GCF_900128455.1 | Bacteroides_mediterraneensis | BQ7480_RS08760 | 2129051..2132638 | 1841856 | BREX-system-P-loop-protein-BrxC-[Bacteroides-mediterraneensis] |
| WP_254795118.1 | Type1.BREX | [illegible] | 1209 | Bacteria>FCB_group>Bacteroidetes/Chlorobi_group>Bacteroidota>Bacteroidia>Bacteroidales | GCF_900142525.1 | Prevotella_ruminicola | BUB86_RS07050 | 46256..49885 | 839 | BREX-system-P-loop-protein-BrxC-[Prevotella-ruminicola] |
| WP_125105113.1 | Type1.BREX | [illegible] | 1217 | Bacteria>FCB_group>Bacteroidetes/Chlorobi_group>Bacteroidota>Bacteroidia>Bacteroidales | GCF_900625065.1 | Prevotella_marseillensis | EJL04_RS12230 | 3058927..3062580 | 2479840 | BREX-system-P-loop-protein-BrxC-[Prevotella-marseillensis] |
| WP_144153273.1 | Type1.BREX | [illegible] | 1194 | Bacteria>FCB_group>Bacteroidetes/Chlorobi_group>Bacteroidota>Bacteroidia>Bacteroidales | GCF_902167755.1 | Prevotella_copri | BIL0145876EF_RS09275 | 107988..111572 | 165179 | BREX-system-P-loop-protein-BrxC-[Prevotella-copri] |
| WP_032543214.1 | Type1.BREX | [illegible] | 1211 | Bacteria>FCB_group>Bacteroidetes/Chlorobi_group>Bacteroidota>Bacteroidia>Bacteroidales | GCF_902364655.1 | Bacteroides_fragilis | PDN68_RS18445 | 100181..103816 | 817 | BREX-system-P-loop-protein-BrxC-[Bacteroides-fragilis] |
| WP_008770354.1 | Type6.BREX | [illegible] | 1219 | Bacteria>FCB_group>Bacteroidetes/Chlorobi_group>Bacteroidota>Bacteroidia>Bacteroidales | GCF_902374095.1 | Bacteroides | BSAG_RS19905 | 423901..427560 | 816 | MULTISPECIES:-BREX-system-P-loop-protein-BrxC-[Bacteroides] |
| WP_060407611.1 | Type1.BREX | [illegible] | 1205 | Bacteria>FCB_group>Bacteroidetes/Chlorobi_group>Bacteroidota>Bacteroidia>Bacteroidales | GCF_902387305.1 | Bacteroides_cellulosilyticus | FYC30_RS03960 | 75039..78656 | 246787 | BREX-system-P-loop-protein-BrxC-[Bacteroides-cellulosilyticus] |
| WP_174646033.1 | Type6.BREX | [illegible] | 1215 | Bacteria>FCB_group>Bacteroidetes/Chlorobi_group>Bacteroidota>Bacteroidia>Bacteroidales | GCF_902825465.1 | Bacteroides_sp._Marseille-P8574 | HUX74_RS00900 | 196439..200086 | 2697504 | BREX-system-P-loop-protein-BrxC-[Bacteroides-sp.-Marseille-P8574] |
| WP_118419043.1 | Type1.BREX | [illegible] | 1202 | Bacteria>FCB_group>Bacteroidetes/Chlorobi_group>Bacteroidota>Bacteroidia>Bacteroidales | GCF_918279245.1 | Bacteroides_ovatus | P0203_RS09380 | 101158..104766 | 28116 | BREX-system-P-loop-protein-BrxC-[Bacteroides-ovatus] |
| WP_117985926.1 | Type1.BREX | [illegible] | 1210 | Bacteria>FCB_group>Bacteroidetes/Chlorobi_group>Bacteroidota>Bacteroidia>Bacteroidales | GCF_934726555.1 | Bacteroides | O1446_RS12920 | 68574..72206 | 816 | MULTISPECIES:-BREX-system-P-loop-protein-BrxC-[Bacteroides] |
| WP_158582848.1 | Type1.BREX | [illegible] | 1210 | Bacteria>FCB_group>Bacteroidetes/Chlorobi_group>Bacteroidota>Bacteroidia>Bacteroidales | GCF_938033175.1 | Parabacteroides_sp._AF17-28 | DWW90_RS15090 | 22936..26568 | 2292241 | BREX-system-P-loop-protein-BrxC-[Parabacteroides-sp.-AF17-28] |
| WP_046149424.1 | Type6.BREX | [illegible] | 1217 | Bacteria>FCB_group>Bacteroidetes/Chlorobi_group>Bacteroidota>Bacteroidia>Bacteroidales | - | Parabacteroides_sp._HGS0025 | HMPREF1212_RS26675 | 2885204..2888857 | 1078087 | BREX-system-P-loop-protein-BrxC-[Parabacteroides-sp.-HGS0025] |
| WP_118139864.1 | Type1.BREX | [illegible] | 1202 | Bacteria>FCB_group>Bacteroidetes/Chlorobi_group>Bacteroidota>Bacteroidia>Bacteroidales | - | Prevotella_copri | DWV60_RS07145 | 31878..35486 | 165179 | BREX-system-P-loop-protein-BrxC-[Prevotella-copri] |
| WP_117695280.1 | Type6.BREX | [illegible] | 1216 | Bacteria>FCB_group>Bacteroidetes/Chlorobi_group>Bacteroidota>Bacteroidia>Bacteroidales | - | Prevotella_copri | DXC61_RS12020 | 26690..30340 | 165179 | BREX-system-P-loop-protein-BrxC-[Prevotella-copri] |
| WP_233339205.1 | Type6.BREX | [illegible] | 1216 | Bacteria>FCB_group>Bacteroidetes/Chlorobi_group>Bacteroidota>Bacteroidia>Bacteroidales | - | Prevotella_copri | LYY06_RS08410 | 34942..38592 | 165179 | BREX-system-P-loop-protein-BrxC-[Prevotella-copri] |
| WP_264908465.1 | Type1.BREX | [illegible] | 1206 | Bacteria>FCB_group>Bacteroidetes/Chlorobi_group>Bacteroidota>Bacteroidia>Bacteroidales | - | Prevotella_copri | ONT12_RS10110 | 2444120..2447740 | 165179 | BREX-system-P-loop-protein-BrxC-[Prevotella-copri] |
| WP_264904325.1 | Type1.BREX | [illegible] | 1206 | Bacteria>FCB_group>Bacteroidetes/Chlorobi_group>Bacteroidota>Bacteroidia>Bacteroidales | - | Prevotella_copri | ONT18_RS04485 | 1079147..1082767 | 165179 | BREX-system-P-loop-protein-BrxC-[Prevotella-copri] |
| WP_073207711.1 | Type1.BREX | [illegible] | 1206 | Bacteria>FCB_group>Bacteroidetes/Chlorobi_group>Bacteroidota>Bacteroidia>Bacteroidales | - | Prevotella_ruminicola | SAMN05216463_10942 | 46265..49885 | 839 | BREX-system-P-loop-protein-BrxC-[Prevotella-ruminicola] |
| WP_181980669.1 | Type1.BREX | [illegible] | 1192 | Bacteria>FCB_group>Bacteroidetes/Chlorobi_group>Bacteroidota>Bacteroidia | GCF_015558535.1 | Bacteroidales | QH473_RS20245 | 26845..30423 | 171549 | MULTISPECIES:-BREX-system-P-loop-protein-BrxC-[Bacteroidales] |
| BBE20443.1 | Type1.BREX | [illegible] | 1173 | Bacteria>FCB_group>Bacteroidetes/Chlorobi_group>Bacteroidota>Bacteroidia>Marinilabiliales | - | Aquipluma_nitroreducens | AQPE_4635 | 5186015..5189536 | 2010828 | hypothetical-protein-AQPE_4635-[Aquipluma-nitroreducens] |
| MCF8362389.1 | Type1.BREX | [illegible] | 1180 | Bacteria>FCB_group>Bacteroidetes/Chlorobi_group>Bacteroidota>Bacteroidia>Marinilabiliales | GCA_021740345.1 | Prolixibacteraceae_bacterium | K9G70_07170 | 11489..15031 | 2053594 | BREX-system-P-loop-protein-BrxC-[Prolixibacteraceae-bacterium] |
| MCK9411439.1 | Type1.BREX | [illegible] | 1173 | Bacteria>FCB_group>Bacteroidetes/Chlorobi_group>Bacteroidota>Bacteroidia>Marinilabiliales | GCA_023229575.1 | Prolixibacteraceae_bacterium | M0Q53_04000 | 25065..28586 | 2053594 | BREX-system-P-loop-protein-BrxC-[Prolixibacteraceae-bacterium] |
| MCU4173933.1 | Type1.BREX | [illegible] | 1167 | Bacteria>FCB_group>Bacteroidetes/Chlorobi_group>Bacteroidota>Bacteroidia>Marinilabiliales | GCA_025499625.1 | Marinilabiliaceae_bacterium_N1Y90 | J1N11_02510 | 10462..13965 | 2816964 | BREX-system-P-loop-protein-BrxC-[Marinilabiliaceae-bacterium-N1Y90] |
| WP_062122677.1 | Type1.BREX | [illegible] | 329 | Bacteria>FCB_group>Bacteroidetes/Chlorobi_group>Bacteroidota>Bacteroidia>Marinilabiliales | GCF_000974365.1 | Geofilum_rubicundum | JCM15548_RS04470 | 156242..157231 | 472113 | hypothetical-protein-[Geofilum-rubicundum] |
| WP_107823954.1 | Type1.BREX | [illegible] | 1168 | Bacteria>FCB_group>Bacteroidetes/Chlorobi_group>Bacteroidota>Bacteroidia>Marinilabiliales | GCF_003046625.1 | Mangrovibacterium_marinum | C8N47_RS17935 | 11546..15052 | 1639118 | BREX-system-P-loop-protein-BrxC-[Mangrovibacterium-marinum] |
| WP_163710191.1 | Type1.BREX | [illegible] | 1168 | Bacteria>FCB_group>Bacteroidetes/Chlorobi_group>Bacteroidota>Bacteroidia>Marinilabiliales | GCF_010882175.1 | Mangrovibacterium_lignilyticum | GJV22_RS06145 | 1519093..1522599 | 2668052 | BREX-system-P-loop-protein-BrxC-[Mangrovibacterium-lignilyticum] |
| WP_250725046.1 | Type1.BREX | [illegible] | 1173 | Bacteria>FCB_group>Bacteroidetes/Chlorobi_group>Bacteroidota>Bacteroidia>Marinilabiliales | GCF_023674465.1 | Alkaliflexus_sp._Ai-910 | M9189_RS04830 | 1155477..1158998 | 2949635 | BREX-system-P-loop-protein-BrxC-[Alkaliflexus-sp.-Ai-910] |
| OJV92258.1 | Type1.BREX | [illegible] | 1200 | Bacteria>FCB_group>Bacteroidetes/Chlorobi_group>Bacteroidota>Bacteroidia>unclassified_Bacteroidia | GCA_001897595.1 | Bacteroidia_bacterium_44-10 | BGO34_14870 | 43076..46678 | 1895721 | hypothetical-protein-BGO34_14870-[Bacteroidia-bacterium-44-10] |
| PJE46487.1 | Type1.BREX | [illegible] | 1169 | Bacteria>FCB_group>Bacteroidetes/Chlorobi_group>Bacteroidota>Chitinophagia>Chitinophagales | GCA_002786355.1 | Sediminibacterium_sp._FEMGT703S | CUR34_07570 | 66784..70293 | 2080766 | MAG:-hypothetical-protein-CUR34_07570-[Sediminibacterium-sp.] |
| QEC55281.1 | Type1.BREX | [illegible] | 1177 | Bacteria>FCB_group>Bacteroidetes/Chlorobi_group>Bacteroidota>Chitinophagia>Chitinophagales | GCA_007970805.1 | Flavisolibacter_ginsenosidimutans | FSB75_04955 | 1196460..1199993 | 661481 | BREX-system-P-loop-protein-BrxC-[Flavisolibacter-ginsenosidimutans] |
| NTS43012.1 | Type1.BREX | [illegible] | 1173 | Bacteria>FCB_group>Bacteroidetes/Chlorobi_group>Bacteroidota>Chitinophagia>Chitinophagales | GCA_013327885.1 | Flavisolibacter_longurius | HRG84_19150 | 28734..32255 | 2740583 | BREX-system-P-loop-protein-BrxC-[Flavisolibacter-longurius] |
| WP_172623061.1 | Type1.BREX | [illegible] | 1174 | Bacteria>FCB_group>Bacteroidetes/Chlorobi_group>Bacteroidota>Chitinophagia>Chitinophagales | GCF_007970805.1 | Flavisolibacter_ginsenosidimutans | FSB75_RS04955 | 1196460..1199984 | 661481 | BREX-system-P-loop-protein-BrxC-[Flavisolibacter-ginsenosidimutans] |
| ADB40822.1 | Type1.BREX | [illegible] | 1186 | Bacteria>FCB_group>Bacteroidetes/Chlorobi_group>Bacteroidota>Cytophagia>Cytophagales | GCA_000024525.1 | Spirosoma_linguale_DSM_74 | Slin_4844 | 5894993..5898553 | 504472 | hypothetical-protein-Slin_4844-[Spirosoma-linguale-DSM-74] |
| UOQ83254.1 | Type1.BREX | [illegible] | 1148 | Bacteria>FCB_group>Bacteroidetes/Chlorobi_group>Bacteroidota>Cytophagia>Cytophagales | GCA_022919455.1 | Hymenobacter_sp._5414T-23 | MUN83_21225 | 87571..91017 | 2932252 | BREX-system-P-loop-protein-BrxC-[Hymenobacter-sp.-5414T-23] |
| WP_028665035.1 | Type1.BREX | [illegible] | 1171 | Bacteria>FCB_group>Bacteroidetes/Chlorobi_group>Bacteroidota>Cytophagia>Cytophagales | GCF_000423565.1 | Runella_zeae | G563_RS0108985 | 25803..29318 | 94255 | BREX-system-P-loop-protein-BrxC-[Runella-zeae] |
| WP_053405044.1 | Type1.BREX | [illegible] | 1179 | Bacteria>FCB_group>Bacteroidetes/Chlorobi_group>Bacteroidota>Cytophagia>Cytophagales | GCF_001274635.1 | Persicobacter_sp._CCB-QB2 | NG22_RS06825 | 1502189..1505728 | 1561025 | BREX-system-P-loop-protein-BrxC-[Persicobacter-sp.-CCB-QB2] |
| WP_073850520.1 | Type1.BREX | [illegible] | 1179 | Bacteria>FCB_group>Bacteroidetes/Chlorobi_group>Bacteroidota>Cytophagia>Cytophagales | GCF_001907195.1 | Pontibacter_flavimaris | A3841_RS01550 | 388287..391826 | 1797110 | BREX-system-P-loop-protein-BrxC-[Pontibacter-flavimaris] |
| WP_146899771.1 | Type1.BREX | [illegible] | 1182 | Bacteria>FCB_group>Bacteroidetes/Chlorobi_group>Bacteroidota>Cytophagia>Cytophagales | GCF_007991635.1 | Adhaeribacter_aerolatus | AAE02nite_RS16235 | 111241..114789 | 670289 | BREX-system-P-loop-protein-BrxC-[Adhaeribacter-aerolatus] |
| WP_168675169.1 | Type1.BREX | [illegible] | 1172 | Bacteria>FCB_group>Bacteroidetes/Chlorobi_group>Bacteroidota>Cytophagia>Cytophagales | GCF_012275535.1 | Hymenobacter_artigasi | HBN54_RS20840 | 55199..58717 | 2719616 | BREX-system-P-loop-protein-BrxC-[Hymenobacter-artigasi] |
| WP_185195977.1 | Type1.BREX | [illegible] | 1179 | Bacteria>FCB_group>Bacteroidetes/Chlorobi_group>Bacteroidota>Cytophagia>Cytophagales | GCF_014212495.1 | Pontibacter_sp._Tf4 | H7F15_RS05560 | 22760..26299 | 2761620 | BREX-system-P-loop-protein-BrxC-[Pontibacter-sp.-Tf4] |
| WP_187455529.1 | Type1.BREX | [illegible] | 1178 | Bacteria>FCB_group>Bacteroidetes/Chlorobi_group>Bacteroidota>Cytophagia>Cytophagales | GCF_014349165.1 | Hymenobacter_sp._BT491 | H9I52_RS21275 | 49708..53244 | 2766779 | BREX-system-P-loop-protein-BrxC-[Hymenobacter-sp.-BT491] |
| WP_188933335.1 | Type1.BREX | [illegible] | 1165 | Bacteria>FCB_group>Bacteroidetes/Chlorobi_group>Bacteroidota>Cytophagia>Cytophagales | GCF_014641595.1 | Dyadobacter_endophyticus | IEY24_RS14735 | 618400..621897 | 1749036 | BREX-system-P-loop-protein-BrxC-[Dyadobacter-endophyticus] |
| WP_191042738.1 | Type1.BREX | [illegible] | 1206 | Bacteria>FCB_group>Bacteroidetes/Chlorobi_group>Bacteroidota>Cytophagia>Cytophagales | GCF_014699035.1 | Spirosoma_validum | IC230_RS29845 | 51060..54680 | 2771355 | BREX-system-P-loop-protein-BrxC-[Spirosoma-validum] |
| WP_191006191.1 | Type6.BREX | [illegible] | 1204 | Bacteria>FCB_group>Bacteroidetes/Chlorobi_group>Bacteroidota>Cytophagia>Cytophagales | GCF_014699115.1 | Hymenobacter_montanus | IC235_RS15945 | 31815..35429 | 2771359 | BREX-system-P-loop-protein-BrxC-[Hymenobacter-montanus] |

| Protein-ID | System | Gene-Neighbourhoods | Length | Taxonomy | Assembly-ID | Strain | Locus-Tag | Genomic-coordinates | TaxID | Fasta-defline |
|---|---|---|---|---|---|---|---|---|---|---|
| WP_204660265.1 | Type6.BREX | <-HTH_3\|\|?->?->CopG/RHH->CR-ATPase5+REase->AbiEi-> BrxE -> BrxA-DNA-binding -> BrxB-BrxF-iSTAND -> DUF6079 *-> N6-METHYLASE -> PglZ -> SIGMA+OB-Fold+AAA-ATPase+HKD-DNase-BrxL ->McrB-AAA->McrC-NTD+REase-><-MORC+DUF3684+REase<-DOC+zhth<-?\|\|HEPN-> | 1216 | Bacteria>FCB_group>Bacteroidetes/Chlorobi_group>Bacteroidota>Cytophagia>Cytophagales | GCF_016894445.1 | Dyadobacter_sandarakinus | HWI92_RS00525 | 139788..143438 | 2747268 | BREX-system-P-loop-protein-BrxC-[Dyadobacter-sandarakinus] |
| WP_205503392.1 | Type1.BREX | <- wHTH+WYL+WCX \|\|SF1-SF2-Helicase->RVT->?-><- SIGMA+OB-Fold+AAA-ATPase+HKD-DNase-BrxL <- PglZ <- N6-METHYLASE <- DUF6079 *<- BrxB-BrxF-iSTAND <- BrxA-DNA-binding <-McrC-NTD+REase<-McrB-AAA\|\|?->DDE_Tnp_4+DDE_Tnp_1_2-><-tRNA\|\|HAD+HAD->Peptidase_M50-> | 1180 | Bacteria>FCB_group>Bacteroidetes/Chlorobi_group>Bacteroidota>Cytophagia>Cytophagales | GCF_016924645.1 | Rufibacter_sp._SYSU_D00308 | JT718_RS14320 | 179098..182640 | 2812556 | BREX-system-P-loop-protein-BrxC-[Rufibacter-sp.-SYSU-D00308] |
| WP_234861365.1 | Type1.BREX | STYKIN-><-?\|\|?->GDResolvase-><-?\|\|TPR+TPR-> BrxA-DNA-binding -> BrxB-BrxF-iSTAND -> DUF6079 *-> N6-METHYLASE ->OLD-ABC+TOPRIM-> PglZ -> SIGMA+OB-Fold+AAA-ATPase+HKD-DNase-BrxL ->TIR->?->?->?->ParB+N6_N4_METHYLASE-> | 1174 | Bacteria>FCB_group>Bacteroidetes/Chlorobi_group>Bacteroidota>Cytophagia>Cytophagales | GCF_021491425.1 | Mariniradius_sp._RY-2 | LOU89_RS09865 | 251856..255380 | 2909237 | BREX-system-P-loop-protein-BrxC-[Mariniradius-sp.-RY-2] |
| WP_241348547.1 | Type1.BREX | <-ParB+N6_N4_METHYLASE<-?<-?<- SIGMA+OB-Fold+AAA-ATPase+HKD-DNase-BrxL <- PglZ <-CR-Kinase+CR-Kinase<- N6-METHYLASE <-?<- DUF6079 *<- BrxB-BrxF-iSTAND <- BrxA-DNA-binding <-TPR+TPR<-MarR-HTH\|\|?->?-><-?<-GDResolvase | 1174 | Bacteria>FCB_group>Bacteroidetes/Chlorobi_group>Bacteroidota>Cytophagia>Cytophagales | GCF_022549775.1 | Belliella_sp._DSM_111904 | MM239_RS12315 | 24692..28216 | 2923435 | BREX-system-P-loop-protein-BrxC-[Belliella-sp.-DSM-111904] |
| WP_243520943.1 | Type6.BREX | Glyoxalase-><-?<-HEPN<-?\|\|?->?-> BrxE -> BrxA-DNA-binding -> BrxB-BrxF-iSTAND -> DUF6079 *-> N6-METHYLASE ->HEPN-> PglZ -> SIGMA+OB-Fold+AAA-ATPase+HKD-DNase-BrxL ->?-> HKD-DNase ->HTH+REase-><-CopG_antitoxin | 1204 | Bacteria>FCB_group>Bacteroidetes/Chlorobi_group>Bacteroidota>Cytophagia>Cytophagales | GCF_022811645.1 | Hymenobacter_monticola | MTP16_RS25545 | 68480..72094 | 1705399 | BREX-system-P-loop-protein-BrxC-[Hymenobacter-monticola] |
| WP_262922299.1 | Type1.BREX | <-pPIWI+pPIWI\|\|HEPN-><-REase<-?\|\|Phage_integrase+Phage_integrase->NUMOD4+HNH->HTH_3-> BrxA-DNA-binding -> BrxB-BrxF-iSTAND -> DUF6079 *-> N6-METHYLASE -p-> N6-METHYLASE -> N6-METHYLASE -p-> PglZ -> SIGMA+OB-Fold+AAA-ATPase+HKD-DNase-BrxL -><-SF1-SF2-Helicase+REase\|\|NtrC-AAA+FIS-HTH->LexA-protease-> | 1164 | Bacteria>FCB_group>Bacteroidetes/Chlorobi_group>Bacteroidota>Cytophagia>Cytophagales | GCF_022919455.1 | Hymenobacter_sp._5414T-23 | MUN83_RS21225 | 87523..91017 | 2932252 | BREX-system-P-loop-protein-BrxC-[Hymenobacter-sp.-5414T-23] |
| WP_235164443.1 | Type6.BREX | <-STYKIN<-?<-?<-?<-McrC-NTD+REase<-HTH+McrB-AAA<- SIGMA+OB-Fold+AAA-ATPase+HKD-DNase-BrxL <- PglZ <- N6-METHYLASE <- DUF6079 *<- BrxB-BrxF-iSTAND <- BrxA-DNA-binding <- BrxE <-AbiEi\|\|NERD+SF1-SF2-Helicase->HTH_21+rve+rve_3->?->wHTH+HNH->REase+N-glycosylase-> | 1216 | Bacteria>FCB_group>Bacteroidetes/Chlorobi_group>Bacteroidota>Cytophagia>Cytophagales | GCF_023821985.2 | Dyadobacter_chenhuakuii | NFI80_RS01415 | 331734..335384 | 2909339 | BREX-system-P-loop-protein-BrxC-[Dyadobacter-chenhuakuii] |
| WP_269011923.1 | Type1.BREX | HTH_3->Couple_hipA+HipA_C-><-?<-HNH+Polbeta<- SIGMA+OB-Fold+AAA-ATPase+HKD-DNase-BrxL <- PglZ <- N6-METHYLASE <- DUF6079 *<- BrxB-BrxF-iSTAND <- BrxA-DNA-binding <-?<-?<-?<-?<-?<- wHTH+WYL+WCX \|\|TPR+TPR+TPR-> | 1166 | Bacteria>FCB_group>Bacteroidetes/Chlorobi_group>Bacteroidota>Cytophagia>Cytophagales | GCF_026938185.1 | Aquirufa_antheringensis | G9H55_RS04755 | 131184..134684 | 2516559 | BREX-system-P-loop-protein-BrxC-[Aquirufa-antheringensis] |
| WP_244679416.1 | Type1.BREX | <-pPIWI+pPIWI\|\|HEPN-><-REase<-?\|\|Phage_integrase+Phage_integrase->NUMOD4+HNH->HTH_3-> BrxA-DNA-binding -> BrxB-BrxF-iSTAND -> DUF6079 *-> N6-METHYLASE -p-> N6-METHYLASE -> N6-METHYLASE -p-> PglZ -> SIGMA+OB-Fold+AAA-ATPase+HKD-DNase-BrxL -><-SF1-SF2-Helicase+REase\|\|NtrC-AAA+FIS-HTH->LexA-protease-> | 1148 | Bacteria>FCB_group>Bacteroidetes/Chlorobi_group>Bacteroidota>Cytophagia>Cytophagales | - | Hymenobacter_sp._5414T-23 | MUN83_21225 | 87571..91017 | 2932252 | BREX-system-P-loop-protein-BrxC-[Hymenobacter-sp.-5414T-23] |
| OIQ21186.1 | Type1.BREX | CHTH+LexA-protease-><-GDResolvase+Recombinase+ZnR<-?\|\|?-><-?\|\| BrxA-DNA-binding -> BrxB-BrxF-iSTAND -> DUF6079 *-> N6-METHYLASE ->SF1-SF2-Helicase->?-> PglZ -> SIGMA+OB-Fold+AAA-ATPase+HKD-DNase-BrxL ->ParB+HNH->URI-DNase->?->GDResolvase-> | 1165 | Bacteria>FCB_group>Bacteroidetes/Chlorobi_group>Bacteroidota>Flavobacteriia>Flavobacteriales | GCA_001874145.1 | Lacinutrix_sp._MedPE-SW | BM549_09430 | 19615..23112 | 1860087 | MAG:-hypothetical-protein-BM549_09430-[Lacinutrix-sp.-MedPE-SW] |
| MAM30277.1 | Type1.BREX | <-ParB+N6_N4_METHYLASE\|\|XIS-HTH-><- SIGMA+OB-Fold+AAA-ATPase+HKD-DNase-BrxL <- PglZ <-TPR+S1<-NERD+SF1-SF2-Helicase<- N6-METHYLASE <-ParB+HNH<- DUF6079 *<- BrxB-BrxF-iSTAND <- BrxA-DNA-binding <-HTH_3\|\|GDResolvase->?->?->ABC-ATPase+SbcC-> N6-METHYLASE -> | 1165 | Bacteria>FCB_group>Bacteroidetes/Chlorobi_group>Bacteroidota>Flavobacteriia>Flavobacteriales | GCA_002693005.1 | Flavobacteriaceae_bacterium | CMC13_14765 | 19860..23357 | 1871037 | hypothetical-protein-[Flavobacteriaceae-bacterium] |
| ROI04325.1 | Type1.BREX | <-HTH_3<-?\|\|?->?-><-?<-?<- SIGMA+OB-Fold+AAA-ATPase+HKD-DNase-BrxL <- PglZ <- N6-METHYLASE <- DUF6079 *<- BrxB-BrxF-iSTAND <- BrxA-DNA-binding <-Schlafen+HATPase+wHTH\|\|?->ChlI+Lon_C+AAA-><-?\|\|?->SLOG-> | 1174 | Bacteria>FCB_group>Bacteroidetes/Chlorobi_group>Bacteroidota>Flavobacteriia>Flavobacteriales | GCA_003729975.1 | Kaistella_haifensis | EGH90_10140 | 55406..58930 | 421525 | BREX-system-P-loop-protein-BrxC-[Kaistella-haifensis] |
| AZB21330.1 | Type1.BREX | <-SLOG<-?\|\|?-><-ChlI+Lon_C+AAA\|\| BrxA-DNA-binding -> BrxB-BrxF-iSTAND -> DUF6079 -p-> DUF6079 -p*-> N6-METHYLASE -> PglZ -> SIGMA+OB-Fold+AAA-ATPase+HKD-DNase-BrxL ->?->?->ZnR->DDE_Tnp_ISL3-> | 808 | Bacteria>FCB_group>Bacteroidetes/Chlorobi_group>Bacteroidota>Flavobacteriia>Flavobacteriales | GCA_003816015.1 | Kaistella_haifensis | EG338_03940 | 799177..801603 | 421525 | BREX-system-P-loop-protein-BrxC-[Kaistella-haifensis] |
| RZJ34503.1 | Type1.BREX | <-GDResolvase\|\|HTH_3-> BrxA-DNA-binding -> BrxB-BrxF-iSTAND -> DUF6079 *->SF1-SF2-Helicase->?-> PglZ -> SIGMA+OB-Fold+AAA-ATPase+HKD-DNase-BrxL -><-URI-DNase\|\|tRNA-><-?<-REC<-AraC-HTH | 1165 | Bacteria>FCB_group>Bacteroidetes/Chlorobi_group>Bacteroidota>Flavobacteriia>Flavobacteriales | GCA_004210845.1 | Flavobacterium_sp. | EOO51_09215 | 4934..8431 | 239 | MAG:-BREX-system-P-loop-protein-BrxC-[Flavobacterium-sp.] |
| MBW7869163.1 | Type1.BREX | HTH_3-> BrxA-DNA-binding -> BrxB-BrxF-iSTAND -> DUF6079 *-> N6-METHYLASE ->?->Schlafen+HATPase+wHTH->KAP-NTPase-> PglZ -> SIGMA+OB-Fold+AAA-ATPase+HKD-DNase-BrxL -> | 1166 | Bacteria>FCB_group>Bacteroidetes/Chlorobi_group>Bacteroidota>Flavobacteriia>Flavobacteriales | GCA_019455205.1 | Brumimicrobium_sp. | H3C31_12675 | 1666..5166 | 2029867 | BREX-system-P-loop-protein-BrxC-[Brumimicrobium-sp.] |
| MBZ0328278.1 | Type1.BREX | <- SIGMA+OB-Fold+AAA-ATPase+HKD-DNase-BrxL <- PglZ <-?<-SF1-SF2-Helicase<-?<- HKD-DNase <- N6-METHYLASE <- DUF6079 *<- BrxB-BrxF-iSTAND <- BrxA-DNA-binding <-HTH_3\|\|?-><-?<-?\|\|?->GDResolvase-> | 1164 | Bacteria>FCB_group>Bacteroidetes/Chlorobi_group>Bacteroidota>Flavobacteriia>Flavobacteriales | GCA_019913785.1 | Altibacter_sp. | K8F54_11775 | 128036..131530 | 2024823 | BREX-system-P-loop-protein-BrxC-[Altibacter-sp.] |
| WP_051224566.1 | Type1.BREX | <-TPR+TPR<-?<- SIGMA+OB-Fold+AAA-ATPase+HKD-DNase-BrxL <- PglZ <-?<-?<- N6-METHYLASE <-?<- DUF6079 *<- BrxB-BrxF-iSTAND <- BrxA-DNA-binding <-HTH_3\|\|?->GDResolvase+Recombinase+ZnR->?-><-?<-Peptidase_M49 | 1168 | Bacteria>FCB_group>Bacteroidetes/Chlorobi_group>Bacteroidota>Flavobacteriia>Flavobacteriales | GCF_000425485.1 | Flavobacterium_tegetincola | H602_RS18955 | 62387..65893 | 150172 | BREX-system-P-loop-protein-BrxC-[Flavobacterium-tegetincola] |
| WP_027137224.1 | Type6.BREX | DarT+Peptidase->MACRODOMAIN->TPR+TPR->?-><-?<- SIGMA+OB-Fold+AAA-ATPase+HKD-DNase-BrxL <- PglZ <-ParB+HNH<- N6-METHYLASE <- DUF6079 *<- BrxB-BrxF-iSTAND <- BrxA-DNA-binding <- BrxE <-?<- wHTH+WYL+WCX \|\|?-><-HEPN<-ARG | 1196 | Bacteria>FCB_group>Bacteroidetes/Chlorobi_group>Bacteroidota>Flavobacteriia>Flavobacteriales | GCF_000425645.1 | Gaetbulibacter_saemankumensis | H611_RS0104475 | 476638..480228 | 311208 | BREX-system-P-loop-protein-BrxC-[Gaetbulibacter-saemankumensis] |
| WP_028123425.1 | Type1.BREX | <-XIS-HTH\|\|?-><-?<-?\|\|HTH_3-> BrxA-DNA-binding -> BrxB-BrxF-iSTAND -> DUF6079 *-> N6-METHYLASE -> PglZ -> SIGMA+OB-Fold+AAA-ATPase+HKD-DNase-BrxL -> | 1188 | Bacteria>FCB_group>Bacteroidetes/Chlorobi_group>Bacteroidota>Flavobacteriia>Flavobacteriales | GCF_000428485.1 | Epilithonimonas_tenax | F564_RS0115480 | 6701..10267 | 191577 | BREX-system-P-loop-protein-BrxC-[Epilithonimonas-tenax] |
| WP_034657463.1 | Type1.BREX | REase->?->HEPN->REase->?-><-?\|\|HTH_3-> BrxA-DNA-binding -> BrxB-BrxF-iSTAND -> DUF6079 *-> N6-METHYLASE ->?->TPR+REase->?->SF1-SF2-Helicase->?->METHYLASE-> PglZ -> SIGMA+OB-Fold+AAA-ATPase+HKD-DNase-BrxL -> | 1160 | Bacteria>FCB_group>Bacteroidetes/Chlorobi_group>Bacteroidota>Flavobacteriia>Flavobacteriales | GCF_000468595.1 | Cellulophaga_baltica | M665_RS0118760 | 90424..93906 | 76594 | BREX-system-P-loop-protein-BrxC-[Cellulophaga-baltica] |
| WP_088323652.1 | Type1.BREX | <-XIS-HTH<-?<-ParB+N6_N4_Mtase<-?<-GDResolvase<-?<- SIGMA+OB-Fold+AAA-ATPase+HKD-DNase-BrxL <- PglZ <- N6-METHYLASE <- DUF6079 *<- BrxB-BrxF-iSTAND <- BrxA-DNA-binding <-HTH_3\|\|?->REase+DUF4268->?->?->?-><-Phage_int_SAM_5+Phage_integrase | 1158 | Bacteria>FCB_group>Bacteroidetes/Chlorobi_group>Bacteroidota>Flavobacteriia>Flavobacteriales | GCF_002201235.1 | Polaribacter_tangerinus | CCS00_RS05195 | 350003..353479 | 1920034 | BREX-system-P-loop-protein-BrxC-[Polaribacter-tangerinus] |
| WP_108879107.1 | Type1.BREX | MoxR-AAA->?-> wHTH+WYL+WCX ->ZnR+CC+REase-> BrxA-DNA-binding -> BrxB-BrxF-iSTAND -> DUF6079 *-> N6-METHYLASE -> PglZ -> SIGMA+OB-Fold+AAA-ATPase+HKD-DNase-BrxL ->?->MORC+DUF3684+REase->SF1-SF2-Helicase+PIN->?-><-AAA-ToxA\|\| wHTH+WYL+WCX -> | 1166 | Bacteria>FCB_group>Bacteroidetes/Chlorobi_group>Bacteroidota>Flavobacteriia>Flavobacteriales | GCF_003095655.1 | Dokdonia_sp._Dokd-P16 | DCS32_RS15465 | 3477371..3480871 | 2173169 | BREX-system-P-loop-protein-BrxC-[Dokdonia-sp.-Dokd-P16] |
| WP_129366655.1 | Type1.BREX | <-HNH<-SbcC<- SIGMA+OB-Fold+AAA-ATPase+HKD-DNase-BrxL <- PglZ <- N6-METHYLASE <- DUF6079 *<- BrxB-BrxF-iSTAND <- BrxA-DNA-binding <-Classical-AAA | 1168 | Bacteria>FCB_group>Bacteroidetes/Chlorobi_group>Bacteroidota>Flavobacteriia>Flavobacteriales | GCF_004121075.1 | Lutibacter_sp._HS1-25 | EC396_RS10485 | 12877..16383 | 2485000 | BREX-system-P-loop-protein-BrxC-[Lutibacter-sp.-HS1-25] |
| WP_132215847.1 | Type1.BREX | <-CARF+REase<-CARF<- HKD-DNase + HKD-DNase <-?<-S1+SF1-SF2-Helicase+S1<- SIGMA+OB-Fold+AAA-ATPase+HKD-DNase-BrxL <- PglZ <-ParB+HNH<- N6-METHYLASE <- DUF6079 *<- BrxB-BrxF-iSTAND <- BrxA-DNA-binding <-ZnR+CC+REase<- wHTH+WYL+WCX <-Arm-DNA-bind_5+Phage_int_SAM_5+Phage_integrase<-ZnR\|\|Phage_integrase+Phage_integrase-><-Zn_Tnp_IS1595+DDE_Tnp_IS1595<-REC+LuxR-HTH | 1169 | Bacteria>FCB_group>Bacteroidetes/Chlorobi_group>Bacteroidota>Flavobacteriia>Flavobacteriales | GCF_004341235.1 | Mariniflexile_fucanivorans | EV196_RS05490 | 443037..446546 | 264023 | BREX-system-P-loop-protein-BrxC-[Mariniflexile-fucanivorans] |
| WP_133546801.1 | Type1.BREX | <-XIS-HTH<-?<-?<-?<-AraC-HTH<-Arm-DNA-bind_5+Phage_int_SAM_5+Phage_integrase\|\|?-> BrxA-DNA-binding -> BrxB-BrxF-iSTAND -> DUF6079 *-> N6-METHYLASE ->REase+DUF5655->NERD+SF1-SF2-Helicase->TPR+S1-> PglZ -> SIGMA+OB-Fold+AAA-ATPase+HKD-DNase-BrxL ->HEPN->?-><-HTH_36 | 1167 | Bacteria>FCB_group>Bacteroidetes/Chlorobi_group>Bacteroidota>Flavobacteriia>Flavobacteriales | GCF_004362395.1 | Salegentibacter_sp._24 | DET49_RS03950 | 90042..93545 | 2183986 | BREX-system-P-loop-protein-BrxC-[Salegentibacter-sp.-24] |
| WP_136564624.1 | Type1.BREX | XIS-HTH->?->AbiEi_4+AbiEi_1->AbiEii+ZnR-> wHTH+WYL+WCX ->ZnR+CC+REase->HTH_3-> BrxA-DNA-binding -> BrxB-BrxF-iSTAND -> DUF6079 *-> N6-METHYLASE -> HKD-DNase ->NERD+SF1-SF2-Helicase->TPR+S1-> PglZ -> SIGMA+OB-Fold+AAA-ATPase+HKD-DNase-BrxL ->?->SMODS-> | 1164 | Bacteria>FCB_group>Bacteroidetes/Chlorobi_group>Bacteroidota>Flavobacteriia>Flavobacteriales | GCF_004916895.1 | Muricauda_alvinocaridis | EZV76_RS00375 | 77026..80520 | 2530200 | BREX-system-P-loop-protein-BrxC-[Muricauda-alvinocaridis] |
| WP_141421442.1 | Type1.BREX | <-HAD+PRTase+PRTase+REase+SF1-SF2-Helicase<-?<- SIGMA+OB-Fold+AAA-ATPase+HKD-DNase-BrxL <- PglZ <-TPR+S1<-NERD+SF1-SF2-Helicase<- N6-METHYLASE <-ParB+HNH<- DUF6079 *<- BrxB-BrxF-iSTAND <- BrxA-DNA-binding <-TPR+TPR<-?<-?<-?<-Phage_int+DNA-glycosylase<-HNH<-OLD-ABC | 1165 | Bacteria>FCB_group>Bacteroidetes/Chlorobi_group>Bacteroidota>Flavobacteriia>Flavobacteriales | GCF_006546625.1 | Haloflavibacter_putidus | FKR84_RS06225 | 21354..24851 | 2576776 | BREX-system-P-loop-protein-BrxC-[Haloflavibacter-putidus] |
| WP_019974718.1 | Type1.BREX | <- SIGMA+OB-Fold+AAA-ATPase+HKD-DNase-BrxL <- PglZ <- N6-METHYLASE <- DUF6079 *<- BrxB-BrxF-iSTAND <- BrxA-DNA-binding <-HTH_3 | 1191 | Bacteria>FCB_group>Bacteroidetes/Chlorobi_group>Bacteroidota>Flavobacteriia>Flavobacteriales | GCF_007990795.1 | Empedobacter_brevis | EB1_RS03060 | 16539..20114 | 247 | BREX-system-P-loop-protein-BrxC-[Empedobacter-brevis] |
| WP_153266434.1 | Type1.BREX | HTH_36->Classical-AAA->TRPR-HTH->STYKinase-like->HTH_3->?->HTH_3-> BrxA-DNA-binding -> BrxB-BrxF-iSTAND -> DUF6079 *->ParB+HNH-> N6-METHYLASE -> PglZ -> SIGMA+OB-Fold+AAA-ATPase+HKD-DNase-BrxL ->HigB->HTH_3->?-><-XIS-HTH | 1157 | Bacteria>FCB_group>Bacteroidetes/Chlorobi_group>Bacteroidota>Flavobacteriia>Flavobacteriales | GCF_009498295.1 | Oceanihabitans_sp._IOP_32 | FEZ18_RS00165 | 31374..34847 | 2529032 | BREX-system-P-loop-protein-BrxC-[Oceanihabitans-sp.-IOP_32] |
| WP_158973850.1 | Type1.BREX | <-ANK+ANK\|\| wHTH+WYL+WCX -> BrxA-DNA-binding -> BrxB-BrxF-iSTAND -> DUF6079 *-> N6-METHYLASE -> PglZ -> SIGMA+OB-Fold+AAA-ATPase+HKD-DNase-BrxL ->?->?->SF1-SF2-Helicase->?-><-D5ATPase | 1170 | Bacteria>FCB_group>Bacteroidetes/Chlorobi_group>Bacteroidota>Flavobacteriia>Flavobacteriales | GCF_009797025.1 | Cellulophaga_sp._L1A9 | GQR94_RS02145 | 525105..528617 | 2686362 | BREX-system-P-loop-protein-BrxC-[Cellulophaga-sp.-L1A9] |
| WP_166386804.1 | Type6.BREX | Arm-DNA-bind_5+Phage_int_SAM_5+Phage_integrase->?->XIS-HTH->?-><-?\|\| BrxE -> BrxA-DNA-binding -> BrxB-BrxF-iSTAND -> DUF6079 *-> N6-METHYLASE -> PglZ -> SIGMA+OB-Fold+AAA-ATPase+HKD-DNase-BrxL -><-TPR+TPR<-?<-SF1-SF2-Helicase | 1195 | Bacteria>FCB_group>Bacteroidetes/Chlorobi_group>Bacteroidota>Flavobacteriia>Flavobacteriales | GCF_011379025.1 | Polaribacter_sp._11A2H | GQS58_RS13590 | 79017..82604 | 2687290 | BREX-system-P-loop-protein-BrxC-[Polaribacter-sp.-11A2H] |
| WP_073117772.1 | Type1.BREX | SLATT_1-><-METHYLASE-p<-?<-SMODS<-?<-?<- SIGMA+OB-Fold+AAA-ATPase+HKD-DNase-BrxL <- PglZ <- N6-METHYLASE <- DUF6079 *<- BrxB-BrxF-iSTAND <- BrxA-DNA-binding <-ZnR+CC+REase<- wHTH+WYL+WCX <-HipA_C<-Couple_hipA<-HTH_3<-?<-Classical-AAA | 1170 | Bacteria>FCB_group>Bacteroidetes/Chlorobi_group>Bacteroidota>Flavobacteriia>Flavobacteriales | GCF_014636515.1 | Wenyingzhuangia_marina | IEV03_RS11465 | 186450..189962 | 1195760 | BREX-system-P-loop-protein-BrxC-[Wenyingzhuangia-marina] |
| WP_193813001.1 | Type6.BREX | <- wHTH+WYL+WCX \|\|ZnR+CC+REase->?-><-?<- SIGMA+OB-Fold+AAA-ATPase+HKD-DNase-BrxL <- PglZ <-ParB+HNH<- N6-METHYLASE <- DUF6079 *<- BrxB-BrxF-iSTAND <- BrxA-DNA-binding <- BrxE <-?<-FGase<-HEPN | 1217 | Bacteria>FCB_group>Bacteroidetes/Chlorobi_group>Bacteroidota>Flavobacteriia>Flavobacteriales | GCF_015191005.1 | Kaistella_flava_(ex_Peng_et_al._2021) | Q73A0000_05090 | 1135416..1139069 | 2038776 | BREX-system-P-loop-protein-BrxC-[Kaistella-flava-(ex-Peng-et-al.-2021)] |
| WP_194850146.1 | Type1.BREX | <-SF1-SF2-Helicase<-OLD-ABC+TOPRIM<- SIGMA+OB-Fold+AAA-ATPase+HKD-DNase-BrxL <- PglZ <-METHYLASE<-?<-SF1-SF2-Helicase<-?<- N6-METHYLASE <- DUF6079 *<- BrxB-BrxF-iSTAND <- BrxA-DNA-binding <-?<-TraN<-?<-TraK<-TrwI<-VirB5<-HerA/FtsK | 1160 | Bacteria>FCB_group>Bacteroidetes/Chlorobi_group>Bacteroidota>Flavobacteriia>Flavobacteriales | GCF_015356755.1 | Nonlabens_antarcticus | IQ282_RS01355 | 76947..80429 | 392714 | BREX-system-P-loop-protein-BrxC-[Nonlabens-antarcticus] |
| WP_199600307.1 | Type1.BREX | Relaxase->VirD4-TraG+VirD4-FtsK->Classical-AAA-><-Peptidase_S8<- SIGMA+OB-Fold+AAA-ATPase+HKD-DNase-BrxL <- PglZ <- N6-METHYLASE <-DEOR-HTH<-Schlafen+HATPase+wHTH<- DUF6079 *<- BrxB-BrxF-iSTAND <- BrxA-DNA-binding <-HTH_3\|\|?-><-?<-TraN<-?<-TraK<-TrwI | 1160 | Bacteria>FCB_group>Bacteroidetes/Chlorobi_group>Bacteroidota>Flavobacteriia>Flavobacteriales | GCF_016469035.1 | Gelidibacter_salicanalis | JEM65_RS13235 | 170248..173730 | 291193 | BREX-system-P-loop-protein-BrxC-[Gelidibacter-salicanalis] |
| WP_203057453.1 | Type1.BREX | <-DUF4011+SF1-SF2-Helicase+REase<- SIGMA+OB-Fold+AAA-ATPase+HKD-DNase-BrxL <-?<-HAD+PRTase+PRTase+REase+SF1-SF2-Helicase<-?<-SIGMA-HTH<- N6-METHYLASE <- DUF6079 *<- BrxB-BrxF-iSTAND <- BrxA-DNA-binding <-?\|\|ZnR+CC+REase->?-><-?<-?<-MACRODOMAIN<-ART-DarT | 1178 | Bacteria>FCB_group>Bacteroidetes/Chlorobi_group>Bacteroidota>Flavobacteriia>Flavobacteriales | GCF_016786255.1 | Robertkochia_sediminum | IS446_RS01110 | 192082..195618 | 2785326 | BREX-system-P-loop-protein-BrxC-[Robertkochia-sediminum] |
| WP_203251454.1 | Type1.BREX | XIS-HTH-><- SIGMA+OB-Fold+AAA-ATPase+HKD-DNase-BrxL <- PglZ <-TPR+S1<-NERD+SF1-SF2-Helicase<- N6-METHYLASE <- N6-METHYLASE <- DUF6079 *<- BrxB-BrxF-iSTAND <- BrxA-DNA-binding <-HTH_3<-REase<-?<-SF1-SF2-Helicase+DUF1998<-?<- wHTH+WYL+WCX <-tRNA<-Glyoxalase_4 | 1160 | Bacteria>FCB_group>Bacteroidetes/Chlorobi_group>Bacteroidota>Flavobacteriia>Flavobacteriales | GCF_016811105.1 | Aurantibacter_crassamenti | JQC67_RS03370 | 810968..814450 | 1837375 | BREX-system-P-loop-protein-BrxC-[Aurantibacter-crassamenti] |
| WP_224137560.1 | Type1.BREX | <-REC<- N6-METHYLASE <-ASCH<-HIRAN<- SIGMA+OB-Fold+AAA-ATPase+HKD-DNase-BrxL <- PglZ <- N6-METHYLASE <- DUF6079 *<- BrxB-BrxF-iSTAND <- BrxA-DNA-binding \|\| wHTH+WYL+WCX ->?->?->?->?->REase-><-Classical-AAA | 1174 | Bacteria>FCB_group>Bacteroidetes/Chlorobi_group>Bacteroidota>Flavobacteriia>Flavobacteriales | GCF_020084865.1 | Candidatus_Kaistella_beijingensis | J4771_RS06115 | 1314125..1317649 | 2820270 | BREX-system-P-loop-protein-BrxC-[Candidatus-Kaistella-beijingensis] |
| WP_224474755.1 | Type1.BREX | <- SIGMA+OB-Fold+AAA-ATPase+HKD-DNase-BrxL <- PglZ <-METHYLASE<-?<-SF1-SF2-Helicase<-?<-REase+DUF5655<- N6-METHYLASE <- DUF6079 *<- BrxB-BrxF-iSTAND <- BrxA-DNA-binding \|\|?->?->?->?->?-><-HNH | 1167 | Bacteria>FCB_group>Bacteroidetes/Chlorobi_group>Bacteroidota>Flavobacteriia>Flavobacteriales | GCF_020164485.1 | Salegentibacter_tibetensis | LB467_RS14420 | 15605..19108 | 2873600 | BREX-system-P-loop-protein-BrxC-[Salegentibacter-tibetensis] |
| WP_226063322.1 | Type1.BREX | SF1-SF2-Helicase->GDResolvase+HIN-HTH->URI-DNase-><-?<-?<-?<- SIGMA+OB-Fold+AAA-ATPase+HKD-DNase-BrxL <- PglZ <- N6-METHYLASE <- DUF6079 *<- BrxB-BrxF-iSTAND <- BrxA-DNA-binding \|\|?->?->REase+35exo-> | 1175 | Bacteria>FCB_group>Bacteroidetes/Chlorobi_group>Bacteroidota>Flavobacteriia>Flavobacteriales | GCF_020410745.1 | Kaistella_sp._GW4-15 | LC814_RS07510 | 1609850..1613377 | 2878534 | BREX-system-P-loop-protein-BrxC-[Kaistella-sp.-GW4-15] |
| WP_232121385.1 | Type1.BREX | <-ArdCN+MPTase<-?<-?\|\|<-TPR+TPR<- SIGMA+OB-Fold+AAA-ATPase+HKD-DNase-BrxL <- PglZ <- N6-METHYLASE <- DUF6079 *<- BrxB-BrxF-iSTAND <- BrxA-DNA-binding <-ZnR+TOPRIM<-?<-ZnR+TOPRIM<-VirD4-TraG<-?<-?<-TraI-VirD2-RCRE | 1176 | Bacteria>FCB_group>Bacteroidetes/Chlorobi_group>Bacteroidota>Flavobacteriia>Flavobacteriales | GCF_021205915.1 | Tenacibaculum_dicentrarchi | LNI89_RS11375 | 20484..24014 | 669041 | BREX-system-P-loop-protein-BrxC-[Tenacibaculum-dicentrarchi] |
| WP_232125551.1 | Type1.BREX | <-ArdCN+MPTase<-?<-?\|\|?-><-TPR+TPR<- SIGMA+OB-Fold+AAA-ATPase+HKD-DNase-BrxL <- PglZ <- N6-METHYLASE <- DUF6079 *<- BrxB-BrxF-iSTAND <- BrxA-DNA-binding <-ZnR+TOPRIM<-?<-ZnR+TOPRIM<-VirD4-TraG<-?<-?<-TraI-VirD2-RCRE | 1176 | Bacteria>FCB_group>Bacteroidetes/Chlorobi_group>Bacteroidota>Flavobacteriia>Flavobacteriales | GCF_021206215.1 | Tenacibaculum_finnmarkense | LNJ08_RS11655 | 18832..22362 | 2781243 | BREX-system-P-loop-protein-BrxC-[Tenacibaculum-finnmarkense] |
| WP_233899778.1 | Type1.BREX | Arm-DNA-bind_5+Phage_int_SAM_5+Phage_integrase->D5ATPase->XIS-HTH->?->D5ATPase-> BrxA-DNA-binding -> BrxB-BrxF-iSTAND -> DUF6079 *-> N6-METHYLASE -> PglZ -> SIGMA+OB-Fold+AAA-ATPase+HKD-DNase-BrxL -><-HipA_C<-Couple_hipA<-HTH_3 | 1176 | Bacteria>FCB_group>Bacteroidetes/Chlorobi_group>Bacteroidota>Flavobacteriia>Flavobacteriales | GCF_021390795.1 | Tenacibaculum_piscium | LOU78_RS02170 | 464400..467930 | 1458515 | BREX-system-P-loop-protein-BrxC-[Tenacibaculum-piscium] |
| WP_232123632.1 | Type1.BREX | ZnR+TOPRIM-> BrxA-DNA-binding -> BrxB-BrxF-iSTAND -> DUF6079 *-> N6-METHYLASE -> PglZ -> SIGMA+OB-Fold+AAA-ATPase+HKD-DNase-BrxL ->TPR+TPR-> | 1176 | Bacteria>FCB_group>Bacteroidetes/Chlorobi_group>Bacteroidota>Flavobacteriia>Flavobacteriales | GCF_022381915.1 | Tenacibaculum_finnmarkense | G1K59_RS11550 | 3814..7344 | 2781243 | BREX-system-P-loop-protein-BrxC-[Tenacibaculum-finnmarkense] |
| WP_193709240.1 | Type1.BREX | TraI-VirD2-RCRE->?->?->VirD4-TraG->?->ZnR+TOPRIM-> BrxA-DNA-binding -> BrxB-BrxF-iSTAND -> DUF6079 *-> N6-METHYLASE -> PglZ -> SIGMA+OB-Fold+AAA-ATPase+HKD-DNase-BrxL ->TPR+TPR-><-?\|\|?->?->ArdCN+MPTase-> | 1176 | Bacteria>FCB_group>Bacteroidetes/Chlorobi_group>Bacteroidota>Flavobacteriia>Flavobacteriales | GCF_022382115.1 | Tenacibaculum_finnmarkense | PJJ26_RS00320 | 69854..73384 | 2781243 | BREX-system-P-loop-protein-BrxC-[Tenacibaculum-finnmarkense] |
| WP_249986967.1 | Type1.BREX | <-DOC+HTH<-ANK+ANK\|\|HTH_3-><-REase<-HNH<-OLD-ABC<- SIGMA+OB-Fold+AAA-ATPase+HKD-DNase-BrxL <- PglZ <- N6-METHYLASE <- DUF6079 *<- BrxB-BrxF-iSTAND <- BrxA-DNA-binding <-?<-?<-?<-?<-?<-ParA-Soj-PloopNTPase | 1176 | Bacteria>FCB_group>Bacteroidetes/Chlorobi_group>Bacteroidota>Flavobacteriia>Flavobacteriales | GCF_023555435.1 | Polaribacter_sp._Z014 | MPF19_RS04515 | 308529..312059 | 2927126 | BREX-system-P-loop-protein-BrxC-[Polaribacter-sp.-Z014] |
| WP_264521759.1 | Type1.BREX | GDResolvase+Recombinase+Zn_ribbon_recom-><-MarR-HTH\|\|?->HTH_3-> BrxA-DNA-binding -> BrxB-BrxF-iSTAND -> DUF6079 *-> N6-METHYLASE ->?-> PglZ -> SIGMA+OB-Fold+AAA-ATPase+HKD-DNase-BrxL ->?-><-?<-HTH_3\|\|?->GDResolvase-> | 1166 | Bacteria>FCB_group>Bacteroidetes/Chlorobi_group>Bacteroidota>Flavobacteriia>Flavobacteriales | GCF_025947145.1 | Flavobacterium_sp._N1994 | OLM53_RS03950 | 842577..846077 | 2986827 | BREX-system-P-loop-protein-BrxC-[Flavobacterium-sp.-N1994] |
| WP_267609146.1 | Type1.BREX | <-Phage_int_SAM_1+Phage_integrase<-?<- SIGMA+OB-Fold+AAA-ATPase+HKD-DNase-BrxL <- PglZ <- N6-METHYLASE <- DUF6079 *<- BrxB-BrxF-iSTAND <- BrxA-DNA-binding \|\|Phage_integrase-><-XIS-HTH<-?<-ParB<-?<-GDResolvase+HIN-HTH | 1173 | Bacteria>FCB_group>Bacteroidetes/Chlorobi_group>Bacteroidota>Flavobacteriia>Flavobacteriales | GCF_026626535.1 | Chryseobacterium_sp._CY353 | OUD28_RS14320 | 10693..14214 | 2997334 | BREX-system-P-loop-protein-BrxC-[Chryseobacterium-sp.-CY353] |
| WP_089756260.1 | Type1.BREX | <-RHS+RHS<-?<-?\|\|NUMOD4+HNH->NUMOD4+HNH-><- SIGMA+OB-Fold+AAA-ATPase+HKD-DNase-BrxL <- PglZ <-ParB+HNH<- N6-METHYLASE <- DUF6079 *<- BrxB-BrxF-iSTAND <- BrxA-DNA-binding <-HTH_3<-HTH_19\|\|SIGMA-HTH-><-REase<-ParB+HNH<-?<-TraI-VirD2-RCRE | 1176 | Bacteria>FCB_group>Bacteroidetes/Chlorobi_group>Bacteroidota>Flavobacteriia>Flavobacteriales | GCF_900100115.1 | Chryseobacterium_soldanellicola | BLT04_RS13255 | 831929..835459 | 311333 | BREX-system-P-loop-protein-BrxC-[Chryseobacterium-soldanellicola] |
| WP_089663492.1 | Type1.BREX | BrxA-DNA-binding -> BrxB-BrxF-iSTAND -> DUF6079 *-> N6-METHYLASE ->?->?->HAD+PRTase+PRTase+REase+SF1-SF2-Helicase->?-> PglZ -> SIGMA+OB-Fold+AAA-ATPase+HKD-DNase-BrxL -> | 1167 | Bacteria>FCB_group>Bacteroidetes/Chlorobi_group>Bacteroidota>Flavobacteriia>Flavobacteriales | GCF_900105035.1 | Gramella_echinicola | BLT93_RS14455 | 3168257..3171760 | 279359 | BREX-system-P-loop-protein-BrxC-[Christiangramia-echinicola] |
| WP_103999283.1 | Type1.BREX | <-GDResolvase+Recombinase+ZnR\|\|?-><-?\|\|?-><-?<-?\|\|HTH_3-> BrxA-DNA-binding -> BrxB-BrxF-iSTAND -> DUF6079 *-> N6-METHYLASE ->ParB+HNH->URI-DNase->RADICAL-SAM->METHYLASE+RNA-Helicase->HTH_3-> PglZ -> SIGMA+OB-Fold+AAA-ATPase+HKD-DNase-BrxL -><-Arm-DNA-bind_5+Phage_int_SAM_5+Phage_integrase | 1170 | Bacteria>FCB_group>Bacteroidetes/Chlorobi_group>Bacteroidota>Flavobacteriia>Flavobacteriales | GCF_900108015.1 | Flavobacterium_urumqiense | C4S35_RS05860 | 195084..198596 | 935224 | BREX-system-P-loop-protein-BrxC-[Flavobacterium-urumqiense] |
| WP_091208763.1 | Type1.BREX | <- SIGMA+OB-Fold+AAA-ATPase+HKD-DNase-BrxL <- PglZ <-TPR+S1<-NERD+SF1-SF2-Helicase<- HKD-DNase <-?<-?<- N6-METHYLASE <-?<- DUF6079 *<- BrxB-BrxF-iSTAND <- BrxA-DNA-binding <-HTH_3\|\|?->HIN-HTH-> | 1172 | Bacteria>FCB_group>Bacteroidetes/Chlorobi_group>Bacteroidota>Flavobacteriia>Flavobacteriales | GCF_900112975.1 | Flavobacterium_xueshanense | BMZ76_RS14825 | 46469..49987 | 935223 | BREX-system-P-loop-protein-BrxC-[Flavobacterium-xueshanense] |
| WP_089819303.1 | Type1.BREX | <-ARG<-?<-HEPN<-?<-?\|\| BrxA-DNA-binding -> BrxB-BrxF-iSTAND -> DUF6079 *-> N6-METHYLASE -> PglZ -> SIGMA+OB-Fold+AAA-ATPase+HKD-DNase-BrxL ->?->?->HTH_Tnp_1->HTH_Tnp_1+HTH_21+rve+rve_3-> | 1174 | Bacteria>FCB_group>Bacteroidetes/Chlorobi_group>Bacteroidota>Flavobacteriia>Flavobacteriales | GCF_900114045.1 | Kaistella_treverensis | BMZ55_RS05400 | 1141726..1145250 | 631455 | BREX-system-P-loop-protein-BrxC-[Kaistella-treverensis] |
| WP_159447525.1 | Type1.BREX | <-HISKIN+REC<-?<-?\|\| wHTH+WYL+WCX ->ZnR+CC+REase-> BrxA-DNA-binding -> BrxB-BrxF-iSTAND -> DUF6079 -p*-> DUF6079 -p-> N6-METHYLASE -p-> N6-METHYLASE ->ParB+HNH->HTH_3->Couple_hipA+HipA_C-> PglZ -> SIGMA+OB-Fold+AAA-ATPase+HKD-DNase-BrxL -> | 719 | Bacteria>FCB_group>Bacteroidetes/Chlorobi_group>Bacteroidota>Flavobacteriia>Flavobacteriales | GCF_900167935.1 | Maribacter_arcticus | B5X55_RS04985 | 472839..474998 | 561365 | BREX-system-P-loop-protein-BrxC-[Maribacter-arcticus] |
| WP_095070899.1 | Type1.BREX | <-AbiEii<-AbiEi_4\|\|XIS-HTH->HTH_36->Classical-AAA-><- wHTH+WYL+WCX \|\| BrxA-DNA-binding -> BrxB-BrxF-iSTAND -> DUF6079 *-> N6-METHYLASE -> PglZ -> SIGMA+OB-Fold+AAA-ATPase+HKD-DNase-BrxL ->ASCH->HIRAN->ASCH->ZnR+CC+REase->ARG-> | 1178 | Bacteria>FCB_group>Bacteroidetes/Chlorobi_group>Bacteroidota>Flavobacteriia>Flavobacteriales | GCF_900187185.1 | Chryseobacterium_taklimakanense | CKV81_RS04675 | 977692..981228 | 536441 | BREX-system-P-loop-protein-BrxC-[Chryseobacterium-taklimakanense] |
| WP_089379946.1 | Type1.BREX | RHH-><- SIGMA+OB-Fold+AAA-ATPase+HKD-DNase-BrxL <- PglZ <-?<-SF1-SF2-Helicase<-?<- wHTH+WYL+WCX -p<- N6-METHYLASE <-REase+DUF5655<- DUF6079 *<- BrxB-BrxF-iSTAND <- BrxA-DNA-binding <-HTH_3<-?<-SF1-Phage_integrase+Phage_integrase\|\|?->Radical_SAM-> | 1168 | Bacteria>FCB_group>Bacteroidetes/Chlorobi_group>Bacteroidota>Flavobacteriia>Flavobacteriales | GCF_900188235.1 | Lutibacter_agarilyticus | SAMN06265371_101285 | 359460..362966 | 1109740 | BREX-system-P-loop-protein-BrxC-[Lutibacter-agarilyticus] |
| WP_213195478.1 | Type1.BREX | TraI-VirD2-RCRE->?->TRPR-HTH->STYKinase-like-> wHTH+WYL+WCX ->ZnR+CC+REase->HIRAN-> BrxA-DNA-binding -> BrxB-BrxF-iSTAND -> DUF6079 *-> N6-METHYLASE ->HTH_3->Couple_hipA+HipA_C-> PglZ -> SIGMA+OB-Fold+AAA-ATPase+HKD-DNase-BrxL ->?->?->?-> wHTH+WYL+WCX -> | 1176 | Bacteria>FCB_group>Bacteroidetes/Chlorobi_group>Bacteroidota>Flavobacteriia>Flavobacteriales | GCF_907163105.1 | Cloacibacterium_caeni | KKQ76_RS01235 | 263131..266661 | 2004710 | BREX-system-P-loop-protein-BrxC-[Cloacibacterium-caeni] |
| WP_271393637.1 | Type1.BREX | <-ParB+HNH<-PIWI<- SIGMA+OB-Fold+AAA-ATPase+HKD-DNase-BrxL <- PglZ <- N6-METHYLASE <- DUF6079 *<- BrxB-BrxF-iSTAND <- BrxA-DNA-binding <-HTH<-?<-?<-XIS-HTH<-XIS-HTH<-?<-Arm-DNA-bind_5+Phage_int_SAM_5+Phage_integrase | 1168 | Bacteria>FCB_group>Bacteroidetes/Chlorobi_group>Bacteroidota>Flavobacteriia>Flavobacteriales | GCF_947492035.1 | Aequorivita_sinensis | PDR58_RS10295 | 126765..130271 | 1382458 | BREX-system-P-loop-protein-BrxC-[Aequorivita-sinensis] |
| WP_271393758.1 | Type1.BREX | <- SIGMA+OB-Fold+AAA-ATPase+HKD-DNase-BrxL <- PglZ <-?<-SF1-SF2-Helicase<-?<- N6-METHYLASE <-ParB+HNH<- DUF6079 *<- BrxB-BrxF-iSTAND <- BrxA-DNA-binding <-?<-STYKinase-like<-TRPR-HTH<-?<-Classical-AAA<-HTH_36<-XIS-HTH | 1165 | Bacteria>FCB_group>Bacteroidetes/Chlorobi_group>Bacteroidota>Flavobacteriia>Flavobacteriales | GCF_947492035.1 | Aequorivita_sinensis | PDR58_RS11110 | 23754..27251 | 1382458 | BREX-system-P-loop-protein-BrxC-[Aequorivita-sinensis] |
| MCB0517804.1 | Type1.BREX | <- SIGMA+OB-Fold+AAA-ATPase+HKD-DNase-BrxL <- PglZ <- N6-METHYLASE <- DUF6079 *<- BrxB-BrxF-iSTAND <- BrxA-DNA-binding <-?<-HNH\|\|REC-><-?<-SF1-SF2-Helicase+RNase+RNase<-?<-RVT | 1174 | Bacteria>FCB_group>Bacteroidetes/Chlorobi_group>Bacteroidota>Saprospiria>Saprospirales | GCA_020436325.1 | Saprospiraceae_bacterium | KDD27_02635 | 9612..13136 | 2202734 | BREX-system-P-loop-protein-BrxC-[Saprospiraceae-bacterium] |
| MCB0524505.1 | Type1.BREX | SF1-SF2-Helicase+RNase+RNase->?->REase+35exo->?->?-> BrxA-DNA-binding -> BrxB-BrxF-iSTAND -> DUF6079 -p-> DUF6079 *-> N6-METHYLASE ->?->McrB-NTD+HNH->Polbeta->?->CR-ATPase5+REase-> PglZ -> SIGMA+OB-Fold+AAA-ATPase+HKD-DNase-BrxL -> | 1094 | Bacteria>FCB_group>Bacteroidetes/Chlorobi_group>Bacteroidota>Saprospiria>Saprospirales | GCA_020436385.1 | Saprospiraceae_bacterium | KDC86_08270 | 14975..18259 | 2202734 | BREX-system-P-loop-protein-BrxC-[Saprospiraceae-bacterium] |
| WP_020534907.1 | Type1.BREX | N6-METHYLASE ->?-><-?\|\|?->?->HTH_3-> BrxA-DNA-binding -> BrxB-BrxF-iSTAND -> DUF6079 *-> N6-METHYLASE ->OLD-ABC->TOPRIM->?->HAD+PRTase+PRTase+REase+SF1-SF2-Helicase->?-> PglZ -> SIGMA+OB-Fold+AAA-ATPase+HKD-DNase-BrxL ->Peptidase-> | 1159 | Bacteria>FCB_group>Bacteroidetes/Chlorobi_group>Bacteroidota>Saprospiria>Saprospirales | GCF_000379805.1 | Lewinella_cohaerens | A3EU_RS29375 | 116498..119977 | 70995 | BREX-system-P-loop-protein-BrxC-[Lewinella-cohaerens] |
| WP_168040578.1 | Type1.BREX | <-REC+LYTTR<-HisKin<-Peptidase_M3\|\|?-><-?\|\| BrxA-DNA-binding -> BrxB-BrxF-iSTAND -> DUF6079 *-> N6-METHYLASE ->HTH+DOC-> PglZ -> SIGMA+OB-Fold+AAA-ATPase+HKD-DNase-BrxL -> | 1211 | Bacteria>FCB_group>Bacteroidetes/Chlorobi_group>Bacteroidota>Saprospiria>Saprospirales | GCF_011927835.1 | Neolewinella_antarctica | GGR27_RS19565 | 12164..15799 | 442734 | BREX-system-P-loop-protein-BrxC-[Neolewinella-antarctica] |
| WP_236977640.1 | Type1.BREX | RNase+DSRBD->?-><-?<-?\|\|tRNA-> BrxA-DNA-binding -> BrxB-BrxF-iSTAND -> DUF6079 *->?-> N6-METHYLASE ->MORC+DUF3684+HNH-> PglZ -> SIGMA+OB-Fold+AAA-ATPase+HKD-DNase-BrxL -><-?\|\|?->?->PIN-> | 1159 | Bacteria>FCB_group>Bacteroidetes/Chlorobi_group>Bacteroidota>Saprospiria>Saprospirales | GCF_021729145.1 | Membranihabitans_sp._C281 | LVD47_RS14080 | 21527..25006 | 2904244 | BREX-system-P-loop-protein-BrxC-[Membranihabitans-maritimus] |
| OYD42847.1 | Type1.BREX | wHTH+WYL+WCX -><-?<-?\|\|?-><-?\|\|sGTP->HTH_3-> BrxA-DNA-binding -> BrxB-BrxF-iSTAND -> DUF6079 *-> N6-METHYLASE ->ParB+HNH-> PglZ -> SIGMA+OB-Fold+AAA-ATPase+HKD-DNase-BrxL -> | 1175 | Bacteria>FCB_group>Bacteroidetes/Chlorobi_group>Bacteroidota>Sphingobacteriia>Sphingobacteriales | GCA_002245855.1 | Sphingobacterium_cellulitidis | CHT99_08530 | 557163..560690 | 1768011 | hypothetical-protein-CHT99_08530-[Sphingobacterium-cellulitidis] |
| WP_157278947.1 | Type1.BREX | Arm-DNA-bind_5+Phage_int_SAM_5+Phage_integrase->?->XIS-HTH->TraJ-RHH->?->?->HTH_3-> BrxA-DNA-binding -> BrxB-BrxF-iSTAND -> DUF6079 *-> N6-METHYLASE ->Schlafen+HATPase+wHTH-> PglZ -> SIGMA+OB-Fold+AAA-ATPase+HKD-DNase-BrxL ->?-><-HTH_21+rve+rve_3 | 1169 | Bacteria>FCB_group>Bacteroidetes/Chlorobi_group>Bacteroidota>Sphingobacteriia>Sphingobacteriales | GCF_000427965.1 | Olivibacter_sitiensis | A37S_RS0120635 | 7227..10736 | 376470 | BREX-system-P-loop-protein-BrxC-[Olivibacter-sitiensis] |
| WP_094282244.1 | Type1.BREX | <- SIGMA+OB-Fold+AAA-ATPase+HKD-DNase-BrxL <- PglZ <-ParB+HNH<- N6-METHYLASE <- N6-METHYLASE -p<- DUF6079 *<- BrxB-BrxF-iSTAND <- BrxA-DNA-binding <-HTH_3<-sGTP\|\|?-><-?\|\|SIGMA-HTH-> | 1175 | Bacteria>FCB_group>Bacteroidetes/Chlorobi_group>Bacteroidota>Sphingobacteriia>Sphingobacteriales | GCF_002245765.1 | Sphingobacterium_cellulitidis | CHU00_RS15775 | 18381..21908 | 1768011 | BREX-system-P-loop-protein-BrxC-[Sphingobacterium-cellulitidis] |
| WP_070561019.1 | Type1.BREX | TPR+TPR+TPR+TPR+TPR->?->ANK->ANK+ANK+ANK-><-?<- SIGMA+OB-Fold+AAA-ATPase+HKD-DNase-BrxL <- PglZ <-HTH+REase<- N6-METHYLASE <- DUF6079 *<- BrxB-BrxF-iSTAND <- BrxA-DNA-binding <-HTH_3<-?\|\|?->XIS-HTH->HTH_36->?->Relaxase-> | 1187 | Bacteria>FCB_group>Bacteroidetes/Chlorobi_group>Bacteroidota>Sphingobacteriia>Sphingobacteriales | GCF_002339225.1 | Sphingobacterium_sp._HMSC13C05 | HMPREF3127_RS01230 | 95716..99279 | 1581095 | BREX-system-P-loop-protein-BrxC-[Sphingobacterium-sp.-HMSC13C05] |
| WP_165305702.1 | Type1.BREX | SLATT_1->?->HEPN+REase->?->?->HTH_3-> BrxA-DNA-binding -> BrxB-BrxF-iSTAND -> DUF6079 *-> N6-METHYLASE ->HTH+REase-> PglZ -> SIGMA+OB-Fold+AAA-ATPase+HKD-DNase-BrxL ->?-><-HTH_3 | 1187 | Bacteria>FCB_group>Bacteroidetes/Chlorobi_group>Bacteroidota>Sphingobacteriia>Sphingobacteriales | GCF_011064245.1 | Sphingobacterium_sp._DR205 | G6053_RS05210 | 1244063..1247626 | 2713573 | BREX-system-P-loop-protein-BrxC-[Sphingobacterium-sp.-DR205] |
| WP_090698254.1 | Type1.BREX | <-HTH_19\|\|?->?-><-?\|\|ANK+ANK-> BrxA-DNA-binding -> BrxB-BrxF-iSTAND -> DUF6079 *-> N6-METHYLASE -> PglZ -> SIGMA+OB-Fold+AAA-ATPase+HKD-DNase-BrxL ->?-><-PHP+SbcC | 1169 | Bacteria>FCB_group>Bacteroidetes/Chlorobi_group>Bacteroidota>Sphingobacteriia>Sphingobacteriales | GCF_900103545.1 | Daejeonella_rubra | BLR27_RS01455 | 316504..320013 | 990371 | BREX-system-P-loop-protein-BrxC-[Daejeonella-rubra] |
| KPP94735.1 | Type1.BREX | SAM-METHYLASE->tRNA-><- SIGMA+OB-Fold+AAA-ATPase+HKD-DNase-BrxL <- PglZ <-Schlafen+HATPase+wHTH<- N6-METHYLASE <-?<- DUF6079 *<- BrxB-BrxF-iSTAND <- BrxA-DNA-binding <-?<-SF1-SF2-Helicase<-?<-?<-DnaB+SMS-KaiC<-HTH_36<-XIS-HTH | 1203 | Bacteria>FCB_group>Bacteroidetes/Chlorobi_group>Bacteroidota>unclassified_Bacteroidota | GCA_001314545.1 | Bacteroidetes_bacterium_HLUCCA01 | HLUCCA01_08750 | 188157..191768 | 1666909 | Brex-phage-defense-system-ATPase-BrxC-[Bacteroidetes-bacterium-HLUCCA01] |
| MBK7566196.1 | Type6.BREX | <-SF1-SF2-Helicase<-MORC+DUF3684+REase<-?<-ParB<-?<-?<- SIGMA+OB-Fold+AAA-ATPase+HKD-DNase-BrxL <- PglZ <- N6-METHYLASE <- DUF6079 *<- BrxB-BrxF-iSTAND <- BrxA-DNA-binding <- BrxE <-HTH<-?<-?<-?<-?<-REC | 1193 | Bacteria>FCB_group>Bacteroidetes/Chlorobi_group>Bacteroidota>unclassified_Bacteroidota | GCA_016706865.1 | Bacteroidota_bacterium | IPI31_00060 | 18734..22315 | 1898104 | BREX-system-P-loop-protein-BrxC-[Bacteroidota-bacterium] |
| MCH7965547.1 | Type1.BREX | <-Aminotran_3\|\|?-><-?\|\|?-><- SIGMA+OB-Fold+AAA-ATPase+HKD-DNase-BrxL <- PglZ <-HTH+REase<- N6-METHYLASE <- DUF6079 *<- BrxB-BrxF-iSTAND <- BrxA-DNA-binding | 1161 | Bacteria>FCB_group>Bacteroidetes/Chlorobi_group>Bacteroidota>unclassified_Bacteroidota | GCA_022564295.1 | Bacteroidota_bacterium | IH852_16610 | 28479..31964 | 1898104 | BREX-system-P-loop-protein-BrxC-[Bacteroidota-bacterium] |
| MA064836.1 | Type1.BREX | Phage_int_SAM_4+Phage_integrase-><-?<- SIGMA+OB-Fold+AAA-ATPase+HKD-DNase-BrxL <- PglZ <-MORC+DUF3684+HNH<-REase+DUF4268<-Trypsin<- N6-METHYLASE <-SIGMA-HTH<- DUF6079 *<- BrxB-BrxF-iSTAND <-REase<-SF1-SF2-Helicase+Classical-AAA<-?<-HNH | 1168 | Bacteria>FCB_group>Bacteroidetes/Chlorobi_group>Balneolota>Balneolia>Balneolales | GCA_002696035.1 | Balneola_sp. | CL666_07525 | 664531..668037 | 2024824 | hypothetical-protein-[Balneola-sp.] |
| MCC5907043.1 | Type1.BREX | <-REase<-?<-?<-?\|\|TPR+TPR-> BrxA-DNA-binding -> BrxB-BrxF-iSTAND -> DUF6079 *->AbiEi->AbiEii-> N6-METHYLASE ->Schlafen+HATPase+wHTH-> PglZ -> SIGMA+OB-Fold+AAA-ATPase+HKD-DNase-BrxL ->REase-><-?\|\|Thymidylate_kin-> | 1167 | Bacteria>FCB_group>Bacteroidetes/Chlorobi_group>Balneolota>Balneolia>Balneolales | GCA_020832285.1 | Balneolaceae_bacterium | JJU13_12585 | 29362..32865 | 2053516 | BREX-system-P-loop-protein-BrxC-[Balneolaceae-bacterium] |
| MCH2449130.1 | Type1.BREX | Phage_int_SAM_4+Phage_integrase->TRPR-HTH->?->HTH_3-> BrxA-DNA-binding -> BrxB-BrxF-iSTAND -> DUF6079 *-> N6-METHYLASE -> PglZ -> SIGMA+OB-Fold+AAA-ATPase+HKD-DNase-BrxL -><-?<-HD\|\|REase->MULE-TRANSPOSASE-> | 1151 | Bacteria>FCB_group>Bacteroidetes/Chlorobi_group>Balneolota>Balneolia>Balneolales | GCA_022450045.1 | Gracilimonas_sp. | MK198_03190 | 295077..298532 | 1974203 | BREX-system-P-loop-protein-BrxC-[Gracilimonas-sp.] |
| WP_237852616.1 | Type1.BREX | <- SIGMA+OB-Fold+AAA-ATPase+HKD-DNase-BrxL <- PglZ \|\|SF1-SF2-Helicase-><-SF1-SF2-Helicase<-?<- N6-METHYLASE <-AbiEii<-AbiEi<- DUF6079 *<- BrxB-BrxF-iSTAND <- BrxA-DNA-binding <-REase\|\|?->?->?->?-><-HEPN | 1164 | Bacteria>FCB_group>Bacteroidetes/Chlorobi_group>Balneolota>Balneolia>Balneolales | GCF_022012475.1 | Rhodohalobacter_sp._WB101 | L6773_RS04280 | 217944..221438 | 2911366 | BREX-system-P-loop-protein-BrxC-[Rhodohalobacter-sp.-WB101] |
| WP_265763895.1 | Type1.BREX | <-HEPN<- SIGMA+OB-Fold+AAA-ATPase+HKD-DNase-BrxL <- PglZ <-?<-Schlafen+HATPase+wHTH<-REase<-?<-?<- N6-METHYLASE <- DUF6079 *<- BrxB-BrxF-iSTAND <- BrxA-DNA-binding <-tRNA\|\|?->?->?-><-?<-?<-ABC-ATPase+SbcC | 1165 | Bacteria>FCB_group>Bacteroidetes/Chlorobi_group>Balneolota>Balneolia>Balneolales | GCF_026229185.1 | Aliifodinibius_sp._1BSP15-2V2 | J6I44_RS00130 | 34405..37902 | 2820877 | BREX-system-P-loop-protein-BrxC-[Aliifodinibius-sp.-1BSP15-2V2] |
| KAA6232806.1 | Type1.BREX | <-McrC-NTD+REase<-McrB-AAA<-HARE-HTH+REase<- SIGMA+OB-Fold+AAA-ATPase+HKD-DNase-BrxL <- PglZ \|\|DDE_Tnp_1_4-><- N6-METHYLASE <-REase<-ABC-ATPase<- DUF6079 *<- BrxB-BrxF-iSTAND <- BrxA-DNA-binding <-XIS-HTH<-?<-?<-?\|\|Nudix_N_2+NUDIX-><-?<-HAMP+PAS+HISKIN | 1204 | Bacteria>FCB_group>Bacteroidetes/Chlorobi_group>Chlorobiota>Chlorobiia>Chlorobiales | GCA_008634005.1 | Chlorobium_phaeovibrioides | FP507_06840 | 1409593..1413207 | 1094 | BREX-system-P-loop-protein-BrxC-[Chlorobium-phaeovibrioides] |
| NMW22669.1 | Type6.BREX | <- SIGMA+OB-Fold+AAA-ATPase+HKD-DNase-BrxL <- PglZ <- N6-METHYLASE <- DUF6079 *<- BrxB-BrxF-iSTAND <- BrxA-DNA-binding <- BrxE | 1210 | Bacteria>FCB_group>Bacteroidetes/Chlorobi_group>Chlorobiota>Chlorobiia>Chlorobiales | GCA_012972835.1 | Chlorobiaceae_bacterium | HKK67_13830 | 33011..36643 | 2067991 | BREX-system-P-loop-protein-BrxC-[Chlorobiaceae-bacterium] |
| WP_204208384.1 | Type1.BREX | ABhydrolase->?->Aminotran_1_2-><-?\|\|DUF772+DDE_Tnp_1_2-> BrxA-DNA-binding -> BrxB-BrxF-iSTAND -> DUF6079 -p*->?->Schlafen+HATPase+wHTH+HTH-> N6-METHYLASE ->DUF4372+DDE_Tnp_1-> N6-METHYLASE -p->ParB+HNH->RVT-> PglZ -> SIGMA+OB-Fold+AAA-ATPase+HKD-DNase-BrxL -> | 997 | Bacteria>FCB_group>Bacteroidetes/Chlorobi_group>Chlorobiota>Chlorobiia>Chlorobiales | GCF_001687065.1 | Prosthecochloris_sp._CIB_2401 | Ptc2401_RS05035 | 1057250..1060243 | 1868325 | BREX-system-P-loop-protein-BrxC-[Prosthecochloris-sp.-CIB-2401] |
| WP_076791024.1 | Type1.BREX | <- SIGMA+OB-Fold+AAA-ATPase+HKD-DNase-BrxL <- PglZ <-DOC+HTH<-?<-?<-Schlafen+HATPase+wHTH+HTH<- DUF6079 *<- BrxB-BrxF-iSTAND <- BrxA-DNA-binding \|\|?->RelE-ParE-><-?<-Peptidase | 1179 | Bacteria>FCB_group>Bacteroidetes/Chlorobi_group>Chlorobiota>Chlorobiia>Chlorobiales | GCF_001975465.1 | Chlorobium_sp._KB01 | BOO20_RS04865 | 12313..15852 | 1917528 | BREX-system-P-loop-protein-BrxC-[Chlorobium-sp.-KB01] |
| WP_192496514.1 | Type1.BREX | <-McrB-AAA<-HARE-HTH+REase<- SIGMA+OB-Fold+AAA-ATPase+HKD-DNase-BrxL <- PglZ <- N6-METHYLASE -p\|\|DDE_Tnp_1_4-><- N6-METHYLASE <-REase<-ABC-ATPase<- DUF6079 *<- BrxB-BrxF-iSTAND <- BrxA-DNA-binding <-XIS-HTH<-?<-?<-?<-?\|\|Nudix_N_2+NUDIX-> | 1203 | Bacteria>FCB_group>Bacteroidetes/Chlorobi_group>Chlorobiota>Chlorobiia>Chlorobiales | GCF_008634005.1 | Chlorobium_phaeovibrioides | FP507_RS07315 | 1409593..1413204 | 1094 | BREX-system-P-loop-protein-BrxC-[Chlorobium-phaeovibrioides] |
| WP_151419733.1 | Type1.BREX | BrxA-DNA-binding -> BrxB-BrxF-iSTAND -> DUF6079 *->REase+iREase->?-> N6-METHYLASE ->HTH+REase->REase-> PglZ -> SIGMA+OB-Fold+AAA-ATPase+HKD-DNase-BrxL ->HTH_Tnp_1->rve-> | 1183 | Bacteria>FCB_group>Bacteroidetes/Chlorobi_group>Chlorobiota>Chlorobiia>Chlorobiales | GCF_008634005.1 | Chlorobium_phaeovibrioides | FP507_RS10470 | 13780..17331 | 1094 | BREX-system-P-loop-protein-BrxC-[Chlorobium-phaeovibrioides] |
| WP_098060820.1 | Type1.BREX | Peptidase->?->?->?-> BrxA-DNA-binding -> BrxB-BrxF-iSTAND -> DUF6079 *-> N6-METHYLASE -> PglZ -> SIGMA+OB-Fold+AAA-ATPase+HKD-DNase-BrxL ->CHTH->Tox-RES->Thioredoxin-> | 1206 | Bacteria>FCB_group>Bacteroidetes/Chlorobi_group>Rhodothermota>Rhodothermia>Rhodothermales | GCF_002554705.1 | Longimonas_halophila | CRI93_RS01465 | 364524..368144 | 1469170 | BREX-system-P-loop-protein-BrxC-[Longimonas-halophila] |
| OQC05381.1 | Type1.BREX | BrxA-DNA-binding -> BrxB-BrxF-iSTAND -> DUF6079 *->OLD-ABC->REase-> N6-METHYLASE -> PglZ -> SIGMA+OB-Fold+AAA-ATPase+HKD-DNase-BrxL ->HEPN-> | 1162 | Bacteria>FCB_group>Candidatus_Cloacimonetes>unclassified_Candidatus_Cloacimonetes | GCA_002070875.1 | Candidatus_Cloacimonetes_bacterium_ADurb.Bin089 | BWX76_00750 | 1812..5300 | 1852836 | MAG:-hypothetical-protein-BWX76_00750-[Candidatus-Cloacimonetes-bacterium-ADurb.Bin089] |
| MCB5271576.1 | Type1.BREX | KAP-NTPase->KAP-glycosylase->?->TatD_DNase-><- SIGMA+OB-Fold+AAA-ATPase+HKD-DNase-BrxL <- PglZ <- N6-METHYLASE <-REase<-OLD-ABC<- DUF6079 *<- BrxB-BrxF-iSTAND <- BrxA-DNA-binding <-NERD+ZnR | 1171 | Bacteria>FCB_group>Candidatus_Cloacimonetes>unclassified_Candidatus_Cloacimonetes | GCA_020532545.1 | Candidatus_Cloacimonetes_bacterium | LHW56_07005 | 18816..22331 | 2030808 | BREX-system-P-loop-protein-BrxC-[Candidatus-Cloacimonadota-bacterium] |
| NQV48959.1 | Type6.BREX | ParB+HNH-p->XIS-HTH-> BrxE -p-> BrxE -p-> BrxA-DNA-binding -> BrxB-BrxF-iSTAND -> DUF6079 *-> N6-METHYLASE -> PglZ -> SIGMA+OB-Fold+AAA-ATPase+HKD-DNase-BrxL ->Phage_int+DNA-glycosylase->?->SbcC-> | 1210 | Bacteria>FCB_group>Candidatus_Marinimicrobia | GCA_013202385.1 | Candidatus_Marinimicrobia_bacterium | HQ507_00570 | 3171..6803 | 2026760 | BREX-system-P-loop-protein-BrxC-[Candidatus-Marinimicrobia-bacterium] |
| MBT4361892.1 | Type1.BREX | DUF6079 *-> N6-METHYLASE -> PglZ -> SIGMA+OB-Fold+AAA-ATPase+HKD-DNase-BrxL ->REase+DUF5655-> | 1148 | Bacteria>FCB_group>Candidatus_Marinimicrobia | GCA_018653465.1 | Candidatus_Marinimicrobia_bacterium | HOD11_13085 | 1..3448 | 2026760 | BREX-system-P-loop-protein-BrxC-[Candidatus-Marinimicrobia-bacterium] |
| NLW30564.1 | Type1.BREX | HNH->?-> BrxA-DNA-binding -> BrxB-BrxF-iSTAND -> DUF6079 *->?->ParB+HNH-> PglZ -> SIGMA+OB-Fold+AAA-ATPase+HKD-DNase-BrxL -> | 1185 | Bacteria>FCB_group>Fibrobacterota>Fibrobacteria>Fibrobacterales>Fibrobacteraceae | GCA_012728995.1 | Fibrobacter_sp. | GXY77_03805 | 4326..7883 | 35828 | BREX-system-P-loop-protein-BrxC-[Fibrobacter-sp.] |
| WP_073424902.1 | Type1.BREX | ABhydrolase->?-><- SIGMA+OB-Fold+AAA-ATPase+HKD-DNase-BrxL <- PglZ <- N6-METHYLASE <- DUF6079 *<- BrxB-BrxF-iSTAND <- BrxA-DNA-binding <-LRR_5<-PIN<-RelB | 1212 | Bacteria>FCB_group>Fibrobacterota>Fibrobacteria>Fibrobacterales>Fibrobacteraceae | GCF_900142945.1 | Fibrobacter_sp._UWB7 | BUC30_RS10995 | 309469..313107 | 1896206 | BREX-system-P-loop-protein-BrxC-[Fibrobacter-sp.-UWB7] |
| WP_240631539.1 | Type6.BREX | <-Tox-RES<-XIS-HTH\|\|HDRnase-><- SIGMA+OB-Fold+AAA-ATPase+HKD-DNase-BrxL <- PglZ <-DOC+HTH<- N6-METHYLASE <-GreA_GreB+SF1-SF2-Helicase<- DUF6079 *<- BrxB-BrxF-iSTAND <- BrxA-DNA-binding <- BrxE <- wHTH+WYL+WCX \|\|MODE-HTH+PBPII-><-HTH_3\|\|LysR-HTH->TRPR-HTH->STYKinase-like-> | 1234 | Bacteria | GCF_003671915.1 | Pseudomonadota | D3M96_RS03425 | 754727..758431 | 1224 | MULTISPECIES:-BREX-system-P-loop-protein-BrxC-[Pseudomonadota] |
| WP_203328719.1 | Type6.BREX | <-TraM<-DUF3262<-?<-?<-SMODS<-?\|\| wHTH+WYL+WCX -> BrxE -> BrxB-BrxF-iSTAND -> DUF6079 *->GreA_GreB+SF1-SF2-Helicase-> N6-METHYLASE -> N6-METHYLASE ->ParB+HNH-> PglZ -> SIGMA+OB-Fold+AAA-ATPase+HKD-DNase-BrxL -><-JAB<-?<-SMODS | 1213 | Bacteria | GCF_016836415.1 | Pseudomonadota | JQ482_RS25705 | 5512214..5515855 | 1224 | MULTISPECIES:-BREX-system-P-loop-protein-BrxC-[Pseudomonadota] |
| WP_240685338.1 | Type6.BREX | SMODS->?->JAB-><- SIGMA+OB-Fold+AAA-ATPase+HKD-DNase-BrxL <- PglZ <-ParB+HNH<- N6-METHYLASE <-GreA_GreB+SF1-SF2-Helicase<- DUF6079 *<- BrxB-BrxF-iSTAND <- BrxA-DNA-binding <- BrxE <- wHTH+WYL+WCX \|\|JetA->JetB->SbcC-> | 1213 | Bacteria | GCF_025461165.1 | Pseudomonadota | JAK52_RS02505 | 287897..291538 | 1224 | MULTISPECIES:-BREX-system-P-loop-protein-BrxC-[Pseudomonadota] |
| WP_172437661.1 | Type1.BREX | DEDD_Tnp_IS110+Transposase_20-><-?\|\|XIS-HTH->TOPRIM->\|\|XIS-HTH-> BrxA-DNA-binding -> BrxB-BrxF-iSTAND -> DUF6079 *->Schlafen+HATPase+wHTH+HTH-> N6-METHYLASE ->HTH+REase-> PglZ -> SIGMA+OB-Fold+AAA-ATPase+HKD-DNase-BrxL ->SbcC+APATPase+SbcC->Calcineurin+Calcineurin-SbcD->SbcC->RNase-> | 1203 | Bacteria>Pseudomonadota>Acidithiobacillia>Acidithiobacillales>Acidithiobacillaceae>Acidithiobacillus | - | Acidithiobacillus_sp._SH | HF670_09010 | 8231..11842 | 187490 | MULTISPECIES:-BREX-system-P-loop-protein-BrxC-[Acidithiobacillus] |
| MBU2839699.1 | Type1.BREX | DEDD_Tnp_IS110+Transposase_20-><-?\|\|XIS-HTH->TOPRIM->\|\|XIS-HTH-> BrxA-DNA-binding -> BrxB-BrxF-iSTAND -> DUF6079 *->Schlafen+HATPase+wHTH+HTH-> N6-METHYLASE ->HTH+REase-> PglZ -> SIGMA+OB-Fold+AAA-ATPase+HKD-DNase-BrxL ->SbcC+APATPase+SbcC->Calcineurin+Calcineurin-SbcD->SbcC->RNase-> | 1203 | Bacteria>Pseudomonadota>Acidithiobacillia>Acidithiobacillales>Acidithiobacillaceae>Acidithiobacillus | GCA_018854655.1 | Acidithiobacillus_thiooxidans | HF670_09010 | 8231..11842 | 930 | BREX-system-P-loop-protein-BrxC-[Acidithiobacillus-thiooxidans] |
| WP_014029717.1 | Type1.BREX | DEDD_Tnp_IS110+Transposase_20-><-DDE_5<-?\|\|HTH_17->Phage_int_SAM_2+Integrase_1-><-SbcC<-Calcineurin+Calcineurin-SbcD\|\| BrxA-DNA-binding -> BrxB-BrxF-iSTAND -> DUF6079 *-> N6-METHYLASE ->Schlafen+HATPase+wHTH-> N6-METHYLASE -p->OLD-ABC->TOPRIM-> PglZ -> SIGMA+OB-Fold+AAA-ATPase+HKD-DNase-BrxL -> | 1179 | Bacteria>Pseudomonadota>Acidithiobacillia>Acidithiobacillales>Acidithiobacillaceae>Acidithiobacillus | GCF_000214095.2 | Acidithiobacillus_ferrivorans | ACIFE_RS11315 | 2362677..2366216 | 160808 | BREX-system-P-loop-protein-BrxC-[Acidithiobacillus-ferrivorans] |
| WP_064218004.1 | Type1.BREX | Arm-DNA-bind_3+Phage_integrase->?->HTH_3->?->?-> BrxA-DNA-binding -> BrxB-BrxF-iSTAND -> DUF6079 *->OLD-ABC->REase-> N6-METHYLASE -> PglZ -> SIGMA+OB-Fold+AAA-ATPase+HKD-DNase-BrxL ->SbcC+SbcC-> | 1187 | Bacteria>Pseudomonadota>Acidithiobacillia>Acidithiobacillales>Acidithiobacillaceae>Acidithiobacillus | GCF_001652185.1 | Acidithiobacillus_ferrooxidans | A4H96_RS01790 | 31399..34962 | 920 | BREX-system-P-loop-protein-BrxC-[Acidithiobacillus-ferrooxidans] |
| WP_163056479.1 | Type1.BREX | XIS-HTH->?->TOPRIM->?->XIS-HTH-> BrxA-DNA-binding -> BrxB-BrxF-iSTAND -> DUF6079 *->OLD-ABC->REase-> N6-METHYLASE -> PglZ -> SIGMA+OB-Fold+AAA-ATPase+HKD-DNase-BrxL ->SbcC+SbcC->Calcineurin+Calcineurin-SbcD->SbcC-> | 1203 | Bacteria>Pseudomonadota>Acidithiobacillia>Acidithiobacillales>Acidithiobacillaceae>Acidithiobacillus | GCF_010577825.1 | Acidithiobacillus_ferrooxidans | X857_RS08730 | 38026..41637 | 920 | BREX-system-P-loop-protein-BrxC-[Acidithiobacillus-ferrooxidans] |
| WP_271780036.1 | Type1.BREX | HTH_17->Phage_int_SAM_2+Integrase_1-><-SbcC<-Calcineurin+Calcineurin-SbcD\|\| BrxA-DNA-binding -> BrxB-BrxF-iSTAND -> DUF6079 *-> N6-METHYLASE ->Schlafen+HATPase+wHTH->SIR2->HerA/FtsK-> N6-METHYLASE -p-> PglZ -> SIGMA+OB-Fold+AAA-ATPase+HKD-DNase-BrxL -> | 1179 | Bacteria>Pseudomonadota>Acidithiobacillia>Acidithiobacillales>Acidithiobacillaceae>Acidithiobacillus | GCF_028215975.1 | Acidithiobacillus_ferriphilus | PJU76_RS00595 | 119550..123089 | 1689834 | BREX-system-P-loop-protein-BrxC-[Acidithiobacillus-ferriphilus] |
| WP_101538034.1 | Type1.BREX | DEDD_Tnp_IS110+Transposase_20-><-?\|\|XIS-HTH->TOPRIM->\|\|XIS-HTH-> BrxA-DNA-binding -> BrxB-BrxF-iSTAND -> DUF6079 *->Schlafen+HATPase+wHTH+HTH-> N6-METHYLASE ->HTH+REase-> PglZ -> SIGMA+OB-Fold+AAA-ATPase+HKD-DNase-BrxL ->SbcC+APATPase+SbcC->Calcineurin+Calcineurin-SbcD->SbcC->RNase-> | 1204 | Bacteria>Pseudomonadota>Acidithiobacillia>Acidithiobacillales>Acidithiobacillaceae | GCF_018854655.1 | Acidithiobacillus | HF670_RS09000 | 8228..11842 | 119977 | MULTISPECIES:-BREX-system-P-loop-protein-BrxC-[Acidithiobacillus] |
| WP_023449856.1 | Type1.BREX | Ntox45->?-><-Couple_hipA+HipA_C<-HTH_3\|\|UDG-conflict-><-ParB+HNH<- SIGMA+OB-Fold+AAA-ATPase+HKD-DNase-BrxL <- PglZ <- N6-METHYLASE <- DUF6079 *<- BrxB-BrxF-iSTAND <- BrxA-DNA-binding <-RelE-ParE<-MetJarc-HTH<-?\|\|AbiEi_4->AbiEii->SWIM-> wHTH+WYL+WCX -> | 1177 | Bacteria>Pseudomonadota>Alphaproteobacteria>Caulobacterales>Caulobacteraceae>Asticcacaulis | GCF_000495835.1 | Asticcacaulis_sp._AC402 | ABAC402_RS31845 | 123615..127148 | 1282361 | BREX-system-P-loop-protein-BrxC-[Asticcacaulis-sp.-AC402] |
| WP_114817812.1 | Type1.BREX | HTH_3-><-?<-ParB+N6_N4_METHYLASE<-?\|\|OLD-ABC+TOPRIM->SF1-SF2-Helicase-><- SIGMA+OB-Fold+AAA-ATPase+HKD-DNase-BrxL <- PglZ <- N6-METHYLASE <- DUF6079 *<- BrxB-BrxF-iSTAND <- BrxA-DNA-binding <- wHTH+WYL+WCX \|\|35Exo+REase-> | 1179 | Bacteria>Pseudomonadota>Alphaproteobacteria>Caulobacterales>Caulobacteraceae>Brevundimonas | GCF_003350205.1 | Brevundimonas_bullata | DXK21_RS10945 | 61535..65074 | 13160 | BREX-system-P-loop-protein-BrxC-[Brevundimonas-bullata] |
| MAS04597.1 | Type1.BREX | XIS-HTH->Phage_integrase->Phage_integrase->TetR-HTH->\|\|?-><-EVE\|\|XIS-HTH-> BrxA-DNA-binding -> BrxB-BrxF-iSTAND -> DUF6079 *-> N6-METHYLASE -> PglZ -> SIGMA+OB-Fold+AAA-ATPase+HKD-DNase-BrxL ->?->SbcC-><-TIR<-TIR<-?<-HIT | 1153 | Bacteria>Pseudomonadota>Alphaproteobacteria>Hyphomicrobiales>Ahrensiaceae>Ahrensia | GCA_002698425.1 | Ahrensia_sp. | CL534_07875 | 15621..19082 | 1872419 | ATP-binding-protein-[Ahrensia-sp.] |
| WP_061974428.1 | Type1.BREX | <-SF1-SF2-Helicase<- HKD-DNase <-DUF6361<-Small-B-barrel+SF1-SF2-Helicase+RdgC+REase<-SF1-SF2-Helicase+REase<-35Exo+REase\|\| wHTH+WYL+WCX -> BrxA-DNA-binding -> BrxB-BrxF-iSTAND -> DUF6079 *-> N6-METHYLASE -> PglZ -> SIGMA+OB-Fold+AAA-ATPase+HKD-DNase-BrxL -><-GreA_GreB+SF1-SF2-Helicase<-?<-?<-REase+N-glycosylase<-PVC-AAA+Classical-AAA<-Ntox49 | 1181 | Bacteria>Pseudomonadota>Alphaproteobacteria>Hyphomicrobiales>Aurantimonadaceae>Aureimonas | GCF_001463725.1 | Aureimonas_sp._AU20 | AUL37_RS03985 | 274964..278509 | 1349819 | BREX-system-P-loop-protein-BrxC-[Aureimonas-sp.-AU20] |
| WP_268883456.1 | Type1.BREX | <- SIGMA+OB-Fold+AAA-ATPase+HKD-DNase-BrxL <- PglZ <- N6-METHYLASE -p<- N6-METHYLASE \|\|?-><-?\|\|HIN-HTH-><- DUF6079 -p<-?<- DUF6079 -p*<- BrxB-BrxF-iSTAND <- BrxA-DNA-binding <-?\|\|<-MULE-TRANSPOSASE\|\|HIN-HTH->?->?-><-?\|\|ST-Hiskin-> | 617 | Bacteria>Pseudomonadota>Alphaproteobacteria>Hyphomicrobiales>Aurantimonadaceae>Jiella | GCF_026898155.1 | Jiella_sp._HL-NP1 | OH818_RS13650 | 2639917..2641770 | 2986949 | BREX-system-P-loop-protein-BrxC-[Jiella-sp.-HL-NP1] |

| Protein-ID | System | Gene-Neighbourhoods | Length | Taxonomy | Assembly-ID | Strain | Locus-Tag | Genomic-coordinates | TaxID | Fasta-define |
|---|---|---|---|---|---|---|---|---|---|---|
| WP_230722685.1 | Type1.BREX | <-ArdCN+MPTase<-DEDD_Tnp_IS110+Transposase_20<-Small-B-barrel+SF1-SF2-Helicase+RdgC+REase<-SF1-SF2-Helicase+REase<-35Exo+REase<-?\|\|wHTH+WYL+WCX->BrxA-DNA-binding->BrxB-BrxF-iSTAND->DUF6079*->OLD-ABC->NERD->N6-METHYLASE->PglZ->SIGMA+OB-Fold+AAA-ATPase+HKD-DNase-BrxL-><-HHH_4+HTH+wHTH+SF1-SF2-Helicase+REase | 1177 | Bacteria>Pseudomonadota>Alphaproteobacteria>Hyphomicrobiales>Aurantimonadaceae>Martelella | GCF_021021415.1 | Martelella_mediterranea | K7H91_RS23870 | 16033..19566 | 293089 | BREX-system-P-loop-protein-BrxC-[Martelella-mediterranea] |
| WP_094540671.1 | Type1.BREX | HTH_3->Recombinase+ZnR-><-SF1-SF2-Helicase<-HKD-DNase+HKD-DNase<-wHTH<-SF1-SF2-Helicase+REase<-SIGMA+OB-Fold+AAA-ATPase+HKD-DNase-BrxL<-PglZ<-N6-METHYLASE<-DUF6079*<-BrxB-BrxF-iSTAND<-BrxA-DNA-binding<-wHTH+WYL+WCX\|\|Calcineurin+Calcineurin-SbcD->SbcC+SbcC-><-MODE-HTH+PBPII\|\|TetR-HTH-><-RHH<-VirB10 | 1179 | Bacteria>Pseudomonadota>Alphaproteobacteria>Hyphomicrobiales>Brucellaceae>Brucella/Ochrobactrum_group | GCF_002252505.1 | Brucella_grignonensis | CEV33_RS11715 | 211751..215290 | 94627 | BREX-system-P-loop-protein-BrxC-[Brucella-grignonensis] |
| WP_113731560.1 | Type1.BREX | VirB8->VirB9->VirB10->RHH-><-TetR-HTH\|\|MODE-HTH+PBPII->wHTH+WYL+WCX->BrxA-DNA-binding->BrxB-BrxF-iSTAND->DUF6079*->N6-METHYLASE->PglZ->SIGMA+OB-Fold+AAA-ATPase+HKD-DNase-BrxL->SF1-SF2-Helicase+REase->SF1-SF2-Helicase+REase->wHTH->HKD-DNase+HKD-DNase->SF1-SF2-Helicase-> | 1179 | Bacteria>Pseudomonadota>Alphaproteobacteria>Hyphomicrobiales>Brucellaceae>Brucella/Ochrobactrum_group | GCF_003261015.1 | Brucella_anthropi | F9L04_RS18285 | 14553..18092 | 529 | BREX-system-P-loop-protein-BrxC-[Brucella-anthropi] |
| WP_151664132.1 | Type1.BREX | ParA-Soj-PloopNTPase->ParB+RepB->ZDNA-HTH+HTH->CHTH-><-SbcC+CCDC158<-Calcineurin-SbcD\|\|wHTH+WYL+WCX->BrxA-DNA-binding->BrxB-BrxF-iSTAND->DUF6079*->N6-METHYLASE->PglZ->SIGMA+OB-Fold+AAA-ATPase+HKD-DNase-BrxL->HTH_3-><-ParB+HNH<-HEPN<-SF1-SF2-Helicase+REase<-SF1-SF2-Helicase+REase<-REase | 1179 | Bacteria>Pseudomonadota>Alphaproteobacteria>Hyphomicrobiales>Brucellaceae>Brucella/Ochrobactrum_group | GCF_008932705.1 | Brucella_anthropi | F9L04_RS20265 | 51555..55094 | 529 | BREX-system-P-loop-protein-BrxC-[Brucella-anthropi] |
| WP_151665645.1 | Type1.BREX | ParB+RepB->ZDNA-HTH+HTH->?->?->LuxR-HTH-><-?<-SIGMA+OB-Fold+AAA-ATPase+HKD-DNase-BrxL<-PglZ<-N6-METHYLASE<-DUF6079*<-BrxB-BrxF-iSTAND<-BrxA-DNA-binding<-wHTH+WYL+WCX\|\|Calcineurin-SbcD->SbcC+CCDC158-> | 1180 | Bacteria>Pseudomonadota>Alphaproteobacteria>Hyphomicrobiales>Brucellaceae>Brucella/Ochrobactrum_group | GCF_008932825.1 | Brucella_anthropi | F9K96_RS24475 | 19243..22785 | 529 | BREX-system-P-loop-protein-BrxC-[Brucella-anthropi] |
| WP_204157651.1 | Type6.BREX | <-TUDOR+SF1-SF2-Helicase+iREase+CTD<-SIGMA+OB-Fold+AAA-ATPase+HKD-DNase-BrxL<-PglZ<-N6-METHYLASE-p<-AAA-ATPase<-rve<-N6-METHYLASE-p<-N6-METHYLASE<-DUF6079*<-BrxB-BrxF-iSTAND<-BrxE<-PhdYeFM_antitox<-TraI-VirD2-RCRE<-TraJ-RHH\|\|?-><-DOC+HTH | 1208 | Bacteria>Pseudomonadota>Alphaproteobacteria>Hyphomicrobiales>Brucellaceae>Brucella/Ochrobactrum_group | GCF_016887905.1 | Brucella_anthropi | JQC79_RS05930 | 1236117..1239743 | 529 | BREX-system-P-loop-protein-BrxC-[Brucella-anthropi] |
| WP_109366634.1 | Type1.BREX | PIN+OST-HTH->DOC+HTH->?-><-SbcC<-SbcC+CCDC158<-Calcineurin-SbcD\|\|wHTH+WYL+WCX->BrxA-DNA-binding->BrxB-BrxF-iSTAND->DUF6079*->N6-METHYLASE->PglZ->SIGMA+OB-Fold+AAA-ATPase+HKD-DNase-BrxL-><-HIN-HTH<-SF1-SF2-Helicase+REase<-SF1-SF2-Helicase+REase<-REase | 1179 | Bacteria>Pseudomonadota>Alphaproteobacteria>Hyphomicrobiales>Brucellaceae>Brucella/Ochrobactrum_group | GCF_900243435.1 | [Ochrobactrum]_soli | OHAE_RS05910 | 16979..20518 | 2448455 | BREX-system-P-loop-protein-BrxC-[[Ochrobactrum] |
| WP_184220677.1 | Type1.BREX | <-ParB\|\|?->?->?-><-?<-SIGMA+OB-Fold+AAA-ATPase+HKD-DNase-BrxL<-PglZ<-N6-METHYLASE<-DUF6079*<-BrxB-BrxF-iSTAND<-BrxA-DNA-binding\|\|?->?->NTP_transferase+MannoseP_isomer->?-><-?\|\|?->SAM-METHYLASE-> | 1179 | Bacteria>Pseudomonadota>Alphaproteobacteria>Hyphomicrobiales>Brucellaceae>Paenochrobactrum | GCF_014205685.1 | Paenochrobactrum_gallinarii | FHS77_RS04490 | 13414..16953 | 643673 | BREX-system-P-loop-protein-BrxC-[Paenochrobactrum-gallinarii] |
| WP_075657534.1 | Type1.BREX | VirB8->VirB9->VirB10->RHH-><-TetR-HTH\|\|MODE-HTH+PBPII->wHTH+WYL+WCX->BrxA-DNA-binding->BrxB-BrxF-iSTAND->DUF6079*->N6-METHYLASE->PglZ->SIGMA+OB-Fold+AAA-ATPase+HKD-DNase-BrxL->SF1-SF2-Helicase+REase->SF1-SF2-Helicase+REase->wHTH->HKD-DNase+HKD-DNase->SF1-SF2-Helicase-> | 1179 | Bacteria>Pseudomonadota>Alphaproteobacteria>Hyphomicrobiales>Brucellaceae>Pseudochrobactrum | GCF_001939785.1 | Pseudochrobactrum_sp._B5 | BKD14_RS17970 | 55973..59512 | 1289478 | BREX-system-P-loop-protein-BrxC-[Pseudochrobactrum-sp.-B5] |
| WP_113944570.1 | Type1.BREX | VirB8->VirB9->VirB10->RHH-><-TetR-HTH\|\|MODE-HTH+PBPII->wHTH+WYL+WCX->BrxA-DNA-binding->BrxB-BrxF-iSTAND->DUF6079*->N6-METHYLASE->PglZ->SIGMA+OB-Fold+AAA-ATPase+HKD-DNase-BrxL->SF1-SF2-Helicase+REase->SF1-SF2-Helicase+REase->wHTH->HKD-DNase+HKD-DNase->SF1-SF2-Helicase-> | 1179 | Bacteria>Pseudomonadota>Alphaproteobacteria>Hyphomicrobiales>Brucellaceae>Pseudochrobactrum | GCF_003314995.1 | Pseudochrobactrum_asaccharolyticum | DFR47_RS11470 | 41745..45284 | 354351 | BREX-system-P-loop-protein-BrxC-[Pseudochrobactrum-asaccharolyticum] |
| MBX3547065.1 | Type1.BREX | <-HIN-HTH<-REase<-SF1-SF2-Helicase<-OLD-ABC+TOPRIM<-AbiEii<-AbiEi_4<-SIGMA+OB-Fold+AAA-ATPase+HKD-DNase-BrxL<-PglZ<-N6-METHYLASE<-DUF6079-p* | 777 | Bacteria>Pseudomonadota>Alphaproteobacteria>Hyphomicrobiales>Chelatococcaceae>Chelatococcus | GCA_019635905.1 | Chelatococcus_sp. | KF818_30290 | 46824..49157 | 1953771 | BREX-system-P-loop-protein-BrxC-[Chelatococcus-sp.] |
| WP_046141798.1 | Type6.BREX | <-XIS-HTH<-?\|\|RAMA->GDResolvase+ZnR->ZnR->?->BrxE->BrxA-DNA-binding->BrxB-BrxF-iSTAND->DUF6079*->N6-METHYLASE->N6-METHYLASE->PglZ->SIGMA+OB-Fold+AAA-ATPase+HKD-DNase-BrxL-><-tRNA<-ADH_N+ADH_zinc_N | 1208 | Bacteria>Pseudomonadota>Alphaproteobacteria>Hyphomicrobiales>Devosiaceae>Devosia | - | Devosia_soli | VW35_RS04250 | 28883..32509 | 361041 | BREX-system-P-loop-protein-BrxC-[Devosia-soli] |
| MBL8596415.1 | Type1.BREX | ANK+ANK->wHTH+WYL+WCX->BrxA-DNA-binding->BrxB-BrxF-iSTAND->DUF6079*->N6-METHYLASE->PglZ->SIGMA+OB-Fold+AAA-ATPase+HKD-DNase-BrxL->?-><-?<-?\|\|HTH_Tnp_1->?->MPTase-> | 1166 | Bacteria>Pseudomonadota>Alphaproteobacteria>Hyphomicrobiales>Devosiaceae>Devosia | GCA_016793625.1 | Devosia_sp. | JNL14_01620 | 65385..68885 | 1871048 | BREX-system-P-loop-protein-BrxC-[Devosia-sp.] |
| WP_240230050.1 | Type1.BREX | <-rve<-REase<-?<-Peptidase<-Classical-AAA\|\|?-><-SIGMA+OB-Fold+AAA-ATPase+HKD-DNase-BrxL<-PglZ<-N6-METHYLASE<-DUF6079*<-BrxB-BrxF-iSTAND<-BrxA-DNA-binding<-wHTH+WYL+WCX<-ANK+ANK\|\|HEPN-><-tRNA\|\|ICLR-HTH-> | 1166 | Bacteria>Pseudomonadota>Alphaproteobacteria>Hyphomicrobiales>Devosiaceae>Devosia | GCF_022429625.1 | Devosia_sp._JXJ_CY_41 | MF606_RS14450 | 2929921..2933421 | 2917991 | BREX-system-P-loop-protein-BrxC-[Devosia-sp.-JXJ-CY-41] |
| WP_203065852.1 | Type1.BREX | VirB10->RHH-><-TetR-HTH\|\|MODE-HTH+PBPII-><-SbcC+SbcC<-Calcineurin+Calcineurin-SbcD\|\|wHTH+WYL+WCX->BrxA-DNA-binding->BrxB-BrxF-iSTAND->DUF6079*->N6-METHYLASE->PglZ->SIGMA+OB-Fold+AAA-ATPase+HKD-DNase-BrxL->OLD-ABC+TOPRIM->SF1-SF2-Helicase-><-?<-Arm-DNA-bind_3+Phage_int_SAM_3+Phage_integrase | 1179 | Bacteria>Pseudomonadota>Alphaproteobacteria>Hyphomicrobiales>Devosiaceae>Paradevosia | GCF_016801025.1 | Paradevosia_shaoguanensis | JNE37_RS07680 | 1630850..1634389 | 1335043 | BREX-system-P-loop-protein-BrxC-[Paradevosia-shaoguanensis] |
| WP_266332120.1 | Type1.BREX | <-SF1-SF2-Helicase<-HKD-DNase+HKD-DNase<-wHTH<-SF1-SF2-Helicase+REase<-SF1-SF2-Helicase+REase<-SIGMA+OB-Fold+AAA-ATPase+HKD-DNase-BrxL<-PglZ<-N6-METHYLASE<-DUF6079*<-BrxB-BrxF-iSTAND<-BrxA-DNA-binding<-wHTH+WYL+WCX<-MODE-HTH+PBPII\|\|TetR-HTH-><-RHH<-VirB10<-VirB9<-VirB8 | 1179 | Bacteria>Pseudomonadota>Alphaproteobacteria>Hyphomicrobiales>Kaistiaceae>Kaistia | GCF_026343955.1 | Kaistia_defluvii | OSH10_RS12655 | 370045..373584 | 410841 | BREX-system-P-loop-protein-BrxC-[Kaistia-defluvii] |
| KFG67132.1 | Type6.BREX | <-AbiGi+seqA+RAMA\|\|BrxE->BrxA-DNA-binding->BrxB-BrxF-iSTAND->DUF6079*->N6-METHYLASE-p->PglZ->SIGMA+OB-Fold+AAA-ATPase+HKD-DNase-BrxL->HNH->HNH->HNH->?-><-HEPN<-Polbeta | 1210 | Bacteria>Pseudomonadota>Alphaproteobacteria>Hyphomicrobiales>Methylobacteriaceae>Microvirga | GCA_000739015.1 | Microvirga_sp._BSC39 | JH26_23850 | 105316..108948 | 1549810 | ATPase-[Microvirga-sp.-BSC39] |
| WP_036357316.1 | Type6.BREX | <-AbiGi+seqA+RAMA<-?\|\|PhdYeFM_antitox->BrxE->BrxA-DNA-binding->BrxB-BrxF-iSTAND->DUF6079*->N6-METHYLASE->PglZ->SIGMA+OB-Fold+AAA-ATPase+HKD-DNase-BrxL->HNH->HNH->HNH->?-><-HEPN<-Polbeta | 1208 | Bacteria>Pseudomonadota>Alphaproteobacteria>Hyphomicrobiales>Methylobacteriaceae>Microvirga | GCF_000739015.1 | Microvirga_sp._BSC39 | MA07_RS22775 | 105322..108948 | 1549810 | BREX-system-P-loop-protein-BrxC-[Microvirga-sp.-BSC39] |
| WP_199050625.1 | Type6.BREX | <-ParB+HNH<-ARG\|\|?-><-?<-SF1-SF2-Helicase+DUF1998<-?\|\|BrxE->BrxA-DNA-binding->BrxB-BrxF-iSTAND->DUF6079*->N6-METHYLASE->N6-METHYLASE-p->PglZ->SIGMA+OB-Fold+AAA-ATPase+HKD-DNase-BrxL-><-Phage_integrase | 1206 | Bacteria>Pseudomonadota>Alphaproteobacteria>Hyphomicrobiales>Methylobacteriaceae>Microvirga | GCF_016427565.1 | Microvirga_splendida | JAO75_RS18515 | 48792..52412 | 2795727 | BREX-system-P-loop-protein-BrxC-[Microvirga-splendida] |
| WP_202057660.1 | Type1.BREX | Phage_integrase+Phage_integrase->REase->DnaK->?->?->?->BrxA-DNA-binding->BrxB-BrxF-iSTAND->DUF6079*->N6-METHYLASE->PglZ->SIGMA+OB-Fold+AAA-ATPase+HKD-DNase-BrxL->?->?->?-><-?<-tRNA<-?<-CDP-OH_P_transf | 1176 | Bacteria>Pseudomonadota>Alphaproteobacteria>Hyphomicrobiales>Methylobacteriaceae>Microvirga | GCF_016743775.1 | Microvirga_aerilata | JKG68_RS07685 | 69600..73130 | 670292 | BREX-system-P-loop-protein-BrxC-[Microvirga-aerilata] |
| WP_036280608.1 | Type1.BREX | HKD-DNase->SF1-SF2-Helicase-><-Small-B-barrel+SF1-SF2-Helicase+RdgC+REase<-SF1-SF2-Helicase+REase<-35Exo+REase\|\|wHTH+WYL+WCX->BrxA-DNA-binding->BrxB-BrxF-iSTAND->DUF6079*->OLD-ABC->REase->N6-METHYLASE->PglZ->SIGMA+OB-Fold+AAA-ATPase+HKD-DNase-BrxL->HIN-HTH->?->LZ_Tnp_IS66+zf-IS66+DDE_Tnp_IS66+DDE_Tnp_IS66_C-> | 1178 | Bacteria>Pseudomonadota>Alphaproteobacteria>Hyphomicrobiales>Methylocystaceae>Methylocystis | GCF_000188155.2 | Methylocystis_sp._ATCC_49242 | MET49242_RS03150 | 256459..259995 | 622637 | BREX-system-P-loop-protein-BrxC-[Methylocystis-sp.-ATCC-49242] |
| WP_253589372.1 | Type6.BREX | <-HIRAN<-BirA-HTH<-SF1-SF2-Helicase+REase<-35Exo+REase<-?\|\|wHTH+WYL+WCX->BrxE->BrxA-DNA-binding->BrxB-BrxF-iSTAND->DUF6079*->N6-METHYLASE->N6-METHYLASE->PglZ->SIGMA+OB-Fold+AAA-ATPase+HKD-DNase-BrxL->?-><-?\|\|?->?->HNH-> | 1208 | Bacteria>Pseudomonadota>Alphaproteobacteria>Hyphomicrobiales>Nitrobacteraceae>Bradyrhizobium | - | Bradyrhizobium_elkanii | J2R96_RS28580 | 6233220..6236846 | 29448 | BREX-system-P-loop-protein-BrxC-[Bradyrhizobium-elkanii] |
| WP_212207485.1 | Type6.BREX | <-PIWI<-SIGMA+OB-Fold+AAA-ATPase+HKD-DNase-BrxL<-PglZ<-ParB+HNH<-N6-METHYLASE-p<-N6-METHYLASE<-DUF6079*<-BrxB-BrxF-iSTAND<-BrxA-DNA-binding<-BrxE<-wHTH+WYL+WCX\|\|35Exo+REase->SF1-SF2-Helicase+REase-> | 1208 | Bacteria>Pseudomonadota>Alphaproteobacteria>Hyphomicrobiales>Nitrobacteraceae>Bradyrhizobium | - | Bradyrhizobium_sp._AUGA_SZCCT0274 | JQ620_09165 | 74241..77867 | 2807670 | BREX-system-P-loop-protein-BrxC-[Bradyrhizobium-sp.-AUGA-SZCCT0274] |
| MBR1240293.1 | Type6.BREX | <-PIWI<-SIGMA+OB-Fold+AAA-ATPase+HKD-DNase-BrxL<-PglZ<-ParB+HNH<-N6-METHYLASE-p<-N6-METHYLASE<-DUF6079*<-BrxB-BrxF-iSTAND<-BrxA-DNA-binding<-BrxE<-wHTH+WYL+WCX\|\|35Exo+REase->SF1-SF2-Helicase+REase-> | 1208 | Bacteria>Pseudomonadota>Alphaproteobacteria>Hyphomicrobiales>Nitrobacteraceae>Bradyrhizobium | GCA_018130325.1 | Bradyrhizobium_sp._AUGA_SZCCT0274 | JQ620_09165 | 74241..77867 | 2807670 | BREX-system-P-loop-protein-BrxC-[Bradyrhizobium-sp.-AUGA-SZCCT0274] |
| WP_130224749.1 | Type1.BREX | <-ArdCN+MPTase<-HEPN<-Small-B-barrel+SF1-SF2-Helicase+RdgC+REase<-35Exo+REase<-?\|\|wHTH+WYL+WCX->BrxA-DNA-binding->BrxB-BrxF-iSTAND->DUF6079*->N6-METHYLASE->PglZ->SIGMA+OB-Fold+AAA-ATPase+HKD-DNase-BrxL->AbiEi_4->AbiEii->OLD-ABC+TOPRIM->SF1-SF2-Helicase->REase-><-ABhydrolase | 1181 | Bacteria>Pseudomonadota>Alphaproteobacteria>Hyphomicrobiales>Nitrobacteraceae>Bradyrhizobium | GCF_004212675.1 | Bradyrhizobium_sp._Leo121 | CWO90_RS07090 | 48469..52014 | 1571195 | BREX-system-P-loop-protein-BrxC-[Bradyrhizobium-sp.-Leo121] |
| WP_157343672.1 | Type1.BREX | ABC-ATPase+SbcC->?->?-><-tRNA<-Small-B-barrel+SF1-SF2-Helicase+RdgC+REase<-SF1-SF2-Helicase+REase<-35Exo+REase\|\|wHTH+WYL+WCX->BrxA-DNA-binding->BrxB-BrxF-iSTAND->DUF6079*->N6-METHYLASE->PglZ->SIGMA+OB-Fold+AAA-ATPase+HKD-DNase-BrxL-><-?<-HTH+REase<-Arm-DNA-bind_3+Phage_integrase<-?\|\|?-><-Peptidase_M24 | 1181 | Bacteria>Pseudomonadota>Alphaproteobacteria>Hyphomicrobiales>Nitrobacteraceae>Bradyrhizobium | GCF_009759865.1 | Bradyrhizobium_pachyrhizi | GPL21_13485 | 267907..271452 | 280333 | BREX-system-P-loop-protein-BrxC-[Bradyrhizobium-pachyrhizi] |
| WP_249144787.1 | Type6.BREX | <-PIWI<-SIGMA+OB-Fold+AAA-ATPase+HKD-DNase-BrxL<-PglZ<-ParB+HNH<-N6-METHYLASE-p<-N6-METHYLASE<-DUF6079*<-BrxB-BrxF-iSTAND<-BrxA-DNA-binding<-BrxE<-wHTH+WYL+WCX\|\|35Exo+REase->SF1-SF2-Helicase+REase-> | 1203 | Bacteria>Pseudomonadota>Alphaproteobacteria>Hyphomicrobiales>Nitrobacteraceae>Bradyrhizobium | GCF_018130325.1 | Bradyrhizobium_sp._AUGA_SZCCT0274 | JQ620_RS09170 | 74241..77852 | 2807670 | BREX-system-P-loop-protein-BrxC-[Bradyrhizobium-sp.-AUGA-SZCCT0274] |
| WP_212056797.1 | Type1.BREX | <-SF1-SF2-Helicase<-MORC+DUF3684+REase<-HHH_4+SF1-SF2-Helicase+REase<-AbiEi_4<-SIGMA+OB-Fold+AAA-ATPase+HKD-DNase-BrxL<-PglZ<-N6-METHYLASE<-DUF6079*<-BrxB-BrxF-iSTAND<-BrxA-DNA-binding<-wHTH+WYL+WCX\|\|35Exo+REase->SF1-SF2-Helicase+REase->Small-B-barrel+SF1-SF2-Helicase+RdgC+REase-><-?\|\|GDResolvase+Recombinase+ZnR-> | 1180 | Bacteria>Pseudomonadota>Alphaproteobacteria>Hyphomicrobiales>Nitrobacteraceae>Bradyrhizobium | GCF_018131045.1 | Bradyrhizobium_elkanii | JQ569_RS15725 | 68040..71582 | 29448 | BREX-system-P-loop-protein-BrxC-[Bradyrhizobium-elkanii] |
| WP_213286691.1 | Type6.BREX | Phage_integrase->MarR-HTH->AbiEii->?->?->BrxE->BrxA-DNA-binding->BrxB-BrxF-iSTAND->DUF6079*->N6-METHYLASE->N6-METHYLASE->ParB+HNH->PglZ->SIGMA+OB-Fold+AAA-ATPase+HKD-DNase-BrxL-><-TPR<-?<-?\|\|wHTH+WYL+WCX-> | 1208 | Bacteria>Pseudomonadota>Alphaproteobacteria>Hyphomicrobiales>Nitrobacteraceae>Bradyrhizobium | GCF_018398875.1 | Bradyrhizobium_sp._sGM-13 | KJQ96_RS11300 | 145012..148638 | 2831781 | BREX-system-P-loop-protein-BrxC-[Bradyrhizobium-sp.-sGM-13] |
| WP_225161486.1 | Type1.BREX | <-ParB+HNH+Schlafen<-?<-SIGMA+OB-Fold+AAA-ATPase+HKD-DNase-BrxL<-PglZ<-N6-METHYLASE<-DUF6079*<-BrxB-BrxF-iSTAND<-BrxA-DNA-binding<-wHTH+WYL+WCX\|\|?->35Exo+REase->SF1-SF2-Helicase+REase->35exo+SF1-SF2-Helicase-><-Phage_integrase | 1183 | Bacteria>Pseudomonadota>Alphaproteobacteria>Hyphomicrobiales>Nitrobacteraceae>Bradyrhizobium | GCF_020194035.1 | Bradyrhizobium_sp._BRP56 | I6F38_19500 | 73459..77010 | 2793819 | BREX-system-P-loop-protein-BrxC-[Bradyrhizobium-sp.-BRP56] |
| WP_247477718.1 | Type6.BREX | BrxE->BrxA-DNA-binding->BrxB-BrxF-iSTAND->DUF6079*->N6-METHYLASE->PglZ->SIGMA+OB-Fold+AAA-ATPase+HKD-DNase-BrxL->SbcC->HNH-><-?<-?\|\|TPR+TPR+TPR+TPR-> | 1206 | Bacteria>Pseudomonadota>Alphaproteobacteria>Hyphomicrobiales>Nitrobacteraceae>Bradyrhizobium | GCF_023072855.1 | Bradyrhizobium_sp._CW9 | IVB57_RS09580 | 204980..208600 | 2782689 | BREX-system-P-loop-protein-BrxC-[Bradyrhizobium-sp.-CW9] |
| WP_259215308.1 | Type6.BREX | DSBH->?->tRNA->BrxE->BrxA-DNA-binding->BrxB-BrxF-iSTAND->DUF6079*->N6-METHYLASE->PglZ->SIGMA+OB-Fold+AAA-ATPase+HKD-DNase-BrxL->SbcC->HNH-><-?<-?<-?<-SIGMA-HTH+HIN-HTH | 1207 | Bacteria>Pseudomonadota>Alphaproteobacteria>Hyphomicrobiales>Nitrobacteraceae>Bradyrhizobium | GCF_024809395.1 | Bradyrhizobium_centrosematis | GGE24_RS17830 | 223880..227503 | 1300039 | BREX-system-P-loop-protein-BrxC-[Bradyrhizobium-centrosematis] |
| WP_260373497.1 | Type1.BREX | <-ParB+HNH+Schlafen<-?<-SIGMA+OB-Fold+AAA-ATPase+HKD-DNase-BrxL<-PglZ<-N6-METHYLASE<-DUF6079*<-BrxB-BrxF-iSTAND<-BrxA-DNA-binding<-wHTH+WYL+WCX\|\|?->35Exo+REase->SF1-SF2-Helicase+REase->35exo+SF1-SF2-Helicase-><-Phage_integrase | 1183 | Bacteria>Pseudomonadota>Alphaproteobacteria>Hyphomicrobiales>Nitrobacteraceae>Bradyrhizobium | GCF_025200925.1 | Bradyrhizobium_sp._CB1015 | N2604_RS01610 | 347161..350712 | 2976822 | BREX-system-P-loop-protein-BrxC-[Bradyrhizobium-sp.-CB1015] |
| WP_110780661.1 | Type1.BREX | <-DOC+HTH<-SF1-SF2-Helicase<-HKD-DNase+HKD-DNase<-wHTH<-SF1-SF2-Helicase+REase<-SF1-SF2-Helicase+REase<-SIGMA+OB-Fold+AAA-ATPase+HKD-DNase-BrxL<-PglZ<-N6-METHYLASE<-DUF6079*<-BrxB-BrxF-iSTAND<-BrxA-DNA-binding<-wHTH+WYL+WCX<-MODE-HTH+PBPII\|\|TetR-HTH-><-RHH<-VirB10<-VirB9<-VirB8 | 1179 | Bacteria>Pseudomonadota>Alphaproteobacteria>Hyphomicrobiales>Nitrobacteraceae>Rhodopseudomonas | GCF_003217325.1 | Rhodopseudomonas_faecalis | BJ122_RS08545 | 91514..95053 | 99655 | BREX-system-P-loop-protein-BrxC-[Rhodopseudomonas-faecalis] |
| MAM94519.1 | Type1.BREX | VirB10->RHH-><-TetR-HTH\|\|MODE-HTH+PBPII-><-SbcC+SbcC<-Calcineurin+Calcineurin-SbcD\|\|wHTH+WYL+WCX->BrxA-DNA-binding->BrxB-BrxF-iSTAND->DUF6079*->N6-METHYLASE->PglZ->SIGMA+OB-Fold+AAA-ATPase+HKD-DNase-BrxL->OLD-ABC+TOPRIM->SF1-SF2-Helicase-><-Arm-DNA-bind_3+Phage_int_SAM_3+Phage_integrase<-?<-?<-MerR-HTH | 1179 | Bacteria>Pseudomonadota>Alphaproteobacteria>Hyphomicrobiales>Parvibaculaceae>Parvibaculum | GCA_002694985.1 | Parvibaculum_sp. | CMI61_07805 | 145242..148781 | 2024848 | hypothetical-protein-[Parvibaculum-sp.] |
| WP_110030326.1 | Type1.BREX | RAMPs->RAMPs->?->RVT->CRISPR_Cas6->?->?<-?\|\|BrxA-DNA-binding->BrxB-BrxF-iSTAND->DUF6079*->N6-METHYLASE->PglZ->SIGMA+OB-Fold+AAA-ATPase+HKD-DNase-BrxL->?->HKD-DNase+HKD-DNase->SF1-SF2-Helicase->MORC+DUF3684+REase->REase+SF1-SF2-Helicase-> | 1173 | Bacteria>Pseudomonadota>Alphaproteobacteria>Hyphomicrobiales>Phyllobacteriaceae>Hoeflea | GCF_003182275.1 | Hoeflea_marina | DFR52_RS02490 | 542434..545955 | 274592 | BREX-system-P-loop-protein-BrxC-[Hoeflea-marina] |
| ESX24129.1 | Type1.BREX | <-SIGMA+OB-Fold+AAA-ATPase+HKD-DNase-BrxL<-PglZ<-N6-METHYLASE<-DUF6079*<-BrxB-BrxF-iSTAND<-BrxA-DNA-binding<-?\|\|REase->HIN-HTH->?->LZ_Tnp_IS66+zf-IS66+DDE_Tnp_IS66+DDE_Tnp_IS66_C-> | 1186 | Bacteria>Pseudomonadota>Alphaproteobacteria>Hyphomicrobiales>Phyllobacteriaceae>Mesorhizobium | GCA_000502755.1 | Mesorhizobium_sp._LSJC264A00 | X767_13205 | 150936..154496 | 1287321 | hypothetical-protein-X767_13205-[Mesorhizobium-sp.-LSJC264A00] |
| ESX84801.1 | Type1.BREX | PIWI->HIN-HTH->HTH_21+rve+rve_3->Y1_Tnp-><-SIGMA+OB-Fold+AAA-ATPase+HKD-DNase-BrxL<-PglZ<-N6-METHYLASE<-NERD<-OLD-ABC<-DUF6079*<-BrxB-BrxF-iSTAND<-BrxA-DNA-binding\|\|?->?-><-HEPN<-HTH_37 | 1177 | Bacteria>Pseudomonadota>Alphaproteobacteria>Hyphomicrobiales>Phyllobacteriaceae>Mesorhizobium | GCA_000503035.1 | Mesorhizobium_sp._LSHC412B00 | X756_23895 | 63877..67410 | 1287285 | hypothetical-protein-X756_23895-[Mesorhizobium-sp.-LSHC412B00] |
| RWC29997.1 | Type1.BREX | SbcC+CCDC158->?->REase+PNTD1+PNTD2->PIWI-><-SIGMA+OB-Fold+AAA-ATPase+HKD-DNase-BrxL<-PglZ<-HNH<-OLD-ABC<-N6-METHYLASE<-DUF6079*<-BrxB-BrxF-iSTAND<-BrxA-DNA-binding\|\|REase-><-?\|\|?-><-?\|\|REase->?->rve-> | 1181 | Bacteria>Pseudomonadota>Alphaproteobacteria>Hyphomicrobiales>Phyllobacteriaceae>Mesorhizobium | GCA_004017395.1 | Mesorhizobium_sp. | EOS27_14215 | 67406..70951 | 1871066 | BREX-system-P-loop-protein-BrxC-[Mesorhizobium-sp.] |
| RWM07956.1 | Type1.BREX | CHTH->Phage_integrase-><-?<-TnsC-ATPase<-rve+Mu-transpos_C<-?\|\|?->BrxA-DNA-binding->BrxB-BrxF-iSTAND->DUF6079*->N6-METHYLASE->PglZ->SIGMA+OB-Fold+AAA-ATPase+HKD-DNase-BrxL->GreA_GreB+SF1-SF2-Helicase-><-wHTH+WYL+WCX\|\|?->?->?-><-HEPN | 1184 | Bacteria>Pseudomonadota>Alphaproteobacteria>Hyphomicrobiales>Phyllobacteriaceae>Mesorhizobium | GCA_004020545.1 | Mesorhizobium_sp. | EOR71_14460 | 11856..15410 | 1871066 | BREX-system-P-loop-protein-BrxC-[Mesorhizobium-sp.] |
| RWD20271.1 | Type1.BREX | AbiEi_4-><-DEDD_Tnp_IS110+Transposase_20<-SIGMA+OB-Fold+AAA-ATPase+HKD-DNase-BrxL<-PglZ<-N6-METHYLASE<-NERD<-OLD-ABC<-DUF6079*<-BrxB-BrxF-iSTAND<-BrxA-DNA-binding<-wHTH+WYL+WCX\|\|35Exo+REase->SF1-SF2-Helicase+REase->Small-B-barrel+SF1-SF2-Helicase+RdgC+REase->NUDIX-><-?\|\|NACHT+SF1-SF2-Helicase+HKD-DNase-> | 1178 | Bacteria>Pseudomonadota>Alphaproteobacteria>Hyphomicrobiales>Phyllobacteriaceae>Mesorhizobium | GCA_004021465.1 | Mesorhizobium_sp. | EOS09_27860 | 18398..21934 | 1871066 | BREX-system-P-loop-protein-BrxC-[Mesorhizobium-sp.] |
| TIL20465.1 | Type6.BREX | BrxE->BrxA-DNA-binding->BrxB-BrxF-iSTAND->DUF6079*->N6-METHYLASE->N6-METHYLASE->ParB+HNH->PglZ->SIGMA+OB-Fold+AAA-ATPase+HKD-DNase-BrxL-> | 1208 | Bacteria>Pseudomonadota>Alphaproteobacteria>Hyphomicrobiales>Phyllobacteriaceae>Mesorhizobium | GCA_004961195.1 | Mesorhizobium_sp. | E5Y88_32415 | 5019..8645 | 1871066 | BREX-system-P-loop-protein-BrxC-[Mesorhizobium-sp.] |
| TIV88924.1 | Type1.BREX | <-35Exo+REase\|\|wHTH+WYL+WCX->BrxA-DNA-binding->BrxB-BrxF-iSTAND->DUF6079*->OLD-ABC->NERD->N6-METHYLASE->PglZ->SIGMA+OB-Fold+AAA-ATPase+HKD-DNase-BrxL-><-HEPN<-RVT<-AbiEi_4 | 1179 | Bacteria>Pseudomonadota>Alphaproteobacteria>Hyphomicrobiales>Phyllobacteriaceae>Mesorhizobium | GCA_004963965.1 | Mesorhizobium_sp. | E5V78_03705 | 4955..8494 | 1871066 | BREX-system-P-loop-protein-BrxC-[Mesorhizobium-sp.] |
| WP_156951459.1 | Type1.BREX | <-SIGMA+OB-Fold+AAA-ATPase+HKD-DNase-BrxL<-PglZ<-N6-METHYLASE<-DUF6079*<-BrxB-BrxF-iSTAND<-BrxA-DNA-binding<-?\|\|REase->HIN-HTH->?->LZ_Tnp_IS66+zf-IS66+DDE_Tnp_IS66+DDE_Tnp_IS66_C-> | 1188 | Bacteria>Pseudomonadota>Alphaproteobacteria>Hyphomicrobiales>Phyllobacteriaceae>Mesorhizobium | GCF_000502755.1 | Mesorhizobium_sp._LSJC264A00 | X768_RS154485 | 150936..154502 | 1287321 | BREX-system-P-loop-protein-BrxC-[Mesorhizobium-sp.-LSJC264A00] |
| WP_156933469.1 | Type1.BREX | REase+PNTD1+PNTD2->PIWI->Y1_Tnp-><-SIGMA+OB-Fold+AAA-ATPase+HKD-DNase-BrxL<-PglZ<-N6-METHYLASE<-NERD<-OLD-ABC<-DUF6079*<-BrxB-BrxF-iSTAND<-BrxA-DNA-binding\|\|?->RVT-><-HEPN<-HTH_37 | 1179 | Bacteria>Pseudomonadota>Alphaproteobacteria>Hyphomicrobiales>Phyllobacteriaceae>Mesorhizobium | GCF_000503035.1 | Mesorhizobium_sp._LSHC412B00 | X757_RS142950 | 63877..67416 | 1287285 | BREX-system-P-loop-protein-BrxC-[Mesorhizobium-sp.-LSHC412B00] |
| WP_128286158.1 | Type1.BREX | ParB+HNH-><-SIGMA+OB-Fold+AAA-ATPase+HKD-DNase-BrxL<-PglZ<-HNH<-OLD-ABC<-N6-METHYLASE<-DUF6079*<-BrxB-BrxF-iSTAND<-BrxA-DNA-binding<-RNaseH | 1181 | Bacteria>Pseudomonadota>Alphaproteobacteria>Hyphomicrobiales>Phyllobacteriaceae>Mesorhizobium | GCF_004023115.1 | Mesorhizobium_sp._M6A.T.Cr.TU.017.01.1.1 | EOB36_RS28010 | 14606..18151 | 2496774 | BREX-system-P-loop-protein-BrxC-[Mesorhizobium-sp.-M6A.T.Cr.TU.017.01.1.1] |
| WP_136091391.1 | Type1.BREX | REase+PNTD1+PNTD2->PIWI->Y1_Tnp-><-SIGMA+OB-Fold+AAA-ATPase+HKD-DNase-BrxL<-PglZ<-HNH<-OLD-ABC<-N6-METHYLASE<-DUF6079*<-BrxB-BrxF-iSTAND<-BrxA-DNA-binding\|\|REase->?-><-?\|\|REase->rve->TnsC-ATPase->TnsC-ATPase-> | 1181 | Bacteria>Pseudomonadota>Alphaproteobacteria>Hyphomicrobiales>Phyllobacteriaceae>Mesorhizobium | GCF_004794715.1 | Mesorhizobium_sp._M8A.F.Ca.ET.207.01.1.1 | EN850_RS12645 | 56072..59617 | 2563968 | BREX-system-P-loop-protein-BrxC-[Mesorhizobium-sp.-M8A.F.Ca.ET.207.01.1.1] |
| WP_140781888.1 | Type6.BREX | <-D5ATPase<-XIS-HTH\|\|MerR-HTH->?->HKD-DNase+HKD-Rease->?->BrxE->BrxA-DNA-binding->BrxB-BrxF-iSTAND->DUF6079*->N6-METHYLASE->PglZ->SIGMA+OB-Fold+AAA-ATPase+HKD-DNase-BrxL-><-?<-?<-CHTH | 1211 | Bacteria>Pseudomonadota>Alphaproteobacteria>Hyphomicrobiales>Phyllobacteriaceae>Mesorhizobium | GCF_006442585.1 | Mesorhizobium_sp._B2-4-8 | FJ947_RS26560 | 105800..109435 | 2589941 | BREX-system-P-loop-protein-BrxC-[Mesorhizobium-sp.-B2-4-8] |
| WP_199202187.1 | Type1.BREX | DOC+HTH-><-?\|\|?-><-Small-B-barrel+SF1-SF2-Helicase+RdgC+REase<-SF1-SF2-Helicase+REase<-35Exo+REase\|\|wHTH+WYL+WCX->BrxA-DNA-binding->BrxB-BrxF-iSTAND->DUF6079*->N6-METHYLASE->PglZ->SIGMA+OB-Fold+AAA-ATPase+HKD-DNase-BrxL->REase+PNTD1+PNTD2->PIWI-><-?<-?\|\|RADICAL-SAM-> | 1185 | Bacteria>Pseudomonadota>Alphaproteobacteria>Hyphomicrobiales>Phyllobacteriaceae>Mesorhizobium | GCF_016756495.1 | Mesorhizobium_sp._113-3-3 | MesoLj113b_71250 | 318356..321913 | 2744516 | BREX-system-P-loop-protein-BrxC-[Mesorhizobium-sp.-113-3-3] |
| WP_092958042.1 | Type1.BREX | <-Aminotran_1_2<-?\|\|?-><-?<-tRNA\|\|Arm-DNA-bind_3+Phage_int_SAM_3+Phage_integrase-><-SF1-SF2-Helicase<-SIGMA+OB-Fold+AAA-ATPase+HKD-DNase-BrxL<-PglZ<-N6-METHYLASE<-DUF6079*<-BrxB-BrxF-iSTAND<-BrxA-DNA-binding<-wHTH+WYL+WCX<-MODE-HTH+PBPII\|\|TetR-HTH-><-RHH<-VirB10<-VirB9<-VirB8 | 1179 | Bacteria>Pseudomonadota>Alphaproteobacteria>Hyphomicrobiales>Phyllobacteriaceae>Mesorhizobium | GCF_020164785.1 | Mesorhizobium_sp._ES1-1 | LB513_RS22695 | 51971..55510 | 2876629 | BREX-system-P-loop-protein-BrxC-[Mesorhizobium-sp.-ES1-1] |
| WP_224719979.1 | Type6.BREX | <-Ntox49<-REase+HTH+35exo+Small-B-barrel+SF1-SF2-Helicase\|\|wHTH+WYL->wHTH+WYL+WCX->?->GreA_GreB+SF1-SF2-Helicase-p->BrxE->BrxA-DNA-binding->BrxB-BrxF-iSTAND->DUF6079*->N6-METHYLASE->N6-METHYLASE->PglZ->SIGMA+OB-Fold+AAA-ATPase+HKD-DNase-BrxL-> | 1208 | Bacteria>Pseudomonadota>Alphaproteobacteria>Hyphomicrobiales>Phyllobacteriaceae>Mesorhizobium | GCF_020164885.1 | Mesorhizobium_sp._CO1-1-7 | LB516_RS16265 | 26347..29973 | 2876632 | BREX-system-P-loop-protein-BrxC-[Mesorhizobium-sp.-CO1-1-7] |
| WP_256745318.1 | Type1.BREX | <-D5ATPase<-XIS-HTH\|\|MerR-HTH->?->?->HKD-DNase+HKD-Rease->?->BrxA-DNA-binding->BrxB-BrxF-iSTAND->DUF6079*->N6-METHYLASE->N6-METHYLASE->PglZ->SIGMA+OB-Fold+AAA-ATPase+HKD-DNase-BrxL-><-?<-?<-CHTH | 1211 | Bacteria>Pseudomonadota>Alphaproteobacteria>Hyphomicrobiales>Phyllobacteriaceae>Mesorhizobium | GCF_024518695.1 | Mesorhizobium_sp._LMG17149 | NP945_RS00755 | 156415..160050 | 2968497 | BREX-system-P-loop-protein-BrxC-[Mesorhizobium-sp.-LMG17149] |
| WP_009450523.1 | Type1.BREX | ACET-><-SF1-SF2-Helicase<-HKD-DNase+HKD-DNase<-wHTH<-SF1-SF2-Helicase+REase<-SF1-SF2-Helicase+REase<-SIGMA+OB-Fold+AAA-ATPase+HKD-DNase-BrxL<-PglZ<-N6-METHYLASE<-DUF6079*<-BrxB-BrxF-iSTAND<-BrxA-DNA-binding<-wHTH+WYL+WCX<-MODE-HTH+PBPII\|\|TetR-HTH-><-RHH<-VirB10<-VirB9<-VirB8 | 1179 | Bacteria>Pseudomonadota>Alphaproteobacteria>Hyphomicrobiales>Phyllobacteriaceae>Nitratireductor | GCF_000300515.1 | Nitratireductor_indicus | NA8A_RS11845 | 137447..140986 | 721133 | BREX-system-P-loop-protein-BrxC-[Nitratireductor-indicus] |
| WP_080921652.1 | Type1.BREX | HEPN->HEPN->?-><-DEDD_Tnp_IS110+Transposase_20\|\|<-?<-HEPN<-SIGMA+OB-Fold+AAA-ATPase+HKD-DNase-BrxL<-PglZ<-N6-METHYLASE<-DUF6079*<-BrxB-BrxF-iSTAND<-BrxA-DNA-binding<-wHTH+WYL+WCX\|\|Calcineurin-SbcD->SbcC+CCDC158->?->HTH_3->Peptidase+SF1-SF2-Helicase+REase-> | 1179 | Bacteria>Pseudomonadota>Alphaproteobacteria>Hyphomicrobiales>Phyllobacteriaceae>Pseudaminobacter | GCF_002075885.1 | Pseudaminobacter_manganicus | BFN67_RS22220 | 17195..20734 | 1873176 | BREX-system-P-loop-protein-BrxC-[Pseudaminobacter-manganicus] |
| WP_150288342.1 | Type1.BREX | Glyco_hydro_53+Phage-tail_3->?->?-><-URI-DNase+RAMA<-SIGMA+OB-Fold+AAA-ATPase+HKD-DNase-BrxL<-PglZ<-N6-METHYLASE<-DUF6079*<-BrxB-BrxF-iSTAND<-BrxA-DNA-binding<-wHTH+WYL+WCX<-XIS-HTH | 1173 | Bacteria>Pseudomonadota>Alphaproteobacteria>Hyphomicrobiales>Rhabdaerophilaceae>Rhabdaerophilum | GCF_008641065.1 | Rhabdaerophilum_calidifontis | FWJ24_RS13435 | 109797..113318 | 2604328 | BREX-system-P-loop-protein-BrxC-[Rhabdaerophilum-calidifontis] |
| WP_154722158.1 | Type1.BREX | <-Classical-AAA<-?<-SMODS<-SF1-SF2-Helicase+REase<-35Exo+REase<-?\|\|wHTH+WYL+WCX->BrxA-DNA-binding->BrxB-BrxF-iSTAND->DUF6079*->N6-METHYLASE->PglZ->SIGMA+OB-Fold+AAA-ATPase+HKD-DNase-BrxL-><-REase<-REase\|\|?->35exo-> | 1181 | Bacteria>Pseudomonadota>Alphaproteobacteria>Hyphomicrobiales>Rhizobiaceae>Ciceribacter | GCF_900492205.1 | Ciceribacter_naphthalenivorans | DYZ14_RS19075 | 12704..16249 | 1118451 | BREX-system-P-loop-protein-BrxC-[Ciceribacter-naphthalenivorans] |
| WP_183807573.1 | Type6.BREX | <-SIGMA+OB-Fold+AAA-ATPase+HKD-DNase-BrxL<-PglZ<-N6-METHYLASE<-N6-METHYLASE<-DUF6079*<-BrxB-BrxF-iSTAND<-BrxE<-PhdYeFM_antitox<-PIN+OST-HTH | 1206 | Bacteria>Pseudomonadota>Alphaproteobacteria>Hyphomicrobiales>Rhizobiaceae>Mycoplana | GCF_014196765.1 | Mycoplana_azooxidifex | GGQ64_RS22460 | 165035..168655 | 1636188 | BREX-system-P-loop-protein-BrxC-[Mycoplana-azooxidifex] |
| WP_225246472.1 | Type1.BREX | <-DDE_Tnp_1_4<-GDResolvase+HIN-HTH<-?<-?<-?<-?\|\|wHTH+WYL+WCX->BrxA-DNA-binding->BrxB-BrxF-iSTAND->DUF6079*->N6-METHYLASE->PglZ->SIGMA+OB-Fold+AAA-ATPase+HKD-DNase-BrxL->?-><-?<-LexA-protease\|\|?-><-?<-HTH_3 | 1175 | Bacteria>Pseudomonadota>Alphaproteobacteria>Hyphomicrobiales>Rhizobiaceae>Rhizobium/Agrobacterium_group | - | Agrobacterium_tumefaciens_complex | HW573_RS24905 | 44719..48246 | 1183400 | MULTISPECIES:-BREX-system-P-loop-protein-BrxC-[Agrobacterium-tumefaciens-complex] |
| WP_174060722.1 | Type1.BREX | DEDD_Tnp_IS110+Transposase_20-><-HTH_3\|\|HTH_3-><-DEDD_Tnp_IS110+Transposase_20\|\|Relaxase+SF1-SF2-Helicase->\|\|EVE+HNH->REase->BrxA-DNA-binding->BrxB-BrxF-iSTAND->DUF6079*->N6-METHYLASE->PglZ->SIGMA+OB-Fold+AAA-ATPase+HKD-DNase-BrxL-><-?<-?<-REase<-MORC+DUF3684+DUF3883\|\|?->\|\|HIN-HTH-> | 1175 | Bacteria>Pseudomonadota>Alphaproteobacteria>Hyphomicrobiales>Rhizobiaceae>Rhizobium/Agrobacterium_group | - | Agrobacterium_tumefaciens | G6M83_RS06120 | 12437..15964 | 358 | BREX-system-P-loop-protein-BrxC-[Agrobacterium-tumefaciens] |
| WP_156535776.1 | Type1.BREX | <-ParB<-ArdCN+MPTase<-Small-B-barrel+SF1-SF2-Helicase+RdgC+REase<-SF1-SF2-Helicase+REase<-35Exo+REase\|\|wHTH+WYL+WCX->BrxA-DNA-binding->BrxB-BrxF-iSTAND->DUF6079*->N6-METHYLASE->PglZ->SIGMA+OB-Fold+AAA-ATPase+HKD-DNase-BrxL->?->Calcineurin+Calcineurin-SbcD->SbcC+CCDC158->?->?-><-HTH+REase | 1181 | Bacteria>Pseudomonadota>Alphaproteobacteria>Hyphomicrobiales>Rhizobiaceae>Rhizobium/Agrobacterium_group | - | Agrobacterium_vitis | GOZ98_21320 | 20620..24165 | 373 | BREX-system-P-loop-protein-BrxC-[Agrobacterium-vitis] |
| PZU76796.1 | Type1.BREX | XIS-HTH->?->?->?->TraJ-RHH->TraI-VirD2-RCRE->?->BrxA-DNA-binding->BrxB-BrxF-iSTAND->DUF6079*->N6-METHYLASE->PglZ->SIGMA+OB-Fold+AAA-ATPase+HKD-DNase-BrxL-><-tRNA<-?\|\|?-><-?\|\|PAS+HISKIN-> | 1188 | Bacteria>Pseudomonadota>Alphaproteobacteria>Hyphomicrobiales>Rhizobiaceae>Rhizobium/Agrobacterium_group | GCA_003248675.1 | Rhizobium_sp. | DI546_06985 | 39524..43090 | 391 | BREX-system-P-loop-protein-BrxC-[Rhizobium-sp.] |
| TBF64186.1 | Type6.BREX | <-XIS-HTH\|\|HTH_19->?->Couple_hipA+HipA_C->CHTH-><-SIGMA+OB-Fold+AAA-ATPase+HKD-DNase-BrxL<-PglZ<-N6-METHYLASE<-N6-METHYLASE<-DUF6079*<-BrxB-BrxF-iSTAND<-BrxA-DNA-binding<-BrxE<-PhdYeFM_antitox<-VirD4-TraG+VirD4-FtsK<-TraI-VirD2-RCRE<-TraJ-RHH<-HTH_36\|\|HIN-HTH-> | 1208 | Bacteria>Pseudomonadota>Alphaproteobacteria>Hyphomicrobiales>Rhizobiaceae>Rhizobium/Agrobacterium_group | GCA_004306065.1 | Rhizobium_leguminosarum | ELG90_15310 | 3002859..3006485 | 384 | BREX-system-P-loop-protein-BrxC-[Rhizobium-leguminosarum] |
| TQN59072.1 | Type1.BREX | <-SF1-SF2-Helicase<-HKD-DNase+HKD-DNase<-wHTH<-SF1-SF2-Helicase+REase<-SIGMA+OB-Fold+AAA-ATPase+HKD-DNase-BrxL<-PglZ<-N6-METHYLASE<-DUF6079*<-BrxB-BrxF-iSTAND<-BrxA-DNA-binding<-wHTH+WYL+WCX<-MODE-HTH+PBPII\|\|TetR-HTH-><-RHH<-VirB10<-VirB9<-VirB8 | 1179 | Bacteria>Pseudomonadota>Alphaproteobacteria>Hyphomicrobiales>Rhizobiaceae>Rhizobium/Agrobacterium_group | GCA_006757705.1 | Agrobacterium_tumefaciens | FLX27_23795 | 22501..26040 | 358 | BREX-system-P-loop-protein-BrxC-[Agrobacterium-tumefaciens] |
| NEH61390.1 | Type1.BREX | <-REase<-Classical-AAA<-Classical-AAA<-REase+ZnR<-ParB\|\|?->?->BrxA-DNA-binding->BrxB-BrxF-iSTAND->DUF6079*->N6-METHYLASE->PglZ->SIGMA+OB-Fold+AAA-ATPase+HKD-DNase-BrxL-><-SbcC<-ABC-3C<-?\|\|NACHT-><-ParB+HNH<-ParB+HNH | 1174 | Bacteria>Pseudomonadota>Alphaproteobacteria>Hyphomicrobiales>Rhizobiaceae>Rhizobium/Agrobacterium_group | GCA_010668215.1 | Rhizobium_leguminosarum | GR207_02220 | 29199..32723 | 384 | BREX-system-P-loop-protein-BrxC-[Rhizobium-leguminosarum] |
| MBA8801280.1 | Type6.BREX | <-SIGMA+OB-Fold+AAA-ATPase+HKD-DNase-BrxL<-PglZ<-N6-METHYLASE<-N6-METHYLASE<-DUF6079*<-BrxB-BrxF-iSTAND<-BrxA-DNA-binding<-BrxE\|\|?-><-wHTH+WYL+WCX\|\|?->35Exo+REase->SMODS-> | 1208 | Bacteria>Pseudomonadota>Alphaproteobacteria>Hyphomicrobiales>Rhizobiaceae>Rhizobium/Agrobacterium_group | GCA_014138475.1 | Agrobacterium_sp._RC10-4-1 | FHW77_005035 | 9105..12731 | 2587039 | hypothetical-protein-[Agrobacterium-sp.-RC10-4-1] |
| MBX5048875.1 | Type6.BREX | TetR-HTH->?-><-SIGMA+OB-Fold+AAA-ATPase+HKD-DNase-BrxL<-PglZ<-ParB+HNH<-N6-METHYLASE<-N6-METHYLASE<-DUF6079*<-BrxB-BrxF-iSTAND<-BrxA-DNA-binding<-BrxE<-tRNA\|\|?->Thioredoxin-> | 1213 | Bacteria>Pseudomonadota>Alphaproteobacteria>Hyphomicrobiales>Rhizobiaceae>Rhizobium/Agrobacterium_group | GCA_019684595.1 | Rhizobium_lentis | HJB64_19910 | 192083..195724 | 1138194 | BREX-system-P-loop-protein-BrxC-[Rhizobium-lentis] |
| MBY3093307.1 | Type1.BREX | Phage_integrase-><-HTH+Phage_integrase+Phage_integrase-><-?<-Classical-AAA<-Classical-AAA<-REase+ZnR<-ParB\|\|BrxA-DNA-binding->BrxB-BrxF-iSTAND->DUF6079*->N6-METHYLASE->PglZ->SIGMA+OB-Fold+AAA-ATPase+HKD-DNase-BrxL->\|\|HIN-HTH->?->LZ_Tnp_IS66+zf-IS66+DDE_Tnp_IS66+DDE_Tnp_IS66_C->?->?-><-SbcC | 1174 | Bacteria>Pseudomonadota>Alphaproteobacteria>Hyphomicrobiales>Rhizobiaceae>Rhizobium/Agrobacterium_group | GCA_019778505.1 | Rhizobium_laguerreae | HF072_21270 | 353189..356713 | 1076926 | BREX-system-P-loop-protein-BrxC-[Rhizobium-laguerreae] |
| MBY5873817.1 | Type1.BREX | DnaB+McrB-AAA->McrC-NTD+REase+ZnR->?->OLD-ABC+TOPRIM->SF1-SF2-Helicase->ParB+HNH+Schlafen-><-SIGMA+OB-Fold+AAA-ATPase+HKD-DNase-BrxL<-PglZ<-N6-METHYLASE<-DUF6079*<-BrxB-BrxF-iSTAND<-BrxA-DNA-binding\|\|ParB->REase+ZnR->?->Classical-AAA->Classical-AAA->REase-> | 1174 | Bacteria>Pseudomonadota>Alphaproteobacteria>Hyphomicrobiales>Rhizobiaceae>Rhizobium/Agrobacterium_group | GCA_019799745.1 | Rhizobium_leguminosarum | HFN53_17500 | 401377..404901 | 384 | BREX-system-P-loop-protein-BrxC-[Rhizobium-leguminosarum] |
| WP_131614693.1 | Type1.BREX | HKD-DNase->SF1-SF2-Helicase->?->?-><-SIGMA+OB-Fold+AAA-ATPase+HKD-DNase-BrxL<-PglZ<-N6-METHYLASE<-NERD<-OLD-ABC<-DUF6079*<-BrxB-BrxF-iSTAND<-BrxA-DNA-binding<-?\|\|ParB->REase+ZnR->Classical-AAA->Classical-AAA->REase-> | 1174 | Bacteria>Pseudomonadota>Alphaproteobacteria>Hyphomicrobiales>Rhizobiaceae>Rhizobium/Agrobacterium_group | GCA_030820635.1 | Rhizobium_leguminosarum | HFN52_RS21030 | 83963..87487 | 384 | BREX-system-P-loop-protein-BrxC-[Rhizobium-leguminosarum] |
| CAD6617282.1 | Type1.BREX | <-XIS-HTH<-Arm-DNA-bind_3+Phage_integrase<-SIGMA+OB-Fold+AAA-ATPase+HKD-DNase-BrxL<-PglZ<-N6-METHYLASE<-REase<-OLD-ABC<-DUF6079*<-BrxB-BrxF-iSTAND<-BrxA-DNA-binding<-wHTH+WYL+WCX\|\|GDResolvase+HIN-HTH-> | 1172 | Bacteria>Pseudomonadota>Alphaproteobacteria>Hyphomicrobiales>Rhizobiaceae>Rhizobium/Agrobacterium_group | GCA_902502895.2 | Rhizobium_sp._Khangiran2 | RKHAN_03345 | 3387102..3390620 | 1435607 | BREX-system-P-loop-protein-BrxC-[Rhizobium-sp.-Khangiran2] |
| WP_027666511.1 | Type1.BREX | <-SbcC<-ABC-3C<-?<-SIGMA+OB-Fold+AAA-ATPase+HKD-DNase-BrxL<-PglZ<-N6-METHYLASE<-DUF6079*<-BrxB-BrxF-iSTAND<-BrxA-DNA-binding<-?<-?\|\|ParB->REase+ZnR->?->Classical-AAA->Classical-AAA-> | 1174 | Bacteria>Pseudomonadota>Alphaproteobacteria>Hyphomicrobiales>Rhizobiaceae>Rhizobium/Agrobacterium_group | GCF_000427945.1 | Rhizobium_leguminosarum | A19Q_RS0117655 | 62384..65908 | 384 | BREX-system-P-loop-protein-BrxC-[Rhizobium-leguminosarum] |
| WP_088677288.1 | Type1.BREX | <-SPOUT\|\|tRNA->tRNA->?->?->?->tRNA->?->Classical-AAA+S2P->HEPN->BrxA-DNA-binding->BrxB-BrxF-iSTAND->DUF6079*->N6-METHYLASE->PglZ->SIGMA+OB-Fold+AAA-ATPase+HKD-DNase-BrxL-><-?\|\|?->?->?->MORC+REase->RNase-> | 1194 | Bacteria>Pseudomonadota>Alphaproteobacteria>Hyphomicrobiales>Rhizobiaceae>Rhizobium/Agrobacterium_group | GCF_002211395.1 | Rhizobium_sp._R339 | ATY76_RS24585 | 353019..356603 | 1764273 | BREX-system-P-loop-protein-BrxC-[Rhizobium-sp.-R339] |
| WP_126920114.1 | Type1.BREX | <-ABC-3C<-?\|\|35exo+SF1-SF2-Helicase-><-HHH_4+HTH+wHTH+SF1-SF2-Helicase+REase\|\|MORC+SF1-SF2-Helicase-><-?<-SIGMA+OB-Fold+AAA-ATPase+HKD-DNase-BrxL<-PglZ<-N6-METHYLASE<-DUF6079*<-BrxB-BrxF-iSTAND<-BrxA-DNA-binding<-nSTAND1+HNH\|\|ParB->REase+ZnR->?->Classical-AAA->Classical-AAA->REase-> | 1175 | Bacteria>Pseudomonadota>Alphaproteobacteria>Hyphomicrobiales>Rhizobiaceae>Rhizobium/Agrobacterium_group | GCF_003985155.1 | Rhizobium_vallis | EFQ99_RS06935 | 372119..375646 | 634290 | BREX-system-P-loop-protein-BrxC-[Rhizobium-vallis] |
| WP_130750054.1 | Type6.BREX | <-XIS-HTH\|\|HTH_19->?->Couple_hipA+HipA_C->CHTH-><-SIGMA+OB-Fold+AAA-ATPase+HKD-DNase-BrxL<-PglZ<-N6-METHYLASE<-N6-METHYLASE<-DUF6079*<-BrxB-BrxF-iSTAND<-BrxA-DNA-binding<-BrxE<-PhdYeFM_antitox<-VirD4-TraG+VirD4-FtsK<-?<-?<-TraJ-RHH<-HTH_36 | 1208 | Bacteria>Pseudomonadota>Alphaproteobacteria>Hyphomicrobiales>Rhizobiaceae>Rhizobium/Agrobacterium_group | GCF_004304245.1 | Rhizobium_leguminosarum | ELI03_RS14940 | 2921080..2924706 | 384 | BREX-system-P-loop-protein-BrxC-[Rhizobium-leguminosarum] |
| WP_130755622.1 | Type1.BREX | 35exo+SF1-SF2-Helicase->MarR-HTH->Polbeta->REase->SbcC-><-Phage_int_SAM_4<-SIGMA+OB-Fold+AAA-ATPase+HKD-DNase-BrxL<-PglZ<-N6-METHYLASE<-DUF6079*<-BrxB-BrxF-iSTAND<-BrxA-DNA-binding<-?\|\|ParB->REase+ZnR->Classical-AAA->Classical-AAA->REase-> | 1174 | Bacteria>Pseudomonadota>Alphaproteobacteria>Hyphomicrobiales>Rhizobiaceae>Rhizobium/Agrobacterium_group | GCF_004304705.1 | Rhizobium_leguminosarum | ELH79_RS02355 | 502725..506249 | 384 | BREX-system-P-loop-protein-BrxC-[Rhizobium-leguminosarum] |
| WP_197729861.1 | Type6.BREX | <-XIS-HTH\|\|HTH_19->?->Couple_hipA+HipA_C->CHTH-><-SIGMA+OB-Fold+AAA-ATPase+HKD-DNase-BrxL<-PglZ<-N6-METHYLASE<-N6-METHYLASE<-DUF6079*<-BrxB-BrxF-iSTAND<-BrxA-DNA-binding-p<-BrxE<-PhdYeFM_antitox<-VirD4-TraG+VirD4-FtsK<-?<-?<-TraJ-RHH<-HTH_36 | 1210 | Bacteria>Pseudomonadota>Alphaproteobacteria>Hyphomicrobiales>Rhizobiaceae>Rhizobium/Agrobacterium_group | GCF_004306065.1 | Rhizobium_leguminosarum | ELG90_RS15310 | 3002859..3006491 | 384 | BREX-system-P-loop-protein-BrxC-[Rhizobium-leguminosarum] |
| WP_131718742.1 | Type1.BREX | URI-DNase->REase->?-><-SbcC+SbcC<-Calcineurin+Calcineurin-SbcD\|\|wHTH+WYL+WCX->BrxA-DNA-binding->BrxB-BrxF-iSTAND->DUF6079*->N6-METHYLASE->PglZ->SIGMA+OB-Fold+AAA-ATPase+HKD-DNase-BrxL-><-?<-GDResolvase<-?<-?<-tRNA\|\|?->SAM-METHYLASE-> | 1178 | Bacteria>Pseudomonadota>Alphaproteobacteria>Hyphomicrobiales>Rhizobiaceae>Rhizobium/Agrobacterium_group | GCF_004330655.1 | Rhizobium_leguminosarum | EOH57_RS05710 | 350884..354420 | 384 | BREX-system-P-loop-protein-BrxC-[Rhizobium-leguminosarum] |
| WP_132661711.1 | Type6.BREX | Glyoxalase+ZnR->HipA_C->?->HKD-DNase+HKD-Rease-><-SIGMA+OB-Fold+AAA-ATPase+HKD-DNase-BrxL<-PglZ<-N6-METHYLASE<-N6-METHYLASE<-DUF6079*<-BrxB-BrxF-iSTAND<-BrxA-DNA-binding<-BrxE<-PhdYeFM_antitox<-wHTH+WYL+WCX\|\|SMODS->?->Classical-AAA->Cap4_nuclease+SAVED-> | 1208 | Bacteria>Pseudomonadota>Alphaproteobacteria>Hyphomicrobiales>Rhizobiaceae>Rhizobium/Agrobacterium_group | GCF_004342325.1 | Rhizobium_azibense | EV130_RS24200 | 158902..162528 | 1136135 | BREX-system-P-loop-protein-BrxC-[Rhizobium-azibense] |
| WP_168337993.1 | Type1.BREX | <-SIGMA+OB-Fold+AAA-ATPase+HKD-DNase-BrxL<-PglZ<-N6-METHYLASE<-DUF6079*<-BrxB-BrxF-iSTAND<-BrxA-DNA-binding\|\|?->ParB->REase+ZnR->?->Classical-AAA->Classical-AAA->REase-> | 1174 | Bacteria>Pseudomonadota>Alphaproteobacteria>Hyphomicrobiales>Rhizobiaceae>Rhizobium/Agrobacterium_group | GCF_012275885.1 | Rhizobium_laguerreae | GFL43_RS28845 | 8439..11963 | 1076926 | BREX-system-P-loop-protein-BrxC-[Rhizobium-laguerreae] |
| WP_168349960.1 | Type1.BREX | <-SIGMA+OB-Fold+AAA-ATPase+HKD-DNase-BrxL<-PglZ<-ParB+HNH<-N6-METHYLASE<-N6-METHYLASE<-DUF6079-p*<-Classical-AAA<-REase+wHTH\|\|?->PIN-><-ParB<-ParB<-GDResolvase+PPP1-HTH\|\|CHTH-> | 190 | Bacteria>Pseudomonadota>Alphaproteobacteria>Hyphomicrobiales>Rhizobiaceae>Rhizobium/Agrobacterium_group | GCF_012276085.1 | Rhizobium_leguminosarum | GFM28_RS29310 | 8641..9213 | 384 | hypothetical-protein-[Rhizobium-leguminosarum] |
| WP_136542766.1 | Type1.BREX | ACET-><-SF1-SF2-Helicase<-HKD-DNase+HKD-DNase<-wHTH<-SF1-SF2-Helicase+REase<-SF1-SF2-Helicase+REase<-SIGMA+OB-Fold+AAA-ATPase+HKD-DNase-BrxL<-PglZ<-N6-METHYLASE<-DUF6079*<-BrxB-BrxF-iSTAND<-BrxA-DNA-binding<-wHTH+WYL+WCX<-MODE-HTH+PBPII\|\|TetR-HTH-><-RHH<-VirB10<-VirB9<-VirB8 | 1179 | Bacteria>Pseudomonadota>Alphaproteobacteria>Hyphomicrobiales>Rhizobiaceae>Rhizobium/Agrobacterium_group | GCF_014202175.1 | Rhizobium_rosettiformans | FAA86_RS19555 | 64899..68438 | 1368430 | BREX-system-P-loop-protein-BrxC-[Rhizobium-rosettiformans] |
| WP_192117159.1 | Type1.BREX | <-REase<-Classical-AAA<-Classical-AAA<-?<-REase+ZnR<-ParB\|\|nSTAND1+HNH->BrxA-DNA-binding->BrxB-BrxF-iSTAND->DUF6079*->N6-METHYLASE->PglZ->SIGMA+OB-Fold+AAA-ATPase+HKD-DNase-BrxL->?->SF1-SF2-Helicase-><-?\|\|\|HTH_Tnp_1+HTH_21+rve+rve_3->?->HKD-DNase+HKD-Rease-> | 1174 | Bacteria>Pseudomonadota>Alphaproteobacteria>Hyphomicrobiales>Rhizobiaceae>Rhizobium/Agrobacterium_group | GCF_014841565.1 | Rhizobium_sp._CFBP_8762 | IFT84_RS03115 | 659482..663006 | 2775279 | BREX-system-P-loop-protein-BrxC-[Rhizobium-sp.-CFBP-8762] |
| WP_215570403.1 | Type1.BREX | HTH_3->?-><-?\|\|LexA-protease->?-><-?<-SIGMA+OB-Fold+AAA-ATPase+HKD-DNase-BrxL<-PglZ<-N6-METHYLASE<-DUF6079*<-BrxB-BrxF-iSTAND<-BrxA-DNA-binding<-wHTH+WYL+WCX\|\|?->?->?->?->GDResolvase+HIN-HTH-><-DUF772+DDE_Tnp_1+DDE_Tnp_1_6 | 1175 | Bacteria>Pseudomonadota>Alphaproteobacteria>Hyphomicrobiales>Rhizobiaceae>Rhizobium/Agrobacterium_group | GCF_018760785.1 | Rhizobium_sp._CSW-27 | KLP42_RS24160 | 30886..34413 | 2839985 | BREX-system-P-loop-protein-BrxC-[Rhizobium-sp.-CSW-27] |
| WP_246638929.1 | Type1.BREX | <-REase<-Classical-AAA<-Classical-AAA<-REase+ZnR<-ParB\|\|?->BrxA-DNA-binding->BrxB-BrxF-iSTAND->DUF6079*->N6-METHYLASE->PglZ->SIGMA+OB-Fold+AAA-ATPase+HKD-DNase-BrxL->\|\|HIN-HTH->?->LZ_Tnp_IS66+zf-IS66+DDE_Tnp_IS66+DDE_Tnp_IS66_C->?->?-><-SbcC | 1170 | Bacteria>Pseudomonadota>Alphaproteobacteria>Hyphomicrobiales>Rhizobiaceae>Rhizobium/Agrobacterium_group | GCF_019778505.1 | Rhizobium_laguerreae | HF072_RS21050 | 353201..356713 | 1076926 | BREX-system-P-loop-protein-BrxC-[Rhizobium-laguerreae] |
| WP_222362949.1 | Type1.BREX | <-SF1-SF2-Helicase<-OLD-ABC+TOPRIM<-SIGMA+OB-Fold+AAA-ATPase+HKD-DNase-BrxL<-PglZ<-N6-METHYLASE<-DUF6079*<-BrxB-BrxF-iSTAND<-BrxA-DNA-binding<-?<-?\|\|ParB->REase+ZnR->?->Classical-AAA->Classical-AAA-> | 1174 | Bacteria>Pseudomonadota>Alphaproteobacteria>Hyphomicrobiales>Rhizobiaceae>Rhizobium/Agrobacterium_group | GCF_019798305.1 | Rhizobium_leguminosarum | HF099_RS19360 | 191490..195014 | 384 | BREX-system-P-loop-protein-BrxC-[Rhizobium-leguminosarum] |
| WP_222342569.1 | Type1.BREX | <-REase<-Classical-AAA<-Classical-AAA\|\|?-><-REase+ZnR<-ParB\|\|BrxA-DNA-binding->BrxB-BrxF-iSTAND->DUF6079*->N6-METHYLASE->PglZ->SIGMA+OB-Fold+AAA-ATPase+HKD-DNase-BrxL-><-SbcC<-ABC-3C | 1174 | Bacteria>Pseudomonadota>Alphaproteobacteria>Hyphomicrobiales>Rhizobiaceae>Rhizobium/Agrobacterium_group | GCF_019799465.1 | Rhizobium_leguminosarum | HF012_RS18165 | 9414..12938 | 384 | BREX-system-P-loop-protein-BrxC-[Rhizobium-leguminosarum] |
| WP_221976415.1 | Type1.BREX | ParB+HNH->?-><-NACHT\|\|?->ABC-3C->SbcC-><-SIGMA+OB-Fold+AAA-ATPase+HKD-DNase-BrxL<-PglZ<-N6-METHYLASE<-DUF6079*<-BrxB-BrxF-iSTAND<-BrxA-DNA-binding<-?\|\|ParB->REase+ZnR-><-?\|\|Classical-AAA->Classical-AAA->REase-> | 1174 | Bacteria>Pseudomonadota>Alphaproteobacteria>Hyphomicrobiales>Rhizobiaceae>Rhizobium/Agrobacterium_group | GCF_021391715.1 | Rhizobium | LZK79_RS04540 | 933670..937194 | 379 | MULTISPECIES:-BREX-system-P-loop-protein-BrxC-[Rhizobium] |
| WP_239619846.1 | Type1.BREX | <-Classical-AAA\|\|?->?-><-REase+ZnR<-REase+ZnR<-ParB<-?\|\|BrxA-DNA-binding->BrxB-BrxF-iSTAND->DUF6079*->N6-METHYLASE->PglZ-p->PglZ->SIGMA+OB-Fold+AAA-ATPase+HKD-DNase-BrxL->?->ABC-3C->SbcC->\|\|HIN-HTH->AAA-ATPase-> | 1174 | Bacteria>Pseudomonadota>Alphaproteobacteria>Hyphomicrobiales>Rhizobiaceae>Rhizobium/Agrobacterium_group | GCF_022354505.1 | Rhizobium_sp._C104 | MF410_RS14985 | 2774142..2777666 | 2917727 | BREX-system-P-loop-protein-BrxC-[Rhizobium-sp.-C104] |
| WP_250810295.1 | Type6.BREX | <-GreA_GreB+SF1-SF2-Helicase<-PIWI<-REase+TIR<-?\|\|wHTH+WYL+WCX->BrxE->BrxA-DNA-binding->BrxB-BrxF-iSTAND->DUF6079*->N6-METHYLASE->PglZ->SIGMA+OB-Fold+AAA-ATPase+HKD-DNase-BrxL-><-?<-ParB+HNH\|\|GDResolvase+HIN-HTH-><-HTH_3 | 1208 | Bacteria>Pseudomonadota>Alphaproteobacteria>Hyphomicrobiales>Rhizobiaceae>Rhizobium/Agrobacterium_group | GCF_023667605.1 | Neorhizobium_tomejilense | L1082_RS18485 | 51181..54807 | 2093828 | BREX-system-P-loop-protein-BrxC-[Neorhizobium-tomejilense] |
| WP_245134299.1 | Type1.BREX | <-Peptidase_M29<-?\|\|?-><-SIGMA+OB-Fold+AAA-ATPase+HKD-DNase-BrxL<-PglZ<-N6-METHYLASE<-DUF6079*<-BrxB-BrxF-iSTAND<-BrxA-DNA-binding<-wHTH+WYL+WCX\|\|35Exo+REase->SF1-SF2-Helicase+REase->Small-B-barrel+SF1-SF2-Helicase+RdgC+REase->?->HKD-DNase->SF1-SF2-Helicase-> | 1177 | Bacteria>Pseudomonadota>Alphaproteobacteria>Hyphomicrobiales>Rhizobiaceae>Rhizobium/Agrobacterium_group | GCF_030369675.1 | Rhizobium_sp._SSM4.3 | MKJ03_RS01550 | 316527..320060 | 2919917 | BREX-system-P-loop-protein-BrxC-[Rhizobium-sp.-SSM4.3] |
| WP_077987199.1 | Type1.BREX | HTH_3->Couple_hipA+HipA_C-><-?\|\|OLD-ABC+TOPRIM->SF1-SF2-Helicase-><-SIGMA+OB-Fold+AAA-ATPase+HKD-DNase-BrxL<-PglZ<-N6-METHYLASE<-DUF6079*<-BrxB-BrxF-iSTAND<-BrxA-DNA-binding<-REase\|\|ParB->REase+ZnR->REase+ZnR->?->Classical-AAA->Classical-AAA-> | 1175 | Bacteria>Pseudomonadota>Alphaproteobacteria>Hyphomicrobiales>Rhizobiaceae>Rhizobium/Agrobacterium_group | GCF_030440405.1 | Rhizobium_pusense | BTE56_RS17195 | 18928..22455 | 648995 | BREX-system-P-loop-protein-BrxC-[Rhizobium-pusense] |
| WP_149748055.1 | Type1.BREX | nSTAND1+HNH->REase->BrxA-DNA-binding->BrxB-BrxF-iSTAND->DUF6079*->N6-METHYLASE->PglZ->SIGMA+OB-Fold+AAA-ATPase+HKD-DNase-BrxL->?->SF1-SF2-Helicase+DUF1998-> | 1175 | Bacteria>Pseudomonadota>Alphaproteobacteria>Hyphomicrobiales>Rhizobiaceae>Rhizobium/Agrobacterium_group | GCF_900156005.1 | Rhizobium_sp._RU35A | F3H95_RS25745 | 3768..7295 | 1907414 | BREX-system-P-loop-protein-BrxC-[Rhizobium-sp.-RU35A] |
| WP_245278427.1 | Type1.BREX | <-REase<-Classical-AAA<-Classical-AAA<-REase+ZnR<-ParB\|\|?->BrxA-DNA-binding->BrxB-BrxF-iSTAND->DUF6079*->N6-METHYLASE->PglZ->SIGMA+OB-Fold+AAA-ATPase+HKD-DNase-BrxL-><-?<-?<-?\|\|?-><-McrC-NTD+REase+ZnR<-DnaB+McrB-AAA | 1170 | Bacteria>Pseudomonadota>Alphaproteobacteria>Hyphomicrobiales>Rhizobiaceae>Rhizobium/Agrobacterium_group | - | Rhizobium_leguminosarum | A3A3_RS0116705 | 192694..196206 | 384 | BREX-system-P-loop-protein-BrxC-[Rhizobium-leguminosarum] |
| WP_131707092.1 | Type1.BREX | <-REase<-Classical-AAA<-Classical-AAA<-REase+ZnR<-ParB\|\|?->BrxA-DNA-binding->BrxB-BrxF-iSTAND->DUF6079*->N6-METHYLASE->PglZ->SIGMA+OB-Fold+AAA-ATPase+HKD-DNase-BrxL-><-?<-?<-SF1-SF2-Helicase<-HKD-DNase<-?<-Peptidase_S8 | 1174 | Bacteria>Pseudomonadota>Alphaproteobacteria>Hyphomicrobiales>Rhizobiaceae>Rhizobium/Agrobacterium_group | - | Rhizobium_leguminosarum | EOH59_RS18255 | 300646..304170 | 384 | BREX-system-P-loop-protein-BrxC-[Rhizobium-leguminosarum] |
| WP_024321352.1 | Type1.BREX | DnaB+McrB-AAA->McrC-NTD+REase+ZnR-><-?\|\|?->?->?-><-SIGMA+OB-Fold+AAA-ATPase+HKD-DNase-BrxL<-PglZ<-N6-METHYLASE<-DUF6079*<-BrxB-BrxF-iSTAND<-BrxA-DNA-binding<-?\|\|ParB->REase+ZnR->Classical-AAA->Classical-AAA->REase-> | 1174 | Bacteria>Pseudomonadota>Alphaproteobacteria>Hyphomicrobiales>Rhizobiaceae>Rhizobium/Agrobacterium_group | - | Rhizobium_leguminosarum | EOJ20_RS04835 | 93871..97395 | 384 | BREX-system-P-loop-protein-BrxC-[Rhizobium-leguminosarum] |
| WP_130784075.1 | Type1.BREX | <-LZ_Tnp_IS66+zf-IS66+DDE_Tnp_IS66+DDE_Tnp_IS66_C<-?<-HIN-HTH<-SIGMA+OB-Fold+AAA-ATPase+HKD-DNase-BrxL<-PglZ<-N6-METHYLASE<-DUF6079*<-BrxB-BrxF-iSTAND<-BrxA-DNA-binding<-wHTH+WYL+WCX\|\|Calcineurin+Calcineurin-SbcD->SbcC+SbcC-><-?<-REase<-URI-DNase | 1178 | Bacteria>Pseudomonadota>Alphaproteobacteria>Hyphomicrobiales>Rhizobiaceae>Rhizobium/Agrobacterium_group | - | Rhizobium_leguminosarum | ELG96_RS28915 | 11184..14720 | 384 | BREX-system-P-loop-protein-BrxC-[Rhizobium-leguminosarum] |
| WP_130764093.1 | Type1.BREX | <-REase<-Classical-AAA<-Classical-AAA\|\|?-><-REase+ZnR<-ParB\|\|<-?\|\|BrxA-DNA-binding->BrxB-BrxF-iSTAND->DUF6079*->N6-METHYLASE->PglZ->SIGMA+OB-Fold+AAA-ATPase+HKD-DNase-BrxL->?->ABC-3C->SbcC->SbcC-><-?<-McrC-NTD+REase+ZnR | 1174 | Bacteria>Pseudomonadota>Alphaproteobacteria>Hyphomicrobiales>Rhizobiaceae>Rhizobium/Agrobacterium_group | - | Rhizobium_leguminosarum | ELI00_RS29085 | 22850..26374 | 384 | BREX-system-P-loop-protein-BrxC-[Rhizobium-leguminosarum] |
| WP_245488667.1 | Type1.BREX | DADA-Ligase->\|\|?->?->SbcC-><-SIGMA+OB-Fold+AAA-ATPase+HKD-DNase-BrxL<-PglZ<-N6-METHYLASE<-DUF6079*<-BrxB-BrxF-iSTAND<-BrxA-DNA-binding<-?\|\|ParB->REase+ZnR->?->Classical-AAA->Classical-AAA->REase-> | 1170 | Bacteria>Pseudomonadota>Alphaproteobacteria>Hyphomicrobiales>Rhizobiaceae>Rhizobium/Agrobacterium_group | - | Rhizobium_leguminosarum | ELI09_RS02630 | 538702..542214 | 384 | BREX-system-P-loop-protein-BrxC-[Rhizobium-leguminosarum] |
| WP_130672003.1 | Type1.BREX | DADA-Ligase->\|\|?->?->SbcC-><-SIGMA+OB-Fold+AAA-ATPase+HKD-DNase-BrxL<-PglZ<-N6-METHYLASE<-DUF6079*<-BrxB-BrxF-iSTAND<-BrxA-DNA-binding<-?\|\|ParB->REase+ZnR->?->Classical-AAA->Classical-AAA->REase-> | 1174 | Bacteria>Pseudomonadota>Alphaproteobacteria>Hyphomicrobiales>Rhizobiaceae>Rhizobium/Agrobacterium_group | - | Rhizobium_leguminosarum | ELI10_RS02625 | 537164..540688 | 384 | BREX-system-P-loop-protein-BrxC-[Rhizobium-leguminosarum] |
| WP_168316672.1 | Type1.BREX | <-Small-B-barrel+SF1-SF2-Helicase+RdgC+REase<-SF1-SF2-Helicase+REase<-35Exo+REase<-?\|\|wHTH+WYL+WCX->BrxA-DNA-binding->BrxB-BrxF-iSTAND->DUF6079*->N6-METHYLASE->PglZ->SIGMA+OB-Fold+AAA-ATPase+HKD-DNase-BrxL-> | 1181 | Bacteria>Pseudomonadota>Alphaproteobacteria>Hyphomicrobiales>Rhizobiaceae>Rhizobium/Agrobacterium_group | - | Rhizobium_leguminosarum | GFL49_RS25835 | 227535..231080 | 384 | BREX-system-P-loop-protein-BrxC-[Rhizobium-leguminosarum] |
| WP_168308740.1 | Type1.BREX | <-ParB<-Small-B-barrel+SF1-SF2-Helicase+RdgC+REase<-SF1-SF2-Helicase+REase<-35Exo+REase<-?\|\|wHTH+WYL+WCX->BrxA-DNA-binding->BrxB-BrxF-iSTAND->DUF6079*->N6-METHYLASE->PglZ->SIGMA+OB-Fold+AAA-ATPase+HKD-DNase-BrxL-><-SIR2<-HTH+REase<-Arm-DNA-bind_3+Phage_integrase\|\|?->Peptidase-> | 1181 | Bacteria>Pseudomonadota>Alphaproteobacteria>Hyphomicrobiales>Rhizobiaceae>Rhizobium/Agrobacterium_group | - | Rhizobium_leguminosarum | GFM13_RS06520 | 103411..106956 | 384 | BREX-system-P-loop-protein-BrxC-[Rhizobium-leguminosarum] |
| WP_246708550.1 | Type1.BREX | <-Classical-AAA<-Classical-AAA\|\|?-><-REase+ZnR<-ParB\|\|?->?->BrxA-DNA-binding->BrxB-BrxF-iSTAND->DUF6079*->N6-METHYLASE->PglZ->SIGMA+OB-Fold+AAA-ATPase+HKD-DNase-BrxL-><-SbcC<-ABC-3C<-?\|\|NACHT-><-ParB+HNH<-ParB+HNH | 1170 | Bacteria>Pseudomonadota>Alphaproteobacteria>Hyphomicrobiales>Rhizobiaceae>Rhizobium/Agrobacterium_group | - | Rhizobium_leguminosarum | GR207_RS02230 | 29211..32723 | 384 | BREX-system-P-loop-protein-BrxC-[Rhizobium-leguminosarum] |
| WP_221145756.1 | Type6.BREX | TetR-HTH->?-><-SIGMA+OB-Fold+AAA-ATPase+HKD-DNase-BrxL<-PglZ<-ParB+HNH<-N6-METHYLASE<-N6-METHYLASE<-DUF6079*<-BrxB-BrxF-iSTAND<-BrxA-DNA-binding<-BrxE<-tRNA\|\|?->Thioredoxin-> | 1213 | Bacteria>Pseudomonadota>Alphaproteobacteria>Hyphomicrobiales>Rhizobiaceae>Rhizobium/Agrobacterium_group | - | Rhizobium_lentis | HJB64_19910 | 192083..195724 | 1138194 | BREX-system-P-loop-protein-BrxC-[Rhizobium-lentis] |
| WP_192067374.1 | Type6.BREX | <-SIGMA+OB-Fold+AAA-ATPase+HKD-DNase-BrxL<-PglZ<-N6-METHYLASE<-N6-METHYLASE<-DUF6079*<-BrxB-BrxF-iSTAND<-BrxA-DNA-binding<-BrxE<-Phage_integrase<-XIS-HTH<-Phage_integrase\|\|REase->wHTH+REase->MPTase-> | 1208 | Bacteria>Pseudomonadota>Alphaproteobacteria>Hyphomicrobiales>Rhizobiaceae>Rhizobium/Agrobacterium_group | - | unclassified_Rhizobium | IFT53_20370 | 8816..12442 | 2613769 | MULTISPECIES:-BREX-system-P-loop-protein-BrxC-[unclassified-Rhizobium] |
| PZU19146.1 | Type1.BREX | HTH_3->?-><-?\|\|LexA-protease->?-><-SIGMA+OB-Fold+AAA-ATPase+HKD-DNase-BrxL<-PglZ<-N6-METHYLASE<-DUF6079*<-BrxB-BrxF-iSTAND<-BrxA-DNA-binding<-wHTH+WYL+WCX\|\|?->?->?->?->GDResolvase+HIN-HTH-> | 1175 | Bacteria>Pseudomonadota>Alphaproteobacteria>Hyphomicrobiales>Rhizobiaceae>Shinella | GCA_003248545.1 | Shinella_sp. | DI589_22170 | 12827..16354 | 1870904 | MAG:-BREX-system-P-loop-protein-BrxC-[Shinella-sp.] |
| KSV61619.1 | Type6.BREX | <-SIGMA+OB-Fold+AAA-ATPase+HKD-DNase-BrxL<-PglZ<-N6-METHYLASE<-DUF6079*<-BrxB-BrxF-iSTAND<-BrxA-DNA-binding<-BrxE<-tRNA\|\|?->?->ACET->ACET-><-?\|\|sGTP-> | 1203 | Bacteria>Pseudomonadota>Alphaproteobacteria>Hyphomicrobiales>Rhizobiaceae>Sinorhizobium/Ensifer_group | GCA_001461715.1 | Sinorhizobium_sp._GL2 | N182_13660 | 115788..119399 | 1358416 | hypothetical-protein-N182_13660-[Sinorhizobium-sp.-GL2] |

| Protein-ID | System | Gene-Neighbourhoods | Length | Taxonomy | Assembly-ID | Strain | Locus-Tag | Genomic-coordinates | TaxID | Fasta-defline |
|---|---|---|---|---|---|---|---|---|---|---|
| WP_196847550.1 | Type1.BREX | [illegible] | 1172 | Bacteria>Pseudomonadota>Alphaproteobacteria>Sphingomonadales>Erythrobacteraceae>Erythrobacter/Porphyrobacter_group | GCF_015767595.1 | Erythrobacter_sp._A30-3 | IT881_RS05450 | 1101811..1105329 | 2785907 | BREX-system-P-loop-protein-BrxC-[Erythrobacter-sp.-A30-3] |
| WP_226699433.1 | Type1.BREX | [illegible] | 1170 | Bacteria>Pseudomonadota>Alphaproteobacteria>Sphingomonadales>Erythrobacteraceae>Qipengyuania | GCF_020524765.1 | Qipengyuania_gaetbuli | LC083_RS04940 | 959412..962924 | 266952 | BREX-system-P-loop-protein-BrxC-[Qipengyuania-gaetbuli] |
| MBB3862674.1 | Type6.BREX | [illegible] | 1208 | Bacteria>Pseudomonadota>Alphaproteobacteria>Sphingomonadales>Sphingomonadaceae>Novosphingobium | GCA_014196055.1 | Novosphingobium_hassiacum | GGQ88_003976 | 14858..18484 | 173676 | hypothetical-protein-[Novosphingobium-hassiacum] |
| WP_246386386.1 | Type6.BREX | [illegible] | 1213 | Bacteria>Pseudomonadota>Alphaproteobacteria>Sphingomonadales>Sphingomonadaceae>Novosphingobium | GCF_014196055.1 | Novosphingobium_hassiacum | GGQ88_RS19485 | 14843..18484 | 173676 | BREX-system-P-loop-protein-BrxC-[Novosphingobium-hassiacum] |
| WP_243992543.1 | Type1.BREX | [illegible] | 1189 | Bacteria>Pseudomonadota>Alphaproteobacteria>Sphingomonadales>Sphingomonadaceae>Novosphingobium | GCF_022832385.1 | Novosphingobium_sp._B2580 | MTR64_RS07820 | 57412..60981 | 2930093 | BREX-system-P-loop-protein-BrxC-[Novosphingobium-sp.-B2580] |
| WP_243995499.1 | Type1.BREX | [illegible] | 1179 | Bacteria>Pseudomonadota>Alphaproteobacteria>Sphingomonadales>Sphingomonadaceae>Novosphingobium | GCF_022832385.1 | Novosphingobium_sp._B2580 | MTR64_RS16290 | 56132..59671 | 2930093 | BREX-system-P-loop-protein-BrxC-[Novosphingobium-sp.-B2580] |
| WP_022673601.1 | Type1.BREX | [illegible] | 1189 | Bacteria>Pseudomonadota>Alphaproteobacteria>Sphingomonadales>Sphingomonadaceae>Novosphingopyxis | GCF_000420305.1 | Novosphingopyxis_baekryungensis | G410_RS0110420 | 254465..258034 | 279369 | BREX-system-P-loop-protein-BrxC-[Novosphingopyxis-baekryungensis] |
| ARR56525.1 | Type1.BREX | [illegible] | 1189 | Bacteria>Pseudomonadota>Alphaproteobacteria>Sphingomonadales>Sphingomonadaceae>Rhizorhabdus | GCA_000715175.2 | Rhizorhabdus_wittichii_DC-6 | HY78_25365 | 5371615..5375184 | 1283312 | hypothetical-protein-HY78_25365-[Rhizorhabdus-wittichii-DC-6] |
| WP_209047668.1 | Type1.BREX | [illegible] | 1179 | Bacteria>Pseudomonadota>Alphaproteobacteria>Sphingomonadales>Sphingomonadaceae>Rhizorhabdus | GCF_029543145.1 | Rhizorhabdus_histidinilytica | P7B04_RS17610 | 19171..22710 | 439228 | MULTISPECIES:-BREX-system-P-loop-protein-BrxC-[Sphingomonadaceae] |
| WP_010336176.1 | Type1.BREX | [illegible] | 1179 | Bacteria>Pseudomonadota>Alphaproteobacteria>Sphingomonadales>Sphingomonadaceae>Sphingobium | GCF_000224695.1 | Sphingobium_yanoikuyae | SYA_RS0104815 | 38393..41932 | 13690 | BREX-system-P-loop-protein-BrxC-[Sphingobium-yanoikuyae] |
| WP_007688323.1 | Type1.BREX | [illegible] | 1189 | Bacteria>Pseudomonadota>Alphaproteobacteria>Sphingomonadales>Sphingomonadaceae>Sphingobium | GCF_000264945.2 | Sphingobium_indicum | SIDU_RS03565 | 700467..704036 | 332055 | BREX-system-P-loop-protein-BrxC-[Sphingobium-indicum] |
| WP_021240096.1 | Type1.BREX | [illegible] | 1177 | Bacteria>Pseudomonadota>Alphaproteobacteria>Sphingomonadales>Sphingomonadaceae>Sphingobium | GCF_000445065.1 | Sphingobium_quisquiliarum | L288_RS41575 | 32905..36438 | 538379 | BREX-system-P-loop-protein-BrxC-[Sphingobium-quisquiliarum] |
| WP_264583946.1 | Type1.BREX | [illegible] | 1179 | Bacteria>Pseudomonadota>Alphaproteobacteria>Sphingomonadales>Sphingomonadaceae>Sphingobium | GCF_025961895.1 | Sphingobium_sp._B7D2B | M2341_RS01365 | 293056..296595 | 2940583 | BREX-system-P-loop-protein-BrxC-[Sphingobium-sp.-B7D2B] |
| NJC05753.1 | Type6.BREX | [illegible] | 1209 | Bacteria>Pseudomonadota>Alphaproteobacteria>Sphingomonadales>Sphingomonadaceae>Sphingomonas | GCA_011927725.1 | Sphingomonas_kaistensis | GGQ97_001546 | 1560422..1564051 | 298708 | hypothetical-protein-[Sphingomonas-kaistensis] |
| WP_066478880.1 | Type1.BREX | [illegible] | 1179 | Bacteria>Pseudomonadota>Alphaproteobacteria>Sphingomonadales>Sphingomonadaceae>Sphingomonas | GCF_001555785.1 | unclassified_Sphingomonas | AX667_RS15610 | 42451..45990 | 196159 | MULTISPECIES:-BREX-system-P-loop-protein-BrxC-[unclassified-Sphingomonas] |
| WP_107968397.1 | Type1.BREX | [illegible] | 1186 | Bacteria>Pseudomonadota>Alphaproteobacteria>Sphingomonadales>Sphingomonadaceae>Sphingomonas | GCF_003050615.1 | Sphingomonas_oleivorans | CLG96_RS13035 | 363272..366832 | 1735121 | BREX-system-P-loop-protein-BrxC-[Sphingomonas-oleivorans] |
| WP_192976950.1 | Type1.BREX | [illegible] | 1177 | Bacteria>Pseudomonadota>Alphaproteobacteria>Sphingomonadales>Sphingomonadaceae>Sphingomonas | GCF_014892665.1 | Sphingomonas_sp._CFBP_13603 | IFR23_RS17335 | 87025..90558 | 2774040 | BREX-system-P-loop-protein-BrxC-[Sphingomonas-sp.-CFBP-13603] |
| WP_089221081.1 | Type1.BREX | [illegible] | 1179 | Bacteria>Pseudomonadota>Alphaproteobacteria>Sphingomonadales>Sphingomonadaceae>Sphingomonas | GCF_900188165.1 | Sphingomonas_laterariae | CHB74_RS18985 | 16083..19622 | 861865 | BREX-system-P-loop-protein-BrxC-[Sphingomonas-laterariae] |
| WP_133494816.1 | Type1.BREX | [illegible] | 1189 | Bacteria>Pseudomonadota>Alphaproteobacteria>Sphingomonadales>Sphingomonadaceae>Stakelama | GCF_014645395.1 | Stakelama_pacifica | IEX20_RS07270 | 305060..308629 | 517720 | BREX-system-P-loop-protein-BrxC-[Stakelama-pacifica] |
| WP_121052748.1 | Type1.BREX | [illegible] | 1179 | Bacteria>Pseudomonadota>Alphaproteobacteria>Sphingomonadales>Sphingosinicellaceae>Sphingosinicella | GCF_003967095.1 | Sphingosinicella_microcystinivorans | SmB9_RS16135 | 3426933..3430472 | 335406 | BREX-system-P-loop-protein-BrxC-[Sphingosinicella-microcystinivorans] |
| OYY96513.1 | Type1.BREX | [illegible] | 1189 | Bacteria>Pseudomonadota>Alphaproteobacteria>Sphingomonadales>unclassified_Sphingomonadales | GCA_002281585.1 | Sphingomonadales_bacterium_28-56-43 | B7Y38_10875 | 3957..7526 | 1970590 | MAG:-hypothetical-protein-B7Y38_10875-[Sphingomonadales-bacterium-28-56-43] |
| OYZ13690.1 | Type5.BREX | [illegible] | 1197 | Bacteria>Pseudomonadota>Alphaproteobacteria>Sphingomonadales>unclassified_Sphingomonadales | GCA_002281615.1 | Sphingomonadales_bacterium_28-64-96 | B7Y35_07475 | 14573..18166 | 1970591 | MAG:-hypothetical-protein-B7Y35_07475-[Sphingomonadales-bacterium-28-64-96] |
| TNE62528.1 | Type1.BREX | [illegible] | 1175 | Bacteria>Pseudomonadota>Alphaproteobacteria>Sphingomonadales>unclassified_Sphingomonadales | GCA_006226385.1 | Sphingomonadales_bacterium | EP341_00225 | 33441..36968 | 1978525 | MAG:-BREX-system-P-loop-protein-BrxC-[Sphingomonadales-bacterium] |
| WP_015740180.1 | Type1.BREX | [illegible] | 1179 | Bacteria>Pseudomonadota>Alphaproteobacteria>Sphingomonadales>Zymomonadaceae>Zymomonas | GCF_000024245.1 | Zymomonas_mobilis | Za10_1374 | 1591366..1594905 | 542 | BREX-system-P-loop-protein-BrxC-[Zymomonas-mobilis] |
| RYG90169.1 | Type6.BREX | [illegible] | 1208 | Bacteria>Pseudomonadota>Alphaproteobacteria>unclassified_Alphaproteobacteria | GCA_004145145.1 | Alphaproteobacteria_bacterium | EON59_00500 | 24262..27888 | 1913988 | BREX-system-P-loop-protein-BrxC-[Alphaproteobacteria-bacterium] |
| NCC22319.1 | Type6.BREX | [illegible] | 1212 | Bacteria>Pseudomonadota>Alphaproteobacteria>unclassified_Alphaproteobacteria | GCA_009929775.1 | Alphaproteobacteria_bacterium | EOM26_07635 | 14398..18036 | 1913988 | BREX-system-P-loop-protein-BrxC-[Alphaproteobacteria-bacterium] |
| MBN9530327.1 | Type1.BREX | [illegible] | 1180 | Bacteria>Pseudomonadota>Alphaproteobacteria>unclassified_Alphaproteobacteria | GCA_017307375.1 | Alphaproteobacteria_bacterium | JOH82_29215 | 188733..192275 | 1913988 | BREX-system-P-loop-protein-BrxC-[Alphaproteobacteria-bacterium] |
| MBQ8347028.1 | Type1.BREX | [illegible] | 1150 | Bacteria>Pseudomonadota>Alphaproteobacteria>unclassified_Alphaproteobacteria | GCA_017462585.1 | Alphaproteobacteria_bacterium | IJY17_05445 | 46698..50150 | 1913988 | BREX-system-P-loop-protein-BrxC-[Alphaproteobacteria-bacterium] |
| MBN8451986.1 | Type6.BREX | [illegible] | 1213 | Bacteria>Pseudomonadota>Betaproteobacteria>Betaproteobacteria_incertae_sedis>Candidatus_Accumulibacter>unclassified_Candidatus_Accumulibacter | GCA_017302415.1 | Candidatus_Accumulibacter_sp. | JOM31_02680 | 99081..102722 | 2053492 | BREX-system-P-loop-protein-BrxC-[Candidatus-Accumulibacter-sp.] |
| WP_042797405.1 | Type6.BREX | [illegible] | 1207 | Bacteria>Pseudomonadota>Betaproteobacteria>Burkholderiales>Alcaligenaceae>Achromobacter | GCF_000783435.2 | Achromobacter_insolitus | MC81_20460 | 4405602..4409225 | 217204 | BREX-system-P-loop-protein-BrxC-[Achromobacter-insolitus] |
| WP_076471941.1 | Type6.BREX | [illegible] | 1207 | Bacteria>Pseudomonadota>Betaproteobacteria>Burkholderiales>Alcaligenaceae>Achromobacter | GCF_001957055.1 | Achromobacter_xylosoxidans | BIZ53_RS00410 | 81330..84953 | 85698 | BREX-system-P-loop-protein-BrxC-[Achromobacter-xylosoxidans] |
| WP_212312045.1 | Type6.BREX | [illegible] | 1217 | Bacteria>Pseudomonadota>Betaproteobacteria>Burkholderiales>Alcaligenaceae>Achromobacter | GCF_018195695.1 | Achromobacter_sp._Marseille-Q0513 | KDH83_RS00260 | 59261..62914 | 2829161 | BREX-system-P-loop-protein-BrxC-[Achromobacter-sp.-Marseille-Q0513] |
| WP_233436978.1 | Type6.BREX | [illegible] | 1234 | Bacteria>Pseudomonadota>Betaproteobacteria>Burkholderiales>Alcaligenaceae>Achromobacter | GCF_022976495.1 | Achromobacter_xylosoxidans | DBA06_RS11680 | 120350..124054 | 85698 | BREX-system-P-loop-protein-BrxC-[Achromobacter-xylosoxidans] |
| WP_259250627.1 | Type6.BREX | [illegible] | 1213 | Bacteria>Pseudomonadota>Betaproteobacteria>Burkholderiales>Alcaligenaceae>Achromobacter | GCF_024806975.1 | Achromobacter_sp._JUb104 | M2361_RS16380 | 989592..993233 | 2940590 | BREX-system-P-loop-protein-BrxC-[Achromobacter-sp.-JUb104] |
| WP_076518288.1 | Type1.BREX | [illegible] | 1181 | Bacteria>Pseudomonadota>Betaproteobacteria>Burkholderiales>Alcaligenaceae>Achromobacter | GCF_900156745.1 | Achromobacter_sp._MFA1_R4 | BXA00_RS09440 | 2091384..2094929 | 1881016 | BREX-system-P-loop-protein-BrxC-[Achromobacter-sp.-MFA1-R4] |
| WP_175212942.1 | Type1.BREX | [illegible] | 1181 | Bacteria>Pseudomonadota>Betaproteobacteria>Burkholderiales>Alcaligenaceae>Achromobacter | GCF_902859965.1 | Achromobacter_aegrifaciens | HV793_RS07505 | 1613364..1616909 | 1287736 | BREX-system-P-loop-protein-BrxC-[Achromobacter-aegrifaciens] |
| WP_180097564.1 | Type6.BREX | [illegible] | 1213 | Bacteria>Pseudomonadota>Betaproteobacteria>Burkholderiales>Alcaligenaceae>Achromobacter | GCF_902859995.1 | Achromobacter_mucicolens | H0520_RS02915 | 616832..620473 | 1389922 | BREX-system-P-loop-protein-BrxC-[Achromobacter-mucicolens] |
| WP_200235209.1 | Type1.BREX | [illegible] | 1190 | Bacteria>Pseudomonadota>Betaproteobacteria>Burkholderiales>Alcaligenaceae>Advenella | GCF_016595155.1 | Advenella_mandrilli | JHL22_RS06635 | 201531..205103 | 2800330 | BREX-system-P-loop-protein-BrxC-[Advenella-mandrilli] |
| AYN22647.1 | Type6.BREX | [illegible] | 1213 | Bacteria>Pseudomonadota>Betaproteobacteria>Burkholderiales>Alcaligenaceae>Alcaligenes | GCA_003671915.1 | Alcaligenes_aquatilis | D3M96_03425 | 754727..758368 | 323284 | BREX-system-P-loop-protein-BrxC-[Alcaligenes-aquatilis] |
| WP_121740016.1 | Type1.BREX | [illegible] | 1190 | Bacteria>Pseudomonadota>Betaproteobacteria>Burkholderiales>Alcaligenaceae>Alcaligenes | GCF_003671915.1 | Alcaligenes_aquatilis | D3M96_RS02075 | 458513..462085 | 323284 | BREX-system-P-loop-protein-BrxC-[Alcaligenes-aquatilis] |
| WP_199265315.1 | Type6.BREX | [illegible] | 1213 | Bacteria>Pseudomonadota>Betaproteobacteria>Burkholderiales>Alcaligenaceae>Alcaligenes | GCF_016446305.1 | Alcaligenes_faecalis | GTU60_RS02720 | 606582..610223 | 511 | BREX-system-P-loop-protein-BrxC-[Alcaligenes-faecalis] |
| CCJ47860.1 | Type1.BREX | [illegible] | 1194 | Bacteria>Pseudomonadota>Betaproteobacteria>Burkholderiales>Alcaligenaceae>Bordetella | GCA_000317935.1 | Bordetella_parapertussis_Bpp5 | BN117_0527 | 555038..558622 | 1208660 | conserved-hypothetical-protein-[Bordetella-parapertussis-Bpp5] |
| WP_164926424.1 | Type1.BREX | [illegible] | 1188 | Bacteria>Pseudomonadota>Betaproteobacteria>Burkholderiales>Alcaligenaceae>Bordetella | GCF_000317935.1 | Bordetella_parapertussis | BN117_RS02695 | 555056..558622 | 519 | BREX-system-P-loop-protein-BrxC-[Bordetella-parapertussis] |
| WP_032972587.1 | Type6.BREX | [illegible] | 1213 | Bacteria>Pseudomonadota>Betaproteobacteria>Burkholderiales>Alcaligenaceae>Bordetella | GCF_006770345.1 | Bordetella_hinzii | CBR67_RS06420 | 1279803..1283444 | 103855 | BREX-system-P-loop-protein-BrxC-[Bordetella-hinzii] |
| WP_165720894.1 | Type1.BREX | [illegible] | 1181 | Bacteria>Pseudomonadota>Betaproteobacteria>Burkholderiales>Alcaligenaceae>Bordetella | GCF_011386965.1 | Bordetella_trematum | G6Z49_RS19625 | 4277130..4280675 | 123899 | BREX-system-P-loop-protein-BrxC-[Bordetella-trematum] |
| WP_068373657.1 | Type1.BREX | [illegible] | 1176 | Bacteria>Pseudomonadota>Betaproteobacteria>Burkholderiales>Alcaligenaceae>Kerstersia | GCF_000988095.1 | Kerstersia_gyiorum | AAV32_RS14310 | 103983..107513 | 206506 | BREX-system-P-loop-protein-BrxC-[Kerstersia-gyiorum] |
| WP_115075260.1 | Type1.BREX | [illegible] | 1187 | Bacteria>Pseudomonadota>Betaproteobacteria>Burkholderiales>Alcaligenaceae>Oligella | GCF_900454255.1 | Oligella_urethralis | DX091_RS10510 | 2175019..2178582 | 90245 | BREX-system-P-loop-protein-BrxC-[Oligella-urethralis] |
| WP_166406662.1 | Type1.BREX | [illegible] | 1190 | Bacteria>Pseudomonadota>Betaproteobacteria>Burkholderiales>Alcaligenaceae>Paenalcaligenes | GCF_011290505.1 | Paenalcaligenes_suwonensis | GY634_RS00685 | 163356..166928 | 1202713 | BREX-system-P-loop-protein-BrxC-[Paenalcaligenes-suwonensis] |
| MCK9506949.1 | Type1.BREX | [illegible] | 1069 | Bacteria>Pseudomonadota>Betaproteobacteria>Burkholderiales>Alcaligenaceae>Pigmentiphaga | GCA_023228545.1 | Pigmentiphaga_sp. | MOQ54_00780 | 41..3250 | 1977564 | BREX-system-P-loop-protein-BrxC-[Pigmentiphaga-sp.] |
| WP_132980217.1 | Type1.BREX | [illegible] | 1181 | Bacteria>Pseudomonadota>Betaproteobacteria>Burkholderiales>Alcaligenaceae>Pigmentiphaga | GCF_004352985.1 | Pigmentiphaga_sp._D-2 | EZY13_RS19720 | 185799..189344 | 1002116 | BREX-system-P-loop-protein-BrxC-[Pigmentiphaga-sp.-D-2] |
| WP_102070953.1 | Type1.BREX | [illegible] | 1181 | Bacteria>Pseudomonadota>Betaproteobacteria>Burkholderiales>Alcaligenaceae>Pollutimonas | GCF_002849615.1 | Pollutimonas_nitritireducens | CR155_RS15505 | 30043..33588 | 2045209 | BREX-system-P-loop-protein-BrxC-[Pollutimonas-nitritireducens] |
| MAO51251.1 | Type6.BREX | [illegible] | 1209 | Bacteria>Pseudomonadota>Betaproteobacteria>Burkholderiales>Alcaligenaceae>Pusillimonas | GCA_002693665.1 | Pusillimonas_sp._(ex_Stolz_et_al._2005) | CML16_10275 | 63709..67338 | 1979962 | ATPase-[Pusillimonas-sp.-(ex-Stolz-et-al.-2005)] |
| NYT45622.1 | Type6.BREX | [illegible] | 1214 | Bacteria>Pseudomonadota>Betaproteobacteria>Burkholderiales>Alcaligenaceae>unclassified_Alcaligenaceae | GCA_013416345.1 | Alcaligenaceae_bacterium | H0A64_12395 | 218321..221965 | 2021368 | BREX-system-P-loop-protein-BrxC-[Alcaligenaceae-bacterium] |
| NYT60502.1 | Type1.BREX | [illegible] | 1188 | Bacteria>Pseudomonadota>Betaproteobacteria>Burkholderiales>Alcaligenaceae>unclassified_Alcaligenaceae | GCA_013416405.1 | Alcaligenaceae_bacterium | H0A65_16390 | 10015..13581 | 2021368 | BREX-system-P-loop-protein-BrxC-[Alcaligenaceae-bacterium] |
| NYT63370.1 | Type1.BREX | [illegible] | 1191 | Bacteria>Pseudomonadota>Betaproteobacteria>Burkholderiales>Alcaligenaceae>unclassified_Alcaligenaceae | GCA_013416445.1 | Alcaligenaceae_bacterium | H0A66_13705 | 63657..67232 | 2021368 | BREX-system-P-loop-protein-BrxC-[Alcaligenaceae-bacterium] |
| NYT66107.1 | Type1.BREX | [illegible] | 1193 | Bacteria>Pseudomonadota>Betaproteobacteria>Burkholderiales>Alcaligenaceae>unclassified_Alcaligenaceae | GCA_013416485.1 | Alcaligenaceae_bacterium | H0A58_09005 | 172517..176098 | 2021368 | BREX-system-P-loop-protein-BrxC-[Alcaligenaceae-bacterium] |
| MCA8264434.1 | Type6.BREX | [illegible] | 1210 | Bacteria>Pseudomonadota>Betaproteobacteria>Burkholderiales>Burkholderiaceae>Burkholderia | GCA_020420005.1 | Burkholderia_multivorans | LGM79_28445 | 10767..14399 | 87883 | BREX-system-P-loop-protein-BrxC-[Burkholderia-multivorans] |
| WP_059944780.1 | Type6.BREX | [illegible] | 1213 | Bacteria>Pseudomonadota>Betaproteobacteria>Burkholderiales>Burkholderiaceae>Burkholderia | GCF_001527205.1 | Burkholderia_territorii | WT00_RS19595 | 122446..126087 | 1503055 | BREX-system-P-loop-protein-BrxC-[Burkholderia-territorii] |
| WP_060189440.1 | Type1.BREX | [illegible] | 1186 | Bacteria>Pseudomonadota>Betaproteobacteria>Burkholderiales>Burkholderiaceae>Burkholderia | GCF_001533155.1 | Burkholderia_diffusa | WL85_RS21680 | 47133..50693 | 488732 | BREX-system-P-loop-protein-BrxC-[Burkholderia-diffusa] |
| WP_124661146.1 | Type6.BREX | [illegible] | 1210 | Bacteria>Pseudomonadota>Betaproteobacteria>Burkholderiales>Burkholderiaceae>Burkholderia | GCF_003854235.1 | Burkholderia_sp._Bp8998 | DIE06_RS09175 | 82371..86003 | 2184557 | BREX-system-P-loop-protein-BrxC-[Burkholderia-sp.-Bp8998] |
| WP_226255239.1 | Type6.BREX | [illegible] | 1208 | Bacteria>Pseudomonadota>Betaproteobacteria>Burkholderiales>Burkholderiaceae>Burkholderia | GCF_020419005.1 | Burkholderia_cepacia | LGM39_RS24930 | 366713..370339 | 292 | BREX-system-P-loop-protein-BrxC-[Burkholderia-cepacia] |
| WP_175041065.1 | Type6.BREX | [illegible] | 1215 | Bacteria>Pseudomonadota>Betaproteobacteria>Burkholderiales>Burkholderiaceae>Burkholderia | GCF_902499225.1 | Burkholderia_contaminans | BLA3211_RS35460 | 10621..14268 | 488447 | BREX-system-P-loop-protein-BrxC-[Burkholderia-contaminans] |
| WP_175704189.1 | Type6.BREX | [illegible] | 1215 | Bacteria>Pseudomonadota>Betaproteobacteria>Burkholderiales>Burkholderiaceae>Burkholderia | GCF_902829775.1 | Burkholderia_ambifaria | HWV86_RS26205 | 192524..196171 | 152480 | BREX-system-P-loop-protein-BrxC-[Burkholderia-ambifaria] |
| WP_186043215.1 | Type6.BREX | [illegible] | 1213 | Bacteria>Pseudomonadota>Betaproteobacteria>Burkholderiales>Burkholderiaceae>Burkholderia | GCF_902830235.1 | Burkholderia_gladioli | H9C96_RS14825 | 15196..18837 | 28095 | BREX-system-P-loop-protein-BrxC-[Burkholderia-gladioli] |
| WP_186236015.1 | Type6.BREX | [illegible] | 1215 | Bacteria>Pseudomonadota>Betaproteobacteria>Burkholderiales>Burkholderiaceae>Burkholderia | GCF_902832475.1 | Burkholderia_gladioli | H9G27_RS00755 | 169921..173568 | 28095 | BREX-system-P-loop-protein-BrxC-[Burkholderia-gladioli] |
| WP_075360437.1 | Type6.BREX | [illegible] | 1210 | Bacteria>Pseudomonadota>Betaproteobacteria>Burkholderiales>Burkholderiaceae>Caballeronia | GCF_001935975.1 | Caballeronia_sordidicola | BSU03_RS34875 | 87497..91129 | 196367 | BREX-system-P-loop-protein-BrxC-[Caballeronia-sordidicola] |
| WP_187608173.1 | Type6.BREX | [illegible] | 1215 | Bacteria>Pseudomonadota>Betaproteobacteria>Burkholderiales>Burkholderiaceae>Caballeronia | GCF_014384845.1 | Caballeronia_sp._EK | IAG25_RS30620 | 154529..158176 | 2767469 | BREX-system-P-loop-protein-BrxC-[Caballeronia-sp.-EK] |
| WP_091908362.1 | Type6.BREX | [illegible] | 1213 | Bacteria>Pseudomonadota>Betaproteobacteria>Burkholderiales>Burkholderiaceae>Chitinasiproducens | GCF_900095735.1 | Chitinasiproducens_palmae | BLU54_RS11860 | 213511..217152 | 1770053 | BREX-system-P-loop-protein-BrxC-[Chitinasiproducens-palmae] |
| WP_013958181.1 | Type6.BREX | [illegible] | 1214 | Bacteria>Pseudomonadota>Betaproteobacteria>Burkholderiales>Burkholderiaceae>Cupriavidus | GCF_000219215.1 | Cupriavidus_necator | CNE_1c33280 | 3480419..3484063 | 106590 | BREX-system-P-loop-protein-BrxC-[Cupriavidus-necator] |
| WP_174424431.1 | Type6.BREX | [illegible] | 1215 | Bacteria>Pseudomonadota>Betaproteobacteria>Burkholderiales>Burkholderiaceae>Cupriavidus | GCF_013337645.1 | Cupriavidus_basilensis | FCJ59_RS11575 | 146108..149755 | 68895 | BREX-system-P-loop-protein-BrxC-[Cupriavidus-basilensis] |
| WP_059038859.1 | Type6.BREX | [illegible] | 1215 | Bacteria>Pseudomonadota>Betaproteobacteria>Burkholderiales>Burkholderiaceae | GCF_001485395.1 | Ralstonia | SAMD00023378_RS10640 | 9198..12845 | 48736 | MULTISPECIES:-BREX-system-P-loop-protein-BrxC-[Ralstonia] |
| WP_220363411.1 | Type6.BREX | [illegible] | 1215 | Bacteria>Pseudomonadota>Betaproteobacteria>Burkholderiales>Burkholderiaceae>Paraburkholderia | GCF_019511605.1 | Paraburkholderia_phenoliruptrix | JNB89_RS20360 | 10910..14557 | 252970 | BREX-system-P-loop-protein-BrxC-[Paraburkholderia-phenoliruptrix] |
| WP_230560751.1 | Type6.BREX | [illegible] | 1215 | Bacteria>Pseudomonadota>Betaproteobacteria>Burkholderiales>Burkholderiaceae>Paraburkholderia | GCF_020881035.1 | Paraburkholderia_sp._MMS20-SJTN17 | LJ655_RS08170 | 67515..71162 | 2886945 | BREX-system-P-loop-protein-BrxC-[Paraburkholderia-sp.-MMS20-SJTN17] |
| WP_167306478.1 | Type1.BREX | [illegible] | 1182 | Bacteria>Pseudomonadota>Betaproteobacteria>Burkholderiales>Burkholderiaceae>Paraburkholderia | GCF_900100565.1 | Paraburkholderia_steynii | BLT65_RS12240 | 289685..293233 | 1245441 | BREX-system-P-loop-protein-BrxC-[Paraburkholderia-steynii] |
| WP_201107905.1 | Type1.BREX | [illegible] | 1179 | Bacteria>Pseudomonadota>Betaproteobacteria>Burkholderiales>Burkholderiaceae>Paraburkholderia | GCF_905220855.1 | Paraburkholderia_domus | KHO36_RS10755 | 51203..54742 | 2793075 | BREX-system-P-loop-protein-BrxC-[Paraburkholderia-domus] |
| WP_183964658.1 | Type6.BREX | [illegible] | 1210 | Bacteria>Pseudomonadota>Betaproteobacteria>Burkholderiales>Burkholderiaceae>Quisquiliibacterium | GCF_014202215.1 | Quisquiliibacterium_transsilvanicum | HNQ70_RS04390 | 209103..212735 | 1549638 | BREX-system-P-loop-protein-BrxC-[Quisquiliibacterium-transsilvanicum] |
| MBP6902893.1 | Type6.BREX | [illegible] | 1219 | Bacteria>Pseudomonadota>Betaproteobacteria>Burkholderiales>Burkholderiaceae>unclassified_Burkholderiaceae | GCA_017992055.1 | Burkholderiaceae_bacterium | KBC73_22560 | 2680..6339 | 2030806 | BREX-system-P-loop-protein-BrxC-[Burkholderiaceae-bacterium] |
| MCP5271536.1 | Type6.BREX | [illegible] | 1219 | Bacteria>Pseudomonadota>Betaproteobacteria>Burkholderiales>Burkholderiaceae>unclassified_Burkholderiaceae | GCA_024235855.1 | Burkholderiaceae_bacterium | H6932_09900 | 152250..155909 | 2030806 | BREX-system-P-loop-protein-BrxC-[Burkholderiaceae-bacterium] |
| WP_143968626.1 | Type6.BREX | [illegible] | 1210 | Bacteria>Pseudomonadota>Betaproteobacteria>Burkholderiales>Burkholderiales_genera_incertae_sedis>Tepidimonas | GCF_007556755.1 | Tepidimonas_fonticaldi | Tfont_RS05070 | 95231..98863 | 1101373 | BREX-system-P-loop-protein-BrxC-[Tepidimonas-fonticaldi] |
| WP_054257718.1 | Type6.BREX | [illegible] | 1215 | Bacteria>Pseudomonadota>Betaproteobacteria>Burkholderiales>Comamonadaceae>Acidovorax | GCF_001298675.1 | Acidovorax_caeni | BN2503_RS16450 | 13891..17538 | 343013 | BREX-system-P-loop-protein-BrxC-[Acidovorax-caeni] |
| WP_057202647.1 | Type6.BREX | [illegible] | 1207 | Bacteria>Pseudomonadota>Betaproteobacteria>Burkholderiales>Comamonadaceae>Acidovorax | GCF_001428645.1 | Acidovorax_sp._Root217 | ASE31_RS15170 | 91296..94919 | 1736492 | BREX-system-P-loop-protein-BrxC-[Acidovorax-sp.-Root217] |
| GAO21740.1 | Type6.BREX | [illegible] | 1213 | Bacteria>Pseudomonadota>Betaproteobacteria>Burkholderiales>Comamonadaceae>Alicycliphilus | GCA_000974645.1 | Alicycliphilus_sp._B1 | ALISP_1560 | 91569..95210 | 1603291 | ATPase-[Alicycliphilus-sp.-B1] |
| WP_034379112.1 | Type6.BREX | [illegible] | 1210 | Bacteria>Pseudomonadota>Betaproteobacteria>Burkholderiales>Comamonadaceae>Comamonas | GCF_000761335.1 | Comamonas_thiooxydans | P245_RS10570 | 85855..89487 | 363952 | BREX-system-P-loop-protein-BrxC-[Comamonas-thiooxydans] |
| KEH09926.1 | Type1.BREX | [illegible] | 1185 | Bacteria>Pseudomonadota>Betaproteobacteria>Burkholderiales>Comamonadaceae>Delftia | GCA_000710165.1 | Delftia_tsuruhatensis | GY14_08945 | 17729..21286 | 180282 | hypothetical-protein-GY14_08945-[Delftia-tsuruhatensis] |
| OJX16153.1 | Type1.BREX | [illegible] | 1181 | Bacteria>Pseudomonadota>Betaproteobacteria>Burkholderiales>Comamonadaceae>Delftia | GCA_001898745.1 | Delftia_sp._67-8 | BGO79_00315 | 75959..79504 | 1895749 | hypothetical-protein-BGO79_00315-[Delftia-sp.-67-8] |
| WP_096915618.1 | Type6.BREX | [illegible] | 1224 | Bacteria>Pseudomonadota>Betaproteobacteria>Burkholderiales>Comamonadaceae>Delftia | GCF_002443055.1 | Delftia_acidovorans | CHL79_RS18790 | 4226898..4230572 | 80866 | BREX-system-P-loop-protein-BrxC-[Delftia-acidovorans] |
| WP_234638190.1 | Type6.BREX | [illegible] | 1219 | Bacteria>Pseudomonadota>Betaproteobacteria>Burkholderiales>Comamonadaceae>Delftia | GCF_903684635.1 | Delftia_tsuruhatensis | MJ845_RS20380 | 81444..85103 | 180282 | BREX-system-P-loop-protein-BrxC-[Delftia-tsuruhatensis] |
| TFI47144.1 | Type1.BREX | [illegible] | 1188 | Bacteria>Pseudomonadota>Betaproteobacteria>Burkholderiales>Comamonadaceae>Diaphorobacter | GCA_004563745.1 | Diaphorobacter_sp._DS2 | E4O93_14250 | 10625..14191 | 1302548 | BREX-system-P-loop-protein-BrxC-[Diaphorobacter-sp.-DS2] |
| WP_047349524.1 | Type6.BREX | [illegible] | 1213 | Bacteria>Pseudomonadota>Betaproteobacteria>Burkholderiales>Comamonadaceae>Diaphorobacter | GCF_001023515.1 | Diaphorobacter_sp._J5-51 | OX89_RS03130 | 14085..17726 | 680496 | BREX-system-P-loop-protein-BrxC-[Diaphorobacter-sp.-J5-51] |
| WP_106685100.1 | Type6.BREX | [illegible] | 1213 | Bacteria>Pseudomonadota>Betaproteobacteria>Burkholderiales>Comamonadaceae>Melaminivora | GCF_003008575.1 | Melaminivora_suipulveris | C6568_RS16475 | 3505730..3509371 | 2109913 | BREX-system-P-loop-protein-BrxC-[Melaminivora-suipulveris] |
| CDS49862.1 | Type6.BREX | [illegible] | 1210 | Bacteria>Pseudomonadota>Betaproteobacteria>Burkholderiales>Comamonadaceae>Polaromonas | - | Polaromonas_sp._CG9_12 | . | 485322..488954 | 1504672 | hypothetical-protein-[Polaromonas-sp.-CG9_12] |
| MBW7834374.1 | Type6.BREX | [illegible] | 1210 | Bacteria>Pseudomonadota>Betaproteobacteria>Burkholderiales>Comamonadaceae>Simplicispira | GCA_019455265.1 | Simplicispira_suum | H3C29_14305 | 45952..49584 | 2109915 | BREX-system-P-loop-protein-BrxC-[Simplicispira-suum] |
| WP_273364231.1 | Type6.BREX | [illegible] | 1210 | Bacteria>Pseudomonadota>Betaproteobacteria>Burkholderiales>Comamonadaceae>Simplicispira | GCF_019455265.1 | Simplicispira_suum | H3C29_RS14325 | 45952..49584 | 2109915 | BREX-system-P-loop-protein-BrxC-[Simplicispira-suum] |
| WP_068673039.1 | Type6.BREX | [illegible] | 1212 | Bacteria>Pseudomonadota>Betaproteobacteria>Burkholderiales>Comamonadaceae>Variovorax | GCF_001541225.1 | unclassified_Variovorax | APY03_RS01000 | 202384..206022 | 663243 | MULTISPECIES:-BREX-system-P-loop-protein-BrxC-[unclassified-Variovorax] |
| WP_218509735.1 | Type6.BREX | [illegible] | 1213 | Bacteria>Pseudomonadota>Betaproteobacteria>Burkholderiales>Comamonadaceae>Variovorax | GCF_019207085.1 | Variovorax_sp._dw_308 | HCU76_RS14255 | 36638..40279 | 2721546 | BREX-system-P-loop-protein-BrxC-[Variovorax-sp.-dw_308] |
| WP_093162784.1 | Type6.BREX | [illegible] | 1212 | Bacteria>Pseudomonadota>Betaproteobacteria>Burkholderiales>Comamonadaceae>Variovorax | GCF_900107745.1 | Variovorax_sp._YR216 | BLW05_RS12080 | 465668..469306 | 1882828 | BREX-system-P-loop-protein-BrxC-[Variovorax-sp.-YR216] |
| WP_186915942.1 | Type1.BREX | [illegible] | 1183 | Bacteria>Pseudomonadota>Betaproteobacteria>Burkholderiales>Oxalobacteraceae | GCF_014284115.1 | Undibacterium | H8K35_RS08325 | 157325..160876 | 401469 | MULTISPECIES:-BREX-system-P-loop-protein-BrxC-[Undibacterium] |
| WP_141835430.1 | Type1.BREX | [illegible] | 1209 | Bacteria>Pseudomonadota>Betaproteobacteria>Burkholderiales>Oxalobacteraceae>Herbaspirillum | - | Herbaspirillum_sp._SJZ107 | FBX97_RS19505 | 315565..319194 | 2572881 | BREX-system-P-loop-protein-BrxC-[Herbaspirillum-sp.-SJZ107] |
| WP_152879050.1 | Type6.BREX | [illegible] | 1215 | Bacteria>Pseudomonadota>Betaproteobacteria>Burkholderiales>Oxalobacteraceae>Massilia_group | GCF_009295655.1 | Duganella_sp._FT27W | F8197_RS00385 | 74619..78266 | 2654636 | BREX-system-P-loop-protein-BrxC-[Duganella-sp.-FT27W] |
| WP_258819858.1 | Type1.BREX | [illegible] | 1186 | Bacteria>Pseudomonadota>Betaproteobacteria>Burkholderiales>Oxalobacteraceae>Massilia_group | GCF_024753285.1 | Massilia_pinisoli | NX784_RS27510 | 42185..45745 | 1772194 | BREX-system-P-loop-protein-BrxC-[Massilia-pinisoli] |
| WP_182219587.1 | Type1.BREX | [illegible] | 1186 | Bacteria>Pseudomonadota>Betaproteobacteria>Burkholderiales>Oxalobacteraceae>Rugamonas | GCF_014042365.1 | Rugamonas_fusca | H3H36_RS18580 | 34208..37768 | 2758568 | BREX-system-P-loop-protein-BrxC-[Rugamonas-fusca] |
| OGB35307.1 | Type6.BREX | [illegible] | 1215 | Bacteria>Pseudomonadota>Betaproteobacteria>Burkholderiales>unclassified_Burkholderiales | GCA_001770935.1 | Burkholderiales_bacterium_RIFCSPLOWO2_12_FULL_61_40 | A3F78_11695 | 46897..50544 | 1797566 | ATPase-[Burkholderiales-bacterium-RIFCSPLOWO2_12_FULL_61_40] |
| MBN8486519.1 | Type1.BREX | [illegible] | 1186 | Bacteria>Pseudomonadota>Betaproteobacteria>Burkholderiales>unclassified_Burkholderiales | GCA_017302485.1 | Burkholderiales_bacterium | JOM20_02125 | 4400..7960 | 1891238 | BREX-system-P-loop-protein-BrxC-[Burkholderiales-bacterium] |
| MCP5275500.1 | Type1.BREX | [illegible] | 1176 | Bacteria>Pseudomonadota>Betaproteobacteria>Burkholderiales>unclassified_Burkholderiales | GCA_024235795.1 | Burkholderiales_bacterium | H6936_11775 | 653459..656989 | 1891238 | BREX-system-P-loop-protein-BrxC-[Burkholderiales-bacterium] |
| WP_194117489.1 | Type6.BREX | [illegible] | 1213 | Bacteria>Pseudomonadota>Betaproteobacteria>Neisseriales>Chromobacteriaceae>Chitinilyticum | GCF_015223195.1 | Chitinilyticum_piscinae | INR99_RS16780 | 39673..43314 | 2866724 | BREX-system-P-loop-protein-BrxC-[Chitinilyticum-piscinae] |
| WP_239907373.1 | Type1.BREX | [illegible] | 1185 | Bacteria>Pseudomonadota>Betaproteobacteria>Neisseriales>Chromobacteriaceae>Laribacter | GCF_022394255.1 | Laribacter_hongkongensis | LH425_05855 | 38844..42401 | 168471 | BREX-system-P-loop-protein-BrxC-[Laribacter-hongkongensis] |

| Protein-ID | System | Gene-Neighbourhoods | Length | Taxonomy | Assembly-ID | Strain | Locus-Tag | Genomic-coordinates | TaxID | Fasta-defline |
|---|---|---|---|---|---|---|---|---|---|---|
| WP_239909460.1 | Type1.BREX | <-CopG/RHH<-?<-?\|\|HTH_3->?->TraI-VirD2-RCRE->wHTH+WYL+WCX->BrxA-DNA-binding->BrxB-BrxF-iSTAND->DUF6079*->N6-METHYLASE->OLD-ABC->REase->SIR2->HerA/FtsK->N6-METHYLASE-p->OLD-ABC->PglZ->SIGMA+OB-Fold+AAA-ATPase+HKD-DNase-BrxL-> | 1186 | Bacteria>Pseudomonadota>Betaproteobacteria>Neisseriales>Chromobacteriaceae>Laribacter | GCF_022394655.1 | Laribacter_hongkongensis | LH423_RS01205 | 99182..102742 | 168471 | BREX-system-P-loop-protein-BrxC-[Laribacter-hongkongensis] |
| WP_088965422.1 | Type6.BREX | <-GTPase-AIG<-?\|\|HTH_3-><-?<-?<-DUF932<-SIGMA+OB-Fold+AAA-ATPase+HKD-DNase-BrxL<-PglZ<-N6-METHYLASE<-DUF6079*<-BrxB-BrxF-iSTAND<-BrxA-DNA-binding<-BrxE<-wHTH+WYL+WCX<-ParB<-ParB<-GDResolvase+PPP1-HTH<-HTH_3 | 1215 | Bacteria>Pseudomonadota>Betaproteobacteria>Neisseriales>Chromobacteriaceae>Vogesella | GCF_900090205.1 | Vogesella_sp._LIG4 | PSELUDRAFT_RS02905 | 629858..633505 | 1192162 | BREX-system-P-loop-protein-BrxC-[Vogesella-sp.-LIG4] |
| WP_153715435.1 | Type6.BREX | <-SIGMA+OB-Fold+AAA-ATPase+HKD-DNase-BrxL<-PglZ<-ParB+HNH<-N6-METHYLASE<-N6-METHYLASE<-N6-METHYLASE<-GreA_GreB+SF1-SF2-Helicase<-DUF6079*<-BrxB-BrxF-iSTAND<-BrxA-DNA-binding<-BrxE<-wHTH+WYL+WCX\|\|?->HNH-> | 1213 | Bacteria>Pseudomonadota>Betaproteobacteria>Neisseriales>Neisseriaceae>Eikenella | GCF_009650905.1 | Eikenella_corrodens | GJ661_RS08680 | 18367..22008 | 539 | BREX-system-P-loop-protein-BrxC-[Eikenella-corrodens] |
| MBK7364915.1 | Type1.BREX | <-HIN-HTH+rve+rve_3\|\|tRNA-><-SIGMA+OB-Fold+AAA-ATPase+HKD-DNase-BrxL<-PglZ<-N6-METHYLASE<-REase<-OLD-ABC<-DUF6079*<-BrxB-BrxF-iSTAND<-BrxA-DNA-binding<-XIS-HTH\|\|ArdA->TRPR-HTH->?-><-DOC+HTH<-HNH<-TraI-VirD2-RCRE | 1187 | Bacteria>Pseudomonadota>Betaproteobacteria>Nitrosomonadales>Nitrosomonadaceae>Nitrosomonas | GCA_016705625.1 | Nitrosomonas_sp. | IPI97_07925 | 1671776..1675339 | 42353 | BREX-system-P-loop-protein-BrxC-[Nitrosomonas-sp.] |
| MBK7352718.1 | Type1.BREX | TraI-VirD2-RCRE->HNH-><-?<-TRPR-HTH<-?<-ArdA\|\|XIS-HTH->BrxA-DNA-binding->BrxB-BrxF-iSTAND->DUF6079-p*->DUF6079-p->?->?->OLD-ABC->N6-METHYLASE->PglZ->SIGMA+OB-Fold+AAA-ATPase+HKD-DNase-BrxL->HIN-HTH->AAA-ATPase-> | 705 | Bacteria>Pseudomonadota>Betaproteobacteria>Nitrosomonadales>Nitrosomonadaceae>Nitrosomonas | GCA_016710135.1 | Nitrosomonas_sp. | IPJ05_03020 | 595038..597155 | 42353 | BREX-system-P-loop-protein-BrxC-[Nitrosomonas-sp.] |
| MBV6446594.1 | Type1.BREX | TraI-VirD2-RCRE->HNH->?-><-Couple_hipA+HipA_C<-TRPR-HTH<-ArdA\|\|XIS-HTH->BrxA-DNA-binding->BrxB-BrxF-iSTAND->DUF6079*->OLD-ABC->REase->N6-METHYLASE->PglZ->SIGMA+OB-Fold+AAA-ATPase+HKD-DNase-BrxL->SbcC+SbcC-><-Peptidase_S8<-BrxB-BrxF-iSTAND\|\|DOC+HTH-> | 1182 | Bacteria>Pseudomonadota>Betaproteobacteria>Nitrosomonadales>Nitrosomonadaceae>Nitrosomonas | GCA_019187415.1 | Nitrosomonas_sp. | FCKEDINB_00187 | 17095..20643 | 42353 | hypothetical-protein-[Nitrosomonas-sp.] |
| MBX3639968.1 | Type1.BREX | <-SIGMA+OB-Fold+AAA-ATPase+HKD-DNase-BrxL<-PglZ<-REase<-N6-METHYLASE<-?<-DUF6079*<-BrxB-BrxF-iSTAND<-BrxA-DNA-binding<-XIS-HTH\|\|HIN-HTH->SIR2-> | 1182 | Bacteria>Pseudomonadota>Betaproteobacteria>Nitrosomonadales>Nitrosomonadaceae>Nitrosomonas | GCA_019635175.1 | Nitrosomonas_sp. | KF888_05565 | 120382..123930 | 42353 | BREX-system-P-loop-protein-BrxC-[Nitrosomonas-sp.] |
| WP_181258202.1 | Type1.BREX | <-REase+TOPRIM<-HEPN+OST-HTH\|\|BrxA-DNA-binding->BrxB-BrxF-iSTAND->DUF6079*->N6-METHYLASE->ParB+HNH->OLD-ABC->?->PglZ->SIGMA+OB-Fold+AAA-ATPase+HKD-DNase-BrxL->SbcC->Calcineurin+Calcineurin-SbcD-><-MULE-TRANSPOSASE | 1179 | Bacteria>Pseudomonadota>Betaproteobacteria>Nitrosomonadales>Nitrosomonadaceae>Nitrosomonas | GCF_003046585.1 | Nitrosomonas_aestuarii | C8R11_107137 | 171439..174978 | 52441 | BREX-system-P-loop-protein-BrxC-[Nitrosomonas-aestuarii] |
| WP_107790392.1 | Type1.BREX | ParB+RepB->?-><-SbcC<-Calcineurin+Calcineurin-SbcD<-SbcC<-SIGMA+OB-Fold+AAA-ATPase+HKD-DNase-BrxL<-PglZ<-N6-METHYLASE<-Schlafen+HATPase+wHTH+HTH<-DUF6079*<-BrxB-BrxF-iSTAND<-BrxA-DNA-binding\|\|?->?->PIN-> | 1180 | Bacteria>Pseudomonadota>Betaproteobacteria>Nitrosomonadales>Nitrosomonadaceae>Nitrosomonas | GCF_003051105.1 | Nitrosomonas_nitrosa | C8R30_RS13785 | 18563..22105 | 52442 | BREX-system-P-loop-protein-BrxC-[Nitrosomonas-nitrosa] |
| WP_204800510.1 | Type1.BREX | <-SbcC<-Calcineurin+Calcineurin-SbcD<-SbcC<-SIGMA+OB-Fold+AAA-ATPase+HKD-DNase-BrxL<-PglZ<-HTH+REase<-?<-Schlafen+HATPase+wHTH+HTH<-DUF6079*<-BrxB-BrxF-iSTAND<-BrxA-DNA-binding<-XIS-HTH<-?<-Arm-DNA-bind_3+Phage_integrase | 1180 | Bacteria>Pseudomonadota>Betaproteobacteria>Nitrosomonadales>Nitrosomonadaceae>Nitrosomonas | GCF_905220645.1 | Nitrosomonas_nitrosa | NUZ5A_RS15530 | 137140..140682 | 52442 | BREX-system-P-loop-protein-BrxC-[Nitrosomonas-nitrosa] |
| MBA0014163.1 | Type6.BREX | <-SIR2<-SIGMA+OB-Fold+AAA-ATPase+HKD-DNase-BrxL<-PglZ<-N6-METHYLASE<-?<-DUF6079*<-BrxB-BrxF-iSTAND<-BrxA-DNA-binding<-BrxE\|\|?->?->HIN-HTH-> | 1210 | Bacteria>Pseudomonadota>Betaproteobacteria>Nitrosomonadales>Nitrosomonadaceae>Nitrosospira | GCA_013450215.1 | Nitrosospira_sp. | HOZ53_00860 | 54117..57749 | 38080 | BREX-system-P-loop-protein-BrxC-[Nitrosospira-sp.] |
| WP_090458040.1 | Type6.BREX | Y1_Tnp-><-?<-?\|\|tRNA->BrxE->BrxA-DNA-binding->BrxB-BrxF-iSTAND->DUF6079*->GreA_GreB+SF1-SF2-Helicase->N6-METHYLASE->N6-METHYLASE->PglZ->SIGMA+OB-Fold+AAA-ATPase+HKD-DNase-BrxL->Glycos_transf_1+Glyco_trans_1_2-><-DADA-Ligase | 1213 | Bacteria>Pseudomonadota>Betaproteobacteria>Nitrosomonadales>Nitrosomonadaceae>Nitrosospira | GCF_900100985.1 | Nitrosospira_sp._Nsp1 | BLP78_RS07395 | 18423..22064 | 136547 | BREX-system-P-loop-protein-BrxC-[Nitrosospira-sp.-Nsp1] |
| SFW23276.1 | Type6.BREX | <-AAA-ATPase<-HIN-HTH\|\|Phage_integrase-><-?<-SIGMA+OB-Fold+AAA-ATPase+HKD-DNase-BrxL<-PglZ<-ParB+HNH<-N6-METHYLASE<-GreA_GreB+SF1-SF2-Helicase<-DUF6079*<-BrxB-BrxF-iSTAND<-BrxA-DNA-binding<-BrxE<-?<-?<-?\|\|?-><-TPR+TPR+TPR+TPR | 1213 | Bacteria>Pseudomonadota>Betaproteobacteria>Nitrosomonadales>Nitrosomonadaceae>Nitrosovibrio | GCA_900119085.1 | Nitrosovibrio_sp._Nv17 | SAMN05216414_10762 | 70156..73797 | 1855339 | hypothetical-protein-SAMN05216414_10762-[Nitrosovibrio-sp.-Nv17] |
| WP_218184829.1 | Type6.BREX | sGTP->Arm-DNA-bind_3+Phage_integrase->Arm-DNA-bind_3+Phage_int_SAM_3+Phage_integrase-><-AAA-ATPase\|\|<-SIGMA+OB-Fold+AAA-ATPase+HKD-DNase-BrxL<-PglZ<-ParB+HNH<-N6-METHYLASE<-GreA_GreB+SF1-SF2-Helicase<-DUF6079*<-BrxB-BrxF-iSTAND<-BrxA-DNA-binding<-BrxE<-?<-?<-?\|\|?-><-TPR+TPR+TPR+TPR | 1234 | Bacteria>Pseudomonadota>Betaproteobacteria>Nitrosomonadales>Nitrosomonadaceae>Nitrosovibrio | GCF_900119085.1 | Nitrosovibrio_sp._Nv17 | BT108_RS05915 | 70156..73860 | 1855339 | BREX-system-P-loop-protein-BrxC-[Nitrosovibrio-sp.-Nv17] |
| WP_121241046.1 | Type6.BREX | RNA_METHYLASE-><-?<-BrxD<-SIGMA+OB-Fold+AAA-ATPase+HKD-DNase-BrxL<-PglZ<-N6-METHYLASE<-DUF6079*<-BrxB-BrxF-iSTAND<-BrxA-DNA-binding<-BrxE<-XIS-HTH<-ArdCN+MPTase<-?<-PadR-HTH<-CopG/RHH | 1202 | Bacteria>Pseudomonadota>Betaproteobacteria>Nitrosomonadales>Sterolibacteriaceae>Sulfurisoma | GCF_003865015.1 | Sulfurisoma_sediminicola | EJE94_RS09880 | 239721..243329 | 1381557 | BREX-system-P-loop-protein-BrxC-[Sulfurisoma-sediminicola] |
| ODU51178.1 | Type6.BREX | <-LysR-HTH\|\|HTH_3-><-MODE-HTH+PBPII\|\|?->wHTH+WYL+WCX->BrxE->BrxA-DNA-binding->BrxB-BrxF-iSTAND->DUF6079*->GreA_GreB+SF1-SF2-Helicase->ParB+HNH->?->DOC+HTH->PglZ->SIGMA+OB-Fold+AAA-ATPase+HKD-DNase-BrxL-><-HDRnase\|\|XIS-HTH->Tox-RES-> | 1213 | Bacteria>Pseudomonadota>Betaproteobacteria>Nitrosomonadales>Thiobacillaceae>Thiobacillus | GCA_001725135.1 | Thiobacillus_sp._SCN_63-374 | ABS92_00645 | 9244..12885 | 1660149 | MAG:-ATPase-[Thiobacillus-sp.-SCN-63-374] |
| MBW8470844.1 | Type6.BREX | <-SIGMA+OB-Fold+AAA-ATPase+HKD-DNase-BrxL<-PglZ<-?<-ParB+HNH<-N6-METHYLASE<-N6-METHYLASE<-DUF6079*<-BrxB-BrxF-iSTAND<-BrxA-DNA-binding<-BrxE\|\|?->?-><-TOPRIM<-?<-XIS-HTH | 1215 | Bacteria>Pseudomonadota>Betaproteobacteria>Nitrosomonadales>Thiobacillaceae>Thiobacillus | GCA_019459125.1 | Thiobacillus_sp. | KOM67_21445 | 16263..19910 | 924 | BREX-system-P-loop-protein-BrxC-[Thiobacillus-sp.] |
| NMG55067.1 | Type1.BREX | <-TraK<-TraE<-TraM<-TraM<-DUF3262<-?\|\|XIS-HTH->BrxA-DNA-binding->BrxB-BrxF-iSTAND->DUF6079*->N6-METHYLASE->OLD-ABC->REase->SIR2->HerA/FtsK->N6-METHYLASE-p->PglZ->SIGMA+OB-Fold+AAA-ATPase+HKD-DNase-BrxL->Phage-HTH-> | 1189 | Bacteria>Pseudomonadota>Betaproteobacteria>Rhodocyclales>Rhodocyclaceae>Aromatoleum | GCA_012910915.1 | Aromatoleum_aromaticum | GPA23_10435 | 14208..17777 | 551760 | BREX-system-P-loop-protein-BrxC-[Aromatoleum-aromaticum] |
| WP_168953997.1 | Type1.BREX | <-TraK<-TraE<-TraM<-TraM<-DUF3262<-?\|\|XIS-HTH->BrxA-DNA-binding->BrxB-BrxF-iSTAND->DUF6079*->N6-METHYLASE->OLD-ABC->REase->SIR2->HerA/FtsK->N6-METHYLASE-p->PglZ->SIGMA+OB-Fold+AAA-ATPase+HKD-DNase-BrxL->Phage-HTH-> | 1188 | Bacteria>Pseudomonadota>Betaproteobacteria>Rhodocyclales>Rhodocyclaceae>Aromatoleum | GCF_023704195.1 | Aromatoleum_aromaticum | L8M16_RS08665 | 1808924..1812490 | 551760 | BREX-system-P-loop-protein-BrxC-[Aromatoleum-aromaticum] |
| MBP6421402.1 | Type1.BREX | SF1-SF2-Helicase-><-SIGMA+OB-Fold+AAA-ATPase+HKD-DNase-BrxL<-PglZ<-ParB+HNH<-DUF488-s<-DUF488-l<-DUF4326<-N6-METHYLASE<-N6-METHYLASE<-DUF6079-p* | 923 | Bacteria>Pseudomonadota>Betaproteobacteria>Rhodocyclales>Rhodocyclaceae>Propionivibrio | GCA_017988215.1 | Propionivibrio_sp. | KA271_00765 | 12654..15427 | 2212460 | BREX-system-P-loop-protein-BrxC-[Propionivibrio-sp.] |
| MBK6907823.1 | Type6.BREX | <-AA_kinase+ACT+ACT_7\|\|?->?-><-BrxD<-SIGMA+OB-Fold+AAA-ATPase+HKD-DNase-BrxL<-PglZ<-DOC+HTH<-DUF6079*<-BrxB-BrxF-iSTAND<-BrxA-DNA-binding<-BrxE<-XIS-HTH<-ArdCN+MPTase<-?<-PadR-HTH<-CopG/RHH | 1202 | Bacteria>Pseudomonadota>Betaproteobacteria>Rhodocyclales>Rhodocyclaceae>unclassified_Rhodocyclaceae | GCA_016702335.1 | Rhodocyclaceae_bacterium | IPH08_12425 | 301349..304957 | 1898103 | BREX-system-P-loop-protein-BrxC-[Rhodocyclaceae-bacterium] |
| MCA3121411.1 | Type6.BREX | <-DNA-glycosylase\|\|HIN-HTH-><-HIN-HTH<-OBFOLD-NUCLEASE<-ParB+HNH<-SIGMA+OB-Fold+AAA-ATPase+HKD-DNase-BrxL<-PglZ<-N6-METHYLASE<-N6-METHYLASE<-DUF6079*<-BrxB-BrxF-iSTAND<-BrxA-DNA-binding<-BrxE<-ANK+ANK<-ANK+ANK<-?<-?<-NUDIX\|\|HD-> | 1210 | Bacteria>Pseudomonadota>Betaproteobacteria>Rhodocyclales>Rhodocyclaceae>unclassified_Rhodocyclaceae | GCA_020248305.1 | Rhodocyclaceae_bacterium | ING90_02100 | 367371..371003 | 1898103 | BREX-system-P-loop-protein-BrxC-[Rhodocyclaceae-bacterium] |
| TVT60730.1 | Type6.BREX | Arm-DNA-bind_3+Phage_integrase-><-LysR-HTH\|\|HTH_3-><-MODE-HTH+PBPII\|\|wHTH+WYL+WCX->BrxE->BrxA-DNA-binding->BrxB-BrxF-iSTAND->DUF6079*->GreA_GreB+SF1-SF2-Helicase->N6-METHYLASE->N6-METHYLASE->PglZ->SIGMA+OB-Fold+AAA-ATPase+HKD-DNase-BrxL->KAP-NTPase->KAP-glycosylase->?->TatD_DNase-> | 1213 | Bacteria>Pseudomonadota>Betaproteobacteria>Rhodocyclales>Zoogloeaceae>Azoarcus | GCA_007713545.1 | Azoarcus_sp._PHD | FHK80_01725 | 174831..178472 | 1628740 | MAG:-BREX-system-P-loop-protein-BrxC-[Azoarcus-sp.-PHD] |
| WP_141017182.1 | Type6.BREX | PhdYeFM_antitox->RelE-ParE-><-DOC+HTH<-Phage_integrase<-?\|\|wHTH+WYL+WCX->BrxE->BrxA-DNA-binding->BrxB-BrxF-iSTAND->DUF6079*->OLD-ABC->PIN->N6-METHYLASE->N6-METHYLASE->?->ParB+HNH->PglZ->SIGMA+OB-Fold+AAA-ATPase+HKD-DNase-BrxL-><-XIS-HTH | 1207 | Bacteria>Pseudomonadota>Betaproteobacteria>Rhodocyclales>Zoogloeaceae>Azoarcus | GCF_006496635.1 | Azoarcus_sp._DD4 | CJ010_RS05935 | 1245819..1249442 | 2027405 | BREX-system-P-loop-protein-BrxC-[Azoarcus-sp.-DD4] |
| NMG49986.1 | Type1.BREX | <-TraE<-TraM<-TraM<-DUF3262<-?\|\|HipA_C->XIS-HTH->BrxA-DNA-binding->BrxB-BrxF-iSTAND->DUF6079*->OLD-ABC->REase->SIR2->HerA/FtsK->N6-METHYLASE-p->PglZ->SIGMA+OB-Fold+AAA-ATPase+HKD-DNase-BrxL->SbcC+SbcC-> | 1189 | Bacteria>Pseudomonadota>Betaproteobacteria>Rhodocyclales>Zoogloeaceae>Parazoarcus | GCF_012910825.1 | Parazoarcus_communis | GO613_17970 | 21745..25314 | 41977 | BREX-system-P-loop-protein-BrxC-[Parazoarcus-communis] |
| WP_168942691.1 | Type1.BREX | <-TraK<-TraE<-TraM<-TraM<-DUF3262<-?\|\|XIS-HTH->BrxA-DNA-binding->BrxB-BrxF-iSTAND->DUF6079*->N6-METHYLASE->OLD-ABC->REase->SIR2->HerA/FtsK->N6-METHYLASE-p->PglZ->SIGMA+OB-Fold+AAA-ATPase+HKD-DNase-BrxL->SbcC+SbcC-> | 1188 | Bacteria>Pseudomonadota>Betaproteobacteria>Rhodocyclales>Zoogloeaceae>Parazoarcus | GCF_012910825.1 | Parazoarcus_communis | GO613_RS17995 | 21748..25314 | 41977 | BREX-system-P-loop-protein-BrxC-[Parazoarcus-communis] |
| MBL8465274.1 | Type6.BREX | ABC-ATPase+SbcC->BrxE->BrxA-DNA-binding->BrxB-BrxF-iSTAND->DUF6079*->MNS+MarR-HTH->N6-METHYLASE->N6-METHYLASE->HEPN->ParA-Soj-PloopNTPase->ORC-CDC6->PglZ->SIGMA+OB-Fold+AAA-ATPase+HKD-DNase-BrxL-> | 1209 | Bacteria>Pseudomonadota>Betaproteobacteria>Rhodocyclales>Zoogloeaceae>Thauera | GCA_016790365.1 | Thauera_sp. | JNL37_10520 | 3981..7610 | 1905334 | BREX-system-P-loop-protein-BrxC-[Thauera-sp.] |
| WP_148578091.1 | Type1.BREX | Arm-DNA-bind_2+Phage_int_SAM_3+Phage_integrase->HDRnase->?-><-HTH_19\|\|?->?->wHTH+WYL+WCX->BrxA-DNA-binding->BrxB-BrxF-iSTAND->DUF6079*->N6-METHYLASE->OLD-ABC->REase->SIR2->HerA/FtsK->N6-METHYLASE-p->HTH+REase->PglZ->SIGMA+OB-Fold+AAA-ATPase+HKD-DNase-BrxL-> | 1186 | Bacteria>Pseudomonadota>Betaproteobacteria>Rhodocyclales>Zoogloeaceae>Zoogloea | GCF_008107625.1 | Zoogloea_oleivorans | ETQ85_RS05700 | 193093..196653 | 1552750 | BREX-system-P-loop-protein-BrxC-[Zoogloea-oleivorans] |
| GHT91750.1 | Type6.BREX | <-SIGMA+OB-Fold+AAA-ATPase+HKD-DNase-BrxL<-PglZ<-DOC+HTH<-TOPRIM<-OLD-ABC<-N6-METHYLASE-p<-N6-METHYLASE<-REase+AIPR<-DUF6079*<-BrxB-BrxF-iSTAND<-BrxA-DNA-binding<-BrxE<-XIS-HTH\|\|HTH_3->Couple_hipA+HipA_C->Phage_integrase->RelE-> | 1209 | Bacteria>Pseudomonadota>Betaproteobacteria>unclassified_Betaproteobacteria | GCA_026000515.1 | Betaproteobacteria_bacterium | FACS1894116_00820 | 42503..46132 | 1891241 | hypothetical-protein-FACS1894116_00820-[Betaproteobacteria-bacterium] |
| AUZ77883.1 | Type1.BREX | <-HIN-HTH<-SIGMA+OB-Fold+AAA-ATPase+HKD-DNase-BrxL<-PglZ<-ParB+HNH<-N6-METHYLASE<-REase<-OLD-ABC<-DUF6079*<-BrxB-BrxF-iSTAND<-BrxA-DNA-binding<-XIS-HTH\|\|<-DOC+HTH<-?<-XIS-HTH<-Arm-DNA-bind_3+Phage_integrase\|\|?-><-Sulfate_transp+STAS | 1183 | Bacteria>Pseudomonadota>Gammaproteobacteria>Aeromonadales>Aeromonadaceae>Aeromonas | GCA_002934425.1 | Aeromonas_sp._ASNIH4 | C2U40_16585 | 3601638..3605189 | 1636609 | BREX-system-P-loop-protein-BrxC-[Aeromonas-sp.-ASNIH4] |
| WP_043557612.1 | Type1.BREX | REase-><-?\|\|?-><-SIGMA+OB-Fold+AAA-ATPase+HKD-DNase-BrxL<-PglZ<-ParB+HNH<-N6-METHYLASE<-REase<-OLD-ABC<-DUF6079*<-BrxB-BrxF-iSTAND<-BrxA-DNA-binding<-XIS-HTH<-?<-AAA+HTH<-?<-XIS-HTH | 1182 | Bacteria>Pseudomonadota>Gammaproteobacteria>Aeromonadales>Aeromonadaceae>Aeromonas | GCF_000819745.1 | Aeromonas_bestiarum | BN1118_RS19650 | 69736..73284 | 105751 | BREX-system-P-loop-protein-BrxC-[Aeromonas-bestiarum] |
| WP_100653505.1 | Type6.BREX | Arm-DNA-bind_3+Phage_integrase->\|\|XIS-HTH->?->TOPRIM-DNAG+D5N+D5ATPase+P4PRIMASE-HTH->HTH_Tnp->BrxE->BrxA-DNA-binding->BrxB-BrxF-iSTAND->DUF6079*->N6-METHYLASE->N6-METHYLASE->PglZ->SIGMA+OB-Fold+AAA-ATPase+HKD-DNase-BrxL-><-?<-?<-?<-?<-HAD+HAD+NUDIX | 1216 | Bacteria>Pseudomonadota>Gammaproteobacteria>Aeromonadales>Aeromonadaceae>Aeromonas | GCF_002803925.1 | Aeromonas_veronii | CVS41_08490 | 1813618..1817268 | 654 | BREX-system-P-loop-protein-BrxC-[Aeromonas-veronii] |
| WP_103244178.1 | Type6.BREX | <-zf-IS66+DDE_Tnp_IS66\|\|?->DnaB->HTH_Tnp-><-DEDD_Tnp_IS110\|\|wHTH+WYL+WCX->BrxE->BrxA-DNA-binding->BrxB-BrxF-iSTAND->DUF6079*->SF1-SF2-Helicase+RNase+RNase->TPR->N6-METHYLASE->N6-METHYLASE->ParB+HNH->PglZ->SIGMA+OB-Fold+AAA-ATPase+HKD-DNase-BrxL-><-Glyoxalase | 1210 | Bacteria>Pseudomonadota>Gammaproteobacteria>Aeromonadales>Aeromonadaceae>Aeromonas | GCF_002902865.1 | Aeromonas_sp._ASNIH5 | C2U30_RS17315 | 3671715..3675347 | 1758179 | BREX-system-P-loop-protein-BrxC-[Aeromonas-sp.-ASNIH5] |
| WP_216948840.1 | Type6.BREX | HTH_Tnp_1+HTH_21+rve+rve_3-><-Arm-DNA-bind_3<-SIGMA+OB-Fold+AAA-ATPase+HKD-DNase-BrxL<-PglZ<-N6-METHYLASE<-N6-METHYLASE<-DUF6079*<-BrxB-BrxF-iSTAND<-BrxA-DNA-binding<-BrxE<-HTH_Tnp<-TOPRIM-DNAG+D5N+D5ATPase+P4PRIMASE-HTH<-?<-XIS-HTH\|\|AAA-ATPase-p-><-Arm-DNA-bind_3+Phage_integrase | 1216 | Bacteria>Pseudomonadota>Gammaproteobacteria>Aeromonadales>Aeromonadaceae>Aeromonas | GCF_019046985.1 | Aeromonas_sp._FDAARGOS_1417 | I6L47_RS05350 | 1228193..1231843 | 2778066 | BREX-system-P-loop-protein-BrxC-[Aeromonas-sp.-FDAARGOS-1417] |
| WP_241417643.1 | Type6.BREX | Arm-DNA-bind_3+Phage_integrase->\|\|XIS-HTH->?->TOPRIM-DNAG+D5N+D5ATPase+P4PRIMASE-HTH->HTH_Tnp->BrxE->BrxA-DNA-binding->BrxB-BrxF-iSTAND->DUF6079*->N6-METHYLASE->N6-METHYLASE->PglZ->SIGMA+OB-Fold+AAA-ATPase+HKD-DNase-BrxL-> | 1216 | Bacteria>Pseudomonadota>Gammaproteobacteria>Aeromonadales>Aeromonadaceae>Aeromonas | GCF_022557195.1 | Aeromonas_hydrophila | MKW86_RS08070 | 1734303..1737953 | 644 | BREX-system-P-loop-protein-BrxC-[Aeromonas-hydrophila] |
| WP_257706959.1 | Type1.BREX | Zn_Tnp_IS1+DDE_Tnp_IS1595-><-SIGMA+OB-Fold+AAA-ATPase+HKD-DNase-BrxL<-PglZ<-ParB+HNH<-N6-METHYLASE<-REase<-OLD-ABC<-DUF6079*<-BrxB-BrxF-iSTAND<-BrxA-DNA-binding<-XIS-HTH | 1182 | Bacteria>Pseudomonadota>Gammaproteobacteria>Aeromonadales>Aeromonadaceae>Aeromonas | GCF_024649325.1 | Aeromonas_caviae | NUK33_RS01495 | 150155..153703 | 648 | BREX-system-P-loop-protein-BrxC-[Aeromonas-caviae] |
| WP_181014924.1 | Type1.BREX | <-HIN-HTH<-SIGMA+OB-Fold+AAA-ATPase+HKD-DNase-BrxL<-PglZ<-ParB+HNH<-N6-METHYLASE<-REase<-OLD-ABC<-DUF6079*<-BrxB-BrxF-iSTAND<-BrxA-DNA-binding<-XIS-HTH\|\|?-><-DOC+HTH<-?<-XIS-HTH\|\|?-><-Arm-DNA-bind_3+Phage_integrase | 1182 | Bacteria>Pseudomonadota>Gammaproteobacteria>Aeromonadales>Aeromonadaceae | GCF_002934425.1 | Aeromonas | C2U40_RS16580 | 3601638..3605186 | 642 | MULTISPECIES:-BREX-system-P-loop-protein-BrxC-[Aeromonas] |
| WP_019002750.1 | Type1.BREX | 35exo->LPD36+METHYLASE+SF1-SF2-Helicase->?->Phage_int_SAM_1+Phage_integrase-><-SIGMA+OB-Fold+AAA-ATPase+HKD-DNase-BrxL<-PglZ<-N6-METHYLASE-p<-N6-METHYLASE<-DUF6079*<-BrxB-BrxF-iSTAND<-BrxA-DNA-binding<-GDResolvase+HIN-HTH<-?\|\|?->?-><-?<-?<-VirB10 | 1210 | Bacteria>Pseudomonadota>Gammaproteobacteria>Aeromonadales>Succinivibrionaceae>Succinimonas | GCF_000378405.1 | Succinimonas_amylolytica | F557_RS20035 | 169312..172944 | 83769 | BREX-system-P-loop-protein-BrxC-[Succinimonas-amylolytica] |
| WP_219938189.1 | Type1.BREX | HIN-HTH->BrxA-DNA-binding->BrxB-BrxF-iSTAND->DUF6079*->N6-METHYLASE->Schlafen+HATPase+wHTH->PglZ->SIGMA+OB-Fold+AAA-ATPase+HKD-DNase-BrxL-> | 1210 | Bacteria>Pseudomonadota>Gammaproteobacteria>Aeromonadales>Succinivibrionaceae>Succinivibrio | GCF_019448115.1 | Succinivibrio_faecicola | J5V48_RS08685 | 2494..6126 | 2820300 | BREX-system-P-loop-protein-BrxC-[Succinivibrio-faecicola] |
| WP_265045927.1 | Type1.BREX | TRPR-HTH-><-HTH_Tnp_1+HTH_21+rve+rve_3\|\|?->NUDIX->BrxA-DNA-binding->BrxB-BrxF-iSTAND->DUF6079*->N6-METHYLASE->PglZ->SIGMA+OB-Fold+AAA-ATPase+HKD-DNase-BrxL->HKD-DNase+DUF3150->McrB-AAA->McrC-NTD+REase->?->SF1-SF2-Helicase+REase-> | 1194 | Bacteria>Pseudomonadota>Gammaproteobacteria>Alteromonadales>Alteromonadaceae>Alkalimarinus | GCF_026016465.1 | Alkalimarinus_sp._A2M4 | NKI27_RS10060 | 2208861..2212445 | 2961619 | BREX-system-P-loop-protein-BrxC-[Alkalimarinus-sp.-A2M4] |
| WP_217358045.1 | Type6.BREX | <-TUDOR+SF1-SF2-Helicase+iREase+CTD\|\|BrxE->BrxA-DNA-binding->BrxB-BrxF-iSTAND-><-DUF772+DDE_Tnp_1\|\|BrxB-BrxF-iSTAND->DUF6079-p*->DUF6079-p->N6-METHYLASE->N6-METHYLASE->PglZ->SIGMA+OB-Fold+AAA-ATPase+HKD-DNase-BrxL-><-NERD+SF1-SF2-Helicase<-?<-SF1-SF2-Helicase<-DrmB | 951 | Bacteria>Pseudomonadota>Gammaproteobacteria>Alteromonadales>Alteromonadaceae>Alteromonas/Salinimonas_group | GCF_002499975.2 | Alteromonas_pelagimontana | CA267_RS01545 | 356718..359573 | 1858656 | BREX-system-P-loop-protein-BrxC-[Alteromonas-pelagimontana] |
| WP_006006589.1 | Type1.BREX | <-PIN<-MazE_antitoxin<-MODE-HTH+PBPII\|\|?->?->?->XIS-HTH->BrxA-DNA-binding->BrxB-BrxF-iSTAND->DUF6079*->OLD-ABC->REase->N6-METHYLASE->PglZ->SIGMA+OB-Fold+AAA-ATPase+HKD-DNase-BrxL-> | 1189 | Bacteria>Pseudomonadota>Gammaproteobacteria>Alteromonadales>Alteromonadaceae>Glaciecola | GCF_000252165.1 | Glaciecola_punicea | GPUN_RS10800 | 142812..146381 | 56804 | BREX-system-P-loop-protein-BrxC-[Glaciecola-punicea] |
| WP_136550441.1 | Type1.BREX | <-SbcC+APATPase+SbcC<-SIGMA+OB-Fold+AAA-ATPase+HKD-DNase-BrxL<-PglZ<-N6-METHYLASE<-REase<-OLD-ABC<-Schlafen+HATPase+wHTH+HTH<-DUF6079*<-BrxB-BrxF-iSTAND<-BrxA-DNA-binding<-XIS-HTH<-MobI | 1193 | Bacteria>Pseudomonadota>Gammaproteobacteria>Alteromonadales>Alteromonadaceae>Hydrocarboniclastica | GCF_004851605.1 | Hydrocarboniclastica_marina | soil367_RS18210 | 4131404..4134985 | 2259620 | BREX-system-P-loop-protein-BrxC-[Hydrocarboniclastica-marina] |
| NQY33786.1 | Type1.BREX | <-REase<-?<-SIGMA+OB-Fold+AAA-ATPase+HKD-DNase-BrxL<-PglZ<-Schlafen+HATPase+wHTH<-PHP+SbcC<-N6-METHYLASE<-NERD<-OLD-ABC<-DUF6079*<-BrxB-BrxF-iSTAND<-BrxA-DNA-binding<-XIS-HTH<-?<-?<-?<-?<-?<-MoxR-AAA | 1200 | Bacteria>Pseudomonadota>Gammaproteobacteria>Alteromonadales>Alteromonadaceae>unclassified_Alteromonadaceae | GCA_013216015.1 | Alteromonadaceae_bacterium | HRT37_02225 | 108462..112064 | 1916082 | BREX-system-P-loop-protein-BrxC-[Alteromonadaceae-bacterium] |
| WP_220104111.1 | Type6.BREX | SbcC+SbcC->?->\|\|?->Arm-DNA-bind_3+Phage_integrase-><-35exo+SF1-SF2-Helicase<-SIGMA+OB-Fold+AAA-ATPase+HKD-DNase-BrxL<-PglZ<-N6-METHYLASE<-N6-METHYLASE<-DUF6079*<-BrxB-BrxF-iSTAND<-BrxA-DNA-binding<-BrxE<-wHTH+WYL+WCX\|\|?->NERD+ZnR-><-Phage_integrase | 1208 | Bacteria>Pseudomonadota>Gammaproteobacteria>Alteromonadales>Alteromonadales_genera_incertae_sedis>Neiella | GCF_019457915.1 | Neiella_holothuriorum | K0504_RS10305 | 24423..28049 | 2870530 | BREX-system-P-loop-protein-BrxC-[Neiella-holothuriorum] |
| WP_130603913.1 | Type1.BREX | NtrC-AAA+FIS-HTH->XIS-HTH->BrxA-DNA-binding->BrxB-BrxF-iSTAND->DUF6079*->N6-METHYLASE->PglZ->SIGMA+OB-Fold+AAA-ATPase+HKD-DNase-BrxL->?->McrB-AAA->McrC-NTD+REase->?->REase-> | 1188 | Bacteria>Pseudomonadota>Gammaproteobacteria>Alteromonadales>Colwelliaceae>Litorilituus | GCF_004295665.1 | Litorilituus_sediminis | EMK97_RS16655 | 3755554..3759120 | 718192 | BREX-system-P-loop-protein-BrxC-[Litorilituus-sediminis] |
| WP_112714387.1 | Type1.BREX | <-SIGMA+OB-Fold+AAA-ATPase+HKD-DNase-BrxL<-PglZ<-ParB+HNH<-N6-METHYLASE<-REase<-OLD-ABC<-DUF6079*<-BrxB-BrxF-iSTAND<-BrxA-DNA-binding<-XIS-HTH\|\|?-><-OrfB_Zn_ribbon<-HTH_OrfB_IS605+OrfB_IS605+OrfB_Zn_ribbon\|\|DEDD_Tnp_IS110+Transposase_20-><-Peptidase_S51 | 1199 | Bacteria>Pseudomonadota>Gammaproteobacteria>Alteromonadales>Moritellaceae>Moritella | GCF_900465055.1 | Moritella_yayanosii | MORIYA_RS08605 | 1893116..1896715 | 69539 | BREX-system-P-loop-protein-BrxC-[Moritella-yayanosii] |
| WP_010607320.1 | Type1.BREX | <-TniQ<-TnsC-ATPase<-?<-REase<-?<-SIGMA+OB-Fold+AAA-ATPase+HKD-DNase-BrxL<-PglZ<-ParB+HNH<-N6-METHYLASE<-DUF6079*<-BrxB-BrxF-iSTAND<-BrxA-DNA-binding<-XIS-HTH<-NtrC-AAA+FIS-HTH<-?<-?<-?<-?\|\|MODE-HTH+PBPII-> | 1188 | Bacteria>Pseudomonadota>Gammaproteobacteria>Alteromonadales>Pseudoalteromonadaceae | GCF_000276645.1 | Pseudoalteromonas | UY7_RS0121200 | 90697..94263 | 53246 | MULTISPECIES:-BREX-system-P-loop-protein-BrxC-[Pseudoalteromonas] |
| WP_024611453.1 | Type1.BREX | <-ArdCN+MPTase<-SIGMA+OB-Fold+AAA-ATPase+HKD-DNase-BrxL<-PglZ<-N6-METHYLASE<-REase<-OLD-ABC<-DUF6079*<-BrxB-BrxF-iSTAND<-BrxA-DNA-binding<-XIS-HTH | 1193 | Bacteria>Pseudomonadota>Gammaproteobacteria>Alteromonadales>Pseudoalteromonadaceae>Pseudoalteromonas | GCF_000498095.1 | Pseudoalteromonas_sp._TB64 | L647_RS0112855 | 131987..135568 | 1938600 | BREX-system-P-loop-protein-BrxC-[Pseudoalteromonas-sp.-TB64] |
| WP_167658112.1 | Type1.BREX | <-rve<-REase<-?<-SIGMA+OB-Fold+AAA-ATPase+HKD-DNase-BrxL<-PglZ<-ParB+HNH<-N6-METHYLASE<-REase<-OLD-ABC<-DUF6079*<-BrxB-BrxF-iSTAND<-BrxA-DNA-binding<-XIS-HTH<-NtrC-AAA+FIS-HTH<-?<-?<-?<-?\|\|MODE-HTH+PBPII-> | 1188 | Bacteria>Pseudomonadota>Gammaproteobacteria>Alteromonadales>Pseudoalteromonadaceae>Pseudoalteromonas | GCF_011806445.1 | Pseudoalteromonas_sp._HF66 | HCZ11_RS07660 | 212811..216377 | 2721559 | BREX-system-P-loop-protein-BrxC-[Pseudoalteromonas-sp.-HF66] |
| WP_200220630.1 | Type1.BREX | <-ArdCN+MPTase<-SIGMA+OB-Fold+AAA-ATPase+HKD-DNase-BrxL<-PglZ<-N6-METHYLASE<-NERD<-OLD-ABC<-DUF6079*<-BrxB-BrxF-iSTAND<-BrxA-DNA-binding<-XIS-HTH<-?<-ADH_N+ADH_zinc_N<-?<-?<-?\|\|MODE-HTH+PBPII-> | 1193 | Bacteria>Pseudomonadota>Gammaproteobacteria>Alteromonadales>Pseudoalteromonadaceae>Pseudoalteromonas | GCF_016598755.1 | Pseudoalteromonas_sp._LC2018020214 | JG479_RS17805 | 301190..304771 | 2799564 | BREX-system-P-loop-protein-BrxC-[Pseudoalteromonas-sp.-LC2018020214] |
| PKG74712.1 | Type1.BREX | <-SIGMA+OB-Fold+AAA-ATPase+HKD-DNase-BrxL<-PglZ<-ParB+HNH<-N6-METHYLASE<-REase<-OLD-ABC<-DUF6079*<-BrxB-BrxF-iSTAND<-BrxA-DNA-binding<-XIS-HTH<-?<-TnsC-ATPase<-rve+Mu-transpos_C<-REase+wHTH | 1191 | Bacteria>Pseudomonadota>Gammaproteobacteria>Alteromonadales>Shewanellaceae>Shewanella | GCA_002836135.1 | Shewanella_sp._GutCb | CXF86_11690 | 122253..125828 | 2058315 | BREX-system-P-loop-protein-BrxC-[Shewanella-sp.-GutCb] |
| MBB1438394.1 | Type1.BREX | <-CHTH\|\|XIS-HTH->BrxA-DNA-binding->BrxB-BrxF-iSTAND->DUF6079*->N6-METHYLASE->PglZ->SIGMA+OB-Fold+AAA-ATPase+HKD-DNase-BrxL-><-?<-MerR-HTH<-?<-HTH_26\|\|Phage_AlpA->Arm-DNA-bind_2+Phage_int_SAM_3+Phage_integrase-> | 1182 | Bacteria>Pseudomonadota>Gammaproteobacteria>Alteromonadales>Shewanellaceae>Shewanella | GCA_014164505.1 | Shewanella_sp._SG41-4 | H5202_06785 | 3542..7090 | 2760976 | BREX-system-P-loop-protein-BrxC-[Shewanella-sp.-SG41-4] |
| WP_257985331.1 | Type1.BREX | <-SIGMA+OB-Fold+AAA-ATPase+HKD-DNase-BrxL<-PglZ<-ParB+HNH<-N6-METHYLASE<-REase<-OLD-ABC<-DUF6079*<-BrxB-BrxF-iSTAND<-BrxA-DNA-binding<-XIS-HTH<-?<-TnsC-ATPase<-rve+Mu-transpos_C<-REase+wHTH | 1193 | Bacteria>Pseudomonadota>Gammaproteobacteria>Alteromonadales>Shewanellaceae>Shewanella | GCF_002836135.1 | Shewanella_sp._GutCb | CXF86_RS11730 | 122253..125834 | 2058315 | BREX-system-P-loop-protein-BrxC-[Shewanella-sp.-GutCb] |
| WP_173496635.1 | Type6.BREX | VIT+VWA->?->DUF3944->?->DDE_Tnp_1_5+DDE_Tnp_1->wHTH+WYL+WCX->BrxE->BrxA-DNA-binding->BrxB-BrxF-iSTAND->DUF6079*->N6-METHYLASE->PglZ->SIGMA+OB-Fold+AAA-ATPase+HKD-DNase-BrxL-><-HNH\|\|?->ANK+ANK+ANK-> | 1211 | Bacteria>Pseudomonadota>Gammaproteobacteria>Alteromonadales>Shewanellaceae>Shewanella | GCF_013283205.1 | Shewanella_sp._ISTPL2 | HUB64_RS06505 | 1443171..1446806 | 2699425 | BREX-system-P-loop-protein-BrxC-[Shewanella-sp.-ISTPL2] |
| WP_259397220.1 | Type1.BREX | <-CHTH\|\|XIS-HTH->BrxA-DNA-binding->BrxB-BrxF-iSTAND->DUF6079*->N6-METHYLASE->PglZ->SIGMA+OB-Fold+AAA-ATPase+HKD-DNase-BrxL-><-?<-MerR-HTH<-?<-HTH_26\|\|Phage_AlpA->Arm-DNA-bind_2+Phage_int_SAM_3+Phage_integrase-> | 1180 | Bacteria>Pseudomonadota>Gammaproteobacteria>Alteromonadales>Shewanellaceae>Shewanella | GCF_014164505.1 | Shewanella_sp._SG41-4 | H5202_RS06785 | 3548..7090 | 2760976 | BREX-system-P-loop-protein-BrxC-[Shewanella-sp.-SG41-4] |
| WP_188922770.1 | Type6.BREX | XIS-HTH->BrxE->BrxA-DNA-binding->BrxB-BrxF-iSTAND->DUF6079*->N6-METHYLASE->PglZ->SIGMA+OB-Fold+AAA-ATPase+HKD-DNase-BrxL-> | 1210 | Bacteria>Pseudomonadota>Gammaproteobacteria>Alteromonadales>Shewanellaceae>Shewanella | GCF_014648295.1 | Shewanella_saliphila | IEY48_RS17840 | 13004..16636 | 2282698 | BREX-system-P-loop-protein-BrxC-[Shewanella-saliphila] |
| WP_219012197.1 | Type6.BREX | Arm-DNA-bind_3+Phage_integrase-><-HTH_3\|\|?-><-SF1-SF2-Helicase<-REase\|\|HNH-><-SIGMA+OB-Fold+AAA-ATPase+HKD-DNase-BrxL<-PglZ<-N6-METHYLASE<-DUF6079*<-BrxB-BrxF-iSTAND<-BrxA-DNA-binding<-BrxE<-XIS-HTH\|\|?->?->DUF2787->Phage_integrase-> | 1211 | Bacteria>Pseudomonadota>Gammaproteobacteria>Alteromonadales>Shewanellaceae>Shewanella | GCF_019303235.1 | Shewanella_algae | KOE59_RS06910 | 134402..138037 | 38313 | BREX-system-P-loop-protein-BrxC-[Shewanella-algae] |
| WP_249554920.1 | Type1.BREX | TetR-HTH->ABhydrolase->?-><-PRTase<-RNasePH\|\|?->Arm-DNA-bind_3+Phage_integrase->BrxA-DNA-binding->BrxB-BrxF-iSTAND->DUF6079*->N6-METHYLASE->PglZ->SIGMA+OB-Fold+AAA-ATPase+HKD-DNase-BrxL->MORC+DUF3684+REase-><-XIS-HTH\|\|?->?->DUF2787->Phage_integrase-> | 1188 | Bacteria>Pseudomonadota>Gammaproteobacteria>Alteromonadales>Shewanellaceae>Shewanella | GCF_023475945.1 | Shewanella_sp._8A | M0413_RS16385 | 25717..29283 | 2943323 | BREX-system-P-loop-protein-BrxC-[Shewanella-sp.-8A] |
| WP_112352819.1 | Type1.BREX | MerR-HTH->?-><-SIGMA+OB-Fold+AAA-ATPase+HKD-DNase-BrxL<-PglZ<-N6-METHYLASE<-REase<-OLD-ABC<-DUF6079*<-BrxB-BrxF-iSTAND<-BrxA-DNA-binding<-XIS-HTH<-HNH+NERD+ZnR<-SAM-METHYLASE+wHTH+HNH<-?\|\|URI-DNase-><-MarR-HTH | 1197 | Bacteria>Pseudomonadota>Gammaproteobacteria>Alteromonadales>Shewanellaceae>Shewanella | GCF_900476435.1 | Shewanella_benthica | SHEWBE_RS12055 | 2634220..2637813 | 43661 | BREX-system-P-loop-protein-BrxC-[Shewanella-benthica] |
| WP_102039083.1 | Type1.BREX | <-SIGMA+OB-Fold+AAA-ATPase+HKD-DNase-BrxL<-PglZ<-ParB+HNH<-N6-METHYLASE<-REase<-OLD-ABC<-DUF6079*<-BrxB-BrxF-iSTAND<-BrxA-DNA-binding<-XIS-HTH<-?<-TnsC-ATPase<-rve+Mu-transpos_C<-REase+wHTH | 1191 | Bacteria>Pseudomonadota>Gammaproteobacteria>Alteromonadales>Shewanellaceae>Shewanella | - | Shewanella_sp._GutCb | CXF86_11690 | 122253..125828 | 2058315 | BREX-system-P-loop-protein-BrxC-[Shewanella-sp.-GutCb] |
| WP_182771402.1 | Type1.BREX | <-CHTH\|\|XIS-HTH->BrxA-DNA-binding->BrxB-BrxF-iSTAND->DUF6079*->N6-METHYLASE->PglZ->SIGMA+OB-Fold+AAA-ATPase+HKD-DNase-BrxL-><-?<-MerR-HTH<-?<-HTH_26\|\|Phage_AlpA->Arm-DNA-bind_2+Phage_int_SAM_3+Phage_integrase-> | 1182 | Bacteria>Pseudomonadota>Gammaproteobacteria>Alteromonadales>Shewanellaceae>Shewanella | - | Shewanella_sp._SG41-4 | H5202_06785 | 3542..7090 | 2760976 | BREX-system-P-loop-protein-BrxC-[Shewanella-sp.-SG41-4] |
| MBV7434872.1 | Type1.BREX | <-SIGMA+OB-Fold+AAA-ATPase+HKD-DNase-BrxL<-PglZ<-ParB+HNH<-N6-METHYLASE<-DUF6079*<-BrxB-BrxF-iSTAND<-BrxA-DNA-binding<-?<-?<-?<-?<-Peptidase_M20 | 1175 | Bacteria>Pseudomonadota>Gammaproteobacteria>Cardiobacteriales>Cardiobacteriaceae>unclassified_Cardiobacteriaceae | GCA_019218315.1 | Cardiobacteriaceae_bacterium_TAE3-ERU3 | KRX19_07510 | 264796..268323 | 2856340 | BREX-system-P-loop-protein-BrxC-[Cardiobacteriaceae-bacterium-TAE3-ERU3] |
| WP_265366287.1 | Type1.BREX | Arm-DNA-bind_3+Phage_integrase-><-XIS-HTH<-AAA+HTH<-XIS-HTH\|\|MobI->XIS-HTH->BrxA-DNA-binding->BrxB-BrxF-iSTAND->DUF6079*->Schlafen+HATPase+wHTH+HTH->OLD-ABC->REase->N6-METHYLASE->ParB+HNH->PglZ->SIGMA+OB-Fold+AAA-ATPase+HKD-DNase-BrxL->SbcC+APATPase+SbcC-> | 1193 | Bacteria>Pseudomonadota>Gammaproteobacteria>Cellvibrionales>Cellvibrionaceae>Marinimicrobium | GCF_026153455.1 | Marinimicrobium_koreense | OOT55_RS13025 | 3024461..3028042 | 306545 | BREX-system-P-loop-protein-BrxC-[Marinimicrobium-sp.-C6131] |
| WP_009576096.1 | Type1.BREX | Calcineurin-><-?<-?<-?<-HD<-SLATT\|\|nSTAND1+HNH->BrxA-DNA-binding->BrxB-BrxF-iSTAND->DUF6079*->N6-METHYLASE->PglZ->SIGMA+OB-Fold+AAA-ATPase+HKD-DNase-BrxL->URI-DNase+RAMA->?->?-><-?<-RADICAL-SAM | 1178 | Bacteria>Pseudomonadota>Gammaproteobacteria>Cellvibrionales>Halieaceae>Aequoribacter | GCF_000204315.1 | Aequoribacter_fuscus | EYZ66_RS04240 | 942285..945821 | 2518989 | BREX-system-P-loop-protein-BrxC-[Aequoribacter-fuscus] |
| WP_084199902.1 | Type1.BREX | CHTH->Calcineurin-><-?<-TnsC-ATPase<-?<-?\|\|XIS-HTH->BrxA-DNA-binding->BrxB-BrxF-iSTAND->DUF6079*->OLD-ABC->REase->N6-METHYLASE->PglZ->SIGMA+OB-Fold+AAA-ATPase+HKD-DNase-BrxL-> | 1184 | Bacteria>Pseudomonadota>Gammaproteobacteria>Cellvibrionales>Halieaceae>Halioglobus | GCF_014652255.1 | Halioglobus_japonicus | C0029_RS05530 | 1195751..1199305 | 930805 | BREX-system-P-loop-protein-BrxC-[Halioglobus-japonicus] |
| WP_148062597.1 | Type1.BREX | XIS-HTH->TOPRIM->AAA+wHTH->?->?->XIS-HTH->BrxA-DNA-binding->BrxB-BrxF-iSTAND->DUF6079*->SF1-SF2-Helicase+PRTase->SLOG+BirA-HTH->N6-METHYLASE->RVT->PglZ->Immank+ANK+ANK->REase->SIGMA+OB-Fold+AAA-ATPase+HKD-DNase-BrxL-> | 1180 | Bacteria>Pseudomonadota>Gammaproteobacteria>Cellvibrionales>Halieaceae>Parahaliea | GCF_008064635.1 | Parahaliea_aestuarii | FVW59_RS02260 | 491803..495345 | 1852021 | BREX-system-P-loop-protein-BrxC-[Parahaliea-aestuarii] |
| WP_036859475.1 | Type1.BREX | Classical-AAA->?->?->?->tRNA->Arm-DNA-bind_3+Phage_integrase->XIS-HTH->BrxA-DNA-binding->BrxB-BrxF-iSTAND->DUF6079*->SF1-SF2-Helicase+PRTase->SLOG+BirA-HTH->N6-METHYLASE->ParB+HNH->PglZ->SIGMA+OB-Fold+AAA-ATPase+HKD-DNase-BrxL->?->XIS-HTH-> | 1187 | Bacteria>Pseudomonadota>Gammaproteobacteria>Cellvibrionales>Porticoccaceae>Porticoccus | GCF_000744735.1 | Porticoccus_hydrocarbonoclasticus | U740_RS05135 | 1073957..1077520 | 1073414 | BREX-system-P-loop-protein-BrxC-[Porticoccus-hydrocarbonoclasticus] |
| WP_273033648.1 | Type1.BREX | <-SIGMA+OB-Fold+AAA-ATPase+HKD-DNase-BrxL<-PglZ<-ParB+HNH<-N6-METHYLASE-p<-HerA/FtsK<-?<-CR-ATPase3<-N6-METHYLASE<-Schlafen+HATPase+wHTH+HTH<-DUF6079*<-BrxB-BrxF-iSTAND<-BrxA-DNA-binding<-XIS-HTH<-MobI\|\|XIS-HTH->AAA+HTH->XIS-HTH-><-Arm-DNA-bind_3+Phage_integrase<-SIGMA-HTH | 1193 | Bacteria>Pseudomonadota>Gammaproteobacteria>Cellvibrionales>Spongiibacteraceae>Spongiibacter | GCF_950061765.1 | Spongiibacter_tropicus | B4C78_RS03795 | 102796..106377 | 454602 | BREX-system-P-loop-protein-BrxC-[Spongiibacter-tropicus] |
| MTW21824.1 | Type1.BREX | <-SAM-METHYLASE<-SIGMA+OB-Fold+AAA-ATPase+HKD-DNase-BrxL<-PglZ<-ParB+HNH<-Schlafen+HATPase+wHTH<-PhdYeFM_antitox<-N6-METHYLASE<-REase<-OLD-ABC<-DUF6079*<-BrxB-BrxF-iSTAND<-BrxA-DNA-binding<-DnaB+HINT+DnaB+HINT | 1173 | Bacteria>Pseudomonadota>Gammaproteobacteria>Chromatiales>Chromatiaceae>Allochromatium | GCA_009720725.1 | Allochromatium_palmeri | GJ668_12065 | 37225..40746 | 231048 | BREX-system-P-loop-protein-BrxC-[Allochromatium-palmeri] |
| WP_186343071.1 | Type1.BREX | <-SAM-METHYLASE<-SIGMA+OB-Fold+AAA-ATPase+HKD-DNase-BrxL<-PglZ<-ParB+HNH<-Schlafen+HATPase+wHTH<-PhdYeFM_antitox<-N6-METHYLASE<-REase<-OLD-ABC<-DUF6079*<-BrxB-BrxF-iSTAND<-BrxA-DNA-binding<-DnaB+HINT+DnaB+HINT | 1172 | Bacteria>Pseudomonadota>Gammaproteobacteria>Chromatiales>Chromatiaceae>Allochromatium | GCF_009720725.1 | Allochromatium_palmeri | GJ668_RS12065 | 37225..40743 | 231048 | BREX-system-P-loop-protein-BrxC-[Allochromatium-palmeri] |
| NEX15944.1 | Type1.BREX | <-NTP_transf<-?<-SIGMA+OB-Fold+AAA-ATPase+HKD-DNase-BrxL<-PglZ-p<-PglZ<-ParB+HNH<-N6-METHYLASE<-REase<-OLD-ABC<-DUF6079*<-BrxB-BrxF-iSTAND<-BrxA-DNA-binding<-?<-?<-ABC-ATPase+SbcC<-?<-?\|\|PALP-> | 1173 | Bacteria>Pseudomonadota>Gammaproteobacteria>Chromatiales>Chromatiaceae>Halochromatium | GCA_010912705.1 | Halochromatium_sp. | C1943_04720 | 67001..70522 | 2049430 | BREX-system-P-loop-protein-BrxC-[Halochromatium-sp.] |
| WP_132582321.1 | Type6.BREX | Phage_integrase->?->HerA/FtsK->TrwC+SF1-SF2-Helicase->?->BrxE->BrxA-DNA-binding->BrxB-BrxF-iSTAND->DUF6079*->N6-METHYLASE->PglZ->SIGMA+OB-Fold+AAA-ATPase+HKD-DNase-BrxL-><-HNH\|\|?->MarR-HTH->AbiEii-> | 1208 | Bacteria>Pseudomonadota>Gammaproteobacteria>Chromatiales>Chromatiaceae>Rheinheimera | GCF_004345025.1 | Rheinheimera_sp._D18 | E0Z06_RS04605 | 954479..958105 | 2545632 | BREX-system-P-loop-protein-BrxC-[Rheinheimera-sp.-D18] |
| WP_120797099.1 | Type1.BREX | <-HTH_3<-DDE_Tnp\|\|?-><-?\|\|?->?->BrxA-DNA-binding->BrxB-BrxF-iSTAND->DUF6079*->N6-METHYLASE-p-><-DDE_Tnp\|\|ParB+HNH->PglZ->?->SIGMA+OB-Fold+AAA-ATPase+HKD-DNase-BrxL-> | 1173 | Bacteria>Pseudomonadota>Gammaproteobacteria>Chromatiales>Chromatiaceae>Thiocapsa | GCF_003634315.1 | Thiocapsa_rosea | BDD21_RS10290 | 2320412..2323933 | 69360 | BREX-system-P-loop-protein-BrxC-[Thiocapsa-rosea] |
| WP_015282553.1 | Type6.BREX | <-SIGMA+OB-Fold+AAA-ATPase+HKD-DNase-BrxL<-PglZ<-PIN<-RHH<-N6-METHYLASE<-?<-SF1-SF2-Helicase<-N6-METHYLASE<-DUF6079*<-BrxB-BrxF-iSTAND<-BrxA-DNA-binding<-BrxE | 1209 | Bacteria>Pseudomonadota>Gammaproteobacteria>Chromatiales>Chromatiaceae>Thioflavicoccus | GCF_000327045.1 | Thioflavicoccus_mobilis | THIMO_RS17980 | 83065..86694 | 80679 | BREX-system-P-loop-protein-BrxC-[Thioflavicoccus-mobilis] |
| WP_058554449.1 | Type6.BREX | TIR->?->?->?->BrxE->BrxA-DNA-binding->BrxB-BrxF-iSTAND->DUF6079*->N6-METHYLASE->?->Schlafen+HATPase+wHTH->N6-METHYLASE->?->PIN->PglZ->SIGMA+OB-Fold+AAA-ATPase+HKD-DNase-BrxL-><-tRNA\|\|HHH_5+SF1-SF2-Helicase-> | 1214 | Bacteria>Pseudomonadota>Gammaproteobacteria>Chromatiales>Chromatiaceae>Thiohalocapsa | GCF_001469165.1 | Thiohalocapsa_sp._ML1 | ML01_RS05745 | 155025..158669 | 1431688 | BREX-system-P-loop-protein-BrxC-[Thiohalocapsa-sp.-ML1] |
| WP_239679777.1 | Type6.BREX | <-Arm-DNA-bind_2+Phage_int_SAM_3+Phage_integrase<-Phage_AlpA\|\|HTH_26-><-SIGMA+OB-Fold+AAA-ATPase+HKD-DNase-BrxL<-PglZ<-DOC+HTH<-ParB+HNH<-N6-METHYLASE<-N6-METHYLASE<-DUF6079*<-BrxB-BrxF-iSTAND<-BrxA-DNA-binding<-BrxE<-tRNA<-?\|\|RADICAL-SAM-> | 1223 | Bacteria>Pseudomonadota>Gammaproteobacteria>Chromatiales>Ectothiorhodospiraceae>Ectothiorhodospira | GCF_022227625.1 | Ectothiorhodospira_variabilis | LPW30_RS00090 | 25011..28682 | 505694 | BREX-system-P-loop-protein-BrxC-[Ectothiorhodospira-variabilis] |
| WP_238639758.1 | Type6.BREX | DUF488->\|\|BrxE->BrxA-DNA-binding->BrxB-BrxF-iSTAND->DUF6079*->N6-METHYLASE->N6-METHYLASE->?->DOC+HTH->PglZ->SIGMA+OB-Fold+AAA-ATPase+HKD-DNase-BrxL->?-><-TRD | 1207 | Bacteria>Pseudomonadota>Gammaproteobacteria>Chromatiales>Ectothiorhodospiraceae>Halorhodospira | GCF_022229325.1 | Halorhodospira_halochloris | LRD17_RS10705 | 4342..7965 | 1052 | BREX-system-P-loop-protein-BrxC-[Halorhodospira-halochloris] |
| WP_253479345.1 | Type6.BREX | GNTR-HTH->tRNA->Phage_integrase-><-REase\|\|ZnR-><-SIGMA+OB-Fold+AAA-ATPase+HKD-DNase-BrxL<-PglZ<-DOC+HTH<-ParB+HNH<-N6-METHYLASE<-DUF6079*<-BrxB-BrxF-iSTAND<-BrxA-DNA-binding<-BrxE\|\|?->?-><-HKD-DNase+SF1-SF2-Helicase | 1216 | Bacteria>Pseudomonadota>Gammaproteobacteria>Chromatiales>Ectothiorhodospiraceae>Natronocella | GCF_024170285.1 | Natronocella_acetinitrilica | J2T57_RS14030 | 326743..330393 | 414046 | BREX-system-P-loop-protein-BrxC-[Natronocella-acetinitrilica] |
| WP_019611713.1 | Type6.BREX | wHTH+WYL+WCX->?->TUDOR+SF1-SF2-Helicase+iREase+CTD->?->?->BrxE->BrxA-DNA-binding->BrxB-BrxF-iSTAND->DUF6079*->N6-METHYLASE-><-?\|\|N6-METHYLASE-p->ParB+HNH->PglZ->SIGMA+OB-Fold+AAA-ATPase+HKD-DNase-BrxL-><-HDRnase<-?<-Arm-DNA-bind_2+Phage_int_SAM_3+Phage_integrase | 1216 | Bacteria>Pseudomonadota>Gammaproteobacteria>Chromatiales>Ectothiorhodospiraceae>Thioalkalivibrio | GCF_000381705.1 | Thioalkalivibrio_sp._AKL7 | G315_RS0100950 | 222198..225848 | 1158155 | BREX-system-P-loop-protein-BrxC-[Thioalkalivibrio-sp.-AKL7] |
| WP_167070220.1 | Type1.BREX | <-wHTH+WYL+WCX\|\|?->ZnR-><-BrxB-BrxF-iSTAND<-GTPase-AIG\|\|XIS-HTH->BrxA-DNA-binding->BrxB-BrxF-iSTAND->DUF6079*->N6-METHYLASE->OLD-ABC->N6-METHYLASE->ParB+HNH->PglZ->SIGMA+OB-Fold+AAA-ATPase+HKD-DNase-BrxL-><-Phage_integrase | 1184 | Bacteria>Pseudomonadota>Gammaproteobacteria>Enterobacterales>Erwiniaceae>Pantoea | GCF_011752565.1 | Pantoea_sp._Ap-967 | F3J45_27210 | 36650..40204 | 2608362 | BREX-system-P-loop-protein-BrxC-[Pantoea-sp.-Ap-967] |
| QIG07578.1 | Type1.BREX | HTH_8->?->XIS-HTH->XIS-HTH->?->?->BrxA-DNA-binding->BrxB-BrxF-iSTAND->DUF6079*->OLD-ABC->REase->N6-METHYLASE->PglZ->SIGMA+OB-Fold+AAA-ATPase+HKD-DNase-BrxL-><-Arm-DNA-bind_3+Phage_integrase | 1183 | Bacteria>Pseudomonadota>Gammaproteobacteria>Enterobacterales>Morganellaceae>Proteus | GCA_011046025.1 | Proteus_sp._ZN5 | GTK47_19170 | 4181966..4185517 | 2697019 | BREX-system-P-loop-protein-BrxC-[Proteus-sp.-ZN5] |
| WP_206535871.1 | Type1.BREX | HTH_8->?->XIS-HTH->XIS-HTH->?->?->BrxA-DNA-binding->BrxB-BrxF-iSTAND->DUF6079*->OLD-ABC->REase->N6-METHYLASE->PglZ->SIGMA+OB-Fold+AAA-ATPase+HKD-DNase-BrxL-><-Arm-DNA-bind_3+Phage_integrase | 1185 | Bacteria>Pseudomonadota>Gammaproteobacteria>Enterobacterales>Morganellaceae>Proteus | GCF_011046025.1 | Proteus_sp._ZN5 | GTK47_RS19230 | 4181960..4185517 | 2697019 | BREX-system-P-loop-protein-BrxC-[Proteus-sp.-ZN5] |
| WP_272670012.1 | Type1.BREX | <-GTPase-AIG<-CopG/RHH\|\|?->XIS-HTH->?->BrxA-DNA-binding->BrxB-BrxF-iSTAND->DUF6079*->OLD-ABC->REase->N6-METHYLASE->PglZ->SIGMA+OB-Fold+AAA-ATPase+HKD-DNase-BrxL-><-HTH_Tnp_1+HTH_21+rve+rve_3 | 1185 | Bacteria>Pseudomonadota>Gammaproteobacteria>Enterobacterales>Morganellaceae>Providencia | GCF_028480605.1 | unclassified_Providencia | PSE59_RS21125 | 8138..11695 | 2633465 | MULTISPECIES:-BREX-system-P-loop-protein-BrxC-[unclassified-Providencia] |
| WP_068809899.1 | Type6.BREX | Arm-DNA-bind_3+Phage_integrase-><-?<-XIS-HTH<-?<-SIGMA+OB-Fold+AAA-ATPase+HKD-DNase-BrxL<-PglZ<-ParB+HNH<-N6-METHYLASE<-N6-METHYLASE<-DUF6079*<-BrxB-BrxF-iSTAND<-BrxA-DNA-binding<-BrxE\|\|?-><-REase+SF1-SF2-Helicase<-DOC+HTH<-TRD+TRD<-N6-METHYLASE<-DOC+HTH | 1214 | Bacteria>Pseudomonadota>Gammaproteobacteria>Gammaproteobacteria_incertae_sedis>Pseudohongiella | GCF_014642795.1 | Pseudohongiella_nitratireducens | IEW51_RS03970 | 206813..210457 | 1768907 | BREX-system-P-loop-protein-BrxC-[Pseudohongiella-nitratireducens] |
| MBT4669056.1 | Type1.BREX | <-MazG<-?<-SIGMA+OB-Fold+AAA-ATPase+HKD-DNase-BrxL<-PglZ<-N6-METHYLASE<-DUF6079*<-BrxB-BrxF-iSTAND<-BrxA-DNA-binding<-Trypsin+PDZ | 1167 | Bacteria>Pseudomonadota>Gammaproteobacteria>Gammaproteobacteria_incertae_sedis>sulfur-oxidizing_symbionts>Candidatus_Ruthia | GCA_018657745.1 | Candidatus_Ruthia_sp. | HN615_06440 | 8657..12160 | 2735566 | BREX-system-P-loop-protein-BrxC-[Candidatus-Ruthturnera-sp.] |
| MCG7867606.1 | Type1.BREX | <-RNase<-SbcC+CCDC158<-Calcineurin-SbcD<-SbcC<-SIGMA+OB-Fold+AAA-ATPase+HKD-DNase-BrxL<-PglZ<-N6-METHYLASE<-NERD<-OLD-ABC<-DUF6079*<-BrxB-BrxF-iSTAND<-BrxA-DNA-binding<-XIS-HTH\|\|HTH->?->?->XIS-HTH->?->TOPRIM-> | 1172 | Bacteria>Pseudomonadota>Gammaproteobacteria>Gammaproteobacteria_incertae_sedis>sulfur-oxidizing_symbionts>Candidatus_Thiodiazotropha | GCA_016842585.1 | Candidatus_Thiodiazotropha_taylori | JAY74_14755 | 23622..27140 | 2792791 | BREX-system-P-loop-protein-BrxC-[Candidatus-Thiodiazotropha-taylori] |
| MCG7876672.1 | Type6.BREX | TraG_N-><-?<-?<-?\|\|wHTH+WYL+WCX->BrxE->BrxA-DNA-binding->BrxB-BrxF-iSTAND->DUF6079*->N6-METHYLASE->ParB+HNH->PglZ->SIGMA+OB-Fold+AAA-ATPase+HKD-DNase-BrxL->Exonuc_V_gamma+REase->SF1-SF2-Helicase+REase->SbcC-> | 1215 | Bacteria>Pseudomonadota>Gammaproteobacteria>Gammaproteobacteria_incertae_sedis>sulfur-oxidizing_symbionts>Candidatus_Thiodiazotropha | GCA_016842655.1 | Candidatus_Thiodiazotropha_taylori | JAY78_10080 | 4978..8625 | 2792791 | BREX-system-P-loop-protein-BrxC-[Candidatus-Thiodiazotropha-taylori] |
| MCG7951159.1 | Type6.BREX | TraH->?->TraG_N-><-?<-?\|\|wHTH+WYL+WCX->BrxE->BrxA-DNA-binding->BrxB-BrxF-iSTAND->DUF6079*->N6-METHYLASE->ParB+HNH->PglZ->SIGMA+OB-Fold+AAA-ATPase+HKD-DNase-BrxL->Exonuc_V_gamma+REase->SF1-SF2-Helicase+REase->SbcC->SbcC->Calcineurin+Calcineurin-SbcD-> | 1215 | Bacteria>Pseudomonadota>Gammaproteobacteria>Gammaproteobacteria_incertae_sedis>sulfur-oxidizing_symbionts>Candidatus_Thiodiazotropha | GCA_016842765.1 | Candidatus_Thiodiazotropha_taylori | JAZ09_08050 | 22496..26143 | 2792791 | BREX-system-P-loop-protein-BrxC-[Candidatus-Thiodiazotropha-taylori] |
| MCG8102611.1 | Type6.BREX | TraH->?->TraG_N-><-?<-?\|\|wHTH+WYL+WCX->BrxE->BrxA-DNA-binding->BrxB-BrxF-iSTAND->DUF6079*->N6-METHYLASE->ParB+HNH->PglZ->SIGMA+OB-Fold+AAA-ATPase+HKD-DNase-BrxL->Exonuc_V_gamma+REase->SF1-SF2-Helicase+REase->SbcC->Calcineurin+Calcineurin-SbcD->SbcC-> | 1215 | Bacteria>Pseudomonadota>Gammaproteobacteria>Gammaproteobacteria_incertae_sedis>sulfur-oxidizing_symbionts>Candidatus_Thiodiazotropha | GCA_016842845.1 | Candidatus_Thiodiazotropha_taylori | JAY70_09490 | 22526..26173 | 2792791 | BREX-system-P-loop-protein-BrxC-[Candidatus-Thiodiazotropha-taylori] |
| MCG8058214.1 | Type6.BREX | <-SbcC<-Calcineurin+Calcineurin-SbcD<-SbcC<-SF1-SF2-Helicase+REase<-Exonuc_V_gamma+REase<-SIGMA+OB-Fold+AAA-ATPase+HKD-DNase-BrxL<-PglZ<-ParB+HNH<-N6-METHYLASE<-DUF6079*<-BrxB-BrxF-iSTAND<-BrxA-DNA-binding<-BrxE<-wHTH+WYL+WCX\|\|?->?-><-TraG_N<-?<-TraH | 1215 | Bacteria>Pseudomonadota>Gammaproteobacteria>Gammaproteobacteria_incertae_sedis>sulfur-oxidizing_symbionts>Candidatus_Thiodiazotropha | GCA_016843165.1 | Candidatus_Thiodiazotropha_taylori | JAZ21_00070 | 27190..30837 | 2792791 | BREX-system-P-loop-protein-BrxC-[Candidatus-Thiodiazotropha-taylori] |
| MCG7871843.1 | Type1.BREX | Phage_integrase-><-?<-SF1-SF2-Helicase<-OLD-ABC+TOPRIM<-SIGMA+OB-Fold+AAA-ATPase+HKD-DNase-BrxL<-PglZ<-REase<-N6-METHYLASE<-DUF6079*<-BrxB-BrxF-iSTAND<-BrxA-DNA-binding<-HTH_26\|\|?-><-Phage_integrase\|\|?-><-?<-?<-ACT | 1174 | Bacteria>Pseudomonadota>Gammaproteobacteria>Gammaproteobacteria_incertae_sedis>sulfur-oxidizing_symbionts>Candidatus_Thiodiazotropha | GCA_016848925.1 | Candidatus_Thiodiazotropha_lotti | JAZ11_07035 | 23024..26548 | 2792787 | BREX-system-P-loop-protein-BrxC-[Candidatus-Thiodiazotropha-lotti] |
| MCG7872379.1 | Type1.BREX | <-SbcC+CCDC158<-Calcineurin\|\|NtrC-AAA+FIS-HTH-><-HTH_3<-SIGMA+OB-Fold+AAA-ATPase+HKD-DNase-BrxL<-PglZ<-RVT<-N6-METHYLASE<-Schlafen+HATPase+wHTH+HTH<-DUF6079*<-BrxB-BrxF-iSTAND<-BrxA-DNA-binding<-XIS-HTH\|\|HNH->?->XIS-HTH->Phage_integrase->Phage_integrase->Phage_integrase-> | 1170 | Bacteria>Pseudomonadota>Gammaproteobacteria>Gammaproteobacteria_incertae_sedis>sulfur-oxidizing_symbionts>Candidatus_Thiodiazotropha | GCA_016848925.1 | Candidatus_Thiodiazotropha_lotti | JAZ11_09800 | 178889..182401 | 2792787 | BREX-system-P-loop-protein-BrxC-[Candidatus-Thiodiazotropha-lotti] |
| WP_207387236.1 | Type1.BREX | <-VirB11+VirB4-FtsK<-?<-?<-Phage_rep\|\|HTH+LexA-protease->XIS-HTH->BrxA-DNA-binding->BrxB-BrxF-iSTAND->DUF6079*->N6-METHYLASE->PglZ->SIGMA+OB-Fold+AAA-ATPase+HKD-DNase-BrxL->N6-METHYLASE->?->?-><-HTH_3\|\|?->Phage_integrase-> | 1194 | Bacteria>Pseudomonadota>Gammaproteobacteria>Legionellales>Legionellaceae>Fluoribacter | GCF_900640105.1 | Fluoribacter_gormanii | E1P13_RS01655 | 70546..74130 | 464 | BREX-system-P-loop-protein-BrxC-[Fluoribacter-gormanii] |
| TIG67286.1 | Type1.BREX | <-HerA/FtsK<-VirB3<-VirB2<-?<-?\|\|HTH+LexA-protease->XIS-HTH->BrxA-DNA-binding->BrxB-BrxF-iSTAND->DUF6079*->?->N6-METHYLASE->PglZ->SIGMA+OB-Fold+AAA-ATPase+HKD-DNase-BrxL-><-REase+RNase<-REase\|\|?-><-AAA-ATPase\|\|DDE_Tnp_1_3-> | 1185 | Bacteria>Pseudomonadota>Gammaproteobacteria>Legionellales>Legionellaceae>Legionella | GCA_004921705.1 | Legionella_pneumophila | JAN10_12325 | 31417..34974 | 446 | BREX-system-P-loop-protein-BrxC-[Legionella-pneumophila] |
| HAT1978996.1 | Type1.BREX | TRD+TRD->REase+SF1-SF2-Helicase->SbcC->REase->REase+RNase-><-SIGMA+OB-Fold+AAA-ATPase+HKD-DNase-BrxL<-PglZ<-N6-METHYLASE<-?<-DUF6079*<-BrxB-BrxF-iSTAND<-BrxA-DNA-binding<-XIS-HTH<-HTH+LexA-protease\|\|Phage_rep->?->?->VirB2->VirB3-> | 1184 | Bacteria>Pseudomonadota>Gammaproteobacteria>Legionellales>Legionellaceae>Legionella | GCA_015895665.1 | Legionella_pneumophila | I8Z83_002252 | 23760..27314 | 446 | BREX-system-P-loop-protein-BrxC-[Legionella-pneumophila] |
| HAT6348316.1 | Type1.BREX | REase+SF1-SF2-Helicase->?->CR-ATPase5+REase->REase->REase+RNase-><-SIGMA+OB-Fold+AAA-ATPase+HKD-DNase-BrxL<-PglZ<-N6-METHYLASE<-?<-DUF6079*<-BrxB-BrxF-iSTAND<-BrxA-DNA-binding<-XIS-HTH<-HTH+LexA-protease | 1185 | Bacteria>Pseudomonadota>Gammaproteobacteria>Legionellales>Legionellaceae>Legionella | GCA_015945075.1 | Legionella_pneumophila | JAJ33_001368 | 159147..162704 | 446 | BREX-system-P-loop-protein-BrxC-[Legionella-pneumophila] |
| HAT6334489.1 | Type1.BREX | <-HerA/FtsK<-VirB3<-VirB2<-?<-?\|\|HTH+LexA-protease->XIS-HTH->BrxA-DNA-binding->BrxB-BrxF-iSTAND->DUF6079*->?->N6-METHYLASE->PglZ->SIGMA+OB-Fold+AAA-ATPase+HKD-DNase-BrxL-><-REase+RNase<-REase\|\|?-><-AAA-ATPase\|\|DDE_Tnp_1_3-> | 1185 | Bacteria>Pseudomonadota>Gammaproteobacteria>Legionellales>Legionellaceae>Legionella | GCA_015946305.1 | Legionella_pneumophila | JAJ38_001743 | 27953..31510 | 446 | BREX-system-P-loop-protein-BrxC-[Legionella-pneumophila] |
| HAT9211453.1 | Type1.BREX | <-XIS-HTH<-?<-Arm-DNA-bind_3+Phage_integrase\|\|REase->REase+RNase-><-SIGMA+OB-Fold+AAA-ATPase+HKD-DNase-BrxL<-PglZ<-N6-METHYLASE<-?<-DUF6079*<-BrxB-BrxF-iSTAND<-BrxA-DNA-binding<-XIS-HTH<-HTH+LexA-protease\|\|?->?->VirB2->VirB3->HerA/FtsK-> | 1185 | Bacteria>Pseudomonadota>Gammaproteobacteria>Legionellales>Legionellaceae>Legionella | GCA_015973865.1 | Legionella_pneumophila_subsp._pneumophila | JBB81_06050 | 33487..37044 | 91891 | BREX-system-P-loop-protein-BrxC-[Legionella-pneumophila-subsp.-pneumophila] |
| HAU0774874.1 | Type1.BREX | HTH+LexA-protease->XIS-HTH->BrxA-DNA-binding->BrxB-BrxF-iSTAND->DUF6079*->?->N6-METHYLASE->PglZ->SIGMA+OB-Fold+AAA-ATPase+HKD-DNase-BrxL-><-REase+RNase<-REase<-?<-REase+SF1-SF2-Helicase<-Schlafen+HATPase+wHTH | 1185 | Bacteria>Pseudomonadota>Gammaproteobacteria>Legionellales>Legionellaceae>Legionella | GCA_015983005.1 | Legionella_pneumophila | JBH09_07400 | 2447..6004 | 446 | BREX-system-P-loop-protein-BrxC-[Legionella-pneumophila] |
| HAU2155475.1 | Type1.BREX | <-HerA/FtsK<-VirB3<-VirB2<-?<-?\|\|HTH+LexA-protease->XIS-HTH->BrxA-DNA-binding->BrxB-BrxF-iSTAND->DUF6079*->?->N6-METHYLASE->PglZ->SIGMA+OB-Fold+AAA-ATPase+HKD-DNase-BrxL-><-REase+RNase<-REase<-SbcC<-REase+SF1-SF2-Helicase<-TRD+TRD | 1185 | Bacteria>Pseudomonadota>Gammaproteobacteria>Legionellales>Legionellaceae>Legionella | GCA_016020315.1 | Legionella_pneumophila | JBK36_03075 | 164784..168341 | 446 | BREX-system-P-loop-protein-BrxC-[Legionella-pneumophila] |
| HAU3861597.1 | Type1.BREX | Arm-DNA-bind_3+Phage_integrase->?->?->?-><-ACET\|\|REase+RNase-><-SIGMA+OB-Fold+AAA-ATPase+HKD-DNase-BrxL<-PglZ<-N6-METHYLASE<-?<-DUF6079*<-BrxB-BrxF-iSTAND<-BrxA-DNA-binding<-XIS-HTH<-HTH+LexA-protease\|\|Phage_rep-> | 1185 | Bacteria>Pseudomonadota>Gammaproteobacteria>Legionellales>Legionellaceae>Legionella | GCA_016062615.1 | Legionella_pneumophila | F7Z80_07285 | 265287..268844 | 446 | BREX-system-P-loop-protein-BrxC-[Legionella-pneumophila] |
| HCJ1122636.1 | Type1.BREX | <-Arm-DNA-bind_3+Phage_integrase\|\|REase->REase+RNase-><-SIGMA+OB-Fold+AAA-ATPase+HKD-DNase-BrxL<-PglZ<-N6-METHYLASE<-?<-DUF6079*<-BrxB-BrxF-iSTAND<-BrxA-DNA-binding<-XIS-HTH<-HTH+LexA-protease\|\|Phage_rep->?->?->VirB2->VirB3-> | 1184 | Bacteria>Pseudomonadota>Gammaproteobacteria>Legionellales>Legionellaceae>Legionella | GCA_024486645.1 | Legionella_pneumophila | NQN52_000248 | 14141..17695 | 446 | BREX-system-P-loop-protein-BrxC-[Legionella-pneumophila] |
| WP_061693758.1 | Type1.BREX | <-DDE_Tnp_1_3\|\|AAA-ATPase-><-?\|\|REase->REase+RNase-><-SIGMA+OB-Fold+AAA-ATPase+HKD-DNase-BrxL<-PglZ<-N6-METHYLASE<-?<-DUF6079*<-BrxB-BrxF-iSTAND<-BrxA-DNA-binding<-XIS-HTH<-HTH+LexA-protease\|\|Phage_rep->?->?->VirB2->VirB3-> | 1184 | Bacteria>Pseudomonadota>Gammaproteobacteria>Legionellales>Legionellaceae>Legionella | GCF_002002625.1 | Legionella_pneumophila | LPM_RS06580 | 16051..19605 | 446 | BREX-system-P-loop-protein-BrxC-[Legionella-pneumophila] |
| WP_192875104.1 | Type1.BREX | <-VirB3<-VirB2<-?<-?<-Phage_rep\|\|HTH+LexA-protease->XIS-HTH->BrxA-DNA-binding->BrxB-BrxF-iSTAND->DUF6079*->?->N6-METHYLASE->PglZ->SIGMA+OB-Fold+AAA-ATPase+HKD-DNase-BrxL-><-REase+RNase<-REase<-?<-REase+SF1-SF2-Helicase<-Schlafen+HATPase+wHTH | 1184 | Bacteria>Pseudomonadota>Gammaproteobacteria>Legionellales>Legionellaceae>Legionella | GCF_003004175.1 | Legionella_pneumophila | CA922_RS00900 | 198017..201571 | 446 | BREX-system-P-loop-protein-BrxC-[Legionella-pneumophila] |
| WP_061721013.1 | Type1.BREX | REase->REase+RNase-><-SIGMA+OB-Fold+AAA-ATPase+HKD-DNase-BrxL<-PglZ<-N6-METHYLASE<-?<-DUF6079*<-BrxB-BrxF-iSTAND<-BrxA-DNA-binding<-XIS-HTH<-HTH+LexA-protease\|\|?-><-AAA-ATPase<-HIN-HTH | 1184 | Bacteria>Pseudomonadota>Gammaproteobacteria>Legionellales>Legionellaceae>Legionella | GCF_900054435.1 | Legionella_pneumophila | A2E62_RS14865 | 12190..15744 | 446 | BREX-system-P-loop-protein-BrxC-[Legionella-pneumophila] |

| Protein-ID | System | Gene-Neighbourhoods | Length | Taxonomy | Assembly-ID | Strain | Locus-Tag | Genomic-coordinates | TaxID | Fasta-define |
|---|---|---|---|---|---|---|---|---|---|---|
| WP_061637858.1 | Type1.BREX | [illegible] | 1184 | Bacteria>Pseudomonadota>Gammaproteobacteria>Legionellales>Legionellaceae>Legionella | GCF_900061645.1 | Legionella_pneumophila | A1L82_RS06820 | 180647..184201 | 446 | BREX-system-P-loop-protein-BrxC-[Legionella-pneumophila] |
| WP_027222687.1 | Type1.BREX | [illegible] | 1184 | Bacteria>Pseudomonadota>Gammaproteobacteria>Legionellales>Legionellaceae>Legionella | GCF_900639945.1 | Legionella_pneumophila | E1076_RS05405 | 246005..249559 | 446 | BREX-system-P-loop-protein-BrxC-[Legionella-pneumophila] |
| WP_165474921.1 | Type1.BREX | [illegible] | 1184 | Bacteria>Pseudomonadota>Gammaproteobacteria>Legionellales>Legionellaceae>Legionella | GCF_900640175.1 | Legionella_sp._W10-070 | E1065_RS06915 | 46772..50326 | 1117709 | BREX-system-P-loop-protein-BrxC-[Legionella-sp.-W10-070] |
| WP_159653266.1 | Type1.BREX | [illegible] | 1184 | Bacteria>Pseudomonadota>Gammaproteobacteria>Methylococcales>Methylococcaceae>Methylicorpusculum | GCF_009828925.2 | Methylicorpusculum_oleiharenae | GO003_RS20515 | 4601128..4604682 | 1338687 | BREX-system-P-loop-protein-BrxC-[Methylicorpusculum-oleiharenae] |
| WP_256360610.1 | Type1.BREX | [illegible] | 1145 | Bacteria>Pseudomonadota>Gammaproteobacteria>Methylococcales>Methylococcaceae>Methylomonas | GCF_001312005.1 | Methylomonas_koyamae | JCM16701_RS06940 | 49293..52730 | 702114 | BREX-system-P-loop-protein-BrxC-[Methylomonas-koyamae] |
| WP_096876853.1 | Type1.BREX | [illegible] | 1180 | Bacteria>Pseudomonadota>Gammaproteobacteria>Methylococcales>Methylococcaceae>Methylomonas | GCF_002441955.1 | Methylomonas_koyamae | MKLM6_RS22285 | 58076..61618 | 702114 | BREX-system-P-loop-protein-BrxC-[Methylomonas-koyamae] |
| WP_221054675.1 | Type1.BREX | [illegible] | 1176 | Bacteria>Pseudomonadota>Gammaproteobacteria>Methylococcales>Methylococcaceae>Methylomonas | GCF_019669905.1 | Methylomonas_koyamae | MKFW12EY_RS21895 | 239254..1338 | 702114 | BREX-system-P-loop-protein-BrxC-[Methylomonas-koyamae] |
| WP_004984466.1 | Type1.BREX | [illegible] | 1190 | Bacteria>Pseudomonadota>Gammaproteobacteria>Moraxellales>Moraxellaceae>Acinetobacter | GCF_000368805.1 | Acinetobacter_johnsonii | F946_RS15710 | 148915..152487 | 40214 | BREX-system-P-loop-protein-BrxC-[Acinetobacter-johnsonii] |
| WP_005162499.1 | Type6.BREX | [illegible] | 1063 | Bacteria>Pseudomonadota>Gammaproteobacteria>Moraxellales>Moraxellaceae>Acinetobacter | GCF_000369425.1 | Acinetobacter_sp._NIPH_284 | F908_RS12580 | 1400419..1403610 | 1217704 | BREX-system-P-loop-protein-BrxC-[Acinetobacter-sp.-NIPH-284] |
| WP_131214141.1 | Type1.BREX | [illegible] | 1190 | Bacteria>Pseudomonadota>Gammaproteobacteria>Moraxellales>Moraxellaceae>Acinetobacter | GCF_004331285.1 | Acinetobacter_sp._ANC_3781 | E0H89_RS01980 | 72363..75935 | 2529835 | BREX-system-P-loop-protein-BrxC-[Acinetobacter-sp.-ANC-3781] |
| WP_140963539.1 | Type6.BREX | [illegible] | 1214 | Bacteria>Pseudomonadota>Gammaproteobacteria>Moraxellales>Moraxellaceae>Acinetobacter | GCF_006494575.1 | Acinetobacter_baumannii | FJU84_RS09090 | 25284..28928 | 470 | BREX-system-P-loop-protein-BrxC-[Acinetobacter-baumannii] |
| WP_151834043.1 | Type1.BREX | [illegible] | 1190 | Bacteria>Pseudomonadota>Gammaproteobacteria>Moraxellales>Moraxellaceae>Acinetobacter | GCF_009012055.1 | Acinetobacter_ursingii | GBX89_RS12550 | 5375..8947 | 108980 | BREX-system-P-loop-protein-BrxC-[Acinetobacter-ursingii] |
| WP_180108075.1 | Type1.BREX | [illegible] | 1190 | Bacteria>Pseudomonadota>Gammaproteobacteria>Moraxellales>Moraxellaceae>Acinetobacter | GCF_013419415.1 | Acinetobacter_sp._YH12085 | FTU28_RS13915 | 49013..52585 | 2601077 | BREX-system-P-loop-protein-BrxC-[Acinetobacter-sp.-YH12085] |
| WP_032068282.1 | Type6.BREX | [illegible] | 1214 | Bacteria>Pseudomonadota>Gammaproteobacteria>Moraxellales>Moraxellaceae>Acinetobacter | GCF_014559915.1 | Acinetobacter_baumannii | HPJ29_RS06280 | 177170..180814 | 470 | BREX-system-P-loop-protein-BrxC-[Acinetobacter-baumannii] |
| WP_216936951.1 | Type1.BREX | [illegible] | 1190 | Bacteria>Pseudomonadota>Gammaproteobacteria>Moraxellales>Moraxellaceae>Acinetobacter | GCF_019038255.1 | unclassified_Acinetobacter | KUG40_RS10660 | 10626..14198 | 196816 | MULTISPECIES:-BREX-system-P-loop-protein-BrxC-[unclassified-Acinetobacter] |
| WP_264453044.1 | Type6.BREX | [illegible] | 1214 | Bacteria>Pseudomonadota>Gammaproteobacteria>Moraxellales>Moraxellaceae>Acinetobacter | GCF_025938795.1 | Acinetobacter_baumannii | OLN40_RS03910 | 58425..62069 | 470 | BREX-system-P-loop-protein-BrxC-[Acinetobacter-baumannii] |
| WP_134244656.1 | Type6.BREX | [illegible] | 1217 | Bacteria>Pseudomonadota>Gammaproteobacteria>Moraxellales>Moraxellaceae>Alkanindiges | GCF_004377485.1 | Alkanindiges_illinoisensis | E2B99_RS09020 | 10631..14284 | 197183 | BREX-system-P-loop-protein-BrxC-[Alkanindiges-illinoisensis] |
| WP_151794804.1 | Type6.BREX | [illegible] | 1214 | Bacteria>Pseudomonadota>Gammaproteobacteria>Moraxellales>Moraxellaceae | GCF_008997035.1 | Acinetobacter | GBP95_RS04055 | 337470..341114 | 469 | MULTISPECIES:-BREX-system-P-loop-protein-BrxC-[Acinetobacter] |
| WP_102077888.1 | Type6.BREX | [illegible] | 1219 | Bacteria>Pseudomonadota>Gammaproteobacteria>Moraxellales>Moraxellaceae>Psychrobacter | GCF_001742065.1 | unclassified_Psychrobacter | CXF69_RS01065 | 108247..111906 | 196806 | MULTISPECIES:-BREX-system-P-loop-protein-BrxC-[unclassified-Psychrobacter] |
| WP_218690652.1 | Type6.BREX | [illegible] | 1213 | Bacteria>Pseudomonadota>Gammaproteobacteria>Moraxellales>Moraxellaceae>Psychrobacter | GCF_019279065.1 | Psychrobacter_sp._BF1 | KZA12_RS14005 | 8359..12000 | 2821147 | BREX-system-P-loop-protein-BrxC-[Psychrobacter-sp.-BF1] |
| WP_238051284.1 | Type6.BREX | [illegible] | 1213 | Bacteria>Pseudomonadota>Gammaproteobacteria>Moraxellales>Moraxellaceae>Psychrobacter | GCF_022129405.1 | Psychrobacter_sp._Ps2 | I3254_RS03805 | 13291..16932 | 2790956 | BREX-system-P-loop-protein-BrxC-[Psychrobacter-sp.-Ps2] |
| WP_201561160.1 | Type6.BREX | [illegible] | 1219 | Bacteria>Pseudomonadota>Gammaproteobacteria>Moraxellales>Moraxellaceae>Psychrobacter | GCF_904846215.1 | Psychrobacter_glacincola | JMW90_RS00425 | 116435..120094 | 56810 | BREX-system-P-loop-protein-BrxC-[Psychrobacter-glacincola] |
| TQC95078.1 | Type1.BREX | [illegible] | 1192 | Bacteria>Pseudomonadota>Gammaproteobacteria>Moraxellales>Moraxellaceae>unclassified_Moraxellaceae | GCA_006546595.1 | Moraxellaceae_bacterium_AER2_44_116 | FK216_14950 | 12152..15730 | 2591038 | BREX-system-P-loop-protein-BrxC-[Moraxellaceae-bacterium-AER2_44_116] |
| MBC7753680.1 | Type1.BREX | [illegible] | 1163 | Bacteria>Pseudomonadota>Gammaproteobacteria>Moraxellales>Moraxellaceae>unclassified_Moraxellaceae | GCA_014378775.1 | Moraxellaceae_bacterium | H7Z71_05545 | 8991..12482 | 1889775 | BREX-system-P-loop-protein-BrxC-[Moraxellaceae-bacterium] |
| MCP5177609.1 | Type1.BREX | [illegible] | 1192 | Bacteria>Pseudomonadota>Gammaproteobacteria>Moraxellales>Moraxellaceae>unclassified_Moraxellaceae | GCA_024234655.1 | Moraxellaceae_bacterium | H6997_08670 | 33703..37281 | 1889775 | BREX-system-P-loop-protein-BrxC-[Moraxellaceae-bacterium] |
| WP_161382580.1 | Type1.BREX | [illegible] | 1193 | Bacteria>Pseudomonadota>Gammaproteobacteria>Oceanospirillales>Alcanivoracaceae>Alcanivorax | - | Alcanivorax_sp._ZXX171 | GN155_RS00640 | 132435..136016 | 2695858 | BREX-system-P-loop-protein-BrxC-[Alcanivorax-sp.-ZXX171] |
| KAF0808603.1 | Type1.BREX | [illegible] | 1165 | Bacteria>Pseudomonadota>Gammaproteobacteria>Oceanospirillales>Alcanivoracaceae>Alcanivorax | GCA_009846755.1 | Alcanivorax_sp._S71-1-4 | A167_02568 | 1..3498 | 1177159 | ATPase,-partial-[Alcanivorax-sp.-S71-1-4] |
| MCH2557814.1 | Type1.BREX | [illegible] | 1193 | Bacteria>Pseudomonadota>Gammaproteobacteria>Oceanospirillales>Alcanivoracaceae>Alcanivorax | GCA_022449455.1 | Alcanivorax_sp. | MK005_11000 | 47715..51296 | 1872427 | BREX-system-P-loop-protein-BrxC-[Alcanivorax-sp.] |
| WP_153499070.1 | Type6.BREX | [illegible] | 1214 | Bacteria>Pseudomonadota>Gammaproteobacteria>Oceanospirillales>Alcanivoracaceae>Alcanivorax | GCF_009601165.1 | Alcanivorax_sediminis | GFN93_RS03795 | 861896..865540 | 2663008 | BREX-system-P-loop-protein-BrxC-[Alcanivorax-sediminis] |
| WP_159558342.1 | Type6.BREX | [illegible] | 1213 | Bacteria>Pseudomonadota>Gammaproteobacteria>Oceanospirillales>Alcanivoracaceae>Alcanivorax | GCF_009846755.1 | Alcanivorax_sp._S71-1-4 | A167_RS05685 | 90732..94373 | 1177159 | BREX-system-P-loop-protein-BrxC-[Alcanivorax-sp.-S71-1-4] |
| WP_159559845.1 | Type1.BREX | [illegible] | 1165 | Bacteria>Pseudomonadota>Gammaproteobacteria>Oceanospirillales>Alcanivoracaceae>Alcanivorax | GCF_009846755.1 | Alcanivorax_sp._S71-1-4 | A167_RS13005 | 1..3498 | 1177159 | BREX-system-P-loop-protein-BrxC,-partial-[Alcanivorax-sp.-S71-1-4] |
| WP_103916700.1 | Type1.BREX | [illegible] | 1193 | Bacteria>Pseudomonadota>Gammaproteobacteria>Oceanospirillales>Alcanivoracaceae>Alcanivorax | GCF_900107995.1 | Alcanivorax_sp._DSM_26293 | C4S63_RS03510 | 780383..783964 | 1798238 | BREX-system-P-loop-protein-BrxC-[Alcanivorax-sp.-DSM-26293] |
| WP_204429962.1 | Type1.BREX | [illegible] | 1193 | Bacteria>Pseudomonadota>Gammaproteobacteria>Oceanospirillales>Alcanivoracaceae>Alloalcanivorax | GCF_016906305.1 | Alloalcanivorax_marinus | IM760_RS12910 | 16365..19946 | 1177169 | BREX-system-P-loop-protein-BrxC-[Alloalcanivorax-marinus] |
| WP_228232948.1 | Type1.BREX | [illegible] | 1193 | Bacteria>Pseudomonadota>Gammaproteobacteria>Oceanospirillales>Alcanivoracaceae>Alloalcanivorax | GCF_020782135.1 | Alloalcanivorax_marinus | LL252_RS03465 | 287759..291340 | 1177169 | BREX-system-P-loop-protein-BrxC-[Alloalcanivorax-marinus] |
| WP_008734404.1 | Type1.BREX | [illegible] | 1192 | Bacteria>Pseudomonadota>Gammaproteobacteria>Oceanospirillales>Alcanivoracaceae>Isoalcanivorax | GCF_000299335.2 | Isoalcanivorax_pacificus | S7S_RS18615 | 4158589..4162167 | 1306787 | BREX-system-P-loop-protein-BrxC-[Isoalcanivorax-pacificus] |
| RLP52821.1 | Type1.BREX | [illegible] | 1193 | Bacteria>Pseudomonadota>Gammaproteobacteria>Oceanospirillales>Alcanivoracaceae>Ketobacter | GCA_003667185.1 | Ketobacter_sp. | D6160_18810 | 15503..19084 | 2083498 | MAG:-BREX-system-P-loop-protein-BrxC-[Ketobacter-sp.] |
| AZZ95337.1 | Type1.BREX | [illegible] | 1183 | Bacteria>Pseudomonadota>Gammaproteobacteria>Oceanospirillales>Hahellaceae>Hahella | GCA_004014695.1 | Hahella_sp._KA22 | ENC22_30655 | 6892256..6895807 | 1628392 | BREX-system-P-loop-protein-BrxC-[Hahella-sp.-KA22] |
| WP_164887423.1 | Type1.BREX | [illegible] | 1182 | Bacteria>Pseudomonadota>Gammaproteobacteria>Oceanospirillales>Hahellaceae>Hahella | GCF_004014695.1 | Hahella_sp._KA22 | EUZ85_RS02365 | 526761..530309 | 1628392 | BREX-system-P-loop-protein-BrxC-[Hahella-sp.-KA22] |
| MBR2514422.1 | Type6.BREX | [illegible] | 1224 | Bacteria>Pseudomonadota>Gammaproteobacteria>Oceanospirillales>Halomonadaceae>Halomonas | GCA_017515565.1 | Halomonas_sp. | IKE45_10465 | 23089..26763 | 1486246 | BREX-system-P-loop-protein-BrxC-[Halomonas-sp.] |
| WP_133481548.1 | Type1.BREX | [illegible] | 1193 | Bacteria>Pseudomonadota>Gammaproteobacteria>Oceanospirillales>Halomonadaceae>Halomonas | GCF_004361885.1 | Halomonas_ventosae | DF068_RS01985 | 17598..21179 | 229007 | BREX-system-P-loop-protein-BrxC-[Halomonas-ventosae] |
| WP_159343441.1 | Type1.BREX | [illegible] | 1193 | Bacteria>Pseudomonadota>Gammaproteobacteria>Oceanospirillales>Halomonadaceae>Halomonas | GCF_009846525.1 | Halomonas_meridiana | CTT34_RS16815 | 3563093..3566674 | 29570 | BREX-system-P-loop-protein-BrxC-[Halomonas-meridiana] |
| WP_161432258.1 | Type6.BREX | [illegible] | 1223 | Bacteria>Pseudomonadota>Gammaproteobacteria>Oceanospirillales>Halomonadaceae>Halomonas | GCF_009902005.1 | Halomonas_alimentaria | GRB96_RS11410 | 2397586..2401257 | 147248 | BREX-system-P-loop-protein-BrxC-[Halomonas-alimentaria] |
| WP_169956446.1 | Type1.BREX | [illegible] | 1193 | Bacteria>Pseudomonadota>Gammaproteobacteria>Oceanospirillales>Halomonadaceae>Halomonas | GCF_013000985.1 | Halomonas_sp._PGE1 | HIR79_RS02110 | 421842..425423 | 2730360 | BREX-system-P-loop-protein-BrxC-[Halomonas-sp.-PGE1] |
| WP_019020105.1 | Type1.BREX | [illegible] | 1193 | Bacteria>Pseudomonadota>Gammaproteobacteria>Oceanospirillales>Halomonadaceae>Halomonas | GCF_014651575.1 | Halomonas_lutea | IE067_RS16235 | 61170..64751 | 453962 | BREX-system-P-loop-protein-BrxC-[Halomonas-lutea] |
| WP_103969332.1 | Type1.BREX | [illegible] | 1193 | Bacteria>Pseudomonadota>Gammaproteobacteria>Oceanospirillales>Halomonadaceae>Halomonas | GCF_900108215.1 | Halomonas_desiderata | C4S71_RS12465 | 87857..91438 | 52021 | BREX-system-P-loop-protein-BrxC-[Halomonas-desiderata] |
| WP_159177832.1 | Type1.BREX | [illegible] | 1193 | Bacteria>Pseudomonadota>Gammaproteobacteria>Oceanospirillales>Halomonadaceae>Halomonas | GCF_902506335.1 | unclassified_Halomonas | JMX33_RS24540 | 5309589..5313170 | 2609666 | MULTISPECIES:-BREX-system-P-loop-protein-BrxC-[unclassified-Halomonas] |
| WP_110669866.1 | Type6.BREX | [illegible] | 1225 | Bacteria>Pseudomonadota>Gammaproteobacteria>Oceanospirillales>Halomonadaceae>Salinicola | GCF_003206155.1 | Salinicola_halophilus | DOM33_RS13680 | 401271..404948 | 184065 | BREX-system-P-loop-protein-BrxC-[Salinicola-halophilus] |
| WP_027854332.1 | Type1.BREX | [illegible] | 1193 | Bacteria>Pseudomonadota>Gammaproteobacteria>Oceanospirillales>Oceanospirillaceae>Marinobacterium | GCF_000428985.1 | Marinobacterium_litorale | G403_RS0112740 | 74962..78543 | 404770 | BREX-system-P-loop-protein-BrxC-[Marinobacterium-litorale] |
| WP_254579406.1 | Type6.BREX | [illegible] | 1212 | Bacteria>Pseudomonadota>Gammaproteobacteria>Oceanospirillales>Oceanospirillaceae>Marinobacterium | GCF_024267675.1 | Marinobacterium_sedimentorum | MRS74_RS06180 | 150386..154024 | 2927804 | BREX-system-P-loop-protein-BrxC-[Marinobacterium-sedimentorum] |
| AWY02455.1 | Type1.BREX | [illegible] | 1197 | Bacteria>Pseudomonadota>Gammaproteobacteria>Oceanospirillales>Oceanospirillaceae>Marinomonas | GCA_003285205.1 | Marinomonas_primoryensis | A8139_14855 | 3171018..3174611 | 178399 | hypothetical-protein-A8139_14855-[Marinomonas-primoryensis] |
| WP_239495510.1 | Type1.BREX | [illegible] | 1199 | Bacteria>Pseudomonadota>Gammaproteobacteria>Oceanospirillales>Oceanospirillaceae>Marinomonas | GCF_003285205.1 | Marinomonas_primoryensis | A8139_RS14980 | 3171018..3174617 | 178399 | BREX-system-P-loop-protein-BrxC-[Marinomonas-primoryensis] |
| WP_205113926.1 | Type1.BREX | [illegible] | 1191 | Bacteria>Pseudomonadota>Gammaproteobacteria>Oceanospirillales>Oceanospirillaceae>Marinomonas | GCF_016906065.1 | Marinomonas_foliarum | JSY38_RS15135 | 3246777..3250352 | 491950 | BREX-system-P-loop-protein-BrxC-[Marinomonas-foliarum] |
| WP_138986811.1 | Type1.BREX | [illegible] | 1190 | Bacteria>Pseudomonadota>Gammaproteobacteria>Oceanospirillales>Oceanospirillaceae>Neptunomonas | GCF_008630635.1 | Neptunomonas_concharum | F0U83_RS12940 | 2741645..2745217 | 1031538 | BREX-system-P-loop-protein-BrxC-[Neptunomonas-concharum] |
| QEW05875.1 | Type6.BREX | [illegible] | 1212 | Bacteria>Pseudomonadota>Gammaproteobacteria>Oceanospirillales>Oceanospirillaceae>Nitrincola | GCA_008727795.1 | Nitrincola_iocasae | F5I99_04885 | 1032629..1036267 | 2614693 | BREX-system-P-loop-protein-BrxC-[Nitrincola-iocasae] |
| WP_225307559.1 | Type6.BREX | [illegible] | 1214 | Bacteria>Pseudomonadota>Gammaproteobacteria>Oceanospirillales>Oceanospirillaceae>Nitrincola | GCF_008727795.1 | Nitrincola_iocasae | F5I99_RS04885 | 1032623..1036267 | 2614693 | BREX-system-P-loop-protein-BrxC-[Nitrincola-iocasae] |
| MAD46040.1 | Type1.BREX | [illegible] | 1189 | Bacteria>Pseudomonadota>Gammaproteobacteria>Oceanospirillales>Oceanospirillaceae>unclassified_Oceanospirillaceae | GCA_002685055.1 | Oceanospirillaceae_bacterium | CMH98_13650 | 9512..13081 | 1899355 | hypothetical-protein-[Oceanospirillaceae-bacterium] |
| MAR01289.1 | Type1.BREX | [illegible] | 1185 | Bacteria>Pseudomonadota>Gammaproteobacteria>Oceanospirillales>Oceanospirillaceae>unclassified_Oceanospirillaceae | GCA_002701445.1 | Oceanospirillaceae_bacterium | CMI00_12230 | 13782..17339 | 1899355 | hypothetical-protein-[Oceanospirillaceae-bacterium] |
| WP_058011517.1 | Type6.BREX | [illegible] | 1213 | Bacteria>Pseudomonadota>Gammaproteobacteria>Pseudomonadales | GCF_901482535.1 | Pseudomonadaceae | FQU16_RS01885 | 279271..282912 | 135621 | MULTISPECIES:-BREX-system-P-loop-protein-BrxC-[Pseudomonadaceae] |
| MBE95557.1 | Type1.BREX | [illegible] | 1193 | Bacteria>Pseudomonadota>Gammaproteobacteria>Pseudomonadales>Marinobacteraceae>Marinobacter | GCA_002714765.1 | Marinobacter_sp. | CMG89_11180 | 696..4277 | 50741 | hypothetical-protein-[Marinobacter-sp.] |
| WP_011784101.1 | Type1.BREX | [illegible] | 1193 | Bacteria>Pseudomonadota>Gammaproteobacteria>Pseudomonadales>Marinobacteraceae>Marinobacter | GCF_000015365.1 | Marinobacter_nauticus | MAQU_RS02805 | 628948..632529 | 2743 | BREX-system-P-loop-protein-BrxC-[Marinobacter-nauticus] |
| WP_007349691.1 | Type5.BREX | [illegible] | 1194 | Bacteria>Pseudomonadota>Gammaproteobacteria>Pseudomonadales>Marinobacteraceae>Marinobacter | GCF_000169375.1 | Marinobacter_sp._ELB17 | MELB17_RS08075 | 35418..39002 | 270374 | BREX-system-P-loop-protein-BrxC-[Marinobacter-sp.-ELB17] |
| WP_018404435.1 | Type1.BREX | [illegible] | 1194 | Bacteria>Pseudomonadota>Gammaproteobacteria>Pseudomonadales>Marinobacteraceae>Marinobacter | GCF_000372805.1 | Marinobacter_gelidimuriae | A3OI_RS0111615 | 2203..5787 | 2739064 | BREX-system-P-loop-protein-BrxC-[Marinobacter-gelidimuriae] |
| WP_022992169.1 | Type1.BREX | [illegible] | 1193 | Bacteria>Pseudomonadota>Gammaproteobacteria>Pseudomonadales>Marinobacteraceae>Marinobacter | GCF_000475355.1 | Marinobacter_sp._C1S70 | Q669_RS10410 | 149187..152768 | 1396859 | BREX-system-P-loop-protein-BrxC-[Marinobacter-sp.-C1S70] |
| WP_023008352.1 | Type1.BREX | [illegible] | 1193 | Bacteria>Pseudomonadota>Gammaproteobacteria>Pseudomonadales>Marinobacteraceae>Marinobacter | GCF_000475375.1 | Marinobacter_sp._EVN1 | Q674_RS26680 | 52773..56354 | 1397532 | BREX-system-P-loop-protein-BrxC-[Marinobacter-sp.-EVN1] |
| WP_104416052.1 | Type1.BREX | [illegible] | 1193 | Bacteria>Pseudomonadota>Gammaproteobacteria>Pseudomonadales>Marinobacteraceae>Marinobacter | GCF_002934485.1 | Marinobacter_persicus | BY455_RS08980 | 65339..68920 | 930118 | BREX-system-P-loop-protein-BrxC-[Marinobacter-persicus] |
| WP_113863603.1 | Type1.BREX | [illegible] | 1193 | Bacteria>Pseudomonadota>Gammaproteobacteria>Pseudomonadales>Marinobacteraceae>Marinobacter | GCF_003315345.1 | Marinobacter_pelagius | DET50_RS14950 | 40642..44223 | 379482 | BREX-system-P-loop-protein-BrxC-[Marinobacter-pelagius] |
| WP_141068072.1 | Type1.BREX | [illegible] | 1193 | Bacteria>Pseudomonadota>Gammaproteobacteria>Pseudomonadales>Marinobacteraceae>Marinobacter | GCF_006516615.1 | Marinobacter_nauticus | FH712_RS17655 | 33566..37147 | 2743 | BREX-system-P-loop-protein-BrxC-[Marinobacter-nauticus] |
| WP_247116152.1 | Type1.BREX | [illegible] | 1189 | Bacteria>Pseudomonadota>Gammaproteobacteria>Pseudomonadales>Marinobacteraceae>Marinobacter | GCF_023016045.1 | Marinobacter_sp._S6332 | MWU62_RS17920 | 19915..23484 | 2926403 | BREX-system-P-loop-protein-BrxC-[Marinobacter-sp.-S6332] |
| WP_092033189.1 | Type1.BREX | [illegible] | 1193 | Bacteria>Pseudomonadota>Gammaproteobacteria>Pseudomonadales>Marinobacteraceae>Marinobacter | GCF_900112835.1 | Marinobacter_sp._DSM_26671 | BM035_RS16335 | 75345..78926 | 1761793 | BREX-system-P-loop-protein-BrxC-[Marinobacter-sp.-DSM-26671] |
| BBJ05051.1 | Type1.BREX | [illegible] | 1193 | Bacteria>Pseudomonadota>Gammaproteobacteria>Pseudomonadales>Marinobacteraceae>Marinobacter | - | Marinobacter_nauticus | YBY_29000 | 3185607..3189188 | 2743 | hypothetical-protein-YBY_29000-[Marinobacter-nauticus] |
| WP_116918937.1 | Type1.BREX | [illegible] | 1193 | Bacteria>Pseudomonadota>Gammaproteobacteria>Pseudomonadales>Marinobacteraceae>Tamilnaduibacter | GCF_003096655.1 | Tamilnaduibacter_salinus | C8D92_104115 | 123558..127139 | 1484056 | BREX-system-P-loop-protein-BrxC-[Tamilnaduibacter-salinus] |
| WP_022966741.1 | Type1.BREX | [illegible] | 1184 | Bacteria>Pseudomonadota>Gammaproteobacteria>Pseudomonadales>Pseudomonadaceae>Denitrificimonas | GCF_000421765.1 | Denitrificimonas_caeni | H591_RS0109630 | 17243..20797 | 521720 | BREX-system-P-loop-protein-BrxC-[Denitrificimonas-caeni] |
| WP_269819144.1 | Type6.BREX | [illegible] | 1219 | Bacteria>Pseudomonadota>Gammaproteobacteria>Pseudomonadales>Pseudomonadaceae>Denitrificimonas | GCF_027498055.1 | Denitrificimonas_caeni | O6P33_RS05150 | 1067699..1071358 | 521720 | BREX-system-P-loop-protein-BrxC-[Denitrificimonas-caeni] |
| WP_019470415.1 | Type6.BREX | [illegible] | 1212 | Bacteria>Pseudomonadota>Gammaproteobacteria>Pseudomonadales>Pseudomonadaceae | GCF_000292285.1 | Pseudomonas | N805_RS25690 | 5809538..5813176 | 286 | MULTISPECIES:-BREX-system-P-loop-protein-BrxC-[Pseudomonas] |
| WP_033898350.1 | Type1.BREX | [illegible] | 1185 | Bacteria>Pseudomonadota>Gammaproteobacteria>Pseudomonadales>Pseudomonadaceae | GCF_000510895.2 | Pseudomonas | GIW75_RS00865 | 8119..11676 | 286 | MULTISPECIES:-BREX-system-P-loop-protein-BrxC-[Pseudomonas] |
| WP_096426782.1 | Type1.BREX | [illegible] | 1185 | Bacteria>Pseudomonadota>Gammaproteobacteria>Pseudomonadales>Pseudomonadaceae | GCF_002356095.1 | Pseudomonas | J3A71_RS03870 | 842777..846334 | 286 | MULTISPECIES:-BREX-system-P-loop-protein-BrxC-[Pseudomonas] |
| WP_105523318.1 | Type1.BREX | [illegible] | 1185 | Bacteria>Pseudomonadota>Gammaproteobacteria>Pseudomonadales>Pseudomonadaceae | GCF_002975295.1 | Pseudomonas | CQ048_RS15780 | 97047..100604 | 286 | MULTISPECIES:-BREX-system-P-loop-protein-BrxC-[Pseudomonas] |
| WP_170037970.1 | Type1.BREX | [illegible] | 1185 | Bacteria>Pseudomonadota>Gammaproteobacteria>Pseudomonadales>Pseudomonadaceae | GCF_012985445.1 | Pseudomonas | HBO05_RS12455 | 196790..200347 | 286 | MULTISPECIES:-BREX-system-P-loop-protein-BrxC-[Pseudomonas] |
| WP_218430053.1 | Type1.BREX | [illegible] | 1189 | Bacteria>Pseudomonadota>Gammaproteobacteria>Pseudomonadales>Pseudomonadaceae | GCF_019219185.1 | Pseudomonas | KW849_RS14820 | 325397..328966 | 286 | MULTISPECIES:-BREX-system-P-loop-protein-BrxC-[Pseudomonas] |
| WP_033986129.1 | Type6.BREX | [illegible] | 1212 | Bacteria>Pseudomonadota>Gammaproteobacteria>Pseudomonadales>Pseudomonadaceae | GCF_020180035.1 | Pseudomonas | AO894_RS16050 | 13512..17150 | 286 | MULTISPECIES:-BREX-system-P-loop-protein-BrxC-[Pseudomonas] |
| WP_060765715.1 | Type1.BREX | [illegible] | 1186 | Bacteria>Pseudomonadota>Gammaproteobacteria>Pseudomonadales>Pseudomonadaceae | GCF_029959765.1 | Pseudomonas | PFL603g_RS03160 | 383988..387548 | 286 | MULTISPECIES:-BREX-system-P-loop-protein-BrxC-[Pseudomonas] |
| WP_104739193.1 | Type1.BREX | [illegible] | 1180 | Bacteria>Pseudomonadota>Gammaproteobacteria>Pseudomonadales>Pseudomonadaceae>Halopseudomonas | GCF_014641295.1 | Halopseudomonas_oceani | IEX06_RS08390 | 366595..370137 | 1708783 | BREX-system-P-loop-protein-BrxC-[Halopseudomonas-oceani] |
| PPB00131.1 | Type1.BREX | [illegible] | 1194 | Bacteria>Pseudomonadota>Gammaproteobacteria>Pseudomonadales>Pseudomonadaceae>Pseudomonas | GCA_002927135.1 | Pseudomonas_aeruginosa | HV84_25295 | 2273516..2277100 | 287 | BREX-system-P-loop-protein-BrxC-[Pseudomonas-aeruginosa] |
| RPD95236.1 | Type1.BREX | [illegible] | 1191 | Bacteria>Pseudomonadota>Gammaproteobacteria>Pseudomonadales>Pseudomonadaceae>Pseudomonas | GCA_003813925.1 | Pseudomonas_monteilii | EGN69_04400 | 958521..962096 | 76759 | BREX-system-P-loop-protein-BrxC-[Pseudomonas-monteilii] |
| MBG3983980.1 | Type6.BREX | [illegible] | 1212 | Bacteria>Pseudomonadota>Gammaproteobacteria>Pseudomonadales>Pseudomonadaceae>Pseudomonas | GCA_015733145.1 | Pseudomonas_aeruginosa | I4919_20885 | 86674..90312 | 287 | BREX-system-P-loop-protein-BrxC-[Pseudomonas-aeruginosa] |
| MBG4530231.1 | Type1.BREX | [illegible] | 1181 | Bacteria>Pseudomonadota>Gammaproteobacteria>Pseudomonadales>Pseudomonadaceae>Pseudomonas | GCA_015734885.1 | Pseudomonas_aeruginosa | I4952_20115 | 27108..30653 | 287 | BREX-system-P-loop-protein-BrxC-[Pseudomonas-aeruginosa] |
| MBG4863325.1 | Type1.BREX | [illegible] | 1184 | Bacteria>Pseudomonadota>Gammaproteobacteria>Pseudomonadales>Pseudomonadaceae>Pseudomonas | GCA_015735985.1 | Pseudomonas_aeruginosa | GHU23_28970 | 1726..5280 | 287 | BREX-system-P-loop-protein-BrxC-[Pseudomonas-aeruginosa] |
| MBG6730505.1 | Type1.BREX | [illegible] | 1184 | Bacteria>Pseudomonadota>Gammaproteobacteria>Pseudomonadales>Pseudomonadaceae>Pseudomonas | GCA_015764285.1 | Pseudomonas_aeruginosa | I5I92_11320 | 4371..7925 | 287 | BREX-system-P-loop-protein-BrxC-[Pseudomonas-aeruginosa] |
| MBH3980671.1 | Type6.BREX | [illegible] | 1213 | Bacteria>Pseudomonadota>Gammaproteobacteria>Pseudomonadales>Pseudomonadaceae>Pseudomonas | GCA_016011075.1 | Pseudomonas_aeruginosa | I5Q70_02570 | 503331..506972 | 287 | BREX-system-P-loop-protein-BrxC-[Pseudomonas-aeruginosa] |
| MBL7230767.1 | Type6.BREX | [illegible] | 1214 | Bacteria>Pseudomonadota>Gammaproteobacteria>Pseudomonadales>Pseudomonadaceae>Pseudomonas | GCA_016785205.1 | Pseudomonas_sp. | JJK56_22660 | 12252..15896 | 306 | BREX-system-P-loop-protein-BrxC-[Pseudomonas-sp.] |
| MBN0299744.1 | Type6.BREX | [illegible] | 1213 | Bacteria>Pseudomonadota>Gammaproteobacteria>Pseudomonadales>Pseudomonadaceae>Pseudomonas | GCA_016922145.1 | Pseudomonas_aeruginosa | JTM78_25120 | 125675..129316 | 287 | BREX-system-P-loop-protein-BrxC-[Pseudomonas-aeruginosa] |
| MBN0536373.1 | Type6.BREX | [illegible] | 1206 | Bacteria>Pseudomonadota>Gammaproteobacteria>Pseudomonadales>Pseudomonadaceae>Pseudomonas | GCA_016922535.1 | Pseudomonas_aeruginosa | JTM59_25840 | 125679..129299 | 287 | BREX-system-P-loop-protein-BrxC-[Pseudomonas-aeruginosa] |
| MBV5724753.1 | Type1.BREX | [illegible] | 1184 | Bacteria>Pseudomonadota>Gammaproteobacteria>Pseudomonadales>Pseudomonadaceae>Pseudomonas | GCA_019166825.1 | Pseudomonas_aeruginosa | KUT37_26485 | 44544..48098 | 287 | BREX-system-P-loop-protein-BrxC-[Pseudomonas-aeruginosa] |
| HBN8119635.1 | Type6.BREX | [illegible] | 1213 | Bacteria>Pseudomonadota>Gammaproteobacteria>Pseudomonadales>Pseudomonadaceae>Pseudomonas | GCA_021586615.1 | Pseudomonas_aeruginosa | L3655_000128 | 111220..114861 | 287 | BREX-system-P-loop-protein-BrxC-[Pseudomonas-aeruginosa] |
| HBO2945375.1 | Type1.BREX | [illegible] | 1184 | Bacteria>Pseudomonadota>Gammaproteobacteria>Pseudomonadales>Pseudomonadaceae>Pseudomonas | GCA_021651875.1 | Pseudomonas_aeruginosa | L4R51_003866 | 17556..21110 | 287 | BREX-system-P-loop-protein-BrxC-[Pseudomonas-aeruginosa] |
| HBO3309314.1 | Type6.BREX | [illegible] | 1188 | Bacteria>Pseudomonadota>Gammaproteobacteria>Pseudomonadales>Pseudomonadaceae>Pseudomonas | GCA_021652015.1 | Pseudomonas_aeruginosa | L4R58_004351 | 30228..33794 | 287 | BREX-system-P-loop-protein-BrxC-[Pseudomonas-aeruginosa] |
| HBO4843314.1 | Type6.BREX | [illegible] | 1207 | Bacteria>Pseudomonadota>Gammaproteobacteria>Pseudomonadales>Pseudomonadaceae>Pseudomonas | GCA_021660595.1 | Pseudomonas_aeruginosa | L4V37_002405 | 61513..65136 | 287 | BREX-system-P-loop-protein-BrxC-[Pseudomonas-aeruginosa] |
| HBO7186446.1 | Type1.BREX | [illegible] | 978 | Bacteria>Pseudomonadota>Gammaproteobacteria>Pseudomonadales>Pseudomonadaceae>Pseudomonas | GCA_021674675.1 | Pseudomonas_aeruginosa | L4968_002122 | 1..2938 | 287 | BREX-system-P-loop-protein-BrxC-[Pseudomonas-aeruginosa] |
| HBP1093904.1 | Type6.BREX | [illegible] | 1213 | Bacteria>Pseudomonadota>Gammaproteobacteria>Pseudomonadales>Pseudomonadaceae>Pseudomonas | GCA_021687815.1 | Pseudomonas_aeruginosa | L5S32_002755 | 180884..184525 | 287 | BREX-system-P-loop-protein-BrxC-[Pseudomonas-aeruginosa] |
| HBP5527123.1 | Type6.BREX | [illegible] | 1212 | Bacteria>Pseudomonadota>Gammaproteobacteria>Pseudomonadales>Pseudomonadaceae>Pseudomonas | GCA_021712095.1 | Pseudomonas_aeruginosa | L6B52_09510 | 5147..8785 | 287 | BREX-system-P-loop-protein-BrxC-[Pseudomonas-aeruginosa] |
| HCE6004670.1 | Type6.BREX | [illegible] | 1213 | Bacteria>Pseudomonadota>Gammaproteobacteria>Pseudomonadales>Pseudomonadaceae>Pseudomonas | GCA_023904815.1 | Pseudomonas_aeruginosa | NHK94_000107 | 121075..124716 | 287 | BREX-system-P-loop-protein-BrxC-[Pseudomonas-aeruginosa] |
| MCO2922446.1 | Type1.BREX | [illegible] | 1189 | Bacteria>Pseudomonadota>Gammaproteobacteria>Pseudomonadales>Pseudomonadaceae>Pseudomonas | GCA_023909145.1 | Pseudomonas_aeruginosa | FA260_04940 | 145579..149148 | 287 | BREX-system-P-loop-protein-BrxC-[Pseudomonas-aeruginosa] |
| HCF1683404.1 | Type6.BREX | [illegible] | 1213 | Bacteria>Pseudomonadota>Gammaproteobacteria>Pseudomonadales>Pseudomonadaceae>Pseudomonas | GCA_023941775.1 | Pseudomonas_aeruginosa | NH867_002024 | 49404..53045 | 287 | BREX-system-P-loop-protein-BrxC-[Pseudomonas-aeruginosa] |
| HCF1762884.1 | Type1.BREX | [illegible] | 1168 | Bacteria>Pseudomonadota>Gammaproteobacteria>Pseudomonadales>Pseudomonadaceae>Pseudomonas | GCA_023941915.1 | Pseudomonas_aeruginosa | NH889_002155 | 1..3508 | 287 | BREX-system-P-loop-protein-BrxC-[Pseudomonas-aeruginosa] |
| HCF5867680.1 | Type1.BREX | [illegible] | 1181 | Bacteria>Pseudomonadota>Gammaproteobacteria>Pseudomonadales>Pseudomonadaceae>Pseudomonas | GCA_023959235.1 | Pseudomonas_aeruginosa | NIH49_002287 | 171515..175060 | 287 | BREX-system-P-loop-protein-BrxC-[Pseudomonas-aeruginosa] |
| MCO7642712.1 | Type6.BREX | [illegible] | 1204 | Bacteria>Pseudomonadota>Gammaproteobacteria>Pseudomonadales>Pseudomonadaceae>Pseudomonas | GCA_024128275.1 | Pseudomonas_sp._S_311-6 | NJI34_38735 | 21195..24809 | 2956727 | BREX-system-P-loop-protein-BrxC-[Pseudomonas-sp.-S-311-6] |

| Protein-ID | System | Gene-Neighbourhoods | Length | Taxonomy | Assembly-ID | Strain | Locus-Tag | Genomic-coordinates | TaxID | Fasta-define |
|---|---|---|---|---|---|---|---|---|---|---|
| WCY64468.1 | Type6.BREX | JAB-><-SIGMA+OB-Fold+AAA-ATPase+HKD-DNase-BrxL<-PglZ<-?<-ParB+HNH<-N6-METHYLASE<-N6-METHYLASE<-GreA_GreB+SF1-SF2-Helicase<-DUF6079*<-BrxB-BrxF-iSTAND<-BrxA-DNA-binding<-BrxE<-wHTH+WYL+WCX\|\|JetA->JetB->SbcC-> | 1213 | Bacteria>Pseudomonadota>Gammaproteobacteria>Pseudomonadales>Pseudomonadaceae>Pseudomonas | GCA_028610105.1 | Pseudomonas_aeruginosa | NHK94_000107 | 121075..124716 | 287 | BREX-system-P-loop-protein-BrxC-[Pseudomonas-aeruginosa] |
| WP_257648015.1 | Type1.BREX | <-SIGMA+OB-Fold+AAA-ATPase+HKD-DNase-BrxL<-PglZ<-N6-METHYLASE-p<-?<-REase<-SbcC-p<-N6-METHYLASE<-DUF6079* | 1073 | Bacteria>Pseudomonadota>Gammaproteobacteria>Pseudomonadales>Pseudomonadaceae>Pseudomonas | GCA_030820635.1 | Pseudomonas_aeruginosa | K3Z95_RS14360 | 10117..13340 | 287 | BREX-system-P-loop-protein-BrxC,-partial-[Pseudomonas-aeruginosa] |
| WP_013692225.1 | Type6.BREX | <-RHH\|\|XIS-HTH->?-><-?\|\|HTH_3->wHTH+WYL+WCX->BrxE->BrxA-DNA-binding->BrxB-BrxF-iSTAND->DUF6079*->N6-METHYLASE->ParB+HNH->PglZ->SIGMA+OB-Fold+AAA-ATPase+HKD-DNase-BrxL-><-HNH<-?<-?<-?<-DUF932 | 1213 | Bacteria>Pseudomonadota>Gammaproteobacteria>Pseudomonadales>Pseudomonadaceae>Pseudomonas | GCF_000194805.1 | Pseudomonas_brassicacearum | PSEBR_RS03955 | 941705..945346 | 930166 | BREX-system-P-loop-protein-BrxC-[Pseudomonas-brassicacearum] |
| WP_007159175.1 | Type6.BREX | PIN+OST-HTH->?-><-SIGMA+OB-Fold+AAA-ATPase+HKD-DNase-BrxL<-PglZ<-ParB+HNH<-N6-METHYLASE<-N6-METHYLASE<-DUF6079*<-BrxB-BrxF-iSTAND<-BrxA-DNA-binding<-BrxE<-SF1-SF2-Helicase<-OLD-ABC+TOPRIM | 1212 | Bacteria>Pseudomonadota>Gammaproteobacteria>Pseudomonadales>Pseudomonadaceae>Pseudomonas | GCF_000236825.1 | Pseudomonas_psychrotolerans | PPL19_RS03910 | 152166..155804 | 237610 | BREX-system-P-loop-protein-BrxC-[Pseudomonas-psychrotolerans] |
| WP_010463595.1 | Type1.BREX | <-SIGMA+OB-Fold+AAA-ATPase+HKD-DNase-BrxL<-PglZ<-ParB+HNH<-N6-METHYLASE-p<-REase<-OLD-ABC<-N6-METHYLASE-p<-DUF6079*<-BrxB-BrxF-iSTAND<-BrxA-DNA-binding<-XIS-HTH<-ARG<-?\|\|wHTH+WYL+WCX-><-sGTP<-sGTP | 1185 | Bacteria>Pseudomonadota>Gammaproteobacteria>Pseudomonadales>Pseudomonadaceae>Pseudomonas | GCF_000257545.3 | Pseudomonas_mandelii | OU5_RS13450 | 3003724..3007281 | 75612 | BREX-system-P-loop-protein-BrxC-[Pseudomonas-mandelii] |
| WP_019485333.1 | Type1.BREX | <-TraE<-TraM<-TraM<-DUF3262<-?\|\|XIS-HTH->BrxA-DNA-binding->BrxB-BrxF-iSTAND->DUF6079*->N6-METHYLASE->OLD-ABC->REase->SIR2->HerA/FtsK->N6-METHYLASE-p->PglZ->SIGMA+OB-Fold+AAA-ATPase+HKD-DNase-BrxL->SbcC+SbcC-> | 1193 | Bacteria>Pseudomonadota>Gammaproteobacteria>Pseudomonadales>Pseudomonadaceae>Pseudomonas | GCF_000290555.2 | Pseudomonas_aeruginosa | E4028_RS01350 | 3998..7579 | 287 | BREX-system-P-loop-protein-BrxC-[Pseudomonas-aeruginosa] |
| WP_017337797.1 | Type6.BREX | MarR-HTH-><-RHH\|\|XIS-HTH-><-HIN-HTH\|\|?->wHTH+WYL+WCX->BrxE->BrxA-DNA-binding->BrxB-BrxF-iSTAND->DUF6079*->N6-METHYLASE->N6-METHYLASE->ParB+HNH->PglZ->SIGMA+OB-Fold+AAA-ATPase+HKD-DNase-BrxL-><-HNH<-?<-?<-DUF932 | 1213 | Bacteria>Pseudomonadota>Gammaproteobacteria>Pseudomonadales>Pseudomonadaceae>Pseudomonas | GCF_000293885.2 | Pseudomonas_fluorescens | B723_RS16820 | 3631858..3635499 | 294 | BREX-system-P-loop-protein-BrxC-[Pseudomonas-fluorescens] |
| WP_042135808.1 | Type1.BREX | <-MarR-HTH<-SIGMA+OB-Fold+AAA-ATPase+HKD-DNase-BrxL<-PglZ<-ParB+HNH<-N6-METHYLASE-p<-RVT<-NERD<-OLD-ABC<-N6-METHYLASE<-DUF6079*<-BrxB-BrxF-iSTAND<-BrxA-DNA-binding<-XIS-HTH<-tRNA<-MerR-HTH | 1184 | Bacteria>Pseudomonadota>Gammaproteobacteria>Pseudomonadales>Pseudomonadaceae>Pseudomonas | GCF_000730625.1 | Pseudomonas_oryzihabitans | POR01S_RS10925 | 121811..125365 | 47885 | BREX-system-P-loop-protein-BrxC-[Pseudomonas-oryzihabitans] |
| WP_034081877.1 | Type1.BREX | MerR-HTH->tRNA->XIS-HTH->BrxA-DNA-binding->BrxB-BrxF-iSTAND->DUF6079*->N6-METHYLASE->OLD-ABC->REase->SIR2->HerA/FtsK->N6-METHYLASE-p->PglZ->SIGMA+OB-Fold+AAA-ATPase+HKD-DNase-BrxL-><-RNase | 1184 | Bacteria>Pseudomonadota>Gammaproteobacteria>Pseudomonadales>Pseudomonadaceae>Pseudomonas | GCF_000793185.1 | Pseudomonas_aeruginosa | NT71_RS08285 | 193065..196619 | 287 | BREX-system-P-loop-protein-BrxC-[Pseudomonas-aeruginosa] |
| WP_044305280.1 | Type6.BREX | JAB-><-SIGMA+OB-Fold+AAA-ATPase+HKD-DNase-BrxL<-PglZ<-?<-ParB+HNH<-N6-METHYLASE<-N6-METHYLASE<-GreA_GreB+SF1-SF2-Helicase<-DUF6079*<-BrxB-BrxF-iSTAND<-BrxA-DNA-binding<-BrxE<-wHTH+WYL+WCX\|\|JetA->JetB->SbcC-> | 1213 | Bacteria>Pseudomonadota>Gammaproteobacteria>Pseudomonadales>Pseudomonadaceae>Pseudomonas | GCF_000793505.1 | Pseudomonas_aeruginosa | NT87_RS20770 | 137629..141270 | 287 | BREX-system-P-loop-protein-BrxC-[Pseudomonas-aeruginosa] |
| WP_041506104.1 | Type1.BREX | DUF4969-><-SIGMA+OB-Fold+AAA-ATPase+HKD-DNase-BrxL<-PglZ<-ParB+HNH<-N6-METHYLASE-p<-?<-N6-METHYLASE<-DUF6079*<-BrxB-BrxF-iSTAND<-BrxA-DNA-binding<-XIS-HTH<-?\|\|wHTH+WYL+WCX->LexA-protease-><-Glyoxalase\|\|?->DUF772+DDE_Tnp_1+DDE_Tnp_1_6-> | 1184 | Bacteria>Pseudomonadota>Gammaproteobacteria>Pseudomonadales>Pseudomonadaceae>Pseudomonas | GCF_000831585.1 | Pseudomonas_plecoglossicida | RK21_RS13525 | 2976546..2980100 | 70775 | BREX-system-P-loop-protein-BrxC-[Pseudomonas-plecoglossicida] |
| WP_044727109.1 | Type1.BREX | RelE-ParE-><-?<-wHTH+WYL+WCX\|\|?->Peptidase_M15->?->XIS-HTH->BrxA-DNA-binding->BrxB-BrxF-iSTAND->DUF6079*->N6-METHYLASE->OLD-ABC->NERD->RVT->N6-METHYLASE-p->PglZ->SIGMA+OB-Fold+AAA-ATPase+HKD-DNase-BrxL-><-Arm-DNA-bind_3+Phage_integrase | 1184 | Bacteria>Pseudomonadota>Gammaproteobacteria>Pseudomonadales>Pseudomonadaceae>Pseudomonas | GCF_000937465.2 | Pseudomonas_aeruginosa | TN45_RS09710 | 568754..572308 | 287 | BREX-system-P-loop-protein-BrxC-[Pseudomonas-aeruginosa] |
| WP_047306816.1 | Type1.BREX | <-SIGMA+OB-Fold+AAA-ATPase+HKD-DNase-BrxL<-PglZ<-HNH<-OLD-ABC<-N6-METHYLASE-p<-RVT<-N6-METHYLASE-p<-DUF6079*<-BrxB-BrxF-iSTAND<-BrxA-DNA-binding<-XIS-HTH<-?\|\|wHTH+WYL+WCX->sGTP->sGTP->sGTP-> | 1185 | Bacteria>Pseudomonadota>Gammaproteobacteria>Pseudomonadales>Pseudomonadaceae>Pseudomonas | GCF_001020715.1 | Pseudomonas_fluorescens | AA061_RS32150 | 16461..20018 | 294 | BREX-system-P-loop-protein-BrxC-[Pseudomonas-fluorescens] |
| WP_048385719.1 | Type1.BREX | <-SIGMA+OB-Fold+AAA-ATPase+HKD-DNase-BrxL<-PglZ<-ParB+HNH<-N6-METHYLASE-p<-REase<-OLD-ABC<-N6-METHYLASE-p<-DUF6079*<-BrxB-BrxF-iSTAND<-BrxA-DNA-binding<-XIS-HTH<-ARG<-?\|\|wHTH+WYL+WCX-><-sGTP<-sGTP | 1185 | Bacteria>Pseudomonadota>Gammaproteobacteria>Pseudomonadales>Pseudomonadaceae>Pseudomonas | GCF_001043065.1 | Pseudomonas_helleri | TU85_RS02975 | 88982..92539 | 1608996 | BREX-system-P-loop-protein-BrxC-[Pseudomonas-helleri] |
| WP_058181876.1 | Type6.BREX | <-SbcC<-JetB<-JetA\|\|wHTH+WYL+WCX->BrxE->BrxA-DNA-binding->BrxB-BrxF-iSTAND->DUF6079*->GreA_GreB+SF1-SF2-Helicase->N6-METHYLASE->N6-METHYLASE->ParB+HNH->?->PglZ->SIGMA+OB-Fold+AAA-ATPase+HKD-DNase-BrxL-><-JAB | 1213 | Bacteria>Pseudomonadota>Gammaproteobacteria>Pseudomonadales>Pseudomonadaceae>Pseudomonas | GCF_001450675.1 | Pseudomonas_aeruginosa | AO933_RS04770 | 131954..135595 | 287 | BREX-system-P-loop-protein-BrxC-[Pseudomonas-aeruginosa] |
| WP_031639867.1 | Type1.BREX | LZ_Tnp_IS66+zf-IS66+DDE_Tnp_IS66+DDE_Tnp_IS66_C-><-SIGMA+OB-Fold+AAA-ATPase+HKD-DNase-BrxL<-PglZ<-N6-METHYLASE-p<-HerA/FtsK<-SIR2<-REase<-OLD-ABC<-N6-METHYLASE<-DUF6079*<-BrxB-BrxF-iSTAND<-BrxA-DNA-binding<-XIS-HTH<-?\|\|?-><-?<-XIS-HTH\|\|CopG/RHH-> | 1184 | Bacteria>Pseudomonadota>Gammaproteobacteria>Pseudomonadales>Pseudomonadaceae>Pseudomonas | GCF_001548335.1 | Pseudomonas_aeruginosa | IOMTU133_RS23320 | 5069423..5072977 | 287 | BREX-system-P-loop-protein-BrxC-[Pseudomonas-aeruginosa] |
| WP_069953193.1 | Type1.BREX | DUF4969-><-SIGMA+OB-Fold+AAA-ATPase+HKD-DNase-BrxL<-PglZ<-N6-METHYLASE-p<-RVT<-REase<-SbcC-p<-N6-METHYLASE<-DUF6079-p* | 828 | Bacteria>Pseudomonadota>Gammaproteobacteria>Pseudomonadales>Pseudomonadaceae>Pseudomonas | GCF_001750225.1 | Pseudomonas_aeruginosa | A7R77_RS00815 | 21562..24049 | 287 | BREX-system-P-loop-protein-BrxC,-partial-[Pseudomonas-aeruginosa] |
| WP_074859114.1 | Type1.BREX | MerR-HTH->tRNA->XIS-HTH->?-><-?<-wHTH+WYL+WCX\|\|?->XIS-HTH->BrxA-DNA-binding->BrxB-BrxF-iSTAND->DUF6079*->N6-METHYLASE->SbcC-p->OLD-ABC->NERD->RVT->N6-METHYLASE-p->PglZ->SIGMA+OB-Fold+AAA-ATPase+HKD-DNase-BrxL-><-DUF4969 | 1184 | Bacteria>Pseudomonadota>Gammaproteobacteria>Pseudomonadales>Pseudomonadaceae>Pseudomonas | GCF_002091815.1 | Pseudomonas_oleovorans | POL01S_RS18595 | 25960..29514 | 301 | BREX-system-P-loop-protein-BrxC-[Pseudomonas-oleovorans] |
| WP_084929168.1 | Type1.BREX | Arm-DNA-bind_3+Phage_integrase-><-SIGMA+OB-Fold+AAA-ATPase+HKD-DNase-BrxL<-PglZ<-N6-METHYLASE-p<-RVT<-REase<-SbcC-p<-N6-METHYLASE<-DUF6079*<-BrxB-BrxF-iSTAND<-BrxA-DNA-binding<-XIS-HTH<-?<-Peptidase_M15<-?\|\|wHTH+WYL+WCX->?-><-RelE-ParE | 1184 | Bacteria>Pseudomonadota>Gammaproteobacteria>Pseudomonadales>Pseudomonadaceae>Pseudomonas | GCF_002094805.1 | Pseudomonas_aeruginosa | B7H20_RS24080 | 14270..17824 | 287 | BREX-system-P-loop-protein-BrxC-[Pseudomonas-aeruginosa] |
| WP_086230732.1 | Type1.BREX | <-wHTH+WYL+WCX\|\|?->Peptidase_M15->Peptidase_M15->?->XIS-HTH->BrxA-DNA-binding->BrxB-BrxF-iSTAND->DUF6079*->N6-METHYLASE->OLD-ABC->NERD->RVT->N6-METHYLASE-p->ParB+HNH->PglZ->SIGMA+OB-Fold+AAA-ATPase+HKD-DNase-BrxL-><-Arm-DNA-bind_3+Phage_integrase | 1184 | Bacteria>Pseudomonadota>Gammaproteobacteria>Pseudomonadales>Pseudomonadaceae>Pseudomonas | GCF_002135805.1 | Pseudomonas_aeruginosa | CAZ03_RS13145 | 128061..131615 | 287 | BREX-system-P-loop-protein-BrxC-[Pseudomonas-aeruginosa] |
| WP_101196767.1 | Type1.BREX | ZnR-><-wHTH+WYL+WCX<-AAA-ToxA+Classical-AAA\|\|?->\|\|XIS-HTH->BrxA-DNA-binding->BrxB-BrxF-iSTAND->DUF6079*->N6-METHYLASE->?->N6-METHYLASE-p->PglZ->SIGMA+OB-Fold+AAA-ATPase+HKD-DNase-BrxL-><-?<-?<-DUF932 | 1180 | Bacteria>Pseudomonadota>Gammaproteobacteria>Pseudomonadales>Pseudomonadaceae>Pseudomonas | GCF_002835905.1 | Pseudomonas_monteilii | CXB65_RS25690 | 173341..176883 | 76759 | BREX-system-P-loop-protein-BrxC-[Pseudomonas-monteilii] |
| WP_101152285.1 | Type6.BREX | CopG/RHH->TrwC+SF1-SF2-Helicase->HNH-><-SIGMA+OB-Fold+AAA-ATPase+HKD-DNase-BrxL<-PglZ<-ParB+HNH<-N6-METHYLASE<-N6-METHYLASE-p<-DUF6079*<-BrxB-BrxF-iSTAND<-BrxA-DNA-binding<-BrxE\|\|?->Arm-DNA-bind_2+Phage_int_SAM_3+Phage_integrase-><-?<-Calcineurin | 1211 | Bacteria>Pseudomonadota>Gammaproteobacteria>Pseudomonadales>Pseudomonadaceae>Pseudomonas | GCF_002836195.1 | Pseudomonas_sp._Choline-3u-10 | CXF92_RS00360 | 72818..76453 | 2058311 | BREX-system-P-loop-protein-BrxC-[Pseudomonas-sp.-Choline-3u-10] |
| WP_102082564.1 | Type1.BREX | DUF4969-><-SIGMA+OB-Fold+AAA-ATPase+HKD-DNase-BrxL<-PglZ<-ParB+HNH<-N6-METHYLASE-p<-RVT<-REase<-SbcC-p<-N6-METHYLASE<-DUF6079*<-BrxB-BrxF-iSTAND<-BrxA-DNA-binding<-XIS-HTH<-?\|\|wHTH+WYL+WCX->LexA-protease->?-><-?<-XIS-HTH | 1184 | Bacteria>Pseudomonadota>Gammaproteobacteria>Pseudomonadales>Pseudomonadaceae>Pseudomonas | GCF_002864915.1 | Pseudomonas_guariconensis | CXG53_RS23735 | 60064..63618 | 1288410 | BREX-system-P-loop-protein-BrxC-[Pseudomonas-guariconensis] |
| WP_102368059.1 | Type1.BREX | <-CopG/RHH<-HTH_Tnp_1+HTH_21+rve+rve_3<-CopG/RHH\|\|XIS-HTH->?->XIS-HTH->BrxA-DNA-binding->BrxB-BrxF-iSTAND->DUF6079*->N6-METHYLASE-p->?->N6-METHYLASE-p->?->PglZ->SIGMA+OB-Fold+AAA-ATPase+HKD-DNase-BrxL-><-HTH_Tnp_1+HTH_21+rve+rve_3 | 1188 | Bacteria>Pseudomonadota>Gammaproteobacteria>Pseudomonadales>Pseudomonadaceae>Pseudomonas | GCF_002874965.1 | Pseudomonas_sp._NC02 | C0058_RS03795 | 845324..848890 | 2067572 | BREX-system-P-loop-protein-BrxC-[Pseudomonas-sp.-NC02] |
| WP_102886089.1 | Type1.BREX | <-SIGMA+OB-Fold+AAA-ATPase+HKD-DNase-BrxL<-PglZ<-N6-METHYLASE-p<-SIR2<-REase<-OLD-ABC<-N6-METHYLASE<-DUF6079*<-BrxB-BrxF-iSTAND<-BrxA-DNA-binding<-?\|\|wHTH+WYL+WCX-p-><-?\|\|sGTP->sGTP->MA+PspA+PspA+sGTP-><-ZnR | 1184 | Bacteria>Pseudomonadota>Gammaproteobacteria>Pseudomonadales>Pseudomonadaceae>Pseudomonas | GCF_002891885.1 | Pseudomonas_putida | CBL13_04161 | 1040459..1044013 | 303 | BREX-system-P-loop-protein-BrxC-[Pseudomonas-putida] |
| WP_116801381.1 | Type1.BREX | DUF4969-><-SIGMA+OB-Fold+AAA-ATPase+HKD-DNase-BrxL<-PglZ<-N6-METHYLASE-p<-RVT<-NERD<-OLD-ABC<-N6-METHYLASE<-DUF6079*<-BrxB-BrxF-iSTAND<-BrxA-DNA-binding<-XIS-HTH<-?\|\|wHTH+WYL+WCX->HEPN-><-SF1-SF2-Helicase | 1184 | Bacteria>Pseudomonadota>Gammaproteobacteria>Pseudomonadales>Pseudomonadaceae>Pseudomonas | GCF_003410555.2 | Pseudomonas_aeruginosa | DZ982_RS28380 | 35413..38967 | 287 | BREX-system-P-loop-protein-BrxC-[Pseudomonas-aeruginosa] |
| WP_121398877.1 | Type1.BREX | Arm-DNA-bind_3+Phage_integrase-><-SIGMA+OB-Fold+AAA-ATPase+HKD-DNase-BrxL<-PglZ<-N6-METHYLASE-p<-HerA/FtsK<-SIR2<-REase<-OLD-ABC<-N6-METHYLASE<-DUF6079*<-BrxB-BrxF-iSTAND<-BrxA-DNA-binding<-XIS-HTH<-?<-Peptidase_M15<-?\|\|wHTH+WYL+WCX->?-><-RelE-ParE | 1180 | Bacteria>Pseudomonadota>Gammaproteobacteria>Pseudomonadales>Pseudomonadaceae>Pseudomonas | GCF_003631245.1 | Pseudomonas_aeruginosa | DQH61_RS02890 | 22908..26450 | 287 | BREX-system-P-loop-protein-BrxC-[Pseudomonas-aeruginosa] |
| WP_128551186.1 | Type1.BREX | Arm-DNA-bind_3+Phage_integrase-><-SIGMA+OB-Fold+AAA-ATPase+HKD-DNase-BrxL<-PglZ<-ParB+HNH<-N6-METHYLASE-p<-RVT<-NERD<-OLD-ABC<-N6-METHYLASE<-DUF6079*<-BrxB-BrxF-iSTAND<-BrxA-DNA-binding<-XIS-HTH<-?<-Peptidase_M15<-Peptidase_M15<-?\|\|wHTH+WYL+WCX-> | 1184 | Bacteria>Pseudomonadota>Gammaproteobacteria>Pseudomonadales>Pseudomonadaceae>Pseudomonas | GCF_003698635.1 | Pseudomonas_aeruginosa | IPC82_RS03385 | 45130..48684 | 287 | BREX-system-P-loop-protein-BrxC-[Pseudomonas-aeruginosa] |
| WP_124126664.1 | Type6.BREX | <-LON-AAA\|\|HTH_19-><-?<-ARG\|\|wHTH+WYL+WCX->BrxE->BrxA-DNA-binding->BrxB-BrxF-iSTAND->DUF6079*->N6-METHYLASE->N6-METHYLASE->?->PglZ->SIGMA+OB-Fold+AAA-ATPase+HKD-DNase-BrxL-><-?<-Arm-DNA-bind_3+Phage_integrase<-sGTP | 1210 | Bacteria>Pseudomonadota>Gammaproteobacteria>Pseudomonadales>Pseudomonadaceae>Pseudomonas | GCF_003836525.1 | Pseudomonas_aeruginosa | IPC776_RS06145 | 34793..38425 | 287 | BREX-system-P-loop-protein-BrxC-[Pseudomonas-aeruginosa] |
| WP_126190092.1 | Type1.BREX | <-RNase<-?<-JetB<-?<-wHTH+WYL+WCX\|\|XIS-HTH->BrxA-DNA-binding->BrxB-BrxF-iSTAND->DUF6079*->N6-METHYLASE->OLD-ABC->NERD->?->N6-METHYLASE-p->PglZ->SIGMA+OB-Fold+AAA-ATPase+HKD-DNase-BrxL-><-DUF4969 | 1185 | Bacteria>Pseudomonadota>Gammaproteobacteria>Pseudomonadales>Pseudomonadaceae>Pseudomonas | GCF_003957495.3 | Pseudomonas_songnenensis | EJA06_RS16615 | 27316..30873 | 1176259 | BREX-system-P-loop-protein-BrxC-[Pseudomonas-songnenensis] |
| WP_128704208.1 | Type1.BREX | <-HTH_20\|\|Arm-DNA-bind_3+Phage_integrase-><-SIGMA+OB-Fold+AAA-ATPase+HKD-DNase-BrxL<-PglZ<-N6-METHYLASE-p<-?<-REase<-SbcC-p<-N6-METHYLASE<-DUF6079*<-BrxB-BrxF-iSTAND<-BrxA-DNA-binding<-XIS-HTH<-?<-Peptidase_M15<-?\|\|wHTH+WYL+WCX-><-?<-PhdYeFM_antitox | 1184 | Bacteria>Pseudomonadota>Gammaproteobacteria>Pseudomonadales>Pseudomonadaceae>Pseudomonas | GCF_003974605.1 | Pseudomonas_aeruginosa | IPC1390_RS17360 | 217632..221186 | 287 | BREX-system-P-loop-protein-BrxC-[Pseudomonas-aeruginosa] |
| WP_126632346.1 | Type6.BREX | <-Small-B-barrel+SF1-SF2-Helicase+RdgC+REase<-35exo+SF1-SF2-Helicase<-XIS-HTH<-SIGMA+OB-Fold+AAA-ATPase+HKD-DNase-BrxL<-PglZ<-ParB+HNH<-N6-METHYLASE<-N6-METHYLASE<-GreA_GreB+SF1-SF2-Helicase<-DUF6079*<-BrxB-BrxF-iSTAND<-BrxA-DNA-binding<-BrxE<-tRNA<-MerR-HTH | 1213 | Bacteria>Pseudomonadota>Gammaproteobacteria>Pseudomonadales>Pseudomonadaceae>Pseudomonas | GCF_003975645.1 | Pseudomonas_aeruginosa | IPC447_RS09565 | 370705..374346 | 287 | BREX-system-P-loop-protein-BrxC-[Pseudomonas-aeruginosa] |
| WP_023094202.1 | Type6.BREX | <-SbcC<-JetB<-JetA\|\|wHTH+WYL+WCX->BrxE->BrxA-DNA-binding->BrxB-BrxF-iSTAND->DUF6079*->GreA_GreB+SF1-SF2-Helicase->N6-METHYLASE->N6-METHYLASE->ParB+HNH->?->PglZ->SIGMA+OB-Fold+AAA-ATPase+HKD-DNase-BrxL-><-JAB | 1213 | Bacteria>Pseudomonadota>Gammaproteobacteria>Pseudomonadales>Pseudomonadaceae>Pseudomonas | GCF_004346985.1 | Pseudomonas_aeruginosa | C9I70_RS03025 | 613762..617403 | 287 | BREX-system-P-loop-protein-BrxC-[Pseudomonas-aeruginosa] |
| WP_134249510.1 | Type6.BREX | <-KAP-glycosylase<-KAP-NTPase<-SIGMA+OB-Fold+AAA-ATPase+HKD-DNase-BrxL<-PglZ<-ParB+HNH<-N6-METHYLASE<-N6-METHYLASE<-GreA_GreB+SF1-SF2-Helicase<-DUF6079*<-BrxB-BrxF-iSTAND<-BrxA-DNA-binding<-BrxE<-wHTH+WYL+WCX\|\|?->SMODS->?->?->DUF3262-> | 1213 | Bacteria>Pseudomonadota>Gammaproteobacteria>Pseudomonadales>Pseudomonadaceae>Pseudomonas | GCF_004374205.1 | Pseudomonas_aeruginosa | IPC115_RS16040 | 120938..124579 | 287 | BREX-system-P-loop-protein-BrxC-[Pseudomonas-aeruginosa] |
| WP_134295820.1 | Type6.BREX | <-KAP-glycosylase<-KAP-NTPase<-SIGMA+OB-Fold+AAA-ATPase+HKD-DNase-BrxL<-PglZ<-ParB+HNH<-N6-METHYLASE<-N6-METHYLASE-p<-N6-METHYLASE-p<-GreA_GreB+SF1-SF2-Helicase<-DUF6079*<-BrxB-BrxF-iSTAND<-BrxA-DNA-binding<-BrxE<-wHTH+WYL+WCX\|\|?->SMODS->?->?->DUF3262-> | 1213 | Bacteria>Pseudomonadota>Gammaproteobacteria>Pseudomonadales>Pseudomonadaceae>Pseudomonas | GCF_004374605.1 | Pseudomonas_aeruginosa | IPC112_RS05880 | 126222..129863 | 287 | BREX-system-P-loop-protein-BrxC-[Pseudomonas-aeruginosa] |
| WP_143505426.1 | Type6.BREX | HAD+HAD->?->?->?->?-><-wHTH+WYL+WCX\|\|BrxE->BrxA-DNA-binding->BrxB-BrxF-iSTAND->DUF6079*->N6-METHYLASE->N6-METHYLASE->ParB+HNH->PglZ->SIGMA+OB-Fold+AAA-ATPase+HKD-DNase-BrxL-><-AAA-ATPase<-HTH+rve<-?<-TrwC+SF1-SF2-Helicase | 1212 | Bacteria>Pseudomonadota>Gammaproteobacteria>Pseudomonadales>Pseudomonadaceae>Pseudomonas | GCF_007049795.1 | Pseudomonas_mendocina | EQ826_RS09455 | 197604..201242 | 300 | BREX-system-P-loop-protein-BrxC-[Pseudomonas-mendocina] |
| WP_150073350.1 | Type1.BREX | XIS-HTH->?->XIS-HTH->BrxA-DNA-binding->BrxB-BrxF-iSTAND->DUF6079*->N6-METHYLASE-p->OLD-ABC->REase->N6-METHYLASE-p->ParB+HNH->PglZ->SIGMA+OB-Fold+AAA-ATPase+HKD-DNase-BrxL-> | 1186 | Bacteria>Pseudomonadota>Gammaproteobacteria>Pseudomonadales>Pseudomonadaceae>Pseudomonas | GCF_008632325.1 | Pseudomonas_lactis | F3K52_RS19340 | 4156..7716 | 1615674 | BREX-system-P-loop-protein-BrxC-[Pseudomonas-lactis] |
| WP_151183841.1 | Type1.BREX | 35exo->AbiEi_3_N+AbiEi_3->AbiEii->?->?->BrxA-DNA-binding->BrxB-BrxF-iSTAND->DUF6079*->N6-METHYLASE->PglZ->SIGMA+OB-Fold+AAA-ATPase+HKD-DNase-BrxL->McrB-AAA->McrC-NTD+REase-><-Peptidase_C39_2<-PVC-AAA+Classical-AAA\|\|wHTH+WYL+WCX-> | 1181 | Bacteria>Pseudomonadota>Gammaproteobacteria>Pseudomonadales>Pseudomonadaceae>Pseudomonas | GCF_008807395.1 | Pseudomonas_sp._C27(2019) | FXF61_RS02795 | 588989..592534 | 2604941 | BREX-system-P-loop-protein-BrxC-[Pseudomonas-sp.-C27(2019)] |
| WP_153329486.1 | Type1.BREX | <-SIGMA+OB-Fold+AAA-ATPase+HKD-DNase-BrxL<-PglZ<-ParB+HNH<-N6-METHYLASE-p<-RVT<-NERD<-OLD-ABC<-SbcC-p<-N6-METHYLASE<-DUF6079*<-BrxB-BrxF-iSTAND<-BrxA-DNA-binding<-XIS-HTH<-?\|\|wHTH+WYL+WCX->LexA-protease-><-?<-XIS-HTH | 1184 | Bacteria>Pseudomonadota>Gammaproteobacteria>Pseudomonadales>Pseudomonadaceae>Pseudomonas | GCF_009600225.1 | Pseudomonas_helleri | GHO39_RS16765 | 34385..37939 | 1608996 | BREX-system-P-loop-protein-BrxC-[Pseudomonas-helleri] |
| WP_153386820.1 | Type1.BREX | Pkinase+Pkinase->?-><-?<-?\|\|XIS-HTH->BrxA-DNA-binding->BrxB-BrxF-iSTAND->DUF6079*->N6-METHYLASE->OLD-ABC->REase->SIR2->HerA/FtsK->N6-METHYLASE-p->PglZ->SIGMA+OB-Fold+AAA-ATPase+HKD-DNase-BrxL-> | 1183 | Bacteria>Pseudomonadota>Gammaproteobacteria>Pseudomonadales>Pseudomonadaceae>Pseudomonas | GCF_009600315.1 | Pseudomonas_helleri | GHN86_RS14975 | 48582..52133 | 1608996 | BREX-system-P-loop-protein-BrxC-[Pseudomonas-helleri] |
| WP_153384138.1 | Type1.BREX | MerR-HTH->tRNA-><-RHH\|\|XIS-HTH->?->XIS-HTH->BrxA-DNA-binding->BrxB-BrxF-iSTAND->DUF6079*->N6-METHYLASE-p->OLD-ABC->REase->N6-METHYLASE-p->ParB+HNH->PglZ->SIGMA+OB-Fold+AAA-ATPase+HKD-DNase-BrxL->\|\|<-?<-wHTH+WYL+WCX | 1186 | Bacteria>Pseudomonadota>Gammaproteobacteria>Pseudomonadales>Pseudomonadaceae>Pseudomonas | GCF_009600565.1 | unclassified_Pseudomonas | GHO24_RS06625 | 35218..38778 | 196821 | MULTISPECIES:-BREX-system-P-loop-protein-BrxC-[unclassified-Pseudomonas] |
| WP_161928857.1 | Type6.BREX | <-Arm-DNA-bind_2+Phage_int_SAM_3+Phage_integrase\|\|?-><-?<-DUF772+DDE_Tnp_1<-wHTH+WYL+WCX\|\|BrxE->BrxA-DNA-binding->BrxB-BrxF-iSTAND->DUF6079*->N6-METHYLASE->N6-METHYLASE->ParB+HNH->PglZ->SIGMA+OB-Fold+AAA-ATPase+HKD-DNase-BrxL->?-><-?<-TrwC+SF1-SF2-Helicase<-CopG/RHH | 1212 | Bacteria>Pseudomonadota>Gammaproteobacteria>Pseudomonadales>Pseudomonadaceae>Pseudomonas | GCF_009932815.1 | Pseudomonas_yangonensis | GYH25_RS19705 | 87173..90811 | 2579922 | BREX-system-P-loop-protein-BrxC-[Pseudomonas-yangonensis] |
| WP_169989748.1 | Type1.BREX | <-SIGMA+OB-Fold+AAA-ATPase+HKD-DNase-BrxL<-PglZ<-N6-METHYLASE-p<-HerA/FtsK<-SIR2<-REase<-OLD-ABC<-N6-METHYLASE<-DUF6079*<-BrxB-BrxF-iSTAND<-BrxA-DNA-binding<-XIS-HTH<-?<-?<-ABC-ATPase+SbcC<-?<-?<-ABC-ATPase+SbcC | 1186 | Bacteria>Pseudomonadota>Gammaproteobacteria>Pseudomonadales>Pseudomonadaceae>Pseudomonas | GCF_012985575.1 | Pseudomonas_sp._WS_5413 | HBO21_RS11490 | 97465..101025 | 2717488 | BREX-system-P-loop-protein-BrxC-[Pseudomonas-sp.-WS-5413] |
| WP_169883492.1 | Type6.BREX | HNH-><-SIGMA+OB-Fold+AAA-ATPase+HKD-DNase-BrxL<-PglZ<-N6-METHYLASE<-N6-METHYLASE<-DUF6079*<-BrxB-BrxF-iSTAND<-BrxE<-wHTH+WYL+WCX<-?<-XIS-HTH\|\|RHH-><-tRNA<-MerR-HTH | 1213 | Bacteria>Pseudomonadota>Gammaproteobacteria>Pseudomonadales>Pseudomonadaceae>Pseudomonas | GCF_012986805.1 | Pseudomonas_fragi | HBN91_RS17750 | 10864..14505 | 296 | BREX-system-P-loop-protein-BrxC-[Pseudomonas-fragi] |
| WP_177031785.1 | Type1.BREX | sGTP-><-wHTH+WYL+WCX\|\|?->ARG->XIS-HTH->BrxA-DNA-binding->BrxB-BrxF-iSTAND->DUF6079*->N6-METHYLASE->?->PglZ->SIGMA+OB-Fold+AAA-ATPase+HKD-DNase-BrxL-><-?<-?<-?<-DUF932 | 1185 | Bacteria>Pseudomonadota>Gammaproteobacteria>Pseudomonadales>Pseudomonadaceae>Pseudomonas | GCF_013386345.1 | Pseudomonas_veronii | HX878_RS08640 | 6154..9711 | 76761 | BREX-system-P-loop-protein-BrxC-[Pseudomonas-veronii] |
| WP_178910878.1 | Type1.BREX | <-SIGMA+OB-Fold+AAA-ATPase+HKD-DNase-BrxL<-PglZ<-N6-METHYLASE-p<-RVT<-NERD<-OLD-ABC<-N6-METHYLASE<-DUF6079*<-BrxB-BrxF-iSTAND<-BrxA-DNA-binding<-XIS-HTH<-?\|\|?->?->\|\|Aminotran_1_2-> | 1186 | Bacteria>Pseudomonadota>Gammaproteobacteria>Pseudomonadales>Pseudomonadaceae>Pseudomonas | GCF_013386495.1 | Pseudomonas_tolaasii | HX799_RS03165 | 122791..126351 | 29442 | BREX-system-P-loop-protein-BrxC-[Pseudomonas-tolaasii] |
| WP_180216146.1 | Type6.BREX | <-DUF3262<-?<-?<-SMODS<-?<-?\|\|wHTH+WYL+WCX->BrxE->BrxB-BrxF-iSTAND->DUF6079*->GreA_GreB+SF1-SF2-Helicase->N6-METHYLASE->N6-METHYLASE->ParB+HNH->PglZ->SIGMA+OB-Fold+AAA-ATPase+HKD-DNase-BrxL-><-JAB | 1213 | Bacteria>Pseudomonadota>Gammaproteobacteria>Pseudomonadales>Pseudomonadaceae>Pseudomonas | GCF_013421145.1 | Pseudomonas_aeruginosa | HOB52_RS17845 | 402112..405753 | 287 | BREX-system-P-loop-protein-BrxC-[Pseudomonas-aeruginosa] |
| WP_180689220.1 | Type1.BREX | ABC-ATPase+SbcC->?->?->?->?->DUF772+DDE_Tnp_1->XIS-HTH->BrxA-DNA-binding->BrxB-BrxF-iSTAND->DUF6079*->N6-METHYLASE->ParB+HNH->PglZ->SIGMA+OB-Fold+AAA-ATPase+HKD-DNase-BrxL-><-Phage_integrase<-Integrase_1<-wHTH<-AAA+wHTH<-Arm-DNA-bind_3+Phage_integrase | 1185 | Bacteria>Pseudomonadota>Gammaproteobacteria>Pseudomonadales>Pseudomonadaceae>Pseudomonas | GCF_013425825.1 | Pseudomonas_putida | HOH12_RS03375 | 759269..762826 | 303 | BREX-system-P-loop-protein-BrxC-[Pseudomonas-putida] |
| WP_181088903.1 | Type6.BREX | TPR->wHTH->Integrase_1->Phage_integrase-><-SIGMA+OB-Fold+AAA-ATPase+HKD-DNase-BrxL<-PglZ<-ParB+HNH<-N6-METHYLASE<-N6-METHYLASE<-DUF6079*<-BrxB-BrxF-iSTAND<-BrxE<-wHTH+WYL+WCX\|\|Peptidase_C39_2->DNA-glycosylase->ParB+HNH-><-?<-wHTH+WYL+WCX | 1214 | Bacteria>Pseudomonadota>Gammaproteobacteria>Pseudomonadales>Pseudomonadaceae>Pseudomonas | GCF_013523035.1 | Pseudomonas_plecoglossicida | G7011_RS14160 | 93028..96672 | 70775 | BREX-system-P-loop-protein-BrxC-[Pseudomonas-plecoglossicida] |
| WP_185818333.1 | Type6.BREX | MerR-HTH->tRNA-><-CopG/RHH\|\|BirA-HTH->?->XIS-HTH-><-HTH_Tnp_1+HTH_21+rve+rve_3\|\|wHTH+WYL+WCX->BrxE->BrxB-BrxF-iSTAND->DUF6079*->N6-METHYLASE->PglZ->SIGMA+OB-Fold+AAA-ATPase+HKD-DNase-BrxL-><-HNH<-?<-?<-?<-DUF932 | 1214 | Bacteria>Pseudomonadota>Gammaproteobacteria>Pseudomonadales>Pseudomonadaceae>Pseudomonas | GCF_014236655.1 | Pseudomonas_kielensis | H7995_RS10625 | 125852..129496 | 2762577 | BREX-system-P-loop-protein-BrxC-[Pseudomonas-kielensis] |
| WP_186598283.1 | Type6.BREX | <-sGTP<-GTPase-AIG<-?<-?\|\|<-CopG/RHH\|\|XIS-HTH->?->BrxE->BrxB-BrxF-iSTAND->DUF6079*->N6-METHYLASE->ParB+HNH->PglZ->SIGMA+OB-Fold+AAA-ATPase+HKD-DNase-BrxL-><-HNH<-DUF4969 | 1212 | Bacteria>Pseudomonadota>Gammaproteobacteria>Pseudomonadales>Pseudomonadaceae>Pseudomonas | GCF_014269085.1 | Pseudomonas_taiwanensis | HU747_RS04800 | 431376..435014 | 470150 | BREX-system-P-loop-protein-BrxC-[Pseudomonas-taiwanensis] |
| WP_198790924.1 | Type1.BREX | <-CopG/RHH\|\|XIS-HTH->?->?->XIS-HTH->BrxA-DNA-binding->BrxB-BrxF-iSTAND->DUF6079*->N6-METHYLASE-p->OLD-ABC->REase->?->N6-METHYLASE-p->ParB+HNH->PglZ->SIGMA+OB-Fold+AAA-ATPase+HKD-DNase-BrxL-> | 1185 | Bacteria>Pseudomonadota>Gammaproteobacteria>Pseudomonadales>Pseudomonadaceae>Pseudomonas | GCF_016405175.1 | Pseudomonas_sp._MF7451 | JFT65_RS10900 | 194530..198087 | 2797538 | BREX-system-P-loop-protein-BrxC-[Pseudomonas-sp.-MF7451] |
| WP_198832884.1 | Type1.BREX | <-SIGMA+OB-Fold+AAA-ATPase+HKD-DNase-BrxL<-PglZ<-?<-N6-METHYLASE<-DUF6079*<-BrxB-BrxF-iSTAND<-BrxA-DNA-binding<-XIS-HTH<-ARG<-?\|\|wHTH+WYL+WCX-><-sGTP | 1185 | Bacteria>Pseudomonadota>Gammaproteobacteria>Pseudomonadales>Pseudomonadaceae>Pseudomonas | GCF_016405445.1 | Pseudomonas_haemolytica | JFT42_RS28230 | 10084..13641 | 2600065 | BREX-system-P-loop-protein-BrxC-[Pseudomonas-haemolytica] |
| WP_198849248.1 | Type1.BREX | wHTH+WYL+WCX-><-SIGMA+OB-Fold+AAA-ATPase+HKD-DNase-BrxL<-HNH<-OLD-ABC<-N6-METHYLASE-p<-HerA/FtsK<-SIR2<-N6-METHYLASE-p<-DUF6079*<-BrxB-BrxF-iSTAND<-BrxA-DNA-binding<-XIS-HTH<-?<-tRNA<-MerR-HTH | 1186 | Bacteria>Pseudomonadota>Gammaproteobacteria>Pseudomonadales>Pseudomonadaceae>Pseudomonas | GCF_016405505.1 | Pseudomonas_sp._MF6772 | JFT60_RS22275 | 24346..27906 | 2797533 | BREX-system-P-loop-protein-BrxC-[Pseudomonas-sp.-MF6772] |
| WP_273507518.1 | Type6.BREX | <-CopG/RHH\|\|CITB-HTH->XIS-HTH->?->wHTH+WYL+WCX->BrxE->BrxA-DNA-binding->BrxB-BrxF-iSTAND->DUF6079*->PglZ->SIGMA+OB-Fold+AAA-ATPase+HKD-DNase-BrxL-><-HNH<-DUF4969 | 1214 | Bacteria>Pseudomonadota>Gammaproteobacteria>Pseudomonadales>Pseudomonadaceae>Pseudomonas | GCF_016785205.1 | Pseudomonas_sp. | JJK56_22660 | 12252..15896 | 306 | BREX-system-P-loop-protein-BrxC-[Pseudomonas-sp.] |
| WP_203328783.1 | Type6.BREX | <-LysR-HTH\|\|HTH_3-><-MODE-HTH+PBPII\|\|wHTH+WYL+WCX->BrxE->BrxA-DNA-binding->BrxB-BrxF-iSTAND->DUF6079*->GreA_GreB+SF1-SF2-Helicase->N6-METHYLASE->N6-METHYLASE->DOC+HTH->PglZ->SIGMA+OB-Fold+AAA-ATPase+HKD-DNase-BrxL-><-HDRnase\|\|XIS-HTH->Tox-RES-> | 1213 | Bacteria>Pseudomonadota>Gammaproteobacteria>Pseudomonadales>Pseudomonadaceae>Pseudomonas | GCF_016836545.1 | Pseudomonas_aeruginosa | JQ483_RS07485 | 1620720..1624361 | 287 | BREX-system-P-loop-protein-BrxC-[Pseudomonas-aeruginosa] |
| WP_209939193.1 | Type6.BREX | <-SbcC<-JetB<-JetA\|\|wHTH+WYL+WCX->BrxE->BrxA-DNA-binding->BrxB-BrxF-iSTAND->DUF6079*->GreA_GreB+SF1-SF2-Helicase->N6-METHYLASE->N6-METHYLASE->ParB+HNH->?->PglZ->SIGMA+OB-Fold+AAA-ATPase+HKD-DNase-BrxL-><-JAB | 1213 | Bacteria>Pseudomonadota>Gammaproteobacteria>Pseudomonadales>Pseudomonadaceae>Pseudomonas | GCF_017838075.1 | Pseudomonas_aeruginosa | JOA89_RS01675 | 341330..344971 | 287 | BREX-system-P-loop-protein-BrxC-[Pseudomonas-aeruginosa] |
| WP_223023201.1 | Type1.BREX | <-REase\|\|DNA_METHYLASE-><-CopG/RHH\|\|XIS-HTH->?->XIS-HTH->BrxA-DNA-binding->BrxB-BrxF-iSTAND->DUF6079*->OLD-ABC->REase->N6-METHYLASE->PglZ->SIGMA+OB-Fold+AAA-ATPase+HKD-DNase-BrxL-><-?<-?<-?\|\|HTH_3-> | 1183 | Bacteria>Pseudomonadota>Gammaproteobacteria>Pseudomonadales>Pseudomonadaceae>Pseudomonas | GCF_019891425.1 | Pseudomonas_carnis | J1G36_06600 | 89435..92986 | 2487355 | BREX-system-P-loop-protein-BrxC-[Pseudomonas-carnis] |
| WP_223717957.1 | Type6.BREX | MerR-HTH->tRNA->BrxE->BrxB-BrxF-iSTAND->DUF6079*->GreA_GreB+SF1-SF2-Helicase->N6-METHYLASE->N6-METHYLASE->ParB+HNH->PglZ->SIGMA+OB-Fold+AAA-ATPase+HKD-DNase-BrxL->XIS-HTH->35exo+SF1-SF2-Helicase->Small-B-barrel+SF1-SF2-Helicase+RdgC+REase-> | 1213 | Bacteria>Pseudomonadota>Gammaproteobacteria>Pseudomonadales>Pseudomonadaceae>Pseudomonas | GCF_020036535.1 | Pseudomonas_aeruginosa | LC626_RS08340 | 482522..486163 | 287 | BREX-system-P-loop-protein-BrxC-[Pseudomonas-aeruginosa] |
| WP_227364598.1 | Type6.BREX | VirD4-TraG->DUF4400-><-SbcC<-SIGMA+OB-Fold+AAA-ATPase+HKD-DNase-BrxL<-PglZ<-ParB+HNH<-N6-METHYLASE<-N6-METHYLASE<-GreA_GreB+SF1-SF2-Helicase<-DUF6079*<-BrxB-BrxF-iSTAND<-BrxE<-wHTH+WYL+WCX\|\|?->SMODS->?->DUF3262->TraM->TraM-> | 1213 | Bacteria>Pseudomonadota>Gammaproteobacteria>Pseudomonadales>Pseudomonadaceae>Pseudomonas | GCF_020639995.1 | Pseudomonas_aeruginosa | LJG07_RS13375 | 107262..110903 | 287 | BREX-system-P-loop-protein-BrxC-[Pseudomonas-aeruginosa] |
| WP_229605965.1 | Type6.BREX | AlcRN+AraC-HTH-><-?\|\|HNH-><-SIGMA+OB-Fold+AAA-ATPase+HKD-DNase-BrxL<-PglZ<-ParB+HNH<-N6-METHYLASE<-N6-METHYLASE<-DUF6079*<-BrxB-BrxF-iSTAND<-BrxE<-?<-ARG<-?\|\|wHTH+WYL+WCX-><-LZ_Tnp_IS481+HTH_32+rve+rve_3<-tRNA<-MerR-HTH | 1212 | Bacteria>Pseudomonadota>Gammaproteobacteria>Pseudomonadales>Pseudomonadaceae>Pseudomonas | GCF_020831405.1 | Pseudomonas_oryzagri | LMJ29_RS13110 | 91421..95059 | 2892443 | BREX-system-P-loop-protein-BrxC-[Pseudomonas-oryzagri] |
| WP_033990403.1 | Type6.BREX | SMODS->?->JAB-><-SIGMA+OB-Fold+AAA-ATPase+HKD-DNase-BrxL<-PglZ<-ParB+HNH<-N6-METHYLASE<-N6-METHYLASE<-GreA_GreB+SF1-SF2-Helicase<-DUF6079*<-BrxB-BrxF-iSTAND<-BrxA-DNA-binding<-BrxE<-wHTH+WYL+WCX\|\|JetA->JetB->SbcC-> | 1213 | Bacteria>Pseudomonadota>Gammaproteobacteria>Pseudomonadales>Pseudomonadaceae>Pseudomonas | GCF_020907765.1 | Pseudomonas_aeruginosa | LOH47_RS10865 | 120991..124632 | 287 | BREX-system-P-loop-protein-BrxC-[Pseudomonas-aeruginosa] |
| WP_196706085.1 | Type1.BREX | <-SIGMA+OB-Fold+AAA-ATPase+HKD-DNase-BrxL<-PglZ<-N6-METHYLASE-p<-HerA/FtsK<-SIR2<-REase<-OLD-ABC<-N6-METHYLASE<-DUF6079*<-BrxB-BrxF-iSTAND<-BrxA-DNA-binding<-XIS-HTH<-?\|\|<-?\|\|sGTP->sGTP->MA+PspA+PspA+sGTP-><-ARG | 1184 | Bacteria>Pseudomonadota>Gammaproteobacteria>Pseudomonadales>Pseudomonadaceae>Pseudomonas | GCF_021460775.1 | Pseudomonas_putida | LH705_RS23300 | 5093270..5096824 | 303 | BREX-system-P-loop-protein-BrxC-[Pseudomonas-putida] |
| WP_236238652.1 | Type1.BREX | <-HIN-HTH<-SIGMA+OB-Fold+AAA-ATPase+HKD-DNase-BrxL<-PglZ<-ParB+HNH<-N6-METHYLASE-p<-REase<-OLD-ABC<-N6-METHYLASE-p<-DUF6079*<-BrxB-BrxF-iSTAND<-BrxA-DNA-binding<-XIS-HTH | 1185 | Bacteria>Pseudomonadota>Gammaproteobacteria>Pseudomonadales>Pseudomonadaceae>Pseudomonas | GCF_021603345.1 | Pseudomonas_rhodesiae | L6E14_RS13730 | 266364..269921 | 76760 | BREX-system-P-loop-protein-BrxC-[Pseudomonas-rhodesiae] |
| WP_236165768.1 | Type6.BREX | DUF932->?->?->HNH-><-SIGMA+OB-Fold+AAA-ATPase+HKD-DNase-BrxL<-PglZ<-ParB+HNH<-N6-METHYLASE<-N6-METHYLASE<-DUF6079*<-BrxB-BrxF-iSTAND<-BrxA-DNA-binding<-BrxE<-?<-XIS-HTH\|\|?->CopG/RHH-> | 1212 | Bacteria>Pseudomonadota>Gammaproteobacteria>Pseudomonadales>Pseudomonadaceae>Pseudomonas | GCF_021603865.1 | Pseudomonas_fulva | L6D10_RS06765 | 41152..44790 | 47880 | BREX-system-P-loop-protein-BrxC-[Pseudomonas-fulva] |
| WP_247325109.1 | Type6.BREX | <-DUF3262<-?<-?<-SMODS<-?\|\|wHTH+WYL+WCX->BrxE->BrxA-DNA-binding->BrxB-BrxF-iSTAND->DUF6079-p*-><-AAA-ATPase<-HIN-HTH\|\|GreA_GreB+SF1-SF2-Helicase-p->N6-METHYLASE->N6-METHYLASE->ParB+HNH->PglZ->SIGMA+OB-Fold+AAA-ATPase+HKD-DNase-BrxL-><-JAB | 371 | Bacteria>Pseudomonadota>Gammaproteobacteria>Pseudomonadales>Pseudomonadaceae>Pseudomonas | GCF_023101305.1 | Pseudomonas_aeruginosa | MX621_RS22735 | 4826578..4827693 | 287 | DUF6079-family-protein-[Pseudomonas-aeruginosa] |
| WP_124171099.1 | Type1.BREX | RelE-ParE-><-?<-wHTH+WYL+WCX\|\|?->Peptidase_M15->?->XIS-HTH->BrxA-DNA-binding->BrxB-BrxF-iSTAND->DUF6079*->N6-METHYLASE->SbcC-p->REase->RVT->N6-METHYLASE-p->PglZ->SIGMA+OB-Fold+AAA-ATPase+HKD-DNase-BrxL-><-Arm-DNA-bind_3+Phage_integrase<-REase+SF1-SF2-Helicase | 1184 | Bacteria>Pseudomonadota>Gammaproteobacteria>Pseudomonadales>Pseudomonadaceae>Pseudomonas | GCF_025697325.1 | Pseudomonas_aeruginosa | IPC368_RS11155 | 506506..510060 | 287 | BREX-system-P-loop-protein-BrxC-[Pseudomonas-aeruginosa] |
| WP_265072431.1 | Type1.BREX | <-SIGMA+OB-Fold+AAA-ATPase+HKD-DNase-BrxL<-PglZ<-ParB+HNH<-N6-METHYLASE-p<-REase<-OLD-ABC<-N6-METHYLASE-p<-DUF6079*<-BrxB-BrxF-iSTAND<-BrxA-DNA-binding<-XIS-HTH\|\|HTH+REase-><-?<-?<-?<-Glyco_hydro_68 | 1186 | Bacteria>Pseudomonadota>Gammaproteobacteria>Pseudomonadales>Pseudomonadaceae>Pseudomonas | GCF_026016265.1 | Pseudomonas_viciae | LOY66_RS25465 | 5798638..5802198 | 2505979 | BREX-system-P-loop-protein-BrxC-[Pseudomonas-viciae] |
| WP_010792139.1 | Type6.BREX | SMODS->?->JAB-><-SIGMA+OB-Fold+AAA-ATPase+HKD-DNase-BrxL<-PglZ<-?<-ParB+HNH<-N6-METHYLASE<-N6-METHYLASE<-DUF6079*<-BrxB-BrxF-iSTAND<-BrxA-DNA-binding<-BrxE<-wHTH+WYL+WCX\|\|JetA->JetB->SbcC-> | 1210 | Bacteria>Pseudomonadota>Gammaproteobacteria>Pseudomonadales>Pseudomonadaceae>Pseudomonas | GCF_026126725.1 | Pseudomonas_aeruginosa | K3Z93_RS02355 | 20406..24038 | 287 | BREX-system-P-loop-protein-BrxC-[Pseudomonas-aeruginosa] |
| WP_265535081.1 | Type1.BREX | <-SIGMA+OB-Fold+AAA-ATPase+HKD-DNase-BrxL<-PglZ<-N6-METHYLASE-p<-HerA/FtsK<-SIR2<-REase<-OLD-ABC<-N6-METHYLASE<-DUF6079*<-BrxB-BrxF-iSTAND<-BrxA-DNA-binding<-XIS-HTH<-?\|\|<-?\|\|sGTP->sGTP->MA+PspA+PspA+sGTP-><-ZnR | 1184 | Bacteria>Pseudomonadota>Gammaproteobacteria>Pseudomonadales>Pseudomonadaceae>Pseudomonas | GCF_026173695.1 | Pseudomonas_saponiphila | ORM81_RS30140 | 14715..18269 | 556534 | BREX-system-P-loop-protein-BrxC-[Pseudomonas-saponiphila] |
| WP_265980027.1 | Type1.BREX | Phage_integrase-><-SIGMA+OB-Fold+AAA-ATPase+HKD-DNase-BrxL<-PglZ<-ParB+HNH<-N6-METHYLASE-p<-?<-NERD<-OLD-ABC<-N6-METHYLASE<-DUF6079*<-BrxB-BrxF-iSTAND<-BrxA-DNA-binding<-XIS-HTH\|\|<-?<-?\|\|wHTH+WYL+WCX-><-JAB | 1184 | Bacteria>Pseudomonadota>Gammaproteobacteria>Pseudomonadales>Pseudomonadaceae>Pseudomonas | GCF_026241005.1 | Pseudomonas_sp._DCB_BG | OO254_RS23990 | 64105..67659 | 2993595 | BREX-system-P-loop-protein-BrxC-[Pseudomonas-sp.-DCB_BG] |
| WP_057380122.1 | Type6.BREX | LZ_Tnp_IS66+zf-IS66+DDE_Tnp_IS66+DDE_Tnp_IS66_C-><-SbcC<-JetB<-JetA\|\|wHTH+WYL+WCX->BrxE->BrxA-DNA-binding->BrxB-BrxF-iSTAND->DUF6079*->GreA_GreB+SF1-SF2-Helicase->N6-METHYLASE->ParB+HNH->?->PglZ->SIGMA+OB-Fold+AAA-ATPase+HKD-DNase-BrxL-><-JAB<-?<-SMODS | 1213 | Bacteria>Pseudomonadota>Gammaproteobacteria>Pseudomonadales>Pseudomonadaceae>Pseudomonas | GCF_026727755.1 | Pseudomonas_aeruginosa | PAC42_RS26725 | 5539866..5543507 | 287 | BREX-system-P-loop-protein-BrxC-[Pseudomonas-aeruginosa] |
| WP_196620444.1 | Type1.BREX | <-XIS-HTH<-SIGMA+OB-Fold+AAA-ATPase+HKD-DNase-BrxL<-PglZ<-N6-METHYLASE-p<-HerA/FtsK<-SIR2<-REase<-OLD-ABC<-N6-METHYLASE<-DUF6079*<-BrxB-BrxF-iSTAND<-BrxA-DNA-binding<-XIS-HTH<-tRNA<-MerR-HTH | 1181 | Bacteria>Pseudomonadota>Gammaproteobacteria>Pseudomonadales>Pseudomonadaceae>Pseudomonas | GCF_027857235.1 | Pseudomonas_aeruginosa | OFL78_RS02500 | 39831..43376 | 287 | BREX-system-P-loop-protein-BrxC-[Pseudomonas-aeruginosa] |
| WP_273916456.1 | Type1.BREX | XIS-HTH->BrxA-DNA-binding->BrxB-BrxF-iSTAND->DUF6079*->N6-METHYLASE->OLD-ABC->NERD->?->?->N6-METHYLASE-p->PglZ->SIGMA+OB-Fold+AAA-ATPase+HKD-DNase-BrxL-> | 1186 | Bacteria>Pseudomonadota>Gammaproteobacteria>Pseudomonadales>Pseudomonadaceae>Pseudomonas | GCF_028656995.1 | Pseudomonas_sp._TNT2022_ID642 | M5G26_RS21745 | 58505..62065 | 2942632 | BREX-system-P-loop-protein-BrxC-[Pseudomonas-sp.-TNT2022-ID642] |
| WP_058409982.1 | Type6.BREX | <-LuxR-HTH<-?<-ZnR+TOPRIM+AAA-helicase+wHTH\|\|XIS-HTH->?->wHTH+WYL+WCX->BrxE->BrxA-DNA-binding->BrxB-BrxF-iSTAND->DUF6079*->N6-METHYLASE->PglZ->SIGMA+OB-Fold+AAA-ATPase+HKD-DNase-BrxL-><-HNH<-?<-?\|\|HTH_3-><-ZnR+ZnR<-DUF4969 | 1212 | Bacteria>Pseudomonadota>Gammaproteobacteria>Pseudomonadales>Pseudomonadaceae>Pseudomonas | GCF_029959515.1 | Pseudomonas_sp._ICMP_19500 | AO239_RS04345 | 8658..12296 | 1208102 | BREX-system-P-loop-protein-BrxC-[Pseudomonas-sp.-ICMP-19500] |
| WP_176947249.1 | Type1.BREX | <-TraK<-TraE<-TraM<-TraM<-DUF3262<-?\|\|XIS-HTH->BrxA-DNA-binding->BrxB-BrxF-iSTAND->DUF6079*->N6-METHYLASE->OLD-ABC->REase->SIR2->HerA/FtsK->N6-METHYLASE-p->PglZ->SIGMA+OB-Fold+AAA-ATPase+HKD-DNase-BrxL->Phage-HTH-> | 1188 | Bacteria>Pseudomonadota>Gammaproteobacteria>Pseudomonadales>Pseudomonadaceae>Pseudomonas | GCF_900100495.1 | Pseudomonas_benzenivorans | BLS63_RS10310 | 86902..90468 | 556533 | BREX-system-P-loop-protein-BrxC-[Pseudomonas-benzenivorans] |
| WP_034038617.1 | Type6.BREX | MerR-HTH->tRNA->BrxE->BrxB-BrxF-iSTAND->DUF6079*->GreA_GreB+SF1-SF2-Helicase->N6-METHYLASE->N6-METHYLASE->ParB+HNH->PglZ->SIGMA+OB-Fold+AAA-ATPase+HKD-DNase-BrxL->XIS-HTH->35exo+SF1-SF2-Helicase->Small-B-barrel+SF1-SF2-Helicase+RdgC+REase-> | 1213 | Bacteria>Pseudomonadota>Gammaproteobacteria>Pseudomonadales>Pseudomonadaceae>Pseudomonas | GCF_900147165.1 | Pseudomonas_aeruginosa | K3Z85_RS02500 | 4652..8293 | 287 | BREX-system-P-loop-protein-BrxC-[Pseudomonas-aeruginosa] |
| WP_076583447.1 | Type1.BREX | DUF4969-><-SIGMA+OB-Fold+AAA-ATPase+HKD-DNase-BrxL<-PglZ<-ParB+HNH<-N6-METHYLASE-p<-RVT<-REase<-OLD-ABC<-N6-METHYLASE<-DUF6079*<-BrxB-BrxF-iSTAND<-BrxA-DNA-binding<-XIS-HTH<-?\|\|wHTH+WYL+WCX-><-?<-?<-KAP-NTPase\|\|LexA-protease-> | 1184 | Bacteria>Pseudomonadota>Gammaproteobacteria>Pseudomonadales>Pseudomonadaceae>Pseudomonas | GCF_900156545.1 | Pseudomonas_alcaligenes | BW984_RS17050 | 273058..276612 | 43263 | BREX-system-P-loop-protein-BrxC-[Pseudomonas-alcaligenes] |
| WP_031684509.1 | Type1.BREX | PhdYeFM_antitox->?-><-wHTH+WYL+WCX\|\|?->Peptidase_M15->?->XIS-HTH->BrxA-DNA-binding->BrxB-BrxF-iSTAND->DUF6079*->N6-METHYLASE->SbcC-p->REase->?->N6-METHYLASE-p->PglZ->SIGMA+OB-Fold+AAA-ATPase+HKD-DNase-BrxL-><-Arm-DNA-bind_3+Phage_integrase\|\|SF1-SF2-Helicase-> | 1184 | Bacteria>Pseudomonadota>Gammaproteobacteria>Pseudomonadales>Pseudomonadaceae>Pseudomonas | GCF_900708485.1 | Pseudomonas_aeruginosa | ID019_RS13300 | 2848972..2852526 | 287 | BREX-system-P-loop-protein-BrxC-[Pseudomonas-aeruginosa] |
| WP_150756947.1 | Type6.BREX | wHTH+WYL+WCX-><-?<-Phage_AlpA<-?<-?\|\|tRNA->BrxE->BrxA-DNA-binding->BrxB-BrxF-iSTAND->DUF6079*->N6-METHYLASE->N6-METHYLASE->N6-METHYLASE->PglZ->SIGMA+OB-Fold+AAA-ATPase+HKD-DNase-BrxL->TUDOR+SF1-SF2-Helicase+iREase+CTD-> | 1210 | Bacteria>Pseudomonadota>Gammaproteobacteria>Pseudomonadales>Pseudomonadaceae>Pseudomonas | GCF_902498135.1 | Pseudomonas_fluorescens | F8N72_RS03340 | 145208..148840 | 294 | BREX-system-P-loop-protein-BrxC-[Pseudomonas-fluorescens] |
| WP_151213669.1 | Type6.BREX | SIR2->?->?-><-?<-TUDOR+SF1-SF2-Helicase+iREase+CTD<-SIGMA+OB-Fold+AAA-ATPase+HKD-DNase-BrxL<-PglZ<-N6-METHYLASE-p<-N6-METHYLASE<-DUF6079*<-BrxB-BrxF-iSTAND<-BrxA-DNA-binding<-BrxE\|\|?->Phage_AlpA->?-><-wHTH+WYL+WCX<-?<-Integrase_1 | 1210 | Bacteria>Pseudomonadota>Gammaproteobacteria>Pseudomonadales>Pseudomonadaceae>Pseudomonas | GCF_911283405.1 | Pseudomonas_fluorescens | F9263_RS07600 | 503497..507129 | 294 | BREX-system-P-loop-protein-BrxC-[Pseudomonas-fluorescens] |
| WP_033944490.1 | Type1.BREX | Arm-DNA-bind_3+Phage_integrase-><-SIGMA+OB-Fold+AAA-ATPase+HKD-DNase-BrxL<-PglZ<-ParB+HNH<-N6-METHYLASE-p<-RVT<-NERD<-OLD-ABC<-N6-METHYLASE<-DUF6079*<-BrxB-BrxF-iSTAND<-BrxA-DNA-binding<-XIS-HTH<-?<-?\|\|?->?->?->HIN-HTH-> | 1184 | Bacteria>Pseudomonadota>Gammaproteobacteria>Pseudomonadales>Pseudomonadaceae>Pseudomonas | GCF_947387505.1 | Pseudomonas_aeruginosa | GPV34_RS03115 | 17071..20625 | 287 | BREX-system-P-loop-protein-BrxC-[Pseudomonas-aeruginosa] |
| WP_095027056.1 | Type6.BREX | <-LZ_Tnp_IS66+zf-IS66+DDE_Tnp_IS66+DDE_Tnp_IS66_C<-?<-HIN-HTH<-CopG/RHH\|\|XIS-HTH->?->wHTH+WYL+WCX->BrxE->BrxB-BrxF-iSTAND->DUF6079*->N6-METHYLASE->ParB+HNH->PglZ->SIGMA+OB-Fold+AAA-ATPase+HKD-DNase-BrxL-><-HNH<-?<-?<-?<-DUF932 | 1213 | Bacteria>Pseudomonadota>Gammaproteobacteria>Pseudomonadales>Pseudomonadaceae | - | Pseudomonas | JFT63_RS16080 | 178886..182527 | 286 | MULTISPECIES:-BREX-system-P-loop-protein-BrxC-[Pseudomonas] |
| WP_121740032.1 | Type6.BREX | <-Tox-RES<-XIS-HTH\|\|HDRnase-><-SIGMA+OB-Fold+AAA-ATPase+HKD-DNase-BrxL<-PglZ<-DOC+HTH<-N6-METHYLASE<-GreA_GreB+SF1-SF2-Helicase<-DUF6079*<-BrxB-BrxF-iSTAND<-BrxA-DNA-binding<-BrxE<-wHTH+WYL+WCX<-?\|\|MODE-HTH+PBPII-><-HTH_3\|\|LysR-HTH->TRPR-HTH-> | 1213 | Bacteria>Pseudomonadota>Gammaproteobacteria>Pseudomonadales>Pseudomonadaceae>Pseudomonas | - | Pseudomonas_aeruginosa | D3M96_03425 | 754727..758368 | 287 | MULTISPECIES:-BREX-system-P-loop-protein-BrxC-[Proteobacteria] |
| WP_150298006.1 | Type6.BREX | <-Arm-DNA-bind_2+Phage_int_SAM_3+Phage_integrase\|\|?-><-?<-RVT\|\|?-><-wHTH+WYL+WCX\|\|BrxE->BrxA-DNA-binding->BrxB-BrxF-iSTAND->DUF6079*->N6-METHYLASE->N6-METHYLASE->ParB+HNH->PglZ->SIGMA+OB-Fold+AAA-ATPase+HKD-DNase-BrxL->?-><-HIN-HTH<-TrwC+SF1-SF2-Helicase<-CopG/RHH | 1212 | Bacteria>Pseudomonadota>Gammaproteobacteria>Pseudomonadales>Pseudomonadaceae>Pseudomonas | - | Pseudomonas_aeruginosa | F3G60_11255 | 510071..513709 | 287 | BREX-system-P-loop-protein-BrxC-[Pseudomonas-aeruginosa] |
| EIU4198817.1 | Type1.BREX | DUF932->?->?->?-><-SIGMA+OB-Fold+AAA-ATPase+HKD-DNase-BrxL<-PglZ<-ParB+HNH<-N6-METHYLASE<-DUF6079*<-BrxB-BrxF-iSTAND<-BrxA-DNA-binding<-XIS-HTH\|\|<-?\|\|sGTP->sGTP->MA+PspA+PspA+sGTP-><-ZnR<-ARG | 1180 | Bacteria>Pseudomonadota>Gammaproteobacteria>Pseudomonadales>Pseudomonadaceae>Pseudomonas | - | Pseudomonas_aeruginosa | L4V49_002912 | 27945..31487 | 287 | BREX-system-P-loop-protein-BrxC-[Pseudomonas-aeruginosa] |
| WP_046164318.1 | Type1.BREX | <-SIGMA+OB-Fold+AAA-ATPase+HKD-DNase-BrxL<-PglZ<-ParB+HNH<-N6-METHYLASE-p<-HerA/FtsK<-SIR2<-REase<-OLD-ABC<-N6-METHYLASE<-DUF6079*<-BrxB-BrxF-iSTAND<-BrxA-DNA-binding<-XIS-HTH<-?\|\|wHTH+WYL+WCX->sGTP->sGTP->sGTP-> | 1186 | Bacteria>Pseudomonadota>Gammaproteobacteria>Pseudomonadales>Pseudomonadaceae>Pseudomonas | - | Pseudomonas_sp._10B238 | RT21_RS21355 | 15645..19205 | 1586417 | BREX-system-P-loop-protein-BrxC-[Pseudomonas-sp.-10B238] |
| WP_223843585.1 | Type6.BREX | JAB-><-wHTH+WYL+WCX\|\|?->?->BrxE->BrxA-DNA-binding->BrxB-BrxF-iSTAND->DUF6079-p->DUF6079-p*->N6-METHYLASE->N6-METHYLASE->?->PglZ->SIGMA+OB-Fold+AAA-ATPase+HKD-DNase-BrxL-><-Phage_integrase<-Integrase_1<-wHTH<-AAA+wHTH | 858 | Bacteria>Pseudomonadota>Gammaproteobacteria>Pseudomonadales>Pseudomonadaceae>Pseudomonas | - | Pseudomonas_sp._CFA | F7661_RS08255 | 1849794..1852370 | 2651048 | BREX-system-P-loop-protein-BrxC-[Pseudomonas-sp.-CFA] |
| WP_223843584.1 | Type6.BREX | JAB-><-wHTH+WYL+WCX\|\|?->?->BrxE->BrxA-DNA-binding->BrxB-BrxF-iSTAND->DUF6079-p*->DUF6079-p->N6-METHYLASE->N6-METHYLASE->?->PglZ->SIGMA+OB-Fold+AAA-ATPase+HKD-DNase-BrxL-><-Phage_integrase<-Integrase_1<-wHTH | 372 | Bacteria>Pseudomonadota>Gammaproteobacteria>Pseudomonadales>Pseudomonadaceae>Pseudomonas | - | Pseudomonas_sp._CFA | F7661_RS27875 | 1848730..1849848 | 2651048 | hypothetical-protein-[Pseudomonas-sp.-CFA] |
| EPL61008.1 | Type1.BREX | <-Phage-HTH<-SIGMA+OB-Fold+AAA-ATPase+HKD-DNase-BrxL<-PglZ<-N6-METHYLASE-p<-HerA/FtsK<-SIR2<-REase<-OLD-ABC<-N6-METHYLASE<-DUF6079*<-BrxB-BrxF-iSTAND<-BrxA-DNA-binding<-XIS-HTH\|\|?->DUF3262->TraM->TraM->TraE->TraK-> | 1188 | Bacteria>Pseudomonadota>Gammaproteobacteria>Pseudomonadales>Pseudomonadaceae>Stutzerimonas | GCA_000416345.1 | Stutzerimonas_stutzeri_B1SMN1 | B382_18428 | 65079..68645 | 1212549 | hypothetical-protein-B382_18428-[Stutzerimonas-stutzeri-B1SMN1] |
| OWG36865.1 | Type1.BREX | <-RNase<-?<-JetB<-?<-wHTH+WYL+WCX\|\|XIS-HTH->BrxA-DNA-binding->BrxB-BrxF-iSTAND->DUF6079*->N6-METHYLASE->ParB+HNH->PglZ->SIGMA+OB-Fold+AAA-ATPase+HKD-DNase-BrxL-><-?<-?<-?<-DUF932 | 1185 | Bacteria>Pseudomonadota>Gammaproteobacteria>Pseudomonadales>Pseudomonadaceae>Stutzerimonas | GCA_002193135.1 | Stutzerimonas_stutzeri | CAQ69_18770 | 20186..23743 | 316 | hypothetical-protein-CAQ69_18770-[Stutzerimonas-stutzeri] |
| PZR87477.1 | Type1.BREX | RHH-><-TetR-HTH\|\|MODE-HTH+PBPII->wHTH+WYL+WCX->BrxA-DNA-binding->BrxB-BrxF-iSTAND->DUF6079*->PglZ->SIGMA+OB-Fold+AAA-ATPase+HKD-DNase-BrxL-><-?<-HAD+HAD | 1179 | Bacteria>Pseudomonadota>Gammaproteobacteria>Pseudomonadales>Pseudomonadaceae>Stutzerimonas | GCA_003243385.1 | Stutzerimonas_stutzeri | DI537_26065 | 4408..7947 | 316 | MAG:-BREX-system-P-loop-protein-BrxC-[Stutzerimonas-stutzeri] |
| MCQ2029875.1 | Type1.BREX | <-sGTP<-sGTP<-sGTP<-wHTH+WYL+WCX\|\|?-><-?\|\|XIS-HTH->BrxA-DNA-binding->BrxB-BrxF-iSTAND->DUF6079*->N6-METHYLASE->OLD-ABC->NERD->RVT->N6-METHYLASE-p->ParB+HNH->PglZ->SIGMA+OB-Fold+AAA-ATPase+HKD-DNase-BrxL->RHH_1-> | 1186 | Bacteria>Pseudomonadota>Gammaproteobacteria>Pseudomonadales>Pseudomonadaceae>Stutzerimonas | GCA_024397535.1 | [Pseudomonas]_zhaodongensis | NA613_09780 | 17231..20791 | 1176257 | BREX-system-P-loop-protein-BrxC-[[Pseudomonas] |
| WP_014819601.1 | Type1.BREX | <-Phage-HTH<-SIGMA+OB-Fold+AAA-ATPase+HKD-DNase-BrxL<-PglZ<-N6-METHYLASE-p<-HerA/FtsK<-SIR2<-REase<-OLD-ABC<-N6-METHYLASE<-DUF6079*<-BrxB-BrxF-iSTAND<-BrxA-DNA-binding<-XIS-HTH\|\|?->DUF3262->TraM->TraM->TraE->TraK-> | 1188 | Bacteria>Pseudomonadota>Gammaproteobacteria>Pseudomonadales>Pseudomonadaceae>Stutzerimonas | GCF_000267545.1 | Stutzerimonas_stutzeri_subgroup | A458_RS28085 | 1395712..1399278 | 578833 | MULTISPECIES:-BREX-system-P-loop-protein-BrxC-[Stutzerimonas-stutzeri-subgroup] |
| WP_023443891.1 | Type6.BREX | <-SbcC<-JetB<-JetA\|\|wHTH+WYL+WCX->BrxE->BrxA-DNA-binding->BrxB-BrxF-iSTAND->DUF6079*->?->GreA_GreB+SF1-SF2-Helicase-p->N6-METHYLASE-><-DUF772+DDE_Tnp_1\|\|N6-METHYLASE->ParB+HNH->PglZ->SIGMA+OB-Fold+AAA-ATPase+HKD-DNase-BrxL-><-JAB | 1213 | Bacteria>Pseudomonadota>Gammaproteobacteria>Pseudomonadales>Pseudomonadaceae>Stutzerimonas | GCF_000495915.1 | Stutzerimonas_stutzeri_group | F753_RS26950 | 57885..61526 | 136846 | MULTISPECIES:-BREX-system-P-loop-protein-BrxC-[Stutzerimonas-stutzeri-group] |
| WP_120997717.1 | Type6.BREX | TrwC+SF1-SF2-Helicase->35exo->?-><-?<-SIGMA+OB-Fold+AAA-ATPase+HKD-DNase-BrxL<-PglZ<-ParB+HNH<-N6-METHYLASE<-N6-METHYLASE<-DUF6079*<-BrxB-BrxF-iSTAND<-BrxA-DNA-binding<-BrxE\|\|?->Arm-DNA-bind_2+Phage_int_SAM_3+Phage_integrase-> | 1212 | Bacteria>Pseudomonadota>Gammaproteobacteria>Pseudomonadales>Pseudomonadaceae>Stutzerimonas | GCF_003640395.1 | [Pseudomonas]_urumqiensis | D7325_RS17370 | 36612..40250 | 638269 | BREX-system-P-loop-protein-BrxC-[[Pseudomonas] |
| WP_125866641.1 | Type6.BREX | JAB-><-SIGMA+OB-Fold+AAA-ATPase+HKD-DNase-BrxL<-PglZ<-?<-ParB+HNH<-N6-METHYLASE<-N6-METHYLASE<-GreA_GreB+SF1-SF2-Helicase<-DUF6079*<-BrxB-BrxF-iSTAND<-BrxA-DNA-binding<-BrxE<-wHTH+WYL+WCX\|\|JetA->JetB->SbcC-> | 1213 | Bacteria>Pseudomonadota>Gammaproteobacteria>Pseudomonadales>Pseudomonadaceae>Stutzerimonas | GCF_003935625.1 | Stutzerimonas_stutzeri | EGJ12_RS17010 | 223453..227094 | 316 | BREX-system-P-loop-protein-BrxC-[Stutzerimonas-stutzeri] |
| WP_212739636.1 | Type6.BREX | <-SbcC<-JetB<-JetA\|\|wHTH+WYL+WCX->BrxE->BrxA-DNA-binding->BrxB-BrxF-iSTAND->DUF6079*->OLD-ABC->PIN-><-HIN-HTH+PAIREDC-HTH\|\|N6-METHYLASE->ParB+HNH->?->PglZ->SIGMA+OB-Fold+AAA-ATPase+HKD-DNase-BrxL-><-JAB | 1207 | Bacteria>Pseudomonadota>Gammaproteobacteria>Pseudomonadales>Pseudomonadaceae>Stutzerimonas | GCF_018316915.2 | Stutzerimonas_balearica | D3P44_RS15315 | 3232724..3236347 | 74829 | BREX-system-P-loop-protein-BrxC-[Stutzerimonas-balearica] |
| WP_255830664.1 | Type1.BREX | <-sGTP<-sGTP<-sGTP<-wHTH+WYL+WCX\|\|?-><-?\|\|XIS-HTH->BrxA-DNA-binding->BrxB-BrxF-iSTAND->DUF6079*->N6-METHYLASE->OLD-ABC->NERD->RVT->N6-METHYLASE-p->ParB+HNH->PglZ->SIGMA+OB-Fold+AAA-ATPase+HKD-DNase-BrxL->RHH_1-> | 1186 | Bacteria>Pseudomonadota>Gammaproteobacteria>Pseudomonadales>Pseudomonadaceae>Stutzerimonas | GCF_024397535.1 | [Pseudomonas]_zhaodongensis | NA613_RS09805 | 17231..20791 | 1176257 | BREX-system-P-loop-protein-BrxC-[[Pseudomonas] |
| WP_238586968.1 | Type1.BREX | <-TraK<-TraE<-TraM<-TraM<-DUF3262<-?\|\|XIS-HTH->BrxA-DNA-binding->BrxB-BrxF-iSTAND->DUF6079-p*->PglZ->SIGMA+OB-Fold+AAA-ATPase+HKD-DNase-BrxL->Phage-HTH-><-DUF4400<-VirD4-TraG | 718 | Bacteria>Pseudomonadota>Gammaproteobacteria>Pseudomonadales>Pseudomonadaceae>Stutzerimonas | - | Stutzerimonas_degradans | . | 57411..59567 | 2968968 | BREX-system-P-loop-protein-BrxC-[Stutzerimonas-degradans] |
| BBJ02099.1 | Type1.BREX | <-TraK<-TraE<-TraM<-TraM<-DUF3262<-?\|\|XIS-HTH->BrxA-DNA-binding->BrxB-BrxF-iSTAND->DUF6079-p*->PglZ->SIGMA+OB-Fold+AAA-ATPase+HKD-DNase-BrxL->Phage-HTH-><-DUF4400<-VirD4-TraG | 718 | Bacteria>Pseudomonadota>Gammaproteobacteria>Pseudomonadales>Pseudomonadaceae>Stutzerimonas | - | Stutzerimonas_degradans | . | 57411..59567 | 2968968 | uncharacterized-protein-sporadically-distributed-in-bacteria-and-archaea-[Stutzerimonas-degradans] |
| WP_020308058.1 | Type1.BREX | <-Phage-HTH<-SIGMA+OB-Fold+AAA-ATPase+HKD-DNase-BrxL<-PglZ<-N6-METHYLASE-p<-HerA/FtsK<-SIR2<-REase<-OLD-ABC<-N6-METHYLASE<-DUF6079*<-BrxB-BrxF-iSTAND<-BrxA-DNA-binding<-XIS-HTH\|\|?->DUF3262->TraM->TraM->TraE->TraK-> | 1188 | Bacteria>Pseudomonadota>Gammaproteobacteria>Pseudomonadales>Pseudomonadaceae>Stutzerimonas | - | Stutzerimonas_stutzeri | B382_18428 | 65079..68645 | 316 | BREX-system-P-loop-protein-BrxC-[Pseudomonas-stutzeri] |
| WP_045633496.1 | Type1.BREX | <-RNase<-?<-JetB<-?<-wHTH+WYL+WCX\|\|XIS-HTH->BrxA-DNA-binding->BrxB-BrxF-iSTAND->DUF6079*->N6-METHYLASE->ParB+HNH->PglZ->SIGMA+OB-Fold+AAA-ATPase+HKD-DNase-BrxL-><-?<-?<-?<-DUF932 | 1185 | Bacteria>Pseudomonadota>Gammaproteobacteria>Pseudomonadales>Pseudomonadaceae>Stutzerimonas | - | Stutzerimonas_stutzeri | CAQ69_18770 | 20186..23743 | 316 | MULTISPECIES:-BREX-system-P-loop-protein-BrxC-[Gammaproteobacteria] |
| WP_041106841.1 | Type6.BREX | <-SbcC<-JetB<-JetA\|\|wHTH+WYL+WCX->BrxE->BrxA-DNA-binding->BrxB-BrxF-iSTAND->DUF6079*->GreA_GreB+SF1-SF2-Helicase->N6-METHYLASE-p->N6-METHYLASE-p->N6-METHYLASE-p->ParB+HNH->?->PglZ->SIGMA+OB-Fold+AAA-ATPase+HKD-DNase-BrxL-><-JAB | 1213 | Bacteria>Pseudomonadota>Gammaproteobacteria>Pseudomonadales>Pseudomonadaceae>Stutzerimonas | - | Stutzerimonas_stutzeri | QX25_RS10015 | 148024..151665 | 316 | BREX-system-P-loop-protein-BrxC-[Stutzerimonas-stutzeri] |
| WP_007146327.1 | Type1.BREX | UbiA-><-HNH<-tRNA<-?<-?<-SIGMA+OB-Fold+AAA-ATPase+HKD-DNase-BrxL<-PglZ<-REase<-ParB+HNH<-N6-METHYLASE<-DUF6079*<-BrxB-BrxF-iSTAND<-BrxA-DNA-binding<-NADAR<-ART-DarT<-?<-?<-Calcineurin | 1179 | Bacteria>Pseudomonadota>Gammaproteobacteria>Thiotrichales>Piscirickettsiaceae>Methylophaga | GCF_000214595.1 | Methylophaga_aminisulfidivorans | MAMP_RS12010 | 87136..90675 | 230105 | BREX-system-P-loop-protein-BrxC-[Methylophaga-aminisulfidivorans] |
| WP_210225661.1 | Type1.BREX | BrxA-DNA-binding->BrxB-BrxF-iSTAND->DUF6079*->ABC-ATPase->?->N6-METHYLASE->N6-METHYLASE-p-><-REase\|\|HATPase+wHTH->PglZ->SIGMA+OB-Fold+AAA-ATPase+HKD-DNase-BrxL-> | 1173 | Bacteria>Pseudomonadota>Gammaproteobacteria>Thiotrichales>Thiotrichaceae>Thiothrix | GCF_017901155.1 | Candidatus_Thiothrix_anitrata | J8380_RS10820 | 2094889..2098410 | 2823902 | BREX-system-P-loop-protein-BrxC-[Candidatus-Thiothrix-anitrata] |

| Protein-ID | System | Gene-Neighbourhoods | Length | Taxonomy | Assembly-ID | Strain | Locus-Tag | Genomic-coordinates | TaxID | Fasta-define |
|---|---|---|---|---|---|---|---|---|---|---|
| WP_130013100.1 | Type1.BREX | MODE-HTH+PBPII->YTCD-HTH-><-HTH_3<-RelE-ParE<-ACET+ACET<-ZnR<-ZnR\|\| BrxA-DNA-binding -> BrxB-BrxF-iSTAND -> DUF6079 *-> PglZ -> SIGMA+OB-Fold+AAA-ATPase+ HKD-DNase -BrxL ->SIR2-><-PRTase | 1214 | Bacteria>Terrabacteria_group>Actinomycetota>Actinomycetes>Bifidobacteriales>Bifidobacteriaceae | GCF_004155015.1 | Bifidobacterium_pseudolongum | PG2054B_RS04700 | 182910..186554 | 1694 | BREX-system-P-loop-protein-BrxC-[Bifidobacterium-pseudolongum] |
| WP_129902658.1 | Type1.BREX | HAD+HAD->MODE-HTH+PBPII->YTCD-HTH-><-HTH_3<-RelE-ParE<-ACET+ACET<-ZnR\|\| BrxA-DNA-binding -> BrxB-BrxF-iSTAND -> DUF6079 *-> N6-METHYLASE -> PglZ -> SIGMA+OB-Fold+AAA-ATPase+ HKD-DNase -BrxL -><-PRTase | 1214 | Bacteria>Terrabacteria_group>Actinomycetota>Actinomycetes>Bifidobacteriales>Bifidobacteriaceae | GCF_004155045.1 | Bifidobacterium_pseudolongum | PG2083B_RS04940 | 90141..93785 | 1694 | BREX-system-P-loop-protein-BrxC-[Bifidobacterium-pseudolongum] |
| WP_129867688.1 | Type1.BREX | HAD+HAD->MODE-HTH+PBPII->YTCD-HTH-><-HTH_3<-RelE-ParE<-ACET+ACET<-ZnR\|\| BrxA-DNA-binding -> BrxB-BrxF-iSTAND -> DUF6079 *-> N6-METHYLASE -> PglZ -> SIGMA+OB-Fold+AAA-ATPase+ HKD-DNase -BrxL -><-PRTase | 1212 | Bacteria>Terrabacteria_group>Actinomycetota>Actinomycetes>Bifidobacteriales>Bifidobacteriaceae | GCF_004155405.1 | Bifidobacterium_pseudolongum | PG102017_RS04060 | 111812..115450 | 1694 | BREX-system-P-loop-protein-BrxC-[Bifidobacterium-pseudolongum] |
| WP_129914280.1 | Type1.BREX | MODE-HTH+PBPII->YTCD-HTH-><-HTH_3<-RelE-ParE<-ACET+ACET<-ZnR<-ZnR\|\| BrxA-DNA-binding -> BrxB-BrxF-iSTAND -> DUF6079 *-> N6-METHYLASE -p-> N6-METHYLASE -> PglZ -> SIGMA+OB-Fold+AAA-ATPase+ HKD-DNase -BrxL ->SIR2-><-PRTase | 1212 | Bacteria>Terrabacteria_group>Actinomycetota>Actinomycetes>Bifidobacteriales>Bifidobacteriaceae | GCF_004155645.1 | Bifidobacterium_pseudolongum | PG2001B_RS04910 | 52519..56157 | 1694 | BREX-system-P-loop-protein-BrxC-[Bifidobacterium-pseudolongum] |
| WP_129869778.1 | Type1.BREX | HAD+HAD->MODE-HTH+PBPII->YTCD-HTH-><-ACET+ACET<-ZnR\|\| BrxA-DNA-binding -> BrxB-BrxF-iSTAND -> DUF6079 *-> N6-METHYLASE -> PglZ -> SIGMA+OB-Fold+AAA-ATPase+ HKD-DNase -BrxL -><-PRTase | 1214 | Bacteria>Terrabacteria_group>Actinomycetota>Actinomycetes>Bifidobacteriales>Bifidobacteriaceae | GCF_004155805.1 | Bifidobacterium_pseudolongum | PG1550B_RS04805 | 1102192..1105836 | 1694 | BREX-system-P-loop-protein-BrxC-[Bifidobacterium-pseudolongum] |
| WP_129900679.1 | Type1.BREX | <-HTH_3<-RelE-ParE<-ACET+ACET<-ZnR\|\| BrxA-DNA-binding -> BrxB-BrxF-iSTAND -> DUF6079 *-> N6-METHYLASE -> PglZ -> SIGMA+OB-Fold+AAA-ATPase+ HKD-DNase -BrxL -><-PRTase | 1214 | Bacteria>Terrabacteria_group>Actinomycetota>Actinomycetes>Bifidobacteriales>Bifidobacteriaceae | GCF_004155855.1 | Bifidobacterium_pseudolongum | PG112206_RS04240 | 4644..8288 | 1694 | BREX-system-P-loop-protein-BrxC-[Bifidobacterium-pseudolongum] |
| WP_129917395.1 | Type1.BREX | BrxA-DNA-binding -> BrxB-BrxF-iSTAND -> DUF6079 *-> N6-METHYLASE -> PglZ -> SIGMA+OB-Fold+AAA-ATPase+ HKD-DNase -BrxL ->SIR2-><-PRTase | 1214 | Bacteria>Terrabacteria_group>Actinomycetota>Actinomycetes>Bifidobacteriales>Bifidobacteriaceae | GCF_004156175.1 | Bifidobacterium_pseudolongum | PG2103B_RS04555 | 1564..5208 | 1694 | BREX-system-P-loop-protein-BrxC-[Bifidobacterium-pseudolongum] |
| WP_131220402.1 | Type1.BREX | HTH_28-> BrxA-DNA-binding -> BrxB-BrxF-iSTAND -> DUF6079 *-> N6-METHYLASE ->?->Schlafen+HATPase+wHTH-> PglZ -> SIGMA+OB-Fold+AAA-ATPase+ HKD-DNase -BrxL ->ZnR+GP->ZnR->ParB+HNH->Sigma-HTH-> | 1219 | Bacteria>Terrabacteria_group>Actinomycetota>Actinomycetes>Bifidobacteriales>Bifidobacteriaceae | GCF_004334345.1 | Bifidobacterium_longum | MCC10040_RS02080 | 100674..104333 | 216816 | BREX-system-P-loop-protein-BrxC-[Bifidobacterium-longum] |
| WP_136510830.1 | Type1.BREX | BrxA-DNA-binding -> BrxB-BrxF-iSTAND -> DUF6079 *-> N6-METHYLASE -> PglZ -> SIGMA+OB-Fold+AAA-ATPase+ HKD-DNase -BrxL ->SIR2-><-PRTase | 1214 | Bacteria>Terrabacteria_group>Actinomycetota>Actinomycetes>Bifidobacteriales>Bifidobacteriaceae | GCF_004801625.1 | Bifidobacterium_pseudolongum | E5991_RS00930 | 3050..6694 | 1694 | BREX-system-P-loop-protein-BrxC-[Bifidobacterium-pseudolongum] |
| WP_140556589.1 | Type1.BREX | <-Cysteine-Desulfurase<-ABC-ATPase+SbcC<-?<-?<-?<- SIGMA+OB-Fold+AAA-ATPase+ HKD-DNase -BrxL <- PglZ <-Schlafen+HATPase+wHTH<- N6-METHYLASE <- DUF6079 *<- BrxB-BrxF-iSTAND <- BrxA-DNA-binding <-?<-Peptidase_M20<-?<-?<-?\|\|Peptidase_A24-> | 1232 | Bacteria>Terrabacteria_group>Actinomycetota>Actinomycetes>Bifidobacteriales>Bifidobacteriaceae | GCF_006408715.1 | Bifidobacterium_sp._UTCIF-39 | EP30_RS04700 | 102584..106282 | 1465359 | BREX-system-P-loop-protein-BrxC-[Bifidobacterium-sp.-UTCIF-39] |
| WP_142421439.1 | Type1.BREX | RNaseH-><-?<- SIGMA+OB-Fold+AAA-ATPase+ HKD-DNase -BrxL <- PglZ <-Schlafen+HATPase+wHTH<-REase<- N6-METHYLASE -p<- N6-METHYLASE <- DUF6079 *<- BrxB-BrxF-iSTAND <- BrxA-DNA-binding \|\|REase->?-><-ACET+ACET\|\|HTH-><-?<-35exo+HRDC | 1218 | Bacteria>Terrabacteria_group>Actinomycetota>Actinomycetes>Bifidobacteriales>Bifidobacteriaceae | GCF_006828885.1 | Bifidobacterium_pseudocatenulatum | GH753_RS08045 | 1981980..1985636 | 28026 | BREX-system-P-loop-protein-BrxC-[Bifidobacterium-pseudocatenulatum] |
| WP_150379196.1 | Type1.BREX | <-SbcD\|\|?-><-ABhydrolase\|\| BrxA-DNA-binding -> BrxB-BrxF-iSTAND -> DUF6079 *->CR-ATPase5+REase->TPR+TPR-> N6-METHYLASE ->HEPN->MazE_antitoxin-> PglZ -> SIGMA+OB-Fold+AAA-ATPase+ HKD-DNase -BrxL ->METHYLASE-> | 1267 | Bacteria>Terrabacteria_group>Actinomycetota>Actinomycetes>Bifidobacteriales>Bifidobacteriaceae | GCF_008698155.1 | Bifidobacterium_myosotis | EM091_RS06000 | 83624..87427 | 1630166 | BREX-system-P-loop-protein-BrxC-[Bifidobacterium-myosotis] |
| WP_150382415.1 | Type1.BREX | ABC-ATPase+SbcC-><-CR-ATPase5+REase\|\|?-><-ZnR\|\|MazE_antitoxin->PemK_toxin-> BrxA-DNA-binding -> BrxB-BrxF-iSTAND -> DUF6079 *-> N6-METHYLASE ->?->?->?->HEPN+RES->MazE_antitoxin->PemK_toxin-> PglZ -> SIGMA+OB-Fold+AAA-ATPase+ HKD-DNase -BrxL -> | 1244 | Bacteria>Terrabacteria_group>Actinomycetota>Actinomycetes>Bifidobacteriales>Bifidobacteriaceae | GCF_008698235.1 | Bifidobacterium_rousetti | CSQ85_RS00640 | 119339..123073 | 2045439 | BREX-system-P-loop-protein-BrxC-[Bifidobacterium-rousetti] |
| WP_151916615.1 | Type1.BREX | HNH->\|\|DOC+HTH-><-sGTP\|\|?->?->?-><-HAD+HAD\|\| BrxA-DNA-binding -> BrxB-BrxF-iSTAND -> DUF6079 *->CR-ATPase5+REase->TPR+TPR-> N6-METHYLASE -><-Tox-RelE<-RelB\|\|CR-ATPase5+REase-> PglZ -> SIGMA+OB-Fold+AAA-ATPase+ HKD-DNase -BrxL ->METHYLASE-> | 1269 | Bacteria>Terrabacteria_group>Actinomycetota>Actinomycetes>Bifidobacteriales>Bifidobacteriaceae | GCF_009078285.1 | Bifidobacterium_jacchi | EHS19_RS04610 | 26586..30395 | 2490545 | BREX-system-P-loop-protein-BrxC-[Bifidobacterium-jacchi] |
| WP_152233646.1 | Type1.BREX | REase+Bacteriocin_IIc->?->?->?->?->?-><-?\|\| BrxA-DNA-binding -> BrxB-BrxF-iSTAND -> DUF6079 *-><-?\|\|CR-ATPase5+REase-> N6-METHYLASE ->?->?->HEPN+RES->Schlafen+HATPase+wHTH-> PglZ -> SIGMA+OB-Fold+AAA-ATPase+ HKD-DNase -BrxL -> | 1235 | Bacteria>Terrabacteria_group>Actinomycetota>Actinomycetes>Bifidobacteriales>Bifidobacteriaceae | GCF_009193355.1 | Bifidobacterium_leontopitheci | F7D09_RS01345 | 123122..126829 | 2650774 | BREX-system-P-loop-protein-BrxC-[Bifidobacterium-leontopitheci] |
| WP_117762529.1 | Type1.BREX | <-ParB+HNH+RAMA<-ZnR<-ZnR+GP<- SIGMA+OB-Fold+AAA-ATPase+ HKD-DNase -BrxL <- PglZ <-Schlafen+HATPase+wHTH<-HEPN<- N6-METHYLASE <- DUF6079 *<- BrxB-BrxF-iSTAND <- BrxA-DNA-binding <-?\|\|?-><-Acyl_transf_3\|\|ACET-> | 1223 | Bacteria>Terrabacteria_group>Actinomycetota>Actinomycetes>Bifidobacteriales>Bifidobacteriaceae | GCF_009832825.1 | Bifidobacterium_adolescentis | DWZ00_RS05085 | 1174565..1178236 | 1680 | BREX-system-P-loop-protein-BrxC-[Bifidobacterium-adolescentis] |
| WP_163196475.1 | Type1.BREX | <- SIGMA+OB-Fold+AAA-ATPase+ HKD-DNase -BrxL <- PglZ <-Schlafen+HATPase+wHTH<-?<- N6-METHYLASE <- DUF6079 *<- BrxB-BrxF-iSTAND <- BrxA-DNA-binding <-ABC-ATPase+SbcC<-?<-ZnR<-?<-Cas_Cas1 | 1248 | Bacteria>Terrabacteria_group>Actinomycetota>Actinomycetes>Bifidobacteriales>Bifidobacteriaceae | GCF_010667645.1 | Bifidobacterium_platyrrhinorum | GFD21_RS03310 | 757948..761694 | 2661628 | BREX-system-P-loop-protein-BrxC-[Bifidobacterium-platyrrhinorum] |
| WP_163113202.1 | Type1.BREX | MULE-TRANSPOSASE-><-ParB+HNH<-ZnR<-ZnR+GP<- SIGMA+OB-Fold+AAA-ATPase+ HKD-DNase -BrxL <- PglZ <-Schlafen+HATPase+wHTH<-SIR2<- N6-METHYLASE <- DUF6079 *<- BrxB-BrxF-iSTAND <- BrxA-DNA-binding \|\|REase-><-HTH+HisKin<-?<-?\|\|?-><-REase | 1218 | Bacteria>Terrabacteria_group>Actinomycetota>Actinomycetes>Bifidobacteriales>Bifidobacteriaceae | GCF_011029205.1 | Bifidobacterium_bifidum | G5T23_RS07350 | 43765..47421 | 1681 | BREX-system-P-loop-protein-BrxC-[Bifidobacterium-bifidum] |
| WP_168445148.1 | Type1.BREX | <-Glyoxalase\|\|DSBH+AraC-HTH+AraC-HTH-><-ZnR<- SIGMA+OB-Fold+AAA-ATPase+ HKD-DNase -BrxL <- PglZ <- PglZ <-Schlafen+HATPase+wHTH<-?<- N6-METHYLASE <- DUF6079 *<- BrxB-BrxF-iSTAND <- BrxA-DNA-binding \|\|REase-><-HEPN<-?\|\|HTH-><-35exo+BRCT<-?<-35exo+HRDC | 1217 | Bacteria>Terrabacteria_group>Actinomycetota>Actinomycetes>Bifidobacteriales>Bifidobacteriaceae | GCF_012310535.1 | Bifidobacterium_dentium | PG133_RS04075 | 130679..134332 | 1689 | BREX-system-P-loop-protein-BrxC-[Bifidobacterium-dentium] |
| WP_169274836.1 | Type1.BREX | <-PIN<-RelB<-?\|\|HTH_3->?->PIN-> BrxA-DNA-binding -> BrxB-BrxF-iSTAND -> DUF6079 *->CR-ATPase5+REase-> N6-METHYLASE ->Schlafen+HATPase+wHTH-> PglZ -> SIGMA+OB-Fold+AAA-ATPase+ HKD-DNase -BrxL ->REase->?-> BrxA-DNA-binding -> | 1226 | Bacteria>Terrabacteria_group>Actinomycetota>Actinomycetes>Bifidobacteriales>Bifidobacteriaceae | GCF_012932365.1 | Bifidobacterium_sp._DSM_109958 | G1C96_RS01295 | 288582..292262 | 2705430 | BREX-system-P-loop-protein-BrxC-[Bifidobacterium-sp.-DSM-109958] |
| WP_193020253.1 | Type1.BREX | HTH+MPTase-><-?<-HEPN<-?<-?\|\|?-> BrxA-DNA-binding -> BrxB-BrxF-iSTAND -> DUF6079 *-> N6-METHYLASE ->REase->Schlafen+wHTH-> PglZ -> SIGMA+OB-Fold+AAA-ATPase+ HKD-DNase -BrxL ->?->?->RNase-> | 1229 | Bacteria>Terrabacteria_group>Actinomycetota>Actinomycetes>Bifidobacteriales>Bifidobacteriaceae | GCF_014898135.1 | Bifidobacterium_longum | BL5934_RS04380 | 976256..979945 | 216816 | BREX-system-P-loop-protein-BrxC-[Bifidobacterium-longum] |
| WP_200406515.1 | Type1.BREX | HTH_28+HTH_28->HTH_Tnp_1+HTH_21+rve+rve_3->HTH_28->DEDD_Tnp_IS110+Transposase_20-> BrxA-DNA-binding -> BrxB-BrxF-iSTAND -> DUF6079 *-> N6-METHYLASE ->?->Schlafen+HATPase+wHTH-> PglZ -> SIGMA+OB-Fold+AAA-ATPase+ HKD-DNase -BrxL ->?-><-?<-?<-ABC-ATPase+SbcC | 1219 | Bacteria>Terrabacteria_group>Actinomycetota>Actinomycetes>Bifidobacteriales>Bifidobacteriaceae | GCF_014898255.1 | Bifidobacterium_longum | BL7055_RS03240 | 790763..794422 | 216816 | BREX-system-P-loop-protein-BrxC-[Bifidobacterium-longum] |
| WP_227411048.1 | Type1.BREX | ACET->ACET+ACET->?->Thioredoxin_3->ArsR-HTH-><-Glycoside_Hydrolase\|\|OLD-ABC-> BrxA-DNA-binding -> BrxB-BrxF-iSTAND -> DUF6079 *-> N6-METHYLASE ->REase->?->Schlafen+HATPase+wHTH-> PglZ -> SIGMA+OB-Fold+AAA-ATPase+ HKD-DNase -BrxL -> | 1242 | Bacteria>Terrabacteria_group>Actinomycetota>Actinomycetes>Bifidobacteriales>Bifidobacteriaceae | GCF_014983005.1 | Bifidobacterium_pullorum | INF27_RS02890 | 183960..187688 | 78448 | BREX-system-P-loop-protein-BrxC-[Bifidobacterium-pullorum] |
| WP_195393981.1 | Type1.BREX | PemK_toxin->?->PolB->HEPN-><-?<-?\|\|?-> BrxA-DNA-binding -> BrxB-BrxF-iSTAND -> DUF6079 *-> N6-METHYLASE ->SIR2->Schlafen+HATPase+wHTH-> PglZ -> SIGMA+OB-Fold+AAA-ATPase+ HKD-DNase -BrxL ->ZnR+GP->ZnR->ParB+HNH+RAMA-> | 1218 | Bacteria>Terrabacteria_group>Actinomycetota>Actinomycetes>Bifidobacteriales>Bifidobacteriaceae | GCF_015553845.1 | Bifidobacterium_adolescentis | I2D91_RS01775 | 423737..427393 | 1680 | BREX-system-P-loop-protein-BrxC-[Bifidobacterium-adolescentis] |
| WP_195451643.1 | Type1.BREX | <-ABhydrolase<-?<-Glyoxalase\|\|DSBH+AraC-HTH+AraC-HTH-><-ZnR<- SIGMA+OB-Fold+AAA-ATPase+ HKD-DNase -BrxL <- PglZ <-Schlafen+HATPase+wHTH<- N6-METHYLASE <- DUF6079 *<- BrxB-BrxF-iSTAND <- BrxA-DNA-binding <-?\|\|ACET-><-?\|\|HTH-><-35exo+BRCT<-35exo+HRDC | 1217 | Bacteria>Terrabacteria_group>Actinomycetota>Actinomycetes>Bifidobacteriales>Bifidobacteriaceae | GCF_015555105.1 | Bifidobacterium_dentium | I2E42_RS03170 | 226129..229782 | 1689 | BREX-system-P-loop-protein-BrxC-[Bifidobacterium-dentium] |
| WP_195482034.1 | Type1.BREX | <-ParB+HNH+RAMA<-ZnR<-ZnR+GP<- SIGMA+OB-Fold+AAA-ATPase+ HKD-DNase -BrxL <- PglZ <-Schlafen+HATPase+wHTH<-SIR2<- N6-METHYLASE <- DUF6079 *<- BrxB-BrxF-iSTAND <- BrxA-DNA-binding <-?\|\|?->?-><-HEPN<-PolB<-PemK_toxin<-RelB | 1218 | Bacteria>Terrabacteria_group>Actinomycetota>Actinomycetes>Bifidobacteriales>Bifidobacteriaceae | GCF_015555595.1 | Bifidobacterium_adolescentis | I2E65_RS01565 | 362593..366249 | 1680 | BREX-system-P-loop-protein-BrxC-[Bifidobacterium-adolescentis] |
| WP_195936831.1 | Type1.BREX | REase-><-?<- SIGMA+OB-Fold+AAA-ATPase+ HKD-DNase -BrxL <- PglZ <-Schlafen+HATPase+wHTH<-?<- N6-METHYLASE <- DUF6079 *<- BrxB-BrxF-iSTAND <- BrxA-DNA-binding <-?<-?<-?<-ABC-ATPase+SbcC<-ZnR | 1218 | Bacteria>Terrabacteria_group>Actinomycetota>Actinomycetes>Bifidobacteriales>Bifidobacteriaceae | GCF_015667555.1 | Bifidobacterium_bifidum | I4U71_RS07235 | 13396..17052 | 1681 | BREX-system-P-loop-protein-BrxC-[Bifidobacterium-bifidum] |
| WP_195938980.1 | Type1.BREX | 35exo+HRDC-><-HTH\|\|ACET+ACET-><-?<-SbcD<-?<-REase\|\| BrxA-DNA-binding -> BrxB-BrxF-iSTAND -> DUF6079 *-> N6-METHYLASE ->SIR2->Schlafen+HATPase+wHTH-> PglZ -> SIGMA+OB-Fold+AAA-ATPase+ HKD-DNase -BrxL ->ZnR+GP->ZnR->ParB+HNH+RAMA-> | 1217 | Bacteria>Terrabacteria_group>Actinomycetota>Actinomycetes>Bifidobacteriales>Bifidobacteriaceae | GCF_015667675.1 | Bifidobacterium_pseudocatenulatum | I4U64_RS06630 | 371599..375252 | 28026 | BREX-system-P-loop-protein-BrxC-[Bifidobacterium-pseudocatenulatum] |
| WP_196336889.1 | Type1.BREX | 35exo+HRDC->35exo+BRCT-><-HTH\|\|?-><-ACET\|\|?-> BrxA-DNA-binding -> BrxB-BrxF-iSTAND -> DUF6079 *-> N6-METHYLASE ->HEPN->?->Schlafen+HATPase+wHTH-> PglZ -> SIGMA+OB-Fold+AAA-ATPase+ HKD-DNase -BrxL ->ZnR-><-DSBH+AraC-HTH+AraC-HTH\|\|Glyoxalase-> | 1217 | Bacteria>Terrabacteria_group>Actinomycetota>Actinomycetes>Bifidobacteriales>Bifidobacteriaceae | GCF_015703835.1 | Bifidobacterium_dentium | IHQ82_RS06050 | 179848..183501 | 1689 | BREX-system-P-loop-protein-BrxC-[Bifidobacterium-dentium] |
| WP_202574831.1 | Type1.BREX | <-Peptidase_S26<-?\|\|ABC-ATPase+SbcC->?-><-?<- SIGMA+OB-Fold+AAA-ATPase+ HKD-DNase -BrxL <- PglZ <-Schlafen+HATPase+wHTH<- N6-METHYLASE <- DUF6079 *<- BrxB-BrxF-iSTAND <- BrxA-DNA-binding <-HTH_28 | 1219 | Bacteria>Terrabacteria_group>Actinomycetota>Actinomycetes>Bifidobacteriales>Bifidobacteriaceae | GCF_016759645.1 | Bifidobacterium_longum | JL737_RS05465 | 24879..28538 | 216816 | BREX-system-P-loop-protein-BrxC-[Bifidobacterium-longum] |
| WP_204464696.1 | Type1.BREX | ACET-><-O6MGT-HTH\|\|Phage_integrase-><-?<- SIGMA+OB-Fold+AAA-ATPase+ HKD-DNase -BrxL <- PglZ <-Schlafen+HATPase+wHTH<- N6-METHYLASE <- DUF6079 *<- BrxB-BrxF-iSTAND <- BrxA-DNA-binding <-?<-?<-?\|\|Glycoside_Hydrolase-><-ArsR-HTH<-Thioredoxin_3 | 1221 | Bacteria>Terrabacteria_group>Actinomycetota>Actinomycetes>Bifidobacteriales>Bifidobacteriaceae | GCF_016899675.1 | Bifidobacterium_pullorum | H6A68_RS03865 | 164290..167955 | 78448 | BREX-system-P-loop-protein-BrxC-[Bifidobacterium-pullorum] |
| WP_204389272.1 | Type1.BREX | <-IstB_IS21\|\|AAA-ATPase-><-CMP-hydrolase<- SIGMA+OB-Fold+AAA-ATPase+ HKD-DNase -BrxL <- PglZ <-Schlafen+HATPase+wHTH<- N6-METHYLASE <- DUF6079 *<- BrxB-BrxF-iSTAND <- BrxA-DNA-binding <-TOPRIM<-OLD-ABC\|\|Glycoside_Hydrolase-><-ACET+ACET<-ACET | 1221 | Bacteria>Terrabacteria_group>Actinomycetota>Actinomycetes>Bifidobacteriales>Bifidobacteriaceae | GCF_016900615.1 | Bifidobacterium_pullorum | H6A14_RS02700 | 186110..189775 | 78448 | BREX-system-P-loop-protein-BrxC-[Bifidobacterium-pullorum] |
| WP_217054570.1 | Type1.BREX | <- SIGMA+OB-Fold+AAA-ATPase+ HKD-DNase -BrxL <- PglZ <-HEPN+RES<-?<-?<-REase<- N6-METHYLASE <- DUF6079 *<- BrxB-BrxF-iSTAND <- BrxA-DNA-binding \|\|?->CR-Kinase-><-CR-ATPase5+REase-p<-REase<-HTH_3 | 1229 | Bacteria>Terrabacteria_group>Actinomycetota>Actinomycetes>Bifidobacteriales>Bifidobacteriaceae | GCF_019041665.1 | Bifidobacterium_longum | KTF47_RS04030 | 15252..18941 | 216816 | BREX-system-P-loop-protein-BrxC-[Bifidobacterium-longum] |
| WP_219100165.1 | Type1.BREX | <-ABhydrolase\|\|?->MACRODOMAIN-><-?<- SIGMA+OB-Fold+AAA-ATPase+ HKD-DNase -BrxL <- PglZ <-?<-?<- N6-METHYLASE <- DUF6079 *<- BrxB-BrxF-iSTAND <- BrxA-DNA-binding \|\|?-><-ABhydrolase | 1253 | Bacteria>Terrabacteria_group>Actinomycetota>Actinomycetes>Bifidobacteriales>Bifidobacteriaceae | GCF_019331695.1 | Bifidobacterium_pluvialisilvae | KIH77_RS05905 | 203323..207084 | 2834436 | BREX-system-P-loop-protein-BrxC-[Bifidobacterium-pluvialisilvae] |
| WP_226586661.1 | Type1.BREX | REC+LuxR-HTH->HISKIN->?->?-><-SbcD<-?<-REase\|\| BrxA-DNA-binding -> BrxB-BrxF-iSTAND -> DUF6079 *-> N6-METHYLASE ->?->Schlafen+HATPase+wHTH-> PglZ -> SIGMA+OB-Fold+AAA-ATPase+ HKD-DNase -BrxL ->SLATT->SMODS->ParB+HNH+RAMA-> | 1219 | Bacteria>Terrabacteria_group>Actinomycetota>Actinomycetes>Bifidobacteriales>Bifidobacteriaceae | GCF_020525765.1 | Bifidobacterium_pseudocatenulatum | PG776_RS01085 | 258368..262027 | 28026 | BREX-system-P-loop-protein-BrxC-[Bifidobacterium-pseudocatenulatum] |
| WP_226650300.1 | Type1.BREX | <- SIGMA+OB-Fold+AAA-ATPase+ HKD-DNase -BrxL <- PglZ <-Schlafen+HATPase+wHTH<-REase<- N6-METHYLASE <- DUF6079 *<- BrxB-BrxF-iSTAND <- BrxA-DNA-binding \|\|REase->?->SbcD-><-?\|\|?-><-?\|\|MerR-HTH-> | 1219 | Bacteria>Terrabacteria_group>Actinomycetota>Actinomycetes>Bifidobacteriales>Bifidobacteriaceae | GCF_020526585.1 | Bifidobacterium_pseudocatenulatum | KZP10_RS05695 | 1065058..1068717 | 28026 | BREX-system-P-loop-protein-BrxC-[Bifidobacterium-pseudocatenulatum] |
| WP_235341612.1 | Type1.BREX | METHYLASE->?->Glycos_transf_4-><-?\|\|?->?-> BrxA-DNA-binding -> BrxB-BrxF-iSTAND -> DUF6079 *-> PglZ -> SIGMA+OB-Fold+AAA-ATPase+ HKD-DNase -BrxL ->?->?->35exo->HAD+S6PP->SF1-SF2-Helicase+RNase-> | 1238 | Bacteria>Terrabacteria_group>Actinomycetota>Actinomycetes>Bifidobacteriales>Bifidobacteriaceae | GCF_021531155.1 | Bifidobacterium_boum | I6E06_RS03585 | 150899..154615 | 78343 | BREX-system-P-loop-protein-BrxC-[Bifidobacterium-boum] |
| WP_237919300.1 | Type1.BREX | ACET+ACET->?-><-?\|\|?-><-SbcD\|\|?-><-?\|\| BrxA-DNA-binding -> BrxB-BrxF-iSTAND -> DUF6079 *-> N6-METHYLASE ->SIR2->Schlafen+HATPase+wHTH-> PglZ -> SIGMA+OB-Fold+AAA-ATPase+ HKD-DNase -BrxL -> | 1219 | Bacteria>Terrabacteria_group>Actinomycetota>Actinomycetes>Bifidobacteriales>Bifidobacteriaceae | GCF_022135125.1 | Bifidobacterium_pseudocatenulatum | LOM88_RS00205 | 52697..56356 | 28026 | BREX-system-P-loop-protein-BrxC-[Bifidobacterium-pseudocatenulatum] |
| WP_237970585.1 | Type1.BREX | HTH_28-> BrxA-DNA-binding -> BrxB-BrxF-iSTAND -> DUF6079 *-> N6-METHYLASE ->?->Schlafen+HATPase+wHTH-> PglZ -> SIGMA+OB-Fold+AAA-ATPase+ HKD-DNase -BrxL ->ZnR+GP->ZnR->ParB+HNH->Sigma-HTH-> | 1218 | Bacteria>Terrabacteria_group>Actinomycetota>Actinomycetes>Bifidobacteriales>Bifidobacteriaceae | GCF_022136875.1 | Bifidobacterium_longum | LOO84_RS04690 | 81950..85606 | 216816 | BREX-system-P-loop-protein-BrxC-[Bifidobacterium-longum] |
| WP_273192946.1 | Type1.BREX | <-SF1-SF2-Helicase\|\|?->?->?-><-SPOUT\|\|ENDO3+NUDIX->?-> BrxA-DNA-binding -> BrxB-BrxF-iSTAND -> DUF6079 *->?-> N6-METHYLASE -><- N6-METHYLASE -p\|\|Phage_int_SAM_3+Phage_integrase-> N6-METHYLASE -p-> PglZ -> SIGMA+OB-Fold+AAA-ATPase+ HKD-DNase -BrxL -> | 1173 | Bacteria>Terrabacteria_group>Actinomycetota>Actinomycetes>Bifidobacteriales>Bifidobacteriaceae | GCF_022487345.1 | Bifidobacterium_tibiigranuli | LKF49_RS06765 | 121598..125119 | 2172043 | BREX-system-P-loop-protein-BrxC-[Bifidobacterium-tibiigranuli] |
| WP_240833140.1 | Type1.BREX | <-Abhydrolase<-ParB+HNH+RAMA<-ZnR<-ZnR+GP<- SIGMA+OB-Fold+AAA-ATPase+ HKD-DNase -BrxL <- PglZ <-Schlafen+HATPase+wHTH<-?<- N6-METHYLASE <- DUF6079 *<- BrxB-BrxF-iSTAND <- BrxA-DNA-binding \|\|REase->?-><-?\|\|SbcD-> | 1217 | Bacteria>Terrabacteria_group>Actinomycetota>Actinomycetes>Bifidobacteriales>Bifidobacteriaceae | GCF_022496395.1 | Bifidobacterium_pseudocatenulatum | LF914_RS07635 | 54595..58248 | 28026 | BREX-system-P-loop-protein-BrxC-[Bifidobacterium-pseudocatenulatum] |
| WP_242075534.1 | Type1.BREX | HTH_28-> BrxA-DNA-binding -> BrxB-BrxF-iSTAND -> DUF6079 *-> N6-METHYLASE ->Schlafen+HATPase+wHTH-> PglZ -> SIGMA+OB-Fold+AAA-ATPase+ HKD-DNase -BrxL ->?-><-?<-?<-ABC-ATPase+SbcC | 1218 | Bacteria>Terrabacteria_group>Actinomycetota>Actinomycetes>Bifidobacteriales>Bifidobacteriaceae | GCF_022643745.1 | Bifidobacterium_longum | G8B15_RS07005 | 1527026..1530682 | 216816 | BREX-system-P-loop-protein-BrxC-[Bifidobacterium-longum] |
| WP_032684176.1 | Type1.BREX | <- SIGMA+OB-Fold+AAA-ATPase+ HKD-DNase -BrxL <- PglZ <-Schlafen+HATPase+wHTH<- N6-METHYLASE -p<-HerA/FtsK<-SIR2<- N6-METHYLASE <- DUF6079 *<- BrxB-BrxF-iSTAND <- BrxA-DNA-binding <-HTH_28 | 1219 | Bacteria>Terrabacteria_group>Actinomycetota>Actinomycetes>Bifidobacteriales>Bifidobacteriaceae | GCF_022701235.1 | Bifidobacterium_longum | LMY38_RS03520 | 757375..761034 | 216816 | BREX-system-P-loop-protein-BrxC-[Bifidobacterium-longum] |
| WP_248110663.1 | Type1.BREX | 35exo+HRDC->?->35exo+BRCT-><-HTH\|\|?-><-ACET\|\|?-> BrxA-DNA-binding -> BrxB-BrxF-iSTAND -> DUF6079 *-> N6-METHYLASE ->?->Schlafen+HATPase+wHTH-> PglZ -> SIGMA+OB-Fold+AAA-ATPase+ HKD-DNase -BrxL ->ZnR-><-DSBH+AraC-HTH+AraC-HTH\|\|Glyoxalase-> | 1219 | Bacteria>Terrabacteria_group>Actinomycetota>Actinomycetes>Bifidobacteriales>Bifidobacteriaceae | GCF_023147995.1 | Bifidobacterium_dentium | KZX63_RS03535 | 164390..168049 | 1689 | BREX-system-P-loop-protein-BrxC-[Bifidobacterium-dentium] |
| WP_269374243.1 | Type1.BREX | <-Sigma-HTH<-ParB+HNH<-ZnR<-ZnR+GP<- SIGMA+OB-Fold+AAA-ATPase+ HKD-DNase -BrxL <- PglZ <-Schlafen+HATPase+wHTH<-SIR2<- N6-METHYLASE <- DUF6079 *<- BrxB-BrxF-iSTAND <- BrxA-DNA-binding | 1219 | Bacteria>Terrabacteria_group>Actinomycetota>Actinomycetes>Bifidobacteriales>Bifidobacteriaceae | GCF_027214485.1 | Bifidobacterium_breve | D4C76_RS09020 | 75244..78903 | 1685 | BREX-system-P-loop-protein-BrxC-[Bifidobacterium-breve] |
| WP_269355165.1 | Type1.BREX | <-Sigma-HTH<-ParB+HNH<-ZnR<-ZnR+GP<- SIGMA+OB-Fold+AAA-ATPase+ HKD-DNase -BrxL <- PglZ <-Schlafen+HATPase+wHTH<-SIR2<- N6-METHYLASE <- DUF6079 *<- BrxB-BrxF-iSTAND <- BrxA-DNA-binding <-HTH_28<-?\|\|tRNA->tRNA->?-><-?<-RNasePH | 1218 | Bacteria>Terrabacteria_group>Actinomycetota>Actinomycetes>Bifidobacteriales>Bifidobacteriaceae | GCF_027215185.1 | Bifidobacterium_breve | D4C27_RS08995 | 67566..71222 | 1685 | BREX-system-P-loop-protein-BrxC-[Bifidobacterium-breve] |
| WP_270294263.1 | Type1.BREX | <- SIGMA+OB-Fold+AAA-ATPase+ HKD-DNase -BrxL <- PglZ <-Schlafen+HATPase+wHTH<-SIR2<- N6-METHYLASE -p<-HerA/FtsK<-SIR2<- N6-METHYLASE <- DUF6079 *<- BrxB-BrxF-iSTAND <- BrxA-DNA-binding \|\|REase->?->SbcD-><-?\|\|?-><-HISKIN<-REC+LuxR-HTH | 1219 | Bacteria>Terrabacteria_group>Actinomycetota>Actinomycetes>Bifidobacteriales>Bifidobacteriaceae | GCF_027659585.1 | Bifidobacterium_adolescentis | PGD90_RS01995 | 459377..463036 | 1680 | BREX-system-P-loop-protein-BrxC-[Bifidobacterium-adolescentis] |
| WP_270267401.1 | Type1.BREX | <- SIGMA+OB-Fold+AAA-ATPase+ HKD-DNase -BrxL <- PglZ <-RNase<-BirA-HTH<-Schlafen+HATPase+wHTH<-SMODS<-SLATT<- N6-METHYLASE <- DUF6079 *<- BrxB-BrxF-iSTAND <- BrxA-DNA-binding \|\|?->CR-Kinase-><-CR-ATPase5+REase-p<- N6-METHYLASE <-RelB | 1229 | Bacteria>Terrabacteria_group>Actinomycetota>Actinomycetes>Bifidobacteriales>Bifidobacteriaceae | GCF_027663845.1 | Bifidobacterium_breve | PGQ90_RS07415 | 209217..212906 | 1685 | BREX-system-P-loop-protein-BrxC-[Bifidobacterium-breve] |
| WP_117779802.1 | Type1.BREX | <- SIGMA+OB-Fold+AAA-ATPase+ HKD-DNase -BrxL <- PglZ <-Schlafen+HATPase+wHTH<-SIR2<- N6-METHYLASE -p<-HerA/FtsK<-SIR2<- N6-METHYLASE <- DUF6079 *<- BrxB-BrxF-iSTAND <- BrxA-DNA-binding \|\|REase->?->?-><-RelE-ParE<-PhdYeFM_antitox\|\|SbcD-> | 1219 | Bacteria>Terrabacteria_group>Actinomycetota>Actinomycetes>Bifidobacteriales>Bifidobacteriaceae | GCF_027664445.1 | Bifidobacterium_adolescentis | PGQ86_RS05915 | 1356414..1360073 | 1680 | BREX-system-P-loop-protein-BrxC-[Bifidobacterium-adolescentis] |
| WP_270271788.1 | Type1.BREX | REase-><-?\|\|?->?->HTH+HisKin-><-REase\|\| BrxA-DNA-binding -> BrxB-BrxF-iSTAND -> DUF6079 *-> N6-METHYLASE ->?->Schlafen+HATPase+wHTH-> PglZ -> SIGMA+OB-Fold+AAA-ATPase+ HKD-DNase -BrxL ->ZnR+GP->ZnR->ParB+HNH-> | 1218 | Bacteria>Terrabacteria_group>Actinomycetota>Actinomycetes>Bifidobacteriales>Bifidobacteriaceae | GCF_027667785.1 | Bifidobacterium_bifidum | PGS22_RS05345 | 127181..130837 | 1681 | BREX-system-P-loop-protein-BrxC-[Bifidobacterium-bifidum] |
| WP_117715835.1 | Type1.BREX | ZnR->?->ABC-ATPase+SbcC->?->?->?-> BrxA-DNA-binding -> BrxB-BrxF-iSTAND -> DUF6079 *-> N6-METHYLASE ->SIR2->Schlafen+HATPase+wHTH-> PglZ -> SIGMA+OB-Fold+AAA-ATPase+ HKD-DNase -BrxL ->?-><-REase\|\|?-><-Glyoxalase | 1217 | Bacteria>Terrabacteria_group>Actinomycetota>Actinomycetes>Bifidobacteriales>Bifidobacteriaceae | GCF_027669765.1 | Bifidobacterium_bifidum | PHA16_RS02375 | 586926..590579 | 1681 | BREX-system-P-loop-protein-BrxC-[Bifidobacterium-bifidum] |
| WP_270200767.1 | Type1.BREX | REC+LuxR-HTH->HISKIN-><-?<-SbcD<-?<-?\|\|Pentapeptide+Pentapeptide+Pentapeptide+Pentapeptide-> BrxA-DNA-binding -> BrxB-BrxF-iSTAND -> DUF6079 -p*-> DUF6079 -p-> N6-METHYLASE ->REase->Schlafen+HATPase+wHTH-> PglZ -> SIGMA+OB-Fold+AAA-ATPase+ HKD-DNase -BrxL ->?-><-RNaseH | 467 | Bacteria>Terrabacteria_group>Actinomycetota>Actinomycetes>Bifidobacteriales>Bifidobacteriaceae | GCF_027671795.1 | Bifidobacterium_pseudocatenulatum | PGT11_RS00265 | 67391..68794 | 28026 | BREX-system-P-loop-protein-BrxC-[Bifidobacterium-pseudocatenulatum] |
| WP_270199955.1 | Type1.BREX | <-SbcD<-?<-?\|\|?->\|\|Pentapeptide+Pentapeptide+Pentapeptide+Pentapeptide-> BrxA-DNA-binding -> BrxB-BrxF-iSTAND -> DUF6079 *-> DUF6079 -p-> N6-METHYLASE ->REase->Schlafen+HATPase+wHTH-> PglZ -> SIGMA+OB-Fold+AAA-ATPase+ HKD-DNase -BrxL ->?-><-RNaseH | 1044 | Bacteria>Terrabacteria_group>Actinomycetota>Actinomycetes>Bifidobacteriales>Bifidobacteriaceae | GCF_027675485.1 | Bifidobacterium_pseudocatenulatum | PGV18_RS06405 | 41954..45088 | 28026 | BREX-system-P-loop-protein-BrxC-[Bifidobacterium-pseudocatenulatum] |
| WP_270271488.1 | Type1.BREX | <-ZnR<-ZnR+GP<- SIGMA+OB-Fold+AAA-ATPase+ HKD-DNase -BrxL <- PglZ <-Schlafen+HATPase+wHTH<-SIR2<- N6-METHYLASE <- DUF6079 *<- BrxB-BrxF-iSTAND <- BrxA-DNA-binding <-Pentapeptide+Pentapeptide+Pentapeptide<-HAD\|\|SbcD-><-?<-ACET+ACET\|\|HTH-><-35exo+BRCT | 1219 | Bacteria>Terrabacteria_group>Actinomycetota>Actinomycetes>Bifidobacteriales>Bifidobacteriaceae | GCF_027679525.1 | Bifidobacterium_adolescentis | PGW19_RS08965 | 15037..18696 | 1680 | BREX-system-P-loop-protein-BrxC-[Bifidobacterium-adolescentis] |
| WP_270447025.1 | Type1.BREX | HISKIN-><-?\|\|<-SbcD<-?<-?\|\|?->\|\|Pentapeptide+Pentapeptide+Pentapeptide+Pentapeptide-> BrxA-DNA-binding -> BrxB-BrxF-iSTAND -> DUF6079 -p*-> DUF6079 -p-> N6-METHYLASE ->SIR2->HerA/FtsK-> N6-METHYLASE -p->SIR2->Schlafen+HATPase+wHTH-> PglZ -> SIGMA+OB-Fold+AAA-ATPase+ HKD-DNase -BrxL -> | 958 | Bacteria>Terrabacteria_group>Actinomycetota>Actinomycetes>Bifidobacteriales>Bifidobacteriaceae | GCF_027682665.1 | Bifidobacterium_pseudocatenulatum | PGX68_RS01375 | 343601..346477 | 28026 | BREX-system-P-loop-protein-BrxC-[Bifidobacterium-pseudocatenulatum] |
| WP_270479593.1 | Type1.BREX | <-ZnR+GP<- SIGMA+OB-Fold+AAA-ATPase+ HKD-DNase -BrxL <- PglZ <-Schlafen+HATPase+wHTH<-SIR2<- N6-METHYLASE -p<-HerA/FtsK<-SIR2<- N6-METHYLASE <- DUF6079 *<- BrxB-BrxF-iSTAND <- BrxA-DNA-binding \|\|?->HEPN-><-?\|\|?->?-><-HEPN<-PolB | 1219 | Bacteria>Terrabacteria_group>Actinomycetota>Actinomycetes>Bifidobacteriales>Bifidobacteriaceae | GCF_027683145.1 | Bifidobacterium_adolescentis | PGX62_RS02115 | 472627..476286 | 1680 | BREX-system-P-loop-protein-BrxC-[Bifidobacterium-adolescentis] |
| WP_117726345.1 | Type1.BREX | ZnR->?->ABC-ATPase+SbcC->?->?->?-> BrxA-DNA-binding -> BrxB-BrxF-iSTAND -> DUF6079 *-> N6-METHYLASE ->?->Schlafen+HATPase+wHTH-> PglZ -> SIGMA+OB-Fold+AAA-ATPase+ HKD-DNase -BrxL ->?-><-REase\|\|?-><-Glyoxalase | 1217 | Bacteria>Terrabacteria_group>Actinomycetota>Actinomycetes>Bifidobacteriales>Bifidobacteriaceae | GCF_027685785.1 | Bifidobacterium_bifidum | PG427_RS02260 | 555349..559002 | 1681 | BREX-system-P-loop-protein-BrxC-[Bifidobacterium-bifidum] |
| WP_270266433.1 | Type1.BREX | <- SIGMA+OB-Fold+AAA-ATPase+ HKD-DNase -BrxL <- PglZ <-Schlafen+HATPase+wHTH<-?<- N6-METHYLASE <- DUF6079 *<- BrxB-BrxF-iSTAND <- BrxA-DNA-binding <-?<-?<-?<-ABC-ATPase+SbcC<-?<-ZnR | 1219 | Bacteria>Terrabacteria_group>Actinomycetota>Actinomycetes>Bifidobacteriales>Bifidobacteriaceae | GCF_027687815.1 | Bifidobacterium_bifidum | PG972_RS08475 | 12443..16102 | 1681 | BREX-system-P-loop-protein-BrxC-[Bifidobacterium-bifidum] |
| WP_117501722.1 | Type1.BREX | REC+LuxR-HTH->HISKIN-><-?\|\|?-><-SbcD<-?<-REase\|\| BrxA-DNA-binding -> BrxB-BrxF-iSTAND -> DUF6079 *-> N6-METHYLASE ->SIR2->Schlafen+HATPase+wHTH-> PglZ -> SIGMA+OB-Fold+AAA-ATPase+ HKD-DNase -BrxL -> | 1219 | Bacteria>Terrabacteria_group>Actinomycetota>Actinomycetes>Bifidobacteriales>Bifidobacteriaceae | GCF_027689935.1 | Bifidobacterium_pseudocatenulatum | PG028_RS01370 | 345834..349493 | 28026 | BREX-system-P-loop-protein-BrxC-[Bifidobacterium-pseudocatenulatum] |
| WP_270558503.1 | Type1.BREX | <-ParB+HNH+RAMA<-ZnR<-ZnR+GP<- SIGMA+OB-Fold+AAA-ATPase+ HKD-DNase -BrxL <- PglZ <-Schlafen+HATPase+wHTH<-REase<- N6-METHYLASE <- DUF6079 *<- BrxB-BrxF-iSTAND <- BrxA-DNA-binding <-?<-?\|\|?-><-Peptidase_M78<-HigA | 1219 | Bacteria>Terrabacteria_group>Actinomycetota>Actinomycetes>Bifidobacteriales>Bifidobacteriaceae | GCF_027690155.1 | Bifidobacterium_adolescentis | PG057_RS05020 | 457971..461630 | 1680 | BREX-system-P-loop-protein-BrxC-[Bifidobacterium-adolescentis] |
| WP_270554212.1 | Type1.BREX | <-SbcD\|\|HAD->PhdYeFM_antitox->RelE-ParE-><-?<-?<-REase\|\| BrxA-DNA-binding -> BrxB-BrxF-iSTAND -> DUF6079 *-> N6-METHYLASE ->REase->Schlafen+HATPase+wHTH-> PglZ -> SIGMA+OB-Fold+AAA-ATPase+ HKD-DNase -BrxL ->ZnR+GP->ZnR->ParB+HNH+RAMA-> | 1219 | Bacteria>Terrabacteria_group>Actinomycetota>Actinomycetes>Bifidobacteriales>Bifidobacteriaceae | GCF_027695195.1 | Bifidobacterium_adolescentis | PG742_RS01120 | 252838..256497 | 1680 | BREX-system-P-loop-protein-BrxC-[Bifidobacterium-adolescentis] |
| WP_270561184.1 | Type1.BREX | 35exo+HRDC->?->35exo+BRCT-><-HTH\|\|?-><-ACET\|\|?-> BrxA-DNA-binding -> BrxB-BrxF-iSTAND -> DUF6079 *->SIR2->HerA/FtsK-> N6-METHYLASE -p->HEPN->?->Schlafen+HATPase+wHTH-> PglZ -> SIGMA+OB-Fold+AAA-ATPase+ HKD-DNase -BrxL ->ZnR-> | 1217 | Bacteria>Terrabacteria_group>Actinomycetota>Actinomycetes>Bifidobacteriales>Bifidobacteriaceae | GCF_027698005.1 | Bifidobacterium_dentium | PG823_RS09640 | 38589..42242 | 1689 | BREX-system-P-loop-protein-BrxC-[Bifidobacterium-dentium] |
| WP_118376243.1 | Type1.BREX | RNaseH-><-?<- SIGMA+OB-Fold+AAA-ATPase+ HKD-DNase -BrxL <- PglZ <-Schlafen+HATPase+wHTH<-SIR2<- N6-METHYLASE <- DUF6079 *<- BrxB-BrxF-iSTAND <- BrxA-DNA-binding \|\|REase->?->?-><-?<-HEPN<-PolB | 1219 | Bacteria>Terrabacteria_group>Actinomycetota>Actinomycetes>Bifidobacteriales>Bifidobacteriaceae | GCF_027698785.1 | Bifidobacterium_pseudocatenulatum | PG820_RS05645 | 117035..120694 | 28026 | BREX-system-P-loop-protein-BrxC-[Bifidobacterium-pseudocatenulatum] |
| WP_271768971.1 | Type1.BREX | <-SbcD<-?<-REase\|\| BrxA-DNA-binding -> BrxB-BrxF-iSTAND -> DUF6079 *-> N6-METHYLASE ->SIR2->Schlafen+HATPase+wHTH-> PglZ -> SIGMA+OB-Fold+AAA-ATPase+ HKD-DNase -BrxL ->ZnR+GP->ZnR->ParB+HNH+RAMA-> | 1219 | Bacteria>Terrabacteria_group>Actinomycetota>Actinomycetes>Bifidobacteriales>Bifidobacteriaceae | GCF_028314485.1 | Bifidobacterium_pseudocatenulatum | PMN34_RS05085 | 218989..222648 | 28026 | BREX-system-P-loop-protein-BrxC-[Bifidobacterium-pseudocatenulatum] |
| WP_116463714.1 | Type1.BREX | HTH_28-> BrxA-DNA-binding -> BrxB-BrxF-iSTAND -> DUF6079 *-> N6-METHYLASE ->?->Schlafen+HATPase+wHTH-> PglZ -> SIGMA+OB-Fold+AAA-ATPase+ HKD-DNase -BrxL ->?-><-?<-?<-ABC-ATPase+SbcC | 1219 | Bacteria>Terrabacteria_group>Actinomycetota>Actinomycetes>Bifidobacteriales>Bifidobacteriaceae | GCF_028898865.1 | Bifidobacterium_longum | PWA56_RS02155 | 487318..490977 | 216816 | BREX-system-P-loop-protein-BrxC-[Bifidobacterium-longum] |
| WP_077383013.1 | Type1.BREX | ABC-ATPase+SbcC->?->?-><-?<- SIGMA+OB-Fold+AAA-ATPase+ HKD-DNase -BrxL <- PglZ <-Schlafen+HATPase+wHTH<-?<- N6-METHYLASE <- DUF6079 *<- BrxB-BrxF-iSTAND <- BrxA-DNA-binding <-HTH_28 | 1219 | Bacteria>Terrabacteria_group>Actinomycetota>Actinomycetes>Bifidobacteriales>Bifidobacteriaceae | GCF_900157155.1 | Bifidobacterium_longum | BQ8631_RS08415 | 45904..49563 | 216816 | BREX-system-P-loop-protein-BrxC-[Bifidobacterium-longum] |
| WP_154050436.1 | Type1.BREX | HAD+S6PP->?->?-><-?<-?<-?<-SIR2\|\| BrxA-DNA-binding -> BrxB-BrxF-iSTAND -> DUF6079 *-> N6-METHYLASE ->SIR2->Schlafen+HATPase+wHTH-> PglZ -> SIGMA+OB-Fold+AAA-ATPase+ HKD-DNase -BrxL ->?-><-ABhydrolase | 1217 | Bacteria>Terrabacteria_group>Actinomycetota>Actinomycetes>Bifidobacteriales>Bifidobacteriaceae | GCF_901212475.1 | Bifidobacterium_longum | FGK88_RS06805 | 85798..89451 | 216816 | BREX-system-P-loop-protein-BrxC-[Bifidobacterium-longum] |
| WP_143724686.1 | Type1.BREX | <- SIGMA+OB-Fold+AAA-ATPase+ HKD-DNase -BrxL <- PglZ <-RNase<-Schlafen+HATPase+wHTH<-SMODS<-SLATT<- N6-METHYLASE <- DUF6079 *<- BrxB-BrxF-iSTAND <- BrxA-DNA-binding <-CR-ATPase5+REase-p<-?\|\|?-><-?<-?\|\|<-?<-MarR_2+ROK-HTH | 1229 | Bacteria>Terrabacteria_group>Actinomycetota>Actinomycetes>Bifidobacteriales>Bifidobacteriaceae | GCF_902167555.1 | Bifidobacterium_longum | FP566_RS07260 | 1570713..1574402 | 216816 | BREX-system-P-loop-protein-BrxC-[Bifidobacterium-longum] |
| WP_143246218.1 | Type1.BREX | <-GDResolvase+HIN-HTH<-?<- SIGMA+OB-Fold+AAA-ATPase+ HKD-DNase -BrxL <- PglZ <-Schlafen+HATPase+wHTH<-SIR2<- N6-METHYLASE <- DUF6079 *<- BrxB-BrxF-iSTAND <- BrxA-DNA-binding | 1218 | Bacteria>Terrabacteria_group>Actinomycetota>Actinomycetes>Bifidobacteriales>Bifidobacteriaceae | GCF_902167575.1 | Bifidobacterium_breve | FPG44_RS02615 | 449270..452926 | 1685 | BREX-system-P-loop-protein-BrxC-[Bifidobacterium-breve] |
| WP_200406352.1 | Type1.BREX | RelB-> N6-METHYLASE ->CR-ATPase5+REase-p-><-CR-Kinase<-?\|\| BrxA-DNA-binding -> BrxB-BrxF-iSTAND -> DUF6079 *->?-> N6-METHYLASE ->SLATT->SMODS->Schlafen-p->Schlafen+wHTH-> PglZ -> SIGMA+OB-Fold+AAA-ATPase+ HKD-DNase -BrxL ->TIR-> | 1229 | Bacteria>Terrabacteria_group>Actinomycetota>Actinomycetes>Bifidobacteriales>Bifidobacteriaceae | GCF_934675445.1 | Bifidobacterium_longum | BL7055_RS05750 | 1406658..1410347 | 216816 | BREX-system-P-loop-protein-BrxC-[Bifidobacterium-longum] |
| WP_245206784.1 | Type1.BREX | <-ZnR<- SIGMA+OB-Fold+AAA-ATPase+ HKD-DNase -BrxL <- PglZ <-Schlafen+HATPase+wHTH<-SIR2<- N6-METHYLASE <-AAA-ATPase<-HTH_7+rve+rve_3<- N6-METHYLASE -p<- DUF6079 *<- DUF6079 -p<- BrxB-BrxF-iSTAND <- BrxA-DNA-binding <-?\|\|ACET-><-?\|\|HTH-><-35exo+BRCT | 1073 | Bacteria>Terrabacteria_group>Actinomycetota>Actinomycetes>Bifidobacteriales>Bifidobacteriaceae | GCF_937997665.1 | Bifidobacterium_dentium | J7M36_RS06065 | 1427241..1430462 | 1689 | BREX-system-P-loop-protein-BrxC-[Bifidobacterium-dentium] |
| WP_250246699.1 | Type1.BREX | MarR_2+ROK-HTH->?->?\|\|?->?-><-?\|\|?->CR-ATPase5+REase-p-> BrxA-DNA-binding -> BrxB-BrxF-iSTAND -> DUF6079 *-> N6-METHYLASE ->SLATT->SMODS->Schlafen+HATPase+wHTH->RNase-> PglZ -> SIGMA+OB-Fold+AAA-ATPase+ HKD-DNase -BrxL -> | 1229 | Bacteria>Terrabacteria_group>Actinomycetota>Actinomycetes>Bifidobacteriales>Bifidobacteriaceae | GCF_939148185.1 | Bifidobacterium_longum | NBW58_RS09955 | 18627..22316 | 216816 | BREX-system-P-loop-protein-BrxC-[Bifidobacterium-longum] |
| WP_196330425.1 | Type1.BREX | 35exo+HRDC->?->35exo+BRCT-><-HTH\|\|?-><-ACET\|\|?-> BrxA-DNA-binding -> BrxB-BrxF-iSTAND -> DUF6079 *-> N6-METHYLASE ->?->Schlafen+HATPase+wHTH-> PglZ -> SIGMA+OB-Fold+AAA-ATPase+ HKD-DNase -BrxL ->ZnR-><-DSBH+AraC-HTH+AraC-HTH\|\|Glyoxalase-> | 1217 | Bacteria>Terrabacteria_group>Actinomycetota>Actinomycetes>Bifidobacteriales>Bifidobacteriaceae | GCF_958433885.1 | Bifidobacterium_dentium | J7M35_RS04855 | 1219062..1222715 | 1689 | BREX-system-P-loop-protein-BrxC-[Bifidobacterium-dentium] |
| WP_085380386.1 | Type1.BREX | <-ParB+HNH+RAMA\|\|MULE-TRANSPOSASE-><-ZnR<-ZnR+GP<- SIGMA+OB-Fold+AAA-ATPase+ HKD-DNase -BrxL <- PglZ <-Schlafen+HATPase+wHTH<-SIR2<- N6-METHYLASE <- DUF6079 *<- BrxB-BrxF-iSTAND <- BrxA-DNA-binding <-Pentapeptide+Pentapeptide+Pentapeptide<-HAD\|\|SbcD-><-?<-ACET+ACET\|\|HTH-><-35exo+BRCT | 1219 | Bacteria>Terrabacteria_group>Actinomycetota>Actinomycetes>Bifidobacteriales>Bifidobacteriaceae | GCF_959020145.1 | Bifidobacterium_adolescentis | LMG10734_RS05600 | 489007..492666 | 1680 | BREX-system-P-loop-protein-BrxC-[Bifidobacterium-adolescentis] |
| WP_196335743.1 | Type1.BREX | <-Glyoxalase\|\|DSBH+AraC-HTH+AraC-HTH-><-ZnR<- SIGMA+OB-Fold+AAA-ATPase+ HKD-DNase -BrxL <- PglZ <-Schlafen+HATPase+wHTH<-HEPN<- N6-METHYLASE <- DUF6079 *<- BrxB-BrxF-iSTAND <- BrxA-DNA-binding \|\|REase-><-HEPN<-?\|\|HTH-><-35exo+BRCT<-35exo+HRDC | 1217 | Bacteria>Terrabacteria_group>Actinomycetota>Actinomycetes>Bifidobacteriales>Bifidobacteriaceae | GCF_959600175.1 | Bifidobacterium_dentium | IHQ76_RS06180 | 228522..232175 | 1689 | BREX-system-P-loop-protein-BrxC-[Bifidobacterium-dentium] |
| WP_096305156.1 | Type5.BREX | <-rve+rve_3\|\|?->?-><-?<-?\|\|Peptidase->XIS-HTH-> BrxA-DNA-binding -> BrxB-BrxF-iSTAND -> DUF6079 *-> N6-METHYLASE -> PglZ -> SIGMA+OB-Fold+AAA-ATPase+ HKD-DNase -BrxL ->SF1-SF2-Helicase-><-AbiEii<-?\|\|?->TUDOR+SF1-SF2-Helicase+iREase+CTD+REase-p-><-HIN-HTH | 1161 | Bacteria>Terrabacteria_group>Actinomycetota>Actinomycetes>Jatrophihabitantales>Jatrophihabitantaceae | GCF_900230215.1 | Jatrophihabitans_sp._GAS493 | CPH63_RS13795 | 2948299..2951784 | 1907575 | BREX-system-P-loop-protein-BrxC-[Jatrophihabitans-sp.-GAS493] |
| EST38900.1 | Type1.BREX | Phage_int_SAM_3+Phage_integrase->DDE_Tnp_1_assoc+DDE_Tnp_1->DDE_Tnp_ISAZ013-><- SIGMA+OB-Fold+AAA-ATPase+ HKD-DNase -BrxL \|\|HTH_29+HTH_32+DDE_3-><- SIGMA+OB-Fold+AAA-ATPase+Lon-Protease-BrxL -p<-?<-HTH_29+HTH_32+DDE_3\|\|zf-ISL3+HTH_Tnp_ISL3-><- DUF6079 -p*<- BrxB-BrxF-iSTAND <- BrxA-DNA-binding \|\|HTH_29+HTH_32+DDE_3-><-DDE_Tnp_1<- BrxA-DNA-binding <-XIS-HTH\|\|HAD+HAD-> | 571 | Bacteria>Terrabacteria_group>Actinomycetota>Actinomycetes>Kitasatosporales>Streptomycetaceae | GCA_000497405.1 | Streptomycetaceae_bacterium_MP113-05 | N566_04930 | 13788..15503 | 1380770 | hypothetical-protein-N566_04930-[Streptomycetaceae-bacterium-MP113-05] |
| MCG7205148.1 | Type1.BREX | <-DUF4096+DDE_Tnp_1+DDE_Tnp_1_2\|\|?->?-><- SIGMA+OB-Fold+AAA-ATPase+ HKD-DNase -BrxL <- PglZ <- N6-METHYLASE <- DUF6079 *<- BrxB-BrxF-iSTAND <- BrxA-DNA-binding <-XIS-HTH\|\| N6-METHYLASE -> | 1161 | Bacteria>Terrabacteria_group>Actinomycetota>Actinomycetes>Kitasatosporales>Streptomycetaceae | GCA_022343545.1 | Streptomyces_arenae | MB828_15195 | 11459..14944 | 29301 | BREX-system-P-loop-protein-BrxC-[Streptomyces-arenae] |
| WP_030399158.1 | Type1.BREX | OLD-ABC+TOPRIM->SF1-SF2-Helicase->?-><-DNA_glycosylase\|\|REase-><- SIGMA+OB-Fold+AAA-ATPase+ HKD-DNase -BrxL <- PglZ <- N6-METHYLASE <- DUF6079 *<- BrxB-BrxF-iSTAND <- BrxA-DNA-binding <-XIS-HTH<-pPIWI+MID+RNaseH<-pPIWI+SF1-SF2-Helicase<-pPIWI+pPIWI<-HTH_3+HTH_3\|\|?-><-DDE_Tnp_1_4 | 1161 | Bacteria>Terrabacteria_group>Actinomycetota>Actinomycetes>Kitasatosporales>Streptomycetaceae | GCF_000719665.1 | unclassified_Streptomyces | ADK87_RS19375 | 27273..30758 | 2593676 | MULTISPECIES:-BREX-system-P-loop-protein-BrxC-[unclassified-Streptomyces] |
| WP_042419815.1 | Type1.BREX | <-VirB6<-?<-TraM<-?\|\|N6-N4-METHYLASE-><-REase+DUF5655\|\|XIS-HTH-> BrxA-DNA-binding -> BrxB-BrxF-iSTAND -> DUF6079 *-> N6-METHYLASE -> PglZ -> SIGMA+OB-Fold+AAA-ATPase+ HKD-DNase -BrxL -><-PIWI<-REase+PNTD1+PNTD2<- HKD-DNase + HKD-DNase | 1161 | Bacteria>Terrabacteria_group>Actinomycetota>Actinomycetes>Kitasatosporales>Streptomycetaceae | GCF_000787855.1 | Streptacidiphilus_anmyonensis | TR50_RS13090 | 101724..105209 | 405782 | BREX-system-P-loop-protein-BrxC-[Streptacidiphilus-anmyonensis] |
| WP_062022136.1 | Type1.BREX | METHYLASE->?->?-><-DNA_glycosylase\|\|REase-><- SIGMA+OB-Fold+AAA-ATPase+ HKD-DNase -BrxL <- PglZ <- N6-METHYLASE <- DUF6079 *<- BrxB-BrxF-iSTAND <- BrxA-DNA-binding <-XIS-HTH<-pPIWI+MID+RNaseH<-pPIWI+SF1-SF2-Helicase<-pPIWI+pPIWI<-HTH_3+HTH_3\|\|REase+DUF5655-><-N6-N4-METHYLASE | 1161 | Bacteria>Terrabacteria_group>Actinomycetota>Actinomycetes>Kitasatosporales>Streptomycetaceae | GCF_001513985.1 | Streptomyces_griseorubiginosus | AQI84_07455 | 596850..600335 | 67304 | BREX-system-P-loop-protein-BrxC-[Streptomyces-griseorubiginosus] |
| WP_129286571.1 | Type1.BREX | PolB->RADICAL-SAM->?->HD->Glyco_trans_1_4-><-?\|\|XIS-HTH-> BrxA-DNA-binding -> BrxB-BrxF-iSTAND -> DUF6079 *-> N6-METHYLASE -> PglZ -> SIGMA+OB-Fold+AAA-ATPase+ HKD-DNase -BrxL -><-Aminotran_1_2\|\|?->\|\|METHYLASE-> | 1163 | Bacteria>Terrabacteria_group>Actinomycetota>Actinomycetes>Kitasatosporales>Streptomycetaceae | GCF_004123705.1 | Streptomyces_sp._GZWMJZ-114 | EJA11_RS17595 | 541597..545088 | 2494734 | BREX-system-P-loop-protein-BrxC-[Streptomyces-sp.-GZWMJZ-114] |
| WP_138352701.1 | Type1.BREX | REase+wHTH->rve->TnsC-ATPase->?->?-><-?<- SIGMA+OB-Fold+AAA-ATPase+ HKD-DNase -BrxL <- PglZ <- N6-METHYLASE <- DUF6079 *<- BrxB-BrxF-iSTAND <- BrxA-DNA-binding <-XIS-HTH<-?\|\|HTH_3->SF1-SF2-Helicase-><-?\|\|?-><-TIR | 1186 | Bacteria>Terrabacteria_group>Actinomycetota>Actinomycetes>Kitasatosporales>Streptomycetaceae | GCF_005860785.2 | Streptomyces_sp._So13.3 | C8250_RS11580 | 2549516..2553076 | 2136173 | BREX-system-P-loop-protein-BrxC-[Streptomyces-sp.-So13.3] |
| WP_161225978.1 | Type1.BREX | <-MarR-HTH<-?<-Hiskin\|\|?->METHYLASE-><-HTH_31+DUF5753<- SIGMA+OB-Fold+AAA-ATPase+ HKD-DNase -BrxL <- PglZ <- N6-METHYLASE <- DUF6079 *<- BrxB-BrxF-iSTAND <-tRNA\|\|CR-Kinase+pi3KINASE-><-?<-?<-Glycos_transf_1+Glyco_trans_1_2<-?<-?<-?\|\|HTH_3-> | 1165 | Bacteria>Terrabacteria_group>Actinomycetota>Actinomycetes>Kitasatosporales>Streptomycetaceae | GCF_009862265.1 | Streptomyces_sp._SID8352 | GTY84_RS10560 | 297618..301115 | 2690338 | BREX-system-P-loop-protein-BrxC-[Streptomyces-sp.-SID8352] |
| WP_164340211.1 | Type1.BREX | <-Pkinase+Pkinase<-ABhydrolase\|\|?-><-?\|\|<-TetR-HTH\|\|?-><- SIGMA+OB-Fold+AAA-ATPase+ HKD-DNase -BrxL <- PglZ <- N6-METHYLASE <- DUF6079 *<- BrxB-BrxF-iSTAND <- BrxA-DNA-binding <-XIS-HTH\|\|Acetyltransf_8->ABC-ATPase->ABC-ATPase-><-?\|\|Peptidase_C25-> | 1163 | Bacteria>Terrabacteria_group>Actinomycetota>Actinomycetes>Kitasatosporales>Streptomycetaceae | GCF_010548045.1 | Streptomyces_anulatus | G3I47_RS13230 | 74640..78131 | 1892 | BREX-system-P-loop-protein-BrxC-[Streptomyces-anulatus] |
| WP_182441002.1 | Type1.BREX | <-PhdYeFM_antitox<-?\|\|?-><-?\|\|?-><-?<- SIGMA+OB-Fold+AAA-ATPase+ HKD-DNase -BrxL <- PglZ <- N6-METHYLASE <- DUF6079 *<- BrxB-BrxF-iSTAND <- BrxA-DNA-binding <-XIS-HTH\|\|REase->ACET-><-?<-HAD+HAD<-PRTase | 1159 | Bacteria>Terrabacteria_group>Actinomycetota>Actinomycetes>Kitasatosporales>Streptomycetaceae | GCF_014084125.1 | Streptacidiphilus_sp._PB12-B1b | GXW83_RS01100 | 242781..246260 | 2705012 | BREX-system-P-loop-protein-BrxC-[Streptacidiphilus-sp.-PB12-B1b] |
| WP_073800314.1 | Type1.BREX | <-Peptidase_C25\|\|?-><-ABC-ATPase+SbcC<-ABC-ATPase<-Acetyltransf_8\|\|XIS-HTH-> BrxA-DNA-binding -> BrxB-BrxF-iSTAND -> DUF6079 *-> N6-METHYLASE -> PglZ -> SIGMA+OB-Fold+AAA-ATPase+ HKD-DNase -BrxL -><-?\|\|TetR-HTH-><-REC+OmpR-HTH | 1163 | Bacteria>Terrabacteria_group>Actinomycetota>Actinomycetes>Kitasatosporales>Streptomycetaceae | GCF_014495705.1 | unclassified_Streptomyces | AMK20_RS03875 | 898838..902329 | 2593676 | MULTISPECIES:-BREX-system-P-loop-protein-BrxC-[unclassified-Streptomyces] |
| WP_265293811.1 | Type1.BREX | Glycos_transf_1+Glyco_trans_1_2->?->HTH_33+DDE_3->?-> BrxA-DNA-binding -> BrxB-BrxF-iSTAND -> DUF6079 *-> N6-METHYLASE -> PglZ -> SIGMA+OB-Fold+AAA-ATPase+ HKD-DNase -BrxL ->HTH_31+DUF5753-><-METHYLASE<-?\|\|Hiskin->?->MarR-HTH-> | 1168 | Bacteria>Terrabacteria_group>Actinomycetota>Actinomycetes>Kitasatosporales>Streptomycetaceae | GCF_026126445.1 | Streptomyces_sp._SHP_1-2 | IBX28_RS14860 | 97274..100780 | 2769489 | BREX-system-P-loop-protein-BrxC-[Streptomyces-sp.-SHP-1-2] |
| WP_266948097.1 | Type1.BREX | <-DUF397<-?\|\|?->\|\|?-><-ACET+ACET\|\|?-><- SIGMA+OB-Fold+AAA-ATPase+ HKD-DNase -BrxL <- PglZ <- N6-METHYLASE <- DUF6079 *<- BrxB-BrxF-iSTAND <- BrxA-DNA-binding <-XIS-HTH\|\|HIN-HTH->?-><-NUDIX<-HTH_31 | 1186 | Bacteria>Terrabacteria_group>Actinomycetota>Actinomycetes>Kitasatosporales>Streptomycetaceae | GCF_026342255.1 | Streptomyces_sp._NBC_00291 | DG914_RS06430 | 1462666..1466226 | 2975704 | BREX-system-P-loop-protein-BrxC-[Streptomyces-sp.-NBC_00291] |
| WP_093695822.1 | Type1.BREX | <-ZDNA-HTH\|\|ICLR-HTH->?->sGTP+sGTP+GTPase-AIG->sGTP-><-DNA_glycosylase<- SIGMA+OB-Fold+AAA-ATPase+ HKD-DNase -BrxL <- PglZ <- N6-METHYLASE <- DUF6079 *<- BrxB-BrxF-iSTAND <- BrxA-DNA-binding <-XIS-HTH<-?<-PRTase<-?\|\|?-><-GNTR-HTH\|\|SKI-> | 1162 | Bacteria>Terrabacteria_group>Actinomycetota>Actinomycetes>Kitasatosporales>Streptomycetaceae | GCF_900105605.1 | Streptomyces_sp._2231.1 | BLV73_RS10010 | 2127135..2130623 | 1855347 | BREX-system-P-loop-protein-BrxC-[Streptomyces-sp.-2231.1] |
| WP_005885100.1 | Type1.BREX | <-ABC-ATPase+SbcC<-?<-?\|\|ACET-><-OLD-ABC<- SIGMA+OB-Fold+AAA-ATPase+ HKD-DNase -BrxL <- PglZ <-HTH+REase<- N6-METHYLASE <- DUF6079 *<- BrxB-BrxF-iSTAND <- BrxA-DNA-binding <-XIS-HTH<-ABhydrolase\|\|?-><-HTH | 1178 | Bacteria>Terrabacteria_group>Actinomycetota>Actinomycetes>Micrococcales>Brevibacteriaceae | GCF_000178455.1 | Brevibacterium_mcbrellneri | HMPREF0183_RS08780 | 50551..54087 | 53363 | BREX-system-P-loop-protein-BrxC-[Brevibacterium-mcbrellneri] |
| WP_125240568.1 | Type1.BREX | <-ACET<-?<-?\|\|ADH_N+ADH_zinc_N-><-HerA/FtsK<-SIR2+SIR2<- SIGMA+OB-Fold+AAA-ATPase+ HKD-DNase -BrxL <- PglZ <- N6-METHYLASE <- DUF6079 *<- BrxB-BrxF-iSTAND <- BrxA-DNA-binding <-?<-?<-?<-?<-?<-?\|\|ICLR-HTH-> | 1170 | Bacteria>Terrabacteria_group>Actinomycetota>Actinomycetes>Micrococcales>Brevibacteriaceae | GCF_003932995.1 | Brevibacterium_aurantiacum | CXR25_RS11695 | 2538315..2541827 | 273384 | BREX-system-P-loop-protein-BrxC-[Brevibacterium-aurantiacum] |
| WP_201670680.1 | Type1.BREX | NUDIX-><-MACRODOMAIN+SIR2\|\|Pkinase+STYKIN-><- SIGMA+OB-Fold+AAA-ATPase+ HKD-DNase -BrxL <- PglZ <- N6-METHYLASE -p<-Phage_int_SAM_3+Phage_integrase\|\| N6-METHYLASE -p-><- N6-METHYLASE <- DUF6079 *<- BrxB-BrxF-iSTAND <- BrxA-DNA-binding <-ADH_N+ADH_zinc_N\|\|?->?->ACET->Aminotran_3->?->wHTH+WYL-> | 1160 | Bacteria>Terrabacteria_group>Actinomycetota>Actinomycetes>Micrococcales>Brevibacteriaceae | GCF_016726245.1 | Brevibacterium_casei | I6I57_RS09610 | 2195903..2199385 | 33889 | BREX-system-P-loop-protein-BrxC-[Brevibacterium-casei] |
| WP_259801020.1 | Type1.BREX | <-VirB11+MuB-ATPase\|\|DNA_METHYLASE-><-HerA/FtsK<-SIR2<- SIGMA+OB-Fold+AAA-ATPase+ HKD-DNase -BrxL <- PglZ <-HTH+REase<- N6-METHYLASE <- DUF6079 *<- BrxB-BrxF-iSTAND <- BrxA-DNA-binding <-XIS-HTH<-ABhydrolase<-?<-MPTase<-HTH\|\|ArdCN+MPTase->Primpol+PriCT1-> | 1173 | Bacteria>Terrabacteria_group>Actinomycetota>Actinomycetes>Micrococcales>Brevibacteriaceae | GCF_025144065.1 | Brevibacterium_casei | M3G43_RS08140 | 29205..32726 | 33889 | BREX-system-P-loop-protein-BrxC-[Brevibacterium-casei] |
| WP_101598836.1 | Type1.BREX | <-Sigma-HTH\|\|NUDIX-><- SIGMA+OB-Fold+AAA-ATPase+ HKD-DNase -BrxL <- PglZ <- N6-METHYLASE <- N6-METHYLASE -p<- DUF6079 *<- BrxB-BrxF-iSTAND <- BrxA-DNA-binding <-ADH_N+ADH_zinc_N\|\|?->?->ACET->Aminotran_3->?->wHTH+WYL-> | 1160 | Bacteria>Terrabacteria_group>Actinomycetota>Actinomycetes>Micrococcales>Brevibacteriaceae | GCF_900169315.1 | Brevibacterium_aurantiacum | BAURA63_03577 | 9324..12806 | 273384 | BREX-system-P-loop-protein-BrxC-[Brevibacterium-aurantiacum] |
| WP_029253807.1 | Type1.BREX | ABC-ATPase+SbcC-><-?<-?<-HEPN<-ParB+HNH<- SIGMA+OB-Fold+AAA-ATPase+ HKD-DNase -BrxL <- PglZ <- N6-METHYLASE <- DUF6079 *<- BrxB-BrxF-iSTAND <- BrxA-DNA-binding <-AbiEii<-AbiEii<-REase | 1160 | Bacteria>Terrabacteria_group>Actinomycetota>Actinomycetes>Micrococcales>Cellulomonadaceae | GCF_000701465.1 | Paraoerskovia_marina | Q343_RS0111945 | 114651..118133 | 545619 | BREX-system-P-loop-protein-BrxC-[Paraoerskovia-marina] |
| WP_130780297.1 | Type1.BREX | <-SPOUT<-AbiEi_4<-?<-?<-PhdYeFM_antitox<-?\|\|XIS-HTH-> BrxA-DNA-binding -> BrxB-BrxF-iSTAND -> DUF6079 *-> N6-METHYLASE -p-> N6-METHYLASE -> PglZ -> SIGMA+OB-Fold+AAA-ATPase+ HKD-DNase -BrxL -><-?<-TRPR-HTH\|\|Phage_int_SAM_1->DDE_Tnp_1_4-> | 1159 | Bacteria>Terrabacteria_group>Actinomycetota>Actinomycetes>Micrococcales>Cellulomonadaceae | GCF_004306155.1 | Cellulomonas_biazotea | CBZ_RS03705 | 43649..47128 | 1709 | BREX-system-P-loop-protein-BrxC-[Cellulomonas-biazotea] |
| WP_010552220.1 | Type1.BREX | <-REase<-ParB+HNH<-ParB+HNH\|\|?-><-MULE-TRANSPOSASE\|\| BrxA-DNA-binding -> BrxB-BrxF-iSTAND -> DUF6079 *-> N6-METHYLASE -> PglZ -> SIGMA+OB-Fold+AAA-ATPase+ HKD-DNase -BrxL ->HIN-HTH-> | 1160 | Bacteria>Terrabacteria_group>Actinomycetota>Actinomycetes>Micrococcales>Dermabacteraceae | GCF_000233655.1 | Brachybacterium_paraconglomeratum | I7Y_RS0116025 | 54770..58252 | 173362 | BREX-system-P-loop-protein-BrxC-[Brachybacterium-paraconglomeratum] |
| WP_114305671.1 | Type1.BREX | <-SF1-SF2-Helicase<-MODE-HTH+PBPII\|\|?-><-?\|\|RNase\|\|?-><-?\|\| BrxA-DNA-binding -> BrxB-BrxF-iSTAND -> DUF6079 *-> N6-METHYLASE -> PglZ -> SIGMA+OB-Fold+AAA-ATPase+ HKD-DNase -BrxL -> | 1160 | Bacteria>Terrabacteria_group>Actinomycetota>Actinomycetes>Micrococcales>Dermabacteraceae | GCF_003335355.1 | Brachybacterium_alimentarium | CIK67_RS09225 | 107631..111113 | 47845 | BREX-system-P-loop-protein-BrxC-[Brachybacterium-alimentarium] |
| WP_249738141.1 | Type1.BREX | <-PTSEIIA1<-?<-HHH_5+SNF2_N+SF1-SF2-Helicase\|\|CASPASE-><- SIGMA+OB-Fold+AAA-ATPase+ HKD-DNase -BrxL <- PglZ <-HTH+REase<- N6-METHYLASE <- DUF6079 *<- BrxB-BrxF-iSTAND <- BrxA-DNA-binding <-SAM-METHYLASE<-?\|\|HIN-HTH->HTH_Tnp_1+HTH_21+rve+rve_3-> | 1173 | Bacteria>Terrabacteria_group>Actinomycetota>Actinomycetes>Micrococcales>Dermabacteraceae | GCF_023509525.1 | Brachybacterium_sp._JHP9 | Bequi_RS11780 | 103104..106625 | 2910770 | BREX-system-P-loop-protein-BrxC-[Brachybacterium-sp.-JHP9] |
| WP_179478704.1 | Type1.BREX | ADH_N+ADH_zinc_N->?->Thymidylate_kin->ABhydrolase->tRNA-><-?<-?<-?\|\|<- SIGMA+OB-Fold+AAA-ATPase+ HKD-DNase -BrxL <- N6-METHYLASE <- DUF6079 *<- BrxB-BrxF-iSTAND <- BrxA-DNA-binding \|\|SF1-SF2-Helicase+REase-p->SF1-SF2-Helicase+REase->REase->?-><-?\|\|MACRODOMAIN+SIR2-> | 1156 | Bacteria>Terrabacteria_group>Actinomycetota>Actinomycetes>Micrococcales>Dermacoccaceae | GCF_013410455.1 | Allobranchiibius_huperziae | HNR15_RS01985 | 409066..412536 | 1874116 | BREX-system-P-loop-protein-BrxC-[Allobranchiibius-huperziae] |
| WP_233763981.1 | Type1.BREX | <-MACRODOMAIN+SIR2\|\|?->?-><-SF1-SF2-Helicase+REase<-SF1-SF2-Helicase+REase\|\| BrxA-DNA-binding -> BrxB-BrxF-iSTAND -> DUF6079 *-> N6-METHYLASE -> PglZ -> SIGMA+OB-Fold+AAA-ATPase+ HKD-DNase -BrxL ->\|\|HTH_Tnp_1+HTH_21+rve+rve_3->\|\|?->?-><-?\|\|?-><-tRNA<-ABhydrolase | 1156 | Bacteria>Terrabacteria_group>Actinomycetota>Actinomycetes>Micrococcales>Dermacoccaceae | GCF_021391335.1 | Allobranchiibius_sp._GilTou73 | LVQ62_RS01295 | 263639..267109 | 2904523 | BREX-system-P-loop-protein-BrxC-[Allobranchiibius-sp.-GilTou73] |
| WP_040162373.1 | Type1.BREX | 35exo->tRNA-><-GDResolvase+Recombinase+ZnR<-?\|\|HIN-HTH->HTH_Tnp_1+HTH_21+rve+rve_3->XIS-HTH-> BrxA-DNA-binding -> BrxB-BrxF-iSTAND -> DUF6079 *-> N6-METHYLASE ->?-> PglZ -> SIGMA+OB-Fold+AAA-ATPase+ HKD-DNase -BrxL ->OLD-ABC+TOPRIM->SF1-SF2-Helicase->GTPase-AIG->MULE-TRANSPOSASE-> | 1178 | Bacteria>Terrabacteria_group>Actinomycetota>Actinomycetes>Micrococcales>Dermatophilaceae | GCF_902375455.1 | Mobilicoccus_massiliensis | FX004_RS16815 | 289663..293199 | 1522310 | BREX-system-P-loop-protein-BrxC-[Mobilicoccus-massiliensis] |
| WP_215519875.1 | Type1.BREX | <-NERD\|\|?->?-><-sGTP<-GTPase-AIG<- SIGMA+OB-Fold+AAA-ATPase+ HKD-DNase -BrxL <- SIGMA+OB-Fold+AAA-ATPase+Lon-Protease-BrxL -p<- PglZ <- N6-METHYLASE <- DUF6079 *<- BrxB-BrxF-iSTAND <- BrxA-DNA-binding <-?<-ZnR+ATPcone | 1156 | Bacteria>Terrabacteria_group>Actinomycetota>Actinomycetes>Micrococcales>Intrasporangiaceae | GCF_018729305.1 | Phycicoccus_mangrovi | KMZ30_RS04765 | 132164..135634 | 2840470 | BREX-system-P-loop-protein-BrxC-[Phycicoccus-mangrovi] |
| WP_265231741.1 | Type1.BREX | <-HD<-DnaB<-DNA_glycosylase<-HNH<-NUDIX\|\|?->XIS-HTH-> BrxA-DNA-binding -> BrxB-BrxF-iSTAND -> DUF6079 *-> N6-METHYLASE -> N6-METHYLASE -p-> PglZ -> N6-METHYLASE -p-> N6-METHYLASE -> SIGMA+OB-Fold+AAA-ATPase+ HKD-DNase -BrxL ->HTH_Tnp_1+HTH_21+rve+rve_3-><-?<-sGTP | 1154 | Bacteria>Terrabacteria_group>Actinomycetota>Actinomycetes>Micrococcales>Intrasporangiaceae | GCF_026114115.1 | Janibacter_hoylei | ON003_RS16090 | 3001704..3005168 | 364298 | BREX-system-P-loop-protein-BrxC-[Janibacter-hoylei] |
| WP_270886872.1 | Type1.BREX | XIS-HTH-> BrxA-DNA-binding -> BrxB-BrxF-iSTAND -> DUF6079 *-> N6-METHYLASE -> PglZ -> SIGMA+OB-Fold+AAA-ATPase+ HKD-DNase -BrxL -><-HIN-HTH<-Phage_int_SAM_3+Phage_integrase<-MarR-HTH<-XIS-HTH<-?\|\|Phage_int_SAM_1+Phage_integrase-> | 1154 | Bacteria>Terrabacteria_group>Actinomycetota>Actinomycetes>Micrococcales>Intrasporangiaceae | GCF_027812955.1 | Pedococcus_sp._5OH_020 | PHA84_RS04350 | 172901..176365 | 2989814 | BREX-system-P-loop-protein-BrxC-[Pedococcus-sp.-5OH_020] |

| Protein-ID | System | Gene-Neighbourhoods | Length | Taxonomy | Assembly-ID | Strain | Locus-Tag | Genomic-coordinates | TaxID | Fasta-defline |
|---|---|---|---|---|---|---|---|---|---|---|
| WP_272461397.1 | Type1.BREX | <-MODE-HTH+PBPII\|\|XIS-HTH->BrxA-DNA-binding->BrxB-BrxF-iSTAND->DUF6079*->N6-METHYLASE->PglZ->SIGMA+OB-Fold+AAA-ATPase+HKD-DNase-BrxL->GTPase-AIG->sGTP->REase->DUF932-><-?<-VirB11+MuB-ATPase | 1154 | Bacteria>Terrabacteria_group>Actinomycetota>Actinomycetes>Micrococcales>Intrasporangiaceae | GCF_028446585.1 | Intrasporangium_calvum | OO014_RS06100 | 11826..15290 | 53358 | BREX-system-P-loop-protein-BrxC-[Intrasporangium-calvum] |
| TFB71537.1 | Type1.BREX | <-URI-DNase<-VirB11+MuB-ATPase<-HEPN+REase<-REase<-SIGMA+OB-Fold+AAA-ATPase+HKD-DNase-BrxL<-PglZ<-N6-METHYLASE<-DUF6079*<-BrxB-BrxF-iSTAND<-BrxA-DNA-binding<-XIS-HTH<-ABhydrolase<-?<-MPTase<-HTH\|\|ArdCN+MPTase->Primpol+PriCT1-> | 1170 | Bacteria>Terrabacteria_group>Actinomycetota>Actinomycetes>Micrococcales>Microbacteriaceae | GCA_004402535.1 | Cryobacterium_sp._HLT2-23 | E3D06_11880 | 40394..43906 | 1259145 | BREX-system-P-loop-protein-BrxC-[Cryobacterium-glaciale] |
| QIM17270.1 | Type1.BREX | <-SIGMA+OB-Fold+AAA-ATPase+HKD-DNase-BrxL<-?<-?<-SIR2<-PglZ<-?<-DUF6079*<-BrxB-BrxF-iSTAND<-BrxA-DNA-binding<-?<-?\|\|TetR-HTH-><-PIN<-PhdYeFM_antitox<-ABC-ATPase+SbcC<-Sulfate_transp+STAS | 1193 | Bacteria>Terrabacteria_group>Actinomycetota>Actinomycetes>Micrococcales>Microbacteriaceae | GCA_011382965.1 | Leucobacter_insecticola | G7067_02015 | 442468..446049 | 2714934 | BREX-system-P-loop-protein-BrxC-[Leucobacter-insecticola] |
| MBX3116430.1 | Type1.BREX | CR-ATPase5+REase-><-?\|\|HTH_6+SIS->?->PolB->BrxA-DNA-binding->BrxB-BrxF-iSTAND->DUF6079*->N6-METHYLASE->OLD-ABC->TOPRIM->PglZ->SIGMA+OB-Fold+AAA-ATPase+HKD-DNase-BrxL->Pkinase+CR-Kinase-><-PIN<-DUF433\|\|REase+HTH+35exo+Small-B-barrel+SF1-SF2-Helicase-> | 1182 | Bacteria>Terrabacteria_group>Actinomycetota>Actinomycetes>Micrococcales>Microbacteriaceae | GCA_019638875.1 | Cryobacterium_sp. | KF808_03680 | 24849..28397 | 1926290 | BREX-system-P-loop-protein-BrxC-[Cryobacterium-sp.] |
| MBX3086848.1 | Type1.BREX | <-VirB11+MuB-ATPase<-?<-SIGMA+OB-Fold+AAA-ATPase+HKD-DNase-BrxL<-PglZ<-N6-METHYLASE<-DUF6079*<-BrxB-BrxF-iSTAND<-BrxA-DNA-binding<-XIS-HTH<-ABhydrolase<-?<-MPTase<-HTH\|\|ArdCN+MPTase->Primpol+PriCT1-> | 1160 | Bacteria>Terrabacteria_group>Actinomycetota>Actinomycetes>Micrococcales>Microbacteriaceae | GCA_019638955.1 | Cryobacterium_sp. | KF742_00030 | 12107..15589 | 1926290 | BREX-system-P-loop-protein-BrxC-[Cryobacterium-sp.] |
| WP_029153784.1 | Type1.BREX | <-REase+SF1-SF2-Helicase<-TRD+TRD\|\|?->?->BrxA-DNA-binding->BrxB-BrxF-iSTAND->DUF6079*->N6-METHYLASE->PglZ->SIGMA+OB-Fold+AAA-ATPase+HKD-DNase-BrxL-><-?\|\|?->?->?->HIN-HTH-> | 1160 | Bacteria>Terrabacteria_group>Actinomycetota>Actinomycetes>Micrococcales>Microbacteriaceae | GCF_000422745.1 | Microbacterium_gubbeenense | G510_RS0111780 | 9571..13053 | 159896 | BREX-system-P-loop-protein-BrxC-[Microbacterium-gubbeenense] |
| WP_028245009.1 | Type1.BREX | <-Thioredoxin\|\|?-><-PALP\|\|tRNA->35exo->tRNA->BrxA-DNA-binding->BrxB-BrxF-iSTAND->DUF6079*->N6-METHYLASE-><-N6-METHYLASE-p\|\|N6-METHYLASE-p->Schlafen+HATPase+wHTH->PglZ->SIGMA+OB-Fold+AAA-ATPase+HKD-DNase-BrxL->SIR2->HerA/FtsK->PhdYeFM_antitox-> | 1170 | Bacteria>Terrabacteria_group>Actinomycetota>Actinomycetes>Micrococcales>Microbacteriaceae | GCF_000423505.1 | Pseudoclavibacter_soli | G557_RS0103810 | 190122..193634 | 452623 | BREX-system-P-loop-protein-BrxC-[Pseudoclavibacter-soli] |
| WP_053098414.1 | Type1.BREX | <-HEPN+SF1-SF2-Helicase+REase\|\|UDG-><-?<-?<-?\|\|BrxA-DNA-binding->BrxB-BrxF-iSTAND->DUF6079*->N6-METHYLASE->PglZ->SIGMA+OB-Fold+AAA-ATPase+HKD-DNase-BrxL->Pkinase->?-><-?<-MarR-HTH\|\|ADH_N+ADH_zinc_N-><-McrC-NTD+REase | 1186 | Bacteria>Terrabacteria_group>Actinomycetota>Actinomycetes>Micrococcales>Microbacteriaceae | GCF_001266755.1 | Microbacterium_sp._CGR1 | AKG07_RS13635 | 2816924..2820484 | 1696072 | BREX-system-P-loop-protein-BrxC-[Microbacterium-sp.-CGR1] |
| WP_053352376.1 | Type1.BREX | <-MODE-HTH+PBPII\|\|?->?->?->?->tRNA->XIS-HTH->BrxA-DNA-binding->BrxB-BrxF-iSTAND->DUF6079*->N6-METHYLASE->PglZ->SIGMA+OB-Fold+AAA-ATPase+HKD-DNase-BrxL-><-?<-?<-?\|\|HNH+SAVED->ParB+HNH-> | 1182 | Bacteria>Terrabacteria_group>Actinomycetota>Actinomycetes>Micrococcales>Microbacteriaceae | GCF_001273845.1 | Leucobacter_musarum | CH42_RS06350 | 60565..64113 | 1930747 | BREX-system-P-loop-protein-BrxC-[Leucobacter-musarum] |
| WP_085605911.1 | Type1.BREX | <-HTH_Tnp_1\|\|DUF6079*->N6-METHYLASE->PglZ->SIGMA+OB-Fold+AAA-ATPase+HKD-DNase-BrxL->?->VirB11+MuB-ATPase-> | 1160 | Bacteria>Terrabacteria_group>Actinomycetota>Actinomycetes>Micrococcales>Microbacteriaceae | GCF_002114135.1 | Microbacterium_sp._LEMMJ01 | B7W94_RS08480 | 394..3876 | 1978350 | BREX-system-P-loop-protein-BrxC-[Microbacterium-sp.-LEMMJ01] |
| WP_104291918.1 | Type1.BREX | McrB-AAA->McrC-NTD+REase->GDResolvase+HIN-HTH-><-?\|\|?->?->XIS-HTH->BrxA-DNA-binding->BrxB-BrxF-iSTAND->DUF6079*->N6-METHYLASE->PglZ->SIGMA+OB-Fold+AAA-ATPase+HKD-DNase-BrxL->REase+HTH+35exo+Small-B-barrel+SF1-SF2-Helicase-><-RNase | 1170 | Bacteria>Terrabacteria_group>Actinomycetota>Actinomycetes>Micrococcales>Microbacteriaceae | GCF_002930935.1 | Rathayibacter_sp._AY1A2 | C5B93_RS04570 | 113948..117460 | 2080520 | BREX-system-P-loop-protein-BrxC-[Rathayibacter-sp.-AY1A2] |
| WP_104323594.1 | Type1.BREX | MarR-HTH->?->?-><-?\|\|?-><-SIGMA+OB-Fold+AAA-ATPase+HKD-DNase-BrxL<-PglZ<-N6-METHYLASE<-DUF6079*<-BrxB-BrxF-iSTAND<-BrxA-DNA-binding | 1160 | Bacteria>Terrabacteria_group>Actinomycetota>Actinomycetes>Micrococcales>Microbacteriaceae | GCF_002931415.1 | unclassified_Rathayibacter | C5C74_RS01635 | 26455..29937 | 2609250 | MULTISPECIES:-BREX-system-P-loop-protein-BrxC-[unclassified-Rathayibacter] |
| WP_104309935.1 | Type1.BREX | TIR-><-?\|\|?-><-TIR<-SIGMA+OB-Fold+AAA-ATPase+HKD-DNase-BrxL<-PglZ<-REase<-N6-METHYLASE<-DUF6079*<-BrxB-BrxF-iSTAND<-BrxA-DNA-binding<-XIS-HTH<-?\|\|GDResolvase+HIN-HTH-><-Acyltransferase<-DSBH<-SIGMA-HTH | 1162 | Bacteria>Terrabacteria_group>Actinomycetota>Actinomycetes>Micrococcales>Microbacteriaceae | GCF_002932095.1 | Rathayibacter_sp._AY1G1 | C5C36_RS14015 | 21701..25189 | 2080564 | BREX-system-P-loop-protein-BrxC-[Rathayibacter-sp.-AY1G1] |
| WP_104347518.1 | Type1.BREX | <-ClpABN-AAA+ClpABC-AAA<-?<-SIGMA+OB-Fold+AAA-ATPase+HKD-DNase-BrxL<-PglZ<-HTH+REase<-N6-METHYLASE<-DUF6079*<-BrxB-BrxF-iSTAND<-BrxA-DNA-binding<-XIS-HTH<-ABhydrolase | 1182 | Bacteria>Terrabacteria_group>Actinomycetota>Actinomycetes>Micrococcales>Microbacteriaceae | GCF_002932695.1 | Rathayibacter_sp._AY1D2 | C5D04_RS07575 | 19648..23196 | 2080543 | BREX-system-P-loop-protein-BrxC-[Rathayibacter-sp.-AY1D2] |
| WP_124973569.1 | Type1.BREX | Aminotran_4-><-?<-NAD_synthase\|\|?-><-tRNA\|\|35exo->tRNA->XIS-HTH->BrxA-DNA-binding->BrxB-BrxF-iSTAND->DUF6079*->N6-METHYLASE->HTH+REase->PglZ->SIGMA+OB-Fold+AAA-ATPase+HKD-DNase-BrxL->SIR2->HerA/FtsK-><-DNA_METHYLASE | 1170 | Bacteria>Terrabacteria_group>Actinomycetota>Actinomycetes>Micrococcales>Microbacteriaceae | GCF_003865375.1 | Gulosibacter_macacae | EG850_RS11335 | 21958..25470 | 2488791 | BREX-system-P-loop-protein-BrxC-[Gulosibacter-macacae] |
| WP_237399358.1 | Type1.BREX | <-Glycos_transf<-Glyco_transf_9<-HAD+HAD\|\|Glyco_transf_9-><-SIGMA-HTH\|\|?->?->tRNA->BrxA-DNA-binding->BrxB-BrxF-iSTAND->DUF6079-p*->N6-METHYLASE-p-><-N6-METHYLASE-p\|\|Phage_int_SAM_3+Phage_integrase->N6-METHYLASE-p->Schlafen+HATPase+wHTH->PglZ->SIGMA+OB-Fold+AAA-ATPase+HKD-DNase-BrxL-> | 878 | Bacteria>Terrabacteria_group>Actinomycetota>Actinomycetes>Micrococcales>Microbacteriaceae | GCF_004011115.1 | Rathayibacter_iranicus | C7V51_RS01465 | 309166..311802 | 59737 | BREX-system-P-loop-protein-BrxC-[Rathayibacter-iranicus] |
| WP_129387833.1 | Type1.BREX | ABhydrolase->?->Peptidase_S15+PepX_C->?-><-?<-?<-Peptidase_S66+Peptidase_S66C\|\|BrxA-DNA-binding->BrxB-BrxF-iSTAND->DUF6079*->N6-METHYLASE-><-N6-METHYLASE-p\|\|Phage_int_SAM_3+Phage_integrase->N6-METHYLASE-p->Schlafen+HATPase+wHTH->PglZ->SIGMA+OB-Fold+AAA-ATPase+HKD-DNase-BrxL-><-PhdYeFM_antitox | 1162 | Bacteria>Terrabacteria_group>Actinomycetota>Actinomycetes>Micrococcales>Microbacteriaceae | GCF_004135285.1 | Microbacterium_protaetiae | ET475_07150 | 1526355..1529843 | 2509458 | BREX-system-P-loop-protein-BrxC-[Microbacterium-protaetiae] |
| WP_130108838.1 | Type1.BREX | <-TetR-HTH\|\|TrwC+SF1-SF2-Helicase+TOPRIM->METHYLASE-><-VirB11+MuB-ATPase<-HerA/FtsK<-SIR2+SIR2<-SIGMA+OB-Fold+AAA-ATPase+HKD-DNase-BrxL<-PglZ<-N6-METHYLASE<-DUF6079*<-BrxB-BrxF-iSTAND<-BrxA-DNA-binding<-XIS-HTH<-ABhydrolase<-?<-MPTase<-HTH\|\|ArdCN+MPTase->Primpol+PriCT1-> | 1160 | Bacteria>Terrabacteria_group>Actinomycetota>Actinomycetes>Micrococcales>Microbacteriaceae | GCF_004208635.1 | Leucobacter_triazinivorans | EVS81_RS01585 | 346722..350204 | 1784719 | BREX-system-P-loop-protein-BrxC-[Leucobacter-triazinivorans] |
| WP_134415956.1 | Type1.BREX | <-HNH<-?<-?<-VirB11+MuB-ATPase<-HerA/FtsK<-SIR2+SIR2<-SIGMA+OB-Fold+AAA-ATPase+HKD-DNase-BrxL<-PglZ<-N6-METHYLASE<-DUF6079*<-BrxB-BrxF-iSTAND<-BrxA-DNA-binding<-XIS-HTH<-ABhydrolase<-?<-MPTase<-HTH\|\|ArdCN+MPTase->Primpol+PriCT1-> | 1170 | Bacteria>Terrabacteria_group>Actinomycetota>Actinomycetes>Micrococcales>Microbacteriaceae | GCF_004402465.1 | Cryobacterium_sp._TMT2-14 | E3028_RS10860 | 27948..31460 | 1259245 | BREX-system-P-loop-protein-BrxC-[Cryobacterium-sp.-TMT2-14] |
| WP_241980122.1 | Type1.BREX | <-URI-DNase<-VirB11+MuB-ATPase<-HEPN+REase<-REase<-SIGMA+OB-Fold+AAA-ATPase+HKD-DNase-BrxL<-PglZ<-N6-METHYLASE<-DUF6079*<-BrxB-BrxF-iSTAND<-BrxA-DNA-binding<-XIS-HTH<-ABhydrolase<-?<-MPTase<-HTH\|\|ArdCN+MPTase->Primpol+PriCT1-> | 1172 | Bacteria>Terrabacteria_group>Actinomycetota>Actinomycetes>Micrococcales>Microbacteriaceae | GCF_004402535.1 | Cryobacterium_sp._HLT2-23 | E3D06_RS11880 | 40394..43912 | 1259145 | BREX-system-P-loop-protein-BrxC-[Cryobacterium-glaciale] |
| WP_134434730.1 | Type1.BREX | <-VirB11+MuB-ATPase<-SbcC<-?<-SIGMA+OB-Fold+AAA-ATPase+HKD-DNase-BrxL<-PglZ<-N6-METHYLASE<-DUF6079*<-BrxB-BrxF-iSTAND<-BrxA-DNA-binding<-XIS-HTH\|\|AraC-HTH+AraC-HTH->ArdCN+MPTase->Primpol+PriCT1->DUF2637+HIN-HTH-> | 1164 | Bacteria>Terrabacteria_group>Actinomycetota>Actinomycetes>Micrococcales>Microbacteriaceae | GCF_004403545.1 | Cryobacterium_sp._TmT2-59 | E3T24_RS13895 | 41136..44630 | 1259264 | BREX-system-P-loop-protein-BrxC-[Cryobacterium-sp.-TmT2-59] |
| WP_146322974.1 | Type1.BREX | DOC+HTH->?->BrxA-DNA-binding->BrxB-BrxF-iSTAND->DUF6079*->N6-METHYLASE-><-N6-METHYLASE-p\|\|Phage_int_SAM_3+Phage_integrase->N6-METHYLASE-p->?->?->PglZ->SIGMA+OB-Fold+AAA-ATPase+HKD-DNase-BrxL->ParB+HNH-> | 1162 | Bacteria>Terrabacteria_group>Actinomycetota>Actinomycetes>Micrococcales>Microbacteriaceae | GCF_007859675.1 | Humibacter_ginsenosidimutans | FPZ11_RS17385 | 3786036..3789524 | 2599293 | BREX-system-P-loop-protein-BrxC-[Humibacter-ginsenosidimutans] |
| WP_200838962.1 | Type1.BREX | <-SIGMA+OB-Fold+AAA-ATPase+HKD-DNase-BrxL<-PglZ<-N6-METHYLASE-p<-Phage_int_SAM_3+Phage_integrase\|\|N6-METHYLASE-p-><-N6-METHYLASE<-DUF6079*<-BrxB-BrxF-iSTAND<-BrxA-DNA-binding<-XIS-HTH\|\|HTH_3-><-XIS-HTH<-tRNA<-ABC-ATPase+ABC-ATPase<-ZnR+TOPRIM+TOPRIM<-HD | 1182 | Bacteria>Terrabacteria_group>Actinomycetota>Actinomycetes>Micrococcales>Microbacteriaceae | GCF_008271365.1 | Microbacterium | FY549_RS13635 | 2819312..2822860 | 33882 | MULTISPECIES:-BREX-system-P-loop-protein-BrxC-[Microbacterium] |
| WP_163473715.1 | Type1.BREX | Aminotran_4->?->tRNA->XIS-HTH->Phage_integrase->TrwC+SF1-SF2-Helicase+TOPRIM->XIS-HTH->BrxA-DNA-binding->BrxB-BrxF-iSTAND->DUF6079*->N6-METHYLASE->REase->PglZ->SIGMA+OB-Fold+AAA-ATPase+HKD-DNase-BrxL->N6-METHYLASE->MORC+HISKIN->?->PIN->PIN-> | 1162 | Bacteria>Terrabacteria_group>Actinomycetota>Actinomycetes>Micrococcales>Microbacteriaceae | GCF_010686725.1 | Galbitalea_soli | HNR17_RS04680 | 978511..981999 | 1268042 | BREX-system-P-loop-protein-BrxC-[Galbitalea-soli] |
| WP_244301194.1 | Type1.BREX | <-SIGMA+OB-Fold+AAA-ATPase+HKD-DNase-BrxL<-?<-?<-HerA/FtsK<-SIR2<-PglZ<-?<-N6-METHYLASE-p<-N6-METHYLASE<-DUF6079*<-BrxB-BrxF-iSTAND<-BrxA-DNA-binding<-?<-?\|\|TetR-HTH-><-PIN<-PhdYeFM_antitox<-ABC-ATPase+SbcC<-Sulfate_transp+STAS | 1194 | Bacteria>Terrabacteria_group>Actinomycetota>Actinomycetes>Micrococcales>Microbacteriaceae | GCF_011382965.1 | Leucobacter_insecticola | G7067_RS02010 | 442468..446052 | 2714934 | BREX-system-P-loop-protein-BrxC-[Leucobacter-insecticola] |
| WP_166866044.1 | Type1.BREX | <-REase<-SbcC<-JetB<-JetA<-SIGMA+OB-Fold+AAA-ATPase+HKD-DNase-BrxL<-PglZ<-N6-METHYLASE<-DUF6079*<-BrxB-BrxF-iSTAND<-BrxA-DNA-binding<-REase<-tRNA<-?<-?<-?\|\|?-><-?<-Peptidase_M28 | 1160 | Bacteria>Terrabacteria_group>Actinomycetota>Actinomycetes>Micrococcales>Microbacteriaceae | GCF_011751865.2 | Salinibacterium_sp._ZJ70 | HCR12_RS10200 | 2144642..2148124 | 2708084 | BREX-system-P-loop-protein-BrxC-[Salinibacterium-sp.-ZJ70] |
| WP_169077984.1 | Type1.BREX | BrxA-DNA-binding->BrxB-BrxF-iSTAND->DUF6079*->N6-METHYLASE->REase->PglZ->SIGMA+OB-Fold+AAA-ATPase+HKD-DNase-BrxL->SIR2->HerA/FtsK-><-Peptidase_M48\|\|?->REase+HTH+35exo+Small-B-barrel+SF1-SF2-Helicase-> | 1162 | Bacteria>Terrabacteria_group>Actinomycetota>Actinomycetes>Micrococcales>Microbacteriaceae | GCF_012845865.1 | Yonghaparkia | HFV16_RS05520 | 206976..210464 | 355929 | MULTISPECIES:-BREX-system-P-loop-protein-BrxC-[Yonghaparkia] |
| WP_121673638.1 | Type1.BREX | Phage_integrase-><-HEPN<-ParB+HNH\|\|VirB11+MuB-ATPase-><-sGTP<-GTPase-AIG<-SIGMA+OB-Fold+AAA-ATPase+HKD-DNase-BrxL<-PglZ<-N6-METHYLASE<-DUF6079*<-BrxB-BrxF-iSTAND<-BrxA-DNA-binding<-XIS-HTH<-?<-?\|\|?->ORC-CDC6->?->N6-METHYLASE-> | 1161 | Bacteria>Terrabacteria_group>Actinomycetota>Actinomycetes>Micrococcales>Microbacteriaceae | GCF_014651755.1 | Mycetocola_manganoxydans | IE057_RS13450 | 51714..55199 | 699879 | BREX-system-P-loop-protein-BrxC-[Mycetocola-manganoxydans] |
| WP_197132648.1 | Type1.BREX | REase-><-?<-sGTP<-GTPase-AIG<-SIGMA+OB-Fold+AAA-ATPase+HKD-DNase-BrxL<-N6-METHYLASE<-N6-METHYLASE<-DUF6079*<-BrxB-BrxF-iSTAND<-BrxA-DNA-binding<-XIS-HTH\|\|Phage_integrase->Phage_integrase->DDE_Tnp_1+DDE_Tnp_1-> | 1171 | Bacteria>Terrabacteria_group>Actinomycetota>Actinomycetes>Micrococcales>Microbacteriaceae | GCF_015864675.1 | Salinibacterium_sp._SWN1162 | I6E58_RS09540 | 2021923..2025438 | 2792053 | BREX-system-P-loop-protein-BrxC-[Salinibacterium-sp.-SWN1162] |
| WP_206395457.1 | Type1.BREX | ParB+HNH-><-ABhydrolase<-?<-SIGMA+OB-Fold+AAA-ATPase+HKD-DNase-BrxL<-PglZ<-N6-METHYLASE-p<-Phage_int_SAM_3+Phage_integrase\|\|N6-METHYLASE-p-><-N6-METHYLASE<-DUF6079*<-BrxB-BrxF-iSTAND<-BrxA-DNA-binding<-tRNA\|\|?->?->?->?->?->?->Pkinase+TPR+TPR-> | 1158 | Bacteria>Terrabacteria_group>Actinomycetota>Actinomycetes>Micrococcales>Microbacteriaceae | GCF_017167395.1 | Pseudoclavibacter_alba | JRG19_RS09065 | 120280..123756 | 272241 | BREX-system-P-loop-protein-BrxC-[Pseudoclavibacter-alba] |
| WP_240240880.1 | Type1.BREX | <-VirB11+MuB-ATPase\|\|DNA_METHYLASE-><-HerA/FtsK<-SIR2+SIR2<-SIGMA+OB-Fold+AAA-ATPase+HKD-DNase-BrxL<-PglZ<-?<-N6-METHYLASE<-DUF6079*<-BrxB-BrxF-iSTAND<-BrxA-DNA-binding<-XIS-HTH<-ABhydrolase<-?<-MPTase<-HTH\|\|ArdCN+MPTase->Primpol+PriCT1-> | 1178 | Bacteria>Terrabacteria_group>Actinomycetota>Actinomycetes>Micrococcales>Microbacteriaceae | GCF_022436485.1 | Agrococcus_sp._ARC_14 | MKD51_RS07980 | 1621402..1624938 | 2919927 | BREX-system-P-loop-protein-BrxC-[Agrococcus-sp.-ARC_14] |
| WP_242086581.1 | Type1.BREX | XIS-HTH->BrxA-DNA-binding->BrxB-BrxF-iSTAND->DUF6079*->N6-METHYLASE-><-N6-METHYLASE-p\|\|Phage_int_SAM_3+Phage_integrase->N6-METHYLASE-p->PglZ->SIGMA+OB-Fold+AAA-ATPase+HKD-DNase-BrxL-> | 1160 | Bacteria>Terrabacteria_group>Actinomycetota>Actinomycetes>Micrococcales>Microbacteriaceae | GCF_022662255.1 | Microbacterium_lacticum | L8F16_RS04085 | 1822..5304 | 33885 | BREX-system-P-loop-protein-BrxC-[Microbacterium-lacticum] |
| WP_156999272.1 | Type1.BREX | <-HEPN<-PolB<-HEPN+SF1-SF2-Helicase+REase\|\|?->CBS+HEPN+HEPN->BrxA-DNA-binding->BrxB-BrxF-iSTAND->DUF6079*->N6-METHYLASE->PglZ->SIGMA+OB-Fold+AAA-ATPase+HKD-DNase-BrxL->sGTP+sGTP+GTPase-AIG->?-><-ParB+HNH\|\|Exonuc_V_gamma+REase->REase->REase-> | 1173 | Bacteria>Terrabacteria_group>Actinomycetota>Actinomycetes>Micrococcales>Microbacteriaceae | GCF_024172065.1 | Agromyces_terreus | BJ978_RS07845 | 1788329..1791850 | 424795 | BREX-system-P-loop-protein-BrxC-[Agromyces-terreus] |
| WP_257523879.1 | Type1.BREX | <-PIN<-URI-DNase\|\|?->?-><-?\|\|XIS-HTH->BrxA-DNA-binding->BrxB-BrxF-iSTAND->DUF6079*->N6-METHYLASE->PglZ->SIGMA+OB-Fold+AAA-ATPase+HKD-DNase-BrxL->CMP-hydrolase-> | 1160 | Bacteria>Terrabacteria_group>Actinomycetota>Actinomycetes>Micrococcales>Microbacteriaceae | GCF_024623505.1 | Microbacterium_sp._zg.B48 | NQ152_RS10060 | 193646..197128 | 2969408 | BREX-system-P-loop-protein-BrxC-[Microbacterium-sp.-zg.B48] |
| WP_259980691.1 | Type1.BREX | MerR-HTH-><-?\|\|N6-METHYLASE-p-><-HerA/FtsK<-SIR2+SIR2<-SIGMA+OB-Fold+AAA-ATPase+HKD-DNase-BrxL<-PglZ<-N6-METHYLASE<-DUF6079*<-BrxB-BrxF-iSTAND<-BrxA-DNA-binding<-XIS-HTH<-DnaK\|\|HTH_3->?-><-DUF6361<-HNH | 1162 | Bacteria>Terrabacteria_group>Actinomycetota>Actinomycetes>Micrococcales>Microbacteriaceae | GCF_025151145.1 | Microbacterium_paraoxydans | M3D59_RS02760 | 52380..55868 | 199592 | BREX-system-P-loop-protein-BrxC-[Microbacterium-paraoxydans] |
| WP_261607502.1 | Type1.BREX | SIR2->ADH_N+ADH_zinc_N-><-?\|\|tRNA->?-><-Phage_int_SAM_3+Phage_integrase<-?<-?\|\|BrxA-DNA-binding->BrxB-BrxF-iSTAND->DUF6079*->N6-METHYLASE->PglZ->SIGMA+OB-Fold+AAA-ATPase+HKD-DNase-BrxL->Pkinase-><-SF1-SF2-Helicase | 1186 | Bacteria>Terrabacteria_group>Actinomycetota>Actinomycetes>Micrococcales>Microbacteriaceae | GCF_025398275.1 | Microbacterium_sp._PMIC_1C1B | N4R40_RS11380 | 57181..60741 | 2978350 | BREX-system-P-loop-protein-BrxC-[Microbacterium-sp.-PMIC_1C1B] |
| WP_263798144.1 | Type1.BREX | <-Phage_int_SAM_3+Phage_integrase\|\|?->BrxA-DNA-binding->BrxB-BrxF-iSTAND->DUF6079*->N6-METHYLASE->PglZ->SIGMA+OB-Fold+AAA-ATPase+HKD-DNase-BrxL->Pkinase-><-?<-REase+HTH+35exo+Small-B-barrel+SF1-SF2-Helicase<-URI-DNase+RAMA\|\|TrwC+SF1-SF2-Helicase->REase-> | 1186 | Bacteria>Terrabacteria_group>Actinomycetota>Actinomycetes>Micrococcales>Microbacteriaceae | GCF_027943945.1 | Microbacterium_terricola | OAU46_RS04855 | 990438..993998 | 344163 | BREX-system-P-loop-protein-BrxC-[Microbacterium-terricola] |
| WP_257508277.1 | Type1.BREX | <-METHYLASE<-?\|\|LACI-HTH+PBPI->?->?->?-><-?\|\|tRNA->\|\|BrxA-DNA-binding->BrxB-BrxF-iSTAND->DUF6079*->N6-METHYLASE->PglZ->SIGMA+OB-Fold+AAA-ATPase+HKD-DNase-BrxL->Pkinase-><-?\|\|?-><-REase+HTH+35exo+Small-B-barrel+SF1-SF2-Helicase<-?\|\|AAA-ATPase-> | 1186 | Bacteria>Terrabacteria_group>Actinomycetota>Actinomycetes>Micrococcales>Microbacteriaceae | GCF_030246865.1 | Microbacterium_sp._zg.B96 | NQ154_RS06000 | 561655..565215 | 2969409 | BREX-system-P-loop-protein-BrxC-[Microbacterium-sp.-zg.B96] |
| WP_257505179.1 | Type1.BREX | <-McrC-NTD+REase<-McrB-AAA<-?\|\|HTH_Tnp_1+HTH_21+rve+rve_3->ACET-><-SIGMA+OB-Fold+AAA-ATPase+HKD-DNase-BrxL<-PglZ<-N6-METHYLASE<-DUF6079*<-BrxB-BrxF-iSTAND<-BrxA-DNA-binding<-HIN-HTH<-HIRAN\|\|REase-><-tRNA\|\|?-><-REase | 1162 | Bacteria>Terrabacteria_group>Actinomycetota>Actinomycetes>Micrococcales>Microbacteriaceae | GCF_030246925.1 | Microbacterium_sp._zg.Y625 | NQ156_RS01310 | 276922..280410 | 2969411 | BREX-system-P-loop-protein-BrxC-[Microbacterium-sp.-zg.Y625] |
| WP_028495577.1 | Type1.BREX | VirB11+MuB-ATPase->?-><-?\|\|XIS-HTH->BrxA-DNA-binding->BrxB-BrxF-iSTAND->DUF6079*->N6-METHYLASE->PglZ->SIGMA+OB-Fold+AAA-ATPase+HKD-DNase-BrxL->?->tRNA->?-><-?<-ADH_N+ADH_zinc_N<-DSBH<-HAD+HAD | 1160 | Bacteria>Terrabacteria_group>Actinomycetota>Actinomycetes>Micrococcales>Microbacteriaceae | GCF_900113885.1 | Microbacterium | BM226_RS05185 | 1039412..1042894 | 33882 | MULTISPECIES:-BREX-system-P-loop-protein-BrxC-[Microbacterium] |
| WP_201614919.1 | Type1.BREX | Aminotran_4-><-?<-Peptidase<-NAD_synthase<-tRNA\|\|35exo->tRNA->BrxA-DNA-binding->BrxB-BrxF-iSTAND->DUF6079*->N6-METHYLASE-><-N6-METHYLASE-p\|\|Phage_int_SAM_3+Phage_integrase->N6-METHYLASE-p->HTH+REase->PglZ->SIGMA+OB-Fold+AAA-ATPase+HKD-DNase-BrxL-> | 1170 | Bacteria>Terrabacteria_group>Actinomycetota>Actinomycetes>Micrococcales>Microbacteriaceae | GCF_904848685.1 | Gulosibacter_hominis | JMX86_RS10045 | 19055..22567 | 2770504 | BREX-system-P-loop-protein-BrxC-[Gulosibacter-hominis] |
| WP_230672563.1 | Type1.BREX | ABC-ATPase+SbcC->?->?-><-?<-?\|\|MACRODOMAIN+SIR2->BrxA-DNA-binding->BrxB-BrxF-iSTAND->DUF6079*->N6-METHYLASE->HTH+REase->HTH+REase->PglZ->SIGMA+OB-Fold+AAA-ATPase+HKD-DNase-BrxL-><-?\|\|?->HEPN+SF1-SF2-Helicase+REase-> | 1176 | Bacteria>Terrabacteria_group>Actinomycetota>Actinomycetes>Micrococcales>Microbacteriaceae | GCF_920939495.1 | Rathayibacter_sp._Leaf248 | LRG51_RS10050 | 2075326..2078856 | 2876555 | BREX-system-P-loop-protein-BrxC-[Rathayibacter-sp.-Leaf248] |
| WP_219885258.1 | Type1.BREX | HIN-HTH-><-SIGMA+OB-Fold+AAA-ATPase+HKD-DNase-BrxL<-PglZ<-N6-METHYLASE<-DUF6079*<-BrxB-BrxF-iSTAND<-XIS-HTH<-?<-?\|\|XIS-HTH->?->HNH-> | 1210 | Bacteria>Terrabacteria_group>Actinomycetota>Actinomycetes>Micrococcales>Micrococcaceae | - | Arthrobacter_sp._PAMC25284 | KY499_09320 | 2016646..2020278 | 2861279 | BREX-system-P-loop-protein-BrxC-[Arthrobacter-sp.-PAMC25284] |
| AFR34584.1 | Type1.BREX | <-SIGMA+OB-Fold+AAA-ATPase+HKD-DNase-BrxL<-PglZ<-N6-METHYLASE<-DUF6079*<-BrxB-BrxF-iSTAND<-BrxA-DNA-binding\|\|ParA-Soj-PloopNTPase->?->?->?->VirB11-> | 1166 | Bacteria>Terrabacteria_group>Actinomycetota>Actinomycetes>Micrococcales>Micrococcaceae | GCA_000294695.2 | Arthrobacter_sp._Rue61a | ARUE_113p00760 | 81229..84729 | 1118963 | putative-ATP-binding-protein-[Arthrobacter-sp.-Rue61a] |
| ERI35655.1 | Type1.BREX | <-Radical-SAM<-?\|\|XIS-HTH->BrxA-DNA-binding->BrxB-BrxF-iSTAND->DUF6079*->N6-METHYLASE->PglZ->SIGMA+OB-Fold+AAA-ATPase+HKD-DNase-BrxL-><-?\|\|URI-DNase->?->?-><-ParB+HNH | 1161 | Bacteria>Terrabacteria_group>Actinomycetota>Actinomycetes>Micrococcales>Micrococcaceae | GCA_000465895.2 | Arthrobacter_sp._AK-YN10 | M707_20590 | 5564..9049 | 1349820 | ATP-binding-protein-[Arthrobacter-sp.-AK-YN10] |
| ALV40633.1 | Type1.BREX | <-HEPN+SF1-SF2-Helicase+REase<-?<-?<-ParB+HNH<-SIGMA+OB-Fold+AAA-ATPase+HKD-DNase-BrxL<-PglZ<-N6-METHYLASE<-DUF6079*<-BrxB-BrxF-iSTAND<-BrxA-DNA-binding<-XIS-HTH<-HKD-DNase+SF1-SF2-Helicase+HTH+HTH+SRA\|\|?->?->?->McrB-AAA->McrC-NTD+REase-> | 1172 | Bacteria>Terrabacteria_group>Actinomycetota>Actinomycetes>Micrococcales>Micrococcaceae | GCA_001484605.1 | Pseudarthrobacter_sulfonivorans | AU252_05160 | 1179330..1182848 | 121292 | hypothetical-protein-AU252_05160-[Pseudarthrobacter-sulfonivorans] |
| AOY70771.1 | Type1.BREX | RNase+DSRBD->?-><-?\|\|XIS-HTH->BrxA-DNA-binding->BrxB-BrxF-iSTAND->DUF6079*->N6-METHYLASE->PglZ->SIGMA+OB-Fold+AAA-ATPase+HKD-DNase-BrxL->?-><-HTH_21+rve+rve_3<-HIN-HTH\|\|?->NERD-> | 1161 | Bacteria>Terrabacteria_group>Actinomycetota>Actinomycetes>Micrococcales>Micrococcaceae | GCA_001854365.1 | Arthrobacter_sp._ZXY-2 | ARZXY2_1218 | 1271114..1274599 | 1806905 | ATP-binding-protein-[Arthrobacter-sp.-ZXY-2] |
| ASN39993.1 | Type1.BREX | <-Peptidase_M16+Peptidase_M16_C<-RNasePH+RNasePH+KH+S1\|\|RNase+PHP-><-?<-TRD<-N6-METHYLASE<-SIGMA+OB-Fold+AAA-ATPase+HKD-DNase-BrxL<-PglZ<-N6-METHYLASE<-DUF6079*<-BrxB-BrxF-iSTAND<-BrxA-DNA-binding<-XIS-HTH\|\|Schlafen-> | 1170 | Bacteria>Terrabacteria_group>Actinomycetota>Actinomycetes>Micrococcales>Micrococcaceae | GCA_002224445.1 | Arthrobacter_sp._7749 | CGQ24_13870 | 3050415..3053927 | 2020130 | ATP-binding-protein-[Arthrobacter-sp.-7749] |
| THE19344.1 | Type1.BREX | <-TIR\|\|XIS-HTH->BrxA-DNA-binding->BrxB-BrxF-iSTAND->DUF6079*->N6-METHYLASE->PglZ->SIGMA+OB-Fold+AAA-ATPase+HKD-DNase-BrxL-><-?\|\|REase->HTH+REase->URI-DNase->DUF6361->HKD-DNase-> | 1166 | Bacteria>Terrabacteria_group>Actinomycetota>Actinomycetes>Micrococcales>Micrococcaceae | GCA_004799725.1 | Kocuria_rosea | E1J17_01420 | 4081..7581 | 1275 | BREX-system-P-loop-protein-BrxC-[Kocuria-rosea] |
| NHQ58332.1 | Type1.BREX | ICLR-HTH-><-?<-?\|\|?-><-?\|\|?->BrxA-DNA-binding->BrxB-BrxF-iSTAND->DUF6079*->N6-METHYLASE->PglZ->SIGMA+OB-Fold+AAA-ATPase+HKD-DNase-BrxL-><-?\|\|?-><-RADICAL-SAM<-?<-ACET<-XIS-HTH | 1174 | Bacteria>Terrabacteria_group>Actinomycetota>Actinomycetes>Micrococcales>Micrococcaceae | GCA_011602345.1 | Micrococcus_luteus | G9B47_07855 | 17274..20798 | 1270 | BREX-system-P-loop-protein-BrxC-[Micrococcus-luteus] |
| MBB4882886.1 | Type1.BREX | 35exo->tRNA-><-Recombinase+ZnR\|\|ABhydrolase->XIS-HTH->BrxA-DNA-binding->BrxB-BrxF-iSTAND->DUF6079*->N6-METHYLASE->HTH+REase->PglZ->SIGMA+OB-Fold+AAA-ATPase+HKD-DNase-BrxL->SIR2->HIN-HTH->HTH_Tnp_1+HTH_21+rve+rve_3-> | 1173 | Bacteria>Terrabacteria_group>Actinomycetota>Actinomycetes>Micrococcales>Micrococcaceae | GCA_014204815.1 | Micrococcus_flavus | BJ976_001237 | 1342727..1346248 | 384602 | hypothetical-protein-[Micrococcus-flavus] |
| MBR9643072.1 | Type1.BREX | <-DNA-glycosylase<-URI-DNase<-MACRODOMAIN+SIR2<-REase\|\|HKD-DNase+SF1-SF2-Helicase+HTH+HTH+SRA-><-SIGMA+OB-Fold+AAA-ATPase+HKD-DNase-BrxL<-PglZ<-N6-METHYLASE<-DUF6079*<-BrxB-BrxF-iSTAND<-BrxA-DNA-binding\|\|McrB-AAA->McrC-NTD+REase->ABhydrolase+ABhydrolase-> | 1173 | Bacteria>Terrabacteria_group>Actinomycetota>Actinomycetes>Micrococcales>Micrococcaceae | GCA_018215265.1 | Paenarthrobacter_sp._GOM3 | IRJ34_14045 | 199455..202976 | 2782567 | BREX-system-P-loop-protein-BrxC-[Paenarthrobacter-sp.-GOM3] |
| QYF88488.1 | Type1.BREX | HIN-HTH-><-SIGMA+OB-Fold+AAA-ATPase+HKD-DNase-BrxL<-PglZ<-N6-METHYLASE<-DUF6079*<-BrxB-BrxF-iSTAND<-XIS-HTH<-?<-?\|\|XIS-HTH->?->HNH-> | 1210 | Bacteria>Terrabacteria_group>Actinomycetota>Actinomycetes>Micrococcales>Micrococcaceae | GCA_019443425.1 | Arthrobacter_sp._PAMC25284 | KY499_09320 | 2016646..2020278 | 2861279 | BREX-system-P-loop-protein-BrxC-[Arthrobacter-sp.-PAMC25284] |
| WP_019309257.1 | Type1.BREX | <-MPTase<-HTH_3\|\|SMODS->?->?-><-SIGMA+OB-Fold+AAA-ATPase+HKD-DNase-BrxL<-PglZ<-N6-METHYLASE<-DUF6079*<-BrxB-BrxF-iSTAND<-BrxA-DNA-binding<-XIS-HTH<-HTH_3<-?<-?<-?<-?\|\|ABhydrolase-> | 1159 | Bacteria>Terrabacteria_group>Actinomycetota>Actinomycetes>Micrococcales>Micrococcaceae | GCF_000214115.1 | Kocuria_rhizophila | GGK_RS0101750 | 388045..391524 | 72000 | BREX-system-P-loop-protein-BrxC-[Kocuria-rhizophila] |
| WP_017834907.1 | Type1.BREX | <-SIGMA+OB-Fold+AAA-ATPase+HKD-DNase-BrxL<-PglZ<-N6-METHYLASE<-DUF6079*<-BrxB-BrxF-iSTAND<-BrxA-DNA-binding<-XIS-HTH<-?<-McrC-NTD+REase<-McrB-AAA<-REase<-?\|\|<-Schlafen+HATPase+wHTH | 1190 | Bacteria>Terrabacteria_group>Actinomycetota>Actinomycetes>Micrococcales>Micrococcaceae | GCF_000349605.1 | Kocuria_sp._UCD-OTCP | H488_RS34905 | 37795..41367 | 1292021 | BREX-system-P-loop-protein-BrxC-[Kocuria-sp.-UCD-OTCP] |
| WP_155853802.1 | Type1.BREX | <-Primpol+PriCT1<-ArdCN+MPTase\|\|HTH->MPTase->?->ABhydrolase->XIS-HTH->BrxA-DNA-binding->BrxB-BrxF-iSTAND->DUF6079*->N6-METHYLASE->PglZ->SIGMA+OB-Fold+AAA-ATPase+HKD-DNase-BrxL->?->VirB11+MuB-ATPase-> | 1170 | Bacteria>Terrabacteria_group>Actinomycetota>Actinomycetes>Micrococcales>Micrococcaceae | GCF_000520595.1 | Arthrobacter_sp._H5 | Y708_RS0109785 | 51778..55290 | 1267973 | BREX-system-P-loop-protein-BrxC-[Arthrobacter-sp.-H5] |
| WP_062292549.1 | Type1.BREX | VirB11+MuB-ATPase->?-><-SIGMA+OB-Fold+AAA-ATPase+HKD-DNase-BrxL<-PglZ<-N6-METHYLASE-p<-Phage_int_SAM_3+Phage_integrase\|\|N6-METHYLASE-p-><-N6-METHYLASE<-DUF6079*<-BrxB-BrxF-iSTAND<-BrxA-DNA-binding<-XIS-HTH<-ArdCN+MPTase<-?<-VirD4-TraG+VirD4-FtsK<-nSTAND3 | 1169 | Bacteria>Terrabacteria_group>Actinomycetota>Actinomycetes>Micrococcales>Micrococcaceae | GCF_001445575.1 | Arthrobacter_alpinus | AS189_RS18910 | 4268679..4272188 | 656366 | BREX-system-P-loop-protein-BrxC-[Arthrobacter-alpinus] |
| WP_099093371.1 | Type1.BREX | <-HEPN+SF1-SF2-Helicase+REase<-?\|\|?-><-ParB+HNH<-SIGMA+OB-Fold+AAA-ATPase+HKD-DNase-BrxL<-PglZ<-N6-METHYLASE<-DUF6079*<-BrxB-BrxF-iSTAND<-BrxA-DNA-binding<-XIS-HTH<-HKD-DNase+SF1-SF2-Helicase+HTH+HTH+SRA\|\|?->?->McrB-AAA->McrC-NTD+REase-> | 1173 | Bacteria>Terrabacteria_group>Actinomycetota>Actinomycetes>Micrococcales>Micrococcaceae | GCF_001484605.1 | Pseudarthrobacter_sulfonivorans | AU252_RS05160 | 1179330..1182851 | 121292 | BREX-system-P-loop-protein-BrxC-[Pseudarthrobacter-sulfonivorans] |
| WP_078028155.1 | Type1.BREX | <-EVE+HNH\|\|?->?->CHTH->?->?-><-SIGMA+OB-Fold+AAA-ATPase+HKD-DNase-BrxL<-PglZ<-N6-METHYLASE<-DUF6079*<-BrxB-BrxF-iSTAND<-BrxA-DNA-binding<-XIS-HTH\|\|?-><-?\|\|?-><-Phage_integrase<-XIS-HTH<-AAA-ATPase | 1176 | Bacteria>Terrabacteria_group>Actinomycetota>Actinomycetes>Micrococcales>Micrococcaceae | GCF_002008315.2 | Arthrobacter_sp._KBS0703 | B1A87_RS03145 | 655510..659040 | 1955698 | BREX-system-P-loop-protein-BrxC-[Arthrobacter-sp.-KBS0703] |
| WP_104138811.1 | Type5.BREX | NUDIX->ARPP2-><-MACRODOMAIN+SIR2<-URI-DNase\|\|HKD-DNase+SF1-SF2-Helicase+HTH+HTH+SRA->XIS-HTH->BrxA-DNA-binding->BrxB-BrxF-iSTAND->DUF6079*->N6-METHYLASE->PglZ->N6-METHYLASE-p->N6-METHYLASE->SIGMA+OB-Fold+AAA-ATPase+HKD-DNase-BrxL->?->ParB+HNH->?->35exo+TUDOR+SF1-SF2-Helicase+iREase+CTD-> | 1173 | Bacteria>Terrabacteria_group>Actinomycetota>Actinomycetes>Micrococcales>Micrococcaceae | GCF_002929275.1 | Arthrobacter_sp._ZGTC131 | CXQ90_RS09900 | 131121..134642 | 2058898 | BREX-system-P-loop-protein-BrxC-[Arthrobacter-sp.-ZGTC131] |
| WP_242630351.1 | Type1.BREX | ParA-Soj-PloopNTPase->?-><-TIR\|\|XIS-HTH->BrxA-DNA-binding->BrxB-BrxF-iSTAND->DUF6079*->N6-METHYLASE->PglZ->SIGMA+OB-Fold+AAA-ATPase+HKD-DNase-BrxL-><-?\|\|REase->HTH+REase->URI-DNase->DUF6361->HKD-DNase-> | 1169 | Bacteria>Terrabacteria_group>Actinomycetota>Actinomycetes>Micrococcales>Micrococcaceae | GCF_004799725.1 | Kocuria_rosea | E1J17_RS01420 | 4072..7581 | 1275 | BREX-system-P-loop-protein-BrxC-[Kocuria-rosea] |
| WP_142936644.1 | Type1.BREX | <-NERD<-?<-PIN<-?<-NAD_binding_10<-SIGMA+OB-Fold+AAA-ATPase+HKD-DNase-BrxL<-PglZ<-N6-METHYLASE<-DUF6079*<-BrxB-BrxF-iSTAND<-BrxA-DNA-binding<-XIS-HTH\|\|?->XIS-HTH->Phage_int_SAM_3->Phage_integrase-> | 1161 | Bacteria>Terrabacteria_group>Actinomycetota>Actinomycetes>Micrococcales>Micrococcaceae | GCF_006964045.1 | Arthrobacter_sp._TS-15 | EU811_RS00880 | 63009..66494 | 2510797 | BREX-system-P-loop-protein-BrxC-[Arthrobacter-sp.-TS-15] |
| WP_208011566.1 | Type1.BREX | ICLR-HTH-><-?<-?\|\|?-><-?\|\|?->BrxA-DNA-binding->BrxB-BrxF-iSTAND->DUF6079*->N6-METHYLASE->PglZ->SIGMA+OB-Fold+AAA-ATPase+HKD-DNase-BrxL-><-?\|\|?-><-RADICAL-SAM<-?<-ACET<-XIS-HTH | 1172 | Bacteria>Terrabacteria_group>Actinomycetota>Actinomycetes>Micrococcales>Micrococcaceae | GCF_011602345.1 | Micrococcus_luteus | G9B47_RS07850 | 17280..20798 | 1270 | BREX-system-P-loop-protein-BrxC-[Micrococcus-luteus] |
| WP_171585597.1 | Type1.BREX | Peptidase_M20-><-Peptidase_M20\|\|DNA_METHYLASE-><-?<-?<-?\|\|?-><-SIGMA+OB-Fold+AAA-ATPase+HKD-DNase-BrxL<-N6-METHYLASE<-DUF6079*<-BrxB-BrxF-iSTAND<-XIS-HTH<-?\|\|?-><-?<-HNH\|\|Peptidase-> | 1171 | Bacteria>Terrabacteria_group>Actinomycetota>Actinomycetes>Micrococcales>Micrococcaceae | GCF_013116745.1 | Arthrobacter_sp._260 | HMI57_RS08915 | 135623..139138 | 2735314 | BREX-system-P-loop-protein-BrxC-[Arthrobacter-sp.-260] |
| WP_135027971.1 | Type1.BREX | 35exo->tRNA-><-Recombinase+ZnR\|\|ABhydrolase->XIS-HTH->BrxA-DNA-binding->BrxB-BrxF-iSTAND->DUF6079*->N6-METHYLASE->HTH+REase->PglZ->SIGMA+OB-Fold+AAA-ATPase+HKD-DNase-BrxL->SIR2->HTH_Tnp_1+HTH_21+rve+rve_3-><-?\|\|?-><-HAD+HAD | 1170 | Bacteria>Terrabacteria_group>Actinomycetota>Actinomycetes>Micrococcales>Micrococcaceae | GCF_014644875.1 | Micrococcus_flavus | E4A47_RS01340 | 286387..289899 | 384602 | BREX-system-P-loop-protein-BrxC-[Micrococcus-flavus] |
| WP_193340403.1 | Type1.BREX | <-SIGMA+OB-Fold+AAA-ATPase+HKD-DNase-BrxL<-PglZ<-REase<-N6-METHYLASE<-DUF6079*<-BrxB-BrxF-iSTAND<-BrxA-DNA-binding<-XIS-HTH\|\|?-><-EVE+REase | 1153 | Bacteria>Terrabacteria_group>Actinomycetota>Actinomycetes>Micrococcales>Micrococcaceae | GCF_014926485.1 | Pseudarthrobacter_sp._AB1 | DAD99_RS03725 | 12045..15506 | 2138309 | BREX-system-P-loop-protein-BrxC-[Pseudarthrobacter-sp.-AB1] |
| WP_258190683.1 | Type1.BREX | HIN-HTH-><-SIGMA+OB-Fold+AAA-ATPase+HKD-DNase-BrxL<-PglZ<-N6-METHYLASE<-DUF6079*<-BrxA-DNA-binding<-XIS-HTH<-?<-?\|\|?->HNH-> | 1170 | Bacteria>Terrabacteria_group>Actinomycetota>Actinomycetes>Micrococcales>Micrococcaceae | GCF_019443425.1 | Arthrobacter_sp._PAMC25284 | KY499_RS09320 | 2016646..2020158 | 2861279 | BREX-system-P-loop-protein-BrxC-[Arthrobacter-sp.-PAMC25284] |
| WP_142938938.1 | Type1.BREX | ParB+HNH-p->ParB+HNH-><-?<-?<-URI-DNase\|\|?-><-SIGMA+OB-Fold+AAA-ATPase+HKD-DNase-BrxL<-PglZ<-N6-METHYLASE<-DUF6079*<-BrxB-BrxF-iSTAND<-BrxA-DNA-binding<-XIS-HTH\|\|?->Radical-SAM-><-METHYLASE<-Phage_integrase+Phage_integrase\|\|Phage_integrase-><-HTH_3 | 1162 | Bacteria>Terrabacteria_group>Actinomycetota>Actinomycetes>Micrococcales>Micrococcaceae | GCF_019976975.1 | unclassified_Arthrobacter | LDN90_RS14640 | 3248673..3252161 | 235627 | MULTISPECIES:-BREX-system-P-loop-protein-BrxC-[unclassified-Arthrobacter] |
| WP_224048794.1 | Type1.BREX | NUDIX->wHTH(SBD)+HNH-><-?\|\|?-><-SF1-SF2-Helicase\|\|XIS-HTH->BrxA-DNA-binding->BrxB-BrxF-iSTAND->DUF6079*->N6-METHYLASE->PglZ->N6-METHYLASE->SIGMA+OB-Fold+AAA-ATPase+HKD-DNase-BrxL-><-ParB+HNH\|\|HEPN+SF1-SF2-Helicase+REase-> | 1173 | Bacteria>Terrabacteria_group>Actinomycetota>Actinomycetes>Micrococcales>Micrococcaceae | GCF_019977335.1 | Arthrobacter_sp._NicSoilB4 | LDD13_RS04105 | 882114..885635 | 2830997 | BREX-system-P-loop-protein-BrxC-[Arthrobacter-sp.-NicSoilB4] |
| WP_226761965.1 | Type1.BREX | LexA-protease->?->?->?->XIS-HTH->BrxA-DNA-binding->BrxB-BrxF-iSTAND->DUF6079*->N6-METHYLASE->PglZ->SIGMA+OB-Fold+AAA-ATPase+HKD-DNase-BrxL-><-?<-?<-?\|\|HEPN->GDResolvase+HIN-HTH-> | 1161 | Bacteria>Terrabacteria_group>Actinomycetota>Actinomycetes>Micrococcales>Micrococcaceae | GCF_020532825.1 | Arthrobacter_sp._SO3 | BJQ90_RS20165 | 19380..22865 | 1897057 | BREX-system-P-loop-protein-BrxC-[Arthrobacter-sp.-SO3] |
| WP_235519134.1 | Type5.BREX | <-MarR-HTH+DTXR-HTH\|\|?-><-HKD-DNase+HKD-Rease<-?\|\|HKD-DNase+SF1-SF2-Helicase+HTH+HTH+SRA-><-SF1-SF2-Helicase<-35exo+TUDOR+SF1-SF2-Helicase+iREase+CTD\|\|BrxA-DNA-binding->BrxB-BrxF-iSTAND->DUF6079*->N6-METHYLASE->PglZ->SIGMA+OB-Fold+AAA-ATPase+HKD-DNase-BrxL-><-?\|\|?-><-REase | 1162 | Bacteria>Terrabacteria_group>Actinomycetota>Actinomycetes>Micrococcales>Micrococcaceae | GCF_021538675.1 | Paenarthrobacter_sp._AR_02 | LRQ04_RS07575 | 297017..300505 | 2899821 | BREX-system-P-loop-protein-BrxC-[Paenarthrobacter-sp.-AR-02] |
| WP_241470046.1 | Type1.BREX | DNA_glycosylase->DNA_glycosylase->?->?-><-SIGMA+OB-Fold+AAA-ATPase+HKD-DNase-BrxL<-PglZ<-N6-METHYLASE<-DUF6079*<-BrxB-BrxF-iSTAND<-BrxA-DNA-binding\|\|?->HTH_26->?-><-ABC-ATPase+SbcC | 1168 | Bacteria>Terrabacteria_group>Actinomycetota>Actinomycetes>Micrococcales>Micrococcaceae | GCF_022565835.1 | Nesterenkonia_sp._DZ6 | LTH93_RS03075 | 681136..684642 | 2901229 | BREX-system-P-loop-protein-BrxC-[Nesterenkonia-sp.-DZ6] |
| WP_241707532.1 | Type1.BREX | REase->ChlI+Lon_C+AAA->?->?->?->XIS-HTH->BrxA-DNA-binding->BrxB-BrxF-iSTAND->DUF6079*->N6-METHYLASE->PglZ->SIGMA+OB-Fold+AAA-ATPase+HKD-DNase-BrxL-><-?<-DNA-glycosylase\|\|?->NACHT-> | 1171 | Bacteria>Terrabacteria_group>Actinomycetota>Actinomycetes>Micrococcales>Micrococcaceae | GCF_022606295.1 | Arthrobacter_bambusae | KNN17_RS03605 | 221894..225409 | 1338426 | BREX-system-P-loop-protein-BrxC-[Arthrobacter-bambusae] |
| WP_247040359.1 | Type1.BREX | HTH_3-><-Phage_integrase\|\|Phage_integrase+Phage_integrase->METHYLASE-><-Radical-SAM<-?\|\|XIS-HTH->BrxA-DNA-binding->BrxB-BrxF-iSTAND->DUF6079*->N6-METHYLASE->PglZ->SIGMA+OB-Fold+AAA-ATPase+HKD-DNase-BrxL-><-?\|\|URI-DNase->?->?-><-ParB+HNH | 1168 | Bacteria>Terrabacteria_group>Actinomycetota>Actinomycetes>Micrococcales>Micrococcaceae | GCF_023062595.1 | Arthrobacter_rhizosphaerae | MU551_RS01475 | 284158..287664 | 2855490 | BREX-system-P-loop-protein-BrxC-[Arthrobacter-rhizosphaerae] |
| WP_248193800.1 | Type1.BREX | <-SIGMA+OB-Fold+AAA-ATPase+HKD-DNase-BrxL<-PglZ<-?<-N6-METHYLASE<-DUF6079*<-BrxB-BrxF-iSTAND<-BrxA-DNA-binding<-XIS-HTH\|\|?-><-HAD+HAD | 1173 | Bacteria>Terrabacteria_group>Actinomycetota>Actinomycetes>Micrococcales>Micrococcaceae | GCF_023147525.1 | Micrococcus_endophyticus | KZX39_RS06505 | 8911..12432 | 455343 | BREX-system-P-loop-protein-BrxC-[Micrococcus-endophyticus] |
| WP_073116219.1 | Type1.BREX | ICLR-HTH-><-?<-?\|\|?-><-?\|\|?->BrxA-DNA-binding->BrxB-BrxF-iSTAND->DUF6079*->N6-METHYLASE->PglZ->SIGMA+OB-Fold+AAA-ATPase+HKD-DNase-BrxL-><-?\|\|?-><-RADICAL-SAM<-?<-ACET<-XIS-HTH | 1174 | Bacteria>Terrabacteria_group>Actinomycetota>Actinomycetes>Micrococcales>Micrococcaceae | GCF_900142795.1 | Micrococcus | BUB74_RS03185 | 94466..97990 | 1269 | MULTISPECIES:-BREX-system-P-loop-protein-BrxC-[Micrococcus] |
| WP_237187176.1 | Type1.BREX | <-HHH_5+SNF2_N+SF1-SF2-Helicase\|\|SWACOS-><-?<-SIGMA+OB-Fold+AAA-ATPase+HKD-DNase-BrxL<-PglZ<-Schlafen+HATPase+wHTH<-N6-METHYLASE<-DUF6079*<-BrxB-BrxF-iSTAND<-BrxA-DNA-binding\|\|?-><-Aminotran_4<-?<-ABC-ATPase+SbcC+ABC-ATPase+SbcC | 1190 | Bacteria>Terrabacteria_group>Actinomycetota>Actinomycetes>Micrococcales>Micrococcaceae | GCF_928381705.1 | Rothia_nasimurium | L6762_RS10435 | 40591..44163 | 85336 | BREX-system-P-loop-protein-BrxC-[Rothia-nasimurium] |
| WP_211713005.1 | Type1.BREX | <-DNA-glycosylase<-URI-DNase<-MACRODOMAIN+SIR2<-REase\|\|HKD-DNase+SF1-SF2-Helicase+HTH+HTH+SRA-><-SIGMA+OB-Fold+AAA-ATPase+HKD-DNase-BrxL<-PglZ<-N6-METHYLASE<-DUF6079*<-BrxB-BrxF-iSTAND<-BrxA-DNA-binding\|\|McrB-AAA->McrC-NTD+REase->ABhydrolase+ABhydrolase-> | 1173 | Bacteria>Terrabacteria_group>Actinomycetota>Actinomycetes>Micrococcales>Micrococcaceae | - | Paenarthrobacter_sp._GOM3 | IRJ34_14045 | 199455..202976 | 2782567 | BREX-system-P-loop-protein-BrxC-[Paenarthrobacter-sp.-GOM3] |
| WP_100350249.1 | Type1.BREX | HEPN+SF1-SF2-Helicase+REase->?-><-Pkinase+CR-Kinase<-SIGMA+OB-Fold+AAA-ATPase+HKD-DNase-BrxL<-PglZ<-N6-METHYLASE-p<-Phage_int_SAM_3+Phage_integrase\|\|N6-METHYLASE-p-><-N6-METHYLASE<-DUF6079*<-BrxB-BrxF-iSTAND<-BrxA-DNA-binding\|\|?->ZNR+RmuC-> | 1179 | Bacteria>Terrabacteria_group>Actinomycetota>Actinomycetes>Micrococcales>Micrococcales_incertae_sedis | GCF_002798015.1 | Luteimicrobium_subarcticum | CLV34_RS10655 | 413171..416710 | 620910 | BREX-system-P-loop-protein-BrxC-[Luteimicrobium-subarcticum] |
| SDF46422.1 | Type1.BREX | ABC-ATPase+SbcC->Cysteine-Desulfurase->?->?->?->BrxA-DNA-binding->BrxB-BrxF-iSTAND->DUF6079*->N6-METHYLASE->PglZ->SIGMA+OB-Fold+AAA-ATPase+HKD-DNase-BrxL->REase->PAIREDC-HTH-> | 1166 | Bacteria>Terrabacteria_group>Actinomycetota>Actinomycetes>Micrococcales>Promicromonosporaceae | GCA_900102445.1 | Cellulosimicrobium_cellulans | SAMN04487781_1443 | 34344..37844 | 1710 | hypothetical-protein-SAMN04487781_1443-[Cellulosimicrobium-cellulans] |
| WP_162290006.1 | Type1.BREX | PIWI-><-HerA/FtsK<-SIR2+SIR2<-SIGMA+OB-Fold+AAA-ATPase+HKD-DNase-BrxL<-PglZ<-N6-METHYLASE<-DUF6079*<-BrxB-BrxF-iSTAND<-BrxA-DNA-binding<-XIS-HTH<-tRNA<-?<-GNTR-HTH\|\|DSBH+AraC-HTH+AraC-HTH-><-HTH_42<-ABhydrolase | 1167 | Bacteria>Terrabacteria_group>Actinomycetota>Actinomycetes>Micrococcales>Promicromonosporaceae | GCF_010287905.1 | Cellulosimicrobium_fucosivorans | GYH36_RS12555 | 12661..16164 | 2850388 | BREX-system-P-loop-protein-BrxC-[Cellulosimicrobium-fucosivorans] |
| WP_253873967.1 | Type1.BREX | <-URI-DNase\|\|?->SIR2->EVE-><-SIGMA+OB-Fold+AAA-ATPase+HKD-DNase-BrxL<-PglZ<-N6-METHYLASE<-DUF6079*<-BrxB-BrxF-iSTAND<-BrxA-DNA-binding\|\|?-><-CHTH\|\|?-><-REase<-REase<-ANK+ANK+ANK<-REase | 1157 | Bacteria>Terrabacteria_group>Actinomycetota>Actinomycetes>Micrococcales>Promicromonosporaceae | GCF_024171995.1 | Promicromonospora_umidemergens | APR04_RS20365 | 11407..14880 | 629679 | BREX-system-P-loop-protein-BrxC-[Promicromonospora-umidemergens] |
| WP_142121348.1 | Type1.BREX | CR-ATPase5+REase-><-REase+SF1-SF2-Helicase-p\|\|BrxA-DNA-binding->BrxB-BrxF-iSTAND->DUF6079*->N6-METHYLASE->?->PglZ->RVT->SIGMA+OB-Fold+AAA-ATPase+HKD-DNase-BrxL-><-?\|\|SF1-SF2-Helicase->wHTH-><-CR-ATPase5+REase | 1170 | Bacteria>Terrabacteria_group>Actinomycetota>Actinomycetes>Micrococcales>Rarobacteraceae | GCF_006716265.1 | Rarobacter_faecitabidus | FB461_RS08150 | 23162..26674 | 13243 | BREX-system-P-loop-protein-BrxC-[Rarobacter-faecitabidus] |
| WP_142113745.1 | Type1.BREX | <-VirB11+MuB-ATPase\|\|DNA_METHYLASE-><-HerA/FtsK<-SIR2<-SIGMA+OB-Fold+AAA-ATPase+HKD-DNase-BrxL<-PglZ<-Schlafen+HATPase+wHTH<-?<-N6-METHYLASE<-DUF6079*<-BrxB-BrxF-iSTAND<-BrxA-DNA-binding<-XIS-HTH<-ABhydrolase<-ABhydrolase<-?<-MPTase<-HTH\|\|ArdCN+MPTase-> | 1173 | Bacteria>Terrabacteria_group>Actinomycetota>Actinomycetes>Micrococcales>Rarobacteraceae | - | Rarobacter_incanus | FB389_RS10090 | 2435944..2439465 | 153494 | BREX-system-P-loop-protein-BrxC-[Rarobacter-incanus] |
| SNU02231.1 | Type1.BREX | <-SbcC<-?<-SIGMA+OB-Fold+AAA-ATPase+HKD-DNase-BrxL<-PglZ<-N6-METHYLASE-p<-Phage_int_SAM_3+Phage_integrase\|\|?->N6-METHYLASE-p-><-N6-METHYLASE<-DUF6079*<-BrxB-BrxF-iSTAND<-BrxA-DNA-binding\|\|SF1-SF2-Helicase+PhospholipaseD-><-?<-?\|\|Peptidase_M23-><-?\|\|TraM-> | 1159 | Bacteria>Terrabacteria_group>Actinomycetota>Actinomycetes>Micrococcales>Ruaniaceae | GCA_900199505.1 | Ruaniaceae_bacterium_KH17 | SAMN06298212_12910 | 14626..18105 | 1945888 | hypothetical-protein-SAMN06298212_12910-[Ruaniaceae-bacterium-KH17] |
| WP_147917199.1 | Type1.BREX | HicA-><-?\|\|?->DNA_glycosylase-><-?<-?<-?\|\|BrxA-DNA-binding->BrxB-BrxF-iSTAND->DUF6079*->N6-METHYLASE->PglZ->SIGMA+OB-Fold+AAA-ATPase+HKD-DNase-BrxL->HTH_3-><-?\|\|?->?->?->ABC-ATPase+SbcC-> | 1152 | Bacteria>Terrabacteria_group>Actinomycetota>Actinomycetes>Micrococcales>Ruaniaceae | GCF_008000995.1 | Ruania_zhangjianzhongii | FU260_RS11560 | 2487086..2490544 | 2603206 | BREX-system-P-loop-protein-BrxC-[Ruania-zhangjianzhongii] |
| WP_163542003.1 | Type1.BREX | <-HIN-HTH\|\|SbcC->?-><-?<-SIGMA+OB-Fold+AAA-ATPase+HKD-DNase-BrxL<-PglZ<-REase<-N6-METHYLASE<-DUF6079*<-BrxB-BrxF-iSTAND<-BrxA-DNA-binding<-XIS-HTH<-?<-?\|\|CN_hydrolase+NAD_synthase->?-><-Pkinase<-RelE-ParE | 1154 | Bacteria>Terrabacteria_group>Actinomycetota>Actinomycetes>Micrococcales>Ruaniaceae | GCF_010994285.1 | Occultella_kanbiaonis | G7B41_RS06055 | 118944..122408 | 2675754 | BREX-system-P-loop-protein-BrxC-[Occultella-kanbiaonis] |
| MCL2802837.1 | Type1.BREX | <-RADICAL-SAM<-sGTP<-?\|\|SWACOS->TPR->?-><-SIGMA+OB-Fold+AAA-ATPase+HKD-DNase-BrxL<-PglZ<-N6-METHYLASE<-DUF6079*<-BrxB-BrxF-iSTAND<-BrxA-DNA-binding\|\|?-><-?\|\|?-><-PhdYeFM_antitox | 1162 | Bacteria>Terrabacteria_group>Actinomycetota>Actinomycetes>Micrococcales>unclassified_Micrococcales | GCA_009784895.1 | Micrococcales_bacterium | FWD29_02600 | 43259..46747 | 2026762 | BREX-system-P-loop-protein-BrxC-[Micrococcales-bacterium] |
| WP_183221872.1 | Type1.BREX | <-STYKIN+Pkinase<-?<-?\|\|?-><-?\|\|BrxA-DNA-binding->BrxB-BrxF-iSTAND->DUF6079*->N6-METHYLASE->PglZ->SIGMA+OB-Fold+AAA-ATPase+HKD-DNase-BrxL-><-GGDEF+EAL<-?\|\|ABhydrolase->?->?->METHYLASE-> | 1164 | Bacteria>Terrabacteria_group>Actinomycetota>Actinomycetes>Micromonosporales>Micromonosporaceae | GCF_014647795.1 | Actinoplanes_campanulatus | IEZ13_RS15515 | 158515..162009 | 113559 | BREX-system-P-loop-protein-BrxC-[Actinoplanes-campanulatus] |
| WP_203833901.1 | Type1.BREX | <-METHYLASE<-?<-?<-ABhydrolase\|\|?->GGDEF+EAL-><-SIGMA+OB-Fold+AAA-ATPase+HKD-DNase-BrxL<-PglZ<-N6-METHYLASE<-DUF6079*<-BrxB-BrxF-iSTAND<-BrxA-DNA-binding\|\|?-><-?\|\|?->?->STYKIN+Pkinase-> | 1164 | Bacteria>Terrabacteria_group>Actinomycetota>Actinomycetes>Micromonosporales>Micromonosporaceae | GCF_016862035.1 | Actinoplanes_campanulatus | Aca09nite_RS24855 | 101506..105000 | 113559 | BREX-system-P-loop-protein-BrxC-[Actinoplanes-campanulatus] |
| WP_188126500.1 | Type1.BREX | <-METHYLASE<-?<-?<-ABhydrolase\|\|GGDEF+EAL-><-SIGMA+OB-Fold+AAA-ATPase+HKD-DNase-BrxL<-PglZ<-N6-METHYLASE<-DUF6079*<-BrxB-BrxF-iSTAND<-BrxA-DNA-binding\|\|?-><-?\|\|?->?->STYKIN+Pkinase-> | 1164 | Bacteria>Terrabacteria_group>Actinomycetota>Actinomycetes>Micromonosporales>Micromonosporaceae | GCF_016862255.1 | Actinoplanes_lobatus | IEZ29_RS29630 | 68673..72167 | 113568 | BREX-system-P-loop-protein-BrxC-[Actinoplanes-lobatus] |
| WP_203928003.1 | Type1.BREX | TPR-><-SIGMA+OB-Fold+AAA-ATPase+HKD-DNase-BrxL<-PglZ<-N6-METHYLASE<-DUF6079*<-BrxB-BrxF-iSTAND<-BrxA-DNA-binding<-APATPase+TPR+TPR+TPR+TPR+TPR+TPR+TPR\|\|NAD_binding_10->?-><-NACHT\|\|SIGMA-HTH-> | 1154 | Bacteria>Terrabacteria_group>Actinomycetota>Actinomycetes>Micromonosporales>Micromonosporaceae | GCF_016863655.1 | Virgisporangium_ochraceum | Voc01_RS14900 | 8921..12385 | 65505 | BREX-system-P-loop-protein-BrxC-[Virgisporangium-ochraceum] |
| WP_260726425.1 | Type1.BREX | <-SIGMA-HTH<-?<-SAM-METHYLASE\|\|?->?->?-><-SIGMA+OB-Fold+AAA-ATPase+HKD-DNase-BrxL<-PglZ<-N6-METHYLASE<-DUF6079*<-BrxB-BrxF-iSTAND<-BrxA-DNA-binding<-XIS-HTH\|\|HAMP+MA+MA->?->SIGMA-HTH-><-Glycos_transf_1+Glyco_trans_1_2<-Glycos_transf_1+Glyco_trans_1_2<-SF1-SF2-Helicase | 1154 | Bacteria>Terrabacteria_group>Actinomycetota>Actinomycetes>Micromonosporales>Micromonosporaceae | GCF_025264685.1 | Dactylosporangium_roseum | Drose_RS01775 | 392440..395904 | 47989 | BREX-system-P-loop-protein-BrxC-[Dactylosporangium-roseum] |
| WP_091673594.1 | Type1.BREX | RADICAL-SAM->?->?-><-HAD\|\|XIS-HTH->Phage_int_SAM_3+Phage_integrase->XIS-HTH->BrxA-DNA-binding->BrxB-BrxF-iSTAND->DUF6079*->N6-METHYLASE->PglZ->SIGMA+OB-Fold+AAA-ATPase+HKD-DNase-BrxL->DNA_glycosylase-><-?<-SF1-SF2-Helicase<-HKD-DNase<-DUF6361<-RNase | 1156 | Bacteria>Terrabacteria_group>Actinomycetota>Actinomycetes>Micromonosporales>Micromonosporaceae | GCF_900089595.1 | Micromonospora_auratinigra | GA0070611_RS28700 | 6259973..6263443 | 261654 | BREX-system-P-loop-protein-BrxC-[Micromonospora-auratinigra] |
| WP_091348932.1 | Type1.BREX | XIS-HTH->BrxA-DNA-binding->BrxB-BrxF-iSTAND->DUF6079*->N6-METHYLASE->PglZ->SIGMA+OB-Fold+AAA-ATPase+HKD-DNase-BrxL->HIN-HTH-><-HTH_32<-HTH_32<-SF1-SF2-Helicase<-HKD-DNase+HKD-Rease | 1153 | Bacteria>Terrabacteria_group>Actinomycetota>Actinomycetes>Micromonosporales>Micromonosporaceae | GCF_900091465.1 | Micromonospora_rhizosphaerae | GA0070624_RS15965 | 3332590..3336051 | 568872 | BREX-system-P-loop-protein-BrxC-[Micromonospora-rhizosphaerae] |
| HJG42479.1 | Type1.BREX | BrxA-DNA-binding->BrxB-BrxF-iSTAND->DUF6079*->N6-METHYLASE->PglZ->SIGMA+OB-Fold+AAA-ATPase+HKD-DNase-BrxL-> | 1176 | Bacteria>Terrabacteria_group>Actinomycetota>Actinomycetes>Mycobacteriales>Corynebacteriaceae | GCA_020743465.1 | Corynebacterium_phoceense | K8V34_00330 | 34206..37736 | 1686286 | BREX-system-P-loop-protein-BrxC-[Corynebacterium-phoceense] |
| WP_102213741.1 | Type1.BREX | XIS-HTH->BrxA-DNA-binding->BrxB-BrxF-iSTAND->DUF6079*->N6-METHYLASE->HTH+REase->PglZ->SIGMA+OB-Fold+AAA-ATPase+HKD-DNase-BrxL-><-ParB+HNH+METHYLASE | 1173 | Bacteria>Terrabacteria_group>Actinomycetota>Actinomycetes>Mycobacteriales>Corynebacteriaceae | GCF_002871715.1 | Corynebacterium_xerosis | CJ204_09375 | 1693..5214 | 1725 | BREX-system-P-loop-protein-BrxC-[Corynebacterium-xerosis] |
| WP_220313057.1 | Type1.BREX | <-REC+OmpR-HTH<-MerR-HTH+TipAS<-?<-SIR2\|\|?->TetR-HTH->?->BrxA-DNA-binding->BrxB-BrxF-iSTAND->DUF6079*->N6-METHYLASE-><-N6-METHYLASE-p\|\|Phage_int_SAM_3+Phage_integrase->N6-METHYLASE-p->PglZ->SIGMA+OB-Fold+AAA-ATPase+HKD-DNase-BrxL-><-?<-?<-ADH_N+ADH_zinc_N | 1160 | Bacteria>Terrabacteria_group>Actinomycetota>Actinomycetes>Mycobacteriales>Corynebacteriaceae | GCF_019551735.1 | Corynebacterium_glutamicum | JJQ73_RS12320 | 2669346..2672828 | 1718 | BREX-system-P-loop-protein-BrxC-[Corynebacterium-glutamicum] |
| WP_259898411.1 | Type1.BREX | <-HIN-HTH<-ParB+HNH<-SIGMA+OB-Fold+AAA-ATPase+HKD-DNase-BrxL<-PglZ<-N6-METHYLASE<-DUF6079*<-BrxB-BrxF-iSTAND<-BrxA-DNA-binding<-XIS-HTH | 1160 | Bacteria>Terrabacteria_group>Actinomycetota>Actinomycetes>Mycobacteriales>Corynebacteriaceae | GCF_025152505.1 | Corynebacterium_sanguinis | M3G46_RS10600 | 10062..13544 | 2594913 | BREX-system-P-loop-protein-BrxC-[Corynebacterium-sanguinis] |
| WP_271034470.1 | Type1.BREX | <-HTH_23+rve+rve_3\|\|?->BrxA-DNA-binding->BrxB-BrxF-iSTAND->DUF6079*->N6-METHYLASE-><-N6-METHYLASE-p\|\|Phage_int_SAM_3+Phage_integrase->N6-METHYLASE-p->PglZ->SIGMA+OB-Fold+AAA-ATPase+HKD-DNase-BrxL-> | 1159 | Bacteria>Terrabacteria_group>Actinomycetota>Actinomycetes>Mycobacteriales>Corynebacteriaceae | GCF_027886475.1 | Corynebacterium_sp._SCR221107 | PAB09_RS02255 | 466456..469935 | 3017361 | BREX-system-P-loop-protein-BrxC-[Corynebacterium-sp.-SCR221107] |
| WP_245802167.1 | Type1.BREX | <-ARG<-SIR2<-SIGMA+OB-Fold+AAA-ATPase+HKD-DNase-BrxL<-PglZ<-Schlafen+HATPase+wHTH<-N6-METHYLASE<-DUF6079*<-BrxB-BrxF-iSTAND<-BrxA-DNA-binding | 1179 | Bacteria>Terrabacteria_group>Actinomycetota>Actinomycetes>Mycobacteriales>Corynebacteriaceae | GCF_900169525.1 | Corynebacterium_pacaense | B7E02_RS05025 | 1051363..1054902 | 1816684 | BREX-system-P-loop-protein-BrxC-[Corynebacterium-pacaense] |
| WP_047036580.1 | Type1.BREX | <-Phage_integrase\|\|ACET-><-HerA/FtsK<-SIR2+SIR2<-SIGMA+OB-Fold+AAA-ATPase+HKD-DNase-BrxL<-PglZ<-N6-METHYLASE<-DUF6079*<-BrxB-BrxF-iSTAND<-BrxA-DNA-binding<-XIS-HTH<-TetR+HTH<-Phage_integrase+Phage_integrase<-Phage_integrase+Phage_integrase<-Phage_int_SAM_1+Phage_integrase<-DNA_glycosylase | 1156 | Bacteria>Terrabacteria_group>Actinomycetota>Actinomycetes>Mycobacteriales | GCF_900141005.1 | Mycobacteriaceae | ABW05_RS31210 | 104572..108042 | 1762 | MULTISPECIES:-BREX-system-P-loop-protein-BrxC-[Mycobacteriaceae] |
| PZT90276.1 | Type1.BREX | HAD+HAD-><-?\|\|XIS-HTH->BrxA-DNA-binding->BrxB-BrxF-iSTAND->DUF6079*->N6-METHYLASE->HTH+REase->PglZ->SIGMA+OB-Fold+AAA-ATPase+HKD-DNase-BrxL->REase-> | 1137 | Bacteria>Terrabacteria_group>Actinomycetota>Actinomycetes>Mycobacteriales>Gordoniaceae | GCA_003248405.1 | Gordonia_sp._(in:_high_G+C_Gram-positive_bacteria) | DI630_31230 | 6915..10328 | 84139 | MAG:-BREX-system-P-loop-protein-BrxC-[Gordonia-sp.-(in:-high-G+C-Gram-positive-bacteria)] |
| WP_042377474.1 | Type1.BREX | TetR-HTH->?-><-?<-SIGMA+OB-Fold+AAA-ATPase+HKD-DNase-BrxL<-PglZ<-N6-METHYLASE<-DUF6079*<-BrxB-BrxF-iSTAND<-BrxA-DNA-binding\|\|Exonuc_V_gamma+REase->SF1-SF2-Helicase+REase-><-?<-DNA_glycosylase<-HNH+HNH | 1161 | Bacteria>Terrabacteria_group>Actinomycetota>Actinomycetes>Mycobacteriales>Gordoniaceae | GCF_000529675.1 | Gordonia_alkanivorans | TY17_RS15305 | 47194..50679 | 84096 | BREX-system-P-loop-protein-BrxC-[Gordonia-alkanivorans] |
| WP_159370828.1 | Type1.BREX | <-DNA_glycosylase\|\|DUF6361->HKD-DNase->SF1-SF2-Helicase-><-SIGMA+OB-Fold+AAA-ATPase+HKD-DNase-BrxL<-PglZ<-N6-METHYLASE<-DUF6079*<-BrxB-BrxF-iSTAND<-BrxA-DNA-binding<-?<-?<-ARG<-?\|\|?-><-SAM-METHYLASE | 1174 | Bacteria>Terrabacteria_group>Actinomycetota>Actinomycetes>Mycobacteriales>Gordoniaceae | GCF_009856645.1 | Gordonia_sp._JH63 | GR168_RS12095 | 2701909..2705433 | 2698900 | BREX-system-P-loop-protein-BrxC-[Gordonia-sp.-JH63] |
| WP_213244950.1 | Type1.BREX | <-SIGMA+OB-Fold+AAA-ATPase+HKD-DNase-BrxL<-PglZ<-HTH+REase<-N6-METHYLASE-p<-?<-Phage_int_SAM_3+Phage_integrase<-Phage_int_SAM_3+Phage_integrase\|\|N6-METHYLASE-p-><-N6-METHYLASE<-DUF6079*<-BrxB-BrxF-iSTAND<-BrxA-DNA-binding | 1174 | Bacteria>Terrabacteria_group>Actinomycetota>Actinomycetes>Mycobacteriales>Gordoniaceae | GCF_009914535.1 | Gordonia_pseudamarae | GII33_RS19265 | 4440551..4444075 | 2831662 | BREX-system-P-loop-protein-BrxC-[Gordonia-pseudamarae] |

| Protein-ID | System | Gene-Neighbourhoods | Length | Taxonomy | Assembly-ID | Strain | Locus-Tag | Genomic-coordinates | TaxID | Fasta-defline |
|---|---|---|---|---|---|---|---|---|---|---|
| WP_223374200.1 | Type1.BREX | REase->sGTP+sGTP+GTPase-AIG->sGTP-><-PHP+SbcC<-?<-SIGMA+OB-Fold+AAA-ATPase+HKD-DNase-BrxL<-PglZ<-N6-METHYLASE<-DUF6079*<-BrxB-BrxF-iSTAND<-BrxA-DNA-binding<-XIS-HTH<-?<-TnsC-ATPase\|\|PAIREDC-HTH-><-rve\|\|HTH_29+HTH_32+DDE_3-><-REase+wHTH | 1154 | Bacteria>Terrabacteria_group>Actinomycetota>Actinomycetes>Mycobacteriales>Gordoniaceae | GCF_019930645.1 | Gordonia_bronchialis | K8093_RS05575 | 1152614..1156078 | 2054 | BREX-system-P-loop-protein-BrxC-[Gordonia-bronchialis] |
| WP_226512485.1 | Type1.BREX | MACRODOMAIN+SIR2-><-?<-?\|\|DUF6361->HKD-DNase->SF1-SF2-Helicase->URI-DNase->BrxA-DNA-binding->BrxB-BrxF-iSTAND->DUF6079-p*->DUF6079-p->XIS-HTH->PIN->N6-METHYLASE-p->N6-METHYLASE->PglZ->?->AbiEi_4+AbiEi_1->SIGMA+OB-Fold+AAA-ATPase+HKD-DNase-BrxL-> | 435 | Bacteria>Terrabacteria_group>Actinomycetota>Actinomycetes>Mycobacteriales>Gordoniaceae | GCF_020520305.1 | Gordonia_sp._WA4-43 | LEL84_RS18910 | 4162672..4163979 | 2878678 | BREX-system-P-loop-protein-BrxC-[Gordonia-sp.-WA4-43] |
| WP_226512486.1 | Type1.BREX | DUF6361->HKD-DNase->SF1-SF2-Helicase->URI-DNase->BrxA-DNA-binding->BrxB-BrxF-iSTAND->DUF6079-p->DUF6079-p*->XIS-HTH->PIN->N6-METHYLASE-p->N6-METHYLASE->PglZ->?->AbiEi_4+AbiEi_1->SIGMA+OB-Fold+AAA-ATPase+HKD-DNase-BrxL-><-ABhydrolase+ABhydrolase | 785 | Bacteria>Terrabacteria_group>Actinomycetota>Actinomycetes>Mycobacteriales>Gordoniaceae | GCF_020520305.1 | Gordonia_sp._WA4-43 | LEL84_RS18915 | 4163880..4166237 | 2878678 | hypothetical-protein-[Gordonia-sp.-WA4-43] |
| WP_246991513.1 | Type1.BREX | <-SIGMA+OB-Fold+AAA-ATPase+HKD-DNase-BrxL<-PglZ<-?<-?<-N6-METHYLASE-p<-Phage_int_SAM_3+Phage_integrase\|\|N6-METHYLASE-p-><-N6-METHYLASE<-DUF6079*<-BrxB-BrxF-iSTAND<-BrxA-DNA-binding<-sGTP<-?\|\|?->O6MGT-HTH-> | 1175 | Bacteria>Terrabacteria_group>Actinomycetota>Actinomycetes>Mycobacteriales>Gordoniaceae | GCF_023017165.1 | Gordonia_alkaliphila | MUG78_01460 | 307013..310540 | 1053547 | BREX-system-P-loop-protein-BrxC-[Gordonia-alkaliphila] |
| OKH76423.1 | Type1.BREX | <-NUDIX\|\|ARG-><-SbcC<-?<-SIGMA+OB-Fold+AAA-ATPase+HKD-DNase-BrxL<-PglZ<-N6-METHYLASE<-DUF6079*<-BrxB-BrxF-iSTAND<-BrxA-DNA-binding<-XIS-HTH | 1156 | Bacteria>Terrabacteria_group>Actinomycetota>Actinomycetes>Mycobacteriales>Mycobacteriaceae | GCA_001905565.1 | Mycobacterium_sp._SWH-M1 | EB72_01455 | 44646..48116 | 1490485 | ATP-binding-protein-[Mycobacterium-sp.-SWH-M1] |
| OKH79002.1 | Type1.BREX | XIS-HTH->BrxA-DNA-binding->BrxB-BrxF-iSTAND->DUF6079*->N6-METHYLASE->PglZ->SIGMA+OB-Fold+AAA-ATPase+HKD-DNase-BrxL->?->SbcC-><-MPTase<-?<-?<-Phage_integrase | 1156 | Bacteria>Terrabacteria_group>Actinomycetota>Actinomycetes>Mycobacteriales>Mycobacteriaceae | GCA_001905655.1 | Mycobacterium_sp._ST-F2 | EB75_25595 | 1846..5316 | 1490484 | ATP-binding-protein-[Mycobacterium-sp.-ST-F2] |
| WP_018600838.1 | Type1.BREX | <-ABhydrolase\|\|?-><-?<-?\|\|?-><-HTH_3\|\|BrxA-DNA-binding->BrxB-BrxF-iSTAND->DUF6079*->N6-METHYLASE->PglZ->SIGMA+OB-Fold+AAA-ATPase+HKD-DNase-BrxL-><-?<-?\|\|?-><-?<-ACT<-SIR2 | 1152 | Bacteria>Terrabacteria_group>Actinomycetota>Actinomycetes>Mycobacteriales>Mycobacteriaceae | GCF_000373905.1 | Mycobacterium_sp._155 | B133_RS0110195 | 2156875..2160333 | 1157943 | BREX-system-P-loop-protein-BrxC-[Mycobacterium-sp.-155] |
| WP_020822248.1 | Type1.BREX | Pkinase+TPR+TPR-><-?\|\|TetR-HTH-><-?\|\|Glyoxalase-><-SIGMA+OB-Fold+AAA-ATPase+HKD-DNase-BrxL<-RVT<-PglZ<-N6-METHYLASE<-DUF6079*<-BrxB-BrxF-iSTAND<-BrxA-DNA-binding<-XIS-HTH<-zf-ISL3+HTH_Tnp_ISL3+DDE_Tnp_ISL3 | 1152 | Bacteria>Terrabacteria_group>Actinomycetota>Actinomycetes>Mycobacteriales>Mycobacteriaceae | GCF_000418535.1 | Mycobacterium_intracellulare | OEM_RS09395 | 2016549..2020007 | 1767 | BREX-system-P-loop-protein-BrxC-[Mycobacterium-intracellulare] |
| WP_036372476.1 | Type5.BREX | <-TRD+TRD<-?<-HTH_Tnp_1+HTH_21+rve+rve_3<-AAA-ATPase<-HEPN+DUF499<-SIR2+SIR2<-SIGMA+OB-Fold+AAA-ATPase+HKD-DNase-BrxL<-PglZ<-N6-METHYLASE<-DUF6079*<-BrxB-BrxF-iSTAND<-BrxA-DNA-binding<-XIS-HTH<-TetR+HTH<-Phage_integrase+Phage_integrase<-Phage_integrase+Phage_integrase<-Phage_integrase<-TUDOR+SF1-SF2-Helicase+iREase+CTD+REase<-XIS-HTH | 1152 | Bacteria>Terrabacteria_group>Actinomycetota>Actinomycetes>Mycobacteriales>Mycobacteriaceae | GCF_000612725.1 | Mycolicibacterium_austroafricanum | BN976_RS17355 | 59223..62681 | 39687 | BREX-system-P-loop-protein-BrxC-[Mycolicibacterium-austroafricanum] |
| WP_052621181.1 | Type1.BREX | <-HerA/FtsK<-?<-?<-SF1-SF2-Helicase+REase<-HerA/FtsK<-SIR2<-SIGMA+OB-Fold+AAA-ATPase+HKD-DNase-BrxL<-PglZ<-N6-METHYLASE<-DUF6079*<-BrxB-BrxF-iSTAND<-BrxA-DNA-binding<-XIS-HTH<-TetR+HTH<-Phage_integrase+Phage_integrase<-Phage_int_SAM_1+Phage_integrase\|\|HEPN-> | 1156 | Bacteria>Terrabacteria_group>Actinomycetota>Actinomycetes>Mycobacteriales>Mycobacteriaceae | GCF_001214625.1 | Mycobacteroides_abscessus | B5319_RS07405 | 1475995..1479465 | 36809 | BREX-system-P-loop-protein-BrxC-[Mycobacteroides-abscessus] |
| WP_065289569.1 | Type1.BREX | DUF5086-><-ANK+ANK<-ZnR\|\|XIS-HTH->BrxA-DNA-binding->BrxB-BrxF-iSTAND->DUF6079*->N6-METHYLASE->PglZ->SIGMA+OB-Fold+AAA-ATPase+HKD-DNase-BrxL-> | 1152 | Bacteria>Terrabacteria_group>Actinomycetota>Actinomycetes>Mycobacteriales>Mycobacteriaceae | GCF_001679965.1 | Mycolicibacter_kumamotonensis | ACT18_RS22010 | 4536..7994 | 354243 | BREX-system-P-loop-protein-BrxC-[Mycolicibacter-kumamotonensis] |
| WP_070953149.1 | Type1.BREX | SAM-METHYLASE-><-SopB+HNH<-?<-?\|\|XIS-HTH->BrxA-DNA-binding->BrxB-BrxF-iSTAND->DUF6079*->N6-METHYLASE->PglZ->SIGMA+OB-Fold+AAA-ATPase+HKD-DNase-BrxL->Acyl_transf_3->?->?-><-ACET | 1156 | Bacteria>Terrabacteria_group>Actinomycetota>Actinomycetes>Mycobacteriales>Mycobacteriaceae | GCF_001853775.1 | Mycobacteroides_chelonae | BKG71_RS10405 | 2100910..2104380 | 1774 | BREX-system-P-loop-protein-BrxC-[Mycobacteroides-chelonae] |
| WP_234813880.1 | Type1.BREX | <-ParB+HNH<-SIGMA+OB-Fold+AAA-ATPase+HKD-DNase-BrxL<-PglZ<-N6-METHYLASE<-DUF6079-p<-DUF6079-p*<-BrxB-BrxF-iSTAND<-BrxA-DNA-binding | 682 | Bacteria>Terrabacteria_group>Actinomycetota>Actinomycetes>Mycobacteriales>Mycobacteriaceae | GCF_002101705.1 | Mycolicibacterium_iranicum | AWC12_RS31570 | 96008..98056 | 912594 | BREX-system-P-loop-protein-BrxC-[Mycolicibacterium-iranicum] |
| WP_119635996.1 | Type1.BREX | <-XIS-HTH<-?<-?<-SF1-SF2-Helicase+REase<-HerA/FtsK<-SIR2+SIR2<-SIGMA+OB-Fold+AAA-ATPase+HKD-DNase-BrxL<-PglZ<-N6-METHYLASE<-DUF6079*<-BrxB-BrxF-iSTAND<-BrxA-DNA-binding<-XIS-HTH<-TetR+HTH<-Phage_integrase+Phage_integrase<-Phage_integrase+Phage_integrase<-Phage_int_SAM_1+Phage_integrase\|\|HEPN-> | 1156 | Bacteria>Terrabacteria_group>Actinomycetota>Actinomycetes>Mycobacteriales>Mycobacteriaceae | GCF_003582985.1 | Mycobacteroides_abscessus | D2E92_RS23075 | 16528..19998 | 36809 | BREX-system-P-loop-protein-BrxC-[Mycobacteroides-abscessus] |
| WP_134206037.1 | Type1.BREX | <-REase<-MORC<-HEPN+DUF499<-METHYLASE-p<-HerA/FtsK<-SIR2+SIR2<-SIGMA+OB-Fold+AAA-ATPase+HKD-DNase-BrxL<-PglZ<-N6-METHYLASE<-DUF6079*<-BrxB-BrxF-iSTAND<-BrxA-DNA-binding<-XIS-HTH<-TetR+HTH<-Phage_integrase+Phage_integrase<-Phage_integrase+Phage_integrase<-Phage_int_SAM_1+Phage_integrase<-REase\|\|DNA_METHYLASE-> | 1155 | Bacteria>Terrabacteria_group>Actinomycetota>Actinomycetes>Mycobacteriales>Mycobacteriaceae | GCF_004366875.1 | Mycobacteroides_salmoniphilum | CCUG62472_RS09145 | 418307..421774 | 404941 | BREX-system-P-loop-protein-BrxC-[Mycobacteroides-salmoniphilum] |
| WP_085074024.1 | Type1.BREX | <-sGTP<-?<-sGTP<-HerA/FtsK<-SIR2+SIR2<-SIGMA+OB-Fold+AAA-ATPase+HKD-DNase-BrxL<-PglZ<-N6-METHYLASE<-DUF6079*<-BrxB-BrxF-iSTAND<-BrxA-DNA-binding<-XIS-HTH | 1154 | Bacteria>Terrabacteria_group>Actinomycetota>Actinomycetes>Mycobacteriales>Mycobacteriaceae | GCF_010723135.1 | Mycobacterium_kubicae | H7H99_RS20140 | 22484..25948 | 120959 | BREX-system-P-loop-protein-BrxC-[Mycobacterium-kubicae] |
| WP_163778779.1 | Type1.BREX | GGDEF+EAL->?-><-Glyco_tran_28_C<-?\|\|?-><-?<-tRNA\|\|XIS-HTH->BrxA-DNA-binding->BrxB-BrxF-iSTAND->DUF6079*->N6-METHYLASE->PglZ->SIGMA+OB-Fold+AAA-ATPase+HKD-DNase-BrxL->ParB+HNH->?->AAA-ATPase->ZnR->?->Phage_integrase-> | 1156 | Bacteria>Terrabacteria_group>Actinomycetota>Actinomycetes>Mycobacteriales>Mycobacteriaceae | GCF_010727945.1 | Mycobacterium_cookii | G6N27_RS18550 | 3937772..3941242 | 1775 | BREX-system-P-loop-protein-BrxC-[Mycobacterium-cookii] |
| WP_085152401.1 | Type1.BREX | <-ABhydrolase+ABhydrolase<-Preader7<-HEPN+SF1-SF2-Helicase+REase<-?<-SIGMA+OB-Fold+AAA-ATPase+HKD-DNase-BrxL<-PglZ<-N6-METHYLASE<-DUF6079*<-BrxB-BrxF-iSTAND<-BrxA-DNA-binding<-XIS-HTH<-HKD-DNase+SF1-SF2-Helicase+HTH+HTH+SRA\|\|35exo+SF1-SF2-Helicase->SF1-SF2-Helicase-> | 1156 | Bacteria>Terrabacteria_group>Actinomycetota>Actinomycetes>Mycobacteriales>Mycobacteriaceae | GCF_010729895.1 | Mycolicibacterium_confluentis | H7K00_RS24370 | 129281..132751 | 28047 | BREX-system-P-loop-protein-BrxC-[Mycolicibacterium-confluentis] |
| WP_196326342.1 | Type1.BREX | ABhydrolase->?-><-tRNA\|\|?-><-?\|\|XIS-HTH->XIS-HTH->BrxA-DNA-binding->BrxB-BrxF-iSTAND->DUF6079*->N6-METHYLASE->PglZ->SIGMA+OB-Fold+AAA-ATPase+HKD-DNase-BrxL->SIR2->HerA/FtsK-><-NUDIX\|\|?->?->HerA/FtsK-> | 1156 | Bacteria>Terrabacteria_group>Actinomycetota>Actinomycetes>Mycobacteriales>Mycobacteriaceae | GCF_015700725.1 | Mycobacteroides_chelonae | HA135_RS11130 | 2312216..2315686 | 1774 | BREX-system-P-loop-protein-BrxC-[Mycobacteroides-chelonae] |
| WP_074292199.1 | Type1.BREX | <-HerA/FtsK<-?<-?\|\|NUDIX-><-HerA/FtsK<-SIR2<-SIGMA+OB-Fold+AAA-ATPase+HKD-DNase-BrxL<-PglZ<-N6-METHYLASE<-DUF6079*<-BrxB-BrxF-iSTAND<-BrxA-DNA-binding<-XIS-HTH<-PIN<-XIS-HTH | 1156 | Bacteria>Terrabacteria_group>Actinomycetota>Actinomycetes>Mycobacteriales>Mycobacteriaceae | GCF_017183935.1 | Mycobacteroides_abscessus | I3U31_RS04170 | 833809..837279 | 36809 | BREX-system-P-loop-protein-BrxC-[Mycobacteroides-abscessus] |
| WP_241204715.1 | Type1.BREX | <-DnaK<-?\|\|?->?-><-HerA/FtsK<-SIR2<-SIGMA+OB-Fold+AAA-ATPase+HKD-DNase-BrxL<-PglZ<-N6-METHYLASE<-DUF6079*<-BrxB-BrxF-iSTAND<-BrxA-DNA-binding<-XIS-HTH\|\|HEPN-><-AAA-ATPase<-HTH_7+rve+rve_3\|\|HEPN-> | 1156 | Bacteria>Terrabacteria_group>Actinomycetota>Actinomycetes>Mycobacteriales>Mycobacteriaceae | GCF_022348085.1 | Mycobacterium_sp._PSTR-4-N | MHN84_RS11260 | 180742..184212 | 2917745 | BREX-system-P-loop-protein-BrxC-[Mycobacterium-sp.-PSTR-4-N] |
| WP_260940163.1 | Type1.BREX | XIS-HTH->BrxA-DNA-binding->BrxB-BrxF-iSTAND->DUF6079*->N6-METHYLASE->PglZ->SIGMA+OB-Fold+AAA-ATPase+HKD-DNase-BrxL->?->ParB+HNH-p->HIN-HTH-><-wHTH\|\|MORC+DUF3684+DUF3883-><-REase+HTH+35exo+Small-B-barrel+SF1-SF2-Helicase | 1156 | Bacteria>Terrabacteria_group>Actinomycetota>Actinomycetes>Mycobacteriales>Mycobacteriaceae | GCF_025331195.1 | Mycolicibacterium_llatzerense | A7G45_RS21390 | 50451..53921 | 280871 | BREX-system-P-loop-protein-BrxC-[Mycolicibacterium-llatzerense] |
| WP_066911619.1 | Type1.BREX | <-UDG-conflict<-HerA/FtsK<-SIR2+SIR2<-SIGMA+OB-Fold+AAA-ATPase+HKD-DNase-BrxL<-PglZ<-N6-METHYLASE<-DUF6079*<-BrxB-BrxF-iSTAND<-BrxA-DNA-binding<-XIS-HTH<-Classical-AAA<-Peptidase_S8 | 1161 | Bacteria>Terrabacteria_group>Actinomycetota>Actinomycetes>Mycobacteriales>Mycobacteriaceae | GCF_025821415.1 | Mycobacterium_interjectum | H7H80_RS25830 | 12521..16006 | 33895 | BREX-system-P-loop-protein-BrxC-[Mycobacterium-interjectum] |
| WP_264033192.1 | Type1.BREX | Classical-AAA->HTH_3-><-SF1-SF2-Helicase+REase<-SF1-SF2-Helicase+REase<-HerA/FtsK<-SIR2<-SIGMA+OB-Fold+AAA-ATPase+HKD-DNase-BrxL<-PglZ<-N6-METHYLASE<-DUF6079*<-BrxB-BrxF-iSTAND | 1156 | Bacteria>Terrabacteria_group>Actinomycetota>Actinomycetes>Mycobacteriales>Mycobacteriaceae | GCF_025821885.1 | Mycolicibacterium_gilvum | H7I56_RS29610 | 57941..61411 | 1804 | BREX-system-P-loop-protein-BrxC-[Mycolicibacterium-gilvum] |
| WP_265123865.1 | Type5.BREX | EVE-><-nSTAND<-SIGMA+OB-Fold+AAA-ATPase+HKD-DNase-BrxL<-HerA/FtsK<-SIR2<-PglZ<-N6-METHYLASE<-DUF6079*<-BrxB-BrxF-iSTAND<-BrxA-DNA-binding<-XIS-HTH\|\|ParB+HNH->?->35exo+TUDOR+SF1-SF2-Helicase+iREase+CTD->SF1-SF2-Helicase->?->ABhydrolase-> | 1159 | Bacteria>Terrabacteria_group>Actinomycetota>Actinomycetes>Mycobacteriales>Mycobacteriaceae | GCF_026041215.1 | Hoyosella_sp._YIM_151337 | ONR57_RS12750 | 17833..21312 | 2992742 | BREX-system-P-loop-protein-BrxC-[Hoyosella-sp.-YIM-151337] |
| WP_074325944.1 | Type1.BREX | <-HEPN\|\|Phage_int_SAM_1+Phage_integrase->Phage_integrase+Phage_integrase->Phage_integrase+Phage_integrase->TetR+HTH->XIS-HTH->BrxA-DNA-binding->BrxB-BrxF-iSTAND->DUF6079*->N6-METHYLASE->PglZ->SIGMA+OB-Fold+AAA-ATPase+HKD-DNase-BrxL->SIR2+SIR2->HerA/FtsK->?->?->XIS-HTH->Phage_int_SAM_3+Phage_integrase-> | 1152 | Bacteria>Terrabacteria_group>Actinomycetota>Actinomycetes>Mycobacteriales>Mycobacteriaceae | GCF_900133325.1 | Mycobacteroides_abscessus | BUU36_RS03865 | 789941..793399 | 36809 | BREX-system-P-loop-protein-BrxC-[Mycobacteroides-abscessus] |
| OFE08162.1 | Type1.BREX | <-Pkinase+TPR+TPR<-?<-?<-?<-?<-?\|\|tRNA-><-?\|\|BrxA-DNA-binding->BrxB-BrxF-iSTAND->DUF6079*->PglZ->SIGMA+OB-Fold+AAA-ATPase+HKD-DNase-BrxL->?->?->?-><-ParB+HNH\|\|?->?->LZ_Tnp_IS481+HTH_32+rve+rve_3-> | 1175 | Bacteria>Terrabacteria_group>Actinomycetota>Actinomycetes>Mycobacteriales>Nocardiaceae | GCA_001757665.1 | Rhodococcus_sp._1139 | A5N83_14165 | 58033..61560 | 1833762 | ATP-binding-protein-[Rhodococcus-sp.-1139] |
| ORL42035.1 | Type1.BREX | <-ANK+ANK<-SIGMA+OB-Fold+AAA-ATPase+HKD-DNase-BrxL<-AbiEii+ZnR<-AbiEi_4+AbiEi_1<-sGTP<-sGTP+sGTP+GTPase-AIG<-PglZ<-N6-METHYLASE<-DUF6079*<-BrxB-BrxF-iSTAND<-BrxA-DNA-binding | 1188 | Bacteria>Terrabacteria_group>Actinomycetota>Actinomycetes>Mycobacteriales>Nocardiaceae | GCA_002094315.1 | Prescottella_equi | A6F59_07450 | 52967..56533 | 43767 | hypothetical-protein-A6F59_07450-[Prescottella-equi] |
| OUS97312.1 | Type5.BREX | <-MULE-TRANSPOSASE<-TPR+TPR+TPR+TPR+TPR+TPR<-?\|\|BrxA-DNA-binding->BrxB-BrxF-iSTAND->DUF6079*->N6-METHYLASE->HTH+REase->PglZ->SIGMA+OB-Fold+AAA-ATPase+HKD-DNase-BrxL-><-DNA_glycosylase\|\|?-><-TUDOR+SF1-SF2-Helicase+iREase+CTD+REase\|\|SF1-SF2-Helicase-> | 1175 | Bacteria>Terrabacteria_group>Actinomycetota>Actinomycetes>Mycobacteriales>Nocardiaceae | GCA_002165735.1 | Rhodococcus_sp._NCIMB_12038 | CA951_02905 | 126594..130121 | 933800 | ATP-binding-protein-[Rhodococcus-sp.-NCIMB-12038] |
| RGP48319.1 | Type1.BREX | <-GDResolvase+HIN-HTH<-SIGMA+OB-Fold+AAA-ATPase+HKD-DNase-BrxL<-PglZ<-N6-METHYLASE<-DUF6079*<-BrxB-BrxF-iSTAND-p | 1175 | Bacteria>Terrabacteria_group>Actinomycetota>Actinomycetes>Mycobacteriales>Nocardiaceae | GCA_003444715.1 | Rhodococcus_erythropolis | AWH04_08905 | 69911..73438 | 1833 | ATP-binding-protein-[Rhodococcus-erythropolis] |
| MBM4589636.1 | Type1.BREX | <-ANK+ANK<-SIGMA+OB-Fold+AAA-ATPase+HKD-DNase-BrxL<-AbiEii+ZnR<-AbiEi_4+AbiEi_1<-sGTP<-sGTP<-sGTP<-PglZ<-N6-METHYLASE<-DUF6079*<-BrxB-BrxF-iSTAND<-BrxA-DNA-binding<-?\|\|?-><-Glyoxalase\|\|?->?->Calcineurin->MalT+LuxR-HTH-> | 1188 | Bacteria>Terrabacteria_group>Actinomycetota>Actinomycetes>Mycobacteriales>Nocardiaceae | GCA_016879295.1 | Prescottella_equi | GS507_15715 | 26092..29658 | 43767 | BREX-system-P-loop-protein-BrxC-[Prescottella-equi] |
| MBM4523030.1 | Type1.BREX | <-MalT+LuxR-HTH<-Calcineurin<-?<-?\|\|Glyoxalase-><-?\|\|?->BrxA-DNA-binding->BrxB-BrxF-iSTAND->DUF6079*->N6-METHYLASE->PglZ->sGTP+sGTP+GTPase-AIG->sGTP->AbiEi_4+AbiEi_1->AbiEii+ZnR->SIGMA+OB-Fold+AAA-ATPase+HKD-DNase-BrxL->ANK+ANK-> | 1188 | Bacteria>Terrabacteria_group>Actinomycetota>Actinomycetes>Mycobacteriales>Nocardiaceae | GCA_016879495.1 | Prescottella_equi | GS442_03230 | 599834..603400 | 43767 | BREX-system-P-loop-protein-BrxC-[Prescottella-equi] |
| MBP1050938.1 | Type1.BREX | <-REase<-?<-?<-ARG\|\|NUDIX->BrxA-DNA-binding->BrxB-BrxF-iSTAND->DUF6079*->N6-METHYLASE->PglZ->SIGMA+OB-Fold+AAA-ATPase+HKD-DNase-BrxL-><-?<-?\|\|GDResolvase+HIN-HTH-><-ZnR\|\|HISKIN-> | 1171 | Bacteria>Terrabacteria_group>Actinomycetota>Actinomycetes>Mycobacteriales>Nocardiaceae | GCA_017831275.1 | Rhodococcus_qingshengii | J6397_12385 | 68679..72194 | 334542 | BREX-system-P-loop-protein-BrxC-[Rhodococcus-qingshengii] |
| MBT2268266.1 | Type1.BREX | <-SIGMA+OB-Fold+AAA-ATPase+HKD-DNase-BrxL<-PglZ<-N6-METHYLASE-p<-N6-METHYLASE-p<-DUF6079*<-BrxB-BrxF-iSTAND<-BrxA-DNA-binding<-?<-TIR-like | 1170 | Bacteria>Terrabacteria_group>Actinomycetota>Actinomycetes>Mycobacteriales>Nocardiaceae | GCA_018615075.1 | Rhodococcus_erythropolis | J7E74_25040 | 11077..14589 | 1833 | BREX-system-P-loop-protein-BrxC-[Rhodococcus-erythropolis] |
| BDU00126.1 | Type1.BREX | <-ABhydrolase\|\|Recombinase+ZnR->BrxA-DNA-binding->BrxB-BrxF-iSTAND->DUF6079*->N6-METHYLASE->?->PglZ->SIGMA+OB-Fold+AAA-ATPase+HKD-DNase-BrxL-><-HTH_21+rve+rve_3\|\|HIN-HTH->HTH_Tnp_1+HTH_21+rve+rve_3-><-HTH_21+rve+rve_3<-HIN-HTH | 1173 | Bacteria>Terrabacteria_group>Actinomycetota>Actinomycetes>Mycobacteriales>Nocardiaceae | GCA_027924405.1 | Nocardia_sp._ON39_IFM12276 | IFM12276_31540 | 3396523..3400044 | 2984338 | hypothetical-protein-IFM12276_31540-[Nocardia-sp.-ON39_IFM12276] |
| WP_020907622.1 | Type1.BREX | <-GDResolvase\|\|?-><-PIN<-?<-GDResolvase+HIN-HTH<-?<-SIGMA+OB-Fold+AAA-ATPase+HKD-DNase-BrxL<-PglZ<-N6-METHYLASE<-DUF6079*<-BrxB-BrxF-iSTAND<-BrxA-DNA-binding<-XIS-HTH<-Phage_integrase | 1168 | Bacteria>Terrabacteria_group>Actinomycetota>Actinomycetes>Mycobacteriales>Nocardiaceae | GCF_000010105.1 | Rhodococcus_erythropolis | RER_RS16440 | 3107254..3110760 | 1833 | BREX-system-P-loop-protein-BrxC-[Rhodococcus-erythropolis] |
| WP_039798161.1 | Type1.BREX | HTH+REase->?->?-><-DDE_Tnp_ISL3<-DDE_Tnp_ISL3<-DEDD_Tnp_IS110+Transposase_20\|\|BrxA-DNA-binding->BrxB-BrxF-iSTAND->DUF6079*->N6-METHYLASE->?->PglZ->SIGMA+OB-Fold+AAA-ATPase+HKD-DNase-BrxL-><-?<-?\|\|NUDIX-><-?<-RADICAL-SAM | 1173 | Bacteria>Terrabacteria_group>Actinomycetota>Actinomycetes>Mycobacteriales>Nocardiaceae | GCF_000308435.1 | Nocardia_araoensis | ON21_RS13315 | 16578..20099 | 228600 | BREX-system-P-loop-protein-BrxC-[Nocardia-araoensis] |
| WP_029256831.1 | Type1.BREX | ParB+HNH-><-?<-?<-SIGMA+OB-Fold+AAA-ATPase+HKD-DNase-BrxL<-PglZ<-N6-METHYLASE<-DUF6079*<-BrxB-BrxF-iSTAND<-BrxA-DNA-binding\|\|?-><-tRNA\|\|?->?->?->?->?->Pkinase+TPR+TPR-> | 1173 | Bacteria>Terrabacteria_group>Actinomycetota>Actinomycetes>Mycobacteriales>Nocardiaceae | GCF_000696675.2 | Rhodococcus_erythropolis | H351_RS02855 | 635383..638904 | 1833 | BREX-system-P-loop-protein-BrxC-[Rhodococcus-erythropolis] |
| WP_032384849.1 | Type1.BREX | <-EVE+REase<-ParB+HNH\|\|SF1-SF2-Helicase+REase->SF1-SF2-Helicase+HTH-><-HerA/FtsK<-SIR2<-SIGMA+OB-Fold+AAA-ATPase+HKD-DNase-BrxL<-PglZ<-N6-METHYLASE<-DUF6079*<-BrxB-BrxF-iSTAND<-BrxA-DNA-binding<-XIS-HTH<-REC+OmpR-HTH<-HisKA_3+HATPase_c\|\|DUF4352->?-><-URI-DNase-> | 1155 | Bacteria>Terrabacteria_group>Actinomycetota>Actinomycetes>Mycobacteriales>Nocardiaceae | GCF_000760875.1 | Rhodococcus_fascians | BH91_RS0103225 | 150682..154149 | 1828 | BREX-system-P-loop-protein-BrxC-[Rhodococcus-fascians] |
| WP_045066308.1 | Type1.BREX | IG+IG-><-?\|\|Acyltransferase->?-><-MORC+DUF3684+REase<-?\|\|BrxA-DNA-binding->BrxB-BrxF-iSTAND->DUF6079*->N6-METHYLASE->PglZ->SIGMA+OB-Fold+AAA-ATPase+HKD-DNase-BrxL->?-><-?<-?<-?\|\|HAD+HAD-> | 1173 | Bacteria>Terrabacteria_group>Actinomycetota>Actinomycetes>Mycobacteriales>Nocardiaceae | GCF_000949305.1 | unclassified_Rhodococcus_(in:_high_G+C_Gram-positive_bacteria) | SZ00_RS11305 | 2127210..2130731 | 192944 | MULTISPECIES:-BREX-system-P-loop-protein-BrxC-[unclassified-Rhodococcus-(in:-high-G+C-Gram-positive-bacteria)] |
| WP_062991956.1 | Type1.BREX | <-MACRODOMAIN<-?\|\|HTH_3->ACET->SWACOS->Cysteine-Desulfurase->Pkinase->BrxA-DNA-binding->BrxB-BrxF-iSTAND->DUF6079*->N6-METHYLASE->?->PglZ->SIGMA+OB-Fold+AAA-ATPase+HKD-DNase-BrxL-><-URI-DNase+RAMA<-?<-AbiEii | 1173 | Bacteria>Terrabacteria_group>Actinomycetota>Actinomycetes>Mycobacteriales>Nocardiaceae | GCF_001613085.1 | Nocardia_salmonicida | NS1_RS36345 | 21477..24998 | 53431 | BREX-system-P-loop-protein-BrxC-[Nocardia-salmonicida] |
| WP_154100274.1 | Type1.BREX | BrxA-DNA-binding->BrxB-BrxF-iSTAND->DUF6079*->N6-METHYLASE->PglZ->SIGMA+OB-Fold+AAA-ATPase+HKD-DNase-BrxL->?-><-?<-RAMA<-?<-?\|\|GDResolvase+HIN-HTH-> | 1172 | Bacteria>Terrabacteria_group>Actinomycetota>Actinomycetes>Mycobacteriales>Nocardiaceae | GCF_001653035.1 | Rhodococcus_sp._YH3-3 | YH3_RS02030 | 410612..414130 | 1803579 | BREX-system-P-loop-protein-BrxC-[Rhodococcus-sp.-YH3-3] |
| WP_070161347.1 | Type1.BREX | <-Pkinase+TPR+TPR<-?<-?<-?<-?<-?\|\|tRNA-><-?\|\|BrxA-DNA-binding->BrxB-BrxF-iSTAND->DUF6079*->N6-METHYLASE-p->N6-METHYLASE->PglZ->SIGMA+OB-Fold+AAA-ATPase+HKD-DNase-BrxL->?->?-><-ParB+HNH | 1175 | Bacteria>Terrabacteria_group>Actinomycetota>Actinomycetes>Mycobacteriales>Nocardiaceae | GCF_001757665.1 | Rhodococcus_sp._1139 | A5N83_RS14165 | 58033..61560 | 1833762 | BREX-system-P-loop-protein-BrxC-[Rhodococcus-sp.-1139] |
| WP_032407000.1 | Type1.BREX | <-DUF4352\|\|HisKA_3+HATPase_c->REC+OmpR-HTH->REase->?->XIS-HTH->BrxA-DNA-binding->BrxB-BrxF-iSTAND->DUF6079*->N6-METHYLASE->PglZ->SIGMA+OB-Fold+AAA-ATPase+HKD-DNase-BrxL->SIR2->HerA/FtsK-><-SF1-SF2-Helicase+HTH<-SF1-SF2-Helicase+REase\|\|ParB+HNH->EVE+REase-> | 1155 | Bacteria>Terrabacteria_group>Actinomycetota>Actinomycetes>Mycobacteriales>Nocardiaceae | GCF_001894785.1 | Rhodococcus_fascians | BI05_RS0109965 | 97728..101195 | 1828 | BREX-system-P-loop-protein-BrxC-[Rhodococcus-fascians] |
| WP_080909365.1 | Type1.BREX | <-GDResolvase+HIN-HTH\|\|?->?->RAMA->?->?-><-SIGMA+OB-Fold+AAA-ATPase+HKD-DNase-BrxL<-PglZ<-N6-METHYLASE<-DUF6079*<-BrxB-BrxF-iSTAND<-BrxA-DNA-binding\|\|?-><-tRNA\|\|?->?->?->?->?->Pkinase+TPR+TPR-> | 1173 | Bacteria>Terrabacteria_group>Actinomycetota>Actinomycetes>Mycobacteriales>Nocardiaceae | GCF_002076325.1 | Rhodococcus_sp._66b | B0E55_RS07060 | 288553..292074 | 1945511 | BREX-system-P-loop-protein-BrxC-[Rhodococcus-sp.-66b] |
| WP_081557823.1 | Type1.BREX | <-PIN<-?<-GDResolvase+HIN-HTH<-?<-SIGMA+OB-Fold+AAA-ATPase+HKD-DNase-BrxL<-PglZ<-N6-METHYLASE-p<-N6-METHYLASE<-DUF6079*<-BrxB-BrxF-iSTAND<-BrxA-DNA-binding<-XIS-HTH<-Phage_integrase | 1168 | Bacteria>Terrabacteria_group>Actinomycetota>Actinomycetes>Mycobacteriales>Nocardiaceae | GCF_002079265.1 | Rhodococcus_sp._BH4 | A0W34_RS13850 | 2965873..2969379 | 1807790 | BREX-system-P-loop-protein-BrxC-[Rhodococcus-sp.-BH4] |
| WP_256973250.1 | Type5.BREX | <-MULE-TRANSPOSASE<-TPR+TPR+TPR+TPR+TPR+TPR\|\|BrxA-DNA-binding->BrxB-BrxF-iSTAND->DUF6079*->N6-METHYLASE->HTH+REase->PglZ->SIGMA+OB-Fold+AAA-ATPase+HKD-DNase-BrxL-><-DNA_glycosylase\|\|?-><-TUDOR+SF1-SF2-Helicase+iREase+CTD+REase\|\|SF1-SF2-Helicase-> | 1213 | Bacteria>Terrabacteria_group>Actinomycetota>Actinomycetes>Mycobacteriales>Nocardiaceae | GCF_002165735.1 | Rhodococcus_sp._NCIMB_12038 | CA951_RS02920 | 126480..130121 | 933800 | BREX-system-P-loop-protein-BrxC-[Rhodococcus-sp.-NCIMB-12038] |
| WP_088896782.1 | Type1.BREX | ParB+HNH->nSTAND1+HNH-><-?\|\|XIS-HTH->BrxA-DNA-binding->BrxB-BrxF-iSTAND->DUF6079*->N6-METHYLASE->PglZ->SIGMA+OB-Fold+AAA-ATPase+HKD-DNase-BrxL->?-><-HTH_21+rve+rve_3 | 1154 | Bacteria>Terrabacteria_group>Actinomycetota>Actinomycetes>Mycobacteriales>Nocardiaceae | GCF_002215235.1 | Rhodococcus_sp._BUPNP1 | B9C99_RS01850 | 433431..436895 | 1432786 | BREX-system-P-loop-protein-BrxC-[Rhodococcus-sp.-BUPNP1] |
| WP_094721084.1 | Type1.BREX | SF1-SF2-Helicase+REase->SF1-SF2-Helicase+HTH-><-N6-METHYLASE\|\|MORC+HISKIN->?->?-><-SIGMA+OB-Fold+AAA-ATPase+HKD-DNase-BrxL<-PglZ<-N6-METHYLASE<-DUF6079*<-BrxB-BrxF-iSTAND<-BrxA-DNA-binding<-HEPN+SF1-SF2-Helicase+REase\|\|HIRAN->?-><-TIR<-HD<-?<-MarR-HTH | 1174 | Bacteria>Terrabacteria_group>Actinomycetota>Actinomycetes>Mycobacteriales>Nocardiaceae | GCF_002259415.1 | Rhodococcus_sp._14-2470-1b | CH293_RS18620 | 24333..27857 | 2023149 | BREX-system-P-loop-protein-BrxC-[Rhodococcus-sp.-14-2470-1b] |
| WP_198387782.1 | Type1.BREX | HEPN+SF1-SF2-Helicase+REase->HTH_36-><-SIGMA+OB-Fold+AAA-ATPase+HKD-DNase-BrxL<-PglZ<-HTH+REase<-N6-METHYLASE-p\|\|<-N6-METHYLASE<-DUF6079*<-BrxB-BrxF-iSTAND<-BrxA-DNA-binding<-DNA_METHYLASE<-DrmB<-SF1-SF2-Helicase<-SF1-SF2-Helicase<-MORC+SF1-SF2-Helicase\|\|ParB+HNH->SF1-SF2-Helicase-> | 1173 | Bacteria>Terrabacteria_group>Actinomycetota>Actinomycetes>Mycobacteriales>Nocardiaceae | GCF_002269365.1 | Rhodococcus_pyridinivorans | CJU69_RS21710 | 4620785..4624306 | 103816 | BREX-system-P-loop-protein-BrxC-[Rhodococcus-pyridinivorans] |
| WP_122191117.1 | Type1.BREX | <-ABhydrolase\|\|?->Glyoxalase-><-?<-SIGMA+OB-Fold+AAA-ATPase+HKD-DNase-BrxL<-PglZ<-?<-N6-METHYLASE<-DUF6079*<-BrxB-BrxF-iSTAND<-BrxA-DNA-binding\|\|?-><-STYKIN<-TPR | 1173 | Bacteria>Terrabacteria_group>Actinomycetota>Actinomycetes>Mycobacteriales>Nocardiaceae | GCF_003696265.1 | Nocardia_stercoris | EBN03_RS27825 | 64799..68320 | 2483361 | BREX-system-P-loop-protein-BrxC-[Nocardia-stercoris] |
| WP_149431764.1 | Type1.BREX | <-SIGMA+OB-Fold+AAA-ATPase+HKD-DNase-BrxL<-AbiEii+ZnR<-AbiEi_4+AbiEi_1<-PglZ<-N6-METHYLASE<-?<-DUF6079*<-BrxB-BrxF-iSTAND<-BrxA-DNA-binding | 1184 | Bacteria>Terrabacteria_group>Actinomycetota>Actinomycetes>Mycobacteriales>Nocardiaceae | GCF_008297975.1 | Rhodococcus_cavernicola | FOY51_RS18630 | 121957..125511 | 2495913 | BREX-system-P-loop-protein-BrxC-[Rhodococcus-cavernicola] |
| WP_225320218.1 | Type1.BREX | <-Pkinase+TPR+TPR<-?<-?<-?<-?<-?\|\|tRNA-><-?\|\|BrxA-DNA-binding->BrxB-BrxF-iSTAND->DUF6079-p*->DUF6079-p->N6-METHYLASE->PglZ->SIGMA+OB-Fold+AAA-ATPase+HKD-DNase-BrxL->?->?-><-AAA+<-rve | 435 | Bacteria>Terrabacteria_group>Actinomycetota>Actinomycetes>Mycobacteriales>Nocardiaceae | GCF_011611865.1 | Rhodococcus_erythropolis | G9444_RS32380 | 46132..47439 | 1833 | BREX-system-P-loop-protein-BrxC-[Rhodococcus-erythropolis] |
| WP_195040827.1 | Type1.BREX | McrB-AAA->McrC-NTD+REase-><-?<-N6-METHYLASE<-?<-?<-SIGMA+OB-Fold+AAA-ATPase+HKD-DNase-BrxL<-PglZ<-N6-METHYLASE<-DUF6079*<-BrxB-BrxF-iSTAND<-BrxA-DNA-binding<-XIS-HTH<-?<-HTH_31+DUF5753 | 1159 | Bacteria>Terrabacteria_group>Actinomycetota>Actinomycetes>Mycobacteriales>Nocardiaceae | GCF_015477975.1 | Nocardia_cyriacigeorgica | IU440_RS19520 | 298293..301772 | 135487 | BREX-system-P-loop-protein-BrxC-[Nocardia-cyriacigeorgica] |
| WP_238424144.1 | Type1.BREX | HTH_3->?-><-?<-HTH_36<-?\|\|BrxA-DNA-binding->BrxB-BrxF-iSTAND->DUF6079*->N6-METHYLASE->HTH+REase->PglZ->SIGMA+OB-Fold+AAA-ATPase+HKD-DNase-BrxL->zf-ISL3+HTH_Tnp_ISL3-><-SF1-SF2-Helicase<-HKD-DNase<-DUF6361\|\|TetR-HTH-> | 1175 | Bacteria>Terrabacteria_group>Actinomycetota>Actinomycetes>Mycobacteriales>Nocardiaceae | GCF_016804345.1 | Rhodococcus_ruber | F1734_RS10025 | 2133745..2137272 | 1830 | BREX-system-P-loop-protein-BrxC-[Rhodococcus-ruber] |
| WP_245219430.1 | Type1.BREX | <-REase<-?<-?\|\|BrxA-DNA-binding->BrxB-BrxF-iSTAND->DUF6079*->N6-METHYLASE->PglZ->SIGMA+OB-Fold+AAA-ATPase+HKD-DNase-BrxL-><-?<-?\|\|GDResolvase+HIN-HTH-><-ZnR\|\|HISKIN-> | 1173 | Bacteria>Terrabacteria_group>Actinomycetota>Actinomycetes>Mycobacteriales>Nocardiaceae | GCF_017831275.1 | Rhodococcus_qingshengii | J6397_RS12385 | 68673..72194 | 334542 | BREX-system-P-loop-protein-BrxC-[Rhodococcus-qingshengii] |
| WP_222635591.1 | Type1.BREX | RAMA->wHTH+REase->HNH->SF1-SF2-Helicase+REase->SF1-SF2-Helicase+HTH->BrxA-DNA-binding->BrxB-BrxF-iSTAND->DUF6079*->N6-METHYLASE->PglZ->SIGMA+OB-Fold+AAA-ATPase+HKD-DNase-BrxL-><-?<-HEPN+SF1-SF2-Helicase+REase\|\|CMP-hydrolase-> | 1188 | Bacteria>Terrabacteria_group>Actinomycetota>Actinomycetes>Mycobacteriales>Nocardiaceae | GCF_019834785.1 | unclassified_Rhodococcus_(in:_high_G+C_Gram-positive_bacteria) | HQ363_RS22195 | 12044..15610 | 192944 | MULTISPECIES:-BREX-system-P-loop-protein-BrxC-[unclassified-Rhodococcus-(in:-high-G+C-Gram-positive-bacteria)] |
| WP_230793610.1 | Type1.BREX | <-GDResolvase+HIN-HTH<-?<-REase<-?<-Polbeta\|\|BrxA-DNA-binding->BrxB-BrxF-iSTAND->DUF6079*->N6-METHYLASE->PglZ->SIGMA+OB-Fold+AAA-ATPase+HKD-DNase-BrxL-><-?<-?\|\|GDResolvase+HIN-HTH->URI-DNase-> | 1173 | Bacteria>Terrabacteria_group>Actinomycetota>Actinomycetes>Mycobacteriales>Nocardiaceae | GCF_021026075.1 | Rhodococcus_qingshengii | LQ424_RS30730 | 10749..14270 | 334542 | BREX-system-P-loop-protein-BrxC-[Rhodococcus-qingshengii] |
| WP_231782753.1 | Type1.BREX | <-Pkinase+TPR+TPR<-?<-?<-?<-?<-?\|\|tRNA-><-?\|\|BrxA-DNA-binding->BrxB-BrxF-iSTAND->DUF6079*->N6-METHYLASE->PglZ->SIGMA+OB-Fold+AAA-ATPase+HKD-DNase-BrxL->?->?->?-><-ParB+HNH\|\|?-><-MULE-TRANSPOSASE | 1173 | Bacteria>Terrabacteria_group>Actinomycetota>Actinomycetes>Mycobacteriales>Nocardiaceae | GCF_021165975.1 | Rhodococcus_qingshengii | LRL17_RS24605 | 5269427..5272948 | 334542 | BREX-system-P-loop-protein-BrxC-[Rhodococcus-qingshengii] |
| WP_233062445.1 | Type1.BREX | <-SAM-METHYLASE<-?<-DSBH<-?\|\|BrxA-DNA-binding->BrxB-BrxF-iSTAND->DUF6079*->N6-METHYLASE->PglZ->SIGMA+OB-Fold+AAA-ATPase+HKD-DNase-BrxL->nSTAND3-> | 1173 | Bacteria>Terrabacteria_group>Actinomycetota>Actinomycetes>Mycobacteriales>Nocardiaceae | GCF_021183645.1 | Nocardia_gipuzkoensis | LTT66_RS33135 | 7242905..7246426 | 2749991 | BREX-system-P-loop-protein-BrxC-[Nocardia-gipuzkoensis] |
| WP_242324362.1 | Type1.BREX | <-SIGMA+OB-Fold+AAA-ATPase+HKD-DNase-BrxL<-AbiEii+ZnR<-AbiEi_4+AbiEi_1<-PglZ<-N6-METHYLASE<-DUF6079*<-BrxB-BrxF-iSTAND<-BrxA-DNA-binding<-RAMA<-?\|\|?-><-REase<-?\|\|TIR-> | 1188 | Bacteria>Terrabacteria_group>Actinomycetota>Actinomycetes>Mycobacteriales>Nocardiaceae | GCF_022655485.1 | Rhodococcus_opacus | MOO23_RS40495 | 105196..108762 | 37919 | BREX-system-P-loop-protein-BrxC-[Rhodococcus-opacus] |
| WP_064077541.1 | Type1.BREX | LuxR-HTH-><-ANK+ANK<-SIGMA+OB-Fold+AAA-ATPase+HKD-DNase-BrxL<-AbiEii+ZnR<-AbiEi_4+AbiEi_1<-sGTP<-PglZ<-N6-METHYLASE<-DUF6079*<-BrxB-BrxF-iSTAND<-BrxA-DNA-binding<-?\|\|?-><-Glyoxalase\|\|?->?->Calcineurin->MalT+LuxR-HTH-> | 1188 | Bacteria>Terrabacteria_group>Actinomycetota>Actinomycetes>Mycobacteriales>Nocardiaceae | GCF_022691665.1 | Prescottella_equi | MPC39_RS12500 | 2698129..2701695 | 43767 | BREX-system-P-loop-protein-BrxC-[Prescottella-equi] |
| WP_242421152.1 | Type1.BREX | SF1-SF2-Helicase+DUF1998->REase-><-DNA_METHYLASE\|\|REase-><-SIGMA+OB-Fold+AAA-ATPase+HKD-DNase-BrxL<-PglZ<-HTH+REase<-DUF6079*<-BrxB-BrxF-iSTAND<-BrxA-DNA-binding<-?\|\|?-><-?\|\|?->RNase-> | 1172 | Bacteria>Terrabacteria_group>Actinomycetota>Actinomycetes>Mycobacteriales>Nocardiaceae | GCF_022691685.1 | Prescottella_equi | MPC38_RS08825 | 1859263..1862781 | 43767 | BREX-system-P-loop-protein-BrxC-[Prescottella-equi] |
| WP_156525037.1 | Type1.BREX | <-GDResolvase+HIN-HTH\|\|?->?-><-?<-Polbeta\|\|BrxA-DNA-binding->BrxB-BrxF-iSTAND->DUF6079*->N6-METHYLASE->PglZ->SIGMA+OB-Fold+AAA-ATPase+HKD-DNase-BrxL-><-REase<-?<-?\|\|GDResolvase+HIN-HTH->Phage_integrase->Phage_integrase-> | 1173 | Bacteria>Terrabacteria_group>Actinomycetota>Actinomycetes>Mycobacteriales>Nocardiaceae | GCF_023221595.1 | Rhodococcus | MO639_RS29930 | 74142..77663 | 1827 | MULTISPECIES:-BREX-system-P-loop-protein-BrxC-[Rhodococcus] |
| WP_253659814.1 | Type1.BREX | <-REase<-SIGMA+OB-Fold+AAA-ATPase+HKD-DNase-BrxL<-PglZ<-N6-METHYLASE<-DUF6079*<-BrxB-BrxF-iSTAND<-BrxA-DNA-binding\|\|DNA_glycosylase->?->REase-><-?<-?<-?\|\|PhdYeFM_antitox-> | 1156 | Bacteria>Terrabacteria_group>Actinomycetota>Actinomycetes>Mycobacteriales>Nocardiaceae | GCF_024171815.1 | Williamsia_maris | LX13_RS03015 | 647894..651364 | 72806 | BREX-system-P-loop-protein-BrxC-[Williamsia-maris] |
| WP_155277929.1 | Type1.BREX | <-Pkinase+TPR+TPR<-?<-?<-?<-?<-?\|\|tRNA-><-?\|\|BrxA-DNA-binding->BrxB-BrxF-iSTAND->DUF6079*->N6-METHYLASE->PglZ->SIGMA+OB-Fold+AAA-ATPase+HKD-DNase-BrxL->?->?-><-ParB+HNH | 1173 | Bacteria>Terrabacteria_group>Actinomycetota>Actinomycetes>Mycobacteriales>Nocardiaceae | GCF_024349805.1 | Rhodococcus | RE2895_RS00215 | 46904..50425 | 1827 | MULTISPECIES:-BREX-system-P-loop-protein-BrxC-[Rhodococcus] |
| WP_256520207.1 | Type1.BREX | <-Pkinase+TPR+TPR<-?<-?<-?<-?<-?\|\|tRNA-><-?\|\|BrxA-DNA-binding->BrxB-BrxF-iSTAND->DUF6079*->N6-METHYLASE->PglZ->SIGMA+OB-Fold+AAA-ATPase+HKD-DNase-BrxL->?->?->?-><-ParB+HNH | 1174 | Bacteria>Terrabacteria_group>Actinomycetota>Actinomycetes>Mycobacteriales>Nocardiaceae | GCF_024498635.1 | Rhodococcus_qingshengii | LRQ08_RS00215 | 46197..49721 | 334542 | BREX-system-P-loop-protein-BrxC-[Rhodococcus-qingshengii] |
| WP_269577378.1 | Type1.BREX | <-Pkinase+TPR+TPR<-?<-?<-?<-?<-?\|\|tRNA-><-?\|\|BrxA-DNA-binding->BrxB-BrxF-iSTAND->DUF6079*->N6-METHYLASE->PglZ->SIGMA+OB-Fold+AAA-ATPase+HKD-DNase-BrxL->?-><-?\|\|?->GDResolvase+HIN-HTH-><-ZnR\|\|HISKIN-> | 1173 | Bacteria>Terrabacteria_group>Actinomycetota>Actinomycetes>Mycobacteriales>Nocardiaceae | GCF_027270845.1 | Rhodococcus_erythropolis | O4216_RS13595 | 117955..121476 | 1833 | BREX-system-P-loop-protein-BrxC-[Rhodococcus-erythropolis] |
| WP_118604627.1 | Type1.BREX | <-GDResolvase+HIN-HTH<-SIGMA+OB-Fold+AAA-ATPase+HKD-DNase-BrxL<-PglZ<-N6-METHYLASE<-DUF6079*<-BrxB-BrxF-iSTAND-p | 1173 | Bacteria>Terrabacteria_group>Actinomycetota>Actinomycetes>Mycobacteriales>Nocardiaceae | - | Rhodococcus_erythropolis | AWH04_RS14655 | 69911..73432 | 1833 | BREX-system-P-loop-protein-BrxC-[Rhodococcus-erythropolis] |
| WP_214887135.1 | Type1.BREX | <-SIGMA+OB-Fold+AAA-ATPase+HKD-DNase-BrxL<-PglZ<-N6-METHYLASE-p<-N6-METHYLASE-p<-DUF6079*<-BrxB-BrxF-iSTAND<-BrxA-DNA-binding<-?<-TIR-like | 1170 | Bacteria>Terrabacteria_group>Actinomycetota>Actinomycetes>Mycobacteriales>Nocardiaceae | - | Rhodococcus_erythropolis | J7E74_25040 | 11077..14589 | 1833 | BREX-system-P-loop-protein-BrxC-[Rhodococcus-erythropolis] |
| TWS24839.1 | Type1.BREX | <-Glyoxalase+Glyoxalase<-TPR\|\|?-><-?<-?<-?<-SIGMA+OB-Fold+AAA-ATPase+HKD-DNase-BrxL<-PglZ<-N6-METHYLASE<-DUF6079*<-BrxB-BrxF-iSTAND<-BrxA-DNA-binding\|\|HTH_3->JetA->JetB->SbcC->?->TetR-HTH-> | 1298 | Bacteria>Terrabacteria_group>Actinomycetota>Actinomycetes>Mycobacteriales>Tsukamurellaceae | GCA_007858445.1 | Tsukamurella_sputi | FK268_06235 | 209185..213081 | 2591848 | BREX-system-P-loop-protein-BrxC-[Tsukamurella-sputi] |
| NKY16937.1 | Type1.BREX | <-TetR-HTH<-?<-SbcC<-JetB<-JetA<-HTH_3\|\|BrxA-DNA-binding->BrxB-BrxF-iSTAND->DUF6079*->N6-METHYLASE->PglZ->SIGMA+OB-Fold+AAA-ATPase+HKD-DNase-BrxL-><-?\|\|<-MoxR-AAA<-?<-?\|\|?->TPR-> | 1182 | Bacteria>Terrabacteria_group>Actinomycetota>Actinomycetes>Mycobacteriales>Tsukamurellaceae | GCA_012396015.1 | Tsukamurella_spumae | HF999_00885 | 181603..185151 | 44753 | BREX-system-P-loop-protein-BrxC-[Tsukamurella-spumae] |
| WP_068569480.1 | Type1.BREX | <-TetR-HTH<-?<-SbcC<-JetB<-JetA<-HTH_3\|\|BrxA-DNA-binding->BrxB-BrxF-iSTAND->DUF6079*->N6-METHYLASE->PglZ->SIGMA+OB-Fold+AAA-ATPase+HKD-DNase-BrxL->HTH+REase->?-><-?<-TraB<-MoxR-AAA | 1182 | Bacteria>Terrabacteria_group>Actinomycetota>Actinomycetes>Mycobacteriales>Tsukamurellaceae | GCF_001575195.1 | Tsukamurella_pseudospumae | AXK60_RS03070 | 654047..657595 | 239498 | BREX-system-P-loop-protein-BrxC-[Tsukamurella-pseudospumae] |
| WP_068743364.1 | Type1.BREX | MoxR-AAA->TraB->?-><-HTH+REase<-HTH+REase<-SIGMA+OB-Fold+AAA-ATPase+HKD-DNase-BrxL<-PglZ<-N6-METHYLASE<-DUF6079*<-BrxB-BrxF-iSTAND<-BrxA-DNA-binding\|\|HTH_3->JetA->JetB->SbcC->?->TetR-HTH-> | 1180 | Bacteria>Terrabacteria_group>Actinomycetota>Actinomycetes>Mycobacteriales>Tsukamurellaceae | GCF_001575205.1 | Tsukamurella_pseudospumae | AXK61_RS01225 | 254752..258294 | 239498 | BREX-system-P-loop-protein-BrxC-[Tsukamurella-pseudospumae] |
| WP_158636061.1 | Type1.BREX | <-Glyoxalase+Glyoxalase<-TPR\|\|?-><-?<-?<-?<-SIGMA+OB-Fold+AAA-ATPase+HKD-DNase-BrxL<-PglZ<-N6-METHYLASE<-DUF6079*<-BrxB-BrxF-iSTAND<-BrxA-DNA-binding\|\|JetA->JetB->SbcC->?->TetR-HTH-> | 1180 | Bacteria>Terrabacteria_group>Actinomycetota>Actinomycetes>Mycobacteriales>Tsukamurellaceae | GCF_007858445.1 | Tsukamurella_sputi | FK268_RS06235 | 209185..212727 | 2591848 | BREX-system-P-loop-protein-BrxC-[Tsukamurella-sputi] |
| WP_246207064.1 | Type1.BREX | <-TetR-HTH<-?<-SbcC<-JetB<-JetA<-HTH_3\|\|BrxA-DNA-binding->BrxB-BrxF-iSTAND->DUF6079*->N6-METHYLASE->SIGMA+OB-Fold+AAA-ATPase+HKD-DNase-BrxL->HTH+REase-><-?<-TraB<-MoxR-AAA | 1179 | Bacteria>Terrabacteria_group>Actinomycetota>Actinomycetes>Mycobacteriales>Tsukamurellaceae | GCF_012396015.1 | Tsukamurella_spumae | HF999_RS00885 | 181612..185151 | 44753 | BREX-system-P-loop-protein-BrxC-[Tsukamurella-spumae] |
| TCC44056.1 | Type1.BREX | ArsR-HTH->?-><-Aminotran_1_2<-?\|\|ABhydrolase->ACET->XIS-HTH->BrxA-DNA-binding->BrxB-BrxF-iSTAND->DUF6079*->N6-METHYLASE->PglZ->SIGMA+OB-Fold+AAA-ATPase+HKD-DNase-BrxL-> | 1159 | Bacteria>Terrabacteria_group>Actinomycetota>Actinomycetes>Propionibacteriales>Kribbellaceae | GCA_004331485.1 | Kribbella_capetownensis | EOH75_37810 | 267915..271394 | 1572659 | BREX-system-P-loop-protein-BrxC-[Kribbella-capetownensis] |
| WP_238163516.1 | Type1.BREX | ArsR-HTH->?-><-Aminotran_1_2<-?\|\|ABhydrolase->ACET->XIS-HTH->BrxA-DNA-binding->BrxB-BrxF-iSTAND->DUF6079*->N6-METHYLASE->PglZ->SIGMA+OB-Fold+AAA-ATPase+HKD-DNase-BrxL-> | 1162 | Bacteria>Terrabacteria_group>Actinomycetota>Actinomycetes>Propionibacteriales>Kribbellaceae | GCF_004331485.1 | Kribbella_capetownensis | EOH75_RS37805 | 267906..271394 | 1572659 | BREX-system-P-loop-protein-BrxC-[Kribbella-capetownensis] |
| WP_010833228.1 | Type1.BREX | <-CHTH<-HerA/FtsK<-SIR2<-SIGMA+OB-Fold+AAA-ATPase+HKD-DNase-BrxL<-PglZ<-N6-METHYLASE<-DUF6079*<-BrxB-BrxF-iSTAND<-BrxA-DNA-binding<-XIS-HTH<-VirB11+MuB-ATPase\|\|?->?->?->?->VirB11-> | 1160 | Bacteria>Terrabacteria_group>Actinomycetota>Actinomycetes>Propionibacteriales>Nocardioidaceae | GCF_000389985.1 | Nocardioides_sp._CF8 | CF8_RS11160 | 2288637..2292119 | 110319 | BREX-system-P-loop-protein-BrxC-[Nocardioides-sp.-CF8] |
| WP_028707702.1 | Type1.BREX | HEPN-><-SIGMA+OB-Fold+AAA-ATPase+HKD-DNase-BrxL<-PglZ<-Schlafen+HATPase+wHTH<-N6-METHYLASE-p<-Phage_int_SAM_3+Phage_integrase\|\|N6-METHYLASE-p->N6-METHYLASE-p-><-N6-METHYLASE<-DUF6079*<-BrxB-BrxF-iSTAND<-BrxA-DNA-binding<-?<-?<-?<-?\|\|?->AbiEi_4-> | 1159 | Bacteria>Terrabacteria_group>Actinomycetota>Actinomycetes>Propionibacteriales>Nocardioidaceae | GCF_000423465.1 | Propionicicella_superfundia | G555_RS0102905 | 100593..104072 | 348582 | BREX-system-P-loop-protein-BrxC-[Propionicicella-superfundia] |
| WP_068111667.1 | Type1.BREX | VirB11+MuB-ATPase->?-><-?\|\|XIS-HTH->BrxA-DNA-binding->BrxB-BrxF-iSTAND->DUF6079*->N6-METHYLASE->REase->PglZ->SIGMA+OB-Fold+AAA-ATPase+HKD-DNase-BrxL->OLD-ABC+TOPRIM->SF1-SF2-Helicase->?->HTH+MPTase->Phage_int_SAM_1+Phage_integrase-> | 1154 | Bacteria>Terrabacteria_group>Actinomycetota>Actinomycetes>Propionibacteriales>Nocardioidaceae | GCF_001653335.1 | Nocardioides_dokdonensis | I601_RS15600 | 3293021..3296485 | 450734 | BREX-system-P-loop-protein-BrxC-[Nocardioides-dokdonensis] |
| WP_107767053.1 | Type1.BREX | LZ_Tnp_IS481+HTH_32+rve+rve_3-><-?<-?\|\|XIS-HTH->BrxA-DNA-binding->BrxB-BrxF-iSTAND->DUF6079*->N6-METHYLASE->PglZ->SIGMA+OB-Fold+AAA-ATPase+HKD-DNase-BrxL-> | 1154 | Bacteria>Terrabacteria_group>Actinomycetota>Actinomycetes>Propionibacteriales>Nocardioidaceae | GCF_003047265.1 | Nocardioides_terrigena | DAE25_RS15680 | 407524..410988 | 424797 | BREX-system-P-loop-protein-BrxC-[Nocardioides-terrigena] |
| WP_116571233.1 | Type1.BREX | VirB11+MuB-ATPase->?-><-?\|\|XIS-HTH->BrxA-DNA-binding->BrxB-BrxF-iSTAND->DUF6079*->N6-METHYLASE->PglZ->SIGMA+OB-Fold+AAA-ATPase+HKD-DNase-BrxL->SIR2->HerA/FtsK->DUF4326->DUF488-1->DUF488-s-> | 1152 | Bacteria>Terrabacteria_group>Actinomycetota>Actinomycetes>Propionibacteriales>Nocardioidaceae | GCF_003076135.1 | Nocardioides_gansuensis | DDE18_RS05460 | 245415..248873 | 2138300 | BREX-system-P-loop-protein-BrxC-[Nocardioides-gansuensis] |
| WP_147685627.1 | Type1.BREX | ACET->?-><-SF1-SF2-Helicase<-SIGMA+OB-Fold+AAA-ATPase+HKD-DNase-BrxL<-PglZ<-N6-METHYLASE<-DUF6079*<-BrxB-BrxF-iSTAND<-BrxA-DNA-binding | 1154 | Bacteria>Terrabacteria_group>Actinomycetota>Actinomycetes>Propionibacteriales>Nocardioidaceae | GCF_008039565.1 | Aeromicrobium_terrae | FHP06_RS08160 | 302941..306405 | 2498846 | BREX-system-P-loop-protein-BrxC-[Aeromicrobium-terrae] |
| WP_148615130.1 | Type1.BREX | XIS-HTH->Phage_int_SAM_3+Phage_integrase-><-REase\|\|?-><-SIGMA+OB-Fold+AAA-ATPase+HKD-DNase-BrxL<-PglZ<-N6-METHYLASE<-DUF6079*<-BrxB-BrxF-iSTAND<-BrxA-DNA-binding<-XIS-HTH | 1154 | Bacteria>Terrabacteria_group>Actinomycetota>Actinomycetes>Propionibacteriales>Nocardioidaceae | GCF_008087085.1 | Nocardioides_rubriscoriae | FZC61_RS15785 | 254232..257696 | 642762 | BREX-system-P-loop-protein-BrxC-[Nocardioides-rubriscoriae] |
| WP_153653111.1 | Type5.BREX | <-HTH_3\|\|?->SF1-SF2-Helicase+REase->SF1-SF2-Helicase+REase->?-><-ParB+HNH<-SIGMA+OB-Fold+AAA-ATPase+HKD-DNase-BrxL<-PglZ<-N6-METHYLASE<-DUF6079*<-BrxB-BrxF-iSTAND<-BrxA-DNA-binding\|\|GTPase-AIG->sGTP-><-?<-SF1-SF2-Helicase<-35exo+TUDOR+SF1-SF2-Helicase+iREase+CTD | 1160 | Bacteria>Terrabacteria_group>Actinomycetota>Actinomycetes>Propionibacteriales>Nocardioidaceae | GCF_009649075.1 | Aeromicrobium_yanjiei | GEV26_RS10980 | 2228820..2232302 | 2662028 | BREX-system-P-loop-protein-BrxC-[Aeromicrobium-yanjiei] |
| WP_166189200.1 | Type1.BREX | XIS-HTH->BrxA-DNA-binding->BrxB-BrxF-iSTAND->DUF6079*->N6-METHYLASE->PglZ->SIGMA+OB-Fold+AAA-ATPase+HKD-DNase-BrxL->REase-><-Phage_int_SAM_3+Phage_integrase<-XIS-HTH | 1154 | Bacteria>Terrabacteria_group>Actinomycetota>Actinomycetes>Propionibacteriales>Nocardioidaceae | GCF_011250565.1 | Nocardioides_sp._IC4_145 | G6553_RS03445 | 61321..64785 | 2714037 | BREX-system-P-loop-protein-BrxC-[Nocardioides-sp.-IC4_145] |
| WP_179652935.1 | Type1.BREX | <-LPD36+METHYLASE+SF1-SF2-Helicase\|\|REase->XIS-HTH->BrxA-DNA-binding->BrxB-BrxF-iSTAND->DUF6079*->N6-METHYLASE->PglZ->SIGMA+OB-Fold+AAA-ATPase+HKD-DNase-BrxL->AAA-ATPase-><-?<-Peptidase_M48 | 1154 | Bacteria>Terrabacteria_group>Actinomycetota>Actinomycetes>Propionibacteriales>Nocardioidaceae | GCF_013408525.1 | Nocardioides_aromaticivorans | BJ993_RS25170 | 157779..161243 | 200618 | BREX-system-P-loop-protein-BrxC-[Nocardioides-aromaticivorans] |
| BAK36822.1 | Type1.BREX | HTH_3->ADH_N+ADH_zinc_N->?->?->?->BrxA-DNA-binding->BrxB-BrxF-iSTAND->DUF6079*->N6-METHYLASE-><-N6-METHYLASE-p\|\|Phage_int_SAM_3+Phage_integrase->N6-METHYLASE-p->Schlafen+HATPase+wHTH->PglZ->SIGMA+OB-Fold+AAA-ATPase+HKD-DNase-BrxL->SIR2-><-DDE_Tnp_1 | 1176 | Bacteria>Terrabacteria_group>Actinomycetota>Actinomycetes>Propionibacteriales>Propionibacteriaceae | GCA_000270245.1 | Microlunatus_phosphovorus_NM-1 | MLP_38080 | 4107257..4110787 | 1032480 | hypothetical-protein-MLP_38080-[Microlunatus-phosphovorus-NM-1] |
| NLI84511.1 | Type1.BREX | HIN-HTH-><-SMODS<-SLATT<-?<-AbiEii<-Schlafen<-?<-SIGMA+OB-Fold+AAA-ATPase+HKD-DNase-BrxL<-PglZ<-N6-METHYLASE<-DUF6079*<-BrxB-BrxF-iSTAND<-BrxA-DNA-binding\|\|?->TIR->?-><-DUF4096+DDE_Tnp_1+DDE_Tnp_1_2 | 1160 | Bacteria>Terrabacteria_group>Actinomycetota>Actinomycetes>Propionibacteriales>Propionibacteriaceae | GCA_012517405.1 | Propionibacterium_sp. | GX440_03735 | 112337..115819 | 1977903 | BREX-system-P-loop-protein-BrxC-[Propionibacterium-sp.] |
| MBK7820961.1 | Type1.BREX | <-DUF2637+HIN-HTH<-ArdCN+MPTase\|\|HTH->MPTase->XIS-HTH->BrxA-DNA-binding->BrxB-BrxF-iSTAND->DUF6079*->N6-METHYLASE->HTH+REase->PglZ->SIGMA+OB-Fold+AAA-ATPase+HKD-DNase-BrxL->OLD-ABC+TOPRIM->SF1-SF2-Helicase->?->DDE_Tnp_1-><-DNA_METHYLASE | 1176 | Bacteria>Terrabacteria_group>Actinomycetota>Actinomycetes>Propionibacteriales>Propionibacteriaceae | GCA_016714275.1 | Tessaracoccus_sp. | IPJ61_07760 | 1693709..1697239 | 1971211 | BREX-system-P-loop-protein-BrxC-[Tessaracoccus-sp.] |
| MCI1746855.1 | Type1.BREX | sGTP->?->?->?->tRNA->tRNA->BrxA-DNA-binding->BrxB-BrxF-iSTAND->DUF6079*->?->N6-METHYLASE-><-N6-METHYLASE-p\|\|Phage_int_SAM_3+Phage_integrase->N6-METHYLASE-p->Schlafen+HATPase+wHTH-p->PglZ->SIGMA+OB-Fold+AAA-ATPase+HKD-DNase-BrxL->Schlafen-> | 1159 | Bacteria>Terrabacteria_group>Actinomycetota>Actinomycetes>Propionibacteriales>Propionibacteriaceae | GCA_022647765.1 | Acidipropionibacterium_sp. | LKI24_01350 | 263130..266609 | 1912218 | BREX-system-P-loop-protein-BrxC-[Acidipropionibacterium-sp.] |
| WP_041792941.1 | Type1.BREX | Glyco_hydro_3+Glyco_hydro_3_C+Fn3-like->ADH_N+ADH_zinc_N->?->HTH_3->ADH_N+ADH_zinc_N->?->?->BrxA-DNA-binding->BrxB-BrxF-iSTAND->DUF6079*->N6-METHYLASE->Phage_int_SAM_3+Phage_integrase->N6-METHYLASE-p->Schlafen+HATPase+wHTH->PglZ->SIGMA+OB-Fold+AAA-ATPase+HKD-DNase-BrxL->SIR2-><-DDE_Tnp_1\|\|HIN-HTH-> | 1173 | Bacteria>Terrabacteria_group>Actinomycetota>Actinomycetes>Propionibacteriales>Propionibacteriaceae | GCF_000270245.1 | Microlunatus_phosphovorus | MLP_RS18400 | 4107266..4110787 | 29405 | BREX-system-P-loop-protein-BrxC-[Microlunatus-phosphovorus] |
| WP_028706416.1 | Type1.BREX | <-SIGMA-HTH+SIGMA-HTH<-SF1-SF2-Helicase+HTH<-SIGMA+OB-Fold+AAA-ATPase+HKD-DNase-BrxL<-PglZ<-Schlafen+HATPase+wHTH-p<-?<-HerA/FtsK<-SIR2<-N6-METHYLASE<-DUF6079*<-BrxB-BrxF-iSTAND<-BrxA-DNA-binding\|\|Glyco_hydro_36N+Melibiase+Glyco_hydro_36C->?->?->?->?-><-XIS-HTH | 1171 | Bacteria>Terrabacteria_group>Actinomycetota>Actinomycetes>Propionibacteriales>Propionibacteriaceae | GCF_000426605.1 | Propionibacterium_acidifaciens | K336_RS0104645 | 45562..49077 | 556499 | BREX-system-P-loop-protein-BrxC-[Propionibacterium-acidifaciens] |
| WP_021796710.1 | Type1.BREX | XIS-HTH-><-?<-?<-?<-Glyco_hydro_36N+Melibiase+Glyco_hydro_36C\|\|BrxA-DNA-binding->BrxB-BrxF-iSTAND->DUF6079*->N6-METHYLASE->REase->?->HATPase+wHTH->PglZ->SIGMA+OB-Fold+AAA-ATPase+HKD-DNase-BrxL->SF1-SF2-Helicase+HTH-> | 1171 | Bacteria>Terrabacteria_group>Actinomycetota>Actinomycetes>Propionibacteriales>Propionibacteriaceae | GCF_000478805.1 | Propionibacterium_acidifaciens | HMPREF0682_RS02920 | 11275..14790 | 556499 | BREX-system-P-loop-protein-BrxC-[Propionibacterium-acidifaciens] |
| WP_048768895.1 | Type1.BREX | <-DUF2637+HIN-HTH<-Primpol+PriCT1<-ArdCN+MPTase<-AraC-HTH\|\|XIS-HTH->BrxA-DNA-binding->BrxB-BrxF-iSTAND->DUF6079*->N6-METHYLASE->REase->?->Schlafen+HATPase+wHTH->PglZ->SIGMA+OB-Fold+AAA-ATPase+HKD-DNase-BrxL->SIR2->HerA/FtsK->VirB11+MuB-ATPase-> | 1170 | Bacteria>Terrabacteria_group>Actinomycetota>Actinomycetes>Propionibacteriales>Propionibacteriaceae | GCF_001050955.1 | Propionibacterium_freudenreichii | PFCIRM1025_RS01850 | 33525..37037 | 1744 | BREX-system-P-loop-protein-BrxC-[Propionibacterium-freudenreichii] |
| WP_055346094.1 | Type1.BREX | <-DUF2637+HIN-HTH<-Primpol+PriCT1<-ArdCN+MPTase<-AraC-HTH\|\|XIS-HTH->BrxA-DNA-binding->BrxB-BrxF-iSTAND->DUF6079*->N6-METHYLASE->REase->?->Schlafen+HATPase+wHTH->PglZ->SIGMA+OB-Fold+AAA-ATPase+HKD-DNase-BrxL->SIR2->HerA/FtsK->VirB11+MuB-ATPase-> | 1170 | Bacteria>Terrabacteria_group>Actinomycetota>Actinomycetes>Propionibacteriales>Propionibacteriaceae | GCF_001369055.1 | Propionibacterium_freudenreichii | PFCIRM516_RS05500 | 48312..51824 | 1744 | BREX-system-P-loop-protein-BrxC-[Propionibacterium-freudenreichii] |
| WP_094364310.1 | Type1.BREX | DNA_METHYLASE-><-sGTP<-GTPase-AIG<-HerA/FtsK<-SIR2+SIR2<-SIGMA+OB-Fold+AAA-ATPase+HKD-DNase-BrxL<-PglZ<-HTH+REase<-N6-METHYLASE<-DUF6079*<-BrxB-BrxF-iSTAND<-BrxA-DNA-binding<-XIS-HTH<-ABhydrolase<-?<-MPTase<-HTH\|\|ArdCN+MPTase->Primpol+PriCT1-> | 1173 | Bacteria>Terrabacteria_group>Actinomycetota>Actinomycetes>Propionibacteriales>Propionibacteriaceae | GCF_002250625.1 | Enemella_dayhoffiae | CGZ93_RS11755 | 175760..179281 | 2016507 | BREX-system-P-loop-protein-BrxC-[Enemella-dayhoffiae] |
| WP_097828824.1 | Type1.BREX | <-ABC-ATPase+SbcC+ABC-ATPase+SbcC<-?<-?<-tRNA\|\|35exo->tRNA->BrxA-DNA-binding->BrxB-BrxF-iSTAND->DUF6079*->N6-METHYLASE-><-N6-METHYLASE-p\|\|Phage_int_SAM_3+Phage_integrase->N6-METHYLASE-p->Schlafen+HATPase+wHTH->PglZ->SIGMA+OB-Fold+AAA-ATPase+HKD-DNase-BrxL-><-?<-HTH_Tnp_1+HTH_21+rve+rve_3 | 1173 | Bacteria>Terrabacteria_group>Actinomycetota>Actinomycetes>Propionibacteriales>Propionibacteriaceae | GCF_004011075.1 | Acidipropionibacterium_jensenii | COZ11_RS09870 | 2230980..2234501 | 1749 | BREX-system-P-loop-protein-BrxC-[Acidipropionibacterium-jensenii] |
| WP_182782739.1 | Type1.BREX | Peptidase_C14->?->?->?->PIN-><-URI-DNase<-TrwC+SF1-SF2-Helicase+TOPRIM\|\|BrxA-DNA-binding->BrxB-BrxF-iSTAND->DUF6079*->N6-METHYLASE->N6-METHYLASE-p->PglZ->SIGMA+OB-Fold+AAA-ATPase+HKD-DNase-BrxL-><-ParB+HNH\|\|NACHT->?->?-><-REase | 1159 | Bacteria>Terrabacteria_group>Actinomycetota>Actinomycetes>Propionibacteriales>Propionibacteriaceae | GCF_014164725.1 | Tessaracoccus_sp._MC1627 | H5399_RS09110 | 126520..129999 | 2760312 | BREX-system-P-loop-protein-BrxC-[Tessaracoccus-sp.-MC1627] |
| WP_182799522.1 | Type1.BREX | <-DOC+HTH<-RelE-ParE<-PhdYeFM_antitox<-Calcineurin<-ABhydrolase\|\|XIS-HTH->BrxA-DNA-binding->BrxB-BrxF-iSTAND->DUF6079*->N6-METHYLASE->?->PglZ->SIGMA+OB-Fold+AAA-ATPase+HKD-DNase-BrxL->OLD-ABC+TOPRIM->SF1-SF2-Helicase->GTPase-AIG->sGTP->SF1-SF2-Helicase-> | 1173 | Bacteria>Terrabacteria_group>Actinomycetota>Actinomycetes>Propionibacteriales>Propionibacteriaceae | GCF_014164775.1 | Tessaracoccus_sp._MC1679 | H5398_RS07140 | 376087..379608 | 2760313 | BREX-system-P-loop-protein-BrxC-[Tessaracoccus-sp.-MC1679] |
| WP_232547416.1 | Type1.BREX | <-OLD-ABC\|\|?->?->?->nSTAND3->nSTAND3->?->BrxA-DNA-binding->BrxB-BrxF-iSTAND->DUF6079*->N6-METHYLASE->PglZ->SIGMA+OB-Fold+AAA-ATPase+HKD-DNase-BrxL->?->VirD4-TraG+VirD4-FtsK->?-><-Peptidase_A24 | 1159 | Bacteria>Terrabacteria_group>Actinomycetota>Actinomycetes>Propionibacteriales>Propionibacteriaceae | GCF_014858005.1 | Propioniciclava_soli | IFY26_RS02480 | 540114..543593 | 2775081 | BREX-system-P-loop-protein-BrxC-[Propioniciclava-soli] |
| WP_204918880.1 | Type1.BREX | <-sGTP<-GTPase-AIG<-SF1-SF2-Helicase<-OLD-ABC+TOPRIM<-SIGMA+OB-Fold+AAA-ATPase+HKD-DNase-BrxL<-PglZ<-?<-N6-METHYLASE<-DUF6079*<-BrxB-BrxF-iSTAND<-BrxA-DNA-binding<-XIS-HTH<-ABhydrolase\|\|Recombinase-><-tRNA<-35exo<-?\|\|tRNA-><-MarR-HTH | 1173 | Bacteria>Terrabacteria_group>Actinomycetota>Actinomycetes>Propionibacteriales>Propionibacteriaceae | GCF_016907535.1 | Microlunatus_panaciterrae | JOE57_RS13940 | 3049076..3052597 | 400768 | BREX-system-P-loop-protein-BrxC-[Microlunatus-panaciterrae] |
| WP_073185198.1 | Type1.BREX | GDResolvase+HIN-HTH->Peptidase_A24-><-?<-VirD4-TraG+VirD4-FtsK<-?<-SIGMA+OB-Fold+AAA-ATPase+HKD-DNase-BrxL<-PglZ<-N6-METHYLASE<-DUF6079*<-BrxB-BrxF-iSTAND<-BrxA-DNA-binding<-nSTAND3 | 1159 | Bacteria>Terrabacteria_group>Actinomycetota>Actinomycetes>Propionibacteriales>Propionibacteriaceae | GCF_900141915.1 | Tessaracoccus_bendigoensis | BUC15_RS00075 | 21322..24801 | 72764 | BREX-system-P-loop-protein-BrxC-[Tessaracoccus-bendigoensis] |
| WP_073189906.1 | Type1.BREX | XIS-HTH->BrxA-DNA-binding->BrxB-BrxF-iSTAND->DUF6079*->N6-METHYLASE->Schlafen+HATPase+wHTH->PglZ->SIGMA+OB-Fold+AAA-ATPase+HKD-DNase-BrxL->SIR2+SIR2->HerA/FtsK-><-?\|\|GTPase-AIG->sGTP-> | 1176 | Bacteria>Terrabacteria_group>Actinomycetota>Actinomycetes>Propionibacteriales>Propionibacteriaceae | GCF_900141915.1 | Tessaracoccus_bendigoensis | BUC15_RS15345 | 1766..5296 | 72764 | BREX-system-P-loop-protein-BrxC-[Tessaracoccus-bendigoensis] |
| WP_130875595.1 | Type1.BREX | <-Peptidase_M20\|\|?-><-SIGMA+OB-Fold+AAA-ATPase+HKD-DNase-BrxL<-PglZ<-Schlafen+HATPase+wHTH<-N6-METHYLASE<-DUF6079*<-BrxB-BrxF-iSTAND<-BrxA-DNA-binding<-?<-HAD<-Trypsin+PDZ\|\|?-><-?<-METHYLASE\|\|ACET-> | 1171 | Bacteria>Terrabacteria_group>Actinomycetota>Actinomycetes>Propionibacteriales>Propionibacteriaceae | GCF_900607225.1 | [Pseudopropionibacterium]_massiliense | EOE39_RS01895 | 450669..454184 | 2220000 | BREX-system-P-loop-protein-BrxC-[[Pseudopropionibacterium] |

<table>
<tr><th>Protein-ID</th><th>System</th><th>Gene-Neighbourhoods</th><th>Length</th><th>Taxonomy</th><th>Assembly-ID</th><th>Strain</th><th>Locus-Tag</th><th>Genomic-coordinates</th><th>TaxID</th><th>Fasta-defline</th></tr>
<tr><td>ECG0705015.1</td><td>Type1.BREX</td><td><-HEPN<-REase+iREase<-?||||YjhX_toxin->SAM-METHYLASE->?->||BrxA-DNA-binding->BrxB-BrxF-iSTAND->DUF6079*->N6-METHYLASE->OLD-ABC->TOPRIM->PglZ->SIGMA+OB-Fold+AAA-ATPase+Lon-Protease-BrxL->HKD-DNase+OGG-like-><-REase<-NACHT+SF1-SF2-Helicase+HKD-DNase<-nSTAND1+HNH</td><td>1213</td><td>Bacteria>Pseudomonadota>Gammaproteobacteria>Enterobacterales>Enterobacteriaceae>Salmonella</td><td>-</td><td>Salmonella_enterica_subsp._salamae</td><td>DL122_04700</td><td>35596..39237</td><td>59202</td><td>BREX-system-P-loop-protein-BrxC-[Salmonella-enterica-subsp.-salamae]</td></tr>
<tr><td>WP_222233110.1</td><td>Type1.BREX</td><td><-HKD-DNase+OGG-like<-SIGMA+OB-Fold+AAA-ATPase+Lon-Protease-BrxL<-PglZ<-TOPRIM<-OLD-ABC<-N6-METHYLASE<-DUF6079*<-BrxB-BrxF-iSTAND<-BrxA-DNA-binding<-?<-?<-?<-?<-PRTase<-DADA-Ligase</td><td>1202</td><td>Bacteria>Pseudomonadota>Gammaproteobacteria>Enterobacterales>Erwiniaceae>Pantoea</td><td>GCF_019795975.1</td><td>Pantoea_sp._DY-17</td><td>K5Y33_RS01225</td><td>252070..255678</td><td>2871490</td><td>BREX-system-P-loop-protein-BrxC-[Pantoea-sp.-DY-17]</td></tr>
<tr><td>WP_060426782.1</td><td>Type1.BREX</td><td><-RNase||?->?->SAD+HNH->HTH+REase->BrxA-DNA-binding->BrxB-BrxF-iSTAND->DUF6079*->N6-METHYLASE->PglZ->SIGMA+OB-Fold+AAA-ATPase+Lon-Protease-BrxL->HKD-DNase+OGG-like-><-?<-?<-tRNA||?->DEOR-HTH+ISOCOT-></td><td>1213</td><td>Bacteria>Pseudomonadota>Gammaproteobacteria>Enterobacterales>Yersiniaceae</td><td>GCF_001537885.1</td><td>Serratia</td><td>AWX06_RS15845</td><td>182718..186359</td><td>613</td><td>MULTISPECIES:-BREX-system-P-loop-protein-BrxC-[Serratia]</td></tr>
<tr><td>WP_112197427.1</td><td>Type1.BREX</td><td><-CopG/RHH||?->XIS-HTH->ZnR->ZnR->?->?->BrxA-DNA-binding->BrxB-BrxF-iSTAND->DUF6079*->N6-METHYLASE->OLD-ABC->TOPRIM->PglZ->SIGMA+OB-Fold+AAA-ATPase+Lon-Protease-BrxL->HKD-DNase+OGG-like-><-Arm-DNA-bind_3+Phage_integrase</td><td>1204</td><td>Bacteria>Pseudomonadota>Gammaproteobacteria>Enterobacterales>Yersiniaceae>Rahnella</td><td>GCF_003263525.1</td><td>Rahnella_sp._NRRL_B-41462</td><td>UC14_RS06670</td><td>31474..35088</td><td>1610579</td><td>BREX-system-P-loop-protein-BrxC-[Rahnella-sp.-NRRL-B-41462]</td></tr>
<tr><td>WP_197785670.1</td><td>Type1.BREX</td><td>Glyco_hydro_1-><-?<-?||?->?->SAD+HNH->HTH+REase->BrxA-DNA-binding->BrxB-BrxF-iSTAND->DUF6079*->N6-METHYLASE->PglZ->SIGMA+OB-Fold+AAA-ATPase+Lon-Protease-BrxL->HKD-DNase+OGG-like-></td><td>1213</td><td>Bacteria>Pseudomonadota>Gammaproteobacteria>Enterobacterales>Yersiniaceae>Serratia</td><td>GCF_016007555.1</td><td>Serratia_ureilytica</td><td>I5P92_RS12860</td><td>24369..28010</td><td>300181</td><td>BREX-system-P-loop-protein-BrxC-[Serratia-ureilytica]</td></tr>
<tr><td>WP_212602139.1</td><td>Type1.BREX</td><td><-AraC-HTH<-?||?->Aminotran_3->wHTH+SAD+HNH-><-?<-REase||BrxA-DNA-binding->BrxB-BrxF-iSTAND->DUF6079*->N6-METHYLASE->PglZ->SIGMA+OB-Fold+AAA-ATPase+Lon-Protease-BrxL->HKD-DNase+OGG-like-><-?<-?<-tRNA||DEOR-HTH+ISOCOT-></td><td>1213</td><td>Bacteria>Pseudomonadota>Gammaproteobacteria>Enterobacterales>Yersiniaceae>Serratia</td><td>GCF_018257315.1</td><td>Serratia_rubidaea</td><td>JK228_RS02460</td><td>34162..37803</td><td>61652</td><td>BREX-system-P-loop-protein-BrxC-[Serratia-rubidaea]</td></tr>
<tr><td>WP_244757920.1</td><td>Type1.BREX</td><td><-HKD-DNase+OGG-like<-SIGMA+OB-Fold+AAA-ATPase+Lon-Protease-BrxL<-PglZ<-N6-METHYLASE<-DUF6079*<-BrxB-BrxF-iSTAND<-BrxA-DNA-binding<-HTH+REase<-SAD+HNH<-?||?->?-><-Glyco_hydro_1</td><td>1213</td><td>Bacteria>Pseudomonadota>Gammaproteobacteria>Enterobacterales>Yersiniaceae>Serratia</td><td>GCF_022870585.1</td><td>Serratia_marcescens</td><td>HW877_RS02460</td><td>535877..539518</td><td>615</td><td>BREX-system-P-loop-protein-BrxC-[Serratia-marcescens]</td></tr>
<tr><td>WP_261139451.1</td><td>Type1.BREX</td><td>DOC+HTH-><-LACI-HTH+PBPI||?->Glyco_hydro_1->?->SAD+HNH->BrxA-DNA-binding->BrxB-BrxF-iSTAND->DUF6079*->PglZ->SIGMA+OB-Fold+AAA-ATPase+Lon-Protease-BrxL->HKD-DNase+OGG-like-><-Couple_hipA+HipA_C<-HTH_3<-?||HIN-HTH-><-HTH_21+rve+rve_3<-rve</td><td>1213</td><td>Bacteria>Pseudomonadota>Gammaproteobacteria>Enterobacterales>Yersiniaceae>Serratia</td><td>GCF_946226675.1</td><td>Serratia_liquefaciens</td><td>N8M82_RS16050</td><td>322435..326076</td><td>614</td><td>BREX-system-P-loop-protein-BrxC-[Serratia-liquefaciens]</td></tr>
<tr><td>WP_261135874.1</td><td>Type1.BREX</td><td>DOC+HTH-><-LACI-HTH+PBPI||?->Glyco_hydro_1->?->SAD+HNH->BrxA-DNA-binding->BrxB-BrxF-iSTAND->DUF6079*->N6-METHYLASE->PglZ->SIGMA+OB-Fold+AAA-ATPase+Lon-Protease-BrxL->HKD-DNase+OGG-like-><-Couple_hipA+HipA_C<-HTH_3</td><td>1213</td><td>Bacteria>Pseudomonadota>Gammaproteobacteria>Enterobacterales>Yersiniaceae>Serratia</td><td>GCF_946226775.1</td><td>Serratia_liquefaciens</td><td>N8M94_RS14775</td><td>322685..326326</td><td>614</td><td>BREX-system-P-loop-protein-BrxC-[Serratia-liquefaciens]</td></tr>
<tr><td>WP_261427666.1</td><td>Type1.BREX</td><td>Glyco_hydro_1-><-?<-?||?->?->SAD+HNH->BrxA-DNA-binding->BrxB-BrxF-iSTAND->DUF6079*->N6-METHYLASE->PglZ->SIGMA+OB-Fold+AAA-ATPase+Lon-Protease-BrxL->HKD-DNase+OGG-like-><-Couple_hipA+HipA_C<-HTH_3</td><td>1213</td><td>Bacteria>Pseudomonadota>Gammaproteobacteria>Enterobacterales>Yersiniaceae>Serratia</td><td>GCF_946227925.1</td><td>Serratia_proteamaculans</td><td>OCH17_RS14295</td><td>357572..361213</td><td>28151</td><td>BREX-system-P-loop-protein-BrxC-[Serratia-proteamaculans]</td></tr>
<tr><td>WP_195869502.1</td><td>Type1.BREX</td><td><-Peptidase_M17_N+Peptidase_M17||?->tRNA->Arm-DNA-bind_3+Phage_integrase-><-HKD-DNase+OGG-like<-SIGMA+OB-Fold+AAA-ATPase+Lon-Protease-BrxL<-PglZ<-TOPRIM<-OLD-ABC<-N6-METHYLASE<-DUF6079*<-BrxB-BrxF-iSTAND<-BrxA-DNA-binding<-?<-?||<-ZnR<-XIS-HTH<-?||OLD-ABC+TOPRIM-></td><td>1202</td><td>Bacteria>Pseudomonadota>Gammaproteobacteria>Enterobacterales>Yersiniaceae>Serratia</td><td>-</td><td>Serratia_liquefaciens</td><td>I1A46_RS14725</td><td>213767..217375</td><td>614</td><td>BREX-system-P-loop-protein-BrxC-[Serratia-liquefaciens]</td></tr>
<tr><td>WP_053011787.1</td><td>Type1.BREX</td><td>XIS-HTH->?->ZnR->XIS-HTH->ZnR->?->HTH+MPTase->||BrxA-DNA-binding->BrxB-BrxF-iSTAND->DUF6079*->N6-METHYLASE->OLD-ABC->TOPRIM->PglZ->SIGMA+OB-Fold+AAA-ATPase+Lon-Protease-BrxL->HKD-DNase+OGG-like-><-Arm-DNA-bind_3+Phage_integrase<-tRNA||?-><-HTH_3</td><td>1202</td><td>Bacteria>Pseudomonadota>Gammaproteobacteria>Enterobacterales>Yersiniaceae>Yersinia</td><td>GCF_001244675.1</td><td>Yersinia_intermedia</td><td>ALC37_RS11575</td><td>459968..463576</td><td>631</td><td>BREX-system-P-loop-protein-BrxC-[Yersinia-intermedia]</td></tr>
<tr><td>WP_044865655.1</td><td>Type1.BREX</td><td><-RHH_antitoxin||?->XIS-HTH->?->ZnR->BrxA-DNA-binding->BrxB-BrxF-iSTAND->DUF6079*->N6-METHYLASE->PglZ->SIGMA+OB-Fold+AAA-ATPase+Lon-Protease-BrxL->HKD-DNase+OGG-like-><-Arm-DNA-bind_3+Phage_integrase</td><td>1213</td><td>Bacteria>Pseudomonadota>Gammaproteobacteria</td><td>GCF_001540705.1</td><td>Enterobacterales</td><td>TJ11_RS03570</td><td>21952..25593</td><td>91347</td><td>MULTISPECIES:-BREX-system-P-loop-protein-BrxC-[Enterobacterales]</td></tr>
<tr><td>WP_031438510.1</td><td>Type1.BREX</td><td>HD->?->?->||Phage_integrase->Phage_integrase->?->wHTH+WYL+WCX->BrxA-DNA-binding->BrxB-BrxF-iSTAND->DUF6079*->HKD-DNase+DUF3150->N6-METHYLASE->OLD-ABC->TOPRIM->DOC+HTH->PglZ->SIGMA+OB-Fold+AAA-ATPase+Lon-Protease-BrxL->SMODS->SLATT-></td><td>1218</td><td>Bacteria>Pseudomonadota>Gammaproteobacteria>Methylococcales>Methylococcaceae>Methylobacter</td><td>GCF_000733835.1</td><td>Methylobacter_tundripaludum</td><td>T449_RS0121480</td><td>4638570..4642226</td><td>173365</td><td>BREX-system-P-loop-protein-BrxC-[Methylobacter-tundripaludum]</td></tr>
<tr><td>WP_136143645.1</td><td>Type1.BREX</td><td>SMODS->?->Classical-AAA->ABhydrolase->?-><-SIGMA+OB-Fold+AAA-ATPase+Lon-Protease-BrxL<-PglZ<-HKD-DNase+DUF3150<-N6-METHYLASE<-DUF6079*<-BrxB-BrxF-iSTAND<-BrxA-DNA-binding<-wHTH+WYL+WCX||DUF4236-><-DOC+HTH<-?<-ParB+HNH<-Phage_integrase+Phage_integrase<-Phage_integrase+Phage_integrase</td><td>1226</td><td>Bacteria>Pseudomonadota>Gammaproteobacteria>Moraxellales>Moraxellaceae>Acinetobacter</td><td>GCA_030820635.1</td><td>Acinetobacter_towneri</td><td>EQG52_RS02450</td><td>14374..18054</td><td>202956</td><td>BREX-system-P-loop-protein-BrxC-[Acinetobacter-towneri]</td></tr>
<tr><td>WP_023013360.1</td><td>Type1.BREX</td><td><-SIGMA+OB-Fold+AAA-ATPase+Lon-Protease-BrxL<-PglZ<-HKD-DNase+DUF3150<-N6-METHYLASE<-DUF6079*<-BrxB-BrxF-iSTAND<-BrxA-DNA-binding<-wHTH+WYL+WCX<-wHTH+HNH<-SF1-SF2-Helicase<-?<-SAM-METHYLASE<-JAB</td><td>1226</td><td>Bacteria>Pseudomonadota>Gammaproteobacteria>Moraxellales>Moraxellaceae>Acinetobacter</td><td>GCF_000475335.1</td><td>Acinetobacter_sp._COS3</td><td>IO68_RS25385</td><td>11690..15370</td><td>1397525</td><td>BREX-system-P-loop-protein-BrxC-[Acinetobacter-sp.-COS3]</td></tr>
<tr><td>WP_039048213.1</td><td>Type1.BREX</td><td><-GDResolvase+HIN-HTH||<-SbcC+ABC-ATPase<-?<-?<-?||<-SIGMA+OB-Fold+AAA-ATPase+Lon-Protease-BrxL<-PglZ<-HKD-DNase+DUF3150<-N6-METHYLASE<-DUF6079*<-BrxB-BrxF-iSTAND<-BrxA-DNA-binding<-wHTH+WYL+WCX||DUF4236-><-CR-ATPase5+REase<-ParB+HNH<-Phage_integrase+Phage_integrase<-Phage_integrase+Phage_integrase</td><td>1226</td><td>Bacteria>Pseudomonadota>Gammaproteobacteria>Moraxellales>Moraxellaceae>Acinetobacter</td><td>GCF_000813495.1</td><td>Acinetobacter_junii</td><td>RP26_RS13815</td><td>17614..21294</td><td>40215</td><td>BREX-system-P-loop-protein-BrxC-[Acinetobacter-junii]</td></tr>
<tr><td>WP_180041457.1</td><td>Type1.BREX</td><td>ParB+HNH-><-?||SAM-METHYLASE->?->SF1-SF2-Helicase->wHTH+HNH->wHTH+WYL+WCX->BrxA-DNA-binding->BrxB-BrxF-iSTAND->DUF6079*->N6-METHYLASE->HKD-DNase+DUF3150->PglZ->SIGMA+OB-Fold+AAA-ATPase+Lon-Protease-BrxL->SF1-SF2-Helicase+REase-></td><td>1221</td><td>Bacteria>Pseudomonadota>Gammaproteobacteria>Moraxellales>Moraxellaceae>Acinetobacter</td><td>GCF_013417925.1</td><td>Acinetobacter_sp._YH12218</td><td>FTV03_RS11390</td><td>13993..17658</td><td>2601152</td><td>BREX-system-P-loop-protein-BrxC-[Acinetobacter-sp.-YH12218]</td></tr>
<tr><td>WP_180131186.1</td><td>Type1.BREX</td><td>Phage_integrase-><-HTH_OrfB_IS605+OrfB_IS605+OrfB_Zn_ribbon||Y1_Tnp->ParB+HNH->CR-ATPase5+REase-><-DUF4236||wHTH+WYL+WCX->BrxA-DNA-binding->BrxB-BrxF-iSTAND->DUF6079*->N6-METHYLASE->HKD-DNase+DUF3150->PglZ->SIGMA+OB-Fold+AAA-ATPase+Lon-Protease-BrxL-></td><td>1226</td><td>Bacteria>Pseudomonadota>Gammaproteobacteria>Moraxellales>Moraxellaceae>Acinetobacter</td><td>GCF_013419565.1</td><td>Acinetobacter_sp._YH12072</td><td>FTU19_RS11080</td><td>35259..38939</td><td>2601068</td><td>BREX-system-P-loop-protein-BrxC-[Acinetobacter-sp.-YH12072]</td></tr>
<tr><td>WP_218698253.1</td><td>Type1.BREX</td><td><-MarR-HTH<-SF1-SF2-Helicase<-STYKIN+CR-Kinase<-?<-SIGMA+OB-Fold+AAA-ATPase+Lon-Protease-BrxL<-PglZ<-HKD-DNase+DUF3150<-ParB+HNH<-N6-METHYLASE<-DUF6079*<-BrxB-BrxF-iSTAND<-BrxA-DNA-binding<-wHTH+WYL+WCX||DUF4236-><-RelE-ParE<-ParD_antitoxin||<-?<-HIN-HTH</td><td>1224</td><td>Bacteria>Pseudomonadota>Gammaproteobacteria>Moraxellales>Moraxellaceae>Acinetobacter</td><td>GCF_019279015.1</td><td>Acinetobacter_harbinensis</td><td>KZA13_RS08660</td><td>34272..37946</td><td>1353941</td><td>BREX-system-P-loop-protein-BrxC-[Acinetobacter-harbinensis]</td></tr>
<tr><td>WP_234115082.1</td><td>Type1.BREX</td><td>Phage_integrase+Phage_integrase->Phage_integrase+Phage_integrase->ParB+HNH->?->DOC+HTH-><-DUF4236||wHTH+WYL+WCX->BrxA-DNA-binding->BrxB-BrxF-iSTAND->DUF6079*->N6-METHYLASE->HKD-DNase+DUF3150->PglZ->SIGMA+OB-Fold+AAA-ATPase+Lon-Protease-BrxL-><-?<-ABhydrolase<-Classical-AAA<-?<-SMODS</td><td>1226</td><td>Bacteria>Pseudomonadota>Gammaproteobacteria>Moraxellales>Moraxellaceae>Acinetobacter</td><td>GCF_021432045.1</td><td>Acinetobacter_towneri</td><td>LZG54_RS14455</td><td>23931..27611</td><td>202956</td><td>BREX-system-P-loop-protein-BrxC-[Acinetobacter-towneri]</td></tr>
<tr><td>WP_234693498.1</td><td>Type1.BREX</td><td>Phage_integrase+Phage_integrase->Phage_integrase+Phage_integrase->ParB+HNH->?->DOC+HTH-><-DUF4236||wHTH+WYL+WCX->BrxA-DNA-binding->BrxB-BrxF-iSTAND->DUF6079*->N6-METHYLASE->HKD-DNase+DUF3150->PglZ->SIGMA+OB-Fold+AAA-ATPase+Lon-Protease-BrxL-><-?<-ABhydrolase<-Classical-AAA<-?<-SMODS</td><td>1226</td><td>Bacteria>Pseudomonadota>Gammaproteobacteria>Moraxellales>Moraxellaceae>Acinetobacter</td><td>GCF_021460555.1</td><td>Acinetobacter_baumannii</td><td>LH676_RS19735</td><td>26787..30467</td><td>470</td><td>BREX-system-P-loop-protein-BrxC-[Acinetobacter-baumannii]</td></tr>
<tr><td>WP_262597505.1</td><td>Type1.BREX</td><td>Zn_Tnp_IS1595+DDE_Tnp_IS1595-><-TIR<-?<-SLOG+BirA-HTH<-SF1-SF2-Helicase+PRTase<-N6-METHYLASE||?-><-?<-HKD-DNase+DUF3150<-DUF6079*<-BrxB-BrxF-iSTAND<-BrxA-DNA-binding<-XIS-HTH||?-><-?||?->GspA-ATPase-><-?||RVT-></td><td>1227</td><td>Bacteria>Pseudomonadota>Gammaproteobacteria>Oceanospirillales>Endozoicomonadaceae>Endozoicomonas</td><td>GCF_025562755.1</td><td>Endozoicomonas_euniceicola</td><td>NX720_RS21900</td><td>5400416..5404099</td><td>1234143</td><td>BREX-system-P-loop-protein-BrxC-[Endozoicomonas-euniceicola]</td></tr>
<tr><td>WP_125460588.1</td><td>Type1.BREX</td><td><-PglZ<-AbiEii<-AbiEi_3_N+AbiEi_3<-SF1-SF2-Helicase<-HKD-DNase+HKD-Rease<-?<-?<-N6-METHYLASE<-AbiGi+seqA+RAMA<-DUF6079*<-BrxB-BrxF-iSTAND<-BrxA-DNA-binding<-DUF772+DDE_Tnp_1||?->Arm-DNA-bind_2+Phage_int_SAM_3+Phage_integrase-></td><td>1220</td><td>Bacteria>Pseudomonadota>Gammaproteobacteria>Pseudomonadales>Pseudomonadaceae>Stutzerimonas</td><td>GCF_003936745.1</td><td>Stutzerimonas_stutzeri</td><td>EGJ43_RS19065</td><td>24827..28489</td><td>316</td><td>BREX-system-P-loop-protein-BrxC-[Stutzerimonas-stutzeri]</td></tr>
<tr><td>MCP5179014.1</td><td>Type1.BREX</td><td><-DUF772+DDE_Tnp_1+DDE_Tnp_1_6<-PglZ<-ParB+HNH<-?<-viral-capsid<-ZnR<-N6-METHYLASE<-HKD-DNase<-sGTP+GTPase-AIG<-DUF6079*<-BrxB-BrxF-iSTAND<-BrxA-DNA-binding<-?||?-><-rve<-rve<-Couple_hipA+HipA_C<-?<-HTH_Tnp_1+HTH_21+rve+rve_3</td><td>1153</td><td>Bacteria>Pseudomonadota>Gammaproteobacteria>Pseudomonadales>unclassified_Pseudomonadales</td><td>-</td><td>Pseudomonadales_bacterium</td><td>H6994_03785</td><td>340860..344321</td><td>1891229</td><td>BREX-system-P-loop-protein-BrxC-[Pseudomonadales-bacterium]</td></tr>
<tr><td>MCF7788313.1</td><td>Type1.BREX</td><td>BrxA-DNA-binding->BrxB-BrxF-iSTAND->DUF6079*->HKD-DNase+DUF3150->MarR-HTH->Polbeta->N6-METHYLASE->?->?->SbcC->PglZ-p->REase-></td><td>1167</td><td>Bacteria>PVC_group>Verrucomicrobiota>Verrucomicrobiae>Verrucomicrobiales>Verrucomicrobiaceae</td><td>GCA_021733925.1</td><td>Prosthecobacter_sp.</td><td>K9N47_19475</td><td>2530..6033</td><td>1965333</td><td>BREX-system-P-loop-protein-BrxC-[Prosthecobacter-sp.]</td></tr>
<tr><td>MCO4261480.1</td><td>Type1.BREX</td><td><-SbcC<-JetB<-JetA<-HEPN<-SF1-SF2-Helicase<-REase<-SIGMA+OB-Fold+AAA-ATPase+Lon-Protease-BrxL<-PglZ<-N6-METHYLASE<-DUF6079*<-BrxB-BrxF-iSTAND<-BrxA-DNA-binding<-HKD-DNase+SF1-SF2-Helicase+HTH+HTH+SRA||URI-DNase->MACRODOMAIN+SIR2->URI-DNase-><-?<-NUDIX</td><td>1151</td><td>Bacteria>Terrabacteria_group>Actinomycetota>Actinomycetes>Micrococcales>Micrococcaceae</td><td>GCA_023913835.1</td><td>Pseudarthrobacter_sp._MDT3-26</td><td>NG697_00745</td><td>166559..170014</td><td>2953651</td><td>BREX-system-P-loop-protein-BrxC-[Pseudarthrobacter-sp.-MDT3-26]</td></tr>
<tr><td>WP_252613231.1</td><td>Type1.BREX</td><td><-SbcC<-JetB<-JetA<-HEPN<-SF1-SF2-Helicase<-REase<-SIGMA+OB-Fold+AAA-ATPase+Lon-Protease-BrxL<-PglZ<-N6-METHYLASE<-DUF6079*<-BrxB-BrxF-iSTAND<-BrxA-DNA-binding<-HKD-DNase+SF1-SF2-Helicase+HTH+HTH+SRA||URI-DNase->MACRODOMAIN+SIR2->URI-DNase-><-?<-NUDIX</td><td>1151</td><td>Bacteria>Terrabacteria_group>Actinomycetota>Actinomycetes>Micrococcales>Micrococcaceae</td><td>-</td><td>Pseudarthrobacter_sp._MDT3-26</td><td>NG697_00745</td><td>166559..170014</td><td>2953651</td><td>BREX-system-P-loop-protein-BrxC-[Pseudarthrobacter-sp.-MDT3-26]</td></tr>
<tr><td>OBF48784.1</td><td>Type6.BREX</td><td>HEPN+SF1-SF2-Helicase+REase->HIRAN-><-HKD-DNase+HKD-DNase<-?||BrxE->BrxA-DNA-binding->BrxB-BrxF-iSTAND->DUF6079*->PglZ->SIGMA+OB-Fold+AAA-ATPase+Lon-Protease-BrxL-><-SF1-SF2-Helicase+REase||?-><-SF1-SF2-Helicase<-SF1-SF2-Helicase+REase</td><td>1143</td><td>Bacteria>Terrabacteria_group>Actinomycetota>Actinomycetes>Mycobacteriales>Mycobacteriaceae</td><td>GCA_001667465.1</td><td>Mycolicibacterium_monacense</td><td>A5778_22445</td><td>215738..219169</td><td>85693</td><td>hypothetical-protein-A5778_22445-[Mycolicibacterium-monacense]</td></tr>
<tr><td>WP_067214457.1</td><td>Type1.BREX</td><td>Calcineurin->SbcC+SbcC->?-><-REase||MACRODOMAIN+SIR2-><-?<-SIGMA+OB-Fold+AAA-ATPase+Lon-Protease-BrxL<-PglZ<-N6-METHYLASE<-DUF6079*<-BrxB-BrxF-iSTAND<-BrxA-DNA-binding||?->HKD-DNase+HKD-DNase-><-Peptidase_M48+TerB<-HIRAN<-?<-HEPN+SF1-SF2-Helicase+REase<-Peptidase_M23+SLT</td><td>1155</td><td>Bacteria>Terrabacteria_group>Actinomycetota>Actinomycetes>Mycobacteriales>Mycobacteriaceae</td><td>GCF_001667065.1</td><td>Mycolicibacterium_celeriflavum</td><td>A5765_RS01610</td><td>25848..29315</td><td>1249101</td><td>BREX-system-P-loop-protein-BrxC-[Mycolicibacterium-celeriflavum]</td></tr>
<tr><td>WP_120314218.1</td><td>Type6.BREX</td><td>HEPN+SF1-SF2-Helicase+REase->HIRAN-><-HKD-DNase+HKD-DNase<-?||BrxE->BrxA-DNA-binding->BrxB-BrxF-iSTAND->DUF6079*->N6-METHYLASE->PglZ->SIGMA+OB-Fold+AAA-ATPase+Lon-Protease-BrxL-><-SF1-SF2-Helicase+REase<-SF1-SF2-Helicase<-SF1-SF2-Helicase+REase<-?<-?<-SbcC+SbcC</td><td>1163</td><td>Bacteria>Terrabacteria_group>Actinomycetota>Actinomycetes>Mycobacteriales>Mycobacteriaceae</td><td>GCF_001667465.1</td><td>Mycolicibacterium_monacense</td><td>A5778_RS22240</td><td>215678..219169</td><td>85693</td><td>BREX-system-P-loop-protein-BrxC-[Mycolicibacterium-monacense]</td></tr>
<tr><td>WP_230497381.1</td><td>Type1.BREX</td><td><-SIR2||?->?->?->BrxA-DNA-binding->BrxB-BrxF-iSTAND->DUF6079*->N6-METHYLASE->PglZ->SIGMA+OB-Fold+AAA-ATPase+Lon-Protease-BrxL-><-HKD-DNase+DUF3150||ZnR+HARE-HTH->?-><-Glyoxalase</td><td>1193</td><td>Bacteria>Terrabacteria_group>Bacillota>Bacilli>Bacillales>Bacillaceae</td><td>GCF_917563885.1</td><td>Pseudoneobacillus_rhizosphaerae</td><td>NEOCIP111885_02886</td><td>60307..63888</td><td>2880968</td><td>BREX-system-P-loop-protein-BrxC-[Pseudoneobacillus-rhizosphaerae]</td></tr>
<tr><td>WP_012369225.1</td><td>Type1.BREX</td><td><-PadR-HTH||?->Phage_holin-><-RNase||?->HKD-DNase+SF1-SF2-Helicase+HTH+HTH+SRA->?->BrxA-DNA-binding->BrxB-BrxF-iSTAND->DUF6079*->N6-METHYLASE->PglZ->SIGMA+OB-Fold+AAA-ATPase+Lon-Protease-BrxL->OLD-ABC->HNH-><-Glyoxalase||LACI-HTH+PBPI-><-HTH_OrfB_IS605+OrfB_IS605+OrfB_Zn_ribbon</td><td>1192</td><td>Bacteria>Terrabacteria_group>Bacillota>Bacilli>Bacillales>Bacillales_incertae_sedis</td><td>GCF_000019905.1</td><td>Exiguobacterium_sibiricum</td><td>Exig_0314</td><td>327084..330662</td><td>332410</td><td>BREX-system-P-loop-protein-BrxC-[Exiguobacterium-sibiricum]</td></tr>
<tr><td>WP_225047865.1</td><td>Type1.BREX</td><td>Thioredoxin->?->?-><-?||ABC-ATPase+SbcC->?->BrxA-DNA-binding->BrxB-BrxF-iSTAND->DUF6079*->N6-METHYLASE-><-Arm-DNA-bind_4+Phage_int_SAM_5+Phage_integrase||PglZ->SIGMA+OB-Fold+AAA-ATPase+Lon-Protease-BrxL->HKD-DNase+SF1-SF2-Helicase+HTH+HTH+SRA-></td><td>1221</td><td>Bacteria>Terrabacteria_group>Bacillota>Bacilli>Lactobacillales>Lactobacillaceae</td><td>GCF_020180945.1</td><td>Lacticaseibacillus_kribbianus</td><td>LE416_RS07415</td><td>6539..10204</td><td>2926292</td><td>BREX-system-P-loop-protein-BrxC-[Lacticaseibacillus-kribbianus]</td></tr>
<tr><td>KJS18733.1</td><td>Type1.BREX</td><td>HNH->?->HEPN->Peptidase->?->HTH_26->BrxA-DNA-binding->BrxB-BrxF-iSTAND->DUF6079*->N6-METHYLASE->?->PglZ->SIGMA+OB-Fold+AAA-ATPase+Lon-Protease-BrxL->HKD-DNase->?->?->Calcineurin->SbcC-></td><td>1210</td><td>Bacteria>Terrabacteria_group>Bacillota>Clostridia>Eubacteriales>Clostridiaceae</td><td>GCA_000961765.1</td><td>Clostridiaceae_bacterium_BRH_c20a</td><td>VR72_21175</td><td>10275..13907</td><td>1629719</td><td>ATPase-[Clostridiaceae-bacterium-BRH_c20a]</td></tr>
<tr><td>MPQ42518.1</td><td>Type1.BREX</td><td><-SIGMA+OB-Fold+AAA-ATPase+Lon-Protease-BrxL<-PglZ<-HTH+HKD-DNase<-Cas4+SF1-SF2-Helicase<-?<-?<-?<-HTH_OrfB_IS605+OrfB_IS605+OrfB_Zn_ribbon<-N6-METHYLASE<-DUF6079-p*<-HTH_OrfB_IS605+OrfB_IS605+OrfB_Zn_ribbon<-BrxB-BrxF-iSTAND<-BrxA-DNA-binding<-Ntox45<-HEPN</td><td>995</td><td>Bacteria>Terrabacteria_group>Bacillota>Clostridia>Eubacteriales>Clostridiaceae</td><td>GCA_009295725.1</td><td>Clostridium_tarantellae</td><td>GBZ86_01880</td><td>24585..27572</td><td>39493</td><td>BREX-system-P-loop-protein-BrxC-[Clostridium-tarantellae]</td></tr>
<tr><td>WP_043662653.1</td><td>Type1.BREX</td><td><-McrC-NTD+REase<-wHTH+McrB-AAA<-SIGMA+OB-Fold+AAA-ATPase+Lon-Protease-BrxL<-?<-PglZ<-HNH<-OLD-ABC<-N6-METHYLASE<-DUF6079*<-BrxB-BrxF-iSTAND<-BrxA-DNA-binding<-HKD-DNase+SF1-SF2-Helicase+HTH+HTH+SRA<-SAM-METHYLASE<-RNase<-HNH+NERD+ZnR<-?<-?<-Amidohydro_3</td><td>1216</td><td>Bacteria>Terrabacteria_group>Bacillota>Clostridia>Eubacteriales>Clostridiaceae</td><td>GCF_000785185.1</td><td>Clostridium_butyricum</td><td>AWN73_RS08010</td><td>18592..22242</td><td>1492</td><td>BREX-system-P-loop-protein-BrxC-[Clostridium-butyricum]</td></tr>
<tr><td>WP_156211829.1</td><td>Type1.BREX</td><td><-wHTH+McrB-AAA<-?<-PglZ<-?<-?<-OLD-ABC<-ParB+HNH<-N6-METHYLASE<-CR-ATPase9+REase<-DUF6079*<-BrxB-BrxF-iSTAND<-BrxA-DNA-binding<-HKD-DNase+SF1-SF2-Helicase+HTH+HTH+SRA<-SAM-METHYLASE<-NERD+HRDC<-Ntox45<-HEPN</td><td>1216</td><td>Bacteria>Terrabacteria_group>Bacillota>Clostridia>Eubacteriales>Clostridiaceae</td><td>GCF_009735365.2</td><td>Clostridium_butyricum</td><td>GND98_RS05145</td><td>54704..58354</td><td>1492</td><td>BREX-system-P-loop-protein-BrxC-[Clostridium-butyricum]</td></tr>
<tr><td>WP_164958329.1</td><td>Type1.BREX</td><td><-HKD-DNase<-N6-METHYLASE<-?<-GDResolvase<-?<-?<-GDResolvase+HIN-HTH<-SIGMA+OB-Fold+AAA-ATPase-BrxL<-PglZ<-DUF6079*<-BrxB-BrxF-iSTAND<-DUF4007<-N6-METHYLASE</td><td>1363</td><td>Bacteria>Terrabacteria_group>Bacillota>Clostridia>Eubacteriales>Clostridiaceae</td><td>GCF_011065935.2</td><td>Clostridium_estertheticum</td><td>G9F72_RS19190</td><td>4031148..4035239</td><td>238834</td><td>hypothetical-protein-[Clostridium-estertheticum]</td></tr>
<tr><td>MCI6150868.1</td><td>Type1.BREX</td><td><-wHTH+WYL<-?<-HTH+MPTase<-ZnR<-SIGMA+OB-Fold+AAA-ATPase+Lon-Protease-BrxL<-PglZ<-HEPN<-HKD-DNase<-N6-METHYLASE<-DUF6079*<-BrxB-BrxF-iSTAND<-BrxA-DNA-binding<-HMG+NLPC+NLPC<-VirB4-FtsK<-?||?-><-TrwI<-TraM<-VirD4-TraG+VirD4-FtsK</td><td>1179</td><td>Bacteria>Terrabacteria_group>Bacillota>Clostridia>Eubacteriales>Eubacteriales_incertae_sedis</td><td>GCA_022772165.1</td><td>Flintibacter_sp.</td><td>MR705_10640</td><td>43131..46670</td><td>1918624</td><td>BREX-system-P-loop-protein-BrxC-[Flintibacter-sp.]</td></tr>
<tr><td>MCC2230802.1</td><td>Type1.BREX</td><td><-HAMP+HISKIN<-REC+OmpR-HTH<-HKD-DNase+HKD-DNase<-HKD-DNase+HKD-DNase<-SIGMA+OB-Fold+AAA-ATPase+Lon-Protease-BrxL<-PglZ<-Schlafen+HATPase+wHTH<-?<-N6-METHYLASE<-DUF6079*<-BrxB-BrxF-iSTAND<-BrxA-DNA-binding<-XIS-HTH<-?<-?<-TraM<-TraJ-RHH<-LPD32+LPD36+N6_Mtase+SF1-SF2-Helicase</td><td>1217</td><td>Bacteria>Terrabacteria_group>Bacillota>Clostridia>Eubacteriales>Lachnospiraceae</td><td>GCA_020687485.1</td><td>Lachnospiraceae_bacterium_CLA-AA-H215</td><td>LKD81_07295</td><td>25894..29547</td><td>2885348</td><td>BREX-system-P-loop-protein-BrxC-[Lachnospiraceae-bacterium-CLA-AA-H215]</td></tr>
<tr><td>WP_090167800.1</td><td>Type1.BREX</td><td><-HKD-DNase<-?<-SIGMA+OB-Fold+AAA-ATPase+Lon-Protease-BrxL<-BrxD-AAA<-?<-PglZ<-HEPN<-N6-METHYLASE<-N6-METHYLASE<-DUF6079*<-BrxB-BrxF-iSTAND<-BrxA-DNA-binding<-HTH_26<-?<-?<-wHTH+WYL+WCX<-HNH<-?<-ARG</td><td>1201</td><td>Bacteria>Terrabacteria_group>Bacillota>Clostridia>Eubacteriales>Lachnospiraceae</td><td>GCF_900100545.1</td><td>Pseudobutyrivibrio_sp._49</td><td>BLP54_RS06100</td><td>185622..189227</td><td>1855344</td><td>BREX-system-P-loop-protein-BrxC-[Pseudobutyrivibrio-sp.-49]</td></tr>
<tr><td>WP_090545210.1</td><td>Type1.BREX</td><td><-HKD-DNase<-?<-SIGMA+OB-Fold+AAA-ATPase+Lon-Protease-BrxL<-BrxD-AAA<-?<-PglZ<-HEPN<-N6-METHYLASE<-N6-METHYLASE<-DUF6079*<-BrxB-BrxF-iSTAND<-BrxA-DNA-binding<-HTH_26<-?<-?<-wHTH+WYL+WCX<-HNH<-?<-ARG</td><td>1201</td><td>Bacteria>Terrabacteria_group>Bacillota>Clostridia>Eubacteriales>Lachnospiraceae</td><td>GCF_900113655.1</td><td>Pseudobutyrivibrio_sp._OR37</td><td>BM097_RS01535</td><td>134498..138103</td><td>1798186</td><td>BREX-system-P-loop-protein-BrxC-[Pseudobutyrivibrio-sp.-OR37]</td></tr>
<tr><td>WP_061213686.1</td><td>Type1.BREX</td><td>RelE-ParE->?->Classical-AAA->CR-ATPase5+REase-><-HEPN||BrxA-DNA-binding->BrxB-BrxF-iSTAND->DUF6079*->N6-METHYLASE->PglZ->SIGMA+OB-Fold+AAA-ATPase+Lon-Protease-BrxL->?->HKD-DNase+SF1-SF2-Helicase+HTH+HTH+SRA-><-PIN<-?||?->RAMPs-></td><td>1199</td><td>Bacteria>Terrabacteria_group>Bacillota>Clostridia>Eubacteriales>Syntrophomonadaceae</td><td>GCF_001570625.1</td><td>Syntrophomonas_wolfei</td><td>AWB54_RS05890</td><td>82988..86587</td><td>863</td><td>BREX-system-P-loop-protein-BrxC-[Syntrophomonas-wolfei]</td></tr>
<tr><td>MBN1450028.1</td><td>Type1.BREX</td><td><-SIGMA+OB-Fold+AAA-ATPase+Lon-Protease-BrxL<-PglZ<-REase+DUF5655<-N6-METHYLASE<-HTH+HKD-DNase<-Pkinase+SF1-SF2-Helicase<-?<-NERD+Pkinase<-DUF6079*<-BrxB-BrxF-iSTAND<-BrxA-DNA-binding</td><td>1209</td><td>Bacteria>Terrabacteria_group>Chloroflexota>Anaerolineae>Anaerolineales>unclassified_Anaerolineales</td><td>GCA_016928525.1</td><td>Anaerolineales_bacterium</td><td>JW963_03360</td><td>18534..22163</td><td>2073117</td><td>BREX-system-P-loop-protein-BrxC-[Anaerolineales-bacterium]</td></tr>
<tr><td>MBI4322558.1</td><td>Type6.BREX</td><td><-DUF6079-p*<-BrxB-BrxF-iSTAND<-BrxA-DNA-binding<-BrxE<-Phage_integrase<-?||?->HKD-DNase+HKD-Rease->REase-></td><td>427</td><td>Bacteria>Terrabacteria_group>Chloroflexota>unclassified_Chloroflexota</td><td>GCA_016210495.1</td><td>Chloroflexota_bacterium</td><td>HY675_29045</td><td>1..1283</td><td>2026724</td><td>BREX-system-P-loop-protein-BrxC-[Chloroflexota-bacterium]</td></tr>
<tr><td>MCL4487040.1</td><td>Type1.BREX</td><td>BrxA-DNA-binding->BrxB-BrxF-iSTAND->Peptidase_M91->N6-METHYLASE->HKD-DNase->DUF6079*->HEPN->PhdYeFM_antitox->PIN->PglZ->?->?->?->MerR-HTH+TipAS-></td><td>1221</td><td>Bacteria>Terrabacteria_group>Chloroflexota>unclassified_Chloroflexota</td><td>GCA_023381515.1</td><td>Chloroflexota_bacterium</td><td>M1570_02770</td><td>10289..13954</td><td>2026724</td><td>BREX-system-P-loop-protein-BrxC-[Chloroflexota-bacterium]</td></tr>
<tr><td>BBD57221.1</td><td>Type1.BREX</td><td><-MerR-HTH||RHH-><-?<-?||?->TPR+TPR+TPR-><-Phage_integrase<-?||BrxB-BrxF-iSTAND->DUF6079*-><-?<-REase||?->PIN->N6-METHYLASE->HEPN->PglZ+Phosphodiest->HKD-DNase->SIGMA+OB-Fold+AAA-ATPase+Lon-Protease-BrxL-></td><td>1176</td><td>Bacteria>Terrabacteria_group>Cyanobacteriota/Melainabacteria_group>Cyanobacteriota>Cyanophyceae>Oscillatoriophycideae</td><td>GCA_003609755.1</td><td>Planktothrix_agardhii_NIES-204</td><td>NIES204_45570</td><td>14633..18163</td><td>282423</td><td>hypothetical-protein-NIES204_45570-[Planktothrix-agardhii-NIES-204]</td></tr>
<tr><td>WP_275522152.1</td><td>Type1.BREX</td><td>ACET->BrxA-DNA-binding-><-SIGMA+OB-Fold+AAA-ATPase+Lon-Protease-BrxL<-HKD-DNase<-PglZ+Phosphodiest<-N6-METHYLASE<-DUF6079*<-BrxB-BrxF-iSTAND<-AbiEii<-AbiEi_4<-?||?-><-TOPRIM||?->RVT+HNH-></td><td>1181</td><td>Bacteria>Terrabacteria_group>Cyanobacteriota/Melainabacteria_group>Cyanobacteriota>Cyanophyceae>Oscillatoriophycideae</td><td>GCF_029167235.1</td><td>Kamptonema_sp._UHCC_0994</td><td>PY364_RS09785</td><td>12149..15694</td><td>3031329</td><td>BREX-system-P-loop-protein-BrxC-[Kamptonema-sp.-UHCC-0994]</td></tr>
<tr><td>WP_254034082.1</td><td>Type1.BREX</td><td>TPR+TPR+TPR+TPR-><-?||BrxA-DNA-binding-><-SIGMA+OB-Fold+AAA-ATPase+Lon-Protease-BrxL<-HKD-DNase<-PglZ+Phosphodiest<-N6-METHYLASE<-PIN<-?<-DUF6079*<-BrxB-BrxF-iSTAND<-?||?->?->Phage_integrase-></td><td>1176</td><td>Bacteria>Terrabacteria_group>Cyanobacteriota/Melainabacteria_group>Cyanobacteriota>Cyanophyceae>Oscillatoriophycideae</td><td>GCF_904830845.1</td><td>Planktothrix_agardhii</td><td>NMG87_RS21740</td><td>24173..27703</td><td>1160</td><td>BREX-system-P-loop-protein-BrxC-[Planktothrix-agardhii]</td></tr>
<tr><td>WP_015862248.1</td><td>Type1.BREX</td><td><-Phage_integrase<-?<-?||ParB+N6_N4_METHYLASE->PHP+SbcC-><-SIGMA+OB-Fold+AAA-ATPase+Lon-Protease-BrxL<-PglZ<-HEPN<-N6-METHYLASE<-DUF6079*<-BrxB-BrxF-iSTAND<-BrxA-DNA-binding<-HKD-DNase<-?<-SbcC<-HAD<-?||?->Radical-SAM-></td><td>1188</td><td>Bacteria>Thermodesulfobacteriota>Desulfovibrionia>Desulfovibrionales>Desulfovibrionaceae>Solidesulfovibrio</td><td>GCF_000010665.1</td><td>Solidesulfovibrio_magneticus</td><td>DMR_RS17110</td><td>4099313..4102879</td><td>184917</td><td>BREX-system-P-loop-protein-BrxC-[Solidesulfovibrio-magneticus]</td></tr>
<tr><td>WP_086519900.1</td><td>Type1.BREX</td><td><-PIN||nSTAND1+HNH->RNaseH->NACHT+SF1-SF2-Helicase+HKD-DNase->REase-><-HKD-DNase+OGG-like<-SIGMA+OB-Fold+AAA-ATPase+Lon-Protease-BrxL<-PglZ<-N6-METHYLASE<-DUF6079*<-BrxB-BrxF-iSTAND<-BrxA-DNA-binding<-?<-ColE7+HNH<-ACET||NUDIX->?-><-DUF2754</td><td>1213</td><td>-</td><td>GCF_900775425.1</td><td>Bacteria</td><td>E7W97_RS22710</td><td>357533..361174</td><td>2</td><td>MULTISPECIES:-BREX-system-P-loop-protein-BrxC-[Bacteria]</td></tr>
</table>

| Protein-ID | System | Gene-Neighbourhoods | Length | Taxonomy | Assembly-ID | Strain | Locus-Tag | Genomic-coordinates | TaxID | Fasta-define |
|---|---|---|---|---|---|---|---|---|---|---|
| -- | | | | | | | | | | |
| -- ParB + HNH -(GmrSD) | | | | | | | | | | |
| -- | | | | | | | | | | |
| WP_103992123.1 | Type5.BREX | [illegible] | 1227 | Archaea>Euryarchaeota>Stenosarchaea_group>Halobacteria>Haloferacales>Haloferacaceae | GCF_004799685.1 | Halobellus_limi | DV707_RS08430 | 1701258..1704941 | 699433 | hypothetical-protein-[Halobellus-limi] |
| WP_103992125.1 | Type5.BREX | [illegible] | 1284 | Archaea>Euryarchaeota>Stenosarchaea_group>Halobacteria>Haloferacales>Haloferacaceae | GCF_004799685.1 | Halobellus_limi | DV707_RS08440 | 1705516..1709370 | 699433 | hypothetical-protein-[Halobellus-limi] |
| WP_076580859.1 | Type6.BREX | [illegible] | 1206 | Archaea>Euryarchaeota>Stenosarchaea_group>Halobacteria>Natrialbales>Natrialbaceae | GCF_001971705.1 | Natronorubrum_daqingense | BW960_RS07735 | 340245..343865 | 588898 | BREX-system-P-loop-protein-BrxC-[Natronorubrum-daqingense] |
| WP_253430473.1 | Type6.BREX | [illegible] | 1206 | Archaea>Euryarchaeota>Stenosarchaea_group>Halobacteria>Natrialbales>Natrialbaceae | GCF_024138145.1 | Halovivax_sp._AGai3-5 | NGM15_RS10380 | 2104102..2107722 | 2953746 | BREX-system-P-loop-protein-BrxC-[Halovivax-sp.-AGai3-5] |
| WP_229395640.1 | Type1.BREX | [illegible] | 1143 | Archaea>Euryarchaeota>Stenosarchaea_group>Methanomicrobia>Methanosarcinales>Methanosarcinaceae | GCF_020804205.1 | Methanosarcina_sp._DH1 | FXW07_RS12025 | 174817..178248 | 2605695 | BREX-system-P-loop-protein-BrxC-[Methanosarcina-sp.-DH1] |
| MBD0371455.1 | Type1.BREX | [illegible] | 1218 | Bacteria>Acidobacteriota>Blastocatellia>Blastocatellales>Pyrinomonadaceae>unclassified_Pyrinomonadaceae | GCA_014534425.1 | Pyrinomonadaceae_bacterium | ICV60_11505 | 29495..33151 | 2283092 | BREX-system-P-loop-protein-BrxC-[Pyrinomonadaceae-bacterium] |
| MBK9963544.1 | Type6.BREX | [illegible] | 1207 | Bacteria>Acidobacteriota>Holophagae>Holophagales>unclassified_Holophagales | GCA_016722225.1 | Holophagales_bacterium | IPP07_01030 | 234150..237773 | 2478486 | BREX-system-P-loop-protein-BrxC-[Holophagales-bacterium] |
| MCG3192298.1 | Type6.BREX | [illegible] | 1183 | Bacteria>Acidobacteriota>Thermoanaerobaculia>unclassified_Thermoanaerobaculia | GCA_022071945.1 | Thermoanaerobaculia_bacterium | DIJKHBIC_01536 | 22288..25839 | 2651171 | hypothetical-protein-[Thermoanaerobaculia-bacterium] |
| MBO7131877.1 | Type1.BREX | [illegible] | 1109 | Bacteria>Bacteria_incertae_sedis>Bacteria_candidate_phyla>Candidatus_Saccharibacteria>unclassified_Candidatus_Saccharibacteria | GCA_017645955.1 | Candidatus_Saccharibacteria_bacterium | J6V85_01240 | 41463..44792 | 2026720 | BREX-system-P-loop-protein-BrxC-[Candidatus-Saccharibacteria-bacterium] |
| WP_146981109.1 | Type6.BREX | [illegible] | 1216 | Bacteria>delta/epsilon_subdivisions>Deltaproteobacteria>Bradymonadales>Lujinxingiaceae>Lujinxingia | GCF_007997015.1 | Lujinxingia_vulgaris | FRC98_RS09600 | 57836..61486 | 2600176 | BREX-system-P-loop-protein-BrxC-[Lujinxingia-vulgaris] |
| WP_251951637.1 | Type5.BREX | [illegible] | 1108 | Bacteria>FCB_group>Bacteroidetes/Chlorobi_group>Bacteroidota>Bacteroidetes_incertae_sedis>Rhodothermaceae | GCF_943908255.1 | Salinibacter_ruber | NGL47_RS11295 | 22718..26044 | 146919 | hypothetical-protein-[Salinibacter-ruber] |
| WP_251951639.1 | Type5.BREX | [illegible] | 1245 | Bacteria>FCB_group>Bacteroidetes/Chlorobi_group>Bacteroidota>Bacteroidetes_incertae_sedis>Rhodothermaceae | GCF_943908255.1 | Salinibacter_ruber | NGL47_RS11305 | 26687..30424 | 146919 | hypothetical-protein-[Salinibacter-ruber] |
| MBS6238336.1 | Type1.BREX | [illegible] | 1205 | Bacteria>FCB_group>Bacteroidetes/Chlorobi_group>Bacteroidota>Bacteroidia>Bacteroidales | GCA_018374845.1 | Bacteroides_sp. | KH897_08200 | 204452..208069 | 29523 | BREX-system-P-loop-protein-BrxC-[Bacteroides-sp.] |
| SHF69941.1 | Type1.BREX | [illegible] | 1182 | Bacteria>FCB_group>Bacteroidetes/Chlorobi_group>Bacteroidota>Bacteroidia>Bacteroidales | GCA_900128905.1 | Bacteroides_luti | SAMN05444405_11263 | 81108..84656 | 1297750 | hypothetical-protein-SAMN05444405_11263-[Bacteroides-luti] |
| SHK68061.1 | Type1.BREX | [illegible] | 1206 | Bacteria>FCB_group>Bacteroidetes/Chlorobi_group>Bacteroidota>Bacteroidia>Bacteroidales | GCA_900142525.1 | Prevotella_ruminicola | SAMN05216463_10942 | 46265..49885 | 839 | hypothetical-protein-SAMN05216463_10942-[Prevotella-ruminicola] |
| WP_073402639.1 | Type1.BREX | [illegible] | 1188 | Bacteria>FCB_group>Bacteroidetes/Chlorobi_group>Bacteroidota>Bacteroidia>Bacteroidales | GCF_900128905.1 | Bacteroides_luti | BUC21_RS11790 | 81108..84674 | 1297750 | BREX-system-P-loop-protein-BrxC-[Bacteroides-luti] |
| WP_254795118.1 | Type1.BREX | [illegible] | 1209 | Bacteria>FCB_group>Bacteroidetes/Chlorobi_group>Bacteroidota>Bacteroidia>Bacteroidales | GCF_900142525.1 | Prevotella_ruminicola | BUB86_RS07050 | 46256..49885 | 839 | BREX-system-P-loop-protein-BrxC-[Prevotella-ruminicola] |
| WP_176134075.1 | Type1.BREX | [illegible] | 1192 | Bacteria>FCB_group>Bacteroidetes/Chlorobi_group>Bacteroidota>Bacteroidia>Bacteroidales | GCF_900168155.1 | Parabacteroides_chartae | B5X57_RS12575 | 48804..52382 | 1037355 | BREX-system-P-loop-protein-BrxC-[Parabacteroides-chartae] |
| WP_073207711.1 | Type1.BREX | [illegible] | 1206 | Bacteria>FCB_group>Bacteroidetes/Chlorobi_group>Bacteroidota>Bacteroidia>Bacteroidales | - | Prevotella_ruminicola | SAMN05216463_10942 | 46265..49885 | 839 | BREX-system-P-loop-protein-BrxC-[Prevotella-ruminicola] |
| WP_075603947.1 | Type1.BREX | [illegible] | 1204 | Bacteria>FCB_group>Bacteroidetes/Chlorobi_group>Bacteroidota>Bacteroidia>Marinilabiliales | GCF_001660705.2 | Saccharicrinis_aurantiacus | A8B35_RS21005 | 65274..68888 | 1849719 | BREX-system-P-loop-protein-BrxC-[Saccharicrinis-aurantiacus] |
| WP_053405044.1 | Type1.BREX | [illegible] | 1179 | Bacteria>FCB_group>Bacteroidetes/Chlorobi_group>Bacteroidota>Cytophagia>Cytophagales | GCF_001274635.1 | Persicobacter_sp._CCB-QB2 | NG22_RS06825 | 1502189..1505728 | 1561025 | BREX-system-P-loop-protein-BrxC-[Persicobacter-sp.-CCB-QB2] |
| OIQ21186.1 | Type1.BREX | [illegible] | 1165 | Bacteria>FCB_group>Bacteroidetes/Chlorobi_group>Bacteroidota>Flavobacteriia>Flavobacteriales | GCA_001874145.1 | Lacinutrix_sp._MedPE-SW | BM549_09430 | 19615..23112 | 1860087 | MAG:-hypothetical-protein-BM549_09430-[Lacinutrix-sp.-MedPE-SW] |
| MAM30277.1 | Type1.BREX | [illegible] | 1165 | Bacteria>FCB_group>Bacteroidetes/Chlorobi_group>Bacteroidota>Flavobacteriia>Flavobacteriales | GCA_002693005.1 | Flavobacteriaceae_bacterium | CMC13_14765 | 19860..23357 | 1871037 | hypothetical-protein-[Flavobacteriaceae-bacterium] |
| GEM55955.1 | Type1.BREX | [illegible] | 1216 | Bacteria>FCB_group>Bacteroidetes/Chlorobi_group>Bacteroidota>Flavobacteriia>Flavobacteriales | GCA_007990815.1 | Flavobacterium_branchiophilum_NBRC_15030_=_ATCC_35035 | FB1_21760 | 14247..17897 | 1218110 | hypothetical-protein-FB1_21760-[Flavobacterium-branchiophilum-NBRC-15030-=-ATCC-35035] |
| WP_027137224.1 | Type6.BREX | [illegible] | 1196 | Bacteria>FCB_group>Bacteroidetes/Chlorobi_group>Bacteroidota>Flavobacteriia>Flavobacteriales | GCF_000425645.1 | Gaetbulibacter_saemankumensis | H611_RS0104475 | 476638..480228 | 311208 | BREX-system-P-loop-protein-BrxC-[Gaetbulibacter-saemankumensis] |
| WP_059014619.1 | Type1.BREX | [illegible] | 1169 | Bacteria>FCB_group>Bacteroidetes/Chlorobi_group>Bacteroidota>Flavobacteriia>Flavobacteriales | GCF_001485515.1 | Flavobacterium_psychrophilum | FPK15_RS07995 | 6803..10312 | 96345 | BREX-system-P-loop-protein-BrxC-[Flavobacterium-psychrophilum] |
| WP_076662677.1 | Type1.BREX | [illegible] | 1201 | Bacteria>FCB_group>Bacteroidetes/Chlorobi_group>Bacteroidota>Flavobacteriia>Flavobacteriales | GCF_001975575.1 | Mangrovimonas_sp._DI_80 | BKM32_RS04295 | 828004..831609 | 1779330 | BREX-system-P-loop-protein-BrxC-[Mangrovimonas-sp.-DI-80] |
| WP_132215847.1 | Type1.BREX | [illegible] | 1169 | Bacteria>FCB_group>Bacteroidetes/Chlorobi_group>Bacteroidota>Flavobacteriia>Flavobacteriales | GCF_004341235.1 | Mariniflexile_fucanivorans | EV196_RS05490 | 443037..446546 | 264023 | BREX-system-P-loop-protein-BrxC-[Mariniflexile-fucanivorans] |
| WP_131908313.1 | Type1.BREX | [illegible] | 1203 | Bacteria>FCB_group>Bacteroidetes/Chlorobi_group>Bacteroidota>Flavobacteriia>Flavobacteriales | GCF_004349085.1 | Flavobacterium_caseinilyticum | EOF89_RS02690 | 598586..602197 | 2541732 | BREX-system-P-loop-protein-BrxC-[Flavobacterium-caseinilyticum] |
| WP_141421442.1 | Type1.BREX | [illegible] | 1165 | Bacteria>FCB_group>Bacteroidetes/Chlorobi_group>Bacteroidota>Flavobacteriia>Flavobacteriales | GCF_006546625.1 | Haloflavibacter_putidus | FKR84_RS06225 | 21354..24851 | 2576776 | BREX-system-P-loop-protein-BrxC-[Haloflavibacter-putidus] |
| WP_089081509.1 | Type1.BREX | [illegible] | 1199 | Bacteria>FCB_group>Bacteroidetes/Chlorobi_group>Bacteroidota>Flavobacteriia>Flavobacteriales | GCF_007990815.1 | Flavobacterium_branchiophilum | FB1_RS11100 | 14247..17846 | 55197 | BREX-system-P-loop-protein-BrxC-[Flavobacterium-branchiophilum] |
| WP_153266434.1 | Type1.BREX | [illegible] | 1157 | Bacteria>FCB_group>Bacteroidetes/Chlorobi_group>Bacteroidota>Flavobacteriia>Flavobacteriales | GCF_009498295.1 | Oceanihabitans_sp._IOP_32 | FEZ18_RS00165 | 31374..34847 | 2529032 | BREX-system-P-loop-protein-BrxC-[Oceanihabitans-sp.-IOP_32] |
| WP_165150638.1 | Type1.BREX | [illegible] | 1198 | Bacteria>FCB_group>Bacteroidetes/Chlorobi_group>Bacteroidota>Flavobacteriia>Flavobacteriales | GCF_011065475.1 | Aequorivita_sp._KMM_9714 | G3A46_RS13570 | 21223..24819 | 2707173 | BREX-system-P-loop-protein-BrxC-[Aequorivita-sp.-KMM-9714] |
| WP_193813001.1 | Type6.BREX | [illegible] | 1217 | Bacteria>FCB_group>Bacteroidetes/Chlorobi_group>Bacteroidota>Flavobacteriia>Flavobacteriales | GCF_015191005.1 | Kaistella_flava_(ex_Peng_et_al._2021) | Q73A0000_05090 | 1135416..1139069 | 2038776 | BREX-system-P-loop-protein-BrxC-[Kaistella-flava-(ex-Peng-et-al.-2021)] |
| WP_198639224.1 | Type1.BREX | [illegible] | 1201 | Bacteria>FCB_group>Bacteroidetes/Chlorobi_group>Bacteroidota>Flavobacteriia>Flavobacteriales | GCF_016236975.1 | Salegentibacter_maritimus | I6U50_RS13470 | 35086..38691 | 2794347 | BREX-system-P-loop-protein-BrxC-[Salegentibacter-maritimus] |
| WP_179314673.1 | Type1.BREX | [illegible] | 1160 | Bacteria>FCB_group>Bacteroidetes/Chlorobi_group>Bacteroidota>Flavobacteriia>Flavobacteriales | GCF_019278425.1 | Winogradskyella | HOI32_RS04080 | 939044..942526 | 286104 | MULTISPECIES:-BREX-system-P-loop-protein-BrxC-[Winogradskyella] |
| WP_244340081.1 | Type1.BREX | [illegible] | 1169 | Bacteria>FCB_group>Bacteroidetes/Chlorobi_group>Bacteroidota>Flavobacteriia>Flavobacteriales | GCF_022836875.1 | Flavobacterium | LZF87_RS14585 | 3214137..3217646 | 237 | MULTISPECIES:-BREX-system-P-loop-protein-BrxC-[Flavobacterium] |
| WP_089756260.1 | Type1.BREX | [illegible] | 1176 | Bacteria>FCB_group>Bacteroidetes/Chlorobi_group>Bacteroidota>Flavobacteriia>Flavobacteriales | GCF_900100115.1 | Chryseobacterium_soldanellicola | BLT04_RS13255 | 831929..835459 | 311333 | BREX-system-P-loop-protein-BrxC-[Chryseobacterium-soldanellicola] |
| WP_103999283.1 | Type1.BREX | [illegible] | 1170 | Bacteria>FCB_group>Bacteroidetes/Chlorobi_group>Bacteroidota>Flavobacteriia>Flavobacteriales | GCF_900108015.1 | Flavobacterium_urumqiense | C4S35_RS05860 | 195084..198596 | 935224 | BREX-system-P-loop-protein-BrxC-[Flavobacterium-urumqiense] |
| WP_090838887.1 | Type1.BREX | [illegible] | 1160 | Bacteria>FCB_group>Bacteroidetes/Chlorobi_group>Bacteroidota>Flavobacteriia>Flavobacteriales | GCF_900114005.1 | Olleya_namhaensis | BMZ09_RS07050 | 330310..333792 | 1144750 | BREX-system-P-loop-protein-BrxC-[Olleya-namhaensis] |
| WP_159447525.1 | Type1.BREX | [illegible] | 719 | Bacteria>FCB_group>Bacteroidetes/Chlorobi_group>Bacteroidota>Flavobacteriia>Flavobacteriales | GCF_900167935.1 | Maribacter_arcticus | B5X55_RS04985 | 472839..474998 | 561365 | BREX-system-P-loop-protein-BrxC-[Maribacter-arcticus] |
| WP_271393758.1 | Type1.BREX | [illegible] | 1165 | Bacteria>FCB_group>Bacteroidetes/Chlorobi_group>Bacteroidota>Flavobacteriia>Flavobacteriales | GCF_947492035.1 | Aequorivita_sinensis | POR58_RS11110 | 23754..27251 | 1382458 | BREX-system-P-loop-protein-BrxC-[Aequorivita-sinensis] |
| WP_272023608.1 | Type1.BREX | [illegible] | 1160 | Bacteria>FCB_group>Bacteroidetes/Chlorobi_group>Bacteroidota>Flavobacteriia>Flavobacteriales | GCF_947496445.1 | Olleya_namhaensis | PQG05_RS12495 | 26845..30327 | 1144750 | BREX-system-P-loop-protein-BrxC-[Olleya-namhaensis] |
| OYD42847.1 | Type1.BREX | [illegible] | 1175 | Bacteria>FCB_group>Bacteroidetes/Chlorobi_group>Bacteroidota>Sphingobacteriia>Sphingobacteriales | GCA_002245855.1 | Sphingobacterium_cellulitidis | CHT99_08530 | 557163..560690 | 1768011 | hypothetical-protein-CHT99_08530-[Sphingobacterium-cellulitidis] |
| WP_094282244.1 | Type1.BREX | [illegible] | 1175 | Bacteria>FCB_group>Bacteroidetes/Chlorobi_group>Bacteroidota>Sphingobacteriia>Sphingobacteriales | GCF_002245765.1 | Sphingobacterium_cellulitidis | CHU00_RS15775 | 18381..21908 | 1768011 | BREX-system-P-loop-protein-BrxC-[Sphingobacterium-cellulitidis] |
| MBS1911241.1 | Type1.BREX | [illegible] | 1145 | Bacteria>FCB_group>Bacteroidetes/Chlorobi_group>Bacteroidota>unclassified_Bacteroidota | GCA_018266795.1 | Bacteroidota_bacterium | JST22_04585 | 55920..59357 | 1898104 | BREX-system-P-loop-protein-BrxC-[Bacteroidota-bacterium] |
| MBV5326252.1 | Type1.BREX | [illegible] | 1179 | Bacteria>FCB_group>Bacteroidetes/Chlorobi_group>Chlorobiota>Chlorobiia>Chlorobiales | GCA_019162945.1 | Chlorobium_sp. | JZU65_01250 | 9464..13003 | 1095 | BREX-system-P-loop-protein-BrxC-[Chlorobium-sp.] |
| WP_204208384.1 | Type1.BREX | [illegible] | 997 | Bacteria>FCB_group>Bacteroidetes/Chlorobi_group>Chlorobiota>Chlorobiia>Chlorobiales | GCF_001687065.1 | Prosthecochloris_sp._CIB_2401 | Ptc2401_RS05035 | 1057250..1060243 | 1868325 | BREX-system-P-loop-protein-BrxC-[Prosthecochloris-sp.-CIB-2401] |
| WP_139626761.1 | Type1.BREX | [illegible] | 1204 | Bacteria>FCB_group>Bacteroidetes/Chlorobi_group>Chlorobiota>Chlorobiia>Chlorobiales | GCF_006265245.1 | Prosthecochloris_vibrioformis | FGF68_RS08505 | 9816..13430 | 1098 | BREX-system-P-loop-protein-BrxC-[Prosthecochloris-vibrioformis] |
| NLW30564.1 | Type1.BREX | [illegible] | 1185 | Bacteria>FCB_group>Fibrobacterota>Fibrobacteria>Fibrobacterales>Fibrobacteraceae | GCA_012728995.1 | Fibrobacter_sp. | GXY77_03805 | 4326..7883 | 35828 | BREX-system-P-loop-protein-BrxC-[Fibrobacter-sp.] |
| SHK42502.1 | Type1.BREX | [illegible] | 1186 | Bacteria>FCB_group>Fibrobacterota>Fibrobacteria>Fibrobacterales>Fibrobacteraceae | GCA_900142505.1 | Fibrobacter_sp._UWH5 | SAMN05720764_101395 | 444346..447906 | 1896211 | hypothetical-protein-SAMN05720764_101395-[Fibrobacter-sp.-UWH5] |
| WP_212668349.1 | Type1.BREX | [illegible] | 1186 | Bacteria>FCB_group>Fibrobacterota>Fibrobacteria>Fibrobacterales>Fibrobacteraceae | GCF_002210355.1 | Fibrobacter_sp._UWH3 | B7993_RS04325 | 153736..157296 | 1964353 | BREX-system-P-loop-protein-BrxC-[Fibrobacter-sp.-UWH3] |
| WP_073283173.1 | Type1.BREX | [illegible] | 1184 | Bacteria>FCB_group>Fibrobacterota>Fibrobacteria>Fibrobacterales>Fibrobacteraceae | GCF_900142465.1 | Fibrobacter_sp._UWH6 | BUB73_RS01535 | 351031..354585 | 1896212 | BREX-system-P-loop-protein-BrxC-[Fibrobacter-sp.-UWH6] |
| WP_073156756.1 | Type1.BREX | [illegible] | 1184 | Bacteria>FCB_group>Fibrobacterota>Fibrobacteria>Fibrobacterales>Fibrobacteraceae | GCF_900142505.1 | Fibrobacter_sp._UWH5 | BUB38_RS01950 | 444352..447906 | 1896211 | BREX-system-P-loop-protein-BrxC-[Fibrobacter-sp.-UWH5] |
| WP_005903411.1 | Type1.BREX | [illegible] | 1250 | Bacteria>Fusobacteriota>Fusobacteriia>Fusobacteriales>Fusobacteriaceae>Fusobacterium | GCF_000178895.1 | Fusobacterium_nucleatum | C4N14_RS08075 | 1667383..1671135 | 851 | BREX-system-P-loop-protein-BrxC-[Fusobacterium-nucleatum] |
| WP_124794840.1 | Type1.BREX | [illegible] | 1250 | Bacteria>Fusobacteriota>Fusobacteriia>Fusobacteriales>Fusobacteriaceae>Fusobacterium | GCF_003859915.1 | Fusobacterium_canifelinum | EII27_RS00780 | 173024..176776 | 285729 | BREX-system-P-loop-protein-BrxC-[Fusobacterium-canifelinum] |
| WP_005980883.1 | Type1.BREX | [illegible] | 1242 | Bacteria>Fusobacteriota>Fusobacteriia>Fusobacteriales>Fusobacteriaceae>Fusobacterium | GCF_902373815.1 | Fusobacterium_ulcerans | FXX38_RS11815 | 320598..324326 | 861 | BREX-system-P-loop-protein-BrxC-[Fusobacterium-ulcerans] |
| WP_008800858.1 | Type1.BREX | [illegible] | 1240 | Bacteria>Fusobacteriota>Fusobacteriia>Fusobacteriales>Fusobacteriaceae>Fusobacterium | GCF_902373895.1 | Fusobacterium_gonidiaformans | FY288_RS06315 | 242664..246386 | 849 | BREX-system-P-loop-protein-BrxC-[Fusobacterium-gonidiaformans] |
| WP_203328719.1 | Type6.BREX | [illegible] | 1213 | Bacteria | GCF_016836415.1 | Pseudomonadota | JQ482_RS25705 | 5512214..5515855 | 1224 | MULTISPECIES:-BREX-system-P-loop-protein-BrxC-[Pseudomonadota] |
| WP_240685338.1 | Type6.BREX | [illegible] | 1213 | Bacteria | GCF_025461165.1 | Pseudomonadota | JAK52_RS02505 | 287897..291538 | 1224 | MULTISPECIES:-BREX-system-P-loop-protein-BrxC-[Pseudomonadota] |
| MBA3548104.1 | Type1.BREX | [illegible] | 1231 | Bacteria>Myxococcota>Polyangia>Nannocystales>Nannocystaceae>Nannocystis | GCA_013812955.1 | Nannocystis_sp. | HOT76_16605 | 30110..33805 | 1962667 | BREX-system-P-loop-protein-BrxC-[Nannocystis-sp.] |
| MBK8262770.1 | Type6.BREX | [illegible] | 1208 | Bacteria>Myxococcota>Polyangia>Nannocystales>Nannocystaceae>Nannocystis | GCA_016712315.1 | Nannocystis_sp. | IPK80_15725 | 176860..180486 | 1962667 | BREX-system-P-loop-protein-BrxC-[Nannocystis-sp.] |
| MCK6512397.1 | Type1.BREX | [illegible] | 1145 | Bacteria>Myxococcota>unclassified_Myxococcota | GCA_023150835.1 | Myxococcota_bacterium | L6R29_20905 | 15899..19336 | 2818507 | BREX-system-P-loop-protein-BrxC-[Myxococcota-bacterium] |
| WP_123104890.1 | Type6.BREX | [illegible] | 1210 | Bacteria>Pseudomonadota>Acidithiobacillia>Acidithiobacillales>Acidithiobacillaceae>Acidithiobacillus | GCF_003721225.1 | Acidithiobacillus_sulfuriphilus | EC580_RS10670 | 12374..16006 | 1867749 | BREX-system-P-loop-protein-BrxC-[Acidithiobacillus-sulfuriphilus] |
| WP_215855738.1 | Type6.BREX | [illegible] | 1210 | Bacteria>Pseudomonadota>Acidithiobacillia>Acidithiobacillales>Acidithiobacillaceae>Acidithiobacillus | GCF_018854635.1 | Acidithiobacillus_ferridurans | HF641_RS00655 | 27295..30927 | 1232575 | BREX-system-P-loop-protein-BrxC-[Acidithiobacillus-ferridurans] |
| WP_023449856.1 | Type1.BREX | [illegible] | 1177 | Bacteria>Pseudomonadota>Alphaproteobacteria>Caulobacterales>Caulobacteraceae>Asticcacaulis | GCF_000495835.1 | Asticcacaulis_sp._AC402 | ABAC402_RS31845 | 123615..127148 | 1282361 | BREX-system-P-loop-protein-BrxC-[Asticcacaulis-sp.-AC402] |
| WP_212207485.1 | Type6.BREX | [illegible] | 1208 | Bacteria>Pseudomonadota>Alphaproteobacteria>Hyphomicrobiales>Nitrobacteraceae>Bradyrhizobium | - | Bradyrhizobium_sp._AUGA_SZCCT0274 | JQ620_09165 | 74241..77867 | 2807670 | BREX-system-P-loop-protein-BrxC-[Bradyrhizobium-sp.-AUGA-SZCCT0274] |
| MBR1240293.1 | Type6.BREX | [illegible] | 1208 | Bacteria>Pseudomonadota>Alphaproteobacteria>Hyphomicrobiales>Nitrobacteraceae>Bradyrhizobium | GCA_018130325.1 | Bradyrhizobium_sp._AUGA_SZCCT0274 | JQ620_09165 | 74241..77867 | 2807670 | BREX-system-P-loop-protein-BrxC-[Bradyrhizobium-sp.-AUGA-SZCCT0274] |
| WP_249144787.1 | Type6.BREX | [illegible] | 1203 | Bacteria>Pseudomonadota>Alphaproteobacteria>Hyphomicrobiales>Nitrobacteraceae>Bradyrhizobium | GCF_018130325.1 | Bradyrhizobium_sp._AUGA_SZCCT0274 | JQ620_RS09170 | 74241..77852 | 2807670 | BREX-system-P-loop-protein-BrxC-[Bradyrhizobium-sp.-AUGA-SZCCT0274] |
| WP_213286691.1 | Type6.BREX | [illegible] | 1208 | Bacteria>Pseudomonadota>Alphaproteobacteria>Hyphomicrobiales>Nitrobacteraceae>Bradyrhizobium | GCF_018398875.1 | Bradyrhizobium_sp._sGM-13 | KJQ96_RS11300 | 145012..148638 | 2831781 | BREX-system-P-loop-protein-BrxC-[Bradyrhizobium-sp.-sGM-13] |
| WP_225161486.1 | Type1.BREX | [illegible] | 1183 | Bacteria>Pseudomonadota>Alphaproteobacteria>Hyphomicrobiales>Nitrobacteraceae>Bradyrhizobium | GCF_020194035.1 | Bradyrhizobium_sp._BRP56 | I6F38_19500 | 73459..77010 | 2793819 | BREX-system-P-loop-protein-BrxC-[Bradyrhizobium-sp.-BRP56] |
| WP_260373497.1 | Type1.BREX | [illegible] | 1183 | Bacteria>Pseudomonadota>Alphaproteobacteria>Hyphomicrobiales>Nitrobacteraceae>Bradyrhizobium | GCF_025200925.1 | Bradyrhizobium_sp._CB1015 | N2604_RS01610 | 347161..350712 | 2976822 | BREX-system-P-loop-protein-BrxC-[Bradyrhizobium-sp.-CB1015] |
| RWD70309.1 | Type6.BREX | [illegible] | 1181 | Bacteria>Pseudomonadota>Alphaproteobacteria>Hyphomicrobiales>Phyllobacteriaceae>Mesorhizobium | GCA_004018245.1 | Mesorhizobium_sp. | EOS37_15285 | 26400..29945 | 1871066 | BREX-system-P-loop-protein-BrxC-[Mesorhizobium-sp.] |
| TIL20465.1 | Type6.BREX | [illegible] | 1208 | Bacteria>Pseudomonadota>Alphaproteobacteria>Hyphomicrobiales>Phyllobacteriaceae>Mesorhizobium | GCA_004961195.1 | Mesorhizobium_sp. | E5Y88_32415 | 5019..8645 | 1871066 | BREX-system-P-loop-protein-BrxC-[Mesorhizobium-sp.] |
| WP_023809232.1 | Type1.BREX | [illegible] | 1150 | Bacteria>Pseudomonadota>Alphaproteobacteria>Hyphomicrobiales>Phyllobacteriaceae>Mesorhizobium | GCF_000502875.1 | Mesorhizobium_sp._L2C084A000 | X735_RS161770 | 861519..864971 | 1287116 | BREX-system-P-loop-protein-BrxC-[Mesorhizobium-sp.-L2C084A000] |
| WP_128286158.1 | Type1.BREX | [illegible] | 1181 | Bacteria>Pseudomonadota>Alphaproteobacteria>Hyphomicrobiales>Phyllobacteriaceae>Mesorhizobium | GCF_004023115.1 | Mesorhizobium_sp._M6A.T.Cr.TU.017.01.1.1 | EOB36_RS28010 | 14606..18151 | 2496774 | BREX-system-P-loop-protein-BrxC-[Mesorhizobium-sp.-M6A.T.Cr.TU.017.01.1.1] |
| MBX5048875.1 | Type6.BREX | [illegible] | 1213 | Bacteria>Pseudomonadota>Alphaproteobacteria>Hyphomicrobiales>Rhizobiaceae>Rhizobium/Agrobacterium_group | GCA_019684595.1 | Rhizobium_lentis | HJB64_19910 | 192083..195724 | 1138194 | BREX-system-P-loop-protein-BrxC-[Rhizobium-lentis] |
| MBY5873817.1 | Type1.BREX | [illegible] | 1174 | Bacteria>Pseudomonadota>Alphaproteobacteria>Hyphomicrobiales>Rhizobiaceae>Rhizobium/Agrobacterium_group | GCA_019799745.1 | Rhizobium_leguminosarum | HFN53_17500 | 401377..404901 | 384 | BREX-system-P-loop-protein-BrxC-[Rhizobium-leguminosarum] |
| WP_168349960.1 | Type1.BREX | [illegible] | 190 | Bacteria>Pseudomonadota>Alphaproteobacteria>Hyphomicrobiales>Rhizobiaceae>Rhizobium/Agrobacterium_group | GCF_012276085.1 | Rhizobium_leguminosarum | GFM28_RS29310 | 8641..9213 | 384 | hypothetical-protein-[Rhizobium-leguminosarum] |
| WP_250810295.1 | Type6.BREX | [illegible] | 1208 | Bacteria>Pseudomonadota>Alphaproteobacteria>Hyphomicrobiales>Rhizobiaceae>Rhizobium/Agrobacterium_group | GCF_023667605.1 | Neorhizobium_tomejilense | L1082_RS18485 | 51181..54807 | 2093828 | BREX-system-P-loop-protein-BrxC-[Neorhizobium-tomejilense] |
| WP_221145756.1 | Type6.BREX | [illegible] | 1213 | Bacteria>Pseudomonadota>Alphaproteobacteria>Hyphomicrobiales>Rhizobiaceae>Rhizobium/Agrobacterium_group | - | Rhizobium_lentis | HJB64_19910 | 192083..195724 | 1138194 | BREX-system-P-loop-protein-BrxC-[Rhizobium-lentis] |
| WP_027288600.1 | Type1.BREX | [illegible] | 1150 | Bacteria>Pseudomonadota>Alphaproteobacteria>Hyphomicrobiales>Rhodovibrionaceae>Rhodovibrio | GCF_000515255.1 | Rhodovibrio_salinarum | RHOSA_RS0110320 | 2222437..2225889 | 1087 | BREX-system-P-loop-protein-BrxC-[Rhodovibrio-salinarum] |
| WP_212538477.1 | Type6.BREX | [illegible] | 1208 | Bacteria>Pseudomonadota>Alphaproteobacteria>Rhodobacterales>Paracoccaceae>Aestuariicoccus | GCF_018224955.1 | Aestuariicoccus_sp._KMU-90 | KB874_RS20730 | 16568..20194 | 2827239 | BREX-system-P-loop-protein-BrxC-[Aestuariicoccus-sp.-KMU-90] |
| WP_274653192.1 | Type1.BREX | [illegible] | 1172 | Bacteria>Pseudomonadota>Alphaproteobacteria>Rhodobacterales>Paracoccaceae>Halocynthiibacter | GCF_028823805.1 | Halocynthiibacter_sp._C4 | OU789_RS03095 | 642215..645733 | 2992758 | BREX-system-P-loop-protein-BrxC-[Halocynthiibacter-sp.-C4] |
| WP_108641830.1 | Type1.BREX | [illegible] | 1173 | Bacteria>Pseudomonadota>Alphaproteobacteria>Rhodobacterales>Paracoccaceae>Thalassorhabdomicrobium | GCF_003072065.1 | Thalassorhabdomicrobium_marinisediminis | DC363_RS14250 | 78584..82105 | 2170577 | BREX-system-P-loop-protein-BrxC-[Thalassorhabdomicrobium-marinisediminis] |
| WP_058289795.1 | Type1.BREX | [illegible] | 1172 | Bacteria>Pseudomonadota>Alphaproteobacteria>Rhodobacterales>Paracoccaceae>Tritonibacter | GCF_001458415.1 | Tritonibacter_multivorans | PGB32_RS12460 | 15411..18929 | 928856 | BREX-system-P-loop-protein-BrxC-[Tritonibacter-multivorans] |
| WP_089279817.1 | Type1.BREX | [illegible] | 1147 | Bacteria>Pseudomonadota>Alphaproteobacteria>Rhodobacterales>Roseobacteraceae>Antarctobacter | GCF_900188425.1 | Antarctobacter_heliothermus | CHB90_RS21205 | 7452..10895 | 74033 | BREX-system-P-loop-protein-BrxC-[Antarctobacter-heliothermus] |
| WP_123792783.1 | Type1.BREX | [illegible] | 1172 | Bacteria>Pseudomonadota>Alphaproteobacteria>Rhodobacterales>Roseobacteraceae>Pacificibacter | GCF_003814915.1 | Pacificibacter_maritimus | EDD53_RS08420 | 394668..398186 | 762213 | BREX-system-P-loop-protein-BrxC-[Pacificibacter-maritimus] |
| MAE92765.1 | Type1.BREX | [illegible] | 1219 | Bacteria>Pseudomonadota>Alphaproteobacteria>Rhodobacterales>Roseobacteraceae>Pelagibaca | GCA_002687285.1 | Pelagibaca_sp. | CMI67_24850 | 4770..8429 | 1979400 | ATPase-[Pelagibaca-sp.] |
| WP_201727547.1 | Type6.BREX | [illegible] | 650 | Bacteria>Pseudomonadota>Alphaproteobacteria>Rhodospirillales>Acetobacteraceae>Acidocella | GCF_907164655.1 | Acidocella_sp._C78 | ACIDOCELLA78_RS32075 | 228724..230676 | 1671486 | BREX-system-P-loop-protein-BrxC-[Acidocella-sp.-C78] |
| WP_201727548.1 | Type6.BREX | [illegible] | 495 | Bacteria>Pseudomonadota>Alphaproteobacteria>Rhodospirillales>Acetobacteraceae>Acidocella | GCF_907164655.1 | Acidocella_sp._C78 | ACIDOCELLA78_RS32100 | 233672..235159 | 1671486 | hypothetical-protein-[Acidocella-sp.-C78] |
| HCW69913.1 | Type6.BREX | [illegible] | 1208 | Bacteria>Pseudomonadota>Alphaproteobacteria>Rhodospirillales>Thalassospiraceae>Thalassospira | GCA_003539675.1 | Thalassospira_lucentensis | DHR80_22445 | 45804..49430 | 168935 | BREX-system-P-loop-protein-BrxC-[Thalassospira-lucentensis] |
| TVR84541.1 | Type6.BREX | [illegible] | 1206 | Bacteria>Pseudomonadota>Alphaproteobacteria>Rhodospirillales>unclassified_Rhodospirillales | GCA_007695045.1 | Rhodospirillales_bacterium | EA405_01435 | 12860..16480 | 2026786 | BREX-system-P-loop-protein-BrxC-[Rhodospirillales-bacterium] |
| WP_193715186.1 | Type6.BREX | [illegible] | 1215 | Bacteria>Pseudomonadota>Alphaproteobacteria>Sneathiellales>Sneathiellaceae>Sneathiella | GCF_015143595.1 | Sneathiella_sp._P13V-1 | GUA87_RS03840 | 775657..779304 | 2697366 | BREX-system-P-loop-protein-BrxC-[Sneathiella-sp.-P13V-1] |
| MBN8436950.1 | Type1.BREX | [illegible] | 1146 | Bacteria>Pseudomonadota>Betaproteobacteria>Betaproteobacteria_incertae_sedis>Candidatus_Accumulibacter>unclassified_Candidatus_Accumulibacter | GCA_017302555.1 | Candidatus_Accumulibacter_sp. | JOL95_02725 | 187324..190764 | 2053492 | BREX-system-P-loop-protein-BrxC-[Candidatus-Accumulibacter-sp.] |
| WP_042797405.1 | Type6.BREX | [illegible] | 1207 | Bacteria>Pseudomonadota>Betaproteobacteria>Burkholderiales>Alcaligenaceae>Achromobacter | GCF_000783435.2 | Achromobacter_insolitus | MC81_20460 | 4405602..4409225 | 217204 | BREX-system-P-loop-protein-BrxC-[Achromobacter-insolitus] |
| WP_050447538.1 | Type1.BREX | [illegible] | 1153 | Bacteria>Pseudomonadota>Betaproteobacteria>Burkholderiales>Alcaligenaceae>Achromobacter | GCF_001189595.1 | Achromobacter_spanius | AFM18_RS14295 | 68240..71701 | 217203 | BREX-system-P-loop-protein-BrxC-[Achromobacter-spanius] |
| WP_063584995.1 | Type1.BREX | [illegible] | 1146 | Bacteria>Pseudomonadota>Betaproteobacteria>Burkholderiales>Alcaligenaceae>Achromobacter | GCF_001637105.1 | Achromobacter_ruhlandii | AOT39_RS22745 | 36149..39589 | 72557 | BREX-system-P-loop-protein-BrxC-[Achromobacter-ruhlandii] |
| WP_076471941.1 | Type6.BREX | [illegible] | 1207 | Bacteria>Pseudomonadota>Betaproteobacteria>Burkholderiales>Alcaligenaceae>Achromobacter | GCF_001957055.1 | Achromobacter_xylosoxidans | BIZ53_RS00410 | 81330..84953 | 85698 | BREX-system-P-loop-protein-BrxC-[Achromobacter-xylosoxidans] |
| WP_212312045.1 | Type6.BREX | [illegible] | 1217 | Bacteria>Pseudomonadota>Betaproteobacteria>Burkholderiales>Alcaligenaceae>Achromobacter | GCF_018195695.1 | Achromobacter_sp._Marseille-Q0513 | KDH83_RS00260 | 59261..62914 | 2829161 | BREX-system-P-loop-protein-BrxC-[Achromobacter-sp.-Marseille-Q0513] |
| WP_233436978.1 | Type6.BREX | [illegible] | 1234 | Bacteria>Pseudomonadota>Betaproteobacteria>Burkholderiales>Alcaligenaceae>Achromobacter | GCF_022976495.1 | Achromobacter_xylosoxidans | DBA06_RS11680 | 120350..124054 | 85698 | BREX-system-P-loop-protein-BrxC-[Achromobacter-xylosoxidans] |
| WP_259250627.1 | Type6.BREX | [illegible] | 1213 | Bacteria>Pseudomonadota>Betaproteobacteria>Burkholderiales>Alcaligenaceae>Achromobacter | GCF_024806975.1 | Achromobacter_sp._JUb104 | M2361_RS16380 | 989592..993233 | 2940590 | BREX-system-P-loop-protein-BrxC-[Achromobacter-sp.-JUb104] |
| WP_180097564.1 | Type6.BREX | [illegible] | 1213 | Bacteria>Pseudomonadota>Betaproteobacteria>Burkholderiales>Alcaligenaceae>Achromobacter | GCF_902859995.1 | Achromobacter_mucicolens | H0520_RS02915 | 616832..620473 | 1389922 | BREX-system-P-loop-protein-BrxC-[Achromobacter-mucicolens] |
| WP_200235209.1 | Type1.BREX | [illegible] | 1190 | Bacteria>Pseudomonadota>Betaproteobacteria>Burkholderiales>Alcaligenaceae>Advenella | GCF_016595155.1 | Advenella_mandrilli | JHL22_RS06635 | 201531..205103 | 2800330 | BREX-system-P-loop-protein-BrxC-[Advenella-mandrilli] |
| WP_109088944.1 | Type1.BREX | [illegible] | 1146 | Bacteria>Pseudomonadota>Betaproteobacteria>Burkholderiales>Alcaligenaceae>Alcaligenes | GCF_003122065.1 | Alcaligenes_faecalis | DF183_RS09485 | 797094..800534 | 511 | BREX-system-P-loop-protein-BrxC-[Alcaligenes-faecalis] |
| WP_199265315.1 | Type6.BREX | [illegible] | 1213 | Bacteria>Pseudomonadota>Betaproteobacteria>Burkholderiales>Alcaligenaceae>Alcaligenes | GCF_016446305.1 | Alcaligenes_faecalis | GTU60_RS02720 | 606582..610223 | 511 | BREX-system-P-loop-protein-BrxC-[Alcaligenes-faecalis] |
| WP_197022064.1 | Type1.BREX | [illegible] | 1150 | Bacteria>Pseudomonadota>Betaproteobacteria>Burkholderiales>Alcaligenaceae>Bordetella | GCF_000518845.1 | Bordetella_petrii | L636_RS0102150 | 157001..160453 | 94624 | BREX-system-P-loop-protein-BrxC-[Bordetella-petrii] |
| WP_032972587.1 | Type6.BREX | [illegible] | 1213 | Bacteria>Pseudomonadota>Betaproteobacteria>Burkholderiales>Alcaligenaceae>Bordetella | GCF_006770345.1 | Bordetella_hinzii | CBR67_RS06420 | 1279803..1283444 | 103855 | BREX-system-P-loop-protein-BrxC-[Bordetella-hinzii] |
| WP_148812759.1 | Type1.BREX | [illegible] | 1153 | Bacteria>Pseudomonadota>Betaproteobacteria>Burkholderiales>Alcaligenaceae>Pigmentiphaga | GCF_008119665.1 | Pigmentiphaga_aceris | FXN63_RS03140 | 736142..739603 | 1940612 | BREX-system-P-loop-protein-BrxC-[Pigmentiphaga-aceris] |
| RIY39966.1 | Type1.BREX | [illegible] | 1145 | Bacteria>Pseudomonadota>Betaproteobacteria>Burkholderiales>Alcaligenaceae>Pusillimonas | GCA_003585905.1 | Pusillimonas_maritima | CJP73_12365 | 2891..6328 | 2026239 | hypothetical-protein-CJP73_12365-[Pusillimonas-maritima] |
| WP_218972830.1 | Type1.BREX | [illegible] | 1146 | Bacteria>Pseudomonadota>Betaproteobacteria>Burkholderiales>Alcaligenaceae>Pusillimonas | GCF_003585905.1 | Pusillimonas_maritima | CJP73_RS12395 | 2888..6328 | 2026239 | BREX-system-P-loop-protein-BrxC-[Pusillimonas-maritima] |
| NYT63370.1 | Type1.BREX | [illegible] | 1191 | Bacteria>Pseudomonadota>Betaproteobacteria>Burkholderiales>Alcaligenaceae>unclassified_Alcaligenaceae | GCA_013416445.1 | Alcaligenaceae_bacterium | H0A66_13705 | 63657..67232 | 2021368 | BREX-system-P-loop-protein-BrxC-[Alcaligenaceae-bacterium] |
| MCA8485082.1 | Type1.BREX | [illegible] | 879 | Bacteria>Pseudomonadota>Betaproteobacteria>Burkholderiales>Burkholderiaceae>Burkholderia | GCA_020421385.1 | Burkholderia_multivorans | LGM85_14180 | 202982..205621 | 87883 | BREX-system-P-loop-protein-BrxC-[Burkholderia-multivorans] |
| MCA8485083.1 | Type1.BREX | [illegible] | 266 | Bacteria>Pseudomonadota>Betaproteobacteria>Burkholderiales>Burkholderiaceae>Burkholderia | GCA_020421385.1 | Burkholderia_multivorans | LGM85_14185 | 205622..206422 | 87883 | hypothetical-protein-[Burkholderia-multivorans] |
| WP_059944780.1 | Type6.BREX | [illegible] | 1213 | Bacteria>Pseudomonadota>Betaproteobacteria>Burkholderiales>Burkholderiaceae>Burkholderia | GCF_001527205.1 | Burkholderia_territorii | WT00_RS19595 | 122446..126087 | 1503055 | BREX-system-P-loop-protein-BrxC-[Burkholderia-territorii] |
| WP_059936887.1 | Type6.BREX | [illegible] | 1213 | Bacteria>Pseudomonadota>Betaproteobacteria>Burkholderiales>Burkholderiaceae>Burkholderia | GCF_001528605.1 | Burkholderia_multivorans | WK33_RS03230 | 12913..16554 | 87883 | BREX-system-P-loop-protein-BrxC-[Burkholderia-multivorans] |
| WP_060225079.1 | Type1.BREX | [illegible] | 1146 | Bacteria>Pseudomonadota>Betaproteobacteria>Burkholderiales>Burkholderiaceae>Burkholderia | GCF_001531525.1 | Burkholderia_cepacia | WL34_RS17275 | 238513..241953 | 292 | BREX-system-P-loop-protein-BrxC-[Burkholderia-cepacia] |
| WP_226255239.1 | Type6.BREX | [illegible] | 1208 | Bacteria>Pseudomonadota>Betaproteobacteria>Burkholderiales>Burkholderiaceae>Burkholderia | GCF_020419005.1 | Burkholderia_cepacia | LGM39_RS24930 | 366713..370339 | 292 | BREX-system-P-loop-protein-BrxC-[Burkholderia-cepacia] |
| WP_186043215.1 | Type6.BREX | [illegible] | 1213 | Bacteria>Pseudomonadota>Betaproteobacteria>Burkholderiales>Burkholderiaceae>Burkholderia | GCF_902830235.1 | Burkholderia_gladioli | H9C96_RS14825 | 15196..18837 | 28095 | BREX-system-P-loop-protein-BrxC-[Burkholderia-gladioli] |
| WP_186236015.1 | Type6.BREX | [illegible] | 1215 | Bacteria>Pseudomonadota>Betaproteobacteria>Burkholderiales>Burkholderiaceae>Burkholderia | GCF_902832475.1 | Burkholderia_gladioli | H9G27_RS00755 | 169921..173568 | 28095 | BREX-system-P-loop-protein-BrxC-[Burkholderia-gladioli] |
| WP_091908362.1 | Type6.BREX | [illegible] | 1213 | Bacteria>Pseudomonadota>Betaproteobacteria>Burkholderiales>Burkholderiaceae>Chitinasiproducens | GCF_900095735.1 | Chitinasiproducens_palmae | BLU54_RS11860 | 213511..217152 | 1770053 | BREX-system-P-loop-protein-BrxC-[Chitinasiproducens-palmae] |
| WP_125094459.1 | Type1.BREX | [illegible] | 1146 | Bacteria>Pseudomonadota>Betaproteobacteria>Burkholderiales>Burkholderiaceae>Lautropia | GCF_003892345.1 | Lautropia_dentalis | EHV23_RS01850 | 480599..484039 | 2490857 | BREX-system-P-loop-protein-BrxC-[Lautropia-dentalis] |
| WP_275272077.1 | Type1.BREX | [illegible] | 1193 | Bacteria>Pseudomonadota>Betaproteobacteria>Burkholderiales>Burkholderiaceae>Limnobacter | GCF_029087225.1 | Limnobacter_sp._P1 | PY990_RS13680 | 12633..16214 | 3031298 | BREX-system-P-loop-protein-BrxC-[Limnobacter-sp.-P1] |
| MCK6421971.1 | Type1.BREX | [illegible] | 1153 | Bacteria>Pseudomonadota>Betaproteobacteria>Burkholderiales>Burkholderiales_genera_incertae_sedis>Aquabacterium | GCA_023150345.1 | Aquabacterium_sp. | L6Q73_13845 | 24265..27726 | 1872578 | BREX-system-P-loop-protein-BrxC-[Aquabacterium-sp.] |
| WP_054257718.1 | Type6.BREX | [illegible] | 1215 | Bacteria>Pseudomonadota>Betaproteobacteria>Burkholderiales>Comamonadaceae>Acidovorax | GCF_001298675.1 | Acidovorax_caeni | BN2503_RS16450 | 13891..17538 | 343013 | BREX-system-P-loop-protein-BrxC-[Acidovorax-caeni] |
| WP_057202647.1 | Type6.BREX | [illegible] | 1207 | Bacteria>Pseudomonadota>Betaproteobacteria>Burkholderiales>Comamonadaceae>Acidovorax | GCF_001428645.1 | Acidovorax_sp._Root217 | ASE31_RS15170 | 91296..94919 | 1736492 | BREX-system-P-loop-protein-BrxC-[Acidovorax-sp.-Root217] |
| WP_086927320.1 | Type1.BREX | [illegible] | 1153 | Bacteria>Pseudomonadota>Betaproteobacteria>Burkholderiales>Comamonadaceae>Acidovorax | GCF_002157125.1 | Acidovorax_carolinensis | CBP36_RS10055 | 2162910..2166371 | 553814 | BREX-system-P-loop-protein-BrxC-[Acidovorax-carolinensis] |
| WP_204247869.1 | Type1.BREX | [illegible] | 590 | Bacteria>Pseudomonadota>Betaproteobacteria>Burkholderiales>Comamonadaceae>Acidovorax | GCF_002157185.1 | Acidovorax_carolinensis | CBP35_RS21920 | 1914645..1916417 | 553814 | BREX-system-P-loop-protein-BrxC-[Acidovorax-carolinensis] |
| WP_034379112.1 | Type6.BREX | [illegible] | 1210 | Bacteria>Pseudomonadota>Betaproteobacteria>Burkholderiales>Comamonadaceae>Comamonas | GCF_000761335.1 | Comamonas_thiooxydans | P245_RS10570 | 85855..89487 | 363952 | BREX-system-P-loop-protein-BrxC-[Comamonas-thiooxydans] |
| KEH09926.1 | Type1.BREX | [illegible] | 1185 | Bacteria>Pseudomonadota>Betaproteobacteria>Burkholderiales>Comamonadaceae>Delftia | GCA_000710165.1 | Delftia_tsuruhatensis | GY14_08945 | 17729..21286 | 180282 | hypothetical-protein-GY14_08945-[Delftia-tsuruhatensis] |

| Protein-ID | System | Gene-Neighbourhoods | Length | Taxonomy | Assembly-ID | Strain | Locus-Tag | Genomic-coordinates | TaxID | Fasta-define |
|---|---|---|---|---|---|---|---|---|---|---|
| WP_101103324.1 | Type1.BREX | [illegible] | 1153 | [illegible] | GCF_002837135.1 | Macromonas_bipunctata | CON76_RS10585 | 3906..7367 | 183670 | [illegible] |
| CDS49862.1 | Type6.BREX | [illegible] | 1210 | [illegible] | - | Polaromonas_sp._CG9_12 | . | 485322..488954 | 1504672 | [illegible] |
| WP_169420224.1 | Type1.BREX | [illegible] | 1153 | [illegible] | GCF_012927085.1 | Ramlibacter_agri | HHL11_RS19385 | 189581..193042 | 2728837 | [illegible] |
| WP_068673039.1 | Type6.BREX | [illegible] | 1212 | [illegible] | GCF_001541225.1 | unclassified_Variovorax | APY03_RS01000 | 202384..206022 | 663243 | [illegible] |
| WP_186915942.1 | Type1.BREX | [illegible] | 1183 | [illegible] | GCF_014284115.1 | Undibacterium | H8K35_RS08325 | 157325..160876 | 401469 | [illegible] |
| WP_141835430.1 | Type6.BREX | [illegible] | 1209 | [illegible] | - | Herbaspirillum_sp._SJZ107 | FBX97_RS19505 | 315565..319194 | 2572881 | [illegible] |
| WP_106758218.1 | Type1.BREX | [illegible] | 1154 | [illegible] | GCF_003011895.2 | Massilia_glaciei | C7C56_RS15200 | 27208..30672 | 1524097 | [illegible] |
| WP_152879050.1 | Type6.BREX | [illegible] | 1215 | [illegible] | GCF_009295655.1 | Duganella_sp._FT27W | F8197_RS00385 | 74619..78266 | 2654636 | [illegible] |
| OGB35307.1 | Type6.BREX | [illegible] | 1215 | [illegible] | GCA_001770935.1 | Burkholderiales_bacterium_RIFCSPLOWO2_12_FULL_61_40 | A3F78_11695 | 46897..50544 | 1797566 | [illegible] |
| MCP5275500.1 | Type1.BREX | [illegible] | 1176 | [illegible] | GCA_024235795.1 | Burkholderiales_bacterium | H6936_11775 | 653459..656989 | 1891238 | [illegible] |
| WP_194117489.1 | Type6.BREX | [illegible] | 1213 | [illegible] | GCF_015223195.1 | Chitinilyticum_piscinae | INR99_RS16780 | 39673..43314 | 2866724 | [illegible] |
| WP_153715435.1 | Type6.BREX | [illegible] | 1213 | [illegible] | GCF_009650905.1 | Eikenella_corrodens | GJ661_RS08680 | 18367..22008 | 539 | [illegible] |
| ROH85757.1 | Type1.BREX | [illegible] | 1146 | [illegible] | GCA_003730115.1 | Pseudomethylobacillus_aquaticus | ED236_08405 | 68439..71879 | 2676064 | [illegible] |
| WP_205125664.1 | Type1.BREX | [illegible] | 1150 | [illegible] | GCF_003730115.1 | Pseudomethylobacillus_aquaticus | ED236_RS08405 | 68439..71891 | 2676064 | [illegible] |
| SDW37048.1 | Type1.BREX | [illegible] | 1145 | [illegible] | GCA_900106545.1 | Nitrosomonas_communis | SAMN05421882_100921 | 53532..56969 | 44574 | [illegible] |
| WP_181258202.1 | Type1.BREX | [illegible] | 1179 | [illegible] | GCF_003046585.1 | Nitrosomonas_aestuarii | C8R11_107137 | 171439..174978 | 52441 | [illegible] |
| WP_211752264.1 | Type1.BREX | [illegible] | 1146 | [illegible] | GCF_900106545.1 | Nitrosomonas_communis | BLV21_RS05435 | 53532..56972 | 44574 | [illegible] |
| SFW23276.1 | Type6.BREX | [illegible] | 1213 | [illegible] | GCA_900119085.1 | Nitrosovibrio_sp._Nv17 | SAMN05216414_10762 | 70156..73797 | 1855339 | [illegible] |
| WP_218184829.1 | Type6.BREX | [illegible] | 1234 | [illegible] | GCF_900119085.1 | Nitrosovibrio_sp._Nv17 | BT108_RS05915 | 70156..73860 | 1855339 | [illegible] |
| MBW8470844.1 | Type6.BREX | [illegible] | 1215 | [illegible] | GCA_019459125.1 | Thiobacillus_sp. | KOM67_21445 | 16263..19910 | 924 | [illegible] |
| MBK9161130.1 | Type1.BREX | [illegible] | 1146 | [illegible] | GCA_016716325.1 | Nitrosomonadales_bacterium | IPM27_06155 | 1292283..1295723 | 2026761 | [illegible] |
| MBP6421402.1 | Type1.BREX | [illegible] | 923 | [illegible] | GCA_017988215.1 | Propionivibrio_sp. | KA271_00765 | 12654..15427 | 2212460 | [illegible] |
| MCA3121411.1 | Type6.BREX | [illegible] | 1210 | [illegible] | GCA_020248305.1 | Rhodocyclaceae_bacterium | ING90_02100 | 367371..371003 | 1898103 | [illegible] |
| WP_141017182.1 | Type6.BREX | [illegible] | 1207 | [illegible] | GCF_006496635.1 | Azoarcus_sp._DD4 | CJ010_RS05935 | 1245819..1249442 | 2027405 | [illegible] |
| WP_148578351.1 | Type1.BREX | [illegible] | 1146 | [illegible] | GCF_008107625.1 | Zoogloea_oleivorans | ETQ85_RS07180 | 242245..245685 | 1552750 | [illegible] |
| AUZ77883.1 | Type1.BREX | [illegible] | 1183 | [illegible] | GCA_002934425.1 | Aeromonas_sp._ASNIH4 | C2U40_16585 | 3601638..3605189 | 1636609 | [illegible] |
| BBR09393.1 | Type1.BREX | [illegible] | 1226 | [illegible] | GCA_014158455.1 | Aeromonas_caviae | WP3S18E02_10540 | 1118346..1122026 | 648 | [illegible] |
| EGX6956813.1 | Type1.BREX | [illegible] | 1218 | [illegible] | GCA_016414985.1 | Aeromonas_hydrophila | JFQ85_000577 | 650804..654460 | 644 | [illegible] |
| MBS4697267.1 | Type1.BREX | [illegible] | 1226 | [illegible] | GCA_018359965.1 | Aeromonas_allosaccharophila | J4G65_17610 | 87511..91191 | 656 | [illegible] |
| BDO08311.1 | Type1.BREX | [illegible] | 1089 | [illegible] | GCA_024347775.1 | Aeromonas_caviae | KAM643c_18840 | 2028863..2032132 | 648 | [illegible] |
| WP_010634561.1 | Type1.BREX | [illegible] | 1157 | [illegible] | GCF_000315195.1 | Aeromonas_dhakensis | AQU_RS10070 | 136801..140274 | 196024 | [illegible] |
| WP_043557612.1 | Type1.BREX | [illegible] | 1182 | [illegible] | GCF_000819745.1 | Aeromonas_bestiarum | BN1118_RS19650 | 69736..73284 | 105751 | [illegible] |
| WP_087756130.1 | Type1.BREX | [illegible] | 1231 | [illegible] | GCF_002180335.1 | Aeromonas_salmonicida | O23A_RS10590 | 2337193..2340888 | 645 | [illegible] |
| WP_095593101.1 | Type1.BREX | [illegible] | 1231 | [illegible] | GCF_002285935.1 | Aeromonas_dhakensis | CK627_RS18865 | 4072109..4075804 | 196024 | [illegible] |
| WP_103244178.1 | Type6.BREX | [illegible] | 1210 | [illegible] | GCF_002902865.1 | Aeromonas_sp._ASNIH5 | C2U30_RS17315 | 3671715..3675347 | 1758179 | [illegible] |
| WP_120415687.1 | Type1.BREX | [illegible] | 1231 | [illegible] | GCF_003611985.1 | Aeromonas_veronii | D6R50_RS16910 | 827539..831234 | 654 | [illegible] |
| WP_139442435.1 | Type5.BREX | [illegible] | 1231 | [illegible] | GCF_006246325.1 | Aeromonas_veronii | CF108_RS03265 | 124178..127873 | 654 | [illegible] |
| WP_159144201.1 | Type1.BREX | [illegible] | 1231 | [illegible] | GCF_009834065.1 | Aeromonas_veronii | GRF56_RS18105 | 3933894..3937589 | 654 | [illegible] |
| WP_171280306.1 | Type1.BREX | [illegible] | 1157 | [illegible] | GCF_013085505.1 | Aeromonas_sp._2692-1 | E5E97_RS12155 | 2531425..2534898 | 2560029 | [illegible] |
| WP_232090117.1 | Type1.BREX | [illegible] | 1281 | [illegible] | GCF_014158455.1 | Aeromonas_caviae | H7X73_RS05535 | 1118181..1122026 | 648 | [illegible] |
| WP_193000848.1 | Type1.BREX | [illegible] | 1226 | [illegible] | GCF_014892695.1 | Aeromonas_simiae | FE240_RS10745 | 2272968..2276648 | 218936 | [illegible] |
| WP_244842130.1 | Type1.BREX | [illegible] | 1281 | [illegible] | GCF_018359965.1 | Aeromonas_allosaccharophila | J4G65_RS17605 | 87511..91356 | 656 | [illegible] |
| WP_203133976.1 | Type1.BREX | [illegible] | 1156 | [illegible] | GCF_019348745.1 | Aeromonas_hydrophila | PVL97_RS11265 | 2527654..2531124 | 644 | [illegible] |
| WP_223931722.1 | Type1.BREX | [illegible] | 1231 | [illegible] | GCF_019972495.1 | Aeromonas_caviae | LDN23_RS02850 | 18313..22008 | 648 | [illegible] |
| WP_241871100.1 | Type1.BREX | [illegible] | 1157 | [illegible] | GCF_022631175.1 | Aeromonas_hydrophila | MOO40_RS17050 | 3752340..3755813 | 644 | [illegible] |
| WP_261925329.1 | Type1.BREX | [illegible] | 1089 | [illegible] | GCF_024347775.1 | Aeromonas_caviae | OCX50_RS09760 | 2028863..2032132 | 648 | [illegible] |
| WP_257706959.1 | Type1.BREX | [illegible] | 1182 | [illegible] | GCF_024649325.1 | Aeromonas_caviae | NUK33_RS01495 | 150155..153703 | 648 | [illegible] |
| WP_270809144.1 | Type1.BREX | [illegible] | 1157 | [illegible] | GCF_027705685.1 | Aeromonas_sp._QDB08 | OKB79_RS12620 | 11744..15217 | 2990480 | [illegible] |
| WP_181014924.1 | Type1.BREX | [illegible] | 1182 | [illegible] | GCF_002934425.1 | Aeromonas | C2U40_RS16580 | 3601638..3605186 | 642 | [illegible] |
| WP_201934227.1 | Type1.BREX | [illegible] | 1231 | [illegible] | GCF_019972675.1 | Aeromonas | MWG22_RS17445 | 1818..5513 | 642 | [illegible] |
| WP_270796656.1 | Type1.BREX | [illegible] | 1157 | [illegible] | GCF_029024985.1 | Aeromonas | OKC40_RS19420 | 7580..11053 | 642 | [illegible] |
| WP_069785631.1 | Type1.BREX | [illegible] | 1157 | [illegible] | GCF_029773025.1 | Aeromonas | QBS29_RS17430 | 3038195..3041668 | 642 | [illegible] |
| MCR4553833.1 | Type1.BREX | [illegible] | 1184 | [illegible] | GCA_024683965.1 | Succinivibrionaceae_bacterium | K5752_02815 | 10310..13864 | 2053620 | [illegible] |
| WP_057786312.1 | Type1.BREX | [illegible] | 1157 | [illegible] | GCF_001441695.1 | Alishewanella_sp._WH16-1 | AAY72_RS06860 | 5881..9354 | 1651088 | [illegible] |
| WP_207611650.1 | Type1.BREX | [illegible] | 1157 | [illegible] | GCF_017368475.1 | Alteromonas_sp._5E99-2 | J3L16_RS04210 | 79358..82831 | 2817683 | [illegible] |
| WP_240919947.1 | Type1.BREX | [illegible] | 1180 | [illegible] | GCF_011378905.1 | Paraglaciecola_sp._20A4 | GQS54_RS13935 | 91436..94978 | 2687288 | [illegible] |
| AWB59509.1 | Type1.BREX | [illegible] | 1232 | [illegible] | GCA_003061515.1 | Colwellia_sp._Arc7-D | DBO93_09830 | 2239405..2243103 | 2161872 | [illegible] |
| WP_239058953.1 | Type1.BREX | [illegible] | 1235 | [illegible] | GCF_003061515.1 | Colwellia_sp._Arc7-D | DBO93_RS09830 | 2239405..2243112 | 2161872 | [illegible] |
| WP_085284213.1 | Type1.BREX | [illegible] | 1243 | [illegible] | GCF_900109795.1 | Colwellia_chukchiensis | B6K83_RS06980 | 108494..112225 | 641665 | [illegible] |
| KXS36208.1 | Type1.BREX | [illegible] | 1150 | [illegible] | GCA_001577655.1 | Idiomarina_sp._T82-3 | AWU56_323 | 88359..91811 | 1794809 | [illegible] |
| WP_112714387.1 | Type1.BREX | [illegible] | 1199 | [illegible] | GCF_900465055.1 | Moritella_yayanosii | MORIYA_RS08605 | 1893116..1896715 | 69539 | [illegible] |
| WP_010607320.1 | Type1.BREX | [illegible] | 1188 | [illegible] | GCF_000276645.1 | Pseudoalteromonas | UY7_RS0121200 | 90697..94263 | 53246 | [illegible] |
| WP_054983666.1 | Type1.BREX | [illegible] | 1159 | [illegible] | GCF_001400005.1 | Pseudoalteromonas_sp._P1-30 | AN395_RS16640 | 61256..64735 | 1723760 | [illegible] |
| WP_064663367.1 | Type1.BREX | [illegible] | 1242 | [illegible] | GCF_001661605.1 | Pseudoalteromonas_sp._MQS005 | A9479_RS10285 | 16730..20458 | 1854052 | [illegible] |
| WP_234000034.1 | Type1.BREX | [illegible] | 1252 | [illegible] | GCF_002964725.1 | Pseudoalteromonas_sp._T1lg76 | C2M35_RS06890 | 48486..52244 | 2077103 | [illegible] |
| WP_130052623.1 | Type1.BREX | [illegible] | 1244 | [illegible] | GCF_004195145.1 | Pseudoalteromonas_sp._MEBiC_03485 | EVU92_RS18430 | 351830..355564 | 2571103 | [illegible] |
| WP_096038338.1 | Type1.BREX | [illegible] | 1159 | [illegible] | GCF_007991235.1 | Pseudoalteromonas_tetraodonis | PTE01_RS15665 | 26152..29631 | 43659 | [illegible] |
| WP_089368387.1 | Type1.BREX | [illegible] | 1229 | [illegible] | GCF_007991255.1 | Pseudoalteromonas_nigrifaciens | PNIG_RS09950 | 2060327..2064016 | 28109 | [illegible] |
| WP_167658112.1 | Type1.BREX | [illegible] | 1188 | [illegible] | GCF_011806445.1 | Pseudoalteromonas_sp._HF66 | HCZ11_RS07660 | 212811..216377 | 2721559 | [illegible] |
| WP_193038371.1 | Type1.BREX | [illegible] | 1159 | [illegible] | GCF_014897925.1 | Pseudoalteromonas_nigrifaciens | INT11_RS00030 | 7982..11461 | 28109 | [illegible] |
| WP_074988793.1 | Type1.BREX | [illegible] | 1242 | [illegible] | GCF_014925285.1 | Pseudoalteromonas_lipolytica | PLIP_RS19215 | 618195..621923 | 570156 | [illegible] |
| WP_037429580.1 | Type1.BREX | [illegible] | 1157 | [illegible] | GCF_000282755.1 | Shewanella | SHEWPOL2_RS15310 | 7136..10609 | 22 | [illegible] |
| PKG74712.1 | Type1.BREX | [illegible] | 1191 | [illegible] | GCA_002836135.1 | Shewanella_sp._GutCb | CXF86_11690 | 122253..125828 | 2058315 | [illegible] |
| HAY94623.1 | Type1.BREX | [illegible] | 1224 | [illegible] | GCA_003486485.1 | Shewanella_sp. | DCY70_12125 | 4933..8607 | 50422 | [illegible] |
| WP_011622483.1 | Type1.BREX | [illegible] | 1157 | [illegible] | GCF_000014685.1 | Shewanella_sp._MR-4 | SHEWMR4_RS09030 | 2019364..2022837 | 60480 | [illegible] |
| WP_011716828.1 | Type1.BREX | [illegible] | 1157 | [illegible] | GCF_000203935.1 | Shewanella_sp._ANA-3 | SHEWANA3_RS09590 | 2144277..2147750 | 94122 | [illegible] |
| WP_006086465.1 | Type1.BREX | [illegible] | 1229 | [illegible] | GCF_000231345.1 | Shewanella_baltica | SBAL625DRAFT_RS15525 | 212954..216643 | 62322 | [illegible] |
| WP_028764576.1 | Type1.BREX | [illegible] | 1157 | [illegible] | GCF_000518705.1 | Shewanella_colwelliana | L876_RS0115155 | 4861..8334 | 23 | [illegible] |
| WP_231570521.1 | Type1.BREX | [illegible] | 941 | [illegible] | GCF_000798835.1 | Shewanella_sp._ZOR0012 | L976_RS20155 | 4209796..4212621 | 1339231 | [illegible] |
| WP_055648628.1 | Type1.BREX | [illegible] | 1157 | [illegible] | GCF_001308045.1 | Shewanella_sp._Sh95 | AEA42_RS19845 | 36695..40168 | 1689868 | [illegible] |
| WP_080916542.1 | Type1.BREX | [illegible] | 1234 | [illegible] | GCF_002075795.1 | Shewanella_japonica | SJ2017_RS16905 | 3936240..3939944 | 93973 | [illegible] |
| WP_257985331.1 | Type1.BREX | [illegible] | 1193 | [illegible] | GCF_002836135.1 | Shewanella_sp._GutCb | CXF86_RS11730 | 122253..125834 | 2058315 | [illegible] |
| WP_108947326.1 | Type1.BREX | [illegible] | 1225 | [illegible] | GCF_003112715.1 | Shewanella_halifaxensis | C6V78_RS24360 | 248833..252510 | 271098 | [illegible] |
| WP_252977707.1 | Type1.BREX | [illegible] | 261 | [illegible] | GCF_013307415.1 | Shewanella_sp._ZOR0012 | HT094_RS07805 | 1783246..1784031 | 1339231 | [illegible] |
| WP_252977708.1 | Type1.BREX | [illegible] | 746 | [illegible] | GCF_013307415.1 | Shewanella_sp._ZOR0012 | HT094_RS07815 | 1785170..1787410 | 1339231 | [illegible] |
| WP_208124435.1 | Type1.BREX | [illegible] | 1157 | [illegible] | GCF_017570045.1 | Shewanella_algae | JKK45_RS07515 | 1709365..1712838 | 38313 | [illegible] |
| WP_101059463.1 | Type1.BREX | [illegible] | 1157 | [illegible] | GCF_017580385.1 | Shewanella_algae | CX652_RS12220 | 2312..5785 | 38313 | [illegible] |
| WP_144191359.1 | Type1.BREX | [illegible] | 1157 | [illegible] | GCF_017580415.1 | Shewanella_algae | I6M45_RS07395 | 495996..499469 | 38313 | [illegible] |
| WP_208223521.1 | Type1.BREX | [illegible] | 1157 | [illegible] | GCF_017580425.1 | Shewanella_algae | I6M46_RS07445 | 1788..5261 | 38313 | [illegible] |
| WP_220062217.1 | Type1.BREX | [illegible] | 1157 | [illegible] | GCF_019457515.1 | Shewanella_sp._FJAT-53870 | KOI73_RS17055 | 3880275..3883748 | 2864208 | [illegible] |
| WP_220768073.1 | Type1.BREX | [illegible] | 1225 | [illegible] | GCF_019654915.1 | Shewanella_sp._MBTL60-007 | K5Q92_RS05655 | 54742..58419 | 2815911 | [illegible] |
| WP_221058246.1 | Type1.BREX | [illegible] | 1199 | [illegible] | GCF_019670685.1 | Shewanella_algae | K5631_RS12960 | 2926333..2929932 | 38313 | [illegible] |
| WP_224020606.1 | Type1.BREX | [illegible] | 1157 | [illegible] | GCF_019973655.1 | Shewanella_xiamenensis | LDO18_RS12250 | 2746465..2749938 | 332186 | [illegible] |
| WP_248084385.1 | Type1.BREX | [illegible] | 1231 | [illegible] | GCF_023159095.1 | unclassified_Shewanella | MOD77_RS02670 | 216752..220447 | 196818 | [illegible] |
| WP_259476695.1 | Type1.BREX | [illegible] | 1225 | [illegible] | GCF_024972795.1 | Shewanella_xiamenensis | K5M76_RS09815 | 2212502..2216179 | 332186 | [illegible] |
| WP_259506670.1 | Type1.BREX | [illegible] | 1157 | [illegible] | GCF_024999425.1 | Shewanella_baltica | G3479_RS14595 | 93624..97097 | 62322 | [illegible] |
| WP_263145911.1 | Type1.BREX | [illegible] | 1224 | [illegible] | GCF_025653395.1 | Shewanella_sp._SW24 | L5M17_RS05810 | 187685..191359 | 2912815 | [illegible] |
| WP_263177453.1 | Type1.BREX | [illegible] | 1225 | [illegible] | GCF_025654415.1 | unclassified_Shewanella | L5M16_RS21425 | 11352..15029 | 196818 | [illegible] |
| WP_264921054.1 | Type1.BREX | [illegible] | 1157 | [illegible] | GCF_026013945.1 | Shewanella_algae | OLL83_RS07695 | 1703466..1706939 | 38313 | [illegible] |
| WP_275482470.1 | Type1.BREX | [illegible] | 1225 | [illegible] | GCF_029167175.1 | Shewanella_sp._A32 | PY479_RS00080 | 16338..20015 | 3031327 | [illegible] |
| WP_102039083.1 | Type1.BREX | [illegible] | 1191 | [illegible] | - | Shewanella_sp._GutCb | CXF86_11690 | 122253..125828 | 2058315 | [illegible] |
| MBV7434872.1 | Type1.BREX | [illegible] | 1175 | [illegible] | GCA_019218315.1 | Cardiobacteriaceae_bacterium_TAE3-ERU3 | KRX19_07510 | 264796..268323 | 2856340 | [illegible] |
| WP_265366287.1 | Type1.BREX | [illegible] | 1193 | [illegible] | GCF_026153455.1 | Marinimicrobium_koreense | OOT55_RS13025 | 3024461..3028042 | 306545 | [illegible] |
| WP_187410606.1 | Type1.BREX | [illegible] | 1146 | [illegible] | GCF_014336955.1 | Saccharophagus_sp._K07 | DWB84_RS14900 | 71821..75261 | 2283636 | [illegible] |
| WP_043319021.1 | Type1.BREX | [illegible] | 1145 | [illegible] | GCF_000708675.1 | Microbulbifer_sp._HZ11 | AU11_RS09640 | 451911..455348 | 1453501 | [illegible] |
| WP_036859475.1 | Type1.BREX | [illegible] | 1187 | [illegible] | GCF_000744735.1 | Porticoccus_hydrocarbonoclasticus | U740_RS05135 | 1073957..1077520 | 1073414 | [illegible] |
| MAT50029.1 | Type6.BREX | [illegible] | 1215 | [illegible] | GCA_002702725.1 | Porticoccaceae_bacterium | CMK32_02455 | 2974..6621 | 2026782 | [illegible] |
| WP_268905395.1 | Type1.BREX | [illegible] | 1220 | [illegible] | GCF_026891035.1 | Dasania_sp._GY-19 | OOV09_RS18305 | 21121..24783 | 2965066 | [illegible] |
| WP_190766547.1 | Type1.BREX | [illegible] | 1192 | [illegible] | GCF_014705705.1 | Spongiibacter_pelagi | IB286_RS13890 | 27960..31538 | 2760804 | [illegible] |
| WP_273033648.1 | Type1.BREX | [illegible] | 1193 | [illegible] | GCF_950061765.1 | Spongiibacter_tropicus | B4C78_RS03795 | 102796..106377 | 454602 | [illegible] |
| MTW21824.1 | Type1.BREX | [illegible] | 1173 | [illegible] | GCA_009720725.1 | Allochromatium_palmeri | GJ668_12065 | 37225..40746 | 231048 | [illegible] |
| WP_186343071.1 | Type1.BREX | [illegible] | 1172 | [illegible] | GCF_009720725.1 | Allochromatium_palmeri | GJ668_RS12065 | 37225..40743 | 231048 | [illegible] |
| NEX15944.1 | Type1.BREX | [illegible] | 1173 | [illegible] | GCA_010912705.1 | Halochromatium_sp. | C1943_04720 | 67001..70522 | 2049430 | [illegible] |
| WP_201222419.1 | Type1.BREX | [illegible] | 1174 | [illegible] | GCF_016653315.1 | Halochromatium_roseum | CCR96_RS19410 | 137258..140782 | 391920 | [illegible] |
| WP_264346732.1 | Type1.BREX | [illegible] | 1157 | [illegible] | GCF_947090785.1 | Rheinheimera_sp. | OM978_RS09960 | 2096767..2100240 | 1869214 | [illegible] |
| WP_120797099.1 | Type1.BREX | [illegible] | 1173 | [illegible] | GCF_003634315.1 | Thiocapsa_rosea | BDD21_RS10290 | 2320412..2323933 | 69360 | [illegible] |
| WP_018716649.1 | Type1.BREX | [illegible] | 1144 | [illegible] | GCF_000374645.1 | Arhodomonas_aquaeolei | C516_RS0101825 | 422257..425691 | 2369 | [illegible] |
| WP_239679777.1 | Type6.BREX | [illegible] | 1223 | [illegible] | GCF_022227625.1 | Ectothiorhodospira_variabilis | LPW30_RS00090 | 25011..28682 | 505694 | [illegible] |
| MBK5942672.1 | Type1.BREX | [illegible] | 1146 | [illegible] | GCA_016653405.1 | Halorhodospira_halophila | CCR79_02095 | 7976..11416 | 1053 | [illegible] |
| WP_242510795.1 | Type1.BREX | [illegible] | 1148 | [illegible] | GCF_016653405.1 | Halorhodospira_halophila | CCR79_RS02135 | 7976..11422 | 1053 | [illegible] |
| WP_253479345.1 | Type6.BREX | [illegible] | 1216 | [illegible] | GCF_024170285.1 | Natronocella_acetinitrilica | J2T57_RS14030 | 326743..330393 | 414046 | [illegible] |
| AGM41053.1 | Type1.BREX | [illegible] | 1145 | [illegible] | GCA_000319575.2 | Spiribacter_salinus_M19-40 | SPISAL_04795 | 979011..982448 | 1260251 | [illegible] |
| KAF0285942.1 | Type1.BREX | [illegible] | 1145 | [illegible] | GCA_009805925.1 | Spiribacter_sp._SSL99 | BA899_01280 | 102096..105533 | 1866884 | [illegible] |
| WP_245539889.1 | Type1.BREX | [illegible] | 1151 | [illegible] | GCF_000319575.2 | Spiribacter_salinus | SPISAL_RS04875 | 979011..982466 | 1335746 | [illegible] |
| WP_154297058.1 | Type1.BREX | [illegible] | 1145 | [illegible] | GCF_009676705.1 | Spiribacter_sp._2438 | GJ672_RS05635 | 1128117..1131554 | 2666185 | [illegible] |
| WP_235013293.1 | Type1.BREX | [illegible] | 1151 | [illegible] | GCF_009805925.1 | Spiribacter_sp._SSL99 | BA899_RS04165 | 102078..105533 | 1866884 | [illegible] |
| WP_019570748.1 | Type1.BREX | [illegible] | 1225 | [illegible] | GCF_000381205.1 | Thioalkalivibrio_sp._ALE11 | F620_RS0112040 | 26049..29726 | 1265494 | [illegible] |
| WP_019611713.1 | Type6.BREX | [illegible] | 1216 | [illegible] | GCF_000381705.1 | Thioalkalivibrio_sp._AKL7 | G315_RS0100950 | 222198..225848 | 1158155 | [illegible] |
| RJT19525.1 | Type1.BREX | [illegible] | 1233 | [illegible] | GCA_003601925.1 | Buttiauxella_izardii | D6029_18525 | 11062..14763 | 82991 | [illegible] |
| WP_244212427.1 | Type1.BREX | [illegible] | 1237 | [illegible] | GCF_003601925.1 | Buttiauxella_izardii | D6029_RS18475 | 11062..14775 | 82991 | [illegible] |
| WP_124023337.1 | Type1.BREX | [illegible] | 1157 | [illegible] | GCF_003818135.1 | Buttiauxella_warmboldiae | EHN07_RS06350 | 61248..64721 | 82993 | [illegible] |
| WP_094760992.1 | Type1.BREX | [illegible] | 1233 | [illegible] | - | Citrobacter_braakii | B9P84_05295 | 62196..65897 | 57706 | [illegible] |

| Protein-ID | System | Gene-Neighbourhoods | Length | Taxonomy | Assembly-ID | Strain | Locus-Tag | Genomic-coordinates | TaxID | Fasta-defline |
|---|---|---|---|---|---|---|---|---|---|---|
| HBK4794038.1 | Type1.BREX | [illegible] | 1233 | Bacteria>Pseudomonadota>Gammaproteobacteria>Enterobacterales>Yersiniaceae>Serratia | GCA_020682225.1 | Serratia_marcescens | LLT15_004323 | 35577..39278 | 615 | BREX-system-P-loop-protein-BrxC-[Serratia-marcescens] |
| WP_218833283.1 | Type1.BREX | [illegible] | 1197 | Bacteria>Pseudomonadota>Gammaproteobacteria>Enterobacterales>Yersiniaceae>Serratia | GCA_013401715.1 | Serratia_fonticola | HZI31_RS15400 | 48979..52572 | 47917 | BREX-system-P-loop-protein-BrxC-[Serratia-fonticola] |
| WP_206196473.1 | Type1.BREX | [illegible] | 1157 | Bacteria>Pseudomonadota>Gammaproteobacteria>Enterobacterales>Yersiniaceae>Serratia | GCF_017152235.1 | Serratia_ureilytica | JY544_RS23375 | 37448..40921 | 300181 | BREX-system-P-loop-protein-BrxC-[Serratia-ureilytica] |
| WP_223536794.1 | Type1.BREX | [illegible] | 1158 | Bacteria>Pseudomonadota>Gammaproteobacteria>Enterobacterales>Yersiniaceae>Serratia | GCF_019968705.1 | unclassified_Serratia_(in:_enterobacteria) | KGP26_RS00385 | 95452..98928 | 2647522 | MULTISPECIES:-BREX-system-P-loop-protein-BrxC-[unclassified-Serratia-(in:-enterobacteria)] |
| WP_260965787.1 | Type1.BREX | [illegible] | 1240 | Bacteria>Pseudomonadota>Gammaproteobacteria>Enterobacterales>Yersiniaceae>Serratia | GCF_025336405.1 | Serratia_marcescens | N7751_RS05105 | 1087639..1091361 | 615 | BREX-system-P-loop-protein-BrxC-[Serratia-marcescens] |
| WP_273866014.1 | Type1.BREX | [illegible] | 1157 | Bacteria>Pseudomonadota>Gammaproteobacteria>Enterobacterales>Yersiniaceae>Serratia | GCF_028616515.1 | Serratia_ureilytica | PVU68_RS17145 | 216123..219596 | 300181 | BREX-system-P-loop-protein-BrxC-[Serratia-ureilytica] |
| WP_262254721.1 | Type1.BREX | [illegible] | 1240 | Bacteria>Pseudomonadota>Gammaproteobacteria>Enterobacterales>Yersiniaceae>Serratia | GCF_946228755.1 | Serratia_proteamaculans | OC575_RS10145 | 2133622..2137344 | 28151 | BREX-system-P-loop-protein-BrxC-[Serratia-proteamaculans] |
| WP_046051668.1 | Type1.BREX | [illegible] | 1157 | Bacteria>Pseudomonadota>Gammaproteobacteria>Enterobacterales>Yersiniaceae>Yersinia | GCF_000968115.1 | Yersinia_enterocolitica | YE5303_RS19630 | 4225906..4229379 | 630 | BREX-system-P-loop-protein-BrxC-[Yersinia-enterocolitica] |
| WP_050073003.1 | Type1.BREX | [illegible] | 1157 | Bacteria>Pseudomonadota>Gammaproteobacteria>Enterobacterales>Yersiniaceae>Yersinia | GCF_001088505.1 | Yersinia_intermedia | AE858_RS05265 | 31386..34859 | 631 | BREX-system-P-loop-protein-BrxC-[Yersinia-intermedia] |
| WP_068809899.1 | Type6.BREX | [illegible] | 1214 | Bacteria>Pseudomonadota>Gammaproteobacteria>Gammaproteobacteria_incertae_sedis>Pseudohongiella | GCF_014642795.1 | Pseudohongiella_nitratireducens | IEW51_RS03970 | 206813..210457 | 1768907 | BREX-system-P-loop-protein-BrxC-[Pseudohongiella-nitratireducens] |
| MCG7876672.1 | Type6.BREX | [illegible] | 1215 | Bacteria>Pseudomonadota>Gammaproteobacteria>Gammaproteobacteria_incertae_sedis>sulfur-oxidizing_symbionts>Candidatus_Thiodiazotropha | GCA_016842655.1 | Candidatus_Thiodiazotropha_taylori | JAY78_10080 | 4978..8625 | 2792791 | BREX-system-P-loop-protein-BrxC-[Candidatus-Thiodiazotropha-taylori] |
| MCG7951159.1 | Type6.BREX | [illegible] | 1215 | Bacteria>Pseudomonadota>Gammaproteobacteria>Gammaproteobacteria_incertae_sedis>sulfur-oxidizing_symbionts>Candidatus_Thiodiazotropha | GCA_016842765.1 | Candidatus_Thiodiazotropha_taylori | JAZ09_08050 | 22496..26143 | 2792791 | BREX-system-P-loop-protein-BrxC-[Candidatus-Thiodiazotropha-taylori] |
| MCG8102611.1 | Type6.BREX | [illegible] | 1215 | Bacteria>Pseudomonadota>Gammaproteobacteria>Gammaproteobacteria_incertae_sedis>sulfur-oxidizing_symbionts>Candidatus_Thiodiazotropha | GCA_016842845.1 | Candidatus_Thiodiazotropha_taylori | JAY70_09490 | 22526..26173 | 2792791 | BREX-system-P-loop-protein-BrxC-[Candidatus-Thiodiazotropha-taylori] |
| MCG8058214.1 | Type6.BREX | [illegible] | 1215 | Bacteria>Pseudomonadota>Gammaproteobacteria>Gammaproteobacteria_incertae_sedis>sulfur-oxidizing_symbionts>Candidatus_Thiodiazotropha | GCA_016843165.1 | Candidatus_Thiodiazotropha_taylori | JAZ21_00070 | 27190..30837 | 2792791 | BREX-system-P-loop-protein-BrxC-[Candidatus-Thiodiazotropha-taylori] |
| WP_223374968.1 | Type1.BREX | [illegible] | 1240 | Bacteria>Pseudomonadota>Gammaproteobacteria | GCF_003051945.2 | Enterobacterales | N5D94_RS08195 | 82368..86090 | 91347 | MULTISPECIES:-BREX-system-P-loop-protein-BrxC-[Enterobacterales] |
| WP_052921406.1 | Type1.BREX | [illegible] | 1225 | Bacteria>Pseudomonadota>Gammaproteobacteria | GCF_003402955.1 | Enterobacterales | DS732_RS04425 | 849664..853341 | 91347 | MULTISPECIES:-BREX-system-P-loop-protein-BrxC-[Enterobacterales] |
| WP_099433763.1 | Type1.BREX | [illegible] | 1233 | Bacteria>Pseudomonadota>Gammaproteobacteria | GCF_014168815.1 | Enterobacterales | H7R56_RS21645 | 4484852..4488553 | 91347 | MULTISPECIES:-BREX-system-P-loop-protein-BrxC-[Enterobacterales] |
| WP_099476278.1 | Type1.BREX | [illegible] | 1226 | Bacteria>Pseudomonadota>Gammaproteobacteria>Moraxellales>Moraxellaceae>Acinetobacter | - | Acinetobacter_pittii | HPG40_RS00205 | 37613..41293 | 48296 | BREX-system-P-loop-protein-BrxC-[Acinetobacter-pittii] |
| WP_019836725.1 | Type1.BREX | [illegible] | 1226 | Bacteria>Pseudomonadota>Gammaproteobacteria>Moraxellales>Moraxellaceae>Acinetobacter | - | Acinetobacter_sp._MDS7A | H048_RS0104285 | 13386..17066 | 1209355 | BREX-system-P-loop-protein-BrxC-[Acinetobacter-sp.-MDS7A] |
| WP_005303500.1 | Type1.BREX | [illegible] | 1226 | Bacteria>Pseudomonadota>Gammaproteobacteria>Moraxellales>Moraxellaceae>Acinetobacter | - | Acinetobacter_sp._NIPH_542 | F886_RS07830 | 464704..468384 | 1217688 | BREX-system-P-loop-protein-BrxC-[Acinetobacter-sp.-NIPH-542] |
| ENX11167.1 | Type1.BREX | [illegible] | 911 | Bacteria>Pseudomonadota>Gammaproteobacteria>Moraxellales>Moraxellaceae>Acinetobacter | GCA_000369645.1 | Acinetobacter_sp._CIP_64.2 | F895_03860 | 13889..16624 | 1217694 | hypothetical-protein-F895_03860-[Acinetobacter-sp.-CIP-64.2] |
| HAV3580163.1 | Type1.BREX | [illegible] | 1226 | Bacteria>Pseudomonadota>Gammaproteobacteria>Moraxellales>Moraxellaceae>Acinetobacter | GCA_016485765.1 | Acinetobacter_baumannii | JIE63_001443 | 180845..184525 | 470 | BREX-system-P-loop-protein-BrxC-[Acinetobacter-baumannii] |
| HAV4979216.1 | Type1.BREX | [illegible] | 1226 | Bacteria>Pseudomonadota>Gammaproteobacteria>Moraxellales>Moraxellaceae>Acinetobacter | GCA_016507565.1 | Acinetobacter_baumannii | JIZ05_14385 | 7251..10931 | 470 | BREX-system-P-loop-protein-BrxC-[Acinetobacter-baumannii] |
| HAV6134262.1 | Type1.BREX | [illegible] | 1226 | Bacteria>Pseudomonadota>Gammaproteobacteria>Moraxellales>Moraxellaceae>Acinetobacter | GCA_016514395.1 | Acinetobacter_baumannii | JI295_12625 | 9155..12835 | 470 | BREX-system-P-loop-protein-BrxC-[Acinetobacter-baumannii] |
| BCZ16231.1 | Type1.BREX | [illegible] | 1142 | Bacteria>Pseudomonadota>Gammaproteobacteria>Moraxellales>Moraxellaceae>Acinetobacter | GCA_019703285.1 | Acinetobacter_baumannii | OCUAc18_37710 | 57364..60792 | 470 | hypothetical-protein-OCUAc18_37710-[Acinetobacter-baumannii] |
| HCA4923662.1 | Type1.BREX | [illegible] | 1226 | Bacteria>Pseudomonadota>Gammaproteobacteria>Moraxellales>Moraxellaceae>Acinetobacter | GCA_022931145.1 | Acinetobacter_baumannii | MW112_001902 | 190856..194536 | 470 | BREX-system-P-loop-protein-BrxC-[Acinetobacter-baumannii] |
| USI88235.1 | Type1.BREX | [illegible] | 1226 | Bacteria>Pseudomonadota>Gammaproteobacteria>Moraxellales>Moraxellaceae>Acinetobacter | GCA_023822225.1 | Acinetobacter_johnsonii | LZ086_18665 | 10226..13906 | 40214 | BREX-system-P-loop-protein-BrxC-[Acinetobacter-johnsonii] |
| WP_005401825.1 | Type1.BREX | [illegible] | 1226 | Bacteria>Pseudomonadota>Gammaproteobacteria>Moraxellales>Moraxellaceae>Acinetobacter | GCF_000162055.1 | Acinetobacter_johnsonii | HMPREF0016_RS15525 | 47644..51324 | 40214 | BREX-system-P-loop-protein-BrxC-[Acinetobacter-johnsonii] |
| WP_010116441.1 | Type1.BREX | [illegible] | 1226 | Bacteria>Pseudomonadota>Gammaproteobacteria>Moraxellales>Moraxellaceae>Acinetobacter | GCF_000214135.1 | Acinetobacter_sp._P8-3-8 | GGG_RS0115320 | 12069..15749 | 1029823 | BREX-system-P-loop-protein-BrxC-[Acinetobacter-sp.-P8-3-8] |
| WP_001021843.1 | Type1.BREX | [illegible] | 1226 | Bacteria>Pseudomonadota>Gammaproteobacteria>Moraxellales>Moraxellaceae>Acinetobacter | GCF_000305235.1 | Acinetobacter_baumannii | ACIN5098_RS09930 | 47279..50959 | 470 | BREX-system-P-loop-protein-BrxC-[Acinetobacter-baumannii] |
| WP_004811070.1 | Type1.BREX | [illegible] | 1225 | Bacteria>Pseudomonadota>Gammaproteobacteria>Moraxellales>Moraxellaceae>Acinetobacter | GCF_000368465.1 | Acinetobacter_schindleri | F965_RS04725 | 393977..397654 | 108981 | BREX-system-P-loop-protein-BrxC-[Acinetobacter-schindleri] |
| WP_005162499.1 | Type6.BREX | [illegible] | 1063 | Bacteria>Pseudomonadota>Gammaproteobacteria>Moraxellales>Moraxellaceae>Acinetobacter | GCF_000369425.1 | Acinetobacter_sp._NIPH_284 | F908_RS12580 | 1400419..1403610 | 1217704 | BREX-system-P-loop-protein-BrxC-[Acinetobacter-sp.-NIPH-284] |
| WP_031946304.1 | Type1.BREX | [illegible] | 1226 | Bacteria>Pseudomonadota>Gammaproteobacteria>Moraxellales>Moraxellaceae>Acinetobacter | GCF_000516575.2 | Acinetobacter_baumannii | UN97_RS12380 | 246428..250108 | 470 | BREX-system-P-loop-protein-BrxC-[Acinetobacter-baumannii] |
| WP_034686569.1 | Type1.BREX | [illegible] | 1226 | Bacteria>Pseudomonadota>Gammaproteobacteria>Moraxellales>Moraxellaceae>Acinetobacter | GCF_000581035.1 | Acinetobacter_sp._869535 | J520_RS12320 | 4406..8086 | 1310621 | BREX-system-P-loop-protein-BrxC-[Acinetobacter-sp.-869535] |
| WP_025095052.1 | Type1.BREX | [illegible] | 1226 | Bacteria>Pseudomonadota>Gammaproteobacteria>Moraxellales>Moraxellaceae>Acinetobacter | GCF_000633005.1 | Acinetobacter_soli | EG58_RS0106955 | 8205..11885 | 487316 | BREX-system-P-loop-protein-BrxC-[Acinetobacter-soli] |
| WP_045544683.1 | Type1.BREX | [illegible] | 1226 | Bacteria>Pseudomonadota>Gammaproteobacteria>Moraxellales>Moraxellaceae>Acinetobacter | GCF_000828935.1 | Acinetobacter_baumannii | IOMTU433_RS19345 | 66852..70532 | 470 | BREX-system-P-loop-protein-BrxC-[Acinetobacter-baumannii] |
| WP_043972752.1 | Type1.BREX | [illegible] | 1226 | Bacteria>Pseudomonadota>Gammaproteobacteria>Moraxellales>Moraxellaceae>Acinetobacter | GCF_000876115.1 | Acinetobacter_sp._NBRC_110496 | Aba26S_RS13795 | 7444..11124 | 1550715 | BREX-system-P-loop-protein-BrxC-[Acinetobacter-sp.-NBRC-110496] |
| WP_032059948.1 | Type1.BREX | [illegible] | 1226 | Bacteria>Pseudomonadota>Gammaproteobacteria>Moraxellales>Moraxellaceae>Acinetobacter | GCF_001077555.2 | Acinetobacter_baumannii | AB895_RS13070 | 2715747..2719427 | 470 | BREX-system-P-loop-protein-BrxC-[Acinetobacter-baumannii] |
| WP_058952662.1 | Type1.BREX | [illegible] | 1220 | Bacteria>Pseudomonadota>Gammaproteobacteria>Moraxellales>Moraxellaceae>Acinetobacter | GCF_001484935.1 | Acinetobacter_johnsonii | EHS42_RS19740 | 21243..24905 | 40214 | BREX-system-P-loop-protein-BrxC-[Acinetobacter-johnsonii] |
| WP_065184483.1 | Type1.BREX | [illegible] | 1226 | Bacteria>Pseudomonadota>Gammaproteobacteria>Moraxellales>Moraxellaceae>Acinetobacter | GCF_001674695.1 | Acinetobacter_baumannii | GKP36_RS06470 | 16105..19785 | 470 | BREX-system-P-loop-protein-BrxC-[Acinetobacter-baumannii] |
| WP_069122561.1 | Type1.BREX | [illegible] | 1226 | Bacteria>Pseudomonadota>Gammaproteobacteria>Moraxellales>Moraxellaceae>Acinetobacter | GCF_001716945.1 | Acinetobacter_pittii | AXH23_RS05625 | 247550..251230 | 48296 | BREX-system-P-loop-protein-BrxC-[Acinetobacter-pittii] |
| WP_075040289.1 | Type1.BREX | [illegible] | 1226 | Bacteria>Pseudomonadota>Gammaproteobacteria>Moraxellales>Moraxellaceae>Acinetobacter | GCF_001917365.1 | Acinetobacter_radioresistens | SA188iCDA_RS03905 | 54372..58052 | 40216 | BREX-system-P-loop-protein-BrxC-[Acinetobacter-radioresistens] |
| WP_086231303.1 | Type1.BREX | [illegible] | 1226 | Bacteria>Pseudomonadota>Gammaproteobacteria>Moraxellales>Moraxellaceae>Acinetobacter | GCF_002137255.1 | Acinetobacter_baumannii | B9Y01_RS02195 | 14367..18047 | 470 | BREX-system-P-loop-protein-BrxC-[Acinetobacter-baumannii] |
| WP_096902445.1 | Type1.BREX | [illegible] | 1222 | Bacteria>Pseudomonadota>Gammaproteobacteria>Moraxellales>Moraxellaceae>Acinetobacter | GCF_002412355.1 | Acinetobacter_sp._YT-02 | CF596_RS10765 | 15331..18999 | 2018564 | BREX-system-P-loop-protein-BrxC-[Acinetobacter-sp.-YT-02] |
| WP_099475573.1 | Type1.BREX | [illegible] | 1226 | Bacteria>Pseudomonadota>Gammaproteobacteria>Moraxellales>Moraxellaceae>Acinetobacter | GCF_002740745.1 | Acinetobacter_pittii | CT134_RS17990 | 28430..32110 | 48296 | BREX-system-P-loop-protein-BrxC-[Acinetobacter-pittii] |
| WP_104442411.1 | Type1.BREX | [illegible] | 1226 | Bacteria>Pseudomonadota>Gammaproteobacteria>Moraxellales>Moraxellaceae>Acinetobacter | GCF_002934965.1 | Acinetobacter_sp._ACNIH2 | C3F34_RS18690 | 72475..76155 | 1758189 | BREX-system-P-loop-protein-BrxC-[Acinetobacter-sp.-ACNIH2] |
| WP_104500994.1 | Type1.BREX | [illegible] | 1226 | Bacteria>Pseudomonadota>Gammaproteobacteria>Moraxellales>Moraxellaceae>Acinetobacter | GCF_002938745.1 | Acinetobacter_indicus | C5505_RS10055 | 19022..22702 | 756892 | BREX-system-P-loop-protein-BrxC-[Acinetobacter-indicus] |
| WP_109291743.1 | Type1.BREX | [illegible] | 1226 | Bacteria>Pseudomonadota>Gammaproteobacteria>Moraxellales>Moraxellaceae>Acinetobacter | GCF_003131095.1 | Acinetobacter_baumannii | DF172_RS10505 | 4712..8392 | 470 | BREX-system-P-loop-protein-BrxC-[Acinetobacter-baumannii] |
| WP_111281982.1 | Type1.BREX | [illegible] | 1226 | Bacteria>Pseudomonadota>Gammaproteobacteria>Moraxellales>Moraxellaceae>Acinetobacter | GCF_003258335.1 | Acinetobacter_radioresistens | NVV25_RS10895 | 13399..17079 | 40216 | BREX-system-P-loop-protein-BrxC-[Acinetobacter-radioresistens] |
| WP_119025213.1 | Type1.BREX | [illegible] | 1226 | Bacteria>Pseudomonadota>Gammaproteobacteria>Moraxellales>Moraxellaceae>Acinetobacter | GCF_003569585.1 | Acinetobacter_soli | D5I70_RS01315 | 8436..12116 | 487316 | BREX-system-P-loop-protein-BrxC-[Acinetobacter-soli] |
| WP_119047908.1 | Type1.BREX | [illegible] | 1226 | Bacteria>Pseudomonadota>Gammaproteobacteria>Moraxellales>Moraxellaceae>Acinetobacter | GCF_003569645.1 | Acinetobacter_baumannii | D5I76_RS05685 | 52847..56527 | 470 | BREX-system-P-loop-protein-BrxC-[Acinetobacter-baumannii] |
| WP_123785432.1 | Type1.BREX | [illegible] | 1226 | Bacteria>Pseudomonadota>Gammaproteobacteria>Moraxellales>Moraxellaceae>Acinetobacter | GCF_003812065.1 | Acinetobacter_baumannii | LH406_RS10280 | 68046..71726 | 470 | BREX-system-P-loop-protein-BrxC-[Acinetobacter-baumannii] |
| WP_131273640.1 | Type1.BREX | [illegible] | 1154 | Bacteria>Pseudomonadota>Gammaproteobacteria>Moraxellales>Moraxellaceae>Acinetobacter | GCF_004331195.1 | Acinetobacter_sp._ANC_4216 | E0H88_RS07250 | 81102..84566 | 2529840 | BREX-system-P-loop-protein-BrxC-[Acinetobacter-sp.-ANC-4216] |
| WP_138000997.1 | Type1.BREX | [illegible] | 1226 | Bacteria>Pseudomonadota>Gammaproteobacteria>Moraxellales>Moraxellaceae>Acinetobacter | GCF_005519305.1 | Acinetobacter_radioresistens | E3H47_RS14125 | 6771..10451 | 40216 | BREX-system-P-loop-protein-BrxC-[Acinetobacter-radioresistens] |
| WP_140948309.1 | Type1.BREX | [illegible] | 1226 | Bacteria>Pseudomonadota>Gammaproteobacteria>Moraxellales>Moraxellaceae>Acinetobacter | GCF_006491975.1 | Acinetobacter_baumannii | FJV17_RS05945 | 24470..28150 | 470 | BREX-system-P-loop-protein-BrxC-[Acinetobacter-baumannii] |
| WP_140963539.1 | Type6.BREX | [illegible] | 1214 | Bacteria>Pseudomonadota>Gammaproteobacteria>Moraxellales>Moraxellaceae>Acinetobacter | GCF_006494575.1 | Acinetobacter_baumannii | FJU84_RS09090 | 25284..28928 | 470 | BREX-system-P-loop-protein-BrxC-[Acinetobacter-baumannii] |
| WP_144583841.1 | Type1.BREX | [illegible] | 1226 | Bacteria>Pseudomonadota>Gammaproteobacteria>Moraxellales>Moraxellaceae>Acinetobacter | GCF_007713425.1 | Acinetobacter_colistiniresistens | FPV60_RS17750 | 3413..7093 | 280145 | BREX-system-P-loop-protein-BrxC-[Acinetobacter-colistiniresistens] |
| WP_153389475.1 | Type1.BREX | [illegible] | 1224 | Bacteria>Pseudomonadota>Gammaproteobacteria>Moraxellales>Moraxellaceae>Acinetobacter | GCF_009601085.1 | Acinetobacter_wanghuae | GHJ48_RS13160 | 17934..21608 | 2662362 | BREX-system-P-loop-protein-BrxC-[Acinetobacter-wanghuae] |
| WP_160255370.1 | Type1.BREX | [illegible] | 1226 | Bacteria>Pseudomonadota>Gammaproteobacteria>Moraxellales>Moraxellaceae>Acinetobacter | GCF_009914395.1 | Acinetobacter_indicus | GA646_RS00530 | 100361..104041 | 756892 | BREX-system-P-loop-protein-BrxC-[Acinetobacter-indicus] |
| WP_163071913.1 | Type1.BREX | [illegible] | 1226 | Bacteria>Pseudomonadota>Gammaproteobacteria>Moraxellales>Moraxellaceae>Acinetobacter | GCF_010499775.1 | Acinetobacter_baumannii | G3N14_RS16470 | 10817..14497 | 470 | BREX-system-P-loop-protein-BrxC-[Acinetobacter-baumannii] |
| WP_168370063.1 | Type1.BREX | [illegible] | 1224 | Bacteria>Pseudomonadota>Gammaproteobacteria>Moraxellales>Moraxellaceae>Acinetobacter | GCF_012278715.1 | Acinetobacter_indicus | HHC64_RS10945 | 2259038..2262712 | 756892 | BREX-system-P-loop-protein-BrxC-[Acinetobacter-indicus] |
| WP_169025139.1 | Type1.BREX | [illegible] | 1226 | Bacteria>Pseudomonadota>Gammaproteobacteria>Moraxellales>Moraxellaceae>Acinetobacter | GCF_012931665.1 | Acinetobacter_baumannii | HIN88_RS18590 | 12016..15696 | 470 | BREX-system-P-loop-protein-BrxC-[Acinetobacter-baumannii] |
| WP_171530296.1 | Type1.BREX | [illegible] | 1226 | Bacteria>Pseudomonadota>Gammaproteobacteria>Moraxellales>Moraxellaceae>Acinetobacter | GCF_013072655.1 | Acinetobacter_defluvii | A7P53_RS01880 | 14398..18078 | 1871111 | BREX-system-P-loop-protein-BrxC-[Acinetobacter-defluvii] |
| WP_174727826.1 | Type1.BREX | [illegible] | 1224 | Bacteria>Pseudomonadota>Gammaproteobacteria>Moraxellales>Moraxellaceae>Acinetobacter | GCF_013344765.1 | Acinetobacter_lactucae | HUN22_RS07895 | 1711..5385 | 1785128 | BREX-system-P-loop-protein-BrxC-[Acinetobacter-lactucae] |
| WP_179994566.1 | Type1.BREX | [illegible] | 1226 | Bacteria>Pseudomonadota>Gammaproteobacteria>Moraxellales>Moraxellaceae>Acinetobacter | GCF_013416255.1 | Acinetobacter_baumannii | HLG75_RS04210 | 863150..866830 | 470 | BREX-system-P-loop-protein-BrxC-[Acinetobacter-baumannii] |
| WP_180045992.1 | Type1.BREX | [illegible] | 1218 | Bacteria>Pseudomonadota>Gammaproteobacteria>Moraxellales>Moraxellaceae>Acinetobacter | GCF_013418865.1 | Acinetobacter_sp._YH12117 | FTU55_RS01900 | 159420..163076 | 2601104 | BREX-system-P-loop-protein-BrxC-[Acinetobacter-sp.-YH12117] |
| WP_185218776.1 | Type1.BREX | [illegible] | 1226 | Bacteria>Pseudomonadota>Gammaproteobacteria>Moraxellales>Moraxellaceae>Acinetobacter | GCF_014218985.1 | Acinetobacter_baumannii | H6W49_RS17735 | 6511..10191 | 470 | BREX-system-P-loop-protein-BrxC-[Acinetobacter-baumannii] |
| WP_032068282.1 | Type6.BREX | [illegible] | 1214 | Bacteria>Pseudomonadota>Gammaproteobacteria>Moraxellales>Moraxellaceae>Acinetobacter | GCF_014559915.1 | Acinetobacter_baumannii | HPJ29_RS06280 | 177170..180814 | 470 | BREX-system-P-loop-protein-BrxC-[Acinetobacter-baumannii] |
| WP_193893631.1 | Type1.BREX | [illegible] | 1226 | Bacteria>Pseudomonadota>Gammaproteobacteria>Moraxellales>Moraxellaceae>Acinetobacter | GCF_015209685.1 | Acinetobacter_albensis | IQR32_RS12760 | 7183..10863 | 1673609 | BREX-system-P-loop-protein-BrxC-[Acinetobacter-albensis] |
| WP_136512845.1 | Type1.BREX | [illegible] | 1226 | Bacteria>Pseudomonadota>Gammaproteobacteria>Moraxellales>Moraxellaceae>Acinetobacter | GCF_015265775.1 | Acinetobacter_baumannii | O9X70_RS06810 | 133723..137403 | 470 | BREX-system-P-loop-protein-BrxC-[Acinetobacter-baumannii] |
| WP_196075355.1 | Type1.BREX | [illegible] | 1204 | Bacteria>Pseudomonadota>Gammaproteobacteria>Moraxellales>Moraxellaceae>Acinetobacter | GCF_015676315.1 | unclassified_Acinetobacter | IMO39_RS19270 | 9799..13413 | 196816 | MULTISPECIES:-BREX-system-P-loop-protein-BrxC-[unclassified-Acinetobacter] |
| WP_198080379.1 | Type1.BREX | [illegible] | 1222 | Bacteria>Pseudomonadota>Gammaproteobacteria>Moraxellales>Moraxellaceae>Acinetobacter | GCF_016082245.1 | Acinetobacter_calcoaceticus | BL242_RS20750 | 15792..19460 | 471 | BREX-system-P-loop-protein-BrxC-[Acinetobacter-calcoaceticus] |
| WP_199939733.1 | Type1.BREX | [illegible] | 1226 | Bacteria>Pseudomonadota>Gammaproteobacteria>Moraxellales>Moraxellaceae>Acinetobacter | GCF_016506315.1 | Acinetobacter_baumannii | I5555_RS13010 | 139911..143591 | 470 | BREX-system-P-loop-protein-BrxC-[Acinetobacter-baumannii] |
| WP_203080037.1 | Type1.BREX | [illegible] | 1226 | Bacteria>Pseudomonadota>Gammaproteobacteria>Moraxellales>Moraxellaceae>Acinetobacter | GCF_016804045.1 | Acinetobacter_pittii | HRJ47_RS04290 | 884929..888609 | 48296 | BREX-system-P-loop-protein-BrxC-[Acinetobacter-pittii] |
| WP_205779680.1 | Type1.BREX | [illegible] | 1226 | Bacteria>Pseudomonadota>Gammaproteobacteria>Moraxellales>Moraxellaceae>Acinetobacter | GCF_017132735.1 | Acinetobacter_indicus | JYB86_RS06365 | 1326455..1330135 | 756892 | BREX-system-P-loop-protein-BrxC-[Acinetobacter-indicus] |
| WP_109426631.1 | Type1.BREX | [illegible] | 1226 | Bacteria>Pseudomonadota>Gammaproteobacteria>Moraxellales>Moraxellaceae>Acinetobacter | GCF_017726575.1 | Acinetobacter_baumannii | J6I88_RS04260 | 873693..877373 | 470 | BREX-system-P-loop-protein-BrxC-[Acinetobacter-baumannii] |
| WP_216174886.1 | Type1.BREX | [illegible] | 1226 | Bacteria>Pseudomonadota>Gammaproteobacteria>Moraxellales>Moraxellaceae>Acinetobacter | GCF_018863255.1 | Acinetobacter_baumannii | KPE97_RS15710 | 83323..87003 | 470 | BREX-system-P-loop-protein-BrxC-[Acinetobacter-baumannii] |
| WP_216116819.1 | Type1.BREX | [illegible] | 1226 | Bacteria>Pseudomonadota>Gammaproteobacteria>Moraxellales>Moraxellaceae>Acinetobacter | GCF_018883565.1 | Acinetobacter_baumannii | KPF69_RS05265 | 35520..39200 | 470 | BREX-system-P-loop-protein-BrxC-[Acinetobacter-baumannii] |
| WP_218698253.1 | Type1.BREX | [illegible] | 1224 | Bacteria>Pseudomonadota>Gammaproteobacteria>Moraxellales>Moraxellaceae>Acinetobacter | GCF_019279015.1 | Acinetobacter_harbinensis | KZA13_RS08660 | 34272..37946 | 1353941 | BREX-system-P-loop-protein-BrxC-[Acinetobacter-harbinensis] |
| WP_221260378.1 | Type1.BREX | [illegible] | 1226 | Bacteria>Pseudomonadota>Gammaproteobacteria>Moraxellales>Moraxellaceae>Acinetobacter | GCF_019702905.1 | Acinetobacter_sp._Tol_5 | TOL5_RS12615 | 2696813..2700493 | 710648 | BREX-system-P-loop-protein-BrxC-[Acinetobacter-sp.-Tol-5] |
| WP_207272916.1 | Type1.BREX | [illegible] | 1226 | Bacteria>Pseudomonadota>Gammaproteobacteria>Moraxellales>Moraxellaceae>Acinetobacter | GCF_019921105.1 | Acinetobacter_baumannii | HLH91_RS14165 | 18027..21707 | 470 | BREX-system-P-loop-protein-BrxC-[Acinetobacter-baumannii] |
| WP_005236280.1 | Type1.BREX | [illegible] | 1226 | Bacteria>Pseudomonadota>Gammaproteobacteria>Moraxellales>Moraxellaceae>Acinetobacter | GCF_020099375.1 | Acinetobacter_variabilis | LA331_RS04655 | 963646..967326 | 70346 | BREX-system-P-loop-protein-BrxC-[Acinetobacter-variabilis] |
| WP_225554349.1 | Type1.BREX | [illegible] | 1226 | Bacteria>Pseudomonadota>Gammaproteobacteria>Moraxellales>Moraxellaceae>Acinetobacter | GCF_020271765.1 | Acinetobacter_pseudolwoffii | LDD52_RS14245 | 193616..1867 | 2053287 | BREX-system-P-loop-protein-BrxC-[Acinetobacter-pseudolwoffii] |
| WP_228414014.1 | Type1.BREX | [illegible] | 1226 | Bacteria>Pseudomonadota>Gammaproteobacteria>Moraxellales>Moraxellaceae>Acinetobacter | GCF_020810655.1 | Acinetobacter_baumannii | NLHDIDDJ_RS06545 | 1352098..1355778 | 470 | BREX-system-P-loop-protein-BrxC-[Acinetobacter-baumannii] |
| WP_233945509.1 | Type1.BREX | [illegible] | 1226 | Bacteria>Pseudomonadota>Gammaproteobacteria>Moraxellales>Moraxellaceae>Acinetobacter | GCF_021391555.1 | Acinetobacter_sp._SH20PTE14 | LXF01_RS17045 | 120215..123895 | 2905879 | BREX-system-P-loop-protein-BrxC-[Acinetobacter-sp.-SH20PTE14] |
| WP_236095241.1 | Type1.BREX | [illegible] | 1226 | Bacteria>Pseudomonadota>Gammaproteobacteria>Moraxellales>Moraxellaceae>Acinetobacter | GCF_021560715.1 | Acinetobacter_sp._SCLZS86 | LZP46_RS14905 | 6362..10042 | 2908637 | BREX-system-P-loop-protein-BrxC-[Acinetobacter-sp.-SCLZS86] |
| WP_236163089.1 | Type1.BREX | [illegible] | 1226 | Bacteria>Pseudomonadota>Gammaproteobacteria>Moraxellales>Moraxellaceae>Acinetobacter | GCF_021602685.1 | unclassified_Acinetobacter | L6B80_RS11000 | 14595..18275 | 196816 | MULTISPECIES:-BREX-system-P-loop-protein-BrxC-[unclassified-Acinetobacter] |
| WP_241338465.1 | Type1.BREX | [illegible] | 1226 | Bacteria>Pseudomonadota>Gammaproteobacteria>Moraxellales>Moraxellaceae>Acinetobacter | GCF_022549435.1 | Acinetobacter_sp._ANC_5659 | MMP64_RS09425 | 122256..125936 | 2923427 | BREX-system-P-loop-protein-BrxC-[Acinetobacter-sp.-ANC-5659] |
| WP_128202440.1 | Type1.BREX | [illegible] | 1226 | Bacteria>Pseudomonadota>Gammaproteobacteria>Moraxellales>Moraxellaceae>Acinetobacter | GCF_023896505.1 | Acinetobacter_baumannii | O9X56_RS14280 | 107190..110870 | 470 | BREX-system-P-loop-protein-BrxC-[Acinetobacter-baumannii] |
| WP_257259040.1 | Type1.BREX | [illegible] | 1226 | Bacteria>Pseudomonadota>Gammaproteobacteria>Moraxellales>Moraxellaceae>Acinetobacter | GCF_024589115.1 | Acinetobacter_baumannii | L2615_RS07865 | 180845..184525 | 470 | BREX-system-P-loop-protein-BrxC-[Acinetobacter-baumannii] |
| WP_257935145.1 | Type1.BREX | [illegible] | 1226 | Bacteria>Pseudomonadota>Gammaproteobacteria>Moraxellales>Moraxellaceae>Acinetobacter | GCF_024705725.1 | Acinetobacter_baumannii | MSH04_RS15115 | 32588..36268 | 470 | BREX-system-P-loop-protein-BrxC-[Acinetobacter-baumannii] |
| WP_238640059.1 | Type1.BREX | [illegible] | 1226 | Bacteria>Pseudomonadota>Gammaproteobacteria>Moraxellales>Moraxellaceae>Acinetobacter | GCF_025264005.1 | Acinetobacter_baumannii | NT399_RS17670 | 10742..14422 | 470 | BREX-system-P-loop-protein-BrxC-[Acinetobacter-baumannii] |
| WP_262579272.1 | Type1.BREX | [illegible] | 1226 | Bacteria>Pseudomonadota>Gammaproteobacteria>Moraxellales>Moraxellaceae>Acinetobacter | GCF_025562355.1 | Acinetobacter_junii | KTH64_RS13010 | 10561..14241 | 40215 | BREX-system-P-loop-protein-BrxC-[Acinetobacter-junii] |
| WP_264453044.1 | Type6.BREX | [illegible] | 1214 | Bacteria>Pseudomonadota>Gammaproteobacteria>Moraxellales>Moraxellaceae>Acinetobacter | GCF_025938795.1 | Acinetobacter_baumannii | OLN40_RS03910 | 58425..62069 | 470 | BREX-system-P-loop-protein-BrxC-[Acinetobacter-baumannii] |
| WP_265466015.1 | Type1.BREX | [illegible] | 1226 | Bacteria>Pseudomonadota>Gammaproteobacteria>Moraxellales>Moraxellaceae>Acinetobacter | GCF_026168575.1 | Acinetobacter_sp._BIT-DXN8 | OKC24_RS16365 | 19397..23077 | 2989714 | BREX-system-P-loop-protein-BrxC-[Acinetobacter-sp.-BIT-DXN8] |
| WP_071225618.1 | Type1.BREX | [illegible] | 1226 | Bacteria>Pseudomonadota>Gammaproteobacteria>Moraxellales>Moraxellaceae>Acinetobacter | GCF_027854895.1 | Acinetobacter_baumannii | MJ031_RS19545 | 43789..47469 | 470 | BREX-system-P-loop-protein-BrxC-[Acinetobacter-baumannii] |
| WP_274579352.1 | Type1.BREX | [illegible] | 1226 | Bacteria>Pseudomonadota>Gammaproteobacteria>Moraxellales>Moraxellaceae>Acinetobacter | GCF_028774645.1 | Acinetobacter_lactucae | MO054_RS17975 | 24360..28040 | 1785128 | BREX-system-P-loop-protein-BrxC-[Acinetobacter-lactucae] |
| WP_275290158.1 | Type1.BREX | [illegible] | 1226 | Bacteria>Pseudomonadota>Gammaproteobacteria>Moraxellales>Moraxellaceae>Acinetobacter | GCF_029100495.1 | Acinetobacter_nosocomialis | JIZ05_14385 | 7251..10931 | 106654 | BREX-system-P-loop-protein-BrxC-[Acinetobacter-nosocomialis] |
| WP_161409476.1 | Type1.BREX | [illegible] | 1226 | Bacteria>Pseudomonadota>Gammaproteobacteria>Moraxellales>Moraxellaceae>Acinetobacter | GCF_030329445.1 | Acinetobacter_haemolyticus | GPS61_RS07925 | 49241..52921 | 29430 | BREX-system-P-loop-protein-BrxC-[Acinetobacter-haemolyticus] |
| WP_005115116.1 | Type1.BREX | [illegible] | 1226 | Bacteria>Pseudomonadota>Gammaproteobacteria>Moraxellales>Moraxellaceae>Acinetobacter | GCF_030413705.1 | Acinetobacter_baumannii | AWM09_RS06130 | 10901..14581 | 470 | BREX-system-P-loop-protein-BrxC-[Acinetobacter-baumannii] |
| WP_002010901.1 | Type1.BREX | [illegible] | 1226 | Bacteria>Pseudomonadota>Gammaproteobacteria>Moraxellales>Moraxellaceae | GCF_000214965.1 | Acinetobacter | ACINNAV18_RS19255 | 65599..69279 | 469 | MULTISPECIES:-BREX-system-P-loop-protein-BrxC-[Acinetobacter] |
| WP_111885889.1 | Type1.BREX | [illegible] | 1226 | Bacteria>Pseudomonadota>Gammaproteobacteria>Moraxellales>Moraxellaceae | GCF_002339435.1 | Acinetobacter | NVV26_RS12530 | 20211..23891 | 469 | MULTISPECIES:-BREX-system-P-loop-protein-BrxC-[Acinetobacter] |
| WP_108675357.1 | Type1.BREX | [illegible] | 1225 | Bacteria>Pseudomonadota>Gammaproteobacteria>Moraxellales>Moraxellaceae | GCF_003072585.2 | Acinetobacter | C0119_RS05575 | 1118908..1122585 | 469 | MULTISPECIES:-BREX-system-P-loop-protein-BrxC-[Acinetobacter] |
| WP_151794804.1 | Type6.BREX | [illegible] | 1214 | Bacteria>Pseudomonadota>Gammaproteobacteria>Moraxellales>Moraxellaceae | GCF_008997035.1 | Acinetobacter | GBP95_RS04055 | 337470..341114 | 469 | MULTISPECIES:-BREX-system-P-loop-protein-BrxC-[Acinetobacter] |
| WP_178319334.1 | Type1.BREX | [illegible] | 1226 | Bacteria>Pseudomonadota>Gammaproteobacteria>Moraxellales>Moraxellaceae | GCF_013390985.1 | Acinetobacter | HX743_RS11980 | 35327..39007 | 469 | MULTISPECIES:-BREX-system-P-loop-protein-BrxC-[Acinetobacter] |
| ABE75386.1 | Type1.BREX | [illegible] | 1201 | Bacteria>Pseudomonadota>Gammaproteobacteria>Moraxellales>Moraxellaceae>Psychrobacter | GCA_000013905.1 | Psychrobacter_cryohalolentis_K5 | Pcryo_1609 | 1979457..1983062 | 335284 | conserved-hypothetical-protein-[Psychrobacter-cryohalolentis-K5] |
| MCG3880112.1 | Type1.BREX | [illegible] | 1154 | Bacteria>Pseudomonadota>Gammaproteobacteria>Moraxellales>Moraxellaceae>Psychrobacter | GCA_022129235.1 | Psychrobacter_sp._Ps6 | I3255_11455 | 48655..52119 | 2790960 | BREX-system-P-loop-protein-BrxC-[Psychrobacter-sp.-Ps6] |
| WP_226939352.1 | Type1.BREX | [illegible] | 1221 | Bacteria>Pseudomonadota>Gammaproteobacteria>Moraxellales>Moraxellaceae>Psychrobacter | GCF_000013905.1 | Psychrobacter_cryohalolentis | CEP87_RS02910 | 676881..680546 | 330922 | BREX-system-P-loop-protein-BrxC-[Psychrobacter-cryohalolentis] |
| WP_261869530.1 | Type1.BREX | [illegible] | 642 | Bacteria>Pseudomonadota>Gammaproteobacteria>Moraxellales>Moraxellaceae>Psychrobacter | GCF_000586435.1 | Psychrobacter_sp._JCM_18901 | TT11_RS05300 | 11820..13748 | 1298609 | BREX-system-P-loop-protein-BrxC-[Psychrobacter-sp.-JCM-18901] |
| WP_089554755.1 | Type1.BREX | [illegible] | 1223 | Bacteria>Pseudomonadota>Gammaproteobacteria>Moraxellales>Moraxellaceae>Psychrobacter | GCF_002224105.1 | Psychrobacter_sp._DAB_AL32B | CAN34_RS05290 | 13543..17214 | 1028414 | BREX-system-P-loop-protein-BrxC-[Psychrobacter-sp.-DAB_AL32B] |
| WP_191691193.1 | Type1.BREX | [illegible] | 1222 | Bacteria>Pseudomonadota>Gammaproteobacteria>Moraxellales>Moraxellaceae>Psychrobacter | GCF_014836505.1 | Psychrobacter_communis | H9653_RS05990 | 24340..28008 | 2762238 | BREX-system-P-loop-protein-BrxC-[Psychrobacter-communis] |
| WP_207991848.1 | Type1.BREX | [illegible] | 1221 | Bacteria>Pseudomonadota>Gammaproteobacteria>Moraxellales>Moraxellaceae>Psychrobacter | GCF_017498085.1 | Psychrobacter_coccoides | J3492_RS09680 | 48612..52277 | 2818440 | BREX-system-P-loop-protein-BrxC-[Psychrobacter-coccoides] |
| WP_218690652.1 | Type6.BREX | [illegible] | 1213 | Bacteria>Pseudomonadota>Gammaproteobacteria>Moraxellales>Moraxellaceae>Psychrobacter | GCF_019279065.1 | Psychrobacter_sp._BF1 | KZA12_RS14005 | 8359..12000 | 2821147 | BREX-system-P-loop-protein-BrxC-[Psychrobacter-sp.-BF1] |
| WP_238038933.1 | Type1.BREX | [illegible] | 1162 | Bacteria>Pseudomonadota>Gammaproteobacteria>Moraxellales>Moraxellaceae>Psychrobacter | GCF_022129235.1 | Psychrobacter_sp._Ps6 | I3255_RS11455 | 48631..52119 | 2790960 | BREX-system-P-loop-protein-BrxC-[Psychrobacter-sp.-Ps6] |
| WP_238051284.1 | Type6.BREX | [illegible] | 1213 | Bacteria>Pseudomonadota>Gammaproteobacteria>Moraxellales>Moraxellaceae>Psychrobacter | GCF_022129405.1 | Psychrobacter_sp._Ps2 | I3254_RS03805 | 13291..16932 | 2790956 | BREX-system-P-loop-protein-BrxC-[Psychrobacter-sp.-Ps2] |
| WP_199774478.1 | Type1.BREX | [illegible] | 1164 | Bacteria>Pseudomonadota>Gammaproteobacteria>Moraxellales>Moraxellaceae>Psychrobacter | GCF_900291405.1 | Psychrobacter_sp._Marseille-P5312 | C6W44_RS10120 | 2338090..2341584 | 2086574 | BREX-system-P-loop-protein-BrxC-[Psychrobacter-sp.-Marseille-P5312] |
| WP_201558729.1 | Type1.BREX | [illegible] | 1201 | Bacteria>Pseudomonadota>Gammaproteobacteria>Moraxellales>Moraxellaceae>Psychrobacter | GCF_904846085.1 | Psychrobacter_immobilis | JMW74_RS07725 | 23770..27375 | 498 | BREX-system-P-loop-protein-BrxC-[Psychrobacter-immobilis] |
| WP_201625912.1 | Type1.BREX | [illegible] | 1146 | Bacteria>Pseudomonadota>Gammaproteobacteria>Moraxellales>Moraxellaceae>Psychrobacter | GCF_904846275.1 | Psychrobacter_immobilis | JMX31_RS11870 | 11441..14881 | 498 | BREX-system-P-loop-protein-BrxC-[Psychrobacter-immobilis] |
| WP_198334111.1 | Type1.BREX | [illegible] | 1210 | Bacteria>Pseudomonadota>Gammaproteobacteria>Moraxellales>Moraxellaceae>Psychrobacter | GCF_904846355.1 | Psychrobacter_namhaensis | JMX36_RS06945 | 21523..25155 | 292734 | BREX-system-P-loop-protein-BrxC-[Psychrobacter-namhaensis] |
| WP_201542137.1 | Type1.BREX | [illegible] | 1210 | Bacteria>Pseudomonadota>Gammaproteobacteria>Moraxellales>Moraxellaceae>Psychrobacter | GCF_904846515.1 | Psychrobacter_sanguinis | LK453_RS13670 | 3248348..3251980 | 861445 | BREX-system-P-loop-protein-BrxC-[Psychrobacter-sanguinis] |
| WP_201579699.1 | Type1.BREX | [illegible] | 1223 | Bacteria>Pseudomonadota>Gammaproteobacteria>Moraxellales>Moraxellaceae>Psychrobacter | GCF_904846705.1 | Psychrobacter_sp._Pi2-52 | JMY00_RS07535 | 19023..22694 | 2774133 | BREX-system-P-loop-protein-BrxC-[Psychrobacter-sp.-Pi2-52] |
| WP_011513935.1 | Type1.BREX | [illegible] | 1201 | Bacteria>Pseudomonadota>Gammaproteobacteria>Moraxellales>Moraxellaceae>Psychrobacter | - | Psychrobacter_cryohalolentis | Pcryo_1609 | 1979457..1983062 | 330922 | BREX-system-P-loop-protein-BrxC-[Psychrobacter-cryohalolentis] |
| WP_010129447.1 | Type1.BREX | [illegible] | 1225 | Bacteria>Pseudomonadota | - | Gammaproteobacteria | MTT88_RS00175 | 38411..42088 | 1236 | MULTISPECIES:-BREX-system-P-loop-protein-BrxC-[Gammaproteobacteria] |
| WP_228349567.1 | Type1.BREX | [illegible] | 1203 | Bacteria>Pseudomonadota>Gammaproteobacteria>Nevskiales>Sinobacteraceae>Flagellatimonas | GCF_016918765.2 | Flagellatimonas_centrodinii | JN531_RS14475 | 2992880..2996491 | 2806210 | BREX-system-P-loop-protein-BrxC-[Flagellatimonas-centrodinii] |
| WP_161384457.1 | Type1.BREX | [illegible] | 1192 | Bacteria>Pseudomonadota>Gammaproteobacteria>Oceanospirillales>Alcanivoracaceae>Alcanivorax | - | Alcanivorax_sp._ZXX171 | GN155_RS10555 | 118232..121810 | 2695858 | BREX-system-P-loop-protein-BrxC-[Alcanivorax-sp.-ZXX171] |
| MBA3979207.1 | Type1.BREX | [illegible] | 1145 | Bacteria>Pseudomonadota>Gammaproteobacteria>Oceanospirillales>Alcanivoracaceae>Alcanivorax | GCA_013823245.1 | Alcanivorax_sp. | C0462_01260 | 7976..11413 | 1872427 | BREX-system-P-loop-protein-BrxC-[Alcanivorax-sp.] |
| WP_153499070.1 | Type6.BREX | [illegible] | 1214 | Bacteria>Pseudomonadota>Gammaproteobacteria>Oceanospirillales>Alcanivoracaceae>Alcanivorax | GCF_009601165.1 | Alcanivorax_sediminis | GFN93_RS03795 | 861896..865540 | 2663008 | BREX-system-P-loop-protein-BrxC-[Alcanivorax-sediminis] |
| WP_103916700.1 | Type1.BREX | [illegible] | 1193 | Bacteria>Pseudomonadota>Gammaproteobacteria>Oceanospirillales>Alcanivoracaceae>Alcanivorax | GCF_900107995.1 | Alcanivorax_sp._DSM_26293 | C4S63_RS03510 | 780383..783964 | 1798238 | BREX-system-P-loop-protein-BrxC-[Alcanivorax-sp.-DSM-26293] |
| WP_101745536.1 | Type1.BREX | [illegible] | 1220 | Bacteria>Pseudomonadota>Gammaproteobacteria>Oceanospirillales>Endozoicomonadaceae>Endozoicomonas | GCF_002864045.1 | Endozoicomonas_acroporae | CX664_RS04280 | 14964..18626 | 1701104 | BREX-system-P-loop-protein-BrxC-[Endozoicomonas-acroporae] |

| Protein-ID | System | Gene-Neighbourhoods | Length | Taxonomy | Assembly-ID | Strain | Locus-Tag | Genomic-coordinates | TaxID | Fasta-defline |
|---|---|---|---|---|---|---|---|---|---|---|
| WP_153766763.1 | Type1.BREX | <- PglZ <- ParB + HNH <-SF1-SF2-Helicase<- N6-METHYLASE <-?<- HNH \|\|?-><-?<- DUF6079 *<- BrxB-BrxF-iSTAND <- BrxA-DNA-binding <-XIS-HTH\|\|HTH_3-><-HTH_3\|\|REase+DDE_Tnp->TnsC-ATPase-> | 1223 | Bacteria>Pseudomonadota>Gammaproteobacteria>Oceanospirillales>Endozoicomonadaceae>Endozoicomonas | GCF_009653635.1 | Endozoicomonas_sp._OPT23 | EOPP23_14760 | 167593..171264 | 2072845 | BREX-system-P-loop-protein-BrxC-[Endozoicomonas-sp.-OPT23] |
| WP_163371285.1 | Type1.BREX | HD-> BrxA-DNA-binding -> BrxB-BrxF-iSTAND -> DUF6079 *-> N6-METHYLASE -> PglZ -> SIGMA+OB-Fold+AAA-ATPase+Lon-Protease-BrxL -> ParB + HNH ->OLD-ABC-> | 1220 | Bacteria>Pseudomonadota>Gammaproteobacteria>Oceanospirillales>Endozoicomonadaceae>Endozoicomonas | GCF_010994335.1 | Endozoicomonas_acroporae | E0230_RS03420 | 3343..7005 | 1701104 | BREX-system-P-loop-protein-BrxC-[Endozoicomonas-acroporae] |
| MCP5206320.1 | Type1.BREX | SF1-SF2-Helicase+REase->SF1-SF2-Helicase+REase-><- PglZ <- ParB + HNH <- HNH <-SbcC<-OLD-ABC<- N6-METHYLASE <- DUF6079 -p*\|\|DUF4372+DDE_Tnp_1-><- DUF6079 -p<- BrxB-BrxF-iSTAND <- [illegible] | 910 | Bacteria>Pseudomonadota>Gammaproteobacteria>Oceanospirillales>Hahellaceae>unclassified_Hahellaceae | GCA_024234475.1 | Hahellaceae_bacterium | H7A01_03865 | 876629..879361 | 2048971 | BREX-system-P-loop-protein-BrxC-[Hahellaceae-bacterium] |
| MCP5206322.1 | Type1.BREX | SF1-SF2-Helicase+REase-><- PglZ <- ParB + HNH <- HNH <-SbcC<-OLD-ABC<- N6-METHYLASE <- DUF6079 -p\|\|DUF4372+DDE_Tnp_1-><- DUF6079 -p*<- BrxB-BrxF-iSTAND <- BrxA-DNA-binding <- [illegible] | 261 | Bacteria>Pseudomonadota>Gammaproteobacteria>Oceanospirillales>Hahellaceae>unclassified_Hahellaceae | GCA_024234475.1 | Hahellaceae_bacterium | H7A01_03875 | 880608..881393 | 2048971 | hypothetical-protein-[Hahellaceae-bacterium] |
| WP_040239271.1 | Type1.BREX | <-sGTP<-?<-?\|\|?->tRNA-> BrxA-DNA-binding -> BrxB-BrxF-iSTAND -> DUF6079 *->OLD-ABC->PIN-> N6-METHYLASE -> ParB + HNH ->?->?->?-> PglZ -> SIGMA+OB-Fold+AAA-ATPase+Lon-Protease-BrxL -> [illegible] | 1144 | Bacteria>Pseudomonadota>Gammaproteobacteria>Oceanospirillales>Halomonadaceae>Chromohalobacter | GCF_000821045.2 | Chromohalobacter_japonicus | BN995_RS00415 | 38131..41565 | 223900 | BREX-system-P-loop-protein-BrxC-[Chromohalobacter-japonicus] |
| PHR03208.1 | Type1.BREX | <- SIGMA+OB-Fold+AAA-ATPase+Lon-Protease-BrxL <- PglZ <- ParB + HNH <-?<-?<-?<- N6-METHYLASE <-Schlafen+HATPase+wHTH<- DUF6079 *<- BrxB-BrxF-iSTAND <- BrxA-DNA-binding \|\|Mob_Pre-> [illegible] | 1144 | Bacteria>Pseudomonadota>Gammaproteobacteria>Oceanospirillales>Halomonadaceae>Halomonas | GCA_002733025.1 | Halomonas_sp. | COB32_05665 | 22826..26260 | 1486246 | hypothetical-protein-COB32_05665-[Halomonas-sp.] |
| NQY75752.1 | Type1.BREX | <-SbcC<-?<-?<- SIGMA+OB-Fold+AAA-ATPase+Lon-Protease-BrxL <- PglZ <- ParB + HNH <-HerA/FtsK<-SIR2<-HTH+REase<- DUF6079 *<- BrxB-BrxF-iSTAND <- BrxA-DNA-binding <-HTH_19<-TnsC- [illegible] | 1227 | Bacteria>Pseudomonadota>Gammaproteobacteria>Oceanospirillales>Halomonadaceae>Halomonas | GCA_013213865.1 | Halomonas_sp. | HRT78_01215 | 164704..168387 | 1486246 | BREX-system-P-loop-protein-BrxC-[Halomonas-sp.] |
| WP_017428908.1 | Type1.BREX | ZnR->?-><-?<- SIGMA+OB-Fold+AAA-ATPase+Lon-Protease-BrxL <- PglZ <- ParB + HNH <-?<- N6-METHYLASE <-RNaseH<- DUF6079 *<- BrxB-BrxF-iSTAND <- BrxA-DNA-binding <-XIS-HTH<- [illegible] | 1247 | Bacteria>Pseudomonadota>Gammaproteobacteria>Oceanospirillales>Halomonadaceae>Halomonas | GCF_000334215.1 | Halomonas_jeotgali | C793_RS0100125 | 27130..30873 | 553386 | BREX-system-P-loop-protein-BrxC-[Halomonas-jeotgali] |
| WP_035558509.1 | Type1.BREX | <-DUF4096+DDE_Tnp_1+DDE_Tnp_1_2<- SIGMA+OB-Fold+AAA-ATPase+Lon-Protease-BrxL <- PglZ <- ParB + HNH <-HerA/FtsK<-SIR2<- N6-METHYLASE <-SLOG+BirA-HTH<-SF1-SF2-Helicase+PRTase<- [illegible] | 1236 | Bacteria>Pseudomonadota>Gammaproteobacteria>Oceanospirillales>Halomonadaceae>Halomonas | GCF_000734975.2 | Halomonas_sp._KO116 | KO116_RS11715 | 2523126..2526836 | 1504981 | BREX-system-P-loop-protein-BrxC-[Halomonas-sp.-KO116] |
| WP_159343441.1 | Type1.BREX | <-Phage_AlpA\|\|HTH_26-><- SIGMA+OB-Fold+AAA-ATPase+HKD-DNase-BrxL <- PglZ <- ParB + HNH <- N6-METHYLASE <-SLOG+BirA-HTH<-SF1-SF2-Helicase+PRTase<-Schlafen+HATPase+wHTH+HTH<- [illegible] | 1193 | Bacteria>Pseudomonadota>Gammaproteobacteria>Oceanospirillales>Halomonadaceae>Halomonas | GCF_009846525.1 | Halomonas_meridiana | CTT34_RS16815 | 3563093..3566674 | 29570 | BREX-system-P-loop-protein-BrxC-[Halomonas-meridiana] |
| WP_161432258.1 | Type6.BREX | Arm-DNA-bind_3+Phage_integrase->?-><- SIGMA+OB-Fold+AAA-ATPase+HKD-DNase-BrxL <- PglZ <-DOC+HTH<- ParB + HNH <- N6-METHYLASE -p\|\|?-><- N6-METHYLASE <- DUF6079 *<- BrxB-BrxF-iSTAND <- [illegible] | 1223 | Bacteria>Pseudomonadota>Gammaproteobacteria>Oceanospirillales>Halomonadaceae>Halomonas | GCF_009902005.1 | Halomonas_alimentaria | GRB96_RS11410 | 2397586..2401257 | 147248 | BREX-system-P-loop-protein-BrxC-[Halomonas-alimentaria] |
| WP_172515133.1 | Type1.BREX | <- PglZ <- ParB + HNH <- N6-METHYLASE -p\|\|<- N6-METHYLASE -p<-HerA/FtsK<-?<-SF1-SF2-Helicase<- N6-METHYLASE <-Schlafen+HATPase+wHTH+HTH<- DUF6079 *<- BrxB-BrxF-iSTAND <- [illegible] | 1193 | Bacteria>Pseudomonadota>Gammaproteobacteria>Oceanospirillales>Halomonadaceae>Halomonas | GCF_011764625.1 | Halomonas_meridiana | HMEPL2_RS11780 | 2496852..2500433 | 29570 | BREX-system-P-loop-protein-BrxC-[Halomonas-meridiana] |
| WP_179915949.1 | Type1.BREX | <-SbcC<-?<-?<- SIGMA+OB-Fold+AAA-ATPase+Lon-Protease-BrxL <- ParB + HNH <-HerA/FtsK<-SIR2<- N6-METHYLASE <-HTH+REase<- DUF6079 *<- BrxB-BrxF-iSTAND <- BrxA-DNA-binding <-HTH_19<- [illegible] | 1227 | Bacteria>Pseudomonadota>Gammaproteobacteria>Oceanospirillales>Halomonadaceae>Halomonas | GCF_013415125.1 | Halomonas_glaciei | HZS80_RS09710 | 42867..46550 | 186761 | BREX-system-P-loop-protein-BrxC-[Halomonas-glaciei] |
| WP_188638083.1 | Type1.BREX | <-TarH+HAMP+MA\|\|ABC-ATPase+SbcC->?->?->?->?->tRNA-> BrxA-DNA-binding -> BrxB-BrxF-iSTAND -> DUF6079 *-> N6-METHYLASE -> ParB + HNH -> OLD-ABC-> HNH -> PglZ -> [illegible] | 1154 | Bacteria>Pseudomonadota>Gammaproteobacteria>Oceanospirillales>Halomonadaceae>Halomonas | GCF_014640815.1 | Halomonas_lutescens | IEW65_RS03380 | 721900..725364 | 1602943 | BREX-system-P-loop-protein-BrxC-[Halomonas-lutescens] |
| WP_019020105.1 | Type1.BREX | <-AlcRN+AraC-HTH<-?<-?<-?\|\|tRNA-><-Phage_integrase\|\|?->XIS-HTH-> BrxA-DNA-binding -> BrxB-BrxF-iSTAND -> DUF6079 *->OLD-ABC->REase-> N6-METHYLASE -> ParB + HNH -> PglZ ->Immank+ANK+ [illegible] | 1193 | Bacteria>Pseudomonadota>Gammaproteobacteria>Oceanospirillales>Halomonadaceae>Halomonas | GCF_014651575.1 | Halomonas_lutea | IE067_RS16235 | 61170..64751 | 453962 | BREX-system-P-loop-protein-BrxC-[Halomonas-lutea] |
| WP_263591023.1 | Type1.BREX | RepB_primase-><- SIGMA+OB-Fold+AAA-ATPase+Lon-Protease-BrxL <- PglZ <- ParB + HNH <-?<-?<-?<- N6-METHYLASE <-Schlafen+HATPase+wHTH<- DUF6079 *<- BrxB-BrxF-iSTAND <- BrxA-DNA-binding [illegible] | 1144 | Bacteria>Pseudomonadota>Gammaproteobacteria>Oceanospirillales>Halomonadaceae>Halomonas | GCF_025725685.1 | Halomonas_sp._M4R1S46 | OCT48_RS01560 | 327087..330521 | 2982692 | BREX-system-P-loop-protein-BrxC-[Halomonas-sp.-M4R1S46] |
| WP_089705394.1 | Type1.BREX | dCache_1+HisKA+HATPase_c-><-GAF+PAS+GGDEF+EAL\|\|?->tRNA->HTH_Tnp_1-><-ZnR+TOPRIM+AAA-helicase+wHTH<-TOPRIM\|\|XIS-HTH-> BrxA-DNA-binding -> BrxB-BrxF-iSTAND -> DUF6079 *->?-> [illegible] | 1189 | Bacteria>Pseudomonadota>Gammaproteobacteria>Oceanospirillales>Halomonadaceae>Halomonas | GCF_900103865.1 | Halomonas_arcis | BLR97_RS09380 | 247223..250792 | 416873 | BREX-system-P-loop-protein-BrxC-[Halomonas-arcis] |
| WP_081193131.1 | Type1.BREX | Peptidase_S49+Peptidase_S49-><-ArsR-HTH\|\|?->ABhydrolase-><-?\|\|?->tRNA-> BrxA-DNA-binding -> BrxB-BrxF-iSTAND -> DUF6079 *-> N6-METHYLASE -> ParB + HNH -> OLD-ABC-> HNH -> PglZ -> [illegible] | 1154 | Bacteria>Pseudomonadota>Gammaproteobacteria>Oceanospirillales>Halomonadaceae>Halomonas | - | Halomonas_sp._BC1 | B7L72_RS05665 | 10416..13880 | 1670448 | BREX-system-P-loop-protein-BrxC-[Halomonas-sp.-BC1] |
| WP_086901154.1 | Type1.BREX | MODE-HTH+PBPII->ACET->Arm-DNA-bind_3+Phage_integrase-><-REase+SF1-SF2-Helicase\|\| BrxA-DNA-binding -> BrxB-BrxF-iSTAND -> DUF6079 *->SF1-SF2-Helicase+PRTase->SLOG+BirA-HTH-> [illegible] | 1237 | Bacteria>Pseudomonadota>Gammaproteobacteria>Oceanospirillales>Halomonadaceae>Kushneria | GCF_002157205.1 | Kushneria_marisflavi | B9H00_RS13900 | 3054889..3058602 | 157779 | BREX-system-P-loop-protein-BrxC-[Kushneria-marisflavi] |
| WP_189514712.1 | Type1.BREX | [illegible] | 1144 | Bacteria>Pseudomonadota>Gammaproteobacteria>Oceanospirillales>Halomonadaceae>Kushneria | GCF_014652795.1 | Kushneria_pakistanensis | IE123_RS02380 | 531699..535133 | 1508770 | BREX-system-P-loop-protein-BrxC-[Kushneria-pakistanensis] |
| WP_110669866.1 | Type6.BREX | [illegible] | 1225 | Bacteria>Pseudomonadota>Gammaproteobacteria>Oceanospirillales>Halomonadaceae>Salinicola | GCF_003206155.1 | Salinicola_halophilus | DOM33_RS13680 | 401271..404948 | 184065 | BREX-system-P-loop-protein-BrxC-[Salinicola-halophilus] |
| WP_261843643.1 | Type1.BREX | [illegible] | 1225 | Bacteria>Pseudomonadota>Gammaproteobacteria>Oceanospirillales>Oceanospirillaceae>Aliamphritea | GCF_024347215.1 | Aliamphritea_ceti | OCU49_RS03510 | 764184..767861 | 1524258 | BREX-system-P-loop-protein-BrxC-[Aliamphritea-ceti] |
| WP_027853089.1 | Type1.BREX | [illegible] | 1154 | Bacteria>Pseudomonadota>Gammaproteobacteria>Oceanospirillales>Oceanospirillaceae>Marinobacterium | GCF_000428985.1 | Marinobacterium_litorale | G403_RS0105385 | 191863..195327 | 404770 | BREX-system-P-loop-protein-BrxC-[Marinobacterium-litorale] |
| WP_181736357.1 | Type1.BREX | [illegible] | 1154 | Bacteria>Pseudomonadota>Gammaproteobacteria>Oceanospirillales>Oceanospirillaceae>Marinobacterium | GCF_013868415.1 | Marinobacterium_sp._3-1745 | H1S06_RS00670 | 30655..34119 | 2756129 | BREX-system-P-loop-protein-BrxC-[Marinobacterium-sp.-3-1745] |
| AWY02455.1 | Type1.BREX | [illegible] | 1197 | Bacteria>Pseudomonadota>Gammaproteobacteria>Oceanospirillales>Oceanospirillaceae>Marinomonas | GCA_003285205.1 | Marinomonas_primoryensis | A8139_14855 | 3171018..3174611 | 178399 | hypothetical-protein-A8139_14855-[Marinomonas-primoryensis] |
| WP_100635909.1 | Type1.BREX | [illegible] | 1157 | Bacteria>Pseudomonadota>Gammaproteobacteria>Oceanospirillales>Oceanospirillaceae>Marinomonas | GCF_002806845.1 | Marinomonas_sp._ef1 | CCS02_RS04315 | 398110..401583 | 2005043 | BREX-system-P-loop-protein-BrxC-[Marinomonas-sp.-ef1] |
| WP_239495510.1 | Type1.BREX | [illegible] | 1199 | Bacteria>Pseudomonadota>Gammaproteobacteria>Oceanospirillales>Oceanospirillaceae>Marinomonas | GCF_003285205.1 | Marinomonas_primoryensis | A8139_RS14980 | 3171018..3174617 | 178399 | BREX-system-P-loop-protein-BrxC-[Marinomonas-primoryensis] |
| QEW05875.1 | Type6.BREX | [illegible] | 1212 | Bacteria>Pseudomonadota>Gammaproteobacteria>Oceanospirillales>Oceanospirillaceae>Nitrincola | GCA_008727795.1 | Nitrincola_iocasae | F5I99_04885 | 1032629..1036267 | 2614693 | BREX-system-P-loop-protein-BrxC-[Nitrincola-iocasae] |
| WP_225307559.1 | Type6.BREX | [illegible] | 1214 | Bacteria>Pseudomonadota>Gammaproteobacteria>Oceanospirillales>Oceanospirillaceae>Nitrincola | GCF_008727795.1 | Nitrincola_iocasae | F5I99_RS04885 | 1032623..1036267 | 2614693 | BREX-system-P-loop-protein-BrxC-[Nitrincola-iocasae] |
| WP_260997651.1 | Type1.BREX | [illegible] | 1156 | Bacteria>Pseudomonadota>Gammaproteobacteria>Oceanospirillales>Oceanospirillaceae>Thalassolituus | GCF_025345565.1 | Thalassolituus_hydrocarbonoclasticus | HUF19_RS16715 | 3723205..3726675 | 2742796 | BREX-system-P-loop-protein-BrxC-[Thalassolituus-hydrocarbonoclasticus] |
| MAS25385.1 | Type1.BREX | [illegible] | 1145 | Bacteria>Pseudomonadota>Gammaproteobacteria>Oceanospirillales>Oceanospirillaceae>unclassified_Oceanospirillaceae | GCA_002698675.1 | Oceanospirillaceae_bacterium | CMI08_10750 | 15208..18645 | 1899355 | hypothetical-protein-[Oceanospirillaceae-bacterium] |
| WP_116688140.1 | Type1.BREX | [illegible] | 1236 | Bacteria>Pseudomonadota>Gammaproteobacteria>Oceanospirillales>Oceanospirillales_incertae_sedis>Pelagibaculum | GCF_003097315.1 | Pelagibaculum_spongiae | DC094_RS16040 | 158438..162148 | 2080658 | BREX-system-P-loop-protein-BrxC-[Pelagibaculum-spongiae] |
| WP_100257386.1 | Type1.BREX | [illegible] | 1215 | Bacteria>Pseudomonadota>Gammaproteobacteria>Oceanospirillales>Saccharospirillaceae>Reinekea | GCF_002795845.1 | Reinekea_forsetii | REIFOR_RS09825 | 2117568..2121215 | 1336806 | BREX-system-P-loop-protein-BrxC-[Reinekea-forsetii] |
| MBR9867112.1 | Type1.BREX | [illegible] | 1225 | Bacteria>Pseudomonadota>Gammaproteobacteria>Oceanospirillales>unclassified_Oceanospirillales | GCA_018223265.1 | Oceanospirillales_bacterium | GYB18_08415 | 21169..24846 | 1914300 | BREX-system-P-loop-protein-BrxC-[Oceanospirillales-bacterium] |
| WP_241468927.1 | Type1.BREX | [illegible] | 1225 | Bacteria>Pseudomonadota>Gammaproteobacteria>Pasteurellales>Pasteurellaceae>Pasteurella | GCF_022559605.1 | Pasteurella_multocida | MMZ64_RS09105 | 1895707..1899384 | 747 | BREX-system-P-loop-protein-BrxC-[Pasteurella-multocida] |
| WP_119518392.1 | Type1.BREX | [illegible] | 1154 | Bacteria>Pseudomonadota>Gammaproteobacteria>Pseudomonadales | GCF_021610005.1 | Pseudomonadaceae | L1776_RS13375 | 2929669..2933133 | 135621 | MULTISPECIES:-BREX-system-P-loop-protein-BrxC-[Pseudomonadaceae] |
| WP_058011517.1 | Type6.BREX | [illegible] | 1213 | Bacteria>Pseudomonadota>Gammaproteobacteria>Pseudomonadales | GCF_901482535.1 | Pseudomonadaceae | FQU16_RS01885 | 279271..282912 | 135621 | MULTISPECIES:-BREX-system-P-loop-protein-BrxC-[Pseudomonadaceae] |
| MBE95557.1 | Type1.BREX | [illegible] | 1193 | Bacteria>Pseudomonadota>Gammaproteobacteria>Pseudomonadales>Marinobacteraceae>Marinobacter | GCA_002714765.1 | Marinobacter_sp. | CMG89_11180 | 696..4277 | 50741 | hypothetical-protein-[Marinobacter-sp.] |
| WP_048384700.1 | Type1.BREX | [illegible] | 1145 | Bacteria>Pseudomonadota>Gammaproteobacteria>Pseudomonadales>Marinobacteraceae>Marinobacter | GCF_001043175.1 | Marinobacter_psychrophilus | ABA45_RS05800 | 1298938..1302375 | 330734 | BREX-system-P-loop-protein-BrxC-[Marinobacter-psychrophilus] |
| WP_071262841.1 | Type1.BREX | [illegible] | 1217 | Bacteria>Pseudomonadota>Gammaproteobacteria>Pseudomonadales>Marinobacteraceae>Marinobacter | GCF_001858325.1 | Marinobacter_sp._AC-23 | BCA33_RS00460 | 104149..107802 | 1879031 | BREX-system-P-loop-protein-BrxC-[Marinobacter-sp.-AC-23] |
| WP_141068072.1 | Type1.BREX | [illegible] | 1193 | Bacteria>Pseudomonadota>Gammaproteobacteria>Pseudomonadales>Marinobacteraceae>Marinobacter | GCF_006516615.1 | Marinobacter_nauticus | FH712_RS17655 | 33566..37147 | 2743 | BREX-system-P-loop-protein-BrxC-[Marinobacter-nauticus] |
| WP_213480159.1 | Type1.BREX | [illegible] | 1145 | Bacteria>Pseudomonadota>Gammaproteobacteria>Pseudomonadales>Marinobacteraceae>Marinobacter | GCF_018424725.1 | Marinobacter_salarius | DYI41_RS19390 | 15297..18734 | 1420917 | BREX-system-P-loop-protein-BrxC-[Marinobacter-salarius] |
| WP_216008134.1 | Type1.BREX | [illegible] | 1217 | Bacteria>Pseudomonadota>Gammaproteobacteria>Pseudomonadales>Marinobacteraceae>Marinobacter | GCF_018860765.1 | Marinobacter_salexigens | KO508_RS09770 | 175903..179556 | 1925763 | BREX-system-P-loop-protein-BrxC-[Marinobacter-salexigens] |
| WP_247116152.1 | Type1.BREX | [illegible] | 1189 | Bacteria>Pseudomonadota>Gammaproteobacteria>Pseudomonadales>Marinobacteraceae>Marinobacter | GCF_023016045.1 | Marinobacter_sp._S6332 | MWU62_RS17920 | 19915..23484 | 2926403 | BREX-system-P-loop-protein-BrxC-[Marinobacter-sp.-S6332] |
| WP_248236652.1 | Type1.BREX | [illegible] | 1145 | Bacteria>Pseudomonadota>Gammaproteobacteria>Pseudomonadales>Marinobacteraceae>Marinobacter | GCF_023156245.1 | Marinobacter_bryozoorum | MLC59_RS17300 | 17294..20731 | 256324 | BREX-system-P-loop-protein-BrxC-[Marinobacter-bryozoorum] |
| WP_249013861.1 | Type1.BREX | [illegible] | 1217 | Bacteria>Pseudomonadota>Gammaproteobacteria>Pseudomonadales>Marinobacteraceae>Marinobacter | GCF_023311895.1 | Marinobacter_sp._M3C | MIH18_RS02775 | 634198..637851 | 2917715 | BREX-system-P-loop-protein-BrxC-[Marinobacter-sp.-M3C] |
| WP_092021229.1 | Type1.BREX | [illegible] | 1219 | Bacteria>Pseudomonadota>Gammaproteobacteria>Pseudomonadales>Marinobacteraceae>Marinobacter | GCF_900114775.1 | Marinobacter_zhejiangensis | BMZ67_RS06955 | 334318..337977 | 488535 | BREX-system-P-loop-protein-BrxC-[Marinobacter-zhejiangensis] |
| WP_019470415.1 | Type6.BREX | [illegible] | 1212 | Bacteria>Pseudomonadota>Gammaproteobacteria>Pseudomonadales>Pseudomonadaceae | GCF_000292285.1 | Pseudomonas | N805_RS25690 | 5809538..5813176 | 286 | MULTISPECIES:-BREX-system-P-loop-protein-BrxC-[Pseudomonas] |
| WP_057024539.1 | Type1.BREX | [illegible] | 1154 | Bacteria>Pseudomonadota>Gammaproteobacteria>Pseudomonadales>Pseudomonadaceae | GCF_002091795.1 | Pseudomonas | TU77_RS22280 | 179418..182882 | 286 | MULTISPECIES:-BREX-system-P-loop-protein-BrxC-[Pseudomonas] |
| WP_096426782.1 | Type1.BREX | [illegible] | 1185 | Bacteria>Pseudomonadota>Gammaproteobacteria>Pseudomonadales>Pseudomonadaceae | GCF_002356095.1 | Pseudomonas | J3A71_RS03870 | 842777..846334 | 286 | MULTISPECIES:-BREX-system-P-loop-protein-BrxC-[Pseudomonas] |
| WP_196184087.1 | Type1.BREX | [illegible] | 1154 | Bacteria>Pseudomonadota>Gammaproteobacteria>Pseudomonadales>Pseudomonadaceae | GCF_015680805.1 | Pseudomonas | IR013_RS15060 | 51443..54907 | 286 | MULTISPECIES:-BREX-system-P-loop-protein-BrxC-[Pseudomonas] |
| WP_033986129.1 | Type6.BREX | [illegible] | 1212 | Bacteria>Pseudomonadota>Gammaproteobacteria>Pseudomonadales>Pseudomonadaceae | GCF_020180035.1 | Pseudomonas | AO894_RS16050 | 13512..17150 | 286 | MULTISPECIES:-BREX-system-P-loop-protein-BrxC-[Pseudomonas] |
| WP_064314265.1 | Type1.BREX | [illegible] | 1154 | Bacteria>Pseudomonadota>Gammaproteobacteria>Pseudomonadales>Pseudomonadaceae | GCF_021601465.1 | Pseudomonas | L6D68_RS06035 | 31411..34875 | 286 | MULTISPECIES:-BREX-system-P-loop-protein-BrxC-[Pseudomonas] |
| WP_021219096.1 | Type1.BREX | [illegible] | 1220 | Bacteria>Pseudomonadota>Gammaproteobacteria>Pseudomonadales>Pseudomonadaceae | GCF_023093875.1 | Pseudomonas | MYO55_RS15275 | 3320590..3324252 | 286 | MULTISPECIES:-BREX-system-P-loop-protein-BrxC-[Pseudomonas] |
| WP_060765715.1 | Type1.BREX | [illegible] | 1186 | Bacteria>Pseudomonadota>Gammaproteobacteria>Pseudomonadales>Pseudomonadaceae | GCF_029959765.1 | Pseudomonas | PFL603g_RS03160 | 383988..387548 | 286 | MULTISPECIES:-BREX-system-P-loop-protein-BrxC-[Pseudomonas] |
| WP_023115268.1 | Type1.BREX | [illegible] | 1154 | Bacteria>Pseudomonadota>Gammaproteobacteria>Pseudomonadales>Pseudomonadaceae | GCF_900707185.1 | Pseudomonas | E4Y87_RS04340 | 6868..10332 | 286 | MULTISPECIES:-BREX-system-P-loop-protein-BrxC-[Pseudomonas] |
| WP_198417420.1 | Type1.BREX | [illegible] | 1159 | Bacteria>Pseudomonadota>Gammaproteobacteria>Pseudomonadales>Pseudomonadaceae | GCF_902329585.1 | Pseudomonas | PPTH39_RS22285 | 114113..117592 | 286 | MULTISPECIES:-BREX-system-P-loop-protein-BrxC-[Pseudomonas] |
| WP_104739193.1 | Type1.BREX | [illegible] | 1180 | Bacteria>Pseudomonadota>Gammaproteobacteria>Pseudomonadales>Pseudomonadaceae>Halopseudomonas | GCF_014641295.1 | Halopseudomonas_oceani | IEX06_RS08390 | 366595..370137 | 1708783 | BREX-system-P-loop-protein-BrxC-[Halopseudomonas-oceani] |
| EQL43544.1 | Type6.BREX | [illegible] | 1213 | Bacteria>Pseudomonadota>Gammaproteobacteria>Pseudomonadales>Pseudomonadaceae>Pseudomonas | GCA_000467675.1 | Pseudomonas_aeruginosa_VRFPA03 | M770_31820 | 18788..22429 | 1350465 | ATPase-[Pseudomonas-aeruginosa-VRFPA03] |
| ARQ76275.1 | Type1.BREX | [illegible] | 1156 | Bacteria>Pseudomonadota>Gammaproteobacteria>Pseudomonadales>Pseudomonadaceae>Pseudomonas | GCA_002128325.1 | Pseudomonas_fragi | B6D87_19590 | 4227178..4230648 | 296 | hypothetical-protein-B6D87_19590-[Pseudomonas-fragi] |
| PAA41060.1 | Type1.BREX | [illegible] | 1156 | Bacteria>Pseudomonadota>Gammaproteobacteria>Pseudomonadales>Pseudomonadaceae>Pseudomonas | GCA_002269445.1 | Pseudomonas_fragi | CJU80_12345 | 107560..111030 | 296 | hypothetical-protein-CJU80_12345-[Pseudomonas-fragi] |
| ATN11139.1 | Type1.BREX | [illegible] | 1222 | Bacteria>Pseudomonadota>Gammaproteobacteria>Pseudomonadales>Pseudomonadaceae>Pseudomonas | GCA_002591235.1 | Pseudomonas_sp._FDAARGOS_380 | CRN80_16460 | 3507093..3510761 | 2018067 | hypothetical-protein-CRN80_16460-[Pseudomonas-sp.-FDAARGOS_380] |
| TKK14312.1 | Type1.BREX | [illegible] | 1222 | Bacteria>Pseudomonadota>Gammaproteobacteria>Pseudomonadales>Pseudomonadaceae>Pseudomonas | GCA_005233695.1 | Pseudomonas_fluorescens | PflCFBP13510_04710 | 495926..499594 | 294 | BREX-system-P-loop-protein-BrxC-[Pseudomonas-fluorescens] |
| MVW77192.1 | Type1.BREX | [illegible] | 1221 | Bacteria>Pseudomonadota>Gammaproteobacteria>Pseudomonadales>Pseudomonadaceae>Pseudomonas | GCA_009763245.1 | Pseudomonas_xionganensis | GJV18_17880 | 24794..28459 | 2654845 | BREX-system-P-loop-protein-BrxC-[Pseudomonas-xionganensis] |
| NLU01326.1 | Type1.BREX | [illegible] | 1222 | Bacteria>Pseudomonadota>Gammaproteobacteria>Pseudomonadales>Pseudomonadaceae>Pseudomonas | GCA_012720005.1 | Pseudomonas_lundensis | GXW88_11975 | 7910..11578 | 86185 | BREX-system-P-loop-protein-BrxC-[Pseudomonas-lundensis] |
| NMY25069.1 | Type1.BREX | [illegible] | 1222 | Bacteria>Pseudomonadota>Gammaproteobacteria>Pseudomonadales>Pseudomonadaceae>Pseudomonas | GCA_012985535.1 | Pseudomonas_sp._WS_5021 | HBO19_03685 | 794090..797758 | 2717490 | BREX-system-P-loop-protein-BrxC-[Pseudomonas-sp.-WS-5021] |
| NWF08290.1 | Type1.BREX | [illegible] | 1222 | Bacteria>Pseudomonadota>Gammaproteobacteria>Pseudomonadales>Pseudomonadaceae>Pseudomonas | GCA_013386885.1 | Pseudomonas_salomonii | HX810_11520 | 509017..512685 | 191391 | BREX-system-P-loop-protein-BrxC-[Pseudomonas-salomonii] |
| MBF9242733.1 | Type1.BREX | [illegible] | 1222 | Bacteria>Pseudomonadota>Gammaproteobacteria>Pseudomonadales>Pseudomonadaceae>Pseudomonas | GCA_015694365.1 | Pseudomonas_syringae_pv._tomato | I4I80_00295 | 46308..49976 | 323 | BREX-system-P-loop-protein-BrxC-[Pseudomonas-syringae-pv.-tomato] |
| MBG3983980.1 | Type6.BREX | [illegible] | 1212 | Bacteria>Pseudomonadota>Gammaproteobacteria>Pseudomonadales>Pseudomonadaceae>Pseudomonas | GCA_015733145.1 | Pseudomonas_aeruginosa | I4919_20885 | 86674..90312 | 287 | BREX-system-P-loop-protein-BrxC-[Pseudomonas-aeruginosa] |
| MBG4373384.1 | Type1.BREX | [illegible] | 1154 | Bacteria>Pseudomonadota>Gammaproteobacteria>Pseudomonadales>Pseudomonadaceae>Pseudomonas | GCA_015734445.1 | Pseudomonas_aeruginosa | I5F50_23085 | 8082..11546 | 287 | BREX-system-P-loop-protein-BrxC-[Pseudomonas-aeruginosa] |
| MBG4395801.1 | Type1.BREX | [illegible] | 1154 | Bacteria>Pseudomonadota>Gammaproteobacteria>Pseudomonadales>Pseudomonadaceae>Pseudomonas | GCA_015734545.1 | Pseudomonas_aeruginosa | GHT73_12640 | 7113..10577 | 287 | BREX-system-P-loop-protein-BrxC-[Pseudomonas-aeruginosa] |
| MBG4463093.1 | Type1.BREX | [illegible] | 1154 | Bacteria>Pseudomonadota>Gammaproteobacteria>Pseudomonadales>Pseudomonadaceae>Pseudomonas | GCA_015734755.1 | Pseudomonas_aeruginosa | GHU50_17045 | 10342..13806 | 287 | BREX-system-P-loop-protein-BrxC-[Pseudomonas-aeruginosa] |
| MBG5216128.1 | Type1.BREX | [illegible] | 1154 | Bacteria>Pseudomonadota>Gammaproteobacteria>Pseudomonadales>Pseudomonadaceae>Pseudomonas | GCA_015737185.1 | Pseudomonas_aeruginosa | GHT75_27125 | 7080..10544 | 287 | BREX-system-P-loop-protein-BrxC-[Pseudomonas-aeruginosa] |
| MBG6382130.1 | Type6.BREX | [illegible] | 1210 | Bacteria>Pseudomonadota>Gammaproteobacteria>Pseudomonadales>Pseudomonadaceae>Pseudomonas | GCA_015762815.1 | Pseudomonas_aeruginosa | I5I12_28760 | 1691..5323 | 287 | BREX-system-P-loop-protein-BrxC-[Pseudomonas-aeruginosa] |
| MBH3980671.1 | Type6.BREX | [illegible] | 1213 | Bacteria>Pseudomonadota>Gammaproteobacteria>Pseudomonadales>Pseudomonadaceae>Pseudomonas | GCA_016011075.1 | Pseudomonas_aeruginosa | I5Q70_02570 | 503331..506972 | 287 | BREX-system-P-loop-protein-BrxC-[Pseudomonas-aeruginosa] |
| MBI8737743.1 | Type1.BREX | [illegible] | 1154 | Bacteria>Pseudomonadota>Gammaproteobacteria>Pseudomonadales>Pseudomonadaceae>Pseudomonas | GCA_016342945.1 | Pseudomonas_aeruginosa | JEZ70_02820 | 55146..58610 | 287 | BREX-system-P-loop-protein-BrxC-[Pseudomonas-aeruginosa] |
| MBM1201834.1 | Type1.BREX | [illegible] | 1154 | Bacteria>Pseudomonadota>Gammaproteobacteria>Pseudomonadales>Pseudomonadaceae>Pseudomonas | GCA_016812215.1 | Pseudomonas_fragi | GYM96_18820 | 91266..94730 | 296 | BREX-system-P-loop-protein-BrxC-[Pseudomonas-fragi] |
| MBN0299744.1 | Type6.BREX | [illegible] | 1213 | Bacteria>Pseudomonadota>Gammaproteobacteria>Pseudomonadales>Pseudomonadaceae>Pseudomonas | GCA_016922145.1 | Pseudomonas_aeruginosa | JTM78_25120 | 125675..129316 | 287 | BREX-system-P-loop-protein-BrxC-[Pseudomonas-aeruginosa] |
| MBN0536373.1 | Type6.BREX | [illegible] | 1206 | Bacteria>Pseudomonadota>Gammaproteobacteria>Pseudomonadales>Pseudomonadaceae>Pseudomonas | GCA_016922535.1 | Pseudomonas_aeruginosa | JTM59_25840 | 125679..129299 | 287 | BREX-system-P-loop-protein-BrxC-[Pseudomonas-aeruginosa] |
| MBV5724753.1 | Type1.BREX | [illegible] | 1184 | Bacteria>Pseudomonadota>Gammaproteobacteria>Pseudomonadales>Pseudomonadaceae>Pseudomonas | GCA_019166825.1 | Pseudomonas_aeruginosa | KUT37_26485 | 44544..48098 | 287 | BREX-system-P-loop-protein-BrxC-[Pseudomonas-aeruginosa] |
| MBX6150345.1 | Type1.BREX | [illegible] | 1154 | Bacteria>Pseudomonadota>Gammaproteobacteria>Pseudomonadales>Pseudomonadaceae>Pseudomonas | GCA_019686265.1 | Pseudomonas_aeruginosa | ISD36_12550 | 248185..251649 | 287 | BREX-system-P-loop-protein-BrxC-[Pseudomonas-aeruginosa] |
| MBX5915342.1 | Type1.BREX | [illegible] | 1154 | Bacteria>Pseudomonadota>Gammaproteobacteria>Pseudomonadales>Pseudomonadaceae>Pseudomonas | GCA_019688355.1 | Pseudomonas_aeruginosa | ISE50_29115 | 6954..10418 | 287 | BREX-system-P-loop-protein-BrxC-[Pseudomonas-aeruginosa] |
| MBY8973665.1 | Type1.BREX | [illegible] | 1222 | Bacteria>Pseudomonadota>Gammaproteobacteria>Pseudomonadales>Pseudomonadaceae>Pseudomonas | GCA_019891455.1 | Pseudomonas_sp._P867 | J1G33_25110 | 13708..17376 | 2816050 | BREX-system-P-loop-protein-BrxC-[Pseudomonas-sp.-P867] |
| HBN8119635.1 | Type6.BREX | [illegible] | 1213 | Bacteria>Pseudomonadota>Gammaproteobacteria>Pseudomonadales>Pseudomonadaceae>Pseudomonas | GCA_021586615.1 | Pseudomonas_aeruginosa | L3655_000128 | 111220..114861 | 287 | BREX-system-P-loop-protein-BrxC-[Pseudomonas-aeruginosa] |
| HBN9708921.1 | Type1.BREX | [illegible] | 1154 | Bacteria>Pseudomonadota>Gammaproteobacteria>Pseudomonadales>Pseudomonadaceae>Pseudomonas | GCA_021591785.1 | Pseudomonas_aeruginosa | L3920_001787 | 275283..278747 | 287 | BREX-system-P-loop-protein-BrxC-[Pseudomonas-aeruginosa] |
| HBO1619179.1 | Type1.BREX | [illegible] | 1154 | Bacteria>Pseudomonadota>Gammaproteobacteria>Pseudomonadales>Pseudomonadaceae>Pseudomonas | GCA_021598085.1 | Pseudomonas_aeruginosa | L4F62_005712 | 16995..20459 | 287 | BREX-system-P-loop-protein-BrxC-[Pseudomonas-aeruginosa] |
| EIU3464059.1 | Type1.BREX | [illegible] | 1154 | Bacteria>Pseudomonadota>Gammaproteobacteria>Pseudomonadales>Pseudomonadaceae>Pseudomonas | GCA_021644165.1 | Pseudomonas_aeruginosa | L4P80_001287 | 38375..41839 | 287 | BREX-system-P-loop-protein-BrxC-[Pseudomonas-aeruginosa] |
| HBO3309314.1 | Type6.BREX | [illegible] | 1188 | Bacteria>Pseudomonadota>Gammaproteobacteria>Pseudomonadales>Pseudomonadaceae>Pseudomonas | GCA_021652015.1 | Pseudomonas_aeruginosa | L4R58_004351 | 30228..33794 | 287 | BREX-system-P-loop-protein-BrxC-[Pseudomonas-aeruginosa] |
| HBO6336105.1 | Type1.BREX | [illegible] | 1154 | Bacteria>Pseudomonadota>Gammaproteobacteria>Pseudomonadales>Pseudomonadaceae>Pseudomonas | GCA_021670735.1 | Pseudomonas_aeruginosa | L4747_005548 | 6948..10412 | 287 | BREX-system-P-loop-protein-BrxC-[Pseudomonas-aeruginosa] |
| HBP6330513.1 | Type1.BREX | [illegible] | 1154 | Bacteria>Pseudomonadota>Gammaproteobacteria>Pseudomonadales>Pseudomonadaceae>Pseudomonas | GCA_021710255.1 | Pseudomonas_aeruginosa | L6A37_21805 | 72354..75818 | 287 | BREX-system-P-loop-protein-BrxC-[Pseudomonas-aeruginosa] |
| HBP5527123.1 | Type6.BREX | [illegible] | 1212 | Bacteria>Pseudomonadota>Gammaproteobacteria>Pseudomonadales>Pseudomonadaceae>Pseudomonas | GCA_021712095.1 | Pseudomonas_aeruginosa | L6B52_09510 | 5147..8785 | 287 | BREX-system-P-loop-protein-BrxC-[Pseudomonas-aeruginosa] |
| MCK9534930.1 | Type1.BREX | [illegible] | 1154 | Bacteria>Pseudomonadota>Gammaproteobacteria>Pseudomonadales>Pseudomonadaceae>Pseudomonas | GCA_023228505.1 | Pseudomonas_sp. | MOQ98_09595 | 5981..9445 | 306 | BREX-system-P-loop-protein-BrxC-[Pseudomonas-sp.] |
| HCE6004670.1 | Type6.BREX | [illegible] | 1213 | Bacteria>Pseudomonadota>Gammaproteobacteria>Pseudomonadales>Pseudomonadaceae>Pseudomonas | GCA_023904815.1 | Pseudomonas_aeruginosa | NHK94_000107 | 121075..124716 | 287 | BREX-system-P-loop-protein-BrxC-[Pseudomonas-aeruginosa] |
| MCO2467177.1 | Type1.BREX | [illegible] | 1146 | Bacteria>Pseudomonadota>Gammaproteobacteria>Pseudomonadales>Pseudomonadaceae>Pseudomonas | GCA_023907445.1 | Pseudomonas_aeruginosa | FA343_15170 | 67465..70905 | 287 | BREX-system-P-loop-protein-BrxC-[Pseudomonas-aeruginosa] |
| HCE7261574.1 | Type1.BREX | [illegible] | 1154 | Bacteria>Pseudomonadota>Gammaproteobacteria>Pseudomonadales>Pseudomonadaceae>Pseudomonas | GCA_023919205.1 | Pseudomonas_aeruginosa | NHQ13_005985 | 6842..10306 | 287 | BREX-system-P-loop-protein-BrxC-[Pseudomonas-aeruginosa] |
| HCF1683404.1 | Type6.BREX | [illegible] | 1213 | Bacteria>Pseudomonadota>Gammaproteobacteria>Pseudomonadales>Pseudomonadaceae>Pseudomonas | GCA_023941775.1 | Pseudomonas_aeruginosa | NH867_002024 | 49404..53045 | 287 | BREX-system-P-loop-protein-BrxC-[Pseudomonas-aeruginosa] |
| HCH0453160.1 | Type1.BREX | [illegible] | 1154 | Bacteria>Pseudomonadota>Gammaproteobacteria>Pseudomonadales>Pseudomonadaceae>Pseudomonas | GCA_024072695.1 | Pseudomonas_aeruginosa | NKO07_001951 | 228071..231535 | 287 | BREX-system-P-loop-protein-BrxC-[Pseudomonas-aeruginosa] |
| BDM24922.1 | Type1.BREX | [illegible] | 1154 | Bacteria>Pseudomonadota>Gammaproteobacteria>Pseudomonadales>Pseudomonadaceae>Pseudomonas | GCA_024349685.1 | Pseudomonas_sp._LRP2-20 | KMS_R46790 | 5328463..5331927 | 2944234 | BREX-system-P-loop-protein-BrxC-[Pseudomonas-sp.-LRP2-20] |
| WCY64468.1 | Type6.BREX | [illegible] | 1213 | Bacteria>Pseudomonadota>Gammaproteobacteria>Pseudomonadales>Pseudomonadaceae>Pseudomonas | GCA_028610105.1 | Pseudomonas_aeruginosa | NHK94_000107 | 121075..124716 | 287 | BREX-system-P-loop-protein-BrxC-[Pseudomonas-aeruginosa] |
| WP_132538218.1 | Type1.BREX | [illegible] | 1154 | Bacteria>Pseudomonadota>Gammaproteobacteria>Pseudomonadales>Pseudomonadaceae>Pseudomonas | GCA_030820635.1 | Pseudomonas_aeruginosa | EQ548_RS01360 | 6947..10411 | 287 | BREX-system-P-loop-protein-BrxC-[Pseudomonas-aeruginosa] |
| WP_230691811.1 | Type1.BREX | [illegible] | 1225 | Bacteria>Pseudomonadota>Gammaproteobacteria>Pseudomonadales>Pseudomonadaceae>Pseudomonas | GCA_030820635.1 | Pseudomonas_syringae_group_genomosp._3 | I4I80_RS00295 | 46299..49976 | 251701 | BREX-system-P-loop-protein-BrxC-[Pseudomonas-syringae-group-genomosp.-3] |
| WP_241090907.1 | Type1.BREX | [illegible] | 1225 | Bacteria>Pseudomonadota>Gammaproteobacteria>Pseudomonadales>Pseudomonadaceae>Pseudomonas | GCA_030820635.1 | Pseudomonas_viridiflava | EE220_RS21960 | 64975..68652 | 33069 | BREX-system-P-loop-protein-BrxC-[Pseudomonas-viridiflava] |
| SBW85146.1 | Type1.BREX | [illegible] | 1154 | Bacteria>Pseudomonadota>Gammaproteobacteria>Pseudomonadales>Pseudomonadaceae>Pseudomonas | GCA_900092355.1 | Pseudomonas_veronii_1YdBTEX2 | PVE_P0103 | 88310..91774 | 1295141 | hypothetical-protein-PVE_P0103-[Pseudomonas-veronii-1YdBTEX2] |
| VVQ23891.1 | Type5.BREX | [illegible] | 1225 | Bacteria>Pseudomonadota>Gammaproteobacteria>Pseudomonadales>Pseudomonadaceae>Pseudomonas | GCA_902498245.1 | Pseudomonas_fluorescens | PS928_05643 | 91300..94977 | 294 | hypothetical-protein-PS928_05643-[Pseudomonas-fluorescens] |
| WP_013692225.1 | Type6.BREX | [illegible] | 1213 | Bacteria>Pseudomonadota>Gammaproteobacteria>Pseudomonadales>Pseudomonadaceae>Pseudomonas | GCF_000194805.1 | Pseudomonas_brassicacearum | PSEBR_RS03955 | 941705..945346 | 930166 | BREX-system-P-loop-protein-BrxC-[Pseudomonas-brassicacearum] |
| WP_007159175.1 | Type6.BREX | [illegible] | 1212 | Bacteria>Pseudomonadota>Gammaproteobacteria>Pseudomonadales>Pseudomonadaceae>Pseudomonas | GCF_000236825.1 | Pseudomonas_psychrotolerans | PPL19_RS03910 | 152166..155804 | 237610 | BREX-system-P-loop-protein-BrxC-[Pseudomonas-psychrotolerans] |
| WP_010463595.1 | Type1.BREX | [illegible] | 1185 | Bacteria>Pseudomonadota>Gammaproteobacteria>Pseudomonadales>Pseudomonadaceae>Pseudomonas | GCF_000257545.3 | Pseudomonas_mandelii | OU5_RS13450 | 3003724..3007281 | 75612 | BREX-system-P-loop-protein-BrxC-[Pseudomonas-mandelii] |
| WP_017337797.1 | Type6.BREX | [illegible] | 1213 | Bacteria>Pseudomonadota>Gammaproteobacteria>Pseudomonadales>Pseudomonadaceae>Pseudomonas | GCF_000293885.2 | Pseudomonas_fluorescens | B723_RS16820 | 3631858..3635499 | 294 | BREX-system-P-loop-protein-BrxC-[Pseudomonas-fluorescens] |
| WP_023113968.1 | Type1.BREX | [illegible] | 1146 | Bacteria>Pseudomonadota>Gammaproteobacteria>Pseudomonadales>Pseudomonadaceae>Pseudomonas | GCF_000481245.1 | Pseudomonas_aeruginosa | Q036_RS16830 | 2151526..2154966 | 287 | BREX-system-P-loop-protein-BrxC-[Pseudomonas-aeruginosa] |
| WP_042135808.1 | Type1.BREX | [illegible] | 1184 | Bacteria>Pseudomonadota>Gammaproteobacteria>Pseudomonadales>Pseudomonadaceae>Pseudomonas | GCF_000730625.1 | Pseudomonas_oryzihabitans | POR01S_RS10925 | 121811..125365 | 47885 | BREX-system-P-loop-protein-BrxC-[Pseudomonas-oryzihabitans] |
| WP_033954069.1 | Type1.BREX | [illegible] | 1154 | Bacteria>Pseudomonadota>Gammaproteobacteria>Pseudomonadales>Pseudomonadaceae>Pseudomonas | GCF_000793385.1 | Pseudomonas_aeruginosa | NT88_RS29575 | 15006..18470 | 287 | BREX-system-P-loop-protein-BrxC-[Pseudomonas-aeruginosa] |
| WP_044305280.1 | Type6.BREX | [illegible] | 1213 | Bacteria>Pseudomonadota>Gammaproteobacteria>Pseudomonadales>Pseudomonadaceae>Pseudomonas | GCF_000793505.1 | Pseudomonas_aeruginosa | NT87_RS20770 | 137629..141270 | 287 | BREX-system-P-loop-protein-BrxC-[Pseudomonas-aeruginosa] |
| WP_202808417.1 | Type1.BREX | [illegible] | 937 | Bacteria>Pseudomonadota>Gammaproteobacteria>Pseudomonadales>Pseudomonadaceae>Pseudomonas | GCF_000794515.1 | Pseudomonas_aeruginosa | NQ84_RS28370 | 16579..19392 | 287 | BREX-system-P-loop-protein-BrxC-[Pseudomonas-aeruginosa] |
| WP_228480203.1 | Type1.BREX | [illegible] | 619 | Bacteria>Pseudomonadota>Gammaproteobacteria>Pseudomonadales>Pseudomonadaceae>Pseudomonas | GCF_000829885.1 | Pseudomonas_aeruginosa | EG09_RS07505 | 1715440..1717299 | 287 | BREX-system-P-loop-protein-BrxC-[Pseudomonas-aeruginosa] |
| WP_041506104.1 | Type1.BREX | [illegible] | 1184 | Bacteria>Pseudomonadota>Gammaproteobacteria>Pseudomonadales>Pseudomonadaceae>Pseudomonas | GCF_000831585.1 | Pseudomonas_plecoglossicida | RK21_RS13525 | 2976546..2980100 | 70775 | BREX-system-P-loop-protein-BrxC-[Pseudomonas-plecoglossicida] |
| WP_047283135.1 | Type1.BREX | [illegible] | 1154 | Bacteria>Pseudomonadota>Gammaproteobacteria>Pseudomonadales>Pseudomonadaceae>Pseudomonas | GCF_001020725.2 | Pseudomonas_lundensis | AA042_RS07115 | 1582722..1586186 | 86185 | BREX-system-P-loop-protein-BrxC-[Pseudomonas-lundensis] |
| WP_048385719.1 | Type1.BREX | [illegible] | 1185 | Bacteria>Pseudomonadota>Gammaproteobacteria>Pseudomonadales>Pseudomonadaceae>Pseudomonas | GCF_001043065.1 | Pseudomonas_helleri | TU85_RS02975 | 88982..92539 | 1608996 | BREX-system-P-loop-protein-BrxC-[Pseudomonas-helleri] |
| WP_057429193.1 | Type1.BREX | [illegible] | 1156 | Bacteria>Pseudomonadota>Gammaproteobacteria>Pseudomonadales>Pseudomonadaceae>Pseudomonas | GCF_001401385.1 | Pseudomonas_syringae_group_genomosp._3 | ALO87_RS04320 | 127987..131457 | 251701 | BREX-system-P-loop-protein-BrxC-[Pseudomonas-syringae-group-genomosp.-3] |
| WP_058181876.1 | Type6.BREX | [illegible] | 1213 | Bacteria>Pseudomonadota>Gammaproteobacteria>Pseudomonadales>Pseudomonadaceae>Pseudomonas | GCF_001450675.1 | Pseudomonas_aeruginosa | AO933_RS04770 | 131954..135595 | 287 | BREX-system-P-loop-protein-BrxC-[Pseudomonas-aeruginosa] |
| WP_058174899.1 | Type1.BREX | [illegible] | 1184 | Bacteria>Pseudomonadota>Gammaproteobacteria>Pseudomonadales>Pseudomonadaceae>Pseudomonas | GCF_001452485.1 | Pseudomonas_aeruginosa | APA89_RS03075 | 652482..656036 | 287 | BREX-system-P-loop-protein-BrxC-[Pseudomonas-aeruginosa] |
| WP_070624003.1 | Type1.BREX | [illegible] | 1154 | Bacteria>Pseudomonadota>Gammaproteobacteria>Pseudomonadales>Pseudomonadaceae>Pseudomonas | GCF_001810465.1 | Pseudomonas_aeruginosa | HMPREF2746_RS21475 | 11855..15319 | 287 | BREX-system-P-loop-protein-BrxC-[Pseudomonas-aeruginosa] |
| WP_079783554.1 | Type1.BREX | [illegible] | 1154 | Bacteria>Pseudomonadota>Gammaproteobacteria>Pseudomonadales>Pseudomonadaceae>Pseudomonas | GCF_002037565.1 | Pseudomonas_sp._LPH1 | BHQ29_RS09350 | 2029984..2033448 | 1898684 | BREX-system-P-loop-protein-BrxC-[Pseudomonas-sp.-LPH1] |
| WP_205881761.1 | Type1.BREX | [illegible] | 1159 | Bacteria>Pseudomonadota>Gammaproteobacteria>Pseudomonadales>Pseudomonadaceae>Pseudomonas | GCF_002128325.1 | Pseudomonas_fragi | B6D87_RS19610 | 4227178..4230657 | 296 | BREX-system-P-loop-protein-BrxC-[Pseudomonas-fragi] |
| WP_086230732.1 | Type1.BREX | [illegible] | 1184 | Bacteria>Pseudomonadota>Gammaproteobacteria>Pseudomonadales>Pseudomonadaceae>Pseudomonas | GCF_002135805.1 | Pseudomonas_aeruginosa | CAZ03_RS13145 | 128061..131615 | 287 | BREX-system-P-loop-protein-BrxC-[Pseudomonas-aeruginosa] |
| WP_095025594.1 | Type1.BREX | [illegible] | 1154 | Bacteria>Pseudomonadota>Gammaproteobacteria>Pseudomonadales>Pseudomonadaceae>Pseudomonas | GCF_002269515.1 | Pseudomonas_fragi | CJU79_RS17655 | 17310..20774 | 296 | BREX-system-P-loop-protein-BrxC-[Pseudomonas-fragi] |
| WP_095626344.1 | Type1.BREX | [illegible] | 1154 | Bacteria>Pseudomonadota>Gammaproteobacteria>Pseudomonadales>Pseudomonadaceae>Pseudomonas | GCF_002287035.1 | Pseudomonas_sp._WN033 | CK507_RS15715 | 130328..133792 | 2032629 | BREX-system-P-loop-protein-BrxC-[Pseudomonas-sp.-WN033] |
| WP_256588486.1 | Type1.BREX | [illegible] | 1225 | Bacteria>Pseudomonadota>Gammaproteobacteria>Pseudomonadales>Pseudomonadaceae>Pseudomonas | GCF_002591235.1 | Pseudomonas_sp._FDAARGOS_380 | CRN80_RS16460 | 3507084..3510761 | 2018067 | BREX-system-P-loop-protein-BrxC-[Pseudomonas-sp.-FDAARGOS_380] |
| WP_101152285.1 | Type6.BREX | [illegible] | 1211 | Bacteria>Pseudomonadota>Gammaproteobacteria>Pseudomonadales>Pseudomonadaceae>Pseudomonas | GCF_002836195.1 | Pseudomonas_sp._Choline-3u-10 | CXF92_RS00360 | 72818..76453 | 2058311 | BREX-system-P-loop-protein-BrxC-[Pseudomonas-sp.-Choline-3u-10] |
| WP_102082564.1 | Type1.BREX | [illegible] | 1184 | Bacteria>Pseudomonadota>Gammaproteobacteria>Pseudomonadales>Pseudomonadaceae>Pseudomonas | GCF_002864915.1 | Pseudomonas_guariconensis | CXG53_RS23735 | 60064..63618 | 1288410 | BREX-system-P-loop-protein-BrxC-[Pseudomonas-guariconensis] |
| WP_102684491.1 | Type1.BREX | [illegible] | 1222 | Bacteria>Pseudomonadota>Gammaproteobacteria>Pseudomonadales>Pseudomonadaceae>Pseudomonas | GCF_002883975.1 | Pseudomonas_sp._FW305-42 | C1X79_RS19065 | 47194..50862 | 2070677 | BREX-system-P-loop-protein-BrxC-[Pseudomonas-sp.-FW305-42] |
| WP_109933363.1 | Type1.BREX | [illegible] | 1154 | Bacteria>Pseudomonadota>Gammaproteobacteria>Pseudomonadales>Pseudomonadaceae>Pseudomonas | GCF_003176655.1 | Pseudomonas_sp._RW407 | DK254_RS08115 | 332506..335970 | 2202894 | BREX-system-P-loop-protein-BrxC-[Pseudomonas-sp.-RW407] |

| Protein-ID | System | Gene-Neighbourhoods | Length | Taxonomy | Assembly-ID | Strain | Locus-Tag | Genomic-coordinates | TaxID | Fasta-defline |
|---|---|---|---|---|---|---|---|---|---|---|
| WP_113603075.1 | Type1.BREX | <-VirD4-TraG+VirB4-FtsK<-HDRnase<-SbcC<- SIGMA+OB-Fold+AAA-ATPase+Lon-Protease-BrxL <- PglZ <- N6-METHYLASE <- ParB + HNH <- DUF6079 *<- BrxB-BrxF-iSTAND <- BrxA-DNA-binding <- wHTH+WYL+WCX <-?<-35exo<-?\|\|LexA-protease->?-><-MobI | 1225 | Bacteria>Pseudomonadota>Gammaproteobacteria>Vibrionales>Vibrionaceae>Vibrio | GCF_003311885.1 | Vibrio_tarriae | DLR65_RS02205 | 60103..63780 | 2014742 | BREX-system-P-loop-protein-BrxC-[Vibrio-tarriae] |
| WP_122015998.1 | Type1.BREX | CR-ATPase4->?-><- SIGMA+OB-Fold+AAA-ATPase+Lon-Protease-BrxL <- PglZ <-?<-?<- ParB + HNH <- N6-METHYLASE <- DUF6079 *<- BrxB-BrxF-iSTAND <- BrxA-DNA-binding <- wHTH+WYL+WCX \|\|SF1-SF2-Helicase->MarR-HTH->HKD-DNase+HKD-DNase->CARF+NtrC-AAA->PIN+OST-HTH-> | 1248 | Bacteria>Pseudomonadota>Gammaproteobacteria>Vibrionales>Vibrionaceae>Vibrio | GCF_003691525.1 | Vibrio_parahaemolyticus | D0871_RS08265 | 1554661..1558407 | 670 | BREX-system-P-loop-protein-BrxC-[Vibrio-parahaemolyticus] |
| WP_130200308.1 | Type1.BREX | AbiEi->viral-protein->SF1-SF2-Helicase+REase->SF1-SF2-Helicase+REase-><-?<-ZetaToxin<- PglZ <- ParB + HNH <- N6-METHYLASE <- DUF6079 *<- BrxB-BrxF-iSTAND <- BrxA-DNA-binding <- wHTH+WYL+WCX <-DDE_Tnp_1_5+DDE_Tnp_1\|\|<-?<-CR-ATPase4\|\|SbcC-> | 1157 | Bacteria>Pseudomonadota>Gammaproteobacteria>Vibrionales>Vibrionaceae>Vibrio | GCF_004214625.1 | Vibrio_vulnificus | D8T51_RS18945 | 13637..17110 | 672 | BREX-system-P-loop-protein-BrxC-[Vibrio-vulnificus] |
| WP_137027181.1 | Type1.BREX | SF1-SF2-Helicase->AbiEi->viral-protein-> wHTH+WYL+WCX -> BrxA-DNA-binding -> BrxB-BrxF-iSTAND -> DUF6079 *-> N6-METHYLASE -> ParB + HNH ->GrpE->?-> PglZ -> SIGMA+OB-Fold+AAA-ATPase+Lon-Protease-BrxL -><-wHTH\|\|DUF4113-> | 1225 | Bacteria>Pseudomonadota>Gammaproteobacteria>Vibrionales>Vibrionaceae>Vibrio | GCF_005145785.1 | Vibrio_kanaloae | FCV44_20755 | 6017..9694 | 170673 | BREX-system-P-loop-protein-BrxC-[Vibrio-kanaloae] |
| WP_140128967.1 | Type1.BREX | AbiEi->viral-protein-> wHTH+WYL+WCX -> BrxA-DNA-binding -> BrxB-BrxF-iSTAND -> DUF6079 *-> N6-METHYLASE -> ParB + HNH ->GrpE->?-> PglZ -> SIGMA+OB-Fold+AAA-ATPase+Lon-Protease-BrxL -><-wHTH<-?<-CR-ATPase4 | 1225 | Bacteria>Pseudomonadota>Gammaproteobacteria>Vibrionales>Vibrionaceae>Vibrio | GCF_006368845.1 | Vibrio_parahaemolyticus | CGJ25_RS23760 | 3581..7258 | 670 | BREX-system-P-loop-protein-BrxC-[Vibrio-parahaemolyticus] |
| WP_140247448.1 | Type1.BREX | AbiEi->viral-protein->SF1-SF2-Helicase+REase->SF1-SF2-Helicase+REase-><-?<-ZetaToxin<- PglZ <- ParB + HNH <- N6-METHYLASE <- DUF6079 *<- BrxB-BrxF-iSTAND <- BrxA-DNA-binding <- wHTH+WYL+WCX \|\|<-?<-CR-ATPase4\|\|PIN+OST-HTH-><-REase+SPF-cons | 1157 | Bacteria>Pseudomonadota>Gammaproteobacteria>Vibrionales>Vibrionaceae>Vibrio | GCF_006371495.1 | Vibrio_parahaemolyticus | CGJ41_RS23680 | 13693..17166 | 670 | BREX-system-P-loop-protein-BrxC-[Vibrio-parahaemolyticus] |
| WP_140344322.1 | Type1.BREX | CR-ATPase4->?->\|\| wHTH+WYL+WCX -> BrxA-DNA-binding -> BrxB-BrxF-iSTAND -> DUF6079 *-> N6-METHYLASE -> ParB + HNH -> PglZ ->SF1-SF2-Helicase-> | 1157 | Bacteria>Pseudomonadota>Gammaproteobacteria>Vibrionales>Vibrionaceae>Vibrio | GCF_006372045.1 | Vibrio_parahaemolyticus | CGJ19_RS21455 | 3824..7297 | 670 | BREX-system-P-loop-protein-BrxC-[Vibrio-parahaemolyticus] |
| WP_140256559.1 | Type1.BREX | MobI-><-?<-LexA-protease\|\|35exo->?-> wHTH+WYL+WCX -> BrxA-DNA-binding -> BrxB-BrxF-iSTAND -> DUF6079 *-> ParB + HNH -> N6-METHYLASE -> PglZ -> SIGMA+OB-Fold+AAA-ATPase+Lon-Protease-BrxL ->SbcC->HDRnase->VirD4-TraG+VirB4-FtsK-> | 1225 | Bacteria>Pseudomonadota>Gammaproteobacteria>Vibrionales>Vibrionaceae>Vibrio | GCF_006372155.1 | Vibrio_parahaemolyticus | D9A28_03775 | 149461..153138 | 670 | BREX-system-P-loop-protein-BrxC-[Vibrio-parahaemolyticus] |
| WP_140272583.1 | Type1.BREX | CR-ATPase4->?-><- SIGMA+OB-Fold+AAA-ATPase+Lon-Protease-BrxL <- PglZ <-?<-?<- ParB + HNH <- N6-METHYLASE <- DUF6079 *<- BrxB-BrxF-iSTAND <- BrxA-DNA-binding <- wHTH+WYL+WCX \|\|SF1-SF2-Helicase->MarR-HTH->HKD-DNase+HKD-DNase->CARF+NtrC-AAA->PIN+OST-HTH-> | 1248 | Bacteria>Pseudomonadota>Gammaproteobacteria>Vibrionales>Vibrionaceae>Vibrio | GCF_006372715.1 | Vibrio_parahaemolyticus | CGI87_RS22525 | 16069..19815 | 670 | BREX-system-P-loop-protein-BrxC-[Vibrio-parahaemolyticus] |
| WP_148513618.1 | Type1.BREX | AbiEi->viral-protein->SF1-SF2-Helicase+REase->SF1-SF2-Helicase+REase-><-TPR<-HEPN+SF1-SF2-Helicase+RNase+RNase<- PglZ <- ParB + HNH <- N6-METHYLASE <- DUF6079 *<- BrxB-BrxF-iSTAND <- BrxA-DNA-binding <- wHTH+WYL+WCX \|\|DDE_Tnp_1_5+DDE_Tnp_1-> | 1157 | Bacteria>Pseudomonadota>Gammaproteobacteria>Vibrionales>Vibrionaceae>Vibrio | GCF_008083315.1 | Vibrio_cholerae | FXF10_RS00340 | 29414..32887 | 666 | BREX-system-P-loop-protein-BrxC-[Vibrio-cholerae] |
| WP_148532770.1 | Type1.BREX | <-VirD4-TraG+VirB4-FtsK<-HDRnase<-SbcC<- SIGMA+OB-Fold+AAA-ATPase+Lon-Protease-BrxL <- PglZ <- N6-METHYLASE <- ParB + HNH <- DUF6079 *<- BrxB-BrxF-iSTAND <- BrxA-DNA-binding <- wHTH+WYL+WCX <-?<-35exo<-?\|\|LexA-protease-> | 1225 | Bacteria>Pseudomonadota>Gammaproteobacteria>Vibrionales>Vibrionaceae>Vibrio | GCF_008084025.1 | Vibrio_cholerae | FXE74_RS08220 | 132887..136564 | 666 | BREX-system-P-loop-protein-BrxC-[Vibrio-cholerae] |
| WP_148490856.1 | Type1.BREX | <-DDE_Tnp_1_5+DDE_Tnp_1\|\|HTH_Tnp_1+HTH_21+rve+rve_3-><-SbcC\|\|CR-ATPase4->?-><-DUF4113\|\| wHTH+WYL+WCX -> BrxA-DNA-binding -> BrxB-BrxF-iSTAND -> DUF6079 *-> N6-METHYLASE -> ParB + HNH -> PglZ ->HEPN+SF1-SF2-Helicase+RNase+RNase->TPR-><-SF1-SF2-Helicase+REase<-SF1-SF2-Helicase+REase<-viral-protein<-AbiEi | 1157 | Bacteria>Pseudomonadota>Gammaproteobacteria>Vibrionales>Vibrionaceae>Vibrio | GCF_008084645.1 | Vibrio_cholerae | FXE54_RS16715 | 26974..30447 | 666 | BREX-system-P-loop-protein-BrxC-[Vibrio-cholerae] |
| WP_148536810.1 | Type1.BREX | <-CARF+NtrC-AAA+FIS-HTH<- SIGMA+OB-Fold+AAA-ATPase+Lon-Protease-BrxL <- PglZ <- N6-METHYLASE <- ParB + HNH <- DUF6079 *<- BrxB-BrxF-iSTAND <- BrxA-DNA-binding <- wHTH+WYL+WCX <-?<-35exo\|\|LexA-protease->?-><-MobI | 1225 | Bacteria>Pseudomonadota>Gammaproteobacteria>Vibrionales>Vibrionaceae>Vibrio | GCF_008085615.1 | Vibrio_cholerae | FXE25_RS13680 | 35779..39456 | 666 | BREX-system-P-loop-protein-BrxC-[Vibrio-cholerae] |
| WP_123161578.1 | Type1.BREX | MobI-><-?<-LexA-protease\|\|35exo->?-> wHTH+WYL+WCX -> BrxA-DNA-binding -> BrxB-BrxF-iSTAND -> DUF6079 *-> ParB + HNH -> N6-METHYLASE -> PglZ -> SIGMA+OB-Fold+AAA-ATPase+Lon-Protease-BrxL ->SbcC->HDRnase->VirD4-TraG+VirB4-FtsK-> | 1225 | Bacteria>Pseudomonadota>Gammaproteobacteria>Vibrionales>Vibrionaceae>Vibrio | GCF_009763665.1 | Vibrio_cholerae | GPY01_RS14950 | 2265324..2269001 | 666 | BREX-system-P-loop-protein-BrxC-[Vibrio-cholerae] |
| WP_172566988.1 | Type1.BREX | CR-ATPase4->?-><-?<-LexA-protease\|\| wHTH+WYL+WCX -> BrxA-DNA-binding -> BrxB-BrxF-iSTAND -> DUF6079 *-> N6-METHYLASE -> ParB + HNH -> PglZ ->ZetaToxin->?-><-SF1-SF2-Helicase+REase<-SF1-SF2-Helicase+REase<-viral-protein<-AbiEi | 1157 | Bacteria>Pseudomonadota>Gammaproteobacteria>Vibrionales>Vibrionaceae>Vibrio | GCF_012274945.1 | Vibrio_fluvialis | HF303_RS09150 | 434217..437690 | 676 | BREX-system-P-loop-protein-BrxC-[Vibrio-fluvialis] |
| WP_169592914.1 | Type1.BREX | MobI-><-Glyoxalase_8<-?<-LexA-protease\|\|35exo->?-> wHTH+WYL+WCX -> BrxA-DNA-binding -> BrxB-BrxF-iSTAND -> DUF6079 *-> ParB + HNH -> N6-METHYLASE -> PglZ -> SIGMA+OB-Fold+AAA-ATPase+Lon-Protease-BrxL ->SbcC->HDRnase->VirD4-TraG+VirB4-FtsK-> | 1225 | Bacteria>Pseudomonadota>Gammaproteobacteria>Vibrionales>Vibrionaceae>Vibrio | GCF_012956215.1 | Vibrio_parahaemolyticus | HKB29_RS26535 | 221026..224703 | 670 | BREX-system-P-loop-protein-BrxC-[Vibrio-parahaemolyticus] |
| WP_171375390.1 | Type1.BREX | <-ZnR<-?\|\|?->SF1-SF2-Helicase->AbiEi->viral-protein-> wHTH+WYL+WCX -> BrxA-DNA-binding -> BrxB-BrxF-iSTAND -> DUF6079 *-> N6-METHYLASE -> ParB + HNH ->GrpE->?-> PglZ -> SIGMA+OB-Fold+AAA-ATPase+Lon-Protease-BrxL -><-TPR<-HEPN+SF1-SF2-Helicase+RNase+RNase\|\|LexA-protease-> | 1225 | Bacteria>Pseudomonadota>Gammaproteobacteria>Vibrionales>Vibrionaceae>Vibrio | GCF_013114495.1 | Vibrio_aestuarianus | F0230_RS00325 | 78960..82637 | 28171 | BREX-system-P-loop-protein-BrxC-[Vibrio-aestuarianus] |
| WP_182291541.1 | Type1.BREX | REase+SPF-cons-><-PIN+OST-HTH\|\|CR-ATPase4->?->\|\| wHTH+WYL+WCX -> BrxA-DNA-binding -> BrxB-BrxF-iSTAND -> DUF6079 *-> N6-METHYLASE -> ParB + HNH -> PglZ ->ZetaToxin->?-><-SF1-SF2-Helicase+REase<-SF1-SF2-Helicase+REase<-viral-protein<-AbiEi | 1157 | Bacteria>Pseudomonadota>Gammaproteobacteria>Vibrionales>Vibrionaceae>Vibrio | GCF_014107515.1 | Vibrio_vulnificus | F6X00_RS18205 | 373031..376504 | 672 | BREX-system-P-loop-protein-BrxC-[Vibrio-vulnificus] |
| WP_238930029.1 | Type1.BREX | HTH_20-><-SbcC\|\|CR-ATPase4->?->\|\| wHTH+WYL+WCX -> BrxA-DNA-binding -> BrxB-BrxF-iSTAND -> DUF6079 -p*-> DUF6079 -p-> N6-METHYLASE -> ParB + HNH -> PglZ ->ZetaToxin->?-><-SF1-SF2-Helicase+REase<-SF1-SF2-Helicase+REase<-Tnp_DNA_bind+DDE_Tnp_1+Dimer_Tnp_Tn5 | 779 | Bacteria>Pseudomonadota>Gammaproteobacteria>Vibrionales>Vibrionaceae>Vibrio | GCF_014525155.1 | Vibrio_vulnificus | GO186_RS04075 | 575963..578302 | 672 | BREX-system-P-loop-protein-BrxC-[Vibrio-vulnificus] |
| WP_238930031.1 | Type1.BREX | HTH_20-><-SbcC\|\|CR-ATPase4->?->\|\| wHTH+WYL+WCX -> BrxA-DNA-binding -> BrxB-BrxF-iSTAND -> DUF6079 -p-> DUF6079 -p*-> N6-METHYLASE -> ParB + HNH -> PglZ ->ZetaToxin->?-><-SF1-SF2-Helicase+REase<-SF1-SF2-Helicase+REase<-Tnp_DNA_bind+DDE_Tnp_1+Dimer_Tnp_Tn5<-AbiEi | 366 | Bacteria>Pseudomonadota>Gammaproteobacteria>Vibrionales>Vibrionaceae>Vibrio | GCF_014525155.1 | Vibrio_vulnificus | GO186_RS25670 | 578332..579432 | 672 | hypothetical-protein-[Vibrio-vulnificus] |
| WP_191116552.1 | Type1.BREX | CR-ATPase4->?-><- SIGMA+OB-Fold+AAA-ATPase+Lon-Protease-BrxL <- PglZ <-?<-?<- ParB + HNH <- N6-METHYLASE <- DUF6079 *<- BrxB-BrxF-iSTAND <- BrxA-DNA-binding <- wHTH+WYL+WCX \|\|SF1-SF2-Helicase->MarR-HTH->HKD-DNase+HKD-DNase->CARF+NtrC-AAA->PIN+OST-HTH-><-AAA-ATPase | 1248 | Bacteria>Pseudomonadota>Gammaproteobacteria>Vibrionales>Vibrionaceae>Vibrio | GCF_014769095.1 | Vibrio_parahaemolyticus | EUV06_RS14935 | 117914..121660 | 670 | BREX-system-P-loop-protein-BrxC-[Vibrio-parahaemolyticus] |
| WP_054579585.1 | Type1.BREX | AbiEi->viral-protein->SF1-SF2-Helicase+REase->SF1-SF2-Helicase+REase-><-?<-ZetaToxin<- PglZ <- ParB + HNH <- N6-METHYLASE <- DUF6079 *<- BrxB-BrxF-iSTAND <- BrxA-DNA-binding <- wHTH+WYL+WCX \|\|<-?<-CR-ATPase4\|\|HEPN-><-sGTP | 1157 | Bacteria>Pseudomonadota>Gammaproteobacteria>Vibrionales>Vibrionaceae>Vibrio | GCF_014769135.1 | Vibrio_alginolyticus | AOR09_RS18625 | 85180..88653 | 663 | BREX-system-P-loop-protein-BrxC-[Vibrio-alginolyticus] |
| WP_206630456.1 | Type1.BREX | GTPase-AIG->GTPase-AIG-><-HEPN\|\|?->?-><- SIGMA+OB-Fold+AAA-ATPase+Lon-Protease-BrxL <- PglZ <- ParB + HNH <- N6-METHYLASE <- DUF6079 *<- BrxB-BrxF-iSTAND <- BrxA-DNA-binding <- wHTH+WYL+WCX <-viral-protein\|\|?->ZnR-><-SF1-SF2-Helicase<-Couple_hipA+HipA_C<-HTH_3 | 1232 | Bacteria>Pseudomonadota>Gammaproteobacteria>Vibrionales>Vibrionaceae>Vibrio | GCF_017291155.1 | Vibrio_vulnificus | JOJ25_RS14690 | 90356..94054 | 672 | BREX-system-P-loop-protein-BrxC-[Vibrio-vulnificus] |
| WP_206946493.1 | Type1.BREX | sGTP-><-HEPN<-?\|\|CR-ATPase4->?->\|\| wHTH+WYL+WCX -> BrxA-DNA-binding -> BrxB-BrxF-iSTAND -> DUF6079 *-> N6-METHYLASE -> ParB + HNH -> PglZ ->ZetaToxin->?-><-SF1-SF2-Helicase+REase<-SF1-SF2-Helicase+REase<-viral-protein<-AbiEi | 1157 | Bacteria>Pseudomonadota>Gammaproteobacteria>Vibrionales>Vibrionaceae>Vibrio | GCF_017313505.1 | Vibrio_sp._Vb2424 | J0679_RS01025 | 227908..231381 | 2816074 | BREX-system-P-loop-protein-BrxC-[Vibrio-sp.-Vb2424] |
| WP_210942219.1 | Type1.BREX | <- wHTH+WYL+WCX \|\|SF1-SF2-Helicase+REase->?->XIS-HTH-> wHTH+WYL+WCX -> BrxA-DNA-binding -> BrxB-BrxF-iSTAND -> DUF6079 *-> N6-METHYLASE -> ParB + HNH -> PglZ ->ZetaToxin->?-><-SF1-SF2-Helicase+REase<-SF1-SF2-Helicase+REase<-Arm-DNA-bind_3+Phage_integrase | 1157 | Bacteria>Pseudomonadota>Gammaproteobacteria>Vibrionales>Vibrionaceae>Vibrio | GCF_018035165.1 | Vibrio_paracholerae | GSG51_RS14590 | 9402..12875 | 650003 | BREX-system-P-loop-protein-BrxC-[Vibrio-paracholerae] |
| WP_213968024.1 | Type1.BREX | <-PIN+OST-HTH<-CARF+NtrC-AAA<-HKD-DNase+HKD-DNase<-MarR-HTH<-SF1-SF2-Helicase\|\| wHTH+WYL+WCX -> BrxA-DNA-binding -> BrxB-BrxF-iSTAND -> DUF6079 *-> N6-METHYLASE -> ParB + HNH ->ZnR+3C-ABC+TPR->?->?-> PglZ -> SIGMA+OB-Fold+AAA-ATPase+Lon-Protease-BrxL -><-?<-CR-ATPase4 | 1248 | Bacteria>Pseudomonadota>Gammaproteobacteria>Vibrionales>Vibrionaceae>Vibrio | GCF_018458215.1 | Vibrio_alginolyticus | J4H41_RS21340 | 12665..16411 | 663 | BREX-system-P-loop-protein-BrxC-[Vibrio-alginolyticus] |
| WP_213970810.1 | Type1.BREX | AbiEi->viral-protein->SF1-SF2-Helicase+REase->SF1-SF2-Helicase+REase-><-?<-ZetaToxin<- PglZ <- ParB + HNH <- N6-METHYLASE <- DUF6079 *<- BrxB-BrxF-iSTAND <- BrxA-DNA-binding <- wHTH+WYL+WCX \|\|DUF4113-><-?<-CR-ATPase4 | 1157 | Bacteria>Pseudomonadota>Gammaproteobacteria>Vibrionales>Vibrionaceae>Vibrio | GCF_018458405.1 | Vibrio_alginolyticus | J4H58_RS14535 | 108443..111916 | 663 | BREX-system-P-loop-protein-BrxC-[Vibrio-alginolyticus] |
| WP_213899843.1 | Type1.BREX | AbiEi->viral-protein->SF1-SF2-Helicase+REase->SF1-SF2-Helicase+REase-><-TPR<-HEPN+SF1-SF2-Helicase+RNase+RNase<- PglZ <- ParB + HNH <- N6-METHYLASE <- DUF6079 *<- BrxB-BrxF-iSTAND <- BrxA-DNA-binding <- wHTH+WYL+WCX \|\|<-?<-CR-ATPase4\|\|SbcC->HEPN-><-HTH_20 | 1157 | Bacteria>Pseudomonadota>Gammaproteobacteria>Vibrionales>Vibrionaceae>Vibrio | GCF_018458525.1 | Vibrio_alginolyticus | J4H25_RS19085 | 22843..26316 | 663 | BREX-system-P-loop-protein-BrxC-[Vibrio-alginolyticus] |
| WP_213874127.1 | Type1.BREX | sGTP-><-HEPN<-?\|\|CR-ATPase4->?->\|\| wHTH+WYL+WCX -> BrxA-DNA-binding -> BrxB-BrxF-iSTAND -> DUF6079 *-> N6-METHYLASE -> ParB + HNH -> PglZ ->ZetaToxin->?-><-SF1-SF2-Helicase+REase<-SF1-SF2-Helicase+REase<-viral-protein<-AbiEi | 1157 | Bacteria>Pseudomonadota>Gammaproteobacteria>Vibrionales>Vibrionaceae>Vibrio | GCF_018458545.1 | Vibrio_alginolyticus | J4H28_RS04435 | 1018248..1021721 | 663 | BREX-system-P-loop-protein-BrxC-[Vibrio-alginolyticus] |
| WP_217555808.1 | Type1.BREX | AbiEi->viral-protein->SF1-SF2-Helicase+REase->SF1-SF2-Helicase+REase-><-?<-ZetaToxin<- PglZ <- ParB + HNH <- N6-METHYLASE <- DUF6079 *<- BrxB-BrxF-iSTAND <- BrxA-DNA-binding <- wHTH+WYL+WCX \|\|LexA-protease-> | 1157 | Bacteria>Pseudomonadota>Gammaproteobacteria>Vibrionales>Vibrionaceae>Vibrio | GCF_019091765.1 | Vibrio_metschnikovii | KVT01_RS15810 | 26769..30242 | 28172 | BREX-system-P-loop-protein-BrxC-[Vibrio-metschnikovii] |
| WP_222133201.1 | Type1.BREX | <-PIN+OST-HTH<-CARF+NtrC-AAA<-HKD-DNase+HKD-DNase<-MarR-HTH<-SF1-SF2-Helicase\|\| wHTH+WYL+WCX -> BrxA-DNA-binding -> BrxB-BrxF-iSTAND -> DUF6079 *-> N6-METHYLASE -> ParB + HNH ->?->?-> PglZ -> SIGMA+OB-Fold+AAA-ATPase+Lon-Protease-BrxL -> | 1248 | Bacteria>Pseudomonadota>Gammaproteobacteria>Vibrionales>Vibrionaceae>Vibrio | GCF_019793245.1 | Vibrio_alginolyticus | K6L11_RS14305 | 27139..30885 | 663 | BREX-system-P-loop-protein-BrxC-[Vibrio-alginolyticus] |
| WP_222134563.1 | Type1.BREX | <-VirD4-TraG+VirB4-FtsK<-HDRnase<-NACHT+HEAT+REase<- SIGMA+OB-Fold+AAA-ATPase+Lon-Protease-BrxL <- PglZ <- N6-METHYLASE <- ParB + HNH <- DUF6079 *<- BrxB-BrxF-iSTAND <- BrxA-DNA-binding <- wHTH+WYL+WCX <-?<-35exo\|\|LexA-protease->?-><-MobI | 1225 | Bacteria>Pseudomonadota>Gammaproteobacteria>Vibrionales>Vibrionaceae>Vibrio | GCF_019793275.1 | Vibrio_cholerae | NB536_RS17220 | 131235..134912 | 666 | BREX-system-P-loop-protein-BrxC-[Vibrio-cholerae] |
| WP_224640811.1 | Type1.BREX | MobI-><-?<-LexA-protease\|\|?->35exo->?-> wHTH+WYL+WCX -> BrxA-DNA-binding -> BrxB-BrxF-iSTAND -> DUF6079 *-> ParB + HNH -> N6-METHYLASE -> PglZ -> SIGMA+OB-Fold+AAA-ATPase+Lon-Protease-BrxL ->NACHT+HEAT+REase->HDRnase->VirD4-TraG+VirB4-FtsK-> | 1225 | Bacteria>Pseudomonadota>Gammaproteobacteria>Vibrionales>Vibrionaceae>Vibrio | GCF_020169625.1 | Vibrio_vulnificus | KUO11_RS09480 | 205814..209491 | 672 | BREX-system-P-loop-protein-BrxC-[Vibrio-vulnificus] |
| WP_225482458.1 | Type1.BREX | <- SIGMA+OB-Fold+AAA-ATPase+Lon-Protease-BrxL <- PglZ \|\|RADICAL-SAM->?-><-?<-?<- ParB + HNH <- N6-METHYLASE <-?<- DUF6079 *<- BrxB-BrxF-iSTAND <- BrxA-DNA-binding <- wHTH+WYL+WCX \|\|?->?-><-viral-protein\|\|?-><-SF1-SF2-Helicase<-Couple_hipA+HipA_C | 1231 | Bacteria>Pseudomonadota>Gammaproteobacteria>Vibrionales>Vibrionaceae>Vibrio | GCF_020251885.1 | Vibrio_vulnificus | JKP23_RS02685 | 250869..254564 | 672 | BREX-system-P-loop-protein-BrxC-[Vibrio-vulnificus] |
| WP_225454267.1 | Type1.BREX | TraJ-RHH->TraI-VirD2-RCRE->REase-><-AbiEi_4\|\|?-> wHTH+WYL+WCX -> BrxA-DNA-binding -> BrxB-BrxF-iSTAND -> DUF6079 *-> N6-METHYLASE -> ParB + HNH ->ANK+ANK+ANK->ZnR+3C-ABC+TPR->?->?-> PglZ -> SIGMA+OB-Fold+AAA-ATPase+Lon-Protease-BrxL -> | 1248 | Bacteria>Pseudomonadota>Gammaproteobacteria>Vibrionales>Vibrionaceae>Vibrio | GCF_020252125.1 | Vibrio_vulnificus | JKP08_RS08925 | 222135..225881 | 672 | BREX-system-P-loop-protein-BrxC-[Vibrio-vulnificus] |
| WP_228096077.1 | Type1.BREX | <-HTH_Tnp_1\|\|SF1-SF2-Helicase+REase->SF1-SF2-Helicase+REase-><-?<-ZetaToxin<- PglZ <- ParB + HNH <- N6-METHYLASE <- DUF6079 *<- BrxB-BrxF-iSTAND <- BrxA-DNA-binding <- wHTH+WYL+WCX \|\|DUF4113-><-?<-CR-ATPase4\|\|SbcC->HEPN-><-HTH_20 | 1157 | Bacteria>Pseudomonadota>Gammaproteobacteria>Vibrionales>Vibrionaceae>Vibrio | GCF_020741035.1 | Vibrio_parahaemolyticus | IB294_RS01075 | 246074..249547 | 670 | BREX-system-P-loop-protein-BrxC-[Vibrio-parahaemolyticus] |
| WP_236110294.1 | Type1.BREX | <-GspA-ATPase\|\|SF1-SF2-Helicase->?->HKD-DNase+HKD-DNase->CARF+NtrC-AAA-> wHTH+WYL+WCX -> BrxA-DNA-binding -> BrxB-BrxF-iSTAND -> DUF6079 *-> N6-METHYLASE -> ParB + HNH -> PglZ ->ZetaToxin->?-><-SF1-SF2-Helicase+REase<-SF1-SF2-Helicase+REase | 1157 | Bacteria>Pseudomonadota>Gammaproteobacteria>Vibrionales>Vibrionaceae>Vibrio | GCF_021568915.1 | Vibrio_sp._McD22-P3 | L2W36_RS13785 | 194928..198401 | 2724880 | BREX-system-P-loop-protein-BrxC-[Vibrio-sp.-McD22-P3] |
| WP_238795743.1 | Type1.BREX | MobI-><-?<-LexA-protease\|\|35exo->?-> wHTH+WYL+WCX -> BrxA-DNA-binding -> BrxB-BrxF-iSTAND -> DUF6079 *-> ParB + HNH -> N6-METHYLASE -> PglZ -> SIGMA+OB-Fold+AAA-ATPase+Lon-Protease-BrxL ->CARF+NtrC-AAA+FIS-HTH->OBFOLD-NUCLEASE-> | 1225 | Bacteria>Pseudomonadota>Gammaproteobacteria>Vibrionales>Vibrionaceae>Vibrio | GCF_022289115.1 | Vibrio_furnissii | K6U56_RS11800 | 143910..147587 | 29494 | BREX-system-P-loop-protein-BrxC-[Vibrio-furnissii] |
| WP_140273790.1 | Type1.BREX | <-AAA-ATPase-p<-HEPN\|\|CR-ATPase4->?->\|\| wHTH+WYL+WCX -> BrxA-DNA-binding -> BrxB-BrxF-iSTAND -> DUF6079 *-> N6-METHYLASE -> ParB + HNH -> PglZ ->ZetaToxin->?-><-SF1-SF2-Helicase+REase<-SF1-SF2-Helicase+REase<-viral-protein<-AbiEi | 1157 | Bacteria>Pseudomonadota>Gammaproteobacteria>Vibrionales>Vibrionaceae>Vibrio | GCF_022291015.1 | Vibrio_parahaemolyticus | K6U15_RS27495 | 32395..35868 | 670 | BREX-system-P-loop-protein-BrxC-[Vibrio-parahaemolyticus] |
| WP_238967171.1 | Type1.BREX | sGTP-><-HEPN<-?\|\|CR-ATPase4->?->\|\| wHTH+WYL+WCX -> BrxA-DNA-binding -> BrxB-BrxF-iSTAND -> DUF6079 *-> N6-METHYLASE -> ParB + HNH -> PglZ ->ZetaToxin->?-><-SF1-SF2-Helicase+REase<-SF1-SF2-Helicase+REase<-viral-protein<-AbiEi | 1157 | Bacteria>Pseudomonadota>Gammaproteobacteria>Vibrionales>Vibrionaceae>Vibrio | GCF_022343145.1 | Vibrio_alginolyticus | K6750_RS21880 | 1365022..1368495 | 663 | BREX-system-P-loop-protein-BrxC-[Vibrio-alginolyticus] |
| WP_243579240.1 | Type1.BREX | <-BrxD<- SIGMA+OB-Fold+AAA-ATPase+Lon-Protease-BrxL <- PglZ <-?<-?<-?<-ZnR+3C-ABC+TPR<-ANK+ANK+ANK<- ParB + HNH <- N6-METHYLASE <- DUF6079 *<- BrxB-BrxF-iSTAND <- BrxA-DNA-binding <- wHTH+WYL+WCX \|\|SF1-SF2-Helicase+HTH->?<-?\|\|?-><-Calcineurin-SbcD<-REase | 1247 | Bacteria>Pseudomonadota>Gammaproteobacteria>Vibrionales>Vibrionaceae>Vibrio | GCF_022812045.1 | Vibrio_splendidus | LTQ02_RS02805 | 621543..625286 | 29497 | BREX-system-P-loop-protein-BrxC-[Vibrio-splendidus] |
| WP_253644856.1 | Type1.BREX | CR-ATPase4->?-><- SIGMA+OB-Fold+AAA-ATPase+Lon-Protease-BrxL <- PglZ <-?<-?<- ParB + HNH <- N6-METHYLASE <- DUF6079 *<- BrxB-BrxF-iSTAND <- BrxA-DNA-binding <- wHTH+WYL+WCX | 1248 | Bacteria>Pseudomonadota>Gammaproteobacteria>Vibrionales>Vibrionaceae>Vibrio | GCF_024181785.1 | Vibrio_sp._Y42_MX_L11 | NL643_RS16680 | 61256..65002 | 2957765 | BREX-system-P-loop-protein-BrxC-[Vibrio-sp.-Y42_MX_L11] |
| WP_253658043.1 | Type1.BREX | <- SIGMA+OB-Fold+AAA-ATPase+Lon-Protease-BrxL <- PglZ <-?<-?<-?<-ZnR+3C-ABC+TPR<- ParB + HNH <- N6-METHYLASE <- DUF6079 *<- BrxB-BrxF-iSTAND <- BrxA-DNA-binding <- wHTH+WYL+WCX \|\|SF1-SF2-Helicase->MarR-HTH->HKD-DNase+HKD-DNase->CARF+NtrC-AAA->PIN+OST-HTH->MORC-> | 1248 | Bacteria>Pseudomonadota>Gammaproteobacteria>Vibrionales>Vibrionaceae>Vibrio | GCF_024181795.1 | unclassified_Vibrio | NL644_RS04175 | 947816..951562 | 2614977 | MULTISPECIES:-BREX-system-P-loop-protein-BrxC-[unclassified-Vibrio] |
| WP_261872676.1 | Type1.BREX | <- wHTH+WYL+WCX \|\|SF1-SF2-Helicase+REase->SF1-SF2-Helicase+REase-><-?<-ZetaToxin<- PglZ <- ParB + HNH <- N6-METHYLASE <- N6-METHYLASE <- DUF6079 *<- BrxB-BrxF-iSTAND <- BrxA-DNA-binding <-tRNA\|\|?-><-?<-TPR+TPR+TPR | 1151 | Bacteria>Pseudomonadota>Gammaproteobacteria>Vibrionales>Vibrionaceae>Vibrio | GCF_024347075.1 | Vibrio_rarus | OCU56_RS07815 | 1705601..1709056 | 413403 | BREX-system-P-loop-protein-BrxC-[Vibrio-rarus] |
| WP_256897643.1 | Type1.BREX | <-TrkH<-REC+OmpR-HTH<-HISKIN<-?<-?<-?\|\| wHTH+WYL+WCX -> BrxA-DNA-binding -> BrxB-BrxF-iSTAND -> DUF6079 *-> N6-METHYLASE -> ParB + HNH ->?->?-> PglZ -> SIGMA+OB-Fold+AAA-ATPase+Lon-Protease-BrxL -><-?<-CR-ATPase4 | 1248 | Bacteria>Pseudomonadota>Gammaproteobacteria>Vibrionales>Vibrionaceae>Vibrio | GCF_024530955.1 | Vibrio_parahaemolyticus | H2C82_RS04830 | 209368..213114 | 670 | BREX-system-P-loop-protein-BrxC-[Vibrio-parahaemolyticus] |
| WP_258495358.1 | Type1.BREX | AbiEi->viral-protein->SF1-SF2-Helicase+REase->SF1-SF2-Helicase+REase-><-?<-ZetaToxin<- PglZ <- ParB + HNH <- N6-METHYLASE <- DUF6079 *<- BrxB-BrxF-iSTAND <- BrxA-DNA-binding <- wHTH+WYL+WCX \|\|<-?<-CR-ATPase4\|\|HEPN-><-sGTP | 1157 | Bacteria>Pseudomonadota>Gammaproteobacteria>Vibrionales>Vibrionaceae>Vibrio | GCF_024744595.1 | Vibrio_alginolyticus | NB477_RS23070 | 13774..17247 | 663 | BREX-system-P-loop-protein-BrxC-[Vibrio-alginolyticus] |
| WP_248354663.1 | Type1.BREX | AbiEi->viral-protein->SF1-SF2-Helicase+REase->SF1-SF2-Helicase+REase-><-TPR<-HEPN+SF1-SF2-Helicase+RNase+RNase<- PglZ <- ParB + HNH <- N6-METHYLASE <- DUF6079 *<- BrxB-BrxF-iSTAND <- BrxA-DNA-binding <- wHTH+WYL+WCX \|\|DDE_Tnp_1_5+DDE_Tnp_1->PIN+OST-HTH-><-?<-HTH_20 | 1157 | Bacteria>Pseudomonadota>Gammaproteobacteria>Vibrionales>Vibrionaceae>Vibrio | GCF_024745175.1 | Vibrio_parahaemolyticus | DDM99_15775 | 19819..23292 | 670 | BREX-system-P-loop-protein-BrxC-[Vibrio-parahaemolyticus] |
| WP_258484886.1 | Type1.BREX | MobI-><-?<-LexA-protease\|\|35exo->?-> wHTH+WYL+WCX -> BrxA-DNA-binding -> BrxB-BrxF-iSTAND -> DUF6079 *-> ParB + HNH -> N6-METHYLASE -> PglZ -> SIGMA+OB-Fold+AAA-ATPase+Lon-Protease-BrxL ->CARF+NtrC-AAA+FIS-HTH->OBFOLD-NUCLEASE-> | 1225 | Bacteria>Pseudomonadota>Gammaproteobacteria>Vibrionales>Vibrionaceae>Vibrio | GCF_024745305.1 | Vibrio_parahaemolyticus | NB589_RS16770 | 223237..226914 | 670 | BREX-system-P-loop-protein-BrxC-[Vibrio-parahaemolyticus] |
| WP_258517485.1 | Type1.BREX | AbiEi->viral-protein-> wHTH+WYL+WCX -> BrxA-DNA-binding -> BrxB-BrxF-iSTAND -> DUF6079 *-> N6-METHYLASE -> ParB + HNH ->GrpE->?-> PglZ -> SIGMA+OB-Fold+AAA-ATPase+Lon-Protease-BrxL -><-wHTH | 1225 | Bacteria>Pseudomonadota>Gammaproteobacteria>Vibrionales>Vibrionaceae>Vibrio | GCF_024745395.1 | Vibrio_parahaemolyticus | NB556_RS01245 | 3462..7139 | 670 | BREX-system-P-loop-protein-BrxC-[Vibrio-parahaemolyticus] |
| WP_264912148.1 | Type1.BREX | <- SIGMA+OB-Fold+AAA-ATPase+Lon-Protease-BrxL <- PglZ <-?<-?<-?<-ZnR+3C-ABC+TPR<- ParB + HNH <- N6-METHYLASE <- DUF6079 *<- BrxB-BrxF-iSTAND <- BrxA-DNA-binding <- wHTH+WYL+WCX \|\|SF1-SF2-Helicase->MarR-HTH->HKD-DNase+HKD-DNase->CARF+NtrC-AAA->PIN+OST-HTH->MORC-> | 1248 | Bacteria>Pseudomonadota>Gammaproteobacteria>Vibrionales>Vibrionaceae>Vibrio | GCF_026000495.1 | Vibrio_sp._STUT-A16 | OO778_RS24845 | 1838056..1841802 | 2976237 | BREX-system-P-loop-protein-BrxC-[Vibrio-sp.-STUT-A16] |
| WP_267365291.1 | Type1.BREX | <-ACET<-ZnR<-?\|\| wHTH+WYL+WCX -> BrxA-DNA-binding -> BrxB-BrxF-iSTAND -> DUF6079 *-> N6-METHYLASE -> ParB + HNH ->ANK+ANK+ANK->ZnR+3C-ABC+TPR->?->?->?->?-> PglZ -> | 1248 | Bacteria>Pseudomonadota>Gammaproteobacteria>Vibrionales>Vibrionaceae>Vibrio | GCF_026537625.1 | Vibrio_parahaemolyticus | NOL37_RS23780 | 19146..22892 | 670 | BREX-system-P-loop-protein-BrxC-[Vibrio-parahaemolyticus] |
| WP_269646266.1 | Type1.BREX | MobI-><-?<-LexA-protease\|\|35exo->?-> wHTH+WYL+WCX -> BrxA-DNA-binding -> BrxB-BrxF-iSTAND -> DUF6079 *-> ParB + HNH -> N6-METHYLASE -> PglZ -> SIGMA+OB-Fold+AAA-ATPase+Lon-Protease-BrxL ->SbcC->HDRnase->VirD4-TraG+VirB4-FtsK-> | 1225 | Bacteria>Pseudomonadota>Gammaproteobacteria>Vibrionales>Vibrionaceae>Vibrio | GCF_027277845.1 | Vibrio_parahaemolyticus | O4N70_RS12810 | 220363..224040 | 670 | BREX-system-P-loop-protein-BrxC-[Vibrio-parahaemolyticus] |
| WP_269665659.1 | Type1.BREX | MobI-><-?<-LexA-protease\|\|35exo->?-> wHTH+WYL+WCX -> BrxA-DNA-binding -> BrxB-BrxF-iSTAND -> DUF6079 *-> ParB + HNH -> N6-METHYLASE -> PglZ -> SIGMA+OB-Fold+AAA-ATPase+Lon-Protease-BrxL ->NACHT+HEAT+REase->HDRnase->VirD4-TraG+VirB4-FtsK-> | 1225 | Bacteria>Pseudomonadota>Gammaproteobacteria>Vibrionales>Vibrionaceae>Vibrio | GCF_027277905.1 | Vibrio_parahaemolyticus | O4M94_RS09585 | 134179..137856 | 670 | BREX-system-P-loop-protein-BrxC-[Vibrio-parahaemolyticus] |
| WP_274020835.1 | Type1.BREX | <- PglZ <-?<-?<-?<-?<-ZnR+3C-ABC+TPR<- ParB + HNH <- N6-METHYLASE -p<- N6-METHYLASE <- DUF6079 *<- BrxB-BrxF-iSTAND <- BrxA-DNA-binding <- wHTH+WYL+WCX \|\|SF1-SF2-Helicase->MarR-HTH->HKD-DNase+HKD-DNase->CARF+NtrC-AAA->PIN+OST-HTH->MORC-> | 1248 | Bacteria>Pseudomonadota>Gammaproteobacteria>Vibrionales>Vibrionaceae>Vibrio | GCF_028617275.1 | Vibrio_parahaemolyticus | O8254_RS13605 | 106668..110414 | 670 | BREX-system-P-loop-protein-BrxC-[Vibrio-parahaemolyticus] |
| WP_069535824.1 | Type1.BREX | REase+SPF-cons-><-PIN+OST-HTH\|\|CR-ATPase4->?->\|\| wHTH+WYL+WCX -> BrxA-DNA-binding -> BrxB-BrxF-iSTAND -> DUF6079 *-> N6-METHYLASE -> ParB + HNH -> PglZ ->ZetaToxin->?-><-SF1-SF2-Helicase+REase<-SF1-SF2-Helicase+REase<-viral-protein<-AbiEi | 1157 | Bacteria>Pseudomonadota>Gammaproteobacteria>Vibrionales>Vibrionaceae>Vibrio | GCF_905331795.1 | Vibrio_parahaemolyticus | LPR62_RS04710 | 409258..412731 | 670 | BREX-system-P-loop-protein-BrxC-[Vibrio-parahaemolyticus] |
| WP_229611425.1 | Type1.BREX | REase+SPF-cons-><-PIN+OST-HTH\|\|CR-ATPase4->?->\|\| wHTH+WYL+WCX -> BrxA-DNA-binding -> BrxB-BrxF-iSTAND -> DUF6079 *-> N6-METHYLASE -> ParB + HNH -> PglZ ->HEPN+SF1-SF2-Helicase+RNase+RNase->TPR-><-SF1-SF2-Helicase+REase<-SF1-SF2-Helicase+REase<-viral-protein<-AbiEi | 1157 | Bacteria>Pseudomonadota>Gammaproteobacteria>Vibrionales>Vibrionaceae>Vibrio | GCF_905331955.1 | Vibrio_parahaemolyticus | LPR99_RS16740 | 18511..21984 | 670 | BREX-system-P-loop-protein-BrxC-[Vibrio-parahaemolyticus] |
| WP_237468615.1 | Type1.BREX | wHTH+WYL+WCX -> BrxA-DNA-binding -> BrxB-BrxF-iSTAND -> DUF6079 *-> N6-METHYLASE -> ParB + HNH -> ParB + HNH -> PglZ -> SIGMA+OB-Fold+AAA-ATPase+Lon-Protease-BrxL -><-Acetate_kinase | 1231 | Bacteria>Pseudomonadota>Gammaproteobacteria>Vibrionales>Vibrionaceae>Vibrio | GCF_921293875.1 | Vibrio_stylophorae | L9P36_RS15955 | 492812..496507 | 659351 | BREX-system-P-loop-protein-BrxC-[Vibrio-stylophorae] |
| WP_168780466.1 | Type1.BREX | <- SIGMA+OB-Fold+AAA-ATPase+Lon-Protease-BrxL <- PglZ <-?<-?<-?<-GrpE<-ANK+ANK+ANK<- ParB + HNH <- N6-METHYLASE <- DUF6079 *<- BrxB-BrxF-iSTAND <- BrxA-DNA-binding <- wHTH+WYL+WCX <-?\|\|AbiEi_4-><-REase<-TraI-VirD2-RCRE<-TraJ-RHH<-SbcC | 1248 | Bacteria>Pseudomonadota>Gammaproteobacteria>Vibrionales>Vibrionaceae>Vibrio | GCF_944738875.1 | Vibrio_aestuarianus | OCX59_RS00150 | 25401..29147 | 28171 | BREX-system-P-loop-protein-BrxC-[Vibrio-aestuarianus] |
| WP_272166069.1 | Type1.BREX | CR-ATPase4->?-><- SIGMA+OB-Fold+AAA-ATPase+Lon-Protease-BrxL <- PglZ <-?<-?<- ParB + HNH <- N6-METHYLASE <- DUF6079 *<- BrxB-BrxF-iSTAND <- BrxA-DNA-binding <- wHTH+WYL+WCX \|\|SF1-SF2-Helicase->MarR-HTH->HKD-DNase+HKD-DNase->CARF+NtrC-AAA->Vrr-REase->HEPN+PAS+DUF4011+SF1-SF2-Helicase+REase-> | 1248 | Bacteria>Pseudomonadota>Gammaproteobacteria>Vibrionales>Vibrionaceae>Vibrio | GCF_947503745.1 | Vibrio_diabolicus | PQF93_RS21015 | 28188..31934 | 50719 | BREX-system-P-loop-protein-BrxC-[Vibrio-diabolicus] |
| WP_142738823.1 | Type1.BREX | MobI-><-?<-LexA-protease\|\|?-> wHTH+WYL+WCX -> BrxA-DNA-binding -> BrxB-BrxF-iSTAND -> DUF6079 *-> ParB + HNH -> N6-METHYLASE ->OLD-ABC->TOPRIM-> PglZ -> SIGMA+OB-Fold+AAA-ATPase+Lon-Protease-BrxL ->SbcC->HDRnase->VirD4-TraG+VirB4-FtsK-> | 1224 | Bacteria>Pseudomonadota>Gammaproteobacteria>Vibrionales>Vibrionaceae>Vibrio | - | Vibrio_cholerae | FLL79_RS01410 | 36024..39698 | 666 | BREX-system-P-loop-protein-BrxC-[Vibrio-cholerae] |
| WP_226420703.1 | Type1.BREX | <-MORC<-PIN+OST-HTH<-CARF+NtrC-AAA<-HKD-DNase+HKD-DNase<-MarR-HTH<-SF1-SF2-Helicase\|\| wHTH+WYL+WCX -> BrxA-DNA-binding -> BrxB-BrxF-iSTAND -> DUF6079 *-> N6-METHYLASE ->?-> ParB + HNH ->ZnR+3C-ABC+TPR->?->?->?->?-> PglZ -> | 1248 | Bacteria>Pseudomonadota>Gammaproteobacteria>Vibrionales>Vibrionaceae>Vibrio | - | Vibrio_sp._E14 | LIX84_RS04085 | 940499..944245 | 2849869 | BREX-system-P-loop-protein-BrxC-[Vibrio-sp.-E14] |
| WP_017462916.1 | Type6.BREX | <-SbcC<-JetB<-JetA\|\| wHTH+WYL+WCX -> BrxE -> BrxA-DNA-binding -> BrxB-BrxF-iSTAND -> DUF6079 *->GreA_GreB+SF1-SF2-Helicase-> N6-METHYLASE -> N6-METHYLASE -> ParB + HNH ->?-> PglZ -> SIGMA+OB-Fold+AAA-ATPase+HKD-DNase-BrxL -><-JAB | 1213 | Bacteria>Pseudomonadota>Gammaproteobacteria>Xanthomonadales>Rhodanobacteraceae>Dyella | GCF_000334915.1 | Dyella_ginsengisoli | C693_RS0114050 | 37588..41229 | 363848 | BREX-system-P-loop-protein-BrxC-[Dyella-ginsengisoli] |
| MBK7210551.1 | Type1.BREX | <-HDRnase<-Arm-DNA-bind_2+Phage_int_SAM_3+Phage_integrase<-Phage_AlpA\|\|HTH_26->SLOG+BirA-HTH-> ParB + HNH -><-?\|\| BrxA-DNA-binding -> BrxB-BrxF-iSTAND -> DUF6079 *->?-> PglZ -> SIGMA+OB-Fold+AAA-ATPase+Lon-Protease-BrxL -><-tRNA\|\|NUDIX->?->?-><-HISKIN+REC+HPT<-REC+GGDEF+EAL | 1144 | Bacteria>Pseudomonadota>Gammaproteobacteria>Xanthomonadales>unclassified_Xanthomonadales | GCA_016707485.1 | Xanthomonadales_bacterium | IPH99_09390 | 1022709..1026143 | 2006849 | BREX-system-P-loop-protein-BrxC-[Xanthomonadales-bacterium] |
| WP_070426722.1 | Type6.BREX | METHYLASE->RNaseH->35exo-><-?\|\|tRNA-> wHTH+WYL+WCX -> BrxE -> BrxA-DNA-binding -> BrxB-BrxF-iSTAND -> DUF6079 *->SF1-SF2-Helicase+RNase+RNase->TPR-> N6-METHYLASE -> N6-METHYLASE -p-> ParB + HNH -> PglZ -> SIGMA+OB-Fold+AAA-ATPase+HKD-DNase-BrxL ->DUF932-> | 1209 | Bacteria>Pseudomonadota>Gammaproteobacteria>Xanthomonadales>Xanthomonadaceae | GCF_900766255.1 | Stenotrophomonas | BIZ42_RS13545 | 3079736..3083365 | 40323 | MULTISPECIES:-BREX-system-P-loop-protein-BrxC-[Stenotrophomonas] |
| WP_162376315.1 | Type1.BREX | <-NTP_transf_3+Hexapep+Hexapep\|\|?-><-?<- SIGMA+OB-Fold+AAA-ATPase+Lon-Protease-BrxL <- PglZ <-Polbeta<-MarR-HTH<- ParB + HNH <- N6-METHYLASE <- DUF6079 *<- BrxB-BrxF-iSTAND <- BrxA-DNA-binding \|\|REase-><-?<-?<-TOPRIM<-XIS-HTH | 1148 | Bacteria>Pseudomonadota>Gammaproteobacteria>Xanthomonadales>Xanthomonadaceae>Pseudoxanthomonas | GCF_010211765.1 | Pseudoxanthomonas_kaohsiungensis | CSC66_RS05120 | 191406..194852 | 283923 | BREX-system-P-loop-protein-BrxC-[Pseudoxanthomonas-kaohsiungensis] |
| WP_271250641.1 | Type1.BREX | <- SIGMA+OB-Fold+AAA-ATPase+Lon-Protease-BrxL <- PglZ <-Polbeta<-MarR-HTH<- ParB + HNH <- N6-METHYLASE <- DUF6079 *<- BrxB-BrxF-iSTAND <- BrxA-DNA-binding \|\|REase-><-?<-?<-?<-TOPRIM<-?<-XIS-HTH | 1148 | Bacteria>Pseudomonadota>Gammaproteobacteria>Xanthomonadales>Xanthomonadaceae>Pseudoxanthomonas | GCF_027941875.1 | Pseudoxanthomonas_mexicana | PE064_RS03375 | 689805..693251 | 128785 | BREX-system-P-loop-protein-BrxC-[Pseudoxanthomonas-mexicana] |
| MBH1702634.1 | Type1.BREX | <-RFC-AAA+RuvB-AAA\|\|tRNA->?->AAA+wHTH-><- SIGMA+OB-Fold+AAA-ATPase+Lon-Protease-BrxL <- PglZ <- HNH <-OLD-ABC <- ParB + HNH <- N6-METHYLASE <- DUF6079 *<- BrxB-BrxF-iSTAND <- BrxA-DNA-binding <- wHTH+WYL+WCX <-?<-?<-?<-DUF932\|\|XIS-HTH-> | 1146 | Bacteria>Pseudomonadota>Gammaproteobacteria>Xanthomonadales>Xanthomonadaceae>Stenotrophomonas | GCA_015998095.1 | Stenotrophomonas_maltophilia | I5U93_05090 | 24102..27542 | 40324 | BREX-system-P-loop-protein-BrxC-[Stenotrophomonas-maltophilia] |
| MBH1753233.1 | Type6.BREX | <-DUF932<- SIGMA+OB-Fold+AAA-ATPase+HKD-DNase-BrxL <- PglZ <- ParB + HNH <- N6-METHYLASE -p<- N6-METHYLASE <-TPR<-SF1-SF2-Helicase+RNase+RNase<- DUF6079 *<- BrxB-BrxF-iSTAND <- BrxA-DNA-binding <- BrxE <- wHTH+WYL+WCX <-tRNA\|\|?-><-35exo<-RNaseH<-METHYLASE | 1209 | Bacteria>Pseudomonadota>Gammaproteobacteria>Xanthomonadales>Xanthomonadaceae>Stenotrophomonas | GCA_015998385.1 | Stenotrophomonas_maltophilia | I5V37_01935 | 25775..29404 | 40324 | BREX-system-P-loop-protein-BrxC-[Stenotrophomonas-maltophilia] |
| MBN5083090.1 | Type6.BREX | <-DUF932<- SIGMA+OB-Fold+AAA-ATPase+HKD-DNase-BrxL <- PglZ <- ParB + HNH <- N6-METHYLASE <- N6-METHYLASE <-PIN<-OLD-ABC<- DUF6079 *<- BrxB-BrxF-iSTAND <- BrxA-DNA-binding <- BrxE <- wHTH+WYL+WCX <-Arm-DNA-bind_3+Phage_integrase<-tRNA<-SIGMA-HTH+SIGMA-HTH+SIGMA-HTH | 1207 | Bacteria>Pseudomonadota>Gammaproteobacteria>Xanthomonadales>Xanthomonadaceae>Stenotrophomonas | GCA_017150495.1 | Stenotrophomonas_maltophilia | JY467_05250 | 61681..65304 | 40324 | BREX-system-P-loop-protein-BrxC-[Stenotrophomonas-maltophilia] |
| WP_004146422.1 | Type6.BREX | <-XIS-HTH\|\|?->Phage_integrase->tRNA->Arm-DNA-bind_3+Phage_integrase-> wHTH+WYL+WCX -> BrxE -> BrxA-DNA-binding -> BrxB-BrxF-iSTAND -> DUF6079 *->OLD-ABC->PIN-> N6-METHYLASE -> N6-METHYLASE -> ParB + HNH -> PglZ -> SIGMA+OB-Fold+AAA-ATPase+HKD-DNase-BrxL ->DUF932-> | 1207 | Bacteria>Pseudomonadota>Gammaproteobacteria>Xanthomonadales>Xanthomonadaceae>Stenotrophomonas | GCF_000355725.1 | Stenotrophomonas_maltophilia | G5A34_RS07915 | 81973..85596 | 40324 | BREX-system-P-loop-protein-BrxC-[Stenotrophomonas-maltophilia] |
| WP_099491466.1 | Type6.BREX | METHYLASE->RNaseH->35exo-><-?\|\|tRNA-> wHTH+WYL+WCX -> BrxE -> BrxA-DNA-binding -> BrxB-BrxF-iSTAND -> DUF6079 *->SF1-SF2-Helicase+RNase+RNase->TPR-> N6-METHYLASE -> N6-METHYLASE -p-> ParB + HNH -> PglZ -> SIGMA+OB-Fold+AAA-ATPase+HKD-DNase-BrxL ->DUF932-> | 1209 | Bacteria>Pseudomonadota>Gammaproteobacteria>Xanthomonadales>Xanthomonadaceae>Stenotrophomonas | GCF_002739275.1 | Stenotrophomonas_maltophilia | BRB91_RS07170 | 53995..57624 | 40324 | BREX-system-P-loop-protein-BrxC-[Stenotrophomonas-maltophilia] |
| WP_125428654.1 | Type1.BREX | <-RFC-AAA+RuvB-AAA\|\|tRNA->?->AAA+wHTH-><- SIGMA+OB-Fold+AAA-ATPase+Lon-Protease-BrxL <- PglZ <- HNH <-OLD-ABC <- ParB + HNH <- N6-METHYLASE <- DUF6079 *<- BrxB-BrxF-iSTAND <- BrxA-DNA-binding <- wHTH+WYL+WCX <-?<-?<-?<-DUF932\|\|XIS-HTH-> | 1146 | Bacteria>Pseudomonadota>Gammaproteobacteria>Xanthomonadales>Xanthomonadaceae>Stenotrophomonas | GCF_003935615.1 | Stenotrophomonas_maltophilia | EGJ89_RS09210 | 127004..130444 | 40324 | BREX-system-P-loop-protein-BrxC-[Stenotrophomonas-maltophilia] |
| WP_164085131.1 | Type6.BREX | METHYLASE->RNaseH->35exo-><-?\|\|tRNA-> wHTH+WYL+WCX -> BrxE -> BrxA-DNA-binding -> BrxB-BrxF-iSTAND -> DUF6079 *->SF1-SF2-Helicase+RNase+RNase->TPR-> N6-METHYLASE -> N6-METHYLASE -p-> ParB + HNH -> | 1209 | Bacteria>Pseudomonadota>Gammaproteobacteria>Xanthomonadales>Xanthomonadaceae>Stenotrophomonas | GCF_010605045.1 | Stenotrophomonas_maltophilia | G5A32_RS08645 | 68360..71989 | 40324 | BREX-system-P-loop-protein-BrxC-[Stenotrophomonas-maltophilia] |
| WP_180875343.1 | Type6.BREX | METHYLASE->RNaseH->35exo-><-?\|\|tRNA-> wHTH+WYL+WCX -> BrxE -> BrxA-DNA-binding -> BrxB-BrxF-iSTAND -> DUF6079 *->SF1-SF2-Helicase+RNase+RNase->TPR-> N6-METHYLASE -> N6-METHYLASE -p-> ParB + HNH -> PglZ -> SIGMA+OB-Fold+AAA-ATPase+HKD-DNase-BrxL ->DUF932-> | 1209 | Bacteria>Pseudomonadota>Gammaproteobacteria>Xanthomonadales>Xanthomonadaceae>Stenotrophomonas | GCF_013463485.1 | Stenotrophomonas_maltophilia | D7U74_RS02160 | 10322..13951 | 40324 | BREX-system-P-loop-protein-BrxC-[Stenotrophomonas-maltophilia] |
| WP_192288052.1 | Type1.BREX | SF1-SF2-Helicase->tRNA->tRNA-> BrxA-DNA-binding -> BrxB-BrxF-iSTAND -> DUF6079 *-> N6-METHYLASE -> ParB + HNH ->MarR-HTH->Polbeta-> PglZ -> SIGMA+OB-Fold+AAA-ATPase+Lon-Protease-BrxL ->?-><-HTH_3\|\|TPR-> | 1148 | Bacteria>Pseudomonadota>Gammaproteobacteria>Xanthomonadales>Xanthomonadaceae>Stenotrophomonas | GCF_014852135.1 | Stenotrophomonas_sp._STM01 | IB227_RS14890 | 150260..153706 | 2769278 | BREX-system-P-loop-protein-BrxC-[Stenotrophomonas-sp.-STM01] |
| WP_049450109.1 | Type6.BREX | <-DUF932<- SIGMA+OB-Fold+AAA-ATPase+HKD-DNase-BrxL <- PglZ <- ParB + HNH <- N6-METHYLASE -p<- N6-METHYLASE <-TPR<-SF1-SF2-Helicase+RNase+RNase<- DUF6079 *<- BrxB-BrxF-iSTAND <- BrxA-DNA-binding <- BrxE <- wHTH+WYL+WCX <-tRNA\|\|?-><-35exo<-RNaseH<-METHYLASE | 1209 | Bacteria>Pseudomonadota>Gammaproteobacteria>Xanthomonadales>Xanthomonadaceae>Stenotrophomonas | GCF_030549525.1 | Stenotrophomonas_maltophilia | MT454_RS13320 | 40759..44388 | 40324 | BREX-system-P-loop-protein-BrxC-[Stenotrophomonas-maltophilia] |
| WP_111722876.1 | Type6.BREX | METHYLASE->RNaseH->35exo-><-?\|\|tRNA-> wHTH+WYL+WCX -> BrxE -> BrxA-DNA-binding -> BrxB-BrxF-iSTAND -> DUF6079 *->SF1-SF2-Helicase+RNase+RNase->TPR-> N6-METHYLASE -> N6-METHYLASE -p-> ParB + HNH -> PglZ -> SIGMA+OB-Fold+AAA-ATPase+HKD-DNase-BrxL ->DUF932-> | 1209 | Bacteria>Pseudomonadota>Gammaproteobacteria>Xanthomonadales>Xanthomonadaceae>Stenotrophomonas | GCF_900475685.1 | Stenotrophomonas_maltophilia | DQN96_RS04535 | 962467..966096 | 40324 | BREX-system-P-loop-protein-BrxC-[Stenotrophomonas-maltophilia] |
| WP_262214630.1 | Type6.BREX | METHYLASE->RNaseH->35exo-><-?\|\|tRNA-> wHTH+WYL+WCX -> BrxE -> BrxA-DNA-binding -> BrxB-BrxF-iSTAND -> DUF6079 *->GreA_GreB+SF1-SF2-Helicase-> N6-METHYLASE -> N6-METHYLASE -> ParB + HNH -> PglZ -> SIGMA+OB-Fold+AAA-ATPase+HKD-DNase-BrxL ->DUF932->?-><-HTH_3 | 1213 | Bacteria>Pseudomonadota>Gammaproteobacteria>Xanthomonadales>Xanthomonadaceae>Stenotrophomonas | - | Stenotrophomonas_maltophilia | JAK45_RS19240 | 75512..79153 | 40324 | BREX-system-P-loop-protein-BrxC-[Stenotrophomonas-maltophilia] |
| VUJ46804.1 | Type6.BREX | <-LexA-protease\|\|?-><-Phage_integrase\|\|Arm-DNA-bind_3+Phage_integrase-> wHTH+WYL+WCX -> BrxE -> BrxA-DNA-binding -> BrxB-BrxF-iSTAND -> DUF6079 *->OLD-ABC->PIN-> N6-METHYLASE -> N6-METHYLASE -> ParB + HNH -> PglZ -> SIGMA+OB-Fold+AAA-ATPase+HKD-DNase-BrxL ->DUF932-> | 1207 | Bacteria>Pseudomonadota>Gammaproteobacteria>Xanthomonadales>Xanthomonadaceae>Stenotrophomonas | - | Stenotrophomonas_maltophilia | N5D45_RS14200 | 24687..28310 | 40324 | hypothetical-protein-NGKLMNPD_00401-[Stenotrophomonas-maltophilia] |
| EGD07603.1 | Type1.BREX | <-RFC-AAA+RuvB-AAA\|\|tRNA->?->AAA+wHTH-><- SIGMA+OB-Fold+AAA-ATPase+Lon-Protease-BrxL <- PglZ <- ParB + HNH <- N6-METHYLASE <- DUF6079 *<- BrxB-BrxF-iSTAND <- BrxA-DNA-binding <- wHTH+WYL+WCX <-DUF772+DDE_Tnp_1\|\|XIS-HTH-> | 1152 | Bacteria>Pseudomonadota>Gammaproteobacteria>Xanthomonadales>Xanthomonadaceae>Xanthomonas | GCA_000192025.2 | Xanthomonas_vesicatoria_ATCC_35937 | XVE_4193 | 13272..16730 | 925775 | hypothetical-protein-XVE_4193-[Xanthomonas-vesicatoria-ATCC-35937] |
| WP_042828568.1 | Type1.BREX | <-RFC-AAA+RuvB-AAA\|\|tRNA->?->AAA+wHTH-><- SIGMA+OB-Fold+AAA-ATPase+Lon-Protease-BrxL <- PglZ <- ParB + HNH <- N6-METHYLASE <- DUF6079 *<- BrxB-BrxF-iSTAND <- BrxA-DNA-binding <- wHTH+WYL+WCX \|\|<-DUF772+DDE_Tnp_1\|\|XIS-HTH-> | 1146 | Bacteria>Pseudomonadota>Gammaproteobacteria>Xanthomonadales>Xanthomonadaceae>Xanthomonas | GCF_000192025.1 | Xanthomonas_vesicatoria | XVE_RS20000 | 13272..16712 | 56460 | BREX-system-P-loop-protein-BrxC-[Xanthomonas-vesicatoria] |
| WP_260816965.1 | Type1.BREX | DnaJ->?->?->?-><-?<-XIS-HTH\|\| wHTH+WYL+WCX -> BrxA-DNA-binding -> BrxB-BrxF-iSTAND -> DUF6079 *-> N6-METHYLASE -> ParB + HNH -> PglZ -> SIGMA+OB-Fold+AAA-ATPase+Lon-Protease-BrxL -><-AAA+wHTH<-?<-tRNA\|\|RFC-AAA+RuvB-AAA->?->HHH_5+RecR+TOPRIM-> | 1146 | Bacteria>Pseudomonadota>Gammaproteobacteria>Xanthomonadales>Xanthomonadaceae>Xanthomonas | GCF_025263745.1 | Xanthomonas_prunicola | MOD45_RS16865 | 3940549..3943989 | 2053930 | BREX-system-P-loop-protein-BrxC-[Xanthomonas-prunicola] |
| WP_000687848.1 | Type1.BREX | <-VirD4-TraG+VirB4-FtsK<-HDRnase<-SbcC<- SIGMA+OB-Fold+AAA-ATPase+Lon-Protease-BrxL <- PglZ <- N6-METHYLASE <- ParB + HNH <- DUF6079 *<- BrxB-BrxF-iSTAND <- BrxA-DNA-binding <- wHTH+WYL+WCX <-?<-35exo<-?\|\|LexA-protease->?->Phage_integrase-> | 1225 | Bacteria>Pseudomonadota | GCF_000022585.1 | Gammaproteobacteria | ALD23_RS14425 | 130597..134274 | 1236 | MULTISPECIES:-BREX-system-P-loop-protein-BrxC-[Gammaproteobacteria] |
| WP_156734107.1 | Type1.BREX | <-LexA-protease\|\|?->35exo->?-> wHTH+WYL+WCX -> BrxA-DNA-binding -> BrxB-BrxF-iSTAND -> DUF6079 *-> ParB + HNH -> N6-METHYLASE -> PglZ -> SIGMA+OB-Fold+AAA-ATPase+Lon-Protease-BrxL ->NACHT+HEAT+REase->HDRnase->VirD4-TraG+VirB4-FtsK-> | 1225 | Bacteria>Pseudomonadota | GCF_009740265.1 | Gammaproteobacteria | GTH23_RS00050 | 5219..8896 | 1236 | MULTISPECIES:-BREX-system-P-loop-protein-BrxC-[Gammaproteobacteria] |
| WP_119567031.1 | Type1.BREX | <-OBFOLD-NUCLEASE<-CARF+NtrC-AAA+FIS-HTH<- SIGMA+OB-Fold+AAA-ATPase+Lon-Protease-BrxL <- PglZ <- N6-METHYLASE <- ParB + HNH <- DUF6079 *<- BrxB-BrxF-iSTAND <- BrxA-DNA-binding <- wHTH+WYL+WCX <-?<-35exo\|\|LexA-protease->?-><-MobI | 1225 | Bacteria>Pseudomonadota | GCF_013602615.1 | Gammaproteobacteria | G9G97_RS25355 | 26642..30319 | 1236 | MULTISPECIES:-BREX-system-P-loop-protein-BrxC-[Gammaproteobacteria] |
| WP_125941663.1 | Type1.BREX | GDResolvase+HIN-HTH-><-?<-?<- SIGMA+OB-Fold+AAA-ATPase+Lon-Protease-BrxL <- PglZ <- ParB + HNH <- N6-METHYLASE -p<-HTH_23+HTH_32+HTH_33+DDE_3<- N6-METHYLASE <- DUF6079 *<- BrxB-BrxF-iSTAND <- BrxA-DNA-binding <- wHTH+WYL+WCX \|\|XIS-HTH->?->?-><-?<-Phage_int_SAM_5+Phage_integrase<-sGTP | 1154 | Bacteria>Pseudomonadota | GCF_019046925.1 | Gammaproteobacteria | I6L32_RS07295 | 1524921..1528385 | 1236 | MULTISPECIES:-BREX-system-P-loop-protein-BrxC-[Gammaproteobacteria] |
| WP_110076276.1 | Type1.BREX | <-VirD4-TraG+VirB4-FtsK<-HDRnase<-SbcC<- SIGMA+OB-Fold+AAA-ATPase+Lon-Protease-BrxL <- PglZ <- N6-METHYLASE <- ParB + HNH <- DUF6079 *<- BrxB-BrxF-iSTAND <- BrxA-DNA-binding <- wHTH+WYL+WCX <-?<-35exo\|\|LexA-protease->\|\|?->?->Pkinase-> | 1225 | Bacteria>Pseudomonadota | GCF_021431945.1 | Gammaproteobacteria | LY388_RS16340 | 2424979..2428656 | 1236 | MULTISPECIES:-BREX-system-P-loop-protein-BrxC-[Gammaproteobacteria] |
| WP_111805008.1 | Type1.BREX | <-VirD4-TraG+VirB4-FtsK<-HDRnase<-SbcC<- SIGMA+OB-Fold+AAA-ATPase+Lon-Protease-BrxL <- PglZ <- N6-METHYLASE <- ParB + HNH <- DUF6079 *<- BrxB-BrxF-iSTAND <- BrxA-DNA-binding <- wHTH+WYL+WCX <-?<-35exo\|\|LexA-protease->?-><-MobI | 1225 | Bacteria>Pseudomonadota | GCF_028482435.1 | Gammaproteobacteria | DR130_RS11140 | 144958..148635 | 1236 | MULTISPECIES:-BREX-system-P-loop-protein-BrxC-[Gammaproteobacteria] |
| PIX55556.1 | Type6.BREX | BrxE -> BrxA-DNA-binding -> BrxB-BrxF-iSTAND -> DUF6079 *-> N6-METHYLASE -> ParB + HNH ->?-> PglZ -> SIGMA+OB-Fold+AAA-ATPase+HKD-DNase-BrxL ->ZNR+RmuC->XIS-HTH-><-HTH_26\|\|Phage_AlpA-> | 1215 | Bacteria>Pseudomonadota>Zetaproteobacteria>unclassified_Zetaproteobacteria | GCA_002782825.1 | Zetaproteobacteria_bacterium_CG_4_10_14_3_um_filter_54_28 | COZ50_02095 | 2235..5882 | 1974114 | BREX-system-P-loop-protein-BrxC-[Zetaproteobacteria-bacterium-CG_4_10_14_3_um_filter_54_28] |
| MBX3380229.1 | Type6.BREX | Pkinase+pi3KINASE->?->?-><-?<- SIGMA+OB-Fold+AAA-ATPase+Lon-Protease-BrxL <- PglZ <- ParB + HNH <- ParB + HNH <- N6-METHYLASE <- DUF6079 *<- BrxB-BrxF-iSTAND <- BrxA-DNA-binding <- BrxE <-?<-Phage_integrase | 1207 | Bacteria>PVC_group>Planctomycetota>Phycisphaerae>Phycisphaerales>Phycisphaeraceae | GCA_019636565.1 | Phycisphaeraceae_bacterium | KF805_09030 | 72021..75644 | 2026777 | BREX-system-P-loop-protein-BrxC-[Phycisphaeraceae-bacterium] |
| WP_145095869.1 | Type1.BREX | <-TPR+TPR<-Peptidase<-?<-?<- SIGMA+OB-Fold+AAA-ATPase+Lon-Protease-BrxL <- PglZ <-RAMA<- N6-METHYLASE <- ParB + HNH <- DUF6079 *<- BrxB-BrxF-iSTAND <- BrxA-DNA-binding <-XIS-HTH\|\|Thermonuclease->OLD-ABC->?->ABhydrolase-> | 1192 | Bacteria>PVC_group>Planctomycetota>Planctomycetia>Pirellulales>Pirellulaceae | GCF_007753095.1 | Rosistilla_carotiformis | Poly24_RS13225 | 3653187..3656765 | 2528017 | BREX-system-P-loop-protein-BrxC-[Rosistilla-carotiformis] |
| WP_207398110.1 | Type1.BREX | <-Phage_int_SAM_1+Phage_integrase<-MerR-HTH<-RFC-AAA<-?<-PIWI<-REase+PNTD1+PNTD2\|\|XIS-HTH-> BrxA-DNA-binding -> BrxB-BrxF-iSTAND -> DUF6079 *-> ParB + HNH -> N6-METHYLASE ->REase->?->REase->?-> PglZ -> SIGMA+OB-Fold+AAA-ATPase+Lon-Protease-BrxL ->KAP-NTPase-> | 1192 | Bacteria>PVC_group>Planctomycetota>Planctomycetia>Pirellulales>Pirellulaceae | GCF_013618625.1 | Bremerella_alba | HOV93_RS18975 | 110238..113816 | 980252 | BREX-system-P-loop-protein-BrxC-[Bremerella-alba] |
| TET34868.1 | Type1.BREX | <- HNH \|\|?-><-?\|\|XIS-HTH->DUF4007->LON-AAA-> N6-METHYLASE -> ParB + HNH ->HEPN-> DUF6079 *-> PglZ ->SIGMA+OB-Fold+AAA-ATPase-BrxL->SbcC->HISKIN->wHTH+WYL->RADICAL-SAM->DEOR-HTH-><-GDResolvase+Recombinase+Zn_ribbon_recom | 1347 | Bacteria>PVC_group>Planctomycetota>unclassified_Planctomycetota | GCA_004376375.1 | Planctomycetota_bacterium | E3J72_12085 | 41890..45933 | 2026780 | MAG:-hypothetical-protein-E3J72_12085-[Planctomycetota-bacterium] |
| TWU09954.1 | Type1.BREX | <-ABhydrolase<-?\|\|?->Calcineurin->SbcC+SbcC->?->XIS-HTH-> BrxA-DNA-binding -> BrxB-BrxF-iSTAND -> DUF6079 *-> ParB + HNH -> N6-METHYLASE ->REase-> PglZ ->SKI->?-> SIGMA+OB-Fold+AAA-ATPase+Lon-Protease-BrxL -> | 1205 | Bacteria>PVC_group>Planctomycetota>unclassified_Planctomycetota | GCA_007860105.1 | Planctomycetes_bacterium_Poly21 | Poly21_52830 | 31284..34901 | 2528014 | hypothetical-protein-Poly21_52830-[Planctomycetes-bacterium-Poly21] |
| MCD6405191.1 | Type1.BREX | <-Schlafen+HATPase+wHTH-p<-PHP+SbcC<- SIGMA+OB-Fold+AAA-ATPase+Lon-Protease-BrxL <- PglZ <- ParB + HNH <- N6-METHYLASE <- DUF6079 *<- BrxB-BrxF-iSTAND <- BrxA-DNA-binding <-XIS-HTH | 1143 | Bacteria>PVC_group>Planctomycetota>unclassified_Planctomycetota | GCA_021160625.1 | Planctomycetota_bacterium | J7M19_05155 | 12563..15994 | 2026780 | BREX-system-P-loop-protein-BrxC-[Planctomycetota-bacterium] |
| WP_215427518.1 | Type1.BREX | ANK+ANK+ANK-> HNH ->HTH_3->Couple_hipA->HipA_C->?->XIS-HTH-> BrxA-DNA-binding -> BrxB-BrxF-iSTAND -> DUF6079 *-> N6-METHYLASE -> PglZ -> SIGMA+OB-Fold+AAA-ATPase+Lon-Protease-BrxL -> HNH -><- ParB + HNH \|\|DUF932->PadR-HTH-> | 1141 | Bacteria>PVC_group>Verrucomicrobiota>Verrucomicrobiae>Verrucomicrobiales>Akkermansiaceae | GCF_018709685.1 | Akkermansia_muciniphila | HHJ06_RS03145 | 88477..91902 | 239935 | BREX-system-P-loop-protein-BrxC-[Akkermansia-muciniphila] |
| MBE7502027.1 | Type6.BREX | HTH_3-><-?<-?<-?<-GNAT<-NERD+ZnR\|\| BrxE -> BrxA-DNA-binding -> BrxB-BrxF-iSTAND -> DUF6079 *-> N6-METHYLASE -><-DDE_Tnp_1\|\| N6-METHYLASE -> ParB + HNH ->OLD-ABC->TOPRIM->DDE_Tnp_1-> PglZ -> SIGMA+OB-Fold+AAA-ATPase+HKD-DNase-BrxL -> | 1231 | Bacteria>PVC_group>Verrucomicrobiota>Verrucomicrobiae>Verrucomicrobiales>unclassified_Verrucomicrobiales | GCA_015075445.1 | Verrucomicrobiales_bacterium | HS113_17385 | 108161..111856 | 2026801 | BREX-system-P-loop-protein-BrxC-[Verrucomicrobiales-bacterium] |
| WP_015675681.1 | Type6.BREX | <-SF1-SF2-Helicase+wHTH+Small-B-barrel+DUF1998<-BrxD-AAA<- PglZ <- N6-METHYLASE -p<-AbiEii<-AbiEi_3_N+AbiEi_3<- ParB + HNH <- N6-METHYLASE <- DUF6079 *<- BrxB-BrxF-iSTAND <- BrxA-DNA-binding <- BrxE <-TPR+TPR<-TPR<-MarR-HTH<- ParB <-ParA-Soj-PloopNTPase | 1203 | Bacteria>Spirochaetota>Spirochaetia>Leptospirales>Leptospiraceae>Leptospira | GCF_000332475.2 | Leptospira_yanagawae | LEP1GSC202_0423 | 26532..30143 | 293069 | BREX-system-P-loop-protein-BrxC-[Leptospira-yanagawae] |
| WP_135627473.1 | Type1.BREX | DUF6079 *-> N6-METHYLASE -> ParB + HNH -> PglZ -> SIGMA+OB-Fold+AAA-ATPase+Lon-Protease-BrxL -> | 1122 | Bacteria>Spirochaetota>Spirochaetia>Leptospirales>Leptospiraceae>Leptospira | GCF_004770835.1 | Leptospira_selangorensis | EHQ82_RS10885 | 1..3369 | 2484982 | BREX-system-P-loop-protein-BrxC,-partial-[Leptospira-selangorensis] |
| WP_238754851.1 | Type6.BREX | ANK->ParA-Soj-PloopNTPase-> ParB ->MarR-HTH->TPR+TPR-> BrxE -> BrxA-DNA-binding -> BrxB-BrxF-iSTAND -> DUF6079 *-> N6-METHYLASE ->?->?->?-> ParB + HNH ->AbiEi_3_N+AbiEi_3->DOC+HTH-> N6-METHYLASE -p-> PglZ -> | 1201 | Bacteria>Spirochaetota>Spirochaetia>Leptospirales>Leptospiraceae>Leptospira | GCF_022267385.1 | Leptospira_bandrabouensis | LFX13_RS16565 | 34041..37646 | 2484903 | BREX-system-P-loop-protein-BrxC-[Leptospira-bandrabouensis] |
| OQY41116.1 | Type1.BREX | REase-> BrxA-DNA-binding -> BrxB-BrxF-iSTAND -> DUF6079 *->OLD-ABC->TOPRIM-> N6-METHYLASE -> ParB + HNH -> ParB + HNH -> SIGMA+OB-Fold+AAA-ATPase+Lon-Protease-BrxL -> | 1207 | Bacteria>Spirochaetota>Spirochaetia>Spirochaetales>Spirochaetaceae>unclassified_Spirochaetaceae | GCA_002084785.1 | Spirochaetaceae_bacterium_4572_7 | B6229_00020 | 2034..5657 | 1972456 | hypothetical-protein-B6229_00020-[Spirochaetaceae-bacterium-4572_7] |
| WP_109093701.1 | Type1.BREX | <-PIN<-PhdYeFM_antitox\|\|?->REase-><- ParB + HNH <- SIGMA+OB-Fold+AAA-ATPase+HKD-DNase-BrxL <-Schlafen+HATPase+wHTH<- N6-METHYLASE <- DUF6079 *<- BrxB-BrxF-iSTAND <- BrxA-DNA-binding \|\|?-><-?<-?<-LACI-HTH+PBPI<-Glyco_hydro_1 | 1170 | Bacteria>Terrabacteria_group>Actinomycetota>Actinomycetes>Actinomycetales>Actinomycetaceae | GCF_003123745.1 | Ancrocorticia_populi | DD236_RS06730 | 275849..279361 | 2175228 | BREX-system-P-loop-protein-BrxC-[Ancrocorticia-populi] |
| QZE24962.1 | Type1.BREX | HTH+MPTase->?->Small-B-barrel+SF1-SF2-Helicase+RdgC+REase-><-?\|\|<-zf-ISL3+HTH_Tnp_ISL3<- ParB + HNH <- SIGMA+OB-Fold+AAA-ATPase+Lon-Protease-BrxL <- PglZ <- N6-METHYLASE <- DUF6079 *<- BrxB-BrxF-iSTAND <- BrxA-DNA-binding <-XIS-HTH<-ABhydrolase<-?<-MPTase<-HTH\|\|ArdCN+MPTase->DUF2637+HIN-HTH-> | 1152 | Bacteria>Terrabacteria_group>Actinomycetota>Actinomycetes>Micrococcales>Brevibacteriaceae | GCA_019720815.1 | Brevibacterium_casei | K4X33_12630 | 2739675..2743133 | 33889 | BREX-system-P-loop-protein-BrxC-[Brevibacterium-casei] |
| WP_186372589.1 | Type1.BREX | <-VirB11+MuB-ATPase\|\|DNA_METHYLASE->DNA_METHYLASE+DCM->DNA_METHYLASE-><- ParB + HNH <- SIGMA+OB-Fold+AAA-ATPase+Lon-Protease-BrxL <- PglZ <- N6-METHYLASE <- DUF6079 *<- BrxB-BrxF-iSTAND <- BrxA-DNA-binding <-XIS-HTH<-ABhydrolase<-?<-MPTase<-HTH\|\|ArdCN+MPTase->Primpol+PriCT1-> | 1152 | Bacteria>Terrabacteria_group>Actinomycetota>Actinomycetes>Micrococcales>Brevibacteriaceae | GCF_002339175.1 | Brevibacterium_casei | B3D11_RS04595 | 80759..84217 | 33889 | BREX-system-P-loop-protein-BrxC-[Brevibacterium-casei] |
| WP_029253807.1 | Type1.BREX | ABC-ATPase+SbcC-><-?<-HEPN<- ParB + HNH <- SIGMA+OB-Fold+AAA-ATPase+HKD-DNase-BrxL <- PglZ <- N6-METHYLASE <- DUF6079 *<- BrxB-BrxF-iSTAND <- BrxA-DNA-binding <-AbiEii<-AbiEii<-REase | 1160 | Bacteria>Terrabacteria_group>Actinomycetota>Actinomycetes>Micrococcales>Cellulomonadaceae | GCF_000701465.1 | Paraoerskovia_marina | Q343_RS0111945 | 114651..118133 | 545619 | BREX-system-P-loop-protein-BrxC-[Paraoerskovia-marina] |
| QAY61399.1 | Type1.BREX | <-VirB11+MuB-ATPase\|\|DNA_METHYLASE-><- ParB + HNH <- SIGMA+OB-Fold+AAA-ATPase+Lon-Protease-BrxL <- PglZ <- N6-METHYLASE <- DUF6079 *<- BrxB-BrxF-iSTAND <- BrxA-DNA-binding <-XIS-HTH<-ABhydrolase<-?<-MPTase<-HTH\|\|ArdCN+MPTase->Primpol+PriCT1-> | 1176 | Bacteria>Terrabacteria_group>Actinomycetota>Actinomycetes>Micrococcales>Microbacteriaceae | GCA_004135285.1 | Microbacterium_protaetiae | ET475_16440 | 3529806..3533336 | 2509458 | BREX-system-P-loop-protein-BrxC-[Microbacterium-protaetiae] |
| MBY6061973.1 | Type1.BREX | TetR-HTH->?-><-?<-Glyoxalase\|\|OLD-ABC+TOPRIM->?-> BrxA-DNA-binding -> BrxB-BrxF-iSTAND -> DUF6079 *-> N6-METHYLASE -> PglZ -> SIGMA+OB-Fold+AAA-ATPase+Lon-Protease-BrxL -><- ParB + HNH \|\|?->MAD-NTD-> | 1152 | Bacteria>Terrabacteria_group>Actinomycetota>Actinomycetes>Micrococcales>Microbacteriaceae | GCA_019801485.1 | Microbacterium_esteraromaticum | KUV70_11015 | 269249..272707 | 57043 | BREX-system-P-loop-protein-BrxC-[Microbacterium-esteraromaticum] |
| WP_053352376.1 | Type1.BREX | <-MODE-HTH+PBPII\|\|?->?->?->tRNA->XIS-HTH-> BrxA-DNA-binding -> BrxB-BrxF-iSTAND -> DUF6079 *-> N6-METHYLASE -> PglZ -> SIGMA+OB-Fold+AAA-ATPase+HKD-DNase-BrxL -><-?<-?<-?\|\| HNH +SAVED-> ParB + HNH -> | 1182 | Bacteria>Terrabacteria_group>Actinomycetota>Actinomycetes>Micrococcales>Microbacteriaceae | GCF_001273845.1 | Leucobacter_musarum | CH42_RS06350 | 60565..64113 | 1930747 | BREX-system-P-loop-protein-BrxC-[Leucobacter-musarum] |
| WP_165310971.1 | Type1.BREX | <-VirB11+MuB-ATPase\|\|DNA_METHYLASE->DNA_METHYLASE+DCM->DNA_METHYLASE-><- ParB + HNH <- SIGMA+OB-Fold+AAA-ATPase+Lon-Protease-BrxL <- PglZ <- N6-METHYLASE <- DUF6079 *<- BrxB-BrxF-iSTAND <- BrxA-DNA-binding <-XIS-HTH<-ABhydrolase<-?<-MPTase<-HTH\|\|ArdCN+MPTase->Primpol+PriCT1-> | 1152 | Bacteria>Terrabacteria_group>Actinomycetota>Actinomycetes>Micrococcales>Microbacteriaceae | GCF_004135285.1 | Microbacterium_protaetiae | ET475_RS16440 | 3529806..3533264 | 2509458 | BREX-system-P-loop-protein-BrxC-[Microbacterium-protaetiae] |
| WP_146322974.1 | Type1.BREX | DOC+HTH->?-> BrxA-DNA-binding -> BrxB-BrxF-iSTAND -> DUF6079 *-> N6-METHYLASE -><- N6-METHYLASE -p\|\|Phage_int_SAM_3+Phage_integrase-> N6-METHYLASE -p->?->?-> PglZ -> SIGMA+OB-Fold+AAA-ATPase+HKD-DNase-BrxL -> ParB + HNH -> | 1162 | Bacteria>Terrabacteria_group>Actinomycetota>Actinomycetes>Micrococcales>Microbacteriaceae | GCF_007859675.1 | Humibacter_ginsenosidimutans | FPZ11_RS17385 | 3786036..3789524 | 2599293 | BREX-system-P-loop-protein-BrxC-[Humibacter-ginsenosidimutans] |
| WP_261393916.1 | Type1.BREX | TetR-HTH->?-><-?<-Glyoxalase\|\|OLD-ABC+TOPRIM->?-> BrxA-DNA-binding -> BrxB-BrxF-iSTAND -> DUF6079 *-> N6-METHYLASE -> PglZ -> SIGMA+OB-Fold+AAA-ATPase+Lon-Protease-BrxL -><- ParB + HNH \|\|?->MAD-NTD-> | 1146 | Bacteria>Terrabacteria_group>Actinomycetota>Actinomycetes>Micrococcales>Microbacteriaceae | GCF_019801485.1 | Microbacterium_esteraromaticum | KUV70_RS11015 | 269267..272707 | 57043 | BREX-system-P-loop-protein-BrxC-[Microbacterium-esteraromaticum] |
| WP_222446684.1 | Type1.BREX | TetR-HTH->?-><-?<-Glyoxalase\|\|OLD-ABC+TOPRIM->?-> BrxA-DNA-binding -> BrxB-BrxF-iSTAND -> DUF6079 *-> N6-METHYLASE -> PglZ -> SIGMA+OB-Fold+AAA-ATPase+Lon-Protease-BrxL -><- ParB + HNH \|\|?->MAD-NTD-> | 1152 | Bacteria>Terrabacteria_group>Actinomycetota>Actinomycetes>Micrococcales>Microbacteriaceae | - | Microbacterium_esteraromaticum | KUV70_11015 | 269249..272707 | 57043 | BREX-system-P-loop-protein-BrxC-[Microbacterium-esteraromaticum] |
| ALV40633.1 | Type1.BREX | <-HEPN+SF1-SF2-Helicase+REase<-?<-?<- ParB + HNH <- SIGMA+OB-Fold+AAA-ATPase+HKD-DNase-BrxL <- PglZ <- N6-METHYLASE <- DUF6079 *<- BrxB-BrxF-iSTAND <- BrxA-DNA-binding <-XIS-HTH<-HKD-DNase+SF1-SF2-Helicase+HTH+HTH+SRA\|\|?->?->?->McrB-AAA->McrC-NTD+REase-> | 1172 | Bacteria>Terrabacteria_group>Actinomycetota>Actinomycetes>Micrococcales>Micrococcaceae | GCA_001484605.1 | Pseudarthrobacter_sulfonivorans | AU252_05160 | 1179330..1182848 | 121292 | hypothetical-protein-AU252_05160-[Pseudarthrobacter-sulfonivorans] |
| BBE21856.1 | Type1.BREX | <-SF1-SF2-Helicase+REase<-SF1-SF2-Helicase+REase-p\|\|?->LZ_Tnp_IS481+HTH_32+rve+rve_3-><-HTH_Tnp_1+HTH_21+rve+rve_3<-HIN-HTH\|\| BrxA-DNA-binding -> BrxB-BrxF-iSTAND -> DUF6079 *-> N6-METHYLASE -> PglZ -> SIGMA+OB-Fold+AAA-ATPase+Lon-Protease-BrxL -> ParB + HNH -><-?<-DDE_Tnp_ISL3<-?\|\|SF1-SF2-Helicase-> | 1144 | Bacteria>Terrabacteria_group>Actinomycetota>Actinomycetes>Micrococcales>Micrococcaceae | GCA_004001285.1 | Arthrobacter_sp._MN05-02 | MN0502_07390 | 781393..784827 | 1571833 | hypothetical-protein-MN0502_07390-[Arthrobacter-sp.-MN05-02] |
| QOD04369.1 | Type1.BREX | NUDIX-><-URI-DNase<-MACRODOMAIN+SIR2<-REase<-?\|\|HKD-DNase+SF1-SF2-Helicase+HTH+HTH+SRA-><- ParB + HNH \|\| BrxA-DNA-binding -> BrxB-BrxF-iSTAND -> DUF6079 *-> N6-METHYLASE -> N6-METHYLASE -> PglZ -> SIGMA+OB-Fold+AAA-ATPase+Lon-Protease-BrxL ->SF1-SF2-Helicase->Sigma-HTH->Sigma-HTH->rve->HTH+rve-> | 1145 | Bacteria>Terrabacteria_group>Actinomycetota>Actinomycetes>Micrococcales>Micrococcaceae | GCA_014764445.1 | Pseudarthrobacter_sp._BIM_B-2242 | IDT60_04725 | 994235..997672 | 2772401 | BREX-system-P-loop-protein-BrxC-[Pseudarthrobacter-sp.-BIM-B-2242] |
| WP_099093371.1 | Type1.BREX | <-HEPN+SF1-SF2-Helicase+REase<-?<-?<- ParB + HNH <- SIGMA+OB-Fold+AAA-ATPase+HKD-DNase-BrxL <- PglZ <- N6-METHYLASE <- DUF6079 *<- BrxB-BrxF-iSTAND <- BrxA-DNA-binding <-XIS-HTH<-HKD-DNase+SF1-SF2-Helicase+HTH+HTH+SRA\|\|?->?->McrB-AAA->McrC-NTD+REase-> | 1173 | Bacteria>Terrabacteria_group>Actinomycetota>Actinomycetes>Micrococcales>Micrococcaceae | GCF_001484605.1 | Pseudarthrobacter_sulfonivorans | AU252_RS05160 | 1179330..1182851 | 121292 | BREX-system-P-loop-protein-BrxC-[Pseudarthrobacter-sulfonivorans] |
| WP_104138811.1 | Type5.BREX | NUDIX->ARPP2-><-MACRODOMAIN+SIR2<-URI-DNase\|\|HKD-DNase+SF1-SF2-Helicase+HTH+HTH+SRA->XIS-HTH-> BrxA-DNA-binding -> BrxB-BrxF-iSTAND -> DUF6079 *-> N6-METHYLASE -> PglZ -> N6-METHYLASE -p-> N6-METHYLASE -> SIGMA+OB-Fold+AAA-ATPase+HKD-DNase-BrxL ->?-> ParB + HNH ->?->35exo+TUDOR+SF1-SF2-Helicase+iREase+CTD-> | 1173 | Bacteria>Terrabacteria_group>Actinomycetota>Actinomycetes>Micrococcales>Micrococcaceae | GCF_002929275.1 | Arthrobacter_sp._ZGTC131 | CXQ90_RS09900 | 131121..134642 | 2058898 | BREX-system-P-loop-protein-BrxC-[Arthrobacter-sp.-ZGTC131] |
| WP_223883883.1 | Type1.BREX | NUDIX-><-URI-DNase<-MACRODOMAIN+SIR2<-REase<-?\|\|HKD-DNase+SF1-SF2-Helicase+HTH+HTH+SRA-><- ParB + HNH \|\| BrxA-DNA-binding -> BrxB-BrxF-iSTAND -> DUF6079 *-> N6-METHYLASE -> N6-METHYLASE -> PglZ -> SIGMA+OB-Fold+AAA-ATPase+Lon-Protease-BrxL ->SF1-SF2-Helicase->Sigma-HTH->HTH+rve->AAA-ATPase-> | 1144 | Bacteria>Terrabacteria_group>Actinomycetota>Actinomycetes>Micrococcales>Micrococcaceae | GCF_014764445.1 | Pseudarthrobacter_sp._BIM_B-2242 | IDT60_RS04725 | 994238..997672 | 2772401 | BREX-system-P-loop-protein-BrxC-[Pseudarthrobacter-sp.-BIM-B-2242] |
| WP_224048794.1 | Type1.BREX | NUDIX-> wHTH(SBD)+ HNH -><-?\|\|?-><-SF1-SF2-Helicase\|\|XIS-HTH-> BrxA-DNA-binding -> BrxB-BrxF-iSTAND -> DUF6079 *-> N6-METHYLASE -> PglZ -> SIGMA+OB-Fold+AAA-ATPase+HKD-DNase-BrxL -><- ParB + HNH \|\|HEPN+SF1-SF2-Helicase+REase-> | 1173 | Bacteria>Terrabacteria_group>Actinomycetota>Actinomycetes>Micrococcales>Micrococcaceae | GCF_019977335.1 | Arthrobacter_sp._NicSoilB4 | LDO13_RS04105 | 882114..885635 | 2830997 | BREX-system-P-loop-protein-BrxC-[Arthrobacter-sp.-NicSoilB4] |
| WP_075958588.1 | Type1.BREX | <-MazG<-nSTAND1+ HNH <-REase+SF1-SF2-Helicase-p<-REase\|\|XIS-HTH-> BrxA-DNA-binding -> BrxB-BrxF-iSTAND -> DUF6079 -p*-> DUF6079 -p-> N6-METHYLASE -> SIGMA+OB-Fold+AAA-ATPase+Lon-Protease-BrxL -> ParB + HNH -><-REase<-SF1-SF2-Helicase+REase<-REase<-REase | 295 | Bacteria>Terrabacteria_group>Actinomycetota>Actinomycetes>Micrococcales>Ornithinimicrobiaceae | GCF_001942405.1 | Ornithinimicrobium_sp._CNJ-824 | BJF81_RS12330 | 99001..99888 | 1904966 | hypothetical-protein-[Ornithinimicrobium-sp.-CNJ-824] |
| WP_075958589.1 | Type1.BREX | <-MazG<-nSTAND1+ HNH <-REase+SF1-SF2-Helicase-p<-REase\|\|XIS-HTH-> BrxA-DNA-binding -> BrxB-BrxF-iSTAND -> DUF6079 -p-> DUF6079 -p*-> N6-METHYLASE -> SIGMA+OB-Fold+AAA-ATPase+Lon-Protease-BrxL -> ParB + HNH -><-REase<-SF1-SF2-Helicase+REase<-REase<-REase | 882 | Bacteria>Terrabacteria_group>Actinomycetota>Actinomycetes>Micrococcales>Ornithinimicrobiaceae | GCF_001942405.1 | Ornithinimicrobium_sp._CNJ-824 | BJF81_RS12335 | 99810..102458 | 1904966 | BREX-system-P-loop-protein-BrxC-[Ornithinimicrobium-sp.-CNJ-824] |
| WP_102213741.1 | Type1.BREX | XIS-HTH-> BrxA-DNA-binding -> BrxB-BrxF-iSTAND -> DUF6079 *-> N6-METHYLASE ->HTH+REase-> PglZ -> SIGMA+OB-Fold+AAA-ATPase+HKD-DNase-BrxL -><- ParB + HNH +METHYLASE | 1173 | Bacteria>Terrabacteria_group>Actinomycetota>Actinomycetes>Mycobacteriales>Corynebacteriaceae | GCF_002871715.1 | Corynebacterium_xerosis | CJ204_09375 | 1693..5214 | 1725 | BREX-system-P-loop-protein-BrxC-[Corynebacterium-xerosis] |
| WP_259898411.1 | Type1.BREX | <-HIN-HTH<- ParB + HNH <- SIGMA+OB-Fold+AAA-ATPase+HKD-DNase-BrxL <- PglZ <- N6-METHYLASE <- DUF6079 *<- BrxB-BrxF-iSTAND <- BrxA-DNA-binding <-XIS-HTH | 1160 | Bacteria>Terrabacteria_group>Actinomycetota>Actinomycetes>Mycobacteriales>Corynebacteriaceae | GCF_025152505.1 | Corynebacterium_sanguinis | M3G46_RS10600 | 10062..13544 | 2594913 | BREX-system-P-loop-protein-BrxC-[Corynebacterium-sanguinis] |
| WP_064441930.1 | Type1.BREX | <- PglZ -p<- N6-METHYLASE <- DUF6079 *<- BrxB-BrxF-iSTAND <- BrxA-DNA-binding \|\| ParB + HNH -> ParB + HNH ->?->DndD->HEPN->?->REase-> | 1148 | Bacteria>Terrabacteria_group>Actinomycetota>Actinomycetes>Mycobacteriales>Mycobacteriaceae | GCF_001653095.1 | Hoyosella_altamirensis | HAL02_RS18845 | 3788..7234 | 616997 | BREX-system-P-loop-protein-BrxC-[Hoyosella-altamirensis] |
| WP_068042721.1 | Type6.BREX | <-Phage_integrase<-?<-ZnR<-AAA-ATPase<- ParB + HNH \|\|?-><-?<- SIGMA+OB-Fold+AAA-ATPase+Lon-Protease-BrxL <- PglZ <- N6-METHYLASE <- DUF6079 *<- BrxB-BrxF-iSTAND <- BrxA-DNA-binding <- BrxE \|\|ABhydrolase+ACYC-> | 1161 | Bacteria>Terrabacteria_group>Actinomycetota>Actinomycetes>Mycobacteriales>Mycobacteriaceae | GCF_001667585.1 | Mycobacterium_sp._E2733 | A5678_RS12510 | 65198..68683 | 1834138 | BREX-system-P-loop-protein-BrxC-[Mycobacterium-sp.-E2733] |
| WP_234813880.1 | Type1.BREX | <- ParB + HNH <- SIGMA+OB-Fold+AAA-ATPase+HKD-DNase-BrxL <- PglZ <- N6-METHYLASE <- DUF6079 -p<- DUF6079 -p*<- BrxB-BrxF-iSTAND <- BrxA-DNA-binding | 682 | Bacteria>Terrabacteria_group>Actinomycetota>Actinomycetes>Mycobacteriales>Mycobacteriaceae | GCF_002101705.1 | Mycolicibacterium_iranicum | AWC12_RS31570 | 96008..98056 | 912594 | BREX-system-P-loop-protein-BrxC-[Mycolicibacterium-iranicum] |
| WP_163778779.1 | Type1.BREX | GGDEF+EAL->?-><-Glyco_tran_28_C<-?\|\|?-><-?<-tRNA\|\|XIS-HTH-> BrxA-DNA-binding -> BrxB-BrxF-iSTAND -> DUF6079 *-> N6-METHYLASE -> PglZ -> SIGMA+OB-Fold+AAA-ATPase+HKD-DNase-BrxL -> ParB + HNH ->?->AAA-ATPase->ZnR->?->Phage_integrase-> | 1156 | Bacteria>Terrabacteria_group>Actinomycetota>Actinomycetes>Mycobacteriales>Mycobacteriaceae | GCF_010727945.1 | Mycobacterium_cookii | G6N27_RS18550 | 3937772..3941242 | 1775 | BREX-system-P-loop-protein-BrxC-[Mycobacterium-cookii] |
| WP_260940163.1 | Type1.BREX | XIS-HTH-> BrxA-DNA-binding -> BrxB-BrxF-iSTAND -> DUF6079 *-> N6-METHYLASE -> PglZ -> SIGMA+OB-Fold+AAA-ATPase+HKD-DNase-BrxL ->?-> ParB + HNH -p->HIN-HTH-><-wHTH\|\|MORC+DUF3684+DUF3883-><-REase+HTH+35exo+Small-B-barrel+SF1-SF2-Helicase | 1156 | Bacteria>Terrabacteria_group>Actinomycetota>Actinomycetes>Mycobacteriales>Mycobacteriaceae | GCF_025331195.1 | Mycolicibacterium_llatzerense | A7G45_RS21390 | 50451..53921 | 280871 | BREX-system-P-loop-protein-BrxC-[Mycolicibacterium-llatzerense] |
| WP_267319448.1 | Type1.BREX | wHTH(SBD)+ HNH -><-HEPN<-DndD\|\|RAMA-><- ParB + HNH <- SIGMA+OB-Fold+AAA-ATPase+Lon-Protease-BrxL <- PglZ <- N6-METHYLASE <- DUF6079 *<- BrxB-BrxF-iSTAND <- BrxA-DNA-binding \|\|DEDD_Tnp_IS110+Transposase_20-> | 1149 | Bacteria>Terrabacteria_group>Actinomycetota>Actinomycetes>Mycobacteriales>Mycobacteriaceae | GCF_026501225.1 | Mycolicibacterium_mucogenicum | DS122_RS24315 | 502363..505812 | 56689 | BREX-system-P-loop-protein-BrxC-[Mycolicibacterium-mucogenicum] |
| OFE08162.1 | Type1.BREX | <-Pkinase+TPR+TPR<-?<-?<-?<-?<-?\|\|tRNA-><-?\|\| BrxA-DNA-binding -> BrxB-BrxF-iSTAND -> DUF6079 *-> N6-METHYLASE -> PglZ -> SIGMA+OB-Fold+AAA-ATPase+HKD-DNase-BrxL ->?->?->?-> ParB + HNH \|\|?->?->LZ_Tnp_IS481+HTH_32+rve+rve_3-> | 1175 | Bacteria>Terrabacteria_group>Actinomycetota>Actinomycetes>Mycobacteriales>Nocardiaceae | GCA_001757665.1 | Rhodococcus_sp._1139 | A5N83_14165 | 58033..61560 | 1833762 | ATP-binding-protein-[Rhodococcus-sp.-1139] |
| MBY6706229.1 | Type1.BREX | <-SF1-SF2-Helicase+DUF1998<-wHTH<-LON-AAA<-REase\|\|?-><-ABhydrolase+ABhydrolase<- HNH \|\| BrxA-DNA-binding -> BrxB-BrxF-iSTAND -> DUF6079 *-> N6-METHYLASE -> PglZ -> SIGMA+OB-Fold+AAA-ATPase+Lon-Protease-BrxL -> ParB + HNH ->\|\|Exonuc_V_gamma+REase->SF1-SF2-Helicase+REase-><-SF1-SF2-Helicase+Classical-AAA<-HD | 1147 | Bacteria>Terrabacteria_group>Actinomycetota>Actinomycetes>Mycobacteriales>Nocardiaceae | GCA_019836145.1 | Rhodococcus_sp._BP-241 | HQ308_05385 | 442671..446114 | 2739441 | BREX-system-P-loop-protein-BrxC-[Rhodococcus-sp.-BP-241] |
| WP_029256831.1 | Type1.BREX | ParB + HNH -><-?<-?<- SIGMA+OB-Fold+AAA-ATPase+HKD-DNase-BrxL <- PglZ <- N6-METHYLASE <- DUF6079 *<- BrxB-BrxF-iSTAND <- BrxA-DNA-binding \|\|?-><-tRNA\|\|?->?->?->?->?->Pkinase+TPR+TPR-> | 1173 | Bacteria>Terrabacteria_group>Actinomycetota>Actinomycetes>Mycobacteriales>Nocardiaceae | GCF_000696675.2 | Rhodococcus_erythropolis | H351_RS02855 | 635383..638904 | 1833 | BREX-system-P-loop-protein-BrxC-[Rhodococcus-erythropolis] |
| WP_070161347.1 | Type1.BREX | <-Pkinase+TPR+TPR<-?<-?<-?<-?<-?\|\|tRNA-><-?\|\| BrxA-DNA-binding -> BrxB-BrxF-iSTAND -> DUF6079 *-> N6-METHYLASE -> N6-METHYLASE -p-> PglZ -> SIGMA+OB-Fold+AAA-ATPase+HKD-DNase-BrxL ->?->?-><- ParB + HNH | 1175 | Bacteria>Terrabacteria_group>Actinomycetota>Actinomycetes>Mycobacteriales>Nocardiaceae | GCF_001757665.1 | Rhodococcus_sp._1139 | A5N83_RS14165 | 58033..61560 | 1833762 | BREX-system-P-loop-protein-BrxC-[Rhodococcus-sp.-1139] |

| Protein-ID | System | Gene-Neighbourhoods | Length | Taxonomy | Assembly-ID | Strain | Locus-Tag | Genomic-coordinates | TaxID | Fasta-defline |
|---|---|---|---|---|---|---|---|---|---|---|
| WP_236656790.1 | Type1.BREX | ASCH->Glyoxalase->ParB+HNH->?->ParB+HNH->BrxA-DNA-binding->BrxB-BrxF-iSTAND->DUF6079*->N6-METHYLASE->nSTAND3-><-MULE-TRANSPOSASE | 1159 | Bacteria>Terrabacteria_group>Bacillota>Bacilli>Lactobacillales>Lactobacillaceae | - | Lactobacillus_helveticus | LHEW6_RS10210 | 11901..15380 | 1587 | BREX-system-P-loop-protein-BrxC-[Lactobacillus-helveticus] |
| WP_168924054.1 | Type1.BREX | McrB-AAA->McrC-NTD+REase->?->BrxA-DNA-binding->BrxB-BrxF-iSTAND->DUF6079*-><-N6-METHYLASE-p<-?\|\|PglZ->SIGMA+OB-Fold+AAA-ATPase+Lon-Protease-BrxL->ParB+HNH-> [illegible] | 1191 | Bacteria>Terrabacteria_group>Bacillota>Bacilli>Lactobacillales>Lactobacillaceae | - | Secundilactobacillus_angelensis | L3V66_04100 | 164372..167947 | 2722706 | BREX-system-P-loop-protein-BrxC-[Secundilactobacillus-angelensis] |
| MCH4057848.1 | Type1.BREX | <-HTH_28+HTH_32\|\|<-?<-ParB+HNH<-SIGMA+OB-Fold+AAA-ATPase+Lon-Protease-BrxL<-PglZ\|\|N6-METHYLASE->Arm-DNA-bind_4+Phage_int_SAM_5+Phage_integrase-><-N6-METHYLASE<-DUF6079 [illegible] | 1214 | Bacteria>Terrabacteria_group>Bacillota>Bacilli>Lactobacillales>unclassified_Lactobacillales | GCA_022483825.1 | Lactobacillaceae_bacterium | LKE89_08070 | 16697..20341 | 2291323 | BREX-system-P-loop-protein-BrxC-[Lactobacillaceae-bacterium] |
| WP_076917462.1 | Type1.BREX | AT-HTH-HYD2->Tox-ART-HYD2->AT-HTH-HYD2-><-SbcC<-ParB+HNH<-SIGMA+OB-Fold+AAA-ATPase+Lon-Protease-BrxL<-PglZ<-N6-METHYLASE<-TIR<-DUF6079*<-BrxB-BrxF-iSTAND<-BrxA-DNA-bi [illegible] | 1178 | Bacteria>Terrabacteria_group>Bacillota>Bacillota_sensu_stricto_incertae_sedis>Negativibacillus | GCF_900148495.1 | Negativibacillus_massiliensis | BQ5375_RS04300 | 340320..343856 | 1871035 | BREX-system-P-loop-protein-BrxC-[Negativibacillus-massiliensis] |
| NFG40378.1 | Type1.BREX | Calcineurin->?->BrxA-DNA-binding->BrxB-BrxF-iSTAND->DUF6079*->N6-METHYLASE->ParB+HNH->OLD-ABC->HNH->PglZ-><-REase\|\|SIGMA+OB-Fold+AAA-ATPase+Lon-Protease-BrxL->Pepti [illegible] | 1215 | Bacteria>Terrabacteria_group>Bacillota>Clostridia>Eubacteriales>Clostridiaceae | GCA_011013445.1 | Clostridium_botulinum | FC789_04035 | 879792..883439 | 1491 | BREX-system-P-loop-protein-BrxC-[Clostridium-botulinum] |
| NFI95403.1 | Type1.BREX | <-Peptidase_M42<-SIGMA+OB-Fold+AAA-ATPase+Lon-Protease-BrxL\|\|REase-><-PglZ<-HNH<-OLD-ABC<-ParB+HNH<-N6-METHYLASE<-DUF6079*<-BrxB-BrxF-iSTAND<-BrxA-DNA-binding [illegible] | 1215 | Bacteria>Terrabacteria_group>Bacillota>Clostridia>Eubacteriales>Clostridiaceae | GCA_011014625.1 | Clostridium_botulinum | FC961_13560 | 15228..18875 | 1491 | BREX-system-P-loop-protein-BrxC-[Clostridium-botulinum] |
| NFO12999.1 | Type1.BREX | Calcineurin->?->BrxA-DNA-binding->BrxB-BrxF-iSTAND->DUF6079*->CR-ATPase9+REase->N6-METHYLASE->ParB+HNH->OLD-ABC->?->PglZ-><-REase\|\|SIGMA+OB-Fold+AAA-ATPase+Lon-Protea [illegible] | 1215 | Bacteria>Terrabacteria_group>Bacillota>Clostridia>Eubacteriales>Clostridiaceae | GCA_011016315.1 | Clostridium_botulinum | FDB34_02025 | 196714..200361 | 1491 | BREX-system-P-loop-protein-BrxC-[Clostridium-botulinum] |
| WP_089984090.1 | Type1.BREX | <-SIGMA-HTH<-?\|\|HNH->?->SbcC->HTH_26->BrxA-DNA-binding->BrxB-BrxF-iSTAND->DUF6079*->N6-METHYLASE->PglZ->SIGMA+OB-Fold+AAA-ATPase+Lon-Protease-BrxL->ParB+HNH->?-><-? [illegible] | 1183 | Bacteria>Terrabacteria_group>Bacillota>Clostridia>Eubacteriales>Clostridiaceae | GCF_001517425.1 | Clostridium_sp._C105KSO13 | BN3456_RS10470 | 25187..28738 | 1776045 | BREX-system-P-loop-protein-BrxC-[Clostridium-sp.-C105KSO13] |
| WP_156211829.1 | Type1.BREX | <-wHTH+McrB-AAA<-?<-PglZ<-?<-?<-OLD-ABC<-ParB+HNH<-N6-METHYLASE<-CR-ATPase9+REase<-DUF6079*<-BrxB-BrxF-iSTAND<-BrxA-DNA-binding<-HKD-DNase+SF1-SF2-Helicase+HTH+HTH+SRA [illegible] | 1216 | Bacteria>Terrabacteria_group>Bacillota>Clostridia>Eubacteriales>Clostridiaceae | GCF_009735365.2 | Clostridium_butyricum | GND98_RS05145 | 54704..58354 | 1492 | BREX-system-P-loop-protein-BrxC-[Clostridium-butyricum] |
| WP_185162822.1 | Type1.BREX | NTP_transf-><-SIGMA+OB-Fold+AAA-ATPase+Lon-Protease-BrxL<-?<-PglZ<-Couple_hipA<-N6-METHYLASE<-?<-DUF6079*<-BrxB-BrxF-iSTAND<-BrxA-DNA-binding\|\|HEPN-><-N6-METHYLASE<-? [illegible] | 1215 | Bacteria>Terrabacteria_group>Bacillota>Clostridia>Eubacteriales>Clostridiaceae | GCF_014212415.1 | Clostridium_algidicarnis | H7D93_RS12595 | 17014..20661 | 37659 | BREX-system-P-loop-protein-BrxC-[Clostridium-algidicarnis] |
| WP_207130137.1 | Type1.BREX | <-SIGMA+OB-Fold+AAA-ATPase+Lon-Protease-BrxL<-PglZ<-AT-HTH-HYD2-2+AT-HTH-HYD2-1<-?<-ParB+HNH<-N6-METHYLASE<-DUF6079*<-BrxB-BrxF-iSTAND<-BrxA-DNA-binding<-ParB [illegible] | 1215 | Bacteria>Terrabacteria_group>Bacillota>Clostridia>Eubacteriales>Clostridiaceae | GCF_017330945.1 | Clostridium_botulinum | EXQ34_RS04235 | 14589..18236 | 1491 | BREX-system-P-loop-protein-BrxC-[Clostridium-botulinum] |
| WP_226104726.1 | Type1.BREX | <-SIGMA+OB-Fold+AAA-ATPase+Lon-Protease-BrxL<-?<-PglZ<-SLOG+BirA-HTH<-SF1-SF2-Helicase+PRTase<-?<-ParB+HNH<-N6-METHYLASE<-?<-DUF6079*<-BrxB-BrxF-iSTAND<-BrxA-DNA-bind [illegible] | 1215 | Bacteria>Terrabacteria_group>Bacillota>Clostridia>Eubacteriales>Clostridiaceae | GCF_020443655.1 | Clostridium_algidicarnis | LGK99_RS11195 | 54237..57884 | 37659 | BREX-system-P-loop-protein-BrxC-[Clostridium-algidicarnis] |
| WP_252235883.1 | Type1.BREX | <-PglZ<-?<-?<-OLD-ABC<-ParB+HNH<-N6-METHYLASE<-?<-Tox-ART-HYD2<-DUF6079*<-BrxB-BrxF-iSTAND<-BrxA-DNA-binding<-?<-Calcineurin | 1215 | Bacteria>Terrabacteria_group>Bacillota>Clostridia>Eubacteriales>Clostridiaceae | GCF_023846115.1 | Clostridium_sp._ZS1 | NG923_RS15480 | 30630..34277 | 2949989 | BREX-system-P-loop-protein-BrxC-[Clostridium-sp.-ZS1] |
| WP_066893479.1 | Type1.BREX | <-SIGMA+OB-Fold+AAA-ATPase+Lon-Protease-BrxL\|\|REase-><-PglZ<-DOC+HTH<-SLOG+BirA-HTH<-SF1-SF2-Helicase+PRTase<-ParB+HNH<-?<-N6-METHYLASE<-DUF6079*<-BrxB-BrxF-iSTAND<-B [illegible] | 1215 | Bacteria>Terrabacteria_group>Bacillota>Clostridia>Eubacteriales>Clostridiaceae | GCF_900086595.1 | Clostridium_nigeriense | BN6774_RS10780 | 2270723..2274370 | 1805470 | BREX-system-P-loop-protein-BrxC-[Clostridium-nigeriense] |
| WP_118637679.1 | Type1.BREX | <-LexA-protease<-MNS+REase<-wHTH+WYL+WCX<-N6-METHYLASE<-HEPN<-ParB+HNH<-SIGMA+OB-Fold+AAA-ATPase+Lon-Protease-BrxL<-PglZ<-N6-METHYLASE<-DUF6079*<-BrxB-BrxF-iSTAND<- [illegible] | 1215 | Bacteria>Terrabacteria_group>Bacillota>Clostridia>Eubacteriales | GCF_027924705.1 | Oscillospiraceae | H6A32_RS13625 | 30472..34119 | 216572 | MULTISPECIES:-BREX-system-P-loop-protein-BrxC-[Oscillospiraceae] |
| TGX97927.1 | Type1.BREX | <-OLD-ABC+TOPRIM-p<-OLD-ABC+TOPRIM-p<-PglZ<-HNH<-OLD-ABC<-N6-METHYLASE<-HEPN<-ParB+HNH<-N6-METHYLASE<-DUF6079*<-BrxB-BrxF-iSTAND<-BrxA-DNA-binding<-HTH_26<-ZnR+ZnR+ [illegible] | 1201 | Bacteria>Terrabacteria_group>Bacillota>Clostridia>Eubacteriales>Lachnospiraceae | GCA_004793585.1 | Lachnospiraceae_bacterium | E5357_10825 | 35909..39514 | 1898203 | BREX-system-P-loop-protein-BrxC-[Lachnospiraceae-bacterium] |
| MBR3171576.1 | Type1.BREX | <-ParB+HNH<-SIGMA+OB-Fold+AAA-ATPase+Lon-Protease-BrxL<-PglZ<-RAMA<-N6-METHYLASE<-DUF6079*<-BrxB-BrxF-iSTAND<-BrxA-DNA-binding<-HTH_26<-OLD-ABC\|\|SMODS->?->?-><-AbiEii [illegible] | 1178 | Bacteria>Terrabacteria_group>Bacillota>Clostridia>Eubacteriales>Lachnospiraceae | GCA_017514845.1 | Lachnospiraceae_bacterium | IKF22_10080 | 15258..18794 | 1898203 | BREX-system-P-loop-protein-BrxC-[Lachnospiraceae-bacterium] |
| MCM1125306.1 | Type1.BREX | MoxR-AAA->LRR_5->Glyco_hydro_3_C+Fn3-like+Glyco_hydro_3->Peptidase_C26->tRNA->tRNA-><-?<-MerR-HTH\|\|HTH_26->BrxA-DNA-binding->BrxB-BrxF-iSTAND->DUF6079*->ParB+HNH->HEPN-> [illegible] | 1205 | Bacteria>Terrabacteria_group>Bacillota>Clostridia>Eubacteriales>Lachnospiraceae | GCA_023663945.1 | Lachnospiraceae_bacterium | NC429_02425 | 28301..31918 | 1898203 | BREX-system-P-loop-protein-BrxC-[Lachnospiraceae-bacterium] |
| SFG07145.1 | Type1.BREX | ArdCN+MPTase-inact+LPD19->TOPRIM+Topoisom_bac->LPD2->TraJ-RHH->?->HTH_26->BrxA-DNA-binding->BrxB-BrxF-iSTAND->DUF6079*->N6-METHYLASE->?->PglZ->AbiGi->SIGMA+OB-Fold+AAA- [illegible] | 1182 | Bacteria>Terrabacteria_group>Bacillota>Clostridia>Eubacteriales>Lachnospiraceae | GCA_900113385.1 | Lachnospiraceae_bacterium_C7 | SAMN04487761_10376 | 78580..82128 | 1792306 | hypothetical-protein-SAMN04487761_10376-[Lachnospiraceae-bacterium-C7] |
| WP_027207205.1 | Type1.BREX | [illegible] | 1196 | Bacteria>Terrabacteria_group>Bacillota>Clostridia>Eubacteriales>Lachnospiraceae | GCF_000423985.1 | Butyrivibrio_fibrisolvens | G591_RS0114720 | 29938..33528 | 831 | BREX-system-P-loop-protein-BrxC-[Butyrivibrio-fibrisolvens] |
| WP_027204781.1 | Type1.BREX | [illegible] | 1181 | Bacteria>Terrabacteria_group>Bacillota>Clostridia>Eubacteriales>Lachnospiraceae | GCF_000424425.1 | Butyrivibrio_fibrisolvens | G624_RS0113140 | 43921..47466 | 831 | BREX-system-P-loop-protein-BrxC-[Butyrivibrio-fibrisolvens] |
| WP_029067206.1 | Type1.BREX | [illegible] | 1182 | Bacteria>Terrabacteria_group>Bacillota>Clostridia>Eubacteriales>Lachnospiraceae | GCF_000687515.1 | Lachnobacterium_bovis | CC91_RS0108435 | 378442..381990 | 140626 | BREX-system-P-loop-protein-BrxC-[Lachnobacterium-bovis] |
| WP_029067213.1 | Type1.BREX | [illegible] | 1182 | Bacteria>Terrabacteria_group>Bacillota>Clostridia>Eubacteriales>Lachnospiraceae | GCF_000687515.1 | Lachnobacterium_bovis | CC91_RS0108490 | 397340..400888 | 140626 | BREX-system-P-loop-protein-BrxC-[Lachnobacterium-bovis] |
| WP_059085682.1 | Type1.BREX | [illegible] | 1196 | Bacteria>Terrabacteria_group>Bacillota>Clostridia>Eubacteriales>Lachnospiraceae | GCF_001487165.1 | Blautia_massiliensis | BN3261_RS05710 | 1028364..1031954 | 1737424 | BREX-system-P-loop-protein-BrxC-[Blautia-massiliensis] |
| WP_114002767.1 | Type1.BREX | [illegible] | 1196 | Bacteria>Terrabacteria_group>Bacillota>Clostridia>Eubacteriales>Lachnospiraceae | GCF_003324155.1 | Blautia_obeum | C4886_RS16030 | 22477..26067 | 40520 | BREX-system-P-loop-protein-BrxC-[Blautia-obeum] |
| WP_195251421.1 | Type1.BREX | [illegible] | 1201 | Bacteria>Terrabacteria_group>Bacillota>Clostridia>Eubacteriales>Lachnospiraceae | GCF_015550485.1 | Coprococcus_comes | I2A60_RS13430 | 4839..8444 | 410072 | BREX-system-P-loop-protein-BrxC-[Coprococcus-comes] |
| WP_118278843.1 | Type1.BREX | [illegible] | 1205 | Bacteria>Terrabacteria_group>Bacillota>Clostridia>Eubacteriales>Lachnospiraceae | GCF_027662525.1 | [Ruminococcus]_lactaris | DW116_RS04365 | 20004..23621 | 46228 | BREX-system-P-loop-protein-BrxC-[[Ruminococcus] |
| WP_271824191.1 | Type1.BREX | [illegible] | 1178 | Bacteria>Terrabacteria_group>Bacillota>Clostridia>Eubacteriales>Lachnospiraceae | GCF_028210655.1 | Enterocloster_clostridioformis | PMX39_RS12925 | 18239..21775 | 1531 | BREX-system-P-loop-protein-BrxC-[Enterocloster-clostridioformis] |
| WP_090815514.1 | Type1.BREX | [illegible] | 1182 | Bacteria>Terrabacteria_group>Bacillota>Clostridia>Eubacteriales>Lachnospiraceae | GCF_900107835.1 | Oribacterium_sp._KHPX15 | BLV86_RS05660 | 48724..52272 | 1855342 | BREX-system-P-loop-protein-BrxC-[Oribacterium-sp.-KHPX15] |
| WP_091825279.1 | Type1.BREX | [illegible] | 1145 | Bacteria>Terrabacteria_group>Bacillota>Clostridia>Eubacteriales>Lachnospiraceae | GCF_900109475.1 | Butyrivibrio_sp._ob235 | BMW61_RS19080 | 17910..21347 | 1761780 | BREX-system-P-loop-protein-BrxC-[Butyrivibrio-sp.-ob235] |
| MCR4638957.1 | Type1.BREX | [illegible] | 1175 | Bacteria>Terrabacteria_group>Bacillota>Clostridia>Eubacteriales>Oscillospiraceae | GCA_024684275.1 | Ruminococcus_sp. | K5677_07000 | 11947..15474 | 41978 | BREX-system-P-loop-protein-BrxC-[Ruminococcus-sp.] |
| UYJ00845.1 | Type1.BREX | [illegible] | 1218 | Bacteria>Terrabacteria_group>Bacillota>Clostridia>Eubacteriales>Oscillospiraceae | GCA_025757765.1 | Faecalibacterium_prausnitzii | OGM65_10405 | 2149952..2153608 | 853 | MAG:-BREX-system-P-loop-protein-BrxC-[Faecalibacterium-prausnitzii] |
| UYJ11714.1 | Type1.BREX | [illegible] | 1182 | Bacteria>Terrabacteria_group>Bacillota>Clostridia>Eubacteriales>Oscillospiraceae | GCA_025757845.1 | Oscillospiraceae_bacterium | OGM78_02700 | 567882..571430 | 2485925 | MAG:-BREX-system-P-loop-protein-BrxC-[Oscillospiraceae-bacterium] |
| WP_024861712.1 | Type1.BREX | [illegible] | 1171 | Bacteria>Terrabacteria_group>Bacillota>Clostridia>Eubacteriales>Oscillospiraceae | GCF_000526795.1 | Ruminococcus_flavefaciens | N774_RS0113310 | 3002422..3005937 | 1265 | BREX-system-P-loop-protein-BrxC-[Ruminococcus-flavefaciens] |
| WP_097782652.1 | Type1.BREX | [illegible] | 1220 | Bacteria>Terrabacteria_group>Bacillota>Clostridia>Eubacteriales>Oscillospiraceae | GCF_002549935.1 | Faecalibacterium_prausnitzii | CGS55_RS03365 | 35885..39547 | 853 | BREX-system-P-loop-protein-BrxC-[Faecalibacterium-prausnitzii] |
| WP_113992229.1 | Type1.BREX | [illegible] | 1218 | Bacteria>Terrabacteria_group>Bacillota>Clostridia>Eubacteriales>Oscillospiraceae | GCF_003324185.1 | Faecalibacterium_prausnitzii | C7J97_RS04695 | 32898..36554 | 853 | BREX-system-P-loop-protein-BrxC-[Faecalibacterium-prausnitzii] |
| WP_117554516.1 | Type1.BREX | [illegible] | 1217 | Bacteria>Terrabacteria_group>Bacillota>Clostridia>Eubacteriales>Oscillospiraceae | GCF_003434135.1 | Faecalibacterium_prausnitzii | DW855_RS10430 | 56056..59709 | 853 | BREX-system-P-loop-protein-BrxC-[Faecalibacterium-prausnitzii] |
| WP_215675194.1 | Type1.BREX | [illegible] | 1182 | Bacteria>Terrabacteria_group>Bacillota>Clostridia>Eubacteriales>Oscillospiraceae | GCF_018784645.1 | Faecalibacterium_prausnitzii | GPK62_RS07855 | 35115..38663 | 853 | BREX-system-P-loop-protein-BrxC-[Faecalibacterium-prausnitzii] |
| WP_235255096.1 | Type1.BREX | [illegible] | 1199 | Bacteria>Terrabacteria_group>Bacillota>Clostridia>Eubacteriales>Oscillospiraceae | GCF_021531255.1 | Fournierella_massiliensis | JQM64_RS00320 | 59255..62854 | 1650663 | BREX-system-P-loop-protein-BrxC-[Fournierella-massiliensis] |
| WP_275252236.1 | Type1.BREX | [illegible] | 1218 | Bacteria>Terrabacteria_group>Bacillota>Clostridia>Eubacteriales>Oscillospiraceae | GCF_029076575.1 | Faecalibacterium_sp._DFI.5.82 | POG09_RS06330 | 132484..136140 | 3031725 | BREX-system-P-loop-protein-BrxC-[Faecalibacterium-sp.-DFI.5.82] |
| WP_158398728.1 | Type1.BREX | [illegible] | 1182 | Bacteria>Terrabacteria_group>Bacillota>Clostridia>Eubacteriales>Oscillospiraceae | GCF_902167845.1 | Faecalibacterium_prausnitzii | FQ260_RS07280 | 12413..15961 | 853 | BREX-system-P-loop-protein-BrxC-[Faecalibacterium-prausnitzii] |
| MBD9283724.1 | Type1.BREX | [illegible] | 1202 | Bacteria>Terrabacteria_group>Bacillota>Clostridia>Eubacteriales>unclassified_Eubacteriales | GCA_014848695.1 | Clostridiales_bacterium | EGR96_02215 | 1987..5595 | 1898207 | BREX-system-P-loop-protein-BrxC-[Clostridiales-bacterium] |
| MCD8148190.1 | Type1.BREX | [illegible] | 773 | Bacteria>Terrabacteria_group>Bacillota>Clostridia>Eubacteriales>unclassified_Eubacteriales | GCA_021203005.1 | Clostridiales_bacterium | LUE92_01205 | 1..2322 | 1898207 | BREX-system-P-loop-protein-BrxC-[Clostridiales-bacterium] |
| WP_271628813.1 | Type1.BREX | [illegible] | 1186 | Bacteria>Terrabacteria_group>Bacillota>Clostridia>Thermoanaerobacterales>Thermoanaerobacterales_Family_III._Incertae_Sedis | GCF_028198085.1 | Caldicellulosiruptor_sp._DIB_104C | PGC14_RS09960 | 2010036..2013596 | 3019889 | BREX-system-P-loop-protein-BrxC-[Caldicellulosiruptor-sp.-DIB-104C] |
| MBQ8616934.1 | Type1.BREX | [illegible] | 1186 | Bacteria>Terrabacteria_group>Bacillota>Clostridia>unclassified_Clostridia | GCA_017472845.1 | Clostridia_bacterium | IJ418_05400 | 5468..9028 | 2044939 | BREX-system-P-loop-protein-BrxC-[Clostridia-bacterium] |
| MBR6253006.1 | Type1.BREX | [illegible] | 1184 | Bacteria>Terrabacteria_group>Bacillota>Clostridia>unclassified_Clostridia | GCA_017539905.1 | Clostridia_bacterium | IKR04_04115 | 193581..197135 | 2044939 | BREX-system-P-loop-protein-BrxC-[Clostridia-bacterium] |
| MCR5783583.1 | Type1.BREX | [illegible] | 1177 | Bacteria>Terrabacteria_group>Bacillota>Clostridia>unclassified_Clostridia | GCA_024699005.1 | Clostridia_bacterium | K6G90_12720 | 47894..51427 | 2044939 | BREX-system-P-loop-protein-BrxC-[Clostridia-bacterium] |
| WP_270396150.1 | Type1.BREX | [illegible] | 1177 | Bacteria>Terrabacteria_group>Bacillota>Erysipelotrichia>Erysipelotrichales>Coprobacillaceae | GCF_027659655.1 | Catenibacterium_mitsuokai | PGO85_RS08450 | 1355..4888 | 100886 | BREX-system-P-loop-protein-BrxC-[Catenibacterium-mitsuokai] |
| NLG80270.1 | Type1.BREX | [illegible] | 1185 | Bacteria>Terrabacteria_group>Bacillota | GCA_012516125.1 | Bacillota_bacterium | GX492_10705 | 20631..24188 | 1879010 | BREX-system-P-loop-protein-BrxC-[Bacillota-bacterium] |
| WP_072743893.1 | Type1.BREX | [illegible] | 1192 | Bacteria>Terrabacteria_group>Bacillota>Tissierellia>Tissierellales>Sporanaerobacteraceae | GCF_900130025.1 | Sporanaerobacter_acetigenes | BUA21_RS05940 | 73351..76929 | 165813 | BREX-system-P-loop-protein-BrxC-[Sporanaerobacter-acetigenes] |
| WP_132026817.1 | Type1.BREX | [illegible] | 1197 | Bacteria>Terrabacteria_group>Bacillota>Tissierellia>Tissierellales>Tepidimicrobiaceae | GCF_016767175.1 | Keratinibaculum_paraultunense | JL105_RS06905 | 1394896..1398489 | 1278232 | BREX-system-P-loop-protein-BrxC-[Keratinibaculum-paraultunense] |
| MBQ6518172.1 | Type1.BREX | [illegible] | 1176 | Bacteria>Terrabacteria_group>Chloroflexota>Anaerolineae>Anaerolineales>Anaerolineaceae | GCA_017433515.1 | Anaerolineaceae_bacterium | IJI14_05585 | 5340..8870 | 2024896 | BREX-system-P-loop-protein-BrxC-[Anaerolineaceae-bacterium] |
| NLF02862.1 | Type1.BREX | [illegible] | 1157 | Bacteria>Terrabacteria_group>Chloroflexota>Anaerolineae>Anaerolineales>unclassified_Anaerolineales | GCA_012514695.1 | Anaerolineales_bacterium | GX601_18015 | 11185..14658 | 2073117 | BREX-system-P-loop-protein-BrxC-[Anaerolineales-bacterium] |
| MCB0100628.1 | Type1.BREX | [illegible] | 1152 | Bacteria>Terrabacteria_group>Chloroflexota>Anaerolineae>Anaerolineales>unclassified_Anaerolineales | GCA_020433725.1 | Anaerolineales_bacterium | KDD74_01185 | 15066..18524 | 2073117 | BREX-system-P-loop-protein-BrxC-[Anaerolineales-bacterium] |
| MBK9096490.1 | Type1.BREX | [illegible] | 900 | Bacteria>Terrabacteria_group>Chloroflexota>Anaerolineae>unclassified_Anaerolineae | GCA_016719785.1 | Anaerolineae_bacterium | IPM84_27780 | 244922..247624 | 2052143 | hypothetical-protein-[Anaerolineae-bacterium] |
| WP_190817045.1 | Type1.BREX | [illegible] | 1234 | Bacteria>Terrabacteria_group>Cyanobacteriota/Melainabacteria_group>Cyanobacteriota>Cyanophyceae>Oscillatoriophycideae | GCF_014698845.1 | Coleofasciculus_sp._FACHB-1120 | H6H02_RS09830 | 171769..175473 | 2692783 | BREX-system-P-loop-protein-BrxC-[Coleofasciculus-sp.-FACHB-1120] |
| WP_156114218.1 | Type1.BREX | [illegible] | 835 | Bacteria>Terrabacteria_group>Cyanobacteriota/Melainabacteria_group>Cyanobacteriota>Cyanophyceae>Pleurocapsales | GCF_000756305.1 | Myxosarcina_sp._GI1 | KV40_RS27950 | 18823..21330 | 1541065 | BREX-system-P-loop-protein-BrxC-[Myxosarcina-sp.-GI1] |
| WP_009625661.1 | Type1.BREX | [illegible] | 1178 | Bacteria>Terrabacteria_group>Cyanobacteriota/Melainabacteria_group>Cyanobacteriota>Cyanophyceae>Pseudanabaenales | GCF_000332215.1 | Pseudanabaena_biceps | PSE7429_RS03435 | 33637..37173 | 927669 | MULTISPECIES:-BREX-system-P-loop-protein-BrxC-[Pseudanabaena] |
| WP_094532951.1 | Type1.BREX | [illegible] | 1181 | Bacteria>Terrabacteria_group>Cyanobacteriota/Melainabacteria_group>Cyanobacteriota>Cyanophyceae>Pseudanabaenales | GCF_002251945.1 | Pseudanabaena_sp._SR411 | B9G53_RS13005 | 14836..18381 | 1980935 | BREX-system-P-loop-protein-BrxC-[Pseudanabaena-sp.-SR411] |
| WP_190797467.1 | Type1.BREX | [illegible] | 1186 | Bacteria>Terrabacteria_group>Cyanobacteriota/Melainabacteria_group>Cyanobacteriota>Cyanophyceae>Pseudanabaenales | GCF_014695875.1 | Leptolyngbya_sp._FACHB-541 | H6G00_RS15685 | 19029..22589 | 2692810 | BREX-system-P-loop-protein-BrxC-[Leptolyngbya-sp.-FACHB-541] |
| MBD1853918.1 | Type1.BREX | [illegible] | 1186 | Bacteria>Terrabacteria_group>Cyanobacteriota/Melainabacteria_group>Cyanobacteriota>unclassified_Cyanobacteriota | GCA_014695325.1 | Cyanobacteria_bacterium_FACHB-502 | H6F87_28810 | 26444..30004 | 2692765 | BREX-system-P-loop-protein-BrxC-[Cyanobacteria-bacterium-FACHB-502] |
| HBT89391.1 | Type1.BREX | [illegible] | 1187 | Bacteria>Thermodesulfobacteriota>Desulfobacteria>Desulfobacterales>Desulfobacteraceae>Desulfobacter | GCA_003510365.1 | Desulfobacter_sp. | DEB50_13295 | 35068..38631 | 2294 | BREX-system-P-loop-protein-BrxC-[Desulfobacter-sp.] |
| WP_035236115.1 | Type1.BREX | [illegible] | 1189 | Bacteria>Thermodesulfobacteriota>Desulfobacteria>Desulfobacterales>Desulfobacteraceae>Desulfobacter | GCF_000745975.1 | Desulfobacter_vibrioformis | Q366_RS02915 | 253933..257502 | 34031 | BREX-system-P-loop-protein-BrxC-[Desulfobacter-vibrioformis] |
| WP_024334643.1 | Type1.BREX | [illegible] | 1144 | Bacteria>Thermodesulfobacteriota>Desulfobacteria>Desulfobacterales>Desulfobacteraceae>Desulfotignum | GCF_000421285.1 | Desulfotignum_balticum | K365_RS0111330 | 2237205..2240639 | 115781 | BREX-system-P-loop-protein-BrxC-[Desulfotignum-balticum] |
| PIP37810.1 | Type1.BREX | [illegible] | 1174 | Bacteria>Thermodesulfobacteriota>Desulfobacteria>Desulfobacterales>unclassified_Desulfobacterales | GCA_002772195.1 | Desulfobacterales_bacterium_CG23_combo_of_CG06-09_8_20_14_all_51_8 | COX19_15630 | 1..3524 | 1973983 | hypothetical-protein-COX19_15630,-partial-[Desulfobacterales-bacterium-CG23_combo_of_CG06-09_8_20_14_all_51_8] |
| MBC8432684.1 | Type6.BREX | [illegible] | 1227 | Bacteria>Thermodesulfobacteriota>Desulfobacteria>Desulfobacterales>unclassified_Desulfobacterales | GCA_014383215.1 | Desulfobacterales_bacterium | H8D96_12300 | 20226..23909 | 2044940 | BREX-system-P-loop-protein-BrxC-[Desulfobacterales-bacterium] |
| KGO34922.1 | Type1.BREX | [illegible] | 1193 | Bacteria>Thermodesulfobacteriota>Desulfobulbia>Desulfobulbales>Desulfobulbaceae>Desulfobulbus | GCA_000769715.1 | Desulfobulbus_sp._Tol-SR | JT06_05465 | 8780..12361 | 1536652 | hypothetical-protein-JT06_05465-[Desulfobulbus-sp.-Tol-SR] |
| WP_092376845.1 | Type1.BREX | [illegible] | 1183 | Bacteria>Thermodesulfobacteriota>Desulfovibrionia>Desulfovibrionales>Desulfomicrobiaceae>Desulfomicrobium | GCF_900114115.1 | Desulfomicrobium_apsheronum | BMZ40_RS13600 | 74396..77947 | 52560 | BREX-system-P-loop-protein-BrxC-[Desulfomicrobium-apsheronum] |
| MBI4803787.1 | Type1.BREX | [illegible] | 1145 | Bacteria>Thermodesulfobacteriota>Desulfovibrionia>Desulfovibrionales>Desulfovibrionaceae>Desulfovibrio | GCA_016208105.1 | Desulfovibrio_sp. | HY795_00975 | 45852..49289 | 885 | BREX-system-P-loop-protein-BrxC-[Desulfovibrio-sp.] |
| MBL3581670.1 | Type1.BREX | [illegible] | 1145 | Bacteria>Thermodesulfobacteriota>Desulfovibrionia>Desulfovibrionales>Desulfovibrionaceae>Oleidesulfovibrio | GCA_016756815.1 | Oleidesulfovibrio_alaskensis | JMJ96_04565 | 108279..111716 | 58180 | BREX-system-P-loop-protein-BrxC-[Oleidesulfovibrio-alaskensis] |
| WP_273192722.1 | Type1.BREX | [illegible] | 1146 | Bacteria>Thermodesulfobacteriota>Desulfovibrionia>Desulfovibrionales>Desulfovibrionaceae>Oleidesulfovibrio | GCF_016756815.1 | Oleidesulfovibrio_alaskensis | JMJ96_RS04575 | 108279..111719 | 58180 | BREX-system-P-loop-protein-BrxC-[Oleidesulfovibrio-alaskensis] |
| AJY71182.1 | Type1.BREX | [illegible] | 1183 | Bacteria>Thermodesulfobacteriota>Desulfuromonadia>Geobacterales>Geobacteraceae>Geobacter | GCA_000961035.1 | Geobacter_sulfurreducens | RW64_17280 | 3775110..3778661 | 35554 | hypothetical-protein-RW64_17280-[Geobacter-sulfurreducens] |
| OPY81323.1 | Type1.BREX | [illegible] | 1144 | Bacteria>Thermodesulfobacteriota>Syntrophia>Syntrophales>Syntrophaceae>Syntrophus | GCA_002067745.1 | Syntrophus_sp._PtaU1.Bin005 | A4E70_01341 | 26340..29774 | 1811712 | MAG:-hypothetical-protein-A4E70_01341-[Syntrophus-sp.-PtaU1.Bin005] |
| WP_103134081.1 | Type1.BREX | [illegible] | 1167 | Bacteria>Thermotogota>Thermotogae>Kosmotogales>Kosmotogaceae>Mesotoga | GCF_002896535.1 | Mesotoga_sp._B105.6.4 | RJ60_RS04180 | 254841..258344 | 1582224 | BREX-system-P-loop-protein-BrxC-[Mesotoga-sp.-B105.6.4] |
| WP_110990429.1 | Type1.BREX | [illegible] | 1163 | Bacteria>Thermotogota>Thermotogae>Kosmotogales>Kosmotogaceae>Mesotoga | GCF_003228205.1 | Mesotoga_sp._TolDC | LH53_RS05220 | 29090..32581 | 1389250 | BREX-system-P-loop-protein-BrxC-[Mesotoga-sp.-TolDC] |
| WP_121509576.1 | Type1.BREX | [illegible] | 1162 | Bacteria>Thermotogota>Thermotogae>Kosmotogales>Kosmotogaceae>Mesotoga | GCF_003664935.1 | Mesotoga_sp._H07pep.5.4 | Y696_RS00810 | 21677..25165 | 1463664 | BREX-system-P-loop-protein-BrxC-[Mesotoga-sp.-H07pep.5.4] |
| MBI1292832.1 | Type1.BREX | [illegible] | 1147 | Bacteria>unclassified_Bacteria | GCA_016125005.1 | bacterium | GC173_16580 | 8915..12358 | 1869227 | BREX-system-P-loop-protein-BrxC-[bacterium] |
| EGR4851343.1 | Type1.BREX | [illegible] | 1225 | - | - | - | C4G37_RS13070 | 147789..151466 | - | DNA-repair-protein-[Vibrio-parahaemolyticus] |
| EGR4808858.1 | Type1.BREX | [illegible] | 1225 | - | - | - | C4G65_RS20915 | 24307..27984 | - | DNA-repair-protein-[Vibrio-parahaemolyticus] |
| EIL2714655.1 | Type1.BREX | [illegible] | 1225 | - | - | - | MOK80_RS10540 | 473799..477476 | - | BREX-system-P-loop-protein-BrxC-[Providencia-rettgeri] |
| -- | | | | | | | | | | |
| -- OLD-ABC->HNH | | | | | | | | | | |
| -- | | | | | | | | | | |
| AGC72568.1 | Type6.BREX | [illegible] | 1181 | Bacteria>environmental_samples | - | uncultured_bacterium_A1Q1_fos_1870 | . | 12880..16425 | 1256554 | hypothetical-protein-[uncultured-bacterium-A1Q1_fos_1870] |
| EFG14417.1 | Type6.BREX | [illegible] | 762 | Bacteria>FCB_group>Bacteroidetes/Chlorobi_group>Bacteroidota>Bacteroidia>Bacteroidales | GCA_000178215.1 | Bacteroides_xylanisolvens_SD_CC_1b | CW3_0497 | 100900..103188 | 702447 | hypothetical-protein-CW3_0497-[Bacteroides-xylanisolvens-SD-CC-1b] |
| EFF57551.1 | Type6.BREX | [illegible] | 706 | Bacteria>FCB_group>Bacteroidetes/Chlorobi_group>Bacteroidota>Bacteroidia>Bacteroidales | GCA_000178295.1 | Bacteroides_xylanisolvens_SD_CC_2a | CW1_2295 | 25339..27459 | 702444 | hypothetical-protein-CW1_2295-[Bacteroides-xylanisolvens-SD-CC-2a] |
| CDM00531.1 | Type6.BREX | [illegible] | 689 | Bacteria>FCB_group>Bacteroidetes/Chlorobi_group>Bacteroidota>Bacteroidia>Bacteroidales | GCA_000577955.1 | Bacteroides_xylanisolvens_SD_CC_2a | BN891_34560 | 156968..159037 | 702444 | hypothetical-protein-BN891_34560-[Bacteroides-xylanisolvens-SD-CC-2a] |
| HAM28416.1 | Type6.BREX | [illegible] | 1218 | Bacteria>FCB_group>Bacteroidetes/Chlorobi_group>Bacteroidota>Bacteroidia>Bacteroidales | GCA_003453655.1 | Prevotella_sp. | DCP79_00675 | 66224..69880 | 59823 | BREX-system-P-loop-protein-BrxC-[Prevotella-sp.] |
| MBQ2852763.1 | Type1.BREX | [illegible] | 1215 | Bacteria>FCB_group>Bacteroidetes/Chlorobi_group>Bacteroidota>Bacteroidia>Bacteroidales | GCA_017409275.1 | Bacteroidales_bacterium | IJE76_05160 | 110084..113731 | 2030927 | hypothetical-protein-[Bacteroidales-bacterium] |
| MCI6342424.1 | Type6.BREX | [illegible] | 1217 | Bacteria>FCB_group>Bacteroidetes/Chlorobi_group>Bacteroidota>Bacteroidia>Bacteroidales | GCA_022777695.1 | Prevotella_sp. | MR788_07720 | 61686..65339 | 59823 | BREX-system-P-loop-protein-BrxC-[Prevotella-sp.] |
| MCI6336457.1 | Type1.BREX | [illegible] | 1215 | Bacteria>FCB_group>Bacteroidetes/Chlorobi_group>Bacteroidota>Bacteroidia>Bacteroidales | GCA_022777825.1 | Prevotella_sp. | MR802_00705 | 5343..8990 | 59823 | BREX-system-P-loop-protein-BrxC-[Prevotella-sp.] |
| WP_232506819.1 | Type6.BREX | [illegible] | 794 | Bacteria>FCB_group>Bacteroidetes/Chlorobi_group>Bacteroidota>Bacteroidia>Bacteroidales | GCF_000178215.1 | Bacteroides_xylanisolvens | CW3_RS0123000 | 100900..103284 | 371601 | hypothetical-protein-[Bacteroides-xylanisolvens] |
| WP_008016782.1 | Type6.BREX | [illegible] | 425 | Bacteria>FCB_group>Bacteroidetes/Chlorobi_group>Bacteroidota>Bacteroidia>Bacteroidales | GCF_000178215.1 | Bacteroides_xylanisolvens | CW3_RS26225 | 103281..104558 | 371601 | DUF6079-family-protein-[Bacteroides-xylanisolvens] |
| WP_004314649.1 | Type6.BREX | [illegible] | 515 | Bacteria>FCB_group>Bacteroidetes/Chlorobi_group>Bacteroidota>Bacteroidia>Bacteroidales | GCF_000178295.1 | Bacteroides_xylanisolvens | CW1_RS26610 | 23799..25346 | 371601 | BREX-system-P-loop-protein-BrxC-[Bacteroides-xylanisolvens] |
| WP_232505074.1 | Type6.BREX | [illegible] | 711 | Bacteria>FCB_group>Bacteroidetes/Chlorobi_group>Bacteroidota>Bacteroidia>Bacteroidales | GCF_000178295.1 | Bacteroides_xylanisolvens | CW1_RS26615 | 25324..27459 | 371601 | hypothetical-protein-[Bacteroides-xylanisolvens] |
| WP_044064719.1 | Type6.BREX | [illegible] | 1217 | Bacteria>FCB_group>Bacteroidetes/Chlorobi_group>Bacteroidota>Bacteroidia>Bacteroidales | GCF_000833995.1 | Prevotella_pectinovora | ST45_RS02505 | 69518..73171 | 1602169 | BREX-system-P-loop-protein-BrxC-[Prevotella-pectinovora] |
| WP_215758559.1 | Type6.BREX | [illegible] | 1217 | Bacteria>FCB_group>Bacteroidetes/Chlorobi_group>Bacteroidota>Bacteroidia>Bacteroidales | GCF_018789675.1 | Hallella_faecis | KM934_00340 | 78197..81850 | 2841596 | BREX-system-P-loop-protein-BrxC-[Hallella-faecis] |
| WP_008770354.1 | Type6.BREX | [illegible] | 1219 | Bacteria>FCB_group>Bacteroidetes/Chlorobi_group>Bacteroidota>Bacteroidia>Bacteroidales | GCF_902374095.1 | Bacteroides | BSAG_RS19905 | 423901..427560 | 816 | MULTISPECIES:-BREX-system-P-loop-protein-BrxC-[Bacteroides] |
| WP_273212250.1 | Type1.BREX | [illegible] | 1173 | Bacteria>FCB_group>Bacteroidetes/Chlorobi_group>Bacteroidota>Cytophagia>Cytophagales | GCF_002483185.1 | Runella_zeae | B4E63_RS12460 | 307458..310979 | 94255 | BREX-system-P-loop-protein-BrxC-[Runella-zeae] |
| WP_159471214.1 | Type1.BREX | [illegible] | 1179 | Bacteria>FCB_group>Bacteroidetes/Chlorobi_group>Bacteroidota>Cytophagia>Cytophagales | GCF_009801115.1 | Dyadobacter_sp._3J3 | FZF08_RS17845 | 89009..92548 | 2606600 | BREX-system-P-loop-protein-BrxC-[Dyadobacter-sp.-3J3] |
| MBN8706499.1 | Type1.BREX | [illegible] | 1173 | Bacteria>FCB_group>Bacteroidetes/Chlorobi_group>Bacteroidota>unclassified_Bacteroidota | GCA_017303995.1 | Bacteroidota_bacterium | J0L62_11545 | 8364..11885 | 1898104 | BREX-system-P-loop-protein-BrxC-[Bacteroidota-bacterium] |
| WP_201422032.1 | Type6.BREX | [illegible] | 1188 | Bacteria>Myxococcota>Myxococcia>Myxococcales>Cystobacterineae>Myxococcaceae | GCF_016698685.1 | Myxococcus_xanthus | JKA73_RS29390 | 7346078..7349644 | 34 | BREX-system-P-loop-protein-BrxC-[Myxococcus-xanthus] |
| WP_267556343.1 | Type6.BREX | [illegible] | 1189 | Bacteria>Myxococcota>Myxococcia>Myxococcales>Cystobacterineae>Myxococcaceae | GCF_026626625.1 | Corallococcus_sp._BB11-1 | OV207_RS17695 | 240885..244454 | 2996783 | BREX-system-P-loop-protein-BrxC-[Corallococcus-sp.-BB11-1] |
| WP_020733584.1 | Type6.BREX | [illegible] | 1182 | Bacteria>Myxococcota>Polyangia>Polyangiales>Polyangiaceae>Sorangium | GCF_000418325.1 | Sorangium_cellulosum | SCE1572_RS08150 | 2251441..2254989 | 56 | BREX-system-P-loop-protein-BrxC-[Sorangium-cellulosum] |
| MCK6536698.1 | Type1.BREX | [illegible] | 1149 | Bacteria>Myxococcota>Polyangia>Polyangiales>Polyangiaceae>unclassified_Polyangiaceae | GCA_023150915.1 | Polyangiaceae_bacterium | L6Q84_27330 | 8687..12136 | 2268199 | BREX-system-P-loop-protein-BrxC-[Polyangiaceae-bacterium] |
| MCK6527600.1 | Type6.BREX | [illegible] | 1184 | Bacteria>Myxococcota>unclassified_Myxococcota | GCA_023150875.1 | Myxococcota_bacterium | L6R50_08565 | 57228..60782 | 2818507 | BREX-system-P-loop-protein-BrxC-[Myxococcota-bacterium] |
| WP_225487034.1 | Type1.BREX | [illegible] | 1146 | Bacteria>Pseudomonadota>Acidithiobacillia>Acidithiobacillales>Acidithiobacillaceae>Acidithiobacillus | - | Acidithiobacillus_ferrooxidans | LDB30_08025 | 1563873..1567313 | 920 | BREX-system-P-loop-protein-BrxC-[Acidithiobacillus-ferrooxidans] |
| RWC29997.1 | Type1.BREX | [illegible] | 1181 | Bacteria>Pseudomonadota>Alphaproteobacteria>Hyphomicrobiales>Phyllobacteriaceae>Mesorhizobium | GCA_004017395.1 | Mesorhizobium_sp. | EOS27_14215 | 67406..70951 | 1871066 | BREX-system-P-loop-protein-BrxC-[Mesorhizobium-sp.] |
| WP_136091391.1 | Type1.BREX | [illegible] | 1181 | Bacteria>Pseudomonadota>Alphaproteobacteria>Hyphomicrobiales>Phyllobacteriaceae>Mesorhizobium | GCF_004794715.1 | Mesorhizobium_sp._M8A.F.Ca.ET.207.01.1.1 | EN850_RS12645 | 56072..59617 | 2563968 | BREX-system-P-loop-protein-BrxC-[Mesorhizobium-sp.-M8A.F.Ca.ET.207.01.1.1] |
| MBF0399382.1 | Type1.BREX | [illegible] | 1144 | Bacteria>Pseudomonadota>Alphaproteobacteria>Magnetococcales>unclassified_Magnetococcales | GCA_015232635.1 | Magnetococcales_bacterium | HQL90_01290 | 18375..21809 | 2026759 | BREX-system-P-loop-protein-BrxC-[Magnetococcales-bacterium] |
| QKS29579.1 | Type6.BREX | [illegible] | 1180 | Bacteria>Pseudomonadota>Betaproteobacteria>Betaproteobacteria_incertae_sedis>Candidatus_Accumulibacter | GCA_013347225.1 | Candidatus_Accumulibacter_similis | HT579_12075 | 2666587..2670129 | 2954388 | MAG:-BREX-system-P-loop-protein-BrxC-[Candidatus-Accumulibacter-similis] |
| MBK8385355.1 | Type1.BREX | [illegible] | 1154 | Bacteria>Pseudomonadota>Betaproteobacteria>Betaproteobacteria_incertae_sedis>Candidatus_Accumulibacter>unclassified_Candidatus_Accumulibacter | GCA_016711725.1 | Candidatus_Accumulibacter_sp. | IPL11_06655 | 113668..117132 | 2053492 | BREX-system-P-loop-protein-BrxC-[Candidatus-Accumulibacter-sp.] |
| MBN8436515.1 | Type1.BREX | [illegible] | 1154 | Bacteria>Pseudomonadota>Betaproteobacteria>Betaproteobacteria_incertae_sedis>Candidatus_Accumulibacter>unclassified_Candidatus_Accumulibacter | GCA_017302555.1 | Candidatus_Accumulibacter_sp. | J0L95_00470 | 105300..108764 | 2053492 | BREX-system-P-loop-protein-BrxC-[Candidatus-Accumulibacter-sp.] |
| WP_043683814.1 | Type1.BREX | [illegible] | 1153 | Bacteria>Pseudomonadota>Betaproteobacteria>Burkholderiales>Alcaligenaceae>Castellaniella | GCF_000612685.1 | Castellaniella_defragrans | BN940_RS13240 | 2967883..2971344 | 75697 | BREX-system-P-loop-protein-BrxC-[Castellaniella-defragrans] |
| WP_085318435.1 | Type1.BREX | [illegible] | 1153 | Bacteria>Pseudomonadota>Betaproteobacteria>Burkholderiales>Alcaligenaceae>Derxia | GCF_002105195.1 | Derxia_lacustris | B9P64_RS23165 | 16077..19538 | 764842 | BREX-system-P-loop-protein-BrxC-[Derxia-lacustris] |
| WP_115063824.1 | Type1.BREX | [illegible] | 1155 | Bacteria>Pseudomonadota>Betaproteobacteria>Burkholderiales>Alcaligenaceae>Oligella | GCF_900454315.1 | Oligella_ureolytica | NCTC11998_02420 | 2456151..2459618 | 90244 | BREX-system-P-loop-protein-BrxC-[Oligella-ureolytica] |
| WP_108330269.1 | Type1.BREX | [illegible] | 1154 | Bacteria>Pseudomonadota>Betaproteobacteria>Burkholderiales>Comamonadaceae>Limnohabitans | GCF_003063435.1 | Limnohabitans_sp._2KL-1 | B9Z47_RS05555 | 224189..227653 | 1100699 | BREX-system-P-loop-protein-BrxC-[Limnohabitans-sp.-2KL-1] |
| WP_108311897.1 | Type1.BREX | [illegible] | 1153 | Bacteria>Pseudomonadota>Betaproteobacteria>Burkholderiales>Comamonadaceae>Limnohabitans | GCF_003063455.1 | Limnohabitans_parvus | B9Z37_RS03875 | 837390..840851 | 540061 | BREX-system-P-loop-protein-BrxC-[Limnohabitans-parvus] |
| MBH2043138.1 | Type1.BREX | [illegible] | 1153 | Bacteria>Pseudomonadota>Betaproteobacteria>Burkholderiales>Comamonadaceae>unclassified_Comamonadaceae | GCA_016000285.1 | Comamonadaceae_bacterium | I8H87_06315 | 1878..5339 | 1871071 | BREX-system-P-loop-protein-BrxC-[Comamonadaceae-bacterium] |
| WP_071078524.1 | Type1.BREX | [illegible] | 1153 | Bacteria>Pseudomonadota>Betaproteobacteria>Burkholderiales>Oxalobacteraceae>Janthinobacterium | GCF_001854915.1 | Janthinobacterium_lividum | AKG95_RS19180 | 51113..54574 | 29581 | BREX-system-P-loop-protein-BrxC-[Janthinobacterium-lividum] |
| WP_046167751.1 | Type1.BREX | [illegible] | 1153 | Bacteria>Pseudomonadota>Betaproteobacteria>Neisseriales>Chromobacteriaceae>Chromobacterium_group | GCF_000971355.1 | Chromobacterium_vaccinii | VL68_RS13365 | 2032..5493 | 1108595 | BREX-system-P-loop-protein-BrxC-[Chromobacterium-vaccinii] |
| WP_096450985.1 | Type6.BREX | [illegible] | 1180 | Bacteria>Pseudomonadota>Betaproteobacteria>Rhodocyclales>Zoogloeaceae>Thauera | GCF_002354895.1 | Thauera_sp._K11 | CCZ27_RS19540 | 4450815..4454357 | 2005884 | BREX-system-P-loop-protein-BrxC-[Thauera-sp.-K11] |

| Protein-ID | System | Gene-Neighbourhoods | Length | Taxonomy | Assembly-ID | Strain | Locus-Tag | Genomic-coordinates | TaxID | Fasta-define |
|---|---|---|---|---|---|---|---|---|---|---|
| EJN6954191.1 | Type1.BREX | <-LexA-protease\|\|?-><-wHTH+WYL+WCX\|\|ARG->?->ZnR->RHS->BrxA-DNA-binding->BrxB-BrxF-iSTAND->DUF6079*->N6-METHYLASE->OLD-ABC->HNH->PglZ->HORMA-><-SF1-SF2-Helicase+REase<-SF1-SF2-Helicase+REase | 1157 | Bacteria>Pseudomonadota>Gammaproteobacteria>Aeromonadales>Aeromonadaceae>Aeromonas | GCA_024506355.1 | Aeromonas_hydrophila | NRD12_000732 | 206979..210452 | 644 | BREX-system-P-loop-protein-BrxC-[Aeromonas-hydrophila] |
| WP_043124789.1 | Type1.BREX | SF1-SF2-Helicase+REase->SF1-SF2-Helicase+REase-><-HORMA<-PglZ<-HNH<-OLD-ABC<-N6-METHYLASE<-DUF6079*<-BrxB-BrxF-iSTAND<-BrxA-DNA-binding<-URI-DNase<-?\|\|LexA-protease-> | 1157 | Bacteria>Pseudomonadota>Gammaproteobacteria>Aeromonadales>Aeromonadaceae>Aeromonas | GCF_000724965.1 | Aeromonas_hydrophila | HR52_RS13935 | 3102820..3106293 | 644 | BREX-system-P-loop-protein-BrxC-[Aeromonas-hydrophila] |
| WP_113721905.1 | Type1.BREX | SF1-SF2-Helicase+REase->SF1-SF2-Helicase+REase-><-HEPN+Sel1+SEC-C<-PglZ<-HNH<-OLD-ABC<-N6-METHYLASE<-DUF6079*<-BrxB-BrxF-iSTAND<-BrxA-DNA-binding<-RHS<-?<-ARG\|\|wHTH+WYL+WCX-><-?\|\|LexA-protease-> | 1157 | Bacteria>Pseudomonadota>Gammaproteobacteria>Aeromonadales>Aeromonadaceae>Aeromonas | GCF_003265515.1 | Aeromonas_salmonicida | CPA05_RS19945 | 20643..24116 | 645 | BREX-system-P-loop-protein-BrxC-[Aeromonas-salmonicida] |
| WP_114523382.1 | Type1.BREX | <-LexA-protease\|\|?->URI-DNase->BrxA-DNA-binding->BrxB-BrxF-iSTAND->DUF6079*->N6-METHYLASE->OLD-ABC->HNH->PglZ-> | 1157 | Bacteria>Pseudomonadota>Gammaproteobacteria>Aeromonadales>Aeromonadaceae>Aeromonas | GCF_003344485.1 | Aeromonas_sp._ARM81 | ASJ36_RS21100 | 4447..7920 | 1747384 | BREX-system-P-loop-protein-BrxC-[Aeromonas-sp.-ARM81] |
| WP_182951579.1 | Type1.BREX | <-LexA-protease\|\|?->REase+N6_METHYLASE->TRD+TRD->REase+SF1-SF2-Helicase->zf-ISL3+HTH_Tnp_ISL3+DDE_Tnp_ISL3->URI-DNase->BrxA-DNA-binding->BrxB-BrxF-iSTAND->DUF6079*->N6-METHYLASE->OLD-ABC->HNH->PglZ->HIN-HTH->HTH_Tnp_1+HTH_21+rve+rve_3->HORMA-><-DDE_Tnp_1<-SF1-SF2-Helicase+REase | 1157 | Bacteria>Pseudomonadota>Gammaproteobacteria>Aeromonadales>Aeromonadaceae>Aeromonas | GCF_014168615.1 | Aeromonas_sp._WP2-W18-CRE-05 | WP2W18C05_13360 | 1426228..1429701 | 2675707 | BREX-system-P-loop-protein-BrxC-[Aeromonas-sp.-WP2-W18-CRE-05] |
| WP_203130882.1 | Type1.BREX | <-wHTH+WYL+WCX\|\|ZnR+ZnR-><-?<-LexA-protease\|\|?->URI-DNase->BrxA-DNA-binding->BrxB-BrxF-iSTAND->DUF6079-p*->DUF6079-p->N6-METHYLASE->OLD-ABC->HNH->PglZ->HORMA-><-SF1-SF2-Helicase+REase<-SF1-SF2-Helicase+REase | 935 | Bacteria>Pseudomonadota>Gammaproteobacteria>Aeromonadales>Aeromonadaceae>Aeromonas | GCF_016805325.1 | Aeromonas_jandaei | GNZ06_RS06060 | 150251..153058 | 650 | BREX-system-P-loop-protein-BrxC-[Aeromonas-jandaei] |
| WP_203130883.1 | Type1.BREX | <-wHTH+WYL+WCX\|\|ZnR+ZnR-><-?<-LexA-protease\|\|?->URI-DNase->BrxA-DNA-binding->BrxB-BrxF-iSTAND->DUF6079-p->DUF6079-p*->N6-METHYLASE->OLD-ABC->HNH->PglZ->HORMA-><-SF1-SF2-Helicase+REase<-SF1-SF2-Helicase+REase | 226 | Bacteria>Pseudomonadota>Gammaproteobacteria>Aeromonadales>Aeromonadaceae>Aeromonas | GCF_016805325.1 | Aeromonas_jandaei | GNZ06_RS06065 | 153058..153738 | 650 | hypothetical-protein-[Aeromonas-jandaei] |
| WP_203133261.1 | Type1.BREX | SF1-SF2-Helicase+REase->SF1-SF2-Helicase+REase-><-HORMA<-PglZ<-HNH<-OLD-ABC<-N6-METHYLASE<-DUF6079*<-BrxB-BrxF-iSTAND<-BrxA-DNA-binding<-URI-DNase<-?\|\|LexA-protease->?-><-ZnR+ZnR\|\|wHTH+WYL+WCX-> | 1157 | Bacteria>Pseudomonadota>Gammaproteobacteria>Aeromonadales>Aeromonadaceae>Aeromonas | GCF_016805365.1 | Aeromonas_jandaei | G4224_RS08640 | 376570..380043 | 650 | BREX-system-P-loop-protein-BrxC-[Aeromonas-jandaei] |
| WP_213387487.1 | Type1.BREX | <-LexA-protease\|\|?-><-wHTH+WYL+WCX\|\|ARG->?->ZnR->RHS->BrxA-DNA-binding->BrxB-BrxF-iSTAND->DUF6079*->N6-METHYLASE->OLD-ABC->HNH->PglZ->HEPN+Sel1+SEC-C->HIN-HTH-><-SF1-SF2-Helicase+REase<-SF1-SF2-Helicase+REase | 1157 | Bacteria>Pseudomonadota>Gammaproteobacteria>Aeromonadales>Aeromonadaceae>Aeromonas | GCF_018359765.1 | Aeromonas_veronii | J4G64_RS11995 | 290568..294041 | 654 | BREX-system-P-loop-protein-BrxC-[Aeromonas-veronii] |
| WP_215805224.1 | Type1.BREX | <-HORMA<-DDE_Tnp_1<-PglZ-p<-PglZ-p<-PglZ<-HNH<-HNH<-OLD-ABC<-N6-METHYLASE<-DUF6079*<-BrxB-BrxF-iSTAND-p<-BrxB-BrxF-iSTAND<-BrxA-DNA-binding\|\|HIN-HTH-><-URI-DNase<-?\|\|LexA-protease-> | 1157 | Bacteria>Pseudomonadota>Gammaproteobacteria>Aeromonadales>Aeromonadaceae>Aeromonas | GCF_018802365.1 | Aeromonas_jandaei | HQ398_RS10760 | 2353585..2357058 | 650 | BREX-system-P-loop-protein-BrxC-[Aeromonas-jandaei] |
| WP_216986928.1 | Type1.BREX | HDRnase-><-?<-LexA-protease\|\|ARG->ZnR->\|\|?->RHS->BrxA-DNA-binding->BrxB-BrxF-iSTAND->DUF6079*->N6-METHYLASE->OLD-ABC->HNH->PglZ->HORMA-><-SF1-SF2-Helicase+REase<-SF1-SF2-Helicase+REase-p<-SF1-SF2-Helicase+REase-p\|\|SAM-METHYLASE-> | 1157 | Bacteria>Pseudomonadota>Gammaproteobacteria>Aeromonadales>Aeromonadaceae>Aeromonas | GCF_019048565.1 | Aeromonas_sp._FDAARGOS_1406 | I6L36_RS00705 | 137070..140543 | 2778055 | BREX-system-P-loop-protein-BrxC-[Aeromonas-sp.-FDAARGOS-1406] |
| WP_216990395.1 | Type1.BREX | DnaJ->SF1-SF2-Helicase+REase->SF1-SF2-Helicase+REase-><-HORMA<-PglZ<-HNH<-OLD-ABC<-N6-METHYLASE<-DUF6079*<-BrxB-BrxF-iSTAND<-BrxA-DNA-binding<-RHS<-?<-?<-RNase<-DUF3584+DUF5401<-JetB<-MukF | 1157 | Bacteria>Pseudomonadota>Gammaproteobacteria>Aeromonadales>Aeromonadaceae>Aeromonas | GCF_019048705.1 | Aeromonas_sp._FDAARGOS_1409 | I6L39_RS17660 | 3801324..3804797 | 2778058 | BREX-system-P-loop-protein-BrxC-[Aeromonas-sp.-FDAARGOS-1409] |
| WP_216992842.1 | Type1.BREX | SF1-SF2-Helicase+REase->SF1-SF2-Helicase+REase-><-HORMA<-PglZ<-HNH<-OLD-ABC<-N6-METHYLASE<-DUF6079*<-BrxB-BrxF-iSTAND<-BrxA-DNA-binding<-?\|\|LexA-protease->HTH+rve->AAA-ATPase->?-><-ZnR+ZnR\|\|wHTH+WYL+WCX-> | 1157 | Bacteria>Pseudomonadota>Gammaproteobacteria>Aeromonadales>Aeromonadaceae>Aeromonas | GCF_019048725.1 | Aeromonas_sp._FDAARGOS_1407 | I6L37_20730 | 4426496..4429969 | 2778056 | BREX-system-P-loop-protein-BrxC-[Aeromonas-sp.-FDAARGOS-1407] |
| WP_223938541.1 | Type1.BREX | <-SAM-METHYLASE<-Tnp_DNA_bind+DDE_Tnp_1+Dimer_Tnp_Tn5\|\|SF1-SF2-Helicase+REase->SF1-SF2-Helicase+REase-><-HORMA<-PglZ<-HNH<-OLD-ABC<-N6-METHYLASE<-DUF6079*<-BrxB-BrxF-iSTAND<-BrxA-DNA-binding<-RHS | 1157 | Bacteria>Pseudomonadota>Gammaproteobacteria>Aeromonadales>Aeromonadaceae>Aeromonas | GCF_019971155.1 | Aeromonas_caviae | LDN24_RS08690 | 33954..37427 | 648 | BREX-system-P-loop-protein-BrxC-[Aeromonas-caviae] |
| WP_235462874.1 | Type1.BREX | XIS-HTH->?->?-><-wHTH+WYL+WCX\|\|ARG->?->RHS->BrxA-DNA-binding->BrxB-BrxF-iSTAND->DUF6079*->N6-METHYLASE->OLD-ABC->HNH->PglZ->HEPN+Sel1+SEC-C-><-SF1-SF2-Helicase+REase-p | 1157 | Bacteria>Pseudomonadota>Gammaproteobacteria>Aeromonadales>Aeromonadaceae>Aeromonas | GCF_021538905.1 | Aeromonas_australiensis | EHZ86_RS12350 | 8704..12177 | 1114880 | BREX-system-P-loop-protein-BrxC-[Aeromonas-australiensis] |
| WP_241325388.1 | Type1.BREX | SF1-SF2-Helicase+REase-p->SF1-SF2-Helicase+REase-p->SF1-SF2-Helicase+REase-p-><-HEPN+Sel1+SEC-C<-PglZ<-HNH<-OLD-ABC<-N6-METHYLASE<-DUF6079*<-BrxB-BrxF-iSTAND<-BrxA-DNA-binding<-?<-?<-?<-ABhydrolase | 1157 | Bacteria>Pseudomonadota>Gammaproteobacteria>Aeromonadales>Aeromonadaceae>Aeromonas | GCF_022549295.1 | Aeromonas_sp._MR7 | MM192_RS09020 | 75622..79095 | 2923419 | BREX-system-P-loop-protein-BrxC-[Aeromonas-sp.-MR7] |
| WP_247824022.1 | Type1.BREX | DOC+HTH-><-?<-LexA-protease\|\|?-><-wHTH+WYL+WCX\|\|ZnR+ZnR->RHS->BrxA-DNA-binding->BrxB-BrxF-iSTAND->DUF6079*->N6-METHYLASE->OLD-ABC->HNH->PglZ->HORMA-><-SF1-SF2-Helicase+REase<-SF1-SF2-Helicase+REase | 1157 | Bacteria>Pseudomonadota>Gammaproteobacteria>Aeromonadales>Aeromonadaceae>Aeromonas | GCF_023100745.1 | Aeromonas_veronii | MYF86_09055 | 1896227..1899700 | 654 | BREX-system-P-loop-protein-BrxC-[Aeromonas-veronii] |
| WP_264787731.1 | Type1.BREX | <-LexA-protease<-?\|\|RHS->ARG->\|\|RHS->BrxA-DNA-binding->BrxB-BrxF-iSTAND->DUF6079*->N6-METHYLASE->OLD-ABC->HNH->PglZ->DDE_Tnp_1->HORMA-><-?<-zf-IS66+DDE_Tnp_IS66<-zf-IS66+DDE_Tnp_IS66 | 1157 | Bacteria>Pseudomonadota>Gammaproteobacteria>Aeromonadales>Aeromonadaceae>Aeromonas | GCF_026000055.1 | Aeromonas_caviae | OPI30_RS10860 | 2213781..2217254 | 648 | BREX-system-P-loop-protein-BrxC-[Aeromonas-caviae] |
| WP_270670318.1 | Type1.BREX | <-HTH+rve\|\|<-HEPN+Sel1+SEC-C<-PglZ<-HNH<-OLD-ABC<-N6-METHYLASE<-DUF6079*<-BrxB-BrxF-iSTAND<-BrxA-DNA-binding<-RHS<-?\|\|<-ZnR<-ARG | 1157 | Bacteria>Pseudomonadota>Gammaproteobacteria>Aeromonadales>Aeromonadaceae>Aeromonas | GCF_027705165.1 | Aeromonas_sp._QDB04 | OKC37_RS13865 | 9577..13050 | 2990477 | BREX-system-P-loop-protein-BrxC-[Aeromonas-sp.-QDB04] |
| WP_270659881.1 | Type1.BREX | SF1-SF2-Helicase+REase->SF1-SF2-Helicase+REase-><-HEPN+Sel1+SEC-C<-PglZ<-HNH<-OLD-ABC<-N6-METHYLASE<-DUF6079*<-BrxB-BrxF-iSTAND<-BrxA-DNA-binding<-URI-DNase<-?\|\|LexA-protease-> | 1157 | Bacteria>Pseudomonadota>Gammaproteobacteria>Aeromonadales>Aeromonadaceae>Aeromonas | GCF_027705965.1 | Aeromonas_sp._Y301-2 | OKB83_RS15925 | 143279..146752 | 2990505 | BREX-system-P-loop-protein-BrxC-[Aeromonas-sp.-Y301-2] |
| GGW57088.1 | Type1.BREX | <-wHTH+WYL+WCX-p\|\|SIR2->?-><-?\|\|HEPN-p->JAB->BrxA-DNA-binding->BrxB-BrxF-iSTAND->DUF6079*->N6-METHYLASE->OLD-ABC->HNH->PglZ->HEPN+Sel1+SEC-C->Immank+ANK+ANK->REase-><-?<-SF1-SF2-Helicase+REase | 1114 | Bacteria>Pseudomonadota>Gammaproteobacteria>Alteromonadales>Alteromonadaceae>Alishewanella | GCA_014652375.1 | Alishewanella_tabrizica | GCM10008111_11430 | 80462..83806 | 671278 | hypothetical-protein-GCM10008111_11430-[Alishewanella-tabrizica] |
| WP_229796961.1 | Type1.BREX | <-wHTH+WYL+WCX-p\|\|SIR2->?-><-?\|\|HEPN-p->JAB->BrxA-DNA-binding->BrxB-BrxF-iSTAND->DUF6079*->N6-METHYLASE->OLD-ABC->HNH->PglZ->HEPN+Sel1+SEC-C->Immank+ANK+ANK->REase-><-?<-SF1-SF2-Helicase+REase | 1125 | Bacteria>Pseudomonadota>Gammaproteobacteria>Alteromonadales>Alteromonadaceae>Alishewanella | GCF_014652375.1 | Alishewanella_tabrizica | IE070_RS05825 | 80429..83806 | 671278 | BREX-system-P-loop-protein-BrxC-[Alishewanella-tabrizica] |
| WP_089368479.1 | Type1.BREX | <-Phage_integrase<-DUF2787<-?\|\|XIS-HTH-><-?<-?\|\|wHTH+WYL+WCX->BrxA-DNA-binding->BrxB-BrxF-iSTAND->DUF6079*->N6-METHYLASE->OLD-ABC->HNH->PglZ->NADAR->rve->AAA-ATPase->ART-DarT->TPR+TPR-> | 1169 | Bacteria>Pseudomonadota>Gammaproteobacteria>Alteromonadales>Pseudoalteromonadaceae>Pseudoalteromonas | GCF_007991255.1 | Pseudoalteromonas_nigrificiens | PNIG_RS10730 | 2243615..2247124 | 28109 | BREX-system-P-loop-protein-BrxC-[Pseudoalteromonas-nigrificiens] |
| WP_182752860.1 | Type1.BREX | <-Phage_integrase<-DUF2787<-?\|\|XIS-HTH-><-?\|\|wHTH+WYL+WCX->BrxA-DNA-binding->BrxB-BrxF-iSTAND->DUF6079*->N6-METHYLASE->OLD-ABC->HNH->PglZ->NADAR->ART-DarT->TPR+TPR->ZetaToxin-> | 1159 | Bacteria>Pseudomonadota>Gammaproteobacteria>Alteromonadales>Pseudoalteromonadaceae>Pseudoalteromonas | GCF_014163875.1 | Pseudoalteromonas_sp._SR43-3 | H5088_RS10925 | 129286..132765 | 2760943 | BREX-system-P-loop-protein-BrxC-[Pseudoalteromonas-sp.-SR43-3] |
| WP_200158841.1 | Type1.BREX | <-SIGMA+OB-Fold+AAA-ATPase+Lon-Protease-BrxL<-PglZ<-HNH<-OLD-ABC<-SF1-SF2-Helicase<-OLD-ABC+TOPRIM<-N6-METHYLASE<-PIN<-OLD-ABC<-DUF6079*<-BrxB-BrxF-iSTAND<-BrxA-DNA-binding<-GDResolvase+HIN-HTH<-ZnR | 1154 | Bacteria>Pseudomonadota>Gammaproteobacteria>Chromatiales>Chromatiaceae>Allochromatium | GCF_016584135.1 | Allochromatium_vinosum | CKO29_RS16480 | 18892..22356 | 1049 | BREX-system-P-loop-protein-BrxC-[Allochromatium-vinosum] |
| WP_200385867.1 | Type1.BREX | DUF6079*->?->PIN-><-?\|\|N6-METHYLASE->OLD-ABC->HNH->PglZ->AbrB-HTH->RHH-> | 1179 | Bacteria>Pseudomonadota>Gammaproteobacteria>Chromatiales>Chromatiaceae>Thiocapsa | GCF_016583985.1 | Thiocapsa_imhoffii | CKO25_RS00005 | 26..3565 | 382777 | BREX-system-P-loop-protein-BrxC-[Thiocapsa-imhoffii] |
| MCC7280699.1 | Type6.BREX | <-SIGMA+OB-Fold+AAA-ATPase+Lon-Protease-BrxL<-PglZ<-HNH<-OLD-ABC<-RHH<-N6-METHYLASE<-Schlafen+HATPase+wHTH+HTH<-DUF6079*<-BrxB-BrxF-iSTAND<-BrxA-DNA-binding<-BrxE | 1181 | Bacteria>Pseudomonadota>Gammaproteobacteria>Chromatiales>Chromatiaceae>unclassified_Chromatiaceae | GCA_020851955.1 | Chromatiaceae_bacterium | IT487_20620 | 13992..17537 | 2049432 | BREX-system-P-loop-protein-BrxC-[Chromatiaceae-bacterium] |
| WP_238620557.1 | Type1.BREX | sGTP-><-35exo+T7DNApol\|\|?->?->BrxA-DNA-binding->BrxB-BrxF-iSTAND->DUF6079*->N6-METHYLASE->OLD-ABC->HNH->OLD-ABC->HNH->PglZ->SIGMA+OB-Fold+AAA-ATPase+Lon-Protease-BrxL-><-Schlafen+HATPase+wHTH<-Schlafen+HATPase+wHTH | 1154 | Bacteria>Pseudomonadota>Gammaproteobacteria>Chromatiales>Ectothiorhodospiraceae>Ectothiorhodospira | GCF_022227785.1 | Ectothiorhodospira_sp._B14B | LQ952_00335 | 78448..81912 | 2899127 | BREX-system-P-loop-protein-BrxC-[Ectothiorhodospira-sp.-B14B] |
| QEP43020.1 | Type1.BREX | <-DDE_Tnp_1<-HTH_Tnp_1+HTH_21+rve+rve_3<-HTH_Tnp_1<-TOPRIM<-OLD-ABC<-RNase<-N6-METHYLASE<-HNH<-OLD-ABC<-DUF6079*<-BrxB-BrxF-iSTAND<-BrxA-DNA-binding<-Phage_holin<-?<-?\|\|GDResolvase+ZnR->?->DDE_Tnp_IS66->HD+SF1-SF2-Helicase-> | 1145 | Bacteria>Pseudomonadota>Gammaproteobacteria>Chromatiales>Ectothiorhodospiraceae>unclassified_Ectothiorhodospiraceae | GCA_008375315.1 | Ectothiorhodospiraceae_bacterium_BW-2 | D5085_07760 | 1613185..1616622 | 947515 | BREX-system-P-loop-protein-BrxC-[Ectothiorhodospiraceae-bacterium-BW-2] |
| WP_132538615.1 | Type6.BREX | <-LexA-protease\|\|?-><-?<-SIGMA+OB-Fold+AAA-ATPase+Lon-Protease-BrxL<-PglZ<-HNH<-OLD-ABC<-N6-METHYLASE<-?<-DUF6079*<-BrxB-BrxF-iSTAND<-BrxA-DNA-binding<-BrxE<-RHH_5\|\|?->?->XIS-HTH->?->ZnR+TOPRIM+AAA-helicase+wHTH-> | 1179 | Bacteria>Pseudomonadota>Gammaproteobacteria>Gammaproteobacteria_incertae_sedis>Candidatus_Competibacteraceae>Plasticicumulans | GCF_004341245.1 | Plasticicumulans_lactativorans | EV699_RS02475 | 34554..38093 | 1133106 | BREX-system-P-loop-protein-BrxC-[Plasticicumulans-lactativorans] |
| WP_132542053.1 | Type1.BREX | <-SIGMA+OB-Fold+AAA-ATPase+Lon-Protease-BrxL<-PglZ<-HNH<-OLD-ABC<-N6-METHYLASE<-HEPN<-PolB<-DUF6079*<-BrxB-BrxF-iSTAND<-BrxA-DNA-binding\|\|ABhydrolase-><-Acyl_transf_3<-?<-?<-?\|\|Peptidase_M3-><-ABhydrolase | 1153 | Bacteria>Pseudomonadota>Gammaproteobacteria>Gammaproteobacteria_incertae_sedis>Candidatus_Competibacteraceae>Plasticicumulans | GCF_004341245.1 | Plasticicumulans_lactativorans | EV699_RS09620 | 184945..188406 | 1133106 | BREX-system-P-loop-protein-BrxC-[Plasticicumulans-lactativorans] |
| WP_071159674.1 | Type1.BREX | <-SIGMA+OB-Fold+AAA-ATPase+Lon-Protease-BrxL<-PglZ<-HNH<-OLD-ABC<-N6-METHYLASE<-DUF6079*<-BrxB-BrxF-iSTAND<-BrxA-DNA-binding<-XIS-HTH\|\|?-><-?<-?<-?<-?<-JAB+sGTP | 1153 | Bacteria>Pseudomonadota>Gammaproteobacteria>Methylococcales>Methylococcaceae>Methylomonas | GCF_001856095.1 | Methylomonas_sp._LWB | BJL95_RS17210 | 210511..213972 | 1905845 | BREX-system-P-loop-protein-BrxC-[Methylomonas-sp.-LWB] |
| WP_262567770.1 | Type1.BREX | <-zf-ISL3+HTH_Tnp_ISL3+DDE_Tnp_ISL3\|\|?-><-Integrase_1<-SIGMA+OB-Fold+AAA-ATPase+Lon-Protease-BrxL<-PglZ<-HNH<-OLD-ABC<-N6-METHYLASE<-DUF6079*<-BrxB-BrxF-iSTAND<-BrxA-DNA-binding<-?<-SF1-SF2-Helicase\|\|?-><-?<-LexA-protease | 1221 | Bacteria>Pseudomonadota>Gammaproteobacteria>Oceanospirillales>Endozoicomonadaceae>Endozoicomonas | GCF_025562715.2 | Endozoicomonas_gorgoniicola | NX722_RS09445 | 2085702..2089367 | 1234144 | BREX-system-P-loop-protein-BrxC-[Endozoicomonas-gorgoniicola] |
| WP_031385055.1 | Type1.BREX | <-PglZ<-HNH<-OLD-ABC<-N6-METHYLASE-p<-HerA/FtsK<-SIR2<-REase<-N6-METHYLASE<-Schlafen+HATPase+wHTH<-DUF6079*<-BrxB-BrxF-iSTAND<-BrxA-DNA-binding<-XIS-HTH<-MobI\|\|XIS-HTH->AAA+HTH->XIS-HTH-><-Arm-DNA-bind_3+Phage_integrase | 1193 | Bacteria>Pseudomonadota>Gammaproteobacteria>Oceanospirillales>Halomonadaceae>Halomonas | GCF_000731955.1 | Halomonas_zincidurans | HALZIN_RS0115275 | 3254805..3258386 | 1178777 | BREX-system-P-loop-protein-BrxC-[Halomonas-zincidurans] |
| WP_107181354.1 | Type1.BREX | <-SIGMA+OB-Fold+AAA-ATPase+Lon-Protease-BrxL<-PglZ<-HNH<-OLD-ABC<-PHP+SbcC<-N6-METHYLASE<-DUF6079*<-BrxB-BrxF-iSTAND<-BrxA-DNA-binding<-XIS-HTH | 1147 | Bacteria>Pseudomonadota>Gammaproteobacteria>Oceanospirillales>Halomonadaceae>Halomonas | GCF_003030865.2 | Halomonas_sp._SL1 | C9J49_RS09500 | 199072..202515 | 2137478 | BREX-system-P-loop-protein-BrxC-[Halomonas-sp.-SL1] |
| WP_087719602.1 | Type1.BREX | <-ICLR-HTH<-?\|\|?->Amidohydro_1->GNTR-HTH->XIS-HTH->BrxA-DNA-binding->BrxB-BrxF-iSTAND->DUF6079*->Schlafen+HATPase+wHTH->N6-METHYLASE->PHP+SbcC->OLD-ABC->HNH->PglZ->SIGMA+OB-Fold+AAA-ATPase+Lon-Protease-BrxL-> | 1145 | Bacteria>Pseudomonadota>Gammaproteobacteria>Oceanospirillales>Halomonadaceae>Salinicola | GCF_002179555.1 | Salinicola_salarius | CCD76_RS11235 | 10059..13496 | 430457 | BREX-system-P-loop-protein-BrxC-[Salinicola-salarius] |
| WP_206838922.1 | Type1.BREX | MODE-HTH+PBPII-><-ADH_N+ADH_zinc_N<-?\|\|?->?->wHTH+WYL+WCX->BrxA-DNA-binding->BrxB-BrxF-iSTAND->DUF6079*->N6-METHYLASE->DUF932->OLD-ABC->TOPRIM->OLD-ABC->HNH->PglZ->SIGMA+OB-Fold+AAA-ATPase+Lon-Protease-BrxL->Integrase_1-> | 1154 | Bacteria>Pseudomonadota>Gammaproteobacteria>Oceanospirillales>Oceanospirillaceae>Marinobacterium | GCF_017310015.1 | Marinobacterium_georgiense | CFI10_RS02405 | 480710..484174 | 48076 | BREX-system-P-loop-protein-BrxC-[Marinobacterium-iners] |
| WP_217336191.1 | Type1.BREX | DUF772+DDE_Tnp_1+DDE_Tnp_1_6->wHTH+WYL+WCX->BrxA-DNA-binding->BrxB-BrxF-iSTAND->DUF6079*->N6-METHYLASE->DUF932->OLD-ABC->HNH->OLD-ABC->HNH->PglZ->SIGMA+OB-Fold+AAA-ATPase+Lon-Protease-BrxL-> | 1154 | Bacteria>Pseudomonadota>Gammaproteobacteria>Oceanospirillales>Oceanospirillaceae>Marinobacterium | GCF_019061305.1 | Marinobacterium_sp._A346 | KTN04_RS15750 | 3266..6730 | 2851016 | BREX-system-P-loop-protein-BrxC-[Marinobacterium-sp.-A346] |
| WP_119395059.1 | Type1.BREX | SF1-SF2-Helicase->HTH+HEPN->McrB-AAA->wHTH+REase+ZnR->HKD-DNase->OLD-ABC->HNH->BrxA-DNA-binding->BrxB-BrxF-iSTAND->DUF6079*->N6-METHYLASE->sGTP->sGTP->?->HerA/FtsK->HNH->?->HNH->PglZ-> | 1155 | Bacteria>Pseudomonadota>Gammaproteobacteria>Oceanospirillales>Saccharospirillaceae>Salinibius | GCF_003577475.1 | Salinibius_halmophilus | DUW71_RS09715 | 145555..149022 | 1853216 | BREX-system-P-loop-protein-BrxC-[Salinibius-halmophilus] |
| MBC7192650.1 | Type1.BREX | <-SIGMA+OB-Fold+AAA-ATPase+Lon-Protease-BrxL<-PglZ<-HNH<-OLD-ABC<-TOPRIM<-OLD-ABC<-HTH+REase<-N6-METHYLASE<-Schlafen+HATPase+wHTH<-DUF6079*<-BrxB-BrxF-iSTAND<-BrxA-DNA-binding<-XIS-HTH\|\|HTH_Tnp_1+HTH_21+rve+rve_3->XIS-HTH->DnaB->?-><-Phage_int_SAM_5+Phage_integrase<-sGTP | 1145 | Bacteria>Pseudomonadota>Gammaproteobacteria>Pseudomonadales>Marinobacteraceae>Marinobacter | GCA_014359475.1 | Marinobacter_sp. | H5U31_09890 | 22634..26071 | 50741 | BREX-system-P-loop-protein-BrxC-[Marinobacter-sp.] |
| WP_007349691.1 | Type5.BREX | <-Glycos_transf_2\|\|tRNA->?->?-><-SIGMA+OB-Fold+AAA-ATPase+HKD-DNase-BrxL<-PglZ<-HNH<-OLD-ABC<-N6-METHYLASE<-Schlafen+HATPase+wHTH+HTH<-DUF6079*<-BrxB-BrxF-iSTAND<-BrxA-DNA-binding<-TUDOR+SF1-SF2-Helicase+iREase+CTD+REase<-TPR | 1194 | Bacteria>Pseudomonadota>Gammaproteobacteria>Pseudomonadales>Marinobacteraceae>Marinobacter | GCF_000169375.1 | Marinobacter_sp._ELB17 | MELB17_RS08075 | 35418..39002 | 270374 | BREX-system-P-loop-protein-BrxC-[Marinobacter-sp.-ELB17] |
| WP_018404435.1 | Type1.BREX | OrfB_Zn_ribbon->BrxA-DNA-binding->BrxB-BrxF-iSTAND->DUF6079*->Schlafen+HATPase+wHTH+HTH->N6-METHYLASE->OLD-ABC->HNH->PglZ->SIGMA+OB-Fold+AAA-ATPase+HKD-DNase-BrxL->N6-METHYLASE-> | 1194 | Bacteria>Pseudomonadota>Gammaproteobacteria>Pseudomonadales>Marinobacteraceae>Marinobacter | GCF_000372805.1 | Marinobacter_gelidimuriae | A3OI_RS0111615 | 2203..5787 | 2739064 | BREX-system-P-loop-protein-BrxC-[Marinobacter-gelidimuriae] |
| WP_033898350.1 | Type1.BREX | <-wHTH+WYL+WCX\|\|?->ARG->XIS-HTH->BrxA-DNA-binding->BrxB-BrxF-iSTAND->DUF6079*->N6-METHYLASE-p->?-><-?\|\|N6-METHYLASE-p->OLD-ABC->HNH->SIGMA+OB-Fold+AAA-ATPase+HKD-DNase-BrxL-> | 1185 | Bacteria>Pseudomonadota>Gammaproteobacteria>Pseudomonadales>Pseudomonadaceae | GCF_000510895.2 | Pseudomonas | GIW75_RS00865 | 8119..11676 | 286 | MULTISPECIES:-BREX-system-P-loop-protein-BrxC-[Pseudomonas] |
| WP_269686934.1 | Type1.BREX | SF1-SF2-Helicase->SF1-SF2-Helicase->?->?-><-RADICAL-SAM<-?<-MazE_antitoxin\|\|wHTH+WYL+WCX->BrxA-DNA-binding->BrxB-BrxF-iSTAND->DUF6079*->N6-METHYLASE->?->?->OLD-ABC->HNH->PglZ->SIGMA+OB-Fold+AAA-ATPase+Lon-Protease-BrxL-><-HTH_3\|\|PadR-HTH-> | 1223 | Bacteria>Pseudomonadota>Gammaproteobacteria>Pseudomonadales>Pseudomonadaceae>Halopseudomonas | GCF_030295685.1 | Halopseudomonas_sp._MFKK-1 | O5048_RS10320 | 2203460..2207131 | 3015994 | BREX-system-P-loop-protein-BrxC-[Halopseudomonas-sp.-MFKK-1] |
| TXH92079.1 | Type1.BREX | <-RelE-ParE<-RHH_1<-PglZ<-HNH<-OLD-ABC<-N6-METHYLASE-p<-NERD<-OLD-ABC<-N6-METHYLASE<-DUF6079*<-BrxB-BrxF-iSTAND<-BrxA-DNA-binding<-XIS-HTH<-?\|\|wHTH+WYL+WCX-> | 1186 | Bacteria>Pseudomonadota>Gammaproteobacteria>Pseudomonadales>Pseudomonadaceae>Pseudomonas | GCA_008015405.1 | Pseudomonas_sp. | E6Q71_01680 | 13269..16829 | 306 | BREX-system-P-loop-protein-BrxC-[Pseudomonas-sp.] |
| MBX5763959.1 | Type1.BREX | Phage_integrase-><-?<-SIGMA+OB-Fold+AAA-ATPase+Lon-Protease-BrxL<-PglZ<-HNH<-OLD-ABC<-?<-?<-N6-METHYLASE<-DUF6079*<-BrxB-BrxF-iSTAND<-BrxA-DNA-binding<-wHTH+WYL+WCX\|\|REase+NACHT->?-><-CHTH<-?\|\|HTH_3-><-SF1-SF2-Helicase | 1223 | Bacteria>Pseudomonadota>Gammaproteobacteria>Pseudomonadales>Pseudomonadaceae>Pseudomonas | GCA_019687095.1 | Pseudomonas_aeruginosa | ISD89_03975 | 862812..866483 | 287 | BREX-system-P-loop-protein-BrxC-[Pseudomonas-aeruginosa] |
| HBN8454461.1 | Type1.BREX | XIS-HTH-><-?<-tRNA\|\|HTH_36->Phage_int_SAM_2+Integrase_1-><-SIGMA+OB-Fold+AAA-ATPase+Lon-Protease-BrxL<-PglZ<-HNH<-OLD-ABC<-N6-METHYLASE<-DUF6079*<-BrxB-BrxF-iSTAND<-BrxA-DNA-binding<-DOC+HTH<-Arm-DNA-bind_3+Phage_integrase<-sGTP | 1146 | Bacteria>Pseudomonadota>Gammaproteobacteria>Pseudomonadales>Pseudomonadaceae>Pseudomonas | GCA_021587585.1 | Pseudomonas_aeruginosa | L3711_005718 | 39513..42953 | 287 | BREX-system-P-loop-protein-BrxC-[Pseudomonas-aeruginosa] |
| HBO3110869.1 | Type1.BREX | Tox-RelE->HTH_3-><-tRNA\|\|HTH_36->Phage_int_SAM_2+Integrase_1-><-SIGMA+OB-Fold+AAA-ATPase+Lon-Protease-BrxL<-PglZ<-HNH<-OLD-ABC<-N6-METHYLASE<-DUF6079*<-BrxB-BrxF-iSTAND<-BrxA-DNA-binding<-DOC+HTH<-Arm-DNA-bind_3+Phage_integrase<-sGTP | 1146 | Bacteria>Pseudomonadota>Gammaproteobacteria>Pseudomonadales>Pseudomonadaceae>Pseudomonas | GCA_021651555.1 | Pseudomonas_aeruginosa | L4R01_001445 | 31462..34902 | 287 | BREX-system-P-loop-protein-BrxC-[Pseudomonas-aeruginosa] |
| HBP6726062.1 | Type1.BREX | SF1-SF2-Helicase-><-HTH_3\|\|?->CHTH-><-?\|\|wHTH+WYL+WCX->BrxA-DNA-binding->BrxB-BrxF-iSTAND->DUF6079*->N6-METHYLASE->?->?->OLD-ABC->HNH->PglZ->SIGMA+OB-Fold+AAA-ATPase+Lon-Protease-BrxL->?-><-Phage_integrase | 1223 | Bacteria>Pseudomonadota>Gammaproteobacteria>Pseudomonadales>Pseudomonadaceae>Pseudomonas | GCA_021711795.1 | Pseudomonas_aeruginosa | L6B36_03955 | 73272..76943 | 287 | BREX-system-P-loop-protein-BrxC-[Pseudomonas-aeruginosa] |
| CAB5551672.1 | Type1.BREX | DUF4969-><-SIGMA+OB-Fold+AAA-ATPase+Lon-Protease-BrxL<-PglZ<-HNH<-OLD-ABC<-?<-DUF932<-N6-METHYLASE<-DUF6079*<-BrxB-BrxF-iSTAND<-BrxA-DNA-binding<-wHTH+WYL+WCX<-?<-XIS-HTH\|\|Arm-DNA-bind_3+Phage_integrase-> | 1154 | Bacteria>Pseudomonadota>Gammaproteobacteria>Pseudomonadales>Pseudomonadaceae>Pseudomonas | GCA_903684265.1 | Pseudomonas_putida | GHA_03446 | 127342..130806 | 303 | Uncharacterised-protein-[Pseudomonas-putida] |
| WP_047306816.1 | Type1.BREX | <-SIGMA+OB-Fold+AAA-ATPase+HKD-DNase-BrxL<-PglZ<-HNH<-OLD-ABC<-N6-METHYLASE-p<-RVT<-N6-METHYLASE-p<-DUF6079*<-BrxB-BrxF-iSTAND<-BrxA-DNA-binding<-XIS-HTH<-?\|\|wHTH+WYL+WCX->sGTP->sGTP->sGTP-> | 1185 | Bacteria>Pseudomonadota>Gammaproteobacteria>Pseudomonadales>Pseudomonadaceae>Pseudomonas | GCF_001020715.1 | Pseudomonas_fluorescens | AA061_RS32150 | 16461..20018 | 294 | BREX-system-P-loop-protein-BrxC-[Pseudomonas-fluorescens] |
| WP_169839752.1 | Type1.BREX | DUF4969-><-SIGMA+OB-Fold+AAA-ATPase+Lon-Protease-BrxL<-PglZ<-HNH<-OLD-ABC<-N6-METHYLASE<-DUF6079*<-BrxB-BrxF-iSTAND<-BrxA-DNA-binding<-wHTH+WYL+WCX<-McrC-NTD+REase+ZnR<-DnaB+McrB-AAA<-Schlafen+HATPase+wHTH<-URI-DNase+RAMA<-?<-XIS-HTH | 1154 | Bacteria>Pseudomonadota>Gammaproteobacteria>Pseudomonadales>Pseudomonadaceae>Pseudomonas | GCF_012986505.1 | Pseudomonas_lundensis | HBN72_RS14625 | 129337..132801 | 86185 | BREX-system-P-loop-protein-BrxC-[Pseudomonas-lundensis] |
| WP_196169752.1 | Type1.BREX | DUF4969-><-SIGMA+OB-Fold+AAA-ATPase+Lon-Protease-BrxL<-PglZ<-HNH<-OLD-ABC<-DUF932<-N6-METHYLASE<-DUF6079*<-BrxB-BrxF-iSTAND<-BrxA-DNA-binding<-wHTH+WYL+WCX<-?<-XIS-HTH\|\|CopG/RHH-><-MODE-HTH+PBPII | 1154 | Bacteria>Pseudomonadota>Gammaproteobacteria>Pseudomonadales>Pseudomonadaceae>Pseudomonas | GCF_015680295.1 | Pseudomonas_guariconensis | IRZ59_RS14965 | 185250..188714 | 1288410 | BREX-system-P-loop-protein-BrxC-[Pseudomonas-guariconensis] |
| WP_198849248.1 | Type1.BREX | wHTH+WYL+WCX-><-SIGMA+OB-Fold+AAA-ATPase+HKD-DNase-BrxL<-HNH<-OLD-ABC<-N6-METHYLASE-p<-HerA/FtsK<-SIR2<-N6-METHYLASE-p<-DUF6079*<-BrxB-BrxF-iSTAND<-BrxA-DNA-binding<-XIS-HTH<-?<-tRNA<-MerR-HTH | 1186 | Bacteria>Pseudomonadota>Gammaproteobacteria>Pseudomonadales>Pseudomonadaceae>Pseudomonas | GCF_016405505.1 | Pseudomonas_sp._MF6772 | JFT60_RS22275 | 24346..27906 | 2797533 | BREX-system-P-loop-protein-BrxC-[Pseudomonas-sp.-MF6772] |
| WP_270945767.1 | Type1.BREX | MerR-HTH->\|\|?->?->DUF772+DDE_Tnp_1-><-?\|\|wHTH+WYL+WCX->BrxA-DNA-binding->BrxB-BrxF-iSTAND->DUF6079*->N6-METHYLASE->?->?->OLD-ABC->HNH->PglZ->SIGMA+OB-Fold+AAA-ATPase+Lon-Protease-BrxL->?->HTH_3-> | 1223 | Bacteria>Pseudomonadota>Gammaproteobacteria>Pseudomonadales>Pseudomonadaceae>Pseudomonas | GCF_027856115.1 | Pseudomonas_aeruginosa | KKQ36_RS16450 | 3486481..3490152 | 287 | BREX-system-P-loop-protein-BrxC-[Pseudomonas-aeruginosa] |
| WP_053526945.1 | Type1.BREX | <-HTH_3<-?<-SIGMA+OB-Fold+AAA-ATPase+Lon-Protease-BrxL<-PglZ<-HNH<-OLD-ABC<-?<-?<-N6-METHYLASE<-DUF6079*<-BrxB-BrxF-iSTAND<-BrxA-DNA-binding<-wHTH+WYL+WCX\|\|?->RVT->?-><-SF1-SF2-Helicase | 1223 | Bacteria>Pseudomonadota>Gammaproteobacteria>Pseudomonadales>Pseudomonadaceae>Stutzerimonas | GCF_001276475.1 | Stutzerimonas_stutzeri | ABW53_RS00980 | 14361..18032 | 316 | BREX-system-P-loop-protein-BrxC-[Stutzerimonas-stutzeri] |
| WP_150138110.1 | Type1.BREX | wHTH+WYL+WCX->BrxA-DNA-binding->BrxB-BrxF-iSTAND->DUF6079*->N6-METHYLASE->OLD-ABC->HNH->PglZ-> | 1159 | Bacteria>Pseudomonadota>Gammaproteobacteria>Vibrionales>Vibrionaceae>Enterovibrio | GCF_008692425.1 | Candidatus_Enterovibrio_escacola | EDM10_RS06800 | 2982..6461 | 1927127 | BREX-system-P-loop-protein-BrxC-[Candidatus-Enterovibrio-escacola] |
| HAS6390651.1 | Type1.BREX | <-HTH\|\|RNase_H-><-DUF2787<-?<-?\|\|XIS-HTH->wHTH+WYL+WCX->BrxA-DNA-binding->BrxB-BrxF-iSTAND->DUF6079*->N6-METHYLASE->OLD-ABC->HNH->PglZ->ZetaToxin-><-SF1-SF2-Helicase+REase<-SF1-SF2-Helicase+REase | 1157 | Bacteria>Pseudomonadota>Gammaproteobacteria>Vibrionales>Vibrionaceae>Vibrio | GCA_015777605.1 | Vibrio_vulnificus | I7251_16610 | 90517..93990 | 672 | BREX-system-P-loop-protein-BrxC-[Vibrio-vulnificus] |
| ODU42543.1 | Type1.BREX | <-SIGMA+OB-Fold+AAA-ATPase+Lon-Protease-BrxL<-PglZ<-Polbeta<-MarR-HTH<-HNH<-OLD-ABC<-N6-METHYLASE<-DUF6079*<-BrxB-BrxF-iSTAND<-BrxA-DNA-binding\|\|?-><-?\|\|HKD-DNase->?->HNH->N6-METHYLASE-> | 1148 | Bacteria>Pseudomonadota>Gammaproteobacteria>Xanthomonadales>Xanthomonadaceae>unclassified_Xanthomonadaceae | GCA_001725645.1 | Xanthomonadaceae_bacterium_SCN_69-123 | ABS96_27585 | 191713..195159 | 1660170 | hypothetical-protein-ABS96_27585-[Xanthomonadaceae-bacterium-SCN-69-123] |
| PIQ12823.1 | Type1.BREX | <-HARE-HTH+REase<-SIGMA+OB-Fold+AAA-ATPase+Lon-Protease-BrxL<-PglZ<-HNH<-OLD-ABC<-N6-METHYLASE<-DUF6079*<-BrxB-BrxF-iSTAND<-BrxA-DNA-binding<-SLOG+BirA-HTH<-SF1-SF2-Helicase+PRTase<-wHTH+WYL+WCX\|\|<-ParB<-ParB<-GDResolvase+PPP1-HTH | 1146 | Bacteria>Pseudomonadota>Hydrogenophilia>Hydrogenophilales>unclassified_Hydrogenophilales | GCA_002770925.1 | Hydrogenophilales_bacterium_CG18_big_fil_WC_8_21_14_2_50_58_12 | CGW70_07940 | 10882..14322 | 1974036 | hypothetical-protein-CGW70_07940-[Hydrogenophilales-bacterium-CG18_big_fil_WC_8_21_14_2_50_58_12] |
| NLF73010.1 | Type1.BREX | XIS-HTH->BrxA-DNA-binding->BrxB-BrxF-iSTAND->DUF6079*->N6-METHYLASE->OLD-ABC->HNH->PglZ->SIGMA+OB-Fold+AAA-ATPase+Lon-Protease-BrxL-> | 1145 | Bacteria>PVC_group>Planctomycetota>Planctomycetia>Pirellulales>Pirellulaceae | GCA_012515235.1 | Candidatus_Anammoximicrobium_sp. | GX575_28570 | 2105..5542 | 2699742 | BREX-system-P-loop-protein-BrxC-[Candidatus-Anammoximicrobium-sp.] |
| MCC5808698.1 | Type1.BREX | Phage_integrase-><-SIGMA+OB-Fold+AAA-ATPase+HKD-DNase-BrxL<-PglZ<-HNH<-OLD-ABC<-N6-METHYLASE<-REase<-OLD-ABC<-DUF6079*<-BrxB-BrxF-iSTAND<-BrxA-DNA-binding<-XIS-HTH | 1183 | Bacteria>PVC_group>Verrucomicrobiota>Opitutae>Opitutales>unclassified_Opitutales | GCA_020831525.1 | Opitutales_bacterium | JJU00_20390 | 12178..15729 | 2026772 | BREX-system-P-loop-protein-BrxC-[Opitutales-bacterium] |
| OPA28160.1 | Type1.BREX | PIN->REase->OLD-ABC->HNH->?->BrxA-DNA-binding->BrxB-BrxF-iSTAND->DUF6079*->N6-METHYLASE->PglZ->SIGMA+OB-Fold+AAA-ATPase+Lon-Protease-BrxL->MarR-HTH->?->?->BZIP+SF1-SF2-Helicase-> | 1190 | Bacteria>Terrabacteria_group>Bacillota>Bacilli>Bacillales>Bacillaceae | GCA_002021465.1 | Bacillus_cereus | BHL47_19745 | 10940..14512 | 1396 | hypothetical-protein-BHL47_19745-[Bacillus-cereus] |
| ATI52289.1 | Type1.BREX | GDResolvase+HIN-HTH->TOPRIM+Topoisom_bac->OLD-ABC->HNH->?->N6-METHYLASE-p->?->BrxA-DNA-binding->BrxB-BrxF-iSTAND->DUF6079*->N6-METHYLASE->PglZ->SIGMA+OB-Fold+AAA-ATPase+Lon-Protease-BrxL->HTH_Tnp_1-> | 1190 | Bacteria>Terrabacteria_group>Bacillota>Bacilli>Bacillales>Bacillaceae | GCA_002504225.1 | Bacillus_cereus | CPZ32_18870 | 3450635..3454207 | 1396 | hypothetical-protein-CPZ32_18870-[Bacillus-cereus] |
| WP_078386197.1 | Type1.BREX | PIN->REase->OLD-ABC->HNH->?->BrxA-DNA-binding->BrxB-BrxF-iSTAND->DUF6079*->N6-METHYLASE->PglZ->SIGMA+OB-Fold+AAA-ATPase+Lon-Protease-BrxL->MarR-HTH->?->?->BZIP+SF1-SF2-Helicase-> | 1190 | Bacteria>Terrabacteria_group>Bacillota>Bacilli>Bacillales>Bacillaceae | GCF_002021465.1 | Bacillus_cereus | BHL47_19745 | 10940..14512 | 1396 | BREX-system-P-loop-protein-BrxC-[Bacillus-cereus] |
| WP_061531636.1 | Type1.BREX | HTH_3-><-ParB\|\|SAD-DUF3427C+SbcC->HNH-><-REase+DUF5655\|\|EVE+HNH->BrxA-DNA-binding->BrxB-BrxF-iSTAND->DUF6079*->N6-METHYLASE->PglZ->SIGMA+OB-Fold+AAA-ATPase+Lon-Protease-BrxL->?-><-DndB<-?\|\|LACI-HTH+PBPI-> | 1195 | Bacteria>Terrabacteria_group>Bacillota>Bacilli>Bacillales>Bacillaceae | GCF_029504955.1 | Bacillus_cereus_group | M2246_RS18875 | 193407..196994 | 86661 | MULTISPECIES:-BREX-system-P-loop-protein-BrxC-[Bacillus] |
| MBN1067552.1 | Type1.BREX | Calcineurin->?->BrxA-DNA-binding->BrxB-BrxF-iSTAND->DUF6079*->CR-ATPase9+REase->N6-METHYLASE->OLD-ABC->HNH->PglZ-><-REase\|\|SIGMA+OB-Fold+AAA-ATPase+Lon-Protease-BrxL->Peptidase_M42->TIR-> | 1215 | Bacteria>Terrabacteria_group>Bacillota>Clostridia>Eubacteriales>Clostridiaceae | GCF_016925955.1 | Clostridium_botulinum | DVW02_06040 | 97050..100697 | 1491 | BREX-system-P-loop-protein-BrxC-[Clostridium-botulinum] |
| WP_043662653.1 | Type1.BREX | <-McrC-NTD+REase<-wHTH+McrB-AAA<-SIGMA+OB-Fold+AAA-ATPase+Lon-Protease-BrxL<-?<-PglZ<-HNH<-OLD-ABC<-N6-METHYLASE<-DUF6079*<-BrxB-BrxF-iSTAND<-BrxA-DNA-binding<-HKD-DNase+SF1-SF2-Helicase+HTH+HTH+SRA<-SAM-METHYLASE<-RNase<-HNH+NERD+ZnR<-?<-?<-Amidohydro_3 | 1216 | Bacteria>Terrabacteria_group>Bacillota>Clostridia>Eubacteriales>Clostridiaceae | GCF_000785185.1 | Clostridium_butyricum | AWN73_RS08010 | 18592..22242 | 1492 | BREX-system-P-loop-protein-BrxC-[Clostridium-butyricum] |
| WP_077851369.1 | Type1.BREX | <-SIGMA+OB-Fold+AAA-ATPase+Lon-Protease-BrxL<-?<-PglZ<-HNH<-OLD-ABC<-N6-METHYLASE<-DUF6079*<-BrxB-BrxF-iSTAND<-BrxA-DNA-binding<-?<-?<-?<-?<-?<-?<-Aminotran_1_2 | 1216 | Bacteria>Terrabacteria_group>Bacillota>Clostridia>Eubacteriales>Clostridiaceae | GCF_002006235.2 | Clostridium_felsineum | CLAUR_RS09645 | 2191072..2194722 | 36839 | BREX-system-P-loop-protein-BrxC-[Clostridium-felsineum] |
| WP_118513467.1 | Type1.BREX | <-Tox-ART-HYD2<-TIR<-PglZ<-Schlafen+HATPase+wHTH<-AbiGi+seqA+RAMA<-N6-METHYLASE<-HNH<-OLD-ABC<-DUF6079*<-BrxB-BrxF-iSTAND<-BrxA-DNA-binding<-HTH_26<-SIGMA-HTH+SIGMA-HTH<-HISKIN<-?<-TPR+TPR+TPR | 1183 | Bacteria>Terrabacteria_group>Bacillota>Clostridia>Eubacteriales>Clostridiaceae | GCF_003477775.1 | Clostridium_sp._AF37-5AT | DW020_RS00895 | 209396..212947 | 2292007 | BREX-system-P-loop-protein-BrxC-[Clostridium-sp.-AF37-5AT] |
| WP_164949146.1 | Type1.BREX | <-HEPN<-SIGMA+OB-Fold+AAA-ATPase+Lon-Protease-BrxL<-PglZ<-HNH<-OLD-ABC<-N6-METHYLASE<-DUF6079*<-BrxB-BrxF-iSTAND<-BrxA-DNA-binding<-?<-?<-CR-ATPase8+REase\|\|?->?-><-?<-?\|\|METHYLASE-> | 1215 | Bacteria>Terrabacteria_group>Bacillota>Clostridia>Eubacteriales>Clostridiaceae | GCF_011065955.2 | Clostridium_sp._FP2 | G9F71_RS17965 | 3736090..3739737 | 2724481 | BREX-system-P-loop-protein-BrxC-[Clostridium-sp.-FP2] |
| WP_171786495.1 | Type1.BREX | <-D5N+D5ATPase<-SIGMA+OB-Fold+AAA-ATPase+Lon-Protease-BrxL<-PglZ<-HNH<-OLD-ABC<-PIWI<-N6-METHYLASE<-DUF6079*<-BrxB-BrxF-iSTAND<-BrxA-DNA-binding<-?<-HTH_3<-?<-?\|\|tRNA-><-RVT | 1216 | Bacteria>Terrabacteria_group>Bacillota>Clostridia>Eubacteriales>Clostridiaceae | GCF_013149405.1 | Clostridium_beijerinckii | BOH39_RS13645 | 3014066..3017716 | 1520 | BREX-system-P-loop-protein-BrxC-[Clostridium-beijerinckii] |
| WP_171780276.1 | Type1.BREX | rve+rve_3->AAA-ATPase-><-D5N+D5ATPase<-SIGMA+OB-Fold+AAA-ATPase+Lon-Protease-BrxL<-PglZ<-HNH<-OLD-ABC<-PIWI<-N6-METHYLASE<-DUF6079*<-BrxB-BrxF-iSTAND<-BrxA-DNA-binding<-?<-?<-HTH_3<-?<-?\|\|tRNA-><-RVT | 1216 | Bacteria>Terrabacteria_group>Bacillota>Clostridia>Eubacteriales>Clostridiaceae | GCF_014230655.1 | Clostridium_beijerinckii | HGI39_RS06505 | 40889..44539 | 1520 | BREX-system-P-loop-protein-BrxC-[Clostridium-beijerinckii] |
| WP_216171751.1 | Type1.BREX | <-SIGMA+OB-Fold+AAA-ATPase+Lon-Protease-BrxL<-PglZ<-HNH<-OLD-ABC<-N6-METHYLASE<-DUF6079*<-BrxB-BrxF-iSTAND<-BrxA-DNA-binding\|\|Phage_int_SAM_3-><-Phage_int_SAM_1+Phage_integrase<-Phage_integrase<-Phage_integrase\|\|Phage_integrase-> | 1215 | Bacteria>Terrabacteria_group>Bacillota>Clostridia>Eubacteriales>Clostridiaceae | GCF_026650905.1 | Clostridium_estertheticum | LL037_RS21145 | 4418214..4421861 | 238834 | BREX-system-P-loop-protein-BrxC-[Clostridium-estertheticum] |
| WP_270487117.1 | Type1.BREX | <-SIGMA+OB-Fold+AAA-ATPase+Lon-Protease-BrxL<-?<-PIN<-OLD-ABC<-PglZ<-HNH<-OLD-ABC<-TIR<-N6-METHYLASE<-DUF6079*<-BrxB-BrxF-iSTAND<-BrxA-DNA-binding<-HTH_26<-PIN<-?<-MACRODOMAIN | 1201 | Bacteria>Terrabacteria_group>Bacillota>Clostridia>Eubacteriales>Clostridiaceae | GCF_027681635.1 | Clostridium_fessum | PGX92_RS02760 | 620693..624298 | 2126740 | BREX-system-P-loop-protein-BrxC-[Clostridium-fessum] |
| WP_115641994.1 | Type1.BREX | NTP_transf->HEPN-><-PglZ<-HNH<-OLD-ABC<-N6-METHYLASE<-?<-DUF6079*<-BrxB-BrxF-iSTAND<-BrxA-DNA-binding\|\|HEPN-><-?<-ABC-ATPase+SbcC<-RADICAL-SAM<-?\|\|DDE_Tnp_IS240+rve_3-> | 1215 | Bacteria>Terrabacteria_group>Bacillota>Clostridia>Eubacteriales>Clostridiaceae | GCF_900461105.1 | Clostridium_putrefaciens | DY168_RS12220 | 2676627..2680274 | 99675 | BREX-system-P-loop-protein-BrxC-[Clostridium-putrefaciens] |
| WP_154780312.1 | Type1.BREX | Peptidase->?->?->MACRODOMAIN->?->ZnR+ZnR+ZnR->HTH_26->BrxA-DNA-binding->BrxB-BrxF-iSTAND->DUF6079*->N6-METHYLASE->RVT->?->OLD-ABC->HNH->?->PglZ->HEPN->SIGMA+OB-Fold+AAA-ATPase+Lon-Protease-BrxL-> | 1201 | Bacteria>Terrabacteria_group>Bacillota>Clostridia>Eubacteriales>Lachnospiraceae | GCF_009707925.1 | Blautia_luti | GKZ57_RS08760 | 74022..77627 | 89014 | BREX-system-P-loop-protein-BrxC-[Blautia-luti] |
| WP_207739085.1 | Type1.BREX | METHYLASE->?->?->?->HTH_26->BrxA-DNA-binding->BrxB-BrxF-iSTAND->DUF6079*->N6-METHYLASE->OLD-ABC->HNH->PglZ->SIGMA+OB-Fold+AAA-ATPase+Lon-Protease-BrxL->?->Arm-DNA-bind_4+Phage_int_SAM_3+Phage_integrase->?->XIS-HTH+XIS-HTH-> | 1173 | Bacteria>Terrabacteria_group>Bacillota>Clostridia>Eubacteriales>Oscillospiraceae | GCF_015558275.1 | Massiliimalia_timonensis | I2G46_RS07880 | 51772..55293 | 1987501 | BREX-system-P-loop-protein-BrxC-[Massiliimalia-timonensis] |
| WP_215659170.1 | Type1.BREX | <-TIR<-SIGMA+OB-Fold+AAA-ATPase+Lon-Protease-BrxL<-PglZ<-DUF4404<-ZnR+ZnR<-N6-METHYLASE<-?<-HNH<-OLD-ABC<-DUF6079*<-BrxB-BrxF-iSTAND<-BrxA-DNA-binding<-HTH_26<-MazE_antitoxin+SpoVT_C<-?\|\|?-><-?<-RADICAL-SAM | 1184 | Bacteria>Terrabacteria_group>Bacillota>Clostridia>Eubacteriales>Oscillospiraceae | GCF_018783905.1 | Ruminococcus_sp._MCC718 | GPK60_RS07265 | 26956..30510 | 2592649 | BREX-system-P-loop-protein-BrxC-[Ruminococcus-sp.-MCC718] |
| CCX63523.1 | Type1.BREX | Phage_integrase-><-RelE-ParE<-MazE_antitoxin<-SIGMA+OB-Fold+AAA-ATPase+Lon-Protease-BrxL<-PglZ<-HNH<-OLD-ABC<-N6-METHYLASE<-DUF6079*<-BrxB-BrxF-iSTAND<-BrxA-DNA-binding<-XIS-HTH\|\|VirD4-TraG+VirD4-FtsK->?->?->ArdCN+MPTase-inact+LPD19->TraJ-RHH-> | 1217 | Bacteria>Terrabacteria_group>Bacillota>environmental_samples | GCA_000431495.1 | Firmicutes_bacterium_CAG:791 | BN785_00943 | 24533..28186 | 1262993 | uncharacterized-protein-BN785_00943-[Firmicutes-bacterium-CAG:791] |
| MBR3457117.1 | Type1.BREX | <-PglZ-p<-HNH<-OLD-ABC<-N6-METHYLASE-p<-?<-PIN<-N6-METHYLASE<-DUF6079*<-BrxB-BrxF-iSTAND-p | 1215 | Bacteria>Terrabacteria_group>Bacillota>Negativicutes>Selenomonadales>Selenomonadaceae | GCA_017509225.1 | Selenomonadaceae_bacterium | IKH16_03040 | 6412..10059 | 2652289 | BREX-system-P-loop-protein-BrxC-[Selenomonadaceae-bacterium] |
| WP_037354735.1 | Type1.BREX | DDE_Tnp_1-><-?<-?<-?\|\|?->BrxA-DNA-binding->BrxB-BrxF-iSTAND->DUF6079*->N6-METHYLASE->OLD-ABC->HNH->?->PglZ->SIGMA+OB-Fold+AAA-ATPase+Lon-Protease-BrxL->HTH_3-> | 1182 | Bacteria>Terrabacteria_group>Bacillota>Negativicutes>Selenomonadales>Selenomonadaceae | GCF_000701565.1 | Selenomonas_sp._FC4001 | T320_RS08130 | 15634..19182 | 1408313 | BREX-system-P-loop-protein-BrxC-[Selenomonas-sp.-FC4001] |
| RGH23082.1 | Type1.BREX | PemK_toxin->?->?->BrxA-DNA-binding->BrxB-BrxF-iSTAND->DUF6079*->N6-METHYLASE->OLD-ABC->HNH->PglZ-><-?\|\|?->?-><-ZnR | 1180 | Bacteria>Terrabacteria_group>Bacillota>unclassified_Bacillota | GCA_003461705.1 | Firmicutes_bacterium_AF12-30 | DWV72_07055 | 48691..52233 | 2292884 | BREX-system-P-loop-protein-BrxC-[Firmicutes-bacterium-AF12-30] |
| MBE9475406.1 | Type1.BREX | BrxB-BrxF-iSTAND-p->BrxA-DNA-binding->BrxB-BrxF-iSTAND->DUF6079-p*->N6-METHYLASE->REase+AIPR->OLD-ABC->HNH->PglZ->SIGMA+OB-Fold+AAA-ATPase+Lon-Protease-BrxL-> | 1145 | Bacteria>Terrabacteria_group>Chloroflexota>unclassified_Chloroflexota | GCA_015222395.1 | Chloroflexota_bacterium | IMY85_10995 | 2370..5807 | 2026724 | BREX-system-P-loop-protein-BrxC-[Chloroflexota-bacterium] |
| MBI4317923.1 | Type1.BREX | Radical_SAM->?->?->?->BrxA-DNA-binding->BrxB-BrxF-iSTAND->DUF6079*->N6-METHYLASE->OLD-ABC->HNH->PglZ->SIGMA+OB-Fold+AAA-ATPase+Lon-Protease-BrxL-> | 1145 | Bacteria>Terrabacteria_group>Chloroflexota>unclassified_Chloroflexota | GCA_016210495.1 | Chloroflexota_bacterium | HY675_05490 | 15525..18962 | 2026724 | BREX-system-P-loop-protein-BrxC-[Chloroflexota-bacterium] |
| MBT9546703.1 | Type1.BREX | Peptidase_M2->BrxA-DNA-binding->BrxB-BrxF-iSTAND->DUF6079*->N6-METHYLASE->RVT->N6-METHYLASE-p->OLD-ABC->HNH->PglZ->SIGMA+OB-Fold+AAA-ATPase+HKD-DNase-BrxL-> | 1179 | Bacteria>Terrabacteria_group>Cyanobacteriota/Melainabacteria_group>Candidatus_Sericytochromatia>unclassified_Candidatus_Sericytochromatia | GCA_018780035.1 | Candidatus_Sericytochromatia_bacterium | IV090_15030 | 94995..98534 | 2762020 | BREX-system-P-loop-protein-BrxC-[Candidatus-Sericytochromatia-bacterium] |
| WP_069067979.1 | Type1.BREX | CHTH->?->?->BrxA-DNA-binding->BrxB-BrxF-iSTAND->DUF6079*->?->N6-METHYLASE->OLD-ABC->HNH->PglZ->SIGMA+OB-Fold+AAA-ATPase+Lon-Protease-BrxL-><-?<-HTH_23 | 1220 | Bacteria>Terrabacteria_group>Cyanobacteriota/Melainabacteria_group>Cyanobacteriota>Cyanophyceae>Nostocales | GCF_001712795.1 | Nostoc_sp._KVJ20 | A4S05_RS00980 | 24778..28440 | 457944 | BREX-system-P-loop-protein-BrxC-[Nostoc-sp.-KVJ20] |
| WP_190429845.1 | Type1.BREX | Pkinase->HTH+HKD-DNase-><-SIGMA+OB-Fold+AAA-ATPase+Lon-Protease-BrxL<-PglZ<-HNH<-OLD-ABC<-N6-METHYLASE<-?<-DUF6079*<-BrxB-BrxF-iSTAND<-BrxA-DNA-binding\|\|ParA-Soj-PloopNTPase->CopG/RHH->?->?->Phage_integrase+Phage_integrase-><-TRPR-HTH | 1222 | Bacteria>Terrabacteria_group>Cyanobacteriota/Melainabacteria_group>Cyanobacteriota>Cyanophyceae>Oscillatoriophycideae | GCF_014695635.1 | Coleofasciculus_sp._FACHB-712 | H6F50_RS18615 | 18748..22416 | 2692789 | BREX-system-P-loop-protein-BrxC-[Coleofasciculus-sp.-FACHB-712] |
| KPQ32454.1 | Type1.BREX | BrxA-DNA-binding->BrxB-BrxF-iSTAND->DUF6079*->OLD-ABC->NERD->N6-METHYLASE->OLD-ABC->HNH->PglZ->SIGMA+OB-Fold+AAA-ATPase+HKD-DNase-BrxL-><-?\|\|DDE_Tnp-> | 1200 | Bacteria>Terrabacteria_group>Cyanobacteriota/Melainabacteria_group>Cyanobacteriota>Cyanophyceae>Pseudanabaenales | GCA_001314865.1 | Phormidesmis_priestleyi_Ana | HLUCCA11_21420 | 1901..5503 | 1666911 | Brex-phage-defense-system-ATPase-BrxC-[Phormidesmis-priestleyi-Ana] |
| HBH74800.1 | Type1.BREX | PIN->?->BrxA-DNA-binding->BrxB-BrxF-iSTAND->DUF6079*->N6-METHYLASE->SAM-METHYLASE->ParA-Soj-PloopNTPase->?->?->PglZ->OLD-ABC->HNH->SIGMA+OB-Fold+AAA-ATPase+Lon-Protease-BrxL-> | 1163 | Bacteria>Terrabacteria_group>Cyanobacteriota/Melainabacteria_group>Cyanobacteriota>Cyanophyceae>Synechococcales | GCA_003508015.1 | Synechococcales_bacterium_UBA10510 | DDY43_15550 | 2736..6227 | 2055795 | BREX-system-P-loop-protein-BrxC-[Synechococcales-bacterium-UBA10510] |
| WP_178365089.1 | Type1.BREX | <-DDE_Tnp\|\|?->?->PRTase->RADICAL-SAM->BrxA-DNA-binding->BrxB-BrxF-iSTAND->DUF6079*->N6-METHYLASE->OLD-ABC->HNH->PglZ->SIGMA+OB-Fold+AAA-ATPase+HKD-DNase-BrxL->tRNA-><-?<-PAS+NtrC-AAA+FIS-HTH\|\|?->REase-> | 1189 | Bacteria>Thermodesulfobacteriota>Desulfobacteria>Desulfobacterales>Desulfobacteraceae>Desulfobacter | GCF_013389365.1 | Desulfobacter_latus | HXW94_RS01255 | 9015..12584 | 2292 | BREX-system-P-loop-protein-BrxC-[Desulfobacter-latus] |
| WP_139446530.1 | Type1.BREX | <-SIGMA+OB-Fold+AAA-ATPase+Lon-Protease-BrxL\|\|DiS_P_DiS+Peptidase_A24-><-?\|\|?-><-SIGMA+OB-Fold+AAA-ATPase+HKD-DNase-BrxL<-PglZ<-HNH<-OLD-ABC<-N6-METHYLASE<-DUF6079*<-BrxB-BrxF-iSTAND<-BrxA-DNA-binding<-tRNA\|\|?->?-><-?<-HDRnase | 1199 | Bacteria>Thermodesulfobacteriota>Desulfobacteria>Desulfobacterales>Desulfobacteraceae>Desulfobotulus | GCF_006175995.1 | Desulfobotulus_mexicanus | FIM25_RS03945 | 130812..134411 | 2586642 | BREX-system-P-loop-protein-BrxC-[Desulfobotulus-mexicanus] |
| RZB38310.1 | Type1.BREX | <-TPR+TPR<-?\|\|?->?->?-><-?\|\|?->BrxA-DNA-binding->BrxB-BrxF-iSTAND->DUF6079*->N6-METHYLASE-p->N6-METHYLASE->OLD-ABC->HNH->PglZ->SIGMA+OB-Fold+AAA-ATPase+HKD-DNase-BrxL-><-?<-?<-ABC-ATPase+SbcC | 1173 | Bacteria>Thermodesulfobacteriota>Desulfobacteria>Desulfobacterales>Desulfobacteraceae>unclassified_Desulfobacteraceae | GCA_004193595.1 | Desulfobacteraceae_bacterium_Eth-SRB2 | SRB2_00058 | 60586..64107 | 2528438 | hypothetical-protein-SRB2_00058-[Desulfobacteraceae-bacterium-Eth-SRB2] |
| WP_155308948.1 | Type1.BREX | VirB10->?->DnaB+HerA/FtsK->TraH->TraG_N-><-?\|\|XIS-HTH->BrxA-DNA-binding->BrxB-BrxF-iSTAND->DUF6079*->OLD-ABC->HNH->N6-METHYLASE->N6-METHYLASE->REase+AIPR->PglZ->SIGMA+OB-Fold+AAA-ATPase+Lon-Protease-BrxL-><-HEPN | 1145 | Bacteria>Thermodesulfobacteriota>Desulfobacteria>Desulfobacterales>Desulfosarcinaceae>Desulfosarcina | GCF_009689005.1 | Desulfosarcina_ovata | GN112_RS03415 | 763655..767092 | 83564 | BREX-system-P-loop-protein-BrxC-[Desulfosarcina-ovata] |
| KGO34873.1 | Type1.BREX | MlaE->ABC-ATPase+SbcC->?-><-?\|\|XIS-HTH->BrxA-DNA-binding->BrxB-BrxF-iSTAND->DUF6079*->OLD-ABC->REase->N6-METHYLASE->?->OLD-ABC->HNH->PglZ->SIGMA+OB-Fold+AAA-ATPase+Lon-Protease-BrxL-> | 1146 | Bacteria>Thermodesulfobacteriota>Desulfobulbia>Desulfobulbales>Desulfobulbaceae>Desulfobulbus | GCA_000769715.1 | Desulfobulbus_sp._Tol-SR | JT06_05895 | 26559..29999 | 1536652 | hypothetical-protein-JT06_05895-[Desulfobulbus-sp.-Tol-SR] |
| WP_066605667.1 | Type1.BREX | HDRnase->?-><-?<-tRNA<-?\|\|?->?->BrxA-DNA-binding->BrxB-BrxF-iSTAND->DUF6079*->N6-METHYLASE->OLD-ABC->HNH->PglZ->SIGMA+OB-Fold+AAA-ATPase+Lon-Protease-BrxL-><-RNA_METHYLASE | 1145 | Bacteria>Thermodesulfobacteriota>Desulfovibrionia>Desulfovibrionales>Desulfomicrobiaceae>Desulfomicrobium | GCF_001553625.1 | Desulfomicrobium_orale | AXF15_RS07915 | 1725501..1728938 | 132132 | BREX-system-P-loop-protein-BrxC-[Desulfomicrobium-orale] |
| MCB6991564.1 | Type1.BREX | <-ACET<-?<-ABC-ATPase+SbcC<-ABC-ATPase+SbcC<-MarR-HTH\|\|?->HTH_26->BrxA-DNA-binding->BrxB-BrxF-iSTAND->DUF6079*->OLD-ABC->HNH->N6-METHYLASE->KAP-NTPase-> | 1183 | Bacteria>unclassified_Bacteria | GCA_020558425.1 | bacterium_210820-DFI.6.38 | LI187_13870 | 12055..15606 | 2883293 | BREX-system-P-loop-protein-BrxC-[bacterium-210820-DFI.6.38] |
| -- | | | | | | | | | | |
| --wHTH(SBD)+HNH | | | | | | | | | | |
| -- | | | | | | | | | | |
| WP_032621388.1 | Type1.BREX | <-SymE_toxin\|\|wHTH(SBD)+HNH-><-SIGMA+OB-Fold+AAA-ATPase+Lon-Protease-BrxL<-PglZ<-TOPRIM<-OLD-ABC<-N6-METHYLASE<-DUF6079*<-BrxB-BrxF-iSTAND<-BrxA-DNA-binding\|\|ZnR-><-tRNA<-?<-?<-?\|\|?->?->ICLR-HTH-> | 1204 | Bacteria>Pseudomonadota>Gammaproteobacteria>Enterobacterales>Enterobacteriaceae>Enterobacter | GCF_023734955.1 | Enterobacter_hormaechei | NVR78_RS04785 | 326171..329785 | 158836 | BREX-system-P-loop-protein-BrxC-[Enterobacter-hormaechei] |
| WP_180264800.1 | Type1.BREX | <-HTH_36<-HTH+HTH<-ParB+BAM<-XIS-HTH<-wHTH<-ZnR\|\|BrxA-DNA-binding->BrxB-BrxF-iSTAND->DUF6079*->N6-METHYLASE->OLD-ABC->TOPRIM->PglZ->SIGMA+OB-Fold+AAA-ATPase+Lon-Protease-BrxL-><-wHTH(SBD)+HNH\|\|SymE_toxin-> | 1204 | Bacteria>Pseudomonadota>Gammaproteobacteria>Enterobacterales>Enterobacteriaceae>Enterobacter | GCF_900558725.1 | Enterobacter_hormaechei | H0848_RS18555 | 27730..31344 | 158836 | BREX-system-P-loop-protein-BrxC-[Enterobacter-hormaechei] |
| WP_063851018.1 | Type1.BREX | <-HTH_36<-HTH+HTH<-ParB+BAM<-XIS-HTH<-wHTH<-ZnR\|\|BrxA-DNA-binding->BrxB-BrxF-iSTAND->DUF6079*->N6-METHYLASE->OLD-ABC->TOPRIM->PglZ->SIGMA+OB-Fold+AAA-ATPase+Lon-Protease-BrxL-><-wHTH(SBD)+HNH\|\|SymE_toxin-> | 1204 | Bacteria>Pseudomonadota>Gammaproteobacteria>Enterobacterales>Enterobacteriaceae>Enterobacter | GCF_902160625.1 | Enterobacter_hormaechei | FN214_RS22220 | 26549..30163 | 158836 | BREX-system-P-loop-protein-BrxC-[Enterobacter-hormaechei] |
| GCK20492.1 | Type1.BREX | <-Fimbrial<-Phage_integrase<-Phage_integrase\|\|?->?->?->wHTH(SBD)+HNH->BrxA-DNA-binding->BrxB-BrxF-iSTAND->DUF6079-p*-> | 621 | Bacteria>Pseudomonadota>Gammaproteobacteria>Enterobacterales>Enterobacteriaceae>Escherichia | GCA_005401165.1 | Escherichia_coli | BvCmsA58A_02120 | 111143..113008 | 562 | ATPase-like-protein-[Escherichia-coli] |
| EEU4096206.1 | Type1.BREX | <-DUF6079-p*<-BrxB-BrxF-iSTAND<-BrxA-DNA-binding\|\|<-wHTH(SBD)+HNH\|\|HNH-><-?<-?<-?\|\|Phage_integrase->Phage_integrase-> | 433 | Bacteria>Pseudomonadota>Gammaproteobacteria>Enterobacterales>Enterobacteriaceae>Escherichia | GCA_011974935.1 | Escherichia_coli | E8H53_004589 | 1..1299 | 562 | BREX-system-P-loop-protein-BrxC-[Escherichia-coli] |
| EEV1146578.1 | Type1.BREX | <-Phage_integrase\|\|?->?->?-><-HNH\|\|wHTH(SBD)+HNH->\|\|BrxA-DNA-binding->BrxB-BrxF-iSTAND->DUF6079*->N6-METHYLASE-> | 1213 | Bacteria>Pseudomonadota>Gammaproteobacteria>Enterobacterales>Enterobacteriaceae>Escherichia | GCA_012013845.1 | Escherichia_coli | EUB49_004354 | 8713..12354 | 562 | BREX-system-P-loop-protein-BrxC-[Escherichia-coli] |
| EEW3089662.1 | Type1.BREX | <-Phage_integrase<-Phage_integrase\|\|?->?->?-><-HNH\|\|wHTH(SBD)+HNH->\|\|BrxA-DNA-binding->BrxB-BrxF-iSTAND->DUF6079*->N6-METHYLASE->PglZ->SIGMA+OB-Fold+AAA-ATPase+Lon-Protease-BrxL->HKD-DNase+OGG-like->\|\|AbiEi_3_N+AbiEi_3->AbiEii-><-DUF6216<-Arm-DNA-bind_3 | 1213 | Bacteria>Pseudomonadota>Gammaproteobacteria>Enterobacterales>Enterobacteriaceae>Escherichia | GCA_012078325.1 | Escherichia_coli | EBA98_002639 | 14697..18338 | 562 | BREX-system-P-loop-protein-BrxC-[Escherichia-coli] |
| EEW3885152.1 | Type1.BREX | <-HNH\|\|wHTH(SBD)+HNH->\|\|BrxA-DNA-binding->BrxB-BrxF-iSTAND->DUF6079*->N6-METHYLASE->PglZ->SIGMA+OB-Fold+AAA-ATPase+Lon-Protease-BrxL->HKD-DNase+OGG-like->\|\|AbiEi_3_N+AbiEi_3->AbiEii-><-DUF6216<-Arm-DNA-bind_3 | 1213 | Bacteria>Pseudomonadota>Gammaproteobacteria>Enterobacterales>Enterobacteriaceae>Escherichia | GCA_012080795.1 | Escherichia_coli | D9Y07_001392 | 7166..10807 | 562 | BREX-system-P-loop-protein-BrxC-[Escherichia-coli] |

| Protein-ID | System | Gene-Neighbourhoods | Length | Taxonomy | Assembly-ID | Strain | Locus-Tag | Genomic-coordinates | TaxID | Fasta-define |
|---|---|---|---|---|---|---|---|---|---|---|
| EEX1719682.1 | Type1.BREX | <-HNH\|\|wHTH(SBD)+HNH->\|\|BrxA-DNA-binding->BrxB-BrxF-iSTAND->DUF6079*->N6-METHYLASE->PglZ->SIGMA+OB-Fold+AAA-ATPase+Lon-Protease-BrxL->HKD-DNase+OGG-like->\|\|AbiEi_3_N+AbiEi_3->AbiEii-><-DUF6216<-Arm-DNA-bind_3 | 1213 | Bacteria>Pseudomonadota>Gammaproteobacteria>Enterobacterales>Enterobacteriaceae>Escherichia | GCA_012115095.1 | Escherichia_coli | D3S79_003407 | 6685..10326 | 562 | BREX-system-P-loop-protein-BrxC-[Escherichia-coli] |
| EEX6733972.1 | Type1.BREX | <-DUF6079*<-BrxB-BrxF-iSTAND<-BrxA-DNA-binding\|\|<-wHTH(SBD)+HNH\|\|HNH-><-?<-?<-?\|\|Phage_integrase->Phage_integrase-> | 1175 | Bacteria>Pseudomonadota>Gammaproteobacteria>Enterobacterales>Enterobacteriaceae>Escherichia | GCA_012135265.1 | Escherichia_coli | L0765_004693 | 1..3524 | 562 | BREX-system-P-loop-protein-BrxC-[Escherichia-coli] |
| EEZ2582139.1 | Type1.BREX | <-DUF6079-p*<-BrxB-BrxF-iSTAND<-BrxA-DNA-binding\|\|<-wHTH(SBD)+HNH\|\|HNH-><-?<-?<-?\|\|Phage_integrase->Phage_integrase-> | 800 | Bacteria>Pseudomonadota>Gammaproteobacteria>Enterobacterales>Enterobacteriaceae>Escherichia | GCA_012201865.1 | Escherichia_coli | DMH54_003958 | 1..2402 | 562 | BREX-system-P-loop-protein-BrxC-[Escherichia-coli] |
| EEZ3215399.1 | Type1.BREX | <-HNH\|\|wHTH(SBD)+HNH->\|\|BrxA-DNA-binding->BrxB-BrxF-iSTAND->DUF6079*->N6-METHYLASE-> | 1213 | Bacteria>Pseudomonadota>Gammaproteobacteria>Enterobacterales>Enterobacteriaceae>Escherichia | GCA_012202545.1 | Escherichia_coli | DKV48_004298 | 6695..10336 | 562 | BREX-system-P-loop-protein-BrxC-[Escherichia-coli] |
| EFC3446201.1 | Type1.BREX | <-DUF6079-p*<-BrxB-BrxF-iSTAND<-BrxA-DNA-binding\|\|<-wHTH(SBD)+HNH\|\|HNH-><-?<-?<-?\|\|Phage_integrase->Phage_integrase-> | 844 | Bacteria>Pseudomonadota>Gammaproteobacteria>Enterobacterales>Enterobacteriaceae>Escherichia | GCA_012346015.1 | Escherichia_coli | CO751_004598 | 1..2534 | 562 | BREX-system-P-loop-protein-BrxC-[Escherichia-coli] |
| EFD3459874.1 | Type1.BREX | <-HNH\|\|wHTH(SBD)+HNH->\|\|BrxA-DNA-binding->BrxB-BrxF-iSTAND->DUF6079*->N6-METHYLASE->PglZ->SIGMA+OB-Fold+AAA-ATPase+Lon-Protease-BrxL-> | 1213 | Bacteria>Pseudomonadota>Gammaproteobacteria>Enterobacterales>Enterobacteriaceae>Escherichia | GCA_012389335.1 | Escherichia_coli | CL211_004302 | 6759..10400 | 562 | BREX-system-P-loop-protein-BrxC-[Escherichia-coli] |
| EFE1965719.1 | Type1.BREX | <-Phage_integrase<-Phage_integrase\|\|?->?->?-><-HNH\|\|wHTH(SBD)+HNH->\|\|BrxA-DNA-binding->BrxB-BrxF-iSTAND->DUF6079*->N6-METHYLASE->PglZ->SIGMA+OB-Fold+AAA-ATPase+Lon-Protease-BrxL->HKD-DNase+OGG-like->\|\|AbiEi_3_N+AbiEi_3->AbiEii-><-DUF6216<-Arm-DNA-bind_3 | 1213 | Bacteria>Pseudomonadota>Gammaproteobacteria>Enterobacterales>Enterobacteriaceae>Escherichia | GCA_012428445.1 | Escherichia_coli | CJV81_002667 | 14696..18337 | 562 | BREX-system-P-loop-protein-BrxC-[Escherichia-coli] |
| EFG4404338.1 | Type1.BREX | <-Phage_integrase<-Phage_integrase\|\|?->?->?-><-HNH\|\|wHTH(SBD)+HNH->\|\|BrxA-DNA-binding->BrxB-BrxF-iSTAND->DUF6079*->N6-METHYLASE->PglZ->SIGMA+OB-Fold+AAA-ATPase+Lon-Protease-BrxL->HKD-DNase+OGG-like->\|\|AbiEi_3_N+AbiEi_3->AbiEii-><-DUF6216<-Arm-DNA-bind_3 | 1213 | Bacteria>Pseudomonadota>Gammaproteobacteria>Enterobacterales>Enterobacteriaceae>Escherichia | GCA_012556825.1 | Escherichia_coli | BM772_002472 | 14697..18338 | 562 | BREX-system-P-loop-protein-BrxC-[Escherichia-coli] |
| EFH0549024.1 | Type1.BREX | <-Phage_integrase<-Phage_integrase\|\|?->?->?-><-HNH\|\|wHTH(SBD)+HNH->\|\|BrxA-DNA-binding->BrxB-BrxF-iSTAND->DUF6079-p*-> | 422 | Bacteria>Pseudomonadota>Gammaproteobacteria>Enterobacterales>Enterobacteriaceae>Escherichia | GCA_012582395.1 | Escherichia_coli | BJE23_003837 | 39992..41258 | 562 | BREX-system-P-loop-protein-BrxC-[Escherichia-coli] |
| EGY5620863.1 | Type1.BREX | <-Phage_integrase<-Phage_integrase\|\|?->?->?-><-HNH\|\|wHTH(SBD)+HNH->\|\|BrxA-DNA-binding->BrxB-BrxF-iSTAND->DUF6079*->N6-METHYLASE->PglZ->SIGMA+OB-Fold+AAA-ATPase+Lon-Protease-BrxL->HKD-DNase+OGG-like->\|\|AbiEi_3_N+AbiEi_3->AbiEii-><-DUF6216<-Arm-DNA-bind_3 | 1213 | Bacteria>Pseudomonadota>Gammaproteobacteria>Enterobacterales>Enterobacteriaceae>Escherichia | GCA_016519195.1 | Escherichia_coli | JI433_002890 | 14696..18337 | 562 | BREX-system-P-loop-protein-BrxC-[Escherichia-coli] |
| EGY7451904.1 | Type1.BREX | <-DUF6079-p*<-BrxB-BrxF-iSTAND<-BrxA-DNA-binding\|\|<-wHTH(SBD)+HNH\|\|HNH-><-?<-?<-?\|\|Phage_integrase->Phage_integrase-> | 968 | Bacteria>Pseudomonadota>Gammaproteobacteria>Enterobacterales>Enterobacteriaceae>Escherichia | GCA_016533805.1 | Escherichia_coli | JI519_004377 | 1..2904 | 562 | BREX-system-P-loop-protein-BrxC-[Escherichia-coli] |
| EGY8948619.1 | Type1.BREX | <-DUF6079-p*<-BrxB-BrxF-iSTAND<-BrxA-DNA-binding\|\|<-wHTH(SBD)+HNH\|\|HNH-><-?<-?<-?\|\|Phage_integrase->Phage_integrase-> | 450 | Bacteria>Pseudomonadota>Gammaproteobacteria>Enterobacterales>Enterobacteriaceae>Escherichia | GCA_016541865.1 | Escherichia_coli | JI613_004635 | 1..1349 | 562 | BREX-system-P-loop-protein-BrxC-[Escherichia-coli] |
| EGY9306082.1 | Type1.BREX | <-HNH\|\|wHTH(SBD)+HNH->\|\|BrxA-DNA-binding->BrxB-BrxF-iSTAND->DUF6079*->N6-METHYLASE->PglZ-> | 1213 | Bacteria>Pseudomonadota>Gammaproteobacteria>Enterobacterales>Enterobacteriaceae>Escherichia | GCA_016543505.1 | Escherichia_coli | HZD96_004640 | 4089..7730 | 562 | BREX-system-P-loop-protein-BrxC-[Escherichia-coli] |
| EGZ1560300.1 | Type1.BREX | <-DUF6079-p*<-BrxB-BrxF-iSTAND<-BrxA-DNA-binding\|\|<-wHTH(SBD)+HNH\|\|HNH-><-?<-?<-?\|\|Phage_integrase->Phage_integrase-> | 444 | Bacteria>Pseudomonadota>Gammaproteobacteria>Enterobacterales>Enterobacteriaceae>Escherichia | GCA_016553465.1 | Escherichia_coli | FAV16_004661 | 1..1332 | 562 | BREX-system-P-loop-protein-BrxC-[Escherichia-coli] |
| EGZ1871551.1 | Type1.BREX | <-DUF6079-p*<-BrxB-BrxF-iSTAND<-BrxA-DNA-binding\|\|<-wHTH(SBD)+HNH\|\|HNH-><-?<-?<-?\|\|Phage_integrase->Phage_integrase-> | 416 | Bacteria>Pseudomonadota>Gammaproteobacteria>Enterobacterales>Enterobacteriaceae>Escherichia | GCA_016554585.1 | Escherichia_coli | EYE54_004341 | 1..1248 | 562 | BREX-system-P-loop-protein-BrxC-[Escherichia-coli] |
| EGZ2189929.1 | Type1.BREX | <-DUF6079-p*<-BrxB-BrxF-iSTAND<-BrxA-DNA-binding\|\|<-wHTH(SBD)+HNH\|\|HNH-><-?<-?<-?\|\|Phage_integrase-> | 628 | Bacteria>Pseudomonadota>Gammaproteobacteria>Enterobacterales>Enterobacteriaceae>Escherichia | GCA_016555885.1 | Escherichia_coli | ED609_004465 | 1..1884 | 562 | BREX-system-P-loop-protein-BrxC-[Escherichia-coli] |
| EGZ2354229.1 | Type1.BREX | <-Phage_integrase<-Phage_integrase\|\|?->?->?-><-HNH\|\|wHTH(SBD)+HNH->\|\|BrxA-DNA-binding->BrxB-BrxF-iSTAND->DUF6079*->N6-METHYLASE->PglZ->SIGMA+OB-Fold+AAA-ATPase+Lon-Protease-BrxL->HKD-DNase+OGG-like->\|\|AbiEi_3_N+AbiEi_3->AbiEii-><-DUF6216<-Arm-DNA-bind_3 | 1213 | Bacteria>Pseudomonadota>Gammaproteobacteria>Enterobacterales>Enterobacteriaceae>Escherichia | GCA_016556665.1 | Escherichia_coli | D7J47_002044 | 14697..18338 | 562 | BREX-system-P-loop-protein-BrxC-[Escherichia-coli] |
| EGZ2777080.1 | Type1.BREX | <-Phage_integrase<-Phage_integrase\|\|?->?->?-><-HNH\|\|wHTH(SBD)+HNH->\|\|BrxA-DNA-binding->BrxB-BrxF-iSTAND->DUF6079*->N6-METHYLASE->PglZ->SIGMA+OB-Fold+AAA-ATPase+Lon-Protease-BrxL->HKD-DNase+OGG-like->\|\|AbiEi_3_N+AbiEi_3->AbiEii-><-DUF6216<-Arm-DNA-bind_3 | 1213 | Bacteria>Pseudomonadota>Gammaproteobacteria>Enterobacterales>Enterobacteriaceae>Escherichia | GCA_016558225.1 | Escherichia_coli | DVO41_002389 | 14697..18338 | 562 | BREX-system-P-loop-protein-BrxC-[Escherichia-coli] |
| EGZ3242782.1 | Type1.BREX | <-Phage_integrase<-Phage_integrase\|\|?->?->?-><-HNH\|\|wHTH(SBD)+HNH->\|\|BrxA-DNA-binding->BrxB-BrxF-iSTAND->DUF6079*->N6-METHYLASE->PglZ->SIGMA+OB-Fold+AAA-ATPase+Lon-Protease-BrxL->HKD-DNase+OGG-like->\|\|AbiEi_3_N+AbiEi_3->AbiEii-><-DUF6216<-Arm-DNA-bind_3 | 1213 | Bacteria>Pseudomonadota>Gammaproteobacteria>Enterobacterales>Enterobacteriaceae>Escherichia | GCA_016560025.1 | Escherichia_coli | C5N70_002404 | 39991..43632 | 562 | BREX-system-P-loop-protein-BrxC-[Escherichia-coli] |
| EGZ5166758.1 | Type1.BREX | <-DUF6079-p*<-BrxB-BrxF-iSTAND<-BrxA-DNA-binding\|\|<-wHTH(SBD)+HNH\|\|HNH-><-?<-?<-?\|\|Phage_integrase->Phage_integrase-> | 624 | Bacteria>Pseudomonadota>Gammaproteobacteria>Enterobacterales>Enterobacteriaceae>Escherichia | GCA_016568175.1 | Escherichia_coli | CDS35_004366 | 1..1874 | 562 | BREX-system-P-loop-protein-BrxC-[Escherichia-coli] |
| EGZ5591347.1 | Type1.BREX | <-DUF6079-p*<-BrxB-BrxF-iSTAND<-BrxA-DNA-binding\|\|<-wHTH(SBD)+HNH\|\|HNH-><-?<-?<-?\|\|Phage_integrase->Phage_integrase-> | 707 | Bacteria>Pseudomonadota>Gammaproteobacteria>Enterobacterales>Enterobacteriaceae>Escherichia | GCA_016569835.1 | Escherichia_coli | BZZ56_004473 | 1..2121 | 562 | BREX-system-P-loop-protein-BrxC-[Escherichia-coli] |
| EGZ5731917.1 | Type1.BREX | <-DUF6079-p*<-BrxB-BrxF-iSTAND<-BrxA-DNA-binding\|\|<-wHTH(SBD)+HNH\|\|HNH-><-?<-?<-?\|\|Phage_integrase->Phage_integrase-> | 885 | Bacteria>Pseudomonadota>Gammaproteobacteria>Enterobacterales>Enterobacteriaceae>Escherichia | GCA_016570515.1 | Escherichia_coli | BL334_004670 | 1..2655 | 562 | BREX-system-P-loop-protein-BrxC-[Escherichia-coli] |
| EGZ5978150.1 | Type1.BREX | <-Phage_integrase<-Phage_integrase\|\|?->?->?-><-HNH\|\|wHTH(SBD)+HNH->\|\|BrxA-DNA-binding->BrxB-BrxF-iSTAND->DUF6079-p*-> | 421 | Bacteria>Pseudomonadota>Gammaproteobacteria>Enterobacterales>Enterobacteriaceae>Escherichia | GCA_016571515.1 | Escherichia_coli | BIB10_004104 | 39992..41255 | 562 | BREX-system-P-loop-protein-BrxC-[Escherichia-coli] |
| EGZ8838308.1 | Type1.BREX | <-Phage_integrase\|\|?->?->?-><-HNH\|\|wHTH(SBD)+HNH->\|\|BrxA-DNA-binding->BrxB-BrxF-iSTAND->DUF6079*->N6-METHYLASE->PglZ->SIGMA+OB-Fold+AAA-ATPase+Lon-Protease-BrxL->HKD-DNase+OGG-like->\|\|AbiEi_3_N+AbiEi_3->AbiEii-><-DUF6216<-Arm-DNA-bind_3 | 1213 | Bacteria>Pseudomonadota>Gammaproteobacteria>Enterobacterales>Enterobacteriaceae>Escherichia | GCA_016590495.1 | Escherichia_coli | AT823_000493 | 8719..12360 | 562 | BREX-system-P-loop-protein-BrxC-[Escherichia-coli] |
| EHA9087556.1 | Type1.BREX | <-Phage_integrase<-Phage_integrase\|\|?->?->?-><-HNH\|\|wHTH(SBD)+HNH->\|\|BrxA-DNA-binding->BrxB-BrxF-iSTAND->DUF6079*->N6-METHYLASE->PglZ->SIGMA+OB-Fold+AAA-ATPase+Lon-Protease-BrxL->HKD-DNase+OGG-like->\|\|AbiEi_3_N+AbiEi_3->AbiEii-><-DUF6216<-Arm-DNA-bind_3 | 1213 | Bacteria>Pseudomonadota>Gammaproteobacteria>Enterobacterales>Enterobacteriaceae>Escherichia | GCA_016846225.1 | Escherichia_coli | D4094_002230 | 14699..18340 | 562 | BREX-system-P-loop-protein-BrxC-[Escherichia-coli] |
| EHJ8640897.1 | Type1.BREX | <-Phage_integrase<-Phage_integrase\|\|?->?->?-><-HNH\|\|wHTH(SBD)+HNH->\|\|BrxA-DNA-binding->BrxB-BrxF-iSTAND->DUF6079*->N6-METHYLASE->PglZ->SIGMA+OB-Fold+AAA-ATPase+Lon-Protease-BrxL->HKD-DNase+OGG-like->\|\|AbiEi_3_N+AbiEi_3->AbiEii->?-><-DUF6216<-Arm-DNA-bind_3 | 1213 | Bacteria>Pseudomonadota>Gammaproteobacteria>Enterobacterales>Enterobacteriaceae>Escherichia | GCA_017945505.1 | Escherichia_coli | KB366_000307 | 330961..334602 | 562 | BREX-system-P-loop-protein-BrxC-[Escherichia-coli] |
| EHL0175571.1 | Type1.BREX | <-DUF6079*<-BrxB-BrxF-iSTAND<-BrxA-DNA-binding\|\|<-wHTH(SBD)+HNH\|\|HNH-><-?<-?<-?\|\|Phage_integrase->Phage_integrase-> | 1016 | Bacteria>Pseudomonadota>Gammaproteobacteria>Enterobacterales>Enterobacteriaceae>Escherichia | GCA_018090545.1 | Escherichia_coli | D7K57_004368 | 1..3050 | 562 | BREX-system-P-loop-protein-BrxC-[Escherichia-coli] |
| EHL0676199.1 | Type1.BREX | <-Phage_integrase<-Phage_integrase\|\|?->?->?-><-HNH\|\|wHTH(SBD)+HNH->\|\|BrxA-DNA-binding->BrxB-BrxF-iSTAND->DUF6079*->N6-METHYLASE-> | 1213 | Bacteria>Pseudomonadota>Gammaproteobacteria>Enterobacterales>Enterobacteriaceae>Escherichia | GCA_018092345.1 | Escherichia_coli | CP667_004253 | 9829..13470 | 562 | BREX-system-P-loop-protein-BrxC-[Escherichia-coli] |
| EHL1134570.1 | Type1.BREX | <-DUF6079-p*<-BrxB-BrxF-iSTAND<-BrxA-DNA-binding\|\|<-wHTH(SBD)+HNH\|\|HNH-><-?<-?<-?\|\|Phage_integrase->Phage_integrase-> | 460 | Bacteria>Pseudomonadota>Gammaproteobacteria>Enterobacterales>Enterobacteriaceae>Escherichia | GCA_018094045.1 | Escherichia_coli | BIB01_004961 | 1..1380 | 562 | BREX-system-P-loop-protein-BrxC-[Escherichia-coli] |
| EHL1162648.1 | Type1.BREX | <-Phage_integrase<-Phage_integrase\|\|?->?->?-><-HNH\|\|wHTH(SBD)+HNH->\|\|BrxA-DNA-binding->BrxB-BrxF-iSTAND->DUF6079*->N6-METHYLASE->PglZ->SIGMA+OB-Fold+AAA-ATPase+Lon-Protease-BrxL->HKD-DNase+OGG-like->\|\|AbiEi_3_N+AbiEi_3->AbiEii->?-><-DUF6216<-Arm-DNA-bind_3 | 1213 | Bacteria>Pseudomonadota>Gammaproteobacteria>Enterobacterales>Enterobacteriaceae>Escherichia | GCA_018094265.1 | Escherichia_coli | BE892_002471 | 14698..18339 | 562 | BREX-system-P-loop-protein-BrxC-[Escherichia-coli] |
| EHN3342070.1 | Type1.BREX | <-Phage_integrase<-Phage_integrase\|\|?->?->?-><-HNH\|\|wHTH(SBD)+HNH->\|\|BrxA-DNA-binding->BrxB-BrxF-iSTAND->DUF6079*->N6-METHYLASE->PglZ->SIGMA+OB-Fold+AAA-ATPase+Lon-Protease-BrxL->HKD-DNase+OGG-like->\|\|AbiEi_3_N+AbiEi_3->AbiEii->?-><-DUF6216<-Arm-DNA-bind_3 | 1213 | Bacteria>Pseudomonadota>Gammaproteobacteria>Enterobacterales>Enterobacteriaceae>Escherichia | GCA_018356405.1 | Escherichia_coli | KIR71_003047 | 14697..18338 | 562 | BREX-system-P-loop-protein-BrxC-[Escherichia-coli] |
| EHY5690081.1 | Type1.BREX | <-Phage_integrase<-Phage_integrase\|\|?->?->?-><-HNH\|\|wHTH(SBD)+HNH->\|\|BrxA-DNA-binding->BrxB-BrxF-iSTAND->DUF6079*->N6-METHYLASE->PglZ->SIGMA+OB-Fold+AAA-ATPase+Lon-Protease-BrxL->HKD-DNase+OGG-like->\|\|AbiEi_3_N+AbiEi_3->AbiEii->?-><-DUF6216<-Arm-DNA-bind_3 | 1213 | Bacteria>Pseudomonadota>Gammaproteobacteria>Enterobacterales>Enterobacteriaceae>Escherichia | GCA_019590565.1 | Escherichia_coli | K4J54_002559 | 14696..18337 | 562 | BREX-system-P-loop-protein-BrxC-[Escherichia-coli] |
| EIB0293606.1 | Type1.BREX | <-DUF6079-p*<-BrxB-BrxF-iSTAND<-BrxA-DNA-binding\|\|<-wHTH(SBD)+HNH\|\|HNH-><-?<-?<-?\|\|Phage_integrase->Phage_integrase-> | 529 | Bacteria>Pseudomonadota>Gammaproteobacteria>Enterobacterales>Enterobacteriaceae>Escherichia | GCA_019848435.1 | Escherichia_coli | K7644_004610 | 1..1589 | 562 | BREX-system-P-loop-protein-BrxC-[Escherichia-coli] |
| EIG6470759.1 | Type1.BREX | <-DUF6079-p*<-BrxB-BrxF-iSTAND<-BrxA-DNA-binding\|\|<-wHTH(SBD)+HNH\|\|HNH-><-?<-?<-?\|\|Phage_integrase->Phage_integrase-> | 634 | Bacteria>Pseudomonadota>Gammaproteobacteria>Enterobacterales>Enterobacteriaceae>Escherichia | GCA_020319175.1 | Escherichia_coli | EG665_005071 | 1..1903 | 562 | BREX-system-P-loop-protein-BrxC-[Escherichia-coli] |
| HBP9919082.1 | Type1.BREX | <-Phage_integrase<-Phage_integrase\|\|?->?->?-><-HNH\|\|wHTH(SBD)+HNH->\|\|BrxA-DNA-binding->BrxB-BrxF-iSTAND->DUF6079*->N6-METHYLASE->PglZ->SIGMA+OB-Fold+AAA-ATPase+Lon-Protease-BrxL->HKD-DNase+OGG-like->\|\|AbiEi_3_N+AbiEi_3->AbiEii->?-><-DUF6216<-Arm-DNA-bind_3 | 1213 | Bacteria>Pseudomonadota>Gammaproteobacteria>Enterobacterales>Enterobacteriaceae>Escherichia | GCA_021766545.1 | Escherichia_coli | L7G70_002290 | 14696..18337 | 562 | BREX-system-P-loop-protein-BrxC-[Escherichia-coli] |
| EJW3212697.1 | Type1.BREX | <-DUF6079-p*<-BrxB-BrxF-iSTAND<-BrxA-DNA-binding\|\|<-wHTH(SBD)+HNH\|\|HNH-><-?<-?<-?\|\|Phage_integrase->Phage_integrase-> | 418 | Bacteria>Pseudomonadota>Gammaproteobacteria>Enterobacterales>Enterobacteriaceae>Escherichia | GCA_025418235.1 | Escherichia_coli | OBZ04_004372 | 1..1254 | 562 | BREX-system-P-loop-protein-BrxC-[Escherichia-coli] |
| WP_109548991.1 | Type1.BREX | <-Fimbrial<-Phage_integrase<-Phage_integrase\|\|?->?->?->\|\|wHTH(SBD)+HNH->\|\|BrxA-DNA-binding->BrxB-BrxF-iSTAND->DUF6079*->N6-METHYLASE->PglZ->SIGMA+OB-Fold+AAA-ATPase+Lon-Protease-BrxL->HKD-DNase+OGG-like->\|\|AbiEi_3_N+AbiEi_3->AbiEii-><-DUF6216 | 1213 | Bacteria>Pseudomonadota>Gammaproteobacteria>Enterobacterales>Enterobacteriaceae>Escherichia | GCF_003145675.1 | Escherichia_coli | B6S34_RS22850 | 50490..54131 | 562 | BREX-system-P-loop-protein-BrxC-[Escherichia-coli] |
| EIY5117082.1 | Type1.BREX | <-Calcineurin<-?<-?<-?<-?<-?\|\|tRNA-><-ZnR\|\|BrxA-DNA-binding->BrxB-BrxF-iSTAND->DUF6079*->N6-METHYLASE->OLD-ABC->TOPRIM->PglZ->SIGMA+OB-Fold+AAA-ATPase+Lon-Protease-BrxL-><-wHTH(SBD)+HNH\|\|MAD-NTD+HNH-><-Glyco_hydro_43+GH43_C2 | 1197 | Bacteria>Pseudomonadota>Gammaproteobacteria>Enterobacterales>Enterobacteriaceae>Klebsiella/Raoultella_group | GCA_022503905.1 | Klebsiella_quasipneumoniae | ML076_004996 | 20031..23624 | 1463165 | BREX-system-P-loop-protein-BrxC-[Klebsiella-quasipneumoniae] |
| EAY5508618.1 | Type1.BREX | <-wHTH\|\|wHTH+WYL+WCX-><-MarR-wHTH<-?<-SIGMA-HTH+REase<-wHTH(SBD)+HNH\|\|?->BrxA-DNA-binding->BrxB-BrxF-iSTAND->DUF6079*->N6-METHYLASE->OLD-ABC->TOPRIM->PglZ->SIGMA+OB-Fold+AAA-ATPase+Lon-Protease-BrxL->HKD-DNase+OGG-like-><-REase<-NACHT+SF1-SF2-Helicase+HKD-DNase<-Arm-DNA-bind_3 | 1204 | Bacteria>Pseudomonadota>Gammaproteobacteria>Enterobacterales>Enterobacteriaceae>Salmonella | GCA_006012725.1 | Salmonella_enterica | NE47_11015 | 49392..53006 | 28901 | BREX-system-P-loop-protein-BrxC-[Salmonella-enterica] |
| EIC8525255.1 | Type1.BREX | <-REase+iREase<-?\|\|YjhX_toxin<-SAM-METHYLASE->?-><-wHTH(SBD)+HNH\|\|?->BrxA-DNA-binding->BrxB-BrxF-iSTAND->DUF6079*->N6-METHYLASE->OLD-ABC->TOPRIM->PglZ->SIGMA+OB-Fold+AAA-ATPase+Lon-Protease-BrxL-><-REase<-NACHT+SF1-SF2-Helicase+HKD-DNase<-Arm-DNA-bind_3 | 1204 | Bacteria>Pseudomonadota>Gammaproteobacteria>Enterobacterales>Enterobacteriaceae>Salmonella | GCA_019993965.1 | Salmonella_enterica | LAR10_002215 | 30447..34061 | 28901 | BREX-system-P-loop-protein-BrxC-[Salmonella-enterica] |
| WP_040031759.1 | Type1.BREX | <-HIN-HTH\|\|DUF4096+DDE_Tnp_1+DDE_Tnp_1_2->?->tRNA->tRNA->Arm-DNA-bind_3+Phage_integrase-><-SIGMA+OB-Fold+AAA-ATPase+Lon-Protease-BrxL<-PglZ<-TOPRIM<-OLD-ABC<-N6-METHYLASE<-DUF6079*<-BrxB-BrxF-iSTAND<-BrxA-DNA-binding<-wHTH(SBD)+HNH<-LuxR-HTH<-?<-TPR | 1202 | Bacteria>Pseudomonadota>Gammaproteobacteria>Enterobacterales>Pectobacteriaceae | GCF_026723725.1 | Pectobacterium | OWC53_RS08230 | 1782306..1785914 | 122277 | MULTISPECIES:-BREX-system-P-loop-protein-BrxC-[Pectobacterium] |
| GKX39595.1 | Type1.BREX | Arm-DNA-bind_3+Phage_integrase-><-SIGMA+OB-Fold+AAA-ATPase+Lon-Protease-BrxL<-PglZ<-TOPRIM<-OLD-ABC<-N6-METHYLASE<-DUF6079*<-BrxB-BrxF-iSTAND<-BrxA-DNA-binding<-wHTH(SBD)+HNH<-LuxR-HTH<-?<-TPR | 1202 | Bacteria>Pseudomonadota>Gammaproteobacteria>Enterobacterales>Pectobacteriaceae>Pectobacterium | GCA_024343445.1 | Pectobacterium_carotovorum_subsp._carotovorum | SOASR014_33340 | 124830..128438 | 555 | hypothetical-protein-SOASR014_33340-[Pectobacterium-carotovorum-subsp.-carotovorum] |
| WP_103199124.1 | Type1.BREX | <-ParA-Soj-PloopNTPase<-XIS-HTH<-MODE-HTH+PBPII\|\|tRNA->tRNA->Arm-DNA-bind_3+Phage_integrase-><-SIGMA+OB-Fold+AAA-ATPase+Lon-Protease-BrxL<-PglZ<-TOPRIM<-OLD-ABC<-N6-METHYLASE<-DUF6079*<-BrxB-BrxF-iSTAND<-BrxA-DNA-binding<-wHTH(SBD)+HNH<-LuxR-HTH | 1202 | Bacteria>Pseudomonadota>Gammaproteobacteria>Enterobacterales>Pectobacteriaceae>Pectobacterium | GCF_002904085.1 | Pectobacterium_odoriferum | BV921_RS15390 | 113976..117584 | 78398 | BREX-system-P-loop-protein-BrxC-[Pectobacterium-odoriferum] |
| WP_116155987.1 | Type1.BREX | LuxR-HTH->wHTH(SBD)+HNH->BrxA-DNA-binding->BrxB-BrxF-iSTAND->DUF6079*->N6-METHYLASE->OLD-ABC->TOPRIM->PglZ->SIGMA+OB-Fold+AAA-ATPase+Lon-Protease-BrxL-><-Arm-DNA-bind_3+Phage_integrase<-tRNA<-tRNA<-?<-URI-DNase+UVR+RNase+HHH<-REC+OmpR-HTH | 1202 | Bacteria>Pseudomonadota>Gammaproteobacteria>Enterobacterales>Pectobacteriaceae>Pectobacterium | GCF_003382585.2 | Pectobacterium_aquaticum | DMB81_RS10735 | 4900..8508 | 2204145 | BREX-system-P-loop-protein-BrxC-[Pectobacterium-aquaticum] |
| WP_181847885.1 | Type1.BREX | REC+OmpR-HTH->?->tRNA->tRNA->tRNA->Arm-DNA-bind_3+Phage_integrase-><-SIGMA+OB-Fold+AAA-ATPase+Lon-Protease-BrxL<-PglZ<-TOPRIM<-OLD-ABC<-N6-METHYLASE<-DUF6079*<-BrxB-BrxF-iSTAND<-BrxA-DNA-binding<-wHTH(SBD)+HNH<-LuxR-HTH<-?<-?<-ZnR<-XIS-HTH | 1204 | Bacteria>Pseudomonadota>Gammaproteobacteria>Enterobacterales>Pectobacteriaceae>Pectobacterium | GCF_013913645.1 | Pectobacterium_aroidearum | H2Y54_RS11285 | 236602..240216 | 1201031 | BREX-system-P-loop-protein-BrxC-[Pectobacterium-aroidearum] |
| HAT4275166.1 | Type1.BREX | <-Glyco_hydro<-?\|\|SbcC->?->wHTH(SBD)+HNH->BrxA-DNA-binding->BrxB-BrxF-iSTAND->DUF6079*->?->N6-METHYLASE->Cas9+HNH+RuvC->Cas_Cas1->CRISPR_Cas2->PglZ->SIGMA+OB-Fold+AAA-ATPase+Lon-Protease-BrxL-><-SH3+SH3+SH3+SH3+SH3+SH3+SH3+SH3 | 1221 | Bacteria>Terrabacteria_group>Bacillota>Clostridia>Eubacteriales>Clostridiaceae | GCA_015933065.1 | Clostridium_perfringens | I9084_000700 | 10095..13760 | 1502 | BREX-system-P-loop-protein-BrxC-[Clostridium-perfringens] |
| EGT3599215.1 | Type1.BREX | <-Glyco_hydro\|\|?-><-?\|\|SbcC->?->wHTH(SBD)+HNH->BrxA-DNA-binding->BrxB-BrxF-iSTAND->DUF6079*->?->N6-METHYLASE->Cas9+HNH+RuvC->Cas_Cas1->CRISPR_Cas2->PglZ->SIGMA+OB-Fold+AAA-ATPase+Lon-Protease-BrxL-><-SH3+SH3+SH3+SH3+SH3+SH3+SH3+SH3 | 1221 | Bacteria>Terrabacteria_group>Bacillota>Clostridia>Eubacteriales>Clostridiaceae | GCA_016068655.1 | Clostridium_perfringens | FT888_03085 | 92913..96578 | 1502 | BREX-system-P-loop-protein-BrxC-[Clostridium-perfringens] |
| EHK2362606.1 | Type1.BREX | SH3+SH3+SH3+SH3+SH3+SH3+SH3+SH3-><-SIGMA+OB-Fold+AAA-ATPase+Lon-Protease-BrxL<-PglZ<-CRISPR_Cas2<-Cas_Cas1<-Cas9+HNH+RuvC<-N6-METHYLASE<-DUF6079*<-BrxB-BrxF-iSTAND<-BrxA-DNA-binding<-wHTH(SBD)+HNH<-SLATT<-SLATT_4<-?<-HTH_OrfB_IS605+OrfB_IS605+OrfB_Zn_ribbon | 1221 | Bacteria>Terrabacteria_group>Bacillota>Clostridia>Eubacteriales>Clostridiaceae | GCA_018011955.1 | Clostridium_perfringens | KCL45_000419 | 449510..453175 | 1502 | BREX-system-P-loop-protein-BrxC-[Clostridium-perfringens] |
| MBS5995886.1 | Type1.BREX | SH3+SH3+SH3+SH3+SH3+SH3+SH3+SH3-><-SIGMA+OB-Fold+AAA-ATPase+Lon-Protease-BrxL<-PglZ<-CRISPR_Cas2<-Cas_Cas1<-Cas9+HNH+RuvC<-N6-METHYLASE<-?<-DUF6079*<-BrxB-BrxF-iSTAND<-BrxA-DNA-binding<-wHTH(SBD)+HNH<-VirB8\|\|?-><-?<-Amidohydro_1 | 1221 | Bacteria>Terrabacteria_group>Bacillota>Clostridia>Eubacteriales>Clostridiaceae | GCA_018372935.1 | Clostridium_perfringens | KIB05_13835 | 52036..55701 | 1502 | BREX-system-P-loop-protein-BrxC-[Clostridium-perfringens] |
| HCG3022573.1 | Type1.BREX | SH3+SH3+SH3+SH3+SH3+SH3+SH3+SH3-><-SIGMA+OB-Fold+AAA-ATPase+Lon-Protease-BrxL<-PglZ<-CRISPR_Cas2<-Cas_Cas1<-Cas9+HNH+RuvC<-N6-METHYLASE<-?<-DUF6079*<-BrxB-BrxF-iSTAND<-BrxA-DNA-binding<-wHTH(SBD)+HNH<-VirB8\|\|?-><-?\|\|Glyco_hydro-><-?<-Amidohydro_1 | 1221 | Bacteria>Terrabacteria_group>Bacillota>Clostridia>Eubacteriales>Clostridiaceae | GCA_024023955.1 | Clostridium_perfringens | NJU98_002507 | 52406..56071 | 1502 | BREX-system-P-loop-protein-BrxC-[Clostridium-perfringens] |
| EJT5935287.1 | Type1.BREX | <-Cas9+HNH+RuvC<-SF1-SF2-Helicase<-MarR-HTH<-?<-HerA/FtsK<-?<-sGTP<-N6-METHYLASE<-?<-DUF6079*<-BrxB-BrxF-iSTAND<-BrxA-DNA-binding<-wHTH(SBD)+HNH<-?<-VirB8\|\|?-><-?\|\|Glyco_hydro-> | 1221 | Bacteria>Terrabacteria_group>Bacillota>Clostridia>Eubacteriales>Clostridiaceae | GCA_025090725.1 | Clostridium_perfringens | N2W52_000012 | 19602..23267 | 1502 | BREX-system-P-loop-protein-BrxC-[Clostridium-perfringens] |
| EJT6164547.1 | Type1.BREX | <-Glyco_hydro\|\|?-><-?\|\|VirB8->?->wHTH(SBD)+HNH->BrxA-DNA-binding->BrxB-BrxF-iSTAND->DUF6079*->?->N6-METHYLASE->Cas9+HNH+RuvC->Cas_Cas1->CRISPR_Cas2->PglZ->SIGMA+OB-Fold+AAA-ATPase+Lon-Protease-BrxL-><-SH3+SH3+SH3+SH3+SH3+SH3+SH3+SH3 | 1221 | Bacteria>Terrabacteria_group>Bacillota>Clostridia>Eubacteriales>Clostridiaceae | GCA_025091785.1 | Clostridium_perfringens | N2W22_000999 | 1099443..1103108 | 1502 | BREX-system-P-loop-protein-BrxC-[Clostridium-perfringens] |
| EJT6169494.1 | Type1.BREX | SH3+SH3+SH3+SH3+SH3+SH3+SH3+SH3-><-SIGMA+OB-Fold+AAA-ATPase+Lon-Protease-BrxL<-PglZ<-CRISPR_Cas2<-Cas_Cas1<-Cas9+HNH+RuvC<-N6-METHYLASE<-?<-DUF6079*<-BrxB-BrxF-iSTAND<-BrxA-DNA-binding<-wHTH(SBD)+HNH<-VirB8\|\|?-><-?\|\|Glyco_hydro-><-?<-Amidohydro_1 | 1221 | Bacteria>Terrabacteria_group>Bacillota>Clostridia>Eubacteriales>Clostridiaceae | GCA_025091845.1 | Clostridium_perfringens | N2W53_000036 | 52483..56148 | 1502 | BREX-system-P-loop-protein-BrxC-[Clostridium-perfringens] |
| EJT6540217.1 | Type1.BREX | SH3+SH3+SH3+SH3+SH3+SH3+SH3+SH3-><-SIGMA+OB-Fold+AAA-ATPase+Lon-Protease-BrxL<-PglZ<-CRISPR_Cas2<-Cas_Cas1<-Cas9+HNH+RuvC<-N6-METHYLASE<-?<-DUF6079*<-BrxB-BrxF-iSTAND<-BrxA-DNA-binding<-wHTH(SBD)+HNH<-VirB8\|\|?-><-?\|\|Glyco_hydro-><-?<-Amidohydro_1 | 1221 | Bacteria>Terrabacteria_group>Bacillota>Clostridia>Eubacteriales>Clostridiaceae | GCA_025095895.1 | Clostridium_perfringens | N2W31_000036 | 52406..56071 | 1502 | BREX-system-P-loop-protein-BrxC-[Clostridium-perfringens] |
| WP_003452047.1 | Type1.BREX | <-Glyco_hydro\|\|?-><-?\|\|VirB8->?->Y1_Tnp->wHTH(SBD)+HNH->BrxA-DNA-binding->BrxB-BrxF-iSTAND->DUF6079*->N6-METHYLASE->HIN-HTH+PAIREDC-HTH->Cas9+HNH+RuvC->Cas_Cas1->CRISPR_Cas2->PglZ->SIGMA+OB-Fold+AAA-ATPase+Lon-Protease-BrxL-><-SH3+SH3+SH3+SH3+SH3+SH3+SH3+SH3 | 1221 | Bacteria>Terrabacteria_group>Bacillota>Clostridia>Eubacteriales>Clostridiaceae | GCF_000171135.1 | Clostridium_perfringens | CPC_RS09775 | 28796..32461 | 1502 | BREX-system-P-loop-protein-BrxC-[Clostridium-perfringens] |
| WP_003473497.1 | Type1.BREX | <-Cas_Cas1<-Cas9+HNH+RuvC<-MarR-HTH<-?<-HerA/FtsK<-?<-sGTP<-N6-METHYLASE<-?<-DUF6079*<-BrxB-BrxF-iSTAND<-BrxA-DNA-binding<-wHTH(SBD)+HNH<-?<-SbcC\|\|?-><-?\|\|Glyco_hydro-> | 1221 | Bacteria>Terrabacteria_group>Bacillota>Clostridia>Eubacteriales>Clostridiaceae | GCF_000172455.1 | Clostridium_perfringens | CJD_RS01695 | 75268..78933 | 1502 | BREX-system-P-loop-protein-BrxC-[Clostridium-perfringens] |
| WP_060670433.1 | Type1.BREX | <-Cas_Cas1<-Cas9+HNH+RuvC<-MarR-HTH<-?<-HerA/FtsK<-?<-sGTP<-N6-METHYLASE<-?<-DUF6079*<-BrxB-BrxF-iSTAND<-BrxA-DNA-binding<-wHTH(SBD)+HNH<-?<-SbcC\|\|?-><-?\|\|Glyco_hydro-> | 1221 | Bacteria>Terrabacteria_group>Bacillota>Clostridia>Eubacteriales>Clostridiaceae | GCF_001304735.1 | Clostridium_perfringens | FORC3_RS07365 | 1728066..1731731 | 1502 | BREX-system-P-loop-protein-BrxC-[Clostridium-perfringens] |
| WP_115613438.1 | Type1.BREX | <-Glyco_hydro\|\|?-><-?\|\|SbcC->?->wHTH(SBD)+HNH->BrxA-DNA-binding->BrxB-BrxF-iSTAND->DUF6079*->?->N6-METHYLASE->Cas9+HNH+RuvC->Cas_Cas1->CRISPR_Cas2->PglZ->SIGMA+OB-Fold+AAA-ATPase+Lon-Protease-BrxL-> | 1221 | Bacteria>Terrabacteria_group>Bacillota>Clostridia>Eubacteriales>Clostridiaceae | GCF_016027155.1 | Clostridium_perfringens | I6G61_RS10175 | 2178205..2181870 | 1502 | BREX-system-P-loop-protein-BrxC-[Clostridium-perfringens] |
| WP_198605367.1 | Type1.BREX | <-Glyco_hydro\|\|?-><-?\|\|SbcC->?->wHTH(SBD)+HNH->BrxA-DNA-binding->BrxB-BrxF-iSTAND->DUF6079*->?->N6-METHYLASE->sGTP->?->HerA/FtsK->?->MarR-HTH->Cas_Cas1->CRISPR_Cas2-> | 1221 | Bacteria>Terrabacteria_group>Bacillota>Clostridia>Eubacteriales>Clostridiaceae | GCF_016236125.1 | Clostridium_perfringens | H8J62_RS02875 | 34894..38559 | 1502 | BREX-system-P-loop-protein-BrxC-[Clostridium-perfringens] |
| WP_198621144.1 | Type1.BREX | <-CRISPR_Cas2<-Cas_Cas1<-MarR-HTH<-?<-HerA/FtsK<-?<-sGTP<-N6-METHYLASE<-?<-DUF6079*<-BrxB-BrxF-iSTAND<-BrxA-DNA-binding<-wHTH(SBD)+HNH<-?<-SbcC\|\|?-><-?\|\|Glyco_hydro-> | 1221 | Bacteria>Terrabacteria_group>Bacillota>Clostridia>Eubacteriales>Clostridiaceae | GCF_016236545.1 | Clostridium_perfringens | H8J92_RS02870 | 79478..83143 | 1502 | BREX-system-P-loop-protein-BrxC-[Clostridium-perfringens] |
| WP_198616415.1 | Type1.BREX | <-Glyco_hydro<-?\|\|VirB8->?->wHTH(SBD)+HNH->BrxA-DNA-binding->BrxB-BrxF-iSTAND->DUF6079*->?->N6-METHYLASE->sGTP->?->HerA/FtsK->?->MarR-HTH->Cas_Cas1->CRISPR_Cas2-> | 1221 | Bacteria>Terrabacteria_group>Bacillota>Clostridia>Eubacteriales>Clostridiaceae | GCF_016236635.1 | Clostridium_perfringens | H8J95_RS08945 | 20074..23739 | 1502 | BREX-system-P-loop-protein-BrxC-[Clostridium-perfringens] |
| WP_164804802.1 | Type1.BREX | Amidohydro_1->?-><-Glyco_hydro\|\|<-?\|\|SbcC->?->wHTH(SBD)+HNH->BrxA-DNA-binding->BrxB-BrxF-iSTAND->DUF6079*->?->N6-METHYLASE->sGTP->?->HerA/FtsK->?->MarR-HTH->Cas_Cas1->CRISPR_Cas2-> | 1221 | Bacteria>Terrabacteria_group>Bacillota>Clostridia>Eubacteriales>Clostridiaceae | GCF_018135545.1 | Clostridium_perfringens | G6Z22_RS06045 | 21931..25596 | 1502 | BREX-system-P-loop-protein-BrxC-[Clostridium-perfringens] |
| WP_198609318.1 | Type1.BREX | SH3+SH3+SH3+SH3+SH3+SH3+SH3+SH3-><-SIGMA+OB-Fold+AAA-ATPase+Lon-Protease-BrxL<-PglZ<-CRISPR_Cas2<-Cas_Cas1<-Cas9+HNH+RuvC<-N6-METHYLASE<-?<-DUF6079*<-BrxB-BrxF-iSTAND<-BrxA-DNA-binding<-wHTH(SBD)+HNH<-?<-SbcC\|\|?-><-?\|\|Glyco_hydro-> | 1221 | Bacteria>Terrabacteria_group>Bacillota>Clostridia>Eubacteriales>Clostridiaceae | GCF_020139275.1 | Clostridium_perfringens | KLF33_RS07605 | 1732833..1736498 | 1502 | BREX-system-P-loop-protein-BrxC-[Clostridium-perfringens] |
| WP_228572337.1 | Type1.BREX | Amidohydro_1->?-><-Glyco_hydro\|\|<-?\|\|VirB8->?->wHTH(SBD)+HNH->BrxA-DNA-binding->BrxB-BrxF-iSTAND->DUF6079*->?->N6-METHYLASE->Cas_Cas1->CRISPR_Cas2->PglZ->SIGMA+OB-Fold+AAA-ATPase+Lon-Protease-BrxL-><-SH3+SH3+SH3+SH3+SH3+SH3+SH3+SH3 | 1221 | Bacteria>Terrabacteria_group>Bacillota>Clostridia>Eubacteriales>Clostridiaceae | GCF_020819925.1 | Clostridium_perfringens | LMS36_RS03155 | 130857..134522 | 1502 | BREX-system-P-loop-protein-BrxC-[Clostridium-perfringens] |
| WP_111744218.1 | Type1.BREX | SbcC->?->wHTH(SBD)+HNH->BrxA-DNA-binding->BrxB-BrxF-iSTAND->DUF6079*->?->N6-METHYLASE->Cas9+HNH+RuvC->Cas_Cas1->CRISPR_Cas2->PglZ->SIGMA+OB-Fold+AAA-ATPase+Lon-Protease-BrxL-> | 1221 | Bacteria>Terrabacteria_group>Bacillota>Clostridia>Eubacteriales>Clostridiaceae | GCF_900475545.1 | Clostridium_perfringens | Cp4442_RS12820 | 4585..8250 | 1502 | BREX-system-P-loop-protein-BrxC-[Clostridium-perfringens] |
| WP_142710734.1 | Type1.BREX | <-Cas9+HNH+RuvC<-SF1-SF2-Helicase<-MarR-HTH<-?<-HerA/FtsK<-?<-sGTP<-N6-METHYLASE<-?<-DUF6079*<-BrxB-BrxF-iSTAND<-BrxA-DNA-binding<-wHTH(SBD)+HNH<-SLATT<-SLATT_4 | 1221 | Bacteria>Terrabacteria_group>Bacillota>Clostridia>Eubacteriales>Clostridiaceae | GCF_902166005.1 | Clostridium_perfringens | FN719_RS05040 | 228419..232084 | 1502 | BREX-system-P-loop-protein-BrxC-[Clostridium-perfringens] |
| WP_142975592.1 | Type1.BREX | SH3+SH3+SH3+SH3+SH3+SH3+SH3+SH3-><-SIGMA+OB-Fold+AAA-ATPase+Lon-Protease-BrxL<-PglZ<-CRISPR_Cas2<-Cas_Cas1<-Cas9+HNH+RuvC<-N6-METHYLASE<-?<-DUF6079*<-BrxB-BrxF-iSTAND<-BrxA-DNA-binding<-wHTH(SBD)+HNH<-?<-SbcC\|\|?-><-?\|\|Glyco_hydro-> | 1221 | Bacteria>Terrabacteria_group>Bacillota>Clostridia>Eubacteriales>Clostridiaceae | GCF_902166025.1 | Clostridium_perfringens | FN715_RS03295 | 751975..755640 | 1502 | BREX-system-P-loop-protein-BrxC-[Clostridium-perfringens] |
| WP_195381616.1 | Type1.BREX | Amidohydro_1->?-><-Glyco_hydro<-?\|\|SbcC->?->wHTH(SBD)+HNH->BrxA-DNA-binding->BrxB-BrxF-iSTAND->DUF6079*->?->N6-METHYLASE->Cas_Cas1->CRISPR_Cas2->PglZ->SIGMA+OB-Fold+AAA-ATPase+Lon-Protease-BrxL-><-SH3+SH3+SH3+SH3+SH3+SH3+SH3+SH3 | 1221 | Bacteria>Terrabacteria_group>Bacillota>Clostridia>Eubacteriales>Clostridiaceae | GCF_951336195.1 | Clostridium_perfringens | H8J86_RS06960 | 66718..70383 | 1502 | BREX-system-P-loop-protein-BrxC-[Clostridium-perfringens] |
| -- | | | | | | | | | | |
| --HNH | | | | | | | | | | |
| -- | | | | | | | | | | |
| MBP7797364.1 | Type6.BREX | <-RNase<-ANK+ANK<-REase<-SbcC<-?<-HNH<-SIGMA+OB-Fold+AAA-ATPase+Lon-Protease-BrxL<-PglZ<-N6-METHYLASE<-DUF6079*<-BrxB-BrxF-iSTAND<-BrxA-DNA-binding<-BrxE\|\|?-><-Phage_integrase<-Phage_integrase<-HerA/FtsK<-?<-CHTH | 1177 | Bacteria>Acidobacteriota>Thermoanaerobaculia>Thermoanaerobaculales>Thermoanaerobaculaceae>unclassified_Thermoanaerobaculaceae | GCA_018000955.1 | Thermoanaerobaculaceae_bacterium | KA072_01920 | 40724..44257 | 2699758 | BREX-system-P-loop-protein-BrxC-[Thermoanaerobaculaceae-bacterium] |
| PKP62156.1 | Type1.BREX | SbcC+SbcC->BrxA-DNA-binding->BrxB-BrxF-iSTAND->DUF6079*->N6-METHYLASE->PglZ->SIGMA+OB-Fold+AAA-ATPase+Lon-Protease-BrxL->HNH->STYKIN->TPR+TPR->TPR-> | 1193 | Bacteria>Atribacterota>unclassified_Atribacterota | GCA_002841055.1 | Candidatus_Atribacteria_bacterium_HGW-Atribacteria-1 | CVT91_00360 | 3425..7006 | 2013676 | BREX-system-P-loop-protein-BrxC-[Candidatus-Atribacteria-bacterium-HGW-Atribacteria-1] |
| MBI2566519.1 | Type1.BREX | NtrC-AAA+FIS-HTH->?->?-><-?<-?\|\|BrxA-DNA-binding->BrxB-BrxF-iSTAND->DUF6079*->?->N6-METHYLASE->HEPN->ParA-Soj-PloopNTPase->?->PglZ->HNH-><-?<-NtrC-AAA+FIS-HTH | 1109 | Bacteria>Bacteria_incertae_sedis>Bacteria_candidate_phyla>Candidatus_Schekmanbacteria | GCA_016188575.1 | Candidatus_Schekmanbacteria_bacterium | HYV63_05760 | 102186..105515 | 2419984 | BREX-system-P-loop-protein-BrxC-[Candidatus-Schekmanbacteria-bacterium] |
| MCM1267126.1 | Type1.BREX | <-SbcC<-?<-HNH<-SIGMA+OB-Fold+AAA-ATPase+Lon-Protease-BrxL<-PglZ<-DOC+HTH<-?<-N6-METHYLASE<-DUF6079*<-BrxB-BrxF-iSTAND<-BrxA-DNA-binding<-?<-?<-?<-CHTH | 1173 | Bacteria>FCB_group>Bacteroidetes/Chlorobi_group>Bacteroidota>Bacteroidia>Bacteroidales | GCA_023665065.1 | Bacteroidales_bacterium | NC302_04400 | 49162..52683 | 2030927 | BREX-system-P-loop-protein-BrxC-[Bacteroidales-bacterium] |
| MCB0517804.1 | Type1.BREX | <-SIGMA+OB-Fold+AAA-ATPase+HKD-DNase-BrxL<-PglZ<-N6-METHYLASE<-DUF6079*<-BrxB-BrxF-iSTAND<-BrxA-DNA-binding<-?<-HNH\|\|REC-><-?<-SF1-SF2-Helicase+RNase+RNase<-?<-RVT | 1174 | Bacteria>FCB_group>Bacteroidetes/Chlorobi_group>Bacteroidota>Saprospiria>Saprospirales | GCA_020436325.1 | Saprospiraceae_bacterium | KDD27_02635 | 9612..13136 | 2202734 | BREX-system-P-loop-protein-BrxC-[Saprospiraceae-bacterium] |
| WP_147932457.1 | Type1.BREX | Calcineurin->RADICAL-SAM->METHYLASE+RNA-Helicase->?->?->DnaJ->BrxA-DNA-binding->BrxB-BrxF-iSTAND->DUF6079*->N6-METHYLASE->?->?->HNH->?->URI-DNase->wHTH+AbiEi_2->AbiEii->PglZ-> | 1198 | Bacteria>FCB_group>Bacteroidetes/Chlorobi_group>Bacteroidota>Saprospiria>Saprospirales | GCF_008014675.1 | Neolewinella_aurantiaca | FUA23_RS19525 | 18155..21751 | 2602767 | BREX-system-P-loop-protein-BrxC-[Neolewinella-aurantiaca] |
| WP_098973854.1 | Type1.BREX | SKI->JAB+sGTP->wHTH+WYL+WCX->?->?->ABC-ATPase+SbcC->?-><-?\|\|DUF6079*->OLD-ABC->TOPRIM->N6-METHYLASE->N6-METHYLASE->MORC+HISKIN->?->HNH->PglZ->SIGMA+OB-Fold+AAA-ATPase+Lon-Protease-BrxL-> | 1250 | Bacteria>Fusobacteriota>Fusobacteriia>Fusobacteriales>Fusobacteriaceae>Fusobacterium | GCF_002591515.1 | Fusobacterium_nucleatum | CBG58_RS00340 | 77641..81393 | 851 | BREX-system-P-loop-protein-BrxC-[Fusobacterium-polymorphum] |
| HCF56393.1 | Type6.BREX | <-Schlafen+HATPase+wHTH<-N6-METHYLASE-p\|\|?->PIN-><-TOPRIM<-OLD-ABC<-DUF6079-p<-APATPase<-DUF6079-p*<-DUF6079-p<-HNH<-BrxB-BrxF-iSTAND<-BrxA-DNA-binding<-BrxA-DNA-binding<-BrxE | 508 | Bacteria>Myxococcota>Myxococcia>Myxococcales>unclassified_Myxococcales | GCA_003534605.1 | Myxococcales_bacterium | DFS52_00150 | 7726..9251 | 2026763 | BREX-system-P-loop-protein-BrxC-[Myxococcales-bacterium] |
| HCF56394.1 | Type6.BREX | <-Schlafen+HATPase+wHTH<-N6-METHYLASE-p\|\|?->PIN-><-TOPRIM<-OLD-ABC<-DUF6079-p<-APATPase<-DUF6079-p*<-DUF6079-p*<-HNH<-BrxB-BrxF-iSTAND<-BrxA-DNA-binding<-BrxA-DNA-binding<-BrxE | 363 | Bacteria>Myxococcota>Myxococcia>Myxococcales>unclassified_Myxococcales | GCA_003534605.1 | Myxococcales_bacterium | DFS52_00155 | 9470..10559 | 2026763 | BREX-system-P-loop-protein-BrxC-[Myxococcales-bacterium] |
| KFG67132.1 | Type6.BREX | <-AbiGi+seqA+RAMA\|\|BrxE->BrxA-DNA-binding->BrxB-BrxF-iSTAND->DUF6079*->N6-METHYLASE-p->PglZ->SIGMA+OB-Fold+AAA-ATPase+HKD-DNase-BrxL->HNH->HNH->HNH->?-><-HEPN<-Polbeta | 1210 | Bacteria>Pseudomonadota>Alphaproteobacteria>Hyphomicrobiales>Methylobacteriaceae>Microvirga | GCA_000739015.1 | Microvirga_sp._BSC39 | JH26_23850 | 105316..108948 | 1549810 | ATPase-[Microvirga-sp.-BSC39] |
| WP_036357316.1 | Type6.BREX | <-AbiGi+seqA+RAMA<-?\|\|PhdYeFM_antitox->BrxE->BrxA-DNA-binding->BrxB-BrxF-iSTAND->DUF6079*->N6-METHYLASE->PglZ->SIGMA+OB-Fold+AAA-ATPase+HKD-DNase-BrxL->HNH->HNH->HNH->?-><-HEPN<-Polbeta | 1208 | Bacteria>Pseudomonadota>Alphaproteobacteria>Hyphomicrobiales>Methylobacteriaceae>Microvirga | GCF_000739015.1 | Microvirga_sp._BSC39 | MA07_RS22775 | 105322..108948 | 1549810 | BREX-system-P-loop-protein-BrxC-[Microvirga-sp.-BSC39] |
| MBK8578271.1 | Type1.BREX | <-SIGMA+OB-Fold+AAA-ATPase+Lon-Protease-BrxL<-PglZ<-N6-METHYLASE<-HNH<-?<-SbcC+NACHT<-DUF6079*<-BrxB-BrxF-iSTAND<-BrxA-DNA-binding<-BrxA-DNA-binding<-Polbeta<-wHTH+AbiEi_2<-?\|\|?-><-wHTH+AbiEi_2 | 1154 | Bacteria>Pseudomonadota>Betaproteobacteria>Betaproteobacteria_incertae_sedis>Candidatus_Accumulibacter>unclassified_Candidatus_Accumulibacter | GCA_016714935.1 | Candidatus_Accumulibacter_sp. | IPN78_08405 | 116670..120134 | 2053492 | BREX-system-P-loop-protein-BrxC-[Candidatus-Accumulibacter-sp.] |
| ABO55019.1 | Type1.BREX | <-HNH<-SIGMA+OB-Fold+AAA-ATPase+Lon-Protease-BrxL<-PglZ<-HTH_29+HTH_32+HTH_33+DDE_3\|\|HIN-HTH+PAIREDC-HTH-><-DUF6079*<-BrxB-BrxF-iSTAND<-BrxA-DNA-binding\|\|?->ANK-><-?\|\|ANK+ANK+ANK->?-><-ANK | 1210 | Bacteria>Pseudomonadota>Betaproteobacteria>Burkholderiales>Burkholderiaceae>Burkholderia | GCA_000016205.1 | Burkholderia_vietnamiensis_G4 | Bcep1808_2017 | 2211669..2215301 | 269482 | ATPase-like-protein-[Burkholderia-vietnamiensis-G4] |
| WP_013697442.1 | Type1.BREX | ZnR+TOPRIM+AAA-helicase+wHTH->?->?->?->SAM-METHYLASE->ACET-><-ANK+ANK\|\|BrxA-DNA-binding->BrxB-BrxF-iSTAND->DUF6079*->N6-METHYLASE->PglZ->SIGMA+OB-Fold+AAA-ATPase+Lon-Protease-BrxL->HNH-><-?\|\|ANK-><-?\|\|wHTH(SBD)+HNH-><-REase | 1223 | Bacteria>Pseudomonadota>Betaproteobacteria>Burkholderiales>Burkholderiaceae>Burkholderia | GCF_000194745.1 | Burkholderia_gladioli | BGLA_RS07070 | 1576041..1579712 | 28095 | BREX-system-P-loop-protein-BrxC-[Burkholderia-gladioli] |
| WP_124831269.1 | Type1.BREX | <-MORC\|\|?-><-HNH<-SIGMA+OB-Fold+AAA-ATPase+Lon-Protease-BrxL<-PglZ<-N6-METHYLASE<-DUF6079*<-BrxB-BrxF-iSTAND<-BrxA-DNA-binding\|\|?-><-?\|\|?-><-?<-?<-REase<-Phage_GPD+Phage_base_V+T6SS_Vgr+DUF2345 | 1200 | Bacteria>Pseudomonadota>Betaproteobacteria>Burkholderiales>Burkholderiaceae>Burkholderia | GCF_003857875.1 | Burkholderia_stagnalis | DF113_RS32940 | 12280..15882 | 1503054 | BREX-system-P-loop-protein-BrxC-[Burkholderia-stagnalis] |
| WP_122833867.1 | Type1.BREX | ANK-><-?<-?<-ANK+ANK\|\|?-><-ANK\|\|BrxA-DNA-binding->BrxB-BrxF-iSTAND->DUF6079*->N6-METHYLASE->PglZ->SIGMA+OB-Fold+AAA-ATPase+Lon-Protease-BrxL->HNH-> | 1223 | Bacteria>Pseudomonadota>Betaproteobacteria>Burkholderiales>Burkholderiaceae>Burkholderia | GCF_900597175.1 | Burkholderia_pseudomallei | EFT75_RS19485 | 195449..199120 | 28450 | BREX-system-P-loop-protein-BrxC-[Burkholderia-pseudomallei] |
| WP_121308231.1 | Type1.BREX | <-MORC<-?\|\|?->?-><-HNH<-SIGMA+OB-Fold+AAA-ATPase+Lon-Protease-BrxL<-PglZ<-N6-METHYLASE<-DUF6079*<-BrxB-BrxF-iSTAND<-BrxA-DNA-binding<-?\|\|?->wHTH+WYL+WCX->?->?-><-?\|\|ANK+ANK-> | 1204 | Bacteria>Pseudomonadota>Betaproteobacteria>Burkholderiales>Burkholderiaceae>Paraburkholderia | GCF_003634175.1 | Paraburkholderia_sp._BL17N1 | BOG82_RS17815 | 3937370..3940984 | 1938798 | BREX-system-P-loop-protein-BrxC-[Paraburkholderia-sp.-BL17N1] |
| WP_201659285.1 | Type1.BREX | SbcC->HTH_3-><-NACHT<-HNH<-SIGMA+OB-Fold+AAA-ATPase+Lon-Protease-BrxL<-PglZ<-N6-METHYLASE<-DUF6079*<-BrxB-BrxF-iSTAND<-BrxA-DNA-binding\|\|PhdYeFM_antitox->RelE-ParE-><-REC<-REC+NtrC-AAA+FIS-HTH\|\|?-><-GGDEF\|\|REC+REC-> | 1218 | Bacteria>Pseudomonadota>Betaproteobacteria>Burkholderiales>Burkholderiaceae>Paraburkholderia | GCF_904848645.1 | Paraburkholderia_sabiae | JMY25_RS35995 | 30018..33674 | 273251 | BREX-system-P-loop-protein-BrxC-[Paraburkholderia-sabiae] |
| WP_094097616.1 | Type1.BREX | <-Peptidase_C39_2<-?\|\|?-><-?\|\|XIS-HTH->Phage_X->BrxA-DNA-binding->BrxB-BrxF-iSTAND->DUF6079*->HNH->HEPN->ParD_antitoxin->RelE-ParE->N6-METHYLASE->OLD-ABC->TOPRIM->ZnR+TOPRIM->PglZ-> | 1214 | Bacteria>Pseudomonadota>Betaproteobacteria>Burkholderiales>Comamonadaceae>Acidovorax | GCF_002157145.1 | Acidovorax_carolinensis | CBP34_RS06480 | 1379414..1383058 | 553814 | BREX-system-P-loop-protein-BrxC-[Acidovorax-carolinensis] |
| WP_086912102.1 | Type1.BREX | HTH_Tnp_1+HTH_21+rve+rve_3-><-PIN<-MazE_antitoxin\|\|?-><-HEPN<-HNH<-DUF6079*<-BrxB-BrxF-iSTAND<-BrxA-DNA-binding<-GNTR-HTH\|\|PBPI-> | 1213 | Bacteria>Pseudomonadota>Betaproteobacteria>Burkholderiales>Comamonadaceae>Acidovorax | GCF_002157165.1 | Acidovorax_carolinensis | CBP33_RS08115 | 1704915..1708556 | 553814 | BREX-system-P-loop-protein-BrxC-[Acidovorax-carolinensis] |
| GAO25248.1 | Type1.BREX | HTH_3-><-SIGMA+OB-Fold+AAA-ATPase+Lon-Protease-BrxL<-PglZ<-ZnR+TOPRIM<-TOPRIM<-OLD-ABC<-N6-METHYLASE<-HEPN<-HNH<-DUF6079*<-BrxB-BrxF-iSTAND<-BrxA-DNA-binding<-HAD+CHTH<-CHTH\|\|?->AAA-ATPase-p->?-><-AAA-ATPase | 1205 | Bacteria>Pseudomonadota>Betaproteobacteria>Burkholderiales>Comamonadaceae>Alicycliphilus | GCA_000974645.1 | Alicycliphilus_sp._B1 | ALISP_5068 | 1538304..1541921 | 1603291 | ATPase-like-protein-[Alicycliphilus-sp.-B1] |
| WP_172207178.1 | Type1.BREX | <-OLD-ABC\|\|<-N6-METHYLASE-p<-RelE-ParE<-ParD_antitoxin<-PIN<-MazE_antitoxin\|\|?-><-HEPN<-HNH<-DUF6079*<-BrxB-BrxF-iSTAND<-BrxA-DNA-binding<-Phage_X<-XIS-HTH\|\|?-><-?\|\|?->Peptidase_C39_2-> | 1213 | Bacteria>Pseudomonadota>Betaproteobacteria>Burkholderiales>Comamonadaceae>Diaphorobacter | GCF_013154915.1 | Diaphorobacter_sp._JS3050 | HND92_RS11875 | 2478331..2481972 | 2735554 | BREX-system-P-loop-protein-BrxC-[Diaphorobacter-sp.-JS3050] |
| MCG9118882.1 | Type1.BREX | <-SIGMA+OB-Fold+AAA-ATPase+Lon-Protease-BrxL<-PglZ<-HNH<-?<-?<-N6-METHYLASE<-?<-DUF6079*<-BrxB-BrxF-iSTAND<-BrxA-DNA-binding<-wHTH+WYL+WCX<-?<-?<-?\|\|?->HTH_Tnp_1+HTH_21+rve+rve_3->Phage_int_SAM_1+Phage_integrase-> | 1221 | Bacteria>Pseudomonadota>Betaproteobacteria>Neisseriales>Chromobacteriaceae>Laribacter | GCA_022394685.1 | Laribacter_hongkongensis | LH456_09760 | 59894..63559 | 168471 | BREX-system-P-loop-protein-BrxC-[Laribacter-hongkongensis] |
| WP_275667170.1 | Type1.BREX | <-SIGMA+OB-Fold+AAA-ATPase+Lon-Protease-BrxL<-PglZ<-HNH<-?<-?<-N6-METHYLASE<-?<-DUF6079*<-BrxB-BrxF-iSTAND<-BrxA-DNA-binding<-wHTH+WYL+WCX<-?<-?<-?\|\|?->HTH_Tnp_1+HTH_21+rve+rve_3->Phage_int_SAM_1+Phage_integrase-> | 1216 | Bacteria>Pseudomonadota>Betaproteobacteria>Neisseriales>Chromobacteriaceae>Laribacter | GCA_022394685.1 | Laribacter_hongkongensis | LH456_RS09740 | 59894..63544 | 168471 | BREX-system-P-loop-protein-BrxC-[Laribacter-hongkongensis] |
| WP_239901116.1 | Type1.BREX | <-SIGMA+OB-Fold+AAA-ATPase+Lon-Protease-BrxL<-PglZ<-HNH<-?<-?<-N6-METHYLASE<-?<-DUF6079*<-BrxB-BrxF-iSTAND<-BrxA-DNA-binding<-wHTH+WYL+WCX<-?<-?<-?\|\|?->HTH_Tnp_1+HTH_21+rve+rve_3->Phage_int_SAM_1+Phage_integrase-> | 1221 | Bacteria>Pseudomonadota>Betaproteobacteria>Neisseriales>Chromobacteriaceae>Laribacter | - | Laribacter_hongkongensis | LH456_09760 | 59894..63559 | 168471 | BREX-system-P-loop-protein-BrxC-[Laribacter-hongkongensis] |
| PZQ93954.1 | Type1.BREX | <-HNH<-DUF6079*<-BrxB-BrxF-iSTAND\|\|HIN-HTH->IstB_IS21-><-HTH_Tnp_1+HTH_21+rve+rve_3 | 1218 | Bacteria>Pseudomonadota>Gammaproteobacteria>Aeromonadales>Aeromonadaceae>Aeromonas | GCA_003243215.1 | Aeromonas_media | DI541_18955 | 3274..6930 | 651 | BREX-system-P-loop-protein-BrxC-[Aeromonas-media] |
| GJA32139.1 | Type1.BREX | <-Peptidase_C39_2<-?<-N6-METHYLASE-p<-HEPN<-N6-METHYLASE-p\|\|DDE_Tnp_1_assoc+DDE_Tnp_1-><-N6-METHYLASE<-?<-HNH<-DUF6079*<-BrxB-BrxF-iSTAND<-BrxA-DNA-binding<-RHS<-?<-Ntox49\|\|wHTH+WYL+WCX-><-XIS-HTH\|\|?-><-HTH_26 | 1227 | Bacteria>Pseudomonadota>Gammaproteobacteria>Aeromonadales>Aeromonadaceae>Aeromonas | GCA_019971155.1 | Aeromonas_caviae | KAM347_17720 | 32751..36434 | 648 | hypothetical-protein-KAM341_18170-[Aeromonas-caviae] |
| WP_059169685.1 | Type1.BREX | <-PglZ<-N6-METHYLASE-p<-HEPN<-RVT<-N6-METHYLASE<-?<-HNH<-TOPRIM<-OLD-ABC<-DUF6079*<-BrxB-BrxF-iSTAND<-BrxA-DNA-binding<-RHS<-ARG<-RHS\|\|LexA-protease->?-><-HDRnase<-Arm-DNA-bind_2+Phage_int_SAM_3+Phage_integrase | 1240 | Bacteria>Pseudomonadota>Gammaproteobacteria>Aeromonadales>Aeromonadaceae>Aeromonas | GCF_001481535.2 | Aeromonas_salmonicida | VO70_RS19205 | 18975..22697 | 645 | BREX-system-P-loop-protein-BrxC-[Aeromonas-salmonicida] |
| WP_210557479.1 | Type1.BREX | DEDD_Tnp_IS110+Transposase_20->\|\|BrxA-DNA-binding->BrxB-BrxF-iSTAND->DUF6079*->HNH->?->N6-METHYLASE->N6-METHYLASE-p->HEPN->N6-METHYLASE-p-> | 1218 | Bacteria>Pseudomonadota>Gammaproteobacteria>Aeromonadales>Aeromonadaceae>Aeromonas | GCF_017939245.1 | Aeromonas_sp._PrichA-15 | J9889_RS17255 | 4362..8018 | 2823360 | BREX-system-P-loop-protein-BrxC-[Aeromonas-sp.-PrichA-15] |
| WP_270805301.1 | Type1.BREX | <-N6-METHYLASE-p<-N6-METHYLASE-p<-N6-METHYLASE<-?<-HNH<-TOPRIM<-OLD-ABC<-DUF6079*<-BrxB-BrxF-iSTAND<-BrxA-DNA-binding<-RHS<-ZnR | 1240 | Bacteria>Pseudomonadota>Gammaproteobacteria>Aeromonadales>Aeromonadaceae>Aeromonas | GCF_027705505.1 | Aeromonas_sp._QDB56 | OKB81_RS14905 | 8131..11853 | 2990495 | BREX-system-P-loop-protein-BrxC-[Aeromonas-sp.-QDB56] |
| WP_182788683.1 | Type1.BREX | HDRnase-><-?<-LexA-protease\|\|RHS->ARG->ZnR->RHS->BrxA-DNA-binding->BrxB-BrxF-iSTAND->DUF6079*->OLD-ABC->TOPRIM->HNH->?->N6-METHYLASE->SIR2->HerA/FtsK->N6-METHYLASE-p-> | 1236 | Bacteria>Pseudomonadota>Gammaproteobacteria>Aeromonadales>Aeromonadaceae | GCF_014161425.1 | Aeromonas | H7K58_RS09750 | 2083810..2087520 | 642 | MULTISPECIES:-BREX-system-P-loop-protein-BrxC-[Aeromonas] |
| WP_223915718.1 | Type1.BREX | <-Peptidase_C39_2<-?<-N6-METHYLASE-p<-HEPN<-N6-METHYLASE-p\|\|DDE_Tnp_1_assoc+DDE_Tnp_1-><-N6-METHYLASE<-?<-HNH<-DUF6079*<-BrxB-BrxF-iSTAND<-BrxA-DNA-binding<-RHS<-?<-Ntox49\|\|wHTH+WYL+WCX-><-XIS-HTH\|\|?-><-HTH_26 | 1218 | Bacteria>Pseudomonadota>Gammaproteobacteria>Aeromonadales>Aeromonadaceae | GCF_019972515.1 | Aeromonas | LDN24_RS09390 | 22833..26489 | 642 | MULTISPECIES:-BREX-system-P-loop-protein-BrxC-[Aeromonas] |
| MBO2546554.1 | Type1.BREX | <-SIGMA+OB-Fold+AAA-ATPase+Lon-Protease-BrxL<-PglZ<-?<-N6-METHYLASE-p<-RVT<-N6-METHYLASE<-?<-HNH<-DUF6079*<-BrxB-BrxF-iSTAND<-BrxA-DNA-binding<-wHTH+WYL+WCX<-?<-?<-XIS-HTH<-XIS-HTH<-?<-Arm-DNA-bind_3+Phage_integrase | 1225 | Bacteria>Pseudomonadota>Gammaproteobacteria>Alteromonadales>Shewanellaceae>Shewanella | GCA_017580215.1 | Shewanella_algae | I6M36_02045 | 117246..120923 | 38313 | BREX-system-P-loop-protein-BrxC-[Shewanella-algae] |
| MCD8561045.1 | Type1.BREX | <-N6-METHYLASE-p<-HEPN<-N6-METHYLASE-p<-N6-METHYLASE<-?<-HNH<-DUF6079*<-BrxB-BrxF-iSTAND<-BrxA-DNA-binding<-wHTH+WYL+WCX<-XIS-HTH | 1231 | Bacteria>Pseudomonadota>Gammaproteobacteria>Alteromonadales>Shewanellaceae>Shewanella | GCA_021209305.1 | Shewanella_xiamenensis | LRY75_19995 | 6155..9850 | 332186 | BREX-system-P-loop-protein-BrxC-[Shewanella-xiamenensis] |
| WP_244159705.1 | Type1.BREX | <-HDRnase<-DDE_Tnp_1<-Arm-DNA-bind_2+Phage_int_SAM_3+Phage_integrase<-Phage_AlpA\|\|HTH_26->wHTH+WYL+WCX->BrxA-DNA-binding->BrxB-BrxF-iSTAND->DUF6079*->HNH->?->N6-METHYLASE->N6-METHYLASE-p->HEPN->N6-METHYLASE-p->MULE-TRANSPOSASE->N6-METHYLASE-p-> | 1254 | Bacteria>Pseudomonadota>Gammaproteobacteria>Alteromonadales>Shewanellaceae>Shewanella | GCF_001723195.1 | Shewanella_xiamenensis | ABT47_RS18205 | 4085889..4089653 | 332186 | BREX-system-P-loop-protein-BrxC-[Shewanella-xiamenensis] |
| WP_069454895.1 | Type1.BREX | <-HDRnase<-Arm-DNA-bind_2+Phage_int_SAM_3+Phage_integrase<-Phage_AlpA\|\|HTH_26->wHTH+WYL+WCX->BrxA-DNA-binding->BrxB-BrxF-iSTAND->DUF6079*->HNH->?->N6-METHYLASE-p->HEPN->N6-METHYLASE-p->MULE-TRANSPOSASE->N6-METHYLASE-p-> | 1240 | Bacteria>Pseudomonadota>Gammaproteobacteria>Alteromonadales>Shewanellaceae>Shewanella | GCF_002074855.1 | Shewanella_xiamenensis | VO00_RS12940 | 4702..8424 | 332186 | BREX-system-P-loop-protein-BrxC-[Shewanella-xiamenensis] |
| WP_107947297.1 | Type1.BREX | <-Phage_integrase<-DUF2787<-DUF2787<-CbeA_antitoxin<-?<-?\|\|wHTH+WYL+WCX->BrxA-DNA-binding->BrxB-BrxF-iSTAND->DUF6079*->HNH->?->N6-METHYLASE->SIR2->HerA/FtsK->N6-METHYLASE-p->PglZ->SIGMA+OB-Fold+AAA-ATPase+Lon-Protease-BrxL-> | 1246 | Bacteria>Pseudomonadota>Gammaproteobacteria>Alteromonadales>Shewanellaceae>Shewanella | GCF_003052765.1 | Shewanella_baltica | C7T95_RS09475 | 2180761..2184501 | 62322 | BREX-system-P-loop-protein-BrxC-[Shewanella-baltica] |
| WP_173294709.1 | Type1.BREX | wHTH(SBD)+HNH->BrxA-DNA-binding->BrxB-BrxF-iSTAND->DUF6079*->N6-METHYLASE->PglZ-><-SF1-SF2-Helicase+REase<-SF1-SF2-Helicase+REase\|\|?-><-PAIREDC-HTH | 1157 | Bacteria>Pseudomonadota>Gammaproteobacteria>Alteromonadales>Shewanellaceae>Shewanella | GCF_013248905.1 | Shewanella_sp._VB17 | HQQ94_RS12350 | 2876249..2879722 | 2739432 | BREX-system-P-loop-protein-BrxC-[Shewanella-sp.-VB17] |
| WP_243891830.1 | Type1.BREX | <-SIGMA+OB-Fold+AAA-ATPase+Lon-Protease-BrxL<-PglZ<-?<-N6-METHYLASE-p<-RVT<-N6-METHYLASE<-?<-HNH<-DUF6079*<-BrxB-BrxF-iSTAND<-BrxA-DNA-binding<-wHTH+WYL+WCX<-?<-?<-XIS-HTH<-?<-Arm-DNA-bind_3+Phage_integrase | 1226 | Bacteria>Pseudomonadota>Gammaproteobacteria>Alteromonadales>Shewanellaceae>Shewanella | GCF_017580215.1 | Shewanella_algae | I6M36_RS02045 | 117246..120926 | 38313 | BREX-system-P-loop-protein-BrxC-[Shewanella-algae] |
| WP_219012197.1 | Type6.BREX | Arm-DNA-bind_3+Phage_integrase-><-HTH_3\|\|?-><-SF1-SF2-Helicase<-REase\|\|HNH-><-SIGMA+OB-Fold+AAA-ATPase+HKD-DNase-BrxL<-PglZ<-N6-METHYLASE<-DUF6079*<-BrxB-BrxF-iSTAND<-BrxA-DNA-binding<-BrxE<-XIS-HTH\|\|?->?->DUF2787->Phage_integrase-> | 1211 | Bacteria>Pseudomonadota>Gammaproteobacteria>Alteromonadales>Shewanellaceae>Shewanella | GCF_019303235.1 | Shewanella_algae | K0E59_RS06910 | 134402..138037 | 38313 | BREX-system-P-loop-protein-BrxC-[Shewanella-algae] |
| WP_263171306.1 | Type1.BREX | <-Phage_integrase<-DUF2787<-DUF2787<-CbeA_antitoxin<-?<-?\|\|wHTH+WYL+WCX->BrxA-DNA-binding->BrxB-BrxF-iSTAND->DUF6079*->HNH->?->N6-METHYLASE->SIR2->HerA/FtsK->N6-METHYLASE-p->PglZ->SIGMA+OB-Fold+AAA-ATPase+Lon-Protease-BrxL-> | 1235 | Bacteria>Pseudomonadota>Gammaproteobacteria>Alteromonadales>Shewanellaceae>Shewanella | GCF_025653795.1 | Shewanella_sp._SM73 | L5M51_RS01900 | 416485..420192 | 2912806 | BREX-system-P-loop-protein-BrxC-[Shewanella-sp.-SM73] |
| WP_263197229.1 | Type1.BREX | <-SIGMA+OB-Fold+AAA-ATPase+Lon-Protease-BrxL<-PglZ<-N6-METHYLASE-p<-HEPN<-N6-METHYLASE-p<-N6-METHYLASE<-?<-HNH<-DUF6079*<-BrxB-BrxF-iSTAND<-BrxA-DNA-binding<-wHTH+WYL+WCX<-HTH_26\|\|Phage_AlpA->Arm-DNA-bind_2+Phage_int_SAM_3+Phage_integrase->HDRnase-> | 1246 | Bacteria>Pseudomonadota>Gammaproteobacteria>Alteromonadales>Shewanellaceae>Shewanella | GCF_025654545.1 | Shewanella_sp._SM20 | L5M18_RS13095 | 42119..45859 | 2912792 | BREX-system-P-loop-protein-BrxC-[Shewanella-sp.-SM20] |
| WP_132582321.1 | Type6.BREX | Phage_integrase->Phage_integrase->?->HerA/FtsK->TrwC+SF1-SF2-Helicase->?->BrxE->BrxA-DNA-binding->BrxB-BrxF-iSTAND->DUF6079*->N6-METHYLASE->PglZ->SIGMA+OB-Fold+AAA-ATPase+HKD-DNase-BrxL-><-HNH\|\|?->MarR-HTH->AbiEii-> | 1208 | Bacteria>Pseudomonadota>Gammaproteobacteria>Chromatiales>Chromatiaceae>Rheinheimera | GCF_004345025.1 | Rheinheimera_sp._D18 | E0Z06_RS04605 | 954479..958105 | 2545632 | BREX-system-P-loop-protein-BrxC-[Rheinheimera-sp.-D18] |
| WP_099719181.1 | Type1.BREX | nSTAND1+HNH->RNaseH->NACHT+SF1-SF2-Helicase+HKD-DNase->REase-><-HKD-DNase+OGG-like<-SIGMA+OB-Fold+AAA-ATPase+Lon-Protease-BrxL<-PglZ<-N6-METHYLASE<-DUF6079*<-BrxB-BrxF-iSTAND<-BrxA-DNA-binding<-?<-HNH<-ACET\|\|?-><-DUF2754 | 1213 | Bacteria>Pseudomonadota>Gammaproteobacteria>Enterobacterales>Enterobacteriaceae>Klebsiella/Raoultella_group | GCF_018137755.1 | Klebsiella_pneumoniae | KFB18_RS20840 | 34480..38121 | 573 | BREX-system-P-loop-protein-BrxC-[Klebsiella-pneumoniae] |
| WP_261139451.1 | Type1.BREX | DOC+HTH-><-LACI-HTH+PBPI\|\|?->Glyco_hydro_1->?->SAD+HNH->BrxA-DNA-binding->BrxB-BrxF-iSTAND->DUF6079*->PglZ->SIGMA+OB-Fold+AAA-ATPase+Lon-Protease-BrxL->HKD-DNase+OGG-like-><-Couple_hipA+HipA_C<-HTH_3<-?\|\|HIN-HTH-><-HTH_21+rve+rve_3<-rve | 1213 | Bacteria>Pseudomonadota>Gammaproteobacteria>Enterobacterales>Yersiniaceae>Serratia | GCF_946226675.1 | Serratia_liquefaciens | N8M82_RS16050 | 322435..326076 | 614 | BREX-system-P-loop-protein-BrxC-[Serratia-liquefaciens] |

<table>
<tr><th>Protein-ID</th><th>System</th><th>Gene-Neighbourhoods</th><th>Length</th><th>Taxonomy</th><th>Assembly-ID</th><th>Strain</th><th>Locus-Tag</th><th>Genomic-coordinates</th><th>TaxID</th><th>Fasta-define</th></tr>
<tr><td>WP_261135874.1</td><td>Type1.BREX</td><td>DOC+HTH-><-LACI-HTH+PBPI||?->Glyco_hydro_1->?-> SAD+ HNH -> BrxA-DNA-binding -> BrxB-BrxF-iSTAND -> DUF6079 *-> N6-METHYLASE -> PglZ -> SIGMA+OB-Fold+AAA-ATPase+Lon-Protease-BrxL ->HKD-DNase+OGG-like-><-Couple_hipA+HipA_C<-HTH_3</td><td>1213</td><td>Bacteria>Pseudomonadota>Gammaproteobacteria>Enterobacterales>Yersiniaceae>Serratia</td><td>GCF_946226775.1</td><td>Serratia_liquefaciens</td><td>N8M94_RS14775</td><td>322685..326326</td><td>614</td><td>BREX-system-P-loop-protein-BrxC-[Serratia-liquefaciens]</td></tr>
<tr><td>WP_261427666.1</td><td>Type1.BREX</td><td>Glyco_hydro_1-><-?<-?||?->?-> SAD+ HNH -> BrxA-DNA-binding -> BrxB-BrxF-iSTAND -> DUF6079 *-> N6-METHYLASE -> PglZ -> SIGMA+OB-Fold+AAA-ATPase+Lon-Protease-BrxL ->HKD-DNase+OGG-like-><-Couple_hipA+HipA_C<-HTH_3</td><td>1213</td><td>Bacteria>Pseudomonadota>Gammaproteobacteria>Enterobacterales>Yersiniaceae>Serratia</td><td>GCF_946227925.1</td><td>Serratia_proteamaculans</td><td>OCH17_RS14295</td><td>357572..361213</td><td>28151</td><td>BREX-system-P-loop-protein-BrxC-[Serratia-proteamaculans]</td></tr>
<tr><td>WP_104852217.1</td><td>Type1.BREX</td><td><-AAA-ATPase<-HIN-HTH||DDE_Tnp_1_3-><-?<-?||HIN-HTH-><- PglZ <- N6-METHYLASE <- DUF6079 *<- BrxB-BrxF-iSTAND <- BrxA-DNA-binding <- wHTH(SBD)+ HNH <-SAM-METHYLASE<-XIS-HTH<-?<-?<-DUF932</td><td>1247</td><td>Bacteria>Pseudomonadota>Gammaproteobacteria>Moraxellales>Moraxellaceae>Acinetobacter</td><td>GCF_002946595.1</td><td>Acinetobacter_sp._SWBY1</td><td>C3Y93_RS04635</td><td>951564..955307</td><td>2079596</td><td>BREX-system-P-loop-protein-BrxC-[Acinetobacter-sp.-SWBY1]</td></tr>
<tr><td>WP_201567145.1</td><td>Type1.BREX</td><td><-REase<- wHTH+WYL+WCX <-?||?->SIR2-><- SIGMA+OB-Fold+AAA-ATPase+Lon-Protease-BrxL <- PglZ <- N6-METHYLASE <- N6-METHYLASE <- DUF6079 *<- BrxB-BrxF-iSTAND <- BrxA-DNA-binding <- wHTH(SBD)+ HNH <-XIS-HTH||35exo-><-?<-HTH_Tnp_1+HTH_21+rve+rve_3</td><td>1221</td><td>Bacteria>Pseudomonadota>Gammaproteobacteria>Moraxellales>Moraxellaceae>Psychrobacter</td><td>GCF_904846115.1</td><td>Psychrobacter_immobilis</td><td>JMW93_RS11045</td><td>31356..35021</td><td>498</td><td>BREX-system-P-loop-protein-BrxC-[Psychrobacter-immobilis]</td></tr>
<tr><td>WP_201591398.1</td><td>Type1.BREX</td><td>Arm-DNA-bind_3+Phage_integrase->Phage_integrase-><- HNH <-?<-?<- SIGMA+OB-Fold+AAA-ATPase+Lon-Protease-BrxL <- PglZ <- N6-METHYLASE <- DUF6079 *<- BrxB-BrxF-iSTAND <- BrxA-DNA-binding <-ZnR||REase->REase-><-?<-?<-GNTR-HTH</td><td>1221</td><td>Bacteria>Pseudomonadota>Gammaproteobacteria>Moraxellales>Moraxellaceae>Psychrobacter</td><td>GCF_904846465.1</td><td>Psychrobacter_pulmonis</td><td>JMX70_RS09025</td><td>249655..253320</td><td>228654</td><td>BREX-system-P-loop-protein-BrxC-[Psychrobacter-pulmonis]</td></tr>
<tr><td>NRB78692.1</td><td>Type1.BREX</td><td><-sGTP<-sGTP<-?<- SIGMA+OB-Fold+AAA-ATPase+Lon-Protease-BrxL <- PglZ <- N6-METHYLASE <- DUF6079 *<- BrxB-BrxF-iSTAND <- BrxA-DNA-binding <- HNH <-?||?->?-><-SF1-SF2-Helicase<-HKD-DNase+HKD-Rease</td><td>1233</td><td>Bacteria>Pseudomonadota>Gammaproteobacteria>Oceanospirillales>Saccharospirillaceae>unclassified_Saccharospirillaceae</td><td>GCA_013214075.1</td><td>Saccharospirillaceae_bacterium</td><td>HRU38_08495</td><td>18482..22183</td><td>2497634</td><td>BREX-system-P-loop-protein-BrxC-[Saccharospirillaceae-bacterium]</td></tr>
<tr><td>KPX73888.1</td><td>Type1.BREX</td><td>PIN+OST-HTH-> HNH ->?->?-> DUF6079 -p*-> PglZ -> SIGMA+OB-Fold+AAA-ATPase+Lon-Protease-BrxL ->HIN-HTH->HTH_Tnp_1+HTH_21+rve+rve_3-></td><td>889</td><td>Bacteria>Pseudomonadota>Gammaproteobacteria>Pseudomonadales>Pseudomonadaceae>Pseudomonas</td><td>GCA_001400855.1</td><td>Pseudomonas_syringae_pv._maculicola</td><td>ALO84_01403</td><td>56181..58850</td><td>59511</td><td>Uncharacterized-protein-ALO84_01403-[Pseudomonas-syringae-pv.-maculicola]</td></tr>
<tr><td>MBL7230767.1</td><td>Type6.BREX</td><td><-CopG/RHH||CITB-HTH->XIS-HTH->?-> wHTH+WYL+WCX -> BrxE -> BrxA-DNA-binding -> BrxB-BrxF-iSTAND -> DUF6079 *-> PglZ -> SIGMA+OB-Fold+AAA-ATPase+HKD-DNase-BrxL -><- HNH <-DUF4969</td><td>1214</td><td>Bacteria>Pseudomonadota>Gammaproteobacteria>Pseudomonadales>Pseudomonadaceae>Pseudomonas</td><td>GCA_016785205.1</td><td>Pseudomonas_sp.</td><td>JJK56_22660</td><td>12252..15896</td><td>306</td><td>BREX-system-P-loop-protein-BrxC-[Pseudomonas-sp.]</td></tr>
<tr><td>WP_052507173.1</td><td>Type1.BREX</td><td>PIN+OST-HTH-> HNH ->?->?->|| BrxB-BrxF-iSTAND -p-> DUF6079 -p*->AbiEii-> PglZ -> SIGMA+OB-Fold+AAA-ATPase+Lon-Protease-BrxL -></td><td>886</td><td>Bacteria>Pseudomonadota>Gammaproteobacteria>Pseudomonadales>Pseudomonadaceae>Pseudomonas</td><td>GCF_000935725.1</td><td>Pseudomonas_syringae_group_genomosp._3</td><td>AC506_RS17885</td><td>8350..11010</td><td>251701</td><td>BREX-system-P-loop-protein-BrxC-[Pseudomonas-syringae-group-genomosp.-3]</td></tr>
<tr><td>WP_169883492.1</td><td>Type6.BREX</td><td>HNH -><- SIGMA+OB-Fold+AAA-ATPase+HKD-DNase-BrxL <- PglZ <- N6-METHYLASE <- N6-METHYLASE <- DUF6079 *<- BrxB-BrxF-iSTAND <- BrxE <- wHTH+WYL+WCX <-?<-XIS-HTH||RHH-><-tRNA<-MerR-HTH</td><td>1213</td><td>Bacteria>Pseudomonadota>Gammaproteobacteria>Pseudomonadales>Pseudomonadaceae>Pseudomonas</td><td>GCF_012986805.1</td><td>Pseudomonas_fragi</td><td>HBN91_RS17750</td><td>10864..14505</td><td>296</td><td>BREX-system-P-loop-protein-BrxC-[Pseudomonas-fragi]</td></tr>
<tr><td>WP_244972677.1</td><td>Type1.BREX</td><td><- SIGMA+OB-Fold+AAA-ATPase+Lon-Protease-BrxL <- PglZ <-AbiEii<-AbiEi_3_N+AbiEi_3<- HNH <- N6-METHYLASE <-DNA-glycosylase<- DUF6079 *<- BrxB-BrxF-iSTAND <- BrxA-DNA-binding <-?|| wHTH+WYL+WCX -><-tRNA<-MerR-HTH</td><td>1225</td><td>Bacteria>Pseudomonadota>Gammaproteobacteria>Pseudomonadales>Pseudomonadaceae>Pseudomonas</td><td>GCF_013386885.1</td><td>Pseudomonas_salomonii</td><td>HX810_RS11525</td><td>509017..512694</td><td>191391</td><td>BREX-system-P-loop-protein-BrxC-[Pseudomonas-salomonii]</td></tr>
<tr><td>WP_273507518.1</td><td>Type6.BREX</td><td><-CopG/RHH||CITB-HTH->XIS-HTH->?-> wHTH+WYL+WCX -> BrxE -> BrxA-DNA-binding -> BrxB-BrxF-iSTAND -> DUF6079 *-> PglZ -> SIGMA+OB-Fold+AAA-ATPase+HKD-DNase-BrxL -><- HNH <-DUF4969</td><td>1214</td><td>Bacteria>Pseudomonadota>Gammaproteobacteria>Pseudomonadales>Pseudomonadaceae>Pseudomonas</td><td>GCF_016785205.1</td><td>Pseudomonas_sp.</td><td>JJK56_22660</td><td>12252..15896</td><td>306</td><td>BREX-system-P-loop-protein-BrxC-[Pseudomonas-sp.]</td></tr>
<tr><td>WP_058409982.1</td><td>Type6.BREX</td><td><-LuxR-HTH<-?<-ZnR+TOPRIM+AAA-helicase+wHTH||XIS-HTH->?-> wHTH+WYL+WCX -> BrxE -> BrxA-DNA-binding -> BrxB-BrxF-iSTAND -> DUF6079 *-> N6-METHYLASE -> PglZ -> SIGMA+OB-Fold+AAA-ATPase+HKD-DNase-BrxL -><- HNH <-?<-?||HTH_3-><-ZnR+ZnR<-DUF4969</td><td>1212</td><td>Bacteria>Pseudomonadota>Gammaproteobacteria>Pseudomonadales>Pseudomonadaceae>Pseudomonas</td><td>GCF_029959515.1</td><td>Pseudomonas_sp._ICMP_19500</td><td>AO239_RS04345</td><td>8658..12296</td><td>1208102</td><td>BREX-system-P-loop-protein-BrxC-[Pseudomonas-sp.-ICMP-19500]</td></tr>
<tr><td>WP_033991202.1</td><td>Type1.BREX</td><td><- HNH <- SIGMA+OB-Fold+AAA-ATPase+Lon-Protease-BrxL <- PglZ <-MNS+REase<-?<- HNH +GCN1<-?<- N6-METHYLASE <- DUF6079 *<- BrxB-BrxF-iSTAND <- BrxA-DNA-binding <-RNase<-?<-JetB<-?<-Peptidase_M15<-?|| wHTH+WYL+WCX -></td><td>1154</td><td>Bacteria>Pseudomonadota>Gammaproteobacteria>Pseudomonadales>Pseudomonadaceae>Pseudomonas</td><td>GCF_929614685.1</td><td>Pseudomonas_aeruginosa</td><td>MJP43_RS27275</td><td>28461..31925</td><td>287</td><td>BREX-system-P-loop-protein-BrxC-[Pseudomonas-aeruginosa]</td></tr>
<tr><td>WP_125870426.1</td><td>Type1.BREX</td><td><-DUF932<-?||NERD->REase-> BrxA-DNA-binding -> BrxB-BrxF-iSTAND -> DUF6079 *-> N6-METHYLASE ->?-> HNH +GCN1->?-> PglZ -> SIGMA+OB-Fold+AAA-ATPase+Lon-Protease-BrxL ->REase->SF1-SF2-Helicase-><-Glycos_transf</td><td>1154</td><td>Bacteria>Pseudomonadota>Gammaproteobacteria>Pseudomonadales>Pseudomonadaceae>Stutzerimonas</td><td>GCF_024199075.1</td><td>Stutzerimonas_stutzeri</td><td>NLG33_RS12680</td><td>207505..210969</td><td>316</td><td>BREX-system-P-loop-protein-BrxC-[Stutzerimonas-stutzeri]</td></tr>
<tr><td>MCJ8311755.1</td><td>Type1.BREX</td><td><-sGTP<-sGTP<-?<- SIGMA+OB-Fold+AAA-ATPase+Lon-Protease-BrxL <- PglZ <- N6-METHYLASE <- DUF6079 *<- BrxB-BrxF-iSTAND <- BrxA-DNA-binding <- HNH <-?||?->?-><-SF1-SF2-Helicase<-HKD-DNase+HKD-Rease</td><td>1233</td><td>Bacteria>Pseudomonadota>Gammaproteobacteria>Pseudomonadales>unclassified_Pseudomonadales</td><td>GCA_022968985.1</td><td>Pseudomonadales_bacterium</td><td>HRU38_08495</td><td>18482..22183</td><td>1891229</td><td>BREX-system-P-loop-protein-BrxC-[Pseudomonadales-bacterium]</td></tr>
<tr><td>MBS1228526.1</td><td>Type1.BREX</td><td>Arm-DNA-bind_3+Phage_integrase-><-Phage_int_SAM_2+Integrase_1||?->?->?->?-> wHTH+WYL+WCX -> BrxA-DNA-binding -> BrxB-BrxF-iSTAND -> DUF6079 *-> N6-METHYLASE -> HNH ->?->HTH+REase->DUF3644-> PglZ -> SIGMA+OB-Fold+AAA-ATPase+Lon-Protease-BrxL ->DOC+HTH-></td><td>1213</td><td>Bacteria>Pseudomonadota>unclassified_Pseudomonadota</td><td>GCA_018262095.1</td><td>Pseudomonadota_bacterium</td><td>H6R17_1803</td><td>75999..79640</td><td>1977087</td><td>ATPase-like-protein-[Pseudomonadota-bacterium]</td></tr>
<tr><td>MBY0113214.1</td><td>Type6.BREX</td><td><-GDResolvase+ZnR<-RAMA<-HARE-HTH||TIR-><- SIGMA+OB-Fold+AAA-ATPase+Lon-Protease-BrxL <- PglZ <- N6-METHYLASE -p<- N6-METHYLASE -p<- N6-METHYLASE <- DUF6079 *<- BrxB-BrxF-iSTAND <- BrxA-DNA-binding <- BrxE <-EVE+ HNH <-?<-LuxR-HTH<- ParB +N6_N4_METHYLASE<-NtrC-AAA</td><td>1179</td><td>Bacteria>PVC_group>Planctomycetota>Phycisphaerae>Phycisphaerales>unclassified_Phycisphaerales</td><td>GCA_019744815.1</td><td>Phycisphaerales_bacterium</td><td>K2Y21_10355</td><td>73058..76597</td><td>2052180</td><td>BREX-system-P-loop-protein-BrxC-[Phycisphaerales-bacterium]</td></tr>
<tr><td>WP_100495666.1</td><td>Type1.BREX</td><td><-ABhydrolase||TPR+TPR+TPR+TPR-> BrxA-DNA-binding -> BrxB-BrxF-iSTAND -> DUF6079 *-><-?<-?<-?|| HNH ->?->SbcC-> N6-METHYLASE ->HEPN->Schlafen+HATPase+wHTH-></td><td>1240</td><td>Bacteria>Terrabacteria_group>Actinomycetota>Actinomycetes>Bifidobacteriales>Bifidobacteriaceae</td><td>GCF_002802865.1</td><td>Bifidobacterium_scaligerum</td><td>CUU80_RS01790</td><td>434556..438278</td><td>2052656</td><td>BREX-system-P-loop-protein-BrxC-[Bifidobacterium-scaligerum]</td></tr>
<tr><td>WP_079914137.1</td><td>Type1.BREX</td><td>REase->?->REase+DUF4268->?-> BrxA-DNA-binding -> BrxB-BrxF-iSTAND -> DUF6079 *-> N6-METHYLASE -> PglZ -> SIGMA+OB-Fold+AAA-ATPase+Lon-Protease-BrxL -> HNH ->?->HTH+MPTase->?->?->Schlafen+HATPase+wHTH-></td><td>1184</td><td>Bacteria>Terrabacteria_group>Bacillota>Bacilli>Bacillales>Paenibacillaceae</td><td>GCF_002042965.1</td><td>Paenibacillus_sp._32352</td><td>B1728_RS30170</td><td>158863..162417</td><td>1969111</td><td>BREX-system-P-loop-protein-BrxC-[Paenibacillus-sp.-32352]</td></tr>
<tr><td>WP_022757624.1</td><td>Type1.BREX</td><td><- HNH +NERD+ZnR||GDResolvase-><-?<-?<- SIGMA+OB-Fold+AAA-ATPase+Lon-Protease-BrxL <- HNH <- PglZ <-?<- N6-METHYLASE <- DUF6079 *<- BrxB-BrxF-iSTAND <- BrxA-DNA-binding <-HTH_26<-?<-TraI-VirD2-RCRE<-TraJ-RHH<-?<-HTH_3||ParA-Soj-PloopNTPase-></td><td>1181</td><td>Bacteria>Terrabacteria_group>Bacillota>Clostridia>Eubacteriales>Lachnospiraceae</td><td>GCF_000420985.1</td><td>Butyrivibrio_fibrisolvens</td><td>G616_RS0108215</td><td>23876..27421</td><td>831</td><td>BREX-system-P-loop-protein-BrxC-[Butyrivibrio-fibrisolvens]</td></tr>
<tr><td>WP_103203921.1</td><td>Type1.BREX</td><td>RHH_8-><-?<-ZnR+TOPRIM||?->SymE_toxin->?-> wHTH(SBD)+ HNH -> BrxA-DNA-binding -> BrxB-BrxF-iSTAND -> DUF6079 *-> N6-METHYLASE -> PglZ ->?-> SIGMA+OB-Fold+AAA-ATPase+Lon-Protease-BrxL ->RAMA->ARG-></td><td>1186</td><td>Bacteria>Terrabacteria_group>Bacillota>Clostridia>Eubacteriales>Lachnospiraceae</td><td>GCF_002904165.1</td><td>Herbinix_hemicellulosilytica</td><td>HHT355_RS13290</td><td>82989..86549</td><td>1564487</td><td>BREX-system-P-loop-protein-BrxC-[Herbinix-hemicellulosilytica]</td></tr>
<tr><td>WP_215631393.1</td><td>Type1.BREX</td><td><-HTH+REase<- wHTH+WYL+WCX <- SIGMA+OB-Fold+AAA-ATPase+Lon-Protease-BrxL <-?<- PglZ <- HNH +CXC3<- N6-METHYLASE <-Four-helical-protein<-wHTH+ZnR<- DUF6079 *<- BrxB-BrxF-iSTAND <- BrxA-DNA-binding <-HTH_26<-HTH_3||?-><-Lysozyme+NLPC<-?<-?<-VirB4-FtsK</td><td>1205</td><td>Bacteria>Terrabacteria_group>Bacillota>Clostridia>Eubacteriales>Lachnospiraceae</td><td>GCF_018785175.1</td><td>Coprococcus_catus</td><td>GPK82_RS01090</td><td>19804..23421</td><td>116085</td><td>BREX-system-P-loop-protein-BrxC-[Coprococcus-catus]</td></tr>
<tr><td>MCC3659975.1</td><td>Type1.BREX</td><td>AraC-HTH+AraC-HTH-><-IONCHANNEL+HISKIN+REC<- SAD+ HNH <-?<- PglZ <-?<- N6-METHYLASE <- DUF6079 *<- BrxB-BrxF-iSTAND <- BrxA-DNA-binding <-HTH_26<-?<-?<-?<-?||?-><-68TM-wHTH+CACHE+GGDEF</td><td>1184</td><td>Bacteria>Terrabacteria_group>Bacillota>Clostridia>Eubacteriales>Oscillospiraceae</td><td>GCA_020740265.1</td><td>Ruminococcus_albus</td><td>LLP35_08975</td><td>12774..16328</td><td>1264</td><td>BREX-system-P-loop-protein-BrxC-[Ruminococcus-albus]</td></tr>
<tr><td>SCI31347.1</td><td>Type1.BREX</td><td>68TM-wHTH+CACHE+GGDEF->?->?->?->?->?->HTH_26-> BrxA-DNA-binding -> BrxB-BrxF-iSTAND -> DUF6079 *-> N6-METHYLASE ->?-> PglZ -> SIGMA+OB-Fold+AAA-ATPase+Lon-Protease-BrxL ->?-> SAD+ HNH ->IONCHANNEL+HISKIN+REC-><-DSBH+AraC-HTH+AraC-HTH</td><td>1184</td><td>Bacteria>Terrabacteria_group>Bacillota>Clostridia>Eubacteriales>Oscillospiraceae</td><td>GCA_900066635.1</td><td>uncultured_Ruminococcus_sp.</td><td>SAMEA3545357_00481</td><td>41099..44653</td><td>165186</td><td>Uncharacterised-protein-[uncultured-Ruminococcus-sp.]</td></tr>
<tr><td>WP_014256029.1</td><td>Type1.BREX</td><td><-Glyoxalase<- N6-METHYLASE <-Peptidase<-?<-?<- SIGMA+OB-Fold+AAA-ATPase+Lon-Protease-BrxL <- PglZ <-Schlafen+HATPase+wHTH<- N6-METHYLASE <- DUF6079 *<- BrxB-BrxF-iSTAND <- BrxA-DNA-binding <- wHTH(SBD)+ HNH <-?<-?<- N6-METHYLASE</td><td>1186</td><td>Bacteria>Terrabacteria_group>Bacillota>Clostridia>Eubacteriales>Oscillospiraceae</td><td>GCF_000237085.1</td><td>Acetivibrio_clariflavus</td><td>CLOCL_RS14360</td><td>3415236..3418796</td><td>288965</td><td>BREX-system-P-loop-protein-BrxC-[Acetivibrio-clariflavus]</td></tr>
<tr><td>WP_027621802.1</td><td>Type1.BREX</td><td><-Glyoxalase<- N6-METHYLASE <-Peptidase<-?<-?<- SIGMA+OB-Fold+AAA-ATPase+Lon-Protease-BrxL <- PglZ <-Schlafen+HATPase+wHTH<- N6-METHYLASE <- DUF6079 *<- BrxB-BrxF-iSTAND <- BrxA-DNA-binding <- wHTH(SBD)+ HNH ||DDE_5+DDE_Tnp_1-><-?<-?<- N6-METHYLASE</td><td>1186</td><td>Bacteria>Terrabacteria_group>Bacillota>Clostridia>Eubacteriales>Oscillospiraceae</td><td>GCF_000519985.1</td><td>Acetivibrio_clariflavus</td><td>CLOCLA_RS0107445</td><td>1195913..1199473</td><td>288965</td><td>BREX-system-P-loop-protein-BrxC-[Acetivibrio-clariflavus]</td></tr>
<tr><td>RJX18832.1</td><td>Type1.BREX</td><td><-Thermonuclease<-?<-?<-PRTase<-?<- HNH <- SIGMA+OB-Fold+AAA-ATPase+Lon-Protease-BrxL <- PglZ <- N6-METHYLASE <- DUF6079 *<- BrxB-BrxF-iSTAND <- BrxA-DNA-binding ||HIN-HTH->AAA-ATPase-></td><td>1147</td><td>Bacteria>Terrabacteria_group>Bacillota>Clostridia>Thermoanaerobacterales>Thermoanaerobacteraceae</td><td>GCA_003604985.1</td><td>Ammonifex_sp.</td><td>C4570_06155</td><td>13420..16863</td><td>1872493</td><td>MAG:-BREX-system-P-loop-protein-BrxC-[Ammonifex-sp.]</td></tr>
<tr><td>MBP3871947.1</td><td>Type1.BREX</td><td><- N6-METHYLASE -p<- PglZ <- DUF6079 *<- BrxB-BrxF-iSTAND <- BrxA-DNA-binding <-?<- HNH <-?<- HNH</td><td>1175</td><td>Bacteria>Terrabacteria_group>Bacillota>Erysipelotrichia>Erysipelotrichales>Erysipelotrichaceae</td><td>GCA_017941505.1</td><td>Faecalicoccus_sp.</td><td>J6E46_13345</td><td>3304..6831</td><td>1971758</td><td>BREX-system-P-loop-protein-BrxC-[Faecalicoccus-sp.]</td></tr>
<tr><td>MCU0565455.1</td><td>Type6.BREX</td><td><-SF1-SF2-Helicase+wHTH+Small-B-barrel+DUF1998<-BrxD-AAA<- PglZ ||?-> HNH -><- N6-METHYLASE <-?<-SKI<- DUF6079 *<- BrxB-BrxF-iSTAND <- BrxA-DNA-binding <- BrxE</td><td>1209</td><td>Bacteria>Terrabacteria_group>Cyanobacteriota/Melainabacteria_group>Cyanobacteriota>Cyanophyceae>Pseudanabaenales</td><td>GCA_025454315.1</td><td>Oculatellaceae_cyanobacterium_Prado106</td><td>MUF49_02535</td><td>23046..26675</td><td>2929566</td><td>BREX-system-P-loop-protein-BrxC-[Oculatellaceae-cyanobacterium-Prado106]</td></tr>
<tr><td>WP_254937144.1</td><td>Type6.BREX</td><td><-SIGMA-HTH||REase-> HNH -><- SIGMA+OB-Fold+AAA-ATPase+Lon-Protease-BrxL <- PglZ <- N6-METHYLASE <- DUF6079 *<- BrxB-BrxF-iSTAND <- BrxA-DNA-binding <- BrxE <-?||Sak4+WYL->?->PemK_toxin-><-?<-SAM-METHYLASE</td><td>1178</td><td>Bacteria>Terrabacteria_group>Cyanobacteriota/Melainabacteria_group>Cyanobacteriota>Cyanophyceae>Synechococcales</td><td>GCF_024345395.1</td><td>Cyanobium_sp._Tous-M-B4</td><td>KBY73_RS11030</td><td>158493..162029</td><td>2823724</td><td>BREX-system-P-loop-protein-BrxC-[Cyanobium-sp.-Tous-M-B4]</td></tr>
<tr><td>WP_221587161.1</td><td>Type1.BREX</td><td><- SIGMA+OB-Fold+AAA-ATPase+Lon-Protease-BrxL <- PglZ ||Herpes_LMP1+REase-><-NADAR<-HTH+ART-DarT<- HNH <- N6-METHYLASE <- DUF6079 *<- BrxB-BrxF-iSTAND <- BrxA-DNA-binding <-CHTH||HKD-DNase+SF1-SF2-Helicase+HTH+HTH+SRA-><-?||?->?->?->Phage_integrase-></td><td>1145</td><td>Bacteria>Terrabacteria_group>Deinococcota>Deinococci>Deinococcales>Deinococcaceae</td><td>-</td><td>Deinococcus_sp._RIT780</td><td>K5D85_00055</td><td>17975..21412</td><td>2870472</td><td>BREX-system-P-loop-protein-BrxC-[Deinococcus-sp.-RIT780]</td></tr>
<tr><td>GGM57141.1</td><td>Type1.BREX</td><td><- SIGMA+OB-Fold+AAA-ATPase+Lon-Protease-BrxL <- PglZ ||Herpes_LMP1+REase-><-NADAR<-fvmX2+ART-DarT<- HNH <- N6-METHYLASE <- DUF6079 *<- BrxB-BrxF-iSTAND <- BrxA-DNA-binding <-CHTH<-?<-?<-?<-REase</td><td>1153</td><td>Bacteria>Terrabacteria_group>Deinococcota>Deinococci>Deinococcales>Deinococcaceae</td><td>GCA_014647995.1</td><td>Deinococcus_arenae</td><td>GCM10008956_36020</td><td>13025..16486</td><td>1452751</td><td>hypothetical-protein-GCM10008956_36020-[Deinococcus-arenae]</td></tr>
<tr><td>MBX8463533.1</td><td>Type1.BREX</td><td><- SIGMA+OB-Fold+AAA-ATPase+Lon-Protease-BrxL <- PglZ ||Herpes_LMP1+REase-><-NADAR<-HTH+ART-DarT<- HNH <- N6-METHYLASE <- DUF6079 *<- BrxB-BrxF-iSTAND <- BrxA-DNA-binding <-CHTH||HKD-DNase+SF1-SF2-Helicase+HTH+HTH+SRA-><-?||?->?->?->Phage_integrase-></td><td>1145</td><td>Bacteria>Terrabacteria_group>Deinococcota>Deinococci>Deinococcales>Deinococcaceae</td><td>GCA_019718715.1</td><td>Deinococcus_sp._RIT780</td><td>K5D85_00055</td><td>17975..21412</td><td>2870472</td><td>BREX-system-P-loop-protein-BrxC-[Deinococcus-sp.-RIT780]</td></tr>
<tr><td>WP_135229998.1</td><td>Type1.BREX</td><td>RNase->?-><-SF1-SF2-Helicase<-REase||<-RNase||?-><-SF1-SF2-Helicase+wHTH+Small-B-barrel+DUF1998|| BrxA-DNA-binding -> BrxB-BrxF-iSTAND -> DUF6079 *->RelE-ParE+SF1-SF2-Helicase->?-> N6-METHYLASE -> HNH ->HTH+ART-DarT->NADAR->REase->PhdYeFM_antitox->PIN-></td><td>1141</td><td>Bacteria>Terrabacteria_group>Deinococcota>Deinococci>Deinococcales>Deinococcaceae</td><td>GCF_004634215.1</td><td>Deinococcus_fonticola</td><td>E5Z01_RS14400</td><td>28954..32379</td><td>2528713</td><td>BREX-system-P-loop-protein-BrxC-[Deinococcus-fonticola]</td></tr>
<tr><td>WP_143721449.1</td><td>Type1.BREX</td><td>SF1-SF2-Helicase+wHTH+Small-B-barrel+DUF1998-><-McrC-NTD+REase<-AAA-ATPase<- SIGMA+OB-Fold+AAA-ATPase+Lon-Protease-BrxL <- PglZ ||?-><- HNH <- N6-METHYLASE <- DUF6079 *<- BrxB-BrxF-iSTAND <- BrxA-DNA-binding <- wHTH+WYL+WCX <-?<-SF1-SF2-Helicase+REase<-Exonuc_V_gamma+REase<-RNase<-?<-MerR-HTH</td><td>1148</td><td>Bacteria>Terrabacteria_group>Deinococcota>Deinococci>Deinococcales>Deinococcaceae</td><td>GCF_007280555.1</td><td>Deinococcus_detaillensis</td><td>FNU79_RS14110</td><td>42957..46403</td><td>2592048</td><td>BREX-system-P-loop-protein-BrxC-[Deinococcus-detaillensis]</td></tr>
<tr><td>WP_189062792.1</td><td>Type5.BREX</td><td><- SIGMA+OB-Fold+AAA-ATPase+Lon-Protease-BrxL <- PglZ ||Herpes_LMP1+REase-><-NADAR<-fvmX2+ART-DarT<- HNH <- N6-METHYLASE <- DUF6079 *<- BrxB-BrxF-iSTAND <- BrxA-DNA-binding <-CHTH<-?<-REase||?->?->?-><-TUDOR+SF1-SF2-Helicase+iREase+CTD+REase</td><td>1145</td><td>Bacteria>Terrabacteria_group>Deinococcota>Deinococci>Deinococcales>Deinococcaceae</td><td>GCF_014647995.1</td><td>Deinococcus_arenae</td><td>IEY66_RS18050</td><td>13025..16462</td><td>1452751</td><td>BREX-system-P-loop-protein-BrxC-[Deinococcus-arenae]</td></tr>
<tr><td>WP_255578705.1</td><td>Type1.BREX</td><td><- SIGMA+OB-Fold+AAA-ATPase+Lon-Protease-BrxL <- PglZ ||Herpes_LMP1+REase-><-NADAR<-ART-DarT<- HNH <- N6-METHYLASE <- DUF6079 *<- BrxB-BrxF-iSTAND <- BrxA-DNA-binding <-CHTH||HKD-DNase+SF1-SF2-Helicase+HTH+HTH+SRA-><-?||?->?->?->Phage_integrase-></td><td>1153</td><td>Bacteria>Terrabacteria_group>Deinococcota>Deinococci>Deinococcales>Deinococcaceae</td><td>GCF_019718715.1</td><td>Deinococcus_sp._RIT780</td><td>K5D85_RS00055</td><td>17975..21436</td><td>2870472</td><td>BREX-system-P-loop-protein-BrxC-[Deinococcus-sp.-RIT780]</td></tr>
<tr><td>WP_092265648.1</td><td>Type1.BREX</td><td><-HIN-HTH<- SIGMA+OB-Fold+AAA-ATPase+Lon-Protease-BrxL <- PglZ <-?<-wHTH||URI-DNase-><- HNH <- N6-METHYLASE <- DUF6079 *<- BrxB-BrxF-iSTAND <- BrxA-DNA-binding <-SF1-SF2-Helicase+REase<-Exonuc_V_gamma+REase<-RNase<-?<-DDE_3</td><td>1160</td><td>Bacteria>Terrabacteria_group>Deinococcota>Deinococci>Deinococcales>Deinococcaceae</td><td>GCF_900109185.1</td><td>Deinococcus_reticulitermitis</td><td>BMY43_RS15290</td><td>11757..15239</td><td>856736</td><td>BREX-system-P-loop-protein-BrxC-[Deinococcus-reticulitermitis]</td></tr>
<tr><td>WP_254867722.1</td><td>Type1.BREX</td><td><-HIN-HTH|| BrxA-DNA-binding -> BrxB-BrxF-iSTAND -> DUF6079 *-> N6-METHYLASE -> PglZ -> SIGMA+OB-Fold+AAA-ATPase+Lon-Protease-BrxL -> HNH ->CR-ATPase-><-?<-?||ABhydrolase-></td><td>1163</td><td>Bacteria>Thermotogota>Thermotogae>Petrotogales>Petrotogaceae>Marinitoga</td><td>GCF_013363285.1</td><td>Marinitoga_sp._1154</td><td>XO12_RS09410</td><td>1913..5404</td><td>1643335</td><td>BREX-system-P-loop-protein-BrxC-[Marinitoga-sp.-1154]</td></tr>
<tr><td>MBU4348826.1</td><td>Type1.BREX</td><td>BrxB-BrxF-iSTAND -> DUF6079 *-> N6-METHYLASE -> N6-METHYLASE -p-> PglZ -> SIGMA+OB-Fold+AAA-ATPase+Lon-Protease-BrxL -> HNH ->?->SF1-SF2-Helicase->?->REase-></td><td>1193</td><td>Bacteria>unclassified_Bacteria</td><td>GCA_018896635.1</td><td>bacterium</td><td>KJ599_00710</td><td>356..3937</td><td>1869227</td><td>BREX-system-P-loop-protein-BrxC-[bacterium]</td></tr>
</table>

<table>
<tr><th>Protein-ID</th><th>System</th><th>Gene-Neighbourhoods</th><th>Length</th><th>Taxonomy</th><th>Assembly-ID</th><th>Strain</th><th>Locus-Tag</th><th>Genomic-coordinates</th><th>TaxID</th><th>Fasta-defline</th></tr>
<tr><td>--</td><td></td><td></td><td></td><td></td><td></td><td></td><td></td><td></td><td></td><td></td></tr>
<tr><td>--PIN-EndoRNase</td><td></td><td></td><td></td><td></td><td></td><td></td><td></td><td></td><td></td><td></td></tr>
<tr><td>--</td><td></td><td></td><td></td><td></td><td></td><td></td><td></td><td></td><td></td><td></td></tr>
<tr><td>MCL2156391.1</td><td>Type1.BREX</td><td>BrxA-DNA-binding->BrxB-BrxF-iSTAND->DUF6079*->Phage_integrase+Phage_integrase-><-N6-METHYLASE||N6-METHYLASE->?->PIN->PglZ->SIGMA+OB-Fold+AAA-ATPase+Lon-Protease-BrxL-></td><td>1194</td><td>Archaea>Euryarchaeota>Methanomada_group>Methanobacteria>Methanobacteriales>Methanobacteriaceae</td><td>GCA_009776495.1</td><td>Methanobrevibacter_sp.</td><td>FWH54_00015</td><td>1416..5000</td><td>66852</td><td>BREX-system-P-loop-protein-BrxC-[Methanobrevibacter-sp.]</td></tr>
<tr><td>ELZ19699.1</td><td>Type1.BREX</td><td><-N6-METHYLASE-p||?->?->?-><-PIN||?-><-REase<-?||?-><-DUF6079*<-BrxB-BrxF-iSTAND<-DUF6079<-BrxA-DNA-binding<-CR-ATPase9+REase<-?<-?||REase->SF1-SF2-Helicase->REase-></td><td>1198</td><td>Archaea>Euryarchaeota>Stenosarchaea_group>Halobacteria>Halobacteriales>Haloarculaceae</td><td>GCA_000337455.1</td><td>Halosimplex_carlsbadense_2-9-1</td><td>C475_22179</td><td>59263..62859</td><td>797114</td><td>hypothetical-protein-C475_22179-[Halosimplex-carlsbadense-2-9-1]</td></tr>
<tr><td>ELZ19701.1</td><td>Type1.BREX</td><td><-PIN||?-><-REase<-?||?-><-DUF6079<-BrxB-BrxF-iSTAND<-DUF6079*<-BrxA-DNA-binding<-CR-ATPase9+REase<-?<-?||REase->SF1-SF2-Helicase->REase->SF1-SF2-Helicase-></td><td>1248</td><td>Archaea>Euryarchaeota>Stenosarchaea_group>Halobacteria>Halobacteriales>Haloarculaceae</td><td>GCA_000337455.1</td><td>Halosimplex_carlsbadense_2-9-1</td><td>C475_22189</td><td>63665..67411</td><td>797114</td><td>hypothetical-protein-C475_22189-[Halosimplex-carlsbadense-2-9-1]</td></tr>
<tr><td>WP_135304847.1</td><td>Type5.BREX</td><td><-REase||?-><-MORC<-TUDOR+SF1-SF2-Helicase+iREase+CTD<-N6-METHYLASE<-PglZ<-N6-METHYLASE<-PIN<-?<-DUF6079*<-BrxB-BrxF-iSTAND<-DUF6079<-BrxA-DNA-binding||?->?->?-><-?||McrB-AAA->wHTH+REase-></td><td>1273</td><td>Archaea>Euryarchaeota>Stenosarchaea_group>Halobacteria>Halobacteriales>Haloarculaceae</td><td>GCF_004681185.1</td><td>Halomicroarcula_amylolytica</td><td>E4S61_RS15390</td><td>199089..202910</td><td>2562280</td><td>hypothetical-protein-[Halomicroarcula-amylolytica]</td></tr>
<tr><td>WP_135304849.1</td><td>Type5.BREX</td><td><-MORC<-TUDOR+SF1-SF2-Helicase+iREase+CTD<-N6-METHYLASE<-PglZ<-N6-METHYLASE<-PIN<-?<-DUF6079<-BrxB-BrxF-iSTAND<-DUF6079*<-BrxA-DNA-binding||?->?->?-><-?||McrB-AAA->wHTH+REase-><-SF1-SF2-Helicase</td><td>1250</td><td>Archaea>Euryarchaeota>Stenosarchaea_group>Halobacteria>Halobacteriales>Haloarculaceae</td><td>GCF_004681185.1</td><td>Halomicroarcula_amylolytica</td><td>E4S61_RS15400</td><td>203494..207246</td><td>2562280</td><td>hypothetical-protein-[Halomicroarcula-amylolytica]</td></tr>
<tr><td>WP_170076589.1</td><td>Type6.BREX</td><td><-SbcC<-Calcineurin<-REase<-?<-Calcineurin-SbcD||BrxE->BrxA-DNA-binding->BrxB-BrxF-iSTAND->DUF6079*->N6-METHYLASE->?->SbcC->PglZ->SIGMA+OB-Fold+AAA-ATPase+Lon-Protease-BrxL->PIN->ARG-></td><td>1214</td><td>Archaea>Euryarchaeota>Stenosarchaea_group>Halobacteria>Haloferacales>Haloferacaceae</td><td>GCF_012726105.1</td><td>Haloferax_alexandrinus</td><td>GOC85_RS10615</td><td>2063512..2067156</td><td>114529</td><td>BREX-system-P-loop-protein-BrxC-[Haloferax-alexandrinus]</td></tr>
<tr><td>WP_188786134.1</td><td>Type5.BREX</td><td><-ABC-ATPase+SbcC<-?<-?||tRNA->tRNA-><-TUDOR+SF1-SF2-Helicase+iREase+CTD<-PglZ<-N6-METHYLASE<-PIN<-?<-?<-DUF6079<-BrxB-BrxF-iSTAND<-DUF6079*<-BrxA-DNA-binding<-Phage_integrase<-REase<-PAS||?-><-DSBH</td><td>1265</td><td>Archaea>Euryarchaeota>Stenosarchaea_group>Halobacteria>Haloferacales>Haloferacaceae</td><td>GCF_014646875.1</td><td>Halobellus_salinus</td><td>IEX72_RS04225</td><td>173567..177364</td><td>931585</td><td>hypothetical-protein-[Halobellus-salinus]</td></tr>
<tr><td>WP_188786136.1</td><td>Type5.BREX</td><td><-TUDOR+SF1-SF2-Helicase+iREase+CTD<-PglZ<-N6-METHYLASE<-PIN<-?<-?<-DUF6079<-BrxB-BrxF-iSTAND<-DUF6079*<-BrxA-DNA-binding<-Phage_integrase<-REase<-PAS||?-><-DSBH<-?<-Sulfatase+Phosphodiest+DUF4976<-NUDIX+DUF429</td><td>1239</td><td>Archaea>Euryarchaeota>Stenosarchaea_group>Halobacteria>Haloferacales>Haloferacaceae</td><td>GCF_014646875.1</td><td>Halobellus_salinus</td><td>IEX72_RS04235</td><td>177948..181667</td><td>931585</td><td>hypothetical-protein-[Halobellus-salinus]</td></tr>
<tr><td>WP_015911586.1</td><td>Type1.BREX</td><td><-N6-METHYLASE-p||MazE_antitoxin->PIN-><-DUF6079*<-BrxB-BrxF-iSTAND<-DUF6079<-BrxA-DNA-binding<-SF1-SF2-Helicase+REase<-SF1-SF2-Helicase+REase</td><td>1270</td><td>Archaea>Euryarchaeota>Stenosarchaea_group>Halobacteria>Haloferacales>Halorubraceae</td><td>GCF_000022205.1</td><td>Halorubrum_lacusprofundi</td><td>CF318_RS14700</td><td>1993..5805</td><td>2247</td><td>hypothetical-protein-[Halorubrum-lacusprofundi]</td></tr>
<tr><td>WP_015911588.1</td><td>Type1.BREX</td><td><-TUDOR+SF1-SF2-Helicase+iREase+CTD<-PglZ<-N6-METHYLASE||DDE_Tnp_1-><-N6-METHYLASE-p||MazE_antitoxin->PIN-><-DUF6079<-BrxB-BrxF-iSTAND<-DUF6079*<-BrxA-DNA-binding<-SF1-SF2-Helicase+REase<-SF1-SF2-Helicase+REase<-?||DDE_Tnp_1-><-?<-MazE_antitoxin</td><td>1239</td><td>Archaea>Euryarchaeota>Stenosarchaea_group>Halobacteria>Haloferacales>Halorubraceae</td><td>GCF_000022205.1</td><td>Halorubrum_lacusprofundi</td><td>CF318_RS14710</td><td>6389..10108</td><td>2247</td><td>hypothetical-protein-[Halorubrum-lacusprofundi]</td></tr>
<tr><td>WP_256418943.1</td><td>Type5.BREX</td><td><-ORC-CDC6+HTH||?-><-TUDOR+SF1-SF2-Helicase+iREase+CTD<-PglZ<-?<-N6-METHYLASE||PIN-><-DUF6079<-BrxB-BrxF-iSTAND<-DUF6079*<-BrxA-DNA-binding||SF1-SF2-Helicase+REase->SF1-SF2-Helicase+REase->ParA-Soj-PloopNTPase-></td><td>1248</td><td>Archaea>Euryarchaeota>Stenosarchaea_group>Halobacteria>Haloferacales>Halorubraceae</td><td>GCF_024494655.1</td><td>Halorubrum_laminariae</td><td>NO887_RS14550</td><td>91131..94877</td><td>1433523</td><td>hypothetical-protein-[Halorubrum-laminariae]</td></tr>
<tr><td>AEN07515.1</td><td>Type5.BREX</td><td>MazE_antitoxin->?-><-DDE_Tnp_1||?->SF1-SF2-Helicase+REase->SF1-SF2-Helicase+REase->?->BrxA-DNA-binding->DUF6079*->BrxB-BrxF-iSTAND->DUF6079-><-PIN<-MazE_antitoxin||N6-METHYLASE->?->PglZ->TUDOR+SF1-SF2-Helicase+iREase+CTD-></td><td>1239</td><td>Archaea>Euryarchaeota>Stenosarchaea_group>Halobacteria>Haloferacales>unclassified_Haloferacales</td><td>GCA_000224475.1</td><td>halophilic_archaeon_DL31</td><td>Halar_0255</td><td>246678..250397</td><td>756883</td><td>hypothetical-protein-Halar_0255-[halophilic-archaeon-DL31]</td></tr>
<tr><td>AEN07517.1</td><td>Type5.BREX</td><td><-DDE_Tnp_1||?->SF1-SF2-Helicase+REase->SF1-SF2-Helicase+REase->?->BrxA-DNA-binding->DUF6079->BrxB-BrxF-iSTAND->DUF6079*-><-PIN<-MazE_antitoxin||N6-METHYLASE->?->PglZ->TUDOR+SF1-SF2-Helicase+iREase+CTD-><-?<-SF1-SF2-Helicase</td><td>1270</td><td>Archaea>Euryarchaeota>Stenosarchaea_group>Halobacteria>Haloferacales>unclassified_Haloferacales</td><td>GCA_000224475.1</td><td>halophilic_archaeon_DL31</td><td>Halar_0257</td><td>250981..254793</td><td>756883</td><td>hypothetical-protein-Halar_0257-[halophilic-archaeon-DL31]</td></tr>
<tr><td>WP_179261303.1</td><td>Type6.BREX</td><td><-HD<-?||ABhydrolase->?-><-SIGMA+OB-Fold+AAA-ATPase+Lon-Protease-BrxL<-PglZ<-N6-METHYLASE<-DUF6079*<-BrxB-BrxF-iSTAND<-BrxA-DNA-binding<-BrxE<-?<-PIN<-MazE_antitoxin</td><td>1214</td><td>Archaea>Euryarchaeota>Stenosarchaea_group>Halobacteria>Natrialbales>Natrialbaceae</td><td>GCF_013402815.2</td><td>Natrinema_halophilum</td><td>HYG82_RS33105</td><td>2537554..2541198</td><td>1699371</td><td>BREX-system-P-loop-protein-BrxC-[Natrinema-halophilum]</td></tr>
<tr><td>MCI0624094.1</td><td>Type1.BREX</td><td><-HHH_5+RecR+TOPRIM<-?<-RFC-AAA+RuvB-AAA<-ncRNA<-PglZ<-PIN<-PhdYeFM_antitox<-REase<-DUF6079*<-BrxB-BrxF-iSTAND<-BrxA-DNA-binding||?->?->?-><-?<-AA_kinase+ACT+ACT_7<-NAD_binding_3+Homoserine_dh<-REC+NtrC-AAA+FIS-HTH<-GAF+PAS+HISKIN</td><td>1197</td><td>Bacteria>Acidobacteriota>unclassified_Acidobacteriota</td><td>GCA_022615935.1</td><td>Acidobacteriota_bacterium</td><td>LO387_20995</td><td>11986..15579</td><td>1978231</td><td>BREX-system-P-loop-protein-BrxC-[Acidobacteriota-bacterium]</td></tr>
<tr><td>PXF57197.1</td><td>Type1.BREX</td><td><-PemK_toxin<-MetJarc-HTH<-PglZ<-TOPRIM<-OLD-ABC<-N6-METHYLASE<-PIN<-Phd-YefM<-DUF6079*<-BrxB-BrxF-iSTAND<-BrxA-DNA-binding<-PIN<-?||?->?->?->?->His_biosynth-></td><td>1215</td><td>Bacteria>delta/epsilon_subdivisions>Deltaproteobacteria>unclassified_Deltaproteobacteria</td><td>GCA_003194495.1</td><td>Deltaproteobacteria_bacterium</td><td>C4B58_10725</td><td>19842..23489</td><td>2026735</td><td>BREX-system-P-loop-protein-BrxC-[Deltaproteobacteria-bacterium]</td></tr>
<tr><td>MBI2378669.1</td><td>Type6.BREX</td><td>PIN-><-?<-PglZ<-Schlafen+HATPase+wHTH<-PglZ-p<-?<-RHH<-PIN<-?<-DUF6079*<-BrxB-BrxF-iSTAND<-BrxA-DNA-binding<-BrxE||PIN->?-><-?<-RADICAL-SAM</td><td>1207</td><td>Bacteria>delta/epsilon_subdivisions>Deltaproteobacteria>unclassified_Deltaproteobacteria</td><td>GCA_016183035.1</td><td>Deltaproteobacteria_bacterium</td><td>HYV07_32040</td><td>22898..26521</td><td>2026735</td><td>BREX-system-P-loop-protein-BrxC-[Deltaproteobacteria-bacterium]</td></tr>
<tr><td>MBI4818899.1</td><td>Type6.BREX</td><td><-PglZ-p<-TOPRIM<-OLD-ABC<-Schlafen+HATPase+wHTH<-?<-Schlafen+HATPase+wHTH<-N6-METHYLASE<-PIN<-?<-DUF6079*<-BrxB-BrxF-iSTAND<-BrxA-DNA-binding<-BrxE<-SF1-SF2-Helicase+wHTH+Small-B-barrel+DUF1998<-BrxD-AAA<-PglZ</td><td>1207</td><td>Bacteria>delta/epsilon_subdivisions>Deltaproteobacteria>unclassified_Deltaproteobacteria</td><td>GCA_016208175.1</td><td>Deltaproteobacteria_bacterium</td><td>HY791_21690</td><td>12157..15780</td><td>2026735</td><td>BREX-system-P-loop-protein-BrxC-[Deltaproteobacteria-bacterium]</td></tr>
<tr><td>MBI5606167.1</td><td>Type1.BREX</td><td><-Schlafen+HATPase+wHTH<-DUF6079*<-BrxB-BrxF-iSTAND<-BrxA-DNA-binding||?-><-PIN<-?||HicA_toxin->HicB_lk_antitox->ACET-></td><td>1147</td><td>Bacteria>delta/epsilon_subdivisions>Deltaproteobacteria>unclassified_Deltaproteobacteria</td><td>GCA_016219225.1</td><td>Deltaproteobacteria_bacterium</td><td>HY879_22775</td><td>11994..15437</td><td>2026735</td><td>BREX-system-P-loop-protein-BrxC-[Deltaproteobacteria-bacterium]</td></tr>
<tr><td>MCD6306396.1</td><td>Type1.BREX</td><td><-N6-METHYLASE-p||PhdYeFM_antitox->PIN-><-DUF6079*<-BrxB-BrxF-iSTAND<-BrxA-DNA-binding||wHTH+WYL+WCX-p-></td><td>1148</td><td>Bacteria>delta/epsilon_subdivisions>Deltaproteobacteria>unclassified_Deltaproteobacteria</td><td>GCA_021159525.1</td><td>Deltaproteobacteria_bacterium</td><td>J7M32_08925</td><td>1514..4960</td><td>2026735</td><td>BREX-system-P-loop-protein-BrxC-[Deltaproteobacteria-bacterium]</td></tr>
<tr><td>WP_259087839.1</td><td>Type1.BREX</td><td>URI-DNase-><-SIGMA+OB-Fold+AAA-ATPase+HKD-DNase-BrxL<-PglZ<-PIN<-XIS-HTH<-?<-?<-N6-METHYLASE<-DUF6079*<-BrxB-BrxF-iSTAND<-BrxA-DNA-binding<-?||Phage_integrase-><-TOPRIM<-?||XIS-HTH->Phage_int_SAM_4+Phage_integrase-><-tRNA||BetaPropeller+ABhydrolase-></td><td>1168</td><td>Bacteria>FCB_group>Bacteroidetes/Chlorobi_group>Bacteroidota>Bacteroidetes_incertae_sedis>Rhodothermaceae</td><td>GCF_024809725.1</td><td>Salinibacter_ruber</td><td>GGQ07_RS07865</td><td>19343..22849</td><td>146919</td><td>BREX-system-P-loop-protein-BrxC-[Salinibacter-ruber]</td></tr>
<tr><td>HAH90337.1</td><td>Type1.BREX</td><td><-D5ATPase+DUF3874<-?||BrxA-DNA-binding->BrxB-BrxF-iSTAND->DUF6079*->HTH+MPTase->PIN->?->?->?->N6-METHYLASE->PglZ->SIGMA+OB-Fold+AAA-ATPase+Lon-Protease-BrxL->HAD+HAD-></td><td>1222</td><td>Bacteria>FCB_group>Bacteroidetes/Chlorobi_group>Bacteroidota>Bacteroidia>Bacteroidales</td><td>GCA_003450425.1</td><td>Prevotella_sp.</td><td>DCL96_01170</td><td>7620..11288</td><td>59823</td><td>BREX-system-P-loop-protein-BrxC-[Prevotella-sp.]</td></tr>
<tr><td>MBR4647494.1</td><td>Type1.BREX</td><td>MNS+CR-ATPase5-><-?||?->?->Calcineurin->SbcC+CCDC158->BrxA-DNA-binding->BrxB-BrxF-iSTAND->DUF6079*->OLD-ABC->PIN->N6-METHYLASE->PglZ->SIGMA+OB-Fold+AAA-ATPase+Lon-Protease-BrxL->?->?->?->Cysteine-Desulfurase-></td><td>1174</td><td>Bacteria>FCB_group>Bacteroidetes/Chlorobi_group>Bacteroidota>Bacteroidia>Bacteroidales</td><td>GCA_017525245.1</td><td>Bacteroidales_bacterium</td><td>IKO75_10320</td><td>20247..23771</td><td>2030927</td><td>BREX-system-P-loop-protein-BrxC-[Bacteroidales-bacterium]</td></tr>
<tr><td>WP_234146092.1</td><td>Type1.BREX</td><td>Arm-DNA-bind_5+Phage_int_SAM_5+Phage_integrase-><-Schlafen+HATPase+wHTH<-SIGMA+OB-Fold+AAA-ATPase+Lon-Protease-BrxL<-PglZ<-N6-METHYLASE<-PIN<-OLD-ABC<-DUF6079*<-BrxB-BrxF-iSTAND<-BrxA-DNA-binding<-HTH_3||?->?->N6-METHYLASE-><-AAA+<-HTH_36<-XIS-HTH</td><td>1180</td><td>Bacteria>FCB_group>Bacteroidetes/Chlorobi_group>Bacteroidota>Bacteroidia>Bacteroidales</td><td>GCF_024760085.1</td><td>Bacteroides_uniformis</td><td>J8872_RS07820</td><td>80453..83995</td><td>820</td><td>BREX-system-P-loop-protein-BrxC-[Bacteroides-uniformis]</td></tr>
<tr><td>WP_264909062.1</td><td>Type1.BREX</td><td><-HAD+HAD<-SIGMA+OB-Fold+AAA-ATPase+Lon-Protease-BrxL<-PglZ<-N6-METHYLASE<-?<-?<-?<-PIN<-HTH+MPTase<-DUF6079*<-BrxB-BrxF-iSTAND<-BrxA-DNA-binding||?->D5ATPase+DUF3874<-?<-CRISPR_Cas2<-Cas_Cas1<-Cas9+HNH+RuvC_III<-HTH+MPTase</td><td>1222</td><td>Bacteria>FCB_group>Bacteroidetes/Chlorobi_group>Bacteroidota>Bacteroidia>Bacteroidales</td><td>-</td><td>Prevotella_copri</td><td>ONT11_RS09740</td><td>2183354..2187022</td><td>165179</td><td>BREX-system-P-loop-protein-BrxC-[Prevotella-copri]</td></tr>
<tr><td>WP_264951499.1</td><td>Type1.BREX</td><td>SLOG+BirA-HTH-><-?||AT-HTH-HYD2->AT-HTH-HYD2-><-HNH<-OLD-ABC<-?||BrxA-DNA-binding->BrxB-BrxF-iSTAND->DUF6079*->HTH+MPTase->PIN->?->?->?->N6-METHYLASE->PglZ->SIGMA+OB-Fold+AAA-ATPase+Lon-Protease-BrxL->HAD+HAD-></td><td>1222</td><td>Bacteria>FCB_group>Bacteroidetes/Chlorobi_group>Bacteroidota>Bacteroidia>Bacteroidales</td><td>-</td><td>Prevotella_copri</td><td>ONT17_RS13510</td><td>3302536..3306204</td><td>165179</td><td>BREX-system-P-loop-protein-BrxC-[Prevotella-copri]</td></tr>
<tr><td>WP_094415998.1</td><td>Type1.BREX</td><td><-PIN<-?<-SIGMA+OB-Fold+AAA-ATPase+Lon-Protease-BrxL<-PglZ<-N6-METHYLASE<-Schlafen+HATPase+wHTH<-DUF6079*<-BrxB-BrxF-iSTAND<-BrxA-DNA-binding<-HTH<-?<-XIS-HTH</td><td>1197</td><td>Bacteria>FCB_group>Bacteroidetes/Chlorobi_group>Bacteroidota>Flavobacteriia>Flavobacteriales</td><td>GCF_002251835.1</td><td>Flavobacterium_cyanobacteriorum</td><td>CHU92_RS12285</td><td>18327..21920</td><td>2022802</td><td>BREX-system-P-loop-protein-BrxC-[Flavobacterium-cyanobacteriorum]</td></tr>
<tr><td>WP_210512160.1</td><td>Type1.BREX</td><td>Phage_int_SAM_5+Phage_integrase->?->?->REase->?->REase->BrxA-DNA-binding->BrxB-BrxF-iSTAND->DUF6079*->N6-METHYLASE-><-AbiEii<-AbiEi_4||XIS-HTH->PIN->PglZ->SIGMA+OB-Fold+AAA-ATPase+Lon-Protease-BrxL-></td><td>1141</td><td>Bacteria>FCB_group>Bacteroidetes/Chlorobi_group>Balneolota>Balneolia>Balneolales</td><td>GCF_017921895.1</td><td>Natronogracilivirga_saccharolytica</td><td>NATSA_RS09880</td><td>86175..89600</td><td>2812953</td><td>BREX-system-P-loop-protein-BrxC-[Natronogracilivirga-saccharolytica]</td></tr>
<tr><td>MCB5286841.1</td><td>Type1.BREX</td><td><-IG+IG||?->?-><-tRNA||?->?->XIS-HTH->BrxA-DNA-binding->BrxB-BrxF-iSTAND->DUF6079*->OLD-ABC->PIN->N6-METHYLASE->BirA-HTH->PglZ->SIGMA+OB-Fold+AAA-ATPase+Lon-Protease-BrxL-></td><td>1145</td><td>Bacteria>FCB_group>Candidatus_Cloacimonetes>unclassified_Candidatus_Cloacimonetes</td><td>GCA_020532145.1</td><td>Candidatus_Cloacimonetes_bacterium</td><td>LHW64_03450</td><td>9843..13280</td><td>2030808</td><td>BREX-system-P-loop-protein-BrxC-[Candidatus-Cloacimonadota-bacterium]</td></tr>
<tr><td>HCF56393.1</td><td>Type6.BREX</td><td><-Schlafen+HATPase+wHTH<-N6-METHYLASE-p||?->PIN-><-TOPRIM<-OLD-ABC<-DUF6079-p<-APATPase<-DUF6079-p*<-DUF6079-p<-HNH<-BrxB-BrxF-iSTAND<-BrxA-DNA-binding<-BrxA-DNA-binding<-BrxE</td><td>508</td><td>Bacteria>Myxococcota>Myxococcia>Myxococcales>unclassified_Myxococcales</td><td>GCA_003534605.1</td><td>Myxococcales_bacterium</td><td>DFS52_00150</td><td>7726..9251</td><td>2026763</td><td>BREX-system-P-loop-protein-BrxC-[Myxococcales-bacterium]</td></tr>
<tr><td>HCF56394.1</td><td>Type6.BREX</td><td><-Schlafen+HATPase+wHTH<-N6-METHYLASE-p||?->PIN-><-TOPRIM<-OLD-ABC<-DUF6079-p<-APATPase<-DUF6079-p<-DUF6079-p*<-HNH<-BrxB-BrxF-iSTAND<-BrxA-DNA-binding<-BrxA-DNA-binding<-BrxE</td><td>363</td><td>Bacteria>Myxococcota>Myxococcia>Myxococcales>unclassified_Myxococcales</td><td>GCA_003534605.1</td><td>Myxococcales_bacterium</td><td>DFS52_00155</td><td>9470..10559</td><td>2026763</td><td>BREX-system-P-loop-protein-BrxC-[Myxococcales-bacterium]</td></tr>
<tr><td>MBK6517532.1</td><td>Type6.BREX</td><td>SF1-SF2-Helicase+REase->SF1-SF2-Helicase+REase->REase-><-TOPRIM<-OLD-ABC<-?||BrxE->BrxA-DNA-binding->BrxB-BrxF-iSTAND->DUF6079*-><-PemK_toxin<-?||N6-METHYLASE->N6-METHYLASE-p->?->?->PIN->PglZ->BrxD-AAA-></td><td>1209</td><td>Bacteria>Myxococcota>Polyangia>Polyangiales>Polyangiaceae>unclassified_Polyangiaceae</td><td>GCA_016704335.1</td><td>Polyangiaceae_bacterium</td><td>IPG04_26305</td><td>282797..286426</td><td>2268199</td><td>BREX-system-P-loop-protein-BrxC-[Polyangiaceae-bacterium]</td></tr>
<tr><td>MBK8937968.1</td><td>Type6.BREX</td><td><-BrxD-AAA<-PglZ<-PIN<-?<-?<-N6-METHYLASE-p<-N6-METHYLASE||?->PemK_toxin-><-DUF6079*<-BrxB-BrxF-iSTAND<-BrxA-DNA-binding<-BrxE||OLD-ABC->TOPRIM-><-REase<-SF1-SF2-Helicase+REase<-SF1-SF2-Helicase<-SF1-SF2-Helicase+REase-p</td><td>1016</td><td>Bacteria>Myxococcota>Polyangia>Polyangiales>Polyangiaceae>unclassified_Polyangiaceae</td><td>GCA_016719945.1</td><td>Polyangiaceae_bacterium</td><td>IPM79_10070</td><td>425737..428787</td><td>2268199</td><td>BREX-system-P-loop-protein-BrxC-[Polyangiaceae-bacterium]</td></tr>
<tr><td>MCB9617769.1</td><td>Type1.BREX</td><td>RAMA->GDResolvase+ZnR-><-SIGMA+OB-Fold+AAA-ATPase+Lon-Protease-BrxL<-PglZ<-Schlafen+HATPase+wHTH<-N6-METHYLASE<-PIN<-DUF6079*<-BrxB-BrxF-iSTAND<-BrxA-DNA-binding<-tRNA||?-><-ADH_N+ADH_zinc_N+METHYLASE</td><td>1156</td><td>Bacteria>Myxococcota>Polyangia>Polyangiales>Sandaracinaceae>Sandaracinus</td><td>GCA_020633155.1</td><td>Sandaracinus_sp.</td><td>H6722_35485</td><td>14055..17525</td><td>2024858</td><td>BREX-system-P-loop-protein-BrxC-[Sandaracinus-sp.]</td></tr>
<tr><td>WP_240530317.1</td><td>Type1.BREX</td><td>N6-METHYLASE->PIN-><-?<-GreA_GreB+SF1-SF2-Helicase<-SF1-SF2-Helicase+REase<-35Exo+REase||wHTH+WYL+WCX->BrxA-DNA-binding->BrxB-BrxF-iSTAND->DUF6079-p*-></td><td>735</td><td>Bacteria>Pseudomonadota>Alphaproteobacteria>Sphingomonadales>Sphingomonadaceae>Novosphingobium</td><td>GCF_002556635.1</td><td>Novosphingobium_sp._PC22D</td><td>B2G71_RS22885</td><td>26272..28479</td><td>1962403</td><td>BREX-system-P-loop-protein-BrxC-[Novosphingobium-sp.-PC22D]</td></tr>
<tr><td>WP_243995499.1</td><td>Type1.BREX</td><td><-PIN<-?<-?<-?<-SIGMA+OB-Fold+AAA-ATPase+HKD-DNase-BrxL<-PglZ<-N6-METHYLASE<-DUF6079*<-BrxB-BrxF-iSTAND<-BrxA-DNA-binding<-wHTH+WYL+WCX||35Exo+REase->SF1-SF2-Helicase+REase->GreA_GreB+SF1-SF2-Helicase->?-><-PIN<-N6-METHYLASE</td><td>1179</td><td>Bacteria>Pseudomonadota>Alphaproteobacteria>Sphingomonadales>Sphingomonadaceae>Novosphingobium</td><td>GCF_022832385.1</td><td>Novosphingobium_sp._B2580</td><td>MTR64_RS16290</td><td>56132..59671</td><td>2930093</td><td>BREX-system-P-loop-protein-BrxC-[Novosphingobium-sp.-B2580]</td></tr>
<tr><td>MBQ2886288.1</td><td>Type1.BREX</td><td><-wHTH+WYL<-REase<-SIGMA+OB-Fold+AAA-ATPase+Lon-Protease-BrxL<-PglZ<-Schlafen+HATPase+wHTH<-N6-METHYLASE<-PIN<-OLD-ABC<-DUF6079*<-BrxB-BrxF-iSTAND<-BrxA-DNA-binding<-HTH_3</td><td>1180</td><td>Bacteria>Pseudomonadota>Alphaproteobacteria>unclassified_Alphaproteobacteria</td><td>GCA_017409945.1</td><td>Alphaproteobacteria_bacterium</td><td>IJE43_21410</td><td>13139..16681</td><td>1913988</td><td>BREX-system-P-loop-protein-BrxC-[Alphaproteobacteria-bacterium]</td></tr>
<tr><td>EXI84705.1</td><td>Type1.BREX</td><td>DDE_Tnp_1-><-?<-MqsA_antitoxin<-HTH<-?<-Arm-DNA-bind_3+Phage_integrase||BrxA-DNA-binding->BrxB-BrxF-iSTAND->DUF6079*->N6-METHYLASE->HTH+REase->PglZ->SIGMA+OB-Fold+AAA-ATPase+Lon-Protease-BrxL->PIN+HTH-></td><td>1204</td><td>Bacteria>Pseudomonadota>Betaproteobacteria>Betaproteobacteria_incertae_sedis>Candidatus_Accumulibacter</td><td>GCA_000585075.1</td><td>Candidatus_Accumulibacter_regalis</td><td>AW11_03760</td><td>6873..10487</td><td>522306</td><td>MAG:-hypothetical-protein-AW11_03760-[Candidatus-Accumulibacter-regalis]</td></tr>
<tr><td>EXI72915.1</td><td>Type1.BREX</td><td><-PIN+OST-HTH<-?||?->?->?-><-?||BrxA-DNA-binding->BrxB-BrxF-iSTAND->DUF6079*->HTH_3->DOC+HTH->N6-METHYLASE-><-PIN<-MazE_antitoxin||SF1-SF2-Helicase+PRTase-><-?||BrxL-></td><td>1155</td><td>Bacteria>Pseudomonadota>Betaproteobacteria>Betaproteobacteria_incertae_sedis>Candidatus_Accumulibacter>unclassified_Candidatus_Accumulibacter</td><td>GCA_000584995.1</td><td>Candidatus_Accumulibacter_sp._SK-11</td><td>AW07_03051</td><td>55405..58872</td><td>1454000</td><td>MAG:-hypothetical-protein-AW07_03051-[Candidatus-Accumulibacter-sp.-SK-11]</td></tr>
<tr><td>HAY27737.1</td><td>Type1.BREX</td><td>BrxA-DNA-binding->DUF6079*->HTH_3->DOC+HTH->N6-METHYLASE-><-PIN<-MazE_antitoxin||SF1-SF2-Helicase+PRTase->SLOG+BirA-HTH-></td><td>1286</td><td>Bacteria>Pseudomonadota>Betaproteobacteria>Betaproteobacteria_incertae_sedis>Candidatus_Accumulibacter>unclassified_Candidatus_Accumulibacter</td><td>GCA_003487685.1</td><td>Candidatus_Accumulibacter_sp.</td><td>DCY47_09605</td><td>6303..10163</td><td>2053492</td><td>BREX-system-P-loop-protein-BrxC-[Candidatus-Accumulibacter-sp.]</td></tr>
<tr><td>WP_076471941.1</td><td>Type6.BREX</td><td><-DUF3262<-?<-?<-?<-JetA||wHTH+WYL+WCX->BrxE->BrxA-DNA-binding->BrxB-BrxF-iSTAND->DUF6079*->OLD-ABC->PIN->N6-METHYLASE->N6-METHYLASE->ParB+HNH->PglZ->SIGMA+OB-Fold+AAA-ATPase+HKD-DNase-BrxL-><-JAB</td><td>1207</td><td>Bacteria>Pseudomonadota>Betaproteobacteria>Burkholderiales>Alcaligenaceae>Achromobacter</td><td>GCF_001957055.1</td><td>Achromobacter_xylosoxidans</td><td>BIZ53_RS00410</td><td>81330..84953</td><td>85698</td><td>BREX-system-P-loop-protein-BrxC-[Achromobacter-xylosoxidans]</td></tr>
<tr><td>WP_250462416.1</td><td>Type6.BREX</td><td><-MODE-HTH+PBPII||?-><-ADH_N+ADH_zinc_N||?-><-?||BrxE->BrxA-DNA-binding->BrxB-BrxF-iSTAND->DUF6079*->Tox-RelA->N6-METHYLASE->PglZ->SIGMA+OB-Fold+AAA-ATPase+Lon-Protease-BrxL->PIN-></td><td>1181</td><td>Bacteria>Pseudomonadota>Betaproteobacteria>Burkholderiales>Burkholderiaceae>Caballeronia</td><td>GCF_023631485.1</td><td>Caballeronia_sp._GAFFF2</td><td>NCT04_RS00125</td><td>29887..33432</td><td>2921741</td><td>BREX-system-P-loop-protein-BrxC-[Caballeronia-sp.-GAFFF2]</td></tr>
<tr><td>WP_108330269.1</td><td>Type1.BREX</td><td><-SIGMA+OB-Fold+AAA-ATPase+Lon-Protease-BrxL<-PglZ<-HNH<-OLD-ABC<-N6-METHYLASE<-REase<-OLD-ABC<-DUF6079*<-BrxB-BrxF-iSTAND<-BrxA-DNA-binding<-PIN<-ZnR+TPR</td><td>1154</td><td>Bacteria>Pseudomonadota>Betaproteobacteria>Burkholderiales>Comamonadaceae>Limnohabitans</td><td>GCF_003063435.1</td><td>Limnohabitans_sp._2KL-1</td><td>B9Z47_RS05555</td><td>224189..227653</td><td>1100699</td><td>BREX-system-P-loop-protein-BrxC-[Limnohabitans-sp.-2KL-1]</td></tr>
<tr><td>WP_108311897.1</td><td>Type1.BREX</td><td><-SF1-SF2-Helicase<-SIGMA+OB-Fold+AAA-ATPase+Lon-Protease-BrxL<-PglZ<-HNH<-OLD-ABC<-N6-METHYLASE<-DUF6079*<-BrxB-BrxF-iSTAND<-BrxA-DNA-binding<-PIN<-ZnR+TPR</td><td>1153</td><td>Bacteria>Pseudomonadota>Betaproteobacteria>Burkholderiales>Comamonadaceae>Limnohabitans</td><td>GCF_003063455.1</td><td>Limnohabitans_parvus</td><td>B9Z37_RS03875</td><td>837390..840851</td><td>540061</td><td>BREX-system-P-loop-protein-BrxC-[Limnohabitans-parvus]</td></tr>
<tr><td>PIQ50876.1</td><td>Type1.BREX</td><td>dCache_1+HisKA+HATPase_c-><-SIGMA+OB-Fold+AAA-ATPase+Lon-Protease-BrxL<-PglZ<-Schlafen+HATPase+wHTH<-SbcC<-N6-METHYLASE<-PIN<-MazE_antitoxin<-DUF6079*<-BrxB-BrxF-iSTAND<-BrxA-DNA-binding<-RHH_5||?->?->SF1-SF2-Helicase+REase->?->RelE-ParE-></td><td>1153</td><td>Bacteria>Pseudomonadota>Betaproteobacteria>Burkholderiales>Comamonadaceae>unclassified_Comamonadaceae</td><td>GCA_002786915.1</td><td>Comamonadaceae_bacterium_CG12_big_fil_rev_8_21_14_0_65_59_15</td><td>COW02_17785</td><td>48153..51614</td><td>1973952</td><td>hypothetical-protein-COW02_17785-[Comamonadaceae-bacterium-CG12_big_fil_rev_8_21_14_0_65_59_15]</td></tr>
<tr><td>ODV10849.1</td><td>Type1.BREX</td><td><-RelE-ParE<-?<-?<-DOC<-MazE_antitoxin||PhdYeFM_antitox->PIN->BrxA-DNA-binding->DUF6079-p*-></td><td>451</td><td>Bacteria>Pseudomonadota>Betaproteobacteria>Nitrosomonadales>Thiobacillaceae>Thiobacillus</td><td>GCA_001725535.1</td><td>Thiobacillus_sp._SCN_64-317</td><td>ABT22_10740</td><td>12066..13421</td><td>1660143</td><td>MAG:-hypothetical-protein-ABT22_10740-[Thiobacillus-sp.-SCN-64-317]</td></tr>
<tr><td>MBN8763784.1</td><td>Type1.BREX</td><td><-PRTase||?->?-><-N6-METHYLASE-p<-DUF6079-p*<-PIN<-PhdYeFM_antitox||MazE_antitoxin->DOC->?->RelE-ParE-></td><td>865</td><td>Bacteria>Pseudomonadota>Betaproteobacteria>Nitrosomonadales>Thiobacillaceae>Thiobacillus</td><td>GCA_017309235.1</td><td>Thiobacillus_sp.</td><td>JOJ02_08935</td><td>6734..9331</td><td>924</td><td>hypothetical-protein-[Thiobacillus-sp.]</td></tr>
<tr><td>WP_011238768.1</td><td>Type1.BREX</td><td><-Peptidase_M48_N+Peptidase_M48||35exo-><-?<-?||?->wHTH+WYL+WCX->BrxA-DNA-binding->BrxB-BrxF-iSTAND->DUF6079*-><-PIN<-RHH_6||?->N6-METHYLASE->OLD-ABC->HNH->OLD-ABC->TOPRIM->OLD-ABC-></td><td>1203</td><td>Bacteria>Pseudomonadota>Betaproteobacteria>Rhodocyclales>Rhodocyclaceae>Aromatoleum</td><td>GCF_000025965.1</td><td>Aromatoleum_aromaticum</td><td>EBN1_RS14750</td><td>3092506..3096117</td><td>551760</td><td>BREX-system-P-loop-protein-BrxC-[Aromatoleum-aromaticum]</td></tr>
<tr><td>WP_141017182.1</td><td>Type6.BREX</td><td>PhdYeFM_antitox->RelE-ParE-><-DOC+HTH<-Phage_integrase<-?||wHTH+WYL+WCX->BrxE->BrxA-DNA-binding->BrxB-BrxF-iSTAND->DUF6079*->OLD-ABC->PIN->N6-METHYLASE->N6-METHYLASE->?->ParB+HNH->PglZ->SIGMA+OB-Fold+AAA-ATPase+HKD-DNase-BrxL-><-XIS-HTH</td><td>1207</td><td>Bacteria>Pseudomonadota>Betaproteobacteria>Rhodocyclales>Zoogloeaceae>Azoarcus</td><td>GCF_006496635.1</td><td>Azoarcus_sp._DD4</td><td>CJ010_RS05935</td><td>1245819..1249442</td><td>2027405</td><td>BREX-system-P-loop-protein-BrxC-[Azoarcus-sp.-DD4]</td></tr>
<tr><td>TXH82461.1</td><td>Type1.BREX</td><td>PhdYeFM_antitox->PIN->BrxA-DNA-binding->BrxB-BrxF-iSTAND->DUF6079*->OLD-ABC->REase->N6-METHYLASE-p-></td><td>1153</td><td>Bacteria>Pseudomonadota>Betaproteobacteria>Rhodocyclales>Zoogloeaceae>Thauera</td><td>GCA_008015255.1</td><td>Thauera_aminoaromatica</td><td>E6Q80_15435</td><td>2474..5935</td><td>164330</td><td>BREX-system-P-loop-protein-BrxC-[Thauera-aminoaromatica]</td></tr>
<tr><td>WP_037984790.1</td><td>Type1.BREX</td><td>wHTH+WYL+WCX->BrxA-DNA-binding->BrxB-BrxF-iSTAND->DUF6079*-><-PIN<-RHH_6||N6-METHYLASE->UDG->REase->?->PglZ-></td><td>1203</td><td>Bacteria>Pseudomonadota>Betaproteobacteria>Rhodocyclales>Zoogloeaceae>Thauera</td><td>GCF_000310145.2</td><td>Thauera_sp._28</td><td>C662_RS14985</td><td>3144..6755</td><td>303682</td><td>BREX-system-P-loop-protein-BrxC-[Thauera-sp.-28]</td></tr>
<tr><td>WP_247724881.1</td><td>Type1.BREX</td><td><-PglZ-p<-REase+AIPR<-N6-METHYLASE<-PIN<-MazE_antitoxin<-DUF6079*<-BrxB-BrxF-iSTAND<-BrxA-DNA-binding<-SPOUT<-RnaseHII</td><td>1153</td><td>Bacteria>Pseudomonadota>Betaproteobacteria>Rhodocyclales>Zoogloeaceae>Thauera</td><td>GCF_023093595.1</td><td>Thauera_aromatica</td><td>MXC99_RS07615</td><td>6533..9994</td><td>59405</td><td>BREX-system-P-loop-protein-BrxC-[Thauera-aromatica]</td></tr>
<tr><td>WP_247725420.1</td><td>Type6.BREX</td><td><-wHTH<-ORC-CDC6<-ParA-Soj-PloopNTPase<-HEPN<-N6-METHYLASE-p<-PIN<-?<-N6-METHYLASE<-MNS+MarR-HTH<-DUF6079*<-BrxB-BrxF-iSTAND<-BrxA-DNA-binding<-BrxE<-?||ABC-ATPase+SbcC+ABC-ATPase+SbcC->HAD-></td><td>1209</td><td>Bacteria>Pseudomonadota>Betaproteobacteria>Rhodocyclales>Zoogloeaceae>Thauera</td><td>GCF_023093595.1</td><td>Thauera_aromatica</td><td>MXC99_RS10610</td><td>105350..108979</td><td>59405</td><td>BREX-system-P-loop-protein-BrxC-[Thauera-aromatica]</td></tr>
<tr><td>WP_247725480.1</td><td>Type1.BREX</td><td><-SIGMA+OB-Fold+AAA-ATPase+Lon-Protease-BrxL<-PglZ<-PIWI<-TOPRIM<-OLD-ABC<-N6-METHYLASE||RHH_6->PIN-><-DUF6079*<-BrxB-BrxF-iSTAND<-BrxA-DNA-binding<-wHTH+WYL+WCX</td><td>1203</td><td>Bacteria>Pseudomonadota>Betaproteobacteria>Rhodocyclales>Zoogloeaceae>Thauera</td><td>GCF_023093595.1</td><td>Thauera_aromatica</td><td>MXC99_RS10910</td><td>53320..56931</td><td>59405</td><td>BREX-system-P-loop-protein-BrxC-[Thauera-aromatica]</td></tr>
<tr><td>MBK6655412.1</td><td>Type1.BREX</td><td>HDRnase->?-><-HTH_19||?->?->PhdYeFM_antitox->PIN->BrxA-DNA-binding->BrxB-BrxF-iSTAND->DUF6079*->OLD-ABC->REase->N6-METHYLASE->ZnR->viral-capsid->?->?->TPR+REase-></td><td>1153</td><td>Bacteria>Pseudomonadota>Betaproteobacteria>Rhodocyclales>Zoogloeaceae>Zoogloea</td><td>GCA_016703645.1</td><td>Zoogloea_sp.</td><td>IPG40_16580</td><td>748210..751671</td><td>49181</td><td>BREX-system-P-loop-protein-BrxC-[Zoogloea-sp.]</td></tr>
<tr><td>MBK7848388.1</td><td>Type1.BREX</td><td>Arm-DNA-bind_2+Phage_int_SAM_3+Phage_integrase->?-><-HTH_19||?->?->PhdYeFM_antitox->PIN->BrxA-DNA-binding->BrxB-BrxF-iSTAND->DUF6079*->REase->ZnR->viral-capsid->?->?->TPR+REase->?->PglZ->SIGMA+OB-Fold+AAA-ATPase+Lon-Protease-BrxL-></td><td>1153</td><td>Bacteria>Pseudomonadota>Betaproteobacteria>Rhodocyclales>Zoogloeaceae>Zoogloea</td><td>GCA_016714095.1</td><td>Zoogloea_sp.</td><td>IPJ73_15250</td><td>34710..38171</td><td>49181</td><td>BREX-system-P-loop-protein-BrxC-[Zoogloea-sp.]</td></tr>
<tr><td>WP_141354347.1</td><td>Type1.BREX</td><td>wHTH+WYL+WCX->BrxA-DNA-binding->BrxB-BrxF-iSTAND->DUF6079*-><-PIN<-RHH_6||N6-METHYLASE-p->HKD-DNase->N6-METHYLASE->HTH+REase->ZnR+TOPRIM->PglZ->SIGMA+OB-Fold+AAA-ATPase+Lon-Protease-BrxL-></td><td>1203</td><td>Bacteria>Pseudomonadota>Betaproteobacteria>Rhodocyclales>Zoogloeaceae>Zoogloea</td><td>GCF_006539865.1</td><td>Zoogloea_ramigera</td><td>ZRA01_RS16820</td><td>2482..6093</td><td>350</td><td>BREX-system-P-loop-protein-BrxC-[Zoogloea-ramigera]</td></tr>
<tr><td>WP_144225094.1</td><td>Type1.BREX</td><td><-Phage_integrase<-DUF2787<-?<-?||XIS-HTH->XIS-HTH->BrxA-DNA-binding->BrxB-BrxF-iSTAND-p->DUF6079*->OLD-ABC->PIN->N6-METHYLASE->PglZ->SIGMA+OB-Fold+AAA-ATPase+Lon-Protease-BrxL->?-><-Arm-DNA-bind_3+Phage_integrase<-?||RNasePH-></td><td>1145</td><td>Bacteria>Pseudomonadota>Gammaproteobacteria>Alteromonadales>Shewanellaceae>Shewanella</td><td>GCF_017580345.1</td><td>Shewanella_algae</td><td>I6M43_RS01845</td><td>49386..52823</td><td>38313</td><td>BREX-system-P-loop-protein-BrxC-[Shewanella-algae]</td></tr>
<tr><td>WP_200158841.1</td><td>Type1.BREX</td><td><-SIGMA+OB-Fold+AAA-ATPase+Lon-Protease-BrxL<-PglZ<-HNH<-OLD-ABC<-SF1-SF2-Helicase<-OLD-ABC+TOPRIM<-N6-METHYLASE<-PIN<-OLD-ABC<-DUF6079*<-BrxB-BrxF-iSTAND<-BrxA-DNA-binding<-GDResolvase+HIN-HTH<-ZnR</td><td>1154</td><td>Bacteria>Pseudomonadota>Gammaproteobacteria>Chromatiales>Chromatiaceae>Allochromatium</td><td>GCF_016584135.1</td><td>Allochromatium_vinosum</td><td>CKO29_RS16480</td><td>18892..22356</td><td>1049</td><td>BREX-system-P-loop-protein-BrxC-[Allochromatium-vinosum]</td></tr>
<tr><td>MBV5272983.1</td><td>Type1.BREX</td><td><-PglZ<-?<-REase+AIPR<-N6-METHYLASE-p<-N6-METHYLASE-p<-PIN<-MazE_antitoxin<-DUF6079*<-BrxB-BrxF-iSTAND<-BrxA-DNA-binding<-Aminotran_3<-FURR-HTH||?-><-?<-?<-?<-Classical-AAA</td><td>1143</td><td>Bacteria>Pseudomonadota>Gammaproteobacteria>Chromatiales>Chromatiaceae>Lamprocystis</td><td>GCA_019163055.1</td><td>Lamprocystis_purpurea</td><td>JZU52_04840</td><td>9550..12981</td><td>61598</td><td>BREX-system-P-loop-protein-BrxC-[Lamprocystis-purpurea]</td></tr>
<tr><td>WP_020507127.1</td><td>Type6.BREX</td><td>XIS-HTH->BrxE->BrxA-DNA-binding->BrxB-BrxF-iSTAND->DUF6079*->Schlafen+HATPase+wHTH+HTH-><-?||N6-METHYLASE->?->PIN->N6-METHYLASE->AbrB-HTH-></td><td>1179</td><td>Bacteria>Pseudomonadota>Gammaproteobacteria>Chromatiales>Chromatiaceae>Lamprocystis</td><td>GCF_000379525.1</td><td>Lamprocystis_purpurea</td><td>A39O_RS0123535</td><td>17517..21056</td><td>61598</td><td>BREX-system-P-loop-protein-BrxC-[Lamprocystis-purpurea]</td></tr>
<tr><td>WP_062271501.1</td><td>Type1.BREX</td><td><-OLD-ABC<-RHH<-SF1-SF2-Helicase<-OLD-ABC+TOPRIM<-N6-METHYLASE<-PIN<-OLD-ABC<-PIN<-?<-DUF6079*<-BrxB-BrxF-iSTAND<-BrxA-DNA-binding<-tRNA<-Aminotran_1_2||SF1-SF2-Helicase-></td><td>1153</td><td>Bacteria>Pseudomonadota>Gammaproteobacteria>Chromatiales>Chromatiaceae>Marichromatium</td><td>GCF_001583505.1</td><td>Marichromatium_gracile</td><td>AY586_RS00820</td><td>23071..26532</td><td>1048</td><td>BREX-system-P-loop-protein-BrxC-[Marichromatium-gracile]</td></tr>
<tr><td>QVL48954.1</td><td>Type1.BREX</td><td><-PglZ<-PIN<-ParD_antitoxin<-wHTH<-TOPRIM<-SbcC<-N6-METHYLASE<-ACET+ACET<-TraJ-RHH<-DUF6079*<-BrxB-BrxF-iSTAND<-BrxA-DNA-binding<-tRNA||RADICAL-SAM->Glycos_transf_2-><-?||?-><-tRNA||NTP_transf+HDRnase-></td><td>1136</td><td>Bacteria>Pseudomonadota>Gammaproteobacteria>Chromatiales>Chromatiaceae>Thiocapsa</td><td>GCA_018399035.1</td><td>Thiocapsa_sp.</td><td>KFB96_26095</td><td>5699625..5703035</td><td>2024551</td><td>MAG:-BREX-system-P-loop-protein-BrxC-[Thiocapsa-sp.]</td></tr>
<tr><td>WP_200385867.1</td><td>Type1.BREX</td><td>DUF6079*->?->PIN-><-?||N6-METHYLASE->OLD-ABC->HNH->PglZ->AbrB-HTH->RHH-></td><td>1179</td><td>Bacteria>Pseudomonadota>Gammaproteobacteria>Chromatiales>Chromatiaceae>Thiocapsa</td><td>GCF_016583985.1</td><td>Thiocapsa_imhoffii</td><td>CKO25_RS00005</td><td>26..3565</td><td>382777</td><td>BREX-system-P-loop-protein-BrxC-[Thiocapsa-imhoffii]</td></tr>
<tr><td>WP_100920354.1</td><td>Type6.BREX</td><td><-HMA+HAD+HAD||?->?->?->BrxE->BrxA-DNA-binding->BrxB-BrxF-iSTAND->?->?||DUF6079*->Schlafen+HATPase+wHTH+HTH-><-?||N6-METHYLASE->?->PIN->N6-METHYLASE->RHH->PIN->RHH-></td><td>1178</td><td>Bacteria>Pseudomonadota>Gammaproteobacteria>Chromatiales>Chromatiaceae>Thiodictyon</td><td>GCF_002813775.1</td><td>Candidatus_Thiodictyon_syntrophicum</td><td>THSYN_17920</td><td>4302311..4305847</td><td>1166950</td><td>BREX-system-P-loop-protein-BrxC-[Candidatus-Thiodictyon-syntrophicum]</td></tr>
<tr><td>WP_100920747.1</td><td>Type1.BREX</td><td>BrxA-DNA-binding->BrxB-BrxF-iSTAND->?->?->DUF6079*->PhdYeFM_antitox->PIN->MqsA_antitoxin->N6-METHYLASE-p->N6-METHYLASE->Transposase_31->OLD-ABC->Polbeta->PglZ-></td><td>1194</td><td>Bacteria>Pseudomonadota>Gammaproteobacteria>Chromatiales>Chromatiaceae>Thiodictyon</td><td>GCF_002813775.1</td><td>Candidatus_Thiodictyon_syntrophicum</td><td>THSYN_RS20415</td><td>4833326..4836910</td><td>1166950</td><td>BREX-system-P-loop-protein-BrxC-[Candidatus-Thiodictyon-syntrophicum]</td></tr>
<tr><td>WP_015282553.1</td><td>Type6.BREX</td><td><-SIGMA+OB-Fold+AAA-ATPase+HKD-DNase-BrxL<-PglZ<-PIN<-RHH<-N6-METHYLASE<-?<-SF1-SF2-Helicase<-N6-METHYLASE<-DUF6079*<-BrxB-BrxF-iSTAND<-BrxA-DNA-binding<-BrxE</td><td>1209</td><td>Bacteria>Pseudomonadota>Gammaproteobacteria>Chromatiales>Chromatiaceae>Thioflavicoccus</td><td>GCF_000327045.1</td><td>Thioflavicoccus_mobilis</td><td>THIMO_RS17980</td><td>83065..86694</td><td>80679</td><td>BREX-system-P-loop-protein-BrxC-[Thioflavicoccus-mobilis]</td></tr>
<tr><td>MCG6940983.1</td><td>Type1.BREX</td><td><-N6-METHYLASE-p<-PIN<-VapB_antitoxin<-DUF6079-p*<-tRNA<-?<-AA_kinase+ACT+ACT_7</td><td>251</td><td>Bacteria>Pseudomonadota>Gammaproteobacteria>Chromatiales>Chromatiaceae>Thiohalocapsa</td><td>GCA_022340605.1</td><td>Thiohalocapsa_sp.</td><td>LJE69_07005</td><td>1995..2750</td><td>2497641</td><td>hypothetical-protein-[Thiohalocapsa-sp.]</td></tr>
<tr><td>WP_058554449.1</td><td>Type6.BREX</td><td>TIR->?->?->?->BrxE->BrxA-DNA-binding->BrxB-BrxF-iSTAND->DUF6079*->N6-METHYLASE->?->Schlafen+HATPase+wHTH->N6-METHYLASE->?->PIN->PglZ->SIGMA+OB-Fold+AAA-ATPase+HKD-DNase-BrxL-><-tRNA||HHH_5+SF1-SF2-Helicase-></td><td>1214</td><td>Bacteria>Pseudomonadota>Gammaproteobacteria>Chromatiales>Chromatiaceae>Thiohalocapsa</td><td>GCF_001469165.1</td><td>Thiohalocapsa_sp._ML1</td><td>ML01_RS05745</td><td>155025..158669</td><td>1431688</td><td>BREX-system-P-loop-protein-BrxC-[Thiohalocapsa-sp.-ML1]</td></tr>
<tr><td>WP_058554877.1</td><td>Type1.BREX</td><td>DDE_Tnp_1->?->?->?->?->AA_kinase+ACT+ACT_7->?->tRNA->BrxA-DNA-binding->BrxB-BrxF-iSTAND->DUF6079*->VapB_antitoxin->VapB_antitoxin->PIN-><-?||N6-METHYLASE-></td><td>1152</td><td>Bacteria>Pseudomonadota>Gammaproteobacteria>Chromatiales>Chromatiaceae>Thiohalocapsa</td><td>GCF_001469165.1</td><td>Thiohalocapsa_sp._ML1</td><td>ML01_RS08025</td><td>156057..159515</td><td>1431688</td><td>BREX-system-P-loop-protein-BrxC-[Thiohalocapsa-sp.-ML1]</td></tr>
<tr><td>WP_200239121.1</td><td>Type1.BREX</td><td><-PglZ<-PIN<-PhdYeFM_antitox<-HicA_toxin<-HicB_lk_antitox<-MazE_antitoxin<-PIN<-MazE_antitoxin<-DUF6079*<-BrxB-BrxF-iSTAND<-BrxA-DNA-binding<-?<-ABhydrolase<-?<-?||?->?-><-REase</td><td>1197</td><td>Bacteria>Pseudomonadota>Gammaproteobacteria>Chromatiales>Chromatiaceae>Thiohalocapsa</td><td>GCF_016583825.1</td><td>Thiohalocapsa_halophila</td><td>CKO31_RS15145</td><td>13443..17036</td><td>69359</td><td>BREX-system-P-loop-protein-BrxC-[Thiohalocapsa-halophila]</td></tr>
<tr><td>WP_164452822.1</td><td>Type6.BREX</td><td><-NtrC-AAA+FIS-HTH||?->?->SF1-SF2-Helicase+DUF1998->GrpE->?->BrxE->BrxA-DNA-binding->BrxB-BrxF-iSTAND->DUF6079*->?->N6-METHYLASE-p->?->PIN->AbrB-HTH->?->N6-METHYLASE->OLD-ABC->TOPRIM-></td><td>1214</td><td>Bacteria>Pseudomonadota>Gammaproteobacteria>Chromatiales>Chromatiaceae>Thiorhodococcus</td><td>GCF_010820565.1</td><td>Thiorhodococcus_minor</td><td>G3446_RS10685</td><td>55754..59398</td><td>57489</td><td>BREX-system-P-loop-protein-BrxC-[Thiorhodococcus-minor]</td></tr>
<tr><td>EIC19850.1</td><td>Type1.BREX</td><td><-SIGMA+OB-Fold+AAA-ATPase+Lon-Protease-BrxL<-PglZ<-RelE-ParE<-PhdYeFM_antitox<-Schlafen+HATPase+wHTH<-N6-METHYLASE<-PIN<-VapB_antitoxin<-DUF6079*<-BrxB-BrxF-iSTAND<-BrxA-DNA-binding<-?<-HISKIN<-?<-?<-?<-Thioredoxin</td><td>1152</td><td>Bacteria>Pseudomonadota>Gammaproteobacteria>Chromatiales>Chromatiaceae>Thiorhodovibrio</td><td>GCA_000228725.3</td><td>Thiorhodovibrio_sp._970</td><td>Thi970DRAFT_03455</td><td>369329..372787</td><td>631362</td><td>hypothetical-protein-Thi970DRAFT_03455-[Thiorhodovibrio-sp.-970]</td></tr>
<tr><td>WP_009149681.1</td><td>Type6.BREX</td><td><-BrxD<-?<-?<-?<-?||BrxE->BrxA-DNA-binding->BrxB-BrxF-iSTAND->DUF6079*->?->RelE-ParE->N6-METHYLASE->?->PIN->N6-METHYLASE-p->?->N6-METHYLASE-p->RHH-></td><td>1208</td><td>Bacteria>Pseudomonadota>Gammaproteobacteria>Chromatiales>Chromatiaceae>Thiorhodovibrio</td><td>GCF_000228725.2</td><td>Thiorhodovibrio_sp._970</td><td>THI970DRAFT_RS14125</td><td>1743457..1747083</td><td>631362</td><td>BREX-system-P-loop-protein-BrxC-[Thiorhodovibrio-sp.-970]</td></tr>
<tr><td>WP_223296802.1</td><td>Type1.BREX</td><td><-SIGMA+OB-Fold+AAA-ATPase+Lon-Protease-BrxL<-PglZ<-RelE-ParE<-PhdYeFM_antitox<-Schlafen+HATPase+wHTH<-N6-METHYLASE<-PIN<-VapB_antitoxin<-DUF6079*<-BrxB-BrxF-iSTAND<-BrxA-DNA-binding<-?<-HISKIN<-?<-?<-Thioredoxin||?->REC+GGDEF+EAL-></td><td>1154</td><td>Bacteria>Pseudomonadota>Gammaproteobacteria>Chromatiales>Chromatiaceae>Thiorhodovibrio</td><td>GCF_000228725.2</td><td>Thiorhodovibrio_sp._970</td><td>THI970DRAFT_RS16355</td><td>369329..372793</td><td>631362</td><td>BREX-system-P-loop-protein-BrxC-[Thiorhodovibrio-sp.-970]</td></tr>
<tr><td>WP_201065314.1</td><td>Type6.BREX</td><td>CR-ATPase5+REase->?-><-?<-HicB<-?||?-><-N6-METHYLASE<-PIN<-wHTH<-DUF6079*<-BrxB-BrxF-iSTAND<-BrxA-DNA-binding<-BrxE||?->?->?->?->?->BrxD-></td><td>1212</td><td>Bacteria>Pseudomonadota>Gammaproteobacteria>Chromatiales>Chromatiaceae>Thiorhodovibrio</td><td>GCF_016653415.1</td><td>Thiorhodovibrio_winogradskyi</td><td>CCR91_RS06145</td><td>56528..60166</td><td>77007</td><td>BREX-system-P-loop-protein-BrxC-[Thiorhodovibrio-winogradskyi]</td></tr>
<tr><td>WP_201066887.1</td><td>Type1.BREX</td><td><-wHTH<-?<-PIN<-?<-?<-N6-METHYLASE<-HicB_lk_antitox<-MazE_antitoxin<-?<-PIN<-DUF6079*<-BrxB-BrxF-iSTAND<-BrxA-DNA-binding<-tRNA<-Sulfate_transp+STAS</td><td>1203</td><td>Bacteria>Pseudomonadota>Gammaproteobacteria>Chromatiales>Chromatiaceae>Thiorhodovibrio</td><td>GCF_016653415.1</td><td>Thiorhodovibrio_winogradskyi</td><td>CCR91_RS10560</td><td>25140..28751</td><td>77007</td><td>BREX-system-P-loop-protein-BrxC-[Thiorhodovibrio-winogradskyi]</td></tr>
<tr><td>TVQ85671.1</td><td>Type6.BREX</td><td><-N6-METHYLASE-p<-PIN<-?<-?<-?<-N6-METHYLASE<-DUF6079*<-BrxB-BrxF-iSTAND<-BrxA-DNA-binding<-BrxE<-?<-Aminotran_1_2<-?<-ABC-ATPase+SbcC</td><td>1212</td><td>Bacteria>Pseudomonadota>Gammaproteobacteria>Chromatiales>Chromatiaceae>unclassified_Chromatiaceae</td><td>GCA_007693535.1</td><td>Chromatiaceae_bacterium</td><td>EA400_15015</td><td>6841..10479</td><td>2049432</td><td>BREX-system-P-loop-protein-BrxC-[Chromatiaceae-bacterium]</td></tr>
<tr><td>MBK8638067.1</td><td>Type1.BREX</td><td>AAA-ATPase->?->ZnR->?->?->?->DUF6079*->PhdYeFM_antitox->AbrB-HTH->PIN->N6-METHYLASE->PIN->?->PglZ->SIGMA+OB-Fold+AAA-ATPase+Lon-Protease-BrxL-><-SIR2</td><td>1067</td><td>Bacteria>Pseudomonadota>Gammaproteobacteria>Chromatiales>Chromatiaceae>unclassified_Chromatiaceae</td><td>GCA_016714645.1</td><td>Chromatiaceae_bacterium</td><td>IPN92_07155</td><td>42706..45909</td><td>2049432</td><td>BREX-system-P-loop-protein-BrxC-[Chromatiaceae-bacterium]</td></tr>
<tr><td>MBK8639641.1</td><td>Type6.BREX</td><td>REC-><-?||BrxE->BrxA-DNA-binding->BrxB-BrxF-iSTAND->DUF6079*->?->N6-METHYLASE->RHH->PIN->PglZ->PolB->HEPN->SIGMA+OB-Fold+AAA-ATPase+Lon-Protease-BrxL-></td><td>1178</td><td>Bacteria>Pseudomonadota>Gammaproteobacteria>Chromatiales>Chromatiaceae>unclassified_Chromatiaceae</td><td>GCA_016714645.1</td><td>Chromatiaceae_bacterium</td><td>IPN92_15725</td><td>846355..849891</td><td>2049432</td><td>BREX-system-P-loop-protein-BrxC-[Chromatiaceae-bacterium]</td></tr>
<tr><td>MBP6733459.1</td><td>Type1.BREX</td><td>BrxA-DNA-binding->BrxB-BrxF-iSTAND->DUF6079*-><-BrxB-BrxF-iSTAND||?->PhdYeFM_antitox->PIN->N6-METHYLASE->HTH+REase->REase->N6-METHYLASE-p-></td><td>1171</td><td>Bacteria>Pseudomonadota>Gammaproteobacteria>Chromatiales>Chromatiaceae>unclassified_Chromatiaceae</td><td>GCA_017990755.1</td><td>Chromatiaceae_bacterium</td><td>KA142_01990</td><td>1791..5306</td><td>2049432</td><td>BREX-system-P-loop-protein-BrxC-[Chromatiaceae-bacterium]</td></tr>
<tr><td>MBP6807296.1</td><td>Type1.BREX</td><td>BrxA-DNA-binding->BrxB-BrxF-iSTAND->DUF6079*->MazE_antitoxin->PIN->N6-METHYLASE-p->AbrB-HTH-></td><td>1194</td><td>Bacteria>Pseudomonadota>Gammaproteobacteria>Chromatiales>Chromatiaceae>unclassified_Chromatiaceae</td><td>GCA_017991215.1</td><td>Chromatiaceae_bacterium</td><td>KA125_03275</td><td>984..4568</td><td>2049432</td><td>BREX-system-P-loop-protein-BrxC-[Chromatiaceae-bacterium]</td></tr>
<tr><td>MCF7978332.1</td><td>Type1.BREX</td><td><-AbrB-HTH<-PIN<-AbrB-HTH<-N6-METHYLASE<-DOC<-MazE_antitoxin<-MqsA_antitoxin<-?<-PIN<-DUF6079*</td><td>1223</td><td>Bacteria>Pseudomonadota>Gammaproteobacteria>Chromatiales>Chromatiaceae>unclassified_Chromatiaceae</td><td>GCA_021736685.1</td><td>Chromatiaceae_bacterium</td><td>K9L82_09985</td><td>9865..13536</td><td>2049432</td><td>BREX-system-P-loop-protein-BrxC-[Chromatiaceae-bacterium]</td></tr>
<tr><td>MCF8016509.1</td><td>Type1.BREX</td><td><-AbrB-HTH<-PIN<-AbrB-HTH<-N6-METHYLASE<-DOC<-MazE_antitoxin<-MqsA_antitoxin<-?<-PIN<-DUF6079-p*</td><td>835</td><td>Bacteria>Pseudomonadota>Gammaproteobacteria>Chromatiales>Chromatiaceae>unclassified_Chromatiaceae</td><td>GCA_021736865.1</td><td>Chromatiaceae_bacterium</td><td>K9L65_12595</td><td>6416..8923</td><td>2049432</td><td>BREX-system-P-loop-protein-BrxC-[Chromatiaceae-bacterium]</td></tr>
<tr><td>MCP5407916.1</td><td>Type1.BREX</td><td>ABhydrolase->?->TraU->?->TraG_N-><-?||BrxA-DNA-binding->BrxB-BrxF-iSTAND->DUF6079*->MazE_antitoxin->PIN->N6-METHYLASE->SIR2->?->PglZ->SIGMA+OB-Fold+AAA-ATPase+Lon-Protease-BrxL->XIS-HTH-></td><td>1194</td><td>Bacteria>Pseudomonadota>Gammaproteobacteria>Chromatiales>Chromatiaceae>unclassified_Chromatiaceae</td><td>GCA_024235335.1</td><td>Chromatiaceae_bacterium</td><td>H6963_01295</td><td>248686..252270</td><td>2049432</td><td>BREX-system-P-loop-protein-BrxC-[Chromatiaceae-bacterium]</td></tr>
<tr><td>MCK7579547.1</td><td>Type1.BREX</td><td><-OLD-ABC+TOPRIM<-?<-?<-N6-METHYLASE<-PIN<-OLD-ABC||?-><-DUF6079-p<-DUF6079-p*<-?<-DUF6079-p<-BrxB-BrxF-iSTAND<-BrxA-DNA-binding-p<-ZnR</td><td>299</td><td>Bacteria>Pseudomonadota>Gammaproteobacteria>Chromatiales>unclassified_Chromatiales</td><td>GCA_023156155.1</td><td>Chromatiales_bacterium</td><td>MZV65_30115</td><td>14491..15390</td><td>2026725</td><td>hypothetical-protein-[Chromatiales-bacterium]</td></tr>
<tr><td>MCK7579549.1</td><td>Type1.BREX</td><td><-N6-METHYLASE<-PIN<-OLD-ABC||?-><-DUF6079-p<-DUF6079-p<-?<-DUF6079-p*<-BrxB-BrxF-iSTAND<-BrxA-DNA-binding-p<-ZnR</td><td>648</td><td>Bacteria>Pseudomonadota>Gammaproteobacteria>Chromatiales>unclassified_Chromatiales</td><td>GCA_023156155.1</td><td>Chromatiales_bacterium</td><td>MZV65_30125</td><td>15595..17541</td><td>2026725</td><td>BREX-system-P-loop-protein-BrxC-[Chromatiales-bacterium]</td></tr>
<tr><td>MBZ4194014.1</td><td>Type1.BREX</td><td><-N6-METHYLASE<-PIN<-HTH+MPTase<-DUF6079*<-BrxB-BrxF-iSTAND<-BrxA-DNA-binding<-?<-TPR</td><td>1138</td><td>Bacteria>Pseudomonadota>Gammaproteobacteria>Gammaproteobacteria_incertae_sedis>Candidatus_Competibacteraceae>Candidatus_Contendobacter</td><td>GCA_020035205.1</td><td>Candidatus_Contendobacter_sp.</td><td>LAE24_06875</td><td>5150..8566</td><td>2651161</td><td>BREX-system-P-loop-protein-BrxC-[Candidatus-Contendobacter-sp.]</td></tr>
<tr><td>MBK8752263.1</td><td>Type1.BREX</td><td><-STYKinase-like<-?<-HTH_37||?-><-RHH_5||?->||BrxA-DNA-binding->BrxB-BrxF-iSTAND->DUF6079*->HTH+MPTase->PIN->N6-METHYLASE->?->PemK_toxin-><-?<-?||PglZ->SIGMA+OB-Fold+AAA-ATPase+Lon-Protease-BrxL-></td><td>1139</td><td>Bacteria>Pseudomonadota>Gammaproteobacteria>Gammaproteobacteria_incertae_sedis>Candidatus_Competibacteraceae>unclassified_Candidatus_Competibacteraceae</td><td>GCA_016710535.1</td><td>Candidatus_Competibacteraceae_bacterium</td><td>IPL99_11760</td><td>9123..12542</td><td>2053538</td><td>BREX-system-P-loop-protein-BrxC-[Candidatus-Competibacteraceae-bacterium]</td></tr>
<tr><td>MCB1769904.1</td><td>Type1.BREX</td><td><-N6-METHYLASE-p<-PIN<-RHH<-MetJarc-HTH<-HicA_toxin<-N6-METHYLASE-p<-PIN<-?<-N6-METHYLASE<-DUF6079-p*</td><td>760</td><td>Bacteria>Pseudomonadota>Gammaproteobacteria>Gammaproteobacteria_incertae_sedis>Candidatus_Competibacteraceae>unclassified_Candidatus_Competibacteraceae</td><td>GCA_020445005.1</td><td>Candidatus_Competibacteraceae_bacterium</td><td>KDJ31_09455</td><td>11685..13969</td><td>2053538</td><td>BREX-system-P-loop-protein-BrxC-[Candidatus-Competibacteraceae-bacterium]</td></tr>
<tr><td>WP_177417917.1</td><td>Type1.BREX</td><td>BrxB-BrxF-iSTAND->DUF6079*->MazE_antitoxin->PIN->N6-METHYLASE->OLD-ABC->TOPRIM->PglZ->SIGMA+OB-Fold+AAA-ATPase+Lon-Protease-BrxL-></td><td>1188</td><td>Bacteria>Pseudomonadota>Gammaproteobacteria>Gammaproteobacteria_incertae_sedis>sulfur-oxidizing_symbionts</td><td>GCF_011947365.2</td><td>endosymbiont_of_Lamellibrachia_barhami</td><td>G9Q73_RS00010</td><td>458..4024</td><td>205975</td><td>BREX-system-P-loop-protein-BrxC-[endosymbiont-of-Lamellibrachia-barhami]</td></tr>
<tr><td>MBS3952424.1</td><td>Type1.BREX</td><td>XIS-HTH->BrxA-DNA-binding->BrxB-BrxF-iSTAND->DUF6079*->OLD-ABC->REase->N6-METHYLASE->PhdYeFM_antitox->PIN->N6-METHYLASE-p->HTH+REase->Schlafen+HATPase+wHTH->SIR2-></td><td>1178</td><td>Bacteria>Pseudomonadota>Gammaproteobacteria>Methylococcales>Methylococcaceae>Methylomicrobium</td><td>GCA_018333785.1</td><td>Methylomicrobium_sp.</td><td>KGZ88_05695</td><td>2093..5629</td><td>2303549</td><td>BREX-system-P-loop-protein-BrxC-[Methylomicrobium-sp.]</td></tr>
<tr><td>WP_024296674.1</td><td>Type1.BREX</td><td>GDResolvase+HIN-HTH->?->BrxA-DNA-binding->BrxB-BrxF-iSTAND->DUF6079*->ParD_antitoxin->PIN->N6-METHYLASE->SbcC->Schlafen+HATPase+wHTH->PglZ->SIGMA+OB-Fold+AAA-ATPase+Lon-Protease-BrxL-></td><td>1202</td><td>Bacteria>Pseudomonadota>Gammaproteobacteria>Methylococcales>Methylococcaceae>Methylomicrobium</td><td>GCF_000527095.1</td><td>Methylomicrobium_lacus</td><td>METLA_RS0100460</td><td>95992..99600</td><td>136992</td><td>BREX-system-P-loop-protein-BrxC-[Methylomicrobium-lacus]</td></tr>
<tr><td>WP_255188315.1</td><td>Type1.BREX</td><td><-PIN<-PIN<-SIGMA+OB-Fold+AAA-ATPase+Lon-Protease-BrxL<-PglZ<-TOPRIM<-OLD-ABC<-N6-METHYLASE<-PIN<-ParD_antitoxin<-DUF6079*<-BrxB-BrxF-iSTAND<-BrxA-DNA-binding||?-><-?<-HIN-HTH||?-><-ACET+ACET<-TraJ-RHH</td><td>1206</td><td>Bacteria>Pseudomonadota>Gammaproteobacteria>Methylococcales>Methylococcaceae>Methylomonas</td><td>GCF_024360925.2</td><td>Methylomonas_rapida</td><td>NM686_RS13235</td><td>2825288..2828908</td><td>2963939</td><td>BREX-system-P-loop-protein-BrxC-[Methylomonas-rapida]</td></tr>
<tr><td>WP_192375581.1</td><td>Type1.BREX</td><td><-KAP-glycosylase<-KAP-NTPase<-PIN<-SIGMA+OB-Fold+AAA-ATPase+Lon-Protease-BrxL<-SIGMA+OB-Fold+AAA-ATPase+Lon-Protease-BrxL<-PglZ<-N6-METHYLASE<-PIN<-ParD_antitoxin<-DUF6079*<-BrxB-BrxF-iSTAND<-BrxA-DNA-binding||RNase-><-?<-GDResolvase+HIN-HTH</td><td>1207</td><td>Bacteria>Pseudomonadota>Gammaproteobacteria>Methylococcales>Methylococcaceae>Methylomonas</td><td>GCF_903064715.1</td><td>Methylomonas_albis</td><td>EBA_RS15675</td><td>3420343..3423966</td><td>1854563</td><td>BREX-system-P-loop-protein-BrxC-[Methylomonas-albis]</td></tr>
<tr><td>WP_080523593.1</td><td>Type1.BREX</td><td><-N6-METHYLASE-p||HTH_Tnp_1+HTH_21+rve+rve_3-><-N6-METHYLASE<-PIN<-?<-ACET+ACET<-BrxB-BrxF-iSTAND<-DUF6079*<-BrxB-BrxF-iSTAND<-BrxA-DNA-binding<-?<-?||GSDH-><-?<-BrxB-BrxF-iSTAND<-?<-PIN</td><td>1206</td><td>Bacteria>Pseudomonadota>Gammaproteobacteria>Methylococcales>Methylococcaceae>Methyloprofundus</td><td>GCF_002072955.1</td><td>Methyloprofundus_sedimenti</td><td>AU255_RS14210</td><td>277762..281382</td><td>1420851</td><td>BREX-system-P-loop-protein-BrxC-[Methyloprofundus-sedimenti]</td></tr>
<tr><td>MCF8007296.1</td><td>Type1.BREX</td><td>BrxA-DNA-binding->BrxB-BrxF-iSTAND->DUF6079-p*->N6-METHYLASE-p->?->PIN->PglZ-p-><-Aminotran_1_2</td><td>781</td><td>Bacteria>Pseudomonadota>Gammaproteobacteria>Methylococcales>Methylococcaceae>Methylovulum</td><td>GCA_021736885.1</td><td>Methylovulum_sp.</td><td>K9K84_07785</td><td>12351..14696</td><td>1916980</td><td>BREX-system-P-loop-protein-BrxC-[Methylovulum-sp.]</td></tr>
<tr><td>WP_088619247.1</td><td>Type1.BREX</td><td><-Glycos_transf<-?<-SIGMA+OB-Fold+AAA-ATPase+Lon-Protease-BrxL<-PglZ<-Schlafen+HATPase+wHTH<-PHP+SbcC<-N6-METHYLASE<-PIN<-ParD_antitoxin<-DUF6079*<-BrxB-BrxF-iSTAND<-BrxA-DNA-binding<-wHTH+WYL+WCX</td><td>1201</td><td>Bacteria>Pseudomonadota>Gammaproteobacteria>Methylococcales>Methylococcaceae>Methylovulum</td><td>GCF_002209385.1</td><td>Methylovulum_psychrotolerans</td><td>CEK71_RS09965</td><td>2189788..2193393</td><td>1704499</td><td>BREX-system-P-loop-protein-BrxC-[Methylovulum-psychrotolerans]</td></tr>
<tr><td>WP_103975698.1</td><td>Type1.BREX</td><td><-SIGMA+OB-Fold+AAA-ATPase+Lon-Protease-BrxL<-PglZ<-Schlafen+HATPase+wHTH<-PHP+SbcC<-?<-?<-N6-METHYLASE<-PIN<-ParD_antitoxin<-DUF6079*<-BrxB-BrxF-iSTAND<-BrxA-DNA-binding<-wHTH+WYL+WCX||?->SAM-METHYLASE->?->SbcC->SAM-METHYLASE->SAM-METHYLASE-></td><td>1201</td><td>Bacteria>Pseudomonadota>Gammaproteobacteria>Methylococcales>Methylococcaceae>Methylovulum</td><td>GCF_002923755.1</td><td>Methylovulum_psychrotolerans</td><td>AADEFJLK_RS21345</td><td>25888..29493</td><td>1704499</td><td>BREX-system-P-loop-protein-BrxC-[Methylovulum-psychrotolerans]</td></tr>
<tr><td>WP_215479507.1</td><td>Type1.BREX</td><td>wHTH+WYL+WCX-p->wHTH+WYL+WCX->BrxA-DNA-binding->BrxB-BrxF-iSTAND->DUF6079*->ParD_antitoxin->PIN->N6-METHYLASE->?->?->PHP+SbcC->Schlafen+HATPase+wHTH->PglZ->SIGMA+OB-Fold+AAA-ATPase+Lon-Protease-BrxL-></td><td>1201</td><td>Bacteria>Pseudomonadota>Gammaproteobacteria>Methylococcales>Methylococcaceae>Methylovulum</td><td>GCF_018722605.1</td><td>Methylovulum_psychrotolerans</td><td>KFZ76_RS21360</td><td>105012..108617</td><td>1704499</td><td>BREX-system-P-loop-protein-BrxC-[Methylovulum-psychrotolerans]</td></tr>
<tr><td>TSA38613.1</td><td>Type1.BREX</td><td>BrxA-DNA-binding->BrxB-BrxF-iSTAND->DUF6079*->PRTase->?->PIN->?->N6-METHYLASE-p->?->?->PIN->N6-METHYLASE-p-></td><td>1218</td><td>Bacteria>Pseudomonadota>Gammaproteobacteria>Methylococcales>Methylococcaceae>unclassified_Methylococcaceae</td><td>GCA_007280895.1</td><td>Methylococcaceae_bacterium</td><td>D4R63_10680</td><td>1521..5177</td><td>1933926</td><td>BREX-system-P-loop-protein-BrxC-[Methylococcaceae-bacterium]</td></tr>
<tr><td>WP_040239271.1</td><td>Type1.BREX</td><td><-sGTP<-?<-?||?->tRNA->BrxA-DNA-binding->BrxB-BrxF-iSTAND->DUF6079*->OLD-ABC->PIN->N6-METHYLASE->ParB+HNH->?->?->PglZ->SIGMA+OB-Fold+AAA-ATPase+Lon-Protease-BrxL-></td><td>1144</td><td>Bacteria>Pseudomonadota>Gammaproteobacteria>Oceanospirillales>Halomonadaceae>Chromohalobacter</td><td>GCF_000821045.2</td><td>Chromohalobacter_japonicus</td><td>BN995_RS00415</td><td>38131..41565</td><td>223900</td><td>BREX-system-P-loop-protein-BrxC-[Chromohalobacter-japonicus]</td></tr>
<tr><td>WP_189514712.1</td><td>Type1.BREX</td><td><-DUF4096+DDE_Tnp_1+DDE_Tnp_1_2<-?||?->DUF932->?->?->?->BrxA-DNA-binding->BrxB-BrxF-iSTAND->DUF6079*->OLD-ABC->PIN->N6-METHYLASE->ParB+HNH->PglZ->SIGMA+OB-Fold+AAA-ATPase+Lon-Protease-BrxL->McrB-AAA->wHTH+REase-></td><td>1144</td><td>Bacteria>Pseudomonadota>Gammaproteobacteria>Oceanospirillales>Halomonadaceae>Kushneria</td><td>GCF_014652795.1</td><td>Kushneria_pakistanensis</td><td>IE123_RS02380</td><td>531699..535133</td><td>1508770</td><td>BREX-system-P-loop-protein-BrxC-[Kushneria-pakistanensis]</td></tr>
<tr><td>WP_163650581.1</td><td>Type1.BREX</td><td><-N6-METHYLASE-p<-PIN<-OLD-ABC<-DUF6079*<-BrxB-BrxF-iSTAND<-BrxA-DNA-binding<-tRNA<-?||?->?->sGTP-></td><td>1144</td><td>Bacteria>Pseudomonadota>Gammaproteobacteria>Oceanospirillales>Halomonadaceae>Modicisalibacter</td><td>GCF_010993675.1</td><td>Modicisalibacter_sp._'Wilcox'</td><td>G7A98_RS15465</td><td>3721..7155</td><td>2679914</td><td>BREX-system-P-loop-protein-BrxC-[Modicisalibacter-sp.-'Wilcox']</td></tr>
<tr><td>WP_224416377.1</td><td>Type1.BREX</td><td><-PglZ<-?<-?<-?<-N6-METHYLASE-p<-HTH_Tnp_1+HTH_21+rve+rve_3<-N6-METHYLASE<-PIN<-OLD-ABC<-DUF6079*<-BrxB-BrxF-iSTAND<-BrxA-DNA-binding<-tRNA<-?||?->?->sGTP-></td><td>1144</td><td>Bacteria>Pseudomonadota>Gammaproteobacteria>Oceanospirillales>Halomonadaceae>Modicisalibacter</td><td>GCF_020147965.1</td><td>Modicisalibacter_tunisiensis</td><td>KGQ90_RS13860</td><td>2977413..2980847</td><td>390637</td><td>BREX-system-P-loop-protein-BrxC-[Modicisalibacter-tunisiensis]</td></tr>
<tr><td>WP_202806908.1</td><td>Type1.BREX</td><td><-PglZ<-TOPRIM<-OLD-ABC<-SbcC<-N6-METHYLASE-p<-N6-METHYLASE<-N6-METHYLASE-p<-PIN<-OLD-ABC<-DUF6079-p*<-DUF6079-p<-BrxB-BrxF-iSTAND<-BrxA-DNA-binding<-XIS-HTH<-?<-HDRnase<-Phage_int_SAM<-Integrase_1||XIS-HTH-></td><td>460</td><td>Bacteria>Pseudomonadota>Gammaproteobacteria>Oceanospirillales>Oceanospirillaceae>Nitrincola</td><td>GCF_000585235.1</td><td>Nitrincola_nitratireducens</td><td>D791_RS20495</td><td>12347..13729</td><td>1229521</td><td>hypothetical-protein-[Nitrincola-nitratireducens]</td></tr>
<tr><td>WP_202806909.1</td><td>Type1.BREX</td><td><-TOPRIM<-OLD-ABC<-SbcC<-N6-METHYLASE-p<-N6-METHYLASE<-N6-METHYLASE-p<-PIN<-OLD-ABC<-DUF6079-p<-DUF6079-p*<-BrxB-BrxF-iSTAND<-BrxA-DNA-binding<-XIS-HTH<-?<-HDRnase<-Phage_int_SAM<-Integrase_1||XIS-HTH->Phage_integrase-></td><td>697</td><td>Bacteria>Pseudomonadota>Gammaproteobacteria>Oceanospirillales>Oceanospirillaceae>Nitrincola</td><td>GCF_000585235.1</td><td>Nitrincola_nitratireducens</td><td>D791_RS20500</td><td>13689..15782</td><td>1229521</td><td>BREX-system-P-loop-protein-BrxC-[Nitrincola-nitratireducens]</td></tr>
<tr><td>HBO4843314.1</td><td>Type6.BREX</td><td><-ACET<-YjhX_toxin||?->?-><-MODE-HTH+PBPII||wHTH+WYL+WCX->BrxE->BrxA-DNA-binding->BrxB-BrxF-iSTAND->DUF6079*->OLD-ABC->PIN->N6-METHYLASE->?->PglZ->SIGMA+OB-Fold+AAA-ATPase+HKD-DNase-BrxL-><-HDRnase||PrlF_antitoxin-></td><td>1207</td><td>Bacteria>Pseudomonadota>Gammaproteobacteria>Pseudomonadales>Pseudomonadaceae>Pseudomonas</td><td>GCA_021660595.1</td><td>Pseudomonas_aeruginosa</td><td>L4V37_002405</td><td>61513..65136</td><td>287</td><td>BREX-system-P-loop-protein-BrxC-[Pseudomonas-aeruginosa]</td></tr>
<tr><td>HFD80332.1</td><td>Type1.BREX</td><td><-SIGMA+OB-Fold+AAA-ATPase+Lon-Protease-BrxL<-PglZ<-Schlafen+HATPase+wHTH<-PHP+SbcC<-N6-METHYLASE<-PIN<-DUF6079*<-BrxB-BrxF-iSTAND<-BrxA-DNA-binding</td><td>1156</td><td>Bacteria>Pseudomonadota>Gammaproteobacteria>unclassified_Gammaproteobacteria</td><td>GCA_011321775.1</td><td>Gammaproteobacteria_bacterium</td><td>ENK05_08090</td><td>12367..15837</td><td>1913989</td><td>BREX-system-P-loop-protein-BrxC-[Gammaproteobacteria-bacterium]</td></tr>
<tr><td>MBT5223355.1</td><td>Type1.BREX</td><td>BrxA-DNA-binding->BrxB-BrxF-iSTAND->DUF6079*->MazE_antitoxin->PIN->N6-METHYLASE->N6-METHYLASE->HEPN->GrpE->?->?->RelE-ParE-></td><td>1192</td><td>Bacteria>Pseudomonadota>Gammaproteobacteria>unclassified_Gammaproteobacteria</td><td>GCA_018661125.1</td><td>Gammaproteobacteria_bacterium</td><td>HOM15_11165</td><td>1377..4955</td><td>1913989</td><td>BREX-system-P-loop-protein-BrxC-[Gammaproteobacteria-bacterium]</td></tr>
<tr><td>MBN5083090.1</td><td>Type6.BREX</td><td><-DUF932<-SIGMA+OB-Fold+AAA-ATPase+HKD-DNase-BrxL<-PglZ<-ParB+HNH<-N6-METHYLASE<-N6-METHYLASE<-PIN<-OLD-ABC<-DUF6079*<-BrxB-BrxF-iSTAND<-BrxA-DNA-binding<-BrxE<-wHTH+WYL+WCX<-Arm-DNA-bind_3+Phage_integrase<-tRNA<-SIGMA-HTH+SIGMA-HTH+SIGMA-HTH</td><td>1207</td><td>Bacteria>Pseudomonadota>Gammaproteobacteria>Xanthomonadales>Xanthomonadaceae>Stenotrophomonas</td><td>GCA_017150495.1</td><td>Stenotrophomonas_maltophilia</td><td>JY467_05250</td><td>61681..65304</td><td>40324</td><td>BREX-system-P-loop-protein-BrxC-[Stenotrophomonas-maltophilia]</td></tr>
<tr><td>WP_004146422.1</td><td>Type6.BREX</td><td><-XIS-HTH||?->Phage_integrase->tRNA->Arm-DNA-bind_3+Phage_integrase->wHTH+WYL+WCX->BrxE->BrxA-DNA-binding->BrxB-BrxF-iSTAND->DUF6079*->OLD-ABC->PIN->N6-METHYLASE->N6-METHYLASE->ParB+HNH->PglZ->SIGMA+OB-Fold+AAA-ATPase+HKD-DNase-BrxL->DUF932-></td><td>1207</td><td>Bacteria>Pseudomonadota>Gammaproteobacteria>Xanthomonadales>Xanthomonadaceae>Stenotrophomonas</td><td>GCF_000355725.1</td><td>Stenotrophomonas_maltophilia</td><td>G5A34_RS07915</td><td>81973..85596</td><td>40324</td><td>BREX-system-P-loop-protein-BrxC-[Stenotrophomonas-maltophilia]</td></tr>
</table>

| Protein-ID | System | Gene-Neighbourhoods | Length | Taxonomy | Assembly-ID | Strain | Locus-Tag | Genomic-coordinates | TaxID | Fasta-defline |
|---|---|---|---|---|---|---|---|---|---|---|
| VUJ46804.1 | Type6.BREX | <-LexA-protease\|\|?-><-Phage_integrase\|\|Arm-DNA-bind_3+Phage_integrase-> wHTH+WYL+WCX -> BrxE -> BrxA-DNA-binding -> BrxB-BrxF-iSTAND -> DUF6079 *-> OLD-ABC-> PIN -> N6-METHYLASE -> N6-METHYLASE ->ParB+HNH-> PglZ -> SIGMA+OB-Fold+AAA-ATPase+HKD-DNase-BrxL ->DUF932-> | 1207 | Bacteria>Pseudomonadota>Gammaproteobacteria>Xanthomonadales>Xanthomonadaceae>Stenotrophomonas | - | Stenotrophomonas_maltophilia | N5D45_RS14200 | 24687..28310 | 40324 | hypothetical-protein-NGKLMNPD_00401-[Stenotrophomonas-maltophilia] |
| WP_142804161.1 | Type6.BREX | TPR+TPR-><-?\|\| BrxE -> BrxA-DNA-binding -> BrxB-BrxF-iSTAND -> DUF6079 *-> N6-METHYLASE ->AbrB-HTH-> PIN -> N6-METHYLASE ->DOC+HTH-> PglZ -> SIGMA+OB-Fold+AAA-ATPase+HKD-DNase-BrxL -> | 1210 | Bacteria>Pseudomonadota>Hydrogenophilia>Hydrogenophilales>Hydrogenophilaceae>Tepidiphilus | GCF_006980785.1 | Tepidiphilus_sp._J10 | EPD81_RS04405 | 897239..900871 | 2502185 | BREX-system-P-loop-protein-BrxC-[Tepidiphilus-sp.-J10] |
| WP_169116165.1 | Type6.BREX | Glyoxalase->?->?->?->?->?-><-?\|\| BrxE -> BrxB-BrxF-iSTAND -> DUF6079 *-> N6-METHYLASE ->AbrB-HTH-> PIN -> N6-METHYLASE ->DOC+HTH-> PglZ -> SIGMA+OB-Fold+AAA-ATPase+HKD-DNase-BrxL -> | 1210 | Bacteria>Pseudomonadota>Hydrogenophilia>Hydrogenophilales>Hydrogenophilaceae>Tepidiphilus | GCF_012911495.1 | Tepidiphilus_baoligensis | GV368_RS08140 | 10579..14211 | 2698687 | BREX-system-P-loop-protein-BrxC-[Tepidiphilus-baoligensis] |
| MCC7337722.1 | Type1.BREX | SF1-SF2-Helicase+wHTH+Small-B-barrel+DUF1998 ->?->?-> PIN ->?-> BrxA-DNA-binding -> BrxB-BrxF-iSTAND -> DUF6079 *->?-> N6-METHYLASE ->REase+TPR+GreAB-C+ PIN ->MORC+DUF3684+REase->SF1-SF2-Helicase->?-> PIN -> PglZ -> SIGMA+OB-Fold+AAA-ATPase+HKD-DNase-BrxL -> | 1171 | Bacteria>PVC_group>Planctomycetota>Planctomycetia>Pirellulales>Pirellulaceae | GCA_020852965.1 | Pirellulaceae_bacterium | IT422_21750 | 7905..11420 | 2699754 | BREX-system-P-loop-protein-BrxC-[Pirellulaceae-bacterium] |
| WP_068129828.1 | Type1.BREX | <- SIGMA+OB-Fold+AAA-ATPase+HKD-DNase-BrxL <-Four-helical-protein<- PglZ <- PIN <-HTH+MPTase<- N6-METHYLASE <- DUF6079 *<- BrxB-BrxF-iSTAND <- BrxA-DNA-binding <-XIS-HTH<-URI-DNase\|\|?-><-Vrr-REase | 1169 | Bacteria>PVC_group>Planctomycetota>Planctomycetia>Pirellulales>Pirellulaceae | GCF_008065135.1 | Roseimaritima_ulvae | UC8_RS27055 | 7584147..7587656 | 980254 | BREX-system-P-loop-protein-BrxC-[Roseimaritima-ulvae] |
| WP_259631594.1 | Type1.BREX | <-SF1-SF2-Helicase<-?<-?<-?\|\|XIS-HTH-> BrxA-DNA-binding -> BrxB-BrxF-iSTAND -> DUF6079 *-> N6-METHYLASE -> PIN ->ParB+HNH->TIR->MarR-HTH->Polbeta-> PglZ -> SIGMA+OB-Fold+AAA-ATPase+Lon-Protease-BrxL -> | 1147 | Bacteria>PVC_group>Planctomycetota>Planctomycetia>Pirellulales>Pirellulaceae | GCF_025060895.1 | Stieleria_sedimenti | NZK35_RS05520 | 42893..46336 | 2976331 | BREX-system-P-loop-protein-BrxC-[Stieleria-sedimenti] |
| HBJ35663.1 | Type1.BREX | <- SIGMA+OB-Fold+AAA-ATPase+HKD-DNase-BrxL <- PglZ <- PIN <-HTH+MPTase\|\|ZDNA-HTH-><-?<-Pkinase+CR-Kinase<- N6-METHYLASE <-Schlafen+HATPase+wHTH<- DUF6079 *<-Four-helical-protein<- BrxB-BrxF-iSTAND <- BrxA-DNA-binding <-XIS-HTH | 1188 | Bacteria>PVC_group>Planctomycetota>Planctomycetia>Planctomycetales>Planctomycetaceae | GCA_003506095.1 | Planctomycetaceae_bacterium | DDZ51_13145 | 16532..20098 | 2026779 | BREX-system-P-loop-protein-BrxC-[Planctomycetaceae-bacterium] |
| MBI3097943.1 | Type1.BREX | <- SIGMA+OB-Fold+AAA-ATPase+Lon-Protease-BrxL <- PglZ <-DOC+HTH<- N6-METHYLASE <- PIN <- DUF6079 *<- BrxB-BrxF-iSTAND <- BrxA-DNA-binding \|\|?->?-><-tRNA\|\|ParA-Soj-PloopNTPase-> | 1104 | Bacteria>PVC_group>Planctomycetota>unclassified_Planctomycetota | GCA_016192525.1 | Planctomycetota_bacterium | HYY93_06795 | 72529..75843 | 2026780 | BREX-system-P-loop-protein-BrxC-[Planctomycetota-bacterium] |
| NBB80453.1 | Type1.BREX | HTH_3-> BrxA-DNA-binding -> BrxB-BrxF-iSTAND -> DUF6079 *-> OLD-ABC-> PIN -> N6-METHYLASE ->OLD-ABC->HNH->ParB+HNH->Schlafen+HATPase+wHTH-> PglZ -> SIGMA+OB-Fold+AAA-ATPase+Lon-Protease-BrxL -> | 1147 | Bacteria>PVC_group>Verrucomicrobiota>unclassified_Verrucomicrobiota | GCA_009908325.1 | Verrucomicrobiota_bacterium | GVY36_13595 | 2227..5670 | 2026799 | BREX-system-P-loop-protein-BrxC-[Verrucomicrobiota-bacterium] |
| MBL9184319.1 | Type1.BREX | DUF932-> BrxA-DNA-binding -> BrxB-BrxF-iSTAND -> DUF6079 *-> PIN ->TIR-> N6-METHYLASE ->?-> PglZ -> SIGMA+OB-Fold+AAA-ATPase+Lon-Protease-BrxL ->REase+PNTD1+PNTD2->PIWI-> | 1153 | Bacteria>PVC_group>Verrucomicrobiota>Verrucomicrobiae>Verrucomicrobiales>Verrucomicrobiaceae | GCA_016795465.1 | Verrucomicrobiaceae_bacterium | JNN17_19395 | 2113..5574 | 2026800 | BREX-system-P-loop-protein-BrxC-[Verrucomicrobiaceae-bacterium] |
| MCB0987052.1 | Type1.BREX | <- N6-METHYLASE -p<- DUF6079 *<- BrxB-BrxF-iSTAND <- BrxA-DNA-binding <-?\|\|?-><-Phage_integrase<- PIN <-HTH+MPTase<-?<-PolB | 1154 | Bacteria>Terrabacteria_group>Actinomycetota>Acidimicrobiia>Acidimicrobiales>unclassified_Acidimicrobiales | GCA_020439505.1 | Acidimicrobiales_bacterium | KDB09_04685 | 1462..4926 | 2201156 | BREX-system-P-loop-protein-BrxC-[Acidimicrobiales-bacterium] |
| WP_169274836.1 | Type1.BREX | <- PIN <-RelB<-?\|\|HTH_3->?-> PIN -> BrxA-DNA-binding -> BrxB-BrxF-iSTAND -> DUF6079 *->CR-ATPase5+REase-> N6-METHYLASE ->Schlafen+HATPase+wHTH-> PglZ -> SIGMA+OB-Fold+AAA-ATPase+HKD-DNase-BrxL ->REase->?-> BrxA-DNA-binding -> | 1226 | Bacteria>Terrabacteria_group>Actinomycetota>Actinomycetes>Bifidobacteriales>Bifidobacteriaceae | GCF_012932365.1 | Bifidobacterium_sp._DSM_109958 | G1C96_RS01295 | 288582..292262 | 2705430 | BREX-system-P-loop-protein-BrxC-[Bifidobacterium-sp.-DSM-109958] |
| WP_226512485.1 | Type1.BREX | MACRODOMAIN+SIR2-><-?<-?\|\|DUF6361->HKD-DNase->SF1-SF2-Helicase->URI-DNase-> BrxA-DNA-binding -> BrxB-BrxF-iSTAND -> DUF6079 -p*-> DUF6079 -p->XIS-HTH-> PIN -> N6-METHYLASE -p-> N6-METHYLASE -> PglZ ->?->AbiEi_4+AbiEi_1-> SIGMA+OB-Fold+AAA-ATPase+HKD-DNase-BrxL -> | 435 | Bacteria>Terrabacteria_group>Actinomycetota>Actinomycetes>Mycobacteriales>Gordoniaceae | GCF_020520305.1 | Gordonia_sp._WA4-43 | LEL84_RS18910 | 4162672..4163979 | 2878678 | BREX-system-P-loop-protein-BrxC-[Gordonia-sp.-WA4-43] |
| WP_226512486.1 | Type1.BREX | DUF6361->HKD-DNase->SF1-SF2-Helicase->URI-DNase-> BrxA-DNA-binding -> BrxB-BrxF-iSTAND -> DUF6079 -p-> DUF6079 -p*->XIS-HTH-> PIN -> N6-METHYLASE -p-> N6-METHYLASE -> PglZ ->?->AbiEi_4+AbiEi_1-> SIGMA+OB-Fold+AAA-ATPase+HKD-DNase-BrxL -><-ABhydrolase+ABhydrolase | 785 | Bacteria>Terrabacteria_group>Actinomycetota>Actinomycetes>Mycobacteriales>Gordoniaceae | GCF_020520305.1 | Gordonia_sp._WA4-43 | LEL84_RS18915 | 4163880..4166237 | 2878678 | hypothetical-protein-[Gordonia-sp.-WA4-43] |
| QIN77519.1 | Type1.BREX | Thermonuclease->DOC+HTH->?->N6_N4_METHYLASE->SF1-SF2-Helicase-> BrxA-DNA-binding -> BrxB-BrxF-iSTAND -> DUF6079 *->?-> N6-METHYLASE ->REase->?-> PIN -> PglZ -> SIGMA+OB-Fold+AAA-ATPase+Lon-Protease-BrxL -><-?<-BirA-HTH | 1167 | Bacteria>Terrabacteria_group>Actinomycetota>Rubrobacteria>Rubrobacterales>Rubrobacteraceae | GCA_011492965.1 | Rubrobacter_marinus | GBA65_02255 | 450674..454177 | 2653852 | BREX-system-P-loop-protein-BrxC-[Rubrobacter-marinus] |
| MBA3426223.1 | Type1.BREX | <- DUF6079 -p*<- BrxB-BrxF-iSTAND <- BrxA-DNA-binding <-?<-?<- PIN <-?<-ParA-Soj-PloopNTPase | 773 | Bacteria>Terrabacteria_group>Actinomycetota>Rubrobacteria>Rubrobacterales>Rubrobacteraceae | GCA_013812245.1 | Rubrobacter_sp. | HOU04_17185 | 1..2321 | 2036010 | BREX-system-P-loop-protein-BrxC-[Rubrobacter-sp.] |
| WP_166395201.1 | Type1.BREX | Thermonuclease->DOC+HTH->?->N6_N4_METHYLASE->SF1-SF2-Helicase-> BrxA-DNA-binding -> BrxB-BrxF-iSTAND -> DUF6079 *->?-> N6-METHYLASE ->REase->?-> PIN -> PglZ -> SIGMA+OB-Fold+AAA-ATPase+Lon-Protease-BrxL -><-?<-BirA-HTH | 1151 | Bacteria>Terrabacteria_group>Actinomycetota>Rubrobacteria>Rubrobacterales>Rubrobacteraceae | GCF_011492965.1 | Rubrobacter_marinus | GBA65_RS02320 | 450722..454177 | 2653852 | BREX-system-P-loop-protein-BrxC-[Rubrobacter-marinus] |
| MBB2481351.1 | Type1.BREX | REase+DUF4268-><-?\|\|ZnR+RHH_7->XIS-HTH->MerR-HTH->?->?-> BrxA-DNA-binding -> BrxB-BrxF-iSTAND -> DUF6079 *-> N6-METHYLASE -> PglZ -> SIGMA+OB-Fold+AAA-ATPase+Lon-Protease-BrxL -> PIN ->HEPN->HEPN->?->MarR-HTH-> | 1187 | Bacteria>Terrabacteria_group>Bacillota>Bacilli>Bacillales>Bacillaceae | GCA_014174455.1 | Bacillus_sp._APMAM | H5P36_14260 | 24010..27573 | 2761103 | BREX-system-P-loop-protein-BrxC-[Bacillus-sp.-APMAM] |
| MBO1513528.1 | Type1.BREX | ZnR+RHH_7->XIS-HTH->MerR-HTH->?-><-?\|\|?-> BrxA-DNA-binding -> BrxB-BrxF-iSTAND -> DUF6079 *-> N6-METHYLASE -> PglZ -> SIGMA+OB-Fold+AAA-ATPase+Lon-Protease-BrxL ->?-> PIN ->HEPN->HEPN->?->MarR-HTH-> | 1187 | Bacteria>Terrabacteria_group>Bacillota>Bacilli>Bacillales>Bacillaceae | GCA_017497975.1 | Metabacillus_bambusae | I7822_17955 | 21833..25396 | 2795218 | BREX-system-P-loop-protein-BrxC-[Metabacillus-bambusae] |
| WP_243459544.1 | Type1.BREX | ZnR+RHH_7->XIS-HTH->MerR-HTH->?-><-?\|\|?-> BrxA-DNA-binding -> BrxB-BrxF-iSTAND -> DUF6079 *-> N6-METHYLASE -> PglZ -> SIGMA+OB-Fold+AAA-ATPase+Lon-Protease-BrxL ->?-> PIN ->HEPN->HEPN->?->MarR-HTH-> | 1182 | Bacteria>Terrabacteria_group>Bacillota>Bacilli>Bacillales>Bacillaceae | GCF_017497975.1 | Metabacillus_bambusae | I7822_RS17965 | 21848..25396 | 2795218 | BREX-system-P-loop-protein-BrxC-[Metabacillus-bambusae] |
| WP_259548277.1 | Type1.BREX | <-MarR-HTH<-?<-?<-HEPN<-HEPN<- PIN <- SIGMA+OB-Fold+AAA-ATPase+Lon-Protease-BrxL <- PglZ <- N6-METHYLASE <- DUF6079 *<- BrxB-BrxF-iSTAND <- BrxA-DNA-binding <-?<-?<-MerR-HTH<-XIS-HTH<-ZnR+RHH_7\|\|?->UDG-> | 1187 | Bacteria>Terrabacteria_group>Bacillota>Bacilli>Bacillales>Bacillaceae | GCF_024971915.1 | Heyndrickxia_oleronia | KXF32_RS03880 | 741080..744643 | 38875 | BREX-system-P-loop-protein-BrxC-[Heyndrickxia-oleronia] |
| CDD75188.1 | Type1.BREX | MACRODOMAIN->?-> PIN ->ZnR+ZnR+ZnR->HTH_26-> BrxA-DNA-binding -> BrxB-BrxF-iSTAND -> DUF6079 *-> OLD-ABC-> PIN -> N6-METHYLASE ->OLD-ABC->NERD->HEPN->Schlafen+HATPase+wHTH-> PglZ ->DUF4365+REase-> | 1200 | Bacteria>Terrabacteria_group>Bacillota>Clostridia>Eubacteriales>Clostridiaceae | GCA_000436175.1 | Clostridium_sp._CAG:62 | BN737_01375 | 69547..73149 | 1262828 | uncharacterized-protein-BN737_01375-[Clostridium-sp.-CAG:62] |
| WP_118744885.1 | Type1.BREX | ACET->REase->?->?->?->?->HTH_26-> BrxA-DNA-binding -> BrxB-BrxF-iSTAND -> DUF6079 *-> OLD-ABC-> PIN -> N6-METHYLASE ->HEPN->?->OLD-ABC->HNH->?->TIR-> | 1203 | Bacteria>Terrabacteria_group>Bacillota>Clostridia>Eubacteriales>Clostridiaceae | GCF_003480745.1 | Clostridium_sp._AM27-31LB | DW721_RS03885 | 61324..64935 | 2293026 | BREX-system-P-loop-protein-BrxC-[Clostridium-sp.-AM27-31LB] |
| WP_173680574.1 | Type1.BREX | <-Cap4_nuclease<-?<-HTH_3\|\|REase-><- SIGMA+OB-Fold+AAA-ATPase+Lon-Protease-BrxL <-?<- PglZ <- PIN <- N6-METHYLASE <- DUF6079 *<- BrxB-BrxF-iSTAND <- BrxA-DNA-binding \|\|PADR-HTH+MarR-HTH-><-D5ATPase<-?<-?<-?<-?\|\|LexA-protease-> | 1216 | Bacteria>Terrabacteria_group>Bacillota>Clostridia>Eubacteriales>Clostridiaceae | GCF_014230485.1 | Clostridium_tetanomorphum | FHX96_RS20165 | 4177507..4181157 | 1553 | BREX-system-P-loop-protein-BrxC-[Clostridium-tetanomorphum] |
| WP_221260245.1 | Type1.BREX | <-HISKIN\|\|RADICAL-SAM->?->?->HEPN->Ntox45->\|\| BrxA-DNA-binding -> BrxB-BrxF-iSTAND -> DUF6079 *-> N6-METHYLASE ->?-> PglZ -><-REase\|\| SIGMA+OB-Fold+AAA-ATPase+Lon-Protease-BrxL ->?-> PIN ->Peptidase_M42-> | 1218 | Bacteria>Terrabacteria_group>Bacillota>Clostridia>Eubacteriales>Clostridiaceae | GCF_019705275.1 | Clostridium_chauvoei | K4H38_08460 | 40416..44072 | 46867 | BREX-system-P-loop-protein-BrxC-[Clostridium-chauvoei] |
| WP_231016614.1 | Type1.BREX | PolB->TOPRIM->?->TrwC+SF1-SF2-Helicase-> BrxA-DNA-binding -> BrxB-BrxF-iSTAND -> DUF6079 *-> N6-METHYLASE ->HTH_3->MPTase-> PIN -> PglZ -> SIGMA+OB-Fold+AAA-ATPase+Lon-Protease-BrxL -><-CopG/RHH | 1189 | Bacteria>Terrabacteria_group>Bacillota>Clostridia>Eubacteriales>Clostridiaceae | GCF_021043395.1 | Clostridium_guangxiense | LQK79_RS11195 | 7642..11211 | 1662055 | BREX-system-P-loop-protein-BrxC-[Clostridium-guangxiense] |
| WP_252217802.1 | Type1.BREX | <-Peptidase_M42<- PIN <-?<- SIGMA+OB-Fold+AAA-ATPase+Lon-Protease-BrxL \|\|REase-><- PglZ <-?<- N6-METHYLASE <- DUF6079 *<- BrxB-BrxF-iSTAND <- BrxA-DNA-binding \|\|<-Ntox45<-HEPN<-HEPN<-?<-?<-RADICAL-SAM\|\|HISKIN-> | 1218 | Bacteria>Terrabacteria_group>Bacillota>Clostridia>Eubacteriales>Clostridiaceae | GCF_023846215.1 | Clostridium_chauvoei | J9878_RS07560 | 18587..22243 | 46867 | BREX-system-P-loop-protein-BrxC-[Clostridium-chauvoei] |
| WP_270487117.1 | Type1.BREX | <- SIGMA+OB-Fold+AAA-ATPase+Lon-Protease-BrxL <-?<- PIN <-SbcC<- PglZ <-HNH<-OLD-ABC<-TIR<- N6-METHYLASE <- DUF6079 *<- BrxB-BrxF-iSTAND <- BrxA-DNA-binding <-HTH_26<- PIN <-?<-MACRODOMAIN | 1201 | Bacteria>Terrabacteria_group>Bacillota>Clostridia>Eubacteriales>Clostridiaceae | GCF_027681635.1 | Clostridium_fessum | PGX92_RS02760 | 620693..624298 | 2126740 | BREX-system-P-loop-protein-BrxC-[Clostridium-fessum] |
| WP_270807575.1 | Type1.BREX | <-LRR_5+LRR_8<- SIGMA+OB-Fold+AAA-ATPase+Lon-Protease-BrxL <-?<- PIN <-SbcC<- PglZ <-HEPN<- N6-METHYLASE <- DUF6079 *<- BrxB-BrxF-iSTAND <- BrxA-DNA-binding <-HTH_26<- PIN <-?<-MACRODOMAIN | 1201 | Bacteria>Terrabacteria_group>Bacillota>Clostridia>Eubacteriales>Clostridiaceae | GCF_027723425.1 | Clostridium_fessum | PG965_RS01510 | 334129..337734 | 2126740 | BREX-system-P-loop-protein-BrxC-[Clostridium-fessum] |
| WP_021876221.1 | Type1.BREX | <-Peptidase_M42<- PIN <-?<- SIGMA+OB-Fold+AAA-ATPase+Lon-Protease-BrxL \|\|REase-><- PglZ <-?<- N6-METHYLASE <- DUF6079 *<- BrxB-BrxF-iSTAND <- BrxA-DNA-binding \|\|<-Ntox45<-HEPN<-?<-?<-RADICAL-SAM\|\|HISKIN-> | 1218 | Bacteria>Terrabacteria_group>Bacillota>Clostridia>Eubacteriales>Clostridiaceae | GCF_900037355.1 | Clostridium_chauvoei | C6H62_RS09310 | 1987009..1990665 | 46867 | BREX-system-P-loop-protein-BrxC-[Clostridium-chauvoei] |
| RGI65884.1 | Type1.BREX | HTH_26-> BrxA-DNA-binding -> BrxB-BrxF-iSTAND -> DUF6079 *-> OLD-ABC-> PIN -> N6-METHYLASE ->TIR->HEPN->Schlafen+HATPase+wHTH-> PglZ ->Tox-ART-HYD3-> SIGMA+OB-Fold+AAA-ATPase+Lon-Protease-BrxL -> | 1196 | Bacteria>Terrabacteria_group>Bacillota>Clostridia>Eubacteriales>Lachnospiraceae | GCA_003436035.1 | [Eubacterium]_rectale | DXD95_13285 | 1619..5209 | 39491 | BREX-system-P-loop-protein-BrxC-[[Eubacterium] |
| HCI18472.1 | Type1.BREX | <- PIN <-?<- SIGMA+OB-Fold+AAA-ATPase+Lon-Protease-BrxL <-OLD-ABC+TOPRIM<- PglZ <-Schlafen+HATPase+wHTH<-REase<- N6-METHYLASE <- DUF6079 -p*<- DUF6079 -p<- BrxB-BrxF-iSTAND <- BrxA-DNA-binding <-HTH_26 | 799 | Bacteria>Terrabacteria_group>Bacillota>Clostridia>Eubacteriales>Lachnospiraceae | GCA_003521585.1 | Lachnospiraceae_bacterium | DFH32_07645 | 14441..16840 | 1898203 | BREX-system-P-loop-protein-BrxC-[Lachnospiraceae-bacterium] |
| HCX41706.1 | Type1.BREX | SKI->?->tRNA->?->?->?->?->HTH_26-> BrxA-DNA-binding -> BrxB-BrxF-iSTAND -> DUF6079 *-> N6-METHYLASE ->REase->Schlafen+HATPase+wHTH-> PglZ ->OLD-ABC+TOPRIM-> SIGMA+OB-Fold+AAA-ATPase+Lon-Protease-BrxL ->?-> PIN ->MACRODOMAIN-> | 1196 | Bacteria>Terrabacteria_group>Bacillota>Clostridia>Eubacteriales>Lachnospiraceae | GCA_003539645.1 | Lachnospiraceae_bacterium | DHH50_05105 | 179559..183149 | 1898203 | BREX-system-P-loop-protein-BrxC-[Lachnospiraceae-bacterium] |
| MBR4345194.1 | Type1.BREX | RelB-> PIN -> BrxA-DNA-binding -> BrxB-BrxF-iSTAND -> DUF6079 *->VbhA+DOC+HTH-> N6-METHYLASE -> N6-METHYLASE -p->HEPN->SbcC+SbcC->Schlafen+HATPase+wHTH-> PglZ -> SIGMA+OB-Fold+AAA-ATPase+Lon-Protease-BrxL ->TIR-> | 1173 | Bacteria>Terrabacteria_group>Bacillota>Clostridia>Eubacteriales>Lachnospiraceae | GCA_017542245.1 | Lachnospiraceae_bacterium | IKP88_21230 | 56947..60468 | 1898203 | BREX-system-P-loop-protein-BrxC-[Lachnospiraceae-bacterium] |
| MCI7121458.1 | Type1.BREX | HATPase+wHTH->?->MACRODOMAIN->?->35exo+ADA-DBD->ZnR+ZnR+ZnR->HTH_26-> BrxA-DNA-binding -> BrxB-BrxF-iSTAND -> DUF6079 *-> OLD-ABC-> PIN -> N6-METHYLASE ->?-> PglZ ->Schlafen+HATPase+wHTH-> SIGMA+OB-Fold+AAA-ATPase+Lon-Protease-BrxL ->LRR_5+LRR_8->Cas9+REC+HNH+Cas9-> | 1201 | Bacteria>Terrabacteria_group>Bacillota>Clostridia>Eubacteriales>Lachnospiraceae | GCA_022775325.1 | [Ruminococcus]_gnavus | MR941_06720 | 10105..13710 | 33038 | BREX-system-P-loop-protein-BrxC-[[Ruminococcus] |
| WP_173865463.1 | Type1.BREX | <-STYKIN<-?<-LRR_5+LRR_8<- SIGMA+OB-Fold+AAA-ATPase+Lon-Protease-BrxL <-Schlafen+HATPase+wHTH<- PglZ <- N6-METHYLASE <- PIN <-OLD-ABC <- DUF6079 *<- BrxB-BrxF-iSTAND <- BrxA-DNA-binding <-HTH_26<-ZnR+ZnR+ZnR<-35exo+ADA-DBD<-?<-MACRODOMAIN<-?<-DOC+zhth | 1201 | Bacteria>Terrabacteria_group>Bacillota>Clostridia>Eubacteriales>Lachnospiraceae | GCF_013300965.1 | Faecalicatena_fissicatena | HFN12_RS00390 | 75728..79333 | 290055 | BREX-system-P-loop-protein-BrxC-[Faecalicatena-fissicatena] |
| WP_270645287.1 | Type1.BREX | HTH_19-><- SIGMA+OB-Fold+AAA-ATPase+Lon-Protease-BrxL <-?<- PglZ <- PIN <-OLD-ABC <-OLD-ABC+TOPRIM<- N6-METHYLASE <- DUF6079 *<- BrxB-BrxF-iSTAND <- BrxA-DNA-binding <-HTH_26<-?<-?<-?<-ArdCN+MPTase-inact+LPD19<-?<-TOPRIM+Topoisom_bac | 1205 | Bacteria>Terrabacteria_group>Bacillota>Clostridia>Eubacteriales>Lachnospiraceae | GCF_027692785.1 | Merdimonas_faecis | PG729_RS15130 | 28205..31822 | 1653435 | BREX-system-P-loop-protein-BrxC-[Merdimonas-faecis] |
| MCI8835830.1 | Type1.BREX | ABC-ATPase->ABC-ATPase+SbcC->?-> BrxA-DNA-binding -> BrxB-BrxF-iSTAND -> DUF6079 *->Schlafen+HATPase+wHTH-> N6-METHYLASE -> OLD-ABC-> PIN -> PglZ -> | 1205 | Bacteria>Terrabacteria_group>Bacillota>Clostridia>Eubacteriales>Oscillospiraceae | GCA_022795995.1 | Ruminococcus_sp. | HFH24_10725 | 31305..34922 | 41978 | BREX-system-P-loop-protein-BrxC-[Ruminococcus-sp.] |
| WP_226924212.1 | Type1.BREX | AraC-HTH-><-?\|\| BrxA-DNA-binding -> BrxB-BrxF-iSTAND -> DUF6079 *-> N6-METHYLASE ->REase-> PIN ->?-> PglZ -> SIGMA+OB-Fold+AAA-ATPase+Lon-Protease-BrxL -> | 1190 | Bacteria>Terrabacteria_group>Bacillota>Clostridia>Eubacteriales>Peptostreptococcaceae | GCF_020537215.1 | Intestinibacter_bartlettii | LIP50_RS08300 | 25949..29521 | 261299 | BREX-system-P-loop-protein-BrxC-[Intestinibacter-bartlettii] |
| MBQ3415902.1 | Type1.BREX | <- SIGMA+OB-Fold+AAA-ATPase+Lon-Protease-BrxL <- PglZ <-?<- PIN <-OLD-ABC <- N6-METHYLASE <-?<- DUF6079 *<- BrxB-BrxF-iSTAND <- BrxA-DNA-binding <-?<-?<-RHH_8 | 1216 | Bacteria>Terrabacteria_group>Bacillota>Clostridia>unclassified_Clostridia | GCA_017404025.1 | Clostridia_bacterium | IJH39_11340 | 11283..14933 | 2044939 | BREX-system-P-loop-protein-BrxC-[Clostridia-bacterium] |
| MBE6098022.1 | Type1.BREX | <- SIGMA+OB-Fold+AAA-ATPase+Lon-Protease-BrxL <- PglZ <-REase<-OLD-ABC<- N6-METHYLASE <- DUF6079 *<- BrxB-BrxF-iSTAND <- BrxA-DNA-binding <- PIN <-?<-Peptidase_M15 | 1220 | Bacteria>Terrabacteria_group>Bacillota>Negativicutes>Selenomonadales>Selenomonadaceae | GCA_015058685.1 | Schwartzia_succinivorans | E7198_09540 | 51689..55351 | 55507 | BREX-system-P-loop-protein-BrxC-[Schwartzia-succinivorans] |
| MBR3457117.1 | Type1.BREX | <- PglZ -p<-HNH<-OLD-ABC<- N6-METHYLASE -p<-?<- PIN <- N6-METHYLASE <- DUF6079 *<- BrxB-BrxF-iSTAND -p | 1215 | Bacteria>Terrabacteria_group>Bacillota>Negativicutes>Selenomonadales>Selenomonadaceae | GCA_017509225.1 | Selenomonadaceae_bacterium | IKH16_03040 | 6412..10059 | 2652289 | BREX-system-P-loop-protein-BrxC-[Selenomonadaceae-bacterium] |
| SEH31389.1 | Type1.BREX | <- PglZ <- PIN <-?<- N6-METHYLASE <- DUF6079 *<- BrxB-BrxF-iSTAND <- BrxA-DNA-binding <-OLD-ABC<-ParB<-ParA-Soj-PloopNTPase<-?<-HTH_3\|\|?->ParA-Soj-PloopNTPase-> | 1188 | Bacteria>Terrabacteria_group>Bacillota>Negativicutes>Selenomonadales>Selenomonadaceae | GCA_900108415.1 | Selenomonas_ruminantium | SAMN05216583_11711 | 10616..14182 | 971 | hypothetical-protein-SAMN05216583_11711-[Selenomonas-ruminantium] |
| WP_273038291.1 | Type1.BREX | <- SIGMA+OB-Fold+AAA-ATPase+Lon-Protease-BrxL <- PglZ <-REase<-OLD-ABC<- N6-METHYLASE <- DUF6079 *<- BrxB-BrxF-iSTAND <- BrxA-DNA-binding <- PIN <-?<-Peptidase_M15 | 1220 | Bacteria>Terrabacteria_group>Bacillota>Negativicutes>Selenomonadales>Selenomonadaceae | GCF_015058685.1 | Schwartzia_succinivorans | E7198_RS09640 | 51689..55351 | 55507 | BREX-system-P-loop-protein-BrxC-[Schwartzia-succinivorans] |
| WP_273698075.1 | Type1.BREX | <-REase<- PglZ <- N6-METHYLASE <- PIN <-OLD-ABC <-HNH<-OLD-ABC<- DUF6079 *<- BrxB-BrxF-iSTAND <- BrxA-DNA-binding <-?<-?<-?<-?<-SKI<-?\|\|O6MGT-HTH-> | 1181 | Bacteria>Terrabacteria_group>Bacillota>Negativicutes>Selenomonadales>Selenomonadaceae | GCF_905371815.1 | Selenomonas_noxia | NW174_RS07850 | 36953..40498 | 135083 | BREX-system-P-loop-protein-BrxC-[Selenomonas-noxia] |
| WP_273079788.1 | Type1.BREX | BrxA-DNA-binding -> BrxB-BrxF-iSTAND -> DUF6079 *->OLD-ABC->HNH-> OLD-ABC-> PIN -> N6-METHYLASE -> PglZ ->SIR2->HerA/FtsK-> | 1187 | Bacteria>Terrabacteria_group>Bacillota>Negativicutes>Selenomonadales>Selenomonadaceae | GCF_905371945.1 | Selenomonas_felix | NW177_RS10645 | 2029..5592 | 1944634 | BREX-system-P-loop-protein-BrxC-[Selenomonas-felix] |
| MBQ6343447.1 | Type1.BREX | BrxA-DNA-binding -> BrxB-BrxF-iSTAND -> DUF6079 *-><-?<-?\|\|PhdYeFM_antitox-> PIN ->?->?->?-> N6-METHYLASE -> | 1175 | Bacteria>Terrabacteria_group>Chloroflexota>Anaerolineae>Anaerolineales>Anaerolineaceae | GCA_017432985.1 | Anaerolineaceae_bacterium | IJI41_10010 | 1911..5438 | 2024896 | BREX-system-P-loop-protein-BrxC-[Anaerolineaceae-bacterium] |
| NJN66674.1 | Type1.BREX | <- N6-METHYLASE -p<- PIN <-CopG/RHH<- N6-METHYLASE -p<- DUF6079 -p<- DUF6079 -p*<- DUF6079 -p<- BrxB-BrxF-iSTAND <- BrxA-DNA-binding <-HTH_3\|\|?->?->?-><-DDE_Tnp\|\|DUF4158-> | 607 | Bacteria>Terrabacteria_group>Chloroflexota>Chloroflexia>Chloroflexales>Chloroflexineae | GCA_012032475.1 | Chloroflexaceae_bacterium | HC884_08105 | 4075..5898 | 2562117 | hypothetical-protein-[Chloroflexaceae-bacterium] |
| WP_097651212.1 | Type1.BREX | <- PglZ <- PIN <-?<- N6-METHYLASE -p<- PIN <-CopG/RHH<-?<-PolB<- N6-METHYLASE <- DUF6079 *<- BrxB-BrxF-iSTAND <- BrxA-DNA-binding <-?\|\|?-><-?<-?<-ParA-Soj-PloopNTPase+ParA-Soj-PloopNTPase | 1204 | Bacteria>Terrabacteria_group>Chloroflexota>Chloroflexia>Chloroflexales>Chloroflexineae | GCF_002532075.1 | Candidatus_Chloroploca_asiatica | A9Q02_RS04750 | 75845..79459 | 1506545 | BREX-system-P-loop-protein-BrxC-[Candidatus-Chloroploca-asiatica] |
| MBI2864205.1 | Type1.BREX | <-ABC-ATPase+SbcC<-ABC-ATPase+SbcC\|\|PolB-><-PhdYeFM_antitox\|\|?-> PIN -> BrxA-DNA-binding -> BrxB-BrxF-iSTAND -> DUF6079 -p*-> | 773 | Bacteria>Terrabacteria_group>Chloroflexota>unclassified_Chloroflexota | GCA_016190625.1 | Chloroflexota_bacterium | HYX94_06540 | 21104..23423 | 2026724 | BREX-system-P-loop-protein-BrxC-[Chloroflexota-bacterium] |
| MBK7180413.1 | Type1.BREX | <-REase<-SF1-SF2-Helicase+DUF1998<-?<- wHTH+WYL+WCX <-?\|\|?-> BrxA-DNA-binding -> BrxB-BrxF-iSTAND -> DUF6079 *-> N6-METHYLASE ->?-><- PIN <-RHH\|\| PglZ -> | 1173 | Bacteria>Terrabacteria_group>Chloroflexota>unclassified_Chloroflexota | GCA_016707865.1 | Chloroflexota_bacterium | IPH82_25035 | 72782..76303 | 2026724 | BREX-system-P-loop-protein-BrxC-[Chloroflexota-bacterium] |
| MCC6629583.1 | Type1.BREX | HIN-HTH-> BrxA-DNA-binding -> BrxB-BrxF-iSTAND -> DUF6079 *->?-> PIN -> N6-METHYLASE ->PolB->HEPN->?-> PglZ -> SIGMA+OB-Fold+AAA-ATPase+Lon-Protease-BrxL -> | 1204 | Bacteria>Terrabacteria_group>Chloroflexota>unclassified_Chloroflexota | GCA_020849725.1 | Chloroflexota_bacterium | IT340_19565 | 31214..34828 | 2026724 | BREX-system-P-loop-protein-BrxC-[Chloroflexota-bacterium] |
| MCL4487040.1 | Type1.BREX | BrxA-DNA-binding -> BrxB-BrxF-iSTAND ->Peptidase_M91-> N6-METHYLASE ->HKD-DNase-> DUF6079 *->HEPN->PhdYeFM_antitox-> PIN -> PglZ ->?->?->?->MerR-HTH+TipAS-> | 1221 | Bacteria>Terrabacteria_group>Chloroflexota>unclassified_Chloroflexota | GCA_023381515.1 | Chloroflexota_bacterium | M1570_02770 | 10289..13954 | 2026724 | BREX-system-P-loop-protein-BrxC-[Chloroflexota-bacterium] |
| WAS06096.1 | Type1.BREX | <- PglZ <- PIN <-PhdYeFM_antitox<-?<- PIN <-PhdYeFM_antitox<- N6-METHYLASE <- PIN <-PhdYeFM_antitox<- DUF6079 *<- BrxB-BrxF-iSTAND <- BrxA-DNA-binding \|\|?-><-NTP_transferase+MannoseP_isomer<-?<-TP_METHYLASE<-NTP_transf_3+Hexapep+Hexapep<-?<-RNase+DSRBD | 1221 | Bacteria>Terrabacteria_group>Cyanobacteriota/Melainabacteria_group>Cyanobacteriota>Cyanophyceae>Gloeomargaritales | GCA_022206215.1 | Gloeomargaritales_cyanobacterium_VI4D9 | LQF76_04175 | 903259..906924 | 2898145 | BREX-system-P-loop-protein-BrxC-[Gloeomargaritales-cyanobacterium-VI4D9] |
| MBW4670333.1 | Type6.BREX | <-Glycos_transf<-TPR\|\|REase-><-Acetate_kinase\|\| BrxE -> BrxA-DNA-binding -> BrxB-BrxF-iSTAND -> DUF6079 *->?-> PIN ->?-><-REase<-?\|\| N6-METHYLASE -> | 1211 | Bacteria>Terrabacteria_group>Cyanobacteriota/Melainabacteria_group>Cyanobacteriota>Cyanophyceae>Nostocales | GCA_019359595.1 | Cyanomargarita_calcarea_GSE-NOS-MK-12-04C | KME60_23685 | 99014..102649 | 2839659 | BREX-system-P-loop-protein-BrxC-[Cyanomargarita-calcarea-GSE-NOS-MK-12-04C] |
| WP_127084959.1 | Type1.BREX | CASPASE->?-><-?<-TOPRIM<-OLD-ABC<-Schlafen+HATPase+wHTH\|\| BrxA-DNA-binding -> BrxB-BrxF-iSTAND -> DUF6079 -p*->?->?-> PIN ->CopG/RHH-> N6-METHYLASE -p->OLD-ABC->HNH-> | 725 | Bacteria>Terrabacteria_group>Cyanobacteriota/Melainabacteria_group>Cyanobacteriota>Cyanophyceae>Nostocales | GCF_003991905.1 | Dulcicalothrix_desertica | DSM106972_RS34665 | 70206..72383 | 32056 | BREX-system-P-loop-protein-BrxC-[Dulcicalothrix-desertica] |
| WP_224095538.1 | Type1.BREX | BrxA-DNA-binding -><- SIGMA+OB-Fold+AAA-ATPase+Lon-Protease-BrxL <- PglZ +Phosphodiest \|\| PIN ->?-><- N6-METHYLASE -p<-HTH_23+HTH_32<- N6-METHYLASE -p<- N6-METHYLASE -p<- DUF6079 *<- BrxB-BrxF-iSTAND <-?<-REase<-?\|\|ArsR-HTH-><-?\|\|?-><-?\|\|HTH_32+DDE_3-> | 1178 | Bacteria>Terrabacteria_group>Cyanobacteriota/Melainabacteria_group>Cyanobacteriota>Cyanophyceae>Nostocales | GCF_019976755.1 | Nostoc_sp._MS1 | NSMS1_RS32170 | 256782..260318 | 2764711 | BREX-system-P-loop-protein-BrxC-[Nostoc-sp.-MS1] |
| WP_227788897.1 | Type1.BREX | DUF6079 *-> PIN ->CopG/RHH->?->?->?-> PglZ -> BrxD-AAA -> SF1-SF2-Helicase+wHTH+Small-B-barrel+DUF1998 ->RHH_9-> | 1169 | Bacteria>Terrabacteria_group>Cyanobacteriota/Melainabacteria_group>Cyanobacteriota>Cyanophyceae>Nostocales | GCF_020702635.1 | Nodularia_sp._LEGE_04288 | IQ240_RS09380 | 506..4015 | 1828639 | BREX-system-P-loop-protein-BrxC-[Nodularia-sp.-LEGE-04288] |
| WP_162398350.1 | Type1.BREX | BrxA-DNA-binding -><- SIGMA+OB-Fold+AAA-ATPase+Lon-Protease-BrxL <- PglZ +Phosphodiest \|\|REase-> PIN -><- N6-METHYLASE \|\|MetJarc-HTH->PemK_toxin-><- DUF6079 *<- BrxB-BrxF-iSTAND \|\|<-tRNA<-?<-?\|\|?->?-><-?\|\|?-><-Peptidase_M28 | 1178 | Bacteria>Terrabacteria_group>Cyanobacteriota/Melainabacteria_group>Cyanobacteriota>Cyanophyceae>Nostocales | - | Nostoc_sp._B(2019) | GS682_RS23175 | 25673..29209 | 2690446 | BREX-system-P-loop-protein-BrxC-[Nostoc-sp.-B(2019)] |
| BBD57221.1 | Type1.BREX | <-MerR-HTH\|\|RHH-><-?<-?\|\|?->TPR+TPR+TPR-><-Phage_integrase<-?\|\| BrxB-BrxF-iSTAND -> DUF6079 *-><-?<-REase\|\|?-> PIN -> N6-METHYLASE ->HEPN-> PglZ +Phosphodiest ->HKD-DNase-> SIGMA+OB-Fold+AAA-ATPase+Lon-Protease-BrxL -> | 1176 | Bacteria>Terrabacteria_group>Cyanobacteriota/Melainabacteria_group>Cyanobacteriota>Cyanophyceae>Oscillatoriophycideae | GCA_003609755.1 | Planktothrix_agardhii_NIES-204 | NIES204_45570 | 14633..18163 | 282423 | hypothetical-protein-NIES204_45570-[Planktothrix-agardhii-NIES-204] |
| WP_235758970.1 | Type1.BREX | <-Pkinase+CR-Kinase\|\| BrxB-BrxF-iSTAND -> DUF6079 *->?-> PIN -> N6-METHYLASE ->?->?-> PglZ +Phosphodiest -> PIN -> SIGMA+OB-Fold+AAA-ATPase+Lon-Protease-BrxL -><- BrxA-DNA-binding | 1176 | Bacteria>Terrabacteria_group>Cyanobacteriota/Melainabacteria_group>Cyanobacteriota>Cyanophyceae>Oscillatoriophycideae | GCA_030820635.1 | Planktothrix_agardhii | L2E81_RS22490 | 2176..5706 | 1160 | BREX-system-P-loop-protein-BrxC-[Planktothrix-agardhii] |
| WP_027255660.1 | Type1.BREX | <-MerR-HTH\|\|?->PAS-><-?<-?<-?\|\|?-> BrxB-BrxF-iSTAND -> DUF6079 *->?-> PIN -> N6-METHYLASE -p-> N6-METHYLASE ->HEPN-> PglZ +Phosphodiest ->HTH_Tnp_IS630+DDE_3-><-DDE_Tnp_1\|\|HKD-DNase-> | 1176 | Bacteria>Terrabacteria_group>Cyanobacteriota/Melainabacteria_group>Cyanobacteriota>Cyanophyceae>Oscillatoriophycideae | GCF_000464825.1 | Planktothrix_agardhii | N316_RS0124710 | 8429..11959 | 1160 | BREX-system-P-loop-protein-BrxC-[Planktothrix-agardhii] |
| WP_254034082.1 | Type1.BREX | TPR+TPR+TPR+TPR-><-?\|\| BrxA-DNA-binding -><- SIGMA+OB-Fold+AAA-ATPase+Lon-Protease-BrxL <-HKD-DNase<- PglZ +Phosphodiest <- N6-METHYLASE <- PIN <-?<- DUF6079 *<- BrxB-BrxF-iSTAND <-?\|\|?->?->Phage_integrase-> | 1176 | Bacteria>Terrabacteria_group>Cyanobacteriota/Melainabacteria_group>Cyanobacteriota>Cyanophyceae>Oscillatoriophycideae | GCF_904830845.1 | Planktothrix_agardhii | NMG87_RS21740 | 24173..27703 | 1160 | BREX-system-P-loop-protein-BrxC-[Planktothrix-agardhii] |
| KPQ33031.1 | Type6.BREX | <- SF1-SF2-Helicase+wHTH+Small-B-barrel+DUF1998 <- BrxD-AAA <- PglZ <- PIN <-?<- N6-METHYLASE <-DUF433<- DUF6079 *<- BrxB-BrxF-iSTAND <- BrxA-DNA-binding <- BrxE <-?<-AAA-ATPase | 1209 | Bacteria>Terrabacteria_group>Cyanobacteriota/Melainabacteria_group>Cyanobacteriota>Cyanophyceae>Pseudanabaenales | GCA_001314865.1 | Phormidesmis_priestleyi_Ana | HLUCCA11_19925 | 26535..30164 | 1666911 | hypothetical-protein-HLUCCA11_19925-[Phormidesmis-priestleyi-Ana] |
| MBW4580237.1 | Type6.BREX | <-ABC-ATPase+SbcC\|\|?->?-><-SAM-METHYLASE<-?\|\|CHTH-> BrxE -> BrxA-DNA-binding -> BrxB-BrxF-iSTAND -> DUF6079 *->?->REase->DUF29->AbrB-HTH-> PIN -> PglZ -> BrxD-AAA -> SF1-SF2-Helicase+wHTH+Small-B-barrel+DUF1998 -> | 1217 | Bacteria>Terrabacteria_group>Cyanobacteriota/Melainabacteria_group>Cyanobacteriota>Cyanophyceae>Pseudanabaenales | GCA_019358895.1 | Tildeniella_nuda_ZEHNDER_1965/U140 | KME42_11780 | 246570..250223 | 2136405 | BREX-system-P-loop-protein-BrxC-[Tildeniella-nuda-ZEHNDER-1965/U140] |
| WP_121969249.1 | Type6.BREX | CHTH->MazG-> BrxE -> BrxA-DNA-binding -> BrxB-BrxF-iSTAND -> DUF6079 *-> N6-METHYLASE ->?-> PIN -> N6-METHYLASE -p->?-> PglZ -> SIGMA+OB-Fold+AAA-ATPase+HKD-DNase-BrxL -> | 1213 | Bacteria>Terrabacteria_group>Cyanobacteriota/Melainabacteria_group>Cyanobacteriota>Cyanophyceae>Pseudanabaenales | GCF_002286735.1 | Leptolyngbya_sp._BC1307 | CKL79_RS08920 | 112124..115765 | 2029589 | BREX-system-P-loop-protein-BrxC-[Leptolyngbya-sp.-BC1307] |
| HBH74800.1 | Type1.BREX | PIN ->?-> BrxA-DNA-binding -> BrxB-BrxF-iSTAND -> DUF6079 *-> N6-METHYLASE ->SAM-METHYLASE->ParA-Soj-PloopNTPase->?->?-> PglZ ->OLD-ABC->HNH-> SIGMA+OB-Fold+AAA-ATPase+Lon-Protease-BrxL -> | 1163 | Bacteria>Terrabacteria_group>Cyanobacteriota/Melainabacteria_group>Cyanobacteriota>Cyanophyceae>Synechococcales | GCA_003508015.1 | Synechococcales_bacterium_UBA10510 | DDY43_15550 | 2736..6227 | 2055795 | BREX-system-P-loop-protein-BrxC-[Synechococcales-bacterium-UBA10510] |
| NJL99823.1 | Type1.BREX | <-SAM-METHYLASE\|\|?-> BrxA-DNA-binding -> BrxB-BrxF-iSTAND -> DUF6079 *-><- PIN <-PhdYeFM_antitox\|\| N6-METHYLASE -p-> N6-METHYLASE -> | 1145 | Bacteria>Terrabacteria_group>Cyanobacteriota/Melainabacteria_group>Cyanobacteriota>Cyanophyceae>Synechococcales | GCA_012031565.1 | Synechococcaceae_cyanobacterium_SM2_3_2 | HC924_13960 | 2310..5747 | 2720471 | BREX-system-P-loop-protein-BrxC-[Synechococcaceae-cyanobacterium-SM2_3_2] |
| WP_094560560.1 | Type1.BREX | <-SIGMA-HTH\|\|HicB_lk_antitox->Unstab_antitox->wHTH+HNH+HIT->EVE+HNH-> BrxA-DNA-binding -> BrxB-BrxF-iSTAND -> DUF6079 *-> N6-METHYLASE -><-?\|\| PglZ ->PhdYeFM_antitox-> PIN -> SIGMA+OB-Fold+AAA-ATPase+HKD-DNase-BrxL ->?-><-SF1-SF2-Helicase<-HEPN+PAS+DUF4011+SF1-SF2-Helicase+REase | 1161 | Bacteria>Terrabacteria_group>Cyanobacteriota/Melainabacteria_group>Cyanobacteriota>Cyanophyceae>Synechococcales | GCF_002252665.1 | Synechococcus_sp._8F6 | CJZ78_RS08320 | 43413..46898 | 2025606 | BREX-system-P-loop-protein-BrxC-[Synechococcus-sp.-8F6] |
| WP_197170384.1 | Type1.BREX | <-Sak4+WYL\|\|?-> PIN -> PIN -><-DUF772+DDE_Tnp_1+DDE_Tnp_1_6<-?\|\|?-> BrxA-DNA-binding -> BrxB-BrxF-iSTAND -> DUF6079 *-> N6-METHYLASE ->GTPase-AIG->sGTP-><-SWACOS+DTXR-HTH\|\| PglZ ->PhdYeFM_antitox-> PIN -> SIGMA+OB-Fold+AAA-ATPase+HKD-DNase-BrxL -><-REase | 1161 | Bacteria>Terrabacteria_group>Cyanobacteriota/Melainabacteria_group>Cyanobacteriota>Cyanophyceae>Synechococcales | GCF_015840915.1 | Synechococcus_sp._CBW1002 | H8F24_RS17545 | 3558842..3562327 | 1353134 | BREX-system-P-loop-protein-BrxC-[Synechococcus-sp.-CBW1002] |
| WP_254967828.1 | Type1.BREX | <- PIN <-MazE_antitoxin\|\|?-><- SIGMA+OB-Fold+AAA-ATPase+HKD-DNase-BrxL <- PIN <-PhdYeFM_antitox<- PglZ <-?<- N6-METHYLASE <- DUF6079 *<- BrxB-BrxF-iSTAND <- BrxA-DNA-binding \|\|DnaJ->?->?->?-><-?\|\|?-><- PIN | 1161 | Bacteria>Terrabacteria_group>Cyanobacteriota/Melainabacteria_group>Cyanobacteriota>Cyanophyceae>Synechococcales | GCF_024346135.1 | Cyanobium_sp._CH-040 | KBY90_RS07090 | 13099..16584 | 2823708 | BREX-system-P-loop-protein-BrxC-[Cyanobium-sp.-CH-040] |
| WP_254970971.1 | Type1.BREX | <- PglZ <-Couple_hipA+HipA_C<-HTH_3<-sGTP<-GTPase-AIG<- N6-METHYLASE <-MNS+REase\|\| PIN -> PIN -><- DUF6079 *<- BrxB-BrxF-iSTAND <- BrxA-DNA-binding \|\|Shikimate_dh_N+SDH_C->?-><-?\|\|?->?-><-Peptidase_M1_N+Peptidase_M1 | 1161 | Bacteria>Terrabacteria_group>Cyanobacteriota/Melainabacteria_group>Cyanobacteriota>Cyanophyceae>Synechococcales | GCF_024346155.1 | Cyanobium_sp._Candia_9D4 | KBZ08_RS02055 | 407526..411011 | 2823707 | BREX-system-P-loop-protein-BrxC-[Cyanobium-sp.-Candia-9D4] |
| WP_255005473.1 | Type1.BREX | PIN -> PIN ->?-><-DOC\|\|?->ArdCN+MPTase-> BrxA-DNA-binding -> BrxB-BrxF-iSTAND -> DUF6079 *-> N6-METHYLASE -> PglZ ->PhdYeFM_antitox-> PIN -> SIGMA+OB-Fold+AAA-ATPase+HKD-DNase-BrxL -><-REase<-TPR<-Peptidase | 1161 | Bacteria>Terrabacteria_group>Cyanobacteriota/Melainabacteria_group>Cyanobacteriota>Cyanophyceae>Synechococcales | GCF_024346315.1 | Cyanobium_sp._ATX_6F1 | KBZ13_RS02100 | 369855..373340 | 2823702 | BREX-system-P-loop-protein-BrxC-[Cyanobium-sp.-ATX-6F1] |
| KPA14790.1 | Type1.BREX | <- N6-METHYLASE <-Schlafen+HATPase+wHTH<- DUF6079 *<- PIN <-?<- BrxB-BrxF-iSTAND <- BrxA-DNA-binding <-?<-?<-HD\|\|?->AAA-ATPase-p-> | 1187 | Bacteria>Thermodesulfobacteriota>Desulfobacteria>Desulfobacterales>Candidatus_Magnetomoraceae>Candidatus_Magnetomorum | GCA_001292585.1 | Candidatus_Magnetomorum_sp._HK-1 | MHK_004993 | 4620..8183 | 1509431 | DNA-repair-ATPase-[Candidatus-Magnetomorum-sp.-HK-1] |
| HJA77858.1 | Type1.BREX | BrxB-BrxF-iSTAND -p-><-?\|\|ACET-> BrxA-DNA-binding -> BrxB-BrxF-iSTAND -> DUF6079 *-> OLD-ABC-> PIN -> N6-METHYLASE ->PHP+SbcC-> PglZ -> SIGMA+OB-Fold+AAA-ATPase+Lon-Protease-BrxL ->SIR2-> | 1145 | Bacteria>Thermodesulfobacteriota>Desulfovibrionia>Desulfovibrionales>Desulfovibrionaceae>Desulfovibrio | GCA_019118745.1 | Candidatus_Desulfovibrio_gallistercoris | H9774_13535 | 7700..11137 | 2838533 | BREX-system-P-loop-protein-BrxC-[Candidatus-Desulfovibrio-gallistercoris] |
| OQA19974.1 | Type1.BREX | BrxA-DNA-binding -> BrxB-BrxF-iSTAND -> DUF6079 *->?->RelE-ParE->VapB_antitoxin-> PIN -> N6-METHYLASE ->REase-> PglZ -> SIGMA+OB-Fold+AAA-ATPase+Lon-Protease-BrxL -> | 1164 | Bacteria>unclassified_Bacteria | GCA_002069435.1 | bacterium_ADurb.Bin363 | BWY64_00492 | 2230..5724 | 1866927 | hypothetical-protein-BWY64_00492-[bacterium-ADurb.Bin363] |
| GBE48762.1 | Type1.BREX | METHYLASE->REC+OmpR-HTH->REC+OmpR-HTH->?->URI-DNase+UVR+RNase+HHH->?->tRNA->tRNA->tRNA->?-> BrxB-BrxF-iSTAND -> DUF6079 *-> PIN -> N6-METHYLASE ->PHP+SbcC->Schlafen+HATPase+wHTH-> PglZ -> SIGMA+OB-Fold+AAA-ATPase+Lon-Protease-BrxL -> | 1155 | Bacteria>unclassified_Bacteria | GCA_002897795.1 | bacterium_BMS3Bbin12 | BMS3Bbin12_01950 | 17892..21359 | 2005735 | hypothetical-protein-BMS3Bbin12_01950-[bacterium-BMS3Bbin12] |

| Protein-ID | System | Gene-Neighbourhoods | Length | Taxonomy | Assembly-ID | Strain | Locus-Tag | Genomic-coordinates | TaxID | Fasta-defline |
|---|---|---|---|---|---|---|---|---|---|---|
| -- | | | | | | | | | | |
| -- TUDOR+SF1-SF2-Helicase+i REase+CTD+REase | | | | | | | | | | |
| -- | | | | | | | | | | |
| WP_007349691.1 | Type5.BREX | <-Glycos_transf_2\|\|tRNA->?->?-><-SIGMA+OB-Fold+AAA-ATPase+HKD-DNase-BrxL<-PglZ<-HNH<-OLD-ABC<-N6-METHYLASE<-Schlafen+HATPase+wHTH+HTH<-DUF6079*<-BrxB-BrxF-iSTAND<-BrxA-DNA-binding<-TUDOR+SF1-SF2-Helicase+i REase+CTD+REase<-TPR | 1194 | Bacteria>Pseudomonadota>Gammaproteobacteria>Pseudomonadales>Marinobacteraceae>Marinobacter | GCF_000169375.1 | Marinobacter_sp._ELB17 | MELB17_RS08075 | 35418..39002 | 270374 | BREX-system-P-loop-protein-BrxC-[Marinobacter-sp.-ELB17] |
| OUS97312.1 | Type5.BREX | <-MULE-TRANSPOSASE<-TPR+TPR+TPR+TPR+TPR+TPR<-?\|\|BrxA-DNA-binding->BrxB-BrxF-iSTAND->DUF6079*->N6-METHYLASE->HTH+REase->PglZ->SIGMA+OB-Fold+AAA-ATPase+HKD-DNase-BrxL-><-DNA_glycosylase\|\|?-><-TUDOR+SF1-SF2-Helicase+i REase+CTD+REase\|\|SF1-SF2-Helicase-> | 1175 | Bacteria>Terrabacteria_group>Actinomycetota>Actinomycetes>Mycobacteriales>Nocardiaceae | GCA_002165735.1 | Rhodococcus_sp._NCIMB_12038 | CA951_02905 | 126594..130121 | 933800 | ATP-binding-protein-[Rhodococcus-sp.-NCIMB-12038] |
| WP_256973250.1 | Type5.BREX | <-MULE-TRANSPOSASE<-TPR+TPR+TPR+TPR+TPR+TPR\|\|BrxA-DNA-binding->BrxB-BrxF-iSTAND->DUF6079*->N6-METHYLASE->HTH+REase->PglZ->SIGMA+OB-Fold+AAA-ATPase+HKD-DNase-BrxL-><-DNA_glycosylase\|\|?-><-TUDOR+SF1-SF2-Helicase+i REase+CTD+REase\|\|SF1-SF2-Helicase-> | 1213 | Bacteria>Terrabacteria_group>Actinomycetota>Actinomycetes>Mycobacteriales>Nocardiaceae | GCF_002165735.1 | Rhodococcus_sp._NCIMB_12038 | CA951_RS02920 | 126480..130121 | 933800 | BREX-system-P-loop-protein-BrxC-[Rhodococcus-sp.-NCIMB-12038] |
| WP_154708534.1 | Type5.BREX | <-REase<-TUDOR+SF1-SF2-Helicase+i REase+CTD+REase<-PglZ<-N6-METHYLASE<-DUF6079*<-BrxB-BrxF-iSTAND<-BrxA-DNA-binding<-?\|\|?->wHTH+HNH->?-><-?<-HTH_36\|\|HTH_3-> | 1187 | Bacteria>Terrabacteria_group>Actinomycetota>Actinomycetes>Propionibacteriales>Propionibacteriaceae | - | Cutibacterium_acnes | CP883_05580 | 10794..14357 | 1747 | BREX-system-P-loop-protein-BrxC-[Cutibacterium-acnes] |
| REB77450.1 | Type5.BREX | <-REase<-TUDOR+SF1-SF2-Helicase+i REase+CTD+REase<-PglZ<-N6-METHYLASE<-DUF6079*<-BrxB-BrxF-iSTAND<-BrxA-DNA-binding<-?\|\|?->wHTH+HNH->?-><-?<-HTH_36\|\|HTH_3-> | 1187 | Bacteria>Terrabacteria_group>Actinomycetota>Actinomycetes>Propionibacteriales>Propionibacteriaceae | GCA_003384685.1 | Cutibacterium_acnes | CP883_05580 | 10794..14357 | 1747 | hypothetical-protein-CP883_05580-[Cutibacterium-acnes] |
| WP_029421470.1 | Type5.BREX | <-ACET<-?<-REase<-TUDOR+SF1-SF2-Helicase+i REase+CTD+REase<-N6-METHYLASE-p<-N6-METHYLASE<-N6-METHYLASE<-PglZ<-DUF6079*<-BrxB-BrxF-iSTAND<-BrxA-DNA-binding<-tRNA<-?<-?<-?<-35exo+T7DNApol | 1205 | Bacteria>Terrabacteria_group>Bacillota>Bacilli>Bacillales>Alicyclobacillaceae | GCF_000702485.1 | Alicyclobacillus_macrosporangiidus | N687_RS0108610 | 1729200..1732817 | 392015 | BREX-system-P-loop-protein-BrxC-[Alicyclobacillus-macrosporangiidus] |
| WP_002116626.1 | Type5.BREX | <-PAS+NtrC-AAA+FIS-HTH\|\|?->?-><-?<-HTH_Tnp_1+HTH_21+rve+rve_3<-HTH_Tnp_1\|\|SF1-SF2-Helicase->BrxA-DNA-binding->BrxB-BrxF-iSTAND->DUF6079*->N6-METHYLASE->PglZ->TUDOR+SF1-SF2-Helicase+i REase+CTD+REase->?->wHTH+WYL+WCX->NUDIX->HKD-DNase+SF1-SF2-Helicase+HTH+HTH+SRA->N6-METHYLASE-> | 1188 | Bacteria>Terrabacteria_group>Bacillota>Bacilli>Bacillales>Bacillaceae | - | Bacillus_cereus | IEQ_RS03950 | 791262..794828 | 1396 | BREX-system-P-loop-protein-BrxC-[Bacillus-cereus] |
| OHR66989.1 | Type5.BREX | GDResolvase+HIN-HTH-><-?<-?<-?\|\|GDResolvase+HIN-HTH->BrxA-DNA-binding->BrxB-BrxF-iSTAND->DUF6079*->N6-METHYLASE->PglZ->TUDOR+SF1-SF2-Helicase+i REase+CTD+REase->?->SF1-SF2-Helicase-><-?\|\|TetR-HTH-> | 1195 | Bacteria>Terrabacteria_group>Bacillota>Bacilli>Bacillales>Bacillaceae | GCA_001838545.1 | Bacillus_sp._HMSC76G11 | HMPREF3291_01045 | 70062..73649 | 1608910 | hypothetical-protein-HMPREF3291_01045-[Bacillus-sp.-HMSC76G11] |
| ASF41331.1 | Type5.BREX | <-REase<-TUDOR+SF1-SF2-Helicase+i REase+CTD+REase<-PglZ<-N6-METHYLASE<-DUF6079*<-BrxB-BrxF-iSTAND<-BrxA-DNA-binding<-?\|\|ABC-ATPase+SbcC->?-><-?<-?<-ZnR | 1188 | Bacteria>Terrabacteria_group>Bacillota>Bacilli>Bacillales>Bacillaceae | GCA_002209425.1 | Halobacillus_halophilus | CEH05_20035 | 4059294..4062860 | 1570 | hypothetical-protein-CEH05_20035-[Halobacillus-halophilus] |
| ASS85704.1 | Type5.BREX | McrC-NTD+REase->?->?->?->HTH_3->?->SF1-SF2-Helicase->BrxA-DNA-binding->BrxB-BrxF-iSTAND->DUF6079*->N6-METHYLASE->PglZ->TUDOR+SF1-SF2-Helicase+i REase+CTD+REase->?-><-DDE_5+DDE_Tnp_2\|\|?->HTH_32+rve+rve_3+Mu-transpos_C->GspA-ATPase-> | 1183 | Bacteria>Terrabacteria_group>Bacillota>Bacilli>Bacillales>Bacillaceae | GCA_002243605.1 | Geobacillus_lituanicus | GLN3_00240 | 44558..48109 | 169283 | hypothetical-protein-GLN3_00240-[Geobacillus-lituanicus] |
| PFG07777.1 | Type5.BREX | <-HKD-DNase+SF1-SF2-Helicase+HTH+HTH+SRA<-NUDIX\|\|wHTH(SBD)+HNH-><-?\|\|HTH_3->?-><-TUDOR+SF1-SF2-Helicase+i REase+CTD+REase<-PglZ<-N6-METHYLASE<-DUF6079*<-BrxB-BrxF-iSTAND<-BrxA-DNA-binding<-ParB+HNH<-?<-HEPN+HTH | 1195 | Bacteria>Terrabacteria_group>Bacillota>Bacilli>Bacillales>Bacillaceae | GCA_002563655.1 | Bacillus_sp._es.034 | ATG71_4686 | 4473829..4477416 | 1761763 | hypothetical-protein-ATG71_4686-[Bacillus-sp.-es.034] |
| PGR68857.1 | Type5.BREX | <-McrC-NTD+REase<-wHTH+McrB-AAA<-?<-wHTH+WYL+WCX<-TUDOR+SF1-SF2-Helicase+i REase+CTD+REase<-PglZ<-N6-METHYLASE<-DUF6079*<-BrxB-BrxF-iSTAND<-BrxA-DNA-binding<-SF1-SF2-Helicase<-?<-NERD+SF1-SF2-Helicase<-REase<-?<-?<-REC+CITB-HTH+ZDNA-HTH | 1191 | Bacteria>Terrabacteria_group>Bacillota>Bacilli>Bacillales>Bacillaceae | GCA_002576445.1 | Bacillus_cereus | COC49_05330 | 39030..42605 | 1396 | hypothetical-protein-COC49_05330-[Bacillus-cereus] |
| PRT22691.1 | Type5.BREX | REC+CITB-HTH+ZDNA-HTH->?->?->REase->NERD+SF1-SF2-Helicase->?->SF1-SF2-Helicase->BrxA-DNA-binding->BrxB-BrxF-iSTAND->DUF6079*->N6-METHYLASE->PglZ->TUDOR+SF1-SF2-Helicase+i REase+CTD+REase->URI-DNase+RAMA->wHTH+WYL+WCX-><-rve | 1188 | Bacteria>Terrabacteria_group>Bacillota>Bacilli>Bacillales>Bacillaceae | GCA_002998815.1 | Bacillus_thuringiensis | C6351_31305 | 13182..16748 | 1428 | BREX-system-P-loop-protein-BrxC-[Bacillus-thuringiensis] |
| MBU9710538.1 | Type5.BREX | RNase->?->?->?->?->ParB->SF1-SF2-Helicase+REase+HTH->BrxA-DNA-binding->BrxB-BrxF-iSTAND->DUF6079*->N6-METHYLASE->PglZ->TUDOR+SF1-SF2-Helicase+i REase+CTD+REase->?->SF1-SF2-Helicase+Z1->?->MazF-><-ABhydrolase | 1188 | Bacteria>Terrabacteria_group>Bacillota>Bacilli>Bacillales>Bacillaceae | GCA_019042215.1 | Evansella_tamaricis | KS419_02125 | 46685..50251 | 2069301 | BREX-system-P-loop-protein-BrxC-[Evansella-tamaricis] |
| WP_019720353.1 | Type5.BREX | BrxA-DNA-binding->BrxB-BrxF-iSTAND->DUF6079*->N6-METHYLASE->PglZ->TUDOR+SF1-SF2-Helicase+i REase+CTD+REase-> | 1187 | Bacteria>Terrabacteria_group>Bacillota>Bacilli>Bacillales>Bacillaceae | GCF_000223155.1 | Weizmannia_coagulans | IOE_RS0101945 | 1303..4866 | 1398 | BREX-system-P-loop-protein-BrxC-[Weizmannia-coagulans] |
| WP_003250543.1 | Type5.BREX | OLD-ABC+TOPRIM->SF1-SF2-Helicase-><-?<-?\|\|HTH->BrxA-DNA-binding->BrxB-BrxF-iSTAND->DUF6079*->N6-METHYLASE->PglZ->TUDOR+SF1-SF2-Helicase+i REase+CTD+REase->HTH_Tnp+HTH_Tnp->?->Phage_integrase-><-Phage_int_SAM_1+Phage_integrase | 1194 | Bacteria>Terrabacteria_group>Bacillota>Bacilli>Bacillales>Bacillaceae | GCF_000258725.1 | Parageobacillus_thermoglucosidasius | GT20_RS09290 | 1817833..1821417 | 1426 | BREX-system-P-loop-protein-BrxC-[Parageobacillus-thermoglucosidasius] |
| WP_223254242.1 | Type5.BREX | <-REase<-TUDOR+SF1-SF2-Helicase+i REase+CTD+REase<-PglZ<-N6-METHYLASE<-DUF6079*<-BrxB-BrxF-iSTAND<-BrxA-DNA-binding<-?\|\|ABC-ATPase+SbcC->?-><-?<-ZnR | 1203 | Bacteria>Terrabacteria_group>Bacillota>Bacilli>Bacillales>Bacillaceae | GCF_000284515.1 | Halobacillus_halophilus | HBHAL_RS19750 | 4059450..4063061 | 1570 | BREX-system-P-loop-protein-BrxC-[Halobacillus-halophilus] |
| WP_026590665.1 | Type5.BREX | <-PAS+NtrC-AAA+FIS-HTH<-NERD<-?<-wHTH+WYL+WCX<-TUDOR+SF1-SF2-Helicase+i REase+CTD+REase<-PglZ<-N6-METHYLASE<-DUF6079*<-BrxB-BrxF-iSTAND<-BrxA-DNA-binding<-SF1-SF2-Helicase\|\|<-ABhydrolase<-REase\|\|?-><-?<-McrB-AAA | 1185 | Bacteria>Terrabacteria_group>Bacillota>Bacilli>Bacillales>Bacillaceae | GCF_000482345.1 | Bacillus_sp._UNC437CL72CviS29 | M014_RS0104560 | 312641..316198 | 1340430 | BREX-system-P-loop-protein-BrxC-[Bacillus-sp.-UNC437CL72CviS29] |
| WP_026908399.1 | Type5.BREX | <-ParB+HNH<-SF1-SF2-Helicase+REase<-?<-?<-?<-REase<-TUDOR+SF1-SF2-Helicase+i REase+CTD+REase<-PglZ<-N6-METHYLASE<-DUF6079*<-BrxB-BrxF-iSTAND<-BrxA-DNA-binding<-?\|\|wHTH->?->wHTH+HNH->?-><-?<-HTH_36 | 1187 | Bacteria>Terrabacteria_group>Bacillota>Bacilli>Bacillales>Bacillaceae | GCF_000482485.1 | Paucisalibacillus_globulus | K332_RS0116125 | 154598..158161 | 351095 | BREX-system-P-loop-protein-BrxC-[Paucisalibacillus-globulus] |
| WP_048718293.1 | Type5.BREX | HTH_36->?-><-?<-wHTH+HNH<-?<-wHTH\|\|?->BrxA-DNA-binding->BrxB-BrxF-iSTAND->DUF6079*->N6-METHYLASE->PglZ->TUDOR+SF1-SF2-Helicase+i REase+CTD+REase->REase->?->TIR->Schlafen+HATPase+wHTH-> | 1187 | Bacteria>Terrabacteria_group>Bacillota>Bacilli>Bacillales>Bacillaceae | GCF_001076885.1 | Bacillus_sp._522_BSPC | ADJ83_RS15690 | 17567..21130 | 1579338 | BREX-system-P-loop-protein-BrxC-[Bacillus-sp.-522_BSPC] |
| WP_061570112.1 | Type5.BREX | ACET-><-?\|\|N6-METHYLASE->?->BrxA-DNA-binding->BrxB-BrxF-iSTAND->DUF6079*->N6-METHYLASE->PglZ->TUDOR+SF1-SF2-Helicase+i REase+CTD+REase->?-><-?\|\|?->?->MerR-HTH-> | 1201 | Bacteria>Terrabacteria_group>Bacillota>Bacilli>Bacillales>Bacillaceae | GCF_001587535.1 | Caldibacillus_debilis | B4135_RS15850 | 22498..26103 | 301148 | BREX-system-P-loop-protein-BrxC-[Caldibacillus-debilis] |
| WP_066159884.1 | Type5.BREX | <-OLD-ABC+TOPRIM<-?<-?<-TUDOR+SF1-SF2-Helicase+i REase+CTD+REase<-PglZ<-N6-METHYLASE<-DUF6079*<-BrxB-BrxF-iSTAND<-BrxA-DNA-binding<-SF1-SF2-Helicase+REase-p<-SF1-SF2-Helicase<-SF1-SF2-Helicase<-ParB<-MerR-HTH | 1205 | Bacteria>Terrabacteria_group>Bacillota>Bacilli>Bacillales>Bacillaceae | GCF_001591945.1 | Halalkalibacter_krulwichiae | BKR01S_RS20150 | 13948..17565 | 199441 | BREX-system-P-loop-protein-BrxC-[Halalkalibacter-krulwichiae] |
| WP_077428558.1 | Type5.BREX | <-SMODS<-?<-?<-SLATT<-ParB+HNH<-?<-TUDOR+SF1-SF2-Helicase+i REase+CTD+REase<-PglZ<-N6-METHYLASE<-DUF6079*<-BrxB-BrxF-iSTAND<-BrxA-DNA-binding<-SF1-SF2-Helicase<-N6-METHYLASE<-HKD-DNase+SF1-SF2-Helicase+HTH+HTH+SRA<-?<-?\|\|?-><-SAM-METHYLASE | 1179 | Bacteria>Terrabacteria_group>Bacillota>Bacilli>Bacillales>Bacillaceae | GCF_001996285.1 | Anoxybacillus_karvacharensis | BO219_RS04435 | 413742..417281 | 1919277 | BREX-system-P-loop-protein-BrxC-[Anoxybacillus-karvacharensis] |
| WP_047753295.1 | Type5.BREX | <-DEDD_Tnp_IS110+Transposase_20<-DDE_3\|\|?->DDE_Tnp_1-><-DDE_5+DDE_Tnp_2<-SAD+REase<-TUDOR+SF1-SF2-Helicase+i REase+CTD+REase<-PglZ<-N6-METHYLASE<-DUF6079*<-BrxB-BrxF-iSTAND<-BrxA-DNA-binding<-SF1-SF2-Helicase\|\|DDE_Tnp_1-><-?<-SF1-SF2-Helicase<-?<-HTH_3<-HigB | 1183 | Bacteria>Terrabacteria_group>Bacillota>Bacilli>Bacillales>Bacillaceae | GCF_002243665.1 | Geobacillus | GT3921_RS13000 | 2562536..2566087 | 129337 | MULTISPECIES:-BREX-system-P-loop-protein-BrxC-[Geobacillus] |
| WP_098359294.1 | Type5.BREX | <-PAS+NtrC-AAA+FIS-HTH<-NERD<-?<-wHTH+WYL+WCX<-TUDOR+SF1-SF2-Helicase+i REase+CTD+REase<-PglZ<-N6-METHYLASE<-DUF6079*<-BrxB-BrxF-iSTAND<-BrxA-DNA-binding<-SF1-SF2-Helicase<-ORC-CDC6<-HEPN<-McrC-NTD+REase<-wHTH+McrB-AAA\|\|ParB-> | 1186 | Bacteria>Terrabacteria_group>Bacillota>Bacilli>Bacillales>Bacillaceae | GCF_002561295.1 | Bacillus_cereus | CN373_RS10800 | 80983..84543 | 1396 | BREX-system-P-loop-protein-BrxC-[Bacillus-cereus] |
| WP_098613368.1 | Type5.BREX | <-McrC-NTD+REase<-wHTH+McrB-AAA<-?<-wHTH+WYL+WCX<-?<-TUDOR+SF1-SF2-Helicase+i REase+CTD+REase<-PglZ<-N6-METHYLASE<-N6-METHYLASE<-DUF6079*<-BrxB-BrxF-iSTAND<-BrxA-DNA-binding<-SF1-SF2-Helicase<-?<-NERD+SF1-SF2-Helicase<-REase<-?<-?<-REC+CITB-HTH+ZDNA-HTH | 1188 | Bacteria>Terrabacteria_group>Bacillota>Bacilli>Bacillales>Bacillaceae | GCF_002567495.1 | Bacillus_cereus | COK05_RS16060 | 61384..64950 | 1396 | BREX-system-P-loop-protein-BrxC-[Bacillus-cereus] |
| WP_098794042.1 | Type5.BREX | <-HKD-DNase<-DrmB<-DrmA\|\|SF1-SF2-Helicase->SF1-SF2-Helicase->DNA_METHYLASE->SF1-SF2-Helicase->BrxA-DNA-binding->BrxB-BrxF-iSTAND->DUF6079*->N6-METHYLASE->PglZ->TUDOR+SF1-SF2-Helicase+i REase+CTD+REase->?->?->?->?-><-?\|\|ACET->TPR-> | 1188 | Bacteria>Terrabacteria_group>Bacillota>Bacilli>Bacillales>Bacillaceae | GCF_002576235.1 | Priestia_megaterium | COA18_RS28020 | 1940977..1944543 | 1404 | BREX-system-P-loop-protein-BrxC-[Priestia-megaterium] |
| WP_098878446.1 | Type5.BREX | <-McrC-NTD+REase<-wHTH+McrB-AAA<-?<-wHTH+WYL+WCX<-?<-TUDOR+SF1-SF2-Helicase+i REase+CTD+REase<-PglZ<-N6-METHYLASE<-DUF6079*<-BrxB-BrxF-iSTAND<-BrxA-DNA-binding<-SF1-SF2-Helicase<-?<-NERD+SF1-SF2-Helicase<-REase<-?<-?<-REC+CITB-HTH+ZDNA-HTH | 1191 | Bacteria>Terrabacteria_group>Bacillota>Bacilli>Bacillales>Bacillaceae | GCF_002576445.1 | Bacillus_cereus | COC49_RS05330 | 39030..42605 | 1396 | BREX-system-P-loop-protein-BrxC-[Bacillus-cereus] |
| WP_100332651.1 | Type5.BREX | ANK->?-><-PolB\|\|<-?\|\|?-><-TUDOR+SF1-SF2-Helicase+i REase+CTD+REase<-PglZ<-N6-METHYLASE<-DUF6079*<-BrxB-BrxF-iSTAND<-BrxA-DNA-binding<-?<-AAA+<-HTH_36\|\|?->?->GDResolvase+Recombinase+ZnR-><-PolB | 1187 | Bacteria>Terrabacteria_group>Bacillota>Bacilli>Bacillales>Bacillaceae | GCF_002797355.1 | Bacillus_xiapuensis | CEF20_RS14915 | 671484..675047 | 2014075 | BREX-system-P-loop-protein-BrxC-[Bacillus-xiapuensis] |
| WP_066188676.1 | Type5.BREX | SPOUT-><-?<-wHTH+HNH<-?<-wHTH\|\|?->BrxA-DNA-binding->BrxB-BrxF-iSTAND->DUF6079*->N6-METHYLASE->PglZ->TUDOR+SF1-SF2-Helicase+i REase+CTD+REase->REase->ZetaToxin->Classical-AAA->Peptidase_S8-> | 1187 | Bacteria>Terrabacteria_group>Bacillota>Bacilli>Bacillales>Bacillaceae | GCF_002835735.1 | Cytobacillus_horneckiae | CWS20_RS08115 | 27127..30690 | 549687 | BREX-system-P-loop-protein-BrxC-[Cytobacillus-horneckiae] |
| WP_106082566.1 | Type5.BREX | REC+CITB-HTH+ZDNA-HTH->?->?->REase->NERD+SF1-SF2-Helicase->?->SF1-SF2-Helicase->BrxA-DNA-binding->BrxB-BrxF-iSTAND->DUF6079*->N6-METHYLASE->PglZ->TUDOR+SF1-SF2-Helicase+i REase+CTD+REase->URI-DNase+RAMA->wHTH+WYL+WCX-><-rve | 1188 | Bacteria>Terrabacteria_group>Bacillota>Bacilli>Bacillales>Bacillaceae | GCF_002998815.1 | Bacillus_thuringiensis | C6351_31305 | 13182..16748 | 1428 | BREX-system-P-loop-protein-BrxC-[Bacillus-thuringiensis] |
| WP_117327463.1 | Type5.BREX | ABC-ATPase+SbcC->?-><-?\|\|Phage_int_SAM_1+Phage_integrase\|\|HIN-HTH-><-TUDOR+SF1-SF2-Helicase+i REase+CTD+REase<-PglZ<-N6-METHYLASE<-DUF6079*<-BrxB-BrxF-iSTAND<-BrxA-DNA-binding<-GDResolvase+HIN-HTH<-Phage_integrase<-Phage_integrase<-NERD+ZnR<-SbcC | 1195 | Bacteria>Terrabacteria_group>Bacillota>Bacilli>Bacillales>Bacillaceae | GCF_003429095.1 | Peribacillus_saganii | D0469_RS14520 | 34367..37954 | 2303992 | BREX-system-P-loop-protein-BrxC-[Peribacillus-saganii] |
| WP_132948756.1 | Type5.BREX | NUDIX->ACET-><-?\|\|McrB-NTD+McrB-AAA->McrC-NTD+REase->?->SF1-SF2-Helicase->BrxA-DNA-binding->BrxB-BrxF-iSTAND->DUF6079*->N6-METHYLASE->PglZ->TUDOR+SF1-SF2-Helicase+i REase+CTD+REase->OLD-ABC+TOPRIM->HIN-HTH-><-?<-?<-ParA-Soj-PloopNTPase | 1183 | Bacteria>Terrabacteria_group>Bacillota>Bacilli>Bacillales>Bacillaceae | GCF_004341205.1 | Thermolongibacillus_altinsuensis | EDD69_RS09635 | 59871..63422 | 575256 | BREX-system-P-loop-protein-BrxC-[Thermolongibacillus-altinsuensis] |
| WP_136831108.1 | Type5.BREX | SPOUT-><-GDResolvase+Recombinase+ZnR<-GDResolvase+Recombinase+ZnR<-?\|\|NUDIX->GDResolvase+HIN-HTH->BrxA-DNA-binding->BrxB-BrxF-iSTAND->DUF6079*->N6-METHYLASE->PglZ->TUDOR+SF1-SF2-Helicase+i REase+CTD+REase-><-PadR-HTH<-?<-?<-HTH_Tnp+HTH_Tnp\|\|HKD-DNase-> | 1195 | Bacteria>Terrabacteria_group>Bacillota>Bacilli>Bacillales>Bacillaceae | GCF_005116465.1 | Robertmurraya_kyonggiensis | FA727_RS11510 | 51719..55306 | 1037680 | BREX-system-P-loop-protein-BrxC-[Robertmurraya-kyonggiensis] |
| WP_144565381.1 | Type5.BREX | <-GTPase-AIG<-GTPase-AIG<-?<-wHTH+WYL+WCX<-URI-DNase+RAMA<-TUDOR+SF1-SF2-Helicase+i REase+CTD+REase<-PglZ<-N6-METHYLASE<-N6-METHYLASE<-DUF6079*<-BrxB-BrxF-iSTAND<-BrxA-DNA-binding<-SF1-SF2-Helicase<-JAB<-SF1-SF2-Helicase+Z1<-?<-REase<-?<-SF1-SF2-Helicase | 1186 | Bacteria>Terrabacteria_group>Bacillota>Bacilli>Bacillales>Bacillaceae | GCF_007681185.1 | Bacillus_mobilis | FS794_RS23165 | 89317..92877 | 2026190 | BREX-system-P-loop-protein-BrxC-[Bacillus-mobilis] |
| WP_144543246.1 | Type5.BREX | Phage_int_SAM_3+Phage_integrase-><-?<-?<-?<-?<-REase<-TUDOR+SF1-SF2-Helicase+i REase+CTD+REase<-PglZ<-N6-METHYLASE<-DUF6079*<-BrxB-BrxF-iSTAND<-BrxA-DNA-binding<-?\|\|wHTH->?->wHTH+HNH->?-><-AAA-ATPase-p | 1192 | Bacteria>Terrabacteria_group>Bacillota>Bacilli>Bacillales>Bacillaceae | GCF_007830235.1 | Cytobacillus_oceanisediminis | IQ19_RS15315 | 231804..235382 | 665099 | BREX-system-P-loop-protein-BrxC-[Cytobacillus-oceanisediminis] |
| WP_093855906.1 | Type5.BREX | <-HNH<-REase<-?<-SF1-SF2-Helicase+REase<-ParB+HNH<-TUDOR+SF1-SF2-Helicase+i REase+CTD+REase<-PglZ<-N6-METHYLASE<-DUF6079*<-BrxB-BrxF-iSTAND<-BrxA-DNA-binding<-?\|\|wHTH->?->wHTH+HNH->?-><-?<-HTH_36 | 1187 | Bacteria>Terrabacteria_group>Bacillota>Bacilli>Bacillales>Bacillaceae | GCF_007989445.1 | Tenuibacillus_multivorans | BLP77_RS04470 | 353994..357557 | 237069 | BREX-system-P-loop-protein-BrxC-[Tenuibacillus-multivorans] |
| WP_155112638.1 | Type5.BREX | <-Peptidase_S8<-Classical-AAA<-?<-REase<-TUDOR+SF1-SF2-Helicase+i REase+CTD+REase<-PglZ<-N6-METHYLASE-p<-N6-METHYLASE<-DUF6079*<-BrxB-BrxF-iSTAND<-BrxA-DNA-binding<-?\|\|wHTH->?->wHTH+HNH->?->AAA-ATPase-> | 1186 | Bacteria>Terrabacteria_group>Bacillota>Bacilli>Bacillales>Bacillaceae | GCF_009711125.1 | Metabacillus_mangrovi | GKZ89_RS11910 | 46455..50015 | 1491830 | BREX-system-P-loop-protein-BrxC-[Metabacillus-mangrovi] |
| WP_167560822.1 | Type5.BREX | <-TUDOR+SF1-SF2-Helicase+i REase+CTD+REase<-PglZ<-N6-METHYLASE<-DUF6079*<-BrxB-BrxF-iSTAND<-BrxA-DNA-binding<-GDResolvase+HIN-HTH\|\|?->Recombinase->GDResolvase->Recombinase+ZnR->?-><-SPOUT | 1192 | Bacteria>Terrabacteria_group>Bacillota>Bacilli>Bacillales>Bacillaceae | GCF_012030395.1 | Bacillus_subtilis | HC742_RS20760 | 3449689..3453267 | 1423 | BREX-system-P-loop-protein-BrxC-[Bacillus-subtilis] |
| WP_184528635.1 | Type5.BREX | <-HEPN<-?<-?<-TUDOR+SF1-SF2-Helicase+i REase+CTD+REase<-PglZ<-N6-METHYLASE<-DUF6079*<-BrxA-DNA-binding<-?\|\|GDResolvase->?->GDResolvase+Zn_ribbon_recom-><-TRAM+METHYLASE | 1192 | Bacteria>Terrabacteria_group>Bacillota>Bacilli>Bacillales>Bacillaceae | GCF_014207535.1 | Bacillus_benzoevorans | HNR53_RS18175 | 67020..70598 | 1456 | BREX-system-P-loop-protein-BrxC-[Bacillus-benzoevorans] |
| WP_188383791.1 | Type5.BREX | <-RVT<-ZnR<-?<-REase<-TUDOR+SF1-SF2-Helicase+i REase+CTD+REase<-PglZ<-N6-METHYLASE<-DUF6079*<-BrxB-BrxF-iSTAND<-BrxA-DNA-binding<-?<-AAA+<-HTH_36\|\|?->?->GDResolvase+Recombinase+ZnR-><-PolB | 1187 | Bacteria>Terrabacteria_group>Bacillota>Bacilli>Bacillales>Bacillaceae | GCF_014637405.1 | Ornithinibacillus_halotolerans | IEV35_RS05945 | 32497..36060 | 1274357 | BREX-system-P-loop-protein-BrxC-[Ornithinibacillus-halotolerans] |
| WP_188386543.1 | Type5.BREX | Pkinase-><-?\|\|?->?->McrC-NTD+REase->SF1-SF2-Helicase->BrxA-DNA-binding->BrxB-BrxF-iSTAND->DUF6079*->N6-METHYLASE->PglZ->TUDOR+SF1-SF2-Helicase+i REase+CTD+REase->wHTH+WYL+WCX-> | 1187 | Bacteria>Terrabacteria_group>Bacillota>Bacilli>Bacillales>Bacillaceae | GCF_014638355.1 | Priestia_taiwanensis | IEV41_RS00670 | 120045..123608 | 1347902 | BREX-system-P-loop-protein-BrxC-[Priestia-taiwanensis] |
| WP_199714161.1 | Type5.BREX | IONCHANNEL+PAS+HISKIN+MORC->REC+CITB-HTH+ZDNA-HTH->?->REase->NERD+SF1-SF2-Helicase->?->SF1-SF2-Helicase->BrxA-DNA-binding->BrxB-BrxF-iSTAND->DUF6079*->N6-METHYLASE->PglZ->TUDOR+SF1-SF2-Helicase+i REase+CTD+REase->?->wHTH+WYL+WCX->?->wHTH+McrB-AAA->McrC-NTD+REase-><-REase+ZnR | 1191 | Bacteria>Terrabacteria_group>Bacillota>Bacilli>Bacillales>Bacillaceae | GCF_016483555.1 | Bacillus_cereus_group | JDS82_RS26190 | 13772..17347 | 86661 | MULTISPECIES:-BREX-system-P-loop-protein-BrxC-[Bacillus-cereus-group] |
| WP_202778095.1 | Type5.BREX | PadR-HTH-><-TatD_DNase<-?<-KAP-glycosylase<-KAP-NTPase<-TUDOR+SF1-SF2-Helicase+i REase+CTD+REase<-PglZ<-N6-METHYLASE<-N6-METHYLASE<-DUF6079*<-BrxB-BrxF-iSTAND<-BrxA-DNA-binding<-GDResolvase+HIN-HTH<-?<-NUDIX\|\|?->GDResolvase+Zn_ribbon_recom->GDResolvase+Zn_ribbon_recom-><-TRAM+METHYLASE | 1192 | Bacteria>Terrabacteria_group>Bacillota>Bacilli>Bacillales>Bacillaceae | GCF_016772275.1 | Heyndrickxia_vini | I5776_RS19095 | 3841015..3844593 | 1476025 | BREX-system-P-loop-protein-BrxC-[Heyndrickxia-vini] |
| WP_205168141.1 | Type5.BREX | XIS-HTH->Phage_integrase->?->?->Beta-lactamase-><-TUDOR+SF1-SF2-Helicase+i REase+CTD+REase<-PglZ<-N6-METHYLASE<-DUF6079*<-BrxB-BrxF-iSTAND<-BrxA-DNA-binding<-GDResolvase+HIN-HTH\|\|?->?-><-XIS-HTH | 1189 | Bacteria>Terrabacteria_group>Bacillota>Bacilli>Bacillales>Bacillaceae | GCF_016908495.1 | Bacillus_pakistanensis | JOC86_RS02215 | 441948..445517 | 992288 | BREX-system-P-loop-protein-BrxC-[Bacillus-pakistanensis] |
| WP_212956143.1 | Type5.BREX | <-sGTP<-McrC-NTD+REase<-wHTH+McrB-AAA<-REase<-?<-wHTH+WYL+WCX<-TUDOR+SF1-SF2-Helicase+i REase+CTD+REase<-PglZ<-N6-METHYLASE<-DUF6079*<-BrxB-BrxF-iSTAND<-BrxA-DNA-binding<-SF1-SF2-Helicase<-JAB<-SF1-SF2-Helicase+Z1<-?<-?<-REC+CITB-HTH+ZDNA-HTH<-IONCHANNEL+PAS+HISKIN+MORC | 1188 | Bacteria>Terrabacteria_group>Bacillota>Bacilli>Bacillales>Bacillaceae | GCF_018332475.1 | Bacillus | KI369_RS12465 | 38598..42164 | 1386 | MULTISPECIES:-BREX-system-P-loop-protein-BrxC-[Bacillus] |
| WP_214723414.1 | Type5.BREX | <-GDResolvase+Recombinase+ZnR\|\|?->?-><-GDResolvase+Recombinase+ZnR<-?\|\|?->GDResolvase+HIN-HTH->BrxA-DNA-binding->BrxB-BrxF-iSTAND->DUF6079*->N6-METHYLASE->PglZ->TUDOR+SF1-SF2-Helicase+i REase+CTD+REase-> | 1189 | Bacteria>Terrabacteria_group>Bacillota>Bacilli>Bacillales>Bacillaceae | GCF_018613175.1 | Bacillus_sp._ISL-47 | J7193_RS03705 | 56511..60080 | 2819130 | BREX-system-P-loop-protein-BrxC-[Bacillus-sp.-ISL-47] |
| WP_215562746.1 | Type5.BREX | IONCHANNEL+PAS+HISKIN+MORC->REC+CITB-HTH+ZDNA-HTH->?->?->NERD+SF1-SF2-Helicase->?->SF1-SF2-Helicase->BrxA-DNA-binding->BrxB-BrxF-iSTAND->DUF6079*->N6-METHYLASE->PglZ->TUDOR+SF1-SF2-Helicase+i REase+CTD+REase->URI-DNase+RAMA->wHTH+WYL+WCX->?->GTPase-AIG->GTPase-AIG->wHTH+McrB-AAA-> | 1185 | Bacteria>Terrabacteria_group>Bacillota>Bacilli>Bacillales>Bacillaceae | GCF_018741585.1 | Bacillus_wiedmannii | EXW48_RS05020 | 1017548..1021105 | 1890302 | BREX-system-P-loop-protein-BrxC-[Bacillus-wiedmannii] |
| WP_221874921.1 | Type5.BREX | <-DnaB+TnsC-ATPase<-?<-?\|\|GDResolvase+HIN-HTH->BrxA-DNA-binding->BrxB-BrxF-iSTAND->DUF6079*->N6-METHYLASE->PglZ->\|\|TUDOR+SF1-SF2-Helicase+i REase+CTD+REase->HNH->?->SbcC-> | 1192 | Bacteria>Terrabacteria_group>Bacillota>Bacilli>Bacillales>Bacillaceae | GCF_019749255.1 | Mesobacillus_maritimus | H0185_RS18235 | 45712..49290 | 1643336 | BREX-system-P-loop-protein-BrxC-[Mesobacillus-maritimus] |
| WP_224936976.1 | Type5.BREX | <-RVT+HEPN\|\|?-><-Pkinase<-?<-REase<-TUDOR+SF1-SF2-Helicase+i REase+CTD+REase<-PglZ<-N6-METHYLASE<-DUF6079*<-BrxB-BrxF-iSTAND<-BrxA-DNA-binding<-?\|\|wHTH->?->wHTH+HNH->?-><-SPOUT | 1187 | Bacteria>Terrabacteria_group>Bacillota>Bacilli>Bacillales>Bacillaceae | GCF_020171845.1 | Cytobacillus_kochii | LCM23_RS10855 | 44257..47820 | 859143 | BREX-system-P-loop-protein-BrxC-[Cytobacillus-kochii] |
| WP_226516206.1 | Type5.BREX | <-Peptidase_S8<-Classical-AAA<-REase<-TUDOR+SF1-SF2-Helicase+i REase+CTD+REase<-PglZ<-N6-METHYLASE<-DUF6079*<-BrxB-BrxF-iSTAND<-BrxA-DNA-binding<-?\|\|?->?->wHTH+HNH->?-><-SPOUT | 1187 | Bacteria>Terrabacteria_group>Bacillota>Bacilli>Bacillales>Bacillaceae | GCF_020519665.1 | Bacillus_shivajii | LGQ02_RS20940 | 4386157..4389720 | 1983719 | BREX-system-P-loop-protein-BrxC-[Bacillus-shivajii] |
| WP_229143299.1 | Type5.BREX | <-REase+ZnR<-McrB-AAA<-?<-REase<-wHTH+WYL+WCX<-URI-DNase+RAMA<-TUDOR+SF1-SF2-Helicase+i REase+CTD+REase<-PglZ<-N6-METHYLASE<-DUF6079*<-BrxB-BrxF-iSTAND<-BrxA-DNA-binding<-SF1-SF2-Helicase<-JAB<-SF1-SF2-Helicase+Z1<-?<-REase<-?<-REC+CITB-HTH+ZDNA-HTH | 1188 | Bacteria>Terrabacteria_group>Bacillota>Bacilli>Bacillales>Bacillaceae | GCF_020686245.1 | Bacillus_paranthracis | LKL84_RS22410 | 92907..96473 | 2026186 | BREX-system-P-loop-protein-BrxC-[Bacillus-paranthracis] |
| WP_048558547.1 | Type5.BREX | IONCHANNEL+PAS+HISKIN+MORC->REC+CITB-HTH+ZDNA-HTH->?->?->SF1-SF2-Helicase+Z1->JAB->SF1-SF2-Helicase->BrxA-DNA-binding->BrxB-BrxF-iSTAND->DUF6079*->N6-METHYLASE->N6-METHYLASE-p->N6-METHYLASE->nSTAND3->PglZ->TUDOR+SF1-SF2-Helicase+i REase+CTD+REase->URI-DNase+RAMA->wHTH+WYL+WCX-> | 1185 | Bacteria>Terrabacteria_group>Bacillota>Bacilli>Bacillales>Bacillaceae | GCF_021655055.1 | Bacillus_cereus | L6497_RS04770 | 943584..947141 | 1396 | BREX-system-P-loop-protein-BrxC-[Bacillus-cereus] |
| WP_200152411.1 | Type5.BREX | IONCHANNEL+PAS+HISKIN+MORC->REC+CITB-HTH+ZDNA-HTH->?->?->SF1-SF2-Helicase+Z1->JAB->SF1-SF2-Helicase->BrxA-DNA-binding->BrxB-BrxF-iSTAND->DUF6079*->N6-METHYLASE->N6-METHYLASE-p->N6-METHYLASE->nSTAND3->PglZ->TUDOR+SF1-SF2-Helicase+i REase+CTD+REase->URI-DNase+RAMA->wHTH+WYL+WCX-> | 1185 | Bacteria>Terrabacteria_group>Bacillota>Bacilli>Bacillales>Bacillaceae | GCF_021655215.1 | Bacillus_cereus | L6484_RS04895 | 983694..987251 | 1396 | BREX-system-P-loop-protein-BrxC-[Bacillus-cereus] |
| WP_242261892.1 | Type5.BREX | <-wHTH+McrB-AAA<-GTPase-AIG<-GTPase-AIG<-?<-wHTH+WYL+WCX<-URI-DNase+RAMA<-TUDOR+SF1-SF2-Helicase+i REase+CTD+REase<-PglZ<-N6-METHYLASE<-DUF6079*<-BrxB-BrxF-iSTAND<-BrxA-DNA-binding<-SF1-SF2-Helicase<-?<-NERD+SF1-SF2-Helicase<-REase<-?<-?<-REC+CITB-HTH+ZDNA-HTH<-IONCHANNEL+PAS+HISKIN+MORC | 1188 | Bacteria>Terrabacteria_group>Bacillota>Bacilli>Bacillales>Bacillaceae | GCF_022662915.1 | Bacillus_cereus_group_sp._BfR-BA-01453 | MR070_RS20010 | 124720..128286 | 2920355 | BREX-system-P-loop-protein-BrxC-[Bacillus-cereus-group-sp.-BfR-BA-01453] |
| WP_242225824.1 | Type5.BREX | <-REase<-McrC-NTD+REase<-wHTH+McrB-AAA<-?<-wHTH+WYL+WCX<-URI-DNase+RAMA<-TUDOR+SF1-SF2-Helicase+i REase+CTD+REase<-PglZ<-N6-METHYLASE<-DUF6079*<-BrxB-BrxF-iSTAND<-BrxA-DNA-binding<-SF1-SF2-Helicase<-?<-NERD+SF1-SF2-Helicase<-REase<-?<-?<-REC+CITB-HTH+ZDNA-HTH | 1185 | Bacteria>Terrabacteria_group>Bacillota>Bacilli>Bacillales>Bacillaceae | GCF_022663655.1 | Bacillus_cereus_group_sp._BfR-BA-01358 | MR078_RS05890 | 38308..41865 | 2920320 | BREX-system-P-loop-protein-BrxC-[Bacillus-cereus-group-sp.-BfR-BA-01358] |
| WP_242320708.1 | Type5.BREX | IONCHANNEL+PAS+HISKIN+MORC->REC+CITB-HTH+ZDNA-HTH->?->REase->NERD+SF1-SF2-Helicase->?->SF1-SF2-Helicase->BrxA-DNA-binding->BrxB-BrxF-iSTAND->DUF6079*->N6-METHYLASE->PglZ->TUDOR+SF1-SF2-Helicase+i REase+CTD+REase->URI-DNase+RAMA->wHTH+WYL+WCX->?->GTPase-AIG->GTPase-AIG->SAD-DUF3427C+SbcC-> | 1186 | Bacteria>Terrabacteria_group>Bacillota>Bacilli>Bacillales>Bacillaceae | GCF_022663735.1 | Bacillus_cereus_group_sp._BfR-BA-01354 | MR162_RS10425 | 194779..198339 | 2920317 | BREX-system-P-loop-protein-BrxC-[Bacillus-cereus-group-sp.-BfR-BA-01354] |
| WP_243571747.1 | Type5.BREX | <-TUDOR+SF1-SF2-Helicase+i REase+CTD+REase<-PglZ<-N6-METHYLASE<-DUF6079*<-BrxB-BrxF-iSTAND<-BrxA-DNA-binding<-GDResolvase+HIN-HTH\|\|GDResolvase->Recombinase+ZnR->?-><-?\|\|?-><-ADH_N+ADH_zinc_N | 1192 | Bacteria>Terrabacteria_group>Bacillota>Bacilli>Bacillales>Bacillaceae | GCF_022814905.1 | Bacillus_stercoris | MSS26_RS04990 | 1003415..1006993 | 2054641 | BREX-system-P-loop-protein-BrxC-[Bacillus-stercoris] |
| WP_251625212.1 | Type5.BREX | MazE_antitoxin+SpoVT_C-><-GDResolvase+HIN-HTH\|\|?->BrxA-DNA-binding->BrxB-BrxF-iSTAND->DUF6079*->N6-METHYLASE->PglZ->TUDOR+SF1-SF2-Helicase+i REase+CTD+REase-><-?\|\|RVT->?-><-?<-LexA-protease | 1192 | Bacteria>Terrabacteria_group>Bacillota>Bacilli>Bacillales>Bacillaceae | GCF_023712125.1 | Metabacillus_idriensis | M4D55_RS07610 | 46091..49669 | 324768 | BREX-system-P-loop-protein-BrxC-[Metabacillus-idriensis] |
| WP_251646840.1 | Type5.BREX | <-Phage_int_SAM_3+Phage_integrase\|\|Glyoxalase->?-><-?<-Arm-DNA-bind_4+Phage_int_SAM_3+Phage_integrase\|\|?->BrxA-DNA-binding->BrxB-BrxF-iSTAND->DUF6079*->N6-METHYLASE->PglZ->TUDOR+SF1-SF2-Helicase+i REase+CTD+REase-> | 1195 | Bacteria>Terrabacteria_group>Bacillota>Bacilli>Bacillales>Bacillaceae | GCF_023714395.1 | Niallia_sp._MER_6 | M3600_17125 | 94088..97675 | 2939567 | BREX-system-P-loop-protein-BrxC-[Niallia-sp.-MER-6] |
| WP_251339432.1 | Type5.BREX | <-HTH_Tnp_1+HTH_21+rve+rve_3<-DndB<-Tox-RelA<-?<-TPR\|\|SLATT<-?<-TUDOR+SF1-SF2-Helicase+i REase+CTD+REase<-PglZ<-N6-METHYLASE<-DUF6079*<-BrxB-BrxF-iSTAND<-BrxA-DNA-binding<-GDResolvase+HIN-HTH\|\|Phage_int_SAM_5+Phage_integrase-> | 1192 | Bacteria>Terrabacteria_group>Bacillota>Bacilli>Bacillales>Bacillaceae | GCF_023714665.1 | Heyndrickxia_oleronia | M3589_RS18030 | 387742..391320 | 38875 | BREX-system-P-loop-protein-BrxC-[Heyndrickxia-oleronia] |
| WP_258922098.1 | Type5.BREX | DDE_Tnp_1_4-><-TUDOR+SF1-SF2-Helicase+i REase+CTD+REase<-PglZ<-N6-METHYLASE<-DUF6079*<-BrxB-BrxF-iSTAND<-BrxA-DNA-binding<-DDE_Tnp_1_4<-?<-?<-?\|\|PadR-HTH-> | 1187 | Bacteria>Terrabacteria_group>Bacillota>Bacilli>Bacillales>Bacillaceae | GCF_024758645.1 | Weizmannia_coagulans | L1879_RS05770 | 1225557..1229120 | 1398 | BREX-system-P-loop-protein-BrxC-[Weizmannia-coagulans] |
| WP_260358530.1 | Type5.BREX | <-DNA_METHYLASE<-SF1-SF2-Helicase<-SF1-SF2-Helicase\|\|DrmA->DrmB->HKD-DNase-><-TUDOR+SF1-SF2-Helicase+i REase+CTD+REase<-PglZ<-N6-METHYLASE<-DUF6079*<-BrxB-BrxF-iSTAND<-BrxA-DNA-binding<-SF1-SF2-Helicase\|\|NUDIX-><-McrC-NTD+REase<-McrB-AAA\|\|SF1-SF2-Helicase-><-REase+SPF-cons | 1177 | Bacteria>Terrabacteria_group>Bacillota>Bacilli>Bacillales>Bacillaceae | GCF_025209795.1 | Peribacillus_frigoritolerans | NOU24_RS18785 | 747765..751298 | 450367 | BREX-system-P-loop-protein-BrxC-[Peribacillus-frigoritolerans] |
| WP_268298773.1 | Type5.BREX | SPOUT-><-Zn_ribbon_recom<-Recombinase<-GDResolvase<-?<-?\|\|GDResolvase+HIN-HTH->BrxA-DNA-binding->BrxB-BrxF-iSTAND->DUF6079*->N6-METHYLASE->PglZ->TUDOR+SF1-SF2-Helicase+i REase+CTD+REase-><-?<-GNTR-HTH+UTRA\|\|?->Glyco_hydro_1-> | 1192 | Bacteria>Terrabacteria_group>Bacillota>Bacilli>Bacillales>Bacillaceae | GCF_026789485.1 | Bacillus_inaquosorum | MOB75_RS13790 | 96491..100069 | 483913 | BREX-system-P-loop-protein-BrxC-[Bacillus-inaquosorum] |
| WP_269897765.1 | Type5.BREX | <-HEPN+HTH<-?<-JAB<-?<-TUDOR+SF1-SF2-Helicase+i REase+CTD+REase<-?<-PglZ<-N6-METHYLASE<-DUF6079*<-BrxB-BrxF-iSTAND<-BrxA-DNA-binding\|\|?->ZnR+TOPRIM+AAA-helicase+wHTH->XIS-HTH->?->Phage_int_SAM_3+Phage_integrase->Arm-DNA-bind_4+Phage_int_SAM_3+Phage_integrase-><-HNH | 1196 | Bacteria>Terrabacteria_group>Bacillota>Bacilli>Bacillales>Bacillaceae | GCF_027557635.1 | Lysinibacillus_sp._JK80 | HXV90_RS18610 | 3707833..3711423 | 2749809 | BREX-system-P-loop-protein-BrxC-[Lysinibacillus-sp.-JK80] |
| WP_231419114.1 | Type5.BREX | TRAM+METHYLASE->?->BrxA-DNA-binding->BrxB-BrxF-iSTAND->DUF6079*->N6-METHYLASE->TUDOR+SF1-SF2-Helicase+i REase+CTD+REase->REase->REase->HNH-><-REase+DUF5655<-SF1-SF2-Helicase\|\|URI-DNase->TIR-> | 1188 | Bacteria>Terrabacteria_group>Bacillota>Bacilli>Bacillales>Bacillaceae | GCF_030166655.1 | Pontibacillus_sp._HN14 | LQ226_RS16640 | 43379..46945 | 2898421 | BREX-system-P-loop-protein-BrxC-[Pontibacillus-sp.-HN14] |
| WP_098147710.1 | Type5.BREX | NERD+SF1-SF2-Helicase->?->SF1-SF2-Helicase->BrxA-DNA-binding->BrxB-BrxF-iSTAND->DUF6079*->N6-METHYLASE->PglZ->TUDOR+SF1-SF2-Helicase+i REase+CTD+REase->wHTH+WYL+WCX->?->wHTH+McrB-AAA->McrC-NTD+REase->REase-> | 1186 | Bacteria>Terrabacteria_group>Bacillota>Bacilli>Bacillales>Bacillaceae | GCF_030676405.1 | Bacillus_wiedmannii | CN685_RS18690 | 6510..10070 | 1890302 | BREX-system-P-loop-protein-BrxC-[Bacillus-wiedmannii] |
| WP_073518919.1 | Type5.BREX | SF1-SF2-Helicase->?->REase->?->SF1-SF2-Helicase+Z1->JAB->SF1-SF2-Helicase->BrxA-DNA-binding->BrxB-BrxF-iSTAND->DUF6079*->N6-METHYLASE->PglZ->TUDOR+SF1-SF2-Helicase+i REase+CTD+REase->wHTH+WYL+WCX->?->GTPase-AIG->GTPase-AIG-><-NACHT | 1185 | Bacteria>Terrabacteria_group>Bacillota>Bacilli>Bacillales>Bacillaceae | GCF_900094815.1 | Bacillus_cereus_group | CCY53_RS04725 | 210220..213777 | 86661 | MULTISPECIES:-BREX-system-P-loop-protein-BrxC-[Bacillus-cereus-group] |
| WP_099159038.1 | Type5.BREX | <-wHTH+HNH<-?<-wHTH<-?<-wHTH\|\|?->BrxA-DNA-binding->BrxB-BrxF-iSTAND->DUF6079*->N6-METHYLASE->PglZ->TUDOR+SF1-SF2-Helicase+i REase+CTD+REase->REase->?->?->?->SF1-SF2-Helicase+REase->ParB+HNH-> | 1187 | Bacteria>Terrabacteria_group>Bacillota>Bacilli>Bacillales>Bacillaceae | GCF_900187325.1 | Virgibacillus_ndiopensis | CS937_RS13675 | 175761..179324 | 2004408 | BREX-system-P-loop-protein-BrxC-[Virgibacillus-ndiopensis] |
| WP_163530739.1 | Type5.BREX | ArdCN+MPTase-inact->TOPRIM->?->BrxA-DNA-binding->BrxB-BrxF-iSTAND->DUF6079*->N6-METHYLASE->PglZ->TUDOR+SF1-SF2-Helicase+i REase+CTD+REase->?->McrB-NTD+HNH->Phage_integrase-><-HEPN<-AAA-ATPase<-HTH_22+rve | 1188 | Bacteria>Terrabacteria_group>Bacillota>Bacilli>Bacillales>Bacillaceae | GCF_902726645.1 | Halobacillus_ihumii | G6R08_RS20435 | 4081488..4085054 | 2686092 | BREX-system-P-loop-protein-BrxC-[Halobacillus-ihumii] |
| WP_026834104.1 | Type5.BREX | <-ABC-ATPase+SbcC<-LYTTR\|\|PAIREDC-HTH+HIN-HTH->?->?->SF1-SF2-Helicase+REase-><-TUDOR+SF1-SF2-Helicase+i REase+CTD+REase<-PglZ<-N6-METHYLASE<-DUF6079*<-BrxB-BrxF-iSTAND<-BrxA-DNA-binding<-HKD-DNase+SF1-SF2-Helicase+HTH+HTH+SRA<-?\|\|RNase-><-HARE-HTH<-?\|\|?->Peptidase-> | 1192 | Bacteria>Terrabacteria_group>Bacillota>Bacilli>Bacillales>Bacillales_incertae_sedis | GCF_000620825.1 | Exiguobacterium_undae | P399_RS0115220 | 2864676..2868254 | 169177 | BREX-system-P-loop-protein-BrxC-[Exiguobacterium-undae] |
| WP_026826561.1 | Type5.BREX | <-HARE-HTH<-Preader17<-HKD-DNase+SF1-SF2-Helicase+HTH+HTH+SRA<-?<-TUDOR+SF1-SF2-Helicase+i REase+CTD+REase<-PglZ<-N6-METHYLASE<-DUF6079*<-BrxB-BrxF-iSTAND<-BrxA-DNA-binding<-?<-?<-?<-?<-?<-HHH+DNA_pol_B+RNase | 1191 | Bacteria>Terrabacteria_group>Bacillota>Bacilli>Bacillales>Bacillales_incertae_sedis | GCF_000620845.1 | Exiguobacterium_marinum | P400_RS0112700 | 2404557..2408132 | 273528 | BREX-system-P-loop-protein-BrxC-[Exiguobacterium-marinum] |
| WP_078145973.1 | Type5.BREX | Trypsin+PDZ->?->SPOUT->?->?-><-PAIREDC-HTH+HIN-HTH\|\|?->BrxA-DNA-binding->BrxB-BrxF-iSTAND->DUF6079*->N6-METHYLASE->PglZ->TUDOR+SF1-SF2-Helicase+i REase+CTD+REase->?-><-AAA-ATPase<-rve | 1187 | Bacteria>Terrabacteria_group>Bacillota>Bacilli>Bacillales>Bacillales_incertae_sedis | GCF_002009685.1 | Exiguobacterium_sp._HVEsp1 | BV910_RS00630 | 122796..126359 | 1934003 | BREX-system-P-loop-protein-BrxC-[Exiguobacterium-sp.-HVEsp1] |
| WP_131508621.1 | Type5.BREX | McrC-NTD+REase-><-NERD+SF1-SF2-Helicase\|\|?->?->?->?-><-wHTH+HNH\|\|BrxA-DNA-binding->BrxB-BrxF-iSTAND->DUF6079*->N6-METHYLASE->PglZ->TUDOR+SF1-SF2-Helicase+i REase+CTD+REase-> | 1200 | Bacteria>Terrabacteria_group>Bacillota>Bacilli>Bacillales>Bacillales_incertae_sedis | GCF_004337165.1 | Exiguobacterium_sp._SHOS1 | EVJ20_RS13610 | 61897..65499 | 2510949 | BREX-system-P-loop-protein-BrxC-[Exiguobacterium-sp.-SHOS1] |
| WP_089525998.1 | Type5.BREX | REase+wHTH-><-zf-IS66\|\|BrxA-DNA-binding->BrxB-BrxF-iSTAND->DUF6079*->N6-METHYLASE->PglZ->TUDOR+SF1-SF2-Helicase+i REase+CTD+REase->Calcineurin->SbcC+SbcC->?->RVT->RNase-> | 1208 | Bacteria>Terrabacteria_group>Bacillota>Bacilli>Bacillales>Paenibacillaceae | GCF_002233675.1 | Paenibacillus_herberti | CGZ75_RS19630 | 18692..22318 | 1619309 | BREX-system-P-loop-protein-BrxC-[Paenibacillus-herberti] |
| WP_272441147.1 | Type5.BREX | <-Phage_int_SAM_5+Phage_integrase\|\|?->?-><-XIS-HTH<-Phage_GPA\|\|BrxA-DNA-binding->BrxB-BrxF-iSTAND->DUF6079*->N6-METHYLASE->ParB+HNH->PglZ->TUDOR+SF1-SF2-Helicase+i REase+CTD+REase->?->DnaB+HINT+DnaB+HINT->ParB+HNH->?-><-XIS-HTH | 1201 | Bacteria>Terrabacteria_group>Bacillota>Bacilli>Bacillales>Paenibacillaceae | GCF_028421645.1 | Aneurinibacillus_sp._Ricciae_BoGa-3 | PP175_RS01545 | 299012..302617 | 3022697 | BREX-system-P-loop-protein-BrxC-[Aneurinibacillus-sp.-Ricciae_BoGa-3] |
| WP_066463066.1 | Type5.BREX | BrxA-DNA-binding->BrxB-BrxF-iSTAND->DUF6079*->N6-METHYLASE->?->PglZ-><-?<-?<-?\|\|TUDOR+SF1-SF2-Helicase+i REase+CTD+REase-> | 1200 | Bacteria>Terrabacteria_group>Bacillota>Bacilli>Bacillales>Planococcaceae | GCF_001700325.1 | Caryophanon_latum | A6K76_RS05925 | 1324..4926 | 33977 | BREX-system-P-loop-protein-BrxC-[Caryophanon-latum] |
| WP_107933601.1 | Type5.BREX | HTH-><-?<-AAA+<-HTH_36<-TUDOR+SF1-SF2-Helicase+i REase+CTD+REase<-PglZ<-N6-METHYLASE<-DUF6079*<-BrxB-BrxF-iSTAND<-BrxA-DNA-binding\|\|?-><-?\|\|?->XIS-HTH->Phage_int_SAM_3+Phage_integrase->Arm-DNA-bind_4+Phage_int_SAM_3+Phage_integrase-> | 1192 | Bacteria>Terrabacteria_group>Bacillota>Bacilli>Bacillales>Planococcaceae | GCF_003049615.1 | Ureibacillus_chungkukjangi | C9J33_RS05745 | 69413..72991 | 1202712 | BREX-system-P-loop-protein-BrxC-[Ureibacillus-chungkukjangi] |
| WP_108587666.1 | Type5.BREX | wHTH+HNH-p->?->MORC+DUF3684+REase->?-><-ParB+HNH\|\|BrxA-DNA-binding->BrxB-BrxF-iSTAND->DUF6079*->N6-METHYLASE->PglZ->TUDOR+SF1-SF2-Helicase+i REase+CTD+REase->GNTR-HTH+UTRA->?->HNH->?->?->ABC-ATPase+SbcC-> | 1191 | Bacteria>Terrabacteria_group>Bacillota>Bacilli>Bacillales>Planococcaceae | GCF_003058165.1 | Paenisporosarcina_sp._OV554 | C8K15_RS17245 | 15921..19496 | 2135694 | BREX-system-P-loop-protein-BrxC-[Paenisporosarcina-sp.-OV554] |
| WP_274307983.1 | Type5.BREX | <-REase<-HEPN+HTH<-JAB<-?<-TUDOR+SF1-SF2-Helicase+i REase+CTD+REase<-?<-PglZ<-N6-METHYLASE<-DUF6079*<-BrxB-BrxF-iSTAND<-BrxA-DNA-binding\|\|XIS-HTH->Phage_int_SAM_3+Phage_integrase->Arm-DNA-bind_4+Phage_int_SAM_3+Phage_integrase-><-?<-?<-RhoN+RhoN+RNase+S1<-ABhydrolase | 1196 | Bacteria>Terrabacteria_group>Bacillota>Bacilli>Bacillales>Planococcaceae | GCF_028747805.1 | Solibacillus_sp._ZS111008 | O7776_RS16115 | 3287543..3291133 | 2912187 | BREX-system-P-loop-protein-BrxC-[Solibacillus-sp.-ZS111008] |
| WP_092984062.1 | Type5.BREX | HTH_7+rve-><-HTH_Tnp_1+HTH_21+rve+rve_3<-HIN-HTH\|\|IstB_IS21-><-?<-TUDOR+SF1-SF2-Helicase+i REase+CTD+REase<-PglZ<-N6-METHYLASE<-DUF6079*<-BrxB-BrxF-iSTAND<-BrxA-DNA-binding<-?<-?\|\|?-><-RelB | 1184 | Bacteria>Terrabacteria_group>Bacillota>Bacilli>Bacillales>Staphylococcaceae | GCF_900101075.1 | Salinicoccus_qingdaonensis | BLR15_RS01740 | 53033..56587 | 576118 | BREX-system-P-loop-protein-BrxC-[Salinicoccus-qingdaonensis] |
| WP_271398742.1 | Type5.BREX | <-ABC-ATPase+SbcC\|\|?->ParB+HNH->HNH->?-><-TUDOR+SF1-SF2-Helicase+i REase+CTD+REase<-PglZ<-N6-METHYLASE<-DUF6079*<-BrxB-BrxF-iSTAND<-BrxA-DNA-binding<-?\|\|SIGMA-HTH+SIGMA-HTH->35exo+wHTH+SF1-SF2-Helicase-> | 1184 | Bacteria>Terrabacteria_group>Bacillota>Bacilli>Bacillales>Staphylococcaceae | GCF_947502135.1 | Salinicoccus_roseus | POR62_RS08110 | 202088..205642 | 45670 | BREX-system-P-loop-protein-BrxC-[Salinicoccus-roseus] |
| KKE80057.1 | Type5.BREX | <-GNTR-HTH\|\|?-><-?\|\|?->?-><-TUDOR+SF1-SF2-Helicase+i REase+CTD+REase<-PglZ<-N6-METHYLASE<-DUF6079*<-BrxB-BrxF-iSTAND<-BrxA-DNA-binding<-?\|\|?->Phage_int_SAM_5+Phage_integrase-><-tRNA<-tRNA<-tRNA<-tRNA<-tRNA<-tRNA<-tRNA<-tRNA<-tRNA<-?\|\|?-><-?<-MarR-HTH | 1187 | Bacteria>Terrabacteria_group>Bacillota>Bacilli>unclassified_Bacilli | GCA_000974285.1 | Bacilli_bacterium_VT-13-104 | WH51_03830 | 66407..69970 | 1637974 | hypothetical-protein-WH51_03830-[Bacilli-bacterium-VT-13-104] |
| WP_164950923.1 | Type5.BREX | <-GDResolvase<-?<-?<-TUDOR+SF1-SF2-Helicase+i REase+CTD+REase<-PglZ<-TOPRIM<-OLD-ABC<-HEPN<-N6-METHYLASE<-DUF6079*<-BrxB-BrxF-iSTAND<-BrxA-DNA-binding<-?<-Calcineurin<-PARG+MACRODOMAIN<-Calcineurin<-HAD<-LRR+LRR | 1191 | Bacteria>Terrabacteria_group>Bacillota>Clostridia>Eubacteriales>Clostridiaceae | GCF_011065955.2 | Clostridium_sp._FP2 | G9F71_RS16320 | 3398773..3402348 | 2724481 | BREX-system-P-loop-protein-BrxC-[Clostridium-sp.-FP2] |
| WP_185649263.1 | Type5.BREX | RADICAL-SAM-><-?\|\|?->?-><-?<-?<-ABhydrolase<-TUDOR+SF1-SF2-Helicase+i REase+CTD+REase<-N6-METHYLASE<-DUF6079*<-BrxB-BrxF-iSTAND<-BrxA-DNA-binding<-HTH_36<-SMS-KaiC<-Peptidase-C39<-?<-?<-HEPN<-DndB | 1189 | Bacteria>Terrabacteria_group>Bacillota>Clostridia>Eubacteriales>Clostridiaceae | GCF_014231265.1 | Clostridium_sp._DJ247 | HGI79_RS03445 | 40571..44140 | 2726188 | BREX-system-P-loop-protein-BrxC-[Clostridium-sp.-DJ247] |
| KJS50653.1 | Type5.BREX | HKD-DNase-><-?<-Polbeta<-TUDOR+SF1-SF2-Helicase+i REase+CTD+REase<-PglZ<-N6-METHYLASE<-DUF6079*<-BrxB-BrxF-iSTAND<-BrxA-DNA-binding<-SF1-SF2-Helicase+REase<-?<-?<-HNH<-OLD-ABC<-?<-HATPase_c | 1187 | Bacteria>Terrabacteria_group>Bacillota>Clostridia>Eubacteriales>Peptococcaceae | GCA_000961805.1 | Peptococcaceae_bacterium_BRH_c23 | VR66_01655 | 19236..22799 | 1629714 | hypothetical-protein-VR66_01655-[Peptococcaceae-bacterium-BRH_c23] |
| WP_146889206.1 | Type5.BREX | <-HNH<-?<-ARSR-HTH<-TUDOR+SF1-SF2-Helicase+i REase+CTD+REase<-?\|\|Peptidase_M48+TerB->SF1-SF2-Helicase+REase->BrxA-DNA-binding->BrxB-BrxF-iSTAND->DUF6079*->N6-METHYLASE->HicB->OLD-ABC->REase->PglZ+Phosphodiest->SIGMA+OB-Fold+AAA-ATPase+Lon-Protease-BrxL->SF1-SF2-Helicase->?->SF1-SF2-Helicase-> | 1186 | Bacteria>Terrabacteria_group>Deinococcota>Deinococci>Deinococcales>Deinococcaceae | GCF_007990775.1 | Deinococcus_cellulosilyticus | DC3_RS23585 | 23663..27223 | 401558 | BREX-system-P-loop-protein-BrxC-[Deinococcus-cellulosilyticus] |
| WP_189062792.1 | Type5.BREX | <-SIGMA+OB-Fold+AAA-ATPase+Lon-Protease-BrxL<-PglZ\|\|Herpes_LMP1+REase-><-NADAR<-fvmX2+ART-DarT<-HNH<-N6-METHYLASE<-DUF6079*<-BrxB-BrxF-iSTAND<-BrxA-DNA-binding<-CHTH<-?<-REase\|\|?->?->?-><-TUDOR+SF1-SF2-Helicase+i REase+CTD+REase | 1145 | Bacteria>Terrabacteria_group>Deinococcota>Deinococci>Deinococcales>Deinococcaceae | GCF_014647995.1 | Deinococcus_arenae | IEY66_RS18050 | 13025..16462 | 1452751 | BREX-system-P-loop-protein-BrxC-[Deinococcus-arenae] |
| -- | | | | | | | | | | |
| --CR-ATPase+REase | | | | | | | | | | |
| -- | | | | | | | | | | |
| ELZ19699.1 | Type1.BREX | <-N6-METHYLASE-p\|\|?->?->?-><-PIN\|\|?-><-REase<-?\|\|?-><-DUF6079*<-BrxB-BrxF-iSTAND<-DUF6079<-BrxA-DNA-binding<-CR-ATPase9+REase<-?<-?\|\|REase->SF1-SF2-Helicase->REase-> | 1198 | Archaea>Euryarchaeota>Stenosarchaea_group>Halobacteria>Halobacteriales>Haloarculaceae | GCA_000337455.1 | Halosimplex_carlsbadense_2-9-1 | C475_22179 | 59263..62859 | 797114 | hypothetical-protein-C475_22179-[Halosimplex-carlsbadense-2-9-1] |
| ELZ19701.1 | Type1.BREX | <-PIN\|\|?-><-?\|\|?-><-REase<-DUF6079*<-BrxB-BrxF-iSTAND<-BrxA-DNA-binding<-CR-ATPase9+REase<-?<-?\|\|REase->SF1-SF2-Helicase->REase-> | 1248 | Archaea>Euryarchaeota>Stenosarchaea_group>Halobacteria>Halobacteriales>Haloarculaceae | GCA_000337455.1 | Halosimplex_carlsbadense_2-9-1 | C475_22189 | 63665..67411 | 797114 | hypothetical-protein-C475_22189-[Halosimplex-carlsbadense-2-9-1] |
| WP_049931199.1 | Type1.BREX | <-PglZ<-N6-METHYLASE<-PIN<-?\|\|?-><-Peptidase_M1<-REase<-?\|\|?-><-DUF6079*<-BrxB-BrxF-iSTAND<-DUF6079<-BrxA-DNA-binding<-CR-ATPase9+REase<-?\|\|REase->SF1-SF2-Helicase->REase->SF1-SF2-Helicase-> | 1272 | Archaea>Euryarchaeota>Stenosarchaea_group>Halobacteria>Halobacteriales>Haloarculaceae | GCF_000337455.1 | Halosimplex_carlsbadense | C475_RS21845 | 59263..63081 | 171164 | hypothetical-protein-[Halosimplex-carlsbadense] |
| WP_049931237.1 | Type1.BREX | <-PIN<-?\|\|?-><-Peptidase_M1<-REase<-?\|\|?-><-DUF6079<-BrxB-BrxF-iSTAND<-DUF6079*<-BrxA-DNA-binding<-CR-ATPase9+REase<-?\|\|REase->SF1-SF2-Helicase->REase->SF1-SF2-Helicase-><-?<-RNase | 1239 | Archaea>Euryarchaeota>Stenosarchaea_group>Halobacteria>Halobacteriales>Haloarculaceae | GCF_000337455.1 | Halosimplex_carlsbadense | C475_RS21855 | 63665..67384 | 171164 | hypothetical-protein-[Halosimplex-carlsbadense] |
| WP_013460546.1 | Type1.BREX | Phage_integrase-><-XIS-HTH<-HTH_36<-?\|\|HTH_3-><-XIS-HTH\|\|BrxA-DNA-binding->BrxB-BrxF-iSTAND->DUF6079*-><-?\|\|N6-METHYLASE->p->?->?->HEPN->BrxD->N6-METHYLASE-p->CR-ATPase5+REase->PglZ-> | 1163 | Bacteria>Campylobacterota>Epsilonproteobacteria>Campylobacterales>Thiovulaceae>Sulfuricurvum | GCF_000183725.1 | Sulfuricurvum_kujiense | SULKU_RS08475 | 1692937..1696428 | 148813 | BREX-system-P-loop-protein-BrxC-[Sulfuricurvum-kujiense] |
| WP_250702187.1 | Type1.BREX | <-GDResolvase+HIN-HTH<-HTH+REase<-SF1-SF2-Helicase\|\|ADH_N+ADH_zinc_N+METHYLASE->?->SPOUT\|\|CR-ATPase5+REase->BrxA-DNA-binding->BrxB-BrxF-iSTAND->DUF6079*->N6-METHYLASE->PglZ->SIGMA+OB-Fold+AAA-ATPase+Lon-Protease-BrxL-><-?<-?<-TraJ-RHH<-ZnR+TOPRIM<-CR-ATPase4<-XIS-HTH | 1215 | Bacteria>FCB_group>Bacteroidetes/Chlorobi_group>Bacteroidota>Bacteroidia>Bacteroidales | GCF_023668595.1 | Bacteroides_ovatus | NCZ26_RS08955 | 97357..101004 | 28116 | BREX-system-P-loop-protein-BrxC-[Bacteroides-ovatus] |
| MCB0524505.1 | Type1.BREX | SF1-SF2-Helicase+RNase+RNase->?->REase+35exo->?->?->BrxA-DNA-binding->BrxB-BrxF-iSTAND->DUF6079-p->DUF6079*->N6-METHYLASE->?->McrB-NTD+HNH->Polbeta->?->CR-ATPase5+REase->PglZ->SIGMA+OB-Fold+AAA-ATPase+HKD-DNase-BrxL-> | 1094 | Bacteria>FCB_group>Bacteroidetes/Chlorobi_group>Bacteroidota>Saprospiria>Saprospirales | GCA_020436385.1 | Saprospiraceae_bacterium | KDC86_08270 | 14975..18259 | 2202734 | BREX-system-P-loop-protein-BrxC-[Saprospiraceae-bacterium] |
| CDA09060.1 | Type1.BREX | <-SIGMA+OB-Fold+AAA-ATPase+Lon-Protease-BrxL<-PglZ<-CR-ATPase8+REase<-OLD-ABC+TOPRIM<-?<-?<-N6-METHYLASE<-TOPRIM<-OLD-ABC<-DUF6079*<-?<-SWIM+DUF3670+SF1-SF2-Helicase<-?<-?<-?<-SF1-SF2-Helicase | 1250 | Bacteria>Fusobacteriota>Fusobacteriia>Fusobacteriales>Fusobacteriaceae>Fusobacterium | GCA_000433695.1 | Fusobacterium_sp._CAG:649 | BN748_00884 | 23068..26820 | 1262900 | putative-uncharacterized-protein-[Fusobacterium-sp.-CAG:649] |
| MCF2639759.1 | Type1.BREX | DUF6079*->N6-METHYLASE->PglZ->BrxA-DNA-binding->BrxB-BrxF-iSTAND->CR-ATPase5+REase->DOC->Calcineurin->HIN-HTH-> | 1244 | Bacteria>Fusobacteriota>Fusobacteriia>Fusobacteriales>Fusobacteriaceae>Fusobacterium | GCA_021531615.1 | Fusobacterium_varium | I6E31_07220 | 5895..9629 | 856 | BREX-system-P-loop-protein-BrxC-[Fusobacterium-varium] |
| WP_008692035.1 | Type1.BREX | DUF3670+SF1-SF2-Helicase->?-><-?<-?\|\|DUF6079*->OLD-ABC+TOPRIM->N6-METHYLASE->?->OLD-ABC+TOPRIM->CR-ATPase8+REase->PglZ->SIGMA+OB-Fold+AAA-ATPase+Lon-Protease-BrxL->BrxA-DNA-binding-> | 1250 | Bacteria>Fusobacteriota>Fusobacteriia>Fusobacteriales>Fusobacteriaceae>Fusobacterium | GCF_000218655.1 | Fusobacterium_nucleatum | HMPREF0401_RS10120 | 557323..561075 | 851 | BREX-system-P-loop-protein-BrxC-[Fusobacterium-animalis] |
| WP_029493951.1 | Type1.BREX | <-BrxB-BrxF-iSTAND<-BrxA-DNA-binding<-SIGMA+OB-Fold+AAA-ATPase+Lon-Protease-BrxL<-PglZ<-CR-ATPase8+REase<-RVT<-N6-METHYLASE<-Schlafen+HATPase+wHTH<-DUF6079*\|\|?->?-><-?<-ABC-ATPase+SbcC<-?<-?<-wHTH+WYL+WCX<-JAB+sGTP<-SKI | 1250 | Bacteria>Fusobacteriota>Fusobacteriia>Fusobacteriales>Fusobacteriaceae>Fusobacterium | GCF_000455885.1 | Fusobacterium_hwasookii | J143_RS0110850 | 68335..72087 | 1583098 | BREX-system-P-loop-protein-BrxC-[Fusobacterium-hwasookii] |
| WP_032838665.1 | Type1.BREX | <-PglZ<-CR-ATPase8+REase<-RVT<-N6-METHYLASE<-Schlafen+HATPase+wHTH<-REase<-CR-ATPase4<-?<-N6-METHYLASE<-DUF6079*\|\|?->?-><-Peptidase_M20<-?\|\|35exo+T7DNApol-> | 1250 | Bacteria>Fusobacteriota>Fusobacteriia>Fusobacteriales>Fusobacteriaceae>Fusobacterium | GCF_000524215.1 | Fusobacterium_sp._CM1 | HMPREF1498_RS02575 | 43304..47056 | 936561 | BREX-system-P-loop-protein-BrxC-[Fusobacterium-sp.-CM1] |
| WP_032836039.1 | Type1.BREX | <-PglZ<-CR-ATPase8+REase<-HEPN<-MORC<-N6-METHYLASE<-Schlafen+HATPase+wHTH<-TOPRIM<-OLD-ABC<-DUF6079*\|\|?-><-?<-ABC-ATPase+SbcC<-?<-?<-wHTH+WYL+WCX<-JAB+sGTP<-SKI | 1250 | Bacteria>Fusobacteriota>Fusobacteriia>Fusobacteriales>Fusobacteriaceae>Fusobacterium | GCF_000524235.1 | Fusobacterium_sp._OBRC1 | HMPREF1501_RS01100 | 63475..67227 | 1032505 | BREX-system-P-loop-protein-BrxC-[Fusobacterium-sp.-OBRC1] |
| WP_100024172.1 | Type1.BREX | <-Peptidase_M20<-?<-?<-?<-RNaseH\|\|Peptidase_M3_N+Peptidase_M3->?->?-><-?\|\|DUF6079*->OLD-ABC+TOPRIM->N6-METHYLASE->HEPN->CR-ATPase8+REase->PglZ->SIGMA+OB-Fold+AAA-ATPase+Lon-Protease-BrxL->BrxA-DNA-binding->BrxB-BrxF-iSTAND-> | 1250 | Bacteria>Fusobacteriota>Fusobacteriia>Fusobacteriales>Fusobacteriaceae>Fusobacterium | GCF_002763625.1 | Fusobacterium_pseudoperiodonticum | CTM72_RS01085 | 199877..203629 | 2663009 | BREX-system-P-loop-protein-BrxC-[Fusobacterium-pseudoperiodonticum] |
| WP_099987862.1 | Type1.BREX | <-BrxA-DNA-binding<-SIGMA+OB-Fold+AAA-ATPase+Lon-Protease-BrxL<-PglZ<-CR-ATPase8+REase<-OLD-ABC+TOPRIM<-?<-N6-METHYLASE<-TOPRIM<-OLD-ABC<-DUF6079*\|\|?->OrfB_IS605+OrfB_Zn_ribbon-><-URI-DNase+SF1-SF2-Helicase<-URI-DNase+SF1-SF2-Helicase<-?<-?<-Peptidase_M3_N+Peptidase_M3\|\|RNaseH-> | 1250 | Bacteria>Fusobacteriota>Fusobacteriia>Fusobacteriales>Fusobacteriaceae>Fusobacterium | GCF_002763695.1 | Fusobacterium_pseudoperiodonticum | CTM74_RS09005 | 1721055..1724807 | 2663009 | BREX-system-P-loop-protein-BrxC-[Fusobacterium-pseudoperiodonticum] |
| WP_238959505.1 | Type1.BREX | <-BrxB-BrxF-iSTAND<-BrxA-DNA-binding<-SIGMA+OB-Fold+AAA-ATPase+Lon-Protease-BrxL<-PglZ<-CR-ATPase8+REase<-HEPN<-MORC<-N6-METHYLASE<-Schlafen+HATPase+wHTH<-DUF6079*\|\|?->?-><-?<-ABC-ATPase+SbcC<-?<-?<-wHTH+WYL+WCX<-JAB+sGTP<-SKI | 1250 | Bacteria>Fusobacteriota>Fusobacteriia>Fusobacteriales>Fusobacteriaceae>Fusobacterium | GCF_022340045.1 | Fusobacterium_nucleatum | K5I20_00280 | 65514..69266 | 851 | BREX-system-P-loop-protein-BrxC-[Fusobacterium-nucleatum] |
| WP_238972693.1 | Type1.BREX | SKI->JAB+sGTP->wHTH+WYL+WCX->?->ABC-ATPase+SbcC->?-><-?<-?\|\|DUF6079*->OLD-ABC+TOPRIM->Schlafen+HATPase+wHTH->N6-METHYLASE->MORC->HEPN->CR-ATPase8+REase->PglZ->SIGMA+OB-Fold+AAA-ATPase+Lon-Protease-BrxL-> | 1250 | Bacteria>Fusobacteriota>Fusobacteriia>Fusobacteriales>Fusobacteriaceae>Fusobacterium | GCF_022340165.1 | Fusobacterium_nucleatum | K5I19_RS02505 | 127932..131684 | 851 | BREX-system-P-loop-protein-BrxC-[Fusobacterium-nucleatum] |

| Protein-ID | System | Gene-Neighbourhoods | Length | Taxonomy | Assembly-ID | Strain | Locus-Tag | Genomic-coordinates | TaxID | Fasta-defline |
|---|---|---|---|---|---|---|---|---|---|---|
| WP_060798221.1 | Type1.BREX | [illegible] | 1250 | Bacteria>Fusobacteriota>Fusobacteriia>Fusobacteriales>Fusobacteriaceae>Fusobacterium | GCF_023383525.1 | Fusobacterium_nucleatum | YWH7054_RS02310 | 21716..25468 | 851 | BREX-system-P-loop-protein-BrxC-[Fusobacterium-nucleatum] |
| WP_022069638.1 | Type1.BREX | [illegible] | 1250 | Bacteria>Fusobacteriota>Fusobacteriia>Fusobacteriales>Fusobacteriaceae>Fusobacterium | GCF_023383545.1 | Fusobacterium_nucleatum | YWH7053_RS01985 | 23867..27619 | 851 | BREX-system-P-loop-protein-BrxC-[Fusobacterium-nucleatum] |
| WP_249524183.1 | Type1.BREX | [illegible] | 1250 | Bacteria>Fusobacteriota>Fusobacteriia>Fusobacteriales>Fusobacteriaceae>Fusobacterium | GCF_023383605.1 | Fusobacterium_nucleatum | YWH7056_RS07045 | 28639..32391 | 851 | BREX-system-P-loop-protein-BrxC-[Fusobacterium-nucleatum] |
| WP_271333596.1 | Type1.BREX | [illegible] | 1250 | Bacteria>Fusobacteriota>Fusobacteriia>Fusobacteriales>Fusobacteriaceae>Fusobacterium | GCF_027947475.1 | Fusobacterium_nucleatum | PGW91_RS03610 | 747072..750824 | 851 | BREX-system-P-loop-protein-BrxC-[Fusobacterium-nucleatum] |
| WP_005980883.1 | Type1.BREX | [illegible] | 1242 | Bacteria>Fusobacteriota>Fusobacteriia>Fusobacteriales>Fusobacteriaceae>Fusobacterium | GCF_902373815.1 | Fusobacterium_ulcerans | FXX38_RS11815 | 320598..324326 | 861 | BREX-system-P-loop-protein-BrxC-[Fusobacterium-ulcerans] |
| WP_008793539.1 | Type1.BREX | [illegible] | 1250 | Bacteria>Fusobacteriota>Fusobacteriia>Fusobacteriales>Fusobacteriaceae>Fusobacterium | GCF_902373915.1 | Fusobacterium_periodonticum | FY302_RS02975 | 615184..618936 | 860 | BREX-system-P-loop-protein-BrxC-[Fusobacterium-periodonticum] |
| NMA29242.1 | Type6.BREX | [illegible] | 1208 | Bacteria>Pseudomonadota>Betaproteobacteria>Burkholderiales>unclassified_Burkholderiales | GCA_012799975.1 | Burkholderiales_bacterium | GX934_15915 | 6559..10185 | 1891238 | BREX-system-P-loop-protein-BrxC-[Burkholderiales-bacterium] |
| WP_109291743.1 | Type1.BREX | [illegible] | 1226 | Bacteria>Pseudomonadota>Gammaproteobacteria>Moraxellales>Moraxellaceae>Acinetobacter | GCF_003131095.1 | Acinetobacter_baumannii | DF172_RS10505 | 4712..8392 | 470 | BREX-system-P-loop-protein-BrxC-[Acinetobacter-baumannii] |
| WP_200291399.1 | Type1.BREX | [illegible] | 1223 | Bacteria>Pseudomonadota>Gammaproteobacteria>Pseudomonadales>Pseudomonadaceae>Stutzerimonas | GCF_016599775.1 | Stutzerimonas_balearica | I6H70_RS19230 | 4083458..4087129 | 74829 | BREX-system-P-loop-protein-BrxC-[Stutzerimonas-balearica] |
| WP_002710590.1 | Type1.BREX | [illegible] | 1209 | Bacteria>Pseudomonadota>Gammaproteobacteria>Thiotrichales>Thiotrichaceae>Thiothrix | GCF_000260135.1 | Thiothrix_nivea | THINI_RS21415 | 4300442..4304071 | 1031 | BREX-system-P-loop-protein-BrxC-[Thiothrix-nivea] |
| MCP5494239.1 | Type1.BREX | [illegible] | 1148 | Bacteria>Spirochaetota>Spirochaetia>Leptospirales>Leptospiraceae>unclassified_Leptospiraceae | GCA_024233865.1 | Leptospiraceae_bacterium | H7A23_06750 | 342984..346430 | 2026757 | BREX-system-P-loop-protein-BrxC-[Leptospiraceae-bacterium] |
| MBR5915376.1 | Type1.BREX | [illegible] | 1170 | Bacteria>Spirochaetota>Spirochaetia>unclassified_Spirochaetia | GCA_017554185.1 | Spirochaetia_bacterium | IKZ57_02880 | 16752..20264 | 2053615 | BREX-system-P-loop-protein-BrxC-[Spirochaetia-bacterium] |
| MBS6134848.1 | Type1.BREX | [illegible] | 1229 | Bacteria>Terrabacteria_group>Actinomycetota>Actinomycetes>Bifidobacteriales>Bifidobacteriaceae | GCA_018374015.1 | Bifidobacterium_longum | KH705_09795 | 10639..14328 | 216816 | BREX-system-P-loop-protein-BrxC-[Bifidobacterium-longum] |
| MCI2117722.1 | Type1.BREX | [illegible] | 1229 | Bacteria>Terrabacteria_group>Actinomycetota>Actinomycetes>Bifidobacteriales>Bifidobacteriaceae | GCA_022653935.1 | Bifidobacterium_breve | LKK17_00135 | 29818..33507 | 1685 | BREX-system-P-loop-protein-BrxC-[Bifidobacterium-breve] |
| WP_051923722.1 | Type1.BREX | [illegible] | 1247 | Bacteria>Terrabacteria_group>Actinomycetota>Actinomycetes>Bifidobacteriales>Bifidobacteriaceae | GCF_000771645.1 | Bifidobacterium_biavatii | OU23_RS06270 | 17800..21543 | 762212 | BREX-system-P-loop-protein-BrxC-[Bifidobacterium-biavatii] |
| WP_052382006.1 | Type1.BREX | [illegible] | 1264 | Bacteria>Terrabacteria_group>Actinomycetota>Actinomycetes>Bifidobacteriales>Bifidobacteriaceae | GCF_000771685.1 | Bifidobacterium_reuteri | OU25_RS00505 | 118725..122519 | 983706 | BREX-system-P-loop-protein-BrxC-[Bifidobacterium-reuteri] |
| WP_050824906.1 | Type1.BREX | [illegible] | 1229 | Bacteria>Terrabacteria_group>Actinomycetota>Actinomycetes>Bifidobacteriales>Bifidobacteriaceae | GCF_001189355.1 | Bifidobacterium_breve | BBRI4_RS05550 | 137028..140717 | 1685 | BREX-system-P-loop-protein-BrxC-[Bifidobacterium-breve] |
| WP_053495781.1 | Type1.BREX | [illegible] | 531 | Bacteria>Terrabacteria_group>Actinomycetota>Actinomycetes>Bifidobacteriales>Bifidobacteriaceae | GCF_001275745.1 | Bifidobacterium_longum | BLOI2_RS03170 | 185000..186595 | 216816 | BREX-system-P-loop-protein-BrxC-[Bifidobacterium-longum] |
| WP_071477881.1 | Type1.BREX | [illegible] | 1229 | Bacteria>Terrabacteria_group>Actinomycetota>Actinomycetes>Bifidobacteriales>Bifidobacteriaceae | GCF_001870755.1 | Bifidobacterium_longum | BFS25_RS03445 | 115725..119414 | 216816 | BREX-system-P-loop-protein-BrxC-[Bifidobacterium-longum] |
| WP_094668334.1 | Type1.BREX | [illegible] | 1270 | Bacteria>Terrabacteria_group>Actinomycetota>Actinomycetes>Bifidobacteriales>Bifidobacteriaceae | GCF_002259745.1 | Bifidobacterium_myosotis | BMYO_RS10065 | 3739..7551 | 1630166 | BREX-system-P-loop-protein-BrxC-[Bifidobacterium-myosotis] |
| WP_109056521.1 | Type1.BREX | [illegible] | 1246 | Bacteria>Terrabacteria_group>Actinomycetota>Actinomycetes>Bifidobacteriales>Bifidobacteriaceae | GCF_003129905.1 | Bifidobacterium_callitrichidarum | DF196_RS03600 | 108404..112144 | 2052941 | BREX-system-P-loop-protein-BrxC-[Bifidobacterium-callitrichidarum] |
| WP_131226808.1 | Type1.BREX | [illegible] | 1229 | Bacteria>Terrabacteria_group>Actinomycetota>Actinomycetes>Bifidobacteriales>Bifidobacteriaceae | GCF_004333385.1 | Bifidobacterium_longum | MCC10066_RS04685 | 5338..9027 | 216816 | BREX-system-P-loop-protein-BrxC-[Bifidobacterium-longum] |
| WP_131210356.1 | Type1.BREX | [illegible] | 442 | Bacteria>Terrabacteria_group>Actinomycetota>Actinomycetes>Bifidobacteriales>Bifidobacteriaceae | GCF_004333565.1 | Bifidobacterium_longum | MCC10048_RS04670 | 1..1326 | 216816 | BREX-system-P-loop-protein-BrxC,-partial-[Bifidobacterium-longum] |
| WP_140489867.1 | Type1.BREX | [illegible] | 1264 | Bacteria>Terrabacteria_group>Actinomycetota>Actinomycetes>Bifidobacteriales>Bifidobacteriaceae | GCF_006408855.1 | Bifidobacterium_sp._UTBIF-68 | BW14_RS07205 | 86138..89932 | 1465262 | BREX-system-P-loop-protein-BrxC-[Bifidobacterium-sp.-UTBIF-68] |
| WP_140558979.1 | Type1.BREX | [illegible] | 1264 | Bacteria>Terrabacteria_group>Actinomycetota>Actinomycetes>Bifidobacteriales>Bifidobacteriaceae | GCF_006408865.1 | Bifidobacterium_sp._UTBIF-56 | BW10_RS07695 | 47972..51766 | 1465261 | BREX-system-P-loop-protein-BrxC-[Bifidobacterium-sp.-UTBIF-56] |
| WP_140463094.1 | Type1.BREX | [illegible] | 1264 | Bacteria>Terrabacteria_group>Actinomycetota>Actinomycetes>Bifidobacteriales>Bifidobacteriaceae | GCF_006408915.1 | unclassified_Bifidobacterium | BW09_RS08245 | 76569..80363 | 2608897 | MULTISPECIES:-BREX-system-P-loop-protein-BrxC-[unclassified-Bifidobacterium] |
| WP_150335795.1 | Type1.BREX | [illegible] | 1264 | Bacteria>Terrabacteria_group>Actinomycetota>Actinomycetes>Bifidobacteriales>Bifidobacteriaceae | GCF_008698125.1 | Bifidobacterium_reuteri | EM092_RS08290 | 2843..6637 | 983706 | BREX-system-P-loop-protein-BrxC-[Bifidobacterium-reuteri] |
| WP_150379196.1 | Type1.BREX | [illegible] | 1267 | Bacteria>Terrabacteria_group>Actinomycetota>Actinomycetes>Bifidobacteriales>Bifidobacteriaceae | GCF_008698155.1 | Bifidobacterium_myosotis | EM091_RS06000 | 83624..87427 | 1630166 | BREX-system-P-loop-protein-BrxC-[Bifidobacterium-myosotis] |
| WP_151916615.1 | Type1.BREX | [illegible] | 1269 | Bacteria>Terrabacteria_group>Actinomycetota>Actinomycetes>Bifidobacteriales>Bifidobacteriaceae | GCF_009078285.1 | Bifidobacterium_jacchi | EHS19_RS04610 | 26586..30395 | 2490545 | BREX-system-P-loop-protein-BrxC-[Bifidobacterium-jacchi] |
| WP_152233646.1 | Type1.BREX | [illegible] | 1235 | Bacteria>Terrabacteria_group>Actinomycetota>Actinomycetes>Bifidobacteriales>Bifidobacteriaceae | GCF_009193355.1 | Bifidobacterium_leontopitheci | F7D09_RS01345 | 123122..126829 | 2650774 | BREX-system-P-loop-protein-BrxC-[Bifidobacterium-leontopitheci] |
| WP_169274836.1 | Type1.BREX | [illegible] | 1226 | Bacteria>Terrabacteria_group>Actinomycetota>Actinomycetes>Bifidobacteriales>Bifidobacteriaceae | GCF_012932365.1 | Bifidobacterium_sp._DSM_109958 | G1C96_RS01295 | 288582..292262 | 2705430 | BREX-system-P-loop-protein-BrxC-[Bifidobacterium-sp.-DSM-109958] |
| WP_077384433.1 | Type1.BREX | [illegible] | 1229 | Bacteria>Terrabacteria_group>Actinomycetota>Actinomycetes>Bifidobacteriales>Bifidobacteriaceae | GCF_017312625.1 | Bifidobacterium_longum | MCC10017_RS04835 | 46545..50234 | 216816 | BREX-system-P-loop-protein-BrxC-[Bifidobacterium-longum] |
| WP_214374600.1 | Type1.BREX | [illegible] | 1249 | Bacteria>Terrabacteria_group>Actinomycetota>Actinomycetes>Bifidobacteriales>Bifidobacteriaceae | GCF_018555335.1 | Bifidobacterium | JS541_RS11260 | 52363..56112 | 1678 | MULTISPECIES:-BREX-system-P-loop-protein-BrxC-[Bifidobacterium] |
| WP_217054570.1 | Type1.BREX | [illegible] | 1229 | Bacteria>Terrabacteria_group>Actinomycetota>Actinomycetes>Bifidobacteriales>Bifidobacteriaceae | GCF_019041665.1 | Bifidobacterium_longum | KTF47_RS04030 | 15252..18941 | 216816 | BREX-system-P-loop-protein-BrxC-[Bifidobacterium-longum] |
| WP_269360645.1 | Type1.BREX | [illegible] | 1229 | Bacteria>Terrabacteria_group>Actinomycetota>Actinomycetes>Bifidobacteriales>Bifidobacteriaceae | GCF_027215095.1 | Bifidobacterium_breve | O4D08_RS00920 | 188382..192071 | 1685 | BREX-system-P-loop-protein-BrxC-[Bifidobacterium-breve] |
| WP_269338054.1 | Type1.BREX | [illegible] | 924 | Bacteria>Terrabacteria_group>Actinomycetota>Actinomycetes>Bifidobacteriales>Bifidobacteriaceae | GCF_027215525.1 | Bifidobacterium_breve | O4C30_RS07905 | 20359..23133 | 1685 | BREX-system-P-loop-protein-BrxC-[Bifidobacterium-breve] |
| WP_270267401.1 | Type1.BREX | [illegible] | 1229 | Bacteria>Terrabacteria_group>Actinomycetota>Actinomycetes>Bifidobacteriales>Bifidobacteriaceae | GCF_027663845.1 | Bifidobacterium_breve | PGQ90_RS07415 | 209217..212906 | 1685 | BREX-system-P-loop-protein-BrxC-[Bifidobacterium-breve] |
| WP_143724686.1 | Type1.BREX | [illegible] | 1229 | Bacteria>Terrabacteria_group>Actinomycetota>Actinomycetes>Bifidobacteriales>Bifidobacteriaceae | GCF_902167555.1 | Bifidobacterium_longum | FP566_RS07260 | 1570713..1574402 | 216816 | BREX-system-P-loop-protein-BrxC-[Bifidobacterium-longum] |
| WP_250246699.1 | Type1.BREX | [illegible] | 1229 | Bacteria>Terrabacteria_group>Actinomycetota>Actinomycetes>Bifidobacteriales>Bifidobacteriaceae | GCF_939148185.1 | Bifidobacterium_longum | NBW58_RS09955 | 18627..22316 | 216816 | BREX-system-P-loop-protein-BrxC-[Bifidobacterium-longum] |
| WP_251163982.1 | Type1.BREX | [illegible] | 1193 | Bacteria>Terrabacteria_group>Actinomycetota>Coriobacteriia>Coriobacteriales>Atopobiaceae | - | Caniella_muris | IM763_04355 | 91958..95539 | 2941502 | BREX-system-P-loop-protein-BrxC-[Caniella-muris] |
| MBF0599289.1 | Type1.BREX | [illegible] | 1193 | Bacteria>Terrabacteria_group>Actinomycetota>Coriobacteriia>Coriobacteriales>Atopobiaceae | GCA_015234195.1 | Atopobiaceae_bacterium_FL090493 | IM763_04355 | 91958..95539 | 2661918 | BREX-system-P-loop-protein-BrxC-[Atopobiaceae-bacterium-FL090493] |
| MBR2835999.1 | Type1.BREX | [illegible] | 1194 | Bacteria>Terrabacteria_group>Actinomycetota>Coriobacteriia>Coriobacteriales>unclassified_Coriobacteriales | GCA_017515595.1 | Coriobacteriales_bacterium | IKE43_09885 | 4830..8414 | 2478480 | BREX-system-P-loop-protein-BrxC-[Coriobacteriales-bacterium] |
| WP_251231669.1 | Type1.BREX | [illegible] | 1213 | Bacteria>Terrabacteria_group>Actinomycetota>Coriobacteriia>Eggerthellales>Eggerthellaceae | GCF_943193145.1 | Adlercreutzia_aquisgranensis | NFR41_RS09570 | 19774..23415 | 2941323 | BREX-system-P-loop-protein-BrxC-[Adlercreutzia-aquisgranensis] |
| WP_251851849.1 | Type1.BREX | [illegible] | 1202 | Bacteria>Terrabacteria_group>Bacillota>Bacilli>Lactobacillales>Enterococcaceae | GCF_023744155.1 | Enterococcus_italicus | NE261_RS07355 | 549628..553236 | 246144 | BREX-system-P-loop-protein-BrxC-[Enterococcus-italicus] |
| NFO12999.1 | Type1.BREX | [illegible] | 1215 | Bacteria>Terrabacteria_group>Bacillota>Clostridia>Eubacteriales>Clostridiaceae | GCA_011016315.1 | Clostridium_botulinum | FDB34_02025 | 196714..200361 | 1491 | BREX-system-P-loop-protein-BrxC-[Clostridium-botulinum] |
| MBN1067552.1 | Type1.BREX | [illegible] | 1215 | Bacteria>Terrabacteria_group>Bacillota>Clostridia>Eubacteriales>Clostridiaceae | GCA_016925955.1 | Clostridium_botulinum | DVW02_06040 | 97050..100697 | 1491 | BREX-system-P-loop-protein-BrxC-[Clostridium-botulinum] |
| WP_039258635.1 | Type1.BREX | [illegible] | 1214 | Bacteria>Terrabacteria_group>Bacillota>Clostridia>Eubacteriales>Clostridiaceae | GCF_000768655.1 | Clostridium_botulinum | ADU88_RS01575 | 72725..76369 | 1491 | BREX-system-P-loop-protein-BrxC-[Clostridium-botulinum] |
| WP_156211829.1 | Type1.BREX | [illegible] | 1216 | Bacteria>Terrabacteria_group>Bacillota>Clostridia>Eubacteriales>Clostridiaceae | GCF_009735365.2 | Clostridium_butyricum | GND98_RS05145 | 54704..58354 | 1492 | BREX-system-P-loop-protein-BrxC-[Clostridium-butyricum] |
| WP_164949146.1 | Type1.BREX | [illegible] | 1215 | Bacteria>Terrabacteria_group>Bacillota>Clostridia>Eubacteriales>Clostridiaceae | GCF_011065955.2 | Clostridium_sp._FP2 | G9F71_RS17965 | 3736090..3739737 | 2724481 | BREX-system-P-loop-protein-BrxC-[Clostridium-sp.-FP2] |
| WP_257411542.1 | Type1.BREX | [illegible] | 1195 | Bacteria>Terrabacteria_group>Bacillota>Clostridia>Eubacteriales>Clostridiaceae | GCF_024622885.1 | Clostridium_sporogenes | NSA52_RS02830 | 6440..10027 | 1509 | BREX-system-P-loop-protein-BrxC-[Clostridium-sporogenes] |
| MBP1566405.1 | Type1.BREX | [illegible] | 1178 | Bacteria>Terrabacteria_group>Bacillota>Clostridia>Eubacteriales>Oscillospiraceae | GCA_017848115.1 | Oscillospiraceae_bacterium | J6A58_11685 | 59197..62733 | 2485925 | BREX-system-P-loop-protein-BrxC-[Oscillospiraceae-bacterium] |
| UYJ11714.1 | Type1.BREX | [illegible] | 1182 | Bacteria>Terrabacteria_group>Bacillota>Clostridia>Eubacteriales>Oscillospiraceae | GCA_025757845.1 | Oscillospiraceae_bacterium | OGM78_02700 | 567882..571430 | 2485925 | MAG:-BREX-system-P-loop-protein-BrxC-[Oscillospiraceae-bacterium] |
| MCR4615485.1 | Type1.BREX | [illegible] | 1161 | Bacteria>Terrabacteria_group>Bacillota>Clostridia>Eubacteriales>unclassified_Eubacteriales | GCA_024684435.1 | Clostridiales_bacterium | K5756_04990 | 82244..85729 | 1898207 | BREX-system-P-loop-protein-BrxC-[Clostridiales-bacterium] |
| MBQ6380318.1 | Type1.BREX | [illegible] | 1174 | Bacteria>Terrabacteria_group>Bacillota>Clostridia>unclassified_Clostridia | GCA_017430965.1 | Clostridia_bacterium | IJJ41_01780 | 27986..31510 | 2044939 | BREX-system-P-loop-protein-BrxC-[Clostridia-bacterium] |
| MBF0120841.1 | Type1.BREX | [illegible] | 1201 | Bacteria>Thermodesulfobacteriota>Desulfobacteria>Desulfobacterales>unclassified_Desulfobacterales | GCA_015228685.1 | Desulfobacterales_bacterium | HQK79_18595 | 27077..30682 | 2044940 | BREX-system-P-loop-protein-BrxC-[Desulfobacterales-bacterium] |
| NQX01743.1 | Type6.BREX | [illegible] | 1210 | Bacteria>unclassified_Bacteria | GCA_013205625.1 | bacterium | HQ447_13895 | 2028..5660 | 1869227 | BREX-system-P-loop-protein-BrxC-[bacterium] |
| -- | | | | | | | | | | |
| -- HTH+REase | | | | | | | | | | |
| -- w HTH+REase | | | | | | | | | | |
| -- HARE-HTH+REase | | | | | | | | | | |
| -- | | | | | | | | | | |
| WP_255191907.1 | Type1.BREX | [illegible] | 1261 | Archaea>Euryarchaeota>Stenosarchaea_group>Halobacteria>Natrialbales>Natrialbaceae | GCF_024362485.1 | Halovivax_sp._WLHSJ27 | NMQ09_RS17725 | 3493167..3496952 | 2963433 | hypothetical-protein-[Halovivax-sp.-WLHSJ27] |
| WP_255191909.1 | Type1.BREX | [illegible] | 1273 | Archaea>Euryarchaeota>Stenosarchaea_group>Halobacteria>Natrialbales>Natrialbaceae | GCF_024362485.1 | Halovivax_sp._WLHSJ27 | NMQ09_RS17735 | 3497545..3501366 | 2963433 | hypothetical-protein-[Halovivax-sp.-WLHSJ27] |
| WP_273738559.1 | Type6.BREX | [illegible] | 1214 | Archaea>Euryarchaeota>Stenosarchaea_group>Halobacteria>Natrialbales>Natrialbaceae | GCF_028618695.1 | Natrinema_soli | PGX04_RS11075 | 6409..10053 | 1930624 | BREX-system-P-loop-protein-BrxC-[Natrinema-soli] |
| WP_004077401.1 | Type1.BREX | [illegible] | 1180 | Archaea>Euryarchaeota>Stenosarchaea_group>Methanomicrobia>Methanomicrobiales>Methanomicrobiaceae | GCF_000243255.1 | Methanoplanus_limicola | METLIM_RS07750 | 1605385..1608927 | 2315 | BREX-system-P-loop-protein-BrxC-[Methanoplanus-limicola] |
| WP_004078487.1 | Type1.BREX | [illegible] | 1155 | Archaea>Euryarchaeota>Stenosarchaea_group>Methanomicrobia>Methanomicrobiales>Methanomicrobiaceae | GCF_000243255.1 | Methanoplanus_limicola | METLIM_RS10810 | 2304713..2308180 | 2315 | BREX-system-P-loop-protein-BrxC-[Methanoplanus-limicola] |
| WP_257742634.1 | Type1.BREX | [illegible] | 1155 | Archaea>Euryarchaeota>Stenosarchaea_group>Methanomicrobia>Methanomicrobiales>Methanomicrobiaceae | GCF_024662215.1 | Methanoplanus_endosymbiosus | L6E24_RS14385 | 3075748..3079215 | 33865 | BREX-system-P-loop-protein-BrxC-[Methanoplanus-endosymbiosus] |
| MBN1165665.1 | Type1.BREX | [illegible] | 1218 | Archaea>Euryarchaeota>Stenosarchaea_group>Methanomicrobia>Methanomicrobiales>Methanospirillaceae | GCA_016926505.1 | Methanospirillaceae_archaeon | JXA44_00885 | 48354..52010 | 2813030 | BREX-system-P-loop-protein-BrxC-[Methanospirillaceae-archaeon] |
| WP_011448873.1 | Type1.BREX | [illegible] | 1178 | Archaea>Euryarchaeota>Stenosarchaea_group>Methanomicrobia>Methanomicrobiales>Methanospirillaceae | GCF_000013445.1 | Methanospirillum_hungatei | MHUN_RS09850 | 2137740..2141276 | 2203 | BREX-system-P-loop-protein-BrxC-[Methanospirillum-hungatei] |
| WP_214418671.1 | Type1.BREX | [illegible] | 1171 | Archaea>Euryarchaeota>Stenosarchaea_group>Methanomicrobia>Methanomicrobiales>Methanospirillaceae | GCF_018502485.1 | Methanospirillum_sp._J.3.6.1-F.2.7.3 | KHC33_RS10925 | 2269426..2272941 | 2834276 | BREX-system-P-loop-protein-BrxC-[Methanospirillum-sp.-J.3.6.1-F.2.7.3] |
| KXS44097.1 | Type1.BREX | [illegible] | 1185 | Archaea>Euryarchaeota>Stenosarchaea_group>Methanomicrobia>Methanosarcinales>Methanosarcinaceae | GCA_001577565.1 | Methanolobus_sp._T82-4 | AWU59_730 | 16022..19579 | 1794908 | MAG:-hypothetical-protein-AWU59_730-[Methanolobus-sp.-T82-4] |
| WP_048162601.1 | Type1.BREX | [illegible] | 1223 | Archaea>Euryarchaeota>Stenosarchaea_group>Methanomicrobia>Methanosarcinales>Methanosarcinaceae | GCF_000979975.1 | Methanosarcina_sp._1.H.A.2.2 | EO93_RS11285 | 63696..67367 | 1483601 | BREX-system-P-loop-protein-BrxC-[Methanosarcina-sp.-1.H.A.2.2] |
| WP_141050577.1 | Type1.BREX | [illegible] | 1162 | Bacteria>Campylobacterota>Epsilonproteobacteria>Campylobacterales>Arcobacteraceae>Aliarcobacter | GCF_006503615.1 | Aliarcobacter_cryaerophilus | AV936_RS08795 | 11744..15232 | 28198 | BREX-system-P-loop-protein-BrxC-[Aliarcobacter-cryaerophilus] |
| RXJ95785.1 | Type1.BREX | [illegible] | 1162 | Bacteria>Campylobacterota>Epsilonproteobacteria>Campylobacterales>Arcobacteraceae>Arcobacter | GCA_004116575.1 | Arcobacter_sp._CECT_9299 | CRU94_03995 | 49858..53346 | 2044506 | hypothetical-protein-CRU94_03995-[Arcobacter-sp.-CECT-9299] |
| WP_129105531.1 | Type1.BREX | [illegible] | 1162 | Bacteria>Campylobacterota>Epsilonproteobacteria>Campylobacterales>Arcobacteraceae>Arcobacter | GCF_004116575.1 | Arcobacter_sp._CECT_9299 | CRU94_RS03995 | 49858..53346 | 2044506 | BREX-system-P-loop-protein-BrxC-[Arcobacter-sp.-CECT-9299] |
| WP_164968523.1 | Type1.BREX | [illegible] | 1160 | Bacteria>Campylobacterota>Epsilonproteobacteria>Campylobacterales>Arcobacteraceae>Arcobacter | GCF_013201725.1 | Arcobacter_defluvii | ADFLV_RS07175 | 1422397..1425879 | 873191 | BREX-system-P-loop-protein-BrxC-[Arcobacter-defluvii] |
| PHS59105.1 | Type1.BREX | [illegible] | 1158 | Bacteria>Campylobacterota>Epsilonproteobacteria>Campylobacterales>Thiovulaceae>Sulfurimonas | GCA_002733155.1 | Sulfurimonas_sp. | COB17_00805 | 100943..104419 | 2022749 | MAG:-hypothetical-protein-COB17_00805-[Sulfurimonas-sp.] |
| MBZ4644075.1 | Type1.BREX | [illegible] | 1168 | Bacteria>Deferribacterota>Deferribacteres>Deferribacterales>Deferribacteraceae>unclassified_Deferribacteraceae | GCA_020054865.1 | Deferribacteraceae_bacterium | JG767_1684 | 7812..11318 | 2053543 | hypothetical-protein-[Deferribacteraceae-bacterium] |
| WP_242820841.1 | Type1.BREX | [illegible] | 1168 | Bacteria>Deferribacterota>Deferribacteres>Deferribacterales>Deferrivibrionaceae>Deferrivibrio | GCF_020480685.1 | Deferrivibrio_essentukiensis | LF845_RS09815 | 29861..33367 | 2880922 | BREX-system-P-loop-protein-BrxC-[Deferrivibrio-essentukiensis] |
| MBF0105032.1 | Type1.BREX | [illegible] | 1166 | Bacteria>delta/epsilon_subdivisions>Deltaproteobacteria>unclassified_Deltaproteobacteria | GCA_015228875.1 | Deltaproteobacteria_bacterium | HQM16_06860 | 10380..13880 | 2026735 | BREX-system-P-loop-protein-BrxC-[Deltaproteobacteria-bacterium] |
| MBF0105437.1 | Type1.BREX | [illegible] | 1155 | Bacteria>delta/epsilon_subdivisions>Deltaproteobacteria>unclassified_Deltaproteobacteria | GCA_015228875.1 | Deltaproteobacteria_bacterium | HQM16_08940 | 21511..24978 | 2026735 | BREX-system-P-loop-protein-BrxC-[Deltaproteobacteria-bacterium] |
| MCP4577326.1 | Type1.BREX | [illegible] | 1196 | Bacteria>delta/epsilon_subdivisions>Deltaproteobacteria>unclassified_Deltaproteobacteria | GCA_024231575.1 | Deltaproteobacteria_bacterium | GY846_13700 | 13452..17042 | 2026735 | BREX-system-P-loop-protein-BrxC-[Deltaproteobacteria-bacterium] |
| NLA16135.1 | Type1.BREX | [illegible] | 1174 | Bacteria>FCB_group>Bacteroidetes/Chlorobi_group>Bacteroidota>Bacteroidia>Bacteroidales | GCA_012510445.1 | Bacteroidales_bacterium | GX877_06335 | 9723..13247 | 2030927 | BREX-system-P-loop-protein-BrxC-[Bacteroidales-bacterium] |
| MBQ9588226.1 | Type1.BREX | [illegible] | 1183 | Bacteria>FCB_group>Bacteroidetes/Chlorobi_group>Bacteroidota>Bacteroidia>Bacteroidales | GCA_017446745.1 | Bacteroidales_bacterium | IJR26_10275 | 19779..23330 | 2030927 | BREX-system-P-loop-protein-BrxC-[Bacteroidales-bacterium] |
| MBW6491485.1 | Type1.BREX | [illegible] | 1144 | Bacteria>FCB_group>Bacteroidetes/Chlorobi_group>Bacteroidota>Bacteroidia>Bacteroidales | GCA_019429275.1 | Lentimicrobium_sp. | KOB15_09875 | 47933..51367 | 2034841 | BREX-system-P-loop-protein-BrxC-[Lentimicrobium-sp.] |
| MCB9017748.1 | Type1.BREX | [illegible] | 1207 | Bacteria>FCB_group>Bacteroidetes/Chlorobi_group>Bacteroidota>Bacteroidia>Bacteroidales | GCA_020636675.1 | Prevotellaceae_bacterium | H6544_03995 | 62697..66320 | 2049047 | BREX-system-P-loop-protein-BrxC-[Prevotellaceae-bacterium] |
| MCC8119638.1 | Type1.BREX | [illegible] | 1189 | Bacteria>FCB_group>Bacteroidetes/Chlorobi_group>Bacteroidota>Bacteroidia>Bacteroidales | GCA_020860515.1 | Bacteroidales_bacterium | LIP09_12950 | 29133..32702 | 2030927 | BREX-system-P-loop-protein-BrxC-[Bacteroidales-bacterium] |
| MCI1742488.1 | Type1.BREX | [illegible] | 1206 | Bacteria>FCB_group>Bacteroidetes/Chlorobi_group>Bacteroidota>Bacteroidia>Bacteroidales | GCA_022647785.1 | Prevotella_sp. | LKI18_09155 | 39395..43015 | 59823 | BREX-system-P-loop-protein-BrxC-[Prevotella-sp.] |
| MCI2087046.1 | Type1.BREX | [illegible] | 1198 | Bacteria>FCB_group>Bacteroidetes/Chlorobi_group>Bacteroidota>Bacteroidia>Bacteroidales | GCA_022653375.1 | Prevotella_sp. | LKK10_02020 | 11419..15015 | 59823 | BREX-system-P-loop-protein-BrxC-[Prevotella-sp.] |
| WP_008760280.1 | Type1.BREX | [illegible] | 1193 | Bacteria>FCB_group>Bacteroidetes/Chlorobi_group>Bacteroidota>Bacteroidia>Bacteroidales | GCF_000162515.1 | Bacteroides_thetaiotaomicron | HMPREF9007_RS20215 | 181866..185447 | 818 | MULTISPECIES:-BREX-system-P-loop-protein-BrxC-[Bacteroides] |
| WP_130041244.1 | Type1.BREX | [illegible] | 1175 | Bacteria>FCB_group>Bacteroidetes/Chlorobi_group>Bacteroidota>Bacteroidia>Bacteroidales | GCF_019733625.1 | Bacteroides_thetaiotaomicron | EAJ10_RS04945 | 124743..128270 | 818 | BREX-system-P-loop-protein-BrxC-[Bacteroides-thetaiotaomicron] |
| WP_225659484.1 | Type1.BREX | [illegible] | 1175 | Bacteria>FCB_group>Bacteroidetes/Chlorobi_group>Bacteroidota>Bacteroidia>Bacteroidales | GCF_020313445.1 | Bacteroides_thetaiotaomicron | KFX61_RS09580 | 34477..38004 | 818 | BREX-system-P-loop-protein-BrxC-[Bacteroides-thetaiotaomicron] |
| WP_258973412.1 | Type1.BREX | [illegible] | 1133 | Bacteria>FCB_group>Bacteroidetes/Chlorobi_group>Bacteroidota>Bacteroidia>Bacteroidales | GCF_024793205.1 | Bacteroides_thetaiotaomicron | NXX32_RS11560 | 3197619..3201020 | 818 | BREX-system-P-loop-protein-BrxC-[Bacteroides-thetaiotaomicron] |
| OSZ81159.1 | Type1.BREX | [illegible] | 1144 | Bacteria>FCB_group>Bacteroidetes/Chlorobi_group>Bacteroidota>Chitinophagia>Chitinophagales | GCA_002127295.1 | Chitinophagaceae_bacterium_IBVUCB2 | CAP36_07965 | 1792100..1795534 | 1985174 | hypothetical-protein-CAP36_07965-[Chitinophagaceae-bacterium-IBVUCB2] |
| MBL7764603.1 | Type1.BREX | [illegible] | 1146 | Bacteria>FCB_group>Bacteroidetes/Chlorobi_group>Bacteroidota>Chitinophagia>Chitinophagales | GCA_016786845.1 | Chitinophagaceae_bacterium | JNJ58_00790 | 47471..50911 | 1869212 | BREX-system-P-loop-protein-BrxC-[Chitinophagaceae-bacterium] |
| MCK9404665.1 | Type1.BREX | [illegible] | 1144 | Bacteria>FCB_group>Bacteroidetes/Chlorobi_group>Bacteroidota>Chitinophagia>Chitinophagales | GCA_023229465.1 | Chitinophagaceae_bacterium | M0Q26_14830 | 11931..15365 | 1869212 | BREX-system-P-loop-protein-BrxC-[Chitinophagaceae-bacterium] |
| WP_072366579.1 | Type1.BREX | [illegible] | 1142 | Bacteria>FCB_group>Bacteroidetes/Chlorobi_group>Bacteroidota>Chitinophagia>Chitinophagales | GCF_900119105.1 | Chitinophaga_sancti | BT118_RS32550 | 2428..5856 | 1004 | BREX-system-P-loop-protein-BrxC-[Chitinophaga-sancti] |
| WP_008625904.1 | Type1.BREX | [illegible] | 1146 | Bacteria>FCB_group>Bacteroidetes/Chlorobi_group>Bacteroidota>Cytophagia>Cytophagales | GCF_000330725.2 | Mariniradius_saccharolyticus | C943_RS07085 | 84904..88344 | 1245591 | BREX-system-P-loop-protein-BrxC-[Mariniradius-saccharolyticus] |
| WP_122934661.1 | Type1.BREX | [illegible] | 1142 | Bacteria>FCB_group>Bacteroidetes/Chlorobi_group>Bacteroidota>Cytophagia>Cytophagales | GCF_003711225.1 | Runella_sp._SP2 | DTQ70_RS30140 | 43566..46994 | 2268026 | BREX-system-P-loop-protein-BrxC-[Runella-sp.-SP2] |
| WP_189584875.1 | Type1.BREX | [illegible] | 1146 | Bacteria>FCB_group>Bacteroidetes/Chlorobi_group>Bacteroidota>Cytophagia>Cytophagales | GCF_014652135.1 | Mongoliitalea_lutea | IE080_RS15850 | 2275..5715 | 849756 | BREX-system-P-loop-protein-BrxC-[Mongoliitalea-lutea] |
| WP_244713422.1 | Type1.BREX | [illegible] | 1145 | Bacteria>FCB_group>Bacteroidetes/Chlorobi_group>Bacteroidota>Cytophagia>Cytophagales | GCF_022919135.1 | Hymenobacter_sp._5116S-27 | MUN80_RS13030 | 3065695..3069132 | 2932249 | BREX-system-P-loop-protein-BrxC-[Hymenobacter-sp.-5116S-27] |
| WP_084121639.1 | Type1.BREX | [illegible] | 1146 | Bacteria>FCB_group>Bacteroidetes/Chlorobi_group>Bacteroidota>Cytophagia>Cytophagales | GCF_900176595.1 | Aquiflexum_balticum | B9A52_RS17175 | 4064780..4068220 | 280473 | BREX-system-P-loop-protein-BrxC-[Aquiflexum-balticum] |
| MCB0520339.1 | Type1.BREX | [illegible] | 1145 | Bacteria>FCB_group>Bacteroidetes/Chlorobi_group>Bacteroidota>Saprospiria>Saprospirales | GCA_020436325.1 | Saprospiraceae_bacterium | KDD27_15480 | 21267..24704 | 2202734 | BREX-system-P-loop-protein-BrxC-[Saprospiraceae-bacterium] |
| WP_013769021.1 | Type1.BREX | [illegible] | 1144 | Bacteria>FCB_group>Bacteroidetes/Chlorobi_group>Bacteroidota>Saprospiria>Saprospirales | GCF_000212735.1 | Haliscomenobacter_hydrossis | HALHY_RS33450 | 27488..30922 | 2350 | BREX-system-P-loop-protein-BrxC-[Haliscomenobacter-hydrossis] |
| WP_070561019.1 | Type1.BREX | [illegible] | 1187 | Bacteria>FCB_group>Bacteroidetes/Chlorobi_group>Bacteroidota>Sphingobacteriia>Sphingobacteriales | GCF_002339225.1 | Sphingobacterium_sp._HMSC13C05 | HMPREF3127_RS01230 | 95716..99279 | 1581095 | BREX-system-P-loop-protein-BrxC-[Sphingobacterium-sp.-HMSC13C05] |
| WP_165305702.1 | Type1.BREX | [illegible] | 1187 | Bacteria>FCB_group>Bacteroidetes/Chlorobi_group>Bacteroidota>Sphingobacteriia>Sphingobacteriales | GCF_011064245.1 | Sphingobacterium_sp._DR205 | G6053_RS05210 | 1244063..1247626 | 2713573 | BREX-system-P-loop-protein-BrxC-[Sphingobacterium-sp.-DR205] |
| KAF0128492.1 | Type1.BREX | [illegible] | 1144 | Bacteria>FCB_group>Bacteroidetes/Chlorobi_group>Bacteroidota>unclassified_Bacteroidota | GCA_009773165.1 | Bacteroidota_bacterium | FD155_3234 | 38823..42257 | 1898104 | hypothetical-protein-FD155_3234-[Bacteroidota-bacterium] |
| NQT60364.1 | Type1.BREX | [illegible] | 1348 | Bacteria>FCB_group>Bacteroidetes/Chlorobi_group>Bacteroidota>unclassified_Bacteroidota | GCA_013202335.1 | Bacteroidota_bacterium | HQ557_15395 | 13554..17600 | 1898104 | hypothetical-protein-[Bacteroidota-bacterium] |
| MCH7965547.1 | Type1.BREX | [illegible] | 1161 | Bacteria>FCB_group>Bacteroidetes/Chlorobi_group>Bacteroidota>unclassified_Bacteroidota | GCA_022564295.1 | Bacteroidota_bacterium | IH852_16610 | 28479..31964 | 1898104 | BREX-system-P-loop-protein-BrxC-[Bacteroidota-bacterium] |
| MCR9155224.1 | Type1.BREX | [illegible] | 1141 | Bacteria>FCB_group>Bacteroidetes/Chlorobi_group>Bacteroidota>unclassified_Bacteroidota | GCA_024742495.1 | Bacteroidota_bacterium | NXI09_14035 | 83336..86761 | 1898104 | BREX-system-P-loop-protein-BrxC-[Bacteroidota-bacterium] |
| KAA6232806.1 | Type1.BREX | [illegible] | 1204 | Bacteria>FCB_group>Bacteroidetes/Chlorobi_group>Chlorobiota>Chlorobiia>Chlorobiales | GCA_008634005.1 | Chlorobium_phaeovibrioides | FP507_06840 | 1409593..1413207 | 1094 | BREX-system-P-loop-protein-BrxC-[Chlorobium-phaeovibrioides] |
| WP_012508724.1 | Type1.BREX | [illegible] | 1211 | Bacteria>FCB_group>Bacteroidetes/Chlorobi_group>Chlorobiota>Chlorobiia>Chlorobiales | GCF_000020645.1 | Pelodictyon_phaeoclathratiforme | PPHA_RS10105 | 2123667..2127302 | 34090 | BREX-system-P-loop-protein-BrxC-[Pelodictyon-phaeoclathratiforme] |
| WP_085659853.1 | Type1.BREX | [illegible] | 1209 | Bacteria>FCB_group>Bacteroidetes/Chlorobi_group>Chlorobiota>Chlorobiia>Chlorobiales | GCF_002113825.1 | Prosthecochloris_sp._HL-130-GSB | B9H02_RS07545 | 1603101..1606730 | 1974213 | BREX-system-P-loop-protein-BrxC-[Prosthecochloris-sp.-HL-130-GSB] |
| WP_192496514.1 | Type1.BREX | [illegible] | 1203 | Bacteria>FCB_group>Bacteroidetes/Chlorobi_group>Chlorobiota>Chlorobiia>Chlorobiales | GCF_008634005.1 | Chlorobium_phaeovibrioides | FP507_RS07315 | 1409593..1413204 | 1094 | BREX-system-P-loop-protein-BrxC-[Chlorobium-phaeovibrioides] |
| MBM3423463.1 | Type1.BREX | [illegible] | 1182 | Bacteria>FCB_group>Bacteroidetes/Chlorobi_group>Chlorobiota>unclassified_Chlorobiota | GCA_016868665.1 | Chlorobiota_bacteirum | FJY09_09520 | 5435..8983 | 2268192 | BREX-system-P-loop-protein-BrxC-[Chlorobiota-bacterium] |
| MBP9037128.1 | Type1.BREX | [illegible] | 1152 | Bacteria>FCB_group>Candidatus_Cloacimonetes>Candidatus_Cloacimonas>unclassified_Candidatus_Cloacimonas | GCA_018057045.1 | Candidatus_Cloacimonas_sp. | KBG38_04995 | 2613..6071 | 2053534 | BREX-system-P-loop-protein-BrxC-[Candidatus-Cloacimonas-sp.] |
| WP_120625208.1 | Type6.BREX | [illegible] | 1203 | Bacteria>Myxococcota>Myxococcia>Myxococcales>Cystobacterineae>Myxococcaceae | GCF_003611735.1 | Corallococcus_sicarius | D7X12_RS10925 | 19643..23254 | 2316726 | BREX-system-P-loop-protein-BrxC-[Corallococcus-sicarius] |
| HCF60732.1 | Type1.BREX | [illegible] | 871 | Bacteria>Myxococcota>Myxococcia>Myxococcales>unclassified_Myxococcales | GCA_003534605.1 | Myxococcales_bacterium | DFS52_22370 | 11512..14124 | 2026763 | BREX-system-P-loop-protein-BrxC-[Myxococcales-bacterium] |

<table>
<tr><th>Protein-ID</th><th>System</th><th>Gene-Neighbourhoods</th><th>Length</th><th>Taxonomy</th><th>Assembly-ID</th><th>Strain</th><th>Locus-Tag</th><th>Genomic-coordinates</th><th>TaxID</th><th>Fasta-define</th></tr>
<tr><td>NUN14061.1</td><td>Type6.BREX</td><td>HTH+ REase -> BrxE -> BrxA-DNA-binding -> BrxB-BrxF-iSTAND -> DUF6079 *->?-> N6-METHYLASE -> PglZ ->BrxD-AAA->SF1-SF2-Helicase+wHTH+Small-B-barrel+DUF1998->Schlafen+HATPase+wHTH-></td><td>1213</td><td>Bacteria>Myxococcota>Myxococcia>Myxococcales>unclassified_Myxococcales</td><td>GCA_013360285.1</td><td>Myxococcales_bacterium</td><td>HUU55_10555</td><td>3445..7086</td><td>2026763</td><td>BREX-system-P-loop-protein-BrxC-[Myxococcales-bacterium]</td></tr>
<tr><td>MBK6463374.1</td><td>Type6.BREX</td><td><-Sulfatase+DUF4976||?->?-><-?||?->?-> BrxE -> BrxA-DNA-binding -> BrxB-BrxF-iSTAND -> DUF6079 *->SbcC-> N6-METHYLASE -p->?-> N6-METHYLASE -p-> N6-METHYLASE -p-> REase -> N6-METHYLASE -p-> HTH+ REase -></td><td>1235</td><td>Bacteria>Myxococcota>Myxococcia>Myxococcales>unclassified_Myxococcales</td><td>GCA_016704525.1</td><td>Myxococcales_bacterium</td><td>IPF92_20555</td><td>135329..139036</td><td>2026763</td><td>BREX-system-P-loop-protein-BrxC-[Myxococcales-bacterium]</td></tr>
<tr><td>MBP9202688.1</td><td>Type6.BREX</td><td><-ChlI+Lon_C+AAA<-HEPN<-SF1-SF2-Helicase+wHTH+Small-B-barrel+35exo<-BrxD-AAA<- PglZ <- HTH+ REase <- N6-METHYLASE <-?<- DUF6079 *<- BrxB-BrxF-iSTAND <- BrxA-DNA-binding <- BrxE</td><td>1203</td><td>Bacteria>Myxococcota>Polyangia>Haliangiales>Kofleriaceae>unclassified_Kofleriaceae</td><td>GCA_018058095.1</td><td>Kofleriaceae_bacterium</td><td>KBG28_01800</td><td>90937..94548</td><td>2212474</td><td>BREX-system-P-loop-protein-BrxC-[Kofleriaceae-bacterium]</td></tr>
<tr><td>MBX9657254.1</td><td>Type1.BREX</td><td><-SF1-SF2-Helicase<-HKD-DNase<-?<-NERD<-HD+HD_assoc<-?<- SIGMA+OB-Fold+AAA-ATPase+Lon-Protease-BrxL <- PglZ <- N6-METHYLASE <- DUF6079 *<- BrxB-BrxF-iSTAND <- BrxA-DNA-binding <- REase +HTH+35exo+Small-B-barrel+SF1-SF2-Helicase<-?<-?<-?<- wHTH+WYL+WCX <-ANK+ANK</td><td>1193</td><td>Bacteria>Nitrospirota>Nitrospiria>Nitrospirales>Nitrospiraceae>unclassified_Nitrospiraceae</td><td>GCA_019748255.1</td><td>Nitrospiraceae_bacterium</td><td>K2X00_01715</td><td>152268..155849</td><td>2026770</td><td>BREX-system-P-loop-protein-BrxC-[Nitrospiraceae-bacterium]</td></tr>
<tr><td>WP_172437661.1</td><td>Type1.BREX</td><td>DEDD_Tnp_IS110+Transposase_20-><-?||XIS-HTH->TOPRIM->||XIS-HTH-> BrxA-DNA-binding -> BrxB-BrxF-iSTAND -> DUF6079 *->Schlafen+HATPase+wHTH+HTH-> N6-METHYLASE -> HTH+ REase -> PglZ -> SIGMA+OB-Fold+AAA-ATPase+HKD-DNase-BrxL ->SbcC+APATPase+SbcC->Calcineurin+Calcineurin-SbcD->SbcC->RNase-></td><td>1203</td><td>Bacteria>Pseudomonadota>Acidithiobacillia>Acidithiobacillales>Acidithiobacillaceae>Acidithiobacillus</td><td>-</td><td>Acidithiobacillus_sp._SH</td><td>HF670_09010</td><td>8231..11842</td><td>187490</td><td>MULTISPECIES:-BREX-system-P-loop-protein-BrxC-[Acidithiobacillus]</td></tr>
<tr><td>MBU2839699.1</td><td>Type1.BREX</td><td>DEDD_Tnp_IS110+Transposase_20-><-?||XIS-HTH->TOPRIM->||XIS-HTH-> BrxA-DNA-binding -> BrxB-BrxF-iSTAND -> DUF6079 *->Schlafen+HATPase+wHTH+HTH-> N6-METHYLASE -> HTH+ REase -> PglZ -> SIGMA+OB-Fold+AAA-ATPase+HKD-DNase-BrxL ->SbcC+APATPase+SbcC->Calcineurin+Calcineurin-SbcD->SbcC->RNase-></td><td>1203</td><td>Bacteria>Pseudomonadota>Acidithiobacillia>Acidithiobacillales>Acidithiobacillaceae>Acidithiobacillus</td><td>GCA_018854655.1</td><td>Acidithiobacillus_thiooxidans</td><td>HF670_09010</td><td>8231..11842</td><td>930</td><td>BREX-system-P-loop-protein-BrxC-[Acidithiobacillus-thiooxidans]</td></tr>
<tr><td>WP_101538034.1</td><td>Type1.BREX</td><td>DEDD_Tnp_IS110+Transposase_20-><-?||XIS-HTH->TOPRIM->||XIS-HTH-> BrxA-DNA-binding -> BrxB-BrxF-iSTAND -> DUF6079 *->Schlafen+HATPase+wHTH+HTH-> N6-METHYLASE -> HTH+ REase -> PglZ -> SIGMA+OB-Fold+AAA-ATPase+HKD-DNase-BrxL ->SbcC+APATPase+SbcC->Calcineurin+Calcineurin-SbcD->SbcC->RNase-></td><td>1204</td><td>Bacteria>Pseudomonadota>Acidithiobacillia>Acidithiobacillales>Acidithiobacillaceae</td><td>GCF_018854655.1</td><td>Acidithiobacillus</td><td>HF670_RS09000</td><td>8228..11842</td><td>119977</td><td>MULTISPECIES:-BREX-system-P-loop-protein-BrxC-[Acidithiobacillus]</td></tr>
<tr><td>WP_164594485.1</td><td>Type6.BREX</td><td><-TIR<- wHTH+WYL+WCX || REase +HTH+35exo+Small-B-barrel+SF1-SF2-Helicase->McrB-AAA-> McrC-NTD+ REase -><- SIGMA+OB-Fold+AAA-ATPase+Lon-Protease-BrxL <- PglZ <- N6-METHYLASE <- N6-METHYLASE <- DUF6079 *<- BrxB-BrxF-iSTAND <- BrxA-DNA-binding <- BrxE ||<-DOC+HTH||PhdYeFM_antitox-><-ZDNA-HTH+HTH<-ParB+RepB<-ParA-Soj-PloopNTPase</td><td>1178</td><td>Bacteria>Pseudomonadota>Alphaproteobacteria>Hyphomicrobiales>Rhizobiaceae>Rhizobium/Agrobacterium_group</td><td>-</td><td>Rhizobium_leguminosarum</td><td>GR223_RS33875</td><td>57482..61018</td><td>384</td><td>BREX-system-P-loop-protein-BrxC-[Rhizobium-leguminosarum]</td></tr>
<tr><td>WP_066770455.1</td><td>Type6.BREX</td><td>McrB-AAA-> McrC-NTD+ REase -><- SIGMA+OB-Fold+AAA-ATPase+Lon-Protease-BrxL <- PglZ <- N6-METHYLASE <- DUF6079 *<- BrxB-BrxF-iSTAND <- BrxA-DNA-binding <- BrxE <-APATPase<-tRNA<-REC+OmpR-HTH||FGase-></td><td>1176</td><td>Bacteria>Pseudomonadota>Alphaproteobacteria>Hyphomonadales>Hyphomonadaceae>Candidatus_Viadribacter</td><td>GCF_001679665.1</td><td>Candidatus_Viadribacter_manganicus</td><td>ATE48_RS09195</td><td>1777065..1780595</td><td>1759059</td><td>BREX-system-P-loop-protein-BrxC-[Candidatus-Viadribacter-manganicus]</td></tr>
<tr><td>WP_009810044.1</td><td>Type1.BREX</td><td><- HARE-HTH + REase <- SIGMA+OB-Fold+AAA-ATPase+HKD-DNase-BrxL <- PglZ <- N6-METHYLASE <- DUF6079 *<- BrxB-BrxF-iSTAND <- BrxA-DNA-binding <- wHTH+WYL+WCX ||?->?->?-><-RNase_T+35exo<-?||DOC+HTH-></td><td>1172</td><td>Bacteria>Pseudomonadota>Alphaproteobacteria>Rhodobacterales>Roseobacteraceae>Roseobacter</td><td>GCF_000152965.1</td><td>Roseobacter_sp._MED193</td><td>MED193_RS12240</td><td>2047846..2051364</td><td>314262</td><td>BREX-system-P-loop-protein-BrxC-[Roseobacter-sp.-MED193]</td></tr>
<tr><td>WP_085595207.1</td><td>Type1.BREX</td><td>XIS-HTH-> BrxA-DNA-binding -> BrxB-BrxF-iSTAND -> DUF6079 *-> OLD-ABC-> REase -> N6-METHYLASE -> HTH+ REase -> PglZ -> SIGMA+OB-Fold+AAA-ATPase+HKD-DNase-BrxL -><-SF1-SF2-Helicase</td><td>1182</td><td>Bacteria>Pseudomonadota>Alphaproteobacteria>Rhodospirillales>Thalassospiraceae>Thalassospira</td><td>GCF_002115825.1</td><td>Thalassospira_sp._MCCC_1A01428</td><td>THS27_RS23975</td><td>1729..5277</td><td>1470575</td><td>BREX-system-P-loop-protein-BrxC-[Thalassospira-sp.-MCCC-1A01428]</td></tr>
<tr><td>EXI84705.1</td><td>Type1.BREX</td><td>DDE_Tnp_1-><-?<-MqsA_antitoxin<-HTH<-?<-Arm-DNA-bind_3+Phage_integrase|| BrxA-DNA-binding -> BrxB-BrxF-iSTAND -> DUF6079 *-> N6-METHYLASE -> HTH+ REase -> PglZ -> SIGMA+OB-Fold+AAA-ATPase+Lon-Protease-BrxL ->PIN+HTH-></td><td>1204</td><td>Bacteria>Pseudomonadota>Betaproteobacteria>Betaproteobacteria_incertae_sedis>Candidatus_Accumulibacter</td><td>GCA_000585075.1</td><td>Candidatus_Accumulibacter_regalis</td><td>AW11_03760</td><td>6873..10487</td><td>522306</td><td>MAG:-hypothetical-protein-AW11_03760-[Candidatus-Accumulibacter-regalis]</td></tr>
<tr><td>WP_169070206.1</td><td>Type1.BREX</td><td>BrxA-DNA-binding -> BrxB-BrxF-iSTAND -> DUF6079 *->OLD-ABC->?->OLD-ABC->TOPRIM-> HTH+ REase ->?-> HTH+ REase ->?-> SIGMA+OB-Fold+AAA-ATPase+Lon-Protease-BrxL -></td><td>1204</td><td>Bacteria>Pseudomonadota>Betaproteobacteria>Betaproteobacteria_incertae_sedis>Candidatus_Accumulibacter</td><td>GCF_012940005.1</td><td>Candidatus_Accumulibacter_contiguus</td><td>E4Q08_RS09460</td><td>34303..37917</td><td>2954381</td><td>BREX-system-P-loop-protein-BrxC-[Candidatus-Accumulibacter-contiguus]</td></tr>
<tr><td>MBK7952886.1</td><td>Type1.BREX</td><td><-TOPRIM<-OLD-ABC<-?<-?<- REase <- HTH+ REase <- N6-METHYLASE <-AbiEii<-AbiEi_4<- DUF6079 *<- BrxB-BrxF-iSTAND <- BrxA-DNA-binding ||?->?-><-tRNA||GGDEF-></td><td>1207</td><td>Bacteria>Pseudomonadota>Betaproteobacteria>Betaproteobacteria_incertae_sedis>Candidatus_Accumulibacter>unclassified_Candidatus_Accumulibacter</td><td>GCA_016713625.1</td><td>Candidatus_Accumulibacter_sp.</td><td>IPK02_02340</td><td>220445..224068</td><td>2053492</td><td>BREX-system-P-loop-protein-BrxC-[Candidatus-Accumulibacter-sp.]</td></tr>
<tr><td>RIY39966.1</td><td>Type1.BREX</td><td>XIS-HTH-> BrxA-DNA-binding -> BrxB-BrxF-iSTAND -> DUF6079 *-> N6-METHYLASE ->PHP+SbcC->ParB+HNH-> PglZ -> SIGMA+OB-Fold+AAA-ATPase+Lon-Protease-BrxL -> HARE-HTH + REase ->?->SF1-SF2-Helicase-></td><td>1145</td><td>Bacteria>Pseudomonadota>Betaproteobacteria>Burkholderiales>Alcaligenaceae>Pusillimonas</td><td>GCA_003585905.1</td><td>Pusillimonas_maritima</td><td>CJP73_12365</td><td>2891..6328</td><td>2026239</td><td>hypothetical-protein-CJP73_12365-[Pusillimonas-maritima]</td></tr>
<tr><td>WP_218972830.1</td><td>Type1.BREX</td><td>XIS-HTH-> BrxA-DNA-binding -> BrxB-BrxF-iSTAND -> DUF6079 *-> N6-METHYLASE ->PHP+SbcC->ParB+HNH-> PglZ -> SIGMA+OB-Fold+AAA-ATPase+Lon-Protease-BrxL -> HARE-HTH + REase ->?->SF1-SF2-Helicase-></td><td>1146</td><td>Bacteria>Pseudomonadota>Betaproteobacteria>Burkholderiales>Alcaligenaceae>Pusillimonas</td><td>GCF_003585905.1</td><td>Pusillimonas_maritima</td><td>CJP73_RS12395</td><td>2888..6328</td><td>2026239</td><td>BREX-system-P-loop-protein-BrxC-[Pusillimonas-maritima]</td></tr>
<tr><td>WP_254334233.1</td><td>Type1.BREX</td><td><-SymE_toxin<-HTH_3<-?<-?<-?||Arm-DNA-bind_3+Phage_integrase+Phage_integr_3-><- SIGMA+OB-Fold+AAA-ATPase+Lon-Protease-BrxL <- PglZ <- N6-METHYLASE <- DUF6079 *<- BrxB-BrxF-iSTAND <- BrxA-DNA-binding <- McrC-NTD+ REase <-LZ_Tnp_IS66+zf-IS66+DDE_Tnp_IS66+DDE_Tnp_IS66_C<-?<-HIN-HTH||HEPN-></td><td>1210</td><td>Bacteria>Pseudomonadota>Betaproteobacteria>Burkholderiales>Burkholderiaceae>Paraburkholderia</td><td>GCF_024206475.1</td><td>Paraburkholderia_sp._CNPSo_3274</td><td>M3I54_RS07285</td><td>43504..47136</td><td>2940932</td><td>BREX-system-P-loop-protein-BrxC-[Paraburkholderia-sp.-CNPSo-3274]</td></tr>
<tr><td>TFI47144.1</td><td>Type1.BREX</td><td><-HTH+rve||?-><- SIGMA+OB-Fold+AAA-ATPase+HKD-DNase-BrxL <- PglZ <- N6-METHYLASE -p<- HTH+ REase <- DUF6079 *<- BrxB-BrxF-iSTAND <- BrxA-DNA-binding <- wHTH+WYL+WCX <-Phage_integrase+Phage_integrase<-?||HIN-HTH-></td><td>1188</td><td>Bacteria>Pseudomonadota>Betaproteobacteria>Burkholderiales>Comamonadaceae>Diaphorobacter</td><td>GCA_004563745.1</td><td>Diaphorobacter_sp._DS2</td><td>E4O93_14250</td><td>10625..14191</td><td>1302548</td><td>BREX-system-P-loop-protein-BrxC-[Diaphorobacter-sp.-DS2]</td></tr>
<tr><td>WP_104799767.1</td><td>Type1.BREX</td><td><- SIGMA+OB-Fold+AAA-ATPase+Lon-Protease-BrxL <- PglZ <-?<-?<- HTH+ REase <-TOPRIM<-OLD-ABC<- N6-METHYLASE <-?<- DUF6079 *<- BrxB-BrxF-iSTAND <- BrxA-DNA-binding <- wHTH+WYL+WCX <-tRNA||?-><-?<-ABhydrolase</td><td>1202</td><td>Bacteria>Pseudomonadota>Betaproteobacteria>Burkholderiales>Comamonadaceae>Limnohabitans</td><td>GCF_002943475.1</td><td>Limnohabitans_sp._TS-CS-82</td><td>C5F52_RS18160</td><td>199496..203104</td><td>2094193</td><td>BREX-system-P-loop-protein-BrxC-[Limnohabitans-sp.-TS-CS-82]</td></tr>
<tr><td>MBG6075367.1</td><td>Type1.BREX</td><td><-SWIM<- SIGMA+OB-Fold+AAA-ATPase+Lon-Protease-BrxL <- PglZ <- HTH+ REase <-DOC+HTH<-?<- N6-METHYLASE <-Couple_hipA+HipA_C<-HTH_3<- DUF6079 -p*||?-><- BrxB-BrxF-iSTAND <- BrxA-DNA-binding <-?<-?<-Phage_integrase||?->?->ABhydrolase+ABhydrolase-></td><td>921</td><td>Bacteria>Pseudomonadota>Betaproteobacteria>Burkholderiales>Comamonadaceae>Polaromonas</td><td>GCA_015752225.1</td><td>Polaromonas_sp._CG_9.11</td><td>IWX85_001192</td><td>30062..32827</td><td>2787730</td><td>hypothetical-protein-[Polaromonas-sp.-CG_9.11]</td></tr>
<tr><td>WP_231379566.1</td><td>Type1.BREX</td><td><-SWIM<- SIGMA+OB-Fold+AAA-ATPase+Lon-Protease-BrxL <- PglZ <- HTH+ REase <-DOC+HTH<-?<- N6-METHYLASE <-Couple_hipA+HipA_C<-HTH_3<- DUF6079 *<- BrxB-BrxF-iSTAND <- BrxA-DNA-binding <-?<-?<-Phage_integrase</td><td>1220</td><td>Bacteria>Pseudomonadota>Betaproteobacteria>Burkholderiales>Comamonadaceae>Polaromonas</td><td>GCF_015752225.1</td><td>Polaromonas_sp._CG_9.11</td><td>IWX85_RS05685</td><td>30062..33724</td><td>2787730</td><td>BREX-system-P-loop-protein-BrxC-[Polaromonas-sp.-CG_9.11]</td></tr>
<tr><td>MBN8486519.1</td><td>Type1.BREX</td><td><-SIR2||?-> wHTH+WYL+WCX -> BrxA-DNA-binding -> BrxB-BrxF-iSTAND -> DUF6079 *-> N6-METHYLASE ->OLD-ABC->NERD->RVT-> N6-METHYLASE -p-> HTH+ REase -> PglZ -> SIGMA+OB-Fold+AAA-ATPase+HKD-DNase-BrxL -><-HISKIN</td><td>1186</td><td>Bacteria>Pseudomonadota>Betaproteobacteria>Burkholderiales>unclassified_Burkholderiales</td><td>GCA_017302485.1</td><td>Burkholderiales_bacterium</td><td>J0M20_02125</td><td>4400..7960</td><td>1891238</td><td>BREX-system-P-loop-protein-BrxC-[Burkholderiales-bacterium]</td></tr>
<tr><td>WP_239907373.1</td><td>Type1.BREX</td><td><- SIGMA+OB-Fold+AAA-ATPase+HKD-DNase-BrxL <- PglZ <- HTH+ REase <- N6-METHYLASE -p<-HerA/FtsK<-SIR2<- REase <-OLD-ABC <- N6-METHYLASE <- DUF6079 *<- BrxB-BrxF-iSTAND <- BrxA-DNA-binding <- wHTH+WYL+WCX <-Phage_integrase+Phage_integrase<-HTH_36<-XIS-HTH<-?<-Arm-DNA-bind_3+Phage_integrase<-sGTP</td><td>1185</td><td>Bacteria>Pseudomonadota>Betaproteobacteria>Neisseriales>Chromobacteriaceae>Laribacter</td><td>GCF_022394255.1</td><td>Laribacter_hongkongensis</td><td>LH425_05855</td><td>38844..42401</td><td>168471</td><td>BREX-system-P-loop-protein-BrxC-[Laribacter-hongkongensis]</td></tr>
<tr><td>WP_013292857.1</td><td>Type1.BREX</td><td>VirD4-TraG+VirB4-FtsK->?->?->Peptidase_M50->?->?-><-?|| BrxA-DNA-binding -> BrxB-BrxF-iSTAND -> DUF6079 *->AbiEi_4->AbiEii-> N6-METHYLASE -> HTH+ REase -> REase ->?->OLD-ABC->TOPRIM->Schlafen+HATPase+wHTH-></td><td>1207</td><td>Bacteria>Pseudomonadota>Betaproteobacteria>Nitrosomonadales>Gallionellaceae>Gallionella</td><td>GCF_000145255.1</td><td>Gallionella_capsiferriformans</td><td>GALF_RS04415</td><td>928777..932400</td><td>370405</td><td>BREX-system-P-loop-protein-BrxC-[Gallionella-capsiferriformans]</td></tr>
<tr><td>ROH85757.1</td><td>Type1.BREX</td><td>Phage_holin->Phage_lysozyme->?-><- HARE-HTH + REase <- SIGMA+OB-Fold+AAA-ATPase+Lon-Protease-BrxL <- PglZ <-ParB+HNH<- N6-METHYLASE <- DUF6079 *<- BrxB-BrxF-iSTAND <- BrxA-DNA-binding <- wHTH+WYL+WCX ||JetA->JetB->SbcC->?-><-PIN<-HTH+MPTase</td><td>1146</td><td>Bacteria>Pseudomonadota>Betaproteobacteria>Nitrosomonadales>Methylophilaceae>Pseudomethylobacillus</td><td>GCA_003730115.1</td><td>Pseudomethylobacillus_aquaticus</td><td>ED236_08405</td><td>68439..71879</td><td>2676064</td><td>BREX-system-P-loop-protein-BrxC-[Pseudomethylobacillus-aquaticus]</td></tr>
<tr><td>WP_205125664.1</td><td>Type1.BREX</td><td>Phage_holin->Phage_lysozyme->?-><- HARE-HTH + REase <- SIGMA+OB-Fold+AAA-ATPase+Lon-Protease-BrxL <- PglZ <-ParB+HNH<- N6-METHYLASE <- DUF6079 *<- BrxB-BrxF-iSTAND <- BrxA-DNA-binding <- wHTH+WYL+WCX ||JetA->JetB->SbcC->?-><-PIN<-HTH+MPTase</td><td>1150</td><td>Bacteria>Pseudomonadota>Betaproteobacteria>Nitrosomonadales>Methylophilaceae>Pseudomethylobacillus</td><td>GCF_003730115.1</td><td>Pseudomethylobacillus_aquaticus</td><td>ED236_RS08405</td><td>68439..71891</td><td>2676064</td><td>BREX-system-P-loop-protein-BrxC-[Pseudomethylobacillus-aquaticus]</td></tr>
<tr><td>MBK7491769.1</td><td>Type1.BREX</td><td>HIN-HTH->?-><-SF1-SF2-Helicase<-?||?->DUF772+DDE_Tnp_1->PIN+OST-HTH-> BrxA-DNA-binding -> BrxB-BrxF-iSTAND -> DUF6079 *-> N6-METHYLASE ->HEPN-> HTH+ REase -> PglZ -> SIGMA+OB-Fold+AAA-ATPase+Lon-Protease-BrxL ->HIN-HTH-></td><td>1203</td><td>Bacteria>Pseudomonadota>Betaproteobacteria>Nitrosomonadales>Nitrosomonadaceae>Nitrosomonas</td><td>GCA_016707165.1</td><td>Nitrosomonas_sp.</td><td>IPI17_07005</td><td>52164..55775</td><td>42353</td><td>BREX-system-P-loop-protein-BrxC-[Nitrosomonas-sp.]</td></tr>
<tr><td>MBX9637025.1</td><td>Type1.BREX</td><td>GDResolvase+HIN-HTH->?-> BrxA-DNA-binding -> BrxB-BrxF-iSTAND -> DUF6079 *-> N6-METHYLASE -> HTH+ REase -> PglZ -> SIGMA+OB-Fold+AAA-ATPase+Lon-Protease-BrxL ->SIR2->HerA/FtsK->Calcineurin-SbcD->SbcC+CCDC158-></td><td>1212</td><td>Bacteria>Pseudomonadota>Betaproteobacteria>Nitrosomonadales>Nitrosomonadaceae>Nitrosomonas</td><td>GCA_019748355.1</td><td>Nitrosomonas_sp.</td><td>K2Q45_05685</td><td>2429..6067</td><td>42353</td><td>BREX-system-P-loop-protein-BrxC-[Nitrosomonas-sp.]</td></tr>
<tr><td>MCB1936159.1</td><td>Type1.BREX</td><td><- HTH+ REase <- N6-METHYLASE <- REase <-ABC-ATPase<- DUF6079 *<- BrxB-BrxF-iSTAND <- BrxA-DNA-binding ||HEPN+OST-HTH-> REase +TOPRIM-></td><td>1174</td><td>Bacteria>Pseudomonadota>Betaproteobacteria>Nitrosomonadales>Nitrosomonadaceae>Nitrosomonas</td><td>GCA_020446045.1</td><td>Nitrosomonas_sp.</td><td>KDF59_09470</td><td>5848..9372</td><td>42353</td><td>BREX-system-P-loop-protein-BrxC-[Nitrosomonas-sp.]</td></tr>
<tr><td>WP_204800510.1</td><td>Type1.BREX</td><td><-SbcC<-Calcineurin+Calcineurin-SbcD<-SbcC<- SIGMA+OB-Fold+AAA-ATPase+HKD-DNase-BrxL <- PglZ <- HTH+ REase <-?<-Schlafen+HATPase+wHTH+HTH<- DUF6079 *<- BrxB-BrxF-iSTAND <- BrxA-DNA-binding <-XIS-HTH<-?<-Arm-DNA-bind_3+Phage_integrase</td><td>1180</td><td>Bacteria>Pseudomonadota>Betaproteobacteria>Nitrosomonadales>Nitrosomonadaceae>Nitrosomonas</td><td>GCF_905220645.1</td><td>Nitrosomonas_nitrosa</td><td>NUZ5A_RS15530</td><td>137140..140682</td><td>52442</td><td>BREX-system-P-loop-protein-BrxC-[Nitrosomonas-nitrosa]</td></tr>
<tr><td>OGS68620.1</td><td>Type1.BREX</td><td>VirD4-TraG+VirB4-FtsK->?->?->Peptidase_M50->?->?-><-?|| BrxA-DNA-binding -> BrxB-BrxF-iSTAND -> DUF6079 *->AbiEi_4->AbiEii-> N6-METHYLASE -> HTH+ REase ->?-> PglZ -> SIGMA+OB-Fold+AAA-ATPase+Lon-Protease-BrxL ->Classical-AAA->Peptidase_S8-></td><td>1207</td><td>Bacteria>Pseudomonadota>Betaproteobacteria>Nitrosomonadales>unclassified_Nitrosomonadales</td><td>GCA_001801025.1</td><td>Gallionellales_bacterium_GWA2_54_124</td><td>A2Z87_08865</td><td>78784..82407</td><td>1798243</td><td>ATPase-[Gallionellales-bacterium-GWA2_54_124]</td></tr>
<tr><td>MBK9161130.1</td><td>Type1.BREX</td><td>Phage_holin->Phage_lysozyme->?-><- HARE-HTH + REase <- SIGMA+OB-Fold+AAA-ATPase+Lon-Protease-BrxL <- PglZ <-ParB+HNH<- N6-METHYLASE <- DUF6079 *<- BrxB-BrxF-iSTAND <- BrxA-DNA-binding <-GreA_GreB+SF1-SF2-Helicase<-SLOG+BirA-HTH<-SF1-SF2-Helicase+PRTase<-XIS-HTH<-?<-ARG</td><td>1146</td><td>Bacteria>Pseudomonadota>Betaproteobacteria>Nitrosomonadales>unclassified_Nitrosomonadales</td><td>GCA_016716325.1</td><td>Nitrosomonadales_bacterium</td><td>IPM27_06155</td><td>1292283..1295723</td><td>2026761</td><td>BREX-system-P-loop-protein-BrxC-[Nitrosomonadales-bacterium]</td></tr>
<tr><td>OHC72318.1</td><td>Type1.BREX</td><td><-Integrase_1|| BrxA-DNA-binding -> BrxB-BrxF-iSTAND -> DUF6079 *-> N6-METHYLASE ->GrpE->?-> HTH+ REase -> PglZ -> SIGMA+OB-Fold+AAA-ATPase+Lon-Protease-BrxL -><-TPR+TPR+TPR+TPR+TPR+TPR+TPR<-CR-Kinase<- REase</td><td>1152</td><td>Bacteria>Pseudomonadota>Betaproteobacteria>Rhodocyclales>unclassified_Rhodocyclales</td><td>GCA_001828935.1</td><td>Rhodocyclales_bacterium_RIFCSPLOWO2_02_FULL_63_24</td><td>A3H93_14585</td><td>39152..42610</td><td>1802018</td><td>MAG:-hypothetical-protein-A3H93_14585-[Rhodocyclales-bacterium-RIFCSPLOWO2_02_FULL_63_24]</td></tr>
<tr><td>WP_108973925.1</td><td>Type1.BREX</td><td>RNase->TPR->NERD+SF1-SF2-Helicase-><-SF1-SF2-Helicase<-MORC+DUF3684+ REase ||DUF772+DDE_Tnp_1+DDE_Tnp_1_6-> wHTH+WYL+WCX -> BrxA-DNA-binding -> BrxB-BrxF-iSTAND -> DUF6079 *-> N6-METHYLASE ->DUF3644-> HTH+ REase ->?-> PglZ ->VapB_antitoxin->PIN-> SIGMA+OB-Fold+AAA-ATPase+Lon-Protease-BrxL -><-Phage_integrase</td><td>1215</td><td>Bacteria>Pseudomonadota>Betaproteobacteria>Rhodocyclales>Zoogloeaceae>Parazoarcus</td><td>GCF_003111665.1</td><td>Parazoarcus_communis</td><td>CEW87_RS14060</td><td>3075229..3078876</td><td>41977</td><td>BREX-system-P-loop-protein-BrxC-[Parazoarcus-communis]</td></tr>
<tr><td>WP_068639549.1</td><td>Type1.BREX</td><td><-HEPN<- HTH+ REase <- N6-METHYLASE <- DUF6079 *<- BrxB-BrxF-iSTAND <- BrxA-DNA-binding <-XIS-HTH</td><td>1188</td><td>Bacteria>Pseudomonadota>Betaproteobacteria>Rhodocyclales>Zoogloeaceae>Thauera</td><td>GCF_001591165.1</td><td>Thauera_butanivorans</td><td>TBU01S_RS21545</td><td>4844..8410</td><td>86174</td><td>BREX-system-P-loop-protein-BrxC-[Thauera-butanivorans]</td></tr>
<tr><td>WP_141354347.1</td><td>Type1.BREX</td><td>wHTH+WYL+WCX -> BrxA-DNA-binding -> BrxB-BrxF-iSTAND -> DUF6079 *-><-PIN<-RHH_6|| N6-METHYLASE -p->HKD-DNase-> N6-METHYLASE -> HTH+ REase ->ZnR+TOPRIM-> PglZ -> SIGMA+OB-Fold+AAA-ATPase+Lon-Protease-BrxL -></td><td>1203</td><td>Bacteria>Pseudomonadota>Betaproteobacteria>Rhodocyclales>Zoogloeaceae>Zoogloea</td><td>GCF_006539865.1</td><td>Zoogloea_ramigera</td><td>ZRA01_RS16820</td><td>2482..6093</td><td>350</td><td>BREX-system-P-loop-protein-BrxC-[Zoogloea-ramigera]</td></tr>
<tr><td>WP_148578091.1</td><td>Type1.BREX</td><td>Arm-DNA-bind_2+Phage_int_SAM_3+Phage_integrase->HDRnase->?-><-HTH_19||?->?-> wHTH+WYL+WCX -> BrxA-DNA-binding -> BrxB-BrxF-iSTAND -> DUF6079 *-> N6-METHYLASE -> OLD-ABC-> REase ->SIR2->HerA/FtsK-> N6-METHYLASE -p-> HTH+ REase -> PglZ -> SIGMA+OB-Fold+AAA-ATPase+HKD-DNase-BrxL -></td><td>1186</td><td>Bacteria>Pseudomonadota>Betaproteobacteria>Rhodocyclales>Zoogloeaceae>Zoogloea</td><td>GCF_008107625.1</td><td>Zoogloea_oleivorans</td><td>ETQ85_RS05700</td><td>193093..196653</td><td>1552750</td><td>BREX-system-P-loop-protein-BrxC-[Zoogloea-oleivorans]</td></tr>
<tr><td>WP_197117904.1</td><td>Type1.BREX</td><td><-MarR-HTH||?-><-?<-?|| BrxA-DNA-binding -> BrxB-BrxF-iSTAND -> DUF6079 *->GP+ REase ->SF1-SF2-Helicase+PRTase->SLOG+BirA-HTH-> N6-METHYLASE ->DnaB+McrB-AAA-> McrC-NTD+ REase +ZnR->ParB+HNH->ZnR+3C-ABC+TPR-></td><td>1229</td><td>Bacteria>Pseudomonadota>Gammaproteobacteria>Alteromonadales>Pseudoalteromonadaceae>Pseudoalteromonas</td><td>GCF_015864765.1</td><td>Pseudoalteromonas_sp._NZS11</td><td>I6E82_RS11050</td><td>319401..323090</td><td>2792049</td><td>BREX-system-P-loop-protein-BrxC-[Pseudoalteromonas-sp.-NZS11]</td></tr>
<tr><td>MBP6733459.1</td><td>Type1.BREX</td><td>BrxA-DNA-binding -> BrxB-BrxF-iSTAND -> DUF6079 *-><- BrxB-BrxF-iSTAND ||?->PhdYeFM_antitox->PIN-> N6-METHYLASE -> HTH+ REase -> REase -> N6-METHYLASE -p-></td><td>1171</td><td>Bacteria>Pseudomonadota>Gammaproteobacteria>Chromatiales>Chromatiaceae>unclassified_Chromatiaceae</td><td>GCA_017990755.1</td><td>Chromatiaceae_bacterium</td><td>KA142_01990</td><td>1791..5306</td><td>2049432</td><td>BREX-system-P-loop-protein-BrxC-[Chromatiaceae-bacterium]</td></tr>
<tr><td>WP_123923007.1</td><td>Type1.BREX</td><td>TOPRIM-DNAG+D5N+D5ATPase+P4PRIMASE-HTH-> BrxB-BrxF-iSTAND ->ACET->?-> BrxA-DNA-binding -> BrxB-BrxF-iSTAND -> DUF6079 *-> N6-METHYLASE ->OLD-ABC->TOPRIM-> PglZ -> SIGMA+OB-Fold+AAA-ATPase+Lon-Protease-BrxL -><- McrC-NTD+ REase ||RVT-><-Glyoxalase</td><td>1204</td><td>Bacteria>Pseudomonadota>Gammaproteobacteria>Enterobacterales>Enterobacteriaceae>Enterobacter</td><td>GCF_003812145.1</td><td>Enterobacter_roggenkampii</td><td>EGY04_RS13705</td><td>2428639..2432253</td><td>1812935</td><td>BREX-system-P-loop-protein-BrxC-[Enterobacter-roggenkampii]</td></tr>
<tr><td>WP_248053563.1</td><td>Type1.BREX</td><td><-ZnR|| BrxA-DNA-binding -> BrxB-BrxF-iSTAND -> DUF6079 *-> N6-METHYLASE ->OLD-ABC->TOPRIM-> PglZ -> SIGMA+OB-Fold+AAA-ATPase+Lon-Protease-BrxL -><- McrC-NTD+ REase <-McrB-AAA||NACHT+SF1-SF2-Helicase+HKD-DNase-><-wHTH+SAD+HNH</td><td>1197</td><td>Bacteria>Pseudomonadota>Gammaproteobacteria>Enterobacterales>Enterobacteriaceae>Enterobacter</td><td>GCF_023154935.1</td><td>Enterobacter_chengduensis</td><td>L8P32_RS20960</td><td>6466..10059</td><td>2494701</td><td>BREX-system-P-loop-protein-BrxC-[Enterobacter-chengduensis]</td></tr>
<tr><td>WP_258247898.1</td><td>Type1.BREX</td><td><-ZnR|| BrxA-DNA-binding -> BrxB-BrxF-iSTAND -> DUF6079 *-> N6-METHYLASE ->OLD-ABC->TOPRIM-> PglZ -> SIGMA+OB-Fold+AAA-ATPase+Lon-Protease-BrxL -><- McrC-NTD+ REase <-McrB-AAA||NACHT+SF1-SF2-Helicase+HKD-DNase-><-wHTH+SAD+HNH</td><td>1204</td><td>Bacteria>Pseudomonadota>Gammaproteobacteria>Enterobacterales>Enterobacteriaceae>Enterobacter</td><td>GCF_024733415.1</td><td>Enterobacter_hormaechei</td><td>NUJ48_RS00150</td><td>29183..32797</td><td>158836</td><td>BREX-system-P-loop-protein-BrxC-[Enterobacter-hormaechei]</td></tr>
<tr><td>WP_060426782.1</td><td>Type1.BREX</td><td><-RNase||?->?->SAD+HNH-> HTH+ REase -> BrxA-DNA-binding -> BrxB-BrxF-iSTAND -> DUF6079 *-> N6-METHYLASE -> PglZ -> SIGMA+OB-Fold+AAA-ATPase+Lon-Protease-BrxL ->HKD-DNase+OGG-like-><-?<-?<-tRNA||?->DEOR-HTH+ISOCOT-></td><td>1213</td><td>Bacteria>Pseudomonadota>Gammaproteobacteria>Enterobacterales>Yersiniaceae</td><td>GCF_001537885.1</td><td>Serratia</td><td>AWX06_RS15845</td><td>182718..186359</td><td>613</td><td>MULTISPECIES:-BREX-system-P-loop-protein-BrxC-[Serratia]</td></tr>
<tr><td>WP_089185250.1</td><td>Type1.BREX</td><td><-Glyoxalase<-?<-?<-?||tRNA->Arm-DNA-bind_3+Phage_integrase-><-?<- SIGMA+OB-Fold+AAA-ATPase+Lon-Protease-BrxL <- PglZ <- N6-METHYLASE <- DUF6079 *<- BrxB-BrxF-iSTAND <- BrxA-DNA-binding <- HTH+ REase <-SAD+HNH<-?||?->?->?-><-Glyco_hydro_1</td><td>1213</td><td>Bacteria>Pseudomonadota>Gammaproteobacteria>Enterobacterales>Yersiniaceae>Serratia</td><td>GCF_002220535.1</td><td>Serratia_marcescens</td><td>BVG90_RS23850</td><td>5002559..5006200</td><td>615</td><td>BREX-system-P-loop-protein-BrxC-[Serratia-marcescens]</td></tr>
<tr><td>WP_197785670.1</td><td>Type1.BREX</td><td>Glyco_hydro_1-><-?<-?<-?||?->?->SAD+HNH-> HTH+ REase -> BrxA-DNA-binding -> BrxB-BrxF-iSTAND -> DUF6079 *-> N6-METHYLASE -> PglZ -> SIGMA+OB-Fold+AAA-ATPase+Lon-Protease-BrxL ->HKD-DNase+OGG-like-></td><td>1213</td><td>Bacteria>Pseudomonadota>Gammaproteobacteria>Enterobacterales>Yersiniaceae>Serratia</td><td>GCF_016007555.1</td><td>Serratia_ureilytica</td><td>I5P92_RS12860</td><td>24369..28010</td><td>300181</td><td>BREX-system-P-loop-protein-BrxC-[Serratia-ureilytica]</td></tr>
<tr><td>WP_222398282.1</td><td>Type1.BREX</td><td>Glyco_hydro_1-><-?<-?<-?||?->?->SAD+HNH-> HTH+ REase -> BrxA-DNA-binding -> BrxB-BrxF-iSTAND -> DUF6079 *-> N6-METHYLASE -> PglZ -> SIGMA+OB-Fold+AAA-ATPase+Lon-Protease-BrxL ->?-><-Arm-DNA-bind_3+Phage_integrase<-tRNA||?->?->?->Glyoxalase-></td><td>1213</td><td>Bacteria>Pseudomonadota>Gammaproteobacteria>Enterobacterales>Yersiniaceae>Serratia</td><td>GCF_019795995.1</td><td>Serratia_marcescens</td><td>L8N14_16510</td><td>93200..96841</td><td>615</td><td>BREX-system-P-loop-protein-BrxC-[Serratia-marcescens]</td></tr>
<tr><td>WP_244757920.1</td><td>Type1.BREX</td><td><-HKD-DNase+OGG-like<- SIGMA+OB-Fold+AAA-ATPase+Lon-Protease-BrxL <- PglZ <- N6-METHYLASE <- DUF6079 *<- BrxB-BrxF-iSTAND <- BrxA-DNA-binding <- HTH+ REase <-SAD+HNH<-?||?->?-><-Glyco_hydro_1</td><td>1213</td><td>Bacteria>Pseudomonadota>Gammaproteobacteria>Enterobacterales>Yersiniaceae>Serratia</td><td>GCF_022870585.1</td><td>Serratia_marcescens</td><td>HW877_RS02460</td><td>535877..539518</td><td>615</td><td>BREX-system-P-loop-protein-BrxC-[Serratia-marcescens]</td></tr>
<tr><td>WP_248566606.1</td><td>Type1.BREX</td><td>SF1-SF2-Helicase+ REase ->?-><- SIGMA+OB-Fold+AAA-ATPase+Lon-Protease-BrxL <- PglZ <-TOPRIM<-OLD-ABC<- HTH+ REase <- N6-METHYLASE <-Schlafen+HATPase+wHTH<- DUF6079 *<- BrxB-BrxF-iSTAND <- BrxA-DNA-binding <-SF1-SF2-Helicase<-HKD-DNase<-DUF6361<-HEPN||?->URI-DNase-></td><td>1145</td><td>Bacteria>Pseudomonadota>Gammaproteobacteria>Gammaproteobacteria_incertae_sedis>Methylonatrum</td><td>GCF_023195885.1</td><td>Methylonatrum_kenyense</td><td>M0534_RS01350</td><td>290769..294206</td><td>455253</td><td>BREX-system-P-loop-protein-BrxC-[Methylonatrum-kenyense]</td></tr>
<tr><td>MAZ66828.1</td><td>Type1.BREX</td><td><- SIGMA+OB-Fold+AAA-ATPase+Lon-Protease-BrxL <- PglZ <-TOPRIM<-OLD-ABC<- N6-METHYLASE <- HTH+ REase <- DUF6079 *<- BrxB-BrxF-iSTAND <- BrxA-DNA-binding <- wHTH+WYL+WCX <-Trypsin+PDZ</td><td>1234</td><td>Bacteria>Pseudomonadota>Gammaproteobacteria>Kangiellales>Kangiellaceae>unclassified_Kangiellaceae</td><td>GCA_002707725.1</td><td>Kangiellaceae_bacterium</td><td>CMF25_06935</td><td>12477..16181</td><td>2026751</td><td>DNA-repair-protein-[Kangiellaceae-bacterium]</td></tr>
<tr><td>MBS3952424.1</td><td>Type1.BREX</td><td>XIS-HTH-> BrxA-DNA-binding -> BrxB-BrxF-iSTAND -> DUF6079 *-> OLD-ABC-> REase -> N6-METHYLASE ->PhdYeFM_antitox->PIN-> N6-METHYLASE -p-> HTH+ REase ->Schlafen+HATPase+wHTH->SIR2-></td><td>1178</td><td>Bacteria>Pseudomonadota>Gammaproteobacteria>Methylococcales>Methylococcaceae>Methylomicrobium</td><td>GCA_018333785.1</td><td>Methylomicrobium_sp.</td><td>KGZ88_05695</td><td>2093..5629</td><td>2303549</td><td>BREX-system-P-loop-protein-BrxC-[Methylomicrobium-sp.]</td></tr>
<tr><td>WP_256360610.1</td><td>Type1.BREX</td><td><- SIGMA+OB-Fold+AAA-ATPase+HKD-DNase-BrxL <- PglZ <- REase <- REase <- HTH+ REase <- N6-METHYLASE -p<- N6-METHYLASE -p<- N6-METHYLASE -p<- N6-METHYLASE -p<- DUF6079 *</td><td>1145</td><td>Bacteria>Pseudomonadota>Gammaproteobacteria>Methylococcales>Methylococcaceae>Methylomonas</td><td>GCF_001312005.1</td><td>Methylomonas_koyamae</td><td>JCM16701_RS06940</td><td>49293..52730</td><td>702114</td><td>BREX-system-P-loop-protein-BrxC-[Methylomonas-koyamae]</td></tr>
<tr><td>WP_096876853.1</td><td>Type1.BREX</td><td>GDResolvase+HIN-HTH->?->XIS-HTH-> BrxA-DNA-binding -> BrxB-BrxF-iSTAND -> DUF6079 *-> OLD-ABC-> REase -> N6-METHYLASE -> HTH+ REase -> PglZ -> SIGMA+OB-Fold+AAA-ATPase+HKD-DNase-BrxL ->Calcineurin-SbcD->SbcC+CCDC158->SbcC-></td><td>1180</td><td>Bacteria>Pseudomonadota>Gammaproteobacteria>Methylococcales>Methylococcaceae>Methylomonas</td><td>GCF_002441955.1</td><td>Methylomonas_koyamae</td><td>MKLM6_RS22285</td><td>58076..61618</td><td>702114</td><td>BREX-system-P-loop-protein-BrxC-[Methylomonas-koyamae]</td></tr>
<tr><td>WP_221054675.1</td><td>Type1.BREX</td><td>DUF6079 *-> N6-METHYLASE -> HTH+ REase -> REase -> PglZ -> SIGMA+OB-Fold+AAA-ATPase+HKD-DNase-BrxL ->Calcineurin-SbcD->SbcC+CCDC158->OLD-ABC+TOPRIM->SF1-SF2-Helicase-></td><td>1176</td><td>Bacteria>Pseudomonadota>Gammaproteobacteria>Methylococcales>Methylococcaceae>Methylomonas</td><td>GCF_019669905.1</td><td>Methylomonas_koyamae</td><td>MKFW12EY_RS21895</td><td>239254..1338</td><td>702114</td><td>BREX-system-P-loop-protein-BrxC-[Methylomonas-koyamae]</td></tr>
<tr><td>MBL1268350.1</td><td>Type1.BREX</td><td><- wHTH+WYL+WCX <-?<-?||JAB->?-> BrxA-DNA-binding -> BrxB-BrxF-iSTAND -> DUF6079 *-> HTH+ REase -> N6-METHYLASE ->?->S1COLD-></td><td>1236</td><td>Bacteria>Pseudomonadota>Gammaproteobacteria>Oceanospirillales>Halomonadaceae>Halomonas</td><td>GCA_002733435.2</td><td>Halomonas_sp.</td><td>COA87_011520</td><td>231341..235051</td><td>1486246</td><td>BREX-system-P-loop-protein-BrxC-[Halomonas-sp.]</td></tr>
<tr><td>NQY75752.1</td><td>Type1.BREX</td><td><-SbcC<-?<-?<- SIGMA+OB-Fold+AAA-ATPase+Lon-Protease-BrxL <- PglZ <-ParB+HNH<-HerA/FtsK<-SIR2<- HTH+ REase <- DUF6079 *<- BrxB-BrxF-iSTAND <- BrxA-DNA-binding <-HTH_19<-TnsC-ATPase<-rve+Mu-transpos_C<- REase +wHTH<-ncRNA<-tRNA<-?||TetR-HTH-></td><td>1227</td><td>Bacteria>Pseudomonadota>Gammaproteobacteria>Oceanospirillales>Halomonadaceae>Halomonas</td><td>GCA_013213865.1</td><td>Halomonas_sp.</td><td>HRT78_01215</td><td>164704..168387</td><td>1486246</td><td>BREX-system-P-loop-protein-BrxC-[Halomonas-sp.]</td></tr>
<tr><td>WP_009095648.1</td><td>Type1.BREX</td><td><-TetR-HTH||?->tRNA->ncRNA->Tn7_Tnp_TnsA_N+rve+Mu-transpos_C->TnsC-ATPase->HTH_19-> BrxA-DNA-binding -> BrxB-BrxF-iSTAND -> DUF6079 *-> HTH+ REase ->Schlafen+HATPase+wHTH-> N6-METHYLASE ->sGTP->?-> N6-METHYLASE -p->NERD+GspA-ATPase->TPR->ParB+HNH-></td><td>1225</td><td>Bacteria>Pseudomonadota>Gammaproteobacteria>Oceanospirillales>Halomonadaceae>Halomonas</td><td>GCF_000236625.2</td><td>Halomonas_sp._GFAJ-1</td><td>BB497_RS11605</td><td>2566793..2570470</td><td>1118153</td><td>BREX-system-P-loop-protein-BrxC-[Halomonas-sp.-GFAJ-1]</td></tr>
<tr><td>WP_027962108.1</td><td>Type1.BREX</td><td>TraG_N-><-?||?-><- SIGMA+OB-Fold+AAA-ATPase+Lon-Protease-BrxL <- PglZ <- HTH+ REase <- N6-METHYLASE <-Schlafen+HATPase+wHTH<- DUF6079 *<- BrxB-BrxF-iSTAND <- BrxA-DNA-binding <-XIS-HTH||?->ARG-><- wHTH+WYL+WCX</td><td>1144</td><td>Bacteria>Pseudomonadota>Gammaproteobacteria>Oceanospirillales>Halomonadaceae>Halomonas</td><td>GCF_000620045.1</td><td>Halomonas_halodenitrificans</td><td>Q358_RS0111245</td><td>19354..22788</td><td>28252</td><td>BREX-system-P-loop-protein-BrxC-[Halomonas-halodenitrificans]</td></tr>
<tr><td>WP_078087405.1</td><td>Type1.BREX</td><td><-TetR-HTH||tRNA->ncRNA-> REase +wHTH->rve+Mu-transpos_C->TnsC-ATPase->HTH_19-> BrxA-DNA-binding -> BrxB-BrxF-iSTAND -> DUF6079 *-> HTH+ REase ->Schlafen+HATPase+wHTH-> N6-METHYLASE ->?->LRR+LRR->Pkinase-> PglZ -> SIGMA+OB-Fold+AAA-ATPase+Lon-Protease-BrxL -></td><td>1225</td><td>Bacteria>Pseudomonadota>Gammaproteobacteria>Oceanospirillales>Halomonadaceae>Halomonas</td><td>GCF_002009255.1</td><td>unclassified_Halomonas</td><td>BV504_RS06350</td><td>1459939..1463616</td><td>2609666</td><td>MULTISPECIES:-BREX-system-P-loop-protein-BrxC-[unclassified-Halomonas]</td></tr>
<tr><td>WP_083023778.1</td><td>Type1.BREX</td><td>REase +wHTH-> REase +wHTH->rve+Mu-transpos_C->TnsC-ATPase->HTH_19-> BrxA-DNA-binding -> BrxB-BrxF-iSTAND -> DUF6079 *-> HTH+ REase -> N6-METHYLASE ->SIR2->HerA/FtsK->ParB+HNH-></td><td>1227</td><td>Bacteria>Pseudomonadota>Gammaproteobacteria>Oceanospirillales>Halomonadaceae>Halomonas</td><td>GCF_002087295.1</td><td>Halomonas_lionensis</td><td>B4967_RS04180</td><td>22810..26493</td><td>1144478</td><td>BREX-system-P-loop-protein-BrxC-[Halomonas-lionensis]</td></tr>
<tr><td>WP_273192082.1</td><td>Type1.BREX</td><td><- wHTH+WYL+WCX <-?<-?||JAB->?-> BrxA-DNA-binding -> BrxB-BrxF-iSTAND -> DUF6079 *-> HTH+ REase -> N6-METHYLASE ->?->S1COLD-></td><td>1236</td><td>Bacteria>Pseudomonadota>Gammaproteobacteria>Oceanospirillales>Halomonadaceae>Halomonas</td><td>GCF_002733435.1</td><td>Halomonas_sp.</td><td>COA87_011520</td><td>231341..235051</td><td>1486246</td><td>BREX-system-P-loop-protein-BrxC-[Halomonas-sp.]</td></tr>
<tr><td>WP_172420975.1</td><td>Type1.BREX</td><td><-TOPRIM<-OLD-ABC<-?<-?<-?<- N6-METHYLASE <-HATPase+wHTH<-Schlafen+HATPase+wHTH-p<- HTH+ REase <- DUF6079 *<- BrxB-BrxF-iSTAND <- BrxA-DNA-binding <-ncRNA<-tRNA<-?||TetR-HTH->?->?->?->ABC-ATPase+SbcC->ABC-ATPase+SbcC-></td><td>1227</td><td>Bacteria>Pseudomonadota>Gammaproteobacteria>Oceanospirillales>Halomonadaceae>Halomonas</td><td>GCF_011399095.1</td><td>Halomonas_hydrothermalis</td><td>HHSLTHF2_RS10320</td><td>2155809..2159492</td><td>115561</td><td>BREX-system-P-loop-protein-BrxC-[Halomonas-hydrothermalis]</td></tr>
<tr><td>WP_179915949.1</td><td>Type1.BREX</td><td><-SbcC<-?<-?<- SIGMA+OB-Fold+AAA-ATPase+Lon-Protease-BrxL <-ParB+HNH<-HerA/FtsK<-SIR2<- N6-METHYLASE <- HTH+ REase <- DUF6079 *<- BrxB-BrxF-iSTAND <- BrxA-DNA-binding <-HTH_19<-TnsC-ATPase<-rve+Mu-transpos_C<- REase +wHTH<-ncRNA<-tRNA<-?||TetR-HTH-></td><td>1227</td><td>Bacteria>Pseudomonadota>Gammaproteobacteria>Oceanospirillales>Halomonadaceae>Halomonas</td><td>GCF_013415125.1</td><td>Halomonas_glaciei</td><td>HZS80_RS09710</td><td>42867..46550</td><td>186761</td><td>BREX-system-P-loop-protein-BrxC-[Halomonas-glaciei]</td></tr>
<tr><td>WP_231664595.1</td><td>Type1.BREX</td><td>GDResolvase+HIN-HTH->?-><- SIGMA+OB-Fold+AAA-ATPase+Lon-Protease-BrxL <- PglZ <-TOPRIM<-OLD-ABC<- HTH+ REase <- N6-METHYLASE <-Schlafen+HATPase+wHTH<- DUF6079 *<- BrxB-BrxF-iSTAND -p<- BrxA-DNA-binding <-XIS-HTH<-MobI</td><td>1153</td><td>Bacteria>Pseudomonadota>Gammaproteobacteria>Oceanospirillales>Halomonadaceae>Halomonas</td><td>GCF_021151495.1</td><td>Halomonas_sp._IOP_31</td><td>K7I12_RS15025</td><td>26514..29975</td><td>2876584</td><td>BREX-system-P-loop-protein-BrxC-[Halomonas-sp.-IOP_31]</td></tr>
<tr><td>WP_110649043.1</td><td>Type1.BREX</td><td><-ICLR-HTH<-?||?->Amidohydro_1->GNTR-HTH->XIS-HTH-> BrxA-DNA-binding -> BrxB-BrxF-iSTAND -> DUF6079 *->Schlafen+HATPase+wHTH-> N6-METHYLASE -> HTH+ REase -><-DDE_Tnp_1_4</td><td>1153</td><td>Bacteria>Pseudomonadota>Gammaproteobacteria>Oceanospirillales>Halomonadaceae>Salinicola</td><td>GCF_003206715.1</td><td>Salinicola_peritrichatus</td><td>DOM57_RS09900</td><td>59337..62798</td><td>1267424</td><td>BREX-system-P-loop-protein-BrxC-[Salinicola-peritrichatus]</td></tr>
<tr><td>WP_036163554.1</td><td>Type1.BREX</td><td><- SIGMA+OB-Fold+AAA-ATPase+Lon-Protease-BrxL <- PglZ <-GP+ REase <- McrC-NTD+ REase <-McrB-AAA<- N6-METHYLASE <-SLOG+BirA-HTH<-SF1-SF2-Helicase+PRTase<- DUF6079 *<- BrxB-BrxF-iSTAND <- BrxA-DNA-binding <-ABhydrolase<-HTH_26||Phage_AlpA->Arm-DNA-bind_2+Phage_int_SAM_3+Phage_integrase->?-><-?<-LexA-protease</td><td>1246</td><td>Bacteria>Pseudomonadota>Gammaproteobacteria>Oceanospirillales>Oceanospirillaceae>Marinomonas</td><td>GCF_000521805.1</td><td>Marinomonas_ushuaiensis</td><td>MUS1_RS14035</td><td>47481..51221</td><td>263818</td><td>BREX-system-P-loop-protein-BrxC-[Marinomonas-ushuaiensis]</td></tr>
<tr><td>MBE0507005.1</td><td>Type1.BREX</td><td>BrxA-DNA-binding -> BrxB-BrxF-iSTAND -> DUF6079 *-> OLD-ABC-> REase -> N6-METHYLASE -> HTH+ REase -> REase -> PglZ -> SIGMA+OB-Fold+AAA-ATPase+Lon-Protease-BrxL -></td><td>1183</td><td>Bacteria>Pseudomonadota>Gammaproteobacteria>Oceanospirillales>Oceanospirillaceae>Marinospirillum</td><td>GCA_014859565.1</td><td>Marinospirillum_sp.</td><td>IBX50_09850</td><td>1716..5267</td><td>2183934</td><td>BREX-system-P-loop-protein-BrxC-[Marinospirillum-sp.]</td></tr>
<tr><td>WP_138986811.1</td><td>Type1.BREX</td><td><- SIGMA+OB-Fold+AAA-ATPase+HKD-DNase-BrxL <- PglZ <- HTH+ REase <- N6-METHYLASE <- DUF6079 *<- BrxB-BrxF-iSTAND <- BrxA-DNA-binding <-XIS-HTH</td><td>1190</td><td>Bacteria>Pseudomonadota>Gammaproteobacteria>Oceanospirillales>Oceanospirillaceae>Neptunomonas</td><td>GCF_008630635.1</td><td>Neptunomonas_concharum</td><td>F0U83_RS12940</td><td>2741645..2745217</td><td>1031538</td><td>BREX-system-P-loop-protein-BrxC-[Neptunomonas-concharum]</td></tr>
<tr><td>MAD46040.1</td><td>Type1.BREX</td><td><- SIGMA+OB-Fold+AAA-ATPase+HKD-DNase-BrxL <- PglZ <- HTH+ REase <- N6-METHYLASE <- DUF6079 *<- BrxB-BrxF-iSTAND <- BrxA-DNA-binding</td><td>1189</td><td>Bacteria>Pseudomonadota>Gammaproteobacteria>Oceanospirillales>Oceanospirillaceae>unclassified_Oceanospirillaceae</td><td>GCA_002685055.1</td><td>Oceanospirillaceae_bacterium</td><td>CMH98_13650</td><td>9512..13081</td><td>1899355</td><td>hypothetical-protein-[Oceanospirillaceae-bacterium]</td></tr>
<tr><td>MBC7192650.1</td><td>Type1.BREX</td><td><- SIGMA+OB-Fold+AAA-ATPase+Lon-Protease-BrxL <- PglZ <-HNH<-OLD-ABC<-TOPRIM<-OLD-ABC<- HTH+ REase <- N6-METHYLASE <-Schlafen+HATPase+wHTH<- DUF6079 *<- BrxB-BrxF-iSTAND <- BrxA-DNA-binding <-XIS-HTH||HTH_Tnp_1+HTH_21+rve+rve_3->XIS-HTH->DnaB->?-><-Phage_int_SAM_5+Phage_integrase<-sGTP</td><td>1145</td><td>Bacteria>Pseudomonadota>Gammaproteobacteria>Pseudomonadales>Marinobacteraceae>Marinobacter</td><td>GCA_014359475.1</td><td>Marinobacter_sp.</td><td>H5U31_09890</td><td>22634..26071</td><td>50741</td><td>BREX-system-P-loop-protein-BrxC-[Marinobacter-sp.]</td></tr>
<tr><td>MCO2467177.1</td><td>Type1.BREX</td><td>Phage_holin->Phage_lysozyme->?-><- HARE-HTH + REase <- SIGMA+OB-Fold+AAA-ATPase+Lon-Protease-BrxL <- PglZ <-ParB+HNH<- N6-METHYLASE <- DUF6079 *<- BrxB-BrxF-iSTAND <- BrxA-DNA-binding <- wHTH+WYL+WCX ||JetA->JetB->SbcC->?->?->TetR-HTH+AbiJ-NTD3+ REase -></td><td>1146</td><td>Bacteria>Pseudomonadota>Gammaproteobacteria>Pseudomonadales>Pseudomonadaceae>Pseudomonas</td><td>GCA_023907445.1</td><td>Pseudomonas_aeruginosa</td><td>FA343_15170</td><td>67465..70905</td><td>287</td><td>BREX-system-P-loop-protein-BrxC-[Pseudomonas-aeruginosa]</td></tr>
<tr><td>WP_023113968.1</td><td>Type1.BREX</td><td>Phage_holin->Phage_lysozyme->?-><- HARE-HTH + REase <- SIGMA+OB-Fold+AAA-ATPase+Lon-Protease-BrxL <- PglZ <-ParB+HNH<- N6-METHYLASE <- DUF6079 *<- BrxB-BrxF-iSTAND <- BrxA-DNA-binding <-SLOG+BirA-HTH<-SF1-SF2-Helicase+PRTase<- wHTH+WYL+WCX ||ARG->?->TetR-HTH+AbiJ-NTD3+ REase -><-AbiGi+seqA+RAMA</td><td>1146</td><td>Bacteria>Pseudomonadota>Gammaproteobacteria>Pseudomonadales>Pseudomonadaceae>Pseudomonas</td><td>GCF_000481245.1</td><td>Pseudomonas_aeruginosa</td><td>Q036_RS16830</td><td>2151526..2154966</td><td>287</td><td>BREX-system-P-loop-protein-BrxC-[Pseudomonas-aeruginosa]</td></tr>
<tr><td>WP_274102063.1</td><td>Type1.BREX</td><td>Arm-DNA-bind_3+Phage_integrase->AAA+wHTH->wHTH->Integrase_1->Phage_integrase-> wHTH+WYL+WCX -> BrxA-DNA-binding -> BrxB-BrxF-iSTAND -> DUF6079 *-> N6-METHYLASE -> PglZ -> SIGMA+OB-Fold+AAA-ATPase+Lon-Protease-BrxL -><- HTH+ REase <-ANK||sGTP->?->sGTP->sGTP-></td><td>1222</td><td>Bacteria>Pseudomonadota>Gammaproteobacteria>Pseudomonadales>Pseudomonadaceae>Pseudomonas</td><td>GCF_028682345.1</td><td>Pseudomonas_putida</td><td>NPS33_RS09465</td><td>85396..89064</td><td>303</td><td>BREX-system-P-loop-protein-BrxC-[Pseudomonas-putida]</td></tr>
<tr><td>MCG8610925.1</td><td>Type1.BREX</td><td><- N6-METHYLASE <- HTH+ REase <- DUF6079 *<- BrxB-BrxF-iSTAND <- BrxA-DNA-binding <- wHTH+WYL+WCX ||ParA-Soj-PloopNTPase-></td><td>1229</td><td>Bacteria>Pseudomonadota>Gammaproteobacteria>Pseudomonadales>unclassified_Pseudomonadales</td><td>GCA_022361325.1</td><td>Pseudomonadales_bacterium</td><td>MI864_10365</td><td>4826..8515</td><td>1891229</td><td>BREX-system-P-loop-protein-BrxC-[Pseudomonadales-bacterium]</td></tr>
<tr><td>MBD3820859.1</td><td>Type1.BREX</td><td>wHTH+WYL+WCX -p-> BrxA-DNA-binding -> BrxB-BrxF-iSTAND -> DUF6079 *-> N6-METHYLASE -> HTH+ REase -> REase -> PglZ -> SIGMA+OB-Fold+AAA-ATPase+HKD-DNase-BrxL -></td><td>1191</td><td>Bacteria>Pseudomonadota>Gammaproteobacteria>Thiotrichales>unclassified_Thiotrichales</td><td>GCA_014764155.1</td><td>Thiotrichales_bacterium</td><td>IE914_01145</td><td>1632..5207</td><td>2026796</td><td>BREX-system-P-loop-protein-BrxC-[Thiotrichales-bacterium]</td></tr>
<tr><td>QFY90721.2</td><td>Type1.BREX</td><td>TRD+TRD->DOC+HTH->?->RNase->SF1-SF2-Helicase-><-rve_3||XIS-HTH-> BrxA-DNA-binding -> BrxB-BrxF-iSTAND -> DUF6079 *->?-> REase -> N6-METHYLASE -> HTH+ REase -> PglZ -> REase -> SIGMA+OB-Fold+AAA-ATPase+HKD-DNase-BrxL ->SbcC->Calcineurin+Calcineurin-SbcD-></td><td>1179</td><td>Bacteria>Pseudomonadota>Gammaproteobacteria>unclassified_Gammaproteobacteria</td><td>GCA_009497875.2</td><td>gamma_proteobacterium_SS-5</td><td>D5125_15330</td><td>2185771..2189310</td><td>947516</td><td>BREX-system-P-loop-protein-BrxC-[gamma-proteobacterium-SS-5]</td></tr>
<tr><td>MBR9924470.1</td><td>Type1.BREX</td><td><-TetR-HTH||?->tRNA->ncRNA-> REase +wHTH->rve->TnsC-ATPase->HTH_19-> BrxA-DNA-binding -> BrxB-BrxF-iSTAND -> DUF6079 *-> HTH+ REase ->Schlafen+HATPase+wHTH-></td><td>1227</td><td>Bacteria>Pseudomonadota>Gammaproteobacteria>unclassified_Gammaproteobacteria</td><td>GCA_018223005.1</td><td>Gammaproteobacteria_bacterium</td><td>GYB28_05690</td><td>375823..379506</td><td>1913989</td><td>BREX-system-P-loop-protein-BrxC-[Gammaproteobacteria-bacterium]</td></tr>
<tr><td>PIQ12823.1</td><td>Type1.BREX</td><td><- HARE-HTH + REase <- SIGMA+OB-Fold+AAA-ATPase+Lon-Protease-BrxL <- PglZ <-HNH<-OLD-ABC<- N6-METHYLASE <- DUF6079 *<- BrxB-BrxF-iSTAND <- BrxA-DNA-binding <-SLOG+BirA-HTH<-SF1-SF2-Helicase+PRTase<- wHTH+WYL+WCX ||<-ParB<-ParB<-GDResolvase+PPP1-HTH</td><td>1146</td><td>Bacteria>Pseudomonadota>Hydrogenophilia>Hydrogenophilales>unclassified_Hydrogenophilales</td><td>GCA_002770925.1</td><td>Hydrogenophilales_bacterium_CG18_big_fil_WC_8_21_14_2_50_58_12</td><td>COW70_07940</td><td>10882..14322</td><td>1974036</td><td>hypothetical-protein-COW70_07940-[Hydrogenophilales-bacterium-CG18_big_fil_WC_8_21_14_2_50_58_12]</td></tr>
<tr><td>PIY00903.1</td><td>Type1.BREX</td><td>Peptidase_M50->?->?-><-ZDNA-HTH<-?|| BrxA-DNA-binding -> BrxB-BrxF-iSTAND -> DUF6079 *->AbiEi_4->AbiEii-> N6-METHYLASE -> HTH+ REase -></td><td>1207</td><td>Bacteria>Pseudomonadota>Hydrogenophilia>Hydrogenophilales>unclassified_Hydrogenophilales</td><td>GCA_002784065.1</td><td>Hydrogenophilales_bacterium_CG_4_10_14_3_um_filter_58_23</td><td>COZ23_06050</td><td>25268..28891</td><td>1974027</td><td>BREX-system-P-loop-protein-BrxC-[Hydrogenophilales-bacterium-CG_4_10_14_3_um_filter_58_23]</td></tr>
<tr><td>MBR6059872.1</td><td>Type1.BREX</td><td><- SIGMA+OB-Fold+AAA-ATPase+Lon-Protease-BrxL <- PglZ <- HTH+ REase <- N6-METHYLASE -p<-?<- N6-METHYLASE <-nSTAND3<-?<- DUF6079 *<- BrxB-BrxF-iSTAND <- BrxA-DNA-binding</td><td>1178</td><td>Bacteria>PVC_group>Lentisphaerota>Lentisphaeria>Victivallales>unclassified_Victivallales</td><td>GCA_017542865.1</td><td>Victivallales_bacterium</td><td>IKP58_17025</td><td>13303..16839</td><td>2212475</td><td>BREX-system-P-loop-protein-BrxC-[Victivallales-bacterium]</td></tr>
<tr><td>OGV31172.1</td><td>Type1.BREX</td><td>XIS-HTH-> BrxA-DNA-binding -> BrxB-BrxF-iSTAND -> DUF6079 *-> HTH+ REase -> N6-METHYLASE -p-></td><td>1201</td><td>Bacteria>PVC_group>Lentisphaerota>unclassified_Lentisphaerota</td><td>GCA_001804785.1</td><td>Lentisphaerae_bacterium_GWF2_45_14</td><td>A2020_01685</td><td>2431..6036</td><td>1798572</td><td>MAG:-hypothetical-protein-A2020_01685-[Lentisphaerae-bacterium-GWF2_45_14]</td></tr>
<tr><td>TVL98972.1</td><td>Type1.BREX</td><td>TPR+TPR-> BrxA-DNA-binding -> BrxB-BrxF-iSTAND -> DUF6079 *-> HTH+ REase -> HTH+ REase -> PglZ -> SIGMA+OB-Fold+AAA-ATPase+Lon-Protease-BrxL -> REase +DUF5655->ParB->SIGMA-HTH-></td><td>1144</td><td>Bacteria>PVC_group>Planctomycetota>Candidatus_Brocadiia>Candidatus_Brocadiales>Candidatus_Brocadiaceae</td><td>GCA_007618135.1</td><td>Candidatus_Brocadia_sp._WS118</td><td>CV087_19775</td><td>5481..8915</td><td>2040625</td><td>MAG:-BREX-system-P-loop-protein-BrxC-[Candidatus-Brocadia-sp.-WS118]</td></tr>
<tr><td>GDY21840.1</td><td>Type6.BREX</td><td>BrxE -> BrxA-DNA-binding -> BrxB-BrxF-iSTAND -> DUF6079 *-> N6-METHYLASE -> HTH+ REase -></td><td>1208</td><td>Bacteria>PVC_group>Verrucomicrobiota>unclassified_Verrucomicrobiota</td><td>GCA_014193395.1</td><td>Verrucomicrobiota_bacterium</td><td>LBMAG56_31870</td><td>2824..6450</td><td>2026799</td><td>hypothetical-protein-LBMAG56_31870-[Verrucomicrobiota-bacterium]</td></tr>
<tr><td>SPE51781.1</td><td>Type1.BREX</td><td><- SIGMA+OB-Fold+AAA-ATPase+Lon-Protease-BrxL <- PglZ -p<- PglZ <- HTH+ REase <- N6-METHYLASE <-?<-SMODS<- DUF6079 *<- BrxB-BrxF-iSTAND <- BrxB-BrxF-iSTAND <- BrxA-DNA-binding</td><td>1149</td><td>Bacteria>PVC_group>Verrucomicrobiota>unclassified_Verrucomicrobiota</td><td>GCA_900290355.1</td><td>Verrucomicrobiota_bacterium</td><td>SBV1_1420061</td><td>72835..76284</td><td>2026799</td><td>putative-P-loop-containing-nucleoside-triphosphate-hydrolase-[Verrucomicrobiota-bacterium]</td></tr>
<tr><td>PAW64461.1</td><td>Type1.BREX</td><td>BrxB-BrxF-iSTAND -> DUF6079 *-> N6-METHYLASE -> HTH+ REase ->Calcineurin-SbcD->SbcC+CCDC158-></td><td>1145</td><td>Bacteria>PVC_group>Verrucomicrobiota>Verrucomicrobiae>unclassified_Verrucomicrobiae</td><td>GCA_002304315.1</td><td>Verrucomicrobiae_bacterium_Tous-C4TDCM</td><td>B9S38_16345</td><td>386..3823</td><td>1982332</td><td>MAG:-hypothetical-protein-B9S38_16345-[Verrucomicrobiae-bacterium-Tous-C4TDCM]</td></tr>
<tr><td>MBL9158389.1</td><td>Type1.BREX</td><td><- SIGMA+OB-Fold+AAA-ATPase+Lon-Protease-BrxL <- PglZ <- HTH+ REase <- N6-METHYLASE -p<-SbcC+ABC-ATPase<- N6-METHYLASE <-TIR<- DUF6079 *<- BrxB-BrxF-iSTAND <- BrxA-DNA-binding ||?->?->?->?->XIS-HTH->RVT-></td><td>1152</td><td>Bacteria>PVC_group>Verrucomicrobiota>Verrucomicrobiae>Verrucomicrobiales>unclassified_Verrucomicrobiales</td><td>GCA_016795385.1</td><td>Verrucomicrobiales_bacterium</td><td>JNJ70_13005</td><td>50985..54443</td><td>2026801</td><td>BREX-system-P-loop-protein-BrxC-[Verrucomicrobiales-bacterium]</td></tr>
<tr><td>MBK6607326.1</td><td>Type6.BREX</td><td>BrxE -> BrxA-DNA-binding -> BrxB-BrxF-iSTAND -> DUF6079 -p*-> DUF6079 -p-> N6-METHYLASE -p-> N6-METHYLASE -p->URI-DNase-> HTH+ REase -> N6-METHYLASE -p->?-> N6-METHYLASE -p-> PglZ -p-></td><td>320</td><td>Bacteria>Spirochaetota>Spirochaetia>Leptospirales>Leptospiraceae>unclassified_Leptospiraceae</td><td>GCA_016703885.1</td><td>Leptospiraceae_bacterium</td><td>IPG24_18070</td><td>61452..62414</td><td>2026757</td><td>hypothetical-protein-[Leptospiraceae-bacterium]</td></tr>
<tr><td>MBK6607327.1</td><td>Type6.BREX</td><td>BrxE -> BrxA-DNA-binding -> BrxB-BrxF-iSTAND -> DUF6079 -p-> DUF6079 -p*-> N6-METHYLASE -p-> N6-METHYLASE -p->URI-DNase-> HTH+ REase -> N6-METHYLASE -p->?-> N6-METHYLASE -p-> PglZ -p-></td><td>877</td><td>Bacteria>Spirochaetota>Spirochaetia>Leptospirales>Leptospiraceae>unclassified_Leptospiraceae</td><td>GCA_016703885.1</td><td>Leptospiraceae_bacterium</td><td>IPG24_18075</td><td>62425..65058</td><td>2026757</td><td>hypothetical-protein-[Leptospiraceae-bacterium]</td></tr>
<tr><td>MCB1192727.1</td><td>Type1.BREX</td><td><-SecA+SF1-SF2-Helicase<-ArdCN|| BrxA-DNA-binding -> BrxB-BrxF-iSTAND -> DUF6079 *-> REase +HTH-> N6-METHYLASE -> PglZ -></td><td>1148</td><td>Bacteria>Spirochaetota>Spirochaetia>Leptospirales>Leptospiraceae>unclassified_Leptospiraceae</td><td>GCA_020440835.1</td><td>Leptospiraceae_bacterium</td><td>KDK90_19935</td><td>18936..22382</td><td>2026757</td><td>BREX-system-P-loop-protein-BrxC-[Leptospiraceae-bacterium]</td></tr>
<tr><td>TXT43486.1</td><td>Type1.BREX</td><td><- SIGMA+OB-Fold+AAA-ATPase+HKD-DNase-BrxL <-?||?-><- HTH+ REase <- N6-METHYLASE -p<- N6-METHYLASE -p<- N6-METHYLASE -p<-Schlafen+HATPase+wHTH-p<- DUF6079 *<- BrxB-BrxF-iSTAND <- BrxA-DNA-binding <-XIS-HTH</td><td>1162</td><td>Bacteria>Spirochaetota>unclassified_Spirochaetota</td><td>GCA_008080685.1</td><td>Spirochaetota_bacterium</td><td>FD137_1824</td><td>16028..19516</td><td>2202144</td><td>hypothetical-protein-FD137_1824-[Spirochaetota-bacterium]</td></tr>
<tr><td>MBL3593011.1</td><td>Type1.BREX</td><td><-CHTH||DUF488-l->DUF488-s->?->SF1-SF2-Helicase->?->XIS-HTH-> BrxA-DNA-binding -> BrxB-BrxF-iSTAND -> DUF6079 *->Schlafen+HATPase+wHTH-> N6-METHYLASE -> PglZ -> SIGMA+OB-Fold+AAA-ATPase+Lon-Protease-BrxL -> HARE-HTH + REase ->?->GGDEF->Aminotran_4->DUF488-></td><td>1149</td><td>Bacteria>Synergistota>Synergistia>Synergistales>Synergistaceae>unclassified_Synergistaceae</td><td>GCA_016756725.1</td><td>Synergistaceae_bacterium</td><td>JMJ93_05810</td><td>374580..378029</td><td>2053624</td><td>BREX-system-P-loop-protein-BrxC-[Synergistaceae-bacterium]</td></tr>
<tr><td>WP_124851247.1</td><td>Type1.BREX</td><td><- SIGMA+OB-Fold+AAA-ATPase+HKD-DNase-BrxL <- PglZ <- HTH+ REase <- N6-METHYLASE -p<-Phage_int_SAM_3+Phage_integrase|| N6-METHYLASE -p-><- N6-METHYLASE <- DUF6079 *<- BrxB-BrxF-iSTAND <- BrxA-DNA-binding <-?||?->?->?->SbcC+SbcC-></td><td>1170</td><td>Bacteria>Terrabacteria_group>Actinomycetota>Actinomycetes>Actinomycetales>Actinomycetaceae</td><td>GCF_003858495.1</td><td>Buchananella_hordeovulneris</td><td>EII12_RS02040</td><td>65131..68643</td><td>52770</td><td>BREX-system-P-loop-protein-BrxC-[Buchananella-hordeovulneris]</td></tr>
<tr><td>WP_004808447.1</td><td>Type1.BREX</td><td><-RNase<-?||?->?->?->?-> HTH+ REase -> BrxA-DNA-binding -> BrxB-BrxF-iSTAND -> DUF6079 *-> N6-METHYLASE -> PglZ -> SIGMA+OB-Fold+AAA-ATPase+HKD-DNase-BrxL -><-?<-?<-ABhydrolase<-DOC+HTH<-?||ZnR-></td><td>1184</td><td>Bacteria>Terrabacteria_group>Actinomycetota>Actinomycetes>Actinomycetales</td><td>GCF_028201595.1</td><td>Actinomycetaceae</td><td>PIG85_RS08980</td><td>1926978..1930532</td><td>2049</td><td>MULTISPECIES:-BREX-system-P-loop-protein-BrxC-[Actinomycetaceae]</td></tr>
<tr><td>WP_005885100.1</td><td>Type1.BREX</td><td><-ABC-ATPase+SbcC<-?<-?||ACET-><-OLD-ABC<- SIGMA+OB-Fold+AAA-ATPase+HKD-DNase-BrxL <- PglZ <- HTH+ REase <- N6-METHYLASE <- DUF6079 *<- BrxB-BrxF-iSTAND <- BrxA-DNA-binding <-XIS-HTH<-ABhydrolase||?-><-HTH</td><td>1178</td><td>Bacteria>Terrabacteria_group>Actinomycetota>Actinomycetes>Micrococcales>Brevibacteriaceae</td><td>GCF_000178455.1</td><td>Brevibacterium_mcbrellneri</td><td>HMPREF0183_RS08780</td><td>50551..54087</td><td>53363</td><td>BREX-system-P-loop-protein-BrxC-[Brevibacterium-mcbrellneri]</td></tr>
<tr><td>WP_259801020.1</td><td>Type1.BREX</td><td><-VirB11+MuB-ATPase||DNA_METHYLASE-><-HerA/FtsK<-SIR2<- SIGMA+OB-Fold+AAA-ATPase+HKD-DNase-BrxL <- PglZ <- HTH+ REase <- N6-METHYLASE <- DUF6079 *<- BrxB-BrxF-iSTAND <- BrxA-DNA-binding <-XIS-HTH<-ABhydrolase<-?<-MPTase<-HTH||ArdCN+MPTase->Primpol+PriCT1-></td><td>1173</td><td>Bacteria>Terrabacteria_group>Actinomycetota>Actinomycetes>Micrococcales>Brevibacteriaceae</td><td>GCF_025144065.1</td><td>Brevibacterium_casei</td><td>M3G43_RS08140</td><td>29205..32726</td><td>33889</td><td>BREX-system-P-loop-protein-BrxC-[Brevibacterium-casei]</td></tr>
<tr><td>WP_249738141.1</td><td>Type1.BREX</td><td><-PTSEIIA1<-?<-HHH_5+SNF2_N+SF1-SF2-Helicase||CASPASE-><- SIGMA+OB-Fold+AAA-ATPase+HKD-DNase-BrxL <- PglZ <- HTH+ REase <- N6-METHYLASE <- DUF6079 *<- BrxB-BrxF-iSTAND <- BrxA-DNA-binding <-SAM-METHYLASE<-?||HIN-HTH->HTH_Tnp_1+HTH_21+rve+rve_3-></td><td>1173</td><td>Bacteria>Terrabacteria_group>Actinomycetota>Actinomycetes>Micrococcales>Dermabacteraceae</td><td>GCF_023509525.1</td><td>Brachybacterium_sp._JHP9</td><td>Bequi_RS11780</td><td>103104..106625</td><td>2910770</td><td>BREX-system-P-loop-protein-BrxC-[Brachybacterium-sp.-JHP9]</td></tr>
<tr><td>WP_104291918.1</td><td>Type1.BREX</td><td>McrB-AAA-> McrC-NTD+ REase ->GDResolvase+HIN-HTH-><-?||?->?->XIS-HTH-> BrxA-DNA-binding -> BrxB-BrxF-iSTAND -> DUF6079 *-> N6-METHYLASE -> PglZ -> SIGMA+OB-Fold+AAA-ATPase+HKD-DNase-BrxL -> REase +HTH+35exo+Small-B-barrel+SF1-SF2-Helicase-><-RNase</td><td>1170</td><td>Bacteria>Terrabacteria_group>Actinomycetota>Actinomycetes>Micrococcales>Microbacteriaceae</td><td>GCF_002930935.1</td><td>Rathayibacter_sp._AY1A2</td><td>C5B93_RS04570</td><td>113948..117460</td><td>2080520</td><td>BREX-system-P-loop-protein-BrxC-[Rathayibacter-sp.-AY1A2]</td></tr>
<tr><td>WP_104347518.1</td><td>Type1.BREX</td><td><-ClpABN-AAA+ClpABC-AAA<-?<- SIGMA+OB-Fold+AAA-ATPase+HKD-DNase-BrxL <- PglZ <- HTH+ REase <- N6-METHYLASE <- DUF6079 *<- BrxB-BrxF-iSTAND <- BrxA-DNA-binding <-XIS-HTH<-ABhydrolase</td><td>1182</td><td>Bacteria>Terrabacteria_group>Actinomycetota>Actinomycetes>Micrococcales>Microbacteriaceae</td><td>GCF_002932695.1</td><td>Rathayibacter_sp._AY1D2</td><td>C5D04_RS07575</td><td>19648..23196</td><td>2080543</td><td>BREX-system-P-loop-protein-BrxC-[Rathayibacter-sp.-AY1D2]</td></tr>
<tr><td>WP_124973569.1</td><td>Type1.BREX</td><td>Aminotran_4-><-?<-NAD_synthase||?-><-tRNA||35exo->tRNA->XIS-HTH-> BrxA-DNA-binding -> BrxB-BrxF-iSTAND -> DUF6079 *-> N6-METHYLASE -> HTH+ REase -> PglZ -> SIGMA+OB-Fold+AAA-ATPase+HKD-DNase-BrxL ->SIR2->HerA/FtsK-><-DNA_METHYLASE</td><td>1170</td><td>Bacteria>Terrabacteria_group>Actinomycetota>Actinomycetes>Micrococcales>Microbacteriaceae</td><td>GCF_003865375.1</td><td>Gulosibacter_macacae</td><td>EG850_RS11335</td><td>21958..25470</td><td>2488791</td><td>BREX-system-P-loop-protein-BrxC-[Gulosibacter-macacae]</td></tr>
<tr><td>WP_207484738.1</td><td>Type1.BREX</td><td><-Primpol+PriCT1<-ArdCN+MPTase||HTH->MPTase->?->ABhydrolase->XIS-HTH-> BrxA-DNA-binding -> BrxB-BrxF-iSTAND -> DUF6079 *-> N6-METHYLASE -><- N6-METHYLASE -p||Phage_int_SAM_3+Phage_integrase-> N6-METHYLASE -p-> HTH+ REase -> PglZ -></td><td>1176</td><td>Bacteria>Terrabacteria_group>Actinomycetota>Actinomycetes>Micrococcales>Microbacteriaceae</td><td>GCF_017355365.1</td><td>Microbacterium_sp._SD291</td><td>IPV10_RS13675</td><td>26562..30092</td><td>2782007</td><td>BREX-system-P-loop-protein-BrxC-[Microbacterium-sp.-SD291]</td></tr>
<tr><td>WP_201614919.1</td><td>Type1.BREX</td><td>Aminotran_4-><-?<-Peptidase<-NAD_synthase<-tRNA||35exo->tRNA-> BrxA-DNA-binding -> BrxB-BrxF-iSTAND -> DUF6079 *-> N6-METHYLASE -><- N6-METHYLASE -p||Phage_int_SAM_3+Phage_integrase-> N6-METHYLASE -p-> HTH+ REase -> PglZ -> SIGMA+OB-Fold+AAA-ATPase+HKD-DNase-BrxL -></td><td>1170</td><td>Bacteria>Terrabacteria_group>Actinomycetota>Actinomycetes>Micrococcales>Microbacteriaceae</td><td>GCF_904848685.1</td><td>Gulosibacter_hominis</td><td>JMX86_RS10045</td><td>19055..22567</td><td>2770504</td><td>BREX-system-P-loop-protein-BrxC-[Gulosibacter-hominis]</td></tr>
<tr><td>WP_230672563.1</td><td>Type1.BREX</td><td>ABC-ATPase+SbcC->?->?-><-?<-?||MACRODOMAIN+SIR2-> BrxA-DNA-binding -> BrxB-BrxF-iSTAND -> DUF6079 *-> N6-METHYLASE -> HTH+ REase -> HTH+ REase -> PglZ -> SIGMA+OB-Fold+AAA-ATPase+HKD-DNase-BrxL -><-?||?->HEPN+ SF1-SF2-Helicase+ REase -></td><td>1176</td><td>Bacteria>Terrabacteria_group>Actinomycetota>Actinomycetes>Micrococcales>Microbacteriaceae</td><td>GCF_920939495.1</td><td>Rathayibacter_sp._Leaf248</td><td>LRG51_RS10050</td><td>2075326..2078856</td><td>2876555</td><td>BREX-system-P-loop-protein-BrxC-[Rathayibacter-sp.-Leaf248]</td></tr>
<tr><td>THE19344.1</td><td>Type1.BREX</td><td><-TIR||XIS-HTH-> BrxA-DNA-binding -> BrxB-BrxF-iSTAND -> DUF6079 *-> N6-METHYLASE -> PglZ -> SIGMA+OB-Fold+AAA-ATPase+HKD-DNase-BrxL -><-?|| REase -> HTH+ REase ->URI-DNase->DUF6361->HKD-DNase-></td><td>1166</td><td>Bacteria>Terrabacteria_group>Actinomycetota>Actinomycetes>Micrococcales>Micrococcaceae</td><td>GCA_004799725.1</td><td>Kocuria_rosea</td><td>E1J17_01420</td><td>4081..7581</td><td>1275</td><td>BREX-system-P-loop-protein-BrxC-[Kocuria-rosea]</td></tr>
<tr><td>MBB4882886.1</td><td>Type1.BREX</td><td>35exo->tRNA-><-Recombinase+ZnR||ABhydrolase->XIS-HTH-> BrxA-DNA-binding -> BrxB-BrxF-iSTAND -> DUF6079 *-> N6-METHYLASE -> HTH+ REase -> PglZ -> SIGMA+OB-Fold+AAA-ATPase+HKD-DNase-BrxL ->SIR2->HIN-HTH->HTH_Tnp_1+HTH_21+rve+rve_3-></td><td>1173</td><td>Bacteria>Terrabacteria_group>Actinomycetota>Actinomycetes>Micrococcales>Micrococcaceae</td><td>GCA_014204815.1</td><td>Micrococcus_flavus</td><td>BJ976_001237</td><td>1342727..1346248</td><td>384602</td><td>hypothetical-protein-[Micrococcus-flavus]</td></tr>
<tr><td>WP_135027971.1</td><td>Type1.BREX</td><td>35exo->tRNA-><-Recombinase+ZnR||ABhydrolase->XIS-HTH-> BrxA-DNA-binding -> BrxB-BrxF-iSTAND -> DUF6079 *-> N6-METHYLASE -> HTH+ REase -> PglZ -> SIGMA+OB-Fold+AAA-ATPase+HKD-DNase-BrxL ->SIR2->HTH_Tnp_1+HTH_21+rve+rve_3-><-?||?-><-HAD+HAD</td><td>1170</td><td>Bacteria>Terrabacteria_group>Actinomycetota>Actinomycetes>Micrococcales>Micrococcaceae</td><td>GCF_014644875.1</td><td>Micrococcus_flavus</td><td>E4A47_RS01340</td><td>286387..289899</td><td>384602</td><td>BREX-system-P-loop-protein-BrxC-[Micrococcus-flavus]</td></tr>
<tr><td>WP_102213741.1</td><td>Type1.BREX</td><td>XIS-HTH-> BrxA-DNA-binding -> BrxB-BrxF-iSTAND -> DUF6079 *-> N6-METHYLASE -> HTH+ REase -> PglZ -> SIGMA+OB-Fold+AAA-ATPase+HKD-DNase-BrxL -><-ParB+HNH+METHYLASE</td><td>1173</td><td>Bacteria>Terrabacteria_group>Actinomycetota>Actinomycetes>Mycobacteriales>Corynebacteriaceae</td><td>GCF_002871715.1</td><td>Corynebacterium_xerosis</td><td>CJ204_09375</td><td>1693..5214</td><td>1725</td><td>BREX-system-P-loop-protein-BrxC-[Corynebacterium-xerosis]</td></tr>
<tr><td>PZT90276.1</td><td>Type1.BREX</td><td>HAD+HAD-><-?||XIS-HTH-> BrxA-DNA-binding -> BrxB-BrxF-iSTAND -> DUF6079 *-> N6-METHYLASE -> HTH+ REase -> PglZ -> SIGMA+OB-Fold+AAA-ATPase+HKD-DNase-BrxL -> REase -></td><td>1137</td><td>Bacteria>Terrabacteria_group>Actinomycetota>Actinomycetes>Mycobacteriales>Gordoniaceae</td><td>GCA_003248405.1</td><td>Gordonia_sp._(in:_high_G+C_Gram-positive_bacteria)</td><td>DI630_31230</td><td>6915..10328</td><td>84139</td><td>MAG:-BREX-system-P-loop-protein-BrxC-[Gordonia-sp.-(in:-high-G+C-Gram-positive-bacteria)]</td></tr>
<tr><td>WP_213244950.1</td><td>Type1.BREX</td><td><- SIGMA+OB-Fold+AAA-ATPase+HKD-DNase-BrxL <- PglZ <- HTH+ REase <- N6-METHYLASE -p<-?<-Phage_int_SAM_3+Phage_integrase<-Phage_int_SAM_3+Phage_integrase|| N6-METHYLASE -p-><- N6-METHYLASE <- DUF6079 *<- BrxB-BrxF-iSTAND <- BrxA-DNA-binding</td><td>1174</td><td>Bacteria>Terrabacteria_group>Actinomycetota>Actinomycetes>Mycobacteriales>Gordoniaceae</td><td>GCF_009914535.1</td><td>Gordonia_pseudamarae</td><td>GII33_RS19265</td><td>4440551..4444075</td><td>2831662</td><td>BREX-system-P-loop-protein-BrxC-[Gordonia-pseudamarae]</td></tr>
<tr><td>GFM17528.1</td><td>Type1.BREX</td><td><- N6-METHYLASE -p<- DUF6079 *<- BrxB-BrxF-iSTAND <- BrxA-DNA-binding <- McrC-NTD+ REase <-McrB-AAA<- HTH+ REase <-HEPN+ SF1-SF2-Helicase+ REase <-?<-Preader7</td><td>1155</td><td>Bacteria>Terrabacteria_group>Actinomycetota>Actinomycetes>Mycobacteriales>Mycobacteriaceae</td><td>GCA_017312405.1</td><td>Mycobacterium_sp._PO1</td><td>PO1_contig-018-2</td><td>841..4308</td><td>1882219</td><td>putative-P-loop-containing-nucleoside-triphospha-te-hydrolase-[Mycobacterium-sp.-PO1]</td></tr>
<tr><td>WP_198387782.1</td><td>Type1.BREX</td><td>HEPN+ SF1-SF2-Helicase+ REase ->HTH_36-><- SIGMA+OB-Fold+AAA-ATPase+HKD-DNase-BrxL <- PglZ <- HTH+ REase <- N6-METHYLASE -p||<- N6-METHYLASE <- DUF6079 *<- BrxB-BrxF-iSTAND <- BrxA-DNA-binding <-DNA_METHYLASE<-DrmB<-SF1-SF2-Helicase<-SF1-SF2-Helicase<-MORC+SF1-SF2-Helicase||ParB+HNH->SF1-SF2-Helicase-></td><td>1173</td><td>Bacteria>Terrabacteria_group>Actinomycetota>Actinomycetes>Mycobacteriales>Nocardiaceae</td><td>GCF_002269365.1</td><td>Rhodococcus_pyridinivorans</td><td>CJU69_RS21710</td><td>4620785..4624306</td><td>103816</td><td>BREX-system-P-loop-protein-BrxC-[Rhodococcus-pyridinivorans]</td></tr>
</table>

<table>
<tr><th>Protein-ID</th><th>System</th><th>Gene-Neighbourhoods</th><th>Length</th><th>Taxonomy</th><th>Assembly-ID</th><th>Strain</th><th>Locus-Tag</th><th>Genomic-coordinates</th><th>TaxID</th><th>Fasta-defline</th></tr>
<tr><td>WP_195040827.1</td><td>Type1.BREX</td><td>McrB-AAA->McrC-NTD+REase-><-?<-N6-METHYLASE<-?<-?<-SIGMA+OB-Fold+AAA-ATPase+HKD-DNase-BrxL<-PglZ<-N6-METHYLASE<-DUF6079*<-BrxB-BrxF-iSTAND<-BrxA-DNA-binding<-XIS-HTH<-?<-HTH_31+DUF5753</td><td>1159</td><td>Bacteria>Terrabacteria_group>Actinomycetota>Actinomycetes>Mycobacteriales>Nocardiaceae</td><td>GCF_015477975.1</td><td>Nocardia_cyriacigeorgica</td><td>IU440_RS19520</td><td>298293..301772</td><td>135487</td><td>BREX-system-P-loop-protein-BrxC-[Nocardia-cyriacigeorgica]</td></tr>
<tr><td>WP_238424144.1</td><td>Type1.BREX</td><td>HTH_3->?-><-?<-HTH_36<-?||BrxA-DNA-binding->BrxB-BrxF-iSTAND->DUF6079*->N6-METHYLASE->HTH+REase->PglZ->SIGMA+OB-Fold+AAA-ATPase+HKD-DNase-BrxL->zf-ISL3+HTH_Tnp_ISL3-><-SF1-SF2-Helicase<-HKD-DNase<-DUF6361||TetR-HTH-></td><td>1175</td><td>Bacteria>Terrabacteria_group>Actinomycetota>Actinomycetes>Mycobacteriales>Nocardiaceae</td><td>GCF_016804345.1</td><td>Rhodococcus_ruber</td><td>F1734_RS10025</td><td>2133745..2137272</td><td>1830</td><td>BREX-system-P-loop-protein-BrxC-[Rhodococcus-ruber]</td></tr>
<tr><td>WP_242421152.1</td><td>Type1.BREX</td><td>SF1-SF2-Helicase+DUF1998->REase-><-DNA_METHYLASE||REase-><-SIGMA+OB-Fold+AAA-ATPase+HKD-DNase-BrxL<-PglZ<-HTH+REase<-N6-METHYLASE<-DUF6079*<-BrxB-BrxF-iSTAND<-BrxA-DNA-binding<-?||?-><-?||?->RNase-></td><td>1172</td><td>Bacteria>Terrabacteria_group>Actinomycetota>Actinomycetes>Mycobacteriales>Nocardiaceae</td><td>GCF_022691685.1</td><td>Prescottella_equi</td><td>MPC38_RS08825</td><td>1859263..1862781</td><td>43767</td><td>BREX-system-P-loop-protein-BrxC-[Prescottella-equi]</td></tr>
<tr><td>WP_068569480.1</td><td>Type1.BREX</td><td><-TetR-HTH<-?<-SbcC<-JetB<-JetA<-HTH_3||BrxA-DNA-binding->BrxB-BrxF-iSTAND->DUF6079*->N6-METHYLASE->PglZ->SIGMA+OB-Fold+AAA-ATPase+HKD-DNase-BrxL->HTH+REase->?-><-?<-TraB<-MoxR-AAA</td><td>1182</td><td>Bacteria>Terrabacteria_group>Actinomycetota>Actinomycetes>Mycobacteriales>Tsukamurellaceae</td><td>GCF_001575195.1</td><td>Tsukamurella_pseudospumae</td><td>AXK60_RS03070</td><td>654047..657595</td><td>239498</td><td>BREX-system-P-loop-protein-BrxC-[Tsukamurella-pseudospumae]</td></tr>
<tr><td>WP_068743364.1</td><td>Type1.BREX</td><td>MoxR-AAA->TraB->?-><-HTH+REase<-HTH+REase<-SIGMA+OB-Fold+AAA-ATPase+HKD-DNase-BrxL<-PglZ<-N6-METHYLASE<-DUF6079*<-BrxB-BrxF-iSTAND<-BrxA-DNA-binding||HTH_3->JetA->JetB->SbcC->?->TetR-HTH-></td><td>1180</td><td>Bacteria>Terrabacteria_group>Actinomycetota>Actinomycetes>Mycobacteriales>Tsukamurellaceae</td><td>GCF_001575205.1</td><td>Tsukamurella_pseudospumae</td><td>AXK61_RS01225</td><td>254752..258294</td><td>239498</td><td>BREX-system-P-loop-protein-BrxC-[Tsukamurella-pseudospumae]</td></tr>
<tr><td>WP_246207064.1</td><td>Type1.BREX</td><td><-TetR-HTH<-?<-SbcC<-JetB<-JetA<-HTH_3||BrxA-DNA-binding->BrxB-BrxF-iSTAND->DUF6079*->N6-METHYLASE->SIGMA+OB-Fold+AAA-ATPase+HKD-DNase-BrxL->HTH+REase-><-?<-TraB<-MoxR-AAA</td><td>1179</td><td>Bacteria>Terrabacteria_group>Actinomycetota>Actinomycetes>Mycobacteriales>Tsukamurellaceae</td><td>GCF_012396015.1</td><td>Tsukamurella_spumae</td><td>HF999_RS00885</td><td>181612..185151</td><td>44753</td><td>BREX-system-P-loop-protein-BrxC-[Tsukamurella-spumae]</td></tr>
<tr><td>MBK7820961.1</td><td>Type1.BREX</td><td><-DUF2637+HIN-HTH<-ArdCN+MPTase||HTH->MPTase->XIS-HTH->BrxA-DNA-binding->BrxB-BrxF-iSTAND->DUF6079*->N6-METHYLASE->HTH+REase->PglZ->SIGMA+OB-Fold+AAA-ATPase+HKD-DNase-BrxL->OLD-ABC+TOPRIM->SF1-SF2-Helicase->?->DDE_Tnp_1-><-DNA_METHYLASE</td><td>1176</td><td>Bacteria>Terrabacteria_group>Actinomycetota>Actinomycetes>Propionibacteriales>Propionibacteriaceae</td><td>GCA_016714275.1</td><td>Tessaracoccus_sp.</td><td>IPJ61_07760</td><td>1693709..1697239</td><td>1971211</td><td>BREX-system-P-loop-protein-BrxC-[Tessaracoccus-sp.]</td></tr>
<tr><td>WP_094364310.1</td><td>Type1.BREX</td><td>DNA_METHYLASE-><-sGTP<-GTPase-AIG<-HerA/FtsK<-SIR2+SIR2<-SIGMA+OB-Fold+AAA-ATPase+HKD-DNase-BrxL<-PglZ<-HTH+REase<-N6-METHYLASE<-DUF6079*<-BrxB-BrxF-iSTAND<-BrxA-DNA-binding<-XIS-HTH<-ABhydrolase<-?<-MPTase<-HTH||ArdCN+MPTase->Primpol+PriCT1-></td><td>1173</td><td>Bacteria>Terrabacteria_group>Actinomycetota>Actinomycetes>Propionibacteriales>Propionibacteriaceae</td><td>GCF_002250625.1</td><td>Enemella_dayhoffiae</td><td>CGZ93_RS11755</td><td>175760..179281</td><td>2016507</td><td>BREX-system-P-loop-protein-BrxC-[Enemella-dayhoffiae]</td></tr>
<tr><td>SFW99921.1</td><td>Type1.BREX</td><td><-DOC+HTH||?->?-><-HTH+REase<-?<-SIGMA+OB-Fold+AAA-ATPase+HKD-DNase-BrxL<-N6-METHYLASE<-PglZ<-DUF6079*<-BrxB-BrxF-iSTAND<-BrxA-DNA-binding<-TraJ-RHH||ParB+HNH->Phage-portal-><-?<-Glyco_hydro_1<-PTSEIIC</td><td>1199</td><td>Bacteria>Terrabacteria_group>Actinomycetota>Coriobacteriia>Coriobacteriales>Atopobiaceae</td><td>GCA_900119625.1</td><td>Olsenella_sp._kh2p3</td><td>SAMN04487823_101110</td><td>126088..129687</td><td>1797112</td><td>hypothetical-protein-SAMN04487823_101110-[Olsenella-sp.-kh2p3]</td></tr>
<tr><td>MBA3474041.1</td><td>Type1.BREX</td><td><-HTH+REase<-PemK_toxin<-?<-METHYLASE<-?<-HEPN<-HEPN<-DUF6079*<-REase<-?<-?<-N6-METHYLASE-p</td><td>1219</td><td>Bacteria>Terrabacteria_group>Actinomycetota>Rubrobacteria>Rubrobacterales>Rubrobacteraceae</td><td>GCA_013813465.1</td><td>Rubrobacter_sp.</td><td>HOT57_12585</td><td>3865..7524</td><td>2036010</td><td>BREX-system-P-loop-protein-BrxC-[Rubrobacter-sp.]</td></tr>
<tr><td>WP_016131320.1</td><td>Type1.BREX</td><td>Phosphodiest->?-><-HTH_Tnp_1+HTH_21+rve+rve_3||?->TOPRIM+Topoisom_bac->wHTH+WYL+WCX-p->BrxA-DNA-binding->BrxB-BrxF-iSTAND->DUF6079*->N6-METHYLASE->PglZ->SIGMA+OB-Fold+AAA-ATPase+Lon-Protease-BrxL->REase+wHTH->rve+Mu-transpos_C->TnsC-ATPase->?-><-HTH_OrfB_IS605||ArsR-HTH-></td><td>1190</td><td>Bacteria>Terrabacteria_group>Bacillota>Bacilli>Bacillales>Bacillaceae</td><td>-</td><td>Bacillus_cereus</td><td>KOW_RS09165</td><td>326636..330208</td><td>1396</td><td>BREX-system-P-loop-protein-BrxC-[Bacillus-cereus]</td></tr>
<tr><td>EOP54853.1</td><td>Type1.BREX</td><td>Phosphodiest->?->?-><-HTH_Tnp_1+HTH_21+rve+rve_3||?->TOPRIM+Topoisom_bac->wHTH+WYL+WCX-p->BrxA-DNA-binding->BrxB-BrxF-iSTAND->DUF6079*->N6-METHYLASE->PglZ->SIGMA+OB-Fold+AAA-ATPase+Lon-Protease-BrxL->REase+wHTH->rve+Mu-transpos_C->TnsC-ATPase->?-><-HTH_OrfB_IS605||ArsR-HTH-></td><td>1193</td><td>Bacteria>Terrabacteria_group>Bacillota>Bacilli>Bacillales>Bacillaceae</td><td>GCA_000399485.1</td><td>Bacillus_cereus_VD136</td><td>IIW_00987</td><td>109394..112975</td><td>1053234</td><td>hypothetical-protein-IIW_00987-[Bacillus-cereus-VD136]</td></tr>
<tr><td>MBT2214695.1</td><td>Type1.BREX</td><td>PTSEIIA1->Cas9+HNH+RuvC->Cas_Cas1->CRISPR_Cas2->BrxA-DNA-binding->BrxB-BrxF-iSTAND->DUF6079*->REase+HTH-><-TetR-HTH</td><td>1340</td><td>Bacteria>Terrabacteria_group>Bacillota>Bacilli>Bacillales>Bacillaceae</td><td>GCA_018603665.1</td><td>Virgibacillus_dakarensis</td><td>KK120_02535</td><td>179341..183363</td><td>1917889</td><td>BREX-system-P-loop-protein-BrxC-[Virgibacillus-dakarensis]</td></tr>
<tr><td>MBW7642206.1</td><td>Type1.BREX</td><td>McrB-NTD+McrB-AAA->McrC-NTD+REase->HTH_3->SF1-SF2-Helicase->BrxA-DNA-binding->BrxB-BrxF-iSTAND->DUF6079*->N6-METHYLASE->PglZ->||REase-><-?||?->?->REase+HTH-></td><td>1188</td><td>Bacteria>Terrabacteria_group>Bacillota>Bacilli>Bacillales>Bacillaceae</td><td>GCA_019448595.1</td><td>Geobacillus_thermoleovorans</td><td>K1I45_03795</td><td>253845..257411</td><td>33941</td><td>BREX-system-P-loop-protein-BrxC-[Geobacillus-thermoleovorans]</td></tr>
<tr><td>UQX55099.1</td><td>Type1.BREX</td><td><-URI-DNase+UVR+UvrC_HhH_N+HHH||NUDIX->?->HTH+McrB-AAA->McrC-NTD+REase->?->BrxA-DNA-binding->BrxB-BrxF-iSTAND->DUF6079*->N6-METHYLASE->N6-METHYLASE-p->PglZ->SIGMA+OB-Fold+AAA-ATPase+Lon-Protease-BrxL->URI-DNase->SbcC->OLD-ABC+TOPRIM->NERD+ZnR-></td><td>1206</td><td>Bacteria>Terrabacteria_group>Bacillota>Bacilli>Bacillales>Bacillaceae</td><td>GCA_023516215.1</td><td>Cytobacillus_oceanisediminis</td><td>M5V91_04885</td><td>909305..912925</td><td>665099</td><td>BREX-system-P-loop-protein-BrxC-[Cytobacillus-oceanisediminis]</td></tr>
<tr><td>WP_033827164.1</td><td>Type1.BREX</td><td>DDE_Tnp_IS240+rve_3-><-?<-DDE_Tnp<-zf-ISL3+HTH_Tnp_ISL3||<-PglZ<-REase+HTH<-N6-METHYLASE<-N6-METHYLASE-p<-N6-METHYLASE-p<-DUF6079*<-BrxB-BrxF-iSTAND<-BrxA-DNA-binding<-REase+HTH<-?<-BrxA-DNA-binding-p<-REase+HTH<-?<-BrxA-DNA-binding<-REase+HTH</td><td>1186</td><td>Bacteria>Terrabacteria_group>Bacillota>Bacilli>Bacillales>Bacillaceae</td><td>GCF_902375605.1</td><td>Bacillus_andreraoultii</td><td>FY465_RS04670</td><td>342053..345613</td><td>1499685</td><td>BREX-system-P-loop-protein-BrxC-[Bacillus-andreraoultii]</td></tr>
<tr><td>WP_102415646.1</td><td>Type1.BREX</td><td>PTSEIIA1->Cas9+HNH+RuvC->Cas_Cas1->CRISPR_Cas2->BrxA-DNA-binding->BrxB-BrxF-iSTAND->DUF6079*->REase+HTH-><-TetR-HTH</td><td>1340</td><td>Bacteria>Terrabacteria_group>Bacillota>Bacilli>Bacillales>Bacillaceae</td><td>-</td><td>Virgibacillus_dakarensis</td><td>KK120_02535</td><td>179341..183363</td><td>1917889</td><td>MULTISPECIES:-BREX-system-P-loop-protein-BrxC-[Terrabacteria-group]</td></tr>
<tr><td>WP_250889608.1</td><td>Type1.BREX</td><td>PTSEIIA1->Cas9+HNH+RuvC->Cas_Cas1->CRISPR_Cas2->BrxA-DNA-binding->BrxB-BrxF-iSTAND->DUF6079*->REase+HTH-><-TetR-HTH</td><td>1335</td><td>Bacteria>Terrabacteria_group>Bacillota>Bacilli>Bacillales>Bacillaceae</td><td>-</td><td>Virgibacillus_dakarensis</td><td>KK120_RS02530</td><td>179356..183363</td><td>1917889</td><td>BREX-system-P-loop-protein-BrxC-[Virgibacillus-dakarensis]</td></tr>
<tr><td>MBT2292692.1</td><td>Type1.BREX</td><td><-MarR-HTH||?->?-><-PadR-HTH<-PadR-HTH<-SIGMA+OB-Fold+AAA-ATPase+Lon-Protease-BrxL<-PglZ<-N6-METHYLASE<-DUF6079*<-BrxB-BrxF-iSTAND<-BrxA-DNA-binding<-?<-wHTH+REase||?-><-HTH_37<-DOC+HTH</td><td>1188</td><td>Bacteria>Terrabacteria_group>Bacillota>Bacilli>Bacillales>Paenibacillaceae</td><td>GCA_018614985.1</td><td>Paenibacillus_albidus</td><td>J7E73_26890</td><td>15469..19035</td><td>2041023</td><td>BREX-system-P-loop-protein-BrxC-[Paenibacillus-albidus]</td></tr>
<tr><td>MCG7380059.1</td><td>Type1.BREX</td><td><-PadR-HTH<-PadR-HTH<-ParA-Soj-PloopNTPase<-?<-SIGMA+OB-Fold+AAA-ATPase+Lon-Protease-BrxL<-PglZ<-N6-METHYLASE<-DUF6079*<-BrxB-BrxF-iSTAND<-BrxA-DNA-binding<-?<-wHTH+REase<-?<-HTH_37<-DOC+HTH</td><td>1187</td><td>Bacteria>Terrabacteria_group>Bacillota>Bacilli>Bacillales>Paenibacillaceae</td><td>GCA_022346845.1</td><td>Paenibacillus_sp._ACRSA</td><td>MH215_23985</td><td>103877..107440</td><td>2918211</td><td>BREX-system-P-loop-protein-BrxC-[Paenibacillus-sp.-ACRSA]</td></tr>
<tr><td>WP_056697878.1</td><td>Type1.BREX</td><td><-MarR-HTH||?->?-><-PadR-HTH<-PadR-HTH<-?<-SIGMA+OB-Fold+AAA-ATPase+Lon-Protease-BrxL<-PglZ<-N6-METHYLASE<-DUF6079*<-BrxB-BrxF-iSTAND<-BrxA-DNA-binding<-?<-wHTH+REase<-?<-HTH_37<-DOC+HTH<-NUDIX</td><td>1188</td><td>Bacteria>Terrabacteria_group>Bacillota>Bacilli>Bacillales>Paenibacillaceae</td><td>GCF_001426865.1</td><td>Paenibacillus_sp._Root52</td><td>ASD24_RS16940</td><td>47514..51080</td><td>1736552</td><td>BREX-system-P-loop-protein-BrxC-[Paenibacillus-sp.-Root52]</td></tr>
<tr><td>WP_155620894.1</td><td>Type1.BREX</td><td><-pPIWI+pPIWI<-?<-?<-RVT<-?<-SIGMA+OB-Fold+AAA-ATPase+Lon-Protease-BrxL<-PglZ<-SIR2<-N6-METHYLASE<-DUF6079*<-BrxB-BrxF-iSTAND<-BrxA-DNA-binding<-McrC-NTD+REase<-McrB-AAA<-?||35exo-><-SF1-SF2-Helicase+REase<-SF1-SF2-Helicase+REase<-wHTH+WYL+WCX</td><td>1206</td><td>Bacteria>Terrabacteria_group>Bacillota>Bacilli>Bacillales>Paenibacillaceae</td><td>GCF_018403165.1</td><td>Paenibacillus_macerans</td><td>KJS71_RS16940</td><td>170315..173935</td><td>44252</td><td>BREX-system-P-loop-protein-BrxC-[Paenibacillus-macerans]</td></tr>
<tr><td>WP_251038338.1</td><td>Type1.BREX</td><td><-MarR-HTH||?->?-><-PadR-HTH<-PadR-HTH<-SIGMA+OB-Fold+AAA-ATPase+Lon-Protease-BrxL<-PglZ<-N6-METHYLASE<-DUF6079*<-BrxB-BrxF-iSTAND<-BrxA-DNA-binding<-?<-wHTH+REase||?-><-HTH_37<-DOC+HTH</td><td>1183</td><td>Bacteria>Terrabacteria_group>Bacillota>Bacilli>Bacillales>Paenibacillaceae</td><td>GCF_018614985.1</td><td>Paenibacillus_albidus</td><td>J7E73_RS26885</td><td>15469..19020</td><td>2041023</td><td>BREX-system-P-loop-protein-BrxC-[Paenibacillus-albidus]</td></tr>
<tr><td>WP_222215805.1</td><td>Type1.BREX</td><td><-MarR-HTH||?->?<-?<-PadR-HTH<-PadR-HTH<-ParA-Soj-PloopNTPase<-SIGMA+OB-Fold+AAA-ATPase+Lon-Protease-BrxL<-PglZ<-N6-METHYLASE<-DUF6079*<-BrxB-BrxF-iSTAND<-BrxA-DNA-binding<-?<-wHTH+REase||?-><-HTH_37<-DOC+HTH||?-><-NUDIX</td><td>1188</td><td>Bacteria>Terrabacteria_group>Bacillota>Bacilli>Bacillales>Paenibacillaceae</td><td>GCF_019793795.1</td><td>Paenibacillus_sp._DR312</td><td>K5K90_RS02600</td><td>595089..598655</td><td>2871175</td><td>BREX-system-P-loop-protein-BrxC-[Paenibacillus-sp.-DR312]</td></tr>
<tr><td>WP_275446318.1</td><td>Type1.BREX</td><td><-PadR-HTH<-PadR-HTH<-ParA-Soj-PloopNTPase<-?<-SIGMA+OB-Fold+AAA-ATPase+Lon-Protease-BrxL<-PglZ<-N6-METHYLASE<-DUF6079*<-BrxB-BrxF-iSTAND<-BrxA-DNA-binding<-?<-wHTH+REase<-?<-HTH_37<-DOC+HTH</td><td>1182</td><td>Bacteria>Terrabacteria_group>Bacillota>Bacilli>Bacillales>Paenibacillaceae</td><td>GCF_022346845.1</td><td>Paenibacillus_sp._ACRSA</td><td>MH215_RS23980</td><td>103877..107425</td><td>2918211</td><td>BREX-system-P-loop-protein-BrxC-[Paenibacillus-sp.-ACRSA]</td></tr>
<tr><td>WP_091013323.1</td><td>Type1.BREX</td><td>DOC+HTH->HTH_37->?->wHTH+REase->?->BrxA-DNA-binding->BrxB-BrxF-iSTAND->DUF6079*->N6-METHYLASE->PglZ->SIGMA+OB-Fold+AAA-ATPase+Lon-Protease-BrxL->ParA-Soj-PloopNTPase->PadR-HTH->PadR-HTH->?->wHTH+WYL-></td><td>1189</td><td>Bacteria>Terrabacteria_group>Bacillota>Bacilli>Bacillales>Paenibacillaceae</td><td>GCF_900095775.1</td><td>Paenibacillus_amylolyticus</td><td>BQ2370_RS05155</td><td>8508..12077</td><td>1451</td><td>BREX-system-P-loop-protein-BrxC-[Paenibacillus-amylolyticus]</td></tr>
<tr><td>WP_215184617.1</td><td>Type1.BREX</td><td><-MarR-HTH||?->?-><-PadR-HTH<-PadR-HTH<-SIGMA+OB-Fold+AAA-ATPase+Lon-Protease-BrxL<-PglZ<-N6-METHYLASE<-DUF6079*<-BrxB-BrxF-iSTAND<-BrxA-DNA-binding<-?<-wHTH+REase||?-><-HTH_37<-DOC+HTH</td><td>1188</td><td>Bacteria>Terrabacteria_group>Bacillota>Bacilli>Bacillales>Paenibacillaceae</td><td>-</td><td>Paenibacillus_albidus</td><td>J7E73_26890</td><td>15469..19035</td><td>2041023</td><td>BREX-system-P-loop-protein-BrxC-[Paenibacillus-albidus]</td></tr>
<tr><td>WP_239293644.1</td><td>Type1.BREX</td><td><-PadR-HTH<-PadR-HTH<-ParA-Soj-PloopNTPase<-?<-SIGMA+OB-Fold+AAA-ATPase+Lon-Protease-BrxL<-PglZ<-N6-METHYLASE<-DUF6079*<-BrxB-BrxF-iSTAND<-BrxA-DNA-binding<-?<-wHTH+REase<-?<-HTH_37<-DOC+HTH</td><td>1187</td><td>Bacteria>Terrabacteria_group>Bacillota>Bacilli>Bacillales>Paenibacillaceae</td><td>-</td><td>Paenibacillus_sp._ACRSA</td><td>MH215_23985</td><td>103877..107440</td><td>2918211</td><td>BREX-system-P-loop-protein-BrxC-[Paenibacillus-sp.-ACRSA]</td></tr>
<tr><td>WP_153832550.1</td><td>Type1.BREX</td><td><-SF1-SF2-Helicase<-OLD-ABC+TOPRIM<-HTH+REase<-?<-SIGMA+OB-Fold+AAA-ATPase+Lon-Protease-BrxL<-PglZ<-N6-METHYLASE<-DUF6079*<-BrxB-BrxF-iSTAND<-BrxA-DNA-binding||Phage_integrase-><-DapB_N+DapB_C</td><td>1185</td><td>Bacteria>Terrabacteria_group>Bacillota>Bacilli>Lactobacillales>Aerococcaceae</td><td>GCF_009659375.1</td><td>Fundicoccus_ignavus</td><td>GF867_RS07800</td><td>14884..18441</td><td>2664442</td><td>BREX-system-P-loop-protein-BrxC-[Fundicoccus-ignavus]</td></tr>
<tr><td>WP_126762085.1</td><td>Type1.BREX</td><td>Peptidase-><-KAP-NTPase<-HTH+REase<-?<-SIGMA+OB-Fold+AAA-ATPase+Lon-Protease-BrxL<-PglZ<-N6-METHYLASE<-DUF6079*<-BrxB-BrxF-iSTAND<-BrxA-DNA-binding<-?<-?<-?<-?<-ABC-ATPase+SbcC<-ABC-ATPase+SbcC</td><td>1186</td><td>Bacteria>Terrabacteria_group>Bacillota>Bacilli>Lactobacillales>Enterococcaceae</td><td>GCF_003987555.1</td><td>Vagococcus_lutrae</td><td>CBF33_RS04670</td><td>25387..28947</td><td>81947</td><td>BREX-system-P-loop-protein-BrxC-[Vagococcus-lutrae]</td></tr>
<tr><td>WP_258233427.1</td><td>Type1.BREX</td><td><-CHTH<-?<-?<-?<-HTH+REase<-SIGMA+OB-Fold+AAA-ATPase+Lon-Protease-BrxL<-PglZ<-N6-METHYLASE<-DUF6079*<-BrxB-BrxF-iSTAND<-BrxA-DNA-binding<-?<-?<-?<-PTSEIIC</td><td>1227</td><td>Bacteria>Terrabacteria_group>Bacillota>Bacilli>Lactobacillales>Enterococcaceae</td><td>GCF_901212505.1</td><td>Enterococcus_avium</td><td>EAVMC1_RS06665</td><td>43716..47399</td><td>33945</td><td>BREX-system-P-loop-protein-BrxC-[Enterococcus-avium]</td></tr>
<tr><td>WP_034992311.1</td><td>Type1.BREX</td><td>GTPase-AIG->?->HTH+REase->BrxA-DNA-binding->BrxB-BrxF-iSTAND->DUF6079*->N6-METHYLASE-></td><td>1228</td><td>Bacteria>Terrabacteria_group>Bacillota>Bacilli>Lactobacillales>Lactobacillaceae</td><td>GCF_000255515.1</td><td>Liquorilactobacillus_vini</td><td>IC26_RS02660</td><td>4237..7923</td><td>238015</td><td>BREX-system-P-loop-protein-BrxC-[Liquorilactobacillus-vini]</td></tr>
<tr><td>WP_009555629.1</td><td>Type1.BREX</td><td>HAD+S6PP->?-><-REase<-?<-HTH+REase<-PglZ<-N6-METHYLASE<-DUF6079*<-BrxB-BrxF-iSTAND<-BrxA-DNA-binding<-?<-?<-HKD-DNase+SF1-SF2-Helicase+HTH+HTH+SRA</td><td>1218</td><td>Bacteria>Terrabacteria_group>Bacillota>Bacilli>Lactobacillales>Lactobacillaceae</td><td>GCF_000317165.1</td><td>Ligilactobacillus_saerimneri</td><td>D271_RS07755</td><td>76703..80359</td><td>228229</td><td>BREX-system-P-loop-protein-BrxC-[Ligilactobacillus-saerimneri]</td></tr>
<tr><td>WP_016383162.1</td><td>Type1.BREX</td><td><-HTH+REase<-SIGMA+OB-Fold+AAA-ATPase+Lon-Protease-BrxL<-PglZ||N6-METHYLASE->Arm-DNA-bind_4+Phage_int_SAM_5+Phage_integrase->zf-ISL3+HTH_Tnp_ISL3+DDE_Tnp_ISL3-><-N6-METHYLASE<-DUF6079*<-BrxB-BrxF-iSTAND<-BrxA-DNA-binding||MazE_antitoxin-><-?<-?<-LACI-HTH+PBPI<-Glyco_hydro_32N+Glyco_hydro_32C<-?<-SAM-METHYLASE</td><td>1215</td><td>Bacteria>Terrabacteria_group>Bacillota>Bacilli>Lactobacillales>Lactobacillaceae</td><td>GCF_000409835.1</td><td>Lacticaseibacillus_paracasei</td><td>J3H89_RS10660</td><td>2208273..2211920</td><td>1597</td><td>BREX-system-P-loop-protein-BrxC-[Lacticaseibacillus-paracasei]</td></tr>
<tr><td>WP_025012991.1</td><td>Type1.BREX</td><td>HTH_6+SIS-><-HTH+REase<-SIGMA+OB-Fold+AAA-ATPase+Lon-Protease-BrxL<-PglZ||N6-METHYLASE->Phage_int_SAM_5+Phage_integrase-><-N6-METHYLASE<-DUF6079*<-BrxB-BrxF-iSTAND<-BrxA-DNA-binding<-?||MazE_antitoxin-><-?<-?<-LACI-HTH+PBPI<-Glyco_hydro_32N+Glyco_hydro_32C</td><td>1214</td><td>Bacteria>Terrabacteria_group>Bacillota>Bacilli>Lactobacillales>Lactobacillaceae</td><td>GCF_000829055.1</td><td>Lacticaseibacillus</td><td>LBCZ_RS09700</td><td>1993700..1997344</td><td>2759736</td><td>MULTISPECIES:-BREX-system-P-loop-protein-BrxC-[Lacticaseibacillus]</td></tr>
<tr><td>WP_010491697.1</td><td>Type1.BREX</td><td>Glyco_hydro_32N+Glyco_hydro_32C->LACI-HTH+PBPI->?->?->?-><-MazE_antitoxin||?->BrxA-DNA-binding->BrxB-BrxF-iSTAND->DUF6079*->N6-METHYLASE-><-Phage_int_SAM_5+Phage_integrase-><-N6-METHYLASE||PglZ->SIGMA+OB-Fold+AAA-ATPase+Lon-Protease-BrxL->HTH+REase->HTH+REase-><-HTH_6+SIS</td><td>1214</td><td>Bacteria>Terrabacteria_group>Bacillota>Bacilli>Lactobacillales>Lactobacillaceae</td><td>GCF_001433745.1</td><td>Lacticaseibacillus_zeae</td><td>FD51_RS09320</td><td>70770..74414</td><td>57037</td><td>BREX-system-P-loop-protein-BrxC-[Lacticaseibacillus-zeae]</td></tr>
<tr><td>WP_056966680.1</td><td>Type1.BREX</td><td>wHTH-><-HTH+REase<-SIGMA+OB-Fold+AAA-ATPase+Lon-Protease-BrxL<-PglZ<-N6-METHYLASE-p||Phage_int_SAM_5+Phage_integrase->N6-METHYLASE-p-><-N6-METHYLASE<-DUF6079*<-BrxB-BrxF-iSTAND<-BrxA-DNA-binding<-PHP+35exo+PHP+ICAD<-?<-S2P+PDZ</td><td>1215</td><td>Bacteria>Terrabacteria_group>Bacillota>Bacilli>Lactobacillales>Lactobacillaceae</td><td>GCF_001435495.1</td><td>Latilactobacillus_curvatus</td><td>LCU_RS04405</td><td>828464..832111</td><td>28038</td><td>BREX-system-P-loop-protein-BrxC-[Latilactobacillus-curvatus]</td></tr>
<tr><td>WP_069468096.1</td><td>Type1.BREX</td><td>S2P+PDZ->?->PHP+35exo+PHP+ICAD->BrxA-DNA-binding->BrxB-BrxF-iSTAND->DUF6079*->N6-METHYLASE-><-N6-METHYLASE-p<-Phage_int_SAM_5+Phage_integrase||N6-METHYLASE->p->PglZ->HTH+REase-><-wHTH</td><td>1215</td><td>Bacteria>Terrabacteria_group>Bacillota>Bacilli>Lactobacillales>Lactobacillaceae</td><td>GCF_001723545.1</td><td>Latilactobacillus_curvatus</td><td>LCW_03905</td><td>751343..754990</td><td>28038</td><td>BREX-system-P-loop-protein-BrxC-[Latilactobacillus-curvatus]</td></tr>
<tr><td>WP_085391474.1</td><td>Type1.BREX</td><td>wHTH-><-HTH+REase<-SIGMA+OB-Fold+AAA-ATPase+Lon-Protease-BrxL<-PglZ<-N6-METHYLASE-p||Phage_int_SAM_5+Phage_integrase->N6-METHYLASE-p-><-N6-METHYLASE<-DUF6079*<-BrxB-BrxF-iSTAND<-BrxA-DNA-binding<-MazG<-PHP+35exo+PHP+ICAD<-?<-S2P+PDZ</td><td>1219</td><td>Bacteria>Terrabacteria_group>Bacillota>Bacilli>Lactobacillales>Lactobacillaceae</td><td>GCF_002105575.1</td><td>Latilactobacillus_sakei</td><td>LP065_RS02285</td><td>462739..466398</td><td>1599</td><td>BREX-system-P-loop-protein-BrxC-[Latilactobacillus-sakei]</td></tr>
<tr><td>WP_096584693.1</td><td>Type1.BREX</td><td>S2P+PDZ->?->PHP+35exo+PHP+ICAD->MazG->BrxA-DNA-binding->BrxB-BrxF-iSTAND->DUF6079*->N6-METHYLASE->AAA-ATPase->HIN-HTH+PAIREDC-HTH->N6-METHYLASE-p->PglZ->SIGMA+OB-Fold+AAA-ATPase+Lon-Protease-BrxL->HTH+REase-><-wHTH<-GNTR-HTH+UTRA</td><td>1215</td><td>Bacteria>Terrabacteria_group>Bacillota>Bacilli>Lactobacillales>Lactobacillaceae</td><td>GCF_002370375.1</td><td>Latilactobacillus_sakei</td><td>CCX78_RS03480</td><td>665145..668792</td><td>1599</td><td>BREX-system-P-loop-protein-BrxC-[Latilactobacillus-sakei]</td></tr>
<tr><td>WP_126133408.1</td><td>Type1.BREX</td><td>wHTH-><-HTH+REase<-SIGMA+OB-Fold+AAA-ATPase+Lon-Protease-BrxL<-PglZ<-N6-METHYLASE-p||Phage_int_SAM_5+Phage_integrase->N6-METHYLASE-p-><-HTH_Tnp_1+HTH_21+rve+rve_3<-N6-METHYLASE<-DUF6079*<-BrxB-BrxF-iSTAND<-BrxA-DNA-binding<-PHP+35exo+PHP+ICAD<-?<-S2P+PDZ</td><td>1219</td><td>Bacteria>Terrabacteria_group>Bacillota>Bacilli>Lactobacillales>Lactobacillaceae</td><td>GCF_003957415.1</td><td>Latilactobacillus_curvatus</td><td>CYK59_RS06580</td><td>1253905..1257564</td><td>28038</td><td>BREX-system-P-loop-protein-BrxC-[Latilactobacillus-curvatus]</td></tr>
<tr><td>WP_126353447.1</td><td>Type1.BREX</td><td><-HTH+REase<-rve<-HTH+REase<-SIGMA+OB-Fold+AAA-ATPase+Lon-Protease-BrxL<-PglZ||N6-METHYLASE->Arm-DNA-bind_4+Phage_int_SAM_5+Phage_integrase-><-N6-METHYLASE<-DUF6079*<-BrxB-BrxF-iSTAND<-BrxA-DNA-binding||MazE_antitoxin-><-?<-?<-LACI-HTH+PBPI<-Glyco_hydro_32N+Glyco_hydro_32C||PAIREDC-HTH+HIN-HTH-><-SAM-METHYLASE</td><td>1212</td><td>Bacteria>Terrabacteria_group>Bacillota>Bacilli>Lactobacillales>Lactobacillaceae</td><td>GCF_003966835.1</td><td>Lacticaseibacillus_paracasei</td><td>EL355_RS11595</td><td>2248100..2251738</td><td>1597</td><td>BREX-system-P-loop-protein-BrxC-[Lacticaseibacillus-paracasei]</td></tr>
<tr><td>WP_150392678.1</td><td>Type1.BREX</td><td><-HTH+REase<-REase<-SIGMA+OB-Fold+AAA-ATPase+Lon-Protease-BrxL<-PglZ||N6-METHYLASE->p->PAIREDC-HTH->Arm-DNA-bind_4+Phage_int_SAM_5+Phage_integrase-><-N6-METHYLASE<-DUF6079*<-BrxB-BrxF-iSTAND<-BrxA-DNA-binding<-SF1-SF2-Helicase||?->?->?->?-><-?<-ABC-ATPase+SbcC</td><td>1241</td><td>Bacteria>Terrabacteria_group>Bacillota>Bacilli>Lactobacillales>Lactobacillaceae</td><td>GCF_008694105.1</td><td>Schleiferilactobacillus_harbinensis</td><td>FMM01_RS11950</td><td>2413874..2417599</td><td>304207</td><td>BREX-system-P-loop-protein-BrxC-[Schleiferilactobacillus-harbinensis]</td></tr>
<tr><td>WP_152261659.1</td><td>Type1.BREX</td><td><-HTH+REase<-SIGMA+OB-Fold+AAA-ATPase+Lon-Protease-BrxL<-PglZ||N6-METHYLASE-><-N6-METHYLASE<-DUF6079*<-BrxB-BrxF-iSTAND<-BrxA-DNA-binding</td><td>1241</td><td>Bacteria>Terrabacteria_group>Bacillota>Bacilli>Lactobacillales>Lactobacillaceae</td><td>GCF_009217765.1</td><td>Schleiferilactobacillus_harbinensis</td><td>D1010_RS17280</td><td>3443993..3447718</td><td>304207</td><td>BREX-system-P-loop-protein-BrxC-[Schleiferilactobacillus-harbinensis]</td></tr>
<tr><td>WP_160528668.1</td><td>Type1.BREX</td><td>BrxA-DNA-binding->BrxB-BrxF-iSTAND->DUF6079*->N6-METHYLASE-><-Arm-DNA-bind_4+Phage_int_SAM_5+Phage_integrase<-N6-METHYLASE||PglZ->SIGMA+OB-Fold+AAA-ATPase+Lon-Protease-BrxL->HTH+REase-></td><td>1217</td><td>Bacteria>Terrabacteria_group>Bacillota>Bacilli>Lactobacillales>Lactobacillaceae</td><td>GCF_009823605.1</td><td>Lacticaseibacillus_paracasei</td><td>GRZ59_RS05150</td><td>1513..5166</td><td>1597</td><td>BREX-system-P-loop-protein-BrxC-[Lacticaseibacillus-paracasei]</td></tr>
<tr><td>WP_162100826.1</td><td>Type1.BREX</td><td>wHTH-><-HTH+REase<-SIGMA+OB-Fold+AAA-ATPase+Lon-Protease-BrxL<-PglZ<-N6-METHYLASE-p||Phage_int_SAM_5+Phage_integrase->N6-METHYLASE-p-><-N6-METHYLASE<-DUF6079*<-BrxB-BrxF-iSTAND<-BrxA-DNA-binding<-MazG<-PHP+35exo+PHP+ICAD<-?<-S2P+PDZ</td><td>1219</td><td>Bacteria>Terrabacteria_group>Bacillota>Bacilli>Lactobacillales>Lactobacillaceae</td><td>GCF_010092945.1</td><td>Latilactobacillus_sakei</td><td>GSF71_RS01930</td><td>394668..398327</td><td>1599</td><td>BREX-system-P-loop-protein-BrxC-[Latilactobacillus-sakei]</td></tr>
<tr><td>WP_179394507.1</td><td>Type1.BREX</td><td><-HTH_3||?->?->SbcC->TOPRIM->BrxA-DNA-binding->BrxB-BrxF-iSTAND->DUF6079*->N6-METHYLASE-><-Arm-DNA-bind_4+Phage_int_SAM_5+Phage_integrase||PglZ->SIGMA+OB-Fold+AAA-ATPase+Lon-Protease-BrxL->HTH+REase->?->ZnR+HARE-HTH->ARG-></td><td>1200</td><td>Bacteria>Terrabacteria_group>Bacillota>Bacilli>Lactobacillales>Lactobacillaceae</td><td>GCF_013407595.1</td><td>Lacticaseibacillus_absianus</td><td>HII33_RS04075</td><td>947851..951453</td><td>2729623</td><td>BREX-system-P-loop-protein-BrxC-[Lacticaseibacillus-absianus]</td></tr>
<tr><td>WP_188891961.1</td><td>Type1.BREX</td><td>ParB+HNH->REase->MULE-TRANSPOSASE->ParB+HNH->BrxA-DNA-binding->BrxB-BrxF-iSTAND->DUF6079*->N6-METHYLASE->nSTAND3->PglZ->?->HTH+REase->TPR+TPR-></td><td>1241</td><td>Bacteria>Terrabacteria_group>Bacillota>Bacilli>Lactobacillales>Lactobacillaceae</td><td>GCF_014656585.1</td><td>Lactobacillus_kefiranofaciens</td><td>ICI50_RS03490</td><td>670073..673798</td><td>267818</td><td>BREX-system-P-loop-protein-BrxC-[Lactobacillus-kefiranofaciens]</td></tr>
<tr><td>WP_251908041.1</td><td>Type1.BREX</td><td>wHTH-><-HTH+REase<-SIGMA+OB-Fold+AAA-ATPase+Lon-Protease-BrxL<-PglZ<-N6-METHYLASE-p||Phage_int_SAM_5+Phage_integrase-><-N6-METHYLASE<-DUF6079*<-BrxB-BrxF-iSTAND<-BrxA-DNA-binding<-MazG<-PHP+35exo+PHP+ICAD<-?<-S2P+PDZ</td><td>1219</td><td>Bacteria>Terrabacteria_group>Bacillota>Bacilli>Lactobacillales>Lactobacillaceae</td><td>GCF_023734335.1</td><td>Latilactobacillus_sakei</td><td>A4W84_RS06360</td><td>1251250..1254909</td><td>1599</td><td>BREX-system-P-loop-protein-BrxC-[Latilactobacillus-sakei]</td></tr>
<tr><td>WP_105300054.1</td><td>Type1.BREX</td><td>S2P+PDZ->?->PHP+35exo+PHP+ICAD->MazG->BrxA-DNA-binding->BrxB-BrxF-iSTAND->DUF6079*->N6-METHYLASE-><-Phage_int_SAM_5+Phage_integrase||N6-METHYLASE-p->PglZ->SIGMA+OB-Fold+AAA-ATPase+Lon-Protease-BrxL->HTH+REase-><-wHTH<-GNTR-HTH+UTRA||Glyco_hydro_1-></td><td>1215</td><td>Bacteria>Terrabacteria_group>Bacillota>Bacilli>Lactobacillales>Lactobacillaceae</td><td>GCF_024022895.1</td><td>Latilactobacillus_sakei</td><td>LAS9267_RS03030</td><td>155179..158826</td><td>1599</td><td>BREX-system-P-loop-protein-BrxC-[Latilactobacillus-sakei]</td></tr>
<tr><td>WP_254681527.1</td><td>Type1.BREX</td><td>S2P+PDZ->?->PHP+35exo+PHP+ICAD->MazG->BrxA-DNA-binding->BrxB-BrxF-iSTAND->DUF6079*->N6-METHYLASE-><-N6-METHYLASE-p<-Phage_int_SAM_5+Phage_integrase||N6-METHYLASE-p->PglZ->SIGMA+OB-Fold+AAA-ATPase+Lon-Protease-BrxL->HTH+REase-><-wHTH<-GNTR-HTH+UTRA</td><td>1219</td><td>Bacteria>Terrabacteria_group>Bacillota>Bacilli>Lactobacillales>Lactobacillaceae</td><td>GCF_024273085.1</td><td>Latilactobacillus_sakei</td><td>LOX63_RS01755</td><td>341729..345388</td><td>1599</td><td>BREX-system-P-loop-protein-BrxC-[Latilactobacillus-sakei]</td></tr>
<tr><td>WP_265678613.1</td><td>Type1.BREX</td><td>wHTH-><-HTH+REase<-HTH+REase<-SIGMA+OB-Fold+AAA-ATPase+Lon-Protease-BrxL<-PglZ<-N6-METHYLASE-p||Phage_int_SAM_5+Phage_integrase-><-N6-METHYLASE<-DUF6079*<-BrxB-BrxF-iSTAND<-BrxA-DNA-binding<-PHP+35exo+PHP+ICAD<-?<-S2P+PDZ</td><td>1215</td><td>Bacteria>Terrabacteria_group>Bacillota>Bacilli>Lactobacillales>Lactobacillaceae</td><td>GCF_026193275.1</td><td>Latilactobacillus_curvatus</td><td>OP752_RS00225</td><td>50217..53864</td><td>28038</td><td>BREX-system-P-loop-protein-BrxC-[Latilactobacillus-curvatus]</td></tr>
<tr><td>WP_270411955.1</td><td>Type1.BREX</td><td><-DDE_Tnp<-?<-HTH+REase<-SIGMA+OB-Fold+AAA-ATPase+Lon-Protease-BrxL<-PglZ<-N6-METHYLASE<-DUF6079*<-BrxB-BrxF-iSTAND<-BrxA-DNA-binding<-HTH_26<-?<-?||HAD-><-?<-?||Peptidase_C69-></td><td>1184</td><td>Bacteria>Terrabacteria_group>Bacillota>Bacilli>Lactobacillales>Lactobacillaceae</td><td>GCF_027672425.1</td><td>Ligilactobacillus_ruminis</td><td>PGT38_RS00055</td><td>14514..18068</td><td>1623</td><td>BREX-system-P-loop-protein-BrxC-[Ligilactobacillus-ruminis]</td></tr>
<tr><td>WP_270447891.1</td><td>Type1.BREX</td><td>ABhydrolase+ABhydrolase-><-?||?->?->?->HTH+REase->BrxA-DNA-binding->BrxB-BrxF-iSTAND->DUF6079*->N6-METHYLASE->PglZ->SIGMA+OB-Fold+AAA-ATPase+Lon-Protease-BrxL->HEPN->HEPN->SF1-SF2-Helicase->?-><-AraC-HTH</td><td>1188</td><td>Bacteria>Terrabacteria_group>Bacillota>Bacilli>Lactobacillales>Lactobacillaceae</td><td>GCF_027681365.1</td><td>Lactobacillus_delbrueckii</td><td>PGX26_RS03415</td><td>69740..73306</td><td>1584</td><td>BREX-system-P-loop-protein-BrxC-[Lactobacillus-delbrueckii]</td></tr>
<tr><td>WP_273725609.1</td><td>Type1.BREX</td><td><-SIGMA+OB-Fold+AAA-ATPase+Lon-Protease-BrxL<-PglZ||N6-METHYLASE->Arm-DNA-bind_4+Phage_int_SAM_5+Phage_integrase-><-N6-METHYLASE<-DUF6079*<-BrxB-BrxF-iSTAND<-BrxA-DNA-binding||?->?-><-McrC-NTD+REase<-ParB+HNH</td><td>1194</td><td>Bacteria>Terrabacteria_group>Bacillota>Bacilli>Lactobacillales>Lactobacillaceae</td><td>GCF_028618945.1</td><td>Leuconostoc_mesenteroides</td><td>PNN51_RS08050</td><td>12149..15733</td><td>1245</td><td>BREX-system-P-loop-protein-BrxC-[Leuconostoc-mesenteroides]</td></tr>
<tr><td>WP_273960123.1</td><td>Type1.BREX</td><td>S2P+PDZ->?->PHP+35exo+PHP+ICAD->PAIREDC-HTH+HIN-HTH->BrxA-DNA-binding->BrxB-BrxF-iSTAND->DUF6079*->N6-METHYLASE-><-N6-METHYLASE-p<-Phage_int_SAM_5+Phage_integrase||N6-METHYLASE-p->PglZ->SIGMA+OB-Fold+AAA-ATPase+Lon-Protease-BrxL->HTH+REase-><-wHTH</td><td>1215</td><td>Bacteria>Terrabacteria_group>Bacillota>Bacilli>Lactobacillales>Lactobacillaceae</td><td>GCF_028622855.1</td><td>Latilactobacillus_curvatus</td><td>PSR33_RS06690</td><td>1305883..1309530</td><td>28038</td><td>BREX-system-P-loop-protein-BrxC-[Latilactobacillus-curvatus]</td></tr>
<tr><td>WP_099947387.1</td><td>Type1.BREX</td><td>GNTR-HTH+UTRA->wHTH-><-HTH+REase<-SIGMA+OB-Fold+AAA-ATPase+Lon-Protease-BrxL<-PglZ<-N6-METHYLASE-p||Phage_int_SAM_5+Phage_integrase->N6-METHYLASE-p-><-N6-METHYLASE<-DUF6079*<-BrxB-BrxF-iSTAND<-BrxA-DNA-binding<-MazG<-PHP+35exo+PHP+ICAD<-?<-S2P+PDZ</td><td>1219</td><td>Bacteria>Terrabacteria_group>Bacillota>Bacilli>Lactobacillales>Lactobacillaceae</td><td>GCF_900234395.1</td><td>Latilactobacillus_sakei</td><td>LSAJ54_RS06515</td><td>1291898..1295557</td><td>1599</td><td>BREX-system-P-loop-protein-BrxC-[Latilactobacillus-sakei]</td></tr>
<tr><td>WP_236911583.1</td><td>Type1.BREX</td><td>Cysteine-Desulfurase->?->AraC-HTH+AraC-HTH->?->?->BrxA-DNA-binding->BrxB-BrxF-iSTAND->DUF6079*->N6-METHYLASE->PglZ->SIGMA+OB-Fold+AAA-ATPase+Lon-Protease-BrxL->HARE-HTH+REase->?->?->RAMA+McrB-AAA->McrC-NTD+REase-></td><td>1193</td><td>Bacteria>Terrabacteria_group>Bacillota>Clostridia>Eubacteriales>Clostridiaceae</td><td>GCF_021655095.1</td><td>Clostridium_sp._Cult1</td><td>C3E89_RS06110</td><td>1240047..1243628</td><td>2079002</td><td>BREX-system-P-loop-protein-BrxC-[Clostridium-sp.-Cult1]</td></tr>
<tr><td>OLA08999.1</td><td>Type1.BREX</td><td><-Phage_int+DNA-glycosylase<-HTH+REase<-SIGMA+OB-Fold+AAA-ATPase+Lon-Protease-BrxL<-PglZ<-AbiGi<-N6-METHYLASE-p<-Schlafen+HATPase+wHTH<-?<-N6-METHYLASE<-DUF6079*<-BrxB-BrxF-iSTAND<-BrxA-DNA-binding<-HTH_26<-?<-HNH+3C-ABC||HTH_36-></td><td>1184</td><td>Bacteria>Terrabacteria_group>Bacillota>Clostridia>Eubacteriales>Eubacteriaceae</td><td>GCA_001916415.1</td><td>Eubacterium_sp._45_250</td><td>BHW21_06160</td><td>165792..169346</td><td>1897004</td><td>MAG:-ATPase-[Eubacterium-sp.-45_250]</td></tr>
<tr><td>EGG92088.1</td><td>Type1.BREX</td><td>Peptidase_M32->DOC+HTH->?->?-><-?<-ABhydrolase||BrxA-DNA-binding->BrxB-BrxF-iSTAND->DUF6079*->EVE+McrB-AAA->McrC-NTD+REase->N6-METHYLASE->RelB->Tox-RelE->PglZ->RADICAL-SAM-></td><td>1217</td><td>Bacteria>Terrabacteria_group>Bacillota>Clostridia>Eubacteriales>Lachnospiraceae</td><td>GCA_000209465.1</td><td>Lachnospiraceae_oral_taxon_107_str._F0167</td><td>HMPREF0491_00115</td><td>36108..39761</td><td>575593</td><td>hypothetical-protein-HMPREF0491_00115-[Lachnospiraceae-oral-taxon-107-str.-F0167]</td></tr>
<tr><td>EDS42584.1</td><td>Type1.BREX</td><td><-SbcC<-SbcC<-Lon_2<-PglZ<-HTH+REase<-N6-METHYLASE<-DUF6079*<-BrxB-BrxF-iSTAND<-BrxA-DNA-binding<-MULE-TRANSPOSASE<-LZ_Tnp_IS66+zf-IS66+DDE_Tnp_IS66+DDE_Tnp_IS66_C<-?<-Classical-AAA</td><td>1182</td><td>Bacteria>Terrabacteria_group>Bacillota>Clostridia>Eubacteriales>Lachnospiraceae</td><td>GCA_000403315.2</td><td>Lachnospiraceae_bacterium_A2</td><td>C810_04454</td><td>133846..137394</td><td>397290</td><td>hypothetical-protein-C810_04454-[Lachnospiraceae-bacterium-A2]</td></tr>
<tr><td>RKW51253.1</td><td>Type1.BREX</td><td><-N6-METHYLASE-p<-McrC-NTD+REase<-EVE+McrB-AAA<-DUF6079*<-BrxB-BrxF-iSTAND<-BrxA-DNA-binding||ABhydrolase-></td><td>1217</td><td>Bacteria>Terrabacteria_group>Bacillota>Clostridia>Eubacteriales>Lachnospiraceae</td><td>GCA_003640285.1</td><td>Lachnospiraceae_bacterium</td><td>D8H95_17195</td><td>4877..8530</td><td>1898203</td><td>BREX-system-P-loop-protein-BrxC-[Lachnospiraceae-bacterium]</td></tr>
<tr><td>MCI6533897.1</td><td>Type1.BREX</td><td><-Arm-DNA-bind_4+Phage_int_SAM_3+Phage_integrase<-HTH<-PemK_toxin<-?<-SIGMA-HTH||HTH_26->?->BrxA-DNA-binding->BrxB-BrxF-iSTAND->DUF6079*->N6-METHYLASE->PglZ->HTH+REase->SIGMA+OB-Fold+AAA-ATPase+Lon-Protease-BrxL->SbcC->DNA_METHYLASE-><-?<-VirB4-FtsK</td><td>1173</td><td>Bacteria>Terrabacteria_group>Bacillota>Clostridia>Eubacteriales>Lachnospiraceae</td><td>GCA_022782135.1</td><td>Lachnospiraceae_bacterium</td><td>MR444_03700</td><td>25882..29403</td><td>1898203</td><td>BREX-system-P-loop-protein-BrxC-[Lachnospiraceae-bacterium]</td></tr>
<tr><td>SFG31088.1</td><td>Type1.BREX</td><td><-SIGMA+OB-Fold+AAA-ATPase+Lon-Protease-BrxL<-PglZ<-HTH+REase<-HNH+ADA-DBD<-?<-N6-METHYLASE<-DUF6079-p*<-DUF6079-p<-BrxB-BrxF-iSTAND<-BrxA-DNA-binding<-HTH_26<-?<-?<-?<-TraI-VirD2-RCRE<-TraI-VirD2-RCRE</td><td>783</td><td>Bacteria>Terrabacteria_group>Bacillota>Clostridia>Eubacteriales>Lachnospiraceae</td><td>GCA_900113415.1</td><td>Oribacterium_sp._WCC10</td><td>SAMN05216356_105117</td><td>149481..151832</td><td>1855343</td><td>hypothetical-protein-SAMN05216356_105117-[Oribacterium-sp.-WCC10]</td></tr>
<tr><td>SFG31099.1</td><td>Type1.BREX</td><td><-SIGMA+OB-Fold+AAA-ATPase+Lon-Protease-BrxL<-PglZ<-HTH+REase<-HNH+ADA-DBD<-?<-N6-METHYLASE<-DUF6079-p<-DUF6079-p*<-BrxB-BrxF-iSTAND<-BrxA-DNA-binding<-HTH_26<-?<-?<-?<-TraI-VirD2-RCRE<-TraI-VirD2-RCRE</td><td>380</td><td>Bacteria>Terrabacteria_group>Bacillota>Clostridia>Eubacteriales>Lachnospiraceae</td><td>GCA_900113415.1</td><td>Oribacterium_sp._WCC10</td><td>SAMN05216356_105118</td><td>151957..153099</td><td>1855343</td><td>hypothetical-protein-SAMN05216356_105118-[Oribacterium-sp.-WCC10]</td></tr>
<tr><td>SFL32270.1</td><td>Type1.BREX</td><td>DDE_Tnp->?->HTH_26->BrxA-DNA-binding->BrxB-BrxF-iSTAND->DUF6079*->N6-METHYLASE->HTH+REase->?->PglZ->SIGMA+OB-Fold+AAA-ATPase+Lon-Protease-BrxL-><-DSBH+AraC-HTH</td><td>1207</td><td>Bacteria>Terrabacteria_group>Bacillota>Clostridia>Eubacteriales>Lachnospiraceae</td><td>GCA_900114405.1</td><td>Lachnospiraceae_bacterium_KH1T2</td><td>SAMN05216390_1196</td><td>2450..6073</td><td>1855374</td><td>hypothetical-protein-SAMN05216390_1196-[Lachnospiraceae-bacterium-KH1T2]</td></tr>
<tr><td>WP_016216871.1</td><td>Type1.BREX</td><td><-HTH||||HTH->||HTH_26->BrxA-DNA-binding->BrxB-BrxF-iSTAND->DUF6079*-><-DEDD_Tnp_IS110+Transposase_20||HEPN->N6-METHYLASE->HTH+REase->Schlafen+HATPase+wHTH->PglZ-></td><td>1182</td><td>Bacteria>Terrabacteria_group>Bacillota>Clostridia>Eubacteriales>Lachnospiraceae</td><td>GCF_000403455.2</td><td>Dorea_sp._5-2</td><td>C817_RS00440</td><td>102069..105617</td><td>1235798</td><td>BREX-system-P-loop-protein-BrxC-[Dorea-sp.-5-2]</td></tr>
<tr><td>WP_173793922.1</td><td>Type1.BREX</td><td><-MazG<-SIGMA-HTH||HNH+3C-ABC->?->?->HTH_26->BrxA-DNA-binding->BrxB-BrxF-iSTAND->DUF6079*->N6-METHYLASE->TIR->PglZ->SIGMA+OB-Fold+AAA-ATPase+Lon-Protease-BrxL->HTH+REase->?->CHTH-></td><td>1184</td><td>Bacteria>Terrabacteria_group>Bacillota>Clostridia>Eubacteriales>Lachnospiraceae</td><td>GCF_013301385.1</td><td>Dorea_formicigenerans</td><td>G4398_RS10070</td><td>34100..37654</td><td>39486</td><td>BREX-system-P-loop-protein-BrxC-[Dorea-formicigenerans]</td></tr>
<tr><td>WP_021651779.1</td><td>Type1.BREX</td><td><-PAS+HISKIN+REC<-?<-ParB<-ParB<-SymE_toxin<-HTH+REase<-SIGMA+OB-Fold+AAA-ATPase+Lon-Protease-BrxL<-PglZ<-N6-METHYLASE<-DUF6079*<-BrxB-BrxF-iSTAND<-BrxA-DNA-binding<-HTH_26<-?<-HNH+3C-ABC||SIGMA-HTH-></td><td>1184</td><td>Bacteria>Terrabacteria_group>Bacillota>Clostridia>Eubacteriales>Lachnospiraceae</td><td>GCF_020735365.1</td><td>Blautia</td><td>LK422_RS11525</td><td>2410424..2413978</td><td>572511</td><td>MULTISPECIES:-BREX-system-P-loop-protein-BrxC-[Blautia]</td></tr>
<tr><td>RGG58361.1</td><td>Type1.BREX</td><td>DSBH+AraC-HTH+AraC-HTH-><-IONCHANNEL+HISKIN+REC<-HTH+REase<-SIGMA+OB-Fold+AAA-ATPase+Lon-Protease-BrxL<-PglZ<-?<-N6-METHYLASE<-DUF6079*<-BrxB-BrxF-iSTAND<-BrxA-DNA-binding<-HTH_26</td><td>1184</td><td>Bacteria>Terrabacteria_group>Bacillota>Clostridia>Eubacteriales>Oscillospiraceae</td><td>GCA_003461215.1</td><td>Ruminococcus_sp._AF19-15</td><td>DWX34_06625</td><td>16920..20474</td><td>2293157</td><td>BREX-system-P-loop-protein-BrxC-[Ruminococcus-sp.-AF19-15]</td></tr>
<tr><td>MCL2053364.1</td><td>Type1.BREX</td><td>TPR+TPR+TPR+TPR->?->TPR+TPR+TPR+TPR+TPR+TPR->BrxA-DNA-binding->BrxB-BrxF-iSTAND->DUF6079*-p*->DUF6079->p->?->SLOG+BirA-HTH->N6-METHYLASE->N6-METHYLASE->p->?->HTH+REase->PglZ->SIGMA+OB-Fold+AAA-ATPase+Lon-Protease-BrxL-></td><td>377</td><td>Bacteria>Terrabacteria_group>Bacillota>Clostridia>Eubacteriales>Oscillospiraceae</td><td>GCA_009777725.1</td><td>Oscillospiraceae_bacterium</td><td>FWG90_02835</td><td>151410..152543</td><td>2485925</td><td>BREX-system-P-loop-protein-BrxC-[Oscillospiraceae-bacterium]</td></tr>
<tr><td>MBQ6611347.1</td><td>Type1.BREX</td><td><-SIGMA+OB-Fold+AAA-ATPase+Lon-Protease-BrxL<-PglZ<-HTH+REase<-N6-METHYLASE<-DUF6079*<-BrxB-BrxF-iSTAND<-BrxA-DNA-binding||SIGMA-HTH->?->MqsA_antitoxin+DUF4065-><-?<-?<-N6-METHYLASE</td><td>1182</td><td>Bacteria>Terrabacteria_group>Bacillota>Clostridia>Eubacteriales>Oscillospiraceae</td><td>GCA_017434435.1</td><td>Oscillospiraceae_bacterium</td><td>IJH70_13030</td><td>9925..13473</td><td>2485925</td><td>BREX-system-P-loop-protein-BrxC-[Oscillospiraceae-bacterium]</td></tr>
<tr><td>UYJ00845.1</td><td>Type1.BREX</td><td><-NUDIX<-?<-HKD-DNase+SF1-SF2-Helicase+HTH+HTH+SRA||XIS-HTH->BrxA-DNA-binding->BrxB-BrxF-iSTAND->DUF6079*->HTH+REase->N6-METHYLASE->TIR->PglZ->SIGMA+OB-Fold+AAA-ATPase+Lon-Protease-BrxL->ParB+HNH->STYKinase-like-></td><td>1218</td><td>Bacteria>Terrabacteria_group>Bacillota>Clostridia>Eubacteriales>Oscillospiraceae</td><td>GCA_025757765.1</td><td>Faecalibacterium_prausnitzii</td><td>OGM65_10405</td><td>2149952..2153608</td><td>853</td><td>MAG:-BREX-system-P-loop-protein-BrxC-[Faecalibacterium-prausnitzii]</td></tr>
<tr><td>SCH98752.1</td><td>Type1.BREX</td><td>68TM-wHTH+CACHE+GGDEF->?->?->?->?->?->HTH_26->BrxA-DNA-binding->BrxB-BrxF-iSTAND->DUF6079*->N6-METHYLASE->?->PglZ->SIGMA+OB-Fold+AAA-ATPase+Lon-Protease-BrxL->HTH+REase->IONCHANNEL+HISKIN+REC->?-><-DSBH+AraC-HTH+AraC-HTH</td><td>1184</td><td>Bacteria>Terrabacteria_group>Bacillota>Clostridia>Eubacteriales>Oscillospiraceae</td><td>GCA_900067025.1</td><td>uncultured_Ruminococcus_sp.</td><td>SAMEA3545436_02036</td><td>58203..61757</td><td>165186</td><td>Uncharacterised-protein-[uncultured-Ruminococcus-sp.]</td></tr>
<tr><td>WP_087287659.1</td><td>Type1.BREX</td><td>Peptidase_M20-><-HTH+REase<-SIGMA+OB-Fold+AAA-ATPase+Lon-Protease-BrxL<-PglZ<-SbcC<-N6-METHYLASE<-DUF6079*<-BrxB-BrxF-iSTAND<-BrxA-DNA-binding<-HTH_26<-AbiEii<-AbiEi_4||TraJ-RHH->RepA_N+HTH->AAA+->XIS-HTH-></td><td>1180</td><td>Bacteria>Terrabacteria_group>Bacillota>Clostridia>Eubacteriales>Oscillospiraceae</td><td>GCF_002160305.1</td><td>Pseudoflavonifractor_sp._An184</td><td>B5F19_RS02685</td><td>31518..35060</td><td>1965576</td><td>BREX-system-P-loop-protein-BrxC-[Pseudoflavonifractor-sp.-An184]</td></tr>
<tr><td>WP_117891999.1</td><td>Type1.BREX</td><td>DSBH+AraC-HTH+AraC-HTH-><-IONCHANNEL+HISKIN+REC<-HTH+REase<-SIGMA+OB-Fold+AAA-ATPase+Lon-Protease-BrxL<-PglZ<-?<-N6-METHYLASE<-DUF6079*<-BrxB-BrxF-iSTAND<-BrxA-DNA-binding<-HTH_26</td><td>1184</td><td>Bacteria>Terrabacteria_group>Bacillota>Clostridia>Eubacteriales>Oscillospiraceae</td><td>GCF_003461215.1</td><td>Ruminococcus_sp._AF19-15</td><td>DWX34_06625</td><td>16920..20474</td><td>2293157</td><td>BREX-system-P-loop-protein-BrxC-[Ruminococcus-sp.-AF19-15]</td></tr>
<tr><td>WP_271847095.1</td><td>Type1.BREX</td><td>URI-DNase+RAMA-><-RelE-ParE<-PhdYeFM_antitox||?->HTH_26->BrxA-DNA-binding->BrxB-BrxF-iSTAND->DUF6079*->N6-METHYLASE->TIR->PglZ->SIGMA+OB-Fold+AAA-ATPase+Lon-Protease-BrxL->HTH+REase->?-><-?||ABC-ATPase+SbcC-></td><td>1184</td><td>Bacteria>Terrabacteria_group>Bacillota>Clostridia>Eubacteriales>Peptostreptococcaceae</td><td>GCF_028326405.1</td><td>Peptostreptococcus_anaerobius</td><td>PN283_RS04660</td><td>12553..16107</td><td>1261</td><td>BREX-system-P-loop-protein-BrxC-[Peptostreptococcus-anaerobius]</td></tr>
<tr><td>MCD8370861.1</td><td>Type1.BREX</td><td>HTH_26->BrxA-DNA-binding->BrxB-BrxF-iSTAND->DUF6079*->N6-METHYLASE->N6-METHYLASE-p->?->HTH+REase->PglZ-p-></td><td>1183</td><td>Bacteria>Terrabacteria_group>Bacillota>Clostridia>Eubacteriales>unclassified_Eubacteriales</td><td>GCA_021205425.1</td><td>Clostridiales_bacterium</td><td>LUC94_11190</td><td>2149..5700</td><td>1898207</td><td>BREX-system-P-loop-protein-BrxC-[Clostridiales-bacterium]</td></tr>
<tr><td>WP_114458231.1</td><td>Type1.BREX</td><td>Peptidase_M42+Peptidase_M28->?->?->?->HEPN+REase+SF1-SF2-Helicase->SF1-SF2-Helicase+REase->?->BrxA-DNA-binding->BrxB-BrxF-iSTAND->DUF6079*->N6-METHYLASE->PglZ->SIGMA+OB-Fold+AAA-ATPase+Lon-Protease-BrxL->?->REase->McrB-AAA->McrC-NTD+REase-></td><td>1191</td><td>Bacteria>Terrabacteria_group>Bacillota>Clostridia>Halanaerobiales>Halanaerobiaceae</td><td>GCF_003337245.1</td><td>Halanaerobium_sp._MA284_MarDTE_T2</td><td>DFR78_RS07540</td><td>27891..31466</td><td>2183913</td><td>BREX-system-P-loop-protein-BrxC-[Halanaerobium-sp.-MA284_MarDTE_T2]</td></tr>
<tr><td>MCR0160131.1</td><td>Type1.BREX</td><td><-XIS-HTH<-?<-HTH+REase<-SIGMA+OB-Fold+AAA-ATPase+Lon-Protease-BrxL<-PglZ<-N6-METHYLASE<-DUF6079*<-BrxB-BrxF-iSTAND<-BrxA-DNA-binding<-HTH_26<-?<-HNH+3C-ABC||SIGMA-HTH->MazG->?->REase-></td><td>1184</td><td>Bacteria>Terrabacteria_group>Bacillota>Erysipelotrichia>Erysipelotrichales>Coprobacillaceae</td><td>GCA_024590065.1</td><td>[Clostridium]_innocuum</td><td>MKC37_06145</td><td>87759..91313</td><td>1522</td><td>BREX-system-P-loop-protein-BrxC-[[Clostridium]</td></tr>
<tr><td>HCR67829.1</td><td>Type1.BREX</td><td>DUF6079*->N6-METHYLASE->?->Schlafen+HATPase+wHTH->PglZ->HTH+REase->SIGMA+OB-Fold+AAA-ATPase+Lon-Protease-BrxL->DUF4365+REase-></td><td>1177</td><td>Bacteria>Terrabacteria_group>Bacillota>Erysipelotrichia>Erysipelotrichales>Erysipelotrichaceae</td><td>GCA_003533525.1</td><td>Erysipelotrichaceae_bacterium</td><td>DIV43_01625</td><td>18..3551</td><td>2049044</td><td>BREX-system-P-loop-protein-BrxC-[Erysipelotrichaceae-bacterium]</td></tr>
<tr><td>GJM58076.1</td><td>Type1.BREX</td><td>SIGMA+OB-Fold+AAA-ATPase+Lon-Protease-BrxL->CR-ATPase5+REase-><-?<-SIGMA+OB-Fold+AAA-ATPase+Lon-Protease-BrxL<-PglZ<-HTH+REase<-HTH+REase<-N6-METHYLASE<-?<-DUF6079*<-BrxB-BrxF-iSTAND<-BrxA-DNA-binding<-HTH_26<-Peptidase_M20||?-><-?<-GspA-ATPase<-HTH_32+rve</td><td>1176</td><td>Bacteria>Terrabacteria_group>Bacillota>Erysipelotrichia>Erysipelotrichales>Erysipelotrichaceae</td><td>GCA_022834735.1</td><td>Erysipelotrichaceae_bacterium_OPF54</td><td>EROP_17690</td><td>1809630..1813160</td><td>2894072</td><td>hypothetical-protein-EROP_17690-[Erysipelotrichaceae-bacterium-OPF54]</td></tr>
<tr><td>MCQ4803584.1</td><td>Type1.BREX</td><td>Phage_int_SAM_3+Phage_integrase->tRNA-><-Arm-DNA-bind_4+Phage_int_SAM_3+Phage_integrase<-Phage_integrase+Phage_integrase<-XIS-HTH<-PemK_toxin<-SIGMA-HTH||HTH_26->BrxA-DNA-binding->BrxB-BrxF-iSTAND->DUF6079*->N6-METHYLASE->?->Schlafen+HATPase+wHTH->PglZ->HTH+REase->SIGMA+OB-Fold+AAA-ATPase+Lon-Protease-BrxL->DUF4365+REase-></td><td>1177</td><td>Bacteria>Terrabacteria_group>Bacillota>Erysipelotrichia>Erysipelotrichales>Erysipelotrichaceae</td><td>GCA_024460475.1</td><td>Holdemanella_sp._MSK.7.32</td><td>NE544_03830</td><td>12806..16339</td><td>2965273</td><td>BREX-system-P-loop-protein-BrxC-[Holdemanella-sp.-MSK.7.32]</td></tr>
<tr><td>WP_006060901.1</td><td>Type1.BREX</td><td><-BirA-HTH<-Schlafen+HATPase+wHTH<-?<-?<-HTH+REase<-?<-SIGMA+OB-Fold+AAA-ATPase+Lon-Protease-BrxL<-PglZ<-N6-METHYLASE<-DUF6079*<-BrxB-BrxF-iSTAND<-BrxA-DNA-binding<-HTH_26<-?<-HNH+3C-ABC||LuxR-HTH->?->SF1-SF2-Helicase-></td><td>1180</td><td>Bacteria>Terrabacteria_group>Bacillota>Erysipelotrichia>Erysipelotrichales>Erysipelotrichaceae</td><td>GCF_000157995.1</td><td>Holdemania_filiformis</td><td>HOLDEFILI_RS15965</td><td>13193..16735</td><td>61171</td><td>BREX-system-P-loop-protein-BrxC-[Holdemania-filiformis]</td></tr>
<tr><td>WP_256268391.1</td><td>Type1.BREX</td><td>Phage_int_SAM_3+Phage_integrase->tRNA-><-Arm-DNA-bind_4+Phage_int_SAM_3+Phage_integrase<-Phage_integrase+Phage_integrase<-XIS-HTH<-PemK_toxin<-SIGMA-HTH||HTH_26->BrxA-DNA-binding->BrxB-BrxF-iSTAND->DUF6079*->N6-METHYLASE->?->Schlafen+HATPase+wHTH->PglZ->HTH+REase->SIGMA+OB-Fold+AAA-ATPase+Lon-Protease-BrxL->DUF4365+REase-></td><td>1177</td><td>Bacteria>Terrabacteria_group>Bacillota>Erysipelotrichia>Erysipelotrichales>Erysipelotrichaceae</td><td>GCF_024460475.1</td><td>Holdemanella_sp._MSK.7.32</td><td>NE544_RS03830</td><td>12806..16339</td><td>2965273</td><td>BREX-system-P-loop-protein-BrxC-[Holdemanella-sp.-MSK.7.32]</td></tr>
<tr><td>WP_270440215.1</td><td>Type1.BREX</td><td><-HTH_Tnp_1+HTH_21+rve+rve_3||PAIREDC-HTH->HTH+REase<-SIGMA+OB-Fold+AAA-ATPase+Lon-Protease-BrxL<-PglZ<-N6-METHYLASE<-DUF6079*<-BrxB-BrxF-iSTAND<-BrxA-DNA-binding<-HTH_26<-?<-HNH+3C-ABC||SIGMA-HTH-></td><td>1171</td><td>Bacteria>Terrabacteria_group>Bacillota>Negativicutes>Acidaminococcales>Acidaminococcaceae</td><td>GCF_027680825.1</td><td>Acidaminococcus_provencensis</td><td>PGX33_RS00725</td><td>148887..152402</td><td>2058289</td><td>BREX-system-P-loop-protein-BrxC-[Acidaminococcus-provencensis]</td></tr>
<tr><td>MBP8598624.1</td><td>Type1.BREX</td><td><-McrC-NTD+REase<-McrB-AAA<-PglZ<-HTH+REase<-N6-METHYLASE<-DUF6079*<-BrxB-BrxF-iSTAND<-BrxA-DNA-binding<-HTH_26<-?<-ACET<-?<-HAMP+HISKIN<-REC+OmpR-HTH<-BirA-HTH</td><td>1194</td><td>Bacteria>Terrabacteria_group>Bacillota>Negativicutes>Selenomonadales>Selenomonadaceae</td><td>GCA_018052705.1</td><td>Selenomonas_sp.</td><td>KBI24_05335</td><td>21839..25423</td><td>2053611</td><td>BREX-system-P-loop-protein-BrxC-[Selenomonas-sp.]</td></tr>
<tr><td>MCH4217192.1</td><td>Type1.BREX</td><td>BrxA-DNA-binding->BrxB-BrxF-iSTAND->DUF6079*->N6-METHYLASE->HTH+REase->SbcC->Schlafen+BirA-HTH->PglZ->SIGMA+OB-Fold+AAA-ATPase+Lon-Protease-BrxL-><-tRNA<-HHH+DNA_pol_B+RNase||Calcineurin-></td><td>1206</td><td>Bacteria>Terrabacteria_group>Bacillota>Negativicutes>Veillonellales>Veillonellaceae</td><td>GCA_022486845.1</td><td>Megasphaera_sp.</td><td>LKF74_01360</td><td>2380..6000</td><td>2023260</td><td>BREX-system-P-loop-protein-BrxC-[Megasphaera-sp.]</td></tr>
<tr><td>MCG8347006.1</td><td>Type1.BREX</td><td><-PolB<-CopG/RHH<-?<-wHTH<-REase+HTH<-N6-METHYLASE<-Y1_Tnp<-DUF6079*<-BrxB-BrxF-iSTAND</td><td>1203</td><td>Bacteria>Terrabacteria_group>Chloroflexota>Chloroflexia>Chloroflexales>unclassified_Chloroflexales</td><td>GCA_022360155.1</td><td>Chloroflexales_bacterium</td><td>MI924_04420</td><td>5729..9340</td><td>2725990</td><td>BREX-system-P-loop-protein-BrxC-[Chloroflexales-bacterium]</td></tr>
<tr><td>OGH97200.1</td><td>Type1.BREX</td><td>ANK-><-?||BrxA-DNA-binding->BrxB-BrxF-iSTAND->DUF6079*->REase->N6-METHYLASE->HTH+REase->PglZ->SIGMA+OB-Fold+AAA-ATPase+Lon-Protease-BrxL->Y2_Tnp-></td><td>1198</td><td>Bacteria>Terrabacteria_group>Cyanobacteriota/Melainabacteria_group>Candidatus_Margulisbacteria>Candidatus_Riflemargulisbacteria</td><td>GCA_001783265.1</td><td>Candidatus_Margulisbacteria_bacterium_GWD2_39_127</td><td>A2X43_08520</td><td>54910..58506</td><td>1817871</td><td>hypothetical-protein-A2X43_08520-[Candidatus-Margulisbacteria-bacterium-GWD2_39_127]</td></tr>
<tr><td>OGI09205.1</td><td>Type1.BREX</td><td><-Y2_Tnp<-SIGMA+OB-Fold+AAA-ATPase+Lon-Protease-BrxL<-PglZ<-HTH+REase<-N6-METHYLASE<-REase<-DUF6079*<-BrxB-BrxF-iSTAND<-BrxA-DNA-binding||?-><-ANK</td><td>1198</td><td>Bacteria>Terrabacteria_group>Cyanobacteriota/Melainabacteria_group>Candidatus_Margulisbacteria>unclassified_Candidatus_Margulisbacteria</td><td>GCA_001784485.1</td><td>Candidatus_Margulisbacteria_bacterium_GWE2_39_32</td><td>A2X41_01355</td><td>17633..21229</td><td>1817872</td><td>hypothetical-protein-A2X41_01355-[Candidatus-Margulisbacteria-bacterium-GWE2_39_32]</td></tr>
<tr><td>PZM83737.1</td><td>Type1.BREX</td><td><-Y2_Tnp<-SIGMA+OB-Fold+AAA-ATPase+Lon-Protease-BrxL<-PglZ<-HTH+REase<-N6-METHYLASE<-REase<-DUF6079*<-BrxB-BrxF-iSTAND<-BrxA-DNA-binding||?-><-ANK</td><td>1198</td><td>Bacteria>Terrabacteria_group>Cyanobacteriota/Melainabacteria_group>Candidatus_Margulisbacteria>unclassified_Candidatus_Margulisbacteria</td><td>GCA_003242895.1</td><td>Candidatus_Margulisbacteria_bacterium</td><td>DKM50_01295</td><td>12966..16562</td><td>2053573</td><td>BREX-system-P-loop-protein-BrxC-[Candidatus-Margulisbacteria-bacterium]</td></tr>
<tr><td>BAZ90345.1</td><td>Type1.BREX</td><td><-SIGMA+OB-Fold+AAA-ATPase+HKD-DNase-BrxL<-PglZ<-HTH+REase<-N6-METHYLASE<-REase<-OLD-ABC<-PIN<-DUF433<-DUF6079*<-BrxB-BrxF-iSTAND||?->?-><-?<-?||?-><-?<-ABC-ATPase+ABC-ATPase<-NTP_transferase+MannoseP_isomer</td><td>1166</td><td>Bacteria>Terrabacteria_group>Cyanobacteriota/Melainabacteria_group>Cyanobacteriota>Cyanophyceae>Nostocales</td><td>GCA_002368135.1</td><td>Raphidiopsis_curvata_NIES-932</td><td>NIES932_18440</td><td>1957862..1961362</td><td>1973482</td><td>hypothetical-protein-NIES932_18440-[Raphidiopsis-curvata-NIES-932]</td></tr>
<tr><td>WP_006277878.1</td><td>Type1.BREX</td><td><-REase||?-><-SIGMA+OB-Fold+AAA-ATPase+HKD-DNase-BrxL<-PglZ<-HTH+REase<-N6-METHYLASE-p<-OLD-ABC<-PIN<-DUF433<-DUF6079*<-BrxB-BrxF-iSTAND||?->?-><-?<-?<-?<-ParA-Soj-PloopNTPase<-?||ACT-></td><td>1166</td><td>Bacteria>Terrabacteria_group>Cyanobacteriota/Melainabacteria_group>Cyanobacteriota>Cyanophyceae>Nostocales</td><td>GCF_000175835.1</td><td>Cylindrospermopsis_raciborskii</td><td>CRC_RS10895</td><td>11801..15301</td><td>77022</td><td>BREX-system-P-loop-protein-BrxC-[Cylindrospermopsis-raciborskii]</td></tr>
<tr><td>WP_057176814.1</td><td>Type1.BREX</td><td><-ACT||?->ParA-Soj-PloopNTPase->?->?->?-><-?<-?||BrxB-BrxF-iSTAND->DUF6079*->DUF433->PIN->OLD-ABC->REase->REase->N6-METHYLASE->HTH+REase->PglZ->SIGMA+OB-Fold+AAA-ATPase+HKD-DNase-BrxL-></td><td>1166</td><td>Bacteria>Terrabacteria_group>Cyanobacteriota/Melainabacteria_group>Cyanobacteriota>Cyanophyceae>Nostocales</td><td>GCF_001432185.1</td><td>Cylindrospermopsis_sp._CR12</td><td>ASL19_RS00210</td><td>43411..46911</td><td>1747196</td><td>BREX-system-P-loop-protein-BrxC-[Cylindrospermopsis-sp.-CR12]</td></tr>
<tr><td>WP_071242349.1</td><td>Type1.BREX</td><td><-ACT||?->ParA-Soj-PloopNTPase->?->?->?-><-?<-?||BrxB-BrxF-iSTAND->DUF6079*->DUF433->PIN->OLD-ABC->REase->N6-METHYLASE->HTH+REase->PglZ->SIGMA+OB-Fold+AAA-ATPase+HKD-DNase-BrxL->REase-></td><td>1166</td><td>Bacteria>Terrabacteria_group>Cyanobacteriota/Melainabacteria_group>Cyanobacteriota>Cyanophyceae>Nostocales</td><td>GCF_001858115.1</td><td>Cylindrospermopsis_raciborskii</td><td>BCV63_RS11865</td><td>45142..48642</td><td>77022</td><td>BREX-system-P-loop-protein-BrxC-[Cylindrospermopsis-raciborskii]</td></tr>
<tr><td>WP_071248143.1</td><td>Type1.BREX</td><td><-REase<-REase<-REase<-REase||?->BrxA-DNA-binding->BrxB-BrxF-iSTAND->DUF6079*->OLD-ABC->REase->N6-METHYLASE->HTH+REase->PglZ->SIGMA+OB-Fold+AAA-ATPase+HKD-DNase-BrxL-><-DsbD<-Thioredoxin</td><td>1172</td><td>Bacteria>Terrabacteria_group>Cyanobacteriota/Melainabacteria_group>Cyanobacteriota>Cyanophyceae>Nostocales</td><td>GCF_001858125.1</td><td>Cylindrospermopsis_raciborskii</td><td>CENA302_RS09780</td><td>17155..20673</td><td>77022</td><td>BREX-system-P-loop-protein-BrxC-[Cylindrospermopsis-raciborskii]</td></tr>
<tr><td>WP_102947438.1</td><td>Type1.BREX</td><td><-SIGMA+OB-Fold+AAA-ATPase+HKD-DNase-BrxL<-PglZ<-HTH+REase<-N6-METHYLASE<-REase<-OLD-ABC<-PIN<-DUF433<-DUF6079*<-BrxB-BrxF-iSTAND||?->?-><-?<-?<-?<-ParA-Soj-PloopNTPase<-?||ACT-></td><td>1166</td><td>Bacteria>Terrabacteria_group>Cyanobacteriota/Melainabacteria_group>Cyanobacteriota>Cyanophyceae>Nostocales</td><td>GCF_002893285.1</td><td>Cylindrospermopsis_raciborskii</td><td>CEP07_RS06090</td><td>15682..19182</td><td>77022</td><td>BREX-system-P-loop-protein-BrxC-[Cylindrospermopsis-raciborskii]</td></tr>
<tr><td>WP_115538370.1</td><td>Type1.BREX</td><td><-HTH+REase<-N6-METHYLASE-p<-DDE_3<-HTH_28+HTH_32+DDE_3<-N6-METHYLASE<-REase<-OLD-ABC<-PIN<-DUF433<-DUF6079*<-BrxB-BrxF-iSTAND||?->?-><-?<-?<-?<-ParA-Soj-PloopNTPase<-?||ACT-></td><td>1166</td><td>Bacteria>Terrabacteria_group>Cyanobacteriota/Melainabacteria_group>Cyanobacteriota>Cyanophyceae>Nostocales</td><td>GCF_003367075.2</td><td>Cylindrospermopsis_raciborskii</td><td>C6N34_008515</td><td>1862557..1866057</td><td>77022</td><td>BREX-system-P-loop-protein-BrxC-[Cylindrospermopsis-raciborskii]</td></tr>
<tr><td>WP_187705577.1</td><td>Type1.BREX</td><td>NTP_transferase+MannoseP_isomer->ABC-ATPase+ABC-ATPase->?-><-?||?->?-><-?<-?||BrxB-BrxF-iSTAND->DUF6079*->DUF433->PIN->OLD-ABC->REase->N6-METHYLASE->HTH+REase->PglZ->SIGMA+OB-Fold+AAA-ATPase+HKD-DNase-BrxL-></td><td>1166</td><td>Bacteria>Terrabacteria_group>Cyanobacteriota/Melainabacteria_group>Cyanobacteriota>Cyanophyceae>Nostocales</td><td>GCF_014489415.1</td><td>Cylindrospermopsis_curvispora</td><td>IAR63_RS13115</td><td>2945663..2949163</td><td>747548</td><td>BREX-system-P-loop-protein-BrxC-[Cylindrospermopsis-curvispora]</td></tr>
<tr><td>WP_211942103.1</td><td>Type1.BREX</td><td><-HTH+REase<-N6-METHYLASE<-REase<-REase<-Y1_Tnp||OrfB_IS605+OrfB_Zn_ribbon-><-OLD-ABC<-PIN<-DUF433<-DUF6079*<-BrxB-BrxF-iSTAND||?->?-><-?<-?<-?<-ParA-Soj-PloopNTPase<-?||ACT-></td><td>1166</td><td>Bacteria>Terrabacteria_group>Cyanobacteriota/Melainabacteria_group>Cyanobacteriota>Cyanophyceae>Nostocales</td><td>GCF_018139025.1</td><td>Cylindrospermopsis_raciborskii</td><td>J9B92_RS02395</td><td>526727..530227</td><td>77022</td><td>BREX-system-P-loop-protein-BrxC-[Cylindrospermopsis-raciborskii]</td></tr>
<tr><td>WP_236308453.1</td><td>Type1.BREX</td><td><-SIGMA+OB-Fold+AAA-ATPase+HKD-DNase-BrxL<-PglZ<-HTH+REase<-N6-METHYLASE<-REase<-OLD-ABC<-PIN<-DUF433<-DUF6079*<-BrxB-BrxF-iSTAND||?->?-><-?<-?<-?<-ParA-Soj-PloopNTPase<-?||ACT-></td><td>1166</td><td>Bacteria>Terrabacteria_group>Cyanobacteriota/Melainabacteria_group>Cyanobacteriota>Cyanophyceae>Nostocales</td><td>GCF_021650815.1</td><td>Cylindrospermopsis_raciborskii</td><td>L3I90_RS00125</td><td>24709..28209</td><td>77022</td><td>BREX-system-P-loop-protein-BrxC-[Cylindrospermopsis-raciborskii]</td></tr>
<tr><td>WP_240869109.1</td><td>Type1.BREX</td><td>ART-DarT-><-SIGMA+OB-Fold+AAA-ATPase+HKD-DNase-BrxL<-PglZ<-HTH+REase<-N6-METHYLASE<-OLD-ABC<-PIN<-DUF433<-DUF6079*<-BrxB-BrxF-iSTAND||?->?-><-?<-?<-?<-ParA-Soj-PloopNTPase<-?||ACT-><-Peptidase_M17_N+Peptidase_M17</td><td>1166</td><td>Bacteria>Terrabacteria_group>Cyanobacteriota/Melainabacteria_group>Cyanobacteriota>Cyanophyceae>Nostocales</td><td>GCF_022496815.1</td><td>Cylindrospermopsis_raciborskii</td><td>GSN00_13735</td><td>14003..17503</td><td>77022</td><td>BREX-system-P-loop-protein-BrxC-[Cylindrospermopsis-raciborskii]</td></tr>
<tr><td>WP_009625661.1</td><td>Type1.BREX</td><td><-CARF+HEPN<-ABhydrolase<-SAM-METHYLASE<-SIGMA+OB-Fold+AAA-ATPase+HKD-DNase-BrxL<-PglZ<-ParB+HNH<-N6-METHYLASE<-NERD<-OLD-ABC<-DUF6079*<-BrxB-BrxF-iSTAND<-BrxA-DNA-binding<-wHTH+REase+ZnR</td><td>1178</td><td>Bacteria>Terrabacteria_group>Cyanobacteriota/Melainabacteria_group>Cyanobacteriota>Cyanophyceae>Pseudanabaenales</td><td>GCF_000332215.1</td><td>Pseudanabaena_biceps</td><td>PSE7429_RS03435</td><td>33637..37173</td><td>927669</td><td>MULTISPECIES:-BREX-system-P-loop-protein-BrxC-[Pseudanabaena]</td></tr>
<tr><td>WP_028946553.1</td><td>Type1.BREX</td><td><-PolB<-ParD_antitoxin||?->PIN-><-CopG/RHH<-N6-METHYLASE||PhdYeFM_antitox->PIN-><-REase+HTH<-DUF6079*<-BrxB-BrxF-iSTAND<-BrxA-DNA-binding<-?<-ParA-Soj-PloopNTPase||HNH->MerR-HTH->?->wHTH-></td><td>1191</td><td>Bacteria>Terrabacteria_group>Cyanobacteriota/Melainabacteria_group>Cyanobacteriota>Cyanophyceae>Synechococcales</td><td>GCF_000478825.2</td><td>Synechocystis_sp._PCC_6714</td><td>D082_RS16385</td><td>87258..90833</td><td>1147</td><td>BREX-system-P-loop-protein-BrxC-[Synechocystis-sp.-PCC-6714]</td></tr>
</table>

| Protein-ID | System | Gene-Neighbourhoods | Length | Taxonomy | Assembly-ID | Strain | Locus-Tag | Genomic-coordinates | TaxID | Fasta-defline |
|---|---|---|---|---|---|---|---|---|---|---|
| WP_188020741.1 | Type1.BREX | <-DUF4236\|\|DOC+HTH->?->?-><-Sak4+WYL\|\|?->?-> BrxA-DNA-binding -> BrxB-BrxF-iSTAND -> DUF6079 *->MNS+REase -> N6-METHYLASE -> PglZ -> SIGMA+OB-Fold+AAA-ATPase+HKD-DNase-BrxL -> HTH+REase -><-ZnR\|\|?-><-SF1-SF2-Helicase<-sGTP+sGTP | 1163 | Bacteria>Terrabacteria_group>Cyanobacteriota/Melainabacteria_group>Cyanobacteriota>Cyanophyceae>Synechococcales | GCF_014279615.1 | Synechococcus_sp._RS9907 | SynRS9907_RS08975 | 1616845..1620336 | 221350 | BREX-system-P-loop-protein-BrxC-[Synechococcus-sp.-RS9907] |
| WP_186471764.1 | Type1.BREX | Peptidase_M28->?->?-><-Sak4+WYL\|\|?->?-> BrxA-DNA-binding -> BrxB-BrxF-iSTAND -> DUF6079 *-> N6-METHYLASE -> N6-METHYLASE -><-SWACOS+DTXR-HTH\|\| PglZ -> SIGMA+OB-Fold+AAA-ATPase+HKD-DNase-BrxL -> HTH+REase -><-ZnR\|\|?-><-SF1-SF2-Helicase | 1163 | Bacteria>Terrabacteria_group>Cyanobacteriota/Melainabacteria_group>Cyanobacteriota>Cyanophyceae>Synechococcales | GCF_014279955.1 | Synechococcus_sp._BMK-MC-1 | SynBMKMC1_RS11005 | 1938101..1941592 | 1442551 | BREX-system-P-loop-protein-BrxC-[Synechococcus-sp.-BMK-MC-1] |
| WP_186513675.1 | Type1.BREX | <-AAA+wHTH+WYL\|\|?->?-> BrxA-DNA-binding -> BrxB-BrxF-iSTAND -> DUF6079 *->MNS+REase -> N6-METHYLASE ->?->HTH_3->Couple_hipA+HipA_C-> PglZ -> SIGMA+OB-Fold+AAA-ATPase+HKD-DNase-BrxL -> HTH+REase -><-ZnR | 1163 | Bacteria>Terrabacteria_group>Cyanobacteriota/Melainabacteria_group>Cyanobacteriota>Cyanophyceae>Synechococcales | GCF_014280115.1 | Synechococcus_sp._A15-44 | SynA1544_02490 | 2002733..2006224 | 1050646 | BREX-system-P-loop-protein-BrxC-[Synechococcus-sp.-A15-44] |
| WP_221091410.1 | Type1.BREX | <- REase <- wHTH+WYL+WCX <- SF1-SF2-Helicase+REase <-Exonuc_V_gamma+REase \|\|?->SF1-SF2-Helicase-> BrxA-DNA-binding -> BrxB-BrxF-iSTAND -> DUF6079 *-> N6-METHYLASE -> McrC-NTD+REase ->HTH+McrB-AAA-> | 1200 | Bacteria>Terrabacteria_group>Deinococcota>Deinococci>Deinococcales>Deinococcaceae | GCF_019693445.1 | Deinococcus_aquaedulcis | KMW22_RS17995 | 15477..19079 | 2840455 | BREX-system-P-loop-protein-BrxC-[Deinococcus-aquaedulcis] |
| WP_207688112.1 | Type1.BREX | <-SF1-SF2-Helicase<-?<-TPR+TPR+TPR+TPR<-SIGMA-HTH+SIGMA-HTH\|\|tRNA->XIS-HTH-> BrxA-DNA-binding -> BrxB-BrxF-iSTAND -> DUF6079 *->?->Schlafen+HATPase+wHTH-> N6-METHYLASE -> HTH+REase ->SbcC-> PglZ -> SIGMA+OB-Fold+AAA-ATPase+Lon-Protease-BrxL -><-HISKIN+REC+HPT | 1185 | Bacteria>Thermodesulfobacteriota>Desulfobacteria>Desulfobacterales>Desulfococcaceae>Desulfonema | GCF_017377355.1 | Desulfonema_limicola | dnl_RS20625 | 4834354..4837911 | 45656 | BREX-system-P-loop-protein-BrxC-[Desulfonema-limicola] |
| MBF0121242.1 | Type1.BREX | <- PglZ <-RelE-ParE<-?<-?<- N6-METHYLASE -p<- HTH+REase <-Schlafen+HATPase+wHTH<- N6-METHYLASE <-Four-helical-protein<- DUF6079 *<- BrxB-BrxF-iSTAND <- BrxA-DNA-binding | 1189 | Bacteria>Thermodesulfobacteriota>Desulfobacteria>Desulfobacterales>unclassified_Desulfobacterales | GCA_015228685.1 | Desulfobacterales_bacterium | HQK79_20605 | 12595..16164 | 2044940 | BREX-system-P-loop-protein-BrxC-[Desulfobacterales-bacterium] |
| WP_205241479.1 | Type1.BREX | <-Arm-DNA-bind_3+Phage_integrase\|\|?->AAA-ATPase->?-><-?<- SIGMA+OB-Fold+AAA-ATPase+Lon-Protease-BrxL <- PglZ <- HTH+REase <- N6-METHYLASE <- DUF6079 *<- BrxB-BrxF-iSTAND <- BrxA-DNA-binding | 1148 | Bacteria>Thermodesulfobacteriota>Desulfobulbia>Desulfobulbales>Desulfobulbaceae>Desulfobulbus | GCF_016918545.1 | Desulfobulbus_alkaliphilus | JWG43_RS03775 | 164535..167981 | 869814 | BREX-system-P-loop-protein-BrxC-[Desulfobulbus-alkaliphilus] |
| MBL3581670.1 | Type1.BREX | <-KAP-glycosylase<-KAP-NTPase<- HARE-HTH + REase <- SIGMA+OB-Fold+AAA-ATPase+Lon-Protease-BrxL <- PglZ <-ParB+HNH<-PHP+SbcC<- N6-METHYLASE <- DUF6079 *<- BrxB-BrxF-iSTAND <- BrxA-DNA-binding <-XIS-HTH<-?<-?<-?<-wHTH | 1145 | Bacteria>Thermodesulfobacteriota>Desulfovibrionia>Desulfovibrionales>Desulfovibrionaceae>Oleidesulfovibrio | GCA_016756815.1 | Oleidesulfovibrio_alaskensis | JMJ96_04565 | 108279..111716 | 58180 | BREX-system-P-loop-protein-BrxC-[Oleidesulfovibrio-alaskensis] |
| WP_273192722.1 | Type1.BREX | <-KAP-glycosylase<-KAP-NTPase<- HARE-HTH + REase <- SIGMA+OB-Fold+AAA-ATPase+Lon-Protease-BrxL <- PglZ <-ParB+HNH<-PHP+SbcC<- N6-METHYLASE <- DUF6079 *<- BrxB-BrxF-iSTAND <- BrxA-DNA-binding <-XIS-HTH<-?<-?<-?<-wHTH | 1146 | Bacteria>Thermodesulfobacteriota>Desulfovibrionia>Desulfovibrionales>Desulfovibrionaceae>Oleidesulfovibrio | GCF_016756815.1 | Oleidesulfovibrio_alaskensis | JMJ96_RS04575 | 108279..111719 | 58180 | BREX-system-P-loop-protein-BrxC-[Oleidesulfovibrio-alaskensis] |
| MCH5276204.1 | Type1.BREX | <- SIGMA+OB-Fold+AAA-ATPase+Lon-Protease-BrxL <- PglZ <-?<- N6-METHYLASE -p<- HTH+REase <- N6-METHYLASE <- REase <-OLD-ABC <- DUF6079 *<- BrxB-BrxF-iSTAND <- BrxA-DNA-binding <-Arm-DNA-bind_3+Phage_integrase<-tRNA<-?<-?<-ABC-ATPase+SbcC | 1145 | Bacteria>Thermodesulfobacteriota>Desulfovibrionia>Desulfovibrionales>Desulfovibrionaceae>unclassified_Desulfovibrionaceae | GCA_022510785.1 | Desulfovibrionaceae_bacterium | J1E80_00045 | 14777..18214 | 2049043 | BREX-system-P-loop-protein-BrxC-[Desulfovibrionaceae-bacterium] |
| WP_216500608.1 | Type6.BREX | Phage_integrase-><-?<- SIGMA+OB-Fold+AAA-ATPase+HKD-DNase-BrxL <- PglZ <- N6-METHYLASE <- DUF6079 *<- BrxB-BrxF-iSTAND <- BrxA-DNA-binding <- BrxE <-MORC+REase <- w HTH+REase +ZnR<-McrB-AAA<-HTH+HEPN<-SF1-SF2-Helicase<-STYKIN+CR-Kinase | 1206 | Bacteria>Thermodesulfobacteriota>Desulfuromonadia>Geobacterales>Geobacteraceae>Geomonas | GCF_018919385.1 | Geomonas_diazotrophica | KOM00_RS19070 | 11745..15365 | 2843197 | BREX-system-P-loop-protein-BrxC-[Geomonas-diazotrophica] |
| OPY81323.1 | Type1.BREX | <-ART-DarT<-SF1-SF2-Helicase<- HARE-HTH <- SIGMA+OB-Fold+AAA-ATPase+Lon-Protease-BrxL <- PglZ <-ParB+HNH<- HTH+REase <- N6-METHYLASE <-HATPase+wHTH<- DUF6079 *<- BrxB-BrxF-iSTAND <- BrxA-DNA-binding | 1144 | Bacteria>Thermodesulfobacteriota>Syntrophia>Syntrophales>Syntrophaceae>Syntrophus | GCA_002067745.1 | Syntrophus_sp._PtaU1.Bin005 | A4E70_01341 | 26340..29774 | 1811712 | MAG:-hypothetical-protein-A4E70_01341-[Syntrophus-sp.-PtaU1.Bin005] |
| CCU85336.1 | Type1.BREX | <-DnaB<-HTH_36<-?\|\| BrxA-DNA-binding -> BrxB-BrxF-iSTAND -> DUF6079 *-> N6-METHYLASE -> PglZ -> SIGMA+OB-Fold+AAA-ATPase+Lon-Protease-BrxL ->?-> REase +DUF4268 ->McrB-AAA-> McrC-NTD+REase -> REase ->AbiEi-> | 1166 | Bacteria>Thermotogota>Thermotogae>Kosmotogales>Kosmotogaceae>Mesotoga | GCA_000367705.1 | Mesotoga_infera | PHOSAC3_150336 | 356643..360143 | 1236046 | hypothetical-protein-PHOSAC3_150336-[Mesotoga-infera] |
| WP_121550387.1 | Type1.BREX | <-AbiEii<-AbiEi<- REase <- McrC-NTD+REase <- REase +DUF4268 <-?<- SIGMA+OB-Fold+AAA-ATPase+Lon-Protease-BrxL <- PglZ <- N6-METHYLASE <- DUF6079 *<- BrxB-BrxF-iSTAND <- BrxA-DNA-binding \|\|?->HTH_36->DnaB-> | 1166 | Bacteria>Thermotogota>Thermotogae>Kosmotogales>Kosmotogaceae>Mesotoga | GCF_003664995.1 | Mesotoga_sp._BH458_6_3_2_1 | Y697_RS03850 | 31631..35131 | 1437446 | BREX-system-P-loop-protein-BrxC-[Mesotoga-sp.-BH458_6_3_2_1] |
| WP_165973936.1 | Type1.BREX | <-RADICAL-SAM<-?\|\|?->?-> BrxA-DNA-binding -> BrxB-BrxF-iSTAND -> DUF6079 -p*-> DUF6079 -p->?-> PglZ -> SIGMA+OB-Fold+AAA-ATPase+Lon-Protease-BrxL -> REase -> HTH+REase ->?->DDE_Tnp_1->SF1-SF2-Helicase-> | 535 | Bacteria>Thermotogota>Thermotogae>Petrotogales>Petrotogaceae>Marinitoga | GCF_004135675.1 | Marinitoga_lauensis | DJ007_RS01225 | 233414..235021 | 2201189 | BREX-system-P-loop-protein-BrxC-[Marinitoga-lauensis] |
| KUO61378.1 | Type1.BREX | <-Schlafen+HATPase+wHTH<- SIGMA+OB-Fold+AAA-ATPase+HKD-DNase-BrxL <- PglZ <- HTH+REase <-?<- N6-METHYLASE <- DUF6079 *<- BrxB-BrxF-iSTAND <- BrxA-DNA-binding <-HTH_Tnp_1+HTH_21+rve+rve_3 | 1157 | Bacteria>unclassified_Bacteria | GCA_001516025.1 | bacterium_BRH_c32 | APF79_11995 | 11772..15245 | 1734400 | hypothetical-protein-APF79_11995-[bacterium-BRH_c32] |
| NBJ82986.1 | Type1.BREX | RADICAL-SAM-><-?<-?\|\|REC+SHELIX+HISKIN-> REase ->?->HTH_26-> BrxA-DNA-binding -> BrxB-BrxF-iSTAND -> DUF6079 *-> N6-METHYLASE -> HTH+REase -> PglZ ->?->?->ZDNA-HTH->AAA+->?->PAS+NtrC-AAA+FIS-HTH-> | 1182 | Bacteria>unclassified_Bacteria | GCA_009917545.1 | bacterium_1XD42-76 | D5271_13855 | 76500..80048 | 2320881 | BREX-system-P-loop-protein-BrxC-[bacterium-1XD42-76] |
| NCP08807.1 | Type1.BREX | HTH_3-><-PIN<-wHTH<-?\|\| CR-ATPase5 + REase -> BrxA-DNA-binding -> BrxB-BrxF-iSTAND -> DUF6079 *->Schlafen+HATPase+wHTH+HTH->?->?-> HTH+REase -> PglZ ->?-> SIGMA+OB-Fold+AAA-ATPase+HKD-DNase-BrxL -> | 1186 | Bacteria>unclassified_Bacteria | GCA_009993525.1 | bacterium | GW860_07900 | 18171..21731 | 1869227 | BREX-system-P-loop-protein-BrxC-[bacterium] |
| -- | | | | | | | | | | |
| -- SF1-SF2-Helicase+REase | | | | | | | | | | |
| -- | | | | | | | | | | |
| MBT4336377.1 | Type1.BREX | <-HIT\|\|CopG/RHH->ZnR->?->DnaB->?-> BrxA-DNA-binding -> BrxB-BrxF-iSTAND -> DUF6079 *-> N6-METHYLASE -> PglZ -> SIGMA+OB-Fold+AAA-ATPase+Lon-Protease-BrxL -> SF1-SF2-Helicase+REase ->?-><-?<-?<-Phage_integrase | 1173 | Archaea>DPANN_group>Candidatus_Woesearchaeota | GCA_018652365.1 | Candidatus_Woesearchaeota_archaeon | HOD66_03370 | 27503..31024 | 2026803 | BREX-system-P-loop-protein-BrxC-[Candidatus-Woesearchaeota-archaeon] |
| MBT4469968.1 | Type1.BREX | Phage_integrase->?->?-><-?<- SF1-SF2-Helicase+REase <- SIGMA+OB-Fold+AAA-ATPase+Lon-Protease-BrxL <- PglZ <- N6-METHYLASE <- DUF6079 *<- BrxB-BrxF-iSTAND <- BrxA-DNA-binding <-?<-DnaB<-?<-ZnR<-CopG/RHH\|\|HIT-> | 1173 | Archaea>DPANN_group>Candidatus_Woesearchaeota | GCA_018656325.1 | Candidatus_Woesearchaeota_archaeon | HOC62_06290 | 58853..62374 | 2026803 | BREX-system-P-loop-protein-BrxC-[Candidatus-Woesearchaeota-archaeon] |
| MBT6744308.1 | Type1.BREX | <-HIT\|\|CopG/RHH->ZnR->?->DnaB->?-> BrxA-DNA-binding -> BrxB-BrxF-iSTAND -> DUF6079 *-> N6-METHYLASE -> PglZ -> SIGMA+OB-Fold+AAA-ATPase+Lon-Protease-BrxL -> SF1-SF2-Helicase+REase ->?-><-?<-?<-Phage_integrase | 1173 | Archaea>DPANN_group>Candidatus_Woesearchaeota | GCA_018696595.1 | Candidatus_Woesearchaeota_archaeon | HOA70_03205 | 27503..31024 | 2026803 | BREX-system-P-loop-protein-BrxC-[Candidatus-Woesearchaeota-archaeon] |
| MBR3112546.1 | Type1.BREX | <-TPR+TPR+TPR+TPR<-?<-OLD-ABC<- SF1-SF2-Helicase+REase <-?<-?<- SIGMA+OB-Fold+AAA-ATPase+Lon-Protease-BrxL <- PglZ <-?<- DUF6079 *<- BrxB-BrxF-iSTAND <- BrxA-DNA-binding <- N6-METHYLASE \|\| N6-METHYLASE -><- N6-METHYLASE -p<- N6-METHYLASE <-Phage_integrase+Phage_integrase\|\| REase +DUF5655-><-ParB+HNH | 1187 | Archaea>Euryarchaeota>Methanomada_group>Methanobacteria>Methanobacteriales>Methanobacteriaceae | GCA_017545145.1 | Methanobrevibacter_sp. | IKH29_02390 | 52350..55913 | 66852 | BREX-system-P-loop-protein-BrxC-[Methanobrevibacter-sp.] |
| WP_121538347.1 | Type5.BREX | <-HTH_20\|\|?->?->ParA-Soj-PloopNTPase->?-><- TUDOR+SF1-SF2-Helicase+i REase +CTD <- PglZ <- N6-METHYLASE \|\|?-><- DUF6079 *<- BrxB-BrxF-iSTAND <- DUF6079 <- BrxA-DNA-binding \|\|Exonuc_V_gamma+REase -> SF1-SF2-Helicase+REase -><- REase \|\|ORC-CDC6+HTH-><-?<- McrC-NTD+REase | 1272 | Archaea>Euryarchaeota>Stenosarchaea_group>Halobacteria>Halobacteriales>Haloarculaceae | GCF_003665885.1 | Haloarcula_sp._Atlit-120R | DVK01_RS19340 | 38493..42311 | 2282135 | hypothetical-protein-[Haloarcula-sp.-Atlit-120R] |
| WP_121538351.1 | Type5.BREX | ParA-Soj-PloopNTPase-><- TUDOR+SF1-SF2-Helicase+i REase +CTD <- PglZ <- N6-METHYLASE \|\|?-><- DUF6079 <- BrxB-BrxF-iSTAND <- DUF6079 *<- BrxA-DNA-binding \|\|Exonuc_V_gamma+REase -> SF1-SF2-Helicase+REase -><- REase \|\|ORC-CDC6+HTH-><-?<- McrC-NTD+REase <-McrB-AAA | 1251 | Archaea>Euryarchaeota>Stenosarchaea_group>Halobacteria>Halobacteriales>Haloarculaceae | GCF_003665885.1 | Haloarcula_sp._Atlit-120R | DVK01_RS19350 | 42895..46650 | 2282135 | hypothetical-protein-[Haloarcula-sp.-Atlit-120R] |
| WP_136689168.1 | Type1.BREX | <-35exo<-35exo<-MNS+REase <-?<-ARG\|\|?->?-> SF1-SF2-Helicase+REase -> BrxA-DNA-binding -> DUF6079 *-> BrxB-BrxF-iSTAND -> DUF6079 ->Zn_Tnp_IS1595+DUF2089+DDE_Tnp_IS1595->SF1-SF2-Helicase->?-><-?\|\|HEPN-> | 1249 | Archaea>Euryarchaeota>Stenosarchaea_group>Halobacteria>Halobacteriales>Haloarculaceae | GCF_004944975.1 | Halorhabdus_amylolytica | E3374_RS11205 | 84563..88312 | 2559573 | hypothetical-protein-[Halorhabdus-amylolytica] |
| WP_136689170.1 | Type1.BREX | <-MNS+REase <-?<-ARG\|\|?->?-> SF1-SF2-Helicase+REase -> BrxA-DNA-binding -> DUF6079 -> BrxB-BrxF-iSTAND -> DUF6079 *->Zn_Tnp_IS1595+DUF2089+DDE_Tnp_IS1595->SF1-SF2-Helicase->?-><-?\|\|HEPN-><-?<-?<-ParA-Soj-PloopNTPase<-Phage_integrase | 1279 | Archaea>Euryarchaeota>Stenosarchaea_group>Halobacteria>Halobacteriales>Haloarculaceae | GCF_004944975.1 | Halorhabdus_amylolytica | E3374_RS11215 | 88926..92765 | 2559573 | hypothetical-protein-[Halorhabdus-amylolytica] |
| WP_152120782.1 | Type1.BREX | BrxB-BrxF-iSTAND -> DUF6079 *-> N6-METHYLASE -> REase -> PglZ -> SIGMA+OB-Fold+AAA-ATPase+Lon-Protease-BrxL ->Exonuc_V_gamma+REase -> SF1-SF2-Helicase+REase ->ParA-Soj-PloopNTPase-> | 1215 | Archaea>Euryarchaeota>Stenosarchaea_group>Halobacteria>Halobacteriales>Haloarculaceae | GCF_009184545.1 | Halosegnis_rubeus | DMP03_RS11345 | 583..4230 | 2212850 | BREX-system-P-loop-protein-BrxC-[Halosegnis-rubeus] |
| WP_188853474.1 | Type1.BREX | <-ABC-ATPase+SbcC+VirB11+ABC-ATPase+SbcC<-?<-Calcineurin<-?<-?\|\|?->?->?-> BrxA-DNA-binding -> DUF6079 *-> BrxB-BrxF-iSTAND -> DUF6079 ->HTH+ICLR-HTH->NTP_transf+wHTH->wHTH-> REase -> N6-METHYLASE -> PglZ -> N6-METHYLASE -> | 1227 | Archaea>Euryarchaeota>Stenosarchaea_group>Halobacteria>Halobacteriales>Haloarculaceae | GCF_014647095.1 | Haloarcula_salaria | IEY10_RS16005 | 45657..49340 | 575195 | hypothetical-protein-[Haloarcula-salaria] |
| WP_188853476.1 | Type1.BREX | <-Calcineurin<-?<-?\|\|?->?->?-> BrxA-DNA-binding -> DUF6079 -> BrxB-BrxF-iSTAND -> DUF6079 *->HTH+ICLR-HTH->NTP_transf+wHTH->wHTH-> REase -> N6-METHYLASE -> PglZ -> N6-METHYLASE -> TUDOR+SF1-SF2-Helicase+i REase +CTD ->Exonuc_V_gamma+REase -> | 1280 | Archaea>Euryarchaeota>Stenosarchaea_group>Halobacteria>Halobacteriales>Haloarculaceae | GCF_014647095.1 | Haloarcula_salaria | IEY10_RS16015 | 49915..53757 | 575195 | hypothetical-protein-[Haloarcula-salaria] |
| WP_135666499.1 | Type1.BREX | <- PglZ <-RelE<-?<- N6-METHYLASE <-TBP+TBP\|\|TBP+TBP->?-><-?<-?<- DUF6079 *<- BrxB-BrxF-iSTAND <- DUF6079 <- BrxA-DNA-binding <- SF1-SF2-Helicase+REase <- SF1-SF2-Helicase+REase <-?<- REase | 1277 | Archaea>Euryarchaeota>Stenosarchaea_group>Halobacteria>Halobacteriales>Haloarculaceae | GCF_900880625.1 | Halorhabdus_rudnickae | BN2694_RS13550 | 37861..41694 | 1775544 | hypothetical-protein-[Halorhabdus-rudnickae] |
| WP_135666504.1 | Type1.BREX | <- N6-METHYLASE <-TBP+TBP\|\|TBP+TBP->?-><-?<-?<- DUF6079 <- BrxB-BrxF-iSTAND <- DUF6079 *<- BrxA-DNA-binding <- SF1-SF2-Helicase+REase <- SF1-SF2-Helicase+REase <-?<- REase | 1248 | Archaea>Euryarchaeota>Stenosarchaea_group>Halobacteria>Halobacteriales>Haloarculaceae | GCF_900880625.1 | Halorhabdus_rudnickae | BN2694_RS13560 | 42308..46054 | 1775544 | hypothetical-protein-[Halorhabdus-rudnickae] |
| WP_058984838.1 | Type5.BREX | <- SF1-SF2-Helicase+REase <-Exonuc_V_gamma+REase <- TUDOR+SF1-SF2-Helicase+i REase +CTD <- REase +METHYLASE<- PglZ <-?<- N6-METHYLASE \|\|viral-protein-><- REase <- DUF6079 *<- BrxB-BrxF-iSTAND <- DUF6079 <- BrxA-DNA-binding \|\|ORC-CDC6->HTH_28->NTP_transf_5->?->?->35exo-> | 1277 | Archaea>Euryarchaeota>Stenosarchaea_group>Halobacteria>Halobacteriales>Halobacteriaceae | GCF_001485535.1 | Halobacterium | AVZ66_RS14305 | 153821..157654 | 2239 | MULTISPECIES:-hypothetical-protein-[Halobacterium] |
| WP_058984840.1 | Type5.BREX | <- TUDOR+SF1-SF2-Helicase+i REase +CTD <- REase +METHYLASE<- PglZ <-?<- N6-METHYLASE \|\|viral-protein-><- REase <- DUF6079 <- BrxB-BrxF-iSTAND <- DUF6079 *<- BrxA-DNA-binding \|\|ORC-CDC6->HTH_28->NTP_transf_5->?->?->35exo-><-?<-HTH_20 | 1227 | Archaea>Euryarchaeota>Stenosarchaea_group>Halobacteria>Halobacteriales>Halobacteriaceae | GCF_001485535.1 | Halobacterium | AVZ66_RS14315 | 158229..161912 | 2239 | MULTISPECIES:-hypothetical-protein-[Halobacterium] |
| WP_123076504.1 | Type1.BREX | <- TUDOR+SF1-SF2-Helicase+i REase +CTD <- PglZ <- N6-METHYLASE \|\|?->?-><-?\|\|?->SF1-SF2-Helicase-><- DUF6079 *<- BrxB-BrxF-iSTAND <- DUF6079 <- BrxA-DNA-binding <-?\|\|?-><- SF1-SF2-Helicase+REase <- SF1-SF2-Helicase+REase | 1277 | Archaea>Euryarchaeota>Stenosarchaea_group>Halobacteria>Halobacteriales>Halobacteriaceae | GCF_003721175.2 | Halarchaeum_sp._CBA1220 | EFA46_RS09075 | 1680640..1684473 | 1853682 | hypothetical-protein-[Halarchaeum-sp.-CBA1220] |
| WP_123076500.1 | Type1.BREX | <- PglZ <- N6-METHYLASE \|\|?->?-><-?\|\|?->SF1-SF2-Helicase-><- DUF6079 <- BrxB-BrxF-iSTAND <- DUF6079 *<- BrxA-DNA-binding <-?\|\|?-><- SF1-SF2-Helicase+REase <- SF1-SF2-Helicase+REase <-?\|\|?-> REase -><-HTH_20 | 1248 | Archaea>Euryarchaeota>Stenosarchaea_group>Halobacteria>Halobacteriales>Halobacteriaceae | GCF_003721175.2 | Halarchaeum_sp._CBA1220 | EFA46_RS09085 | 1685087..1688833 | 1853682 | hypothetical-protein-[Halarchaeum-sp.-CBA1220] |
| WP_233265034.1 | Type1.BREX | <-NTP_transf\|\|?->?-><-PIWI<- SF1-SF2-Helicase+REase <- SF1-SF2-Helicase+REase <-SF1-SF2-Helicase+wHTH+Small-B-barrel+DUF1998<- REase +METHYLASE\|\| BrxA-DNA-binding -> DUF6079 -p*-> | 962 | Archaea>Euryarchaeota>Stenosarchaea_group>Halobacteria>Halobacteriales>Halobacteriaceae | GCF_009791395.1 | Halomarina_oriensis | GQS65_RS04705 | 84531..87418 | 671145 | hypothetical-protein,-partial-[Halomarina-oriensis] |
| WP_205252856.1 | Type1.BREX | <-Exonuc_V_gamma+REase <- TUDOR+SF1-SF2-Helicase+i REase +CTD <-HEPN<- PglZ <-?<- N6-METHYLASE \|\|?->?-><- REase <- DUF6079 *<- BrxB-BrxF-iSTAND <- DUF6079 <- BrxA-DNA-binding \|\|ORC-CDC6->ORC-CDC6+HTH-><-?\|\|MarR-HTH-> | 1280 | Archaea>Euryarchaeota>Stenosarchaea_group>Halobacteria>Halobacteriales>Halobacteriaceae | GCF_016917835.2 | Halobacterium_sp._BOL4-2 | JRZ79_RS24035 | 52137..55979 | 2810537 | hypothetical-protein-[Halobacterium-sp.-BOL4-2] |
| WP_205252782.1 | Type1.BREX | <-HEPN<- PglZ <-?<- N6-METHYLASE \|\|?->?-><- REase <- DUF6079 <- BrxB-BrxF-iSTAND <- DUF6079 *<- BrxA-DNA-binding \|\|ORC-CDC6->ORC-CDC6+HTH-><-?\|\|MarR-HTH-><-?<-?\|\|HTH_28->NTP_transf_5-> | 1227 | Archaea>Euryarchaeota>Stenosarchaea_group>Halobacteria>Halobacteriales>Halobacteriaceae | GCF_016917835.2 | Halobacterium_sp._BOL4-2 | JRZ79_RS24045 | 56554..60237 | 2810537 | hypothetical-protein-[Halobacterium-sp.-BOL4-2] |
| WP_232700617.1 | Type5.BREX | <- TUDOR+SF1-SF2-Helicase+i REase +CTD \|\|TBP+TBP-><- PglZ <-?<- N6-METHYLASE <-?<- DUF6079 *<- BrxB-BrxF-iSTAND <- DUF6079 <- BrxA-DNA-binding <- SF1-SF2-Helicase+REase <- SF1-SF2-Helicase+REase \|\|?->?->?->ABC-ATPase+SbcC-> | 1277 | Archaea>Euryarchaeota>Stenosarchaea_group>Halobacteria>Halobacteriales>Halobacteriaceae | GCF_021249345.1 | Halobacterium_sp._Gai3-17 | LT965_RS09790 | 1831133..1834966 | 2902623 | hypothetical-protein-[Halobacterium-sp.-Gai3-17] |
| WP_232700619.1 | Type5.BREX | <- TUDOR+SF1-SF2-Helicase+i REase +CTD \|\|TBP+TBP-><- PglZ <-?<- N6-METHYLASE <-?<- DUF6079 <- BrxB-BrxF-iSTAND <- DUF6079 *<- BrxA-DNA-binding <- SF1-SF2-Helicase+REase <- SF1-SF2-Helicase+REase \|\|?->?->?->ABC-ATPase+SbcC->ABC-ATPase+SbcC-> | 1249 | Archaea>Euryarchaeota>Stenosarchaea_group>Halobacteria>Halobacteriales>Halobacteriaceae | GCF_021249345.1 | Halobacterium_sp._Gai3-17 | LT965_RS09800 | 1835580..1839329 | 2902623 | hypothetical-protein-[Halobacterium-sp.-Gai3-17] |
| WP_247994824.1 | Type5.BREX | ORC-CDC6+HTH-><-?<-?<-ABC-ATPase+SbcC\|\| SF1-SF2-Helicase+REase -> SF1-SF2-Helicase+REase -><- TUDOR+SF1-SF2-Helicase+i REase +CTD <- PglZ <- N6-METHYLASE <- DUF6079 *<- BrxB-BrxF-iSTAND <- DUF6079 <- BrxA-DNA-binding <-?<-?\|\|?-><-PIWI | 1274 | Archaea>Euryarchaeota>Stenosarchaea_group>Halobacteria>Halobacteriales>Halobacteriaceae | GCF_023115355.1 | Halocatena_salina | MW046_RS14210 | 287042..290866 | 2934340 | hypothetical-protein-[Halocatena-salina] |
| WP_247994826.1 | Type5.BREX | <-ABC-ATPase+SbcC\|\| SF1-SF2-Helicase+REase -> SF1-SF2-Helicase+REase -><- TUDOR+SF1-SF2-Helicase+i REase +CTD <- PglZ <- N6-METHYLASE <- DUF6079 <- BrxB-BrxF-iSTAND <- DUF6079 *<- BrxA-DNA-binding <-?<-?\|\|?-><-PIWI | 1247 | Archaea>Euryarchaeota>Stenosarchaea_group>Halobacteria>Halobacteriales>Halobacteriaceae | GCF_023115355.1 | Halocatena_salina | MW046_RS14220 | 291483..295226 | 2934340 | hypothetical-protein-[Halocatena-salina] |
| WP_254547347.1 | Type6.BREX | <-URI-DNase\|\|?->ArdCN+MPTase-inact->?-> BrxE -> BrxA-DNA-binding -> BrxB-BrxF-iSTAND -> DUF6079 *-> N6-METHYLASE -><-Zn_Tnp_IS1595+DDE_Tnp_IS1595<-?\|\| PglZ -> SIGMA+OB-Fold+AAA-ATPase+Lon-Protease-BrxL -><- SF1-SF2-Helicase+REase <- SF1-SF2-Helicase+REase <-ORC-CDC6+HTH | 1206 | Archaea>Euryarchaeota>Stenosarchaea_group>Halobacteria>Halobacteriales>Halobacteriaceae | GCF_024228315.1 | Halomarina_sp._BND7 | NKI68_RS22870 | 57828..61448 | 2961599 | BREX-system-P-loop-protein-BrxC-[Halomarina-sp.-BND7] |
| WP_267638585.1 | Type5.BREX | Phage_integrase-><-?<-?<-ARG\|\|?-><-?<-Preader+URI-DNase\|\| BrxA-DNA-binding -> DUF6079 *-> BrxB-BrxF-iSTAND -> DUF6079 -> N6-METHYLASE -> PglZ -> TUDOR+SF1-SF2-Helicase+i REase +CTD -><- SF1-SF2-Helicase+REase <-Exonuc_V_gamma+REase \|\|HAD->HKD-DNase+HKD-Rease-> | 1251 | Archaea>Euryarchaeota>Stenosarchaea_group>Halobacteria>Halobacteriales>Halobacteriaceae | GCF_026586665.1 | Halovenus_rubra | N5098_RS14425 | 45600..49355 | 869890 | hypothetical-protein-[Halovenus-rubra] |
| WP_244705671.1 | Type5.BREX | <- SF1-SF2-Helicase+REase <- SF1-SF2-Helicase+REase <- TUDOR+SF1-SF2-Helicase+i REase +CTD <- PglZ <- N6-METHYLASE \|\|TBP+TBP-><- DUF6079 *<- BrxB-BrxF-iSTAND <- DUF6079 -p\|\|?-><- BrxA-DNA-binding \|\|?-><-?<-Glyoxalase | 1269 | Archaea>Euryarchaeota>Stenosarchaea_group>Halobacteria>Halobacteriales>Halococcaceae | GCF_022870485.1 | Halococcus_dombrowskii | MUK72_RS16110 | 375497..379306 | 179637 | hypothetical-protein-[Halococcus-dombrowskii] |
| WP_244705675.1 | Type5.BREX | <- SF1-SF2-Helicase+REase <- SF1-SF2-Helicase+REase <- TUDOR+SF1-SF2-Helicase+i REase +CTD <- PglZ <- N6-METHYLASE \|\|TBP+TBP-><- DUF6079 <- BrxB-BrxF-iSTAND <- DUF6079 -p*\|\|?-><- BrxA-DNA-binding \|\|?-><-?<-Glyoxalase\|\|?->?->AbiEi_4+AbiEi_1->AbiEii+ZnR-> | 848 | Archaea>Euryarchaeota>Stenosarchaea_group>Halobacteria>Halobacteriales>Halococcaceae | GCF_022870485.1 | Halococcus_dombrowskii | MUK72_RS16120 | 379890..382436 | 179637 | hypothetical-protein-[Halococcus-dombrowskii] |
| WP_122106903.1 | Type1.BREX | <-DDE_Tnp_1<-?<-?\|\|MarR-HTH-><-?<-ParA-Soj-PloopNTPase<- SF1-SF2-Helicase+REase <-Exonuc_V_gamma+REase \|\| BrxA-DNA-binding -> DUF6079 *-> BrxB-BrxF-iSTAND -> DUF6079 -><-?<-?\|\|?-> N6-METHYLASE -p-> | 1248 | Archaea>Euryarchaeota>Stenosarchaea_group>Halobacteria>Haloferacales>Haloferacaceae | GCF_003697845.1 | Halobellus_captivus | EAO78_RS16655 | 16343..20089 | 2592614 | hypothetical-protein-[Halobellus-captivus] |
| WP_122106905.1 | Type1.BREX | MarR-HTH-><-?<-ParA-Soj-PloopNTPase<- SF1-SF2-Helicase+REase <-Exonuc_V_gamma+REase \|\| BrxA-DNA-binding -> DUF6079 -> BrxB-BrxF-iSTAND -> DUF6079 *-><-?<-?\|\|?-> N6-METHYLASE -p-> | 1279 | Archaea>Euryarchaeota>Stenosarchaea_group>Halobacteria>Haloferacales>Haloferacaceae | GCF_003697845.1 | Halobellus_captivus | EAO78_RS16665 | 20703..24542 | 2592614 | hypothetical-protein-[Halobellus-captivus] |
| TQQ78814.1 | Type5.BREX | <-Polbeta<-HTH_20<-?<-ParA-Soj-PloopNTPase<- SF1-SF2-Helicase+REase <-Exonuc_V_gamma+REase <- PglZ <- DUF6079 <- BrxB-BrxF-iSTAND <- DUF6079 *<- BrxA-DNA-binding \|\| N6-METHYLASE -> TUDOR+SF1-SF2-Helicase+i REase +CTD ->ORC-CDC6->?-><- REase | 1233 | Archaea>Euryarchaeota>Stenosarchaea_group>Halobacteria>Haloferacales>Halorubraceae | GCA_006861665.1 | Halonotius_terrestris | EGH24_12695 | 42218..45919 | 2487750 | hypothetical-protein-EGH24_12695-[Halonotius-terrestris] |
| WP_015911586.1 | Type1.BREX | <- N6-METHYLASE -p\|\|MazE_antitoxin->PIN-><- DUF6079 *<- BrxB-BrxF-iSTAND <- DUF6079 <- BrxA-DNA-binding <- SF1-SF2-Helicase+REase <- SF1-SF2-Helicase+REase | 1270 | Archaea>Euryarchaeota>Stenosarchaea_group>Halobacteria>Haloferacales>Halorubraceae | GCF_000022205.1 | Halorubrum_lacusprofundi | CF318_RS14700 | 1993..5805 | 2247 | hypothetical-protein-[Halorubrum-lacusprofundi] |
| WP_015911588.1 | Type1.BREX | <- TUDOR+SF1-SF2-Helicase+i REase +CTD <- PglZ <- N6-METHYLASE \|\|DDE_Tnp_1-><- N6-METHYLASE -p\|\|MazE_antitoxin->PIN-><- DUF6079 <- BrxB-BrxF-iSTAND <- DUF6079 *<- BrxA-DNA-binding <- SF1-SF2-Helicase+REase <- SF1-SF2-Helicase+REase <-?\|\|DDE_Tnp_1-><-?<-MazE_antitoxin | 1239 | Archaea>Euryarchaeota>Stenosarchaea_group>Halobacteria>Haloferacales>Halorubraceae | GCF_000022205.1 | Halorubrum_lacusprofundi | CF318_RS14710 | 6389..10108 | 2247 | hypothetical-protein-[Halorubrum-lacusprofundi] |
| WP_096390020.1 | Type6.BREX | <-Calcineurin<-GDResolvase+HIN-HTH<- REase <-?<-Calcineurin-SbcD\|\| BrxE -> BrxA-DNA-binding -> BrxB-BrxF-iSTAND -> DUF6079 *-> REase -> N6-METHYLASE -> REase -> PglZ -> SIGMA+OB-Fold+AAA-ATPase+Lon-Protease-BrxL ->Exonuc_V_gamma+REase -><-DDE_Tnp_1\|\| SF1-SF2-Helicase+REase -> SF1-SF2-Helicase+REase -> | 1214 | Archaea>Euryarchaeota>Stenosarchaea_group>Halobacteria>Haloferacales>Halorubraceae | GCF_002355635.1 | Halopenitus_persicus | CPZ00_RS05685 | 1169858..1173502 | 1048396 | BREX-system-P-loop-protein-BrxC-[Halopenitus-persicus] |
| WP_136591377.1 | Type5.BREX | <-HTH_28\|\|ParA-Soj-PloopNTPase->?-><-Phage_integrase\|\|SF1-SF2-Helicase-> SF1-SF2-Helicase+REase -> SF1-SF2-Helicase+REase -> BrxA-DNA-binding -> DUF6079 *-> BrxB-BrxF-iSTAND -> DUF6079 -><-?\|\| N6-METHYLASE ->?-> PglZ -> TUDOR+SF1-SF2-Helicase+i REase +CTD ->?-><-OrfB_IS605+OrfB_Zn_ribbon | 1258 | Archaea>Euryarchaeota>Stenosarchaea_group>Halobacteria>Haloferacales>Halorubraceae | GCF_004916935.1 | Salinigranum_halophilum | E6C73_RS15450 | 221772..225548 | 2565931 | hypothetical-protein-[Salinigranum-halophilum] |
| WP_136591379.1 | Type5.BREX | ParA-Soj-PloopNTPase->?-><-Phage_integrase\|\|SF1-SF2-Helicase-> SF1-SF2-Helicase+REase -> SF1-SF2-Helicase+REase -> BrxA-DNA-binding -> DUF6079 -> BrxB-BrxF-iSTAND -> DUF6079 *-><-?\|\| N6-METHYLASE ->?-> PglZ -> TUDOR+SF1-SF2-Helicase+i REase +CTD ->?-><-OrfB_IS605+OrfB_Zn_ribbon | 1269 | Archaea>Euryarchaeota>Stenosarchaea_group>Halobacteria>Haloferacales>Halorubraceae | GCF_004916935.1 | Salinigranum_halophilum | E6C73_RS15460 | 226132..229941 | 2565931 | hypothetical-protein-[Salinigranum-halophilum] |
| WP_142980503.1 | Type5.BREX | <-Phage_int_SAM_5<-?<-Polbeta<-HTH_20<-?<-ParA-Soj-PloopNTPase<- SF1-SF2-Helicase+REase <-Exonuc_V_gamma+REase <- PglZ <- DUF6079 *<- BrxB-BrxF-iSTAND <- DUF6079 <- BrxA-DNA-binding \|\| N6-METHYLASE -> TUDOR+SF1-SF2-Helicase+i REase +CTD ->ORC-CDC6->?-><- REase | 1230 | Archaea>Euryarchaeota>Stenosarchaea_group>Halobacteria>Haloferacales>Halorubraceae | GCF_006861665.1 | Halonotius_terrestris | EGH24_RS12690 | 37910..41602 | 2487750 | hypothetical-protein-[Halonotius-terrestris] |
| WP_221886744.1 | Type5.BREX | <-Polbeta<-HTH_20<-?<-ParA-Soj-PloopNTPase<- SF1-SF2-Helicase+REase <-Exonuc_V_gamma+REase <- PglZ <- DUF6079 <- BrxB-BrxF-iSTAND <- DUF6079 *<- BrxA-DNA-binding \|\| N6-METHYLASE -> TUDOR+SF1-SF2-Helicase+i REase +CTD ->ORC-CDC6->?-><- REase | 1230 | Archaea>Euryarchaeota>Stenosarchaea_group>Halobacteria>Haloferacales>Halorubraceae | GCF_006861665.1 | Halonotius_terrestris | EGH24_RS12700 | 42218..45910 | 2487750 | hypothetical-protein-[Halonotius-terrestris] |
| AEN07515.1 | Type5.BREX | MazE_antitoxin->?-><-DDE_Tnp_1\|\|?-> SF1-SF2-Helicase+REase -> SF1-SF2-Helicase+REase ->?-> BrxA-DNA-binding -> DUF6079 *-> BrxB-BrxF-iSTAND -> DUF6079 -><-PIN<-MazE_antitoxin\|\| N6-METHYLASE ->?-> PglZ -> TUDOR+SF1-SF2-Helicase+i REase +CTD -> | 1239 | Archaea>Euryarchaeota>Stenosarchaea_group>Halobacteria>Haloferacales>unclassified_Haloferacales | GCA_000224475.1 | halophilic_archaeon_DL31 | Halar_0255 | 246678..250397 | 756883 | hypothetical-protein-Halar_0255-[halophilic-archaeon-DL31] |
| AEN07517.1 | Type5.BREX | <-DDE_Tnp_1\|\|?-> SF1-SF2-Helicase+REase -> SF1-SF2-Helicase+REase ->?-> BrxA-DNA-binding -> DUF6079 -> BrxB-BrxF-iSTAND -> DUF6079 *-><-PIN<-MazE_antitoxin\|\| N6-METHYLASE ->?-> PglZ -> TUDOR+SF1-SF2-Helicase+i REase +CTD -><-SF1-SF2-Helicase | 1270 | Archaea>Euryarchaeota>Stenosarchaea_group>Halobacteria>Haloferacales>unclassified_Haloferacales | GCA_000224475.1 | halophilic_archaeon_DL31 | Halar_0257 | 250981..254793 | 756883 | hypothetical-protein-Halar_0257-[halophilic-archaeon-DL31] |
| WP_049967095.1 | Type5.BREX | HTH_20->?-><-35exo<-?<-35exo\|\|?-><-ORC-CDC6\|\| BrxA-DNA-binding -> BrxB-BrxF-iSTAND -> DUF6079 *-><-?\|\|?-><-OLD-ABC+TOPRIM<-ZnR+REase \|\| PglZ -> TUDOR+SF1-SF2-Helicase+i REase +CTD -> SF1-SF2-Helicase+REase -><-DDE_Tnp_IS240 | 1277 | Archaea>Euryarchaeota>Stenosarchaea_group>Halobacteria>Natrialbales>Natrialbaceae | GCF_000710605.1 | Natrinema_thermotolerans | DVR14_24745 | 889875..893708 | 121872 | hypothetical-protein-[Natrinema-thermotolerans] |
| WP_049967097.1 | Type5.BREX | <- TUDOR+SF1-SF2-Helicase+i REase +CTD <- REase +METHYLASE<- PglZ \|\|ZnR+REase ->OLD-ABC+TOPRIM-><- N6-METHYLASE \|\|?-><- DUF6079 <- BrxB-BrxF-iSTAND <- DUF6079 *<- BrxA-DNA-binding \|\|ORC-CDC6-><-?\|\|?->?->?->35exo-><-?<-HTH_20 | 1227 | Archaea>Euryarchaeota>Stenosarchaea_group>Halobacteria>Natrialbales>Natrialbaceae | GCF_000710605.1 | Natrinema_thermotolerans | HL44_RS19275 | 148133..151816 | 121872 | hypothetical-protein-[Natrinema-thermotolerans] |
| MBK9154354.1 | Type1.BREX | Phage_integrase->DUF488-><-?<-?<-?\|\|HTH_3->Peptidase_M78-> BrxA-DNA-binding -> BrxB-BrxF-iSTAND -> DUF6079 -p*->?-> DUF6079 -p->AIPR->?-> PglZ -> PglZ ->Exonuc_V_gamma+REase ->SF1-SF2-Helicase->SF1-SF2-Helicase-> | 481 | Bacteria>Acidobacteriota>Blastocatellia>Chloracidobacterium>unclassified_Chloracidobacterium | GCA_016716365.1 | Chloracidobacterium_sp. | IPM25_09080 | 169835..171280 | 2099375 | BREX-system-P-loop-protein-BrxC-[Chloracidobacterium-sp.] |
| WP_203057453.1 | Type1.BREX | <-DUF4011+ SF1-SF2-Helicase+REase <- SIGMA+OB-Fold+AAA-ATPase+HKD-DNase-BrxL <-?<-HAD+PRTase+PRTase+ REase +SF1-SF2-Helicase <-?<-SIGMA-HTH<- N6-METHYLASE <- DUF6079 *<- BrxB-BrxF-iSTAND <- BrxA-DNA-binding <-?\|\|ZnR+CC+REase ->?-><-?<-?<-MACRODOMAIN<-ART-DarT | 1178 | Bacteria>FCB_group>Bacteroidetes/Chlorobi_group>Bacteroidota>Flavobacteriia>Flavobacteriales | GCF_016786255.1 | Robertkochia_sediminum | IS446_RS01110 | 192082..195618 | 2785326 | BREX-system-P-loop-protein-BrxC-[Robertkochia-sediminum] |
| WP_044220103.1 | Type1.BREX | <-HTH_3\|\|?->?->?-><-NtrC-AAA+FIS-HTH\|\| SF1-SF2-Helicase+REase ->RNase-> BrxA-DNA-binding -> BrxB-BrxF-iSTAND -> DUF6079 *-> N6-METHYLASE ->?-> REase ->HTH+rve->AAA-ATPase->HTH_3->DDE_Tnp_1_4-><-?\|\|SbcC-> | 1186 | Bacteria>FCB_group>Bacteroidetes/Chlorobi_group>Bacteroidota>Saprospiria>Saprospirales | GCF_000759025.1 | Phaeodactylibacter_xiamenensis | IX84_RS10880 | 11358..14918 | 1524460 | BREX-system-P-loop-protein-BrxC-[Phaeodactylibacter-xiamenensis] |
| WP_147165921.1 | Type1.BREX | <-HTH_3<- REase \|\|?->PIN-><-?<-HTH_3<-HigB<- REase <- N6-METHYLASE <- DUF6079 *<- BrxB-BrxF-iSTAND <- BrxA-DNA-binding <-RNase<- SF1-SF2-Helicase+REase \|\|NtrC-AAA+FIS-HTH-><-?<-?<-?\|\|HTH_3-> | 1186 | Bacteria>FCB_group>Bacteroidetes/Chlorobi_group>Bacteroidota>Saprospiria>Saprospirales | GCF_007993045.1 | Phaeodactylibacter_luteus | FRY97_RS02895 | 87702..91262 | 1564516 | BREX-system-P-loop-protein-BrxC-[Phaeodactylibacter-luteus] |
| WP_094540671.1 | Type1.BREX | HTH_3->Recombinase+ZnR-><-SF1-SF2-Helicase<-HKD-DNase+HKD-DNase<-wHTH<- SF1-SF2-Helicase+REase <- SIGMA+OB-Fold+AAA-ATPase+HKD-DNase-BrxL <- PglZ <- N6-METHYLASE <- DUF6079 *<- BrxB-BrxF-iSTAND <- BrxA-DNA-binding <- wHTH+WYL+WCX \|\|Calcineurin+Calcineurin-SbcD->SbcC+SbcC-><-MODE-HTH+PBPII\|\|TetR-HTH-><-RHH<-VirB10 | 1179 | Bacteria>Pseudomonadota>Alphaproteobacteria>Hyphomicrobiales>Brucellaceae>Brucella/Ochrobactrum_group | GCF_002252505.1 | Brucella_grignonensis | CEV33_RS11715 | 211751..215290 | 94627 | BREX-system-P-loop-protein-BrxC-[Brucella-grignonensis] |
| WP_113731560.1 | Type1.BREX | VirB8->VirB9->VirB10->RHH-><-TetR-HTH\|\|MODE-HTH+PBPII-> wHTH+WYL+WCX -> BrxA-DNA-binding -> BrxB-BrxF-iSTAND -> DUF6079 *-> N6-METHYLASE -> PglZ -> SIGMA+OB-Fold+AAA-ATPase+HKD-DNase-BrxL -> SF1-SF2-Helicase+REase -> SF1-SF2-Helicase+REase ->wHTH->HKD-DNase+HKD-DNase->SF1-SF2-Helicase-> | 1179 | Bacteria>Pseudomonadota>Alphaproteobacteria>Hyphomicrobiales>Brucellaceae>Brucella/Ochrobactrum_group | GCF_003261015.1 | Brucella_anthropi | F9L04_RS18285 | 14553..18092 | 529 | BREX-system-P-loop-protein-BrxC-[Brucella-anthropi] |
| WP_075657534.1 | Type1.BREX | VirB8->VirB9->VirB10->RHH-><-TetR-HTH\|\|MODE-HTH+PBPII-> wHTH+WYL+WCX -> BrxA-DNA-binding -> BrxB-BrxF-iSTAND -> DUF6079 *-> N6-METHYLASE -> PglZ -> SIGMA+OB-Fold+AAA-ATPase+HKD-DNase-BrxL -> SF1-SF2-Helicase+REase -> SF1-SF2-Helicase+REase ->wHTH->HKD-DNase+HKD-DNase->SF1-SF2-Helicase-> | 1179 | Bacteria>Pseudomonadota>Alphaproteobacteria>Hyphomicrobiales>Brucellaceae>Pseudochrobactrum | GCF_001939785.1 | Pseudochrobactrum_sp._B5 | BKD14_RS17970 | 55973..59512 | 1289478 | BREX-system-P-loop-protein-BrxC-[Pseudochrobactrum-sp.-B5] |
| WP_113944570.1 | Type1.BREX | VirB8->VirB9->VirB10->RHH-><-TetR-HTH\|\|MODE-HTH+PBPII-> wHTH+WYL+WCX -> BrxA-DNA-binding -> BrxB-BrxF-iSTAND -> DUF6079 *-> N6-METHYLASE -> PglZ -> SIGMA+OB-Fold+AAA-ATPase+HKD-DNase-BrxL -> SF1-SF2-Helicase+REase -> SF1-SF2-Helicase+REase ->wHTH->HKD-DNase+HKD-DNase->SF1-SF2-Helicase-> | 1179 | Bacteria>Pseudomonadota>Alphaproteobacteria>Hyphomicrobiales>Brucellaceae>Pseudochrobactrum | GCF_003314995.1 | Pseudochrobactrum_asaccharolyticum | DFR47_RS11470 | 41745..45284 | 354351 | BREX-system-P-loop-protein-BrxC-[Pseudochrobactrum-asaccharolyticum] |
| WP_266332120.1 | Type1.BREX | <-SF1-SF2-Helicase<-HKD-DNase+HKD-DNase<-wHTH<- SF1-SF2-Helicase+REase <- SF1-SF2-Helicase+REase <- SIGMA+OB-Fold+AAA-ATPase+HKD-DNase-BrxL <- PglZ <- N6-METHYLASE <- DUF6079 *<- BrxB-BrxF-iSTAND <- BrxA-DNA-binding <- wHTH+WYL+WCX <-MODE-HTH+PBPII\|\|TetR-HTH-><-RHH<-VirB10<-VirB9<-VirB8 | 1179 | Bacteria>Pseudomonadota>Alphaproteobacteria>Hyphomicrobiales>Kaistiaceae>Kaistia | GCF_026343955.1 | Kaistia_defluvii | OSH10_RS12655 | 370045..373584 | 410841 | BREX-system-P-loop-protein-BrxC-[Kaistia-defluvii] |
| WP_110780661.1 | Type1.BREX | <-DOC+HTH<-SF1-SF2-Helicase<-HKD-DNase+HKD-DNase<-wHTH<- SF1-SF2-Helicase+REase <- SF1-SF2-Helicase+REase <- SIGMA+OB-Fold+AAA-ATPase+HKD-DNase-BrxL <- PglZ <- N6-METHYLASE <- DUF6079 *<- BrxB-BrxF-iSTAND <- BrxA-DNA-binding <- wHTH+WYL+WCX <-MODE-HTH+PBPII\|\|TetR-HTH-><-RHH<-VirB10<-VirB9<-VirB8 | 1179 | Bacteria>Pseudomonadota>Alphaproteobacteria>Hyphomicrobiales>Nitrobacteraceae>Rhodopseudomonas | GCF_003217325.1 | Rhodopseudomonas_faecalis | BJ122_RS08545 | 91514..95053 | 99655 | BREX-system-P-loop-protein-BrxC-[Rhodopseudomonas-faecalis] |
| WP_009450523.1 | Type1.BREX | ACET-><-SF1-SF2-Helicase<-HKD-DNase+HKD-DNase<-wHTH<- SF1-SF2-Helicase+REase <- SF1-SF2-Helicase+REase <- SIGMA+OB-Fold+AAA-ATPase+HKD-DNase-BrxL <- PglZ <- N6-METHYLASE <- DUF6079 *<- BrxB-BrxF-iSTAND <- BrxA-DNA-binding <- wHTH+WYL+WCX <-MODE-HTH+PBPII\|\|TetR-HTH-><-RHH<-VirB10<-VirB9<-VirB8 | 1179 | Bacteria>Pseudomonadota>Alphaproteobacteria>Hyphomicrobiales>Phyllobacteriaceae>Nitratireductor | GCF_000300515.1 | Nitratireductor_indicus | NA8A_RS11845 | 137447..140986 | 721133 | BREX-system-P-loop-protein-BrxC-[Nitratireductor-indicus] |
| TQN59072.1 | Type1.BREX | <-SF1-SF2-Helicase<-HKD-DNase+HKD-DNase<-wHTH<- SF1-SF2-Helicase+REase <- SIGMA+OB-Fold+AAA-ATPase+HKD-DNase-BrxL <- PglZ <- N6-METHYLASE <- DUF6079 *<- BrxB-BrxF-iSTAND <- BrxA-DNA-binding <- wHTH+WYL+WCX <-MODE-HTH+PBPII\|\|TetR-HTH-><-RHH<-VirB10<-VirB9<-VirB8 | 1179 | Bacteria>Pseudomonadota>Alphaproteobacteria>Hyphomicrobiales>Rhizobiaceae>Rhizobium/Agrobacterium_group | GCA_006757705.1 | Agrobacterium_tumefaciens | FLX27_23795 | 22501..26040 | 358 | BREX-system-P-loop-protein-BrxC-[Agrobacterium-tumefaciens] |
| WP_136542766.1 | Type1.BREX | ACET-><-SF1-SF2-Helicase<-HKD-DNase+HKD-DNase<-wHTH<- SF1-SF2-Helicase+REase <- SF1-SF2-Helicase+REase <- SIGMA+OB-Fold+AAA-ATPase+HKD-DNase-BrxL <- PglZ <- N6-METHYLASE <- DUF6079 *<- BrxB-BrxF-iSTAND <- BrxA-DNA-binding <- wHTH+WYL+WCX <-MODE-HTH+PBPII\|\|TetR-HTH-><-RHH<-VirB10<-VirB9<-VirB8 | 1179 | Bacteria>Pseudomonadota>Alphaproteobacteria>Hyphomicrobiales>Rhizobiaceae>Rhizobium/Agrobacterium_group | GCF_014202175.1 | Rhizobium_rosettiformans | FAA86_RS19555 | 64899..68438 | 1368430 | BREX-system-P-loop-protein-BrxC-[Rhizobium-rosettiformans] |
| WP_217361198.1 | Type1.BREX | <-HATPase_c\|\|Arm-DNA-bind_3+Phage_integrase->LON-AAA->?->?-><-?\|\| wHTH+WYL+WCX -> BrxA-DNA-binding -> BrxB-BrxF-iSTAND -> DUF6079 *-> N6-METHYLASE -> PglZ -> SIGMA+OB-Fold+AAA-ATPase+HKD-DNase-BrxL ->?->HEPN+PAS+DUF4011+ SF1-SF2-Helicase+REase ->ParB+HNH+Schlafen-> | 1168 | Bacteria>Pseudomonadota>Alphaproteobacteria>Rhodobacterales>Paracoccaceae>Aliiroseovarius | GCF_013283115.1 | unclassified_Aliiroseovarius | XMV209_RS00060 | 11614..15120 | 2623558 | MULTISPECIES:-BREX-system-P-loop-protein-BrxC-[unclassified-Aliiroseovarius] |
| WP_131389735.1 | Type1.BREX | VirB8->VirB9->VirB10->RHH-><-TetR-HTH\|\|MODE-HTH+PBPII-> wHTH+WYL+WCX -> BrxA-DNA-binding -> BrxB-BrxF-iSTAND -> DUF6079 *-> N6-METHYLASE -> PglZ -> SIGMA+OB-Fold+AAA-ATPase+HKD-DNase-BrxL -> SF1-SF2-Helicase+REase -> SF1-SF2-Helicase+REase ->wHTH->HKD-DNase+HKD-DNase->SF1-SF2-Helicase-> | 1179 | Bacteria>Pseudomonadota>Alphaproteobacteria>Rhodobacterales>Paracoccaceae>Haematobacter | GCF_004328555.1 | Haematobacter_massiliensis | HmaOT1_RS15910 | 65338..68877 | 195105 | BREX-system-P-loop-protein-BrxC-[Haematobacter-massiliensis] |
| APO88218.1 | Type1.BREX | <-HEPN+PAS+DUF4011+ SF1-SF2-Helicase+REase <-?<- SIGMA+OB-Fold+AAA-ATPase+HKD-DNase-BrxL <- PglZ <- N6-METHYLASE <- DUF6079 *<- BrxB-BrxF-iSTAND <- BrxA-DNA-binding <- wHTH+WYL+WCX \|\|?-><-LON-AAA\|\|<-tRNA\|\|PRTase->?->REC+OmpR-HTH->HPT+HISKIN+CheW+REC-> | 1166 | Bacteria>Pseudomonadota>Alphaproteobacteria>Rhodobacterales>Paracoccaceae>Marivivens_group | GCA_001908835.1 | Marivivens_sp._JLT3646 | BSK21_01620 | 313072..316572 | 1920883 | hypothetical-protein-BSK21_01620-[Marivivens-sp.-JLT3646] |
| WP_199257821.1 | Type1.BREX | GDResolvase+Recombinase+ZnR-><-SF1-SF2-Helicase<-HKD-DNase+HKD-DNase<-wHTH<- SF1-SF2-Helicase+REase <- SF1-SF2-Helicase+REase <- SIGMA+OB-Fold+AAA-ATPase+HKD-DNase-BrxL <- PglZ <- N6-METHYLASE <- DUF6079 *<- BrxB-BrxF-iSTAND <- BrxA-DNA-binding <- wHTH+WYL+WCX <-MODE-HTH+PBPII\|\|TetR-HTH-><-RHH<-VirB10<-VirB9<-VirB8 | 1179 | Bacteria>Pseudomonadota>Alphaproteobacteria>Rhodobacterales>Paracoccaceae>Paracoccus | GCF_016446475.1 | Paracoccus_binzhouensis | I9P22_RS02450 | 23714..27253 | 2796149 | BREX-system-P-loop-protein-BrxC-[Paracoccus-binzhouensis] |
| MAD36284.1 | Type1.BREX | DUF6079 -p*-> N6-METHYLASE -> PglZ -> SIGMA+OB-Fold+AAA-ATPase+Lon-Protease-BrxL ->Exonuc_V_gamma+REase -> SF1-SF2-Helicase+REase ->HKD-DNase+OGG-like-> | 971 | Bacteria>Pseudomonadota>Alphaproteobacteria>Rhodospirillales>Geminicoccaceae>Tistrella | GCA_002684955.1 | Tistrella_sp. | CMO28_06235 | 72..2987 | 2024861 | hypothetical-protein-[Tistrella-sp.] |
| MBA78722.1 | Type1.BREX | DUF6079 -p*-> N6-METHYLASE -> PglZ -> SIGMA+OB-Fold+AAA-ATPase+Lon-Protease-BrxL ->Exonuc_V_gamma+REase -> SF1-SF2-Helicase+REase ->HKD-DNase+OGG-like-> | 971 | Bacteria>Pseudomonadota>Alphaproteobacteria>Rhodospirillales>Geminicoccaceae>Tistrella | GCA_002709065.1 | Tistrella_sp. | CMO30_25940 | 72..2987 | 2024861 | hypothetical-protein-[Tistrella-sp.] |
| WP_086466635.1 | Type1.BREX | ParB-><-?<-HKD-DNase+OGG-like<- SF1-SF2-Helicase+REase <-Exonuc_V_gamma+REase <- SIGMA+OB-Fold+AAA-ATPase+Lon-Protease-BrxL <- PglZ <- N6-METHYLASE <- DUF6079 *<- BrxB-BrxF-iSTAND <- BrxA-DNA-binding \|\|Classical-AAA->Peptidase_S8-><-?<-AlcRN+AraC-HTH+AraC-HTH\|\|?->Thermonuclease->PAPS-reduct-> | 1149 | Bacteria>Pseudomonadota>Alphaproteobacteria>Rhodospirillales>Thalassobaculaceae>Oceanibaculum | GCF_002148795.1 | Oceanibaculum_nanhaiense | BKM74_RS15615 | 60819..64268 | 1909734 | BREX-system-P-loop-protein-BrxC-[Oceanibaculum-nanhaiense] |
| MBL4843424.1 | Type1.BREX | <-SF1-SF2-Helicase<-HKD-DNase+HKD-DNase<-wHTH<- SF1-SF2-Helicase+REase <- SF1-SF2-Helicase+REase <- SIGMA+OB-Fold+AAA-ATPase+HKD-DNase-BrxL <- PglZ <- N6-METHYLASE <- DUF6079 *<- BrxB-BrxF-iSTAND <- BrxA-DNA-binding <- wHTH+WYL+WCX \|\|Calcineurin+Calcineurin-SbcD->SbcC+SbcC-><-MODE-HTH+PBPII\|\|TetR-HTH-><-RHH\|\|HTH_Tnp_1+HTH_21+rve+rve_3-> | 1179 | Bacteria>Pseudomonadota>Alphaproteobacteria>Rhodospirillales>Thalassospiraceae>Thalassospira | GCA_016765055.1 | Thalassospira_sp. | JKY47_21635 | 20483..24022 | 1912094 | BREX-system-P-loop-protein-BrxC-[Thalassospira-sp.] |
| WP_010336176.1 | Type1.BREX | VirB8->VirB9->VirB10->RHH-><-TetR-HTH\|\|MODE-HTH+PBPII-> wHTH+WYL+WCX -> BrxA-DNA-binding -> BrxB-BrxF-iSTAND -> DUF6079 *-> N6-METHYLASE -> PglZ -> SIGMA+OB-Fold+AAA-ATPase+HKD-DNase-BrxL -> SF1-SF2-Helicase+REase -> SF1-SF2-Helicase+REase ->wHTH->HKD-DNase+HKD-DNase->SF1-SF2-Helicase-> | 1179 | Bacteria>Pseudomonadota>Alphaproteobacteria>Sphingomonadales>Sphingomonadaceae>Sphingobium | GCF_000224695.1 | Sphingobium_yanoikuyae | SYA_RS0104815 | 38393..41932 | 13690 | BREX-system-P-loop-protein-BrxC-[Sphingobium-yanoikuyae] |
| WP_066478880.1 | Type1.BREX | Peptidase_S26->DUF736->?->TraI-VirD2-RCRE->\|\|MODE-HTH+PBPII-> wHTH+WYL+WCX -> BrxA-DNA-binding -> BrxB-BrxF-iSTAND -> DUF6079 *-> N6-METHYLASE -> PglZ -> SIGMA+OB-Fold+AAA-ATPase+HKD-DNase-BrxL -> SF1-SF2-Helicase+REase -> SF1-SF2-Helicase+REase ->wHTH->HKD-DNase+HKD-DNase->SF1-SF2-Helicase-> | 1179 | Bacteria>Pseudomonadota>Alphaproteobacteria>Sphingomonadales>Sphingomonadaceae>Sphingomonas | GCF_001555785.1 | unclassified_Sphingomonas | AX667_RS15610 | 42451..45990 | 196159 | MULTISPECIES:-BREX-system-P-loop-protein-BrxC-[unclassified-Sphingomonas] |
| WP_015740180.1 | Type1.BREX | VirB10->RHH-><-TetR-HTH\|\|MODE-HTH+PBPII-><-SbcC+SbcC<-Calcineurin+Calcineurin-SbcD\|\| wHTH+WYL+WCX -> BrxA-DNA-binding -> BrxB-BrxF-iSTAND -> DUF6079 *-> N6-METHYLASE -> PglZ -> SIGMA+OB-Fold+AAA-ATPase+HKD-DNase-BrxL -> SF1-SF2-Helicase+REase ->?->?-><-MODE-HTH+PBPII<-ArdCN\|\|GNTR-HTH+Aminotran_1_2-> | 1179 | Bacteria>Pseudomonadota>Alphaproteobacteria>Sphingomonadales>Zymomonadaceae>Zymomonas | GCF_000024245.1 | Zymomonas_mobilis | Za10_1374 | 1591366..1594905 | 542 | BREX-system-P-loop-protein-BrxC-[Zymomonas-mobilis] |
| MCK9506949.1 | Type1.BREX | DUF6079 *-> N6-METHYLASE -> PglZ -> SIGMA+OB-Fold+AAA-ATPase+HKD-DNase-BrxL -> SF1-SF2-Helicase+REase -> SF1-SF2-Helicase+REase ->wHTH->HKD-DNase+HKD-DNase->SF1-SF2-Helicase-> | 1069 | Bacteria>Pseudomonadota>Betaproteobacteria>Burkholderiales>Alcaligenaceae>Pigmentiphaga | GCA_023228545.1 | Pigmentiphaga_sp. | MOQ54_00780 | 41..3250 | 1977564 | BREX-system-P-loop-protein-BrxC-[Pigmentiphaga-sp.] |
| MBP7846551.1 | Type1.BREX | <-TrwC+SF1-SF2-Helicase<- SF1-SF2-Helicase+REase <- SIGMA+OB-Fold+AAA-ATPase+Lon-Protease-BrxL <- PglZ <-TOPRIM<-OLD-ABC<- N6-METHYLASE -p<- N6-METHYLASE <- DUF6079 *<- BrxB-BrxF-iSTAND <- BrxA-DNA-binding <-SWIM+DUF3670+SF1-SF2-Helicase<-?<-SWIM+DUF3670+SF1-SF2-Helicase | 1193 | Bacteria>Pseudomonadota>Betaproteobacteria>Burkholderiales>unclassified_Burkholderiales | GCA_018001735.1 | Burkholderiales_bacterium | KAZ94_00285 | 68005..71586 | 1891238 | BREX-system-P-loop-protein-BrxC-[Burkholderiales-bacterium] |
| WP_010634561.1 | Type1.BREX | DUF2787->?->?->?-> SF1-SF2-Helicase+REase -> SF1-SF2-Helicase+REase -><- PglZ <-ParB+HNH<- N6-METHYLASE <- DUF6079 *<- BrxB-BrxF-iSTAND <- BrxA-DNA-binding <-RHS<-ZnR<-?<-ARG\|\| wHTH+WYL+WCX -><-?\|\|LexA-protease-> | 1157 | Bacteria>Pseudomonadota>Gammaproteobacteria>Aeromonadales>Aeromonadaceae>Aeromonas | GCF_000315195.1 | Aeromonas_dhakensis | AQU_RS10070 | 136801..140274 | 196024 | BREX-system-P-loop-protein-BrxC-[Aeromonas-dhakensis] |
| WP_014998461.1 | Type1.BREX | <-Phage_integrase<-?\|\|XIS-HTH-> SF1-SF2-Helicase+REase -> SF1-SF2-Helicase+REase -><- PglZ <-TOPRIM<-OLD-ABC<- N6-METHYLASE <- DUF6079 *<- BrxB-BrxF-iSTAND <- BrxA-DNA-binding <- wHTH+WYL+WCX <-Arm-DNA-bind_3+Phage_integrase<-tRNA<-tRNA<-tRNA<-?<-URI-DNase+UVR+RNase+HHH<-REC+OmpR-HTH<-DSBH<-Aminotran_1_2 | 1157 | Bacteria>Pseudomonadota>Gammaproteobacteria>Alteromonadales>Alteromonadaceae>Alteromonas/Salinimonas_group | - | Alteromonas_macleodii | AMBLS11_RS09820 | 2297331..2300804 | 28108 | BREX-system-P-loop-protein-BrxC-[Alteromonas-macleodii] |
| WP_100972015.1 | Type1.BREX | SF1-SF2-Helicase+REase -> SF1-SF2-Helicase+REase -p->rve->AAA-ATPase-> SF1-SF2-Helicase+REase -p-><- PglZ <-TOPRIM<-OLD-ABC<- N6-METHYLASE <- DUF6079 *<- BrxB-BrxF-iSTAND <- BrxA-DNA-binding <- wHTH+WYL+WCX \|\|?->GTPase-AIG->GTPase-AIG->?->?-><-?<-GAF+GGDEF | 1157 | Bacteria>Pseudomonadota>Gammaproteobacteria>Alteromonadales>Alteromonadaceae>Alteromonas/Salinimonas_group | GCF_002831605.1 | Alteromonas_sp._MB-3u-76 | CW735_RS10480 | 2338685..2342158 | 2058133 | BREX-system-P-loop-protein-BrxC-[Alteromonas-sp.-MB-3u-76] |
| WP_207611650.1 | Type1.BREX | <-Peptidase_M20<-?<-?\|\|?->tRNA->Arm-DNA-bind_3+Phage_integrase-> wHTH+WYL+WCX -> BrxA-DNA-binding -> BrxB-BrxF-iSTAND -> DUF6079 *-> N6-METHYLASE ->ParB+HNH-> PglZ -><- SF1-SF2-Helicase+REase <- SF1-SF2-Helicase+REase \|\|?->Phage_integrase-> | 1157 | Bacteria>Pseudomonadota>Gammaproteobacteria>Alteromonadales>Alteromonadaceae>Alteromonas/Salinimonas_group | GCF_017368475.1 | Alteromonas_sp._5E99-2 | J3L16_RS04210 | 79358..82831 | 2817683 | BREX-system-P-loop-protein-BrxC-[Alteromonas-sp.-5E99-2] |
| WP_173294709.1 | Type1.BREX | wHTH(SBD)+HNH-> BrxA-DNA-binding -> BrxB-BrxF-iSTAND -> DUF6079 *-> N6-METHYLASE -> PglZ -><- SF1-SF2-Helicase+REase <- SF1-SF2-Helicase+REase \|\|?-><-PAIREDC-HTH | 1157 | Bacteria>Pseudomonadota>Gammaproteobacteria>Alteromonadales>Shewanellaceae>Shewanella | GCF_013248905.1 | Shewanella_sp._VB17 | HQQ94_RS12350 | 2876249..2879722 | 2739432 | BREX-system-P-loop-protein-BrxC-[Shewanella-sp.-VB17] |
| WP_076500079.1 | Type1.BREX | SF1-SF2-Helicase+REase -> SF1-SF2-Helicase+REase -><- REase +SF1-SF2-Helicase <- REase <- PglZ <-TOPRIM<-OLD-ABC<- N6-METHYLASE <- DUF6079 *<- BrxB-BrxF-iSTAND <- BrxA-DNA-binding \|\|?->?->CbeA_antitoxin->DUF2787->DUF2787->Phage_integrase-> | 1157 | Bacteria>Pseudomonadota>Gammaproteobacteria>Alteromonadales>Shewanellaceae>Shewanella | GCF_900156405.1 | Shewanella_morhuae | BW947_RS12490 | 20495..23968 | 365591 | BREX-system-P-loop-protein-BrxC-[Shewanella-morhuae] |
| WP_019531408.1 | Type1.BREX | <- DUF6079 *<- BrxB-BrxF-iSTAND <- BrxA-DNA-binding <- SF1-SF2-Helicase+REase <-Exonuc_V_gamma+REase <-MULE-TRANSPOSASE<-?<-TraJ-RHH\|\|?->Peptidase_M50+PDZ_2+Lon_C-> | 1156 | Bacteria>Pseudomonadota>Gammaproteobacteria>Cellvibrionales>Spongiibacteraceae>Dasania | GCF_000373485.1 | Dasania_marina | B067_RS0117580 | 1458..4928 | 471499 | BREX-system-P-loop-protein-BrxC-[Dasania-marina] |
| WP_124023337.1 | Type1.BREX | <-sGTP<-wHTH+WYL<-NACHT+SF1-SF2-Helicase+HKD-DNase\|\| SF1-SF2-Helicase+REase -> SF1-SF2-Helicase+REase -><- PglZ <-ParB+HNH<- N6-METHYLASE <- DUF6079 *<- BrxB-BrxF-iSTAND <- BrxA-DNA-binding <- wHTH+WYL+WCX <-Arm-DNA-bind_3+Phage_integrase\|\|ParA-Soj-PloopNTPase->?->DnaB+TnsC-ATPase->ParB+BAM->HTH+HTH-> | 1157 | Bacteria>Pseudomonadota>Gammaproteobacteria>Enterobacterales>Enterobacteriaceae>Buttiauxella | GCF_003818135.1 | Buttiauxella_warmboldiae | EHN07_RS06350 | 61248..64721 | 82993 | BREX-system-P-loop-protein-BrxC-[Buttiauxella-warmboldiae] |
| HAT7499716.1 | Type1.BREX | <-Peptidase_M17_N+Peptidase_M17\|\|?->?-><-HerA/FtsK+A32<-?\|\|tRNA->Arm-DNA-bind_3+Phage_integrase-> wHTH+WYL+WCX -> BrxA-DNA-binding -> BrxB-BrxF-iSTAND -> DUF6079 *-> N6-METHYLASE ->ParB+HNH-> PglZ -><- SF1-SF2-Helicase+REase <- SF1-SF2-Helicase+REase \|\|NACHT+SF1-SF2-Helicase+HKD-DNase-><-?<-Glyoxalase<-ZnR | 1157 | Bacteria>Pseudomonadota>Gammaproteobacteria>Enterobacterales>Enterobacteriaceae>Citrobacter | GCA_015956985.1 | Citrobacter_freundii | JAV71_002653 | 147091..150564 | 546 | BREX-system-P-loop-protein-BrxC-[Citrobacter-freundii] |
| WP_200054816.1 | Type1.BREX | <-Peptidase_M17_N+Peptidase_M17\|\|?->?-><-HerA/FtsK+A32<-?\|\|tRNA->Arm-DNA-bind_3+Phage_integrase-> wHTH+WYL+WCX -> BrxA-DNA-binding -> BrxB-BrxF-iSTAND -> DUF6079 *-> N6-METHYLASE ->ParB+HNH-> PglZ -><- SF1-SF2-Helicase+REase <- SF1-SF2-Helicase+REase \|\|NACHT+SF1-SF2-Helicase+HKD-DNase-><-?<-ZnR<-XIS-HTH | 1157 | Bacteria>Pseudomonadota>Gammaproteobacteria>Enterobacterales>Enterobacteriaceae>Citrobacter | GCF_016504925.1 | Citrobacter_freundii | GHT45_RS09240 | 237227..240700 | 546 | BREX-system-P-loop-protein-BrxC-[Citrobacter-freundii] |
| WP_161575571.1 | Type1.BREX | <-Peptidase_M17_N+Peptidase_M17\|\|?->?->?->?-><-?\|\|tRNA->Arm-DNA-bind_3+Phage_integrase-> BrxA-DNA-binding -> BrxB-BrxF-iSTAND -> DUF6079 *-> N6-METHYLASE ->ParB+HNH-> PglZ -><- SF1-SF2-Helicase+REase <- SF1-SF2-Helicase+REase <-?<-?<-XIS-HTH | 1157 | Bacteria>Pseudomonadota>Gammaproteobacteria>Enterobacterales>Enterobacteriaceae>Cronobacter | GCF_009938355.1 | Cronobacter_sakazakii | EHJ08_RS11755 | 205127..208600 | 28141 | BREX-system-P-loop-protein-BrxC-[Cronobacter-sakazakii] |
| WP_126800556.1 | Type1.BREX | SF1-SF2-Helicase+REase -> SF1-SF2-Helicase+REase -><- PglZ <-ParB+HNH<- N6-METHYLASE <- DUF6079 *<- BrxB-BrxF-iSTAND <- BrxA-DNA-binding <-Arm-DNA-bind_3+Phage_integrase<-tRNA<-?<-?<-?\|\|?->?->ICLR-HTH-> | 1157 | Bacteria>Pseudomonadota>Gammaproteobacteria>Enterobacterales>Enterobacteriaceae>Enterobacter | GCF_003986875.1 | Enterobacter_sp._WCHEn090032 | EKN83_RS06700 | 53011..56484 | 2497435 | BREX-system-P-loop-protein-BrxC-[Enterobacter-sp.-WCHEn090032] |
| WP_129694025.1 | Type1.BREX | <- REase <-?<-HTH_Tnp_1+HTH_21+rve+rve_3\|\| SF1-SF2-Helicase+REase -> SF1-SF2-Helicase+REase -><- PglZ <-ParB+HNH<- N6-METHYLASE <- DUF6079 *<- BrxB-BrxF-iSTAND <- BrxA-DNA-binding <-tRNA<-?<-?<-?\|\|?->?->ICLR-HTH-><-Glyco_hydro_43+GH43_C2 | 1157 | Bacteria>Pseudomonadota>Gammaproteobacteria>Enterobacterales>Enterobacteriaceae>Enterobacter | GCF_004138605.1 | Enterobacter_roggenkampii | D8675_RS06390 | 1085829..1089302 | 1812935 | BREX-system-P-loop-protein-BrxC-[Enterobacter-roggenkampii] |
| WP_164474334.1 | Type1.BREX | SF1-SF2-Helicase+REase -> SF1-SF2-Helicase+REase -><- PglZ <-ParB+HNH<- N6-METHYLASE <- N6-METHYLASE <- DUF6079 *<- BrxB-BrxF-iSTAND <- BrxA-DNA-binding <-Arm-DNA-bind_3+Phage_integrase | 1157 | Bacteria>Pseudomonadota>Gammaproteobacteria>Enterobacterales>Enterobacteriaceae>Enterobacter | GCF_011045335.1 | Enterobacter_hormaechei | G5C52_RS10105 | 2135134..2138607 | 158836 | BREX-system-P-loop-protein-BrxC-[Enterobacter-hormaechei] |
| WP_211449384.1 | Type1.BREX | <-SymE_toxin\|\|wHTH(SBD)+HNH-><-NACHT+SF1-SF2-Helicase+HKD-DNase\|\| SF1-SF2-Helicase+REase -> SF1-SF2-Helicase+REase -><- PglZ <-ParB+HNH<- N6-METHYLASE <- DUF6079 *<- BrxB-BrxF-iSTAND <- BrxA-DNA-binding <- wHTH+WYL+WCX \|\|ParA-Soj-PloopNTPase->?->DnaB+TnsC-ATPase->ParB+BAM->HTH+HTH->HTH_36-> | 1157 | Bacteria>Pseudomonadota>Gammaproteobacteria>Enterobacterales>Enterobacteriaceae>Enterobacter | GCF_018299605.1 | Enterobacter_mori | GUY16_RS21845 | 58984..62457 | 539813 | BREX-system-P-loop-protein-BrxC-[Enterobacter-mori] |
| POU04217.1 | Type1.BREX | HNH-><-?\|\|?->XIS-HTH-> SF1-SF2-Helicase+REase -> SF1-SF2-Helicase+REase -><- PglZ <-ParB+HNH<- N6-METHYLASE <- DUF6079 *<- BrxB-BrxF-iSTAND <- BrxA-DNA-binding <-Arm-DNA-bind_3+Phage_integrase<-tRNA\|\|wHTH(SBD)+HNH->SAM-METHYLASE-> | 1157 | Bacteria>Pseudomonadota>Gammaproteobacteria>Enterobacterales>Enterobacteriaceae>Escherichia | GCA_002918705.1 | Escherichia_sp._ESNIH1 | C3369_02380 | 500495..503968 | 1985876 | BREX-system-P-loop-protein-BrxC-[Escherichia-sp.-ESNIH1] |
| HBQ4939450.1 | Type1.BREX | XIS-HTH-> SF1-SF2-Helicase+REase -> SF1-SF2-Helicase+REase -><- PglZ <-ParB+HNH<- N6-METHYLASE <- DUF6079 *<- BrxB-BrxF-iSTAND <- BrxA-DNA-binding <-Arm-DNA-bind_3+Phage_integrase | 1157 | Bacteria>Pseudomonadota>Gammaproteobacteria>Enterobacterales>Enterobacteriaceae>Klebsiella/Raoultella_group | GCA_021861305.1 | Klebsiella_pneumoniae | L7366_003904 | 52372..55845 | 573 | BREX-system-P-loop-protein-BrxC-[Klebsiella-pneumoniae] |
| HBS5818882.1 | Type1.BREX | Arm-DNA-bind_3+Phage_integrase-> BrxA-DNA-binding -> BrxB-BrxF-iSTAND -> DUF6079 *-> N6-METHYLASE ->ParB+HNH-> PglZ -><- SF1-SF2-Helicase+REase <- SF1-SF2-Helicase+REase <-XIS-HTH | 1157 | Bacteria>Pseudomonadota>Gammaproteobacteria>Enterobacterales>Enterobacteriaceae>Klebsiella/Raoultella_group | GCA_021971955.1 | Klebsiella_pneumoniae | MAK71_003676 | 3132..6605 | 573 | BREX-system-P-loop-protein-BrxC-[Klebsiella-pneumoniae] |
| WP_096812545.1 | Type1.BREX | Arm-DNA-bind_3+Phage_integrase-> BrxA-DNA-binding -> BrxB-BrxF-iSTAND -> DUF6079 *-> N6-METHYLASE ->ParB+HNH-> PglZ -><- SF1-SF2-Helicase+REase <- SF1-SF2-Helicase+REase <-XIS-HTH | 1157 | Bacteria>Pseudomonadota>Gammaproteobacteria>Enterobacterales>Enterobacteriaceae>Klebsiella/Raoultella_group | GCF_009907625.1 | Klebsiella_pneumoniae | GV742_RS00800 | 3194..6667 | 573 | BREX-system-P-loop-protein-BrxC-[Klebsiella-pneumoniae] |

| Protein-ID | System | Gene-Neighbourhoods | Length | Taxonomy | Assembly-ID | Strain | Locus-Tag | Genomic-coordinates | TaxID | Fasta-defline |
|---|---|---|---|---|---|---|---|---|---|---|
| WP_023329155.1 | Type1.BREX | [illegible] | 1157 | Bacteria>Pseudomonadota>Gammaproteobacteria>Enterobacterales>Enterobacteriaceae>Klebsiella/Raoultella_group | GCF_021264605.1 | Klebsiella_oxytoca | LVW29_RS13105 | 76515..79988 | 571 | BREX-system-P-loop-protein-BrxC-[Klebsiella-oxytoca] |
| ECJ2539588.1 | Type1.BREX | [illegible] | 1157 | Bacteria>Pseudomonadota>Gammaproteobacteria>Enterobacterales>Enterobacteriaceae>Salmonella | GCA_007808255.1 | Salmonella_enterica_subsp._salamae | FNI64_03990 | 384975..388448 | 59202 | BREX-system-P-loop-protein-BrxC-[Salmonella-enterica-subsp.-salamae] |
| WP_158149594.1 | Type1.BREX | [illegible] | 1157 | Bacteria>Pseudomonadota>Gammaproteobacteria>Enterobacterales>Erwiniaceae>Pantoea | GCF_009765475.1 | Pantoea_agglomerans | GSQ69_RS16285 | 3495115..3498588 | 549 | BREX-system-P-loop-protein-BrxC-[Pantoea-agglomerans] |
| WP_187492467.1 | Type1.BREX | [illegible] | 1157 | Bacteria>Pseudomonadota>Gammaproteobacteria>Enterobacterales>Erwiniaceae>Pantoea | GCF_014353895.1 | Pantoea_agglomerans | IBU46_RS01740 | 23077..26550 | 549 | BREX-system-P-loop-protein-BrxC-[Pantoea-agglomerans] |
| WP_191919036.1 | Type1.BREX | [illegible] | 1157 | Bacteria>Pseudomonadota>Gammaproteobacteria>Enterobacterales>Erwiniaceae>Pantoea | GCF_014838975.1 | Pantoea_agglomerans | IFU26_RS12730 | 289507..292980 | 549 | BREX-system-P-loop-protein-BrxC-[Pantoea-agglomerans] |
| WP_272844016.1 | Type1.BREX | [illegible] | 1157 | Bacteria>Pseudomonadota>Gammaproteobacteria>Enterobacterales>Erwiniaceae>Pantoea | GCF_028555165.1 | Pantoea_ananatis | SZ66_RS15300 | 342532..346005 | 553 | BREX-system-P-loop-protein-BrxC-[Pantoea-ananatis] |
| WP_117259785.1 | Type1.BREX | [illegible] | 1157 | Bacteria>Pseudomonadota>Gammaproteobacteria>Enterobacterales | GCF_013395115.1 | Enterobacteriaceae | HYG96_RS13000 | 239421..242894 | 543 | MULTISPECIES:-BREX-system-P-loop-protein-BrxC-[Enterobacteriaceae] |
| WP_224243997.1 | Type1.BREX | [illegible] | 1157 | Bacteria>Pseudomonadota>Gammaproteobacteria>Enterobacterales | GCF_023752605.1 | Enterobacteriaceae | M8S83_RS11470 | 342574..346047 | 543 | MULTISPECIES:-BREX-system-P-loop-protein-BrxC-[Enterobacteriaceae] |
| WP_045345932.1 | Type1.BREX | [illegible] | 1157 | Bacteria>Pseudomonadota>Gammaproteobacteria>Enterobacterales | GCF_902160145.1 | Enterobacteriaceae | FN179_RS14130 | 343492..346965 | 543 | MULTISPECIES:-BREX-system-P-loop-protein-BrxC-[Enterobacteriaceae] |
| GKV76958.1 | Type1.BREX | [illegible] | 1157 | Bacteria>Pseudomonadota>Gammaproteobacteria>Enterobacterales>Pectobacteriaceae>Pectobacterium | GCA_024342785.1 | Pectobacterium_carotovorum_subsp._carotovorum | PEC106568_21320 | 196499..199972 | 555 | hypothetical-protein-PEC106568_21320-[Pectobacterium-carotovorum-subsp.-carotovorum] |
| WP_211024717.1 | Type1.BREX | [illegible] | 1157 | Bacteria>Pseudomonadota>Gammaproteobacteria>Enterobacterales>Pectobacteriaceae>Pectobacterium | GCF_018094745.1 | Pectobacterium_versatile | F9U42_RS09295 | 89992..93465 | 2488639 | BREX-system-P-loop-protein-BrxC-[Pectobacterium-versatile] |
| WP_249681848.1 | Type1.BREX | [illegible] | 1157 | Bacteria>Pseudomonadota>Gammaproteobacteria>Enterobacterales>Pectobacteriaceae>Pectobacterium | - | Pectobacterium_polaris | EXT50_03115 | 82893..86366 | 2042057 | BREX-system-P-loop-protein-BrxC-[Pectobacterium-polaris] |
| WP_232770162.1 | Type1.BREX | [illegible] | 1157 | Bacteria>Pseudomonadota>Gammaproteobacteria>Enterobacterales>Yersiniaceae>Rahnella | GCF_021276285.1 | Rahnella_victoriana | J9880_RS04875 | 1051789..1055262 | 1510570 | BREX-system-P-loop-protein-BrxC-[Rahnella-victoriana] |
| WP_266057125.1 | Type1.BREX | [illegible] | 1157 | Bacteria>Pseudomonadota>Gammaproteobacteria>Enterobacterales>Yersiniaceae>Rahnella | GCF_026241995.1 | Rahnella_perminowiae | ORG37_RS00070 | 21089..24562 | 2816244 | BREX-system-P-loop-protein-BrxC-[Rahnella-perminowiae] |
| NYA44684.1 | Type1.BREX | [illegible] | 1158 | Bacteria>Pseudomonadota>Gammaproteobacteria>Enterobacterales>Yersiniaceae>Serratia | GCA_013401715.1 | Serratia_fonticola | HZI31_15400 | 48979..52455 | 47917 | BREX-system-P-loop-protein-BrxC-[Serratia-fonticola] |
| WP_218833283.1 | Type1.BREX | [illegible] | 1197 | Bacteria>Pseudomonadota>Gammaproteobacteria>Enterobacterales>Yersiniaceae>Serratia | GCF_013401715.1 | Serratia_fonticola | HZI31_RS15400 | 48979..52572 | 47917 | BREX-system-P-loop-protein-BrxC-[Serratia-fonticola] |
| WP_206196473.1 | Type1.BREX | [illegible] | 1157 | Bacteria>Pseudomonadota>Gammaproteobacteria>Enterobacterales>Yersiniaceae>Serratia | GCF_017152235.1 | Serratia_ureilytica | JY544_RS23375 | 37448..40921 | 300181 | BREX-system-P-loop-protein-BrxC-[Serratia-ureilytica] |
| WP_223536794.1 | Type1.BREX | [illegible] | 1158 | Bacteria>Pseudomonadota>Gammaproteobacteria>Enterobacterales>Yersiniaceae>Serratia | GCF_019968705.1 | unclassified_Serratia_(in:_enterobacteria) | KGP26_RS00385 | 95452..98928 | 2647522 | MULTISPECIES:-BREX-system-P-loop-protein-BrxC-[unclassified-Serratia-(in:-enterobacteria)] |
| WP_273866014.1 | Type1.BREX | [illegible] | 1157 | Bacteria>Pseudomonadota>Gammaproteobacteria>Enterobacterales>Yersiniaceae>Serratia | GCF_028616515.1 | Serratia_ureilytica | PVU68_RS17145 | 216123..219596 | 300181 | BREX-system-P-loop-protein-BrxC-[Serratia-ureilytica] |
| WP_046051668.1 | Type1.BREX | [illegible] | 1157 | Bacteria>Pseudomonadota>Gammaproteobacteria>Enterobacterales>Yersiniaceae>Yersinia | GCF_000968115.1 | Yersinia_enterocolitica | YE5303_RS19630 | 4225906..4229379 | 630 | BREX-system-P-loop-protein-BrxC-[Yersinia-enterocolitica] |
| MCG7876672.1 | Type6.BREX | [illegible] | 1215 | Bacteria>Pseudomonadota>Gammaproteobacteria>Gammaproteobacteria_incertae_sedis>sulfur-oxidizing_symbionts>Candidatus_Thiodiazotropha | GCA_016842655.1 | Candidatus_Thiodiazotropha_taylori | JAY78_10080 | 4978..8625 | 2792791 | BREX-system-P-loop-protein-BrxC-[Candidatus-Thiodiazotropha-taylori] |
| MCG7951159.1 | Type6.BREX | [illegible] | 1215 | Bacteria>Pseudomonadota>Gammaproteobacteria>Gammaproteobacteria_incertae_sedis>sulfur-oxidizing_symbionts>Candidatus_Thiodiazotropha | GCA_016842765.1 | Candidatus_Thiodiazotropha_taylori | JAZ09_08050 | 22496..26143 | 2792791 | BREX-system-P-loop-protein-BrxC-[Candidatus-Thiodiazotropha-taylori] |
| MCG8102611.1 | Type6.BREX | [illegible] | 1215 | Bacteria>Pseudomonadota>Gammaproteobacteria>Gammaproteobacteria_incertae_sedis>sulfur-oxidizing_symbionts>Candidatus_Thiodiazotropha | GCA_016842845.1 | Candidatus_Thiodiazotropha_taylori | JAY70_09490 | 22526..26173 | 2792791 | BREX-system-P-loop-protein-BrxC-[Candidatus-Thiodiazotropha-taylori] |
| MCG8058214.1 | Type6.BREX | [illegible] | 1215 | Bacteria>Pseudomonadota>Gammaproteobacteria>Gammaproteobacteria_incertae_sedis>sulfur-oxidizing_symbionts>Candidatus_Thiodiazotropha | GCA_016843165.1 | Candidatus_Thiodiazotropha_taylori | JAZ21_00070 | 27190..30837 | 2792791 | BREX-system-P-loop-protein-BrxC-[Candidatus-Thiodiazotropha-taylori] |
| WP_252316424.1 | Type1.BREX | [illegible] | 1212 | Bacteria>Pseudomonadota>Gammaproteobacteria>Moraxellales>Moraxellaceae>Acinetobacter | - | Acinetobacter_sp._225588 | J687_RS15310 | 12712..16350 | 1310788 | BREX-system-P-loop-protein-BrxC-[Acinetobacter-sp.-225588] |
| WP_005303500.1 | Type1.BREX | [illegible] | 1226 | Bacteria>Pseudomonadota>Gammaproteobacteria>Moraxellales>Moraxellaceae>Acinetobacter | - | Acinetobacter_sp._NIPH_542 | F886_RS07830 | 464704..468384 | 1217688 | BREX-system-P-loop-protein-BrxC-[Acinetobacter-sp.-NIPH-542] |
| CAM84763.1 | Type1.BREX | [illegible] | 1229 | Bacteria>Pseudomonadota>Gammaproteobacteria>Moraxellales>Moraxellaceae>Acinetobacter | GCA_000069245.1 | Acinetobacter_baumannii_AYE | p3ABAYE0070 | 63476..67165 | 509173 | putative-ATPase-like-protein-[Acinetobacter-baumannii-AYE] |
| EXS00102.1 | Type1.BREX | [illegible] | 1190 | Bacteria>Pseudomonadota>Gammaproteobacteria>Moraxellales>Moraxellaceae>Acinetobacter | GCA_000588415.1 | Acinetobacter_sp._225588 | J687_2205 | 12778..16350 | 1310788 | hypothetical-protein-J687_2205-[Acinetobacter-sp.-225588] |
| HAV3580163.1 | Type1.BREX | [illegible] | 1226 | Bacteria>Pseudomonadota>Gammaproteobacteria>Moraxellales>Moraxellaceae>Acinetobacter | GCA_016485765.1 | Acinetobacter_baumannii | JIE63_001443 | 180845..184525 | 470 | BREX-system-P-loop-protein-BrxC-[Acinetobacter-baumannii] |
| HAV6134262.1 | Type1.BREX | [illegible] | 1226 | Bacteria>Pseudomonadota>Gammaproteobacteria>Moraxellales>Moraxellaceae>Acinetobacter | GCA_016514395.1 | Acinetobacter_baumannii | JI295_12625 | 9155..12835 | 470 | BREX-system-P-loop-protein-BrxC-[Acinetobacter-baumannii] |
| WP_001021844.1 | Type1.BREX | [illegible] | 1226 | Bacteria>Pseudomonadota>Gammaproteobacteria>Moraxellales>Moraxellaceae>Acinetobacter | GCF_000069245.1 | Acinetobacter_calcoaceticus/baumannii_complex | ABAYE_RS00380 | 63476..67156 | 909768 | MULTISPECIES:-BREX-system-P-loop-protein-BrxC-[Acinetobacter-calcoaceticus/baumannii-complex] |
| WP_001021843.1 | Type1.BREX | [illegible] | 1226 | Bacteria>Pseudomonadota>Gammaproteobacteria>Moraxellales>Moraxellaceae>Acinetobacter | GCF_000305235.1 | Acinetobacter_baumannii | ACIN5098_RS09930 | 47279..50959 | 470 | BREX-system-P-loop-protein-BrxC-[Acinetobacter-baumannii] |
| WP_004811070.1 | Type1.BREX | [illegible] | 1225 | Bacteria>Pseudomonadota>Gammaproteobacteria>Moraxellales>Moraxellaceae>Acinetobacter | GCF_000368465.1 | Acinetobacter_schindleri | F965_RS04725 | 393977..397654 | 108981 | BREX-system-P-loop-protein-BrxC-[Acinetobacter-schindleri] |
| WP_005247073.1 | Type1.BREX | [illegible] | 1226 | Bacteria>Pseudomonadota>Gammaproteobacteria>Moraxellales>Moraxellaceae>Acinetobacter | GCF_000369665.1 | Acinetobacter_sp._CIP_51.11 | F894_RS08220 | 945028..948708 | 1144670 | BREX-system-P-loop-protein-BrxC-[Acinetobacter-sp.-CIP-51.11] |
| WP_031946304.1 | Type1.BREX | [illegible] | 1226 | Bacteria>Pseudomonadota>Gammaproteobacteria>Moraxellales>Moraxellaceae>Acinetobacter | GCF_000516575.2 | Acinetobacter_baumannii | UN97_RS12380 | 246428..250108 | 470 | BREX-system-P-loop-protein-BrxC-[Acinetobacter-baumannii] |
| WP_025095052.1 | Type1.BREX | [illegible] | 1226 | Bacteria>Pseudomonadota>Gammaproteobacteria>Moraxellales>Moraxellaceae>Acinetobacter | GCF_000633005.1 | Acinetobacter_soli | EG58_RS0106955 | 8205..11885 | 487316 | BREX-system-P-loop-protein-BrxC-[Acinetobacter-soli] |
| WP_043972752.1 | Type1.BREX | [illegible] | 1226 | Bacteria>Pseudomonadota>Gammaproteobacteria>Moraxellales>Moraxellaceae>Acinetobacter | GCF_000876115.1 | Acinetobacter_sp._NBRC_110496 | Aba26S_RS13795 | 7444..11124 | 1550715 | BREX-system-P-loop-protein-BrxC-[Acinetobacter-sp.-NBRC-110496] |
| WP_075040289.1 | Type1.BREX | [illegible] | 1226 | Bacteria>Pseudomonadota>Gammaproteobacteria>Moraxellales>Moraxellaceae>Acinetobacter | GCF_001917365.1 | Acinetobacter_radioresistens | SA188iCDA_RS03905 | 54372..58052 | 40216 | BREX-system-P-loop-protein-BrxC-[Acinetobacter-radioresistens] |
| WP_111281982.1 | Type1.BREX | [illegible] | 1226 | Bacteria>Pseudomonadota>Gammaproteobacteria>Moraxellales>Moraxellaceae>Acinetobacter | GCF_003258335.1 | Acinetobacter_radioresistens | NVV25_RS10895 | 13399..17079 | 40216 | BREX-system-P-loop-protein-BrxC-[Acinetobacter-radioresistens] |
| WP_119025213.1 | Type1.BREX | [illegible] | 1226 | Bacteria>Pseudomonadota>Gammaproteobacteria>Moraxellales>Moraxellaceae>Acinetobacter | GCF_003569585.1 | Acinetobacter_soli | D5I70_RS01315 | 8436..12116 | 487316 | BREX-system-P-loop-protein-BrxC-[Acinetobacter-soli] |
| WP_119047908.1 | Type1.BREX | [illegible] | 1226 | Bacteria>Pseudomonadota>Gammaproteobacteria>Moraxellales>Moraxellaceae>Acinetobacter | GCF_003569645.1 | Acinetobacter_baumannii | D5I76_RS05685 | 52847..56527 | 470 | BREX-system-P-loop-protein-BrxC-[Acinetobacter-baumannii] |
| WP_121523424.1 | Type1.BREX | [illegible] | 1226 | Bacteria>Pseudomonadota>Gammaproteobacteria>Moraxellales>Moraxellaceae>Acinetobacter | GCF_003664645.1 | Acinetobacter_chengduensis | D9K81_RS12845 | 89350..93030 | 2420890 | BREX-system-P-loop-protein-BrxC-[Acinetobacter-chengduensis] |
| WP_140948309.1 | Type1.BREX | [illegible] | 1226 | Bacteria>Pseudomonadota>Gammaproteobacteria>Moraxellales>Moraxellaceae>Acinetobacter | GCF_006491975.1 | Acinetobacter_baumannii | FJV17_RS05945 | 24470..28150 | 470 | BREX-system-P-loop-protein-BrxC-[Acinetobacter-baumannii] |
| WP_151732002.1 | Type1.BREX | [illegible] | 1226 | Bacteria>Pseudomonadota>Gammaproteobacteria>Moraxellales>Moraxellaceae>Acinetobacter | GCF_008982325.1 | Acinetobacter_ursingii | GBJ63_RS14825 | 20101..23781 | 108980 | BREX-system-P-loop-protein-BrxC-[Acinetobacter-ursingii] |
| WP_153569296.1 | Type1.BREX | [illegible] | 1226 | Bacteria>Pseudomonadota>Gammaproteobacteria>Moraxellales>Moraxellaceae>Acinetobacter | GCF_009645555.1 | Acinetobacter_nosocomialis | F4T96_RS05690 | 68625..72305 | 106654 | BREX-system-P-loop-protein-BrxC-[Acinetobacter-nosocomialis] |
| WP_168396480.1 | Type1.BREX | [illegible] | 1224 | Bacteria>Pseudomonadota>Gammaproteobacteria>Moraxellales>Moraxellaceae>Acinetobacter | GCF_012366875.1 | Acinetobacter_indicus | HG607_RS04385 | 142310..145984 | 756892 | BREX-system-P-loop-protein-BrxC-[Acinetobacter-indicus] |
| WP_179994566.1 | Type1.BREX | [illegible] | 1226 | Bacteria>Pseudomonadota>Gammaproteobacteria>Moraxellales>Moraxellaceae>Acinetobacter | GCF_013416255.1 | Acinetobacter_baumannii | HLG75_RS04210 | 863150..866830 | 470 | BREX-system-P-loop-protein-BrxC-[Acinetobacter-baumannii] |
| WP_180041457.1 | Type1.BREX | [illegible] | 1221 | Bacteria>Pseudomonadota>Gammaproteobacteria>Moraxellales>Moraxellaceae>Acinetobacter | GCF_013417925.1 | Acinetobacter_sp._YH12218 | FTV03_RS11390 | 13993..17658 | 2601152 | BREX-system-P-loop-protein-BrxC-[Acinetobacter-sp.-YH12218] |
| WP_180174851.1 | Type1.BREX | [illegible] | 1226 | Bacteria>Pseudomonadota>Gammaproteobacteria>Moraxellales>Moraxellaceae>Acinetobacter | GCF_013420235.1 | Acinetobacter_sp._YH01022 | FTT87_RS10805 | 36696..40376 | 2601036 | BREX-system-P-loop-protein-BrxC-[Acinetobacter-sp.-YH01022] |
| WP_182925313.1 | Type1.BREX | [illegible] | 1226 | Bacteria>Pseudomonadota>Gammaproteobacteria>Moraxellales>Moraxellaceae>Acinetobacter | GCF_014169655.1 | Acinetobacter_baumannii | H7R41_RS13110 | 2778553..2782233 | 470 | BREX-system-P-loop-protein-BrxC-[Acinetobacter-baumannii] |
| WP_198080379.1 | Type1.BREX | [illegible] | 1222 | Bacteria>Pseudomonadota>Gammaproteobacteria>Moraxellales>Moraxellaceae>Acinetobacter | GCF_016082245.1 | Acinetobacter_calcoaceticus | BL242_RS20750 | 15792..19460 | 471 | BREX-system-P-loop-protein-BrxC-[Acinetobacter-calcoaceticus] |
| WP_032001825.1 | Type1.BREX | [illegible] | 1226 | Bacteria>Pseudomonadota>Gammaproteobacteria>Moraxellales>Moraxellaceae>Acinetobacter | GCF_016502765.1 | Acinetobacter_calcoaceticus/baumannii_complex | I6M92_RS14775 | 66667..70347 | 909768 | MULTISPECIES:-BREX-system-P-loop-protein-BrxC-[Acinetobacter-calcoaceticus/baumannii-complex] |
| WP_203080037.1 | Type1.BREX | [illegible] | 1226 | Bacteria>Pseudomonadota>Gammaproteobacteria>Moraxellales>Moraxellaceae>Acinetobacter | GCF_016804045.1 | Acinetobacter_pittii | HRJ47_RS04290 | 884929..888609 | 48296 | BREX-system-P-loop-protein-BrxC-[Acinetobacter-pittii] |
| WP_204086201.1 | Type1.BREX | [illegible] | 1226 | Bacteria>Pseudomonadota>Gammaproteobacteria>Moraxellales>Moraxellaceae>Acinetobacter | GCF_016889825.1 | Acinetobacter_pittii | I6J44_RS19355 | 45796..49476 | 48296 | BREX-system-P-loop-protein-BrxC-[Acinetobacter-pittii] |
| WP_205779680.1 | Type1.BREX | [illegible] | 1226 | Bacteria>Pseudomonadota>Gammaproteobacteria>Moraxellales>Moraxellaceae>Acinetobacter | GCF_017132735.1 | Acinetobacter_indicus | JYB86_RS06365 | 1326455..1330135 | 756892 | BREX-system-P-loop-protein-BrxC-[Acinetobacter-indicus] |
| WP_206896797.1 | Type1.BREX | [illegible] | 1226 | Bacteria>Pseudomonadota>Gammaproteobacteria>Moraxellales>Moraxellaceae>Acinetobacter | GCF_017309445.1 | Acinetobacter_indicus | JOW33_RS13795 | 2887588..2891268 | 756892 | BREX-system-P-loop-protein-BrxC-[Acinetobacter-indicus] |
| WP_109426631.1 | Type1.BREX | [illegible] | 1226 | Bacteria>Pseudomonadota>Gammaproteobacteria>Moraxellales>Moraxellaceae>Acinetobacter | GCF_017726575.1 | Acinetobacter_baumannii | J6I88_RS04260 | 873693..877373 | 470 | BREX-system-P-loop-protein-BrxC-[Acinetobacter-baumannii] |
| WP_212272673.1 | Type1.BREX | [illegible] | 1226 | Bacteria>Pseudomonadota>Gammaproteobacteria>Moraxellales>Moraxellaceae>Acinetobacter | GCF_018139165.1 | Acinetobacter_nosocomialis | KC103_RS15085 | 69521..73201 | 106654 | BREX-system-P-loop-protein-BrxC-[Acinetobacter-nosocomialis] |
| WP_216174886.1 | Type1.BREX | [illegible] | 1226 | Bacteria>Pseudomonadota>Gammaproteobacteria>Moraxellales>Moraxellaceae>Acinetobacter | GCF_018863255.1 | Acinetobacter_baumannii | KPE97_RS15710 | 83323..87003 | 470 | BREX-system-P-loop-protein-BrxC-[Acinetobacter-baumannii] |
| WP_216116819.1 | Type1.BREX | [illegible] | 1226 | Bacteria>Pseudomonadota>Gammaproteobacteria>Moraxellales>Moraxellaceae>Acinetobacter | GCF_018883565.1 | Acinetobacter_baumannii | KPF69_RS05265 | 35520..39200 | 470 | BREX-system-P-loop-protein-BrxC-[Acinetobacter-baumannii] |
| WP_207272916.1 | Type1.BREX | [illegible] | 1226 | Bacteria>Pseudomonadota>Gammaproteobacteria>Moraxellales>Moraxellaceae>Acinetobacter | GCF_019921105.1 | Acinetobacter_baumannii | HLH91_RS14165 | 18027..21707 | 470 | BREX-system-P-loop-protein-BrxC-[Acinetobacter-baumannii] |
| WP_005236280.1 | Type1.BREX | [illegible] | 1226 | Bacteria>Pseudomonadota>Gammaproteobacteria>Moraxellales>Moraxellaceae>Acinetobacter | GCF_020099375.1 | Acinetobacter_variabilis | LA331_RS04655 | 963646..967326 | 70346 | BREX-system-P-loop-protein-BrxC-[Acinetobacter-variabilis] |
| WP_233945509.1 | Type1.BREX | [illegible] | 1226 | Bacteria>Pseudomonadota>Gammaproteobacteria>Moraxellales>Moraxellaceae>Acinetobacter | GCF_021391555.1 | Acinetobacter_sp._SH20PTE14 | LXF01_RS17045 | 120215..123895 | 2905879 | BREX-system-P-loop-protein-BrxC-[Acinetobacter-sp.-SH20PTE14] |
| WP_128202440.1 | Type1.BREX | [illegible] | 1226 | Bacteria>Pseudomonadota>Gammaproteobacteria>Moraxellales>Moraxellaceae>Acinetobacter | GCF_023896505.1 | Acinetobacter_baumannii | O9X56_RS14280 | 107190..110870 | 470 | BREX-system-P-loop-protein-BrxC-[Acinetobacter-baumannii] |
| WP_257259040.1 | Type1.BREX | [illegible] | 1226 | Bacteria>Pseudomonadota>Gammaproteobacteria>Moraxellales>Moraxellaceae>Acinetobacter | GCF_024589115.1 | Acinetobacter_baumannii | L2615_RS07865 | 180845..184525 | 470 | BREX-system-P-loop-protein-BrxC-[Acinetobacter-baumannii] |
| WP_257935145.1 | Type1.BREX | [illegible] | 1226 | Bacteria>Pseudomonadota>Gammaproteobacteria>Moraxellales>Moraxellaceae>Acinetobacter | GCF_024705725.1 | Acinetobacter_baumannii | MSH04_RS15115 | 32588..36268 | 470 | BREX-system-P-loop-protein-BrxC-[Acinetobacter-baumannii] |
| WP_265466015.1 | Type1.BREX | [illegible] | 1226 | Bacteria>Pseudomonadota>Gammaproteobacteria>Moraxellales>Moraxellaceae>Acinetobacter | GCF_026168575.1 | Acinetobacter_sp._BIT-DXN8 | OKC24_RS16365 | 19397..23077 | 2989714 | BREX-system-P-loop-protein-BrxC-[Acinetobacter-sp.-BIT-DXN8] |
| WP_274579352.1 | Type1.BREX | [illegible] | 1226 | Bacteria>Pseudomonadota>Gammaproteobacteria>Moraxellales>Moraxellaceae>Acinetobacter | GCF_028774645.1 | Acinetobacter_lactucae | MOO54_RS17975 | 24360..28040 | 1785128 | BREX-system-P-loop-protein-BrxC-[Acinetobacter-lactucae] |
| WP_002010901.1 | Type1.BREX | [illegible] | 1226 | Bacteria>Pseudomonadota>Gammaproteobacteria>Moraxellales>Moraxellaceae | GCF_000214965.1 | Acinetobacter | ACINNAV18_RS19255 | 65599..69279 | 469 | MULTISPECIES:-BREX-system-P-loop-protein-BrxC-[Acinetobacter] |
| WP_152630429.1 | Type1.BREX | [illegible] | 1226 | Bacteria>Pseudomonadota>Gammaproteobacteria>Moraxellales>Moraxellaceae | GCF_009014155.1 | Acinetobacter | QJ110_RS14020 | 22608..26288 | 469 | MULTISPECIES:-BREX-system-P-loop-protein-BrxC-[Acinetobacter] |
| MCP5206320.1 | Type1.BREX | [illegible] | 910 | Bacteria>Pseudomonadota>Gammaproteobacteria>Oceanospirillales>Hahellaceae>unclassified_Hahellaceae | GCA_024234475.1 | Hahellaceae_bacterium | H7A01_03865 | 876629..879361 | 2048971 | BREX-system-P-loop-protein-BrxC-[Hahellaceae-bacterium] |
| MCP5206322.1 | Type1.BREX | [illegible] | 261 | Bacteria>Pseudomonadota>Gammaproteobacteria>Oceanospirillales>Hahellaceae>unclassified_Hahellaceae | GCA_024234475.1 | Hahellaceae_bacterium | H7A01_03875 | 880608..881393 | 2048971 | hypothetical-protein-[Hahellaceae-bacterium] |
| MBJ2272566.1 | Type5.BREX | [illegible] | 1224 | Bacteria>Pseudomonadota>Gammaproteobacteria>Pseudomonadales>Pseudomonadaceae>Pseudomonas | GCA_016405625.1 | Pseudomonas_haemolytica | JFT41_06185 | 268773..272447 | 2600065 | BREX-system-P-loop-protein-BrxC-[Pseudomonas-haemolytica] |
| WP_274151131.1 | Type1.BREX | [illegible] | 1223 | Bacteria>Pseudomonadota>Gammaproteobacteria>Pseudomonadales>Pseudomonadaceae>Pseudomonas | GCA_028682495.1 | Pseudomonas_sp._25571 | NP527_RS02645 | 592231..595902 | 2967216 | BREX-system-P-loop-protein-BrxC-[Pseudomonas-sp.-25571] |
| QOQ70156.1 | Type1.BREX | [illegible] | 1154 | Bacteria>Pseudomonadota>Gammaproteobacteria>Vibrionales>Vibrionaceae>Photobacterium | GCA_014930975.1 | Photobacterium_damselae_subsp._damselae | IL982_10195 | 2142038..2145502 | 85581 | BREX-system-P-loop-protein-BrxC-[Photobacterium-damselae-subsp.-damselae] |
| WP_232092144.1 | Type1.BREX | [illegible] | 1153 | Bacteria>Pseudomonadota>Gammaproteobacteria>Vibrionales>Vibrionaceae>Photobacterium | GCA_014930975.1 | Photobacterium_damselae | IL982_RS10195 | 2142038..2145499 | 38293 | BREX-system-P-loop-protein-BrxC-[Photobacterium-damselae] |
| WP_077485085.1 | Type1.BREX | [illegible] | 1157 | Bacteria>Pseudomonadota>Gammaproteobacteria>Vibrionales>Vibrionaceae>Salinivibrio | GCF_001995745.1 | Salinivibrio_sp._ML323 | BZG13_RS03555 | 125341..128814 | 1909474 | BREX-system-P-loop-protein-BrxC-[Salinivibrio-sp.-ML323] |
| ODY54465.1 | Type1.BREX | [illegible] | 1154 | Bacteria>Pseudomonadota>Gammaproteobacteria>Vibrionales>Vibrionaceae>Vibrio | GCA_001727265.1 | Vibrio_parahaemolyticus | BBM25_06240 | 3424..6888 | 670 | hypothetical-protein-BBM25_06240-[Vibrio-parahaemolyticus] |
| EGR0286978.1 | Type1.BREX | [illegible] | 1154 | Bacteria>Pseudomonadota>Gammaproteobacteria>Vibrionales>Vibrionaceae>Vibrio | GCA_015789595.1 | Vibrio_cholerae | FK206_11700 | 85283..88747 | 666 | BREX-system-P-loop-protein-BrxC-[Vibrio-cholerae] |
| EGR1144248.1 | Type1.BREX | [illegible] | 1154 | Bacteria>Pseudomonadota>Gammaproteobacteria>Vibrionales>Vibrionaceae>Vibrio | GCA_015798095.1 | Vibrio_parahaemolyticus | EC562_15505 | 41063..44527 | 670 | BREX-system-P-loop-protein-BrxC-[Vibrio-parahaemolyticus] |
| EGR1548466.1 | Type1.BREX | [illegible] | 1154 | Bacteria>Pseudomonadota>Gammaproteobacteria>Vibrionales>Vibrionaceae>Vibrio | GCA_015798755.1 | Vibrio_parahaemolyticus | D6Y11_16950 | 161246..164710 | 670 | BREX-system-P-loop-protein-BrxC-[Vibrio-parahaemolyticus] |
| EGR2430327.1 | Type1.BREX | [illegible] | 1154 | Bacteria>Pseudomonadota>Gammaproteobacteria>Vibrionales>Vibrionaceae>Vibrio | GCA_015802095.1 | Vibrio_cholerae | DYB83_10140 | 10866..14330 | 666 | BREX-system-P-loop-protein-BrxC-[Vibrio-cholerae] |
| EGR3302280.1 | Type1.BREX | [illegible] | 1154 | Bacteria>Pseudomonadota>Gammaproteobacteria>Vibrionales>Vibrionaceae>Vibrio | GCA_015805535.1 | Vibrio_parahaemolyticus | DMJ21_13130 | 8098..11562 | 670 | BREX-system-P-loop-protein-BrxC-[Vibrio-parahaemolyticus] |
| EJG0981293.1 | Type1.BREX | [illegible] | 1154 | Bacteria>Pseudomonadota>Gammaproteobacteria>Vibrionales>Vibrionaceae>Vibrio | GCA_015813375.2 | Vibrio_parahaemolyticus | C4G85_RS03500 | 776707..780171 | 670 | BREX-system-P-loop-protein-BrxC-[Vibrio-parahaemolyticus] |
| EHH2507631.1 | Type1.BREX | [illegible] | 1154 | Bacteria>Pseudomonadota>Gammaproteobacteria>Vibrionales>Vibrionaceae>Vibrio | GCA_017637525.1 | Vibrio_parahaemolyticus | J7X10_003794 | 109543..113007 | 670 | BREX-system-P-loop-protein-BrxC-[Vibrio-parahaemolyticus] |
| EJC1210117.1 | Type1.BREX | [illegible] | 1154 | Bacteria>Pseudomonadota>Gammaproteobacteria>Vibrionales>Vibrionaceae>Vibrio | GCA_022997325.1 | Vibrio_parahaemolyticus | MX350_002594 | 5689..9153 | 670 | BREX-system-P-loop-protein-BrxC-[Vibrio-parahaemolyticus] |
| EJL6578344.1 | Type1.BREX | [illegible] | 1154 | Bacteria>Pseudomonadota>Gammaproteobacteria>Vibrionales>Vibrionaceae>Vibrio | GCA_024217525.1 | Vibrio_cholerae | NMS50_002630 | 71340..74804 | 666 | BREX-system-P-loop-protein-BrxC-[Vibrio-cholerae] |
| HCM0920431.1 | Type1.BREX | [illegible] | 1154 | Bacteria>Pseudomonadota>Gammaproteobacteria>Vibrionales>Vibrionaceae>Vibrio | GCA_025068375.1 | Vibrio_parahaemolyticus | N2828_003369 | 93414..96878 | 670 | BREX-system-P-loop-protein-BrxC-[Vibrio-parahaemolyticus] |
| WP_253277773.1 | Type1.BREX | [illegible] | 1153 | Bacteria>Pseudomonadota>Gammaproteobacteria>Vibrionales>Vibrionaceae>Vibrio | GCF_000959745.1 | Vibrio_vulnificus | WG98_RS00450 | 91740..95201 | 672 | BREX-system-P-loop-protein-BrxC-[Vibrio-vulnificus] |
| WP_061065627.1 | Type1.BREX | [illegible] | 1157 | Bacteria>Pseudomonadota>Gammaproteobacteria>Vibrionales>Vibrionaceae>Vibrio | GCF_001558435.2 | Vibrio_harveyi | AL538_RS13470 | 2675377..2678850 | 669 | BREX-system-P-loop-protein-BrxC-[Vibrio-harveyi] |
| WP_241782177.1 | Type1.BREX | [illegible] | 1153 | Bacteria>Pseudomonadota>Gammaproteobacteria>Vibrionales>Vibrionaceae>Vibrio | GCF_001608855.1 | Vibrio_parahaemolyticus | AWQ17_RS02375 | 93824..97285 | 670 | BREX-system-P-loop-protein-BrxC-[Vibrio-parahaemolyticus] |
| WP_256093930.1 | Type1.BREX | [illegible] | 1153 | Bacteria>Pseudomonadota>Gammaproteobacteria>Vibrionales>Vibrionaceae>Vibrio | GCF_001727265.1 | Vibrio_parahaemolyticus | BBM25_RS06240 | 3427..6888 | 670 | BREX-system-P-loop-protein-BrxC-[Vibrio-parahaemolyticus] |
| WP_235868160.1 | Type1.BREX | [illegible] | 1153 | Bacteria>Pseudomonadota>Gammaproteobacteria>Vibrionales>Vibrionaceae>Vibrio | GCF_003311905.1 | Vibrio_paracholerae | DLR66_RS03590 | 70731..74192 | 650003 | BREX-system-P-loop-protein-BrxC-[Vibrio-paracholerae] |
| WP_253254707.1 | Type1.BREX | [illegible] | 1153 | Bacteria>Pseudomonadota>Gammaproteobacteria>Vibrionales>Vibrionaceae>Vibrio | GCF_030238805.1 | Vibrio_parahaemolyticus | CSR03_RS22350 | 121618..125079 | 670 | BREX-system-P-loop-protein-BrxC-[Vibrio-parahaemolyticus] |
| WP_069490026.1 | Type1.BREX | [illegible] | 1154 | Bacteria>Pseudomonadota>Gammaproteobacteria>Vibrionales>Vibrionaceae>Vibrio | - | Vibrio_parahaemolyticus | BBM25_06240 | 3424..6888 | 670 | BREX-system-P-loop-protein-BrxC-[Vibrio-parahaemolyticus] |
| WP_254664663.1 | Type1.BREX | [illegible] | 1153 | Bacteria>Pseudomonadota>Gammaproteobacteria>Vibrionales>Vibrionaceae>Vibrio | - | Vibrio_parahaemolyticus | BED52_RS16125 | 3457..6918 | 670 | BREX-system-P-loop-protein-BrxC-[Vibrio-parahaemolyticus] |
| WP_047508417.1 | Type1.BREX | [illegible] | 1154 | Bacteria>Pseudomonadota>Gammaproteobacteria>Vibrionales>Vibrionaceae>Vibrio | - | Vibrio_parahaemolyticus | DMJ21_13130 | 8098..11562 | 670 | BREX-system-P-loop-protein-BrxC-[Vibrio-parahaemolyticus] |
| WP_062854373.1 | Type1.BREX | [illegible] | 1154 | Bacteria>Pseudomonadota>Gammaproteobacteria>Vibrionales>Vibrionaceae>Vibrio | - | Vibrio_parahaemolyticus | LZX80_001782 | 93413..96877 | 670 | BREX-system-P-loop-protein-BrxC-[Vibrio-parahaemolyticus] |
| WP_069709528.1 | Type1.BREX | [illegible] | 1179 | Bacteria>Pseudomonadota | GCF_001742225.1 | Alphaproteobacteria | BES08_RS24075 | 1524283..1527822 | 28211 | MULTISPECIES:-BREX-system-P-loop-protein-BrxC-[Alphaproteobacteria] |
| WP_066700609.1 | Type1.BREX | [illegible] | 1179 | Bacteria>Pseudomonadota | GCF_003967075.1 | Alphaproteobacteria | SAMIE_RS17570 | 3625371..3628910 | 28211 | MULTISPECIES:-BREX-system-P-loop-protein-BrxC-[Alphaproteobacteria] |
| MBK8482432.1 | Type1.BREX | [illegible] | 896 | Bacteria>Pseudomonadota>unclassified_Pseudomonadota | GCA_016711215.1 | Pseudomonadota_bacterium | IPL40_14915 | 141091..143781 | 1977087 | hypothetical-protein-[Pseudomonadota-bacterium] |
| WP_165367554.1 | Type1.BREX | [illegible] | 1157 | Bacteria>Terrabacteria_group>Actinomycetota>Actinomycetes>Jiangellales>Jiangellaceae | GCF_004138495.1 | Phytoactinopolyspora_endophytica | ELR45_RS07100 | 275319..278792 | 1642495 | BREX-system-P-loop-protein-BrxC-[Phytoactinopolyspora-endophytica] |
| WP_005267613.1 | Type1.BREX | [illegible] | 1151 | Bacteria>Terrabacteria_group>Actinomycetota>Actinomycetes>Micrococcales>Micrococcaceae | GCF_000328305.2 | Arthrobacter_crystallopoietes | D477_RS04285 | 11236..14691 | 37928 | BREX-system-P-loop-protein-BrxC-[Arthrobacter-crystallopoietes] |
| WP_104161527.1 | Type1.BREX | [illegible] | 1151 | Bacteria>Terrabacteria_group>Actinomycetota>Actinomycetes>Micrococcales>Micrococcaceae | GCF_002929255.1 | Arthrobacter_sp._ZGTC212 | CXQ89_RS12670 | 85850..89305 | 2058899 | BREX-system-P-loop-protein-BrxC-[Arthrobacter-sp.-ZGTC212] |
| WP_036470026.1 | Type5.BREX | [illegible] | 1155 | Bacteria>Terrabacteria_group>Actinomycetota>Actinomycetes>Mycobacteriales>Mycobacteriaceae | GCF_000689255.1 | Mycobacterium_triplex | BN973_RS18575 | 3994507..3997974 | 47839 | BREX-system-P-loop-protein-BrxC-[Mycobacterium-triplex] |
| WP_085152401.1 | Type1.BREX | [illegible] | 1156 | Bacteria>Terrabacteria_group>Actinomycetota>Actinomycetes>Mycobacteriales>Mycobacteriaceae | GCF_010729895.1 | Mycolicibacterium_confluentis | H7K00_RS24370 | 129281..132751 | 28047 | BREX-system-P-loop-protein-BrxC-[Mycolicibacterium-confluentis] |

| Protein-ID | System | Gene-Neighbourhoods | Length | Taxonomy | Assembly-ID | Strain | Locus-Tag | Genomic-coordinates | TaxID | Fasta-defline |
|---|---|---|---|---|---|---|---|---|---|---|
| WP_094721084.1 | Type1.BREX | [illegible] | 1174 | Bacteria>Terrabacteria_group>Actinomycetota>Actinomycetes>Mycobacteriales>Nocardiaceae | GCF_002259415.1 | Rhodococcus_sp._14-2470-1b | CH293_RS18620 | 24333..27857 | 2023149 | BREX-system-P-loop-protein-BrxC-[Rhodococcus-sp.-14-2470-1b] |
| WP_222635591.1 | Type1.BREX | [illegible] | 1188 | Bacteria>Terrabacteria_group>Actinomycetota>Actinomycetes>Mycobacteriales>Nocardiaceae | GCF_019834785.1 | unclassified_Rhodococcus_(in:_high_G+C_Gram-positive_bacteria) | HQ363_RS22195 | 12044..15610 | 192944 | MULTISPECIES:-BREX-system-P-loop-protein-BrxC-[unclassified-Rhodococcus-(in:-high-G+C-Gram-positive-bacteria)] |
| WP_136846141.1 | Type1.BREX | [illegible] | 1192 | Bacteria>Terrabacteria_group>Actinomycetota>Coriobacteriia>Coriobacteriales>Coriobacteriaceae | GCF_948466405.1 | Parvibacter_caecicola | E5982_RS08750 | 41748..45326 | 747645 | BREX-system-P-loop-protein-BrxC-[Parvibacter-caecicola] |
| PKF86734.1 | Type1.BREX | [illegible] | 1193 | Bacteria>Terrabacteria_group>Bacillota>Bacilli>Bacillales>Bacillaceae | GCA_002835675.1 | Bacillus_sp._BA3 | CW306_21620 | 276915..280496 | 2057910 | BREX-system-P-loop-protein-BrxC-[Bacillus-sp.-BA3] |
| WP_197035844.1 | Type6.BREX | [illegible] | 1207 | Bacteria>Terrabacteria_group>Bacillota>Bacilli>Bacillales>Bacillaceae | GCF_001878675.1 | Virgibacillus_halodenitrificans | BME96_RS06225 | 1179792..1183415 | 1482 | BREX-system-P-loop-protein-BrxC-[Virgibacillus-halodenitrificans] |
| WP_209482153.1 | Type1.BREX | [illegible] | 1187 | Bacteria>Terrabacteria_group>Bacillota>Bacilli>Bacillales>Bacillaceae | GCF_017873675.1 | Virgibacillus_litoralis | J2Z82_RS17835 | 25252..28815 | 578221 | BREX-system-P-loop-protein-BrxC-[Virgibacillus-litoralis] |
| WP_028107090.1 | Type1.BREX | [illegible] | 1192 | Bacteria>Terrabacteria_group>Bacillota>Bacilli>Bacillales>Bacillales_incertae_sedis | GCF_000620805.1 | Exiguobacterium_undae | P397_RS0115435 | 2856472..2860050 | 169177 | BREX-system-P-loop-protein-BrxC-[Exiguobacterium-undae] |
| WP_088074350.1 | Type1.BREX | [illegible] | 1205 | Bacteria>Terrabacteria_group>Bacillota>Bacilli>Bacillales | GCF_019042735.1 | Bacillaceae | KS407_RS09130 | 3322..6939 | 186817 | MULTISPECIES:-BREX-system-P-loop-protein-BrxC-[Bacillaceae] |
| WP_036689483.1 | Type1.BREX | [illegible] | 1187 | Bacteria>Terrabacteria_group>Bacillota>Bacilli>Bacillales>Paenibacillaceae | GCF_000517925.1 | Paenibacillus | C171_RS32010 | 162271..165834 | 44249 | MULTISPECIES:-BREX-system-P-loop-protein-BrxC-[Paenibacillus] |
| WP_060645783.1 | Type1.BREX | [illegible] | 1187 | Bacteria>Terrabacteria_group>Bacillota>Bacilli>Bacillales>Paenibacillaceae | GCF_001427935.1 | Paenibacillus_sp._Soil724D2 | ASG85_RS07160 | 104661..108224 | 1736392 | BREX-system-P-loop-protein-BrxC-[Paenibacillus-sp.-Soil724D2] |
| WP_053168317.1 | Type1.BREX | [illegible] | 1190 | Bacteria>Terrabacteria_group>Bacillota>Bacilli>Bacillales>Planococcaceae | GCF_001270075.1 | Planococcus_glaciei | AC739_RS18785 | 11378..14950 | 459472 | BREX-system-P-loop-protein-BrxC-[Planococcus-glaciei] |
| WP_068488209.1 | Type1.BREX | [illegible] | 1190 | Bacteria>Terrabacteria_group>Bacillota>Bacilli>Bacillales>Planococcaceae | GCF_001592825.1 | Planococcus_maritimus | AY633_RS09895 | 276418..279990 | 192421 | BREX-system-P-loop-protein-BrxC-[Planococcus-maritimus] |
| WP_108306485.1 | Type1.BREX | [illegible] | 1204 | Bacteria>Terrabacteria_group>Bacillota>Bacilli>Bacillales>Planococcaceae | GCF_003057615.1 | Metalysinibacillus_jejuensis | DDE80_RS04455 | 46948..50562 | 914327 | BREX-system-P-loop-protein-BrxC-[Metalysinibacillus-jejuensis] |
| WP_216971036.1 | Type1.BREX | [illegible] | 1193 | Bacteria>Terrabacteria_group>Bacillota>Bacilli>Bacillales>Planococcaceae | GCF_019042655.1 | unclassified_Planococcus_(in:_firmicutes) | KTG14_RS11250 | 8641..12222 | 2662419 | MULTISPECIES:-BREX-system-P-loop-protein-BrxC-[unclassified-Planococcus-(in:-firmicutes)] |
| WP_223643511.1 | Type1.BREX | [illegible] | 1207 | Bacteria>Terrabacteria_group>Bacillota>Bacilli>Bacillales>Planococcaceae | GCF_020023385.1 | Planococcus_sp._4-30 | K8U60_RS18515 | 8957..12580 | 2874583 | BREX-system-P-loop-protein-BrxC-[Planococcus-sp.-4-30] |
| WP_010024517.1 | Type1.BREX | [illegible] | 1189 | Bacteria>Terrabacteria_group>Bacillota>Bacilli>Bacillales>Sporolactobacillaceae | GCF_000222445.2 | Sporolactobacillus_inulinus | SINU_RS07430 | 4602..8171 | 2078 | BREX-system-P-loop-protein-BrxC-[Sporolactobacillus-inulinus] |
| WP_108025903.1 | Type1.BREX | [illegible] | 1166 | Bacteria>Terrabacteria_group>Bacillota>Bacilli>Bacillales>Thermoactinomycetaceae | GCF_003054245.1 | Melghirimyces_profundicolus | C8P63_RS16380 | 15718..19218 | 1242148 | BREX-system-P-loop-protein-BrxC-[Melghirimyces-profundicolus] |
| WP_068559440.1 | Type1.BREX | [illegible] | 1207 | Bacteria>Terrabacteria_group>Bacillota>Bacilli>Lactobacillales>Carnobacteriaceae | GCF_900067165.1 | Trichococcus | BUA33_RS02210 | 213489..217112 | 82802 | MULTISPECIES:-BREX-system-P-loop-protein-BrxC-[Trichococcus] |
| MBR6404277.1 | Type1.BREX | [illegible] | 1160 | Bacteria>Terrabacteria_group>Bacillota>Clostridia>Eubacteriales>Eubacteriaceae | GCA_017539165.1 | Eubacterium_sp. | IKS48_12920 | 16986..20468 | 142586 | BREX-system-P-loop-protein-BrxC-[Eubacterium-sp.] |
| MCR5727513.1 | Type1.BREX | [illegible] | 1173 | Bacteria>Terrabacteria_group>Bacillota>Clostridia>Eubacteriales>Lachnospiraceae | GCA_024699435.1 | Lachnospiraceae_bacterium | K6G24_08610 | 2669..6190 | 1898203 | BREX-system-P-loop-protein-BrxC-[Lachnospiraceae-bacterium] |
| WP_022770173.1 | Type1.BREX | [illegible] | 1202 | Bacteria>Terrabacteria_group>Bacillota>Clostridia>Eubacteriales>Lachnospiraceae | GCF_000421405.1 | Butyrivibrio_sp._NC2007 | G632_RS0114760 | 79363..82971 | 1280683 | BREX-system-P-loop-protein-BrxC-[Butyrivibrio-sp.-NC2007] |
| WP_089716529.1 | Type1.BREX | [illegible] | 1190 | Bacteria>Terrabacteria_group>Bacillota>Clostridia>Halanaerobiales>Halanaerobiaceae | GCF_900100055.1 | Halanaerobium_congolense | BMY48_RS04305 | 58895..62467 | 54121 | BREX-system-P-loop-protein-BrxC-[Halanaerobium-congolense] |
| WP_018248942.1 | Type1.BREX | [illegible] | 1197 | Bacteria>Terrabacteria_group>Bacillota>Clostridia>Halanaerobiales>Halobacteroidaceae | GCF_000379025.1 | Orenia_marismortui | OREMA_RS0108995 | 158007..161600 | 46469 | BREX-system-P-loop-protein-BrxC-[Orenia-marismortui] |
| MCL4488347.1 | Type6.BREX | [illegible] | 1220 | Bacteria>Terrabacteria_group>Chloroflexota>unclassified_Chloroflexota | GCA_023381515.1 | Chloroflexota_bacterium | M1570_09495 | 91471..95133 | 2026724 | BREX-system-P-loop-protein-BrxC-[Chloroflexota-bacterium] |
| MCL5952139.1 | Type6.BREX | [illegible] | 1220 | Bacteria>Terrabacteria_group>Chloroflexota>unclassified_Chloroflexota | GCA_023489485.1 | Chloroflexota_bacterium | M1132_10550 | 55663..59325 | 2026724 | BREX-system-P-loop-protein-BrxC-[Chloroflexota-bacterium] |
| WP_092265648.1 | Type1.BREX | [illegible] | 1160 | Bacteria>Terrabacteria_group>Deinococcota>Deinococci>Deinococcales>Deinococcaceae | GCF_900109185.1 | Deinococcus_reticulitermitis | BMY43_RS15290 | 11757..15239 | 856736 | BREX-system-P-loop-protein-BrxC-[Deinococcus-reticulitermitis] |
| MBI1293571.1 | Type1.BREX | [illegible] | 1173 | Bacteria>unclassified_Bacteria | GCA_016124945.1 | bacterium | GC175_01265 | 40562..44083 | 1869227 | BREX-system-P-loop-protein-BrxC-[bacterium] |
| -- | | | | | | | | | | |
| -- REase +DUF4268 | | | | | | | | | | |
| -- | | | | | | | | | | |
| WP_100909051.1 | Type1.BREX | [illegible] | 1197 | Archaea>Euryarchaeota>Methanomada_group>Methanobacteria>Methanobacteriales>Methanobacteriaceae | GCF_002813695.1 | Methanobacterium_subterraneum | BK009_RS03450 | 727019..730612 | 59277 | BREX-system-P-loop-protein-BrxC-[Methanobacterium-subterraneum] |
| MAO064836.1 | Type1.BREX | [illegible] | 1168 | Bacteria>FCB_group>Bacteroidetes/Chlorobi_group>Balneolota>Balneolia>Balneolales | GCA_002696035.1 | Balneola_sp. | CL666_07525 | 664531..668037 | 2024824 | hypothetical-protein-[Balneola-sp.] |
| QSO48381.1 | Type1.BREX | [illegible] | 1175 | Bacteria>Terrabacteria_group>Bacillota>Bacilli>Bacillales>Alicyclobacillaceae | GCA_017298635.1 | Alicyclobacillus_mengziensis | JZ786_05180 | 1119199..1122726 | 2931921 | BREX-system-P-loop-protein-BrxC-[Alicyclobacillus-mengziensis] |
| WP_021297234.1 | Type1.BREX | [illegible] | 1177 | Bacteria>Terrabacteria_group>Bacillota>Bacilli>Bacillales>Alicyclobacillaceae | GCF_000444055.1 | Alicyclobacillus_acidoterrestris | N007_RS35530 | 2450..5983 | 1450 | BREX-system-P-loop-protein-BrxC-[Alicyclobacillus-acidoterrestris] |
| WP_067929227.1 | Type1.BREX | [illegible] | 1178 | Bacteria>Terrabacteria_group>Bacillota>Bacilli>Bacillales>Alicyclobacillaceae | GCF_001552255.1 | Alicyclobacillus_shizuokensis | ASH01S_RS14820 | 6987..10523 | 392014 | BREX-system-P-loop-protein-BrxC-[Alicyclobacillus-shizuokensis] |
| WP_206657716.1 | Type1.BREX | [illegible] | 1175 | Bacteria>Terrabacteria_group>Bacillota>Bacilli>Bacillales>Alicyclobacillaceae | GCF_017298635.1 | Alicyclobacillus_mengziensis | JZ786_RS05180 | 1119199..1122726 | 2931921 | BREX-system-P-loop-protein-BrxC-[Alicyclobacillus-mengziensis] |
| WP_119547238.1 | Type1.BREX | [illegible] | 1193 | Bacteria>Terrabacteria_group>Bacillota>Bacilli>Bacillales>Bacillaceae | GCF_003581585.1 | Bacillus_salacetis | D3H55_RS12405 | 3906..7487 | 2315464 | BREX-system-P-loop-protein-BrxC-[Bacillus-salacetis] |
| WP_216889200.1 | Type1.BREX | [illegible] | 1197 | Bacteria>Terrabacteria_group>Bacillota>Bacilli>Bacillales>Bacillaceae | GCF_019036865.1 | Bacillus_pumilus | KM878_RS13750 | 41263..44856 | 1408 | BREX-system-P-loop-protein-BrxC-[Bacillus-pumilus] |
| WP_240257149.1 | Type1.BREX | [illegible] | 1195 | Bacteria>Terrabacteria_group>Bacillota>Bacilli>Bacillales>Bacillaceae | GCF_022427965.1 | Fredinandcohnia_sp._SECRCQ15 | MJG50_RS17985 | 30135..33722 | 2918902 | BREX-system-P-loop-protein-BrxC-[Fredinandcohnia-sp.-SECRCQ15] |
| WP_175638347.1 | Type1.BREX | [illegible] | 1193 | Bacteria>Terrabacteria_group>Bacillota>Bacilli>Bacillales>Bacillaceae | GCF_903166545.1 | Metabacillus_schmidteae | HWV59_RS06330 | 1297576..1301157 | 2730405 | BREX-system-P-loop-protein-BrxC-[Metabacillus-schmidteae] |
| MXO77664.1 | Type1.BREX | [illegible] | 1188 | Bacteria>Terrabacteria_group>Bacillota>Bacilli>Bacillales>Paenibacillaceae | GCA_009827695.1 | Paenibacillus_sp._OT2-17 | GRP75_07340 | 6853..10419 | 2691605 | BREX-system-P-loop-protein-BrxC-[Paenibacillus-sp.-OT2-17] |
| NUU75083.1 | Type1.BREX | [illegible] | 1187 | Bacteria>Terrabacteria_group>Bacillota>Bacilli>Bacillales>Paenibacillaceae | GCA_013359935.1 | Paenibacillus_xylanilyticus | HP552_07495 | 27888..31451 | 248903 | BREX-system-P-loop-protein-BrxC-[Paenibacillus-xylanilyticus] |
| MBO2945633.1 | Type1.BREX | [illegible] | 1188 | Bacteria>Terrabacteria_group>Bacillota>Bacilli>Bacillales>Paenibacillaceae | GCA_017582705.1 | Paenibacillus_sp._F411 | JJQ72_16765 | 27941..31507 | 2820239 | BREX-system-P-loop-protein-BrxC-[Paenibacillus-sp.-F411] |
| MBP1153944.1 | Type1.BREX | [illegible] | 1188 | Bacteria>Terrabacteria_group>Bacillota>Bacilli>Bacillales>Paenibacillaceae | GCA_017832315.1 | Paenibacillus_sp._PvP091 | JOE36_000751 | 753497..757063 | 2806590 | hypothetical-protein-[Paenibacillus-sp.-PvP091] |
| WP_095288526.1 | Type1.BREX | [illegible] | 1188 | Bacteria>Terrabacteria_group>Bacillota>Bacilli>Bacillales>Paenibacillaceae | GCF_002272715.1 | Paenibacillus_sp._7516 | CHI14_RS11220 | 25623..29189 | 2022549 | BREX-system-P-loop-protein-BrxC-[Paenibacillus-sp.-7516] |
| WP_153976927.1 | Type1.BREX | [illegible] | 1188 | Bacteria>Terrabacteria_group>Bacillota>Bacilli>Bacillales>Paenibacillaceae | GCF_009664365.1 | Paenibacillus_xylanilyticus | F4V51_RS03900 | 837268..840834 | 248903 | BREX-system-P-loop-protein-BrxC-[Paenibacillus-xylanilyticus] |
| WP_202404033.1 | Type1.BREX | [illegible] | 1183 | Bacteria>Terrabacteria_group>Bacillota>Bacilli>Bacillales>Paenibacillaceae | GCF_009827695.1 | Paenibacillus_sp._OT2-17 | GRP75_RS07325 | 6853..10404 | 2691605 | BREX-system-P-loop-protein-BrxC-[Paenibacillus-sp.-OT2-17] |
| WP_217506352.1 | Type1.BREX | [illegible] | 1182 | Bacteria>Terrabacteria_group>Bacillota>Bacilli>Bacillales>Paenibacillaceae | GCF_013359935.1 | Paenibacillus_xylanilyticus | HP552_RS07485 | 27903..31451 | 248903 | BREX-system-P-loop-protein-BrxC-[Paenibacillus-xylanilyticus] |
| WP_243896166.1 | Type1.BREX | [illegible] | 1183 | Bacteria>Terrabacteria_group>Bacillota>Bacilli>Bacillales>Paenibacillaceae | GCF_017582705.1 | Paenibacillus_sp._F411 | JJQ72_RS16770 | 27941..31492 | 2820239 | BREX-system-P-loop-protein-BrxC-[Paenibacillus-sp.-F411] |
| WP_245247302.1 | Type1.BREX | [illegible] | 1183 | Bacteria>Terrabacteria_group>Bacillota>Bacilli>Bacillales>Paenibacillaceae | GCF_017832315.1 | unclassified_Paenibacillus | JOE36_RS03880 | 753512..757063 | 185978 | MULTISPECIES:-BREX-system-P-loop-protein-BrxC-[unclassified-Paenibacillus] |
| WP_274359496.1 | Type1.BREX | [illegible] | 1188 | Bacteria>Terrabacteria_group>Bacillota>Bacilli>Bacillales>Paenibacillaceae | GCF_028747665.1 | Paenibacillus_urinalis | PUW23_RS03705 | 829690..833256 | 521520 | BREX-system-P-loop-protein-BrxC-[Paenibacillus-urinalis] |
| WP_251582630.1 | Type1.BREX | [illegible] | 1188 | Bacteria>Terrabacteria_group>Bacillota>Bacilli>Bacillales>Paenibacillaceae | - | Paenibacillus_sp._MER_78 | M3625_RS10220 | 1040527..1044093 | 2939571 | BREX-system-P-loop-protein-BrxC-[Paenibacillus-sp.-MER-78] |
| WP_027725313.1 | Type1.BREX | [illegible] | 1183 | Bacteria>Terrabacteria_group>Bacillota>Bacilli>Bacillales>Sporolactobacillaceae | GCF_000430685.1 | Tuberibacillus_calidus | H532_RS0109650 | 26446..29997 | 340097 | BREX-system-P-loop-protein-BrxC-[Tuberibacillus-calidus] |
| WP_013777644.1 | Type1.BREX | [illegible] | 1188 | Bacteria>Terrabacteria_group>Bacillota>Clostridia>Thermosediminibacterales>Tepidanaerobacteraceae | GCF_000213235.1 | Tepidanaerobacter_acetatoxydans | TEPRE1_RS02650 | 565756..569322 | 499229 | BREX-system-P-loop-protein-BrxC-[Tepidanaerobacter-acetatoxydans] |
| MBD8958802.1 | Type1.BREX | [illegible] | 1182 | Bacteria>Terrabacteria_group>Bacillota>Erysipelotrichia>Erysipelotrichales>Erysipelotrichaceae | GCA_014847315.1 | Holdemanella_biformis | EGQ83_10165 | 17990..21538 | 1735 | BREX-system-P-loop-protein-BrxC-[Holdemanella-biformis] |
| WP_019892646.1 | Type1.BREX | [illegible] | 1175 | Bacteria>Terrabacteria_group>Bacillota>Erysipelotrichia>Erysipelotrichales>Erysipelotrichaceae | GCF_000384195.1 | Allobaculum_stercoricanis | ALLST_RS0103345 | 705410..708937 | 174709 | BREX-system-P-loop-protein-BrxC-[Allobaculum-stercoricanis] |
| WP_186999074.1 | Type1.BREX | [illegible] | 1182 | Bacteria>Terrabacteria_group>Bacillota>Erysipelotrichia>Erysipelotrichales>Erysipelotrichaceae | GCF_014306155.1 | Holdemanella_hominis | H8911_RS06490 | 3688..7236 | 2764327 | BREX-system-P-loop-protein-BrxC-[Holdemanella-hominis] |
| WP_169698881.1 | Type1.BREX | [illegible] | 1164 | Bacteria>Thermotogota>Thermotogae>Kosmotogales>Kosmotogaceae>Mesotoga | GCF_900157305.1 | Mesotoga_infera | MESINF_RS05385 | 1186645..1190139 | 1236046 | BREX-system-P-loop-protein-BrxC-[Mesotoga-infera] |
| -- | | | | | | | | | | |
| -- REase +RNase | | | | | | | | | | |
| -- | | | | | | | | | | |
| MCB0345181.1 | Type6.BREX | [illegible] | 811 | Bacteria>Bdellovibrionota>Bdellovibrionia>Bdellovibrionales>unclassified_Bdellovibrionales | GCA_020434585.1 | Bdellovibrionales_bacterium | KDD66_08680 | 10999..13432 | 2053517 | BREX-system-P-loop-protein-BrxC-[Bdellovibrionales-bacterium] |
| WP_119843039.1 | Type1.BREX | [illegible] | 1201 | Bacteria>FCB_group>Bacteroidetes/Chlorobi_group>Bacteroidota>Bacteroidetes_incertae_sedis>Rhodothermaceae | GCF_002290345.1 | Salinibacter_ruber | B7W84_RS15115 | 76596..80201 | 146919 | BREX-system-P-loop-protein-BrxC-[Salinibacter-ruber] |
| WP_103016193.1 | Type5.BREX | [illegible] | 1108 | Bacteria>FCB_group>Bacteroidetes/Chlorobi_group>Bacteroidota>Bacteroidetes_incertae_sedis>Rhodothermaceae | GCF_002894575.1 | Salinibacter_ruber | B7R92_RS05385 | 33133..36459 | 146919 | hypothetical-protein-[Salinibacter-ruber] |
| WP_146031917.1 | Type5.BREX | [illegible] | 1245 | Bacteria>FCB_group>Bacteroidetes/Chlorobi_group>Bacteroidota>Bacteroidetes_incertae_sedis>Rhodothermaceae | GCF_002894575.1 | Salinibacter_ruber | B7R92_RS05395 | 37102..40839 | 146919 | hypothetical-protein-[Salinibacter-ruber] |
| WP_259165690.1 | Type5.BREX | [illegible] | 1245 | Bacteria>FCB_group>Bacteroidetes/Chlorobi_group>Bacteroidota>Bacteroidetes_incertae_sedis>Rhodothermaceae | GCF_024806515.1 | Salinibacter_ruber | GGP90_RS03930 | 969591..973328 | 146919 | hypothetical-protein-[Salinibacter-ruber] |
| WP_259182568.1 | Type5.BREX | [illegible] | 1248 | Bacteria>FCB_group>Bacteroidetes/Chlorobi_group>Bacteroidota>Bacteroidetes_incertae_sedis>Rhodothermaceae | GCF_024806795.1 | Salinibacter_ruber | GGQ19_RS12625 | 57748..61494 | 146919 | hypothetical-protein-[Salinibacter-ruber] |
| WP_259125819.1 | Type5.BREX | [illegible] | 1245 | Bacteria>FCB_group>Bacteroidetes/Chlorobi_group>Bacteroidota>Bacteroidetes_incertae_sedis>Rhodothermaceae | GCF_024806955.1 | Salinibacter_ruber | GGQ05_001713 | 83114..86851 | 146919 | hypothetical-protein-[Salinibacter-ruber] |
| WP_259220531.1 | Type5.BREX | [illegible] | 1245 | Bacteria>FCB_group>Bacteroidetes/Chlorobi_group>Bacteroidota>Bacteroidetes_incertae_sedis>Rhodothermaceae | GCF_024807065.1 | Salinibacter_ruber | GGP94_RS13020 | 49040..52777 | 146919 | hypothetical-protein-[Salinibacter-ruber] |
| WP_259253366.1 | Type5.BREX | [illegible] | 1245 | Bacteria>FCB_group>Bacteroidetes/Chlorobi_group>Bacteroidota>Bacteroidetes_incertae_sedis>Rhodothermaceae | GCF_024807415.1 | Salinibacter_ruber | GGP63_RS14335 | 56454..60191 | 146919 | hypothetical-protein-[Salinibacter-ruber] |
| WP_259130005.1 | Type6.BREX | [illegible] | 1199 | Bacteria>FCB_group>Bacteroidetes/Chlorobi_group>Bacteroidota>Bacteroidetes_incertae_sedis>Rhodothermaceae | GCF_024807455.1 | Salinibacter_ruber | GGQ04_003012 | 22826..26425 | 146919 | BREX-system-P-loop-protein-BrxC-[Salinibacter-ruber] |
| WP_259113946.1 | Type5.BREX | [illegible] | 1245 | Bacteria>FCB_group>Bacteroidetes/Chlorobi_group>Bacteroidota>Bacteroidetes_incertae_sedis>Rhodothermaceae | GCF_024807465.1 | Salinibacter_ruber | GGP53_RS11770 | 37254..40991 | 146919 | hypothetical-protein-[Salinibacter-ruber] |
| WP_259127211.1 | Type5.BREX | [illegible] | 1255 | Bacteria>FCB_group>Bacteroidetes/Chlorobi_group>Bacteroidota>Bacteroidetes_incertae_sedis>Rhodothermaceae | GCF_024807525.1 | Salinibacter_ruber | GGP44_RS04645 | 1121352..1125119 | 146919 | hypothetical-protein-[Salinibacter-ruber] |
| WP_259123903.1 | Type5.BREX | [illegible] | 1245 | Bacteria>FCB_group>Bacteroidetes/Chlorobi_group>Bacteroidota>Bacteroidetes_incertae_sedis>Rhodothermaceae | GCF_024808095.1 | Salinibacter_ruber | GGP68_RS06295 | 37891..41628 | 146919 | hypothetical-protein-[Salinibacter-ruber] |
| WP_259086338.1 | Type5.BREX | [illegible] | 1245 | Bacteria>FCB_group>Bacteroidetes/Chlorobi_group>Bacteroidota>Bacteroidetes_incertae_sedis>Rhodothermaceae | GCF_024808115.1 | Salinibacter_ruber | GGP80_RS14460 | 1984..5721 | 146919 | hypothetical-protein-[Salinibacter-ruber] |
| WP_259107773.1 | Type5.BREX | [illegible] | 1245 | Bacteria>FCB_group>Bacteroidetes/Chlorobi_group>Bacteroidota>Bacteroidetes_incertae_sedis>Rhodothermaceae | GCF_024808295.1 | Salinibacter_ruber | GGP92_RS13390 | 54298..58035 | 146919 | hypothetical-protein-[Salinibacter-ruber] |
| WP_259048220.1 | Type5.BREX | [illegible] | 1245 | Bacteria>FCB_group>Bacteroidetes/Chlorobi_group>Bacteroidota>Bacteroidetes_incertae_sedis>Rhodothermaceae | GCF_024808395.1 | Salinibacter_ruber | GGP52_RS13845 | 3637..7374 | 146919 | hypothetical-protein-[Salinibacter-ruber] |
| WP_259120983.1 | Type5.BREX | [illegible] | 1245 | Bacteria>FCB_group>Bacteroidetes/Chlorobi_group>Bacteroidota>Bacteroidetes_incertae_sedis>Rhodothermaceae | GCF_024808535.1 | Salinibacter_ruber | GGP96_RS14190 | 56803..60540 | 146919 | hypothetical-protein-[Salinibacter-ruber] |
| WP_259122105.1 | Type5.BREX | [illegible] | 1245 | Bacteria>FCB_group>Bacteroidetes/Chlorobi_group>Bacteroidota>Bacteroidetes_incertae_sedis>Rhodothermaceae | GCF_024809285.1 | Salinibacter_ruber | GGQ10_RS14855 | 72088..75825 | 146919 | hypothetical-protein-[Salinibacter-ruber] |
| WP_259134094.1 | Type5.BREX | [illegible] | 1245 | Bacteria>FCB_group>Bacteroidetes/Chlorobi_group>Bacteroidota>Bacteroidetes_incertae_sedis>Rhodothermaceae | GCF_024810035.1 | Salinibacter_ruber | GGP48_RS11375 | 70595..74332 | 146919 | hypothetical-protein-[Salinibacter-ruber] |
| WP_251951637.1 | Type5.BREX | [illegible] | 1108 | Bacteria>FCB_group>Bacteroidetes/Chlorobi_group>Bacteroidota>Bacteroidetes_incertae_sedis>Rhodothermaceae | GCF_943908255.1 | Salinibacter_ruber | NGL47_RS11295 | 22718..26044 | 146919 | hypothetical-protein-[Salinibacter-ruber] |
| WP_251951639.1 | Type5.BREX | [illegible] | 1245 | Bacteria>FCB_group>Bacteroidetes/Chlorobi_group>Bacteroidota>Bacteroidetes_incertae_sedis>Rhodothermaceae | GCF_943908255.1 | Salinibacter_ruber | NGL47_RS11305 | 26687..30424 | 146919 | hypothetical-protein-[Salinibacter-ruber] |
| PJE46487.1 | Type1.BREX | [illegible] | 1169 | Bacteria>FCB_group>Bacteroidetes/Chlorobi_group>Bacteroidota>Chitinophagia>Chitinophagales | GCA_002786355.1 | Sediminibacterium_sp._FEMGT703S | CUR34_07570 | 66784..70293 | 2080766 | MAG:-hypothetical-protein-CUR34_07570-[Sediminibacterium-sp.] |
| MBP6234856.1 | Type1.BREX | [illegible] | 1152 | Bacteria>FCB_group>Bacteroidetes/Chlorobi_group>Bacteroidota>Saprospiria>Saprospirales | GCA_017986015.1 | Saprospiraceae_bacterium | KA536_01890 | 136824..140282 | 2202734 | BREX-system-P-loop-protein-BrxC-[Saprospiraceae-bacterium] |
| RYG90169.1 | Type6.BREX | [illegible] | 1208 | Bacteria>Pseudomonadota>Alphaproteobacteria>unclassified_Alphaproteobacteria | GCA_004145145.1 | Alphaproteobacteria_bacterium | EON59_00500 | 24262..27888 | 1913988 | BREX-system-P-loop-protein-BrxC-[Alphaproteobacteria-bacterium] |
| HAT1978996.1 | Type1.BREX | [illegible] | 1184 | Bacteria>Pseudomonadota>Gammaproteobacteria>Legionellales>Legionellaceae>Legionella | GCA_015895665.1 | Legionella_pneumophila | I8Z83_002252 | 23760..27314 | 446 | BREX-system-P-loop-protein-BrxC-[Legionella-pneumophila] |
| HAT9211453.1 | Type1.BREX | [illegible] | 1185 | Bacteria>Pseudomonadota>Gammaproteobacteria>Legionellales>Legionellaceae>Legionella | GCA_015973865.1 | Legionella_pneumophila_subsp._pneumophila | JBB81_06050 | 33487..37044 | 91891 | BREX-system-P-loop-protein-BrxC-[Legionella-pneumophila-subsp.-pneumophila] |
| HAU3861597.1 | Type1.BREX | [illegible] | 1185 | Bacteria>Pseudomonadota>Gammaproteobacteria>Legionellales>Legionellaceae>Legionella | GCA_016062615.1 | Legionella_pneumophila | F7Z80_07285 | 265287..268844 | 446 | BREX-system-P-loop-protein-BrxC-[Legionella-pneumophila] |
| WP_061721013.1 | Type1.BREX | [illegible] | 1184 | Bacteria>Pseudomonadota>Gammaproteobacteria>Legionellales>Legionellaceae>Legionella | GCF_900054435.1 | Legionella_pneumophila | A2E62_RS14865 | 12190..15744 | 446 | BREX-system-P-loop-protein-BrxC-[Legionella-pneumophila] |
| -- | | | | | | | | | | |
| -- REase +AIPR | | | | | | | | | | |
| -- | | | | | | | | | | |
| MBI4904787.1 | Type1.BREX | [illegible] | 1144 | Bacteria>Acidobacteriota>unclassified_Acidobacteriota | GCA_016213775.1 | Acidobacteriota_bacterium | HY820_14195 | 24664..28098 | 1978231 | BREX-system-P-loop-protein-BrxC-[Acidobacteriota-bacterium] |
| MBF0098005.1 | Type1.BREX | [illegible] | 1144 | Bacteria>Pseudomonadota>Alphaproteobacteria>Magnetococcales>unclassified_Magnetococcales | GCA_015228825.1 | Magnetococcales_bacterium | HQM05_11780 | 14372..17806 | 2026759 | BREX-system-P-loop-protein-BrxC-[Magnetococcales-bacterium] |
| WP_137104425.1 | Type1.BREX | [illegible] | 1148 | Bacteria>Pseudomonadota>Alphaproteobacteria>Rhodospirillales>Azospirillaceae>Azospirillum | GCF_005222185.1 | Azospirillum_argentinense | D3867_15345 | 176394..179840 | 2970906 | BREX-system-P-loop-protein-BrxC-[Azospirillum-argentinense] |
| WP_089417825.1 | Type1.BREX | [illegible] | 1153 | Bacteria>Pseudomonadota>Betaproteobacteria>Neisseriales>Neisseriaceae>Vitreoscilla | GCF_002222655.1 | Vitreoscilla_filiformis | VITFI_RS15935 | 3397543..3401004 | 63 | BREX-system-P-loop-protein-BrxC-[Vitreoscilla-filiformis] |
| WP_043743640.1 | Type1.BREX | [illegible] | 1153 | Bacteria>Pseudomonadota>Betaproteobacteria>Rhodocyclales>Zoogloeaceae>Thauera | GCF_000831325.1 | Thauera_sp._SWB20 | PO78_RS10565 | 31733..35194 | 1572758 | BREX-system-P-loop-protein-BrxC-[Thauera-sp.-SWB20] |
| WP_068803137.1 | Type1.BREX | [illegible] | 1153 | Bacteria>Pseudomonadota>Betaproteobacteria>Rhodocyclales>Zoogloeaceae>Thauera | GCF_001696715.1 | Thauera_phenolivorans | BB334_RS00200 | 55015..58476 | 1792543 | BREX-system-P-loop-protein-BrxC-[Thauera-phenolivorans] |
| WP_247724881.1 | Type1.BREX | [illegible] | 1153 | Bacteria>Pseudomonadota>Betaproteobacteria>Rhodocyclales>Zoogloeaceae>Thauera | GCF_023093595.1 | Thauera_aromatica | MXC99_RS07615 | 6533..9994 | 59405 | BREX-system-P-loop-protein-BrxC-[Thauera-aromatica] |
| GHT91750.1 | Type6.BREX | [illegible] | 1209 | Bacteria>Pseudomonadota>Betaproteobacteria>unclassified_Betaproteobacteria | GCA_026000515.1 | Betaproteobacteria_bacterium | FACS1894116_00820 | 42503..46132 | 1891241 | hypothetical-protein-FACS1894116_00820-[Betaproteobacteria-bacterium] |
| MBV5272983.1 | Type1.BREX | [illegible] | 1143 | Bacteria>Pseudomonadota>Gammaproteobacteria>Chromatiales>Chromatiaceae>Lamprocystis | GCA_019163055.1 | Lamprocystis_purpurea | JZU52_04840 | 9550..12981 | 61598 | BREX-system-P-loop-protein-BrxC-[Lamprocystis-purpurea] |
| MBK8754669.1 | Type6.BREX | [illegible] | 1209 | Bacteria>Pseudomonadota>Gammaproteobacteria>Gammaproteobacteria_incertae_sedis>Candidatus_Competibacteraceae>unclassified_Candidatus_Competibacteraceae | GCA_016710535.1 | Candidatus_Competibacteraceae_bacterium | IPL99_24865 | 32033..35662 | 2053538 | BREX-system-P-loop-protein-BrxC-[Candidatus-Competibacteraceae-bacterium] |
| WP_159653266.1 | Type1.BREX | [illegible] | 1184 | Bacteria>Pseudomonadota>Gammaproteobacteria>Methylococcales>Methylococcaceae>Methylicorpusculum | GCF_009828925.2 | Methylicorpusculum_oleiharenae | GO003_RS20515 | 4601128..4604682 | 1338687 | BREX-system-P-loop-protein-BrxC-[Methylicorpusculum-oleiharenae] |
| MBE0434438.1 | Type1.BREX | [illegible] | 1177 | Bacteria>Pseudomonadota>Gammaproteobacteria>Methylococcales>Methylococcaceae>Methylomicrobium | GCA_014859425.1 | Methylomicrobium_sp. | IBX56_01390 | 2720..6253 | 2303549 | BREX-system-P-loop-protein-BrxC-[Methylomicrobium-sp.] |
| WP_180058343.1 | Type1.BREX | [illegible] | 1223 | Bacteria>Pseudomonadota>Gammaproteobacteria>Moraxellales>Moraxellaceae>Acinetobacter | GCF_013417785.1 | Acinetobacter_sp._YH12227 | FTV09_RS11090 | 15051..18722 | 2601158 | BREX-system-P-loop-protein-BrxC-[Acinetobacter-sp.-YH12227] |
| WP_133481656.1 | Type1.BREX | [illegible] | 1144 | Bacteria>Pseudomonadota>Gammaproteobacteria>Oceanospirillales>Halomonadaceae>Halomonas | GCF_004361885.1 | Halomonas_ventosae | DFO68_RS02360 | 102047..105481 | 229007 | BREX-system-P-loop-protein-BrxC-[Halomonas-ventosae] |
| WP_018403235.1 | Type1.BREX | [illegible] | 1145 | Bacteria>Pseudomonadota>Gammaproteobacteria>Pseudomonadales>Marinobacteraceae>Marinobacter | GCF_000372805.1 | Marinobacter_gelidimuriae | A3OI_RS0105075 | 19872..23309 | 2739064 | BREX-system-P-loop-protein-BrxC-[Marinobacter-gelidimuriae] |
| WP_077575502.1 | Type6.BREX | [illegible] | 1211 | Bacteria>Pseudomonadota>Gammaproteobacteria>Pseudomonadales>Pseudomonadaceae>Pseudomonas | GCF_002003595.1 | Pseudomonas_aeruginosa | BWR00_RS05555 | 37630..41265 | 287 | BREX-system-P-loop-protein-BrxC-[Pseudomonas-aeruginosa] |

| Protein-ID | System | Gene-Neighbourhoods | Length | Taxonomy | Assembly-ID | Strain | Locus-Tag | Genomic-coordinates | TaxID | Fasta-define |
|---|---|---|---|---|---|---|---|---|---|---|
| KAA3613611.1 | Type1.BREX | [illegible] | 1159 | [illegible] | GCA_008501795.1 | Calditrichota_bacterium | DWQ03_12295 | 106728..110207 | 2212469 | [illegible] |
| WP_187475233.1 | Type1.BREX | [illegible] | 1166 | [illegible] | GCF_014352915.1 | Aliarcobacter_cryaerophilus | HO034_RS05640 | 1060903..1064403 | 28198 | [illegible] |
| WP_152054592.1 | Type1.BREX | [illegible] | 1162 | [illegible] | GCF_902500815.1 | Aliarcobacter_butzleri | GDI74_RS00980 | 25690..29178 | 28197 | [illegible] |
| MBF0239771.1 | Type1.BREX | [illegible] | 1178 | [illegible] | GCA_015231535.1 | SAR324_cluster_bacterium | HQM12_18890 | 8068..11604 | 2024889 | [illegible] |
| TDB40153.1 | Type1.BREX | [illegible] | 1180 | [illegible] | GCA_004348915.1 | Deltaproteobacteria_bacterium | D9V46_00535 | 112112..115654 | 2026735 | [illegible] |
| MBC8466717.1 | Type1.BREX | [illegible] | 1172 | [illegible] | GCA_014384065.1 | Deltaproteobacteria_bacterium | H8D55_02650 | 10326..13844 | 2026735 | [illegible] |
| MBI4519050.1 | Type6.BREX | [illegible] | 1218 | [illegible] | GCA_016210005.1 | Deltaproteobacteria_bacterium | HY699_24940 | 63374..67030 | 2026735 | [illegible] |
| MBK9368670.1 | Type6.BREX | [illegible] | 1184 | [illegible] | GCA_016719515.1 | Deltaproteobacteria_bacterium | IPN01_20575 | 101681..105235 | 2026735 | [illegible] |
| MBW1967272.1 | Type1.BREX | [illegible] | 1154 | [illegible] | GCA_019308155.1 | Deltaproteobacteria_bacterium | JRI48_07780 | 2430..5894 | 2026735 | [illegible] |
| MCC6503799.1 | Type1.BREX | [illegible] | 1181 | [illegible] | GCA_020846435.1 | Deltaproteobacteria_bacterium | IT362_11790 | 16699..20244 | 2026735 | [illegible] |
| MCD6532985.1 | Type1.BREX | [illegible] | 691 | [illegible] | GCA_021163445.1 | Deltaproteobacteria_bacterium | J7L25_02755 | 20972..23047 | 2026735 | [illegible] |
| WP_259094322.1 | Type1.BREX | [illegible] | 1143 | [illegible] | GCF_024809025.1 | Salinibacter_ruber | GGP87_RS12315 | 32840..36271 | 146919 | [illegible] |
| WP_251951689.1 | Type1.BREX | [illegible] | 1143 | [illegible] | GCF_943908255.1 | Salinibacter_ruber | NGL47_RS12295 | 24332..27763 | 146919 | [illegible] |
| RHC75801.1 | Type1.BREX | [illegible] | 691 | [illegible] | GCA_003468465.1 | Bacteroides_uniformis | DW831_02855 | 19890..21965 | 820 | [illegible] |
| HBQ57053.1 | Type1.BREX | [illegible] | 1176 | [illegible] | GCA_003511405.1 | Porphyromonadaceae_bacterium | DD786_07890 | 28635..32165 | 2049046 | [illegible] |
| ROT18783.1 | Type1.BREX | [illegible] | 1223 | [illegible] | GCA_003833095.1 | Muribaculaceae_bacterium_Isolate-114_(HZI) | EEL53_11930 | 25441..29112 | 2486475 | [illegible] |
| KAB5317657.1 | Type1.BREX | [illegible] | 1196 | [illegible] | GCA_009025145.1 | Bacteroides_stercoris | F9949_11590 | 61674..65264 | 46506 | [illegible] |
| NBJ07681.1 | Type1.BREX | [illegible] | 1223 | [illegible] | GCA_009911575.1 | Alistipes_sp._Z76 | D1647_16085 | 15947..19618 | 2304565 | [illegible] |
| MBD5357714.1 | Type1.BREX | [illegible] | 1203 | [illegible] | GCA_014802265.1 | Bacteroides_sp. | HDR88_12035 | 22916..26527 | 29523 | [illegible] |
| MBQ6178640.1 | Type1.BREX | [illegible] | 1202 | [illegible] | GCA_017429185.1 | Bacteroidales_bacterium | IJK32_03880 | 18867..22475 | 2030927 | [illegible] |
| MBQ6577354.1 | Type1.BREX | [illegible] | 744 | [illegible] | GCA_017438445.1 | Bacteroidales_bacterium | IJL91_06365 | 8459..10695 | 2030927 | [illegible] |
| MBV1818013.1 | Type1.BREX | [illegible] | 1195 | [illegible] | GCA_019084065.1 | Bacteroidales_bacterium_MSK.15.36 | KUA25_08040 | 15312..18899 | 2850325 | [illegible] |
| WP_042520323.1 | Type1.BREX | [illegible] | 1204 | [illegible] | GCF_000833975.1 | Prevotella_pectinovora | ST44_RS13175 | 10747..14361 | 1602169 | [illegible] |
| WP_044073670.1 | Type1.BREX | [illegible] | 1206 | [illegible] | GCF_000834015.1 | Prevotella_pectinovora | ST42_RS12125 | 1627..5247 | 1602169 | [illegible] |
| WP_094389547.1 | Type1.BREX | [illegible] | 1228 | [illegible] | GCF_002251345.1 | Prevotella_sp._P4-119 | CIK89_RS00860 | 20388..24074 | 2024218 | [illegible] |
| WP_204506195.1 | Type1.BREX | [illegible] | 1187 | [illegible] | GCF_016902095.1 | Phocaeicola_plebeius | H6A37_RS09755 | 27157..30720 | 310297 | [illegible] |
| WP_138333738.1 | Type6.BREX | [illegible] | 1224 | [illegible] | GCF_019412725.1 | Bacteroides | GPL18_RS15350 | 126249..129923 | 816 | [illegible] |
| WP_181980669.1 | Type1.BREX | [illegible] | 1192 | [illegible] | GCF_015558535.1 | Bacteroidales | QH473_RS20245 | 26845..30423 | 171549 | [illegible] |
| WP_046151576.1 | Type1.BREX | [illegible] | 1184 | [illegible] | GCF_900102645.1 | Bacteroidales | BLR14_RS08960 | 69320..72874 | 171549 | [illegible] |
| BBE18573.1 | Type1.BREX | [illegible] | 1211 | [illegible] | - | Aquipluma_nitroreducens | AQPE_2736 | 3049074..3052709 | 2010828 | [illegible] |
| PLW98642.1 | Type6.BREX | [illegible] | 249 | [illegible] | GCA_002869405.1 | Marinilabiliales_bacterium | C0593_05050 | 18447..19193 | 2053303 | [illegible] |
| WP_273212250.1 | Type1.BREX | [illegible] | 1173 | [illegible] | GCF_002483185.1 | Runella_zeae | B4E63_RS12460 | 307458..310979 | 94255 | [illegible] |
| WP_100629834.1 | Type1.BREX | [illegible] | 553 | [illegible] | GCF_002807035.1 | Algoriphagus_formosus | CDH51_RS17530 | 460328..461989 | 2007308 | [illegible] |
| WP_159471214.1 | Type1.BREX | [illegible] | 1179 | [illegible] | GCF_009801115.1 | Dyadobacter_sp._3J3 | FZF08_RS17845 | 89009..92548 | 2606600 | [illegible] |
| WP_183972373.1 | Type1.BREX | [illegible] | 1202 | [illegible] | GCF_014195535.1 | Runella_defluvii | FHS57_RS08095 | 111516..115124 | 370973 | [illegible] |
| WP_205503392.1 | Type1.BREX | [illegible] | 1180 | [illegible] | GCF_016924645.1 | Rufibacter_sp._SYSU_D00308 | JT718_RS14320 | 179098..182640 | 2812556 | [illegible] |
| WP_255036052.1 | Type1.BREX | [illegible] | 1199 | [illegible] | GCF_024343775.1 | Lacihabitans_soyangensis | EGI31_RS04905 | 50707..54306 | 869394 | [illegible] |
| MAL23063.1 | Type1.BREX | [illegible] | 1157 | [illegible] | GCA_002694765.1 | Xanthomarina_sp. | CMP04_07455 | 107327..110800 | 1931211 | [illegible] |
| WP_034657463.1 | Type1.BREX | [illegible] | 1160 | [illegible] | GCF_000468595.1 | Cellulophaga_baltica | M665_RS0118760 | 90424..93906 | 76594 | [illegible] |
| WP_108879107.1 | Type1.BREX | [illegible] | 1166 | [illegible] | GCF_003095655.1 | Dokdonia_sp._Dokd-P16 | DCS32_RS15465 | 3477371..3480871 | 2173169 | [illegible] |
| WP_132215847.1 | Type1.BREX | [illegible] | 1169 | [illegible] | GCF_004341235.1 | Mariniflexile_fucanivorans | EV196_RS05490 | 443037..446546 | 264023 | [illegible] |
| WP_133546801.1 | Type1.BREX | [illegible] | 1167 | [illegible] | GCF_004362395.1 | Salegentibacter_sp._24 | DET49_RS03950 | 90042..93545 | 2183986 | [illegible] |
| WP_073117772.1 | Type1.BREX | [illegible] | 1170 | [illegible] | GCF_014636515.1 | Wenyingzhuangia_marina | IEV03_RS11465 | 186450..189962 | 1195760 | [illegible] |
| WP_224474755.1 | Type1.BREX | [illegible] | 1167 | [illegible] | GCF_020164485.1 | Salegentibacter_tibetensis | LB467_RS14420 | 15605..19108 | 2873600 | [illegible] |
| WP_159447525.1 | Type1.BREX | [illegible] | 719 | [illegible] | GCF_900167935.1 | Maribacter_arcticus | B5X55_RS04985 | 472839..474998 | 561365 | [illegible] |
| WP_089379946.1 | Type1.BREX | [illegible] | 1168 | [illegible] | GCF_900188235.1 | Lutibacter_agarilyticus | SAMN06265371_101285 | 359460..362966 | 1109740 | [illegible] |
| WP_089376855.1 | Type1.BREX | [illegible] | 1165 | [illegible] | GCF_900188355.1 | Lutibacter_flavus | CHB63_RS02565 | 583517..587014 | 691689 | [illegible] |
| WP_272023608.1 | Type1.BREX | [illegible] | 1160 | [illegible] | GCF_947496445.1 | Olleya_namhaensis | PQG05_RS12495 | 26845..30327 | 1144750 | [illegible] |
| MBK7107567.1 | Type1.BREX | [illegible] | 1197 | [illegible] | GCA_016708205.1 | Bacteroidota_bacterium | IPH61_00035 | 12776..16369 | 1898104 | [illegible] |
| MBL0279470.1 | Type1.BREX | [illegible] | 1197 | [illegible] | GCA_016720495.1 | Bacteroidota_bacterium | IPQ11_03885 | 937205..940798 | 1898104 | [illegible] |
| MCB0536651.1 | Type1.BREX | [illegible] | 1197 | [illegible] | GCA_020436205.1 | Bacteroidota_bacterium | KDE33_03895 | 1012..4605 | 1898104 | [illegible] |
| MCC5907043.1 | Type1.BREX | [illegible] | 1167 | [illegible] | GCA_020832285.1 | Balneolaceae_bacterium | JJU13_12585 | 29362..32865 | 2053516 | [illegible] |
| WP_210512160.1 | Type1.BREX | [illegible] | 1141 | [illegible] | GCF_017921895.1 | Natronogracilivirga_saccharolytica | NATSA_RS09880 | 86175..89600 | 2812953 | [illegible] |
| WP_237852616.1 | Type1.BREX | [illegible] | 1164 | [illegible] | GCF_022012475.1 | Rhodohalobacter_sp._WB101 | L6773_RS04280 | 217944..221438 | 2911366 | [illegible] |
| WP_265763895.1 | Type1.BREX | [illegible] | 1165 | [illegible] | GCF_026229185.1 | Aliifodinibius_sp._1BSP15-2V2 | J6I44_RS00130 | 34405..37902 | 2820877 | [illegible] |
| MCK6605711.1 | Type1.BREX | [illegible] | 1179 | [illegible] | GCA_023151225.1 | Ignavibacteriaceae_bacterium | L6Q59_12515 | 25884..29423 | 2651163 | [illegible] |
| OQC05381.1 | Type1.BREX | [illegible] | 1162 | [illegible] | GCA_002070875.1 | Candidatus_Cloacimonetes_bacterium_ADurb.Bin089 | BWX76_00750 | 1812..5300 | 1852836 | [illegible] |
| MCB5271576.1 | Type1.BREX | [illegible] | 1171 | [illegible] | GCA_020532545.1 | Candidatus_Cloacimonetes_bacterium | LHW56_07005 | 18816..22331 | 2030808 | [illegible] |
| MBT4361892.1 | Type1.BREX | [illegible] | 1148 | [illegible] | GCA_018653465.1 | Candidatus_Marinimicrobia_bacterium | HOD11_13085 | 1..3448 | 2026760 | [illegible] |
| WP_212668349.1 | Type1.BREX | [illegible] | 1186 | [illegible] | GCF_002210355.1 | Fibrobacter_sp._UWH3 | B7993_RS04325 | 153736..157296 | 1964353 | [illegible] |
| WP_073283173.1 | Type1.BREX | [illegible] | 1184 | [illegible] | GCF_900142465.1 | Fibrobacter_sp._UWH6 | BUB73_RS01535 | 351031..354585 | 1896212 | [illegible] |
| WP_021768239.1 | Type1.BREX | [illegible] | 1193 | [illegible] | GCF_000469525.1 | Leptotrichia_sp._oral_taxon_225 | HMPREF9108_RS00960 | 38226..41807 | 671213 | [illegible] |
| WP_026745399.1 | Type1.BREX | [illegible] | 1193 | [illegible] | GCF_007990525.1 | Leptotrichia_hofstadii | FVE77_RS00755 | 171012..174593 | 157688 | [illegible] |
| MCA3011281.1 | Type1.BREX | [illegible] | 1152 | [illegible] | GCA_020247085.1 | Myxococcaceae_bacterium | INH41_02665 | 329..3787 | 2484252 | [illegible] |
| MCK6514475.1 | Type6.BREX | [illegible] | 1083 | [illegible] | GCA_023150775.1 | Myxococcota_bacterium | L6R46_05405 | 17071..20323 | 2818507 | [illegible] |
| MBX3319023.1 | Type1.BREX | [illegible] | 1193 | [illegible] | GCA_019636895.1 | Nitrospira_sp. | KF890_04020 | 66850..70431 | 70125 | [illegible] |
| MCC6543788.1 | Type1.BREX | [illegible] | 1181 | [illegible] | GCA_020846775.1 | Nitrospirota_bacterium | IT392_04715 | 12437..15982 | 2026887 | [illegible] |
| WP_064218004.1 | Type1.BREX | [illegible] | 1187 | [illegible] | GCF_001652185.1 | Acidithiobacillus_ferrooxidans | A4H96_RS01790 | 31399..34962 | 920 | [illegible] |
| WP_163056479.1 | Type1.BREX | [illegible] | 1203 | [illegible] | GCF_010577825.1 | Acidithiobacillus_ferrooxidans | X857_RS08730 | 38026..41637 | 920 | [illegible] |
| WP_009764060.1 | Type1.BREX | [illegible] | 1149 | [illegible] | GCF_000262405.1 | Microvirga_lotononidis | MICLODRAFT_RS22145 | 1808535..1811984 | 864069 | [illegible] |
| WP_036280608.1 | Type1.BREX | [illegible] | 1178 | [illegible] | GCF_000188155.2 | Methylocystis_sp._ATCC_49242 | MET49242_RS03150 | 256459..259995 | 622637 | [illegible] |
| RWE30575.1 | Type1.BREX | [illegible] | 1034 | [illegible] | GCA_004017865.1 | Mesorhizobium_sp. | EOS77_19540 | 17552..20653 | 1871066 | [illegible] |
| WP_199202187.1 | Type1.BREX | [illegible] | 1185 | [illegible] | GCF_016756495.1 | Mesorhizobium_sp._113-3-3 | MesoLj113b_71250 | 318356..321913 | 2744516 | [illegible] |
| WP_154722158.1 | Type1.BREX | [illegible] | 1181 | [illegible] | GCF_900492205.1 | Ciceribacter_naphthalenivorans | DYZ14_RS19075 | 12704..16249 | 1118451 | [illegible] |
| WP_174060722.1 | Type1.BREX | [illegible] | 1175 | [illegible] | - | Agrobacterium_tumefaciens | G6M83_RS06120 | 12437..15964 | 358 | [illegible] |
| CAD6617282.1 | Type1.BREX | [illegible] | 1172 | [illegible] | GCA_902502895.2 | Rhizobium_sp._Khangiran2 | RKHAN_03345 | 3387102..3390620 | 1435607 | [illegible] |
| WP_077987199.1 | Type1.BREX | [illegible] | 1175 | [illegible] | GCF_030440405.1 | Rhizobium_pusense | BTE56_RS17195 | 18928..22455 | 648995 | [illegible] |
| WP_149748055.1 | Type1.BREX | [illegible] | 1175 | [illegible] | GCF_900156005.1 | Rhizobium_sp._RU35A | F3H95_RS25745 | 3768..7295 | 1907414 | [illegible] |
| WP_004434288.1 | Type1.BREX | [illegible] | 1177 | [illegible] | GCF_000236945.1 | Sinorhizobium_meliloti | SM0020_RS29775 | 166866..170399 | 382 | [illegible] |
| WP_037424884.1 | Type1.BREX | [illegible] | 1181 | [illegible] | GCF_000261005.1 | Sinorhizobium_sp._CCBAU_05631 | SS05631_RS30325 | 18322..21867 | 794846 | [illegible] |
| WP_074444907.1 | Type1.BREX | [illegible] | 1148 | [illegible] | GCF_900094735.1 | Saliniramus_fredricksonii | GA0071312_RS10335 | 795252..798698 | 1653334 | [illegible] |
| WP_199258165.1 | Type1.BREX | [illegible] | 1173 | [illegible] | GCF_016446475.1 | Paracoccus_binzhouensis | I9P22_RS04365 | 3344..6865 | 2796149 | [illegible] |
| MBD1206697.1 | Type1.BREX | [illegible] | 1181 | [illegible] | GCA_014654775.1 | Paracoccaceae_bacterium | H9533_21535 | 9124..12669 | 1904441 | [illegible] |
| WP_106262543.1 | Type1.BREX | [illegible] | 1172 | [illegible] | GCF_003003355.1 | Donghicola_tyrosinivorans | CLV74_RS02110 | 436162..439680 | 1652492 | [illegible] |
| WP_274591702.1 | Type1.BREX | [illegible] | 1173 | [illegible] | GCF_028807375.1 | Marinovum_sp._PR37 | PVW49_RS22970 | 3287..6808 | 3028382 | [illegible] |
| WP_123792783.1 | Type1.BREX | [illegible] | 1172 | [illegible] | GCF_003814915.1 | Pacificibacter_maritimus | EDD53_RS08420 | 394668..398186 | 762213 | [illegible] |
| MAE92765.1 | Type1.BREX | [illegible] | 1219 | [illegible] | GCA_002687285.1 | Pelagibaca_sp. | CMI67_24850 | 4770..8429 | 1979400 | [illegible] |
| WP_150282504.1 | Type1.BREX | [illegible] | 1173 | [illegible] | GCF_008692285.1 | Salipiger_aestuarii | AL036_RS18780 | 13068..16589 | 568098 | [illegible] |
| WP_200477164.1 | Type6.BREX | [illegible] | 1208 | [illegible] | GCF_016622105.1 | Azospirillum_brasilense | GAY29_RS06575 | 31945..35571 | 192 | [illegible] |
| WP_022673601.1 | Type1.BREX | [illegible] | 1189 | [illegible] | GCF_000420305.1 | Novosphingopyxis_baekryungensis | G410_RS0110420 | 254465..258034 | 279369 | [illegible] |
| WP_021240096.1 | Type1.BREX | [illegible] | 1177 | [illegible] | GCF_000445065.1 | Sphingobium_quisquiliarum | L288_RS41575 | 32905..36438 | 538379 | [illegible] |
| WP_192976950.1 | Type1.BREX | [illegible] | 1177 | [illegible] | GCF_014892665.1 | Sphingomonas_sp._CFBP_13603 | IFR23_RS17335 | 87025..90558 | 2774040 | [illegible] |
| TNE62528.1 | Type1.BREX | [illegible] | 1175 | [illegible] | GCA_006226385.1 | Sphingomonadales_bacterium | EP341_00225 | 33441..36968 | 1978525 | [illegible] |
| MBQ2886288.1 | Type1.BREX | [illegible] | 1180 | [illegible] | GCA_017409945.1 | Alphaproteobacteria_bacterium | IJE43_21410 | 13139..16681 | 1913988 | [illegible] |
| HCN68765.1 | Type1.BREX | [illegible] | 1154 | [illegible] | GCA_003535635.1 | Candidatus_Accumulibacter_sp. | DIT03_10970 | 31798..35262 | 2053492 | [illegible] |
| WP_076518288.1 | Type1.BREX | [illegible] | 1181 | [illegible] | GCF_900156745.1 | Achromobacter_sp._MFA1_R4 | BXA00_RS09440 | 2091384..2094929 | 1881016 | [illegible] |
| WP_121740016.1 | Type1.BREX | [illegible] | 1190 | [illegible] | GCF_003671915.1 | Alcaligenes_aquatilis | D3M96_RS02075 | 458513..462085 | 323284 | [illegible] |
| WP_068373657.1 | Type1.BREX | [illegible] | 1176 | [illegible] | GCF_000988095.1 | Kerstersia_gyiorum | AAV32_RS14310 | 103983..107513 | 206506 | [illegible] |
| WP_115075260.1 | Type1.BREX | [illegible] | 1187 | [illegible] | GCF_900454255.1 | Oligella_urethralis | DX091_RS10510 | 2175019..2178582 | 90245 | [illegible] |
| WP_132980217.1 | Type1.BREX | [illegible] | 1181 | [illegible] | GCF_004352985.1 | Pigmentiphaga_sp._D-2 | EZY13_RS19720 | 185799..189344 | 1002116 | [illegible] |
| WP_124831269.1 | Type1.BREX | [illegible] | 1200 | [illegible] | GCF_003857875.1 | Burkholderia_stagnalis | DF113_RS32940 | 12280..15882 | 1503054 | [illegible] |
| WP_167306478.1 | Type1.BREX | [illegible] | 1182 | [illegible] | GCF_900100565.1 | Paraburkholderia_steynii | BLT65_RS12240 | 289685..293233 | 1245441 | [illegible] |
| MBN6206497.1 | Type1.BREX | [illegible] | 1366 | [illegible] | GCA_017162415.1 | Ralstonia_pickettii | JYK21_08530 | 22216..26316 | 329 | [illegible] |
| MCK6425125.1 | Type1.BREX | [illegible] | 1081 | [illegible] | GCA_023150375.1 | Burkholderiaceae_bacterium | L6Q75_08530 | 1698..4943 | 2030806 | [illegible] |
| OJX16153.1 | Type1.BREX | [illegible] | 1181 | [illegible] | GCA_001898745.1 | Delftia_sp._67-8 | BGO79_00315 | 75959..79504 | 1895749 | [illegible] |
| WP_108330269.1 | Type1.BREX | [illegible] | 1154 | [illegible] | GCF_003063435.1 | Limnohabitans_sp._2KL-1 | B9Z47_RS05555 | 224189..227653 | 1100699 | [illegible] |
| WP_169420224.1 | Type1.BREX | [illegible] | 1153 | [illegible] | GCF_012927085.1 | Ramlibacter_agri | HHL11_RS19385 | 189581..193042 | 2728837 | [illegible] |
| MBP9058688.1 | Type1.BREX | [illegible] | 1153 | [illegible] | GCA_018057375.1 | Rhodoferax_sp. | KBF98_00065 | 15882..19343 | 50421 | [illegible] |
| WP_186915942.1 | Type1.BREX | [illegible] | 1183 | [illegible] | GCF_014284115.1 | Undibacterium | H8K35_RS08325 | 157325..160876 | 401469 | [illegible] |
| WP_219136384.1 | Type1.BREX | [illegible] | 1153 | [illegible] | GCF_900095235.1 | Janthinobacterium_sp._UMAB-60 | K0F86_RS22385 | 15275..18736 | 1365365 | [illegible] |
| WP_258819858.1 | Type1.BREX | [illegible] | 1186 | [illegible] | GCF_024753285.1 | Massilia_pinisoli | NX784_RS27510 | 42185..45745 | 1772194 | [illegible] |
| MCP5275500.1 | Type1.BREX | [illegible] | 1176 | [illegible] | GCA_024235795.1 | Burkholderiales_bacterium | H6936_11775 | 653459..656989 | 1891238 | [illegible] |
| MCP5275800.1 | Type1.BREX | [illegible] | 1177 | [illegible] | GCA_024235795.1 | Burkholderiales_bacterium | H6936_13310 | 1016712..1020245 | 1891238 | [illegible] |
| MBK7364915.1 | Type1.BREX | [illegible] | 1187 | [illegible] | GCA_016705625.1 | Nitrosomonas_sp. | IPI97_07925 | 1671776..1675339 | 42353 | [illegible] |
| MBV6446594.1 | Type1.BREX | [illegible] | 1182 | [illegible] | GCA_019187415.1 | Nitrosomonas_sp. | FCKEOINB_00187 | 17095..20643 | 42353 | [illegible] |
| MBX3639968.1 | Type1.BREX | [illegible] | 1182 | [illegible] | GCA_019635175.1 | Nitrosomonas_sp. | KF888_05565 | 120382..123930 | 42353 | [illegible] |
| WP_205230204.1 | Type1.BREX | [illegible] | 1153 | [illegible] | GCF_905120365.1 | Azospira_oryzae | JYJ48_RS05820 | 17001..20462 | 146939 | [illegible] |
| TXH82461.1 | Type1.BREX | [illegible] | 1153 | [illegible] | GCA_008015255.1 | Thauera_aminoaromatica | E6Q80_15435 | 2474..5935 | 164330 | [illegible] |
| WP_012584348.1 | Type5.BREX | [illegible] | 1203 | [illegible] | GCF_000021765.1 | Thauera_sp._MZ1T | TMZ1T_RS01390 | 330563..334174 | 85643 | [illegible] |
| WP_107221110.1 | Type1.BREX | [illegible] | 1153 | [illegible] | GCF_003030465.1 | Thauera_aromatica | Tharo_RS10350 | 2186270..2189731 | 59405 | [illegible] |
| WP_247737217.1 | Type1.BREX | [illegible] | 1153 | [illegible] | - | Thauera_aromatica | MX653_RS17175 | 1332..4793 | 59405 | [illegible] |
| MBK6655412.1 | Type1.BREX | [illegible] | 1153 | [illegible] | GCA_016703645.1 | Zoogloea_sp. | IPG40_16580 | 748210..751671 | 49181 | [illegible] |
| MBK7848388.1 | Type1.BREX | [illegible] | 1153 | [illegible] | GCA_016714095.1 | Zoogloea_sp. | IPJ73_15250 | 34710..38171 | 49181 | [illegible] |

| Protein-ID | System | Gene-Neighbourhoods | Length | Taxonomy | Assembly-ID | Strain | Locus-Tag | Genomic-coordinates | TaxID | Fasta-defline |
|---|---|---|---|---|---|---|---|---|---|---|
| AUZ77883.1 | Type1.BREX | <-HIN-HTH<- SIGMA+OB-Fold+AAA-ATPase+HKD-DNase-BrxL <- PglZ <-ParB+HNH<- N6-METHYLASE <- REase <-OLD-ABC <- DUF6079 *<- BrxB-BrxF-iSTAND <- BrxA-DNA-binding <-XIS-HTH\|\|<-DOC [illegible] | 1183 | Bacteria>Pseudomonadota>Gammaproteobacteria>Aeromonadales>Aeromonadaceae>Aeromonas | GCA_002934425.1 | Aeromonas_sp._ASNIH4 | C2U40_16585 | 3601638..3605189 | 1636609 | BREX-system-P-loop-protein-BrxC-[Aeromonas-sp.-ASNIH4] |
| WP_043557612.1 | Type1.BREX | REase -><-?\|\|?-><- SIGMA+OB-Fold+AAA-ATPase+HKD-DNase-BrxL <- PglZ <-ParB+HNH<- N6-METHYLASE <- REase <-OLD-ABC <- DUF6079 *<- BrxB-BrxF-iSTAND <- BrxA-DNA-binding <-XIS-HTH<-? [illegible] | 1182 | Bacteria>Pseudomonadota>Gammaproteobacteria>Aeromonadales>Aeromonadaceae>Aeromonas | GCF_000819745.1 | Aeromonas_bestiarum | BN1118_RS19650 | 69736..73284 | 105751 | BREX-system-P-loop-protein-BrxC-[Aeromonas-bestiarum] |
| WP_257706959.1 | Type1.BREX | Zn_Tnp_IS1+DDE_Tnp_IS1595-><- SIGMA+OB-Fold+AAA-ATPase+HKD-DNase-BrxL <- PglZ <-ParB+HNH<- N6-METHYLASE <- REase <-OLD-ABC <- DUF6079 *<- BrxB-BrxF-iSTAND <- BrxA-DNA-binding [illegible] | 1182 | Bacteria>Pseudomonadota>Gammaproteobacteria>Aeromonadales>Aeromonadaceae>Aeromonas | GCF_024649325.1 | Aeromonas_caviae | NUK33_RS01495 | 150155..153703 | 648 | BREX-system-P-loop-protein-BrxC-[Aeromonas-caviae] |
| WP_270661186.1 | Type1.BREX | XIS-HTH->?->DOC+HTH-><-?\|\|XIS-HTH-> BrxA-DNA-binding -> BrxB-BrxF-iSTAND -> DUF6079 *-> OLD-ABC-> REase -> N6-METHYLASE ->ParB-> | 1182 | Bacteria>Pseudomonadota>Gammaproteobacteria>Aeromonadales>Aeromonadaceae>Aeromonas | GCF_027705205.1 | unclassified_Aeromonas | OKC30_RS15245 | 6003..9551 | 257493 | MULTISPECIES:-BREX-system-P-loop-protein-BrxC-[unclassified-Aeromonas] |
| WP_181014924.1 | Type1.BREX | <-HIN-HTH<- SIGMA+OB-Fold+AAA-ATPase+HKD-DNase-BrxL <- PglZ <-ParB+HNH<- N6-METHYLASE <- REase <-OLD-ABC <- DUF6079 *<- BrxB-BrxF-iSTAND <- BrxA-DNA-binding <-XIS-HTH\|\|?-><-DOC [illegible] | 1182 | Bacteria>Pseudomonadota>Gammaproteobacteria>Aeromonadales>Aeromonadaceae | GCF_002934425.1 | Aeromonas | C2U40_RS16580 | 3601638..3605186 | 642 | MULTISPECIES:-BREX-system-P-loop-protein-BrxC-[Aeromonas] |
| WP_006006589.1 | Type1.BREX | <-PIN<-MazE_antitoxin<-MODE-HTH+PBPII\|\|?->?->?->XIS-HTH-> BrxA-DNA-binding -> BrxB-BrxF-iSTAND -> DUF6079 *-> OLD-ABC-> REase -> N6-METHYLASE -> PglZ -> [illegible] | 1189 | Bacteria>Pseudomonadota>Gammaproteobacteria>Alteromonadales>Alteromonadaceae>Glaciecola | GCF_000252165.1 | Glaciecola_punicea | GPUN_RS10800 | 142812..146381 | 56804 | BREX-system-P-loop-protein-BrxC-[Glaciecola-punicea] |
| WP_136550441.1 | Type1.BREX | <-SbcC+APATPase+SbcC<- SIGMA+OB-Fold+AAA-ATPase+HKD-DNase-BrxL <- PglZ <- N6-METHYLASE <- REase <-OLD-ABC <-Schlafen+HATPase+wHTH+HTH<- DUF6079 *<- BrxB-BrxF-iSTAND <- [illegible] | 1193 | Bacteria>Pseudomonadota>Gammaproteobacteria>Alteromonadales>Alteromonadaceae>Hydrocarboniclastica | GCF_004851605.1 | Hydrocarboniclastica_marina | soil367_RS18210 | 4131404..4134985 | 2259620 | BREX-system-P-loop-protein-BrxC-[Hydrocarboniclastica-marina] |
| ASP49913.1 | Type1.BREX | Trypsin-><-LZ_Tnp_IS66+zf-IS66+DDE_Tnp_IS66+DDE_Tnp_IS66_C<-HIN-HTH<-SF1-SF2-Helicase<-PadR-HTH\|\|HTH_3-> wHTH+WYL+WCX -> BrxA-DNA-binding -> BrxB-BrxF-iSTAND -> DUF6079 *-> [illegible] | 1229 | Bacteria>Pseudomonadota>Gammaproteobacteria>Alteromonadales>Colwelliaceae>Cognaticolwellia | GCA_002076895.1 | Cognaticolwellia_beringensis | B5D82_06290 | 1500021..1503710 | 1967665 | DNA-repair-protein-[Cognaticolwellia-beringensis] |
| WP_245807568.1 | Type1.BREX | <-LZ_Tnp_IS66+zf-IS66+DDE_Tnp_IS66_C<-?<-HIN-HTH<-SF1-SF2-Helicase<-PadR-HTH\|\|HTH_3-> wHTH+WYL+WCX -> BrxA-DNA-binding -> BrxB-BrxF-iSTAND -> DUF6079 *-> [illegible] | 1232 | Bacteria>Pseudomonadota>Gammaproteobacteria>Alteromonadales>Colwelliaceae>Cognaticolwellia | GCF_002076895.1 | Cognaticolwellia_beringensis | B5D82_RS06290 | 1500012..1503710 | 1967665 | BREX-system-P-loop-protein-BrxC-[Cognaticolwellia-beringensis] |
| WP_258240870.1 | Type1.BREX | <-RNasePH<-HTH_20\|\|?->?->DOC->XIS-HTH-> wHTH+WYL+WCX -> BrxA-DNA-binding -> BrxB-BrxF-iSTAND -> DUF6079 *-> N6-METHYLASE ->PHP+SbcC-> PglZ -> [illegible] | 1150 | Bacteria>Pseudomonadota>Gammaproteobacteria>Alteromonadales>Idiomarinaceae>Pseudidiomarina | GCF_024723895.1 | Pseudidiomarina_homiensis | NV376_RS09715 | 149869..153321 | 364198 | BREX-system-P-loop-protein-BrxC-[Pseudidiomarina-homiensis] |
| WP_112714387.1 | Type1.BREX | <- SIGMA+OB-Fold+AAA-ATPase+HKD-DNase-BrxL <- PglZ <-ParB+HNH<- N6-METHYLASE <- REase <-OLD-ABC <- DUF6079 *<- BrxB-BrxF-iSTAND <- BrxA-DNA-binding <-XIS-HTH\|\|?-><- [illegible] | 1199 | Bacteria>Pseudomonadota>Gammaproteobacteria>Alteromonadales>Moritellaceae>Moritella | GCF_900465055.1 | Moritella_yayanosii | MORIYA_RS08605 | 1893116..1896715 | 69539 | BREX-system-P-loop-protein-BrxC-[Moritella-yayanosii] |
| WP_010607320.1 | Type1.BREX | <-TniQ<-TnsC-ATPase<-?<- REase <-?<- SIGMA+OB-Fold+AAA-ATPase+HKD-DNase-BrxL <- PglZ <-ParB+HNH<- N6-METHYLASE <- DUF6079 *<- BrxB-BrxF-iSTAND <- BrxA-DNA-binding <-XIS-HTH<- [illegible] | 1188 | Bacteria>Pseudomonadota>Gammaproteobacteria>Alteromonadales>Pseudoalteromonadaceae | GCF_000276645.1 | Pseudoalteromonas | UY7_RS0121200 | 90697..94263 | 53246 | MULTISPECIES:-BREX-system-P-loop-protein-BrxC-[Pseudoalteromonas] |
| WP_024611453.1 | Type1.BREX | <-ArdCN+MPTase<- SIGMA+OB-Fold+AAA-ATPase+HKD-DNase-BrxL <- PglZ <- N6-METHYLASE <- REase <-OLD-ABC <- DUF6079 *<- BrxB-BrxF-iSTAND <- BrxA-DNA-binding <-XIS-HTH | 1193 | Bacteria>Pseudomonadota>Gammaproteobacteria>Alteromonadales>Pseudoalteromonadaceae>Pseudoalteromonas | GCF_000498095.1 | Pseudoalteromonas_sp._TB64 | L647_RS0112855 | 131987..135568 | 1938600 | BREX-system-P-loop-protein-BrxC-[Pseudoalteromonas-sp.-TB64] |
| WP_089368387.1 | Type1.BREX | LZ_Tnp_IS66+zf-IS66+DDE_Tnp_IS66+DDE_Tnp_IS66_C-><- SIGMA+OB-Fold+AAA-ATPase+Lon-Protease-BrxL <- PglZ <- REase +NACHT<-?<-RADICAL-SAM<- N6-METHYLASE <-ParB+HNH<- DUF6079 *<- [illegible] | 1229 | Bacteria>Pseudomonadota>Gammaproteobacteria>Alteromonadales>Pseudoalteromonadaceae>Pseudoalteromonas | GCF_007991255.1 | Pseudoalteromonas_nigrifaciens | PNIG_RS09950 | 2060327..2064016 | 28109 | BREX-system-P-loop-protein-BrxC-[Pseudoalteromonas-nigrifaciens] |
| WP_167658112.1 | Type1.BREX | <-rve<- REase <-?<- SIGMA+OB-Fold+AAA-ATPase+HKD-DNase-BrxL <- PglZ <-ParB+HNH<- N6-METHYLASE <- REase <-OLD-ABC <- DUF6079 *<- BrxB-BrxF-iSTAND <- BrxA-DNA-binding <-XIS-HTH<- [illegible] | 1188 | Bacteria>Pseudomonadota>Gammaproteobacteria>Alteromonadales>Pseudoalteromonadaceae>Pseudoalteromonas | GCF_011806445.1 | Pseudoalteromonas_sp._HF66 | HCZ11_RS07660 | 212811..216377 | 2721559 | BREX-system-P-loop-protein-BrxC-[Pseudoalteromonas-sp.-HF66] |
| PKG74712.1 | Type1.BREX | <- SIGMA+OB-Fold+AAA-ATPase+HKD-DNase-BrxL <- PglZ <-ParB+HNH<- N6-METHYLASE <- REase <-OLD-ABC <- DUF6079 *<- BrxB-BrxF-iSTAND <- BrxA-DNA-binding <-XIS-HTH<-?<-TnsC-ATPase<- [illegible] | 1191 | Bacteria>Pseudomonadota>Gammaproteobacteria>Alteromonadales>Shewanellaceae>Shewanella | GCA_002836135.1 | Shewanella_sp._GutCb | CXF86_11690 | 122253..125828 | 2058315 | BREX-system-P-loop-protein-BrxC-[Shewanella-sp.-GutCb] |
| VEE64276.1 | Type1.BREX | <-Cas10\|\|Cas_NE0113-><-?<-HDRnase<-Arm-DNA-bind_2+Phage_int_SAM_3+Phage_integrase<-Phage_AlpA\|\| wHTH+WYL+WCX -> BrxA-DNA-binding -> BrxB-BrxF-iSTAND -> DUF6079 *-> [illegible] | 1268 | Bacteria>Pseudomonadota>Gammaproteobacteria>Alteromonadales>Shewanellaceae>Shewanella | GCA_900636665.1 | Shewanella_putrefaciens | NCTC12093_04119 | 4646571..4650377 | 24 | Uncharacterised-protein-[Shewanella-putrefaciens] |
| WP_257985331.1 | Type1.BREX | [illegible] | 1193 | Bacteria>Pseudomonadota>Gammaproteobacteria>Alteromonadales>Shewanellaceae>Shewanella | GCF_002836135.1 | Shewanella_sp._GutCb | CXF86_RS11730 | 122253..125834 | 2058315 | BREX-system-P-loop-protein-BrxC-[Shewanella-sp.-GutCb] |
| WP_259573703.1 | Type1.BREX | [illegible] | 1240 | Bacteria>Pseudomonadota>Gammaproteobacteria>Alteromonadales>Shewanellaceae>Shewanella | GCF_024999165.1 | Shewanella_baltica | G7A68_RS06675 | 25637..29359 | 62322 | BREX-system-P-loop-protein-BrxC-[Shewanella-baltica] |
| WP_112352819.1 | Type1.BREX | [illegible] | 1197 | Bacteria>Pseudomonadota>Gammaproteobacteria>Alteromonadales>Shewanellaceae>Shewanella | GCF_900476435.1 | Shewanella_benthica | SHEWBE_RS12055 | 2634220..2637813 | 43661 | BREX-system-P-loop-protein-BrxC-[Shewanella-benthica] |
| WP_126513302.1 | Type1.BREX | [illegible] | 1254 | Bacteria>Pseudomonadota>Gammaproteobacteria>Alteromonadales>Shewanellaceae>Shewanella | - | Shewanella_putrefaciens | EL161_RS20440 | 4646613..4650377 | 24 | BREX-system-P-loop-protein-BrxC-[Shewanella-putrefaciens] |
| WP_102039083.1 | Type1.BREX | [illegible] | 1191 | Bacteria>Pseudomonadota>Gammaproteobacteria>Alteromonadales>Shewanellaceae>Shewanella | - | Shewanella_sp._GutCb | CXF86_11690 | 122253..125828 | 2058315 | BREX-system-P-loop-protein-BrxC-[Shewanella-sp.-GutCb] |
| WP_265366287.1 | Type1.BREX | [illegible] | 1193 | Bacteria>Pseudomonadota>Gammaproteobacteria>Cellvibrionales>Cellvibrionaceae>Marinimicrobium | GCF_026153455.1 | Marinimicrobium_koreense | OOT55_RS13025 | 3024461..3028042 | 306545 | BREX-system-P-loop-protein-BrxC-[Marinimicrobium-sp.-C6131] |
| WP_084199902.1 | Type1.BREX | [illegible] | 1184 | Bacteria>Pseudomonadota>Gammaproteobacteria>Cellvibrionales>Halieaceae>Halioglobus | GCF_014652255.1 | Halioglobus_japonicus | COO29_RS05530 | 1195751..1199305 | 930805 | BREX-system-P-loop-protein-BrxC-[Halioglobus-japonicus] |
| WP_148062597.1 | Type1.BREX | [illegible] | 1180 | Bacteria>Pseudomonadota>Gammaproteobacteria>Cellvibrionales>Halieaceae>Parahaliea | GCF_008064635.1 | Parahaliea_aestuarii | FVW59_RS02260 | 491803..495345 | 1852021 | BREX-system-P-loop-protein-BrxC-[Parahaliea-aestuarii] |
| MTW21824.1 | Type1.BREX | [illegible] | 1173 | Bacteria>Pseudomonadota>Gammaproteobacteria>Chromatiales>Chromatiaceae>Allochromatium | GCA_009720725.1 | Allochromatium_palmeri | GJ668_12065 | 37225..40746 | 231048 | BREX-system-P-loop-protein-BrxC-[Allochromatium-palmeri] |
| WP_186343071.1 | Type1.BREX | [illegible] | 1172 | Bacteria>Pseudomonadota>Gammaproteobacteria>Chromatiales>Chromatiaceae>Allochromatium | GCF_009720725.1 | Allochromatium_palmeri | GJ668_RS12065 | 37225..40743 | 231048 | BREX-system-P-loop-protein-BrxC-[Allochromatium-palmeri] |
| NEX15944.1 | Type1.BREX | [illegible] | 1173 | Bacteria>Pseudomonadota>Gammaproteobacteria>Chromatiales>Chromatiaceae>Halochromatium | GCA_010912705.1 | Halochromatium_sp. | C1943_04720 | 67001..70522 | 2049430 | BREX-system-P-loop-protein-BrxC-[Halochromatium-sp.] |
| WP_201222419.1 | Type1.BREX | [illegible] | 1174 | Bacteria>Pseudomonadota>Gammaproteobacteria>Chromatiales>Chromatiaceae>Halochromatium | GCF_016653315.1 | Halochromatium_roseum | CCR96_RS19410 | 137258..140782 | 391920 | BREX-system-P-loop-protein-BrxC-[Halochromatium-roseum] |
| WP_200250838.1 | Type1.BREX | [illegible] | 1173 | Bacteria>Pseudomonadota>Gammaproteobacteria>Chromatiales>Chromatiaceae>Lamprobacter | GCF_016583905.1 | Lamprobacter_modestohalophilus | CKO42_RS24315 | 8724..12245 | 1064514 | BREX-system-P-loop-protein-BrxC-[Lamprobacter-modestohalophilus] |
| WP_244406389.1 | Type1.BREX | [illegible] | 1041 | Bacteria>Pseudomonadota>Gammaproteobacteria>Chromatiales>Ectothiorhodospiraceae>Thioalkalivibrio | GCF_000025545.1 | Thioalkalivibrio_sp._K90mix | TK90_RS05670 | 1196495..1199620 | 396595 | BREX-system-P-loop-protein-BrxC-[Thioalkalivibrio-sp.-K90mix] |
| MCB1864346.1 | Type1.BREX | [illegible] | 1147 | Bacteria>Pseudomonadota>Gammaproteobacteria>Chromatiales>unclassified_Chromatiales | GCA_020445605.1 | Chromatiales_bacterium | KDG50_02880 | 27912..31355 | 2026725 | BREX-system-P-loop-protein-BrxC-[Chromatiales-bacterium] |
| RFF27207.1 | Type1.BREX | [illegible] | 1166 | Bacteria>Pseudomonadota>Gammaproteobacteria>Chromatiales>Wenzhouxiangellaceae>Wenzhouxiangella | GCA_003410035.1 | Wenzhouxiangella_sp._15181 | DZK25_09525 | 145045..148545 | 2301224 | BREX-system-P-loop-protein-BrxC-[Wenzhouxiangella-sp.-15181] |
| WP_181918413.1 | Type1.BREX | [illegible] | 1149 | Bacteria>Pseudomonadota>Gammaproteobacteria>Chromatiales>Wenzhouxiangellaceae>Wenzhouxiangella | GCF_003410035.1 | unclassified_Wenzhouxiangella | DZK25_RS09530 | 145045..148494 | 2613841 | MULTISPECIES:-BREX-system-P-loop-protein-BrxC-[unclassified-Wenzhouxiangella] |
| WP_230863208.1 | Type1.BREX | [illegible] | 1240 | Bacteria>Pseudomonadota>Gammaproteobacteria>Enterobacterales>Enterobacteriaceae>Enterobacter | GCF_015704905.1 | Enterobacter_cloacae | IOR24_RS17915 | 3774454..3778176 | 550 | BREX-system-P-loop-protein-BrxC-[Enterobacter-cloacae] |
| HAO0324358.1 | Type1.BREX | [illegible] | 1213 | Bacteria>Pseudomonadota>Gammaproteobacteria>Enterobacterales>Enterobacteriaceae>Escherichia | GCA_014771515.1 | Escherichia_coli | IGZ64_004245 | 6773..10414 | 562 | BREX-system-P-loop-protein-BrxC-[Escherichia-coli] |
| HBR4497442.1 | Type1.BREX | [illegible] | 1213 | Bacteria>Pseudomonadota>Gammaproteobacteria>Enterobacterales>Enterobacteriaceae>Klebsiella/Raoultella_group | GCA_021926505.1 | Klebsiella_pneumoniae | L9V75_004938 | 11616..15257 | 573 | BREX-system-P-loop-protein-BrxC-[Klebsiella-pneumoniae] |
| EBU6515329.1 | Type1.BREX | [illegible] | 1213 | Bacteria>Pseudomonadota>Gammaproteobacteria>Enterobacterales>Enterobacteriaceae>Salmonella | GCA_007137465.1 | Salmonella_enterica_subsp._enterica_serovar_Typhimurium | DKJ21_01470 | 4814..8455 | 90371 | BREX-system-P-loop-protein-BrxC-[Salmonella-enterica-subsp.-enterica-serovar-Typhimurium] |
| WP_094887773.1 | Type1.BREX | [illegible] | 1213 | Bacteria>Pseudomonadota>Gammaproteobacteria>Enterobacterales>Enterobacteriaceae>Salmonella | GCF_002265975.1 | Salmonella_enterica | CCD43_RS23255 | 12854..16495 | 28901 | BREX-system-P-loop-protein-BrxC-[Salmonella-enterica] |
| QIG07578.1 | Type1.BREX | [illegible] | 1183 | Bacteria>Pseudomonadota>Gammaproteobacteria>Enterobacterales>Morganellaceae>Proteus | GCA_011046025.1 | Proteus_sp._ZN5 | GTK47_19170 | 4181966..4185517 | 2697019 | BREX-system-P-loop-protein-BrxC-[Proteus-sp.-ZN5] |
| WP_206535871.1 | Type1.BREX | [illegible] | 1185 | Bacteria>Pseudomonadota>Gammaproteobacteria>Enterobacterales>Morganellaceae>Proteus | GCF_011046025.1 | Proteus_sp._ZN5 | GTK47_RS19230 | 4181960..4185517 | 2697019 | BREX-system-P-loop-protein-BrxC-[Proteus-sp.-ZN5] |
| WP_272670012.1 | Type1.BREX | [illegible] | 1185 | Bacteria>Pseudomonadota>Gammaproteobacteria>Enterobacterales>Morganellaceae>Providencia | GCF_028480605.1 | unclassified_Providencia | PSE59_RS21125 | 8138..11695 | 2633465 | MULTISPECIES:-BREX-system-P-loop-protein-BrxC-[unclassified-Providencia] |
| MCG7871843.1 | Type1.BREX | [illegible] | 1174 | Bacteria>Pseudomonadota>Gammaproteobacteria>Gammaproteobacteria_incertae_sedis>sulfur-oxidizing_symbionts>Candidatus_Thiodiazotropha | GCA_016848925.1 | Candidatus_Thiodiazotropha_lotti | JAZ11_07035 | 23024..26548 | 2792787 | BREX-system-P-loop-protein-BrxC-[Candidatus-Thiodiazotropha-lotti] |
| MCF7971355.1 | Type1.BREX | [illegible] | 1151 | Bacteria>Pseudomonadota>Gammaproteobacteria>Methylococcales>Methylococcaceae>unclassified_Methylococcaceae | GCA_021736705.1 | Methylococcaceae_bacterium | K9L22_09360 | 3141..6596 | 1933926 | BREX-system-P-loop-protein-BrxC-[Methylococcaceae-bacterium] |
| WP_004984466.1 | Type1.BREX | [illegible] | 1190 | Bacteria>Pseudomonadota>Gammaproteobacteria>Moraxellales>Moraxellaceae>Acinetobacter | GCF_000368805.1 | Acinetobacter_johnsonii | F946_RS15710 | 148915..152487 | 40214 | BREX-system-P-loop-protein-BrxC-[Acinetobacter-johnsonii] |
| WP_131214141.1 | Type1.BREX | [illegible] | 1190 | Bacteria>Pseudomonadota>Gammaproteobacteria>Moraxellales>Moraxellaceae>Acinetobacter | GCF_004331285.1 | Acinetobacter_sp._ANC_3781 | E0H89_RS01980 | 72363..75935 | 2529835 | BREX-system-P-loop-protein-BrxC-[Acinetobacter-sp.-ANC-3781] |
| WP_151834043.1 | Type1.BREX | [illegible] | 1190 | Bacteria>Pseudomonadota>Gammaproteobacteria>Moraxellales>Moraxellaceae>Acinetobacter | GCF_009012055.1 | Acinetobacter_ursingii | GBX89_RS12550 | 5375..8947 | 108980 | BREX-system-P-loop-protein-BrxC-[Acinetobacter-ursingii] |
| WP_180108075.1 | Type1.BREX | [illegible] | 1190 | Bacteria>Pseudomonadota>Gammaproteobacteria>Moraxellales>Moraxellaceae>Acinetobacter | GCF_013419415.1 | Acinetobacter_sp._YH12085 | FTU28_RS13915 | 49013..52585 | 2601077 | BREX-system-P-loop-protein-BrxC-[Acinetobacter-sp.-YH12085] |
| WP_216936951.1 | Type1.BREX | [illegible] | 1190 | Bacteria>Pseudomonadota>Gammaproteobacteria>Moraxellales>Moraxellaceae>Acinetobacter | GCF_019038255.1 | unclassified_Acinetobacter | KUG40_RS10660 | 10626..14198 | 196816 | MULTISPECIES:-BREX-system-P-loop-protein-BrxC-[unclassified-Acinetobacter] |
| WP_102077888.1 | Type6.BREX | [illegible] | 1219 | Bacteria>Pseudomonadota>Gammaproteobacteria>Moraxellales>Moraxellaceae>Psychrobacter | GCF_001742065.1 | unclassified_Psychrobacter | CXF69_RS01065 | 108247..111906 | 196806 | MULTISPECIES:-BREX-system-P-loop-protein-BrxC-[unclassified-Psychrobacter] |
| MBL0229728.1 | Type1.BREX | [illegible] | 1147 | Bacteria>Pseudomonadota>Gammaproteobacteria>Moraxellales>Moraxellaceae>unclassified_Moraxellaceae | GCA_016721165.1 | Moraxellaceae_bacterium | IPP76_02595 | 6399..9842 | 1889775 | BREX-system-P-loop-protein-BrxC-[Moraxellaceae-bacterium] |
| MCP5177609.1 | Type1.BREX | [illegible] | 1192 | Bacteria>Pseudomonadota>Gammaproteobacteria>Moraxellales>Moraxellaceae>unclassified_Moraxellaceae | GCA_024234655.1 | Moraxellaceae_bacterium | H6997_08670 | 33703..37281 | 1889775 | BREX-system-P-loop-protein-BrxC-[Moraxellaceae-bacterium] |
| MAD69915.1 | Type1.BREX | [illegible] | 1193 | Bacteria>Pseudomonadota>Gammaproteobacteria>Oceanospirillales>Alcanivoracaceae>Alcanivorax | GCA_002685555.1 | Alcanivorax_sp. | CL538_05260 | 26266..29847 | 1872427 | hypothetical-protein-[Alcanivorax-sp.] |
| HIK75733.1 | Type1.BREX | [illegible] | 1193 | Bacteria>Pseudomonadota>Gammaproteobacteria>Oceanospirillales>Alcanivoracaceae>Alcanivorax | GCA_015660495.1 | Alcanivorax_sp. | EYG00_13690 | 18356..21937 | 1872427 | BREX-system-P-loop-protein-BrxC-[Alcanivorax-sp.] |
| MCH2557814.1 | Type1.BREX | [illegible] | 1193 | Bacteria>Pseudomonadota>Gammaproteobacteria>Oceanospirillales>Alcanivoracaceae>Alcanivorax | GCA_022449455.1 | Alcanivorax_sp. | MK005_11000 | 47715..51296 | 1872427 | BREX-system-P-loop-protein-BrxC-[Alcanivorax-sp.] |
| WP_103916700.1 | Type1.BREX | [illegible] | 1193 | Bacteria>Pseudomonadota>Gammaproteobacteria>Oceanospirillales>Alcanivoracaceae>Alcanivorax | GCF_900107995.1 | Alcanivorax_sp._DSM_26293 | C4S63_RS03510 | 780383..783964 | 1798238 | BREX-system-P-loop-protein-BrxC-[Alcanivorax-sp.-DSM-26293] |
| WP_194855579.1 | Type1.BREX | [illegible] | 1192 | Bacteria>Pseudomonadota>Gammaproteobacteria>Oceanospirillales>Alcanivoracaceae>Alloalcanivorax | GCF_015356855.1 | Alloalcanivorax_venustensis | ISO4_RS06450 | 133547..137125 | 172371 | BREX-system-P-loop-protein-BrxC-[Alloalcanivorax-venustensis] |
| WP_228232948.1 | Type1.BREX | [illegible] | 1193 | Bacteria>Pseudomonadota>Gammaproteobacteria>Oceanospirillales>Alcanivoracaceae>Alloalcanivorax | GCF_020782135.1 | Alloalcanivorax_marinus | LL252_RS03465 | 287759..291340 | 1177169 | BREX-system-P-loop-protein-BrxC-[Alloalcanivorax-marinus] |
| WP_008734404.1 | Type1.BREX | [illegible] | 1192 | Bacteria>Pseudomonadota>Gammaproteobacteria>Oceanospirillales>Alcanivoracaceae>Isoalcanivorax | GCF_000299335.2 | Isoalcanivorax_pacificus | S7S_RS18615 | 4158589..4162167 | 1306787 | BREX-system-P-loop-protein-BrxC-[Isoalcanivorax-pacificus] |
| WP_257265108.1 | Type1.BREX | [illegible] | 1183 | Bacteria>Pseudomonadota>Gammaproteobacteria>Oceanospirillales>Endozoicomonadaceae>Endozoicomonas | GCF_024606365.1 | Endozoicomonas_sp._ONNA2 | KDK14_RS10760 | 1444..4995 | 2828741 | BREX-system-P-loop-protein-BrxC-[Endozoicomonas-sp.-ONNA2] |
| WP_164887423.1 | Type1.BREX | [illegible] | 1182 | Bacteria>Pseudomonadota>Gammaproteobacteria>Oceanospirillales>Hahellaceae>Hahella | GCF_004014695.1 | Hahella_sp._KA22 | EUZ85_RS02365 | 526761..530309 | 1628392 | BREX-system-P-loop-protein-BrxC-[Hahella-sp.-KA22] |
| MCP5213002.1 | Type1.BREX | [illegible] | 1218 | Bacteria>Pseudomonadota>Gammaproteobacteria>Oceanospirillales>Hahellaceae>unclassified_Hahellaceae | GCA_024234575.1 | Hahellaceae_bacterium | H6998_18960 | 8319..11975 | 2048971 | BREX-system-P-loop-protein-BrxC-[Hahellaceae-bacterium] |
| WP_031385055.1 | Type1.BREX | [illegible] | 1193 | Bacteria>Pseudomonadota>Gammaproteobacteria>Oceanospirillales>Halomonadaceae>Halomonas | GCF_000731955.1 | Halomonas_zincidurans | HALZIN_RS0115275 | 3254805..3258386 | 1178777 | BREX-system-P-loop-protein-BrxC-[Halomonas-zincidurans] |
| WP_019020105.1 | Type1.BREX | [illegible] | 1193 | Bacteria>Pseudomonadota>Gammaproteobacteria>Oceanospirillales>Halomonadaceae>Halomonas | GCF_014651575.1 | Halomonas_lutea | IE067_RS16235 | 61170..64751 | 453962 | BREX-system-P-loop-protein-BrxC-[Halomonas-lutea] |
| WP_239296660.1 | Type1.BREX | [illegible] | 1193 | Bacteria>Pseudomonadota>Gammaproteobacteria>Oceanospirillales>Halomonadaceae>Halomonas | GCF_022348215.1 | unclassified_Halomonas | MHM35_RS10400 | 116133..119714 | 2609666 | MULTISPECIES:-BREX-system-P-loop-protein-BrxC-[unclassified-Halomonas] |
| WP_027854332.1 | Type1.BREX | [illegible] | 1193 | Bacteria>Pseudomonadota>Gammaproteobacteria>Oceanospirillales>Oceanospirillaceae>Marinobacterium | GCF_000428985.1 | Marinobacterium_litorale | G403_RS0112740 | 74962..78543 | 404770 | BREX-system-P-loop-protein-BrxC-[Marinobacterium-litorale] |
| AWY02455.1 | Type1.BREX | [illegible] | 1197 | Bacteria>Pseudomonadota>Gammaproteobacteria>Oceanospirillales>Oceanospirillaceae>Marinomonas | GCA_003285205.1 | Marinomonas_primoryensis | A8139_14855 | 3171018..3174611 | 178399 | hypothetical-protein-A8139_14855-[Marinomonas-primoryensis] |
| WP_239495510.1 | Type1.BREX | [illegible] | 1199 | Bacteria>Pseudomonadota>Gammaproteobacteria>Oceanospirillales>Oceanospirillaceae>Marinomonas | GCF_003285205.1 | Marinomonas_primoryensis | A8139_RS14980 | 3171018..3174617 | 178399 | BREX-system-P-loop-protein-BrxC-[Marinomonas-primoryensis] |
| WP_205113926.1 | Type1.BREX | [illegible] | 1191 | Bacteria>Pseudomonadota>Gammaproteobacteria>Oceanospirillales>Oceanospirillaceae>Marinomonas | GCF_016906065.1 | Marinomonas_foliarum | JSY38_RS15135 | 3246777..3250352 | 491950 | BREX-system-P-loop-protein-BrxC-[Marinomonas-foliarum] |
| MAR01289.1 | Type1.BREX | [illegible] | 1185 | Bacteria>Pseudomonadota>Gammaproteobacteria>Oceanospirillales>Oceanospirillaceae>unclassified_Oceanospirillaceae | GCA_002701445.1 | Oceanospirillaceae_bacterium | CMI00_12230 | 13782..17339 | 1899355 | hypothetical-protein-[Oceanospirillaceae-bacterium] |
| TVT30739.1 | Type1.BREX | [illegible] | 1145 | Bacteria>Pseudomonadota>Gammaproteobacteria>Pseudomonadales>Marinobacteraceae>Marinobacter | GCA_007713625.1 | Marinobacter_vinifirmus | FHK81_16390 | 18158..21595 | 355591 | BREX-system-P-loop-protein-BrxC-[Marinobacter-vinifirmus] |
| WP_022988850.1 | Type1.BREX | [illegible] | 1193 | Bacteria>Pseudomonadota>Gammaproteobacteria>Pseudomonadales>Marinobacteraceae>Marinobacter | GCF_000475255.1 | Marinobacter_sp._ES-1 | Q668_RS06800 | 88554..92135 | 1396858 | BREX-system-P-loop-protein-BrxC-[Marinobacter-sp.-ES-1] |
| WP_022990203.1 | Type1.BREX | [illegible] | 1189 | Bacteria>Pseudomonadota>Gammaproteobacteria>Pseudomonadales>Marinobacteraceae>Marinobacter | GCF_000475255.1 | Marinobacter_sp._ES-1 | Q668_RS13780 | 2583..6152 | 1396858 | BREX-system-P-loop-protein-BrxC-[Marinobacter-sp.-ES-1] |
| WP_023011723.1 | Type1.BREX | [illegible] | 1193 | Bacteria>Pseudomonadota>Gammaproteobacteria>Pseudomonadales>Marinobacteraceae>Marinobacter | GCF_000475315.1 | Marinobacter_sp._EN3 | Q675_RS32885 | 62164..65745 | 1397533 | BREX-system-P-loop-protein-BrxC-[Marinobacter-sp.-EN3] |
| WP_141068072.1 | Type1.BREX | [illegible] | 1193 | Bacteria>Pseudomonadota>Gammaproteobacteria>Pseudomonadales>Marinobacteraceae>Marinobacter | GCF_006516615.1 | Marinobacter_nauticus | FH712_RS17655 | 33566..37147 | 2743 | BREX-system-P-loop-protein-BrxC-[Marinobacter-nauticus] |
| WP_273135111.1 | Type1.BREX | [illegible] | 1145 | Bacteria>Pseudomonadota>Gammaproteobacteria>Pseudomonadales>Marinobacteraceae>Marinobacter | GCF_007713625.1 | Marinobacter_vinifirmus | FHK81_RS16410 | 18158..21595 | 355591 | BREX-system-P-loop-protein-BrxC-[Marinobacter-vinifirmus] |
| WP_247116152.1 | Type1.BREX | [illegible] | 1189 | Bacteria>Pseudomonadota>Gammaproteobacteria>Pseudomonadales>Marinobacteraceae>Marinobacter | GCF_023016045.1 | Marinobacter_sp._S6332 | MWU62_RS17920 | 19915..23484 | 2926403 | BREX-system-P-loop-protein-BrxC-[Marinobacter-sp.-S6332] |
| WP_092033189.1 | Type1.BREX | [illegible] | 1193 | Bacteria>Pseudomonadota>Gammaproteobacteria>Pseudomonadales>Marinobacteraceae>Marinobacter | GCF_900112835.1 | Marinobacter_sp._DSM_26671 | BM035_RS16335 | 75345..78926 | 1761793 | BREX-system-P-loop-protein-BrxC-[Marinobacter-sp.-DSM-26671] |
| WP_092021229.1 | Type1.BREX | [illegible] | 1219 | Bacteria>Pseudomonadota>Gammaproteobacteria>Pseudomonadales>Marinobacteraceae>Marinobacter | GCF_900114775.1 | Marinobacter_zhejiangensis | BMZ67_RS06955 | 334318..337977 | 488535 | BREX-system-P-loop-protein-BrxC-[Marinobacter-zhejiangensis] |
| WP_116918937.1 | Type1.BREX | [illegible] | 1193 | Bacteria>Pseudomonadota>Gammaproteobacteria>Pseudomonadales>Marinobacteraceae>Tamilnaduibacter | GCF_003096655.1 | Tamilnaduibacter_salinus | C8D92_104115 | 123558..127139 | 1484056 | BREX-system-P-loop-protein-BrxC-[Tamilnaduibacter-salinus] |
| WP_105523318.1 | Type1.BREX | [illegible] | 1185 | Bacteria>Pseudomonadota>Gammaproteobacteria>Pseudomonadales>Pseudomonadaceae | GCF_002975295.1 | Pseudomonas | CQ048_RS15780 | 97047..100604 | 286 | MULTISPECIES:-BREX-system-P-loop-protein-BrxC-[Pseudomonas] |
| WP_176516884.1 | Type1.BREX | [illegible] | 1223 | Bacteria>Pseudomonadota>Gammaproteobacteria>Pseudomonadales>Pseudomonadaceae | GCF_013373895.1 | Pseudomonas | HXS67_RS28935 | 22447..26118 | 286 | MULTISPECIES:-BREX-system-P-loop-protein-BrxC-[Pseudomonas] |
| WP_060765715.1 | Type1.BREX | [illegible] | 1186 | Bacteria>Pseudomonadota>Gammaproteobacteria>Pseudomonadales>Pseudomonadaceae | GCF_029959765.1 | Pseudomonas | PFL603g_RS03160 | 383988..387548 | 286 | MULTISPECIES:-BREX-system-P-loop-protein-BrxC-[Pseudomonas] |
| WP_023115268.1 | Type1.BREX | [illegible] | 1154 | Bacteria>Pseudomonadota>Gammaproteobacteria>Pseudomonadales>Pseudomonadaceae | GCF_900707185.1 | Pseudomonas | E4Y87_RS04340 | 6868..10332 | 286 | MULTISPECIES:-BREX-system-P-loop-protein-BrxC-[Pseudomonas] |
| MBG5216128.1 | Type1.BREX | [illegible] | 1154 | Bacteria>Pseudomonadota>Gammaproteobacteria>Pseudomonadales>Pseudomonadaceae>Pseudomonas | GCA_015737185.1 | Pseudomonas_aeruginosa | GHT75_27125 | 7080..10544 | 287 | BREX-system-P-loop-protein-BrxC-[Pseudomonas-aeruginosa] |
| MBG6730505.1 | Type1.BREX | [illegible] | 1184 | Bacteria>Pseudomonadota>Gammaproteobacteria>Pseudomonadales>Pseudomonadaceae>Pseudomonas | GCA_015764285.1 | Pseudomonas_aeruginosa | I5I92_11320 | 4371..7925 | 287 | BREX-system-P-loop-protein-BrxC-[Pseudomonas-aeruginosa] |
| MBX5915342.1 | Type1.BREX | [illegible] | 1154 | Bacteria>Pseudomonadota>Gammaproteobacteria>Pseudomonadales>Pseudomonadaceae>Pseudomonas | GCA_019688355.1 | Pseudomonas_aeruginosa | ISE50_29115 | 6954..10418 | 287 | BREX-system-P-loop-protein-BrxC-[Pseudomonas-aeruginosa] |
| HBO3135817.1 | Type1.BREX | [illegible] | 1184 | Bacteria>Pseudomonadota>Gammaproteobacteria>Pseudomonadales>Pseudomonadaceae>Pseudomonas | GCA_021651475.1 | Pseudomonas_aeruginosa | L4Q96_001363 | 40663..44217 | 287 | BREX-system-P-loop-protein-BrxC-[Pseudomonas-aeruginosa] |
| HBO2945375.1 | Type1.BREX | [illegible] | 1184 | Bacteria>Pseudomonadota>Gammaproteobacteria>Pseudomonadales>Pseudomonadaceae>Pseudomonas | GCA_021651875.1 | Pseudomonas_aeruginosa | L4R51_003866 | 17556..21110 | 287 | BREX-system-P-loop-protein-BrxC-[Pseudomonas-aeruginosa] |
| HBO6336105.1 | Type1.BREX | [illegible] | 1154 | Bacteria>Pseudomonadota>Gammaproteobacteria>Pseudomonadales>Pseudomonadaceae>Pseudomonas | GCA_021670735.1 | Pseudomonas_aeruginosa | L4747_005548 | 6948..10412 | 287 | BREX-system-P-loop-protein-BrxC-[Pseudomonas-aeruginosa] |
| HBO7186446.1 | Type1.BREX | [illegible] | 978 | Bacteria>Pseudomonadota>Gammaproteobacteria>Pseudomonadales>Pseudomonadaceae>Pseudomonas | GCA_021674675.1 | Pseudomonas_aeruginosa | L4968_002122 | 1..2938 | 287 | BREX-system-P-loop-protein-BrxC-[Pseudomonas-aeruginosa] |
| HCE7261574.1 | Type1.BREX | [illegible] | 1154 | Bacteria>Pseudomonadota>Gammaproteobacteria>Pseudomonadales>Pseudomonadaceae>Pseudomonas | GCA_023919205.1 | Pseudomonas_aeruginosa | NHQ13_005985 | 6842..10306 | 287 | BREX-system-P-loop-protein-BrxC-[Pseudomonas-aeruginosa] |
| HCF1762884.1 | Type1.BREX | [illegible] | 1168 | Bacteria>Pseudomonadota>Gammaproteobacteria>Pseudomonadales>Pseudomonadaceae>Pseudomonas | GCA_023941915.1 | Pseudomonas_aeruginosa | NH889_002155 | 1..3508 | 287 | BREX-system-P-loop-protein-BrxC-[Pseudomonas-aeruginosa] |
| HCF5867680.1 | Type1.BREX | [illegible] | 1181 | Bacteria>Pseudomonadota>Gammaproteobacteria>Pseudomonadales>Pseudomonadaceae>Pseudomonas | GCA_023959235.1 | Pseudomonas_aeruginosa | NIH49_002287 | 171515..175060 | 287 | BREX-system-P-loop-protein-BrxC-[Pseudomonas-aeruginosa] |
| WP_132538218.1 | Type1.BREX | [illegible] | 1154 | Bacteria>Pseudomonadota>Gammaproteobacteria>Pseudomonadales>Pseudomonadaceae>Pseudomonas | GCA_030820635.1 | Pseudomonas_aeruginosa | EQ548_RS01360 | 6947..10411 | 287 | BREX-system-P-loop-protein-BrxC-[Pseudomonas-aeruginosa] |
| WP_257648015.1 | Type1.BREX | [illegible] | 1073 | Bacteria>Pseudomonadota>Gammaproteobacteria>Pseudomonadales>Pseudomonadaceae>Pseudomonas | GCA_030820635.1 | Pseudomonas_aeruginosa | K3Z95_RS14360 | 10117..13340 | 287 | BREX-system-P-loop-protein-BrxC,-partial-[Pseudomonas-aeruginosa] |
| WP_010463595.1 | Type1.BREX | [illegible] | 1185 | Bacteria>Pseudomonadota>Gammaproteobacteria>Pseudomonadales>Pseudomonadaceae>Pseudomonas | GCF_000257545.3 | Pseudomonas_mandelii | OU5_RS13450 | 3003724..3007281 | 75612 | BREX-system-P-loop-protein-BrxC-[Pseudomonas-mandelii] |
| WP_033954069.1 | Type1.BREX | [illegible] | 1154 | Bacteria>Pseudomonadota>Gammaproteobacteria>Pseudomonadales>Pseudomonadaceae>Pseudomonas | GCF_000793385.1 | Pseudomonas_aeruginosa | NT88_RS29575 | 15006..18470 | 287 | BREX-system-P-loop-protein-BrxC-[Pseudomonas-aeruginosa] |
| WP_048385719.1 | Type1.BREX | [illegible] | 1185 | Bacteria>Pseudomonadota>Gammaproteobacteria>Pseudomonadales>Pseudomonadaceae>Pseudomonas | GCF_001043065.1 | Pseudomonas_helleri | TU85_RS02975 | 88982..92539 | 1608996 | BREX-system-P-loop-protein-BrxC-[Pseudomonas-helleri] |
| WP_116805555.1 | Type1.BREX | [illegible] | 1154 | Bacteria>Pseudomonadota>Gammaproteobacteria>Pseudomonadales>Pseudomonadaceae>Pseudomonas | GCF_003412045.2 | Pseudomonas_aeruginosa | L4E78_005908 | 8239..11703 | 287 | BREX-system-P-loop-protein-BrxC-[Pseudomonas-aeruginosa] |
| WP_128704208.1 | Type1.BREX | [illegible] | 1184 | Bacteria>Pseudomonadota>Gammaproteobacteria>Pseudomonadales>Pseudomonadaceae>Pseudomonas | GCF_003974605.1 | Pseudomonas_aeruginosa | IPC1390_RS17360 | 217632..221186 | 287 | BREX-system-P-loop-protein-BrxC-[Pseudomonas-aeruginosa] |
| WP_134242485.1 | Type1.BREX | [illegible] | 1154 | Bacteria>Pseudomonadota>Gammaproteobacteria>Pseudomonadales>Pseudomonadaceae>Pseudomonas | GCF_004370105.1 | Pseudomonas_aeruginosa | IPC1592_RS30135 | 15141..18605 | 287 | BREX-system-P-loop-protein-BrxC-[Pseudomonas-aeruginosa] |
| WP_202876546.1 | Type1.BREX | [illegible] | 531 | Bacteria>Pseudomonadota>Gammaproteobacteria>Pseudomonadales>Pseudomonadaceae>Pseudomonas | GCF_004373095.1 | Pseudomonas_aeruginosa | IPC174_RS30040 | 16693..18288 | 287 | BREX-system-P-loop-protein-BrxC-[Pseudomonas-aeruginosa] |
| WP_202876713.1 | Type1.BREX | [illegible] | 539 | Bacteria>Pseudomonadota>Gammaproteobacteria>Pseudomonadales>Pseudomonadaceae>Pseudomonas | GCF_004373095.1 | Pseudomonas_aeruginosa | IPC175_RS30340 | 15284..16903 | 287 | hypothetical-protein-[Pseudomonas-aeruginosa] |
| WP_134282343.1 | Type1.BREX | [illegible] | 1154 | Bacteria>Pseudomonadota>Gammaproteobacteria>Pseudomonadales>Pseudomonadaceae>Pseudomonas | GCF_004373575.1 | Pseudomonas_aeruginosa | IPC135_RS30160 | 15285..18749 | 287 | BREX-system-P-loop-protein-BrxC-[Pseudomonas-aeruginosa] |
| WP_202876545.1 | Type1.BREX | [illegible] | 624 | Bacteria>Pseudomonadota>Gammaproteobacteria>Pseudomonadales>Pseudomonadaceae>Pseudomonas | GCF_004373665.1 | Pseudomonas_aeruginosa | IPC174_RS30035 | 14826..16700 | 287 | hypothetical-protein-[Pseudomonas-aeruginosa] |
| WP_150073350.1 | Type1.BREX | [illegible] | 1186 | Bacteria>Pseudomonadota>Gammaproteobacteria>Pseudomonadales>Pseudomonadaceae>Pseudomonas | GCF_008632325.1 | Pseudomonas_lactis | F3K52_RS19340 | 4156..7716 | 1615674 | BREX-system-P-loop-protein-BrxC-[Pseudomonas-lactis] |
| WP_152745526.1 | Type1.BREX | [illegible] | 1183 | Bacteria>Pseudomonadota>Gammaproteobacteria>Pseudomonadales>Pseudomonadaceae>Pseudomonas | GCF_009296165.1 | Pseudomonas_kitaguniensis | F0169_RS03300 | 21652..25203 | 2607908 | BREX-system-P-loop-protein-BrxC-[Pseudomonas-kitaguniensis] |
| WP_153384138.1 | Type1.BREX | [illegible] | 1186 | Bacteria>Pseudomonadota>Gammaproteobacteria>Pseudomonadales>Pseudomonadaceae>Pseudomonas | GCF_009600565.1 | unclassified_Pseudomonas | GHO24_RS06625 | 35218..38778 | 196821 | MULTISPECIES:-BREX-system-P-loop-protein-BrxC-[unclassified-Pseudomonas] |
| WP_177095197.1 | Type1.BREX | [illegible] | 1185 | Bacteria>Pseudomonadota>Gammaproteobacteria>Pseudomonadales>Pseudomonadaceae>Pseudomonas | GCF_013387065.1 | Pseudomonas_reactans | HX796_RS11550 | 4461..8018 | 117680 | BREX-system-P-loop-protein-BrxC-[Pseudomonas-reactans] |
| WP_196476665.1 | Type1.BREX | [illegible] | 1154 | Bacteria>Pseudomonadota>Gammaproteobacteria>Pseudomonadales>Pseudomonadaceae>Pseudomonas | GCF_015712105.2 | Pseudomonas_sp._LMG_31766 | H3221_RS24535 | 30166..33630 | 2758695 | BREX-system-P-loop-protein-BrxC-[Pseudomonas-chaetocerotis] |
| WP_198790924.1 | Type1.BREX | [illegible] | 1185 | Bacteria>Pseudomonadota>Gammaproteobacteria>Pseudomonadales>Pseudomonadaceae>Pseudomonas | GCF_016405175.1 | Pseudomonas_sp._MF7451 | JFT65_RS10900 | 194530..198087 | 2797538 | BREX-system-P-loop-protein-BrxC-[Pseudomonas-sp.-MF7451] |
| WP_223023201.1 | Type1.BREX | [illegible] | 1183 | Bacteria>Pseudomonadota>Gammaproteobacteria>Pseudomonadales>Pseudomonadaceae>Pseudomonas | GCF_019891425.1 | Pseudomonas_carnis | J1G36_06600 | 89435..92986 | 2487355 | BREX-system-P-loop-protein-BrxC-[Pseudomonas-carnis] |
| WP_236238652.1 | Type1.BREX | [illegible] | 1185 | Bacteria>Pseudomonadota>Gammaproteobacteria>Pseudomonadales>Pseudomonadaceae>Pseudomonas | GCF_021603345.1 | Pseudomonas_rhodesiae | L6E14_RS13730 | 266364..269921 | 76760 | BREX-system-P-loop-protein-BrxC-[Pseudomonas-rhodesiae] |
| WP_260092499.1 | Type1.BREX | [illegible] | 1154 | Bacteria>Pseudomonadota>Gammaproteobacteria>Pseudomonadales>Pseudomonadaceae>Pseudomonas | GCF_025152895.1 | Pseudomonas_aeruginosa | MNU23_RS16440 | 110369..113833 | 287 | BREX-system-P-loop-protein-BrxC-[Pseudomonas-aeruginosa] |
| WP_124171099.1 | Type1.BREX | [illegible] | 1184 | Bacteria>Pseudomonadota>Gammaproteobacteria>Pseudomonadales>Pseudomonadaceae>Pseudomonas | GCF_025697325.1 | Pseudomonas_aeruginosa | IPC368_RS11155 | 506506..510060 | 287 | BREX-system-P-loop-protein-BrxC-[Pseudomonas-aeruginosa] |
| WP_265072431.1 | Type1.BREX | [illegible] | 1186 | Bacteria>Pseudomonadota>Gammaproteobacteria>Pseudomonadales>Pseudomonadaceae>Pseudomonas | GCF_026016265.1 | Pseudomonas_viciae | LOY66_RS25465 | 5798638..5802198 | 2505979 | BREX-system-P-loop-protein-BrxC-[Pseudomonas-viciae] |
| WP_090192550.1 | Type1.BREX | [illegible] | 1153 | Bacteria>Pseudomonadota>Gammaproteobacteria>Pseudomonadales>Pseudomonadaceae>Pseudomonas | GCF_900105995.1 | Pseudomonas_pohangensis | BLT89_RS00315 | 68625..72086 | 364197 | BREX-system-P-loop-protein-BrxC-[Pseudomonas-pohangensis] |

<table>
<tr><th>Protein-ID</th><th>System</th><th>Gene-Neighbourhoods</th><th>Length</th><th>Taxonomy</th><th>Assembly-ID</th><th>Strain</th><th>Locus-Tag</th><th>Genomic-coordinates</th><th>TaxID</th><th>Fasta-defline</th></tr>
<tr><td>WP_031684509.1</td><td>Type1.BREX</td><td>PhdYeFM_antitox->?-><- wHTH+WYL+WCX ||?->Peptidase_M15->?->XIS-HTH-> BrxA-DNA-binding -> BrxB-BrxF-iSTAND -> DUF6079 *-> N6-METHYLASE ->SbcC-p-> REase ->?-> N6-METHYLASE -p-> PglZ -> SIGMA+OB-Fold+AAA-ATPase+HKD-DNase-BrxL -><-Arm-DNA-bind_3+Phage_integrase||SF1-SF2-Helicase-></td><td>1184</td><td>Bacteria>Pseudomonadota>Gammaproteobacteria>Pseudomonadales>Pseudomonadaceae>Pseudomonas</td><td>GCF_900708485.1</td><td>Pseudomonas_aeruginosa</td><td>ID019_RS13300</td><td>2848972..2852526</td><td>287</td><td>BREX-system-P-loop-protein-BrxC-[Pseudomonas-aeruginosa]</td></tr>
<tr><td>WP_033991202.1</td><td>Type1.BREX</td><td><-HNH<- SIGMA+OB-Fold+AAA-ATPase+Lon-Protease-BrxL <- PglZ <-MNS+ REase <-?<-HNH+GCN1<-?<- N6-METHYLASE <- DUF6079 *<- BrxB-BrxF-iSTAND <- BrxA-DNA-binding <-RNase<-?<-JetB<-?<-Peptidase_M15<-?|| wHTH+WYL+WCX -></td><td>1154</td><td>Bacteria>Pseudomonadota>Gammaproteobacteria>Pseudomonadales>Pseudomonadaceae>Pseudomonas</td><td>GCF_929614685.1</td><td>Pseudomonas_aeruginosa</td><td>MJP43_RS27275</td><td>28461..31925</td><td>287</td><td>BREX-system-P-loop-protein-BrxC-[Pseudomonas-aeruginosa]</td></tr>
<tr><td>EIU5588796.1</td><td>Type1.BREX</td><td>PhdYeFM_antitox->RelE-ParE-><-?<- wHTH+WYL+WCX ||?->Peptidase_M15->?-> BrxA-DNA-binding -> BrxB-BrxF-iSTAND -> DUF6079 *-> N6-METHYLASE ->?->HNH+GCN1->?->ParB+HNH-> PglZ -> SIGMA+OB-Fold+AAA-ATPase+Lon-Protease-BrxL -> REase ->SF1-SF2-Helicase-></td><td>1154</td><td>Bacteria>Pseudomonadota>Gammaproteobacteria>Pseudomonadales>Pseudomonadaceae>Pseudomonas</td><td>-</td><td>Pseudomonas_aeruginosa</td><td>L4689_001050</td><td>82930..86394</td><td>287</td><td>BREX-system-P-loop-protein-BrxC-[Pseudomonas-aeruginosa]</td></tr>
<tr><td>WP_198809634.1</td><td>Type5.BREX</td><td><- TUDOR+SF1-SF2-Helicase+i REase +CTD <-HHH_4+ SF1-SF2-Helicase+ REase <-?||?-> wHTH+WYL+WCX -> BrxA-DNA-binding -> BrxB-BrxF-iSTAND -> DUF6079 *->?->?-> N6-METHYLASE ->GP+ REase -> PglZ -> SIGMA+OB-Fold+AAA-ATPase+Lon-Protease-BrxL -><-tRNA<-MerR-HTH</td><td>1224</td><td>Bacteria>Pseudomonadota>Gammaproteobacteria>Pseudomonadales>Pseudomonadaceae>Pseudomonas</td><td>-</td><td>Pseudomonas_shahriarae</td><td>NEE03_RS00560</td><td>109508..113182</td><td>2745512</td><td>BREX-system-P-loop-protein-BrxC-[Pseudomonas-shahriarae]</td></tr>
<tr><td>MCQ4256509.1</td><td>Type1.BREX</td><td><-SF1-SF2-Helicase<- REase <- SIGMA+OB-Fold+AAA-ATPase+Lon-Protease-BrxL <- PglZ <-ParB+HNH<-?<-MarR-HTH<-MNS+ORC-CDC6<- N6-METHYLASE <- DUF6079 *<- BrxB-BrxF-iSTAND <- BrxA-DNA-binding <- REase <-NERD||?->DUF932-></td><td>1154</td><td>Bacteria>Pseudomonadota>Gammaproteobacteria>Pseudomonadales>Pseudomonadaceae>Stutzerimonas</td><td>GCA_024448895.1</td><td>Stutzerimonas_stutzeri</td><td>NA619_00645</td><td>147157..150621</td><td>316</td><td>BREX-system-P-loop-protein-BrxC-[Stutzerimonas-stutzeri]</td></tr>
<tr><td>WP_218421912.1</td><td>Type1.BREX</td><td><- SIGMA+OB-Fold+AAA-ATPase+Lon-Protease-BrxL <- PglZ <- REase <- N6-METHYLASE -p<-?<-HerA/FtsK<-SIR2<- N6-METHYLASE <- DUF6079 *<- BrxB-BrxF-iSTAND <- BrxA-DNA-binding <- wHTH+WYL+WCX ||?->MORC->RVT-></td><td>1223</td><td>Bacteria>Pseudomonadota>Gammaproteobacteria>Pseudomonadales>Pseudomonadaceae>Stutzerimonas</td><td>GCF_019203165.1</td><td>Stutzerimonas_stutzeri</td><td>KUD99_RS18220</td><td>3895806..3899477</td><td>316</td><td>BREX-system-P-loop-protein-BrxC-[Stutzerimonas-stutzeri]</td></tr>
<tr><td>WP_125870426.1</td><td>Type1.BREX</td><td><-DUF932<-?||NERD-> REase -> BrxA-DNA-binding -> BrxB-BrxF-iSTAND -> DUF6079 *-> N6-METHYLASE ->?->HNH+GCN1->?-> PglZ -> SIGMA+OB-Fold+AAA-ATPase+Lon-Protease-BrxL -> REase ->SF1-SF2-Helicase-><-Glycos_transf</td><td>1154</td><td>Bacteria>Pseudomonadota>Gammaproteobacteria>Pseudomonadales>Pseudomonadaceae>Stutzerimonas</td><td>GCF_024199075.1</td><td>Stutzerimonas_stutzeri</td><td>NLG33_RS12680</td><td>207505..210969</td><td>316</td><td>BREX-system-P-loop-protein-BrxC-[Stutzerimonas-stutzeri]</td></tr>
<tr><td>WP_255958979.1</td><td>Type1.BREX</td><td><-SF1-SF2-Helicase<- REase <- SIGMA+OB-Fold+AAA-ATPase+Lon-Protease-BrxL <- PglZ <-ParB+HNH<-?<-MarR-HTH<-MNS+ORC-CDC6<- N6-METHYLASE <- DUF6079 *<- BrxB-BrxF-iSTAND <- BrxA-DNA-binding <- REase <-NERD||?->DUF932-></td><td>1154</td><td>Bacteria>Pseudomonadota>Gammaproteobacteria>Pseudomonadales>Pseudomonadaceae>Stutzerimonas</td><td>GCF_024448895.1</td><td>Stutzerimonas_stutzeri</td><td>NA619_RS00645</td><td>147157..150621</td><td>316</td><td>BREX-system-P-loop-protein-BrxC-[Stutzerimonas-stutzeri]</td></tr>
<tr><td>WP_269263428.1</td><td>Type1.BREX</td><td><-HerA/FtsK<-SIR2<- N6-METHYLASE -p<-GP+ REase <- N6-METHYLASE <-DOC+HTH<-?<-?<- DUF6079 *<- BrxB-BrxF-iSTAND <- BrxA-DNA-binding ||Integrase_1->Phage_int_SAM-><-?||DSBH->?-><-Phage_integrase||DUF772+DDE_Tnp_1-></td><td>1223</td><td>Bacteria>Pseudomonadota>Gammaproteobacteria>Pseudomonadales>Pseudomonadaceae>Stutzerimonas</td><td>GCF_027158265.1</td><td>Stutzerimonas_balearica</td><td>D3V18_RS18735</td><td>22059..25730</td><td>74829</td><td>BREX-system-P-loop-protein-BrxC-[Stutzerimonas-balearica]</td></tr>
<tr><td>MAQ23221.1</td><td>Type1.BREX</td><td><-Calcineurin+Calcineurin-SbcD<- PglZ <-ParB+HNH<- N6-METHYLASE <- REase <-OLD-ABC <-Schlafen+HATPase+wHTH+HTH<- DUF6079 *<- BrxB-BrxF-iSTAND <- BrxA-DNA-binding <-XIS-HTH<-MobI||XIS-HTH->AAA+HTH->XIS-HTH-><-Arm-DNA-bind_3+Phage_integrase</td><td>1193</td><td>Bacteria>Pseudomonadota>Gammaproteobacteria>Pseudomonadales>unclassified_Pseudomonadales</td><td>GCA_002701315.1</td><td>Pseudomonadales_bacterium</td><td>CMK83_03295</td><td>18015..21596</td><td>1891229</td><td>hypothetical-protein-[Pseudomonadales-bacterium]</td></tr>
<tr><td>MCP5344826.1</td><td>Type1.BREX</td><td><-PALP<- SIGMA+OB-Fold+AAA-ATPase+Lon-Protease-BrxL <- PglZ <-PHP+SbcC<- N6-METHYLASE <- REase <-OLD-ABC <- DUF6079 -p*</td><td>834</td><td>Bacteria>Pseudomonadota>Gammaproteobacteria>Pseudomonadales>unclassified_Pseudomonadales</td><td>GCA_024234345.1</td><td>Pseudomonadales_bacterium</td><td>H7A05_09410</td><td>50993..53498</td><td>1891229</td><td>BREX-system-P-loop-protein-BrxC-[Pseudomonadales-bacterium]</td></tr>
<tr><td>WP_007146327.1</td><td>Type1.BREX</td><td>UbiA-><-HNH<-tRNA<-?<-?<- SIGMA+OB-Fold+AAA-ATPase+HKD-DNase-BrxL <- PglZ <- REase <-ParB+HNH<- N6-METHYLASE <- DUF6079 *<- BrxB-BrxF-iSTAND <- BrxA-DNA-binding <-NADAR<-ART-DarT<-?<-?<-Calcineurin</td><td>1179</td><td>Bacteria>Pseudomonadota>Gammaproteobacteria>Thiotrichales>Piscirickettsiaceae>Methylophaga</td><td>GCF_000214595.1</td><td>Methylophaga_aminisulfidivorans</td><td>MAMP_RS12010</td><td>87136..90675</td><td>230105</td><td>BREX-system-P-loop-protein-BrxC-[Methylophaga-aminisulfidivorans]</td></tr>
<tr><td>CAA6806547.1</td><td>Type1.BREX</td><td><-ParB+HNH<- N6-METHYLASE <- N6-METHYLASE -p<-?<- REase <-ABC-ATPase<- DUF6079 -p*</td><td>849</td><td>Bacteria>Pseudomonadota>Gammaproteobacteria>Thiotrichales>Thiotrichaceae>environmental_samples</td><td>-</td><td>uncultured_Thiotrichaceae_bacterium</td><td>HELGO_WM31114</td><td>6431..8980</td><td>298394</td><td>FIG00607287:-hypothetical-protein,-partial-[uncultured-Thiotrichaceae-bacterium]</td></tr>
<tr><td>WP_210225661.1</td><td>Type1.BREX</td><td>BrxA-DNA-binding -> BrxB-BrxF-iSTAND -> DUF6079 *->ABC-ATPase->?-> N6-METHYLASE -> N6-METHYLASE -p-><- REase ||HATPase+wHTH-> PglZ -> SIGMA+OB-Fold+AAA-ATPase+HKD-DNase-BrxL -></td><td>1173</td><td>Bacteria>Pseudomonadota>Gammaproteobacteria>Thiotrichales>Thiotrichaceae>Thiothrix</td><td>GCF_017901155.1</td><td>Candidatus_Thiothrix_anitrata</td><td>J8380_RS10820</td><td>2094889..2098410</td><td>2823902</td><td>BREX-system-P-loop-protein-BrxC-[Candidatus-Thiothrix-anitrata]</td></tr>
<tr><td>PCJ88241.1</td><td>Type1.BREX</td><td><-RelE-ParE<-MetJarc-HTH||?-><-HKD-DNase+DUF3150<- SIGMA+OB-Fold+AAA-ATPase+HKD-DNase-BrxL <- PglZ <- N6-METHYLASE <- REase <-OLD-ABC <- DUF6079 *<- BrxB-BrxF-iSTAND <- BrxA-DNA-binding <-XIS-HTH<-?<-?||RHH_5->?-><-HISKIN</td><td>1177</td><td>Bacteria>Pseudomonadota>Gammaproteobacteria>Thiotrichales>Thiotrichaceae>unclassified_Thiotrichaceae</td><td>GCA_002746365.1</td><td>Thiotrichaceae_bacterium</td><td>COA54_02955</td><td>15853..19386</td><td>2030882</td><td>hypothetical-protein-COA54_02955-[Thiotrichaceae-bacterium]</td></tr>
<tr><td>HIJ32929.1</td><td>Type1.BREX</td><td>BrxA-DNA-binding -> BrxB-BrxF-iSTAND -> DUF6079 *-> OLD-ABC-> REase -> N6-METHYLASE ->p-></td><td>1200</td><td>Bacteria>Pseudomonadota>Gammaproteobacteria>unclassified_Gammaproteobacteria</td><td>GCA_013349805.1</td><td>Gammaproteobacteria_bacterium</td><td>HPP65_00765</td><td>772..4374</td><td>1913989</td><td>BREX-system-P-loop-protein-BrxC-[Gammaproteobacteria-bacterium]</td></tr>
<tr><td>MBT4196271.1</td><td>Type1.BREX</td><td><- SIGMA+OB-Fold+AAA-ATPase+HKD-DNase-BrxL <- REase <-Immank+ANK+ANK<- PglZ <- N6-METHYLASE <-SLOG+BirA-HTH<-SF1-SF2-Helicase+PRTase<- DUF6079 *<- BrxB-BrxF-iSTAND <- BrxA-DNA-binding <-HTH_26||?->?-> REase ->rve+Mu-transpos_C->TnsC-ATPase->ZnR-></td><td>1178</td><td>Bacteria>Pseudomonadota>Gammaproteobacteria>unclassified_Gammaproteobacteria</td><td>GCA_018651435.1</td><td>Gammaproteobacteria_bacterium</td><td>HOE12_18240</td><td>31044..34580</td><td>1913989</td><td>BREX-system-P-loop-protein-BrxC-[Gammaproteobacteria-bacterium]</td></tr>
<tr><td>WP_245395138.1</td><td>Type1.BREX</td><td>LON-AAA->Calcineurin->SbcD->Calcineurin-><-?||?->|| BrxA-DNA-binding -> DUF6079 -p-><- REase <-OLD-ABC <- DUF6079 *<-HDRnase||<-Arm-DNA-bind_2+Phage_int_SAM_3+Phage_integrase<-Phage_AlpA||HTH_26-><-?<-tRNA||RADICAL-SAM->?->DDE_Tnp_1-></td><td>1115</td><td>Bacteria>Pseudomonadota>Gammaproteobacteria>unclassified_Gammaproteobacteria</td><td>GCF_016097405.1</td><td>methane-oxidizing_endosymbiont_of_Gigantopelta_aegis</td><td>JEU20_RS06805</td><td>1356880..1360227</td><td>2794938</td><td>BREX-system-P-loop-protein-BrxC-[methane-oxidizing-endosymbiont-of-Gigantopelta-aegis]</td></tr>
<tr><td>WP_006647488.1</td><td>Type1.BREX</td><td>HTH_Tnp_1+HTH_21+rve+rve_3-><- SIGMA+OB-Fold+AAA-ATPase+HKD-DNase-BrxL <- PglZ <-ParB+HNH<- N6-METHYLASE <- REase <-OLD-ABC <-HTH_Tnp_1+HTH_21+rve+rve_3<-OLD-ABC<- DUF6079 *<- BrxB-BrxF-iSTAND <- BrxA-DNA-binding <-XIS-HTH<-?<-TnsC-ATPase||HTH_Tnp_1+HTH_21+rve+rve_3-><-rve+Mu-transpos_C<- REase +wHTH</td><td>1193</td><td>Bacteria>Pseudomonadota>Gammaproteobacteria>Vibrionales>Vibrionaceae>Photobacterium</td><td>GCA_000153325.1</td><td>Photobacterium_sp._SKA34</td><td>SKA34_RS21260</td><td>87534..91115</td><td>121723</td><td>BREX-system-P-loop-protein-BrxC-[Photobacterium-sp.-SKA34]</td></tr>
<tr><td>ASG04983.1</td><td>Type1.BREX</td><td><-DeoR-HTH+ISOCOT<-DeoR-HTH+ISOCOT<-DUF2787<-?<-?||XIS-HTH->XIS-HTH-> BrxA-DNA-binding -> BrxB-BrxF-iSTAND -> DUF6079 *-> OLD-ABC-> REase -> N6-METHYLASE -> PglZ -> SIGMA+OB-Fold+AAA-ATPase+HKD-DNase-BrxL -><-Arm-DNA-bind_3+Phage_integrase</td><td>1185</td><td>Bacteria>Pseudomonadota>Gammaproteobacteria>Vibrionales>Vibrionaceae>Vibrio</td><td>GCA_002211985.1</td><td>Vibrio_anguillarum</td><td>CEJ46_04105</td><td>862780..866337</td><td>55601</td><td>hypothetical-protein-CEJ46_04105-[Vibrio-anguillarum]</td></tr>
<tr><td>NAZ69809.1</td><td>Type1.BREX</td><td><- SIGMA+OB-Fold+AAA-ATPase+HKD-DNase-BrxL <- PglZ <- N6-METHYLASE <- REase <-OLD-ABC <- DUF6079 *<- BrxB-BrxF-iSTAND <- BrxA-DNA-binding <-XIS-HTH<-HNH<-?<-XIS-HTH||?->DDE_Tnp_1_5+DDE_Tnp_1-></td><td>1185</td><td>Bacteria>Pseudomonadota>Gammaproteobacteria>Vibrionales>Vibrionaceae>Vibrio</td><td>GCA_009906175.1</td><td>Vibrio_toranzoniae</td><td>GL180_08045</td><td>10464..14021</td><td>1194427</td><td>BREX-system-P-loop-protein-BrxC-[Vibrio-toranzoniae]</td></tr>
<tr><td>EGQ8448095.1</td><td>Type1.BREX</td><td><- SIGMA+OB-Fold+AAA-ATPase+HKD-DNase-BrxL <- PglZ <- N6-METHYLASE <- REase <-OLD-ABC <- DUF6079 *<- BrxB-BrxF-iSTAND <- BrxA-DNA-binding <-XIS-HTH<-HNH<-XIS-HTH||?->?->DUF2787-> wHTH+WYL+WCX -></td><td>1185</td><td>Bacteria>Pseudomonadota>Gammaproteobacteria>Vibrionales>Vibrionaceae>Vibrio</td><td>GCA_015775815.1</td><td>Vibrio_alginolyticus</td><td>GTP59_14930</td><td>64166..67723</td><td>663</td><td>BREX-system-P-loop-protein-BrxC-[Vibrio-alginolyticus]</td></tr>
<tr><td>MBH9740666.1</td><td>Type1.BREX</td><td>SF1-SF2-Helicase+HTH->MukF->JetB->XIS-HTH-> BrxA-DNA-binding -> BrxB-BrxF-iSTAND -> DUF6079 *-> OLD-ABC-> REase -> N6-METHYLASE -> PglZ -> SIGMA+OB-Fold+AAA-ATPase+HKD-DNase-BrxL -></td><td>1185</td><td>Bacteria>Pseudomonadota>Gammaproteobacteria>Vibrionales>Vibrionaceae>Vibrio</td><td>GCA_016094345.1</td><td>Vibrio_navarrensis</td><td>CJ419_13180</td><td>152917..156474</td><td>29495</td><td>hypothetical-protein-[Vibrio-navarrensis]</td></tr>
<tr><td>MBT0011870.1</td><td>Type1.BREX</td><td><- SIGMA+OB-Fold+AAA-ATPase+HKD-DNase-BrxL <- PglZ <- N6-METHYLASE <- REase <-OLD-ABC <- DUF6079 *<- BrxB-BrxF-iSTAND <- BrxA-DNA-binding <-XIS-HTH<-HNH<-XIS-HTH||?->?->DUF2787-> wHTH+WYL+WCX -></td><td>1187</td><td>Bacteria>Pseudomonadota>Gammaproteobacteria>Vibrionales>Vibrionaceae>Vibrio</td><td>GCA_018457755.1</td><td>Vibrio_alginolyticus</td><td>N7I24_002738</td><td>64007..67570</td><td>663</td><td>BREX-system-P-loop-protein-BrxC-[Vibrio-alginolyticus]</td></tr>
<tr><td>EIZ1010574.1</td><td>Type1.BREX</td><td>HTH_3->Couple_hipA+HipA_C->SAM-METHYLASE->HKD-DNase+SF1-SF2-Helicase+HTH+HTH+SRA-><- SIGMA+OB-Fold+AAA-ATPase+HKD-DNase-BrxL <- PglZ <- N6-METHYLASE <- REase <-OLD-ABC <- DUF6079 *<- BrxB-BrxF-iSTAND <- BrxA-DNA-binding <-XIS-HTH<-RNase<-DUF3584+DUF5401<-JetB<-MukF|| wHTH+WYL+WCX -><-TPR</td><td>1187</td><td>Bacteria>Pseudomonadota>Gammaproteobacteria>Vibrionales>Vibrionaceae>Vibrio</td><td>GCA_022609845.1</td><td>Vibrio_vulnificus</td><td>MGS09_002154</td><td>288384..291947</td><td>672</td><td>BREX-system-P-loop-protein-BrxC-[Vibrio-vulnificus]</td></tr>
<tr><td>HCG7393723.1</td><td>Type1.BREX</td><td>McrB-AAA-> McrC-NTD+ REase ->?->HNH->?-><-HEPN||XIS-HTH-> BrxA-DNA-binding -> BrxB-BrxF-iSTAND -> DUF6079 *-> OLD-ABC-> REase -> N6-METHYLASE -> PglZ -> SIGMA+OB-Fold+AAA-ATPase+HKD-DNase-BrxL ->Phage_integrase-><-ParB<-GDResolvase+HIN-HTH||OLD-ABC-></td><td>1187</td><td>Bacteria>Pseudomonadota>Gammaproteobacteria>Vibrionales>Vibrionaceae>Vibrio</td><td>GCA_024044075.1</td><td>Vibrio_parahaemolyticus</td><td>NJ383_004129</td><td>13694..17257</td><td>670</td><td>BREX-system-P-loop-protein-BrxC-[Vibrio-parahaemolyticus]</td></tr>
<tr><td>EJV5950193.1</td><td>Type1.BREX</td><td><- SIGMA+OB-Fold+AAA-ATPase+HKD-DNase-BrxL <- PglZ <- N6-METHYLASE <- REase <-OLD-ABC <- DUF6079 *<- BrxB-BrxF-iSTAND <- BrxA-DNA-binding <-XIS-HTH<-HNH<-XIS-HTH||?->?->DUF2787-> wHTH+WYL+WCX -></td><td>1187</td><td>Bacteria>Pseudomonadota>Gammaproteobacteria>Vibrionales>Vibrionaceae>Vibrio</td><td>GCA_025320405.1</td><td>Vibrio_alginolyticus</td><td>N7I24_002738</td><td>64007..67570</td><td>663</td><td>BREX-system-P-loop-protein-BrxC-[Vibrio-alginolyticus]</td></tr>
<tr><td>GLT15886.1</td><td>Type1.BREX</td><td><- wHTH+WYL+WCX ||MukF->JetB->DUF3584+DUF5401->RNase->XIS-HTH-> BrxA-DNA-binding -> BrxB-BrxF-iSTAND -> DUF6079 *-> OLD-ABC-> REase -> N6-METHYLASE -> PglZ -> SIGMA+OB-Fold+AAA-ATPase+HKD-DNase-BrxL ->?->HTH_3->Couple_hipA+HipA_C-></td><td>1186</td><td>Bacteria>Pseudomonadota>Gammaproteobacteria>Vibrionales>Vibrionaceae>Vibrio</td><td>GCA_030161215.1</td><td>Vibrio_algivorus</td><td>GCM10007931_28610</td><td>496776..500336</td><td>1667024</td><td>hypothetical-protein-GCM10007931_28610-[Vibrio-algivorus]</td></tr>
<tr><td>WP_198300681.1</td><td>Type1.BREX</td><td><-DeoR-HTH+ISOCOT<-DeoR-HTH+ISOCOT<-?<-?||XIS-HTH->XIS-HTH-> BrxA-DNA-binding -> BrxB-BrxF-iSTAND -> DUF6079 *-> OLD-ABC-> REase -> N6-METHYLASE -> PglZ -> SIGMA+OB-Fold+AAA-ATPase+HKD-DNase-BrxL -><-Arm-DNA-bind_3+Phage_integrase</td><td>1187</td><td>Bacteria>Pseudomonadota>Gammaproteobacteria>Vibrionales>Vibrionaceae>Vibrio</td><td>GCF_002211985.1</td><td>Vibrio_anguillarum</td><td>CEJ46_RS04105</td><td>862774..866337</td><td>55601</td><td>BREX-system-P-loop-protein-BrxC-[Vibrio-anguillarum]</td></tr>
<tr><td>WP_089123599.1</td><td>Type1.BREX</td><td><-Couple_hipA+HipA_C<-HTH_3<-?<- SIGMA+OB-Fold+AAA-ATPase+HKD-DNase-BrxL <- PglZ <- N6-METHYLASE <- REase <-OLD-ABC <- DUF6079 *<- BrxB-BrxF-iSTAND <- BrxA-DNA-binding <-XIS-HTH<-RNase<-DUF3584+DUF5401<-JetB<-MukF|| wHTH+WYL+WCX -></td><td>1184</td><td>Bacteria>Pseudomonadota>Gammaproteobacteria>Vibrionales>Vibrionaceae>Vibrio</td><td>GCF_002218005.2</td><td>Vibrio_algivorus</td><td>AVR_RS07075</td><td>1502603..1506157</td><td>1667024</td><td>BREX-system-P-loop-protein-BrxC-[Vibrio-algivorus]</td></tr>
<tr><td>WP_202601719.1</td><td>Type1.BREX</td><td><- SIGMA+OB-Fold+AAA-ATPase+HKD-DNase-BrxL <- PglZ <- N6-METHYLASE <- REase <-OLD-ABC <- DUF6079 *<- BrxB-BrxF-iSTAND <- BrxA-DNA-binding <-XIS-HTH<-HNH<-SF1-SF2-Helicase+HTH<-SF1-SF2-Helicase<-?<-XIS-HTH</td><td>1187</td><td>Bacteria>Pseudomonadota>Gammaproteobacteria>Vibrionales>Vibrionaceae>Vibrio</td><td>GCF_009906175.1</td><td>Vibrio_toranzoniae</td><td>GL180_RS08040</td><td>10464..14027</td><td>1194427</td><td>BREX-system-P-loop-protein-BrxC-[Vibrio-toranzoniae]</td></tr>
<tr><td>WP_198106729.1</td><td>Type1.BREX</td><td>SF1-SF2-Helicase+HTH->MukF->JetB->DUF3584+DUF5401->RNase->XIS-HTH-> BrxA-DNA-binding -> BrxB-BrxF-iSTAND -> DUF6079 *-> OLD-ABC-> REase -> N6-METHYLASE -> PglZ -> SIGMA+OB-Fold+AAA-ATPase+HKD-DNase-BrxL ->Phage_integrase-></td><td>1187</td><td>Bacteria>Pseudomonadota>Gammaproteobacteria>Vibrionales>Vibrionaceae>Vibrio</td><td>GCF_016094345.1</td><td>Vibrio_navarrensis</td><td>CJ419_RS12980</td><td>152911..156474</td><td>29495</td><td>BREX-system-P-loop-protein-BrxC-[Vibrio-navarrensis]</td></tr>
<tr><td>WP_250156418.1</td><td>Type1.BREX</td><td><- wHTH+WYL+WCX <-DUF2787<-?<-?||XIS-HTH->HNH->XIS-HTH-> BrxA-DNA-binding -> BrxB-BrxF-iSTAND -> DUF6079 *-> OLD-ABC-> REase -> N6-METHYLASE -> PglZ -> SIGMA+OB-Fold+AAA-ATPase+HKD-DNase-BrxL -></td><td>1185</td><td>Bacteria>Pseudomonadota>Gammaproteobacteria>Vibrionales>Vibrionaceae>Vibrio</td><td>GCF_018457845.1</td><td>Vibrio_alginolyticus</td><td>J4H60_RS12980</td><td>321648..325205</td><td>663</td><td>BREX-system-P-loop-protein-BrxC-[Vibrio-alginolyticus]</td></tr>
<tr><td>WP_228097414.1</td><td>Type6.BREX</td><td>ParB-><-?<-Phage_integrase<-HNH<-OLD-ABC<- REase <- SIGMA+OB-Fold+AAA-ATPase+HKD-DNase-BrxL <- PglZ <- N6-METHYLASE <- DUF6079 *<- BrxB-BrxF-iSTAND <- BrxA-DNA-binding <- BrxE <-RNase<-DUF3584+DUF5401<-JetB<-MukF|| wHTH+WYL+WCX -><-ANK+ANK</td><td>1208</td><td>Bacteria>Pseudomonadota>Gammaproteobacteria>Vibrionales>Vibrionaceae>Vibrio</td><td>GCF_020741135.1</td><td>Vibrio_parahaemolyticus</td><td>IB300_RS08390</td><td>552360..555986</td><td>670</td><td>BREX-system-P-loop-protein-BrxC-[Vibrio-parahaemolyticus]</td></tr>
<tr><td>WP_267363047.1</td><td>Type1.BREX</td><td><-XIS-HTH<-Arm-DNA-bind_3+Phage_int_SAM_3+Phage_integrase||?->?->||?-><-HEPN||XIS-HTH-> BrxA-DNA-binding -> BrxB-BrxF-iSTAND -> DUF6079 *-> OLD-ABC-> REase -> N6-METHYLASE -> PglZ -> SIGMA+OB-Fold+AAA-ATPase+HKD-DNase-BrxL -></td><td>1187</td><td>Bacteria>Pseudomonadota>Gammaproteobacteria>Vibrionales>Vibrionaceae>Vibrio</td><td>GCF_026537625.1</td><td>Vibrio_parahaemolyticus</td><td>NOL37_RS24175</td><td>9284..12847</td><td>670</td><td>BREX-system-P-loop-protein-BrxC-[Vibrio-parahaemolyticus]</td></tr>
<tr><td>WP_268670710.1</td><td>Type1.BREX</td><td>MODE-HTH+PBPII-><-?||XIS-HTH->?->SF1-SF2-Helicase+HTH->HNH->XIS-HTH-> BrxA-DNA-binding -> BrxB-BrxF-iSTAND -> DUF6079 *-> OLD-ABC-> REase -> N6-METHYLASE -> PglZ -> SIGMA+OB-Fold+AAA-ATPase+HKD-DNase-BrxL -><-PALP<-?<-MODE-HTH+PBPII</td><td>1185</td><td>Bacteria>Pseudomonadota>Gammaproteobacteria>Vibrionales>Vibrionaceae>Vibrio</td><td>GCF_026798035.1</td><td>Vibrio_coralliirubri</td><td>OTK49_RS06670</td><td>229636..233193</td><td>1516159</td><td>BREX-system-P-loop-protein-BrxC-[Vibrio-coralliirubri]</td></tr>
<tr><td>WP_274674022.1</td><td>Type1.BREX</td><td><-HAMP+GGDEF<-?<- SIGMA+OB-Fold+AAA-ATPase+HKD-DNase-BrxL <- PglZ <- N6-METHYLASE <- REase <-OLD-ABC <- DUF6079 *<- BrxB-BrxF-iSTAND <- BrxA-DNA-binding <-XIS-HTH</td><td>1187</td><td>Bacteria>Pseudomonadota>Gammaproteobacteria>Vibrionales>Vibrionaceae>Vibrio</td><td>GCF_028857845.1</td><td>Vibrio_aestuarianus</td><td>L9W73_RS08980</td><td>42250..45813</td><td>28171</td><td>BREX-system-P-loop-protein-BrxC-[Vibrio-aestuarianus]</td></tr>
<tr><td>WP_213868350.1</td><td>Type1.BREX</td><td><- SIGMA+OB-Fold+AAA-ATPase+HKD-DNase-BrxL <- PglZ <- N6-METHYLASE <- REase <-OLD-ABC <- DUF6079 *<- BrxB-BrxF-iSTAND <- BrxA-DNA-binding <-XIS-HTH<-HNH<-XIS-HTH||?->?->DUF2787-> wHTH+WYL+WCX -></td><td>1187</td><td>Bacteria>Pseudomonadota>Gammaproteobacteria>Vibrionales>Vibrionaceae>Vibrio</td><td>-</td><td>Vibrio_alginolyticus</td><td>N7I24_002738</td><td>64007..67570</td><td>663</td><td>BREX-system-P-loop-protein-BrxC-[Vibrio-alginolyticus]</td></tr>
<tr><td>WP_225763940.1</td><td>Type1.BREX</td><td><-TRPR-HTH||?->PRTase->HTH_Tnp_1+HTH_21+rve+rve_3->XIS-HTH-> BrxA-DNA-binding -> BrxB-BrxF-iSTAND -> DUF6079 *->?-> N6-METHYLASE -> OLD-ABC-> REase ->?-> N6-METHYLASE -p-> PglZ -> SIGMA+OB-Fold+AAA-ATPase+HKD-DNase-BrxL -></td><td>1181</td><td>Bacteria>Pseudomonadota>Gammaproteobacteria>Xanthomonadales>Xanthomonadaceae>Stenotrophomonas</td><td>GCF_916618915.1</td><td>Stenotrophomonas_sp._Marseille-Q4652</td><td>LG380_RS05370</td><td>1171221..1174766</td><td>2866595</td><td>BREX-system-P-loop-protein-BrxC-[Stenotrophomonas-sp.-Marseille-Q4652]</td></tr>
<tr><td>RYD14825.1</td><td>Type1.BREX</td><td><-35exo<-SF1-SF2-Helicase+wHTH+Small-B-barrel+DUF1998||HTH_3->Couple_hipA+HipA_C-> BrxA-DNA-binding -> BrxB-BrxF-iSTAND -> DUF6079 *-> N6-METHYLASE -> PglZ -> SIGMA+OB-Fold+AAA-ATPase+Lon-Protease-BrxL -> REase -><-tRNA||NUDIX->?->?-><-REC+GGDEF+EAL</td><td>1207</td><td>Bacteria>Pseudomonadota>Gammaproteobacteria>Xanthomonadales>Xanthomonadaceae>unclassified_Xanthomonadaceae</td><td>GCA_004143945.1</td><td>Xanthomonadaceae_bacterium</td><td>EOP90_11180</td><td>76143..79766</td><td>1926873</td><td>BREX-system-P-loop-protein-BrxC-[Xanthomonadaceae-bacterium]</td></tr>
<tr><td>WP_218196661.1</td><td>Type1.BREX</td><td>Arm-DNA-bind_3+Phage_integrase-><- SIGMA+OB-Fold+AAA-ATPase+HKD-DNase-BrxL <- PglZ <- N6-METHYLASE -p<-?<- REase <-OLD-ABC <- N6-METHYLASE <-?<- DUF6079 *<- BrxB-BrxF-iSTAND <- BrxA-DNA-binding <-XIS-HTH<-Phage_int_SAM_2+Integrase_1<-Phage_rep</td><td>1182</td><td>Bacteria>Pseudomonadota>Gammaproteobacteria>Xanthomonadales>Xanthomonadaceae>Xanthomonas</td><td>GCF_019193005.1</td><td>Xanthomonas_euvesicatoria</td><td>KV691_RS15985</td><td>182541..186089</td><td>456327</td><td>BREX-system-P-loop-protein-BrxC-[Xanthomonas-euvesicatoria]</td></tr>
<tr><td>MBS0349076.1</td><td>Type1.BREX</td><td>RHH_5-> BrxA-DNA-binding -> BrxB-BrxF-iSTAND -> DUF6079 *->?-> N6-METHYLASE ->PHP+SbcC->SF1-SF2-Helicase+PRTase->SLOG+BirA-HTH-> PglZ -> SIGMA+OB-Fold+AAA-ATPase+Lon-Protease-BrxL -> REase +TIR->PIWI-></td><td>1153</td><td>Bacteria>Pseudomonadota>unclassified_Pseudomonadota</td><td>GCA_018241665.1</td><td>Pseudomonadota_bacterium</td><td>JSR69_21735</td><td>2072..5533</td><td>1977087</td><td>BREX-system-P-loop-protein-BrxC-[Pseudomonadota-bacterium]</td></tr>
<tr><td>MBU0673651.1</td><td>Type1.BREX</td><td>RADICAL-SAM-><-?||?-> BrxA-DNA-binding -> BrxB-BrxF-iSTAND -> DUF6079 *->Schlafen+HATPase+wHTH->?-> N6-METHYLASE -> REase ->ParB+HNH->OLD-ABC->?-> PglZ -> SIGMA+OB-Fold+AAA-ATPase+HKD-DNase-BrxL -></td><td>1167</td><td>Bacteria>Pseudomonadota>unclassified_Pseudomonadota</td><td>GCA_018814305.1</td><td>Pseudomonadota_bacterium</td><td>KJ950_03300</td><td>48865..52368</td><td>1977087</td><td>BREX-system-P-loop-protein-BrxC-[Pseudomonadota-bacterium]</td></tr>
<tr><td>MBU4235493.1</td><td>Type1.BREX</td><td><- N6-METHYLASE <- REase <-OLD-ABC <-Schlafen+HATPase+wHTH<- DUF6079 *<- BrxB-BrxF-iSTAND <- BrxA-DNA-binding <-XIS-HTH</td><td>1147</td><td>Bacteria>Pseudomonadota>unclassified_Pseudomonadota</td><td>GCA_018897885.1</td><td>Pseudomonadota_bacterium</td><td>KKI01_04080</td><td>7059..10502</td><td>1977087</td><td>BREX-system-P-loop-protein-BrxC-[Pseudomonadota-bacterium]</td></tr>
<tr><td>WP_053322564.1</td><td>Type1.BREX</td><td>HNH->?->?->LON-AAA->Calcineurin->?-> BrxA-DNA-binding -> BrxB-BrxF-iSTAND -> DUF6079 *-> N6-METHYLASE -p->Schlafen+HATPase+wHTH->HEPN-> REase -> N6-METHYLASE -p->HerA/FtsK-> PglZ -> SIGMA+OB-Fold+AAA-ATPase+HKD-DNase-BrxL -></td><td>1181</td><td>Bacteria>Pseudomonadota>Zetaproteobacteria>Mariprofundales>Mariprofundaceae>Mariprofundus</td><td>GCF_001273615.1</td><td>Mariprofundus_ferrooxydans</td><td>AL013_RS02410</td><td>88551..92096</td><td>314344</td><td>BREX-system-P-loop-protein-BrxC-[Mariprofundus-ferrooxydans]</td></tr>
<tr><td>MCL4743298.1</td><td>Type1.BREX</td><td><-Phage_integrase<-XIS-HTH<-Primpol+D5ATPase<-XIS-HTH<-?<-?|| BrxA-DNA-binding -> BrxB-BrxF-iSTAND -> DUF6079 *-> N6-METHYLASE -> PglZ -> SIGMA+OB-Fold+AAA-ATPase+Lon-Protease-BrxL -><-Phage_integrase<- REase <-Phage_integrase</td><td>1150</td><td>Bacteria>PVC_group>Planctomycetota>Phycisphaerae>Phycisphaerales>unclassified_Phycisphaerales</td><td>GCA_023384195.1</td><td>Phycisphaerales_bacterium</td><td>KJZ54_13970</td><td>8587..12039</td><td>2052180</td><td>BREX-system-P-loop-protein-BrxC-[Phycisphaerales-bacterium]</td></tr>
<tr><td>WP_145350724.1</td><td>Type1.BREX</td><td><-SIR2<-?<- SIGMA+OB-Fold+AAA-ATPase+HKD-DNase-BrxL <- PglZ <- REase +TPR+GreAB-C+PIN<- N6-METHYLASE <-Schlafen+HATPase+wHTH<- REase <-OLD-ABC <- DUF6079 *<- BrxB-BrxF-iSTAND <- BrxA-DNA-binding <-?||TrwC+SF1-SF2-Helicase->?-><-?<-?<-ArdCN+MPTase</td><td>1169</td><td>Bacteria>PVC_group>Planctomycetota>Planctomycetia>Pirellulales>Pirellulaceae</td><td>GCF_007741495.1</td><td>Roseimaritima_multifibrata</td><td>FF011L_RS05780</td><td>1542279..1545788</td><td>1930274</td><td>BREX-system-P-loop-protein-BrxC-[Roseimaritima-multifibrata]</td></tr>
<tr><td>WP_184301662.1</td><td>Type1.BREX</td><td><-nSTAND1+HNH<-URI-DNase+RAMA<- SIGMA+OB-Fold+AAA-ATPase+HKD-DNase-BrxL <- PglZ <- REase +TPR+GreAB-C+PIN<- N6-METHYLASE <-?<-ABC-ATPase<- DUF6079 *<- BrxB-BrxF-iSTAND <- BrxA-DNA-binding <-XIS-HTH||OLD-ABC+TOPRIM->SF1-SF2-Helicase-></td><td>1169</td><td>Bacteria>PVC_group>Planctomycetota>Planctomycetia>Pirellulales>Pirellulaceae</td><td>GCF_014192335.1</td><td>Rhodopirellula_rubra</td><td>FHS27_RS03210</td><td>253910..257419</td><td>980271</td><td>BREX-system-P-loop-protein-BrxC-[Rhodopirellula-rubra]</td></tr>
<tr><td>GIX04293.1</td><td>Type1.BREX</td><td><- SIGMA+OB-Fold+AAA-ATPase+Lon-Protease-BrxL <- PglZ <- N6-METHYLASE <- REase <- DUF6079 *<- BrxB-BrxF-iSTAND <- BrxA-DNA-binding <-XIS-HTH<-SF1-SF2-Helicase</td><td>1221</td><td>Bacteria>PVC_group>Planctomycetota>Planctomycetia>Planctomycetales>Planctomycetaceae</td><td>GCA_026004855.1</td><td>Planctomycetaceae_bacterium</td><td>KatS3mg114_0162</td><td>186824..190489</td><td>2026779</td><td>MAG:-hypothetical-protein-KatS3mg114_0162-[Planctomycetaceae-bacterium]</td></tr>
<tr><td>MCA9231627.1</td><td>Type6.BREX</td><td><- REase <- SIGMA+OB-Fold+AAA-ATPase+Lon-Protease-BrxL <- PglZ <- N6-METHYLASE <- DUF6079 *<- BrxB-BrxF-iSTAND <- BrxA-DNA-binding <- BrxE <-?|| REase -></td><td>1178</td><td>Bacteria>PVC_group>Planctomycetota>Planctomycetia>Planctomycetales>unclassified_Planctomycetales</td><td>GCA_020427155.1</td><td>Planctomycetales_bacterium</td><td>KDA57_13330</td><td>9427..12963</td><td>2053591</td><td>BREX-system-P-loop-protein-BrxC-[Planctomycetales-bacterium]</td></tr>
<tr><td>TWU09954.1</td><td>Type1.BREX</td><td><-ABhydrolase<-?||?->Calcineurin->SbcC+SbcC->?->XIS-HTH-> BrxA-DNA-binding -> BrxB-BrxF-iSTAND -> DUF6079 *->?->ParB+HNH-> N6-METHYLASE -> REase -> PglZ ->?->SKI->?-> SIGMA+OB-Fold+AAA-ATPase+Lon-Protease-BrxL -></td><td>1205</td><td>Bacteria>PVC_group>Planctomycetota>unclassified_Planctomycetota</td><td>GCA_007860105.1</td><td>Planctomycetes_bacterium_Poly21</td><td>Poly21_52830</td><td>31284..34901</td><td>2528014</td><td>hypothetical-protein-Poly21_52830-[Planctomycetes-bacterium-Poly21]</td></tr>
<tr><td>AHF94674.1</td><td>Type1.BREX</td><td><- PglZ <-TOPRIM<-OLD-ABC<-PIWI<- REase +PNTD1+PNTD2<- N6-METHYLASE -p<-DOC+HTH<- N6-METHYLASE <-?<- DUF6079 *<- BrxB-BrxF-iSTAND <- BrxA-DNA-binding <-?||TrwC+SF1-SF2-Helicase->?->?->ClpABC-AAA-></td><td>1248</td><td>Bacteria>PVC_group>Verrucomicrobiota>Opitutae>Opitutales>Opitutaceae</td><td>GCA_000242935.3</td><td>Opitutaceae_bacterium_TAV5</td><td>OPIT5_23275</td><td>5485357..5489103</td><td>794903</td><td>hypothetical-protein-OPIT5_23275-[Opitutaceae-bacterium-TAV5]</td></tr>
<tr><td>RRK00196.1</td><td>Type6.BREX</td><td><-DNA_METHYLASE<- PglZ -p<- PglZ <- PglZ <-?<-OLD-ABC<-ParB<- DUF6079 *<- BrxB-BrxF-iSTAND <- BrxA-DNA-binding <- BrxE <- REase <-?<-HEPN</td><td>1207</td><td>Bacteria>PVC_group>Verrucomicrobiota>Opitutae>Opitutales>Opitutaceae</td><td>GCA_001653815.2</td><td>Opitutaceae_bacterium_TAV3</td><td>Ga0100230_019740</td><td>2814873..2818496</td><td>278958</td><td>BREX-system-P-loop-protein-BrxC-[Opitutaceae-bacterium-TAV3]</td></tr>
<tr><td>RRJ96051.1</td><td>Type6.BREX</td><td><-TrwC+SF1-SF2-Helicase||?-><-DNA_METHYLASE<- SIGMA+OB-Fold+AAA-ATPase+HKD-DNase-BrxL <- PglZ <-?<-OLD-ABC<-ParB<- DUF6079 *<- BrxB-BrxF-iSTAND <- BrxA-DNA-binding <- BrxE <- REase <-?<-Phage_int_SAM_1+Phage_integrase<-HEPN<-ZnR+ATPcone</td><td>1206</td><td>Bacteria>PVC_group>Verrucomicrobiota>Opitutae>Opitutales>Opitutaceae</td><td>GCA_001653825.2</td><td>Opitutaceae_bacterium_TAV4</td><td>Ga0100231_019065</td><td>2732930..2736550</td><td>278959</td><td>BREX-system-P-loop-protein-BrxC-[Opitutaceae-bacterium-TAV4]</td></tr>
<tr><td>MBC7388090.1</td><td>Type1.BREX</td><td>DUF6079 -p*-> REase -> N6-METHYLASE ->?-> PglZ -> SIGMA+OB-Fold+AAA-ATPase+Lon-Protease-BrxL -></td><td>828</td><td>Bacteria>PVC_group>Verrucomicrobiota>Opitutae>Opitutales>Opitutaceae</td><td>GCA_014376535.1</td><td>Opitutaceae_bacterium</td><td>H7329_02660</td><td>1..2487</td><td>2006848</td><td>hypothetical-protein-[Opitutaceae-bacterium]</td></tr>
<tr><td>MCC5808698.1</td><td>Type1.BREX</td><td>Phage_integrase-><- SIGMA+OB-Fold+AAA-ATPase+HKD-DNase-BrxL <- PglZ <-HNH<-OLD-ABC<- N6-METHYLASE <- REase <-OLD-ABC <- DUF6079 *<- BrxB-BrxF-iSTAND <- BrxA-DNA-binding <-XIS-HTH</td><td>1183</td><td>Bacteria>PVC_group>Verrucomicrobiota>Opitutae>Opitutales>unclassified_Opitutales</td><td>GCA_020831525.1</td><td>Opitutales_bacterium</td><td>JJU00_20390</td><td>12178..15729</td><td>2026772</td><td>BREX-system-P-loop-protein-BrxC-[Opitutales-bacterium]</td></tr>
<tr><td>HCF96075.1</td><td>Type1.BREX</td><td><- N6-METHYLASE -p<- N6-METHYLASE <- REase <-OLD-ABC <- DUF6079 -p<- DUF6079 -p*<- DUF6079 -p<- DUF6079 -p<- DUF6079 -p<- BrxB-BrxF-iSTAND <- BrxA-DNA-binding</td><td>268</td><td>Bacteria>PVC_group>Verrucomicrobiota>unclassified_Verrucomicrobiota</td><td>GCA_003534995.1</td><td>Verrucomicrobiota_bacterium</td><td>DEW46_13535</td><td>6770..7576</td><td>2026799</td><td>BREX-system-P-loop-protein-BrxC-[Verrucomicrobiota-bacterium]</td></tr>
<tr><td>HCF96078.1</td><td>Type1.BREX</td><td><- N6-METHYLASE -p<- N6-METHYLASE <- REase <-OLD-ABC <- DUF6079 -p<- DUF6079 -p<- DUF6079 -p<- DUF6079 -p<- DUF6079 -p*<- BrxB-BrxF-iSTAND <- BrxA-DNA-binding</td><td>531</td><td>Bacteria>PVC_group>Verrucomicrobiota>unclassified_Verrucomicrobiota</td><td>GCA_003534995.1</td><td>Verrucomicrobiota_bacterium</td><td>DEW46_13550</td><td>8732..10325</td><td>2026799</td><td>BREX-system-P-loop-protein-BrxC-[Verrucomicrobiota-bacterium]</td></tr>
<tr><td>MBL9184319.1</td><td>Type1.BREX</td><td>DUF932-> BrxA-DNA-binding -> BrxB-BrxF-iSTAND -> DUF6079 *->PIN->TIR-> N6-METHYLASE ->?-> PglZ -> SIGMA+OB-Fold+AAA-ATPase+Lon-Protease-BrxL -> REase +PNTD1+PNTD2->PIWI-></td><td>1153</td><td>Bacteria>PVC_group>Verrucomicrobiota>Verrucomicrobiae>Verrucomicrobiales>Verrucomicrobiaceae</td><td>GCA_016795465.1</td><td>Verrucomicrobiaceae_bacterium</td><td>JNN17_19395</td><td>2113..5574</td><td>2026800</td><td>BREX-system-P-loop-protein-BrxC-[Verrucomicrobiaceae-bacterium]</td></tr>
<tr><td>MCF7788313.1</td><td>Type1.BREX</td><td>BrxA-DNA-binding -> BrxB-BrxF-iSTAND -> DUF6079 *->HKD-DNase+DUF3150->MarR-HTH->Polbeta-> N6-METHYLASE ->?->?->SbcC-> PglZ -p-> REase -></td><td>1167</td><td>Bacteria>PVC_group>Verrucomicrobiota>Verrucomicrobiae>Verrucomicrobiales>Verrucomicrobiaceae</td><td>GCA_021733925.1</td><td>Prosthecobacter_sp.</td><td>K9N47_19475</td><td>2530..6033</td><td>1965333</td><td>BREX-system-P-loop-protein-BrxC-[Prosthecobacter-sp.]</td></tr>
<tr><td>WP_114628929.1</td><td>Type1.BREX</td><td><-Thymidylate_kin||ParB+HNH->ParB+HNH-><-?<-D5N+D5ATPase||XIS-HTH-> BrxA-DNA-binding -> BrxB-BrxF-iSTAND -> DUF6079 *-> OLD-ABC-> REase -> N6-METHYLASE -> SIGMA+OB-Fold+AAA-ATPase+HKD-DNase-BrxL -><-AbiEii<-AbiEi_3_N+AbiEi_3</td><td>1173</td><td>Bacteria>Spirochaetota>Spirochaetia>Spirochaetales>Spirochaetaceae>Oceanispirochaeta</td><td>GCF_013166645.2</td><td>unclassified_Oceanispirochaeta</td><td>HNV09_RS06050</td><td>178318..181839</td><td>2635722</td><td>MULTISPECIES:-BREX-system-P-loop-protein-BrxC-[unclassified-Oceanispirochaeta]</td></tr>
<tr><td>OQY41116.1</td><td>Type1.BREX</td><td>REase -> BrxA-DNA-binding -> BrxB-BrxF-iSTAND -> DUF6079 *->OLD-ABC->TOPRIM-> N6-METHYLASE ->ParB+HNH->ParB+HNH-> SIGMA+OB-Fold+AAA-ATPase+Lon-Protease-BrxL -></td><td>1207</td><td>Bacteria>Spirochaetota>Spirochaetia>Spirochaetales>Spirochaetaceae>unclassified_Spirochaetaceae</td><td>GCA_002084785.1</td><td>Spirochaetaceae_bacterium_4572_7</td><td>B6229_00020</td><td>2034..5657</td><td>1972456</td><td>hypothetical-protein-B6229_00020-[Spirochaetaceae-bacterium-4572_7]</td></tr>
<tr><td>GHV28822.1</td><td>Type1.BREX</td><td><-HicA<-?<- REase <-?<-?<- N6-METHYLASE <- N6-METHYLASE -p<-NERD<-OLD-ABC<- DUF6079 *<- BrxB-BrxF-iSTAND <- BrxA-DNA-binding</td><td>1195</td><td>Bacteria>Synergistota>Synergistia>Synergistales>unclassified_Synergistales</td><td>GCA_026002695.1</td><td>Synergistales_bacterium</td><td>FACS1894167_06880</td><td>20211..23798</td><td>2268201</td><td>hypothetical-protein-FACS1894167_06880-[Synergistales-bacterium]</td></tr>
<tr><td>WP_247595282.1</td><td>Type1.BREX</td><td>ADH_N+ADH_zinc_N->TetR-HTH->ABC-ATPase+SbcC->?->?-><- SIGMA+OB-Fold+AAA-ATPase+HKD-DNase-BrxL <- PglZ <-Schlafen+HATPase+wHTH<- REase <- N6-METHYLASE <- DUF6079 *<- BrxB-BrxF-iSTAND <- BrxA-DNA-binding ||?->PhoH->?->?->sGTP->Peptidase_S9-></td><td>1174</td><td>Bacteria>Terrabacteria_group>Actinomycetota>Actinomycetes>Actinomycetales>Actinomycetaceae</td><td>GCF_004796555.1</td><td>Actinomyces_procaprae</td><td>FAQ94_RS07530</td><td>102106..105630</td><td>2560010</td><td>BREX-system-P-loop-protein-BrxC-[Actinomyces-procaprae]</td></tr>
<tr><td>WP_126382192.1</td><td>Type1.BREX</td><td><- SIGMA+OB-Fold+AAA-ATPase+HKD-DNase-BrxL <- PglZ <-Schlafen+HATPase+wHTH<- REase <- N6-METHYLASE <- DUF6079 *<- BrxB-BrxF-iSTAND <- BrxA-DNA-binding <-SIR2<-?<-?<-?||GNTR-HTH+UTRA-></td><td>1170</td><td>Bacteria>Terrabacteria_group>Actinomycetota>Actinomycetes>Actinomycetales>Actinomycetaceae</td><td>GCF_900637165.1</td><td>Actinomyces_howellii</td><td>GYB74_RS04555</td><td>58123..61635</td><td>52771</td><td>BREX-system-P-loop-protein-BrxC-[Actinomyces-howellii]</td></tr>
<tr><td>MBE5064977.1</td><td>Type1.BREX</td><td>ACET->ACET+ACET->?->Thioredoxin_3->?-><-Glycoside_Hydrolase||OLD-ABC-> BrxA-DNA-binding -> BrxB-BrxF-iSTAND -> DUF6079 *-> N6-METHYLASE -> REase ->?->Schlafen+HATPase+wHTH-> PglZ -> SIGMA+OB-Fold+AAA-ATPase+HKD-DNase-BrxL -></td><td>1221</td><td>Bacteria>Terrabacteria_group>Actinomycetota>Actinomycetes>Bifidobacteriales>Bifidobacteriaceae</td><td>GCA_014983005.1</td><td>Bifidobacterium_pullorum_subsp._saeculare</td><td>INF27_02890</td><td>184023..187688</td><td>78257</td><td>BREX-system-P-loop-protein-BrxC-[Bifidobacterium-pullorum-subsp.-saeculare]</td></tr>
<tr><td>MBS6717590.1</td><td>Type1.BREX</td><td>HTH_28-> BrxA-DNA-binding -> BrxB-BrxF-iSTAND -> DUF6079 *-> REase ->Schlafen+HATPase+wHTH-> PglZ -> SIGMA+OB-Fold+AAA-ATPase+HKD-DNase-BrxL ->SLATT->SMODS-></td><td>1218</td><td>Bacteria>Terrabacteria_group>Actinomycetota>Actinomycetes>Bifidobacteriales>Bifidobacteriaceae</td><td>GCA_018378875.1</td><td>Bifidobacterium_longum</td><td>KH250_10240</td><td>6375..10031</td><td>216816</td><td>BREX-system-P-loop-protein-BrxC-[Bifidobacterium-longum]</td></tr>
<tr><td>WP_003819016.1</td><td>Type1.BREX</td><td>ZnR->?->ABC-ATPase+SbcC->?->?->?-> BrxA-DNA-binding -> BrxB-BrxF-iSTAND -> DUF6079 *-> N6-METHYLASE -> REase ->Schlafen+HATPase+wHTH-> PglZ -> SIGMA+OB-Fold+AAA-ATPase+HKD-DNase-BrxL ->?-><- REase ||?-><-Glyoxalase</td><td>1217</td><td>Bacteria>Terrabacteria_group>Actinomycetota>Actinomycetes>Bifidobacteriales>Bifidobacteriaceae</td><td>GCF_000300215.1</td><td>Bifidobacterium_bifidum</td><td>IF1010_RS06520</td><td>82392..86045</td><td>1681</td><td>BREX-system-P-loop-protein-BrxC-[Bifidobacterium-bifidum]</td></tr>
<tr><td>WP_046999386.1</td><td>Type1.BREX</td><td>REase -><-ZnR<-ZnR+GP<- SIGMA+OB-Fold+AAA-ATPase+HKD-DNase-BrxL <- PglZ <-Schlafen+HATPase+wHTH<- REase <- N6-METHYLASE <- DUF6079 *<- BrxB-BrxF-iSTAND <- BrxA-DNA-binding <-Pentapeptide+Pentapeptide+Pentapeptide<-HAD||SbcD-><-?||HTH-><-35exo+BRCT</td><td>1217</td><td>Bacteria>Terrabacteria_group>Actinomycetota>Actinomycetes>Bifidobacteriales>Bifidobacteriaceae</td><td>GCF_001010915.1</td><td>Bifidobacterium_adolescentis</td><td>AAX71_RS05490</td><td>58570..62223</td><td>1680</td><td>BREX-system-P-loop-protein-BrxC-[Bifidobacterium-adolescentis]</td></tr>
<tr><td>WP_065445635.1</td><td>Type1.BREX</td><td>HTH_28-> BrxA-DNA-binding -> BrxB-BrxF-iSTAND -> DUF6079 *-> N6-METHYLASE -> REase ->Schlafen+HATPase+wHTH-> PglZ -> SIGMA+OB-Fold+AAA-ATPase+HKD-DNase-BrxL ->ZnR+GP->ZnR->ParB+HNH->Sigma-HTH-></td><td>1219</td><td>Bacteria>Terrabacteria_group>Actinomycetota>Actinomycetes>Bifidobacteriales>Bifidobacteriaceae</td><td>GCF_001686165.1</td><td>Bifidobacterium_longum</td><td>MCC10077_RS10215</td><td>2512..6171</td><td>216816</td><td>BREX-system-P-loop-protein-BrxC-[Bifidobacterium-longum]</td></tr>
<tr><td>WP_118279738.1</td><td>Type1.BREX</td><td>REase -><-?<- SIGMA+OB-Fold+AAA-ATPase+HKD-DNase-BrxL <- PglZ <-Schlafen+HATPase+wHTH<- REase <- N6-METHYLASE <- DUF6079 *<- BrxB-BrxF-iSTAND <- BrxA-DNA-binding || REase ->?->SbcD-><-?||?-><-HISKIN<-REC+LuxR-HTH</td><td>1219</td><td>Bacteria>Terrabacteria_group>Actinomycetota>Actinomycetes>Bifidobacteriales>Bifidobacteriaceae</td><td>GCF_003472685.1</td><td>Bifidobacterium_pseudocatenulatum</td><td>DW102_RS06345</td><td>114145..117804</td><td>28026</td><td>BREX-system-P-loop-protein-BrxC-[Bifidobacterium-pseudocatenulatum]</td></tr>
<tr><td>WP_126016404.1</td><td>Type1.BREX</td><td><-ParB+HNH+RAMA<-ParB+HNH<-ZnR<-ZnR+GP<- SIGMA+OB-Fold+AAA-ATPase+HKD-DNase-BrxL <- PglZ <-Schlafen+HATPase+wHTH<- REase <- N6-METHYLASE <- DUF6079 *<- BrxB-BrxF-iSTAND <- BrxA-DNA-binding <-?||SbcD-><-ACET+ACET||HTH-><-?<-35exo+HRDC</td><td>1217</td><td>Bacteria>Terrabacteria_group>Actinomycetota>Actinomycetes>Bifidobacteriales>Bifidobacteriaceae</td><td>GCF_003952825.1</td><td>Bifidobacterium_pseudocatenulatum</td><td>CWS99_RS05770</td><td>1339669..1343322</td><td>28026</td><td>BREX-system-P-loop-protein-BrxC-[Bifidobacterium-pseudocatenulatum]</td></tr>
<tr><td>WP_142421439.1</td><td>Type1.BREX</td><td>RNaseH-><-?<- SIGMA+OB-Fold+AAA-ATPase+HKD-DNase-BrxL <- PglZ <-Schlafen+HATPase+wHTH<- REase <- N6-METHYLASE -p<- N6-METHYLASE <- DUF6079 *<- BrxB-BrxF-iSTAND <- BrxA-DNA-binding || REase ->?-><-ACET+ACET||HTH-><-?<-35exo+HRDC</td><td>1218</td><td>Bacteria>Terrabacteria_group>Actinomycetota>Actinomycetes>Bifidobacteriales>Bifidobacteriaceae</td><td>GCF_006828885.1</td><td>Bifidobacterium_pseudocatenulatum</td><td>GH753_RS08045</td><td>1981980..1985636</td><td>28026</td><td>BREX-system-P-loop-protein-BrxC-[Bifidobacterium-pseudocatenulatum]</td></tr>
<tr><td>WP_193020253.1</td><td>Type1.BREX</td><td>HTH+MPTase-><-?<-HEPN<-?<-?||?-> BrxA-DNA-binding -> BrxB-BrxF-iSTAND -> DUF6079 *-> N6-METHYLASE -> REase ->Schlafen+wHTH-> PglZ -> SIGMA+OB-Fold+AAA-ATPase+HKD-DNase-BrxL ->?->?->RNase-></td><td>1229</td><td>Bacteria>Terrabacteria_group>Actinomycetota>Actinomycetes>Bifidobacteriales>Bifidobacteriaceae</td><td>GCF_014898135.1</td><td>Bifidobacterium_longum</td><td>BL5934_RS04380</td><td>976256..979945</td><td>216816</td><td>BREX-system-P-loop-protein-BrxC-[Bifidobacterium-longum]</td></tr>
<tr><td>WP_227411048.1</td><td>Type1.BREX</td><td>ACET->ACET+ACET->?->Thioredoxin_3->ArsR-HTH-><-Glycoside_Hydrolase||OLD-ABC-> BrxA-DNA-binding -> BrxB-BrxF-iSTAND -> DUF6079 *-> N6-METHYLASE -> REase ->?->Schlafen+HATPase+wHTH-> PglZ -> SIGMA+OB-Fold+AAA-ATPase+HKD-DNase-BrxL -></td><td>1242</td><td>Bacteria>Terrabacteria_group>Actinomycetota>Actinomycetes>Bifidobacteriales>Bifidobacteriaceae</td><td>GCF_014983005.1</td><td>Bifidobacterium_pullorum</td><td>INF27_RS02890</td><td>183960..187688</td><td>78448</td><td>BREX-system-P-loop-protein-BrxC-[Bifidobacterium-pullorum]</td></tr>
<tr><td>WP_195506526.1</td><td>Type1.BREX</td><td><-Schlafen+HATPase+wHTH<- REase <- N6-METHYLASE <- DUF6079 *<- BrxB-BrxF-iSTAND <- BrxA-DNA-binding <-Pentapeptide+Pentapeptide+Pentapeptide||<-?||?->?->SbcD-></td><td>1218</td><td>Bacteria>Terrabacteria_group>Actinomycetota>Actinomycetes>Bifidobacteriales>Bifidobacteriaceae</td><td>GCF_015556885.1</td><td>Bifidobacterium_pseudocatenulatum</td><td>I2F64_RS01000</td><td>6102..9758</td><td>28026</td><td>BREX-system-P-loop-protein-BrxC-[Bifidobacterium-pseudocatenulatum]</td></tr>
<tr><td>WP_226650300.1</td><td>Type1.BREX</td><td><- SIGMA+OB-Fold+AAA-ATPase+HKD-DNase-BrxL <- PglZ <-Schlafen+HATPase+wHTH<- REase <- N6-METHYLASE <- DUF6079 *<- BrxB-BrxF-iSTAND <- BrxA-DNA-binding || REase ->?->SbcD-><-?||?-><-?||MerR-HTH-></td><td>1219</td><td>Bacteria>Terrabacteria_group>Actinomycetota>Actinomycetes>Bifidobacteriales>Bifidobacteriaceae</td><td>GCF_020526585.1</td><td>Bifidobacterium_pseudocatenulatum</td><td>KZP10_RS05695</td><td>1065058..1068717</td><td>28026</td><td>BREX-system-P-loop-protein-BrxC-[Bifidobacterium-pseudocatenulatum]</td></tr>
<tr><td>WP_117715835.1</td><td>Type1.BREX</td><td>ZnR->?->ABC-ATPase+SbcC->?->?->?-> BrxA-DNA-binding -> BrxB-BrxF-iSTAND -> DUF6079 *-> N6-METHYLASE ->SIR2->Schlafen+HATPase+wHTH-> PglZ -> SIGMA+OB-Fold+AAA-ATPase+HKD-DNase-BrxL ->?-><- REase ||?-><-Glyoxalase</td><td>1217</td><td>Bacteria>Terrabacteria_group>Actinomycetota>Actinomycetes>Bifidobacteriales>Bifidobacteriaceae</td><td>GCF_027669765.1</td><td>Bifidobacterium_bifidum</td><td>PHA16_RS02375</td><td>586926..590579</td><td>1681</td><td>BREX-system-P-loop-protein-BrxC-[Bifidobacterium-bifidum]</td></tr>
<tr><td>WP_270200767.1</td><td>Type1.BREX</td><td>REC+LuxR-HTH->HISKIN-><-?<-SbcD<-?<-?||Pentapeptide+Pentapeptide+Pentapeptide+Pentapeptide-> BrxA-DNA-binding -> BrxB-BrxF-iSTAND -> DUF6079 -p*-> DUF6079 -p-> N6-METHYLASE -> REase ->Schlafen+HATPase+wHTH-> PglZ -> SIGMA+OB-Fold+AAA-ATPase+HKD-DNase-BrxL ->?-><-RNaseH</td><td>467</td><td>Bacteria>Terrabacteria_group>Actinomycetota>Actinomycetes>Bifidobacteriales>Bifidobacteriaceae</td><td>GCF_027671795.1</td><td>Bifidobacterium_pseudocatenulatum</td><td>PGT11_RS00265</td><td>67391..68794</td><td>28026</td><td>BREX-system-P-loop-protein-BrxC-[Bifidobacterium-pseudocatenulatum]</td></tr>
<tr><td>WP_270199955.1</td><td>Type1.BREX</td><td><-SbcD<-?<-?||?->||Pentapeptide+Pentapeptide+Pentapeptide+Pentapeptide-> BrxA-DNA-binding -> BrxB-BrxF-iSTAND -> DUF6079 -p-> N6-METHYLASE -> REase ->Schlafen+HATPase+wHTH-> PglZ -> SIGMA+OB-Fold+AAA-ATPase+HKD-DNase-BrxL ->?-><-RNaseH</td><td>1044</td><td>Bacteria>Terrabacteria_group>Actinomycetota>Actinomycetes>Bifidobacteriales>Bifidobacteriaceae</td><td>GCF_027675485.1</td><td>Bifidobacterium_pseudocatenulatum</td><td>PGV18_RS06405</td><td>41954..45088</td><td>28026</td><td>BREX-system-P-loop-protein-BrxC-[Bifidobacterium-pseudocatenulatum]</td></tr>
<tr><td>WP_117726345.1</td><td>Type1.BREX</td><td>ZnR->?->ABC-ATPase+SbcC->?->?->?-> BrxA-DNA-binding -> BrxB-BrxF-iSTAND -> DUF6079 *-> N6-METHYLASE ->?->Schlafen+HATPase+wHTH-> PglZ -> SIGMA+OB-Fold+AAA-ATPase+HKD-DNase-BrxL ->?-><- REase ||?-><-Glyoxalase</td><td>1217</td><td>Bacteria>Terrabacteria_group>Actinomycetota>Actinomycetes>Bifidobacteriales>Bifidobacteriaceae</td><td>GCF_027685785.1</td><td>Bifidobacterium_bifidum</td><td>PG427_RS02260</td><td>555349..559002</td><td>1681</td><td>BREX-system-P-loop-protein-BrxC-[Bifidobacterium-bifidum]</td></tr>
<tr><td>WP_270558503.1</td><td>Type1.BREX</td><td><-ParB+HNH+RAMA<-ParB+HNH<-ZnR<-ZnR+GP<- SIGMA+OB-Fold+AAA-ATPase+HKD-DNase-BrxL <- PglZ <-Schlafen+HATPase+wHTH<- REase <- N6-METHYLASE <- DUF6079 *<- BrxB-BrxF-iSTAND <- BrxA-DNA-binding <-?<-?||?-><-Peptidase_M78<-HigA</td><td>1219</td><td>Bacteria>Terrabacteria_group>Actinomycetota>Actinomycetes>Bifidobacteriales>Bifidobacteriaceae</td><td>GCF_027690155.1</td><td>Bifidobacterium_adolescentis</td><td>PG057_RS05020</td><td>457971..461630</td><td>1680</td><td>BREX-system-P-loop-protein-BrxC-[Bifidobacterium-adolescentis]</td></tr>
<tr><td>WP_270554212.1</td><td>Type1.BREX</td><td><-SbcD||HAD->PhdYeFM_antitox->RelE-ParE-><-?<-?<- REase || BrxA-DNA-binding -> BrxB-BrxF-iSTAND -> DUF6079 *-> N6-METHYLASE -> REase ->Schlafen+HATPase+wHTH-> PglZ -> SIGMA+OB-Fold+AAA-ATPase+HKD-DNase-BrxL ->ZnR+GP->ZnR->ParB+HNH+RAMA-></td><td>1219</td><td>Bacteria>Terrabacteria_group>Actinomycetota>Actinomycetes>Bifidobacteriales>Bifidobacteriaceae</td><td>GCF_027695195.1</td><td>Bifidobacterium_adolescentis</td><td>PG742_RS01120</td><td>252838..256497</td><td>1680</td><td>BREX-system-P-loop-protein-BrxC-[Bifidobacterium-adolescentis]</td></tr>
<tr><td>WP_196335743.1</td><td>Type1.BREX</td><td><-Glyoxalase||DSBH+AraC-HTH+AraC-HTH-><-ZnR<- SIGMA+OB-Fold+AAA-ATPase+HKD-DNase-BrxL <- PglZ <-Schlafen+HATPase+wHTH<- REase <- N6-METHYLASE <- DUF6079 *<- BrxB-BrxF-iSTAND <- BrxA-DNA-binding || REase -><-HEPN<-?||HTH-><-35exo+BRCT<-35exo+HRDC</td><td>1217</td><td>Bacteria>Terrabacteria_group>Actinomycetota>Actinomycetes>Bifidobacteriales>Bifidobacteriaceae</td><td>GCF_959600175.1</td><td>Bifidobacterium_dentium</td><td>IHQ76_RS06180</td><td>228522..232175</td><td>1689</td><td>BREX-system-P-loop-protein-BrxC-[Bifidobacterium-dentium]</td></tr>
<tr><td>WP_030399158.1</td><td>Type1.BREX</td><td>OLD-ABC+TOPRIM->SF1-SF2-Helicase->?-><-DNA_glycosylase|| REase -><- SIGMA+OB-Fold+AAA-ATPase+HKD-DNase-BrxL <- PglZ <- N6-METHYLASE <- DUF6079 *<- BrxB-BrxF-iSTAND <- BrxA-DNA-binding <-XIS-HTH<-pPIWI+MID+RNaseH<-pPIWI+SF1-SF2-Helicase<-pPIWI+pPIWI<-HTH_3+HTH_3||?-><-DDE_Tnp_1_4</td><td>1161</td><td>Bacteria>Terrabacteria_group>Actinomycetota>Actinomycetes>Kitasatosporales>Streptomycetaceae</td><td>GCF_000719665.1</td><td>unclassified_Streptomyces</td><td>ADK87_RS19375</td><td>27273..30758</td><td>2593676</td><td>MULTISPECIES:-BREX-system-P-loop-protein-BrxC-[unclassified-Streptomyces]</td></tr>
<tr><td>WP_062022136.1</td><td>Type1.BREX</td><td>METHYLASE->?->?-><-DNA_glycosylase|| REase -><- SIGMA+OB-Fold+AAA-ATPase+HKD-DNase-BrxL <- PglZ <- N6-METHYLASE <- DUF6079 *<- BrxB-BrxF-iSTAND <- BrxA-DNA-binding <-XIS-HTH<-pPIWI+MID+RNaseH<-pPIWI+SF1-SF2-Helicase<-pPIWI+pPIWI<-HTH_3+HTH_3|| REase +DUF5655-><-N6-N4-METHYLASE</td><td>1161</td><td>Bacteria>Terrabacteria_group>Actinomycetota>Actinomycetes>Kitasatosporales>Streptomycetaceae</td><td>GCF_001513985.1</td><td>Streptomyces_griseorubiginosus</td><td>AQI84_07455</td><td>596850..600335</td><td>67304</td><td>BREX-system-P-loop-protein-BrxC-[Streptomyces-griseorubiginosus]</td></tr>
<tr><td>TFB71537.1</td><td>Type1.BREX</td><td><-URI-DNase<-VirB11+MuB-ATPase<-HEPN+ REase <- REase <- SIGMA+OB-Fold+AAA-ATPase+HKD-DNase-BrxL <- PglZ <- N6-METHYLASE <- DUF6079 *<- BrxB-BrxF-iSTAND <- BrxA-DNA-binding <-XIS-HTH<-ABhydrolase<-?<-MPTase<-HTH||ArdCN+MPTase->Primpol+PriCT1-></td><td>1170</td><td>Bacteria>Terrabacteria_group>Actinomycetota>Actinomycetes>Micrococcales>Microbacteriaceae</td><td>GCA_004402535.1</td><td>Cryobacterium_sp._HLT2-23</td><td>E3006_11880</td><td>40394..43906</td><td>1259145</td><td>BREX-system-P-loop-protein-BrxC-[Cryobacterium-glaciale]</td></tr>
<tr><td>NYD70997.1</td><td>Type1.BREX</td><td>LZ_Tnp_IS481+HTH_32+rve+rve_3->?-><-?||?-> BrxA-DNA-binding -> BrxB-BrxF-iSTAND -> DUF6079 *-> N6-METHYLASE -> PglZ -> SIGMA+OB-Fold+AAA-ATPase+Lon-Protease-BrxL -><- REase <-SF1-SF2-Helicase<- REase <-YTCD-HTH</td><td>1151</td><td>Bacteria>Terrabacteria_group>Actinomycetota>Actinomycetes>Micrococcales>Microbacteriaceae</td><td>GCA_013409865.1</td><td>Herbiconiux_flava</td><td>BJ984_002155</td><td>2240959..2244414</td><td>881268</td><td>hypothetical-protein-[Herbiconiux-flava]</td></tr>
<tr><td>WP_104309935.1</td><td>Type1.BREX</td><td>TIR-><-?||?-><-TIR<- SIGMA+OB-Fold+AAA-ATPase+HKD-DNase-BrxL <- PglZ <- REase <- N6-METHYLASE <- DUF6079 *<- BrxB-BrxF-iSTAND <- BrxA-DNA-binding <-XIS-HTH<-?||GDResolvase+HIN-HTH-><-Acyltransferase<-DSBH<-SIGMA-HTH</td><td>1162</td><td>Bacteria>Terrabacteria_group>Actinomycetota>Actinomycetes>Micrococcales>Microbacteriaceae</td><td>GCF_002932095.1</td><td>Rathayibacter_sp._AY1G1</td><td>C5C36_RS14015</td><td>21701..25189</td><td>2080564</td><td>BREX-system-P-loop-protein-BrxC-[Rathayibacter-sp.-AY1G1]</td></tr>
<tr><td>WP_241980122.1</td><td>Type1.BREX</td><td><-URI-DNase<-VirB11+MuB-ATPase<-HEPN+ REase <- REase <- SIGMA+OB-Fold+AAA-ATPase+HKD-DNase-BrxL <- PglZ <- N6-METHYLASE <- DUF6079 *<- BrxB-BrxF-iSTAND <- BrxA-DNA-binding <-XIS-HTH<-ABhydrolase<-?<-MPTase<-HTH||ArdCN+MPTase->Primpol+PriCT1-></td><td>1172</td><td>Bacteria>Terrabacteria_group>Actinomycetota>Actinomycetes>Micrococcales>Microbacteriaceae</td><td>GCF_004402535.1</td><td>Cryobacterium_sp._HLT2-23</td><td>E3006_RS11880</td><td>40394..43912</td><td>1259145</td><td>BREX-system-P-loop-protein-BrxC-[Cryobacterium-glaciale]</td></tr>
<tr><td>WP_163473715.1</td><td>Type1.BREX</td><td>Aminotran_4->?->tRNA->XIS-HTH->Phage_integrase->TrwC+SF1-SF2-Helicase+TOPRIM->XIS-HTH-> BrxA-DNA-binding -> BrxB-BrxF-iSTAND -> DUF6079 *-> N6-METHYLASE -> REase -> PglZ -> SIGMA+OB-Fold+AAA-ATPase+HKD-DNase-BrxL -> N6-METHYLASE ->MORC+HISKIN->?->PIN->PIN-></td><td>1162</td><td>Bacteria>Terrabacteria_group>Actinomycetota>Actinomycetes>Micrococcales>Microbacteriaceae</td><td>GCF_010686725.1</td><td>Galbitalea_soli</td><td>HNR17_RS04680</td><td>978511..981999</td><td>1268042</td><td>BREX-system-P-loop-protein-BrxC-[Galbitalea-soli]</td></tr>
<tr><td>WP_166866044.1</td><td>Type1.BREX</td><td><- REase <-SbcC<-JetB<-JetA<- SIGMA+OB-Fold+AAA-ATPase+HKD-DNase-BrxL <- PglZ <- N6-METHYLASE <- DUF6079 *<- BrxB-BrxF-iSTAND <- BrxA-DNA-binding <- REase <-tRNA<-?<-?<-?||?-><-?<-Peptidase_M28</td><td>1160</td><td>Bacteria>Terrabacteria_group>Actinomycetota>Actinomycetes>Micrococcales>Microbacteriaceae</td><td>GCF_011751865.2</td><td>Salinibacterium_sp._ZJ70</td><td>HCR12_RS10200</td><td>2144642..2148124</td><td>2708084</td><td>BREX-system-P-loop-protein-BrxC-[Salinibacterium-sp.-ZJ70]</td></tr>
<tr><td>WP_169077984.1</td><td>Type1.BREX</td><td>BrxA-DNA-binding -> BrxB-BrxF-iSTAND -> DUF6079 *-> N6-METHYLASE -> REase -> PglZ -> SIGMA+OB-Fold+AAA-ATPase+HKD-DNase-BrxL ->SIR2->HerA/FtsK-><-Peptidase_M48||?-> REase +HTH+35exo+Small-B-barrel+SF1-SF2-Helicase-></td><td>1162</td><td>Bacteria>Terrabacteria_group>Actinomycetota>Actinomycetes>Micrococcales>Microbacteriaceae</td><td>GCF_012845865.1</td><td>Yonghaparkia</td><td>HFV16_RS05520</td><td>206976..210464</td><td>355929</td><td>MULTISPECIES:-BREX-system-P-loop-protein-BrxC-[Yonghaparkia]</td></tr>
<tr><td>WP_179548032.1</td><td>Type1.BREX</td><td>YTCD-HTH-> REase ->SF1-SF2-Helicase-> REase -><- SIGMA+OB-Fold+AAA-ATPase+Lon-Protease-BrxL <- PglZ <- N6-METHYLASE <- DUF6079 *<- BrxB-BrxF-iSTAND <- BrxA-DNA-binding</td><td>1151</td><td>Bacteria>Terrabacteria_group>Actinomycetota>Actinomycetes>Micrococcales>Microbacteriaceae</td><td>GCF_027922025.1</td><td>Herbiconiux_flava</td><td>PKE81_RS17675</td><td>1519620..1523075</td><td>881268</td><td>BREX-system-P-loop-protein-BrxC-[Herbiconiux-flava]</td></tr>
<tr><td>MCO4261480.1</td><td>Type1.BREX</td><td><-SbcC<-JetB<-JetA<-HEPN<-SF1-SF2-Helicase<- REase <- SIGMA+OB-Fold+AAA-ATPase+Lon-Protease-BrxL <- PglZ <- N6-METHYLASE <- DUF6079 *<- BrxB-BrxF-iSTAND <- BrxA-DNA-binding <-HKD-DNase+SF1-SF2-Helicase+HTH+HTH+SRA||URI-DNase->MACRODOMAIN+SIR2->URI-DNase-><-?<-NUDIX</td><td>1151</td><td>Bacteria>Terrabacteria_group>Actinomycetota>Actinomycetes>Micrococcales>Micrococcaceae</td><td>GCA_023913835.1</td><td>Pseudarthrobacter_sp._MDT3-26</td><td>NG697_00745</td><td>166559..170014</td><td>2953651</td><td>BREX-system-P-loop-protein-BrxC-[Pseudarthrobacter-sp.-MDT3-26]</td></tr>
<tr><td>WP_193340403.1</td><td>Type1.BREX</td><td><- SIGMA+OB-Fold+AAA-ATPase+HKD-DNase-BrxL <- PglZ <- REase <- N6-METHYLASE <- DUF6079 *<- BrxB-BrxF-iSTAND <- BrxA-DNA-binding <-XIS-HTH||?-><-EVE+ REase</td><td>1153</td><td>Bacteria>Terrabacteria_group>Actinomycetota>Actinomycetes>Micrococcales>Micrococcaceae</td><td>GCF_014926485.1</td><td>Pseudarthrobacter_sp._AB1</td><td>DAD99_RS03725</td><td>12045..15506</td><td>2138309</td><td>BREX-system-P-loop-protein-BrxC-[Pseudarthrobacter-sp.-AB1]</td></tr>
<tr><td>WP_252613231.1</td><td>Type1.BREX</td><td><-SbcC<-JetB<-JetA<-HEPN<-SF1-SF2-Helicase<- REase <- SIGMA+OB-Fold+AAA-ATPase+Lon-Protease-BrxL <- PglZ <- N6-METHYLASE <- DUF6079 *<- BrxB-BrxF-iSTAND <- BrxA-DNA-binding <-HKD-DNase+SF1-SF2-Helicase+HTH+HTH+SRA||URI-DNase->MACRODOMAIN+SIR2->URI-DNase-><-?<-NUDIX</td><td>1151</td><td>Bacteria>Terrabacteria_group>Actinomycetota>Actinomycetes>Micrococcales>Micrococcaceae</td><td>-</td><td>Pseudarthrobacter_sp._MDT3-26</td><td>NG697_00745</td><td>166559..170014</td><td>2953651</td><td>BREX-system-P-loop-protein-BrxC-[Pseudarthrobacter-sp.-MDT3-26]</td></tr>
<tr><td>SDF46422.1</td><td>Type1.BREX</td><td>ABC-ATPase+SbcC->Cysteine-Desulfurase->?->?->?-> BrxA-DNA-binding -> BrxB-BrxF-iSTAND -> DUF6079 *-> N6-METHYLASE -> PglZ -> SIGMA+OB-Fold+AAA-ATPase+HKD-DNase-BrxL -> REase ->PAIREDC-HTH-></td><td>1166</td><td>Bacteria>Terrabacteria_group>Actinomycetota>Actinomycetes>Micrococcales>Promicromonosporaceae</td><td>GCA_900102445.1</td><td>Cellulosimicrobium_cellulans</td><td>SAMN04487781_1443</td><td>34344..37844</td><td>1710</td><td>hypothetical-protein-SAMN04487781_1443-[Cellulosimicrobium-cellulans]</td></tr>
<tr><td>WP_163542003.1</td><td>Type1.BREX</td><td><-HIN-HTH||SbcC->?-><-?<- SIGMA+OB-Fold+AAA-ATPase+HKD-DNase-BrxL <- PglZ <- REase <- N6-METHYLASE <- DUF6079 *<- BrxB-BrxF-iSTAND <- BrxA-DNA-binding <-XIS-HTH<-?<-?||CN_hydrolase+NAD_synthase->?-><-Pkinase<-RelE-ParE</td><td>1154</td><td>Bacteria>Terrabacteria_group>Actinomycetota>Actinomycetes>Micrococcales>Ruaniaceae</td><td>GCF_010994285.1</td><td>Occultella_kanbiaonis</td><td>G7B41_RS06055</td><td>118944..122408</td><td>2675754</td><td>BREX-system-P-loop-protein-BrxC-[Occultella-kanbiaonis]</td></tr>
<tr><td>WP_208468634.1</td><td>Type6.BREX</td><td>XIS-HTH->URI-DNase-> REase -> BrxE -> BrxA-DNA-binding -> BrxB-BrxF-iSTAND -> DUF6079 *->Schlafen+HATPase+wHTH-> N6-METHYLASE -> N6-METHYLASE -> PglZ -> SIGMA+OB-Fold+AAA-ATPase+Lon-Protease-BrxL -><-DDE_Tnp_1_assoc+DDE_Tnp_1</td><td>1183</td><td>Bacteria>Terrabacteria_group>Actinomycetota>Actinomycetes>Micromonosporales>Micromonosporaceae</td><td>GCF_017592555.1</td><td>Actinoplanes_flavus</td><td>J5X75_RS18430</td><td>129103..132654</td><td>2820290</td><td>BREX-system-P-loop-protein-BrxC-[Actinoplanes-flavus]</td></tr>
<tr><td>MBP2454134.1</td><td>Type6.BREX</td><td><-McrB-AAA<-Preader7<-HEPN+ SF1-SF2-Helicase+ REase <-?|| BrxE -p-> BrxA-DNA-binding -> BrxB-BrxF-iSTAND -> DUF6079 -p->zf-ISL3+HTH_Tnp_ISL3+DDE_Tnp_ISL3-> DUF6079 -p*-> N6-METHYLASE -p-><- REase <- N6-METHYLASE <-?<-TetR+HTH<-Phage_integrase+Phage_integrase<-Phage_integrase+Phage_integrase<-Phage_int_SAM_1+Phage_integrase<-DNA_glycosylase</td><td>870</td><td>Bacteria>Terrabacteria_group>Actinomycetota>Actinomycetes>Mycobacteriales>Mycobacteriaceae</td><td>GCA_017876775.1</td><td>Mycolicibacterium_lutetiense</td><td>JOF57_004047</td><td>2230868..2233480</td><td>1641992</td><td>hypothetical-protein-[Mycolicibacterium-lutetiense]</td></tr>
<tr><td>WP_253659814.1</td><td>Type1.BREX</td><td><- REase <- SIGMA+OB-Fold+AAA-ATPase+HKD-DNase-BrxL <- PglZ <- N6-METHYLASE <- DUF6079 *<- BrxB-BrxF-iSTAND <- BrxA-DNA-binding ||DNA_glycosylase->?-> REase -><-?<-?<-?||PhdYeFM_antitox-></td><td>1156</td><td>Bacteria>Terrabacteria_group>Actinomycetota>Actinomycetes>Mycobacteriales>Nocardiaceae</td><td>GCF_024171815.1</td><td>Williamsia_maris</td><td>LX13_RS03015</td><td>647894..651364</td><td>72806</td><td>BREX-system-P-loop-protein-BrxC-[Williamsia-maris]</td></tr>
<tr><td>WP_068111667.1</td><td>Type1.BREX</td><td>VirB11+MuB-ATPase->?-><-?||XIS-HTH-> BrxA-DNA-binding -> BrxB-BrxF-iSTAND -> DUF6079 *-> N6-METHYLASE -> REase -> PglZ -> SIGMA+OB-Fold+AAA-ATPase+HKD-DNase-BrxL ->OLD-ABC+TOPRIM->SF1-SF2-Helicase->?->HTH+MPTase->Phage_int_SAM_1+Phage_integrase-></td><td>1154</td><td>Bacteria>Terrabacteria_group>Actinomycetota>Actinomycetes>Propionibacteriales>Nocardioidaceae</td><td>GCF_001653335.1</td><td>Nocardioides_dokdonensis</td><td>I601_RS15600</td><td>3293021..3296485</td><td>450734</td><td>BREX-system-P-loop-protein-BrxC-[Nocardioides-dokdonensis]</td></tr>
<tr><td>WP_166189200.1</td><td>Type1.BREX</td><td>XIS-HTH-> BrxA-DNA-binding -> BrxB-BrxF-iSTAND -> DUF6079 *-> N6-METHYLASE -> PglZ -> SIGMA+OB-Fold+AAA-ATPase+HKD-DNase-BrxL -> REase -><-Phage_int_SAM_3+Phage_integrase<-XIS-HTH</td><td>1154</td><td>Bacteria>Terrabacteria_group>Actinomycetota>Actinomycetes>Propionibacteriales>Nocardioidaceae</td><td>GCF_011250565.1</td><td>Nocardioides_sp._IC4_145</td><td>G6553_RS03445</td><td>61321..64785</td><td>2714037</td><td>BREX-system-P-loop-protein-BrxC-[Nocardioides-sp.-IC4_145]</td></tr>
<tr><td>NLI84511.1</td><td>Type1.BREX</td><td>HIN-HTH-><-SMODS<-SLATT<-?<-AbiEii<- REase <- SIGMA+OB-Fold+AAA-ATPase+HKD-DNase-BrxL <- PglZ <- N6-METHYLASE <- DUF6079 *<- BrxB-BrxF-iSTAND <- BrxA-DNA-binding ||?->TIR->?-><-DUF4096+DDE_Tnp_1_2</td><td>1160</td><td>Bacteria>Terrabacteria_group>Actinomycetota>Actinomycetes>Propionibacteriales>Propionibacteriaceae</td><td>GCA_012517405.1</td><td>Propionibacterium_sp.</td><td>GX440_03735</td><td>112337..115819</td><td>1977903</td><td>BREX-system-P-loop-protein-BrxC-[Propionibacterium-sp.]</td></tr>
<tr><td>WP_021796710.1</td><td>Type1.BREX</td><td>XIS-HTH-><-?<-?<-?<-Glyco_hydro_36N+Melibiase+Glyco_hydro_36C|| BrxA-DNA-binding -> BrxB-BrxF-iSTAND -> DUF6079 *-> N6-METHYLASE -> REase ->?->HATPase+wHTH-> PglZ -> SIGMA+OB-Fold+AAA-ATPase+HKD-DNase-BrxL ->SF1-SF2-Helicase+HTH-></td><td>1171</td><td>Bacteria>Terrabacteria_group>Actinomycetota>Actinomycetes>Propionibacteriales>Propionibacteriaceae</td><td>GCF_000478805.1</td><td>Propionibacterium_acidifaciens</td><td>HMPREF0682_RS02920</td><td>11275..14790</td><td>556499</td><td>BREX-system-P-loop-protein-BrxC-[Propionibacterium-acidifaciens]</td></tr>
<tr><td>WP_048768895.1</td><td>Type1.BREX</td><td><-DUF2637+HIN-HTH<-Primpol+PriCT1<-ArdCN+MPTase<-AraC-HTH||XIS-HTH-> BrxA-DNA-binding -> BrxB-BrxF-iSTAND -> DUF6079 *-> N6-METHYLASE -> REase ->?->Schlafen+HATPase+wHTH-> PglZ -> SIGMA+OB-Fold+AAA-ATPase+HKD-DNase-BrxL ->SIR2->HerA/FtsK->VirB11+MuB-ATPase-></td><td>1170</td><td>Bacteria>Terrabacteria_group>Actinomycetota>Actinomycetes>Propionibacteriales>Propionibacteriaceae</td><td>GCF_001050955.1</td><td>Propionibacterium_freudenreichii</td><td>PFCIRM1025_RS01850</td><td>33525..37037</td><td>1744</td><td>BREX-system-P-loop-protein-BrxC-[Propionibacterium-freudenreichii]</td></tr>
<tr><td>WP_055346094.1</td><td>Type1.BREX</td><td><-DUF2637+HIN-HTH<-Primpol+PriCT1<-ArdCN+MPTase<-AraC-HTH||XIS-HTH-> BrxA-DNA-binding -> BrxB-BrxF-iSTAND -> DUF6079 *-> N6-METHYLASE -> REase ->?->Schlafen+HATPase+wHTH-> PglZ -> SIGMA+OB-Fold+AAA-ATPase+HKD-DNase-BrxL ->SIR2->HerA/FtsK->VirB11+MuB-ATPase-></td><td>1170</td><td>Bacteria>Terrabacteria_group>Actinomycetota>Actinomycetes>Propionibacteriales>Propionibacteriaceae</td><td>GCF_001369055.1</td><td>Propionibacterium_freudenreichii</td><td>PFCIRM516_RS05500</td><td>48312..51824</td><td>1744</td><td>BREX-system-P-loop-protein-BrxC-[Propionibacterium-freudenreichii]</td></tr>
<tr><td>WP_142096818.1</td><td>Type1.BREX</td><td>MODE-HTH+PBPII-><-?||?->?-> REase -> BrxA-DNA-binding -> BrxB-BrxF-iSTAND -> DUF6079 *-> N6-METHYLASE -> PglZ ->PolB->?-> SIGMA+OB-Fold+AAA-ATPase+Lon-Protease-BrxL ->HTH_31->?-><-DUF4352||HHH_5+SNF2_N+SF1-SF2-Helicase-></td><td>1155</td><td>Bacteria>Terrabacteria_group>Actinomycetota>Actinomycetes>Pseudonocardiales>Pseudonocardiaceae</td><td>GCF_006717045.1</td><td>Pseudonocardia_cypriaca</td><td>FB388_RS03550</td><td>743160..746627</td><td>882449</td><td>BREX-system-P-loop-protein-BrxC-[Pseudonocardia-cypriaca]</td></tr>
<tr><td>WP_154435944.1</td><td>Type1.BREX</td><td>ACET->ParB+HNH-><-?<-TraI-VirD2-RCRE<-TraJ-RHH||TraJ-RHH-> BrxA-DNA-binding -> BrxB-BrxF-iSTAND -> DUF6079 *->?-> N6-METHYLASE ->?-><- REase <-XIS-HTH<-Phage_int_SAM_5+Phage_integrase||HTH_3->?->Schlafen+HATPase+wHTH-></td><td>1196</td><td>Bacteria>Terrabacteria_group>Actinomycetota>Coriobacteriia>Coriobacteriales>Atopobiaceae</td><td>GCF_009695875.1</td><td>Olsenella_porci</td><td>FYJ68_RS09105</td><td>10364..13954</td><td>2652279</td><td>BREX-system-P-loop-protein-BrxC-[Olsenella-porci]</td></tr>
<tr><td>RRF95308.1</td><td>Type1.BREX</td><td><-TraI-VirD2-RCRE<-TraJ-RHH||TraJ-RHH-> BrxA-DNA-binding -> BrxB-BrxF-iSTAND -> DUF6079 *-> PglZ -> N6-METHYLASE ->?-> SIGMA+OB-Fold+AAA-ATPase+HKD-DNase-BrxL -><- REase <-XIS-HTH<-Phage_int_SAM_5+Phage_integrase</td><td>1196</td><td>Bacteria>Terrabacteria_group>Actinomycetota>Coriobacteriia>Coriobacteriales>Coriobacteriaceae</td><td>GCA_003862165.1</td><td>Coriobacteriaceae_bacterium</td><td>DUD31_01665</td><td>3961..7551</td><td>2011094</td><td>BREX-system-P-loop-protein-BrxC-[Coriobacteriaceae-bacterium]</td></tr>
<tr><td>WP_204559883.1</td><td>Type1.BREX</td><td>Cysteine-Desulfurase->?-> SIGMA+OB-Fold+AAA-ATPase+HKD-DNase-BrxL <- PglZ <-Schlafen+HATPase+wHTH<- REase <- N6-METHYLASE <-HEPN<- DUF6079 *<- BrxB-BrxF-iSTAND <- BrxA-DNA-binding <-TraJ-RHH</td><td>1171</td><td>Bacteria>Terrabacteria_group>Actinomycetota>Coriobacteriia>Coriobacteriales>Coriobacteriaceae</td><td>GCF_016902615.1</td><td>Collinsella_tanakaei</td><td>H6A21_RS01790</td><td>55502..59017</td><td>626935</td><td>BREX-system-P-loop-protein-BrxC-[Collinsella-tanakaei]</td></tr>
<tr><td>QIN77519.1</td><td>Type1.BREX</td><td>Thermonuclease->DOC+HTH->?->N6-N4-METHYLASE->SF1-SF2-Helicase-> BrxA-DNA-binding -> BrxB-BrxF-iSTAND -> DUF6079 *->?-> N6-METHYLASE -> REase ->?->PIN-> PglZ -> SIGMA+OB-Fold+AAA-ATPase+Lon-Protease-BrxL -><-?<-BirA-HTH</td><td>1167</td><td>Bacteria>Terrabacteria_group>Actinomycetota>Rubrobacteria>Rubrobacterales>Rubrobacteraceae</td><td>GCA_011492965.1</td><td>Rubrobacter_marinus</td><td>GBA65_02255</td><td>450674..454177</td><td>2653852</td><td>BREX-system-P-loop-protein-BrxC-[Rubrobacter-marinus]</td></tr>
<tr><td>WP_166395201.1</td><td>Type1.BREX</td><td>Thermonuclease->DOC+HTH->?->N6-N4-METHYLASE->SF1-SF2-Helicase-> BrxA-DNA-binding -> BrxB-BrxF-iSTAND -> DUF6079 *->?-> N6-METHYLASE -> REase ->?->PIN-> PglZ -> SIGMA+OB-Fold+AAA-ATPase+Lon-Protease-BrxL -><-?<-BirA-HTH</td><td>1151</td><td>Bacteria>Terrabacteria_group>Actinomycetota>Rubrobacteria>Rubrobacterales>Rubrobacteraceae</td><td>GCF_011492965.1</td><td>Rubrobacter_marinus</td><td>GBA65_RS02320</td><td>450722..454177</td><td>2653852</td><td>BREX-system-P-loop-protein-BrxC-[Rubrobacter-marinus]</td></tr>
<tr><td>APC47779.1</td><td>Type6.BREX</td><td><-TetR-HTH||ABC-ATPase+SbcC->?->HEPN+PAS+DUF4011+ SF1-SF2-Helicase+ REase -> REase -> BrxE -> BrxA-DNA-binding -> BrxB-BrxF-iSTAND -> DUF6079 *-> N6-METHYLASE -> PglZ ->BrxD-AAA->SF1-SF2-Helicase+wHTH+Small-B-barrel+DUF1998->ParB+HNH->NUDIX->?-><-HKD-DNase+SF1-SF2-Helicase+HTH+HTH+SRA</td><td>1225</td><td>Bacteria>Terrabacteria_group>Bacillota>Bacilli>Bacillales>Bacillaceae</td><td>GCA_001878675.1</td><td>Virgibacillus_halodenitrificans</td><td>BME96_06155</td><td>1179738..1183415</td><td>1482</td><td>hypothetical-protein-BME96_06155-[Virgibacillus-halodenitrificans]</td></tr>
<tr><td>GGJ55411.1</td><td>Type1.BREX</td><td><-Phage_int_SAM_1+Phage_integrase||?-><-SF1-SF2-Helicase+wHTH+Small-B-barrel+DUF1998<-BrxD-AAA<- PglZ <- DUF6079 *<- BrxB-BrxF-iSTAND <- BrxA-DNA-binding <- N6-METHYLASE <- REase <-?<-?<-SF1-SF2-Helicase<-OLD-ABC+TOPRIM</td><td>1365</td><td>Bacteria>Terrabacteria_group>Bacillota>Bacilli>Bacillales>Bacillaceae</td><td>GCA_014647615.1</td><td>Virgibacillus_kapii</td><td>GCM10007111_17110</td><td>601764..605861</td><td>1638645</td><td>hypothetical-protein-GCM10007111_17110-[Virgibacillus-kapii]</td></tr>
<tr><td>MCI1589808.1</td><td>Type1.BREX</td><td>ParD_antitoxin-><- REase <- SIGMA+OB-Fold+AAA-ATPase+Lon-Protease-BrxL <- PglZ <-?<-?<-HEPN<- N6-METHYLASE <- DUF6079 *<- BrxB-BrxF-iSTAND <- BrxA-DNA-binding <-?<-?<-HTH_31<-TRAM+METHYLASE||MerR-HTH-></td><td>1190</td><td>Bacteria>Terrabacteria_group>Bacillota>Bacilli>Bacillales>Bacillaceae</td><td>GCA_022646685.1</td><td>Heyndrickxia_oleronia</td><td>LKH79_04555</td><td>33937..37509</td><td>38875</td><td>BREX-system-P-loop-protein-BrxC-[Heyndrickxia-oleronia]</td></tr>
<tr><td>MCI1592662.1</td><td>Type1.BREX</td><td><-HTH_3<-?||?->ParD_antitoxin-><- REase <- SIGMA+OB-Fold+AAA-ATPase+Lon-Protease-BrxL <- PglZ <-AbiGi+seqA+RAMA<- N6-METHYLASE <- DUF6079 *<- BrxB-BrxF-iSTAND <- BrxA-DNA-binding <-?<-?<-HTH_31<-HTH+HTH+SF1-SF2-Helicase</td><td>1190</td><td>Bacteria>Terrabacteria_group>Bacillota>Bacilli>Bacillales>Bacillaceae</td><td>GCA_022646685.1</td><td>Heyndrickxia_oleronia</td><td>LKH79_19270</td><td>20507..24079</td><td>38875</td><td>BREX-system-P-loop-protein-BrxC-[Heyndrickxia-oleronia]</td></tr>
<tr><td>WP_233275777.1</td><td>Type1.BREX</td><td><-wHTH+HNH<-?<-wHTH||?-> BrxA-DNA-binding -> BrxB-BrxF-iSTAND -> DUF6079 *-> N6-METHYLASE -> PglZ -> SIGMA+OB-Fold+AAA-ATPase+Lon-Protease-BrxL -> REase ->ParB+HNH->?-><-SH3</td><td>1184</td><td>Bacteria>Terrabacteria_group>Bacillota>Bacilli>Bacillales>Bacillaceae</td><td>GCF_000429685.1</td><td>Salibacterium_aidingense</td><td>G475_RS0113415</td><td>15614..19168</td><td>384933</td><td>BREX-system-P-loop-protein-BrxC-[Salibacterium-aidingense]</td></tr>
<tr><td>WP_026673780.1</td><td>Type1.BREX</td><td>ParD_antitoxin-><- SIGMA+OB-Fold+AAA-ATPase+Lon-Protease-BrxL <- PglZ <- REase <-OLD-ABC <- N6-METHYLASE <- DUF6079 *<- BrxB-BrxF-iSTAND <- BrxA-DNA-binding <-?<-?<-HTH_31</td><td>1188</td><td>Bacteria>Terrabacteria_group>Bacillota>Bacilli>Bacillales>Bacillaceae</td><td>GCF_000621445.1</td><td>Alkalihalobacillus_bogoriensis</td><td>T323_RS0113555</td><td>41147..44713</td><td>246272</td><td>BREX-system-P-loop-protein-BrxC-[Alkalihalobacillus-bogoriensis]</td></tr>
<tr><td>WP_053997117.1</td><td>Type1.BREX</td><td><-LZ_Tnp_IS66+zf-IS66+DDE_Tnp_IS66+DDE_Tnp_IS66_C<-?<-HIN-HTH||HTH_31-><-MULE-TRANSPOSASE||?->?-> BrxA-DNA-binding -> BrxB-BrxF-iSTAND -> DUF6079 *-> N6-METHYLASE ->RVT->RVT-> PglZ -> SIGMA+OB-Fold+AAA-ATPase+Lon-Protease-BrxL -> REase ->primpol+D5N+D5ATPase-><-HTH_58+RHH</td><td>1190</td><td>Bacteria>Terrabacteria_group>Bacillota>Bacilli>Bacillales>Bacillaceae</td><td>GCF_001281525.1</td><td>Lysinibacillus_macroides</td><td>ADM90_RS22630</td><td>144729..148301</td><td>33935</td><td>BREX-system-P-loop-protein-BrxC-[Lysinibacillus-macroides]</td></tr>
<tr><td>WP_062353349.1</td><td>Type1.BREX</td><td>HTH+HTH+SF1-SF2-Helicase->HTH_31->?->?-> BrxA-DNA-binding -> BrxB-BrxF-iSTAND -> DUF6079 *-> N6-METHYLASE ->HEPN-> PglZ -> SIGMA+OB-Fold+AAA-ATPase+Lon-Protease-BrxL -> REase -> REase -><-HTH_Tnp_1+HTH_21+rve+rve_3||?->Schlafen+HATPase+wHTH-></td><td>1190</td><td>Bacteria>Terrabacteria_group>Bacillota>Bacilli>Bacillales>Bacillaceae</td><td>GCF_001375515.1</td><td>Bacillus_kwashiorkori</td><td>BN1242_RS07000</td><td>23447..27019</td><td>1522318</td><td>BREX-system-P-loop-protein-BrxC-[Bacillus-kwashiorkori]</td></tr>
<tr><td>WP_058950067.1</td><td>Type1.BREX</td><td>IstB_IS21-><-SF1-SF2-Helicase+wHTH+Small-B-barrel+DUF1998<-BrxD-AAA<- PglZ <- DUF6079 *<- BrxB-BrxF-iSTAND <- BrxA-DNA-binding <- N6-METHYLASE <- REase <-?<-?<-?<-SF1-SF2-Helicase<-OLD-ABC+TOPRIM</td><td>1366</td><td>Bacteria>Terrabacteria_group>Bacillota>Bacilli>Bacillales>Bacillaceae</td><td>GCF_001485235.1</td><td>Oceanobacillus_picturae</td><td>OPHB3_RS08850</td><td>6149..10249</td><td>171693</td><td>hypothetical-protein-[Oceanobacillus-picturae]</td></tr>
<tr><td>WP_071743491.1</td><td>Type1.BREX</td><td>IONCHANNEL+PAS+HISKIN+MORC->REC+CITB-HTH+ZDNA-HTH->?-> BrxA-DNA-binding -> BrxB-BrxF-iSTAND -> DUF6079 *-> N6-METHYLASE -> PglZ -> SIGMA+OB-Fold+AAA-ATPase+Lon-Protease-BrxL ->?->?-> REase ->ABhydrolase-></td><td>1195</td><td>Bacteria>Terrabacteria_group>Bacillota>Bacilli>Bacillales>Bacillaceae</td><td>GCF_001884125.1</td><td>Bacillus</td><td>BAU22_RS23055</td><td>7102..10689</td><td>1386</td><td>MULTISPECIES:-BREX-system-P-loop-protein-BrxC-[Bacillus]</td></tr>
</table>

| Protein-ID | System | Gene-Neighbourhoods | Length | Taxonomy | Assembly-ID | Strain | Locus-Tag | Genomic-coordinates | TaxID | Fasta-defline |
|---|---|---|---|---|---|---|---|---|---|---|
| WP_121661845.1 | Type1.BREX | HAD+HAD-><-SF1-SF2-Helicase+wHTH+Small-B-barrel+DUF1998<-BrxD-AAA<- PglZ <- DUF6079 *<- BrxB-BrxF-iSTAND <- BrxA-DNA-binding <- N6-METHYLASE <- REase <-?<- SF1-SF2-Helicase+REase -p<- SF1-SF2-Helicase+REase -p<-SF1-SF2-Helicase<-wHTH+HNH | 1365 | Bacteria>Terrabacteria_group>Bacillota>Bacilli>Bacillales>Bacillaceae | GCF_003667825.1 | Metabacillus_litoralis | D9842_RS06665 | 1400368..1404465 | 152268 | hypothetical-protein-[Metabacillus-litoralis] |
| WP_126293910.1 | Type1.BREX | TRAM+METHYLASE-><-?\|\|HTH_31->?->?-> BrxA-DNA-binding -> BrxB-BrxF-iSTAND -> DUF6079 *-> N6-METHYLASE ->HEPN-> PglZ -> SIGMA+OB-Fold+AAA-ATPase+Lon-Protease-BrxL -> REase -><-ParD_antitoxin<-?\|\|?->HTH_3-> | 1190 | Bacteria>Terrabacteria_group>Bacillota>Bacilli>Bacillales>Bacillaceae | GCF_003966145.1 | Lysinibacillus_telephonicus | EKG35_RS07955 | 47066..50638 | 1714840 | BREX-system-P-loop-protein-BrxC-[Lysinibacillus-telephonicus] |
| WP_137744598.1 | Type1.BREX | HNH+NERD+ZnR->?-><-HTH_3\|\|GDResolvase->HTH_31->?->?-> BrxA-DNA-binding -> BrxB-BrxF-iSTAND -> DUF6079 *-> N6-METHYLASE ->?->RVT-> PglZ -> SIGMA+OB-Fold+AAA-ATPase+Lon-Protease-BrxL -> REase -><-ParD_antitoxin | 1190 | Bacteria>Terrabacteria_group>Bacillota>Bacilli>Bacillales>Bacillaceae | GCF_005502275.1 | Robertmurraya_siralis | FCJ93_RS20760 | 22138..25710 | 77777 | BREX-system-P-loop-protein-BrxC-[Robertmurraya-siralis] |
| WP_163243257.1 | Type1.BREX | BrxA-DNA-binding -> BrxB-BrxF-iSTAND -> DUF6079 *-> N6-METHYLASE ->RVT-> PglZ -> SIGMA+OB-Fold+AAA-ATPase+Lon-Protease-BrxL -> REase -><-HTH_58+RHH | 1190 | Bacteria>Terrabacteria_group>Bacillota>Bacilli>Bacillales>Bacillaceae | GCF_011008855.1 | Bacillus_aquiflavi | H1Z61_RS15490 | 1670..5242 | 2672567 | BREX-system-P-loop-protein-BrxC-[Bacillus-aquiflavi] |
| WP_171510262.1 | Type1.BREX | <-Phage_int_SAM_1+Phage_integrase\|\|?-><-SF1-SF2-Helicase+wHTH+Small-B-barrel+DUF1998<-BrxD-AAA<- PglZ <- DUF6079 *<- BrxB-BrxF-iSTAND <- BrxA-DNA-binding <- N6-METHYLASE <- REase <-?<-?<-SF1-SF2-Helicase<-OLD-ABC+TOPRIM | 1366 | Bacteria>Terrabacteria_group>Bacillota>Bacilli>Bacillales>Bacillaceae | GCF_013138395.1 | Niallia_circulans | HLK66_RS25080 | 5165831..5169931 | 1397 | hypothetical-protein-[Niallia-circulans] |
| WP_210367870.1 | Type1.BREX | <-HTH_Tnp\|\|?->?-> BrxA-DNA-binding -> BrxB-BrxF-iSTAND -> DUF6079 *-> N6-METHYLASE ->?->RVT-> PglZ -> SIGMA+OB-Fold+AAA-ATPase+Lon-Protease-BrxL -> REase ->ZnR+NACHT-> | 1190 | Bacteria>Terrabacteria_group>Bacillota>Bacilli>Bacillales>Bacillaceae | GCF_017912555.1 | Bacillus_sp._REN3 | JJQ88_RS20340 | 2531..6103 | 2802440 | BREX-system-P-loop-protein-BrxC-[Bacillus-sp.-REN3] |
| WP_213108269.1 | Type1.BREX | <-EVE+McrB-AAA<-SbcC<-?<-HNH<-TRPR-HTH<- SIGMA+OB-Fold+AAA-ATPase+Lon-Protease-BrxL <- PglZ <- REase <- N6-METHYLASE <- DUF6079 *<- BrxB-BrxF-iSTAND <- BrxA-DNA-binding <-?<-NERD | 1190 | Bacteria>Terrabacteria_group>Bacillota>Bacilli>Bacillales>Bacillaceae | GCF_018343555.1 | Bacillus_sp._FJAT-49731 | KHA89_RS14830 | 208852..212424 | 2833584 | BREX-system-P-loop-protein-BrxC-[Bacillus-sp.-FJAT-49731] |
| WP_223087052.1 | Type1.BREX | HTH_58+RHH-><- REase <- SIGMA+OB-Fold+AAA-ATPase+Lon-Protease-BrxL <- PglZ <-RVT<- N6-METHYLASE <- DUF6079 *<- BrxB-BrxF-iSTAND <- BrxA-DNA-binding <-?<-?<-SPOUT<-?<-Trypsin+PDZ | 1190 | Bacteria>Terrabacteria_group>Bacillota>Bacilli>Bacillales>Bacillaceae | GCF_019915265.1 | Bacillus_aquiflavi | K6959_RS16340 | 3377801..3381373 | 2672567 | BREX-system-P-loop-protein-BrxC-[Bacillus-aquiflavi] |
| WP_229142420.1 | Type1.BREX | <-Four-helical-protein+Four-helical-protein<-ZnR<-MarR-HTH<-primpol+D5N+D5ATPase<-HTH_31<- SIGMA+OB-Fold+AAA-ATPase+Lon-Protease-BrxL <- PglZ <- REase <- N6-METHYLASE <- DUF6079 *<- BrxB-BrxF-iSTAND <- BrxA-DNA-binding <-?<-HTH\|\|HTH_58+RHH-><-ANK | 1192 | Bacteria>Terrabacteria_group>Bacillota>Bacilli>Bacillales>Bacillaceae | GCF_020731355.1 | Bacillus_sp._REN16 | LKW28_RS21250 | 18532..22110 | 2887296 | BREX-system-P-loop-protein-BrxC-[Bacillus-sp.-REN16] |
| WP_236787490.1 | Type1.BREX | FURR-HTH-><-?\|\|rRNA->rRNA->rRNA->tRNA->tRNA->tRNA->tRNA->tRNA->tRNA->tRNA->tRNA->tRNA->tRNA->tRNA->tRNA->tRNA->STYKinase-like->HEPN-> REase -> N6-METHYLASE -> BrxA-DNA-binding -> BrxB-BrxF-iSTAND -> DUF6079 *-> PglZ ->BrxD-AAA->SF1-SF2-Helicase+wHTH+Small-B-barrel+DUF1998-> | 1364 | Bacteria>Terrabacteria_group>Bacillota>Bacilli>Bacillales>Bacillaceae | GCF_021654895.1 | Bacillus_cereus | L6489_RS02850 | 541168..545262 | 1396 | hypothetical-protein-[Bacillus-cereus] |
| WP_257351442.1 | Type1.BREX | REase ->?->SbcC-><-URI-DNase<- REase <- REase <- SIGMA+OB-Fold+AAA-ATPase+Lon-Protease-BrxL <- PglZ <- N6-METHYLASE <- DUF6079 *<- BrxB-BrxF-iSTAND <- BrxA-DNA-binding <-?\|\|wHTH+HNH->?->?-><-HTH_3 | 1188 | Bacteria>Terrabacteria_group>Bacillota>Bacilli>Bacillales>Bacillaceae | GCF_024609785.1 | Pseudalkalibacillus_decolorationis | NON16_RS21340 | 32931..36497 | 163879 | BREX-system-P-loop-protein-BrxC-[Pseudalkalibacillus-decolorationis] |
| WP_110928088.1 | Type1.BREX | AAA+HTH->N6-N4-METHYLASE-><-?\|\|?-> BrxA-DNA-binding -> BrxB-BrxF-iSTAND -> DUF6079 *-> N6-METHYLASE -> N6-METHYLASE -> PglZ -> SIGMA+OB-Fold+AAA-ATPase+Lon-Protease-BrxL -> REase -><-YTCD-HTH\|\|?->?->MarR-HTH-> | 1197 | Bacteria>Terrabacteria_group>Bacillota>Bacilli>Bacillales>Bacillaceae | GCF_900098925.1 | Bacillus_massiliglaciei | BQ4305_RS12315 | 67463..71056 | 1816693 | BREX-system-P-loop-protein-BrxC-[Bacillus-massiliglaciei] |
| WP_088051628.1 | Type1.BREX | <-Phage_integrase<-Phage_integrase<-Phage_int_SAM_1+Phage_integrase\|\|HTH+rve->AAA-ATPase->?-><-SF1-SF2-Helicase+wHTH+Small-B-barrel+DUF1998<-BrxD-AAA<- PglZ <- DUF6079 *<- BrxB-BrxF-iSTAND <- BrxA-DNA-binding <-?<-HTH_Tnp_1+HTH_21+rve+rve_3<- N6-METHYLASE <- REase | 1366 | Bacteria>Terrabacteria_group>Bacillota>Bacilli>Bacillales>Bacillaceae | GCF_900155625.1 | Virgibacillus_dakarensis | CZ382_RS12920 | 682519..686619 | 1917889 | hypothetical-protein-[Virgibacillus-dakarensis] |
| WP_042357731.1 | Type1.BREX | <-VirB4-FtsK<-HTH_3\|\|?->ParD_antitoxin-><- REase <- SIGMA+OB-Fold+AAA-ATPase+Lon-Protease-BrxL <- PglZ <-AbiGi+seqA+RAMA<- N6-METHYLASE <- DUF6079 *<- BrxB-BrxF-iSTAND <- BrxA-DNA-binding <-?<-?<-HTH_31 | 1190 | Bacteria>Terrabacteria_group>Bacillota>Bacilli>Bacillales>Bacillaceae | GCF_902375395.1 | Bacillus_rubiinfantis | FY453_RS20345 | 135883..139455 | 1499680 | BREX-system-P-loop-protein-BrxC-[Bacillus-rubiinfantis] |
| WP_147675231.1 | Type1.BREX | <-HTH_Tnp_1+HTH_21+rve+rve_3<-HTH_Tnp_1<-?\|\|?-><-?<-?<-HTH+TPR+TPR+TPR+TPR+TPR<-HTH+TPR+TPR+TPR+TPR+TPR<- N6-METHYLASE <- DUF6079 *<- BrxB-BrxF-iSTAND <- BrxA-DNA-binding <- REase <-?\|\|?->Peptidase_M20-> | 1260 | Bacteria>Terrabacteria_group>Bacillota>Bacilli>Bacillales>Bacillaceae | GCF_902375755.1 | Numidum_massiliense | BN1247_RS10940 | 2280999..2284781 | 1522315 | BREX-system-P-loop-protein-BrxC-[Numidum-massiliense] |
| WP_263326060.1 | Type6.BREX | GDResolvase+Recombinase->AAA-ATPase-> REase -> BrxE -> BrxA-DNA-binding -> BrxB-BrxF-iSTAND -> DUF6079 *-> N6-METHYLASE -> PglZ -> SIGMA+OB-Fold+AAA-ATPase+HKD-DNase-BrxL ->Trypsin->GDResolvase+HIN-HTH->ParB+HNH->HEPN-> | 1207 | Bacteria>Terrabacteria_group>Bacillota>Bacilli>Bacillales>Bacillaceae | GCF_946151075.1 | Neobacillus_sp._Marseille-Q6967 | OGZ30_RS15230 | 466591..470214 | 2963105 | BREX-system-P-loop-protein-BrxC-[Neobacillus-sp.-Marseille-Q6967] |
| WP_099517056.1 | Type1.BREX | <-SIGMA-HTH<- PglZ <- REase <-OLD-ABC <- N6-METHYLASE <- DUF6079 *<- BrxB-BrxF-iSTAND <- BrxA-DNA-binding <-HTH_26<-?<-Tox-RES<-TRAM+METHYLASE<-?<-?<-TRPR-HTH | 1196 | Bacteria>Terrabacteria_group>Bacillota>Bacilli>Bacillales>Paenibacillaceae | GCF_002741035.1 | Paenibacillus_sp._BIHB4019 | BBD42_RS03740 | 863001..866591 | 1870819 | BREX-system-P-loop-protein-BrxC-[Paenibacillus-sp.-BIHB-4019] |
| MBO0589396.1 | Type1.BREX | <- N6-METHYLASE <- DUF6079 *<- BrxB-BrxF-iSTAND <- BrxA-DNA-binding <-?<-?<-?<-?<- REase <-SSB | 1191 | Bacteria>Terrabacteria_group>Bacillota>Bacilli>Bacillales>Planococcaceae | GCA_017347095.1 | Sporosarcina_sp._E16_8 | I2484_19050 | 2440..6015 | 2789295 | BREX-system-P-loop-protein-BrxC-[Sporosarcina-sp.-E16_8] |
| WP_252990012.1 | Type1.BREX | <-GDResolvase+Recombinase+ZnR<-GDResolvase+Recombinase+ZnR<-?\|\|HTH_31->?->?->?-> BrxA-DNA-binding -> BrxB-BrxF-iSTAND -> DUF6079 *-> N6-METHYLASE ->RVT-> PglZ -> SIGMA+OB-Fold+AAA-ATPase+Lon-Protease-BrxL -> REase ->?-><-HTH_58+RHH\|\|?->RNase-> | 1185 | Bacteria>Terrabacteria_group>Bacillota>Bacilli>Bacillales>Planococcaceae | GCF_000220335.1 | Sporosarcina_newyorkensis | HMPREF9372_RS16970 | 828586..832143 | 759851 | BREX-system-P-loop-protein-BrxC-[Sporosarcina-newyorkensis] |
| WP_083033314.1 | Type1.BREX | <-RVT<- REase <- SIGMA+OB-Fold+AAA-ATPase+Lon-Protease-BrxL <- PglZ <-?<-?<-?<- N6-METHYLASE <- N6-METHYLASE <- DUF6079 *<- BrxB-BrxF-iSTAND <- BrxA-DNA-binding <-?\|\|?->?->?->?-><-DOC+HTH<-RelE-ParE | 1190 | Bacteria>Terrabacteria_group>Bacillota>Bacilli>Bacillales>Planococcaceae | GCF_002082015.1 | Sporosarcina_ureae | SporoP17a_RS05210 | 1053755..1057327 | 1571 | BREX-system-P-loop-protein-BrxC-[Sporosarcina-ureae] |
| WP_125903398.1 | Type1.BREX | <-HAD\|\|?->?-><-?\|\|?->?-><-SF1-SF2-Helicase+wHTH+Small-B-barrel+DUF1998<-BrxD-AAA<- PglZ <- DUF6079 *<- BrxB-BrxF-iSTAND <- BrxA-DNA-binding <- N6-METHYLASE <- REase <-?<-?<-SbcC\|\|OLD-ABC-p->SF1-SF2-Helicase-> | 1366 | Bacteria>Terrabacteria_group>Bacillota>Bacilli>Bacillales>Planococcaceae | GCF_003944745.1 | Bhargavaea_beijingensis | EJA12_RS02395 | 145968..150068 | 426756 | hypothetical-protein-[Bhargavaea-beijingensis] |
| WP_208650578.1 | Type1.BREX | <-HTH_Tnp<-?\|\|?-><-?<- REase <- SIGMA+OB-Fold+AAA-ATPase+Lon-Protease-BrxL <- PglZ <-RelE-ParE<-?<- N6-METHYLASE <- DUF6079 *<- BrxB-BrxF-iSTAND <- BrxA-DNA-binding \|\|?->?-><-?<-HIN-HTH | 1191 | Bacteria>Terrabacteria_group>Bacillota>Bacilli>Bacillales>Planococcaceae | GCF_004331915.1 | Ureibacillus_thermophilus | DKZ56_RS14305 | 2835227..2838802 | 367743 | BREX-system-P-loop-protein-BrxC-[Ureibacillus-thermophilus] |
| WP_004894779.1 | Type1.BREX | MarR-HTH+DTXR-HTH->ABC-ATPase+SbcC->?-><-MODE-HTH+PBPII\|\|RFC-AAA->?->\|\|?-> BrxA-DNA-binding -> BrxB-BrxF-iSTAND -> DUF6079 *-> N6-METHYLASE ->nSTAND3->HEPN-p-> REase -> PglZ ->TetR-HTH->?->?->ABC-ATPase-> | 1243 | Bacteria>Terrabacteria_group>Bacillota>Bacilli>Lactobacillales>Lactobacillaceae | GCF_001433975.1 | Lactobacillus_johnsonii | FC22_RS00455 | 98654..102385 | 33959 | BREX-system-P-loop-protein-BrxC-[Lactobacillus-johnsonii] |
| WP_117285422.1 | Type1.BREX | MarR-HTH+DTXR-HTH->ABC-ATPase+SbcC->?-><-MODE-HTH+PBPII\|\|RFC-AAA->?->\|\|?-> BrxA-DNA-binding -> BrxB-BrxF-iSTAND -> DUF6079 *-> N6-METHYLASE ->nSTAND3->HEPN+ REase -> PglZ ->TetR-HTH->?->?->ABC-ATPase-> | 1243 | Bacteria>Terrabacteria_group>Bacillota>Bacilli>Lactobacillales>Lactobacillaceae | GCF_003428395.1 | Lactobacillus_johnsonii | DOY49_RS01485 | 293356..297087 | 33959 | BREX-system-P-loop-protein-BrxC-[Lactobacillus-johnsonii] |
| WP_113532397.1 | Type1.BREX | YTCD-HTH-><-?<-?<-TetR-HTH<-TetR-HTH<- PglZ <-HEPN+ REase <-nSTAND3<- N6-METHYLASE <- DUF6079 *<- BrxB-BrxF-iSTAND <- BrxA-DNA-binding <-?<-?<-?<-NUDIX<-HAD+HAD | 1239 | Bacteria>Terrabacteria_group>Bacillota>Bacilli>Lactobacillales>Lactobacillaceae | GCF_003584685.1 | Lactobacillus_paragasseri | LpgJCM5343_RS05125 | 1058950..1062669 | 2107999 | BREX-system-P-loop-protein-BrxC-[Lactobacillus-paragasseri] |
| MCG0730673.1 | Type1.BREX | <-HIN-HTH\|\| BrxA-DNA-binding -> BrxB-BrxF-iSTAND -> DUF6079 *-> N6-METHYLASE -><-Phage_integrase\|\| PglZ -> SIGMA+OB-Fold+AAA-ATPase+Lon-Protease-BrxL -p-> REase ->GDResolvase+HIN-HTH-> | 1213 | Bacteria>Terrabacteria_group>Bacillota>Bacilli>Lactobacillales>Lactobacillaceae | - | Lactiplantibacillus_plantarum | IMAU60171_02928 | 2439..6080 | 1590 | BREX-system-P-loop-protein-BrxC-[Lactiplantibacillus-plantarum] |
| WP_168924054.1 | Type1.BREX | McrB-AAA-> McrC-NTD+REase ->?-> BrxA-DNA-binding -> BrxB-BrxF-iSTAND -> DUF6079 *-><- N6-METHYLASE -p<-?\|\| PglZ -> SIGMA+OB-Fold+AAA-ATPase+Lon-Protease-BrxL ->ParB+HNH-> | 1191 | Bacteria>Terrabacteria_group>Bacillota>Bacilli>Lactobacillales>Lactobacillaceae | - | Secundilactobacillus_angelensis | L3V66_04100 | 164372..167947 | 2722706 | BREX-system-P-loop-protein-BrxC-[Secundilactobacillus-angelensis] |
| MBR6516599.1 | Type1.BREX | N6-METHYLASE -p-> BrxA-DNA-binding -> BrxB-BrxF-iSTAND -> DUF6079 *->?-> PglZ -> SIGMA+OB-Fold+AAA-ATPase+Lon-Protease-BrxL -> REase -> | 1196 | Bacteria>Terrabacteria_group>Bacillota>Bacilli>unclassified_Bacilli | GCA_017537305.1 | Bacilli_bacterium | IKT40_07050 | 1703..5293 | 1903720 | BREX-system-P-loop-protein-BrxC-[Bacilli-bacterium] |
| CDA58916.1 | Type1.BREX | <-NACHT<- SIGMA+OB-Fold+AAA-ATPase+Lon-Protease-BrxL <- PglZ <-?<- N6-METHYLASE <- DUF6079 *<- BrxB-BrxF-iSTAND <- BrxA-DNA-binding <- REase +N-glycosylase<-35exo+wHTH+SF1-SF2-Helicase+HTH\|\|?-><-?<-?<-MerR-HTH<-ASCH | 1178 | Bacteria>Terrabacteria_group>Bacillota>Clostridia>Eubacteriales>Clostridiaceae | GCA_000435175.1 | Clostridium_sp._CAG:245 | BN559_00263 | 10316..13852 | 1262784 | putative-uncharacterized-protein-[Clostridium-sp.-CAG:245] |
| ODM26889.1 | Type1.BREX | <- REase +HTH\|\|?-><-?<-Schlafen+ REase <- SIGMA+OB-Fold+AAA-ATPase+Lon-Protease-BrxL <-?<- PglZ <- N6-METHYLASE <- DUF6079 *<- BrxB-BrxF-iSTAND <- BrxA-DNA-binding <-?<-?<-?<-?<-?<-SpoOE-like | 1186 | Bacteria>Terrabacteria_group>Bacillota>Clostridia>Eubacteriales>Clostridiaceae | GCA_001717745.1 | Clostridium_sp._Bc-iso-3 | A7W90_12080 | 2831354..2834914 | 1848158 | hypothetical-protein-A7W90_12080-[Clostridium-sp.-Bc-iso-3] |
| NFG40378.1 | Type1.BREX | Calcineurin->?-> BrxA-DNA-binding -> BrxB-BrxF-iSTAND -> DUF6079 *-> N6-METHYLASE ->ParB+HNH->OLD-ABC->HNH-> PglZ -><- REase \|\| SIGMA+OB-Fold+AAA-ATPase+Lon-Protease-BrxL ->Peptidase_M42->TIR-> | 1215 | Bacteria>Terrabacteria_group>Bacillota>Clostridia>Eubacteriales>Clostridiaceae | GCA_011013445.1 | Clostridium_botulinum | FC789_04035 | 879792..883439 | 1491 | BREX-system-P-loop-protein-BrxC-[Clostridium-botulinum] |
| NFI95403.1 | Type1.BREX | <-Peptidase_M42<- SIGMA+OB-Fold+AAA-ATPase+Lon-Protease-BrxL \|\| REase -><- PglZ <-HNH<-OLD-ABC<-ParB+HNH<- N6-METHYLASE <- DUF6079 *<- BrxB-BrxF-iSTAND <- BrxA-DNA-binding | 1215 | Bacteria>Terrabacteria_group>Bacillota>Clostridia>Eubacteriales>Clostridiaceae | GCA_011014625.1 | Clostridium_botulinum | FC961_13560 | 15228..18875 | 1491 | BREX-system-P-loop-protein-BrxC-[Clostridium-botulinum] |
| MCI8983480.1 | Type1.BREX | ABC-ATPase+SbcC->Sulfatase+DUF4976->dCache_1+HAMP+HisKin+HATPase_c->REC+AraC-HTH+AraC-HTH->Sulfatase-> REase -> BrxA-DNA-binding -> BrxB-BrxF-iSTAND -> DUF6079 *-> N6-METHYLASE ->?-> N6-METHYLASE -p->?->?-> PglZ -> | 1182 | Bacteria>Terrabacteria_group>Bacillota>Clostridia>Eubacteriales>Clostridiaceae | GCA_022796695.1 | Hungatella_sp. | HFG78_13950 | 16181..19729 | 2613924 | BREX-system-P-loop-protein-BrxC-[Hungatella-sp.] |
| WP_078025149.1 | Type1.BREX | <-Phage_int_SAM_1+Phage_integrase<-DndB<-?<-ZnR+ HARE-HTH <-CarD+HTH<-?<-SF1-SF2-Helicase+wHTH+Small-B-barrel+DUF1998<-BrxD-AAA<- PglZ <- DUF6079 *<- BrxB-BrxF-iSTAND <- BrxA-DNA-binding <- N6-METHYLASE <- REase <-RHH_8<-?<-?\|\|?-><-primpol+D5N+D5ATPase | 1380 | Bacteria>Terrabacteria_group>Bacillota>Clostridia>Eubacteriales>Clostridiaceae | GCF_002008265.1 | Clostridium_tepidum | BS637_RS12490 | 20116..24258 | 1962263 | hypothetical-protein-[Clostridium-tepidum] |
| WP_027639925.1 | Type1.BREX | wHTH+HNH->?-> REase -> BrxA-DNA-binding -> BrxB-BrxF-iSTAND -> DUF6079 *->?-> N6-METHYLASE ->Cas9+HNH+RuvC->Cas_Cas1->CRISPR_Cas2-> PglZ -><- REase \|\| SIGMA+OB-Fold+AAA-ATPase+Lon-Protease-BrxL ->ARG-> | 1214 | Bacteria>Terrabacteria_group>Bacillota>Clostridia>Eubacteriales>Clostridiaceae | GCF_012844035.1 | Clostridium_cadaveris | HF846_RS03710 | 105265..108909 | 1529 | BREX-system-P-loop-protein-BrxC-[Clostridium-cadaveris] |
| WP_178312104.1 | Type1.BREX | <-ARG<-?<-DUF4326<-DUF488-s<-DUF488-1\|\| REase -><-?<- N6-METHYLASE <- DUF6079 *<- BrxB-BrxF-iSTAND <- BrxA-DNA-binding <- REase <-?<-wHTH+HNH | 1218 | Bacteria>Terrabacteria_group>Bacillota>Clostridia>Eubacteriales>Clostridiaceae | GCF_013390975.1 | Clostridium_cadaveris | HXZ52_RS14490 | 15645..19301 | 1529 | BREX-system-P-loop-protein-BrxC-[Clostridium-cadaveris] |
| WP_173680574.1 | Type1.BREX | <-Cap4_nuclease<-?<-HTH_3\|\| REase -><- SIGMA+OB-Fold+AAA-ATPase+Lon-Protease-BrxL <-?<- PglZ <-PIN<- N6-METHYLASE <- DUF6079 *<- BrxB-BrxF-iSTAND <- BrxA-DNA-binding \|\|PADR-HTH+MarR-HTH-><-D5ATPase<-?<-?<-?<-?\|\|LexA-protease-> | 1216 | Bacteria>Terrabacteria_group>Bacillota>Clostridia>Eubacteriales>Clostridiaceae | GCF_014230485.1 | Clostridium_tetanomorphum | FHX96_RS20165 | 4177507..4181157 | 1553 | BREX-system-P-loop-protein-BrxC-[Clostridium-tetanomorphum] |
| WP_209702448.1 | Type1.BREX | BrxA-DNA-binding ->HTH-> DUF6079 *-> PglZ ->BrxD-AAA->SF1-SF2-Helicase+wHTH+Small-B-barrel+DUF1998->?->BrxL-> REase -><-HTH_3<-Phage_integrase\|\|PAS+GGDEF+EAL-> | 1377 | Bacteria>Terrabacteria_group>Bacillota>Clostridia>Eubacteriales>Clostridiaceae | GCF_017874415.1 | Clostridium_algifaecis | J2Z42_RS09735 | 1889..6022 | 1472040 | hypothetical-protein-[Clostridium-algifaecis] |
| WP_217303321.1 | Type1.BREX | OLD-ABC-><-DOC<-Phd-YefM<- SIGMA+OB-Fold+AAA-ATPase+Lon-Protease-BrxL <- PglZ <- REase <-OLD-ABC <-?<- N6-METHYLASE <- DUF6079 *<- BrxB-BrxF-iSTAND <- BrxA-DNA-binding <-tRNA<-?<-?<-?<-HTH_3<-HTH_OrfB_IS605+OrfB_IS605+OrfB_Zn_ribbon | 1188 | Bacteria>Terrabacteria_group>Bacillota>Clostridia>Eubacteriales>Clostridiaceae | GCF_019056555.1 | Clostridium_sp._001 | B5S50_RS05665 | 1268220..1271786 | 1970093 | BREX-system-P-loop-protein-BrxC-[Clostridium-sp.-001] |
| WP_221260245.1 | Type1.BREX | <-HISKIN\|\|RADICAL-SAM->?->?->HEPN->Ntox45->\|\| BrxA-DNA-binding -> BrxB-BrxF-iSTAND -> DUF6079 *-> N6-METHYLASE ->?-> PglZ -><- REase \|\| SIGMA+OB-Fold+AAA-ATPase+Lon-Protease-BrxL ->?->PIN->Peptidase_M42-> | 1218 | Bacteria>Terrabacteria_group>Bacillota>Clostridia>Eubacteriales>Clostridiaceae | GCF_019705275.1 | Clostridium_chauvoei | K4H38_08460 | 40416..44072 | 46867 | BREX-system-P-loop-protein-BrxC-[Clostridium-chauvoei] |
| WP_238021483.1 | Type1.BREX | OLD-ABC+TOPRIM->SF1-SF2-Helicase-> REase ->?-> BrxA-DNA-binding -> BrxB-BrxF-iSTAND -> DUF6079 *-> N6-METHYLASE -> PglZ -> SIGMA+OB-Fold+AAA-ATPase+Lon-Protease-BrxL -p-> SIGMA+OB-Fold+AAA-ATPase+Lon-Protease-BrxL -p->SF1-SF2-Helicase+RdgC+ REase ->?->?->D5N+D5ATPase-> | 1195 | Bacteria>Terrabacteria_group>Bacillota>Clostridia>Eubacteriales>Clostridiaceae | GCF_022133285.1 | Clostridium_cochlearium | L0O71_RS07755 | 6541..10128 | 1494 | BREX-system-P-loop-protein-BrxC-[Clostridium-cochlearium] |
| WP_252217802.1 | Type1.BREX | <-Peptidase_M42<-PIN<-?<- SIGMA+OB-Fold+AAA-ATPase+Lon-Protease-BrxL \|\| REase -><- PglZ <-?<- N6-METHYLASE <- DUF6079 *<- BrxB-BrxF-iSTAND <- BrxA-DNA-binding \|\|<-Ntox45<-HEPN<-HEPN<-?<-?<-RADICAL-SAM\|\|HISKIN-> | 1218 | Bacteria>Terrabacteria_group>Bacillota>Clostridia>Eubacteriales>Clostridiaceae | GCF_023846215.1 | Clostridium_chauvoei | J9878_RS07560 | 18587..22243 | 46867 | BREX-system-P-loop-protein-BrxC-[Clostridium-chauvoei] |
| WP_021876221.1 | Type1.BREX | <-Peptidase_M42<-PIN<-?<- SIGMA+OB-Fold+AAA-ATPase+Lon-Protease-BrxL \|\| REase -><- PglZ <-?<- N6-METHYLASE <- DUF6079 *<- BrxB-BrxF-iSTAND <- BrxA-DNA-binding \|\|<-Ntox45<-HEPN<-?<-?<-RADICAL-SAM\|\|HISKIN-> | 1218 | Bacteria>Terrabacteria_group>Bacillota>Clostridia>Eubacteriales>Clostridiaceae | GCF_900037355.1 | Clostridium_chauvoei | C6H62_RS09310 | 1987009..1990665 | 46867 | BREX-system-P-loop-protein-BrxC-[Clostridium-chauvoei] |
| WP_066893479.1 | Type1.BREX | <- SIGMA+OB-Fold+AAA-ATPase+Lon-Protease-BrxL \|\| REase -><- PglZ <-DOC+HTH<-SLOG+BirA-HTH<-SF1-SF2-Helicase+PRTase<-ParB+HNH<-?<- N6-METHYLASE <- DUF6079 *<- BrxB-BrxF-iSTAND <- BrxA-DNA-binding <-?<-SF1-SF2-Helicase\|\|?->?-><-?<-ABC-ATPase+SbcC | 1215 | Bacteria>Terrabacteria_group>Bacillota>Clostridia>Eubacteriales>Clostridiaceae | GCF_900086595.1 | Clostridium_nigeriense | BN6774_RS10780 | 2270723..2274370 | 1805470 | BREX-system-P-loop-protein-BrxC-[Clostridium-nigeriense] |
| WP_143148632.1 | Type1.BREX | <-N6-N4-METHYLASE<-SF1-SF2-Helicase+wHTH+Small-B-barrel+DUF1998<-BrxD-AAA<- PglZ <- DUF6079 *<-HTH<- BrxA-DNA-binding <- N6-METHYLASE <- REase <-HKD-DNase+SF1-SF2-Helicase+HTH+HTH+SRA | 1377 | Bacteria>Terrabacteria_group>Bacillota>Clostridia>Eubacteriales>Clostridiaceae | GCF_900142075.1 | Clostridium_amylolyticum | BUA82_RS13560 | 29831..33964 | 1121298 | hypothetical-protein-[Clostridium-amylolyticum] |
| WP_073026825.1 | Type1.BREX | <-TIR<-HNH<-SF1-SF2-Helicase+wHTH+Small-B-barrel+DUF1998<-BrxD-AAA<- PglZ <- DUF6079 *<- BrxB-BrxF-iSTAND <- BrxA-DNA-binding <- N6-METHYLASE <- REase <-?<-?<-DndB<-GGDEF\|\|HTH_3-> | 1381 | Bacteria>Terrabacteria_group>Bacillota>Clostridia>Eubacteriales>Clostridiaceae | GCF_900142105.1 | Lutispora_thermophila | BUB48_RS13880 | 40274..44419 | 288966 | hypothetical-protein-[Lutispora-thermophila] |
| EOS24741.1 | Type1.BREX | ACET->DDE_Tnp->DDE_5+DDE_Tnp_1->HTH_26-> BrxA-DNA-binding -> BrxB-BrxF-iSTAND -> DUF6079 *-> REase ->HicA-> N6-METHYLASE ->MORC+ REase ->?->HNH->HTH_28+HTH_32+DDE_3->DDE_3-> | 1175 | Bacteria>Terrabacteria_group>Bacillota>Clostridia>Eubacteriales>Lachnospiraceae | GCA_000403215.2 | Lachnospiraceae_bacterium_A4 | C804_05106 | 19579..23106 | 397291 | hypothetical-protein-C804_05106-[Lachnospiraceae-bacterium-A4] |
| EOS28360.1 | Type1.BREX | <- REase <- SIGMA+OB-Fold+AAA-ATPase+Lon-Protease-BrxL <- PglZ <-?<-?<-DEDD_Tnp_IS110+Transposase_20<-?<-ORC-CDC6+ZnR<- N6-METHYLASE <- DUF6079 *<-?<- BrxB-BrxF-iSTAND <- BrxA-DNA-binding <-HTH_26\|\|HTH_28+HTH_32+DDE_3-><-?<-TraI-VirD2-RCRE<-TraJ-RHH<-CopG/RHH | 1224 | Bacteria>Terrabacteria_group>Bacillota>Clostridia>Eubacteriales>Lachnospiraceae | GCA_000403275.2 | Lachnospiraceae_bacterium_28-4 | C807_03898 | 219781..223455 | 397287 | hypothetical-protein-C807_03898-[Lachnospiraceae-bacterium-28-4] |
| EOS37735.1 | Type1.BREX | <-HTH_26<-?<- PglZ -p<-OLD-ABC<-ParB+HNH<-NACHT<- N6-METHYLASE -p<- N6-METHYLASE -p<-DUF1016+ REase <- DUF6079 *<- BrxB-BrxF-iSTAND <- BrxA-DNA-binding <-?<-?<-SAM-METHYLASE<-HISKIN+MORC+REC+OmpR-HTH\|\|ABC-ATPase+SbcC-> | 1218 | Bacteria>Terrabacteria_group>Bacillota>Clostridia>Eubacteriales>Lachnospiraceae | GCA_000403295.2 | Lachnospiraceae_bacterium_M18-1 | C808_03715 | 724875..728531 | 1235792 | hypothetical-protein-C808_03715-[Lachnospiraceae-bacterium-M18-1] |
| ONI39098.1 | Type1.BREX | <-ABC-ATPase+SbcC<-HISKIN<-REC+OmpR-HTH<- PglZ <- N6-METHYLASE -p<- REase <- N6-METHYLASE <- N6-METHYLASE <- DUF6079 *<- BrxB-BrxF-iSTAND <- BrxA-DNA-binding | 1197 | Bacteria>Terrabacteria_group>Bacillota>Clostridia>Eubacteriales>Lachnospiraceae | GCA_001983455.1 | Epulopiscium_sp._SCG-B11WGA-EpuloA1 | AN396_09505 | 82489..86082 | 1710701 | ATPase-[Epulopiscium-sp.-SCG-B11WGA-EpuloA1] |
| HAB87400.1 | Type1.BREX | ZnR+ZnR+ZnR->?->HTH_26-> BrxA-DNA-binding -> BrxB-BrxF-iSTAND -> DUF6079 *-> N6-METHYLASE -> REase ->Schlafen+HATPase+wHTH-> PglZ ->Peptidase_U49-> SIGMA+OB-Fold+AAA-ATPase+Lon-Protease-BrxL ->?->LRR_5+LRR_8-> | 1196 | Bacteria>Terrabacteria_group>Bacillota>Clostridia>Eubacteriales>Lachnospiraceae | GCA_003448155.1 | Coprococcus_sp. | DCE60_00030 | 3490..7080 | 2049024 | BREX-system-P-loop-protein-BrxC-[Coprococcus-sp.] |
| HCI18472.1 | Type1.BREX | <-PIN<-?<- SIGMA+OB-Fold+AAA-ATPase+Lon-Protease-BrxL <-OLD-ABC+TOPRIM<- PglZ <-Schlafen+HATPase+wHTH<- REase <- N6-METHYLASE <- DUF6079 -p*<- DUF6079 -p<- BrxB-BrxF-iSTAND <- BrxA-DNA-binding <-HTH_26 | 799 | Bacteria>Terrabacteria_group>Bacillota>Clostridia>Eubacteriales>Lachnospiraceae | GCA_003521585.1 | Lachnospiraceae_bacterium | DFH32_07645 | 14441..16840 | 1898203 | BREX-system-P-loop-protein-BrxC-[Lachnospiraceae-bacterium] |
| HCX41706.1 | Type1.BREX | SKI->?->tRNA->?->?->HTH_26-> BrxA-DNA-binding -> BrxB-BrxF-iSTAND -> DUF6079 *-> N6-METHYLASE -> REase ->Schlafen+HATPase+wHTH-> PglZ ->OLD-ABC+TOPRIM-> SIGMA+OB-Fold+AAA-ATPase+Lon-Protease-BrxL ->?->PIN->MACRODOMAIN-> | 1196 | Bacteria>Terrabacteria_group>Bacillota>Clostridia>Eubacteriales>Lachnospiraceae | GCA_003539645.1 | Lachnospiraceae_bacterium | DHH50_05105 | 179559..183149 | 1898203 | BREX-system-P-loop-protein-BrxC-[Lachnospiraceae-bacterium] |
| TGY98248.1 | Type1.BREX | <-MarR-HTH<-?<-RhoN+RhoN+RNase+S1<-?<-?<- PglZ <- REase <-OLD-ABC <- N6-METHYLASE <- DUF6079 *<- BrxB-BrxF-iSTAND <- BrxA-DNA-binding <-?<-PRTase | 1218 | Bacteria>Terrabacteria_group>Bacillota>Clostridia>Eubacteriales>Lachnospiraceae | GCA_004793545.1 | Petralouisia_muris | E5329_00220 | 47888..51544 | 3032872 | BREX-system-P-loop-protein-BrxC-[Petralouisia-muris] |
| TYL56933.1 | Type1.BREX | HTH_26-> BrxA-DNA-binding -> BrxB-BrxF-iSTAND -> DUF6079 *-> N6-METHYLASE -> REase ->Schlafen+HATPase+wHTH-> PglZ ->?->HEPN-> SIGMA+OB-Fold+AAA-ATPase+Lon-Protease-BrxL ->?->LRR_5+LRR_8-> | 1196 | Bacteria>Terrabacteria_group>Bacillota>Clostridia>Eubacteriales>Lachnospiraceae | GCA_008123415.1 | [Eubacterium]_rectale | FYL31_13935 | 1744..5334 | 39491 | BREX-system-P-loop-protein-BrxC-[[Eubacterium] |
| MBR4606065.1 | Type1.BREX | Phage_integrase-> REase -> REase ->?-> BrxA-DNA-binding -> BrxB-BrxF-iSTAND -> DUF6079 *-> N6-METHYLASE -p-> | 1191 | Bacteria>Terrabacteria_group>Bacillota>Clostridia>Eubacteriales>Lachnospiraceae | GCA_017536085.1 | Lachnospiraceae_bacterium | IKO41_07545 | 5884..9459 | 1898203 | BREX-system-P-loop-protein-BrxC-[Lachnospiraceae-bacterium] |
| MCD7743820.1 | Type1.BREX | HTH_42->ABhydrolase+ABhydrolase->?->?->?->HTH_26-> BrxA-DNA-binding -> BrxB-BrxF-iSTAND -> DUF6079 *-> N6-METHYLASE -> OLD-ABC-> REase -> PglZ ->?-> SIGMA+OB-Fold+AAA-ATPase+Lon-Protease-BrxL ->MACRODOMAIN-> | 1187 | Bacteria>Terrabacteria_group>Bacillota>Clostridia>Eubacteriales>Lachnospiraceae | GCA_021199475.1 | Lachnospiraceae_bacterium | LUI13_00760 | 15415..18978 | 1898203 | BREX-system-P-loop-protein-BrxC-[Lachnospiraceae-bacterium] |
| WP_075720910.1 | Type1.BREX | <-LRR_5+LRR_8<-?<-?<- SIGMA+OB-Fold+AAA-ATPase+Lon-Protease-BrxL <- PglZ <-Schlafen+HATPase+wHTH<- REase <- N6-METHYLASE <- DUF6079 *<- BrxB-BrxF-iSTAND <- BrxA-DNA-binding <-HTH_26<-ZnR+ZnR+ZnR<-HicA_toxin<-?<-?<-MACRODOMAIN<-35exo+ADA-DBD | 1196 | Bacteria>Terrabacteria_group>Bacillota>Clostridia>Eubacteriales>Lachnospiraceae | GCF_001940225.1 | Roseburia_sp._499 | BIV20_RS10280 | 367523..371113 | 1261634 | BREX-system-P-loop-protein-BrxC-[Roseburia-sp.-499] |
| WP_118390632.1 | Type1.BREX | wHTH+WYL+WCX ->HTH_26-> BrxA-DNA-binding -> BrxB-BrxF-iSTAND -> DUF6079 *-> N6-METHYLASE -> REase ->Schlafen+HATPase+wHTH-> PglZ ->Pkinase-> SIGMA+OB-Fold+AAA-ATPase+Lon-Protease-BrxL ->?->SF1-SF2-Helicase+TUDOR->MPTase-> | 1196 | Bacteria>Terrabacteria_group>Bacillota>Clostridia>Eubacteriales>Lachnospiraceae | GCF_003474935.1 | Lachnospira_eligens | DWZ79_RS02870 | 260992..264582 | 39485 | BREX-system-P-loop-protein-BrxC-[Lachnospira-eligens] |
| WP_135987576.1 | Type1.BREX | <-MarR-HTH<-?<-RhoN+RhoN+RNase+S1<-?<-?<- PglZ <- REase <-OLD-ABC <- N6-METHYLASE <- DUF6079 *<- BrxB-BrxF-iSTAND <- BrxA-DNA-binding <-?<-PRTase | 1218 | Bacteria>Terrabacteria_group>Bacillota>Clostridia>Eubacteriales>Lachnospiraceae | GCF_004793545.1 | Petralouisia_muris | E5329_RS00220 | 47888..51544 | 3032872 | BREX-system-P-loop-protein-BrxC-[Petralouisia-muris] |
| WP_173753518.1 | Type1.BREX | wHTH+WYL+WCX ->?->?->?->HTH_26-> BrxA-DNA-binding -> BrxB-BrxF-iSTAND -> DUF6079 *-> N6-METHYLASE ->DUF4365+ REase -> PglZ ->HEPN+SH3-like+SF1-SF2-Helicase+wHTH->MORC+DUF3684+HNH->SF1-SF2-Helicase->PIN-> | 1203 | Bacteria>Terrabacteria_group>Bacillota>Clostridia>Eubacteriales>Lachnospiraceae | GCF_013301805.1 | Blautia | G5A92_RS05405 | 158405..162016 | 572511 | MULTISPECIES:-BREX-system-P-loop-protein-BrxC-[Blautia] |
| WP_128132381.1 | Type1.BREX | MerR-HTH-><-?<- REase <-?<- SIGMA+OB-Fold+AAA-ATPase+Lon-Protease-BrxL <- PglZ <-HEPN<- N6-METHYLASE <- DUF6079 *<- BrxB-BrxF-iSTAND <- BrxA-DNA-binding <-HTH_26<-HTH\|\|?-><-?<-TraJ-RHH<-SF1-SF2-Helicase | 1182 | Bacteria>Terrabacteria_group>Bacillota>Clostridia>Eubacteriales>Lachnospiraceae | GCF_015547745.1 | Enterocloster_bolteae | I2D45_RS23920 | 73790..77338 | 208479 | BREX-system-P-loop-protein-BrxC-[Enterocloster-bolteae] |
| WP_216429335.1 | Type1.BREX | HTH_26-> BrxA-DNA-binding -> BrxB-BrxF-iSTAND -> DUF6079 *-> N6-METHYLASE ->RAMA->HEPN-> PglZ -> SIGMA+OB-Fold+AAA-ATPase+Lon-Protease-BrxL ->?-> REase ->?-><-HTH_3 | 1182 | Bacteria>Terrabacteria_group>Bacillota>Clostridia>Eubacteriales>Lachnospiraceae | GCF_018918145.1 | Anaerotignum_sp._MSJ-24 | KQI66_RS10650 | 65878..69426 | 2841521 | BREX-system-P-loop-protein-BrxC-[Anaerotignum-sp.-MSJ-24] |
| WP_227178521.1 | Type1.BREX | RADICAL-SAM->?->?-><-Peptidase<-?\|\|HTH->HTH_26-> BrxA-DNA-binding -> BrxB-BrxF-iSTAND -> DUF6079 *-> N6-METHYLASE ->Schlafen+HATPase+wHTH-> PglZ ->?-> SIGMA+OB-Fold+AAA-ATPase+Lon-Protease-BrxL ->?-> REase ->?-><-MPTase | 1182 | Bacteria>Terrabacteria_group>Bacillota>Clostridia>Eubacteriales>Lachnospiraceae | GCF_020555685.1 | Enterocloster_bolteae | LIO18_RS10100 | 522012..525560 | 208479 | BREX-system-P-loop-protein-BrxC-[Enterocloster-bolteae] |
| WP_055287913.1 | Type1.BREX | <-SF1-SF2-Helicase+TUDOR<-?<- SIGMA+OB-Fold+AAA-ATPase+Lon-Protease-BrxL <-OLD-ABC+TOPRIM<- PglZ <-Schlafen+HATPase+wHTH<- REase <- N6-METHYLASE <- DUF6079 *<- BrxB-BrxF-iSTAND <- BrxA-DNA-binding <-HTH_26<- wHTH+WYL+WCX | 1196 | Bacteria>Terrabacteria_group>Bacillota>Clostridia>Eubacteriales>Lachnospiraceae | GCF_902362785.1 | Lachnospira_eligens | FXW70_RS12185 | 167515..171105 | 39485 | BREX-system-P-loop-protein-BrxC-[Lachnospira-eligens] |
| MBD5149892.1 | Type1.BREX | ArsR-HTH->HTH_3-><-?<-?<-NTP_transf_3\|\|?->HTH_26-> BrxA-DNA-binding -> BrxB-BrxF-iSTAND -> DUF6079 *-> N6-METHYLASE -> OLD-ABC-> REase -> PglZ ->DDE_Tnp_1_assoc->?->?->ABC-ATPase+SbcC+ABC-ATPase+SbcC->HTH-> | 1183 | Bacteria>Terrabacteria_group>Bacillota>Clostridia>Eubacteriales>Oscillospiraceae | GCA_014799845.1 | Oscillibacter_sp. | HDT18_05880 | 26304..29855 | 1945593 | BREX-system-P-loop-protein-BrxC-[Oscillibacter-sp.] |
| MBQ9664285.1 | Type1.BREX | REase -><- SIGMA+OB-Fold+AAA-ATPase+Lon-Protease-BrxL <- PglZ <-?<- N6-METHYLASE <- DUF6079 *<- BrxB-BrxF-iSTAND <- BrxA-DNA-binding | 1198 | Bacteria>Terrabacteria_group>Bacillota>Clostridia>Eubacteriales>Oscillospiraceae | GCA_017458065.1 | Oscillospiraceae_bacterium | IJV40_14155 | 25902..29498 | 2485925 | BREX-system-P-loop-protein-BrxC-[Oscillospiraceae-bacterium] |
| WP_010247739.1 | Type1.BREX | <-Herpes_LMP1+ REase <- SIGMA+OB-Fold+AAA-ATPase+Lon-Protease-BrxL <- PglZ <- REase <-OLD-ABC <- N6-METHYLASE <- DUF6079 *<- BrxB-BrxF-iSTAND <- BrxA-DNA-binding <-?<-?<-NUDIX+DUF429<-?<-DDE_Tnp_IS66+DDE_Tnp_IS66_C<-?<-HIN-HTH<-ParB+HNH | 1183 | Bacteria>Terrabacteria_group>Bacillota>Clostridia>Eubacteriales>Oscillospiraceae | GCF_000179595.2 | Acetivibrio_cellulolyticus | ACECE_RS0212925 | 179923..183474 | 35830 | BREX-system-P-loop-protein-BrxC-[Acetivibrio-cellulolyticus] |
| WP_094549793.1 | Type1.BREX | <-ZnR<-ARG<-?<-TIR<-?<-HNH<-SF1-SF2-Helicase+wHTH+Small-B-barrel+DUF1998<-BrxD-AAA<- PglZ <- DUF6079 *<- BrxB-BrxF-iSTAND <- BrxA-DNA-binding <- N6-METHYLASE <- REase <-?<-?<-?<-?<-DndB | 1381 | Bacteria>Terrabacteria_group>Bacillota>Clostridia>Eubacteriales>Oscillospiraceae | GCF_002252565.1 | Petroclostridium_xylanilyticum | CIB29_RS11425 | 318791..322936 | 1792311 | hypothetical-protein-[Petroclostridium-xylanilyticum] |
| WP_137698823.1 | Type1.BREX | <-PhdYeFM_antitox<-?<-?<-?<-?<- REase <- SIGMA+OB-Fold+AAA-ATPase+Lon-Protease-BrxL <- PglZ <-?<- N6-METHYLASE <- DUF6079 *<- BrxB-BrxF-iSTAND <- BrxA-DNA-binding <-?<-?<-?<-?<-Phage_int_SAM_5+Phage_integrase | 1191 | Bacteria>Terrabacteria_group>Bacillota>Clostridia>Eubacteriales>Oscillospiraceae | GCF_005473905.2 | Ruminiclostridium_herbifermentans | EHE19_RS13505 | 3333193..3336768 | 2488810 | BREX-system-P-loop-protein-BrxC-[Ruminiclostridium-herbifermentans] |
| WP_120485839.1 | Type1.BREX | <- SIGMA+OB-Fold+AAA-ATPase+Lon-Protease-BrxL <- PglZ <- REase <-OLD-ABC <- N6-METHYLASE <- DUF6079 *<- BrxB-BrxF-iSTAND <- BrxA-DNA-binding <-HTH_26<-JetC<-?<-HNH\|\|HTH_3->PemK_toxin->ParB-> | 1178 | Bacteria>Terrabacteria_group>Bacillota>Clostridia>Eubacteriales>Oscillospiraceae | GCF_009917605.1 | unclassified_Anaerotruncus | D5269_RS12205 | 11128..14664 | 2641626 | MULTISPECIES:-BREX-system-P-loop-protein-BrxC-[unclassified-Anaerotruncus] |
| WP_204901129.1 | Type1.BREX | SF1-SF2-Helicase-><- SIGMA+OB-Fold+AAA-ATPase+Lon-Protease-BrxL <- PglZ <- REase <-OLD-ABC <-?<-Tox-RES<- N6-METHYLASE <-SIR2<- DUF6079 *<- BrxB-BrxF-iSTAND <- BrxA-DNA-binding <-SIGMA-HTH | 1179 | Bacteria>Terrabacteria_group>Bacillota>Clostridia>Eubacteriales>Oscillospiraceae | GCF_016902575.1 | Pseudoflavonifractor_capillosus | H9X86_RS08600 | 19493..23032 | 106588 | BREX-system-P-loop-protein-BrxC-[Pseudoflavonifractor-capillosus] |
| WP_118142807.1 | Type1.BREX | wHTH+WYL+WCX ->?->?->?->HTH_26-> BrxA-DNA-binding -> BrxB-BrxF-iSTAND -> DUF6079 *-> N6-METHYLASE ->DUF4365+ REase -> PglZ ->HEPN+SH3-like+SF1-SF2-Helicase+wHTH->MORC+DUF3684+HNH->SF1-SF2-Helicase->PIN->?->CHTH-> | 1203 | Bacteria>Terrabacteria_group>Bacillota>Clostridia>Eubacteriales>Oscillospiraceae | - | Ruminococcus_sp._OM07-7 | DXC06_07755 | 161101..164712 | 2293234 | BREX-system-P-loop-protein-BrxC-[Ruminococcus-sp.-OM07-7] |
| WP_057535321.1 | Type1.BREX | TetR-HTH->?-><-?\|\| BrxA-DNA-binding -> BrxB-BrxF-iSTAND -> DUF6079 *-> N6-METHYLASE ->?->HTH_3->MPTase->?->?-> PglZ -> REase +DUF5655-> | 1185 | Bacteria>Terrabacteria_group>Bacillota>Clostridia>Eubacteriales>Peptostreptococcaceae | GCF_001372455.1 | Paeniclostridium_sordellii | R26833_RS12520 | 91086..94643 | 1505 | BREX-system-P-loop-protein-BrxC-[Paeniclostridium-sordellii] |
| WP_099191952.1 | Type1.BREX | DNA_METHYLASE->?->HHH+DNA_pol_B+RNase->TRAM+METHYLASE->NERD+HRDC->?-> REase -> BrxA-DNA-binding -> BrxB-BrxF-iSTAND -> DUF6079 *-> N6-METHYLASE ->OLD-ABC->TOPRIM-> SF1-SF2-Helicase+ REase +HTH-> OLD-ABC-> REase -> PglZ -> SIGMA+OB-Fold+AAA-ATPase+Lon-Protease-BrxL -> | 1190 | Bacteria>Terrabacteria_group>Bacillota>Clostridia>Eubacteriales>Peptostreptococcaceae | GCF_002724055.1 | Tepidibacter_mesophilus | Tmeso_RS13165 | 34071..37643 | 655607 | BREX-system-P-loop-protein-BrxC-[Tepidibacter-mesophilus] |
| WP_226924212.1 | Type1.BREX | AraC-HTH-><-?\|\| BrxA-DNA-binding -> BrxB-BrxF-iSTAND -> DUF6079 *-> N6-METHYLASE -> REase ->PIN->?-> PglZ -> SIGMA+OB-Fold+AAA-ATPase+Lon-Protease-BrxL -> | 1190 | Bacteria>Terrabacteria_group>Bacillota>Clostridia>Eubacteriales>Peptostreptococcaceae | GCF_020537215.1 | Intestinibacter_bartlettii | LIP50_RS08300 | 25949..29521 | 261299 | BREX-system-P-loop-protein-BrxC-[Intestinibacter-bartlettii] |
| MBP2018669.1 | Type6.BREX | <- REase <- SIGMA+OB-Fold+AAA-ATPase+Lon-Protease-BrxL <- PglZ <- N6-METHYLASE <- N6-METHYLASE <- DUF6079 *<- BrxB-BrxF-iSTAND <- BrxA-DNA-binding <- BrxE \|\| McrC-NTD+ REase ->McrB-AAA-><-SF1-SF2-Helicase+wHTH+Small-B-barrel+DUF1998<-?<-?\|\|SF1-SF2-Helicase-> | 1199 | Bacteria>Terrabacteria_group>Bacillota>Clostridia>Eubacteriales>Symbiobacteriaceae | GCA_017874315.1 | Symbiobacterium_terraclitae | J2Z79_002084 | 10592..14191 | 557451 | hypothetical-protein-[Symbiobacterium-terraclitae] |
| WP_245302581.1 | Type1.BREX | <- REase <- SIGMA+OB-Fold+AAA-ATPase+Lon-Protease-BrxL <- PglZ <- N6-METHYLASE <- N6-METHYLASE <- DUF6079 *<- BrxB-BrxF-iSTAND <- BrxA-DNA-binding \|\| McrC-NTD+ REase ->McrB-AAA-><-SF1-SF2-Helicase+wHTH+Small-B-barrel+DUF1998<-?<-?\|\|SF1-SF2-Helicase-> | 1198 | Bacteria>Terrabacteria_group>Bacillota>Clostridia>Eubacteriales>Symbiobacteriaceae | GCF_017874315.1 | Symbiobacterium_terraclitae | J2Z79_RS10300 | 10592..14188 | 557451 | BREX-system-P-loop-protein-BrxC-[Symbiobacterium-terraclitae] |
| WP_011640409.1 | Type1.BREX | Classical-AAA-> CR-ATPase5 + REase ->AbiEii-><-?\|\| BrxA-DNA-binding -> BrxB-BrxF-iSTAND -> DUF6079 *-> N6-METHYLASE -p->?-> REase -> N6-METHYLASE -p->HTH_Tnp_1+HTH_21+rve+rve_3-> N6-METHYLASE -p-> PglZ -> SIGMA+OB-Fold+AAA-ATPase+Lon-Protease-BrxL -> | 1199 | Bacteria>Terrabacteria_group>Bacillota>Clostridia>Eubacteriales>Syntrophomonadaceae | GCF_000014725.1 | Syntrophomonas_wolfei | SWOL_RS05025 | 1120940..1124539 | 863 | BREX-system-P-loop-protein-BrxC-[Syntrophomonas-wolfei] |
| WP_061215054.1 | Type1.BREX | PolB->HEPN->HicB_1k_antitox-> BrxA-DNA-binding -> BrxB-BrxF-iSTAND -> DUF6079 *-> N6-METHYLASE -> PglZ -> SIGMA+OB-Fold+AAA-ATPase+Lon-Protease-BrxL -> REase -> | 1184 | Bacteria>Terrabacteria_group>Bacillota>Clostridia>Eubacteriales>Syntrophomonadaceae | GCF_001570625.1 | Syntrophomonas_wolfei | AWB54_RS13115 | 26224..29778 | 863 | BREX-system-P-loop-protein-BrxC-[Syntrophomonas-wolfei] |
| WP_129723506.1 | Type1.BREX | <-HTH_3\|\|GGDEF->?-> REase -> N6-METHYLASE -> BrxA-DNA-binding -> BrxB-BrxF-iSTAND -> DUF6079 *-> PglZ ->BrxD-AAA->SF1-SF2-Helicase+wHTH+Small-B-barrel+DUF1998->HNH->ZnR->?->SF1-SF2-Helicase-> | 1381 | Bacteria>Terrabacteria_group>Bacillota>Clostridia>Eubacteriales>Xylanivirgaceae | GCF_004138105.1 | Xylanivirga_thermophila | EJN67_RS06315 | 20747..24892 | 2496273 | hypothetical-protein-[Xylanivirga-thermophila] |
| WP_258359412.1 | Type1.BREX | <-XIS-HTH\|\|?->?->Phage_int+DNA-glycosylase-> REase ->?->?-> BrxA-DNA-binding -> BrxB-BrxF-iSTAND -> DUF6079 *-> N6-METHYLASE -> PglZ -> SIGMA+OB-Fold+AAA-ATPase+Lon-Protease-BrxL ->GspA-ATPase->?->Schlafen+PIWI->DEDD_Tnp_IS110+Transposase_20-> REase +EVE-> REase -> | 1148 | Bacteria>Terrabacteria_group>Bacillota>Clostridia>Thermoanaerobacterales>Thermoanaerobacteraceae | GCF_024733845.1 | Moorella_sp._SLA38 | NWF31_RS05005 | 117585..121031 | 2972948 | BREX-system-P-loop-protein-BrxC-[Moorella-sp.-SLA38] |
| WP_094397497.1 | Type1.BREX | HTH_32+rve+rve_3+Mu-transpos_C->GspA-ATPase->?->?->?-> REase -> N6-METHYLASE -> BrxA-DNA-binding -> BrxB-BrxF-iSTAND -> DUF6079 *-> PglZ ->BrxD-AAA->SF1-SF2-Helicase+wHTH+Small-B-barrel+DUF1998->HNH->TIR->?->ARG-> | 1381 | Bacteria>Terrabacteria_group>Bacillota>Clostridia>Thermoanaerobacterales>Thermoanaerobacterales_Family_III._Incertae_Sedis | GCF_002250075.1 | Thermoanaerobacterium_thermosaccharolyticum | Thert_RS08605 | 1721072..1725217 | 1517 | hypothetical-protein-[Thermoanaerobacterium-thermosaccharolyticum] |
| HHW04166.1 | Type1.BREX | <-SF1-SF2-Helicase<-OLD-ABC+TOPRIM<-?\|\|Classical-AAA->Peptidase_S8-><-?<- REase +DUF5655<- SIGMA+OB-Fold+AAA-ATPase+Lon-Protease-BrxL <- PglZ <- DUF6079 *<- BrxB-BrxF-iSTAND <- BrxA-DNA-binding <-TraJ-RHH | 1186 | Bacteria>Terrabacteria_group>Bacillota>Clostridia>Thermoanaerobacterales>unclassified_Thermoanaerobacterales | GCA_012840135.1 | Thermoanaerobacterales_bacterium | GXX35_15425 | 56222..59782 | 2304039 | BREX-system-P-loop-protein-BrxC-[Thermoanaerobacterales-bacterium] |
| MBQ9252308.1 | Type1.BREX | TOPRIM->AbiEi->?->AbiEii-> REase -> BrxA-DNA-binding -> BrxB-BrxF-iSTAND -> DUF6079 *-> N6-METHYLASE ->Tox-ART-HYD2-> PglZ -> SIGMA+OB-Fold+AAA-ATPase+Lon-Protease-BrxL ->?-><-Amidohydro_1\|\|Glyco_hydro_4+Glyco_hydro_4C->?->URI-DNase+UVR+UvrC_HhH_N+HHH-> | 1218 | Bacteria>Terrabacteria_group>Bacillota>Clostridia>unclassified_Clostridia | GCA_017470145.1 | Clostridia_bacterium | IJ188_06715 | 56969..60625 | 2044939 | BREX-system-P-loop-protein-BrxC-[Clostridia-bacterium] |
| EHM91311.1 | Type1.BREX | <-HTH_3\|\|HTH->Arm-DNA-bind_4+Phage_int_SAM_3+Phage_integrase->TraJ-RHH->?->?->GDResolvase+HIN-HTH-> BrxA-DNA-binding -p-> BrxB-BrxF-iSTAND -> DUF6079 -p*-><-Integrase_DNA+Phage_integrase<-?<-ABC-ATPase+SbcC<-?\|\| DUF6079 -p-> REase -> N6-METHYLASE ->METHYLASE->RADICAL-SAM-> | 917 | Bacteria>Terrabacteria_group>Bacillota>Erysipelotrichia>Erysipelotrichales>Coprobacillaceae | GCA_000239735.1 | Coprobacillus_sp._3_3_56FAA | HMPREF1021_02117 | 35070..37823 | 665941 | hypothetical-protein-HMPREF1021_02117-[Coprobacillus-sp.-3_3_56FAA] |
| WP_008791974.1 | Type1.BREX | <-Peptidase<-?\|\|HTH-><-HTH_38\|\|HTH_26-> BrxA-DNA-binding -> BrxB-BrxF-iSTAND -> DUF6079 *-> N6-METHYLASE -p-> N6-METHYLASE ->AbiGi-> PglZ -> SIGMA+OB-Fold+AAA-ATPase+Lon-Protease-BrxL ->?-> REase -> | 1182 | Bacteria>Terrabacteria_group>Bacillota>Erysipelotrichia>Erysipelotrichales>Coprobacillaceae | GCF_020097255.1 | Thomasclavelia_ramosa | HMPREF0978_01713 | 101921..105469 | 1547 | BREX-system-P-loop-protein-BrxC-[Thomasclavelia-ramosa] |
| WP_270396150.1 | Type1.BREX | BrxA-DNA-binding -> BrxB-BrxF-iSTAND -> DUF6079 *-> N6-METHYLASE ->ParB+HNH->HEPN+HEPN-> PglZ ->SbcC-> SIGMA+OB-Fold+AAA-ATPase+Lon-Protease-BrxL -> REase -> | 1177 | Bacteria>Terrabacteria_group>Bacillota>Erysipelotrichia>Erysipelotrichales>Coprobacillaceae | GCF_027659655.1 | Catenibacterium_mitsuokai | PGO85_RS08450 | 1355..4888 | 100886 | BREX-system-P-loop-protein-BrxC-[Catenibacterium-mitsuokai] |
| NLG80270.1 | Type1.BREX | ABC-ATPase+SbcC->ABC-ATPase+SbcC-><- REase +NACHT<- PglZ <- N6-METHYLASE <- DUF6079 *<- BrxB-BrxF-iSTAND <- BrxA-DNA-binding <- SIGMA+OB-Fold+AAA-ATPase+Lon-Protease-BrxL <-ParB+HNH<-AAA-ATPase-p<-DEDD_Tnp_IS110+Transposase_20\|\|?->DDE_Tnp_ISL3-> | 1185 | Bacteria>Terrabacteria_group>Bacillota | GCA_012516125.1 | Bacillota_bacterium | GX492_10705 | 20631..24188 | 1879010 | BREX-system-P-loop-protein-BrxC-[Bacillota-bacterium] |
| NPV28413.1 | Type1.BREX | NAD_synthase->?->?->?-> REase -> BrxA-DNA-binding -> BrxB-BrxF-iSTAND -> DUF6079 *-> N6-METHYLASE ->PHP+SbcC-> PglZ -> SIGMA+OB-Fold+AAA-ATPase+Lon-Protease-BrxL ->?->35exo+T7DNApol-> | 1208 | Bacteria>Terrabacteria_group>Bacillota | GCA_013177945.1 | Bacillota_bacterium | HPY58_01910 | 11153..14779 | 1879010 | BREX-system-P-loop-protein-BrxC-[Bacillota-bacterium] |
| MBP2637786.1 | Type1.BREX | HTH_26-> BrxA-DNA-binding -> BrxB-BrxF-iSTAND -> DUF6079 *-> N6-METHYLASE ->?->?-> PglZ -> SIGMA+OB-Fold+AAA-ATPase+Lon-Protease-BrxL -> REase +DUF5655-> | 1214 | Bacteria>Terrabacteria_group>Bacillota | GCA_017889705.1 | Bacillota_bacterium | H6Q72_3693 | 16334..19978 | 1879010 | hypothetical-protein-[Bacillota-bacterium] |
| MBS5426410.1 | Type1.BREX | <-Schlafen+HATPase+wHTH<-?\|\|?-><- SIGMA+OB-Fold+AAA-ATPase+Lon-Protease-BrxL <- PglZ <- REase <-OLD-ABC <- N6-METHYLASE <-Schlafen+HATPase+wHTH<- DUF6079 *<- BrxB-BrxF-iSTAND <- BrxA-DNA-binding <-zf-ISL3+HTH_Tnp_ISL3+DDE_Tnp_ISL3 | 1207 | Bacteria>Terrabacteria_group>Bacillota>Negativicutes>Acidaminococcales>Acidaminococcaceae | GCA_018368575.1 | Phascolarctobacterium_succinatutens | KHX39_04785 | 71877..75500 | 626940 | BREX-system-P-loop-protein-BrxC-[Phascolarctobacterium-succinatutens] |
| MBE6098022.1 | Type1.BREX | <- SIGMA+OB-Fold+AAA-ATPase+Lon-Protease-BrxL <- PglZ <- REase <-OLD-ABC <- N6-METHYLASE <- DUF6079 *<- BrxB-BrxF-iSTAND <- BrxA-DNA-binding <-PIN<-?<-Peptidase_M15 | 1220 | Bacteria>Terrabacteria_group>Bacillota>Negativicutes>Selenomonadales>Selenomonadaceae | GCA_015058685.1 | Schwartzia_succinivorans | E7198_09540 | 51689..55351 | 55507 | BREX-system-P-loop-protein-BrxC-[Schwartzia-succinivorans] |
| WP_273038291.1 | Type1.BREX | <- SIGMA+OB-Fold+AAA-ATPase+Lon-Protease-BrxL <- PglZ <- REase <-OLD-ABC <- N6-METHYLASE <- DUF6079 *<- BrxB-BrxF-iSTAND <- BrxA-DNA-binding <-PIN<-?<-Peptidase_M15 | 1220 | Bacteria>Terrabacteria_group>Bacillota>Negativicutes>Selenomonadales>Selenomonadaceae | GCF_015058685.1 | Schwartzia_succinivorans | E7198_RS09640 | 51689..55351 | 55507 | BREX-system-P-loop-protein-BrxC-[Schwartzia-succinivorans] |
| WP_273698075.1 | Type1.BREX | <- REase <- PglZ <- N6-METHYLASE <-PIN<-OLD-ABC<-HNH<-OLD-ABC<- DUF6079 *<- BrxB-BrxF-iSTAND <- BrxA-DNA-binding <-?<-?<-?<-?<-SKI<-?\|\|O6MGT-HTH-> | 1181 | Bacteria>Terrabacteria_group>Bacillota>Negativicutes>Selenomonadales>Selenomonadaceae | GCF_905371815.1 | Selenomonas_noxia | NW174_RS07850 | 36953..40498 | 135083 | BREX-system-P-loop-protein-BrxC-[Selenomonas-noxia] |
| WP_154440327.1 | Type1.BREX | primpol+D5N+D5ATPase-><-?\|\|?->?->RHH_8-> REase -> N6-METHYLASE -> BrxA-DNA-binding -> BrxB-BrxF-iSTAND -> DUF6079 *-> PglZ ->BrxD-AAA->SF1-SF2-Helicase+wHTH+Small-B-barrel+DUF1998->?->CarD+HTH->ZnR+ HARE-HTH -> | 1380 | Bacteria>Terrabacteria_group>Bacillota>Tissierellia>Tissierellales>Tissierellaceae | GCF_018917845.1 | Tissierella_pigra | KQI41_RS04690 | 949565..953707 | 2607614 | hypothetical-protein-[Tissierella-pigra] |
| WP_073050525.1 | Type1.BREX | <-RNase<-NERD<-OLD-ABC<- REase <- SIGMA+OB-Fold+AAA-ATPase+Lon-Protease-BrxL <- PglZ <-HEPN<- N6-METHYLASE <-?<- DUF6079 *<- BrxB-BrxF-iSTAND <- BrxA-DNA-binding <-HTH_26 | 1214 | Bacteria>Terrabacteria_group>Bacillota>Tissierellia>Tissierellia_incertae_sedis>Dethiosulfatibacter | GCF_900142005.1 | Dethiosulfatibacter_aminovorans | BUB05_RS15460 | 41174..44818 | 332095 | BREX-system-P-loop-protein-BrxC-[Dethiosulfatibacter-aminovorans] |
| OUL09467.1 | Type1.BREX | <- SF1-SF2-Helicase+ REase <- SF1-SF2-Helicase+ REase <-?<- SIGMA+OB-Fold+AAA-ATPase+Lon-Protease-BrxL <- PglZ <- REase <- N6-METHYLASE <- DUF6079 *<- BrxB-BrxF-iSTAND <- BrxA-DNA-binding <-HTH_26<-?<-?<- HTH+ REase <-?<-?<-Glyco_hydro_1 | 1207 | Bacteria>Terrabacteria_group>Bacillota>Tissierellia>Tissierellia_incertae_sedis>Sedimentibacter | GCA_002155105.1 | Sedimentibacter_sp._SX930 | B0533_04695 | 278529..282152 | 1955060 | ATPase-[Sedimentibacter-sp.-SX930] |
| MCC6190561.1 | Type6.BREX | MPTase->?->HerA/FtsK->HEPN-> REase ->?-> BrxE -> BrxA-DNA-binding -> BrxB-BrxF-iSTAND -> DUF6079 *-> N6-METHYLASE ->SF1-SF2-Helicase->HTH+HEPN->McrB-AAA-> REase -> PglZ -> SIGMA+OB-Fold+AAA-ATPase+Lon-Protease-BrxL ->?->Phage_int+DNA-glycosylase-> | 1208 | Bacteria>Terrabacteria_group>Chloroflexota>Anaerolineae>Anaerolineales>unclassified_Anaerolineales | - | Anaerolineales_bacterium | IT318_16150 | 14120..17746 | 2073117 | BREX-system-P-loop-protein-BrxC-[Anaerolineales-bacterium] |
| MBN1450028.1 | Type1.BREX | <- SIGMA+OB-Fold+AAA-ATPase+Lon-Protease-BrxL <- PglZ <- REase +DUF5655<- N6-METHYLASE <-HTH+HKD-DNase<-Pkinase+SF1-SF2-Helicase<-?<-NERD+Pkinase<- DUF6079 *<- BrxB-BrxF-iSTAND <- BrxA-DNA-binding | 1209 | Bacteria>Terrabacteria_group>Chloroflexota>Anaerolineae>Anaerolineales>unclassified_Anaerolineales | GCA_016928525.1 | Anaerolineales_bacterium | JW963_03360 | 18534..22163 | 2073117 | BREX-system-P-loop-protein-BrxC-[Anaerolineales-bacterium] |
| MBK9096490.1 | Type1.BREX | <- SIGMA+OB-Fold+AAA-ATPase+Lon-Protease-BrxL <- PglZ <-ParB+HNH<-HEPN+ REase <-?<- N6-METHYLASE <-TOPRIM<-OLD-ABC<- DUF6079 -p*<- DUF6079 -p<- BrxB-BrxF-iSTAND <- BrxA-DNA-binding <-?\|\|?-> N6-METHYLASE -p->RVT-> | 900 | Bacteria>Terrabacteria_group>Chloroflexota>Anaerolineae>unclassified_Anaerolineae | GCA_016719785.1 | Anaerolineae_bacterium | IPM84_27780 | 244922..247624 | 2052143 | hypothetical-protein-[Anaerolineae-bacterium] |
| MCL4299606.1 | Type1.BREX | <-DnaJ<-?<- N6-METHYLASE <- N6-METHYLASE <-?<-?<-?<- REase <- DUF6079 *<- BrxB-BrxF-iSTAND <- BrxA-DNA-binding <-MPTase<-HTH_3\|\|?->?->SMODS->?->JAB-> | 1231 | Bacteria>Terrabacteria_group>Chloroflexota>Anaerolineae>unclassified_Anaerolineae | GCA_023382535.1 | Anaerolineae_bacterium | KJ077_27970 | 96973..100668 | 2052143 | BREX-system-P-loop-protein-BrxC-[Anaerolineae-bacterium] |
| KPV54581.1 | Type1.BREX | <- N6-METHYLASE <- REase <- DUF6079 -p*<- DUF6079 -p<-HEPN<- BrxB-BrxF-iSTAND <- BrxA-DNA-binding <-XIS-HTH | 802 | Bacteria>Terrabacteria_group>Chloroflexota>Chloroflexia>Chloroflexales>Roseiflexineae | GCA_001399705.1 | Kouleothrix_aurantiaca | SE17_02870 | 3869..6277 | 186479 | hypothetical-protein-SE17_02870-[Kouleothrix-aurantiaca] |
| MBE0430956.1 | Type1.BREX | XIS-HTH-> BrxA-DNA-binding -> BrxB-BrxF-iSTAND -> DUF6079 *->Schlafen+HATPase+wHTH+HTH-> OLD-ABC-> REase -> N6-METHYLASE -> PglZ -p-> | 1193 | Bacteria>Terrabacteria_group>Chloroflexota>Dehalococcoidia>unclassified_Dehalococcoidia | GCA_014859205.1 | Dehalococcoidia_bacterium | IBX67_03920 | 8033..11614 | 2026734 | BREX-system-P-loop-protein-BrxC-[Dehalococcoidia-bacterium] |
| WP_201375473.1 | Type1.BREX | <- SIGMA+OB-Fold+AAA-ATPase+Lon-Protease-BrxL <- PglZ <-?<- N6-METHYLASE <- DUF6079 *<- BrxB-BrxF-iSTAND <- BrxA-DNA-binding <-?<-?\|\| REase -><-?\|\|HKD-DNase->?->SF1-SF2-Helicase-> | 1216 | Bacteria>Terrabacteria_group>Chloroflexota>Ktedonobacteria>Ktedonobacterales>Ktedonobacteraceae | GCF_016587375.1 | Ktedonobacter_robiniae | KSB_77480 | 1130085..1133735 | 2778365 | BREX-system-P-loop-protein-BrxC-[Ktedonobacter-robiniae] |
| KAA3643964.1 | Type1.BREX | <-ParA-Soj-PloopNTPase\|\|GspA-ATPase->?->?->?->?-> BrxA-DNA-binding -> BrxB-BrxF-iSTAND -> DUF6079 *-> N6-METHYLASE -> PglZ -> SIGMA+OB-Fold+AAA-ATPase+Lon-Protease-BrxL -> REase -><-?<-NTP_transf_5<-REC | 1152 | Bacteria>Terrabacteria_group>Chloroflexota>unclassified_Chloroflexota | GCA_008501685.1 | Chloroflexota_bacterium | DWQ07_17765 | 184634..188092 | 2026724 | BREX-system-P-loop-protein-BrxC-[Chloroflexota-bacterium] |
| NWJ94851.1 | Type1.BREX | <- SIGMA+OB-Fold+AAA-ATPase+Lon-Protease-BrxL <- PglZ <-?<- REase <-?<- N6-METHYLASE <-?<-?<- DUF6079 *<- BrxB-BrxF-iSTAND <- BrxA-DNA-binding <-XIS-HTH<-?<-XIS-HTH\|\|?->MoxR-AAA->VWA+vWA-L+ CR-ATPase9 ->CR-ATPase-> | 1218 | Bacteria>Terrabacteria_group>Chloroflexota>unclassified_Chloroflexota | GCA_013390945.1 | Chloroflexota_bacterium | HXX20_03630 | 11914..15570 | 2026724 | BREX-system-P-loop-protein-BrxC-[Chloroflexota-bacterium] |
| MBX9724057.1 | Type1.BREX | <- SIGMA+OB-Fold+AAA-ATPase+HKD-DNase-BrxL \|\| BrxA-DNA-binding -> BrxB-BrxF-iSTAND -> DUF6079 *-> OLD-ABC-> REase -> N6-METHYLASE -> PglZ -p-> | 1175 | Bacteria>Terrabacteria_group>Cyanobacteriota/Melainabacteria_group>Candidatus_Melainabacteria>Candidatus_Obscuribacterales>unclassified_Candidatus_Obscuribacterales | GCA_019747345.1 | Candidatus_Obscuribacterales_bacterium | K2X81_21805 | 3793..7320 | 2364082 | BREX-system-P-loop-protein-BrxC-[Candidatus-Obscuribacterales-bacterium] |
| WP_190963244.1 | Type6.BREX | HTH_26->?->?->?->?->HEPN-> BrxE -> BrxA-DNA-binding -> BrxB-BrxF-iSTAND -> DUF6079 *-> N6-METHYLASE ->MORC+DUF3684->MORC+DUF3684+ REase -> PglZ ->BrxD-AAA->SF1-SF2-Helicase+wHTH+Small-B-barrel+DUF1998-> | 1207 | Bacteria>Terrabacteria_group>Cyanobacteriota/Melainabacteria_group>Cyanobacteriota>Cyanophyceae>Nostocales | - | Desmonostoc_muscorum | H6G91_22885 | 8366..11989 | 1179 | BREX-system-P-loop-protein-BrxC-[Desmonostoc-muscorum] |
| MBU7587299.1 | Type1.BREX | ParA-Soj-PloopNTPase->?-> BrxA-DNA-binding -> BrxB-BrxF-iSTAND -> DUF6079 *-> REase -> N6-METHYLASE -> | 1221 | Bacteria>Terrabacteria_group>Cyanobacteriota/Melainabacteria_group>Cyanobacteriota>Cyanophyceae>Nostocales | GCA_018993265.1 | Nostoc_sp._TH1S01 | KAF91_31390 | 3365..7030 | 442893 | BREX-system-P-loop-protein-BrxC-[Nostoc-sp.-TH1S01] |
| MBW4670333.1 | Type6.BREX | <-Glycos_transf<-TPR\|\| REase -><-Acetate_kinase\|\| BrxE -> BrxA-DNA-binding -> BrxB-BrxF-iSTAND -> DUF6079 *->?->PIN->?-><- REase <-?\|\| N6-METHYLASE -> | 1211 | Bacteria>Terrabacteria_group>Cyanobacteriota/Melainabacteria_group>Cyanobacteriota>Cyanophyceae>Nostocales | GCA_019359595.1 | Cyanomargarita_calcarea_GSE-NOS-MK-12-04C | KME60_23685 | 99014..102649 | 2839659 | BREX-system-P-loop-protein-BrxC-[Cyanomargarita-calcarea-GSE-NOS-MK-12-04C] |
| WP_012411904.1 | Type1.BREX | REase -><-?<-?<-?\|\|?->?<-?\|\|tRNA-><-Cas12k\|\|MerR-HTH-> BrxB-BrxF-iSTAND -> DUF6079 *-> N6-METHYLASE -><-?<- REase \|\| PglZ +Phosphodiest -> SIGMA+OB-Fold+AAA-ATPase+Lon-Protease-BrxL -><- BrxA-DNA-binding <-RHH\|\|?->CHTH-> | 1179 | Bacteria>Terrabacteria_group>Cyanobacteriota/Melainabacteria_group>Cyanobacteriota>Cyanophyceae>Nostocales | GCF_000020025.1 | Nostoc_punctiforme | NPUN_RS28630 | 6987385..6990924 | 272131 | BREX-system-P-loop-protein-BrxC-[Nostoc-punctiforme] |
| WP_141302954.1 | Type1.BREX | <- PglZ \|\|DDE_Tnp_IS1-><- N6-METHYLASE -p<- N6-METHYLASE -p<- N6-METHYLASE -p<- REase <-OLD-ABC <-PIN<-DUF433<- DUF6079 *<- BrxB-BrxF-iSTAND \|\|?->?-><-?<-?<-?<-ParA-Soj-PloopNTPase<-?\|\|ACT-> | 1166 | Bacteria>Terrabacteria_group>Cyanobacteriota/Melainabacteria_group>Cyanobacteriota>Cyanophyceae>Nostocales | GCF_006523545.1 | Cylindrospermopsis_raciborskii | FIV49_RS01515 | 324659..328159 | 77022 | BREX-system-P-loop-protein-BrxC-[Cylindrospermopsis-raciborskii] |
| WP_194144969.1 | Type1.BREX | BrxA-DNA-binding -><- SIGMA+OB-Fold+AAA-ATPase+Lon-Protease-BrxL <- PglZ +Phosphodiest \|\| REase ->?-><- N6-METHYLASE <- DUF6079 *<- BrxB-BrxF-iSTAND <-?\|\|?->CHTH-> | 1179 | Bacteria>Terrabacteria_group>Cyanobacteriota/Melainabacteria_group>Cyanobacteriota>Cyanophyceae>Nostocales | GCF_015206985.1 | Nostoc_sp._LEGE_12447 | IQ274_RS34165 | 10352..13891 | 1828640 | BREX-system-P-loop-protein-BrxC-[Nostoc-sp.-LEGE-12447] |
| WP_235116163.1 | Type1.BREX | <-Glyco_hydro_63\|\| BrxA-DNA-binding -><- SIGMA+OB-Fold+AAA-ATPase+Lon-Protease-BrxL <- PglZ +Phosphodiest \|\| REase ->?-><- N6-METHYLASE \|\|MetJarc-HTH->PemK_toxin-> DUF6079 *<- BrxB-BrxF-iSTAND <-MerR-HTH | 1180 | Bacteria>Terrabacteria_group>Cyanobacteriota/Melainabacteria_group>Cyanobacteriota>Cyanophyceae>Nostocales | GCF_015207005.2 | Desmonostoc_muscorum | IQ276_RS32480 | 7902408..7905950 | 1179 | BREX-system-P-loop-protein-BrxC-[Desmonostoc-muscorum] |
| WP_224095538.1 | Type1.BREX | BrxA-DNA-binding -><- SIGMA+OB-Fold+AAA-ATPase+Lon-Protease-BrxL <- PglZ +Phosphodiest \|\|PIN->?-><-HTH_23+HTH_32<- N6-METHYLASE -p<- N6-METHYLASE -p<- DUF6079 *<- BrxB-BrxF-iSTAND <-?<- REase <-?\|\|ArsR-HTH-><-?\|\|?-><-?\|\|HTH_32+DDE_3-> | 1178 | Bacteria>Terrabacteria_group>Cyanobacteriota/Melainabacteria_group>Cyanobacteriota>Cyanophyceae>Nostocales | GCF_019976755.1 | Nostoc_sp._MS1 | NSMS1_RS32170 | 256782..260318 | 2764711 | BREX-system-P-loop-protein-BrxC-[Nostoc-sp.-MS1] |
| WP_229553232.1 | Type1.BREX | ZDNA-HTH->rve->?->TnsC-ATPase->?-> BrxA-DNA-binding -> BrxB-BrxF-iSTAND -> DUF6079 *-> REase -> N6-METHYLASE -> N6-METHYLASE -p-> | 1231 | Bacteria>Terrabacteria_group>Cyanobacteriota/Melainabacteria_group>Cyanobacteriota>Cyanophyceae>Nostocales | GCF_020829625.1 | Nostoc_sp._XA010 | LC608_RS09180 | 421109..424804 | 2780407 | BREX-system-P-loop-protein-BrxC-[Nostoc-sp.-XA010] |
| WP_162398350.1 | Type1.BREX | BrxA-DNA-binding -><- SIGMA+OB-Fold+AAA-ATPase+Lon-Protease-BrxL <- PglZ +Phosphodiest \|\| REase ->PIN-><- N6-METHYLASE \|\|MetJarc-HTH->PemK_toxin-> DUF6079 *<- BrxB-BrxF-iSTAND \|\|<-tRNA<-?<-?\|\|?->?-><-?\|\|?-><-Peptidase_M28 | 1178 | Bacteria>Terrabacteria_group>Cyanobacteriota/Melainabacteria_group>Cyanobacteriota>Cyanophyceae>Nostocales | - | Nostoc_sp._B(2019) | GS682_RS23175 | 25673..29209 | 2690446 | BREX-system-P-loop-protein-BrxC-[Nostoc-sp.-B(2019)] |
| CCI01315.1 | Type1.BREX | Pkinase+TPR+TPR+TPR-> BrxB-BrxF-iSTAND -> DUF6079 *-> OLD-ABC-> REase -> N6-METHYLASE -> REase ->?-> PglZ -><-SPOUT\|\|Peptidase_S51-> | 1188 | Bacteria>Terrabacteria_group>Cyanobacteriota/Melainabacteria_group>Cyanobacteriota>Cyanophyceae>Oscillatoriophycideae | GCA_000312185.1 | Microcystis_aeruginosa_PCC_9443 | MICAC_1830011 | 77203..80769 | 1160281 | conserved-hypothetical-protein-[Microcystis-aeruginosa-PCC-9443] |
| REJ45948.1 | Type1.BREX | Pkinase+TPR+TPR+TPR-> BrxB-BrxF-iSTAND -> DUF6079 *-> OLD-ABC-> REase -> N6-METHYLASE ->DOC+HTH-> PglZ -><-SPOUT\|\|Peptidase_S51->?-><-DDE_5+DDE_Tnp_1 | 1168 | Bacteria>Terrabacteria_group>Cyanobacteriota/Melainabacteria_group>Cyanobacteriota>Cyanophyceae>Oscillatoriophycideae | GCA_003389435.1 | Microcystis_flos-aquae_DF17 | DWQ53_12245 | 7047..10553 | 2060470 | BREX-system-P-loop-protein-BrxC-[Microcystis-flos-aquae-DF17] |
| BBD57221.1 | Type1.BREX | <-MerR-HTH\|\|RHH-><-?<-?\|\|?->TPR+TPR+TPR-><-Phage_integrase<-?\|\| BrxB-BrxF-iSTAND -> DUF6079 *-><-?<- REase \|\|?->PIN-> N6-METHYLASE ->HEPN-> PglZ +Phosphodiest ->HKD-DNase-> SIGMA+OB-Fold+AAA-ATPase+Lon-Protease-BrxL -> | 1176 | Bacteria>Terrabacteria_group>Cyanobacteriota/Melainabacteria_group>Cyanobacteriota>Cyanophyceae>Oscillatoriophycideae | GCA_003609755.1 | Planktothrix_agardhii_NIES-204 | NIES204_45570 | 14633..18163 | 282423 | hypothetical-protein-NIES204_45570-[Planktothrix-agardhii-NIES-204] |
| TRU20104.1 | Type1.BREX | OrfB_IS605+OrfB_Zn_ribbon-><-Pkinase<-Peptidase_M16+Peptidase_M16_C\|\|ncRNA->?->?-><-?\|\|?->tRNA->Pkinase+TPR+TPR+TPR-> BrxB-BrxF-iSTAND -> DUF6079 *-> OLD-ABC-> REase -> N6-METHYLASE ->DOC+HTH-> PglZ -><-SPOUT\|\|Peptidase_S51->?->DUF488-1-> | 1168 | Bacteria>Terrabacteria_group>Cyanobacteriota/Melainabacteria_group>Cyanobacteriota>Cyanophyceae>Oscillatoriophycideae | GCA_007095795.1 | Microcystis_aeruginosa_Ma_MB_S_20031200_S102D | EWV79_18805 | 31642..35148 | 2486255 | BREX-system-P-loop-protein-BrxC-[Microcystis-aeruginosa-Ma_MB_S_20031200_S102D] |
| NCR78979.1 | Type1.BREX | <-Peptidase_S51\|\|SPOUT-><- PglZ <-?<- REase <-?<- N6-METHYLASE <- REase <-OLD-ABC <- DUF6079 *<- BrxB-BrxF-iSTAND <-Pkinase+TPR+TPR+TPR<-tRNA<-HTH_Tnp_1 | 1168 | Bacteria>Terrabacteria_group>Cyanobacteriota/Melainabacteria_group>Cyanobacteriota>Cyanophyceae>Oscillatoriophycideae | GCA_009996105.1 | Microcystis_aeruginosa_K13-10 | GPI91_02970 | 22657..26163 | 2685970 | BREX-system-P-loop-protein-BrxC-[Microcystis-aeruginosa-K13-10] |
| NCS23184.1 | Type1.BREX | <-Peptidase_S51\|\|SPOUT-><- PglZ <-DOC+HTH<-?<- N6-METHYLASE <- REase <-OLD-ABC <- DUF6079 *<- BrxB-BrxF-iSTAND <-Pkinase+TPR+TPR+TPR<-tRNA<-?\|\|?-><-?<-?<-ncRNA\|\|Peptidase_M16+Peptidase_M16_C->Pkinase-><-OrfB_IS605+OrfB_Zn_ribbon | 1168 | Bacteria>Terrabacteria_group>Cyanobacteriota/Melainabacteria_group>Cyanobacteriota>Cyanophyceae>Oscillatoriophycideae | GCA_009996485.1 | Microcystis_aeruginosa_BS13-02 | GPI96_01080 | 43062..46568 | 2685952 | BREX-system-P-loop-protein-BrxC-[Microcystis-aeruginosa-BS13-02] |

| Protein-ID | System | Gene-Neighbourhoods | Length | Taxonomy | Assembly-ID | Strain | Locus-Tag | Genomic-coordinates | TaxID | Fasta-define |
|---|---|---|---|---|---|---|---|---|---|---|
| MBE5230557.1 | Type1.BREX | Pkinase+TPR+TPR+TPR-> BrxB-BrxF-iSTAND -> DUF6079 *-> OLD-ABC-> REase -> N6-METHYLASE -> | 1168 | Bacteria>Terrabacteria_group>Cyanobacteriota/Melainabacteria_group>Cyanobacteriota>Cyanophyceae>Oscillatoriophycideae | GCA_014946735.1 | Microcystis_aeruginosa_PMC_728.11 | INR81_16200 | 47803..51309 | 2316423 | BREX-system-P-loop-protein-BrxC-[Microcystis-aeruginosa-PMC-728.11] |
| MCA2504292.1 | Type1.BREX | <-DUF488-l<-?<-Peptidase_S51\|\|SPOUT-><- PglZ <-DOC+HTH<- N6-METHYLASE <- REase <-OLD-ABC <- DUF6079 *<- BrxB-BrxF-iSTAND <-Pkinase+TPR+TPR+TPR | 1168 | Bacteria>Terrabacteria_group>Cyanobacteriota/Melainabacteria_group>Cyanobacteriota>Cyanophyceae>Oscillatoriophycideae | GCA_020244565.1 | Microcystis_sp._M62BS1 | IM561_02895 | 20537..24043 | 2771204 | BREX-system-P-loop-protein-BrxC-[Microcystis-sp.-M62BS1] |
| WP_004266780.1 | Type1.BREX | <-Peptidase_S51\|\|SPOUT-><- PglZ <-?<- REase <- N6-METHYLASE -p<- N6-METHYLASE <- REase <-OLD-ABC <- DUF6079 *<- BrxB-BrxF-iSTAND <-Pkinase+TPR+TPR+TPR<-tRNA<-?<-?\|\|?-><-?<-?<-ncRNA<-?<-DADA-Ligase | 1168 | Bacteria>Terrabacteria_group>Cyanobacteriota/Melainabacteria_group>Cyanobacteriota>Cyanophyceae>Oscillatoriophycideae | GCF_000312165.1 | Microcystis_aeruginosa | MICAB_RS21670 | 36173..39679 | 1126 | BREX-system-P-loop-protein-BrxC-[Microcystis-aeruginosa] |
| WP_043996480.1 | Type1.BREX | Pkinase+TPR+TPR+TPR-> BrxB-BrxF-iSTAND -> DUF6079 *-> OLD-ABC-> REase -> N6-METHYLASE -> REase ->?-> PglZ -><-SPOUT\|\|Peptidase_S51-> | 1168 | Bacteria>Terrabacteria_group>Cyanobacteriota/Melainabacteria_group>Cyanobacteriota>Cyanophyceae>Oscillatoriophycideae | GCF_000312185.1 | Microcystis_aeruginosa | MICAC_RS06610 | 77263..80769 | 1126 | BREX-system-P-loop-protein-BrxC-[Microcystis-aeruginosa] |
| WP_002790046.1 | Type1.BREX | PIN-><-HicB_lk_antitox<-?\|\|?-><- N6-METHYLASE -p<- REase <-OLD-ABC <- DUF6079 -p<- DUF6079 -p*<- DUF6079 -p<- BrxB-BrxF-iSTAND <-?<-Pkinase+TPR+TPR+TPR | 532 | Bacteria>Terrabacteria_group>Cyanobacteriota/Melainabacteria_group>Cyanobacteriota>Cyanophyceae>Oscillatoriophycideae | GCF_000312225.1 | Microcystis_aeruginosa | MICAF_RS30295 | 136943..138541 | 1126 | BREX-system-P-loop-protein-BrxC-[Microcystis-aeruginosa] |
| WP_193959346.1 | Type1.BREX | <- PglZ <-?<- REase <- N6-METHYLASE <- REase <-OLD-ABC <- DUF6079 *<- BrxB-BrxF-iSTAND | 1168 | Bacteria>Terrabacteria_group>Cyanobacteriota/Melainabacteria_group>Cyanobacteriota>Cyanophyceae>Oscillatoriophycideae | GCF_015207185.1 | Microcystis_sp._LEGE_08355 | IQ242_RS05955 | 8581..12087 | 1828687 | BREX-system-P-loop-protein-BrxC-[Microcystis-sp.-LEGE-08355] |
| WP_271989206.1 | Type1.BREX | <-Peptidase_S51\|\|SPOUT-><- PglZ <-?<- REase <- N6-METHYLASE <- REase <-OLD-ABC <- DUF6079 *<- BrxB-BrxF-iSTAND <-Pkinase+TPR+TPR+TPR | 1168 | Bacteria>Terrabacteria_group>Cyanobacteriota/Melainabacteria_group>Cyanobacteriota>Cyanophyceae>Oscillatoriophycideae | GCF_028330905.1 | Microcystis_aeruginosa | PN502_RS07790 | 14255..17761 | 1126 | BREX-system-P-loop-protein-BrxC-[Microcystis-aeruginosa] |
| WP_272076547.1 | Type1.BREX | <- REase <-Pkinase<-Peptidase_M16+Peptidase_M16_C\|\|ncRNA->?->?-><-?\|\|?->tRNA->Pkinase+TPR+TPR+TPR-> BrxB-BrxF-iSTAND -> DUF6079 *-> OLD-ABC-> REase -> N6-METHYLASE ->DOC+HTH-> PglZ -><-SPOUT\|\|Peptidase_S51->?->DUF488-l-> | 1168 | Bacteria>Terrabacteria_group>Cyanobacteriota/Melainabacteria_group>Cyanobacteriota>Cyanophyceae>Oscillatoriophycideae | GCF_028331105.1 | Microcystis_aeruginosa | PN487_RS04630 | 38433..41939 | 1126 | BREX-system-P-loop-protein-BrxC-[Microcystis-aeruginosa] |
| MBW4580237.1 | Type6.BREX | <-ABC-ATPase+SbcC\|\|?->?-><-SAM-METHYLASE<-?\|\|CHTH-> BrxE -> BrxA-DNA-binding -> BrxB-BrxF-iSTAND -> DUF6079 *->?-> REase ->DUF29->AbrB-HTH->PIN-> PglZ ->BrxD-AAA->SF1-SF2-Helicase+wHTH+Small-B-barrel+DUF1998-> | 1217 | Bacteria>Terrabacteria_group>Cyanobacteriota/Melainabacteria_group>Cyanobacteriota>Cyanophyceae>Pseudanabaenales | GCA_019358895.1 | Tildeniella_nuda_ZEHNDER_1965/U140 | KME42_11780 | 246570..250223 | 2136405 | BREX-system-P-loop-protein-BrxC-[Tildeniella-nuda-ZEHNDER-1965/U140] |
| MCL1489589.1 | Type6.BREX | HTH_32+DDE_3->?->?-><- N6-METHYLASE -p<-DDE_Tnp_1_assoc+DDE_Tnp_1<- N6-METHYLASE -p<-?<- BrxB-BrxF-iSTAND <- REase <- DUF6079 *<- BrxB-BrxF-iSTAND <- BrxA-DNA-binding <- BrxE \|\|ParA-Soj-PloopNTPase->ParB-><-NACHT<-PrimPol | 1227 | Bacteria>Terrabacteria_group>Cyanobacteriota/Melainabacteria_group>Cyanobacteriota>Cyanophyceae>Pseudanabaenales | GCA_023336205.1 | Pseudanabaena_sp._Salubria-1 | M1G31_02380 | 45126..48809 | 2938918 | BREX-system-P-loop-protein-BrxC-[Pseudanabaena-sp.-Salubria-1] |
| WP_009625711.1 | Type6.BREX | <-SF1-SF2-Helicase+wHTH+Small-B-barrel+DUF1998<-BrxD-AAA<- PglZ \|\|PhdYeFM_antitox-> REase ->?-><- N6-METHYLASE <-ACET<- BrxB-BrxF-iSTAND <- DUF6079 *<- BrxB-BrxF-iSTAND <- BrxA-DNA-binding <- BrxE \|\|ParA-Soj-PloopNTPase->ParB-> | 1222 | Bacteria>Terrabacteria_group>Cyanobacteriota/Melainabacteria_group>Cyanobacteriota>Cyanophyceae>Pseudanabaenales | GCF_000332215.1 | Pseudanabaena_biceps | PSE7429_RS03680 | 23473..27141 | 927669 | MULTISPECIES:-BREX-system-P-loop-protein-BrxC-[Pseudanabaena] |
| WP_068818822.1 | Type1.BREX | Phage_int_SAM_1->Phage_integrase-> BrxB-BrxF-iSTAND -> DUF6079 *->MazE_antitoxin->PIN->Polbeta-> N6-METHYLASE ->?-> REase -> PglZ +Phosphodiest -> SIGMA+OB-Fold+AAA-ATPase+Lon-Protease-BrxL -><- BrxA-DNA-binding | 1182 | Bacteria>Terrabacteria_group>Cyanobacteriota/Melainabacteria_group>Cyanobacteriota>Cyanophyceae>Pseudanabaenales | GCF_001650195.1 | Phormidesmis_priestleyi | A8707_RS19220 | 3171..6719 | 268141 | BREX-system-P-loop-protein-BrxC-[Phormidesmis-priestleyi] |
| WP_094536005.1 | Type6.BREX | TOPRIM+SF1-SF2-Helicase->NACHT-><-ParB<-ParA-Soj-PloopNTPase\|\| BrxE -> BrxA-DNA-binding -> BrxB-BrxF-iSTAND -> DUF6079 *-> REase -> REase -> N6-METHYLASE -> | 1222 | Bacteria>Terrabacteria_group>Cyanobacteriota/Melainabacteria_group>Cyanobacteriota>Cyanophyceae>Pseudanabaenales | GCF_002251945.1 | Pseudanabaena_sp._SR411 | B9G53_RS24225 | 12197..15865 | 1980935 | BREX-system-P-loop-protein-BrxC-[Pseudanabaena-sp.-SR411] |
| WP_169613736.1 | Type1.BREX | Phage_integrase+Phage_integrase-><-?\|\|?->?-><-HTH\|\|?-> BrxA-DNA-binding -> BrxB-BrxF-iSTAND -> DUF6079 *-> N6-METHYLASE ->OLD-ABC->HNH-> REase ->?-> PglZ -> SIGMA+OB-Fold+AAA-ATPase+Lon-Protease-BrxL ->AAA-ATPase-> | 1150 | Bacteria>Terrabacteria_group>Cyanobacteriota/Melainabacteria_group>Cyanobacteriota>Cyanophyceae>Pseudanabaenales | GCF_012911975.1 | Nodosilinea_sp._P-1105 | E1H13_RS12425 | 20381..23833 | 2546229 | BREX-system-P-loop-protein-BrxC-[Nodosilinea-sp.-P-1105] |
| WP_193855895.1 | Type1.BREX | <-Acyltransferase<-ACET<-SF1-SF2-Helicase<-?<-TRD\|\| BrxA-DNA-binding -> BrxB-BrxF-iSTAND -> DUF6079 *-> OLD-ABC-> REase -> N6-METHYLASE -> | 1185 | Bacteria>Terrabacteria_group>Cyanobacteriota/Melainabacteria_group>Cyanobacteriota>Cyanophyceae>Pseudanabaenales | GCF_015207105.1 | cf._Phormidesmis_sp._LEGE_11477 | IQ256_RS12585 | 42701..46258 | 1828680 | BREX-system-P-loop-protein-BrxC-[cf.-Phormidesmis-sp.-LEGE-11477] |
| WP_194058630.1 | Type6.BREX | pPIWI+pPIWI->pPIWI+SF1-SF2-Helicase->pPIWI-><-SF1-SF2-Helicase+wHTH+Small-B-barrel+DUF1998<-BrxD-AAA<- PglZ <-?<- REase <- N6-METHYLASE <- DUF6079 *<- BrxB-BrxF-iSTAND <- BrxA-DNA-binding <- BrxE <-CHTH\|\|?-><-pPIWI<-?<- N6-METHYLASE | 1220 | Bacteria>Terrabacteria_group>Cyanobacteriota/Melainabacteria_group>Cyanobacteriota>Cyanophyceae>Pseudanabaenales | GCF_015207525.1 | Nodosilinea_sp._LEGE_06152 | IQ265_RS24920 | 20188..23850 | 2777966 | BREX-system-P-loop-protein-BrxC-[Nodosilinea-sp.-LEGE-06152] |
| MCE2837751.1 | Type1.BREX | <- BrxB-BrxF-iSTAND -p<-?\|\|?->SAM-METHYLASE->DDE_Tnp_1-><-?\|\| BrxA-DNA-binding -> BrxB-BrxF-iSTAND -> DUF6079 -p*-> DUF6079 -p->MNS+ REase ->MNS+ REase ->?->PIN-><-PIN\|\|?->PIN->HTH_3-> | 600 | Bacteria>Terrabacteria_group>Cyanobacteriota/Melainabacteria_group>Cyanobacteriota>Cyanophyceae>Synechococcales | GCA_021298515.1 | Cyanobium_sp._49614_E6 | LW834_12445 | 8564..10363 | 2904989 | BREX-system-P-loop-protein-BrxC-[Cyanobium-sp.-49614_E6] |
| WP_007100811.1 | Type1.BREX | AAA-ATPase-p-><-SIGMA-HTH<-?<-?<- SIGMA+OB-Fold+AAA-ATPase+Lon-Protease-BrxL <- PglZ <- N6-METHYLASE <- N6-METHYLASE <- DUF6079 *<- BrxB-BrxF-iSTAND <- BrxA-DNA-binding <- REase <-?<-?<-?<-?\|\|McrB-AAA-> | 1161 | Bacteria>Terrabacteria_group>Cyanobacteriota/Melainabacteria_group>Cyanobacteriota>Cyanophyceae>Synechococcales | GCF_000153065.1 | unclassified_Synechococcus | RS9917_RS03965 | 839955..843440 | 2626047 | MULTISPECIES:-BREX-system-P-loop-protein-BrxC-[unclassified-Synechococcus] |
| WP_186696561.1 | Type1.BREX | HIN-HTH->HAD+CHTH->?->?->ArdCN+MPTase-> BrxA-DNA-binding -> BrxB-BrxF-iSTAND -> DUF6079 *-> N6-METHYLASE -> PglZ -> SIGMA+OB-Fold+AAA-ATPase+HKD-DNase-BrxL -><- REase \|\|DUF4236+TPR-><-ABhydrolase<-PIN | 1164 | Bacteria>Terrabacteria_group>Cyanobacteriota/Melainabacteria_group>Cyanobacteriota>Cyanophyceae>Synechococcales | GCF_014280235.1 | Cyanobium_sp._NS01 | CyaNS01_RS07595 | 1454207..1457701 | 261284 | BREX-system-P-loop-protein-BrxC-[Cyanobium-sp.-NS01] |
| WP_255087766.1 | Type1.BREX | REase -><- SIGMA+OB-Fold+AAA-ATPase+HKD-DNase-BrxL <- PglZ <-sGTP<-GTPase-AIG<- N6-METHYLASE <-MNS+ REase <- DUF6079 *<- BrxB-BrxF-iSTAND <- BrxA-DNA-binding | 1161 | Bacteria>Terrabacteria_group>Cyanobacteriota/Melainabacteria_group>Cyanobacteriota>Cyanophyceae>Synechococcales | GCF_024345415.1 | unclassified_Synechococcus | KBY92_RS03480 | 114966..118451 | 2626047 | MULTISPECIES:-BREX-system-P-loop-protein-BrxC-[unclassified-Synechococcus] |
| WP_254955669.1 | Type1.BREX | <- N6-METHYLASE <-HTH_3<-PIN<-?\|\|PIN->PIN-><-PIN<-?<-MNS+ REase <- DUF6079 *<- BrxB-BrxF-iSTAND <- BrxA-DNA-binding \|\|Shikimate_dh_N+SDH_C->?-><-?\|\|?->?-><-Peptidase_M1_N+Peptidase_M1 | 1161 | Bacteria>Terrabacteria_group>Cyanobacteriota/Melainabacteria_group>Cyanobacteriota>Cyanophyceae>Synechococcales | GCF_024346055.1 | unclassified_Cyanobium | KBZ12_RS00475 | 78178..81663 | 2627006 | MULTISPECIES:-BREX-system-P-loop-protein-BrxC-[unclassified-Cyanobium] |
| WP_254970971.1 | Type1.BREX | <- PglZ <-Couple_hipA+HipA_C<-HTH_3<-sGTP<-GTPase-AIG<- N6-METHYLASE <-MNS+ REase \|\|PIN->PIN-><- DUF6079 *<- BrxB-BrxF-iSTAND <- BrxA-DNA-binding \|\|Shikimate_dh_N+SDH_C->?-><-?\|\|?->?-><-Peptidase_M1_N+Peptidase_M1 | 1161 | Bacteria>Terrabacteria_group>Cyanobacteriota/Melainabacteria_group>Cyanobacteriota>Cyanophyceae>Synechococcales | GCF_024346155.1 | Cyanobium_sp._Candia_9D4 | KBZ08_RS02055 | 407526..411011 | 2823707 | BREX-system-P-loop-protein-BrxC-[Cyanobium-sp.-Candia-9D4] |
| WP_255125212.1 | Type1.BREX | <- PglZ -p<-Couple_hipA+HipA_C<-HTH_3<- N6-METHYLASE <-MNS+ REase <- DUF6079 *<- BrxB-BrxF-iSTAND <- BrxA-DNA-binding | 1165 | Bacteria>Terrabacteria_group>Cyanobacteriota/Melainabacteria_group>Cyanobacteriota>Cyanophyceae>Synechococcales | GCF_024346215.1 | Synechococcus_lacustris | KBY79_RS12110 | 7241..10738 | 2116544 | BREX-system-P-loop-protein-BrxC-[Synechococcus-lacustris] |
| WP_255005473.1 | Type1.BREX | PIN->PIN->?-><-DOC\|\|?->ArdCN+MPTase-> BrxA-DNA-binding -> BrxB-BrxF-iSTAND -> DUF6079 *-> N6-METHYLASE -> PglZ ->PhdYeFM_antitox->PIN-> SIGMA+OB-Fold+AAA-ATPase+HKD-DNase-BrxL -><- REase <-TPR<-Peptidase | 1161 | Bacteria>Terrabacteria_group>Cyanobacteriota/Melainabacteria_group>Cyanobacteriota>Cyanophyceae>Synechococcales | GCF_024346315.1 | Cyanobium_sp._ATX_6F1 | KBZ13_RS02100 | 369855..373340 | 2823702 | BREX-system-P-loop-protein-BrxC-[Cyanobium-sp.-ATX-6F1] |
| WP_255017269.1 | Type1.BREX | <-Sak4+WYL\|\|?-> REase ->?-> BrxA-DNA-binding -> BrxB-BrxF-iSTAND -> DUF6079 *-> N6-METHYLASE ->PRTase-> PglZ -> SIGMA+OB-Fold+AAA-ATPase+Lon-Protease-BrxL ->?->?->?->SIGMA-HTH-> | 1162 | Bacteria>Terrabacteria_group>Cyanobacteriota/Melainabacteria_group>Cyanobacteriota>Cyanophyceae>Synechococcales | GCF_024346475.1 | Cyanobium_sp._A2C-AMD | KBZ17_RS12415 | 5448..8936 | 2823695 | BREX-system-P-loop-protein-BrxC-[Cyanobium-sp.-A2C-AMD] |
| MBU6345615.1 | Type1.BREX | <- N6-METHYLASE -p<- REase <-OLD-ABC <-PIN<-DUF433<- DUF6079 *<- BrxB-BrxF-iSTAND \|\|?->?-><-?<-?<-?<-ParA-Soj-PloopNTPase<-?\|\|ACT-> | 1166 | Bacteria>Terrabacteria_group>Cyanobacteriota/Melainabacteria_group>Cyanobacteriota>unclassified_Cyanobacteriota | GCA_018969285.1 | Cyanobacteria_bacterium_REEB494 | KGR70_09985 | 4368..7868 | 2800107 | BREX-system-P-loop-protein-BrxC-[Cyanobacteria-bacterium-REEB494] |
| WP_104992429.1 | Type1.BREX | HTH_32+DDE_3->OrfB_Zn_ribbon-><-ASCH\|\|PhdYeFM_antitox->RelE-ParE->OrfB_Zn_ribbon-> BrxA-DNA-binding -> BrxB-BrxF-iSTAND -> DUF6079 *->RelE-ParE+SF1-SF2-Helicase-> N6-METHYLASE ->HNH->HTH+ART-DarT->HTH_Tnp_1+HTH_21+rve+rve_3->NADAR-> REase -> PglZ -> SIGMA+OB-Fold+AAA-ATPase+Lon-Protease-BrxL -> | 1140 | Bacteria>Terrabacteria_group>Deinococcota>Deinococci>Deinococcales>Deinococcaceae | GCF_002953415.1 | Deinococcus_sp._NW-56 | C3K08_RS15840 | 30048..33470 | 2080419 | BREX-system-P-loop-protein-BrxC-[Deinococcus-sp.-NW-56] |
| WP_028324416.1 | Type1.BREX | CHTH-> N6-METHYLASE ->Schlafen+HATPase+wHTH-> REase +SF1-SF2-Helicase ->DOC+HTH-> BrxA-DNA-binding -> BrxB-BrxF-iSTAND -> DUF6079 *-> OLD-ABC-> REase -> N6-METHYLASE ->OLD-ABC->TOPRIM-> PglZ -> SIGMA+OB-Fold+AAA-ATPase+Lon-Protease-BrxL ->SbcC->PIN-> | 1145 | Bacteria>Thermodesulfobacteriota>Desulfobacteria>Desulfobacterales>Desulfatirhabdiaceae>Desulfatirhabdium | GCF_000429925.1 | Desulfatirhabdium_butyrativorans | G492_RS0109380 | 84085..87522 | 340467 | BREX-system-P-loop-protein-BrxC-[Desulfatirhabdium-butyrativorans] |
| MBF0232702.1 | Type1.BREX | <- N6-METHYLASE <- REase <-OLD-ABC <- DUF6079 *<- BrxB-BrxF-iSTAND <- BrxA-DNA-binding | 1220 | Bacteria>Thermodesulfobacteriota>Desulfobacteria>Desulfobacterales>Desulfobacteraceae>Desulfamplus | GCA_015231525.1 | Desulfamplus_sp. | HQK65_06645 | 4455..8117 | 2006184 | BREX-system-P-loop-protein-BrxC-[Desulfamplus-sp.] |
| WP_080799293.1 | Type1.BREX | <-HEPN<-PolB<- N6-METHYLASE -p<- N6-METHYLASE <-MNS+ REase <-HEPN<-ParA-Soj-PloopNTPase<-CR-ATPase<-ParA-Soj-PloopNTPase<- DUF6079 *<- BrxB-BrxF-iSTAND <- BrxA-DNA-binding <-tRNA\|\|Peptidase_M20->?->?->?->?->?->PAS+PAS+PAS+HISKIN-> | 1193 | Bacteria>Thermodesulfobacteriota>Desulfobacteria>Desulfobacterales>Desulfobacteraceae>Desulfamplus | GCF_900170035.1 | Desulfamplus_magnetovallimortis | MTBBW1_RS03710 | 949012..952593 | 1246637 | BREX-system-P-loop-protein-BrxC-[Desulfamplus-magnetovallimortis] |
| WP_163354568.1 | Type1.BREX | <-35exo\|\|GDResolvase+HIN-HTH-><-?<-HHH_5+SF1-SF2-Helicase\|\|XIS-HTH-> BrxA-DNA-binding -> BrxB-BrxF-iSTAND -> DUF6079 *-> OLD-ABC-> REase -> N6-METHYLASE -> PglZ -> SIGMA+OB-Fold+AAA-ATPase+HKD-DNase-BrxL -><-Zn_Tnp_IS91+Y2_Tnp | 1187 | Bacteria>Thermodesulfobacteriota>Desulfobacteria>Desulfobacterales>Desulfobacteraceae>Desulfobacter | - | Desulfobacter_hydrogenophilus | G3N28_17830 | 17749..21312 | 2291 | BREX-system-P-loop-protein-BrxC-[Desulfobacter-hydrogenophilus] |
| NDY73873.1 | Type1.BREX | <-35exo\|\|GDResolvase+HIN-HTH-><-?<-HHH_5+SF1-SF2-Helicase\|\|XIS-HTH-> BrxA-DNA-binding -> BrxB-BrxF-iSTAND -> DUF6079 *-> OLD-ABC-> REase -> N6-METHYLASE -> PglZ -> SIGMA+OB-Fold+AAA-ATPase+HKD-DNase-BrxL -><-Zn_Tnp_IS91+Y2_Tnp | 1187 | Bacteria>Thermodesulfobacteriota>Desulfobacteria>Desulfobacterales>Desulfobacteraceae>Desulfobacter | GCA_010499545.1 | Desulfobacter_hydrogenophilus | G3N28_17830 | 17749..21312 | 2291 | BREX-system-P-loop-protein-BrxC-[Desulfobacter-hydrogenophilus] |
| WP_035236115.1 | Type1.BREX | <- SIGMA+OB-Fold+AAA-ATPase+HKD-DNase-BrxL <- PglZ <-ParB+HNH<- N6-METHYLASE <- REase <-OLD-ABC <- DUF6079 *<- BrxB-BrxF-iSTAND <- BrxA-DNA-binding <-XIS-HTH\|\|?->?->EAL->S1+GTPase-AIG-> | 1189 | Bacteria>Thermodesulfobacteriota>Desulfobacteria>Desulfobacterales>Desulfobacteraceae>Desulfobacter | GCF_000745975.1 | Desulfobacter_vibrioformis | Q366_RS02915 | 253933..257502 | 34031 | BREX-system-P-loop-protein-BrxC-[Desulfobacter-vibrioformis] |
| WP_111959628.1 | Type1.BREX | Zn_Tnp_IS91+Y2_Tnp-><- SIGMA+OB-Fold+AAA-ATPase+HKD-DNase-BrxL <- PglZ <- N6-METHYLASE <- REase <-OLD-ABC <- DUF6079 *<- BrxB-BrxF-iSTAND <- BrxA-DNA-binding <-XIS-HTH\|\|HHH_5+SF1-SF2-Helicase->?-><-GDResolvase+HIN-HTH\|\|35exo-> | 1188 | Bacteria>Thermodesulfobacteriota>Desulfobacteria>Desulfobacterales>Desulfobacteraceae>Desulfobacter | GCF_004319545.1 | Desulfobacter_hydrogenophilus | DO021_RS19095 | 11943..15509 | 2291 | BREX-system-P-loop-protein-BrxC-[Desulfobacter-hydrogenophilus] |
| MBL6994747.1 | Type1.BREX | BrxA-DNA-binding -> BrxB-BrxF-iSTAND -> DUF6079 *-> N6-METHYLASE ->RVT-> N6-METHYLASE -p->?-> PglZ -> SIGMA+OB-Fold+AAA-ATPase+Lon-Protease-BrxL -> REase +nSTAND3->?->SbcC-> | 1186 | Bacteria>Thermodesulfobacteriota>Desulfobacteria>Desulfobacterales>Desulfobacteraceae>Desulfobacula | GCA_016785005.1 | Desulfobacula_sp. | ISR66_12690 | 4271..7831 | 2593537 | BREX-system-P-loop-protein-BrxC-[Desulfobacula-sp.] |
| MCD4719518.1 | Type6.BREX | BrxE -> BrxA-DNA-binding -> BrxB-BrxF-iSTAND -> DUF6079 *-> OLD-ABC-> REase -> N6-METHYLASE -> PglZ -> | 1208 | Bacteria>Thermodesulfobacteriota>Desulfobacteria>Desulfobacterales>Desulfobacteraceae>Desulfobacula | GCA_021107415.1 | Desulfobacula_sp. | K8S13_06615 | 2783..6409 | 2593537 | BREX-system-P-loop-protein-BrxC-[Desulfobacula-sp.] |
| MBL7173047.1 | Type1.BREX | <- wHTH+WYL+WCX <-AbiEi_4\|\|?-> BrxA-DNA-binding -> BrxB-BrxF-iSTAND -> DUF6079 *-> OLD-ABC-> REase -> N6-METHYLASE -p-> | 1172 | Bacteria>Thermodesulfobacteriota>Desulfobacteria>Desulfobacterales>Desulfobacteraceae>unclassified_Desulfobacteraceae | GCA_016782905.1 | Desulfobacteraceae_bacterium | ISS68_09440 | 5839..9357 | 2049433 | BREX-system-P-loop-protein-BrxC-[Desulfobacteraceae-bacterium] |
| CBX26795.1 | Type6.BREX | <-rve<- SIGMA+OB-Fold+AAA-ATPase+HKD-DNase-BrxL <- PglZ <-Schlafen+HATPase+wHTH<-DrHyd<- N6-METHYLASE <- REase <- REase <-OLD-ABC <- DUF6079 *<- BrxB-BrxF-iSTAND <- BrxA-DNA-binding <- BrxE \|\|URI-DNase-><-tRNA<-?\|\|?-><-PRTase<-DnaJ | 1208 | Bacteria>Thermodesulfobacteriota>Desulfobacteria>Desulfobacterales>Desulfobacteriaceae>Desulfobacterium | - | uncultured_Desulfobacterium_sp. | N47_A08240 | 98686..102312 | 201089 | hypothetical-protein-N47_A08240-[uncultured-Desulfobacterium-sp.] |
| MBC8421197.1 | Type1.BREX | <-AbiEii<-AbiEi_4\|\|?-> BrxA-DNA-binding -> BrxB-BrxF-iSTAND -> DUF6079 *-> OLD-ABC-> REase -> N6-METHYLASE -> | 1172 | Bacteria>Thermodesulfobacteriota>Desulfobacteria>Desulfobacterales>unclassified_Desulfobacterales | GCA_014382905.1 | Desulfobacterales_bacterium | H8E10_21680 | 4515..8033 | 2044940 | BREX-system-P-loop-protein-BrxC-[Desulfobacterales-bacterium] |
| KGO34873.1 | Type1.BREX | MlaE->ABC-ATPase+SbcC->?-><-?\|\|XIS-HTH-> BrxA-DNA-binding -> BrxB-BrxF-iSTAND -> DUF6079 *-> OLD-ABC-> REase -> N6-METHYLASE ->?->OLD-ABC->HNH-> PglZ -> SIGMA+OB-Fold+AAA-ATPase+Lon-Protease-BrxL -> | 1146 | Bacteria>Thermodesulfobacteriota>Desulfobulbia>Desulfobulbales>Desulfobulbaceae>Desulfobulbus | GCA_000769715.1 | Desulfobulbus_sp._Tol-SR | JT06_05895 | 26559..29999 | 1536652 | hypothetical-protein-JT06_05895-[Desulfobulbus-sp.-Tol-SR] |
| WP_205243275.1 | Type1.BREX | <- SIGMA+OB-Fold+AAA-ATPase+Lon-Protease-BrxL <- REase <- PglZ <-Schlafen+HATPase+wHTH<- N6-METHYLASE <-?<- DUF6079 *<- REase <- BrxB-BrxF-iSTAND <- BrxA-DNA-binding <-?\|\| wHTH+WYL+WCX -> | 1162 | Bacteria>Thermodesulfobacteriota>Desulfobulbia>Desulfobulbales>Desulfobulbaceae>Desulfobulbus | GCF_016918545.1 | Desulfobulbus_alkaliphilus | JWG43_RS12510 | 49790..53278 | 869814 | BREX-system-P-loop-protein-BrxC-[Desulfobulbus-alkaliphilus] |
| WP_205228200.1 | Type1.BREX | <-ParB+HNH<-SbcC<-Calcineurin+Calcineurin-SbcD<-SbcC+SbcC<- SIGMA+OB-Fold+AAA-ATPase+HKD-DNase-BrxL <- PglZ <- N6-METHYLASE <- REase <-ABC-ATPase<- DUF6079 *<- BrxB-BrxF-iSTAND <- BrxA-DNA-binding <-XIS-HTH<-?<-?<-?<-?<-wHTH | 1182 | Bacteria>Thermodesulfobacteriota>Desulfobulbia>Desulfobulbales>Desulfobulbaceae>Desulfobulbus | GCF_016919065.1 | Desulfobulbus_rhabdoformis | JWJ90_RS15890 | 30219..33767 | 34032 | BREX-system-P-loop-protein-BrxC-[Desulfobulbus-rhabdoformis] |
| WP_028584971.1 | Type1.BREX | <-ZnR<- SIGMA+OB-Fold+AAA-ATPase+HKD-DNase-BrxL <- PglZ <-PadR-HTH<-Schlafen+HATPase+wHTH<- N6-METHYLASE <- REase <-OLD-ABC <- DUF6079 *<- BrxB-BrxF-iSTAND <- BrxA-DNA-binding <-?<-VirB11<-?<-PCMT+METHYLASE | 1201 | Bacteria>Thermodesulfobacteriota>Desulfobulbia>Desulfobulbales>Desulfobulbaceae>Desulfogranum | GCF_000429965.1 | Desulfogranum_mediterraneum | G494_RS0113505 | 11738..15343 | 160661 | BREX-system-P-loop-protein-BrxC-[Desulfogranum-mediterraneum] |
| MBA3007526.1 | Type1.BREX | Phage_integrase->XIS-HTH-> BrxA-DNA-binding -> BrxB-BrxF-iSTAND -> DUF6079 *->Schlafen+HATPase+wHTH-> OLD-ABC-> REase -> N6-METHYLASE ->Schlafen+HATPase+wHTH-> PglZ -> SIGMA+OB-Fold+AAA-ATPase+Lon-Protease-BrxL -><-?\|\|35exo+SF1-SF2-Helicase-> | 1147 | Bacteria>Thermodesulfobacteriota>Desulfobulbia>Desulfobulbales>Desulfocapsaceae>Desulfocapsa | GCA_013792135.1 | Desulfocapsa_sp. | FP810_13320 | 53855..57298 | 2066021 | BREX-system-P-loop-protein-BrxC-[Desulfocapsa-sp.] |
| TKB07808.1 | Type1.BREX | <-HEPN<-SbcC<-Calcineurin+Calcineurin-SbcD<-SbcC+SbcC<- SIGMA+OB-Fold+AAA-ATPase+HKD-DNase-BrxL <- PglZ <- N6-METHYLASE <- REase <-OLD-ABC <- DUF6079 *<- BrxB-BrxF-iSTAND <- BrxA-DNA-binding <-XIS-HTH<-?<-?<-?<-?<-wHTH | 1183 | Bacteria>Thermodesulfobacteriota>Desulfobulbia>Desulfobulbales>Desulfocapsaceae>Desulforhopalus | GCA_005116575.1 | Desulforhopalus_sp._IMCC35007 | FCL48_15775 | 80281..83832 | 2569543 | BREX-system-P-loop-protein-BrxC-[Desulforhopalus-sp.-IMCC35007] |
| WP_169309248.1 | Type1.BREX | <-HEPN<-SbcC<-Calcineurin+Calcineurin-SbcD<-SbcC+SbcC<- SIGMA+OB-Fold+AAA-ATPase+HKD-DNase-BrxL <- PglZ <- N6-METHYLASE <- REase <-OLD-ABC <- DUF6079 *<- BrxB-BrxF-iSTAND <- BrxA-DNA-binding <-XIS-HTH<-?<-?<-?<-?<-wHTH | 1182 | Bacteria>Thermodesulfobacteriota>Desulfobulbia>Desulfobulbales>Desulfocapsaceae>Desulforhopalus | GCF_005116575.1 | Desulforhopalus_sp._IMCC35007 | FCL48_RS15780 | 80281..83829 | 2569543 | BREX-system-P-loop-protein-BrxC-[Desulforhopalus-sp.-IMCC35007] |
| WP_027370868.1 | Type1.BREX | <-PIN<-MazE_antitoxin\|\|?->RHH->PIN->?->GGDEF+EAL->tRNA-> BrxA-DNA-binding -> BrxB-BrxF-iSTAND -> DUF6079 *-> N6-METHYLASE ->?->Schlafen+HATPase+wHTH-> PglZ ->?->HicB_lk_antitox-> REase -> SIGMA+OB-Fold+AAA-ATPase+Lon-Protease-BrxL ->Pkinase+Pkinase-> | 1190 | Bacteria>Thermodesulfobacteriota>Desulfovibrionia>Desulfovibrionales>Desulfohalobiaceae>Desulfovermiculus | GCF_000620765.1 | Desulfovermiculus_halophilus | N902_RS0110295 | 13006..16578 | 339722 | BREX-system-P-loop-protein-BrxC-[Desulfovermiculus-halophilus] |
| WP_035067990.1 | Type1.BREX | <-35exo<-?<-SbcC+SbcC<- SIGMA+OB-Fold+AAA-ATPase+HKD-DNase-BrxL <- PglZ <- N6-METHYLASE <- REase <-OLD-ABC <- DUF6079 *<- BrxB-BrxF-iSTAND <- BrxA-DNA-binding <- wHTH+WYL+WCX \|\|RHH_1->RelE-ParE-><-?\|\|Phd-YefM->PIN-> | 1182 | Bacteria>Thermodesulfobacteriota>Desulfovibrionia>Desulfovibrionales>Desulfovibrionaceae>Cupidesulfovibrio | GCF_000504305.1 | Cupidesulfovibrio_termitidis | DESTE_RS12450 | 3095808..3099356 | 42252 | BREX-system-P-loop-protein-BrxC-[Cupidesulfovibrio-termitidis] |
| AAS96500.1 | Type1.BREX | <-GGDEF<-ZnR+HTH_Tnp_ISL3+DDE_Tnp_ISL3<- SIGMA+OB-Fold+AAA-ATPase+HKD-DNase-BrxL <- PglZ <- REase <- N6-METHYLASE <- REase <-OLD-ABC <- DUF6079 *<- BrxB-BrxF-iSTAND \|\|?-><- wHTH+WYL+WCX -p\|\|HIN-HTH->HIN-HTH-><-?\|\|?->35exo-> | 1183 | Bacteria>Thermodesulfobacteriota>Desulfovibrionia>Desulfovibrionales>Desulfovibrionaceae>Desulfovibrio | GCA_000195755.1 | Desulfovibrio_vulgaris_str._Hildenborough | DVU_2025 | 2106300..2109851 | 882 | conserved-hypothetical-protein-[Desulfovibrio-vulgaris-str.-Hildenborough] |
| WP_014524353.1 | Type1.BREX | <-GGDEF<-ZnR+HTH_Tnp_ISL3+DDE_Tnp_ISL3<-SbcC<- SIGMA+OB-Fold+AAA-ATPase+HKD-DNase-BrxL <- PglZ <- REase <- N6-METHYLASE <- REase <-OLD-ABC <- DUF6079 *<- BrxB-BrxF-iSTAND <- BrxA-DNA-binding <- wHTH+WYL+WCX -p\|\|HIN-HTH-><- wHTH+WYL+WCX \|\|?->35exo->?-><-RelE-ParE | 1182 | Bacteria>Thermodesulfobacteriota>Desulfovibrionia>Desulfovibrionales>Desulfovibrionaceae>Desulfovibrio | GCF_000195755.1 | Desulfovibrio_vulgaris | DVU_RS09565 | 2106300..2109848 | 881 | BREX-system-P-loop-protein-BrxC-[Desulfovibrio-vulgaris] |
| WP_043638458.1 | Type1.BREX | NTP_transferase-><-?\|\|GAF+GAF+HISKIN-><-?<-?\|\|ParB+N6_N4_METHYLASE->?-> BrxA-DNA-binding -> BrxB-BrxF-iSTAND -> DUF6079 *-> OLD-ABC-> REase -> N6-METHYLASE -> PglZ -> SIGMA+OB-Fold+AAA-ATPase+HKD-DNase-BrxL -> | 1182 | Bacteria>Thermodesulfobacteriota>Desulfovibrionia>Desulfovibrionales>Desulfovibrionaceae>Desulfovibrio | GCF_000801335.2 | Desulfovibrio_sp._TomC | NY78_RS16925 | 95476..99024 | 1562888 | BREX-system-P-loop-protein-BrxC-[Desulfovibrio-sp.-TomC] |
| MBE0574417.1 | Type1.BREX | <-VirD4-TraG+VirD4-FtsK<-?<-?\|\|XIS-HTH-> BrxA-DNA-binding -> BrxB-BrxF-iSTAND -> DUF6079 *-> OLD-ABC-> REase -> N6-METHYLASE -> REase -> PglZ -> SIGMA+OB-Fold+AAA-ATPase+HKD-DNase-BrxL ->SbcC+APATPase+SbcC->Calcineurin+Calcineurin-SbcD->SbcC-> | 1183 | Bacteria>Thermodesulfobacteriota>Desulfuromonadia>Desulfuromonadales>unclassified_Desulfuromonadales | GCA_014860225.1 | Desulfuromonadales_bacterium | IH613_00815 | 71489..75040 | 2099678 | BREX-system-P-loop-protein-BrxC-[Desulfuromonadales-bacterium] |
| MBW6509585.1 | Type1.BREX | Amidohydro_1->?->?-><-Four-helical-protein\|\|XIS-HTH-> BrxA-DNA-binding -> BrxB-BrxF-iSTAND -> DUF6079 *->Schlafen+HATPase+wHTH+HTH-> OLD-ABC-> REase -> N6-METHYLASE -> REase -> | 1178 | Bacteria>Thermodesulfobacteriota>Desulfuromonadia>Desulfuromonadales>unclassified_Desulfuromonadales | GCA_019429255.1 | Desulfuromonadales_bacterium | K0A94_08605 | 43514..47050 | 2099678 | BREX-system-P-loop-protein-BrxC-[Desulfuromonadales-bacterium] |
| WP_207162791.1 | Type1.BREX | <-Peptidase_C14<-TIR<- SIGMA+OB-Fold+AAA-ATPase+HKD-DNase-BrxL <- PglZ <-TOPRIM<-OLD-ABC<- N6-METHYLASE <- REase <-OLD-ABC <- DUF6079 *<- BrxB-BrxF-iSTAND <- BrxA-DNA-binding <-AbiEi_4+AbiEi_1<-ParB<-ParB<-GDResolvase+PPP1-HTH\|\|?-><-Arm-DNA-bind_3+Phage_integrase | 1175 | Bacteria>Thermodesulfobacteriota>Desulfuromonadia>Geobacterales>Geobacteraceae>Geobacter | GCF_017338855.1 | Geobacter_benzoatilyticus | JZM60_RS12610 | 2741421..2744948 | 2815309 | BREX-system-P-loop-protein-BrxC-[Geobacter-benzoatilyticus] |
| MBJ6802709.1 | Type1.BREX | BrxA-DNA-binding -> BrxB-BrxF-iSTAND -> DUF6079 *-> OLD-ABC-> REase -> N6-METHYLASE -> PglZ -> SIGMA+OB-Fold+AAA-ATPase+HKD-DNase-BrxL ->?-><-?<-NERD+ZnR<-wHTH+SAD+HNH | 1183 | Bacteria>Thermodesulfobacteriota>Desulfuromonadia>Geobacterales>Geobacteraceae>Geomonas | GCA_016458235.1 | Geomonas_propionica | JFN90_21475 | 2703..6254 | 2798582 | BREX-system-P-loop-protein-BrxC-[Geomonas-propionica] |
| WP_236019113.1 | Type1.BREX | BrxA-DNA-binding -> BrxB-BrxF-iSTAND -> DUF6079 *-> OLD-ABC-> REase -> N6-METHYLASE -> PglZ -> SIGMA+OB-Fold+AAA-ATPase+HKD-DNase-BrxL ->?-><-?<-NERD+ZnR<-wHTH+SAD+HNH | 1191 | Bacteria>Thermodesulfobacteriota>Desulfuromonadia>Geobacterales>Geobacteraceae>Geomonas | GCF_016458235.1 | Geomonas_propionica | JFN90_RS21475 | 2679..6254 | 2798582 | BREX-system-P-loop-protein-BrxC-[Geomonas-propionica] |
| OQA19974.1 | Type1.BREX | BrxA-DNA-binding -> BrxB-BrxF-iSTAND -> DUF6079 *->?->RelE-ParE->VapB_antitoxin->PIN-> N6-METHYLASE -> REase -> PglZ -> SIGMA+OB-Fold+AAA-ATPase+Lon-Protease-BrxL -> | 1164 | Bacteria>unclassified_Bacteria | GCA_002069435.1 | bacterium_ADurb.Bin363 | BWY64_00492 | 2230..5724 | 1866927 | hypothetical-protein-BWY64_00492-[bacterium-ADurb.Bin363] |
| TRZ93676.1 | Type1.BREX | <- REase <-HerA/FtsK<-SIR2<- REase <- SIGMA+OB-Fold+AAA-ATPase+Lon-Protease-BrxL <- PglZ <- N6-METHYLASE <- DUF6079 *<- BrxB-BrxF-iSTAND <- BrxA-DNA-binding <-XIS-HTH<-?<- REase <-?<- REase +35exo | 1181 | Bacteria>unclassified_Bacteria | GCA_007280235.1 | bacterium | D4R78_07780 | 14482..18027 | 1869227 | BREX-system-P-loop-protein-BrxC-[bacterium] |
| MBS2037451.1 | Type6.BREX | BrxE -> BrxA-DNA-binding -> BrxB-BrxF-iSTAND -> DUF6079 *-><- REase \|\|<- REase <- REase \|\|?-><-?<-?<-?\|\|MerR-HTH-><-MODE-HTH+PBPII | 1202 | Bacteria>unclassified_Bacteria | GCA_018266105.1 | bacterium | JST97_20850 | 5122..8730 | 1869227 | BREX-system-P-loop-protein-BrxC-[bacterium] |
| MCE5247410.1 | Type1.BREX | <- REase <-?<- SIGMA+OB-Fold+AAA-ATPase+Lon-Protease-BrxL <- PglZ <- N6-METHYLASE \|\|?-><- DUF6079 *<- BrxB-BrxF-iSTAND <- BrxA-DNA-binding | 1148 | Bacteria>unclassified_Bacteria | GCA_021372775.1 | bacterium | LLG88_10905 | 14182..17628 | 1869227 | BREX-system-P-loop-protein-BrxC-[bacterium] |
| MCP4549289.1 | Type6.BREX | <-XIS-HTH<-?\|\| BrxE -> BrxA-DNA-binding -> BrxB-BrxF-iSTAND -> N6-METHYLASE ->?-> REase -> DUF6079 *->HEPN-> N6-METHYLASE -><-?\|\|?-> N6-METHYLASE ->AbiEi_4+AbiEi_1->AbiEii+ZnR-> | 1242 | Bacteria>unclassified_Bacteria | GCA_024231955.1 | bacterium | GY835_22785 | 11465..15193 | 1869227 | BREX-system-P-loop-protein-BrxC-[bacterium] |

| Protein-ID | System | Gene-Neighbourhoods | Length | Taxonomy | Assembly-ID | Strain | Locus-Tag | Genomic-coordinates | TaxID | Fasta-define |
|---|---|---|---|---|---|---|---|---|---|---|
| MCC8119638.1 | Type1.BREX | <-PglZ<-Schlafen+HATPase+wHTH<-?<-ParB+HNH<-AAA-ATPase-p<-Schlafen+HATPase+wHTH<-?<-N6-METHYLASE<-HTH+REase<-DUF6079*<-BrxB-BrxF-iSTAND<-BrxA-DNA-binding<-tRNA<-35exo+T7DNApol | 1189 | Bacteria>FCB_group>Bacteroidetes/Chlorobi_group>Bacteroidota>Bacteroidia>Bacteroidales | GCA_020860515.1 | Bacteroidales_bacterium | LIP09_12950 | 29133..32702 | 2030927 | BREX-system-P-loop-protein-BrxC-[Bacteroidales-bacterium] |
| MCF0219894.1 | Type1.BREX | CR-Kinase+Classical-AAA->?-><-TPR<-SIGMA+OB-Fold+AAA-ATPase+Lon-Protease-BrxL<-PglZ<-?<-Schlafen+HATPase+wHTH<-N6-METHYLASE<-DUF6079*<-BrxB-BrxF-iSTAND<-BrxA-DNA-binding<-?<-XIS-HTH<-?<-Phage_int_SAM_5+Phage_integrase\|\|LRR_5+LRR_5+LRR_5-> | 1215 | Bacteria>FCB_group>Bacteroidetes/Chlorobi_group>Bacteroidota>Bacteroidia>Bacteroidales | GCA_021443165.1 | Muribaculaceae_bacterium | HUK14_08940 | 85831..89478 | 2498093 | BREX-system-P-loop-protein-BrxC-[Muribaculaceae-bacterium] |
| MCF2581772.1 | Type1.BREX | <-SIGMA+OB-Fold+AAA-ATPase+Lon-Protease-BrxL<-PglZ<-Schlafen+HATPase+wHTH<-N6-METHYLASE<-SLOG+BirA-HTH<-SF1-SF2-Helicase+PRTase<-HNH<-OLD-ABC<-DUF6079*<-BrxB-BrxF-iSTAND<-BrxA-DNA-binding | 1182 | Bacteria>FCB_group>Bacteroidetes/Chlorobi_group>Bacteroidota>Bacteroidia>Bacteroidales | GCA_021530995.1 | Bacteroides_caecigallinarum | I6E24_07160 | 15567..19115 | 1411144 | BREX-system-P-loop-protein-BrxC-[Bacteroides-caecigallinarum] |
| MCH4018978.1 | Type1.BREX | XIS-HTH->?->AAA+-><-?\|\|wHTH+WYL+WCX->HTH_3->BrxA-DNA-binding->BrxB-BrxF-iSTAND->DUF6079*->N6-METHYLASE->Schlafen+HATPase+wHTH->SIR2->PglZ->SIGMA+OB-Fold+AAA-ATPase+HKD-DNase-BrxL->SIR2+SIR2->ZnR->DDE_Tnp_ISL3->Tox-RelE-> | 1198 | Bacteria>FCB_group>Bacteroidetes/Chlorobi_group>Bacteroidota>Bacteroidia>Bacteroidales | GCA_022483425.1 | Prevotella_sp. | LKE74_11130 | 2793807..2797403 | 59823 | BREX-system-P-loop-protein-BrxC-[Prevotella-sp.] |
| MCI1742488.1 | Type1.BREX | <-HTH_31\|\|tRNA-><-?<-SIR2+SIR2<-SIGMA+OB-Fold+AAA-ATPase+HKD-DNase-BrxL<-PglZ<-HTH+REase<-Schlafen+HATPase+wHTH<-N6-METHYLASE<-DUF6079*<-BrxB-BrxF-iSTAND<-BrxA-DNA-binding<-HTH_3<-wHTH+WYL+WCX\|\|?-><-AAA+<-?<-XIS-HTH | 1206 | Bacteria>FCB_group>Bacteroidetes/Chlorobi_group>Bacteroidota>Bacteroidia>Bacteroidales | GCA_022647785.1 | Prevotella_sp. | LKI18_09155 | 39395..43015 | 59823 | BREX-system-P-loop-protein-BrxC-[Prevotella-sp.] |
| MCI2087046.1 | Type1.BREX | <-SIGMA+OB-Fold+AAA-ATPase+HKD-DNase-BrxL<-PglZ<-HTH+REase<-Schlafen+HATPase+wHTH<-N6-METHYLASE<-DUF6079*<-BrxB-BrxF-iSTAND<-BrxA-DNA-binding<-HTH_3<-wHTH+WYL+WCX\|\|?-><-AAA+<-?<-XIS-HTH | 1198 | Bacteria>FCB_group>Bacteroidetes/Chlorobi_group>Bacteroidota>Bacteroidia>Bacteroidales | GCA_022653375.1 | Prevotella_sp. | LKK10_02020 | 11419..15015 | 59823 | BREX-system-P-loop-protein-BrxC-[Prevotella-sp.] |
| MCI7017346.1 | Type1.BREX | <-SIGMA+OB-Fold+AAA-ATPase+HKD-DNase-BrxL<-PglZ<-?<-Schlafen+HATPase+wHTH<-N6-METHYLASE<-AT-HTH-HYD2<-Tox-ART-HYD2<-AT-HTH-HYD2<-DUF6079*<-BrxB-BrxF-iSTAND<-BrxA-DNA-binding<-wHTH+WYL+WCX | 1189 | Bacteria>FCB_group>Bacteroidetes/Chlorobi_group>Bacteroidota>Bacteroidia>Bacteroidales | GCA_022775925.1 | Prevotella_sp. | MR923_09655 | 21723..25292 | 59823 | BREX-system-P-loop-protein-BrxC-[Prevotella-sp.] |
| MCI6403340.1 | Type1.BREX | AAA-ATPase-p-><-?\|\|BrxA-DNA-binding->BrxB-BrxF-iSTAND->DUF6079*->Schlafen+HATPase+wHTH->N6-METHYLASE-> | 1217 | Bacteria>FCB_group>Bacteroidetes/Chlorobi_group>Bacteroidota>Bacteroidia>Bacteroidales | GCA_022777225.1 | Prevotella_sp. | MR820_04565 | 20272..23925 | 59823 | BREX-system-P-loop-protein-BrxC-[Prevotella-sp.] |
| MCQ2071598.1 | Type1.BREX | <-TOPRIM<-?<-?\|\|Peptidase_M23->BrxA-DNA-binding->BrxB-BrxF-iSTAND->DUF6079*->N6-METHYLASE->Schlafen+HATPase+wHTH->PglZ->SIGMA+OB-Fold+AAA-ATPase+HKD-DNase-BrxL->XIS-HTH->wHTH+WYL+WCX->?->HHH_5+SF1-SF2-Helicase-> | 1190 | Bacteria>FCB_group>Bacteroidetes/Chlorobi_group>Bacteroidota>Bacteroidia>Bacteroidales | GCA_024398655.1 | Bacteroidaceae_bacterium | MJY96_00535 | 107887..111459 | 2212467 | BREX-system-P-loop-protein-BrxC-[Bacteroidaceae-bacterium] |
| SCH71957.1 | Type1.BREX | <-SIGMA+OB-Fold+AAA-ATPase+Lon-Protease-BrxL<-PglZ<-Schlafen+HATPase+wHTH<-N6-METHYLASE<-SLOG+BirA-HTH<-SF1-SF2-Helicase+PRTase<-HNH<-OLD-ABC<-DUF6079*<-BrxB-BrxF-iSTAND<-BrxA-DNA-binding | 1179 | Bacteria>FCB_group>Bacteroidetes/Chlorobi_group>Bacteroidota>Bacteroidia>Bacteroidales | GCA_900066445.1 | uncultured_Bacteroides_sp. | SAMEA3545309_01215 | 56504..60043 | 162156 | Uncharacterised-protein-[uncultured-Bacteroides-sp.] |
| SHF69941.1 | Type1.BREX | <-RADICAL-SAM<-?\|\|tRNA-><-SIGMA+OB-Fold+AAA-ATPase+Lon-Protease-BrxL<-PglZ<-Schlafen+HATPase+wHTH<-ParB+HNH<-N6-METHYLASE<-?<-DUF6079*<-BrxB-BrxF-iSTAND<-BrxA-DNA-binding | 1182 | Bacteria>FCB_group>Bacteroidetes/Chlorobi_group>Bacteroidota>Bacteroidia>Bacteroidales | GCA_900128905.1 | Bacteroides_luti | SAMN05444405_11263 | 81108..84656 | 1297750 | hypothetical-protein-SAMN05444405_11263-[Bacteroides-luti] |
| WP_002561973.1 | Type1.BREX | 35exo+SF1-SF2-Helicase-><-DDE_Tnp\|\|<-SIGMA+OB-Fold+AAA-ATPase+HKD-DNase-BrxL<-PglZ<-Schlafen+HATPase+wHTH<-SF1-SF2-Helicase<-OLD-ABC+TOPRIM<-N6-METHYLASE<-DUF6079*<-BrxB-BrxF-iSTAND<-BrxA-DNA-binding | 1202 | Bacteria>FCB_group>Bacteroidetes/Chlorobi_group>Bacteroidota>Bacteroidia>Bacteroidales | GCF_000382465.1 | Bacteroides_sp._HPS0048 | HMPREF1214_RS19625 | 483033..486641 | 1078089 | BREX-system-P-loop-protein-BrxC-[Bacteroides-sp.-HPS0048] |
| WP_024993540.1 | Type1.BREX | <-wHTH+WYL+WCX<-wHTH+WYL+WCX<-SIGMA+OB-Fold+AAA-ATPase+HKD-DNase-BrxL<-PglZ<-Schlafen+HATPase+wHTH<-N6-METHYLASE<-DUF6079*<-BrxB-BrxF-iSTAND<-BrxA-DNA-binding<-HTH_3 | 1210 | Bacteria>FCB_group>Bacteroidetes/Chlorobi_group>Bacteroidota>Bacteroidia>Bacteroidales | GCF_000613805.1 | Phocaeicola_paurosaccharolyticus | JCM15092_RS04015 | 53285..56917 | 732242 | BREX-system-P-loop-protein-BrxC-[Phocaeicola-paurosaccharolyticus] |
| WP_122342696.1 | Type1.BREX | Arm-DNA-bind_5+Phage_int_SAM_5+Phage_integrase->?->AbrB-HTH-><-HIN-HTH+rve+rve_3<-KAP-NTPase<-SIGMA+OB-Fold+AAA-ATPase+Lon-Protease-BrxL<-PglZ<-Schlafen+HATPase+wHTH<-N6-METHYLASE<-DUF6079*<-BrxB-BrxF-iSTAND<-BrxA-DNA-binding<-HTH_3\|\|?->XIS-HTH->XIS-HTH->Arm-DNA-bind_5+Phage_int_SAM_5+Phage_integrase->XIS-HTH->XIS-HTH-> | 1211 | Bacteria>FCB_group>Bacteroidetes/Chlorobi_group>Bacteroidota>Bacteroidia>Bacteroidales | GCF_003475265.1 | Parabacteroides_merdae | DWZ81_RS06115 | 100594..104229 | 46503 | BREX-system-P-loop-protein-BrxC-[Parabacteroides-merdae] |
| WP_135993587.1 | Type1.BREX | <-SF1-SF2-Helicase<-SIGMA+OB-Fold+AAA-ATPase+HKD-DNase-BrxL<-PglZ<-Schlafen+HATPase+wHTH<-SIR2<-N6-METHYLASE<-DUF6079*<-BrxB-BrxF-iSTAND<-BrxA-DNA-binding<-HTH_3<-?\|\|Phage_int_SAM_1+Phage_integrase->Phage_integrase->Phage_int_SAM_1+Phage_integrase-><-Peptidase_M23 | 1221 | Bacteria>FCB_group>Bacteroidetes/Chlorobi_group>Bacteroidota>Bacteroidia>Bacteroidales | GCF_004793655.1 | Muribaculum_intestinale | E5330_RS10970 | 18051..21716 | 1796646 | BREX-system-P-loop-protein-BrxC-[Muribaculum-intestinale] |
| WP_172774764.1 | Type1.BREX | OLD-ABC->TOPRIM->XIS-HTH-><-?<-?<-?\|\|HTH_3->BrxA-DNA-binding->BrxB-BrxF-iSTAND->DUF6079*->N6-METHYLASE->OLD-ABC+TOPRIM->SF1-SF2-Helicase->Schlafen+HATPase+wHTH->PglZ->SIGMA+OB-Fold+AAA-ATPase+HKD-DNase-BrxL->NADAR->RADICAL-SAM->HIRAN-> | 1217 | Bacteria>FCB_group>Bacteroidetes/Chlorobi_group>Bacteroidota>Bacteroidia>Bacteroidales | GCF_012976275.1 | Phocaeicola_vulgatus | HKQ53_RS16515 | 79396..83049 | 821 | BREX-system-P-loop-protein-BrxC-[Phocaeicola-vulgatus] |
| WP_181213825.1 | Type1.BREX | TOPRIM->?->TOPRIM->?-><-KAP-NTPase<-SIGMA+OB-Fold+AAA-ATPase+Lon-Protease-BrxL<-PglZ<-Schlafen+HATPase+wHTH<-N6-METHYLASE<-DUF6079*<-BrxB-BrxF-iSTAND<-BrxA-DNA-binding<-HTH_3\|\|?->?-><-XIS-HTH\|\|?->?->wHTH+WYL+WCX-> | 1212 | Bacteria>FCB_group>Bacteroidetes/Chlorobi_group>Bacteroidota>Bacteroidia>Bacteroidales | GCF_013618865.1 | Phocaeicola_faecicola | H3001_RS00990 | 112732..116370 | 2739389 | BREX-system-P-loop-protein-BrxC-[Phocaeicola-faecicola] |
| WP_115498725.1 | Type1.BREX | <-SIGMA+OB-Fold+AAA-ATPase+HKD-DNase-BrxL<-PglZ<-Schlafen+HATPase+wHTH<-N6-METHYLASE<-DUF6079*<-BrxB-BrxF-iSTAND<-BrxA-DNA-binding\|\|?->?->?->?-><-tRNA\|\|Cysteine-Desulfurase->?-><-ACET | 1211 | Bacteria>FCB_group>Bacteroidetes/Chlorobi_group>Bacteroidota>Bacteroidia>Bacteroidales | GCF_014385205.1 | Parabacteroides_acidifaciens | H8784_RS05825 | 88295..91930 | 2290935 | BREX-system-P-loop-protein-BrxC-[Parabacteroides-acidifaciens] |
| WP_004319896.1 | Type1.BREX | BrxA-DNA-binding->BrxB-BrxF-iSTAND->DUF6079*->N6-METHYLASE->SIR2+SIR2->Schlafen+HATPase+wHTH->PglZ->SIGMA+OB-Fold+AAA-ATPase+HKD-DNase-BrxL-><-35exo+SF1-SF2-Helicase\|\|BACON+Peptidase_M64-> | 1202 | Bacteria>FCB_group>Bacteroidetes/Chlorobi_group>Bacteroidota>Bacteroidia>Bacteroidales | GCF_015548155.1 | Bacteroides | P0193_RS10880 | 44218..47826 | 816 | MULTISPECIES:-BREX-system-P-loop-protein-BrxC-[Bacteroides] |
| WP_118403962.1 | Type1.BREX | <-Phage_int_SAM_5+Phage_integrase\|\|STYKinase-like->?-><-HEPN\|\|HTH_3->BrxA-DNA-binding->BrxB-BrxF-iSTAND->DUF6079*->N6-METHYLASE->Schlafen+HATPase+wHTH->PglZ->SIGMA+OB-Fold+AAA-ATPase+HKD-DNase-BrxL->?->URI-DNase+RAMA-> | 1211 | Bacteria>FCB_group>Bacteroidetes/Chlorobi_group>Bacteroidota>Bacteroidia>Bacteroidales | GCF_015553065.1 | Bacteroides_cellulosilyticus | I2F01_RS18130 | 73980..77615 | 246787 | BREX-system-P-loop-protein-BrxC-[Bacteroides-cellulosilyticus] |
| WP_195571580.1 | Type1.BREX | BrxA-DNA-binding->BrxB-BrxF-iSTAND->DUF6079*->N6-METHYLASE->SIR2+SIR2->Schlafen+HATPase+wHTH->PglZ->SIGMA+OB-Fold+AAA-ATPase+HKD-DNase-BrxL-><-35exo+SF1-SF2-Helicase\|\|?->SIGMA-HTH-> | 1202 | Bacteria>FCB_group>Bacteroidetes/Chlorobi_group>Bacteroidota>Bacteroidia>Bacteroidales | GCF_015557365.1 | Bacteroides_nordii | I2G99_RS18035 | 12742..16350 | 291645 | BREX-system-P-loop-protein-BrxC-[Bacteroides-nordii] |
| WP_203049164.1 | Type6.BREX | <-TOPRIM<-NUDIX<-?<-DUF4369+Thioredoxin_8\|\|RNasePH+RNasePH+KH+S1->HTH_3->BrxE->BrxA-DNA-binding->BrxB-BrxF-iSTAND->DUF6079*->N6-METHYLASE->Schlafen+HATPase+wHTH->PglZ->SIGMA+OB-Fold+AAA-ATPase+HKD-DNase-BrxL->SLATT_4-> | 1216 | Bacteria>FCB_group>Bacteroidetes/Chlorobi_group>Bacteroidota>Bacteroidia>Bacteroidales | GCF_016803255.1 | Prevotella_copri | F8Q89_RS01960 | 71962..75612 | 165179 | BREX-system-P-loop-protein-BrxC-[Prevotella-copri] |
| WP_204430319.1 | Type1.BREX | <-NADAR<-SIGMA+OB-Fold+AAA-ATPase+HKD-DNase-BrxL<-PglZ<-Schlafen+HATPase+wHTH<-SIR2<-N6-METHYLASE<-DUF6079*<-BrxB-BrxF-iSTAND<-BrxA-DNA-binding<-HTH_3\|\|?->?->?->?-><-XIS-HTH<-TOPRIM | 1217 | Bacteria>FCB_group>Bacteroidetes/Chlorobi_group>Bacteroidota>Bacteroidia>Bacteroidales | GCF_016899805.1 | Bacteroides_gallinaceum | H6B13_RS12495 | 13542..17195 | 1462571 | BREX-system-P-loop-protein-BrxC-[Bacteroides-gallinaceum] |
| WP_205111283.1 | Type1.BREX | OLD-ABC->XIS-HTH-><-?<-?<-?\|\|HTH_3->BrxA-DNA-binding->BrxB-BrxF-iSTAND->DUF6079*->N6-METHYLASE->AT-HTH-HYD2->Tox-ART-HYD2->AT-HTH-HYD2->?->N6-METHYLASE-p->SIR2+SIR2->Schlafen+HATPase+wHTH->PglZ-> | 1217 | Bacteria>FCB_group>Bacteroidetes/Chlorobi_group>Bacteroidota>Bacteroidia>Bacteroidales | GCF_016899955.1 | Marseilla_massiliensis | H6B30_RS12995 | 19775..23428 | 1841864 | BREX-system-P-loop-protein-BrxC-[Marseilla-massiliensis] |
| WP_204506195.1 | Type1.BREX | HTH_3->BrxA-DNA-binding->BrxB-BrxF-iSTAND->DUF6079*->N6-METHYLASE->Schlafen+HATPase+wHTH->PglZ->SIGMA+OB-Fold+AAA-ATPase+HKD-DNase-BrxL->?->?->?->?->REase-> | 1187 | Bacteria>FCB_group>Bacteroidetes/Chlorobi_group>Bacteroidota>Bacteroidia>Bacteroidales | GCF_016902095.1 | Phocaeicola_plebeius | H6A37_RS09755 | 27157..30720 | 310297 | BREX-system-P-loop-protein-BrxC-[Phocaeicola-plebeius] |
| WP_204401091.1 | Type1.BREX | Calcineurin+Calcineurin-conf->?->?->?->?->?->BrxA-DNA-binding->BrxB-BrxF-iSTAND->DUF6079*->OLD-ABC->HNH->SF1-SF2-Helicase+PRTase->SLOG+BirA-HTH->N6-METHYLASE->Schlafen+HATPase+wHTH->PglZ->SIGMA+OB-Fold+AAA-ATPase+Lon-Protease-BrxL->SbcC-> | 1179 | Bacteria>FCB_group>Bacteroidetes/Chlorobi_group>Bacteroidota>Bacteroidia>Bacteroidales | GCF_016902515.1 | Bacteroides_caecigallinarum | H6A30_RS00985 | 221918..225457 | 1411144 | BREX-system-P-loop-protein-BrxC-[Bacteroides-caecigallinarum] |
| WP_217718390.1 | Type1.BREX | TOPRIM->XIS-HTH-><-?<-?<-?<-?\|\|HTH_3->BrxA-DNA-binding->BrxB-BrxF-iSTAND->DUF6079*->N6-METHYLASE->OLD-ABC+TOPRIM->SF1-SF2-Helicase->Schlafen+HATPase+wHTH->PglZ->SIGMA+OB-Fold+AAA-ATPase+HKD-DNase-BrxL->NADAR->RADICAL-SAM->HIRAN-> | 1217 | Bacteria>FCB_group>Bacteroidetes/Chlorobi_group>Bacteroidota>Bacteroidia>Bacteroidales | GCF_019127825.1 | Bacteroides_uniformis | KSX11_RS12350 | 118043..121696 | 820 | BREX-system-P-loop-protein-BrxC-[Bacteroides-uniformis] |
| WP_217725669.1 | Type1.BREX | <-BACON+Peptidase_M64\|\|35exo+SF1-SF2-Helicase-><-SIGMA+OB-Fold+AAA-ATPase+HKD-DNase-BrxL<-PglZ<-Schlafen+HATPase+wHTH<-SIR2+SIR2<-N6-METHYLASE<-DUF6079*<-BrxB-BrxF-iSTAND<-BrxA-DNA-binding | 1202 | Bacteria>FCB_group>Bacteroidetes/Chlorobi_group>Bacteroidota>Bacteroidia>Bacteroidales | GCF_019128915.1 | unclassified_Bacteroides | KSX87_RS10005 | 229853..233461 | 2646097 | MULTISPECIES:-BREX-system-P-loop-protein-BrxC-[unclassified-Bacteroides] |
| WP_115473019.1 | Type1.BREX | <-OLD-ABC\|\|?-><-Phage_int_SAM_5+Phage_integrase\|\|?->?-><-HEPN\|\|HTH_3->BrxA-DNA-binding->BrxB-BrxF-iSTAND->DUF6079*->N6-METHYLASE->Schlafen+HATPase+wHTH->PglZ->SIGMA+OB-Fold+AAA-ATPase+HKD-DNase-BrxL->?-><-MULE-TRANSPOSASE\|\|DOC+HTH-> | 1211 | Bacteria>FCB_group>Bacteroidetes/Chlorobi_group>Bacteroidota>Bacteroidia>Bacteroidales | GCF_019733435.1 | Bacteroides_fragilis | G7060_RS07670 | 242068..245703 | 817 | BREX-system-P-loop-protein-BrxC-[Bacteroides-fragilis] |
| WP_120489887.1 | Type1.BREX | BrxA-DNA-binding->BrxB-BrxF-iSTAND->DUF6079*->N6-METHYLASE->SIR2->Schlafen+HATPase+wHTH->PglZ->SIGMA+OB-Fold+AAA-ATPase+HKD-DNase-BrxL-><-35exo+SF1-SF2-Helicase\|\|BACON+Peptidase_M64-> | 1202 | Bacteria>FCB_group>Bacteroidetes/Chlorobi_group>Bacteroidota>Bacteroidia>Bacteroidales | GCF_019733535.1 | Bacteroides_ovatus | G7086_RS03160 | 85646..89254 | 28116 | BREX-system-P-loop-protein-BrxC-[Bacteroides-ovatus] |
| WP_060386337.1 | Type1.BREX | <-SF1-SF2-Helicase<-?\|\|?->AbrB-HTH-><-KAP-NTPase<-SIGMA+OB-Fold+AAA-ATPase+Lon-Protease-BrxL<-PglZ<-Schlafen+HATPase+wHTH<-N6-METHYLASE<-DUF6079*<-BrxB-BrxF-iSTAND<-BrxA-DNA-binding<-HTH_3\|\|?->XIS-HTH->XIS-HTH-><-DEDD_Tnp_IS110+Transposase_20\|\|?->Trypsin-> | 1213 | Bacteria>FCB_group>Bacteroidetes/Chlorobi_group>Bacteroidota>Bacteroidia>Bacteroidales | GCF_020091485.1 | Bacteroides_stercoris | K6V31_RS06895 | 1552414..1556055 | 46506 | BREX-system-P-loop-protein-BrxC-[Bacteroides-stercoris] |
| WP_227088012.1 | Type1.BREX | ArdCN+MPTase->TOPRIM->?->TOPRIM->?->?->HTH_3->BrxA-DNA-binding->BrxB-BrxF-iSTAND->DUF6079*->N6-METHYLASE->Schlafen+HATPase+wHTH->PglZ->SIGMA+OB-Fold+AAA-ATPase+Lon-Protease-BrxL->RNaseH-><-?\|\|?->?-><-XIS-HTH | 1176 | Bacteria>FCB_group>Bacteroidetes/Chlorobi_group>Bacteroidota>Bacteroidia>Bacteroidales | GCF_020560105.1 | Bacteroides_cellulosilyticus | LI190_RS20855 | 41329..44859 | 246787 | BREX-system-P-loop-protein-BrxC-[Bacteroides-cellulosilyticus] |
| WP_234117060.1 | Type1.BREX | <-BACON+Peptidase_M64\|\|35exo+SF1-SF2-Helicase-><-SIGMA+OB-Fold+AAA-ATPase+HKD-DNase-BrxL<-PglZ<-Schlafen+HATPase+wHTH<-SIR2+SIR2<-N6-METHYLASE<-DUF6079*<-BrxB-BrxF-iSTAND<-BrxA-DNA-binding | 1202 | Bacteria>FCB_group>Bacteroidetes/Chlorobi_group>Bacteroidota>Bacteroidia>Bacteroidales | GCF_021410745.1 | Bacteroides_ovatus | K0G75_RS22190 | 19952..23560 | 28116 | BREX-system-P-loop-protein-BrxC-[Bacteroides-ovatus] |
| WP_235327427.1 | Type1.BREX | <-SIGMA+OB-Fold+AAA-ATPase+Lon-Protease-BrxL<-PglZ<-Schlafen+HATPase+wHTH<-N6-METHYLASE<-SLOG+BirA-HTH<-SF1-SF2-Helicase+PRTase<-HNH<-OLD-ABC<-DUF6079*<-BrxB-BrxF-iSTAND<-BrxA-DNA-binding | 1179 | Bacteria>FCB_group>Bacteroidetes/Chlorobi_group>Bacteroidota>Bacteroidia>Bacteroidales | GCF_021530995.1 | Bacteroides_caecigallinarum | I6E24_RS07160 | 15567..19106 | 1411144 | BREX-system-P-loop-protein-BrxC-[Bacteroides-caecigallinarum] |
| WP_235417293.1 | Type1.BREX | BrxA-DNA-binding->BrxB-BrxF-iSTAND->DUF6079*->N6-METHYLASE->Schlafen+HATPase+wHTH->PglZ->SIGMA+OB-Fold+AAA-ATPase+Lon-Protease-BrxL->?->REase+Bacteriocin_IIc-> | 1176 | Bacteria>FCB_group>Bacteroidetes/Chlorobi_group>Bacteroidota>Bacteroidia>Bacteroidales | GCF_021531515.1 | Prevotella_stercorea | I6E41_RS10105 | 1347..4877 | 363265 | BREX-system-P-loop-protein-BrxC-[Prevotella-stercorea] |
| WP_249093422.1 | Type1.BREX | TOPRIM->XIS-HTH-><-?<-?<-?<-?\|\|HTH_3->BrxA-DNA-binding->BrxB-BrxF-iSTAND->DUF6079*->N6-METHYLASE->SIR2->Schlafen+HATPase+wHTH->PglZ->SIGMA+OB-Fold+AAA-ATPase+HKD-DNase-BrxL->NADAR->RADICAL-SAM->HIRAN->HHH_5+SF1-SF2-Helicase-> | 1217 | Bacteria>FCB_group>Bacteroidetes/Chlorobi_group>Bacteroidota>Bacteroidia>Bacteroidales | GCF_023337025.1 | Mediterranea_sp._ET5 | M3087_RS02205 | 173446..177099 | 2939419 | BREX-system-P-loop-protein-BrxC-[Mediterranea-sp.-ET5] |
| WP_252760927.1 | Type1.BREX | DDE_Tnp_ISL3->XIS-HTH->?->AAA+-><-?\|\|wHTH+WYL+WCX->HTH_3->BrxA-DNA-binding->BrxB-BrxF-iSTAND->DUF6079*->N6-METHYLASE->Schlafen+HATPase+wHTH->PglZ->SIGMA+OB-Fold+AAA-ATPase+HKD-DNase-BrxL->SIR2+SIR2->?->?-><-tRNA\|\|HTH_31-> | 1206 | Bacteria>FCB_group>Bacteroidetes/Chlorobi_group>Bacteroidota>Bacteroidia>Bacteroidales | GCF_024054555.1 | Prevotella_cerevisiae | NG821_RS06905 | 6531..10151 | 2053716 | BREX-system-P-loop-protein-BrxC-[Prevotella-cerevisiae] |
| WP_258327305.1 | Type1.BREX | <-HHH_5+SF1-SF2-Helicase<-HIRAN<-RADICAL-SAM<-NADAR<-SIGMA+OB-Fold+AAA-ATPase+HKD-DNase-BrxL<-PglZ<-Schlafen+HATPase+wHTH<-SIR2<-N6-METHYLASE<-DUF6079*<-BrxB-BrxF-iSTAND<-BrxA-DNA-binding<-HTH_3\|\|?->?->?->?-><-XIS-HTH<-TOPRIM | 1217 | Bacteria>FCB_group>Bacteroidetes/Chlorobi_group>Bacteroidota>Bacteroidia>Bacteroidales | GCF_024733785.1 | Barnesiella_sp._ET7 | NW211_RS06370 | 166743..170396 | 2972460 | BREX-system-P-loop-protein-BrxC-[Barnesiella-sp.-ET7] |
| WP_234146092.1 | Type1.BREX | Arm-DNA-bind_5+Phage_int_SAM_5+Phage_integrase-><-Schlafen+HATPase+wHTH<-SIGMA+OB-Fold+AAA-ATPase+Lon-Protease-BrxL<-PglZ<-N6-METHYLASE<-PIN<-OLD-ABC<-DUF6079*<-BrxB-BrxF-iSTAND<-BrxA-DNA-binding<-HTH_3\|\|?->?->N6-METHYLASE-><-AAA+<-HTH_36<-XIS-HTH | 1180 | Bacteria>FCB_group>Bacteroidetes/Chlorobi_group>Bacteroidota>Bacteroidia>Bacteroidales | GCF_024760085.1 | Bacteroides_uniformis | J8872_RS07820 | 80453..83995 | 820 | BREX-system-P-loop-protein-BrxC-[Bacteroides-uniformis] |
| WP_259011403.1 | Type1.BREX | BrxA-DNA-binding->BrxB-BrxF-iSTAND->DUF6079*->N6-METHYLASE->ParB->OLD-ABC->?->Schlafen+HATPase+wHTH->PglZ->SIGMA+OB-Fold+AAA-ATPase+HKD-DNase-BrxL-><-35exo+SF1-SF2-Helicase | 1202 | Bacteria>FCB_group>Bacteroidetes/Chlorobi_group>Bacteroidota>Bacteroidia>Bacteroidales | GCF_024791585.1 | Bacteroides_sp._BFG-638 | NXX54_RS14145 | 3701947..3705555 | 2972765 | BREX-system-P-loop-protein-BrxC-[Bacteroides-sp.-BFG-638] |
| WP_258979859.1 | Type1.BREX | BrxA-DNA-binding->BrxB-BrxF-iSTAND->DUF6079-p*->DUF6079-p->N6-METHYLASE->ParB->OLD-ABC->?->Schlafen+HATPase+wHTH->PglZ->SIGMA+OB-Fold+AAA-ATPase+HKD-DNase-BrxL-> | 780 | Bacteria>FCB_group>Bacteroidetes/Chlorobi_group>Bacteroidota>Bacteroidia>Bacteroidales | GCF_024793605.1 | Bacteroides_sp._BFG-637 | NXX38_RS14295 | 3701972..3704314 | 2972764 | BREX-system-P-loop-protein-BrxC-[Bacteroides-sp.-BFG-637] |
| WP_009000708.1 | Type1.BREX | BrxA-DNA-binding->BrxB-BrxF-iSTAND->DUF6079*->N6-METHYLASE->?->?->Schlafen+HATPase+wHTH->PglZ->SIGMA+OB-Fold+AAA-ATPase+HKD-DNase-BrxL-><-35exo+SF1-SF2-Helicase\|\|BACON+Peptidase_M64-> | 1202 | Bacteria>FCB_group>Bacteroidetes/Chlorobi_group>Bacteroidota>Bacteroidia>Bacteroidales | GCF_025147085.1 | Bacteroides | NQ505_RS12870 | 3648371..3651979 | 816 | MULTISPECIES:-BREX-system-P-loop-protein-BrxC-[Bacteroides] |
| WP_262301578.1 | Type1.BREX | <-SIGMA+OB-Fold+AAA-ATPase+Lon-Protease-BrxL<-PglZ<-Schlafen+HATPase+wHTH<-N6-METHYLASE<-DUF6079*<-BrxB-BrxF-iSTAND<-BrxA-DNA-binding<-REase | 1218 | Bacteria>FCB_group>Bacteroidetes/Chlorobi_group>Bacteroidota>Bacteroidia>Bacteroidales | GCF_025460925.1 | Prevotella_hominis | OCW41_RS08900 | 100342..103998 | 2518605 | BREX-system-P-loop-protein-BrxC-[Prevotella-hominis] |
| WP_195661285.1 | Type1.BREX | BrxA-DNA-binding->BrxB-BrxF-iSTAND->DUF6079*->N6-METHYLASE->SIR2+SIR2->Schlafen+HATPase+wHTH->PglZ->SIGMA+OB-Fold+AAA-ATPase+HKD-DNase-BrxL-> | 1202 | Bacteria>FCB_group>Bacteroidetes/Chlorobi_group>Bacteroidota>Bacteroidia>Bacteroidales | GCF_025914175.1 | Bacteroides_nordii | KQP55_RS20540 | 5436855..5440463 | 291645 | BREX-system-P-loop-protein-BrxC-[Bacteroides-nordii] |
| WP_264965401.1 | Type1.BREX | ABC-ATPase+ABC-ATPase+ABC-ATPase+SbcC->Peptidase_M13_N+Peptidase_M13-><-SIGMA+OB-Fold+AAA-ATPase+HKD-DNase-BrxL<-PglZ<-N6-METHYLASE-p<-Schlafen+HATPase+wHTH<-N6-METHYLASE<-DUF6079*<-BrxB-BrxF-iSTAND<-BrxA-DNA-binding<-?<-?\|\|?-><-?<-?<-ABC-ATPase+SbcC | 1194 | Bacteria>FCB_group>Bacteroidetes/Chlorobi_group>Bacteroidota>Bacteroidia>Bacteroidales | GCF_026015685.1 | Prevotella_copri | ONT16_RS03155 | 729578..733162 | 165179 | BREX-system-P-loop-protein-BrxC-[Prevotella-copri] |
| WP_271704578.1 | Type1.BREX | HTH_3->MORC+DUF3684+REase->SF1-SF2-Helicase+PIN->HTH_3->BrxA-DNA-binding->BrxB-BrxF-iSTAND->DUF6079*->N6-METHYLASE->SIR2+SIR2->Schlafen+HATPase+wHTH->PglZ->SIGMA+OB-Fold+AAA-ATPase+HKD-DNase-BrxL-> | 1209 | Bacteria>FCB_group>Bacteroidetes/Chlorobi_group>Bacteroidota>Bacteroidia>Bacteroidales | GCF_028200085.1 | Bacteroides_xylanisolvens | PL493_RS08910 | 20513..24142 | 371601 | BREX-system-P-loop-protein-BrxC-[Bacteroides-xylanisolvens] |
| WP_274224871.1 | Type1.BREX | Arm-DNA-bind_5+Phage_int_SAM_5+Phage_integrase->?->AbrB-HTH-><-KAP-NTPase<-SIGMA+OB-Fold+AAA-ATPase+Lon-Protease-BrxL<-PglZ<-Schlafen+HATPase+wHTH<-N6-METHYLASE<-DUF6079*<-BrxB-BrxF-iSTAND<-BrxA-DNA-binding<-HTH_3\|\|?->XIS-HTH->XIS-HTH-> | 1213 | Bacteria>FCB_group>Bacteroidetes/Chlorobi_group>Bacteroidota>Bacteroidia>Bacteroidales | GCF_028743275.1 | Bacteroides_fragilis | PT255_RS11125 | 2684967..2688608 | 817 | BREX-system-P-loop-protein-BrxC-[Bacteroides-fragilis] |
| WP_032560932.1 | Type1.BREX | SF1-SF2-Helicase->?-><-DOC+HTH<-URI-DNase+RAMA<-?<-SIGMA+OB-Fold+AAA-ATPase+HKD-DNase-BrxL<-PglZ<-Schlafen+HATPase+wHTH<-N6-METHYLASE<-DUF6079*<-BrxB-BrxF-iSTAND<-BrxA-DNA-binding<-HTH_3\|\|HEPN-><-?<-?<-STYKinase-like\|\|Phage_int_SAM_5+Phage_integrase-> | 1211 | Bacteria>FCB_group>Bacteroidetes/Chlorobi_group>Bacteroidota>Bacteroidia>Bacteroidales | GCF_029872615.1 | Bacteroides_fragilis | M068_RS14675 | 124538..128173 | 817 | BREX-system-P-loop-protein-BrxC-[Bacteroides-fragilis] |
| WP_154043608.1 | Type1.BREX | <-SbcC<-SIGMA+OB-Fold+AAA-ATPase+Lon-Protease-BrxL<-PglZ<-Schlafen+HATPase+wHTH<-N6-METHYLASE<-SLOG+BirA-HTH<-SF1-SF2-Helicase+PRTase<-HNH<-OLD-ABC<-DUF6079*<-BrxB-BrxF-iSTAND<-BrxA-DNA-binding<-?<-?<-?<-?<-?<-Calcineurin+Calcineurin-conf | 1179 | Bacteria>FCB_group>Bacteroidetes/Chlorobi_group>Bacteroidota>Bacteroidia>Bacteroidales | GCF_900066445.1 | Bacteroides_cellulolyticus | OCV82_RS05980 | 56504..60043 | 2981780 | BREX-system-P-loop-protein-BrxC-[Bacteroides-cellulolyticus] |
| WP_074557116.1 | Type1.BREX | <-HAD+PRTase+PRTase+REase+SF1-SF2-Helicase<-?<-TPR+TPR<-?<-Schlafen+HATPase+wHTH<-TrwF\|\|BrxA-DNA-binding->BrxB-BrxF-iSTAND->DUF6079*->N6-METHYLASE->SIR2->Schlafen+HATPase+wHTH->PglZ->SIGMA+OB-Fold+AAA-ATPase+HKD-DNase-BrxL-><-35exo+SF1-SF2-Helicase\|\|BACON+Peptidase_M64-> | 1202 | Bacteria>FCB_group>Bacteroidetes/Chlorobi_group>Bacteroidota>Bacteroidia>Bacteroidales | GCF_900102645.1 | Bacteroides_ovatus | BLR14_RS05840 | 106864..110472 | 28116 | BREX-system-P-loop-protein-BrxC-[Bacteroides-ovatus] |
| WP_073402639.1 | Type1.BREX | <-RADICAL-SAM<-?\|\|tRNA-><-SIGMA+OB-Fold+AAA-ATPase+Lon-Protease-BrxL<-PglZ<-Schlafen+HATPase+wHTH<-ParB+HNH<-N6-METHYLASE<-?<-DUF6079*<-BrxB-BrxF-iSTAND<-BrxA-DNA-binding | 1188 | Bacteria>FCB_group>Bacteroidetes/Chlorobi_group>Bacteroidota>Bacteroidia>Bacteroidales | GCF_900128905.1 | Bacteroides_luti | BUC21_RS11790 | 81108..84674 | 1297750 | BREX-system-P-loop-protein-BrxC-[Bacteroides-luti] |
| WP_077197246.1 | Type1.BREX | DDE_Tnp_IS66+DDE_Tnp_IS66_C-><-?<-?<-?<-?<-?\|\|HTH_3->BrxA-DNA-binding->BrxB-BrxF-iSTAND->DUF6079*->N6-METHYLASE->Schlafen+HATPase+wHTH->PglZ->SIGMA+OB-Fold+AAA-ATPase+Lon-Protease-BrxL->?->\|\|HTH_38+rve-><-?\|\|DDE_Tnp_1->HTH_3-> | 1176 | Bacteria>FCB_group>Bacteroidetes/Chlorobi_group>Bacteroidota>Bacteroidia>Bacteroidales | GCF_900155655.1 | Prevotella_ihumii | CZ367_RS13300 | 3061778..3065308 | 1917878 | BREX-system-P-loop-protein-BrxC-[Prevotella-ihumii] |
| WP_130829424.1 | Type1.BREX | <-SIGMA+OB-Fold+AAA-ATPase+Lon-Protease-BrxL<-PglZ<-Schlafen+HATPase+wHTH<-Schlafen<-N6-METHYLASE<-DUF6079*<-BrxB-BrxF-iSTAND<-BrxA-DNA-binding<-HTH_3<-?\|\|?-><-PTSEIIA1\|\|?->PRTase+MutS-> | 1153 | Bacteria>FCB_group>Bacteroidetes/Chlorobi_group>Bacteroidota>Bacteroidia>Bacteroidales | GCF_900607215.1 | Prevotella | ETF27_RS09605 | 9675..13136 | 838 | MULTISPECIES:-BREX-system-P-loop-protein-BrxC-[Prevotella] |
| WP_125105113.1 | Type1.BREX | <-HHH_5+SF1-SF2-Helicase<-HIRAN<-RADICAL-SAM<-NADAR<-SIGMA+OB-Fold+AAA-ATPase+HKD-DNase-BrxL<-PglZ<-Schlafen+HATPase+wHTH<-SIR2<-N6-METHYLASE<-DUF6079*<-BrxB-BrxF-iSTAND<-BrxA-DNA-binding<-HTH_3\|\|?->?->?->?-><-XIS-HTH<-TOPRIM | 1217 | Bacteria>FCB_group>Bacteroidetes/Chlorobi_group>Bacteroidota>Bacteroidia>Bacteroidales | GCF_900625065.1 | Prevotella_marseillensis | EJL04_RS12230 | 3058927..3062580 | 2479840 | BREX-system-P-loop-protein-BrxC-[Prevotella-marseillensis] |
| WP_032543214.1 | Type1.BREX | <-METHYLASE<-ACET<-DOC+HTH<-URI-DNase+RAMA<-?<-SIGMA+OB-Fold+AAA-ATPase+HKD-DNase-BrxL<-PglZ<-Schlafen+HATPase+wHTH<-N6-METHYLASE<-DUF6079*<-BrxB-BrxF-iSTAND<-BrxA-DNA-binding<-HTH_3<-?<-?<-TraJ-RHH<-ZnR+TOPRIM<-CR-ATPase4<-XIS-HTH | 1211 | Bacteria>FCB_group>Bacteroidetes/Chlorobi_group>Bacteroidota>Bacteroidia>Bacteroidales | GCF_902364655.1 | Bacteroides_fragilis | PDN68_RS18445 | 100181..103816 | 817 | BREX-system-P-loop-protein-BrxC-[Bacteroides-fragilis] |
| WP_060407611.1 | Type1.BREX | <-SIGMA+OB-Fold+AAA-ATPase+HKD-DNase-BrxL<-PglZ<-Schlafen+HATPase+wHTH<-SIR2<-N6-METHYLASE-p<-N6-METHYLASE<-DUF6079*<-BrxB-BrxF-iSTAND<-BrxA-DNA-binding<-HTH_3<-?<-?<-?\|\|SIGMA-HTH-> | 1205 | Bacteria>FCB_group>Bacteroidetes/Chlorobi_group>Bacteroidota>Bacteroidia>Bacteroidales | GCF_902387305.1 | Bacteroides_cellulosilyticus | FYC30_RS03960 | 75039..78656 | 246787 | BREX-system-P-loop-protein-BrxC-[Bacteroides-cellulosilyticus] |
| WP_118419043.1 | Type1.BREX | BrxA-DNA-binding->BrxB-BrxF-iSTAND->DUF6079*->N6-METHYLASE->SIR2+SIR2->Schlafen+HATPase+wHTH->PglZ->SIGMA+OB-Fold+AAA-ATPase+HKD-DNase-BrxL-><-35exo+SF1-SF2-Helicase\|\|BACON+Peptidase_M64-> | 1202 | Bacteria>FCB_group>Bacteroidetes/Chlorobi_group>Bacteroidota>Bacteroidia>Bacteroidales | GCF_918279245.1 | Bacteroides_ovatus | P0203_RS09380 | 101158..104766 | 28116 | BREX-system-P-loop-protein-BrxC-[Bacteroides-ovatus] |
| WP_117985926.1 | Type1.BREX | XIS-HTH->SMS-KaiC-><-SF1-SF2-Helicase<-SIGMA+OB-Fold+AAA-ATPase+HKD-DNase-BrxL<-PglZ<-Schlafen+HATPase+wHTH<-SF1-SF2-Helicase<-OLD-ABC+TOPRIM<-N6-METHYLASE<-DUF6079*<-BrxB-BrxF-iSTAND<-BrxA-DNA-binding<-HTH_3<-AbiEi_4+AbiEi_1<-STYKinase-like<-TRPR-HTH\|\|SbcC->DEDD_Tnp_IS110+Transposase_20-> | 1210 | Bacteria>FCB_group>Bacteroidetes/Chlorobi_group>Bacteroidota>Bacteroidia>Bacteroidales | GCF_934726555.1 | Bacteroides | O1446_RS12920 | 68574..72206 | 816 | MULTISPECIES:-BREX-system-P-loop-protein-BrxC-[Bacteroides] |
| WP_158582848.1 | Type1.BREX | ACET-><-?<-Cysteine-Desulfurase\|\|tRNA-><-?<-?<-?\|\|BrxA-DNA-binding->BrxB-BrxF-iSTAND->DUF6079*->N6-METHYLASE->Schlafen+HATPase+wHTH->PglZ->SIGMA+OB-Fold+AAA-ATPase+HKD-DNase-BrxL-><-?\|\|?->PolB->HEPN-> | 1210 | Bacteria>FCB_group>Bacteroidetes/Chlorobi_group>Bacteroidota>Bacteroidia>Bacteroidales | GCF_938033175.1 | Parabacteroides_sp._AF17-28 | DWW90_RS15090 | 22936..26568 | 2292241 | BREX-system-P-loop-protein-BrxC-[Parabacteroides-sp.-AF17-28] |
| WP_117695280.1 | Type6.BREX | <-SLATT_4<-SIGMA+OB-Fold+AAA-ATPase+HKD-DNase-BrxL<-PglZ-p<-PglZ<-Schlafen+HATPase+wHTH<-N6-METHYLASE<-DUF6079*<-BrxB-BrxF-iSTAND<-BrxA-DNA-binding<-BrxE<-HTH_3<-RNasePH+RNasePH+KH+S1\|\|DUF4369+Thioredoxin_8->?->NUDIX->TOPRIM-> | 1216 | Bacteria>FCB_group>Bacteroidetes/Chlorobi_group>Bacteroidota>Bacteroidia>Bacteroidales | - | Prevotella_copri | DXC61_RS12020 | 26690..30340 | 165179 | BREX-system-P-loop-protein-BrxC-[Prevotella-copri] |
| WP_233339205.1 | Type6.BREX | <-SLATT_4<-SIGMA+OB-Fold+AAA-ATPase+HKD-DNase-BrxL<-PglZ<-Schlafen+HATPase+wHTH<-N6-METHYLASE<-DUF6079*<-BrxB-BrxF-iSTAND<-BrxA-DNA-binding<-BrxE<-HTH_3<-RNasePH+RNasePH+KH+S1\|\|DUF4369+Thioredoxin_8->?->NUDIX->TOPRIM-> | 1216 | Bacteria>FCB_group>Bacteroidetes/Chlorobi_group>Bacteroidota>Bacteroidia>Bacteroidales | - | Prevotella_copri | LYY06_RS08410 | 34942..38592 | 165179 | BREX-system-P-loop-protein-BrxC-[Prevotella-copri] |
| WP_057098610.1 | Type1.BREX | <-KAP-NTPase<-SIGMA+OB-Fold+AAA-ATPase+Lon-Protease-BrxL<-PglZ<-Schlafen+HATPase+wHTH<-N6-METHYLASE<-DUF6079*<-BrxB-BrxF-iSTAND<-BrxA-DNA-binding<-HTH_3\|\|XIS-HTH->XIS-HTH-><-?<-?<-ABhydrolase | 1212 | Bacteria>FCB_group>Bacteroidetes/Chlorobi_group>Bacteroidota>Bacteroidia | GCF_017309675.2 | Bacteroidales | DWX60_RS07020 | 13179..16817 | 171549 | MULTISPECIES:-BREX-system-P-loop-protein-BrxC-[Bacteroidales] |
| WP_046151576.1 | Type1.BREX | <-wHTH+WYL+WCX<-ParB+HNH<-SIR2<-SIGMA+OB-Fold+AAA-ATPase+Lon-Protease-BrxL<-PglZ<-?<-Schlafen+HATPase+wHTH<-REase<-N6-METHYLASE<-DUF6079*<-BrxB-BrxF-iSTAND<-BrxA-DNA-binding<-HTH_3\|\|SIR2->HerA/FtsK-><-Phage_lysozyme<-VirD4-TraG+VirD4-FtsK | 1184 | Bacteria>FCB_group>Bacteroidetes/Chlorobi_group>Bacteroidota>Bacteroidia | GCF_900102645.1 | Bacteroidales | BLR14_RS08960 | 69320..72874 | 171549 | MULTISPECIES:-BREX-system-P-loop-protein-BrxC-[Bacteroidales] |
| MCF8362389.1 | Type1.BREX | <-SIGMA+OB-Fold+AAA-ATPase+HKD-DNase-BrxL<-PglZ<-PglZ<-Schlafen+HATPase+wHTH<-N6-METHYLASE<-N6-METHYLASE-p<-DUF6079*<-BrxB-BrxF-iSTAND<-BrxA-DNA-binding<-HTH_3<-Schlafen+HATPase+wHTH-p | 1180 | Bacteria>FCB_group>Bacteroidetes/Chlorobi_group>Bacteroidota>Bacteroidia>Marinilabiliales | GCA_021740345.1 | Prolixibacteraceae_bacterium | K9G70_07170 | 11489..15031 | 2053594 | BREX-system-P-loop-protein-BrxC-[Prolixibacteraceae-bacterium] |
| MCK9411439.1 | Type1.BREX | Peptidase_A8->Peptidase_A8-><-wHTH+WYL+WCX\|\|?->SMODS->?->HTH_3->BrxA-DNA-binding->BrxB-BrxF-iSTAND->DUF6079*->N6-METHYLASE->Schlafen+HATPase+wHTH->?->PglZ->AbiEi_4->SIGMA+OB-Fold+AAA-ATPase+HKD-DNase-BrxL->?->REC+LYTTR-> | 1173 | Bacteria>FCB_group>Bacteroidetes/Chlorobi_group>Bacteroidota>Bacteroidia>Marinilabiliales | GCA_023229575.1 | Prolixibacteraceae_bacterium | M0Q53_04000 | 25065..28586 | 2053594 | BREX-system-P-loop-protein-BrxC-[Prolixibacteraceae-bacterium] |
| WP_062122677.1 | Type1.BREX | ABhydrolase->tRNA-><-SIGMA+OB-Fold+AAA-ATPase+HKD-DNase-BrxL<-PglZ<-Schlafen+HATPase+wHTH<-N6-METHYLASE<-DUF6079-p<-DUF499<-DUF6079-p*<-BrxB-BrxF-iSTAND<-BrxA-DNA-binding<-HTH_3\|\|?-><-?<-?<-?<-SIGMA-HTH | 329 | Bacteria>FCB_group>Bacteroidetes/Chlorobi_group>Bacteroidota>Bacteroidia>Marinilabiliales | GCF_000974365.1 | Geofilum_rubicundum | JCM15548_RS04470 | 156242..157231 | 472113 | hypothetical-protein-[Geofilum-rubicundum] |
| WP_107823954.1 | Type1.BREX | <-STYKinase-like<-Couple_hipA<-HTH_3<-SIGMA+OB-Fold+AAA-ATPase+HKD-DNase-BrxL<-PglZ<-N6-METHYLASE<-Schlafen+HATPase+wHTH<-DUF6079*<-BrxB-BrxF-iSTAND<-BrxA-DNA-binding<-HTH_3 | 1168 | Bacteria>FCB_group>Bacteroidetes/Chlorobi_group>Bacteroidota>Bacteroidia>Marinilabiliales | GCF_003046625.1 | Mangrovibacterium_marinum | C8N47_RS17935 | 11546..15052 | 1639118 | BREX-system-P-loop-protein-BrxC-[Mangrovibacterium-marinum] |
| WP_163710191.1 | Type1.BREX | HTH_3->BrxA-DNA-binding->BrxB-BrxF-iSTAND->DUF6079*->Schlafen+HATPase+wHTH->SbcC->?->N6-METHYLASE->SF1-SF2-Helicase->PglZ->SIGMA+OB-Fold+AAA-ATPase+HKD-DNase-BrxL-> | 1168 | Bacteria>FCB_group>Bacteroidetes/Chlorobi_group>Bacteroidota>Bacteroidia>Marinilabiliales | GCF_010882175.1 | Mangrovibacterium_lignilyticum | GJV22_RS06145 | 1519093..1522599 | 2668052 | BREX-system-P-loop-protein-BrxC-[Mangrovibacterium-lignilyticum] |
| WP_013929909.1 | Type1.BREX | Peptidase_S51->?-><-SIGMA+OB-Fold+AAA-ATPase+Lon-Protease-BrxL<-PglZ<-Schlafen+HATPase+wHTH<-HipA_C<-Couple_hipA<-HTH_3<-N6-METHYLASE<-DUF6079*<-BrxB-BrxF-iSTAND<-BrxA-DNA-binding<-DDE_Tnp<-?<-DDE_Tnp_1_assoc<-MULE-TRANSPOSASE<-SF1-SF2-Helicase<-?<-HEPN+DUF499 | 1173 | Bacteria>FCB_group>Bacteroidetes/Chlorobi_group>Bacteroidota>Cytophagia>Cytophagales | GCF_000218895.1 | Runella_slithyformis | RUNSL_RS20935 | 4933430..4936951 | 106 | BREX-system-P-loop-protein-BrxC-[Runella-slithyformis] |
| WP_273212250.1 | Type1.BREX | <-AbiEii<-wHTH+AbiEi_2<-SIGMA+OB-Fold+AAA-ATPase+Lon-Protease-BrxL<-PglZ<-Schlafen+HATPase+wHTH<-REase<-N6-METHYLASE<-HNH<-OLD-ABC<-DUF6079*<-BrxB-BrxF-iSTAND<-BrxA-DNA-binding<-?<-?<-D5ATPase<-XIS-HTH<-?<-Arm-DNA-bind_5+Phage_int_SAM_5+Phage_integrase | 1173 | Bacteria>FCB_group>Bacteroidetes/Chlorobi_group>Bacteroidota>Cytophagia>Cytophagales | GCF_002483185.1 | Runella_zeae | B4E63_RS12460 | 307458..310979 | 94255 | BREX-system-P-loop-protein-BrxC-[Runella-zeae] |
| WP_138994213.1 | Type1.BREX | TraN->?->?->?->?->GDResolvase+HIN-HTH->BrxA-DNA-binding->BrxB-BrxF-iSTAND->DUF6079*->N6-METHYLASE->Schlafen+HATPase+wHTH->PglZ->SIGMA+OB-Fold+AAA-ATPase+Lon-Protease-BrxL->MORC+DUF3684+REase->SF1-SF2-Helicase->HEPN+PAS+DUF4011+SF1-SF2-Helicase+REase-><-SF1-SF2-Helicase+wHTH+Small-B-barrel+DUF1998<-BrxD-AAA | 1214 | Bacteria>FCB_group>Bacteroidetes/Chlorobi_group>Bacteroidota>Cytophagia>Cytophagales | GCF_006149005.1 | Larkinella_sp._C7 | FH871_RS25315 | 15156..18800 | 2576607 | BREX-system-P-loop-protein-BrxC-[Larkinella-sp.-C7] |
| WP_255036052.1 | Type1.BREX | XIS-HTH->D5ATPase->?->?->?->?->BrxA-DNA-binding->BrxB-BrxF-iSTAND->DUF6079*->Schlafen+HATPase+wHTH->N6-METHYLASE->?->PglZ->SIGMA+OB-Fold+AAA-ATPase+Lon-Protease-BrxL->REase+DUF5655->?->?->SF1-SF2-Helicase-> | 1199 | Bacteria>FCB_group>Bacteroidetes/Chlorobi_group>Bacteroidota>Cytophagia>Cytophagales | GCF_024343775.1 | Lacihabitans_soyangensis | EGI31_RS04905 | 50707..54306 | 869394 | BREX-system-P-loop-protein-BrxC-[Lacihabitans-soyangensis] |
| MAL23063.1 | Type1.BREX | <-METHYLASE<-?<-SF1-SF2-Helicase<-?<-?<-REase+DUF5655<-N6-METHYLASE<-?<-Schlafen+HATPase+wHTH<-DUF6079*<-BrxB-BrxF-iSTAND<-BrxA-DNA-binding<-HTH_3<-?<-HipA_C<-Couple_hipA<-HTH_3 | 1157 | Bacteria>FCB_group>Bacteroidetes/Chlorobi_group>Bacteroidota>Flavobacteriia>Flavobacteriales | GCA_002694765.1 | Xanthomarina_sp. | CMP04_07455 | 107327..110800 | 1931211 | hypothetical-protein-[Xanthomarina-sp.] |
| ROI04325.1 | Type1.BREX | <-HTH_3<-?\|\|?->?-><-?<-?<-SIGMA+OB-Fold+AAA-ATPase+HKD-DNase-BrxL<-PglZ<-N6-METHYLASE<-DUF6079*<-BrxB-BrxF-iSTAND<-BrxA-DNA-binding<-Schlafen+HATPase+wHTH\|\|?->ChlI+Lon_C+AAA-><-?\|\|?->SLOG-> | 1174 | Bacteria>FCB_group>Bacteroidetes/Chlorobi_group>Bacteroidota>Flavobacteriia>Flavobacteriales | GCA_003729975.1 | Kaistella_haifensis | EGH90_10140 | 55406..58930 | 421525 | BREX-system-P-loop-protein-BrxC-[Kaistella-haifensis] |
| NJY61426.1 | Type1.BREX | Arm-DNA-bind_5+Phage_int_SAM_5+Phage_integrase-><-?<-SIGMA+OB-Fold+AAA-ATPase+Lon-Protease-BrxL<-PglZ<-Schlafen+HATPase+wHTH<-N6-METHYLASE<-DUF6079*<-BrxB-BrxF-iSTAND<-BrxA-DNA-binding<-ANK+ANK<-?<-TraI-VirD2-RCRE<-TraJ-RHH<-ZnR+TOPRIM<-D5ATPase<-XIS-HTH | 1202 | Bacteria>FCB_group>Bacteroidetes/Chlorobi_group>Bacteroidota>Flavobacteriia>Flavobacteriales | GCA_012038135.1 | Salinimicrobium_nanhaiense | HC174_01475 | 311288..314896 | 2722701 | BREX-system-P-loop-protein-BrxC-[Salinimicrobium-nanhaiense] |
| MBQ0768324.1 | Type1.BREX | <-N6-METHYLASE<-Schlafen+HATPase+wHTH<-DUF6079*<-BrxB-BrxF-iSTAND<-BrxA-DNA-binding | 1160 | Bacteria>FCB_group>Bacteroidetes/Chlorobi_group>Bacteroidota>Flavobacteriia>Flavobacteriales | GCA_018067115.1 | Bizionia_sp. | KBT58_03470 | 3917..7399 | 1954480 | BREX-system-P-loop-protein-BrxC-[Bizionia-sp.] |
| MBW7869163.1 | Type1.BREX | HTH_3->BrxA-DNA-binding->BrxB-BrxF-iSTAND->DUF6079*->N6-METHYLASE->?->Schlafen+HATPase+wHTH->KAP-NTPase->PglZ->SIGMA+OB-Fold+AAA-ATPase+HKD-DNase-BrxL-> | 1166 | Bacteria>FCB_group>Bacteroidetes/Chlorobi_group>Bacteroidota>Flavobacteriia>Flavobacteriales | GCA_019455205.1 | Brumimicrobium_sp. | H3C31_12675 | 1666..5166 | 2029867 | BREX-system-P-loop-protein-BrxC-[Brumimicrobium-sp.] |
| WP_026451333.1 | Type1.BREX | VirB5->TrwI->TraK->?->TraN->?->HTH_3->BrxA-DNA-binding->BrxB-BrxF-iSTAND->DUF6079*->Schlafen+HATPase+wHTH->?->N6-METHYLASE->?->?->SF1-SF2-Helicase->?->METHYLASE->PglZ-> | 1160 | Bacteria>FCB_group>Bacteroidetes/Chlorobi_group>Bacteroidota>Flavobacteriia>Flavobacteriales | GCF_000429125.1 | Aequorivita_capsosiphonis | G440_RS0110860 | 108295..111777 | 487317 | BREX-system-P-loop-protein-BrxC-[Aequorivita-capsosiphonis] |
| WP_034059321.1 | Type1.BREX | <-SAM-METHYLASE<-?<-PglZ<-METHYLASE<-?<-SF1-SF2-Helicase<-?<-N6-METHYLASE<-Schlafen+HATPase+wHTH<-DUF6079*<-BrxB-BrxF-iSTAND<-BrxA-DNA-binding<-HTH_3<-?<-TPR+TPR+TPR\|\|ABhydrolase->SIGMA-HTH-> | 1160 | Bacteria>FCB_group>Bacteroidetes/Chlorobi_group>Bacteroidota>Flavobacteriia>Flavobacteriales | GCF_000797445.1 | Lacinutrix_jangbogonensis | RH82_RS07100 | 67712..71194 | 1469557 | BREX-system-P-loop-protein-BrxC-[Lacinutrix-jangbogonensis] |
| WP_094415998.1 | Type1.BREX | <-PIN<-?<-SIGMA+OB-Fold+AAA-ATPase+Lon-Protease-BrxL<-PglZ<-N6-METHYLASE<-Schlafen+HATPase+wHTH<-DUF6079*<-BrxB-BrxF-iSTAND<-BrxA-DNA-binding<-HTH<-?<-XIS-HTH | 1197 | Bacteria>FCB_group>Bacteroidetes/Chlorobi_group>Bacteroidota>Flavobacteriia>Flavobacteriales | GCF_002251835.1 | Flavobacterium_cyanobacteriorum | CHU92_RS12285 | 18327..21920 | 2022802 | BREX-system-P-loop-protein-BrxC-[Flavobacterium-cyanobacteriorum] |
| WP_129016554.1 | Type1.BREX | <-SIGMA+OB-Fold+AAA-ATPase+Lon-Protease-BrxL<-PglZ<-Schlafen+HATPase+wHTH<-N6-METHYLASE<-DUF6079*<-BrxB-BrxF-iSTAND<-BrxA-DNA-binding\|\|?-><-ACET<-SIGMA-HTH | 1202 | Bacteria>FCB_group>Bacteroidetes/Chlorobi_group>Bacteroidota>Flavobacteriia>Flavobacteriales | GCF_004115975.1 | Gelidibacter_gilvus | ESZ48_RS06605 | 20961..24569 | 59602 | BREX-system-P-loop-protein-BrxC-[Gelidibacter-gilvus] |
| WP_093139951.1 | Type1.BREX | <-SF1-SF2-Helicase<-SIGMA+OB-Fold+AAA-ATPase+Lon-Protease-BrxL<-PglZ<-Schlafen+HATPase+wHTH<-?<-N6-METHYLASE<-DUF6079*<-BrxB-BrxF-iSTAND<-BrxA-DNA-binding\|\|?-><-?<-?<-?\|\|?->?->HisKin-> | 1202 | Bacteria>FCB_group>Bacteroidetes/Chlorobi_group>Bacteroidota>Flavobacteriia>Flavobacteriales | GCF_014651275.1 | Ulvibacter_litoralis | IE050_RS03690 | 820645..824253 | 227084 | BREX-system-P-loop-protein-BrxC-[Ulvibacter-litoralis] |
| WP_199600307.1 | Type1.BREX | Relaxase->VirD4-TraG+VirD4-FtsK->Classical-AAA-><-Peptidase_S8<-SIGMA+OB-Fold+AAA-ATPase+HKD-DNase-BrxL<-PglZ<-N6-METHYLASE<-DEOR-HTH<-Schlafen+HATPase+wHTH<-DUF6079*<-BrxB-BrxF-iSTAND<-BrxA-DNA-binding<-HTH_3\|\|?-><-?<-TraN<-?<-TraK<-TrwI | 1160 | Bacteria>FCB_group>Bacteroidetes/Chlorobi_group>Bacteroidota>Flavobacteriia>Flavobacteriales | GCF_016469035.1 | Gelidibacter_salicanalis | JEM65_RS13235 | 170248..173730 | 291193 | BREX-system-P-loop-protein-BrxC-[Gelidibacter-salicanalis] |
| WP_157278947.1 | Type1.BREX | Arm-DNA-bind_5+Phage_int_SAM_5+Phage_integrase->?->XIS-HTH->TraJ-RHH->?->?->HTH_3->BrxA-DNA-binding->BrxB-BrxF-iSTAND->DUF6079*->N6-METHYLASE->Schlafen+HATPase+wHTH->PglZ->SIGMA+OB-Fold+AAA-ATPase+HKD-DNase-BrxL->?-><-HTH_21+rve+rve_3 | 1169 | Bacteria>FCB_group>Bacteroidetes/Chlorobi_group>Bacteroidota>Sphingobacteriia>Sphingobacteriales | GCF_000427965.1 | Olivibacter_sitiensis | A37S_RS0120635 | 7227..10736 | 376470 | BREX-system-P-loop-protein-BrxC-[Olivibacter-sitiensis] |
| KPP94735.1 | Type1.BREX | SAM-METHYLASE->tRNA-><-SIGMA+OB-Fold+AAA-ATPase+HKD-DNase-BrxL<-PglZ<-Schlafen+HATPase+wHTH<-N6-METHYLASE<-?<-DUF6079*<-BrxB-BrxF-iSTAND<-BrxA-DNA-binding<-?<-SF1-SF2-Helicase<-?<-?<-DnaB+SMS-KaiC<-HTH_36<-XIS-HTH | 1203 | Bacteria>FCB_group>Bacteroidetes/Chlorobi_group>Bacteroidota>unclassified_Bacteroidota | GCA_001314545.1 | Bacteroidetes_bacterium_HLUCCA01 | HLUCCA01_08750 | 188157..191768 | 1666909 | Brex-phage-defense-system-ATPase-BrxC-[Bacteroidetes-bacterium-HLUCCA01] |
| KAF0128492.1 | Type1.BREX | <-SIGMA+OB-Fold+AAA-ATPase+Lon-Protease-BrxL<-PglZ<-wHTH+REase<-McrB-AAA<-Schlafen+HATPase+wHTH<-N6-METHYLASE<-DUF6079*<-BrxB-BrxF-iSTAND<-BrxA-DNA-binding | 1144 | Bacteria>FCB_group>Bacteroidetes/Chlorobi_group>Bacteroidota>unclassified_Bacteroidota | GCA_009773165.1 | Bacteroidota_bacterium | FD155_3234 | 38823..42257 | 1898104 | hypothetical-protein-FD155_3234-[Bacteroidota-bacterium] |
| NUQ82570.1 | Type1.BREX | <-GspA-ATPase<-TrkH<-?<-?\|\|?->BrxA-DNA-binding->BrxB-BrxF-iSTAND->DUF6079*->N6-METHYLASE->Schlafen+HATPase+wHTH->PglZ->SIGMA+OB-Fold+AAA-ATPase+Lon-Protease-BrxL->DSBH+AraC-HTH+AraC-HTH->?->?->?-><-ABhydrolase | 1197 | Bacteria>FCB_group>Bacteroidetes/Chlorobi_group>Bacteroidota>unclassified_Bacteroidota | GCA_013360915.1 | Bacteroidota_bacterium | HUU10_13230 | 47380..50973 | 1898104 | BREX-system-P-loop-protein-BrxC-[Bacteroidota-bacterium] |
| MBN8706499.1 | Type1.BREX | Phage_int_SAM_4+Phage_integrase->35exo+URI-DNase->tRNA->HTH->BrxA-DNA-binding->BrxB-BrxF-iSTAND->DUF6079*->OLD-ABC->HNH->N6-METHYLASE->?->Schlafen+HATPase+wHTH->PglZ->SIGMA+OB-Fold+AAA-ATPase+Lon-Protease-BrxL->SbcC-> | 1173 | Bacteria>FCB_group>Bacteroidetes/Chlorobi_group>Bacteroidota>unclassified_Bacteroidota | GCA_017303995.1 | Bacteroidota_bacterium | J0L62_11545 | 8364..11885 | 1898104 | BREX-system-P-loop-protein-BrxC-[Bacteroidota-bacterium] |
| MCC5907043.1 | Type1.BREX | <-REase<-?<-?<-?\|\|TPR+TPR->BrxA-DNA-binding->BrxB-BrxF-iSTAND->DUF6079*->AbiEi->AbiEii->N6-METHYLASE->Schlafen+HATPase+wHTH->PglZ->SIGMA+OB-Fold+AAA-ATPase+HKD-DNase-BrxL->REase-><-?\|\|Thymidylate_kin-> | 1167 | Bacteria>FCB_group>Bacteroidetes/Chlorobi_group>Balneolota>Balneolia>Balneolales | GCA_020832285.1 | Balneolaceae_bacterium | JJU13_12585 | 29362..32865 | 2053516 | BREX-system-P-loop-protein-BrxC-[Balneolaceae-bacterium] |
| WP_265763895.1 | Type1.BREX | <-HEPN<-SIGMA+OB-Fold+AAA-ATPase+HKD-DNase-BrxL<-PglZ<-?<-Schlafen+HATPase+wHTH<-REase<-?<-N6-METHYLASE<-DUF6079*<-BrxB-BrxF-iSTAND<-BrxA-DNA-binding<-tRNA\|\|?->?->?<-?<-?<-ABC-ATPase+SbcC | 1165 | Bacteria>FCB_group>Bacteroidetes/Chlorobi_group>Balneolota>Balneolia>Balneolales | GCF_026229185.1 | Aliifodinibius_sp._1BSP15-2V2 | J6I44_RS00130 | 34405..37902 | 2820877 | BREX-system-P-loop-protein-BrxC-[Aliifodinibius-sp.-1BSP15-2V2] |
| ANT65175.1 | Type1.BREX | <-HTH_21+rve+rve_3<-HTH_Tnp_1\|\|DUF772+DDE_Tnp_1_2-><-SIGMA+OB-Fold+AAA-ATPase+Lon-Protease-BrxL<-PglZ<-Schlafen+HATPase+wHTH<-SbcC<-N6-METHYLASE<-Schlafen+HATPase+wHTH<-DUF6079*<-BrxB-BrxF-iSTAND<-BrxA-DNA-binding<-XIS-HTH<-RVT<-RVT\|\|AAA-ATPase-p-><-?\|\|?-><-Thioredoxin | 1144 | Bacteria>FCB_group>Bacteroidetes/Chlorobi_group>Chlorobiota>Chlorobiia>Chlorobiales | GCA_001687065.1 | Prosthecochloris_sp._CIB_2401 | Ptc2401_01411 | 1430885..1434319 | 1868325 | hypothetical-protein-Ptc2401_01411-[Prosthecochloris-sp.-CIB-2401] |
| TNJ34090.1 | Type1.BREX | HIN-HTH->HTH_Tnp_1+HTH_21+rve+rve_3-><-SIGMA+OB-Fold+AAA-ATPase+Lon-Protease-BrxL<-PglZ<-TOPRIM<-ParB+SbcC<-N6-METHYLASE<-Schlafen+HATPase+wHTH<-DUF6079*<-BrxB-BrxF-iSTAND<-BrxA-DNA-binding<-XIS-HTH<-RVT<-RVT | 1144 | Bacteria>FCB_group>Bacteroidetes/Chlorobi_group>Chlorobiota>Chlorobiia>Chlorobiales | GCA_006265245.1 | Prosthecochloris_vibrioformis | FGF68_10440 | 14740..18174 | 1098 | BREX-system-P-loop-protein-BrxC-[Prosthecochloris-vibrioformis] |
| NMW20007.1 | Type1.BREX | DUF6079*->Schlafen+HATPase+wHTH->HTH->N6-METHYLASE-p->N6-METHYLASE->Schlafen+HATPase+wHTH-p-> | 1160 | Bacteria>FCB_group>Bacteroidetes/Chlorobi_group>Chlorobiota>Chlorobiia>Chlorobiales | GCA_012972825.1 | Chlorobiaceae_bacterium | HKK66_13560 | 1..3483 | 2067991 | BREX-system-P-loop-protein-BrxC-[Chlorobiaceae-bacterium] |
| MBV5326244.1 | Type1.BREX | BrxA-DNA-binding->BrxB-BrxF-iSTAND->DUF6079*->Schlafen+HATPase+wHTH->OLD-ABC->REase-> | 1179 | Bacteria>FCB_group>Bacteroidetes/Chlorobi_group>Chlorobiota>Chlorobiia>Chlorobiales | GCA_019162945.1 | Chlorobium_sp. | JZU65_01205 | 2432..5971 | 1095 | BREX-system-P-loop-protein-BrxC-[Chlorobium-sp.] |

| Protein-ID | System | Gene-Neighbourhoods | Length | Taxonomy | Assembly-ID | Strain | Locus-Tag | Genomic-coordinates | TaxID | Fasta-defline |
|---|---|---|---|---|---|---|---|---|---|---|
| MBV5326252.1 | Type1.BREX | <-ParB+HNH<-N6-METHYLASE-p<-RVT<-Schlafen+HATPase+wHTH+HTH<-DUF6079*<-BrxB-BrxF-iSTAND<-BrxA-DNA-binding<-XIS-HTH<-?<-tRNA<-HslUClpX-AAA<-?<-?<-?<-His_Phos_1 | 1179 | Bacteria>FCB_group>Bacteroidetes/Chlorobi_group>Chlorobiota>Chlorobiia>Chlorobiales | GCA_019162945.1 | Chlorobium_sp. | JZU65_01250 | 9464..13003 | 1095 | BREX-system-P-loop-protein-BrxC-[Chlorobium-sp.] |
| MBV5329669.1 | Type1.BREX | <-N6-METHYLASE-p<-Schlafen+HATPase+wHTH<-DUF6079*<-BrxB-BrxF-iSTAND<-BrxA-DNA-binding<-XIS-HTH<-HTH_3\|\|XIS-HTH->XIS-HTH->SMS-KaiC-> | 1150 | Bacteria>FCB_group>Bacteroidetes/Chlorobi_group>Chlorobiota>Chlorobiia>Chlorobiales | GCA_019162945.1 | Chlorobium_sp. | JZU65_18935 | 2439..5891 | 1095 | BREX-system-P-loop-protein-BrxC-[Chlorobium-sp.] |
| MCF8215288.1 | Type1.BREX | HslUClpX-AAA->tRNA->MNS+MarR-HTH->?-><-METHYLASE\|\|?->XIS-HTH->BrxA-DNA-binding->BrxB-BrxF-iSTAND->DUF6079*->Schlafen+HATPase+wHTH->N6-METHYLASE->SbcC->Schlafen+HATPase+wHTH->PglZ->SIGMA+OB-Fold+AAA-ATPase+Lon-Protease-BrxL->RVT-><-Calcineurin\|\|RADICAL-SAM-> | 1145 | Bacteria>FCB_group>Bacteroidetes/Chlorobi_group>Chlorobiota>Chlorobiia>Chlorobiales | GCA_021738525.1 | Chlorobium_sp. | K9I96_00250 | 58948..62385 | 1095 | BREX-system-P-loop-protein-BrxC-[Chlorobium-sp.] |
| WP_204208384.1 | Type1.BREX | ABhydrolase->?->Aminotran_1_2-><-?\|\|DUF772+DDE_Tnp_1_2->BrxA-DNA-binding->BrxB-BrxF-iSTAND->DUF6079-p*->?->Schlafen+HATPase+wHTH+HTH->N6-METHYLASE->DUF4372+DDE_Tnp_1->N6-METHYLASE-p->ParB+HNH->RVT->PglZ->SIGMA+OB-Fold+AAA-ATPase+HKD-DNase-BrxL-> | 997 | Bacteria>FCB_group>Bacteroidetes/Chlorobi_group>Chlorobiota>Chlorobiia>Chlorobiales | GCF_001687065.1 | Prosthecochloris_sp._CIB_2401 | Ptc2401_RS05035 | 1057250..1060243 | 1868325 | BREX-system-P-loop-protein-BrxC-[Prosthecochloris-sp.-CIB-2401] |
| WP_204208337.1 | Type1.BREX | <-HTH_21+rve+rve_3<-HTH_Tnp_1\|\|DUF772+DDE_Tnp_1_2-><-SIGMA+OB-Fold+AAA-ATPase+Lon-Protease-BrxL<-PglZ<-Schlafen+HATPase+wHTH<-SbcC<-N6-METHYLASE<-Schlafen+HATPase+wHTH<-DUF6079*<-BrxB-BrxF-iSTAND<-BrxA-DNA-binding<-XIS-HTH\|\|AAA-ATPase-p->\|\|?-><-Thioredoxin | 1145 | Bacteria>FCB_group>Bacteroidetes/Chlorobi_group>Chlorobiota>Chlorobiia>Chlorobiales | GCF_001687065.1 | Prosthecochloris_sp._CIB_2401 | Ptc2401_RS06780 | 1430885..1434322 | 1868325 | BREX-system-P-loop-protein-BrxC-[Prosthecochloris-sp.-CIB-2401] |
| WP_076791024.1 | Type1.BREX | <-SIGMA+OB-Fold+AAA-ATPase+HKD-DNase-BrxL<-PglZ<-DOC+HTH<-?<-?<-Schlafen+HATPase+wHTH+HTH<-DUF6079*<-BrxB-BrxF-iSTAND<-BrxA-DNA-binding\|\|?->RelE-ParE-><-?<-Peptidase | 1179 | Bacteria>FCB_group>Bacteroidetes/Chlorobi_group>Chlorobiota>Chlorobiia>Chlorobiales | GCF_001975465.1 | Chlorobium_sp._KB01 | BOO20_RS04865 | 12313..15852 | 1917528 | BREX-system-P-loop-protein-BrxC-[Chlorobium-sp.-KB01] |
| WP_139626761.1 | Type1.BREX | <-PglZ<-ParB+HNH<-N6-METHYLASE<-N6-METHYLASE-p<-Schlafen+HATPase+wHTH+HTH<-DUF6079*<-BrxB-BrxF-iSTAND<-BrxA-DNA-binding\|\|<-Aminotran_1_2<-?<-ABhydrolase | 1204 | Bacteria>FCB_group>Bacteroidetes/Chlorobi_group>Chlorobiota>Chlorobiia>Chlorobiales | GCF_006265245.1 | Prosthecochloris_vibrioformis | FGF68_RS08505 | 9816..13430 | 1098 | BREX-system-P-loop-protein-BrxC-[Prosthecochloris-vibrioformis] |
| WP_211340437.1 | Type1.BREX | HIN-HTH->HTH_Tnp_1+HTH_21+rve+rve_3-><-GDResolvase<-SIGMA+OB-Fold+AAA-ATPase+Lon-Protease-BrxL<-PglZ<-TOPRIM<-ParB+SbcC<-N6-METHYLASE<-Schlafen+HATPase+wHTH<-DUF6079*<-BrxB-BrxF-iSTAND<-BrxA-DNA-binding<-XIS-HTH | 1145 | Bacteria>FCB_group>Bacteroidetes/Chlorobi_group>Chlorobiota>Chlorobiia>Chlorobiales | GCF_006265245.1 | Prosthecochloris_vibrioformis | FGF68_RS10435 | 14740..18177 | 1098 | BREX-system-P-loop-protein-BrxC-[Prosthecochloris-vibrioformis] |
| MBM3423463.1 | Type1.BREX | HTH+REase->BrxA-DNA-binding->BrxB-BrxF-iSTAND->DUF6079*->Schlafen+HATPase+wHTH+HTH-> | 1182 | Bacteria>FCB_group>Bacteroidetes/Chlorobi_group>Chlorobiota>unclassified_Chlorobiota | GCA_016868665.1 | Chlorobiota_bacteirum | FJY09_09520 | 5435..8983 | 2268192 | BREX-system-P-loop-protein-BrxC-[Chlorobiota-bacterium] |
| QOJ30044.1 | Type1.BREX | SF1-SF2-Helicase+PRTase-><-?<-?<-PGK<-?\|\|BrxA-DNA-binding->BrxB-BrxF-iSTAND->DUF6079*->Schlafen+HATPase+wHTH->N6-METHYLASE->PglZ->SIGMA+OB-Fold+AAA-ATPase+Lon-Protease-BrxL-><-?<-?<-ABC-ATPase+SbcC | 1198 | Bacteria>FCB_group>Bacteroidetes/Chlorobi_group>Ignavibacteriota>Ignavibacteria>Ignavibacteriales | GCA_015709675.1 | Ignavibacteriales_bacterium | HRU80_14675 | 3719207..3722803 | 2049428 | BREX-system-P-loop-protein-BrxC-[Ignavibacteriales-bacterium] |
| MBV6512081.1 | Type1.BREX | SF1-SF2-Helicase+PRTase-><-?<-?<-PGK<-?\|\|BrxA-DNA-binding->BrxB-BrxF-iSTAND->DUF6079*->Schlafen+HATPase+wHTH->N6-METHYLASE->PglZ->SIGMA+OB-Fold+AAA-ATPase+Lon-Protease-BrxL-><-?<-?<-ABC-ATPase+SbcC | 1198 | Bacteria>FCB_group>Bacteroidetes/Chlorobi_group>Ignavibacteriota>Ignavibacteria>Ignavibacteriales | GCA_019187025.1 | Ignavibacteriaceae_bacterium | FMNOHCHN_01571 | 145578..149174 | 2651163 | hypothetical-protein-[Ignavibacteriaceae-bacterium] |
| RKY59330.1 | Type1.BREX | SIGMA+OB-Fold+AAA-ATPase+Lon-Protease-BrxL->BrxA-DNA-binding->BrxB-BrxF-iSTAND->DUF6079*->N6-METHYLASE->Schlafen+HATPase+wHTH->N6-METHYLASE-> | 1192 | Bacteria>FCB_group>Candidatus_Latescibacteria>unclassified_Candidatus_Latescibacteria | GCA_003645885.1 | Candidatus_Latescibacteria_bacterium | DRP94_03915 | 4031..7609 | 2053570 | BREX-system-P-loop-protein-BrxC-[Candidatus-Latescibacteria-bacterium] |
| OWV23454.1 | Type1.BREX | DNA_pol3_beta+DNA_pol3_beta_2+DNA_pol3_beta_3->?-><-?<-?<-?<-SIGMA+OB-Fold+AAA-ATPase+Lon-Protease-BrxL<-PglZ<-Schlafen+HATPase+wHTH<-N6-METHYLASE<-DUF6079*<-BrxB-BrxF-iSTAND<-BrxA-DNA-binding<-?<-MODE-HTH+PBPII\|\|?->?->Phos_pyr_kin->RADICAL-SAM-> | 1174 | Bacteria>FCB_group>Fibrobacterota>Fibrobacteria>Fibrobacterales>Fibrobacteraceae | GCA_002210425.1 | Fibrobacter_sp._UWB2 | B7982_07645 | 365376..368900 | 1964358 | ATPase-[Fibrobacter-sp.-UWB2] |
| NLW30641.1 | Type1.BREX | BrxA-DNA-binding->BrxB-BrxF-iSTAND->DUF6079*->Schlafen+HATPase+wHTH+HTH->N6-METHYLASE->PglZ->SIGMA+OB-Fold+AAA-ATPase+Lon-Protease-BrxL->ArdCN+MPTase-> | 1213 | Bacteria>FCB_group>Fibrobacterota>Fibrobacteria>Fibrobacterales>Fibrobacteraceae | GCA_012728995.1 | Fibrobacter_sp. | GXY77_04195 | 2375..6016 | 35828 | BREX-system-P-loop-protein-BrxC-[Fibrobacter-sp.] |
| SHK42502.1 | Type1.BREX | <-VirE<-?<-XIS-HTH\|\|?->?->?->BrxA-DNA-binding->BrxB-BrxF-iSTAND->DUF6079*->ParB+HNH->N6-METHYLASE->Schlafen+HATPase+wHTH->PglZ->SIGMA+OB-Fold+AAA-ATPase+Lon-Protease-BrxL->TIR->?->REase->Phage_int_SAM_1+Phage_integrase-> | 1186 | Bacteria>FCB_group>Fibrobacterota>Fibrobacteria>Fibrobacterales>Fibrobacteraceae | GCA_900142505.1 | Fibrobacter_sp._UWH5 | SAMN05720764_101395 | 444346..447906 | 1896211 | hypothetical-protein-SAMN05720764_101395-[Fibrobacter-sp.-UWH5] |
| WP_212668349.1 | Type1.BREX | <-VirE<-?<-XIS-HTH\|\|?->?->?->BrxA-DNA-binding->BrxB-BrxF-iSTAND->DUF6079*->ParB+HNH->N6-METHYLASE->Schlafen+HATPase+wHTH->PglZ->SIGMA+OB-Fold+AAA-ATPase+Lon-Protease-BrxL->REase->REase->Phage_int_SAM_1+Phage_integrase-> | 1186 | Bacteria>FCB_group>Fibrobacterota>Fibrobacteria>Fibrobacterales>Fibrobacteraceae | GCF_002210355.1 | Fibrobacter_sp._UWH3 | B7993_RS04325 | 153736..157296 | 1964353 | BREX-system-P-loop-protein-BrxC-[Fibrobacter-sp.-UWH3] |
| WP_198953245.1 | Type1.BREX | DNA_pol3_beta+DNA_pol3_beta_2+DNA_pol3_beta_3->?-><-?<-?<-?<-SIGMA+OB-Fold+AAA-ATPase+Lon-Protease-BrxL<-PglZ<-Schlafen+HATPase+wHTH<-N6-METHYLASE<-DUF6079*<-BrxB-BrxF-iSTAND<-BrxA-DNA-binding<-?<-MODE-HTH+PBPII\|\|?->?->Phos_pyr_kin->RADICAL-SAM-> | 1176 | Bacteria>FCB_group>Fibrobacterota>Fibrobacteria>Fibrobacterales>Fibrobacteraceae | GCF_002210425.1 | Fibrobacter_sp._UWB2 | B7982_RS07630 | 365376..368906 | 1964358 | BREX-system-P-loop-protein-BrxC-[Fibrobacter-sp.-UWB2] |
| WP_073283173.1 | Type1.BREX | <-VirE<-?<-XIS-HTH\|\|?->?->?->BrxA-DNA-binding->BrxB-BrxF-iSTAND->DUF6079*->ParB+HNH->N6-METHYLASE->Schlafen+HATPase+wHTH->PglZ->SIGMA+OB-Fold+AAA-ATPase+Lon-Protease-BrxL->REase->REase->Phage_int_SAM_1+Phage_integrase-> | 1184 | Bacteria>FCB_group>Fibrobacterota>Fibrobacteria>Fibrobacterales>Fibrobacteraceae | GCF_900142465.1 | Fibrobacter_sp._UWH6 | BUB73_RS01535 | 351031..354585 | 1896212 | BREX-system-P-loop-protein-BrxC-[Fibrobacter-sp.-UWH6] |
| WP_217650997.1 | Type1.BREX | <-AbiEii<-AbiEi_4<-?\|\|?-><-MORC+REase\|\|<-PIN<-Phd-YefM\|\|BrxA-DNA-binding->BrxB-BrxF-iSTAND->DUF6079*->SbcC->SbcC->N6-METHYLASE->Schlafen+HATPase+wHTH->PglZ->SIGMA+OB-Fold+AAA-ATPase+Lon-Protease-BrxL->RNase-> | 1176 | Bacteria>FCB_group>Fibrobacterota>Fibrobacteria>Fibrobacterales>Fibrobacteraceae | GCF_900142475.1 | Fibrobacter_sp._UWH4 | BUA93_RS10220 | 85922..89452 | 1896210 | BREX-system-P-loop-protein-BrxC-[Fibrobacter-sp.-UWH4] |
| WP_073156756.1 | Type1.BREX | <-VirE<-?<-XIS-HTH\|\|?->?->?->BrxA-DNA-binding->BrxB-BrxF-iSTAND->DUF6079*->ParB+HNH->N6-METHYLASE->Schlafen+HATPase+wHTH->PglZ->SIGMA+OB-Fold+AAA-ATPase+Lon-Protease-BrxL->TIR->?->REase->Phage_int_SAM_1+Phage_integrase-> | 1184 | Bacteria>FCB_group>Fibrobacterota>Fibrobacteria>Fibrobacterales>Fibrobacteraceae | GCF_900142505.1 | Fibrobacter_sp._UWH5 | BUB38_RS01950 | 444352..447906 | 1896211 | BREX-system-P-loop-protein-BrxC-[Fibrobacter-sp.-UWH5] |
| ALF25292.1 | Type1.BREX | SKI->JAB+sGTP->wHTH+WYL+WCX->?->?->ABC-ATPase+SbcC->?-><-?<-?\|\|DUF6079*->OLD-ABC->TOPRIM->Schlafen+HATPase+wHTH->N6-METHYLASE->KAP-NTPase->HNH->SbcC->SIR2->HerA/FtsK-> | 1250 | Bacteria>Fusobacteriota>Fusobacteriia>Fusobacteriales>Fusobacteriaceae>Fusobacterium | GCA_001296185.1 | Fusobacterium_nucleatum_subsp._nucleatum | RN95_02150 | 455192..458944 | 76856 | cobalamin-adenosyltransferase-[Fusobacterium-nucleatum-subsp.-nucleatum] |
| WP_005903411.1 | Type1.BREX | <-PglZ<-ParB+HNH+RAMA<-OLD-ABC+TOPRIM<-N6-METHYLASE<-Schlafen+HATPase+wHTH<-?<-RNase+PHP<-?<-OLD-ABC<-DUF6079*\|\|?->?-><-?<-ABC-ATPase+SbcC<-?<-?<-wHTH+WYL+WCX<-JAB+sGTP<-SKI | 1250 | Bacteria>Fusobacteriota>Fusobacteriia>Fusobacteriales>Fusobacteriaceae>Fusobacterium | GCF_000178895.1 | Fusobacterium_nucleatum | C4N14_RS08075 | 1667383..1671135 | 851 | BREX-system-P-loop-protein-BrxC-[Fusobacterium-nucleatum] |
| WP_029493951.1 | Type1.BREX | <-BrxB-BrxF-iSTAND<-BrxA-DNA-binding<-SIGMA+OB-Fold+AAA-ATPase+Lon-Protease-BrxL<-PglZ<-CR-ATPase8+REase<-RVT<-N6-METHYLASE<-Schlafen+HATPase+wHTH<-DUF6079*\|\|?->?-><-?<-ABC-ATPase+SbcC<-?<-?<-wHTH+WYL+WCX<-JAB+sGTP<-SKI | 1250 | Bacteria>Fusobacteriota>Fusobacteriia>Fusobacteriales>Fusobacteriaceae>Fusobacterium | GCF_000455885.1 | Fusobacterium_hwasookii | J143_RS0110850 | 68335..72087 | 1583098 | BREX-system-P-loop-protein-BrxC-[Fusobacterium-hwasookii] |
| WP_032838665.1 | Type1.BREX | <-PglZ<-CR-ATPase8+REase<-RVT<-N6-METHYLASE<-Schlafen+HATPase+wHTH<-REase<-CR-ATPase4<-?<-N6-METHYLASE<-DUF6079*\|\|?->?-><-Peptidase_M20<-?\|\|35exo+T7DNApol-> | 1250 | Bacteria>Fusobacteriota>Fusobacteriia>Fusobacteriales>Fusobacteriaceae>Fusobacterium | GCF_000524215.1 | Fusobacterium_sp._CM1 | HMPREF1498_RS02575 | 43304..47056 | 936561 | BREX-system-P-loop-protein-BrxC-[Fusobacterium-sp.-CM1] |
| WP_032836039.1 | Type1.BREX | <-PglZ<-CR-ATPase8+REase<-HEPN<-HEPN<-MORC<-N6-METHYLASE<-Schlafen+HATPase+wHTH<-TOPRIM<-OLD-ABC<-DUF6079*\|\|?-><-?<-ABC-ATPase+SbcC<-?<-?<-wHTH+WYL+WCX<-JAB+sGTP<-SKI | 1250 | Bacteria>Fusobacteriota>Fusobacteriia>Fusobacteriales>Fusobacteriaceae>Fusobacterium | GCF_000524235.1 | Fusobacterium_sp._OBRC1 | HMPREF1501_RS01100 | 63475..67227 | 1032505 | BREX-system-P-loop-protein-BrxC-[Fusobacterium-sp.-OBRC1] |
| WP_032846252.1 | Type1.BREX | <-BrxA-DNA-binding<-SIGMA+OB-Fold+AAA-ATPase+Lon-Protease-BrxL<-PglZ<-nSTAND<-?<-N6-METHYLASE<-Schlafen+HATPase+wHTH<-TOPRIM<-OLD-ABC<-DUF6079*\|\|?->?->?-><-?<-ABC-ATPase+SbcC<-?<-?<-wHTH+WYL+WCX<-JAB+sGTP | 1250 | Bacteria>Fusobacteriota>Fusobacteriia>Fusobacteriales>Fusobacteriaceae>Fusobacterium | GCF_000524395.1 | Fusobacterium_sp._CM22 | HMPREF1500_RS09190 | 124439..128191 | 936563 | BREX-system-P-loop-protein-BrxC-[Fusobacterium-sp.-CM22] |
| WP_070766192.1 | Type1.BREX | SKI->JAB+sGTP->wHTH+WYL+WCX->?->?->ABC-ATPase+SbcC->?-><-?\|\|DUF6079*->OLD-ABC->TOPRIM->Schlafen+HATPase+wHTH->N6-METHYLASE->N6-METHYLASE->MORC+HISKIN->MORC+HISKIN->?->HNH-> | 1250 | Bacteria>Fusobacteriota>Fusobacteriia>Fusobacteriales>Fusobacteriaceae>Fusobacterium | GCF_001815715.1 | Fusobacterium_sp._HMSC064B11 | HMPREF3051_RS04300 | 251838..255590 | 1739543 | BREX-system-P-loop-protein-BrxC-[Fusobacterium-sp.-HMSC064B11] |
| WP_158411765.1 | Type1.BREX | <-35exo+T7DNApol\|\|?->Peptidase_M20-><-?<-?\|\|DUF6079*->Schlafen+HATPase+wHTH->Schlafen+HATPase+wHTH->N6-METHYLASE->KAP-NTPase->KAP-NTPase->HNH->SbcC->SIR2->HerA/FtsK-> | 1250 | Bacteria>Fusobacteriota>Fusobacteriia>Fusobacteriales>Fusobacteriaceae>Fusobacterium | GCF_002573475.1 | Fusobacterium_nucleatum | RN90_RS06260 | 1289254..1293006 | 851 | BREX-system-P-loop-protein-BrxC-[Fusobacterium-animalis] |
| WP_098997105.1 | Type1.BREX | <-NADAR<-ART-DarT<-N6-METHYLASE<-Schlafen+HATPase+wHTH<-Schlafen+HATPase+wHTH<-PHP+SbcC<-OLD-ABC<-DUF6079*\|\|?-><-?<-ABC-ATPase+SbcC<-?<-?<-wHTH+WYL+WCX<-JAB+sGTP<-SKI | 1250 | Bacteria>Fusobacteriota>Fusobacteriia>Fusobacteriales>Fusobacteriaceae>Fusobacterium | GCF_002591465.1 | Fusobacterium_nucleatum | CBG56_RS03320 | 143954..147706 | 851 | MULTISPECIES:-BREX-system-P-loop-protein-BrxC-[Fusobacterium] |
| WP_099989146.1 | Type1.BREX | <-BrxA-DNA-binding<-SIGMA+OB-Fold+AAA-ATPase+Lon-Protease-BrxL<-PglZ<-Peptidase_S8<-Classical-AAA<-CR-ATPase8+REase<-RVT<-N6-METHYLASE<-Schlafen+HATPase+wHTH<-DUF6079*\|\|?-><-?<-?<-Peptidase_M3_N+Peptidase_M3\|\|RNaseH->?->?->?->Peptidase_M20-> | 1250 | Bacteria>Fusobacteriota>Fusobacteriia>Fusobacteriales>Fusobacteriaceae>Fusobacterium | GCF_002763735.1 | Fusobacterium_pseudoperiodonticum | CTM78_RS04805 | 952520..956272 | 2663009 | BREX-system-P-loop-protein-BrxC-[Fusobacterium-pseudoperiodonticum] |
| WP_220302801.1 | Type1.BREX | SKI->JAB+sGTP->wHTH+WYL+WCX->?->?->ABC-ATPase+SbcC->?-><-?<-?\|\|DUF6079*->OLD-ABC->TOPRIM->Schlafen+HATPase+wHTH->N6-METHYLASE->KAP-NTPase->HNH->SbcC->SIR2->HerA/FtsK-> | 1250 | Bacteria>Fusobacteriota>Fusobacteriia>Fusobacteriales>Fusobacteriaceae>Fusobacterium | GCF_019552045.1 | Fusobacterium_nucleatum | JY397_RS08300 | 1731485..1735237 | 851 | BREX-system-P-loop-protein-BrxC-[Fusobacterium-polymorphum] |
| WP_220282655.1 | Type1.BREX | <-BrxB-BrxF-iSTAND<-BrxA-DNA-binding<-SIGMA+OB-Fold+AAA-ATPase+Lon-Protease-BrxL<-PglZ<-RVT<-N6-METHYLASE<-Schlafen+HATPase+wHTH<-DUF6079*\|\|?->?->?-><-?<-ABC-ATPase+SbcC<-?<-?<-wHTH+WYL+WCX<-JAB+sGTP | 1250 | Bacteria>Fusobacteriota>Fusobacteriia>Fusobacteriales>Fusobacteriaceae>Fusobacterium | GCF_019552065.1 | Fusobacterium_nucleatum | JY402_RS06730 | 1356729..1360481 | 851 | BREX-system-P-loop-protein-BrxC-[Fusobacterium-polymorphum] |
| WP_220297607.1 | Type1.BREX | <-PAS+NtrC-AAA+FIS-HTH<-BrxB-BrxF-iSTAND<-BrxA-DNA-binding<-SIGMA+OB-Fold+AAA-ATPase+Lon-Protease-BrxL<-PglZ<-RVT<-N6-METHYLASE<-Schlafen+HATPase+wHTH<-Schlafen+HATPase+wHTH<-DUF6079*\|\|?->?-><-Peptidase_M20<-?\|\|35exo+T7DNApol-> | 1250 | Bacteria>Fusobacteriota>Fusobacteriia>Fusobacteriales>Fusobacteriaceae>Fusobacterium | GCF_019552085.1 | Fusobacterium_nucleatum | JY398_RS09460 | 1985091..1988843 | 851 | BREX-system-P-loop-protein-BrxC-[Fusobacterium-animalis] |
| WP_238959505.1 | Type1.BREX | <-BrxB-BrxF-iSTAND<-BrxA-DNA-binding<-SIGMA+OB-Fold+AAA-ATPase+Lon-Protease-BrxL<-PglZ<-CR-ATPase8+REase<-HEPN<-MORC<-N6-METHYLASE<-Schlafen+HATPase+wHTH<-DUF6079*\|\|?-><-?<-ABC-ATPase+SbcC<-?<-?<-wHTH+WYL+WCX<-JAB+sGTP<-SKI | 1250 | Bacteria>Fusobacteriota>Fusobacteriia>Fusobacteriales>Fusobacteriaceae>Fusobacterium | GCF_022340045.1 | Fusobacterium_nucleatum | K5I20_00280 | 65514..69266 | 851 | BREX-system-P-loop-protein-BrxC-[Fusobacterium-nucleatum] |
| WP_238972693.1 | Type1.BREX | SKI->JAB+sGTP->wHTH+WYL+WCX->?->?->ABC-ATPase+SbcC->?-><-?<-?\|\|DUF6079*->OLD-ABC->TOPRIM->Schlafen+HATPase+wHTH->N6-METHYLASE->MORC->HEPN->CR-ATPase8+REase->PglZ->SIGMA+OB-Fold+AAA-ATPase+Lon-Protease-BrxL-> | 1250 | Bacteria>Fusobacteriota>Fusobacteriia>Fusobacteriales>Fusobacteriaceae>Fusobacterium | GCF_022340165.1 | Fusobacterium_nucleatum | K5I19_RS02505 | 127932..131684 | 851 | BREX-system-P-loop-protein-BrxC-[Fusobacterium-nucleatum] |
| WP_271333596.1 | Type1.BREX | <-PglZ<-CR-ATPase8+REase<-?<-N6-METHYLASE<-Schlafen+HATPase+wHTH<-REase<-CR-ATPase4<-?<-N6-METHYLASE<-DUF6079*\|\|?-><-?<-ABC-ATPase+SbcC<-?<-?<-wHTH+WYL+WCX<-JAB+sGTP<-SKI | 1250 | Bacteria>Fusobacteriota>Fusobacteriia>Fusobacteriales>Fusobacteriaceae>Fusobacterium | GCF_027947475.1 | Fusobacterium_nucleatum | PGW91_RS03610 | 747072..750824 | 851 | BREX-system-P-loop-protein-BrxC-[Fusobacterium-nucleatum] |
| WP_274179672.1 | Type1.BREX | <-KAP-NTPase<-N6-METHYLASE<-Schlafen+HATPase+wHTH<-Schlafen+HATPase+wHTH<-Schlafen+HATPase+wHTH<-REase<-CR-ATPase4<-?<-N6-METHYLASE<-DUF6079*\|\|?-><-?<-ABC-ATPase+SbcC<-?<-?<-wHTH+WYL+WCX<-JAB+sGTP<-SKI | 1250 | Bacteria>Fusobacteriota>Fusobacteriia>Fusobacteriales>Fusobacteriaceae>Fusobacterium | GCF_028735995.1 | Fusobacterium_nucleatum | PSC67_RS07260 | 1410490..1414242 | 851 | BREX-system-P-loop-protein-BrxC-[Fusobacterium-nucleatum] |
| WP_008800858.1 | Type1.BREX | PrmA+METHYLASE->?->?->METHYLASE->?->DUF6079*->Schlafen+HATPase+wHTH->N6-METHYLASE->ParB+HNH+RAMA->PglZ->BrxA-DNA-binding->BrxB-BrxF-iSTAND-> | 1240 | Bacteria>Fusobacteriota>Fusobacteriia>Fusobacteriales>Fusobacteriaceae>Fusobacterium | GCF_902373895.1 | Fusobacterium_gonidiaformans | FY288_RS06315 | 242664..246386 | 849 | BREX-system-P-loop-protein-BrxC-[Fusobacterium-gonidiaformans] |
| WP_008793539.1 | Type1.BREX | <-RNaseH\|\|Peptidase_M3_N+Peptidase_M3->?-><-?<-HTH_OrfB_IS605+OrfB_IS605+OrfB_Zn_ribbon\|\|DUF6079*->OLD-ABC->RNase+PHP->Schlafen+HATPase+wHTH->N6-METHYLASE->HEPN->CR-ATPase8+REase->PglZ->SIGMA+OB-Fold+AAA-ATPase+Lon-Protease-BrxL->BrxA-DNA-binding-> | 1250 | Bacteria>Fusobacteriota>Fusobacteriia>Fusobacteriales>Fusobacteriaceae>Fusobacterium | GCF_902373915.1 | Fusobacterium_periodonticum | FY302_RS02975 | 615184..618936 | 860 | BREX-system-P-loop-protein-BrxC-[Fusobacterium-periodonticum] |
| HCF56393.1 | Type6.BREX | <-Schlafen+HATPase+wHTH<-N6-METHYLASE-p\|\|?->PIN-><-TOPRIM<-OLD-ABC<-DUF6079-p<-APATPase<-DUF6079-p<-DUF6079-p*<-HNH<-BrxB-BrxF-iSTAND<-BrxA-DNA-binding<-BrxA-DNA-binding<-BrxE | 508 | Bacteria>Myxococcota>Myxococcia>Myxococcales>unclassified_Myxococcales | GCA_003534605.1 | Myxococcales_bacterium | DFS52_00150 | 7726..9251 | 2026763 | BREX-system-P-loop-protein-BrxC-[Myxococcales-bacterium] |
| HCF56394.1 | Type6.BREX | <-Schlafen+HATPase+wHTH<-N6-METHYLASE-p\|\|?->PIN-><-TOPRIM<-OLD-ABC<-DUF6079-p<-APATPase<-DUF6079-p<-DUF6079-p*<-HNH<-BrxB-BrxF-iSTAND<-BrxA-DNA-binding<-BrxA-DNA-binding<-BrxE | 363 | Bacteria>Myxococcota>Myxococcia>Myxococcales>unclassified_Myxococcales | GCA_003534605.1 | Myxococcales_bacterium | DFS52_00155 | 9470..10559 | 2026763 | BREX-system-P-loop-protein-BrxC-[Myxococcales-bacterium] |
| WP_020733584.1 | Type6.BREX | ABC-ATPase+SbcC-><-SIGMA+OB-Fold+AAA-ATPase+Lon-Protease-BrxL<-PglZ<-HNH<-OLD-ABC<-N6-METHYLASE<-N6-METHYLASE<-Schlafen+HATPase+wHTH<-DUF6079*<-BrxB-BrxF-iSTAND<-BrxA-DNA-binding<-BrxE<-XIS-HTH\|\|?->?->ACET-> | 1182 | Bacteria>Myxococcota>Polyangia>Polyangiales>Polyangiaceae>Sorangium | GCF_000418325.1 | Sorangium_cellulosum | SCE1572_RS08150 | 2251441..2254989 | 56 | BREX-system-P-loop-protein-BrxC-[Sorangium-cellulosum] |
| WP_129354457.1 | Type6.BREX | <-Glyco_hydro_15<-?\|\|?-><-SIGMA+OB-Fold+AAA-ATPase+Lon-Protease-BrxL<-PglZ<-N6-METHYLASE<-Schlafen+HATPase+wHTH<-DUF6079*<-BrxB-BrxF-iSTAND<-BrxA-DNA-binding<-BrxE<-REC+FIS-HTH<-HISKIN\|\|Methyltransf_33->FGS-><-SAM-METHYLASE | 1180 | Bacteria>Myxococcota>Polyangia>Polyangiales>Polyangiaceae>Sorangium | GCF_004135735.1 | Sorangium_cellulosum | SOCEGT47_RS35865 | 9457770..9461312 | 56 | BREX-system-P-loop-protein-BrxC-[Sorangium-cellulosum] |
| MCL2723928.1 | Type6.BREX | Glyoxalase->?->?-><-SIGMA+OB-Fold+AAA-ATPase+Lon-Protease-BrxL<-PglZ<-Schlafen+HATPase+wHTH<-SbcC<-N6-METHYLASE<-N6-METHYLASE<-DUF6079*<-BrxB-BrxF-iSTAND<-BrxA-DNA-binding<-BrxE<-?<-?\|\|?-><-?<-?\|\|Pkinase+Pkinase-> | 1179 | Bacteria>Myxococcota>Polyangia>Polyangiales>Polyangiaceae>unclassified_Polyangiaceae | GCA_009781005.1 | Polyangiaceae_bacterium | FWD69_05765 | 18977..22516 | 2268199 | BREX-system-P-loop-protein-BrxC-[Polyangiaceae-bacterium] |
| MCB9617769.1 | Type1.BREX | RAMA->GDResolvase+ZnR-><-SIGMA+OB-Fold+AAA-ATPase+Lon-Protease-BrxL<-PglZ<-Schlafen+HATPase+wHTH<-N6-METHYLASE<-PIN<-DUF6079*<-BrxB-BrxF-iSTAND<-BrxA-DNA-binding<-tRNA\|\|?-><-ADH_N+ADH_zinc_N+METHYLASE | 1156 | Bacteria>Myxococcota>Polyangia>Polyangiales>Sandaracinaceae>Sandaracinus | GCA_020633155.1 | Sandaracinus_sp. | H6722_35485 | 14055..17525 | 2024858 | BREX-system-P-loop-protein-BrxC-[Sandaracinus-sp.] |
| MCK6514475.1 | Type6.BREX | <-SF1-SF2-Helicase<-HamA<-SIGMA+OB-Fold+AAA-ATPase+Lon-Protease-BrxL<-PglZ<-OLD-ABC<-REase<-N6-METHYLASE-p<-N6-METHYLASE<-Schlafen+HATPase+wHTH<-DUF6079*<-DUF6079-p<-BrxB-BrxF-iSTAND<-BrxA-DNA-binding<-BrxE<-DNA-glycosylase\|\|RADICAL-SAM-> | 1083 | Bacteria>Myxococcota>unclassified_Myxococcota | GCA_023150775.1 | Myxococcota_bacterium | L6R46_05405 | 17071..20323 | 2818507 | BREX-system-P-loop-protein-BrxC-[Myxococcota-bacterium] |
| MBI5192441.1 | Type1.BREX | <-MORC+HISKIN<-TPR+TPR+TPR+TPR<-REC+NtrC-AAA+FIS-HTH<-SIGMA+OB-Fold+AAA-ATPase+Lon-Protease-BrxL<-PglZ<-Schlafen+HATPase+wHTH<-PHP+SbcC<-?<-Schlafen+HATPase+wHTH<-DUF6079*<-BrxB-BrxF-iSTAND<-BrxA-DNA-binding\|\|HTH_8->wHTH+WYL-> | 1181 | Bacteria>Nitrospirota>unclassified_Nitrospirota | GCA_016212215.1 | Nitrospirota_bacterium | HZA08_03235 | 28914..32459 | 2026887 | BREX-system-P-loop-protein-BrxC-[Nitrospirota-bacterium] |
| MCC6543788.1 | Type1.BREX | <-N6-METHYLASE-p<-Schlafen+HATPase+wHTH<-REase<-SIGMA+OB-Fold+AAA-ATPase+Lon-Protease-BrxL<-PglZ<-Schlafen+HATPase+wHTH<-N6-METHYLASE<-DUF6079*<-BrxB-BrxF-iSTAND<-BrxA-DNA-binding<-XIS-HTH | 1181 | Bacteria>Nitrospirota>unclassified_Nitrospirota | GCA_020846775.1 | Nitrospirota_bacterium | IT392_04715 | 12437..15982 | 2026887 | BREX-system-P-loop-protein-BrxC-[Nitrospirota-bacterium] |
| WP_206385353.1 | Type1.BREX | Integrase_1-><-SbcC<-Calcineurin+Calcineurin-SbcD\|\|BrxA-DNA-binding->BrxB-BrxF-iSTAND->DUF6079*->N6-METHYLASE->Schlafen+HATPase+wHTH->SIR2->HerA/FtsK-> | 1179 | Bacteria>Pseudomonadota>Acidithiobacillia>Acidithiobacillales>Acidithiobacillaceae>Acidithiobacillus | - | Acidithiobacillus_sp._MC6.1 | JKG47_07195 | 11507..15046 | 2801580 | BREX-system-P-loop-protein-BrxC-[Acidithiobacillus-sp.-MC6.1] |
| WP_172437661.1 | Type1.BREX | DEDD_Tnp_IS110+Transposase_20-><-?\|\|XIS-HTH->TOPRIM->\|\|XIS-HTH->BrxA-DNA-binding->BrxB-BrxF-iSTAND->DUF6079*->Schlafen+HATPase+wHTH+HTH->N6-METHYLASE->HTH+REase->PglZ->SIGMA+OB-Fold+AAA-ATPase+HKD-DNase-BrxL->SbcC+APATPase+SbcC->Calcineurin+Calcineurin-SbcD->SbcC->RNase-> | 1203 | Bacteria>Pseudomonadota>Acidithiobacillia>Acidithiobacillales>Acidithiobacillaceae>Acidithiobacillus | - | Acidithiobacillus_sp._SH | HF670_09010 | 8231..11842 | 187490 | MULTISPECIES:-BREX-system-P-loop-protein-BrxC-[Acidithiobacillus] |
| MBU2839699.1 | Type1.BREX | DEDD_Tnp_IS110+Transposase_20-><-?\|\|XIS-HTH->TOPRIM->\|\|XIS-HTH->BrxA-DNA-binding->BrxB-BrxF-iSTAND->DUF6079*->Schlafen+HATPase+wHTH+HTH->N6-METHYLASE->HTH+REase->PglZ->SIGMA+OB-Fold+AAA-ATPase+HKD-DNase-BrxL->SbcC+APATPase+SbcC->Calcineurin+Calcineurin-SbcD->SbcC->RNase-> | 1203 | Bacteria>Pseudomonadota>Acidithiobacillia>Acidithiobacillales>Acidithiobacillaceae>Acidithiobacillus | GCA_018854655.1 | Acidithiobacillus_thiooxidans | HF670_09010 | 8231..11842 | 930 | BREX-system-P-loop-protein-BrxC-[Acidithiobacillus-thiooxidans] |
| WP_014029717.1 | Type1.BREX | DEDD_Tnp_IS110+Transposase_20-><-DDE_5<-?\|\|HTH_17->Phage_int_SAM_2+Integrase_1-><-SbcC<-Calcineurin+Calcineurin-SbcD\|\|BrxA-DNA-binding->BrxB-BrxF-iSTAND->DUF6079*->N6-METHYLASE->Schlafen+HATPase+wHTH->N6-METHYLASE-p->OLD-ABC->TOPRIM->PglZ->SIGMA+OB-Fold+AAA-ATPase+HKD-DNase-BrxL-> | 1179 | Bacteria>Pseudomonadota>Acidithiobacillia>Acidithiobacillales>Acidithiobacillaceae>Acidithiobacillus | GCF_000214095.2 | Acidithiobacillus_ferrivorans | ACIFE_RS11315 | 2362677..2366216 | 160808 | BREX-system-P-loop-protein-BrxC-[Acidithiobacillus-ferrivorans] |
| WP_054609107.1 | Type1.BREX | Arm-DNA-bind_3+Phage_integrase-><-AAA-ATPase<-rve\|\|RVT-><-PglZ<-N6-METHYLASE-p<-N6-METHYLASE-p<-Schlafen+HATPase+wHTH<-N6-METHYLASE-p<-DUF6079*<-BrxB-BrxF-iSTAND<-BrxA-DNA-binding\|\|Calcineurin+Calcineurin-SbcD->SbcC-><-Phage_int_SAM_2+Integrase_1\|\|?->?-><-?\|\|REase+wHTH-> | 1180 | Bacteria>Pseudomonadota>Acidithiobacillia>Acidithiobacillales>Acidithiobacillaceae>Acidithiobacillus | GCF_020887015.1 | Acidithiobacillus_ferrooxidans | EC577_RS04340 | 17685..21227 | 920 | BREX-system-P-loop-protein-BrxC-[Acidithiobacillus-ferrooxidans] |
| WP_271780036.1 | Type1.BREX | HTH_17->Phage_int_SAM_2+Integrase_1-><-SbcC<-Calcineurin+Calcineurin-SbcD\|\|BrxA-DNA-binding->BrxB-BrxF-iSTAND->DUF6079*->N6-METHYLASE->Schlafen+HATPase+wHTH->SIR2->HerA/FtsK->N6-METHYLASE-p->PglZ->SIGMA+OB-Fold+AAA-ATPase+HKD-DNase-BrxL-> | 1179 | Bacteria>Pseudomonadota>Acidithiobacillia>Acidithiobacillales>Acidithiobacillaceae>Acidithiobacillus | GCF_028215975.1 | Acidithiobacillus_ferriphilus | PJU76_RS00595 | 119550..123089 | 1689834 | BREX-system-P-loop-protein-BrxC-[Acidithiobacillus-ferriphilus] |
| WP_101538034.1 | Type1.BREX | DEDD_Tnp_IS110+Transposase_20-><-?\|\|XIS-HTH->TOPRIM->\|\|XIS-HTH->BrxA-DNA-binding->BrxB-BrxF-iSTAND->DUF6079*->Schlafen+HATPase+wHTH+HTH->N6-METHYLASE->HTH+REase->PglZ->SIGMA+OB-Fold+AAA-ATPase+HKD-DNase-BrxL->SbcC+APATPase+SbcC->Calcineurin+Calcineurin-SbcD->SbcC->RNase-> | 1204 | Bacteria>Pseudomonadota>Acidithiobacillia>Acidithiobacillales>Acidithiobacillaceae | GCF_018854655.1 | Acidithiobacillus | HF670_RS09000 | 8228..11842 | 119977 | MULTISPECIES:-BREX-system-P-loop-protein-BrxC-[Acidithiobacillus] |
| WP_225161486.1 | Type1.BREX | <-ParB+HNH+Schlafen<-?<-SIGMA+OB-Fold+AAA-ATPase+HKD-DNase-BrxL<-PglZ<-N6-METHYLASE<-DUF6079*<-BrxB-BrxF-iSTAND<-BrxA-DNA-binding<-wHTH+WYL+WCX\|\|?->35Exo+REase->SF1-SF2-Helicase+REase->35exo+SF1-SF2-Helicase-><-Phage_integrase | 1183 | Bacteria>Pseudomonadota>Alphaproteobacteria>Hyphomicrobiales>Nitrobacteraceae>Bradyrhizobium | GCF_020194035.1 | Bradyrhizobium_sp._BRP56 | I6F38_19500 | 73459..77010 | 2793819 | BREX-system-P-loop-protein-BrxC-[Bradyrhizobium-sp.-BRP56] |
| WP_260373497.1 | Type1.BREX | <-ParB+HNH+Schlafen<-?<-SIGMA+OB-Fold+AAA-ATPase+HKD-DNase-BrxL<-PglZ<-N6-METHYLASE<-DUF6079*<-BrxB-BrxF-iSTAND<-BrxA-DNA-binding<-wHTH+WYL+WCX\|\|?->35Exo+REase->SF1-SF2-Helicase+REase->35exo+SF1-SF2-Helicase-><-Phage_integrase | 1183 | Bacteria>Pseudomonadota>Alphaproteobacteria>Hyphomicrobiales>Nitrobacteraceae>Bradyrhizobium | GCF_025200925.1 | Bradyrhizobium_sp._CB1015 | N2604_RS01610 | 347161..350712 | 2976822 | BREX-system-P-loop-protein-BrxC-[Bradyrhizobium-sp.-CB1015] |
| MBY5873817.1 | Type1.BREX | DnaB+McrB-AAA->wHTH+REase+ZnR->?->OLD-ABC+TOPRIM->SF1-SF2-Helicase->ParB+HNH+Schlafen-><-SIGMA+OB-Fold+AAA-ATPase+HKD-DNase-BrxL<-PglZ<-N6-METHYLASE<-DUF6079*<-BrxB-BrxF-iSTAND<-BrxA-DNA-binding\|\|ParB->REase+ZnR->?->Classical-AAA->Classical-AAA->REase-> | 1174 | Bacteria>Pseudomonadota>Alphaproteobacteria>Hyphomicrobiales>Rhizobiaceae>Rhizobium/Agrobacterium_group | GCA_019799745.1 | Rhizobium_leguminosarum | HFN53_17500 | 401377..404901 | 384 | BREX-system-P-loop-protein-BrxC-[Rhizobium-leguminosarum] |
| MBF0098005.1 | Type1.BREX | <-PglZ<-Schlafen+HATPase+wHTH<-PHP+SbcC<-HNH<-OLD-ABC<-REase+AIPR<-N6-METHYLASE<-HicA_toxin<-HicB_lk_antitox<-DUF6079*<-BrxB-BrxF-iSTAND<-BrxA-DNA-binding<-Radical_SAM | 1144 | Bacteria>Pseudomonadota>Alphaproteobacteria>Magnetococcales>unclassified_Magnetococcales | GCA_015228825.1 | Magnetococcales_bacterium | HQM05_11780 | 14372..17806 | 2026759 | BREX-system-P-loop-protein-BrxC-[Magnetococcales-bacterium] |
| WP_274653192.1 | Type1.BREX | <-ParB+HNH+Schlafen<-?<-SIGMA+OB-Fold+AAA-ATPase+HKD-DNase-BrxL<-PglZ<-N6-METHYLASE<-N6-METHYLASE<-NERD<-OLD-ABC<-DUF6079*<-BrxB-BrxF-iSTAND<-BrxA-DNA-binding<-wHTH+WYL+WCX\|\|ArdCN+MPTase->?->?->FurR-HTH->?->HIN-HTH+TraJ-RHH-> | 1172 | Bacteria>Pseudomonadota>Alphaproteobacteria>Rhodobacterales>Paracoccaceae>Halocynthiibacter | GCF_028823805.1 | Halocynthiibacter_sp._C4 | OU789_RS03095 | 642215..645733 | 2992758 | BREX-system-P-loop-protein-BrxC-[Halocynthiibacter-sp.-C4] |
| WP_108641830.1 | Type1.BREX | <-Schlafen<-?<-ParB+HNH+Schlafen<-?<-SIGMA+OB-Fold+AAA-ATPase+HKD-DNase-BrxL<-PglZ<-N6-METHYLASE<-DUF6079*<-BrxB-BrxF-iSTAND<-BrxA-DNA-binding<-wHTH+WYL+WCX\|\|?->Thermonuclease->TrbM-><-?<-HTH_3 | 1173 | Bacteria>Pseudomonadota>Alphaproteobacteria>Rhodobacterales>Paracoccaceae>Thalassorhabdomicrobium | GCF_003072065.1 | Thalassorhabdomicrobium_marinisediminis | DC363_RS14250 | 78584..82105 | 2170577 | BREX-system-P-loop-protein-BrxC-[Thalassorhabdomicrobium-marinisediminis] |
| WP_058289795.1 | Type1.BREX | <-TraI-VirD2-RCRE<-HIN-HTH+TraJ-RHH<-?<-FurR-HTH<-?<-ArdCN+MPTase\|\|wHTH+WYL+WCX->BrxA-DNA-binding->BrxB-BrxF-iSTAND->DUF6079*->OLD-ABC->NERD->N6-METHYLASE->PglZ->SIGMA+OB-Fold+AAA-ATPase+HKD-DNase-BrxL->ParB+HNH+Schlafen-><-?<-Couple_hipA+HipA_C<-HTH_3 | 1172 | Bacteria>Pseudomonadota>Alphaproteobacteria>Rhodobacterales>Paracoccaceae>Tritonibacter | GCF_001458415.1 | Tritonibacter_multivorans | PGB32_RS12460 | 15411..18929 | 928856 | BREX-system-P-loop-protein-BrxC-[Tritonibacter-multivorans] |
| WP_123792783.1 | Type1.BREX | <-ParB+HNH+Schlafen<-SIGMA+OB-Fold+AAA-ATPase+HKD-DNase-BrxL<-PglZ<-N6-METHYLASE<-REase<-OLD-ABC<-N6-METHYLASE<-DUF6079*<-BrxB-BrxF-iSTAND<-BrxA-DNA-binding<-HIN-HTH<-HTH_36<-?<-?<-Arm-DNA-bind_3+Phage_integrase | 1172 | Bacteria>Pseudomonadota>Alphaproteobacteria>Rhodobacterales>Roseobacteraceae>Pacificibacter | GCF_003814915.1 | Pacificibacter_maritimus | EDD53_RS08420 | 394668..398186 | 762213 | BREX-system-P-loop-protein-BrxC-[Pacificibacter-maritimus] |
| TNE62528.1 | Type1.BREX | REase+wHTH->rve->TnsC-ATPase->LON-AAA->?-><-Schlafen+REase<-SIGMA+OB-Fold+AAA-ATPase+HKD-DNase-BrxL<-PglZ<-N6-METHYLASE<-DUF6079*<-BrxB-BrxF-iSTAND<-BrxA-DNA-binding<-wHTH+WYL+WCX\|\|REase+wHTH->HTH_28+rve+Mu-transpos_C-> | 1175 | Bacteria>Pseudomonadota>Alphaproteobacteria>Sphingomonadales>unclassified_Sphingomonadales | GCA_006226385.1 | Sphingomonadales_bacterium | EP341_00225 | 33441..36968 | 1978525 | MAG:-BREX-system-P-loop-protein-BrxC-[Sphingomonadales-bacterium] |
| MBQ2886288.1 | Type1.BREX | <-wHTH+WYL<-REase<-SIGMA+OB-Fold+AAA-ATPase+Lon-Protease-BrxL<-PglZ<-Schlafen+HATPase+wHTH<-N6-METHYLASE<-PIN<-DUF6079*<-BrxB-BrxF-iSTAND<-BrxA-DNA-binding<-HTH_3 | 1180 | Bacteria>Pseudomonadota>Alphaproteobacteria>unclassified_Alphaproteobacteria | GCA_017409945.1 | Alphaproteobacteria_bacterium | IJE43_21410 | 13139..16681 | 1913988 | BREX-system-P-loop-protein-BrxC-[Alphaproteobacteria-bacterium] |
| WP_200235209.1 | Type1.BREX | GDResolvase+HIN-HTH->?->?-><-SIGMA+OB-Fold+AAA-ATPase+HKD-DNase-BrxL<-PglZ<-Schlafen+HATPase+wHTH<-N6-METHYLASE-p<-RVT<-N6-METHYLASE<-DUF6079*<-BrxB-BrxF-iSTAND<-BrxA-DNA-binding<-XIS-HTH<-wHTH+HNH-p\|\|N6-METHYLASE->?->LexA-protease->?->ParB+HNH-> | 1190 | Bacteria>Pseudomonadota>Betaproteobacteria>Burkholderiales>Alcaligenaceae>Advenella | GCF_016595155.1 | Advenella_mandrilli | JHL22_RS06635 | 201531..205103 | 2800330 | BREX-system-P-loop-protein-BrxC-[Advenella-mandrilli] |
| WP_121740016.1 | Type1.BREX | Phage_integrase-><-RHH_1\|\|DDC+HTH->HTH_36->DUF772+DDE_Tnp_1_6-><-CHTH\|\|XIS-HTH->BrxA-DNA-binding->BrxB-BrxF-iSTAND->DUF6079*->N6-METHYLASE->?->RVT->N6-METHYLASE-p->Schlafen+HATPase+wHTH->PglZ->SIGMA+OB-Fold+AAA-ATPase+HKD-DNase-BrxL->REase-> | 1190 | Bacteria>Pseudomonadota>Betaproteobacteria>Burkholderiales>Alcaligenaceae>Alcaligenes | GCF_003671915.1 | Alcaligenes_aquatilis | D3M96_RS02075 | 458513..462085 | 323284 | BREX-system-P-loop-protein-BrxC-[Alcaligenes-aquatilis] |
| WP_066458434.1 | Type1.BREX | XIS-HTH->BrxA-DNA-binding->BrxB-BrxF-iSTAND->DUF6079*->Schlafen+HATPase+wHTH+HTH->N6-METHYLASE->OLD-ABC-> | 1196 | Bacteria>Pseudomonadota>Betaproteobacteria>Burkholderiales>Alcaligenaceae>Castellaniella | GCF_001592225.1 | Castellaniella_caeni | CUB94_RS15500 | 2114..5704 | 266123 | BREX-system-P-loop-protein-BrxC-[Castellaniella-caeni] |
| WP_166406662.1 | Type1.BREX | Integrase_1-><-?\|\|SbcC->GDResolvase+HIN-HTH-><-SIGMA+OB-Fold+AAA-ATPase+HKD-DNase-BrxL<-PglZ<-Schlafen+HATPase+wHTH<-N6-METHYLASE<-DUF6079*<-BrxB-BrxF-iSTAND<-BrxA-DNA-binding<-XIS-HTH\|\|Arm-DNA-bind_3+Phage_integrase+Phage_integr_3->?->?->?-><-?<-REase+SF1-SF2-Helicase | 1190 | Bacteria>Pseudomonadota>Betaproteobacteria>Burkholderiales>Alcaligenaceae>Paenalcaligenes | GCF_011290505.1 | Paenalcaligenes_suwonensis | GY634_RS00685 | 163356..166928 | 1202713 | BREX-system-P-loop-protein-BrxC-[Paenalcaligenes-suwonensis] |
| NYT60502.1 | Type1.BREX | <-SIGMA+OB-Fold+AAA-ATPase+HKD-DNase-BrxL<-PglZ<-N6-METHYLASE-p<-Schlafen+HATPase+wHTH<-N6-METHYLASE<-DUF6079*<-BrxB-BrxF-iSTAND<-BrxA-DNA-binding<-XIS-HTH\|\|?-><-tRNA\|\|?-><-RelE-ParE<-CHTH<-?<-RRXRR+HNH | 1188 | Bacteria>Pseudomonadota>Betaproteobacteria>Burkholderiales>Alcaligenaceae>unclassified_Alcaligenaceae | GCA_013416405.1 | Alcaligenaceae_bacterium | HOA65_16390 | 10015..13581 | 2021368 | BREX-system-P-loop-protein-BrxC-[Alcaligenaceae-bacterium] |
| NYT60768.1 | Type1.BREX | <-Phage_AlpA\|\|YdaS_antitoxin-><-SIGMA+OB-Fold+AAA-ATPase+Lon-Protease-BrxL<-PglZ<-Schlafen+HATPase+wHTH<-SbcC<-TOPRIM<-OLD-ABC<-N6-METHYLASE<-DUF6079*<-BrxB-BrxF-iSTAND<-BrxA-DNA-binding<-wHTH+WYL+WCX\|\|SbcC-> | 1146 | Bacteria>Pseudomonadota>Betaproteobacteria>Burkholderiales>Alcaligenaceae>unclassified_Alcaligenaceae | GCA_013416445.1 | Alcaligenaceae_bacterium | HOA66_00285 | 70051..73491 | 2021368 | BREX-system-P-loop-protein-BrxC-[Alcaligenaceae-bacterium] |
| MBK8767853.1 | Type1.BREX | <-AAA-ATPase<-rve\|\|?-><-Preader17<-SAM-METHYLASE<-?<-CHTH\|\|BrxA-DNA-binding->BrxB-BrxF-iSTAND->DUF6079*->N6-METHYLASE->Schlafen+HATPase+wHTH->Schlafen+HATPase+wHTH->PglZ->PglZ->SIGMA+OB-Fold+AAA-ATPase+Lon-Protease-BrxL-> | 1148 | Bacteria>Pseudomonadota>Betaproteobacteria>Burkholderiales>Burkholderiaceae>unclassified_Burkholderiaceae | GCA_016710525.1 | Burkholderiaceae_bacterium | IPM01_25890 | 21329..24775 | 2030806 | BREX-system-P-loop-protein-BrxC-[Burkholderiaceae-bacterium] |
| OJX16153.1 | Type1.BREX | <-SIGMA+OB-Fold+AAA-ATPase+HKD-DNase-BrxL<-PglZ<-Schlafen+HATPase+wHTH<-N6-METHYLASE-p<-?<-REase<-OLD-ABC<-N6-METHYLASE<-?<-DUF6079*<-BrxB-BrxF-iSTAND<-BrxA-DNA-binding<-Phage_int_SAM_2+Integrase_1<-Phage_rep\|\|CNMP-><-PRTase\|\|CBS+CBS->HAD+HAD-> | 1181 | Bacteria>Pseudomonadota>Betaproteobacteria>Burkholderiales>Comamonadaceae>Delftia | GCA_001898745.1 | Delftia_sp._67-8 | BGO79_00315 | 75959..79504 | 1895749 | hypothetical-protein-BGO79_00315-[Delftia-sp.-67-8] |
| WP_108373352.1 | Type1.BREX | <-TOPRIM<-OLD-ABC<-?<-TOPRIM<-OLD-ABC<-TOPRIM<-OLD-ABC<-N6-METHYLASE<-Schlafen+HATPase+wHTH<-DUF6079*<-BrxB-BrxF-iSTAND<-BrxA-DNA-binding<-wHTH+WYL+WCX\|\|?->?->?-><-?<-?<-SPOUT | 1209 | Bacteria>Pseudomonadota>Betaproteobacteria>Burkholderiales>Comamonadaceae>Limnohabitans | GCF_003063315.1 | Limnohabitans_sp._T6-5 | B9Z51_RS07570 | 412563..416192 | 1100724 | BREX-system-P-loop-protein-BrxC-[Limnohabitans-sp.-T6-5] |
| PIQ50876.1 | Type1.BREX | dCache_1+HisKA+HATPase_c-><-SIGMA+OB-Fold+AAA-ATPase+Lon-Protease-BrxL<-PglZ<-Schlafen+HATPase+wHTH<-SbcC<-N6-METHYLASE<-PIN<-MazE_antitoxin<-DUF6079*<-BrxB-BrxF-iSTAND<-BrxA-DNA-binding<-RHH_5\|\|?->?->SF1-SF2-Helicase+REase->?->RelE-ParE-> | 1153 | Bacteria>Pseudomonadota>Betaproteobacteria>Burkholderiales>Comamonadaceae>unclassified_Comamonadaceae | GCA_002786915.1 | Comamonadaceae_bacterium_CG12_big_fil_rev_8_21_14_0_65_59_15 | COW02_17785 | 48153..51614 | 1973952 | hypothetical-protein-COW02_17785-[Comamonadaceae-bacterium-CG12_big_fil_rev_8_21_14_0_65_59_15] |
| RIK89376.1 | Type1.BREX | AAA-ATPase->HIN-HTH-><-?<-?<-?<-?\|\|GNTR-HTH+Aminotran_1_2->BrxA-DNA-binding->BrxB-BrxF-iSTAND->DUF6079*->Schlafen+HATPase+wHTH->ParA-Soj-PloopNTPase->PglZ->SIGMA+OB-Fold+AAA-ATPase+Lon-Protease-BrxL-><-?<-ABC-ATPase+SbcC<-ABC-ATPase+SbcC | 1146 | Bacteria>Pseudomonadota>Betaproteobacteria>Burkholderiales>unclassified_Burkholderiales | GCA_003577305.1 | Burkholderiales_bacterium | DCC70_08090 | 13043..16483 | 1891238 | BREX-system-P-loop-protein-BrxC-[Burkholderiales-bacterium] |
| MCP5252671.1 | Type1.BREX | <-HNH<-TraI-VirD2-RCRE<-TraJ-RHH<-XIS-HTH<-?\|\|DEDD_Tnp_IS110+Transposase_20-><-ParA\|\|AAA-ATPase->BrxB-BrxF-iSTAND->DUF6079*->Schlafen+HATPase+wHTH-><-?<-HTH_3<-Tox-RelE<-Arm-DNA-bind_3<-?\|\|HKD-DNase+HKD-DNase-> | 1144 | Bacteria>Pseudomonadota>Betaproteobacteria>Burkholderiales>unclassified_Burkholderiales | GCA_024235735.1 | Burkholderiales_bacterium | H6939_13335 | 16624..20058 | 1891238 | BREX-system-P-loop-protein-BrxC-[Burkholderiales-bacterium] |
| MCP5292583.1 | Type1.BREX | <-HKD-DNase+HKD-DNase\|\|?->Arm-DNA-bind_3->Tox-RelE->HTH_3->?-><-Schlafen+HATPase+wHTH<-DUF6079*<-BrxB-BrxF-iSTAND<-AAA-ATPase-p | 1144 | Bacteria>Pseudomonadota>Betaproteobacteria>Burkholderiales>unclassified_Burkholderiales | GCA_024235775.1 | Burkholderiales_bacterium | H6938_08650 | 46881..50315 | 1891238 | BREX-system-P-loop-protein-BrxC-[Burkholderiales-bacterium] |
| SDW37048.1 | Type1.BREX | <-SIGMA+OB-Fold+AAA-ATPase+Lon-Protease-BrxL<-PglZ<-ParB+HNH+PHP+SbcC<-N6-METHYLASE<-Schlafen+HATPase+wHTH<-Schlafen+HATPase+wHTH<-DUF6079*<-BrxB-BrxF-iSTAND<-BrxA-DNA-binding\|\|JetA->JetB->SbcC->REase-> | 1145 | Bacteria>Pseudomonadota>Betaproteobacteria>Nitrosomonadales>Nitrosomonadaceae>Nitrosomonas | GCA_900106545.1 | Nitrosomonas_communis | SAMN05421882_100921 | 53532..56969 | 44574 | hypothetical-protein-SAMN05421882_100921-[Nitrosomonas-communis] |
| WP_107790392.1 | Type1.BREX | ParB+RepB->?-><-SbcC<-Calcineurin+Calcineurin-SbcD<-SbcC<-SIGMA+OB-Fold+AAA-ATPase+HKD-DNase-BrxL<-PglZ<-N6-METHYLASE<-Schlafen+HATPase+wHTH+HTH<-DUF6079*<-BrxB-BrxF-iSTAND<-BrxA-DNA-binding\|\|?->?->PIN-> | 1180 | Bacteria>Pseudomonadota>Betaproteobacteria>Nitrosomonadales>Nitrosomonadaceae>Nitrosomonas | GCF_003051105.1 | Nitrosomonas_nitrosa | C8R30_RS13785 | 18563..22105 | 52442 | BREX-system-P-loop-protein-BrxC-[Nitrosomonas-nitrosa] |
| WP_211752264.1 | Type1.BREX | <-SIGMA+OB-Fold+AAA-ATPase+Lon-Protease-BrxL<-PglZ<-ParB+HNH<-PHP+SbcC<-N6-METHYLASE<-Schlafen+HATPase+wHTH<-Schlafen+HATPase+wHTH<-DUF6079*<-BrxB-BrxF-iSTAND<-BrxA-DNA-binding\|\|JetA->JetB->SbcC->REase->?-><-?<-ABC-ATPase+SbcC | 1146 | Bacteria>Pseudomonadota>Betaproteobacteria>Nitrosomonadales>Nitrosomonadaceae>Nitrosomonas | GCF_900106545.1 | Nitrosomonas_communis | BLV21_RS05435 | 53532..56972 | 44574 | BREX-system-P-loop-protein-BrxC-[Nitrosomonas-communis] |
| WP_204800510.1 | Type1.BREX | <-SbcC<-Calcineurin+Calcineurin-SbcD<-SbcC<-SIGMA+OB-Fold+AAA-ATPase+HKD-DNase-BrxL<-PglZ<-HTH+REase<-?<-Schlafen+HATPase+wHTH+HTH<-DUF6079*<-BrxB-BrxF-iSTAND<-BrxA-DNA-binding<-XIS-HTH<-?<-Arm-DNA-bind_3+Phage_integrase | 1180 | Bacteria>Pseudomonadota>Betaproteobacteria>Nitrosomonadales>Nitrosomonadaceae>Nitrosomonas | GCF_905220645.1 | Nitrosomonas_nitrosa | NUZ5A_RS15530 | 137140..140682 | 52442 | BREX-system-P-loop-protein-BrxC-[Nitrosomonas-nitrosa] |
| WP_219938189.1 | Type1.BREX | HIN-HTH->BrxA-DNA-binding->BrxB-BrxF-iSTAND->DUF6079*->N6-METHYLASE->Schlafen+HATPase+wHTH->PglZ->SIGMA+OB-Fold+AAA-ATPase+HKD-DNase-BrxL-> | 1210 | Bacteria>Pseudomonadota>Gammaproteobacteria>Aeromonadales>Succinivibrionaceae>Succinivibrio | GCF_019448115.1 | Succinivibrio_faecicola | J5V48_RS08685 | 2494..6126 | 2820300 | BREX-system-P-loop-protein-BrxC-[Succinivibrio-faecicola] |
| WP_136550441.1 | Type1.BREX | <-SbcC+APATPase+SbcC<-SIGMA+OB-Fold+AAA-ATPase+HKD-DNase-BrxL<-PglZ<-N6-METHYLASE<-REase<-OLD-ABC<-Schlafen+HATPase+wHTH+HTH<-DUF6079*<-BrxB-BrxF-iSTAND<-BrxA-DNA-binding<-XIS-HTH<-MobI | 1193 | Bacteria>Pseudomonadota>Gammaproteobacteria>Alteromonadales>Alteromonadaceae>Hydrocarboniclastica | GCF_004851605.1 | Hydrocarboniclastica_marina | soil367_RS18210 | 4131404..4134985 | 2259620 | BREX-system-P-loop-protein-BrxC-[Hydrocarboniclastica-marina] |
| NQY33786.1 | Type1.BREX | <-REase<-?<-SIGMA+OB-Fold+AAA-ATPase+HKD-DNase-BrxL<-PglZ<-Schlafen+HATPase+wHTH<-PHP+SbcC<-N6-METHYLASE<-NERD<-OLD-ABC<-DUF6079*<-BrxB-BrxF-iSTAND<-BrxA-DNA-binding<-XIS-HTH<-?<-?<-?<-?<-?<-MoxR-AAA | 1200 | Bacteria>Pseudomonadota>Gammaproteobacteria>Alteromonadales>Alteromonadaceae>unclassified_Alteromonadaceae | GCA_013216015.1 | Alteromonadaceae_bacterium | HRT37_02225 | 108462..112064 | 1916082 | BREX-system-P-loop-protein-BrxC-[Alteromonadaceae-bacterium] |
| WP_234000034.1 | Type1.BREX | <-SIGMA+OB-Fold+AAA-ATPase+Lon-Protease-BrxL<-PglZ<-?<-?<-ParB+HNH<-SF1-SF2-Helicase<-?<-N6-METHYLASE<-Schlafen+HATPase+wHTH<-DUF6079*<-BrxB-BrxF-iSTAND<-BrxA-DNA-binding<-wHTH+WYL+WCX<-?<-?\|\|?-><-Peptidase_S8<-?<-REC+LuxR-HTH | 1252 | Bacteria>Pseudomonadota>Gammaproteobacteria>Alteromonadales>Pseudoalteromonadaceae>Pseudoalteromonas | GCF_002964725.1 | Pseudoalteromonas_sp._T1lg76 | C2M35_RS06890 | 48486..52244 | 2077103 | BREX-system-P-loop-protein-BrxC-[Pseudoalteromonas-sp.-T1lg76] |
| WP_233998422.1 | Type1.BREX | <-PglZ<-?<-?<-SF1-SF2-Helicase<-HEPN<-N6-METHYLASE<-N6-METHYLASE-p<-HATPase+wHTH<-Schlafen+HATPase+wHTH<-DUF6079*<-BrxB-BrxF-iSTAND<-BrxA-DNA-binding<-wHTH+WYL+WCX<-?<-?\|\|?-><-?\|\|?-><-SIR2 | 1252 | Bacteria>Pseudomonadota>Gammaproteobacteria>Alteromonadales>Pseudoalteromonadaceae>Pseudoalteromonas | GCF_002964785.1 | Pseudoalteromonas_sp._T1lg48 | C2M29_RS12220 | 655771..659529 | 2077100 | BREX-system-P-loop-protein-BrxC-[Pseudoalteromonas-sp.-T1lg48] |
| WP_265366287.1 | Type1.BREX | Arm-DNA-bind_3+Phage_integrase-><-XIS-HTH<-AAA+HTH<-XIS-HTH\|\|MobI->XIS-HTH->BrxA-DNA-binding->BrxB-BrxF-iSTAND->DUF6079*->Schlafen+HATPase+wHTH+HTH->OLD-ABC->REase->N6-METHYLASE->ParB+HNH->PglZ->SIGMA+OB-Fold+AAA-ATPase+HKD-DNase-BrxL->SbcC+APATPase+SbcC-> | 1193 | Bacteria>Pseudomonadota>Gammaproteobacteria>Cellvibrionales>Cellvibrionaceae>Marinimicrobium | GCF_026153455.1 | Marinimicrobium_koreense | OOT55_RS13025 | 3024461..3028042 | 306545 | BREX-system-P-loop-protein-BrxC-[Marinimicrobium-sp.-C6131] |
| WP_043319021.1 | Type1.BREX | <-TetR-HTH<-?\|\|?-><-SIGMA+OB-Fold+AAA-ATPase+Lon-Protease-BrxL<-PglZ<-ParB+HNH<-N6-METHYLASE<-Schlafen+HATPase+wHTH<-DUF6079*<-BrxB-BrxF-iSTAND<-BrxA-DNA-binding<-tRNA<-?<-?<-?<-?<-Classical-AAA<-SAM-METHYLASE | 1145 | Bacteria>Pseudomonadota>Gammaproteobacteria>Cellvibrionales>Microbulbiferaceae>Microbulbifer | GCF_000708675.1 | Microbulbifer_sp._HZ11 | AU11_RS09640 | 451911..455348 | 1453501 | BREX-system-P-loop-protein-BrxC-[Microbulbifer-sp.-HZ11] |
| MAT50826.1 | Type1.BREX | <-N6-METHYLASE<-SLOG+BirA-HTH<-SF1-SF2-Helicase+PRTase<-Schlafen+HATPase+wHTH+HTH<-DUF6079*<-BrxB-BrxF-iSTAND<-BrxA-DNA-binding<-XIS-HTH\|\|?->?-><-GDResolvase | 1182 | Bacteria>Pseudomonadota>Gammaproteobacteria>Cellvibrionales>Porticoccaceae>unclassified_Porticoccaceae | GCA_002702725.1 | Porticoccaceae_bacterium | CMK32_06560 | 7929..11477 | 2026782 | hypothetical-protein-[Porticoccaceae-bacterium] |
| WP_268905395.1 | Type1.BREX | Phage_integrase-><-?<-SMODS<-SLATT<-SIGMA+OB-Fold+AAA-ATPase+Lon-Protease-BrxL<-PglZ<-ParB+HNH<-N6-METHYLASE<-Schlafen+HATPase+wHTH<-DUF6079*<-BrxB-BrxF-iSTAND<-BrxA-DNA-binding<-wHTH+WYL+WCX\|\|GGDEF+EAL-><-HTH_3 | 1220 | Bacteria>Pseudomonadota>Gammaproteobacteria>Cellvibrionales>Spongiibacteraceae>Dasania | GCF_026891035.1 | Dasania_sp._GY-19 | OOV09_RS18305 | 21121..24783 | 2965066 | BREX-system-P-loop-protein-BrxC-[Dasania-sp.-GY-19] |
| WP_273033648.1 | Type1.BREX | <-SIGMA+OB-Fold+AAA-ATPase+HKD-DNase-BrxL<-PglZ<-ParB+HNH<-N6-METHYLASE-p<-HerA/FtsK<-?<-CR-ATPase3<-N6-METHYLASE<-Schlafen+HATPase+wHTH+HTH<-DUF6079*<-BrxB-BrxF-iSTAND<-BrxA-DNA-binding<-XIS-HTH<-MobI\|\|XIS-HTH->AAA+HTH->XIS-HTH-><-Arm-DNA-bind_3+Phage_integrase<-SIGMA-HTH | 1193 | Bacteria>Pseudomonadota>Gammaproteobacteria>Cellvibrionales>Spongiibacteraceae>Spongiibacter | GCF_950061765.1 | Spongiibacter_tropicus | B4C78_RS03795 | 102796..106377 | 454602 | BREX-system-P-loop-protein-BrxC-[Spongiibacter-tropicus] |
| WP_201222419.1 | Type1.BREX | <-PCMT+METHYLASE<-?<-?<-?\|\|?->tRNA->?->BrxA-DNA-binding->BrxB-BrxF-iSTAND->DUF6079*->OLD-ABC->REase->N6-METHYLASE->Schlafen+HATPase+wHTH->ParB+HNH->PglZ->REase->MIT_C-> | 1174 | Bacteria>Pseudomonadota>Gammaproteobacteria>Chromatiales>Chromatiaceae>Halochromatium | GCF_016653315.1 | Halochromatium_roseum | CCR96_RS19410 | 137258..140782 | 391920 | BREX-system-P-loop-protein-BrxC-[Halochromatium-roseum] |
| WP_020505452.1 | Type1.BREX | RNase-><-?<-?\|\|BrxA-DNA-binding->BrxB-BrxF-iSTAND->DUF6079*-><-?\|\|N6-METHYLASE->Schlafen+HATPase+wHTH->PglZ->SIGMA+OB-Fold+AAA-ATPase+Lon-Protease-BrxL-> | 1146 | Bacteria>Pseudomonadota>Gammaproteobacteria>Chromatiales>Chromatiaceae>Lamprocystis | GCF_000379525.1 | Lamprocystis_purpurea | A39O_RS0114850 | 122427..125867 | 61598 | BREX-system-P-loop-protein-BrxC-[Lamprocystis-purpurea] |
| WP_020507127.1 | Type6.BREX | XIS-HTH->BrxE->BrxA-DNA-binding->BrxB-BrxF-iSTAND->DUF6079*->Schlafen+HATPase+wHTH+HTH-><-?\|\|N6-METHYLASE->?->PIN->N6-METHYLASE->AbrB-HTH-> | 1179 | Bacteria>Pseudomonadota>Gammaproteobacteria>Chromatiales>Chromatiaceae>Lamprocystis | GCF_000379525.1 | Lamprocystis_purpurea | A39O_RS0123535 | 17517..21056 | 61598 | BREX-system-P-loop-protein-BrxC-[Lamprocystis-purpurea] |

| Protein-ID | System | Gene-Neighbourhoods | Length | Taxonomy | Assembly-ID | Strain | Locus-Tag | Genomic-coordinates | TaxID | Fasta-define |
|---|---|---|---|---|---|---|---|---|---|---|
| WP_014777950.1 | Type6.BREX | <-DUF4096+DDE_Tnp_1+DDE_Tnp_1_2<-SF1-SF2-Helicase+wHTH+Small-B-barrel+DUF1998-p<-BrxD-AAA<-PglZ<-Schlafen+HATPase+wHTH<-PHP+SbcC<-N6-METHYLASE<-DOC<-MazE_antitoxin<-DUF6079*<-BrxB-BrxF-iSTAND<-BrxA-DNA-binding<-BrxE<-?<-?<-?<-?<-MoxR-AAA | 1206 | Bacteria>Pseudomonadota>Gammaproteobacteria>Chromatiales>Chromatiaceae>Thiocystis | GCF_000227745.2 | Thiocystis_violascens | THIVI_RS07160 | 1619300..1622920 | 73141 | BREX-system-P-loop-protein-BrxC-[Thiocystis-violascens] |
| WP_200373981.1 | Type6.BREX | N6-METHYLASE->?->BrxE->BrxA-DNA-binding->BrxB-BrxF-iSTAND->DUF6079*->MNS+REase->N6-METHYLASE->SbcC->Schlafen+HATPase+wHTH->PglZ->BrxD-AAA->SF1-SF2-Helicase+wHTH+Small-B-barrel+DUF1998-><-ParA | 1206 | Bacteria>Pseudomonadota>Gammaproteobacteria>Chromatiales>Chromatiaceae>Thiocystis | GCF_016583575.1 | Thiocystis_violacea | CKO27_RS06060 | 39556..43176 | 13725 | BREX-system-P-loop-protein-BrxC-[Thiocystis-violacea] |
| WP_100920354.1 | Type6.BREX | <-HMA+HAD+HAD\|\|?->?->?->BrxE->BrxA-DNA-binding->BrxB-BrxF-iSTAND->?->\|\|DUF6079*->Schlafen+HATPase+wHTH+HTH-><-?\|\|N6-METHYLASE->?->PIN->N6-METHYLASE->RHH->PIN->RHH-> | 1178 | Bacteria>Pseudomonadota>Gammaproteobacteria>Chromatiales>Chromatiaceae>Thiodictyon | GCF_002813775.1 | Candidatus_Thiodictyon_syntrophicum | THSYN_17920 | 4302311..4305847 | 1166950 | BREX-system-P-loop-protein-BrxC-[Candidatus-Thiodictyon-syntrophicum] |
| MBK1629582.1 | Type1.BREX | <-SbcC<-?<-RFC-AAA+DNAA-HTH\|\|?->RNaseP->?->sGTP->BrxA-DNA-binding->BrxB-BrxF-iSTAND->DUF6079*->MazE_antitoxin->PemK_toxin->DOC+HTH->N6-METHYLASE-p->PHP+SbcC->Schlafen+HATPase+wHTH->PglZ->SIGMA+OB-Fold+AAA-ATPase+Lon-Protease-BrxL-> | 1152 | Bacteria>Pseudomonadota>Gammaproteobacteria>Chromatiales>Chromatiaceae>Thiohalocapsa | GCA_016583825.1 | Thiohalocapsa_halophila | CKO31_02265 | 100295..103753 | 69359 | hypothetical-protein-[Thiohalocapsa-halophila] |
| WP_058554449.1 | Type6.BREX | TIR->?->?->?->BrxE->BrxA-DNA-binding->BrxB-BrxF-iSTAND->DUF6079*->N6-METHYLASE->?->Schlafen+HATPase+wHTH->N6-METHYLASE->?->PIN->PglZ->SIGMA+OB-Fold+AAA-ATPase+HKD-DNase-BrxL-><-tRNA\|\|HHH_5+SF1-SF2-Helicase-> | 1214 | Bacteria>Pseudomonadota>Gammaproteobacteria>Chromatiales>Chromatiaceae>Thiohalocapsa | GCF_001469165.1 | Thiohalocapsa_sp._ML1 | ML01_RS05745 | 155025..158669 | 1431688 | BREX-system-P-loop-protein-BrxC-[Thiohalocapsa-sp.-ML1] |
| EIC19850.1 | Type1.BREX | <-SIGMA+OB-Fold+AAA-ATPase+Lon-Protease-BrxL<-PglZ<-RelE-ParE<-PhdYeFM_antitox<-Schlafen+HATPase+wHTH<-N6-METHYLASE<-PIN<-VapB_antitoxin<-DUF6079*<-BrxB-BrxF-iSTAND<-BrxA-DNA-binding<-?<-HISKIN<-?<-?<-?<-Thioredoxin | 1152 | Bacteria>Pseudomonadota>Gammaproteobacteria>Chromatiales>Chromatiaceae>Thiorhodovibrio | GCA_000228725.3 | Thiorhodovibrio_sp._970 | Thi970DRAFT_03455 | 369329..372787 | 631362 | hypothetical-protein-Thi970DRAFT_03455-[Thiorhodovibrio-sp.-970] |
| WP_223296802.1 | Type1.BREX | <-SIGMA+OB-Fold+AAA-ATPase+Lon-Protease-BrxL<-PglZ<-RelE-ParE<-PhdYeFM_antitox<-Schlafen+HATPase+wHTH<-N6-METHYLASE<-PIN<-VapB_antitoxin<-DUF6079*<-BrxB-BrxF-iSTAND<-BrxA-DNA-binding<-?<-HISKIN<-?<-?<-Thioredoxin\|\|?->REC+GGDEF+EAL-> | 1154 | Bacteria>Pseudomonadota>Gammaproteobacteria>Chromatiales>Chromatiaceae>Thiorhodovibrio | GCF_000228725.2 | Thiorhodovibrio_sp._970 | THI970DRAFT_RS16355 | 369329..372793 | 631362 | BREX-system-P-loop-protein-BrxC-[Thiorhodovibrio-sp.-970] |
| MBP6733683.1 | Type6.BREX | BrxE->BrxA-DNA-binding->BrxB-BrxF-iSTAND->DUF6079*->Schlafen+HATPase+wHTH+HTH->N6-METHYLASE-p-> | 1181 | Bacteria>Pseudomonadota>Gammaproteobacteria>Chromatiales>Chromatiaceae>unclassified_Chromatiaceae | GCA_017990755.1 | Chromatiaceae_bacterium | KA142_03150 | 2562..6107 | 2049432 | BREX-system-P-loop-protein-BrxC-[Chromatiaceae-bacterium] |
| MCC7280699.1 | Type6.BREX | <-SIGMA+OB-Fold+AAA-ATPase+Lon-Protease-BrxL<-PglZ<-HNH<-OLD-ABC<-RHH<-N6-METHYLASE<-Schlafen+HATPase+wHTH+HTH<-DUF6079*<-BrxB-BrxF-iSTAND<-BrxA-DNA-binding<-BrxE | 1181 | Bacteria>Pseudomonadota>Gammaproteobacteria>Chromatiales>Chromatiaceae>unclassified_Chromatiaceae | GCA_020851955.1 | Chromatiaceae_bacterium | IT487_20620 | 13992..17537 | 2049432 | BREX-system-P-loop-protein-BrxC-[Chromatiaceae-bacterium] |
| WP_018716649.1 | Type1.BREX | HerA/FtsK->Relaxase+SF1-SF2-Helicase->?->XIS-HTH->BrxA-DNA-binding->BrxB-BrxF-iSTAND->DUF6079*->Schlafen+HATPase+wHTH->N6-METHYLASE->ParB+HNH->PglZ->SIGMA+OB-Fold+AAA-ATPase+Lon-Protease-BrxL-> | 1144 | Bacteria>Pseudomonadota>Gammaproteobacteria>Chromatiales>Ectothiorhodospiraceae>Arhodomonas | GCF_000374645.1 | Arhodomonas_aquaeolei | C516_RS0101825 | 422257..425691 | 2369 | BREX-system-P-loop-protein-BrxC-[Arhodomonas-aquaeolei] |
| WP_063464066.1 | Type1.BREX | ZnR->RADICAL-SAM->?->?->?->GGDEF->XIS-HTH->BrxA-DNA-binding->BrxB-BrxF-iSTAND->DUF6079*->Schlafen+HATPase+wHTH+HTH->N6-METHYLASE-p->OLD-ABC->REase->SIR2->HerA/FtsK->N6-METHYLASE-p->OLD-ABC->HNH-> | 1189 | Bacteria>Pseudomonadota>Gammaproteobacteria>Chromatiales>Ectothiorhodospiraceae>Ectothiorhodospira | GCF_001632845.1 | Ectothiorhodospira_sp._BSL-9 | ECTOBSL9_RS04500 | 914827..918396 | 1442136 | BREX-system-P-loop-protein-BrxC-[Ectothiorhodospira-sp.-BSL-9] |
| WP_238631208.1 | Type1.BREX | <-PglZ<-TOPRIM<-OLD-ABC<-N6-METHYLASE-p<-HerA/FtsK<-?<-CR-ATPase3<-N6-METHYLASE<-Schlafen+HATPase+wHTH+HTH<-DUF6079*<-BrxB-BrxF-iSTAND<-BrxA-DNA-binding<-XIS-HTH<-?<-?<-?<-RADICAL-SAM<-ZnR\|\|SWACOS-> | 1189 | Bacteria>Pseudomonadota>Gammaproteobacteria>Chromatiales>Ectothiorhodospiraceae>Ectothiorhodospira | GCF_022227695.1 | Ectothiorhodospira_sp._9905 | LPW28_RS03070 | 43824..47393 | 2897387 | BREX-system-P-loop-protein-BrxC-[Ectothiorhodospira-sp.-9905] |
| WP_238620557.1 | Type1.BREX | sGTP-><-35exo+T7DNApol\|\|?->?->BrxA-DNA-binding->BrxB-BrxF-iSTAND->DUF6079*->N6-METHYLASE->OLD-ABC->HNH->OLD-ABC->HNH->PglZ->SIGMA+OB-Fold+AAA-ATPase+Lon-Protease-BrxL-><-Schlafen+HATPase+wHTH<-Schlafen+HATPase+wHTH | 1154 | Bacteria>Pseudomonadota>Gammaproteobacteria>Chromatiales>Ectothiorhodospiraceae>Ectothiorhodospira | GCF_022227785.1 | Ectothiorhodospira_sp._B14B | LQ952_00335 | 78448..81912 | 2899127 | BREX-system-P-loop-protein-BrxC-[Ectothiorhodospira-sp.-B14B] |
| TQE98798.1 | Type1.BREX | BrxA-DNA-binding->BrxB-BrxF-iSTAND->DUF6079*->Schlafen+HATPase+wHTH->N6-METHYLASE-p-> | 1193 | Bacteria>Pseudomonadota>Gammaproteobacteria>Chromatiales>Ectothiorhodospiraceae>Spiribacter | GCA_006569195.1 | Spiribacter_salinus | FKY71_11930 | 1820..5401 | 1335746 | BREX-system-P-loop-protein-BrxC-[Spiribacter-salinus] |
| WP_079805580.1 | Type1.BREX | SLOG->HAD-><-SIGMA+OB-Fold+AAA-ATPase+Lon-Protease-BrxL<-PglZ<-N6-METHYLASE-p<-RVT<-HEPN<-N6-METHYLASE<-ParB+HNH<-DUF6079*<-BrxB-BrxF-iSTAND<-BrxA-DNA-binding\|\|Schlafen+HATPase+wHTH-><-?<-?<-XIS-HTH | 1240 | Bacteria>Pseudomonadota>Gammaproteobacteria>Enterobacterales>Enterobacteriaceae>Salmonella | GCF_016903075.1 | Salmonella_enterica | AJH70_RS01330 | 291078..294800 | 28901 | BREX-system-P-loop-protein-BrxC-[Salmonella-enterica] |
| WP_248566606.1 | Type1.BREX | SF1-SF2-Helicase+REase->?-><-SIGMA+OB-Fold+AAA-ATPase+Lon-Protease-BrxL<-PglZ<-TOPRIM<-OLD-ABC<-HTH+REase<-N6-METHYLASE<-Schlafen+HATPase+wHTH<-DUF6079*<-BrxB-BrxF-iSTAND<-BrxA-DNA-binding<-SF1-SF2-Helicase<-HKD-DNase<-DUF6361<-HEPN\|\|?->URI-DNase-> | 1145 | Bacteria>Pseudomonadota>Gammaproteobacteria>Gammaproteobacteria_incertae_sedis>Methylonatrum | GCF_023195885.1 | Methylonatrum_kenyense | M0534_RS01350 | 290769..294206 | 455253 | BREX-system-P-loop-protein-BrxC-[Methylonatrum-kenyense] |
| WP_029131813.1 | Type1.BREX | TraU->TraH->?->TraG_N-><-?<-?<-HTH\|\|BrxA-DNA-binding->BrxB-BrxF-iSTAND->DUF6079*->N6-METHYLASE->PHP+SbcC->Schlafen+HATPase+wHTH->PglZ->SIGMA+OB-Fold+AAA-ATPase+Lon-Protease-BrxL-> | 1147 | Bacteria>Pseudomonadota>Gammaproteobacteria>Gammaproteobacteria_incertae_sedis>Sedimenticola | GCF_000428045.1 | Sedimenticola_selenatireducens | A3GO_RS0101005 | 196433..199876 | 191960 | BREX-system-P-loop-protein-BrxC-[Sedimenticola-selenatireducens] |
| MCG8047516.1 | Type1.BREX | FtsK+VirB4-FtsK+A32->?->TraU->TraH->?->TraG_N-><-?\|\|BrxA-DNA-binding->BrxB-BrxF-iSTAND->DUF6079*->N6-METHYLASE->SbcC->Schlafen+HATPase+wHTH->PglZ->PglZ->SIGMA+OB-Fold+AAA-ATPase+Lon-Protease-BrxL->REase+TIR->PIWI->TPR-> | 1147 | Bacteria>Pseudomonadota>Gammaproteobacteria>Gammaproteobacteria_incertae_sedis>sulfur-oxidizing_symbionts>Candidatus_Thiodiazotropha | GCA_016843075.1 | Candidatus_Thiodiazotropha_taylori | JAY66_17825 | 23755..27198 | 2792791 | BREX-system-P-loop-protein-BrxC-[Candidatus-Thiodiazotropha-taylori] |
| MCG7872379.1 | Type1.BREX | <-SbcC+CCDC158<-Calcineurin\|\|NtrC-AAA+FIS-HTH-><-HTH_3<-SIGMA+OB-Fold+AAA-ATPase+HKD-DNase-BrxL<-PglZ<-RVT<-N6-METHYLASE<-Schlafen+HATPase+wHTH+HTH<-DUF6079*<-BrxB-BrxF-iSTAND<-BrxA-DNA-binding<-XIS-HTH\|\|HNH->?->XIS-HTH->Phage_integrase->Phage_integrase->Phage_integrase-> | 1170 | Bacteria>Pseudomonadota>Gammaproteobacteria>Gammaproteobacteria_incertae_sedis>sulfur-oxidizing_symbionts>Candidatus_Thiodiazotropha | GCA_016848925.1 | Candidatus_Thiodiazotropha_lotti | JAZ11_09800 | 178889..182401 | 2792787 | BREX-system-P-loop-protein-BrxC-[Candidatus-Thiodiazotropha-lotti] |
| MCG7872685.1 | Type1.BREX | TraU->TraH->?->TraG_N->HTH_26->BrxA-DNA-binding->BrxB-BrxF-iSTAND->DUF6079*->N6-METHYLASE->SbcC->Schlafen+HATPase+wHTH->PglZ->SIGMA+OB-Fold+AAA-ATPase+Lon-Protease-BrxL-><-TatD_DNase<-?<-KAP-glycosylase<-KAP-NTPase | 1151 | Bacteria>Pseudomonadota>Gammaproteobacteria>Gammaproteobacteria_incertae_sedis>sulfur-oxidizing_symbionts>Candidatus_Thiodiazotropha | GCA_016848925.1 | Candidatus_Thiodiazotropha_lotti | JAZ11_11345 | 100935..104390 | 2792787 | BREX-system-P-loop-protein-BrxC-[Candidatus-Thiodiazotropha-lotti] |
| WP_174625954.1 | Type1.BREX | N6-METHYLASE->?->DNA-glycosylase->XIS-HTH->BrxA-DNA-binding->BrxB-BrxF-iSTAND->DUF6079*->N6-METHYLASE->PHP+SbcC->Schlafen+HATPase+wHTH->PglZ-><-Phage_integrase<-?\|\|SSB-> | 1149 | Bacteria>Pseudomonadota>Gammaproteobacteria>Methylococcales>Methylococcaceae>Methylobacter | GCF_902806695.1 | Candidatus_Methylobacter_favarea | HUX64_RS09295 | 14302..17751 | 2707345 | BREX-system-P-loop-protein-BrxC-[Candidatus-Methylobacter-favarea] |
| WP_024296674.1 | Type1.BREX | GDResolvase+HIN-HTH->?->BrxA-DNA-binding->BrxB-BrxF-iSTAND->DUF6079*->ParD_antitoxin->PIN->N6-METHYLASE->SbcC->Schlafen+HATPase+wHTH->PglZ->SIGMA+OB-Fold+AAA-ATPase+Lon-Protease-BrxL-> | 1202 | Bacteria>Pseudomonadota>Gammaproteobacteria>Methylococcales>Methylococcaceae>Methylomicrobium | GCF_000527095.1 | Methylomicrobium_lacus | METLA_RS0100460 | 95992..99600 | 136992 | BREX-system-P-loop-protein-BrxC-[Methylomicrobium-lacus] |
| WP_088619247.1 | Type1.BREX | <-Glycos_transf<-?<-SIGMA+OB-Fold+AAA-ATPase+Lon-Protease-BrxL<-PglZ<-Schlafen+HATPase+wHTH<-PHP+SbcC<-N6-METHYLASE<-PIN<-ParD_antitoxin<-DUF6079*<-BrxB-BrxF-iSTAND<-BrxA-DNA-binding<-wHTH+WYL+WCX | 1201 | Bacteria>Pseudomonadota>Gammaproteobacteria>Methylococcales>Methylococcaceae>Methylovulum | GCF_002209385.1 | Methylovulum_psychrotolerans | CEK71_RS09965 | 2189788..2193393 | 1704499 | BREX-system-P-loop-protein-BrxC-[Methylovulum-psychrotolerans] |
| WP_103975698.1 | Type1.BREX | <-SIGMA+OB-Fold+AAA-ATPase+Lon-Protease-BrxL<-PglZ<-Schlafen+HATPase+wHTH<-PHP+SbcC<-?<-?<-N6-METHYLASE<-PIN<-ParD_antitoxin<-DUF6079*<-BrxB-BrxF-iSTAND<-BrxA-DNA-binding<-wHTH+WYL+WCX\|\|?->SAM-METHYLASE->?->SbcC->SAM-METHYLASE->SAM-METHYLASE-> | 1201 | Bacteria>Pseudomonadota>Gammaproteobacteria>Methylococcales>Methylococcaceae>Methylovulum | GCF_002923755.1 | Methylovulum_psychrotolerans | AADEFJLK_RS21345 | 25888..29493 | 1704499 | BREX-system-P-loop-protein-BrxC-[Methylovulum-psychrotolerans] |
| WP_215479507.1 | Type1.BREX | wHTH+WYL+WCX-p->wHTH+WYL+WCX->BrxA-DNA-binding->BrxB-BrxF-iSTAND->DUF6079*->ParD_antitoxin->PIN->N6-METHYLASE->?->?->PHP+SbcC->Schlafen+HATPase+wHTH->PglZ->SIGMA+OB-Fold+AAA-ATPase+Lon-Protease-BrxL-> | 1201 | Bacteria>Pseudomonadota>Gammaproteobacteria>Methylococcales>Methylococcaceae>Methylovulum | GCF_018722605.1 | Methylovulum_psychrotolerans | KFZ76_RS21360 | 105012..108617 | 1704499 | BREX-system-P-loop-protein-BrxC-[Methylovulum-psychrotolerans] |
| WP_241878421.1 | Type1.BREX | <-MIT_C<-MIT_C<-SIGMA+OB-Fold+AAA-ATPase+Lon-Protease-BrxL<-PglZ<-N6-METHYLASE-p<-N6-METHYLASE-p<-N6-METHYLASE-p<-N6-METHYLASE-p<-?<-DUF6079-p*<-DUF6079-p<-?<-BrxB-BrxF-iSTAND<-BrxA-DNA-binding\|\|?-><-Schlafen+HATPase+wHTH-p<-?<-Schlafen+HATPase+wHTH-p<-Schlafen | 709 | Bacteria>Pseudomonadota>Gammaproteobacteria>Moraxellales>Moraxellaceae>Psychrobacter | GCF_022631235.2 | Psychrobacter_sp._PraFG1 | MN210_RS12710 | 2340484..2342613 | 2565531 | BREX-system-P-loop-protein-BrxC-[Psychrobacter-sp.-PraFG1] |
| WP_063140235.1 | Type1.BREX | <-PRTase<-RNasePH\|\|?->Arm-DNA-bind_3+Phage_integrase-><-XIS-HTH\|\|MobI->XIS-HTH->BrxA-DNA-binding->BrxB-BrxF-iSTAND->DUF6079*->Schlafen+HATPase+wHTH->HIN-HTH-> | 1145 | Bacteria>Pseudomonadota>Gammaproteobacteria>Oceanospirillales>Alcanivoracaceae>Alcanivorax | - | Alcanivorax_sp._KX64203 | A3Q32_03160 | 117678..121115 | 1817793 | BREX-system-P-loop-protein-BrxC-[Alcanivorax-sp.-KX64203] |
| WP_161382580.1 | Type1.BREX | Phage_integrase-><-XIS-HTH<-AAA+HTH<-XIS-HTH\|\|MobI->XIS-HTH->BrxA-DNA-binding->BrxB-BrxF-iSTAND->DUF6079*->Schlafen+HATPase+wHTH+HTH->N6-METHYLASE->SF1-SF2-Helicase->?->HerA/FtsK->N6-METHYLASE-p->PglZ->SIGMA+OB-Fold+AAA-ATPase+HKD-DNase-BrxL->OLD-ABC+TOPRIM-> | 1193 | Bacteria>Pseudomonadota>Gammaproteobacteria>Oceanospirillales>Alcanivoracaceae>Alcanivorax | - | Alcanivorax_sp._ZXX171 | GN155_RS00640 | 132435..136016 | 2695858 | BREX-system-P-loop-protein-BrxC-[Alcanivorax-sp.-ZXX171] |
| MAD69915.1 | Type1.BREX | CHTH-><-?\|\|wHTH+WYL+WCX-><-?<-wHTH+WYL+WCX\|\|?->XIS-HTH->BrxA-DNA-binding->BrxB-BrxF-iSTAND->DUF6079*->Schlafen+HATPase+wHTH+HTH->N6-METHYLASE->REase->SIR2->HerA/FtsK->N6-METHYLASE-p->OLD-ABC->TOPRIM->PglZ-> | 1193 | Bacteria>Pseudomonadota>Gammaproteobacteria>Oceanospirillales>Alcanivoracaceae>Alcanivorax | GCA_002685555.1 | Alcanivorax_sp. | CL538_05260 | 26266..29847 | 1872427 | hypothetical-protein-[Alcanivorax-sp.] |
| MBA3979207.1 | Type1.BREX | PhdYeFM_antitox->?-><-Trypsin+PDZ<-Mrr-N\|\|wHTH+WYL+WCX->XIS-HTH->BrxA-DNA-binding->BrxB-BrxF-iSTAND->DUF6079*->Schlafen+HATPase+wHTH->N6-METHYLASE->PHP+SbcC->ParB+HNH->PglZ->SIGMA+OB-Fold+AAA-ATPase+Lon-Protease-BrxL-><-?<-SbcC+SidE<-JetB | 1145 | Bacteria>Pseudomonadota>Gammaproteobacteria>Oceanospirillales>Alcanivoracaceae>Alcanivorax | GCA_013823245.1 | Alcanivorax_sp. | C0462_01260 | 7976..11413 | 1872427 | BREX-system-P-loop-protein-BrxC-[Alcanivorax-sp.] |
| HIK75733.1 | Type1.BREX | <-PglZ<-TOPRIM<-OLD-ABC<-N6-METHYLASE<-HerA/FtsK<-SIR2<-REase<-N6-METHYLASE<-Schlafen+HATPase+wHTH+HTH<-DUF6079*<-BrxB-BrxF-iSTAND<-BrxA-DNA-binding<-XIS-HTH<-?\|\|wHTH+WYL+WCX->?-><-wHTH+WYL+WCX\|\|?-><-CHTH | 1193 | Bacteria>Pseudomonadota>Gammaproteobacteria>Oceanospirillales>Alcanivoracaceae>Alcanivorax | GCA_015660495.1 | Alcanivorax_sp. | EYG00_13690 | 18356..21937 | 1872427 | BREX-system-P-loop-protein-BrxC-[Alcanivorax-sp.] |
| MCH2557814.1 | Type1.BREX | Arm-DNA-bind_3+Phage_integrase-><-XIS-HTH<-AAA+HTH<-XIS-HTH\|\|MobI->XIS-HTH->BrxA-DNA-binding->BrxB-BrxF-iSTAND->DUF6079*->Schlafen+HATPase+wHTH+HTH->OLD-ABC->REase->N6-METHYLASE->PglZ->SIGMA+OB-Fold+AAA-ATPase+HKD-DNase-BrxL->SbcC+SbcC->Calcineurin+Calcineurin-SbcD->SbcC-> | 1193 | Bacteria>Pseudomonadota>Gammaproteobacteria>Oceanospirillales>Alcanivoracaceae>Alcanivorax | GCA_022449455.1 | Alcanivorax_sp. | MK005_11000 | 47715..51296 | 1872427 | BREX-system-P-loop-protein-BrxC-[Alcanivorax-sp.] |
| WP_103916700.1 | Type1.BREX | <-SIGMA+OB-Fold+AAA-ATPase+HKD-DNase-BrxL<-?<-PglZ<-ParB+HNH<-N6-METHYLASE<-REase<-OLD-ABC<-Schlafen+HATPase+wHTH+HTH<-DUF6079*<-BrxB-BrxF-iSTAND<-BrxA-DNA-binding<-XIS-HTH<-?\|\|wHTH+WYL+WCX-><-XIS-HTH\|\|HIN-HTH-> | 1193 | Bacteria>Pseudomonadota>Gammaproteobacteria>Oceanospirillales>Alcanivoracaceae>Alcanivorax | GCF_900107995.1 | Alcanivorax_sp._DSM_26293 | C4S63_RS03510 | 780383..783964 | 1798238 | BREX-system-P-loop-protein-BrxC-[Alcanivorax-sp.-DSM-26293] |
| WP_204429962.1 | Type1.BREX | <-Calcineurin+Calcineurin-SbcD<-SbcC<-SIGMA+OB-Fold+AAA-ATPase+HKD-DNase-BrxL<-PglZ<-N6-METHYLASE-p<-HerA/FtsK<-RVT<-N6-METHYLASE<-Schlafen+HATPase+wHTH+HTH<-DUF6079*<-BrxB-BrxF-iSTAND<-BrxA-DNA-binding<-XIS-HTH<-MobI\|\|XIS-HTH->AAA+HTH->XIS-HTH-><-Phage_integrase | 1193 | Bacteria>Pseudomonadota>Gammaproteobacteria>Oceanospirillales>Alcanivoracaceae>Alloalcanivorax | GCF_016906305.1 | Alloalcanivorax_marinus | IM760_RS12910 | 16365..19946 | 1177169 | BREX-system-P-loop-protein-BrxC-[Alloalcanivorax-marinus] |
| WP_228232948.1 | Type1.BREX | <-SbcC<-Calcineurin+Calcineurin-SbcD<-SbcC+SbcC<-SIGMA+OB-Fold+AAA-ATPase+HKD-DNase-BrxL<-PglZ<-N6-METHYLASE<-REase<-OLD-ABC<-Schlafen+HATPase+wHTH+HTH<-DUF6079*<-BrxB-BrxF-iSTAND<-BrxA-DNA-binding<-XIS-HTH<-MobI\|\|XIS-HTH->AAA+HTH->XIS-HTH-><-Phage_integrase | 1193 | Bacteria>Pseudomonadota>Gammaproteobacteria>Oceanospirillales>Alcanivoracaceae>Alloalcanivorax | GCF_020782135.1 | Alloalcanivorax_marinus | LL252_RS03465 | 287759..291340 | 1177169 | BREX-system-P-loop-protein-BrxC-[Alloalcanivorax-marinus] |
| WP_262731202.1 | Type1.BREX | <-OLD-ABC<-N6-METHYLASE<-Schlafen+HATPase+wHTH<-DUF6079*<-BrxB-BrxF-iSTAND<-BrxA-DNA-binding<-XIS-HTH<-XIS-HTH\|\|?->?-><-HTH\|\|IG+IG-> | 1145 | Bacteria>Pseudomonadota>Gammaproteobacteria>Oceanospirillales>Alcanivoracaceae>Alloalcanivorax | GCF_025532125.1 | Alloalcanivorax_marinus | B27N_RS13560 | 6066..9503 | 1177169 | BREX-system-P-loop-protein-BrxC-[Alloalcanivorax-marinus] |
| WP_008734404.1 | Type1.BREX | <-SIGMA+OB-Fold+AAA-ATPase+HKD-DNase-BrxL<-PglZ<-N6-METHYLASE<-REase<-OLD-ABC<-Schlafen+HATPase+wHTH+HTH<-DUF6079*<-BrxB-BrxF-iSTAND<-BrxA-DNA-binding<-XIS-HTH<-sGTP<-?<-RNaseP | 1192 | Bacteria>Pseudomonadota>Gammaproteobacteria>Oceanospirillales>Alcanivoracaceae>Isoalcanivorax | GCF_000299335.2 | Isoalcanivorax_pacificus | S7S_RS18615 | 4158589..4162167 | 1306787 | BREX-system-P-loop-protein-BrxC-[Isoalcanivorax-pacificus] |
| RLP52821.1 | Type1.BREX | Arm-DNA-bind_3+Phage_integrase-><-XIS-HTH<-AAA+HTH<-XIS-HTH\|\|MobI->XIS-HTH->BrxA-DNA-binding->BrxB-BrxF-iSTAND->DUF6079*->Schlafen+HATPase+wHTH+HTH->OLD-ABC->NERD->N6-METHYLASE->PglZ->SIGMA+OB-Fold+AAA-ATPase+HKD-DNase-BrxL-><-?<-GDResolvase+HIN-HTH | 1193 | Bacteria>Pseudomonadota>Gammaproteobacteria>Oceanospirillales>Alcanivoracaceae>Ketobacter | GCA_003667185.1 | Ketobacter_sp. | D6160_18810 | 15503..19084 | 2083498 | MAG:-BREX-system-P-loop-protein-BrxC-[Ketobacter-sp.] |
| WP_062359357.1 | Type1.BREX | <-N6-METHYLASE<-Schlafen+HATPase+wHTH<-DUF6079*<-BrxB-BrxF-iSTAND<-BrxA-DNA-binding<-ncRNA<-tRNA<-?\|\|TetR-HTH->?->?->?->ABC-ATPase+SbcC->ABC-ATPase+SbcC-> | 1225 | Bacteria>Pseudomonadota>Gammaproteobacteria>Oceanospirillales>Halomonadaceae | GCF_020782315.1 | Halomonas | TK80_RS02225 | 5404..9081 | 2745 | MULTISPECIES:-BREX-system-P-loop-protein-BrxC-[Halomonas] |
| WP_041158482.1 | Type1.BREX | <-PglZ<-TOPRIM<-OLD-ABC<-ParB+HNH<-TPR<-NERD+GspA-ATPase<-N6-METHYLASE-p<-N6-METHYLASE<-Schlafen+HATPase+wHTH<-DUF6079*<-BrxB-BrxF-iSTAND<-BrxA-DNA-binding<-HTH_19<-TnsC-ATPase<-rve+Mu-transpos_C<-REase+wHTH<-ncRNA<-tRNA<-?\|\|TetR-HTH-> | 1227 | Bacteria>Pseudomonadota>Gammaproteobacteria>Oceanospirillales>Halomonadaceae | GCF_900099955.1 | Halomonas | BLR10_RS07715 | 76580..80263 | 2745 | MULTISPECIES:-BREX-system-P-loop-protein-BrxC-[Halomonas] |
| PHR03208.1 | Type1.BREX | <-SIGMA+OB-Fold+AAA-ATPase+Lon-Protease-BrxL<-PglZ<-ParB+HNH<-?<-?<-?<-N6-METHYLASE<-Schlafen+HATPase+wHTH<-DUF6079*<-BrxB-BrxF-iSTAND<-BrxA-DNA-binding\|\|Mob_Pre->XIS-HTH-><-tRNA\|\|?->?->?-><-TarH+HAMP+MA | 1144 | Bacteria>Pseudomonadota>Gammaproteobacteria>Oceanospirillales>Halomonadaceae>Halomonas | GCA_002733025.1 | Halomonas_sp. | COB32_05665 | 22826..26260 | 1486246 | hypothetical-protein-COB32_05665-[Halomonas-sp.] |
| PHR03139.1 | Type1.BREX | <-ABC-ATPase+SbcC<-ABC-ATPase+SbcC<-?<-?<-?<-TetR-HTH\|\|?->tRNA->ncRNA->BrxA-DNA-binding->BrxB-BrxF-iSTAND->DUF6079*->Phd-YefM->PIN->Schlafen+HATPase+wHTH->N6-METHYLASE-> | 1227 | Bacteria>Pseudomonadota>Gammaproteobacteria>Oceanospirillales>Halomonadaceae>Halomonas | GCA_002733025.1 | Halomonas_sp. | COB32_06215 | 64469..68152 | 1486246 | ATPase-[Halomonas-sp.] |
| HBQ05713.1 | Type1.BREX | <-N6-METHYLASE-p<-N6-METHYLASE-p<-Schlafen+HATPase+wHTH<-PIN<-Phd-YefM<-DUF6079-p<-DUF6079-p*<-DUF6079-p<-BrxB-BrxF-iSTAND<-BrxA-DNA-binding | 642 | Bacteria>Pseudomonadota>Gammaproteobacteria>Oceanospirillales>Halomonadaceae>Halomonas | GCA_003529085.1 | Halomonas_sp. | DD676_03555 | 4448..6375 | 1486246 | BREX-system-P-loop-protein-BrxC-[Halomonas-sp.] |
| WP_008956851.1 | Type1.BREX | <-GP+REase<-?<-NADAR<-ART-DarT<-N6-METHYLASE-p<-N6-METHYLASE<-Schlafen+HATPase+wHTH<-PIN<-Phd-YefM<-DUF6079*<-BrxB-BrxF-iSTAND<-BrxA-DNA-binding<-tRNA<-?<-?<-?\|\|VirB11->?->DiS_P_DiS+Peptidase_A24-> | 1230 | Bacteria>Pseudomonadota>Gammaproteobacteria>Oceanospirillales>Halomonadaceae>Halomonas | GCF_000235725.1 | Halomonas_sp._HAL1 | HAL1_RS04490 | 19261..22953 | 550984 | BREX-system-P-loop-protein-BrxC-[Halomonas-sp.-HAL1] |
| WP_009095648.1 | Type1.BREX | <-TetR-HTH\|\|?->tRNA->ncRNA->Tn7_Tnp_TnsA_N+rve+Mu-transpos_C->TnsC-ATPase->HTH_19->BrxA-DNA-binding->BrxB-BrxF-iSTAND->DUF6079*->HTH+REase->Schlafen+HATPase+wHTH->N6-METHYLASE->sGTP->?->N6-METHYLASE-p->NERD+GspA-ATPase->TPR->ParB+HNH-> | 1225 | Bacteria>Pseudomonadota>Gammaproteobacteria>Oceanospirillales>Halomonadaceae>Halomonas | GCF_000236625.2 | Halomonas_sp._GFAJ-1 | BB497_RS11605 | 2566793..2570470 | 1118153 | BREX-system-P-loop-protein-BrxC-[Halomonas-sp.-GFAJ-1] |
| WP_027962108.1 | Type1.BREX | TraG_N-><-?\|\|?-><-SIGMA+OB-Fold+AAA-ATPase+Lon-Protease-BrxL<-PglZ<-HTH+REase<-N6-METHYLASE<-Schlafen+HATPase+wHTH<-DUF6079*<-BrxB-BrxF-iSTAND<-BrxA-DNA-binding<-XIS-HTH\|\|?->ARG-><-wHTH+WYL+WCX | 1144 | Bacteria>Pseudomonadota>Gammaproteobacteria>Oceanospirillales>Halomonadaceae>Halomonas | GCF_000620045.1 | Halomonas_halodenitrificans | Q358_RS0111245 | 19354..22788 | 28252 | BREX-system-P-loop-protein-BrxC-[Halomonas-halodenitrificans] |
| WP_031385055.1 | Type1.BREX | <-PglZ<-HNH<-OLD-ABC<-N6-METHYLASE-p<-HerA/FtsK<-SIR2<-REase<-N6-METHYLASE<-Schlafen+HATPase+wHTH+HTH<-DUF6079*<-BrxB-BrxF-iSTAND<-BrxA-DNA-binding<-XIS-HTH<-MobI\|\|XIS-HTH->AAA+HTH->XIS-HTH-><-Arm-DNA-bind_3+Phage_integrase | 1193 | Bacteria>Pseudomonadota>Gammaproteobacteria>Oceanospirillales>Halomonadaceae>Halomonas | GCF_000731955.1 | Halomonas_zincidurans | HALZIN_RS0115275 | 3254805..3258386 | 1178777 | BREX-system-P-loop-protein-BrxC-[Halomonas-zincidurans] |
| WP_253272941.1 | Type1.BREX | <-N6-METHYLASE-p<-N6-METHYLASE-p<-Schlafen+HATPase+wHTH+HTH<-DUF6079-p<-DUF6079-p*<-BrxB-BrxF-iSTAND<-BrxA-DNA-binding<-XIS-HTH\|\|?-><-?<-SIR2\|\|ICLR-HTH-> | 525 | Bacteria>Pseudomonadota>Gammaproteobacteria>Oceanospirillales>Halomonadaceae>Halomonas | GCF_001039575.1 | Halomonas_sp._PR-M31 | CH74_RS19345 | 4877..6454 | 1471202 | BREX-system-P-loop-protein-BrxC-[Halomonas-sp.-PR-M31] |
| WP_062374360.1 | Type1.BREX | <-ABC-ATPase+SbcC<-ABC-ATPase+SbcC<-?<-?<-?<-TetR-HTH\|\|?->tRNA->ncRNA->BrxA-DNA-binding->BrxB-BrxF-iSTAND->DUF6079*->Phd-YefM->PIN->Schlafen+HATPase+wHTH->N6-METHYLASE->N6-METHYLASE-p->HEPN->OLD-ABC->TOPRIM-> | 1227 | Bacteria>Pseudomonadota>Gammaproteobacteria>Oceanospirillales>Halomonadaceae>Halomonas | GCF_001596805.1 | Halomonas_sp._KX33721 | A3220_RS08340 | 63501..67184 | 1819251 | BREX-system-P-loop-protein-BrxC-[Halomonas-sp.-KX33721] |
| WP_078087405.1 | Type1.BREX | <-TetR-HTH\|\|tRNA->ncRNA->REase+wHTH->rve+Mu-transpos_C->TnsC-ATPase->HTH_19->BrxA-DNA-binding->BrxB-BrxF-iSTAND->DUF6079*->HTH+REase->Schlafen+HATPase+wHTH->N6-METHYLASE->?->LRR+LRR->Pkinase->PglZ->SIGMA+OB-Fold+AAA-ATPase+Lon-Protease-BrxL-> | 1225 | Bacteria>Pseudomonadota>Gammaproteobacteria>Oceanospirillales>Halomonadaceae>Halomonas | GCF_002009255.1 | unclassified_Halomonas | BV504_RS06350 | 1459939..1463616 | 2609666 | MULTISPECIES:-BREX-system-P-loop-protein-BrxC-[unclassified-Halomonas] |
| WP_101147206.1 | Type1.BREX | <-DiS_P_DiS+Peptidase_A24<-?<-VirB11\|\|?->?->?->tRNA->BrxA-DNA-binding->BrxB-BrxF-iSTAND->DUF6079*->Phd-YefM->PIN->Schlafen+HATPase+wHTH->N6-METHYLASE->HEPN-><-?\|\|OLD-ABC->TOPRIM-> | 1230 | Bacteria>Pseudomonadota>Gammaproteobacteria>Oceanospirillales>Halomonadaceae>Halomonas | GCF_002836035.1 | Halomonas_sp._MES3-P3E | CXF87_RS15340 | 100201..103893 | 2058321 | BREX-system-P-loop-protein-BrxC-[Halomonas-sp.-MES3-P3E] |
| WP_125748596.1 | Type1.BREX | <-ABC-ATPase+SbcC<-ABC-ATPase+SbcC<-?<-?<-?<-TetR-HTH\|\|?->tRNA->ncRNA->BrxA-DNA-binding->BrxB-BrxF-iSTAND->DUF6079*->Schlafen+HATPase+wHTH->N6-METHYLASE->?-><-?\|\|PglZ->SIGMA+OB-Fold+AAA-ATPase+Lon-Protease-BrxL->?->DUF488-> | 1227 | Bacteria>Pseudomonadota>Gammaproteobacteria>Oceanospirillales>Halomonadaceae>Halomonas | GCF_003950215.1 | Halomonas_venusta | DDR56_RS08110 | 77879..81562 | 44935 | BREX-system-P-loop-protein-BrxC-[Halomonas-venusta] |
| WP_133481548.1 | Type1.BREX | <-SIGMA+OB-Fold+AAA-ATPase+HKD-DNase-BrxL<-PglZ<-TOPRIM<-OLD-ABC<-HNH<-OLD-ABC<-N6-METHYLASE<-Schlafen+HATPase+wHTH+HTH<-DUF6079*<-BrxB-BrxF-iSTAND<-BrxA-DNA-binding<-XIS-HTH<-MobI\|\|XIS-HTH->AAA+HTH->XIS-HTH-><-Arm-DNA-bind_3+Phage_integrase | 1193 | Bacteria>Pseudomonadota>Gammaproteobacteria>Oceanospirillales>Halomonadaceae>Halomonas | GCF_004361885.1 | Halomonas_ventosae | DFO68_RS01985 | 17598..21179 | 229007 | BREX-system-P-loop-protein-BrxC-[Halomonas-ventosae] |
| WP_133481656.1 | Type1.BREX | MoxR-AAA->?->?-><-SIGMA+OB-Fold+AAA-ATPase+Lon-Protease-BrxL<-PglZ<-REase+AIPR<-N6-METHYLASE<-Schlafen+HATPase+wHTH<-DUF6079*<-BrxB-BrxF-iSTAND<-BrxA-DNA-binding<-XIS-HTH<-?<-?<-Peptidase_S51 | 1144 | Bacteria>Pseudomonadota>Gammaproteobacteria>Oceanospirillales>Halomonadaceae>Halomonas | GCF_004361885.1 | Halomonas_ventosae | DFO68_RS02360 | 102047..105481 | 229007 | BREX-system-P-loop-protein-BrxC-[Halomonas-ventosae] |
| WP_074210658.1 | Type1.BREX | <-ABC-ATPase+SbcC<-ABC-ATPase+SbcC<-?<-?<-?<-TetR-HTH\|\|?->tRNA->ncRNA->BrxA-DNA-binding->BrxB-BrxF-iSTAND->DUF6079*->Schlafen+HATPase+wHTH->N6-METHYLASE-> | 1227 | Bacteria>Pseudomonadota>Gammaproteobacteria>Oceanospirillales>Halomonadaceae>Halomonas | GCF_006540125.1 | Halomonas_meridiana | HME01_RS04155 | 107071..110754 | 29570 | BREX-system-P-loop-protein-BrxC-[Halomonas-meridiana] |
| WP_159343441.1 | Type1.BREX | <-Phage_AlpA\|\|HTH_26-><-SIGMA+OB-Fold+AAA-ATPase+HKD-DNase-BrxL<-PglZ<-ParB+HNH<-N6-METHYLASE<-SLOG+BirA-HTH<-SF1-SF2-Helicase+PRTase<-Schlafen+HATPase+wHTH+HTH<-DUF6079*<-BrxB-BrxF-iSTAND<-BrxA-DNA-binding<-tRNA\|\|?->?->?-><-AraC-HTH+AraC-HTH<-HTH_1+PBP_like | 1193 | Bacteria>Pseudomonadota>Gammaproteobacteria>Oceanospirillales>Halomonadaceae>Halomonas | GCF_009846525.1 | Halomonas_meridiana | CTT34_RS16815 | 3563093..3566674 | 29570 | BREX-system-P-loop-protein-BrxC-[Halomonas-meridiana] |
| WP_163558376.1 | Type1.BREX | Mrr-N-><-SIGMA+OB-Fold+AAA-ATPase+Lon-Protease-BrxL<-PglZ<-?<-?<-?<-N6-METHYLASE\|\|?-><-Schlafen+HATPase+wHTH<-DUF6079*<-BrxB-BrxF-iSTAND<-BrxA-DNA-binding<-XIS-HTH\|\|?->?->?-><-?<-LexA-protease | 1144 | Bacteria>Pseudomonadota>Gammaproteobacteria>Oceanospirillales>Halomonadaceae>Halomonas | GCF_010994375.1 | Halomonas_sp._NO4 | FEF15_RS06635 | 15883..19317 | 2484813 | BREX-system-P-loop-protein-BrxC-[Halomonas-sp.-NO4] |
| WP_172514441.1 | Type1.BREX | <-TetR-HTH\|\|?->?->tRNA->ncRNA->ABhydrolase->\|\|BrxA-DNA-binding->BrxB-BrxF-iSTAND->DUF6079*->Phd-YefM->PIN->Schlafen+HATPase+wHTH->N6-METHYLASE->ORC-CDC6->?->?->OLD-ABC->TOPRIM-> | 1227 | Bacteria>Pseudomonadota>Gammaproteobacteria>Oceanospirillales>Halomonadaceae>Halomonas | GCF_011764625.1 | Halomonas_meridiana | HMEPL2_RS01760 | 381650..385333 | 29570 | BREX-system-P-loop-protein-BrxC-[Halomonas-meridiana] |
| WP_172515133.1 | Type1.BREX | <-PglZ<-ParB+HNH<-N6-METHYLASE-p\|\|<-N6-METHYLASE-p<-HerA/FtsK<-?<-SF1-SF2-Helicase<-N6-METHYLASE<-Schlafen+HATPase+wHTH+HTH<-DUF6079*<-BrxB-BrxF-iSTAND<-BrxA-DNA-binding<-XIS-HTH<-MobI\|\|XIS-HTH->AAA+HTH->XIS-HTH->SIGMA-HTH+HIN-HTH-> | 1193 | Bacteria>Pseudomonadota>Gammaproteobacteria>Oceanospirillales>Halomonadaceae>Halomonas | GCF_011764625.1 | Halomonas_meridiana | HMEPL2_RS11780 | 2496852..2500433 | 29570 | BREX-system-P-loop-protein-BrxC-[Halomonas-meridiana] |
| WP_153844139.1 | Type1.BREX | <-PglZ<-Pkinase<-REase<-?<-NADAR<-ART-DarT<-?<-N6-METHYLASE<-Schlafen+HATPase+wHTH<-DUF6079*<-BrxB-BrxF-iSTAND<-BrxA-DNA-binding<-HTH_19<-TnsC-ATPase<-rve\|\|DUF772+DDE_Tnp_1-><-rve+Mu-transpos_C<-REase+wHTH\|\|ZnR+HTH_Tnp_ISL3+DDE_Tnp_ISL3-> | 1227 | Bacteria>Pseudomonadota>Gammaproteobacteria>Oceanospirillales>Halomonadaceae>Halomonas | GCF_012427705.1 | Halomonas_piezotolerans | GYM47_RS09720 | 2110719..2114402 | 2609667 | BREX-system-P-loop-protein-BrxC-[Halomonas-piezotolerans] |
| WP_169956446.1 | Type1.BREX | XIS-HTH->BrxA-DNA-binding->BrxB-BrxF-iSTAND->DUF6079*->Schlafen+HATPase+wHTH+HTH->N6-METHYLASE->OLD-ABC->TOPRIM->PglZ->SIGMA+OB-Fold+AAA-ATPase+HKD-DNase-BrxL-><-REC+OmpR-HTH | 1193 | Bacteria>Pseudomonadota>Gammaproteobacteria>Oceanospirillales>Halomonadaceae>Halomonas | GCF_013000985.1 | Halomonas_sp._PGE1 | HIR79_RS02110 | 421842..425423 | 2730360 | BREX-system-P-loop-protein-BrxC-[Halomonas-sp.-PGE1] |
| WP_180091726.1 | Type1.BREX | <-REase<-ParB+HNH<-TPR<-NERD+GspA-ATPase<-N6-METHYLASE-p<-?<-sGTP<-N6-METHYLASE<-Schlafen+HATPase+wHTH<-DUF6079*<-BrxB-BrxF-iSTAND<-?<-JAB\|\|?->wHTH+WYL+WCX->?->?-><-?<-CN_hydrolase+NAD_synthase | 1227 | Bacteria>Pseudomonadota>Gammaproteobacteria>Oceanospirillales>Halomonadaceae>Halomonas | GCF_013416325.1 | Halomonas_sedimenti | HZU72_RS10350 | 63822..67505 | 2729618 | BREX-system-P-loop-protein-BrxC-[Halomonas-sedimenti] |
| WP_183333607.1 | Type1.BREX | <-HIN-HTH\|\|XIS-HTH->BrxA-DNA-binding->BrxB-BrxF-iSTAND->DUF6079*->Schlafen+HATPase+wHTH->N6-METHYLASE->OLD-ABC->NERD->PglZ->SIGMA+OB-Fold+AAA-ATPase+Lon-Protease-BrxL-><-?\|\|?-><-TraG_N | 1144 | Bacteria>Pseudomonadota>Gammaproteobacteria>Oceanospirillales>Halomonadaceae>Halomonas | GCF_014193375.1 | Halomonas_campaniensis | BDK63_RS15585 | 2230..5664 | 213554 | BREX-system-P-loop-protein-BrxC-[Halomonas-campaniensis] |
| WP_192538995.1 | Type1.BREX | <-TetR-HTH\|\|?->tRNA->ncRNA->REase+wHTH->rve->TnsC-ATPase->HTH_19->BrxA-DNA-binding->BrxB-BrxF-iSTAND->DUF6079*->Schlafen+HATPase+wHTH->N6-METHYLASE-> | 1227 | Bacteria>Pseudomonadota>Gammaproteobacteria>Oceanospirillales>Halomonadaceae>Halomonas | GCF_014861465.1 | Halomonas_sp._FME20 | EI547_RS13715 | 35008..38691 | 2487703 | BREX-system-P-loop-protein-BrxC-[Halomonas-sp.-FME20] |
| WP_193059740.1 | Type1.BREX | TnsC-ATPase->HTH_19->BrxA-DNA-binding->BrxB-BrxF-iSTAND->DUF6079*->Schlafen+HATPase+wHTH->N6-METHYLASE->?->?->OLD-ABC->TOPRIM->PglZ->SIGMA+OB-Fold+AAA-ATPase+Lon-Protease-BrxL-> | 1227 | Bacteria>Pseudomonadota>Gammaproteobacteria>Oceanospirillales>Halomonadaceae>Halomonas | GCF_014897995.1 | unclassified_Halomonas | INT28_RS17780 | 3434..7117 | 2609666 | MULTISPECIES:-BREX-system-P-loop-protein-BrxC-[unclassified-Halomonas] |
| WP_218314905.1 | Type1.BREX | <-ABC-ATPase+SbcC<-ABC-ATPase+SbcC<-?<-?<-?<-TetR-HTH\|\|tRNA->ncRNA->BrxA-DNA-binding->BrxB-BrxF-iSTAND->DUF6079*->Schlafen+HATPase+wHTH->N6-METHYLASE->?->?->OLD-ABC->TOPRIM->PglZ->SIGMA+OB-Fold+AAA-ATPase+Lon-Protease-BrxL-> | 1227 | Bacteria>Pseudomonadota>Gammaproteobacteria>Oceanospirillales>Halomonadaceae>Halomonas | GCF_019211725.1 | Halomonas_sp._18071143 | KVY02_RS07715 | 1661242..1664925 | 2855441 | BREX-system-P-loop-protein-BrxC-[Halomonas-sp.-18071143] |
| WP_231664595.1 | Type1.BREX | GDResolvase+HIN-HTH->?-><-SIGMA+OB-Fold+AAA-ATPase+Lon-Protease-BrxL<-PglZ<-TOPRIM<-OLD-ABC<-HTH+REase<-N6-METHYLASE<-Schlafen+HATPase+wHTH<-DUF6079*<-BrxB-BrxF-iSTAND-p<-BrxA-DNA-binding<-XIS-HTH<-MobI | 1153 | Bacteria>Pseudomonadota>Gammaproteobacteria>Oceanospirillales>Halomonadaceae>Halomonas | GCF_021151495.1 | Halomonas_sp._IOP_31 | K7I12_RS15025 | 26514..29975 | 2876584 | BREX-system-P-loop-protein-BrxC-[Halomonas-sp.-IOP_31] |
| WP_239296660.1 | Type1.BREX | Phage_integrase-><-XIS-HTH<-AAA+HTH<-XIS-HTH\|\|MobI->XIS-HTH->BrxA-DNA-binding->BrxB-BrxF-iSTAND->DUF6079*->Schlafen+HATPase+wHTH+HTH->N6-METHYLASE->REase->SIR2->HerA/FtsK->N6-METHYLASE-p->OLD-ABC->TOPRIM->PglZ-> | 1193 | Bacteria>Pseudomonadota>Gammaproteobacteria>Oceanospirillales>Halomonadaceae>Halomonas | GCF_022348215.1 | unclassified_Halomonas | MHM35_RS10400 | 116133..119714 | 2609666 | MULTISPECIES:-BREX-system-P-loop-protein-BrxC-[unclassified-Halomonas] |
| WP_253557196.1 | Type1.BREX | <-N6-METHYLASE<-Schlafen+HATPase+wHTH<-DUF6079*<-BrxB-BrxF-iSTAND<-BrxA-DNA-binding<-ncRNA<-tRNA<-?\|\|TetR-HTH->?->?->?->ABC-ATPase+SbcC->ABC-ATPase+SbcC-> | 1225 | Bacteria>Pseudomonadota>Gammaproteobacteria>Oceanospirillales>Halomonadaceae>Halomonas | GCF_024158065.1 | unclassified_Halomonas | BIS08_RS08535 | 5439..9116 | 2609666 | MULTISPECIES:-BREX-system-P-loop-protein-BrxC-[unclassified-Halomonas] |
| WP_263591023.1 | Type1.BREX | RepB_primase-><-SIGMA+OB-Fold+AAA-ATPase+Lon-Protease-BrxL<-PglZ<-ParB+HNH<-?<-?<-?<-N6-METHYLASE<-Schlafen+HATPase+wHTH<-DUF6079*<-BrxB-BrxF-iSTAND<-BrxA-DNA-binding<-tRNA<-?\|\|?->?->sGTP-> | 1144 | Bacteria>Pseudomonadota>Gammaproteobacteria>Oceanospirillales>Halomonadaceae>Halomonas | GCF_025725685.1 | Halomonas_sp._M4R1S46 | OCT48_RS01560 | 327087..330521 | 2982692 | BREX-system-P-loop-protein-BrxC-[Halomonas-sp.-M4R1S46] |
| WP_103969332.1 | Type1.BREX | HicA_toxin->HicB_lk_antitox-><-?<-?<-?\|\|XIS-HTH->BrxA-DNA-binding->BrxB-BrxF-iSTAND->DUF6079*->Schlafen+HATPase+wHTH+HTH->SF1-SF2-Helicase+PRTase->SLOG+BirA-HTH->N6-METHYLASE->PglZ->SIGMA+OB-Fold+AAA-ATPase+HKD-DNase-BrxL-><-REC+OmpR-HTH | 1193 | Bacteria>Pseudomonadota>Gammaproteobacteria>Oceanospirillales>Halomonadaceae>Halomonas | GCF_900108215.1 | Halomonas_desiderata | C4S71_RS12465 | 87857..91438 | 52021 | BREX-system-P-loop-protein-BrxC-[Halomonas-desiderata] |
| WP_089675693.1 | Type1.BREX | <-SIGMA+OB-Fold+AAA-ATPase+Lon-Protease-BrxL<-PglZ<-DOC+HTH<-?<-?<-N6-METHYLASE<-Schlafen+HATPase+wHTH<-PIN<-Phd-YefM<-DUF6079*<-BrxB-BrxF-iSTAND<-BrxA-DNA-binding<-ncRNA<-tRNA<-?\|\|TetR-HTH->?->?->?->ABC-ATPase+SbcC->ABC-ATPase+SbcC-> | 1227 | Bacteria>Pseudomonadota>Gammaproteobacteria>Oceanospirillales>Halomonadaceae>Halomonas | GCF_900110265.1 | Halomonas_aquamarina | BMY49_RS12195 | 13754..17437 | 77097 | BREX-system-P-loop-protein-BrxC-[Halomonas-aquamarina] |
| WP_079550430.1 | Type1.BREX | <-TetR-HTH\|\|?->tRNA->ncRNA-><-?\|\|BrxA-DNA-binding->BrxB-BrxF-iSTAND->DUF6079*->Schlafen+HATPase+wHTH-p->HATPase_c->N6-METHYLASE->OLD-ABC->TOPRIM->?->DOC+HTH->PglZ->SIGMA+OB-Fold+AAA-ATPase+Lon-Protease-BrxL-> | 1240 | Bacteria>Pseudomonadota>Gammaproteobacteria>Oceanospirillales>Halomonadaceae>Halomonas | GCF_900142895.1 | Halomonas_subglaciescola | B5495_RS00605 | 132715..136437 | 29571 | BREX-system-P-loop-protein-BrxC-[Halomonas-subglaciescola] |
| WP_159177832.1 | Type1.BREX | <-SF1-SF2-Helicase<-OLD-ABC+TOPRIM<-SIGMA+OB-Fold+AAA-ATPase+HKD-DNase-BrxL<-PglZ<-N6-METHYLASE<-SLOG+BirA-HTH<-SF1-SF2-Helicase+PRTase<-Schlafen+HATPase+wHTH+HTH<-DUF6079*<-BrxB-BrxF-iSTAND<-BrxA-DNA-binding<-XIS-HTH<-MobI<-?\|\|XIS-HTH->AAA+HTH->XIS-HTH-><-Arm-DNA-bind_3+Phage_integrase | 1193 | Bacteria>Pseudomonadota>Gammaproteobacteria>Oceanospirillales>Halomonadaceae>Halomonas | GCF_902506335.1 | unclassified_Halomonas | JMX33_RS24540 | 5309589..5313170 | 2609666 | MULTISPECIES:-BREX-system-P-loop-protein-BrxC-[unclassified-Halomonas] |
| WP_179915499.1 | Type1.BREX | <-N6-METHYLASE<-Schlafen+HATPase+wHTH<-DUF6079*<-BrxB-BrxF-iSTAND<-BrxA-DNA-binding<-Aminotran_3<-?\|\|GNTR-HTH+Aminotran_1_2-> | 1144 | Bacteria>Pseudomonadota>Gammaproteobacteria>Oceanospirillales>Halomonadaceae>Halomonas | GCF_947331475.1 | Halomonas_glaciei | HZS80_RS06355 | 4022..7456 | 186761 | BREX-system-P-loop-protein-BrxC-[Halomonas-glaciei] |
| WP_254893719.1 | Type1.BREX | <-ABC-ATPase+SbcC<-ABC-ATPase+SbcC<-?<-?<-?<-?<-TetR-HTH\|\|tRNA->ncRNA->BrxA-DNA-binding->BrxB-BrxF-iSTAND->DUF6079*->Schlafen+HATPase+wHTH->N6-METHYLASE->?->?->OLD-ABC->TOPRIM->MULE-TRANSPOSASE-> | 1210 | Bacteria>Pseudomonadota>Gammaproteobacteria>Oceanospirillales>Halomonadaceae>Halomonas | - | Halomonas_sp._BC1 | B7L72_RS00095 | 19375..23007 | 1670448 | BREX-system-P-loop-protein-BrxC-[Halomonas-sp.-BC1] |
| WP_087719602.1 | Type1.BREX | <-ICLR-HTH<-?\|\|?->Amidohydro_1->GNTR-HTH->XIS-HTH->BrxA-DNA-binding->BrxB-BrxF-iSTAND->DUF6079*->Schlafen+HATPase+wHTH->N6-METHYLASE->PHP+SbcC->OLD-ABC->HNH->PglZ->SIGMA+OB-Fold+AAA-ATPase+Lon-Protease-BrxL-> | 1145 | Bacteria>Pseudomonadota>Gammaproteobacteria>Oceanospirillales>Halomonadaceae>Salinicola | GCF_002179555.1 | Salinicola_salarius | CCD76_RS11235 | 10059..13496 | 430457 | BREX-system-P-loop-protein-BrxC-[Salinicola-salarius] |
| WP_110649043.1 | Type1.BREX | <-ICLR-HTH<-?\|\|?->Amidohydro_1->GNTR-HTH->XIS-HTH->BrxA-DNA-binding->BrxB-BrxF-iSTAND->DUF6079*->Schlafen+HATPase+wHTH->N6-METHYLASE->HTH+REase-><-DDE_Tnp_1_4 | 1153 | Bacteria>Pseudomonadota>Gammaproteobacteria>Oceanospirillales>Halomonadaceae>Salinicola | GCF_003206715.1 | Salinicola_peritrichatus | DOM57_RS09900 | 59337..62798 | 1267424 | BREX-system-P-loop-protein-BrxC-[Salinicola-peritrichatus] |
| WP_027854332.1 | Type1.BREX | GDResolvase+HIN-HTH->?->?-><-SIGMA+OB-Fold+AAA-ATPase+HKD-DNase-BrxL<-PglZ<-N6-METHYLASE<-REase<-OLD-ABC<-Schlafen+HATPase+wHTH+HTH<-DUF6079*<-BrxB-BrxF-iSTAND<-BrxA-DNA-binding<-XIS-HTH<-MobI\|\|XIS-HTH->AAA+HTH->XIS-HTH-><-Arm-DNA-bind_3+Phage_integrase | 1193 | Bacteria>Pseudomonadota>Gammaproteobacteria>Oceanospirillales>Oceanospirillaceae>Marinobacterium | GCF_000428985.1 | Marinobacterium_litorale | G403_RS0112740 | 74962..78543 | 404770 | BREX-system-P-loop-protein-BrxC-[Marinobacterium-litorale] |
| MAS25385.1 | Type1.BREX | <-SIGMA+OB-Fold+AAA-ATPase+Lon-Protease-BrxL<-PglZ<-ParB+HNH<-PHP+SbcC<-N6-METHYLASE<-Schlafen+HATPase+wHTH<-DUF6079*<-BrxB-BrxF-iSTAND<-BrxA-DNA-binding<-XIS-HTH | 1145 | Bacteria>Pseudomonadota>Gammaproteobacteria>Oceanospirillales>Oceanospirillaceae>unclassified_Oceanospirillaceae | GCA_002698675.1 | Oceanospirillaceae_bacterium | CMI08_10750 | 15208..18645 | 1899355 | hypothetical-protein-[Oceanospirillaceae-bacterium] |
| KXS53697.1 | Type1.BREX | <-HNH<-?<-OLD-ABC<-N6-METHYLASE<-HTH+HTH<-Schlafen+HATPase+wHTH<-DUF6079-p\|\|?-><-DUF6079-p*<-BrxB-BrxF-iSTAND<-BrxA-DNA-binding<-XIS-HTH<-MobI\|\|XIS-HTH->AAA+HTH->XIS-HTH-><-Arm-DNA-bind_3+Phage_integrase | 539 | Bacteria>Pseudomonadota>Gammaproteobacteria>Pseudomonadales>Marinobacteraceae>Marinobacter | GCA_001577575.1 | Marinobacter_sp._T13-3 | AWU57_1918 | 9032..10651 | 1794906 | hypothetical-protein-AWU57_1918-[Marinobacter-sp.-T13-3] |
| MAH31343.1 | Type1.BREX | <-Schlafen+HATPase+wHTH<-DUF6079*<-BrxB-BrxF-iSTAND<-BrxA-DNA-binding<-XIS-HTH<-MobI\|\|XIS-HTH->AAA+HTH->XIS-HTH-><-Arm-DNA-bind_3+Phage_integrase | 1193 | Bacteria>Pseudomonadota>Gammaproteobacteria>Pseudomonadales>Marinobacteraceae>Marinobacter | GCA_002690365.1 | Marinobacter_sp. | CMG78_04665 | 1378..4959 | 50741 | hypothetical-protein-[Marinobacter-sp.] |
| MBE95557.1 | Type1.BREX | BrxB-BrxF-iSTAND->DUF6079*->Schlafen+HATPase+wHTH+HTH->SF1-SF2-Helicase+PRTase->SLOG+BirA-HTH->N6-METHYLASE->ParB+HNH->PglZ->SIGMA+OB-Fold+AAA-ATPase+HKD-DNase-BrxL-><-?<-IstB_IS21 | 1193 | Bacteria>Pseudomonadota>Gammaproteobacteria>Pseudomonadales>Marinobacteraceae>Marinobacter | GCA_002714765.1 | Marinobacter_sp. | CMG89_11180 | 696..4277 | 50741 | hypothetical-protein-[Marinobacter-sp.] |
| TVT30739.1 | Type1.BREX | GDResolvase+HIN-HTH-><-?<-REase<-SIGMA+OB-Fold+AAA-ATPase+Lon-Protease-BrxL<-PglZ<-Schlafen+HATPase+wHTH<-PHP+SbcC<-N6-METHYLASE<-Schlafen+HATPase+wHTH<-DUF6079*<-BrxB-BrxF-iSTAND<-BrxA-DNA-binding<-XIS-HTH\|\|XIS-HTH->APATPase->?-><-Phage_int_SAM_5+Phage_integrase | 1145 | Bacteria>Pseudomonadota>Gammaproteobacteria>Pseudomonadales>Marinobacteraceae>Marinobacter | GCA_007713625.1 | Marinobacter_vinifirmus | FHK81_16390 | 18158..21595 | 355591 | BREX-system-P-loop-protein-BrxC-[Marinobacter-vinifirmus] |
| MBC7192650.1 | Type1.BREX | <-SIGMA+OB-Fold+AAA-ATPase+Lon-Protease-BrxL<-PglZ<-HNH<-OLD-ABC<-TOPRIM<-OLD-ABC<-HTH+REase<-N6-METHYLASE<-Schlafen+HATPase+wHTH<-DUF6079*<-BrxB-BrxF-iSTAND<-BrxA-DNA-binding<-XIS-HTH\|\|HTH_Tnp_1+HTH_21+rve+rve_3->XIS-HTH->DnaB->?-><-Phage_int_SAM_5+Phage_integrase<-sGTP | 1145 | Bacteria>Pseudomonadota>Gammaproteobacteria>Pseudomonadales>Marinobacteraceae>Marinobacter | GCA_014359475.1 | Marinobacter_sp. | H5U31_09890 | 22634..26071 | 50741 | BREX-system-P-loop-protein-BrxC-[Marinobacter-sp.] |
| WP_011784101.1 | Type1.BREX | Arm-DNA-bind_3+Phage_integrase-><-XIS-HTH<-AAA+HTH<-XIS-HTH\|\|MobI->XIS-HTH->BrxA-DNA-binding->BrxB-BrxF-iSTAND->DUF6079*->Schlafen+HATPase+wHTH+HTH->SF1-SF2-Helicase+PRTase->SLOG+BirA-HTH->N6-METHYLASE->OLD-ABC->TOPRIM->PglZ->SIGMA+OB-Fold+AAA-ATPase+HKD-DNase-BrxL->SbcC-> | 1193 | Bacteria>Pseudomonadota>Gammaproteobacteria>Pseudomonadales>Marinobacteraceae>Marinobacter | GCF_000015365.1 | Marinobacter_nauticus | MAQU_RS02805 | 628948..632529 | 2743 | BREX-system-P-loop-protein-BrxC-[Marinobacter-nauticus] |
| WP_007349110.1 | Type1.BREX | <-Phage_int_SAM_1+Phage_integrase<-REase+N6_Mtase<-HTH_26<-TetR-HTH<-Phage_integrase+Phage_integrase<-Phage_integrase+Phage_integrase<-Phage_integrase<-N6-METHYLASE-p<-Schlafen+HATPase+wHTH<-DUF6079*<-BrxB-BrxF-iSTAND<-BrxA-DNA-binding<-XIS-HTH\|\|Arm-DNA-bind_3->?->DDE_Tnp_1_4-><-?<-?<-DDE_Tnp_IS66 | 1145 | Bacteria>Pseudomonadota>Gammaproteobacteria>Pseudomonadales>Marinobacteraceae>Marinobacter | GCF_000169375.1 | Marinobacter_sp._ELB17 | MELB17_RS05280 | 142493..145930 | 270374 | BREX-system-P-loop-protein-BrxC-[Marinobacter-sp.-ELB17] |
| WP_007349691.1 | Type5.BREX | <-Glycos_transf_2\|\|tRNA->?->?-><-SIGMA+OB-Fold+AAA-ATPase+HKD-DNase-BrxL<-PglZ<-HNH<-OLD-ABC<-N6-METHYLASE<-Schlafen+HATPase+wHTH+HTH<-DUF6079*<-BrxB-BrxF-iSTAND<-BrxA-DNA-binding<-TUDOR+SF1-SF2-Helicase+iREase+CTD+REase<-TPR | 1194 | Bacteria>Pseudomonadota>Gammaproteobacteria>Pseudomonadales>Marinobacteraceae>Marinobacter | GCF_000169375.1 | Marinobacter_sp._ELB17 | MELB17_RS08075 | 35418..39002 | 270374 | BREX-system-P-loop-protein-BrxC-[Marinobacter-sp.-ELB17] |
| WP_008174228.1 | Type1.BREX | Arm-DNA-bind_3+Phage_integrase-><-XIS-HTH<-AAA+HTH<-XIS-HTH\|\|MobI->XIS-HTH->BrxA-DNA-binding->BrxB-BrxF-iSTAND->DUF6079*->Schlafen+HATPase+wHTH+HTH->N6-METHYLASE->OLD-ABC->TOPRIM-> | 1193 | Bacteria>Pseudomonadota>Gammaproteobacteria>Pseudomonadales>Marinobacteraceae>Marinobacter | GCF_000235625.1 | Marinobacter_manganoxydans | KYE_13690 | 42273..45854 | 1150997 | BREX-system-P-loop-protein-BrxC-[Marinobacter-manganoxydans] |
| WP_018403235.1 | Type1.BREX | CHTH->GDResolvase->ZnR-><-GDResolvase+HIN-HTH<-SIGMA+OB-Fold+AAA-ATPase+Lon-Protease-BrxL<-PglZ<-REase+AIPR<-N6-METHYLASE<-Schlafen+HATPase+wHTH<-DUF6079*<-BrxB-BrxF-iSTAND<-BrxA-DNA-binding<-XIS-HTH<-tRNA<-HAD+HAD<-HDRnase<-?<-?<-DNA_METHYLASE | 1145 | Bacteria>Pseudomonadota>Gammaproteobacteria>Pseudomonadales>Marinobacteraceae>Marinobacter | GCF_000372805.1 | Marinobacter_gelidimuriae | A3OI_RS0105075 | 19872..23309 | 2739064 | BREX-system-P-loop-protein-BrxC-[Marinobacter-gelidimuriae] |
| WP_018404435.1 | Type1.BREX | OrfB_Zn_ribbon->BrxA-DNA-binding->BrxB-BrxF-iSTAND->DUF6079*->Schlafen+HATPase+wHTH+HTH->N6-METHYLASE->OLD-ABC->HNH->PglZ->SIGMA+OB-Fold+AAA-ATPase+HKD-DNase-BrxL->N6-METHYLASE-> | 1194 | Bacteria>Pseudomonadota>Gammaproteobacteria>Pseudomonadales>Marinobacteraceae>Marinobacter | GCF_000372805.1 | Marinobacter_gelidimuriae | A3OI_RS0111615 | 2203..5787 | 2739064 | BREX-system-P-loop-protein-BrxC-[Marinobacter-gelidimuriae] |
| WP_022988850.1 | Type1.BREX | <-PglZ<-TOPRIM<-OLD-ABC<-HNH<-OLD-ABC<-N6-METHYLASE<-REase<-OLD-ABC<-Schlafen+HATPase+wHTH+HTH<-DUF6079*<-BrxB-BrxF-iSTAND<-BrxA-DNA-binding<-XIS-HTH<-MobI | 1193 | Bacteria>Pseudomonadota>Gammaproteobacteria>Pseudomonadales>Marinobacteraceae>Marinobacter | GCF_000475255.1 | Marinobacter_sp._ES-1 | Q668_RS06800 | 88554..92135 | 1396858 | BREX-system-P-loop-protein-BrxC-[Marinobacter-sp.-ES-1] |
| WP_022990203.1 | Type1.BREX | <-DnaB<-XIS-HTH\|\|XIS-HTH->BrxA-DNA-binding->BrxB-BrxF-iSTAND->DUF6079*->Schlafen+HATPase+wHTH+HTH->N6-METHYLASE->REase->SIR2->HerA/FtsK->N6-METHYLASE-p->PglZ->Immank+ANK+ANK->REase-> | 1189 | Bacteria>Pseudomonadota>Gammaproteobacteria>Pseudomonadales>Marinobacteraceae>Marinobacter | GCF_000475255.1 | Marinobacter_sp._ES-1 | Q668_RS13780 | 2583..6152 | 1396858 | BREX-system-P-loop-protein-BrxC-[Marinobacter-sp.-ES-1] |
| WP_023011723.1 | Type1.BREX | <-OLD-ABC<-N6-METHYLASE-p<-HerA/FtsK<-SIR2<-REase<-N6-METHYLASE<-SLOG+BirA-HTH<-SF1-SF2-Helicase+PRTase<-Schlafen+HATPase+wHTH+HTH<-DUF6079*<-BrxB-BrxF-iSTAND<-BrxA-DNA-binding<-XIS-HTH<-MobI\|\|XIS-HTH->AAA+HTH->XIS-HTH-><-Arm-DNA-bind_3+Phage_integrase | 1193 | Bacteria>Pseudomonadota>Gammaproteobacteria>Pseudomonadales>Marinobacteraceae>Marinobacter | GCF_000475315.1 | Marinobacter_sp._EN3 | Q675_RS32885 | 62164..65745 | 1397533 | BREX-system-P-loop-protein-BrxC-[Marinobacter-sp.-EN3] |

| Protein-ID | System | Gene-Neighbourhoods | Length | Taxonomy | Assembly-ID | Strain | Locus-Tag | Genomic-coordinates | TaxID | Fasta-defline |
|---|---|---|---|---|---|---|---|---|---|---|
| WP_195393981.1 | Type1.BREX | PemK_toxin->?->PolB->HEPN-><-?<-?\|\|?->BrxA-DNA-binding->BrxB-BrxF-iSTAND->DUF6079*->N6-METHYLASE->SIR2->Schlafen+HATPase+wHTH->PglZ->SIGMA+OB-Fold+AAA-ATPase+HKD-DNase-BrxL->ZnR+GP->ZnR->ParB+HNH+RAMA-> | 1218 | Bacteria>Terrabacteria_group>Actinomycetota>Actinomycetes>Bifidobacteriales>Bifidobacteriaceae | GCF_015553845.1 | Bifidobacterium_adolescentis | I2D91_RS01775 | 423737..427393 | 1680 | BREX-system-P-loop-protein-BrxC-[Bifidobacterium-adolescentis] |
| WP_195451643.1 | Type1.BREX | <-ABhydrolase<-?<-Glyoxalase\|\|DSBH+AraC-HTH+AraC-HTH-><-ZnR<-SIGMA+OB-Fold+AAA-ATPase+HKD-DNase-BrxL<-PglZ<-Schlafen+HATPase+wHTH<-N6-METHYLASE<-DUF6079*<-BrxB-BrxF-iSTAND<-BrxA-DNA-binding<-?\|\|ACET-><-?\|\|HTH-><-35exo+BRCT<-35exo+HRDC | 1217 | Bacteria>Terrabacteria_group>Actinomycetota>Actinomycetes>Bifidobacteriales>Bifidobacteriaceae | GCF_015555105.1 | Bifidobacterium_dentium | I2E42_RS03170 | 226129..229782 | 1689 | BREX-system-P-loop-protein-BrxC-[Bifidobacterium-dentium] |
| WP_195482034.1 | Type1.BREX | <-ParB+HNH+RAMA<-ZnR<-ZnR+GP<-SIGMA+OB-Fold+AAA-ATPase+HKD-DNase-BrxL<-PglZ<-Schlafen+HATPase+wHTH<-SIR2<-N6-METHYLASE<-DUF6079*<-BrxB-BrxF-iSTAND<-BrxA-DNA-binding<-?\|\|?->?-><-HEPN<-PolB<-PemK_toxin<-RelB | 1218 | Bacteria>Terrabacteria_group>Actinomycetota>Actinomycetes>Bifidobacteriales>Bifidobacteriaceae | GCF_015555595.1 | Bifidobacterium_adolescentis | I2E65_RS01565 | 362593..366249 | 1680 | BREX-system-P-loop-protein-BrxC-[Bifidobacterium-adolescentis] |
| WP_195936831.1 | Type1.BREX | REase-><-?<-SIGMA+OB-Fold+AAA-ATPase+HKD-DNase-BrxL<-PglZ<-Schlafen+HATPase+wHTH<-?<-N6-METHYLASE<-DUF6079*<-BrxB-BrxF-iSTAND<-BrxA-DNA-binding<-?<-?<-?<-ABC-ATPase+SbcC<-ZnR | 1218 | Bacteria>Terrabacteria_group>Actinomycetota>Actinomycetes>Bifidobacteriales>Bifidobacteriaceae | GCF_015667555.1 | Bifidobacterium_bifidum | I4U71_RS07235 | 13396..17052 | 1681 | BREX-system-P-loop-protein-BrxC-[Bifidobacterium-bifidum] |
| WP_195938980.1 | Type1.BREX | 35exo+HRDC-><-HTH\|\|ACET+ACET-><-?<-SbcD<-?<-REase\|\|BrxA-DNA-binding->BrxB-BrxF-iSTAND->DUF6079*->N6-METHYLASE->SIR2->Schlafen+HATPase+wHTH->PglZ->SIGMA+OB-Fold+AAA-ATPase+HKD-DNase-BrxL->ZnR+GP->ZnR->ParB+HNH+RAMA-> | 1217 | Bacteria>Terrabacteria_group>Actinomycetota>Actinomycetes>Bifidobacteriales>Bifidobacteriaceae | GCF_015667675.1 | Bifidobacterium_pseudocatenulatum | I4U64_RS06630 | 371599..375252 | 28026 | BREX-system-P-loop-protein-BrxC-[Bifidobacterium-pseudocatenulatum] |
| WP_196336889.1 | Type1.BREX | 35exo+HRDC->35exo+BRCT-><-HTH\|\|?-><-ACET\|\|?->BrxA-DNA-binding->BrxB-BrxF-iSTAND->DUF6079*->N6-METHYLASE->HEPN->?->Schlafen+HATPase+wHTH->PglZ->SIGMA+OB-Fold+AAA-ATPase+HKD-DNase-BrxL->ZnR-><-DSBH+AraC-HTH+AraC-HTH\|\|Glyoxalase-> | 1217 | Bacteria>Terrabacteria_group>Actinomycetota>Actinomycetes>Bifidobacteriales>Bifidobacteriaceae | GCF_015703835.1 | Bifidobacterium_dentium | IHQ82_RS06050 | 179848..183501 | 1689 | BREX-system-P-loop-protein-BrxC-[Bifidobacterium-dentium] |
| WP_202574831.1 | Type1.BREX | <-Peptidase_S26<-?\|\|ABC-ATPase+SbcC->?-><-?<-SIGMA+OB-Fold+AAA-ATPase+HKD-DNase-BrxL<-PglZ<-Schlafen+HATPase+wHTH<-N6-METHYLASE<-DUF6079*<-BrxB-BrxF-iSTAND<-BrxA-DNA-binding<-HTH_28 | 1219 | Bacteria>Terrabacteria_group>Actinomycetota>Actinomycetes>Bifidobacteriales>Bifidobacteriaceae | GCF_016759645.1 | Bifidobacterium_longum | JL737_RS05465 | 24879..28538 | 216816 | BREX-system-P-loop-protein-BrxC-[Bifidobacterium-longum] |
| WP_204464696.1 | Type1.BREX | ACET-><-O6MGT-HTH\|\|Phage_integrase-><-?<-SIGMA+OB-Fold+AAA-ATPase+HKD-DNase-BrxL<-PglZ<-Schlafen+HATPase+wHTH<-N6-METHYLASE<-DUF6079*<-BrxB-BrxF-iSTAND<-BrxA-DNA-binding<-?<-?<-?\|\|Glycoside_Hydrolase-><-ArsR-HTH<-Thioredoxin_3 | 1221 | Bacteria>Terrabacteria_group>Actinomycetota>Actinomycetes>Bifidobacteriales>Bifidobacteriaceae | GCF_016899675.1 | Bifidobacterium_pullorum | H6A68_RS03865 | 164290..167955 | 78448 | BREX-system-P-loop-protein-BrxC-[Bifidobacterium-pullorum] |
| WP_204389272.1 | Type1.BREX | <-IstB_IS21\|\|AAA-ATPase-><-CMP-hydrolase<-SIGMA+OB-Fold+AAA-ATPase+HKD-DNase-BrxL<-PglZ<-Schlafen+HATPase+wHTH<-N6-METHYLASE<-DUF6079*<-BrxB-BrxF-iSTAND<-BrxA-DNA-binding<-TOPRIM<-OLD-ABC\|\|Glycoside_Hydrolase-><-ACET+ACET<-ACET | 1221 | Bacteria>Terrabacteria_group>Actinomycetota>Actinomycetes>Bifidobacteriales>Bifidobacteriaceae | GCF_016900615.1 | Bifidobacterium_pullorum | H6A14_RS02700 | 186110..189775 | 78448 | BREX-system-P-loop-protein-BrxC-[Bifidobacterium-pullorum] |
| WP_214374600.1 | Type1.BREX | ZnR->?->ABC-ATPase+SbcC-><-?\|\|?-><-ABhydrolase\|\|BrxA-DNA-binding->BrxB-BrxF-iSTAND->DUF6079*->CR-ATPase5+REase->N6-METHYLASE->?->?->HEPN+RES->Schlafen+HATPase+wHTH->PglZ->MIT_C->METHYLASE-> | 1249 | Bacteria>Terrabacteria_group>Actinomycetota>Actinomycetes>Bifidobacteriales>Bifidobacteriaceae | GCF_018555335.1 | Bifidobacterium | JS541_RS11260 | 52363..56112 | 1678 | MULTISPECIES:-BREX-system-P-loop-protein-BrxC-[Bifidobacterium] |
| WP_226586661.1 | Type1.BREX | REC+LuxR-HTH->HISKIN->?->?-><-SbcD<-?<-REase\|\|BrxA-DNA-binding->BrxB-BrxF-iSTAND->DUF6079*->N6-METHYLASE->?->Schlafen+HATPase+wHTH->PglZ->SIGMA+OB-Fold+AAA-ATPase+HKD-DNase-BrxL->SLATT->SMODS->ParB+HNH+RAMA-> | 1219 | Bacteria>Terrabacteria_group>Actinomycetota>Actinomycetes>Bifidobacteriales>Bifidobacteriaceae | GCF_020525765.1 | Bifidobacterium_pseudocatenulatum | PG776_RS01085 | 258368..262027 | 28026 | BREX-system-P-loop-protein-BrxC-[Bifidobacterium-pseudocatenulatum] |
| WP_226650300.1 | Type1.BREX | <-SIGMA+OB-Fold+AAA-ATPase+HKD-DNase-BrxL<-PglZ<-Schlafen+HATPase+wHTH<-REase<-N6-METHYLASE<-DUF6079*<-BrxB-BrxF-iSTAND<-BrxA-DNA-binding\|\|REase->?->SbcD-><-?\|\|?-><-?\|\|MerR-HTH-> | 1219 | Bacteria>Terrabacteria_group>Actinomycetota>Actinomycetes>Bifidobacteriales>Bifidobacteriaceae | GCF_020526585.1 | Bifidobacterium_pseudocatenulatum | KZP10_RS05695 | 1065058..1068717 | 28026 | BREX-system-P-loop-protein-BrxC-[Bifidobacterium-pseudocatenulatum] |
| WP_237919300.1 | Type1.BREX | ACET+ACET->?-><-?\|\|?-><-SbcD\|\|?-><-?\|\|BrxA-DNA-binding->BrxB-BrxF-iSTAND->DUF6079*->N6-METHYLASE->SIR2->Schlafen+HATPase+wHTH->PglZ->SIGMA+OB-Fold+AAA-ATPase+HKD-DNase-BrxL-> | 1219 | Bacteria>Terrabacteria_group>Actinomycetota>Actinomycetes>Bifidobacteriales>Bifidobacteriaceae | GCF_022135125.1 | Bifidobacterium_pseudocatenulatum | LOM88_RS00205 | 52697..56356 | 28026 | BREX-system-P-loop-protein-BrxC-[Bifidobacterium-pseudocatenulatum] |
| WP_237970585.1 | Type1.BREX | HTH_28->BrxA-DNA-binding->BrxB-BrxF-iSTAND->DUF6079*->N6-METHYLASE->?->Schlafen+HATPase+wHTH->PglZ->SIGMA+OB-Fold+AAA-ATPase+HKD-DNase-BrxL->ZnR+GP->ZnR->ParB+HNH->Sigma-HTH-> | 1218 | Bacteria>Terrabacteria_group>Actinomycetota>Actinomycetes>Bifidobacteriales>Bifidobacteriaceae | GCF_022136875.1 | Bifidobacterium_longum | LOO84_RS04690 | 81950..85606 | 216816 | BREX-system-P-loop-protein-BrxC-[Bifidobacterium-longum] |
| WP_240833140.1 | Type1.BREX | <-ABhydrolase<-ParB+HNH+RAMA<-ZnR<-ZnR+GP<-SIGMA+OB-Fold+AAA-ATPase+HKD-DNase-BrxL<-PglZ<-Schlafen+HATPase+wHTH<-?<-N6-METHYLASE<-DUF6079*<-BrxB-BrxF-iSTAND<-BrxA-DNA-binding\|\|REase->?-><-?\|\|SbcD-> | 1217 | Bacteria>Terrabacteria_group>Actinomycetota>Actinomycetes>Bifidobacteriales>Bifidobacteriaceae | GCF_022496395.1 | Bifidobacterium_pseudocatenulatum | LF914_RS07635 | 54595..58248 | 28026 | BREX-system-P-loop-protein-BrxC-[Bifidobacterium-pseudocatenulatum] |
| WP_242075534.1 | Type1.BREX | HTH_28->BrxA-DNA-binding->BrxB-BrxF-iSTAND->DUF6079*->N6-METHYLASE->Schlafen+HATPase+wHTH->PglZ->SIGMA+OB-Fold+AAA-ATPase+HKD-DNase-BrxL->?-><-?<-?<-ABC-ATPase+SbcC | 1218 | Bacteria>Terrabacteria_group>Actinomycetota>Actinomycetes>Bifidobacteriales>Bifidobacteriaceae | GCF_022643745.1 | Bifidobacterium_longum | G8B15_RS07005 | 1527026..1530682 | 216816 | BREX-system-P-loop-protein-BrxC-[Bifidobacterium-longum] |
| WP_032684176.1 | Type1.BREX | <-SIGMA+OB-Fold+AAA-ATPase+HKD-DNase-BrxL<-PglZ<-Schlafen+HATPase+wHTH<-N6-METHYLASE-p<-HerA/FtsK<-SIR2<-N6-METHYLASE<-DUF6079*<-BrxB-BrxF-iSTAND<-BrxA-DNA-binding<-HTH_28 | 1219 | Bacteria>Terrabacteria_group>Actinomycetota>Actinomycetes>Bifidobacteriales>Bifidobacteriaceae | GCF_022701235.1 | Bifidobacterium_longum | LMY38_RS03520 | 757375..761034 | 216816 | BREX-system-P-loop-protein-BrxC-[Bifidobacterium-longum] |
| WP_248110663.1 | Type1.BREX | 35exo+HRDC->?->35exo+BRCT-><-HTH\|\|?-><-ACET\|\|?->BrxA-DNA-binding->BrxB-BrxF-iSTAND->DUF6079*->N6-METHYLASE->?->Schlafen+HATPase+wHTH->PglZ->SIGMA+OB-Fold+AAA-ATPase+HKD-DNase-BrxL->ZnR-><-DSBH+AraC-HTH+AraC-HTH\|\|Glyoxalase-> | 1219 | Bacteria>Terrabacteria_group>Actinomycetota>Actinomycetes>Bifidobacteriales>Bifidobacteriaceae | GCF_023147995.1 | Bifidobacterium_dentium | KZX63_RS03535 | 164390..168049 | 1689 | BREX-system-P-loop-protein-BrxC-[Bifidobacterium-dentium] |
| WP_269374243.1 | Type1.BREX | <-Sigma-HTH<-ParB+HNH<-ZnR<-ZnR+GP<-SIGMA+OB-Fold+AAA-ATPase+HKD-DNase-BrxL<-PglZ<-Schlafen+HATPase+wHTH<-SIR2<-N6-METHYLASE<-DUF6079*<-BrxB-BrxF-iSTAND<-BrxA-DNA-binding | 1219 | Bacteria>Terrabacteria_group>Actinomycetota>Actinomycetes>Bifidobacteriales>Bifidobacteriaceae | GCF_027214485.1 | Bifidobacterium_breve | O4C76_RS09020 | 75244..78903 | 1685 | BREX-system-P-loop-protein-BrxC-[Bifidobacterium-breve] |
| WP_269355165.1 | Type1.BREX | <-Sigma-HTH<-ParB+HNH<-ZnR<-ZnR+GP<-SIGMA+OB-Fold+AAA-ATPase+HKD-DNase-BrxL<-PglZ<-Schlafen+HATPase+wHTH<-SIR2<-N6-METHYLASE<-DUF6079*<-BrxB-BrxF-iSTAND<-BrxA-DNA-binding<-HTH_28<-?\|\|tRNA->tRNA->?-><-?<-RNasePH | 1218 | Bacteria>Terrabacteria_group>Actinomycetota>Actinomycetes>Bifidobacteriales>Bifidobacteriaceae | GCF_027215185.1 | Bifidobacterium_breve | O4C27_RS08995 | 67566..71222 | 1685 | BREX-system-P-loop-protein-BrxC-[Bifidobacterium-breve] |
| WP_270294263.1 | Type1.BREX | <-SIGMA+OB-Fold+AAA-ATPase+HKD-DNase-BrxL<-PglZ<-Schlafen+HATPase+wHTH<-SIR2<-N6-METHYLASE-p<-HerA/FtsK<-SIR2<-N6-METHYLASE<-DUF6079*<-BrxB-BrxF-iSTAND<-BrxA-DNA-binding\|\|REase->?->SbcD-><-?\|\|?-><-HISKIN<-REC+LuxR-HTH | 1219 | Bacteria>Terrabacteria_group>Actinomycetota>Actinomycetes>Bifidobacteriales>Bifidobacteriaceae | GCF_027659585.1 | Bifidobacterium_adolescentis | PG090_RS01995 | 459377..463036 | 1680 | BREX-system-P-loop-protein-BrxC-[Bifidobacterium-adolescentis] |
| WP_117779802.1 | Type1.BREX | <-SIGMA+OB-Fold+AAA-ATPase+HKD-DNase-BrxL<-PglZ<-Schlafen+HATPase+wHTH<-SIR2<-N6-METHYLASE-p<-HerA/FtsK<-SIR2<-N6-METHYLASE<-DUF6079*<-BrxB-BrxF-iSTAND<-BrxA-DNA-binding\|\|REase->?->?-><-RelE-ParE<-PhdYeFM_antitox\|\|SbcD-> | 1219 | Bacteria>Terrabacteria_group>Actinomycetota>Actinomycetes>Bifidobacteriales>Bifidobacteriaceae | GCF_027664445.1 | Bifidobacterium_adolescentis | PGQ86_RS05915 | 1356414..1360073 | 1680 | BREX-system-P-loop-protein-BrxC-[Bifidobacterium-adolescentis] |
| WP_270271788.1 | Type1.BREX | REase-><-?\|\|?->?->HTH+HisKin-><-REase\|\|BrxA-DNA-binding->BrxB-BrxF-iSTAND->DUF6079*->N6-METHYLASE->?->Schlafen+HATPase+wHTH->PglZ->SIGMA+OB-Fold+AAA-ATPase+HKD-DNase-BrxL->ZnR+GP->ZnR->ParB+HNH-> | 1218 | Bacteria>Terrabacteria_group>Actinomycetota>Actinomycetes>Bifidobacteriales>Bifidobacteriaceae | GCF_027667785.1 | Bifidobacterium_bifidum | PGS22_RS05345 | 127181..130837 | 1681 | BREX-system-P-loop-protein-BrxC-[Bifidobacterium-bifidum] |
| WP_117715835.1 | Type1.BREX | ZnR->?->ABC-ATPase+SbcC->?->?->?->BrxA-DNA-binding->BrxB-BrxF-iSTAND->DUF6079*->N6-METHYLASE->SIR2->Schlafen+HATPase+wHTH->PglZ->SIGMA+OB-Fold+AAA-ATPase+HKD-DNase-BrxL->?-><-REase\|\|?-><-Glyoxalase | 1217 | Bacteria>Terrabacteria_group>Actinomycetota>Actinomycetes>Bifidobacteriales>Bifidobacteriaceae | GCF_027669765.1 | Bifidobacterium_bifidum | PHA16_RS02375 | 586926..590579 | 1681 | BREX-system-P-loop-protein-BrxC-[Bifidobacterium-bifidum] |
| WP_270200767.1 | Type1.BREX | REC+LuxR-HTH->HISKIN-><-?<-SbcD<-?<-?\|\|Pentapeptide+Pentapeptide+Pentapeptide+Pentapeptide->BrxA-DNA-binding->BrxB-BrxF-iSTAND->DUF6079-p*->DUF6079-p->N6-METHYLASE->REase->Schlafen+HATPase+wHTH->PglZ->SIGMA+OB-Fold+AAA-ATPase+HKD-DNase-BrxL->?-><-RNaseH | 467 | Bacteria>Terrabacteria_group>Actinomycetota>Actinomycetes>Bifidobacteriales>Bifidobacteriaceae | GCF_027671795.1 | Bifidobacterium_pseudocatenulatum | PGT11_RS00265 | 67391..68794 | 28026 | BREX-system-P-loop-protein-BrxC-[Bifidobacterium-pseudocatenulatum] |
| WP_270199955.1 | Type1.BREX | <-SbcD<-?<-?\|\|?->\|\|Pentapeptide+Pentapeptide+Pentapeptide+Pentapeptide->BrxA-DNA-binding->BrxB-BrxF-iSTAND->DUF6079*->DUF6079-p->N6-METHYLASE->REase->Schlafen+HATPase+wHTH->PglZ->SIGMA+OB-Fold+AAA-ATPase+HKD-DNase-BrxL->?-><-RNaseH | 1044 | Bacteria>Terrabacteria_group>Actinomycetota>Actinomycetes>Bifidobacteriales>Bifidobacteriaceae | GCF_027675485.1 | Bifidobacterium_pseudocatenulatum | PGV18_RS06405 | 41954..45088 | 28026 | BREX-system-P-loop-protein-BrxC-[Bifidobacterium-pseudocatenulatum] |
| WP_270271488.1 | Type1.BREX | <-ZnR<-ZnR+GP<-SIGMA+OB-Fold+AAA-ATPase+HKD-DNase-BrxL<-PglZ<-Schlafen+HATPase+wHTH<-SIR2<-N6-METHYLASE<-DUF6079*<-BrxB-BrxF-iSTAND<-BrxA-DNA-binding<-Pentapeptide+Pentapeptide+Pentapeptide<-HAD\|\|SbcD-><-?<-ACET+ACET\|\|HTH-><-35exo+BRCT | 1219 | Bacteria>Terrabacteria_group>Actinomycetota>Actinomycetes>Bifidobacteriales>Bifidobacteriaceae | GCF_027679525.1 | Bifidobacterium_adolescentis | PGW19_RS08965 | 15037..18696 | 1680 | BREX-system-P-loop-protein-BrxC-[Bifidobacterium-adolescentis] |
| WP_270447025.1 | Type1.BREX | HISKIN-><-?\|\|<-SbcD<-?<-?\|\|?->\|\|Pentapeptide+Pentapeptide+Pentapeptide+Pentapeptide->BrxA-DNA-binding->BrxB-BrxF-iSTAND->DUF6079-p*->DUF6079-p->N6-METHYLASE->SIR2->HerA/FtsK->N6-METHYLASE-p->SIR2->Schlafen+HATPase+wHTH->PglZ->SIGMA+OB-Fold+AAA-ATPase+HKD-DNase-BrxL-> | 958 | Bacteria>Terrabacteria_group>Actinomycetota>Actinomycetes>Bifidobacteriales>Bifidobacteriaceae | GCF_027682665.1 | Bifidobacterium_pseudocatenulatum | PGX68_RS01375 | 343601..346477 | 28026 | BREX-system-P-loop-protein-BrxC-[Bifidobacterium-pseudocatenulatum] |
| WP_270479593.1 | Type1.BREX | <-ZnR+GP<-SIGMA+OB-Fold+AAA-ATPase+HKD-DNase-BrxL<-PglZ<-Schlafen+HATPase+wHTH<-SIR2<-N6-METHYLASE-p<-HerA/FtsK<-SIR2<-N6-METHYLASE<-DUF6079*<-BrxB-BrxF-iSTAND<-BrxA-DNA-binding\|\|?->HEPN-><-?\|\|?->?-><-HEPN<-PolB | 1219 | Bacteria>Terrabacteria_group>Actinomycetota>Actinomycetes>Bifidobacteriales>Bifidobacteriaceae | GCF_027683145.1 | Bifidobacterium_adolescentis | PGX62_RS02115 | 472627..476286 | 1680 | BREX-system-P-loop-protein-BrxC-[Bifidobacterium-adolescentis] |
| WP_117726345.1 | Type1.BREX | ZnR->?->ABC-ATPase+SbcC->?->?->?->BrxA-DNA-binding->BrxB-BrxF-iSTAND->DUF6079*->N6-METHYLASE->?->Schlafen+HATPase+wHTH->PglZ->SIGMA+OB-Fold+AAA-ATPase+HKD-DNase-BrxL->?-><-REase\|\|?-><-Glyoxalase | 1217 | Bacteria>Terrabacteria_group>Actinomycetota>Actinomycetes>Bifidobacteriales>Bifidobacteriaceae | GCF_027685785.1 | Bifidobacterium_bifidum | PG427_RS02260 | 555349..559002 | 1681 | BREX-system-P-loop-protein-BrxC-[Bifidobacterium-bifidum] |
| WP_270266433.1 | Type1.BREX | <-SIGMA+OB-Fold+AAA-ATPase+HKD-DNase-BrxL<-PglZ<-Schlafen+HATPase+wHTH<-?<-N6-METHYLASE<-DUF6079*<-BrxB-BrxF-iSTAND<-BrxA-DNA-binding<-?<-?<-?<-ABC-ATPase+SbcC<-?<-ZnR | 1219 | Bacteria>Terrabacteria_group>Actinomycetota>Actinomycetes>Bifidobacteriales>Bifidobacteriaceae | GCF_027687815.1 | Bifidobacterium_bifidum | PG972_RS08475 | 12443..16102 | 1681 | BREX-system-P-loop-protein-BrxC-[Bifidobacterium-bifidum] |
| WP_117501722.1 | Type1.BREX | REC+LuxR-HTH->HISKIN-><-?\|\|?-><-SbcD<-?<-REase\|\|BrxA-DNA-binding->BrxB-BrxF-iSTAND->DUF6079*->N6-METHYLASE->SIR2->Schlafen+HATPase+wHTH->PglZ->SIGMA+OB-Fold+AAA-ATPase+HKD-DNase-BrxL-> | 1219 | Bacteria>Terrabacteria_group>Actinomycetota>Actinomycetes>Bifidobacteriales>Bifidobacteriaceae | GCF_027689935.1 | Bifidobacterium_pseudocatenulatum | PG028_RS01370 | 345834..349493 | 28026 | BREX-system-P-loop-protein-BrxC-[Bifidobacterium-pseudocatenulatum] |
| WP_270558503.1 | Type1.BREX | <-ParB+HNH+RAMA<-ZnR<-ZnR+GP<-SIGMA+OB-Fold+AAA-ATPase+HKD-DNase-BrxL<-PglZ<-Schlafen+HATPase+wHTH<-REase<-N6-METHYLASE<-DUF6079*<-BrxB-BrxF-iSTAND<-BrxA-DNA-binding<-?<-?\|\|?-><-Peptidase_M78<-HigA | 1219 | Bacteria>Terrabacteria_group>Actinomycetota>Actinomycetes>Bifidobacteriales>Bifidobacteriaceae | GCF_027690155.1 | Bifidobacterium_adolescentis | PG057_RS05020 | 457971..461630 | 1680 | BREX-system-P-loop-protein-BrxC-[Bifidobacterium-adolescentis] |
| WP_270554212.1 | Type1.BREX | <-SbcD\|\|HAD->PhdYeFM_antitox->RelE-ParE-><-?<-?<-REase\|\|BrxA-DNA-binding->BrxB-BrxF-iSTAND->DUF6079*->N6-METHYLASE->REase->Schlafen+HATPase+wHTH->PglZ->SIGMA+OB-Fold+AAA-ATPase+HKD-DNase-BrxL->ZnR+GP->ZnR->ParB+HNH+RAMA-> | 1219 | Bacteria>Terrabacteria_group>Actinomycetota>Actinomycetes>Bifidobacteriales>Bifidobacteriaceae | GCF_027695195.1 | Bifidobacterium_adolescentis | PG742_RS01120 | 252838..256497 | 1680 | BREX-system-P-loop-protein-BrxC-[Bifidobacterium-adolescentis] |
| WP_270561184.1 | Type1.BREX | 35exo+HRDC->?->35exo+BRCT-><-HTH\|\|?-><-ACET\|\|?->BrxA-DNA-binding->BrxB-BrxF-iSTAND->DUF6079*->SIR2->HerA/FtsK->N6-METHYLASE-p->HEPN->?->Schlafen+HATPase+wHTH->PglZ->SIGMA+OB-Fold+AAA-ATPase+HKD-DNase-BrxL->ZnR-> | 1217 | Bacteria>Terrabacteria_group>Actinomycetota>Actinomycetes>Bifidobacteriales>Bifidobacteriaceae | GCF_027698005.1 | Bifidobacterium_dentium | PG823_RS09640 | 38589..42242 | 1689 | BREX-system-P-loop-protein-BrxC-[Bifidobacterium-dentium] |
| WP_118376243.1 | Type1.BREX | RNaseH-><-?<-SIGMA+OB-Fold+AAA-ATPase+HKD-DNase-BrxL<-PglZ<-Schlafen+HATPase+wHTH<-SIR2<-N6-METHYLASE<-DUF6079*<-BrxB-BrxF-iSTAND<-BrxA-DNA-binding\|\|REase->?->?-><-?<-HEPN<-PolB | 1219 | Bacteria>Terrabacteria_group>Actinomycetota>Actinomycetes>Bifidobacteriales>Bifidobacteriaceae | GCF_027698785.1 | Bifidobacterium_pseudocatenulatum | PG820_RS05645 | 117035..120694 | 28026 | BREX-system-P-loop-protein-BrxC-[Bifidobacterium-pseudocatenulatum] |
| WP_271768971.1 | Type1.BREX | <-SbcD<-?<-REase\|\|BrxA-DNA-binding->BrxB-BrxF-iSTAND->DUF6079*->N6-METHYLASE->SIR2->Schlafen+HATPase+wHTH->PglZ->SIGMA+OB-Fold+AAA-ATPase+HKD-DNase-BrxL->ZnR+GP->ZnR->ParB+HNH+RAMA-> | 1219 | Bacteria>Terrabacteria_group>Actinomycetota>Actinomycetes>Bifidobacteriales>Bifidobacteriaceae | GCF_028314485.1 | Bifidobacterium_pseudocatenulatum | PMN34_RS05085 | 218989..222648 | 28026 | BREX-system-P-loop-protein-BrxC-[Bifidobacterium-pseudocatenulatum] |
| WP_116463714.1 | Type1.BREX | HTH_28->BrxA-DNA-binding->BrxB-BrxF-iSTAND->DUF6079*->N6-METHYLASE->?->Schlafen+HATPase+wHTH->PglZ->SIGMA+OB-Fold+AAA-ATPase+HKD-DNase-BrxL->?-><-?<-?<-ABC-ATPase+SbcC | 1219 | Bacteria>Terrabacteria_group>Actinomycetota>Actinomycetes>Bifidobacteriales>Bifidobacteriaceae | GCF_028898865.1 | Bifidobacterium_longum | PWA56_RS02155 | 487318..490977 | 216816 | BREX-system-P-loop-protein-BrxC-[Bifidobacterium-longum] |
| WP_077383013.1 | Type1.BREX | ABC-ATPase+SbcC->?->?-><-?<-SIGMA+OB-Fold+AAA-ATPase+HKD-DNase-BrxL<-PglZ<-Schlafen+HATPase+wHTH<-?<-N6-METHYLASE<-DUF6079*<-BrxB-BrxF-iSTAND<-BrxA-DNA-binding<-HTH_28 | 1219 | Bacteria>Terrabacteria_group>Actinomycetota>Actinomycetes>Bifidobacteriales>Bifidobacteriaceae | GCF_900157155.1 | Bifidobacterium_longum | BQ8631_RS08415 | 45904..49563 | 216816 | BREX-system-P-loop-protein-BrxC-[Bifidobacterium-longum] |
| WP_154050436.1 | Type1.BREX | HAD+S6PP->?->?-><-?<-?<-?<-SIR2\|\|BrxA-DNA-binding->BrxB-BrxF-iSTAND->DUF6079*->N6-METHYLASE->SIR2->Schlafen+HATPase+wHTH->PglZ->SIGMA+OB-Fold+AAA-ATPase+HKD-DNase-BrxL->?-><-ABhydrolase | 1217 | Bacteria>Terrabacteria_group>Actinomycetota>Actinomycetes>Bifidobacteriales>Bifidobacteriaceae | GCF_901212475.1 | Bifidobacterium_longum | FGK88_RS06805 | 85798..89451 | 216816 | BREX-system-P-loop-protein-BrxC-[Bifidobacterium-longum] |
| WP_143246218.1 | Type1.BREX | <-GDResolvase+HIN-HTH<-?<-SIGMA+OB-Fold+AAA-ATPase+HKD-DNase-BrxL<-PglZ<-Schlafen+HATPase+wHTH<-SIR2<-N6-METHYLASE<-DUF6079*<-BrxB-BrxF-iSTAND<-BrxA-DNA-binding | 1218 | Bacteria>Terrabacteria_group>Actinomycetota>Actinomycetes>Bifidobacteriales>Bifidobacteriaceae | GCF_902167575.1 | Bifidobacterium_breve | FPG44_RS02615 | 449270..452926 | 1685 | BREX-system-P-loop-protein-BrxC-[Bifidobacterium-breve] |
| WP_200406352.1 | Type1.BREX | RelB->N6-METHYLASE->CR-ATPase5+REase-p-><-CR-Kinase<-?\|\|BrxA-DNA-binding->BrxB-BrxF-iSTAND->DUF6079*->?->N6-METHYLASE->SLATT->SMODS->Schlafen-p->Schlafen+wHTH->PglZ->SIGMA+OB-Fold+AAA-ATPase+HKD-DNase-BrxL->TIR-> | 1229 | Bacteria>Terrabacteria_group>Actinomycetota>Actinomycetes>Bifidobacteriales>Bifidobacteriaceae | GCF_934675445.1 | Bifidobacterium_longum | BL7055_RS05750 | 1406658..1410347 | 216816 | BREX-system-P-loop-protein-BrxC-[Bifidobacterium-longum] |
| WP_245206784.1 | Type1.BREX | <-ZnR<-SIGMA+OB-Fold+AAA-ATPase+HKD-DNase-BrxL<-PglZ<-Schlafen+HATPase+wHTH<-SIR2<-N6-METHYLASE<-AAA-ATPase<-HTH_7+rve+rve_3<-N6-METHYLASE-p<-DUF6079*<-DUF6079-p<-BrxB-BrxF-iSTAND<-BrxA-DNA-binding<-?\|\|ACET-><-?\|\|HTH-><-35exo+BRCT | 1073 | Bacteria>Terrabacteria_group>Actinomycetota>Actinomycetes>Bifidobacteriales>Bifidobacteriaceae | GCF_937997665.1 | Bifidobacterium_dentium | J7M36_RS06065 | 1427241..1430462 | 1689 | BREX-system-P-loop-protein-BrxC-[Bifidobacterium-dentium] |
| WP_196330425.1 | Type1.BREX | 35exo+HRDC->?->35exo+BRCT-><-HTH\|\|?-><-ACET\|\|?->BrxA-DNA-binding->BrxB-BrxF-iSTAND->DUF6079*->N6-METHYLASE->?->Schlafen+HATPase+wHTH->PglZ->SIGMA+OB-Fold+AAA-ATPase+HKD-DNase-BrxL->ZnR-><-DSBH+AraC-HTH+AraC-HTH\|\|Glyoxalase-> | 1217 | Bacteria>Terrabacteria_group>Actinomycetota>Actinomycetes>Bifidobacteriales>Bifidobacteriaceae | GCF_958433885.1 | Bifidobacterium_dentium | J7M35_RS04855 | 1219062..1222715 | 1689 | BREX-system-P-loop-protein-BrxC-[Bifidobacterium-dentium] |
| WP_085380386.1 | Type1.BREX | <-ParB+HNH+RAMA\|\|MULE-TRANSPOSASE-><-ZnR<-ZnR+GP<-SIGMA+OB-Fold+AAA-ATPase+HKD-DNase-BrxL<-PglZ<-Schlafen+HATPase+wHTH<-SIR2<-N6-METHYLASE<-DUF6079*<-BrxB-BrxF-iSTAND<-BrxA-DNA-binding<-Pentapeptide+Pentapeptide+Pentapeptide<-HAD\|\|SbcD-><-?<-ACET+ACET\|\|HTH-><-35exo+BRCT | 1219 | Bacteria>Terrabacteria_group>Actinomycetota>Actinomycetes>Bifidobacteriales>Bifidobacteriaceae | GCF_959020145.1 | Bifidobacterium_adolescentis | LMG10734_RS05600 | 489007..492666 | 1680 | BREX-system-P-loop-protein-BrxC-[Bifidobacterium-adolescentis] |
| WP_196335743.1 | Type1.BREX | <-Glyoxalase\|\|DSBH+AraC-HTH+AraC-HTH-><-ZnR<-SIGMA+OB-Fold+AAA-ATPase+HKD-DNase-BrxL<-PglZ<-Schlafen+HATPase+wHTH<-REase<-N6-METHYLASE<-DUF6079*<-BrxB-BrxF-iSTAND<-BrxA-DNA-binding\|\|REase-><-HEPN<-?\|\|HTH-><-35exo+BRCT<-35exo+HRDC | 1217 | Bacteria>Terrabacteria_group>Actinomycetota>Actinomycetes>Bifidobacteriales>Bifidobacteriaceae | GCF_959600175.1 | Bifidobacterium_dentium | IHQ76_RS06180 | 228522..232175 | 1689 | BREX-system-P-loop-protein-BrxC-[Bifidobacterium-dentium] |
| SBW20208.1 | Type6.BREX | BrxE->BrxA-DNA-binding->BrxB-BrxF-iSTAND->DUF6079*->Schlafen+HATPase+wHTH->N6-METHYLASE->N6-METHYLASE-> | 1180 | Bacteria>Terrabacteria_group>Actinomycetota>Actinomycetes>Frankiales>Frankiaceae | - | Candidatus_Frankia_californiensis | FDG2_1623 | 2594..6136 | 1839754 | hypothetical-protein-FDG2_1623-[Candidatus-Frankia-californiensis] |
| WP_028245009.1 | Type1.BREX | <-Thioredoxin\|\|?-><-PALP\|\|tRNA->35exo->tRNA->BrxA-DNA-binding->BrxB-BrxF-iSTAND->DUF6079*->N6-METHYLASE-><-N6-METHYLASE-p\|\|N6-METHYLASE-p->Schlafen+HATPase+wHTH->PglZ->SIGMA+OB-Fold+AAA-ATPase+HKD-DNase-BrxL->SIR2->HerA/FtsK->PhdYeFM_antitox-> | 1170 | Bacteria>Terrabacteria_group>Actinomycetota>Actinomycetes>Micrococcales>Microbacteriaceae | GCF_000423505.1 | Pseudoclavibacter_soli | G557_RS0103810 | 190122..193634 | 452623 | BREX-system-P-loop-protein-BrxC-[Pseudoclavibacter-soli] |
| WP_237399358.1 | Type1.BREX | <-Glycos_transf<-Glyco_transf_9-><-HAD+HAD\|\|Glyco_transf_9-><-SIGMA-HTH\|\|?->?->tRNA->BrxA-DNA-binding->BrxB-BrxF-iSTAND->DUF6079-p*->N6-METHYLASE-p-><-N6-METHYLASE-p\|\|Phage_int_SAM_3+Phage_integrase->N6-METHYLASE-p->Schlafen+HATPase+wHTH->PglZ->SIGMA+OB-Fold+AAA-ATPase+HKD-DNase-BrxL-> | 878 | Bacteria>Terrabacteria_group>Actinomycetota>Actinomycetes>Micrococcales>Microbacteriaceae | GCF_004011115.1 | Rathayibacter_iranicus | C7V51_RS01465 | 309166..311802 | 59737 | BREX-system-P-loop-protein-BrxC-[Rathayibacter-iranicus] |
| WP_129387833.1 | Type1.BREX | ABhydrolase->?->Peptidase_S15+PepX_C->?-><-?<-?<-Peptidase_S66+Peptidase_S66C\|\|BrxA-DNA-binding->BrxB-BrxF-iSTAND->DUF6079*->N6-METHYLASE-><-N6-METHYLASE-p\|\|Phage_int_SAM_3+Phage_integrase->N6-METHYLASE-p->Schlafen+HATPase+wHTH->PglZ->SIGMA+OB-Fold+AAA-ATPase+HKD-DNase-BrxL-><-PhdYeFM_antitox | 1162 | Bacteria>Terrabacteria_group>Actinomycetota>Actinomycetes>Micrococcales>Microbacteriaceae | GCF_004135285.1 | Microbacterium_protaetiae | ET475_07150 | 1526355..1529843 | 2509458 | BREX-system-P-loop-protein-BrxC-[Microbacterium-protaetiae] |
| WP_237187176.1 | Type1.BREX | <-HHH_5+SNF2_N+SF1-SF2-Helicase\|\|SWACOS-><-?<-SIGMA+OB-Fold+AAA-ATPase+HKD-DNase-BrxL<-PglZ<-Schlafen+HATPase+wHTH<-N6-METHYLASE<-DUF6079*<-BrxB-BrxF-iSTAND<-BrxA-DNA-binding\|\|?-><-Aminotran_4<-?<-ABC-ATPase+SbcC+ABC-ATPase+SbcC | 1190 | Bacteria>Terrabacteria_group>Actinomycetota>Actinomycetes>Micrococcales>Micrococcaceae | GCF_928381705.1 | Rothia_nasimurium | L6762_RS10435 | 40591..44163 | 85336 | BREX-system-P-loop-protein-BrxC-[Rothia-nasimurium] |
| WP_142113745.1 | Type1.BREX | <-VirB11+MuB-ATPase\|\|DNA_METHYLASE-><-HerA/FtsK<-SIR2<-SIGMA+OB-Fold+AAA-ATPase+HKD-DNase-BrxL<-PglZ<-Schlafen+HATPase+wHTH<-?<-N6-METHYLASE<-DUF6079*<-BrxB-BrxF-iSTAND<-BrxA-DNA-binding<-XIS-HTH<-ABhydrolase<-ABhydrolase<-?<-MPTase<-HTH\|\|ArdCN+MPTase-> | 1173 | Bacteria>Terrabacteria_group>Actinomycetota>Actinomycetes>Micrococcales>Rarobacteraceae | - | Rarobacter_incanus | FB389_RS10090 | 2435944..2439465 | 153494 | BREX-system-P-loop-protein-BrxC-[Rarobacter-incanus] |
| WP_208468634.1 | Type6.BREX | XIS-HTH->URI-DNase->REase->BrxE->BrxA-DNA-binding->BrxB-BrxF-iSTAND->DUF6079*->Schlafen+HATPase+wHTH->N6-METHYLASE->N6-METHYLASE->PglZ->SIGMA+OB-Fold+AAA-ATPase+Lon-Protease-BrxL-><-DDE_Tnp_1_assoc+DDE_Tnp_1 | 1183 | Bacteria>Terrabacteria_group>Actinomycetota>Actinomycetes>Micromonosporales>Micromonosporaceae | GCF_017592555.1 | Actinoplanes_flavus | J5X75_RS18430 | 129103..132654 | 2820290 | BREX-system-P-loop-protein-BrxC-[Actinoplanes-flavus] |
| WP_245802167.1 | Type1.BREX | <-ARG<-SIR2<-SIGMA+OB-Fold+AAA-ATPase+HKD-DNase-BrxL<-PglZ<-Schlafen+HATPase+wHTH<-N6-METHYLASE<-DUF6079*<-BrxB-BrxF-iSTAND<-BrxA-DNA-binding | 1179 | Bacteria>Terrabacteria_group>Actinomycetota>Actinomycetes>Mycobacteriales>Corynebacteriaceae | GCF_900169525.1 | Corynebacterium_pacaense | B7E02_RS05025 | 1051363..1054902 | 1816684 | BREX-system-P-loop-protein-BrxC-[Corynebacterium-pacaense] |
| WP_028707702.1 | Type1.BREX | HEPN-><-SIGMA+OB-Fold+AAA-ATPase+HKD-DNase-BrxL<-PglZ<-Schlafen+HATPase+wHTH<-N6-METHYLASE-p<-Phage_int_SAM_3+Phage_integrase\|\|N6-METHYLASE-p->N6-METHYLASE-p-><-N6-METHYLASE<-DUF6079*<-BrxB-BrxF-iSTAND<-BrxA-DNA-binding<-?<-?<-?<-?\|\|?->AbiEi_4-> | 1159 | Bacteria>Terrabacteria_group>Actinomycetota>Actinomycetes>Propionibacteriales>Nocardioidaceae | GCF_000423465.1 | Propionicicella_superfundia | G555_RS0102905 | 100593..104072 | 348582 | BREX-system-P-loop-protein-BrxC-[Propionicicella-superfundia] |
| BAK36822.1 | Type1.BREX | HTH_3->ADH_N+ADH_zinc_N->?->?->?->BrxA-DNA-binding->BrxB-BrxF-iSTAND->DUF6079*->N6-METHYLASE-><-N6-METHYLASE-p\|\|Phage_int_SAM_3+Phage_integrase->N6-METHYLASE-p->Schlafen+HATPase+wHTH->PglZ->SIGMA+OB-Fold+AAA-ATPase+HKD-DNase-BrxL->SIR2-><-DDE_Tnp_1 | 1176 | Bacteria>Terrabacteria_group>Actinomycetota>Actinomycetes>Propionibacteriales>Propionibacteriaceae | GCA_000270245.1 | Microlunatus_phosphovorus_NM-1 | MLP_38080 | 4107257..4110787 | 1032480 | hypothetical-protein-MLP_38080-[Microlunatus-phosphovorus-NM-1] |
| MCI1746855.1 | Type1.BREX | sGTP->?->?->?->tRNA->tRNA->BrxA-DNA-binding->BrxB-BrxF-iSTAND->DUF6079*->?->N6-METHYLASE-><-N6-METHYLASE-p\|\|Phage_int_SAM_3+Phage_integrase->N6-METHYLASE-p->Schlafen+HATPase+wHTH-p->PglZ->SIGMA+OB-Fold+AAA-ATPase+HKD-DNase-BrxL->Schlafen-> | 1159 | Bacteria>Terrabacteria_group>Actinomycetota>Actinomycetes>Propionibacteriales>Propionibacteriaceae | GCA_022647765.1 | Acidipropionibacterium_sp. | LKI24_01350 | 263130..266609 | 1912218 | BREX-system-P-loop-protein-BrxC-[Acidipropionibacterium-sp.] |
| WP_041792941.1 | Type1.BREX | Glyco_hydro_3+Glyco_hydro_3_C+Fn3-like->ADH_N+ADH_zinc_N->?->HTH_3->ADH_N+ADH_zinc_N->?->?->BrxA-DNA-binding->BrxB-BrxF-iSTAND->DUF6079*->N6-METHYLASE->Phage_int_SAM_3+Phage_integrase->N6-METHYLASE-p->Schlafen+HATPase+wHTH->PglZ->SIGMA+OB-Fold+AAA-ATPase+HKD-DNase-BrxL->SIR2-><-DDE_Tnp_1\|\|HIN-HTH-> | 1173 | Bacteria>Terrabacteria_group>Actinomycetota>Actinomycetes>Propionibacteriales>Propionibacteriaceae | GCF_000270245.1 | Microlunatus_phosphovorus | MLP_RS18400 | 4107266..4110787 | 29405 | BREX-system-P-loop-protein-BrxC-[Microlunatus-phosphovorus] |
| WP_028706416.1 | Type1.BREX | <-SIGMA-HTH+SIGMA-HTH<-SF1-SF2-Helicase+HTH<-SIGMA+OB-Fold+AAA-ATPase+HKD-DNase-BrxL<-PglZ<-Schlafen+HATPase+wHTH-p<-?<-HerA/FtsK<-SIR2<-N6-METHYLASE<-DUF6079*<-BrxB-BrxF-iSTAND<-BrxA-DNA-binding\|\|Glyco_hydro_36N+Melibiase+Glyco_hydro_36C->?->?->?->?-><-XIS-HTH | 1171 | Bacteria>Terrabacteria_group>Actinomycetota>Actinomycetes>Propionibacteriales>Propionibacteriaceae | GCF_000426605.1 | Propionibacterium_acidifaciens | K336_RS0104645 | 45562..49077 | 556499 | BREX-system-P-loop-protein-BrxC-[Propionibacterium-acidifaciens] |
| WP_048768895.1 | Type1.BREX | <-DUF2637+HIN-HTH<-Primpol+PriCT1<-ArdCN+MPTase<-AraC-HTH\|\|XIS-HTH->BrxA-DNA-binding->BrxB-BrxF-iSTAND->DUF6079*->N6-METHYLASE->REase->?->Schlafen+HATPase+wHTH->PglZ->SIGMA+OB-Fold+AAA-ATPase+HKD-DNase-BrxL->SIR2->HerA/FtsK->VirB11+MuB-ATPase-> | 1170 | Bacteria>Terrabacteria_group>Actinomycetota>Actinomycetes>Propionibacteriales>Propionibacteriaceae | GCF_001050955.1 | Propionibacterium_freudenreichii | PFCIRM1025_RS01850 | 33525..37037 | 1744 | BREX-system-P-loop-protein-BrxC-[Propionibacterium-freudenreichii] |
| WP_055346094.1 | Type1.BREX | <-DUF2637+HIN-HTH<-Primpol+PriCT1<-ArdCN+MPTase<-AraC-HTH\|\|XIS-HTH->BrxA-DNA-binding->BrxB-BrxF-iSTAND->DUF6079*->N6-METHYLASE->REase->?->Schlafen+HATPase+wHTH->PglZ->SIGMA+OB-Fold+AAA-ATPase+HKD-DNase-BrxL->SIR2->HerA/FtsK->VirB11+MuB-ATPase-> | 1170 | Bacteria>Terrabacteria_group>Actinomycetota>Actinomycetes>Propionibacteriales>Propionibacteriaceae | GCF_001369055.1 | Propionibacterium_freudenreichii | PFCIRM516_RS05500 | 48312..51824 | 1744 | BREX-system-P-loop-protein-BrxC-[Propionibacterium-freudenreichii] |
| WP_097828824.1 | Type1.BREX | <-ABC-ATPase+SbcC+ABC-ATPase+SbcC<-?<-?<-tRNA\|\|35exo->tRNA->BrxA-DNA-binding->BrxB-BrxF-iSTAND->DUF6079*->N6-METHYLASE-><-N6-METHYLASE-p\|\|Phage_int_SAM_3+Phage_integrase->N6-METHYLASE-p->Schlafen+HATPase+wHTH->PglZ->SIGMA+OB-Fold+AAA-ATPase+HKD-DNase-BrxL-><-?<-HTH_Tnp_1+HTH_21+rve+rve_3 | 1173 | Bacteria>Terrabacteria_group>Actinomycetota>Actinomycetes>Propionibacteriales>Propionibacteriaceae | GCF_004011075.1 | Acidipropionibacterium_jensenii | COZ11_RS09870 | 2230980..2234501 | 1749 | BREX-system-P-loop-protein-BrxC-[Acidipropionibacterium-jensenii] |
| WP_073189906.1 | Type1.BREX | XIS-HTH->BrxA-DNA-binding->BrxB-BrxF-iSTAND->DUF6079*->N6-METHYLASE->Schlafen+HATPase+wHTH->PglZ->SIGMA+OB-Fold+AAA-ATPase+HKD-DNase-BrxL->SIR2->SIR2->HerA/FtsK-><-?\|\|GTPase-AIG->sGTP-> | 1176 | Bacteria>Terrabacteria_group>Actinomycetota>Actinomycetes>Propionibacteriales>Propionibacteriaceae | GCF_900141915.1 | Tessaracoccus_bendigoensis | BUC15_RS15345 | 1766..5296 | 72764 | BREX-system-P-loop-protein-BrxC-[Tessaracoccus-bendigoensis] |
| WP_130875595.1 | Type1.BREX | <-Peptidase_M20\|\|?-><-SIGMA+OB-Fold+AAA-ATPase+HKD-DNase-BrxL<-PglZ<-Schlafen+HATPase+wHTH<-N6-METHYLASE<-DUF6079*<-BrxB-BrxF-iSTAND<-BrxA-DNA-binding<-?<-HAD<-Trypsin+PDZ\|\|?-><-?<-METHYLASE\|\|ACET-> | 1171 | Bacteria>Terrabacteria_group>Actinomycetota>Actinomycetes>Propionibacteriales>Propionibacteriaceae | GCF_900607225.1 | [Pseudopropionibacterium]_massiliense | EOE39_RS01895 | 450669..454184 | 2220000 | BREX-system-P-loop-protein-BrxC-[[Pseudopropionibacterium] |
| MCA0378127.1 | Type1.BREX | TraJ-RHH->?-><-?<-MACRODOMAIN\|\|MerR-HTH->MerR-HTH->XIS-HTH->BrxA-DNA-binding->BrxB-BrxF-iSTAND->DUF6079*->N6-METHYLASE-><-N6-METHYLASE-p\|\|Phage_int_SAM_3+Phage_integrase->N6-METHYLASE-p->Schlafen+HATPase+wHTH->PglZ->RVT->SIGMA+OB-Fold+AAA-ATPase+HKD-DNase-BrxL-><-SMODS | 1183 | Bacteria>Terrabacteria_group>Actinomycetota>Actinomycetes>unclassified_Actinomycetes | GCA_020161315.1 | Actinomycetia_bacterium | LCH36_01625 | 63135..66686 | 1883427 | BREX-system-P-loop-protein-BrxC-[Actinomycetia-bacterium] |
| WP_204537521.1 | Type1.BREX | Cysteine-Desulfurase->?-><-SIGMA+OB-Fold+AAA-ATPase+HKD-DNase-BrxL<-PglZ<-Schlafen+HATPase+wHTH<-N6-METHYLASE<-HEPN<-DUF6079*<-BrxB-BrxF-iSTAND<-BrxA-DNA-binding<-TraJ-RHH | 1171 | Bacteria>Terrabacteria_group>Actinomycetota>Coriobacteriia>Coriobacteriales>Coriobacteriaceae | - | Collinsella_tanakaei | H6A33_RS00145 | 33492..37007 | 626935 | BREX-system-P-loop-protein-BrxC-[Collinsella-tanakaei] |
| WP_195443573.1 | Type1.BREX | <-HTH_3\|\|HTH->Arm-DNA-bind_4+Phage_int_SAM_3+Phage_integrase->TraJ-RHH-><-SF1-SF2-Helicase+HamA<-SIGMA+OB-Fold+AAA-ATPase+Lon-Protease-BrxL<-Schlafen+HATPase+wHTH<-N6-METHYLASE<-DUF6079*<-BrxB-BrxF-iSTAND<-BrxA-DNA-binding<-TraJ-RHH\|\|?->?->ParA-Soj-PloopNTPase->ParB->TraJ-RHH->TraI-VirD2-RCRE-> | 1199 | Bacteria>Terrabacteria_group>Actinomycetota>Coriobacteriia>Coriobacteriales>Coriobacteriaceae | GCF_015555045.1 | Collinsella_aerofaciens | I2E45_RS00495 | 74218..77817 | 74426 | BREX-system-P-loop-protein-BrxC-[Collinsella-aerofaciens] |
| WP_204559883.1 | Type1.BREX | Cysteine-Desulfurase->?-><-SIGMA+OB-Fold+AAA-ATPase+HKD-DNase-BrxL<-PglZ<-Schlafen+HATPase+wHTH<-REase<-N6-METHYLASE<-HEPN<-DUF6079*<-BrxB-BrxF-iSTAND<-BrxA-DNA-binding<-TraJ-RHH | 1171 | Bacteria>Terrabacteria_group>Actinomycetota>Coriobacteriia>Coriobacteriales>Coriobacteriaceae | GCF_016902615.1 | Collinsella_tanakaei | H6A21_RS01790 | 55502..59017 | 626935 | BREX-system-P-loop-protein-BrxC-[Collinsella-tanakaei] |
| WP_273384065.1 | Type1.BREX | HAD+HAD-><-LACI-HTH+PBPI\|\|?-><-NUDIX<-?\|\|BrxA-DNA-binding->BrxB-BrxF-iSTAND->DUF6079*->N6-METHYLASE->XIS-HTH-><-?\|\|N6-METHYLASE-p->Schlafen+HATPase+wHTH->?->HEPN+ParB->SF1-SF2-Helicase->PglZ-> | 1183 | Bacteria>Terrabacteria_group>Actinomycetota>Coriobacteriia>Coriobacteriales>Coriobacteriaceae | GCF_944320695.1 | Enorma_phocaeensis | PUQ31_RS06660 | 14198..17749 | 1871019 | BREX-system-P-loop-protein-BrxC-[Enorma-phocaeensis] |
| WP_195347157.1 | Type1.BREX | <-HTH_3\|\|HTH->Arm-DNA-bind_4+Phage_int_SAM_3+Phage_integrase->TraJ-RHH-><-HEPN<-SIGMA+OB-Fold+AAA-ATPase+Lon-Protease-BrxL<-PglZ<-Schlafen+HATPase+wHTH<-N6-METHYLASE<-DUF6079*<-BrxB-BrxF-iSTAND<-BrxA-DNA-binding<-TraJ-RHH\|\|?->?->ParA-Soj-PloopNTPase->ParB->TraJ-RHH->TraI-VirD2-RCRE-> | 1201 | Bacteria>Terrabacteria_group>Actinomycetota>Coriobacteriia>Eggerthellales>Eggerthellaceae | GCF_028209285.1 | Eggerthella_lenta | PMV52_RS07270 | 251002..254607 | 84112 | unnamed-protein-product-[Eggerthella-lenta] |
| WP_227070741.1 | Type1.BREX | <-TraI-VirD2-RCRE<-TraJ-RHH<-ParB<-ParA-Soj-PloopNTPase<-?<-?\|\|TraJ-RHH->BrxA-DNA-binding->BrxB-BrxF-iSTAND->DUF6079*->N6-METHYLASE->Schlafen+HATPase+wHTH->SIGMA+OB-Fold+AAA-ATPase+Lon-Protease-BrxL->HEPN-><-TraJ-RHH<-Arm-DNA-bind_4+Phage_int_SAM_3+Phage_integrase<-HTH\|\|HTH_3-> | 1199 | Bacteria>Terrabacteria_group>Actinomycetota>Coriobacteriia>Eggerthellales | GCF_015551405.1 | Eggerthellaceae | I2D50_RS06370 | 44195..47794 | 1643826 | MULTISPECIES:-BREX-system-P-loop-protein-BrxC-[Eggerthellaceae] |
| NCO65434.1 | Type1.BREX | STAS->ST-Hiskin->tRNA->BrxA-DNA-binding->BrxB-BrxF-iSTAND->DUF6079*->Schlafen+HATPase+wHTH+HTH->N6-METHYLASE-p-> | 1161 | Bacteria>Terrabacteria_group>Actinomycetota>Thermoleophilia>Solirubrobacterales>Solirubrobacteraceae | GCA_009993245.1 | Solirubrobacter_sp. | GW874_02325 | 6745..10230 | 1932788 | BREX-system-P-loop-protein-BrxC-[Solirubrobacter-sp.] |
| WP_237075536.1 | Type1.BREX | ParB+HNH+Schlafen->?->?->BrxA-DNA-binding->BrxB-BrxF-iSTAND->DUF6079*->N6-METHYLASE->SIGMA+OB-Fold+AAA-ATPase+Lon-Protease-BrxL-><-RVT<-SLATT | 1192 | Bacteria>Terrabacteria_group>Bacillota>Bacilli>Bacillales>Bacillaceae | GCF_021864615.1 | Priestia_megaterium | L4R70_RS26975 | 37253..40831 | 1404 | BREX-system-P-loop-protein-BrxC-[Priestia-megaterium] |
| WP_124068694.1 | Type1.BREX | BrxA-DNA-binding->BrxB-BrxF-iSTAND->DUF6079*->N6-METHYLASE->Schlafen+HATPase+wHTH->PglZ-><-?\|\|TPR-><-?<-TetR-HTH\|\|?->TPR-> | 1184 | Bacteria>Terrabacteria_group>Bacillota>Bacilli>Bacillales>Planococcaceae | GCF_900609045.1 | Filibacter_tadaridae | FILTAD_RS01265 | 2469..6023 | 2483811 | BREX-system-P-loop-protein-BrxC-[Filibacter-tadaridae] |
| MBK0347250.1 | Type1.BREX | HTH_36->?->HTH_26->BrxA-DNA-binding->BrxB-BrxF-iSTAND->DUF6079*->Schlafen+HATPase+wHTH->N6-METHYLASE->HEPN->PglZ->wHTH+WYL+WCX->?->ParA-Soj-PloopNTPase-><-DEOR-HTH | 1198 | Bacteria>Terrabacteria_group>Bacillota>Bacilli>Lactobacillales>Aerococcaceae | GCA_016522115.1 | Aerococcaceae_bacterium_zg-1578 | JDW15_01195 | 55878..59474 | 2796350 | BREX-system-P-loop-protein-BrxC-[Aerococcaceae-bacterium-zg-1578] |
| WP_137638544.1 | Type1.BREX | SF1-SF2-Helicase->ParB->ParB-><-DEDD_Tnp_IS110+Transposase_20\|\|Cas9+HNH+RuvC->Cas_Cas1->CRISPR_Cas2->BrxA-DNA-binding->BrxB-BrxF-iSTAND->DUF6079*->N6-METHYLASE->PglZ->Schlafen+HATPase+wHTH-> | 1204 | Bacteria>Terrabacteria_group>Bacillota>Bacilli>Lactobacillales>Lactobacillaceae | GCF_005405165.1 | Loigolactobacillus_binensis | FFM78_RS11390 | 22588..26202 | 2559922 | BREX-system-P-loop-protein-BrxC-[Loigolactobacillus-binensis] |
| WP_231545262.1 | Type1.BREX | Glyco_hydro_36N+Melibiase+Glyco_hydro_36C->?-><-SIGMA-HTH\|\|HNH+3C-ABC->HNH-p->?->?->BrxA-DNA-binding->BrxB-BrxF-iSTAND->DUF6079*->N6-METHYLASE->Schlafen+HATPase+wHTH->PglZ->SIGMA+OB-Fold+AAA-ATPase+Lon-Protease-BrxL->?->GGDEF+PAS-><-ABhydrolase\|\|ABhydrolase->ABhydrolase+ABhydrolase-> | 1171 | Bacteria>Terrabacteria_group>Bacillota>Bacilli>Lactobacillales>Lactobacillaceae | GCF_021134235.1 | Lactobacillus_delbrueckii | LOB65_RS04175 | 29230..32745 | 1584 | BREX-system-P-loop-protein-BrxC-[Lactobacillus-delbrueckii] |
| WP_236150042.1 | Type1.BREX | <-ABhydrolase+ABhydrolase<-ABhydrolase\|\|ABhydrolase-><-GGDEF+PAS<-?<-SIGMA+OB-Fold+AAA-ATPase+Lon-Protease-BrxL<-PglZ<-Schlafen+HATPase+wHTH<-N6-METHYLASE<-DUF6079*<-BrxB-BrxF-iSTAND<-BrxA-DNA-binding<-?<-?<-HNH-p<-HNH+3C-ABC\|\|SIGMA-HTH-><-?<-Glyco_hydro_36N+Melibiase+Glyco_hydro_36C | 1171 | Bacteria>Terrabacteria_group>Bacillota>Bacilli>Lactobacillales>Lactobacillaceae | GCF_021600205.1 | Lactobacillus_delbrueckii | L5979_RS00560 | 22704..26219 | 1584 | BREX-system-P-loop-protein-BrxC-[Lactobacillus-delbrueckii] |
| HIZ73445.1 | Type1.BREX | <-Mob_Pre<-?\|\|?->BrxA-DNA-binding->BrxB-BrxF-iSTAND->DUF6079*->N6-METHYLASE->?->N6-METHYLASE-p->PglZ->SIGMA+OB-Fold+AAA-ATPase+Lon-Protease-BrxL->Schlafen+HATPase+wHTH-> | 1196 | Bacteria>Terrabacteria_group>Bacillota>Clostridia>Clostridia_incertae_sedis>Candidatus_Gallimonas | GCA_019116755.1 | Candidatus_Gallimonas_intestinavium | H9964_07670 | 4171..7761 | 2838603 | BREX-system-P-loop-protein-BrxC-[Candidatus-Gallimonas-intestinavium] |
| WP_118669593.1 | Type1.BREX | <-SIGMA+OB-Fold+AAA-ATPase+Lon-Protease-BrxL<-OLD-ABC+TOPRIM<-PglZ<-Schlafen+HATPase+wHTH<-HEPN<-TIR<-N6-METHYLASE<-DUF6079*<-BrxB-BrxF-iSTAND<-BrxA-DNA-binding<-HTH_26 | 1196 | Bacteria>Terrabacteria_group>Bacillota>Clostridia>Eubacteriales>Clostridiaceae | - | Clostridium_sp._TF08-15 | DXC82_15795 | 32814..36404 | 2293052 | BREX-system-P-loop-protein-BrxC-[Clostridium-sp.-TF08-15] |
| CDD75188.1 | Type1.BREX | MACRODOMAIN->?->PIN->ZnR+ZnR+ZnR->HTH_26->BrxA-DNA-binding->BrxB-BrxF-iSTAND->DUF6079*->OLD-ABC->PIN->N6-METHYLASE->OLD-ABC->NERD->HEPN->Schlafen+HATPase+wHTH->PglZ->DUF4365+REase-> | 1200 | Bacteria>Terrabacteria_group>Bacillota>Clostridia>Eubacteriales>Clostridiaceae | GCA_000436175.1 | Clostridium_sp._CAG:62 | BN737_01375 | 69547..73149 | 1262828 | uncharacterized-protein-BN737_01375-[Clostridium-sp.-CAG:62] |
| ODM26889.1 | Type1.BREX | <-REase+HTH\|\|?-><-?<-Schlafen+REase<-SIGMA+OB-Fold+AAA-ATPase+Lon-Protease-BrxL<-?<-PglZ<-N6-METHYLASE<-DUF6079*<-BrxB-BrxF-iSTAND<-BrxA-DNA-binding<-?<-?<-?<-?<-?<-?<-SpoOE-like | 1186 | Bacteria>Terrabacteria_group>Bacillota>Clostridia>Eubacteriales>Clostridiaceae | GCA_001717745.1 | Clostridium_sp._Bc-iso-3 | A7W90_12080 | 2831354..2834914 | 1848158 | hypothetical-protein-A7W90_12080-[Clostridium-sp.-Bc-iso-3] |
| RGF28780.1 | Type1.BREX | <-OLD-ABC+TOPRIM<-PglZ<-Schlafen+HATPase+wHTH<-HEPN<-N6-METHYLASE<-DUF6079*<-BrxB-BrxF-iSTAND<-BrxA-DNA-binding<-HTH_26 | 1196 | Bacteria>Terrabacteria_group>Bacillota>Clostridia>Eubacteriales>Clostridiaceae | GCA_003435925.1 | Clostridium_sp._AF46-9NS | DW081_14990 | 34449..38039 | 2293020 | BREX-system-P-loop-protein-BrxC-[Clostridium-sp.-AF46-9NS] |
| RHU61449.1 | Type1.BREX | <-SIGMA+OB-Fold+AAA-ATPase+Lon-Protease-BrxL<-OLD-ABC+TOPRIM<-PglZ<-Schlafen+HATPase+wHTH<-HEPN<-TIR<-N6-METHYLASE<-DUF6079*<-BrxB-BrxF-iSTAND<-BrxA-DNA-binding<-HTH_26 | 1196 | Bacteria>Terrabacteria_group>Bacillota>Clostridia>Eubacteriales>Clostridiaceae | GCA_003481275.1 | Clostridium_sp._TF08-15 | DXC82_15795 | 32814..36404 | 2293052 | BREX-system-P-loop-protein-BrxC-[Clostridium-sp.-TF08-15] |
| NBH35981.1 | Type1.BREX | <-SIR2<-TetR-HTH<-?<-?<-PglZ<-Schlafen+HATPase+wHTH<-DUF4404<-ZnR+ZnR<-N6-METHYLASE<-DUF6079*<-BrxB-BrxF-iSTAND<-BrxA-DNA-binding<-HTH_26\|\|<-HDRnase<-?<-DSBH+AraC-HTH+AraC-HTH | 1182 | Bacteria>Terrabacteria_group>Bacillota>Clostridia>Eubacteriales>Clostridiaceae | GCA_009911065.1 | Clostridiaceae_bacterium | D3Z58_21210 | 19493..23041 | 1898204 | BREX-system-P-loop-protein-BrxC-[Clostridiaceae-bacterium] |
| NBH17641.1 | Type1.BREX | HDRnase->\|\|HTH_26->BrxA-DNA-binding->BrxB-BrxF-iSTAND->DUF6079*->N6-METHYLASE->ZnR+ZnR->DUF4404->Schlafen+HATPase+wHTH->PglZ->?->TetR-HTH->SIR2->ACT-> | 1182 | Bacteria>Terrabacteria_group>Bacillota>Clostridia>Eubacteriales>Clostridiaceae | GCA_009911105.1 | Clostridiaceae_bacterium | D3Z55_09200 | 40728..44276 | 1898204 | BREX-system-P-loop-protein-BrxC-[Clostridiaceae-bacterium] |
| NBH73079.1 | Type1.BREX | Phage_integrase->HTH_26->BrxA-DNA-binding->BrxB-BrxF-iSTAND->DUF6079*->N6-METHYLASE->Schlafen+HATPase+wHTH->PglZ->SIGMA+OB-Fold+AAA-ATPase+Lon-Protease-BrxL-><-ACET<-REase+35exo<-Schlafen+HATPase+wHTH | 1212 | Bacteria>Terrabacteria_group>Bacillota>Clostridia>Eubacteriales>Clostridiaceae | GCA_009911255.1 | Clostridiaceae_bacterium | D3Z51_13830 | 3930..7568 | 1898204 | BREX-system-P-loop-protein-BrxC-[Clostridiaceae-bacterium] |
| NBI69485.1 | Type1.BREX | <-Peptidase_S11+PBP5_C\|\|?-><-?<-?<-GspA-ATPase<-?\|\|<-PglZ<-Schlafen+HATPase+wHTH<-N6-METHYLASE<-DUF6079*<-BrxB-BrxF-iSTAND<-BrxA-DNA-binding<-HTH_26<-?<-ABhydrolase<-GNTR-HTH<-?<-GNTR-HTH+UTRA | 1212 | Bacteria>Terrabacteria_group>Bacillota>Clostridia>Eubacteriales>Clostridiaceae | GCA_009911305.1 | Clostridiaceae_bacterium | D3Z50_00070 | 19913..23551 | 1898204 | BREX-system-P-loop-protein-BrxC-[Clostridiaceae-bacterium] |
| MBS5468798.1 | Type1.BREX | ACT->?-><-MODE-HTH+PBPII\|\|?->?->HTH_26->BrxA-DNA-binding->BrxB-BrxF-iSTAND->DUF6079*->?->N6-METHYLASE->ZnR+ZnR->DUF4404->Schlafen+HATPase+wHTH->PglZ->SIGMA+OB-Fold+AAA-ATPase+Lon-Protease-BrxL->?->ABC-ATPase+SbcC+ABC-ATPase+SbcC-> | 1178 | Bacteria>Terrabacteria_group>Bacillota>Clostridia>Eubacteriales>Clostridiaceae | GCA_018368355.1 | Clostridium_sp. | KHX69_08270 | 35146..38682 | 1506 | BREX-system-P-loop-protein-BrxC-[Clostridium-sp.] |
| MCI8726399.1 | Type1.BREX | <-HKD-DNase+HKD-DNase<-?<-?\|\|?-><-ClpABN-AAA+ClpABC-AAA<-PglZ<-Schlafen+HATPase+wHTH<-N6-METHYLASE<-DUF6079*<-BrxB-BrxF-iSTAND<-BrxA-DNA-binding<-HTH_26 | 1179 | Bacteria>Terrabacteria_group>Bacillota>Clostridia>Eubacteriales>Clostridiaceae | GCA_022794715.1 | Hungatella_sp. | HFG66_09835 | 51517..55056 | 2613924 | BREX-system-P-loop-protein-BrxC-[Hungatella-sp.] |
| WP_103227376.1 | Type1.BREX | <-SIGMA+OB-Fold+AAA-ATPase+Lon-Protease-BrxL<-PglZ<-Schlafen+HATPase+wHTH<-N6-METHYLASE<-DUF6079*<-BrxB-BrxF-iSTAND<-BrxA-DNA-binding<-HTH_26<-?<-?\|\|?->GNTR-HTH-> | 1215 | Bacteria>Terrabacteria_group>Bacillota>Clostridia>Eubacteriales>Clostridiaceae | GCF_002899675.1 | Clostridium_sp._chh4-2 | C0033_RS08180 | 119935..123582 | 2067550 | BREX-system-P-loop-protein-BrxC-[Clostridium-sp.-chh4-2] |
| WP_118320912.1 | Type1.BREX | TPR+TPR+TPR->?->HISKIN->SIGMA-HTH+SIGMA-HTH->HTH_26->BrxA-DNA-binding->BrxB-BrxF-iSTAND->DUF6079*->TIR->N6-METHYLASE->AbiGi->Schlafen+HATPase+wHTH->PglZ->SIGMA+OB-Fold+AAA-ATPase+Lon-Protease-BrxL->TIR->AT-HTH-HYD2-2-> | 1183 | Bacteria>Terrabacteria_group>Bacillota>Clostridia>Eubacteriales>Clostridiaceae | GCF_003473525.1 | Clostridium_sp._AM22-16AC | DW260_RS00800 | 177692..181243 | 2292009 | BREX-system-P-loop-protein-BrxC-[Clostridium-sp.-AM22-16AC] |
| WP_118513467.1 | Type1.BREX | <-Tox-ART-HYD2<-TIR<-PglZ<-Schlafen+HATPase+wHTH<-AbiGi+seqA+RAMA<-N6-METHYLASE<-HNH<-OLD-ABC<-DUF6079*<-BrxB-BrxF-iSTAND<-BrxA-DNA-binding<-HTH_26<-SIGMA-HTH+SIGMA-HTH<-HISKIN<-?<-TPR+TPR+TPR | 1183 | Bacteria>Terrabacteria_group>Bacillota>Clostridia>Eubacteriales>Clostridiaceae | GCF_003477775.1 | Clostridium_sp._AF37-5AT | DW020_RS00895 | 209396..212947 | 2292007 | BREX-system-P-loop-protein-BrxC-[Clostridium-sp.-AF37-5AT] |
| WP_118731364.1 | Type1.BREX | PALP->?->RVT->HTH_26->BrxA-DNA-binding->BrxB-BrxF-iSTAND->DUF6079*->N6-METHYLASE->HEPN->Schlafen+HATPase+wHTH->PglZ->OLD-ABC+TOPRIM->SIGMA+OB-Fold+AAA-ATPase+Lon-Protease-BrxL-> | 1196 | Bacteria>Terrabacteria_group>Bacillota>Clostridia>Eubacteriales>Clostridiaceae | GCF_003480225.1 | Clostridium_sp._AM34-9AC | DW835_RS14805 | 20266..23856 | 2293030 | BREX-system-P-loop-protein-BrxC-[Clostridium-sp.-AM34-9AC] |

| Protein-ID | System | Gene-Neighbourhoods | Length | Taxonomy | Assembly-ID | Strain | Locus-Tag | Genomic-coordinates | TaxID | Fasta-defline |
|---|---|---|---|---|---|---|---|---|---|---|
| WP_118740431.1 | Type1.BREX | <-OLD-ABC+TOPRIM<- SIGMA+OB-Fold+AAA-ATPase+Lon-Protease-BrxL <- PglZ <- Schlafen +HATPase+wHTH <-DUF4404<-ZnR+ZnR<- N6-METHYLASE <-TIR<-?<- DUF6079 *<- BrxB-BrxF-iSTAND <- [illegible] | 1182 | Bacteria>Terrabacteria_group>Bacillota>Clostridia>Eubacteriales>Clostridiaceae | GCF_003480345.1 | Clostridium_sp._AM32-2 | DW807_RS02880 | 180384..183932 | 2293029 | BREX-system-P-loop-protein-BrxC-[Clostridium-sp.-AM32-2] |
| WP_118740660.1 | Type1.BREX | <- SIGMA+OB-Fold+AAA-ATPase+Lon-Protease-BrxL <- PglZ <- Schlafen +HATPase+wHTH <-AbiGi+seqA+RAMA<-?<- DUF6079 *<- BrxB-BrxF-iSTAND <- BrxA-DNA-binding <-HTH_26 | 1182 | Bacteria>Terrabacteria_group>Bacillota>Clostridia>Eubacteriales>Clostridiaceae | GCF_003480345.1 | Clostridium_sp._AM32-2 | DW807_RS07265 | 17813..21361 | 2293029 | BREX-system-P-loop-protein-BrxC-[Clostridium-sp.-AM32-2] |
| WP_118742502.1 | Type1.BREX | 35exo+T7DNApol->?->?->?->?->HTH_26-> BrxA-DNA-binding -> BrxB-BrxF-iSTAND -> DUF6079 *-> N6-METHYLASE ->HEPN-> Schlafen +HATPase+wHTH -> PglZ -> SIGMA+OB-Fold+AAA-ATPase+Lon-Prot[illegible] | 1182 | Bacteria>Terrabacteria_group>Bacillota>Clostridia>Eubacteriales>Clostridiaceae | GCF_003480665.1 | Clostridium_sp._AM28-20LB | DW739_RS01385 | 291245..294793 | 2293027 | BREX-system-P-loop-protein-BrxC-[Clostridium-sp.-AM28-20LB] |
| WP_118670884.1 | Type1.BREX | wHTH+WYL+WCX ->?->?->?->?->?-> BrxA-DNA-binding -> BrxB-BrxF-iSTAND -> DUF6079 *-> N6-METHYLASE ->?->?-> Schlafen +HATPase+wHTH -> PglZ ->?-> SIGMA+OB-Fold+AAA-ATPase+Lon-Protea[illegible] | 1196 | Bacteria>Terrabacteria_group>Bacillota>Clostridia>Eubacteriales>Clostridiaceae | GCF_003481335.1 | Clostridium_sp._OM08-29 | DXC24_RS05235 | 319406..322996 | 2293049 | BREX-system-P-loop-protein-BrxC-[Clostridium-sp.-OM08-29] |
| WP_195540766.1 | Type1.BREX | <-HNH<-ClpABC-AAA<-?<- SIGMA+OB-Fold+AAA-ATPase+Lon-Protease-BrxL <- PglZ <- Schlafen +HATPase+wHTH <-HEPN<- N6-METHYLASE <-?<- DUF6079 *<- BrxB-BrxF-iSTAND <- BrxA-DNA-binding [illegible] | 1182 | Bacteria>Terrabacteria_group>Bacillota>Clostridia>Eubacteriales>Clostridiaceae | GCF_015558125.1 | Clostridium_sp._D43t1_170807_D5 | I2G39_RS02195 | 465793..469341 | 2787150 | BREX-system-P-loop-protein-BrxC-[Clostridium-sp.-D43t1_170807_D5] |
| WP_216137994.1 | Type1.BREX | MqsA_antitoxin+DUF4065->?->HTH_26-> BrxA-DNA-binding -> BrxB-BrxF-iSTAND -> DUF6079 *-> Schlafen +HATPase+wHTH -p->?-> Schlafen +HATPase+wHTH -p-> N6-METHYLASE -> PglZ -> SIGMA+O[illegible] | 1203 | Bacteria>Terrabacteria_group>Bacillota>Clostridia>Eubacteriales>Clostridiaceae | GCF_018861855.1 | Clostridium_estertheticum | KPL49_RS19700 | 51196..54807 | 238834 | BREX-system-P-loop-protein-BrxC-[Clostridium-estertheticum] |
| WP_216574650.1 | Type1.BREX | <-ABhydrolase\|\|IG+IG->?->MazE_antitoxin+SpoVT_C->HTH_26-> BrxA-DNA-binding -> BrxB-BrxF-iSTAND -> DUF6079 *-> N6-METHYLASE ->HEPN-> Schlafen +HATPase+wHTH -> PglZ -> SIGMA+OB-Fo[illegible] | 1182 | Bacteria>Terrabacteria_group>Bacillota>Clostridia>Eubacteriales>Clostridiaceae | GCF_018918265.1 | Falcatimonas_sp._MSJ-15 | KQI85_RS03175 | 188313..191861 | 2841515 | BREX-system-P-loop-protein-BrxC-[Falcatimonas-sp.-MSJ-15] |
| WP_073026962.1 | Type1.BREX | <-HTH_3\|\|GGDEF->?->SpoOE-like->DndB->?-> BrxA-DNA-binding -> BrxB-BrxF-iSTAND -> DUF6079 *-> N6-METHYLASE -> Schlafen +HATPase+wHTH -> PglZ -> SIGMA+OB-Fold+AAA-ATPase+Lon-Protea[illegible] | 1186 | Bacteria>Terrabacteria_group>Bacillota>Clostridia>Eubacteriales>Clostridiaceae | GCF_900142105.1 | Lutispora_thermophila | BUB48_RS14275 | 12515..16075 | 288966 | BREX-system-P-loop-protein-BrxC-[Lutispora-thermophila] |
| WP_055656468.1 | Type1.BREX | <-HD+SF1-SF2-Helicase<-Peptidase<- SIGMA+OB-Fold+AAA-ATPase+Lon-Protease-BrxL <- PglZ <- Schlafen +HATPase+wHTH <- N6-METHYLASE <- DUF6079 *<- BrxB-BrxF-iSTAND <- BrxA-DNA-bindi[illegible] | 1220 | Bacteria>Terrabacteria_group>Bacillota>Clostridia>Eubacteriales>Clostridiaceae | GCF_902362405.1 | Hungatella_hathewayi | FY393_RS15415 | 235272..238934 | 154046 | BREX-system-P-loop-protein-BrxC-[Hungatella-hathewayi] |
| WP_117466987.1 | Type1.BREX | <-OLD-ABC+TOPRIM<- PglZ <- Schlafen +HATPase+wHTH <-HEPN<- N6-METHYLASE <- DUF6079 *<- BrxB-BrxF-iSTAND <- BrxA-DNA-binding <-HTH_26 | 1196 | Bacteria>Terrabacteria_group>Bacillota>Clostridia>Eubacteriales>Clostridiaceae | - | unclassified_Clostridium | DW081_14990 | 34449..38039 | 2614128 | MULTISPECIES:-BREX-system-P-loop-protein-BrxC-[unclassified-Clostridium] |
| WP_117821980.1 | Type1.BREX | MACRODOMAIN->?->ZnR+ZnR+ZnR->?-><-?<-HTH_38\|\|HTH_26-> BrxA-DNA-binding -> BrxB-BrxF-iSTAND -> DUF6079 *-> N6-METHYLASE -> N6-METHYLASE -p->?-> Schlafen +HATPase+wHTH -> PglZ ->OL[illegible] | 1196 | Bacteria>Terrabacteria_group>Bacillota>Clostridia>Eubacteriales>Clostridiaceae | - | unclassified_Clostridium | DW959_RS12365 | 44943..48533 | 2614128 | MULTISPECIES:-BREX-system-P-loop-protein-BrxC-[unclassified-Clostridium] |
| OLA08999.1 | Type1.BREX | <-Phage_int+DNA-glycosylase<-HTH+REase<- SIGMA+OB-Fold+AAA-ATPase+Lon-Protease-BrxL <- PglZ <-AbiGi<- N6-METHYLASE -p<- Schlafen +HATPase+wHTH <-?<- N6-METHYLASE <- DUF6079 *<- [illegible] | 1184 | Bacteria>Terrabacteria_group>Bacillota>Clostridia>Eubacteriales>Eubacteriaceae | GCA_001916415.1 | Eubacterium_sp._45_250 | BHW21_06160 | 165792..169346 | 1897004 | MAG:-ATPase-[Eubacterium-sp.-45_250] |
| MBS6586983.1 | Type1.BREX | <-GATase_6<-?<-?\|\|NUDIX-><-?\|\| Schlafen +HATPase+wHTH -><- SIGMA+OB-Fold+AAA-ATPase+Lon-Protease-BrxL <- PglZ <- N6-METHYLASE <- DUF6079 *<- BrxB-BrxF-iSTAND <- BrxA-DNA-binding [illegible] | 1209 | Bacteria>Terrabacteria_group>Bacillota>Clostridia>Eubacteriales>Eubacteriaceae | GCA_018377805.1 | Eubacterium_sp. | KH345_12625 | 21110..24739 | 142586 | BREX-system-P-loop-protein-BrxC-[Eubacterium-sp.] |
| MCH4073094.1 | Type1.BREX | <-ClpABC-AAA<-?<-HEPN<-HEPN<- SIGMA+OB-Fold+AAA-ATPase+Lon-Protease-BrxL <- PglZ <- Schlafen +HATPase+wHTH <- Schlafen +HATPase+wHTH <- N6-METHYLASE <- DUF6079 *<- BrxB-BrxF-iST[illegible] | 1171 | Bacteria>Terrabacteria_group>Bacillota>Clostridia>Eubacteriales>Eubacteriaceae | GCA_022483875.1 | Pseudoramibacter_sp. | LKF02_09135 | 1854333..1857848 | 2034862 | BREX-system-P-loop-protein-BrxC-[Pseudoramibacter-sp.] |
| MCI5510134.1 | Type1.BREX | CopG/RHH->?->TraJ[illegible] | 1182 | Bacteria>Terrabacteria_group>Bacillota>Clostridia>Eubacteriales>Eubacteriaceae | GCA_022764165.1 | Eubacterium_sp. | MR436_06790 | 9461..13009 | 142586 | BREX-system-P-loop-protein-BrxC-[Eubacterium-sp.] |
| WP_055289382.1 | Type1.BREX | HTH-><-?<-PHP+Sbc[illegible] | 1183 | Bacteria>Terrabacteria_group>Bacillota>Clostridia>Eubacteriales>Eubacteriaceae | GCF_001406295.1 | Eubacterium_ramulus | ARA08_RS03325 | 204677..208228 | 39490 | BREX-system-P-loop-protein-BrxC-[Eubacterium-ramulus] |
| WP_186842510.1 | Type1.BREX | Aminotran_1_2->?-[illegible] | 1222 | Bacteria>Terrabacteria_group>Bacillota>Clostridia>Eubacteriales>Eubacteriaceae | GCF_014284475.1 | Acetobacterium_fimetarium | GH808_RS09300 | 27272..30940 | 52691 | BREX-system-P-loop-protein-BrxC-[Acetobacterium-fimetarium] |
| WP_087184149.1 | Type1.BREX | TraJ-RHH->?->?-><[illegible] | 1178 | Bacteria>Terrabacteria_group>Bacillota>Clostridia>Eubacteriales>Eubacteriales_incertae_sedis | GCF_002159185.1 | Gemmiger_sp._An87 | B5G38_RS07650 | 33911..37447 | 1965662 | BREX-system-P-loop-protein-BrxC-[Gemmiger-sp.-An87] |
| WP_055219000.1 | Type1.BREX | ZnR->?->HTH_26->[illegible] | 1182 | Bacteria>Terrabacteria_group>Bacillota>Clostridia>Eubacteriales | GCF_013300755.1 | Lachnospiraceae | G5A93_RS10915 | 4301..7849 | 186803 | MULTISPECIES:-BREX-system-P-loop-protein-BrxC-[Lachnospiraceae] |
| WP_117739785.1 | Type1.BREX | HTH_36->?->CopG/R[illegible] | 1183 | Bacteria>Terrabacteria_group>Bacillota>Clostridia>Eubacteriales | GCF_013301605.1 | Lachnospiraceae | G4434_RS12220 | 38820..42371 | 186803 | MULTISPECIES:-BREX-system-P-loop-protein-BrxC-[Lachnospiraceae] |
| WP_138373965.1 | Type1.BREX | <-SF1-SF2-Helicas[illegible] | 1179 | Bacteria>Terrabacteria_group>Bacillota>Clostridia>Eubacteriales | GCF_015554965.1 | Oscillospiraceae | I2E95_RS16325 | 14974..18513 | 216572 | MULTISPECIES:-BREX-system-P-loop-protein-BrxC-[Oscillospiraceae] |
| WP_118699091.1 | Type1.BREX | HTH_36->?->CopG/R[illegible] | 1182 | Bacteria>Terrabacteria_group>Bacillota>Clostridia>Eubacteriales | GCF_020560115.1 | Lachnospiraceae | LI126_RS00410 | 82443..85991 | 186803 | MULTISPECIES:-BREX-system-P-loop-protein-BrxC-[Lachnospiraceae] |
| WP_118447467.1 | Type1.BREX | CopG/RHH->?->TraJ[illegible] | 1182 | Bacteria>Terrabacteria_group>Bacillota>Clostridia>Eubacteriales | GCF_022900215.1 | Lachnospiraceae | MUY38_RS08320 | 63562..67110 | 186803 | MULTISPECIES:-BREX-system-P-loop-protein-BrxC-[Lachnospiraceae] |
| WP_117761890.1 | Type1.BREX | <-Phage_integrase[illegible] | 1182 | Bacteria>Terrabacteria_group>Bacillota>Clostridia>Eubacteriales | GCF_902364755.1 | Lachnospiraceae | FXX70_RS12975 | 76873..80421 | 186803 | MULTISPECIES:-BREX-system-P-loop-protein-BrxC-[Lachnospiraceae] |
| WP_256171760.1 | Type1.BREX | HTH_36->?->CopG/R[illegible] | 1182 | Bacteria>Terrabacteria_group>Bacillota>Clostridia>Eubacteriales>Lachnospiraceae | - | Dorea_longicatena | NE607_07100 | 21358..24906 | 88431 | BREX-system-P-loop-protein-BrxC-[Dorea-longicatena] |
| EFV19659.1 | Type1.BREX | wHTH+WYL+WCX ->?-[illegible] | 1177 | Bacteria>Terrabacteria_group>Bacillota>Clostridia>Eubacteriales>Lachnospiraceae | GCA_000185545.1 | Lachnospiraceae_bacterium_8_1_57FAA | HMPREF1026_01169 | 31500..35033 | 665951 | hypothetical-protein-HMPREF1026_01169-[Lachnospiraceae-bacterium-8_1_57FAA] |
| EGN43144.1 | Type1.BREX | <-TIR<-?<- PglZ <-[illegible] | 1182 | Bacteria>Terrabacteria_group>Bacillota>Clostridia>Eubacteriales>Lachnospiraceae | GCA_000218445.1 | Lachnospiraceae_bacterium_1_1_57FAA | HMPREF0990_02349 | 46029..49577 | 658081 | hypothetical-protein-HMPREF0990_02349-[Lachnospiraceae-bacterium-1_1_57FAA] |
| EHO53370.1 | Type1.BREX | Peptidase_M32->TP[illegible] | 1186 | Bacteria>Terrabacteria_group>Bacillota>Clostridia>Eubacteriales>Lachnospiraceae | GCA_000242315.1 | Lachnospiraceae_bacterium_oral_taxon_082_str._F0431 | HMPREF9099_00896 | 82439..85999 | 861454 | hypothetical-protein-HMPREF9099_00896-[Lachnospiraceae-bacterium-oral-taxon-082-str.-F0431] |
| EOS37361.1 | Type1.BREX | <- PglZ <- Schlafe[illegible] | 1182 | Bacteria>Terrabacteria_group>Bacillota>Clostridia>Eubacteriales>Lachnospiraceae | GCA_000403295.2 | Lachnospiraceae_bacterium_M18-1 | C808_03339 | 122087..125635 | 1235792 | hypothetical-protein-C808_03339-[Lachnospiraceae-bacterium-M18-1] |
| CUO05580.1 | Type1.BREX | <- SIGMA+OB-Fold+[illegible] | 1182 | Bacteria>Terrabacteria_group>Bacillota>Clostridia>Eubacteriales>Lachnospiraceae | GCA_001405875.1 | Roseburia_hominis | ERS852391_01284 | 73560..77108 | 301301 | Uncharacterised-protein-[Roseburia-hominis] |
| CUM83721.1 | Type1.BREX | <- SIGMA+OB-Fold+[illegible] | 1182 | Bacteria>Terrabacteria_group>Bacillota>Clostridia>Eubacteriales>Lachnospiraceae | GCA_001406375.1 | [Eubacterium]_rectale | ERS852580_00755 | 13381..16929 | 39491 | Uncharacterised-protein-[[Eubacterium] |
| RGI65884.1 | Type1.BREX | HTH_26-> BrxA-DNA-[illegible] | 1196 | Bacteria>Terrabacteria_group>Bacillota>Clostridia>Eubacteriales>Lachnospiraceae | GCA_003436035.1 | [Eubacterium]_rectale | DXD95_13285 | 1619..5209 | 39491 | BREX-system-P-loop-protein-BrxC-[[Eubacterium] |
| HAB87400.1 | Type1.BREX | ZnR+ZnR+ZnR->?->H[illegible] | 1196 | Bacteria>Terrabacteria_group>Bacillota>Clostridia>Eubacteriales>Lachnospiraceae | GCA_003448155.1 | Coprococcus_sp. | DCE60_00030 | 3490..7080 | 2049024 | BREX-system-P-loop-protein-BrxC-[Coprococcus-sp.] |
| RHA07177.1 | Type1.BREX | <-TIR<- SIGMA+OB-[illegible] | 1182 | Bacteria>Terrabacteria_group>Bacillota>Clostridia>Eubacteriales>Lachnospiraceae | GCA_003466745.1 | [Eubacterium]_rectale | DW951_01530 | 328230..331778 | 39491 | BREX-system-P-loop-protein-BrxC-[[Eubacterium] |
| RGZ95621.1 | Type1.BREX | HTH_26-> BrxB-Brx[illegible] | 1182 | Bacteria>Terrabacteria_group>Bacillota>Clostridia>Eubacteriales>Lachnospiraceae | GCA_003467085.1 | [Eubacterium]_rectale | DW967_00020 | 2042..5590 | 39491 | BREX-system-P-loop-protein-BrxC-[[Eubacterium] |
| RHD97433.1 | Type1.BREX | HTH_26-> BrxA-DNA-[illegible] | 1182 | Bacteria>Terrabacteria_group>Bacillota>Clostridia>Eubacteriales>Lachnospiraceae | GCA_003468965.1 | [Eubacterium]_rectale | DW775_02150 | 1680..5228 | 39491 | BREX-system-P-loop-protein-BrxC-[[Eubacterium] |
| HBA06127.1 | Type1.BREX | MNS+REase->TOPRIM[illegible] | 1182 | Bacteria>Terrabacteria_group>Bacillota>Clostridia>Eubacteriales>Lachnospiraceae | GCA_003491515.1 | Roseburia_sp. | DC452_03320 | 12130..15678 | 2049040 | BREX-system-P-loop-protein-BrxC-[Roseburia-sp.] |
| HCI18472.1 | Type1.BREX | <-PIN<-?<- SIGMA+[illegible] | 799 | Bacteria>Terrabacteria_group>Bacillota>Clostridia>Eubacteriales>Lachnospiraceae | GCA_003521585.1 | Lachnospiraceae_bacterium | DFH32_07645 | 14441..16840 | 1898203 | BREX-system-P-loop-protein-BrxC-[Lachnospiraceae-bacterium] |
| HCS67503.1 | Type1.BREX | <- PglZ <- Schlafe[illegible] | 1189 | Bacteria>Terrabacteria_group>Bacillota>Clostridia>Eubacteriales>Lachnospiraceae | GCA_003531095.1 | Oribacterium_sp. | DIW34_04725 | 7705..11274 | 1969407 | BREX-system-P-loop-protein-BrxC-[Oribacterium-sp.] |
| HCX41706.1 | Type1.BREX | SKI->?->tRNA->?->[illegible] | 1196 | Bacteria>Terrabacteria_group>Bacillota>Clostridia>Eubacteriales>Lachnospiraceae | GCA_003539645.1 | Lachnospiraceae_bacterium | DHH50_05105 | 179559..183149 | 1898203 | BREX-system-P-loop-protein-BrxC-[Lachnospiraceae-bacterium] |
| HCU04414.1 | Type1.BREX | <- SIGMA+OB-Fold+[illegible] | 1182 | Bacteria>Terrabacteria_group>Bacillota>Clostridia>Eubacteriales>Lachnospiraceae | GCA_003543095.1 | Roseburia_sp. | DF667_12405 | 43487..47035 | 2049040 | BREX-system-P-loop-protein-BrxC-[Roseburia-sp.] |
| RKW57361.1 | Type1.BREX | Schlafen +HATPase[illegible] | 1217 | Bacteria>Terrabacteria_group>Bacillota>Clostridia>Eubacteriales>Lachnospiraceae | GCA_003640285.1 | Lachnospiraceae_bacterium | D8H95_03165 | 3356..7009 | 1898203 | BREX-system-P-loop-protein-BrxC-[Lachnospiraceae-bacterium] |
| TYL56933.1 | Type1.BREX | HTH_26-> BrxA-DNA-[illegible] | 1196 | Bacteria>Terrabacteria_group>Bacillota>Clostridia>Eubacteriales>Lachnospiraceae | GCA_008123415.1 | [Eubacterium]_rectale | FYL31_13935 | 1744..5334 | 39491 | BREX-system-P-loop-protein-BrxC-[[Eubacterium] |
| QNM03383.1 | Type1.BREX | GNTR-HTH+Aminotra[illegible] | 1215 | Bacteria>Terrabacteria_group>Bacillota>Clostridia>Eubacteriales>Lachnospiraceae | GCA_014337175.1 | Simiaoa_sunii | H9Q77_04470 | 956158..959805 | 2763672 | BREX-system-P-loop-protein-BrxC-[Simiaoa-sunii] |
| MBN2952027.1 | Type1.BREX | ZnR->HTH_26-> Brx[illegible] | 1182 | Bacteria>Terrabacteria_group>Bacillota>Clostridia>Eubacteriales>Lachnospiraceae | GCA_016939215.1 | Fusicatenibacter_saccharivorans | JTJ23_00185 | 3299..6847 | 1150298 | BREX-system-P-loop-protein-BrxC-[Fusicatenibacter-saccharivorans] |
| MBQ9134971.1 | Type1.BREX | <-REase+iREase<-?[illegible] | 1213 | Bacteria>Terrabacteria_group>Bacillota>Clostridia>Eubacteriales>Lachnospiraceae | GCA_017463605.1 | Lachnospiraceae_bacterium | IJX66_02610 | 29830..33471 | 1898203 | BREX-system-P-loop-protein-BrxC-[Lachnospiraceae-bacterium] |
| MBR4345194.1 | Type1.BREX | RelB->PIN-> BrxA-[illegible] | 1173 | Bacteria>Terrabacteria_group>Bacillota>Clostridia>Eubacteriales>Lachnospiraceae | GCA_017542245.1 | Lachnospiraceae_bacterium | IKP88_21230 | 56947..60468 | 1898203 | BREX-system-P-loop-protein-BrxC-[Lachnospiraceae-bacterium] |
| MBP3802955.1 | Type1.BREX | RelB-><-?\|\|HTH_26[illegible] | 1204 | Bacteria>Terrabacteria_group>Bacillota>Clostridia>Eubacteriales>Lachnospiraceae | GCA_017940955.1 | Oribacterium_sp. | J6I76_03520 | 9410..13024 | 1969407 | BREX-system-P-loop-protein-BrxC-[Oribacterium-sp.] |
| HIV92250.1 | Type1.BREX | TrkH->?->CBS->URI[illegible] | 1180 | Bacteria>Terrabacteria_group>Bacillota>Clostridia>Eubacteriales>Lachnospiraceae | GCA_019113175.1 | Candidatus_Eisenbergiella_stercoravium | H9744_11330 | 86529..90071 | 2838556 | BREX-system-P-loop-protein-BrxC-[Candidatus-Eisenbergiella-stercoravium] |
| HJC65843.1 | Type1.BREX | <-HTH+REase<-?<-?[illegible] | 1180 | Bacteria>Terrabacteria_group>Bacillota>Clostridia>Eubacteriales>Lachnospiraceae | GCA_019119955.1 | Candidatus_Enterocloster_excrementigallinarum | H9931_03865 | 69183..72725 | 2838558 | BREX-system-P-loop-protein-BrxC-[Candidatus-Enterocloster-excrementigallinarum] |
| MCB7506159.1 | Type1.BREX | <-ClpABN-AAA+ClpA[illegible] | 1182 | Bacteria>Terrabacteria_group>Bacillota>Clostridia>Eubacteriales>Lachnospiraceae | GCA_020553125.1 | Blautia_sp._MSK20_18 | LI091_01875 | 117354..120902 | 2883186 | BREX-system-P-loop-protein-BrxC-[Blautia-sp.-MSK20_18] |
| MCB7110231.1 | Type1.BREX | METHYLASE->?->ZnR[illegible] | 1182 | Bacteria>Terrabacteria_group>Bacillota>Clostridia>Eubacteriales>Lachnospiraceae | GCA_020554685.1 | [Eubacterium]_rectale | LON19_09275 | 43133..46681 | 39491 | BREX-system-P-loop-protein-BrxC-[[Eubacterium] |
| MCC2229908.1 | Type1.BREX | HTH_36->?->CopG/R[illegible] | 1182 | Bacteria>Terrabacteria_group>Bacillota>Clostridia>Eubacteriales>Lachnospiraceae | GCA_020687485.1 | Lachnospiraceae_bacterium_CLA-AA-H215 | LKD81_02670 | 26793..30341 | 2885348 | BREX-system-P-loop-protein-BrxC-[Lachnospiraceae-bacterium-CLA-AA-H215] |
| MCC2230802.1 | Type1.BREX | <-HAMP+HISKIN<-RE[illegible] | 1217 | Bacteria>Terrabacteria_group>Bacillota>Clostridia>Eubacteriales>Lachnospiraceae | GCA_020687485.1 | Lachnospiraceae_bacterium_CLA-AA-H215 | LKD81_07295 | 25894..29547 | 2885348 | BREX-system-P-loop-protein-BrxC-[Lachnospiraceae-bacterium-CLA-AA-H215] |
| MCD7763957.1 | Type1.BREX | HTH_26-> BrxA-DNA-[illegible] | 1215 | Bacteria>Terrabacteria_group>Bacillota>Clostridia>Eubacteriales>Lachnospiraceae | GCA_021199655.1 | Lachnospiraceae_bacterium | LUI14_12340 | 45356..49003 | 1898203 | BREX-system-P-loop-protein-BrxC-[Lachnospiraceae-bacterium] |
| MCD8362902.1 | Type1.BREX | NTP_transf+HDRnas[illegible] | 1212 | Bacteria>Terrabacteria_group>Bacillota>Clostridia>Eubacteriales>Lachnospiraceae | GCA_021205385.1 | Lachnospiraceae_bacterium | LUC98_08085 | 19577..23215 | 1898203 | BREX-system-P-loop-protein-BrxC-[Lachnospiraceae-bacterium] |
| MCH3986384.1 | Type1.BREX | TOPRIM+Topoisom_b[illegible] | 1223 | Bacteria>Terrabacteria_group>Bacillota>Clostridia>Eubacteriales>Lachnospiraceae | GCA_022483185.1 | Lachnospiraceae_bacterium | LKE69_00320 | 66250..69921 | 1898203 | BREX-system-P-loop-protein-BrxC-[Lachnospiraceae-bacterium] |
| MCI7121458.1 | Type1.BREX | HATPase+wHTH->?->[illegible] | 1201 | Bacteria>Terrabacteria_group>Bacillota>Clostridia>Eubacteriales>Lachnospiraceae | GCA_022775325.1 | [Ruminococcus]_gnavus | MR941_06720 | 10105..13710 | 33038 | BREX-system-P-loop-protein-BrxC-[[Ruminococcus] |
| MCI6977806.1 | Type1.BREX | sGTP->?->?->?->?-[illegible] | 1182 | Bacteria>Terrabacteria_group>Bacillota>Clostridia>Eubacteriales>Lachnospiraceae | GCA_022775645.1 | Lachnospiraceae_bacterium | MR930_04145 | 182006..185554 | 1898203 | BREX-system-P-loop-protein-BrxC-[Lachnospiraceae-bacterium] |
| MCI9239442.1 | Type1.BREX | HTH_26-> BrxA-DNA-[illegible] | 1182 | Bacteria>Terrabacteria_group>Bacillota>Clostridia>Eubacteriales>Lachnospiraceae | GCA_022798835.1 | Dorea_sp. | HFH07_13900 | 2922..6470 | 2040332 | BREX-system-P-loop-protein-BrxC-[Dorea-sp.] |
| MCQ4892848.1 | Type1.BREX | HTH_36->?->CopG/R[illegible] | 1182 | Bacteria>Terrabacteria_group>Bacillota>Clostridia>Eubacteriales>Lachnospiraceae | GCA_024461355.1 | Dorea_longicatena | NE607_07100 | 21358..24906 | 88431 | BREX-system-P-loop-protein-BrxC-[Dorea-longicatena] |
| SCG99927.1 | Type1.BREX | <- PglZ <- Schlafe[illegible] | 1182 | Bacteria>Terrabacteria_group>Bacillota>Clostridia>Eubacteriales>Lachnospiraceae | GCA_900066205.1 | uncultured_Blautia_sp. | SAMEA3545256_00122 | 140025..143573 | 765821 | Uncharacterised-protein-[uncultured-Blautia-sp.] |
| SEQ84346.1 | Type1.BREX | Classical-AAA->MA[illegible] | 1212 | Bacteria>Terrabacteria_group>Bacillota>Clostridia>Eubacteriales>Lachnospiraceae | GCA_900110635.1 | Lachnospiraceae_bacterium_NE2001 | SAMN02910369_02521 | 19203..22841 | 1520823 | hypothetical-protein-SAMN02910369_02521-[Lachnospiraceae-bacterium-NE2001] |
| SEP83303.1 | Type1.BREX | DUF6079 -p*-> N6-[illegible] | 728 | Bacteria>Terrabacteria_group>Bacillota>Clostridia>Eubacteriales>Lachnospiraceae | GCA_900110975.1 | Lachnospiraceae_bacterium_RM5 | SAMN02910289_00625 | 328..2514 | 1200721 | hypothetical-protein-SAMN02910289_00625-[Lachnospiraceae-bacterium-RM5] |
| WP_009536434.1 | Type1.BREX | <-SF1-SF2-Helicas[illegible] | 1217 | Bacteria>Terrabacteria_group>Bacillota>Clostridia>Eubacteriales>Lachnospiraceae | GCF_000238075.1 | Oribacterium_asaccharolyticum | HMPREF9624_00544 | 316049..319702 | 1501332 | BREX-system-P-loop-protein-BrxC-[Oribacterium-asaccharolyticum] |
| WP_002578404.1 | Type1.BREX | HTH-><-SF1-SF2-He[illegible] | 1182 | Bacteria>Terrabacteria_group>Bacillota>Clostridia>Eubacteriales>Lachnospiraceae | GCF_000371725.1 | Enterocloster_bolteae | HMPREF1095_RS19895 | 1377442..1380990 | 208479 | BREX-system-P-loop-protein-BrxC-[Enterocloster-bolteae] |
| WP_016216871.1 | Type1.BREX | <-HTH\|\|\|HTH->\|\|H[illegible] | 1182 | Bacteria>Terrabacteria_group>Bacillota>Clostridia>Eubacteriales>Lachnospiraceae | GCF_000403455.2 | Dorea_sp._5-2 | C817_RS00440 | 102069..105617 | 1235798 | BREX-system-P-loop-protein-BrxC-[Dorea-sp.-5-2] |
| WP_055268215.1 | Type1.BREX | ZnR->HTH_26-> Brx[illegible] | 1182 | Bacteria>Terrabacteria_group>Bacillota>Clostridia>Eubacteriales>Lachnospiraceae | GCF_001406335.1 | Fusicatenibacter_saccharivorans | ARA47_RS16350 | 4112..7660 | 1150298 | BREX-system-P-loop-protein-BrxC-[Fusicatenibacter-saccharivorans] |
| WP_060932093.1 | Type1.BREX | Peptidase_M32->TP[illegible] | 1215 | Bacteria>Terrabacteria_group>Bacillota>Clostridia>Eubacteriales>Lachnospiraceae | GCF_001552975.1 | Lachnoanaerobaculum_saburreum | HMPREF1866_RS12095 | 13224..16871 | 467210 | BREX-system-P-loop-protein-BrxC-[Lachnoanaerobaculum-saburreum] |
| WP_075720910.1 | Type1.BREX | <-LRR_5+LRR_8<-?<[illegible] | 1196 | Bacteria>Terrabacteria_group>Bacillota>Clostridia>Eubacteriales>Lachnospiraceae | GCF_001940225.1 | Roseburia_sp._499 | BIV20_RS10280 | 367523..371113 | 1261634 | BREX-system-P-loop-protein-BrxC-[Roseburia-sp.-499] |
| WP_075672058.1 | Type1.BREX | N-OB+HDRnase->TRA[illegible] | 1182 | Bacteria>Terrabacteria_group>Bacillota>Clostridia>Eubacteriales>Lachnospiraceae | GCF_001940245.1 | Anaerostipes_sp._992a | BHF69_RS12275 | 110494..114042 | 1261637 | BREX-system-P-loop-protein-BrxC-[Anaerostipes-sp.-992a] |
| WP_087163144.1 | Type1.BREX | <-HTH_Tnp_1<-SF1-[illegible] | 1179 | Bacteria>Terrabacteria_group>Bacillota>Clostridia>Eubacteriales>Lachnospiraceae | GCF_002160525.1 | Lachnoclostridium_sp._An169 | B5F07_RS20950 | 16171..19710 | 1965569 | BREX-system-P-loop-protein-BrxC-[Lachnoclostridium-sp.-An169] |
| WP_087273661.1 | Type1.BREX | DNA_METHYLASE->Pa[illegible] | 1212 | Bacteria>Terrabacteria_group>Bacillota>Clostridia>Eubacteriales>Lachnospiraceae | GCF_002161645.1 | Blautia_sp._An46 | B5G00_RS13700 | 83455..87093 | 1965636 | BREX-system-P-loop-protein-BrxC-[Blautia-sp.-An46] |
| WP_118348289.1 | Type1.BREX | <- SIGMA+OB-Fold+[illegible] | 1182 | Bacteria>Terrabacteria_group>Bacillota>Clostridia>Eubacteriales>Lachnospiraceae | GCF_003458885.1 | Coprococcus_sp._AF21-14LB | DWX73_RS07815 | 11153..14701 | 2292231 | BREX-system-P-loop-protein-BrxC-[Coprococcus-sp.-AF21-14LB] |
| WP_118051545.1 | Type1.BREX | <-Phage_int_SAM_5[illegible] | 1183 | Bacteria>Terrabacteria_group>Bacillota>Clostridia>Eubacteriales>Lachnospiraceae | GCF_003464645.1 | Blautia_obeum | DWW07_RS08215 | 33645..37196 | 40520 | BREX-system-P-loop-protein-BrxC-[Blautia-obeum] |
| WP_118012961.1 | Type1.BREX | <-Phage_int_SAM_5[illegible] | 1183 | Bacteria>Terrabacteria_group>Bacillota>Clostridia>Eubacteriales>Lachnospiraceae | GCF_003467455.1 | Blautia_obeum | DW880_RS12300 | 36915..40466 | 40520 | BREX-system-P-loop-protein-BrxC-[Blautia-obeum] |
| WP_118198114.1 | Type1.BREX | <- SIGMA+OB-Fold+[illegible] | 1183 | Bacteria>Terrabacteria_group>Bacillota>Clostridia>Eubacteriales>Lachnospiraceae | GCF_003471125.1 | Blautia_obeum | DW272_RS15845 | 24197..27748 | 40520 | BREX-system-P-loop-protein-BrxC-[Blautia-obeum] |
| WP_118390632.1 | Type1.BREX | wHTH+WYL+WCX ->HT[illegible] | 1196 | Bacteria>Terrabacteria_group>Bacillota>Clostridia>Eubacteriales>Lachnospiraceae | GCF_003474935.1 | Lachnospira_eligens | DWZ79_RS02870 | 260992..264582 | 39485 | BREX-system-P-loop-protein-BrxC-[Lachnospira-eligens] |
| WP_118601699.1 | Type1.BREX | HTH_Tnp_1->?->ZnR[illegible] | 1182 | Bacteria>Terrabacteria_group>Bacillota>Clostridia>Eubacteriales>Lachnospiraceae | GCF_003479155.1 | Blautia_sp._AF19-1 | DWX26_RS14805 | 4595..8143 | 2292960 | BREX-system-P-loop-protein-BrxC-[Blautia-sp.-AF19-1] |
| WP_118742111.1 | Type1.BREX | ZnR->?->HTH_26->[illegible] | 1182 | Bacteria>Terrabacteria_group>Bacillota>Clostridia>Eubacteriales>Lachnospiraceae | GCF_003480245.1 | Blautia_sp._AM42-2 | DW915_10705 | 4262..7810 | 2292976 | BREX-system-P-loop-protein-BrxC-[Blautia-sp.-AM42-2] |
| WP_124952173.1 | Type1.BREX | ABhydrolase->?->?[illegible] | 1215 | Bacteria>Terrabacteria_group>Bacillota>Clostridia>Eubacteriales>Lachnospiraceae | GCF_003862485.1 | Lachnoanaerobaculum_orale | EHW90_RS07145 | 1507422..1511069 | 979627 | BREX-system-P-loop-protein-BrxC-[Lachnoanaerobaculum-orale] |
| WP_168932969.1 | Type1.BREX | <-OLD-ABC+TOPRIM<[illegible] | 1182 | Bacteria>Terrabacteria_group>Bacillota>Clostridia>Eubacteriales>Lachnospiraceae | GCF_012843825.1 | Dorea_formicigenerans | HF855_RS01350 | 86312..89860 | 39486 | BREX-system-P-loop-protein-BrxC-[Dorea-formicigenerans] |
| WP_173900038.1 | Type1.BREX | <- SIGMA+OB-Fold+[illegible] | 1183 | Bacteria>Terrabacteria_group>Bacillota>Clostridia>Eubacteriales>Lachnospiraceae | GCF_013299585.1 | Blautia_obeum | G4467_RS13410 | 30685..34236 | 40520 | BREX-system-P-loop-protein-BrxC-[Blautia-obeum] |
| WP_173865463.1 | Type1.BREX | <-STYKIN<-?<-LRR_[illegible] | 1201 | Bacteria>Terrabacteria_group>Bacillota>Clostridia>Eubacteriales>Lachnospiraceae | GCF_013300965.1 | Faecalicatena_fissicatena | HFN12_RS00390 | 75728..79333 | 290055 | BREX-system-P-loop-protein-BrxC-[Faecalicatena-fissicatena] |
| WP_173813806.1 | Type1.BREX | ZnR->?->HTH_26->[illegible] | 1182 | Bacteria>Terrabacteria_group>Bacillota>Clostridia>Eubacteriales>Lachnospiraceae | GCF_013301105.1 | Fusicatenibacter_saccharivorans | G5B05_RS14750 | 4301..7849 | 1150298 | BREX-system-P-loop-protein-BrxC-[Fusicatenibacter-saccharivorans] |
| WP_173794823.1 | Type1.BREX | CopG/RHH->?->TraJ[illegible] | 1183 | Bacteria>Terrabacteria_group>Bacillota>Clostridia>Eubacteriales>Lachnospiraceae | GCF_013301435.1 | Coprococcus_eutactus | G4458_RS07820 | 91077..94628 | 33043 | BREX-system-P-loop-protein-BrxC-[Coprococcus-eutactus] |
| WP_173770663.1 | Type1.BREX | HTH-><-PHP+SbcC<-[illegible] | 1182 | Bacteria>Terrabacteria_group>Bacillota>Clostridia>Eubacteriales>Lachnospiraceae | GCF_013302375.1 | Blautia_faecis | G5A01_RS01800 | 135200..138748 | 871665 | BREX-system-P-loop-protein-BrxC-[Blautia-faecis] |
| WP_002585535.1 | Type1.BREX | <-ClpABC-AAA<-?<-[illegible] | 1178 | Bacteria>Terrabacteria_group>Bacillota>Clostridia>Eubacteriales>Lachnospiraceae | GCF_013304275.1 | Enterocloster_clostridioformis | HMPREF1088_RS04980 | 1098149..1101685 | 1531 | BREX-system-P-loop-protein-BrxC-[Enterocloster-clostridioformis] |
| WP_181475440.1 | Type1.BREX | REase->?->Phage_i[illegible] | 1182 | Bacteria>Terrabacteria_group>Bacillota>Clostridia>Eubacteriales>Lachnospiraceae | GCF_013694215.1 | Sellimonas_intestinalis | H1A12_RS00250 | 51099..54647 | 1653434 | BREX-system-P-loop-protein-BrxC-[Sellimonas-intestinalis] |
| WP_249326641.1 | Type1.BREX | GNTR-HTH+Aminotra[illegible] | 1215 | Bacteria>Terrabacteria_group>Bacillota>Clostridia>Eubacteriales>Lachnospiraceae | GCF_014337175.1 | Simiaoa_sunii | H9Q77_RS04465 | 956158..959805 | 2763672 | BREX-system-P-loop-protein-BrxC-[Simiaoa-sunii] |
| WP_195441337.1 | Type1.BREX | CopG/RHH->?->TraJ[illegible] | 1182 | Bacteria>Terrabacteria_group>Bacillota>Clostridia>Eubacteriales>Lachnospiraceae | GCF_015554785.1 | Roseburia_intestinalis | I2F09_RS06795 | 71540..75088 | 166486 | BREX-system-P-loop-protein-BrxC-[Roseburia-intestinalis] |
| WP_138345918.1 | Type1.BREX | <-SIGMA-HTH+SIGMA[illegible] | 1182 | Bacteria>Terrabacteria_group>Bacillota>Clostridia>Eubacteriales>Lachnospiraceae | GCF_015669935.1 | Roseburia | FGQ84_RS06860 | 104625..108173 | 841 | MULTISPECIES:-BREX-system-P-loop-protein-BrxC-[Roseburia] |
| WP_215631370.1 | Type1.BREX | GTPase-AIG->?->MA[illegible] | 1201 | Bacteria>Terrabacteria_group>Bacillota>Clostridia>Eubacteriales>Lachnospiraceae | GCF_018785175.1 | Coprococcus_catus | GPK82_RS00920 | 42842..46447 | 116085 | BREX-system-P-loop-protein-BrxC-[Coprococcus-catus] |
| WP_252210442.1 | Type1.BREX | <-TIR<- SIGMA+OB-[illegible] | 1099 | Bacteria>Terrabacteria_group>Bacillota>Clostridia>Eubacteriales>Lachnospiraceae | GCF_018785625.1 | Blautia_faecis | GPK86_RS03045 | 13256..16555 | 871665 | BREX-system-P-loop-protein-BrxC-[Blautia-faecis] |

| Protein-ID | System | Gene-Neighbourhoods | Length | Taxonomy | Assembly-ID | Strain | Locus-Tag | Genomic-coordinates | TaxID | Fasta-defline |
|---|---|---|---|---|---|---|---|---|---|---|
| -- | | | | | | | | | | |
| --OLD-ABC->TOPRIM | | | | | | | | | | |
| --(PARIS-like) | | | | | | | | | | |
| -- | | | | | | | | | | |
| WP_015789649.1 | Type1.BREX | <-REase<-?<-?\|\|DOC+zhth-><-BirA-HTH<-?\|\|?->BrxA-DNA-binding->DUF6079*->BrxB-BrxF-iSTAND->DUF6079-><-?<-?\|\|?-><-OLD-ABC+TOPRIM\|\|N6-METHYLASE->TBP+TBP->PglZ-> | 1236 | Archaea>Euryarchaeota>Stenosarchaea_group>Halobacteria>Halobacteriales>Haloarculaceae | GCF_000023945.1 | Halorhabdus_utahensis | HUTA_RS09330 | 1919731..1923441 | 146826 | hypothetical-protein-[Halorhabdus-utahensis] |
| WP_015789651.1 | Type1.BREX | DOC+zhth-><-BirA-HTH<-?\|\|?->BrxA-DNA-binding->DUF6079->BrxB-BrxF-iSTAND->DUF6079*-><-?<-?\|\|?-><-OLD-ABC+TOPRIM\|\|N6-METHYLASE->TBP+TBP->PglZ->HEPN->TUDOR+SF1-SF2-Helicase+iREase+CTD-> | 1280 | Archaea>Euryarchaeota>Stenosarchaea_group>Halobacteria>Halobacteriales>Haloarculaceae | GCF_000023945.1 | Halorhabdus_utahensis | HUTA_RS09340 | 1924016..1927858 | 146826 | hypothetical-protein-[Halorhabdus-utahensis] |
| WP_049892750.1 | Type1.BREX | <-N6-METHYLASE\|\|OLD-ABC+TOPRIM-><-DDE_Tnp_1\|\|?-><-?<-NTP_transf+wHTH<-NTP_transf+wHTH<-CopG/RHH<-HTH+ICLR-HTH<-DUF6079*<-BrxB-BrxF-iSTAND<-DUF6079<-BrxA-DNA-binding\|\|DDE_Tnp_1->\|\|DDE_Tnp_1->BirA-HTH->PIN->Phage_int_SAM_5->CopG/RHH-> | 1280 | Archaea>Euryarchaeota>Stenosarchaea_group>Halobacteria>Halobacteriales>Halobacteriaceae | GCF_000069025.1 | Halobacterium_salinarum | OE_RS13050 | 230465..234307 | 2242 | hypothetical-protein-[Halobacterium-salinarum] |
| WP_049892751.1 | Type1.BREX | <-DDE_Tnp_1\|\|?-><-?<-NTP_transf+wHTH<-NTP_transf+wHTH<-CopG/RHH<-HTH+ICLR-HTH<-DUF6079<-BrxB-BrxF-iSTAND<-DUF6079*<-BrxA-DNA-binding\|\|DDE_Tnp_1->\|\|DDE_Tnp_1->BirA-HTH->PIN->Phage_int_SAM_5->CopG/RHH-><-?<-Classical-AAA | 1227 | Archaea>Euryarchaeota>Stenosarchaea_group>Halobacteria>Halobacteriales>Halobacteriaceae | GCF_000069025.1 | Halobacterium_salinarum | OE_RS13060 | 234882..238565 | 2242 | hypothetical-protein-[Halobacterium-salinarum] |
| WP_247995979.1 | Type5.BREX | <-ARG<-wHTH+REase<-McrB-AAA<-SF1-SF2-Helicase+REase<-SF1-SF2-Helicase+REase\|\|BrxA-DNA-binding->DUF6079*->BrxB-BrxF-iSTAND->DUF6079->REase->N6-METHYLASE-><-OLD-ABC+TOPRIM\|\|PglZ->TUDOR+SF1-SF2-Helicase+iREase+CTD->?->BrxD-AAA-> | 1225 | Archaea>Euryarchaeota>Stenosarchaea_group>Halobacteria>Halobacteriales>Halobacteriaceae | GCF_023115355.1 | Halocatena_salina | MW046_RS18845 | 37719..41396 | 2934340 | hypothetical-protein-[Halocatena-salina] |
| WP_247995981.1 | Type5.BREX | <-ARG<-wHTH+REase<-McrB-AAA<-SF1-SF2-Helicase+REase<-SF1-SF2-Helicase+REase\|\|BrxA-DNA-binding->DUF6079->BrxB-BrxF-iSTAND->DUF6079*->REase->N6-METHYLASE-><-OLD-ABC+TOPRIM\|\|PglZ->TUDOR+SF1-SF2-Helicase+iREase+CTD->?->BrxD-AAA->SF1-SF2-Helicase+wHTH+Small-B-barrel+DUF1998-><-CARF+HTH | 1278 | Archaea>Euryarchaeota>Stenosarchaea_group>Halobacteria>Halobacteriales>Halobacteriaceae | GCF_023115355.1 | Halocatena_salina | MW046_RS18855 | 41971..45807 | 2934340 | hypothetical-protein-[Halocatena-salina] |
| WP_103992123.1 | Type5.BREX | <-Calcineurin<-HKD-DNase\|\|CARF+HTH->?-><-?<-?<-?\|\|ParB+HNH->BrxA-DNA-binding->DUF6079*->BrxB-BrxF-iSTAND->DUF6079-><-OLD-ABC+TOPRIM\|\|N6-METHYLASE-><-REase\|\|PglZ->REase+METHYLASE->TUDOR+SF1-SF2-Helicase+iREase+CTD-> | 1227 | Archaea>Euryarchaeota>Stenosarchaea_group>Halobacteria>Haloferacales>Haloferacaceae | GCF_004799685.1 | Halobellus_limi | DV707_RS08430 | 1701258..1704941 | 699433 | hypothetical-protein-[Halobellus-limi] |
| WP_103992125.1 | Type5.BREX | CARF+HTH->?-><-?<-?<-?\|\|ParB+HNH->BrxA-DNA-binding->DUF6079->BrxB-BrxF-iSTAND->DUF6079*-><-OLD-ABC+TOPRIM\|\|N6-METHYLASE-><-REase\|\|PglZ->REase+METHYLASE->TUDOR+SF1-SF2-Helicase+iREase+CTD-> | 1284 | Archaea>Euryarchaeota>Stenosarchaea_group>Halobacteria>Haloferacales>Haloferacaceae | GCF_004799685.1 | Halobellus_limi | DV707_RS08440 | 1705516..1709370 | 699433 | hypothetical-protein-[Halobellus-limi] |
| ELZ39021.1 | Type1.BREX | OrfB_IS605+OrfB_Zn_ribbon->DDE_Tnp_1->?-><-ORC-CDC6\|\|BrxA-DNA-binding->DUF6079->BrxB-BrxF-iSTAND->DUF6079*->REase-><-viral-protein\|\|N6-METHYLASE-><-OLD-ABC+TOPRIM<-ZnR+REase | 1280 | Archaea>Euryarchaeota>Stenosarchaea_group>Halobacteria>Haloferacales>Halorubraceae | GCA_000336875.1 | Halorubrum_californiense_DSM_19288 | C463_17695 | 23296..27138 | 1227465 | hypothetical-protein-C463_17695-[Halorubrum-californiense-DSM-19288] |
| SFP79187.1 | Type1.BREX | <-PglZ\|\|ZnR+REase->OLD-ABC+TOPRIM-><-N6-METHYLASE\|\|viral-protein-><-REase\|\|?-><-DUF6079<-BrxB-BrxF-iSTAND<-DUF6079*<-BrxA-DNA-binding\|\|?->ORC-CDC6-><-?\|\|?->?->?->ORC-CDC6+HTH-> | 1232 | Archaea>Euryarchaeota>Stenosarchaea_group>Halobacteria>Haloferacales>Halorubraceae | GCA_900115675.1 | Halolamina_pelagica | SAMN05216277_10870 | 87547..91245 | 699431 | hypothetical-protein-SAMN05216277_10870-[Halolamina-pelagica] |
| WP_008446367.1 | Type1.BREX | OrfB_IS605+OrfB_Zn_ribbon->\|\|DDE_Tnp_1->?-><-ORC-CDC6\|\|BrxA-DNA-binding->DUF6079*->BrxB-BrxF-iSTAND->DUF6079->REase-><-viral-protein\|\|N6-METHYLASE-><-OLD-ABC+TOPRIM<-ZnR+REase | 1227 | Archaea>Euryarchaeota>Stenosarchaea_group>Halobacteria>Haloferacales>Halorubraceae | GCF_000336875.1 | Halorubrum_californiense | C463_RS17190 | 19038..22721 | 416585 | hypothetical-protein-[Halorubrum-californiense] |
| WP_239638934.1 | Type1.BREX | OrfB_IS605+OrfB_Zn_ribbon->\|\|DDE_Tnp_1->?-><-ORC-CDC6\|\|BrxA-DNA-binding->DUF6079->BrxB-BrxF-iSTAND->DUF6079*->REase-><-viral-protein\|\|N6-METHYLASE-><-OLD-ABC+TOPRIM<-ZnR+REase | 1235 | Archaea>Euryarchaeota>Stenosarchaea_group>Halobacteria>Haloferacales>Halorubraceae | GCF_000336875.1 | Halorubrum_californiense | C463_RS17200 | 23431..27138 | 416585 | hypothetical-protein-[Halorubrum-californiense] |
| WP_157746008.1 | Type5.BREX | DUF6079*->?->N6-METHYLASE-><-OLD-ABC+TOPRIM<-ZnR+REase\|\|PglZ->REase+METHYLASE->TUDOR+SF1-SF2-Helicase+iREase+CTD-> | 1280 | Archaea>Euryarchaeota>Stenosarchaea_group>Halobacteria>Haloferacales>Halorubraceae | GCF_002355655.1 | Halorubrum_trapanicum | CPZ01_RS15580 | 1415231..3842 | 29284 | hypothetical-protein-[Halorubrum-trapanicum] |
| WP_074878712.1 | Type5.BREX | <-TUDOR+SF1-SF2-Helicase+iREase+CTD<-REase+METHYLASE<-PglZ\|\|ZnR+REase->OLD-ABC+TOPRIM-><-N6-METHYLASE\|\|viral-protein-><-REase\|\|?-><-DUF6079*<-BrxB-BrxF-iSTAND<-DUF6079<-BrxA-DNA-binding\|\|ORC-CDC6-> | 1279 | Archaea>Euryarchaeota>Stenosarchaea_group>Halobacteria>Haloferacales>Halorubraceae | GCF_900115675.1 | Halolamina | BMZ83_RS11295 | 83133..86972 | 1075397 | MULTISPECIES:-hypothetical-protein-[Halolamina] |
| WP_074878715.1 | Type5.BREX | <-PglZ\|\|ZnR+REase->OLD-ABC+TOPRIM-><-N6-METHYLASE\|\|viral-protein-><-REase\|\|?-><-DUF6079<-BrxB-BrxF-iSTAND<-DUF6079*<-BrxA-DNA-binding\|\|ORC-CDC6-><-?\|\|?->?->?->ORC-CDC6+HTH-> | 1227 | Archaea>Euryarchaeota>Stenosarchaea_group>Halobacteria>Haloferacales>Halorubraceae | GCF_900115675.1 | Halolamina | BMZ83_RS11305 | 87547..91230 | 1075397 | MULTISPECIES:-hypothetical-protein-[Halolamina] |
| WP_049967095.1 | Type5.BREX | HTH_20->?-><-35exo<-?<-35exo\|\|?-><-ORC-CDC6\|\|BrxA-DNA-binding->BrxB-BrxF-iSTAND->DUF6079*-><-?\|\|?-><-OLD-ABC+TOPRIM<-ZnR+REase\|\|PglZ->TUDOR+SF1-SF2-Helicase+iREase+CTD->SF1-SF2-Helicase+REase-><-DDE_Tnp_IS240 | 1277 | Archaea>Euryarchaeota>Stenosarchaea_group>Halobacteria>Natrialbales>Natrialbaceae | GCF_000710605.1 | Natrinema_thermotolerans | DVR14_24745 | 889875..893708 | 121872 | hypothetical-protein-[Natrinema-thermotolerans] |
| WP_049967097.1 | Type5.BREX | <-TUDOR+SF1-SF2-Helicase+iREase+CTD<-REase+METHYLASE<-PglZ\|\|ZnR+REase->OLD-ABC+TOPRIM-><-N6-METHYLASE\|\|?-><-DUF6079<-BrxB-BrxF-iSTAND<-DUF6079*<-BrxA-DNA-binding\|\|ORC-CDC6-><-?\|\|?->?->?->35exo-><-?<-HTH_20 | 1227 | Archaea>Euryarchaeota>Stenosarchaea_group>Halobacteria>Natrialbales>Natrialbaceae | GCF_000710605.1 | Natrinema_thermotolerans | HL44_RS19275 | 148133..151816 | 121872 | hypothetical-protein-[Natrinema-thermotolerans] |
| WP_253488285.1 | Type1.BREX | <-SIGMA+OB-Fold+AAA-ATPase+Lon-Protease-BrxL<-PglZ<-TOPRIM<-OLD-ABC<-N6-METHYLASE<-DUF6079*<-BrxB-BrxF-iSTAND<-BrxA-DNA-binding<-35exo | 1155 | Archaea>Euryarchaeota>Stenosarchaea_group>Methanomicrobia>Methanomicrobiales>Methanocalculaceae | GCF_024170505.1 | Methanocalculus_alkaliphilus | J2T58_RS06410 | 9528..12995 | 768730 | BREX-system-P-loop-protein-BrxC-[Methanocalculus-alkaliphilus] |
| ABD40832.1 | Type1.BREX | <-N6-METHYLASE<-?<-SIGMA+OB-Fold+AAA-ATPase+Lon-Protease-BrxL<-PglZ<-TOPRIM<-OLD-ABC<-N6-METHYLASE<-RHH<-DUF6079*<-BrxB-BrxF-iSTAND<-BrxA-DNA-binding<-?<-?<-Glyco_hydro | 1166 | Archaea>Euryarchaeota>Stenosarchaea_group>Methanomicrobia>Methanomicrobiales>Methanospirillaceae | GCA_000013445.1 | Methanospirillum_hungatei_JF-1 | Mhun_1083 | 1223800..1227300 | 323259 | hypothetical-protein-Mhun_1083-[Methanospirillum-hungatei-JF-1] |
| WP_052288823.1 | Type1.BREX | <-SIGMA+OB-Fold+AAA-ATPase+Lon-Protease-BrxL<-PglZ<-TOPRIM<-OLD-ABC<-N6-METHYLASE<-?<-RHH<-DUF6079*<-BrxB-BrxF-iSTAND<-BrxA-DNA-binding<-?<-?<-?<-Glyco_hydro | 1159 | Archaea>Euryarchaeota>Stenosarchaea_group>Methanomicrobia>Methanomicrobiales>Methanospirillaceae | GCF_000013445.1 | Methanospirillum_hungatei | MHUN_RS05665 | 1223800..1227279 | 2203 | BREX-system-P-loop-protein-BrxC-[Methanospirillum-hungatei] |
| WP_011448873.1 | Type1.BREX | CR-ATPase8+REase-><-TPR+TPR<-TPR<-TOPRIM<-OLD-ABC<-?\|\|BrxB-BrxF-iSTAND->DUF6079*->N6-METHYLASE->?->?->?->HATPase+wHTH->HTH+REase->N6-METHYLASE-p->?<-DDE_Tnp\|\|N6-METHYLASE-p-> | 1178 | Archaea>Euryarchaeota>Stenosarchaea_group>Methanomicrobia>Methanomicrobiales>Methanospirillaceae | GCF_000013445.1 | Methanospirillum_hungatei | MHUN_RS09850 | 2137740..2141276 | 2203 | BREX-system-P-loop-protein-BrxC-[Methanospirillum-hungatei] |
| KXS44097.1 | Type1.BREX | <-SIGMA+OB-Fold+AAA-ATPase+Lon-Protease-BrxL<-PglZ<-TOPRIM<-OLD-ABC<-?<-N6-METHYLASE-p<-Schlafen+HATPase+wHTH<-N6-METHYLASE<-DUF6079*<-BrxB-BrxF-iSTAND<-wHTH+REase<-EVE+McrB-AAA<-?\|\|REase->MNS+REase->RADICAL-SAM-> | 1185 | Archaea>Euryarchaeota>Stenosarchaea_group>Methanomicrobia>Methanosarcinales>Methanosarcinaceae | GCA_001577565.1 | Methanolobus_sp._T82-4 | AWU59_730 | 16022..19579 | 1794908 | MAG:-hypothetical-protein-AWU59_730-[Methanolobus-sp.-T82-4] |
| NPE28183.1 | Type1.BREX | <-SIGMA+OB-Fold+AAA-ATPase+Lon-Protease-BrxL<-PglZ<-TOPRIM<-OLD-ABC<-TIR<-N6-METHYLASE-p<-Schlafen+HATPase+wHTH<-N6-METHYLASE<-DUF6079*<-BrxB-BrxF-iSTAND<-URI-DNase<-?<-SF1-SF2-Helicase+REase\|\|<-?\|\|?->HNH-> | 1185 | Archaea>Euryarchaeota>Stenosarchaea_group>Methanomicrobia>Methanosarcinales>Methanosarcinaceae | GCA_013166765.1 | Methanococcoides_sp._SA1 | HNV12_09470 | 12693..16250 | 2735871 | BREX-system-P-loop-protein-BrxC-[Methanococcoides-sp.-SA1] |
| HII01702.1 | Type1.BREX | <-SF1-SF2-Helicase<-?<-OLD-ABC+TOPRIM<-?<-N6-METHYLASE<-DUF6079*<-BrxB-BrxF-iSTAND\|\|?->SAM-METHYLASE->?->MNS-><-?\|\|HAD+HAD-><-TPR+TPR+TPR+TPR<-REC+OmpR-HTH | 1196 | Archaea>Euryarchaeota>Stenosarchaea_group>Methanomicrobia>Methanosarcinales>Methanosarcinaceae | GCA_013329415.1 | Methanosarcinaceae_archaeon | HA351_08670 | 9336..12926 | 1945595 | BREX-system-P-loop-protein-BrxC-[Methanosarcinaceae-archaeon] |
| WP_270096529.1 | Type1.BREX | SF1-SF2-Helicase+REase->HNH-><-SIGMA+OB-Fold+AAA-ATPase+Lon-Protease-BrxL<-PglZ<-TOPRIM<-OLD-ABC<-HD<-N6-METHYLASE-p<-N6-METHYLASE-p<-DUF6079*<-BrxB-BrxF-iSTAND<-REase<-?<-?<-RNase\|\|Classical-AAA->SF1-SF2-Helicase-> | 1190 | Archaea>Euryarchaeota>Stenosarchaea_group>Methanomicrobia>Methanosarcinales>Methanosarcinaceae | GCF_027594145.1 | Methanococcoides_alaskense | PBV53_RS07490 | 737341..740913 | 325778 | BREX-system-P-loop-protein-BrxC-[Methanococcoides-alaskense] |
| WP_091932407.1 | Type1.BREX | SF1-SF2-Helicase+REase->?-><-SIGMA+OB-Fold+AAA-ATPase+Lon-Protease-BrxL<-PglZ<-TOPRIM<-OLD-ABC<-N6-METHYLASE-p<-Schlafen+HATPase+wHTH<-N6-METHYLASE<-DUF6079*<-BrxB-BrxF-iSTAND\|\|wHTH+REase->McrB-AAA->SF1-SF2-Helicase-><-DDE_Tnp_1+DDE_Tnp_1_2 | 1184 | Archaea>Euryarchaeota>Stenosarchaea_group>Methanomicrobia>Methanosarcinales>Methanosarcinaceae | GCF_900114835.1 | Methanolobus_profundi | BM008_RS01885 | 368008..371562 | 487685 | BREX-system-P-loop-protein-BrxC-[Methanolobus-profundi] |
| WP_172363658.1 | Type1.BREX | <-SIGMA+OB-Fold+AAA-ATPase+Lon-Protease-BrxL<-PglZ<-TOPRIM<-OLD-ABC<-TIR<-N6-METHYLASE-p<-Schlafen+HATPase+wHTH<-N6-METHYLASE<-DUF6079*<-BrxB-BrxF-iSTAND<-URI-DNase<-?<-SF1-SF2-Helicase+REase\|\|<-?\|\|?->HNH-> | 1185 | Archaea>Euryarchaeota>Stenosarchaea_group>Methanomicrobia>Methanosarcinales>Methanosarcinaceae | - | Methanococcoides_sp._SA1 | HNV12_09470 | 12693..16250 | 2735871 | BREX-system-P-loop-protein-BrxC-[Methanococcoides-sp.-SA1] |
| MBK9963544.1 | Type6.BREX | HTH+MPTase-><-SF1-SF2-Helicase+wHTH+Small-B-barrel+DUF1998<-ParB+HNH<-PglZ<-N6-METHYLASE<-OLD-ABC+TOPRIM<-ASCH<-PRTase<-?<-DUF6079*<-BrxB-BrxF-iSTAND<-BrxA-DNA-binding<-BrxE\|\|?-><-NUDIX<-SAM-METHYLASE | 1207 | Bacteria>Acidobacteriota>Holophagae>Holophagales>unclassified_Holophagales | GCA_016722225.1 | Holophagales_bacterium | IPP07_01030 | 234150..237773 | 2478486 | BREX-system-P-loop-protein-BrxC-[Holophagales-bacterium] |
| MCG3192298.1 | Type6.BREX | <-DUF932<-ParB+HNH<-?<-SIGMA+OB-Fold+AAA-ATPase+Lon-Protease-BrxL<-PglZ<-TOPRIM<-OLD-ABC<-N6-METHYLASE<-DUF6079*<-BrxB-BrxF-iSTAND<-BrxA-DNA-binding<-BrxE<-?<-?<-?<-HTH+LexA-protease | 1183 | Bacteria>Acidobacteriota>Thermoanaerobaculia>unclassified_Thermoanaerobaculia | GCA_022071945.1 | Thermoanaerobaculia_bacterium | DIJKHBIC_01536 | 22288..25839 | 2651171 | hypothetical-protein-[Thermoanaerobaculia-bacterium] |
| MYN66269.1 | Type1.BREX | BrxA-DNA-binding->BrxB-BrxF-iSTAND->DUF6079*->OLD-ABC->TOPRIM->Schlafen+HATPase+wHTH->N6-METHYLASE->?->?->SbcC->PHP+SbcC->Schlafen+HATPase+wHTH-> | 1283 | Bacteria>Acidobacteriota>unclassified_Acidobacteriota | GCA_009861545.1 | Acidobacteriota_bacterium | F4X11_14760 | 2943..6794 | 1978231 | BREX-system-P-loop-protein-BrxC-[Acidobacteriota-bacterium] |
| MCC6130024.1 | Type6.BREX | HTH+LexA-protease->?->?->?->BrxE->BrxA-DNA-binding->BrxB-BrxF-iSTAND->DUF6079*->N6-METHYLASE->N6-METHYLASE-p->OLD-ABC->TOPRIM->PglZ->SIGMA+OB-Fold+AAA-ATPase+Lon-Protease-BrxL->?->REase->REase-> | 1183 | Bacteria>Acidobacteriota>unclassified_Acidobacteriota | GCA_020845555.1 | Acidobacteriota_bacterium | IT186_08845 | 15940..19491 | 1978231 | BREX-system-P-loop-protein-BrxC-[Acidobacteriota-bacterium] |
| MBF0500206.1 | Type1.BREX | IG+IG-><-ANK<-?<-ANK+ANK+ANK<-?\|\|EVE->BrxA-DNA-binding->BrxB-BrxF-iSTAND->DUF6079*->N6-METHYLASE->OLD-ABC->TOPRIM->N6-METHYLASE->PglZ->SIGMA+OB-Fold+AAA-ATPase+Lon-Protease-BrxL->DDE_Tnp_IS66-><-SbcC | 1142 | Bacteria>Bacteria_incertae_sedis>Bacteria_candidate_phyla>Candidatus_Riflebacteria>unclassified_Candidatus_Riflebacteria | GCA_015233155.1 | Candidatus_Riflebacteria_bacterium | HQM09_08735 | 64284..67712 | 2053586 | BREX-system-P-loop-protein-BrxC-[Candidatus-Riflebacteria-bacterium] |
| EFK06105.1 | Type1.BREX | <-Acyltransferase<-?<-?<-SIGMA+OB-Fold+AAA-ATPase+Lon-Protease-BrxL<-PglZ<-TOPRIM<-OLD-ABC<-N6-METHYLASE<-HEPN<-DUF6079*<-BrxB-BrxF-iSTAND-p\|\|?-><-BrxA-DNA-binding\|\|?->?->?-><-DnaJ | 1215 | Bacteria>delta/epsilon_subdivisions>Deltaproteobacteria>unclassified_Deltaproteobacteria | GCA_000179315.1 | delta_proteobacterium_NaphS2 | NPH_2628 | 46790..50437 | 88274 | conserved-hypothetical-protein-[delta-proteobacterium-NaphS2] |
| PJB45699.1 | Type6.BREX | <-SF1-SF2-Helicase+wHTH+Small-B-barrel+DUF1998-p<-PglZ<-N6-METHYLASE-p<-N6-METHYLASE<-TOPRIM<-OLD-ABC<-DUF6079*<-BrxB-BrxF-iSTAND<-BrxA-DNA-binding<-BrxE\|\|DSBH-> | 1207 | Bacteria>delta/epsilon_subdivisions>Deltaproteobacteria>unclassified_Deltaproteobacteria | GCA_002789955.1 | Deltaproteobacteria_bacterium_CG_4_9_14_3_um_filter_63_12 | CO108_06840 | 12981..16604 | 1973967 | BREX-system-P-loop-protein-BrxC-[Deltaproteobacteria-bacterium-CG_4_9_14_3_um_filter_63_12] |
| PXF57197.1 | Type1.BREX | <-PemK_toxin<-MetJarc-HTH<-PglZ<-TOPRIM<-OLD-ABC<-N6-METHYLASE<-PIN<-Phd-YefM<-DUF6079*<-BrxB-BrxF-iSTAND<-BrxA-DNA-binding<-PIN<-?\|\|?->?->?->His_biosynth-> | 1215 | Bacteria>delta/epsilon_subdivisions>Deltaproteobacteria>unclassified_Deltaproteobacteria | GCA_003194495.1 | Deltaproteobacteria_bacterium | C4B58_10725 | 19842..23489 | 2026735 | BREX-system-P-loop-protein-BrxC-[Deltaproteobacteria-bacterium] |
| MBI4818899.1 | Type6.BREX | <-PglZ-p<-TOPRIM<-OLD-ABC<-Schlafen+HATPase+wHTH<-?<-Schlafen+HATPase+wHTH<-N6-METHYLASE<-PIN<-?<-DUF6079*<-BrxB-BrxF-iSTAND<-BrxA-DNA-binding<-BrxE<-SF1-SF2-Helicase+wHTH+Small-B-barrel+DUF1998<-BrxD-AAA<-PglZ | 1207 | Bacteria>delta/epsilon_subdivisions>Deltaproteobacteria>unclassified_Deltaproteobacteria | GCA_016208175.1 | Deltaproteobacteria_bacterium | HY791_21690 | 12157..15780 | 2026735 | BREX-system-P-loop-protein-BrxC-[Deltaproteobacteria-bacterium] |
| MBI4706185.1 | Type6.BREX | XIS-HTH->BrxE->BrxA-DNA-binding->BrxB-BrxF-iSTAND->DUF6079*->N6-METHYLASE-p->OLD-ABC->TOPRIM->PglZ->SIGMA+OB-Fold+AAA-ATPase+Lon-Protease-BrxL->MORC+DUF3684+DUF3883-><-?<-REase+SF1-SF2-Helicase<-Schlafen+HATPase+wHTH | 1236 | Bacteria>delta/epsilon_subdivisions>Deltaproteobacteria>unclassified_Deltaproteobacteria | GCA_016209145.1 | Deltaproteobacteria_bacterium | HY744_34245 | 2867..6577 | 2026735 | BREX-system-P-loop-protein-BrxC-[Deltaproteobacteria-bacterium] |
| MBN1905588.1 | Type1.BREX | PAS+NtrC-AAA+FIS-HTH-><-SIGMA+OB-Fold+AAA-ATPase+Lon-Protease-BrxL<-PglZ<-N6-METHYLASE<-N6-METHYLASE-p<-TOPRIM<-OLD-ABC<-DUF6079*<-BrxA-DNA-binding | 1190 | Bacteria>delta/epsilon_subdivisions>Deltaproteobacteria>unclassified_Deltaproteobacteria | GCA_016931115.1 | Deltaproteobacteria_bacterium | JW927_10865 | 38502..42074 | 2026735 | BREX-system-P-loop-protein-BrxC-[Deltaproteobacteria-bacterium] |
| MBT3255199.1 | Type1.BREX | <-wHTH+WYL+WCX<-?\|\|?->DUF932->BrxA-DNA-binding->BrxB-BrxF-iSTAND->DUF6079*->N6-METHYLASE->OLD-ABC->TOPRIM->PglZ->SIGMA+OB-Fold+AAA-ATPase+Lon-Protease-BrxL->?->?-><-SAVED<-JAB | 1216 | Bacteria>delta/epsilon_subdivisions>Deltaproteobacteria>unclassified_Deltaproteobacteria | GCA_018644785.1 | Deltaproteobacteria_bacterium | HN366_01900 | 5574..9224 | 2026735 | BREX-system-P-loop-protein-BrxC-[Deltaproteobacteria-bacterium] |
| MCC6811534.1 | Type6.BREX | STYKinase-like->?->HTH_32+DDE_3->HTH_32+DDE_3->Four-helical-protein->SF1-SF2-Helicase->BrxE->BrxA-DNA-binding->BrxB-BrxF-iSTAND->DUF6079*->OLD-ABC->TOPRIM->N6-METHYLASE->OLD-ABC->TOPRIM->PglZ->BrxD-AAA->SF1-SF2-Helicase+wHTH+Small-B-barrel+DUF1998->Peptidase_C26-> | 1210 | Bacteria>delta/epsilon_subdivisions>Deltaproteobacteria>unclassified_Deltaproteobacteria | GCA_020848545.1 | Deltaproteobacteria_bacterium | IT381_29155 | 76632..80264 | 2026735 | BREX-system-P-loop-protein-BrxC-[Deltaproteobacteria-bacterium] |
| WP_259165690.1 | Type5.BREX | <-SbcC<-Calcineurin-SbcD<-OLD-ABC+TOPRIM<-TUDOR+SF1-SF2-Helicase+iREase+CTD<-PglZ<-N6-METHYLASE<-DUF6079<-BrxB-BrxF-iSTAND<-DUF6079*<-BrxA-DNA-binding<-REase+RNase<-viral-protein<-?<-ZnR | 1245 | Bacteria>FCB_group>Bacteroidetes/Chlorobi_group>Bacteroidota>Bacteroidetes_incertae_sedis>Rhodothermaceae | GCF_024806515.1 | Salinibacter_ruber | GGP90_RS03930 | 969591..973328 | 146919 | hypothetical-protein-[Salinibacter-ruber] |
| WP_259253366.1 | Type5.BREX | <-HD<-SbcC<-Calcineurin-SbcD<-OLD-ABC+TOPRIM<-TUDOR+SF1-SF2-Helicase+iREase+CTD<-PglZ<-N6-METHYLASE<-DUF6079<-BrxB-BrxF-iSTAND<-DUF6079*<-BrxA-DNA-binding<-REase+RNase<-?<-viral-protein<-?<-?<-?<-Phage_holin_2_4 | 1245 | Bacteria>FCB_group>Bacteroidetes/Chlorobi_group>Bacteroidota>Bacteroidetes_incertae_sedis>Rhodothermaceae | GCF_024807415.1 | Salinibacter_ruber | GGP63_RS14335 | 56454..60191 | 146919 | hypothetical-protein-[Salinibacter-ruber] |
| WP_259129819.1 | Type5.BREX | TraJ-RHH->RHH->Phage_holin_2_4->?->REase+RNase->BrxA-DNA-binding->DUF6079*->BrxB-BrxF-iSTAND->DUF6079->N6-METHYLASE->PglZ->TUDOR+SF1-SF2-Helicase+iREase+CTD->OLD-ABC+TOPRIM->Calcineurin-SbcD->SbcC-> | 1245 | Bacteria>FCB_group>Bacteroidetes/Chlorobi_group>Bacteroidota>Bacteroidetes_incertae_sedis>Rhodothermaceae | GCF_024807455.1 | Salinibacter_ruber | GGQ04_RS13395 | 34758..38495 | 146919 | hypothetical-protein-[Salinibacter-ruber] |
| WP_259127211.1 | Type5.BREX | <-HD<-SbcC<-Calcineurin-SbcD<-OLD-ABC+TOPRIM<-TUDOR+SF1-SF2-Helicase+iREase+CTD<-PglZ<-N6-METHYLASE<-DUF6079<-BrxB-BrxF-iSTAND<-DUF6079*<-BrxA-DNA-binding<-REase+RNase<-viral-protein<-?<-DDE_Tnp_1\|\|?-><-?<-35exo<-viral-protein | 1255 | Bacteria>FCB_group>Bacteroidetes/Chlorobi_group>Bacteroidota>Bacteroidetes_incertae_sedis>Rhodothermaceae | GCF_024807525.1 | Salinibacter_ruber | GGP44_RS04645 | 1121352..1125119 | 146919 | hypothetical-protein-[Salinibacter-ruber] |
| WP_259086338.1 | Type5.BREX | REase+RNase->BrxA-DNA-binding->DUF6079*->BrxB-BrxF-iSTAND->DUF6079->N6-METHYLASE->PglZ->?->TUDOR+SF1-SF2-Helicase+iREase+CTD->OLD-ABC+TOPRIM->Calcineurin-SbcD->SbcC-> | 1245 | Bacteria>FCB_group>Bacteroidetes/Chlorobi_group>Bacteroidota>Bacteroidetes_incertae_sedis>Rhodothermaceae | GCF_024808115.1 | Salinibacter_ruber | GGP80_RS14460 | 1984..5721 | 146919 | hypothetical-protein-[Salinibacter-ruber] |
| WP_259107773.1 | Type5.BREX | <-DDE_Tnp_1\|\|?->viral-protein-><-?<-HORMA<-URI-DNase\|\|?->REase+RNase->BrxA-DNA-binding->DUF6079*->BrxB-BrxF-iSTAND->DUF6079->N6-METHYLASE->PglZ->TUDOR+SF1-SF2-Helicase+iREase+CTD->OLD-ABC+TOPRIM->Calcineurin-SbcD->SbcC->HD-> | 1245 | Bacteria>FCB_group>Bacteroidetes/Chlorobi_group>Bacteroidota>Bacteroidetes_incertae_sedis>Rhodothermaceae | GCF_024808295.1 | Salinibacter_ruber | GGP92_RS13390 | 54298..58035 | 146919 | hypothetical-protein-[Salinibacter-ruber] |
| WP_259048220.1 | Type5.BREX | viral-protein->REase+RNase->BrxA-DNA-binding->DUF6079*->BrxB-BrxF-iSTAND->DUF6079->N6-METHYLASE->PglZ->TUDOR+SF1-SF2-Helicase+iREase+CTD->OLD-ABC+TOPRIM->Calcineurin-SbcD->SbcC->HD-> | 1245 | Bacteria>FCB_group>Bacteroidetes/Chlorobi_group>Bacteroidota>Bacteroidetes_incertae_sedis>Rhodothermaceae | GCF_024808395.1 | Salinibacter_ruber | GGP52_RS13845 | 3637..7374 | 146919 | hypothetical-protein-[Salinibacter-ruber] |
| WP_259099618.1 | Type1.BREX | <-HD<-SbcC<-Calcineurin-SbcD<-SIGMA+OB-Fold+AAA-ATPase+Lon-Protease-BrxL<-PglZ<-OLD-ABC+TOPRIM<-N6-METHYLASE<-DOC+HTH<-DUF6079*<-BrxB-BrxF-iSTAND<-BrxA-DNA-binding\|\|REase-><-?<-viral-protein | 1143 | Bacteria>FCB_group>Bacteroidetes/Chlorobi_group>Bacteroidota>Bacteroidetes_incertae_sedis>Rhodothermaceae | GCF_024808405.1 | Salinibacter_ruber | GGP77_002965 | 30959..34390 | 146919 | BREX-system-P-loop-protein-BrxC-[Salinibacter-ruber] |
| WP_259120983.1 | Type5.BREX | <-HTH_3+Peptidase_S24\|\|TraJ-RHH->RHH->?->viral-protein->REase+RNase->BrxA-DNA-binding->DUF6079*->BrxB-BrxF-iSTAND->DUF6079->N6-METHYLASE->PglZ->TUDOR+SF1-SF2-Helicase+iREase+CTD->OLD-ABC+TOPRIM->Calcineurin-SbcD->SbcC->HD-> | 1245 | Bacteria>FCB_group>Bacteroidetes/Chlorobi_group>Bacteroidota>Bacteroidetes_incertae_sedis>Rhodothermaceae | GCF_024808535.1 | Salinibacter_ruber | GGP96_RS14190 | 56803..60540 | 146919 | hypothetical-protein-[Salinibacter-ruber] |
| WP_259122105.1 | Type5.BREX | TraJ-RHH->RHH->Phage_holin_2_4-><-?<-CHTH\|\|?->viral-protein->REase+RNase->BrxA-DNA-binding->DUF6079*->BrxB-BrxF-iSTAND->DUF6079->N6-METHYLASE->PglZ->TUDOR+SF1-SF2-Helicase+iREase+CTD->OLD-ABC+TOPRIM->Calcineurin-SbcD->SbcC->HD-> | 1245 | Bacteria>FCB_group>Bacteroidetes/Chlorobi_group>Bacteroidota>Bacteroidetes_incertae_sedis>Rhodothermaceae | GCF_024809285.1 | Salinibacter_ruber | GGQ10_RS14855 | 72088..75825 | 146919 | hypothetical-protein-[Salinibacter-ruber] |
| WP_259195582.1 | Type5.BREX | Phage_integrase->Phage_integrase->TetR-HTH-><-HTH_3\|\|?->Phage_int_SAM_1+Phage_integrase-><-REase\|\|BrxA-DNA-binding->BrxB-BrxF-iSTAND->DUF6079*->DOC+HTH->N6-METHYLASE->OLD-ABC+TOPRIM->PglZ->SIGMA+OB-Fold+AAA-ATPase+Lon-Protease-BrxL->Calcineurin-SbcD->SbcC+SbcC->REase+DUF4268-> | 1144 | Bacteria>FCB_group>Bacteroidetes/Chlorobi_group>Bacteroidota>Bacteroidetes_incertae_sedis>Rhodothermaceae | GCF_024809635.1 | Salinibacter_ruber | GGP50_RS04130 | 10424..13858 | 146919 | BREX-system-P-loop-protein-BrxC-[Salinibacter-ruber] |
| WP_251951639.1 | Type5.BREX | <-OLD-ABC+TOPRIM<-TUDOR+SF1-SF2-Helicase+iREase+CTD<-?<-DDE_Tnp_1<-PglZ<-ParB+HNH<-N6-METHYLASE<-DUF6079<-BrxB-BrxF-iSTAND<-DUF6079*<-BrxA-DNA-binding<-REase+RNase<-viral-protein<-?<-RHH<-TraJ-RHH\|\|HTH_3+Peptidase_S24-> | 1245 | Bacteria>FCB_group>Bacteroidetes/Chlorobi_group>Bacteroidota>Bacteroidetes_incertae_sedis>Rhodothermaceae | GCF_943908255.1 | Salinibacter_ruber | NGL47_RS11305 | 26687..30424 | 146919 | hypothetical-protein-[Salinibacter-ruber] |
| WP_204433169.1 | Type1.BREX | BrxA-DNA-binding->BrxB-BrxF-iSTAND->DUF6079*->OLD-ABC->TOPRIM->N6-METHYLASE-> | 1195 | Bacteria>FCB_group>Bacteroidetes/Chlorobi_group>Bacteroidota>Bacteroidia>Bacteroidales | - | Bacteroides_mediterraneensis | H6B28_RS04775 | 126749..130336 | 1841856 | BREX-system-P-loop-protein-BrxC-[Bacteroides-mediterraneensis] |
| CDD50103.1 | Type1.BREX | OLD-ABC->?->?-><-?<-?<-?<-?\|\|BrxA-DNA-binding->BrxB-BrxF-iSTAND->DUF6079*->SbcC->OLD-ABC-p->TOPRIM->N6-METHYLASE->N6-METHYLASE-p-> | 1195 | Bacteria>FCB_group>Bacteroidetes/Chlorobi_group>Bacteroidota>Bacteroidia>Bacteroidales | GCA_000435415.1 | Bacteroides_sp._CAG:875 | BN800_01988 | 17775..21362 | 1262752 | putative-uncharacterized-protein-[Bacteroides-sp.-CAG:875] |
| CDD33290.1 | Type1.BREX | <-TOPRIM<-OLD-ABC<-DUF6079*<-BrxB-BrxF-iSTAND<-BrxA-DNA-binding<-HTH_3<-Phage_holin<-HD | 1195 | Bacteria>FCB_group>Bacteroidetes/Chlorobi_group>Bacteroidota>Bacteroidia>Bacteroidales | GCA_000436055.1 | Bacteroides_sp._CAG:714 | BN762_00981 | 3639..7226 | 1262749 | putative-uncharacterized-protein-[Bacteroides-sp.-CAG:714] |
| HAH91106.1 | Type1.BREX | <-HIRAN<-RADICAL-SAM<-ARG<-?<-?\|\|HTH_3->wHTH+WYL+WCX->BrxA-DNA-binding->BrxB-BrxF-iSTAND->DUF6079*->OLD-ABC->TOPRIM->?->SF1-SF2-Helicase->PglZ->SIGMA+OB-Fold+AAA-ATPase+HKD-DNase-BrxL->McrB-AAA->wHTH+REase-> | 1202 | Bacteria>FCB_group>Bacteroidetes/Chlorobi_group>Bacteroidota>Bacteroidia>Bacteroidales | GCA_003450425.1 | Prevotella_sp. | DCL96_05230 | 7787..11395 | 59823 | BREX-system-P-loop-protein-BrxC-[Prevotella-sp.] |
| MBD5181931.1 | Type1.BREX | <-Acyltransferase<-?<-?\|\|HISKIN+MORC->REC+OmpR-HTH->BrxA-DNA-binding->BrxB-BrxF-iSTAND->DUF6079*->N6-METHYLASE->OLD-ABC->TOPRIM->PglZ->SIGMA+OB-Fold+AAA-ATPase+HKD-DNase-BrxL-> | 1193 | Bacteria>FCB_group>Bacteroidetes/Chlorobi_group>Bacteroidota>Bacteroidia>Bacteroidales | GCA_014800235.1 | Bacteroidales_bacterium | HDS98_05210 | 20716..24297 | 2030927 | BREX-system-P-loop-protein-BrxC-[Bacteroidales-bacterium] |
| MBN2915288.1 | Type1.BREX | <-wHTH+REase<-McrB-AAA<-SIGMA+OB-Fold+AAA-ATPase+HKD-DNase-BrxL<-PglZ<-SIR2+SIR2<-N6-METHYLASE<-TOPRIM<-OLD-ABC<-DUF6079-p* | 971 | Bacteria>FCB_group>Bacteroidetes/Chlorobi_group>Bacteroidota>Bacteroidia>Bacteroidales | GCA_016939035.1 | Prevotella_sp. | JTJ26_04215 | 25012..27927 | 59823 | BREX-system-P-loop-protein-BrxC-[Prevotella-sp.] |
| HJF82219.1 | Type1.BREX | <-SF1-SF2-Helicase-p<-HTH_3<-?<-?\|\|BrxA-DNA-binding->BrxB-BrxF-iSTAND->DUF6079*->OLD-ABC->TOPRIM->N6-METHYLASE->SIR2+SIR2->PglZ->SIGMA+OB-Fold+AAA-ATPase+HKD-DNase-BrxL-> | 1195 | Bacteria>FCB_group>Bacteroidetes/Chlorobi_group>Bacteroidota>Bacteroidia>Bacteroidales | GCA_020742805.1 | Phocaeicola_plebeius | K8V40_11345 | 4107..7694 | 310297 | BREX-system-P-loop-protein-BrxC-[Phocaeicola-plebeius] |
| MCI6195664.1 | Type1.BREX | <-PglZ<-SIR2+SIR2<-Peptidase_C39_2<-?<-SIR2<-?<-N6-METHYLASE<-TOPRIM<-OLD-ABC<-DUF6079*<-BrxB-BrxF-iSTAND<-BrxA-DNA-binding<-SF1-SF2-Helicase<-?\|\|CR-ATPase5+REase-><-ARG<-?<-?\|\|SbcC-> | 1190 | Bacteria>FCB_group>Bacteroidetes/Chlorobi_group>Bacteroidota>Bacteroidia>Bacteroidales | GCA_022773545.1 | Prevotella_sp. | MR670_00305 | 49581..53153 | 59823 | BREX-system-P-loop-protein-BrxC-[Prevotella-sp.] |
| MCI6336758.1 | Type1.BREX | DUF6079-p*->OLD-ABC->TOPRIM->N6-METHYLASE->?->SIR2->ACET+TPR->?->Peptidase_C39_2-> | 873 | Bacteria>FCB_group>Bacteroidetes/Chlorobi_group>Bacteroidota>Bacteroidia>Bacteroidales | GCA_022777825.1 | Prevotella_sp. | MR802_02260 | 1..2624 | 59823 | BREX-system-P-loop-protein-BrxC-[Prevotella-sp.] |
| SCI35540.1 | Type1.BREX | <-N6-METHYLASE<-TOPRIM<-OLD-ABC<-DUF6079*<-BrxB-BrxF-iSTAND<-BrxA-DNA-binding | 1195 | Bacteria>FCB_group>Bacteroidetes/Chlorobi_group>Bacteroidota>Bacteroidia>Bacteroidales | GCA_900066455.1 | uncultured_Bacteroides_sp. | OCV81_RS14585 | 3983..7570 | 162156 | Uncharacterised-protein-[uncultured-Bacteroides-sp.] |
| WP_094881847.1 | Type1.BREX | <-TOPRIM<-OLD-ABC<-DUF6079*<-BrxB-BrxF-iSTAND<-BrxA-DNA-binding<-wHTH+WYL+WCX<-HTH_3 | 1206 | Bacteria>FCB_group>Bacteroidetes/Chlorobi_group>Bacteroidota>Bacteroidia>Bacteroidales | GCF_002265625.1 | Prevotella_sp._885 | CHL74_RS14190 | 4100..7720 | 2022527 | BREX-system-P-loop-protein-BrxC-[Prevotella-sp.-885] |
| WP_118441918.1 | Type1.BREX | HTH_Tnp_1+HTH_21+rve+rve_3-><-?<-?\|\|BrxA-DNA-binding->BrxB-BrxF-iSTAND->DUF6079*->OLD-ABC->TOPRIM->N6-METHYLASE->?->PglZ->SIGMA+OB-Fold+AAA-ATPase+HKD-DNase-BrxL->?->?->Peptidase_M28-> | 1195 | Bacteria>FCB_group>Bacteroidetes/Chlorobi_group>Bacteroidota>Bacteroidia>Bacteroidales | GCF_003474535.1 | Phocaeicola_plebeius | DW041_RS13375 | 4564..8151 | 310297 | BREX-system-P-loop-protein-BrxC-[Phocaeicola-plebeius] |
| WP_217745553.1 | Type1.BREX | <-HIRAN<-RADICAL-SAM<-ARG<-?<-?\|\|HTH_3->wHTH+WYL+WCX->BrxA-DNA-binding->BrxB-BrxF-iSTAND->DUF6079*->OLD-ABC->TOPRIM->N6-METHYLASE->SIR2+SIR2->PglZ->SIGMA+OB-Fold+AAA-ATPase+HKD-DNase-BrxL->McrB-AAA->wHTH+REase-> | 1206 | Bacteria>FCB_group>Bacteroidetes/Chlorobi_group>Bacteroidota>Bacteroidia>Bacteroidales | GCF_019127365.1 | Prevotella_copri | KSW85_RS06175 | 41108..44728 | 165179 | BREX-system-P-loop-protein-BrxC-[Prevotella-copri] |
| WP_148330685.1 | Type1.BREX | <-N6-METHYLASE<-TOPRIM<-OLD-ABC<-DUF6079*<-BrxB-BrxF-iSTAND<-BrxA-DNA-binding | 1195 | Bacteria>FCB_group>Bacteroidetes/Chlorobi_group>Bacteroidota>Bacteroidia>Bacteroidales | GCF_025567125.1 | Phocaeicola_fibrisolvens | OCV81_RS14585 | 3983..7570 | 2981793 | BREX-system-P-loop-protein-BrxC-[Phocaeicola-fibrisolvens] |
| WP_072542576.1 | Type1.BREX | <-AAA-ATPase<-HTH+rve<-?<-SIGMA+OB-Fold+AAA-ATPase+HKD-DNase-BrxL<-PglZ<-SIR2+SIR2<-N6-METHYLASE<-TOPRIM<-OLD-ABC<-DUF6079*<-BrxB-BrxF-iSTAND<-BrxA-DNA-binding\|\|?->?->?->?-><-?<-?<-OLD-ABC | 1195 | Bacteria>FCB_group>Bacteroidetes/Chlorobi_group>Bacteroidota>Bacteroidia>Bacteroidales | GCF_900128455.1 | Bacteroides_mediterraneensis | BQ7480_RS08760 | 2129051..2132638 | 1841856 | BREX-system-P-loop-protein-BrxC-[Bacteroides-mediterraneensis] |
| WP_118139864.1 | Type1.BREX | <-HIRAN<-RADICAL-SAM<-ARG<-?<-?\|\|HTH_3->wHTH+WYL+WCX->BrxA-DNA-binding->BrxB-BrxF-iSTAND->DUF6079*->OLD-ABC->TOPRIM->N6-METHYLASE->N6-METHYLASE-p->?->SF1-SF2-Helicase->PglZ->SIGMA+OB-Fold+AAA-ATPase+HKD-DNase-BrxL->McrB-AAA-> | 1202 | Bacteria>FCB_group>Bacteroidetes/Chlorobi_group>Bacteroidota>Bacteroidia>Bacteroidales | - | Prevotella_copri | DWV60_RS07145 | 31878..35486 | 165179 | BREX-system-P-loop-protein-BrxC-[Prevotella-copri] |
| WP_188169955.1 | Type1.BREX | <-OLD-ABC<-Arm-DNA-bind_5+Phage_int_SAM_5+Phage_integrase<-MACRODOMAIN<-ART-DarT<-SF1-SF2-Helicase<-SIGMA+OB-Fold+AAA-ATPase+Lon-Protease-BrxL<-PglZ<-OLD-ABC+TOPRIM<-N6-METHYLASE<-DUF6079*<-BrxB-BrxF-iSTAND<-BrxA-DNA-binding<-HTH_3<-?<-?<-Pkinase\|\|Calcineurin-> | 1216 | Bacteria>FCB_group>Bacteroidetes/Chlorobi_group>Bacteroidota>Cytophagia>Cytophagales | GCF_014534745.1 | Flammeovirga_sp._EKP202 | IC221_RS16125 | 83755..87405 | 2770592 | BREX-system-P-loop-protein-BrxC-[Flammeovirga-sp.-EKP202] |
| WP_201982535.1 | Type1.BREX | HEPN->CHTH->TOPRIM->Sp-><-ParB<-OLD-ABC+TOPRIM<-SIGMA+OB-Fold+AAA-ATPase+Lon-Protease-BrxL<-PglZ<-N6-METHYLASE<-DUF6079*<-BrxB-BrxF-iSTAND<-BrxA-DNA-binding<-?\|\|REase+PNTD1+PNTD2->PIWI-><-?<-Phage_integrase+Phage_integrase<-ACET\|\|35exo-> | 1143 | Bacteria>FCB_group>Bacteroidetes/Chlorobi_group>Bacteroidota>Cytophagia>Cytophagales | GCF_016734815.1 | Hymenobacter_rubidus | JKL48_RS11490 | 20898..24329 | 1441626 | BREX-system-P-loop-protein-BrxC-[Hymenobacter-rubidus] |
| WP_234861365.1 | Type1.BREX | STYKIN-><-?\|\|?->GDResolvase-><-?\|\|TPR+TPR->BrxA-DNA-binding->BrxB-BrxF-iSTAND->DUF6079*->N6-METHYLASE->OLD-ABC+TOPRIM->PglZ->SIGMA+OB-Fold+AAA-ATPase+HKD-DNase-BrxL->TIR->?->?->?->ParB+N6_N4_METHYLASE-> | 1174 | Bacteria>FCB_group>Bacteroidetes/Chlorobi_group>Bacteroidota>Cytophagia>Cytophagales | GCF_021491425.1 | Mariniradius_sp._RY-2 | LOU89_RS09865 | 251856..255380 | 2909237 | BREX-system-P-loop-protein-BrxC-[Mariniradius-sp.-RY-2] |
| MCB9053313.1 | Type1.BREX | <-PglZ<-TOPRIM<-OLD-ABC<-?<-NACHT<-?<-REase<-TPR+S1<-NERD+SF1-SF2-Helicase<-DUF6079*<-BrxB-BrxF-iSTAND<-BrxA-DNA-binding<-?<-RNase<-SF1-SF2-Helicase+REase\|\|NtrC-AAA+FIS-HTH->?->OLD-ABC->TOPRIM | 1180 | Bacteria>FCB_group>Bacteroidetes/Chlorobi_group>Bacteroidota>Saprospiria>Saprospirales | GCA_020636405.1 | Lewinellaceae_bacterium | H6556_28080 | 325530..329072 | 2760807 | BREX-system-P-loop-protein-BrxC-[Lewinellaceae-bacterium] |
| WP_020534907.1 | Type1.BREX | N6-METHYLASE->?-><-?<-?\|\|?->?->HTH_3->BrxA-DNA-binding->BrxB-BrxF-iSTAND->DUF6079*->N6-METHYLASE->OLD-ABC->TOPRIM->?->HAD+PRTase+PRTase+REase+SF1-SF2-Helicase->?->PglZ->SIGMA+OB-Fold+AAA-ATPase+HKD-DNase-BrxL->Peptidase-> | 1159 | Bacteria>FCB_group>Bacteroidetes/Chlorobi_group>Bacteroidota>Saprospiria>Saprospirales | GCF_000379805.1 | Lewinella_cohaerens | A3EU_RS29375 | 116498..119977 | 70995 | BREX-system-P-loop-protein-BrxC-[Lewinella-cohaerens] |
| TNJ34090.1 | Type1.BREX | HIN-HTH->HTH_Tnp_1+HTH_21+rve+rve_3-><-SIGMA+OB-Fold+AAA-ATPase+Lon-Protease-BrxL<-PglZ<-TOPRIM<-OLD-ABC<-N6-METHYLASE<-Schlafen+HATPase+wHTH<-DUF6079*<-BrxB-BrxF-iSTAND<-BrxA-DNA-binding<-XIS-HTH<-RVT<-RVT | 1144 | Bacteria>FCB_group>Bacteroidetes/Chlorobi_group>Chlorobiota>Chlorobiia>Chlorobiales | GCA_006265245.1 | Prosthecochloris_vibrioformis | FGF68_10440 | 14740..18174 | 1098 | BREX-system-P-loop-protein-BrxC-[Prosthecochloris-vibrioformis] |
| MBV5328349.1 | Type1.BREX | <-SIGMA+OB-Fold+AAA-ATPase+Lon-Protease-BrxL<-PglZ<-TOPRIM<-OLD-ABC<-N6-METHYLASE<-DUF6079*<-BrxB-BrxF-iSTAND<-BrxA-DNA-binding<-XIS-HTH | 1208 | Bacteria>FCB_group>Bacteroidetes/Chlorobi_group>Chlorobiota>Chlorobiia>Chlorobiales | GCA_019162945.1 | Chlorobium_sp. | JZU65_12040 | 10298..13924 | 1095 | BREX-system-P-loop-protein-BrxC-[Chlorobium-sp.] |
| WP_211340437.1 | Type1.BREX | HIN-HTH->HTH_Tnp_1+HTH_21+rve+rve_3-><-GDResolvase<-SIGMA+OB-Fold+AAA-ATPase+Lon-Protease-BrxL<-PglZ<-TOPRIM<-ParB+SbcC<-N6-METHYLASE<-Schlafen+HATPase+wHTH<-DUF6079*<-BrxB-BrxF-iSTAND<-BrxA-DNA-binding<-XIS-HTH | 1145 | Bacteria>FCB_group>Bacteroidetes/Chlorobi_group>Chlorobiota>Chlorobiia>Chlorobiales | GCF_006265245.1 | Prosthecochloris_vibrioformis | FGF68_RS10435 | 14740..18177 | 1098 | BREX-system-P-loop-protein-BrxC-[Prosthecochloris-vibrioformis] |
| MCB5285398.1 | Type6.BREX | <-DNA_METHYLASE<-SIGMA+OB-Fold+AAA-ATPase+Lon-Protease-BrxL<-TOPRIM<-OLD-ABC<-HTH_29+HTH_32+HTH_33+DDE_3<-PglZ<-TOPRIM<-OLD-ABC<-N6-METHYLASE<-DUF6079*<-BrxB-BrxF-iSTAND<-BrxA-DNA-binding<-BrxE<-CHTH | 1227 | Bacteria>FCB_group>Candidatus_Cloacimonetes>unclassified_Candidatus_Cloacimonetes | GCA_020532085.1 | Candidatus_Cloacimonetes_bacterium | LHW45_07405 | 38085..41768 | 2030808 | BREX-system-P-loop-protein-BrxC-[Candidatus-Cloacimonadota-bacterium] |
| MBX6362419.1 | Type6.BREX | Peptidase-><-?\|\|BrxE->BrxA-DNA-binding->BrxB-BrxF-iSTAND->DUF6079*->N6-METHYLASE->HKD-DNase->?->N6-METHYLASE-p->OLD-ABC->TOPRIM->?->N6-METHYLASE->PglZ-> | 1201 | Bacteria>FCB_group>Gemmatimonadota>unclassified_Gemmatimonadota | GCA_019689775.1 | Gemmatimonadota_bacterium | IRZ00_01015 | 11407..15012 | 2026742 | BREX-system-P-loop-protein-BrxC-[Gemmatimonadota-bacterium] |
| CDA09060.1 | Type1.BREX | <-SIGMA+OB-Fold+AAA-ATPase+Lon-Protease-BrxL<-PglZ<-CR-ATPase8+REase<-OLD-ABC+TOPRIM<-?<-?<-N6-METHYLASE<-TOPRIM<-OLD-ABC<-DUF6079*<-?<-SWIM+DUF3670+SF1-SF2-Helicase<-?<-?<-?<-SF1-SF2-Helicase | 1250 | Bacteria>Fusobacteriota>Fusobacteriia>Fusobacteriales>Fusobacteriaceae>Fusobacterium | GCA_000433695.1 | Fusobacterium_sp._CAG:649 | BN748_00884 | 23068..26820 | 1262900 | putative-uncharacterized-protein-[Fusobacterium-sp.-CAG:649] |
| ALF25292.1 | Type1.BREX | SKI->JAB+sGTP->wHTH+WYL+WCX->?->?->ABC-ATPase+SbcC->?-><-?\|\|DUF6079*->OLD-ABC->TOPRIM->Schlafen+HATPase+wHTH->N6-METHYLASE->KAP-NTPase->HNH->SbcC->SIR2->HerA/FtsK-> | 1250 | Bacteria>Fusobacteriota>Fusobacteriia>Fusobacteriales>Fusobacteriaceae>Fusobacterium | GCA_001296185.1 | Fusobacterium_nucleatum_subsp._nucleatum | RN95_02150 | 455192..458944 | 76856 | cobalamin-adenosyltransferase-[Fusobacterium-nucleatum-subsp.-nucleatum] |
| WP_008797395.1 | Type1.BREX | <-HerA/FtsK<-SIR2<-SbcC<-HNH<-KAP-NTPase<-KAP-NTPase<-N6-METHYLASE<-TOPRIM<-OLD-ABC<-DUF6079*\|\|?->?-><-?<-ABC-ATPase+SbcC<-?<-?<-wHTH+WYL+WCX<-JAB+sGTP<-SKI | 1249 | Bacteria>Fusobacteriota>Fusobacteriia>Fusobacteriales>Fusobacteriaceae>Fusobacterium | GCF_000163915.2 | Fusobacterium_nucleatum | HMPREF0405_RS05640 | 1192926..1196675 | 851 | BREX-system-P-loop-protein-BrxC-[Fusobacterium-vincentii] |
| WP_005903411.1 | Type1.BREX | <-PglZ<-ParB+HNH+RAMA<-OLD-ABC+TOPRIM<-N6-METHYLASE<-Schlafen+HATPase+wHTH<-?<-RNase+PHP<-?<-OLD-ABC<-DUF6079*\|\|?->?-><-?<-ABC-ATPase+SbcC<-?<-?<-wHTH+WYL+WCX<-JAB+sGTP<-SKI | 1250 | Bacteria>Fusobacteriota>Fusobacteriia>Fusobacteriales>Fusobacteriaceae>Fusobacterium | GCF_000178895.1 | Fusobacterium_nucleatum | C4N14_RS08075 | 1667383..1671135 | 851 | BREX-system-P-loop-protein-BrxC-[Fusobacterium-nucleatum] |
| WP_008692035.1 | Type1.BREX | DUF3670+SF1-SF2-Helicase->?-><-?<-?<-?\|\|DUF6079*->OLD-ABC->TOPRIM->N6-METHYLASE->?->OLD-ABC+TOPRIM->CR-ATPase8+REase->PglZ->SIGMA+OB-Fold+AAA-ATPase+Lon-Protease-BrxL->BrxA-DNA-binding-> | 1250 | Bacteria>Fusobacteriota>Fusobacteriia>Fusobacteriales>Fusobacteriaceae>Fusobacterium | GCF_000218655.1 | Fusobacterium_nucleatum | HMPREF0401_RS10120 | 557323..561075 | 851 | BREX-system-P-loop-protein-BrxC-[Fusobacterium-animalis] |
| WP_008700610.1 | Type1.BREX | SWIM+DUF3670+SF1-SF2-Helicase->?-><-?<-?<-?<-?\|\|DUF6079*->OLD-ABC->TOPRIM->KAP-NTPase->HNH->SbcC->SIR2->HerA/FtsK->CR-ATPase8+REase->Classical-AAA-> | 1250 | Bacteria>Fusobacteriota>Fusobacteriia>Fusobacteriales>Fusobacteriaceae>Fusobacterium | GCF_000234075.2 | Fusobacterium_nucleatum | FSDG_RS04805 | 1011636..1015388 | 851 | MULTISPECIES:-BREX-system-P-loop-protein-BrxC-[Fusobacterium] |
| WP_029598126.1 | Type1.BREX | SWIM+DUF3670+SF1-SF2-Helicase->?-><-?<-?<-?\|\|DUF6079*->OLD-ABC->TOPRIM->N6-METHYLASE->KAP-NTPase->HNH->SbcC->SIR2->HerA/FtsK-> | 1250 | Bacteria>Fusobacteriota>Fusobacteriia>Fusobacteriales>Fusobacteriaceae>Fusobacterium | GCF_000455985.1 | Fusobacterium_nucleatum | J141_RS0104310 | 22455..26207 | 851 | BREX-system-P-loop-protein-BrxC-[Fusobacterium-animalis] |
| WP_023036666.1 | Type1.BREX | <-HerA/FtsK<-SIR2<-SbcC<-HNH<-KAP-NTPase<-N6-METHYLASE<-TOPRIM<-?<-OLD-ABC<-DUF6079*\|\|?-><-?<-ABC-ATPase+SbcC<-?<-?<-wHTH+WYL+WCX<-JAB+sGTP<-SKI | 1250 | Bacteria>Fusobacteriota>Fusobacteriia>Fusobacteriales>Fusobacteriaceae>Fusobacterium | GCF_000479185.1 | Fusobacterium_nucleatum | HMPREF1767_RS09525 | 33414..37166 | 851 | BREX-system-P-loop-protein-BrxC-[Fusobacterium-nucleatum] |
| WP_032836039.1 | Type1.BREX | <-PglZ<-CR-ATPase8+REase<-HEPN<-MORC<-N6-METHYLASE<-Schlafen+HATPase+wHTH<-TOPRIM<-OLD-ABC<-DUF6079*\|\|?-><-?<-ABC-ATPase+SbcC<-?<-?<-wHTH+WYL+WCX<-JAB+sGTP<-SKI | 1250 | Bacteria>Fusobacteriota>Fusobacteriia>Fusobacteriales>Fusobacteriaceae>Fusobacterium | GCF_000524235.1 | Fusobacterium_sp._OBRC1 | HMPREF1501_RS01100 | 63475..67227 | 1032505 | BREX-system-P-loop-protein-BrxC-[Fusobacterium-sp.-OBRC1] |
| WP_032846252.1 | Type1.BREX | <-BrxA-DNA-binding<-SIGMA+OB-Fold+AAA-ATPase+Lon-Protease-BrxL<-PglZ<-nSTAND<-?<-N6-METHYLASE<-Schlafen+HATPase+wHTH<-TOPRIM<-OLD-ABC<-DUF6079*\|\|?->?->?-><-?<-ABC-ATPase+SbcC<-?<-?<-wHTH+WYL+WCX<-JAB+sGTP | 1250 | Bacteria>Fusobacteriota>Fusobacteriia>Fusobacteriales>Fusobacteriaceae>Fusobacterium | GCF_000524395.1 | Fusobacterium_sp._CM22 | HMPREF1500_RS09190 | 124439..128191 | 936563 | BREX-system-P-loop-protein-BrxC-[Fusobacterium-sp.-CM22] |
| WP_060676307.1 | Type1.BREX | <-HerA/FtsK<-SIR2<-?<-SbcC<-HNH<-KAP-NTPase<-N6-METHYLASE<-TOPRIM<-OLD-ABC<-DUF6079*\|\|?->?<-Peptidase_M20<-?\|\|35exo+T7DNApol-> | 1250 | Bacteria>Fusobacteriota>Fusobacteriia>Fusobacteriales>Fusobacteriaceae>Fusobacterium | GCF_001296085.1 | Fusobacterium_nucleatum | RN98_RS07270 | 1461443..1465195 | 851 | BREX-system-P-loop-protein-BrxC-[Fusobacterium-animalis] |
| WP_070766192.1 | Type1.BREX | SKI->JAB+sGTP->wHTH+WYL+WCX->?->?->ABC-ATPase+SbcC->?-><-?\|\|DUF6079*->OLD-ABC->TOPRIM->Schlafen+HATPase+wHTH->N6-METHYLASE->N6-METHYLASE->MORC+HISKIN->MORC+HISKIN->?->HNH-> | 1250 | Bacteria>Fusobacteriota>Fusobacteriia>Fusobacteriales>Fusobacteriaceae>Fusobacterium | GCF_001815715.1 | Fusobacterium_sp._HMSC064B11 | HMPREF3051_RS04300 | 251838..255590 | 1739543 | BREX-system-P-loop-protein-BrxC-[Fusobacterium-sp.-HMSC064B11] |
| WP_088387741.1 | Type1.BREX | SKI->JAB+sGTP->wHTH+WYL+WCX->?->?->ABC-ATPase+SbcC->?-><-?\|\|DUF6079*->OLD-ABC->TOPRIM->N6-METHYLASE->?->nSTAND->CR-ATPase8+REase->ParB+HNH+RAMA->Classical-AAA->Peptidase_S8-> | 1250 | Bacteria>Fusobacteriota>Fusobacteriia>Fusobacteriales>Fusobacteriaceae>Fusobacterium | GCF_002204435.1 | Fusobacterium_nucleatum | CA839_RS02150 | 467099..470851 | 851 | BREX-system-P-loop-protein-BrxC-[Fusobacterium-polymorphum] |
| WP_098702328.1 | Type1.BREX | SKI->JAB+sGTP->wHTH+WYL+WCX->?->?->ABC-ATPase+SbcC->?-><-?\|\|DUF6079*->OLD-ABC->TOPRIM->N6-METHYLASE->HEPN->CR-ATPase8+REase->Classical-AAA->Peptidase_S8->PglZ->SIGMA+OB-Fold+AAA-ATPase+Lon-Protease-BrxL-> | 1250 | Bacteria>Fusobacteriota>Fusobacteriia>Fusobacteriales>Fusobacteriaceae>Fusobacterium | GCF_002573625.1 | Fusobacterium_nucleatum | RN96_RS03470 | 669523..673275 | 851 | BREX-system-P-loop-protein-BrxC-[Fusobacterium-polymorphum] |
| WP_098973854.1 | Type1.BREX | SKI->JAB+sGTP->wHTH+WYL+WCX->?->?->ABC-ATPase+SbcC->?-><-?\|\|DUF6079*->OLD-ABC->TOPRIM->N6-METHYLASE->MORC+HISKIN->?->HNH->PglZ->SIGMA+OB-Fold+AAA-ATPase+Lon-Protease-BrxL-> | 1250 | Bacteria>Fusobacteriota>Fusobacteriia>Fusobacteriales>Fusobacteriaceae>Fusobacterium | GCF_002591515.1 | Fusobacterium_nucleatum | CBG58_RS00340 | 77641..81393 | 851 | BREX-system-P-loop-protein-BrxC-[Fusobacterium-polymorphum] |
| WP_099972121.1 | Type1.BREX | <-PglZ<-Peptidase_S8<-Classical-AAA<-ParB+HNH+RAMA<-nSTAND<-?<-N6-METHYLASE<-TOPRIM<-OLD-ABC<-DUF6079*\|\|?-><-?<-?<-Peptidase_M3_N+Peptidase_M3\|\|RNaseH->?->?->?->Peptidase_M20-> | 1250 | Bacteria>Fusobacteriota>Fusobacteriia>Fusobacteriales>Fusobacteriaceae>Fusobacterium | GCF_002761935.1 | Fusobacterium_pseudoperiodonticum | CTM69_RS07785 | 481250..485002 | 2663009 | BREX-system-P-loop-protein-BrxC-[Fusobacterium-pseudoperiodonticum] |
| WP_099959237.1 | Type1.BREX | <-SIGMA+OB-Fold+AAA-ATPase+Lon-Protease-BrxL<-PglZ<-Peptidase_S8<-Classical-AAA<-CR-ATPase8+REase<-HEPN<-N6-METHYLASE<-TOPRIM<-OLD-ABC<-DUF6079*\|\|?-><-?<-?<-Peptidase_M3_N+Peptidase_M3\|\|RNaseH->?->?->?->Peptidase_M20-> | 1250 | Bacteria>Fusobacteriota>Fusobacteriia>Fusobacteriales>Fusobacteriaceae>Fusobacterium | GCF_002761955.1 | Fusobacterium_pseudoperiodonticum | CTM71_RS09955 | 1967192..1970944 | 2663009 | BREX-system-P-loop-protein-BrxC-[Fusobacterium-pseudoperiodonticum] |
| WP_158410489.1 | Type1.BREX | <-TOPRIM<-OLD-ABC<-DUF6079*\|\|?->?-><-Peptidase_M20<-?\|\|35exo+T7DNApol-> | 1250 | Bacteria>Fusobacteriota>Fusobacteriia>Fusobacteriales>Fusobacteriaceae>Fusobacterium | GCF_002762015.1 | Fusobacterium_nucleatum | CI111_RS06870 | 6549..10301 | 851 | BREX-system-P-loop-protein-BrxC-[Fusobacterium-animalis] |
| WP_100024172.1 | Type1.BREX | <-Peptidase_M20<-?<-?<-?<-RNaseH\|\|Peptidase_M3_N+Peptidase_M3->?->?->?-><-?\|\|DUF6079*->OLD-ABC->TOPRIM->N6-METHYLASE->HEPN->CR-ATPase8+REase->PglZ->SIGMA+OB-Fold+AAA-ATPase+Lon-Protease-BrxL->BrxA-DNA-binding->BrxB-BrxF-iSTAND-> | 1250 | Bacteria>Fusobacteriota>Fusobacteriia>Fusobacteriales>Fusobacteriaceae>Fusobacterium | GCF_002763625.1 | Fusobacterium_pseudoperiodonticum | CTM72_RS01085 | 199877..203629 | 2663009 | BREX-system-P-loop-protein-BrxC-[Fusobacterium-pseudoperiodonticum] |
| WP_099987862.1 | Type1.BREX | <-BrxA-DNA-binding<-SIGMA+OB-Fold+AAA-ATPase+Lon-Protease-BrxL<-PglZ<-CR-ATPase8+REase<-OLD-ABC+TOPRIM<-?<-?<-N6-METHYLASE<-TOPRIM<-OLD-ABC<-DUF6079*\|\|?->OrfB_IS605+OrfB_Zn_ribbon-><-URI-DNase+SF1-SF2-Helicase<-URI-DNase+SF1-SF2-Helicase<-?<-?<-Peptidase_M3_N+Peptidase_M3\|\|RNaseH-> | 1250 | Bacteria>Fusobacteriota>Fusobacteriia>Fusobacteriales>Fusobacteriaceae>Fusobacterium | GCF_002763695.1 | Fusobacterium_pseudoperiodonticum | CTM74_RS09005 | 1721055..1724807 | 2663009 | BREX-system-P-loop-protein-BrxC-[Fusobacterium-pseudoperiodonticum] |
| WP_124794840.1 | Type1.BREX | wHTH+WYL+WCX->?->?->ABC-ATPase+SbcC->?-><-?<-?<-?<-?\|\|DUF6079*->OLD-ABC->TOPRIM->N6-METHYLASE->?->OLD-ABC+TOPRIM->CR-ATPase8+REase->ParB+HNH+RAMA->PglZ->SIGMA+OB-Fold+AAA-ATPase+Lon-Protease-BrxL-> | 1250 | Bacteria>Fusobacteriota>Fusobacteriia>Fusobacteriales>Fusobacteriaceae>Fusobacterium | GCF_003859915.1 | Fusobacterium_canifelinum | EII27_RS00780 | 173024..176776 | 285729 | BREX-system-P-loop-protein-BrxC-[Fusobacterium-canifelinum] |
| WP_220302801.1 | Type1.BREX | SKI->JAB+sGTP->wHTH+WYL+WCX->?->?->ABC-ATPase+SbcC->?-><-?\|\|DUF6079*->OLD-ABC->TOPRIM->Schlafen+HATPase+wHTH->N6-METHYLASE->KAP-NTPase->HNH->SbcC->SIR2->HerA/FtsK-> | 1250 | Bacteria>Fusobacteriota>Fusobacteriia>Fusobacteriales>Fusobacteriaceae>Fusobacterium | GCF_019552045.1 | Fusobacterium_nucleatum | JY397_RS08300 | 1731485..1735237 | 851 | BREX-system-P-loop-protein-BrxC-[Fusobacterium-polymorphum] |
| WP_220308657.1 | Type1.BREX | <-PglZ<-Peptidase_S8<-Classical-AAA<-ParB+HNH+RAMA<-OLD-ABC+TOPRIM<-?<-N6-METHYLASE<-TOPRIM<-OLD-ABC<-DUF6079*\|\|?->?-><-Peptidase_M20<-?\|\|35exo+T7DNApol-> | 1250 | Bacteria>Fusobacteriota>Fusobacteriia>Fusobacteriales>Fusobacteriaceae>Fusobacterium | GCF_019552125.1 | Fusobacterium_nucleatum | JY401_RS04550 | 954810..958562 | 851 | BREX-system-P-loop-protein-BrxC-[Fusobacterium-animalis] |
| WP_238972693.1 | Type1.BREX | SKI->JAB+sGTP->wHTH+WYL+WCX->?->?->ABC-ATPase+SbcC->?-><-?\|\|DUF6079*->OLD-ABC->TOPRIM->Schlafen+HATPase+wHTH->N6-METHYLASE->MORC->HEPN->CR-ATPase8+REase->PglZ->SIGMA+OB-Fold+AAA-ATPase+Lon-Protease-BrxL-> | 1250 | Bacteria>Fusobacteriota>Fusobacteriia>Fusobacteriales>Fusobacteriaceae>Fusobacterium | GCF_022340165.1 | Fusobacterium_nucleatum | K5I19_RS02505 | 127932..131684 | 851 | BREX-system-P-loop-protein-BrxC-[Fusobacterium-nucleatum] |

<table>
<tr><th>Protein-ID</th><th>System</th><th>Gene-Neighbourhoods</th><th>Length</th><th>Taxonomy</th><th>Assembly-ID</th><th>Strain</th><th>Locus-Tag</th><th>Genomic-coordinates</th><th>TaxID</th><th>Fasta-defline</th></tr>
<tr><td>WP_060798221.1</td><td>Type1.BREX</td><td><- BrxA-DNA-binding <- SIGMA+OB-Fold+AAA-ATPase+Lon-Protease-BrxL <- PglZ <-CR-ATPase8+REase<-HEPN<-MORC<- N6-METHYLASE <- TOPRIM <- OLD-ABC <- DUF6079 *<-?<-SWIM+DUF3670+SF1-SF2-Helicase<-?<-?<-?<-SF1-SF2-Helicase</td><td>1250</td><td>Bacteria>Fusobacteriota>Fusobacteriia>Fusobacteriales>Fusobacteriaceae>Fusobacterium</td><td>GCF_023383525.1</td><td>Fusobacterium_nucleatum</td><td>YWH7054_RS02310</td><td>21716..25468</td><td>851</td><td>BREX-system-P-loop-protein-BrxC-[Fusobacterium-nucleatum]</td></tr>
<tr><td>WP_022069638.1</td><td>Type1.BREX</td><td><- BrxA-DNA-binding <- SIGMA+OB-Fold+AAA-ATPase+Lon-Protease-BrxL <- PglZ <-CR-ATPase8+REase<- OLD-ABC + TOPRIM <-?<- N6-METHYLASE <- TOPRIM <- OLD-ABC <- DUF6079 *</td><td>1250</td><td>Bacteria>Fusobacteriota>Fusobacteriia>Fusobacteriales>Fusobacteriaceae>Fusobacterium</td><td>GCF_023383545.1</td><td>Fusobacterium_nucleatum</td><td>YWH7053_RS01985</td><td>23867..27619</td><td>851</td><td>BREX-system-P-loop-protein-BrxC-[Fusobacterium-nucleatum]</td></tr>
<tr><td>WP_249524183.1</td><td>Type1.BREX</td><td><- BrxA-DNA-binding <- SIGMA+OB-Fold+AAA-ATPase+Lon-Protease-BrxL <- PglZ <-CR-ATPase8+REase<-HEPN<-MORC<- N6-METHYLASE <- TOPRIM <- OLD-ABC <- DUF6079 *<-?<-SWIM+DUF3670+SF1-SF2-Helicase<-?<-?<-?<-SF1-SF2-Helicase</td><td>1250</td><td>Bacteria>Fusobacteriota>Fusobacteriia>Fusobacteriales>Fusobacteriaceae>Fusobacterium</td><td>GCF_023383605.1</td><td>Fusobacterium_nucleatum</td><td>YWH7056_RS07045</td><td>28639..32391</td><td>851</td><td>BREX-system-P-loop-protein-BrxC-[Fusobacterium-nucleatum]</td></tr>
<tr><td>WP_249526950.1</td><td>Type1.BREX</td><td><- BrxB-BrxF-iSTAND <- BrxA-DNA-binding <- SIGMA+OB-Fold+AAA-ATPase+Lon-Protease-BrxL <- PglZ <-HEPN<-MORC<- N6-METHYLASE <- TOPRIM <- OLD-ABC <- DUF6079 *<-?<-SWIM+DUF3670+SF1-SF2-Helicase<-?<-?<-?<-SF1-SF2-Helicase</td><td>1250</td><td>Bacteria>Fusobacteriota>Fusobacteriia>Fusobacteriales>Fusobacteriaceae>Fusobacterium</td><td>GCF_023383645.1</td><td>Fusobacterium_nucleatum</td><td>YWH7199_RS06535</td><td>22791..26543</td><td>851</td><td>BREX-system-P-loop-protein-BrxC-[Fusobacterium-nucleatum]</td></tr>
<tr><td>WP_023036012.1</td><td>Type1.BREX</td><td><-SbcC<-HNH<-KAP-NTPase<- N6-METHYLASE <- TOPRIM <- OLD-ABC <- DUF6079 *||?->?-><-?<-ABC-ATPase+SbcC<-?<-?<- wHTH+WYL+WCX <-JAB+sGTP<-SKI</td><td>1249</td><td>Bacteria>Fusobacteriota>Fusobacteriia>Fusobacteriales>Fusobacteriaceae>Fusobacterium</td><td>GCF_024140205.1</td><td>Fusobacterium_nucleatum</td><td>BKN39_RS05495</td><td>10931..14680</td><td>851</td><td>MULTISPECIES:-BREX-system-P-loop-protein-BrxC-[Fusobacterium]</td></tr>
<tr><td>WP_265152268.1</td><td>Type1.BREX</td><td><-RADICAL-SAM<-Peptidase_M42<-?||RNaseH->?-><-?<-?<-?<-?|| DUF6079 *-> OLD-ABC -> TOPRIM -> N6-METHYLASE ->?->nSTAND->CR-ATPase-></td><td>1250</td><td>Bacteria>Fusobacteriota>Fusobacteriia>Fusobacteriales>Fusobacteriaceae>Fusobacterium</td><td>GCF_026089295.1</td><td>Fusobacterium_simiae</td><td>OCK72_RS06710</td><td>29932..33684</td><td>855</td><td>BREX-system-P-loop-protein-BrxC-[Fusobacterium-simiae]</td></tr>
<tr><td>HCF56393.1</td><td>Type6.BREX</td><td><-Schlafen+HATPase+wHTH<- N6-METHYLASE -p||?->PIN-><- TOPRIM <- OLD-ABC <- DUF6079 -p<-APATPase<- DUF6079 -p*<- DUF6079 -p<-HNH<- BrxB-BrxF-iSTAND <- BrxA-DNA-binding <- BrxA-DNA-binding <- BrxE</td><td>508</td><td>Bacteria>Myxococcota>Myxococcia>Myxococcales>unclassified_Myxococcales</td><td>GCA_003534605.1</td><td>Myxococcales_bacterium</td><td>DFS52_00150</td><td>7726..9251</td><td>2026763</td><td>BREX-system-P-loop-protein-BrxC-[Myxococcales-bacterium]</td></tr>
<tr><td>HCF56394.1</td><td>Type6.BREX</td><td><-Schlafen+HATPase+wHTH<- N6-METHYLASE -p||?->PIN-><- TOPRIM <- OLD-ABC <- DUF6079 -p<-APATPase<- DUF6079 -p<- DUF6079 -p*<-HNH<- BrxB-BrxF-iSTAND <- BrxA-DNA-binding <- BrxA-DNA-binding <- BrxE</td><td>363</td><td>Bacteria>Myxococcota>Myxococcia>Myxococcales>unclassified_Myxococcales</td><td>GCA_003534605.1</td><td>Myxococcales_bacterium</td><td>DFS52_00155</td><td>9470..10559</td><td>2026763</td><td>BREX-system-P-loop-protein-BrxC-[Myxococcales-bacterium]</td></tr>
<tr><td>MBK8262770.1</td><td>Type6.BREX</td><td><-DnaJ||?-><-?|| BrxE -> BrxA-DNA-binding -> BrxB-BrxF-iSTAND -> DUF6079 *-> OLD-ABC -> TOPRIM -> N6-METHYLASE -> PglZ -> BrxD-AAA ->ParB+HNH->SF1-SF2-Helicase+wHTH+Small-B-barrel+35exo-><-HEPN</td><td>1208</td><td>Bacteria>Myxococcota>Polyangia>Nannocystales>Nannocystaceae>Nannocystis</td><td>GCA_016712315.1</td><td>Nannocystis_sp.</td><td>IPK80_15725</td><td>176860..180486</td><td>1962667</td><td>BREX-system-P-loop-protein-BrxC-[Nannocystis-sp.]</td></tr>
<tr><td>MBK6517532.1</td><td>Type6.BREX</td><td>SF1-SF2-Helicase+REase->SF1-SF2-Helicase+REase->REase-><- TOPRIM <- OLD-ABC <-?|| BrxE -> BrxA-DNA-binding -> BrxB-BrxF-iSTAND -> DUF6079 *-><-PemK_toxin<-?|| N6-METHYLASE -> N6-METHYLASE -p->?->?->PIN-> PglZ -> BrxD-AAA -></td><td>1209</td><td>Bacteria>Myxococcota>Polyangia>Polyangiales>Polyangiaceae>unclassified_Polyangiaceae</td><td>GCA_016704335.1</td><td>Polyangiaceae_bacterium</td><td>IPG04_26305</td><td>282797..286426</td><td>2268199</td><td>BREX-system-P-loop-protein-BrxC-[Polyangiaceae-bacterium]</td></tr>
<tr><td>MBK8937968.1</td><td>Type6.BREX</td><td><- BrxD-AAA <- PglZ <-PIN<-?<-?<- N6-METHYLASE -p<- N6-METHYLASE ||?->PemK_toxin-><- DUF6079 *<- BrxB-BrxF-iSTAND <- BrxA-DNA-binding <- BrxE || OLD-ABC -> TOPRIM -><-REase<-SF1-SF2-Helicase+REase<-SF1-SF2-Helicase<-SF1-SF2-Helicase+REase-p</td><td>1016</td><td>Bacteria>Myxococcota>Polyangia>Polyangiales>Polyangiaceae>unclassified_Polyangiaceae</td><td>GCA_016719945.1</td><td>Polyangiaceae_bacterium</td><td>IPM79_10070</td><td>425737..428787</td><td>2268199</td><td>BREX-system-P-loop-protein-BrxC-[Polyangiaceae-bacterium]</td></tr>
<tr><td>MBN2191668.1</td><td>Type1.BREX</td><td><-TrkH<-?||?->?-><-Schlafen+HATPase+wHTH-p<-wHTH<- TOPRIM <- OLD-ABC <- N6-METHYLASE <- DUF6079 -p*</td><td>869</td><td>Bacteria>Myxococcota>Polyangia>Polyangiales>Polyangiaceae>unclassified_Polyangiaceae</td><td>GCA_016933275.1</td><td>Polyangiaceae_bacterium</td><td>JW751_02535</td><td>24054..26663</td><td>2268199</td><td>BREX-system-P-loop-protein-BrxC-[Polyangiaceae-bacterium]</td></tr>
<tr><td>MCK6575226.1</td><td>Type6.BREX</td><td><- PglZ -p<- TOPRIM <- OLD-ABC <- N6-METHYLASE <-?<- OLD-ABC <- DUF6079 *<- BrxB-BrxF-iSTAND <- BrxA-DNA-binding <- BrxE ||MPTase->?->?-><-REase</td><td>1247</td><td>Bacteria>Myxococcota>unclassified_Myxococcota</td><td>GCA_023150895.1</td><td>Myxococcota_bacterium</td><td>L6V77_29520</td><td>8524..12267</td><td>2818507</td><td>BREX-system-P-loop-protein-BrxC-[Myxococcota-bacterium]</td></tr>
<tr><td>MCU0671310.1</td><td>Type6.BREX</td><td><- TOPRIM <- OLD-ABC <- DUF6079 *<- BrxB-BrxF-iSTAND <- BrxA-DNA-binding <- BrxE ||?-><-?||?->XIS-HTH-><-HTH_3</td><td>1203</td><td>Bacteria>Myxococcota>unclassified_Myxococcota</td><td>GCA_025453555.1</td><td>Myxococcota_bacterium</td><td>MUE69_00810</td><td>1555..5166</td><td>2818507</td><td>BREX-system-P-loop-protein-BrxC-[Myxococcota-bacterium]</td></tr>
<tr><td>NOS82785.1</td><td>Type1.BREX</td><td><-SbcC+CCDC158<-Calcineurin+Calcineurin-SbcD<- OLD-ABC + TOPRIM <- SIGMA+OB-Fold+AAA-ATPase+Lon-Protease-BrxL <- PglZ <- N6-METHYLASE <- DUF6079 *<- BrxB-BrxF-iSTAND <- BrxA-DNA-binding ||?->LexA-protease->?-><-Phage_int_SAM_1+Phage_integrase</td><td>1196</td><td>Bacteria>Nitrospirota>Nitrospiria>Nitrospirales>Nitrospiraceae>Nitrospira</td><td>GCA_013140535.1</td><td>Nitrospira_sp.</td><td>HOP32_14505</td><td>14147..17737</td><td>70125</td><td>BREX-system-P-loop-protein-BrxC-[Nitrospira-sp.]</td></tr>
<tr><td>MCC6543975.1</td><td>Type1.BREX</td><td>BrnT_toxin->CopG_antitoxin->CopG_antitoxin-><-PIN<-DUF433<-?||?-> BrxA-DNA-binding -> BrxB-BrxF-iSTAND -> DUF6079 *-> OLD-ABC -> TOPRIM -> N6-METHYLASE -> PglZ -> SIGMA+OB-Fold+AAA-ATPase+Lon-Protease-BrxL ->TPR+TPR+TPR+TPR->?-><-PolB</td><td>1143</td><td>Bacteria>Nitrospirota>unclassified_Nitrospirota</td><td>GCA_020846775.1</td><td>Nitrospirota_bacterium</td><td>IT392_05655</td><td>165915..169346</td><td>2026887</td><td>BREX-system-P-loop-protein-BrxC-[Nitrospirota-bacterium]</td></tr>
<tr><td>WP_014029717.1</td><td>Type1.BREX</td><td>DEDD_Tnp_IS110+Transposase_20-><-DDE_5<-?||HTH_17->Phage_int_SAM_2+Integrase_1-><-SbcC<-Calcineurin+Calcineurin-SbcD|| BrxA-DNA-binding -> BrxB-BrxF-iSTAND -> DUF6079 *-> N6-METHYLASE ->Schlafen+HATPase+wHTH-> N6-METHYLASE -p-> OLD-ABC -> TOPRIM -> PglZ -> SIGMA+OB-Fold+AAA-ATPase+HKD-DNase-BrxL -></td><td>1179</td><td>Bacteria>Pseudomonadota>Acidithiobacillia>Acidithiobacillales>Acidithiobacillaceae>Acidithiobacillus</td><td>GCF_000214095.2</td><td>Acidithiobacillus_ferrivorans</td><td>ACIFE_RS11315</td><td>2362677..2366216</td><td>160808</td><td>BREX-system-P-loop-protein-BrxC-[Acidithiobacillus-ferrivorans]</td></tr>
<tr><td>WP_123104890.1</td><td>Type6.BREX</td><td>SIGMA-HTH->tRNA-><- SIGMA+OB-Fold+AAA-ATPase+Lon-Protease-BrxL <- PglZ <-ParB+HNH<- TOPRIM <- OLD-ABC <- N6-METHYLASE <- N6-METHYLASE <- DUF6079 *<- BrxB-BrxF-iSTAND <- BrxE ||<-?<-XIS-HTH||?->?-> TOPRIM -></td><td>1210</td><td>Bacteria>Pseudomonadota>Acidithiobacillia>Acidithiobacillales>Acidithiobacillaceae>Acidithiobacillus</td><td>GCF_003721225.1</td><td>Acidithiobacillus_sulfuriphilus</td><td>EC580_RS10670</td><td>12374..16006</td><td>1867749</td><td>BREX-system-P-loop-protein-BrxC-[Acidithiobacillus-sulfuriphilus]</td></tr>
<tr><td>VXB71403.1</td><td>Type1.BREX</td><td>SF1-SF2-Helicase-><-?<-DUF4096+DDE_Tnp_1+DDE_Tnp_1_2<-DUF4096+DDE_Tnp_1_2<-?||?-> BrxA-DNA-binding -> BrxB-BrxF-iSTAND -> DUF6079 *-> N6-METHYLASE -> OLD-ABC -> TOPRIM -> PglZ -></td><td>1238</td><td>Bacteria>Pseudomonadota>Alphaproteobacteria>Hyphomicrobiales>Hyphomicrobiales_incertae_sedis>Enhydrobacter</td><td>GCA_902506225.1</td><td>Enhydrobacter_sp._8BJ</td><td>ENHYD8BJ_80433</td><td>457912..461628</td><td>2653137</td><td>conserved-hypothetical-protein-[Enhydrobacter-sp.-8BJ]</td></tr>
<tr><td>WP_271692167.1</td><td>Type1.BREX</td><td>ACET-><-?<-?<-?<-?||MULE-TRANSPOSASE-><-?||MULE-TRANSPOSASE-><- N6-METHYLASE -p<- DUF6079 *<- BrxB-BrxF-iSTAND <- BrxA-DNA-binding <- wHTH+WYL+WCX <-?|| OLD-ABC + TOPRIM ->?-><-LON-AAA</td><td>1174</td><td>Bacteria>Pseudomonadota>Alphaproteobacteria>Rhodobacterales>Roseobacteraceae>Sulfitobacter</td><td>GCF_028201715.1</td><td>Sulfitobacter_faviae</td><td>PL335_RS12830</td><td>2584668..2588192</td><td>1775881</td><td>BREX-system-P-loop-protein-BrxC-[Sulfitobacter-faviae]</td></tr>
<tr><td>WP_137104425.1</td><td>Type1.BREX</td><td><-wHTH+REase<-wHTH+McrB-AAA<- SIGMA+OB-Fold+AAA-ATPase+Lon-Protease-BrxL <- PglZ <-REase+AIPR<- N6-METHYLASE <- TOPRIM <- OLD-ABC <- DUF6079 *<- BrxB-BrxF-iSTAND <- BrxA-DNA-binding ||?->?-><-?<-?<-MODE-HTH+PBPII</td><td>1148</td><td>Bacteria>Pseudomonadota>Alphaproteobacteria>Rhodospirillales>Azospirillaceae>Azospirillum</td><td>GCF_005222185.1</td><td>Azospirillum_argentinense</td><td>D3867_15345</td><td>176394..179840</td><td>2970906</td><td>BREX-system-P-loop-protein-BrxC-[Azospirillum-argentinense]</td></tr>
<tr><td>RDE49751.1</td><td>Type1.BREX</td><td><-PIN+HTH<- SIGMA+OB-Fold+AAA-ATPase+Lon-Protease-BrxL <- PglZ <- TOPRIM <- OLD-ABC <- DUF6079 *<- BrxB-BrxF-iSTAND <- BrxA-DNA-binding ||?->?-><-?<-?||?-><-?<-Zn_Tnp_IS91+Y2_Tnp</td><td>1071</td><td>Bacteria>Pseudomonadota>Betaproteobacteria>Betaproteobacteria_incertae_sedis>Candidatus_Accumulibacter</td><td>GCA_003332265.1</td><td>Candidatus_Accumulibacter_meliphilus</td><td>DVS81_14920</td><td>17642..20857</td><td>2211374</td><td>MAG:-BREX-system-P-loop-protein-BrxC-[Candidatus-Accumulibacter-meliphilus]</td></tr>
<tr><td>WP_169070206.1</td><td>Type1.BREX</td><td>BrxA-DNA-binding -> BrxB-BrxF-iSTAND -> DUF6079 *-> OLD-ABC ->?-> OLD-ABC -> TOPRIM ->HTH+REase->?->HTH+REase->?-> SIGMA+OB-Fold+AAA-ATPase+Lon-Protease-BrxL -></td><td>1204</td><td>Bacteria>Pseudomonadota>Betaproteobacteria>Betaproteobacteria_incertae_sedis>Candidatus_Accumulibacter</td><td>GCF_012940005.1</td><td>Candidatus_Accumulibacter_contiguus</td><td>E4Q08_RS09460</td><td>34303..37917</td><td>2954381</td><td>BREX-system-P-loop-protein-BrxC-[Candidatus-Accumulibacter-contiguus]</td></tr>
<tr><td>MBN8451986.1</td><td>Type6.BREX</td><td><-KAP-glycosylase<-KAP-NTPase<- SIGMA+OB-Fold+AAA-ATPase+HKD-DNase-BrxL <- PglZ <- TOPRIM <- OLD-ABC <- N6-METHYLASE <- N6-METHYLASE <-GreA_GreB+SF1-SF2-Helicase<- DUF6079 *<- BrxB-BrxF-iSTAND <- BrxA-DNA-binding <- BrxE <- wHTH+WYL+WCX ||?->SMODS->MODE-HTH+PBPII-><-HTH_3||LysR-HTH-></td><td>1213</td><td>Bacteria>Pseudomonadota>Betaproteobacteria>Betaproteobacteria_incertae_sedis>Candidatus_Accumulibacter>unclassified_Candidatus_Accumulibacter</td><td>GCA_017302415.1</td><td>Candidatus_Accumulibacter_sp.</td><td>J0M31_02680</td><td>99081..102722</td><td>2053492</td><td>BREX-system-P-loop-protein-BrxC-[Candidatus-Accumulibacter-sp.]</td></tr>
<tr><td>WP_115063824.1</td><td>Type1.BREX</td><td>Phage_AlpA->?->TraJ-RHH-><-REase+NACHT<-?<-RADICAL-SAM<-DUF772+DDE_Tnp_1|| BrxA-DNA-binding -> BrxB-BrxF-iSTAND -> DUF6079 *-> N6-METHYLASE ->DUF3644-> OLD-ABC -> TOPRIM -> OLD-ABC ->HNH-> PglZ -> SIGMA+OB-Fold+AAA-ATPase+Lon-Protease-BrxL -><-Y1_Tnp</td><td>1155</td><td>Bacteria>Pseudomonadota>Betaproteobacteria>Burkholderiales>Alcaligenaceae>Oligella</td><td>GCF_900454315.1</td><td>Oligella_ureolytica</td><td>NCTC11998_02420</td><td>2456151..2459618</td><td>90244</td><td>BREX-system-P-loop-protein-BrxC-[Oligella-ureolytica]</td></tr>
<tr><td>NYT60768.1</td><td>Type1.BREX</td><td><-Phage_AlpA||YdaS_antitoxin-><- SIGMA+OB-Fold+AAA-ATPase+Lon-Protease-BrxL <- PglZ <-Schlafen+HATPase+wHTH<-SbcC<- TOPRIM <- OLD-ABC <- N6-METHYLASE <- DUF6079 *<- BrxB-BrxF-iSTAND <- BrxA-DNA-binding <- wHTH+WYL+WCX ||SbcC-></td><td>1146</td><td>Bacteria>Pseudomonadota>Betaproteobacteria>Burkholderiales>Alcaligenaceae>unclassified_Alcaligenaceae</td><td>GCA_013416445.1</td><td>Alcaligenaceae_bacterium</td><td>HOA66_00285</td><td>70051..73491</td><td>2021368</td><td>BREX-system-P-loop-protein-BrxC-[Alcaligenaceae-bacterium]</td></tr>
<tr><td>WP_059936887.1</td><td>Type6.BREX</td><td><-HIN-HTH<- PglZ <-ParB+HNH<- N6-METHYLASE <- OLD-ABC + TOPRIM <- N6-METHYLASE <-GreA_GreB+SF1-SF2-Helicase<- DUF6079 *<- BrxB-BrxF-iSTAND <- BrxA-DNA-binding <- BrxE <- wHTH+WYL+WCX ||ARG->?-><-HTH_19||LON-AAA-></td><td>1213</td><td>Bacteria>Pseudomonadota>Betaproteobacteria>Burkholderiales>Burkholderiaceae>Burkholderia</td><td>GCF_001528605.1</td><td>Burkholderia_multivorans</td><td>WK33_RS03230</td><td>12913..16554</td><td>87883</td><td>BREX-system-P-loop-protein-BrxC-[Burkholderia-multivorans]</td></tr>
<tr><td>WP_243738604.1</td><td>Type1.BREX</td><td><-Phage_integrase<-?||TRPR-HTH-> BrxA-DNA-binding -> BrxB-BrxF-iSTAND -> DUF6079 *-> N6-METHYLASE -> OLD-ABC -> TOPRIM -> PglZ -> SIGMA+OB-Fold+AAA-ATPase+Lon-Protease-BrxL -><-?<-?<-HISKIN</td><td>1224</td><td>Bacteria>Pseudomonadota>Betaproteobacteria>Burkholderiales>Burkholderiales_genera_incertae_sedis>Aquabacterium</td><td>GCF_004362855.1</td><td>Aquabacterium_commune</td><td>EV672_RS09250</td><td>184197..187871</td><td>70586</td><td>BREX-system-P-loop-protein-BrxC-[Aquabacterium-commune]</td></tr>
<tr><td>WP_143968626.1</td><td>Type6.BREX</td><td>PglZ -> SIGMA+OB-Fold+AAA-ATPase+HKD-DNase-BrxL ->||Phage_integrase->Phage_integrase->Phage_integrase->?-> BrxE -> BrxA-DNA-binding -> BrxB-BrxF-iSTAND -> DUF6079 *-> N6-METHYLASE -> OLD-ABC + TOPRIM -> N6-METHYLASE -> PglZ -> SIGMA+OB-Fold+AAA-ATPase+HKD-DNase-BrxL -></td><td>1210</td><td>Bacteria>Pseudomonadota>Betaproteobacteria>Burkholderiales>Burkholderiales_genera_incertae_sedis>Tepidimonas</td><td>GCF_007556755.1</td><td>Tepidimonas_fonticaldi</td><td>Tfont_RS05070</td><td>95231..98863</td><td>1101373</td><td>BREX-system-P-loop-protein-BrxC-[Tepidimonas-fonticaldi]</td></tr>
<tr><td>WP_008906812.1</td><td>Type1.BREX</td><td>XIS-HTH->?->?-><-?<-?||?-> BrxA-DNA-binding -> BrxB-BrxF-iSTAND -> DUF6079 *->ParD_antitoxin->RelE-ParE-> N6-METHYLASE -> OLD-ABC -> TOPRIM -> PglZ -> SIGMA+OB-Fold+AAA-ATPase+Lon-Protease-BrxL -><-CHTH<-RelE</td><td>1213</td><td>Bacteria>Pseudomonadota>Betaproteobacteria>Burkholderiales>Comamonadaceae>Acidovorax</td><td>GCF_000238595.1</td><td>Acidovorax_sp._NO-1</td><td>KYG_RS20325</td><td>5938..9579</td><td>512030</td><td>BREX-system-P-loop-protein-BrxC-[Acidovorax-sp.-NO-1]</td></tr>
<tr><td>WP_094097616.1</td><td>Type1.BREX</td><td><-Peptidase_C39_2<-?||?-><-?||XIS-HTH->Phage_X-> BrxA-DNA-binding -> BrxB-BrxF-iSTAND -> DUF6079 *->HNH->HEPN->ParD_antitoxin->RelE-ParE-> N6-METHYLASE -> OLD-ABC -> TOPRIM ->ZnR+ TOPRIM -> PglZ -></td><td>1214</td><td>Bacteria>Pseudomonadota>Betaproteobacteria>Burkholderiales>Comamonadaceae>Acidovorax</td><td>GCF_002157145.1</td><td>Acidovorax_carolinensis</td><td>CBP34_RS06480</td><td>1379414..1383058</td><td>553814</td><td>BREX-system-P-loop-protein-BrxC-[Acidovorax-carolinensis]</td></tr>
<tr><td>GAO25248.1</td><td>Type1.BREX</td><td>HTH_3-><- SIGMA+OB-Fold+AAA-ATPase+Lon-Protease-BrxL <- PglZ <-ZnR+ TOPRIM <- TOPRIM <- OLD-ABC <- N6-METHYLASE <-HEPN<-HNH<- DUF6079 *<- BrxB-BrxF-iSTAND <- BrxA-DNA-binding <-HAD+CHTH<-CHTH||?->AAA-ATPase-p->?-><-AAA-ATPase</td><td>1205</td><td>Bacteria>Pseudomonadota>Betaproteobacteria>Burkholderiales>Comamonadaceae>Alicycliphilus</td><td>GCA_000974645.1</td><td>Alicycliphilus_sp._B1</td><td>ALISP_5068</td><td>1538304..1541921</td><td>1603291</td><td>ATPase-like-protein-[Alicycliphilus-sp.-B1]</td></tr>
<tr><td>WP_054075976.1</td><td>Type1.BREX</td><td><-HNH<-?<- wHTH+WYL+WCX <-?<-?<-?||HTH_19-> BrxA-DNA-binding -> BrxB-BrxF-iSTAND -> DUF6079 *-> N6-METHYLASE -> OLD-ABC -> TOPRIM -></td><td>1209</td><td>Bacteria>Pseudomonadota>Betaproteobacteria>Burkholderiales>Comamonadaceae</td><td>GCF_001294075.1</td><td>Comamonas</td><td>KF712C_RS25790</td><td>15422..19051</td><td>283</td><td>MULTISPECIES:-BREX-system-P-loop-protein-BrxC-[Comamonas]</td></tr>
<tr><td>WP_104799767.1</td><td>Type1.BREX</td><td><- SIGMA+OB-Fold+AAA-ATPase+Lon-Protease-BrxL <- PglZ <-?<-?<-HTH+REase<- TOPRIM <- OLD-ABC <- N6-METHYLASE <-?<- DUF6079 *<- BrxB-BrxF-iSTAND <- BrxA-DNA-binding <- wHTH+WYL+WCX <-tRNA||?-><-?<-ABhydrolase</td><td>1202</td><td>Bacteria>Pseudomonadota>Betaproteobacteria>Burkholderiales>Comamonadaceae>Limnohabitans</td><td>GCF_002943475.1</td><td>Limnohabitans_sp._TS-CS-82</td><td>C5F52_RS18160</td><td>199496..203104</td><td>2094193</td><td>BREX-system-P-loop-protein-BrxC-[Limnohabitans-sp.-TS-CS-82]</td></tr>
<tr><td>WP_108373352.1</td><td>Type1.BREX</td><td><- TOPRIM <- OLD-ABC <-?<- TOPRIM <- OLD-ABC <- TOPRIM <- OLD-ABC <- N6-METHYLASE <-Schlafen+HATPase+wHTH<- DUF6079 *<- BrxB-BrxF-iSTAND <- BrxA-DNA-binding <- wHTH+WYL+WCX ||?->?->?-><-?<-?<-SPOUT</td><td>1209</td><td>Bacteria>Pseudomonadota>Betaproteobacteria>Burkholderiales>Comamonadaceae>Limnohabitans</td><td>GCF_003063315.1</td><td>Limnohabitans_sp._T6-5</td><td>B9Z51_RS07570</td><td>412563..416192</td><td>1100724</td><td>BREX-system-P-loop-protein-BrxC-[Limnohabitans-sp.-T6-5]</td></tr>
<tr><td>WP_096234088.1</td><td>Type1.BREX</td><td>HTH_3->?->?->?-> BrxA-DNA-binding -> BrxB-BrxF-iSTAND -> DUF6079 *-> N6-METHYLASE -> OLD-ABC -> TOPRIM -> PglZ -> SIGMA+OB-Fold+AAA-ATPase+Lon-Protease-BrxL ->HEPN-><-TPR</td><td>1204</td><td>Bacteria>Pseudomonadota>Betaproteobacteria>Burkholderiales>Oxalobacteraceae>Janthinobacterium</td><td>GCF_002327145.1</td><td>Janthinobacterium_svalbardensis</td><td>CNX70_RS06880</td><td>1564323..1567937</td><td>368607</td><td>BREX-system-P-loop-protein-BrxC-[Janthinobacterium-svalbardensis]</td></tr>
<tr><td>MBP7846551.1</td><td>Type1.BREX</td><td><-TrwC+SF1-SF2-Helicase<-SF1-SF2-Helicase+REase<- SIGMA+OB-Fold+AAA-ATPase+Lon-Protease-BrxL <- PglZ <- TOPRIM <- OLD-ABC <- N6-METHYLASE -p<- N6-METHYLASE <- DUF6079 *<- BrxB-BrxF-iSTAND <- BrxA-DNA-binding <-SWIM+DUF3670+SF1-SF2-Helicase<-?<-SWIM+DUF3670+SF1-SF2-Helicase</td><td>1193</td><td>Bacteria>Pseudomonadota>Betaproteobacteria>Burkholderiales>unclassified_Burkholderiales</td><td>GCA_018001735.1</td><td>Burkholderiales_bacterium</td><td>KAZ94_00285</td><td>68005..71586</td><td>1891238</td><td>BREX-system-P-loop-protein-BrxC-[Burkholderiales-bacterium]</td></tr>
<tr><td>MBX9598966.1</td><td>Type1.BREX</td><td>BrxA-DNA-binding -> BrxB-BrxF-iSTAND -> DUF6079 *-> N6-METHYLASE -> TOPRIM -> OLD-ABC -> TOPRIM -> PglZ -> SIGMA+OB-Fold+AAA-ATPase+Lon-Protease-BrxL ->?->Radical_SAM->PIN-></td><td>1193</td><td>Bacteria>Pseudomonadota>Betaproteobacteria>Burkholderiales>unclassified_Burkholderiales</td><td>GCA_019748155.1</td><td>Burkholderiales_bacterium</td><td>K2X04_10405</td><td>1415..4996</td><td>1891238</td><td>BREX-system-P-loop-protein-BrxC-[Burkholderiales-bacterium]</td></tr>
<tr><td>WP_047248551.1</td><td>Type1.BREX</td><td>SbcC-><- SIGMA+OB-Fold+AAA-ATPase+Lon-Protease-BrxL <- PglZ <- PglZ -p<- TOPRIM <- OLD-ABC <- N6-METHYLASE <- DUF6079 *<- BrxB-BrxF-iSTAND <- BrxA-DNA-binding <- wHTH+WYL+WCX ||DUF932->?->?->?-><-HTH_19</td><td>1228</td><td>Bacteria>Pseudomonadota>Betaproteobacteria>Neisseriales>Chromobacteriaceae>Chromobacterium_group</td><td>GCF_001020585.1</td><td>Chromobacterium_subtsugae</td><td>AA094_RS04310</td><td>55231..58917</td><td>251747</td><td>BREX-system-P-loop-protein-BrxC-[Chromobacterium-subtsugae]</td></tr>
<tr><td>MCG9052529.1</td><td>Type1.BREX</td><td>HIN-HTH-><- SIGMA+OB-Fold+AAA-ATPase+Lon-Protease-BrxL <- PglZ <- TOPRIM <- OLD-ABC <- N6-METHYLASE <-DOC+HTH<-Couple_hipA+HipA_C<-TRPR-HTH<- DUF6079 *<- BrxB-BrxF-iSTAND <- BrxA-DNA-binding <- wHTH+WYL+WCX -p<-?<-XIS-HTH||?-><-Arm-DNA-bind_3+Phage_integrase</td><td>1224</td><td>Bacteria>Pseudomonadota>Betaproteobacteria>Neisseriales>Chromobacteriaceae>Laribacter</td><td>GCA_022394235.1</td><td>Laribacter_hongkongensis</td><td>LH447_05335</td><td>18538..22212</td><td>168471</td><td>BREX-system-P-loop-protein-BrxC-[Laribacter-hongkongensis]</td></tr>
<tr><td>MCG9059785.1</td><td>Type1.BREX</td><td>DDE_Tnp_1-><-?||ARG-><- SIGMA+OB-Fold+AAA-ATPase+Lon-Protease-BrxL <- PglZ <- TOPRIM <- OLD-ABC <- N6-METHYLASE <-HKD-DNase<- DUF6079 *<- BrxB-BrxF-iSTAND <- BrxA-DNA-binding <- wHTH+WYL+WCX ||||?->DOC-><-DUF772+DDE_Tnp_1||?->||AAA-ATPase-><-CRISPR_Cas6</td><td>1217</td><td>Bacteria>Pseudomonadota>Betaproteobacteria>Neisseriales>Chromobacteriaceae>Laribacter</td><td>GCA_022394335.1</td><td>Laribacter_hongkongensis</td><td>LH452_12755</td><td>30373..34026</td><td>168471</td><td>BREX-system-P-loop-protein-BrxC-[Laribacter-hongkongensis]</td></tr>
<tr><td>WP_275663441.1</td><td>Type1.BREX</td><td>wHTH+WYL+WCX -p-> BrxA-DNA-binding -> BrxB-BrxF-iSTAND -> DUF6079 *->TRPR-HTH->Couple_hipA+HipA_C->DOC+HTH-> N6-METHYLASE -> OLD-ABC -> TOPRIM -> PglZ -> SIGMA+OB-Fold+AAA-ATPase+Lon-Protease-BrxL -><-HIN-HTH</td><td>1229</td><td>Bacteria>Pseudomonadota>Betaproteobacteria>Neisseriales>Chromobacteriaceae>Laribacter</td><td>GCF_022394235.1</td><td>Laribacter_hongkongensis</td><td>LH442_RS13325</td><td>2333..6022</td><td>168471</td><td>BREX-system-P-loop-protein-BrxC-[Laribacter-hongkongensis]</td></tr>
<tr><td>WP_239898705.1</td><td>Type1.BREX</td><td>DDE_Tnp_1-><-?||ARG-><- SIGMA+OB-Fold+AAA-ATPase+Lon-Protease-BrxL <- PglZ <- TOPRIM <- OLD-ABC <- N6-METHYLASE <-HKD-DNase<- DUF6079 *<- BrxB-BrxF-iSTAND <- BrxA-DNA-binding <- wHTH+WYL+WCX ||||?->DOC-><-DUF772+DDE_Tnp_1||?->||AAA-ATPase-><-Cas_Cas1</td><td>1222</td><td>Bacteria>Pseudomonadota>Betaproteobacteria>Neisseriales>Chromobacteriaceae>Laribacter</td><td>GCF_022394335.1</td><td>Laribacter_hongkongensis</td><td>LH452_RS12745</td><td>30373..34041</td><td>168471</td><td>BREX-system-P-loop-protein-BrxC-[Laribacter-hongkongensis]</td></tr>
<tr><td>WP_239900332.1</td><td>Type1.BREX</td><td>HIN-HTH-><- SIGMA+OB-Fold+AAA-ATPase+Lon-Protease-BrxL <- PglZ <- TOPRIM <- OLD-ABC <- N6-METHYLASE <-DOC+HTH<-Couple_hipA+HipA_C<-TRPR-HTH<- DUF6079 *<- BrxB-BrxF-iSTAND <- BrxA-DNA-binding <- wHTH+WYL+WCX -p<-?<-XIS-HTH||?-><-Arm-DNA-bind_3+Phage_integrase</td><td>1224</td><td>Bacteria>Pseudomonadota>Betaproteobacteria>Neisseriales>Chromobacteriaceae>Laribacter</td><td>-</td><td>Laribacter_hongkongensis</td><td>LH447_05335</td><td>18538..22212</td><td>168471</td><td>BREX-system-P-loop-protein-BrxC-[Laribacter-hongkongensis]</td></tr>
<tr><td>WP_239891840.1</td><td>Type1.BREX</td><td>DDE_Tnp_1-><-?||ARG-><- SIGMA+OB-Fold+AAA-ATPase+Lon-Protease-BrxL <- PglZ <- TOPRIM <- OLD-ABC <- N6-METHYLASE <-HKD-DNase<- DUF6079 *<- BrxB-BrxF-iSTAND <- BrxA-DNA-binding <- wHTH+WYL+WCX ||||?->DOC-><-DUF772+DDE_Tnp_1||?->||AAA-ATPase-><-CRISPR_Cas6</td><td>1217</td><td>Bacteria>Pseudomonadota>Betaproteobacteria>Neisseriales>Chromobacteriaceae>Laribacter</td><td>-</td><td>Laribacter_hongkongensis</td><td>LH452_12755</td><td>30373..34026</td><td>168471</td><td>BREX-system-P-loop-protein-BrxC-[Laribacter-hongkongensis]</td></tr>
<tr><td>WP_239900962.1</td><td>Type1.BREX</td><td>TOPRIM +Primase-><- SIGMA+OB-Fold+AAA-ATPase+Lon-Protease-BrxL <- PglZ <- TOPRIM <- OLD-ABC <- N6-METHYLASE <-DOC+HTH<-Couple_hipA+HipA_C<-TRPR-HTH<- DUF6079 *<- BrxB-BrxF-iSTAND <- BrxA-DNA-binding <- wHTH+WYL+WCX <-?||?->35exo->?-><-Calcineurin-><-Arm-DNA-bind_3+Phage_integrase</td><td>1224</td><td>Bacteria>Pseudomonadota>Betaproteobacteria>Neisseriales>Chromobacteriaceae>Laribacter</td><td>-</td><td>Laribacter_hongkongensis</td><td>LH454_02040</td><td>72475..76149</td><td>168471</td><td>BREX-system-P-loop-protein-BrxC-[Laribacter-hongkongensis]</td></tr>
<tr><td>WP_013292857.1</td><td>Type1.BREX</td><td>VirD4-TraG+VirB4-FtsK->?->?->Peptidase_M50->?->?-><-?|| BrxA-DNA-binding -> BrxB-BrxF-iSTAND -> DUF6079 *->AbiEi_4->AbiEii-> N6-METHYLASE ->HTH+REase->REase->?-> OLD-ABC -> TOPRIM ->Schlafen+HATPase+wHTH-></td><td>1207</td><td>Bacteria>Pseudomonadota>Betaproteobacteria>Nitrosomonadales>Gallionellaceae>Gallionella</td><td>GCF_000145255.1</td><td>Gallionella_capsiferriformans</td><td>GALF_RS04415</td><td>928777..932400</td><td>370405</td><td>BREX-system-P-loop-protein-BrxC-[Gallionella-capsiferriformans]</td></tr>
<tr><td>TXI44178.1</td><td>Type1.BREX</td><td><-EVE+HNH<- wHTH+WYL+WCX <-REase+HTH+35exo+Small-B-barrel+SF1-SF2-Helicase||?-> BrxA-DNA-binding -> BrxB-BrxF-iSTAND -> DUF6079 *-> N6-METHYLASE ->SH3-> OLD-ABC + TOPRIM ->ZNR+RmuC->DOC+HTH-> PglZ -> SIGMA+OB-Fold+AAA-ATPase+Lon-Protease-BrxL -></td><td>1198</td><td>Bacteria>Pseudomonadota>Betaproteobacteria>Nitrosomonadales>Methylophilaceae>Methylophilus</td><td>GCA_008015755.1</td><td>Methylophilus_sp.</td><td>E6Q52_09965</td><td>12844..16440</td><td>29541</td><td>MAG:-BREX-system-P-loop-protein-BrxC-[Methylophilus-sp.]</td></tr>
<tr><td>NOT17080.1</td><td>Type1.BREX</td><td><- PglZ <-SF1-SF2-Helicase<-REase<-HTH+REase<- TOPRIM <- OLD-ABC <- N6-METHYLASE <-AbiEii<-AbiEi_4<- DUF6079 *<- BrxB-BrxF-iSTAND <- BrxA-DNA-binding <- wHTH+WYL+WCX <-SbcC+CCDC158<-SbcC+CCDC158<-Calcineurin+Calcineurin-SbcD<-LZ_Tnp_IS66+zf-IS66+DDE_Tnp_IS66+DDE_Tnp_IS66_C<-TnpB_IS66<-HIN-HTH</td><td>1202</td><td>Bacteria>Pseudomonadota>Betaproteobacteria>Nitrosomonadales>Sulfuricellaceae>Sulfuriferula</td><td>GCA_013140765.1</td><td>Sulfuriferula_sp.</td><td>HOP20_03285</td><td>21073..24681</td><td>2025307</td><td>BREX-system-P-loop-protein-BrxC-[Sulfuriferula-sp.]</td></tr>
<tr><td>MCK0506222.1</td><td>Type1.BREX</td><td><-METHYLASE<- SIGMA+OB-Fold+AAA-ATPase+Lon-Protease-BrxL <- PglZ <-PIWI<- OLD-ABC <- TOPRIM <- OLD-ABC <- N6-METHYLASE ||RHH_6-><- DUF6079 *<- BrxB-BrxF-iSTAND <- BrxA-DNA-binding <- wHTH+WYL+WCX ||?->?-><-?<-35exo||Peptidase_M48_N+Peptidase_M48->S1+GTPase-AIG-></td><td>1186</td><td>Bacteria>Pseudomonadota>Betaproteobacteria>Rhodocyclales>Rhodocyclaceae>Aromatoleum</td><td>GCA_012910705.2</td><td>Aromatoleum_anaerobium</td><td>GD606_004945</td><td>104237..107797</td><td>182180</td><td>BREX-system-P-loop-protein-BrxC-[Aromatoleum-anaerobium]</td></tr>
<tr><td>WP_011238768.1</td><td>Type1.BREX</td><td><-Peptidase_M48_N+Peptidase_M48||35exo-><-?<-?<-?||?-> wHTH+WYL+WCX -> BrxA-DNA-binding -> BrxB-BrxF-iSTAND -> DUF6079 *-><-PIN<-RHH_6||?-> N6-METHYLASE -> OLD-ABC ->HNH-> OLD-ABC -> TOPRIM -> OLD-ABC -></td><td>1203</td><td>Bacteria>Pseudomonadota>Betaproteobacteria>Rhodocyclales>Rhodocyclaceae>Aromatoleum</td><td>GCF_000025965.1</td><td>Aromatoleum_aromaticum</td><td>EBN1_RS14750</td><td>3092506..3096117</td><td>551760</td><td>BREX-system-P-loop-protein-BrxC-[Aromatoleum-aromaticum]</td></tr>
<tr><td>WP_169117250.1</td><td>Type1.BREX</td><td><-METHYLASE<- SIGMA+OB-Fold+AAA-ATPase+Lon-Protease-BrxL <- PglZ <-PIWI<- OLD-ABC <- TOPRIM <- OLD-ABC <- N6-METHYLASE ||RHH_6-><- DUF6079 *<- BrxB-BrxF-iSTAND <- BrxA-DNA-binding <- wHTH+WYL+WCX ||?->?-><-?<-35exo||Peptidase_M48_N+Peptidase_M48->S1+GTPase-AIG-></td><td>1203</td><td>Bacteria>Pseudomonadota>Betaproteobacteria>Rhodocyclales>Rhodocyclaceae>Aromatoleum</td><td>GCF_012910705.2</td><td>Aromatoleum_anaerobium</td><td>GD606_RS04945</td><td>104237..107848</td><td>182180</td><td>BREX-system-P-loop-protein-BrxC-[Aromatoleum-anaerobium]</td></tr>
<tr><td>TRZ91915.1</td><td>Type6.BREX</td><td><- PglZ <- TOPRIM <- OLD-ABC <-ParB+HNH<- TOPRIM <- OLD-ABC <- N6-METHYLASE <- DUF6079 *<- BrxB-BrxF-iSTAND <- BrxA-DNA-binding <- BrxE <-XIS-HTH<-?<-?<-?|| wHTH+WYL+WCX -></td><td>1212</td><td>Bacteria>Pseudomonadota>Betaproteobacteria>Rhodocyclales>Rhodocyclaceae>unclassified_Rhodocyclaceae</td><td>GCA_007280105.1</td><td>Rhodocyclaceae_bacterium</td><td>D4R84_13335</td><td>10594..14232</td><td>1898103</td><td>BREX-system-P-loop-protein-BrxC-[Rhodocyclaceae-bacterium]</td></tr>
<tr><td>CAH1748211.1</td><td>Type1.BREX</td><td><-DUF3584+DUF5401<-JetB<-MukF<-?||Terminase_5+HTH_33+DDE_3-><-?|| wHTH+WYL+WCX -> BrxA-DNA-binding -> BrxB-BrxF-iSTAND -> DUF6079 *->HKD-DNase-> N6-METHYLASE ->TIR->SbcC+NACHT-> OLD-ABC -> TOPRIM ->PIWI-> PglZ -> SIGMA+OB-Fold+AAA-ATPase+Lon-Protease-BrxL -></td><td>1203</td><td>Bacteria>Pseudomonadota>Betaproteobacteria>Rhodocyclales>Zoogloeaceae>Thauera</td><td>GCA_932126575.1</td><td>Thauera_sp._PIV-1</td><td>C4PIVTH_3071</td><td>3140142..3143753</td><td>204461</td><td>ATPase-like-protein-[Thauera-sp.-PIV-1]</td></tr>
<tr><td>WP_247725480.1</td><td>Type1.BREX</td><td><- SIGMA+OB-Fold+AAA-ATPase+Lon-Protease-BrxL <- PglZ <-PIWI<- TOPRIM <- OLD-ABC <- N6-METHYLASE ||RHH_6->PIN-><- DUF6079 *<- BrxB-BrxF-iSTAND <- BrxA-DNA-binding <- wHTH+WYL+WCX</td><td>1203</td><td>Bacteria>Pseudomonadota>Betaproteobacteria>Rhodocyclales>Zoogloeaceae>Thauera</td><td>GCF_023093595.1</td><td>Thauera_aromatica</td><td>MXC99_RS10910</td><td>53320..56931</td><td>59405</td><td>BREX-system-P-loop-protein-BrxC-[Thauera-aromatica]</td></tr>
<tr><td>MBI3372694.1</td><td>Type1.BREX</td><td><-AAA-ATPase||?->?-><-?|| BrxA-DNA-binding -> BrxB-BrxF-iSTAND -> DUF6079 *-> OLD-ABC -> TOPRIM ->?->?->SbcC-> N6-METHYLASE ->ArdCN+MPTase-> PglZ -> SIGMA+OB-Fold+AAA-ATPase+Lon-Protease-BrxL -></td><td>1148</td><td>Bacteria>Pseudomonadota>Betaproteobacteria>unclassified_Betaproteobacteria</td><td>GCA_016195915.1</td><td>Betaproteobacteria_bacterium</td><td>HY017_13195</td><td>52819..56265</td><td>1891241</td><td>BREX-system-P-loop-protein-BrxC-[Betaproteobacteria-bacterium]</td></tr>
<tr><td>GHT91750.1</td><td>Type6.BREX</td><td><- SIGMA+OB-Fold+AAA-ATPase+HKD-DNase-BrxL <- PglZ <-DOC+HTH<- TOPRIM <- OLD-ABC <- N6-METHYLASE -p<- N6-METHYLASE <-REase+AIPR<- DUF6079 *<- BrxB-BrxF-iSTAND <- BrxA-DNA-binding <- BrxE <-XIS-HTH||HTH_3->Couple_hipA+HipA_C->Phage_integrase->RelE-></td><td>1209</td><td>Bacteria>Pseudomonadota>Betaproteobacteria>unclassified_Betaproteobacteria</td><td>GCA_026000515.1</td><td>Betaproteobacteria_bacterium</td><td>FACS1894116_00820</td><td>42503..46132</td><td>1891241</td><td>hypothetical-protein-FACS1894116_00820-[Betaproteobacteria-bacterium]</td></tr>
<tr><td>WP_219254923.1</td><td>Type1.BREX</td><td><- PglZ <- TOPRIM <- OLD-ABC <- N6-METHYLASE -p<-HerA/FtsK<-SIR2<- N6-METHYLASE <- TOPRIM <- OLD-ABC <- DUF6079 *<- BrxB-BrxF-iSTAND <- BrxA-DNA-binding <-RHS<-?|| wHTH+WYL+WCX -><-?||LexA-protease->?-><-DOC+HTH</td><td>1226</td><td>Bacteria>Pseudomonadota>Gammaproteobacteria>Aeromonadales>Aeromonadaceae>Aeromonas</td><td>-</td><td>Aeromonas_hydrophila</td><td>GPV00_14065</td><td>286470..290150</td><td>644</td><td>BREX-system-P-loop-protein-BrxC-[Aeromonas-hydrophila]</td></tr>
<tr><td>BBR09393.1</td><td>Type1.BREX</td><td>Arm-DNA-bind_3+Phage_integrase-><-?||?->?-> wHTH+WYL+WCX ||?->RHS-> BrxA-DNA-binding -> BrxB-BrxF-iSTAND -> DUF6079 *-> OLD-ABC -> TOPRIM ->ParB+HNH-> N6-METHYLASE -> OLD-ABC -> TOPRIM -> PglZ -> SIGMA+OB-Fold+AAA-ATPase+Lon-Protease-BrxL -></td><td>1226</td><td>Bacteria>Pseudomonadota>Gammaproteobacteria>Aeromonadales>Aeromonadaceae>Aeromonas</td><td>GCA_014158455.1</td><td>Aeromonas_caviae</td><td>WP3S18E02_10540</td><td>1118346..1122026</td><td>648</td><td>hypothetical-protein-WP3S18E02_10540-[Aeromonas-caviae]</td></tr>
<tr><td>HAU4888102.1</td><td>Type1.BREX</td><td><- N6-METHYLASE -p<- TOPRIM <- OLD-ABC <- DUF6079 *<- BrxB-BrxF-iSTAND <- BrxA-DNA-binding <-RHS<-ZnR+ZnR|| wHTH+WYL+WCX -><-?||LexA-protease->?-><-DOC+HTH</td><td>1231</td><td>Bacteria>Pseudomonadota>Gammaproteobacteria>Aeromonadales>Aeromonadaceae>Aeromonas</td><td>GCA_016067875.1</td><td>Aeromonas_hydrophila</td><td>OD804_RS12725</td><td>3923..7618</td><td>644</td><td>BREX-system-P-loop-protein-BrxC-[Aeromonas-hydrophila]</td></tr>
<tr><td>MBS4697267.1</td><td>Type1.BREX</td><td><- SIGMA+OB-Fold+AAA-ATPase+Lon-Protease-BrxL <- PglZ <- TOPRIM <- OLD-ABC <- N6-METHYLASE <-ParB+HNH<- TOPRIM <- OLD-ABC <- DUF6079 *<- BrxB-BrxF-iSTAND <- BrxA-DNA-binding <-RHS</td><td>1226</td><td>Bacteria>Pseudomonadota>Gammaproteobacteria>Aeromonadales>Aeromonadaceae>Aeromonas</td><td>GCA_018359965.1</td><td>Aeromonas_allosaccharophila</td><td>J4G65_17610</td><td>87511..91191</td><td>656</td><td>BREX-system-P-loop-protein-BrxC-[Aeromonas-allosaccharophila]</td></tr>
<tr><td>MBW3845327.1</td><td>Type1.BREX</td><td><- PglZ <- TOPRIM <- OLD-ABC <- N6-METHYLASE -p<-HerA/FtsK<-SIR2<- N6-METHYLASE <- TOPRIM <- OLD-ABC <- DUF6079 *<- BrxB-BrxF-iSTAND <- BrxA-DNA-binding <-RHS<-?|| wHTH+WYL+WCX -><-?||LexA-protease->?-><-DOC+HTH</td><td>1226</td><td>Bacteria>Pseudomonadota>Gammaproteobacteria>Aeromonadales>Aeromonadaceae>Aeromonas</td><td>GCA_019348915.1</td><td>Aeromonas_hydrophila</td><td>GPV00_14065</td><td>286470..290150</td><td>644</td><td>BREX-system-P-loop-protein-BrxC-[Aeromonas-hydrophila]</td></tr>
<tr><td>WP_042011559.1</td><td>Type1.BREX</td><td><- SIGMA+OB-Fold+AAA-ATPase+Lon-Protease-BrxL <- PglZ <- TOPRIM <- OLD-ABC <-?<-DUF3944<- N6-METHYLASE <- TOPRIM <- OLD-ABC <- DUF6079 *<- BrxB-BrxF-iSTAND <- BrxA-DNA-binding <-RHS<-HKD-DNase<-Tnp_DNA_bind+DDE_Tnp_1+Dimer_Tnp_Tn5</td><td>1239</td><td>Bacteria>Pseudomonadota>Gammaproteobacteria>Aeromonadales>Aeromonadaceae>Aeromonas</td><td>GCF_000819885.1</td><td>Aeromonas_fluvialis</td><td>BN1125_RS07645</td><td>79425..83144</td><td>591962</td><td>BREX-system-P-loop-protein-BrxC-[Aeromonas-fluvialis]</td></tr>
<tr><td>WP_059169685.1</td><td>Type1.BREX</td><td><- PglZ <- N6-METHYLASE -p<-HEPN<-RVT<- N6-METHYLASE <-?<-HNH<- TOPRIM <- OLD-ABC <- DUF6079 *<- BrxB-BrxF-iSTAND <- BrxA-DNA-binding <-RHS<-ARG<-RHS||LexA-protease->?-><-HDRnase<-Arm-DNA-bind_2+Phage_int_SAM_3+Phage_integrase</td><td>1240</td><td>Bacteria>Pseudomonadota>Gammaproteobacteria>Aeromonadales>Aeromonadaceae>Aeromonas</td><td>GCF_001481535.2</td><td>Aeromonas_salmonicida</td><td>VO70_RS19205</td><td>18975..22697</td><td>645</td><td>BREX-system-P-loop-protein-BrxC-[Aeromonas-salmonicida]</td></tr>
<tr><td>WP_087756130.1</td><td>Type1.BREX</td><td><- SIGMA+OB-Fold+AAA-ATPase+Lon-Protease-BrxL <- PglZ <-?<-DUF3944<- N6-METHYLASE -p<- N6-METHYLASE <-ParB+HNH<- TOPRIM <- OLD-ABC <- DUF6079 *<- BrxB-BrxF-iSTAND <- BrxA-DNA-binding <-RHS||ZnR->HEPN-><-?|| wHTH+WYL+WCX -><-sGTP</td><td>1231</td><td>Bacteria>Pseudomonadota>Gammaproteobacteria>Aeromonadales>Aeromonadaceae>Aeromonas</td><td>GCF_002180335.1</td><td>Aeromonas_salmonicida</td><td>O23A_RS10590</td><td>2337193..2340888</td><td>645</td><td>BREX-system-P-loop-protein-BrxC-[Aeromonas-salmonicida]</td></tr>
<tr><td>WP_101617899.1</td><td>Type1.BREX</td><td><-TetR-HTH<-Phage_integrase+Phage_integrase<-Phage_integrase+Phage_integrase<-Phage_integrase+Phage_integrase<-LexA-protease||?->RHS-> BrxA-DNA-binding -> BrxB-BrxF-iSTAND -> DUF6079 *-> OLD-ABC -> TOPRIM -> N6-METHYLASE -> N6-METHYLASE -p->HEPN-> N6-METHYLASE -p-><-AAA-ATPase<-HTH+rve||DUF3944-></td><td>1231</td><td>Bacteria>Pseudomonadota>Gammaproteobacteria>Aeromonadales>Aeromonadaceae>Aeromonas</td><td>GCF_002850695.3</td><td>Aeromonas_hydrophila</td><td>C1A23_RS07645</td><td>1215070..1218765</td><td>644</td><td>BREX-system-P-loop-protein-BrxC-[Aeromonas-hydrophila]</td></tr>
<tr><td>WP_123173885.1</td><td>Type1.BREX</td><td>DOC+HTH-><-?<-LexA-protease||?-><- wHTH+WYL+WCX ||ZnR+ZnR->RHS-> BrxA-DNA-binding -> BrxB-BrxF-iSTAND -> DUF6079 *-> OLD-ABC -> TOPRIM -> N6-METHYLASE -> N6-METHYLASE ->DUF3944->?-><-HTH_Tnp_1+HTH_21+rve+rve_3</td><td>1231</td><td>Bacteria>Pseudomonadota>Gammaproteobacteria>Aeromonadales>Aeromonadaceae>Aeromonas</td><td>GCF_003722175.1</td><td>Aeromonas_veronii</td><td>EFI48_RS19250</td><td>4187979..4191674</td><td>654</td><td>BREX-system-P-loop-protein-BrxC-[Aeromonas-veronii]</td></tr>
<tr><td>WP_128297388.1</td><td>Type1.BREX</td><td>RHS-> BrxA-DNA-binding -> BrxB-BrxF-iSTAND -> DUF6079 *-> OLD-ABC -> TOPRIM -> N6-METHYLASE -></td><td>1225</td><td>Bacteria>Pseudomonadota>Gammaproteobacteria>Aeromonadales>Aeromonadaceae>Aeromonas</td><td>GCF_004024195.1</td><td>Aeromonas_caviae</td><td>DN612_RS19420</td><td>2423..6100</td><td>648</td><td>BREX-system-P-loop-protein-BrxC-[Aeromonas-caviae]</td></tr>
<tr><td>WP_240039238.1</td><td>Type1.BREX</td><td><- PglZ <- TOPRIM <- OLD-ABC <- N6-METHYLASE -p<-HerA/FtsK<-SIR2<- N6-METHYLASE <- TOPRIM <- OLD-ABC <- DUF6079 *<- BrxB-BrxF-iSTAND <- BrxA-DNA-binding <-RHS<-?|| wHTH+WYL+WCX -><-?||LexA-protease->?-><-DOC+HTH</td><td>1280</td><td>Bacteria>Pseudomonadota>Gammaproteobacteria>Aeromonadales>Aeromonadaceae>Aeromonas</td><td>GCF_006243275.1</td><td>Aeromonas_hydrophila</td><td>CF138_RS14425</td><td>159803..163645</td><td>644</td><td>BREX-system-P-loop-protein-BrxC-[Aeromonas-hydrophila]</td></tr>
<tr><td>WP_160837717.1</td><td>Type1.BREX</td><td><- N6-METHYLASE -p<-TIR<-HEPN<- N6-METHYLASE <- TOPRIM <- OLD-ABC <- DUF6079 *<- BrxB-BrxF-iSTAND <- BrxA-DNA-binding <-RHS<-ZnR<-ARG<-RHS||LexA-protease->?-><-HDRnase</td><td>1231</td><td>Bacteria>Pseudomonadota>Gammaproteobacteria>Aeromonadales>Aeromonadaceae>Aeromonas</td><td>GCF_009831085.1</td><td>Aeromonas_caviae</td><td>GR302_RS08560</td><td>7464..11159</td><td>648</td><td>BREX-system-P-loop-protein-BrxC-[Aeromonas-caviae]</td></tr>
<tr><td>WP_159144201.1</td><td>Type1.BREX</td><td><- SIGMA+OB-Fold+AAA-ATPase+Lon-Protease-BrxL <- PglZ <- TOPRIM <- OLD-ABC <-?<-DUF3944<- N6-METHYLASE <- N6-METHYLASE <-ParB+HNH<- DUF6079 *<- BrxB-BrxF-iSTAND <- BrxA-DNA-binding <-RHS<-?||LexA-protease->?-><-RADICAL-SAM||ANK+ANK+ANK-><-ZnR+ZnR</td><td>1231</td><td>Bacteria>Pseudomonadota>Gammaproteobacteria>Aeromonadales>Aeromonadaceae>Aeromonas</td><td>GCF_009834065.1</td><td>Aeromonas_veronii</td><td>GRF56_RS18105</td><td>3933894..3937589</td><td>654</td><td>BREX-system-P-loop-protein-BrxC-[Aeromonas-veronii]</td></tr>
<tr><td>WP_232090117.1</td><td>Type1.BREX</td><td>XIS-HTH->?->?-><- wHTH+WYL+WCX ||?->RHS-> BrxA-DNA-binding -> BrxB-BrxF-iSTAND -> DUF6079 *-> OLD-ABC -> TOPRIM ->ParB+HNH-> N6-METHYLASE -> OLD-ABC -> TOPRIM -> PglZ -> SIGMA+OB-Fold+AAA-ATPase+Lon-Protease-BrxL -></td><td>1281</td><td>Bacteria>Pseudomonadota>Gammaproteobacteria>Aeromonadales>Aeromonadaceae>Aeromonas</td><td>GCF_014158455.1</td><td>Aeromonas_caviae</td><td>H7X73_RS05535</td><td>1118181..1122026</td><td>648</td><td>BREX-system-P-loop-protein-BrxC-[Aeromonas-caviae]</td></tr>
<tr><td>WP_190283730.1</td><td>Type1.BREX</td><td><- PglZ <-?<-DUF3944<- N6-METHYLASE -p<-HerA/FtsK<-SIR2<- N6-METHYLASE <- TOPRIM <- OLD-ABC <- DUF6079 *<- BrxB-BrxF-iSTAND <- BrxA-DNA-binding <-RHS<-?<-DDE_Tnp_1||?->HIN-HTH->ADH_N+ADH_zinc_N_2-></td><td>1226</td><td>Bacteria>Pseudomonadota>Gammaproteobacteria>Aeromonadales>Aeromonadaceae>Aeromonas</td><td>GCF_014679585.1</td><td>Aeromonas_hydrophila</td><td>IFA22_RS17960</td><td>3928101..3931781</td><td>644</td><td>BREX-system-P-loop-protein-BrxC-[Aeromonas-hydrophila]</td></tr>
<tr><td>WP_193000848.1</td><td>Type1.BREX</td><td><-DUF3944<- N6-METHYLASE -p<-HerA/FtsK<-SIR2<- N6-METHYLASE <-ParB+HNH<- TOPRIM <- OLD-ABC <- DUF6079 *<- BrxB-BrxF-iSTAND <- BrxA-DNA-binding <-RHS||HTH_Tnp_1+HTH_21+rve+rve_3-><-?<-?<-?<-HAMP+GGDEF+EAL</td><td>1226</td><td>Bacteria>Pseudomonadota>Gammaproteobacteria>Aeromonadales>Aeromonadaceae>Aeromonas</td><td>GCF_014892695.1</td><td>Aeromonas_simiae</td><td>FE240_RS10745</td><td>2272968..2276648</td><td>218936</td><td>BREX-system-P-loop-protein-BrxC-[Aeromonas-simiae]</td></tr>
<tr><td>WP_201944276.1</td><td>Type1.BREX</td><td><- SIGMA+OB-Fold+AAA-ATPase+Lon-Protease-BrxL <- PglZ <-?<-DUF3944<- N6-METHYLASE -p<- N6-METHYLASE <- TOPRIM <- OLD-ABC <- DUF6079 *<- BrxB-BrxF-iSTAND <- BrxA-DNA-binding <-RHS<-?||LexA-protease->?->PBPB+PBPB+PAS+PAS+HISKIN+REC+HPT->REC+EAL-></td><td>1231</td><td>Bacteria>Pseudomonadota>Gammaproteobacteria>Aeromonadales>Aeromonadaceae>Aeromonas</td><td>GCF_016729595.1</td><td>Aeromonas_veronii</td><td>JD494_RS16545</td><td>22676..26371</td><td>654</td><td>BREX-system-P-loop-protein-BrxC-[Aeromonas-veronii]</td></tr>
<tr><td>WP_201932781.1</td><td>Type1.BREX</td><td><- SIGMA+OB-Fold+AAA-ATPase+Lon-Protease-BrxL <- PglZ <-?<-DUF3944<- N6-METHYLASE -p<-RVT<- N6-METHYLASE <- TOPRIM <- OLD-ABC <- DUF6079 *<- BrxB-BrxF-iSTAND <- BrxA-DNA-binding <-RHS<-?|| wHTH+WYL+WCX -><-?||LexA-protease->Phage_integrase->Phage_integrase+Phage_integrase-></td><td>1226</td><td>Bacteria>Pseudomonadota>Gammaproteobacteria>Aeromonadales>Aeromonadaceae>Aeromonas</td><td>GCF_016729935.1</td><td>Aeromonas_caviae</td><td>JD546_RS01995</td><td>64762..68442</td><td>648</td><td>BREX-system-P-loop-protein-BrxC-[Aeromonas-caviae]</td></tr>
<tr><td>WP_202047008.1</td><td>Type1.BREX</td><td><- SIGMA+OB-Fold+AAA-ATPase+Lon-Protease-BrxL <- PglZ <- TOPRIM <- OLD-ABC <- N6-METHYLASE <- TOPRIM <- OLD-ABC <- DUF6079 *<- BrxB-BrxF-iSTAND <- BrxA-DNA-binding <-RHS<-?|| wHTH+WYL+WCX -><-?||LexA-protease->?-><-DOC+HTH</td><td>1226</td><td>Bacteria>Pseudomonadota>Gammaproteobacteria>Aeromonadales>Aeromonadaceae>Aeromonas</td><td>GCF_016730065.1</td><td>Aeromonas_media</td><td>JD497_RS12375</td><td>531279..534959</td><td>651</td><td>BREX-system-P-loop-protein-BrxC-[Aeromonas-media]</td></tr>
<tr><td>WP_244842130.1</td><td>Type1.BREX</td><td><- SIGMA+OB-Fold+AAA-ATPase+Lon-Protease-BrxL <- PglZ <- TOPRIM <- OLD-ABC <- N6-METHYLASE <-ParB+HNH<- TOPRIM <- OLD-ABC <- DUF6079 *<- BrxB-BrxF-iSTAND <- BrxA-DNA-binding <-RHS</td><td>1281</td><td>Bacteria>Pseudomonadota>Gammaproteobacteria>Aeromonadales>Aeromonadaceae>Aeromonas</td><td>GCF_018359965.1</td><td>Aeromonas_allosaccharophila</td><td>J4G65_RS17605</td><td>87511..91356</td><td>656</td><td>BREX-system-P-loop-protein-BrxC-[Aeromonas-allosaccharophila]</td></tr>
<tr><td>WP_257621860.1</td><td>Type1.BREX</td><td><- PglZ <- TOPRIM <- OLD-ABC <- N6-METHYLASE -p<-HerA/FtsK<-SIR2<- N6-METHYLASE <- TOPRIM <- OLD-ABC <- DUF6079 *<- BrxB-BrxF-iSTAND <- BrxA-DNA-binding <-RHS<-?|| wHTH+WYL+WCX -><-?||LexA-protease->?-><-DOC+HTH</td><td>1281</td><td>Bacteria>Pseudomonadota>Gammaproteobacteria>Aeromonadales>Aeromonadaceae>Aeromonas</td><td>GCF_019348915.1</td><td>Aeromonas_hydrophila</td><td>GPV00_RS14000</td><td>286470..290315</td><td>644</td><td>BREX-system-P-loop-protein-BrxC-[Aeromonas-hydrophila]</td></tr>
<tr><td>WP_234578020.1</td><td>Type1.BREX</td><td><- SIGMA+OB-Fold+AAA-ATPase+Lon-Protease-BrxL <- PglZ <-?<-DUF3944<- N6-METHYLASE -p<-HEPN<- N6-METHYLASE <- TOPRIM <- OLD-ABC <- DUF6079 *<- BrxB-BrxF-iSTAND <- BrxA-DNA-binding <-RHS<-ZnR+ZnR|| wHTH+WYL+WCX -><-?||LexA-protease->?-><-SF1-SF2-Helicase</td><td>1231</td><td>Bacteria>Pseudomonadota>Gammaproteobacteria>Aeromonadales>Aeromonadaceae>Aeromonas</td><td>GCF_021440865.1</td><td>Aeromonas_salmonicida</td><td>LZ632_RS15270</td><td>23400..27095</td><td>645</td><td>BREX-system-P-loop-protein-BrxC-[Aeromonas-salmonicida]</td></tr>
<tr><td>WP_236274804.1</td><td>Type1.BREX</td><td><-zf-IS66+DDE_Tnp_IS66||NTP_transferase-><-zf-IS66+DDE_Tnp_IS66<-?<-DUF3944<- N6-METHYLASE -p<- N6-METHYLASE <- TOPRIM <- OLD-ABC <- DUF6079 *<- BrxB-BrxF-iSTAND <- BrxA-DNA-binding <-RHS<-?||LexA-protease->?-><-DOC+HTH<-?||HIN-HTH-></td><td>1231</td><td>Bacteria>Pseudomonadota>Gammaproteobacteria>Aeromonadales>Aeromonadaceae>Aeromonas</td><td>GCF_021609985.1</td><td>Aeromonas_caviae</td><td>L1871_RS17025</td><td>3638140..3641835</td><td>648</td><td>BREX-system-P-loop-protein-BrxC-[Aeromonas-caviae]</td></tr>
<tr><td>WP_139392567.1</td><td>Type1.BREX</td><td><-Phage_integrase+Phage_integrase<-Phage_integrase<-LexA-protease||?-><- wHTH+WYL+WCX ||?->RHS-> BrxA-DNA-binding -> BrxB-BrxF-iSTAND -> DUF6079 *-> OLD-ABC -> TOPRIM -> N6-METHYLASE ->RVT->HEPN-> N6-METHYLASE -p->DUF3944->?-> PglZ -></td><td>1225</td><td>Bacteria>Pseudomonadota>Gammaproteobacteria>Aeromonadales>Aeromonadaceae>Aeromonas</td><td>GCF_021654355.1</td><td>Aeromonas_hydrophila</td><td>L6424_RS03515</td><td>743150..746827</td><td>644</td><td>BREX-system-P-loop-protein-BrxC-[Aeromonas-hydrophila]</td></tr>
<tr><td>WP_252657151.1</td><td>Type1.BREX</td><td><-LexA-protease||?-><- wHTH+WYL+WCX ||ZnR+ZnR->RHS-> BrxA-DNA-binding -> BrxB-BrxF-iSTAND -> DUF6079 *-> OLD-ABC -> TOPRIM -> N6-METHYLASE -> N6-METHYLASE -p->DUF3944->?-> PglZ -> SIGMA+OB-Fold+AAA-ATPase+Lon-Protease-BrxL -></td><td>1231</td><td>Bacteria>Pseudomonadota>Gammaproteobacteria>Aeromonadales>Aeromonadaceae>Aeromonas</td><td>GCF_023958325.1</td><td>Aeromonas_veronii</td><td>NG896_RS17505</td><td>21137..24832</td><td>654</td><td>BREX-system-P-loop-protein-BrxC-[Aeromonas-veronii]</td></tr>
<tr><td>WP_257688091.1</td><td>Type1.BREX</td><td><- N6-METHYLASE -p<- TOPRIM <- OLD-ABC <- DUF6079 *<- BrxB-BrxF-iSTAND <- BrxA-DNA-binding <-RHS<-ZnR+ZnR|| wHTH+WYL+WCX -><-?||LexA-protease-></td><td>1231</td><td>Bacteria>Pseudomonadota>Gammaproteobacteria>Aeromonadales>Aeromonadaceae>Aeromonas</td><td>GCF_024649335.1</td><td>Aeromonas_veronii</td><td>NUK45_RS19510</td><td>3671..7366</td><td>654</td><td>BREX-system-P-loop-protein-BrxC-[Aeromonas-veronii]</td></tr>
<tr><td>WP_257712004.1</td><td>Type1.BREX</td><td><- PglZ <- TOPRIM <- OLD-ABC <- N6-METHYLASE -p<-HEPN<-RVT<- N6-METHYLASE <- TOPRIM <- OLD-ABC <- DUF6079 *<- BrxB-BrxF-iSTAND <- BrxA-DNA-binding <-RHS<-?|| wHTH+WYL+WCX -></td><td>1281</td><td>Bacteria>Pseudomonadota>Gammaproteobacteria>Aeromonadales>Aeromonadaceae>Aeromonas</td><td>GCF_024649525.1</td><td>Aeromonas_caviae</td><td>NUK31_RS14230</td><td>17564..21409</td><td>648</td><td>BREX-system-P-loop-protein-BrxC-[Aeromonas-caviae]</td></tr>
<tr><td>WP_265455065.1</td><td>Type1.BREX</td><td><- SIGMA+OB-Fold+AAA-ATPase+Lon-Protease-BrxL <- PglZ <-?<-DUF3944<- N6-METHYLASE <- TOPRIM <- OLD-ABC <- DUF6079 *<- BrxB-BrxF-iSTAND <- BrxA-DNA-binding <-RHS<-?||LexA-protease-></td><td>1231</td><td>Bacteria>Pseudomonadota>Gammaproteobacteria>Aeromonadales>Aeromonadaceae>Aeromonas</td><td>GCF_026174125.1</td><td>Aeromonas_salmonicida</td><td>ORM31_RS04470</td><td>132358..136053</td><td>645</td><td>BREX-system-P-loop-protein-BrxC-[Aeromonas-salmonicida]</td></tr>
<tr><td>WP_265776110.1</td><td>Type1.BREX</td><td><- N6-METHYLASE -p<- TOPRIM <- OLD-ABC <- DUF6079 *<- BrxB-BrxF-iSTAND <- BrxA-DNA-binding <-RHS<-ZnR+ZnR|| wHTH+WYL+WCX -><-?||LexA-protease->?-><-DOC+HTH</td><td>1231</td><td>Bacteria>Pseudomonadota>Gammaproteobacteria>Aeromonadales>Aeromonadaceae>Aeromonas</td><td>GCF_026230455.1</td><td>Aeromonas_veronii</td><td>OD804_RS12725</td><td>3923..7618</td><td>654</td><td>BREX-system-P-loop-protein-BrxC-[Aeromonas-veronii]</td></tr>
<tr><td>WP_265783524.1</td><td>Type1.BREX</td><td><- TOPRIM <- OLD-ABC <- N6-METHYLASE -p<-TIR<-HEPN<- N6-METHYLASE <-HEPN<- TOPRIM <- OLD-ABC <- DUF6079 *<- BrxB-BrxF-iSTAND <- BrxA-DNA-binding <-RHS<-?||LexA-protease->?->PBPB+PBPB+PAS+PAS+HISKIN+REC+HPT->REC+EAL-></td><td>1235</td><td>Bacteria>Pseudomonadota>Gammaproteobacteria>Aeromonadales>Aeromonadaceae>Aeromonas</td><td>GCF_026230585.1</td><td>Aeromonas_veronii</td><td>OD810_RS03320</td><td>107717..111424</td><td>654</td><td>BREX-system-P-loop-protein-BrxC-[Aeromonas-veronii]</td></tr>
<tr><td>WP_270698108.1</td><td>Type1.BREX</td><td><-LexA-protease||?-><- wHTH+WYL+WCX ||?->RHS-> BrxA-DNA-binding -> BrxB-BrxF-iSTAND -> DUF6079 *-> OLD-ABC -> TOPRIM -> N6-METHYLASE ->RVT->HEPN-> N6-METHYLASE -> OLD-ABC -> TOPRIM -> PglZ -></td><td>1280</td><td>Bacteria>Pseudomonadota>Gammaproteobacteria>Aeromonadales>Aeromonadaceae>Aeromonas</td><td>GCF_027705185.1</td><td>Aeromonas_sp._QDB05</td><td>OKC32_RS09690</td><td>4766..8608</td><td>2990478</td><td>BREX-system-P-loop-protein-BrxC-[Aeromonas-sp.-QDB05]</td></tr>
<tr><td>WP_270793601.1</td><td>Type1.BREX</td><td>RHS-> BrxA-DNA-binding -> BrxB-BrxF-iSTAND -> DUF6079 *-> OLD-ABC -> TOPRIM -> N6-METHYLASE -> N6-METHYLASE -p-> OLD-ABC -> TOPRIM -> PglZ -> SIGMA+OB-Fold+AAA-ATPase+Lon-Protease-BrxL -></td><td>1226</td><td>Bacteria>Pseudomonadota>Gammaproteobacteria>Aeromonadales>Aeromonadaceae>Aeromonas</td><td>GCF_027705385.1</td><td>unclassified_Aeromonas</td><td>OKB76_RS13245</td><td>1893..5573</td><td>257493</td><td>MULTISPECIES:-BREX-system-P-loop-protein-BrxC-[unclassified-Aeromonas]</td></tr>
<tr><td>WP_270805301.1</td><td>Type1.BREX</td><td><- N6-METHYLASE -p<- N6-METHYLASE <-?<-HNH<- TOPRIM <- OLD-ABC <- DUF6079 *<- BrxB-BrxF-iSTAND <- BrxA-DNA-binding <-RHS<-ZnR</td><td>1240</td><td>Bacteria>Pseudomonadota>Gammaproteobacteria>Aeromonadales>Aeromonadaceae>Aeromonas</td><td>GCF_027705505.1</td><td>Aeromonas_sp._QDB56</td><td>OKB81_RS14905</td><td>8131..11853</td><td>2990495</td><td>BREX-system-P-loop-protein-BrxC-[Aeromonas-sp.-QDB56]</td></tr>
<tr><td>WP_275056847.1</td><td>Type1.BREX</td><td><-zf-IS66+DDE_Tnp_IS66<-?<-DUF3944<- N6-METHYLASE -p<-HerA/FtsK<-SIR2<- N6-METHYLASE <- TOPRIM <- OLD-ABC <- DUF6079 *<- BrxB-BrxF-iSTAND <- BrxA-DNA-binding <-RHS<-?||LexA-protease->?-><-DOC+HTH<-?<-XIS-HTH</td><td>1231</td><td>Bacteria>Pseudomonadota>Gammaproteobacteria>Aeromonadales>Aeromonadaceae>Aeromonas</td><td>GCF_029024985.1</td><td>Aeromonas_allosaccharophila</td><td>PYU98_RS18280</td><td>3929296..3932991</td><td>656</td><td>BREX-system-P-loop-protein-BrxC-[Aeromonas-allosaccharophila]</td></tr>
<tr><td>WP_076362312.1</td><td>Type1.BREX</td><td>SF1-SF2-Helicase-><-?<-LexA-protease||?-><- wHTH+WYL+WCX ||ZnR+ZnR->RHS-> BrxA-DNA-binding -> BrxB-BrxF-iSTAND -> DUF6079 *-> OLD-ABC -> TOPRIM -> N6-METHYLASE -> N6-METHYLASE -p-> N6-METHYLASE -p->DUF3944->?-> PglZ -> SIGMA+OB-Fold+AAA-ATPase+Lon-Protease-BrxL -></td><td>1231</td><td>Bacteria>Pseudomonadota>Gammaproteobacteria>Aeromonadales>Aeromonadaceae>Aeromonas</td><td>GCF_900155975.1</td><td>Aeromonas_hydrophila</td><td>BW963_RS17620</td><td>135140..138835</td><td>644</td><td>BREX-system-P-loop-protein-BrxC-[Aeromonas-hydrophila]</td></tr>
<tr><td>WP_050664996.1</td><td>Type1.BREX</td><td><-LexA-protease||?-><- wHTH+WYL+WCX ||ARG->?->ZnR->RHS-> BrxA-DNA-binding -> BrxB-BrxF-iSTAND -> DUF6079 *-> OLD-ABC -> TOPRIM -> N6-METHYLASE ->DUF3944->?-> PglZ -> SIGMA+OB-Fold+AAA-ATPase+Lon-Protease-BrxL -></td><td>1231</td><td>Bacteria>Pseudomonadota>Gammaproteobacteria>Aeromonadales>Aeromonadaceae</td><td>GCF_000820105.1</td><td>Aeromonas</td><td>BN1133_RS02615</td><td>38531..42226</td><td>642</td><td>MULTISPECIES:-BREX-system-P-loop-protein-BrxC-[Aeromonas]</td></tr>
<tr><td>WP_182788683.1</td><td>Type1.BREX</td><td>HDRnase-><-?<-LexA-protease||RHS->ARG->ZnR->RHS-> BrxA-DNA-binding -> BrxB-BrxF-iSTAND -> DUF6079 *-> OLD-ABC -> TOPRIM ->HNH->?-> N6-METHYLASE ->SIR2->HerA/FtsK-> N6-METHYLASE -p-></td><td>1236</td><td>Bacteria>Pseudomonadota>Gammaproteobacteria>Aeromonadales>Aeromonadaceae</td><td>GCF_014161425.1</td><td>Aeromonas</td><td>H7K58_RS09750</td><td>2083810..2087520</td><td>642</td><td>MULTISPECIES:-BREX-system-P-loop-protein-BrxC-[Aeromonas]</td></tr>
<tr><td>WP_171273737.1</td><td>Type1.BREX</td><td>Arm-DNA-bind_2+Phage_int_SAM_3+Phage_integrase->HDRnase-><-?||RHS->ARG->ZnR->RHS-> BrxA-DNA-binding -> BrxB-BrxF-iSTAND -> DUF6079 *-> OLD-ABC -> TOPRIM -> N6-METHYLASE ->HEPN->TIR-> N6-METHYLASE -p->DUF3944->?->|| PglZ -></td><td>1231</td><td>Bacteria>Pseudomonadota>Gammaproteobacteria>Aeromonadales>Aeromonadaceae</td><td>GCF_021654375.1</td><td>Aeromonas</td><td>E4185_RS15845</td><td>3355159..3358854</td><td>642</td><td>MULTISPECIES:-BREX-system-P-loop-protein-BrxC-[Aeromonas]</td></tr>
<tr><td>WP_234605103.1</td><td>Type1.BREX</td><td><- PglZ <-?<-DUF3944<- N6-METHYLASE -p<-TIR<- N6-METHYLASE -p<- N6-METHYLASE <- TOPRIM <- OLD-ABC <- DUF6079 *<- BrxB-BrxF-iSTAND <- BrxA-DNA-binding <-RHS<-ZnR+ZnR|| wHTH+WYL+WCX -><-?||LexA-protease->HTH_Tnp_1+HTH_21+rve+rve_3-><-DOC+HTH</td><td>1231</td><td>Bacteria>Pseudomonadota>Gammaproteobacteria>Aeromonadales>Aeromonadaceae</td><td>GCF_026173065.1</td><td>Aeromonas</td><td>ORM39_RS00835</td><td>21547..25242</td><td>642</td><td>BREX-system-P-loop-protein-BrxC-[Aeromonas-salmonicida]</td></tr>
<tr><td>WP_163133170.1</td><td>Type1.BREX</td><td>GDResolvase+HIN-HTH->SF1-SF2-Helicase+REase->SF1-SF2-Helicase+REase-><-SLOG+BirA-HTH<-SF1-SF2-Helicase+PRTase<- PglZ <- TOPRIM <- OLD-ABC <- N6-METHYLASE <- DUF6079 *<- BrxB-BrxF-iSTAND <- BrxA-DNA-binding <-XIS-HTH<-SAM-METHYLASE+wHTH+HNH||?-><-?||?-><-Arm-DNA-bind_3+Phage_integrase</td><td>1160</td><td>Bacteria>Pseudomonadota>Gammaproteobacteria>Alteromonadales>Alteromonadaceae>Agarivorans</td><td>GCF_900537085.1</td><td>Agarivorans_sp._Alg241-V36</td><td>G6R11_RS11270</td><td>302012..305494</td><td>2305992</td><td>BREX-system-P-loop-protein-BrxC-[Agarivorans-sp.-Alg241-V36]</td></tr>
<tr><td>WP_014998461.1</td><td>Type1.BREX</td><td><-Phage_integrase<-?||XIS-HTH->SF1-SF2-Helicase+REase->SF1-SF2-Helicase+REase-><- PglZ <- TOPRIM <- OLD-ABC <- N6-METHYLASE <- DUF6079 *<- BrxB-BrxF-iSTAND <- BrxA-DNA-binding <- wHTH+WYL+WCX <-Arm-DNA-bind_3+Phage_integrase<-tRNA<-tRNA<-?<-URI-DNase+UVR+RNase+HHH<-REC+OmpR-HTH<-DSBH<-Aminotran_1_2</td><td>1157</td><td>Bacteria>Pseudomonadota>Gammaproteobacteria>Alteromonadales>Alteromonadaceae>Alteromonas/Salinimonas_group</td><td>-</td><td>Alteromonas_macleodii</td><td>AMBLS11_RS09820</td><td>2297331..2300804</td><td>28108</td><td>BREX-system-P-loop-protein-BrxC-[Alteromonas-macleodii]</td></tr>
<tr><td>WP_100972015.1</td><td>Type1.BREX</td><td>SF1-SF2-Helicase+REase->SF1-SF2-Helicase+REase-p->rve->AAA-ATPase->SF1-SF2-Helicase+REase-p-><- PglZ <- TOPRIM <- OLD-ABC <- N6-METHYLASE <- DUF6079 *<- BrxB-BrxF-iSTAND <- BrxA-DNA-binding <- wHTH+WYL+WCX ||?->GTPase-AIG->GTPase-AIG->?->?-><-GAF+GGDEF</td><td>1157</td><td>Bacteria>Pseudomonadota>Gammaproteobacteria>Alteromonadales>Alteromonadaceae>Alteromonas/Salinimonas_group</td><td>GCF_002831605.1</td><td>Alteromonas_sp._MB-3u-76</td><td>CW735_RS10480</td><td>2338685..2342158</td><td>2058133</td><td>BREX-system-P-loop-protein-BrxC-[Alteromonas-sp.-MB-3u-76]</td></tr>
<tr><td>WP_146799747.1</td><td>Type1.BREX</td><td>TraI-VirD2-RCRE->SF1-SF2-Helicase+REase->SF1-SF2-Helicase+REase-><-?<-ZetaToxin<- PglZ <- TOPRIM <- OLD-ABC <- N6-METHYLASE <- DUF6079 *<- BrxB-BrxF-iSTAND <- BrxA-DNA-binding <- wHTH+WYL+WCX <-SF1-SF2-Helicase</td><td>1165</td><td>Bacteria>Pseudomonadota>Gammaproteobacteria>Alteromonadales>Colwelliaceae>Colwellia</td><td>GCF_007954305.1</td><td>Colwellia_hornerae</td><td>ESZ26_RS11985</td><td>115858..119355</td><td>89402</td><td>BREX-system-P-loop-protein-BrxC-[Colwellia-hornerae]</td></tr>
<tr><td>WP_130146693.1</td><td>Type1.BREX</td><td>DeoR-HTH+ISOCOT->?-><- SIGMA+OB-Fold+AAA-ATPase+Lon-Protease-BrxL <- PglZ <- N6-METHYLASE <- TOPRIM <- OLD-ABC <- DUF6079 *<- BrxB-BrxF-iSTAND <- BrxA-DNA-binding <- wHTH+WYL+WCX ||HTH_Tnp_1+HTH_21+rve+rve_3-></td><td>1225</td><td>Bacteria>Pseudomonadota>Gammaproteobacteria>Alteromonadales>Pseudoalteromonadaceae>Pseudoalteromonas</td><td>GCF_004208955.1</td><td>Pseudoalteromonas_sp._CO325X</td><td>EXT46_RS08650</td><td>70759..74436</td><td>1777262</td><td>BREX-system-P-loop-protein-BrxC-[Pseudoalteromonas-sp.-CO325X]</td></tr>
<tr><td>WP_165723736.1</td><td>Type1.BREX</td><td>REC+LuxR-HTH-><-PAS+SHELIX+PAS+PAS+PAS<-sGTP+sGTP||?-><- SIGMA+OB-Fold+AAA-ATPase+Lon-Protease-BrxL <- PglZ <- N6-METHYLASE <- TOPRIM <- OLD-ABC <- DUF6079 *<- BrxB-BrxF-iSTAND <- BrxA-DNA-binding <- wHTH+WYL+WCX <-XIS-HTH||?->DUF2787->Phage_integrase-></td><td>1225</td><td>Bacteria>Pseudomonadota>Gammaproteobacteria>Alteromonadales>Pseudoalteromonadaceae>Pseudoalteromonas</td><td>GCF_943789085.1</td><td>Pseudoalteromonas_sp._Z1A2</td><td>QDP19_RS17640</td><td>3065370..3069047</td><td>2686350</td><td>MULTISPECIES:-BREX-system-P-loop-protein-BrxC-[Pseudoalteromonas]</td></tr>
<tr><td>WP_006086465.1</td><td>Type1.BREX</td><td><-TraI-VirD2-RCRE||?-><-?||?-> wHTH+WYL+WCX ->Peptidase_C47-> wHTH+WYL+WCX -> BrxA-DNA-binding -> BrxB-BrxF-iSTAND -> DUF6079 *->ParB+HNH-> OLD-ABC -> TOPRIM -> N6-METHYLASE -> PglZ -> SIGMA+OB-Fold+AAA-ATPase+Lon-Protease-BrxL ->SF1-SF2-Helicase->?-><-DDE_Tnp_1_assoc+DDE_Tnp_1</td><td>1229</td><td>Bacteria>Pseudomonadota>Gammaproteobacteria>Alteromonadales>Shewanellaceae>Shewanella</td><td>GCF_000231345.1</td><td>Shewanella_baltica</td><td>SBAL625DRAFT_RS15525</td><td>212954..216643</td><td>62322</td><td>BREX-system-P-loop-protein-BrxC-[Shewanella-baltica]</td></tr>
</table>

| Protein-ID | System | Gene-Neighbourhoods | Length | Taxonomy | Assembly-ID | Strain | Locus-Tag | Genomic-coordinates | TaxID | Fasta-defline |
|---|---|---|---|---|---|---|---|---|---|---|
| WP_080250285.1 | Type1.BREX | <-HEPN<-?\|\|YjhX_toxin->SAM-METHYLASE->?-> BrxA-DNA-binding -> BrxB-BrxF-iSTAND -> DUF6079 *-> N6-METHYLASE -> OLD-ABC -> TOPRIM -> PglZ -> SIGMA+OB-Fold+AAA-ATPase+Lon-Protease-BrxL ->HKD-DNase+HKD-DNase-><-REase<-NACHT+SF1-SF2-Helicase+HKD-DNase<-Arm-DNA-bind_3 | 1213 | Bacteria>Pseudomonadota>Gammaproteobacteria>Enterobacterales>Enterobacteriaceae>Salmonella | GCF_002066965.1 | Salmonella_enterica | RZ13_RS02295 | 164882..168523 | 28901 | BREX-system-P-loop-protein-BrxC-[Salmonella-enterica] |
| WP_111769566.1 | Type1.BREX | NACHT+SF1-SF2-Helicase+HKD-DNase->REase-><-?<- SIGMA+OB-Fold+AAA-ATPase+Lon-Protease-BrxL <- PglZ <- TOPRIM <- OLD-ABC <- N6-METHYLASE <- DUF6079 *<- BrxB-BrxF-iSTAND <- BrxA-DNA-binding <-?<-SAM-METHYLASE<-YjhX_toxin\|\|?->HEPN-> | 1213 | Bacteria>Pseudomonadota>Gammaproteobacteria>Enterobacterales>Enterobacteriaceae>Salmonella | GCF_002106425.1 | Salmonella_enterica | R537_RS06030 | 154441..158082 | 28901 | BREX-system-P-loop-protein-BrxC-[Salmonella-enterica] |
| WP_086805524.1 | Type1.BREX | NACHT+SF1-SF2-Helicase+HKD-DNase->REase-><-HKD-DNase+HKD-DNase<- SIGMA+OB-Fold+AAA-ATPase+Lon-Protease-BrxL <- PglZ <- TOPRIM <- OLD-ABC <- N6-METHYLASE <- DUF6079 *<- BrxB-BrxF-iSTAND <- BrxA-DNA-binding \|\|ZnR-><-?\|\|SIGMA-HTH+REase->?->MarR-wHTH-><- wHTH+WYL+WCX \|\|wHTH-> | 1204 | Bacteria>Pseudomonadota>Gammaproteobacteria>Enterobacterales>Enterobacteriaceae>Salmonella | GCF_002156225.1 | Salmonella_enterica | CCL33_RS10730 | 52147..55761 | 28901 | BREX-system-P-loop-protein-BrxC-[Salmonella-enterica] |
| WP_065348243.1 | Type1.BREX | <-HEPN<-?\|\|YjhX_toxin->SAM-METHYLASE->?-> BrxA-DNA-binding -> BrxB-BrxF-iSTAND -> DUF6079 *-> N6-METHYLASE -> OLD-ABC -> TOPRIM -> PglZ -> SIGMA+OB-Fold+AAA-ATPase+Lon-Protease-BrxL ->?-><-REase<-NACHT+SF1-SF2-Helicase+HKD-DNase | 1213 | Bacteria>Pseudomonadota>Gammaproteobacteria>Enterobacterales>Enterobacteriaceae>Salmonella | GCF_002196535.1 | Salmonella_enterica | GX95_RS20905 | 4055067..4058708 | 28901 | BREX-system-P-loop-protein-BrxC-[Salmonella-enterica] |
| WP_088546528.1 | Type1.BREX | <-GntR-HTH++TrmB+ZDNA-HTH\|\|?-><-?<-HEPN<-?\|\|YjhX_toxin->?-> BrxA-DNA-binding -> BrxB-BrxF-iSTAND -> DUF6079 *-> N6-METHYLASE -> OLD-ABC -> TOPRIM -> PglZ -> SIGMA+OB-Fold+AAA-ATPase+Lon-Protease-BrxL ->?-><-REase<-NACHT+SF1-SF2-Helicase+HKD-DNase | 1213 | Bacteria>Pseudomonadota>Gammaproteobacteria>Enterobacterales>Enterobacteriaceae>Salmonella | GCF_002208255.1 | Salmonella_enterica | LFZ35_RS21090 | 4061970..4065611 | 28901 | BREX-system-P-loop-protein-BrxC-[Salmonella-enterica] |
| WP_088759215.1 | Type1.BREX | <-MukB<-MukE<-wHTH\|\| wHTH+WYL+WCX -><-MarR-wHTH<-SIGMA-HTH+REase\|\|?-> BrxA-DNA-binding -> BrxB-BrxF-iSTAND -> DUF6079 *-> N6-METHYLASE -> OLD-ABC -> TOPRIM -> PglZ -> SIGMA+OB-Fold+AAA-ATPase+Lon-Protease-BrxL -><-REase<-NACHT+SF1-SF2-Helicase+HKD-DNase<-Arm-DNA-bind_3<- TOPRIM -DNAG+D5N+D5ATPase+P4PRIMASE-HTH | 1204 | Bacteria>Pseudomonadota>Gammaproteobacteria>Enterobacterales>Enterobacteriaceae>Salmonella | GCF_002211845.1 | Salmonella_enterica | CE137_RS21050 | 4095775..4099389 | 28901 | BREX-system-P-loop-protein-BrxC-[Salmonella-enterica] |
| WP_080090697.1 | Type1.BREX | <-HEPN<-REase+iREase<-?\|\|YjhX_toxin->SAM-METHYLASE->?-> BrxA-DNA-binding -> BrxB-BrxF-iSTAND -> DUF6079 *-> N6-METHYLASE -> OLD-ABC -> TOPRIM -> PglZ -> SIGMA+OB-Fold+AAA-ATPase+Lon-Protease-BrxL -><-REase<-Couple_hipA+HipA_C<-HTH_3 | 1213 | Bacteria>Pseudomonadota>Gammaproteobacteria>Enterobacterales>Enterobacteriaceae>Salmonella | GCF_002234755.2 | Salmonella_enterica | CHD02_RS05505 | 1125404..1129045 | 28901 | BREX-system-P-loop-protein-BrxC-[Salmonella-enterica] |
| WP_053445538.1 | Type1.BREX | <-HEPN<-?\|\|YjhX_toxin->SAM-METHYLASE->?-> BrxA-DNA-binding -> BrxB-BrxF-iSTAND -> DUF6079 *-> N6-METHYLASE -> OLD-ABC -> TOPRIM -> PglZ -> SIGMA+OB-Fold+AAA-ATPase+Lon-Protease-BrxL -><-REase | 1213 | Bacteria>Pseudomonadota>Gammaproteobacteria>Enterobacterales>Enterobacteriaceae>Salmonella | GCF_002238275.1 | Salmonella_enterica | F1E41_RS17160 | 150695..154336 | 28901 | BREX-system-P-loop-protein-BrxC-[Salmonella-enterica] |
| WP_094797764.1 | Type1.BREX | <-HEPN<-REase+iREase<-?\|\|YjhX_toxin->SAM-METHYLASE->?-> BrxA-DNA-binding -> BrxB-BrxF-iSTAND -> DUF6079 *-> N6-METHYLASE -> OLD-ABC -> TOPRIM -> PglZ -> SIGMA+OB-Fold+AAA-ATPase+Lon-Protease-BrxL -><-REase<-NACHT+SF1-SF2-Helicase+HKD-DNase<-Arm-DNA-bind_3 | 1213 | Bacteria>Pseudomonadota>Gammaproteobacteria>Enterobacterales>Enterobacteriaceae>Salmonella | GCF_002260565.1 | Salmonella_enterica | CFB03_RS03285 | 15872..19513 | 28901 | BREX-system-P-loop-protein-BrxC-[Salmonella-enterica] |
| WP_094887773.1 | Type1.BREX | REase-><- SIGMA+OB-Fold+AAA-ATPase+Lon-Protease-BrxL <- PglZ <- TOPRIM <- OLD-ABC <- N6-METHYLASE <- DUF6079 *<- BrxB-BrxF-iSTAND <- BrxA-DNA-binding | 1213 | Bacteria>Pseudomonadota>Gammaproteobacteria>Enterobacterales>Enterobacteriaceae>Salmonella | GCF_002265975.1 | Salmonella_enterica | CCO43_RS23255 | 12854..16495 | 28901 | BREX-system-P-loop-protein-BrxC-[Salmonella-enterica] |
| WP_094964979.1 | Type1.BREX | <-HEPN<-?\|\|YjhX_toxin->SAM-METHYLASE->?-> BrxA-DNA-binding -> BrxB-BrxF-iSTAND -> DUF6079 *-> N6-METHYLASE -> OLD-ABC -> TOPRIM -> PglZ -> SIGMA+OB-Fold+AAA-ATPase+Lon-Protease-BrxL ->\|\|<-HTH_Tnp_1+HTH_21+rve+rve_3\|\|PIN-><-HIN-HTH | 1213 | Bacteria>Pseudomonadota>Gammaproteobacteria>Enterobacterales>Enterobacteriaceae>Salmonella | GCF_002266055.1 | Salmonella_enterica | CCO49_RS14335 | 399067..402708 | 28901 | BREX-system-P-loop-protein-BrxC-[Salmonella-enterica] |
| WP_094946975.1 | Type1.BREX | <-HEPN<-?\|\|YjhX_toxin->SAM-METHYLASE->?-> BrxA-DNA-binding -> BrxB-BrxF-iSTAND -> DUF6079 *-> N6-METHYLASE -> OLD-ABC -> TOPRIM -> PglZ -> SIGMA+OB-Fold+AAA-ATPase+Lon-Protease-BrxL -><-REase<-NACHT+SF1-SF2-Helicase+HKD-DNase | 1213 | Bacteria>Pseudomonadota>Gammaproteobacteria>Enterobacterales>Enterobacteriaceae>Salmonella | GCF_002266315.1 | Salmonella_enterica | CCO36_RS01880 | 387042..390683 | 28901 | BREX-system-P-loop-protein-BrxC-[Salmonella-enterica] |
| WP_097369131.1 | Type1.BREX | <-HEPN<-REase+iREase<-?\|\|YjhX_toxin->SAM-METHYLASE->?-> BrxA-DNA-binding -> BrxB-BrxF-iSTAND -> DUF6079 *-> N6-METHYLASE -> OLD-ABC -> TOPRIM -> PglZ -> SIGMA+OB-Fold+AAA-ATPase+Lon-Protease-BrxL -><-REase<-NACHT+SF1-SF2-Helicase+HKD-DNase<-Arm-DNA-bind_3 | 1213 | Bacteria>Pseudomonadota>Gammaproteobacteria>Enterobacterales>Enterobacteriaceae>Salmonella | GCF_002507575.1 | Salmonella_enterica | CBI65_RS08825 | 170892..174533 | 28901 | BREX-system-P-loop-protein-BrxC-[Salmonella-enterica] |
| WP_097368036.1 | Type1.BREX | <-HEPN<-REase+iREase<-?\|\|YjhX_toxin->SAM-METHYLASE->?-> BrxA-DNA-binding -> BrxB-BrxF-iSTAND -> DUF6079 *-> N6-METHYLASE -> OLD-ABC -> TOPRIM -> PglZ -> SIGMA+OB-Fold+AAA-ATPase+Lon-Protease-BrxL -><-REase<-NACHT+SF1-SF2-Helicase+HKD-DNase<-Arm-DNA-bind_3 | 1213 | Bacteria>Pseudomonadota>Gammaproteobacteria>Enterobacterales>Enterobacteriaceae>Salmonella | GCF_002507625.1 | Salmonella_enterica | CBI66_RS11180 | 171059..174700 | 28901 | BREX-system-P-loop-protein-BrxC-[Salmonella-enterica] |
| WP_097375567.1 | Type1.BREX | <-HEPN<-REase+iREase<-?\|\|YjhX_toxin->SAM-METHYLASE->?-> BrxA-DNA-binding -> BrxB-BrxF-iSTAND -> DUF6079 *-> N6-METHYLASE -> OLD-ABC -> TOPRIM -> PglZ -> SIGMA+OB-Fold+AAA-ATPase+Lon-Protease-BrxL -><-REase<-NACHT+SF1-SF2-Helicase+HKD-DNase<-Arm-DNA-bind_3 | 1213 | Bacteria>Pseudomonadota>Gammaproteobacteria>Enterobacterales>Enterobacteriaceae>Salmonella | GCF_002507635.1 | Salmonella_enterica | CBI67_RS08835 | 170640..174281 | 28901 | BREX-system-P-loop-protein-BrxC-[Salmonella-enterica] |
| WP_107326618.1 | Type1.BREX | Arm-DNA-bind_3->NACHT+SF1-SF2-Helicase+HKD-DNase->REase-><- SIGMA+OB-Fold+AAA-ATPase+Lon-Protease-BrxL <- PglZ <- TOPRIM <- OLD-ABC <- N6-METHYLASE <- DUF6079 *<- BrxB-BrxF-iSTAND <- BrxA-DNA-binding <-?<-SAM-METHYLASE<-YjhX_toxin\|\|?->REase+iREase->HEPN-> | 1213 | Bacteria>Pseudomonadota>Gammaproteobacteria>Enterobacterales>Enterobacteriaceae>Salmonella | GCF_003032235.1 | Salmonella_enterica | C5I04_RS23495 | 346379..350020 | 28901 | BREX-system-P-loop-protein-BrxC-[Salmonella-enterica] |
| WP_116574272.1 | Type1.BREX | TOPRIM -DNAG+D5N+D5ATPase+P4PRIMASE-HTH->Arm-DNA-bind_3+Phage_integrase-><-?<- SIGMA+OB-Fold+AAA-ATPase+Lon-Protease-BrxL <- PglZ <- N6-METHYLASE <- DUF6079 *<- BrxB-BrxF-iSTAND <- BrxA-DNA-binding <-?<-SAM-METHYLASE<-YjhX_toxin\|\|?->HEPN-> | 1213 | Bacteria>Pseudomonadota>Gammaproteobacteria>Enterobacterales>Enterobacteriaceae>Salmonella | GCF_003077035.1 | Salmonella_enterica | C4874_RS10725 | 43609..47250 | 28901 | BREX-system-P-loop-protein-BrxC-[Salmonella-enterica] |
| WP_058114204.1 | Type1.BREX | <-GntR-HTH++TrmB+ZDNA-HTH\|\|<-?<-HEPN<-REase+iREase<-?\|\|YjhX_toxin->?->\|\| BrxA-DNA-binding -> BrxB-BrxF-iSTAND -> DUF6079 *-> N6-METHYLASE -> OLD-ABC -> TOPRIM -> PglZ -> SIGMA+OB-Fold+AAA-ATPase+Lon-Protease-BrxL ->?->\|\|<-DEDD_Tnp_IS110+Transposase_20\|\|<-tRNA\|\|?-><-SKI | 1213 | Bacteria>Pseudomonadota>Gammaproteobacteria>Enterobacterales>Enterobacteriaceae>Salmonella | GCF_003324795.1 | Salmonella_enterica | DOE60_RS06770 | 1364315..1367956 | 28901 | BREX-system-P-loop-protein-BrxC-[Salmonella-enterica] |
| WP_114051212.1 | Type1.BREX | HTH_3->Couple_hipA+HipA_C->REase-><- SIGMA+OB-Fold+AAA-ATPase+Lon-Protease-BrxL <- PglZ <- TOPRIM <- OLD-ABC <- DUF6079 *<- BrxB-BrxF-iSTAND <- BrxA-DNA-binding <-SAM-METHYLASE<-YjhX_toxin\|\|?->?-><-?\|\|GntR-HTH++TrmB+ZDNA-HTH-> | 1213 | Bacteria>Pseudomonadota>Gammaproteobacteria>Enterobacterales>Enterobacteriaceae>Salmonella | GCF_003325275.1 | Salmonella_enterica | CHE40_RS23180 | 4543503..4547144 | 28901 | BREX-system-P-loop-protein-BrxC-[Salmonella-enterica] |
| WP_140237021.1 | Type1.BREX | BrxA-DNA-binding -> BrxB-BrxF-iSTAND -> DUF6079 *-> N6-METHYLASE -> OLD-ABC -> TOPRIM -> PglZ -> SIGMA+OB-Fold+AAA-ATPase+Lon-Protease-BrxL -><-REase<-Couple_hipA+HipA_C<-HTH_3 | 1213 | Bacteria>Pseudomonadota>Gammaproteobacteria>Enterobacterales>Enterobacteriaceae>Salmonella | GCF_006379925.2 | Salmonella_enterica | FG677_RS21910 | 2844..6485 | 28901 | BREX-system-P-loop-protein-BrxC-[Salmonella-enterica] |
| WP_080166838.1 | Type1.BREX | <-HEPN<-?\|\|YjhX_toxin->SAM-METHYLASE->?-> BrxA-DNA-binding -> BrxB-BrxF-iSTAND -> DUF6079 *-> N6-METHYLASE -> OLD-ABC -> TOPRIM -> PglZ -> SIGMA+OB-Fold+AAA-ATPase+Lon-Protease-BrxL ->?-><-REase<-NACHT+SF1-SF2-Helicase+HKD-DNase | 1213 | Bacteria>Pseudomonadota>Gammaproteobacteria>Enterobacterales>Enterobacteriaceae>Salmonella | GCF_012273475.1 | Salmonella_enterica | FOC17_RS02975 | 512521..516162 | 28901 | BREX-system-P-loop-protein-BrxC-[Salmonella-enterica] |
| WP_181299515.1 | Type1.BREX | <-HEPN<-?\|\|\|\|YjhX_toxin->SAM-METHYLASE->?->?-> BrxA-DNA-binding -> BrxB-BrxF-iSTAND -> DUF6079 *-> N6-METHYLASE -> OLD-ABC -> TOPRIM -> PglZ -> SIGMA+OB-Fold+AAA-ATPase+Lon-Protease-BrxL -><-REase<-Couple_hipA+HipA_C<-HTH_3 | 1213 | Bacteria>Pseudomonadota>Gammaproteobacteria>Enterobacterales>Enterobacteriaceae>Salmonella | GCF_013624525.1 | Salmonella_enterica | D6413_08715 | 52330..55971 | 28901 | BREX-system-P-loop-protein-BrxC-[Salmonella-enterica] |
| WP_080213758.1 | Type1.BREX | <-GntR-HTH++TrmB+ZDNA-HTH\|\|?-><-?<-?\|\|YjhX_toxin->SAM-METHYLASE->?-> BrxA-DNA-binding -> BrxB-BrxF-iSTAND -> DUF6079 *-> N6-METHYLASE -> OLD-ABC -> TOPRIM -> PglZ -> SIGMA+OB-Fold+AAA-ATPase+Lon-Protease-BrxL ->?-><-REase | 1213 | Bacteria>Pseudomonadota>Gammaproteobacteria>Enterobacterales>Enterobacteriaceae>Salmonella | GCF_015180625.1 | Salmonella_enterica | AS04_RS02250 | 44431..48072 | 28901 | BREX-system-P-loop-protein-BrxC-[Salmonella-enterica] |
| WP_079788274.1 | Type1.BREX | <-HEPN<-REase+iREase<-?\|\|YjhX_toxin->SAM-METHYLASE->?-> BrxA-DNA-binding -> BrxB-BrxF-iSTAND -> DUF6079 *-> N6-METHYLASE -> OLD-ABC -> TOPRIM -> PglZ -> SIGMA+OB-Fold+AAA-ATPase+Lon-Protease-BrxL -><-REase<-NACHT+SF1-SF2-Helicase+HKD-DNase<-Arm-DNA-bind_3 | 1213 | Bacteria>Pseudomonadota>Gammaproteobacteria>Enterobacterales>Enterobacteriaceae>Salmonella | GCF_016772815.1 | Salmonella_enterica | JM978_RS19965 | 4115108..4118749 | 28901 | BREX-system-P-loop-protein-BrxC-[Salmonella-enterica] |
| WP_210138875.1 | Type1.BREX | SKI-><-?\|\|tRNA->Arm-DNA-bind_3->NACHT+SF1-SF2-Helicase+HKD-DNase->REase-><- SIGMA+OB-Fold+AAA-ATPase+Lon-Protease-BrxL <- TOPRIM <- OLD-ABC <- N6-METHYLASE <- DUF6079 *<- BrxB-BrxF-iSTAND <- BrxA-DNA-binding <-?<-SAM-METHYLASE<-YjhX_toxin\|\|?->REase+iREase->HEPN-> | 1213 | Bacteria>Pseudomonadota>Gammaproteobacteria>Enterobacterales>Enterobacteriaceae>Salmonella | GCF_017893505.1 | Salmonella_enterica | JJB80_RS22605 | 4685033..4688674 | 28901 | BREX-system-P-loop-protein-BrxC-[Salmonella-enterica] |
| WP_241575978.1 | Type1.BREX | Arm-DNA-bind_3->NACHT+SF1-SF2-Helicase+HKD-DNase->REase-><- SIGMA+OB-Fold+AAA-ATPase+Lon-Protease-BrxL <- PglZ <- TOPRIM <- OLD-ABC <- N6-METHYLASE <- DUF6079 *<- BrxB-BrxF-iSTAND <- BrxA-DNA-binding <-?<-SAM-METHYLASE<-YjhX_toxin\|\|?->REase+iREase->HEPN-> | 1213 | Bacteria>Pseudomonadota>Gammaproteobacteria>Enterobacterales>Enterobacteriaceae>Salmonella | GCF_022603615.1 | Salmonella_enterica | K5800_RS10145 | 108176..111817 | 28901 | BREX-system-P-loop-protein-BrxC-[Salmonella-enterica] |
| WP_247258912.1 | Type1.BREX | NACHT+SF1-SF2-Helicase+HKD-DNase->REase-><-HKD-DNase+HKD-DNase<- SIGMA+OB-Fold+AAA-ATPase+Lon-Protease-BrxL <- PglZ <- TOPRIM <- OLD-ABC <- N6-METHYLASE <- DUF6079 *<- BrxB-BrxF-iSTAND <- BrxA-DNA-binding \|\|ZnR-><-?\|\|<-?<-SAM-METHYLASE<-YjhX_toxin\|\|?->HEPN-> | 1204 | Bacteria>Pseudomonadota>Gammaproteobacteria>Enterobacterales>Enterobacteriaceae>Salmonella | GCF_023007725.1 | Salmonella_enterica | MA804_RS14880 | 357716..361330 | 28901 | BREX-system-P-loop-protein-BrxC-[Salmonella-enterica] |
| WP_247223752.1 | Type1.BREX | <-HEPN<-REase+iREase<-?\|\|YjhX_toxin->SAM-METHYLASE->?-> BrxA-DNA-binding -> BrxB-BrxF-iSTAND -> DUF6079 *-> N6-METHYLASE -> OLD-ABC -> TOPRIM -> PglZ -> SIGMA+OB-Fold+AAA-ATPase+Lon-Protease-BrxL -><-REase<-NACHT+SF1-SF2-Helicase+HKD-DNase<-Arm-DNA-bind_3 | 1213 | Bacteria>Pseudomonadota>Gammaproteobacteria>Enterobacterales>Enterobacteriaceae>Salmonella | GCF_023016865.1 | Salmonella_sp._L-S2638 | MVT37_RS15260 | 170948..174589 | 2933299 | BREX-system-P-loop-protein-BrxC-[Salmonella-sp.-L-S2638] |
| WP_198899489.1 | Type1.BREX | <-HEPN<-REase+iREase<-?\|\|YjhX_toxin->SAM-METHYLASE->?-> BrxA-DNA-binding -> BrxB-BrxF-iSTAND -> DUF6079 *-> N6-METHYLASE -> OLD-ABC -> TOPRIM -> PglZ -> SIGMA+OB-Fold+AAA-ATPase+Lon-Protease-BrxL -><-REase<-NACHT+SF1-SF2-Helicase+HKD-DNase<-Arm-DNA-bind_3 | 1213 | Bacteria>Pseudomonadota>Gammaproteobacteria>Enterobacterales>Enterobacteriaceae>Salmonella | GCF_023017625.1 | Salmonella_sp._14ESS1484 | MVA85_RS11375 | 170949..174590 | 2933314 | BREX-system-P-loop-protein-BrxC-[Salmonella-sp.-14ESS1484] |
| WP_254499948.1 | Type1.BREX | Arm-DNA-bind_3->NACHT+SF1-SF2-Helicase+HKD-DNase->REase-><- SIGMA+OB-Fold+AAA-ATPase+Lon-Protease-BrxL <- PglZ <- TOPRIM <- OLD-ABC <- N6-METHYLASE <- DUF6079 *<- BrxB-BrxF-iSTAND <- BrxA-DNA-binding <-?<-SAM-METHYLASE<-YjhX_toxin\|\|?->REase+iREase->HEPN-> | 1213 | Bacteria>Pseudomonadota>Gammaproteobacteria>Enterobacterales>Enterobacteriaceae>Salmonella | GCF_024261625.1 | Salmonella_enterica | NNJ27_RS08465 | 108147..111788 | 28901 | BREX-system-P-loop-protein-BrxC-[Salmonella-enterica] |
| WP_255223204.1 | Type1.BREX | <-HEPN<-REase+iREase<-?<-PAIREDC-HTH+HIN-HTH\|\|YjhX_toxin->SAM-METHYLASE->?-> BrxA-DNA-binding -> BrxB-BrxF-iSTAND -> DUF6079 *-> N6-METHYLASE -> OLD-ABC -> TOPRIM -> PglZ -> SIGMA+OB-Fold+AAA-ATPase+Lon-Protease-BrxL -><-REase<-NACHT+SF1-SF2-Helicase+HKD-DNase<-Arm-DNA-bind_3 | 1213 | Bacteria>Pseudomonadota>Gammaproteobacteria>Enterobacterales>Enterobacteriaceae>Salmonella | GCF_024364985.1 | Salmonella_enterica | NMU37_RS20780 | 4227543..4231184 | 28901 | BREX-system-P-loop-protein-BrxC-[Salmonella-enterica] |
| WP_255929194.1 | Type1.BREX | <-HEPN<-?\|\|YjhX_toxin->SAM-METHYLASE->?->\|\|?-><-ZnR\|\| BrxA-DNA-binding -> BrxB-BrxF-iSTAND -> DUF6079 *-> N6-METHYLASE -> OLD-ABC -> TOPRIM -> PglZ -> SIGMA+OB-Fold+AAA-ATPase+Lon-Protease-BrxL ->HKD-DNase+HKD-DNase-><-REase<-NACHT+SF1-SF2-Helicase+HKD-DNase | 1204 | Bacteria>Pseudomonadota>Gammaproteobacteria>Enterobacterales>Enterobacteriaceae>Salmonella | GCF_024442275.1 | Salmonella_enterica | NOQ95_RS21560 | 4448475..4452089 | 28901 | BREX-system-P-loop-protein-BrxC-[Salmonella-enterica] |
| WP_080092317.1 | Type1.BREX | <-HEPN<-?\|\|YjhX_toxin->SAM-METHYLASE->?-> BrxA-DNA-binding -> BrxB-BrxF-iSTAND -> DUF6079 *-> N6-METHYLASE -> OLD-ABC -> TOPRIM -> PglZ -> SIGMA+OB-Fold+AAA-ATPase+Lon-Protease-BrxL ->?-><-REase<-NACHT+SF1-SF2-Helicase+HKD-DNase | 1213 | Bacteria>Pseudomonadota>Gammaproteobacteria>Enterobacterales>Enterobacteriaceae>Salmonella | GCF_025594515.1 | Salmonella_enterica | J3T40_RS14905 | 380423..384064 | 28901 | BREX-system-P-loop-protein-BrxC-[Salmonella-enterica] |
| WP_267272757.1 | Type1.BREX | <- SIGMA+OB-Fold+AAA-ATPase+Lon-Protease-BrxL <- PglZ <- TOPRIM <- OLD-ABC <- N6-METHYLASE <- DUF6079 *<- BrxB-BrxF-iSTAND <- BrxA-DNA-binding <-?<-SAM-METHYLASE<-YjhX_toxin\|\|?->HEPN-> | 1213 | Bacteria>Pseudomonadota>Gammaproteobacteria>Enterobacterales>Enterobacteriaceae>Salmonella | GCF_026460795.1 | Salmonella_enterica | OWO74_RS14195 | 11359..15000 | 28901 | BREX-system-P-loop-protein-BrxC-[Salmonella-enterica] |
| WP_097173203.1 | Type1.BREX | Arm-DNA-bind_3->NACHT+SF1-SF2-Helicase+HKD-DNase->REase-><- SIGMA+OB-Fold+AAA-ATPase+Lon-Protease-BrxL <- PglZ <- TOPRIM <- OLD-ABC <- N6-METHYLASE <- DUF6079 *<- BrxB-BrxF-iSTAND <- BrxA-DNA-binding <-?<-SAM-METHYLASE<-YjhX_toxin\|\|?->REase+iREase->HEPN-> | 1213 | Bacteria>Pseudomonadota>Gammaproteobacteria>Enterobacterales>Enterobacteriaceae>Salmonella | GCF_900003195.1 | Salmonella_enterica | CRP09_RS12230 | 81403..85044 | 28901 | BREX-system-P-loop-protein-BrxC-[Salmonella-enterica] |
| WP_094914772.1 | Type1.BREX | <-HEPN<-?\|\|YjhX_toxin->SAM-METHYLASE->?-> BrxA-DNA-binding -> BrxB-BrxF-iSTAND -> DUF6079 *-> N6-METHYLASE -> OLD-ABC -> TOPRIM -> PglZ -> SIGMA+OB-Fold+AAA-ATPase+Lon-Protease-BrxL ->?-><-REase<-NACHT+SF1-SF2-Helicase+HKD-DNase<-HTH_Tnp_1+HTH_21+rve+rve_3 | 1213 | Bacteria>Pseudomonadota>Gammaproteobacteria>Enterobacterales>Enterobacteriaceae>Salmonella | GCF_903930985.1 | Salmonella_enterica | DQN83_RS20180 | 3946680..3950321 | 28901 | BREX-system-P-loop-protein-BrxC-[Salmonella-enterica] |
| WP_088524188.1 | Type1.BREX | NACHT+SF1-SF2-Helicase+HKD-DNase->REase-><-HKD-DNase+HKD-DNase<- SIGMA+OB-Fold+AAA-ATPase+Lon-Protease-BrxL <- PglZ <- TOPRIM <- OLD-ABC <- N6-METHYLASE -p<- DUF6079 *<- BrxB-BrxF-iSTAND <- BrxA-DNA-binding \|\|ZnR->?-><-?\|\|SIGMA-HTH+REase->?->MarR-wHTH-><- wHTH+WYL+WCX | 1204 | Bacteria>Pseudomonadota>Gammaproteobacteria>Enterobacterales>Enterobacteriaceae>Salmonella | GCF_903931065.1 | Salmonella_enterica | LFZ40_19410 | 3810154..3813768 | 28901 | BREX-system-P-loop-protein-BrxC-[Salmonella-enterica] |
| WP_183072251.1 | Type1.BREX | <-HEPN<-REase+iREase<-?\|\|YjhX_toxin->SAM-METHYLASE->?-> BrxA-DNA-binding -> BrxB-BrxF-iSTAND -> DUF6079 *-> N6-METHYLASE -> OLD-ABC -> TOPRIM -> PglZ -> SIGMA+OB-Fold+AAA-ATPase+Lon-Protease-BrxL -><-REase<-Couple_hipA+HipA_C<-HTH_3 | 1213 | Bacteria>Pseudomonadota>Gammaproteobacteria>Enterobacterales>Enterobacteriaceae>Salmonella | GCF_903931105.1 | Salmonella_enterica | H7T17_RS07610 | 54171..57812 | 28901 | BREX-system-P-loop-protein-BrxC-[Salmonella-enterica] |
| WP_058685106.1 | Type1.BREX | Arm-DNA-bind_3->NACHT+SF1-SF2-Helicase+HKD-DNase->REase-><- SIGMA+OB-Fold+AAA-ATPase+Lon-Protease-BrxL <- PglZ <- TOPRIM <- OLD-ABC <- N6-METHYLASE <- DUF6079 *<- BrxB-BrxF-iSTAND <- BrxA-DNA-binding <-?<-SAM-METHYLASE<-YjhX_toxin\|\|?->REase+iREase->HEPN-> | 1213 | Bacteria>Pseudomonadota>Gammaproteobacteria>Enterobacterales>Enterobacteriaceae>Salmonella | GCF_940677185.1 | Salmonella_enterica | NBW86_RS10200 | 108141..111782 | 28901 | BREX-system-P-loop-protein-BrxC-[Salmonella-enterica] |
| WP_080205562.1 | Type1.BREX | <-HEPN<-REase+iREase<-?\|\|YjhX_toxin->SAM-METHYLASE->?-> BrxA-DNA-binding -> BrxB-BrxF-iSTAND -> DUF6079 *-> N6-METHYLASE -> OLD-ABC -> TOPRIM -> PglZ -> SIGMA+OB-Fold+AAA-ATPase+Lon-Protease-BrxL -><-REase<-Couple_hipA+HipA_C<-HTH_3 | 1213 | Bacteria>Pseudomonadota>Gammaproteobacteria>Enterobacterales>Enterobacteriaceae>Salmonella | GCF_943114325.1 | Salmonella_enterica | OCU44_RS00985 | 212044..215685 | 28901 | BREX-system-P-loop-protein-BrxC-[Salmonella-enterica] |
| WP_039590210.1 | Type1.BREX | <-HEPN<-REase+iREase<-?\|\|YjhX_toxin->SAM-METHYLASE->?-> BrxA-DNA-binding -> BrxB-BrxF-iSTAND -> DUF6079 *-> N6-METHYLASE -> OLD-ABC -> TOPRIM -> PglZ -> SIGMA+OB-Fold+AAA-ATPase+Lon-Protease-BrxL -><-REase<-Couple_hipA+HipA_C<-HTH_3 | 1213 | Bacteria>Pseudomonadota>Gammaproteobacteria>Enterobacterales>Enterobacteriaceae>Salmonella | GCF_943119195.1 | Salmonella_enterica | OCU40_RS05645 | 1155326..1158967 | 28901 | BREX-system-P-loop-protein-BrxC-[Salmonella-enterica] |
| EJF5860346.1 | Type1.BREX | <-HEPN<-REase+iREase<-?\|\|YjhX_toxin->SAM-METHYLASE->?-> BrxA-DNA-binding -> BrxB-BrxF-iSTAND -> DUF6079 *-> N6-METHYLASE -> OLD-ABC -> TOPRIM -> PglZ -> SIGMA+OB-Fold+AAA-ATPase+Lon-Protease-BrxL -><-REase<-NACHT+SF1-SF2-Helicase+HKD-DNase<-Arm-DNA-bind_3 | 1213 | Bacteria>Pseudomonadota>Gammaproteobacteria>Enterobacterales>Enterobacteriaceae>Salmonella | - | Salmonella_enterica | M8X04_001881 | 170514..174155 | 28901 | BREX-system-P-loop-protein-BrxC-[Salmonella-enterica] |
| ECG0705015.1 | Type1.BREX | <-HEPN<-REase+iREase<-?\|\|\|\|YjhX_toxin->SAM-METHYLASE->?->\|\| BrxA-DNA-binding -> BrxB-BrxF-iSTAND -> DUF6079 *-> N6-METHYLASE -> OLD-ABC -> TOPRIM -> PglZ -> SIGMA+OB-Fold+AAA-ATPase+Lon-Protease-BrxL ->HKD-DNase+HKD-DNase-><-REase<-NACHT+SF1-SF2-Helicase+HKD-DNase<-nSTAND1+HNH | 1213 | Bacteria>Pseudomonadota>Gammaproteobacteria>Enterobacterales>Enterobacteriaceae>Salmonella | - | Salmonella_enterica_subsp._salamae | DL122_04700 | 35596..39237 | 59202 | BREX-system-P-loop-protein-BrxC-[Salmonella-enterica-subsp.-salamae] |
| WP_007890763.1 | Type1.BREX | ABhydrolase+METHYLASE->?->?->?->?->?-><-?\|\| BrxA-DNA-binding -> BrxB-BrxF-iSTAND -> DUF6079 *-> N6-METHYLASE -> OLD-ABC -> TOPRIM -> PglZ -> SIGMA+OB-Fold+AAA-ATPase+Lon-Protease-BrxL -><-tRNA<-?\|\|?->?-><- OLD-ABC | 1202 | Bacteria>Pseudomonadota>Gammaproteobacteria>Enterobacterales>Erwiniaceae>Pantoea | GCF_000282675.1 | Pantoea_sp._GM01 | PMI17_RS16905 | 182397..186005 | 1144320 | BREX-system-P-loop-protein-BrxC-[Pantoea-sp.-GM01] |
| WP_137241123.1 | Type1.BREX | <-SymE_toxin<-CHTH<-RelE\|\| BrxA-DNA-binding -> BrxB-BrxF-iSTAND -> DUF6079 *-> N6-METHYLASE -> OLD-ABC -> TOPRIM -> PglZ -> SIGMA+OB-Fold+AAA-ATPase+Lon-Protease-BrxL -> | 1202 | Bacteria>Pseudomonadota>Gammaproteobacteria>Enterobacterales>Erwiniaceae>Pantoea | GCF_005233495.1 | Pantoea_agglomerans | PagCFBP13505_RS08740 | 40890..44498 | 549 | BREX-system-P-loop-protein-BrxC-[Pantoea-agglomerans] |
| WP_154209858.1 | Type1.BREX | <- SIGMA+OB-Fold+AAA-ATPase+Lon-Protease-BrxL <- PglZ <- TOPRIM <- OLD-ABC <- N6-METHYLASE <- DUF6079 *<- BrxB-BrxF-iSTAND <- BrxA-DNA-binding \|\|SymE_toxin-> | 1202 | Bacteria>Pseudomonadota>Gammaproteobacteria>Enterobacterales>Erwiniaceae>Pantoea | GCF_009668105.1 | Pantoea_agglomerans | GJV05_RS04780 | 299589..303197 | 549 | BREX-system-P-loop-protein-BrxC-[Pantoea-agglomerans] |
| WP_222233110.1 | Type1.BREX | <-HKD-DNase+HKD-DNase<- SIGMA+OB-Fold+AAA-ATPase+Lon-Protease-BrxL <- PglZ <- TOPRIM <- OLD-ABC <- N6-METHYLASE <- DUF6079 *<- BrxB-BrxF-iSTAND <- BrxA-DNA-binding <-?<-?<-?<-?<-PRTase<-DADA-Ligase | 1202 | Bacteria>Pseudomonadota>Gammaproteobacteria>Enterobacterales>Erwiniaceae>Pantoea | GCF_019795975.1 | Pantoea_sp._DY-17 | K5Y33_RS01225 | 252070..255678 | 2871490 | BREX-system-P-loop-protein-BrxC-[Pantoea-sp.-DY-17] |
| WP_245115608.1 | Type1.BREX | <-SymE_toxin<-CHTH\|\|ColE7+HNH->MarR-HTH-><-wHTH+REase<-McrB-AAA\|\| BrxA-DNA-binding -> BrxB-BrxF-iSTAND -> DUF6079 *-> N6-METHYLASE -> OLD-ABC -> TOPRIM -> PglZ -> SIGMA+OB-Fold+AAA-ATPase+Lon-Protease-BrxL -> | 1204 | Bacteria>Pseudomonadota>Gammaproteobacteria>Enterobacterales>Erwiniaceae>Pantoea | GCF_022966105.1 | Pantoea_agglomerans | LZ609_RS16055 | 2632558..2636172 | 549 | BREX-system-P-loop-protein-BrxC-[Pantoea-agglomerans] |
| WP_115625931.1 | Type1.BREX | <-HKD-DNase+HKD-DNase\|\|HIN-HTH->\|\|?->LZ_Tnp_IS66+zf-IS66+DDE_Tnp_IS66+DDE_Tnp_IS66_C->\|\|RVT->HTH_32+rve-><-ZnR\|\| BrxA-DNA-binding -> BrxB-BrxF-iSTAND -> DUF6079 *-> N6-METHYLASE -> OLD-ABC -> TOPRIM -> PglZ -> SIGMA+OB-Fold+AAA-ATPase+Lon-Protease-BrxL -><-?<-DUF772+DDE_Tnp_1<-XIS-HTH\|\|<-Phage_Cox | 1204 | Bacteria>Pseudomonadota>Gammaproteobacteria>Enterobacterales | GCF_013635755.1 | Enterobacteriaceae | HV027_RS18510 | 3797222..3800836 | 543 | MULTISPECIES:-BREX-system-P-loop-protein-BrxC-[Enterobacteriaceae] |
| WP_222691531.1 | Type1.BREX | BrxA-DNA-binding -> BrxB-BrxF-iSTAND -> DUF6079 *-> N6-METHYLASE -> OLD-ABC -> TOPRIM -> PglZ -> SIGMA+OB-Fold+AAA-ATPase+Lon-Protease-BrxL ->SF1-SF2-Helicase+HTH-><-Arm-DNA-bind_3+Phage_integrase\|\|Phage_integrase-> | 1204 | Bacteria>Pseudomonadota>Gammaproteobacteria>Enterobacterales | GCF_019841255.1 | Enterobacteriaceae | K7A35_RS15405 | 1428..5042 | 543 | MULTISPECIES:-BREX-system-P-loop-protein-BrxC-[Enterobacteriaceae] |
| WP_117108864.1 | Type1.BREX | <-METHYLASE\|\|SKI-><-?\|\|tRNA-><- SIGMA+OB-Fold+AAA-ATPase+Lon-Protease-BrxL <- PglZ <- TOPRIM <- OLD-ABC <- N6-METHYLASE <- DUF6079 *<- BrxB-BrxF-iSTAND <- BrxA-DNA-binding \|\|<-DEDD_Tnp_IS110+Transposase_20\|\|HTH_58->Phage_integrase->Arm-DNA-bind_3+Phage_integrase->XIS-HTH->DnaB-> | 1189 | Bacteria>Pseudomonadota>Gammaproteobacteria>Enterobacterales | GCF_903684385.1 | Enterobacteriaceae | MJ801_RS10050 | 169597..173166 | 543 | MULTISPECIES:-BREX-system-P-loop-protein-BrxC-[Enterobacteriaceae] |
| WP_226228481.1 | Type1.BREX | GNTR-HTH+Aminotran_1_2->?-><-wHTH+REase<-McrB-AAA\|\| BrxA-DNA-binding -> BrxB-BrxF-iSTAND -> DUF6079 *-> N6-METHYLASE -> OLD-ABC -> TOPRIM -> PglZ -> SIGMA+OB-Fold+AAA-ATPase+Lon-Protease-BrxL ->SF1-SF2-Helicase+HTH-> | 1202 | Bacteria>Pseudomonadota>Gammaproteobacteria>Enterobacterales>Hafniaceae>Hafnia | GCF_020450285.1 | Hafnia_paralvei | LA328_RS17860 | 3833328..3836936 | 546367 | BREX-system-P-loop-protein-BrxC-[Hafnia-paralvei] |
| WP_241543889.1 | Type1.BREX | <-HTH_37<-?\|\|GDResolvase+HIN-HTH-> wHTH+WYL+WCX -> BrxA-DNA-binding -> BrxB-BrxF-iSTAND -> DUF6079 *-> OLD-ABC -> TOPRIM -> N6-METHYLASE -> PglZ -> SIGMA+OB-Fold+AAA-ATPase+Lon-Protease-BrxL -> | 1225 | Bacteria>Pseudomonadota>Gammaproteobacteria>Enterobacterales>Morganellaceae>Moellerella | GCF_022591895.1 | Moellerella_wisconsensis | MNY64_RS16755 | 120814..124491 | 158849 | BREX-system-P-loop-protein-BrxC-[Moellerella-wisconsensis] |
| WP_241501907.1 | Type1.BREX | <-HTH_37<-?\|\|GDResolvase+HIN-HTH-> wHTH+WYL+WCX -> BrxA-DNA-binding -> BrxB-BrxF-iSTAND -> DUF6079 *-> OLD-ABC -> TOPRIM -> N6-METHYLASE -> PglZ -> SIGMA+OB-Fold+AAA-ATPase+Lon-Protease-BrxL -> | 1225 | Bacteria>Pseudomonadota>Gammaproteobacteria>Enterobacterales>Morganellaceae>Moellerella | GCF_022592395.1 | Moellerella_wisconsensis | MNY66_RS17045 | 197034..200711 | 158849 | BREX-system-P-loop-protein-BrxC-[Moellerella-wisconsensis] |
| HAT2164885.1 | Type1.BREX | <-MULE-TRANSPOSASE<-?<-LexA-protease\|\|?->35exo->?-> wHTH+WYL+WCX -> BrxA-DNA-binding -> BrxB-BrxF-iSTAND -> DUF6079 *->ParB+HNH-> N6-METHYLASE -> OLD-ABC -> TOPRIM -> PglZ -> SIGMA+OB-Fold+AAA-ATPase+Lon-Protease-BrxL ->SbcC-> | 1225 | Bacteria>Pseudomonadota>Gammaproteobacteria>Enterobacterales>Morganellaceae>Proteus | GCA_015896365.1 | Proteus_mirabilis | I8052_002910 | 10079..13756 | 584 | BREX-system-P-loop-protein-BrxC-[Proteus-mirabilis] |
| MBI6417160.1 | Type1.BREX | <-CARF+NtrC-AAA+FIS-HTH<- SIGMA+OB-Fold+AAA-ATPase+Lon-Protease-BrxL <- PglZ <- TOPRIM <- OLD-ABC <- N6-METHYLASE <-ParB+HNH<- DUF6079 *<- BrxB-BrxF-iSTAND <- BrxA-DNA-binding <- wHTH+WYL+WCX <-XIS-HTH<-?\|\|?->?->wHTH+WYL-> | 1225 | Bacteria>Pseudomonadota>Gammaproteobacteria>Enterobacterales>Morganellaceae>Proteus | GCA_016238675.1 | Proteus_mirabilis | JEP37_15655 | 69055..72732 | 584 | BREX-system-P-loop-protein-BrxC-[Proteus-mirabilis] |
| WP_210849185.1 | Type1.BREX | <-OBFOLD-NUCLEASE<-CARF+NtrC-AAA+FIS-HTH<- SIGMA+OB-Fold+AAA-ATPase+Lon-Protease-BrxL <- PglZ <- TOPRIM <- OLD-ABC <- N6-METHYLASE <-ParB+HNH<- DUF6079 *<- BrxB-BrxF-iSTAND <- BrxA-DNA-binding <-XIS-HTH<-?\|\|?->?->wHTH+WYL-> | 1225 | Bacteria>Pseudomonadota>Gammaproteobacteria>Enterobacterales>Morganellaceae>Proteus | GCF_022349905.1 | Proteus_mirabilis | MJ636_RS02915 | 47390..51067 | 584 | BREX-system-P-loop-protein-BrxC-[Proteus-mirabilis] |
| WP_209036130.1 | Type1.BREX | <-wHTH+WYL<-?<-?\|\|?->XIS-HTH-> BrxA-DNA-binding -> BrxB-BrxF-iSTAND -> DUF6079 *-> OLD-ABC -> TOPRIM -> N6-METHYLASE -> PglZ -> SIGMA+OB-Fold+AAA-ATPase+Lon-Protease-BrxL ->SbcC-><-ArdA<-?<-Arm-DNA-bind_3+Phage_integrase | 1225 | Bacteria>Pseudomonadota>Gammaproteobacteria>Enterobacterales>Morganellaceae>Proteus | GCF_029542405.1 | Proteus_mirabilis | IGB04_RS13130 | 82611..86288 | 584 | BREX-system-P-loop-protein-BrxC-[Proteus-mirabilis] |
| WP_137022260.1 | Type1.BREX | <-GTPase-AIG+sGTP<-CopG/RHH\|\|?->XIS-HTH-> BrxA-DNA-binding -> BrxB-BrxF-iSTAND -> DUF6079 *-> N6-METHYLASE -> OLD-ABC -> TOPRIM -> PglZ -> SIGMA+OB-Fold+AAA-ATPase+Lon-Protease-BrxL -><-Arm-DNA-bind_3+Phage_integrase<-tRNA\|\|?->?->Peptidase_C80-> | 1204 | Bacteria>Pseudomonadota>Gammaproteobacteria>Enterobacterales>Morganellaceae>Providencia | GCF_005157325.1 | Providencia_heimbachae | C9446_RS17890 | 3853703..3857317 | 333962 | BREX-system-P-loop-protein-BrxC-[Providencia-heimbachae] |
| WP_265497329.1 | Type1.BREX | <-HTH_37<-?\|\|GDResolvase+HIN-HTH-> wHTH+WYL+WCX -> BrxA-DNA-binding -> BrxB-BrxF-iSTAND -> DUF6079 *-> OLD-ABC -> TOPRIM -> N6-METHYLASE -> PglZ -> SIGMA+OB-Fold+AAA-ATPase+Lon-Protease-BrxL -> | 1225 | Bacteria>Pseudomonadota>Gammaproteobacteria>Enterobacterales>Morganellaceae>Providencia | GCF_026173365.1 | Providencia_rustigianii | ORM28_RS01805 | 9364..13041 | 158850 | BREX-system-P-loop-protein-BrxC-[Providencia-rustigianii] |
| WP_272668877.1 | Type1.BREX | XIS-HTH->HEPN+SF1-SF2-Helicase+RNase+RNase->TPR->\|\| wHTH+WYL+WCX -> BrxA-DNA-binding -> BrxB-BrxF-iSTAND -> DUF6079 *->ParB+HNH-> N6-METHYLASE -> OLD-ABC -> TOPRIM -> PglZ -> SIGMA+OB-Fold+AAA-ATPase+Lon-Protease-BrxL ->CARF+NtrC-AAA+FIS-HTH->OBFOLD-NUCLEASE-> | 1225 | Bacteria>Pseudomonadota>Gammaproteobacteria>Enterobacterales>Morganellaceae>Providencia | GCF_028481005.1 | Providencia_sp._PROV151 | PSE52_RS00990 | 220703..224380 | 2949861 | BREX-system-P-loop-protein-BrxC-[Providencia-sp.-PROV151] |
| WP_045967689.1 | Type1.BREX | <-ACET<- BrxB-BrxF-iSTAND <- SIGMA+OB-Fold+AAA-ATPase+Lon-Protease-BrxL <- PglZ <- TOPRIM <- OLD-ABC <- N6-METHYLASE <- DUF6079 *<- BrxB-BrxF-iSTAND <- BrxA-DNA-binding <-DDE_3 | 1212 | Bacteria>Pseudomonadota>Gammaproteobacteria>Enterobacterales>Morganellaceae>Xenorhabdus | GCF_000968195.1 | Xenorhabdus_doucetiae | LY16_RS12985 | 24233..27871 | 351671 | BREX-system-P-loop-protein-BrxC-[Xenorhabdus-doucetiae] |
| WP_040031759.1 | Type1.BREX | <-HIN-HTH\|\|DUF4096+DDE_Tnp_1+DDE_Tnp_1_2->?->tRNA->tRNA->Arm-DNA-bind_3+Phage_integrase-><- SIGMA+OB-Fold+AAA-ATPase+Lon-Protease-BrxL <- PglZ <- TOPRIM <- OLD-ABC <- N6-METHYLASE <- DUF6079 *<- BrxB-BrxF-iSTAND <- BrxA-DNA-binding <-wHTH+SAD+HNH<-LuxR-HTH<-?<-TPR | 1202 | Bacteria>Pseudomonadota>Gammaproteobacteria>Enterobacterales>Pectobacteriaceae | GCF_026723725.1 | Pectobacterium | OWC53_RS08230 | 1782306..1785914 | 122277 | MULTISPECIES:-BREX-system-P-loop-protein-BrxC-[Pectobacterium] |
| GKX39595.1 | Type1.BREX | Arm-DNA-bind_3+Phage_integrase-><- SIGMA+OB-Fold+AAA-ATPase+Lon-Protease-BrxL <- PglZ <- TOPRIM <- OLD-ABC <- N6-METHYLASE <- DUF6079 *<- BrxB-BrxF-iSTAND <- BrxA-DNA-binding <-wHTH+SAD+HNH<-LuxR-HTH<-?<-TPR | 1202 | Bacteria>Pseudomonadota>Gammaproteobacteria>Enterobacterales>Pectobacteriaceae>Pectobacterium | GCA_024343445.1 | Pectobacterium_carotovorum_subsp._carotovorum | SOASR014_33340 | 124830..128438 | 555 | hypothetical-protein-SOASR014_33340-[Pectobacterium-carotovorum-subsp.-carotovorum] |
| WP_005974434.1 | Type1.BREX | REC+OmpR-HTH->URI-DNase+UVR+RNase+HHH->?->tRNA->tRNA->tRNA->Arm-DNA-bind_3+Phage_integrase-><- SIGMA+OB-Fold+AAA-ATPase+Lon-Protease-BrxL <- PglZ <- TOPRIM <- OLD-ABC <- N6-METHYLASE <- DUF6079 *<- BrxB-BrxF-iSTAND <- BrxA-DNA-binding <-LuxR-HTH<-?<-ZnR<-XIS-HTH<-?\|\|CopG/RHH->CopG/RHH-> | 1204 | Bacteria>Pseudomonadota>Gammaproteobacteria>Enterobacterales>Pectobacteriaceae>Pectobacterium | GCF_000291725.1 | Pectobacterium_wasabiae | Y17_RS17690 | 22336..25950 | 55208 | BREX-system-P-loop-protein-BrxC-[Pectobacterium-wasabiae] |
| WP_103160989.1 | Type1.BREX | LuxR-HTH-> BrxA-DNA-binding -> BrxB-BrxF-iSTAND -> DUF6079 *-> N6-METHYLASE -> OLD-ABC -> TOPRIM -> PglZ -> SIGMA+OB-Fold+AAA-ATPase+Lon-Protease-BrxL -> | 1202 | Bacteria>Pseudomonadota>Gammaproteobacteria>Enterobacterales>Pectobacteriaceae>Pectobacterium | GCF_002904045.1 | Pectobacterium_odoriferum | BVY05_RS19610 | 74039..77647 | 78398 | BREX-system-P-loop-protein-BrxC-[Pectobacterium-odoriferum] |
| WP_103199124.1 | Type1.BREX | <-ParA-Soj-PloopNTPase<-XIS-HTH<-MODE-HTH+PBPII\|\|tRNA->tRNA->Arm-DNA-bind_3+Phage_integrase-><- SIGMA+OB-Fold+AAA-ATPase+Lon-Protease-BrxL <- PglZ <- TOPRIM <- OLD-ABC <- N6-METHYLASE <- DUF6079 *<- BrxB-BrxF-iSTAND <- BrxA-DNA-binding <-wHTH+SAD+HNH<-LuxR-HTH | 1202 | Bacteria>Pseudomonadota>Gammaproteobacteria>Enterobacterales>Pectobacteriaceae>Pectobacterium | GCF_002904085.1 | Pectobacterium_odoriferum | BV921_RS15390 | 113976..117584 | 78398 | BREX-system-P-loop-protein-BrxC-[Pectobacterium-odoriferum] |
| WP_116155987.1 | Type1.BREX | LuxR-HTH->wHTH+SAD+HNH-> BrxA-DNA-binding -> BrxB-BrxF-iSTAND -> DUF6079 *-> N6-METHYLASE -> OLD-ABC -> TOPRIM -> PglZ -> SIGMA+OB-Fold+AAA-ATPase+Lon-Protease-BrxL -><-Arm-DNA-bind_3+Phage_integrase<-tRNA<-tRNA<-?<-URI-DNase+UVR+RNase+HHH<-REC+OmpR-HTH | 1202 | Bacteria>Pseudomonadota>Gammaproteobacteria>Enterobacterales>Pectobacteriaceae>Pectobacterium | GCF_003382585.2 | Pectobacterium_aquaticum | DMB81_RS10735 | 4900..8508 | 2204145 | BREX-system-P-loop-protein-BrxC-[Pectobacterium-aquaticum] |
| WP_181847885.1 | Type1.BREX | REC+OmpR-HTH->?->tRNA->tRNA->Arm-DNA-bind_3+Phage_integrase-><- SIGMA+OB-Fold+AAA-ATPase+Lon-Protease-BrxL <- PglZ <- TOPRIM <- OLD-ABC <- N6-METHYLASE <- DUF6079 *<- BrxB-BrxF-iSTAND <- BrxA-DNA-binding <-wHTH+SAD+HNH<-LuxR-HTH<-?<-?<-ZnR<-XIS-HTH | 1204 | Bacteria>Pseudomonadota>Gammaproteobacteria>Enterobacterales>Pectobacteriaceae>Pectobacterium | GCF_013913645.1 | Pectobacterium_aroidearum | H2Y54_RS11285 | 236602..240216 | 1201031 | BREX-system-P-loop-protein-BrxC-[Pectobacterium-aroidearum] |
| WP_138775748.1 | Type1.BREX | ZnR->XIS-HTH->ZnR->?->?->HTH+MPTase-><-?\|\| BrxA-DNA-binding -> BrxB-BrxF-iSTAND -> DUF6079 *-> N6-METHYLASE -> OLD-ABC -> TOPRIM -> PglZ -> SIGMA+OB-Fold+AAA-ATPase+Lon-Protease-BrxL -><-Arm-DNA-bind_3+Phage_integrase<-tRNA<-?<-?\|\|Peptidase_M17_N+Peptidase_M17-> | 1202 | Bacteria>Pseudomonadota>Gammaproteobacteria>Enterobacterales>Yersiniaceae | GCF_902168625.1 | Yersinia | FFE93_018510 | 3893626..3897234 | 629 | MULTISPECIES:-BREX-system-P-loop-protein-BrxC-[Yersinia] |
| WP_112197427.1 | Type1.BREX | <-CopG/RHH\|\|?->XIS-HTH->ZnR->ZnR->?->?-> BrxA-DNA-binding -> BrxB-BrxF-iSTAND -> DUF6079 *-> N6-METHYLASE -> OLD-ABC -> TOPRIM -> PglZ -> SIGMA+OB-Fold+AAA-ATPase+Lon-Protease-BrxL ->HKD-DNase+HKD-DNase-><-Arm-DNA-bind_3+Phage_integrase | 1204 | Bacteria>Pseudomonadota>Gammaproteobacteria>Enterobacterales>Yersiniaceae>Rahnella | GCF_003263525.1 | Rahnella_sp._NRRL_B-41462 | UC14_RS06670 | 31474..35088 | 1610579 | BREX-system-P-loop-protein-BrxC-[Rahnella-sp.-NRRL-B-41462] |
| HBE9089660.1 | Type1.BREX | HTH+McrB-AAA->wHTH+REase-><-?<-?<-?<-wHTH+WYL<-nSTAND1+GTPase-AIG+sGTP\|\| BrxA-DNA-binding -> BrxB-BrxF-iSTAND -> DUF6079 *-> N6-METHYLASE -> OLD-ABC -> TOPRIM -> PglZ -> SIGMA+OB-Fold+AAA-ATPase+Lon-Protease-BrxL -><-?<-AAA+wHTH<-XIS-HTH | 1204 | Bacteria>Pseudomonadota>Gammaproteobacteria>Enterobacterales>Yersiniaceae>Serratia | GCA_018627075.1 | Serratia_fonticola | KNG84_001684 | 34672..38286 | 47917 | BREX-system-P-loop-protein-BrxC-[Serratia-fonticola] |
| WP_195869502.1 | Type1.BREX | <-Peptidase_M17_N+Peptidase_M17\|\|?->tRNA->Arm-DNA-bind_3+Phage_integrase-><-HKD-DNase+HKD-DNase<- SIGMA+OB-Fold+AAA-ATPase+Lon-Protease-BrxL <- PglZ <- TOPRIM <- OLD-ABC <- N6-METHYLASE <- DUF6079 *<- BrxB-BrxF-iSTAND <- BrxA-DNA-binding <-?<-?\|\|<-ZnR<-XIS-HTH<-?\|\| OLD-ABC + TOPRIM -> | 1202 | Bacteria>Pseudomonadota>Gammaproteobacteria>Enterobacterales>Yersiniaceae>Serratia | - | Serratia_liquefaciens | I1A46_RS14725 | 213767..217375 | 614 | BREX-system-P-loop-protein-BrxC-[Serratia-liquefaciens] |
| WP_004722527.1 | Type1.BREX | <-GTPase-AIG<-CopG/RHH\|\|?->XIS-HTH->?->HTH+MPTase->\|\| BrxA-DNA-binding -> BrxB-BrxF-iSTAND -> DUF6079 *-> N6-METHYLASE -> OLD-ABC -> TOPRIM -> PglZ -> SIGMA+OB-Fold+AAA-ATPase+Lon-Protease-BrxL -><-Arm-DNA-bind_3+Phage_integrase | 1202 | Bacteria>Pseudomonadota>Gammaproteobacteria>Enterobacterales>Yersiniaceae>Yersinia | GCF_000173755.1 | Yersinia_ruckeri | YRUCK0001_RS14080 | 15308..18916 | 29486 | BREX-system-P-loop-protein-BrxC-[Yersinia-ruckeri] |
| WP_050124850.1 | Type1.BREX | DUF4007->?->\|\|?-><-nSTAND1+HNH\|\| BrxA-DNA-binding -> BrxB-BrxF-iSTAND -> DUF6079 *-> N6-METHYLASE -> OLD-ABC -> TOPRIM -> PglZ -> SIGMA+OB-Fold+AAA-ATPase+Lon-Protease-BrxL -><-Arm-DNA-bind_3+Phage_integrase<-tRNA<-?<-?\|\|Peptidase_M17_N+Peptidase_M17-> | 1202 | Bacteria>Pseudomonadota>Gammaproteobacteria>Enterobacterales>Yersiniaceae>Yersinia | GCF_001115185.1 | Yersinia_aleksiciae | AEB18_RS00865 | 216657..220265 | 263819 | BREX-system-P-loop-protein-BrxC-[Yersinia-aleksiciae] |
| WP_053011787.1 | Type1.BREX | XIS-HTH->?->ZnR->XIS-HTH->ZnR->?->HTH+MPTase->\|\| BrxA-DNA-binding -> BrxB-BrxF-iSTAND -> DUF6079 *-> N6-METHYLASE -> OLD-ABC -> TOPRIM -> PglZ -> SIGMA+OB-Fold+AAA-ATPase+Lon-Protease-BrxL ->HKD-DNase+HKD-DNase-><-Arm-DNA-bind_3+Phage_integrase<-tRNA\|\|?->HTH_3 | 1202 | Bacteria>Pseudomonadota>Gammaproteobacteria>Enterobacterales>Yersiniaceae>Yersinia | GCF_001244675.1 | Yersinia_intermedia | ALC37_RS11575 | 459968..463576 | 631 | BREX-system-P-loop-protein-BrxC-[Yersinia-intermedia] |
| WP_128451204.1 | Type1.BREX | Phage_Cox->XIS-HTH->AAA+wHTH->?-><- SIGMA+OB-Fold+AAA-ATPase+Lon-Protease-BrxL <- PglZ <- TOPRIM <- OLD-ABC <- N6-METHYLASE <- DUF6079 *<- BrxB-BrxF-iSTAND <- BrxA-DNA-binding \|\|nSTAND1+GTPase-AIG+sGTP->DUF932->wHTH+WYL->?->?->DUF905->ArdB-> | 1202 | Bacteria>Pseudomonadota>Gammaproteobacteria>Enterobacterales>Yersiniaceae>Yersinia | GCF_004100405.1 | Yersinia_sp._2105_StPb_PI | EQP49_RS07200 | 292011..295619 | 2507058 | BREX-system-P-loop-protein-BrxC-[Yersinia-sp.-2105-StPb-PI] |
| WP_145931270.1 | Type1.BREX | XIS-HTH->ZnR->?->HTH+MPTase->\|\| BrxA-DNA-binding -> BrxB-BrxF-iSTAND -> DUF6079 *-> N6-METHYLASE -> OLD-ABC -> TOPRIM -> PglZ -> SIGMA+OB-Fold+AAA-ATPase+Lon-Protease-BrxL -><-Arm-DNA-bind_3+Phage_integrase<-tRNA<-?<-?\|\|Peptidase_M17_N+Peptidase_M17-> | 1202 | Bacteria>Pseudomonadota>Gammaproteobacteria>Enterobacterales>Yersiniaceae>Yersinia | GCF_902168565.1 | Yersinia_bercovieri | FS908_RS01775 | 53708..57316 | 634 | BREX-system-P-loop-protein-BrxC-[Yersinia-bercovieri] |
| WP_145532653.1 | Type1.BREX | XIS-HTH->ZnR->?->?->HTH+MPTase->\|\| BrxA-DNA-binding -> BrxB-BrxF-iSTAND -> DUF6079 *-> N6-METHYLASE -> OLD-ABC -> TOPRIM -> PglZ -> SIGMA+OB-Fold+AAA-ATPase+Lon-Protease-BrxL -><-Arm-DNA-bind_3+Phage_integrase<-tRNA<-?<-?\|\|Peptidase_M17_N+Peptidase_M17-> | 1202 | Bacteria>Pseudomonadota>Gammaproteobacteria>Enterobacterales>Yersiniaceae>Yersinia | GCF_902170345.1 | Yersinia_alsatica | FTB07_RS01240 | 22698..26306 | 2890317 | BREX-system-P-loop-protein-BrxC-[Yersinia-alsatica] |
| WP_145580439.1 | Type1.BREX | XIS-HTH->?->HTH+MPTase->\|\| BrxA-DNA-binding -> BrxB-BrxF-iSTAND -> DUF6079 *-> N6-METHYLASE -> OLD-ABC -> TOPRIM -> PglZ -> SIGMA+OB-Fold+AAA-ATPase+Lon-Protease-BrxL -><-Arm-DNA-bind_3+Phage_integrase<-tRNA<-?<-?\|\|Peptidase_M17_N+Peptidase_M17-> | 1202 | Bacteria>Pseudomonadota>Gammaproteobacteria>Enterobacterales>Yersiniaceae>Yersinia | GCF_902170795.1 | Yersinia_mollaretii | FTB33_RS16505 | 6738..10346 | 33060 | BREX-system-P-loop-protein-BrxC-[Yersinia-mollaretii] |
| WP_132538615.1 | Type6.BREX | <-LexA-protease\|\|?-><-?<- SIGMA+OB-Fold+AAA-ATPase+Lon-Protease-BrxL <- PglZ <-HNH<- OLD-ABC <- N6-METHYLASE <-?<- DUF6079 *<- BrxB-BrxF-iSTAND <- BrxA-DNA-binding <- BrxE <-RHH_5\|\|?->?->XIS-HTH->?->ZnR+ TOPRIM +AAA-helicase+wHTH-> | 1179 | Bacteria>Pseudomonadota>Gammaproteobacteria>Gammaproteobacteria_incertae_sedis>Candidatus_Competibacteraceae>Plasticicumulans | GCF_004341245.1 | Plasticicumulans_lactativorans | EV699_RS02475 | 34554..38093 | 1133106 | BREX-system-P-loop-protein-BrxC-[Plasticicumulans-lactativorans] |
| MBK8754669.1 | Type6.BREX | ACET-><-CR-ATPase<-?\|\|?->ParB+HNH->SF1-SF2-Helicase-> BrxE -> BrxA-DNA-binding -> BrxB-BrxF-iSTAND -> DUF6079 *->REase+AIPR-> N6-METHYLASE ->HTH_Tnp_4+DDE_Tnp_4->DDE_Tnp->?-> OLD-ABC -> TOPRIM -> | 1209 | Bacteria>Pseudomonadota>Gammaproteobacteria>Gammaproteobacteria_incertae_sedis>Candidatus_Competibacteraceae>unclassified_Candidatus_Competibacteraceae | GCA_016710535.1 | Candidatus_Competibacteraceae_bacterium | IPL99_24865 | 32033..35662 | 2053538 | BREX-system-P-loop-protein-BrxC-[Candidatus-Competibacteraceae-bacterium] |
| WP_248566606.1 | Type1.BREX | SF1-SF2-Helicase+REase->?-><- SIGMA+OB-Fold+AAA-ATPase+Lon-Protease-BrxL <- PglZ <- TOPRIM <- OLD-ABC <-HTH+REase<- N6-METHYLASE <-Schlafen+HATPase+wHTH<- DUF6079 *<- BrxB-BrxF-iSTAND <- BrxA-DNA-binding <-SF1-SF2-Helicase<-HKD-DNase<-DUF6361<-HEPN\|\|?->URI-DNase-> | 1145 | Bacteria>Pseudomonadota>Gammaproteobacteria>Gammaproteobacteria_incertae_sedis>Methylonatrum | GCF_023195885.1 | Methylonatrum_kenyense | M0534_RS01350 | 290769..294206 | 455253 | BREX-system-P-loop-protein-BrxC-[Methylonatrum-kenyense] |
| WP_177417917.1 | Type1.BREX | BrxB-BrxF-iSTAND -> DUF6079 *->MazE_antitoxin->PIN-> N6-METHYLASE -> OLD-ABC -> TOPRIM -> PglZ -> SIGMA+OB-Fold+AAA-ATPase+Lon-Protease-BrxL -> | 1188 | Bacteria>Pseudomonadota>Gammaproteobacteria>Gammaproteobacteria_incertae_sedis>sulfur-oxidizing_symbionts | GCF_011947365.2 | endosymbiont_of_Lamellibrachia_barhami | G9Q73_RS00010 | 458..4024 | 205975 | BREX-system-P-loop-protein-BrxC-[endosymbiont-of-Lamellibrachia-barhami] |
| WP_112020499.1 | Type1.BREX | <-DUF905<-?<-?<-?<-wHTH+WYL<-DUF932<-nSTAND1+GTPase-AIG+sGTP\|\| BrxA-DNA-binding -> BrxB-BrxF-iSTAND -> DUF6079 *-> N6-METHYLASE -> OLD-ABC -> TOPRIM -> PglZ -> SIGMA+OB-Fold+AAA-ATPase+Lon-Protease-BrxL -> | 1204 | Bacteria>Pseudomonadota>Gammaproteobacteria | GCF_013927185.1 | Enterobacterales | DP180_RS12665 | 36252..39866 | 91347 | MULTISPECIES:-BREX-system-P-loop-protein-BrxC-[Enterobacterales] |
| MAZ66828.1 | Type1.BREX | <- SIGMA+OB-Fold+AAA-ATPase+Lon-Protease-BrxL <- PglZ <- TOPRIM <- OLD-ABC <- N6-METHYLASE <-HTH+REase<- DUF6079 *<- BrxB-BrxF-iSTAND <- BrxA-DNA-binding <- wHTH+WYL+WCX <-Trypsin+PDZ | 1234 | Bacteria>Pseudomonadota>Gammaproteobacteria>Kangiellales>Kangiellaceae>unclassified_Kangiellaceae | GCA_002707725.1 | Kangiellaceae_bacterium | CMF25_06935 | 12477..16181 | 2026751 | DNA-repair-protein-[Kangiellaceae-bacterium] |
| WP_031438510.1 | Type1.BREX | HD->?->?->\|\|Phage_integrase->Phage_integrase-> wHTH+WYL+WCX -> BrxA-DNA-binding -> BrxB-BrxF-iSTAND -> DUF6079 *->HKD-DNase+DUF3150-> N6-METHYLASE -> OLD-ABC -> TOPRIM ->DOC+HTH-> PglZ -> SIGMA+OB-Fold+AAA-ATPase+Lon-Protease-BrxL ->SMODS->SLATT-> | 1218 | Bacteria>Pseudomonadota>Gammaproteobacteria>Methylococcales>Methylococcaceae>Methylobacter | GCF_000733835.1 | Methylobacter_tundripaludum | T449_RS0121480 | 4638570..4642226 | 173365 | BREX-system-P-loop-protein-BrxC-[Methylobacter-tundripaludum] |
| WP_262965888.1 | Type1.BREX | SIGMA+OB-Fold+AAA-ATPase+Lon-Protease-BrxL ->?-><-DDE_3\|\|?-> wHTH+WYL+WCX -> BrxA-DNA-binding -> BrxB-BrxF-iSTAND -> DUF6079 *->Calcineurin->SbcC+SbcC-> N6-METHYLASE -> OLD-ABC -> TOPRIM -> PglZ -> SIGMA+OB-Fold+AAA-ATPase+Lon-Protease-BrxL ->?->ANK+ANK+ANK-> | 1212 | Bacteria>Pseudomonadota>Gammaproteobacteria>Methylococcales>Methylococcaceae>Methylobacter | GCF_025583945.1 | Methylobacter_psychrophilus | N7974_RS12230 | 10707..14345 | 96941 | BREX-system-P-loop-protein-BrxC-[Methylobacter-psychrophilus] |
| MBS3964846.1 | Type1.BREX | HTH_Tnp_1+HTH_21+rve+rve_3-> BrxA-DNA-binding -> BrxB-BrxF-iSTAND -> DUF6079 *->Calcineurin->SbcC+SbcC->SIR2-> N6-METHYLASE -> OLD-ABC -> TOPRIM ->DOC+HTH-> PglZ -> SIGMA+OB-Fold+AAA-ATPase+Lon-Protease-BrxL -> | 1209 | Bacteria>Pseudomonadota>Gammaproteobacteria>Methylococcales>Methylococcaceae>Methylomonas | GCA_018333915.1 | Methylomonas_sp. | KGZ80_10205 | 1968..5597 | 418 | BREX-system-P-loop-protein-BrxC-[Methylomonas-sp.] |
| WP_255188315.1 | Type1.BREX | <-PIN<-PIN<- SIGMA+OB-Fold+AAA-ATPase+Lon-Protease-BrxL <- PglZ <- TOPRIM <- OLD-ABC <- N6-METHYLASE <-PIN<-ParD_antitoxin<- DUF6079 *<- BrxB-BrxF-iSTAND <- BrxA-DNA-binding \|\|?-><-?<-HIN-HTH\|\|?-><-ACET+ACET<-TraJ-RHH | 1206 | Bacteria>Pseudomonadota>Gammaproteobacteria>Methylococcales>Methylococcaceae>Methylomonas | GCF_024360925.2 | Methylomonas_rapida | NM686_RS13235 | 2825288..2828908 | 2963939 | BREX-system-P-loop-protein-BrxC-[Methylomonas-rapida] |
| WP_255189595.1 | Type1.BREX | <-DDE_Tnp_1_4\|\|?-><-zf-ISL3+HTH_Tnp_ISL3+DDE_Tnp_ISL3\|\|?->DDE_Tnp_1_4->Phage_integrase+Phage_integrase->35exo-> wHTH+WYL+WCX -> BrxB-BrxF-iSTAND -> DUF6079 *->Calcineurin->SbcC+SbcC-> N6-METHYLASE -p->zf-ISL3+HTH_Tnp_ISL3+DDE_Tnp_ISL3-> N6-METHYLASE -> OLD-ABC -> TOPRIM ->DOC+HTH-> PglZ -> | 1209 | Bacteria>Pseudomonadota>Gammaproteobacteria>Methylococcales>Methylococcaceae>Methylomonas | GCF_024360925.2 | Methylomonas_rapida | NM686_RS20105 | 4241783..4245412 | 2963939 | BREX-system-P-loop-protein-BrxC-[Methylomonas-rapida] |
| MBE0471466.1 | Type1.BREX | <- N6-METHYLASE -p\|\|SbcC-> TOPRIM <-?<- DUF6079 *<- BrxB-BrxF-iSTAND <- BrxA-DNA-binding | 1214 | Bacteria>Pseudomonadota>Gammaproteobacteria>Methylococcales>Methylococcaceae>Methyloprofundus | GCA_014859465.1 | Methyloprofundus_sp. | IBX55_18395 | 2520..6164 | 2020875 | BREX-system-P-loop-protein-BrxC-[Methyloprofundus-sp.] |
| CAM84763.1 | Type1.BREX | <- OLD-ABC + TOPRIM <-HIN-HTH<-SF1-SF2-Helicase+REase+HTH<- SIGMA+OB-Fold+AAA-ATPase+Lon-Protease-BrxL <- PglZ <-?<- N6-METHYLASE <- DUF6079 *<- BrxB-BrxF-iSTAND <- BrxA-DNA-binding <- wHTH+WYL+WCX \|\|<-?\|\|DDE_Tnp_1_5+DDE_Tnp_1-><-?<-?<-?\|\|TRPR-HTH-> | 1229 | Bacteria>Pseudomonadota>Gammaproteobacteria>Moraxellales>Moraxellaceae>Acinetobacter | GCA_000069245.1 | Acinetobacter_baumannii_AYE | p3ABAYE0070 | 63476..67165 | 509173 | putative-ATPase-like-protein-[Acinetobacter-baumannii-AYE] |
| EXS00102.1 | Type1.BREX | <-DDE_Tnp_1_5+DDE_Tnp_1\|\|?->DOC+HTH-> wHTH+WYL+WCX -> BrxA-DNA-binding -> BrxB-BrxF-iSTAND -> DUF6079 *-> N6-METHYLASE ->?-> PglZ -> SIGMA+OB-Fold+AAA-ATPase+Lon-Protease-BrxL ->SF1-SF2-Helicase+REase->HIN-HTH->HTH_Tnp_1+HTH_21+rve+rve_3-> OLD-ABC + TOPRIM -><-PhdYeFM_antitox | 1190 | Bacteria>Pseudomonadota>Gammaproteobacteria>Moraxellales>Moraxellaceae>Acinetobacter | GCA_000588415.1 | Acinetobacter_sp._225588 | J687_2205 | 12778..16350 | 1310788 | hypothetical-protein-J687_2205-[Acinetobacter-sp.-225588] |
| WP_001021844.1 | Type1.BREX | PhdYeFM_antitox->< OLD-ABC + TOPRIM <-SF1-SF2-Helicase+REase<- SIGMA+OB-Fold+AAA-ATPase+Lon-Protease-BrxL <- PglZ <-?<- N6-METHYLASE <- DUF6079 *<- BrxB-BrxF-iSTAND <- BrxA-DNA-binding <- wHTH+WYL+WCX \|\|<-DOC+HTH<-?\|\|DDE_Tnp_1_5+DDE_Tnp_1-> | 1226 | Bacteria>Pseudomonadota>Gammaproteobacteria>Moraxellales>Moraxellaceae>Acinetobacter | GCF_000069245.1 | Acinetobacter_calcoaceticus/baumannii_complex | ABAYE_RS00380 | 63476..67156 | 909768 | MULTISPECIES:-BREX-system-P-loop-protein-BrxC-[Acinetobacter-calcoaceticus/baumannii-complex] |
| WP_087528528.1 | Type1.BREX | <-NADAR<-ART-DarT\|\|TPR+TPR-><-?<- SIGMA+OB-Fold+AAA-ATPase+Lon-Protease-BrxL <- PglZ <- TOPRIM <- OLD-ABC <- N6-METHYLASE <- DUF6079 *<- BrxB-BrxF-iSTAND <- BrxA-DNA-binding <- wHTH+WYL+WCX \|\|DUF4236-><-CR-ATPase5+REase<-ParB+HNH<-HIN-HTH+PAIREDC-HTH\|\|<-Zn_Tnp_IS91+Y2_Tnp | 1226 | Bacteria>Pseudomonadota>Gammaproteobacteria>Moraxellales>Moraxellaceae>Acinetobacter | GCF_002165305.2 | Acinetobacter_sp._WCHA55 | CDG62_RS00815 | 35375..39055 | 2004646 | BREX-system-P-loop-protein-BrxC-[Acinetobacter-sp.-WCHA55] |
| WP_153569296.1 | Type1.BREX | Amidohydro_1->35exo->tRNA-><-?<-DDE_Tnp_1_5+DDE_Tnp_1\|\|?->DOC+HTH->\|\| wHTH+WYL+WCX -> BrxA-DNA-binding -> BrxB-BrxF-iSTAND -> DUF6079 *-> N6-METHYLASE ->?-> PglZ -> SIGMA+OB-Fold+AAA-ATPase+Lon-Protease-BrxL ->SF1-SF2-Helicase+REase-> OLD-ABC + TOPRIM -><-PhdYeFM_antitox | 1226 | Bacteria>Pseudomonadota>Gammaproteobacteria>Moraxellales>Moraxellaceae>Acinetobacter | GCF_009645555.1 | Acinetobacter_nosocomialis | F4T96_RS02690 | 68625..72305 | 106654 | BREX-system-P-loop-protein-BrxC-[Acinetobacter-nosocomialis] |
| WP_032001825.1 | Type1.BREX | PhdYeFM_antitox->< OLD-ABC + TOPRIM <-SF1-SF2-Helicase+REase<- SIGMA+OB-Fold+AAA-ATPase+Lon-Protease-BrxL <- PglZ <-?<- N6-METHYLASE <- DUF6079 *<- BrxB-BrxF-iSTAND <- BrxA-DNA-binding <- wHTH+WYL+WCX \|\|<-DOC+HTH<-?\|\|DDE_Tnp_1_5+DDE_Tnp_1-> | 1226 | Bacteria>Pseudomonadota>Gammaproteobacteria>Moraxellales>Moraxellaceae>Acinetobacter | GCF_016502765.1 | Acinetobacter_calcoaceticus/baumannii_complex | I6M92_RS14775 | 66667..70347 | 909768 | MULTISPECIES:-BREX-system-P-loop-protein-BrxC-[Acinetobacter-calcoaceticus/baumannii-complex] |
| WP_204086201.1 | Type1.BREX | PhdYeFM_antitox->< OLD-ABC + TOPRIM <-SF1-SF2-Helicase+REase<- SIGMA+OB-Fold+AAA-ATPase+Lon-Protease-BrxL <- PglZ <-?<- N6-METHYLASE <- DUF6079 *<- BrxB-BrxF-iSTAND <- BrxA-DNA-binding <- wHTH+WYL+WCX \|\|<-DOC+HTH<-?\|\|DDE_Tnp_1_5+DDE_Tnp_1-> | 1226 | Bacteria>Pseudomonadota>Gammaproteobacteria>Moraxellales>Moraxellaceae>Acinetobacter | GCF_016889825.1 | Acinetobacter_pittii | I6J44_RS19355 | 45796..49476 | 48296 | BREX-system-P-loop-protein-BrxC-[Acinetobacter-pittii] |
| WP_212272673.1 | Type1.BREX | <-DDE_Tnp_1_5+DDE_Tnp_1\|\|?->DOC+HTH->\|\| wHTH+WYL+WCX -> BrxA-DNA-binding -> BrxB-BrxF-iSTAND -> DUF6079 *-> N6-METHYLASE ->?-> PglZ -> SIGMA+OB-Fold+AAA-ATPase+Lon-Protease-BrxL ->SF1-SF2-Helicase+REase-> OLD-ABC + TOPRIM -> | 1226 | Bacteria>Pseudomonadota>Gammaproteobacteria>Moraxellales>Moraxellaceae>Acinetobacter | GCF_018139165.1 | Acinetobacter_nosocomialis | KC103_RS15085 | 69521..73201 | 106654 | BREX-system-P-loop-protein-BrxC-[Acinetobacter-nosocomialis] |
| WP_274579352.1 | Type1.BREX | <-DDE_Tnp_1_5+DDE_Tnp_1\|\|?->DOC+HTH-><-DUF4236\|\| wHTH+WYL+WCX -> BrxA-DNA-binding -> BrxB-BrxF-iSTAND -> DUF6079 *-> N6-METHYLASE ->ParB+HNH-> PglZ -> SIGMA+OB-Fold+AAA-ATPase+Lon-Protease-BrxL ->SF1-SF2-Helicase+REase-> OLD-ABC + TOPRIM -p-> | 1226 | Bacteria>Pseudomonadota>Gammaproteobacteria>Moraxellales>Moraxellaceae>Acinetobacter | GCF_028774645.1 | Acinetobacter_lactucae | MOO54_RS17975 | 24360..28040 | 1785128 | BREX-system-P-loop-protein-BrxC-[Acinetobacter-lactucae] |
| WP_002120185.1 | Type1.BREX | PhdYeFM_antitox->< OLD-ABC + TOPRIM \|\|HIN-HTH+PAIREDC-HTH-><- OLD-ABC + TOPRIM -p<- SIGMA+OB-Fold+AAA-ATPase+Lon-Protease-BrxL <- PglZ <-?<- N6-METHYLASE <- DUF6079 *<- BrxB-BrxF-iSTAND <- BrxA-DNA-binding <- wHTH+WYL+WCX \|\|<-DOC+HTH<-?\|\|DDE_Tnp_1_5+DDE_Tnp_1-> | 1226 | Bacteria>Pseudomonadota>Gammaproteobacteria>Moraxellales>Moraxellaceae | - | Acinetobacter | LG653_RS19250 | 36203..39883 | 469 | MULTISPECIES:-BREX-system-P-loop-protein-BrxC-[Acinetobacter] |
| WP_107866915.1 | Type1.BREX | <- SIGMA+OB-Fold+AAA-ATPase+Lon-Protease-BrxL <- PglZ <- TOPRIM <-?<- OLD-ABC <- N6-METHYLASE <-Couple_hipA+HipA_C<-HTH_3<- DUF6079 *<- BrxB-BrxF-iSTAND <- BrxA-DNA-binding <- wHTH+WYL+WCX | 1216 | Bacteria>Pseudomonadota>Gammaproteobacteria>Moraxellales>Moraxellaceae>Agitococcus | GCF_003051055.1 | Agitococcus_lubricus | C8N29_RS15175 | 21839..25489 | 1077255 | BREX-system-P-loop-protein-BrxC-[Agitococcus-lubricus] |
| WP_108675357.1 | Type1.BREX | Arm-DNA-bind_3+Phage_integrase-><-?\|\|?->ZnR+ZnR->?->?->XIS-HTH-> BrxA-DNA-binding -> BrxB-BrxF-iSTAND -> DUF6079 *->ParB+HNH-> OLD-ABC -> TOPRIM -> N6-METHYLASE -> PglZ -> SIGMA+OB-Fold+AAA-ATPase+Lon-Protease-BrxL ->\|\|?->?->Thioredoxin_3-> | 1225 | Bacteria>Pseudomonadota>Gammaproteobacteria>Moraxellales>Moraxellaceae | GCF_003072585.2 | Acinetobacter | C0119_RS05575 | 1118908..1122585 | 469 | MULTISPECIES:-BREX-system-P-loop-protein-BrxC-[Acinetobacter] |

| Protein-ID | System | Gene-Neighbourhoods | Length | Taxonomy | Assembly-ID | Strain | Locus-Tag | Genomic-coordinates | TaxID | Fasta-defline |
|---|---|---|---|---|---|---|---|---|---|---|
| OLF41388.1 | Type1.BREX | Cas_Cas1->?-><-REase\|\|BrxA-DNA-binding->BrxB-BrxF-iSTAND->DUF6079*->N6-METHYLASE->DUF3644->OLD-ABC->TOPRIM->DUF932-> | 1154 | Bacteria>Pseudomonadota>Gammaproteobacteria>Moraxellales>Moraxellaceae>Psychrobacter | GCA_001921755.1 | Psychrobacter_sp._Rd_27.2 | BTV99_04785 | 263905..267369 | 1926479 | hypothetical-protein-BTV99_04785-[Psychrobacter-sp.-Rd-27.2] |
| WP_218607358.1 | Type1.BREX | Cas_Cas1->?-><-REase\|\|BrxA-DNA-binding->BrxB-BrxF-iSTAND->DUF6079*->N6-METHYLASE->DUF3644->OLD-ABC->TOPRIM->DUF932->ParB+HNH-> | 1162 | Bacteria>Pseudomonadota>Gammaproteobacteria>Moraxellales>Moraxellaceae>Psychrobacter | GCF_001921755.1 | Psychrobacter_sp._Rd_27.2 | BTV99_RS04780 | 263881..267369 | 1926479 | BREX-system-P-loop-protein-BrxC-[Psychrobacter-sp.-Rd-27.2] |
| WP_198336379.1 | Type1.BREX | Arm-DNA-bind_3+Phage_integrase-><-PglZ<-N6-METHYLASE<-DUF6079*<-BrxB-BrxF-iSTAND<-BrxA-DNA-binding\|\|SbcC->TOPRIM-><-?<-?<-TPR+TPR+TPR+TPR | 1201 | Bacteria>Pseudomonadota>Gammaproteobacteria>Moraxellales>Moraxellaceae>Psychrobacter | GCF_904845945.1 | Psychrobacter_celer | KF936_RS04990 | 111156..114761 | 306572 | BREX-system-P-loop-protein-BrxC-[Psychrobacter-celer] |
| WP_201558729.1 | Type1.BREX | Arm-DNA-bind_3+Phage_integrase-><-?<-?<-ParB+HNH<-PglZ<-N6-METHYLASE<-DUF6079*<-BrxB-BrxF-iSTAND<-BrxA-DNA-binding\|\|SbcC->TOPRIM-><-XIS-HTH<-?<-?<-DUF932 | 1201 | Bacteria>Pseudomonadota>Gammaproteobacteria>Moraxellales>Moraxellaceae>Psychrobacter | GCF_904846085.1 | Psychrobacter_immobilis | JMW74_RS07725 | 23770..27375 | 498 | BREX-system-P-loop-protein-BrxC-[Psychrobacter-immobilis] |
| TQC95078.1 | Type1.BREX | CbiA->?-><-SIGMA+OB-Fold+AAA-ATPase+HKD-DNase-BrxL<-PglZ<-TOPRIM<-OLD-ABC<-N6-METHYLASE<-DUF6079*<-BrxB-BrxF-iSTAND<-BrxA-DNA-binding\|\|DDE_Tnp_1_5+DDE_Tnp_1-> | 1192 | Bacteria>Pseudomonadota>Gammaproteobacteria>Moraxellales>Moraxellaceae>unclassified_Moraxellaceae | GCA_006546595.1 | Moraxellaceae_bacterium_AER2_44_116 | FK216_14950 | 12152..15730 | 2591038 | BREX-system-P-loop-protein-BrxC-[Moraxellaceae-bacterium-AER2_44_116] |
| MCP5177609.1 | Type1.BREX | DOC+HTH->TRD-><-DUF4096+DDE_Tnp_1+DDE_Tnp_1_2\|\|REase->?->?->?->BrxA-DNA-binding->BrxB-BrxF-iSTAND->DUF6079*->N6-METHYLASE->OLD-ABC->TOPRIM->PglZ->SIGMA+OB-Fold+AAA-ATPase+HKD-DNase-BrxL-><-?<-ParB+RepB<-ParA-Soj-PloopNTPase | 1192 | Bacteria>Pseudomonadota>Gammaproteobacteria>Moraxellales>Moraxellaceae>unclassified_Moraxellaceae | GCA_024234655.1 | Moraxellaceae_bacterium | H6997_08670 | 33703..37281 | 1889775 | BREX-system-P-loop-protein-BrxC-[Moraxellaceae-bacterium] |
| WP_161382580.1 | Type1.BREX | Phage_integrase-><-XIS-HTH<-AAA+HTH<-XIS-HTH\|\|MobI->XIS-HTH->BrxA-DNA-binding->BrxB-BrxF-iSTAND->DUF6079*->Schlafen+HATPase+wHTH+HTH->N6-METHYLASE->SF1-SF2-Helicase->?->HerA/FtsK->N6-METHYLASE-p->PglZ->SIGMA+OB-Fold+AAA-ATPase+HKD-DNase-BrxL->OLD-ABC+TOPRIM-> | 1193 | Bacteria>Pseudomonadota>Gammaproteobacteria>Oceanospirillales>Alcanivoracaceae>Alcanivorax | - | Alcanivorax_sp._ZXX171 | GN155_RS00640 | 132435..136016 | 2695858 | BREX-system-P-loop-protein-BrxC-[Alcanivorax-sp.-ZXX171] |
| MAD69915.1 | Type1.BREX | CHTH-><-?\|\|wHTH+WYL+WCX-><-?<-wHTH+WYL+WCX\|\|?->XIS-HTH->BrxA-DNA-binding->BrxB-BrxF-iSTAND->DUF6079*->Schlafen+HATPase+wHTH+HTH->N6-METHYLASE->REase->SIR2->HerA/FtsK->N6-METHYLASE-p->OLD-ABC->TOPRIM->PglZ-> | 1193 | Bacteria>Pseudomonadota>Gammaproteobacteria>Oceanospirillales>Alcanivoracaceae>Alcanivorax | GCA_002685555.1 | Alcanivorax_sp. | CL538_05260 | 26266..29847 | 1872427 | hypothetical-protein-[Alcanivorax-sp.] |
| HIK75733.1 | Type1.BREX | <-PglZ<-TOPRIM<-OLD-ABC<-N6-METHYLASE<-HerA/FtsK<-SIR2<-REase<-N6-METHYLASE<-Schlafen+HATPase+wHTH+HTH<-DUF6079*<-BrxB-BrxF-iSTAND<-BrxA-DNA-binding<-XIS-HTH<-?\|\|wHTH+WYL+WCX-><-wHTH+WYL+WCX\|\|?-><-CHTH | 1193 | Bacteria>Pseudomonadota>Gammaproteobacteria>Oceanospirillales>Alcanivoracaceae>Alcanivorax | GCA_015660495.1 | Alcanivorax_sp. | EYG00_13690 | 18356..21937 | 1872427 | BREX-system-P-loop-protein-BrxC-[Alcanivorax-sp.] |
| WP_272975234.1 | Type1.BREX | HTH_3->?-><-SIGMA+OB-Fold+AAA-ATPase+Lon-Protease-BrxL<-PglZ<-?<-ZNR+RmuC<-OLD-ABC+TOPRIM<-N6-METHYLASE<-DUF6079*<-BrxB-BrxF-iSTAND<-BrxA-DNA-binding<-?\|\|wHTH+WYL+WCX-><-?<-ZnR+TOPRIM+AAA-helicase+wHTH<-XIS-HTH<-?<-Arm-DNA-bind_3+Phage_integrase | 1192 | Bacteria>Pseudomonadota>Gammaproteobacteria>Oceanospirillales>Alcanivoracaceae>Alcanivorax | GCF_002354495.1 | Alcanivorax_jadensis | B2Z32_RS11825 | 47111..50689 | 64988 | BREX-system-P-loop-protein-BrxC-[Alcanivorax-jadensis] |
| MBE8233358.1 | Type1.BREX | XIS-HTH->BrxA-DNA-binding->BrxB-BrxF-iSTAND->DUF6079*->N6-METHYLASE->OLD-ABC->TOPRIM->N6-METHYLASE->PglZ-> | 1234 | Bacteria>Pseudomonadota>Gammaproteobacteria>Oceanospirillales>Endozoicomonadaceae>unclassified_Endozoicomonadaceae | GCA_015163765.1 | Endozoicomonadaceae_bacterium | HAW67_06440 | 1819..5523 | 2590577 | BREX-system-P-loop-protein-BrxC-[Endozoicomonadaceae-bacterium] |
| WP_041158482.1 | Type1.BREX | <-PglZ<-TOPRIM<-OLD-ABC<-ParB+HNH<-TPR<-NERD+GspA-ATPase<-N6-METHYLASE-p<-N6-METHYLASE<-Schlafen+HATPase+wHTH<-DUF6079*<-BrxB-BrxF-iSTAND<-BrxA-DNA-binding<-HTH_19<-TnsC-ATPase<-rve+Mu-transpos_C<-REase+wHTH<-ncRNA<-tRNA<-?\|\|TetR-HTH-> | 1227 | Bacteria>Pseudomonadota>Gammaproteobacteria>Oceanospirillales>Halomonadaceae | GCF_900099955.1 | Halomonas | BLR10_RS07715 | 76580..80263 | 2745 | MULTISPECIES:-BREX-system-P-loop-protein-BrxC-[Halomonas] |
| WP_062374360.1 | Type1.BREX | <-ABC-ATPase+SbcC<-ABC-ATPase+SbcC<-?<-?<-?<-TetR-HTH\|\|?->tRNA->ncRNA->BrxA-DNA-binding->BrxB-BrxF-iSTAND->DUF6079*->Phd-YefM->PIN->Schlafen+HATPase+wHTH->N6-METHYLASE->N6-METHYLASE-p->HEPN->OLD-ABC->TOPRIM-> | 1227 | Bacteria>Pseudomonadota>Gammaproteobacteria>Oceanospirillales>Halomonadaceae>Halomonas | GCF_001596805.1 | Halomonas_sp._KX33721 | A3220_RS08340 | 63501..67184 | 1819251 | BREX-system-P-loop-protein-BrxC-[Halomonas-sp.-KX33721] |
| WP_101147206.1 | Type1.BREX | <-DiS_P_DiS+Peptidase_A24<-?<-VirB11\|\|?->?->?->tRNA->BrxA-DNA-binding->BrxB-BrxF-iSTAND->DUF6079*->Phd-YefM->PIN->Schlafen+HATPase+wHTH->N6-METHYLASE->HEPN-><-?\|\|OLD-ABC->TOPRIM-> | 1230 | Bacteria>Pseudomonadota>Gammaproteobacteria>Oceanospirillales>Halomonadaceae>Halomonas | GCF_002836035.1 | Halomonas_sp._MES3-P3E | CXF87_RS15340 | 100201..103893 | 2058321 | BREX-system-P-loop-protein-BrxC-[Halomonas-sp.-MES3-P3E] |
| WP_133481548.1 | Type1.BREX | <-SIGMA+OB-Fold+AAA-ATPase+HKD-DNase-BrxL<-PglZ<-TOPRIM<-OLD-ABC<-HNH<-OLD-ABC<-N6-METHYLASE<-Schlafen+HATPase+wHTH+HTH<-DUF6079*<-BrxB-BrxF-iSTAND<-BrxA-DNA-binding<-XIS-HTH<-MobI\|\|XIS-HTH->AAA+HTH->XIS-HTH-><-Arm-DNA-bind_3+Phage_integrase | 1193 | Bacteria>Pseudomonadota>Gammaproteobacteria>Oceanospirillales>Halomonadaceae>Halomonas | GCF_004361885.1 | Halomonas_ventosae | DFO68_RS01985 | 17598..21179 | 229007 | BREX-system-P-loop-protein-BrxC-[Halomonas-ventosae] |
| WP_172420975.1 | Type1.BREX | <-TOPRIM<-OLD-ABC<-?<-?<-?<-N6-METHYLASE<-HATPase+wHTH<-Schlafen+HATPase+wHTH-p<-HTH+REase<-DUF6079*<-BrxB-BrxF-iSTAND<-BrxA-DNA-binding<-ncRNA<-tRNA<-?\|\|TetR-HTH->?->?->?->ABC-ATPase+SbcC->ABC-ATPase+SbcC-> | 1227 | Bacteria>Pseudomonadota>Gammaproteobacteria>Oceanospirillales>Halomonadaceae>Halomonas | GCF_011399095.1 | Halomonas_hydrothermalis | HHSLTHF2_RS10320 | 2155809..2159492 | 115561 | BREX-system-P-loop-protein-BrxC-[Halomonas-hydrothermalis] |
| WP_169956446.1 | Type1.BREX | XIS-HTH->BrxA-DNA-binding->BrxB-BrxF-iSTAND->DUF6079*->Schlafen+HATPase+wHTH+HTH->N6-METHYLASE->OLD-ABC->TOPRIM->PglZ->SIGMA+OB-Fold+AAA-ATPase+HKD-DNase-BrxL-><-REC+OmpR-HTH | 1193 | Bacteria>Pseudomonadota>Gammaproteobacteria>Oceanospirillales>Halomonadaceae>Halomonas | GCF_013000985.1 | Halomonas_sp._PGE1 | HIR79_RS02110 | 421842..425423 | 2730360 | BREX-system-P-loop-protein-BrxC-[Halomonas-sp.-PGE1] |
| WP_193059740.1 | Type1.BREX | TnsC-ATPase->HTH_19->BrxA-DNA-binding->BrxB-BrxF-iSTAND->DUF6079*->Schlafen+HATPase+wHTH->N6-METHYLASE->?->?->OLD-ABC->TOPRIM->PglZ->SIGMA+OB-Fold+AAA-ATPase+Lon-Protease-BrxL-> | 1227 | Bacteria>Pseudomonadota>Gammaproteobacteria>Oceanospirillales>Halomonadaceae>Halomonas | GCF_014897995.1 | unclassified_Halomonas | INT28_RS17780 | 3434..7117 | 2609666 | MULTISPECIES:-BREX-system-P-loop-protein-BrxC-[unclassified-Halomonas] |
| WP_218314905.1 | Type1.BREX | <-ABC-ATPase+SbcC<-ABC-ATPase+SbcC<-?<-?<-?<-TetR-HTH\|\|tRNA->ncRNA->BrxA-DNA-binding->BrxB-BrxF-iSTAND->DUF6079*->Schlafen+HATPase+wHTH->N6-METHYLASE->?->?->OLD-ABC->TOPRIM->PglZ->SIGMA+OB-Fold+AAA-ATPase+Lon-Protease-BrxL-> | 1227 | Bacteria>Pseudomonadota>Gammaproteobacteria>Oceanospirillales>Halomonadaceae>Halomonas | GCF_019211725.1 | Halomonas_sp._18071143 | KVY02_RS07715 | 1661242..1664925 | 2855441 | BREX-system-P-loop-protein-BrxC-[Halomonas-sp.-18071143] |
| WP_231664595.1 | Type1.BREX | GDResolvase+HIN-HTH->?-><-SIGMA+OB-Fold+AAA-ATPase+Lon-Protease-BrxL<-PglZ<-TOPRIM<-OLD-ABC<-HTH+REase<-N6-METHYLASE<-Schlafen+HATPase+wHTH<-DUF6079*<-BrxB-BrxF-iSTAND-p<-BrxA-DNA-binding<-XIS-HTH<-MobI | 1153 | Bacteria>Pseudomonadota>Gammaproteobacteria>Oceanospirillales>Halomonadaceae>Halomonas | GCF_021151495.1 | Halomonas_sp._IOP_31 | K7I12_RS15025 | 26514..29975 | 2876584 | BREX-system-P-loop-protein-BrxC-[Halomonas-sp.-IOP_31] |
| WP_239296660.1 | Type1.BREX | Phage_integrase-><-XIS-HTH<-AAA+HTH<-XIS-HTH\|\|MobI->XIS-HTH->BrxA-DNA-binding->BrxB-BrxF-iSTAND->DUF6079*->Schlafen+HATPase+wHTH+HTH->N6-METHYLASE->REase->SIR2->HerA/FtsK->N6-METHYLASE-p->OLD-ABC->TOPRIM->PglZ-> | 1193 | Bacteria>Pseudomonadota>Gammaproteobacteria>Oceanospirillales>Halomonadaceae>Halomonas | GCF_022348215.1 | unclassified_Halomonas | MHM35_RS10400 | 116133..119714 | 2609666 | MULTISPECIES:-BREX-system-P-loop-protein-BrxC-[unclassified-Halomonas] |
| WP_079550430.1 | Type1.BREX | <-TetR-HTH\|\|?->tRNA->ncRNA-><-?\|\|BrxA-DNA-binding->BrxB-BrxF-iSTAND->DUF6079*->Schlafen+HATPase+wHTH-p->HATPase_c->N6-METHYLASE->OLD-ABC->TOPRIM->?->DOC+HTH->PglZ->SIGMA+OB-Fold+AAA-ATPase+Lon-Protease-BrxL-> | 1240 | Bacteria>Pseudomonadota>Gammaproteobacteria>Oceanospirillales>Halomonadaceae>Halomonas | GCF_900142895.1 | Halomonas_subglaciescola | B5495_RS00605 | 132715..136437 | 29571 | BREX-system-P-loop-protein-BrxC-[Halomonas-subglaciescola] |
| WP_213229339.1 | Type1.BREX | <-OLD-ABC+TOPRIM<-CHTH<-?<-SIGMA+OB-Fold+AAA-ATPase+Lon-Protease-BrxL<-PglZ<-?<-NADAR<-ART-DarT<-N6-METHYLASE<-DUF6079*<-BrxB-BrxF-iSTAND<-BrxA-DNA-binding<-?<-Phage_integrase<-?<-Phage_integrase | 1241 | Bacteria>Pseudomonadota>Gammaproteobacteria>Oceanospirillales>Halomonadaceae>Halomonas | - | Halomonas_litopenaei | DYI26_RS12555 | 49628..53353 | 2109328 | BREX-system-P-loop-protein-BrxC-[Halomonas-litopenaei] |
| WP_254893719.1 | Type1.BREX | <-ABC-ATPase+SbcC<-ABC-ATPase+SbcC<-?<-?<-?<-TetR-HTH\|\|tRNA->ncRNA->BrxA-DNA-binding->BrxB-BrxF-iSTAND->DUF6079*->Schlafen+HATPase+wHTH->N6-METHYLASE->?->?->OLD-ABC->TOPRIM->MULE-TRANSPOSASE-> | 1210 | Bacteria>Pseudomonadota>Gammaproteobacteria>Oceanospirillales>Halomonadaceae>Halomonas | - | Halomonas_sp._BC1 | B7L72_RS00095 | 19375..23007 | 1670448 | BREX-system-P-loop-protein-BrxC-[Halomonas-sp.-BC1] |
| WP_027853089.1 | Type1.BREX | <-SIGMA+OB-Fold+AAA-ATPase+Lon-Protease-BrxL<-PglZ<-HNH<-OLD-ABC<-TOPRIM<-OLD-ABC<-ParB+HNH<-N6-METHYLASE<-DUF6079*<-BrxB-BrxF-iSTAND<-BrxA-DNA-binding<-wHTH+WYL+WCX<-TIR<-?<-?<-wHTH+WYL+WCX | 1154 | Bacteria>Pseudomonadota>Gammaproteobacteria>Oceanospirillales>Oceanospirillaceae>Marinobacterium | GCF_000428985.1 | Marinobacterium_litorale | G403_RS0105385 | 191863..195327 | 404770 | BREX-system-P-loop-protein-BrxC-[Marinobacterium-litorale] |
| WP_206838922.1 | Type1.BREX | MODE-HTH+PBPII-><-ADH_N+ADH_zinc_N<-?\|\|?->?->wHTH+WYL+WCX->BrxA-DNA-binding->BrxB-BrxF-iSTAND->DUF6079*->N6-METHYLASE->DUF932->OLD-ABC->TOPRIM->OLD-ABC->HNH->PglZ->SIGMA+OB-Fold+AAA-ATPase+Lon-Protease-BrxL->Integrase_1-> | 1154 | Bacteria>Pseudomonadota>Gammaproteobacteria>Oceanospirillales>Oceanospirillaceae>Marinobacterium | GCF_017310015.1 | Marinobacterium_georgiense | CFI10_RS02405 | 480710..484174 | 48076 | BREX-system-P-loop-protein-BrxC-[Marinobacterium-iners] |
| WP_254579406.1 | Type6.BREX | wHTH+REase+ZnR-><-SbcC<-?->?->RNase->wHTH+WYL+WCX->BrxE->BrxA-DNA-binding->BrxB-BrxF-iSTAND->DUF6079*->N6-METHYLASE->OLD-ABC->TOPRIM->PglZ->SIGMA+OB-Fold+AAA-ATPase+HKD-DNase-BrxL->UDG-conflict->REase-><-CARF+NtrC-AAA+FIS-HTH | 1212 | Bacteria>Pseudomonadota>Gammaproteobacteria>Oceanospirillales>Oceanospirillaceae>Marinobacterium | GCF_024267675.1 | Marinobacterium_sedimentorum | MRS74_RS06180 | 150386..154024 | 2927804 | BREX-system-P-loop-protein-BrxC-[Marinobacterium-sedimentorum] |
| WP_226752736.1 | Type1.BREX | <-SIGMA+OB-Fold+AAA-ATPase+Lon-Protease-BrxL<-PglZ<-Peptidase_S8<-Classical-AAA<-SbcC<-N6-METHYLASE<-TOPRIM<-OLD-ABC<-DUF6079*<-BrxB-BrxF-iSTAND<-BrxA-DNA-binding<-HTH_3\|\|wHTH+WYL->SF1-SF2-Helicase->Phage_integrase-><-CACHE+GGDEF | 1237 | Bacteria>Pseudomonadota>Gammaproteobacteria>Oceanospirillales>Oceanospirillaceae>Marinomonas | GCF_020532605.1 | Marinomonas_algarum | LG368_RS00225 | 46423..50136 | 2883105 | BREX-system-P-loop-protein-BrxC-[Marinomonas-algarum] |
| WP_067206359.1 | Type1.BREX | <-VirD4-TraG+VirB4-FtsK<-HDRnase<-SbcC<-SIGMA+OB-Fold+AAA-ATPase+Lon-Protease-BrxL<-PglZ<-N6-METHYLASE<-TOPRIM<-OLD-ABC<-DUF6079*<-BrxB-BrxF-iSTAND<-BrxA-DNA-binding<-wHTH+WYL+WCX<-?<-35exo\|\|LexA-protease->?-><-MobI | 1225 | Bacteria>Pseudomonadota>Gammaproteobacteria>Oceanospirillales>Oceanospirillaceae>Marinomonas | GCF_900089755.1 | Marinomonas_aquimarina | MAQ5080_RS04935 | 83930..87607 | 295068 | BREX-system-P-loop-protein-BrxC-[Marinomonas-aquimarina] |
| QEW05875.1 | Type6.BREX | XIS-HTH->?->CHTH->?->SF1-SF2-Helicase->BrxE->BrxA-DNA-binding->BrxB-BrxF-iSTAND->DUF6079*->N6-METHYLASE->OLD-ABC->TOPRIM->ParB+HNH->PglZ->SIGMA+OB-Fold+AAA-ATPase+HKD-DNase-BrxL->?-><-?\|\|HTH_Tnp_1-> | 1212 | Bacteria>Pseudomonadota>Gammaproteobacteria>Oceanospirillales>Oceanospirillaceae>Nitrincola | GCA_008727795.1 | Nitrincola_iocasae | F5I99_04885 | 1032629..1036267 | 2614693 | BREX-system-P-loop-protein-BrxC-[Nitrincola-iocasae] |
| WP_202806908.1 | Type1.BREX | <-PglZ<-TOPRIM<-OLD-ABC<-SbcC<-N6-METHYLASE-p<-N6-METHYLASE<-N6-METHYLASE-p<-PIN<-OLD-ABC<-DUF6079-p*<-DUF6079-p<-BrxB-BrxF-iSTAND<-BrxA-DNA-binding<-XIS-HTH<-?<-HDRnase<-Phage_int_SAM<-Integrase_1\|\|XIS-HTH-> | 460 | Bacteria>Pseudomonadota>Gammaproteobacteria>Oceanospirillales>Oceanospirillaceae>Nitrincola | GCF_000585235.1 | Nitrincola_nitratireducens | D791_RS20495 | 12347..13729 | 1229521 | hypothetical-protein-[Nitrincola-nitratireducens] |
| WP_202806909.1 | Type1.BREX | <-TOPRIM<-OLD-ABC<-SbcC<-N6-METHYLASE-p<-N6-METHYLASE<-N6-METHYLASE-p<-PIN<-OLD-ABC<-DUF6079-p<-DUF6079-p*<-BrxB-BrxF-iSTAND<-BrxA-DNA-binding<-XIS-HTH<-?<-HDRnase<-Phage_int_SAM<-Integrase_1\|\|XIS-HTH->Phage_integrase-> | 697 | Bacteria>Pseudomonadota>Gammaproteobacteria>Oceanospirillales>Oceanospirillaceae>Nitrincola | GCF_000585235.1 | Nitrincola_nitratireducens | D791_RS20500 | 13689..15782 | 1229521 | BREX-system-P-loop-protein-BrxC-[Nitrincola-nitratireducens] |
| WP_225307559.1 | Type6.BREX | XIS-HTH->?->CHTH->?->SF1-SF2-Helicase->BrxE->BrxA-DNA-binding->BrxB-BrxF-iSTAND->DUF6079*->N6-METHYLASE->OLD-ABC->TOPRIM->ParB+HNH->PglZ->SIGMA+OB-Fold+AAA-ATPase+HKD-DNase-BrxL->?->?-><-?\|\|PAIREDC-HTH+HIN-HTH-> | 1214 | Bacteria>Pseudomonadota>Gammaproteobacteria>Oceanospirillales>Oceanospirillaceae>Nitrincola | GCF_008727795.1 | Nitrincola_iocasae | F5I99_RS04885 | 1032623..1036267 | 2614693 | BREX-system-P-loop-protein-BrxC-[Nitrincola-iocasae] |
| WP_260997651.1 | Type1.BREX | <-Arm-DNA-bind_2+Phage_int_SAM_3+Phage_integrase<-Phage_AlpA\|\|HTH_26-><-SIGMA+OB-Fold+AAA-ATPase+Lon-Protease-BrxL<-PglZ<-TOPRIM<-OLD-ABC<-ParB+HNH<-N6-METHYLASE<-DUF6079*<-BrxB-BrxF-iSTAND<-BrxA-DNA-binding<-wHTH+WYL+WCX | 1156 | Bacteria>Pseudomonadota>Gammaproteobacteria>Oceanospirillales>Oceanospirillaceae>Thalassolituus | GCF_025345565.1 | Thalassolituus_hydrocarbonoclasticus | HUF19_RS16715 | 3723205..3726675 | 2742796 | BREX-system-P-loop-protein-BrxC-[Thalassolituus-hydrocarbonoclasticus] |
| MBM94539.1 | Type1.BREX | CRISPR_Cas6->?-><-SIGMA+OB-Fold+AAA-ATPase+Lon-Protease-BrxL<-PglZ<-TOPRIM<-OLD-ABC<-N6-METHYLASE<-TOPRIM<-OLD-ABC<-DUF6079*<-BrxB-BrxF-iSTAND<-BrxA-DNA-binding\|\|?-><-?<-?<-Arm-DNA-bind_2+Phage_int_SAM_3+Phage_integrase<-Phage_AlpA\|\|HTH_26-> | 1225 | Bacteria>Pseudomonadota>Gammaproteobacteria>Oceanospirillales>Oceanospirillaceae>unclassified_Oceanospirillaceae | GCA_002723385.1 | Oceanospirillaceae_bacterium | CMI09_01635 | 47843..51520 | 1899355 | DNA-repair-protein-[Oceanospirillaceae-bacterium] |
| WP_116688140.1 | Type1.BREX | DDE_Tnp-><-HTH_3\|\|?-><-?\|\|?->DUF4096+DDE_Tnp_1+DDE_Tnp_1_2->HTH_26->BrxA-DNA-binding->BrxB-BrxF-iSTAND->DUF6079*->ParB+HNH->N6-METHYLASE->OLD-ABC->TOPRIM->OLD-ABC->TOPRIM->PglZ->SIGMA+OB-Fold+AAA-ATPase+Lon-Protease-BrxL-><-ANK+ANK | 1236 | Bacteria>Pseudomonadota>Gammaproteobacteria>Oceanospirillales>Oceanospirillales_incertae_sedis>Pelagibaculum | GCF_003097315.1 | Pelagibaculum_spongiae | DC094_RS16040 | 158438..162148 | 2080658 | BREX-system-P-loop-protein-BrxC-[Pelagibaculum-spongiae] |
| MBR9867112.1 | Type1.BREX | <-SbcC<-SIGMA+OB-Fold+AAA-ATPase+Lon-Protease-BrxL<-PglZ<-TOPRIM<-OLD-ABC<-N6-METHYLASE<-ParB+HNH<-DUF6079*<-BrxB-BrxF-iSTAND<-BrxA-DNA-binding<-wHTH+WYL+WCX<-?<-?<-SLOG+BirA-HTH<-SF1-SF2-Helicase+PRTase\|\|DnaJ-> | 1225 | Bacteria>Pseudomonadota>Gammaproteobacteria>Oceanospirillales>unclassified_Oceanospirillales | GCA_018223265.1 | Oceanospirillales_bacterium | GYB18_08415 | 21169..24846 | 1914300 | BREX-system-P-loop-protein-BrxC-[Oceanospirillales-bacterium] |
| KXS54417.1 | Type1.BREX | BrxA-DNA-binding->BrxB-BrxF-iSTAND->DUF6079*->DOC+HTH->N6-METHYLASE->OLD-ABC+TOPRIM->OLD-ABC-> | 1221 | Bacteria>Pseudomonadota>Gammaproteobacteria>Pseudomonadales>Marinobacteraceae>Marinobacter | GCA_001577575.1 | Marinobacter_sp._T13-3 | AWU57_1199 | 1913..5578 | 1794906 | hypothetical-protein-AWU57_1199-[Marinobacter-sp.-T13-3] |
| MBC7192650.1 | Type1.BREX | <-SIGMA+OB-Fold+AAA-ATPase+Lon-Protease-BrxL<-PglZ<-HNH<-OLD-ABC<-TOPRIM<-OLD-ABC<-HTH+REase<-N6-METHYLASE<-Schlafen+HATPase+wHTH<-DUF6079*<-BrxB-BrxF-iSTAND<-BrxA-DNA-binding<-XIS-HTH\|\|HTH_Tnp_1+HTH_21+rve+rve_3->XIS-HTH->DnaB->?-><-Phage_int_SAM_5+Phage_integrase<-sGTP | 1145 | Bacteria>Pseudomonadota>Gammaproteobacteria>Pseudomonadales>Marinobacteraceae>Marinobacter | GCA_014359475.1 | Marinobacter_sp. | H5U31_09890 | 22634..26071 | 50741 | BREX-system-P-loop-protein-BrxC-[Marinobacter-sp.] |
| WP_011784101.1 | Type1.BREX | Arm-DNA-bind_3+Phage_integrase-><-XIS-HTH<-AAA+HTH<-XIS-HTH\|\|MobI->XIS-HTH->BrxA-DNA-binding->BrxB-BrxF-iSTAND->DUF6079*->Schlafen+HATPase+wHTH+HTH->SF1-SF2-Helicase+PRTase->SLOG+BirA-HTH->N6-METHYLASE->OLD-ABC->TOPRIM->PglZ->SIGMA+OB-Fold+AAA-ATPase+HKD-DNase-BrxL->SbcC-> | 1193 | Bacteria>Pseudomonadota>Gammaproteobacteria>Pseudomonadales>Marinobacteraceae>Marinobacter | GCF_000015365.1 | Marinobacter_nauticus | MAQU_RS02805 | 628948..632529 | 2743 | BREX-system-P-loop-protein-BrxC-[Marinobacter-nauticus] |
| WP_008174228.1 | Type1.BREX | Arm-DNA-bind_3+Phage_integrase-><-XIS-HTH<-AAA+HTH<-XIS-HTH\|\|MobI->XIS-HTH->BrxA-DNA-binding->BrxB-BrxF-iSTAND->DUF6079*->Schlafen+HATPase+wHTH+HTH->N6-METHYLASE->OLD-ABC->TOPRIM-> | 1193 | Bacteria>Pseudomonadota>Gammaproteobacteria>Pseudomonadales>Marinobacteraceae>Marinobacter | GCF_000235625.1 | Marinobacter_manganoxydans | KYE_13690 | 42273..45854 | 1150997 | BREX-system-P-loop-protein-BrxC-[Marinobacter-manganoxydans] |
| WP_022988850.1 | Type1.BREX | <-PglZ<-TOPRIM<-OLD-ABC<-HNH<-OLD-ABC<-N6-METHYLASE<-REase<-OLD-ABC<-Schlafen+HATPase+wHTH+HTH<-DUF6079*<-BrxB-BrxF-iSTAND<-BrxA-DNA-binding<-XIS-HTH<-MobI | 1193 | Bacteria>Pseudomonadota>Gammaproteobacteria>Pseudomonadales>Marinobacteraceae>Marinobacter | GCF_000475255.1 | Marinobacter_sp._ES-1 | Q668_RS06800 | 88554..92135 | 1396858 | BREX-system-P-loop-protein-BrxC-[Marinobacter-sp.-ES-1] |
| WP_022992169.1 | Type1.BREX | Arm-DNA-bind_3+Phage_integrase-><-XIS-HTH<-AAA+HTH<-XIS-HTH\|\|MobI->XIS-HTH->BrxA-DNA-binding->BrxB-BrxF-iSTAND->DUF6079*->Schlafen+HATPase+wHTH+HTH->N6-METHYLASE->OLD-ABC->HNH->OLD-ABC->TOPRIM->PglZ->SIGMA+OB-Fold+AAA-ATPase+HKD-DNase-BrxL-> | 1193 | Bacteria>Pseudomonadota>Gammaproteobacteria>Pseudomonadales>Marinobacteraceae>Marinobacter | GCF_000475355.1 | Marinobacter_sp._C1S70 | Q669_RS10410 | 149187..152768 | 1396859 | BREX-system-P-loop-protein-BrxC-[Marinobacter-sp.-C1S70] |
| WP_023008352.1 | Type1.BREX | <-SbcC<-SIGMA+OB-Fold+AAA-ATPase+HKD-DNase-BrxL<-PglZ<-TOPRIM<-OLD-ABC<-HNH<-OLD-ABC<-N6-METHYLASE<-Schlafen+HATPase+wHTH+HTH<-DUF6079*<-BrxB-BrxF-iSTAND<-BrxA-DNA-binding<-XIS-HTH<-MobI\|\|XIS-HTH->AAA+HTH->XIS-HTH-><-Arm-DNA-bind_3+Phage_integrase | 1193 | Bacteria>Pseudomonadota>Gammaproteobacteria>Pseudomonadales>Marinobacteraceae>Marinobacter | GCF_000475375.1 | Marinobacter_sp._EVN1 | Q674_RS26680 | 52773..56354 | 1397532 | BREX-system-P-loop-protein-BrxC-[Marinobacter-sp.-EVN1] |
| WP_104416052.1 | Type1.BREX | Arm-DNA-bind_3+Phage_integrase-><-XIS-HTH<-AAA+HTH<-XIS-HTH\|\|MobI->XIS-HTH->BrxA-DNA-binding->BrxB-BrxF-iSTAND->DUF6079*->Schlafen+HATPase+wHTH+HTH->N6-METHYLASE->OLD-ABC->HNH->OLD-ABC->TOPRIM->PglZ->SIGMA+OB-Fold+AAA-ATPase+HKD-DNase-BrxL-> | 1193 | Bacteria>Pseudomonadota>Gammaproteobacteria>Pseudomonadales>Marinobacteraceae>Marinobacter | GCF_002934485.1 | Marinobacter_persicus | BY455_RS08980 | 65339..68920 | 930118 | BREX-system-P-loop-protein-BrxC-[Marinobacter-persicus] |
| WP_113863603.1 | Type1.BREX | Arm-DNA-bind_3+Phage_integrase-><-XIS-HTH<-AAA+HTH<-XIS-HTH\|\|MobI->XIS-HTH->BrxA-DNA-binding->BrxB-BrxF-iSTAND->DUF6079*->Schlafen+HATPase+wHTH+HTH->N6-METHYLASE->OLD-ABC->HNH->OLD-ABC->TOPRIM->PglZ->SIGMA+OB-Fold+AAA-ATPase+HKD-DNase-BrxL-> | 1193 | Bacteria>Pseudomonadota>Gammaproteobacteria>Pseudomonadales>Marinobacteraceae>Marinobacter | GCF_003315345.1 | Marinobacter_pelagius | DET50_RS14950 | 40642..44223 | 379482 | BREX-system-P-loop-protein-BrxC-[Marinobacter-pelagius] |
| WP_092021229.1 | Type1.BREX | DUF932->?->?-><-HTH_3\|\|?->?->wHTH+WYL+WCX->BrxA-DNA-binding->BrxB-BrxF-iSTAND->DUF6079*->N6-METHYLASE->ParB+HNH->OLD-ABC->TOPRIM->PglZ->SIGMA+OB-Fold+AAA-ATPase+Lon-Protease-BrxL->REase+PNTD1+PNTD2->PIWI->TOPRIM-> | 1219 | Bacteria>Pseudomonadota>Gammaproteobacteria>Pseudomonadales>Marinobacteraceae>Marinobacter | GCF_900114775.1 | Marinobacter_zhejiangensis | BMZ67_RS06955 | 334318..337977 | 488535 | BREX-system-P-loop-protein-BrxC-[Marinobacter-zhejiangensis] |
| BBJ05051.1 | Type1.BREX | Arm-DNA-bind_3+Phage_integrase-><-XIS-HTH<-AAA+HTH<-XIS-HTH\|\|MobI->XIS-HTH->BrxA-DNA-binding->BrxB-BrxF-iSTAND->DUF6079*->Schlafen+HATPase+wHTH+HTH->N6-METHYLASE->RVT->HerA/FtsK->N6-METHYLASE-p->OLD-ABC->TOPRIM->PglZ->SIGMA+OB-Fold+AAA-ATPase+HKD-DNase-BrxL-> | 1193 | Bacteria>Pseudomonadota>Gammaproteobacteria>Pseudomonadales>Marinobacteraceae>Marinobacter | - | Marinobacter_nauticus | YBY_29000 | 3185607..3189188 | 2743 | hypothetical-protein-YBY_29000-[Marinobacter-nauticus] |
| WP_152390158.1 | Type1.BREX | VirB10->?-><-SIGMA+OB-Fold+AAA-ATPase+Lon-Protease-BrxL<-PglZ<-TOPRIM<-OLD-ABC<-N6-METHYLASE<-Schlafen+HATPase+wHTH<-DUF6079*<-BrxB-BrxF-iSTAND<-BrxA-DNA-binding<-wHTH+WYL+WCX<-Tox-RES<-CHTH\|\|?->?->DnaB+HerA/FtsK-> | 1224 | Bacteria>Pseudomonadota>Gammaproteobacteria>Pseudomonadales>Pseudomonadaceae>Azotobacter_group | GCF_009363155.1 | Azotobacter_salinestris | GCU53_RS24425 | 150496..154170 | 69964 | BREX-system-P-loop-protein-BrxC-[Azotobacter-salinestris] |
| WP_269819144.1 | Type6.BREX | <-SAM-METHYLASE<-N6-METHYLASE<-?\|\|?->METHYLASE-><-?\|\|tRNA->tRNA->BrxE->BrxA-DNA-binding->BrxB-BrxF-iSTAND->DUF6079*->N6-METHYLASE->PglZ->SIGMA+OB-Fold+AAA-ATPase+HKD-DNase-BrxL-><-OLD-ABC+TOPRIM<-HTH_26\|\|Phage_AlpA->Arm-DNA-bind_2+Phage_int_SAM_3+Phage_integrase->HDRnase-><-Phage_integrase | 1219 | Bacteria>Pseudomonadota>Gammaproteobacteria>Pseudomonadales>Pseudomonadaceae>Denitrificimonas | GCF_027498055.1 | Denitrificimonas_caeni | O6P33_RS05150 | 1067699..1071358 | 521720 | BREX-system-P-loop-protein-BrxC-[Denitrificimonas-caeni] |
| ARQ76275.1 | Type1.BREX | DUF4969-><-SIGMA+OB-Fold+AAA-ATPase+Lon-Protease-BrxL<-PglZ<-TOPRIM<-OLD-ABC<-ParB+HNH<-N6-METHYLASE<-DUF6079*<-BrxB-BrxF-iSTAND<-BrxA-DNA-binding<-?<-XIS-HTH\|\|?-><-?<-?<-?\|\|Aminotran_1_2-> | 1156 | Bacteria>Pseudomonadota>Gammaproteobacteria>Pseudomonadales>Pseudomonadaceae>Pseudomonas | GCA_002128325.1 | Pseudomonas_fragi | B6D87_19590 | 4227178..4230648 | 296 | hypothetical-protein-B6D87_19590-[Pseudomonas-fragi] |
| MBB59705.1 | Type1.BREX | <-DUF3584+DUF5401<-MukF<-SIGMA+OB-Fold+AAA-ATPase+Lon-Protease-BrxL<-PglZ<-TOPRIM<-OLD-ABC<-ParB+HNH<-DUF932<-N6-METHYLASE<-DUF6079*<-BrxB-BrxF-iSTAND<-BrxA-DNA-binding<-wHTH+WYL+WCX<-?<-ARG\|\|wHTH+WYL+WCX->LexA-protease-> | 1154 | Bacteria>Pseudomonadota>Gammaproteobacteria>Pseudomonadales>Pseudomonadaceae>Pseudomonas | GCA_002709345.1 | Pseudomonas_sp. | CML00_00850 | 39770..43234 | 306 | hypothetical-protein-[Pseudomonas-sp.] |
| MBG4863325.1 | Type1.BREX | <-HIN-HTH\|\|XIS-HTH->BrxA-DNA-binding->BrxB-BrxF-iSTAND->DUF6079*->N6-METHYLASE->OLD-ABC->NERD->N6-METHYLASE-p->OLD-ABC->TOPRIM->PglZ->SIGMA+OB-Fold+AAA-ATPase+HKD-DNase-BrxL->HIN-HTH-> | 1184 | Bacteria>Pseudomonadota>Gammaproteobacteria>Pseudomonadales>Pseudomonadaceae>Pseudomonas | GCA_015735985.1 | Pseudomonas_aeruginosa | GHU23_28970 | 1726..5280 | 287 | BREX-system-P-loop-protein-BrxC-[Pseudomonas-aeruginosa] |
| MBG6730505.1 | Type1.BREX | <-LexA-protease<-wHTH+WYL+WCX\|\|?->XIS-HTH->BrxA-DNA-binding->BrxB-BrxF-iSTAND->DUF6079*->N6-METHYLASE->SbcC-p->REase->N6-METHYLASE-p->OLD-ABC->TOPRIM->PglZ->SIGMA+OB-Fold+AAA-ATPase+HKD-DNase-BrxL-><-DUF4969 | 1184 | Bacteria>Pseudomonadota>Gammaproteobacteria>Pseudomonadales>Pseudomonadaceae>Pseudomonas | GCA_015764285.1 | Pseudomonas_aeruginosa | I5I92_11320 | 4371..7925 | 287 | BREX-system-P-loop-protein-BrxC-[Pseudomonas-aeruginosa] |
| HBO1619179.1 | Type1.BREX | <-DUF3584+DUF5401<-JetB<-MukF<-SIGMA+OB-Fold+AAA-ATPase+Lon-Protease-BrxL<-PglZ<-TOPRIM<-OLD-ABC<-ParB+HNH<-N6-METHYLASE<-DUF6079*<-BrxB-BrxF-iSTAND<-BrxA-DNA-binding<-wHTH+WYL+WCX<-?<-ARG\|\|wHTH+WYL+WCX->?-><-NUDIX | 1154 | Bacteria>Pseudomonadota>Gammaproteobacteria>Pseudomonadales>Pseudomonadaceae>Pseudomonas | GCA_021598085.1 | Pseudomonas_aeruginosa | L4F62_005712 | 16995..20459 | 287 | BREX-system-P-loop-protein-BrxC-[Pseudomonas-aeruginosa] |
| HBO7921132.1 | Type1.BREX | ParB+HNH-><-LexA-protease<-wHTH+WYL+WCX\|\|?->XIS-HTH->BrxA-DNA-binding->BrxB-BrxF-iSTAND->DUF6079*->N6-METHYLASE->OLD-ABC->REase->SIR2->HerA/FtsK->N6-METHYLASE->OLD-ABC->TOPRIM->PglZ-> | 1184 | Bacteria>Pseudomonadota>Gammaproteobacteria>Pseudomonadales>Pseudomonadaceae>Pseudomonas | GCA_021678155.1 | Pseudomonas_aeruginosa | L5C37_005426 | 13663..17217 | 287 | BREX-system-P-loop-protein-BrxC-[Pseudomonas-aeruginosa] |
| MCK9534930.1 | Type1.BREX | HTH_Tnp_1->?->LZ_Tnp_IS66+zf-IS66+DDE_Tnp_IS66_C-><-?\|\|?->wHTH+WYL+WCX->BrxA-DNA-binding->BrxB-BrxF-iSTAND->DUF6079*->N6-METHYLASE->ParB+HNH->OLD-ABC->TOPRIM->OLD-ABC->HNH->PglZ->SIGMA+OB-Fold+AAA-ATPase+Lon-Protease-BrxL-> | 1154 | Bacteria>Pseudomonadota>Gammaproteobacteria>Pseudomonadales>Pseudomonadaceae>Pseudomonas | GCA_023228505.1 | Pseudomonas_sp. | M0Q98_09595 | 5981..9445 | 306 | BREX-system-P-loop-protein-BrxC-[Pseudomonas-sp.] |
| CAB5576233.1 | Type1.BREX | wHTH+WYL+WCX->BrxA-DNA-binding->BrxB-BrxF-iSTAND->DUF6079*->?->?->DOC+HTH->N6-METHYLASE->OLD-ABC->TOPRIM->TIR-><-ZnR | 1223 | Bacteria>Pseudomonadota>Gammaproteobacteria>Pseudomonadales>Pseudomonadaceae>Pseudomonas | GCA_903684465.1 | Pseudomonas_putida | GHA_00144 | 159878..163549 | 303 | ATPase-involved-in-DNA-repair-[Pseudomonas-putida] |
| WP_047283135.1 | Type1.BREX | DUF4969-><-SIGMA+OB-Fold+AAA-ATPase+Lon-Protease-BrxL<-PglZ<-TOPRIM<-OLD-ABC<-ParB+HNH<-N6-METHYLASE<-DUF6079*<-BrxB-BrxF-iSTAND<-BrxA-DNA-binding<-wHTH+WYL+WCX<-?<-XIS-HTH<-BirA-HTH\|\|<-Pkinase+CR-Kinase | 1154 | Bacteria>Pseudomonadota>Gammaproteobacteria>Pseudomonadales>Pseudomonadaceae>Pseudomonas | GCF_001020725.2 | Pseudomonas_lundensis | AA042_RS07115 | 1582722..1586186 | 86185 | BREX-system-P-loop-protein-BrxC-[Pseudomonas-lundensis] |
| WP_057391032.1 | Type1.BREX | <-wHTH+WYL+WCX\|\|ARG->?->wHTH+WYL+WCX->BrxA-DNA-binding->BrxB-BrxF-iSTAND->DUF6079*->N6-METHYLASE->DUF932->ParB+HNH->OLD-ABC->TOPRIM->PglZ->SIGMA+OB-Fold+AAA-ATPase+Lon-Protease-BrxL->MukF->JetB-> | 1154 | Bacteria>Pseudomonadota>Gammaproteobacteria>Pseudomonadales>Pseudomonadaceae>Pseudomonas | GCF_001374115.1 | Pseudomonas_aeruginosa | AYU10_RS17350 | 11442..14906 | 287 | BREX-system-P-loop-protein-BrxC-[Pseudomonas-aeruginosa] |
| WP_059319043.1 | Type1.BREX | <-JetB<-MukF<-SIGMA+OB-Fold+AAA-ATPase+Lon-Protease-BrxL<-PglZ<-TOPRIM<-OLD-ABC<-ParB+HNH<-DUF932<-N6-METHYLASE<-DUF6079*<-BrxB-BrxF-iSTAND<-BrxA-DNA-binding<-wHTH+WYL+WCX<-?<-ARG\|\|wHTH+WYL+WCX->?-><-NUDIX | 1154 | Bacteria>Pseudomonadota>Gammaproteobacteria>Pseudomonadales>Pseudomonadaceae>Pseudomonas | GCF_001516265.1 | Pseudomonas_aeruginosa | HV84_RS13920 | 329043..332507 | 287 | BREX-system-P-loop-protein-BrxC-[Pseudomonas-aeruginosa] |
| WP_205881761.1 | Type1.BREX | DUF4969-><-SIGMA+OB-Fold+AAA-ATPase+Lon-Protease-BrxL<-PglZ<-TOPRIM<-OLD-ABC<-ParB+HNH<-N6-METHYLASE<-DUF6079*<-BrxB-BrxF-iSTAND<-BrxA-DNA-binding<-wHTH+WYL+WCX<-?<-XIS-HTH | 1159 | Bacteria>Pseudomonadota>Gammaproteobacteria>Pseudomonadales>Pseudomonadaceae>Pseudomonas | GCF_002128325.1 | Pseudomonas_fragi | B6D87_RS19610 | 4227178..4230657 | 296 | BREX-system-P-loop-protein-BrxC-[Pseudomonas-fragi] |
| WP_106118416.1 | Type1.BREX | <-ARG<-?\|\|wHTH+WYL+WCX->wHTH+WYL+WCX->BrxA-DNA-binding->BrxB-BrxF-iSTAND->DUF6079*->OLD-ABC->TOPRIM->N6-METHYLASE->N6-METHYLASE-p-> | 1223 | Bacteria>Pseudomonadota>Gammaproteobacteria>Pseudomonadales>Pseudomonadaceae>Pseudomonas | GCF_003001555.1 | Pseudomonas_fluorescens | C7A10_RS23710 | 130630..134301 | 294 | BREX-system-P-loop-protein-BrxC-[Pseudomonas-fluorescens] |
| WP_124127279.1 | Type1.BREX | SF1-SF2-Helicase-><-HTH_3\|\|?->CHTH-><-?<-REase+NACHT\|\|wHTH+WYL+WCX->BrxA-DNA-binding->BrxB-BrxF-iSTAND->DUF6079*->N6-METHYLASE->OLD-ABC->TOPRIM->TIR->PglZ->SIGMA+OB-Fold+AAA-ATPase+Lon-Protease-BrxL->?->?-><-Phage_integrase | 1223 | Bacteria>Pseudomonadota>Gammaproteobacteria>Pseudomonadales>Pseudomonadaceae>Pseudomonas | GCF_003836525.1 | Pseudomonas_aeruginosa | IPC776_RS25325 | 73240..76911 | 287 | BREX-system-P-loop-protein-BrxC-[Pseudomonas-aeruginosa] |
| WP_161892243.1 | Type1.BREX | MerR-HTH->tRNA->?-><-CopG/RHH\|\|XIS-HTH->?->wHTH+WYL+WCX-p->wHTH+WYL+WCX->BrxA-DNA-binding->BrxB-BrxF-iSTAND->DUF6079*->N6-METHYLASE->DUF932->ParB+HNH->OLD-ABC->TOPRIM->PglZ->SIGMA+OB-Fold+AAA-ATPase+Lon-Protease-BrxL-><-DUF4969 | 1154 | Bacteria>Pseudomonadota>Gammaproteobacteria>Pseudomonadales>Pseudomonadaceae>Pseudomonas | GCF_009932375.1 | Pseudomonas_juntendi | GYA94_RS10075 | 73827..77291 | 2666183 | BREX-system-P-loop-protein-BrxC-[Pseudomonas-juntendi] |
| WP_094987826.1 | Type1.BREX | DUF4969-><-SIGMA+OB-Fold+AAA-ATPase+Lon-Protease-BrxL<-PglZ<-TOPRIM<-OLD-ABC<-ParB+HNH<-N6-METHYLASE<-DUF6079*<-BrxB-BrxF-iSTAND<-BrxA-DNA-binding<-wHTH+WYL+WCX<-wHTH+REase+ZnR<-DnaB+McrB-AAA<-Schlafen+HATPase+wHTH<-URI-DNase+RAMA<-?<-XIS-HTH | 1154 | Bacteria>Pseudomonadota>Gammaproteobacteria>Pseudomonadales>Pseudomonadaceae>Pseudomonas | GCF_012986325.1 | Pseudomonas_lundensis | HBN80_RS14475 | 101799..105263 | 86185 | BREX-system-P-loop-protein-BrxC-[Pseudomonas-lundensis] |
| WP_206408616.1 | Type1.BREX | ARG->?->wHTH+WYL+WCX->BrxA-DNA-binding->BrxB-BrxF-iSTAND->DUF6079*->N6-METHYLASE->OLD-ABC->HNH->ParB+HNH->OLD-ABC->TOPRIM->PglZ->SIGMA+OB-Fold+AAA-ATPase+Lon-Protease-BrxL->MukF-> | 1154 | Bacteria>Pseudomonadota>Gammaproteobacteria>Pseudomonadales>Pseudomonadaceae>Pseudomonas | GCF_017168035.1 | Pseudomonas_oleovorans | ACP89_RS12320 | 3896..7360 | 301 | BREX-system-P-loop-protein-BrxC-[Pseudomonas-oleovorans] |
| WP_213936039.1 | Type1.BREX | DUF4969-><-SIGMA+OB-Fold+AAA-ATPase+Lon-Protease-BrxL<-PglZ<-TOPRIM<-OLD-ABC<-ParB+HNH<-N6-METHYLASE<-DUF6079*<-BrxB-BrxF-iSTAND<-BrxA-DNA-binding<-wHTH+WYL+WCX<-?<-XIS-HTH<-BirA-HTH\|\|CopG/RHH-><-Pkinase+CR-Kinase | 1157 | Bacteria>Pseudomonadota>Gammaproteobacteria>Pseudomonadales>Pseudomonadaceae>Pseudomonas | GCF_018449005.1 | Pseudomonas_lundensis | KIY13_RS15530 | 3400509..3403982 | 86185 | BREX-system-P-loop-protein-BrxC-[Pseudomonas-lundensis] |
| WP_222018162.1 | Type1.BREX | N6-METHYLASE->?-><-SIGMA+OB-Fold+AAA-ATPase+Lon-Protease-BrxL<-PglZ\|\|?-><-N6-METHYLASE<-TOPRIM<-OLD-ABC<-DUF6079*<-BrxB-BrxF-iSTAND<-BrxA-DNA-binding<-wHTH+WYL+WCX<-?<-?<-?<-DUF932 | 1223 | Bacteria>Pseudomonadota>Gammaproteobacteria>Pseudomonadales>Pseudomonadaceae>Pseudomonas | GCF_019775695.1 | Pseudomonas_sp._AO-1 | KVQ82_RS04465 | 959176..962847 | 2855434 | BREX-system-P-loop-protein-BrxC-[Pseudomonas-sp.-AO-1] |
| WP_236310112.1 | Type1.BREX | <-HTH+rve<-?\|\|XIS-HTH->?->wHTH+WYL+WCX->BrxA-DNA-binding->BrxB-BrxF-iSTAND->DUF6079*->N6-METHYLASE->DUF932->ParB+HNH->OLD-ABC->TOPRIM->PglZ->SIGMA+OB-Fold+AAA-ATPase+Lon-Protease-BrxL-><-DUF4969 | 1154 | Bacteria>Pseudomonadota>Gammaproteobacteria>Pseudomonadales>Pseudomonadaceae>Pseudomonas | GCF_021605645.1 | Pseudomonas_gessardii | GIW56_RS19725 | 10406..13870 | 78544 | BREX-system-P-loop-protein-BrxC-[Pseudomonas-gessardii] |
| WP_261753794.1 | Type1.BREX | <-SIGMA+OB-Fold+AAA-ATPase+Lon-Protease-BrxL<-PglZ\|\|ZnR-><-TOPRIM<-OLD-ABC<-N6-METHYLASE<-DOC+HTH<-DUF6079*<-BrxB-BrxF-iSTAND<-BrxA-DNA-binding<-wHTH+WYL+WCX<-?<-Phage_AlpA<-?<-Arm-DNA-bind_3+Phage_integrase | 1225 | Bacteria>Pseudomonadota>Gammaproteobacteria>Pseudomonadales>Pseudomonadaceae>Pseudomonas | GCF_025397875.1 | Pseudomonas_veronii | MN113_RS13530 | 508981..512658 | 76761 | BREX-system-P-loop-protein-BrxC-[Pseudomonas-veronii] |
| WP_023127678.1 | Type1.BREX | NUDIX-><-?<-wHTH+WYL+WCX\|\|ARG->?->wHTH+WYL+WCX->BrxA-DNA-binding->BrxB-BrxF-iSTAND->DUF6079*->N6-METHYLASE->DUF932->ParB+HNH->OLD-ABC->TOPRIM->PglZ->SIGMA+OB-Fold+AAA-ATPase+Lon-Protease-BrxL->MukF->JetB-> | 1154 | Bacteria>Pseudomonadota>Gammaproteobacteria>Pseudomonadales>Pseudomonadaceae>Pseudomonas | GCF_025696785.1 | Pseudomonas_aeruginosa | KUC66_RS22970 | 76351..79815 | 287 | BREX-system-P-loop-protein-BrxC-[Pseudomonas-aeruginosa] |
| WP_273917578.1 | Type1.BREX | <-SIGMA+OB-Fold+AAA-ATPase+Lon-Protease-BrxL<-PglZ<-HEPN<-TPR<-NERD+GspA-ATPase<-N6-METHYLASE<-N6-METHYLASE<-TOPRIM<-OLD-ABC<-DUF6079*<-BrxB-BrxF-iSTAND<-BrxA-DNA-binding<-wHTH+WYL+WCX<-wHTH+WYL+WCX\|\|?->ARG-> | 1223 | Bacteria>Pseudomonadota>Gammaproteobacteria>Pseudomonadales>Pseudomonadaceae>Pseudomonas | GCF_028656895.1 | Pseudomonas_sp._TNT2022_ID1044 | M5G20_RS01570 | 347595..351266 | 2942636 | BREX-system-P-loop-protein-BrxC-[Pseudomonas-sp.-TNT2022-ID1044] |
| WP_150549236.1 | Type1.BREX | <-DUF3584+DUF5401<-JetB<-MukF<-SIGMA+OB-Fold+AAA-ATPase+Lon-Protease-BrxL<-PglZ<-TOPRIM<-OLD-ABC<-ParB+HNH<-N6-METHYLASE<-DUF6079*<-BrxB-BrxF-iSTAND<-BrxA-DNA-binding<-wHTH+WYL+WCX<-?<-?<-?<-XIS-HTH\|\|?-><-tRNA<-MerR-HTH | 1154 | Bacteria>Pseudomonadota>Gammaproteobacteria>Pseudomonadales>Pseudomonadaceae>Pseudomonas | GCF_900682675.2 | Pseudomonas_marincola | PMYSY11_RS11215 | 2416219..2419683 | 437900 | BREX-system-P-loop-protein-BrxC-[Pseudomonas-marincola] |
| WP_229999483.1 | Type1.BREX | XIS-HTH->?->URI-DNase+RAMA->Schlafen+HATPase+wHTH->DnaB+McrB-AAA->wHTH+REase+ZnR->wHTH+WYL+WCX->BrxA-DNA-binding->BrxB-BrxF-iSTAND->DUF6079*->N6-METHYLASE->ParB+HNH->OLD-ABC->TOPRIM->PglZ->SIGMA+OB-Fold+AAA-ATPase+Lon-Protease-BrxL-><-DUF4969 | 1156 | Bacteria>Pseudomonadota>Gammaproteobacteria>Pseudomonadales>Pseudomonadaceae>Pseudomonas | GCF_918697895.1 | Pseudomonas_carnis | LQ224_RS08485 | 110382..113852 | 2487355 | BREX-system-P-loop-protein-BrxC-[Pseudomonas-carnis] |
| WP_240043383.1 | Type1.BREX | <-Thymidylat_kin<-?\|\|ZnR-><-?<-?<-TOPRIM<-OLD-ABC<-N6-METHYLASE<-DOC+HTH<-DUF6079*<-BrxB-BrxF-iSTAND<-BrxA-DNA-binding<-wHTH+WYL+WCX | 1225 | Bacteria>Pseudomonadota>Gammaproteobacteria>Pseudomonadales>Pseudomonadaceae>Pseudomonas | - | Pseudomonas_viridiflava | EFA20_RS09810 | 85838..89515 | 33069 | BREX-system-P-loop-protein-BrxC-[Pseudomonas-viridiflava] |
| MCQ4319996.1 | Type1.BREX | Phage_integrase->?->?-><-?<-?\|\|wHTH+WYL+WCX->BrxA-DNA-binding->BrxB-BrxF-iSTAND->DUF6079*->OLD-ABC->TOPRIM->N6-METHYLASE->sGTP->?->PglZ->SIGMA+OB-Fold+AAA-ATPase+Lon-Protease-BrxL-> | 1223 | Bacteria>Pseudomonadota>Gammaproteobacteria>Pseudomonadales>Pseudomonadaceae>Stutzerimonas | GCA_024448935.1 | Stutzerimonas_stutzeri | NA637_07215 | 17580..21251 | 316 | BREX-system-P-loop-protein-BrxC-[Stutzerimonas-stutzeri] |
| WP_095461351.1 | Type1.BREX | <-wHTH+WYL+WCX-p\|\|?->wHTH+WYL+WCX->BrxA-DNA-binding->BrxB-BrxF-iSTAND->DUF6079*->N6-METHYLASE->DUF932->ParB+HNH->OLD-ABC->TOPRIM->PglZ->SIGMA+OB-Fold+AAA-ATPase+Lon-Protease-BrxL->MukF->JetB-> | 1154 | Bacteria>Pseudomonadota>Gammaproteobacteria>Pseudomonadales>Pseudomonadaceae>Stutzerimonas | - | Stutzerimonas_stutzeri | BV581_21185 | 4303..7767 | 316 | BREX-system-P-loop-protein-BrxC-[Stutzerimonas-stutzeri] |
| WP_053105412.1 | Type1.BREX | METHYLASE->?->?-><-?<-?<-?\|\|wHTH+WYL+WCX->BrxA-DNA-binding->BrxB-BrxF-iSTAND->DUF6079*->N6-METHYLASE->ParB+HNH->OLD-ABC->TOPRIM->HEPN->PglZ->SIGMA+OB-Fold+AAA-ATPase+Lon-Protease-BrxL->Arm-DNA-bind_3+Phage_integrase-><-SIGMA+OB-Fold+AAA-ATPase+Lon-Protease-BrxL | 1155 | Bacteria>Pseudomonadota>Gammaproteobacteria>Pseudomonadales>Pseudomonadaceae>Thiopseudomonas | GCF_001267275.1 | Thiopseudomonas_alkaliphila | AKN90_RS00615 | 123321..126788 | 1697053 | BREX-system-P-loop-protein-BrxC-[Thiopseudomonas-alkaliphila] |
| MAB42111.1 | Type1.BREX | <-SIGMA+OB-Fold+AAA-ATPase+Lon-Protease-BrxL<-PglZ<-TOPRIM<-OLD-ABC<-ParB+HNH<-N6-METHYLASE<-DUF6079*<-BrxB-BrxF-iSTAND<-BrxA-DNA-binding\|\|wHTH+WYL+WCX-> | 1154 | Bacteria>Pseudomonadota>Gammaproteobacteria>Pseudomonadales>unclassified_Pseudomonadales | GCA_002684155.1 | Pseudomonadales_bacterium | CMX77_07490 | 13688..17152 | 1891229 | hypothetical-protein-[Pseudomonadales-bacterium] |
| MCP5320940.1 | Type1.BREX | <-SbcC+CCDC158<-Calcineurin-SbcD<-PglZ<-ParA-Soj-PloopNTPase<-Schlafen+HATPase+wHTH<-Schlafen+HATPase+wHTH<-N6-METHYLASE<-TOPRIM<-OLD-ABC<-DUF6079*<-BrxB-BrxF-iSTAND<-BrxA-DNA-binding\|\|MerR-HTH->DUF772+DDE_Tnp_1+DDE_Tnp_1_6->?->?-><-SF1-SF2-Helicase<-SF1-SF2-Helicase<-AbiEii+ZnR | 1148 | Bacteria>Pseudomonadota>Gammaproteobacteria>Pseudomonadales>unclassified_Pseudomonadales | GCA_024234165.1 | Pseudomonadales_bacterium | H7A12_08980 | 738054..741500 | 1891229 | BREX-system-P-loop-protein-BrxC-[Pseudomonadales-bacterium] |
| WP_022954802.1 | Type1.BREX | CR-ATPase5+REase-><-?\|\|wHTH+WYL+WCX->BrxA-DNA-binding->BrxB-BrxF-iSTAND->DUF6079*->N6-METHYLASE->OLD-ABC->TOPRIM->PglZ->SIGMA+OB-Fold+AAA-ATPase+Lon-Protease-BrxL-> | 1222 | Bacteria>Pseudomonadota>Gammaproteobacteria>Thiotrichales>Thiotrichaceae>Leucothrix | GCF_000419525.1 | Leucothrix_mucor | LEUMU_RS0123780 | 65328..68996 | 45248 | BREX-system-P-loop-protein-BrxC-[Leucothrix-mucor] |
| OQX14377.1 | Type1.BREX | <-TRPR-HTH<-?<-DUF3670+SF1-SF2-Helicase\|\|?-><-SIGMA+OB-Fold+AAA-ATPase+Lon-Protease-BrxL<-PglZ<-TOPRIM<-OLD-ABC<-N6-METHYLASE<-DUF6079*<-BrxB-BrxF-iSTAND<-BrxA-DNA-binding<-TIR<-?<-?\|\|XIS-HTH-> | 1145 | Bacteria>Pseudomonadota>Gammaproteobacteria>Thiotrichales>Thiotrichaceae>Thiothrix | GCA_002083875.1 | Thiothrix_lacustris | BWK73_09690 | 20114..23551 | 525917 | hypothetical-protein-BWK73_09690-[Thiothrix-lacustris] |
| QLQ31990.1 | Type1.BREX | DrsE-><-?\|\|REase->?->Arm-DNA-bind_3+Phage_integrase-><-HTH+DOC<-?<-LexA-protease\|\|BrxA-DNA-binding->BrxB-BrxF-iSTAND->DUF6079*->AbiEi_4+AbiEi_1->AbiEii->OLD-ABC->OLD-ABC->TOPRIM-><-?\|\|N6-METHYLASE->N6-METHYLASE-p->HIN-HTH-> | 1209 | Bacteria>Pseudomonadota>Gammaproteobacteria>Thiotrichales>Thiotrichaceae>Thiothrix | GCA_013693955.1 | Candidatus_Thiothrix_singaporensis | HZT40_10770 | 2133663..2137292 | 2799669 | BREX-system-P-loop-protein-BrxC-[Candidatus-Thiothrix-singaporensis] |
| WP_002710590.1 | Type1.BREX | HAMP+PAS+HISKIN->REC+NtrC-AAA+FIS-HTH->BrxA-DNA-binding->BrxB-BrxF-iSTAND->DUF6079*->AbiEi_4+AbiEi_1->AbiEii->OLD-ABC->TOPRIM->N6-METHYLASE->CR-ATPase8+REase->PglZ->SIGMA+OB-Fold+AAA-ATPase+Lon-Protease-BrxL-> | 1209 | Bacteria>Pseudomonadota>Gammaproteobacteria>Thiotrichales>Thiotrichaceae>Thiothrix | GCF_000260135.1 | Thiothrix_nivea | THINI_RS21415 | 4300442..4304071 | 1031 | BREX-system-P-loop-protein-BrxC-[Thiothrix-nivea] |
| WP_210226002.1 | Type6.BREX | <-PAS\|\|?->?->BrxE->BrxA-DNA-binding->BrxB-BrxF-iSTAND->DUF6079*->N6-METHYLASE->OLD-ABC->TOPRIM->PglZ->SIGMA+OB-Fold+AAA-ATPase+HKD-DNase-BrxL-><-?<-?<-?<-SAM-METHYLASE | 1210 | Bacteria>Pseudomonadota>Gammaproteobacteria>Thiotrichales>Thiotrichaceae>Thiothrix | GCF_017901155.1 | Candidatus_Thiothrix_anitrata | J8380_RS12830 | 2543544..2547176 | 2823902 | BREX-system-P-loop-protein-BrxC-[Candidatus-Thiothrix-anitrata] |
| NLY14765.1 | Type1.BREX | <-SIGMA+OB-Fold+AAA-ATPase+Lon-Protease-BrxL<-PglZ<-HEPN<-TOPRIM<-OLD-ABC<-ParB+HNH<-N6-METHYLASE<-DUF6079-p* | 438 | Bacteria>Pseudomonadota>Gammaproteobacteria>unclassified_Gammaproteobacteria | GCA_012728415.1 | Gammaproteobacteria_bacterium | GXZ05_00075 | 12795..14111 | 1913989 | hypothetical-protein-[Gammaproteobacteria-bacterium] |
| NPU94904.1 | Type6.BREX | <-DnaK\|\|35exo-><-SIGMA+OB-Fold+AAA-ATPase+HKD-DNase-BrxL<-PglZ<-TOPRIM<-OLD-ABC<-N6-METHYLASE<-N6-METHYLASE<-GreA_GreB+SF1-SF2-Helicase<-DUF6079*<-BrxB-BrxF-iSTAND<-BrxA-DNA-binding<-BrxE<-wHTH+WYL+WCX\|\|REase->rve->TnsC-ATPase-> | 1213 | Bacteria>Pseudomonadota>Gammaproteobacteria>unclassified_Gammaproteobacteria | GCA_013178375.1 | Gammaproteobacteria_bacterium | HPY82_23615 | 18761..22402 | 1913989 | BREX-system-P-loop-protein-BrxC-[Gammaproteobacteria-bacterium] |
| MCP4410534.1 | Type1.BREX | HTH_26->BrxA-DNA-binding->BrxB-BrxF-iSTAND->DUF6079-p*->OLD-ABC->TOPRIM->Schlafen+HATPase+wHTH->PglZ->SIGMA+OB-Fold+AAA-ATPase+Lon-Protease-BrxL-> | 791 | Bacteria>Pseudomonadota>Gammaproteobacteria>unclassified_Gammaproteobacteria | GCA_024222475.1 | Gammaproteobacteria_bacterium | GY807_22880 | 5382..7757 | 1913989 | BREX-system-P-loop-protein-BrxC-[Gammaproteobacteria-bacterium] |
| MCP5451255.1 | Type1.BREX | <-ParB+HTH<-ParA-Soj-PloopNTPase\|\|?->BrxA-DNA-binding->BrxB-BrxF-iSTAND->DUF6079*->ParD_antitoxin->RelE-ParE->N6-METHYLASE->OLD-ABC->TOPRIM->REase+AIPR->PglZ->SIGMA+OB-Fold+AAA-ATPase+Lon-Protease-BrxL-><-Thermonuclease | 1154 | Bacteria>Pseudomonadota>Gammaproteobacteria>unclassified_Gammaproteobacteria | GCA_024235115.1 | Gammaproteobacteria_bacterium | H6972_12070 | 41829..45293 | 1913989 | BREX-system-P-loop-protein-BrxC-[Gammaproteobacteria-bacterium] |
| WP_190395653.1 | Type1.BREX | <-Couple_hipA+HipA_C<-HTH_3\|\|?-><-RNase<-PALP\|\|wHTH+WYL+WCX->BrxA-DNA-binding->BrxB-BrxF-iSTAND->DUF6079*->N6-METHYLASE->OLD-ABC->TOPRIM->PglZ->HEPN+SF1-SF2-Helicase+RNase+RNase->TPR-><-SF1-SF2-Helicase+REase<-SF1-SF2-Helicase+REase<-Peptidase_M32 | 1157 | Bacteria>Pseudomonadota>Gammaproteobacteria>Vibrionales>Vibrionaceae>Aliivibrio | GCF_014694495.1 | Aliivibrio_sp._S10_S31 | HC728_RS13680 | 57939..61412 | 2720224 | BREX-system-P-loop-protein-BrxC-[Aliivibrio-sp.-S10_S31] |
| WP_199414880.1 | Type1.BREX | Peptidase_M32->SF1-SF2-Helicase+REase->SF1-SF2-Helicase+REase-><-TPR<-HEPN+SF1-SF2-Helicase+RNase+RNase<-PglZ<-TOPRIM<-OLD-ABC<-N6-METHYLASE<-DUF6079*<-BrxB-BrxF-iSTAND<-BrxA-DNA-binding<-wHTH+WYL+WCX\|\|PALP->RNase-><-?\|\|HTH_3->Couple_hipA+HipA_C-><-HNH | 1157 | Bacteria>Pseudomonadota>Gammaproteobacteria>Vibrionales>Vibrionaceae>Aliivibrio | GCF_023983475.1 | Aliivibrio_fischeri | AVFI_RS07680 | 1646065..1649538 | 668 | BREX-system-P-loop-protein-BrxC-[Aliivibrio-fischeri] |
| WP_103201396.1 | Type1.BREX | <-VirD4-TraG+VirB4-FtsK<-HDRnase<-SbcC<-SIGMA+OB-Fold+AAA-ATPase+Lon-Protease-BrxL<-PglZ<-N6-METHYLASE<-TOPRIM<-OLD-ABC<-DUF6079*<-BrxB-BrxF-iSTAND<-BrxA-DNA-binding<-wHTH+WYL+WCX<-?<-35exo<-?\|\|LexA-protease->?-><-MobI | 1225 | Bacteria>Pseudomonadota>Gammaproteobacteria>Vibrionales>Vibrionaceae | GCF_003047085.1 | Vibrio | DA442_RS13700 | 2691122..2694799 | 662 | MULTISPECIES:-BREX-system-P-loop-protein-BrxC-[Vibrio] |
| QOQ70156.1 | Type1.BREX | SF1-SF2-Helicase+REase->SF1-SF2-Helicase+REase-><-PglZ<-TOPRIM<-OLD-ABC<-N6-METHYLASE<-DUF6079*<-BrxB-BrxF-iSTAND<-BrxA-DNA-binding<-wHTH+WYL+WCX<-XIS-HTH<-?<-REase+SF1-SF2-Helicase\|\|SAM-METHYLASE->HKD-DNase+SF1-SF2-Helicase+HTH+HTH+SRA-> | 1154 | Bacteria>Pseudomonadota>Gammaproteobacteria>Vibrionales>Vibrionaceae>Photobacterium | GCA_014930975.1 | Photobacterium_damselae_subsp._damselae | IL982_10195 | 2142038..2145502 | 85581 | BREX-system-P-loop-protein-BrxC-[Photobacterium-damselae-subsp.-damselae] |
| WP_176285150.1 | Type1.BREX | <-SF1-SF2-Helicase\|\|?-><-sGTP+sGTP\|\|XIS-HTH->?->ANK+ANK->BrxA-DNA-binding->BrxB-BrxF-iSTAND->DUF6079*->N6-METHYLASE->OLD-ABC->TOPRIM->PglZ->ZetaToxin->?-><-SF1-SF2-Helicase+REase<-SF1-SF2-Helicase+REase | 1157 | Bacteria>Pseudomonadota>Gammaproteobacteria>Vibrionales>Vibrionaceae>Photobacterium | GCF_013371045.1 | Photobacterium_damselae | HU831_RS17605 | 57170..60643 | 38293 | BREX-system-P-loop-protein-BrxC-[Photobacterium-damselae] |
| WP_232092144.1 | Type1.BREX | SF1-SF2-Helicase+REase->SF1-SF2-Helicase+REase-><-PglZ<-TOPRIM<-OLD-ABC<-N6-METHYLASE<-DUF6079*<-BrxB-BrxF-iSTAND<-BrxA-DNA-binding<-wHTH+WYL+WCX<-XIS-HTH<-?<-REase+SF1-SF2-Helicase\|\|SAM-METHYLASE-><-?\|\|HTH_26-> | 1153 | Bacteria>Pseudomonadota>Gammaproteobacteria>Vibrionales>Vibrionaceae>Photobacterium | GCF_014930975.1 | Photobacterium_damselae | IL982_RS10195 | 2142038..2145499 | 38293 | BREX-system-P-loop-protein-BrxC-[Photobacterium-damselae] |
| WP_101770358.1 | Type1.BREX | <-SIGMA+OB-Fold+AAA-ATPase+Lon-Protease-BrxL<-PglZ<-N6-METHYLASE<-DUF6079<-TOPRIM<-OLD-ABC<-DUF6079*<-BrxB-BrxF-iSTAND<-BrxA-DNA-binding<-Recombinase<-?<-Arm-DNA-bind_2+Phage_int_SAM_3+Phage_integrase<-Phage_AlpA\|\|HTH_26-> | 1223 | Bacteria>Pseudomonadota>Gammaproteobacteria>Vibrionales>Vibrionaceae>Photobacterium | GCF_027923345.1 | Photobacterium_carnosum | CIK00_20185 | 17344..21015 | 2023717 | BREX-system-P-loop-protein-BrxC-[Photobacterium-carnosum] |
| CAQ34953.1 | Type1.BREX | HIN-HTH->HTH_Tnp_1+HTH_21+rve+rve_3-><-Phage_integrase<-LexA-protease\|\|35exo->?->wHTH+WYL+WCX->BrxA-DNA-binding->BrxB-BrxF-iSTAND->DUF6079*->OLD-ABC->TOPRIM->N6-METHYLASE-p-><-DDE_Tnp_1\|\|PglZ->SIGMA+OB-Fold+AAA-ATPase+Lon-Protease-BrxL->SbcC->HDRnase->VirD4-TraG+VirB4-FtsK-> | 1225 | Bacteria>Pseudomonadota>Gammaproteobacteria>Vibrionales>Vibrionaceae>Photobacterium | - | Photobacterium_damselae_subsp._piscicida | . | 21165..24842 | 38294 | hypothetical-protein-[Photobacterium-damselae-subsp.-piscicida] |
| WP_077485085.1 | Type1.BREX | Glyco_hydro_4+Glyco_hydro_4C-><-wHTH+WYL+WCX\|\|SF1-SF2-Helicase+REase->?->wHTH+WYL+WCX->BrxA-DNA-binding->BrxB-BrxF-iSTAND->DUF6079*->N6-METHYLASE->OLD-ABC->TOPRIM->PglZ-><-SF1-SF2-Helicase+REase<-SF1-SF2-Helicase+REase\|\|ZnR-> | 1157 | Bacteria>Pseudomonadota>Gammaproteobacteria>Vibrionales>Vibrionaceae>Salinivibrio | GCF_001995745.1 | Salinivibrio_sp._ML323 | BZG13_RS03555 | 125341..128814 | 1909474 | BREX-system-P-loop-protein-BrxC-[Salinivibrio-sp.-ML323] |
| TNF18455.1 | Type1.BREX | Phage_holin->Phage_lysozyme->XIS-HTH->BrxA-DNA-binding->BrxB-BrxF-iSTAND->DUF6079*->N6-METHYLASE->OLD-ABC->TOPRIM->PglZ->SIGMA+OB-Fold+AAA-ATPase+Lon-Protease-BrxL-> | 1145 | Bacteria>Pseudomonadota>Gammaproteobacteria>Vibrionales>Vibrionaceae>unclassified_Vibrionaceae | GCA_006226665.1 | Vibrionaceae_bacterium | EP325_05025 | 51838..55275 | 1889773 | BREX-system-P-loop-protein-BrxC-[Vibrionaceae-bacterium] |
| KOY19349.1 | Type1.BREX | <-GDResolvase+HIN-HTH\|\|?->Arm-DNA-bind_3+Phage_integrase->SF1-SF2-Helicase+REase->SF1-SF2-Helicase+REase-><-PglZ<-TOPRIM<-OLD-ABC<-N6-METHYLASE<-DUF6079*<-BrxB-BrxF-iSTAND<-BrxA-DNA-binding<-wHTH+WYL+WCX<-XIS-HTH\|\|HTH_Tnp_1-> | 1154 | Bacteria>Pseudomonadota>Gammaproteobacteria>Vibrionales>Vibrionaceae>Vibrio | GCA_001280635.1 | Vibrio_parahaemolyticus | ACX06_21960 | 167502..170966 | 670 | hypothetical-protein-ACX06_21960-[Vibrio-parahaemolyticus] |
| ODY54465.1 | Type1.BREX | <-HTH_Tnp_1\|\|XIS-HTH->wHTH+WYL+WCX->BrxA-DNA-binding->BrxB-BrxF-iSTAND->DUF6079*->N6-METHYLASE->OLD-ABC->TOPRIM->PglZ-><-SF1-SF2-Helicase+REase<-SF1-SF2-Helicase+REase<-Arm-DNA-bind_3+Phage_integrase | 1154 | Bacteria>Pseudomonadota>Gammaproteobacteria>Vibrionales>Vibrionaceae>Vibrio | GCA_001727265.1 | Vibrio_parahaemolyticus | BBM25_06240 | 3424..6888 | 670 | hypothetical-protein-BBM25_06240-[Vibrio-parahaemolyticus] |
| MBE3824873.1 | Type1.BREX | <-VirD4-TraG+VirB4-FtsK<-HDRnase<-SbcC<-SIGMA+OB-Fold+AAA-ATPase+Lon-Protease-BrxL<-PglZ<-TOPRIM<-OLD-ABC<-N6-METHYLASE<-ParB+HNH<-DUF6079*<-BrxB-BrxF-iSTAND<-BrxA-DNA-binding<-wHTH+WYL+WCX<-?<-35exo\|\|LexA-protease->?-><-MobI | 1225 | Bacteria>Pseudomonadota>Gammaproteobacteria>Vibrionales>Vibrionaceae>Vibrio | GCA_014918955.1 | Vibrio_parahaemolyticus | HJ179_16470 | 255963..259640 | 670 | BREX-system-P-loop-protein-BrxC-[Vibrio-parahaemolyticus] |
| MBE3881970.1 | Type1.BREX | MobI-><-?<-LexA-protease\|\|35exo->?->wHTH+WYL+WCX->BrxA-DNA-binding->BrxB-BrxF-iSTAND->DUF6079*->OLD-ABC->TOPRIM->N6-METHYLASE->PglZ->SIGMA+OB-Fold+AAA-ATPase+Lon-Protease-BrxL->NACHT+HEAT+REase->HDRnase->VirD4-TraG+VirB4-FtsK-> | 1225 | Bacteria>Pseudomonadota>Gammaproteobacteria>Vibrionales>Vibrionaceae>Vibrio | GCA_014919165.1 | Vibrio_parahaemolyticus | HJ168_09865 | 226253..229930 | 670 | BREX-system-P-loop-protein-BrxC-[Vibrio-parahaemolyticus] |

<table>
<tr><th>Protein-ID</th><th>System</th><th>Gene-Neighbourhoods</th><th>Length</th><th>Taxonomy</th><th>Assembly-ID</th><th>Strain</th><th>Locus-Tag</th><th>Genomic-coordinates</th><th>TaxID</th><th>Fasta-defline</th></tr>
<tr><td>HAS6259595.1</td><td>Type1.BREX</td><td><-VirD4-TraG+VirB4-FtsK<-HDRnase<-NACHT+HEAT+REase<- SIGMA+OB-Fold+AAA-ATPase+Lon-Protease-BrxL <- PglZ <- TOPRIM <- OLD-ABC <- N6-METHYLASE <-ParB+HNH<- DUF6079 *<- BrxB-BrxF-iSTAND <- BrxA-DNA-binding <- wHTH+WYL+WCX <-?<-35exo||LexA-protease->?-><-MobI</td><td>1225</td><td>Bacteria>Pseudomonadota>Gammaproteobacteria>Vibrionales>Vibrionaceae>Vibrio</td><td>GCA_015777345.1</td><td>Vibrio_vulnificus</td><td>I7226_03370</td><td>263913..267590</td><td>672</td><td>BREX-system-P-loop-protein-BrxC-[Vibrio-vulnificus]</td></tr>
<tr><td>EGQ8649255.1</td><td>Type1.BREX</td><td><-VirD4-TraG+VirB4-FtsK<-HDRnase<-SbcC<- SIGMA+OB-Fold+AAA-ATPase+Lon-Protease-BrxL <- PglZ <- N6-METHYLASE <- TOPRIM <- OLD-ABC <- DUF6079 *<- BrxB-BrxF-iSTAND <- BrxA-DNA-binding <- wHTH+WYL+WCX <-?<-35exo<-?||LexA-protease-></td><td>1225</td><td>Bacteria>Pseudomonadota>Gammaproteobacteria>Vibrionales>Vibrionaceae>Vibrio</td><td>GCA_015781355.1</td><td>Vibrio_cholerae</td><td>GNS53_08430</td><td>57189..60866</td><td>666</td><td>BREX-system-P-loop-protein-BrxC-[Vibrio-cholerae]</td></tr>
<tr><td>EGR0286978.1</td><td>Type1.BREX</td><td>Arm-DNA-bind_3+Phage_integrase->SF1-SF2-Helicase+REase->SF1-SF2-Helicase+REase-><- PglZ <- TOPRIM <- OLD-ABC <- N6-METHYLASE <- DUF6079 *<- BrxB-BrxF-iSTAND <- BrxA-DNA-binding <- wHTH+WYL+WCX <-XIS-HTH||?->?->DUF2787->HTH-> wHTH+WYL+WCX -></td><td>1154</td><td>Bacteria>Pseudomonadota>Gammaproteobacteria>Vibrionales>Vibrionaceae>Vibrio</td><td>GCA_015789595.1</td><td>Vibrio_cholerae</td><td>FK206_11700</td><td>85283..88747</td><td>666</td><td>BREX-system-P-loop-protein-BrxC-[Vibrio-cholerae]</td></tr>
<tr><td>EGR1144248.1</td><td>Type1.BREX</td><td>SF1-SF2-Helicase+REase->SF1-SF2-Helicase+REase-><- PglZ <- TOPRIM <- OLD-ABC <- N6-METHYLASE <- DUF6079 *<- BrxB-BrxF-iSTAND <- BrxA-DNA-binding <- wHTH+WYL+WCX <-XIS-HTH||?->?->DUF2787->||HTH-> wHTH+WYL+WCX -></td><td>1154</td><td>Bacteria>Pseudomonadota>Gammaproteobacteria>Vibrionales>Vibrionaceae>Vibrio</td><td>GCA_015798095.1</td><td>Vibrio_parahaemolyticus</td><td>EC562_15505</td><td>41063..44527</td><td>670</td><td>BREX-system-P-loop-protein-BrxC-[Vibrio-parahaemolyticus]</td></tr>
<tr><td>EGR1548466.1</td><td>Type1.BREX</td><td>Arm-DNA-bind_3+Phage_integrase->SF1-SF2-Helicase+REase->SF1-SF2-Helicase+REase-><- PglZ <- TOPRIM <- OLD-ABC <- N6-METHYLASE <- DUF6079 *<- BrxB-BrxF-iSTAND <- BrxA-DNA-binding <- wHTH+WYL+WCX <-XIS-HTH</td><td>1154</td><td>Bacteria>Pseudomonadota>Gammaproteobacteria>Vibrionales>Vibrionaceae>Vibrio</td><td>GCA_015798755.1</td><td>Vibrio_parahaemolyticus</td><td>D6Y11_16950</td><td>161246..164710</td><td>670</td><td>BREX-system-P-loop-protein-BrxC-[Vibrio-parahaemolyticus]</td></tr>
<tr><td>EGR2430327.1</td><td>Type1.BREX</td><td><- wHTH+WYL+WCX <-HTH<-DUF2787<-?<-?||XIS-HTH-> wHTH+WYL+WCX -> BrxA-DNA-binding -> BrxB-BrxF-iSTAND -> DUF6079 *-> N6-METHYLASE -> OLD-ABC -> TOPRIM -> PglZ -><-SF1-SF2-Helicase+REase<-SF1-SF2-Helicase+REase<-Arm-DNA-bind_3+Phage_integrase</td><td>1154</td><td>Bacteria>Pseudomonadota>Gammaproteobacteria>Vibrionales>Vibrionaceae>Vibrio</td><td>GCA_015802095.1</td><td>Vibrio_cholerae</td><td>DYB83_10140</td><td>10866..14330</td><td>666</td><td>BREX-system-P-loop-protein-BrxC-[Vibrio-cholerae]</td></tr>
<tr><td>EGR3302280.1</td><td>Type1.BREX</td><td>XIS-HTH-> wHTH+WYL+WCX -> BrxA-DNA-binding -> BrxB-BrxF-iSTAND -> DUF6079 *-> N6-METHYLASE -> OLD-ABC -> TOPRIM -> PglZ -><-SF1-SF2-Helicase+REase<-SF1-SF2-Helicase+REase<-Arm-DNA-bind_3+Phage_integrase<-?||GDResolvase+HIN-HTH-></td><td>1154</td><td>Bacteria>Pseudomonadota>Gammaproteobacteria>Vibrionales>Vibrionaceae>Vibrio</td><td>GCA_015805535.1</td><td>Vibrio_parahaemolyticus</td><td>DMJ21_13130</td><td>8098..11562</td><td>670</td><td>BREX-system-P-loop-protein-BrxC-[Vibrio-parahaemolyticus]</td></tr>
<tr><td>EJG0981293.1</td><td>Type1.BREX</td><td>Arm-DNA-bind_3+Phage_integrase->SF1-SF2-Helicase+REase->SF1-SF2-Helicase+REase-><- PglZ <- TOPRIM <- OLD-ABC <- N6-METHYLASE <- DUF6079 *<- BrxB-BrxF-iSTAND <- BrxA-DNA-binding <- wHTH+WYL+WCX <-XIS-HTH</td><td>1154</td><td>Bacteria>Pseudomonadota>Gammaproteobacteria>Vibrionales>Vibrionaceae>Vibrio</td><td>GCA_015813375.2</td><td>Vibrio_parahaemolyticus</td><td>C4G85_RS03500</td><td>776707..780171</td><td>670</td><td>BREX-system-P-loop-protein-BrxC-[Vibrio-parahaemolyticus]</td></tr>
<tr><td>EJG0936611.1</td><td>Type1.BREX</td><td><-GDResolvase+HIN-HTH<-?||Arm-DNA-bind_3+Phage_integrase->SF1-SF2-Helicase+REase->SF1-SF2-Helicase+REase-><- PglZ <- TOPRIM <- OLD-ABC <- N6-METHYLASE <- DUF6079 *<- BrxB-BrxF-iSTAND <- BrxA-DNA-binding <- wHTH+WYL+WCX ||?->?->DUF2787->?->||<-?||MODE-HTH+PBPII-></td><td>1154</td><td>Bacteria>Pseudomonadota>Gammaproteobacteria>Vibrionales>Vibrionaceae>Vibrio</td><td>GCA_015814185.2</td><td>Vibrio_parahaemolyticus</td><td>C4G97_RS14675</td><td>118301..121765</td><td>670</td><td>BREX-system-P-loop-protein-BrxC-[Vibrio-parahaemolyticus]</td></tr>
<tr><td>EHH2507631.1</td><td>Type1.BREX</td><td>Arm-DNA-bind_3+Phage_integrase->SF1-SF2-Helicase+REase->SF1-SF2-Helicase+REase-><- PglZ <- TOPRIM <- OLD-ABC <- N6-METHYLASE <- DUF6079 *<- BrxB-BrxF-iSTAND <- BrxA-DNA-binding <- wHTH+WYL+WCX <-XIS-HTH</td><td>1154</td><td>Bacteria>Pseudomonadota>Gammaproteobacteria>Vibrionales>Vibrionaceae>Vibrio</td><td>GCA_017637525.1</td><td>Vibrio_parahaemolyticus</td><td>J7X10_003794</td><td>109543..113007</td><td>670</td><td>BREX-system-P-loop-protein-BrxC-[Vibrio-parahaemolyticus]</td></tr>
<tr><td>EII2403787.1</td><td>Type1.BREX</td><td><-GDResolvase+HIN-HTH<-?||Arm-DNA-bind_3+Phage_integrase->SF1-SF2-Helicase+REase->SF1-SF2-Helicase+REase-><- PglZ <- TOPRIM <- OLD-ABC <- N6-METHYLASE <- DUF6079 *<- BrxB-BrxF-iSTAND <- BrxA-DNA-binding <- wHTH+WYL+WCX <-XIS-HTH</td><td>1154</td><td>Bacteria>Pseudomonadota>Gammaproteobacteria>Vibrionales>Vibrionaceae>Vibrio</td><td>GCA_020407245.1</td><td>Vibrio_parahaemolyticus</td><td>LG159_002760</td><td>116849..120313</td><td>670</td><td>BREX-system-P-loop-protein-BrxC-[Vibrio-parahaemolyticus]</td></tr>
<tr><td>EII3725635.1</td><td>Type1.BREX</td><td><-VirD4-TraG+VirB4-FtsK<-HDRnase<-SbcC<- SIGMA+OB-Fold+AAA-ATPase+Lon-Protease-BrxL <- PglZ <- TOPRIM <- OLD-ABC <- N6-METHYLASE <-ParB+HNH<- DUF6079 *<- BrxB-BrxF-iSTAND <- BrxA-DNA-binding <- wHTH+WYL+WCX <-?<-35exo<-?||LexA-protease->?-><-MobI</td><td>1225</td><td>Bacteria>Pseudomonadota>Gammaproteobacteria>Vibrionales>Vibrionaceae>Vibrio</td><td>GCA_020414365.1</td><td>Vibrio_cholerae</td><td>LG468_000348</td><td>129198..132875</td><td>666</td><td>BREX-system-P-loop-protein-BrxC-[Vibrio-cholerae]</td></tr>
<tr><td>EJB8347940.1</td><td>Type1.BREX</td><td><-VirD4-TraG+VirB4-FtsK<-HDRnase<-NACHT+HEAT+REase<- SIGMA+OB-Fold+AAA-ATPase+Lon-Protease-BrxL <- PglZ <- N6-METHYLASE <- TOPRIM <- OLD-ABC <- DUF6079 *<- BrxB-BrxF-iSTAND <- BrxA-DNA-binding <- wHTH+WYL+WCX <-?<-35exo||LexA-protease->?-><-MobI</td><td>1225</td><td>Bacteria>Pseudomonadota>Gammaproteobacteria>Vibrionales>Vibrionaceae>Vibrio</td><td>GCA_022966375.1</td><td>Vibrio_cholerae</td><td>MW328_000282</td><td>40595..44272</td><td>666</td><td>BREX-system-P-loop-protein-BrxC-[Vibrio-cholerae]</td></tr>
<tr><td>EJB8581804.1</td><td>Type1.BREX</td><td><-VirD4-TraG+VirB4-FtsK<-HDRnase<-SbcC<- SIGMA+OB-Fold+AAA-ATPase+Lon-Protease-BrxL <- PglZ <- N6-METHYLASE <- TOPRIM <- OLD-ABC <- DUF6079 *<- BrxB-BrxF-iSTAND <- BrxA-DNA-binding <- wHTH+WYL+WCX <-?<-35exo||LexA-protease->?->Glyoxalase_8-><-MobI</td><td>1229</td><td>Bacteria>Pseudomonadota>Gammaproteobacteria>Vibrionales>Vibrionaceae>Vibrio</td><td>GCA_022967455.1</td><td>Vibrio_cholerae</td><td>MW361_003123</td><td>34970..38659</td><td>666</td><td>BREX-system-P-loop-protein-BrxC-[Vibrio-cholerae]</td></tr>
<tr><td>EJC1210117.1</td><td>Type1.BREX</td><td><-DUF2787<-?<-?||XIS-HTH-> wHTH+WYL+WCX -> BrxA-DNA-binding -> BrxB-BrxF-iSTAND -> DUF6079 *-> N6-METHYLASE -> OLD-ABC -> TOPRIM -> PglZ -><-SF1-SF2-Helicase+REase<-SF1-SF2-Helicase+REase<-Arm-DNA-bind_3+Phage_integrase||?->GDResolvase+HIN-HTH-></td><td>1154</td><td>Bacteria>Pseudomonadota>Gammaproteobacteria>Vibrionales>Vibrionaceae>Vibrio</td><td>GCA_022997325.1</td><td>Vibrio_parahaemolyticus</td><td>MX350_002594</td><td>5689..9153</td><td>670</td><td>BREX-system-P-loop-protein-BrxC-[Vibrio-parahaemolyticus]</td></tr>
<tr><td>EJE4207362.1</td><td>Type1.BREX</td><td><-VirD4-TraG+VirB4-FtsK<-HDRnase<-SbcC<- SIGMA+OB-Fold+AAA-ATPase+Lon-Protease-BrxL <- PglZ <- N6-METHYLASE <- TOPRIM <- OLD-ABC <- DUF6079 *<- BrxB-BrxF-iSTAND <- BrxA-DNA-binding <- wHTH+WYL+WCX <-?<-35exo||LexA-protease->?-><-MobI</td><td>1225</td><td>Bacteria>Pseudomonadota>Gammaproteobacteria>Vibrionales>Vibrionaceae>Vibrio</td><td>GCA_023313315.1</td><td>Vibrio_parahaemolyticus</td><td>M3899_001773</td><td>267109..270786</td><td>670</td><td>BREX-system-P-loop-protein-BrxC-[Vibrio-parahaemolyticus]</td></tr>
<tr><td>HCE3362112.1</td><td>Type1.BREX</td><td><-VirD4-TraG+VirB4-FtsK<-HDRnase<-SbcC<- SIGMA+OB-Fold+AAA-ATPase+Lon-Protease-BrxL <- PglZ <- N6-METHYLASE <- TOPRIM <- OLD-ABC <- DUF6079 *<- BrxB-BrxF-iSTAND <- BrxA-DNA-binding <- wHTH+WYL+WCX <-?<-35exo<-?||LexA-protease->?->Glyoxalase_8-></td><td>1225</td><td>Bacteria>Pseudomonadota>Gammaproteobacteria>Vibrionales>Vibrionaceae>Vibrio</td><td>GCA_023821665.1</td><td>Vibrio_parahaemolyticus</td><td>NG589_000244</td><td>272459..276136</td><td>670</td><td>BREX-system-P-loop-protein-BrxC-[Vibrio-parahaemolyticus]</td></tr>
<tr><td>EJK2098984.1</td><td>Type1.BREX</td><td><-VirD4-TraG+VirB4-FtsK<-HDRnase<-SbcC<- SIGMA+OB-Fold+AAA-ATPase+Lon-Protease-BrxL <- PglZ <- N6-METHYLASE <- TOPRIM <- OLD-ABC <- DUF6079 *<- BrxB-BrxF-iSTAND <- BrxA-DNA-binding <- wHTH+WYL+WCX <-?<-HTH_23+HTH_32<-35exo||LexA-protease-></td><td>1225</td><td>Bacteria>Pseudomonadota>Gammaproteobacteria>Vibrionales>Vibrionaceae>Vibrio</td><td>GCA_024103575.1</td><td>Vibrio_cholerae</td><td>NK421_001793</td><td>214069..217746</td><td>666</td><td>BREX-system-P-loop-protein-BrxC-[Vibrio-cholerae]</td></tr>
<tr><td>EJL6578344.1</td><td>Type1.BREX</td><td>Arm-DNA-bind_3+Phage_integrase->SF1-SF2-Helicase+REase->SF1-SF2-Helicase+REase-><- PglZ <- TOPRIM <- OLD-ABC <- N6-METHYLASE <- DUF6079 *<- BrxB-BrxF-iSTAND <- BrxA-DNA-binding <- wHTH+WYL+WCX <-XIS-HTH||?->?->DUF2787->GTPase-AIG-></td><td>1154</td><td>Bacteria>Pseudomonadota>Gammaproteobacteria>Vibrionales>Vibrionaceae>Vibrio</td><td>GCA_024217525.1</td><td>Vibrio_cholerae</td><td>NMS50_002630</td><td>71340..74804</td><td>666</td><td>BREX-system-P-loop-protein-BrxC-[Vibrio-cholerae]</td></tr>
<tr><td>HCM0920431.1</td><td>Type1.BREX</td><td><- wHTH+WYL+WCX <-HTH||<-DUF2787<-?<-?||XIS-HTH-> wHTH+WYL+WCX -> BrxA-DNA-binding -> BrxB-BrxF-iSTAND -> DUF6079 *-> N6-METHYLASE -> OLD-ABC -> TOPRIM -> PglZ -><-SF1-SF2-Helicase+REase<-SF1-SF2-Helicase+REase</td><td>1154</td><td>Bacteria>Pseudomonadota>Gammaproteobacteria>Vibrionales>Vibrionaceae>Vibrio</td><td>GCA_025068375.1</td><td>Vibrio_parahaemolyticus</td><td>N2828_003369</td><td>93414..96878</td><td>670</td><td>BREX-system-P-loop-protein-BrxC-[Vibrio-parahaemolyticus]</td></tr>
<tr><td>WP_017044128.1</td><td>Type1.BREX</td><td>HTH_19-><-ParE<-MobI<-MobI<- SIGMA+OB-Fold+AAA-ATPase+Lon-Protease-BrxL <- PglZ <- N6-METHYLASE <- TOPRIM <- OLD-ABC <- DUF6079 *<- BrxB-BrxF-iSTAND <- BrxA-DNA-binding <- wHTH+WYL+WCX ||XIS-HTH->AAA+HTH->XIS-HTH-><-Phage_int_SAM_5+Phage_integrase<-sGTP</td><td>1225</td><td>Bacteria>Pseudomonadota>Gammaproteobacteria>Vibrionales>Vibrionaceae>Vibrio</td><td>GCF_000287075.2</td><td>Vibrio_ordalii</td><td>A1QQ_08605</td><td>37533..41210</td><td>28174</td><td>BREX-system-P-loop-protein-BrxC-[Vibrio-ordalii]</td></tr>
<tr><td>WP_032482144.1</td><td>Type1.BREX</td><td><-LexA-protease||35exo->?-> wHTH+WYL+WCX -> BrxA-DNA-binding -> BrxB-BrxF-iSTAND -> DUF6079 *->ParB+HNH-> N6-METHYLASE -> OLD-ABC -> TOPRIM -> PglZ -> SIGMA+OB-Fold+AAA-ATPase+Lon-Protease-BrxL ->SbcC->HDRnase->VirD4-TraG+VirB4-FtsK-></td><td>1225</td><td>Bacteria>Pseudomonadota>Gammaproteobacteria>Vibrionales>Vibrionaceae>Vibrio</td><td>GCF_000736875.1</td><td>Vibrio_cholerae</td><td>DN38_RS05545</td><td>8784..12461</td><td>666</td><td>BREX-system-P-loop-protein-BrxC-[Vibrio-cholerae]</td></tr>
<tr><td>WP_253277773.1</td><td>Type1.BREX</td><td><- wHTH+WYL+WCX <-DUF2787<-?<-?|| wHTH+WYL+WCX -> BrxA-DNA-binding -> BrxB-BrxF-iSTAND -> DUF6079 *-> N6-METHYLASE -> OLD-ABC -> TOPRIM -> PglZ -><-SF1-SF2-Helicase+REase<-SF1-SF2-Helicase+REase<-Arm-DNA-bind_3+Phage_integrase</td><td>1153</td><td>Bacteria>Pseudomonadota>Gammaproteobacteria>Vibrionales>Vibrionaceae>Vibrio</td><td>GCF_000959745.1</td><td>Vibrio_vulnificus</td><td>WG98_RS00450</td><td>91740..95201</td><td>672</td><td>BREX-system-P-loop-protein-BrxC-[Vibrio-vulnificus]</td></tr>
<tr><td>WP_241772680.1</td><td>Type1.BREX</td><td><-GDResolvase+HIN-HTH||?->Arm-DNA-bind_3+Phage_integrase->SF1-SF2-Helicase+REase->SF1-SF2-Helicase+REase-><- PglZ <- TOPRIM <- OLD-ABC <- N6-METHYLASE <- DUF6079 *<- BrxB-BrxF-iSTAND <- BrxA-DNA-binding <- wHTH+WYL+WCX <-XIS-HTH</td><td>1153</td><td>Bacteria>Pseudomonadota>Gammaproteobacteria>Vibrionales>Vibrionaceae>Vibrio</td><td>GCF_001280635.1</td><td>Vibrio_parahaemolyticus</td><td>ACX06_RS21960</td><td>167502..170963</td><td>670</td><td>BREX-system-P-loop-protein-BrxC-[Vibrio-parahaemolyticus]</td></tr>
<tr><td>WP_054541287.1</td><td>Type1.BREX</td><td>BPD_transp_2->?->?->SF1-SF2-Helicase+HTH->HNH->XIS-HTH-> BrxA-DNA-binding -> BrxB-BrxF-iSTAND -> DUF6079 *-> N6-METHYLASE -> OLD-ABC -> TOPRIM -> N6-METHYLASE -> PglZ -> SIGMA+OB-Fold+AAA-ATPase+Lon-Protease-BrxL -><-?||?-><-MODE-HTH+PBPII</td><td>1235</td><td>Bacteria>Pseudomonadota>Gammaproteobacteria>Vibrionales>Vibrionaceae>Vibrio</td><td>GCF_001306215.1</td><td>Vibrio_splendidus</td><td>AN167_RS02930</td><td>103766..107473</td><td>29497</td><td>BREX-system-P-loop-protein-BrxC-[Vibrio-splendidus]</td></tr>
<tr><td>WP_241782177.1</td><td>Type1.BREX</td><td><-HTH||RNase-><-DUF2787<-?<-?||XIS-HTH-> wHTH+WYL+WCX -> BrxA-DNA-binding -> BrxB-BrxF-iSTAND -> DUF6079 *-> N6-METHYLASE -> OLD-ABC -> TOPRIM -> PglZ -><-SF1-SF2-Helicase+REase<-SF1-SF2-Helicase+REase||?->SF1-SF2-Helicase-><-Mob_Pre</td><td>1153</td><td>Bacteria>Pseudomonadota>Gammaproteobacteria>Vibrionales>Vibrionaceae>Vibrio</td><td>GCF_001608855.1</td><td>Vibrio_parahaemolyticus</td><td>AWQ17_RS02375</td><td>93824..97285</td><td>670</td><td>BREX-system-P-loop-protein-BrxC-[Vibrio-parahaemolyticus]</td></tr>
<tr><td>WP_256093930.1</td><td>Type1.BREX</td><td>XIS-HTH-> wHTH+WYL+WCX -> BrxA-DNA-binding -> BrxB-BrxF-iSTAND -> DUF6079 *-> N6-METHYLASE -> OLD-ABC -> TOPRIM -> PglZ -><-SF1-SF2-Helicase+REase<-SF1-SF2-Helicase+REase<-Arm-DNA-bind_3+Phage_integrase</td><td>1153</td><td>Bacteria>Pseudomonadota>Gammaproteobacteria>Vibrionales>Vibrionaceae>Vibrio</td><td>GCF_001727265.1</td><td>Vibrio_parahaemolyticus</td><td>BBM25_RS06240</td><td>3427..6888</td><td>670</td><td>BREX-system-P-loop-protein-BrxC-[Vibrio-parahaemolyticus]</td></tr>
<tr><td>WP_139834421.1</td><td>Type1.BREX</td><td>HTH_19-><-ParE<-MobI<-MobI<- SIGMA+OB-Fold+AAA-ATPase+Lon-Protease-BrxL <- PglZ <- N6-METHYLASE <- TOPRIM <- OLD-ABC <- DUF6079 *</td><td>1085</td><td>Bacteria>Pseudomonadota>Gammaproteobacteria>Vibrionales>Vibrionaceae>Vibrio</td><td>GCF_002114605.1</td><td>Vibrio_parahaemolyticus</td><td>B5C31_RS21495</td><td>50871..54130</td><td>670</td><td>BREX-system-P-loop-protein-BrxC,-partial-[Vibrio-parahaemolyticus]</td></tr>
<tr><td>WP_102376359.1</td><td>Type1.BREX</td><td>ParB-><- SIGMA+OB-Fold+AAA-ATPase+Lon-Protease-BrxL <- PglZ <- TOPRIM <- OLD-ABC <-?<-?<- N6-METHYLASE <-ParB+HNH<- DUF6079 *<- BrxB-BrxF-iSTAND <- BrxA-DNA-binding <- wHTH+WYL+WCX <-?<-XIS-HTH<-Arm-DNA-bind_3+Phage_integrase</td><td>1233</td><td>Bacteria>Pseudomonadota>Gammaproteobacteria>Vibrionales>Vibrionaceae>Vibrio</td><td>GCF_002876115.1</td><td>Vibrio_breoganii</td><td>BCT61_RS03015</td><td>196035..199736</td><td>553239</td><td>BREX-system-P-loop-protein-BrxC-[Vibrio-breoganii]</td></tr>
<tr><td>WP_104969987.1</td><td>Type1.BREX</td><td>sGTP->Phage_int_SAM_5+Phage_integrase-><-XIS-HTH<-AAA+HTH<-XIS-HTH|| wHTH+WYL+WCX -> BrxA-DNA-binding -> BrxB-BrxF-iSTAND -> DUF6079 *-> OLD-ABC -> TOPRIM -> N6-METHYLASE -> PglZ -> SIGMA+OB-Fold+AAA-ATPase+Lon-Protease-BrxL ->MobI->MobI-></td><td>1225</td><td>Bacteria>Pseudomonadota>Gammaproteobacteria>Vibrionales>Vibrionaceae>Vibrio</td><td>GCF_002953335.1</td><td>Vibrio_diabolicus</td><td>AL552_RS16285</td><td>3102231..3105908</td><td>50719</td><td>BREX-system-P-loop-protein-BrxC-[Vibrio-diabolicus]</td></tr>
<tr><td>WP_108187618.1</td><td>Type1.BREX</td><td><- SIGMA+OB-Fold+AAA-ATPase+Lon-Protease-BrxL <- PglZ <- TOPRIM <- OLD-ABC <-?<-?<-?<- N6-METHYLASE <-ParB+HNH<- DUF6079 *<- BrxB-BrxF-iSTAND <- BrxA-DNA-binding <- wHTH+WYL+WCX ||SF1-SF2-Helicase->MarR-HTH->HKD-DNase+HKD-DNase->CARF+NtrC-AAA->PIN+OST-HTH->HIN-HTH-></td><td>1233</td><td>Bacteria>Pseudomonadota>Gammaproteobacteria>Vibrionales>Vibrionaceae>Vibrio</td><td>GCF_003049965.1</td><td>Vibrio_splendidus</td><td>CWD07_RS09240</td><td>50503..54204</td><td>29497</td><td>BREX-system-P-loop-protein-BrxC-[Vibrio-splendidus]</td></tr>
<tr><td>WP_235868160.1</td><td>Type1.BREX</td><td>Arm-DNA-bind_3+Phage_integrase->SF1-SF2-Helicase+REase->SF1-SF2-Helicase+REase-><- PglZ <- TOPRIM <- OLD-ABC <- N6-METHYLASE <- DUF6079 *<- BrxB-BrxF-iSTAND <- BrxA-DNA-binding <- wHTH+WYL+WCX <-XIS-HTH||?->?->DUF2787->HTH-> wHTH+WYL+WCX -></td><td>1153</td><td>Bacteria>Pseudomonadota>Gammaproteobacteria>Vibrionales>Vibrionaceae>Vibrio</td><td>GCF_003311905.1</td><td>Vibrio_paracholerae</td><td>DLR66_RS03590</td><td>70731..74192</td><td>650003</td><td>BREX-system-P-loop-protein-BrxC-[Vibrio-paracholerae]</td></tr>
<tr><td>WP_116869874.1</td><td>Type1.BREX</td><td>sGTP->Phage_int_SAM_5+Phage_integrase-><-XIS-HTH<-AAA+HTH<-XIS-HTH|| wHTH+WYL+WCX -> BrxA-DNA-binding -> BrxB-BrxF-iSTAND -> DUF6079 *-> OLD-ABC -> TOPRIM -> N6-METHYLASE -> PglZ -> SIGMA+OB-Fold+AAA-ATPase+Lon-Protease-BrxL ->MobI->MobI->ParE-><-HTH_19</td><td>1225</td><td>Bacteria>Pseudomonadota>Gammaproteobacteria>Vibrionales>Vibrionaceae>Vibrio</td><td>GCF_003408655.1</td><td>Vibrio_splendidus</td><td>D0583_RS01655</td><td>344855..348532</td><td>29497</td><td>BREX-system-P-loop-protein-BrxC-[Vibrio-splendidus]</td></tr>
<tr><td>WP_170894125.1</td><td>Type1.BREX</td><td>HTH_19-><-ParE<-MobI<-MobI<- SIGMA+OB-Fold+AAA-ATPase+Lon-Protease-BrxL <- PglZ <- N6-METHYLASE <- TOPRIM <- OLD-ABC <- DUF6079 *<- BrxB-BrxF-iSTAND <- BrxA-DNA-binding <- wHTH+WYL+WCX ||XIS-HTH->AAA+HTH->XIS-HTH-><-Phage_int_SAM_5+Phage_integrase<-sGTP</td><td>1225</td><td>Bacteria>Pseudomonadota>Gammaproteobacteria>Vibrionales>Vibrionaceae>Vibrio</td><td>GCF_013048535.1</td><td>unclassified_Vibrio</td><td>FKN13_RS12350</td><td>70010..73687</td><td>2614977</td><td>MULTISPECIES:-BREX-system-P-loop-protein-BrxC-[unclassified-Vibrio]</td></tr>
<tr><td>WP_191786376.1</td><td>Type1.BREX</td><td>HTH_19-><-ParE<-MobI<-MobI<- SIGMA+OB-Fold+AAA-ATPase+Lon-Protease-BrxL <- PglZ <- N6-METHYLASE <- TOPRIM <- OLD-ABC <- DUF6079 *<- BrxB-BrxF-iSTAND <- BrxA-DNA-binding <- wHTH+WYL+WCX ||XIS-HTH->AAA+HTH->XIS-HTH-><-Phage_int_SAM_5+Phage_integrase<-sGTP</td><td>1225</td><td>Bacteria>Pseudomonadota>Gammaproteobacteria>Vibrionales>Vibrionaceae>Vibrio</td><td>GCF_014836305.1</td><td>Vibrio_parahaemolyticus</td><td>G3220_RS19475</td><td>66422..70099</td><td>670</td><td>BREX-system-P-loop-protein-BrxC-[Vibrio-parahaemolyticus]</td></tr>
<tr><td>WP_210447442.1</td><td>Type1.BREX</td><td>XIS-HTH-><-DUF2787<-?<-?||XIS-HTH->?-> wHTH+WYL+WCX -> BrxA-DNA-binding -> BrxB-BrxF-iSTAND -> DUF6079 *-> N6-METHYLASE -> OLD-ABC -> TOPRIM -> PglZ ->SF1-SF2-Helicase+PRTase->SLOG+BirA-HTH->?->URI-DNase->sGTP-></td><td>1157</td><td>Bacteria>Pseudomonadota>Gammaproteobacteria>Vibrionales>Vibrionaceae>Vibrio</td><td>GCF_017917465.1</td><td>Vibrio_crassostreae</td><td>KB624_RS13660</td><td>42369..45842</td><td>246167</td><td>BREX-system-P-loop-protein-BrxC-[Vibrio-crassostreae]</td></tr>
<tr><td>WP_238812493.1</td><td>Type1.BREX</td><td><-CopG/RHH||<-?<-DUF2787<-?<-?|| wHTH+WYL+WCX -> BrxA-DNA-binding -> BrxB-BrxF-iSTAND -> DUF6079 *-> N6-METHYLASE -> OLD-ABC -> TOPRIM -> PglZ -></td><td>1153</td><td>Bacteria>Pseudomonadota>Gammaproteobacteria>Vibrionales>Vibrionaceae>Vibrio</td><td>GCF_022290755.1</td><td>Vibrio_parahaemolyticus</td><td>K6U44_RS14230</td><td>7820..11281</td><td>670</td><td>BREX-system-P-loop-protein-BrxC-[Vibrio-parahaemolyticus]</td></tr>
<tr><td>WP_261872478.1</td><td>Type1.BREX</td><td><-ABC-ATPase+SbcC<-?<-?||?-><-?||Peptidase_M32-> wHTH+WYL+WCX -> BrxA-DNA-binding -> BrxB-BrxF-iSTAND -> DUF6079 *-> N6-METHYLASE -> OLD-ABC -> TOPRIM -> PglZ -> SIGMA+OB-Fold+AAA-ATPase+Lon-Protease-BrxL -><-HIN-HTH</td><td>1239</td><td>Bacteria>Pseudomonadota>Gammaproteobacteria>Vibrionales>Vibrionaceae>Vibrio</td><td>GCF_024347075.1</td><td>Vibrio_rarus</td><td>OCU56_RS06785</td><td>1441704..1445423</td><td>413403</td><td>BREX-system-P-loop-protein-BrxC-[Vibrio-rarus]</td></tr>
<tr><td>WP_274056407.1</td><td>Type1.BREX</td><td>HTH_19-><-ParE<-MobI<-MobI<- SIGMA+OB-Fold+AAA-ATPase+Lon-Protease-BrxL <- PglZ <- N6-METHYLASE <- TOPRIM <- OLD-ABC <- DUF6079 *<- BrxB-BrxF-iSTAND <- BrxA-DNA-binding <- wHTH+WYL+WCX ||XIS-HTH->AAA+HTH->XIS-HTH-><-Phage_int_SAM_5+Phage_integrase<-sGTP</td><td>1225</td><td>Bacteria>Pseudomonadota>Gammaproteobacteria>Vibrionales>Vibrionaceae>Vibrio</td><td>GCF_028616275.1</td><td>Vibrio_parahaemolyticus</td><td>O8051_RS20180</td><td>64443..68120</td><td>670</td><td>BREX-system-P-loop-protein-BrxC-[Vibrio-parahaemolyticus]</td></tr>
<tr><td>WP_274695775.1</td><td>Type1.BREX</td><td>sGTP->Phage_int_SAM_5+Phage_integrase-><-XIS-HTH<-AAA+HTH<-XIS-HTH|| wHTH+WYL+WCX -> BrxA-DNA-binding -> BrxB-BrxF-iSTAND -> DUF6079 *-> OLD-ABC -> TOPRIM -> N6-METHYLASE -> PglZ -> SIGMA+OB-Fold+AAA-ATPase+Lon-Protease-BrxL ->MobI->MobI->ParE->ParE-></td><td>1225</td><td>Bacteria>Pseudomonadota>Gammaproteobacteria>Vibrionales>Vibrionaceae>Vibrio</td><td>GCF_028858205.1</td><td>Vibrio_aestuarianus</td><td>MEG05_RS14945</td><td>3088024..3091701</td><td>28171</td><td>BREX-system-P-loop-protein-BrxC-[Vibrio-aestuarianus]</td></tr>
<tr><td>WP_253254707.1</td><td>Type1.BREX</td><td><- wHTH+WYL+WCX <-?<-DUF2787<-?<-?|| wHTH+WYL+WCX -> BrxA-DNA-binding -> BrxB-BrxF-iSTAND -> DUF6079 *-> N6-METHYLASE -> OLD-ABC -> TOPRIM -> PglZ -><-SF1-SF2-Helicase+REase<-SF1-SF2-Helicase+REase<-Arm-DNA-bind_3+Phage_integrase</td><td>1153</td><td>Bacteria>Pseudomonadota>Gammaproteobacteria>Vibrionales>Vibrionaceae>Vibrio</td><td>GCF_030238805.1</td><td>Vibrio_parahaemolyticus</td><td>CSR03_RS22350</td><td>121618..125079</td><td>670</td><td>BREX-system-P-loop-protein-BrxC-[Vibrio-parahaemolyticus]</td></tr>
<tr><td>WP_241766700.1</td><td>Type1.BREX</td><td><-GDResolvase+HIN-HTH||?->Arm-DNA-bind_3+Phage_integrase->SF1-SF2-Helicase+REase->SF1-SF2-Helicase+REase-><- PglZ <- TOPRIM <- OLD-ABC <- N6-METHYLASE <- DUF6079 *<- BrxB-BrxF-iSTAND <- BrxA-DNA-binding <- wHTH+WYL+WCX <-XIS-HTH||?->?->?-><-XIS-HTH<-ANK+ANK</td><td>1153</td><td>Bacteria>Pseudomonadota>Gammaproteobacteria>Vibrionales>Vibrionaceae>Vibrio</td><td>GCF_948193335.1</td><td>Vibrio_parahaemolyticus</td><td>PO77_RS19450</td><td>167850..171311</td><td>670</td><td>BREX-system-P-loop-protein-BrxC-[Vibrio-parahaemolyticus]</td></tr>
<tr><td>AAL59733.1</td><td>Type1.BREX</td><td>DUF4158-><-Phage_integrase<-?<-LexA-protease||35exo->?-> wHTH+WYL+WCX -> BrxA-DNA-binding -> BrxB-BrxF-iSTAND -> DUF6079 -p*-> DUF6079 -p-> OLD-ABC -> OLD-ABC -> TOPRIM -> N6-METHYLASE -p-> N6-METHYLASE -p-> N6-METHYLASE -p->?-> PglZ -></td><td>631</td><td>Bacteria>Pseudomonadota>Gammaproteobacteria>Vibrionales>Vibrionaceae>Vibrio</td><td>-</td><td>Vibrio_cholerae</td><td>.</td><td>28378..30273</td><td>666</td><td>unknown-[Vibrio-cholerae]</td></tr>
<tr><td>WP_142738823.1</td><td>Type1.BREX</td><td>MobI-><-?<-LexA-protease||?-> wHTH+WYL+WCX -> BrxA-DNA-binding -> BrxB-BrxF-iSTAND -> DUF6079 *->ParB+HNH-> N6-METHYLASE -> OLD-ABC -> TOPRIM -> PglZ -> SIGMA+OB-Fold+AAA-ATPase+Lon-Protease-BrxL ->SbcC->HDRnase->VirD4-TraG+VirB4-FtsK-></td><td>1224</td><td>Bacteria>Pseudomonadota>Gammaproteobacteria>Vibrionales>Vibrionaceae>Vibrio</td><td>-</td><td>Vibrio_cholerae</td><td>FLL79_RS01410</td><td>36024..39698</td><td>666</td><td>BREX-system-P-loop-protein-BrxC-[Vibrio-cholerae]</td></tr>
<tr><td>WP_069490026.1</td><td>Type1.BREX</td><td><-HTH_Tnp_1||XIS-HTH-> wHTH+WYL+WCX -> BrxA-DNA-binding -> BrxB-BrxF-iSTAND -> DUF6079 *-> N6-METHYLASE -> OLD-ABC -> TOPRIM -> PglZ -><-SF1-SF2-Helicase+REase<-SF1-SF2-Helicase+REase<-Arm-DNA-bind_3+Phage_integrase</td><td>1154</td><td>Bacteria>Pseudomonadota>Gammaproteobacteria>Vibrionales>Vibrionaceae>Vibrio</td><td>-</td><td>Vibrio_parahaemolyticus</td><td>BBM25_06240</td><td>3424..6888</td><td>670</td><td>BREX-system-P-loop-protein-BrxC-[Vibrio-parahaemolyticus]</td></tr>
<tr><td>WP_254664663.1</td><td>Type1.BREX</td><td>XIS-HTH-> wHTH+WYL+WCX -> BrxA-DNA-binding -> BrxB-BrxF-iSTAND -> DUF6079 *-> N6-METHYLASE -> OLD-ABC -> TOPRIM -> PglZ -><-SF1-SF2-Helicase+REase<-SF1-SF2-Helicase+REase<-Arm-DNA-bind_3+Phage_integrase</td><td>1153</td><td>Bacteria>Pseudomonadota>Gammaproteobacteria>Vibrionales>Vibrionaceae>Vibrio</td><td>-</td><td>Vibrio_parahaemolyticus</td><td>BED52_RS16125</td><td>3457..6918</td><td>670</td><td>BREX-system-P-loop-protein-BrxC-[Vibrio-parahaemolyticus]</td></tr>
<tr><td>WP_047508417.1</td><td>Type1.BREX</td><td>XIS-HTH-> wHTH+WYL+WCX -> BrxA-DNA-binding -> BrxB-BrxF-iSTAND -> DUF6079 *-> N6-METHYLASE -> OLD-ABC -> TOPRIM -> PglZ -><-SF1-SF2-Helicase+REase<-SF1-SF2-Helicase+REase<-Arm-DNA-bind_3+Phage_integrase<-?||GDResolvase+HIN-HTH-></td><td>1154</td><td>Bacteria>Pseudomonadota>Gammaproteobacteria>Vibrionales>Vibrionaceae>Vibrio</td><td>-</td><td>Vibrio_parahaemolyticus</td><td>DMJ21_13130</td><td>8098..11562</td><td>670</td><td>BREX-system-P-loop-protein-BrxC-[Vibrio-parahaemolyticus]</td></tr>
<tr><td>WP_062854373.1</td><td>Type1.BREX</td><td><- wHTH+WYL+WCX <-HTH||<-DUF2787<-?<-?||XIS-HTH-> wHTH+WYL+WCX -> BrxA-DNA-binding -> BrxB-BrxF-iSTAND -> DUF6079 *-> N6-METHYLASE -> OLD-ABC -> TOPRIM -> PglZ -><-SF1-SF2-Helicase+REase<-SF1-SF2-Helicase+REase</td><td>1154</td><td>Bacteria>Pseudomonadota>Gammaproteobacteria>Vibrionales>Vibrionaceae>Vibrio</td><td>-</td><td>Vibrio_parahaemolyticus</td><td>LZX80_001782</td><td>93413..96877</td><td>670</td><td>BREX-system-P-loop-protein-BrxC-[Vibrio-parahaemolyticus]</td></tr>
<tr><td>WP_000687849.1</td><td>Type1.BREX</td><td><-LexA-protease||?->35exo->?-> wHTH+WYL+WCX -> BrxA-DNA-binding -> BrxB-BrxF-iSTAND -> DUF6079 *-> OLD-ABC -> TOPRIM -> N6-METHYLASE -> PglZ -> SIGMA+OB-Fold+AAA-ATPase+Lon-Protease-BrxL ->SbcC->HDRnase->VirD4-TraG+VirB4-FtsK-></td><td>1225</td><td>Bacteria>Pseudomonadota</td><td>GCF_000299535.2</td><td>Gammaproteobacteria</td><td>B934_RS0100870</td><td>5534..9211</td><td>1236</td><td>MULTISPECIES:-BREX-system-P-loop-protein-BrxC-[Gammaproteobacteria]</td></tr>
<tr><td>MBK8482432.1</td><td>Type1.BREX</td><td><- SIGMA+OB-Fold+AAA-ATPase+Lon-Protease-BrxL <- PglZ <-?<- N6-METHYLASE -p<-DOC+HTH<- N6-METHYLASE -p||?-><-SF1-SF2-Helicase+REase-p<- OLD-ABC + TOPRIM <- DUF6079 -p*<- DUF6079 -p<- BrxB-BrxF-iSTAND <- BrxA-DNA-binding <-XIS-HTH<-?<-GDResolvase+ZnR<-GDResolvase<-RAMA</td><td>896</td><td>Bacteria>Pseudomonadota>unclassified_Pseudomonadota</td><td>GCA_016711215.1</td><td>Pseudomonadota_bacterium</td><td>IPL40_14915</td><td>141091..143781</td><td>1977087</td><td>hypothetical-protein-[Pseudomonadota-bacterium]</td></tr>
<tr><td>NPU98919.1</td><td>Type6.BREX</td><td>CHTH-> BrxE -> BrxA-DNA-binding -> BrxB-BrxF-iSTAND -> DUF6079 *->?->?-> OLD-ABC -> TOPRIM -> PglZ -> SIGMA+OB-Fold+AAA-ATPase+Lon-Protease-BrxL ->DNA_METHYLASE->SF1-SF2-Helicase+REase->REase-></td><td>1224</td><td>Bacteria>PVC_group>Candidatus_Omnitrophica>unclassified_Candidatus_Omnitrophica</td><td>GCA_013177805.1</td><td>Candidatus_Omnitrophica_bacterium</td><td>HPY51_17110</td><td>7762..11436</td><td>2035772</td><td>BREX-system-P-loop-protein-BrxC-[Candidatus-Omnitrophota-bacterium]</td></tr>
<tr><td>MCF7790541.1</td><td>Type1.BREX</td><td><-HEPN<- SIGMA+OB-Fold+AAA-ATPase+HKD-DNase-BrxL <- PglZ <- TOPRIM <- OLD-ABC <-?<-?<-?<- DUF6079 *<- BrxB-BrxF-iSTAND <- BrxA-DNA-binding <- wHTH+WYL+WCX <-?||?-><-?<-?||S2P+PDZ-></td><td>1176</td><td>Bacteria>PVC_group>Lentisphaerota>Lentisphaeria>Victivallales>unclassified_Victivallales</td><td>GCA_021733985.1</td><td>Victivallales_bacterium</td><td>K9M56_00960</td><td>14578..18108</td><td>2212475</td><td>BREX-system-P-loop-protein-BrxC-[Victivallales-bacterium]</td></tr>
<tr><td>MBP7937491.1</td><td>Type1.BREX</td><td>Peptidase->Phage_int_SAM_1+Phage_integrase-><- SIGMA+OB-Fold+AAA-ATPase+Lon-Protease-BrxL <- PglZ <- TOPRIM <- OLD-ABC <- N6-METHYLASE <- DUF6079 *<- BrxB-BrxF-iSTAND <- BrxA-DNA-binding ||Polbeta-></td><td>1149</td><td>Bacteria>PVC_group>Planctomycetota>Phycisphaerae>unclassified_Phycisphaerae</td><td>GCA_018003015.1</td><td>Phycisphaerae_bacterium</td><td>KA354_22855</td><td>13009..16458</td><td>2026778</td><td>BREX-system-P-loop-protein-BrxC-[Phycisphaerae-bacterium]</td></tr>
<tr><td>WP_145167965.1</td><td>Type6.BREX</td><td><-Tox-RES||HIN-HTH->ParA-Soj-PloopNTPase-><- SIGMA+OB-Fold+AAA-ATPase+Lon-Protease-BrxL <- PglZ <- N6-METHYLASE <- TOPRIM <- OLD-ABC <- DUF6079 *<- BrxB-BrxF-iSTAND <- BrxA-DNA-binding <- BrxE <-XIS-HTH<-ASCH||?-><-DEOR-HTH<-D5N+D5ATPase</td><td>1219</td><td>Bacteria>PVC_group>Planctomycetota>Planctomycetia>Pirellulales>Pirellulaceae</td><td>GCF_007741535.1</td><td>Rubripirellula_lacrimiformis</td><td>K227x_RS02725</td><td>780099..783758</td><td>1930273</td><td>BREX-system-P-loop-protein-BrxC-[Rubripirellula-lacrimiformis]</td></tr>
<tr><td>HCS52258.1</td><td>Type1.BREX</td><td><- SIGMA+OB-Fold+AAA-ATPase+Lon-Protease-BrxL <- PglZ <- TOPRIM <-SbcC<- N6-METHYLASE <- N6-METHYLASE -p<- DUF6079 -p*<- DUF6079 -p<- BrxB-BrxF-iSTAND <- BrxA-DNA-binding <-XIS-HTH||DDE_Tnp_IS66->DDE_Tnp_1_assoc+DDE_Tnp_1->?-><-ParB</td><td>340</td><td>Bacteria>PVC_group>Planctomycetota>Planctomycetia>Planctomycetales>Planctomycetaceae</td><td>GCA_003531135.1</td><td>Planctomycetaceae_bacterium</td><td>DIW81_11810</td><td>12228..13250</td><td>2026779</td><td>BREX-system-P-loop-protein-BrxC-[Planctomycetaceae-bacterium]</td></tr>
<tr><td>HCS52259.1</td><td>Type1.BREX</td><td><- SIGMA+OB-Fold+AAA-ATPase+Lon-Protease-BrxL <- PglZ <- TOPRIM <-SbcC<- N6-METHYLASE <- N6-METHYLASE -p<- DUF6079 -p<- DUF6079 -p*<- BrxB-BrxF-iSTAND <- BrxA-DNA-binding <-XIS-HTH||DDE_Tnp_IS66->DDE_Tnp_1_assoc+DDE_Tnp_1->?-><-ParB<-ParB</td><td>393</td><td>Bacteria>PVC_group>Planctomycetota>Planctomycetia>Planctomycetales>Planctomycetaceae</td><td>GCA_003531135.1</td><td>Planctomycetaceae_bacterium</td><td>DIW81_11815</td><td>13400..14582</td><td>2026779</td><td>BREX-system-P-loop-protein-BrxC-[Planctomycetaceae-bacterium]</td></tr>
<tr><td>HCS52260.1</td><td>Type1.BREX</td><td><- SIGMA+OB-Fold+AAA-ATPase+Lon-Protease-BrxL <- PglZ <- TOPRIM <-SbcC<- N6-METHYLASE <- N6-METHYLASE -p<- DUF6079 -p<- DUF6079 -p*<- BrxB-BrxF-iSTAND <- BrxA-DNA-binding <-XIS-HTH||DDE_Tnp_IS66->DDE_Tnp_1_assoc+DDE_Tnp_1->?-><-ParB<-ParB<-GDResolvase+PPP1-HTH</td><td>395</td><td>Bacteria>PVC_group>Planctomycetota>Planctomycetia>Planctomycetales>Planctomycetaceae</td><td>GCA_003531135.1</td><td>Planctomycetaceae_bacterium</td><td>DIW81_11820</td><td>14698..15884</td><td>2026779</td><td>BREX-system-P-loop-protein-BrxC-[Planctomycetaceae-bacterium]</td></tr>
<tr><td>MBA4188926.1</td><td>Type1.BREX</td><td><-Sulfatase+DUF4994||GDResolvase+HIN-HTH-><-HTH_3||?->?-><-?|| BrxA-DNA-binding -> BrxB-BrxF-iSTAND -> DUF6079 *-> TOPRIM -> PglZ -> SIGMA+OB-Fold+AAA-ATPase+Lon-Protease-BrxL -><-?<-?<-GDResolvase<-Sulfatase||PAS+NtrC-AAA+FIS-HTH-></td><td>1142</td><td>Bacteria>PVC_group>Planctomycetota>Planctomycetia>Planctomycetales>Planctomycetaceae</td><td>GCA_013823135.1</td><td>Planctomycetaceae_bacterium</td><td>C0467_13090</td><td>97209..100637</td><td>2026779</td><td>BREX-system-P-loop-protein-BrxC-[Planctomycetaceae-bacterium]</td></tr>
<tr><td>WP_013111330.1</td><td>Type6.BREX</td><td><-DUF4096+DDE_Tnp_1+DDE_Tnp_1_2<-?<-?<-DUF4096+DDE_Tnp_1+DDE_Tnp_1_2<- SF1-SF2-Helicase+wHTH+Small-B-barrel+DUF1998 <- BrxD-AAA <- PglZ <- N6-METHYLASE <- DUF6079 *<- BrxB-BrxF-iSTAND <- BrxA-DNA-binding <- BrxE <- TOPRIM -DNAG+T7DNApol<-AAA+HTH<-?<-?||?->XIS-HTH-></td><td>1208</td><td>Bacteria>PVC_group>Planctomycetota>Planctomycetia>Planctomycetales>Planctomycetaceae</td><td>GCF_000092105.1</td><td>Planctopirus_limnophila</td><td>PLIM_RS15820</td><td>3989938..3993564</td><td>120</td><td>BREX-system-P-loop-protein-BrxC-[Planctopirus-limnophila]</td></tr>
<tr><td>HBI47071.1</td><td>Type1.BREX</td><td><-HTH_3<-?<- TOPRIM +Primase<-?<- TOPRIM <-XIS-HTH<-?|| BrxA-DNA-binding -> BrxB-BrxF-iSTAND -> DUF6079 *->?-> N6-METHYLASE -> OLD-ABC -> TOPRIM -> PglZ -> SIGMA+OB-Fold+AAA-ATPase+Lon-Protease-BrxL -><-?<-HTH_Tnp</td><td>1147</td><td>Bacteria>PVC_group>Planctomycetota>Planctomycetia>Planctomycetales>unclassified_Planctomycetales</td><td>GCA_003508275.1</td><td>Planctomycetales_bacterium</td><td>DDY78_30090</td><td>20346..23789</td><td>2053591</td><td>BREX-system-P-loop-protein-BrxC-[Planctomycetales-bacterium]</td></tr>
<tr><td>MCA9126638.1</td><td>Type1.BREX</td><td>SKI->?-><-?<-?<-SMODS|| BrxA-DNA-binding -> BrxB-BrxF-iSTAND -> DUF6079 *->Schlafen+HATPase+wHTH-> N6-METHYLASE ->?->DOC+HTH-> OLD-ABC -> TOPRIM -> PglZ -> SIGMA+OB-Fold+AAA-ATPase+Lon-Protease-BrxL ->ANK+ANK+ANK-></td><td>1149</td><td>Bacteria>PVC_group>Planctomycetota>Planctomycetia>Planctomycetales>unclassified_Planctomycetales</td><td>GCA_020426625.1</td><td>Planctomycetales_bacterium</td><td>KDB22_06125</td><td>7854..11303</td><td>2053591</td><td>BREX-system-P-loop-protein-BrxC-[Planctomycetales-bacterium]</td></tr>
<tr><td>OQA03646.1</td><td>Type1.BREX</td><td>HTH_7+rve+rve_3-><-?||?-> BrxA-DNA-binding -> BrxB-BrxF-iSTAND -> DUF6079 *-> N6-METHYLASE -> OLD-ABC -> TOPRIM -> PglZ -> SIGMA+OB-Fold+AAA-ATPase+Lon-Protease-BrxL -></td><td>1145</td><td>Bacteria>PVC_group>Planctomycetota>unclassified_Planctomycetota</td><td>GCA_002068635.1</td><td>Planctomycetes_bacterium_ADurb.Bin401</td><td>BWY69_00331</td><td>3976..7413</td><td>1852904</td><td>hypothetical-protein-BWY69_00331-[Planctomycetes-bacterium-ADurb.Bin401]</td></tr>
<tr><td>RIK67118.1</td><td>Type1.BREX</td><td><-XIS-HTH<-MPTase<-?<-?<-HTH_3||ACET->?<-?|| BrxA-DNA-binding -> BrxB-BrxF-iSTAND -> DUF6079 *-> N6-METHYLASE -> OLD-ABC -> TOPRIM -> PglZ -> SIGMA+OB-Fold+AAA-ATPase+Lon-Protease-BrxL -><-Phage_int_SAM_1+Phage_integrase||?-><-SIGMA-HTH</td><td>1149</td><td>Bacteria>PVC_group>Planctomycetota>unclassified_Planctomycetota</td><td>GCA_003576905.1</td><td>Planctomycetota_bacterium</td><td>DCC65_07820</td><td>141257..144706</td><td>2026780</td><td>MAG:-BREX-system-P-loop-protein-BrxC-[Planctomycetota-bacterium]</td></tr>
<tr><td>MBI4586093.1</td><td>Type1.BREX</td><td>Peptidase_M28->PALP-><-Pkinase+CR-Kinase||?-><- SIGMA+OB-Fold+AAA-ATPase+HKD-DNase-BrxL <- PglZ <-Schlafen+HATPase+wHTH+HTH<- N6-METHYLASE <- DUF6079 *<- BrxB-BrxF-iSTAND <- BrxA-DNA-binding || OLD-ABC -> TOPRIM ->TetR-HTH->?->ABC-ATPase+SbcC-></td><td>1226</td><td>Bacteria>PVC_group>Planctomycetota>unclassified_Planctomycetota</td><td>GCA_016209605.1</td><td>Planctomycetota_bacterium</td><td>HY717_18950</td><td>18652..22332</td><td>2026780</td><td>BREX-system-P-loop-protein-BrxC-[Planctomycetota-bacterium]</td></tr>
<tr><td>MBW3538721.1</td><td>Type1.BREX</td><td><- SIGMA+OB-Fold+AAA-ATPase+Lon-Protease-BrxL <- PglZ <- TOPRIM <- OLD-ABC <-?<- N6-METHYLASE <- N6-METHYLASE -p||?-><- DUF6079 *</td><td>1230</td><td>Bacteria>PVC_group>Planctomycetota>unclassified_Planctomycetota</td><td>GCA_019347905.1</td><td>Planctomycetota_bacterium</td><td>KY476_00475</td><td>36012..39705</td><td>2026780</td><td>BREX-system-P-loop-protein-BrxC-[Planctomycetota-bacterium]</td></tr>
<tr><td>AHF94674.1</td><td>Type1.BREX</td><td><- PglZ <- TOPRIM <- OLD-ABC <-PIWI<-REase+PNTD1+PNTD2<- N6-METHYLASE -p<-DOC+HTH<- N6-METHYLASE <-?<- DUF6079 *<- BrxB-BrxF-iSTAND <- BrxA-DNA-binding <-?||TrwC+SF1-SF2-Helicase->?->?->ClpABC-AAA-></td><td>1248</td><td>Bacteria>PVC_group>Verrucomicrobiota>Opitutae>Opitutales>Opitutaceae</td><td>GCA_000242935.3</td><td>Opitutaceae_bacterium_TAV5</td><td>OPIT5_23275</td><td>5485357..5489103</td><td>794903</td><td>hypothetical-protein-OPIT5_23275-[Opitutaceae-bacterium-TAV5]</td></tr>
<tr><td>MBI4624648.1</td><td>Type1.BREX</td><td><-PadR-HTH||?->?->?->HTH_Tnp-> BrxA-DNA-binding -> BrxB-BrxF-iSTAND -> DUF6079 *-> OLD-ABC -> TOPRIM -></td><td>1155</td><td>Bacteria>PVC_group>Verrucomicrobiota>unclassified_Verrucomicrobiota</td><td>GCA_016209255.1</td><td>Verrucomicrobiota_bacterium</td><td>HY736_15695</td><td>9824..13291</td><td>2026799</td><td>BREX-system-P-loop-protein-BrxC-[Verrucomicrobiota-bacterium]</td></tr>
<tr><td>MBM3880197.1</td><td>Type1.BREX</td><td>XIS-HTH-> BrxA-DNA-binding -> BrxB-BrxF-iSTAND -> DUF6079 *-> N6-METHYLASE ->SbcC->PIN->?->DOC+HTH-> N6-METHYLASE -p-> OLD-ABC -> TOPRIM -> PglZ -></td><td>1246</td><td>Bacteria>PVC_group>Verrucomicrobiota>unclassified_Verrucomicrobiota</td><td>GCA_016871715.1</td><td>Verrucomicrobiota_bacterium</td><td>FJ387_10840</td><td>14385..18125</td><td>2026799</td><td>BREX-system-P-loop-protein-BrxC-[Verrucomicrobiota-bacterium]</td></tr>
<tr><td>MBE7502027.1</td><td>Type6.BREX</td><td>HTH_3-><-?<-?<-?<-GNAT<-NERD+ZnR|| BrxE -> BrxA-DNA-binding -> BrxB-BrxF-iSTAND -> DUF6079 *-> N6-METHYLASE -><-DDE_Tnp_1|| N6-METHYLASE ->ParB+HNH-> OLD-ABC -> TOPRIM ->DDE_Tnp_1-> PglZ -> SIGMA+OB-Fold+AAA-ATPase+HKD-DNase-BrxL -></td><td>1231</td><td>Bacteria>PVC_group>Verrucomicrobiota>Verrucomicrobiae>Verrucomicrobiales>unclassified_Verrucomicrobiales</td><td>GCA_015075445.1</td><td>Verrucomicrobiales_bacterium</td><td>HS113_17385</td><td>108161..111856</td><td>2026801</td><td>BREX-system-P-loop-protein-BrxC-[Verrucomicrobiales-bacterium]</td></tr>
<tr><td>MCI0536692.1</td><td>Type1.BREX</td><td>TIR-><-?|| BrxA-DNA-binding -> BrxB-BrxF-iSTAND -> DUF6079 *-> OLD-ABC -> TOPRIM -> N6-METHYLASE -p-></td><td>1149</td><td>Bacteria>PVC_group>Verrucomicrobiota>Verrucomicrobiae>Verrucomicrobiales>unclassified_Verrucomicrobiales</td><td>GCA_022615925.1</td><td>Verrucomicrobiales_bacterium</td><td>LOZ50_15855</td><td>3424..6873</td><td>2026801</td><td>BREX-system-P-loop-protein-BrxC-[Verrucomicrobiales-bacterium]</td></tr>
<tr><td>OQY41116.1</td><td>Type1.BREX</td><td>REase-> BrxA-DNA-binding -> BrxB-BrxF-iSTAND -> DUF6079 *-> OLD-ABC -> TOPRIM -> N6-METHYLASE ->ParB+HNH->ParB+HNH-> SIGMA+OB-Fold+AAA-ATPase+Lon-Protease-BrxL -></td><td>1207</td><td>Bacteria>Spirochaetota>Spirochaetia>Spirochaetales>Spirochaetaceae>unclassified_Spirochaetaceae</td><td>GCA_002084785.1</td><td>Spirochaetaceae_bacterium_4572_7</td><td>B6229_00020</td><td>2034..5657</td><td>1972456</td><td>hypothetical-protein-B6229_00020-[Spirochaetaceae-bacterium-4572_7]</td></tr>
<tr><td>AOS47539.1</td><td>Type1.BREX</td><td><-LACI-HTH+PBPI<-?<-?||?->DOC+HTH-> BrxA-DNA-binding -> BrxB-BrxF-iSTAND -> DUF6079 *-> N6-METHYLASE -> OLD-ABC -> TOPRIM -> PglZ -> SIGMA+OB-Fold+AAA-ATPase+HKD-DNase-BrxL -><-ABC-3<-ABC-ATPase+SbcC</td><td>1176</td><td>Bacteria>Terrabacteria_group>Actinomycetota>Actinomycetes>Actinomycetales>Actinomycetaceae</td><td>GCA_001746855.1</td><td>Pauljensenia_hongkongensis</td><td>BH719_06520</td><td>1584234..1587764</td><td>178339</td><td>hypothetical-protein-BH719_06520-[Pauljensenia-hongkongensis]</td></tr>
<tr><td>OFQ22233.1</td><td>Type1.BREX</td><td><-ABC-ATPase+SbcC+ABC-ATPase+SbcC<-?<- SIGMA+OB-Fold+AAA-ATPase+HKD-DNase-BrxL <- PglZ <- TOPRIM <- OLD-ABC <- N6-METHYLASE <- DUF6079 *<- BrxB-BrxF-iSTAND <- BrxA-DNA-binding</td><td>1173</td><td>Bacteria>Terrabacteria_group>Actinomycetota>Actinomycetes>Actinomycetales>Actinomycetaceae</td><td>GCA_001813985.1</td><td>Actinomyces_sp._HMSC062G12</td><td>HMPREF2946_04765</td><td>260177..263698</td><td>1739450</td><td>hypothetical-protein-HMPREF2946_04765-[Actinomyces-sp.-HMSC062G12]</td></tr>
<tr><td>MBF0967807.1</td><td>Type1.BREX</td><td>BrxB-BrxF-iSTAND -> DUF6079 *-> N6-METHYLASE -> OLD-ABC -> TOPRIM -> PglZ -> SIGMA+OB-Fold+AAA-ATPase+HKD-DNase-BrxL -><-?||?->ABC-ATPase+SbcC+ABC-ATPase+SbcC-></td><td>1169</td><td>Bacteria>Terrabacteria_group>Actinomycetota>Actinomycetes>Actinomycetales>Actinomycetaceae</td><td>GCA_015259445.1</td><td>Actinomyces_sp.</td><td>HXK08_00790</td><td>718..4227</td><td>29317</td><td>BREX-system-P-loop-protein-BrxC-[Actinomyces-sp.]</td></tr>
<tr><td>MBS6968676.1</td><td>Type1.BREX</td><td>DEOR-HTH+ISOCOT->?-><- SIGMA+OB-Fold+AAA-ATPase+HKD-DNase-BrxL <- PglZ <- TOPRIM <- OLD-ABC <- N6-METHYLASE <- DUF6079 *<- BrxB-BrxF-iSTAND <- BrxA-DNA-binding</td><td>1169</td><td>Bacteria>Terrabacteria_group>Actinomycetota>Actinomycetes>Actinomycetales>Actinomycetaceae</td><td>GCA_018381115.1</td><td>Actinomyces_sp.</td><td>KH140_06985</td><td>13628..17137</td><td>29317</td><td>BREX-system-P-loop-protein-BrxC-[Actinomyces-sp.]</td></tr>
<tr><td>WP_173647185.1</td><td>Type1.BREX</td><td><-LACI-HTH+PBPI<-?<-?||?->DOC+HTH-> BrxA-DNA-binding -> BrxB-BrxF-iSTAND -> DUF6079 *-> N6-METHYLASE -> OLD-ABC -> TOPRIM -> PglZ -> SIGMA+OB-Fold+AAA-ATPase+HKD-DNase-BrxL ->?-><-ABC-3<-ABC-ATPase+SbcC</td><td>1174</td><td>Bacteria>Terrabacteria_group>Actinomycetota>Actinomycetes>Actinomycetales>Actinomycetaceae</td><td>GCF_001746855.1</td><td>Pauljensenia_hongkongensis</td><td>BH719_RS06520</td><td>1584240..1587764</td><td>178339</td><td>BREX-system-P-loop-protein-BrxC-[Pauljensenia-hongkongensis]</td></tr>
<tr><td>PJM72971.1</td><td>Type1.BREX</td><td>MACRODOMAIN-><-?<- SIGMA+OB-Fold+AAA-ATPase+HKD-DNase-BrxL <- PglZ <-?|| OLD-ABC -> TOPRIM -><- N6-METHYLASE <- DUF6079 *<- BrxB-BrxF-iSTAND <- BrxA-DNA-binding ||?->?-><-?<-?<-JetB</td><td>1243</td><td>Bacteria>Terrabacteria_group>Actinomycetota>Actinomycetes>Bifidobacteriales>Bifidobacteriaceae</td><td>GCA_002802875.1</td><td>Bifidobacterium_primatium</td><td>CS006_06875</td><td>170776..174507</td><td>2045438</td><td>hypothetical-protein-CS006_06875-[Bifidobacterium-primatium]</td></tr>
<tr><td>WP_033509374.1</td><td>Type1.BREX</td><td><- SIGMA+OB-Fold+AAA-ATPase+HKD-DNase-BrxL <- PglZ <-Schlafen+HATPase+wHTH<- N6-METHYLASE -p<-HerA/FtsK<-SIR2<- N6-METHYLASE <- DUF6079 *<- BrxB-BrxF-iSTAND <- BrxA-DNA-binding <- TOPRIM <- OLD-ABC ||Glycoside_Hydrolase-><-?<-Thioredoxin_3<-?<-ACET+ACET</td><td>1221</td><td>Bacteria>Terrabacteria_group>Actinomycetota>Actinomycetes>Bifidobacteriales>Bifidobacteriaceae</td><td>GCF_000770965.1</td><td>Bifidobacterium_pullorum</td><td>OU28_RS04650</td><td>22611..26276</td><td>78448</td><td>BREX-system-P-loop-protein-BrxC-[Bifidobacterium-pullorum]</td></tr>
<tr><td>WP_072727063.1</td><td>Type1.BREX</td><td><- OLD-ABC + TOPRIM <- N6-METHYLASE <- N6-METHYLASE -p<- DUF6079 *<- BrxB-BrxF-iSTAND <- BrxA-DNA-binding ||?->NTP_transf->?-><-CR-ATPase5+REase</td><td>1208</td><td>Bacteria>Terrabacteria_group>Actinomycetota>Actinomycetes>Bifidobacteriales>Bifidobacteriaceae</td><td>GCF_001895165.1</td><td>Bifidobacterium_lemurum</td><td>BRTL_RS11125</td><td>5981..9607</td><td>1603886</td><td>BREX-system-P-loop-protein-BrxC-[Bifidobacterium-lemurum]</td></tr>
<tr><td>WP_100511074.1</td><td>Type1.BREX</td><td>MACRODOMAIN-><-?<- SIGMA+OB-Fold+AAA-ATPase+HKD-DNase-BrxL <- PglZ <-?|| OLD-ABC -> TOPRIM -><- N6-METHYLASE <- DUF6079 *<- BrxB-BrxF-iSTAND <- BrxA-DNA-binding ||?->?-><-?<-?<-JetB</td><td>1251</td><td>Bacteria>Terrabacteria_group>Actinomycetota>Actinomycetes>Bifidobacteriales>Bifidobacteriaceae</td><td>GCF_002802875.1</td><td>Bifidobacterium_primatium</td><td>CS006_RS06855</td><td>170776..174531</td><td>2045438</td><td>BREX-system-P-loop-protein-BrxC-[Bifidobacterium-primatium]</td></tr>
<tr><td>WP_115955260.1</td><td>Type1.BREX</td><td><- OLD-ABC + TOPRIM <-ZnR<-ZnR+GP<- SIGMA+OB-Fold+AAA-ATPase+HKD-DNase-BrxL <- PglZ <-Schlafen+HATPase+wHTH<-?<-HEPN<- N6-METHYLASE <- DUF6079 *<- BrxB-BrxF-iSTAND <- BrxA-DNA-binding <-?<-?<-?<-ABC-ATPase+SbcC<-?<-ZnR</td><td>1217</td><td>Bacteria>Terrabacteria_group>Actinomycetota>Actinomycetes>Bifidobacteriales>Bifidobacteriaceae</td><td>GCF_003390735.1</td><td>Bifidobacterium_bifidum</td><td>CJD48_RS06210</td><td>1417204..1420857</td><td>1681</td><td>BREX-system-P-loop-protein-BrxC-[Bifidobacterium-bifidum]</td></tr>
<tr><td>WP_204389272.1</td><td>Type1.BREX</td><td><-IstB_IS21||AAA-ATPase-><-CMP-hydrolase<- SIGMA+OB-Fold+AAA-ATPase+HKD-DNase-BrxL <- PglZ <-Schlafen+HATPase+wHTH<- N6-METHYLASE <- DUF6079 *<- BrxB-BrxF-iSTAND <- BrxA-DNA-binding <- TOPRIM <- OLD-ABC ||Glycoside_Hydrolase-><-ACET+ACET<-ACET</td><td>1221</td><td>Bacteria>Terrabacteria_group>Actinomycetota>Actinomycetes>Bifidobacteriales>Bifidobacteriaceae</td><td>GCF_016900615.1</td><td>Bifidobacterium_pullorum</td><td>H6A14_RS02700</td><td>186110..189775</td><td>78448</td><td>BREX-system-P-loop-protein-BrxC-[Bifidobacterium-pullorum]</td></tr>
<tr><td>MBX3116430.1</td><td>Type1.BREX</td><td>CR-ATPase5+REase-><-?||HTH_6+SIS->?->PolB-> BrxA-DNA-binding -> BrxB-BrxF-iSTAND -> DUF6079 *-> N6-METHYLASE -> OLD-ABC -> TOPRIM -> PglZ -> SIGMA+OB-Fold+AAA-ATPase+HKD-DNase-BrxL ->Pkinase+CR-Kinase-><-PIN<-DUF433||REase+HTH+35exo+Small-B-barrel+SF1-SF2-Helicase-></td><td>1182</td><td>Bacteria>Terrabacteria_group>Actinomycetota>Actinomycetes>Micrococcales>Microbacteriaceae</td><td>GCA_019638875.1</td><td>Cryobacterium_sp.</td><td>KF808_03680</td><td>24849..28397</td><td>1926290</td><td>BREX-system-P-loop-protein-BrxC-[Cryobacterium-sp.]</td></tr>
<tr><td>MBY6061973.1</td><td>Type1.BREX</td><td>TetR-HTH->?-><-?<-Glyoxalase|| OLD-ABC + TOPRIM ->?-> BrxA-DNA-binding -> BrxB-BrxF-iSTAND -> DUF6079 *-> N6-METHYLASE -> PglZ -> SIGMA+OB-Fold+AAA-ATPase+Lon-Protease-BrxL -><-ParB+HNH||?->MAD-NTD-></td><td>1152</td><td>Bacteria>Terrabacteria_group>Actinomycetota>Actinomycetes>Micrococcales>Microbacteriaceae</td><td>GCA_019801485.1</td><td>Microbacterium_esteraromaticum</td><td>KUV70_11015</td><td>269249..272707</td><td>57043</td><td>BREX-system-P-loop-protein-BrxC-[Microbacterium-esteraromaticum]</td></tr>
<tr><td>WP_261393916.1</td><td>Type1.BREX</td><td>TetR-HTH->?-><-?<-Glyoxalase|| OLD-ABC + TOPRIM ->?-> BrxA-DNA-binding -> BrxB-BrxF-iSTAND -> DUF6079 *-> N6-METHYLASE -> PglZ -> SIGMA+OB-Fold+AAA-ATPase+Lon-Protease-BrxL -><-ParB+HNH||?->MAD-NTD-></td><td>1146</td><td>Bacteria>Terrabacteria_group>Actinomycetota>Actinomycetes>Micrococcales>Microbacteriaceae</td><td>GCF_019801485.1</td><td>Microbacterium_esteraromaticum</td><td>KUV70_RS11015</td><td>269267..272707</td><td>57043</td><td>BREX-system-P-loop-protein-BrxC-[Microbacterium-esteraromaticum]</td></tr>
<tr><td>WP_222446684.1</td><td>Type1.BREX</td><td>TetR-HTH->?-><-?<-Glyoxalase|| OLD-ABC + TOPRIM ->?-> BrxA-DNA-binding -> BrxB-BrxF-iSTAND -> DUF6079 *-> N6-METHYLASE -> PglZ -> SIGMA+OB-Fold+AAA-ATPase+Lon-Protease-BrxL -><-ParB+HNH||?->MAD-NTD-></td><td>1152</td><td>Bacteria>Terrabacteria_group>Actinomycetota>Actinomycetes>Micrococcales>Microbacteriaceae</td><td>-</td><td>Microbacterium_esteraromaticum</td><td>KUV70_11015</td><td>269249..272707</td><td>57043</td><td>BREX-system-P-loop-protein-BrxC-[Microbacterium-esteraromaticum]</td></tr>
<tr><td>KUK48423.1</td><td>Type6.BREX</td><td>CHTH-> BrxE -> BrxA-DNA-binding -> BrxB-BrxF-iSTAND -> DUF6079 *-> N6-METHYLASE -> OLD-ABC + TOPRIM -> PglZ -> PglZ -> SIGMA+OB-Fold+AAA-ATPase+Lon-Protease-BrxL ->DNA_METHYLASE->SF1-SF2-Helicase+REase->REase-></td><td>1227</td><td>Bacteria>Terrabacteria_group>Actinomycetota>Actinomycetes>unclassified_Actinomycetes</td><td>GCA_001508275.1</td><td>Actinobacteria_bacterium_66_15</td><td>XD74_0976</td><td>7706..11389</td><td>1635289</td><td>Uncharacterized-protein-XD74_0976-[Actinobacteria-bacterium-66_15]</td></tr>
<tr><td>MBN1458231.1</td><td>Type1.BREX</td><td><-HEPN|| SF1-SF2-Helicase+wHTH+Small-B-barrel+DUF1998 -><-?<-Phage_int_SAM_3+Phage_integrase<-HTH_3||?->?-> BrxA-DNA-binding -> BrxB-BrxF-iSTAND -> DUF6079 *-> N6-METHYLASE -> OLD-ABC + TOPRIM -> N6-METHYLASE -> PglZ -> SIGMA+OB-Fold+AAA-ATPase+Lon-Protease-BrxL -></td><td>1222</td><td>Bacteria>Terrabacteria_group>Armatimonadota>unclassified_Armatimonadota</td><td>GCA_016928475.1</td><td>Armatimonadota_bacterium</td><td>JXA57_01760</td><td>13903..17571</td><td>2033014</td><td>BREX-system-P-loop-protein-BrxC-[Armatimonadota-bacterium]</td></tr>
<tr><td>WP_028784627.1</td><td>Type6.BREX</td><td>ArdCN+MPTase-inact-> TOPRIM -> BrxE -> BrxA-DNA-binding -> BrxB-BrxF-iSTAND -> DUF6079 *-> N6-METHYLASE -> PglZ ->ArsR-HTH->?->?->PNPase-></td><td>1207</td><td>Bacteria>Terrabacteria_group>Bacillota>Bacilli>Bacillales>Bacillaceae</td><td>GCF_000496835.1</td><td>Thalassobacillus_devorans</td><td>P855_RS0116640</td><td>86998..90621</td><td>279813</td><td>BREX-system-P-loop-protein-BrxC-[Thalassobacillus-devorans]</td></tr>
<tr><td>WP_066159884.1</td><td>Type5.BREX</td><td><- OLD-ABC + TOPRIM <-?<-?<- TUDOR+SF1-SF2-Helicase+iREase+CTD +REase<- PglZ <- N6-METHYLASE <- DUF6079 *<- BrxB-BrxF-iSTAND <- BrxA-DNA-binding <-SF1-SF2-Helicase+REase-p<-SF1-SF2-Helicase<-SF1-SF2-Helicase<-ParB<-MerR-HTH</td><td>1205</td><td>Bacteria>Terrabacteria_group>Bacillota>Bacilli>Bacillales>Bacillaceae</td><td>GCF_001591945.1</td><td>Halalkalibacter_krulwichiae</td><td>BKR01S_RS20150</td><td>13948..17565</td><td>199441</td><td>BREX-system-P-loop-protein-BrxC-[Halalkalibacter-krulwichiae]</td></tr>
<tr><td>WP_132948756.1</td><td>Type5.BREX</td><td>NUDIX-><-ACET-><-?||McrB-NTD+McrB-AAA->wHTH+REase->?->SF1-SF2-Helicase-> BrxA-DNA-binding -> BrxB-BrxF-iSTAND -> DUF6079 *-> N6-METHYLASE -> PglZ -> TUDOR+SF1-SF2-Helicase+iREase+CTD +REase-> OLD-ABC + TOPRIM ->HIN-HTH-><-?<-?<-ParA-Soj-PloopNTPase</td><td>1183</td><td>Bacteria>Terrabacteria_group>Bacillota>Bacilli>Bacillales>Bacillaceae</td><td>GCF_004341205.1</td><td>Thermolongibacillus_altinsuensis</td><td>EDD69_RS09635</td><td>59871..63422</td><td>575256</td><td>BREX-system-P-loop-protein-BrxC-[Thermolongibacillus-altinsuensis]</td></tr>
<tr><td>WP_163530739.1</td><td>Type5.BREX</td><td>ArdCN+MPTase-inact-> TOPRIM ->?-> BrxA-DNA-binding -> BrxB-BrxF-iSTAND -> DUF6079 *-> N6-METHYLASE -> PglZ -> TUDOR+SF1-SF2-Helicase+iREase+CTD +REase->?->McrB-NTD+HNH->Phage_integrase-><-HEPN<-AAA-ATPase<-HTH_22+rve</td><td>1188</td><td>Bacteria>Terrabacteria_group>Bacillota>Bacilli>Bacillales>Bacillaceae</td><td>GCF_902726645.1</td><td>Halobacillus_ihumii</td><td>G6R08_RS20435</td><td>4081488..4085054</td><td>2686092</td><td>BREX-system-P-loop-protein-BrxC-[Halobacillus-ihumii]</td></tr>
<tr><td>WP_077588117.1</td><td>Type1.BREX</td><td><-XIS-HTH<-?||?->SLATT->RVT->?-> BrxA-DNA-binding -> BrxB-BrxF-iSTAND -> DUF6079 *-> N6-METHYLASE -> PglZ -> SIGMA+OB-Fold+AAA-ATPase+Lon-Protease-BrxL -> OLD-ABC -> TOPRIM ->HAD+PRTase+PRTase+REase+SF1-SF2-Helicase->?->HIRAN-></td><td>1190</td><td>Bacteria>Terrabacteria_group>Bacillota>Bacilli>Bacillales>Planococcaceae</td><td>GCF_001999905.1</td><td>Planococcus_lenghuensis</td><td>B0X71_03360</td><td>617590..621162</td><td>2213202</td><td>BREX-system-P-loop-protein-BrxC-[Planococcus-lenghuensis]</td></tr>
<tr><td>MBD8998786.1</td><td>Type1.BREX</td><td><- wHTH+WYL+WCX <-TIR<-?<- SIGMA+OB-Fold+AAA-ATPase+Lon-Protease-BrxL <-?<- PglZ <- OLD-ABC + TOPRIM <- N6-METHYLASE <-TIR<- DUF6079 *<- BrxB-BrxF-iSTAND <- BrxA-DNA-binding <-HTH_26<-?<-?<-?||<-Classical-AAA<-PRTase</td><td>1203</td><td>Bacteria>Terrabacteria_group>Bacillota>Bacilli>Lactobacillales>Lactobacillaceae</td><td>GCA_014847545.1</td><td>Ligilactobacillus_ruminis</td><td>EGQ92_00800</td><td>15462..19073</td><td>1623</td><td>BREX-system-P-loop-protein-BrxC-[Ligilactobacillus-ruminis]</td></tr>
<tr><td>WP_087716717.1</td><td>Type1.BREX</td><td><-TetR-HTH||Cas9+HNH+RuvC->Cas_Cas1->CRISPR_Cas2-><- PglZ || N6-METHYLASE -><- N6-METHYLASE <- DUF6079 *<- BrxB-BrxF-iSTAND <- BrxA-DNA-binding <- TOPRIM <-SbcC</td><td>1221</td><td>Bacteria>Terrabacteria_group>Bacillota>Bacilli>Lactobacillales>Lactobacillaceae</td><td>GCF_002179515.1</td><td>Levilactobacillus_brevis</td><td>CA846_RS04700</td><td>148535..152200</td><td>1580</td><td>BREX-system-P-loop-protein-BrxC-[Levilactobacillus-brevis]</td></tr>
<tr><td>WP_127848553.1</td><td>Type1.BREX</td><td><-ClpABN-AAA+SF1-SF2-Helicase<-?<-?<- SIGMA+OB-Fold+AAA-ATPase+Lon-Protease-BrxL <- PglZ || N6-METHYLASE ->Arm-DNA-bind_4+Phage_int_SAM_5+Phage_integrase-><- N6-METHYLASE <- DUF6079 *<- BrxB-BrxF-iSTAND <- BrxA-DNA-binding <-?<- TOPRIM <-SbcC</td><td>1214</td><td>Bacteria>Terrabacteria_group>Bacillota>Bacilli>Lactobacillales>Lactobacillaceae</td><td>GCF_004010095.1</td><td>Lacticaseibacillus_hulanensis</td><td>EKN00_RS02415</td><td>61527..65171</td><td>2493111</td><td>BREX-system-P-loop-protein-BrxC-[Lacticaseibacillus-hulanensis]</td></tr>
<tr><td>WP_179394507.1</td><td>Type1.BREX</td><td><-HTH_3||?->?->SbcC-> TOPRIM -> BrxA-DNA-binding -> BrxB-BrxF-iSTAND -> DUF6079 *-> N6-METHYLASE -><-Arm-DNA-bind_4+Phage_int_SAM_5+Phage_integrase|| PglZ -> SIGMA+OB-Fold+AAA-ATPase+Lon-Protease-BrxL ->HTH+REase->?->ZnR+HARE-HTH->ARG-></td><td>1200</td><td>Bacteria>Terrabacteria_group>Bacillota>Bacilli>Lactobacillales>Lactobacillaceae</td><td>GCF_013407595.1</td><td>Lacticaseibacillus_absianus</td><td>HII33_RS04075</td><td>947851..951453</td><td>2729623</td><td>BREX-system-P-loop-protein-BrxC-[Lacticaseibacillus-absianus]</td></tr>
<tr><td>WP_058211096.1</td><td>Type1.BREX</td><td><- PglZ <- TOPRIM <- OLD-ABC <-?<-?<- N6-METHYLASE <- DUF6079 *<- BrxB-BrxF-iSTAND <- BrxA-DNA-binding <-XIS-HTH<-?<-?<-?<-REase</td><td>1179</td><td>Bacteria>Terrabacteria_group>Bacillota>Bacilli>Lactobacillales>Streptococcaceae</td><td>GCF_001456725.1</td><td>Lactococcus_lactis</td><td>KF134_RS08810</td><td>324861..328400</td><td>1358</td><td>BREX-system-P-loop-protein-BrxC-[Lactococcus-lactis]</td></tr>
<tr><td>WP_191672696.1</td><td>Type1.BREX</td><td>Glyoxalase-><-?<-?||?->XIS-HTH-> BrxA-DNA-binding -> BrxB-BrxF-iSTAND -> DUF6079 *-> N6-METHYLASE ->?->?-> OLD-ABC -> TOPRIM -> PglZ -></td><td>1179</td><td>Bacteria>Terrabacteria_group>Bacillota>Bacilli>Lactobacillales>Streptococcaceae</td><td>GCF_014830225.1</td><td>Lactococcus_petauri</td><td>DXX96_RS03180</td><td>80359..83898</td><td>1940789</td><td>BREX-system-P-loop-protein-BrxC-[Lactococcus-petauri]</td></tr>
<tr><td>WP_251420892.1</td><td>Type1.BREX</td><td><- PglZ <- TOPRIM <- OLD-ABC <-?<- N6-METHYLASE <- DUF6079 *<- BrxB-BrxF-iSTAND <- BrxA-DNA-binding <-XIS-HTH<-?<-?<-?<-Aminotran_1_2<-?<-35exo</td><td>1179</td><td>Bacteria>Terrabacteria_group>Bacillota>Bacilli>Lactobacillales>Streptococcaceae</td><td>GCF_943192955.1</td><td>Lactococcus</td><td>NFR64_RS02430</td><td>42877..46416</td><td>1357</td><td>MULTISPECIES:-BREX-system-P-loop-protein-BrxC-[Lactococcus]</td></tr>
<tr><td>WP_118669593.1</td><td>Type1.BREX</td><td><- SIGMA+OB-Fold+AAA-ATPase+Lon-Protease-BrxL <- OLD-ABC + TOPRIM <- PglZ <-Schlafen+HATPase+wHTH<-HEPN<-TIR<- N6-METHYLASE <- DUF6079 *<- BrxB-BrxF-iSTAND <- BrxA-DNA-binding <-HTH_26</td><td>1196</td><td>Bacteria>Terrabacteria_group>Bacillota>Clostridia>Eubacteriales>Clostridiaceae</td><td>-</td><td>Clostridium_sp._TF08-15</td><td>DXC82_15795</td><td>32814..36404</td><td>2293052</td><td>BREX-system-P-loop-protein-BrxC-[Clostridium-sp.-TF08-15]</td></tr>
<tr><td>RGF28780.1</td><td>Type1.BREX</td><td><- OLD-ABC + TOPRIM <- PglZ <-Schlafen+HATPase+wHTH<-HEPN<- N6-METHYLASE <- DUF6079 *<- BrxB-BrxF-iSTAND <- BrxA-DNA-binding <-HTH_26</td><td>1196</td><td>Bacteria>Terrabacteria_group>Bacillota>Clostridia>Eubacteriales>Clostridiaceae</td><td>GCA_003435925.1</td><td>Clostridium_sp._AF46-9NS</td><td>DW081_14990</td><td>34449..38039</td><td>2293020</td><td>BREX-system-P-loop-protein-BrxC-[Clostridium-sp.-AF46-9NS]</td></tr>
<tr><td>RHU61449.1</td><td>Type1.BREX</td><td><- SIGMA+OB-Fold+AAA-ATPase+Lon-Protease-BrxL <- OLD-ABC + TOPRIM <- PglZ <-Schlafen+HATPase+wHTH<-HEPN<-TIR<- N6-METHYLASE <- DUF6079 *<- BrxB-BrxF-iSTAND <- BrxA-DNA-binding <-HTH_26</td><td>1196</td><td>Bacteria>Terrabacteria_group>Bacillota>Clostridia>Eubacteriales>Clostridiaceae</td><td>GCA_003481275.1</td><td>Clostridium_sp._TF08-15</td><td>DXC82_15795</td><td>32814..36404</td><td>2293052</td><td>BREX-system-P-loop-protein-BrxC-[Clostridium-sp.-TF08-15]</td></tr>
<tr><td>NLE27040.1</td><td>Type1.BREX</td><td><-MORC+REase-p<-IG+IG<-Phage_integrase|| BrxA-DNA-binding -> BrxB-BrxF-iSTAND -> DUF6079 *->?-> N6-METHYLASE ->ArdCN+MPTase-> PglZ -> SIGMA+OB-Fold+AAA-ATPase+Lon-Protease-BrxL -> OLD-ABC + TOPRIM ->?->DUF488->SIR2-></td><td>1147</td><td>Bacteria>Terrabacteria_group>Bacillota>Clostridia>Eubacteriales>Clostridiaceae</td><td>GCA_012514215.1</td><td>Clostridiaceae_bacterium</td><td>GX625_17200</td><td>20680..24123</td><td>1898204</td><td>BREX-system-P-loop-protein-BrxC-[Clostridiaceae-bacterium]</td></tr>
</table>

| Protein-ID | System | Gene-Neighbourhoods | Length | Taxonomy | Assembly-ID | Strain | Locus-Tag | Genomic-coordinates | TaxID | Fasta-defline |
|---|---|---|---|---|---|---|---|---|---|---|
| WP_035381181.1 | Type1.BREX | HTH_Tnp_1+HTH_21+rve+rve_3->rve-><-?\|\| BrxB-BrxF-iSTAND -> DUF6079 *-> N6-METHYLASE -> OLD-ABC ->?-> TOPRIM -> PglZ -> [illegible] | 1193 | Bacteria>Terrabacteria_group>Bacillota>Clostridia>Eubacteriales>Clostridiaceae | GCF_000601455.1 | Fervidicella_metallireducens | Q428_RS11710 | 3478..7059 | 655338 | BREX-system-P-loop-protein-BrxC-[Fervidicella-metallireducens] |
| WP_118731364.1 | Type1.BREX | PALP->?->RVT->HTH_26-> BrxA-DNA-binding -> BrxB-BrxF-iSTAND -> DUF6079 *-> N6-METHYLASE ->HEPN->Schlafen+HATPase+wHTH-> PglZ -> OLD-ABC + TOPRIM -> [illegible] | 1196 | Bacteria>Terrabacteria_group>Bacillota>Clostridia>Eubacteriales>Clostridiaceae | GCF_003480225.1 | Clostridium_sp._AM34-9AC | DW835_RS14805 | 20266..23856 | 2293030 | BREX-system-P-loop-protein-BrxC-[Clostridium-sp.-AM34-9AC] |
| WP_118740431.1 | Type1.BREX | <- OLD-ABC + TOPRIM <- SIGMA+OB-Fold+AAA-ATPase+Lon-Protease-BrxL <- PglZ <-Schlafen+HATPase+wHTH<-DUF4404<-ZnR+ZnR<- N6-METHYLASE <-TIR<-?<- DUF6079 *<- BrxB-BrxF-iSTAND <- [illegible] | 1182 | Bacteria>Terrabacteria_group>Bacillota>Clostridia>Eubacteriales>Clostridiaceae | GCF_003480345.1 | Clostridium_sp._AM32-2 | DW807_RS02880 | 180384..183932 | 2293029 | BREX-system-P-loop-protein-BrxC-[Clostridium-sp.-AM32-2] |
| WP_164950923.1 | Type5.BREX | <-GDResolvase<-?<-?<- TUDOR+SF1-SF2-Helicase+iREase+CTD +REase<- PglZ <- TOPRIM <- OLD-ABC <-HEPN<- N6-METHYLASE <- DUF6079 *<- BrxB-BrxF-iSTAND <- BrxA-DNA-binding <-?<- [illegible] | 1191 | Bacteria>Terrabacteria_group>Bacillota>Clostridia>Eubacteriales>Clostridiaceae | GCF_011065955.2 | Clostridium_sp._FP2 | G9F71_RS16320 | 3398773..3402348 | 2724481 | BREX-system-P-loop-protein-BrxC-[Clostridium-sp.-FP2] |
| WP_072897339.1 | Type1.BREX | <-ABhydrolase<-AraC-HTH<-?<-?<-?<-Y1_Tnp<- PglZ <- N6-METHYLASE <- OLD-ABC + TOPRIM <- DUF6079 *<- BrxB-BrxF-iSTAND <- BrxA-DNA-binding <-Peptidase_M20 | 1218 | Bacteria>Terrabacteria_group>Bacillota>Clostridia>Eubacteriales>Clostridiaceae | GCF_900129365.1 | Clostridium_fallax | BUA70_RS11140 | 23714..27370 | 1533 | BREX-system-P-loop-protein-BrxC-[Clostridium-fallax] |
| WP_117466987.1 | Type1.BREX | <- OLD-ABC + TOPRIM <- PglZ <-Schlafen+HATPase+wHTH<-HEPN<- N6-METHYLASE <- DUF6079 *<- BrxB-BrxF-iSTAND <- BrxA-DNA-binding <-HTH_26 | 1196 | Bacteria>Terrabacteria_group>Bacillota>Clostridia>Eubacteriales>Clostridiaceae | - | unclassified_Clostridium | DW081_14990 | 34449..38039 | 2614128 | MULTISPECIES:-BREX-system-P-loop-protein-BrxC-[unclassified-Clostridium] |
| WP_117821980.1 | Type1.BREX | MACRODOMAIN->?->ZnR+ZnR+ZnR->?-><-?<-HTH_38\|\|HTH_26-> BrxA-DNA-binding -> BrxB-BrxF-iSTAND -> DUF6079 *-> N6-METHYLASE -> N6-METHYLASE -p->?->Schlafen+HATPase+wHTH-> PglZ -> [illegible] | 1196 | Bacteria>Terrabacteria_group>Bacillota>Clostridia>Eubacteriales>Clostridiaceae | - | unclassified_Clostridium | DW959_RS12365 | 44943..48533 | 2614128 | MULTISPECIES:-BREX-system-P-loop-protein-BrxC-[unclassified-Clostridium] |
| KGP76494.1 | Type1.BREX | sGTP->?->?-> BrxA-DNA-binding -> BrxB-BrxF-iSTAND -> DUF6079 *-> N6-METHYLASE -> OLD-ABC ->PIN->AbiGi-> OLD-ABC -> TOPRIM -> PglZ -> SIGMA+OB-Fold+AAA-ATPase+Lon-Protease-BrxL -> [illegible] | 1189 | Bacteria>Terrabacteria_group>Bacillota>Clostridia>Eubacteriales>Desulfitobacteriaceae | GCA_000770645.1 | Desulfosporosinus_sp._Tol-M | JT05_04435 | 4696..8265 | 1536651 | hypothetical-protein-JT05_04435-[Desulfosporosinus-sp.-Tol-M] |
| WP_091728844.1 | Type1.BREX | <- SIGMA+OB-Fold+AAA-ATPase+Lon-Protease-BrxL <- PglZ <- TOPRIM <- OLD-ABC <- N6-METHYLASE <- DUF6079 *<- BrxB-BrxF-iSTAND <- BrxA-DNA-binding \|\|?->?-><-?<-SF1-SF2-Helicase<- [illegible] | 1193 | Bacteria>Terrabacteria_group>Bacillota>Clostridia>Eubacteriales>Eubacteriales_incertae_sedis | GCF_900107485.1 | Proteiniborus_ethanoligenes | BLV37_RS06315 | 24858..28439 | 415015 | BREX-system-P-loop-protein-BrxC-[Proteiniborus-ethanoligenes] |
| RGC74026.1 | Type1.BREX | Peptidase_M20->MACRODOMAIN->?->?->?->HTH_26-> BrxA-DNA-binding -> BrxB-BrxF-iSTAND -> DUF6079 *-> N6-METHYLASE -> PglZ -> OLD-ABC + TOPRIM -> [illegible] | 1203 | Bacteria>Terrabacteria_group>Bacillota>Clostridia>Eubacteriales>Lachnospiraceae | GCA_003434385.1 | Lachnospiraceae_bacterium_AM23-2LB | DW655_01895 | 389201..392812 | 2302964 | BREX-system-P-loop-protein-BrxC-[Lachnospiraceae-bacterium-AM23-2LB] |
| HCI18472.1 | Type1.BREX | <-PIN<-?<- SIGMA+OB-Fold+AAA-ATPase+Lon-Protease-BrxL <- OLD-ABC + TOPRIM <- PglZ <-Schlafen+HATPase+wHTH<-REase<- N6-METHYLASE <- DUF6079 -p*<- DUF6079 -p<- BrxB-BrxF-iSTAND <- [illegible] | 799 | Bacteria>Terrabacteria_group>Bacillota>Clostridia>Eubacteriales>Lachnospiraceae | GCA_003521585.1 | Lachnospiraceae_bacterium | DFH32_07645 | 14441..16840 | 1898203 | BREX-system-P-loop-protein-BrxC-[Lachnospiraceae-bacterium] |
| HCX41706.1 | Type1.BREX | SKI->?->tRNA->?->?->?->?->HTH_26-> BrxA-DNA-binding -> BrxB-BrxF-iSTAND -> DUF6079 *-> N6-METHYLASE ->REase->Schlafen+HATPase+wHTH-> PglZ -> OLD-ABC + TOPRIM -> [illegible] | 1196 | Bacteria>Terrabacteria_group>Bacillota>Clostridia>Eubacteriales>Lachnospiraceae | GCA_003539645.1 | Lachnospiraceae_bacterium | DHH50_05105 | 179559..183149 | 1898203 | BREX-system-P-loop-protein-BrxC-[Lachnospiraceae-bacterium] |
| TGX97927.1 | Type1.BREX | <- OLD-ABC + TOPRIM -p<- OLD-ABC + TOPRIM -p<- PglZ <-HNH<- OLD-ABC <- N6-METHYLASE <-HEPN<-ParB+HNH<- N6-METHYLASE <- DUF6079 *<- BrxB-BrxF-iSTAND <- BrxA-DNA-binding <-HTH_26<- [illegible] | 1201 | Bacteria>Terrabacteria_group>Bacillota>Clostridia>Eubacteriales>Lachnospiraceae | GCA_004793585.1 | Lachnospiraceae_bacterium | E5357_10825 | 35909..39514 | 1898203 | BREX-system-P-loop-protein-BrxC-[Lachnospiraceae-bacterium] |
| NBJ03627.1 | Type1.BREX | <- PglZ <- TOPRIM <- OLD-ABC <- N6-METHYLASE <- DUF6079 *<- BrxB-BrxF-iSTAND <- BrxA-DNA-binding | 1205 | Bacteria>Terrabacteria_group>Bacillota>Clostridia>Eubacteriales>Lachnospiraceae | GCA_009911175.1 | Lachnospiraceae_bacterium | D3Z62_26960 | 10883..14500 | 1898203 | BREX-system-P-loop-protein-BrxC-[Lachnospiraceae-bacterium] |
| MBR4608207.1 | Type1.BREX | <-REase<-Phage_integrase<-Phage_integrase\|\|?->MobA_MobL->?->?-> BrxA-DNA-binding -> BrxB-BrxF-iSTAND -> DUF6079 *-> N6-METHYLASE -p-> N6-METHYLASE -> OLD-ABC -> TOPRIM -> PglZ -> [illegible] | 1155 | Bacteria>Terrabacteria_group>Bacillota>Clostridia>Eubacteriales>Lachnospiraceae | GCA_017536085.1 | Lachnospiraceae_bacterium | IK041_18520 | 18020..21487 | 1898203 | BREX-system-P-loop-protein-BrxC-[Lachnospiraceae-bacterium] |
| WP_168932969.1 | Type1.BREX | <- OLD-ABC + TOPRIM <- SIGMA+OB-Fold+AAA-ATPase+Lon-Protease-BrxL <- PglZ <-Schlafen+HATPase+wHTH<-DUF4404<-ZnR+ZnR<- N6-METHYLASE <-TIR<-?<- DUF6079 *<- BrxB-BrxF-iSTAND <- [illegible] | 1182 | Bacteria>Terrabacteria_group>Bacillota>Clostridia>Eubacteriales>Lachnospiraceae | GCF_012843825.1 | Dorea_formicigenerans | HF855_RS01350 | 86312..89860 | 39486 | BREX-system-P-loop-protein-BrxC-[Dorea-formicigenerans] |
| WP_195609804.1 | Type1.BREX | Peptidase_M20->MACRODOMAIN->?->?->?->?->HTH_26-> BrxA-DNA-binding -> BrxB-BrxF-iSTAND -> DUF6079 *-> N6-METHYLASE -> PglZ -> OLD-ABC + TOPRIM -> [illegible] | 1203 | Bacteria>Terrabacteria_group>Bacillota>Clostridia>Eubacteriales>Lachnospiraceae | GCF_015560805.1 | Mediterraneibacter_glycyrrhizinilyticus | I2H87_RS09315 | 27635..31246 | 342942 | BREX-system-P-loop-protein-BrxC-[Mediterraneibacter-glycyrrhizinilyticus] |
| WP_227165162.1 | Type1.BREX | MACRODOMAIN->?-> [illegible] | 1201 | Bacteria>Terrabacteria_group>Bacillota>Clostridia>Eubacteriales>Lachnospiraceae | GCF_020560115.1 | Mediterraneibacter_sp._210702-DFI.5.30 | LI126_RS08100 | 75769..79374 | 2883232 | BREX-system-P-loop-protein-BrxC-[Mediterraneibacter-sp.-210702-DFI.5.30] |
| WP_270389656.1 | Type1.BREX | MACRODOMAIN->?->?-> [illegible] | 1201 | Bacteria>Terrabacteria_group>Bacillota>Clostridia>Eubacteriales>Lachnospiraceae | GCF_027661925.1 | Blautia_massiliensis | PGQ23_RS05910 | 13798..17403 | 1737424 | BREX-system-P-loop-protein-BrxC-[Blautia-massiliensis] |
| WP_270645287.1 | Type1.BREX | HTH_19-><- [illegible] | 1205 | Bacteria>Terrabacteria_group>Bacillota>Clostridia>Eubacteriales>Lachnospiraceae | GCF_027692785.1 | Merdimonas_faecis | PG729_RS15130 | 28205..31822 | 1653435 | BREX-system-P-loop-protein-BrxC-[Merdimonas-faecis] |
| WP_073386276.1 | Type1.BREX | OLD-ABC + TOPRIM -> [illegible] | 1206 | Bacteria>Terrabacteria_group>Bacillota>Clostridia>Eubacteriales>Lachnospiraceae | GCF_900129945.1 | Butyrivibrio_fibrisolvens | BUC27_RS06005 | 31476..35096 | 831 | BREX-system-P-loop-protein-BrxC-[Butyrivibrio-fibrisolvens] |
| WP_055287913.1 | Type1.BREX | [illegible] | 1196 | Bacteria>Terrabacteria_group>Bacillota>Clostridia>Eubacteriales>Lachnospiraceae | GCF_902362785.1 | Lachnospira_eligens | FXW70_RS12185 | 167515..171105 | 39485 | BREX-system-P-loop-protein-BrxC-[Lachnospira-eligens] |
| WP_120485415.1 | Type1.BREX | MACRODOMAIN->?-> [illegible] | 1201 | Bacteria>Terrabacteria_group>Bacillota>Clostridia>Eubacteriales>Oscillospiraceae | - | Anaerotruncus_sp._1XD22-93 | D3Z44_12640 | 10997..14602 | 2320082 | BREX-system-P-loop-protein-BrxC-[Anaerotruncus-sp.-1XD22-93] |
| MBS3950095.1 | Type1.BREX | OLD-ABC + TOPRIM -p-> [illegible] | 1183 | Bacteria>Terrabacteria_group>Bacillota>Clostridia>Eubacteriales>Peptococcaceae | GCA_018333435.1 | Peptococcaceae_bacterium | KGZ53_05495 | 2469..6020 | 2052179 | BREX-system-P-loop-protein-BrxC-[Peptococcaceae-bacterium] |
| WP_099191952.1 | Type1.BREX | DNA_METHYLASE->?-> [illegible] | 1190 | Bacteria>Terrabacteria_group>Bacillota>Clostridia>Eubacteriales>Peptostreptococcaceae | GCF_002724055.1 | Tepidibacter_mesophilus | Tmeso_RS13165 | 34071..37643 | 655607 | BREX-system-P-loop-protein-BrxC-[Tepidibacter-mesophilus] |
| WP_166083468.1 | Type1.BREX | [illegible] | 1192 | Bacteria>Terrabacteria_group>Bacillota>Erysipelotrichia>Erysipelotrichales>Erysipelotrichaceae | GCF_011421495.1 | Erysipelothrix_anatis | GON62_RS10715 | 11860..15438 | 2683713 | BREX-system-P-loop-protein-BrxC-[Erysipelothrix-anatis] |
| NLN06909.1 | Type6.BREX | BrxE -> [illegible] | 1222 | Bacteria>Terrabacteria_group>Bacillota | GCA_012522075.1 | Bacillota_bacterium | GX167_04740 | 2708..6376 | 1879010 | BREX-system-P-loop-protein-BrxC-[Bacillota-bacterium] |
| MBU4541895.1 | Type1.BREX | <- PglZ -p<- TOPRIM [illegible] | 1187 | Bacteria>Terrabacteria_group>Bacillota | GCA_018903255.1 | Bacillota_bacterium | KJ774_11820 | 7195..10758 | 1879010 | BREX-system-P-loop-protein-BrxC-[Bacillota-bacterium] |
| WP_072743893.1 | Type1.BREX | HTH_26->?->DOC+HTH-> [illegible] | 1192 | Bacteria>Terrabacteria_group>Bacillota>Tissierellia>Tissierellales>Sporanaerobacteraceae | GCF_900130025.1 | Sporanaerobacter_acetigenes | BUA21_RS05940 | 73351..76929 | 165813 | BREX-system-P-loop-protein-BrxC-[Sporanaerobacter-acetigenes] |
| WP_054870024.1 | Type1.BREX | [illegible] | 1187 | Bacteria>Terrabacteria_group>Bacillota>Tissierellia>Tissierellales>Thermohalobacteraceae | GCF_001316435.1 | Caloranaerobacter_sp._TR13 | TR13x_RS01045 | 223070..226633 | 1302151 | BREX-system-P-loop-protein-BrxC-[Caloranaerobacter-sp.-TR13] |
| MBQ6502300.1 | Type1.BREX | DOC+HTH-><-?<-?\|\|?-> [illegible] | 1194 | Bacteria>Terrabacteria_group>Chloroflexota>Anaerolineae>Anaerolineales>Anaerolineaceae | GCA_017432685.1 | Flexilinea_sp. | IJI57_00125 | 8965..12549 | 2017390 | BREX-system-P-loop-protein-BrxC-[Flexilinea-sp.] |
| WP_062419291.1 | Type6.BREX | [illegible] | 1206 | Bacteria>Terrabacteria_group>Chloroflexota>Anaerolineae>Anaerolineales>Anaerolineaceae | GCA_001050255.2 | Levilinea_saccharolytica | LSAC_RS18205 | 3303565..3307185 | 229921 | BREX-system-P-loop-protein-BrxC-[Levilinea-saccharolytica] |
| MBK9093475.1 | Type1.BREX | RNase->?->?->?-> [illegible] | 1151 | Bacteria>Terrabacteria_group>Chloroflexota>Anaerolineae>unclassified_Anaerolineae | GCA_016719785.1 | Anaerolineae_bacterium | IPM84_12015 | 171966..175421 | 2052143 | BREX-system-P-loop-protein-BrxC-[Anaerolineae-bacterium] |
| MBK9096490.1 | Type1.BREX | <- [illegible] | 900 | Bacteria>Terrabacteria_group>Chloroflexota>Anaerolineae>unclassified_Anaerolineae | GCA_016719785.1 | Anaerolineae_bacterium | IPM84_27780 | 244922..247624 | 2052143 | hypothetical-protein-[Anaerolineae-bacterium] |
| NUQ36643.1 | Type1.BREX | <- N6-METHYLASE -p<- [illegible] | 1152 | Bacteria>Terrabacteria_group>Chloroflexota>Caldilineae>Caldilineales>unclassified_Caldilineales | GCA_013360575.1 | Caldilineales_bacterium | HUU23_02375 | 3098..6556 | 2306055 | BREX-system-P-loop-protein-BrxC-[Caldilineales-bacterium] |
| HHY55161.1 | Type1.BREX | <-ACET<- PglZ <-?<- [illegible] | 1149 | Bacteria>Terrabacteria_group>Chloroflexota>unclassified_Chloroflexota | GCA_012842785.1 | Chloroflexota_bacterium | GYA08_06960 | 29927..33376 | 2026724 | BREX-system-P-loop-protein-BrxC-[Chloroflexota-bacterium] |
| MBM3127996.1 | Type1.BREX | XIS-HTH-><- [illegible] | 1153 | Bacteria>Terrabacteria_group>Chloroflexota>unclassified_Chloroflexota | GCA_016865935.1 | Chloroflexota_bacterium | FJ009_05090 | 25630..29091 | 2026724 | BREX-system-P-loop-protein-BrxC-[Chloroflexota-bacterium] |
| WP_127084959.1 | Type1.BREX | CASPASE->?-><-?<- [illegible] | 725 | Bacteria>Terrabacteria_group>Cyanobacteriota/Melainabacteria_group>Cyanobacteriota>Cyanophyceae>Nostocales | GCF_003991905.1 | Dulcicalothrix_desertica | DSM106972_RS34665 | 70206..72383 | 32056 | BREX-system-P-loop-protein-BrxC-[Dulcicalothrix-desertica] |
| MCA6571981.1 | Type1.BREX | BrxB-BrxF-iSTAND -> [illegible] | 1202 | Bacteria>Terrabacteria_group>Cyanobacteriota/Melainabacteria_group>Cyanobacteriota>Cyanophyceae>Pseudanabaenales | GCA_020404105.1 | Pseudanabaena_sp._M53BS1SP1A06MG | IM536_21850 | 2730..6338 | 2771239 | BREX-system-P-loop-protein-BrxC-[Pseudanabaena-sp.-M53BS1SP1A06MG] |
| MCA6593937.1 | Type1.BREX | <- [illegible] | 813 | Bacteria>Terrabacteria_group>Cyanobacteriota/Melainabacteria_group>Cyanobacteriota>Cyanophyceae>Pseudanabaenales | GCA_020404945.1 | Pseudanabaena_sp._M38BS1SP1A06MG | IM535_17950 | 20684..23126 | 2771238 | BREX-system-P-loop-protein-BrxC-[Pseudanabaena-sp.-M38BS1SP1A06MG] |
| MCA6597699.1 | Type1.BREX | BrxB-BrxF-iSTAND -> [illegible] | 1202 | Bacteria>Terrabacteria_group>Cyanobacteriota/Melainabacteria_group>Cyanobacteriota>Cyanophyceae>Pseudanabaenales | GCA_020405005.1 | Pseudanabaena_sp._M046S1SP1A06QC | IM539_15710 | 2728..6336 | 2771236 | BREX-system-P-loop-protein-BrxC-[Pseudanabaena-sp.-M046S1SP1A06QC] |
| MCF8140154.1 | Type6.BREX | <-AAA+wHTH+WYL\|\|?-> [illegible] | 1215 | Bacteria>Terrabacteria_group>Cyanobacteriota/Melainabacteria_group>Cyanobacteriota>Cyanophyceae>Synechococcales | GCA_021737825.1 | Cyanobium_usitatum_Tobar12.5m-G36 | K9J75_03480 | 41417..45064 | 2874848 | BREX-system-P-loop-protein-BrxC-[Cyanobium-usitatum-Tobar12.5m-G36] |
| WP_073476316.1 | Type1.BREX | <-MoxR-AAA\|\| [illegible] | 1215 | Bacteria>Thermodesulfobacteriota>Desulfobacteria>Desulfobacterales>Desulfatibacillaceae>Desulfatibacillum | GCF_900142135.1 | Desulfatibacillum_alkenivorans | BUE18_RS12840 | 56211..59858 | 259354 | BREX-system-P-loop-protein-BrxC-[Desulfatibacillum-alkenivorans] |
| WP_028324416.1 | Type1.BREX | CHTH-> N6-METHYLASE [illegible] | 1145 | Bacteria>Thermodesulfobacteriota>Desulfobacteria>Desulfobacterales>Desulfatirhabdiaceae>Desulfatirhabdium | GCF_000429925.1 | Desulfatirhabdium_butyrativorans | G492_RS0109380 | 84085..87522 | 340467 | BREX-system-P-loop-protein-BrxC-[Desulfatirhabdium-butyrativorans] |
| MBF0468020.1 | Type1.BREX | <- N6-METHYLASE <- [illegible] | 1187 | Bacteria>Thermodesulfobacteriota>Desulfobacteria>Desulfobacterales>Desulfobacteraceae>Desulfamplus | GCA_015232935.1 | Desulfamplus_sp. | HQK61_03910 | 4881..8444 | 2006184 | BREX-system-P-loop-protein-BrxC-[Desulfamplus-sp.] |
| MCF8089785.1 | Type1.BREX | OLD-ABC ->REase-> [illegible] | 1350 | Bacteria>Thermodesulfobacteriota>Desulfobacteria>Desulfobacterales>Desulfobacteraceae>Desulfotignum | GCA_021737465.1 | Desulfotignum_sp. | K9L23_17210 | 8544..12596 | 2049434 | hypothetical-protein-[Desulfotignum-sp.] |
| MBC8432684.1 | Type6.BREX | <-REase<- [illegible] | 1227 | Bacteria>Thermodesulfobacteriota>Desulfobacteria>Desulfobacterales>unclassified_Desulfobacterales | GCA_014383215.1 | Desulfobacterales_bacterium | H8D96_12300 | 20226..23909 | 2044940 | BREX-system-P-loop-protein-BrxC-[Desulfobacterales-bacterium] |
| MBF0101465.1 | Type1.BREX | <-Glyco_hydro_19<- [illegible] | 1145 | Bacteria>Thermodesulfobacteriota>Desulfobacteria>Desulfobacterales>unclassified_Desulfobacterales | GCA_015228675.1 | Desulfobacterales_bacterium | HQK77_11205 | 12812..16249 | 2044940 | BREX-system-P-loop-protein-BrxC-[Desulfobacterales-bacterium] |
| MCG2778948.1 | Type1.BREX | DUF6079 -p*->HEPN-> [illegible] | 757 | Bacteria>Thermodesulfobacteriota>Desulfobacteria>Desulfobacterales>unclassified_Desulfobacterales | GCA_022013615.1 | Desulfobacterales_bacterium | L6406_25005 | 1..2276 | 2044940 | hypothetical-protein-[Desulfobacterales-bacterium] |
| WP_013164351.1 | Type1.BREX | <- [illegible] | 1145 | Bacteria>Thermodesulfobacteriota>Desulfobulbia>Desulfobulbales>Desulfobulbaceae>Desulfurivibrio | GCF_000092205.1 | Desulfurivibrio_alkaliphilus | DAAHT2_RS10985 | 2524530..2527967 | 427923 | BREX-system-P-loop-protein-BrxC-[Desulfurivibrio-alkaliphilus] |
| BDV01246.1 | Type6.BREX | <-DTXR-HTH\|\|?-> [illegible] | 1210 | Bacteria>Thermodesulfobacteriota>Desulfovibrionia>Desulfovibrionales>Desulfomicrobiaceae>Thermodesulfomicrobium | GCA_027925145.1 | Thermodesulfomicrobium_sp._WS | TDMWS_13310 | 1427384..1431016 | 3004129 | hypothetical-protein-TDMWS_13310-[Thermodesulfomicrobium-sp.-WS] |
| WP_008869235.1 | Type1.BREX | <-Y1_Tnp<-ARG<-?<- [illegible] | 1147 | Bacteria>Thermodesulfobacteriota>Desulfovibrionia>Desulfovibrionales>Desulfonatronovibrionaceae>Desulfonatronospira | GCF_000174435.1 | Desulfonatronospira_thiodismutans | DTHIO_RS04855 | 1069759..1073202 | 488939 | BREX-system-P-loop-protein-BrxC-[Desulfonatronospira-thiodismutans] |
| MCI7405014.1 | Type1.BREX | <-35exo<-?\|\|?->?-> [illegible] | 1199 | Bacteria>Thermodesulfobacteriota>Desulfovibrionia>Desulfovibrionales>Desulfovibrionaceae>Desulfovibrio | GCA_022783735.1 | Desulfovibrio_piger | MSS64_01040 | 35161..38760 | 901 | BREX-system-P-loop-protein-BrxC-[Desulfovibrio-piger] |
| PLX83270.1 | Type1.BREX | <-STYKinase-like\|\| [illegible] | 1175 | Bacteria>Thermodesulfobacteriota>Desulfuromonadia>Desulfuromonadales>Desulfuromonadaceae>Desulfuromonas | GCA_002868845.1 | Desulfuromonas_sp. | C0617_12115 | 157475..161002 | 892 | MAG:-BREX-system-P-loop-protein-BrxC-[Desulfuromonas-sp.] |
| WP_207162791.1 | Type1.BREX | <-Peptidase_C14<- [illegible] | 1175 | Bacteria>Thermodesulfobacteriota>Desulfuromonadia>Geobacterales>Geobacteraceae>Geobacter | GCF_017338855.1 | Geobacter_benzoatilyticus | JZM60_RS12610 | 2741421..2744948 | 2815309 | BREX-system-P-loop-protein-BrxC-[Geobacter-benzoatilyticus] |
| OPY81706.1 | Type1.BREX | HIN-HTH-> [illegible] | 1200 | Bacteria>Thermodesulfobacteriota>Syntrophia>Syntrophales>Syntrophaceae>Syntrophus | GCA_002067745.1 | Syntrophus_sp._PtaU1.Bin005 | A4E70_01072 | 12725..16327 | 1811712 | MAG:-hypothetical-protein-A4E70_01072-[Syntrophus-sp.-PtaU1.Bin005] |
| PIV05610.1 | Type1.BREX | HerA/FtsK-> [illegible] | 1346 | Bacteria>Thermodesulfobacteriota>Syntrophobacteria>Syntrophobacterales>unclassified_Syntrophobacterales | GCA_002779905.1 | Syntrophobacterales_bacterium_CG03_land_8_20_14_0_80_58_14 | COS57_07000 | 12627..16667 | 1974100 | hypothetical-protein-COS57_07000-[Syntrophobacterales-bacterium-CG03_land_8_20_14_0_80_58_14] |
| OPY73251.1 | Type6.BREX | Phage_integrase-><- [illegible] | 1224 | Bacteria>Thermodesulfobacteriota>Syntrophorhabdia>Syntrophorhabdales>Syntrophorhabdaceae>Syntrophorhabdus | GCA_002067415.1 | Syntrophorhabdus_sp._PtaU1.Bin002 | A4E62_00567 | 11466..15140 | 1811702 | MAG:-hypothetical-protein-A4E62_00567-[Syntrophorhabdus-sp.-PtaU1.Bin002] |
| MBD5384945.1 | Type1.BREX | <-Acyltransferase<- [illegible] | 843 | Bacteria>unclassified_Bacteria | GCA_014802625.1 | bacterium | HDR69_00765 | 93850..96381 | 1869227 | BREX-system-P-loop-protein-BrxC-[bacterium] |
| MCR9098968.1 | Type1.BREX | <- PglZ \|\|?->ParB+ [illegible] | 1186 | Bacteria>unclassified_Bacteria | GCA_024742755.1 | bacterium | NXI25_03390 | 27983..31543 | 1869227 | BREX-system-P-loop-protein-BrxC-[bacterium] |
| --<br>-- OLD-ABC + TOPRIM ->SF2-Helicase<br>--(GABIJA-like)<br>-- | | | | | | | | | | |
| MBR1368475.1 | Type1.BREX | <-HTH_Tnp_1\|\| [illegible] | 1155 | Archaea>Euryarchaeota>Stenosarchaea_group>Methanomicrobia>Methanomicrobiales>Methanocalculaceae | GCA_018132105.1 | Methanocalculus_chunghsingensis | RJ53_02730 | 5881..9348 | 156457 | hypothetical-protein-[Methanocalculus-chunghsingensis] |
| WP_246495968.1 | Type1.BREX | OLD-ABC + TOPRIM -> [illegible] | 1148 | Archaea>Euryarchaeota>Stenosarchaea_group>Methanomicrobia>Methanomicrobiales>Methanocalculaceae | GCF_018132105.1 | Methanocalculus_chunghsingensis | RJ53_RS02795 | 5902..9348 | 156457 | BREX-system-P-loop-protein-BrxC-[Methanocalculus-chunghsingensis] |
| WP_048126494.1 | Type1.BREX | <-DDE_Tnp_1<- [illegible] | 1211 | Archaea>Euryarchaeota>Stenosarchaea_group>Methanomicrobia>Methanosarcinales>Methanosarcinaceae | GCF_000970265.1 | Methanosarcina_lacustris | MSLAZ_RS10160 | 2470398..2474033 | 170861 | BREX-system-P-loop-protein-BrxC-[Methanosarcina-lacustris] |
| WP_244603415.1 | Type1.BREX | <-SIR2<- [illegible] | 1181 | Archaea>Euryarchaeota>Stenosarchaea_group>Methanomicrobia>Methanosarcinales>Methanosarcinaceae | GCF_900774285.1 | Methanococcoides_sp._NM1 | E7X56_RS05470 | 155089..158634 | 1201013 | BREX-system-P-loop-protein-BrxC-[Methanococcoides-sp.-NM1] |
| WP_048171216.1 | Type1.BREX | Glyco_hydro_130-> [illegible] | 1204 | Archaea>Euryarchaeota>Stenosarchaea_group>Methanomicrobia>Methanosarcinales>Methanosarcinaceae | - | Methanosarcina_sp._2.H.A.1B.4 | ED92_RS09675 | 12629..16243 | 1483600 | BREX-system-P-loop-protein-BrxC-[Methanosarcina-sp.-2.H.A.1B.4] |
| QNO42144.1 | Type1.BREX | <- [illegible] | 488 | Archaea>Euryarchaeota>Stenosarchaea_group>Methanomicrobia>Methanosarcinales>unclassified_Methanosarcinales | - | Methanosarcinales_archaeon_ANME-2c_ERB4 | CAOPPJJI_00010 | 16805..18271 | 2759911 | hypothetical-protein-CAOPPJJI_00010-[Methanosarcinales-archaeon-ANME-2c-ERB4] |
| WP_226069221.1 | Type1.BREX | Phage_integrase-> [illegible] | 1201 | Bacteria>Campylobacterota>Epsilonproteobacteria>Campylobacterales>Thiovulaceae>Sulfurimonas | GCF_020410885.1 | Sulfurimonas_sp._SWIR-19 | LCX93_RS05490 | 1055837..1059442 | 2878390 | BREX-system-P-loop-protein-BrxC-[Sulfurimonas-sp.-SWIR-19] |
| MCC7070407.1 | Type6.BREX | VirB6->?->?-> [illegible] | 1209 | Bacteria>delta/epsilon_subdivisions>Deltaproteobacteria>unclassified_Deltaproteobacteria | GCA_020850995.1 | Deltaproteobacteria_bacterium | IT383_03735 | 19295..22924 | 2026735 | BREX-system-P-loop-protein-BrxC-[Deltaproteobacteria-bacterium] |
| WP_103016193.1 | Type5.BREX | <-SbcC<- [illegible] | 1108 | Bacteria>FCB_group>Bacteroidetes/Chlorobi_group>Bacteroidota>Bacteroidetes_incertae_sedis>Rhodothermaceae | GCF_002894575.1 | Salinibacter_ruber | B7R92_RS05385 | 33133..36459 | 146919 | hypothetical-protein-[Salinibacter-ruber] |
| WP_146031917.1 | Type5.BREX | <- [illegible] | 1245 | Bacteria>FCB_group>Bacteroidetes/Chlorobi_group>Bacteroidota>Bacteroidetes_incertae_sedis>Rhodothermaceae | GCF_002894575.1 | Salinibacter_ruber | B7R92_RS05395 | 37102..40839 | 146919 | hypothetical-protein-[Salinibacter-ruber] |
| WP_259182568.1 | Type5.BREX | <-PIN\|\| [illegible] | 1248 | Bacteria>FCB_group>Bacteroidetes/Chlorobi_group>Bacteroidota>Bacteroidetes_incertae_sedis>Rhodothermaceae | GCF_024806795.1 | Salinibacter_ruber | GGQ19_RS12625 | 57748..61494 | 146919 | hypothetical-protein-[Salinibacter-ruber] |
| WP_259220531.1 | Type5.BREX | <-SbcC<- [illegible] | 1245 | Bacteria>FCB_group>Bacteroidetes/Chlorobi_group>Bacteroidota>Bacteroidetes_incertae_sedis>Rhodothermaceae | GCF_024807065.1 | Salinibacter_ruber | GGP94_RS13020 | 49040..52777 | 146919 | hypothetical-protein-[Salinibacter-ruber] |
| WP_259084851.1 | Type1.BREX | <-REase+DUF4268<- [illegible] | 1143 | Bacteria>FCB_group>Bacteroidetes/Chlorobi_group>Bacteroidota>Bacteroidetes_incertae_sedis>Rhodothermaceae | GCF_024808015.1 | Salinibacter_ruber | GGP69_RS05925 | 149309..152740 | 146919 | BREX-system-P-loop-protein-BrxC-[Salinibacter-ruber] |
| WP_259094322.1 | Type1.BREX | <-HD<-SbcC+SbcC<- [illegible] | 1143 | Bacteria>FCB_group>Bacteroidetes/Chlorobi_group>Bacteroidota>Bacteroidetes_incertae_sedis>Rhodothermaceae | GCF_024809025.1 | Salinibacter_ruber | GGP87_RS12315 | 32840..36271 | 146919 | BREX-system-P-loop-protein-BrxC-[Salinibacter-ruber] |
| WP_251951637.1 | Type5.BREX | <- [illegible] | 1108 | Bacteria>FCB_group>Bacteroidetes/Chlorobi_group>Bacteroidota>Bacteroidetes_incertae_sedis>Rhodothermaceae | GCF_943908255.1 | Salinibacter_ruber | NGL47_RS11295 | 22718..26044 | 146919 | hypothetical-protein-[Salinibacter-ruber] |
| WP_251951639.1 | Type5.BREX | <- OLD-ABC + TOPRIM <- [illegible] | 1245 | Bacteria>FCB_group>Bacteroidetes/Chlorobi_group>Bacteroidota>Bacteroidetes_incertae_sedis>Rhodothermaceae | GCF_943908255.1 | Salinibacter_ruber | NGL47_RS11305 | 26687..30424 | 146919 | hypothetical-protein-[Salinibacter-ruber] |
| WP_002561973.1 | Type1.BREX | [illegible] | 1202 | Bacteria>FCB_group>Bacteroidetes/Chlorobi_group>Bacteroidota>Bacteroidia>Bacteroidales | GCF_000382465.1 | Bacteroides_sp._HPS0048 | HMPREF1214_RS19625 | 483033..486641 | 1078089 | BREX-system-P-loop-protein-BrxC-[Bacteroides-sp.-HPS0048] |
| WP_172774764.1 | Type1.BREX | OLD-ABC -> TOPRIM -> [illegible] | 1217 | Bacteria>FCB_group>Bacteroidetes/Chlorobi_group>Bacteroidota>Bacteroidia>Bacteroidales | GCF_012976275.1 | Phocaeicola_vulgatus | HKQ53_RS16515 | 79396..83049 | 821 | BREX-system-P-loop-protein-BrxC-[Phocaeicola-vulgatus] |
| WP_217718390.1 | Type1.BREX | TOPRIM ->XIS-HTH-><- [illegible] | 1217 | Bacteria>FCB_group>Bacteroidetes/Chlorobi_group>Bacteroidota>Bacteroidia>Bacteroidales | GCF_019127825.1 | Bacteroides_uniformis | KSX11_RS12350 | 118043..121696 | 820 | BREX-system-P-loop-protein-BrxC-[Bacteroides-uniformis] |
| WP_176134075.1 | Type1.BREX | <-Phage_holin\|\|?-> [illegible] | 1192 | Bacteria>FCB_group>Bacteroidetes/Chlorobi_group>Bacteroidota>Bacteroidia>Bacteroidales | GCF_900168155.1 | Parabacteroides_chartae | B5X57_RS12575 | 48804..52382 | 1037355 | BREX-system-P-loop-protein-BrxC-[Parabacteroides-chartae] |
| WP_117985926.1 | Type1.BREX | XIS-HTH->SMS-KaiC-> [illegible] | 1210 | Bacteria>FCB_group>Bacteroidetes/Chlorobi_group>Bacteroidota>Bacteroidia>Bacteroidales | GCF_934726555.1 | Bacteroides | O1446_RS12920 | 68574..72206 | 816 | MULTISPECIES:-BREX-system-P-loop-protein-BrxC-[Bacteroides] |
| WP_194850146.1 | Type1.BREX | <- SF1-SF2-Helicase [illegible] | 1160 | Bacteria>FCB_group>Bacteroidetes/Chlorobi_group>Bacteroidota>Flavobacteriia>Flavobacteriales | GCF_015356755.1 | Nonlabens_antarcticus | IQ282_RS01355 | 76947..80429 | 392714 | BREX-system-P-loop-protein-BrxC-[Nonlabens-antarcticus] |
| NUM53512.1 | Type1.BREX | <- SF1-SF2-Helicase [illegible] | 1149 | Bacteria>FCB_group>Candidatus_Hydrogenedentes>unclassified_Candidatus_Hydrogenedentes | GCA_013359265.1 | Candidatus_Hydrogenedentes_bacterium | HUU46_07705 | 154693..158142 | 2030809 | BREX-system-P-loop-protein-BrxC-[Candidatus-Hydrogenedentes-bacterium] |
| VTX48853.1 | Type1.BREX | <-RelE-ParE<-RelB<- [illegible] | 1192 | Bacteria>Fusobacteriota>Fusobacteriia>Fusobacteriales>Leptotrichiaceae>Leptotrichia | GCA_901873485.1 | uncultured_Leptotrichia_sp. | FPIDKHFM_01499 | 24244..27822 | 159271 | Uncharacterised-protein-[uncultured-Leptotrichia-sp.] |
| WP_114817812.1 | Type1.BREX | HTH_3-><-?<- [illegible] | 1179 | Bacteria>Pseudomonadota>Alphaproteobacteria>Caulobacterales>Caulobacteraceae>Brevundimonas | GCF_003350205.1 | Brevundimonas_bullata | DXK21_RS10945 | 61535..65074 | 13160 | BREX-system-P-loop-protein-BrxC-[Brevundimonas-bullata] |
| MBX3547065.1 | Type1.BREX | <-HIN-HTH<-REase<- [illegible] | 777 | Bacteria>Pseudomonadota>Alphaproteobacteria>Hyphomicrobiales>Chelatococcaceae>Chelatococcus | GCA_019635905.1 | Chelatococcus_sp. | KF818_30290 | 46824..49157 | 1953771 | BREX-system-P-loop-protein-BrxC-[Chelatococcus-sp.] |
| WP_203065852.1 | Type1.BREX | VirB10->RHH-><- [illegible] | 1179 | Bacteria>Pseudomonadota>Alphaproteobacteria>Hyphomicrobiales>Devosiaceae>Paradevosia | GCF_016801025.1 | Paradevosia_shaoguanensis | JNE37_RS07680 | 1630850..1634389 | 1335043 | BREX-system-P-loop-protein-BrxC-[Paradevosia-shaoguanensis] |
| WP_130224749.1 | Type1.BREX | <-ArdCN+MPTase<- [illegible] | 1181 | Bacteria>Pseudomonadota>Alphaproteobacteria>Hyphomicrobiales>Nitrobacteraceae>Bradyrhizobium | GCF_004212675.1 | Bradyrhizobium_sp._Leo121 | CW090_RS07090 | 48469..52014 | 1571195 | BREX-system-P-loop-protein-BrxC-[Bradyrhizobium-sp.-Leo121] |
| MAM94519.1 | Type1.BREX | VirB10->RHH-><- [illegible] | 1179 | Bacteria>Pseudomonadota>Alphaproteobacteria>Hyphomicrobiales>Parvibaculaceae>Parvibaculum | GCA_002694985.1 | Parvibaculum_sp. | CMI61_07805 | 145242..148781 | 2024848 | hypothetical-protein-[Parvibaculum-sp.] |
| WP_222362949.1 | Type1.BREX | <- SF1-SF2-Helicase [illegible] | 1174 | Bacteria>Pseudomonadota>Alphaproteobacteria>Hyphomicrobiales>Rhizobiaceae>Rhizobium/Agrobacterium_group | GCF_019798305.1 | Rhizobium_leguminosarum | HFO99_RS19360 | 191490..195014 | 384 | BREX-system-P-loop-protein-BrxC-[Rhizobium-leguminosarum] |
| WP_077987199.1 | Type1.BREX | HTH_3->Couple_hipA+ [illegible] | 1175 | Bacteria>Pseudomonadota>Alphaproteobacteria>Hyphomicrobiales>Rhizobiaceae>Rhizobium/Agrobacterium_group | GCF_030440405.1 | Rhizobium_pusense | BTE56_RS17195 | 18928..22455 | 648995 | BREX-system-P-loop-protein-BrxC-[Rhizobium-pusense] |

| Protein-ID | System | Gene-Neighbourhoods | Length | Taxonomy | Assembly-ID | Strain | Locus-Tag | Genomic-coordinates | TaxID | Fasta-defline |
|---|---|---|---|---|---|---|---|---|---|---|
| WP_076528450.1 | Type1.BREX | Arm-DNA-bind_3+Phage_int_SAM_3+Phage_integrase-><-SF1-SF2-Helicase<-OLD-ABC + TOPRIM <- SIGMA+OB-Fold+AAA-ATPase+HKD-DNase-BrxL <- PglZ <- N6-METHYLASE <- DUF6079 *<- BrxB-BrxF-iSTAND <- BrxA-DNA-binding <- wHTH+WYL+WCX \|\|Calcineurin+Calcineurin-SbcD->SbcC+SbcC-><-MODE-HTH+PBPII\|\|TetR-HTH-><-RHH<-VirB10 | 1179 | Bacteria>Pseudomonadota>Alphaproteobacteria>Rhodobacterales>Paracoccaceae>Gemmobacter | GCF_014636395.1 | Gemmobacter_megaterium | IEV79_RS02805 | 604470..608009 | 1086013 | BREX-system-P-loop-protein-BrxC-[Gemmobacter-megaterium] |
| HCN68765.1 | Type1.BREX | HTH_3-><-?<-?\|\|?->wHTH+AbiEi_2->Polbeta->REase-> BrxA-DNA-binding -> BrxB-BrxF-iSTAND -> DUF6079 *->HEPN-> N6-METHYLASE -> OLD-ABC + TOPRIM ->SF1-SF2-Helicase -> N6-METHYLASE -> PglZ -> SIGMA+OB-Fold+AAA-ATPase+Lon-Protease-BrxL -> | 1154 | Bacteria>Pseudomonadota>Betaproteobacteria>Betaproteobacteria_incertae_sedis>Candidatus_Accumulibacter>unclassified_Candidatus_Accumulibacter | GCA_003535635.1 | Candidatus_Accumulibacter_sp. | DIT03_10970 | 31798..35262 | 2053492 | BREX-system-P-loop-protein-BrxC-[Candidatus-Accumulibacter-sp.] |
| WP_059567309.1 | Type1.BREX | <- SIGMA+OB-Fold+AAA-ATPase+Lon-Protease-BrxL <- PglZ <- SF1-SF2-Helicase<- OLD-ABC + TOPRIM <- N6-METHYLASE <- DUF6079 *<- BrxB-BrxF-iSTAND <- BrxA-DNA-binding <-RHH_5 | 1153 | Bacteria>Pseudomonadota>Betaproteobacteria>Burkholderiales>Burkholderiaceae>Burkholderia | GCF_001523965.1 | Burkholderia_cepacia_complex | C6T69_RS26905 | 12049..15510 | 87882 | MULTISPECIES:-BREX-system-P-loop-protein-BrxC-[Burkholderia-cepacia-complex] |
| WP_269509108.1 | Type1.BREX | <-MerR-HTH<-TnsC-ATPase<-HTH_Tnp+DDE_Tnp<-REase<-?\|\|MORC->RHH_5-> BrxA-DNA-binding -> BrxB-BrxF-iSTAND -> DUF6079 *-> N6-METHYLASE -> OLD-ABC + TOPRIM ->SF1-SF2-Helicase -> PglZ -> SIGMA+OB-Fold+AAA-ATPase+Lon-Protease-BrxL -><-MORC+DNA_gyrase+ TOPRIM <-?<-RFC-AAA+DNAA-HTH | 1153 | Bacteria>Pseudomonadota>Betaproteobacteria>Burkholderiales>Burkholderiaceae>Burkholderia | GCF_027257875.1 | Burkholderia_sp._IMCC1007 | O1Q03_RS33565 | 11993..15454 | 3004104 | BREX-system-P-loop-protein-BrxC-[Burkholderia-sp.-IMCC1007] |
| WP_186109347.1 | Type1.BREX | RFC-AAA+DNAA-HTH->?->MORC+DNA_gyrase+ TOPRIM -><- SIGMA+OB-Fold+AAA-ATPase+Lon-Protease-BrxL <- PglZ <- SF1-SF2-Helicase<- OLD-ABC + TOPRIM <- N6-METHYLASE <- DUF6079 *<- BrxB-BrxF-iSTAND <- BrxA-DNA-binding <-RHH_5\|\|<-?\|\|DUF4096+DDE_Tnp_1+DDE_Tnp_1_2-> | 1153 | Bacteria>Pseudomonadota>Betaproteobacteria>Burkholderiales>Burkholderiaceae>Burkholderia | GCF_902831275.1 | Burkholderia_gladioli | H9D39_RS35055 | 68649..72110 | 28095 | BREX-system-P-loop-protein-BrxC-[Burkholderia-gladioli] |
| WP_186089005.1 | Type1.BREX | RFC-AAA+DNAA-HTH->?->MORC+DNA_gyrase+ TOPRIM -><- SIGMA+OB-Fold+AAA-ATPase+Lon-Protease-BrxL <- PglZ <- SF1-SF2-Helicase<- OLD-ABC + TOPRIM <- N6-METHYLASE <- DUF6079 *<- BrxB-BrxF-iSTAND <- BrxA-DNA-binding <-RHH_5\|\|?->?->?->?->ABhydrolase-> | 1153 | Bacteria>Pseudomonadota>Betaproteobacteria>Burkholderiales>Burkholderiaceae>Burkholderia | GCF_902831595.1 | Burkholderia_gladioli | H9D22_RS09670 | 586813..590274 | 28095 | BREX-system-P-loop-protein-BrxC-[Burkholderia-gladioli] |
| WP_186146941.1 | Type1.BREX | REC+LuxR-HTH-><-ABhydrolase<-?<-?<-?<-?\|\|RHH_5-> BrxA-DNA-binding -> BrxB-BrxF-iSTAND -> DUF6079 *-> N6-METHYLASE -> OLD-ABC + TOPRIM ->SF1-SF2-Helicase -> PglZ -> SIGMA+OB-Fold+AAA-ATPase+Lon-Protease-BrxL -><-MORC+DNA_gyrase+ TOPRIM <-?<-RFC-AAA+DNAA-HTH | 1153 | Bacteria>Pseudomonadota>Betaproteobacteria>Burkholderiales>Burkholderiaceae>Burkholderia | GCF_902831935.1 | Burkholderia_gladioli | H9E05_RS08890 | 201279..204740 | 28095 | BREX-system-P-loop-protein-BrxC-[Burkholderia-gladioli] |
| WP_186177578.1 | Type1.BREX | RFC-AAA+DNAA-HTH->?->MORC+DNA_gyrase+ TOPRIM -><- SIGMA+OB-Fold+AAA-ATPase+Lon-Protease-BrxL <- PglZ <- SF1-SF2-Helicase<- OLD-ABC + TOPRIM <- N6-METHYLASE <- DUF6079 *<- BrxB-BrxF-iSTAND <- BrxA-DNA-binding <-RHH_5\|\|?->?->?->?->ABhydrolase-><-REC+LuxR-HTH | 1153 | Bacteria>Pseudomonadota>Betaproteobacteria>Burkholderiales>Burkholderiaceae>Burkholderia | GCF_902832085.1 | Burkholderia_gladioli | H9E56_RS05565 | 447000..450461 | 28095 | BREX-system-P-loop-protein-BrxC-[Burkholderia-gladioli] |
| WP_087048072.1 | Type1.BREX | <- SIGMA+OB-Fold+AAA-ATPase+Lon-Protease-BrxL <- PglZ <- SF1-SF2-Helicase<- OLD-ABC + TOPRIM <- N6-METHYLASE <- DUF6079 *<- BrxB-BrxF-iSTAND <- BrxA-DNA-binding <-RHH_5<-?\|\|?-><-?\|\|?-><-SF1-SF2-Helicase | 1153 | Bacteria>Pseudomonadota>Betaproteobacteria>Burkholderiales>Burkholderiaceae>Caballeronia | GCF_001545075.1 | Caballeronia_pteroechthonis | AWB83_04706 | 123490..126951 | 1777144 | BREX-system-P-loop-protein-BrxC-[Caballeronia-pteroechthonis] |
| WP_250522809.1 | Type1.BREX | <-HTH+MPTase<-?<- SIGMA+OB-Fold+AAA-ATPase+Lon-Protease-BrxL <- PglZ <- SF1-SF2-Helicase<- OLD-ABC + TOPRIM <- N6-METHYLASE <- DUF6079 *<- BrxB-BrxF-iSTAND <- BrxA-DNA-binding <-RHH_5 | 1153 | Bacteria>Pseudomonadota>Betaproteobacteria>Burkholderiales>Burkholderiaceae>Caballeronia | GCF_023631205.1 | unclassified_Caballeronia | NCS65_RS00835 | 183401..186862 | 2646786 | MULTISPECIES:-BREX-system-P-loop-protein-BrxC-[unclassified-Caballeronia] |
| WP_086927320.1 | Type1.BREX | <- SIGMA+OB-Fold+AAA-ATPase+Lon-Protease-BrxL <- PglZ <-ParB+HNH<- SF1-SF2-Helicase<- OLD-ABC + TOPRIM <- N6-METHYLASE <-HEPN<- DUF6079 *<- BrxB-BrxF-iSTAND <- BrxA-DNA-binding <-RHH_5<-?<-XIS-HTH<-tRNA<-tRNA<-35exo<-?<-?<-DnaJ | 1153 | Bacteria>Pseudomonadota>Betaproteobacteria>Burkholderiales>Comamonadaceae>Acidovorax | GCF_002157125.1 | Acidovorax_carolinensis | CBP36_RS10055 | 2162910..2166371 | 553814 | BREX-system-P-loop-protein-BrxC-[Acidovorax-carolinensis] |
| WP_204247869.1 | Type1.BREX | DnaJ->?->?->tRNA->tRNA->XIS-HTH->?->RHH_5-> BrxA-DNA-binding -> BrxB-BrxF-iSTAND -> DUF6079 -p*-> DUF6079 -p->HEPN-> N6-METHYLASE -> OLD-ABC + TOPRIM ->SF1-SF2-Helicase ->ParB+HNH-> PglZ -> SIGMA+OB-Fold+AAA-ATPase+Lon-Protease-BrxL -> | 590 | Bacteria>Pseudomonadota>Betaproteobacteria>Burkholderiales>Comamonadaceae>Acidovorax | GCF_002157185.1 | Acidovorax_carolinensis | CBP35_RS21920 | 1914645..1916417 | 553814 | BREX-system-P-loop-protein-BrxC-[Acidovorax-carolinensis] |
| WP_101103324.1 | Type1.BREX | HTH_Tnp_1+HTH_21+rve+rve_3->RHH_5-> BrxA-DNA-binding -> BrxB-BrxF-iSTAND -> DUF6079 *->HEPN-> N6-METHYLASE -> OLD-ABC + TOPRIM ->SF1-SF2-Helicase -> OLD-ABC ->HNH->ParB+HNH-> PglZ -> SIGMA+OB-Fold+AAA-ATPase+Lon-Protease-BrxL -> | 1153 | Bacteria>Pseudomonadota>Betaproteobacteria>Burkholderiales>Comamonadaceae>Macromonas | GCF_002837135.1 | Macromonas_bipunctata | CON76_RS10585 | 3906..7367 | 183670 | BREX-system-P-loop-protein-BrxC-[Macromonas-bipunctata] |
| WP_239902638.1 | Type1.BREX | ABC-ATPase+ABC-ATPase+ABC-ATPase+SbcC->ABC-ATPase+SbcC-><-?\|\|RHH_5-> BrxA-DNA-binding -> BrxB-BrxF-iSTAND -> DUF6079 *-> N6-METHYLASE -> OLD-ABC + TOPRIM ->SF1-SF2-Helicase -> PglZ -> SIGMA+OB-Fold+AAA-ATPase+Lon-Protease-BrxL ->HTH_19-><-?<-Phage_integrase+Phage_integrase | 1153 | Bacteria>Pseudomonadota>Betaproteobacteria>Neisseriales>Chromobacteriaceae>Laribacter | GCF_022394475.1 | Laribacter_hongkongensis | LH449_RS00385 | 78525..81986 | 168471 | BREX-system-P-loop-protein-BrxC-[Laribacter-hongkongensis] |
| MBT9520412.1 | Type1.BREX | <- SF1-SF2-Helicase<- OLD-ABC + TOPRIM <-MORC<- SIGMA+OB-Fold+AAA-ATPase+Lon-Protease-BrxL <- PglZ <-MarR-HTH<- N6-METHYLASE <-AbiEii<-AbiEi_4<- DUF6079 *<- BrxB-BrxF-iSTAND <- BrxA-DNA-binding <- wHTH+WYL+WCX <-REase<-Phage_integrase<-XIS-HTH<-HTH_3 | 1208 | Bacteria>Pseudomonadota>Betaproteobacteria>Rhodocyclales>Azonexaceae>Dechloromonas | GCA_018780195.1 | Dechloromonas_sp. | IV101_05910 | 54359..57985 | 1917218 | BREX-system-P-loop-protein-BrxC-[Dechloromonas-sp.] |
| WP_205230204.1 | Type1.BREX | SbcC->?-><- SIGMA+OB-Fold+AAA-ATPase+Lon-Protease-BrxL <- PglZ <- SF1-SF2-Helicase<- OLD-ABC + TOPRIM <- N6-METHYLASE <-REase<- OLD-ABC <- DUF6079 *<- BrxB-BrxF-iSTAND <- BrxA-DNA-binding <-RHH_5<-HTH_37\|\|Tn7_Tnp_TnsA_N+Tn7_Tnp_TnsA_C+rve+Mu-transpos_C->TnsC-ATPase-> | 1153 | Bacteria>Pseudomonadota>Betaproteobacteria>Rhodocyclales>Rhodocyclaceae>Azospira | GCF_905120365.1 | Azospira_oryzae | JYJ48_RS05820 | 17001..20462 | 146939 | BREX-system-P-loop-protein-BrxC-[Azospira-oryzae] |
| WP_200158841.1 | Type1.BREX | <- SIGMA+OB-Fold+AAA-ATPase+Lon-Protease-BrxL <- PglZ <-HNH<- OLD-ABC <- SF1-SF2-Helicase<- OLD-ABC + TOPRIM <- N6-METHYLASE <-PIN<- OLD-ABC <- DUF6079 *<- BrxB-BrxF-iSTAND <- BrxA-DNA-binding <-GDResolvase+HIN-HTH<-ZnR | 1154 | Bacteria>Pseudomonadota>Gammaproteobacteria>Chromatiales>Chromatiaceae>Allochromatium | GCF_016584135.1 | Allochromatium_vinosum | CK029_RS16480 | 18892..22356 | 1049 | BREX-system-P-loop-protein-BrxC-[Allochromatium-vinosum] |
| WP_200346460.1 | Type1.BREX | CR-ATPase5+REase->MazE_antitoxin->PemK_toxin->VapB_antitoxin-> BrxA-DNA-binding -> BrxB-BrxF-iSTAND -> DUF6079 *->RHH_6-> N6-METHYLASE -> OLD-ABC + TOPRIM ->SF1-SF2-Helicase ->Schlafen+HATPase+wHTH->MerR-HTH+GDResolvase->HTH_OrfB_IS605+OrfB_IS605+OrfB_Zn_ribbon->HTH_3->PhdYeFM_antitox-> | 1152 | Bacteria>Pseudomonadota>Gammaproteobacteria>Chromatiales>Chromatiaceae>Halochromatium | GCF_016584085.1 | Halochromatium_glycolicum | CK040_RS12030 | 27931..31389 | 85075 | BREX-system-P-loop-protein-BrxC-[Halochromatium-glycolicum] |
| WP_201220083.1 | Type1.BREX | <- OLD-ABC <- SF1-SF2-Helicase<- OLD-ABC + TOPRIM <- N6-METHYLASE <-VapB_antitoxin<-PemK_toxin<-MazE_antitoxin<-PemK_toxin<-RHH<- DUF6079 *<- BrxB-BrxF-iSTAND <- BrxA-DNA-binding <-?<-?<-?\|\|ABhydrolase->MlaE->ABC-ATPase+SbcC-> | 1154 | Bacteria>Pseudomonadota>Gammaproteobacteria>Chromatiales>Chromatiaceae>Halochromatium | GCF_016653315.1 | Halochromatium_roseum | CCR96_RS10495 | 53560..57024 | 391920 | BREX-system-P-loop-protein-BrxC-[Halochromatium-roseum] |
| WP_062271501.1 | Type1.BREX | <- OLD-ABC <-RHH<- SF1-SF2-Helicase<- OLD-ABC + TOPRIM <- N6-METHYLASE <-PIN<- OLD-ABC <-PIN<-?<- DUF6079 *<- BrxB-BrxF-iSTAND <- BrxA-DNA-binding <-tRNA<-Aminotran_1_2\|\|SF1-SF2-Helicase-> | 1153 | Bacteria>Pseudomonadota>Gammaproteobacteria>Chromatiales>Chromatiaceae>Marichromatium | GCF_001583505.1 | Marichromatium_gracile | AY586_RS00820 | 23071..26532 | 1048 | BREX-system-P-loop-protein-BrxC-[Marichromatium-gracile] |
| KAB0815149.1 | Type1.BREX | <-wHTH+HNH\|\|tRNA->Arm-DNA-bind_3+Phage_integrase-><-SF1-SF2-Helicase\|\| OLD-ABC + TOPRIM ->SF1-SF2-Helicase -><- SIGMA+OB-Fold+AAA-ATPase+Lon-Protease-BrxL <- PglZ <- N6-METHYLASE <-ParB+HNH<- DUF6079 *<- BrxB-BrxF-iSTAND <- BrxA-DNA-binding \|\|TIR-><-?<-XIS-HTH\|\|?->nSTAND1+GTPase-AIG+sGTP->wHTH+WYL-> | 1233 | Bacteria>Pseudomonadota>Gammaproteobacteria>Enterobacterales>Enterobacteriaceae>Cronobacter | GCA_008805175.1 | Cronobacter_sakazakii | FZI24_17905 | 37972..41673 | 28141 | BREX-system-P-loop-protein-BrxC-[Cronobacter-sakazakii] |
| WP_239984508.1 | Type1.BREX | <-wHTH+WYL<-GTPase-AIG<-?\|\|XIS-HTH->?-><-TIR\|\| BrxA-DNA-binding -> BrxB-BrxF-iSTAND -> DUF6079 *->ParB+HNH-> N6-METHYLASE -> PglZ -> SIGMA+OB-Fold+AAA-ATPase+Lon-Protease-BrxL -><- SF1-SF2-Helicase<- OLD-ABC + TOPRIM \|\|SF1-SF2-Helicase-><-?<-HIN-HTH | 1240 | Bacteria>Pseudomonadota>Gammaproteobacteria>Enterobacterales>Enterobacteriaceae>Cronobacter | GCF_002942265.1 | Cronobacter_sakazakii | C3D67_RS10955 | 271432..275154 | 28141 | BREX-system-P-loop-protein-BrxC-[Cronobacter-sakazakii] |
| WP_182062523.1 | Type1.BREX | OLD-ABC + TOPRIM ->SF1-SF2-Helicase -><-?<-?<-?<-wHTH+WYL<-nSTAND1+GTPase-AIG+sGTP\|\| BrxA-DNA-binding -> BrxB-BrxF-iSTAND -> DUF6079 *-> N6-METHYLASE -> OLD-ABC -> TOPRIM -> PglZ -> SIGMA+OB-Fold+AAA-ATPase+Lon-Protease-BrxL -><-?<-AAA+wHTH<-XIS-HTH<-Phage_Cox | 1202 | Bacteria>Pseudomonadota>Gammaproteobacteria>Enterobacterales>Enterobacteriaceae>Enterobacter | GCF_013927005.1 | Enterobacter_sp._RHBSTW-00175 | HV107_RS06435 | 1368672..1372280 | 2742639 | BREX-system-P-loop-protein-BrxC-[Enterobacter-sp.-RHBSTW-00175] |
| MCG7871843.1 | Type1.BREX | Phage_integrase->?->?<- SF1-SF2-Helicase<- OLD-ABC + TOPRIM <- SIGMA+OB-Fold+AAA-ATPase+HKD-DNase-BrxL <- PglZ <-REase<- N6-METHYLASE <- DUF6079 *<- BrxB-BrxF-iSTAND <- BrxA-DNA-binding <-HTH_26\|\|?-><-Phage_integrase\|\|?-><-?<-?<-ACT | 1174 | Bacteria>Pseudomonadota>Gammaproteobacteria>Gammaproteobacteria_incertae_sedis>sulfur-oxidizing_symbionts>Candidatus_Thiodiazotropha | GCA_016848925.1 | Candidatus_Thiodiazotropha_lotti | JAZ11_07035 | 23024..26548 | 2792787 | BREX-system-P-loop-protein-BrxC-[Candidatus-Thiodiazotropha-lotti] |
| WP_159177832.1 | Type1.BREX | <- SF1-SF2-Helicase<- OLD-ABC + TOPRIM <- SIGMA+OB-Fold+AAA-ATPase+HKD-DNase-BrxL <- PglZ <- N6-METHYLASE <-SLOG+BirA-HTH<-SF1-SF2-Helicase+PRTase<-Schlafen+HATPase+wHTH+HTH<- DUF6079 *<- BrxB-BrxF-iSTAND <- BrxA-DNA-binding <-XIS-HTH<-MobI<-?\|\|XIS-HTH->AAA+HTH->XIS-HTH-><-Arm-DNA-bind_3+Phage_integrase | 1193 | Bacteria>Pseudomonadota>Gammaproteobacteria>Oceanospirillales>Halomonadaceae>Halomonas | GCF_902506335.1 | unclassified_Halomonas | JMX33_RS24540 | 5309589..5313170 | 2609666 | MULTISPECIES:-BREX-system-P-loop-protein-BrxC-[unclassified-Halomonas] |
| ERV97546.1 | Type1.BREX | LZ_Tnp_IS66+zf-IS66+DDE_Tnp_IS66+DDE_Tnp_IS66_C->?->DDE_Tnp_1-><-?<- N6-METHYLASE <-DOC+HTH<-?<-?<- DUF6079 *<- BrxB-BrxF-iSTAND <- BrxA-DNA-binding <- wHTH+WYL+WCX <-?\|\| OLD-ABC + TOPRIM ->SF1-SF2-Helicase ->SF1-SF2-Helicase->?-><-Phage_integrase | 1223 | Bacteria>Pseudomonadota>Gammaproteobacteria>Pseudomonadales>Pseudomonadaceae>Pseudomonas | GCA_000481225.1 | Pseudomonas_aeruginosa_BWHPSA024 | Q037_05156 | 13246..16917 | 1402524 | hypothetical-protein-Q037_05156-[Pseudomonas-aeruginosa-BWHPSA024] |
| MBM2633269.1 | Type1.BREX | <-ABC-ATPase+SbcC<-?<-?<-MODE-HTH+PBPII\|\|<-?<- N6-METHYLASE <-DOC+HTH<-?<-?<- DUF6079 *<- BrxB-BrxF-iSTAND <- BrxA-DNA-binding <- wHTH+WYL+WCX \|\| OLD-ABC + TOPRIM ->SF1-SF2-Helicase ->?->SF1-SF2-Helicase-><-Phage_integrase | 1223 | Bacteria>Pseudomonadota>Gammaproteobacteria>Pseudomonadales>Pseudomonadaceae>Pseudomonas | GCA_016836445.1 | Pseudomonas_aeruginosa | JQ480_07705 | 1618759..1622430 | 287 | BREX-system-P-loop-protein-BrxC-[Pseudomonas-aeruginosa] |
| WP_007159175.1 | Type6.BREX | PIN+OST-HTH->?-><- SIGMA+OB-Fold+AAA-ATPase+HKD-DNase-BrxL <- PglZ <-ParB+HNH<- N6-METHYLASE <- N6-METHYLASE <- DUF6079 *<- BrxB-BrxF-iSTAND <- BrxA-DNA-binding <- BrxE <- SF1-SF2-Helicase<- OLD-ABC + TOPRIM | 1212 | Bacteria>Pseudomonadota>Gammaproteobacteria>Pseudomonadales>Pseudomonadaceae>Pseudomonas | GCF_000236825.1 | Pseudomonas_psychrotolerans | PPL19_RS03910 | 152166..155804 | 237610 | BREX-system-P-loop-protein-BrxC-[Pseudomonas-psychrotolerans] |
| WP_228382933.1 | Type1.BREX | DDE_Tnp_IS66+DDE_Tnp_IS66_C-><-MODE-HTH+PBPII\|\|?->LZ_Tnp_IS66+zf-IS66+DDE_Tnp_IS66+DDE_Tnp_IS66_C-><-?<- N6-METHYLASE <-DOC+HTH<-?<-?<- DUF6079 *<- BrxB-BrxF-iSTAND <- BrxA-DNA-binding <- wHTH+WYL+WCX \|\| OLD-ABC + TOPRIM ->SF1-SF2-Helicase ->?->SF1-SF2-Helicase-><-Phage_integrase | 1225 | Bacteria>Pseudomonadota>Gammaproteobacteria>Pseudomonadales>Pseudomonadaceae>Pseudomonas | GCF_000481225.1 | Pseudomonas_aeruginosa | Q037_RS32345 | 325963..329640 | 287 | BREX-system-P-loop-protein-BrxC-[Pseudomonas-aeruginosa] |
| WP_239000709.1 | Type1.BREX | <-ABC-ATPase+SbcC<-?<-?<-MODE-HTH+PBPII\|\|<-?<- N6-METHYLASE <-DOC+HTH<-?<-?<- DUF6079 *<- BrxB-BrxF-iSTAND <- BrxA-DNA-binding <- wHTH+WYL+WCX \|\| OLD-ABC + TOPRIM ->SF1-SF2-Helicase ->?->SF1-SF2-Helicase-><-Phage_integrase | 1225 | Bacteria>Pseudomonadota>Gammaproteobacteria>Pseudomonadales>Pseudomonadaceae>Pseudomonas | GCF_016836415.1 | Pseudomonas_aeruginosa | JQ482_RS07395 | 1563823..1567500 | 287 | BREX-system-P-loop-protein-BrxC-[Pseudomonas-aeruginosa] |
| WP_176605727.1 | Type1.BREX | <-MODE-HTH+PBPII\|\|<-?<- N6-METHYLASE <-DOC+HTH<-?<-?<- DUF6079 *<- BrxB-BrxF-iSTAND <- BrxA-DNA-binding <- wHTH+WYL+WCX \|\| OLD-ABC + TOPRIM ->SF1-SF2-Helicase ->?->SF1-SF2-Helicase-><-Phage_integrase | 1225 | Bacteria>Pseudomonadota>Gammaproteobacteria>Pseudomonadales>Pseudomonadaceae>Pseudomonas | GCF_900708415.1 | Pseudomonas_aeruginosa | BUG32_RS23730 | 10964..14641 | 287 | BREX-system-P-loop-protein-BrxC-[Pseudomonas-aeruginosa] |
| EIU3152871.1 | Type1.BREX | Phage_integrase->DDE_Tnp_1-><-SF1-SF2-Helicase<-?<- SF1-SF2-Helicase<- OLD-ABC + TOPRIM \|\| wHTH+WYL+WCX -> BrxA-DNA-binding -> BrxB-BrxF-iSTAND -> DUF6079 *->?->?->DOC+HTH-> N6-METHYLASE -p-> | 1223 | Bacteria>Pseudomonadota>Gammaproteobacteria>Pseudomonadales>Pseudomonadaceae>Pseudomonas | - | Pseudomonas_aeruginosa | L4P27_006182 | 20932..24603 | 287 | BREX-system-P-loop-protein-BrxC-[Pseudomonas-aeruginosa] |
| WP_023124092.1 | Type1.BREX | Phage_integrase->DDE_Tnp_1-><-SF1-SF2-Helicase<-?<- SF1-SF2-Helicase<- OLD-ABC + TOPRIM \|\| wHTH+WYL+WCX -> BrxA-DNA-binding -> BrxB-BrxF-iSTAND -> DUF6079 *->?->?->DOC+HTH-> N6-METHYLASE -p-> | 1223 | Bacteria>Pseudomonadota>Gammaproteobacteria>Pseudomonadales>Pseudomonadaceae>Pseudomonas | - | Pseudomonas_aeruginosa | L4P27_006182 | 20932..24603 | 287 | BREX-system-P-loop-protein-BrxC-[Pseudomonas-aeruginosa] |
| WP_207395558.1 | Type1.BREX | BrxB-BrxF-iSTAND ->N6-N4-METHYLASE->Peptidase->?->?-><- SIGMA+OB-Fold+AAA-ATPase+Lon-Protease-BrxL <- PglZ <- N6-METHYLASE <- DUF6079 *<- BrxB-BrxF-iSTAND <- BrxA-DNA-binding \|\|HTH-> OLD-ABC + TOPRIM ->SF1-SF2-Helicase ->ArdCN+MPTase->?->?->RFC-AAA-> | 1152 | Bacteria>PVC_group>Planctomycetota>Planctomycetia>Pirellulales>Pirellulaceae | GCF_013618625.1 | Bremerella_alba | HOV93_RS05900 | 45922..49380 | 980252 | BREX-system-P-loop-protein-BrxC-[Bremerella-alba] |
| WP_184301662.1 | Type1.BREX | <-nSTAND1+HNH<-URI-DNase+RAMA<- SIGMA+OB-Fold+AAA-ATPase+HKD-DNase-BrxL <- PglZ <-REase+TPR+GreAB-C+PIN<- N6-METHYLASE <-?<-ABC-ATPase<- DUF6079 *<- BrxB-BrxF-iSTAND <- BrxA-DNA-binding <-XIS-HTH\|\| OLD-ABC + TOPRIM ->SF1-SF2-Helicase -> | 1169 | Bacteria>PVC_group>Planctomycetota>Planctomycetia>Pirellulales>Pirellulaceae | GCF_014192335.1 | Rhodopirellula_rubra | FHS27_RS03210 | 253910..257419 | 980271 | BREX-system-P-loop-protein-BrxC-[Rhodopirellula-rubra] |
| WP_040162373.1 | Type1.BREX | 35exo->tRNA-><-GDResolvase+Recombinase+ZnR<-?\|\|HIN-HTH->HTH_Tnp_1+HTH_21+rve+rve_3->XIS-HTH-> BrxA-DNA-binding -> BrxB-BrxF-iSTAND -> DUF6079 *-> N6-METHYLASE ->?-> PglZ -> SIGMA+OB-Fold+AAA-ATPase+HKD-DNase-BrxL -> OLD-ABC + TOPRIM ->SF1-SF2-Helicase ->GTPase-AIG->MULE-TRANSPOSASE-> | 1178 | Bacteria>Terrabacteria_group>Actinomycetota>Actinomycetes>Micrococcales>Dermatophilaceae | GCF_902375455.1 | Mobilicoccus_massiliensis | FX004_RS16815 | 289663..293199 | 1522310 | BREX-system-P-loop-protein-BrxC-[Mobilicoccus-massiliensis] |
| WP_068111667.1 | Type1.BREX | VirB11+MuB-ATPase->?-><-?\|\|XIS-HTH-> BrxA-DNA-binding -> BrxB-BrxF-iSTAND -> DUF6079 *-> N6-METHYLASE ->REase-> PglZ -> SIGMA+OB-Fold+AAA-ATPase+HKD-DNase-BrxL -> OLD-ABC + TOPRIM ->SF1-SF2-Helicase ->?->HTH+MPTase->Phage_int_SAM_1+Phage_integrase-> | 1154 | Bacteria>Terrabacteria_group>Actinomycetota>Actinomycetes>Propionibacteriales>Nocardioidaceae | GCF_001653335.1 | Nocardioides_dokdonensis | I601_RS15600 | 3293021..3296485 | 450734 | BREX-system-P-loop-protein-BrxC-[Nocardioides-dokdonensis] |
| MBK7820961.1 | Type1.BREX | <-DUF2637+HIN-HTH<-ArdCN+MPTase\|\|HTH->MPTase->XIS-HTH-> BrxA-DNA-binding -> BrxB-BrxF-iSTAND -> DUF6079 *-> N6-METHYLASE ->HTH+REase-> PglZ -> SIGMA+OB-Fold+AAA-ATPase+HKD-DNase-BrxL -> OLD-ABC + TOPRIM ->SF1-SF2-Helicase ->?->DDE_Tnp_1-><-DNA_METHYLASE | 1176 | Bacteria>Terrabacteria_group>Actinomycetota>Actinomycetes>Propionibacteriales>Propionibacteriaceae | GCA_016714275.1 | Tessaracoccus_sp. | IPJ61_07760 | 1693709..1697239 | 1971211 | BREX-system-P-loop-protein-BrxC-[Tessaracoccus-sp.] |
| WP_182799522.1 | Type1.BREX | <-DOC+HTH<-RelE-ParE<-PhdYeFM_antitox<-Calcineurin<-ABhydrolase\|\|XIS-HTH-> BrxA-DNA-binding -> BrxB-BrxF-iSTAND -> DUF6079 *-> N6-METHYLASE ->?-> PglZ -> SIGMA+OB-Fold+AAA-ATPase+HKD-DNase-BrxL -> OLD-ABC + TOPRIM ->SF1-SF2-Helicase ->GTPase-AIG->sGTP->SF1-SF2-Helicase-> | 1173 | Bacteria>Terrabacteria_group>Actinomycetota>Actinomycetes>Propionibacteriales>Propionibacteriaceae | GCF_014164775.1 | Tessaracoccus_sp._MC1679 | H5398_RS07140 | 376087..379608 | 2760313 | BREX-system-P-loop-protein-BrxC-[Tessaracoccus-sp.-MC1679] |
| WP_204918880.1 | Type1.BREX | <-sGTP<-GTPase-AIG<- SF1-SF2-Helicase<- OLD-ABC + TOPRIM <- SIGMA+OB-Fold+AAA-ATPase+HKD-DNase-BrxL <- PglZ <-?<- N6-METHYLASE <- DUF6079 *<- BrxB-BrxF-iSTAND <- BrxA-DNA-binding <-XIS-HTH<-ABhydrolase\|\|Recombinase-><-tRNA<-35exo<-?\|\|tRNA-><-MarR-HTH | 1173 | Bacteria>Terrabacteria_group>Actinomycetota>Actinomycetes>Propionibacteriales>Propionibacteriaceae | GCF_016907535.1 | Microlunatus_panaciterrae | JOE57_RS13940 | 3049076..3052597 | 400768 | BREX-system-P-loop-protein-BrxC-[Microlunatus-panaciterrae] |
| MBM6989329.1 | Type1.BREX | <-TraI-VirD2-RCRE<-TraJ-RHH\|\|TraJ-RHH-> BrxA-DNA-binding -> BrxB-BrxF-iSTAND -> DUF6079 -p-> DUF6079 *-> N6-METHYLASE -> PglZ -> SIGMA+OB-Fold+AAA-ATPase+Lon-Protease-BrxL -> OLD-ABC + TOPRIM ->SF1-SF2-Helicase -><-REase<-?\|\|?-><-HEPN | 1040 | Bacteria>Terrabacteria_group>Actinomycetota>Coriobacteriia>Coriobacteriales>Atopobiaceae | GCA_016901295.1 | Parafannyhessea_umbonata | I3I99_06995 | 31435..34557 | 604330 | BREX-system-P-loop-protein-BrxC-[Parafannyhessea-umbonata] |
| WP_259420301.1 | Type1.BREX | <-Phage_CRI+Phage_X<-ParB+HNH<- SIGMA+OB-Fold+AAA-ATPase+Lon-Protease-BrxL <- PglZ <-?<- SF1-SF2-Helicase<- OLD-ABC + TOPRIM <-SbcC<- N6-METHYLASE <- DUF6079 *<- BrxB-BrxF-iSTAND <- BrxA-DNA-binding <-?\|\|XIS-HTH->Arm-DNA-bind_4+Phage_int_SAM_3+Phage_integrase-><-?<-?\|\|CR-Kinase-><-GDResolvase+HIN-HTH | 1191 | Bacteria>Terrabacteria_group>Bacillota>Bacilli>Bacillales>Bacillaceae | GCF_024919535.1 | Lysinibacillus_sp._A4 | NY607_RS20580 | 136984..140559 | 2976269 | BREX-system-P-loop-protein-BrxC-[Lysinibacillus-sp.-A4] |
| WP_068559440.1 | Type1.BREX | <-OrfB_IS605+OrfB_Zn_ribbon<-SF1-SF2-Helicase+REase<-SF1-SF2-Helicase+REase<- SF1-SF2-Helicase<- OLD-ABC + TOPRIM <- SIGMA+OB-Fold+AAA-ATPase+Lon-Protease-BrxL <- PglZ <-REase<- N6-METHYLASE <- DUF6079 *<- BrxB-BrxF-iSTAND <- BrxA-DNA-binding <-HTH_26<-?<-?<-?<-?<-REase\|\|DDE_Tnp_IS240+rve_3-> | 1207 | Bacteria>Terrabacteria_group>Bacillota>Bacilli>Lactobacillales>Carnobacteriaceae | GCF_900067165.1 | Trichococcus | BUA33_RS02210 | 213489..217112 | 82802 | MULTISPECIES:-BREX-system-P-loop-protein-BrxC-[Trichococcus] |
| WP_118650255.1 | Type1.BREX | TPR+TPR+TPR->?->HISKIN->SIGMA-HTH+SIGMA-HTH->HTH_26-> BrxA-DNA-binding -> BrxB-BrxF-iSTAND -> DUF6079 *-> N6-METHYLASE ->AT-HTH-HYD2->Tox-ART-HYD2->AT-HTH-HYD2-> PglZ -> SIGMA+OB-Fold+AAA-ATPase+Lon-Protease-BrxL -> OLD-ABC + TOPRIM ->SF1-SF2-Helicase -> | 1182 | Bacteria>Terrabacteria_group>Bacillota>Clostridia>Eubacteriales>Clostridiaceae | GCF_003482245.1 | Clostridium_sp._OF09-10 | DXA91_RS01310 | 45088..48636 | 2292309 | BREX-system-P-loop-protein-BrxC-[Clostridium-sp.-OF09-10] |
| MCI5510134.1 | Type1.BREX | CopG/RHH->?->TraJ-RHH->TraI-VirD2-RCRE->?->HTH_26-> BrxA-DNA-binding -> BrxB-BrxF-iSTAND -> DUF6079 *->?-> N6-METHYLASE ->HEPN->Schlafen+HATPase+wHTH-> PglZ -> SIGMA+OB-Fold+AAA-ATPase+Lon-Protease-BrxL -> OLD-ABC + TOPRIM ->SF1-SF2-Helicase -><-HTH | 1182 | Bacteria>Terrabacteria_group>Bacillota>Clostridia>Eubacteriales>Eubacteriaceae | GCA_022764165.1 | Eubacterium_sp. | MR436_06790 | 9461..13009 | 142586 | BREX-system-P-loop-protein-BrxC-[Eubacterium-sp.] |
| WP_055219000.1 | Type1.BREX | ZnR->?->HTH_26-> BrxA-DNA-binding -> BrxB-BrxF-iSTAND -> DUF6079 *-> N6-METHYLASE ->AbiGi+seqA+RAMA->Schlafen+HATPase+wHTH-> PglZ -> SIGMA+OB-Fold+AAA-ATPase+Lon-Protease-BrxL -> OLD-ABC + TOPRIM ->SF1-SF2-Helicase ->Phage_integrase-> | 1182 | Bacteria>Terrabacteria_group>Bacillota>Clostridia>Eubacteriales | GCF_013300755.1 | Lachnospiraceae | G5A93_RS10915 | 4301..7849 | 186803 | MULTISPECIES:-BREX-system-P-loop-protein-BrxC-[Lachnospiraceae] |
| WP_138373965.1 | Type1.BREX | <- SF1-SF2-Helicase<- OLD-ABC + TOPRIM <- SIGMA+OB-Fold+AAA-ATPase+Lon-Protease-BrxL <- PglZ <-Schlafen+HATPase+wHTH<-DUF4404<-ZnR+ZnR<- N6-METHYLASE <-?<- DUF6079 *<- BrxB-BrxF-iSTAND <- BrxA-DNA-binding <-HTH_26<-ParB+HNH | 1179 | Bacteria>Terrabacteria_group>Bacillota>Clostridia>Eubacteriales | GCF_015554965.1 | Oscillospiraceae | I2E95_RS16325 | 14974..18513 | 216572 | MULTISPECIES:-BREX-system-P-loop-protein-BrxC-[Oscillospiraceae] |
| WP_256171760.1 | Type1.BREX | HTH_36->?->CopG/RHH->?->TraJ-RHH->TraI-VirD2-RCRE->?->HTH_26-> BrxA-DNA-binding -> BrxB-BrxF-iSTAND -> DUF6079 *->?-> N6-METHYLASE ->HEPN->Schlafen+HATPase+wHTH-> PglZ -> SIGMA+OB-Fold+AAA-ATPase+Lon-Protease-BrxL -> OLD-ABC + TOPRIM ->SF1-SF2-Helicase -><-HTH | 1182 | Bacteria>Terrabacteria_group>Bacillota>Clostridia>Eubacteriales>Lachnospiraceae | - | Dorea_longicatena | NE607_07100 | 21358..24906 | 88431 | BREX-system-P-loop-protein-BrxC-[Dorea-longicatena] |
| MCC2229908.1 | Type1.BREX | HTH_36->?->CopG/RHH->?->TraJ-RHH->TraI-VirD2-RCRE->?->HTH_26-> BrxA-DNA-binding -> BrxB-BrxF-iSTAND -> DUF6079 *->?-> N6-METHYLASE ->HEPN->Schlafen+HATPase+wHTH-> PglZ -> SIGMA+OB-Fold+AAA-ATPase+Lon-Protease-BrxL -> OLD-ABC + TOPRIM ->SF1-SF2-Helicase -><-HTH | 1182 | Bacteria>Terrabacteria_group>Bacillota>Clostridia>Eubacteriales>Lachnospiraceae | GCA_020687485.1 | Lachnospiraceae_bacterium_CLA-AA-H215 | LKD81_02670 | 26793..30341 | 2885348 | BREX-system-P-loop-protein-BrxC-[Lachnospiraceae-bacterium-CLA-AA-H215] |
| MCQ4892848.1 | Type1.BREX | HTH_36->?->CopG/RHH->?->TraJ-RHH->TraI-VirD2-RCRE->?->HTH_26-> BrxA-DNA-binding -> BrxB-BrxF-iSTAND -> DUF6079 *->?-> N6-METHYLASE ->HEPN->Schlafen+HATPase+wHTH-> PglZ -> SIGMA+OB-Fold+AAA-ATPase+Lon-Protease-BrxL -> OLD-ABC + TOPRIM ->SF1-SF2-Helicase -><-HTH | 1182 | Bacteria>Terrabacteria_group>Bacillota>Clostridia>Eubacteriales>Lachnospiraceae | GCA_024461355.1 | Dorea_longicatena | NE607_07100 | 21358..24906 | 88431 | BREX-system-P-loop-protein-BrxC-[Dorea-longicatena] |
| WP_002578404.1 | Type1.BREX | HTH-><- SF1-SF2-Helicase<- OLD-ABC + TOPRIM <- SIGMA+OB-Fold+AAA-ATPase+Lon-Protease-BrxL <- PglZ <-Schlafen+HATPase+wHTH<-HEPN<- N6-METHYLASE <-?<- DUF6079 *<- BrxB-BrxF-iSTAND <- BrxA-DNA-binding <-HTH_26<-HTH_36<-TraI-VirD2-RCRE<-TraJ-RHH<-CopG/RHH<-?<-HTH_36 | 1182 | Bacteria>Terrabacteria_group>Bacillota>Clostridia>Eubacteriales>Lachnospiraceae | GCF_000371725.1 | Enterocloster_bolteae | HMPREF1095_RS19895 | 1377442..1380990 | 208479 | BREX-system-P-loop-protein-BrxC-[Enterocloster-bolteae] |
| WP_026495918.1 | Type1.BREX | HTH_3->Resolvase+Recombinase->?->?->ParB->ParB->HTH_26-> BrxA-DNA-binding -> BrxB-BrxF-iSTAND -> DUF6079 *-> N6-METHYLASE ->?->METHYLASE-> PglZ -> SIGMA+OB-Fold+AAA-ATPase+Lon-Protease-BrxL -> OLD-ABC + TOPRIM ->SF1-SF2-Helicase -> | 1181 | Bacteria>Terrabacteria_group>Bacillota>Clostridia>Eubacteriales>Lachnospiraceae | GCF_000424265.1 | Butyrivibrio_sp._WCD3002 | G608_RS0110170 | 160583..164128 | 1280676 | BREX-system-P-loop-protein-BrxC-[Butyrivibrio-sp.-WCD3002] |
| WP_055268215.1 | Type1.BREX | ZnR->HTH_26-> BrxA-DNA-binding -> BrxB-BrxF-iSTAND -> DUF6079 *-> N6-METHYLASE ->AT-HTH-HYD2->Schlafen+HATPase+wHTH-> PglZ -> SIGMA+OB-Fold+AAA-ATPase+Lon-Protease-BrxL -> OLD-ABC + TOPRIM ->SF1-SF2-Helicase ->Phage_integrase-> | 1182 | Bacteria>Terrabacteria_group>Bacillota>Clostridia>Eubacteriales>Lachnospiraceae | GCF_001406335.1 | Fusicatenibacter_saccharivorans | ARA47_RS16350 | 4112..7660 | 1150298 | BREX-system-P-loop-protein-BrxC-[Fusicatenibacter-saccharivorans] |
| WP_087163144.1 | Type1.BREX | <-HTH_Tnp_1<- SF1-SF2-Helicase<- OLD-ABC + TOPRIM <- SIGMA+OB-Fold+AAA-ATPase+Lon-Protease-BrxL <- PglZ <-Schlafen+HATPase+wHTH<-SIR2<- N6-METHYLASE <-?<- DUF6079 *<- BrxB-BrxF-iSTAND <- BrxA-DNA-binding <-HTH_26\|\|HIN-HTH->?->LZ_Tnp_IS66+zf-IS66+DDE_Tnp_IS66+DDE_Tnp_IS66_C-> | 1179 | Bacteria>Terrabacteria_group>Bacillota>Clostridia>Eubacteriales>Lachnospiraceae | GCF_002160525.1 | Lachnoclostridium_sp._An169 | B5F07_RS20950 | 16171..19710 | 1965569 | BREX-system-P-loop-protein-BrxC-[Lachnoclostridium-sp.-An169] |
| WP_118601699.1 | Type1.BREX | HTH_Tnp_1->?->ZnR->?->HTH_26-> BrxA-DNA-binding -> BrxB-BrxF-iSTAND -> DUF6079 *-> N6-METHYLASE ->AT-HTH-HYD2->Schlafen+HATPase+wHTH-> PglZ -> SIGMA+OB-Fold+AAA-ATPase+Lon-Protease-BrxL -> OLD-ABC + TOPRIM ->SF1-SF2-Helicase ->Phage_integrase-> | 1182 | Bacteria>Terrabacteria_group>Bacillota>Clostridia>Eubacteriales>Lachnospiraceae | GCF_003479155.1 | Blautia_sp._AF19-1 | DWX26_RS14805 | 4595..8143 | 2292960 | BREX-system-P-loop-protein-BrxC-[Blautia-sp.-AF19-1] |
| WP_118742111.1 | Type1.BREX | ZnR->?->HTH_26-> BrxA-DNA-binding -> BrxB-BrxF-iSTAND -> DUF6079 *-> N6-METHYLASE ->RAMA->HEPN->Schlafen+HATPase+wHTH-> PglZ -> SIGMA+OB-Fold+AAA-ATPase+Lon-Protease-BrxL -> OLD-ABC + TOPRIM ->SF1-SF2-Helicase -> | 1182 | Bacteria>Terrabacteria_group>Bacillota>Clostridia>Eubacteriales>Lachnospiraceae | GCF_003480245.1 | Blautia_sp._AM42-2 | DW915_10705 | 4262..7810 | 2292976 | BREX-system-P-loop-protein-BrxC-[Blautia-sp.-AM42-2] |
| WP_173813806.1 | Type1.BREX | ZnR->?->HTH_26-> BrxA-DNA-binding -> BrxB-BrxF-iSTAND -> DUF6079 *-> N6-METHYLASE ->AbiGi+seqA+RAMA->Schlafen+HATPase+wHTH-> PglZ -> SIGMA+OB-Fold+AAA-ATPase+Lon-Protease-BrxL -> OLD-ABC + TOPRIM ->SF1-SF2-Helicase ->Phage_integrase->Phage_integrase-> | 1182 | Bacteria>Terrabacteria_group>Bacillota>Clostridia>Eubacteriales>Lachnospiraceae | GCF_013301105.1 | Fusicatenibacter_saccharivorans | G5B05_RS14750 | 4301..7849 | 1150298 | BREX-system-P-loop-protein-BrxC-[Fusicatenibacter-saccharivorans] |
| WP_238033441.1 | Type1.BREX | <-Phage_integrase<-Phage_integrase<- SF1-SF2-Helicase<- OLD-ABC + TOPRIM <- SIGMA+OB-Fold+AAA-ATPase+Lon-Protease-BrxL <- PglZ <-Schlafen+HATPase+wHTH<-AbiGi+seqA+RAMA<- N6-METHYLASE <- DUF6079 *<- BrxB-BrxF-iSTAND <- BrxA-DNA-binding <-HTH_26<-?<-ZnR | 1182 | Bacteria>Terrabacteria_group>Bacillota>Clostridia>Eubacteriales>Lachnospiraceae | GCF_022136435.1 | Fusicatenibacter_saccharivorans | LON21_RS13045 | 78196..81744 | 1150298 | BREX-system-P-loop-protein-BrxC-[Fusicatenibacter-saccharivorans] |
| WP_270422341.1 | Type1.BREX | CopG/RHH->?->TraJ-RHH->TraI-VirD2-RCRE->?->HTH_26-> BrxA-DNA-binding -> BrxB-BrxF-iSTAND -> DUF6079 *->?-> N6-METHYLASE ->HEPN->Schlafen+HATPase+wHTH-> PglZ -> SIGMA+OB-Fold+AAA-ATPase+Lon-Protease-BrxL -> OLD-ABC + TOPRIM ->SF1-SF2-Helicase -><-HTH | 1182 | Bacteria>Terrabacteria_group>Bacillota>Clostridia>Eubacteriales>Lachnospiraceae | GCF_027661365.1 | Fusicatenibacter_saccharivorans | PGQ16_RS03035 | 254369..257917 | 1150298 | BREX-system-P-loop-protein-BrxC-[Fusicatenibacter-saccharivorans] |
| WP_119247667.1 | Type1.BREX | HTH_36->?->CopG/RHH->?->TraJ-RHH->TraI-VirD2-RCRE->?->HTH_26-> BrxA-DNA-binding -> BrxB-BrxF-iSTAND -> DUF6079 *->?-> N6-METHYLASE ->HEPN->Schlafen+HATPase+wHTH-> PglZ -> SIGMA+OB-Fold+AAA-ATPase+Lon-Protease-BrxL -> OLD-ABC + TOPRIM ->SF1-SF2-Helicase -><-HTH | 1182 | Bacteria>Terrabacteria_group>Bacillota>Clostridia>Eubacteriales>Lachnospiraceae | GCF_027674485.1 | Dorea_longicatena | PGV47_RS07575 | 41898..45446 | 88431 | BREX-system-P-loop-protein-BrxC-[Dorea-longicatena] |
| MCQ4886543.1 | Type1.BREX | <-MORC+REase<-STYKinase-like<- SF1-SF2-Helicase<- OLD-ABC + TOPRIM <- SIGMA+OB-Fold+AAA-ATPase+Lon-Protease-BrxL <- PglZ <-NERD<- N6-METHYLASE <-?<- DUF6079 *<- BrxB-BrxF-iSTAND <- BrxA-DNA-binding <-XIS-HTH\|\|HKD-DNase+SF1-SF2-Helicase+HTH+HTH+SRA->?->NUDIX->DUF932-> | 1218 | Bacteria>Terrabacteria_group>Bacillota>Clostridia>Eubacteriales>Oscillospiraceae | GCA_024460915.1 | Faecalibacterium_prausnitzii | NE589_05495 | 127412..131068 | 853 | BREX-system-P-loop-protein-BrxC-[Faecalibacterium-prausnitzii] |
| WP_256266343.1 | Type1.BREX | <-MORC+REase<-STYKinase-like<- SF1-SF2-Helicase<- OLD-ABC + TOPRIM <- SIGMA+OB-Fold+AAA-ATPase+Lon-Protease-BrxL <- PglZ <-NERD<- N6-METHYLASE <-?<- DUF6079 *<- BrxB-BrxF-iSTAND <- BrxA-DNA-binding <-XIS-HTH\|\|HKD-DNase+SF1-SF2-Helicase+HTH+HTH+SRA->?->NUDIX->DUF932-> | 1218 | Bacteria>Terrabacteria_group>Bacillota>Clostridia>Eubacteriales>Oscillospiraceae | GCF_024460915.1 | Faecalibacterium_prausnitzii | NE589_RS05495 | 127412..131068 | 853 | BREX-system-P-loop-protein-BrxC-[Faecalibacterium-prausnitzii] |
| WP_005488038.1 | Type1.BREX | HTH_3->?->?->REase+HTH->?->wHTH+wHTH-> BrxA-DNA-binding -> DUF6079 *->?->?->?->?->SWIM+DUF3670+SF1-SF2-Helicase-> OLD-ABC + TOPRIM ->SF1-SF2-Helicase ->DUF499-> | 1306 | Bacteria>Terrabacteria_group>Bacillota>Clostridia>Halanaerobiales>Halanaerobiaceae | GCF_000350165.1 | Halanaerobium_saccharolyticum | HSACCH_RS04005 | 779901..783821 | 43595 | DUF6079-family-protein-[Halanaerobium-saccharolyticum] |
| QRN85655.1 | Type1.BREX | APATPase+FtsK+VirB4->?->SMS-KaiC->HTH_36->HTH_26-> BrxA-DNA-binding -> BrxB-BrxF-iSTAND -> DUF6079 *->?-> N6-METHYLASE -> PglZ -> SIGMA+OB-Fold+AAA-ATPase+Lon-Protease-BrxL ->?->?-> OLD-ABC + TOPRIM ->SF1-SF2-Helicase -><-MPTase | 1213 | Bacteria>Terrabacteria_group>Bacillota>Clostridia>unclassified_Clostridia | GCA_016887505.1 | Clostridia_bacterium | JR334_12015 | 2538815..2542456 | 2044939 | BREX-system-P-loop-protein-BrxC-[Clostridia-bacterium] |
| MCH4019936.1 | Type1.BREX | Schlafen+HATPase+wHTH->Schlafen+HATPase+wHTH->?->?->GDResolvase+HIN-HTH->HTH_26-> BrxA-DNA-binding -> BrxB-BrxF-iSTAND ->?-> DUF6079 *-> N6-METHYLASE ->Schlafen+HATPase+wHTH-> PglZ -> SIGMA+OB-Fold+AAA-ATPase+Lon-Protease-BrxL -> OLD-ABC + TOPRIM ->SF1-SF2-Helicase -> | 1223 | Bacteria>Terrabacteria_group>Bacillota>Erysipelotrichia>Erysipelotrichales>Erysipelotrichaceae | GCA_022483385.1 | Erysipelotrichaceae_bacterium | LKE61_03575 | 742879..746550 | 2049044 | BREX-system-P-loop-protein-BrxC-[Erysipelotrichaceae-bacterium] |

| Protein-ID | System | Gene-Neighbourhoods | Length | Taxonomy | Assembly-ID | Strain | Locus-Tag | Genomic-coordinates | TaxID | Fasta-defline |
|---|---|---|---|---|---|---|---|---|---|---|
| -- | | | | | | | | | | |
| -- HKD-DNase | | | | | | | | | | |
| -- | | | | | | | | | | |
| MCB1051856.1 | Type2.Pgl.like.BREX | N6-METHYLASE -p-> DUF6079 *-> PglZ -><-RelE-ParE<-HEPN\|\| BrxD-AAA -> SF1-SF2-Helicase+wHTH+Small-B-barrel+DUF1998 -> SF1-SF2-Helicase+ HKD-DNase -> | 1210 | Bacteria>Acidobacteriota>unclassified_Acidobacteriota | GCA_020439765.1 | Acidobacteriota_bacterium | KDC71_14760 | 1153..4785 | 1978231 | phage-resistance-protein-[Acidobacteriota-bacterium] |
| MCB9398403.1 | Type2.Pgl.like.BREX | SLOG+BirA-HTH->?-><- SF1-SF2-Helicase+ HKD-DNase <- SF1-SF2-Helicase+wHTH+Small-B-barrel+DUF1998 <- BrxD-AAA \|\|HEPN->RelE-ParE-><- PglZ <- DUF6079 *<- N6-METHYLASE <- PglW <-?\|\|<-?<-?\|\|?->Y1_Tnp-> | 1210 | Bacteria>Acidobacteriota>unclassified_Acidobacteriota | GCA_020637335.1 | Acidobacteriota_bacterium | H6510_11360 | 13924..17556 | 1978231 | phage-resistance-protein-[Acidobacteriota-bacterium] |
| WP_037951095.1 | Type2.Pgl.like.BREX | <-TOPRIM<-OLD-ABC\|\| N6-METHYLASE -><- SF1-SF2-Helicase+ HKD-DNase \|\| PglW -> N6-METHYLASE -><-ST-Hiskin\|\|HTH_31+DUF5753->DUF397-> DUF6079 *-> PglZ -><-CBS+HEPN<-ParB+HNH\|\| BrxD-AAA -> SF1-SF2-Helicase+wHTH+Small-B-barrel+DUF1998 ->EVE+SAD+REase-><-?\|\|REase+SF1-SF2-Helicase->REase+SF1-SF2-Helicase-> | 1320 | Bacteria>Terrabacteria_group>Actinomycetota>Actinomycetes>Kitasatosporales>Streptomycetaceae | GCF_000590515.1 | Streptomyces_sp._PRh5 | Z951_RS11140 | 96158..100120 | 1158056 | hypothetical-protein-[Streptomyces-sp.-PRh5] |
| WP_123562585.1 | Type2.Pgl.like.BREX | <-REase+SF1-SF2-Helicase\|\|REase+SF1-SF2-Helicase->ParB+HNH->CBS+HEPN-> PglW -> N6-METHYLASE ->PhdYeFM_antitox->PIN-> DUF6079 *-> PglZ -> BrxD-AAA -> SF1-SF2-Helicase+wHTH+Small-B-barrel+DUF1998 -><- SF1-SF2-Helicase+ HKD-DNase <-?\|\|<-LZ_Tnp_IS481+HTH_32+rve+rve_3\|\|N6_N4_METHYLASE-> | 1307 | Bacteria>Terrabacteria_group>Actinomycetota>Actinomycetes>Kitasatosporales>Streptomycetaceae | GCF_003751605.1 | Kitasatospora_cineracea | EDD39_RS31525 | 1175476..1179399 | 88074 | phage-resistance-protein-[Kitasatospora-cineracea] |
| WP_179084887.1 | Type2.Pgl.like.BREX | AA_permease_2->TRAM+SAM-methylase-> SF1-SF2-Helicase+ HKD-DNase -><- SF1-SF2-Helicase+wHTH+Small-B-barrel+DUF1998 <- BrxD-AAA \|\|ParB+HNH->CBS+HEPN-><- PglZ <- DUF6079 *<-?<-DUF2171\|\|HTH_31+DUF5753-><- N6-METHYLASE <- PglW \|\|DUF4011+SF1-SF2-Helicase+REase->?-><-Pkinase\|\|REase-> | 1304 | Bacteria>Terrabacteria_group>Actinomycetota>Actinomycetes>Kitasatosporales>Streptomycetaceae | GCF_013394065.1 | Streptomyces_rectiverticillatus | FGW37_RS24660 | 5638152..5642066 | 173860 | DUF6079 -family-protein-[Streptomyces-rectiverticillatus] |
| WP_150527865.1 | Type2.Pgl.like.BREX | SF1-SF2-Helicase+ HKD-DNase -> PglW -> N6-METHYLASE -><-DUF397\|\|DUF1995->DUF2171->REase+SF1-SF2-Helicase-> DUF6079 *-> PglZ ->DndB-> BrxD-AAA -> SF1-SF2-Helicase+wHTH+Small-B-barrel+DUF1998 -><-?<-TRAM+SAM-methylase<-AA_permease_2 | 1301 | Bacteria>Terrabacteria_group>Actinomycetota>Actinomycetes>Kitasatosporales>Streptomycetaceae | GCF_014656155.1 | Streptomyces_vinaceus | IE310_RS19410 | 98676..102581 | 1960 | phage-resistance-protein-[Streptomyces-vinaceus] |
| WP_217245869.1 | Type2.Pgl.like.BREX | <-ORC-CDC6<-ParA-Soj-PloopNTPase<-HEPN\|\| N6-METHYLASE -><-DUF397<-HTH_31+DUF5753\|\|ST-Hiskin-><-?<- SF1-SF2-Helicase+ HKD-DNase \|\| DUF6079 *-> PglZ -> BrxD-AAA -> SF1-SF2-Helicase+wHTH+Small-B-barrel+DUF1998 -><-?\|\|?->?-><-DUF397 | 1323 | Bacteria>Terrabacteria_group>Actinomycetota>Actinomycetes>Kitasatosporales>Streptomycetaceae | GCF_019049125.1 | Streptomyces_sp._AC602_WCS936 | KUM03_RS05570 | 51050..55021 | 2823685 | PglY-protein-[Streptomyces-sp.-AC602_WCS936] |
| WP_179779345.1 | Type2.Pgl.like.BREX | HD->?-><-?<-?\|\|?-><- SF1-SF2-Helicase+wHTH+Small-B-barrel+DUF1998 <- PglZ <- DUF6079 *<- HKD-DNase \|\|?-><-SLATT_1\|\|HTH_3-><-ParB<-TOPRIM<-OLD-ABC<-?<- N6-METHYLASE | 1251 | Bacteria>Terrabacteria_group>Actinomycetota>Actinomycetes>Micromonosporales>Micromonosporaceae | GCF_013410645.1 | Micromonospora_jinlongensis | HNR22_RS05305 | 1251485..1255240 | 1287877 | phage-resistance-protein-[Micromonospora-jinlongensis] |
| WP_043526153.1 | Type2.Pgl.like.BREX | SF1-SF2-Helicase+ HKD-DNase -><- SF1-SF2-Helicase+wHTH+Small-B-barrel+DUF1998 <- BrxD-AAA <- PglZ <- DUF6079 *<- N6-METHYLASE \|\|ARG-><-REase<- PglW \|\|HerA/FtsK-> | 1242 | Bacteria>Terrabacteria_group>Actinomycetota>Actinomycetes>Micromonosporales>Micromonosporaceae | GCF_016862455.1 | Actinoplanes_utahensis | Aut01nite_RS42125 | 11222..14950 | 1869 | hypothetical-protein-[Actinoplanes-utahensis] |
| WP_328811366.1 | Type2.Pgl.like.BREX | SF1-SF2-Helicase+REase->SF1-SF2-Helicase->?-> PglZ -><- SF1-SF2-Helicase+wHTH+Small-B-barrel+DUF1998 <- BrxD-AAA <- PglZ <- DUF6079 *<- N6-METHYLASE <- PglW <- HKD-DNase \|\|McrB-AAA-ATPase->DUF2357+REase-><-CBS+HEPN+HEPN<-HEPN+SF1-SF2-Helicase+REase | 1245 | Bacteria>Terrabacteria_group>Actinomycetota>Actinomycetes>Mycobacteriales | GCF_036163995.1 | Rhodococcus_sp._NBC_00294 | OHB23_RS09005 | 1915935..1919672 | 2976004 | DUF6079 -family-protein-[Rhodococcus-sp.-NBC_00294] |
| WP_068057216.1 | Type2.Pgl.like.BREX | <-TRD+TRD<- N6-METHYLASE \|\|?->?->?->DndB-><-?\|\| PglW -> N6-METHYLASE -> DUF6079 *-> PglZ ->ParB+HNH->CBS+HEPN-> BrxD-AAA -> SF1-SF2-Helicase+wHTH+Small-B-barrel+DUF1998 -> SF1-SF2-Helicase+ HKD-DNase -><-REase+SF1-SF2-Helicase | 1303 | Bacteria>Terrabacteria_group>Actinomycetota>Actinomycetes>Mycobacteriales>Nocardiaceae | GCF_001613365.1 | Nocardia_xishanensis | NX1_RS10700 | 31578..35489 | 238964 | DUF6079 -family-protein-[Nocardia-xishanensis] |
| WP_113691957.1 | Type2.Pgl.like.BREX | <-SIGMA-HTH<-Peptidase_M20\|\|tRNA-><-REase<-AAA-ATPase<- SF1-SF2-Helicase+ HKD-DNase <-HTH_Tnp_4+DDE_Tnp_4<- SF1-SF2-Helicase+wHTH+Small-B-barrel+DUF1998 <- BrxD-AAA <- PglZ <- DUF6079 *<- N6-METHYLASE <- PglW <-SF1-SF2-Helicase+REase<-?<- BrxA-DNA-binding \|\|REase+SF1-SF2-Helicase-><-YTCD-HTH\|\|ABhydrolase-> | 1264 | Bacteria>Terrabacteria_group>Actinomycetota>Actinomycetes>Pseudonocardiales>Pseudonocardiaceae | GCF_003312875.1 | Amycolatopsis_albispora | A4R43_RS09335 | 2006879..2010673 | 1804986 | PglY-protein-[Amycolatopsis-albispora] |

| Protein-ID | System | Gene-Neighbourhoods | Length | Taxonomy | Assembly-ID | Strain | Locus-Tag | Genomic-coordinates | TaxID | Fasta-define |
|---|---|---|---|---|---|---|---|---|---|---|
| WP_319723082.1 | Type2.Pgl.like.BREX | AA_permease_2->TRAM+SAM-methylase+METHYLASE-><-SF1-SF2-Helicase+wHTH+Small-B-barrel+DUF1998<-BrxD-AAA<-AIPR<-PglZ<-DUF6079*<-REase+SF1-SF2-Helicase<-?<-?\|\|HTH_31+DUF5753->DUF397-><-N6-METHYLASE<-PglW | 1301 | Bacteria>Terrabacteria_group>Actinomycetota>Actinomycetes>Kitasatosporales | GCF_033794505.1 | Streptomyces_sp._MB09-01 | PV721_RS11725 | 18486..22391 | 3028666 | phage-resistance-protein-[Streptomyces-sp.-MB09-01] |
| WP_324980990.1 | Type2.Pgl.like.BREX | PglW->?-><-?\|\|?->?->?->DUF6079*-> | 1308 | Bacteria>Terrabacteria_group>Actinomycetota>Actinomycetes>Kitasatosporales | GCF_035687565.1 | Streptomyces_sp. | VFG16_RS18990 | 19668..23594 | 1931 | phage-resistance-protein-[Streptomyces-sp.] |
| WP_326435021.1 | Type2.Pgl.like.BREX | PglW->?->?-><-?<-?\|\|?->DUF6079*-> | 1323 | Bacteria>Terrabacteria_group>Actinomycetota>Actinomycetes>Kitasatosporales | GCF_035828615.1 | Streptomyces_sp._H27-D2 | QMK28_RS01135 | 61413..65384 | 3046304 | PglY-protein-[Streptomyces-sp.-H27-D2] |
| WP_326661331.1 | Type2.Pgl.like.BREX | <-OLD-ABC\|\|PglW-><-?\|\|HTH_31+DUF5753->DUF397->N6-METHYLASE->DUF6079*->PglZ->BrxD-AAA->SF1-SF2-Helicase+wHTH+Small-B-barrel+DUF1998-> | 1295 | Bacteria>Terrabacteria_group>Actinomycetota>Actinomycetes>Kitasatosporales | GCF_035917155.1 | Streptomyces_sp._NBC_00385 | OG959_RS23215 | 5186083..5189970 | 2975733 | phage-resistance-protein-[Streptomyces-sp.-NBC_00385] |
| WP_326723507.1 | Type2.Pgl.like.BREX | REase->?->ncRNA-><-SF1-SF2-Helicase+wHTH+Small-B-barrel+DUF1998<-BrxD-AAA\|\|ParB+HNH->CBS+HEPN-><-PglZ<-DUF6079*<-?<-DUF397<-HTH_31+DUF5753\|\|ST-Hiskin-><-N6-METHYLASE<-PglW\|\|SAD(SRA)+wHTH+HNH-> | 1308 | Bacteria>Terrabacteria_group>Actinomycetota>Actinomycetes>Kitasatosporales | GCF_035917175.1 | Streptomyces_sp._NBC_00243 | OHT59_RS33045 | 7427318..7431244 | 2975688 | phage-resistance-protein-[Streptomyces-sp.-NBC_00243] |
| WP_326795582.1 | Type2.Pgl.like.BREX | <-MerR-HTH<-?\|\|?-><-?\|\|ncRNA-><-PglZ<-DUF6079*<-N6-METHYLASE<-PglW\|\|BrxD-AAA->SF1-SF2-Helicase+wHTH+Small-B-barrel+DUF1998->N6-METHYLASE-> | 1284 | Bacteria>Terrabacteria_group>Actinomycetota>Actinomycetes>Kitasatosporales | GCF_035917395.1 | Streptomyces_sp._NBC_01808 | OG946_RS08875 | 2094020..2097874 | 2975947 | DUF6079-family-protein-[Streptomyces-sp.-NBC_01808] |
| WP_326818697.1 | Type2.Pgl.like.BREX | <-pPIWI+MID+RNaseH<-pPIWI+SFII-RAD3<-pPIWI+pPIWI<-SF1-SF2-Helicase+wHTH+Small-B-barrel+DUF1998<-BrxD-AAA<-PglZ<-DUF6079*<-N6-METHYLASE<-PglW\|\|REase+DUF5655-><-?<-?<-?<-DCM | 1297 | Bacteria>Terrabacteria_group>Actinomycetota>Actinomycetes>Kitasatosporales | GCF_035917875.1 | unclassified_Streptomyces | OHA26_RS39625 | 8793533..8797426 | 2593676 | MULTISPECIES:-phage-resistance-protein-[unclassified-Streptomyces] |
| WP_326752420.1 | Type2.Pgl.like.BREX | PglW->N6-METHYLASE-><-ST-Hiskin\|\|HTH_31+DUF5753->DUF397->?->?->DUF6079*->PglZ-><-CBS+HEPN<-ParB+HNH\|\|BrxD-AAA->SF1-SF2-Helicase+wHTH+Small-B-barrel+DUF1998-><-REase | 1310 | Bacteria>Terrabacteria_group>Actinomycetota>Actinomycetes>Kitasatosporales | GCF_035918035.1 | Streptomyces_hirsutus | OIE73_RS11080 | 2415560..2419492 | 35620 | phage-resistance-protein-[Streptomyces-hirsutus] |
| WP_326669072.1 | Type2.Pgl.like.BREX | PglW-><-ORC-CDC6\|\|N6-METHYLASE-><-DUF397<-HTH_31+DUF5753\|\|ST-Hiskin-><-?\|\|DUF6079*->PglZ->BrxD-AAA->SF1-SF2-Helicase+wHTH+Small-B-barrel+DUF1998-><-REase+SF1-SF2-Helicase<-ParB+HNH | 1301 | Bacteria>Terrabacteria_group>Actinomycetota>Actinomycetes>Kitasatosporales | GCF_035918175.1 | Streptomyces_canus | OG925_RS34790 | 7576793..7580698 | 58343 | PglY-protein-[Streptomyces-canus] |
| WP_328677725.1 | Type2.Pgl.like.BREX | <-REase<-?\|\|PglW->N6-METHYLASE-><-DUF397<-HTH_31+DUF5753\|\|REase+SF1-SF2-Helicase->DUF6079*->PglZ->AIPR->BrxD-AAA->SF1-SF2-Helicase+wHTH+Small-B-barrel+DUF1998-><-?<-TRAM+SAM-methylase+METHYLASE<-AA_permease_2 | 1295 | Bacteria>Terrabacteria_group>Actinomycetota>Actinomycetes>Kitasatosporales | GCF_036118775.1 | Streptomyces_sp._NBC_00343 | OHA62_RS11625 | 2752223..2756110 | 2975719 | phage-resistance-protein-[Streptomyces-sp.-NBC_00343] |
| WP_328775746.1 | Type2.Pgl.like.BREX | <-REase+SF1-SF2-Helicase\|\|PglW->N6-METHYLASE-><-DUF397<-HTH_31+DUF5753\|\|?->?->DUF6079*->PglZ-><-CBS+HEPN<-ParB+HNH\|\|BrxD-AAA->SF1-SF2-Helicase+wHTH+Small-B-barrel+DUF1998-><-TRAM+SAM-methylase+METHYLASE<-AA_permease_2 | 1299 | Bacteria>Terrabacteria_group>Actinomycetota>Actinomycetes>Kitasatosporales | GCF_036175285.1 | Streptomyces_goshikiensis | OHU17_RS09805 | 2191702..2195601 | 1942 | phage-resistance-protein-[Streptomyces-goshikiensis] |
| WP_328764025.1 | Type2.Pgl.like.BREX | <-GDResolvase\|\|?->HTH_29+HTH_32+DDE_3-><-SF1-SF2-Helicase+wHTH+Small-B-barrel+DUF1998<-BrxD-AAA<-PglZ<-DUF6079*<-N6-METHYLASE<-PglW\|\|DUF6731-> | 1295 | Bacteria>Terrabacteria_group>Actinomycetota>Actinomycetes>Kitasatosporales | GCF_036178565.1 | Streptomyces_sp._NBC_00272 | OG827_RS28355 | 6050650..6054537 | 2975698 | PglY-protein-[Streptomyces-sp.-NBC_00272] |
| WP_326712555.1 | Type2.Pgl.like.BREX | HTH-><-?\|\|PglW->HEPN->ParA-Soj-PloopNTPase->BrxD-AAA->N6-METHYLASE->DUF6079*->PglZ->BrxD-AAA->SF1-SF2-Helicase+wHTH+Small-B-barrel+DUF1998->HTH-><-?\|\|?-><-ACET | 1292 | Bacteria>Terrabacteria_group>Actinomycetota>Actinomycetes>Kitasatosporales | GCF_036178905.1 | unclassified_Streptomyces | OHA28_RS28115 | 6167427..6171305 | 2593676 | MULTISPECIES:-hypothetical-protein-[unclassified-Streptomyces] |
| WP_328846446.1 | Type2.Pgl.like.BREX | <-REase+SF1-SF2-Helicase<-?<-SF1-SF2-Helicase+wHTH+Small-B-barrel+DUF1998<-BrxD-AAA\|\|ParB+HNH->CBS+HEPN-><-PglZ<-DUF6079*<-?<-DUF397<-HTH_31+DUF5753\|\|ST-Hiskin-><-N6-METHYLASE<-PglW | 1326 | Bacteria>Terrabacteria_group>Actinomycetota>Actinomycetes>Kitasatosporales | GCF_036182465.1 | Streptomyces_sp._NBC_00258 | OG718_RS39415 | 8843524..8847504 | 2903642 | phage-resistance-protein-[Streptomyces-sp.-NBC_00258] |
| WP_328932270.1 | Type2.Pgl.like.BREX | <-REase<-?\|\|PglW->N6-METHYLASE-><-DUF397<-HTH_31+DUF5753\|\|REase+SF1-SF2-Helicase->DUF6079*->PglZ->AIPR->BrxD-AAA->SF1-SF2-Helicase+wHTH+Small-B-barrel+DUF1998-><-?<-TRAM+SAM-methylase+METHYLASE<-AA_permease_2 | 1295 | Bacteria>Terrabacteria_group>Actinomycetota>Actinomycetes>Kitasatosporales | GCF_036196325.1 | Streptomyces_sp._NBC_00228 | OG537_RS11480 | 2681580..2685467 | 2903637 | phage-resistance-protein-[Streptomyces-sp.-NBC_00228] |
| WP_328934070.1 | Type2.Pgl.like.BREX | <-DUF6079*<-?<-PglW | 1304 | Bacteria>Terrabacteria_group>Actinomycetota>Actinomycetes>Kitasatosporales | GCF_036196325.1 | Streptomyces_sp._NBC_00228 | OG537_RS29365 | 6628887..6632801 | 2903637 | PglY-protein-[Streptomyces-sp.-NBC_00228] |
| WP_328937459.1 | Type2.Pgl.like.BREX | PglW->N6-METHYLASE-><-DUF397<-HTH_31+DUF5753\|\|<-?<-?\|\|DUF6079*->PglZ->BrxD-AAA->SF1-SF2-Helicase+wHTH+Small-B-barrel+DUF1998-><-?\|\|?->HKD-DNase-> | 1298 | Bacteria>Terrabacteria_group>Actinomycetota>Actinomycetes>Kitasatosporales | GCF_036204065.1 | Streptomyces_tauricus | OG288_RS12455 | 3035680..3039576 | 68274 | phage-resistance-protein-[Streptomyces-tauricus] |
| WP_329283861.1 | Type2.Pgl.like.BREX | AA_permease_2->TRAM+SAM-methylase+METHYLASE-><-YjhX_toxin<-MORC+REase\|\|REase+DUF5655->PglW->N6-METHYLASE->DUF6079*->PglZ->BrxD-AAA->SF1-SF2-Helicase+wHTH+Small-B-barrel+DUF1998-><-?\|\|NUDIX-> | 1297 | Bacteria>Terrabacteria_group>Actinomycetota>Actinomycetes>Kitasatosporales | GCF_036226665.1 | Streptomyces_sp._NBC_00691 | OG392_RS27675 | 6136672..6140565 | 2903671 | phage-resistance-protein-[Streptomyces-sp.-NBC_00691] |
| WP_329024165.1 | Type2.Pgl.like.BREX | PglW->?-><-?<-?\|\|?->?->DUF6079*-> | 1309 | Bacteria>Terrabacteria_group>Actinomycetota>Actinomycetes>Kitasatosporales | GCF_036226685.1 | Streptomyces_sp._NBC_00690 | OID54_RS28780 | 6564231..6568160 | 2975808 | phage-resistance-protein-[Streptomyces-sp.-NBC_00690] |
| WP_329299357.1 | Type2.Pgl.like.BREX | PglW->N6-METHYLASE-><-ST-Hiskin\|\|HTH_31+DUF5753->DUF397->DUF6079*->PglZ-><-CBS+HEPN<-ParB+HNH\|\|BrxD-AAA->SF1-SF2-Helicase+wHTH+Small-B-barrel+DUF1998-><-SAD(SRA)+wHTH+HNH | 1322 | Bacteria>Terrabacteria_group>Actinomycetota>Actinomycetes>Kitasatosporales | GCF_036226925.1 | Streptomyces_sp._NBC_00659 | OG410_RS13480 | 3111600..3115568 | 2903669 | phage-resistance-protein-[Streptomyces-sp.-NBC_00659] |
| WP_329359101.1 | Type2.Pgl.like.BREX | <-DUF4096+DDE_Tnp_1+DDE_Tnp_1_2<-?\|\|PglW->N6-METHYLASE->DUF6079*->PglZ->BrxD-AAA->SF1-SF2-Helicase+wHTH+Small-B-barrel+DUF1998-> | 1296 | Bacteria>Terrabacteria_group>Actinomycetota>Actinomycetes>Kitasatosporales | GCF_036227165.1 | Streptomyces_sp._NBC_01483 | OG318_RS52040 | 11321104..11324994 | 2903883 | PglY-protein-[Streptomyces-sp.-NBC_01483] |
| WP_329248535.1 | Type2.Pgl.like.BREX | Phd-YefM->RelE-ParE+Tox-RelE->PglW->N6-METHYLASE-><-DUF397<-HTH_31+DUF5753\|\|REase+SF1-SF2-Helicase->DUF6079*->PglZ->AIPR->BrxD-AAA->SF1-SF2-Helicase+wHTH+Small-B-barrel+DUF1998-><-ncRNA\|\|?-><-?<-TetR-HTH | 1296 | Bacteria>Terrabacteria_group>Actinomycetota>Actinomycetes>Kitasatosporales | GCF_036227225.1 | Streptomyces_sp._NBC_01478 | OG223_RS16420 | 3653394..3657284 | 2903882 | DUF6079-family-protein-[Streptomyces-sp.-NBC_01478] |
| WP_329173286.1 | Type2.Pgl.like.BREX | PglW->N6-METHYLASE->DUF6079*->PglZ->BrxD-AAA->SF1-SF2-Helicase+wHTH+Small-B-barrel+DUF1998-> | 1299 | Bacteria>Terrabacteria_group>Actinomycetota>Actinomycetes>Kitasatosporales | GCF_036227245.1 | Streptomyces_sp._NBC_01477 | OHA86_RS06555 | 1702839..1706738 | 2976015 | phage-resistance-protein-[Streptomyces-sp.-NBC_01477] |
| WP_329144525.1 | Type2.Pgl.like.BREX | PglW->N6-METHYLASE-><-ST-Hiskin\|\|HTH_31+DUF5753->DUF397->?->DUF6079*->PglZ-><-CBS+HEPN<-ParB+HNH\|\|BrxD-AAA->SF1-SF2-Helicase+wHTH+Small-B-barrel+DUF1998-><-ncRNA\|\|MerR-HTH-> | 1326 | Bacteria>Terrabacteria_group>Actinomycetota>Actinomycetes>Kitasatosporales | GCF_036227585.1 | Streptomyces_niveus | OG275_RS09080 | 2069272..2073252 | 193462 | phage-resistance-protein-[Streptomyces-niveus] |
| WP_329194497.1 | Type2.Pgl.like.BREX | PglW->N6-METHYLASE-><-ST-Hiskin\|\|HTH_31+DUF5753->DUF397->Phd-YefM->PIN->DUF6079*->PglZ-><-CBS+HEPN<-ParB+HNH\|\|BrxD-AAA->SF1-SF2-Helicase+wHTH+Small-B-barrel+DUF1998-><-SAD(SRA)+wHTH+HNH | 1309 | Bacteria>Terrabacteria_group>Actinomycetota>Actinomycetes>Kitasatosporales | GCF_036228565.1 | Streptomyces_sp._NBC_01435 | OG769_RS10315 | 2474520..2478449 | 2903865 | phage-resistance-protein-[Streptomyces-sp.-NBC_01435] |
| WP_330354472.1 | Type2.Pgl.like.BREX | PglW->HEPN->ParA-Soj-PloopNTPase->BrxD-AAA->N6-METHYLASE->DUF6079*->PglZ->BrxD-AAA->SF1-SF2-Helicase+wHTH+Small-B-barrel+DUF1998-><-?<-SAD+REase | 1305 | Bacteria>Terrabacteria_group>Actinomycetota>Actinomycetes>Kitasatosporales | GCF_036345055.1 | Streptomyces_chartreusis | OG997_RS17120 | 3862934..3866851 | 1969 | hypothetical-protein-[Streptomyces-chartreusis] |
| WP_330344465.1 | Type2.Pgl.like.BREX | PglW->N6-METHYLASE-><-ST-Hiskin\|\|HTH_31+DUF5753->DUF397->?->DUF6079*->PglZ-><-CBS+HEPN<-ParB+HNH\|\|BrxD-AAA->SF1-SF2-Helicase+wHTH+Small-B-barrel+DUF1998-> | 1308 | Bacteria>Terrabacteria_group>Actinomycetota>Actinomycetes>Kitasatosporales | GCF_036346085.1 | Streptomyces_longwoodensis | OHA09_RS12455 | 3070861..3074787 | 68231 | phage-resistance-protein-[Streptomyces-longwoodensis] |
| WP_330333133.1 | Type2.Pgl.like.BREX | AA_permease_2->TRAM+SAM-methylase+METHYLASE-><-SF1-SF2-Helicase+wHTH+Small-B-barrel+DUF1998<-BrxD-AAA<-AIPR<-PglZ<-DUF6079*<-?<-?\|\|HTH_31+DUF5753->DUF397-><-N6-METHYLASE<-PglW\|\|REase+SF1-SF2-Helicase-> | 1298 | Bacteria>Terrabacteria_group>Actinomycetota>Actinomycetes>Kitasatosporales | GCF_036346295.1 | Streptomyces_sp._NBC_00536 | OHS33_RS27775 | 6379336..6383232 | 2975769 | phage-resistance-protein-[Streptomyces-sp.-NBC_00536] |
| WP_330811472.1 | Type2.Pgl.like.BREX | <-SAD+REase<-SF1-SF2-Helicase+wHTH+Small-B-barrel+DUF1998<-BrxD-AAA\|\|ParB+HNH->CBS+HEPN-><-PglZ<-DUF6079*<-DUF397<-HTH_31+DUF5753\|\|ST-Hiskin-><-N6-METHYLASE<-PglW\|\|REase+SF1-SF2-Helicase->HNH-> | 1320 | Bacteria>Terrabacteria_group>Actinomycetota>Actinomycetes>Kitasatosporales | GCF_036352935.1 | Streptomyces | V2J94_RS25890 | 112923..116885 | 1883 | phage-resistance-protein-[Streptomyces-sp.-DSM-41524] |
| WP_331447533.1 | Type2.Pgl.like.BREX | <-N6-METHYLASE\|\|PglW->N6-METHYLASE->DUF6079*->PglZ->PglZ->BrxD-AAA->SF1-SF2-Helicase+wHTH+Small-B-barrel+DUF1998-><-REase\|\|?->Phage_integrase-> | 1297 | Bacteria>Terrabacteria_group>Actinomycetota>Actinomycetes>Kitasatosporales | GCF_036541015.1 | Streptomyces_xanthochromogenes | V3O60_RS08980 | 2066752..2070645 | 67384 | phage-resistance-protein-[Streptomyces-xanthochromogenes] |
| WP_334580654.1 | Type2.Pgl.like.BREX | HEPN+VirB11+NACHT+FGS->?->?-><-?<-DUF6731\|\|PglW->N6-METHYLASE->DUF6079*-><-DEDD_Tnp_IS110+Transposase_20\|\|PglZ->BrxD-AAA->SF1-SF2-Helicase+wHTH+Small-B-barrel+DUF1998-><-?\|\|?-><-Phage_int_SAM_3+Phage_integrase | 1312 | Bacteria>Terrabacteria_group>Actinomycetota>Actinomycetes>Kitasatosporales | GCF_036898445.1 | Streptomyces_sp._B21-108 | QBA54_RS35325 | 7740309..7744247 | 3039419 | PglY-protein-[Streptomyces-sp.-B21-108] |
| WP_334662749.1 | Type2.Pgl.like.BREX | HTH-><-?\|\|PglW->HEPN->ParA-Soj-PloopNTPase->BrxD-AAA->N6-METHYLASE->\|\|DUF6079*->PglZ->BrxD-AAA->SF1-SF2-Helicase+wHTH+Small-B-barrel+DUF1998-><-SF1-SF2-Helicase<-HTH_Tnp_1+HTH_21+rve+rve_3 | 1347 | Bacteria>Terrabacteria_group>Actinomycetota>Actinomycetes>Kitasatosporales | GCF_036919415.1 | Streptomyces_cyaneofuscatus | VVF06_RS00805 | 19993..24036 | 66883 | hypothetical-protein-[Streptomyces-cyaneofuscatus] |
| WP_336050190.1 | Type2.Pgl.like.BREX | <-SF1-SF2-Helicase+wHTH+Small-B-barrel+DUF1998<-BrxD-AAA\|\|ParB+HNH->CBS+HEPN-><-PglZ<-DUF6079*<-DUF397<-HTH_31+DUF5753\|\|ST-Hiskin-><-N6-METHYLASE<-PglW | 1317 | Bacteria>Terrabacteria_group>Actinomycetota>Actinomycetes>Kitasatosporales | GCF_037034225.1 | Streptomyces_sp._CA2R101 | WA779_RS15055 | 59886..63839 | 3120152 | phage-resistance-protein-[Streptomyces-sp.-CA2R101] |
| WP_337304542.1 | Type2.Pgl.like.BREX | DUF4096+DDE_Tnp_1+DDE_Tnp_1_2-><-?<-?\|\|?-><-SF1-SF2-Helicase+wHTH+Small-B-barrel+DUF1998<-BrxD-AAA<-PglZ<-DUF6079*<-ST-Hiskin\|\|HTH_31+DUF5753->DUF397-><-N6-METHYLASE<-PglW\|\|?-><-ACET | 1298 | Bacteria>Terrabacteria_group>Actinomycetota>Actinomycetes>Kitasatosporales | GCF_037204065.1 | unclassified_Streptomyces | V9L26_RS11245 | 111223..115119 | 2593676 | MULTISPECIES:-phage-resistance-protein-[unclassified-Streptomyces] |
| WP_279949630.1 | Type2.Pgl.like.BREX | REase+SF1-SF2-Helicase-><-?\|\|HTH_31+DUF5753->DUF397->?->PglW->N6-METHYLASE->DUF6079*->PglZ-><-?\|\|BrxD-AAA->SF1-SF2-Helicase+wHTH+Small-B-barrel+DUF1998-> | 1295 | Bacteria>Terrabacteria_group>Actinomycetota>Actinomycetes>Kitasatosporales | GCF_039540175.1 | Streptomyces_chitinivorans | ACG5V6_RS01050 | 76981..80868 | 1257027 | phage-resistance-protein-[Streptomyces-chitinivorans] |
| WP_184852634.1 | Type2.Pgl.like.BREX | <-Pkinase+TPR\|\|<-?\|\|?->PglW->N6-METHYLASE->DUF6079*->PglZ-><-DUF397<-HTH_31+DUF5753\|\|ST-Hiskin-><-CBS+HEPN<-ParB+HNH | 1298 | Bacteria>Terrabacteria_group>Actinomycetota>Actinomycetes>Kitasatosporales | GCF_039540655.1 | Streptomyces_collinus | HNR72_RS32100 | 7190212..7194108 | 42684 | phage-resistance-protein-[Streptomyces-collinus] |
| WP_330466073.1 | Type2.Pgl.like.BREX | PglW->N6-METHYLASE-><-ST-Hiskin\|\|HTH_31+DUF5753->DUF397->?->DUF6079*->PglZ-><-CBS+HEPN<-ParB+HNH\|\|BrxD-AAA->SF1-SF2-Helicase+wHTH+Small-B-barrel+DUF1998-> | 1308 | Bacteria>Terrabacteria_group>Actinomycetota>Actinomycetes>Kitasatosporales | GCF_045161325.1 | Streptomyces_longwoodensis | OG547_RS11280 | 2752021..2755947 | 68231 | phage-resistance-protein-[Streptomyces-longwoodensis] |
| WP_326746002.1 | Type2.Pgl.like.BREX | <-pPIWI+SFII-RAD3<-?<-pPIWI+pPIWI<-SF1-SF2-Helicase+wHTH+Small-B-barrel+DUF1998<-BrxD-AAA<-PglZ<-DUF6079*<-N6-METHYLASE<-PglW\|\|REase+DUF5655-><-?<-?<-?<-DCM | 1297 | Bacteria>Terrabacteria_group>Actinomycetota>Actinomycetes>Kitasatosporales | GCF_045206205.1 | Streptomyces_sp._NBC_01760 | OG807_RS36730 | 8213139..8217032 | 2975931 | phage-resistance-protein-[Streptomyces-sp.-NBC_01760] |
| WP_282203631.1 | Type2.Pgl.like.BREX | <-SF1-SF2-Helicase+wHTH+Small-B-barrel+DUF1998<-BrxD-AAA<-PglZ<-DUF6079*<-PglW<-?\|\|?->N6-METHYLASE->N6-METHYLASE-><-REase+SF1-SF2-Helicase | 1346 | Bacteria>Terrabacteria_group>Actinomycetota>Actinomycetes>Kitasatosporales | GCF_948472415.1 | Kitasatospora_fiedleri | QMQ26_RS31720 | 7019286..7023326 | 2991545 | phage-resistance-protein-[Kitasatospora-fiedleri] |
| WTC44582.1 | Type2.Pgl.like.BREX | AA_permease_2->TRAM+SAM-methylase+METHYLASE-><-SF1-SF2-Helicase+wHTH+Small-B-barrel+DUF1998<-BrxD-AAA<-PglZ<-DUF6079*\|\|HTH_31+DUF5753->DUF397-><-N6-METHYLASE<-PglW\|\|REase->?-><-CASPASE+SF1-SF2-Helicase-> | 1298 | Bacteria>Terrabacteria_group>Actinomycetota>Actinomycetes>Kitasatosporales | - | Streptomyces_albidoflavus | OH810_24955 | 5684021..5687917 | 1886 | phage-resistance-protein-[Streptomyces-albidoflavus] |
| EST37408.1 | Type2.Pgl.like.BREX | <-BrxD-AAA-p<-DndB<-PglZ<-DUF6079*<-REase+SF1-SF2-Helicase<-DUF2171<-DUF1995\|\|HTH_31+DUF5753-><-N6-METHYLASE<-DDE_Tnp_1<-PglW\|\|?->REase-> | 1295 | Bacteria>Terrabacteria_group>Actinomycetota>Actinomycetes>Kitasatosporales>Streptomycetaceae | GCA_000497405.1 | Streptomycetaceae_bacterium_MP113-05 | N566_13090 | 5298..9185 | 1380770 | hypothetical-protein-N566_13090-[Streptomycetaceae-bacterium-MP113-05] |
| KMS82322.1 | Type2.Pgl.like.BREX | Methyltransf_21->His_Phos_1-><-?\|\|?->?-><-SF1-SF2-Helicase+wHTH+Small-B-barrel+DUF1998<-BrxD-AAA<-PglZ<-DUF6079*<-ST-Hiskin\|\|HTH_31+DUF5753->DUF397-><-N6-METHYLASE\|\|HEPN->ParA-Soj-PloopNTPase->ORC-CDC6-><-PglW | 1321 | Bacteria>Terrabacteria_group>Actinomycetota>Actinomycetes>Kitasatosporales>Streptomycetaceae | GCA_001047335.1 | Streptomyces_regensis | ACZ91_59210 | 21770..25735 | 68263 | PglY-protein-[Streptomyces-regensis] |
| KPC88168.1 | Type2.Pgl.like.BREX | REase+SF1-SF2-Helicase->REase+SF1-SF2-Helicase->?->PglW->N6-METHYLASE->?->HTH_31+DUF5753->DUF397->?->DUF6079*->PglZ-><-CBS+HEPN<-ParB+HNH<-?\|\|BrxD-AAA->SF1-SF2-Helicase+wHTH+Small-B-barrel+DUF1998-><-DUF4011+SF1-SF2-Helicase+REase\|\|ACET-> | 1295 | Bacteria>Terrabacteria_group>Actinomycetota>Actinomycetes>Kitasatosporales>Streptomycetaceae | GCA_001294395.1 | Streptomyces_sp._NRRL_WC-3753 | ADL35_06445 | 142151..146038 | 1609104 | bacteriophage-resistance-protein-[Streptomyces-sp.-NRRL-WC-3753] |
| AMM07779.1 | Type2.Pgl.like.BREX | <-REase\|\|PglW->N6-METHYLASE-><-DUF397<-HTH_31+DUF5753\|\|ST-Hiskin->DUF6079*->PglZ->BrxD-AAA->SF1-SF2-Helicase+wHTH+Small-B-barrel+DUF1998-><-TRAM+SAM-methylase+METHYLASE<-AA_permease_2 | 1295 | Bacteria>Terrabacteria_group>Actinomycetota>Actinomycetes>Kitasatosporales>Streptomycetaceae | GCA_001577385.1 | Streptomyces_albidoflavus | Salbus254_1235 | 1445801..1449688 | 1886 | bacteriophage-resistance-protein-[Streptomyces-albidoflavus] |
| ARF59786.1 | Type2.Pgl.like.BREX | REase+SF1-SF2-Helicase-><-REase+SF1-SF2-Helicase<-SF1-SF2-Helicase+wHTH+Small-B-barrel+DUF1998<-BrxD-AAA\|\|RNaseH-><-PglZ<-DUF6079*\|\|HTH_31+DUF5753->DUF397-><-N6-METHYLASE<-PglW<-NAD_binding_10<-?\|\|MarR-HTH-> | 1299 | Bacteria>Terrabacteria_group>Actinomycetota>Actinomycetes>Kitasatosporales>Streptomycetaceae | GCA_002082195.1 | Streptomyces_gilvosporeus | B1H19_30120 | 6750768..6754667 | 553510 | phage-resistance-protein-[Streptomyces-gilvosporeus] |
| ROR38155.1 | Type2.Pgl.like.BREX | REase+SF1-SF2-Helicase->ParB+HNH->CBS+HEPN->PglW->N6-METHYLASE->Phd-YefM->PIN->DUF6079*->PglZ->BrxD-AAA->SF1-SF2-Helicase+wHTH+Small-B-barrel+DUF1998-><-CASPASE+SF1-SF2-Helicase<-HTH_21+rve+rve_2<-HIN-HTH | 1320 | Bacteria>Terrabacteria_group>Actinomycetota>Actinomycetes>Kitasatosporales>Streptomycetaceae | GCA_003751605.1 | Kitasatospora_cineracea | EDD39_6327 | 1175437..1179399 | 88074 | hypothetical-protein-EDD39_6327-[Kitasatospora-cineracea] |
| RPK54894.1 | Type2.Pgl.like.BREX | AA_permease_2->TRAM+SAM-methylase+METHYLASE-><-SF1-SF2-Helicase+wHTH+Small-B-barrel+DUF1998<-BrxD-AAA<-PglZ<-DUF6079*<-ST-Hiskin\|\|HTH_31+DUF5753->DUF397-><-N6-METHYLASE<-PglW<-?<-BrxA-DNA-binding | 1295 | Bacteria>Terrabacteria_group>Actinomycetota>Actinomycetes>Kitasatosporales>Streptomycetaceae | GCA_003846265.1 | Streptomyces_sp._ADI96-15 | EES44_29045 | 154353..158240 | 1522761 | hypothetical-protein-EES44_29045-[Streptomyces-sp.-ADI96-15] |
| TKA04731.1 | Type2.Pgl.like.BREX | transposase->?-><-?\|\|?->EVE+REase-p->\|\|PglW->N6-METHYLASE->DUF6079*->PglZ->BrxD-AAA->SF1-SF2-Helicase+wHTH+Small-B-barrel+DUF1998-><-?<-DUF2171\|\|HTH_3-><-HAD+HAD | 1316 | Bacteria>Terrabacteria_group>Actinomycetota>Actinomycetes>Kitasatosporales>Streptomycetaceae | GCA_005047355.1 | Actinacidiphila_oryziradicis | FCI23_34795 | 22737..26687 | 2571141 | PglY-protein-[Actinacidiphila-oryziradicis] |
| GDY52272.1 | Type2.Pgl.like.BREX | DUF4096+DDE_Tnp_1+DDE_Tnp_1_2-><-?\|\|DUF4096+DDE_Tnp_1+DDE_Tnp_1_2->DUF4096+DDE_Tnp_1+DDE_Tnp_1_2-><-DUF2637+HTH\|\|HTH_32->rve_3-><-PglZ<-DUF6079-p<-DUF6079-p*<-DUF6079-p<-N6-METHYLASE<-N6-METHYLASE-p<-PglW\|\|BrxD-AAA->SF1-SF2-Helicase+wHTH+Small-B-barrel+DUF1998-p->SF1-SF2-Helicase+wHTH+Small-B-barrel+DUF1998-><-?<-DUF2637+HTH | 606 | Bacteria>Terrabacteria_group>Actinomycetota>Actinomycetes>Kitasatosporales>Streptomycetaceae | GCA_005405945.1 | Streptomyces_violaceusniger | SVIO_028950 | 3096806..3098626 | 68280 | hypothetical-protein-SVIO_028950-[Streptomyces-violaceusniger] |
| GDY52273.1 | Type2.Pgl.like.BREX | DUF4096+DDE_Tnp_1+DDE_Tnp_1_2->DUF4096+DDE_Tnp_1+DDE_Tnp_1_2-><-DUF2637+HTH\|\|HTH_32->rve_3-><-PglZ<-DUF6079-p<-DUF6079-p<-DUF6079-p*<-N6-METHYLASE<-N6-METHYLASE-p<-PglW\|\|BrxD-AAA->SF1-SF2-Helicase+wHTH+Small-B-barrel+DUF1998-p->SF1-SF2-Helicase+wHTH+Small-B-barrel+DUF1998-><-?<-DUF2637+HTH<-Phage_int_SAM_4+Phage_integrase | 222 | Bacteria>Terrabacteria_group>Actinomycetota>Actinomycetes>Kitasatosporales>Streptomycetaceae | GCA_005405945.1 | Streptomyces_violaceusniger | SVIO_028960 | 3098551..3099219 | 68280 | hypothetical-protein-SVIO_028960-[Streptomyces-violaceusniger] |
| BCK67994.1 | Type2.Pgl.like.BREX | PglW->N6-METHYLASE->N6-METHYLASE-p-><-DUF397<-HTH_31+DUF5753\|\|ST-Hiskin-><-PIN<-CopG/RHH\|\|DUF6079*->PglZ->REase->BrxD-AAA->SF1-SF2-Helicase+wHTH+Small-B-barrel+DUF1998-><-?<-TRAM+SAM-methylase<-?<-AA_permease_2 | 1298 | Bacteria>Terrabacteria_group>Actinomycetota>Actinomycetes>Kitasatosporales>Streptomycetaceae | GCA_009811615.2 | Streptomyces_libani_subsp._rufus | Srufu_019470 | 2037295..2041191 | 249582 | hypothetical-protein-Srufu_019470-[Streptomyces-libani-subsp.-rufus] |
| NEE18804.1 | Type2.Pgl.like.BREX | REase+SF1-SF2-Helicase->REase+SF1-SF2-Helicase<-?\|\|TRD+TRD->REase+N6_Mtase-><-wHTH+SAD+HNH\|\|PglW->N6-METHYLASE->MNS+REase->DUF6079*->PglZ-><-CBS+HEPN<-ParB+HNH\|\|BrxD-AAA->SF1-SF2-Helicase+wHTH+Small-B-barrel+DUF1998-><-ncRNA<-DSBH+AraC-HTH+AraC-HTH\|\|MerR-HTH-> | 1307 | Bacteria>Terrabacteria_group>Actinomycetota>Actinomycetes>Kitasatosporales>Streptomycetaceae | GCA_010550285.1 | Streptomyces_sp._SID7499 | G3M58_71575 | 24154..28077 | 2706086 | type-II-toxin-antitoxin-system-VapC-family-toxin-[Streptomyces-sp.-SID7499] |
| NED01599.1 | Type2.Pgl.like.BREX | ROK-HTH-><-?\|\|sGTP-><-RNaseH<-SF1-SF2-Helicase+wHTH+Small-B-barrel+DUF1998<-BrxD-AAA\|\|PglW->N6-METHYLASE->DUF6079*->PglZ-><-MetJarc-HTH<-?\|\|FAD_binding_9+SIP+Methyltransf_2->?->?->?->ABhydrolase->ABC-ATPase-> | 1299 | Bacteria>Terrabacteria_group>Actinomycetota>Actinomycetes>Kitasatosporales>Streptomycetaceae | GCA_010550305.1 | Streptomyces_sp._SID6648 | G3I55_08000 | 34752..38651 | 2706084 | phage-resistance-protein-[Streptomyces-sp.-SID6648] |
| NEE38448.1 | Type2.Pgl.like.BREX | <-Peptidase_C14+SF1-SF2-Helicase<-?<-REase\|\|PglW->N6-METHYLASE-><-DUF397<-HTH_31+DUF5753\|\|ST-Hiskin->DUF6079*->PglZ->BrxD-AAA->SF1-SF2-Helicase+wHTH+Small-B-barrel+DUF1998-><-?<-HIN-HTH<-?<-ABhydrolase | 1298 | Bacteria>Terrabacteria_group>Actinomycetota>Actinomycetes>Kitasatosporales>Streptomycetaceae | GCA_010550345.1 | Streptomyces_sp._SID7982 | G3M53_75280 | 35734..39630 | 2706094 | phage-resistance-protein-[Streptomyces-sp.-SID7982] |
| NUP22768.1 | Type2.Pgl.like.BREX | <-McrC-NTD+REase<-RAMA+McrB-AAA<-URI-DNase\|\|?->RNaseH-><-SF1-SF2-Helicase+wHTH+Small-B-barrel+DUF1998<-BrxD-AAA<-PglZ<-DUF6079*\|\|BrxA-DNA-binding->?-><-ST-Hiskin\|\|HTH_31+DUF5753->DUF397-><-N6-METHYLASE\|\|HTH+MPTase->PIN-><-PglW | 1306 | Bacteria>Terrabacteria_group>Actinomycetota>Actinomycetes>Kitasatosporales>Streptomycetaceae | GCA_013361465.1 | Streptomyces_sp. | HOZ81_43260 | 19809..23729 | 1931 | PglY-protein-[Streptomyces-sp.] |
| MBB5815397.1 | Type2.Pgl.like.BREX | <-Pkinase+Pkinase\|\|REase+SF1-SF2-Helicase-p->REase+SF1-SF2-Helicase-><-?\|\|?->PglW->N6-METHYLASE->DUF6079*->PglZ-><-DUF397<-HTH_31+DUF5753\|\|ST-Hiskin-><-CBS+CBS+HEPN<-ParB+HNH<-Phage_holin | 1298 | Bacteria>Terrabacteria_group>Actinomycetota>Actinomycetes>Kitasatosporales>Streptomycetaceae | GCA_014204745.1 | Streptomyces_collinus | HNR72_006425 | 7190212..7194108 | 42684 | hypothetical-protein-[Streptomyces-collinus] |
| MBD0735466.1 | Type2.Pgl.like.BREX | <-REC+OmpR-HTH<-HisKA_3+HATPase_c<-?<-?\|\|Peptidase_M4+Peptidase_M4_C->?-><-SF1-SF2-Helicase+wHTH+Small-B-barrel+DUF1998<-BrxD-AAA<-PglZ<-DUF6079*\|\|BrxA-DNA-binding->?-><-ST-Hiskin\|\|HTH_31+DUF5753->DUF397-><-N6-METHYLASE\|\|<-PglW\|\|REase+SF1-SF2-Helicase-> | 1301 | Bacteria>Terrabacteria_group>Actinomycetota>Actinomycetes>Kitasatosporales>Streptomycetaceae | GCA_014596445.1 | Streptomyces_sp._CBMA29 | BGM09_02935 | 64585..68490 | 1896314 | PglY-protein-[Streptomyces-sp.-CBMA29] |
| GGX84370.1 | Type2.Pgl.like.BREX | <-ST-Hiskin\|\|HTH_31+DUF5753->DUF397-><-RAMA\|\|?-><-SF1-SF2-Helicase+wHTH+Small-B-barrel+DUF1998<-BrxD-AAA<-PglZ<-DUF6079-p<-DUF6079-p*<-N6-METHYLASE-p\|\|?-><-N6-METHYLASE<-ORC-CDC6<-ParA-Soj-PloopNTPase<-PglW | 874 | Bacteria>Terrabacteria_group>Actinomycetota>Actinomycetes>Kitasatosporales>Streptomycetaceae | GCA_014650655.1 | Streptomyces_anandii_JCM_4720 | GCM10010510_31730 | 222719..225343 | 1306180 | hypothetical-protein-GCM10010510_31730-[Streptomyces-anandii-JCM-4720] |
| MBT2380833.1 | Type2.Pgl.like.BREX | <-N6-METHYLASE<-DrHyd\|\|SLATT->?-><-SF1-SF2-Helicase+wHTH+Small-B-barrel+DUF1998<-BrxD-AAA<-PglZ<-DUF6079*\|\|<-N6-METHYLASE<-BrxD-AAA<-ParA-Soj-PloopNTPase<-HEPN<-PglW\|\|REase+DUF5655-><-TRD+TRD | 1314 | Bacteria>Terrabacteria_group>Actinomycetota>Actinomycetes>Kitasatosporales>Streptomycetaceae | GCA_018614515.1 | Streptomyces_sp._ISL-111 | J7E90_26860 | 114837..118781 | 2819175 | hypothetical-protein-[Streptomyces-sp.-ISL-111] |
| MCC8482294.1 | Type2.Pgl.like.BREX | PglW->N6-METHYLASE-><-DUF397<-HTH_31+DUF5753\|\|DUF1995->DUF2171->REase+SF1-SF2-Helicase->DUF6079*->PglZ-><-RNaseH\|\|BrxD-AAA->SF1-SF2-Helicase+wHTH+Small-B-barrel+DUF1998-><-GNTR-HTH | 1299 | Bacteria>Terrabacteria_group>Actinomycetota>Actinomycetes>Kitasatosporales>Streptomycetaceae | GCA_020881045.1 | Streptomyces_globisporus | LMJ41_31040 | 42048..45947 | 1908 | phage-resistance-protein-[Streptomyces-globisporus] |
| MCP9962035.1 | Type2.Pgl.like.BREX | AraC-HTH->?-><-?<-PglZ<-DUF6079*<-ST-Hiskin\|\|HTH_31+DUF5753->DUF397-><-N6-METHYLASE<-N6-METHYLASE\|\|?-><-?<-HEPN<-PglW-p | 1300 | Bacteria>Terrabacteria_group>Actinomycetota>Actinomycetes>Kitasatosporales>Streptomycetaceae | GCA_024349105.1 | Streptomyces_somaliensis | LUX05_12805 | 2765612..2769514 | 78355 | hypothetical-protein-[Streptomyces-somaliensis] |
| MCP9974853.1 | Type2.Pgl.like.BREX | AraC-HTH->TPR+TPR+TPR+TPR+TPR+TPR-><-?<-BrxD-AAA<-DUF6079-p*<-ST-Hiskin\|\|HTH_31+DUF5753->DUF397-><-N6-METHYLASE\|\|?->?-><-ParA-Soj-PloopNTPase<-HEPN<-PglW-p | 761 | Bacteria>Terrabacteria_group>Actinomycetota>Actinomycetes>Kitasatosporales>Streptomycetaceae | GCA_024349205.1 | Streptomyces_somaliensis | LUX02_14085 | 3014227..3016512 | 78355 | hypothetical-protein-[Streptomyces-somaliensis] |
| MCQ4043243.1 | Type2.Pgl.like.BREX | <-SF1-SF2-Helicase+wHTH+Small-B-barrel+DUF1998<-BrxD-AAA\|\|ParB+HNH->CBS+HEPN-><-PglZ<-DUF6079*<-ST-Hiskin\|\|HTH_31+DUF5753->?-><-N6-METHYLASE<-PglW<-REase+SF1-SF2-Helicase | 1314 | Bacteria>Terrabacteria_group>Actinomycetota>Actinomycetes>Kitasatosporales>Streptomycetaceae | GCA_024436055.1 | Streptomyces_rubrisoli | NON19_14685 | 110508..114452 | 1387313 | phage-resistance-protein-[Streptomyces-rubrisoli] |
| MCQ4207771.1 | Type2.Pgl.like.BREX | AA_permease_2->TRAM+SAM-methylase+METHYLASE-><-SF1-SF2-Helicase+wHTH+Small-B-barrel+DUF1998<-BrxD-AAA<-AIPR<-PglZ<-DUF6079*<-REase+SF1-SF2-Helicase<-DUF2171<-DUF1995\|\|DUF397-><-N6-METHYLASE<-PglW | 1296 | Bacteria>Terrabacteria_group>Actinomycetota>Actinomycetes>Kitasatosporales>Streptomycetaceae | GCA_024448165.1 | Streptomyces_longispororuber | M4J07_005562 | 11824..15714 | 68230 | DUF6079-family-protein-[Streptomyces-longispororuber] |
| WDV31004.1 | Type2.Pgl.like.BREX | BrxA-DNA-binding->?->PglW->?->?->N6-METHYLASE->?->N6-METHYLASE-p-><-DUF397\|\|DUF6079*->PglZ-><-HTH_Tnp_4+DDE_Tnp_4\|\|HTH_Tnp_4+DDE_Tnp_4-><-CR-ATPase9<-?<-?<-?\|\|GNTR-HTH+UTRA-> | 1272 | Bacteria>Terrabacteria_group>Actinomycetota>Actinomycetes>Kitasatosporales>Streptomycetaceae | GCA_028884895.1 | Streptomyces_sp._AD16 | OIM90_05400 | 1179426..1183244 | 1816220 | phage-resistance-protein-[Streptomyces-sp.-AD16] |
| WP_173872026.1 | Type2.Pgl.like.BREX | <-JetB<-DUF2397\|\|PglW-><-BrxD-AAA<-ParA-Soj-PloopNTPase<-HEPN\|\|N6-METHYLASE-><-DUF397\|\|DUF6079*->PglZ->BrxD-AAA->SF1-SF2-Helicase+wHTH+Small-B-barrel+DUF1998-><-Beta-lactamase<-?<-?\|\|?-><-His_Phos_1<-Methyltransf_21 | 1321 | Bacteria>Terrabacteria_group>Actinomycetota>Actinomycetes>Kitasatosporales>Streptomycetaceae | GCA_030820635.1 | Streptomyces_albus | FM076_18390 | 4166115..4170080 | 1888 | PglY-protein-[Streptomyces-albus] |
| WP_012378658.1 | Type2.Pgl.like.BREX | PglW->N6-METHYLASE-><-DUF397<-HTH_31+DUF5753\|\|DUF1995->DUF2171->REase+SF1-SF2-Helicase->DUF6079*->PglZ-><-RNaseH\|\|BrxD-AAA->SF1-SF2-Helicase+wHTH+Small-B-barrel+DUF1998-><-GNTR-HTH+Aminotran_1_2\|\|?-><-?<-?<-ABC-ATPase+SbcC | 1299 | Bacteria>Terrabacteria_group>Actinomycetota>Actinomycetes>Kitasatosporales>Streptomycetaceae | GCF_000010605.1 | Streptomyces | SGR_RS07860 | 1889546..1893445 | 1883 | MULTISPECIES:-hypothetical-protein-[Streptomyces] |
| WP_015660833.1 | Type2.Pgl.like.BREX | <-wHTH+SAD+HNH\|\|PglW->N6-METHYLASE-><-ST-Hiskin\|\|DUF397->?-><-PIN<-MetJarc-HTH\|\|DUF6079*->PglZ-><-CBS+CBS+HEPN<-ParB+HNH\|\|BrxD-AAA->SF1-SF2-Helicase+wHTH+Small-B-barrel+DUF1998-><-?\|\|PadR-wHTH->JetB-> | 1294 | Bacteria>Terrabacteria_group>Actinomycetota>Actinomycetes>Kitasatosporales>Streptomycetaceae | GCF_000349325.1 | Streptomyces_davaonensis | BN159_RS30305 | 6862049..6865933 | 348043 | hypothetical-protein-[Streptomyces-davaonensis] |
| WP_015662322.1 | Type2.Pgl.like.BREX | REase+DUF5655-><-pPIWI+MID+RNaseH<-pPIWI+SF1-SF2-Helicase<-pPIWI+pPIWI<-HTH_3+HTH_3<-HTH_3+HTH_3<-SF1-SF2-Helicase+wHTH+Small-B-barrel+DUF1998<-BrxD-AAA<-PglZ<-DUF6079*<-N6-METHYLASE<-PglW\|\|?->?->?-><-?<-?<-?<-DnaB+DnaB | 1300 | Bacteria>Terrabacteria_group>Actinomycetota>Actinomycetes>Kitasatosporales>Streptomycetaceae | GCF_000349325.1 | Streptomyces_davaonensis | BN159_RS37770 | 8561539..8565441 | 348043 | hypothetical-protein-[Streptomyces-davaonensis] |
| WP_018530191.1 | Type2.Pgl.like.BREX | sGTP->MetJarc-HTH->PIN-><-ANK+ANK<-?<-SF1-SF2-Helicase+wHTH+Small-B-barrel+DUF1998<-BrxD-AAA<-PglZ<-DUF6079*<-ST-Hiskin\|\|HTH_31+DUF5753->DUF397-><-N6-METHYLASE<-ORC-CDC6<-ParA-Soj-PloopNTPase<-HEPN<-PglW<-REase+SF1-SF2-Helicase | 1308 | Bacteria>Terrabacteria_group>Actinomycetota>Actinomycetes>Kitasatosporales>Streptomycetaceae | GCF_000373565.1 | unclassified_Streptomyces | B072_RS0112005 | 248212..252138 | 2593676 | MULTISPECIES:-hypothetical-protein-[unclassified-Streptomyces] |
| WP_020115998.1 | Type2.Pgl.like.BREX | PAS+GAF+HISKIN+REC+GAF+GAF+PP2C-><-SF1-SF2-Helicase+wHTH+Small-B-barrel+DUF1998<-BrxD-AAA<-PglZ<-DUF6079*\|\|?-><-ST-Hiskin\|\|HTH_31+DUF5753->DUF397-><-N6-METHYLASE\|\|HEPN->ParA-Soj-PloopNTPase->ORC-CDC6-><-PglW | 1301 | Bacteria>Terrabacteria_group>Actinomycetota>Actinomycetes>Kitasatosporales>Streptomycetaceae | GCF_000383615.1 | Streptomyces_canus | H293_RS0100895 | 199491..203396 | 58343 | hypothetical-protein-[Streptomyces-canus] |
| WP_020140010.1 | Type2.Pgl.like.BREX | REase->REase+SF1-SF2-Helicase-><-SF1-SF2-Helicase+wHTH+Small-B-barrel+DUF1998<-BrxD-AAA<-PglZ<-DUF6079*\|\|?-><-ST-Hiskin\|\|HTH_31+DUF5753->DUF397-><-N6-METHYLASE\|\|HEPN->ParA-Soj-PloopNTPase->ORC-CDC6-><-PglW | 1301 | Bacteria>Terrabacteria_group>Actinomycetota>Actinomycetes>Kitasatosporales>Streptomycetaceae | GCF_000383655.1 | Streptomyces_sp._351MFTsu5.1 | H295_RS0134960 | 91744..95649 | 1172180 | hypothetical-protein-[Streptomyces-sp.-351MFTsu5.1] |
| WP_020868709.1 | Type2.Pgl.like.BREX | PglW->HEPN->ParA-Soj-PloopNTPase->ORC-CDC6->N6-METHYLASE-><-?<-?\|\|HTH_31+DUF5753->DUF397->DUF6079*->PglZ->BrxD-AAA->SF1-SF2-Helicase+wHTH+Small-B-barrel+DUF1998->XIS-HTH-><-sGTP\|\|?-><-ROK-HTH | 1306 | Bacteria>Terrabacteria_group>Actinomycetota>Actinomycetes>Kitasatosporales>Streptomycetaceae | GCF_000418455.1 | Streptomyces_rapamycinicus | M271_RS19160 | 4730013..4733933 | 1226757 | hypothetical-protein-[Streptomyces-rapamycinicus] |
| WP_023416579.1 | Type2.Pgl.like.BREX | ADH_N+ADH_zinc_N->BrxA-DNA-binding->?->PglW->N6-METHYLASE-><-DUF397<-HTH_31+DUF5753\|\|DUF6079*->PglZ->BrxD-AAA->SF1-SF2-Helicase+wHTH+Small-B-barrel+DUF1998-><-TRAM+SAM-methylase<-AA_permease_2 | 1298 | Bacteria>Terrabacteria_group>Actinomycetota>Actinomycetes>Kitasatosporales>Streptomycetaceae | GCF_000495755.1 | unclassified_Streptomyces | B591_RS05565 | 1334058..1337954 | 2593676 | MULTISPECIES:-hypothetical-protein-[unclassified-Streptomyces] |
| WP_037951095.1 | Type2.Pgl.like.BREX | <-TOPRIM<-OLD-ABC\|\|N6-METHYLASE-><-SF1-SF2-Helicase+HKD-DNase\|\|PglW->N6-METHYLASE-><-ST-Hiskin\|\|HTH_31+DUF5753->DUF397->DUF6079*->PglZ-><-CBS+HEPN<-ParB+HNH\|\|BrxD-AAA->SF1-SF2-Helicase+wHTH+Small-B-barrel+DUF1998->EVE+SAD+REase-><-?\|\|REase+SF1-SF2-Helicase->REase+SF1-SF2-Helicase-> | 1320 | Bacteria>Terrabacteria_group>Actinomycetota>Actinomycetes>Kitasatosporales>Streptomycetaceae | GCF_000590515.1 | Streptomyces_sp._PRh5 | Z951_RS11140 | 96158..100120 | 1158056 | hypothetical-protein-[Streptomyces-sp.-PRh5] |
| WP_033825778.1 | Type2.Pgl.like.BREX | <-DSBH\|\|?->?-><-?\|\|PglW->N6-METHYLASE->DUF6079*->PglZ->BrxD-AAA->SF1-SF2-Helicase+wHTH+Small-B-barrel+DUF1998->pPIWI+pPIWI->pPIWI+SF1-SF2-Helicase->pPIWI+MID+RNaseH-><-?<-Phage_integrase | 1312 | Bacteria>Terrabacteria_group>Actinomycetota>Actinomycetes>Kitasatosporales>Streptomycetaceae | GCF_000710405.2 | unclassified_Kitasatospora | BI06_RS11260 | 47992..51930 | 2633591 | MULTISPECIES:-DUF6079-family-protein-[unclassified-Kitasatospora] |
| WP_030321491.1 | Type2.Pgl.like.BREX | <-ABhydrolase\|\|SIGMA-HTH-><-?\|\|REase+SF1-SF2-Helicase->?-><-REase+SF1-SF2-Helicase<-SF1-SF2-Helicase+wHTH+Small-B-barrel+DUF1998<-BrxD-AAA<-PglZ<-DUF6079*\|\|SF1-SF2-Helicase+HKD-DNase-><-ST-Hiskin\|\|HTH_31+DUF5753->DUF397-><-N6-METHYLASE\|\|HEPN->ParA-Soj-PloopNTPase->ORC-CDC6-><-PglW | 1318 | Bacteria>Terrabacteria_group>Actinomycetota>Actinomycetes>Kitasatosporales>Streptomycetaceae | GCF_000717595.1 | Streptomyces_flavochromogenes | IF00_RS0124625 | 51178..55134 | 68199 | hypothetical-protein-[Streptomyces-flavochromogenes] |
| WP_030293901.1 | Type2.Pgl.like.BREX | <-HTH+HTH\|\|?->ST-Hiskin->?->\|\|?-><-SF1-SF2-Helicase+wHTH+Small-B-barrel+DUF1998<-BrxD-AAA<-DndB<-PglZ<-DUF6079*<-REase+SF1-SF2-Helicase<-DUF2171<-DUF1995\|\|HTH_31+DUF5753->DUF397-><-N6-METHYLASE<-PglW<-Pkinase+Pkinase<-Pkinase | 1300 | Bacteria>Terrabacteria_group>Actinomycetota>Actinomycetes>Kitasatosporales>Streptomycetaceae | GCF_000717655.1 | Streptomyces_katrae | IF09_RS0109400 | 71835..75737 | 68223 | hypothetical-protein-[Streptomyces-katrae] |
| WP_030145801.1 | Type2.Pgl.like.BREX | MORC+REase-><-AAA-ATPase<-XIS-HTH\|\|MarR-HTH->Phage_int_SAM_3+Phage_integrase-><-XIS-HTH<-SF1-SF2-Helicase+wHTH+Small-B-barrel+DUF1998<-BrxD-AAA<-PglZ<-DUF6079*<-ST-Hiskin\|\|HTH_31+DUF5753->DUF397-><-N6-METHYLASE<-ORC-CDC6<-ParA-Soj-PloopNTPase<-HEPN<-PglW<-SWACOS+TPR+TPR+LuxR-HTH | 1303 | Bacteria>Terrabacteria_group>Actinomycetota>Actinomycetes>Kitasatosporales>Streptomycetaceae | GCF_000717995.1 | Streptomyces_violaceorubidus | IE94_RS0103125 | 657978..661889 | 284042 | hypothetical-protein-[Streptomyces-violaceorubidus] |
| WP_030698798.1 | Type2.Pgl.like.BREX | <-ST-Hiskin\|\|HTH_3->?->DUF4096+DDE_Tnp_1+DDE_Tnp_1_2-><-SF1-SF2-Helicase+wHTH+Small-B-barrel+DUF1998<-BrxD-AAA<-PglZ<-DUF6079*<-ST-Hiskin\|\|HTH_31+DUF5753->DUF397-><-N6-METHYLASE\|\|HEPN->ParA-Soj-PloopNTPase->ORC-CDC6-><-PglW<-Peptidase_C14 | 1324 | Bacteria>Terrabacteria_group>Actinomycetota>Actinomycetes>Kitasatosporales>Streptomycetaceae | GCF_000719495.1 | Streptomyces_albidoflavus | IF80_RS0123230 | 18564..22538 | 1886 | hypothetical-protein-[Streptomyces-albidoflavus] |
| WP_030881234.1 | Type2.Pgl.like.BREX | MetJarc-HTH->PIN-><-sGTP\|\|GDResolvase+Recombinase+Zn_ribbon_recom->?->sGTP+sGTP+GTPase-AIG->sGTP->PglW->N6-METHYLASE->DUF6079*->PglZ->BrxD-AAA->SF1-SF2-Helicase+wHTH+Small-B-barrel+DUF1998-><-N6-N4-METHYLASE\|\|?->TraM->?->VirB6-> | 1298 | Bacteria>Terrabacteria_group>Actinomycetota>Actinomycetes>Kitasatosporales>Streptomycetaceae | GCF_000720405.1 | Streptomyces_sp._NRRL_B-24051 | IG70_RS0127770 | 73848..77744 | 1463832 | hypothetical-protein-[Streptomyces-sp.-NRRL-B-24051] |
| WP_063767425.1 | Type2.Pgl.like.BREX | <-SF1-SF2-Helicase+wHTH+Small-B-barrel+DUF1998<-BrxD-AAA<-PglZ<-DUF6079*<-?<-?\|\|GntR-HTH-><-N6-METHYLASE<-ORC-CDC6<-ParA-Soj-PloopNTPase<-HEPN<-PglW | 1353 | Bacteria>Terrabacteria_group>Actinomycetota>Actinomycetes>Kitasatosporales>Streptomycetaceae | GCF_000721305.1 | Streptomyces_sp._NRRL_S-1868 | IG92_RS0121360 | 18400..22461 | 1463892 | hypothetical-protein-[Streptomyces-sp.-NRRL-S-1868] |
| WP_037930213.1 | Type2.Pgl.like.BREX | PglW->?-><-ORC-CDC6<-ParA-Soj-PloopNTPase<-HEPN\|\|N6-METHYLASE-><-DUF397<-HTH_31+DUF5753\|\|?->DUF6079*->PglZ->BrxD-AAA->SF1-SF2-Helicase+wHTH+Small-B-barrel+DUF1998->NACHT-><-?<-?<-?\|\|?-><-LACI-HTH+PBPI-> | 1323 | Bacteria>Terrabacteria_group>Actinomycetota>Actinomycetes>Kitasatosporales>Streptomycetaceae | GCF_000725125.1 | Streptomyces_toyocaensis | BU52_RS07635 | 197363..201334 | 55952 | hypothetical-protein-[Streptomyces-toyocaensis] |
| WP_031178323.1 | Type2.Pgl.like.BREX | ACET-><-ABhydrolase<-?<-ABhydrolase\|\|HisKA_3+HATPase_c->REC+OmpR-HTH-><-SF1-SF2-Helicase+wHTH+Small-B-barrel+DUF1998<-BrxD-AAA<-PglZ<-DUF6079*<-?<-?\|\|GntR-HTH-><-N6-METHYLASE<-ORC-CDC6<-ParA-Soj-PloopNTPase<-HEPN<-PglW | 1313 | Bacteria>Terrabacteria_group>Actinomycetota>Actinomycetes>Kitasatosporales>Streptomycetaceae | GCF_000725635.1 | Streptomyces_sp._NRRL_B-3253 | IO21_RS0123010 | 108342..112283 | 1463837 | hypothetical-protein-[Streptomyces-sp.-NRRL-B-3253] |
| WP_031126955.1 | Type2.Pgl.like.BREX | GNTR-HTH->HKD-DNase+LuxR-HTH-><-SF1-SF2-Helicase+wHTH+Small-B-barrel+DUF1998<-BrxD-AAA\|\|RNaseH-><-PglZ<-DUF6079*<-REase+SF1-SF2-Helicase<-DUF2171<-DUF1995\|\|HTH_31+DUF5753->DUF397-><-N6-METHYLASE<-PglW | 1299 | Bacteria>Terrabacteria_group>Actinomycetota>Actinomycetes>Kitasatosporales>Streptomycetaceae | GCF_000725705.1 | Streptomyces | AOK13_RS15160 | 17218..21117 | 1883 | MULTISPECIES:-hypothetical-protein-[Streptomyces] |
| WP_043500398.1 | Type2.Pgl.like.BREX | <-N6-METHYLASE\|\|OLD-ABC->TOPRIM-><-?<-SF1-SF2-Helicase+wHTH+Small-B-barrel+DUF1998<-BrxD-AAA<-PglZ<-DUF6079*<-N6-METHYLASE<-PglW<-DUF397\|\|ST-Hiskin->?>SF1-SF2-Helicase-><-?\|\|HTH+McrB-AAA->McrC-NTD+REase-> | 1294 | Bacteria>Terrabacteria_group>Actinomycetota>Actinomycetes>Kitasatosporales>Streptomycetaceae | GCF_000761215.1 | Streptomyces_glaucescens | SGLAU_RS10295 | 2386733..2390617 | 1907 | hypothetical-protein-[Streptomyces-glaucescens] |
| WP_042366886.1 | Type2.Pgl.like.BREX | <-XIS-HTH\|\|HTH_3->Phage_int_SAM_3+Phage_integrase-><-DDE_Tnp_1<-RVT<-REase<-SF1-SF2-Helicase+wHTH+Small-B-barrel+DUF1998<-BrxD-AAA<-PglZ<-DUF6079*<-N6-METHYLASE<-ORC-CDC6<-ParA-Soj-PloopNTPase<-HEPN<-PglW\|\|?-><-tRNA\|\|?-><-?<-SF1-SF2-Helicase | 1311 | Bacteria>Terrabacteria_group>Actinomycetota>Actinomycetes>Kitasatosporales>Streptomycetaceae | GCF_000787815.1 | Streptacidiphilus_neutrinimicus | TR48_RS14550 | 88859..92794 | 105420 | hypothetical-protein-[Streptacidiphilus-neutrinimicus] |
| WP_052192753.1 | Type2.Pgl.like.BREX | ADH_N+ADH_zinc_N->BrxA-DNA-binding->?->PglW->N6-METHYLASE-><-DUF397<-HTH_31+DUF5753\|\|DUF6079*->PglZ->BrxD-AAA->SF1-SF2-Helicase+wHTH+Small-B-barrel+DUF1998-><-TRAM+SAM-methylase+METHYLASE<-AA_permease_2 | 1298 | Bacteria>Terrabacteria_group>Actinomycetota>Actinomycetes>Kitasatosporales>Streptomycetaceae | GCF_000797385.1 | Streptomyces_sp._CNQ431 | PH39_RS06250 | 109132..113028 | 1571532 | hypothetical-protein-[Streptomyces-sp.-CNQ431] |
| WP_086083008.1 | Type2.Pgl.like.BREX | <-ADH_N+ADH_zinc_N_2<-TetR-HTH<-?\|\|?-><-SF1-SF2-Helicase+wHTH+Small-B-barrel+DUF1998<-BrxD-AAA\|\|Phage_holin-><-PglZ<-DUF6079*<-NUDIX<-?<-DUF397<-HTH_31+DUF5753\|\|ST-Hiskin-><-N6-METHYLASE<-PglW<-SIGMA-HTH\|\|MerR-HTH-> | 1302 | Bacteria>Terrabacteria_group>Actinomycetota>Actinomycetes>Kitasatosporales>Streptomycetaceae | GCF_000802245.2 | Streptomyces_pluripotens | LK06_RS01830 | 426923..430831 | 1355015 | phage-resistance-protein-[Streptomyces-pluripotens] |
| WP_044379847.1 | Type2.Pgl.like.BREX | TRAM+SAM-methylase->REase+SF1-SF2-Helicase-><-?<-REase+SF1-SF2-Helicase<-SF1-SF2-Helicase+wHTH+Small-B-barrel+DUF1998<-BrxD-AAA\|\|ParB+HNH->CBS+HEPN-><-PglZ<-DUF6079*<-DUF2171<-DUF1995\|\|DUF397-><-N6-METHYLASE<-PglW\|\|?->JetB->SbcCD_C->DUF3323+TOPRIM-> | 1298 | Bacteria>Terrabacteria_group>Actinomycetota>Actinomycetes>Kitasatosporales>Streptomycetaceae | GCF_000935185.3 | Streptomyces_noursei | SE95_RS29285 | 6702907..6706803 | 1971 | hypothetical-protein-[Streptomyces-noursei] |
| WP_047015461.1 | Type2.Pgl.like.BREX | RhoN+RNase+S1-><-MerR-HTH<-?\|\|DSBH+AraC-HTH+AraC-HTH->ncRNA-><-PglZ<-DUF6079*<-N6-METHYLASE<-PglW\|\|BrxD-AAA->SF1-SF2-Helicase+wHTH+Small-B-barrel+DUF1998->MNS+REase-> | 1287 | Bacteria>Terrabacteria_group>Actinomycetota>Actinomycetes>Kitasatosporales>Streptomycetaceae | GCF_001011035.1 | Streptomyces_sp._CNQ-509 | AA958_RS07555 | 1821426..1825289 | 444103 | DUF6079-family-protein-[Streptomyces-sp.-CNQ-509] |
| WP_047139866.1 | Type2.Pgl.like.BREX | <-Phage_integrase<-HEPN+ZNR<-?<-?<-?<-?<-SF1-SF2-Helicase+wHTH+Small-B-barrel+DUF1998<-BrxD-AAA<-PglZ<-DUF6079*\|\|HTH_31+DUF5753->DUF397-><-N6-METHYLASE<-PglW\|\|<-REase<-?\|\|REase+SF1-SF2-Helicase-><-?<-REase+SF1-SF2-Helicase | 1298 | Bacteria>Terrabacteria_group>Actinomycetota>Actinomycetes>Kitasatosporales>Streptomycetaceae | GCF_001014595.1 | Streptomyces_sp._KE1 | WQ59_RS08655 | 50789..54685 | 1638939 | hypothetical-protein-[Streptomyces-sp.-KE1] |
| WP_234319629.1 | Type2.Pgl.like.BREX | HTH_26->DUF772+DDE_Tnp_1_6-><-?\|\|HisKA_3+HATPase_c->REC+OmpR-HTH-><-SF1-SF2-Helicase+wHTH+Small-B-barrel+DUF1998<-BrxD-AAA<-PglZ<-DUF6079*<-?<-?\|\|GntR-HTH-><-N6-METHYLASE<-ORC-CDC6<-ParA-Soj-PloopNTPase<-HEPN<-PglW | 1343 | Bacteria>Terrabacteria_group>Actinomycetota>Actinomycetes>Kitasatosporales>Streptomycetaceae | GCF_001014595.1 | Streptomyces_sp._KE1 | WQ59_RS14830 | 95422..99453 | 1638939 | hypothetical-protein-[Streptomyces-sp.-KE1] |
| WP_048572280.1 | Type2.Pgl.like.BREX | ABC-ATPase+SbcC->REase+SF1-SF2-Helicase->?-><-?\|\|PglW->N6-METHYLASE->DUF6079*->PglZ-><-DUF397<-HTH_31+DUF5753\|\|ST-Hiskin-><-CBS+CBS+HEPN<-ParB+HNH<-RVT+ArdCN+MPTase\|\|BrxD-AAA->SF1-SF2-Helicase+wHTH+Small-B-barrel+DUF1998-> | 1294 | Bacteria>Terrabacteria_group>Actinomycetota>Actinomycetes>Kitasatosporales>Streptomycetaceae | GCF_001047315.1 | Streptomyces_leeuwenhoekii | ACH49_RS07740 | 40979..44863 | 1437453 | hypothetical-protein-[Streptomyces-leeuwenhoekii] |
| WP_053623542.1 | Type2.Pgl.like.BREX | PglW-><-ORC-CDC6<-ParA-Soj-PloopNTPase<-HEPN\|\|N6-METHYLASE-><-DUF397\|\|<-?<-?<-BrxA-DNA-binding\|\|DUF6079*->PglZ->BrxD-AAA->SF1-SF2-Helicase+wHTH+Small-B-barrel+DUF1998->PhdYeFM_antitox->RelE-ParE->?-><-MarR-HTH | 1308 | Bacteria>Terrabacteria_group>Actinomycetota>Actinomycetes>Kitasatosporales>Streptomycetaceae | GCF_001279425.1 | Streptomyces_sp._NRRL_F-2295 | ADK88_RS11800 | 69838..73764 | 1519477 | hypothetical-protein-[Streptomyces-sp.-NRRL-F-2295] |
| WP_055507306.1 | Type2.Pgl.like.BREX | TetR-HTH-><-STAS<-LuxR-HTH\|\|?->RADICAL-SAM-><-?<-?\|\|PglW->N6-METHYLASE->DUF6079*->PglZ->BrxD-AAA->SF1-SF2-Helicase+wHTH+Small-B-barrel+DUF1998-><-tRNA<-TRAM+SAM-methylase+METHYLASE<-AA_permease_2 | 1304 | Bacteria>Terrabacteria_group>Actinomycetota>Actinomycetes>Kitasatosporales>Streptomycetaceae | GCF_001418335.1 | Streptomyces_aurantiacus | AOK01_RS00950 | 70246..74160 | 47760 | hypothetical-protein-[Streptomyces-aurantiacus] |
| WP_058042191.1 | Type2.Pgl.like.BREX | AA_permease_2->TRAM+SAM-methylase->REase+DUF5655-><-REase<-?<-SF1-SF2-Helicase+wHTH+Small-B-barrel+DUF1998<-BrxD-AAA\|\|REase-><-PglZ<-DUF6079*<-N6-METHYLASE<-PglW\|\|REase+N6_METHYLASE->REase-><-DnaB+DnaB<-DNA_METHYLASE<-VirD4-TraG+VirD4-FtsK | 1299 | Bacteria>Terrabacteria_group>Actinomycetota>Actinomycetes>Kitasatosporales>Streptomycetaceae | GCF_001445655.1 | Streptomyces_roseifaciens | AS857_RS06470 | 89914..93813 | 1488406 | hypothetical-protein-[Streptomyces-roseifaciens] |
| WP_067237065.1 | Type2.Pgl.like.BREX | <-EVE+SAD+REase\|\|PglW-><-ORC-CDC6<-ParA-Soj-PloopNTPase<-HEPN\|\|N6-METHYLASE->DUF397->DUF6079*->PglZ->BrxD-AAA->SF1-SF2-Helicase+wHTH+Small-B-barrel+DUF1998->?->?->HTH_31+DUF5753->ST-Hiskin->SIGMA-HTH-> | 1325 | Bacteria>Terrabacteria_group>Actinomycetota>Actinomycetes>Kitasatosporales>Streptomycetaceae | GCF_001514125.1 | Streptomyces_longwoodensis | AQJ30_RS22670 | 258338..262315 | 68231 | PglY-protein-[Streptomyces-longwoodensis] |
| WP_061915750.1 | Type2.Pgl.like.BREX | <-35exo<-?<-?\|\|GDResolvase+HIN-HTH-><-XIS-HTH\|\|REase+DUF5655-><-SF1-SF2-Helicase+wHTH+Small-B-barrel+DUF1998<-BrxD-AAA<-PglZ<-DUF6079*<-N6-METHYLASE<-PglW<-?<-DnaB+DnaB<-DNA_METHYLASE<-PAPS-reduct | 1299 | Bacteria>Terrabacteria_group>Actinomycetota>Actinomycetes>Kitasatosporales>Streptomycetaceae | GCF_001514215.1 | Streptomyces_bungoensis | AQJ66_RS02630 | 666355..670254 | 285568 | hypothetical-protein-[Streptomyces-bungoensis] |
| WP_060944831.1 | Type2.Pgl.like.BREX | <-REase+SF1-SF2-Helicase<-?<-REase+SF1-SF2-Helicase<-SF1-SF2-Helicase+wHTH+Small-B-barrel+DUF1998<-BrxD-AAA\|\|ParB+HNH->CBS+HEPN-><-PglZ<-DUF6079*<-DUF397<-HTH_31+DUF5753\|\|ST-Hiskin-><-N6-METHYLASE<-PglW\|\|REase+SF1-SF2-Helicase->EVE+HNH-><-N6-METHYLASE\|\|OLD-ABC-> | 1320 | Bacteria>Terrabacteria_group>Actinomycetota>Actinomycetes>Kitasatosporales>Streptomycetaceae | GCF_001553455.1 | Streptomyces_hygroscopicus | AXW62_RS14140 | 42881..46843 | 1912 | hypothetical-protein-[Streptomyces-hygroscopicus] |
| WP_062010201.1 | Type2.Pgl.like.BREX | N6-METHYLASE-p-><-SF1-SF2-Helicase+HKD-DNase\|\|PglW->N6-METHYLASE-><-ST-Hiskin\|\|HTH_31+DUF5753->DUF397->DUF6079*->PglZ-><-CBS+HEPN<-ParB+HNH\|\|BrxD-AAA->SF1-SF2-Helicase+wHTH+Small-B-barrel+DUF1998->REase+SF1-SF2-Helicase->REase+SF1-SF2-Helicase-><-?<-NTP_transf_9 | 1320 | Bacteria>Terrabacteria_group>Actinomycetota>Actinomycetes>Kitasatosporales>Streptomycetaceae | GCF_001570645.1 | Streptomyces_hygroscopicus | AXW65_RS13440 | 81555..85517 | 1912 | hypothetical-protein-[Streptomyces-hygroscopicus] |
| WP_061404090.1 | Type2.Pgl.like.BREX | <-Peptidase_C14+SF1-SF2-Helicase<-?<-REase\|\|PglW->N6-METHYLASE-><-DUF397<-HTH_31+DUF5753\|\|DUF6079*->PglZ->BrxD-AAA->SF1-SF2-Helicase+wHTH+Small-B-barrel+DUF1998-><-TRAM+SAM-methylase<-AA_permease_2 | 1298 | Bacteria>Terrabacteria_group>Actinomycetota>Actinomycetes>Kitasatosporales>Streptomycetaceae | GCF_001577385.1 | Streptomyces_albidoflavus | Salbus254_RS06075 | 1445792..1449688 | 1886 | hypothetical-protein-[Streptomyces-albidoflavus] |
| WP_062926533.1 | Type2.Pgl.like.BREX | TRD+TRD->REase+N6_Mtase-><-wHTH+SAD+HNH\|\|PglW->N6-METHYLASE-><-ST-Hiskin\|\|HTH_31+DUF5753->DUF397->MNS+REase->DUF6079*->PglZ-><-CBS+HEPN<-ParB+HNH\|\|BrxD-AAA->SF1-SF2-Helicase+wHTH+Small-B-barrel+DUF1998-><-ncRNA\|\|?->?-><-DSBH+AraC-HTH+AraC-HTH<-REase | 1309 | Bacteria>Terrabacteria_group>Actinomycetota>Actinomycetes>Kitasatosporales>Streptomycetaceae | GCF_001611795.1 | Streptomyces_qaidamensis | A4E84_RS11860 | 2715868..2719797 | 1783515 | hypothetical-protein-[Streptomyces-qaidamensis] |
| WP_063727790.1 | Type2.Pgl.like.BREX | PglW-><-PIN<-HTH+MPTase\|\|N6-METHYLASE-><-DUF397<-HTH_31+DUF5753\|\|ST-Hiskin->?-><-?\|\|DUF6079*->PglZ->BrxD-AAA->SF1-SF2-Helicase+wHTH+Small-B-barrel+DUF1998-><-RNaseH | 1299 | Bacteria>Terrabacteria_group>Actinomycetota>Actinomycetes>Kitasatosporales>Streptomycetaceae | GCF_001650215.1 | Streptomyces_sp._RTd22 | A7B19_RS30860 | 7245544..7249443 | 1841249 | hypothetical-protein-[Streptomyces-sp.-RTd22] |
| WP_067397487.1 | Type2.Pgl.like.BREX | REase->?->SF1-SF2-Helicase-><-SF1-SF2-Helicase+wHTH+Small-B-barrel+DUF1998<-BrxD-AAA<-?<-PglZ<-DUF6079*<-N6-METHYLASE<-PglW<-DUF397<-HTH_31+DUF5753<-REase+SF1-SF2-Helicase<-?<-CR-ATPase9<-?<-35exo+SF1-SF2-Helicase | 1298 | Bacteria>Terrabacteria_group>Actinomycetota>Actinomycetes>Kitasatosporales>Streptomycetaceae | GCF_001653115.1 | Streptomyces_sp._F-3 | STXM2123_RS19035 | 36552..40448 | 1840095 | phage-resistance-protein-[Streptomyces-sp.-F-3] |
| WP_067441133.1 | Type2.Pgl.like.BREX | AbiEii->ANK+ANK-><-Phage_int_SAM_3+Phage_integrase<-XIS-HTH<-Phage_integrase<-REase+N6_METHYLASE\|\|PglW->N6-METHYLASE->DUF6079*->PglZ->BrxD-AAA->SF1-SF2-Helicase+wHTH+Small-B-barrel+DUF1998->?\|\|PIWI->pPIWI+SF1-SF2-Helicase->pPIWI+MID+RNaseH-><-TRAM+SAM-methylase<-AA_permease_2 | 1297 | Bacteria>Terrabacteria_group>Actinomycetota>Actinomycetes>Kitasatosporales>Streptomycetaceae | GCF_001685355.1 | Streptomyces_lincolnensis | SLCG_RS11545 | 2586949..2590842 | 1915 | phage-resistance-protein-[Streptomyces-lincolnensis] |
| WP_071373158.1 | Type2.Pgl.like.BREX | <-RNaseH<-SF1-SF2-Helicase+wHTH+Small-B-barrel+DUF1998<-BrxD-AAA\|\|PglW->N6-METHYLASE->DUF6079*->PglZ-> | 1279 | Bacteria>Terrabacteria_group>Actinomycetota>Actinomycetes>Kitasatosporales>Streptomycetaceae | GCF_001866675.1 | Streptomyces_sp._MUSC_14 | BIV25_RS51055 | 161897..165736 | 1354889 | DUF6079-family-protein-[Streptomyces-sp.-MUSC-14] |
| WP_073727340.1 | Type2.Pgl.like.BREX | <-SF1-SF2-Helicase+HKD-DNase\|\|PglW->N6-METHYLASE-><-PIN<-ST-Hiskin\|\|HTH_31+DUF5753->DUF397->PhdYeFM_antitox->PIN->DUF6079*->PglZ-><-CBS+CBS+HEPN<-ParB+HNH\|\|BrxD-AAA->SF1-SF2-Helicase+wHTH+Small-B-barrel+DUF1998-><-?\|\|TPR-> | 1307 | Bacteria>Terrabacteria_group>Actinomycetota>Actinomycetes>Kitasatosporales>Streptomycetaceae | GCF_001905125.1 | Streptomyces_sp._TSRI0281 | A6A29_RS36835 | 505881..509804 | 1718998 | phage-resistance-protein-[Streptomyces-sp.-TSRI0281] |
| WP_073931399.1 | Type2.Pgl.like.BREX | REase+SF1-SF2-Helicase->PglW->N6-METHYLASE-><-ST-Hiskin\|\|HTH_31+DUF5753->?->?->?->?->DUF6079*->PglZ-><-CBS+HEPN<-ParB+HNH<-?\|\|BrxD-AAA->SF1-SF2-Helicase+wHTH+Small-B-barrel+DUF1998-><-Phage_holin<-DUF4011+SF1-SF2-Helicase+REase<-SF1-SF2-Helicase | 1293 | Bacteria>Terrabacteria_group>Actinomycetota>Actinomycetes>Kitasatosporales>Streptomycetaceae | GCF_001905725.1 | Streptomyces_sp._CB02400 | AMK33_04750 | 1152826..1156707 | 1703944 | phage-resistance-protein-[Streptomyces-sp.-CB02400] |
| WP_073897500.1 | Type2.Pgl.like.BREX | PglW->N6-METHYLASE-><-DUF397<-HTH_31+DUF5753\|\|DUF1995->DUF2171->REase+SF1-SF2-Helicase->DUF6079*->PglZ->DndB->BrxD-AAA->SF1-SF2-Helicase+wHTH+Small-B-barrel+DUF1998-><-MORC+DUF3684+DUF3883 | 1294 | Bacteria>Terrabacteria_group>Actinomycetota>Actinomycetes>Kitasatosporales>Streptomycetaceae | GCF_001905855.1 | Streptomyces_sp._CB01883 | AMK32_RS27770 | 364608..368492 | 1703943 | phage-resistance-protein-[Streptomyces-sp.-CB01883] |
| WP_077193671.1 | Type2.Pgl.like.BREX | HIN-HTH-><-?\|\|REase+SF1-SF2-Helicase-><-?<-REase+SF1-SF2-Helicase<-SF1-SF2-Helicase+wHTH+Small-B-barrel+DUF1998<-BrxD-AAA\|\|RNaseH-><-PglZ<-DUF6079*\|\|HTH_31+DUF5753->DUF397-><-N6-METHYLASE<-PglW\|\|?->REC+OmpR-HTH->HerA/FtsK-> | 1301 | Bacteria>Terrabacteria_group>Actinomycetota>Actinomycetes>Kitasatosporales>Streptomycetaceae | GCF_001984445.1 | Streptomyces_lydicus | BXT88_RS28535 | 6380732..6384637 | 47763 | phage-resistance-protein-[Streptomyces-lydicus] |
| WP_078633283.1 | Type2.Pgl.like.BREX | <-SF1-SF2-Helicase+wHTH+Small-B-barrel+DUF1998<-BrxD-AAA\|\|ParB+HNH->CBS+HEPN->Phage_holin-><-PglZ<-DUF6079*\|\|MetJarc-HTH->PIN-><-?<-DUF397<-HTH_31+DUF5753\|\|ST-Hiskin-><-N6-METHYLASE<-PglW\|\|wHTH+SAD+HNH-> | 1295 | Bacteria>Terrabacteria_group>Actinomycetota>Actinomycetes>Kitasatosporales>Streptomycetaceae | GCF_002019855.1 | Streptomyces_antibioticus | HCX60_RS12060 | 2686146..2690033 | 1890 | phage-resistance-protein-[Streptomyces-antibioticus] |
| WP_081522440.1 | Type2.Pgl.like.BREX | <-SAD(SRA)+wHTH+HNH\|\|PglW->N6-METHYLASE-><-REase<-ST-Hiskin\|\|HTH_31+DUF5753->DUF397->DUF6079*->PglZ-><-CBS+HEPN<-ParB+HNH\|\|BrxD-AAA->SF1-SF2-Helicase+wHTH+Small-B-barrel+DUF1998-> | 1312 | Bacteria>Terrabacteria_group>Actinomycetota>Actinomycetes>Kitasatosporales>Streptomycetaceae | GCF_002080455.1 | Streptomyces_sp._Sge12 | B6R96_RS11890 | 2759625..2763563 | 1972846 | phage-resistance-protein-[Streptomyces-sp.-Sge12] |
| WP_107426180.1 | Type2.Pgl.like.BREX | TRAM+SAM-methylase-><-?\|\|REase+SF1-SF2-Helicase-><-?<-REase+SF1-SF2-Helicase<-SF1-SF2-Helicase+wHTH+Small-B-barrel+DUF1998<-BrxD-AAA\|\|RNaseH-><-PglZ<-DUF6079*\|\|HTH_31+DUF5753->DUF397-><-N6-METHYLASE<-PglW<-NAD_binding_10<-?\|\|<-STYKIN+Pkinase | 1301 | Bacteria>Terrabacteria_group>Actinomycetota>Actinomycetes>Kitasatosporales>Streptomycetaceae | GCF_002082195.1 | Streptomyces_gilvosporeus | B1H19_RS30135 | 6750768..6754673 | 553510 | phage-resistance-protein-[Streptomyces-gilvosporeus] |
| WP_084898312.1 | Type2.Pgl.like.BREX | <-SF1-SF2-Helicase+wHTH+Small-B-barrel+DUF1998<-BrxD-AAA<-PglZ<-DUF6079*<-N6-METHYLASE\|\|HEPN->ParA-Soj-PloopNTPase->BrxD-AAA-><-PglW\|\|DUF2397-><-N6-METHYLASE | 1326 | Bacteria>Terrabacteria_group>Actinomycetota>Actinomycetes>Kitasatosporales>Streptomycetaceae | GCF_002099185.1 | Streptomyces_sp._CB03238 | BKD26_RS14940 | 390423..394403 | 1907777 | PglY-protein-[Streptomyces-sp.-CB03238] |
| WP_087772686.1 | Type2.Pgl.like.BREX | <-REase\|\|PglW->N6-METHYLASE-><-DUF397<-HTH_31+DUF5753\|\|ST-Hiskin->MerR-HTH->OrfB_Zn_ribbon->DUF6079*->PglZ->BrxD-AAA->SF1-SF2-Helicase+wHTH+Small-B-barrel+DUF1998->?->?->?->?->HEPN+ZNR->Phage_integrase-> | 1298 | Bacteria>Terrabacteria_group>Actinomycetota>Actinomycetes>Kitasatosporales>Streptomycetaceae | GCF_002188355.1 | Streptomyces_sp._CS227 | B9W68_RS05745 | 1341091..1344987 | 1982763 | phage-resistance-protein-[Streptomyces-sp.-CS227] |
| WP_096622839.1 | Type2.Pgl.like.BREX | <-DUF3323+TOPRIM<-?<-JetB<-PadR-wHTH\|\|PglW-><-ORC-CDC6<-ParA-Soj-PloopNTPase<-HEPN\|\|N6-METHYLASE->DUF6079*->PglZ->BrxD-AAA->SF1-SF2-Helicase+wHTH+Small-B-barrel+DUF1998-><-?<-?\|\|?-><-His_Phos_1<-Methyltransf_21 | 1325 | Bacteria>Terrabacteria_group>Actinomycetota>Actinomycetes>Kitasatosporales>Streptomycetaceae | GCF_002382885.1 | Streptomyces_sp._WZ.A104 | CIB93_RS00460 | 105119..109096 | 2023771 | PglY-protein-[Streptomyces-sp.-WZ.A104] |
| WP_218923825.1 | Type2.Pgl.like.BREX | PglW->HEPN->ParA-Soj-PloopNTPase->ORC-CDC6->?->?->?->N6-METHYLASE-><-GntR-HTH\|\|DUF6079*->PglZ->BrxD-AAA->SF1-SF2-Helicase+wHTH+Small-B-barrel+DUF1998-><-HHH_4+HTH+wHTH+SF1-SF2-Helicase+REase\|\|?-><-?\|\|SIGMA-HTH+PR-1->ACET-><-ACET | 1348 | Bacteria>Terrabacteria_group>Actinomycetota>Actinomycetes>Kitasatosporales>Streptomycetaceae | GCF_002551015.1 | unclassified_Streptomyces | CR491_RS01065 | 17513..21559 | 2593676 | MULTISPECIES:-hypothetical-protein-[unclassified-Streptomyces] |
| WP_097880367.1 | Type2.Pgl.like.BREX | PglW->N6-METHYLASE-><-DUF397<-HTH_31+DUF5753\|\|DUF1995->DUF2171->REase+SF1-SF2-Helicase->DUF6079*->PglZ-><-RNaseH\|\|BrxD-AAA->SF1-SF2-Helicase+wHTH+Small-B-barrel+DUF1998-><-GNTR-HTH+Aminotran_1_2\|\|?-><-?<-?\|\|GNTR-HTH-> | 1299 | Bacteria>Terrabacteria_group>Actinomycetota>Actinomycetes>Kitasatosporales>Streptomycetaceae | GCF_002551135.1 | Streptomyces_sp._st140 | CR469_RS01835 | 68816..72715 | 1828052 | phage-resistance-protein-[Streptomyces-sp.-st140] |

| Protein-ID | System | Gene-Neighbourhoods | Length | Taxonomy | Assembly-ID | Strain | Locus-Tag | Genomic-coordinates | TaxID | Fasta-define |
|---|---|---|---|---|---|---|---|---|---|---|
| WP_098891635.1 | Type2.Pgl.like.BREX | PglW->N6-METHYLASE-><-DUF397<-HTH_31+DUF5753\|\|DUF1995->DUF2171->REase+SF1-SF2-Helicase->DUF6079*->PglZ-><-RNaseH\|\|BrxD-AAA->SF1-SF2-Helicase+wHTH+Small-B-barrel+DUF1998-><-GNTR-HTH+Aminotran_1_2\|\|?-><-?<-?<-ABC-ATPase+SbcC | 1299 | Bacteria>Terrabacteria_group>Actinomycetota>Actinomycetes>Kitasatosporales>Streptomycetaceae | GCF_002551295.1 | Streptomyces_sp._or3 | CR480_RS10855 | 74126..78025 | 1828020 | phage-resistance-protein-[Streptomyces-sp.-or3] |
| WP_097873672.1 | Type2.Pgl.like.BREX | <-ANK+ANK<-Phage_int_SAM_3+Phage_integrase<-XIS-HTH<-Phage_integrase<-HTH_Tnp_4+DDE_Tnp_4\|\|?->PglW->N6-METHYLASE->DUF6079*->PglZ->BrxD-AAA->SF1-SF2-Helicase+wHTH+Small-B-barrel+DUF1998-><-?<-BrxA-DNA-binding<-TRAM+SAM-methylase<-AA_permease_2 | 1300 | Bacteria>Terrabacteria_group>Actinomycetota>Actinomycetes>Kitasatosporales>Streptomycetaceae | GCF_002551335.1 | Streptomyces_sp._ms184 | CR468_RS02325 | 19903..23805 | 1827974 | phage-resistance-protein-[Streptomyces-sp.-ms184] |
| WP_099217838.1 | Type2.Pgl.like.BREX | RADICAL-SAM->?->SAM-METHYLASE->?->?->REase-><-SF1-SF2-Helicase+wHTH+Small-B-barrel+DUF1998<-PglZ<-DUF6079*<-N6-METHYLASE<-PglW\|\|REase+N6_METHYLASE->HTH_Tnp_4-> | 1299 | Bacteria>Terrabacteria_group>Actinomycetota>Actinomycetes>Kitasatosporales>Streptomycetaceae | GCF_002705725.1 | unclassified_Streptomyces | GTY74_RS33690 | 48529..52428 | 2593676 | MULTISPECIES:-phage-resistance-protein-[unclassified-Streptomyces] |
| WP_100203497.1 | Type2.Pgl.like.BREX | NUDIX-><-?<-?<-?<-SF1-SF2-Helicase+wHTH+Small-B-barrel+DUF1998<-BrxD-AAA\|\|ParB+HNH->CBS+CBS+HEPN-><-PglZ<-DUF6079*<-?<-?\|\|?-><-N6-METHYLASE<-PglW<-?<-DUF397<-HTH_31+DUF5753 | 1295 | Bacteria>Terrabacteria_group>Actinomycetota>Actinomycetes>Kitasatosporales>Streptomycetaceae | GCF_002794255.1 | Streptomyces_carminius | CUT44_RS21120 | 73830..77717 | 2665496 | phage-resistance-protein-[Streptomyces-carminius] |
| WP_101279159.1 | Type2.Pgl.like.BREX | REase+SF1-SF2-Helicase->?-><-REase+SF1-SF2-Helicase\|\|?->REase->\|\|PglW->N6-METHYLASE-><-DUF397<-HTH_31+DUF5753\|\|DUF6079*->PglZ->BrxD-AAA->SF1-SF2-Helicase+wHTH+Small-B-barrel+DUF1998-><-TRAM+SAM-methylase<-AA_permease_2 | 1298 | Bacteria>Terrabacteria_group>Actinomycetota>Actinomycetes>Kitasatosporales>Streptomycetaceae | GCF_002844215.1 | Streptomyces_sp._EAG2 | CWE27_RS16835 | 41462..45358 | 2056495 | phage-resistance-protein-[Streptomyces-sp.-EAG2] |
| WP_102639909.1 | Type2.Pgl.like.BREX | <-HISKIN\|\|TRPR-HTH->tRNA->NTP_transf_3+Hexapep+Hexapep->?->?-><-SF1-SF2-Helicase+wHTH+Small-B-barrel+DUF1998<-BrxD-AAA<-PglZ<-DUF6079*<-N6-METHYLASE<-DUF397<-HTH_31+DUF5753\|\|?->DUF2171-><-PglW\|\|OLD-ABC->TOPRIM-> | 1302 | Bacteria>Terrabacteria_group>Actinomycetota>Actinomycetes>Kitasatosporales>Streptomycetaceae | GCF_002879675.3 | Streptomyces_sp._SMS_SU21 | C1C94_RS24615 | 1664504..1668412 | 2069440 | phage-resistance-protein-[Streptomyces-sp.-SMS_SU21] |
| WP_103503695.1 | Type2.Pgl.like.BREX | <-STYKIN<-?<-?<-Peptidase_C14+SF1-SF2-Helicase<-?\|\|PglW->N6-METHYLASE-><-DUF397<-HTH_31+DUF5753\|\|DUF6079*->PglZ->BrxD-AAA->SF1-SF2-Helicase+wHTH+Small-B-barrel+DUF1998->?-><-TRAM+SAM-methylase | 1297 | Bacteria>Terrabacteria_group>Actinomycetota>Actinomycetes>Kitasatosporales>Streptomycetaceae | GCF_002910755.1 | Streptomyces_sp._SM14 | B9S65_RS21040 | 21301..25194 | 1736045 | phage-resistance-protein-[Streptomyces-sp.-SM14] |
| WP_103506852.1 | Type2.Pgl.like.BREX | REC+OmpR-HTH->?-><-?<-?\|\|AA_permease_2->TRAM+SAM-methylase-><-SF1-SF2-Helicase+wHTH+Small-B-barrel+DUF1998<-BrxD-AAA<-PglZ<-DUF6079*\|\|HTH_31+DUF5753->DUF397-><-N6-METHYLASE<-PglW\|\|REase->?->Peptidase_C14+SF1-SF2-Helicase-> | 1298 | Bacteria>Terrabacteria_group>Actinomycetota>Actinomycetes>Kitasatosporales>Streptomycetaceae | GCF_002910815.1 | Streptomyces_sp._FM008 | B9S63_RS32625 | 27016..30912 | 1983802 | phage-resistance-protein-[Streptomyces-sp.-FM008] |
| WP_103529453.1 | Type2.Pgl.like.BREX | <-STYKIN<-?<-?<-Peptidase_C14+SF1-SF2-Helicase<-?\|\|PglW->N6-METHYLASE-><-DUF397<-HTH_31+DUF5753\|\|DUF6079*->PglZ->BrxD-AAA->SF1-SF2-Helicase+wHTH+Small-B-barrel+DUF1998->?-><-TRAM+SAM-methylase | 1296 | Bacteria>Terrabacteria_group>Actinomycetota>Actinomycetes>Kitasatosporales>Streptomycetaceae | GCF_002910855.1 | Streptomyces_sp._SM12 | B9S59_RS23865 | 21242..25132 | 1071602 | phage-resistance-protein-[Streptomyces-sp.-SM12] |
| WP_099498686.1 | Type2.Pgl.like.BREX | <-REase+DUF5655\|\|DDE_5-><-RNase\|\|YTCD-HTH->DDE_5-><-HTH_29+HTH_32+DDE_3<-DDE_5\|\|PglW->N6-METHYLASE->DUF6079*->PglZ->BrxD-AAA->SF1-SF2-Helicase+wHTH+Small-B-barrel+DUF1998->pPIWI+pPIWI->pPIWI+SF1-SF2-Helicase->pPIWI+MID+RNaseH-><-SF1-SF2-Helicase<-OLD-ABC+TOPRIM\|\|ZnR+REase-> | 1297 | Bacteria>Terrabacteria_group>Actinomycetota>Actinomycetes>Kitasatosporales>Streptomycetaceae | GCF_002946835.1 | Streptomyces | C4B68_RS07930 | 1717869..1721762 | 1883 | MULTISPECIES:-phage-resistance-protein-[Streptomyces] |
| WP_099498944.1 | Type2.Pgl.like.BREX | PglW->N6-METHYLASE-><-DUF397<-HTH_31+DUF5753\|\|DUF1995->DUF2171->PhdYeFM_antitox->DUF6079*->PglZ->BrxD-AAA->SF1-SF2-Helicase+wHTH+Small-B-barrel+DUF1998->?->?-><-REC+OmpR-HTH | 1303 | Bacteria>Terrabacteria_group>Actinomycetota>Actinomycetes>Kitasatosporales>Streptomycetaceae | GCF_002946835.1 | Streptomyces | C4B68_RS09525 | 2094134..2098045 | 1883 | MULTISPECIES:-phage-resistance-protein-[Streptomyces] |
| WP_099504556.1 | Type2.Pgl.like.BREX | HHH_5+SF1-SF2-Helicase->SAM-METHYLASE->HEPN+VirB11+NACHT+FGS->?->SLATT-><-?<-?\|\|PglW->N6-METHYLASE->DUF6079*->PglZ->BrxD-AAA->SF1-SF2-Helicase+wHTH+Small-B-barrel+DUF1998-><-?\|\|HEPN-><-Phage_integrase\|\|HTH_3->OLD-ABC+TOPRIM->SF1-SF2-Helicase-> | 1312 | Bacteria>Terrabacteria_group>Actinomycetota>Actinomycetes>Kitasatosporales>Streptomycetaceae | GCF_002946835.1 | Streptomyces | C4B68_RS28490 | 6170305..6174243 | 1883 | MULTISPECIES:-PglY-protein-[Streptomyces] |
| WP_105973292.1 | Type2.Pgl.like.BREX | ABhydrolase->?-><-?\|\|?-><-SF1-SF2-Helicase+wHTH+Small-B-barrel+DUF1998<-BrxD-AAA\|\|ParB+HNH->CBS+HEPN-><-PglZ<-DUF6079*<-DUF397<-HTH_31+DUF5753\|\|ST-Hiskin-><-N6-METHYLASE<-PglW\|\|?-><-TOPRIM<-OLD-ABC\|\|N6-METHYLASE-> | 1328 | Bacteria>Terrabacteria_group>Actinomycetota>Actinomycetes>Kitasatosporales>Streptomycetaceae | GCF_002954775.1 | Streptomyces_geranii | CW359_RS29840 | 34991..38977 | 2058923 | phage-resistance-protein-[Streptomyces-geranii] |
| WP_109284035.1 | Type2.Pgl.like.BREX | <-REase+SF1-SF2-Helicase\|\|?-><-?\|\|PglW->N6-METHYLASE-><-DUF397<-HTH_31+DUF5753\|\|DUF2171->?->DUF6079*->PglZ-><-CBS+HEPN<-ParB+HNH\|\|BrxD-AAA->SF1-SF2-Helicase+wHTH+Small-B-barrel+DUF1998-><-TRAM+SAM-methylase<-AA_permease_2 | 1304 | Bacteria>Terrabacteria_group>Actinomycetota>Actinomycetes>Kitasatosporales>Streptomycetaceae | GCF_003121295.1 | Streptomyces_orinoci | CV755_RS31530 | 49510..53424 | 67339 | phage-resistance-protein-[Streptomyces-orinoci] |
| WP_114054613.1 | Type2.Pgl.like.BREX | <-Pkinase+Pkinase<-SF1-SF2-Helicase+wHTH+Small-B-barrel+DUF1998<-BrxD-AAA\|\|ParB+HNH->CBS+HEPN-><-PglZ<-DUF6079*<-DUF2171<-DUF1995\|\|HTH_31+DUF5753->DUF397-><-N6-METHYLASE<-PglW\|\|REase+SF1-SF2-Helicase-><-?<-REase+SF1-SF2-Helicase | 1298 | Bacteria>Terrabacteria_group>Actinomycetota>Actinomycetes>Kitasatosporales>Streptomycetaceae | GCF_003325375.1 | Streptomyces_globosus | C0216_RS08100 | 1744779..1748675 | 68209 | phage-resistance-protein-[Streptomyces-globosus] |
| WP_128502442.1 | Type2.Pgl.like.BREX | <-SF1-SF2-Helicase+wHTH+Small-B-barrel+DUF1998<-BrxD-AAA<-PglZ<-DUF6079*<-N6-METHYLASE<-PglW\|\|?->?->?-><-?<-SAM-METHYLASE | 1304 | Bacteria>Terrabacteria_group>Actinomycetota>Actinomycetes>Kitasatosporales>Streptomycetaceae | GCF_003389455.1 | Streptomyces_inhibens | DY245_RS00990 | 15167..19081 | 2293571 | PglY-protein-[Streptomyces-inhibens] |
| WP_120685184.1 | Type2.Pgl.like.BREX | <-SF1-SF2-Helicase+wHTH+Small-B-barrel+DUF1998<-BrxD-AAA<-PglZ<-DUF6079*<-N6-METHYLASE<-PglW | 1300 | Bacteria>Terrabacteria_group>Actinomycetota>Actinomycetes>Kitasatosporales>Streptomycetaceae | GCF_003626535.1 | Streptomyces_hoynatensis | D7294_RS31160 | 7577..11479 | 1141874 | phage-resistance-protein-[Streptomyces-hoynatensis] |
| WP_121887751.1 | Type2.Pgl.like.BREX | REase+SF1-SF2-Helicase-><-SAD(SRA)+wHTH+HNH\|\|PglW->N6-METHYLASE->?-><-?<-?\|\|?->APATPase+TPR+TPR+TPR->DUF6079*->PglZ-><-ParB+HNH\|\|BrxD-AAA->SF1-SF2-Helicase+wHTH+Small-B-barrel+DUF1998->?->?->?-><-REC+OmpR-HTH<-ST-Hiskin | 1295 | Bacteria>Terrabacteria_group>Actinomycetota>Actinomycetes>Kitasatosporales>Streptomycetaceae | GCF_003688995.1 | Streptomyces_shenzhenensis | CTZ28_RS03710 | 158817..162704 | 943815 | phage-resistance-protein-[Streptomyces-shenzhenensis] |
| WP_123562585.1 | Type2.Pgl.like.BREX | <-REase+SF1-SF2-Helicase\|\|REase+SF1-SF2-Helicase->ParB+HNH->CBS+HEPN->PglW->N6-METHYLASE->PhdYeFM_antitox->PIN->DUF6079*->PglZ->BrxD-AAA->SF1-SF2-Helicase+wHTH+Small-B-barrel+DUF1998-><-SF1-SF2-Helicase+HKD-DNase<-?\|\|<-LZ_Tnp_IS481+HTH_32+rve+rve_3\|\|N6-N4-METHYLASE-> | 1307 | Bacteria>Terrabacteria_group>Actinomycetota>Actinomycetes>Kitasatosporales>Streptomycetaceae | GCF_003751605.1 | Kitasatospora_cineracea | EDD39_RS31525 | 1175476..1179399 | 88074 | phage-resistance-protein-[Kitasatospora-cineracea] |
| WP_123459797.1 | Type2.Pgl.like.BREX | NTP_transf_3+Hexapep+Hexapep->?->?->?->?-><-SF1-SF2-Helicase+wHTH+Small-B-barrel+DUF1998<-BrxD-AAA<-PglZ<-DUF6079*<-DUF397<-HTH_31+DUF5753\|\|?->DUF2171-><-PglW\|\|OLD-ABC->TOPRIM-> | 1295 | Bacteria>Terrabacteria_group>Actinomycetota>Actinomycetes>Kitasatosporales>Streptomycetaceae | GCF_003752275.1 | Streptomyces_sp._CEV_2-1 | EDD95_RS00765 | 172440..176327 | 2485153 | phage-resistance-protein-[Streptomyces-sp.-CEV-2-1] |
| WP_124273797.1 | Type2.Pgl.like.BREX | AA_permease_2->TRAM+SAM-methylase+METHYLASE-><-SF1-SF2-Helicase+wHTH+Small-B-barrel+DUF1998<-BrxD-AAA<-PglZ<-DUF6079*\|\|HTH_31+DUF5753->DUF397-><-N6-METHYLASE<-PglW<-?<-BrxA-DNA-binding<-ADH_N+ADH_zinc_N | 1298 | Bacteria>Terrabacteria_group>Actinomycetota>Actinomycetes>Kitasatosporales>Streptomycetaceae | GCF_003846265.1 | Streptomyces_sp._ADI96-15 | EES44_RS29825 | 154353..158249 | 1522761 | phage-resistance-protein-[Streptomyces-sp.-ADI96-15] |
| WP_125502694.1 | Type2.Pgl.like.BREX | <-SF1-SF2-Helicase+wHTH+Small-B-barrel+DUF1998<-BrxD-AAA\|\|Phage_holin->ParB+HNH->CBS+CBS+HEPN-><-PglZ<-DUF6079*<-APATPase+TPR+TPR+TPR<-?\|\|?->?-><-DUF397<-HTH_31+DUF5753\|\|ST-Hiskin-><-N6-METHYLASE<-PglW | 1295 | Bacteria>Terrabacteria_group>Actinomycetota>Actinomycetes>Kitasatosporales>Streptomycetaceae | GCF_003945525.1 | unclassified_Streptomyces | DLM49_RS04285 | 990903..994790 | 2593676 | MULTISPECIES:-phage-resistance-protein-[unclassified-Streptomyces] |
| WP_125530906.1 | Type2.Pgl.like.BREX | <-pPIWI+pPIWI<-REase\|\|Pkinase+Pkinase-><-?\|\|?->?->XIS-HTH->PglW->N6-METHYLASE->DUF6079*->PglZ->BrxD-AAA->SF1-SF2-Helicase+wHTH+Small-B-barrel+DUF1998-><-TRAM+SAM-methylase<-AA_permease_2 | 1298 | Bacteria>Terrabacteria_group>Actinomycetota>Actinomycetes>Kitasatosporales>Streptomycetaceae | GCF_003947585.1 | Streptomyces_sp._WAC_06725 | DMH15_RS04680 | 32673..36569 | 2203209 | phage-resistance-protein-[Streptomyces-sp.-WAC-06725] |
| WP_260614935.1 | Type2.Pgl.like.BREX | REase+SF1-SF2-Helicase->?-><-REase+SF1-SF2-Helicase\|\|PglW->N6-METHYLASE-><-DUF397<-HTH_31+DUF5753\|\|DUF1995->DUF2171->DUF6079*-> | 1241 | Bacteria>Terrabacteria_group>Actinomycetota>Actinomycetes>Kitasatosporales>Streptomycetaceae | GCF_003949955.1 | Streptomyces_sp._WAC05292 | EF903_RS07655 | 26077..29800 | 2487418 | phage-resistance-protein,-partial-[Streptomyces-sp.-WAC05292] |
| WP_128462336.1 | Type2.Pgl.like.BREX | ADH_N+ADH_zinc_N->BrxA-DNA-binding->?->PglW->N6-METHYLASE-><-DUF397<-HTH_31+DUF5753\|\|DUF6079*->PglZ-> | 1298 | Bacteria>Terrabacteria_group>Actinomycetota>Actinomycetes>Kitasatosporales>Streptomycetaceae | GCF_004078575.1 | Streptomyces_albidoflavus | EQK42_RS04975 | 17321..21217 | 1886 | phage-resistance-protein-[Streptomyces-albidoflavus] |
| WP_136222946.1 | Type2.Pgl.like.BREX | <-TUDOR+SF1-SF2-Helicase+iREase+CTD+REase\|\|?->XIS-HTH->Phage_int_SAM_3+Phage_integrase-><-DUF4011+SF1-SF2-Helicase+REase<-SF1-SF2-Helicase+wHTH+Small-B-barrel+DUF1998<-BrxD-AAA<-DndB<-PglZ<-DUF6079*<-REase+SF1-SF2-Helicase<-DUF2171<-DUF1995\|\|HTH_31+DUF5753->DUF397-><-N6-METHYLASE<-PglW<-SF1-SF2-Helicase+HKD-DNase | 1301 | Bacteria>Terrabacteria_group>Actinomycetota>Actinomycetes>Kitasatosporales>Streptomycetaceae | GCF_004794975.1 | Streptomyces_sp._A1547 | E6W17_RS04085 | 285079..288984 | 2563105 | phage-resistance-protein-[Streptomyces-sp.-A1547] |
| WP_136181970.1 | Type2.Pgl.like.BREX | <-REase<-?\|\|PglW->N6-METHYLASE-><-DUF397<-HTH_31+DUF5753\|\|DUF1995->DUF2171->REase+SF1-SF2-Helicase->DUF6079*->PglZ->DndB->BrxD-AAA->SF1-SF2-Helicase+wHTH+Small-B-barrel+DUF1998->REase-><-LZ_Tnp_IS481+HTH_32+rve+rve_3\|\|HTH_Tnp_1+HTH_21+rve+rve_3-> | 1295 | Bacteria>Terrabacteria_group>Actinomycetota>Actinomycetes>Kitasatosporales>Streptomycetaceae | GCF_004795255.1 | Streptomyces_sp._RKND-216 | E4198_RS04240 | 941855..945742 | 2562581 | phage-resistance-protein-[Streptomyces-sp.-RKND-216] |
| WP_136728164.1 | Type2.Pgl.like.BREX | <-METHYLASE<-?\|\|transposase->?->\|\|?->EVE+REase-p-><-?\|\|PglW->N6-METHYLASE->DUF6079*->PglZ->BrxD-AAA->SF1-SF2-Helicase+wHTH+Small-B-barrel+DUF1998-><-?<-?<-DUF2171\|\|HTH_3-><-HAD+HAD | 1316 | Bacteria>Terrabacteria_group>Actinomycetota>Actinomycetes>Kitasatosporales>Streptomycetaceae | GCF_005047355.1 | Actinacidiphila_oryziradicis | FCI23_RS62555 | 22737..26687 | 2571141 | PglY-protein-[Actinacidiphila-oryziradicis] |
| WP_137230275.1 | Type2.Pgl.like.BREX | ABC-ATPase+SbcC->?->?-><-?\|\|GNTR-HTH+Aminotran_1_2-><-SF1-SF2-Helicase+wHTH+Small-B-barrel+DUF1998<-BrxD-AAA\|\|RNaseH-><-PglZ<-DUF6079*<-REase+SF1-SF2-Helicase<-DUF2171<-DUF1995\|\|HTH_31+DUF5753->DUF397-><-N6-METHYLASE<-PglW | 1299 | Bacteria>Terrabacteria_group>Actinomycetota>Actinomycetes>Kitasatosporales>Streptomycetaceae | GCF_005234185.1 | Streptomyces_sp._BPSDS2 | FD916_RS06870 | 197367..201266 | 2571021 | phage-resistance-protein-[Streptomyces-sp.-BPSDS2] |
| WP_137992471.1 | Type2.Pgl.like.BREX | REC+OmpR-HTH-><-?\|\|NTP_transf_3->?->?-><-?<-SF1-SF2-Helicase+wHTH+Small-B-barrel+DUF1998<-BrxD-AAA<-PglZ<-DUF6079*<-?<-PIN\|\|?-><-OLD-ABC+TOPRIM\|\|DUF397-><-N6-METHYLASE<-PglW\|\|?->REase+SF1-SF2-Helicase-> | 1286 | Bacteria>Terrabacteria_group>Actinomycetota>Actinomycetes>Kitasatosporales>Streptomycetaceae | GCF_005517195.1 | Streptomyces_exfoliatus | FDM97_RS23450 | 5014702..5018562 | 1905 | PglY-protein-[Streptomyces-exfoliatus] |
| WP_144316319.1 | Type2.Pgl.like.BREX | PglW-><-ORC-CDC6<-ParA-Soj-PloopNTPase<-HEPN\|\|N6-METHYLASE-><-DUF397<-HTH_31+DUF5753\|\|ST-Hiskin-><-?\|\|DUF6079*->PglZ->BrxD-AAA->SF1-SF2-Helicase+wHTH+Small-B-barrel+DUF1998-><-HTH_Tnp_1+HTH_21+rve+rve_3\|\|RAMA+McrB-AAA->McrC-NTD+REase-><-DDE_Tnp_IS240<-ABC-ATPase<-PCMT+METHYLASE | 1300 | Bacteria>Terrabacteria_group>Actinomycetota>Actinomycetes>Kitasatosporales>Streptomycetaceae | GCF_007113425.2 | Streptomyces_sp._RLB3-17 | FNV62_RS53960 | 11911719..11915621 | 2594455 | PglY-protein-[Streptomyces-sp.-RLB3-17] |
| WP_143606190.1 | Type2.Pgl.like.BREX | 35exo->AAA-ATPase->?->NUDIX->ABC-ATPase+SbcC-><-SF1-SF2-Helicase+wHTH+Small-B-barrel+DUF1998<-BrxD-AAA<-PglZ<-DUF6079*\|\|?-><-ST-Hiskin\|\|HTH_31+DUF5753->DUF397-><-N6-METHYLASE\|\|HEPN->ParA-Soj-PloopNTPase->ORC-CDC6-><-PglW | 1300 | Bacteria>Terrabacteria_group>Actinomycetota>Actinomycetes>Kitasatosporales>Streptomycetaceae | GCF_007113545.2 | unclassified_Streptomyces | FNV68_RS00625 | 125740..129642 | 2593676 | MULTISPECIES:-PglY-protein-[unclassified-Streptomyces] |
| WP_150231687.1 | Type2.Pgl.like.BREX | <-HAMP+HISKIN<-?<-?\|\|Peptidase_M4+Peptidase_M4_C-><-SF1-SF2-Helicase+wHTH+Small-B-barrel+DUF1998<-BrxD-AAA\|\|ParB+HNH->CBS+HEPN-><-PglZ<-DUF6079*<-DUF397<-HTH_31+DUF5753\|\|ST-Hiskin->REase-><-N6-METHYLASE<-PglW\|\|SAD(SRA)+wHTH+HNH-><-tRNA\|\|REC+OmpR-HTH-> | 1310 | Bacteria>Terrabacteria_group>Actinomycetota>Actinomycetes>Kitasatosporales>Streptomycetaceae | GCF_008634015.1 | Streptomyces_filamentosus | CP979_RS24300 | 5261397..5265329 | 67294 | phage-resistance-protein-[Streptomyces-filamentosus] |
| WP_150950778.1 | Type2.Pgl.like.BREX | <-SF1-SF2-Helicase+wHTH+Small-B-barrel+DUF1998<-BrxD-AAA\|\|ParB+HNH->CBS+HEPN-><-PglZ<-DUF6079*<-PIN<-HTH+MPTase<-DUF397<-HTH_31+DUF5753\|\|ST-Hiskin-><-N6-METHYLASE<-PglW\|\|Phage_integrase-> | 1308 | Bacteria>Terrabacteria_group>Actinomycetota>Actinomycetes>Kitasatosporales>Streptomycetaceae | GCF_008806595.1 | Streptomyces_luteolifulvus | F7R91_RS21840 | 77489..81415 | 2615112 | phage-resistance-protein-[Streptomyces-luteolifulvus] |
| WP_134111864.1 | Type2.Pgl.like.BREX | PglW->N6-METHYLASE-><-ST-Hiskin\|\|HTH_31+DUF5753->DUF397->?->?->?->DUF6079*->PglZ-><-CBS+HEPN<-ParB+HNH\|\|BrxD-AAA->SF1-SF2-Helicase+wHTH+Small-B-barrel+DUF1998-><-SF1-SF2-Helicase\|\|ACET-> | 1294 | Bacteria>Terrabacteria_group>Actinomycetota>Actinomycetes>Kitasatosporales>Streptomycetaceae | GCF_008931305.1 | Streptomyces_coelicolor | FQ762_RS34260 | 7363057..7366941 | 1902 | phage-resistance-protein-[Streptomyces-coelicolor] |
| WP_077964595.1 | Type2.Pgl.like.BREX | <-HTH_Tnp_4+DDE_Tnp_4<-N6-METHYLASE\|\|OLD-ABC->TOPRIM-><-?<-SF1-SF2-Helicase+wHTH+Small-B-barrel+DUF1998<-BrxD-AAA<-PglZ<-DUF6079*<-N6-METHYLASE<-PglW<-?\|\|HTH_3->?->SF1-SF2-Helicase-><-?<-TnsC-ATPase | 1296 | Bacteria>Terrabacteria_group>Actinomycetota>Actinomycetes>Kitasatosporales>Streptomycetaceae | GCF_009296025.1 | Streptomyces_tsukubensis | GBW32_RS05195 | 1168905..1172795 | 83656 | hypothetical-protein-[Streptomyces-tsukubensis] |
| WP_153520868.1 | Type2.Pgl.like.BREX | CASPASE+RNA-Helicase+REase->PglW->N6-METHYLASE-><-DUF397<-HTH_31+DUF5753\|\|DUF1995->DUF2171->DUF6079*->PglZ->SF1-SF2-Helicase+wHTH+Small-B-barrel+DUF1998->?-><-DUF3323+TOPRIM<-SbcCD_C<-JetB<-?<-DUF397<-HTH_31+DUF5753 | 1297 | Bacteria>Terrabacteria_group>Actinomycetota>Actinomycetes>Kitasatosporales>Streptomycetaceae | GCF_009600885.1 | Streptomyces_jumonjinensis | FF041_RS02770 | 20638..24531 | 1945 | phage-resistance-protein-[Streptomyces-jumonjinensis] |
| WP_155072369.1 | Type2.Pgl.like.BREX | ABhydrolase->\|\|?->PglW->N6-METHYLASE-><-ST-Hiskin\|\|HTH_31+DUF5753->DUF397->?->SAM-METHYLASE->DUF6079*->PglZ-><-DUF397<-ARG\|\|BrxD-AAA->SF1-SF2-Helicase+wHTH+Small-B-barrel+DUF1998-> | 1296 | Bacteria>Terrabacteria_group>Actinomycetota>Actinomycetes>Kitasatosporales>Streptomycetaceae | GCF_009709575.1 | Streptomyces_taklimakanensis | FOL17_RS21690 | 4932691..4936581 | 2569853 | phage-resistance-protein-[Streptomyces-taklimakanensis] |
| WP_156694667.1 | Type2.Pgl.like.BREX | REase->ncRNA->PglW->N6-METHYLASE-><-ST-Hiskin\|\|HTH_31+DUF5753->DUF397->?-><-?<-RelE-ParE<-PhdYeFM_antitox\|\|DUF6079*->PglZ-><-Phage_holin<-CBS+CBS+HEPN<-ParB+HNH\|\|BrxD-AAA->SF1-SF2-Helicase+wHTH+Small-B-barrel+DUF1998-> | 1295 | Bacteria>Terrabacteria_group>Actinomycetota>Actinomycetes>Kitasatosporales>Streptomycetaceae | GCF_009739905.1 | Streptomyces_ficellus | EIZ62_RS23460 | 5231851..5235738 | 1977088 | phage-resistance-protein-[Streptomyces-ficellus] |
| WP_190140957.1 | Type2.Pgl.like.BREX | <-HSP70+PIN<-GrpE<-?<-REase\|\|?-><-?<-?\|\|PglW->N6-METHYLASE->DUF6079*->PglZ->BrxD-AAA->SF1-SF2-Helicase+wHTH+Small-B-barrel+DUF1998->?->?->?-><-tRNA\|\|Peptidase_M20-> | 1298 | Bacteria>Terrabacteria_group>Actinomycetota>Actinomycetes>Kitasatosporales>Streptomycetaceae | GCF_009811575.1 | Streptomyces_glebosus | Sgleb_RS29750 | 6831668..6835564 | 249580 | PglY-protein-[Streptomyces-glebosus] |
| WP_159742973.1 | Type2.Pgl.like.BREX | PglW->N6-METHYLASE->DUF6079*->PglZ->BrxD-AAA->SF1-SF2-Helicase+wHTH+Small-B-barrel+DUF1998->NUDIX-><-rRNA<-rRNA<-rRNA<-TRAM+SAM-methylase<-AA_permease_2 | 1299 | Bacteria>Terrabacteria_group>Actinomycetota>Actinomycetes>Kitasatosporales>Streptomycetaceae | GCF_009811635.1 | Streptomyces_tubercidicus | Stube_RS06795 | 1610444..1614343 | 47759 | phage-resistance-protein-[Streptomyces-tubercidicus] |
| WP_161069245.1 | Type2.Pgl.like.BREX | <-SF1-SF2-Helicase+wHTH+Small-B-barrel+DUF1998\|\|ParB+HNH->CBS+HEPN-><-PglZ<-DUF6079*<-MNS+REase<-DUF397<-HTH_31+DUF5753\|\|ST-Hiskin-><-N6-METHYLASE<-PglW\|\|SAD(SRA)+wHTH+HNH-> | 1308 | Bacteria>Terrabacteria_group>Actinomycetota>Actinomycetes>Kitasatosporales>Streptomycetaceae | GCF_009827025.1 | Streptomyces_sp._HUCO-GS316 | GR925_RS33060 | 14814..18740 | 2692198 | phage-resistance-protein-[Streptomyces-sp.-HUCO-GS316] |
| WP_111382238.1 | Type2.Pgl.like.BREX | <-NAD_binding_10\|\|ABC-ATPase+SbcC->ABC-ATPase+SbcC->BrxA-DNA-binding->?-><-SF1-SF2-Helicase+wHTH+Small-B-barrel+DUF1998<-BrxD-AAA<-PglZ<-DUF6079*<-N6-METHYLASE<-PglW<-?\|\|HTH_Tnp_4+DDE_Tnp_4->Phage_integrase->XIS-HTH->Phage_int_SAM_3+Phage_integrase->ANK+ANK-><-APATPase+TPR+TPR+TPR+TPR | 1300 | Bacteria>Terrabacteria_group>Actinomycetota>Actinomycetes>Kitasatosporales>Streptomycetaceae | GCF_009862055.1 | Streptomyces_sp._SID8358 | GTW52_RS08645 | 18687..22589 | 2690342 | phage-resistance-protein-[Streptomyces-sp.-SID8358] |
| WP_161168814.1 | Type2.Pgl.like.BREX | CBS+CBS+HEPN-><-PglZ<-DUF6079*<-?<-DUF397<-HTH_31+DUF5753\|\|ST-Hiskin-><-N6-METHYLASE<-PglW\|\|SF1-SF2-Helicase+HKD-DNase->?-><-REase+SF1-SF2-Helicase | 1296 | Bacteria>Terrabacteria_group>Actinomycetota>Actinomycetes>Kitasatosporales>Streptomycetaceae | GCF_009862125.1 | Streptomyces_sp._SID5910 | GTW67_RS35430 | 5133..9023 | 2690312 | phage-resistance-protein-[Streptomyces-sp.-SID5910] |
| WP_011031061.1 | Type2.Pgl.like.BREX | PglW->N6-METHYLASE-><-ST-Hiskin\|\|HTH_31+DUF5753->DUF397->?->?->?->DUF6079*->PglZ-><-CBS+HEPN<-ParB+HNH\|\|BrxD-AAA->SF1-SF2-Helicase+wHTH+Small-B-barrel+DUF1998-><-SF1-SF2-Helicase\|\|ACET-> | 1294 | Bacteria>Terrabacteria_group>Actinomycetota>Actinomycetes>Kitasatosporales>Streptomycetaceae | GCF_009862445.1 | Streptomyces_sp._SID7813 | GTY87_RS33150 | 7362943..7366827 | 2690330 | hypothetical-protein-[Streptomyces-sp.-SID7813] |
| WP_052855836.1 | Type2.Pgl.like.BREX | <-ST-Hiskin<-NUDIX<-?<-?\|\|?->?-><-?<-?<-DUF6079*<-N6-METHYLASE<-PglW\|\|BrxD-AAA->SF1-SF2-Helicase+wHTH+Small-B-barrel+DUF1998->RNaseH-><-sGTP<-ROK-HTH | 1280 | Bacteria>Terrabacteria_group>Actinomycetota>Actinomycetes>Kitasatosporales>Streptomycetaceae | GCF_009862505.1 | Streptomyces_sp._SID7805 | GTY53_RS24335 | 41453..45295 | 2690328 | DUF6079-family-protein-[Streptomyces-sp.-SID7805] |
| WP_164183062.1 | Type2.Pgl.like.BREX | AA_permease_2->TRAM+SAM-methylase-><-SF1-SF2-Helicase+wHTH+Small-B-barrel+DUF1998<-BrxD-AAA<-PglZ<-DUF6079*\|\|HTH_31+DUF5753->DUF397-><-N6-METHYLASE<-PglW<-?<-BrxA-DNA-binding<-ADH_N+ADH_zinc_N | 1298 | Bacteria>Terrabacteria_group>Actinomycetota>Actinomycetes>Kitasatosporales>Streptomycetaceae | GCF_010523235.1 | Streptomyces_sp._604F | C3K23_RS25915 | 5696613..5700509 | 1476754 | phage-resistance-protein-[Streptomyces-sp.-604F] |
| WP_164407618.1 | Type2.Pgl.like.BREX | <-REase+SF1-SF2-Helicase\|\|PglW-><-?\|\|N6-METHYLASE-><-DUF397<-HTH_31+DUF5753\|\|ST-Hiskin-><-?<-BrxA-DNA-binding\|\|DUF6079*->PglZ->BrxD-AAA->SF1-SF2-Helicase+wHTH+Small-B-barrel+DUF1998-><-RNaseH\|\|ParB+HNH-><-SWACOS+LuxR-HTH\|\|REC+OmpR-HTH-> | 1306 | Bacteria>Terrabacteria_group>Actinomycetota>Actinomycetes>Kitasatosporales>Streptomycetaceae | GCF_010548275.1 | Streptomyces_sp._SID12488 | G3I56_05390 | 276309..280229 | 2706040 | PglY-protein-[Streptomyces-sp.-SID12488] |
| WP_164615103.1 | Type2.Pgl.like.BREX | ABC-ATPase+SbcC->ABC-ATPase+SbcC->?->?-><-?\|\|GNTR-HTH+Aminotran_1_2-><-SF1-SF2-Helicase+wHTH+Small-B-barrel+DUF1998\|\|RNaseH-><-PglZ<-DUF6079*<-REase+SF1-SF2-Helicase<-DUF1995\|\|DUF397-><-N6-METHYLASE<-PglW\|\|?->?->MoxR-AAA->REase+SF1-SF2-Helicase-> | 1299 | Bacteria>Terrabacteria_group>Actinomycetota>Actinomycetes>Kitasatosporales>Streptomycetaceae | GCF_010548505.1 | Streptomyces_sp._SID14515 | G3I54_RS15510 | 49251..53150 | 2706074 | phage-resistance-protein-[Streptomyces-sp.-SID14515] |
| WP_164373105.1 | Type2.Pgl.like.BREX | JetB->DUF3323+TOPRIM->SAD(SRA)+wHTH+HNH-><-SF1-SF2-Helicase+wHTH+Small-B-barrel+DUF1998\|\|ParB+HNH->CBS+HEPN-><-PglZ<-DUF6079*<-DUF397<-HTH_31+DUF5753\|\|ST-Hiskin-><-N6-METHYLASE<-PglW\|\|?-><-SWACOS+LuxR-HTH\|\|REC+OmpR-HTH-> | 1322 | Bacteria>Terrabacteria_group>Actinomycetota>Actinomycetes>Kitasatosporales>Streptomycetaceae | GCF_010548515.1 | Streptomyces_sp._SID14446 | G3I62_RS24970 | 24029..27997 | 2706072 | phage-resistance-protein-[Streptomyces-sp.-SID14446] |
| WP_165234047.1 | Type2.Pgl.like.BREX | DUF5086->DUF5086->Recombinase+ZnR->DUF2637+HTH->PglW-><-ORC-CDC6<-ParA-Soj-PloopNTPase<-HEPN\|\|N6-METHYLASE->DUF6079*->PglZ->SF1-SF2-Helicase+wHTH+Small-B-barrel+DUF1998->SWACOS-><-ACET<-HTH_3<-GNTR-HTH | 1325 | Bacteria>Terrabacteria_group>Actinomycetota>Actinomycetes>Kitasatosporales>Streptomycetaceae | GCF_011044975.1 | Streptomyces_coryli | G5C51_RS08080 | 27992..31969 | 1128680 | PglY-protein-[Streptomyces-coryli] |
| WP_193454094.1 | Type2.Pgl.like.BREX | CR-ATPase9->CR-ATPase9->\|\|HTH_Tnp_4+DDE_Tnp_4-><-SF1-SF2-Helicase+wHTH+Small-B-barrel+DUF1998<-BrxD-AAA<-PglZ<-DUF6079*\|\|HTH_31+DUF5753->DUF397-><-N6-METHYLASE<-PglW\|\|REase->?->Peptidase_C14+SF1-SF2-Helicase-> | 1298 | Bacteria>Terrabacteria_group>Actinomycetota>Actinomycetes>Kitasatosporales>Streptomycetaceae | GCF_011170105.1 | Streptomyces_rutgersensis | Srut_RS22845 | 716645..720541 | 53451 | phage-resistance-protein-[Streptomyces-rutgersensis] |
| WP_167901571.1 | Type2.Pgl.like.BREX | PglW->?-><-?<-?\|\|?->DUF6079*-> | 1297 | Bacteria>Terrabacteria_group>Actinomycetota>Actinomycetes>Kitasatosporales>Streptomycetaceae | GCF_011882225.1 | Streptomyces_sp._NEAU-H3 | HC023_RS01900 | 16923..20816 | 2720636 | phage-resistance-protein-[Streptomyces-sp.-NEAU-H3] |
| WP_167967748.1 | Type2.Pgl.like.BREX | SF1-SF2-Helicase-><-SF1-SF2-Helicase+REase+HTH<-?\|\|PglW->N6-METHYLASE-><-DUF397<-HTH_31+DUF5753<-MetJarc-HTH\|\|DUF6079*->PglZ->BrxD-AAA->SF1-SF2-Helicase+wHTH+Small-B-barrel+DUF1998-><-RHH_1\|\|<-?<-TRAM+SAM-methylase | 1298 | Bacteria>Terrabacteria_group>Actinomycetota>Actinomycetes>Kitasatosporales>Streptomycetaceae | GCF_012033735.1 | Streptomyces_lonarensis | HCN56_RS02350 | 22960..26856 | 700599 | phage-resistance-protein-[Streptomyces-lonarensis] |
| WP_167931758.1 | Type2.Pgl.like.BREX | TRAM+SAM-methylase-><-PIN<-MetJarc-HTH\|\|RHH_1->PIN-><-SF1-SF2-Helicase+wHTH+Small-B-barrel+DUF1998<-PglZ<-DUF6079*\|\|MetJarc-HTH->HTH_31+DUF5753->DUF397-><-N6-METHYLASE<-PglW\|\|?->?->DUF3323+TOPRIM-> | 1298 | Bacteria>Terrabacteria_group>Actinomycetota>Actinomycetes>Kitasatosporales>Streptomycetaceae | GCF_012034385.1 | Streptomyces_spiramenti | HCJ92_RS02745 | 20478..24374 | 2720606 | phage-resistance-protein-[Streptomyces-spiramenti] |
| WP_169086808.1 | Type2.Pgl.like.BREX | <-HTH+REase<-SF1-SF2-Helicase+REase+HTH\|\|<-wHTH+SAD+HNH\|\|PglW->N6-METHYLASE-><-REase<-ST-Hiskin\|\|HTH_31+DUF5753->DUF397->DUF6079*->PglZ-><-CBS+HEPN<-ParB+HNH\|\|BrxD-AAA->SF1-SF2-Helicase+wHTH+Small-B-barrel+DUF1998-><-Peptidase_M4+Peptidase_M4_C | 1309 | Bacteria>Terrabacteria_group>Actinomycetota>Actinomycetes>Kitasatosporales>Streptomycetaceae | GCF_012927225.1 | unclassified_Streptomyces | HHL18_RS29280 | 65110..69039 | 2593676 | MULTISPECIES:-phage-resistance-protein-[unclassified-Streptomyces] |
| WP_171109616.1 | Type2.Pgl.like.BREX | <-HTH_3\|\|HTH_31+DUF5753->DUF397-><-REase\|\|SF1-SF2-Helicase->PIN-><-SF1-SF2-Helicase+wHTH+Small-B-barrel+DUF1998<-BrxD-AAA<-PglZ<-DUF6079*<-ST-Hiskin\|\|HTH_31+DUF5753->DUF397-><-N6-METHYLASE\|\|HEPN->ParA-Soj-PloopNTPase->BrxD-AAA-><-PglW<-Peptidase_C14 | 1326 | Bacteria>Terrabacteria_group>Actinomycetota>Actinomycetes>Kitasatosporales>Streptomycetaceae | GCF_013009255.1 | Streptomyces | HJD23_RS18045 | 4041978..4045958 | 1883 | MULTISPECIES:-PglY-protein-[Streptomyces] |
| WP_171141282.1 | Type2.Pgl.like.BREX | APATPase+TPR+TPR+TPR->APATPase+TPR+TPR+TPR-><-REase<-SF1-SF2-Helicase+wHTH+Small-B-barrel+DUF1998<-BrxD-AAA<-PglZ<-DUF6079*<-N6-METHYLASE\|\|HEPN->ParA-Soj-PloopNTPase->ORC-CDC6-><-PglW<-Peptidase_C14<-?<-BrxA-DNA-binding | 1320 | Bacteria>Terrabacteria_group>Actinomycetota>Actinomycetes>Kitasatosporales>Streptomycetaceae | GCF_013044505.1 | Streptomyces_sp._S3(2020) | HLK59_RS11655 | 32079..36041 | 2732044 | PglY-protein-[Streptomyces-sp.-S3(2020)] |
| WP_172387270.1 | Type2.Pgl.like.BREX | HTH_31->?->HTH_3->?-><-HEPN+Polbeta\|\|REase-><-SF1-SF2-Helicase+wHTH+Small-B-barrel+DUF1998<-BrxD-AAA<-PglZ<-DUF6079*<-N6-METHYLASE<-PglW\|\|REase+N6_METHYLASE-><-pPIWI+MID+RNaseH-p | 1299 | Bacteria>Terrabacteria_group>Actinomycetota>Actinomycetes>Kitasatosporales>Streptomycetaceae | GCF_013177615.1 | Streptomyces_sp._MNP-20 | HC073_RS37710 | 26479..30378 | 2721165 | phage-resistance-protein-[Streptomyces-sp.-MNP-20] |
| WP_175410415.1 | Type2.Pgl.like.BREX | <-JetB<-?\|\|PglW->N6-METHYLASE-><-REase<-ST-Hiskin<-ST-Hiskin\|\|HTH_31+DUF5753->DUF397->DUF6079*->PglZ-><-CBS+HEPN<-ParB+HNH\|\|BrxD-AAA->SF1-SF2-Helicase+wHTH+Small-B-barrel+DUF1998->?->SIGMA-HTH->ACET-> | 1323 | Bacteria>Terrabacteria_group>Actinomycetota>Actinomycetes>Kitasatosporales>Streptomycetaceae | GCF_013362995.1 | Streptomyces_sp._TRM64462 | HTZ82_RS17995 | 58952..62923 | 2741726 | phage-resistance-protein-[Streptomyces-sp.-TRM64462] |
| WP_175458172.1 | Type2.Pgl.like.BREX | REase+SF1-SF2-Helicase->?-><-REase+SF1-SF2-Helicase\|\|?->REase->\|\|PglW->N6-METHYLASE-><-DUF397<-HTH_31+DUF5753\|\|DUF6079*->PglZ->SF1-SF2-Helicase+wHTH+Small-B-barrel+DUF1998-><-TRAM+SAM-methylase<-AA_permease_2 | 1298 | Bacteria>Terrabacteria_group>Actinomycetota>Actinomycetes>Kitasatosporales>Streptomycetaceae | GCF_013363465.1 | Streptomyces | G6W59_RS10675 | 242164..246060 | 1883 | MULTISPECIES:-phage-resistance-protein-[Streptomyces] |
| WP_176117073.1 | Type2.Pgl.like.BREX | PglW->N6-METHYLASE-><-ST-Hiskin\|\|HTH_31+DUF5753->DUF397->?->?->DUF6079*->PglZ-><-CBS+HEPN<-ParB+HNH\|\|BrxD-AAA->SF1-SF2-Helicase+wHTH+Small-B-barrel+DUF1998-> | 1309 | Bacteria>Terrabacteria_group>Actinomycetota>Actinomycetes>Kitasatosporales>Streptomycetaceae | GCF_013364175.1 | Streptomyces_sp._NA02536 | HUT14_RS10185 | 2278747..2282676 | 2742133 | phage-resistance-protein-[Streptomyces-sp.-NA02536] |
| WP_179084887.1 | Type2.Pgl.like.BREX | AA_permease_2->TRAM+SAM-methylase->SF1-SF2-Helicase+HKD-DNase-><-SF1-SF2-Helicase+wHTH+Small-B-barrel+DUF1998<-BrxD-AAA\|\|ParB+HNH->CBS+HEPN-><-PglZ<-DUF6079*<-?<-DUF2171\|\|HTH_31+DUF5753-><-N6-METHYLASE<-PglW\|\|DUF4011+SF1-SF2-Helicase+REase->?-><-Pkinase\|\|REase-> | 1304 | Bacteria>Terrabacteria_group>Actinomycetota>Actinomycetes>Kitasatosporales>Streptomycetaceae | GCF_013394065.1 | Streptomyces_rectiverticillatus | FGW37_RS24660 | 5638152..5642066 | 173860 | DUF6079-family-protein-[Streptomyces-rectiverticillatus] |
| WP_179181139.1 | Type2.Pgl.like.BREX | <-DUF4096+DDE_Tnp_1+DDE_Tnp_1_2\|\|REase+SF1-SF2-Helicase->?-><-REase+SF1-SF2-Helicase<-SF1-SF2-Helicase+wHTH+Small-B-barrel+DUF1998<-BrxD-AAA<-PglZ<-DUF6079*\|\|SF1-SF2-Helicase+HKD-DNase-><-ST-Hiskin\|\|DUF397-><-N6-METHYLASE\|\|HEPN->ParA-Soj-PloopNTPase->ORC-CDC6-><-PglW<-REase | 1318 | Bacteria>Terrabacteria_group>Actinomycetota>Actinomycetes>Kitasatosporales>Streptomycetaceae | GCF_013401535.1 | Streptomyces_sp._CB04723 | HXS80_RS07185 | 1703905..1707861 | 2748873 | PglY-protein-[Streptomyces-sp.-CB04723] |
| WP_181660536.1 | Type2.Pgl.like.BREX | TetR-HTH-><-?\|\|PglW->N6-METHYLASE->DUF6079*->PglZ->BrxD-AAA->SF1-SF2-Helicase+wHTH+Small-B-barrel+DUF1998-><-?<-ParB+HNH<-?<-HTH_36 | 1304 | Bacteria>Terrabacteria_group>Actinomycetota>Actinomycetes>Kitasatosporales>Streptomycetaceae | GCF_013778455.1 | Streptomyces_himalayensis | H1D24_RS28240 | 43695..47609 | 2820085 | PglY-protein-[Streptomyces-himalayensis] |
| WP_191852498.1 | Type2.Pgl.like.BREX | Peptidase_C14->PglW-><-BrxD-AAA<-ParA-Soj-PloopNTPase<-HEPN\|\|N6-METHYLASE-><-DUF397<-HTH_31+DUF5753\|\|ST-Hiskin->DUF6079*->PglZ->BrxD-AAA->SF1-SF2-Helicase+wHTH+Small-B-barrel+DUF1998->HAD+HAD-><-HTH_3\|\|?->RADICAL-SAM->Radical_SAM-> | 1324 | Bacteria>Terrabacteria_group>Actinomycetota>Actinomycetes>Kitasatosporales>Streptomycetaceae | GCF_013913525.1 | Streptomyces_griseoaurantiacus | H1X69_RS09375 | 152668..156642 | 68213 | PglY-protein-[Streptomyces-griseoaurantiacus] |
| WP_161173771.1 | Type2.Pgl.like.BREX | <-HTH+McrB-AAA\|\|?-><-SF1-SF2-Helicase<-?<-ST-Hiskin\|\|HTH_31+DUF5753->DUF397->PglW->N6-METHYLASE->DUF6079*->PglZ->SF1-SF2-Helicase+wHTH+Small-B-barrel+DUF1998->?-><-TOPRIM<-OLD-ABC\|\|N6-METHYLASE->\|\|Hiskin->?->MarR-HTH-> | 1294 | Bacteria>Terrabacteria_group>Actinomycetota>Actinomycetes>Kitasatosporales>Streptomycetaceae | GCF_014138545.1 | Streptomyces | BJ962_RS25945 | 2051930..2055814 | 1883 | MULTISPECIES:-phage-resistance-protein-[Streptomyces] |
| WP_184574795.1 | Type2.Pgl.like.BREX | PglW->HEPN->ParA-Soj-PloopNTPase->BrxD-AAA->N6-METHYLASE->DUF6079*->PglZ->BrxD-AAA->SF1-SF2-Helicase+wHTH+Small-B-barrel+DUF1998-><-CASPASE+SF1-SF2-Helicase | 1305 | Bacteria>Terrabacteria_group>Actinomycetota>Actinomycetes>Kitasatosporales>Streptomycetaceae | GCF_014203645.1 | Streptomyces_zagrosensis | FHS42_RS22770 | 242905..246822 | 1042984 | hypothetical-protein-[Streptomyces-zagrosensis] |
| WP_187063990.1 | Type2.Pgl.like.BREX | METHYLASE-><-?\|\|?->ParB+HNH+METHYLASE-><-PIN<-RHH_1<-?<-tRNA\|\|PglW-p->N6-METHYLASE->DUF6079*->PglZ->BrxD-AAA->SF1-SF2-Helicase+wHTH+Small-B-barrel+DUF1998-><-TRAM+SAM-methylase<-AA_permease_2<-?<-?<-?<-MarR-HTH | 1297 | Bacteria>Terrabacteria_group>Actinomycetota>Actinomycetes>Kitasatosporales>Streptomycetaceae | GCF_014295035.1 | Streptomyces_buecherae | H7H31_RS06855 | 1887405..1891298 | 2763006 | phage-resistance-protein-[Streptomyces-buecherae] |
| WP_223186495.1 | Type2.Pgl.like.BREX | <-REC+OmpR-HTH<-HisKA_3+HATPase_c<-?<-?\|\|Peptidase_M4+Peptidase_M4_C->?-><-SF1-SF2-Helicase+wHTH+Small-B-barrel+DUF1998<-PglZ<-DUF6079*\|\|BrxA-DNA-binding->?-><-ST-Hiskin\|\|HTH_31+DUF5753->DUF397-><-N6-METHYLASE\|\|HTH+MPTase-><-PglW\|\|REase+SF1-SF2-Helicase-> | 1324 | Bacteria>Terrabacteria_group>Actinomycetota>Actinomycetes>Kitasatosporales>Streptomycetaceae | GCF_014596445.1 | Streptomyces_sp._CBMA29 | BGM09_RS08230 | 64585..68559 | 1896314 | PglY-protein-[Streptomyces-sp.-CBMA29] |
| WP_188302216.1 | Type2.Pgl.like.BREX | <-XIS-HTH\|\|HTH->Phage_integrase-><-?<-SF1-SF2-Helicase+wHTH+Small-B-barrel+DUF1998<-BrxD-AAA<-PglZ<-DUF6079*<-N6-METHYLASE<-PglW\|\|?->\|\|?->?-><-HEPN\|\|PAIREDC-HTH-> | 1312 | Bacteria>Terrabacteria_group>Actinomycetota>Actinomycetes>Kitasatosporales>Streptomycetaceae | GCF_014596465.1 | Streptomyces_sp._CBMA123 | BG452_RS01300 | 274066..278004 | 1896313 | DUF6079-family-protein-[Streptomyces-sp.-CBMA123] |
| WP_189191867.1 | Type2.Pgl.like.BREX | <-TRD+TRD<-N6-METHYLASE<-?\|\|PglW-><-?<-ST-Hiskin\|\|HTH_31+DUF5753->DUF397->MNS+REase->DUF6079*->PglZ-><-CBS+CBS+HEPN<-ParB+HNH<-?\|\|BrxD-AAA->SF1-SF2-Helicase+wHTH+Small-B-barrel+DUF1998->wHTH+WYL->HTH_Tnp-> | 1308 | Bacteria>Terrabacteria_group>Actinomycetota>Actinomycetes>Kitasatosporales>Streptomycetaceae | GCF_014645835.1 | Streptomyces_albiflavescens | IEY95_RS43770 | 44051..47977 | 1623582 | phage-resistance-protein-[Streptomyces-albiflavescens] |
| WP_189215567.1 | Type2.Pgl.like.BREX | <-STAS<-GAF+PP2C<-STAS<-PAS+GAF+HISKIN+REC+GAF+GAF+PP2C\|\|STAS-><-SF1-SF2-Helicase+wHTH+Small-B-barrel+DUF1998<-BrxD-AAA\|\|RNaseH-><-PglZ<-DUF6079*\|\|NACHT+68TM-wHTH->?-><-DUF1995\|\|HTH_31+DUF5753->DUF397-><-N6-METHYLASE<-PglW | 1309 | Bacteria>Terrabacteria_group>Actinomycetota>Actinomycetes>Kitasatosporales>Streptomycetaceae | GCF_014648075.1 | Streptomyces_ruber | IEZ04_RS05700 | 261607..265536 | 83378 | phage-resistance-protein-[Streptomyces-ruber] |
| WP_189273365.1 | Type2.Pgl.like.BREX | PglW->N6-METHYLASE-><-PIN<-PhdYeFM_antitox<-ST-Hiskin\|\|HTH_31+DUF5753->DUF397->PhdYeFM_antitox->PIN->DUF6079*->PglZ-><-CBS+HEPN<-ParB+HNH\|\|BrxD-AAA->SF1-SF2-Helicase+wHTH+Small-B-barrel+DUF1998->Peptidase-S8+Peptidase-S8-> | 1310 | Bacteria>Terrabacteria_group>Actinomycetota>Actinomycetes>Kitasatosporales>Streptomycetaceae | GCF_014648655.1 | Streptomyces_atratus | GCM10010207_01540 | 179323..183255 | 1893 | phage-resistance-protein-[Streptomyces-atratus] |
| WP_189233278.1 | Type2.Pgl.like.BREX | <-TOPRIM<-OLD-ABC\|\|PglW-><-DUF2171<-?\|\|HTH_31+DUF5753->DUF397-><-N6-METHYLASE->DUF6079*->PglZ->BrxD-AAA->SF1-SF2-Helicase+wHTH+Small-B-barrel+DUF1998-><-?<-?<-?<-NTP_transf_3+Hexapep+Hexapep<-tRNA\|\|HISKIN-> | 1302 | Bacteria>Terrabacteria_group>Actinomycetota>Actinomycetes>Kitasatosporales>Streptomycetaceae | GCF_014648815.1 | Streptomyces_flaveolus | IEZ05_RS29325 | 142464..146372 | 67297 | phage-resistance-protein-[Streptomyces-flaveolus] |
| WP_189925600.1 | Type2.Pgl.like.BREX | VirD4-TraG+VirD4-FtsK->DCM->DnaB+DnaB->?->PglW->N6-METHYLASE->DUF6079*->PglZ->BrxD-AAA->SF1-SF2-Helicase+wHTH+Small-B-barrel+DUF1998-><-NUDIX<-REase+DUF5655\|\|XIS-HTH->SMODS->?->HNH+SAVED-> | 1299 | Bacteria>Terrabacteria_group>Actinomycetota>Actinomycetes>Kitasatosporales>Streptomycetaceae | GCF_014649075.1 | Streptomyces_asoensis | Saso_RS30340 | 1677261..1681160 | 249586 | phage-resistance-protein-[Streptomyces-asoensis] |
| WP_189531547.1 | Type2.Pgl.like.BREX | <-SAD(SRA)+wHTH+HNH\|\|?-><-?\|\|PglW->N6-METHYLASE-><-ST-Hiskin\|\|HTH_31+DUF5753->DUF397->?->DUF6079*->PglZ-><-CBS+CBS+HEPN<-ParB+HNH<-Phage_holin\|\|BrxD-AAA->SF1-SF2-Helicase+wHTH+Small-B-barrel+DUF1998-><-?\|\|?->Sensor+HisKA_3+HATPase_c-> | 1298 | Bacteria>Terrabacteria_group>Actinomycetota>Actinomycetes>Kitasatosporales>Streptomycetaceae | GCF_014649335.1 | Streptomyces_roseolilacinus | IE106_RS08455 | 240309..244205 | 66904 | phage-resistance-protein-[Streptomyces-roseolilacinus] |
| WP_190076975.1 | Type2.Pgl.like.BREX | Calcineurin->?->tRNA-><-?\|\|<-SF1-SF2-Helicase+wHTH+Small-B-barrel+DUF1998<-BrxD-AAA<-PglZ<-DUF6079*<-ST-Hiskin<-N6-METHYLASE\|\|HEPN->ParA-Soj-PloopNTPase->BrxD-AAA-><-PglW\|\|DUF2397-> | 1321 | Bacteria>Terrabacteria_group>Actinomycetota>Actinomycetes>Kitasatosporales>Streptomycetaceae | GCF_014649475.1 | Streptomyces_daghestanicus | IE186_RS08935 | 129191..133156 | 66885 | PglY-protein-[Streptomyces-daghestanicus] |
| WP_189596065.1 | Type2.Pgl.like.BREX | TPR+TPR+TPR-><-?<-?<-?<-SF1-SF2-Helicase+wHTH+Small-B-barrel+DUF1998<-BrxD-AAA<-DndB<-PglZ<-DUF6079*<-REase+SF1-SF2-Helicase<-DUF2171<-DUF1995\|\|HTH_31+DUF5753->DUF397-><-N6-METHYLASE<-PglW | 1294 | Bacteria>Terrabacteria_group>Actinomycetota>Actinomycetes>Kitasatosporales>Streptomycetaceae | GCF_014649515.1 | Streptomyces_pilosus | IE141_RS07070 | 323618..327502 | 28893 | phage-resistance-protein-[Streptomyces-pilosus] |
| WP_189553697.1 | Type2.Pgl.like.BREX | <-REase+DUF5655<-HTH+MPTase<-?<-SF1-SF2-Helicase+wHTH+Small-B-barrel+DUF1998<-BrxD-AAA<-PglZ<-DUF6079*<-N6-METHYLASE<-PglW<-TRAM+SAM-methylase<-AA_permease_2\|\|?->?-><-?<-REC+OmpR-HTH | 1297 | Bacteria>Terrabacteria_group>Actinomycetota>Actinomycetes>Kitasatosporales>Streptomycetaceae | GCF_014649735.1 | Streptomyces_lavendofoliae | IE109_RS25595 | 27723..31616 | 67314 | phage-resistance-protein-[Streptomyces-lavendofoliae] |
| WP_189965530.1 | Type2.Pgl.like.BREX | XIS-HTH->?-><-SF1-SF2-Helicase+wHTH+Small-B-barrel+DUF1998<-BrxD-AAA<-PglZ<-DUF6079*<-N6-METHYLASE-p<-N6-METHYLASE<-PglW\|\|?->?-><-?\|\|REase+SF1-SF2-Helicase-><-PIN<-MetJarc-HTH | 1299 | Bacteria>Terrabacteria_group>Actinomycetota>Actinomycetes>Kitasatosporales>Streptomycetaceae | GCF_014649955.1 | Streptomyces_violascens | IE206_RS24480 | 114519..118418 | 67381 | PglY-protein-[Streptomyces-violascens] |
| WP_189744979.1 | Type2.Pgl.like.BREX | REase+SF1-SF2-Helicase-><-SAD(SRA)+wHTH+HNH\|\|PglW->N6-METHYLASE-><-ST-Hiskin\|\|HTH_31+DUF5753->DUF397->?->DUF6079*->PglZ-><-CBS+CBS+HEPN<-ParB+HNH<-Phage_holin\|\|BrxD-AAA->SF1-SF2-Helicase+wHTH+Small-B-barrel+DUF1998-><-Phage_holin\|\|REase->DUF4258->ACET-> | 1295 | Bacteria>Terrabacteria_group>Actinomycetota>Actinomycetes>Kitasatosporales>Streptomycetaceae | GCF_014650435.1 | Streptomyces_tendae | IE258_RS18980 | 415118..419005 | 1932 | phage-resistance-protein-[Streptomyces-tendae] |
| WP_189823142.1 | Type2.Pgl.like.BREX | LACI-HTH+PBPI->\|\|RNaseH-><-SF1-SF2-Helicase+wHTH+Small-B-barrel+DUF1998<-BrxD-AAA<-PglZ<-DUF6079*\|\|BrxA-DNA-binding->?-><-ST-Hiskin\|\|HTH_31+DUF5753->DUF397-><-N6-METHYLASE\|\|SMODS->Schlafen+HATPase+wHTH-><-PglW | 1310 | Bacteria>Terrabacteria_group>Actinomycetota>Actinomycetes>Kitasatosporales>Streptomycetaceae | GCF_014650515.1 | Streptomyces_finlayi | IE282_RS09950 | 502304..506236 | 67296 | PglY-protein-[Streptomyces-finlayi] |
| WP_189852660.1 | Type2.Pgl.like.BREX | <-SbcCD_C<-JetB<-?\|\|PglW->N6-METHYLASE-><-DUF397<-HTH_31+DUF5753\|\|DUF1995->DUF2171->DUF6079*->PglZ-><-CBS+HEPN<-ParB+HNH\|\|BrxD-AAA->SF1-SF2-Helicase+wHTH+Small-B-barrel+DUF1998->REase+SF1-SF2-Helicase->?-><-TRAM+SAM-methylase<-AA_permease_2 | 1297 | Bacteria>Terrabacteria_group>Actinomycetota>Actinomycetes>Kitasatosporales>Streptomycetaceae | GCF_014650895.1 | Streptomyces_omiyaensis | IE287_RS31550 | 67641..71534 | 68247 | phage-resistance-protein-[Streptomyces-omiyaensis] |
| WP_190048743.1 | Type2.Pgl.like.BREX | DUF4011+SF1-SF2-Helicase+REase->PglW->N6-METHYLASE->?-><-SAM-METHYLASE<-?\|\|<-RelE-ParE<-PhdYeFM_antitox\|\|DUF6079*->PglZ-><-Phage_holin<-CBS+CBS+HEPN<-ParB+HNH\|\|BrxD-AAA->SF1-SF2-Helicase+wHTH+Small-B-barrel+DUF1998-><-?<-ParA-Soj-PloopNTPase | 1295 | Bacteria>Terrabacteria_group>Actinomycetota>Actinomycetes>Kitasatosporales>Streptomycetaceae | GCF_014651035.1 | Streptomyces_lomondensis | IE255_RS04370 | 1003302..1007189 | 68229 | phage-resistance-protein-[Streptomyces-lomondensis] |
| WP_190127345.1 | Type2.Pgl.like.BREX | AA_permease_2->TRAM+SAM-methylase-><-?<-PIN<-MetJarc-HTH\|\|PglW->N6-METHYLASE->DUF6079*->PglZ->BrxD-AAA->SF1-SF2-Helicase+wHTH+Small-B-barrel+DUF1998->?->?->\|\|?->?->Sigma70_r2+Sigma70_r4_2-> | 1300 | Bacteria>Terrabacteria_group>Actinomycetota>Actinomycetes>Kitasatosporales>Streptomycetaceae | GCF_014654785.1 | Streptomyces_mashuensis | IE305_RS00650 | 166112..170014 | 33904 | phage-resistance-protein-[Streptomyces-mashuensis] |
| WP_190043128.1 | Type2.Pgl.like.BREX | METHYLASE->PRTase->HAD-><-?\|\|NUDIX-><-SF1-SF2-Helicase+wHTH+Small-B-barrel+DUF1998<-BrxD-AAA<-PglZ<-DUF6079*<-N6-METHYLASE<-PglW\|\|HDRnase-><-NUDIX<-HTH+HKD-DNase\|\|HAD+HAD-> | 1297 | Bacteria>Terrabacteria_group>Actinomycetota>Actinomycetes>Kitasatosporales>Streptomycetaceae | GCF_014654895.1 | Streptomyces_filamentosus | IE326_RS25570 | 1478299..1482192 | 67294 | phage-resistance-protein-[Streptomyces-filamentosus] |
| WP_190135917.1 | Type2.Pgl.like.BREX | <-Peptidase_C14+SF1-SF2-Helicase<-?<-REase\|\|PglW->N6-METHYLASE-><-DUF397\|\|ST-Hiskin->DUF6079*->PglZ->BrxD-AAA->SF1-SF2-Helicase+wHTH+Small-B-barrel+DUF1998-><-?\|\|ZnR+REase-><-DUF4158+DDE_Tnp_Tn3 | 1298 | Bacteria>Terrabacteria_group>Actinomycetota>Actinomycetes>Kitasatosporales>Streptomycetaceae | GCF_014656095.1 | Streptomyces_longispororuber | IE330_RS11995 | 43958..47854 | 68230 | phage-resistance-protein-[Streptomyces-longispororuber] |
| WP_150527865.1 | Type2.Pgl.like.BREX | SF1-SF2-Helicase+HKD-DNase->PglW->N6-METHYLASE-><-DUF397\|\|DUF1995->DUF2171->REase+SF1-SF2-Helicase->DUF6079*->PglZ->DndB->BrxD-AAA->SF1-SF2-Helicase+wHTH+Small-B-barrel+DUF1998-><-?<-TRAM+SAM-methylase<-AA_permease_2 | 1301 | Bacteria>Terrabacteria_group>Actinomycetota>Actinomycetes>Kitasatosporales>Streptomycetaceae | GCF_014656155.1 | Streptomyces_vinaceus | IE310_RS19410 | 98676..102581 | 1960 | phage-resistance-protein-[Streptomyces-vinaceus] |
| WP_189832564.1 | Type2.Pgl.like.BREX | TRAM+SAM-methylase->REase+SF1-SF2-Helicase-><-?<-REase+SF1-SF2-Helicase<-SF1-SF2-Helicase+wHTH+Small-B-barrel+DUF1998<-BrxD-AAA\|\|ParB+HNH->CBS+HEPN-><-PglZ<-DUF6079*<-DUF2171<-DUF1995\|\|HTH_31+DUF5753->DUF397-><-N6-METHYLASE<-PglW\|\|?->JetB->SbcCD_C-> | 1298 | Bacteria>Terrabacteria_group>Actinomycetota>Actinomycetes>Kitasatosporales>Streptomycetaceae | GCF_014656215.1 | Streptomyces_zaomyceticus | IE311_RS17210 | 206117..210013 | 68286 | phage-resistance-protein-[Streptomyces-zaomyceticus] |
| WP_189999228.1 | Type2.Pgl.like.BREX | ACET->?->?-><-?<-?\|\|tRNA->REase-><-SF1-SF2-Helicase+wHTH+Small-B-barrel+DUF1998<-BrxD-AAA<-PglZ<-DUF6079*<-N6-METHYLASE<-PglW\|\|REase+N6_METHYLASE-> | 1299 | Bacteria>Terrabacteria_group>Actinomycetota>Actinomycetes>Kitasatosporales>Streptomycetaceae | GCF_014656255.1 | Streptomyces_rubradiris | IE323_RS35215 | 23790..27689 | 285531 | phage-resistance-protein-[Streptomyces-rubradiris] |
| WP_190849997.1 | Type2.Pgl.like.BREX | TetR-HTH-><-STAS<-LuxR-HTH\|\|?->RADICAL-SAM-><-?<-DUF6731\|\|PglW->N6-METHYLASE->DUF6079*->PglZ->BrxD-AAA->SF1-SF2-Helicase+wHTH+Small-B-barrel+DUF1998-><-tRNA<-TRAM+SAM-methylase+METHYLASE<-AA_permease_2 | 1304 | Bacteria>Terrabacteria_group>Actinomycetota>Actinomycetes>Kitasatosporales>Streptomycetaceae | GCF_014701115.1 | Streptomyces_aurantiacus | IGS71_RS09715 | 2395233..2399147 | 47760 | PglY-protein-[Streptomyces-aurantiacus] |
| WP_194046780.1 | Type2.Pgl.like.BREX | DSBH+AraC-HTH+AraC-HTH->ncRNA->?-><-SF1-SF2-Helicase<-?<-ST-Hiskin\|\|HTH_31+DUF5753->DUF397->PglW->N6-METHYLASE->DUF6079*->PglZ->BrxD-AAA->SF1-SF2-Helicase+wHTH+Small-B-barrel+DUF1998->?->EVE+REase->REase+SF1-SF2-Helicase-> | 1301 | Bacteria>Terrabacteria_group>Actinomycetota>Actinomycetes>Kitasatosporales>Streptomycetaceae | GCF_015160855.1 | Streptomyces_ferrugineus | IM697_RS10270 | 2254443..2258348 | 1413221 | phage-resistance-protein-[Streptomyces-ferrugineus] |
| WP_196191504.1 | Type2.Pgl.like.BREX | Peptidase_C14->PglW-><-BrxD-AAA<-ParA-Soj-PloopNTPase<-HEPN\|\|N6-METHYLASE-><-DUF397<-HTH_31+DUF5753\|\|ST-Hiskin->DUF6079*->PglZ->SF1-SF2-Helicase+wHTH+Small-B-barrel+DUF1998-><-?\|\|Phage_integrase->Phage_integrase-><-REase\|\|ST-Hiskin-> | 1324 | Bacteria>Terrabacteria_group>Actinomycetota>Actinomycetes>Kitasatosporales>Streptomycetaceae | GCF_015680775.2 | Streptomyces_sp._BRB081 | ICN76_RS15730 | 3658471..3662445 | 2769544 | PglY-protein-[Streptomyces-sp.-BRB081] |
| WP_197923435.1 | Type2.Pgl.like.BREX | PglW->N6-METHYLASE-><-ST-Hiskin\|\|HTH_31+DUF5753->DUF397->?-><-?<-?\|\|DUF6079*->PglZ-><-CBS+HEPN<-ParB+HNH<-?\|\|BrxD-AAA->SF1-SF2-Helicase+wHTH+Small-B-barrel+DUF1998->ACET-> | 1235 | Bacteria>Terrabacteria_group>Actinomycetota>Actinomycetes>Kitasatosporales>Streptomycetaceae | GCF_016017415.1 | Streptomyces_sp._HB-N217 | I3J14_RS01390 | 16464..20171 | 2792016 | phage-resistance-protein-[Streptomyces-sp.-HB-N217] |
| WP_201851327.1 | Type2.Pgl.like.BREX | <-N6-METHYLASE<-?\|\|PglW->N6-METHYLASE-><-ST-Hiskin\|\|HTH_31+DUF5753->DUF397->DUF397->DUF6079*->PglZ-><-CBS+HEPN<-ParB+HNH\|\|SF1-SF2-Helicase+wHTH+Small-B-barrel+DUF1998->EVE+SAD+REase->\|\|REase+SF1-SF2-Helicase->?->REase+SF1-SF2-Helicase-><-DUF4011+SF1-SF2-Helicase+REase | 1319 | Bacteria>Terrabacteria_group>Actinomycetota>Actinomycetes>Kitasatosporales>Streptomycetaceae | GCF_016741935.1 | Streptomyces_endocoffeicus | JK364_RS14425 | 428407..432366 | 2898945 | phage-resistance-protein-[Streptomyces-endocoffeicus] |
| WP_129245459.1 | Type2.Pgl.like.BREX | TRAM+SAM-methylase-><-ANK+ANK<-?<-BrxD-AAA<-?<-DUF6079*<-?<-PglW\|\|?->?->REase+SF1-SF2-Helicase-> | 1307 | Bacteria>Terrabacteria_group>Actinomycetota>Actinomycetes>Kitasatosporales>Streptomycetaceae | GCF_016876065.1 | Streptomyces_sioyaensis | EST54_RS05070 | 11707..15630 | 67364 | PglY-protein-[Streptomyces-sioyaensis] |
| WP_205378071.1 | Type2.Pgl.like.BREX | PolB->RADICAL-SAM->?->HD->Glyco_trans_1_4-><-?\|\|NUDIX->\|\|PglW-><-HTH_23+HTH_32+HTH_33\|\|DUF6079*->PglZ->BrxD-AAA->SF1-SF2-Helicase+wHTH+Small-B-barrel+DUF1998->?->NUDIX-><-?\|\|<-DUF3323+TOPRIM<-SbcC/MukB | 1297 | Bacteria>Terrabacteria_group>Actinomycetota>Actinomycetes>Kitasatosporales>Streptomycetaceae | GCF_016919245.1 | Streptomyces_zhihengii | JE024_RS36170 | 1980886..1984779 | 1818004 | phage-resistance-protein-[Streptomyces-zhihengii] |
| WP_207248053.1 | Type2.Pgl.like.BREX | <-SIGMA-HTH<-?<-BrxD-AAA\|\|ParB+HNH->CBS+HEPN-><-PglZ<-DUF6079*<-?<-?\|\|DUF397-><-?<-PglW | 1302 | Bacteria>Terrabacteria_group>Actinomycetota>Actinomycetes>Kitasatosporales>Streptomycetaceae | GCF_017356985.1 | Streptomyces | J1792_RS22685 | 920556..924464 | 1883 | MULTISPECIES:-phage-resistance-protein-[Streptomyces] |
| WP_209214544.1 | Type2.Pgl.like.BREX | <-TOPRIM<-OLD-ABC\|\|N6-METHYLASE->PglW->N6-METHYLASE-><-ST-Hiskin\|\|HTH_31+DUF5753->DUF397->DUF6079*->PglZ-><-CBS+HEPN<-ParB+HNH\|\|SF1-SF2-Helicase+wHTH+Small-B-barrel+DUF1998-> | 1314 | Bacteria>Terrabacteria_group>Actinomycetota>Actinomycetes>Kitasatosporales>Streptomycetaceae | GCF_017676345.1 | Streptomyces_smyrnaeus | JW613_RS33010 | 30286..34230 | 1387713 | phage-resistance-protein-[Streptomyces-smyrnaeus] |
| WP_209240964.1 | Type2.Pgl.like.BREX | <-SF1-SF2-Helicase+HKD-DNase\|\|PglW->N6-METHYLASE-><-ST-Hiskin\|\|HTH_31+DUF5753->DUF397->DUF6079*-> | 1229 | Bacteria>Terrabacteria_group>Actinomycetota>Actinomycetes>Kitasatosporales>Streptomycetaceae | GCF_017676365.1 | Streptomyces_oryzae | ITI46_RS19580 | 22640..26326 | 1434886 | phage-resistance-protein,-partial-[Streptomyces-oryzae] |
| WP_209340089.1 | Type2.Pgl.like.BREX | PglW-><-ORC-CDC6<-ParA-Soj-PloopNTPase<-HEPN\|\|N6-METHYLASE-><-DUF397<-HTH_31+DUF5753<-?<-BrxA-DNA-binding\|\|DUF6079*->PglZ->BrxD-AAA->SF1-SF2-Helicase+wHTH+Small-B-barrel+DUF1998->?-><-REC+OmpR-HTH<-HISKIN<-?<-SIGMA-HTH<-ACET | 1324 | Bacteria>Terrabacteria_group>Actinomycetota>Actinomycetes>Kitasatosporales>Streptomycetaceae | GCF_017813245.1 | Streptomyces_montanisoli | JFN87_RS12560 | 26746..30720 | 2798581 | PglY-protein-[Streptomyces-montanisoli] |
| WP_210930997.1 | Type2.Pgl.like.BREX | ABC-ATPase+SbcC->ABC-ATPase+SbcC->?->?-><-?\|\|GNTR-HTH+Aminotran_1_2-><-SF1-SF2-Helicase+wHTH+Small-B-barrel+DUF1998\|\|RNaseH-><-PglZ<-DUF6079*<-REase+SF1-SF2-Helicase<-DUF1995\|\|HTH_31+DUF5753->DUF397-><-N6-METHYLASE<-PglW\|\|?->?->STYKIN-> | 1299 | Bacteria>Terrabacteria_group>Actinomycetota>Actinomycetes>Kitasatosporales>Streptomycetaceae | GCF_018070665.1 | Streptomyces_sp._404i | KBY46_RS00835 | 204983..208882 | 2824902 | phage-resistance-protein-[Streptomyces-sp.-404i] |
| WP_211038996.1 | Type2.Pgl.like.BREX | PglW->N6-METHYLASE-><-DUF397<-HTH_31+DUF5753\|\|ST-Hiskin->TraJ-RHH->PIN-><-PIN<-CopG/RHH\|\|DUF6079*->PglZ->SF1-SF2-Helicase+wHTH+Small-B-barrel+DUF1998-><-?\|\|?->CNMP+CRP-HTH->?->?->?-><-TRAM+SAM-methylase | 1297 | Bacteria>Terrabacteria_group>Actinomycetota>Actinomycetes>Kitasatosporales>Streptomycetaceae | GCF_018101125.2 | Streptomyces_tardus | JGS22_RS25655 | 96725..100618 | 2780544 | phage-resistance-protein-[Streptomyces-tardus] |
| WP_212726653.1 | Type2.Pgl.like.BREX | <-Phage_integrase<-NUDIX\|\|pPIWI+pPIWI->pPIWI+SF1-SF2-Helicase->pPIWI+MID+RNaseH-><-MULE-TRANSPOSASE<-SF1-SF2-Helicase+wHTH+Small-B-barrel+DUF1998<-BrxD-AAA<-PglZ<-DUF6079*<-N6-METHYLASE<-PglW<-PIWI<-REase+PNTD1+PNTD2\|\|URI-DNase->DUF4096+DDE_Tnp_1+DDE_Tnp_1_2->HTH-><-DNA-glycosylase\|\|XIS-HTH-> | 1299 | Bacteria>Terrabacteria_group>Actinomycetota>Actinomycetes>Kitasatosporales>Streptomycetaceae | GCF_018310535.1 | Streptomyces_mirabilis | SMIR_RS05040 | 1089234..1093133 | 68239 | phage-resistance-protein-[Streptomyces-mirabilis] |
| WP_212908905.1 | Type2.Pgl.like.BREX | SWIM-><-DUF3323+TOPRIM\|\|XIS-HTH->REase+SF1-SF2-Helicase-><-SF1-SF2-Helicase+wHTH+Small-B-barrel+DUF1998<-BrxD-AAA<-PglZ<-DUF6079*<-DUF2171<-DUF1995\|\|HTH_31+DUF5753->DUF397-><-N6-METHYLASE<-PglW\|\|ParB+HNH->CBS+HEPN-> | 1296 | Bacteria>Terrabacteria_group>Actinomycetota>Actinomycetes>Kitasatosporales>Streptomycetaceae | GCF_018327685.1 | Streptomyces_sp._TS71-3 | Sm713_RS07750 | 1878907..1882797 | 2733862 | phage-resistance-protein-[Streptomyces-sp.-TS71-3] |
| WP_215129314.1 | Type2.Pgl.like.BREX | <-REase+SF1-SF2-Helicase\|\|PglW-><-?\|\|N6-METHYLASE-><-DUF397<-HTH_31+DUF5753\|\|ST-Hiskin-><-?\|\|?->DUF6079*->PglZ->SF1-SF2-Helicase+wHTH+Small-B-barrel+DUF1998-><-RNaseH+RNaseH\|\|REase->?->Peptidase_C14+SF1-SF2-Helicase-><-STYKIN | 1301 | Bacteria>Terrabacteria_group>Actinomycetota>Actinomycetes>Kitasatosporales>Streptomycetaceae | GCF_018614035.1 | Streptomyces_sp._ISL-96 | J7E96_RS09030 | 27274..31179 | 2819191 | PglY-protein-[Streptomyces-sp.-ISL-96] |
| WP_214957746.1 | Type2.Pgl.like.BREX | <-CBS+HEPN<-ParB+HNH\|\|PglW->N6-METHYLASE-><-DUF397\|\|DUF1995->DUF2171->Schlafen+HATPase+wHTH->ParA-Soj-PloopNTPase->DUF6079*->PglZ->BrxD-AAA->SF1-SF2-Helicase+wHTH+Small-B-barrel+DUF1998->?->?->CASPASE+RNA-Helicase+REase-><-DDE_Tnp_1_assoc+DDE_Tnp_1 | 1299 | Bacteria>Terrabacteria_group>Actinomycetota>Actinomycetes>Kitasatosporales>Streptomycetaceae | GCF_018614195.1 | Streptomyces_sp._ISL-43 | J7F03_RS38005 | 20163..24062 | 2819183 | DUF6079-family-protein-[Streptomyces-sp.-ISL-43] |
| WP_215167460.1 | Type2.Pgl.like.BREX | <-METHYLASE\|\|?->?->?-><-SF1-SF2-Helicase+wHTH+Small-B-barrel+DUF1998<-BrxD-AAA<-PglZ<-DUF6079*<-TOPRIM<-OLD-ABC<-ST-Hiskin\|\|HTH_31+DUF5753-><-N6-METHYLASE<-?<-TOPRIM<-OLD-ABC<-PglW | 1307 | Bacteria>Terrabacteria_group>Actinomycetota>Actinomycetes>Kitasatosporales>Streptomycetaceae | GCF_018614415.1 | Streptomyces_sp._ISL-12 | J7I94_RS14400 | 76255..80178 | 2819177 | hypothetical-protein-[Streptomyces-sp.-ISL-12] |
| WP_251074221.1 | Type2.Pgl.like.BREX | <-N6-METHYLASE<-DrHyd\|\|SLATT->?-><-SF1-SF2-Helicase+wHTH+Small-B-barrel+DUF1998<-BrxD-AAA<-PglZ<-DUF6079*\|\|DUF397-><-N6-METHYLASE<-BrxD-AAA<-ParA-Soj-PloopNTPase<-HEPN<-PglW\|\|REase+DUF5655-> | 1338 | Bacteria>Terrabacteria_group>Actinomycetota>Actinomycetes>Kitasatosporales>Streptomycetaceae | GCF_018614515.1 | Streptomyces_sp._ISL-111 | J7E90_RS26865 | 114837..118853 | 2819175 | hypothetical-protein-[Streptomyces-sp.-ISL-111] |
| WP_215084214.1 | Type2.Pgl.like.BREX | TetR-HTH->?->PglW->N6-METHYLASE-><-DUF397<-HTH_31+DUF5753\|\|DUF2171->?->DUF6079*->PglZ-><-CBS+HEPN<-ParB+HNH\|\|BrxD-AAA->SF1-SF2-Helicase+wHTH+Small-B-barrel+DUF1998->SF1-SF2-Helicase+HKD-DNase->HKD-DNase->REase+SF1-SF2-Helicase->\|\|ArsR-HTH-> | 1302 | Bacteria>Terrabacteria_group>Actinomycetota>Actinomycetes>Kitasatosporales>Streptomycetaceae | GCF_018614525.1 | Streptomyces_sp._ISL-11 | J7E86_RS24340 | 24343..28251 | 2819174 | phage-resistance-protein-[Streptomyces-sp.-ISL-11] |
| WP_214971885.1 | Type2.Pgl.like.BREX | Phosphodiest->Sensor+HisKA_3+HATPase_c->REC+LuxR-HTH-><-TetR-HTH\|\|?->?-><-SF1-SF2-Helicase+wHTH+Small-B-barrel+DUF1998<-BrxD-AAA<-PglZ<-DUF6079*<-N6-METHYLASE<-DUF397<-HTH_31+DUF5753\|\|?->DUF2171-><-PglW\|\|OLD-ABC->TOPRIM-><-REase | 1309 | Bacteria>Terrabacteria_group>Actinomycetota>Actinomycetes>Kitasatosporales>Streptomycetaceae | GCF_018614615.1 | Streptomyces_sp._ISL-100 | J7E89_RS10735 | 38743..42672 | 2819173 | DUF6079-family-protein-[Streptomyces-sp.-ISL-100] |
| WP_216339501.1 | Type2.Pgl.like.BREX | DUF4011+SF1-SF2-Helicase+REase->\|\|?-><-EVE+SAD+REase<-SF1-SF2-Helicase+wHTH+Small-B-barrel+DUF1998<-BrxD-AAA\|\|ParB+HNH->CBS+HEPN-><-PglZ<-DUF6079*<-DUF397<-HTH_31+DUF5753\|\|ST-Hiskin-><-N6-METHYLASE<-PglW\|\|REase+SF1-SF2-Helicase->EVE+HNH-><-N6-METHYLASE\|\|OLD-ABC-> | 1320 | Bacteria>Terrabacteria_group>Actinomycetota>Actinomycetes>Kitasatosporales>Streptomycetaceae | GCF_018883605.1 | Streptomyces_sp._4503 | KN815_RS01825 | 18413..22375 | 2842201 | phage-resistance-protein-[Streptomyces-sp.-4503] |
| WP_217199264.1 | Type2.Pgl.like.BREX | MarR-HTH->?->?->?->AA_permease_2->TRAM+SAM-methylase-><-SF1-SF2-Helicase+wHTH+Small-B-barrel+DUF1998<-BrxD-AAA<-PglZ<-DUF6079*<-N6-METHYLASE<-PglW-p\|\|tRNA->?->?->RHH_1->PIN-><-ParB+HNH+METHYLASE | 1297 | Bacteria>Terrabacteria_group>Actinomycetota>Actinomycetes>Kitasatosporales>Streptomycetaceae | GCF_019049085.1 | Streptomyces_buecherae | KUM04_RS25325 | 52290..56183 | 2763006 | phage-resistance-protein-[Streptomyces-buecherae] |
| WP_218197629.1 | Type2.Pgl.like.BREX | HTH->XIS-HTH->Phage_integrase-><-?<-SF1-SF2-Helicase+wHTH+Small-B-barrel+DUF1998<-BrxD-AAA<-PglZ<-DUF6079*<-N6-METHYLASE<-PglW | 1313 | Bacteria>Terrabacteria_group>Actinomycetota>Actinomycetes>Kitasatosporales>Streptomycetaceae | GCF_019192735.1 | Kitasatospora_aureofaciens | KV557_RS01080 | 237396..241337 | 1894 | DUF6079-family-protein-[Kitasatospora-aureofaciens] |
| WP_219664043.1 | Type2.Pgl.like.BREX | Peptidase->?->Glyco_tran_28_C->PglW->N6-METHYLASE-><-DUF397<-HTH_31+DUF5753\|\|DUF1995->REase+SF1-SF2-Helicase->DUF6079*->PglZ->DndB->BrxD-AAA->SF1-SF2-Helicase+wHTH+Small-B-barrel+DUF1998-> | 1299 | Bacteria>Terrabacteria_group>Actinomycetota>Actinomycetes>Kitasatosporales>Streptomycetaceae | GCF_019400985.1 | Streptomyces_bambusae | GPJ59_RS00105 | 29061..32960 | 1550616 | DUF6079-family-protein-[Streptomyces-bambusae] |
| WP_222995822.1 | Type2.Pgl.like.BREX | pPIWI+SF1-SF2-Helicase->pPIWI+MID+RNaseH->?->SF1-SF2-Helicase->HNH+PNTD1+PNTD2->PIWI->\|\|REase+DUF5655-><-SF1-SF2-Helicase+wHTH+Small-B-barrel+DUF1998<-PglZ<-DUF6079*<-N6-METHYLASE<-PglW<-?<-DnaB+DnaB\|\|?-><-?<-Peptidase_M15<-NUDIX<-NUDIX | 1299 | Bacteria>Terrabacteria_group>Actinomycetota>Actinomycetes>Kitasatosporales>Streptomycetaceae | GCF_019890645.1 | Streptomyces_sp._SP2-10 | K7864_RS12155 | 58567..62466 | 2873385 | phage-resistance-protein-[Streptomyces-sp.-SP2-10] |
| WP_223735561.1 | Type2.Pgl.like.BREX | <-REC+OmpR-HTH<-HAMP+HISKIN<-?\|\|ACET->?-><-SF1-SF2-Helicase+wHTH+Small-B-barrel+DUF1998<-PglZ<-DUF6079*<-ST-Hiskin\|\|<-N6-METHYLASE<-PglW<-ANK+ANK<-?\|\|ACET+PIN+ACET->ASCH->\|\|Phage_integrase-> | 1298 | Bacteria>Terrabacteria_group>Actinomycetota>Actinomycetes>Kitasatosporales>Streptomycetaceae | GCF_020024005.1 | Streptomyces_purpurogeneiscleroticus | CCS38_RS25010 | 26602..30498 | 68259 | phage-resistance-protein-[Streptomyces-purpurogeneiscleroticus] |
| WP_223780477.1 | Type2.Pgl.like.BREX | ROK-HTH-><-?\|\|sGTP-><-SF1-SF2-Helicase+wHTH+Small-B-barrel+DUF1998<-BrxD-AAA<-PglZ<-DUF6079*<-ST-Hiskin\|\|HTH_31+DUF5753->DUF397-><-N6-METHYLASE<-ORC-CDC6<-ParA-Soj-PloopNTPase<-HEPN<-PglW<-TetR-HTH | 1303 | Bacteria>Terrabacteria_group>Actinomycetota>Actinomycetes>Kitasatosporales>Streptomycetaceae | GCF_020034615.1 | Streptomyces_sp._A144 | K5X85_RS23165 | 5194666..5198577 | 2871487 | hypothetical-protein-[Streptomyces-sp.-A144] |

| Protein-ID | System | Gene-Neighbourhoods | Length | Taxonomy | Assembly-ID | Strain | Locus-Tag | Genomic-coordinates | TaxID | Fasta-defline |
|---|---|---|---|---|---|---|---|---|---|---|
| WP_258081613.1 | Type2.Pgl.like.BREX | HTH_3->?-><- SF1-SF2-Helicase+wHTH+Small-B-barrel+DUF1998 <- BrxD-AAA <- PglZ <- DUF6079 *<- N6-METHYLASE <- PglW <-?<-?\|\| N6-METHYLASE ->TRD+TRD->Pkinase+TPR+TPR-> | 1329 | Bacteria>Terrabacteria_group>Actinomycetota>Actinomycetes>Mycobacteriales>Nocardiaceae | GCF_002933535.1 | Nocardia_nova | C5E45_RS23115 | 140922..144911 | 37330 | DUF6079-family-protein-[Nocardia-nova] |
| WP_119699787.1 | Type2.Pgl.like.BREX | His_Phos_1->?->STYKinase-like->?-> PglW -> N6-METHYLASE -><-HEPN+REase\|\| DUF6079 *-> PglZ -> BrxD-AAA ->HHH_4+HTH+wHTH+SF1-SF2-Helicase+REase-><-HicB\|\| SF1-SF2-Helicase+wHTH+Small-B-barrel+DUF1998 -><-?\|\|ZnR-><-SbcC/MukB | 1235 | Bacteria>Terrabacteria_group>Actinomycetota>Actinomycetes>Mycobacteriales>Nocardiaceae | GCF_003586525.1 | Rhodococcus_ruber | YT1_RS14630 | 2910213..2913920 | 1830 | DUF6079-family-protein-[Rhodococcus-ruber] |
| WP_127946806.1 | Type2.Pgl.like.BREX | TetR-HTH-> PglW -> N6-METHYLASE ->ParB+HNH-> DUF6079 *-> PglZ -> BrxD-AAA ->TraJ-RHH->DOC-> SF1-SF2-Helicase+wHTH+Small-B-barrel+DUF1998 -><-GDResolvase+Recombinase | 1232 | Bacteria>Terrabacteria_group>Actinomycetota>Actinomycetes>Mycobacteriales>Nocardiaceae | GCF_004011835.1 | Rhodococcus_spongiicola | EF834_RS08540 | 73116..76814 | 2487352 | DUF6079-family-protein-[Rhodococcus-spongiicola] |
| WP_242496869.1 | Type2.Pgl.like.BREX | <-HEPN+RES<-HNH<-?<-MORC\|\| PglW -> N6-METHYLASE -> DUF6079 *-> PglZ -> BrxD-AAA -> SF1-SF2-Helicase+wHTH+Small-B-barrel+DUF1998 ->URI-DNase+URI-DNase->MAD-NTD+EVE+McrB-AAA->McrC-NTD+REase->ANK+ANK->SF1-SF2-Helicase->ParB+HNH-> | 1242 | Bacteria>Terrabacteria_group>Actinomycetota>Actinomycetes>Mycobacteriales>Nocardiaceae | GCF_004153645.1 | Rhodococcus_sp._Q1 | EQ874_RS21100 | 15779..19507 | 2508718 | DUF6079-family-protein-[Rhodococcus-sp.-Q1] |
| WP_135046364.1 | Type2.Pgl.like.BREX | <-CASPASE\|\|SF1-SF2-Helicase+REase->SF1-SF2-Helicase+HTH->wHTH+REase->ANK+ANK-><-ParB+HNH<- SF1-SF2-Helicase+wHTH+Small-B-barrel+DUF1998 <- BrxD-AAA <- PglZ <- DUF6079 *<- N6-METHYLASE <- PglW \|\|?->?-><-?\|\|MODE-HTH+PBPII-><-?<-?<-ABC-ATPase+SbcC | 1217 | Bacteria>Terrabacteria_group>Actinomycetota>Actinomycetes>Mycobacteriales>Nocardiaceae | GCF_004563845.1 | Rhodococcus_sp._1R11 | E4P29_RS16905 | 125769..129422 | 2559614 | phage-resistance-protein-[Rhodococcus-sp.-1R11] |
| WP_137797778.1 | Type2.Pgl.like.BREX | ParA-Soj-PloopNTPase->HHH_4->SIGMA-HTH+SIGMA-HTH-><-nSTAND1+HNH<-REase+SF1-SF2-Helicase<-ParB+HNH+Schlafen<- SF1-SF2-Helicase+wHTH+Small-B-barrel+DUF1998 <- BrxD-AAA <- PglZ <- DUF6079 *\|\|HEPN-><-RADICAL-SAM<-HKD-DNase<- N6-METHYLASE <- PglW <-?<-MORC\|\|HHH_5+SNF2_N+SF1-SF2-Helicase-><-wHTH | 1264 | Bacteria>Terrabacteria_group>Actinomycetota>Actinomycetes>Mycobacteriales>Nocardiaceae | GCF_005484805.1 | Rhodococcus_sp._SGAir0479 | E7742_RS04110 | 877804..881598 | 2567884 | DUF6079-family-protein-[Rhodococcus-sp.-SGAir0479] |
| WP_227833967.1 | Type2.Pgl.like.BREX | PglW -> N6-METHYLASE -> DUF6079 *-> PglZ -> BrxD-AAA -> SF1-SF2-Helicase+wHTH+Small-B-barrel+DUF1998 -><-?\|\|?->?->APATPase+TPR+TPR+TPR+TPR->?->METHYLASE-> | 1284 | Bacteria>Terrabacteria_group>Actinomycetota>Actinomycetes>Mycobacteriales>Nocardiaceae | GCF_009604405.1 | Nocardia_macrotermitis | NRB20_RS28450 | 166856..170710 | 2585198 | phage-resistance-protein-[Nocardia-macrotermitis] |
| WP_202400078.1 | Type2.Pgl.like.BREX | <-HNH<-?<-HATPase\|\| PglW -> N6-METHYLASE -> DUF6079 *-> PglZ -> BrxD-AAA -> SF1-SF2-Helicase+wHTH+Small-B-barrel+DUF1998 ->URI-DNase+URI-DNase->McrB-AAA-ATPase->McrC-NTD+REase->ANK+ANK-> | 1242 | Bacteria>Terrabacteria_group>Actinomycetota>Actinomycetes>Mycobacteriales>Nocardiaceae | GCF_009831075.1 | Rhodococcus_rhodochrous | GSM98_RS14610 | 113607..117335 | 1829 | DUF6079-family-protein-[Rhodococcus-rhodochrous] |
| WP_168579650.1 | Type2.Pgl.like.BREX | His_Phos_1->?->STYKinase-like->?-> PglW -> N6-METHYLASE ->CR-ATPase-> DUF6079 *-> PglZ -> BrxD-AAA ->?-><-HicB<-HicA\|\| SF1-SF2-Helicase+wHTH+Small-B-barrel+DUF1998 -><-HTH_36\|\|ZnR-><-SbcC/MukB | 1237 | Bacteria>Terrabacteria_group>Actinomycetota>Actinomycetes>Mycobacteriales>Nocardiaceae | GCF_012272955.1 | Rhodococcus_sp._DMU1 | HFP48_RS14070 | 3003998..3007711 | 2722825 | DUF6079-family-protein-[Rhodococcus-sp.-DMU1] |
| WP_206038827.1 | Type2.Pgl.like.BREX | <-MAD-NTD+EVE+McrB-AAA<-URI-DNase<- SF1-SF2-Helicase+wHTH+Small-B-barrel+DUF1998 <- BrxD-AAA <- PglZ <- DUF6079 *<- N6-METHYLASE <- PglW <-REase+SF1-SF2-Helicase | 1228 | Bacteria>Terrabacteria_group>Actinomycetota>Actinomycetes>Mycobacteriales>Nocardiaceae | GCF_012844225.1 | Rhodococcus_sp._105337 | HF886_RS14710 | 10453..14139 | 2725310 | DUF6079-family-protein-[Rhodococcus-sp.-105337] |
| WP_249343668.1 | Type2.Pgl.like.BREX | <-MPTase<-?<-ZetaToxin\|\| PglW -p-><-DDE_Tnp_1\|\| PglW -> N6-METHYLASE -> DUF6079 *-> PglZ -> BrxD-AAA -> SF1-SF2-Helicase+wHTH+Small-B-barrel+DUF1998 ->HEPN+RES->CHTH-><-?<-?<-?<-ParA-Soj-PloopNTPase | 1271 | Bacteria>Terrabacteria_group>Actinomycetota>Actinomycetes>Mycobacteriales>Nocardiaceae | GCF_012971805.1 | Rhodococcus_opacus | HJ581_RS11030 | 86597..90412 | 37919 | DUF6079-family-protein-[Rhodococcus-opacus] |
| WP_188543681.1 | Type2.Pgl.like.BREX | DUF6361->HKD-DNase->D6+SF1-SF2-Helicase->?-><- SF1-SF2-Helicase+wHTH+Small-B-barrel+DUF1998 <- BrxD-AAA <- PglZ <- DUF6079 *<- N6-METHYLASE <- PglW \|\| PglZ -><- BrxD-AAA <-HKD-DNase | 1212 | Bacteria>Terrabacteria_group>Actinomycetota>Actinomycetes>Mycobacteriales>Nocardiaceae | GCF_014635345.1 | Rhodococcus_trifolii | IEW02_RS05775 | 1232248..1235886 | 908250 | phage-resistance-protein-[Rhodococcus-trifolii] |
| WP_063000546.1 | Type2.Pgl.like.BREX | DUF3323+TOPRIM->HEPN+SF1-SF2-Helicase+REase->?-><-REase<-?\|\| PglW -> N6-METHYLASE -> DUF6079 *-> PglZ -> BrxD-AAA -> SF1-SF2-Helicase+wHTH+Small-B-barrel+DUF1998 -><-DUF397<-HTH_31+DUF5753\|\|?->?->?->HKD-DNase-> | 1249 | Bacteria>Terrabacteria_group>Actinomycetota>Actinomycetes>Mycobacteriales>Nocardiaceae | GCF_014645735.1 | Nocardia_jinanensis | IEY68_RS31650 | 40816..44565 | 382504 | DUF6079-family-protein-[Nocardia-jinanensis] |
| WP_229684222.1 | Type2.Pgl.like.BREX | <-NUDIX\|\|?->?-><-?<-?<-?<-Pkinase+TPR+TPR\|\| PglW -> N6-METHYLASE -> DUF6079 *-> PglZ -> BrxD-AAA -> SF1-SF2-Helicase+wHTH+Small-B-barrel+DUF1998 ->DndD->HEPN->?-><-?<-?<-PIN | 1364 | Bacteria>Terrabacteria_group>Actinomycetota>Actinomycetes>Mycobacteriales>Nocardiaceae | GCF_014646135.1 | Nocardia_camponoti | IEX78_RS21925 | 193544..197638 | 1616106 | phage-resistance-protein-[Nocardia-camponoti] |
| WP_257233264.1 | Type2.Pgl.like.BREX | SIGMA-HTH->MORC->?->ParA-Soj-PloopNTPase->HHH_4->wHTH+REase-><- SF1-SF2-Helicase+wHTH+Small-B-barrel+DUF1998 <- BrxD-AAA <- PglZ <- DUF6079 *<- N6-METHYLASE <- PglW <-DNA_glycosylase\|\|EVE+HNH-><-nSTAND1+HNH<-REase+SF1-SF2-Helicase | 1296 | Bacteria>Terrabacteria_group>Actinomycetota>Actinomycetes>Mycobacteriales>Nocardiaceae | GCF_015034605.1 | Rhodococcus_qingshengii | IM699_RS24685 | 5335189..5339079 | 334542 | DUF6079-family-protein-[Rhodococcus-qingshengii] |
| WP_228538510.1 | Type2.Pgl.like.BREX | PglW -> N6-METHYLASE -> DUF6079 *-> PglZ -> BrxD-AAA -> SF1-SF2-Helicase+wHTH+Small-B-barrel+DUF1998 -><-SF1-SF2-Helicase | 1278 | Bacteria>Terrabacteria_group>Actinomycetota>Actinomycetes>Mycobacteriales>Nocardiaceae | GCF_015355755.1 | Nocardia_sp._XZ_19_385 | IBX22_RS05375 | 1166777..1170613 | 2769488 | phage-resistance-protein-[Nocardia-sp.-XZ_19_385] |
| WP_228824462.1 | Type2.Pgl.like.BREX | SF1-SF2-Helicase-><- SF1-SF2-Helicase+wHTH+Small-B-barrel+DUF1998 <- BrxD-AAA <- PglZ <- DUF6079 *<- N6-METHYLASE <- PglW <-?\|\|?->?->?-><-?<-TetR-HTH | 1278 | Bacteria>Terrabacteria_group>Actinomycetota>Actinomycetes>Mycobacteriales>Nocardiaceae | GCF_015477355.1 | Nocardia_blacklockiae | IU476_RS27875 | 116199..120035 | 480036 | phage-resistance-protein-[Nocardia-blacklockiae] |
| WP_195024438.1 | Type2.Pgl.like.BREX | HTH_31+DUF5753->?->?->?-><- SF1-SF2-Helicase+wHTH+Small-B-barrel+DUF1998 <-CBS+HEPN<-ParB+HNH<- PglZ <- DUF6079 *<- N6-METHYLASE <- PglW <-?\|\| N6-METHYLASE ->Pkinase+TPR+TPR-><-SF1-SF2-Helicase | 1300 | Bacteria>Terrabacteria_group>Actinomycetota>Actinomycetes>Mycobacteriales>Nocardiaceae | GCF_015477405.1 | Nocardia_farcinica | IU473_RS11255 | 584605..588507 | 37329 | DUF6079-family-protein-[Nocardia-farcinica] |
| WP_228782161.1 | Type2.Pgl.like.BREX | HTH_3->STYKIN->?-> PglW -> N6-METHYLASE -> DUF6079 *-> PglZ -> SF1-SF2-Helicase+wHTH+Small-B-barrel+DUF1998 -><-DUF3323+TOPRIM<-?<-?<-PadR-wHTH<-?<-ABhydrolase | 1245 | Bacteria>Terrabacteria_group>Actinomycetota>Actinomycetes>Mycobacteriales>Nocardiaceae | GCF_015477435.1 | Nocardia_abscessus | IU470_RS12230 | 97395..101132 | 120957 | DUF6079-family-protein-[Nocardia-abscessus] |
| WP_228838699.1 | Type2.Pgl.like.BREX | DUF2397->?->?->DUF3323+TOPRIM-><- SF1-SF2-Helicase+wHTH+Small-B-barrel+DUF1998 <- BrxD-AAA <- PglZ <- DUF6079 *<- N6-METHYLASE <- PglW <-HTH_31+DUF5753\|\|?->?-><-?<-Abhydrolase_3<-SF1-SF2-Helicase | 1245 | Bacteria>Terrabacteria_group>Actinomycetota>Actinomycetes>Mycobacteriales>Nocardiaceae | GCF_015477605.1 | Nocardia_amamiensis | IU459_RS29160 | 22411..26148 | 404578 | DUF6079-family-protein-[Nocardia-amamiensis] |
| WP_195000848.1 | Type2.Pgl.like.BREX | <-SF1-SF2-Helicase<-Pkinase+TPR+TPR<-TRD+TRD<- N6-METHYLASE \|\|?-> PglW -> N6-METHYLASE -> DUF6079 *-> PglZ -><-DOC+HTH\|\|?->?->HTH_3-> | 1316 | Bacteria>Terrabacteria_group>Actinomycetota>Actinomycetes>Mycobacteriales>Nocardiaceae | GCF_015477835.1 | Nocardia_higoensis | IU449_RS05565 | 1242547..1246497 | 228599 | DUF6079-family-protein-[Nocardia-higoensis] |
| WP_228801228.1 | Type2.Pgl.like.BREX | <- N6-METHYLASE <-?<-?\|\| PglW -> N6-METHYLASE -> DUF6079 *-> PglZ -> BrxD-AAA -> SF1-SF2-Helicase+wHTH+Small-B-barrel+DUF1998 -><-?<-DUF397<-HTH_31+DUF5753 | 1271 | Bacteria>Terrabacteria_group>Actinomycetota>Actinomycetes>Mycobacteriales>Nocardiaceae | GCF_015477865.1 | Nocardia_cyriacigeorgica | IU409_RS23450 | 21336..25151 | 135487 | DUF6079-family-protein-[Nocardia-cyriacigeorgica] |
| WP_195065211.1 | Type2.Pgl.like.BREX | <-DUF4011+SF1-SF2-Helicase+REase\|\|DOC+HTH->?->SF1-SF2-Helicase->?->?->STYKIN-> PglW -> N6-METHYLASE -> DUF6079 *-> PglZ -> SF1-SF2-Helicase+wHTH+Small-B-barrel+DUF1998 ->?->?-><-HTH_3 | 1285 | Bacteria>Terrabacteria_group>Actinomycetota>Actinomycetes>Mycobacteriales>Nocardiaceae | GCF_015478245.1 | Nocardia_abscessus | IU428_RS07225 | 143380..147237 | 120957 | phage-resistance-protein-[Nocardia-abscessus] |
| WP_195027567.1 | Type2.Pgl.like.BREX | <- SF1-SF2-Helicase+wHTH+Small-B-barrel+DUF1998 <- BrxD-AAA <-CBS+HEPN<-ParB+HNH<- PglZ <- DUF6079 *<-?<- PglW <-?\|\| N6-METHYLASE ->?->Pkinase+TPR-> | 1300 | Bacteria>Terrabacteria_group>Actinomycetota>Actinomycetes>Mycobacteriales>Nocardiaceae | GCF_015478425.1 | Nocardia_farcinica | AB8M53_RS07390 | 596656..600558 | 37329 | DUF6079-family-protein-[Nocardia-farcinica] |
| WP_072634863.1 | Type2.Pgl.like.BREX | <-HTH_Tnp_1+HTH_21+rve+rve_3\|\|<-?\|\|?-><- SF1-SF2-Helicase+wHTH+Small-B-barrel+DUF1998 <- BrxD-AAA <- PglZ <- DUF6079 *<- N6-METHYLASE <- PglW \|\|DUF4352->\|\|DDE_Tnp_1-><-TUDOR+SF1-SF2-Helicase+iREase+CTD+REase\|\|HHH_5+SNF2_N+SF1-SF2-Helicase->?-><-wHTH\|\|wHTH+SAD+HNH-p-> | 1243 | Bacteria>Terrabacteria_group>Actinomycetota>Actinomycetes>Mycobacteriales>Nocardiaceae | GCF_015654185.1 | Rhodococcus_sp._M8 | ISO16_RS27035 | 5936618..5940349 | 1925550 | DUF6079-family-protein-[Rhodococcus-sp.-M8] |
| WP_203010703.1 | Type2.Pgl.like.BREX | His_Phos_1->?->STYKinase-like->?-> PglW -> N6-METHYLASE -><-DDE_Tnp_1<-?\|\| DUF6079 *-> PglZ -> BrxD-AAA ->?-><-HicB<-HicA\|\| SF1-SF2-Helicase+wHTH+Small-B-barrel+DUF1998 -> | 1235 | Bacteria>Terrabacteria_group>Actinomycetota>Actinomycetes>Mycobacteriales>Nocardiaceae | GCF_016804345.1 | Rhodococcus_ruber | F1734_RS14140 | 3038038..3041745 | 1830 | DUF6079-family-protein-[Rhodococcus-ruber] |
| WP_254206127.1 | Type2.Pgl.like.BREX | PglW -> N6-METHYLASE -> DUF6079 *-> PglZ -> BrxD-AAA -> SF1-SF2-Helicase+wHTH+Small-B-barrel+DUF1998 -><-?<-HTH_3 | 1278 | Bacteria>Terrabacteria_group>Actinomycetota>Actinomycetes>Mycobacteriales>Nocardiaceae | GCF_019038435.1 | Nocardia_sp._ncl2 | JZ791_RS22515 | 9851..13687 | 2815723 | phage-resistance-protein-[Nocardia-sp.-ncl2] |
| WP_219130349.1 | Type2.Pgl.like.BREX | STYKinase-like->?-><-SF1-SF2-Helicase\|\| PglW -> N6-METHYLASE -><-HEPN+REase\|\| DUF6079 *-> PglZ -> BrxD-AAA ->SF1-SF2-Helicase-> SF1-SF2-Helicase+wHTH+Small-B-barrel+DUF1998 -> | 1235 | Bacteria>Terrabacteria_group>Actinomycetota>Actinomycetes>Mycobacteriales>Nocardiaceae | GCF_019317345.1 | Rhodococcus_sp._LW-XY12 | KXC42_RS11380 | 2418472..2422179 | 2856851 | DUF6079-family-protein-[Rhodococcus-sp.-LW-XY12] |
| WP_084848434.1 | Type2.Pgl.like.BREX | PglW -> N6-METHYLASE ->McrB-AAA->McrC-NTD+REase->EVE+REase->?-> DUF6079 *-> PglZ -> BrxD-AAA -> SF1-SF2-Helicase+wHTH+Small-B-barrel+DUF1998 ->?->DDE_Tnp_1_2->REase+SF1-SF2-Helicase->nSTAND1+HNH-> | 1264 | Bacteria>Terrabacteria_group>Actinomycetota>Actinomycetes>Mycobacteriales>Nocardiaceae | GCF_019703975.1 | Prescottella_equi | A5N78_RS19000 | 135247..139041 | 43767 | DUF6079-family-protein-[Prescottella-equi] |
| WP_222677599.1 | Type2.Pgl.like.BREX | <-HISKIN\|\|DUF4352->?->MORC+DUF3684+REase-><-EVE+REase\|\|SF1-SF2-Helicase+REase->SF1-SF2-Helicase+HTH-><- SF1-SF2-Helicase+wHTH+Small-B-barrel+DUF1998 <- PglZ <- DUF6079 *<- N6-METHYLASE <- PglW \|\|wHTH-><- BrxD-AAA \|\|McrB-AAA->McrC-NTD+REase->MORC->REase-> | 1213 | Bacteria>Terrabacteria_group>Actinomycetota>Actinomycetes>Mycobacteriales>Nocardiaceae | GCF_019834505.1 | Rhodococcus_corynebacterioides | HQ609_RS09015 | 112218..115859 | 53972 | phage-resistance-protein-[Rhodococcus-corynebacterioides] |
| WP_222635286.1 | Type2.Pgl.like.BREX | PglW -> N6-METHYLASE -> DUF6079 *-> PglZ -> BrxD-AAA ->SF1-SF2-Helicase-> SF1-SF2-Helicase+wHTH+Small-B-barrel+DUF1998 -><-GDResolvase+HIN-HTH<-TetR-HTH | 1237 | Bacteria>Terrabacteria_group>Actinomycetota>Actinomycetes>Mycobacteriales>Nocardiaceae | GCF_019834785.1 | unclassified_Rhodococcus_(in:_high_G+C_Gram-positive_bacteria) | HQ360_RS18085 | 40159..43872 | 192944 | MULTISPECIES:-DUF6079-family-protein-[unclassified-Rhodococcus-(in:-high-G+C-Gram-positive-bacteria)] |
| WP_222632854.1 | Type2.Pgl.like.BREX | <-SF1-SF2-Helicase+REase<-Exonuc_V_gamma+REase<-?\|\|OLD-ABC+TOPRIM-><- SF1-SF2-Helicase+wHTH+Small-B-barrel+DUF1998 <- BrxD-AAA <- PglZ <- DUF6079 *<- N6-METHYLASE <- PglW \|\|DUF6361->HKD-DNase->SF1-SF2-Helicase->HNH->ABhydrolase+ABhydrolase-><-?\|\|REase-> | 1293 | Bacteria>Terrabacteria_group>Actinomycetota>Actinomycetes>Mycobacteriales>Nocardiaceae | GCF_019836125.1 | unclassified_Rhodococcus_(in:_high_G+C_Gram-positive_bacteria) | HQ302_RS17065 | 242661..246542 | 192944 | MULTISPECIES:-DUF6079-family-protein-[unclassified-Rhodococcus-(in:-high-G+C-Gram-positive-bacteria)] |
| WP_252173979.1 | Type2.Pgl.like.BREX | <-ParB+HNH<-SF1-SF2-Helicase<-ANK+ANK<-McrC-NTD+REase<-MAD-NTD+EVE+McrB-AAA<-URI-DNase+URI-DNase<- SF1-SF2-Helicase+wHTH+Small-B-barrel+DUF1998 <- BrxD-AAA <- PglZ <- DUF6079 *<- N6-METHYLASE <- PglW \|\|?->HNH->?->?-><-HTH_36<-nSTAND1+HNH<-REase+SF1-SF2-Helicase | 1242 | Bacteria>Terrabacteria_group>Actinomycetota>Actinomycetes>Mycobacteriales>Nocardiaceae | GCF_023822085.1 | Rhodococcus_pyridinivorans | LLA01_RS07335 | 1548248..1551976 | 103816 | DUF6079-family-protein-[Rhodococcus-pyridinivorans] |
| WP_269419422.1 | Type2.Pgl.like.BREX | HD-><-?\|\|CR-ATPase9+Schlafen-> PglW -> N6-METHYLASE -> DUF6079 *-> PglZ -> BrxD-AAA -> SF1-SF2-Helicase+wHTH+Small-B-barrel+DUF1998 -> | 1219 | Bacteria>Terrabacteria_group>Actinomycetota>Actinomycetes>Mycobacteriales>Nocardiaceae | GCF_027213855.1 | Rhodococcus_yunnanensis | O4H54_RS22340 | 14314..17973 | 278209 | phage-resistance-protein-[Rhodococcus-yunnanensis] |
| WP_274752681.1 | Type2.Pgl.like.BREX | <- SF1-SF2-Helicase+wHTH+Small-B-barrel+DUF1998 <- BrxD-AAA <- PglZ <- DUF6079 *<- N6-METHYLASE <- PglW | 1269 | Bacteria>Terrabacteria_group>Actinomycetota>Actinomycetes>Mycobacteriales>Nocardiaceae | GCF_028863315.1 | Nocardia_gipuzkoensis | PWJ90_RS35990 | 6594..10403 | 2749991 | hypothetical-protein-[Nocardia-gipuzkoensis] |
| WP_218474947.1 | Type2.Pgl.like.BREX | <-Phage_integrase<-?<- SF1-SF2-Helicase+wHTH+Small-B-barrel+DUF1998 <- BrxD-AAA <- PglZ <- DUF6079 *<- N6-METHYLASE <- PglW \|\|HTH-><-tRNA\|\|?-><-tRNA<-35exo | 1263 | Bacteria>Terrabacteria_group>Actinomycetota>Actinomycetes>Mycobacteriales>Nocardiaceae | GCF_039543325.1 | Nocardia_iowensis | ACGFNT_RS21195 | 377182..380973 | 204891 | phage-resistance-protein-[Nocardia-iowensis] |
| WP_068365410.1 | Type2.Pgl.like.BREX | SF1-SF2-Helicase-><-?\|\|SF1-SF2-Helicase+REase->SF1-SF2-Helicase+HTH->wHTH+REase->wHTH-><- SF1-SF2-Helicase+wHTH+Small-B-barrel+DUF1998 <- BrxD-AAA <- PglZ <- DUF6079 *<- N6-METHYLASE <- PglW \|\|?-><-TetR-HTH\|\|?-><-?<-?<-SF1-SF2-Helicase<-HKD-DNase+HKD-Rease | 1246 | Bacteria>Terrabacteria_group>Actinomycetota>Actinomycetes>Mycobacteriales>Nocardiaceae | GCF_900111805.1 | Rhodococcus_kroppenstedtii | BMZ18_RS12920 | 141320..145060 | 293050 | DUF6079-family-protein-[Rhodococcus-kroppenstedtii] |
| WP_076482662.1 | Type2.Pgl.like.BREX | <-NUDIX<-McrB-AAA<-McrB-AAA\|\|SF1-SF2-Helicase+REase+HTH->SF1-SF2-Helicase+HTH->\|\|<- SF1-SF2-Helicase+wHTH+Small-B-barrel+DUF1998 <- BrxD-AAA <- PglZ <- DUF6079 *<-ParB+HNH<- N6-METHYLASE <- PglW <-REase+SF1-SF2-Helicase<-DNA-glycosylase<-SF1-SF2-Helicase<-HKD-DNase+HKD-Rease<-?<-ABhydrolase+ABhydrolase | 1235 | Bacteria>Terrabacteria_group>Actinomycetota>Actinomycetes>Mycobacteriales>Nocardiaceae | GCF_900156495.1 | Williamsia_sterculiae | BW971_RS19020 | 25213..28920 | 1344003 | DUF6079-family-protein-[Williamsia-sterculiae] |
| WP_097245973.1 | Type2.Pgl.like.BREX | NUDIX-><-?<-RADICAL-SAM<-APATPase\|\|HTH_3->STYKIN-><- PglZ <- DUF6079 *<- N6-METHYLASE <- PglW <-DndB<-?<-?<-?\|\| N6-METHYLASE ->TRD+TRD->Pkinase+TPR+TPR-> | 1301 | Bacteria>Terrabacteria_group>Actinomycetota>Actinomycetes>Mycobacteriales>Nocardiaceae | GCF_900215415.1 | Nocardia_amikacinitolerans | CRP45_RS17645 | 723031..726936 | 756689 | phage-resistance-protein-[Nocardia-amikacinitolerans] |
| WP_104380333.1 | Type2.Pgl.like.BREX | HTH->?-><- SF1-SF2-Helicase+wHTH+Small-B-barrel+DUF1998 <- BrxD-AAA <- PglZ <- DUF6079 *<- N6-METHYLASE <- PglW <-?<-?\|\| N6-METHYLASE -> N6-METHYLASE ->Pkinase+TPR-> | 1300 | Bacteria>Terrabacteria_group>Actinomycetota>Actinomycetes>Mycobacteriales>Nocardiaceae | - | Nocardia_nova | C5E45_23110 | 140922..144824 | 37330 | DUF6079-family-protein-[Nocardia-nova] |
| KQU58166.1 | Type2.Pgl.like.BREX | AAA-ATPase-><-?<-REase<-MORC\|\| PglW -> N6-METHYLASE -> DUF6079 *-> PglZ -> BrxD-AAA -> SF1-SF2-Helicase+wHTH+Small-B-barrel+DUF1998 -><-ANK+ANK<-?<-SbcC | 1248 | Bacteria>Terrabacteria_group>Actinomycetota>Actinomycetes>Mycobacteriales>Nocardiaceae | - | Rhodococcus_sp._Leaf278 | ASG84_16105 | 127927..131673 | 1736319 | phage-resistance-protein-[Rhodococcus-sp.-Leaf278] |
| WP_068569102.1 | Type2.Pgl.like.BREX | <-sGTP<-?\|\| PglW -> N6-METHYLASE ->?-> DUF6079 *-> PglZ -> BrxD-AAA -><-?\|\| SF1-SF2-Helicase+wHTH+Small-B-barrel+DUF1998 -><-nSTAND1+HNH<-REase+SF1-SF2-Helicase\|\|DUF4352->ParD_antitoxin->SKI-> | 1251 | Bacteria>Terrabacteria_group>Actinomycetota>Actinomycetes>Mycobacteriales>Tsukamurellaceae | GCF_001575195.1 | Tsukamurella_pseudospumae | AXK60_RS00825 | 181660..185415 | 239498 | DUF6079-family-protein-[Tsukamurella-pseudospumae] |
| WP_224646689.1 | Type2.Pgl.like.BREX | <-MMR_HSR1<-?\|\| PglW -> N6-METHYLASE ->HEPN-> DUF6079 *-> PglZ -> BrxD-AAA -> SF1-SF2-Helicase+wHTH+Small-B-barrel+DUF1998 -><-HNH<-REase+SF1-SF2-Helicase\|\|DUF4352->ParD_antitoxin-> | 1249 | Bacteria>Terrabacteria_group>Actinomycetota>Actinomycetes>Mycobacteriales>Tsukamurellaceae | GCF_020169415.1 | Tsukamurella_sp._M9C | LB823_RS09350 | 44778..48527 | 2877520 | DUF6079-family-protein-[Tsukamurella-sp.-M9C] |
| GGL99858.1 | Type2.Pgl.like.BREX | <- DUF6079 *<-?\|\|?->?-><-?<- PglW | 1250 | Bacteria>Terrabacteria_group>Actinomycetota>Actinomycetes>Nakamurellales | GCA_014646215.1 | Nakamurella_endophytica | GCM10011594_19780 | 571447..575199 | 1748367 | hypothetical-protein-GCM10011594_19780-[Nakamurella-endophytica] |
| WP_188941325.1 | Type2.Pgl.like.BREX | SF1-SF2-Helicase+REase->?->?-><-?<-?<-?<- SF1-SF2-Helicase+wHTH+Small-B-barrel+DUF1998 <- BrxD-AAA <- PglZ <- DUF6079 *\|\|Phd-YefM->RelE-ParE-><- N6-METHYLASE <- PglW <-?\|\|?-><-?<-?\|\|Glycos_transf-> | 1250 | Bacteria>Terrabacteria_group>Actinomycetota>Actinomycetes>Nakamurellales>Nakamurellaceae | GCF_014646215.1 | Nakamurella_endophytica | IEY37_RS09760 | 571447..575199 | 1748367 | DUF6079-family-protein-[Nakamurella-endophytica] |
| WP_179789934.1 | Type2.Pgl.like.BREX | <-MULE-TRANSPOSASE<-35exo+BRCT<-SF1-SF2-Helicase<- SF1-SF2-Helicase+wHTH+Small-B-barrel+DUF1998 <- PglZ <- DUF6079 *<- N6-METHYLASE <- PglW \|\|CASPASE-><-ParB+HNH<-tRNA\|\|?-><-?<-?<-?<-ABC-ATPase+SbcC | 1219 | Bacteria>Terrabacteria_group>Actinomycetota>Actinomycetes>Propionibacteriales>Actinopolymorphaceae | GCF_013407855.1 | Actinopolymorpha_rutila | F4554_RS25530 | 5730929..5734588 | 446787 | DUF6079-family-protein-[Actinopolymorpha-rutila] |
| WP_192748454.1 | Type2.Pgl.like.BREX | ABhydrolase->?->HKD-DNase-><-HTH_Tnp_1+HTH_21+rve+rve_3\|\|HKD-DNase+SF1-SF2-Helicase+HTH+HTH+SRA->ArdCN-><- SF1-SF2-Helicase+wHTH+Small-B-barrel+DUF1998 <- PglZ <- DUF6079 *<- N6-METHYLASE <- PglW \|\|?->?->?->STYKIN-><-?<-SF1-SF2-Helicase<-HKD-DNase | 1234 | Bacteria>Terrabacteria_group>Actinomycetota>Actinomycetes>Propionibacteriales>Actinopolymorphaceae | GCF_014873875.1 | Actinopolymorpha_pittospori | HEB94_RS02555 | 543266..546970 | 648752 | DUF6079-family-protein-[Actinopolymorpha-pittospori] |
| WP_092652883.1 | Type2.Pgl.like.BREX | Peptidase_M48-><-?\|\|tRNA->ParB+HNH-><-CASPASE\|\| PglW -> N6-METHYLASE -> DUF6079 *-> PglZ -> BrxD-AAA -> SF1-SF2-Helicase+wHTH+Small-B-barrel+DUF1998 -><-STYKIN | 1221 | Bacteria>Terrabacteria_group>Actinomycetota>Actinomycetes>Propionibacteriales>Actinopolymorphaceae | GCF_900104745.1 | Actinopolymorpha_singaporensis | BLU27_RS10730 | 2355351..2359016 | 117157 | hypothetical-protein-[Actinopolymorpha-singaporensis] |
| WP_139239151.1 | Type2.Pgl.like.BREX | ParB+HNH-><-CASPASE\|\| PglW -> N6-METHYLASE -> DUF6079 *-> PglZ -> BrxD-AAA -> SF1-SF2-Helicase+wHTH+Small-B-barrel+DUF1998 ->SF1-SF2-Helicase->35exo+BRCT-> | 1219 | Bacteria>Terrabacteria_group>Actinomycetota>Actinomycetes>Propionibacteriales>Actinopolymorphaceae | GCF_900113145.1 | Actinopolymorpha_cephalotaxi | BM325_RS25875 | 20933..24592 | 504797 | phage-resistance-protein-[Actinopolymorpha-cephalotaxi] |
| NYG54950.1 | Type2.Pgl.like.BREX | <-REase\|\|MORC->?->?-> PglW -> N6-METHYLASE -> DUF6079 *-> PglZ ->?->?->?->SF1-SF2-Helicase->?->SF1-SF2-Helicase+DUF1998-> | 1220 | Bacteria>Terrabacteria_group>Actinomycetota>Actinomycetes>Propionibacteriales | GCA_013409425.1 | Nocardioides_perillae | BJ989_001254 | 1361369..1365031 | 1119534 | hypothetical-protein-[Nocardioides-perillae] |
| MBB1516007.1 | Type2.Pgl.like.BREX | <- SF1-SF2-Helicase+wHTH+Small-B-barrel+DUF1998 <- BrxD-AAA <-?<-HTH<-?<- PglZ <- DUF6079 *<- N6-METHYLASE <- PglW <-?<-?\|\|?->?-><-ParB+HNH | 1247 | Bacteria>Terrabacteria_group>Actinomycetota>Actinomycetes>Propionibacteriales | GCA_014164775.1 | Tessaracoccus_sp._MC1679 | H5398_08510 | 279146..282889 | 2760313 | phage-resistance-protein-[Tessaracoccus-sp.-MC1679] |
| MEA5119840.1 | Type2.Pgl.like.BREX | <- SF1-SF2-Helicase+wHTH+Small-B-barrel+DUF1998 <- BrxD-AAA <-?<- PglZ <- DUF6079 *<- N6-METHYLASE <- PglW | 1265 | Bacteria>Terrabacteria_group>Actinomycetota>Actinomycetes>Propionibacteriales | GCA_034933485.1 | Propionibacterium_sp. | VB040_01040 | 50756..54553 | 1977903 | hypothetical-protein-[Propionibacterium-sp.] |
| HQA31350.1 | Type2.Pgl.like.BREX | PglW ->ParB+HNH-> N6-METHYLASE -> DUF6079 *-> PglZ -> BrxD-AAA -> SF1-SF2-Helicase+wHTH+Small-B-barrel+DUF1998 -> | 1248 | Bacteria>Terrabacteria_group>Actinomycetota>Actinomycetes>Propionibacteriales | GCA_035412375.1 | Propioniciclava_tarda | PKX10_08015 | 10082..13828 | 433330 | hypothetical-protein-[Propioniciclava-tarda] |
| GAA1393352.1 | Type2.Pgl.like.BREX | PglW ->?-> DUF6079 *-> | 1247 | Bacteria>Terrabacteria_group>Actinomycetota>Actinomycetes>Propionibacteriales | GCA_039526385.1 | Luteococcus_peritonei | GCM10009599_16100 | 160074..163817 | 88874 | hypothetical-protein-GCM10009599_16100-[Luteococcus-peritonei] |
| GAA1089909.1 | Type2.Pgl.like.BREX | REase->SF1-SF2-Helicase-><-?<-?<- SF1-SF2-Helicase+wHTH+Small-B-barrel+DUF1998 <- BrxD-AAA <- PglZ <- DUF6079 *<- N6-METHYLASE <- PglW <-?<-tRNA<-His_Phos_1 | 1261 | Bacteria>Terrabacteria_group>Actinomycetota>Actinomycetes>Propionibacteriales | GCA_039528155.1 | Nocardioides_dubius | GCM10009668_00430 | 62597..66382 | 317019 | hypothetical-protein-GCM10009668_00430-[Nocardioides-dubius] |
| SDL43696.1 | Type2.Pgl.like.BREX | PglW -> N6-METHYLASE -> DUF6079 *-> PglZ ->HEPN+SNF+SF1-SF2-Helicase->?-> BrxD-AAA -> SF1-SF2-Helicase+wHTH+Small-B-barrel+DUF1998 -> | 1240 | Bacteria>Terrabacteria_group>Actinomycetota>Actinomycetes>Propionibacteriales | GCA_900103275.1 | Tessaracoccus_oleiagri | SAMN04488242_1594 | 616502..620224 | 686624 | hypothetical-protein-SAMN04488242_1594-[Tessaracoccus-oleiagri] |
| WP_221025642.1 | Type2.Pgl.like.BREX | <-REase\|\|MORC->SF1-SF2-Helicase->REase-> PglW -> N6-METHYLASE -> DUF6079 *-> PglZ -><-?<-?<-ParB+HNH<-ParB+HNH | 1223 | Bacteria>Terrabacteria_group>Actinomycetota>Actinomycetes>Propionibacteriales | GCF_019679455.1 | Nocardioides_jiangsuensis | K1X13_RS14055 | 1162868..1166539 | 2866161 | DUF6079-family-protein-[Nocardioides-jiangsuensis] |
| WP_281145638.1 | Type2.Pgl.like.BREX | PglW ->?-> DUF6079 *-> | 1282 | Bacteria>Terrabacteria_group>Actinomycetota>Actinomycetes>Propionibacteriales | GCF_029917005.1 | Tessaracoccus_sp._T21 | QH948_RS04195 | 925172..929020 | 3041766 | hypothetical-protein-[Tessaracoccus-sp.-T21] |
| WP_300083660.1 | Type2.Pgl.like.BREX | <- PglZ <- DUF6079 *<- PglW <- PglW <-SF1-SF2-Helicase | 1239 | Bacteria>Terrabacteria_group>Actinomycetota>Actinomycetes>Propionibacteriales | GCF_029977985.1 | Propioniciclava_sp. | Q1K57_RS13985 | 1803..5522 | 2038686 | hypothetical-protein-[Propioniciclava-sp.] |
| WP_286267699.1 | Type2.Pgl.like.BREX | METHYLASE->?->?->tRNA-> PglW -> N6-METHYLASE -> DUF6079 *-> PglZ ->?-> BrxD-AAA -> SF1-SF2-Helicase+wHTH+Small-B-barrel+DUF1998 -> | 1265 | Bacteria>Terrabacteria_group>Actinomycetota>Actinomycetes>Propionibacteriales | GCF_030297015.1 | Brooklawnia_propionicigenes | QUE25_RS03735 | 818664..822461 | 3041175 | hypothetical-protein-[Brooklawnia-sp.-SH051] |
| WP_131296141.1 | Type2.Pgl.like.BREX | TPR+TPR->McrB-AAA->McrC-NTD+REase->?->?->?-><- SF1-SF2-Helicase+wHTH+Small-B-barrel+DUF1998 <- BrxD-AAA <- PglZ <- DUF6079 *<- N6-METHYLASE <- PglW <-?<-?\|\|?-><-?<-HTH_3 | 1261 | Bacteria>Terrabacteria_group>Actinomycetota>Actinomycetes>Propionibacteriales>Kribbellaceae | GCF_004331435.1 | Kribbella_sindirgiensis | EOH50_RS38770 | 82473..86258 | 1124744 | DUF6079-family-protein-[Kribbella-sindirgiensis] |
| GGD24638.1 | Type2.Pgl.like.BREX | <- SF1-SF2-Helicase+wHTH+Small-B-barrel+DUF1998 <- BrxD-AAA <- PglZ <- DUF6079 *<- N6-METHYLASE <- PglW <-?<-?<-PIN\|\|APATPase+TPR+TPR+TPR+TPR+TPR-><-SF1-SF2-Helicase+REase | 1227 | Bacteria>Terrabacteria_group>Actinomycetota>Actinomycetes>Propionibacteriales>Nocardioidaceae | GCA_014635145.1 | Nocardioides_daphniae | GCM10007231_24780 | 66469..70152 | 402297 | hypothetical-protein-GCM10007231_24780-[Nocardioides-daphniae] |
| MBY9075953.1 | Type2.Pgl.like.BREX | DNA_METHYLASE-><-REase\|\|MORC->SF1-SF2-Helicase->REase-> PglW -> N6-METHYLASE -> DUF6079 *-> PglZ -><-?<-?<-ParB+HNH<-ParB+HNH | 1223 | Bacteria>Terrabacteria_group>Actinomycetota>Actinomycetes>Propionibacteriales>Nocardioidaceae | GCA_019679455.1 | Nocardioides_sp._WL0053 | K1X13_14055 | 1162868..1166539 | 2866161 | DUF6079-family-protein-[Nocardioides-sp.-WL0053] |
| WP_056862668.1 | Type2.Pgl.like.BREX | Phage_int_SAM_3+Phage_integrase-><-ParB+HNH\|\|?->VirB11+MuB-ATPase->?->?-> PglW -> N6-METHYLASE -> DUF6079 *-> PglZ -> BrxD-AAA -> SF1-SF2-Helicase+wHTH+Small-B-barrel+DUF1998 -><-?<-?<-?<-SF1-SF2-Helicase+HTH\|\|?->FGase-> | 1261 | Bacteria>Terrabacteria_group>Actinomycetota>Actinomycetes>Propionibacteriales>Nocardioidaceae | GCF_001422955.1 | Nocardioides_sp._Leaf285 | ASF47_RS04185 | 904317..908102 | 1736322 | DUF6079-family-protein-[Nocardioides-sp.-Leaf285] |
| WP_056707023.1 | Type2.Pgl.like.BREX | 35exo+TUDOR+SF1-SF2-Helicase+iREase+CTD->SF1-SF2-Helicase->?->?->?->?-><- SF1-SF2-Helicase+wHTH+Small-B-barrel+DUF1998 <- BrxD-AAA <- PglZ <- DUF6079 *<- N6-METHYLASE <- PglW <-?<-MACRODOMAIN+SIR2<-HNH<-SbcC<-?\|\|TUDOR+SF1-SF2-Helicase+iREase+CTD+REase->HTH_3-> | 1260 | Bacteria>Terrabacteria_group>Actinomycetota>Actinomycetes>Propionibacteriales>Nocardioidaceae | GCF_001427545.1 | unclassified_Nocardioides | ASD81_RS04170 | 894964..898746 | 2615069 | MULTISPECIES:-DUF6079-family-protein-[unclassified-Nocardioides] |
| WP_135832439.1 | Type2.Pgl.like.BREX | <-APATPase+TPR+TPR+TPR+TPR+TPR\|\|PIN->?->?-> PglW -> PglW -p-><- N6-METHYLASE -p-> N6-METHYLASE -> DUF6079 *-> PglZ -> PglZ -p-> BrxD-AAA -> SF1-SF2-Helicase+wHTH+Small-B-barrel+DUF1998 -> | 1164 | Bacteria>Terrabacteria_group>Actinomycetota>Actinomycetes>Propionibacteriales>Nocardioidaceae | GCF_004777465.1 | Nocardioides_daphniae | E2C04_RS09760 | 1990153..1993647 | 402297 | hypothetical-protein-[Nocardioides-daphniae] |
| WP_168929655.1 | Type2.Pgl.like.BREX | <-ARG<-?<-SIGMA-HTH<-REase<-SF1-SF2-Helicase+DUF1998<-?\|\|HEPN+SF1-SF2-Helicase+REase-><-?<- PglZ <- DUF6079 *<- N6-METHYLASE <- PglW <-REase<-SF1-SF2-Helicase<-?<-MORC\|\|REase->wHTH-><-DNA_METHYLASE | 1236 | Bacteria>Terrabacteria_group>Actinomycetota>Actinomycetes>Propionibacteriales>Nocardioidaceae | GCF_004919085.1 | Nocardioides_sp._GY_10127 | E8D37_RS19755 | 682983..686693 | 2569762 | phage-resistance-protein-[Nocardioides-sp.-GY-10127] |
| WP_166390126.1 | Type2.Pgl.like.BREX | <-ZnR<-?<-LuxR-HTH<-?<-?<-Acyl_transf_3+SGNH\|\|?-> PglW -> N6-METHYLASE -> DUF6079 *-> PglZ ->TPR+TPR+TPR+TPR+TPR+TPR-> BrxD-AAA -> SF1-SF2-Helicase+wHTH+Small-B-barrel+DUF1998 -><-MarR-HTH<-?<-ParB+HNH<-TRD+TRD<-HTH_Tnp | 1218 | Bacteria>Terrabacteria_group>Actinomycetota>Actinomycetes>Propionibacteriales>Nocardioidaceae | GCF_011421535.1 | Nocardioides_ochotonae | HBO44_RS08040 | 518625..522281 | 2685869 | hypothetical-protein-[Nocardioides-ochotonae] |
| WP_179517479.1 | Type2.Pgl.like.BREX | DNA_METHYLASE-><-wHTH<-REase\|\|MORC->SF1-SF2-Helicase->REase-> PglW -> N6-METHYLASE -> DUF6079 *-> PglZ ->?->?->SF1-SF2-Helicase->?->SF1-SF2-Helicase+DUF1998->CASPASE-><-NUDIX | 1220 | Bacteria>Terrabacteria_group>Actinomycetota>Actinomycetes>Propionibacteriales>Nocardioidaceae | GCF_013409425.1 | Nocardioides_perillae | BJ989_RS06365 | 1361369..1365031 | 1119534 | phage-resistance-protein-[Nocardioides-perillae] |
| WP_188421964.1 | Type2.Pgl.like.BREX | <- SF1-SF2-Helicase+wHTH+Small-B-barrel+DUF1998 <- BrxD-AAA <- PglZ <- DUF6079 *<- N6-METHYLASE <- PglW <-?<-?<-PIN\|\|APATPase+TPR+TPR+TPR+TPR+TPR-><-?<-SF1-SF2-Helicase<-35exo+TUDOR+SF1-SF2-Helicase+iREase+CTD | 1223 | Bacteria>Terrabacteria_group>Actinomycetota>Actinomycetes>Propionibacteriales>Nocardioidaceae | GCF_014635145.1 | Nocardioides_daphniae | IEV66_RS12415 | 66469..70140 | 402297 | hypothetical-protein-[Nocardioides-daphniae] |
| WP_193638512.1 | Type2.Pgl.like.BREX | PglW -> N6-METHYLASE -> DUF6079 *-> PglZ -> BrxD-AAA -> SF1-SF2-Helicase+wHTH+Small-B-barrel+DUF1998 -> | 1238 | Bacteria>Terrabacteria_group>Actinomycetota>Actinomycetes>Propionibacteriales>Nocardioidaceae | GCF_015070385.1 | Nocardioides_malaquae | IEQ44_RS10970 | 8267..11983 | 2773426 | DUF6079-family-protein-[Nocardioides-malaquae] |
| WP_225915232.1 | Type2.Pgl.like.BREX | <-REase\|\|MORC->SF1-SF2-Helicase->REase-> PglW -> N6-METHYLASE -> DUF6079 *-> PglZ -><-?<-?<-ParB+HNH<-ParB+HNH | 1221 | Bacteria>Terrabacteria_group>Actinomycetota>Actinomycetes>Propionibacteriales>Nocardioidaceae | GCF_019679455.1 | Nocardioides_sp._WL0053 | K1X13_RS14055 | 1162874..1166539 | 2866161 | DUF6079-family-protein-[Nocardioides-sp.-WL0053] |
| WP_264701740.1 | Type2.Pgl.like.BREX | <-Calcineurin<-XIS-HTH<-TrwC+SF1-SF2-Helicase+TOPRIM<-?<-?<- SF1-SF2-Helicase+wHTH+Small-B-barrel+DUF1998 <- BrxD-AAA <- PglZ <- DUF6079 *<- N6-METHYLASE <- PglW \|\|SF1-SF2-Helicase+HTH-><-AbiEi_4<-?<-?\|\|REC+OmpR-HTH-> | 1240 | Bacteria>Terrabacteria_group>Actinomycetota>Actinomycetes>Propionibacteriales>Nocardioidaceae | GCF_025972045.1 | Micropruina_sp._KQZ13P-5 | OIE40_RS08295 | 24265..27987 | 2986940 | DUF6079-family-protein-[Micropruina-sp.-KQZ13P-5] |
| NLI49793.1 | Type2.Pgl.like.BREX | 35exo->tRNA-><-ArdCN+MPTase<-?\|\|?->?->?-> PglW -> N6-METHYLASE -> DUF6079 *-> PglZ ->MNS+MarR-HTH-> BrxD-AAA -> | 1241 | Bacteria>Terrabacteria_group>Actinomycetota>Actinomycetes>Propionibacteriales>Propionibacteriaceae | GCA_012517965.1 | Propionibacterium_sp. | GX412_05150 | 15390..19115 | 1977903 | phage-resistance-protein-[Propionibacterium-sp.] |
| NNG21072.1 | Type2.Pgl.like.BREX | PRTase->?->?->ABhydrolase-> PglW -> N6-METHYLASE -> DUF6079 *-> PglZ -> BrxD-AAA -> SF1-SF2-Helicase+wHTH+Small-B-barrel+DUF1998 -><-?\|\|?-><-?<-Pkinase | 1242 | Bacteria>Terrabacteria_group>Actinomycetota>Actinomycetes>Propionibacteriales>Propionibacteriaceae | GCA_013002685.1 | Naumannella_sp._ID2617S | HJ590_16175 | 146853..150581 | 1656893 | phage-resistance-protein-[Naumannella-sp.-ID2617S] |
| WP_013863744.1 | Type2.Pgl.like.BREX | <-HAD+HAD<-?<-?<-Aminotran_1_2<-?<-?<- SF1-SF2-Helicase+wHTH+Small-B-barrel+DUF1998 <- BrxD-AAA <- PglZ <- DUF6079 *<- N6-METHYLASE <- PglW | 1239 | Bacteria>Terrabacteria_group>Actinomycetota>Actinomycetes>Propionibacteriales>Propionibacteriaceae | GCF_000270245.1 | Microlunatus_phosphovorus | MLP_RS13845 | 3081945..3085664 | 29405 | DUF6079-family-protein-[Microlunatus-phosphovorus] |
| WP_133626269.1 | Type2.Pgl.like.BREX | <-SIGMA+OB-Fold+AAA-ATPase+Lon-Protease-BrxL<-SIR2\|\|Glyco_hydro_15->PRTase->?->?->ABhydrolase-> PglW -> N6-METHYLASE -> DUF6079 *-> PglZ -> BrxD-AAA -> SF1-SF2-Helicase+wHTH+Small-B-barrel+DUF1998 -><-?\|\|<-Pkinase<-tRNA<-tRNA<-FGS | 1247 | Bacteria>Terrabacteria_group>Actinomycetota>Actinomycetes>Propionibacteriales>Propionibacteriaceae | GCF_004362375.1 | Enemella_evansiae | C8D81_RS13920 | 550694..554437 | 2016499 | phage-resistance-protein-[Enemella-evansiae] |
| WP_182817604.1 | Type2.Pgl.like.BREX | Peptidase_M48-><- SF1-SF2-Helicase+wHTH+Small-B-barrel+DUF1998 <- BrxD-AAA \|\|Schlafen+HATPase+wHTH-><- PglZ <- DUF6079 *<-?<- PglW <-McrC-NTD+REase<-McrB-AAA<-URI-DNase\|\|XIS-HTH-> | 1251 | Bacteria>Terrabacteria_group>Actinomycetota>Actinomycetes>Propionibacteriales>Propionibacteriaceae | GCF_014164675.1 | Propioniciclava_sp._MC1683 | H5397_RS11645 | 19536..23291 | 2760309 | phage-resistance-protein-[Propioniciclava-sp.-MC1683] |
| WP_182799709.1 | Type2.Pgl.like.BREX | viral-protein->SF1-SF2-Helicase+DUF1998->?->?-><- SF1-SF2-Helicase+wHTH+Small-B-barrel+DUF1998 <- BrxD-AAA <-?<-HTH_3<-HigB<- DUF6079 *<- N6-METHYLASE <- PglW <-?<-?\|\|?->?-><-ParB+HNH<-HEPN\|\|METHYLASE-> | 1247 | Bacteria>Terrabacteria_group>Actinomycetota>Actinomycetes>Propionibacteriales>Propionibacteriaceae | GCF_014164775.1 | Tessaracoccus_sp._MC1679 | H5398_RS08510 | 279146..282889 | 2760313 | DUF6079-family-protein-[Tessaracoccus-sp.-MC1679] |
| WP_188333772.1 | Type2.Pgl.like.BREX | <-Glyco_hydro_1<-?<-?<-?\|\|SF1-SF2-Helicase-p-><- SF1-SF2-Helicase+wHTH+Small-B-barrel+DUF1998 <- BrxD-AAA <-MNS+MarR-HTH<- PglZ <- DUF6079 *<- N6-METHYLASE <- PglW <-HD\|\|?-><-Couple_hipA+HipA_C<-HTH_3<-DEOR-HTH\|\|JetA->JetB-> | 1233 | Bacteria>Terrabacteria_group>Actinomycetota>Actinomycetes>Propionibacteriales>Propionibacteriaceae | GCF_017569205.1 | Propioniciclava_sp._MC1595 | J4NO2_RS00605 | 137322..141023 | 2760308 | phage-resistance-protein-[Propioniciclava-sp.-MC1595] |
| WP_257478499.1 | Type2.Pgl.like.BREX | <-ArsR-HTH\|\|PTSEIIA1->HIT->?-><- SF1-SF2-Helicase+wHTH+Small-B-barrel+DUF1998 <- BrxD-AAA <- PglZ <- DUF6079 *<- N6-METHYLASE <- PglW \|\|TIR-><-?<-?<-KAP-NTPase\|\|?-><-VirB11+MuB-ATPase\|\|GDResolvase+Recombinase+Zn_ribbon_recom-> | 1236 | Bacteria>Terrabacteria_group>Actinomycetota>Actinomycetes>Propionibacteriales>Propionibacteriaceae | GCF_024623345.1 | Acidipropionibacterium_jensenii | NVR01_RS09600 | 147995..151705 | 1749 | DUF6079-family-protein-[Acidipropionibacterium-jensenii] |
| WP_093250694.1 | Type2.Pgl.like.BREX | SIGMA-HTH-><-?<-?\|\|REase-> PglW -> N6-METHYLASE -> DUF6079 *-> PglZ ->HEPN+SH3-like+SF1-SF2-Helicase+wHTH->MORC+DUF3684+HNH-> BrxD-AAA -> SF1-SF2-Helicase+wHTH+Small-B-barrel+DUF1998 -><-?\|\|?-><-HTH_Tnp_1+HTH_21+rve+rve_3\|\|MULE-TRANSPOSASE-> | 1240 | Bacteria>Terrabacteria_group>Actinomycetota>Actinomycetes>Propionibacteriales>Propionibacteriaceae | GCF_900103275.1 | Tessaracoccus_oleiagri | BLR46_RS07920 | 616502..620224 | 686624 | DUF6079-family-protein-[Tessaracoccus-oleiagri] |
| WP_073188979.1 | Type2.Pgl.like.BREX | <-RAMPs<-?<-RAMPs<-?<-?\|\| PglW -> N6-METHYLASE -> DUF6079 *-> PglZ ->HigB->HTH_3->?-> BrxD-AAA -> SF1-SF2-Helicase+wHTH+Small-B-barrel+DUF1998 -><-GDResolvase\|\|CASPASE-> | 1235 | Bacteria>Terrabacteria_group>Actinomycetota>Actinomycetes>Propionibacteriales>Propionibacteriaceae | GCF_900141915.1 | Tessaracoccus_bendigoensis | BUC15_RS12935 | 18375..22082 | 72764 | DUF6079-family-protein-[Tessaracoccus-bendigoensis] |
| CAM04375.1 | Type2.Pgl.like.BREX | <-HTH_21+rve+rve_2<-HIN-HTH<-REase+SAVED\|\|rve_3-><- SF1-SF2-Helicase+wHTH+Small-B-barrel+DUF1998 <- BrxD-AAA <- PglZ <- DUF6079 *<- N6-METHYLASE <- PglW \|\|SF1-SF2-Helicase+REase->HTH_29+HTH_32+DDE_3-><-tRNA\|\|?-><-?<-ABhydrolase | 1273 | Bacteria>Terrabacteria_group>Actinomycetota>Actinomycetes>Pseudonocardiales | GCA_000062885.1 | Saccharopolyspora_erythraea_NRRL_2338 | SACE_5129 | 5720356..5724177 | 405948 | bacteriophage-(phiC31)-resistance-gene-PglY-[Saccharopolyspora-erythraea-NRRL-2338] |
| EQD84160.1 | Type2.Pgl.like.BREX | <-REase+SAVED\|\|rve_3-><- SF1-SF2-Helicase+wHTH+Small-B-barrel+DUF1998 <- BrxD-AAA <- PglZ <- DUF6079 *<- N6-METHYLASE <- PglW \|\|SF1-SF2-Helicase+REase->HTH_29+HTH_32+DDE_3-> | 1273 | Bacteria>Terrabacteria_group>Actinomycetota>Actinomycetes>Pseudonocardiales | GCA_000448385.1 | Saccharopolyspora_erythraea_D | N599_21475 | 12158..15979 | 1382595 | hypothetical-protein-N599_21475-[Saccharopolyspora-erythraea-D] |
| KFU81956.1 | Type2.Pgl.like.BREX | <-SF1-SF2-Helicase\|\|wHTH+SAD+HNH->?->?-> PglW -> N6-METHYLASE -> DUF6079 *-> PglZ -> BrxD-AAA -> SF1-SF2-Helicase+wHTH+Small-B-barrel+DUF1998 ->REase+SF1-SF2-Helicase->?->HKD-DNase-> | 1273 | Bacteria>Terrabacteria_group>Actinomycetota>Actinomycetes>Pseudonocardiales | GCA_000749465.3 | Amycolatopsis_lurida_NRRL_2430 | BB31_06335 | 65008..68829 | 1460371 | hypothetical-protein-BB31_06335-[Amycolatopsis-lurida-NRRL-2430] |
| ODC03654.1 | Type2.Pgl.like.BREX | <-REase+SF1-SF2-Helicase\|\|REase+SF1-SF2-Helicase-><-?<- SF1-SF2-Helicase+wHTH+Small-B-barrel+DUF1998 <- BrxD-AAA <- PglZ <- DUF6079 *<- N6-METHYLASE <-?<-Classical-AAA\|\|HIN-HTH-><-?<-?<- PglW | 1234 | Bacteria>Terrabacteria_group>Actinomycetota>Actinomycetes>Pseudonocardiales | GCA_001995215.1 | Amycolatopsis_azurea_DSM_43854 | BO293_25575 | 80450..84154 | 1238180 | PglY-protein-[Amycolatopsis-azurea-DSM-43854] |
| MBB5890596.1 | Type2.Pgl.like.BREX | PglW -> N6-METHYLASE -> DUF6079 *-> PglZ -><-CBS+HEPN<-ParB+HNH\|\| BrxD-AAA -> SF1-SF2-Helicase+wHTH+Small-B-barrel+DUF1998 -> | 1308 | Bacteria>Terrabacteria_group>Actinomycetota>Actinomycetes>Pseudonocardiales | GCA_014203355.1 | Kutzneria_kofuensis | BJ998_001792 | 1945314..1949240 | 103725 | hypothetical-protein-[Kutzneria-kofuensis] |
| QRK88134.1 | Type2.Pgl.like.BREX | <-REase+SAVED\|\|?->rve_3-><- SF1-SF2-Helicase+wHTH+Small-B-barrel+DUF1998 <- BrxD-AAA <- PglZ <- DUF6079 *<- N6-METHYLASE <- PglW \|\|SF1-SF2-Helicase+REase->?->HTH_29+HTH_32+DDE_3-> | 1273 | Bacteria>Terrabacteria_group>Actinomycetota>Actinomycetes>Pseudonocardiales | GCA_016859185.1 | Saccharopolyspora_erythraea | JQX30_25960 | 5815005..5818826 | 1836 | PglY-protein-[Saccharopolyspora-erythraea] |
| GLZ41161.1 | Type2.Pgl.like.BREX | <- SF1-SF2-Helicase+wHTH+Small-B-barrel+DUF1998 <- BrxD-AAA <- PglZ <- DUF6079 *<- N6-METHYLASE <- PglW <-STYKinase-like<-ParB+HNH | 1262 | Bacteria>Terrabacteria_group>Actinomycetota>Actinomycetes>Pseudonocardiales | GCA_030269645.1 | Actinokineospora_sp._NBRC_105648 | Acsp05_47850 | 11746..15534 | 3032206 | hypothetical-protein-Acsp05_47850-[Actinokineospora-sp.-NBRC-105648] |
| GAA0926818.1 | Type2.Pgl.like.BREX | <-HNH<-SF1-SF2-Helicase\|\| PglW -> N6-METHYLASE ->?->NACHT+REase-DUF3883->ParB+HNH-> DUF6079 *-> PglZ -> BrxD-AAA -> SF1-SF2-Helicase+wHTH+Small-B-barrel+DUF1998 -> | 1228 | Bacteria>Terrabacteria_group>Actinomycetota>Actinomycetes>Pseudonocardiales | GCA_039525585.1 | Pseudonocardia_zijingensis | GCM10009559_12190 | 45144..48830 | 153376 | hypothetical-protein-GCM10009559_12190-[Pseudonocardia-zijingensis] |
| GAA1837104.1 | Type2.Pgl.like.BREX | <-ParB+HNH<- SF1-SF2-Helicase+wHTH+Small-B-barrel+DUF1998 <- BrxD-AAA <- PglZ <- DUF6079 *<- N6-METHYLASE <- PglW <-ParB+HNH\|\|?->CASPASE+SF1-SF2-Helicase-> | 1229 | Bacteria>Terrabacteria_group>Actinomycetota>Actinomycetes>Pseudonocardiales | GCA_039530145.1 | Pseudonocardia_carboxydivorans | GCM10009772_13250 | 35431..39120 | 415010 | hypothetical-protein-GCM10009772_13250-[Pseudonocardia-carboxydivorans] |
| GAA4535316.1 | Type2.Pgl.like.BREX | <- SF1-SF2-Helicase+wHTH+Small-B-barrel+DUF1998 <- BrxD-AAA <- PglZ <- DUF6079 *<- N6-METHYLASE <- PglW | 1204 | Bacteria>Terrabacteria_group>Actinomycetota>Actinomycetes>Pseudonocardiales | GCA_039541715.1 | Pseudonocardia_xishanensis | GCM10023175_00740 | 72067..75681 | 630995 | hypothetical-protein-GCM10023175_00740-[Pseudonocardia-xishanensis] |
| GAA4855310.1 | Type2.Pgl.like.BREX | <- SF1-SF2-Helicase+wHTH+Small-B-barrel+DUF1998 <- BrxD-AAA <- PglZ <- DUF6079 *<- N6-METHYLASE <- PglW | 1258 | Bacteria>Terrabacteria_group>Actinomycetota>Actinomycetes>Pseudonocardiales | GCA_039544495.1 | Saccharopolyspora_rosea | GCM10025787_44210 | 6730..10506 | 524884 | hypothetical-protein-GCM10025787_44210-[Saccharopolyspora-rosea] |
| GAA4991782.1 | Type2.Pgl.like.BREX | ParB+HNH-> PglW ->DOC-> N6-METHYLASE -> DUF6079 *-> PglZ -> BrxD-AAA -> SF1-SF2-Helicase+wHTH+Small-B-barrel+DUF1998 -><-SF1-SF2-Helicase+REase\|\|?-><-?\|\|ACET-> | 1247 | Bacteria>Terrabacteria_group>Actinomycetota>Actinomycetes>Pseudonocardiales | GCA_039544595.1 | Pseudonocardia_tropica | GCM10025792_45750 | 212546..216289 | 681289 | hypothetical-protein-GCM10025792_45750-[Pseudonocardia-tropica] |
| WFG45383.1 | Type2.Pgl.like.BREX | ParB+HNH-> PglW ->DOC-> N6-METHYLASE -> DUF6079 *-> PglZ -> BrxD-AAA -> SF1-SF2-Helicase+wHTH+Small-B-barrel+DUF1998 ->ParB+HNH-> | 1247 | Bacteria>Terrabacteria_group>Actinomycetota>Actinomycetes>Pseudonocardiales | GCA_045159825.1 | Pseudonocardia_alni | PaSha_20215 | 4112617..4116360 | 33907 | PglY-protein-[Pseudonocardia-alni] |
| WP_199444075.1 | Type2.Pgl.like.BREX | <-REase+SF1-SF2-Helicase<-wHTH+SAD+HNH<-REase+SF1-SF2-Helicase<- SF1-SF2-Helicase+wHTH+Small-B-barrel+DUF1998 <- BrxD-AAA <- PglZ <- DUF6079 *<- N6-METHYLASE <- PglW <-HTH_31+DUF5753\|\|CHTH+SAM-methylase-><-?<-CHTH+SAM-methylase<-HTH | 1288 | Bacteria>Terrabacteria_group>Actinomycetota>Actinomycetes>Pseudonocardiales | GCF_016464305.1 | Umezawaea_beigongshangensis | IMZ35_RS27315 | 28668..32534 | 2780383 | PglY-protein-[Umezawaea-beigongshangensis] |
| WP_308011205.1 | Type2.Pgl.like.BREX | Phage_integrase-><-Phage_integrase<-AbrB\|\|?-> PglW -> N6-METHYLASE -> DUF6079 *-> PglZ -> BrxD-AAA -> SF1-SF2-Helicase+wHTH+Small-B-barrel+DUF1998 -><-SF1-SF2-Helicase\|\|?-><-?<-MazG | 1267 | Bacteria>Terrabacteria_group>Actinomycetota>Actinomycetes>Pseudonocardiales | GCF_020177455.1 | unclassified_Saccharopolyspora | LCD33_RS01215 | 29832..33635 | 2646250 | MULTISPECIES:-PglY-protein-[unclassified-Saccharopolyspora] |
| WP_285505841.1 | Type2.Pgl.like.BREX | <- SF1-SF2-Helicase+wHTH+Small-B-barrel+DUF1998 <- BrxD-AAA <- PglZ <- DUF6079 *<- N6-METHYLASE <- PglW <-STYKinase-like<-ParB+HNH | 1262 | Bacteria>Terrabacteria_group>Actinomycetota>Actinomycetes>Pseudonocardiales | GCF_030269645.1 | Actinokineospora_sp._NBRC_105648 | QSK30_RS23520 | 11746..15534 | 3032206 | PglY-protein-[Actinokineospora-sp.-NBRC-105648] |
| WP_285588125.1 | Type2.Pgl.like.BREX | ABhydrolase->HTH_29+HTH_32+DDE_3-><-?<-?<- SF1-SF2-Helicase+wHTH+Small-B-barrel+DUF1998 <- BrxD-AAA <- PglZ <- DUF6079 *<-?<- N6-METHYLASE <- PglW \|\|SNF+SF1-SF2-Helicase->SF1-SF2-Helicase-><-REase<-SF1-SF2-Helicase | 1221 | Bacteria>Terrabacteria_group>Actinomycetota>Actinomycetes>Pseudonocardiales | GCF_030269685.1 | Actinomycetospora_sp._NBRC_106378 | QSK00_RS04795 | 77910..81575 | 3032208 | hypothetical-protein-[Actinomycetospora-sp.-NBRC-106378] |
| WP_308676743.1 | Type2.Pgl.like.BREX | SF1-SF2-Helicase-><- SF1-SF2-Helicase+wHTH+Small-B-barrel+DUF1998 <- BrxD-AAA <- PglZ <- DUF6079 *<- N6-METHYLASE <- PglW <-?<-?\|\|REase+SF1-SF2-Helicase-> | 1262 | Bacteria>Terrabacteria_group>Actinomycetota>Actinomycetes>Pseudonocardiales | GCF_031082405.1 | Amycolatopsis_sp._A133 | Q5425_RS35455 | 68266..72054 | 3064472 | PglY-protein-[Amycolatopsis-sp.-A133] |
| WP_315933119.1 | Type2.Pgl.like.BREX | HIN-HTH->?-><-PIN<-XIS-HTH\|\|?-> PglW -> N6-METHYLASE -> DUF6079 *-> PglZ -> BrxD-AAA -> SF1-SF2-Helicase+wHTH+Small-B-barrel+DUF1998 -><-?<-HTH | 1223 | Bacteria>Terrabacteria_group>Actinomycetota>Actinomycetes>Pseudonocardiales | GCF_032460545.1 | Umezawaea_sp._Da_62-37 | RM788_RS19500 | 4346679..4350350 | 3075927 | hypothetical-protein-[Umezawaea-sp.-Da-62-37] |
| WP_329045578.1 | Type2.Pgl.like.BREX | <- DUF6079 *<-?<- PglW | 1263 | Bacteria>Terrabacteria_group>Actinomycetota>Actinomycetes>Pseudonocardiales | GCF_036227105.1 | Amycolatopsis_sp._NBC_01488 | OG738_RS28905 | 6084944..6088735 | 2903563 | PglY-protein-[Amycolatopsis-sp.-NBC_01488] |
| WP_329787392.1 | Type2.Pgl.like.BREX | SAM-methylase->HisKA_3+HATPase_c-><-?<-?<- SF1-SF2-Helicase+wHTH+Small-B-barrel+DUF1998 <- BrxD-AAA <- PglZ <- DUF6079 *<-?<- N6-METHYLASE <- PglW \|\|REase+SF1-SF2-Helicase->REase+SF1-SF2-Helicase-> | 1250 | Bacteria>Terrabacteria_group>Actinomycetota>Actinomycetes>Pseudonocardiales | GCF_036321065.1 | Lentzea_sp._DG1S-22 | V1227_RS21575 | 4717613..4721365 | 3108822 | PglY-protein-[Lentzea-sp.-DG1S-22] |
| WP_184672148.1 | Type2.Pgl.like.BREX | <-HTH<-?\|\|?-><-?\|\| PglW -> N6-METHYLASE -> DUF6079 *-> PglZ -> BrxD-AAA -> SF1-SF2-Helicase+wHTH+Small-B-barrel+DUF1998 -><-SAM-methylase<-?\|\| PglZ ->METHYLASE-> | 1272 | Bacteria>Terrabacteria_group>Actinomycetota>Actinomycetes>Pseudonocardiales | GCF_039537345.1 | Saccharothrix_violaceirubra | F4559_RS23655 | 5188653..5192471 | 413306 | PglY-protein-[Saccharothrix-violaceirubra] |
| CTQ94910.1 | Type2.Pgl.like.BREX | <- SF1-SF2-Helicase+wHTH+Small-B-barrel+DUF1998 <- BrxD-AAA <- PglZ <- DUF6079 *<- N6-METHYLASE <- PglW <-REase | 1227 | Bacteria>Terrabacteria_group>Actinomycetota>Actinomycetes>Pseudonocardiales | - | Kibdelosporangium_sp._MJ126-NF4 | . | 7089226..7092909 | 703222 | Bacteriophage-(PhiC31)-resistance-gene-pglY-[Kibdelosporangium-sp.-MJ126-NF4] |
| WP_030106731.1 | Type2.Pgl.like.BREX | ABC-ATPase+SbcC-><- SF1-SF2-Helicase+wHTH+Small-B-barrel+DUF1998 <- BrxD-AAA -p<- BrxD-AAA -p<- PglZ <- DUF6079 -p<- DUF6079 -p*<- N6-METHYLASE -p<- N6-METHYLASE <- PglW <-METHYLASE<-METHYLASE<-?<-?<-?<-ABC-ATPase+SbcC | 896 | Bacteria>Terrabacteria_group>Actinomycetota>Actinomycetes>Pseudonocardiales>Pseudonocardiaceae | - | Actinoalloteichus_caeruleus | HY74_RS31825 | 75989..78677 | 2893586 | hypothetical-protein,-partial-[Actinoalloteichus-caeruleus] |
| ONI70891.1 | Type2.Pgl.like.BREX | HTH_31+DUF5753->DUF397->?-> PglW -> N6-METHYLASE -> DUF6079 *-> PglZ -> BrxD-AAA -> SF1-SF2-Helicase+wHTH+Small-B-barrel+DUF1998 ->?-><-GNTR-HTH | 1287 | Bacteria>Terrabacteria_group>Actinomycetota>Actinomycetes>Pseudonocardiales>Pseudonocardiaceae | GCA_001984155.1 | Actinosynnema_sp._ALI-1.44 | ALI144C_50850 | 853553..857416 | 1933779 | PglY-protein-[Actinosynnema-sp.-ALI-1.44] |
| RJQ78833.1 | Type2.Pgl.like.BREX | TetR-HTH-><-?<-Peptidase_M43+Peptidase_M10<-Peptidase_M43+Peptidase_M10<-SF1-SF2-Helicase+HKD-DNase\|\|SF1-SF2-Helicase->REase-> PglW -> N6-METHYLASE -> DUF6079 *-> PglZ -> BrxD-AAA -> SF1-SF2-Helicase+wHTH+Small-B-barrel+DUF1998 -> | 1241 | Bacteria>Terrabacteria_group>Actinomycetota>Actinomycetes>Pseudonocardiales>Pseudonocardiaceae | GCA_003600225.1 | Pseudonocardiaceae_bacterium_YIM_PH_21723 | D5S17_12215 | 80514..84239 | 2340915 | PglY-protein-[Pseudonocardiaceae-bacterium-YIM-PH-21723] |
| RSM79384.1 | Type2.Pgl.like.BREX | wHTH+SAD+HNH-><-REase+SF1-SF2-Helicase<-?\|\| PglW -> N6-METHYLASE -><-CHTH\|\| DUF6079 *-> PglZ -> BrxD-AAA -> SF1-SF2-Helicase+wHTH+Small-B-barrel+DUF1998 -><-tRNA\|\|REC+OmpR-HTH-> | 1140 | Bacteria>Terrabacteria_group>Actinomycetota>Actinomycetes>Pseudonocardiales>Pseudonocardiaceae | GCA_003947235.1 | Kibdelosporangium_aridum | DMH04_32060 | 79921..83343 | 2030 | phage-resistance-protein-[Kibdelosporangium-aridum] |
| MBB4967435.1 | Type2.Pgl.like.BREX | <-ABC-ATPase+SbcC<-HTH_26<-?\|\|?-><-?\|\| PglW -> N6-METHYLASE -> DUF6079 *-> PglZ -> BrxD-AAA -> SF1-SF2-Helicase+wHTH+Small-B-barrel+DUF1998 -><-SAM-METHYLASE<-?\|\|Phosphodiest->ADH_N+ADH_zinc_N-> | 1272 | Bacteria>Terrabacteria_group>Actinomycetota>Actinomycetes>Pseudonocardiales>Pseudonocardiaceae | GCA_014203755.1 | Saccharothrix_violaceirubra | F4559_004794 | 5188653..5192471 | 413306 | hypothetical-protein-[Saccharothrix-violaceirubra] |
| MCE7001655.1 | Type2.Pgl.like.BREX | REase->Mrr-N+REase-><-?<-?<-?<-?\|\|DDE_Tnp_1_4-><- DUF6079 *<- N6-METHYLASE -p<-DDE_Tnp_4<-DDE_Tnp_4\|\|DDE_Tnp_1_assoc+DDE_Tnp_1-><- N6-METHYLASE -p<- PglW | 1125 | Bacteria>Terrabacteria_group>Actinomycetota>Actinomycetes>Pseudonocardiales>Pseudonocardiaceae | GCA_021390435.3 | Kibdelosporangium_philippinense | LWC34_02185 | 453095..456472 | 211113 | DUF6079-family-protein-[Kibdelosporangium-philippinense] |
| SDG04215.1 | Type2.Pgl.like.BREX | MODE-HTH+PBPII-><- SF1-SF2-Helicase+wHTH+Small-B-barrel+DUF1998 <- BrxD-AAA <- PglZ <- DUF6079 *<- N6-METHYLASE -p<- PglW <-ParB+HNH\|\|YbaB_DNA_bd-> | 1266 | Bacteria>Terrabacteria_group>Actinomycetota>Actinomycetes>Pseudonocardiales>Pseudonocardiaceae | GCA_900100955.1 | Lentzea_fradiae | SAMN05216553_10535 | 41765..45565 | 200378 | hypothetical-protein-SAMN05216553_10535-[Lentzea-fradiae] |
| WP_012796025.1 | Type2.Pgl.like.BREX | PglW ->?->?->?->?->?-><-RHH<-?\|\| N6-METHYLASE -> DUF6079 *-> PglZ -> BrxD-AAA -> SF1-SF2-Helicase+wHTH+Small-B-barrel+DUF1998 ->?->?->?->?->?-><-FAP+Peptidase_S11 | 1285 | Bacteria>Terrabacteria_group>Actinomycetota>Actinomycetes>Pseudonocardiales>Pseudonocardiaceae | GCF_000023865.1 | Saccharomonospora_viridis | SVIR_RS02555 | 520806..524663 | 1852 | hypothetical-protein-[Saccharomonospora-viridis] |
| WP_009942486.1 | Type2.Pgl.like.BREX | <-REase+SAVED\|\|?->rve_3-><- SF1-SF2-Helicase+wHTH+Small-B-barrel+DUF1998 <- BrxD-AAA <- PglZ <- DUF6079 *<- N6-METHYLASE <- PglW \|\|SF1-SF2-Helicase+REase->?->HTH_29+HTH_32+DDE_3-> | 1273 | Bacteria>Terrabacteria_group>Actinomycetota>Actinomycetes>Pseudonocardiales>Pseudonocardiaceae | GCF_000062885.1 | Saccharopolyspora_erythraea | HP400_RS18960 | 4310719..4314540 | 1836 | hypothetical-protein-[Saccharopolyspora-erythraea] |
| WP_005443481.1 | Type2.Pgl.like.BREX | PolB-> PglW ->?->?-><-RHH<-?\|\| N6-METHYLASE -> DUF6079 *-> PglZ -> BrxD-AAA -> SF1-SF2-Helicase+wHTH+Small-B-barrel+DUF1998 ->?-><-?\|\|?->?->?-><-Peptidase_S11 | 1295 | Bacteria>Terrabacteria_group>Actinomycetota>Actinomycetes>Pseudonocardiales>Pseudonocardiaceae | GCF_000231055.2 | Saccharomonospora_azurea | SACAZDRAFT_RS16200 | 3524892..3528779 | 40988 | hypothetical-protein-[Saccharomonospora-azurea] |
| WP_005461571.1 | Type2.Pgl.like.BREX | PglW ->?->?-><-RHH\|\|DUF4096+DDE_Tnp_1+DDE_Tnp_1_2->?-><-RHH<-?\|\| N6-METHYLASE -> DUF6079 *-> PglZ -> BrxD-AAA -> SF1-SF2-Helicase+wHTH+Small-B-barrel+DUF1998 -><-lipoprotein\|\|?->?->?->?-><-FAP+Peptidase_S11 | 1295 | Bacteria>Terrabacteria_group>Actinomycetota>Actinomycetes>Pseudonocardiales>Pseudonocardiaceae | GCF_000243395.2 | Saccharomonospora_glauca | SACGLDRAFT_RS02735 | 552674..556561 | 40990 | hypothetical-protein-[Saccharomonospora-glauca] |
| WP_005453484.1 | Type2.Pgl.like.BREX | <-SF1-SF2-Helicase+REase+HTH\|\| PglW ->?->?->?-><-RHH<-?\|\|MerR-HTH-> N6-METHYLASE -> DUF6079 *-> PglZ -> BrxD-AAA -> SF1-SF2-Helicase+wHTH+Small-B-barrel+DUF1998 -><-lipoprotein\|\|?->?->?->?-><-FAP+Peptidase_S11 | 1293 | Bacteria>Terrabacteria_group>Actinomycetota>Actinomycetes>Pseudonocardiales>Pseudonocardiaceae | GCF_000244975.1 | Saccharomonospora_cyanea | SACCYDRAFT_RS03075 | 615748..619629 | 40989 | hypothetical-protein-[Saccharomonospora-cyanea] |
| WP_006237865.1 | Type2.Pgl.like.BREX | FAP+Peptidase_S11-><-?<-?<-?<-?\|\|lipoprotein-><- SF1-SF2-Helicase+wHTH+Small-B-barrel+DUF1998 <- BrxD-AAA <- PglZ <- DUF6079 *<- N6-METHYLASE \|\|?->RHH-><-?<-?<-?<- PglW | 1336 | Bacteria>Terrabacteria_group>Actinomycetota>Actinomycetes>Pseudonocardiales>Pseudonocardiaceae | GCF_000258175.1 | Saccharomonospora_xinjiangensis | SACXIDRAFT_RS07210 | 1702574..1706584 | 75294 | hypothetical-protein-[Saccharomonospora-xinjiangensis] |

| Protein-ID | System | Gene-Neighbourhoods | Length | Taxonomy | Assembly-ID | Strain | Locus-Tag | Genomic-coordinates | TaxID | Fasta-defline |
|---|---|---|---|---|---|---|---|---|---|---|
| MBN5005304.1 | Type3.MTaseII.containing.BREX | SIGMA-HTH+SIGMA-HTH+SIGMA-HTH->tRNA->Arm-DNA-bind_3+Phage_integrase-> wHTH+WYL+WCX -> BrxB-BrxF-iSTAND -> DUF6079 *-> N6-N4-METHYLASE -> OLD-ABC-> REase -> TUDOR+SF1-SF2-Helicase+i REase +CTD -> PglZ -> BrxA-DNA-binding ->ParB+HNH->?->DUF932-> | 1258 | Bacteria>Pseudomonadota>Gammaproteobacteria>Xanthomonadales>Xanthomonadaceae>Stenotrophomonas | GCA_017149675.1 | Stenotrophomonas_maltophilia | JY447_14725 | 10675..14451 | 40324 | hypothetical-protein-[Stenotrophomonas-maltophilia] |
| WP_135965894.1 | Type3.MTaseII.containing.BREX | METHYLASE->RNaseH->35exo-><-?\|\|tRNA-> wHTH+WYL+WCX -> BrxB-BrxF-iSTAND -> DUF6079 *->?-> N6-N4-METHYLASE -> OLD-ABC-> REase -> TUDOR+SF1-SF2-Helicase+i REase +CTD -> PglZ -> BrxA-DNA-binding ->ParB+HNH->DUF932-> | 1258 | Bacteria>Pseudomonadota>Gammaproteobacteria>Xanthomonadales>Xanthomonadaceae>Stenotrophomonas | GCF_029839415.1 | Stenotrophomonas_maltophilia | N5C11_RS09555 | 28442..32218 | 40324 | MULTISPECIES:- DUF6079 -family-protein-[Stenotrophomonas] |
| WP_100449517.1 | Type3.MTaseII.containing.BREX | METHYLASE->RNaseH->35exo-><-?\|\|tRNA-> wHTH+WYL+WCX -> BrxB-BrxF-iSTAND -> DUF6079 *->?-> N6-N4-METHYLASE -> OLD-ABC-> REase -> TUDOR+SF1-SF2-Helicase+i REase +CTD -> PglZ -> BrxA-DNA-binding ->ParB+HNH->DUF932-> | 1258 | Bacteria>Pseudomonadota>Gammaproteobacteria>Xanthomonadales>Xanthomonadaceae>Stenotrophomonas | - | Stenotrophomonas_maltophilia | B9Y88_RS03200 | 89921..93697 | 40324 | DUF6079 -family-protein-[Stenotrophomonas-maltophilia] |
| MBK7497323.1 | Type3.MTaseII.containing.BREX | <- PglZ <- TUDOR+SF1-SF2-Helicase+i REase +CTD <-MORC<-Schlafen+HATPase+wHTH<-SF1-SF2-Helicase<-SF1-SF2-Helicase<- DUF6079 *<- REase <- N6-N4-METHYLASE -p | 1241 | Bacteria>PVC_group>Candidatus_Omnitrophica>unclassified_Candidatus_Omnitrophica | GCA_016706995.1 | Candidatus_Omnitrophica_bacterium | IPI28_18700 | 38924..42649 | 2035772 | ATP-binding-protein-[Candidatus-Omnitrophota-bacterium] |
| WP_096895081.1 | Type3.MTaseII.containing.BREX | Aminotran_3->?->?->TPR+TPR+TPR+TPR-><-?\|\| BrxB-BrxF-iSTAND -> DUF6079 *-> N6-N4-METHYLASE -> REase ->Schlafen+HATPase+wHTH-> TUDOR+SF1-SF2-Helicase+i REase +CTD -> PglZ -> BrxA-DNA-binding -><-?\|\|RADICAL-SAM-> | 1248 | Bacteria>PVC_group>Planctomycetota>Candidatus_Brocadiia>Candidatus_Brocadiales>Candidatus_Scalinduaceae | GCF_002443295.1 | Candidatus_Scalindua_japonica | SCALIN_RS13975 | 114317..118063 | 1284222 | DUF6079 -family-protein-[Candidatus-Scalindua-japonica] |
| WP_146535598.1 | Type3.MTaseII.containing.BREX | <-ABhydrolase<-SF1-SF2-Helicase<-PIWI<- REase +PNTD1+PNTD2\|\|XIS-HTH-> BrxB-BrxF-iSTAND -> DUF6079 *->HHH_4+HTH+wHTH+ SF1-SF2-Helicase+ REase -> N6-N4-METHYLASE -> TUDOR+SF1-SF2-Helicase+i REase +CTD -> PglZ -> BrxA-DNA-binding ->HEPN->DrHyd-> | 1234 | Bacteria>PVC_group>Planctomycetota>Planctomycetia>Pirellulales>Pirellulaceae | GCF_007860175.1 | Rubripirellula_reticaptiva | Poly59_RS18475 | 382484..386188 | 2528013 | DUF6079 -family-protein-[Rubripirellula-reticaptiva] |
| WP_150079789.1 | Type3.MTaseII.containing.BREX | <-ArdCN+MPTase\|\|?-><-?<-SMODS<-?\|\|HTH_3->Peptidase_M78->?-> BrxB-BrxF-iSTAND -> DUF6079 *->HHH_4+HTH+wHTH+ SF1-SF2-Helicase+ REase -> N6-N4-METHYLASE -> TUDOR+SF1-SF2-Helicase+i REase +CTD -> PglZ -> BrxA-DNA-binding -> | 1243 | Bacteria>PVC_group>Planctomycetota>Planctomycetia>Pirellulales>Pirellulaceae | GCF_008629675.1 | Roseiconus_nitratireducens | FYK55_RS27325 | 17824..21555 | 2605748 | DUF6079 -family-protein-[Roseiconus-nitratireducens] |
| ADY58758.1 | Type3.MTaseII.containing.BREX | <-HEPN<-?<-HEPN<- BrxA-DNA-binding <- PglZ <- TUDOR+SF1-SF2-Helicase+i REase +CTD <- N6-N4-METHYLASE <-HHH_4+HTH+wHTH+ SF1-SF2-Helicase+ REase <- DUF6079 *<- BrxB-BrxF-iSTAND \|\|HTH->MPTase->?->ZnR-><-?\|\|ArdCN+MPTase-> | 1251 | Bacteria>PVC_group>Planctomycetota>Planctomycetia>Planctomycetales>Planctomycetaceae | GCA_000165715.3 | Rubinisphaera_brasiliensis_DSM_5305 | Plabr_1142 | 1331346..1335101 | 756272 | hypothetical-protein-Plabr_1142-[Rubinisphaera-brasiliensis-DSM-5305] |
| WP_041397812.1 | Type3.MTaseII.containing.BREX | <-HEPN<-?<-HEPN<- BrxA-DNA-binding <- PglZ <- TUDOR+SF1-SF2-Helicase+i REase +CTD <- N6-N4-METHYLASE <-HHH_4+HTH+wHTH+ SF1-SF2-Helicase+ REase <- DUF6079 *<- BrxB-BrxF-iSTAND \|\|HTH->MPTase->?->ZnR-><-?\|\|ArdCN+MPTase-> | 1232 | Bacteria>PVC_group>Planctomycetota>Planctomycetia>Planctomycetales>Planctomycetaceae | GCF_000165715.2 | Rubinisphaera_brasiliensis | PLABR_RS05750 | 1331346..1335044 | 119 | DUF6079 -family-protein-[Rubinisphaera-brasiliensis] |
| RLT19656.1 | Type3.MTaseII.containing.BREX | <-SF1-SF2-Helicase\|\|?-><- BrxA-DNA-binding <- PglZ <- TUDOR+SF1-SF2-Helicase+i REase +CTD <- N6-N4-METHYLASE <-HHH_4+HTH+wHTH+ SF1-SF2-Helicase+ REase <- DUF6079 -p* | 987 | Bacteria>PVC_group>Planctomycetota>unclassified_Planctomycetota | GCA_003672035.1 | Planctomycetota_bacterium | DWI28_02425 | 16452..19415 | 2026780 | MAG:-ATP-binding-protein-[Planctomycetota-bacterium] |
| WP_012462666.1 | Type3.MTaseII.containing.BREX | <-SF1-SF2-Helicase<-?<- N6-N4-METHYLASE <- BrxA-DNA-binding <- PglZ <- TUDOR+SF1-SF2-Helicase+i REase +CTD <-PHP+SbcC<- N6-N4-METHYLASE <- REase <- DUF6079 *<- BrxB-BrxF-iSTAND <-XIS-HTH<-?\|\|?-><-tRNA\|\|SAM-METHYLASE->nSTAND1+SbcC+ABC-ATPase+SbcC-> | 1243 | Bacteria>PVC_group>Verrucomicrobiota>Methylacidiphilae>Methylacidiphilales>Methylacidiphilaceae | GCF_000019665.1 | Methylacidiphilum_infernorum | MINF_RS01440 | 329568..333299 | 511746 | DUF6079 -family-protein-[Methylacidiphilum-infernorum] |
| HCF94153.1 | Type3.MTaseII.containing.BREX | XIS-HTH-> BrxB-BrxF-iSTAND -> DUF6079 *->Schlafen+HATPase+wHTH+HTH-> N6-N4-METHYLASE ->?->OLD-ABC->PIN->?-> TUDOR+SF1-SF2-Helicase+i REase +CTD -> REase -> PglZ -> | 1288 | Bacteria>PVC_group>Verrucomicrobiota>unclassified_Verrucomicrobiota | GCA_003534995.1 | Verrucomicrobiota_bacterium | DEW46_03760 | 1029..4895 | 2026799 | ATPase-[Verrucomicrobiota-bacterium] |
| MCF7763181.1 | Type3.MTaseII.containing.BREX | <-RVT<- BrxA-DNA-binding <- PglZ <- TUDOR+SF1-SF2-Helicase+i REase +CTD <- REase +AIPR<-?<- N6-N4-METHYLASE <- DUF6079 *<- BrxB-BrxF-iSTAND <-?<-?\|\|HTH_3-><-TPR+TPR+TPR+TPR<-?\|\|?->?->TraJ-RHH-> | 1257 | Bacteria>PVC_group>Verrucomicrobiota>unclassified_Verrucomicrobiota | GCA_021733945.1 | Verrucomicrobiota_bacterium | K9N62_05860 | 12732..16505 | 2026799 | DUF6079 -family-protein-[Verrucomicrobiota-bacterium] |
| MBX3742594.1 | Type3.MTaseII.containing.BREX | TrwC+SF1-SF2-Helicase->?->?->HTH+MPTase-> BrxB-BrxF-iSTAND -> DUF6079 *->Schlafen+HATPase+wHTH-> REase -> N6-N4-METHYLASE ->nSTAND1+HNH->TIR-> TUDOR+SF1-SF2-Helicase+i REase +CTD -> PglZ -> BrxA-DNA-binding ->XIS-HTH-> | 1243 | Bacteria>PVC_group>Verrucomicrobiota>Verrucomicrobiae>Verrucomicrobiales>Akkermansiaceae | GCA_019634595.1 | Akkermansiaceae_bacterium | KF712_16525 | 314704..318435 | 2562705 | hypothetical-protein-[Akkermansiaceae-bacterium] |
| MCD8271875.1 | Type3.MTaseII.containing.BREX | ANK+ANK+ANK->HNH-> REase -><- BrxA-DNA-binding <- PglZ <- TUDOR+SF1-SF2-Helicase+i REase +CTD <-HEPN<- N6-N4-METHYLASE <- DUF6079 *<- BrxB-BrxF-iSTAND \|\|DUF932->PadR-HTH->?->?-><-SAD+HNH<-STYKinase-like<-TRPR-HTH | 1242 | Bacteria>PVC_group>Verrucomicrobiota>Verrucomicrobiae>Verrucomicrobiales>Akkermansiaceae | GCA_021204505.1 | Akkermansia_sp. | LUC46_04005 | 29201..32929 | 1872421 | DUF6079 -family-protein-[Akkermansia-sp.] |
| WP_102758537.1 | Type3.MTaseII.containing.BREX | TRPR-HTH->STYKinase-like->HEPN-><-?<-?<-?<-PadR-HTH<-DUF932\|\| BrxB-BrxF-iSTAND -> DUF6079 *-> N6-N4-METHYLASE ->HEPN-> TUDOR+SF1-SF2-Helicase+i REase +CTD -> PglZ -> BrxA-DNA-binding -><- REase <-HNH<-PIN<-HTH+MPTase | 1242 | Bacteria>PVC_group>Verrucomicrobiota>Verrucomicrobiae>Verrucomicrobiales>Akkermansiaceae | GCF_001917295.1 | Akkermansia_muciniphila | CXU07_RS02965 | 669523..673251 | 239935 | DUF6079 -family-protein-[Akkermansia-muciniphila] |
| WP_215833375.1 | Type3.MTaseII.containing.BREX | STYKinase-like->SKI->?-><-?<-?<-?<-PadR-HTH<-DUF932\|\| BrxB-BrxF-iSTAND -> DUF6079 *-> N6-N4-METHYLASE ->HEPN-> TUDOR+SF1-SF2-Helicase+i REase +CTD -> PglZ -> BrxA-DNA-binding -><- REase <-HNH | 1242 | Bacteria>PVC_group>Verrucomicrobiota>Verrucomicrobiae>Verrucomicrobiales>Akkermansiaceae | GCF_018847255.1 | Akkermansia_muciniphila | J5W48_RS06490 | 1515271..1518999 | 239935 | DUF6079 -family-protein-[Akkermansia-muciniphila] |
| WP_010252922.1 | Type3.MTaseII.containing.BREX | <- HTH+ REase <-DUF499<-?<-METHYLASE-p<- TUDOR+SF1-SF2-Helicase+i REase +CTD + REase \|\|?-> CR-ATPase5 + REase ->Phage_integrase-> BrxB-BrxF-iSTAND -> DUF6079 *-> N6-N4-METHYLASE ->?->OLD-ABC+TOPRIM->SF1-SF2-Helicase->Schlafen+HATPase+wHTH-> TUDOR+SF1-SF2-Helicase+i REase +CTD ->?-> PglZ -> BrxA-DNA-binding -> | 1224 | Bacteria>Spirochaetota>Spirochaetia>Spirochaetales>Treponemataceae>Treponema | GCF_000297095.1 | Treponema_primitia | TPRIMZ1_RS18025 | 17306..20980 | 88058 | DUF6079 -family-protein-[Treponema-primitia] |
| OQB94579.1 | Type3.MTaseII.containing.BREX | <-HTH_Tnp_1<- BrxA-DNA-binding <- PglZ <- TUDOR+SF1-SF2-Helicase+i REase +CTD <-HHH_4+ SF1-SF2-Helicase+ REase <- N6-N4-METHYLASE <-Schlafen+HATPase+wHTH+HTH<- DUF6079 *<- BrxB-BrxF-iSTAND \|\|?->HEPN->NTP_transferase+MannoseP_isomer->TetR-HTH-> | 1246 | Bacteria>Spirochaetota>unclassified_Spirochaetota | GCA_002070755.1 | Spirochaetes_bacterium_ADurb.Bin110 | BWX81_02438 | 21902..25642 | 1852908 | hypothetical-protein-BWX81_02438-[Spirochaetes-bacterium-ADurb.Bin110] |
| MCK9435717.1 | Type3.MTaseII.containing.BREX | PemK_toxin->?-> CR-ATPase5 + REase ->?->HTH_3-> BrxB-BrxF-iSTAND -> DUF6079 *->HHH_4+ SF1-SF2-Helicase+ REase -> N6-N4-METHYLASE -> TUDOR+SF1-SF2-Helicase+i REase +CTD -> | 1219 | Bacteria>Synergistota>Synergistia>Synergistales>Synergistaceae>unclassified_Synergistaceae | GCA_023229345.1 | Synergistaceae_bacterium | MOQ12_00750 | 4305..7964 | 2053624 | DUF6079 -family-protein-[Synergistaceae-bacterium] |
| WP_181742021.1 | Type3.MTaseII.containing.BREX | <- BrxA-DNA-binding <- PglZ <- N6-N4-METHYLASE <- DUF6079 *<- BrxB-BrxF-iSTAND <- REase <- SF1-SF2-Helicase+ REase -p | 1225 | Bacteria>Terrabacteria_group>Bacillota>Bacilli>Bacillales>Thermoactinomycetaceae | GCF_013868055.1 | Thermoactinomyces_mirandus | H2C83_RS15065 | 6174..9851 | 2756294 | DUF6079 -family-protein-[Thermoactinomyces-mirandus] |
| MBT8908682.1 | Type3.MTaseII.containing.BREX | NUDIX->?-><-AraC-HTH+O6MGT-HTH<-NERD+SF1-SF2-Helicase<-?<- BrxA-DNA-binding -p<- PglZ <- TUDOR+SF1-SF2-Helicase+i REase +CTD <- N6-N4-METHYLASE <- DUF6079 -p*<- BrxB-BrxF-iSTAND <-?<- SF1-SF2-Helicase+ REase +HTH<-?<-?\|\|?-><-HAD+HAD<-SIGMA-HTH+SIGMA-HTH<-SIGMA-HTH+SIGMA-HTH+SIGMA-HTH | 941 | Bacteria>Terrabacteria_group>Bacillota>Bacilli>Lactobacillales>Lactobacillaceae | GCA_018720765.1 | Lactobacillus_delbrueckii_subsp._bulgaricus | BTI62_02310 | 20564..23389 | 1585 | hypothetical-protein-[Lactobacillus-delbrueckii-subsp.-bulgaricus] |
| MCI9500566.1 | Type3.MTaseII.containing.BREX | <- REase <- BrxA-DNA-binding <- PglZ <- TUDOR+SF1-SF2-Helicase+i REase +CTD <- N6-N4-METHYLASE <-?<- DUF6079 -p*<- BrxB-BrxF-iSTAND <-RADICAL-SAM<-METHYLASE<-HTH_26\|\|MPTase->?-><-?<-SymE_toxin<-TRAM+METHYLASE | 941 | Bacteria>Terrabacteria_group>Bacillota>Clostridia>Eubacteriales>Clostridiaceae | GCA_022800895.1 | Hungatella_sp. | HFG76_00915 | 32675..35500 | 2613924 | hypothetical-protein-[Hungatella-sp.] |
| WP_068882975.1 | Type3.MTaseII.containing.BREX | <-AAA-ATPase<-HTH+rve<-?<- REase <-Preader17<- BrxA-DNA-binding <- PglZ <- TUDOR+SF1-SF2-Helicase+i REase +CTD <- N6-N4-METHYLASE <- DUF6079 *<- BrxB-BrxF-iSTAND <-SIGMA-HTH<-Radical_SAM<-?<-Peptidase_U32<-RADICAL-SAM | 1202 | Bacteria>Terrabacteria_group>Bacillota>Clostridia>Eubacteriales>Desulfitobacteriaceae | GCF_001707455.1 | Dehalobacter_sp._TeCB1 | A7D23_05610 | 38739..42347 | 1843715 | DUF6079 -family-protein-[Dehalobacter-sp.-TeCB1] |
| OPX93595.1 | Type3.MTaseII.containing.BREX | <-ZnR+Fer2_BFD<-?<-ZnR+ HARE-HTH <- REase +i REase <- BrxA-DNA-binding <- PglZ <- TUDOR+SF1-SF2-Helicase+i REase +CTD <-SbcC<- N6-N4-METHYLASE <- DUF6079 *<- BrxB-BrxF-iSTAND <-HTH_26<-?<-?<-HTH_36<-VirD4-TraG+VirD4-FtsK<-ZnR+TOPRIM | 1229 | Bacteria>Terrabacteria_group>Bacillota>Clostridia>Eubacteriales>Desulfotomaculaceae | GCA_002067205.1 | Pelotomaculum_sp._PtaB.Bin104 | A4E53_00238 | 25374..29063 | 1811693 | MAG:-hypothetical-protein-A4E53_00238-[Pelotomaculum-sp.-PtaB.Bin104] |
| WP_134213462.1 | Type3.MTaseII.containing.BREX | SMODS->METHYLASE->?->?->ncRNA-> BrxB-BrxF-iSTAND -> DUF6079 *-> N6-N4-METHYLASE ->HEPN->Polbeta-> PglZ -> BrxA-DNA-binding -> SF1-SF2-Helicase+ REase -p->?-><- wHTH+WYL+WCX | 1225 | Bacteria>Terrabacteria_group>Bacillota>Clostridia>Eubacteriales>Desulfotomaculaceae | GCF_004369225.1 | Pelotomaculum_propionicicum | Pmgp_RS07945 | 51068..54745 | 258475 | DUF6079 -family-protein-[Pelotomaculum-propionicicum] |
| PSR33881.1 | Type3.MTaseII.containing.BREX | HISKIN->?-><-?<-Calcineurin\|\|?-> REase ->HEPN+PAS+DUF4011+ SF1-SF2-Helicase+ REase -> BrxB-BrxF-iSTAND -> DUF6079 *-> N6-N4-METHYLASE -> TUDOR+SF1-SF2-Helicase+i REase +CTD -> PglZ -> BrxA-DNA-binding ->?->?->?->SF1-SF2-Helicase->SF1-SF2-Helicase-> | 1224 | Bacteria>Terrabacteria_group>Bacillota>Clostridia>Eubacteriales>Eubacteriales_incertae_sedis | GCA_003023695.1 | Sulfobacillus_benefaciens | C7B46_07770 | 13431..17105 | 453960 | hypothetical-protein-C7B46_07770-[Sulfobacillus-benefaciens] |
| MST58790.1 | Type3.MTaseII.containing.BREX | <-LexA-protease<-SbcC<- wHTH+WYL+WCX <-HNH<- REase <- BrxA-DNA-binding <- PglZ <- TUDOR+SF1-SF2-Helicase+i REase +CTD <- N6-N4-METHYLASE <- DUF6079 -p*<- BrxB-BrxF-iSTAND <-XIS-HTH<-RNase<- SF1-SF2-Helicase+ REase <-HTH_3<-?<-?<-TraJ-RHH<-LPD2 | 943 | Bacteria>Terrabacteria_group>Bacillota>Clostridia>Eubacteriales>Lachnospiraceae | GCA_009695995.1 | Waltera_intestinalis | FYJ59_11185 | 33716..36547 | 2606635 | hypothetical-protein-[Waltera-intestinalis] |
| HBR02120.1 | Type3.MTaseII.containing.BREX | DDE_Tnp_1->?->?-><-?<- BrxA-DNA-binding -p<- PglZ <- TUDOR+SF1-SF2-Helicase+i REase +CTD <- REase <- N6-N4-METHYLASE <- DUF6079 -p*<- DUF6079 -p<- BrxB-BrxF-iSTAND \|\|Trypsin+PDZ-> | 942 | Bacteria>Terrabacteria_group>Bacillota>Clostridia>Eubacteriales>Oscillospiraceae | GCA_003535195.1 | Ruminiclostridium_sp. | DD738_05890 | 21181..24009 | 2053608 | hypothetical-protein-[Ruminiclostridium-sp.] |
| WP_004621790.1 | Type3.MTaseII.containing.BREX | SIGMA-HTH+SIGMA-HTH+SIGMA-HTH->tRNA->tRNA-><-?\|\|METHYLASE->?->ncRNA->Phage_integrase->?->SbcC-><-SbcC\|\| BrxB-BrxF-iSTAND -> DUF6079 *-> N6-N4-METHYLASE -> TUDOR+SF1-SF2-Helicase+i REase +CTD -> PglZ -> BrxA-DNA-binding -><- REase \|\|HHH_4+ SF1-SF2-Helicase+ REase ->URI-DNase-> | 1222 | Bacteria>Terrabacteria_group>Bacillota>Clostridia>Eubacteriales>Oscillospiraceae | GCF_000175795.2 | Ruminiclostridium_papyrosolvens | P0092_RS08120 | 1816637..1820305 | 29362 | DUF6079 -family-protein-[Ruminiclostridium-papyrosolvens] |
| KFD41493.1 | Type3.MTaseII.containing.BREX | <-RAMPs<-HD<-SF1-SF2-Helicase<- N6-N4-METHYLASE <- REase <-N6-METHYLASE<- DUF6079 -p* | 112 | Bacteria>Terrabacteria_group>Bacillota>Clostridia>Eubacteriales>Peptococcaceae | GCA_000681355.2 | Peptococcaceae_bacterium_SCADC1_2_3 | DK28_0208715 | 20340..20678 | 1487582 | hypothetical-protein-DK28_0208715-[Peptococcaceae-bacterium-SCADC1_2_3] |
| KFI36120.1 | Type3.MTaseII.containing.BREX | <-RAMPs<-HD<-SF1-SF2-Helicase<- N6-N4-METHYLASE <- REase <-N6-METHYLASE<- DUF6079 -p*<- BrxB-BrxF-iSTAND \|\|GNTR-HTH+UTRA->?->HNH->?->ABC-ATPase+SbcC-><-?<-?<-DSBH+DSBH | 808 | Bacteria>Terrabacteria_group>Bacillota>Clostridia>Eubacteriales>Peptococcaceae | GCA_000740805.1 | Peptococcaceae_bacterium_SCADC1_2_3 | HY00_07510 | 43328..45754 | 1487582 | hypothetical-protein-HY00_07510-[Peptococcaceae-bacterium-SCADC1_2_3] |
| NHM28431.1 | Type3.MTaseII.containing.BREX | <-OLD-ABC<-ParB<- BrxA-DNA-binding <- PglZ <-?<- REase <- N6-N4-METHYLASE <- DUF6079 *<- BrxB-BrxF-iSTAND | 1226 | Bacteria>Terrabacteria_group>Bacillota>Clostridia>Eubacteriales>Peptococcaceae | GCA_011393055.1 | Desulfofundulus_sp._TPOSR | G7K71_15925 | 30486..34166 | 2714340 | ATP-binding-protein-[Desulfofundulus-sp.-TPOSR] |
| WP_028308210.1 | Type3.MTaseII.containing.BREX | CopG/RHH->RHH_8->?-><-DEDD_Tnp_IS110+Transposase_20\|\|ZnR+TOPRIM->VirD4-TraG+VirD4-FtsK->HTH_36->HTH_26-> BrxB-BrxF-iSTAND -> DUF6079 *-> N6-N4-METHYLASE ->?-> TUDOR+SF1-SF2-Helicase+i REase +CTD -> PglZ -> BrxA-DNA-binding -> REase +i REase ->Preader+URI-DNase->ZnR+ HARE-HTH -> | 1229 | Bacteria>Terrabacteria_group>Bacillota>Clostridia>Eubacteriales>Peptococcaceae | GCF_000620305.1 | Desulfitibacter_alkalitolerans | K364_RS0111915 | 37253..40942 | 264641 | DUF6079 -family-protein-[Desulfitibacter-alkalitolerans] |
| RKL62727.1 | Type3.MTaseII.containing.BREX | <-HEPN<-PolB<-?<-DUF4236<-HEPN<-ParB+HNH<- BrxA-DNA-binding <- PglZ <- N6-N4-METHYLASE <- DUF6079 *<- BrxB-BrxF-iSTAND <- REase +HTH | 1223 | Bacteria>Terrabacteria_group>Bacillota>Clostridia>Thermoanaerobacterales>Thermoanaerobacteraceae | GCA_003615265.1 | Thermoanaerobacteraceae_bacterium_SP2 | DXT63_09740 | 11701..15372 | 2292448 | exonuclease-SbcC-[Thermoanaerobacteraceae-bacterium-SP2] |
| WP_043883710.1 | Type3.MTaseII.containing.BREX | DsbD->?-><-?\|\|?->tRNA->Phage_integrase->PhdYeFM_antitox->PIN->XIS-HTH->\|\| BrxB-BrxF-iSTAND -> DUF6079 *-> N6-N4-METHYLASE -> TUDOR+SF1-SF2-Helicase+i REase +CTD -> PglZ -> BrxA-DNA-binding -> SF1-SF2-Helicase+ REase +HTH->ParB+HNH-> | 1235 | Bacteria>Terrabacteria_group>Bacillota>Clostridia>Thermoanaerobacterales>Thermoanaerobacteraceae | GCF_000156275.2 | Caldanaerobacter_subterraneus | CDSM653_RS04915 | 16153..19860 | 911092 | DUF6079 -family-protein-[Caldanaerobacter-subterraneus] |
| MCF6096371.1 | Type3.MTaseII.containing.BREX | <-HTH_3<-RelE<-ParB+HNH<-DUF4011+ SF1-SF2-Helicase+ REase <- BrxA-DNA-binding <- PglZ <- TUDOR+SF1-SF2-Helicase+i REase +CTD <- N6-N4-METHYLASE <- DUF6079 *<- BrxB-BrxF-iSTAND \|\|DEOR-HTH+ISOCOT->?->?->?-><-?<-?<-?<-Glycos_transf | 1232 | Bacteria>Terrabacteria_group>Bacillota>Clostridia>Thermosediminibacterales>Thermosediminibacteraceae | GCA_021608145.1 | Thermovorax_subterraneus | L1766_05030 | 56890..60588 | 496831 | DUF6079 -family-protein-[Thermovorax-subterraneus] |
| WP_021167322.1 | Type3.MTaseII.containing.BREX | <-DOC+HTH<-?\|\|ArsR-HTH->?->?->DOC+HTH->HD-><- CR-ATPase5 + REase \|\| BrxB-BrxF-iSTAND -> DUF6079 *-> N6-N4-METHYLASE ->SbcC->?-> PglZ -> BrxA-DNA-binding -> REase ->\|\|GDResolvase+Recombinase+ZnR->SIGMA-HTH+SIGMA-HTH-> | 1242 | Bacteria>Terrabacteria_group>Bacillota>Negativicutes>Selenomonadales>Sporomusaceae | GCF_000445445.1 | Sporomusa_ovata | SOV_RS04400 | 941244..944972 | 2378 | DUF6079 -family-protein-[Sporomusa-ovata] |
| WP_075063463.1 | Type3.MTaseII.containing.BREX | RAMPs->?->?->CRISPR_Cas6->RVT->Cas_Cas1->?->HTH_3-> BrxB-BrxF-iSTAND -> DUF6079 *->HKD-DNase-> N6-N4-METHYLASE -> REase -> PglZ -> BrxA-DNA-binding -> | 1258 | Bacteria>Terrabacteria_group>Chloroflexota>Anaerolineae>Anaerolineales>Anaerolineaceae | GCF_001306115.1 | Ornatilinea_apprima | ADN00_RS13180 | 17619..21395 | 1134406 | DUF6079 -family-protein-[Ornatilinea-apprima] |
| MCB0191463.1 | Type3.MTaseII.containing.BREX | <-GDResolvase+HIN-HTH\|\| BrxB-BrxF-iSTAND -> DUF6079 *->?-> CR-ATPase9 -> N6-N4-METHYLASE -> TUDOR+SF1-SF2-Helicase+i REase +CTD -> REase ->?->?->MORC+DUF3684+ REase ->SF1-SF2-Helicase-> | 1225 | Bacteria>Terrabacteria_group>Chloroflexota>Anaerolineae>unclassified_Anaerolineae | GCA_020433805.1 | Anaerolineae_bacterium | KDJ65_05920 | 1466..5143 | 2052143 | hypothetical-protein-[Anaerolineae-bacterium] |
| MCK6629895.1 | Type3.MTaseII.containing.BREX | Glyoxalase->?->AraC-HTH->?->?-> BrxB-BrxF-iSTAND -> DUF6079 *-> N6-N4-METHYLASE -> REase -> PglZ -> BrxA-DNA-binding -> | 1241 | Bacteria>Terrabacteria_group>Chloroflexota>Anaerolineae>unclassified_Anaerolineae | GCA_023151115.1 | Anaerolineae_bacterium | L6R45_32560 | 7534..11259 | 2052143 | DUF6079 -family-protein-[Anaerolineae-bacterium] |
| MCQ3973388.1 | Type3.MTaseII.containing.BREX | ABhydrolase->SF1-SF2-Helicase-><- REase <-PhdYeFM_antitox<- BrxA-DNA-binding <- PglZ <- REase <- N6-N4-METHYLASE <-?<- DUF6079 *<- BrxB-BrxF-iSTAND <-XIS-HTH<-?<-?<-wHTH<-?<-?<-AraC-HTH | 1241 | Bacteria>Terrabacteria_group>Chloroflexota>Anaerolineae>unclassified_Anaerolineae | GCA_024434315.1 | Anaerolineae_bacterium | DPW09_08095 | 27820..31545 | 2052143 | ATP-binding-protein-[Anaerolineae-bacterium] |
| MCR4407612.1 | Type3.MTaseII.containing.BREX | <-PhdYeFM_antitox<-?<-?<-?<-?<-?<- N6-N4-METHYLASE -p<-NTP_transf_5<-?<- DUF6079 -p*<- BrxB-BrxF-iSTAND <- REase <-?\|\|?-><-RelE-ParE<-Phd-YefM<-Phage_int_SAM_4+Phage_integrase\|\|MazE_antitoxin->PIN-> | 600 | Bacteria>Terrabacteria_group>Chloroflexota>Anaerolineae>unclassified_Anaerolineae | GCA_024653325.1 | Anaerolineae_bacterium | NUW24_11945 | 13415..15217 | 2052143 | DUF6079 -family-protein-[Anaerolineae-bacterium] |
| MBR8836037.1 | Type3.MTaseII.containing.BREX | <-ENDO3+NUDIX\|\|ABhydrolase->?-><- REase <-TraJ-RHH\|\|Exonuc_V_gamma+ REase -> SF1-SF2-Helicase+ REase -> BrxB-BrxF-iSTAND -> DUF6079 -p*-> N6-N4-METHYLASE ->Schlafen+HATPase+wHTH-p->OLD-ABC-> TUDOR+SF1-SF2-Helicase+i REase +CTD -> PglZ -> BrxA-DNA-binding -><-TOPRIM<-OLD-ABC | 493 | Bacteria>Terrabacteria_group>Cyanobacteriota/Melainabacteria_group>Cyanobacteriota>Cyanophyceae>Nostocales | GCA_016903675.1 | Stigonema_ocellatum_SAG_48.90_=_DSM_106950 | DSM106950_18940 | 32709..34190 | 221294 | hypothetical-protein-[Stigonema-ocellatum-SAG-48.90-=-DSM-106950] |
| WP_096597413.1 | Type3.MTaseII.containing.BREX | <-REC+OmpR-HTH\|\|DnaJ-><-?\|\|tRNA-><-Cas12k<-ParB+HNH<-CHTH<-CHTH\|\|Exonuc_V_gamma+ REase -> SF1-SF2-Helicase+ REase -> DUF6079 *-> N6-N4-METHYLASE -> TUDOR+SF1-SF2-Helicase+i REase +CTD -> PglZ -> BrxA-DNA-binding -><-?<-TnsC-ATPase<-HTH_28+HTH_32+rve+Mu-transpos_C\|\|?->SF1-SF2-Helicase-> | 1236 | Bacteria>Terrabacteria_group>Cyanobacteriota/Melainabacteria_group>Cyanobacteriota>Cyanophyceae>Nostocales | GCF_002368195.1 | Calothrix_sp._NIES-2100 | CA734_RS02095 | 551989..555699 | 1954172 | DUF6079 -family-protein-[Calothrix-sp.-NIES-2100] |
| WP_096683471.1 | Type3.MTaseII.containing.BREX | <-RHH_9<- BrxA-DNA-binding <- PglZ <- TUDOR+SF1-SF2-Helicase+i REase +CTD <-HEPN<- N6-N4-METHYLASE <- N6-N4-METHYLASE <- DUF6079 *<- SF1-SF2-Helicase+ REase <-Exonuc_V_gamma+ REase <-MetJarc-HTH\|\|ArdA-> | 1236 | Bacteria>Terrabacteria_group>Cyanobacteriota/Melainabacteria_group>Cyanobacteriota>Cyanophyceae>Nostocales | GCF_002368215.1 | Nostoc_sp._NIES-2111 | CA737_RS32345 | 59866..63576 | 1973475 | DUF6079 -family-protein-[Nostoc-sp.-NIES-2111] |
| WP_168540454.1 | Type3.MTaseII.containing.BREX | <- BrxA-DNA-binding <- TUDOR+SF1-SF2-Helicase+i REase +CTD <-NERD<-OLD-ABC<- N6-N4-METHYLASE <- DUF6079 *<- SF1-SF2-Helicase+ REase | 1238 | Bacteria>Terrabacteria_group>Cyanobacteriota/Melainabacteria_group>Cyanobacteriota>Cyanophyceae>Nostocales | GCF_009712085.1 | Anabaena_sp._UHCC_0253 | FJR38_26290 | 50119..53835 | 2590019 | DUF6079 -family-protein-[Anabaena-sp.-UHCC-0253] |
| WP_190937388.1 | Type3.MTaseII.containing.BREX | MORC+DUF3684->MORC+DUF3684+ REase ->SF1-SF2-Helicase-><- BrxA-DNA-binding <- TUDOR+SF1-SF2-Helicase+i REase +CTD <-HEPN<- N6-N4-METHYLASE <- DUF6079 *<- SF1-SF2-Helicase+ REase <-Exonuc_V_gamma+ REase | 1237 | Bacteria>Terrabacteria_group>Cyanobacteriota/Melainabacteria_group>Cyanobacteriota>Cyanophyceae>Nostocales | GCF_014698055.1 | Nostoc_sp._FACHB-133 | H6G86_RS30115 | 33879..37592 | 2692835 | DUF6079 -family-protein-[Nostoc-sp.-FACHB-133] |
| WP_255117825.1 | Type3.MTaseII.containing.BREX | REase ->HNH-><- BrxA-DNA-binding <- PglZ <- TUDOR+SF1-SF2-Helicase+i REase +CTD <- N6-N4-METHYLASE <- DUF6079 *<- BrxB-BrxF-iSTAND <-TPR<-Peptidase<-?\|\|?->?-><-?<-Pkinase+S1COLD<-Pkinase | 1242 | Bacteria>Terrabacteria_group>Cyanobacteriota/Melainabacteria_group>Cyanobacteriota>Cyanophyceae>Synechococcales | GCF_024345835.1 | Synechococcus_sp._HJ21-Hayes | KBZ14_RS08140 | 86825..90553 | 2823736 | DUF6079 -family-protein-[Synechococcus-sp.-HJ21-Hayes] |
| WP_254950517.1 | Type3.MTaseII.containing.BREX | Peptidase->TPR-> BrxB-BrxF-iSTAND -> DUF6079 *-> N6-N4-METHYLASE -> TUDOR+SF1-SF2-Helicase+i REase +CTD -> PglZ -> BrxA-DNA-binding -><-HNH<- REase \|\|?->SIGMA-HTH-> | 1233 | Bacteria>Terrabacteria_group>Cyanobacteriota/Melainabacteria_group>Cyanobacteriota>Cyanophyceae>Synechococcales | GCF_024345955.1 | unclassified_Cyanobium | KBZ34_RS03540 | 162386..166087 | 2627006 | MULTISPECIES:- DUF6079 -family-protein-[unclassified-Cyanobium] |
| WP_073478764.1 | Type3.MTaseII.containing.BREX | REase +DUF5655->Phage_integrase->MerR-HTH->HTH_3->XIS-HTH->TOPRIM->?-> BrxB-BrxF-iSTAND -> DUF6079 *->Schlafen+HATPase+wHTH-> N6-N4-METHYLASE -> REase +AIPR-> TUDOR+SF1-SF2-Helicase+i REase +CTD -> PglZ -> BrxA-DNA-binding -> | 1242 | Bacteria>Thermodesulfobacteriota>Desulfobacteria>Desulfobacterales>Desulfatibacillaceae>Desulfatibacillum | GCF_900142135.1 | Desulfatibacillum_alkenivorans | BUE18_RS24405 | 18194..21922 | 259354 | DUF6079 -family-protein-[Desulfatibacillum-alkenivorans] |
| MCP3942933.1 | Type3.MTaseII.containing.BREX | <- DUF6079 *<- BrxB-BrxF-iSTAND <- BrxA-DNA-binding <-?\|\| N6-N4-METHYLASE -> SF1-SF2-Helicase+ REase -><-SbcC\|\|?-><-Phage_holin | 1204 | Bacteria>Thermodesulfobacteriota>Desulfobacteria>Desulfobacterales>Desulfobacteraceae>unclassified_Desulfobacteraceae | GCA_024226935.1 | Desulfobacteraceae_bacterium | GY710_15805 | 766..4380 | 2049433 | BREX-system-P-loop-protein-BrxC-[Desulfobacteraceae-bacterium] |
| HBH32876.1 | Type3.MTaseII.containing.BREX | DUF6079 -p*->Schlafen+HATPase+wHTH-> N6-N4-METHYLASE -> REase ->?-> PglZ -> BrxA-DNA-binding ->DEOR-HTH-> | 155 | Bacteria>Thermodesulfobacteriota>Desulfobulbia>Desulfobulbales>Desulfocapsaceae>Desulfofustis | GCA_003499505.1 | Desulfofustis_sp. | DDX81_13445 | 1..470 | 2053546 | ATPase-[Desulfofustis-sp.] |
| MBL0380707.1 | Type3.MTaseII.containing.BREX | <-TPR<-?<-DEOR-HTH<- BrxA-DNA-binding <- PglZ <-?<- REase <- N6-N4-METHYLASE <-Schlafen+HATPase+wHTH<- DUF6079 *<- BrxB-BrxF-iSTAND <-XIS-HTH<-tRNA<-?<-Cysteine-Desulfurase<-ABhydrolase\|\|?->?->?->MlaE-> | 1242 | Bacteria>Thermodesulfobacteriota>Desulfobulbia>Desulfobulbales>Desulfocapsaceae>Desulfofustis | GCA_016722945.1 | Desulfofustis_sp._PB-SRB1 | JKP90_13820 | 384202..387930 | 1385624 | ATP-binding-protein-[Desulfofustis-sp.-PB-SRB1] |
| WP_205217413.1 | Type3.MTaseII.containing.BREX | XIS-HTH-> BrxB-BrxF-iSTAND -> DUF6079 *-> N6-N4-METHYLASE -> REase ->Schlafen+HATPase+wHTH-> TUDOR+SF1-SF2-Helicase+i REase +CTD -> PglZ -> BrxA-DNA-binding -><-DTXR-HTH\|\|?->SWIM+DUF3670+SF1-SF2-Helicase-> | 1249 | Bacteria>Thermodesulfobacteriota>Desulfobulbia>Desulfobulbales>Desulfocapsaceae>Desulforhopalus | GCF_016918505.1 | Desulforhopalus_vacuolatus | JWG39_RS00650 | 155250..158999 | 40414 | DUF6079 -family-protein-[Desulforhopalus-vacuolatus] |
| WP_258932099.1 | Type3.MTaseII.containing.BREX | <- REase <- TUDOR+SF1-SF2-Helicase+i REase +CTD <-?<-PIN<-OLD-ABC<- N6-N4-METHYLASE <-?<-DOC+HTH<-Four-helical-protein<- DUF6079 -p*<- DUF6079 -p<-?\|\|?->DDE_3-><- DUF6079 -p<- BrxB-BrxF-iSTAND <-XIS-HTH<-TPR | 174 | Bacteria>Thermodesulfobacteriota>Syntrophobacteria>Syntrophobacterales>Syntrophobacteraceae>Desulfoferrobacter | GCF_022603395.1 | Desulfoferrobacter_suflitae | K1V27_RS20480 | 64243..64767 | 2865782 | DUF6079 -family-protein-[Desulfoferrobacter-suflitae] |

| Protein-ID | System | Gene-Neighbourhoods | Length | Taxonomy | Assembly-ID | Strain | Locus-Tag | Genomic-coordinates | TaxID | Fasta-define |
|---|---|---|---|---|---|---|---|---|---|---|
| MCF7967519.1 | Type3.MTaseII.containing.BREX | HigB_toxin->HTH_3-><-?\|\|XIS-HTH->BrxB-BrxF-iSTAND->DUF6079*->Schlafen+HATPase+wHTH->N6-N4-METHYLASE-> | 1246 | Bacteria>Pseudomonadota>Gammaproteobacteria>Methylococcales>Methylococcaceae>Methylobacter | GCA_021736725.1 | Methylobacter_tundripaludum | K9L79_18560 | 2994..6734 | 173365 | DUF6079-family-protein-[Methylobacter-tundripaludum] |
| WP_174627171.1 | Type3.MTaseII.containing.BREX | <-BrxA-DNA-binding<-PglZ<-TUDOR+SF1-SF2-Helicase+iREase+CTD<-HKD-DNase<-SF1-SF2-Helicase<-N6-N4-METHYLASE<-Schlafen+HATPase+wHTH<-DUF6079*<-BrxB-BrxF-iSTAND | 1240 | Bacteria>Pseudomonadota>Gammaproteobacteria>Methylococcales>Methylococcaceae>Methylobacter | GCF_902806695.1 | Candidatus_Methylobacter_favarea | HUX64_RS15760 | 14501..18223 | 2707345 | DUF6079-family-protein-[Candidatus-Methylobacter-favarea] |
| WP_218812148.1 | Type3.MTaseII.containing.BREX | Integrase_1->Phage_integrase-><-BrxA-DNA-binding<-PglZ<-TUDOR+SF1-SF2-Helicase+iREase+CTD<-Schlafen+HATPase+wHTH<-N6-N4-METHYLASE<-DUF6079*<-BrxB-BrxF-iSTAND\|\|?->HerA/FtsK->?->N6-N4-METHYLASE-><-Arm-DNA-bind_3+Phage_integrase<-sGTP | 1241 | Bacteria>Pseudomonadota>Gammaproteobacteria>Methylococcales>Methylococcaceae>Methylococcus | GCF_019285535.1 | Methylococcus_capsulatus | KW112_RS09060 | 1958527..1962252 | 414 | DUF6079-family-protein-[Methylococcus-capsulatus] |
| WP_218800686.1 | Type3.MTaseII.containing.BREX | RNaseP->?->?->sGTP->Arm-DNA-bind_3+Phage_integrase-><-N6-N4-METHYLASE<-HerA/FtsK<-?\|\|BrxB-BrxF-iSTAND->DUF6079*->OLD-ABC->TOPRIM->N6-N4-METHYLASE->Schlafen+HATPase+wHTH->TUDOR+SF1-SF2-Helicase+iREase+CTD->PglZ->BrxA-DNA-binding-><-Phage_integrase<-Integrase_1 | 1241 | Bacteria>Pseudomonadota>Gammaproteobacteria>Methylococcales>Methylococcaceae>Methylococcus | GCF_019285575.1 | Methylococcus_capsulatus | KW113_RS00555 | 116161..119886 | 414 | DUF6079-family-protein-[Methylococcus-capsulatus] |
| WP_249385185.1 | Type3.MTaseII.containing.BREX | Integrase_1->Phage_integrase-><-BrxA-DNA-binding<-PglZ<-TUDOR+SF1-SF2-Helicase+iREase+CTD<-Schlafen+HATPase+wHTH<-N6-N4-METHYLASE<-DUF6079*<-BrxB-BrxF-iSTAND\|\|?->HerA/FtsK->N6-N4-METHYLASE-><-Arm-DNA-bind_3+Phage_integrase<-sGTP<-?<-?<-RNaseP | 1241 | Bacteria>Pseudomonadota>Gammaproteobacteria>Methylococcales>Methylococcaceae>Methylococcus | GCF_023370095.1 | Methylococcus_capsulatus | M3M30_RS10805 | 2341456..2345181 | 414 | DUF6079-family-protein-[Methylococcus-capsulatus] |
| PZN70748.1 | Type3.MTaseII.containing.BREX | <-BrxA-DNA-binding<-PglZ<-?<-TUDOR+SF1-SF2-Helicase+iREase+CTD<-?<-Schlafen+HATPase+wHTH<-PHP+SbcC<-N6-N4-METHYLASE<-Schlafen+HATPase+wHTH<-DUF6079*<-XIS-HTH | 1249 | Bacteria>Pseudomonadota>Gammaproteobacteria>Methylococcales>Methylococcales_incertae_sedis>Candidatus_Methylumidiphilus | GCA_003242955.1 | Candidatus_Methylumidiphilus_alinenensis | DM484_28025 | 24021..27770 | 2202197 | ATPase-[Candidatus-Methylumidiphilus-alinenensis] |
| MCH2557387.1 | Type3.MTaseII.containing.BREX | <-BrxA-DNA-binding<-PglZ<-TUDOR+SF1-SF2-Helicase+iREase+CTD<-Schlafen+HATPase+wHTH<-N6-N4-METHYLASE<-HerA/FtsK<-SIR2<-OLD-ABC+TOPRIM<-DUF6079*<-BrxB-BrxF-iSTAND<-wHTH+WYL+WCX<-TraG_N<-?<-TraH<-TraU<-?<-?<-Thioredoxin_4 | 1241 | Bacteria>Pseudomonadota>Gammaproteobacteria>Oceanospirillales>Alcanivoracaceae>Alcanivorax | GCA_022449455.1 | Alcanivorax_sp. | MK005_08810 | 144603..148328 | 1872427 | DUF6079-family-protein-[Alcanivorax-sp.] |
| WP_008928712.1 | Type3.MTaseII.containing.BREX | <-BrxA-DNA-binding<-PglZ<-TUDOR+SF1-SF2-Helicase+iREase+CTD<-Schlafen+HATPase+wHTH<-N6-N4-METHYLASE<-HerA/FtsK<-SIR2<-OLD-ABC+TOPRIM<-DUF6079*<-BrxB-BrxF-iSTAND<-wHTH+WYL+WCX\|\|ARG->?->?-><-TraG_N<-?<-TraH<-TraU | 1241 | Bacteria>Pseudomonadota>Gammaproteobacteria>Oceanospirillales>Alcanivoracaceae>Alcanivorax | GCF_000300995.1 | Alcanivorax_hongdengensis | A11A3_RS07680 | 81332..85057 | 519051 | DUF6079-family-protein-[Alcanivorax-hongdengensis] |
| MCP5207854.1 | Type3.MTaseII.containing.BREX | NTP_transf_3+Hexapep+Hexapep->?->REase+wHTH->?->TnsC-ATPase->?->?->XIS-HTH->BrxB-BrxF-iSTAND->DUF6079*->Schlafen+HATPase+wHTH->N6-N4-METHYLASE->TUDOR+SF1-SF2-Helicase+iREase+CTD->PglZ->BrxA-DNA-binding-><-REase+SF1-SF2-Helicase<-?<-SF1-SF2-Helicase<-DrmB | 1243 | Bacteria>Pseudomonadota>Gammaproteobacteria>Oceanospirillales>Hahellaceae>unclassified_Hahellaceae | GCA_024234475.1 | Hahellaceae_bacterium | H7A01_11660 | 134013..137744 | 2048971 | ATP-binding-protein-[Hahellaceae-bacterium] |
| WP_136248423.1 | Type3.MTaseII.containing.BREX | <-DsbC+DsbD+Thioredoxin_7<-?\|\|XIS-HTH->?->DUF4236->Mrr-N->wHTH+WYL+WCX->BrxB-BrxF-iSTAND->DUF6079*->N6-N4-METHYLASE->Schlafen+HATPase+wHTH->TUDOR+SF1-SF2-Helicase+iREase+CTD->PglZ->BrxA-DNA-binding-> | 1241 | Bacteria>Pseudomonadota>Gammaproteobacteria>Oceanospirillales>Halomonadaceae>Halomonas | GCF_004798905.1 | Halomonas_borealis | ETC07_RS10250 | 48079..51804 | 2508710 | DUF6079-family-protein-[Halomonas-borealis] |
| WP_197566989.1 | Type3.MTaseII.containing.BREX | UDG->?->DUF932->?->?->?->?->wHTH+WYL+WCX->BrxB-BrxF-iSTAND->DUF6079*->N6-N4-METHYLASE->Schlafen+HATPase+wHTH->TUDOR+SF1-SF2-Helicase+iREase+CTD->PglZ->BrxA-DNA-binding-><-?<-?<-XIS-HTH<-CITB-HTH | 1241 | Bacteria>Pseudomonadota>Gammaproteobacteria>Oceanospirillales>Halomonadaceae>Halomonas | GCF_014931605.1 | Halomonas_sp._MCCC_1A13316 | HN052_RS20455 | 4389739..4393464 | 2733487 | DUF6079-family-protein-[Halomonas-sp.-MCCC-1A13316] |
| WP_095610363.1 | Type3.MTaseII.containing.BREX | HIN-HTH->HIN-HTH-><-?\|\|?-><-BrxA-DNA-binding<-PglZ<-TUDOR+SF1-SF2-Helicase+iREase+CTD<-Schlafen+HATPase+wHTH<-N6-N4-METHYLASE<-DUF6079*<-BrxB-BrxF-iSTAND<-wHTH+WYL+WCX\|\|Integrase_1->Phage_int_SAM-> | 1241 | Bacteria>Pseudomonadota>Gammaproteobacteria>Pseudomonadales>Marinobacteraceae>Tamilnaduibacter | GCF_002287355.1 | Tamilnaduibacter_salinus | CF392_RS04965 | 53834..57559 | 1484056 | DUF6079-family-protein-[Tamilnaduibacter-salinus] |
| RRI70507.1 | Type3.MTaseII.containing.BREX | <-ACET<-YjhX_toxin\|\|?->?-><-LysR-HTH\|\|HTH_3-><-MODE-HTH+PBPII\|\|wHTH+WYL+WCX->BrxB-BrxF-iSTAND->DUF6079*->SIR2->HerA/FtsK->N6-N4-METHYLASE->OLD-ABC->TOPRIM->Schlafen+HATPase+wHTH->TUDOR+SF1-SF2-Helicase+iREase+CTD->PglZ->BrxA-DNA-binding-> | 1246 | Bacteria>Pseudomonadota>Gammaproteobacteria>Pseudomonadales>Pseudomonadaceae>Pseudomonas | GCA_003863585.1 | Pseudomonas_aeruginosa | EIM10_18005 | 61537..65277 | 287 | ATP-binding-protein-[Pseudomonas-aeruginosa] |
| MBA1269669.1 | Type3.MTaseII.containing.BREX | <-BrxA-DNA-binding<-PglZ<-TUDOR+SF1-SF2-Helicase+iREase+CTD<-Schlafen+HATPase+wHTH<-N6-N4-METHYLASE<-HEPN<-MORC<-TOPRIM<-OLD-ABC<-DUF6079*<-BrxB-BrxF-iSTAND<-wHTH+WYL+WCX\|\|MODE-HTH-><-HIN-HTH\|\|<-HTH_3\|\|LysR-HTH->?->O6MGT-HTH-> | 1242 | Bacteria>Pseudomonadota>Gammaproteobacteria>Pseudomonadales>Pseudomonadaceae>Pseudomonas | GCA_013522615.1 | Pseudomonas_carnis | G7029_19575 | 130618..134346 | 2487355 | ATP-binding-protein-[Pseudomonas-carnis] |
| MBO3071654.1 | Type3.MTaseII.containing.BREX | HDRnase-><-PspB<-BrxA-DNA-binding<-PglZ<-TUDOR+SF1-SF2-Helicase+iREase+CTD<-Schlafen+HATPase+wHTH<-N6-N4-METHYLASE<-HEPN<-MORC<-DUF6079*<-BrxB-BrxF-iSTAND<-wHTH+WYL+WCX\|\|MODE-HTH+PBPII-><-?<-?\|\|YjhX_toxin->ACET-><-Glyoxalase | 1246 | Bacteria>Pseudomonadota>Gammaproteobacteria>Pseudomonadales>Pseudomonadaceae>Pseudomonas | GCA_017582945.1 | Pseudomonas_aeruginosa | J4H77_25770 | 188215..191955 | 287 | ATP-binding-protein-[Pseudomonas-aeruginosa] |
| MBX6030772.1 | Type3.MTaseII.containing.BREX | <-BrxA-DNA-binding<-PglZ<-TUDOR+SF1-SF2-Helicase+iREase+CTD<-Schlafen+HATPase+wHTH<-TOPRIM<-OLD-ABC<-N6-N4-METHYLASE<-?<-HEPN<-DUF6079*<-BrxB-BrxF-iSTAND<-wHTH+WYL+WCX<-TrbL<-DOC+HTH<-?<-ArdA<-?<-?<-MODE-HTH+PBPII | 1241 | Bacteria>Pseudomonadota>Gammaproteobacteria>Pseudomonadales>Pseudomonadaceae>Pseudomonas | GCA_019688625.1 | Pseudomonas_aeruginosa | ISE67_08735 | 287568..291293 | 287 | ATP-binding-protein-[Pseudomonas-aeruginosa] |
| MCO1977065.1 | Type3.MTaseII.containing.BREX | <-PspB<-BrxA-DNA-binding<-PglZ<-TUDOR+SF1-SF2-Helicase+iREase+CTD<-Schlafen+HATPase+wHTH<-N6-N4-METHYLASE<-?<-HerA/FtsK<-SIR2<-DUF6079*<-BrxB-BrxF-iSTAND<-wHTH+WYL+WCX\|\|MODE-HTH+PBPII->HTH_3->Classical-AAA->Peptidase_S8-><-HTH_3\|\|LysR-HTH-> | 1242 | Bacteria>Pseudomonadota>Gammaproteobacteria>Pseudomonadales>Pseudomonadaceae>Pseudomonas | GCA_023905865.1 | Pseudomonas_aeruginosa | FA422_11970 | 22597..26325 | 287 | ATP-binding-protein-[Pseudomonas-aeruginosa] |
| MCO3958051.1 | Type3.MTaseII.containing.BREX | <-SIR2<-BrxA-DNA-binding<-PglZ<-TUDOR+SF1-SF2-Helicase+iREase+CTD<-Schlafen+HATPase+wHTH<-N6-N4-METHYLASE<-?<-HerA/FtsK<-SIR2<-DUF6079*<-BrxB-BrxF-iSTAND<-wHTH+WYL+WCX\|\|MODE-HTH+PBPII->HTH_3->Classical-AAA->Peptidase_S8-><-HTH_3\|\|LysR-HTH-> | 1242 | Bacteria>Pseudomonadota>Gammaproteobacteria>Pseudomonadales>Pseudomonadaceae>Pseudomonas | GCA_023912735.1 | Pseudomonas_aeruginosa | FA108_20890 | 55174..58902 | 287 | ATP-binding-protein-[Pseudomonas-aeruginosa] |
| HCF1890561.1 | Type3.MTaseII.containing.BREX | DOC+HTH->HTH_36->TraJ-RHH->VirB5->?->TrbL->wHTH+WYL+WCX->BrxB-BrxF-iSTAND->DUF6079*->OLD-ABC+TOPRIM->PadR-HTH->N6-N4-METHYLASE->HKD-DNase->Schlafen+HATPase+wHTH->TUDOR+SF1-SF2-Helicase+iREase+CTD->PglZ->BrxA-DNA-binding-> | 1241 | Bacteria>Pseudomonadota>Gammaproteobacteria>Pseudomonadales>Pseudomonadaceae>Pseudomonas | GCA_023942415.1 | Pseudomonas_aeruginosa | NH920_001195 | 99271..102996 | 287 | ATP-binding-protein-[Pseudomonas-aeruginosa] |
| HCF2077872.1 | Type3.MTaseII.containing.BREX | DUF4969->ZnR+ZnR-><-HTH_3\|\|?->?->HNH->wHTH+WYL+WCX->BrxB-BrxF-iSTAND->DUF6079*->OLD-ABC->PIN->MORC->Schlafen+HATPase+wHTH->N6-N4-METHYLASE->TUDOR+SF1-SF2-Helicase+iREase+CTD->PglZ->BrxA-DNA-binding-> | 1241 | Bacteria>Pseudomonadota>Gammaproteobacteria>Pseudomonadales>Pseudomonadaceae>Pseudomonas | GCA_023942995.1 | Pseudomonas_aeruginosa | NH997_004587 | 20722..24447 | 287 | ATP-binding-protein-[Pseudomonas-aeruginosa] |
| HCG0728340.1 | Type3.MTaseII.containing.BREX | <-LysR-HTH\|\|HTH_3-><-Peptidase_S8<-Classical-AAA<-HTH_3<-MODE-HTH+PBPII\|\|wHTH+WYL+WCX->BrxB-BrxF-iSTAND->DUF6079*->SIR2->HerA/FtsK->?->?->N6-N4-METHYLASE->Schlafen+HATPase+wHTH->TUDOR+SF1-SF2-Helicase+iREase+CTD->PglZ->BrxA-DNA-binding-> | 1246 | Bacteria>Pseudomonadota>Gammaproteobacteria>Pseudomonadales>Pseudomonadaceae>Pseudomonas | GCA_024002515.1 | Pseudomonas_aeruginosa | NJJ48_005263 | 12717..16457 | 287 | ATP-binding-protein-[Pseudomonas-aeruginosa] |
| WP_033991628.1 | Type3.MTaseII.containing.BREX | <-Phos_pyr_kin<-ACET<-?<-?\|\|GNTR-HTH+Aminotran_1_2-><-LysR-HTH<-?\|\|wHTH+WYL+WCX->BrxB-BrxF-iSTAND->DUF6079*->MORC->?->?->N6-N4-METHYLASE->Schlafen+HATPase+wHTH->TUDOR+SF1-SF2-Helicase+iREase+CTD->PglZ->BrxA-DNA-binding->SIR2-> | 1246 | Bacteria>Pseudomonadota>Gammaproteobacteria>Pseudomonadales>Pseudomonadaceae>Pseudomonas | GCF_000795045.1 | Pseudomonas_aeruginosa | NR30_RS22645 | 72439..76179 | 287 | DUF6079-family-protein-[Pseudomonas-aeruginosa] |
| WP_050481652.1 | Type3.MTaseII.containing.BREX | HDRnase-><-PspB<-BrxA-DNA-binding<-PglZ<-TUDOR+SF1-SF2-Helicase+iREase+CTD<-Schlafen+HATPase+wHTH<-N6-N4-METHYLASE<-HEPN<-MORC<-DUF6079*<-BrxB-BrxF-iSTAND<-wHTH+WYL+WCX\|\|MODE-HTH+PBPII-><-?<-?\|\|YjhX_toxin->ACET-><-Glyoxalase | 1242 | Bacteria>Pseudomonadota>Gammaproteobacteria>Pseudomonadales>Pseudomonadaceae>Pseudomonas | GCF_001181125.1 | Pseudomonas_aeruginosa | J4H77_RS25765 | 188215..191943 | 287 | DUF6079-family-protein-[Pseudomonas-aeruginosa] |
| WP_249920774.1 | Type3.MTaseII.containing.BREX | <-BrxA-DNA-binding<-PglZ<-TUDOR+SF1-SF2-Helicase+iREase+CTD<-N6-N4-METHYLASE<-Schlafen+HATPase+wHTH<-MORC<-PIN<-OLD-ABC<-DUF6079*<-BrxB-BrxF-iSTAND<-wHTH+WYL+WCX<-HNH<-?<-?\|\|HTH_3-><-ZnR+ZnR<-DUF4969 | 1243 | Bacteria>Pseudomonadota>Gammaproteobacteria>Pseudomonadales>Pseudomonadaceae>Pseudomonas | GCF_001765455.1 | Pseudomonas_aeruginosa | BAW40_RS11025 | 50364..54095 | 287 | DUF6079-family-protein-[Pseudomonas-aeruginosa] |
| WP_070341174.1 | Type3.MTaseII.containing.BREX | <-BrxA-DNA-binding<-PglZ<-TUDOR+SF1-SF2-Helicase+iREase+CTD<-N6-N4-METHYLASE<-Schlafen+HATPase+wHTH<-MORC<-PIN<-OLD-ABC<-DUF6079*<-BrxB-BrxF-iSTAND<-wHTH+WYL+WCX<-HNH<-?<-?\|\|HTH_3-><-ZnR+ZnR<-DUF4969 | 1241 | Bacteria>Pseudomonadota>Gammaproteobacteria>Pseudomonadales>Pseudomonadaceae>Pseudomonas | GCF_001765485.1 | Pseudomonas_aeruginosa | BAW37_RS11780 | 29962..33687 | 287 | DUF6079-family-protein-[Pseudomonas-aeruginosa] |
| WP_070331909.1 | Type3.MTaseII.containing.BREX | <-BrxA-DNA-binding<-PglZ<-TUDOR+SF1-SF2-Helicase+iREase+CTD<-N6-N4-METHYLASE<-Schlafen+HATPase+wHTH<-MORC<-PIN<-OLD-ABC<-DUF6079*<-BrxB-BrxF-iSTAND<-wHTH+WYL+WCX<-HNH<-?<-?\|\|HTH_3-><-ZnR+ZnR<-DUF4969 | 1241 | Bacteria>Pseudomonadota>Gammaproteobacteria>Pseudomonadales>Pseudomonadaceae>Pseudomonas | GCF_001766035.1 | Pseudomonas_aeruginosa | BAW72_RS11835 | 79269..82994 | 287 | DUF6079-family-protein-[Pseudomonas-aeruginosa] |
| WP_196496122.1 | Type3.MTaseII.containing.BREX | HDRnase-><-SIR2<-BrxA-DNA-binding<-PglZ<-TUDOR+SF1-SF2-Helicase+iREase+CTD<-Schlafen+HATPase+wHTH<-N6-N4-METHYLASE<-HEPN<-MORC<-DUF6079*<-BrxB-BrxF-iSTAND<-wHTH+WYL+WCX\|\|MODE-HTH+PBPII->HTH_3->Classical-AAA->Peptidase_S8-><-HTH_3\|\|LysR-HTH-> | 1246 | Bacteria>Pseudomonadota>Gammaproteobacteria>Pseudomonadales>Pseudomonadaceae>Pseudomonas | GCF_002242855.1 | Pseudomonas_aeruginosa | ACG91_RS23775 | 44073..47813 | 287 | DUF6079-family-protein-[Pseudomonas-aeruginosa] |
| WP_140786438.1 | Type3.MTaseII.containing.BREX | NUDIX+NAAR1+NAAR2->HARE-HTH+REase-><-BrxA-DNA-binding<-PglZ<-TUDOR+SF1-SF2-Helicase+iREase+CTD<-N6-N4-METHYLASE<-Schlafen+HATPase+wHTH<-MORC<-DUF6079*<-BrxB-BrxF-iSTAND<-wHTH+WYL+WCX<-HNH<-Arm-DNA-bind_3+Phage_integrase\|\|SF1-SF2-Helicase->AIPR->N6-N4-METHYLASE->SF1-SF2-Helicase+REase-><-URI-DNase+RAMA | 1241 | Bacteria>Pseudomonadota>Gammaproteobacteria>Pseudomonadales>Pseudomonadaceae>Pseudomonas | GCF_003626935.1 | Pseudomonas_aeruginosa | D7H65_RS11285 | 2305878..2309603 | 287 | DUF6079-family-protein-[Pseudomonas-aeruginosa] |
| WP_124182728.1 | Type3.MTaseII.containing.BREX | <-BrxA-DNA-binding<-PglZ<-TUDOR+SF1-SF2-Helicase+iREase+CTD<-N6-N4-METHYLASE<-Schlafen+HATPase+wHTH<-DUF6079*<-BrxB-BrxF-iSTAND<-wHTH+WYL+WCX<-HNH<-?<-?\|\|HTH_3-><-ZnR+ZnR<-DUF4969 | 1241 | Bacteria>Pseudomonadota>Gammaproteobacteria>Pseudomonadales>Pseudomonadaceae>Pseudomonas | GCF_003839865.1 | Pseudomonas_aeruginosa | IPC226_RS26580 | 200303..204028 | 287 | DUF6079-family-protein-[Pseudomonas-aeruginosa] |
| WP_124984784.1 | Type3.MTaseII.containing.BREX | ZnR+ZnR+Pkinase+SF1-SF2-Helicase->FUSC->SymE_toxin->MazG->REase+SF1-SF2-Helicase->Arm-DNA-bind_3+Phage_integrase->HNH->wHTH+WYL+WCX->BrxB-BrxF-iSTAND->DUF6079*->Schlafen+HATPase+wHTH->N6-N4-METHYLASE->TUDOR+SF1-SF2-Helicase+iREase+CTD->PglZ->BrxA-DNA-binding-><-HARE-HTH+REase<-NUDIX+NAAR1+NAAR2<-?<-Peptidase_M15 | 1241 | Bacteria>Pseudomonadota>Gammaproteobacteria>Pseudomonadales>Pseudomonadaceae>Pseudomonas | GCF_003863685.1 | Pseudomonas_aeruginosa | EIM22_RS03570 | 276481..280206 | 287 | DUF6079-family-protein-[Pseudomonas-aeruginosa] |
| WP_124988190.1 | Type3.MTaseII.containing.BREX | <-PglZ<-TUDOR+SF1-SF2-Helicase+iREase+CTD<-Schlafen+HATPase+wHTH<-TOPRIM<-OLD-ABC<-N6-N4-METHYLASE<-HerA/FtsK<-SIR2<-DUF6079*<-BrxB-BrxF-iSTAND<-wHTH+WYL+WCX\|\|MODE-HTH+PBPII-><-HTH_3\|\|LysR-HTH-><-?<-?\|\|YjhX_toxin->ACET-> | 1246 | Bacteria>Pseudomonadota>Gammaproteobacteria>Pseudomonadales>Pseudomonadaceae>Pseudomonas | GCF_003863815.1 | Pseudomonas_aeruginosa | EIM11_RS16920 | 77381..81121 | 287 | DUF6079-family-protein-[Pseudomonas-aeruginosa] |
| WP_209007133.1 | Type3.MTaseII.containing.BREX | <-ParA-Soj-PloopNTPase<-METHYLASE<-?<-?\|\|Phage_rep->HTH_3->Phage_int_SAM_2+Integrase_1->XIS-HTH->BrxB-BrxF-iSTAND->DUF6079*->SIR2->HerA/FtsK->?->N6-N4-METHYLASE->Schlafen+HATPase+wHTH->TUDOR+SF1-SF2-Helicase+iREase+CTD->PglZ->BrxA-DNA-binding->PspB-> | 1246 | Bacteria>Pseudomonadota>Gammaproteobacteria>Pseudomonadales>Pseudomonadaceae>Pseudomonas | GCF_013276295.1 | Pseudomonas_aeruginosa | GWK58_RS24945 | 23431..27171 | 287 | DUF6079-family-protein-[Pseudomonas-aeruginosa] |
| WP_181127557.1 | Type3.MTaseII.containing.BREX | <-BrxA-DNA-binding<-PglZ<-TUDOR+SF1-SF2-Helicase+iREase+CTD<-Schlafen+HATPase+wHTH<-N6-N4-METHYLASE<-HEPN<-MORC<-TOPRIM<-OLD-ABC<-DUF6079*<-BrxB-BrxF-iSTAND<-wHTH+WYL+WCX\|\|<-HIN-HTH\|\|<-HTH_3\|\|LysR-HTH->?->O6MGT-HTH-> | 1242 | Bacteria>Pseudomonadota>Gammaproteobacteria>Pseudomonadales>Pseudomonadaceae>Pseudomonas | GCF_013522615.1 | Pseudomonas_carnis | G7029_RS19485 | 130618..134346 | 2487355 | DUF6079-family-protein-[Pseudomonas-carnis] |
| WP_188867720.1 | Type3.MTaseII.containing.BREX | XIS-HTH->?-><-BrxA-DNA-binding<-PglZ<-TUDOR+SF1-SF2-Helicase+iREase+CTD<-?<-N6-N4-METHYLASE<-Schlafen+HATPase+wHTH<-MORC<-DUF6079*<-BrxB-BrxF-iSTAND<-wHTH+WYL+WCX<-HNH<-?\|\|CR-ATPase-> | 1241 | Bacteria>Pseudomonadota>Gammaproteobacteria>Pseudomonadales>Pseudomonadaceae>Pseudomonas | GCF_014646755.1 | Pseudomonas_asuensis | IEX86_RS19080 | 192626..196351 | 1825787 | DUF6079-family-protein-[Pseudomonas-asuensis] |
| WP_228292836.1 | Type3.MTaseII.containing.BREX | <-BrxA-DNA-binding<-PglZ<-TUDOR+SF1-SF2-Helicase+iREase+CTD<-N6-N4-METHYLASE<-Schlafen+HATPase+wHTH<-MORC<-PIN<-OLD-ABC<-DUF6079*<-BrxB-BrxF-iSTAND<-wHTH+WYL+WCX<-HNH<-?<-?\|\|HTH_3-><-ZnR+ZnR<-DUF4969 | 1243 | Bacteria>Pseudomonadota>Gammaproteobacteria>Pseudomonadales>Pseudomonadaceae>Pseudomonas | GCF_017741985.1 | Pseudomonas_aeruginosa | BAW68_RS10935 | 136915..140646 | 287 | DUF6079-family-protein-[Pseudomonas-aeruginosa] |
| WP_218424333.1 | Type3.MTaseII.containing.BREX | HNH-><-BrxA-DNA-binding<-PglZ<-TUDOR+SF1-SF2-Helicase+iREase+CTD<-Schlafen+HATPase+wHTH<-N6-N4-METHYLASE<-DUF6079*<-BrxB-BrxF-iSTAND<-tRNA<-MerR-HTH | 1244 | Bacteria>Pseudomonadota>Gammaproteobacteria>Pseudomonadales>Pseudomonadaceae>Pseudomonas | GCF_019218965.1 | Pseudomonas_sp._PDM31 | KW839_RS11070 | 28010..31744 | 2854778 | DUF6079-family-protein-[Pseudomonas-sp.-PDM31] |
| WP_220803981.1 | Type3.MTaseII.containing.BREX | <-BrxA-DNA-binding<-PglZ<-TUDOR+SF1-SF2-Helicase+iREase+CTD<-N6-N4-METHYLASE<-Schlafen+HATPase+wHTH<-MORC<-DUF6079*<-BrxB-BrxF-iSTAND<-wHTH+WYL+WCX<-TrbL<-?<-VirB5<-TraJ-RHH<-HTH_36<-DOC+HTH | 1241 | Bacteria>Pseudomonadota>Gammaproteobacteria>Pseudomonadales>Pseudomonadaceae>Pseudomonas | GCF_019656335.1 | Pseudomonas_sp._NCCP-436 | NCCP436_RS10650 | 99901..103626 | 2842481 | DUF6079-family-protein-[Pseudomonas-sp.-NCCP-436] |
| WP_033990875.1 | Type3.MTaseII.containing.BREX | <-BrxA-DNA-binding<-PglZ<-TUDOR+SF1-SF2-Helicase+iREase+CTD<-Schlafen+HATPase+wHTH<-N6-N4-METHYLASE<-?<-?<-HerA/FtsK<-SIR2<-DUF6079*<-BrxB-BrxF-iSTAND<-wHTH+WYL+WCX\|\|?->LysR-HTH-><-GNTR-HTH+Aminotran_1_2\|\|?->?->ACET->Phos_pyr_kin+Amdase-> | 1246 | Bacteria>Pseudomonadota>Gammaproteobacteria>Pseudomonadales>Pseudomonadaceae>Pseudomonas | GCF_020990445.1 | Pseudomonas_aeruginosa | LO755_RS25575 | 5366903..5370643 | 287 | DUF6079-family-protein-[Pseudomonas-aeruginosa] |
| WP_239610961.1 | Type3.MTaseII.containing.BREX | DUF4969->HNH-><-BrxA-DNA-binding<-PglZ<-TUDOR+SF1-SF2-Helicase+iREase+CTD<-Schlafen+HATPase+wHTH<-N6-N4-METHYLASE<-MORC<-DUF6079*<-BrxB-BrxF-iSTAND<-wHTH+WYL+WCX<-NERD\|\|REase+ZnR-><-?<-XIS-HTH<-BirA-HTH\|\|CopG/RHH-><-Phage_integrase | 1250 | Bacteria>Pseudomonadota>Gammaproteobacteria>Pseudomonadales>Pseudomonadaceae>Pseudomonas | GCF_022355455.1 | Pseudomonas_sp._KSR10 | MCB86_RS11370 | 524669..528421 | 2916654 | DUF6079-family-protein-[Pseudomonas-sp.-KSR10] |
| WP_247258974.1 | Type3.MTaseII.containing.BREX | <-TUDOR+SF1-SF2-Helicase+iREase+CTD<-Schlafen+HATPase+wHTH<-TOPRIM<-OLD-ABC<-N6-N4-METHYLASE<-?<-?<-HerA/FtsK<-SIR2<-DUF6079*<-BrxB-BrxF-iSTAND<-wHTH+WYL+WCX\|\|MODE-HTH+PBPII-><-HTH_3\|\|LysR-HTH-><-?<-?\|\|YjhX_toxin->ACET-> | 1246 | Bacteria>Pseudomonadota>Gammaproteobacteria>Pseudomonadales>Pseudomonadaceae>Pseudomonas | GCF_023060325.1 | Pseudomonas_aeruginosa | MX104_RS19120 | 73167..76907 | 287 | DUF6079-family-protein-[Pseudomonas-aeruginosa] |
| WP_031631068.1 | Type3.MTaseII.containing.BREX | <-BrxA-DNA-binding<-PglZ<-TUDOR+SF1-SF2-Helicase+iREase+CTD<-N6-N4-METHYLASE<-Schlafen+HATPase+wHTH<-MORC<-PIN<-OLD-ABC<-DUF6079*<-BrxB-BrxF-iSTAND<-wHTH+WYL+WCX<-HNH<-?<-?\|\|HTH_3-><-ZnR+ZnR<-DUF4969 | 1241 | Bacteria>Pseudomonadota>Gammaproteobacteria>Pseudomonadales>Pseudomonadaceae>Pseudomonas | GCF_900707525.1 | Pseudomonas_aeruginosa | E4Z30_RS22025 | 15967..19692 | 287 | DUF6079-family-protein-[Pseudomonas-aeruginosa] |
| WP_077565984.1 | Type3.MTaseII.containing.BREX | <-LysR-HTH\|\|HTH_3-><-MODE-HTH+PBPII\|\|wHTH+WYL+WCX->BrxB-BrxF-iSTAND->DUF6079*->OLD-ABC+TOPRIM->HEPN->N6-N4-METHYLASE->Schlafen+HATPase+wHTH->TUDOR+SF1-SF2-Helicase+iREase+CTD->PglZ->BrxA-DNA-binding->PspB-><-HDRnase | 1248 | Bacteria>Pseudomonadota>Gammaproteobacteria>Pseudomonadales>Pseudomonadaceae>Pseudomonas | - | Pseudomonas_aeruginosa | BM548_RS18915 | 99013..102759 | 287 | DUF6079-family-protein-[Pseudomonas-aeruginosa] |
| EIU1316754.1 | Type3.MTaseII.containing.BREX | <-Phos_pyr_kin+Amdase<-ACET<-?<-?\|\|GNTR-HTH+Aminotran_1_2-><-LysR-HTH<-?\|\|wHTH+WYL+WCX->BrxB-BrxF-iSTAND->DUF6079*->SIR2->HerA/FtsK->?->?->N6-N4-METHYLASE->Schlafen+HATPase+wHTH->TUDOR+SF1-SF2-Helicase+iREase+CTD->PglZ->BrxA-DNA-binding-> | 1250 | Bacteria>Pseudomonadota>Gammaproteobacteria>Pseudomonadales>Pseudomonadaceae>Pseudomonas | - | Pseudomonas_aeruginosa | L3560_003636 | 71973..75725 | 287 | ATP-binding-protein-[Pseudomonas-aeruginosa] |
| MBL1294161.1 | Type3.MTaseII.containing.BREX | <-SbcC+CCDC158<-Calcineurin-SbcD<-BrxA-DNA-binding<-PglZ<-TUDOR+SF1-SF2-Helicase+iREase+CTD<-N6-N4-METHYLASE<-Schlafen+HATPase+wHTH<-DUF6079*<-BrxB-BrxF-iSTAND<-XIS-HTH<-HTH_3\|\|?-><-?<-?<-SMODS<-SAVED | 1247 | Bacteria>Pseudomonadota>Gammaproteobacteria>Thiotrichales>unclassified_Thiotrichales | GCA_002401375.2 | Thiotrichales_bacterium | CDB61_009840 | 15508..19251 | 2026796 | hypothetical-protein-[Thiotrichales-bacterium] |
| PKM04365.1 | Type3.MTaseII.containing.BREX | <-DSBH<-tRNA<-?<-BrxA-DNA-binding<-PglZ<-TUDOR+SF1-SF2-Helicase+iREase+CTD<-N6-N4-METHYLASE<-Schlafen+HATPase+wHTH<-MORC<-DUF6079*<-BrxB-BrxF-iSTAND<-wHTH+WYL+WCX<-TrbL<-?<-VirB5<-TraJ-RHH<-DOC+HTH\|\|?-><-Phage_integrase | 1241 | Bacteria>Pseudomonadota>Gammaproteobacteria>unclassified_Gammaproteobacteria | GCA_002839965.1 | Gammaproteobacteria_bacterium_HGW-Gammaproteobacteria-6 | CVV16_05345 | 31508..35233 | 2013802 | ATPase-[Gammaproteobacteria-bacterium-HGW-Gammaproteobacteria-6] |
| MBU0782760.1 | Type3.MTaseII.containing.BREX | <-Phage_integrase<-TrwC+SF1-SF2-Helicase<-HerA/FtsK<-?<-SIR2<-?<-?\|\|wHTH+WYL+WCX->BrxB-BrxF-iSTAND->DUF6079*->N6-N4-METHYLASE->Schlafen+HATPase+wHTH->TUDOR+SF1-SF2-Helicase+iREase+CTD->PglZ->BrxA-DNA-binding->REase-><-?<-DOC+HTH | 1246 | Bacteria>Pseudomonadota>Gammaproteobacteria>unclassified_Gammaproteobacteria | GCA_018815355.1 | Gammaproteobacteria_bacterium | KJ979_02525 | 62574..66314 | 1913989 | ATP-binding-protein-[Gammaproteobacteria-bacterium] |
| MBU1223667.1 | Type3.MTaseII.containing.BREX | <-Thermonuclease\|\|?-><-AbiEi_4\|\|Phage_int_SAM_1+Phage_integrase-><-BrxA-DNA-binding<-PglZ<-TUDOR+SF1-SF2-Helicase+iREase+CTD<-Schlafen+HATPase+wHTH<-N6-N4-METHYLASE<-DUF6079*<-BrxB-BrxF-iSTAND<-SF1-SF2-Helicase<-?<-METHYLASE-p<-?<-?<-DNA_METHYLASE | 1241 | Bacteria>Pseudomonadota>Gammaproteobacteria>unclassified_Gammaproteobacteria | GCA_018819475.1 | Gammaproteobacteria_bacterium | KKI05_07425 | 77391..81116 | 1913989 | ATP-binding-protein-[Gammaproteobacteria-bacterium] |
| MBL8299346.1 | Type3.MTaseII.containing.BREX | <-N6-N4-METHYLASE<-HEPN<-Schlafen+HATPase+wHTH<-DUF6079*<-BrxB-BrxF-iSTAND<-XIS-HTH<-HTH+REase<-?<-?<-?<-RADICAL_SAM | 1246 | Bacteria>Pseudomonadota>Gammaproteobacteria>Xanthomonadales>Rhodanobacteraceae>unclassified_Rhodanobacteraceae | GCA_016790075.1 | Rhodanobacteraceae_bacterium | JNN30_13490 | 4880..8620 | 2052182 | hypothetical-protein-[Rhodanobacteraceae-bacterium] |
| WP_075676612.1 | Type3.MTaseII.containing.BREX | <-DUF932<-ParB+HNH<-BrxA-DNA-binding<-PglZ<-TUDOR+SF1-SF2-Helicase+iREase+CTD<-HTH+REase<-N6-N4-METHYLASE<-HEPN<-Schlafen+HATPase+wHTH<-DUF6079*<-BrxB-BrxF-iSTAND<-wHTH+WYL+WCX<-tRNA\|\|?-><-35exo<-RNaseH<-METHYLASE | 1246 | Bacteria>Pseudomonadota>Gammaproteobacteria>Xanthomonadales>Xanthomonadaceae | GCF_001939825.1 | Stenotrophomonas | BKE09_RS06610 | 1401977..1405717 | 40323 | MULTISPECIES:-DUF6079-family-protein-[Stenotrophomonas] |
| WP_033836879.1 | Type3.MTaseII.containing.BREX | <-Peptidase_M24<-?<-BrxA-DNA-binding<-PglZ<-TUDOR+SF1-SF2-Helicase+iREase+CTD<-N6-N4-METHYLASE<-MORC<-Schlafen+HATPase+wHTH<-DUF6079*<-BrxB-BrxF-iSTAND<-XIS-HTH<-Arm-DNA-bind_3+Phage_integrase<-tRNA<-?<-SIGMA-HTH+SIGMA-HTH+SIGMA-HTH<-?\|\|?-><-ncRNA<-ACET | 1248 | Bacteria>Pseudomonadota>Gammaproteobacteria>Xanthomonadales>Xanthomonadaceae>Xanthomonas | GCF_000454525.1 | Xanthomonas_citri | XMLX20_RS19135 | 4482061..4485807 | 346 | DUF6079-family-protein-[Xanthomonas-citri] |
| WP_230434266.1 | Type3.MTaseII.containing.BREX | <-BrxA-DNA-binding<-PglZ<-TUDOR+SF1-SF2-Helicase+iREase+CTD<-N6-N4-METHYLASE<-MORC<-Schlafen+HATPase+wHTH<-TOPRIM<-OLD-ABC<-DUF6079*<-BrxB-BrxF-iSTAND<-XIS-HTH<-Phage_int_SAM_2+Integrase_1<-Phage_rep | 1246 | Bacteria>Pseudomonadota>Gammaproteobacteria>Xanthomonadales>Xanthomonadaceae>Xanthomonas | GCF_020990895.1 | Xanthomonas_melonis | JWH05_RS03165 | 14918..18658 | 56456 | DUF6079-family-protein-[Xanthomonas-melonis] |
| WP_065846576.1 | Type3.MTaseII.containing.BREX | VirB8->VirB9->RHH-><-MODE-HTH+PBPII<-?<-LuxR-HTH<-SIGMA-HTH\|\|XIS-HTH->BrxB-BrxF-iSTAND->DUF6079*->OLD-ABC->TOPRIM->Schlafen+HATPase+wHTH->MORC->N6-N4-METHYLASE->TUDOR+SF1-SF2-Helicase+iREase+CTD->PglZ->BrxA-DNA-binding->Phage_AlpA-> | 1246 | Bacteria>Pseudomonadota | GCF_011762225.1 | Alphaproteobacteria | FHS82_RS15730 | 23668..27408 | 28211 | MULTISPECIES:-DUF6079-family-protein-[Alphaproteobacteria] |
| WP_119335620.1 | Type3.MTaseII.containing.BREX | <-BrxA-DNA-binding<-PglZ\|\|DUF772+DDE_Tnp_1-><-TOPRIM<-OLD-ABC<-N6-N4-METHYLASE<-Schlafen+HATPase+wHTH<-DUF6079*<-BrxB-BrxF-iSTAND\|\|?->?-><-?<-?<-ACET+ACET<-RNA_METHYLASE\|\|tRNA->ArsR-HTH-> | 1246 | Bacteria>Pseudomonadota>Hydrogenophilia>Hydrogenophilales>Hydrogenophilaceae>Hydrogenophilus | GCF_003574215.1 | Hydrogenophilus_thermoluteolus | HPTL_1674 | 1775492..1779232 | 297 | DUF6079-family-protein-[Hydrogenophilus-thermoluteolus] |
| WP_217125923.1 | Type3.MTaseII.containing.BREX | <-Thioredoxin_3<-?<-?<-ArsR-HTH<-tRNA\|\|ACET+ACET->?-><-?<-?\|\|BrxB-BrxF-iSTAND->DUF6079*->Schlafen+HATPase+wHTH->N6-N4-METHYLASE->TUDOR+SF1-SF2-Helicase+iREase+CTD->PglZ->BrxA-DNA-binding-><-?\|\|?-><-?\|\|sGTP-> | 1243 | Bacteria>Pseudomonadota>Hydrogenophilia>Hydrogenophilales>Hydrogenophilaceae>Hydrogenophilus | GCF_019049855.1 | Hydrogenophilus_thiooxidans | KUL93_RS04625 | 87236..90967 | 2820326 | DUF6079-family-protein-[Hydrogenophilus-thiooxidans] |
| WP_142810711.1 | Type3.MTaseII.containing.BREX | HAMP+HISKIN-><-EAL\|\|MerR-HTH->XIS-HTH->XIS-HTH+XIS-HTH->PrlF_antitoxin->?->XIS-HTH->BrxB-BrxF-iSTAND->DUF6079*->?->N6-N4-METHYLASE->Schlafen+HATPase+wHTH->TUDOR+SF1-SF2-Helicase+iREase+CTD->PglZ->BrxA-DNA-binding-><-Phage_lysozyme<-Phage_holin | 1241 | Bacteria>Pseudomonadota>Hydrogenophilia>Hydrogenophilales>Hydrogenophilaceae>Tepidiphilus | GCF_006980705.1 | Tepidiphilus_olei | EPD80_RS07330 | 156841..160566 | 2502184 | DUF6079-family-protein-[Tepidiphilus-olei] |
| MBK6849133.1 | Type3.MTaseII.containing.BREX | <-MPTase<-HTH\|\|?->RAMA->?->?->XIS-HTH->BrxB-BrxF-iSTAND->DUF6079*->TUDOR+SF1-SF2-Helicase+iREase+CTD->PglZ->?->BrxA-DNA-binding->glycosylase->HATPase+wHTH->Schlafen+HATPase+wHTH->DOC+HTH->GDResolvase-> | 1232 | Bacteria>Pseudomonadota>unclassified_Pseudomonadota | GCA_016702545.1 | Pseudomonadota_bacterium | IPG96_16905 | 43585..47283 | 1977087 | ATP-binding-protein-[Pseudomonadota-bacterium] |
| MBS0577369.1 | Type3.MTaseII.containing.BREX | XIS-HTH->BrxB-BrxF-iSTAND->DUF6079*->Schlafen+HATPase+wHTH->HEPN->N6-N4-METHYLASE->TUDOR+SF1-SF2-Helicase+iREase+CTD->PglZ->BrxA-DNA-binding->PspB-> | 1246 | Bacteria>Pseudomonadota>unclassified_Pseudomonadota | GCA_018240435.1 | Pseudomonadota_bacterium | JSS45_13265 | 2298..6038 | 1977087 | ATP-binding-protein-[Pseudomonadota-bacterium] |
| MCA0325062.1 | Type3.MTaseII.containing.BREX | <-TUDOR+SF1-SF2-Helicase+iREase+CTD<-N6-N4-METHYLASE<-HEPN<-Schlafen+HATPase+wHTH<-DUF6079*<-BrxB-BrxF-iSTAND<-XIS-HTH<-Calcineurin-SbcD<-?\|\|CopG/RHH->?->?->SIR2-> | 1246 | Bacteria>Pseudomonadota>unclassified_Pseudomonadota | GCA_020161215.1 | Pseudomonadota_bacterium | LCH89_05600 | 7182..10922 | 1977087 | DUF6079-family-protein-[Pseudomonadota-bacterium] |
| MBK7497323.1 | Type3.MTaseII.containing.BREX | <-PglZ<-TUDOR+SF1-SF2-Helicase+iREase+CTD<-MORC<-Schlafen+HATPase+wHTH<-SF1-SF2-Helicase<-SF1-SF2-Helicase<-DUF6079*<-REase<-N6-N4-METHYLASE-p | 1241 | Bacteria>PVC_group>Candidatus_Omnitrophica>unclassified_Candidatus_Omnitrophica | GCA_016706995.1 | Candidatus_Omnitrophica_bacterium | IPI28_18700 | 38924..42649 | 2035772 | ATP-binding-protein-[Candidatus-Omnitrophota-bacterium] |
| NLF62180.1 | Type3.MTaseII.containing.BREX | <-IG+IG\|\|BrxB-BrxF-iSTAND->DUF6079*->OLD-ABC->TOPRIM->?->?->Schlafen+HATPase+wHTH+HTH->N6-N4-METHYLASE->SbcC->PIN-> | 1251 | Bacteria>PVC_group>Lentisphaerota>unclassified_Lentisphaerota | GCA_012515215.1 | Lentisphaerota_bacterium | GX574_13635 | 31199..34954 | 1932692 | ATP-binding-protein-[Lentisphaerota-bacterium] |
| MBT3379341.1 | Type3.MTaseII.containing.BREX | ZnR->?->?-><-?<-Radical_SAM\|\|Schlafen+HATPase+wHTH->XIS-HTH->BrxB-BrxF-iSTAND->DUF6079*->Schlafen+HATPase+wHTH->N6-N4-METHYLASE->OLD-ABC+TOPRIM->SF1-SF2-Helicase->TUDOR+SF1-SF2-Helicase+iREase+CTD->PglZ->BrxA-DNA-binding->TPR+HEPN->KAP-NTPase-> | 1252 | Bacteria>PVC_group>Lentisphaerota>unclassified_Lentisphaerota | GCA_018648355.1 | Lentisphaerota_bacterium | HN406_27320 | 7239..10997 | 1932692 | ATP-binding-protein-[Lentisphaerota-bacterium] |
| MBT4818762.1 | Type3.MTaseII.containing.BREX | <-N6-N4-METHYLASE-p<-SF1-SF2-Helicase<-OLD-ABC+TOPRIM<-N6-N4-METHYLASE<-Schlafen+HATPase+wHTH<-DUF6079-p* | 298 | Bacteria>PVC_group>Lentisphaerota>unclassified_Lentisphaerota | GCA_018664235.1 | Lentisphaerota_bacterium | HON70_23840 | 8568..9465 | 1932692 | ATP-binding-protein-[Lentisphaerota-bacterium] |
| MBT5606091.1 | Type3.MTaseII.containing.BREX | ZnR->?->?-><-?<-Radical_SAM\|\|Schlafen+HATPase+wHTH->XIS-HTH->BrxB-BrxF-iSTAND->DUF6079-p*-> | 722 | Bacteria>PVC_group>Lentisphaerota>unclassified_Lentisphaerota | GCA_018666605.1 | Lentisphaerota_bacterium | HOJ57_09130 | 7357..9522 | 1932692 | ATP-binding-protein-[Lentisphaerota-bacterium] |
| MBT6049514.1 | Type3.MTaseII.containing.BREX | <-BrxA-DNA-binding<-PglZ<-TUDOR+SF1-SF2-Helicase+iREase+CTD<-Schlafen+HATPase+wHTH<-N6-N4-METHYLASE<-Schlafen+HATPase+wHTH<-DUF6079*<-BrxB-BrxF-iSTAND | 1248 | Bacteria>PVC_group>Planctomycetota>Candidatus_Brocadiia>Candidatus_Brocadiales>Candidatus_Scalinduaceae | GCA_018697755.1 | Candidatus_Scalindua_sp. | HOG49_22170 | 17519..21265 | 2268189 | ATP-binding-protein-[Candidatus-Scalindua-sp.] |
| WP_096895081.1 | Type3.MTaseII.containing.BREX | Aminotran_3->?->?->TPR+TPR+TPR+TPR-><-?\|\|BrxB-BrxF-iSTAND->DUF6079*->N6-N4-METHYLASE->REase->Schlafen+HATPase+wHTH->TUDOR+SF1-SF2-Helicase+iREase+CTD->PglZ->BrxA-DNA-binding-><-?\|\|RADICAL-SAM-> | 1248 | Bacteria>PVC_group>Planctomycetota>Candidatus_Brocadiia>Candidatus_Brocadiales>Candidatus_Scalinduaceae | GCF_002443295.1 | Candidatus_Scalindua_japonica | SCALIN_RS13975 | 114317..118063 | 1284222 | DUF6079-family-protein-[Candidatus-Scalindua-japonica] |
| NQV34261.1 | Type3.MTaseII.containing.BREX | <-SbcC<-N6-N4-METHYLASE<-Schlafen+HATPase+wHTH<-DUF6079*<-BrxB-BrxF-iSTAND<-XIS-HTH\|\|XIS-HTH-><-?<-GDResolvase+HIN-HTH | 1244 | Bacteria>PVC_group>Planctomycetota>Phycisphaerae>Phycisphaerales>Phycisphaeraceae | GCA_013203945.1 | Phycisphaeraceae_bacterium | HQ515_16320 | 6009..9743 | 2026777 | ATP-binding-protein-[Phycisphaeraceae-bacterium] |
| WP_145380206.1 | Type3.MTaseII.containing.BREX | <-D5N+D5ATPase<-?<-?<-BrxA-DNA-binding<-PglZ<-TUDOR+SF1-SF2-Helicase+iREase+CTD<-Schlafen+HATPase+wHTH<-N6-N4-METHYLASE<-DUF6079*<-BrxB-BrxF-iSTAND<-XIS-HTH<-?\|\|wHTH+SAD+HNH-><-Methyltransf_25<-?<-Phage_integrase | 1234 | Bacteria>PVC_group>Planctomycetota>Planctomycetia>Planctomycetales>Planctomycetaceae | GCF_007747995.1 | Symmachiella_dynata | Mal52_RS28010 | 7362136..7365840 | 2527995 | DUF6079-family-protein-[Symmachiella-dynata] |
| OQZ06364.1 | Type3.MTaseII.containing.BREX | <-Schlafen+HATPase+wHTH<-DUF6079*<-BrxB-BrxF-iSTAND<-?\|\|METHYLASE+SF1-SF2-Helicase-> | 1231 | Bacteria>PVC_group>Planctomycetota>unclassified_Planctomycetota | GCA_002050205.1 | Planctomycetes_bacterium_UTPLA1 | B6D36_05375 | 2828..6523 | 1961112 | ATPase-[Planctomycetes-bacterium-UTPLA1] |
| MBI4882201.1 | Type3.MTaseII.containing.BREX | <-N6-N4-METHYLASE<-Schlafen+HATPase+wHTH<-DUF6079*<-BrxB-BrxF-iSTAND<-?\|\|Glyco_transf_9->Sulfatase-><-REC+OmpR-HTH<-?<-REC+OmpR-HTH<-dCache_1+HisKA+HATPase_c | 1237 | Bacteria>PVC_group>Planctomycetota>unclassified_Planctomycetota | GCA_016213895.1 | Planctomycetota_bacterium | HY812_21445 | 2712..6425 | 2026780 | ATP-binding-protein-[Planctomycetota-bacterium] |
| MBL9185851.1 | Type3.MTaseII.containing.BREX | <-HerA/FtsK<-?<-?<-BrxA-DNA-binding<-PglZ<-TUDOR+SF1-SF2-Helicase+iREase+CTD<-N6-N4-METHYLASE<-MORC<-Schlafen+HATPase+wHTH<-DUF6079*<-BrxB-BrxF-iSTAND<-?<-?<-?<-TOPRIM-DNAG+T7DNApol<-BirA-HTH\|\|?-><-DNA_METHYLASE<-HerA/FtsK | 1236 | Bacteria>PVC_group>Verrucomicrobiota>Opitutae>Opitutales>Opitutaceae | GCA_016794785.1 | Opitutaceae_bacterium | JNK23_00090 | 29545..33255 | 2006848 | hypothetical-protein-[Opitutaceae-bacterium] |
| HCF94153.1 | Type3.MTaseII.containing.BREX | XIS-HTH->BrxB-BrxF-iSTAND->DUF6079*->Schlafen+HATPase+wHTH+HTH->N6-N4-METHYLASE->?->OLD-ABC->PIN->?->TUDOR+SF1-SF2-Helicase+iREase+CTD->REase->PglZ-> | 1288 | Bacteria>PVC_group>Verrucomicrobiota>unclassified_Verrucomicrobiota | GCA_003534995.1 | Verrucomicrobiota_bacterium | DEW46_03760 | 1029..4895 | 2026799 | ATPase-[Verrucomicrobiota-bacterium] |
| MBI4023511.1 | Type3.MTaseII.containing.BREX | ANK-><-Glycos_transf<-DUF6079*<-N6-N4-METHYLASE<-Schlafen+HATPase+wHTH<-BrxB-BrxF-iSTAND<-BrxA-DNA-binding<-SIGMA+OB-Fold+AAA-ATPase+Lon-Protease-BrxL<-PglZ | 1371 | Bacteria>PVC_group>Verrucomicrobiota>unclassified_Verrucomicrobiota | GCA_016198745.1 | Verrucomicrobiota_bacterium | HY360_00925 | 8259..12374 | 2026799 | hypothetical-protein-[Verrucomicrobiota-bacterium] |
| MBX3742594.1 | Type3.MTaseII.containing.BREX | TrwC+SF1-SF2-Helicase->?->?->HTH+MPTase->BrxB-BrxF-iSTAND->DUF6079*->Schlafen+HATPase+wHTH->REase->N6-N4-METHYLASE->nSTAND1+HNH->TIR->TUDOR+SF1-SF2-Helicase+iREase+CTD->PglZ->BrxA-DNA-binding->XIS-HTH-> | 1243 | Bacteria>PVC_group>Verrucomicrobiota>Verrucomicrobiae>Verrucomicrobiales>Akkermansiaceae | GCA_019634595.1 | Akkermansiaceae_bacterium | KF712_16525 | 314704..318435 | 2562705 | hypothetical-protein-[Akkermansiaceae-bacterium] |
| RFU95875.1 | Type3.MTaseII.containing.BREX | <-TrkH<-?<-?<-?\|\|tRNA-><-AbiEii<-AbiEi_4\|\|XIS-HTH->BrxB-BrxF-iSTAND->DUF6079*->N6-N4-METHYLASE->?->?->?->Schlafen+HATPase+wHTH->TUDOR+SF1-SF2-Helicase+iREase+CTD->PglZ->BrxA-DNA-binding->?->REC+LuxR-HTH-> | 1246 | Bacteria>Spirochaetota>Spirochaetia>Spirochaetales>Sphaerochaetaceae>Sphaerochaeta | GCA_003429665.1 | Sphaerochaeta_halotolerans | DYP60_02410 | 179528..183268 | 2293840 | ATP-binding-protein-[Sphaerochaeta-halotolerans] |
| WP_206363090.1 | Type3.MTaseII.containing.BREX | <-TrkH<-?\|\|tRNA-><-AbiEii<-AbiEi_4\|\|?->XIS-HTH->?->BrxB-BrxF-iSTAND->DUF6079*->N6-N4-METHYLASE->?->?->?->Schlafen+HATPase+wHTH->TUDOR+SF1-SF2-Helicase+iREase+CTD->PglZ->BrxA-DNA-binding->?->REC+LuxR-HTH-> | 1255 | Bacteria>Spirochaetota>Spirochaetia>Spirochaetales>Sphaerochaetaceae>Sphaerochaeta | GCF_003429665.1 | Sphaerochaeta_halotolerans | DYP60_RS02405 | 179501..183268 | 2293840 | DUF6079-family-protein-[Sphaerochaeta-halotolerans] |
| MBO7485543.1 | Type3.MTaseII.containing.BREX | <-Tox-RelE\|\|tRNA-><-BrxA-DNA-binding<-PglZ<-TUDOR+SF1-SF2-Helicase+iREase+CTD<-Schlafen+HATPase+wHTH<-?<-N6-N4-METHYLASE<-TOPRIM<-OLD-ABC<-DUF6079*<-BrxB-BrxF-iSTAND\|\|wHTH+WYL+WCX->AAA-ATPase->?->?->?->35exo+T7DNApol-> | 1226 | Bacteria>Spirochaetota>Spirochaetia>Spirochaetales>Spirochaetaceae>unclassified_Spirochaetaceae | GCA_017649985.1 | Spirochaetaceae_bacterium | J6T84_05730 | 14536..18216 | 1898206 | hypothetical-protein-[Spirochaetaceae-bacterium] |
| WP_010252922.1 | Type3.MTaseII.containing.BREX | <-HTH+REase<-DUF499<-?<-METHYLASE-p<-TUDOR+SF1-SF2-Helicase+iREase+CTD+REase\|\|?->CR-ATPase5+REase->Phage_integrase->BrxB-BrxF-iSTAND->DUF6079*->N6-N4-METHYLASE->?->OLD-ABC+TOPRIM->SF1-SF2-Helicase->Schlafen+HATPase+wHTH->TUDOR+SF1-SF2-Helicase+iREase+CTD->?->PglZ->BrxA-DNA-binding-> | 1224 | Bacteria>Spirochaetota>Spirochaetia>Spirochaetales>Treponemataceae>Treponema | GCF_000297095.1 | Treponema_primitia | TPRIMZ1_RS18025 | 17306..20980 | 88058 | DUF6079-family-protein-[Treponema-primitia] |
| GHT84073.1 | Type3.MTaseII.containing.BREX | <-N6-N4-METHYLASE<-TIR<-Schlafen+HATPase+wHTH<-OLD-ABC<-DUF6079-p*\|\|DUF6079-p->OLD-ABC->Schlafen+HATPase+wHTH->TIR->N6-N4-METHYLASE-> | 177 | Bacteria>Spirochaetota>Spirochaetia>unclassified_Spirochaetia | GCA_026002105.1 | Spirochaetia_bacterium | FACS1894137_06670 | 4966..5499 | 2053615 | hypothetical-protein-FACS1894137_06670-[Spirochaetia-bacterium] |
| GHT84077.1 | Type3.MTaseII.containing.BREX | <-N6-N4-METHYLASE<-TIR<-Schlafen+HATPase+wHTH<-OLD-ABC<-DUF6079-p\|\|DUF6079-p*->OLD-ABC->Schlafen+HATPase+wHTH->TIR->N6-N4-METHYLASE-> | 180 | Bacteria>Spirochaetota>Spirochaetia>unclassified_Spirochaetia | GCA_026002105.1 | Spirochaetia_bacterium | FACS1894137_06680 | 5766..6308 | 2053615 | hypothetical-protein-FACS1894137_06680-[Spirochaetia-bacterium] |
| OQB94579.1 | Type3.MTaseII.containing.BREX | <-HTH_Tnp_1<-BrxA-DNA-binding<-PglZ<-TUDOR+SF1-SF2-Helicase+iREase+CTD<-HHH_4+SF1-SF2-Helicase+REase<-N6-N4-METHYLASE<-Schlafen+HATPase+wHTH+HTH<-DUF6079*<-BrxB-BrxF-iSTAND\|\|?->HEPN->NTP_transferase+MannoseP_isomer->TetR-HTH-> | 1246 | Bacteria>Spirochaetota>unclassified_Spirochaetota | GCA_002070755.1 | Spirochaetes_bacterium_ADurb.Bin110 | BWX81_02438 | 21902..25642 | 1852908 | hypothetical-protein-BWX81_02438-[Spirochaetes-bacterium-ADurb.Bin110] |
| WP_005662440.1 | Type3.MTaseII.containing.BREX | <-HicB_lk_antitox<-BrxA-DNA-binding<-PglZ<-TUDOR+SF1-SF2-Helicase+iREase+CTD<-PADR-HTH<-DUF772+DDE_Tnp_1<-Schlafen+HATPase+wHTH<-N6-N4-METHYLASE<-Schlafen+HATPase+wHTH+HTH<-DUF6079*<-BrxB-BrxF-iSTAND<-?<-?\|\|?-><-PAS+GGDEF+EAL<-Aminotran_4<-?<-?<-PALP | 1247 | Bacteria>Synergistota>Synergistia>Synergistales>Dethiosulfovibrionaceae>Dethiosulfovibrio | GCF_000172975.1 | Dethiosulfovibrio_peptidovorans | DPEP_RS12015 | 2485235..2488978 | 47055 | DUF6079-family-protein-[Dethiosulfovibrio-peptidovorans] |
| WP_236100997.1 | Type3.MTaseII.containing.BREX | Aminotran_4->PAS+GGDEF+EAL-><-?\|\|?->\|\|?->?->BrxB-BrxF-iSTAND->DUF6079*->N6-N4-METHYLASE->Schlafen+HATPase+wHTH->TUDOR+SF1-SF2-Helicase+iREase+CTD->PglZ->BrxA-DNA-binding-> | 1246 | Bacteria>Synergistota>Synergistia>Synergistales>Dethiosulfovibrionaceae>Dethiosulfovibrio | GCF_021568795.1 | Dethiosulfovibrio_faecalis | L2W58_RS00505 | 84364..88104 | 2720018 | DUF6079-family-protein-[Dethiosulfovibrio-faecalis] |

| Protein-ID | System | Gene-Neighbourhoods | Length | Taxonomy | Assembly-ID | Strain | Locus-Tag | Genomic-coordinates | TaxID | Fasta-defline |
|---|---|---|---|---|---|---|---|---|---|---|
| -- | | | | | | | | | | |
| --HEPN+DUF499 | | | | | | | | | | |
| -- | | | | | | | | | | |
| WP_283168466.1 | DUF499_1.RM.like.systems | NurA->HerA/FtsK->DUF4338-><-HEPN+DUF499*<-METHYLASE-p | 1093 | Archaea>Euryarchaeota>Stenosarchaea_group>Methanomicrobia>Methanocellales | GCF_030019315.1 | Methanocella_conradii | QMC99_RS12775 | 4236..7517 | 1175444 | DUF499-domain-containing-protein-[Methanocella-conradii] |
| ABD41477.1 | DUF499_1.RM.like.systems | OLD-ABC->OLD-ABC->REase->METHYLASE->OLD-ABC->TOPRIM->HEPN+DUF499*-> | 1094 | Archaea>Euryarchaeota>Stenosarchaea_group>Methanomicrobia>Methanomicrobiales | GCA_000013445.1 | Methanospirillum_hungatei_JF-1 | Mhun_1756 | 1984858..1988142 | 323259 | conserved-hypothetical-protein-[Methanospirillum-hungatei-JF-1] |
| KUL01096.1 | DUF499_1.RM.like.systems | METHYLASE-p->HEPN+DUF499*-> | 1106 | Archaea>Euryarchaeota>Stenosarchaea_group>Methanomicrobia>Methanomicrobiales | GCA_001508695.1 | Methanomicrobiales_archaeon_53_19 | XE11_2241 | 598..3918 | 1641394 | Uncharacterized-protein-XE11_2241-[Methanomicrobiales-archaeon-53_19] |
| WP_204222940.1 | DUF499_1.RM.like.systems | OLD-ABC->OLD-ABC->REase->METHYLASE->OLD-ABC->TOPRIM->HEPN+DUF499*-> | 1082 | Archaea>Euryarchaeota>Stenosarchaea_group>Methanomicrobia>Methanomicrobiales | GCF_000013445.1 | Methanospirillum_hungatei | MHUN_RS09170 | 1984894..1988142 | 2203 | DUF499-domain-containing-protein-[Methanospirillum-hungatei] |
| WP_167816002.1 | DUF499_1.RM.like.systems | Schlafen+HATPase+HTH->?->Schlafen+HATPase+HTH->METHYLASE->OLD-ABC->TOPRIM->HEPN+DUF499*-> | 756 | Archaea>Euryarchaeota>Stenosarchaea_group>Methanomicrobia>Methanomicrobiales | GCF_012062385.1 | Methanocorpusculum_sp._GPch4 | EYA77_RS04465 | 15957..18225 | 2527877 | Swt1-family-HEPN-domain-containing-protein,-partial-[Methanocorpusculum-sp.-GPch4] |
| KXS42417.1 | DUF499_1.RM.like.systems | <-HEPN+DUF499*<-METHYLASE<-MORC+DUF3684<-TUDOR+SF1-SF2-Helicase+iREase+CTD+REase | 1129 | Archaea>Euryarchaeota>Stenosarchaea_group>Methanomicrobia>Methanosarcinales | GCA_001577565.1 | Methanolobus_sp._T82-4 | AWU59_1648 | 511..3900 | 1794908 | MAG:-ATPase-(AAA+-superfamily)-like-protein-[Methanolobus-sp.-T82-4] |
| MCG7852630.1 | DUF499_1.RM.like.systems | METHYLASE->HEPN+DUF499*-> | 1119 | Archaea>Euryarchaeota>Stenosarchaea_group>Methanomicrobia>Methanosarcinales | GCA_022353215.1 | Methanosarcinaceae_archaeon | MI092_08920 | 2388..5747 | 1945595 | Swt1-family-HEPN-domain-containing-protein-[Methanosarcinaceae-archaeon] |
| WP_048043068.1 | DUF499_1.RM.like.systems | TUDOR+SF1-SF2-Helicase+iREase+CTD+REase->METHYLASE->HEPN+DUF499*-><-ZnR<-tRNA<-?<-?\|\|RelE-ParE-><-PRTase | 1126 | Archaea>Euryarchaeota>Stenosarchaea_group>Methanomicrobia>Methanosarcinales | GCF_000970185.1 | Methanosarcina_mazei | MSMAP_RS05770 | 1314759..1318139 | 2209 | Swt1-family-HEPN-domain-containing-protein-[Methanosarcina-mazei] |
| WP_048105728.1 | DUF499_1.RM.like.systems | PRTase-><-RelE-ParE\|\|?->?->tRNA-><-HEPN+DUF499*<-METHYLASE<-TUDOR+SF1-SF2-Helicase+iREase+CTD+REase<-?<-?<-Primpol+D5N+D5ATPase | 1126 | Archaea>Euryarchaeota>Stenosarchaea_group>Methanomicrobia>Methanosarcinales | GCF_000970305.1 | Methanosarcina_barkeri | MSBR3_RS00635 | 148480..151860 | 2208 | Swt1-family-HEPN-domain-containing-protein-[Methanosarcina-barkeri] |
| WP_048037397.1 | DUF499_1.RM.like.systems | PRTase-><-RelE-ParE\|\|?->?->tRNA-><-HEPN+DUF499*<-METHYLASE<-TUDOR+SF1-SF2-Helicase+iREase+CTD+REase<-?<-?<-Primpol+D5N+D5ATPase | 1126 | Archaea>Euryarchaeota>Stenosarchaea_group>Methanomicrobia>Methanosarcinales | GCF_000978965.1 | Methanosarcina_mazei | DU68_RS02805 | 48841..52221 | 2209 | Swt1-family-HEPN-domain-containing-protein-[Methanosarcina-mazei] |
| WP_048048703.1 | DUF499_1.RM.like.systems | ParA-Soj-PloopNTPase->?->?-><-HEPN+DUF499*<-METHYLASE<-TUDOR+SF1-SF2-Helicase+iREase+CTD+REase<-Peptidase_C47 | 1113 | Archaea>Euryarchaeota>Stenosarchaea_group>Methanomicrobia>Methanosarcinales | GCF_000979015.1 | Methanosarcina_mazei | DU63_RS12210 | 161635..164976 | 2209 | DUF499-domain-containing-protein-[Methanosarcina-mazei] |
| WP_048047296.1 | DUF499_1.RM.like.systems | <-HEPN+DUF499*<-METHYLASE<-TUDOR+SF1-SF2-Helicase+iREase+CTD+REase<-?<-?<-Primpol+D5N+D5ATPase | 513 | Archaea>Euryarchaeota>Stenosarchaea_group>Methanomicrobia>Methanosarcinales | GCF_000979035.1 | Methanosarcina_mazei | DU66_RS08905 | 1..1541 | 2209 | Swt1-family-HEPN-domain-containing-protein,-partial-[Methanosarcina-mazei] |
| WP_048045029.1 | DUF499_1.RM.like.systems | Primpol+D5N+D5ATPase->?->?->TUDOR+SF1-SF2-Helicase+iREase+CTD+REase->METHYLASE->HEPN+DUF499*-><-?<-?<-HNH | 1126 | Archaea>Euryarchaeota>Stenosarchaea_group>Methanomicrobia>Methanosarcinales | GCF_000979105.1 | Methanosarcina_mazei | DU65_RS07520 | 78194..81574 | 2209 | Swt1-family-HEPN-domain-containing-protein-[Methanosarcina-mazei] |
| WP_048047674.1 | DUF499_1.RM.like.systems | Primpol+D5N+D5ATPase->?->?->TUDOR+SF1-SF2-Helicase+iREase+CTD+REase->METHYLASE->HEPN+DUF499*-><-?<-tRNA<-?<-?\|\|RelE-ParE-><-PRTase | 1126 | Archaea>Euryarchaeota>Stenosarchaea_group>Methanomicrobia>Methanosarcinales | GCF_000979995.1 | Methanosarcina_mazei | DU74_RS10215 | 73839..77219 | 2209 | Swt1-family-HEPN-domain-containing-protein-[Methanosarcina-mazei] |
| MBC7080549.1 | DUF499_1.RM.like.systems | TUDOR+SF1-SF2-Helicase+iREase+CTD+REase->METHYLASE->RNase->HEPN+DUF499*-> | 1102 | Archaea>Euryarchaeota>Stenosarchaea_group>Methanomicrobia>Methanotrichales | GCA_014361255.1 | Methanothrix_sp. | H5T42_08820 | 8191..11499 | 90426 | ATP-binding-protein-[Methanothrix-sp.] |
| MBK7385730.1 | DUF499_1.RM.like.systems | Methyltransf-><-HEPN+DUF499*<-TOPRIM<-OLD-ABC<-METHYLASE\|\|?->?-><-TUDOR+SF1-SF2-Helicase+iREase+CTD+REase | 1119 | Archaea>Euryarchaeota>Stenosarchaea_group>Methanomicrobia>Methanotrichales | GCA_016706325.1 | Methanothrix_sp. | IPI63_02955 | 568305..571664 | 90426 | DUF499-domain-containing-protein-[Methanothrix-sp.] |
| MCX6676562.1 | DUF499_1.RM.like.systems | TUDOR+SF1-SF2-Helicase+iREase+CTD+REase-><-?<-?<-?\|\|METHYLASE->OLD-ABC->HEPN+DUF499*-> | 1111 | Archaea>Euryarchaeota>Stenosarchaea_group>Methanomicrobia>Methanotrichales | GCA_026394565.1 | Methanothrix_sp. | NTU95_01275 | 32506..35841 | 90426 | MAG:-DUF499-domain-containing-protein-[Methanothrix-sp.] |
| MDD4161340.1 | DUF499_1.RM.like.systems | ParB+HNH+Schlafen-><-DEDD_Tnp_IS110+Transposase_20<-?\|\|TUDOR+SF1-SF2-Helicase+iREase+CTD+REase->METHYLASE->HEPN+DUF499*->?-><-SF1-SF2-Helicase-p | 1125 | Archaea>Euryarchaeota>Stenosarchaea_group>Methanomicrobia>Methanotrichales | GCA_028702095.1 | Methanothrix_sp. | PHW87_02550 | 13922..17299 | 90426 | MAG:-Swt1-family-HEPN-domain-containing-protein-[Methanothrix-sp.] |
| WP_273278519.1 | DUF499_1.RM.like.systems | Primpol->?->ArdCN+MPTase->?->?-><-REase<-HEPN+DUF499*<-TOPRIM<-OLD-ABC<-PolB<-METHYLASE<-HEPN<-TUDOR+SF1-SF2-Helicase+iREase+CTD+REase | 1113 | Archaea>Euryarchaeota>Stenosarchaea_group>Methanomicrobia>Methanotrichales | GCF_913757535.1 | Methanothrix_soehngenii | NW278_RS08460 | 1661104..1664445 | 2223 | DUF499-domain-containing-protein-[Methanothrix-soehngenii] |
| OQA55746.1 | DUF499_1.RM.like.systems | METHYLASE->?->OLD-ABC->?->TOPRIM->HEPN+DUF499*-> | 1082 | Archaea>Euryarchaeota>unclassified_Euryarchaeota | GCA_002069575.1 | Euryarchaeota_archaeon_ADurb.Bin294 | BWY45_02073 | 4487..7735 | 1852878 | hypothetical-protein-BWY45_02073-[Euryarchaeota-archaeon-ADurb.Bin294] |
| MCL2643280.1 | DUF499_1.RM.like.systems | <-DDE_Tnp_ISAZ013<-HEPN+DUF499*<-TUDOR+SF1-SF2-Helicase+iREase+CTD+REase | 1134 | Archaea>TACK_group>Candidatus_Bathyarchaeota>Candidatus_Bathyarchaeia>Candidatus_Bathyarchaeia_incertae_sedis | GCA_009781395.1 | Candidatus_Termiticorpusculum_sp. | FWD52_07200 | 1651..5055 | 2994959 | DUF499-domain-containing-protein-[Candidatus-Bathyarchaeota-archaeon] |
| TSA50211.1 | DUF499_1.RM.like.systems | SF1-SF2-Helicase->SF1-SF2-Helicase->METHYLASE->?->HEPN+DUF499*-> | 607 | Archaea>unclassified_Archaea | GCA_007281275.1 | archaeon | D4R47_02600 | 5332..7152 | 1906665 | ATP-binding-protein,-partial-[archaeon] |
| MBA3712812.1 | DUF499_1.RM.like.systems | Schlafen+HATPase+HTH->?->METHYLASE->DUF433->PIN->HEPN+DUF499*->TIR+PNTD2->PIWI->?->Phage_integrase-> | 1118 | Bacteria>Acidobacteriota>Blastocatellia>Blastocatellales>Pyrinomonadaceae | GCA_013816505.1 | Pyrinomonadaceae_bacterium | HOW76_10240 | 5538..8894 | 2283092 | ATP-binding-protein-[Pyrinomonadaceae-bacterium] |
| QUV94961.1 | DUF499_1.RM.like.systems | ABhydrolase-><-?\|\|TUDOR+SF1-SF2-Helicase+iREase+CTD+REase->METHYLASE->HEPN+DUF499*->?->Response_reg+LuxR-HTH->CarboxypepD-reg-IG->BetaPropeller+ABhydrolase-> | 1109 | Bacteria>Acidobacteriota>Blastocatellia>Chloracidobacterium>Chloracidobacterium_aggregatum | GCA_018304765.1 | Chloracidobacterium_sp._N | J8C05_05820 | 1401260..1404589 | 2821540 | DUF499-domain-containing-protein-[Chloracidobacterium-sp.-N] |
| QUW01026.1 | DUF499_1.RM.like.systems | ABhydrolase-><-?\|\|TUDOR+SF1-SF2-Helicase+iREase+CTD+REase->METHYLASE->HEPN+DUF499*->?->Response_reg+LuxR-HTH->CarboxypepD-reg-IG->BetaPropeller+ABhydrolase-> | 1109 | Bacteria>Acidobacteriota>Blastocatellia>Chloracidobacterium>Chloracidobacterium_aggregatum | GCA_018304805.1 | Chloracidobacterium_sp._MS_40/45 | J8C02_05835 | 1402639..1405968 | 2821542 | DUF499-domain-containing-protein-[Chloracidobacterium-sp.-MS-40/45] |
| AEP11956.1 | DUF499_1.RM.like.systems | ABhydrolase-><-?\|\|TUDOR+SF1-SF2-Helicase+iREase+CTD+REase->METHYLASE->HEPN+DUF499*->?->Response_reg+LuxR-HTH->CarboxypepD-reg-IG->BetaPropeller+ABhydrolase-> | 1108 | Bacteria>Acidobacteriota>Blastocatellia>Chloracidobacterium>Chloracidobacterium_thermophilum | GCA_000226295.1 | Chloracidobacterium_thermophilum_B | Cabther_A1202 | 1425690..1429016 | 981222 | hypothetical-protein-Cabther_A1202-[Chloracidobacterium-thermophilum-B] |
| WP_041569618.1 | DUF499_1.RM.like.systems | ABhydrolase-><-?\|\|TUDOR+SF1-SF2-Helicase+iREase+CTD+REase->METHYLASE->HEPN+DUF499*->?->Response_reg+LuxR-HTH->CarboxypepD-reg-IG->BetaPropeller+ABhydrolase-> | 1110 | Bacteria>Acidobacteriota>Blastocatellia>Chloracidobacterium | GCF_000226295.1 | Chloracidobacterium_thermophilum | CABTHER_RS05910 | 1425684..1429016 | 458033 | Swt1-family-HEPN-domain-containing-protein-[Chloracidobacterium-thermophilum] |
| WP_058867444.1 | DUF499_1.RM.like.systems | TUDOR+SF1-SF2-Helicase+iREase+CTD+REase->PrlF_antitoxin->METHYLASE->PolB->HEPN->HEPN+DUF499*->?->?->?->?->NUDIX-> | 1098 | Bacteria>Acidobacteriota>Blastocatellia>Chloracidobacterium | GCF_001482755.1 | Chloracidobacterium_thermophilum | AVR57_RS08855 | 227701..230997 | 458033 | Swt1-family-HEPN-domain-containing-protein-[Chloracidobacterium-thermophilum] |
| WP_211431037.1 | DUF499_1.RM.like.systems | ABhydrolase-><-?\|\|TUDOR+SF1-SF2-Helicase+iREase+CTD+REase->METHYLASE->HEPN+DUF499*->?->Response_reg+LuxR-HTH->CarboxypepD-reg-IG->BetaPropeller+ABhydrolase-> | 1100 | Bacteria>Acidobacteriota>Blastocatellia>Chloracidobacterium | GCF_018304725.1 | Chloracidobacterium_aggregatum | J8C07_RS06025 | 1402377..1405679 | 2851959 | Swt1-family-HEPN-domain-containing-protein-[Chloracidobacterium-aggregatum] |
| WP_246840661.1 | DUF499_1.RM.like.systems | ABhydrolase-><-?\|\|TUDOR+SF1-SF2-Helicase+iREase+CTD+REase->METHYLASE->HEPN+DUF499*->?->Response_reg+LuxR-HTH->CarboxypepD-reg-IG->BetaPropeller+ABhydrolase-> | 1107 | Bacteria>Acidobacteriota>Blastocatellia>Chloracidobacterium | GCF_018304765.1 | Chloracidobacterium_aggregatum | J8C05_RS05820 | 1401266..1404589 | 2851959 | Swt1-family-HEPN-domain-containing-protein-[Chloracidobacterium-aggregatum] |
| WP_246604568.1 | DUF499_1.RM.like.systems | ABhydrolase-><-?\|\|TUDOR+SF1-SF2-Helicase+iREase+CTD+REase->METHYLASE->HEPN+DUF499*->?->Response_reg+LuxR-HTH->CarboxypepD-reg-IG->BetaPropeller+ABhydrolase-> | 1107 | Bacteria>Acidobacteriota>Blastocatellia>Chloracidobacterium | GCF_018304805.1 | Chloracidobacterium_aggregatum | J8C02_RS05835 | 1402645..1405968 | 2851959 | Swt1-family-HEPN-domain-containing-protein-[Chloracidobacterium-aggregatum] |
| WP_211430024.1 | DUF499_1.RM.like.systems | Glyoxalase->?->?->?->TUDOR+SF1-SF2-Helicase+iREase+CTD+REase->METHYLASE->HEPN+DUF499*->?-><-?<-Glycos_trans_3N+Glycos_transf_3\|\|HTH+WYL->PIWI-> | 1106 | Bacteria>Acidobacteriota>Blastocatellia>Chloracidobacterium | GCF_018304825.1 | Chloracidobacterium_validum | J8C06_RS13915 | 660771..664091 | 2821543 | Swt1-family-HEPN-domain-containing-protein-[Chloracidobacterium-validum] |
| MBK9214625.1 | DUF499_1.RM.like.systems | HTH->?->GDResolvase+Recombinase+ZnR-><-?<-SF1-SF2-Helicase+DUF1998+HKD-DNase+HKD-DNase<-?<-HEPN+DUF499*<-?<-METHYLASE<-ArdCN+MPTase<-TUDOR+SF1-SF2-Helicase+iREase+CTD+REase<-?\|\|HTH-> | 1119 | Bacteria>Acidobacteriota>Blastocatellia>Chloracidobacterium>unclassified_Chloracidobacterium | GCA_016715795.1 | Chloracidobacterium_sp. | IPM59_03360 | 738232..741591 | 2099375 | DUF499-domain-containing-protein-[Chloracidobacterium-sp.] |
| MBK9173225.1 | DUF499_1.RM.like.systems | <-ZnR+TOPRIM+D5ATPase<-XIS-HTH<-?<-HEPN+DUF499*<-?<-?<-METHYLASE<-?<-?<-ParB+HNH | 1118 | Bacteria>Acidobacteriota>Blastocatellia>Chloracidobacterium>unclassified_Chloracidobacterium | GCA_016716305.1 | Chloracidobacterium_sp. | IPM28_09505 | 1672158..1675514 | 2099375 | DUF499-domain-containing-protein-[Chloracidobacterium-sp.] |
| MCS7080483.1 | DUF499_1.RM.like.systems | METHYLASE-p->?->SF1-SF2-Helicase->?->HKD-DNase->DDE_Tnp_IS1->HEPN+DUF499*-> | 691 | Bacteria>Acidobacteriota>Blastocatellia>Chloracidobacterium>unclassified_Chloracidobacterium | GCA_025057975.1 | Chloracidobacterium_sp. | NZ585_10625 | 170541..172613 | 2099375 | DUF499-domain-containing-protein-[Chloracidobacterium-sp.] |
| MCS7081031.1 | DUF499_1.RM.like.systems | HEPN+DUF499*-><-METHYLASE-p\|\|?-><-?<-?<-?<-SAM-methylase | 1113 | Bacteria>Acidobacteriota>Blastocatellia>Chloracidobacterium>unclassified_Chloracidobacterium | GCA_025057975.1 | Chloracidobacterium_sp. | NZ585_13400 | 246..3587 | 2099375 | Swt1-family-HEPN-domain-containing-protein-[Chloracidobacterium-sp.] |
| WP_211431305.1 | DUF499_1.RM.like.systems | ABhydrolase-><-?\|\|TUDOR+SF1-SF2-Helicase+iREase+CTD+REase->METHYLASE->HEPN+DUF499*->?->Response_reg+LuxR-HTH->CarboxypepD-reg-IG->BetaPropeller+ABhydrolase-> | 1108 | Bacteria>Acidobacteriota>Blastocatellia>Chloracidobacterium>unclassified_Chloracidobacterium | GCF_018304685.1 | Chloracidobacterium_sp._D | J8C01_RS05850 | 1398986..1402312 | 2821536 | Swt1-family-HEPN-domain-containing-protein-[Chloracidobacterium-sp.-D] |
| MBL8148954.1 | DUF499_1.RM.like.systems | VirB11+MoxR-AAA->?->Radical-SAM->Radical-SAM->?-><-HEPN+DUF499*\|\|REase+HTH-> | 1094 | Bacteria>Acidobacteriota>Blastocatellia>unclassified_Blastocatellia | GCA_016789295.1 | Blastocatellia_bacterium | JNN15_03400 | 5699..8983 | 2052146 | DUF499-domain-containing-protein-[Blastocatellia-bacterium] |
| MBO0720791.1 | DUF499_1.RM.like.systems | <-HEPN+DUF499*<-METHYLASE-p | 1119 | Bacteria>Acidobacteriota>Blastocatellia>unclassified_Blastocatellia | GCA_017353965.1 | Blastocatellia_bacterium | J2P41_08210 | 1861..5220 | 2052146 | DUF499-domain-containing-protein-[Blastocatellia-bacterium] |
| MBX7223403.1 | DUF499_1.RM.like.systems | <-HEPN+DUF499*<-METHYLASE<-NUDIX<-TUDOR+SF1-SF2-Helicase+iREase+CTD+REase | 1127 | Bacteria>Acidobacteriota>Blastocatellia>unclassified_Blastocatellia | GCA_019695185.1 | Blastocatellia_bacterium | K1Y36_26050 | 326..3709 | 2052146 | DUF499-domain-containing-protein-[Blastocatellia-bacterium] |
| MYA06552.1 | DUF499_1.RM.like.systems | <-Radical-SAM+TRAM<-?<-?\|\|?->TUDOR+SF1-SF2-Helicase+iREase+CTD+REase->HEPN+DUF499*-> | 1134 | Bacteria>Acidobacteriota>Holophagae>Holophagales>unclassified_Holophagales | GCA_009838785.1 | Holophagales_bacterium | F4Y32_01005 | 72319..75723 | 2478486 | ATP-binding-protein-[Holophagales-bacterium] |
| QQR73412.1 | DUF499_1.RM.like.systems | STAS+HisKin->?->?->METHYLASE->OLD-ABC->TOPRIM->HEPN+DUF499*-><-?<-Aminotran_1_2<-?<-?\|\|ACET-> | 1117 | Bacteria>Acidobacteriota>Holophagae>Holophagales>unclassified_Holophagales | GCA_016699405.1 | Holophagales_bacterium | IPJ17_18310 | 4435335..4438688 | 2478486 | DUF499-domain-containing-protein-[Holophagales-bacterium] |
| MBE0656246.1 | DUF499_1.RM.like.systems | <-HEPN+DUF499*<-METHYLASE<-sGTP<-HKD-DNase<-?<-TUDOR+SF1-SF2-Helicase+iREase+CTD+REase | 1127 | Bacteria>Acidobacteriota>Terriglobia>Bryobacterales>Bryobacteraceae | GCA_014860455.1 | Bryobacteraceae_bacterium | IH602_01075 | 1306..4689 | 2212468 | DUF499-domain-containing-protein-[Bryobacteraceae-bacterium] |
| MBZ2178820.1 | DUF499_1.RM.like.systems | TUDOR+SF1-SF2-Helicase+iREase+CTD+REase->?->METHYLASE->HEPN+DUF499*->MerR-HTH->?->KAP-NTPase->?->Cluster2725_2clades->Terminase-> | 1118 | Bacteria>Acidobacteriota>Terriglobia>Bryobacterales>Bryobacteraceae | GCA_019931245.1 | Bryobacteraceae_bacterium_CoA2_C42 | K7J47_14040 | 14682..18038 | 2967959 | DUF499-domain-containing-protein-[Acidobacteria-bacterium] |
| MCL6508118.1 | DUF499_1.RM.like.systems | <-HEPN+DUF499*<-METHYLASE-p | 671 | Bacteria>Acidobacteriota>Terriglobia>Bryobacterales>Bryobacteraceae | GCA_023511775.1 | Bryobacteraceae_bacterium | K6T59_13960 | 1..2014 | 2212468 | DUF499-domain-containing-protein-[Bryobacteraceae-bacterium] |
| SPE35562.1 | DUF499_1.RM.like.systems | METHYLASE->METHYLASE-p-><-?\|\|ParB->PolB->HEPN->HEPN+DUF499*->DUF499-p-> | 906 | Bacteria>Acidobacteriota>Terriglobia>Bryobacterales>Solibacteraceae | GCA_900290325.1 | Candidatus_Sulfopaludibacter_sp._SbA6 | SBA6_470068 | 62270..64990 | 2043166 | conserved-hypothetical-protein-[Candidatus-Sulfopaludibacter-sp.-SbA6] |
| MBK9166737.1 | DUF499_1.RM.like.systems | Schlafen+HATPase+HTH->METHYLASE-p-><-rve+rve_3<-HTH_Tnp_1\|\|METHYLASE-p->ZDNA-HTH->HEPN+DUF499*->?->?->ABhydrolase-> | 1121 | Bacteria>Acidobacteriota>Terriglobia>Bryobacterales>unclassified_Bryobacterales | GCA_016716385.1 | Bryobacterales_bacterium | IPM24_04655 | 396638..400003 | 2026791 | DUF499-domain-containing-protein-[Bryobacterales-bacterium] |
| MCL4784604.1 | DUF499_1.RM.like.systems | <-HTH\|\|?->?->?-><-HEPN+DUF499*<-METHYLASE<-?<-TUDOR+SF1-SF2-Helicase+iREase+CTD+REase | 1109 | Bacteria>Acidobacteriota>Terriglobia>Bryobacterales>unclassified_Bryobacterales | GCA_023384735.1 | Bryobacterales_bacterium | KJZ70_16335 | 4638..7967 | 2026791 | DUF499-domain-containing-protein-[Bryobacterales-bacterium] |
| MCU1329937.1 | DUF499_1.RM.like.systems | <-HEPN+DUF499*<-METHYLASE<-TUDOR+SF1-SF2-Helicase+iREase+CTD+REase | 1116 | Bacteria>Acidobacteriota>Terriglobia>Bryobacterales>unclassified_Bryobacterales | GCA_025462945.1 | Bryobacterales_bacterium | JWN34_5307 | 1570..4920 | 2026791 | ATPase-superfamily-[Bryobacterales-bacterium] |
| MCY4596713.1 | DUF499_1.RM.like.systems | TUDOR+SF1-SF2-Helicase+iREase+CTD+REase->HEPN+DUF499*->?-><-METHYLASE-p | 1110 | Bacteria>Acidobacteriota>Terriglobia>Bryobacterales>unclassified_Bryobacterales | GCA_026713455.1 | Bryobacterales_bacterium | OXC19_18175 | 3853..7185 | 2026791 | MAG:-Swt1-family-HEPN-domain-containing-protein-[Bryobacterales-bacterium] |
| MDE0101184.1 | DUF499_1.RM.like.systems | PadR-HTH-><-?\|\|N6_MTase-><-HEPN+DUF499*<-HEPN<-Polbeta<-METHYLASE<-HEPN<-TUDOR+SF1-SF2-Helicase+iREase+CTD+REase\|\|REase-> | 1110 | Bacteria>Acidobacteriota>Terriglobia>Bryobacterales>unclassified_Bryobacterales | GCA_028819235.1 | Bryobacterales_bacterium | OXN89_02285 | 6150..9482 | 2026791 | MAG:-Swt1-family-HEPN-domain-containing-protein-[Bryobacterales-bacterium] |
| MBP7799276.1 | DUF499_1.RM.like.systems | <-HEPN+DUF499*<-PIN<-OLD-ABC<-METHYLASE<-TOPRIM<-OLD-ABC | 1114 | Bacteria>Acidobacteriota>Thermoanaerobaculia>Thermoanaerobaculales>Thermoanaerobaculaceae | GCA_018000955.1 | Thermoanaerobaculaceae_bacterium | KA072_11695 | 22806..26150 | 2699758 | DUF499-domain-containing-protein-[Thermoanaerobaculaceae-bacterium] |
| WP_038049904.1 | DUF499_1.RM.like.systems | Radical-SAM-><-?<-?<-ORC-CDC6+REase\|\|?-><-HEPN+DUF499*<-METHYLASE<-MORC+DUF3684<-TUDOR+SF1-SF2-Helicase+iREase+CTD+REase\|\|ORC-CDC6+REase->?->DUF433-> | 1117 | Bacteria>Acidobacteriota>Thermoanaerobaculia>Thermoanaerobaculales>Thermoanaerobaculaceae | GCF_000687145.1 | Thermoanaerobaculum_aquaticum | EG19_RS09435 | 122506..125859 | 1312852 | Swt1-family-HEPN-domain-containing-protein-[Thermoanaerobaculum-aquaticum] |
| KAB2964430.1 | DUF499_1.RM.like.systems | REase->TUDOR+SF1-SF2-Helicase+iREase+CTD+REase->ParB+HNH->HEPN->METHYLASE->?->HEPN+DUF499*->HNH-> | 1109 | Bacteria>Acidobacteriota>Thermoanaerobaculia>unclassified_Thermoanaerobaculia | GCA_009026165.1 | Thermoanaerobaculia_bacterium | F9K16_04350 | 12106..15435 | 2651171 | MAG:-ATP-binding-protein-[Thermoanaerobaculia-bacterium] |
| MCG3192277.1 | DUF499_1.RM.like.systems | TUDOR+SF1-SF2-Helicase+iREase+CTD+REase->OLD-ABC->TOPRIM->METHYLASE->PHP+SbcC->Schlafen+HATPase+HTH->HEPN+DUF499*->?->?->PAIREDC-HTH-> | 1114 | Bacteria>Acidobacteriota>Thermoanaerobaculia>unclassified_Thermoanaerobaculia | GCA_022071945.1 | Thermoanaerobaculia_bacterium | DIJKHBIC_01515 | 140917..144261 | 2651171 | hypothetical-protein-[Thermoanaerobaculia-bacterium] |
| MCK6683888.1 | DUF499_1.RM.like.systems | TUDOR+SF1-SF2-Helicase+iREase+CTD+REase->?->?->MACRODOMAIN->METHYLASE->HEPN+DUF499*->?->OLD-ABC+TOPRIM->DUF932->?->ASCH+DUF3780-> | 1093 | Bacteria>Acidobacteriota>Thermoanaerobaculia>unclassified_Thermoanaerobaculia | GCA_023151375.1 | Thermoanaerobaculia_bacterium | L6R30_15920 | 12714..15995 | 2651171 | DUF499-domain-containing-protein-[Thermoanaerobaculia-bacterium] |
| ODS52398.1 | DUF499_1.RM.like.systems | <-MethyltransfD12<-ParB<-HEPN+DUF499*<-METHYLASE<-Schlafen+HATPase+HTH<-TUDOR+SF1-SF2-Helicase+iREase+CTD+REase<-?\|\|?->Peptidase_S41-> | 1114 | Bacteria>Acidobacteriota>unclassified_Acidobacteriota | GCA_001724025.1 | Acidobacteria_bacterium_SCN_69-37 | ABS36_17605 | 152750..156094 | 1660160 | AAA+-family-ATPase-[Acidobacteria-bacterium-SCN-69-37] |
| PYQ27742.1 | DUF499_1.RM.like.systems | ABhydrolase-><-?<-HTH\|\|?-><-?<-HEPN+DUF499*\|\|HD->DUF4065-><-METHYLASE<-TUDOR+SF1-SF2-Helicase+iREase+CTD+REase | 1105 | Bacteria>Acidobacteriota>unclassified_Acidobacteriota | GCA_003223645.1 | Acidobacteriota_bacterium | DMF56_19255 | 118621..121938 | 1978231 | AAA+-family-ATPase-[Acidobacteria-bacterium] |
| RIK10616.1 | DUF499_1.RM.like.systems | DOC->?->MPTase->METHYLASE->HEPN+DUF499*-><-?<-Schlafen<-?<-tRNA<-tRNA<-?\|\|tRNA-><-METHYLASE-p | 1113 | Bacteria>Acidobacteriota>unclassified_Acidobacteriota | GCA_003577105.1 | Acidobacteriota_bacterium | DCC49_02885 | 59433..62774 | 1978231 | AAA+-family-ATPase-[Acidobacteria-bacterium] |
| RMH20600.1 | DUF499_1.RM.like.systems | METHYLASE-p->DUF433->PIN->HEPN+DUF499*-> | 1096 | Bacteria>Acidobacteriota>unclassified_Acidobacteriota | GCA_003697015.1 | Acidobacteriota_bacterium | D6696_07825 | 1571..4858 | 1978231 | ATP-binding-protein,-partial-[Acidobacteria-bacterium] |
| TAM79789.1 | DUF499_1.RM.like.systems | PadR-HTH->?->?-><-?<-HEPN+DUF499*<-DUF433<-METHYLASE<-?<-TUDOR+SF1-SF2-Helicase+iREase+CTD+REase<-MPTase<-HTH | 1104 | Bacteria>Acidobacteriota>unclassified_Acidobacteriota | GCA_004298155.1 | Acidobacteriota_bacterium | EPN47_18525 | 398399..401713 | 1978231 | DUF499-domain-containing-protein-[Acidobacteria-bacterium] |
| MXW70175.1 | DUF499_1.RM.like.systems | TUDOR+SF1-SF2-Helicase+iREase+CTD+REase->HEPN+DUF499*-> | 1106 | Bacteria>Acidobacteriota>unclassified_Acidobacteriota | GCA_009835985.1 | Acidobacteriota_bacterium | F4Z74_01830 | 14208..17528 | 1978231 | ATP-binding-protein-[Acidobacteria-bacterium] |
| MXW37283.1 | DUF499_1.RM.like.systems | TUDOR+SF1-SF2-Helicase+iREase+CTD+REase-p->HEPN+DUF499*-> | 1106 | Bacteria>Acidobacteriota>unclassified_Acidobacteriota | GCA_009836105.1 | Acidobacteriota_bacterium | F4Z65_03330 | 485..3805 | 1978231 | ATP-binding-protein-[Acidobacteria-bacterium] |
| MYD85827.1 | DUF499_1.RM.like.systems | <-HEPN+DUF499*<-?<-?<-METHYLASE | 1107 | Bacteria>Acidobacteriota>unclassified_Acidobacteriota | GCA_009839145.1 | Acidobacteriota_bacterium | F4Y14_06400 | 201..3524 | 1978231 | ATP-binding-protein-[Acidobacteria-bacterium] |
| MYD70260.1 | DUF499_1.RM.like.systems | <-HEPN+DUF499*<-TUDOR+SF1-SF2-Helicase+iREase+CTD+REase\|\|Peptidase-> | 1104 | Bacteria>Acidobacteriota>unclassified_Acidobacteriota | GCA_009841675.1 | Acidobacteriota_bacterium | F4W89_05855 | 745..4059 | 1978231 | ATP-binding-protein-[Acidobacteria-bacterium] |
| HHN74473.1 | DUF499_1.RM.like.systems | <-MORC+DUF3684+REase<-HEPN+DUF499* | 1091 | Bacteria>Acidobacteriota>unclassified_Acidobacteriota | GCA_011374785.1 | Acidobacteriota_bacterium | ENK10_04515 | 4666..7941 | 1978231 | ATP-binding-protein-[Acidobacteria-bacterium] |
| NLT66835.1 | DUF499_1.RM.like.systems | METHYLASE-p->HEPN+DUF499*-> | 772 | Bacteria>Acidobacteriota>unclassified_Acidobacteriota | GCA_012719355.1 | Acidobacteriota_bacterium | GXX84_09545 | 1125..3439 | 1978231 | ATP-binding-protein-[Acidobacteria-bacterium] |
| NMD09846.1 | DUF499_1.RM.like.systems | <-HEPN+DUF499*<-METHYLASE-p | 545 | Bacteria>Acidobacteriota>unclassified_Acidobacteriota | GCA_012797375.1 | Acidobacteriota_bacterium | GYA74_01585 | 1..1635 | 1978231 | ATP-binding-protein-[Acidobacteriota-bacterium] |
| NOT26039.1 | DUF499_1.RM.like.systems | <-METHYLASE-p<-?<-?<-?\|\|?->?-><-HEPN+DUF499* | 1117 | Bacteria>Acidobacteriota>unclassified_Acidobacteriota | GCA_013140775.1 | Acidobacteriota_bacterium | HOP16_08040 | 9763..13116 | 1978231 | ATP-binding-protein-[Acidobacteria-bacterium] |
| MBA4124439.1 | DUF499_1.RM.like.systems | DDE_Tnp_1_assoc+DDE_Tnp_1->CHTH->?->TUDOR+SF1-SF2-Helicase+iREase+CTD+REase->ParB+HNH->METHYLASE->HEPN+DUF499*->?->?->BirA-HTH-><-35exo<-Phage_integrase | 1091 | Bacteria>Acidobacteriota>unclassified_Acidobacteriota | GCA_013820745.1 | Acidobacteriota_bacterium | HOX72_18505 | 11663..14938 | 1978231 | DUF499-domain-containing-protein-[Acidobacteria-bacterium] |
| MBI1792208.1 | DUF499_1.RM.like.systems | TUDOR+SF1-SF2-Helicase+iREase+CTD+REase->Schlafen+HATPase+HTH->METHYLASE->PolB->HEPN+DUF499*->OLD-ABC+TOPRIM-> | 1093 | Bacteria>Acidobacteriota>unclassified_Acidobacteriota | GCA_016178945.1 | Acidobacteriota_bacterium | HYR60_32190 | 8610..11891 | 1978231 | ATP-binding-protein-[Acidobacteria-bacterium] |
| MBI4902187.1 | DUF499_1.RM.like.systems | TUDOR+SF1-SF2-Helicase+iREase+CTD+REase->ParB+HNH->METHYLASE->OLD-ABC->TOPRIM->HEPN+DUF499*->Cluster2725_2clades->Terminase-> | 1115 | Bacteria>Acidobacteriota>unclassified_Acidobacteriota | GCA_016213775.1 | Acidobacteriota_bacterium | HY820_01040 | 17603..20950 | 1978231 | ATP-binding-protein-[Acidobacteria-bacterium] |
| MBM3792128.1 | DUF499_1.RM.like.systems | METHYLASE-p->HEPN+DUF499*-> | 464 | Bacteria>Acidobacteriota>unclassified_Acidobacteriota | GCA_016871075.1 | Acidobacteriota_bacterium | FJW35_17505 | 483..1875 | 1978231 | ATP-binding-protein-[Acidobacteriota-bacterium] |
| MBP7569108.1 | DUF499_1.RM.like.systems | Cas_Cmr3->?->Cas_Cmr5->?->CARF+HEPN->HEPN+DUF499*->Schlafen-> | 1117 | Bacteria>Acidobacteriota>unclassified_Acidobacteriota | GCA_017998655.1 | Acidobacteriota_bacterium | KBA95_03500 | 22125..25478 | 1978231 | DUF499-domain-containing-protein-[Acidobacteria-bacterium] |
| MBP7777875.1 | DUF499_1.RM.like.systems | TUDOR+SF1-SF2-Helicase+iREase+CTD+REase->DUF932->METHYLASE->HEPN+DUF499*->?->Pkinase->ParB+HNH->REase->?->HicB_lk_antitox-> | 1114 | Bacteria>Acidobacteriota>unclassified_Acidobacteriota | GCA_018000455.1 | Acidobacteriota_bacterium | KA371_12150 | 120064..123408 | 1978231 | DUF499-domain-containing-protein-[Acidobacteria-bacterium] |
| MBS1816795.1 | DUF499_1.RM.like.systems | <-XIS-HTH<-?\|\|HTH+PolB->TUDOR+SF1-SF2-Helicase+iREase+CTD+REase->METHYLASE->?->HEPN+DUF499*->?-><-OLD-ABC+TOPRIM<-?<-?<-AbiEi_4\|\|REase-> | 1111 | Bacteria>Acidobacteriota>unclassified_Acidobacteriota | GCA_018268955.1 | Acidobacteriota_bacterium | JSU08_02600 | 186783..190118 | 1978231 | DUF499-domain-containing-protein-[Acidobacteria-bacterium] |
| MBS1819352.1 | DUF499_1.RM.like.systems | <-SF1-SF2-Helicase<-OLD-ABC+TOPRIM<-HEPN+DUF499*<-METHYLASE-p | 1115 | Bacteria>Acidobacteriota>unclassified_Acidobacteriota | GCA_018268955.1 | Acidobacteriota_bacterium | JSU08_15565 | 76293..79640 | 1978231 | DUF499-domain-containing-protein-[Acidobacteria-bacterium] |
| MBS1794111.1 | DUF499_1.RM.like.systems | <-Phage_integrase\|\|TUDOR+SF1-SF2-Helicase+iREase+CTD+REase->?->METHYLASE->?->HEPN+DUF499*->?->?->REase->?->SF1-SF2-Helicase-> | 1100 | Bacteria>Acidobacteriota>unclassified_Acidobacteriota | GCA_018269075.1 | Acidobacteriota_bacterium | JSS81_09670 | 12622..15924 | 1978231 | DUF499-domain-containing-protein-[Acidobacteria-bacterium] |
| MCC6131982.1 | DUF499_1.RM.like.systems | <-HEPN+DUF499*<-Schlafen+HATPase+HTH<-PHP+SbcC<-METHYLASE<-TOPRIM<-OLD-ABC<-TUDOR+SF1-SF2-Helicase+iREase+CTD+REase | 1114 | Bacteria>Acidobacteriota>unclassified_Acidobacteriota | GCA_020845555.1 | Acidobacteriota_bacterium | IT186_18830 | 2559..5903 | 1978231 | DUF499-domain-containing-protein-[Acidobacteria-bacterium] |
| MCI0388810.1 | DUF499_1.RM.like.systems | HTH+PolB->HEPN->HEPN+DUF499*-><-Peptidase | 1116 | Bacteria>Acidobacteriota>unclassified_Acidobacteriota | GCA_022614935.1 | Acidobacteriota_bacterium | MOB07_08610 | 4163..7513 | 1978231 | Swt1-family-HEPN-domain-containing-protein-[Acidobacteria-bacterium] |
| MCI0622286.1 | DUF499_1.RM.like.systems | <-REase<-HEPN+DUF499*<-?<-DUF488-l<-?<-METHYLASE<-HEPN<-METHYLASE-p | 1117 | Bacteria>Acidobacteriota>unclassified_Acidobacteriota | GCA_022615935.1 | Acidobacteriota_bacterium | L0387_11575 | 924..4277 | 1978231 | DUF499-domain-containing-protein-[Acidobacteria-bacterium] |
| MCI0623777.1 | DUF499_1.RM.like.systems | HTH-><-HEPN+DUF499*<-METHYLASE<-PIN<-?<-STAS+HisKin<-TUDOR+SF1-SF2-Helicase+iREase+CTD+REase<-XIS-HTH | 1110 | Bacteria>Acidobacteriota>unclassified_Acidobacteriota | GCA_022615935.1 | Acidobacteriota_bacterium | L0387_19325 | 508..3840 | 1978231 | DUF499-domain-containing-protein-[Acidobacteria-bacterium] |
| MCL5262739.1 | DUF499_1.RM.like.systems | <-HEPN+DUF499*<-METHYLASE<-?<-TUDOR+SF1-SF2-Helicase+iREase+CTD+REase-p<-REase<-HKD-DNase+SF1-SF2-Helicase+TRD | 1103 | Bacteria>Acidobacteriota>unclassified_Acidobacteriota | GCA_023480755.1 | Acidobacteriota_bacterium | M1568_00545 | 1..3309 | 1978231 | Swt1-family-HEPN-domain-containing-protein-[Acidobacteria-bacterium] |
| MCY3930234.1 | DUF499_1.RM.like.systems | <-Phage_int_SAM_3+Phage_integrase<-Phage_int_SAM_3+Phage_integrase\|\|METHYLASE-><-HEPN+DUF499*<-TUDOR+SF1-SF2-Helicase+iREase+CTD+REase<-?<-OLD-ABC-p | 1112 | Bacteria>Acidobacteriota>unclassified_Acidobacteriota | GCA_026707325.1 | Acidobacteriota_bacterium | OXH70_00800 | 52234..55572 | 1978231 | MAG:-DUF499-domain-containing-protein-[Acidobacteria-bacterium] |
| MDD8029960.1 | DUF499_1.RM.like.systems | METHYLASE->HEPN+DUF499*-><-?<-?<-Cysteine_desulfurase\|\|Peptidase_M24-> | 1114 | Bacteria>Acidobacteriota>unclassified_Acidobacteriota | GCA_028749155.1 | Acidobacteriota_bacterium | PHE62_10480 | 3051..6395 | 1978231 | MAG:-Swt1-family-HEPN-domain-containing-protein-[Acidobacteriota-bacterium] |
| MDE2851190.1 | DUF499_1.RM.like.systems | METHYLASE->?-><-HEPN+DUF499*<-TUDOR+SF1-SF2-Helicase+iREase+CTD+REase<-?<-?<-?<-?<-MqsA_antitoxin+HTH_3 | 1104 | Bacteria>Acidobacteriota>unclassified_Acidobacteriota | GCA_028874635.1 | Acidobacteriota_bacterium | OXP74_11315 | 2020..5334 | 1978231 | MAG:-Swt1-family-HEPN-domain-containing-protein-[Acidobacteriota-bacterium] |
| MDE2692299.1 | DUF499_1.RM.like.systems | <-METHYLASE\|\|Phage_integrase->Phage_int_SAM_3+Phage_integrase->HEPN->METHYLASE->?-><-HEPN+DUF499*<-TUDOR+SF1-SF2-Helicase+iREase+CTD+REase<-?\|\|Arm-DNA-bind_3+Phage_integrase-> | 1104 | Bacteria>Acidobacteriota>unclassified_Acidobacteriota | GCA_028875725.1 | Acidobacteriota_bacterium | OXI49_17510 | 451971..455285 | 1978231 | MAG:-Swt1-family-HEPN-domain-containing-protein-[Acidobacteriota-bacterium] |
| MDE3262363.1 | DUF499_1.RM.like.systems | <-ArdCN+MPTase<-?<-?<-XIS-HTH<-?\|\|TUDOR+SF1-SF2-Helicase+iREase+CTD+REase->HEPN+DUF499*-> | 1106 | Bacteria>Acidobacteriota>unclassified_Acidobacteriota | GCA_028877265.1 | Acidobacteriota_bacterium | OYL41_10335 | 11539..14859 | 1978231 | MAG:-Swt1-family-HEPN-domain-containing-protein-[Acidobacteriota-bacterium] |
| MCL4821144.1 | DUF499_1.RM.like.systems | STYKinase-><-HEPN+DUF499*<-?<-ORC-CDC6<-ParA-Soj-PloopNTPase<-HEPN<-METHYLASE<-PIN | 1116 | Bacteria>Acidobacteriota>Vicinamibacteria>unclassified_Vicinamibacteria | GCA_023384615.1 | Vicinamibacteria_bacterium | KJ067_18565 | 2540..5890 | 2793154 | DUF499-domain-containing-protein-[Vicinamibacteria-bacterium] |
| MCL4821456.1 | DUF499_1.RM.like.systems | TUDOR+SF1-SF2-Helicase+iREase+CTD+REase-p->XIS-HTH+RVT->METHYLASE->OLD-ABC->PIN->HEPN+DUF499*->SF1-SF2-Helicase-><-?<-?\|\|Pkinase-> | 1115 | Bacteria>Acidobacteriota>Vicinamibacteria>unclassified_Vicinamibacteria | GCA_023384615.1 | Vicinamibacteria_bacterium | KJ067_20160 | 8492..11839 | 2793154 | DUF499-domain-containing-protein-[Vicinamibacteria-bacterium] |
| NSW76333.1 | DUF499_1.RM.like.systems | METHYLASE->HEPN+DUF499*->PolB-> | 1120 | Bacteria>Atribacterota>Atribacteria>unclassified_Atribacteria | GCA_013314795.1 | Candidatus_Atribacteria_bacterium | HPY68_06090 | 2996..6358 | 2053509 | ATP-binding-protein-[Candidatus-Atribacteria-bacterium] |
| MBE3120494.1 | DUF499_1.RM.like.systems | <-KAP-NTPase<-HEPN+DUF499*<-METHYLASE<-Schlafen+HATPase+HTH | 1116 | Bacteria>Atribacterota>Atribacteria>unclassified_Atribacteria | GCA_014894575.1 | Candidatus_Atribacteria_bacterium | IMZ50_17295 | 4085..7435 | 2053509 | DUF499-domain-containing-protein-[Candidatus-Atribacteria-bacterium] |
| KAA0227462.1 | DUF499_1.RM.like.systems | TUDOR+SF1-SF2-Helicase+iREase+CTD+REase->ArdCN+MPTase-><-PIN<-?\|\|HEPN+DUF499*-><-?<-STYKIN<-?<-?\|\|?->HAD-> | 1113 | Bacteria>Bacteria_incertae_sedis>Bacteria_candidate_phyla>candidate_division_KSB1>unclassified_candidate_division_KSB1 | GCA_008363195.1 | candidate_division_KSB1_bacterium | EDS67_15790 | 88921..92262 | 2172550 | ATP-binding-protein-[candidate-division-KSB1-bacterium] |
| PWB47246.1 | DUF499_1.RM.like.systems | TUDOR+SF1-SF2-Helicase+iREase+CTD+REase->Schlafen+HATPase+HTH->METHYLASE->HEPN+DUF499*->?->?-><-?<-REase<-?<-CopG_antitoxin | 1102 | Bacteria>Bacteria_incertae_sedis>Bacteria_candidate_phyla>candidate_division_NC10>unclassified_candidate_division_NC10 | GCA_003104975.1 | candidate_division_NC10_bacterium | C3F12_04520 | 184009..187317 | 2072417 | AAA+-family-ATPase-[candidate-division-NC10-bacterium] |
| MDE2058883.1 | DUF499_1.RM.like.systems | <-HEPN+DUF499*<-?<-METHYLASE<-TOPRIM<-OLD-ABC<-ORC-CDC6+REase<-TUDOR+SF1-SF2-Helicase+iREase+CTD+REase | 1122 | Bacteria>Bacteria_incertae_sedis>Bacteria_candidate_phyla>candidate_division_NC10>unclassified_candidate_division_NC10 | GCA_028864815.1 | candidate_division_NC10_bacterium | KGJ27_04040 | 4506..7874 | 2072417 | MAG:-DUF499-domain-containing-protein-[candidate-division-NC10-bacterium] |
| MCX6842110.1 | DUF499_1.RM.like.systems | NTP_transf->TUDOR+SF1-SF2-Helicase+iREase+CTD+REase->Schlafen+HATPase+HTH->?->METHYLASE-p->METHYLASE->HEPN+DUF499*-> | 1030 | Bacteria>Bacteria_incertae_sedis>Bacteria_candidate_phyla>candidate_division_WOR-3 | GCA_026397595.1 | candidate_division_WOR-3_bacterium | NTX53_07520 | 11691..14783 | 2052148 | MAG:-Swt1-family-HEPN-domain-containing-protein-[candidate-division-WOR-3-bacterium] |
| OQB57508.1 | DUF499_1.RM.like.systems | TUDOR+SF1-SF2-Helicase+iREase+CTD+REase->Schlafen+HATPase+HTH->METHYLASE->HEPN+DUF499*-><-?<-?<-Cysteine_desulfurase\|\|Peptidase_M24-> | 1114 | Bacteria>Bacteria_incertae_sedis>Bacteria_candidate_phyla>Candidatus_Aminicenantes>unclassified_Candidatus_Aminicenantes | GCA_002070465.1 | Candidatus_Aminicenantes_bacterium_ADurb.Bin147 | BWX98_01441 | 17398..20742 | 1852831 | hypothetical-protein-BWX98_01441-[Candidatus-Aminicenantes-bacterium-ADurb.Bin147] |
| QAA75938.1 | DUF499_1.RM.like.systems | <-RNaseH<-?<-?<-?<-?<-?<-HEPN+DUF499*<-?<-METHYLASE<-TOPRIM<-OLD-ABC<-TUDOR+SF1-SF2-Helicase+iREase+CTD+REase | 1113 | Bacteria>Bacteria_incertae_sedis>Bacteria_candidate_phyla>Candidatus_Bipolaricaulota>Bipolaricaulis | GCA_004102645.1 | Candidatus_Bipolaricaulis_sibiricus | BIP78_0170 | 158925..162266 | 2501609 | hypothetical-protein-BIP78_0170-[Candidatus-Bipolaricaulis-sibiricus] |
| MDD5263468.1 | DUF499_1.RM.like.systems | <-HEPN+DUF499*\|\|?-><-OLD-ABC<-METHYLASE<-Schlafen+HATPase+HTH<-TUDOR+SF1-SF2-Helicase+iREase+CTD+REase-p | 1114 | Bacteria>Bacteria_incertae_sedis>Bacteria_candidate_phyla>Candidatus_Bipolaricaulota>Bipolaricaulis | GCA_028713635.1 | Candidatus_Bipolaricaulis_sp. | PHU43_01355 | 30296..33640 | 2814139 | MAG:-Swt1-family-HEPN-domain-containing-protein-[Candidatus-Bipolaricaulis-sp.] |
| WP_122031531.1 | DUF499_1.RM.like.systems | TUDOR+SF1-SF2-Helicase+iREase+CTD+REase->ORC-CDC6+REase->METHYLASE->OLD-ABC->PIN->HEPN+DUF499*->Radical-SAM+TRAM->PfkB->?->35exo+SF1-SF2-Helicase->?-><-NTP_transf_2 | 1110 | Bacteria>Bacteria_incertae_sedis>Bacteria_candidate_phyla>Candidatus_Bipolaricaulota>Bipolaricaulis | GCF_900465355.1 | Candidatus_Bipolaricaulis_anaerobius | BARAN1_RS05370 | 1116813..1120145 | 2026885 | Swt1-family-HEPN-domain-containing-protein-[Candidatus-Bipolaricaulis-anaerobius] |
| KUK26942.1 | DUF499_1.RM.like.systems | <-DUF499-p<-HEPN+DUF499*<-PolB-NTase<-METHYLASE-p<-HEPN<-PolB<-METHYLASE-p<-PIN | 493 | Bacteria>Bacteria_incertae_sedis>Bacteria_candidate_phyla>Candidatus_Bipolaricaulota>unclassified_Candidatus_Bipolaricaulota | GCA_001508155.1 | Acetothermia_bacterium_64_32 | XD60_0831 | 38456..39937 | 1635286 | Uncharacterized-protein-XD60_0831-[Acetothermia-bacterium-64_32] |
| HAF70679.1 | DUF499_1.RM.like.systems | <-DUF499-p<-HEPN+DUF499*<-PolB-NTase<-METHYLASE-p<-HEPN<-PolB<-METHYLASE-p<-PIN | 515 | Bacteria>Bacteria_incertae_sedis>Bacteria_candidate_phyla>Candidatus_Bipolaricaulota>unclassified_Candidatus_Bipolaricaulota | GCA_003445465.1 | Candidatus_Acetothermia_bacterium | DCL37_04935 | 22341..23888 | 2053493 | hypothetical-protein-[Candidatus-Acetothermia-bacterium] |
| HDN19517.1 | DUF499_1.RM.like.systems | TUDOR+SF1-SF2-Helicase+iREase+CTD+REase->OLD-ABC+TOPRIM->METHYLASE->PolB->HEPN+DUF499*->MazE_antitoxin->PIN-> | 1111 | Bacteria>Bacteria_incertae_sedis>Bacteria_candidate_phyla>Candidatus_Bipolaricaulota>unclassified_Candidatus_Bipolaricaulota | GCA_011042375.1 | Candidatus_Acetothermia_bacterium | ENF46_00270 | 10022..13357 | 2053493 | ATP-binding-protein-[Candidatus-Acetothermia-bacterium] |
| MBK8229368.1 | DUF499_1.RM.like.systems | <-PIN<-HTH+MPTase\|\|?->?->HTH+UbE1->TUDOR+SF1-SF2-Helicase+iREase+CTD+REase->HEPN+DUF499*->PAIREDC-HTH->DDE_3->DUF499-p->?->?->LZ_Tnp_IS66+zf-IS66+DDE_Tnp_IS66+DDE_Tnp_IS66_C-> | 799 | Bacteria>Bacteria_incertae_sedis>Bacteria_candidate_phyla>Candidatus_Eisenbacteria>unclassified_Candidatus_Eisenbacteria | GCA_016712505.1 | Candidatus_Eisenbacteria_bacterium | IPK72_02030 | 519396..521795 | 2212470 | DUF499-domain-containing-protein-[Candidatus-Eisenbacteria-bacterium] |
| MXV83700.1 | DUF499_1.RM.like.systems | <-URI-DNase<-?<-METHYLASE<-REase<-REase<-REase<-HEPN+DUF499*<-PIN | 1088 | Bacteria>Bacteria_incertae_sedis>Bacteria_candidate_phyla>Candidatus_Poribacteria>unclassified_Candidatus_Poribacteria | GCA_009835485.1 | Candidatus_Poribacteria_bacterium | F4Z98_10065 | 15224..18490 | 2026781 | ATP-binding-protein-[Candidatus-Poribacteria-bacterium] |
| MYB95133.1 | DUF499_1.RM.like.systems | <-METHYLASE<-REase+HTH<-REase+HTH<-HEPN+DUF499*<-?<-?<-SF1-SF2-Helicase+REase<-ABC-ATPase+SbcC | 1108 | Bacteria>Bacteria_incertae_sedis>Bacteria_candidate_phyla>Candidatus_Poribacteria>unclassified_Candidatus_Poribacteria | GCA_009838805.1 | Candidatus_Poribacteria_bacterium | F4Y43_14870 | 2954..6280 | 2026781 | ATP-binding-protein-[Candidatus-Poribacteria-bacterium] |
| MYC78293.1 | DUF499_1.RM.like.systems | <-HEPN<-URI-DNase<-?<-REase<-REase<-REase<-HEPN+DUF499*<-PIN<-HTH+MPTase<-?<-?<-?<-SF1-SF2-Helicase+REase | 1093 | Bacteria>Bacteria_incertae_sedis>Bacteria_candidate_phyla>Candidatus_Poribacteria>unclassified_Candidatus_Poribacteria | GCA_009841255.1 | Candidatus_Poribacteria_bacterium | F4X10_21230 | 47302..50583 | 2026781 | ATP-binding-protein-[Candidatus-Poribacteria-bacterium] |
| MYK17049.1 | DUF499_1.RM.like.systems | HEPN+DUF499*->REase+HTH->REase->METHYLASE->URI-DNase-> | 1090 | Bacteria>Bacteria_incertae_sedis>Bacteria_candidate_phyla>Candidatus_Poribacteria>unclassified_Candidatus_Poribacteria | GCA_009845505.1 | Candidatus_Poribacteria_bacterium | F4055_02625 | 1958..5230 | 2026781 | ATP-binding-protein-[Candidatus-Poribacteria-bacterium] |
| MBI5115316.1 | DUF499_1.RM.like.systems | <-SF1-SF2-Helicase<-?<-HEPN+REase<-?<-PIN<-HTH+MPTase<-HEPN+DUF499*<-METHYLASE-p | 1118 | Bacteria>Bacteria_incertae_sedis>Bacteria_candidate_phyla>Candidatus_Poribacteria>unclassified_Candidatus_Poribacteria | GCA_016215705.1 | Candidatus_Poribacteria_bacterium | HZA56_02480 | 5349..8705 | 2026781 | ATP-binding-protein-[Candidatus-Poribacteria-bacterium] |
| MCY3724179.1 | DUF499_1.RM.like.systems | PIN->HEPN+DUF499*->REase+HTH->REase->METHYLASE->?->URI-DNase-> | 1093 | Bacteria>Bacteria_incertae_sedis>Bacteria_candidate_phyla>Candidatus_Poribacteria>unclassified_Candidatus_Poribacteria | GCA_026705955.1 | Candidatus_Poribacteria_bacterium | OXG97_18330 | 699..3980 | 2026781 | MAG:-DUF499-domain-containing-protein-[Candidatus-Poribacteria-bacterium] |
| MDE0020779.1 | DUF499_1.RM.like.systems | <-METHYLASE<-HEPN+DUF499*<-?<-?<-?<-SIGMA-HTH | 1121 | Bacteria>Bacteria_incertae_sedis>Bacteria_candidate_phyla>Candidatus_Poribacteria>unclassified_Candidatus_Poribacteria | GCA_028818695.1 | Candidatus_Poribacteria_bacterium | OXT69_05310 | 103742..107107 | 2026781 | MAG:-DUF499-domain-containing-protein-[Candidatus-Poribacteria-bacterium] |
| MDE0014209.1 | DUF499_1.RM.like.systems | HTH+MPTase->PIN->HEPN+DUF499*->REase+HTH->REase->METHYLASE->URI-DNase-> | 1095 | Bacteria>Bacteria_incertae_sedis>Bacteria_candidate_phyla>Candidatus_Poribacteria>unclassified_Candidatus_Poribacteria | GCA_028818705.1 | Candidatus_Poribacteria_bacterium | OXU36_23900 | 11254..14541 | 2026781 | MAG:-DUF499-domain-containing-protein-[Candidatus-Poribacteria-bacterium] |
| MDE0088638.1 | DUF499_1.RM.like.systems | ABC-ATPase+SbcC->ABC-ATPase+SbcC->?-><-URI-DNase<-METHYLASE<-HEPN+DUF499*<-Aldedh | 1101 | Bacteria>Bacteria_incertae_sedis>Bacteria_candidate_phyla>Candidatus_Poribacteria>unclassified_Candidatus_Poribacteria | GCA_028818995.1 | Candidatus_Poribacteria_bacterium | OXU23_23155 | 9286..12591 | 2026781 | MAG:-hypothetical-protein-OXU23_23155-[Candidatus-Poribacteria-bacterium] |
| MDE0423419.1 | DUF499_1.RM.like.systems | SF1-SF2-Helicase+REase->?->?->?->?->?->HEPN+DUF499*->REase+HTH->REase->METHYLASE->URI-DNase-> | 1090 | Bacteria>Bacteria_incertae_sedis>Bacteria_candidate_phyla>Candidatus_Poribacteria>unclassified_Candidatus_Poribacteria | GCA_028820845.1 | Candidatus_Poribacteria_bacterium | OXN25_00975 | 9456..12728 | 2026781 | MAG:-DUF499-domain-containing-protein-[Candidatus-Poribacteria-bacterium] |
| MDE0468570.1 | DUF499_1.RM.like.systems | <-URI-DNase<-METHYLASE<-?<-HEPN+DUF499* | 1092 | Bacteria>Bacteria_incertae_sedis>Bacteria_candidate_phyla>Candidatus_Poribacteria>unclassified_Candidatus_Poribacteria | GCA_028821605.1 | Candidatus_Poribacteria_bacterium | OYL97_16085 | 27558..30836 | 2026781 | MAG:-DUF499-domain-containing-protein-[Candidatus-Poribacteria-bacterium] |
| MDE0681304.1 | DUF499_1.RM.like.systems | HTH+MPTase->PIN->HEPN+DUF499*->METHYLASE-> | 1094 | Bacteria>Bacteria_incertae_sedis>Bacteria_candidate_phyla>Candidatus_Poribacteria>unclassified_Candidatus_Poribacteria | GCA_028824575.1 | Candidatus_Poribacteria_bacterium | OXI63_00195 | 3088..6372 | 2026781 | MAG:-DUF499-domain-containing-protein-[Candidatus-Poribacteria-bacterium] |
| MBI2524457.1 | DUF499_1.RM.like.systems | REase->MORC+DUF3684->METHYLASE->?->?->HEPN+DUF499*->?->?->?->XIS-HTH->Phage_int_SAM_1+Phage_integrase-> | 1121 | Bacteria>Bacteria_incertae_sedis>Bacteria_candidate_phyla>Candidatus_Rokubacteria>unclassified_Candidatus_Rokubacteria | GCA_016188005.1 | Candidatus_Rokubacteria_bacterium | HYV93_00585 | 34529..37894 | 2053607 | ATP-binding-protein-[Candidatus-Rokubacteria-bacterium] |
| MBU0999702.1 | DUF499_1.RM.like.systems | METHYLASE-p->HEPN+DUF499*-> | 1122 | Bacteria>Bacteria_incertae_sedis>Bacteria_candidate_phyla>Patescibacteria_group>unclassified_Patescibacteria_group | GCA_018816985.1 | Patescibacteria_group_bacterium | KKG24_05405 | 385..3750 | 2052139 | DUF499-domain-containing-protein-[Patescibacteria-group-bacterium] |
| HBM97560.1 | DUF499_1.RM.like.systems | <-HEPN+DUF499*<-METHYLASE | 1146 | Bacteria>Bacteria_incertae_sedis>Bacteria_candidate_phyla>unclassified_Bacteria_candidate_phyla | GCA_003504095.1 | bacterium_UBP9_UBA11836 | DD394_08835 | 303..3743 | 2060920 | hypothetical-protein-[bacterium-UBP9_UBA11836] |

| Protein-ID | System | Gene-Neighbourhoods | Length | Taxonomy | Assembly-ID | Strain | Locus-Tag | Genomic-coordinates | TaxID | Fasta-define |
|---|---|---|---|---|---|---|---|---|---|---|
| WP_142699382.1 | DUF499_1.RM.like.systems | HTH-><-?\|\|?->TUDOR+SF1-SF2-Helicase+iREase+CTD+REase->METHYLASE->HEPN+DUF499*->REase-><-ASCH | 1099 | Bacteria>Bdellovibrionota>Bdellovibrionia>Bdellovibrionales>Pseudobdellovibrionaceae | GCF_006874645.1 | Bdellovibrio_sp._ZAP7 | DOM22_RS05315 | 1082808..1086107 | 2231053 | Swt1-family-HEPN-domain-containing-protein-[Bdellovibrio-sp.-ZAP7] |
| MBK8201825.1 | DUF499_1.RM.like.systems | <-HTH\|\|HTH->?->?->HEPN+DUF499*->DUF499->MarR-HTH-> | 351 | Bacteria>Bdellovibrionota>Bdellovibrionia>Bdellovibrionales>unclassified_Bdellovibrionales | GCA_016712575.1 | Bdellovibrionales_bacterium | IPK68_05755 | 72390..73445 | 2053517 | hypothetical-protein-[Bdellovibrionales-bacterium] |
| MBK8201826.1 | DUF499_1.RM.like.systems | <-HTH\|\|HTH->?->?->HEPN+DUF499->DUF499*->MarR-HTH-> | 741 | Bacteria>Bdellovibrionota>Bdellovibrionia>Bdellovibrionales>unclassified_Bdellovibrionales | GCA_016712575.1 | Bdellovibrionales_bacterium | IPK68_05760 | 73468..75693 | 2053517 | ATP-binding-protein-[Bdellovibrionales-bacterium] |
| MCB0334043.1 | DUF499_1.RM.like.systems | METHYLASE-p->HEPN+DUF499*-> | 570 | Bacteria>Bdellovibrionota>Bdellovibrionia>Bdellovibrionales>unclassified_Bdellovibrionales | GCA_020434525.1 | Bdellovibrionales_bacterium | KDD55_11125 | 601..2311 | 2053517 | DUF499-domain-containing-protein-[Bdellovibrionales-bacterium] |
| MBN1883904.1 | DUF499_1.RM.like.systems | Peptidase_M48->TUDOR+SF1-SF2-Helicase+iREase+CTD+REase->Schlafen+HATPase+HTH->METHYLASE->OLD-ABC->TOPRIM->HEPN+DUF499*->PIN+OST-HTH-><-?<-?<-REase<-?\|\|DOC+HTH-> | 1113 | Bacteria>Candidatus_Krumholzibacteriota>unclassified_Candidatus_Krumholzibacteriota | GCA_016931295.1 | Candidatus_Krumholzibacteriota_bacterium | JW876_00100 | 28696..32037 | 2725313 | DUF499-domain-containing-protein-[Candidatus-Krumholzibacteriota-bacterium] |
| OGQ97428.1 | DUF499_1.RM.like.systems | REase->?->?->?-><-?<-SF1-SF2-Helicase<-HEPN+DUF499*<-METHYLASE<-?<-?<-ParB+HNH<-TOPRIM<-OLD-ABC | 1111 | Bacteria>delta/epsilon_subdivisions>Deltaproteobacteria>unclassified_Deltaproteobacteria | GCA_001799275.1 | Deltaproteobacteria_bacterium_RIFOXYD12_FULL_57_12 | A2521_08565 | 7962..11297 | 1797906 | AAA+-family-ATPase-[Deltaproteobacteria-bacterium-RIFOXYD12_FULL_57_12] |
| OQB73667.1 | DUF499_1.RM.like.systems | Classical-AAA->Peptidase_S8-><-HEPN+DUF499* | 1112 | Bacteria>delta/epsilon_subdivisions>Deltaproteobacteria>unclassified_Deltaproteobacteria | GCA_002070575.1 | Deltaproteobacteria_bacterium_ADurb.Bin135 | BWX92_03289 | 4255..7593 | 1852867 | hypothetical-protein-BWX92_03289-[Deltaproteobacteria-bacterium-ADurb.Bin135] |
| PJB39408.1 | DUF499_1.RM.like.systems | <-DUF499-p<-?<-HEPN+DUF499*<-Schlafen+HATPase+HTH<-PHP+SbcC<-METHYLASE-p | 1117 | Bacteria>delta/epsilon_subdivisions>Deltaproteobacteria>unclassified_Deltaproteobacteria | GCA_002789955.1 | Deltaproteobacteria_bacterium_CG_4_9_14_3_um_filter_63_12 | CO108_17305 | 2362..5715 | 1973967 | AAA+-family-ATPase-[Deltaproteobacteria-bacterium-CG_4_9_14_3_um_filter_63_12] |
| MBI4801911.1 | DUF499_1.RM.like.systems | STAS+HisKin->?->PIN->OLD-ABC->HNH->METHYLASE->HEPN+DUF499*->Glycos_trans_3N+Glycos_transf_3->Radical-SAM->Radical-SAM->?->?->RADICAL-SAM-> | 1132 | Bacteria>Elusimicrobiota>unclassified_Elusimicrobiota | GCA_016208085.1 | Elusimicrobiota_bacterium | HY796_05240 | 17514..20912 | 2030800 | ATP-binding-protein-[Elusimicrobia-bacterium] |
| MBI5243147.1 | DUF499_1.RM.like.systems | <-ORC-CDC6+REase\|\|?->PIN-><-TPR\|\|?-><-HEPN+DUF499*<-METHYLASE<-PIN<-?<-STAS+HisKin<-TUDOR+SF1-SF2-Helicase+iREase+CTD+REase | 1133 | Bacteria>Elusimicrobiota>unclassified_Elusimicrobiota | GCA_016212885.1 | Elusimicrobiota_bacterium | HY922_05595 | 9033..12434 | 2030800 | ATP-binding-protein-[Elusimicrobia-bacterium] |
| MCX5796464.1 | DUF499_1.RM.like.systems | TUDOR+SF1-SF2-Helicase+iREase+CTD+REase->STAS+HisKin->?->PIN->METHYLASE->OLD-ABC+TOPRIM->HEPN+DUF499*-> | 1133 | Bacteria>Elusimicrobiota>unclassified_Elusimicrobiota | GCA_026388095.1 | Elusimicrobiota_bacterium | NTY77_13305 | 62095..65496 | 2030800 | MAG:-Swt1-family-HEPN-domain-containing-protein-[Elusimicrobia-bacterium] |
| CAG1064741.1 | DUF499_1.RM.like.systems | <-DOC<-MazE_antitoxin<-?<-TUDOR+SF1-SF2-Helicase+iREase+CTD<-?<-HEPN+DUF499*<-METHYLASE<-ArdCN+MPTase<-Schlafen+HATPase+HTH<-ArdCN+MPTase<-TUDOR+SF1-SF2-Helicase+iREase+CTD+REase<-HTH_26 | 1109 | Bacteria>environmental_samples | GCA_905339405.1 | uncultured_bacterium | BAC1_00306 | 123828..127157 | 77133 | hypothetical-protein-BAC1_00306-[uncultured-bacterium] |
| BAH89290.1 | DUF499_1.RM.like.systems | <-UbE1+JAB<-SMODS\|\|HTH+WYL->TUDOR+SF1-SF2-Helicase+iREase+CTD+REase->TUDOR+SF1-SF2-Helicase+iREase+CTD+REase-p->METHYLASE->HEPN+DUF499*->?-><-MODE-HTH+PBPII\|\|TetR-HTH-><-?\|\|?->MODE-HTH+PBPII-> | 1099 | Bacteria>environmental_samples | - | uncultured_bacterium | . | 18704..22003 | 77133 | conserved-hypothetical-protein-[uncultured-bacterium] |
| ART37960.1 | DUF499_1.RM.like.systems | <-PemK_toxin<-?\|\|Phage_integrase-><-HEPN+DUF499* | 1098 | Bacteria>environmental_samples | - | uncultured_bacterium | . | 22115..25411 | 77133 | F28-[uncultured-bacterium] |
| CDD51558.1 | DUF499_1.RM.like.systems | <-XIS-HTH<-?<-Phage_integrase\|\|TIR->TIR-><-?<-HEPN+DUF499*<-METHYLASE<-REase+ParB<-TUDOR+SF1-SF2-Helicase+iREase+CTD+REase<-HTH_3<-DnaJ | 1098 | Bacteria>FCB_group>Bacteroidota/Chlorobiota_group>Bacteroidota>Bacteroidia | GCA_000435415.1 | Bacteroides_sp._CAG:875 | BN800_02603 | 61429..64725 | 1262752 | putative-uncharacterized-protein-[Bacteroides-sp.-CAG:875] |
| CDD33920.1 | DUF499_1.RM.like.systems | <-Peptidase_M3<-?\|\|?-><-?<-HEPN+DUF499*<-METHYLASE<-REase+ParB<-TUDOR+SF1-SF2-Helicase+iREase+CTD+REase<-Radical-SAM | 1098 | Bacteria>FCB_group>Bacteroidota/Chlorobiota_group>Bacteroidota>Bacteroidia | GCA_000436055.1 | Bacteroides_sp._CAG:714 | BN762_01576 | 11019..14315 | 1262749 | putative-uncharacterized-protein-[Bacteroides-sp.-CAG:714] |
| EXY75216.1 | DUF499_1.RM.like.systems | <-ACET<-Calcineurin<-ZDNA-HTH<-DOC+HTH<-HEPN+DUF499*<-METHYLASE-p | 1101 | Bacteria>FCB_group>Bacteroidota/Chlorobiota_group>Bacteroidota>Bacteroidia | GCA_000598365.1 | Bacteroides_fragilis_str._3988T(B)14 | M124_1038 | 75233..78538 | 1339315 | hypothetical-protein-M124_1038-[Bacteroides-fragilis-str.-3988T(B)14] |
| KXT50696.1 | DUF499_1.RM.like.systems | DEDD_Tnp_IS110->TUDOR+SF1-SF2-Helicase+iREase+CTD+REase->METHYLASE->?->HEPN+DUF499*-> | 1098 | Bacteria>FCB_group>Bacteroidota/Chlorobiota_group>Bacteroidota>Bacteroidia | GCA_001578575.1 | Bacteroides_ovatus | HMPREF2532_00822 | 114886..118182 | 28116 | hypothetical-protein-HMPREF2532_00822-[Bacteroides-ovatus] |
| HBF05518.1 | DUF499_1.RM.like.systems | <-ORC-CDC6+REase<-HEPN+DUF499*<-METHYLASE-p | 1108 | Bacteria>FCB_group>Bacteroidota/Chlorobiota_group>Bacteroidota>Bacteroidia | GCA_003501465.1 | Candidatus_Segatella_violae | DDW28_05315 | 5202..8528 | 1947177 | hypothetical-protein-[Prevotella-sp.] |
| KAB4460058.1 | DUF499_1.RM.like.systems | <-URI-DNase+SF1-SF2-Helicase<-?\|\|sGTP->TUDOR+SF1-SF2-Helicase+iREase+CTD+REase->METHYLASE->HEPN+DUF499*->DOC+HTH-><-HKD-DNase<-SF1-SF2-Helicase<-SF1-SF2-Helicase<-?<-NUDIX | 1099 | Bacteria>FCB_group>Bacteroidota/Chlorobiota_group>Bacteroidota>Bacteroidia | GCA_009021605.1 | Bacteroides_thetaiotaomicron | GAN98_17895 | 29744..33043 | 818 | ATP-binding-protein-[Bacteroides-thetaiotaomicron] |
| MCL2726873.1 | DUF499_1.RM.like.systems | <-ABhydrolase\|\|REase->Peptidase-><-Peptidase_M24<-Peptidase<-?<-HEPN+DUF499*<-TOPRIM<-OLD-ABC<-METHYLASE<-TUDOR+SF1-SF2-Helicase+iREase+CTD+REase<-LysE | 1124 | Bacteria>FCB_group>Bacteroidota/Chlorobiota_group>Bacteroidota>Bacteroidia | GCA_009781275.1 | Bacteroidales_bacterium | FWD56_00650 | 13721..17095 | 2030927 | DUF499-domain-containing-protein-[Bacteroidales-bacterium] |
| NCB25654.1 | DUF499_1.RM.like.systems | TUDOR+SF1-SF2-Helicase+iREase+CTD+REase->METHYLASE->HEPN+DUF499*->REase->Pkinase-> | 1134 | Bacteria>FCB_group>Bacteroidota/Chlorobiota_group>Bacteroidota>Bacteroidia | GCA_009929165.1 | Bacteroidia_bacterium | EOM62_09315 | 4747..8151 | 2044936 | ATP-binding-protein-[Bacteroidia-bacterium] |
| NLB25730.1 | DUF499_1.RM.like.systems | <-HEPN+DUF499*<-METHYLASE-p | 1096 | Bacteria>FCB_group>Bacteroidota/Chlorobiota_group>Bacteroidota>Bacteroidia | GCA_012511595.1 | Bacteroidales_bacterium | GX820_03445 | 1391..4681 | 2030927 | ATP-binding-protein-[Bacteroidales-bacterium] |
| MBE6276390.1 | DUF499_1.RM.like.systems | SbcC->TOPRIM->HicB->TIR->TIR-><-?<-HEPN+DUF499*<-METHYLASE<-TUDOR+SF1-SF2-Helicase+iREase+CTD+REase<-HTH_3<-XIS-HTH | 1095 | Bacteria>FCB_group>Bacteroidota/Chlorobiota_group>Bacteroidota>Bacteroidia | GCA_015060735.1 | Bacteroides_sp. | E7096_08645 | 4627..7914 | 29523 | ATP-binding-protein-[Bacteroides-sp.] |
| MBN1185441.1 | DUF499_1.RM.like.systems | METHYLASE-p->HEPN+DUF499*->TPR+TPR->?->MazG->REase+SF1-SF2-Helicase-> | 1076 | Bacteria>FCB_group>Bacteroidota/Chlorobiota_group>Bacteroidota>Bacteroidia | GCA_016926235.1 | Bacteroidales_bacterium | JXB49_24360 | 1099..4329 | 2030927 | DUF499-domain-containing-protein-[Bacteroidales-bacterium] |
| MBQ8713170.1 | DUF499_1.RM.like.systems | METHYLASE-p->HEPN+DUF499*-> | 1120 | Bacteria>FCB_group>Bacteroidota/Chlorobiota_group>Bacteroidota>Bacteroidia | GCA_017471565.1 | Prevotella_sp. | IJ551_10195 | 948..4310 | 59823 | DUF499-domain-containing-protein-[Prevotella-sp.] |
| MBR1880072.1 | DUF499_1.RM.like.systems | TUDOR+SF1-SF2-Helicase+iREase+CTD+REase->METHYLASE-><-PIN<-?\|\|HEPN+DUF499*-> | 593 | Bacteria>FCB_group>Bacteroidota/Chlorobiota_group>Bacteroidota>Bacteroidia | GCA_017478995.1 | Prevotella_sp. | IJ804_04915 | 6915..8693 | 59823 | DUF499-domain-containing-protein-[Prevotella-sp.] |
| MBO5419352.1 | DUF499_1.RM.like.systems | XIS-HTH->?->?-><-?<-?\|\|ORC-CDC6+REase-><-HEPN+DUF499*<-METHYLASE-p | 1102 | Bacteria>FCB_group>Bacteroidota/Chlorobiota_group>Bacteroidota>Bacteroidia | GCA_017625725.1 | Bacteroidales_bacterium | J6A22_04620 | 7056..10364 | 2030927 | DUF499-domain-containing-protein-[Bacteroidales-bacterium] |
| MBO6254120.1 | DUF499_1.RM.like.systems | <-HEPN+DUF499*<-METHYLASE-p | 1108 | Bacteria>FCB_group>Bacteroidota/Chlorobiota_group>Bacteroidota>Bacteroidia | GCA_017635295.1 | Bacteroidaceae_bacterium | J6O49_10775 | 248..3574 | 2212467 | DUF499-domain-containing-protein-[Bacteroidaceae-bacterium] |
| MBO7596490.1 | DUF499_1.RM.like.systems | <-HEPN+DUF499*<-METHYLASE | 1105 | Bacteria>FCB_group>Bacteroidota/Chlorobiota_group>Bacteroidota>Bacteroidia | GCA_017651345.1 | Bacteroidales_bacterium | J6T70_05520 | 1703..5020 | 2030927 | DUF499-domain-containing-protein-[Bacteroidales-bacterium] |
| MBP6872709.1 | DUF499_1.RM.like.systems | TUDOR+SF1-SF2-Helicase+iREase+CTD+REase->AbiEi_1->AbiEii->ParB+HNH->METHYLASE->HEPN+DUF499*->?->SbcC->?-><-?<-REase+SF1-SF2-Helicase<-MazG | 1092 | Bacteria>FCB_group>Bacteroidota/Chlorobiota_group>Bacteroidota>Bacteroidia | GCA_017991835.1 | Bacteroidales_bacterium | KBC43_11940 | 58567..61845 | 2030927 | ATP-binding-protein-[Bacteroidales-bacterium] |
| MBP7851039.1 | DUF499_1.RM.like.systems | METHYLASE-p->HEPN+DUF499*-> | 1097 | Bacteria>FCB_group>Bacteroidota/Chlorobiota_group>Bacteroidota>Bacteroidia | GCA_018001695.1 | Lentimicrobiaceae_bacterium | KA053_12270 | 826..4119 | 2026756 | DUF499-domain-containing-protein-[Lentimicrobiaceae-bacterium] |
| MBW6537009.1 | DUF499_1.RM.like.systems | <-HEPN+DUF499*<-METHYLASE<-AbiEii<-wHTH+AbiEi<-TUDOR+SF1-SF2-Helicase+iREase+CTD+REase<-TUDOR+SF1-SF2-Helicase+iREase+CTD+REase-p | 696 | Bacteria>FCB_group>Bacteroidota/Chlorobiota_group>Bacteroidota>Bacteroidia | GCA_019429205.1 | Mariniphaga_sp. | KOB11_18520 | 1..2089 | 1954475 | DUF499-domain-containing-protein-[Mariniphaga-sp.] |
| MCC8143016.1 | DUF499_1.RM.like.systems | <-AraC-HTH+SHS2+SHS2<-Peptidase_M28<-?<-BirA-HTH<-Schlafen<-HEPN+DUF499*<-METHYLASE | 1102 | Bacteria>FCB_group>Bacteroidota/Chlorobiota_group>Bacteroidota>Bacteroidia | GCA_020860665.1 | Tannerellaceae_bacterium | LIO97_03250 | 4810..8118 | 2785930 | DUF499-domain-containing-protein-[Tannerellaceae-bacterium] |
| MCD8261448.1 | DUF499_1.RM.like.systems | <-STYKIN<-Schlafen<-HEPN+DUF499*<-METHYLASE-p<-METHYLASE-p | 1102 | Bacteria>FCB_group>Bacteroidota/Chlorobiota_group>Bacteroidota>Bacteroidia | GCA_021204375.1 | Bacteroides_sp. | LUD15_08080 | 1305..4613 | 29523 | DUF499-domain-containing-protein-[Bacteroides-sp.] |
| MCR5642603.1 | DUF499_1.RM.like.systems | <-Peptidase<-DOC<-?<-HEPN+DUF499*<-METHYLASE<-TUDOR+SF1-SF2-Helicase+iREase+CTD+REase | 1106 | Bacteria>FCB_group>Bacteroidota/Chlorobiota_group>Bacteroidota>Bacteroidia | GCA_024690905.1 | Prevotella_sp. | K6G32_04600 | 31632..34952 | 59823 | DUF499-domain-containing-protein-[Prevotella-sp.] |
| MCS2505372.1 | DUF499_1.RM.like.systems | TUDOR+SF1-SF2-Helicase+iREase+CTD+REase->TUDOR+SF1-SF2-Helicase+iREase+CTD+REase-p->METHYLASE-p->METHYLASE-p->METHYLASE-p->HEPN-p->HEPN+DUF499*->DUF499-><-?<-HicB<-TOPRIM<-SbcC\|\|BirA-HTH-> | 434 | Bacteria>FCB_group>Bacteroidota/Chlorobiota_group>Bacteroidota>Bacteroidia | GCA_024788865.1 | Phocaeicola_vulgatus | NXW48_05305 | 1159689..1160993 | 821 | DUF499-domain-containing-protein-[Phocaeicola-vulgatus] |
| MCS2505373.1 | DUF499_1.RM.like.systems | TUDOR+SF1-SF2-Helicase+iREase+CTD+REase-p->METHYLASE-p->METHYLASE-p->METHYLASE-p->HEPN-p->HEPN+DUF499->DUF499*-><-?<-HicB<-TOPRIM<-SbcC\|\|BirA-HTH-><-Phage_int_SAM_5+Phage_integrase | 640 | Bacteria>FCB_group>Bacteroidota/Chlorobiota_group>Bacteroidota>Bacteroidia | GCA_024788865.1 | Phocaeicola_vulgatus | NXW48_05310 | 1160902..1162824 | 821 | ATP-binding-protein-[Phocaeicola-vulgatus] |
| MCS2502386.1 | DUF499_1.RM.like.systems | <-HEPN+DUF499*<-METHYLASE<-METHYLASE-p<-TUDOR+SF1-SF2-Helicase+iREase+CTD+REase<-DEDD_Tnp_IS110 | 1026 | Bacteria>FCB_group>Bacteroidota/Chlorobiota_group>Bacteroidota>Bacteroidia | GCA_024788905.1 | Bacteroides_ovatus | NXV80_13315 | 3514916..3517996 | 28116 | DUF499-domain-containing-protein-[Bacteroides-ovatus] |
| MCS3108285.1 | DUF499_1.RM.like.systems | <-ACET<-Calcineurin<-ZDNA-HTH<-DOC+HTH<-DUF499-p<-HEPN+DUF499*<-METHYLASE<-TUDOR+SF1-SF2-Helicase+iREase+CTD+REase<-TUDOR+SF1-SF2-Helicase+iREase+CTD+REase-p<-HTH_3<-?<-TraJ-RHH | 722 | Bacteria>FCB_group>Bacteroidota/Chlorobiota_group>Bacteroidota>Bacteroidia | GCA_024792645.1 | Bacteroides_fragilis | NXV07_08905 | 2189665..2191833 | 817 | DUF499-domain-containing-protein-[Bacteroides-fragilis] |
| MDD4108176.1 | DUF499_1.RM.like.systems | <-HEPN+DUF499*<-?<-METHYLASE | 1002 | Bacteria>FCB_group>Bacteroidota/Chlorobiota_group>Bacteroidota>Bacteroidia | GCA_028701215.1 | Prolixibacteraceae_bacterium | PHH93_05600 | 1..3006 | 2053594 | MAG:-DUF499-domain-containing-protein,-partial-[Prolixibacteraceae-bacterium] |
| MDD4460430.1 | DUF499_1.RM.like.systems | METHYLASE-p->HEPN+DUF499*-> | 1132 | Bacteria>FCB_group>Bacteroidota/Chlorobiota_group>Bacteroidota>Bacteroidia | GCA_028705295.1 | Proteiniphilum_sp. | PHQ24_11635 | 346..3744 | 1926877 | MAG:-DUF499-domain-containing-protein-[Proteiniphilum-sp.] |
| MDD7697305.1 | DUF499_1.RM.like.systems | XIS-HTH->?->?->HTH_3->TUDOR+SF1-SF2-Helicase+iREase+CTD+REase->METHYLASE->HEPN+DUF499*-><-?<-HicB<-TOPRIM<-SbcC\|\|BirA-HTH-><-Phage_int_SAM_5+Phage_integrase | 1096 | Bacteria>FCB_group>Bacteroidota/Chlorobiota_group>Bacteroidota>Bacteroidia | GCA_028727985.1 | Phocaeicola_vulgatus | PUJ58_17085 | 44225..47515 | 821 | MAG:-DUF499-domain-containing-protein-[Phocaeicola-vulgatus] |
| WP_018709397.1 | DUF499_1.RM.like.systems | REase+SF1-SF2-Helicase->?->Radical-SAM->TUDOR+SF1-SF2-Helicase+iREase+CTD+REase->REase+ParB->METHYLASE->HEPN+DUF499*->?-><-?\|\|?->Peptidase_M3-> | 1098 | Bacteria>FCB_group>Bacteroidota/Chlorobiota_group>Bacteroidota>Bacteroidia | GCF_000374585.1 | Phocaeicola_barnesiae | C510_RS0100610 | 147306..150602 | 376804 | DUF499-domain-containing-protein-[Phocaeicola-barnesiae] |
| WP_016268236.1 | DUF499_1.RM.like.systems | <-URI-DNase+SF1-SF2-Helicase<-?<-?\|\|sGTP->TUDOR+SF1-SF2-Helicase+iREase+CTD+REase->METHYLASE->HEPN+DUF499*->DOC+HTH->Glyoxalase+Glyoxalase-><-HKD-DNase<-SF1-SF2-Helicase<-SF1-SF2-Helicase | 1099 | Bacteria>FCB_group>Bacteroidota/Chlorobiota_group>Bacteroidota>Bacteroidia | GCF_000403155.2 | Bacteroides_thetaiotaomicron | C799_RS12880 | 2627963..2631262 | 818 | DUF499-domain-containing-protein-[Bacteroides-thetaiotaomicron] |
| WP_032587868.1 | DUF499_1.RM.like.systems | <-ACET<-Calcineurin<-ZDNA-HTH<-DOC+HTH<-HEPN+DUF499* | 1099 | Bacteria>FCB_group>Bacteroidota/Chlorobiota_group>Bacteroidota>Bacteroidia | GCF_000598365.1 | Bacteroides_fragilis | M084_RS16910 | 33544..36843 | 817 | DUF499-domain-containing-protein-[Bacteroides-fragilis] |
| WP_038554549.1 | DUF499_1.RM.like.systems | TUDOR+SF1-SF2-Helicase+iREase+CTD+REase->?->OLD-ABC->?->METHYLASE->HEPN+DUF499*->DUF6361->HKD-DNase->SF1-SF2-Helicase-><-?<-Primpol | 1096 | Bacteria>FCB_group>Bacteroidota/Chlorobiota_group>Bacteroidota>Bacteroidia | GCF_000626635.1 | Draconibacterium_orientale | BMW53_RS05405 | 116803..120093 | 1168034 | DUF499-domain-containing-protein-[Draconibacterium-orientale] |
| WP_032540897.1 | DUF499_1.RM.like.systems | <-HEPN+DUF499*<-METHYLASE-p<-METHYLASE<-TUDOR+SF1-SF2-Helicase+iREase+CTD+REase<-HTH_3<-DEDD_Tnp_IS110 | 1096 | Bacteria>FCB_group>Bacteroidota/Chlorobiota_group>Bacteroidota>Bacteroidia | GCF_000724815.2 | Bacteroides | IB64_RS10255 | 2443318..2446608 | 816 | DUF499-domain-containing-protein-[Bacteroides-fragilis] |
| WP_032953489.1 | DUF499_1.RM.like.systems | XIS-HTH->?->?->HTH_3->TUDOR+SF1-SF2-Helicase+iREase+CTD+REase->METHYLASE->HEPN+DUF499*-><-?<-HicB | 1096 | Bacteria>FCB_group>Bacteroidota/Chlorobiota_group>Bacteroidota>Bacteroidia | GCF_001405155.1 | Bacteroidaceae | KSZ98_RS20820 | 10462..13752 | 815 | MULTISPECIES:-DUF499-domain-containing-protein-[Bacteroidaceae] |
| WP_087396459.1 | DUF499_1.RM.like.systems | REase->?-><-?<-DOC+HTH<-HEPN+DUF499*<-METHYLASE<-REase+ParB<-TUDOR+SF1-SF2-Helicase+iREase+CTD+REase<-DOC<-HTH_3<-FAD-NAD-dep-oxidoreductase | 1097 | Bacteria>FCB_group>Bacteroidota/Chlorobiota_group>Bacteroidota>Bacteroidia | GCF_002161565.1 | Bacteroides_sp._An51A | B5G04_RS06890 | 141402..144695 | 1965640 | DUF499-domain-containing-protein-[Bacteroides-sp.-An51A] |
| WP_278365558.1 | DUF499_1.RM.like.systems | <-Peptidase_M3<-?\|\|?-><-?<-HEPN+DUF499*<-METHYLASE<-REase+ParB<-TUDOR+SF1-SF2-Helicase+iREase+CTD+REase<-Radical-SAM<-?<-SF1-SF2-Helicase | 1098 | Bacteria>FCB_group>Bacteroidota/Chlorobiota_group>Bacteroidota>Bacteroidia | GCF_002359135.1 | Phocaeicola_barnesiae | B3R84_RS06870 | 398812..402108 | 376804 | DUF499-domain-containing-protein-[Phocaeicola-barnesiae] |
| WP_117747735.1 | DUF499_1.RM.like.systems | <-HEPN+DUF499*<-METHYLASE<-REase+ParB<-TUDOR+SF1-SF2-Helicase+iREase+CTD+REase<-HTH_3<-DnaJ | 1098 | Bacteria>FCB_group>Bacteroidota/Chlorobiota_group>Bacteroidota>Bacteroidia | GCF_003438135.1 | Phocaeicola_plebeius | DXC17_RS06660 | 1179..4475 | 310297 | DUF499-domain-containing-protein-[Phocaeicola-plebeius] |
| WP_151851142.1 | DUF499_1.RM.like.systems | <-BirA-HTH\|\|SbcC->TOPRIM->HicB->?->?-><-HEPN+DUF499*<-METHYLASE<-TUDOR+SF1-SF2-Helicase+iREase+CTD+REase<-HTH_3<-?<-?<-XIS-HTH | 1096 | Bacteria>FCB_group>Bacteroidota/Chlorobiota_group>Bacteroidota>Bacteroidia | GCF_009025535.1 | Phocaeicola_vulgatus | F9001_RS04885 | 42984..46274 | 821 | DUF499-domain-containing-protein-[Phocaeicola-vulgatus] |
| WP_195204977.1 | DUF499_1.RM.like.systems | DnaJ->HTH_3->TUDOR+SF1-SF2-Helicase+iREase+CTD+REase->REase+ParB->METHYLASE-p->METHYLASE->HEPN+DUF499*-> | 1098 | Bacteria>FCB_group>Bacteroidota/Chlorobiota_group>Bacteroidota>Bacteroidia | GCF_015546785.1 | Phocaeicola_plebeius | I1Q75_RS16665 | 24617..27913 | 310297 | DUF499-domain-containing-protein-[Phocaeicola-plebeius] |
| WP_195365726.1 | DUF499_1.RM.like.systems | <-HEPN+DUF499*<-METHYLASE<-TUDOR+SF1-SF2-Helicase+iREase+CTD+REase<-DEDD_Tnp_IS110<-?<-?\|\|HTH-> | 1096 | Bacteria>FCB_group>Bacteroidota/Chlorobiota_group>Bacteroidota>Bacteroidia | GCF_015550085.1 | Bacteroides_ovatus | PO368_RS10195 | 12026..15316 | 28116 | DUF499-domain-containing-protein-[Bacteroides-ovatus] |
| WP_195425521.1 | DUF499_1.RM.like.systems | STYKinase-like->SF1-SF2-Helicase-><-?<-Glyoxalase+Glyoxalase<-DOC+HTH<-HEPN+DUF499*<-METHYLASE<-TUDOR+SF1-SF2-Helicase+iREase+CTD+REase<-HTH_3<-?<-?<-TraJ-RHH | 1099 | Bacteria>FCB_group>Bacteroidota/Chlorobiota_group>Bacteroidota>Bacteroidia | GCF_015551145.1 | Bacteroides_cellulosilyticus | PQH05_RS04750 | 125017..128316 | 246787 | DUF499-domain-containing-protein-[Bacteroides-cellulosilyticus] |
| WP_196069848.1 | DUF499_1.RM.like.systems | <-REase+SF1-SF2-Helicase<-MazG<-METHYLASE-p<-ABhydrolase<-ACET<-DOC+HTH<-HEPN+DUF499*<-METHYLASE<-TUDOR+SF1-SF2-Helicase+iREase+CTD+REase\|\|?->HTH_3->?->Phage_int_SAM_5+Phage_integrase-> | 1099 | Bacteria>FCB_group>Bacteroidota/Chlorobiota_group>Bacteroidota>Bacteroidia | GCF_015670455.1 | Bacteroides_thetaiotaomicron | PO078_RS08820 | 41190..44489 | 818 | DUF499-domain-containing-protein-[Bacteroides-thetaiotaomicron] |
| WP_220575703.1 | DUF499_1.RM.like.systems | DEDD_Tnp_IS110->HTH_3->TUDOR+SF1-SF2-Helicase+iREase+CTD+REase->METHYLASE-p->METHYLASE->HEPN+DUF499*-> | 1096 | Bacteria>FCB_group>Bacteroidota/Chlorobiota_group>Bacteroidota>Bacteroidia | GCF_019633785.1 | Bacteroides | ISX59_RS04600 | 376325..379615 | 816 | MULTISPECIES:-DUF499-domain-containing-protein-[Bacteroides] |
| WP_229532986.1 | DUF499_1.RM.like.systems | <-SF1-SF2-Helicase<-MAD-NTD+HNH<-SF1-SF2-Helicase+DUF1998<-?<-HEPN+DUF499*<-METHYLASE<-TUDOR+SF1-SF2-Helicase+iREase+CTD+REase<-WYL+WYL-C<-HTH_3 | 1108 | Bacteria>FCB_group>Bacteroidota/Chlorobiota_group>Bacteroidota>Bacteroidia | GCF_020827855.1 | Phocaeicola_vulgatus | HZY60_RS19310 | 19399..22725 | 821 | DUF499-domain-containing-protein-[Phocaeicola-vulgatus] |
| WP_234147452.1 | DUF499_1.RM.like.systems | sGTP->TUDOR+SF1-SF2-Helicase+iREase+CTD+REase->METHYLASE->HEPN+DUF499*->DOC+HTH->Glyoxalase+Glyoxalase-><-HKD-DNase<-SF1-SF2-Helicase<-SF1-SF2-Helicase | 1099 | Bacteria>FCB_group>Bacteroidota/Chlorobiota_group>Bacteroidota>Bacteroidia | GCF_021410605.1 | Bacteroides_thetaiotaomicron | KOG45_RS08855 | 122149..125448 | 818 | DUF499-domain-containing-protein-[Bacteroides-thetaiotaomicron] |
| WP_235327793.1 | DUF499_1.RM.like.systems | SbcC->TOPRIM->HicB->TIR->TIR-><-?<-HEPN+DUF499*<-METHYLASE<-TUDOR+SF1-SF2-Helicase+iREase+CTD+REase<-HTH_3<-?<-?<-tRNA<-Peptidase_S41 | 1094 | Bacteria>FCB_group>Bacteroidota/Chlorobiota_group>Bacteroidota>Bacteroidia | GCF_021530995.1 | Bacteroides_caecigallinarum | I6E24_RS08290 | 4207..7491 | 1411144 | DUF499-domain-containing-protein-[Bacteroides-caecigallinarum] |
| WP_235301263.1 | DUF499_1.RM.like.systems | TUDOR+SF1-SF2-Helicase+iREase+CTD+REase->REase+ParB->METHYLASE->HEPN+DUF499*->?-><-?\|\|?->Peptidase_M3-> | 1098 | Bacteria>FCB_group>Bacteroidota/Chlorobiota_group>Bacteroidota>Bacteroidia | GCF_021531035.1 | Phocaeicola_barnesiae | I6E10_RS09100 | 8385..11681 | 376804 | DUF499-domain-containing-protein-[Phocaeicola-barnesiae] |
| WP_235386280.1 | DUF499_1.RM.like.systems | METHYLASE->HEPN+DUF499*->?->SF1-SF2-Helicase+DUF1998->MAD-NTD+HNH->SF1-SF2-Helicase-> | 1119 | Bacteria>FCB_group>Bacteroidota/Chlorobiota_group>Bacteroidota>Bacteroidia | GCF_021531665.1 | Prevotellamassilia_timonensis | I6E40_RS06760 | 3178..6537 | 1852370 | DUF499-domain-containing-protein-[Prevotellamassilia-timonensis] |
| WP_250712821.1 | DUF499_1.RM.like.systems | SF1-SF2-Helicase->?->DEDD_Tnp_IS110->HTH_3->TUDOR+SF1-SF2-Helicase+iREase+CTD+REase->METHYLASE->HEPN+DUF499*-> | 1096 | Bacteria>FCB_group>Bacteroidota/Chlorobiota_group>Bacteroidota>Bacteroidia | GCF_023654475.1 | Bacteroides_fragilis | J8K93_RS06720 | 1584260..1587550 | 817 | DUF499-domain-containing-protein-[Bacteroides-fragilis] |
| WP_118201547.1 | DUF499_1.RM.like.systems | <-Aminotran_1_2<-?\|\|?->?->TUDOR+SF1-SF2-Helicase+iREase+CTD+REase->METHYLASE->HEPN+DUF499*-> | 1122 | Bacteria>FCB_group>Bacteroidota/Chlorobiota_group>Bacteroidota>Bacteroidia | GCF_024330865.1 | Segatella_copri | NNC52_RS13920 | 15349..18717 | 165179 | DUF499-domain-containing-protein-[Prevotella-copri] |
| WP_258968658.1 | DUF499_1.RM.like.systems | XIS-HTH->?->?->HTH_3->TUDOR+SF1-SF2-Helicase+iREase+CTD+REase->METHYLASE->HEPN+DUF499*-><-?<-HicB<-TOPRIM<-SbcC\|\|BirA-HTH-><-Phage_int_SAM_5+Phage_integrase | 1096 | Bacteria>FCB_group>Bacteroidota/Chlorobiota_group>Bacteroidota>Bacteroidia | GCF_024790655.1 | Phocaeicola_vulgatus | NXV64_RS08565 | 2221353..2224643 | 821 | DUF499-domain-containing-protein-[Phocaeicola-vulgatus] |
| WP_055289369.1 | DUF499_1.RM.like.systems | REC-><-METHYLASE-p<-ABhydrolase<-ACET<-DOC+HTH<-HEPN+DUF499*<-METHYLASE<-TUDOR+SF1-SF2-Helicase+iREase+CTD+REase | 1099 | Bacteria>FCB_group>Bacteroidota/Chlorobiota_group>Bacteroidota>Bacteroidia | GCF_025914195.1 | Bacteroidales | KSY52_RS08740 | 147903..151202 | 171549 | MULTISPECIES:-DUF499-domain-containing-protein-[Bacteroidales] |
| WP_118211914.1 | DUF499_1.RM.like.systems | PrmC_N+Methyltransf_25->?->PRTase->?->tRNA->tRNA-><-HEPN+DUF499*<-METHYLASE<-REase+ParB<-TUDOR+SF1-SF2-Helicase+iREase+CTD+REase-p<-TUDOR+SF1-SF2-Helicase+iREase+CTD+REase-p<-HTH_3<-DnaJ | 1098 | Bacteria>FCB_group>Bacteroidota/Chlorobiota_group>Bacteroidota>Bacteroidia | GCF_027661405.1 | Phocaeicola_plebeius | DWY45_RS04060 | 51608..54904 | 310297 | DUF499-domain-containing-protein-[Phocaeicola-plebeius] |
| WP_118026126.1 | DUF499_1.RM.like.systems | DnaJ->HTH_3->TUDOR+SF1-SF2-Helicase+iREase+CTD+REase->REase+ParB->METHYLASE->HEPN+DUF499*-><-tRNA<-tRNA<-?<-?<-PRTase<-PRTase<-?<-PrmC_N+Methyltransf_25 | 1098 | Bacteria>FCB_group>Bacteroidota/Chlorobiota_group>Bacteroidota>Bacteroidia | GCF_027686295.1 | Phocaeicola_plebeius | PGZ43_RS01830 | 134398..137694 | 310297 | DUF499-domain-containing-protein-[Phocaeicola-plebeius] |
| WP_270668664.1 | DUF499_1.RM.like.systems | <-BirA-HTH\|\|SbcC->TOPRIM->HicB->?-><-?<-HEPN+DUF499*<-METHYLASE<-TUDOR+SF1-SF2-Helicase+iREase+CTD+REase<-HTH_3<-?<-?<-XIS-HTH | 1096 | Bacteria>FCB_group>Bacteroidota/Chlorobiota_group>Bacteroidota>Bacteroidia | GCF_027686505.1 | Phocaeicola_vulgatus | PG001_RS15345 | 5861..9151 | 821 | DUF499-domain-containing-protein-[Phocaeicola-vulgatus] |
| WP_071137164.1 | DUF499_1.RM.like.systems | PAIREDC-HTH-><-SF1-SF2-Helicase+DUF1998<-?<-HEPN+DUF499*<-METHYLASE<-TUDOR+SF1-SF2-Helicase+iREase+CTD+REase<-DDE_Tnp_1_3<-WYL+WYL-C<-HTH_3 | 1108 | Bacteria>FCB_group>Bacteroidota/Chlorobiota_group>Bacteroidota>Bacteroidia | GCF_900095795.1 | Petrimonas_mucosa | ING2E5A_RS09235 | 2277768..2281094 | 1642646 | DUF499-domain-containing-protein-[Petrimonas-mucosa] |
| WP_074635742.1 | DUF499_1.RM.like.systems | DEDD_Tnp_IS110->HTH_3->TUDOR+SF1-SF2-Helicase+iREase+CTD+REase->METHYLASE->HEPN+DUF499*->Schlafen+HATPase+HTH->?->HTH_3->DNA_methylase->MORC-> | 1096 | Bacteria>FCB_group>Bacteroidota/Chlorobiota_group>Bacteroidota>Bacteroidia | GCF_900100465.1 | Bacteroides_ovatus | BLS71_RS01410 | 96540..99830 | 28116 | DUF499-domain-containing-protein-[Bacteroides-ovatus] |
| WP_032849356.1 | DUF499_1.RM.like.systems | <-HEPN+DUF499*<-METHYLASE<-TUDOR+SF1-SF2-Helicase+iREase+CTD+REase<-DEDD_Tnp_IS110<-?<-?\|\|HTH-> | 1096 | Bacteria>FCB_group>Bacteroidota/Chlorobiota_group>Bacteroidota>Bacteroidia | GCF_901212435.1 | Bacteroides | PO242_RS12935 | 72725..76015 | 816 | MULTISPECIES:-DUF499-domain-containing-protein-[Bacteroides] |
| WP_101543458.1 | DUF499_1.RM.like.systems | <-HEPN+DUF499*<-METHYLASE<-TUDOR+SF1-SF2-Helicase+iREase+CTD+REase<-HTH_3<-DDE_Tnp_1<-HAD+HAD | 1096 | Bacteria>FCB_group>Bacteroidota/Chlorobiota_group>Bacteroidota>Bacteroidia | GCF_902385525.1 | Bacteroidaceae | C1A09_RS08785 | 351382..354672 | 815 | MULTISPECIES:-DUF499-domain-containing-protein-[Bacteroidaceae] |
| WP_118170513.1 | DUF499_1.RM.like.systems | <-HEPN+DUF499*<-METHYLASE<-TUDOR+SF1-SF2-Helicase+iREase+CTD+REase<-HTH_3<-XIS-HTH | 1096 | Bacteria>FCB_group>Bacteroidota/Chlorobiota_group>Bacteroidota>Bacteroidia | GCF_934726555.1 | Bacteroidaceae | QV389_RS11685 | 2128..5418 | 815 | MULTISPECIES:-DUF499-domain-containing-protein-[Bacteroidaceae] |
| WP_118432131.1 | DUF499_1.RM.like.systems | PrmC_N+Methyltransf_25->?->PRTase->PRTase->?->?->tRNA->tRNA-><-HEPN+DUF499*<-METHYLASE<-REase+ParB<-TUDOR+SF1-SF2-Helicase+iREase+CTD+REase<-HTH_3<-DnaJ | 1098 | Bacteria>FCB_group>Bacteroidota/Chlorobiota_group>Bacteroidota>Bacteroidia | GCF_958447975.1 | Phocaeicola_plebeius | PGY45_RS13550 | 13557..16853 | 310297 | DUF499-domain-containing-protein-[Phocaeicola-plebeius] |
| WP_118338055.1 | DUF499_1.RM.like.systems | <-HEPN+DUF499*<-METHYLASE<-REase+ParB<-TUDOR+SF1-SF2-Helicase+iREase+CTD+REase<-HTH_3<-DnaJ | 1098 | Bacteria>FCB_group>Bacteroidota/Chlorobiota_group>Bacteroidota>Bacteroidia | GCF_959025615.1 | Phocaeicola_plebeius | PG845_RS15420 | 1179..4475 | 310297 | DUF499-domain-containing-protein-[Phocaeicola-plebeius] |
| WP_284525582.1 | DUF499_1.RM.like.systems | <-DUF499-p<-DUF499-p<-HEPN+DUF499*<-METHYLASE-p<-METHYLASE<-REase+ParB<-?<-TUDOR+SF1-SF2-Helicase+iREase+CTD+REase<-HTH_3 | 518 | Bacteria>FCB_group>Bacteroidota/Chlorobiota_group>Bacteroidota>Bacteroidia | - | Phocaeicola_plebeius | DW982_RS17975 | 3045..4601 | 310297 | DUF499-domain-containing-protein-[Phocaeicola-plebeius] |
| MBB1285247.1 | DUF499_1.RM.like.systems | <-HEPN+DUF499*<-METHYLASE<-AbiEii<-wHTH+AbiEi<-TUDOR+SF1-SF2-Helicase+iREase+CTD+REase<-?\|\|MerR-HTH-> | 1093 | Bacteria>FCB_group>Bacteroidota/Chlorobiota_group>Bacteroidota>Chitinophagia | GCA_014163555.1 | Flavisolibacter_longurius | HRH25_12755 | 53623..56904 | 2740583 | DUF499-domain-containing-protein-[Flavisolibacter-longurius] |
| MBN8676639.1 | DUF499_1.RM.like.systems | REase+35exo+Beta-Barrel+SF1-SF2-Helicase->?->?->TUDOR+SF1-SF2-Helicase+iREase+CTD+REase->ParB+HNH->METHYLASE->HEPN+DUF499*->REase-> | 1087 | Bacteria>FCB_group>Bacteroidota/Chlorobiota_group>Bacteroidota>Chitinophagia | GCA_017303475.1 | Chitinophagales_bacterium | JOM29_00350 | 102288..105551 | 2448779 | DUF499-domain-containing-protein-[Chitinophagales-bacterium] |
| MBW7891560.1 | DUF499_1.RM.like.systems | TUDOR+SF1-SF2-Helicase+iREase+CTD+REase-p->AbiEi_1->METHYLASE->HEPN+DUF499*-> | 917 | Bacteria>FCB_group>Bacteroidota/Chlorobiota_group>Bacteroidota>Chitinophagia | GCA_019454915.1 | Chitinophagaceae_bacterium | H3C48_11165 | 5923..8672 | 1869212 | ATP-binding-protein-[Chitinophagaceae-bacterium] |
| MCB9047412.1 | DUF499_1.RM.like.systems | <-REase+35exo<-?<-WYL+WYL-C<-ParB+HNH<-?<-HEPN+DUF499*<-METHYLASE<-TUDOR+SF1-SF2-Helicase+iREase+CTD+REase\|\|HKD-DNase->wHTH+SAD+HNH-> | 1086 | Bacteria>FCB_group>Bacteroidota/Chlorobiota_group>Bacteroidota>Chitinophagia | GCA_020636495.1 | Chitinophagales_bacterium | H6550_14865 | 3305993..3309253 | 2448779 | DUF499-domain-containing-protein-[Chitinophagales-bacterium] |
| MCG9899089.1 | DUF499_1.RM.like.systems | <-REase<-?\|\|?-><-?<-RVT<-?<-HEPN+DUF499*<-METHYLASE<-AbiEii<-AbiEi_1<-TUDOR+SF1-SF2-Helicase+iREase+CTD+REase | 1108 | Bacteria>FCB_group>Bacteroidota/Chlorobiota_group>Bacteroidota>Chitinophagia | GCA_022402565.1 | Hydrotalea_sp. | MH132_03730 | 9399..12725 | 2881279 | DUF499-domain-containing-protein-[Hydrotalea-sp.] |
| MBC7408026.1 | DUF499_1.RM.like.systems | AbiEii->METHYLASE->HEPN+DUF499*-> | 559 | Bacteria>FCB_group>Bacteroidota/Chlorobiota_group>Bacteroidota>Cytophagia | GCA_014376355.1 | Arcicella_sp. | H7339_06525 | 3538..5214 | 1872628 | DUF499-domain-containing-protein-[Arcicella-sp.] |
| MBK7652070.1 | DUF499_1.RM.like.systems | HTH->REase->METHYLASE-p->HEPN+DUF499*->DUF499-p->DUF499-p-> | 498 | Bacteria>FCB_group>Bacteroidota/Chlorobiota_group>Bacteroidota>Cytophagia | GCA_016709845.1 | Flammeovirgaceae_bacterium | IPJ20_16990 | 23879..25375 | 2026740 | DUF499-domain-containing-protein-[Flammeovirgaceae-bacterium] |
| WP_013929916.1 | DUF499_1.RM.like.systems | <-DDE_Tnp_1<-BirA-HTH<-DDE_Tnp_1_assoc+DDE_Tnp_1<-MULE-TRANSPOSASE<-SF1-SF2-Helicase<-MORC+DUF3684+REase<-HEPN+DUF499*<-METHYLASE-p\|\|DDE_Tnp_1-><-METHYLASE<-REase<-TUDOR+SF1-SF2-Helicase+iREase+CTD+REase<-AbiEi_4 | 1094 | Bacteria>FCB_group>Bacteroidota/Chlorobiota_group>Bacteroidota>Cytophagia | GCF_000218895.1 | Runella_slithyformis | RUNSL_RS20985 | 4951748..4955032 | 106 | DUF499-domain-containing-protein-[Runella-slithyformis] |
| WP_027303393.1 | DUF499_1.RM.like.systems | MNS+REase->CHTH->?->?-><-?\|\|MORC+HisKin+REase-><-HEPN+DUF499*<-wHTH+REase<-METHYLASE\|\|?-><-TUDOR+SF1-SF2-Helicase+iREase+CTD+REase<-PAIREDC-HTH\|\|Phage_integrase-> | 1100 | Bacteria>FCB_group>Bacteroidota/Chlorobiota_group>Bacteroidota>Cytophagia | GCF_000383955.1 | Rudanella_lutea | RUDLU_RS0125935 | 41364..44666 | 451374 | DUF499-domain-containing-protein-[Rudanella-lutea] |
| WP_070725327.1 | DUF499_1.RM.like.systems | <-SF1-SF2-Helicase+REase<-?\|\|TUDOR+SF1-SF2-Helicase+iREase+CTD+REase->OLD-ABC->HNH->METHYLASE->HEPN+DUF499*-><-HTH | 1110 | Bacteria>FCB_group>Bacteroidota/Chlorobiota_group>Bacteroidota>Cytophagia | GCF_001816145.1 | Hymenobacter_lapidarius | BEN47_RS03945 | 71126..74458 | 1908237 | DUF499-domain-containing-protein-[Hymenobacter-lapidarius] |
| WP_106140933.1 | DUF499_1.RM.like.systems | TUDOR+SF1-SF2-Helicase+iREase+CTD+REase->wHTH+AbiEi->AbiEii->METHYLASE->HEPN+DUF499*->?->SF1-SF2-Helicase+DUF1998+HKD-DNase+HKD-DNase-> | 1123 | Bacteria>FCB_group>Bacteroidota/Chlorobiota_group>Bacteroidota>Cytophagia | GCF_003002825.1 | Spirosoma_oryzae | CLV58_RS26865 | 12572..15943 | 1469603 | DUF499-domain-containing-protein-[Spirosoma-oryzae] |
| WP_114065477.1 | DUF499_1.RM.like.systems | <-TUDOR+SF1-SF2-Helicase+iREase+CTD<-HEPN+DUF499*<-?<-OLD-ABC<-ParB+HNH<-METHYLASE<-REase<-TUDOR+SF1-SF2-Helicase+iREase+CTD+REase | 1094 | Bacteria>FCB_group>Bacteroidota/Chlorobiota_group>Bacteroidota>Cytophagia | GCF_003325355.1 | Runella_rosea | DR864_RS02565 | 579082..582366 | 2259595 | DUF499-domain-containing-protein-[Runella-rosea] |
| WP_130024109.1 | DUF499_1.RM.like.systems | <-HTH<-XIS-HTH<-?<-HEPN+DUF499*<-METHYLASE<-AbiEii<-wHTH+AbiEi<-REase<-TUDOR+SF1-SF2-Helicase+iREase+CTD+REase | 1096 | Bacteria>FCB_group>Bacteroidota/Chlorobiota_group>Bacteroidota>Cytophagia | GCF_004168285.1 | Emticicia_agri | EWM59_RS25780 | 4303..7593 | 2492393 | DUF499-domain-containing-protein-[Emticicia-agri] |
| WP_209329595.1 | DUF499_1.RM.like.systems | <-BirA-HTH<-?<-?<-HEPN+DUF499*<-AAA-ATPase+REase<-METHYLASE<-ParB+HNH<-TUDOR+SF1-SF2-Helicase+iREase+CTD+REase | 1097 | Bacteria>FCB_group>Bacteroidota/Chlorobiota_group>Bacteroidota>Cytophagia | GCF_017811155.1 | Lunatimonas_salinarum | IAQ59_RS04505 | 160980..164273 | 1774590 | DUF499-domain-containing-protein-[Lunatimonas-salinarum] |
| WP_163384119.1 | DUF499_1.RM.like.systems | HEPN->?->?->?->Peptidase-><-HEPN+DUF499*<-METHYLASE<-TUDOR+SF1-SF2-Helicase+iREase+CTD+REase<-HTH<-?<-?<-XIS-HTH | 1103 | Bacteria>FCB_group>Bacteroidota/Chlorobiota_group>Bacteroidota>Cytophagia | GCF_030409895.1 | Cyclobacterium_jeungdonense | QWZ15_RS05330 | 512550..515861 | 708087 | Swt1-family-HEPN-domain-containing-protein-[Cyclobacterium-jeungdonense] |
| WP_092673506.1 | DUF499_1.RM.like.systems | HIN-HTH->HTH_21+rve+rve_3-><-SIR2\|\|?->?-><-HEPN+DUF499*<-METHYLASE<-TUDOR+SF1-SF2-Helicase+iREase+CTD+REase<-HTH_3<-?<-URI-DNase<-URI-DNase | 1128 | Bacteria>FCB_group>Bacteroidota/Chlorobiota_group>Bacteroidota>Cytophagia | GCF_900115775.1 | Hymenobacter_arizonensis | BMZ66_RS12440 | 170708..174094 | 1227077 | DUF499-domain-containing-protein-[Siccationidurans-arizonensis] |
| MAM23120.1 | DUF499_1.RM.like.systems | HTH+MPTase->PIN->HTH_3->TUDOR+SF1-SF2-Helicase+iREase+CTD+REase->METHYLASE->HEPN+DUF499*-> | 869 | Bacteria>FCB_group>Bacteroidota/Chlorobiota_group>Bacteroidota>Flavobacteriia | GCA_002695265.1 | Croceibacter_sp. | CL817_05250 | 9003..11611 | 2024829 | hypothetical-protein-[Croceibacter-sp.] |
| HCE56000.1 | DUF499_1.RM.like.systems | <-PIN<-HEPN+DUF499*<-METHYLASE<-AbiEii<-AbiEi_1<-TUDOR+SF1-SF2-Helicase+iREase+CTD+REase<-AbiEii<-AbiEi_4 | 1089 | Bacteria>FCB_group>Bacteroidota/Chlorobiota_group>Bacteroidota>Flavobacteriia | GCA_003516285.1 | Lutibacter_sp. | DER05_13810 | 23309..26578 | 1925666 | hypothetical-protein-[Lutibacter-sp.] |
| NEW78740.1 | DUF499_1.RM.like.systems | HKD-DNase->?-><-HEPN<-HKD-DNase<-HEPN+DUF499*<-METHYLASE<-AbiEii<-AbiEi_1<-TUDOR+SF1-SF2-Helicase+iREase+CTD+REase | 1089 | Bacteria>FCB_group>Bacteroidota/Chlorobiota_group>Bacteroidota>Flavobacteriia | GCA_010906935.1 | Gelidibacter_sp. | GZ086_04810 | 7309..10578 | 2018083 | ATP-binding-protein-[Gelidibacter-sp.] |
| MBX7052699.1 | DUF499_1.RM.like.systems | MazG->REase+SF1-SF2-Helicase->TUDOR+SF1-SF2-Helicase+iREase+CTD+REase->OLD-ABC->TOPRIM->METHYLASE->HEPN+DUF499*-> | 1088 | Bacteria>FCB_group>Bacteroidota/Chlorobiota_group>Bacteroidota>Flavobacteriia | GCA_019694805.1 | Flavobacteriales_bacterium | K1X54_11735 | 22264..25530 | 2021391 | DUF499-domain-containing-protein-[Flavobacteriales-bacterium] |
| WP_143387496.1 | DUF499_1.RM.like.systems | TUDOR+SF1-SF2-Helicase+iREase+CTD+REase->AbiEi_1->AbiEii->METHYLASE->HEPN+DUF499*->?->?->?->PIN->?-><-WYL+WYL-C | 1088 | Bacteria>FCB_group>Bacteroidota/Chlorobiota_group>Bacteroidota>Flavobacteriia | GCF_007097285.1 | Flavobacterium_gawalongense | FNW12_RS09775 | 80798..84064 | 2594432 | DUF499-domain-containing-protein-[Flavobacterium-gawalongense] |
| EJF54826.1 | DUF499_1.RM.like.systems | TUDOR+SF1-SF2-Helicase+iREase+CTD+REase->DUF4096+DDE_Tnp_1+DDE_Tnp_1_2->HEPN+DUF499*-> | 1104 | Bacteria>FCB_group>Bacteroidota/Chlorobiota_group>Bacteroidota>Saprospiria | GCA_000275825.1 | Saprospira_grandis_DSM_2844 | SapgrDRAFT_3181 | 3964674..3967988 | 694433 | putative-ATPase-(AAA+-superfamily)-[Saprospira-grandis-DSM-2844] |
| NUQ24578.1 | DUF499_1.RM.like.systems | TUDOR+SF1-SF2-Helicase+iREase+CTD+REase->OLD-ABC->HNH->METHYLASE->AAA-ATPase+REase->HEPN+DUF499*->?->PolB->HEPN-><-ORC-CDC6+REase | 1095 | Bacteria>FCB_group>Bacteroidota/Chlorobiota_group>Bacteroidota>Saprospiria | GCA_013360405.1 | Saprospiraceae_bacterium | HUU34_11550 | 44219..47506 | 2202734 | DUF499-domain-containing-protein-[Saprospiraceae-bacterium] |
| MBK6932297.1 | DUF499_1.RM.like.systems | <-Glyco_hydro_1\|\|?-><-DUF499-p<-HEPN+DUF499*<-URI-DNase<-METHYLASE-p<-?<-METHYLASE-p<-HNH | 951 | Bacteria>FCB_group>Bacteroidota/Chlorobiota_group>Bacteroidota>Saprospiria | GCA_016709195.1 | Saprospirales_bacterium | IPH12_16140 | 92714..95569 | 2026790 | DUF499-domain-containing-protein-[Saprospirales-bacterium] |
| MBK8877301.1 | DUF499_1.RM.like.systems | TUDOR+SF1-SF2-Helicase+iREase+CTD+REase->HEPN+DUF499*->?->?-><-?<-SMODS | 1105 | Bacteria>FCB_group>Bacteroidota/Chlorobiota_group>Bacteroidota>Saprospiria | GCA_016715005.1 | Haliscomenobacter_sp. | IPN74_01745 | 433020..436337 | 2717303 | DUF499-domain-containing-protein-[Haliscomenobacter-sp.] |
| MBK8920115.1 | DUF499_1.RM.like.systems | <-Glyco_hydro_1<-?<-HEPN+DUF499*<-URI-DNase<-METHYLASE-p<-METHYLASE-p<-METHYLASE-p<-TOPRIM<-OLD-ABC | 1088 | Bacteria>FCB_group>Bacteroidota/Chlorobiota_group>Bacteroidota>Saprospiria | GCA_016719895.1 | Saprospirales_bacterium | IPM81_01165 | 280177..283443 | 2026790 | DUF499-domain-containing-protein-[Saprospirales-bacterium] |
| MCS7037545.1 | DUF499_1.RM.like.systems | TUDOR+SF1-SF2-Helicase+iREase+CTD+REase-p->METHYLASE->HEPN+DUF499*-> | 1108 | Bacteria>FCB_group>Bacteroidota/Chlorobiota_group>Bacteroidota>Saprospiria | GCA_025056735.1 | Saprospiraceae_bacterium | NZM41_12835 | 4336..7662 | 2202734 | DUF499-domain-containing-protein-[Saprospiraceae-bacterium] |
| WP_051018640.1 | DUF499_1.RM.like.systems | TUDOR+SF1-SF2-Helicase+iREase+CTD+REase->METHYLASE-p->?->DDE_Tnp_1+DDE_Tnp_1_2->METHYLASE->HEPN+DUF499*->?->?->?->?-><-?<-Peptidase | 1105 | Bacteria>FCB_group>Bacteroidota/Chlorobiota_group>Bacteroidota>Saprospiria | GCF_000275825.1 | Saprospira_grandis | SAPGRDRAFT_RS17610 | 3964671..3967988 | 1008 | DUF499-domain-containing-protein-[Saprospira-grandis] |
| WP_222581795.1 | DUF499_1.RM.like.systems | <-HIN-HTH\|\|METHYLASE->HEPN+DUF499*-> | 1089 | Bacteria>FCB_group>Bacteroidota/Chlorobiota_group>Bacteroidota>Saprospiria | GCF_019802045.1 | Membranihabitans_marinus | KUV50_RS18995 | 2282..5551 | 1227546 | DUF499-domain-containing-protein-[Membranihabitans-marinus] |
| WP_270099212.1 | DUF499_1.RM.like.systems | Peptidase-><-?<-?<-?<-HEPN+DUF499*<-METHYLASE<-TUDOR+SF1-SF2-Helicase+iREase+CTD+REase\|\|?->?->Y1_Tnp-> | 1105 | Bacteria>FCB_group>Bacteroidota/Chlorobiota_group>Bacteroidota>Saprospiria | GCF_027594745.1 | Saprospira_grandis | OP864_RS16310 | 4154293..4157610 | 1008 | DUF499-domain-containing-protein-[Saprospira-grandis] |
| OWY19366.1 | DUF499_1.RM.like.systems | <-HEPN+DUF499*<-METHYLASE<-TUDOR+SF1-SF2-Helicase+iREase+CTD+REase | 1103 | Bacteria>FCB_group>Bacteroidota/Chlorobiota_group>Bacteroidota>Sphingobacteriia | GCA_002212985.1 | Sphingobacteriales_bacterium_TSM_CSS | B6N25_14540 | 46408..49719 | 1972511 | hypothetical-protein-B6N25_14540-[Sphingobacteriales-bacterium-TSM_CSS] |
| MDB5032646.1 | DUF499_1.RM.like.systems | Phage_int_SAM_5+Phage_integrase->DEDD_Tnp_IS110+Transposase_20-><-ABhydrolase<-HEPN+DUF499*<-METHYLASE<-TUDOR+SF1-SF2-Helicase+iREase+CTD+REase<-?\|\|SMODS->TIR-> | 1091 | Bacteria>FCB_group>Bacteroidota/Chlorobiota_group>Bacteroidota>Sphingobacteriia | GCA_028287845.1 | Mucilaginibacter_sp. | JWP71_3367 | 6008..9283 | 1882438 | MAG:-hypothetical-protein-JWP71_3367-[Mucilaginibacter-sp.] |
| WP_090973307.1 | DUF499_1.RM.like.systems | TUDOR+SF1-SF2-Helicase+iREase+CTD+REase->METHYLASE->?->DOC->HEPN+DUF499*->?->PIN->tRNA->?->Synaptojanin-> | 1100 | Bacteria>FCB_group>Bacteroidota/Chlorobiota_group>Bacteroidota>Sphingobacteriia | GCF_900112315.1 | Parapedobacter_composti | BM030_RS09250 | 52170..55472 | 623281 | DUF499-domain-containing-protein-[Parapedobacter-composti] |
| RLD44971.1 | DUF499_1.RM.like.systems | <-HEPN+DUF499*<-METHYLASE-p<-METHYLASE-p | 726 | Bacteria>FCB_group>Bacteroidota/Chlorobiota_group>Bacteroidota>unclassified_Bacteroidota | GCA_003650385.1 | Bacteroidota_bacterium | DRI88_08990 | 1..2178 | 1898104 | AAA+-family-ATPase,-partial-[Bacteroidetes-bacterium] |
| MBT3751042.1 | DUF499_1.RM.like.systems | METHYLASE-p->HEPN+DUF499*->?->?->GDResolvase+Recombinase+ZnR-> | 1103 | Bacteria>FCB_group>Bacteroidota/Chlorobiota_group>Bacteroidota>unclassified_Bacteroidota | GCA_018653945.1 | Bacteroidota_bacterium | HOG34_18835 | 220..3531 | 1898104 | ATP-binding-protein-[Bacteroidetes-bacterium] |
| MCB0540282.1 | DUF499_1.RM.like.systems | METHYLASE-p->HEPN+DUF499*-> | 769 | Bacteria>FCB_group>Bacteroidota/Chlorobiota_group>Bacteroidota>unclassified_Bacteroidota | GCA_020436205.1 | Bacteroidota_bacterium | KDE33_22385 | 287..2592 | 1898104 | DUF499-domain-containing-protein-[Bacteroidetes-bacterium] |
| MCL5268276.1 | DUF499_1.RM.like.systems | HNH->?-><-?<-REase<-REase<-TIR<-HEPN+DUF499*<-METHYLASE<-DrHyd+BirA-HTH<-TUDOR+SF1-SF2-Helicase+iREase+CTD+REase<-?<-?<-METHYLASE-p | 1111 | Bacteria>FCB_group>Bacteroidota/Chlorobiota_group>Bacteroidota>unclassified_Bacteroidota | GCA_023480735.1 | Bacteroidota_bacterium | M1469_09265 | 8457..11792 | 1898104 | Swt1-family-HEPN-domain-containing-protein-[Bacteroidetes-bacterium] |

| Protein-ID | System | Gene-Neighbourhoods | Length | Taxonomy | Assembly-ID | Strain | Locus-Tag | Genomic-coordinates | TaxID | Fasta-define |
|---|---|---|---|---|---|---|---|---|---|---|
| MCY3629603.1 | DUF499_1.RM.like.systems | <-MNS\|\|TUDOR+ SF1-SF2-Helicase+iREase+CTD+REase->HEPN+DUF499*->Phage_integrase->Phage_int_SAM_3+Phage_integrase-> | 1099 | Bacteria>FCB_group>Bacteroidota/Chlorobiota_group>Bacteroidota>unclassified_Bacteroidota | GCA_026705375.1 | Bacteroidota_bacterium | OXG94_06260 | 16273..19572 | 1898104 | MAG:-DUF499-domain-containing-protein-[Bacteroidetes-bacterium] |
| MCY3999495.1 | DUF499_1.RM.like.systems | REase->?->?->?->?->TUDOR+ SF1-SF2-Helicase+iREase+CTD+REase->HEPN+DUF499*->Phage_int_SAM_3+Phage_integrase->Phage_int_SAM_3+Phage_integrase->METHYLASE-p-> | 1095 | Bacteria>FCB_group>Bacteroidota/Chlorobiota_group>Bacteroidota>unclassified_Bacteroidota | GCA_026708685.1 | Bacteroidota_bacterium | OXF84_01635 | 21572..24859 | 1898104 | MAG:-DUF499-domain-containing-protein-[Bacteroidetes-bacterium] |
| MCY4223999.1 | DUF499_1.RM.like.systems | <-Phage_int_SAM_3+Phage_integrase<-Phage_integrase<-HEPN+DUF499* | 1027 | Bacteria>FCB_group>Bacteroidota/Chlorobiota_group>Bacteroidota>unclassified_Bacteroidota | GCA_026709435.1 | Bacteroidota_bacterium | OXF06_04095 | 2186..5267 | 1898104 | MAG:-DUF499-domain-containing-protein,-partial-[Bacteroidetes-bacterium] |
| MDE2834954.1 | DUF499_1.RM.like.systems | MNS-><-?\|\|TUDOR+ SF1-SF2-Helicase+iREase+CTD+REase->HEPN+DUF499*->Phage_integrase->Phage_int_SAM_3+Phage_integrase-> | 1086 | Bacteria>FCB_group>Bacteroidota/Chlorobiota_group>Bacteroidota>unclassified_Bacteroidota | GCA_028874735.1 | Bacteroidota_bacterium | OXM02_10600 | 6360..9620 | 1898104 | MAG:-Swt1-family-HEPN-domain-containing-protein-[Bacteroidota-bacterium] |
| MDE2771369.1 | DUF499_1.RM.like.systems | <-DnaJ<-?<-?\|\|?->HEPN+DUF499*->Phage_integrase->Phage_integrase-> | 1094 | Bacteria>FCB_group>Bacteroidota/Chlorobiota_group>Bacteroidota>unclassified_Bacteroidota | GCA_028875125.1 | Bacteroidota_bacterium | OXI44_09415 | 24188..27472 | 1898104 | MAG:-DUF499-domain-containing-protein-[Bacteroidota-bacterium] |
| MDE2731786.1 | DUF499_1.RM.like.systems | <-METHYLASE-p<-Phage_integrase\|\|?-><-HEPN+DUF499*<-TUDOR+ SF1-SF2-Helicase+iREase+CTD+REase<-MNS | 1086 | Bacteria>FCB_group>Bacteroidota/Chlorobiota_group>Bacteroidota>unclassified_Bacteroidota | GCA_028875295.1 | Bacteroidota_bacterium | OXI38_10375 | 1957..5217 | 1898104 | MAG:-Swt1-family-HEPN-domain-containing-protein-[Bacteroidota-bacterium] |
| WP_210512925.1 | DUF499_1.RM.like.systems | Ntox49->?->tRNA-><-XIS-HTH\|\|TUDOR+ SF1-SF2-Helicase+iREase+CTD+REase->METHYLASE->HEPN+DUF499*-> | 1089 | Bacteria>FCB_group>Bacteroidota/Chlorobiota_group>Balneolota>Balneolia | GCF_017921895.1 | Natronogracilivirga_saccharolytica | NATSA_RS12455 | 139567..142836 | 2812953 | DUF499-domain-containing-protein-[Natronogracilivirgula-saccharolytica] |
| GIV55345.1 | DUF499_1.RM.like.systems | <-LexA-HTH+LexA-protease\|\|?->Methyltransf_5-><-MORC+DUF3684<-HEPN+DUF499*<-METHYLASE<-TUDOR+ SF1-SF2-Helicase+iREase+CTD+REase | 1140 | Bacteria>FCB_group>Bacteroidota/Chlorobiota_group>Candidatus_Kapabacteria>unclassified_Candidatus_Kapabacteria | GCA_026003495.1 | Candidatus_Kapabacteria_bacterium | KatS3mg040_0113 | 136007..139429 | 2447898 | MAG:-hypothetical-protein-KatS3mg040_0113-[Candidatus-Kapabacteria-bacterium] |
| MBL6956529.1 | DUF499_1.RM.like.systems | <-HTH+MPTase<-Tox-RelE<-?<-HEPN+DUF499*<-METHYLASE<-wHTH+REase<-TUDOR+ SF1-SF2-Helicase+iREase+CTD+REase | 1115 | Bacteria>FCB_group>Bacteroidota/Chlorobiota_group>Chlorobiota>Chlorobiia | GCA_016776555.1 | Chlorobium_phaeobacteroides | ISR54_06890 | 24108..27455 | 1096 | DUF499-domain-containing-protein-[Chlorobium-phaeobacteroides] |
| WP_059138717.1 | DUF499_1.RM.like.systems | RelE-ParE->TUDOR+ SF1-SF2-Helicase+iREase+CTD+REase->wHTH+REase->METHYLASE->HEPN+DUF499*->?->?->?->TraJ-RHH-> | 1115 | Bacteria>FCB_group>Bacteroidota/Chlorobiota_group>Chlorobiota>Chlorobiia | GCF_001509575.1 | Chlorobium_limicola | ASB62_RS03910 | 18289..21636 | 1092 | Swt1-family-HEPN-domain-containing-protein-[Chlorobium-limicola] |
| ARA93108.1 | DUF499_1.RM.like.systems | <-RVT<-MORC<-MethyltransfD12<-?<-DUF4011+SF1-SF2-Helicase+REase<-HEPN+DUF499*<-METHYLASE<-MORC+DUF3684<-TUDOR+ SF1-SF2-Helicase+iREase+CTD+REase<-?\|\|PIN-> | 1112 | Bacteria>FCB_group>Bacteroidota/Chlorobiota_group>Rhodothermota>Rhodothermia | GCA_001518995.2 | Rhodothermaceae_bacterium_RA | AWN76_007995 | 1928774..1932112 | 1779382 | AAA+-family-ATPase-[Rhodothermaceae-bacterium-RA] |
| MXW14654.1 | DUF499_1.RM.like.systems | <-MNS\|\|TUDOR+ SF1-SF2-Helicase+iREase+CTD+REase->HEPN+DUF499*->Phage_integrase->Phage_int_SAM_3+Phage_integrase-> | 1099 | Bacteria>FCB_group>Bacteroidota/Chlorobiota_group>Rhodothermota>Rhodothermia | GCA_009835915.1 | Rhodothermaceae_bacterium | F4Z76_07000 | 16270..19569 | 2026787 | ATP-binding-protein-[Rhodothermaceae-bacterium] |
| MXW34066.1 | DUF499_1.RM.like.systems | <-MNS\|\|TUDOR+ SF1-SF2-Helicase+iREase+CTD+REase->HEPN+DUF499*->Phage_integrase->Phage_int_SAM_3+Phage_integrase-> | 1099 | Bacteria>FCB_group>Bacteroidota/Chlorobiota_group>Rhodothermota>Rhodothermia | GCA_009836215.1 | Rhodothermaceae_bacterium | F4Z62_13320 | 16280..19579 | 2026787 | ATP-binding-protein-[Rhodothermaceae-bacterium] |
| WP_210373541.1 | DUF499_1.RM.like.systems | Peptidase_M28-><-?\|\|?->?->SIGMA-HTH->?-><-HEPN+DUF499*<-METHYLASE<-?<-TUDOR+ SF1-SF2-Helicase+iREase+CTD+REase | 1114 | Bacteria>FCB_group>Bacteroidota/Chlorobiota_group>Rhodothermota>Rhodothermia | GCF_017908595.1 | Rhodothermus_bifroesti | J8E65_RS01005 | 262538..265882 | 2823335 | Swt1-family-HEPN-domain-containing-protein-[Rhodothermus-bifroesti] |
| MBW7874866.1 | DUF499_1.RM.like.systems | <-XIS-HTH\|\|METHYLASE->HTH+MPTase->?->Classical-AAA->HEPN+DUF499*->?->?->Polbeta->HEPN-> | 1093 | Bacteria>FCB_group>Candidatus_Cloacimonadota>unclassified_Candidatus_Cloacimonadota | GCA_019454875.1 | Candidatus_Cloacimonadota_bacterium | H3C47_02625 | 20403..23684 | 2030808 | ATP-binding-protein-[Candidatus-Cloacimonetes-bacterium] |
| MBP7719545.1 | DUF499_1.RM.like.systems | TUDOR+ SF1-SF2-Helicase+iREase+CTD+REase->METHYLASE->OLD-ABC->TOPRIM->HEPN+DUF499*-> | 1111 | Bacteria>FCB_group>Candidatus_Fermentibacteria>Candidatus_Fermentibacteria_(class)>Candidatus_Fermentibacterales | GCA_017999735.1 | Candidatus_Fermentibacter_sp. | KA108_02790 | 14441..17776 | 2779163 | DUF499-domain-containing-protein-[Candidatus-Fermentibacter-sp.] |
| MCC6694832.1 | DUF499_1.RM.like.systems | TUDOR+ SF1-SF2-Helicase+iREase+CTD+REase->REase->HEPN->METHYLASE-p->?->METHYLASE->HEPN+DUF499*->?->?->ArdCN+MPTase-><-Arm-DNA-bind_3+Phage_integrase | 1102 | Bacteria>FCB_group>Candidatus_Hydrogenedentes>unclassified_Candidatus_Hydrogenedentes | GCA_020849535.1 | Candidatus_Hydrogenedentes_bacterium | IT365_04280 | 13299..16607 | 2030809 | DUF499-domain-containing-protein-[Candidatus-Hydrogenedentes-bacterium] |
| MBI2502153.1 | DUF499_1.RM.like.systems | MACRODOMAIN->METHYLASE->?->DUF488-1->?->HEPN+DUF499*->HNH->?->DDE_Tnp_1_4-> | 1125 | Bacteria>FCB_group>Candidatus_Latescibacteria>unclassified_Candidatus_Latescibacteria | GCA_016187725.1 | Candidatus_Latescibacteria_bacterium | HYW07_02835 | 5709..9086 | 2053570 | ATP-binding-protein-[Candidatus-Latescibacteria-bacterium] |
| MCF7823046.1 | DUF499_1.RM.like.systems | <-MazG<-?<-?<-MORC+HISKIN<-METHYLASE<-HEPN+DUF499*<-METHYLASE<-MORC+HisKin+DUF3684<-TUDOR+ SF1-SF2-Helicase+iREase+CTD+REase<-XIS-HTH<-?<-Schlafen+HATPase+HTH | 1111 | Bacteria>FCB_group>Candidatus_Marinimicrobia>unclassified_Candidatus_Marinimicrobia | GCA_021734325.1 | Candidatus_Marinimicrobia_bacterium | K9N35_02645 | 72966..76301 | 2026760 | DUF499-domain-containing-protein-[Candidatus-Marinimicrobia-bacterium] |
| MCF7837530.1 | DUF499_1.RM.like.systems | Phage_integrase-><-ZnR+TOPRIM+D5ATPase<-XIS-HTH\|\|HTH->XIS-HTH-><-HEPN+DUF499*<-OLD-ABC+TOPRIM<-METHYLASE-p | 1110 | Bacteria>FCB_group>Candidatus_Marinimicrobia>unclassified_Candidatus_Marinimicrobia | GCA_021734645.1 | Candidatus_Marinimicrobia_bacterium | K9N49_02780 | 12805..16137 | 2026760 | DUF499-domain-containing-protein-[Candidatus-Marinimicrobia-bacterium] |
| MBN1984134.1 | DUF499_1.RM.like.systems | ABC-ATPase+SbcC->?->?-><-?<-HEPN+DUF499*<-METHYLASE<-DrHyd+BirA-HTH<-TUDOR+ SF1-SF2-Helicase+iREase+CTD+REase-p | 1112 | Bacteria>FCB_group>Fibrobacterota>Chitinivibrionia>Chitinivibrionales | GCA_016931815.1 | Chitinivibrionales_bacterium | JW795_21570 | 16556..19894 | 2663014 | DUF499-domain-containing-protein-[Chitinivibrionales-bacterium] |
| MCZ0935996.1 | DUF499_1.RM.like.systems | SIGMA-HTH->?->?->OLD-ABC+TOPRIM->SF1-SF2-Helicase->REase-><-HEPN+DUF499*<-METHYLASE-p\|\|Zn_Tnp_IS1595+DDE_Tnp_IS1595-><-?\|\|METHYLASE+HNH->?-><-METHYLASE | 1098 | Bacteria>FCB_group>Gemmatimonadota>Gemmatimonadetes>Candidatus_Palauibacterales | GCA_026914155.1 | Candidatus_Palauibacter_rhopaloidicola | OXJ54_12525 | 22300..25596 | 3056738 | MAG:-DUF499-domain-containing-protein-[Gemmatimonadetes-bacterium] |
| MDE2877868.1 | DUF499_1.RM.like.systems | DNA_methylase->REase->DNA_methylase->SF1-SF2-Helicase+REase->OLD-ABC->METHYLASE->HEPN+DUF499*->?->?->?->MULE-TRANSPOSASE-> | 1098 | Bacteria>FCB_group>Gemmatimonadota>Gemmatimonadetes>Candidatus_Palauibacterales | GCA_028874535.1 | Candidatus_Palauibacter_soopunensis | OXO55_03070 | 20960..24256 | 3056739 | MAG:-DUF499-domain-containing-protein-[Gemmatimonadota-bacterium] |
| MXX77944.1 | DUF499_1.RM.like.systems | OLD-ABC->TOPRIM->HEPN+DUF499*-><-RNase<-?<-REase | 1102 | Bacteria>FCB_group>Gemmatimonadota>Gemmatimonadetes>Gemmatimonadales | GCA_009836785.1 | Gemmatimonadales_bacterium | F4Z33_02945 | 1211..4519 | 2448054 | ATP-binding-protein-[Gemmatimonadales-bacterium] |
| MXX78159.1 | DUF499_1.RM.like.systems | <-HEPN+DUF499*<-METHYLASE<-TUDOR+ SF1-SF2-Helicase+iREase+CTD+REase<-BirA-HTH | 1138 | Bacteria>FCB_group>Gemmatimonadota>Gemmatimonadetes>Gemmatimonadales | GCA_009836785.1 | Gemmatimonadales_bacterium | F4Z33_04085 | 250..3666 | 2448054 | ATP-binding-protein-[Gemmatimonadales-bacterium] |
| MBV6522081.1 | DUF499_1.RM.like.systems | TUDOR+ SF1-SF2-Helicase+iREase+CTD+REase->ORC-CDC6+REase->METHYLASE->ORC-CDC6+REase->HEPN+DUF499*-><-Pkinase | 1116 | Bacteria>FCB_group>Gemmatimonadota>Gemmatimonadetes>Gemmatimonadales | GCA_019186945.1 | Gemmatimonadaceae_bacterium | MNPFHGCM_02226 | 129597..132947 | 2026741 | hypothetical-protein-[Gemmatimonadaceae-bacterium] |
| MBX6330503.1 | DUF499_1.RM.like.systems | HEPN-p->Primpol+D5N+D5ATPase->HEPN+DUF499*->Glyoxalase->?->?->HTH-> | 1110 | Bacteria>FCB_group>Gemmatimonadota>Gemmatimonadetes>Gemmatimonadales | GCA_019689955.1 | Gemmatimonadaceae_bacterium | IRY91_01515 | 6551..9883 | 2026741 | DUF499-domain-containing-protein-[Gemmatimonadaceae-bacterium] |
| MCC7134741.1 | DUF499_1.RM.like.systems | <-LuxR-HTH\|\|TUDOR+ SF1-SF2-Helicase+iREase+CTD+REase->?->METHYLASE->HEPN+DUF499*-> | 765 | Bacteria>FCB_group>Gemmatimonadota>Gemmatimonadetes>Gemmatimonadales | GCA_020851315.1 | Gemmatimonadales_bacterium | IT352_18930 | 11948..14243 | 2448054 | DUF499-domain-containing-protein-[Gemmatimonadales-bacterium] |
| MCC7194161.1 | DUF499_1.RM.like.systems | METHYLASE-p->ORC-CDC6+REase->HEPN+DUF499*-><-?\|\|SIGMA-HTH-> | 1104 | Bacteria>FCB_group>Gemmatimonadota>Gemmatimonadetes>Gemmatimonadales | GCA_020851455.1 | Gemmatimonadaceae_bacterium | IT356_01265 | 1930..5244 | 2026741 | DUF499-domain-containing-protein-[Gemmatimonadaceae-bacterium] |
| MCR4340303.1 | DUF499_1.RM.like.systems | <-HEPN+DUF499*<-METHYLASE-p | 1109 | Bacteria>FCB_group>Gemmatimonadota>Gemmatimonadetes>Gemmatimonadales | GCA_024654005.1 | Gemmatimonadaceae_bacterium | NUW01_10510 | 510..3839 | 2026741 | DUF499-domain-containing-protein-[Gemmatimonadaceae-bacterium] |
| ODT03617.1 | DUF499_1.RM.like.systems | TUDOR+ SF1-SF2-Helicase+iREase+CTD+REase->Schlafen+HATPase+HTH->METHYLASE->HEPN+DUF499*->Primpol+D5N+D5ATPase->PAIREDC-HTH->ORC-CDC6+REase-> | 1109 | Bacteria>FCB_group>Gemmatimonadota>unclassified_Gemmatimonadota | GCA_001724275.1 | Gemmatimonadetes_bacterium_SCN_70-22 | ABS52_08400 | 10589..13918 | 1660163 | AAA+-family-ATPase-[Gemmatimonadetes-bacterium-SCN-70-22] |
| MBU10399.1 | DUF499_1.RM.like.systems | METHYLASE->HEPN+DUF499*-><-HIN-HTH | 1090 | Bacteria>FCB_group>Gemmatimonadota>unclassified_Gemmatimonadota | GCA_002730265.1 | Gemmatimonadota_bacterium | CME13_20820 | 6712..9984 | 2026742 | AAA+-family-ATPase-[Gemmatimonadetes-bacterium] |
| MXW81679.1 | DUF499_1.RM.like.systems | <-TUDOR+ SF1-SF2-Helicase+iREase+CTD+REase<-HEPN+DUF499*\|\|REase+HTH->REase+HTH-><-METHYLASE | 1112 | Bacteria>FCB_group>Gemmatimonadota>unclassified_Gemmatimonadota | GCA_009836325.1 | Gemmatimonadota_bacterium | F4Z57_22370 | 10515..13853 | 2026742 | ATP-binding-protein-[Gemmatimonadetes-bacterium] |
| MXZ76449.1 | DUF499_1.RM.like.systems | Synaptojanin->TUDOR+ SF1-SF2-Helicase+iREase+CTD+REase->OLD-ABC->METHYLASE->HEPN+DUF499*->ParB+HNH-><-?<-MazE_antitoxin | 1113 | Bacteria>FCB_group>Gemmatimonadota>unclassified_Gemmatimonadota | GCA_009837395.1 | Gemmatimonadota_bacterium | F4Z03_15245 | 64974..68315 | 2026742 | ATP-binding-protein-[Gemmatimonadetes-bacterium] |
| MYC72435.1 | DUF499_1.RM.like.systems | TUDOR+ SF1-SF2-Helicase+iREase+CTD+REase-><-?<-ParB<-DNA_methylase\|\|METHYLASE->HEPN+DUF499*-> | 1131 | Bacteria>FCB_group>Gemmatimonadota>unclassified_Gemmatimonadota | GCA_009841115.1 | Gemmatimonadota_bacterium | F4X17_17170 | 10634..14029 | 2026742 | ATP-binding-protein-[Gemmatimonadetes-bacterium] |
| MYE70134.1 | DUF499_1.RM.like.systems | TUDOR+ SF1-SF2-Helicase+iREase+CTD+REase-p->METHYLASE->HEPN+DUF499*-> | 1129 | Bacteria>FCB_group>Gemmatimonadota>unclassified_Gemmatimonadota | GCA_009841855.1 | Gemmatimonadota_bacterium | F4237_08820 | 3222..6610 | 2026742 | ATP-binding-protein-[Gemmatimonadetes-bacterium] |
| MYH20375.1 | DUF499_1.RM.like.systems | TUDOR+ SF1-SF2-Helicase+iREase+CTD+REase->ParB+HNH->METHYLASE->HEPN+DUF499*->?->TPR-><-Peptidase-S8<-Classical-AAA<-Peptidase_M1 | 1108 | Bacteria>FCB_group>Gemmatimonadota>unclassified_Gemmatimonadota | GCA_009844015.1 | Gemmatimonadota_bacterium | F4132_14865 | 91560..94886 | 2026742 | ATP-binding-protein-[Gemmatimonadetes-bacterium] |
| MYL00076.1 | DUF499_1.RM.like.systems | TUDOR+ SF1-SF2-Helicase+iREase+CTD+REase->ParB+HNH-p->METHYLASE->HEPN+DUF499*->?->TPR-><-Peptidase-S8 | 1108 | Bacteria>FCB_group>Gemmatimonadota>unclassified_Gemmatimonadota | GCA_009846305.1 | Gemmatimonadota_bacterium | F4014_15170 | 26347..29673 | 2026742 | ATP-binding-protein-[Gemmatimonadetes-bacterium] |
| MBT7594687.1 | DUF499_1.RM.like.systems | <-HEPN+DUF499*<-METHYLASE-p | 1031 | Bacteria>FCB_group>Gemmatimonadota>unclassified_Gemmatimonadota | GCA_018675915.1 | Gemmatimonadota_bacterium | HN559_07235 | 1..3094 | 2026742 | ATP-binding-protein-[Gemmatimonadetes-bacterium] |
| MCY4572509.1 | DUF499_1.RM.like.systems | SF1-SF2-Helicase-><-?<-HEPN+DUF499*<-OLD-ABC<-METHYLASE | 1114 | Bacteria>FCB_group>Gemmatimonadota>unclassified_Gemmatimonadota | GCA_026713345.1 | Gemmatimonadota_bacterium | OXF01_06870 | 2905..6249 | 2026742 | MAG:-Swt1-family-HEPN-domain-containing-protein-[Gemmatimonadetes-bacterium] |
| MDE2984847.1 | DUF499_1.RM.like.systems | TUDOR+ SF1-SF2-Helicase+iREase+CTD+REase->METHYLASE->METHYLASE-p->HEPN+DUF499*-> | 1104 | Bacteria>FCB_group>Gemmatimonadota>unclassified_Gemmatimonadota | GCA_028873845.1 | Gemmatimonadota_bacterium | OXU69_09080 | 7213..10527 | 2026742 | MAG:-DUF499-domain-containing-protein-[Gemmatimonadota-bacterium] |
| MDE2875355.1 | DUF499_1.RM.like.systems | TUDOR+ SF1-SF2-Helicase+iREase+CTD+REase->METHYLASE->HEPN+DUF499*-><-?\|\|?->?-><-HerA/FtsK<-?\|\|DUF429-> | 1104 | Bacteria>FCB_group>Gemmatimonadota>unclassified_Gemmatimonadota | GCA_028874445.1 | Gemmatimonadota_bacterium | OXQ93_07945 | 7211..10525 | 2026742 | MAG:-DUF499-domain-containing-protein-[Gemmatimonadota-bacterium] |
| MDE2733147.1 | DUF499_1.RM.like.systems | <-TUDOR+ SF1-SF2-Helicase+iREase+CTD+REase<-HEPN+DUF499*\|\|REase->?-><-?<-METHYLASE<-REase<-OLD-ABC | 1112 | Bacteria>FCB_group>Gemmatimonadota>unclassified_Gemmatimonadota | GCA_028875335.1 | Gemmatimonadota_bacterium | OXI72_02050 | 10955..14293 | 2026742 | MAG:-Swt1-family-HEPN-domain-containing-protein-[Gemmatimonadota-bacterium] |
| MDE2654306.1 | DUF499_1.RM.like.systems | <-REC<-HEPN+DUF499*<-TOPRIM<-OLD-ABC<-METHYLASE-p | 1116 | Bacteria>FCB_group>Gemmatimonadota>unclassified_Gemmatimonadota | GCA_028875795.1 | Gemmatimonadota_bacterium | OXI71_10900 | 83236..86586 | 2026742 | MAG:-Swt1-family-HEPN-domain-containing-protein-[Gemmatimonadota-bacterium] |
| NMB74747.1 | DUF499_1.RM.like.systems | TUDOR+ SF1-SF2-Helicase+iREase+CTD+REase->METHYLASE->HEPN+DUF499*->?->?-><-?<-?<-?\|\|HAMP+ACYC-> | 1111 | Bacteria>Myxococcota>Myxococcia>Myxococcales>unclassified_Myxococcales | GCA_012798435.1 | Myxococcales_bacterium | GYA21_06410 | 6654..9989 | 2026763 | ATP-binding-protein-[Myxococcales-bacterium] |
| MBI3184674.1 | DUF499_1.RM.like.systems | ParB->?->XIS-HTH-><-REase<-DNA_methylase<-MORC+REase<-HEPN+DUF499*<-METHYLASE<-?<-TUDOR+ SF1-SF2-Helicase+iREase+CTD+REase<-XIS-HTH\|\|AbiEi_1->AbiEii-> | 1116 | Bacteria>Myxococcota>Myxococcia>Myxococcales>unclassified_Myxococcales | GCA_016191815.1 | Myxococcales_bacterium | HYZ28_21265 | 61053..64403 | 2026763 | ATP-binding-protein-[Myxococcales-bacterium] |
| MBK7583793.1 | DUF499_1.RM.like.systems | <-HTH\|\|?->REase->TUDOR+ SF1-SF2-Helicase+iREase+CTD+REase->METHYLASE->HEPN+DUF499*->REase-> | 1105 | Bacteria>Myxococcota>Myxococcia>Myxococcales>unclassified_Myxococcales | GCA_016706225.1 | Myxococcales_bacterium | IPI67_26815 | 461421..464738 | 2026763 | DUF499-domain-containing-protein-[Myxococcales-bacterium] |
| MBL9045676.1 | DUF499_1.RM.like.systems | TUDOR+ SF1-SF2-Helicase+iREase+CTD+REase->METHYLASE->OLD-ABC->TOPRIM->HEPN+DUF499*-><-?<-HTH | 1115 | Bacteria>Myxococcota>Myxococcia>Myxococcales>unclassified_Myxococcales | GCA_016794465.1 | Myxococcales_bacterium | JNM83_28965 | 9575..12922 | 2026763 | DUF499-domain-containing-protein-[Myxococcales-bacterium] |
| MCB9507837.1 | DUF499_1.RM.like.systems | REase-><-Schlafen+HATPase+HTH<-SF1-SF2-Helicase+HRDC<-?<-NERD+SF1-SF2-Helicase<-HEPN+DUF499*<-METHYLASE<-?<-TUDOR+ SF1-SF2-Helicase+iREase+CTD+REase\|\|DNA-glycosylase->?->HTH-> | 1103 | Bacteria>Myxococcota>Myxococcia>Myxococcales>unclassified_Myxococcales | GCA_020633615.1 | Myxococcales_bacterium | H6697_09260 | 18345..21656 | 2026763 | ATP-binding-protein-[Myxococcales-bacterium] |
| MCC6334307.1 | DUF499_1.RM.like.systems | <-REase\|\|?-><-?<-?<-REase+CTD6<-CHTH<-HEPN+DUF499*<-METHYLASE<-TUDOR+ SF1-SF2-Helicase+iREase+CTD+REase | 1100 | Bacteria>Myxococcota>Myxococcia>Myxococcales>unclassified_Myxococcales | GCA_020846875.1 | Myxococcales_bacterium | IT380_10010 | 27471..30773 | 2026763 | DUF499-domain-containing-protein-[Myxococcales-bacterium] |
| MAG32379.1 | DUF499_1.RM.like.systems | METHYLASE-p->OLD-ABC->Schlafen+HATPase+HTH->HEPN+DUF499*->?-><-METHYLASE-p<-TOPRIM<-OLD-ABC\|\|?->METHYLASE-p-> | 1115 | Bacteria>Myxococcota>Myxococcia>unclassified_Myxococcia | GCA_002688255.1 | Deltaproteobacteria_bacterium | CL908_15970 | 4485..7832 | 2026735 | AAA+-family-ATPase-[Deltaproteobacteria-bacterium] |
| PXF55798.1 | DUF499_1.RM.like.systems | TUDOR+ SF1-SF2-Helicase+iREase+CTD+REase->Schlafen+HATPase+HTH->?->METHYLASE-p-><-MULE-TRANSPOSASE\|\|METHYLASE-p->HEPN+DUF499*->MULE-TRANSPOSASE->?-><-LD-peptidase | 1108 | Bacteria>Myxococcota>Myxococcia>unclassified_Myxococcia | GCA_003194485.1 | Deltaproteobacteria_bacterium | C4B57_02005 | 126317..129643 | 2026735 | AAA+-family-ATPase-[Deltaproteobacteria-bacterium] |
| PXF60387.1 | DUF499_1.RM.like.systems | PAIREDC-HTH->Schlafen+HATPase+HTH->METHYLASE->METHYLASE-p->HEPN+DUF499*-> | 1108 | Bacteria>Myxococcota>Myxococcia>unclassified_Myxococcia | GCA_003194495.1 | Deltaproteobacteria_bacterium | C4B58_00710 | 172327..175653 | 2026735 | AAA+-family-ATPase-[Deltaproteobacteria-bacterium] |
| RME24627.1 | DUF499_1.RM.like.systems | <-HEPN+DUF499*<-SbcC<-METHYLASE-p | 1118 | Bacteria>Myxococcota>Myxococcia>unclassified_Myxococcia | GCA_003694325.1 | Deltaproteobacteria_bacterium | D6798_10765 | 99..3455 | 2026735 | ATP-binding-protein-[Deltaproteobacteria-bacterium] |
| TRZ75307.1 | DUF499_1.RM.like.systems | METHYLASE->OLD-ABC->TOPRIM->HEPN+DUF499*-> | 362 | Bacteria>Myxococcota>Myxococcia>unclassified_Myxococcia | GCA_007279985.1 | Deltaproteobacteria_bacterium | D4R93_05190 | 4209..5295 | 2026735 | ATP-binding-protein,-partial-[Deltaproteobacteria-bacterium] |
| NOZ87479.1 | DUF499_1.RM.like.systems | METHYLASE->?->Peptidase_C47->HEPN+DUF499*->MethyltransfD12->HerA/FtsK-> | 1103 | Bacteria>Myxococcota>Myxococcia>unclassified_Myxococcia | GCA_013152945.1 | Deltaproteobacteria_bacterium | GXP49_14650 | 6485..9796 | 2026735 | ATP-binding-protein-[Deltaproteobacteria-bacterium] |
| MBI2357704.1 | DUF499_1.RM.like.systems | <-HEPN+DUF499*<-HEPN<-HEPN<-PolB<-Schlafen+HATPase+HTH | 747 | Bacteria>Myxococcota>Myxococcia>unclassified_Myxococcia | GCA_016183075.1 | Deltaproteobacteria_bacterium | HYV04_02095 | 1..2242 | 2026735 | ATP-binding-protein-[Deltaproteobacteria-bacterium] |
| MBI3073658.1 | DUF499_1.RM.like.systems | <-HTH_3\|\|TUDOR+ SF1-SF2-Helicase+iREase+CTD+REase->?->METHYLASE-><-PIN<-MetJarc-HTH\|\|HEPN+DUF499*->sGTP->?->Phage_integrase-> | 1105 | Bacteria>Myxococcota>Myxococcia>unclassified_Myxococcia | GCA_016192705.1 | Deltaproteobacteria_bacterium | HYY84_16220 | 33822..37139 | 2026735 | ATP-binding-protein-[Deltaproteobacteria-bacterium] |
| MBI3249564.1 | DUF499_1.RM.like.systems | METHYLASE->REase->HEPN+DUF499*-><-HicB_lk_antitox\|\|Phage_integrase-><-tRNA<-ABhydrolase | 1110 | Bacteria>Myxococcota>Myxococcia>unclassified_Myxococcia | GCA_016197285.1 | Deltaproteobacteria_bacterium | HYZ50_23930 | 2669..6001 | 2026735 | ATP-binding-protein-[Deltaproteobacteria-bacterium] |
| MBI3768225.1 | DUF499_1.RM.like.systems | <-HNH\|\|HTH_3->TUDOR+ SF1-SF2-Helicase+iREase+CTD+REase->?->?->METHYLASE->HEPN+DUF499*->?->?-><-REase | 1104 | Bacteria>Myxococcota>Myxococcia>unclassified_Myxococcia | GCA_016200545.1 | Deltaproteobacteria_bacterium | HY271_06975 | 73636..76950 | 2026735 | ATP-binding-protein-[Deltaproteobacteria-bacterium] |
| MBI4517435.1 | DUF499_1.RM.like.systems | TUDOR+ SF1-SF2-Helicase+iREase+CTD+REase-><-?\|\|?->PolB->HEPN->HEPN+DUF499*-><-?<-CHTH<-PolB<-?<-Aminotran_4 | 1154 | Bacteria>Myxococcota>Myxococcia>unclassified_Myxococcia | GCA_016210005.1 | Deltaproteobacteria_bacterium | HY699_16655 | 22012..25476 | 2026735 | ATP-binding-protein-[Deltaproteobacteria-bacterium] |
| MBI4488650.1 | DUF499_1.RM.like.systems | HEPN->NTP_transf_2->HEPN+DUF499*->?->?->DUF4527-> | 1115 | Bacteria>Myxococcota>Myxococcia>unclassified_Myxococcia | GCA_016210105.1 | Deltaproteobacteria_bacterium | HY694_06155 | 1154..4501 | 2026735 | ATP-binding-protein-[Deltaproteobacteria-bacterium] |
| MBI4527695.1 | DUF499_1.RM.like.systems | TUDOR+ SF1-SF2-Helicase+iREase+CTD+REase->PolB->HEPN->HEPN->HEPN+DUF499*->ABhydrolase-> | 1116 | Bacteria>Myxococcota>Myxococcia>unclassified_Myxococcia | GCA_016210115.1 | Deltaproteobacteria_bacterium | HY695_28195 | 8921..12271 | 2026735 | DUF499-domain-containing-protein-[Deltaproteobacteria-bacterium] |
| MBI5504530.1 | DUF499_1.RM.like.systems | MazE_antitoxin->DOC-><-HNH<-?<-?<-HEPN+DUF499*<-METHYLASE<-?<-ParB+HNH<-TUDOR+ SF1-SF2-Helicase+iREase+CTD+REase<-?\|\|HIN-HTH-> | 1116 | Bacteria>Myxococcota>Myxococcia>unclassified_Myxococcia | GCA_016218925.1 | Deltaproteobacteria_bacterium | HY899_06990 | 27440..30790 | 2026735 | ATP-binding-protein-[Deltaproteobacteria-bacterium] |
| MBK8015115.1 | DUF499_1.RM.like.systems | DEDD_Tnp_IS110+Transposase_20->Y2_Tnp-><-REase<-?<-HEPN+DUF499*<-METHYLASE<-TUDOR+ SF1-SF2-Helicase+iREase+CTD+REase<-?<-GDResolvase+Recombinase+ZnR<-HTH | 1099 | Bacteria>Myxococcota>Myxococcia>unclassified_Myxococcia | GCA_016713425.1 | Deltaproteobacteria_bacterium | IPK13_27690 | 7179186..7182485 | 2026735 | DUF499-domain-containing-protein-[Deltaproteobacteria-bacterium] |
| MBM4343558.1 | DUF499_1.RM.like.systems | MPTase->?->NUDIX-><-DUF499-p<-HEPN+DUF499*<-METHYLASE\|\|?->?->\|\|CHTH-> | 454 | Bacteria>Myxococcota>Myxococcia>unclassified_Myxococcia | GCA_016875335.1 | Deltaproteobacteria_bacterium | FJ100_09290 | 14865..16229 | 2026735 | hypothetical-protein-[Deltaproteobacteria-bacterium] |
| MBN1382687.1 | DUF499_1.RM.like.systems | <-HNH<-ZDNA-HTH<-?<-MetJarc-HTH<-?<-HEPN+DUF499*<-METHYLASE<-?<-MqsA_antitoxin+HTH_3<-MqsR_toxin<-TUDOR+ SF1-SF2-Helicase+iREase+CTD+REase | 1117 | Bacteria>Myxococcota>Myxococcia>unclassified_Myxococcia | GCA_016928025.1 | Deltaproteobacteria_bacterium | JXA41_13530 | 29023..32376 | 2026735 | DUF499-domain-containing-protein-[Deltaproteobacteria-bacterium] |
| MBN2299509.1 | DUF499_1.RM.like.systems | <-HEPN+DUF499*<-METHYLASE<-HKD-DNase<-Cluster1090_5clades<-DrHyd+BirA-HTH<-TUDOR+ SF1-SF2-Helicase+iREase+CTD+REase | 1097 | Bacteria>Myxococcota>Myxococcia>unclassified_Myxococcia | GCA_016934135.1 | Deltaproteobacteria_bacterium | JXM72_12985 | 2480..5773 | 2026735 | DUF499-domain-containing-protein-[Deltaproteobacteria-bacterium] |
| MBV8135224.1 | DUF499_1.RM.like.systems | <-HEPN+DUF499*<-METHYLASE<-TUDOR+ SF1-SF2-Helicase+iREase+CTD+REase-p | 478 | Bacteria>Myxococcota>Myxococcia>unclassified_Myxococcia | GCA_019237115.1 | Deltaproteobacteria_bacterium | JO121_06220 | 1..1435 | 2026735 | DUF499-domain-containing-protein-[Deltaproteobacteria-bacterium] |
| MBW1777804.1 | DUF499_1.RM.like.systems | METHYLASE->HEPN+DUF499*-> | 1113 | Bacteria>Myxococcota>Myxococcia>unclassified_Myxococcia | GCA_019306905.1 | Deltaproteobacteria_bacterium | JRJ54_09460 | 3651..6992 | 2026735 | DUF499-domain-containing-protein-[Deltaproteobacteria-bacterium] |
| MBW1922080.1 | DUF499_1.RM.like.systems | ORC-CDC6+REase->TUDOR+ SF1-SF2-Helicase+iREase+CTD+REase-p->Schlafen+HATPase+HTH->METHYLASE->PolB->HEPN->HEPN+DUF499*-> | 1062 | Bacteria>Myxococcota>Myxococcia>unclassified_Myxococcia | GCA_019307945.1 | Deltaproteobacteria_bacterium | JRJ35_01275 | 87250..90435 | 2026735 | DUF499-domain-containing-protein-[Deltaproteobacteria-bacterium] |
| MBW1949838.1 | DUF499_1.RM.like.systems | TUDOR+ SF1-SF2-Helicase+iREase+CTD+REase-p->Schlafen+HATPase+HTH->?->METHYLASE->PolB->HEPN->HEPN+DUF499*-> | 1092 | Bacteria>Myxococcota>Myxococcia>unclassified_Myxococcia | GCA_019308065.1 | Deltaproteobacteria_bacterium | JRJ18_09875 | 33770..37048 | 2026735 | DUF499-domain-containing-protein-[Deltaproteobacteria-bacterium] |
| MBW1956070.1 | DUF499_1.RM.like.systems | <-Radical-SAM<-?<-HEPN+DUF499*<-METHYLASE<-?<-MqsA_antitoxin+HTH_3<-MqsR_toxin<-TUDOR+ SF1-SF2-Helicase+iREase+CTD+REase | 1113 | Bacteria>Myxococcota>Myxococcia>unclassified_Myxococcia | GCA_019308105.1 | Deltaproteobacteria_bacterium | JRI83_10180 | 2201..5542 | 2026735 | DUF499-domain-containing-protein-[Deltaproteobacteria-bacterium] |
| MBW2050448.1 | DUF499_1.RM.like.systems | <-HEPN+DUF499*<-HEPN<-PolB<-METHYLASE-p<-METHYLASE-p | 362 | Bacteria>Myxococcota>Myxococcia>unclassified_Myxococcia | GCA_019308945.1 | Deltaproteobacteria_bacterium | JRJ09_18250 | 1..1087 | 2026735 | DUF499-domain-containing-protein-[Deltaproteobacteria-bacterium] |
| MBW2077703.1 | DUF499_1.RM.like.systems | TUDOR+ SF1-SF2-Helicase+iREase+CTD+REase-p->Schlafen+HATPase+HTH->METHYLASE->URI-DNase->PolB->HEPN->HEPN+DUF499*-> | 1093 | Bacteria>Myxococcota>Myxococcia>unclassified_Myxococcia | GCA_019309065.1 | Deltaproteobacteria_bacterium | JRI71_09200 | 7259..10540 | 2026735 | DUF499-domain-containing-protein-[Deltaproteobacteria-bacterium] |
| MBW2131484.1 | DUF499_1.RM.like.systems | TUDOR+ SF1-SF2-Helicase+iREase+CTD+REase->MqsR_toxin->MqsA_antitoxin+HTH_3->?->METHYLASE->HEPN+DUF499*->?->Radical-SAM-> | 1113 | Bacteria>Myxococcota>Myxococcia>unclassified_Myxococcia | GCA_019309485.1 | Deltaproteobacteria_bacterium | JRG88_02955 | 150894..154235 | 2026735 | DUF499-domain-containing-protein-[Deltaproteobacteria-bacterium] |
| MBW2310290.1 | DUF499_1.RM.like.systems | TUDOR+ SF1-SF2-Helicase+iREase+CTD+REase->MORC+HisKin+DUF3684->METHYLASE->URI-DNase->PolB->HEPN->HEPN+DUF499*-> | 1093 | Bacteria>Myxococcota>Myxococcia>unclassified_Myxococcia | GCA_019310505.1 | Deltaproteobacteria_bacterium | JRF35_04345 | 28703..31984 | 2026735 | DUF499-domain-containing-protein-[Deltaproteobacteria-bacterium] |
| MCC6750891.1 | DUF499_1.RM.like.systems | ParB-><-SIR2<-HEPN+DUF499*<-METHYLASE<-TUDOR+ SF1-SF2-Helicase+iREase+CTD+REase\|\|PIN-> | 1103 | Bacteria>Myxococcota>Myxococcia>unclassified_Myxococcia | GCA_020848905.1 | Deltaproteobacteria_bacterium | IT371_24750 | 4486..7797 | 2026735 | DUF499-domain-containing-protein-[Deltaproteobacteria-bacterium] |
| MCC7383915.1 | DUF499_1.RM.like.systems | <-SIGMA-HTH<-Peptidase_S8<-Classical-AAA<-HEPN+DUF499*<-METHYLASE<-TUDOR+ SF1-SF2-Helicase+iREase+CTD+REase | 1093 | Bacteria>Myxococcota>Myxococcia>unclassified_Myxococcia | GCA_020850435.1 | Deltaproteobacteria_bacterium | IT384_18880 | 106091..109372 | 2026735 | DUF499-domain-containing-protein-[Deltaproteobacteria-bacterium] |
| MCY4442678.1 | DUF499_1.RM.like.systems | AbiEi_1-><-HEPN+DUF499*<-METHYLASE<-MORC+DUF3684<-TUDOR+ SF1-SF2-Helicase+iREase+CTD+REase<-Cysteine_desulfurase\|\|35exo-> | 1115 | Bacteria>Myxococcota>Myxococcia>unclassified_Myxococcia | GCA_026710965.1 | Deltaproteobacteria_bacterium | OXE53_21030 | 4124..7471 | 2026735 | MAG:-Swt1-family-HEPN-domain-containing-protein-[Deltaproteobacteria-bacterium] |
| MCY4488956.1 | DUF499_1.RM.like.systems | DDE_Tnp_1->BrnT_toxin->HEPN+DUF499*-><-?<-tRNA\|\|?-><-DnaB<-Aminotran_3<-Aminotran_1_2 | 1099 | Bacteria>Myxococcota>Myxococcia>unclassified_Myxococcia | GCA_026712905.1 | Deltaproteobacteria_bacterium | OXF11_17810 | 2694..5993 | 2026735 | MAG:-DUF499-domain-containing-protein-[Deltaproteobacteria-bacterium] |
| MDD3553583.1 | DUF499_1.RM.like.systems | XIS-HTH->TUDOR+ SF1-SF2-Helicase+iREase+CTD+REase->METHYLASE->OLD-ABC->TOPRIM->HEPN+DUF499*-> | 1108 | Bacteria>Myxococcota>Myxococcia>unclassified_Myxococcia | GCA_028695735.1 | Deltaproteobacteria_bacterium | PHC35_03575 | 35937..39263 | 2026735 | MAG:-Swt1-family-HEPN-domain-containing-protein-[Deltaproteobacteria-bacterium] |
| MDD9852708.1 | DUF499_1.RM.like.systems | TUDOR+ SF1-SF2-Helicase+iREase+CTD+REase->?->?->METHYLASE->OLD-ABC->PIN->HEPN+DUF499*-> | 632 | Bacteria>Myxococcota>Myxococcia>unclassified_Myxococcia | GCA_028817165.1 | Deltaproteobacteria_bacterium | OXU78_02000 | 21524..23422 | 2026735 | MAG:-Swt1-family-HEPN-domain-containing-protein-[Deltaproteobacteria-bacterium] |
| MDE0031020.1 | DUF499_1.RM.like.systems | DUF433->PIN->Trypsin-><-Trypsin\|\|Phage_integrase-><-HEPN+DUF499*<-BrnT_toxin<-BrnT_toxin<-METHYLASE-p | 1098 | Bacteria>Myxococcota>Myxococcia>unclassified_Myxococcia | GCA_028818775.1 | Deltaproteobacteria_bacterium | OXU42_16660 | 47371..50667 | 2026735 | MAG:-DUF499-domain-containing-protein-[Deltaproteobacteria-bacterium] |
| MDE0344143.1 | DUF499_1.RM.like.systems | <-Arm-DNA-bind_3+Phage_integrase<-?<-HNH<-HEPN+DUF499*<-?\|\|Phage_integrase->METHYLASE-p-> | 1111 | Bacteria>Myxococcota>Myxococcia>unclassified_Myxococcia | GCA_028821055.1 | Deltaproteobacteria_bacterium | OXK82_13420 | 9792..13127 | 2026735 | MAG:-Swt1-family-HEPN-domain-containing-protein-[Deltaproteobacteria-bacterium] |
| MCC6527279.1 | DUF499_1.RM.like.systems | METHYLASE-p->HEPN+DUF499*->LZ_Tnp_IS66+zf-IS66+DDE_Tnp_IS66+DDE_Tnp_IS66_C-> | 1095 | Bacteria>Myxococcota>Polyangia>Polyangiales>Polyangiaceae | GCA_020848455.1 | Polyangiaceae_bacterium | IT373_31815 | 1148..4435 | 2268199 | DUF499-domain-containing-protein-[Polyangiaceae-bacterium] |
| MCS6898143.1 | DUF499_1.RM.like.systems | METHYLASE->HEPN+DUF499*->DUF499-p-><-?<-?<-FURR-HTH | 1001 | Bacteria>Myxococcota>Polyangia>Polyangiales>Polyangiaceae | GCA_025057345.1 | Polyangiaceae_bacterium | NZX77_00010 | 2850..5855 | 2268199 | Swt1-family-HEPN-domain-containing-protein-[Polyangiaceae-bacterium] |
| MAT27035.1 | DUF499_1.RM.like.systems | HEPN->MORC+HisKin+DUF3684-><-?\|\|TUDOR+ SF1-SF2-Helicase+iREase+CTD+REase->HEPN->METHYLASE->HEPN+DUF499*->MORC+HisKin+REase->?->METHYLASE-p-> | 1093 | Bacteria>Myxococcota>Polyangia>Polyangiales>Sandaracinaceae | GCA_002699025.1 | Sandaracinus_sp. | CMN29_19140 | 67210..70491 | 2024858 | AAA+-family-ATPase-[Sandaracinus-sp.] |
| MBX3269805.1 | DUF499_1.RM.like.systems | PIN->TUDOR+ SF1-SF2-Helicase+iREase+CTD+REase->?->HEPN->METHYLASE->METHYLASE->HEPN+DUF499*->?-><-HIN-HTH | 1090 | Bacteria>Myxococcota>Polyangia>Polyangiales>Sandaracinaceae | GCA_019637095.1 | Sandaracinaceae_bacterium | KF729_06070 | 123033..126305 | 2510337 | DUF499-domain-containing-protein-[Sandaracinaceae-bacterium] |
| MBP7125459.1 | DUF499_1.RM.like.systems | TUDOR+ SF1-SF2-Helicase+iREase+CTD+REase->Schlafen+HATPase+HTH->METHYLASE->OLD-ABC->TOPRIM->HEPN+DUF499*-> | 1039 | Bacteria>Myxococcota>unclassified_Myxococcota | GCA_017994695.1 | Myxococcota_bacterium | KBD49_03725 | 64170..67286 | 2818507 | DUF499-domain-containing-protein-[Myxococcota-bacterium] |
| MBY0401387.1 | DUF499_1.RM.like.systems | TPR->GspA-ATPase-><-Mbetalac\|\|tRNA->?->?-><-HEPN+DUF499*<-METHYLASE-p | 1114 | Bacteria>Myxococcota>unclassified_Myxococcota | GCA_019751155.1 | Myxococcota_bacterium | K2X89_13900 | 4687..8031 | 2818507 | DUF499-domain-containing-protein-[Myxococcota-bacterium] |
| MCG3173785.1 | DUF499_1.RM.like.systems | <-HEPN+DUF499*<-HEPN+HEPN<-RAMPs<-RAMPs<-?<-METHYLASE | 1117 | Bacteria>Myxococcota>unclassified_Myxococcota | GCA_022072125.1 | Myxococcota_bacterium | GMKNLPBB_01987 | 84461..87814 | 2818507 | hypothetical-protein-[Myxococcota-bacterium] |
| MCS6914091.1 | DUF499_1.RM.like.systems | Cas_Cmr3->?->Cas_Cmr5->?->CARF+HEPN->HEPN+DUF499*->Radical-SAM-> | 1114 | Bacteria>Myxococcota>unclassified_Myxococcota | GCA_025057315.1 | Myxococcota_bacterium | NZ890_12730 | 8654..11998 | 2818507 | DUF499-domain-containing-protein-[Myxococcota-bacterium] |
| MDE0208007.1 | DUF499_1.RM.like.systems | <-HEPN+DUF499*<-Schlafen+HATPase+HTH<-METHYLASE<-TUDOR+ SF1-SF2-Helicase+iREase+CTD+REase<-BrnA_antitoxin | 1111 | Bacteria>Nitrospinae/Tectomicrobia_group>Candidatus_Tectomicrobia>unclassified_Candidatus_Tectomicrobia | GCA_028819935.1 | Candidatus_Tectomicrobia_bacterium | OXP66_18475 | 1406..4741 | 2528274 | MAG:-Swt1-family-HEPN-domain-containing-protein-[Candidatus-Tectomicrobia-bacterium] |
| MYA94924.1 | DUF499_1.RM.like.systems | ACET-><-?\|\|?->tRNA-><-HEPN+DUF499*<-METHYLASE<-?<-?<-?<-?\|\|Peptidase-> | 1086 | Bacteria>Nitrospinae/Tectomicrobia_group>Nitrospinota>unclassified_Nitrospinota | GCA_009839615.1 | Nitrospinota_bacterium | F4X91_00725 | 11133..14393 | 2026769 | ATP-binding-protein-[Nitrospinae-bacterium] |
| MBZ0154723.1 | DUF499_1.RM.like.systems | PIN->OLD-ABC->HNH->METHYLASE->?->RelE-ParE->HEPN+DUF499*-> | 1114 | Bacteria>Nitrospirota>Nitrospiria>Nitrospirales>Nitrospiraceae | - | Candidatus_Nitrobium_versatile | K8I29_00740 | 55978..59322 | 2884831 | DUF499-domain-containing-protein-[Alphaproteobacteria-bacterium] |
| NGZ08749.1 | DUF499_1.RM.like.systems | Glycos_transf_4->TUDOR+ SF1-SF2-Helicase+iREase+CTD+REase->ParB+HNH->METHYLASE->HEPN+DUF499*-> | 1121 | Bacteria>Nitrospirota>Nitrospiria>Nitrospirales>Nitrospiraceae | GCA_011090425.1 | Nitrospira_sp._LK70 | CV088_05085 | 57956..61321 | 2039809 | ATP-binding-protein-[Nitrospira-sp.-LK70] |
| MBP6261842.1 | DUF499_1.RM.like.systems | <-XIS-HTH\|\|TUDOR+ SF1-SF2-Helicase+iREase+CTD+REase->AbiGi->ParB+HNH->METHYLASE->HEPN+DUF499*->HEPN+CBS->?-><-?\|\|HIN-HTH-> | 1121 | Bacteria>Nitrospirota>Nitrospiria>Nitrospirales>Nitrospiraceae | GCA_017986555.1 | Nitrospira_sp. | KA426_01230 | 11559..14924 | 70125 | DUF499-domain-containing-protein-[Nitrospira-sp.] |
| MBS0184610.1 | DUF499_1.RM.like.systems | TUDOR+ SF1-SF2-Helicase+iREase+CTD+REase-p->TUDOR+ SF1-SF2-Helicase+iREase+CTD+REase-p->ParB+HNH->METHYLASE->METHYLASE-p->HEPN->HEPN+DUF499*-> | 1121 | Bacteria>Nitrospirota>Nitrospiria>Nitrospirales>Nitrospiraceae | GCA_018242685.1 | Nitrospira_sp. | JSS39_19650 | 30690..34055 | 70125 | DUF499-domain-containing-protein-[Nitrospira-sp.] |
| MBS0151038.1 | DUF499_1.RM.like.systems | TUDOR+ SF1-SF2-Helicase+iREase+CTD+REase->REase->METHYLASE->?->DUF488-1->?->HEPN+DUF499*->?-><-?\|\|?-><-?<-SMODS | 1121 | Bacteria>Nitrospirota>Nitrospiria>Nitrospirales>Nitrospiraceae | GCA_018242725.1 | Nitrospira_sp. | JSR31_08860 | 34905..38270 | 70125 | DUF499-domain-containing-protein-[Nitrospira-sp.] |
| MBX3234602.1 | DUF499_1.RM.like.systems | ArdCN+MPTase->?->METHYLASE->?->DUF488-1->?->HEPN+DUF499*->?->sGTP-> | 1121 | Bacteria>Nitrospirota>Nitrospiria>Nitrospirales>Nitrospiraceae | GCA_019637075.1 | Nitrospiraceae_bacterium | KF814_00500 | 97696..101061 | 2026770 | DUF499-domain-containing-protein-[Nitrospiraceae-bacterium] |
| MBZ0154906.1 | DUF499_1.RM.like.systems | TIR->PAIREDC-HTH+HIN-HTH->DDE_3-><-SF1-SF2-Helicase<-HKD-DNase<-?<-HEPN+DUF499*<-METHYLASE<-?<-?<-Schlafen+HATPase+HTH\|\|REase-> | 1108 | Bacteria>Nitrospirota>Nitrospiria>Nitrospirales>Nitrospiraceae | GCA_019912415.1 | Candidatus_Nitrobium_versatile | K8I29_01665 | 56123..59449 | 2884831 | DUF499-domain-containing-protein-[Alphaproteobacteria-bacterium] |
| MCS6288012.1 | DUF499_1.RM.like.systems | <-XIS-HTH\|\|TUDOR+ SF1-SF2-Helicase+iREase+CTD+REase->ParB+HNH->METHYLASE->HEPN+DUF499*->RelE-ParE->HTH->SF1-SF2-Helicase-> | 1121 | Bacteria>Nitrospirota>Nitrospiria>Nitrospirales>Nitrospiraceae | GCA_024998625.1 | Nitrospira_sp. | H8K10_03430 | 260842..264207 | 70125 | ATP-binding-protein-[Nitrospira-sp.] |
| MDA8090379.1 | DUF499_1.RM.like.systems | ACET-><-?<-?<-REase<-HEPN+DUF499*<-METHYLASE<-TIR+PNTD2<-TUDOR+ SF1-SF2-Helicase+iREase+CTD+REase<-?\|\|Tox-RelE->HTH_3-> | 1107 | Bacteria>Nitrospirota>Nitrospiria>Nitrospirales>Nitrospiraceae | GCA_028064195.1 | Nitrospiraceae_bacterium | MOZ61_09200 | 18029..21352 | 2026770 | MAG:-Swt1-family-HEPN-domain-containing-protein-[Nitrospiraceae-bacterium] |
| WP_090899592.1 | DUF499_1.RM.like.systems | PIN->TUDOR+ SF1-SF2-Helicase+iREase+CTD+REase->STAS+HisKin->?->PIN->METHYLASE->HEPN+DUF499*->?->KAP-NTPase->?->PAPS_reduct-> | 1117 | Bacteria>Nitrospirota>Nitrospiria>Nitrospirales>Nitrospiraceae | GCF_001458775.1 | Candidatus_Nitrospira_nitrificans | COMA2_RS14290 | 32189..35542 | 1742973 | Swt1-family-HEPN-domain-containing-protein-[Candidatus-Nitrospira-nitrificans] |
| PIX82647.1 | DUF499_1.RM.like.systems | METHYLASE->OLD-ABC->TOPRIM->HEPN+DUF499*-> | 561 | Bacteria>Nitrospirota>unclassified_Nitrospirota | GCA_002783945.1 | Nitrospirae_bacterium_CG_4_10_14_3_um_filter_70_108 | COZ33_09565 | 4155..5838 | 1974054 | AAA+-family-ATPase,-partial-[Nitrospirae-bacterium-CG_4_10_14_3_um_filter_70_108] |
| KAF0183094.1 | DUF499_1.RM.like.systems | MODE-HTH+PBPII->?->?->?->?-><-HEPN+DUF499*<-METHYLASE<-?<-MqsA_antitoxin+HTH_3<-MqsR_toxin<-TUDOR+ SF1-SF2-Helicase+iREase+CTD+REase | 1120 | Bacteria>Nitrospirota>unclassified_Nitrospirota | GCA_009773415.1 | Nitrospirota_bacterium | FD164_823 | 11555..14917 | 2026887 | ATPase-(AAA+-superfamily)-like-protein-[Nitrospirae-bacterium] |
| KJV40259.1 | DUF499_1.RM.like.systems | ParG_RHH-><-MODE-HTH+PBPII<-HEPN+DUF499*<-METHYLASE<-MORC+DUF3684<-YtH<-TUDOR+ SF1-SF2-Helicase+iREase+CTD+REase<-HTH+WYL\|\|SF1-SF2-Helicase+REase-> | 1098 | Bacteria>Pseudomonadota>Alphaproteobacteria>Caulobacterales>Caulobacteraceae | GCA_000964075.1 | Brevundimonas_sp._KM4 | VH88_11615 | 37954..41250 | 1628191 | ATPase-AAA-[Brevundimonas-sp.-KM4] |
| KAF0185697.1 | DUF499_1.RM.like.systems | MazE_antitoxin->PIN->TUDOR+ SF1-SF2-Helicase+iREase+CTD+REase->METHYLASE->HEPN+DUF499*-><-ParB+HNH\|\|?->?->?->Phage_AlpA-><-Arm-DNA-bind_3+Phage_integrase | 1077 | Bacteria>Pseudomonadota>Alphaproteobacteria>Caulobacterales>Caulobacteraceae | GCA_009773335.1 | Caulobacteraceae_bacterium | FD160_506 | 42091..45324 | 1978230 | ATPase--like-protein-[Caulobacteraceae-bacterium] |
| QLQ13550.1 | DUF499_1.RM.like.systems | VirD4-TraG+VirD4-FtsK->CopG/RHH->VirB11+VirB4-FtsK->?->?-><-MODE-HTH+PBPII<-HEPN+DUF499*<-METHYLASE<-MORC+DUF3684<-HTH+WYL\|\|SF1-SF2-Helicase+REase->SF1-SF2-Helicase+REase->SF1-SF2-helicase+RdgC+Beta-barrel+REase-> | 1096 | Bacteria>Pseudomonadota>Alphaproteobacteria>Caulobacterales>Caulobacteraceae | GCA_013693835.1 | Brevundimonas_sp. | HZY74_10140 | 2044155..2047445 | 1871086 | MAG:-ATP-binding-protein-[Brevundimonas-sp.] |
| MBB3889803.1 | DUF499_1.RM.like.systems | <-SF1-SF2-Helicase+REase\|\|HTH+WYL->TUDOR+ SF1-SF2-Helicase+iREase+CTD+REase->REase->METHYLASE->HEPN+DUF499*->MODE-HTH+PBPII-><-ParG_RHH | 1098 | Bacteria>Pseudomonadota>Alphaproteobacteria>Caulobacterales>Caulobacteraceae | GCA_014196295.1 | Phenylobacterium_haematophilum | GGQ61_000500 | 500754..504050 | 98513 | hypothetical-protein-[Phenylobacterium-haematophilum] |
| MBL8553710.1 | DUF499_1.RM.like.systems | Primpol+PriCT1-><-?<-?\|\|HTH_3->TIR+PNTD2->PIWI-><-HEPN+DUF499*<-METHYLASE-p | 1114 | Bacteria>Pseudomonadota>Alphaproteobacteria>Caulobacterales>Caulobacteraceae | GCA_016793285.1 | Phenylobacterium_sp. | JNL41_05490 | 46797..50141 | 1871053 | DUF499-domain-containing-protein-[Phenylobacterium-sp.] |
| MCA3722145.1 | DUF499_1.RM.like.systems | <-METHYLASE-p\|\|?->?-><-PIWI<-TIR+PNTD2<-HEPN+DUF499*<-METHYLASE-p | 1098 | Bacteria>Pseudomonadota>Alphaproteobacteria>Caulobacterales>Caulobacteraceae | GCA_020253645.1 | Phenylobacterium_sp. | INE98_13845 | 11536..14832 | 1871053 | DUF499-domain-containing-protein-[Phenylobacterium-sp.] |
| MCA6347246.1 | DUF499_1.RM.like.systems | <-HEPN+DUF499*<-METHYLASE<-TUDOR+ SF1-SF2-Helicase+iREase+CTD+REase<-XIS-HTH<-Pkinase | 1101 | Bacteria>Pseudomonadota>Alphaproteobacteria>Caulobacterales>Caulobacteraceae | GCA_020402685.1 | Phenylobacterium_sp. | IM625_01145 | 33818..37123 | 1871053 | DUF499-domain-containing-protein-[Phenylobacterium-sp.] |
| MDB5438829.1 | DUF499_1.RM.like.systems | METHYLASE->HEPN+DUF499*-> | 1126 | Bacteria>Pseudomonadota>Alphaproteobacteria>Caulobacterales>Caulobacteraceae | GCA_028291105.1 | Caulobacteraceae_bacterium | JWM33_1256 | 2767..6147 | 1978230 | MAG:-superfamily-helicase,-family-[Caulobacteraceae-bacterium] |
| WEK58847.1 | DUF499_1.RM.like.systems | <-SF1-SF2-Helicase+REase<-SF1-SF2-Helicase+REase\|\|HTH+WYL->TUDOR+ SF1-SF2-Helicase+iREase+CTD+REase->MORC+DUF3684->METHYLASE->HEPN+DUF499*-><-?<-?<-ACYC+ACYC<-CNMP+TIR\|\|HTH+WYL-> | 1085 | Bacteria>Pseudomonadota>Alphaproteobacteria>Caulobacterales>Caulobacteraceae | GCA_029203145.1 | Brevundimonas_sp. | POY52_04750 | 987163..990420 | 1871086 | MAG:-DUF499-domain-containing-protein-[Brevundimonas-sp.] |
| WP_039244289.1 | DUF499_1.RM.like.systems | <-SF1-SF2-Helicase+REase<-?\|\|HTH+WYL->TUDOR+ SF1-SF2-Helicase+iREase+CTD+REase->?->METHYLASE->HEPN+DUF499*->MODE-HTH+PBPII-><-ParG_RHH | 1098 | Bacteria>Pseudomonadota>Alphaproteobacteria>Caulobacterales>Caulobacteraceae | GCF_000813765.1 | Brevundimonas_nasdae | RM53_RS03375 | 209717..213013 | 172043 | DUF499-domain-containing-protein-[Brevundimonas-nasdae] |
| WP_231576665.1 | DUF499_1.RM.like.systems | ParG_RHH-><-MODE-HTH+PBPII<-HEPN+DUF499*<-METHYLASE<-MORC+DUF3684<-YtH<-TUDOR+ SF1-SF2-Helicase+iREase+CTD+REase<-HTH+WYL\|\|SF1-SF2-Helicase+REase-> | 1086 | Bacteria>Pseudomonadota>Alphaproteobacteria>Caulobacterales>Caulobacteraceae | GCF_000964075.1 | Brevundimonas_sp._KM4 | VH88_RS11620 | 37954..41214 | 1628191 | DUF499-domain-containing-protein-[Brevundimonas-sp.-KM4] |

| Protein-ID | System | Gene-Neighbourhoods | Length | Taxonomy | Assembly-ID | Strain | Locus-Tag | Genomic-coordinates | TaxID | Fasta-defline |
|---|---|---|---|---|---|---|---|---|---|---|
| WP_091738393.1 | DUF499_1.RM.like.systems | <-SF1-SF2-Helicase+REase<-?\|\|HTH+WYL->TUDOR+ SF1-SF2-Helicase+iREase+CTD+REase->ASCH->METHYLASE->HEPN+DUF499*->MODE-HTH+PBPII-><-ParG_RHH | 1099 | Bacteria>Pseudomonadota>Alphaproteobacteria>Caulobacterales>Caulobacteraceae | GCF_001375595.1 | Phenylobacterium_immobile | BN1313_RS07235 | 1497979..1501278 | 21 | DUF499-domain-containing-protein-[Phenylobacterium-immobile] |
| WP_246370526.1 | DUF499_1.RM.like.systems | <-SF1-SF2-Helicase+REase<-SF1-SF2-Helicase+REase\|\|HTH+WYL->TUDOR+ SF1-SF2-Helicase+iREase+CTD+REase->REase->METHYLASE->HEPN+DUF499*->MODE-HTH+PBPII-><-ParG_RHH | 1086 | Bacteria>Pseudomonadota>Alphaproteobacteria>Caulobacterales>Caulobacteraceae | GCF_014196295.1 | Phenylobacterium_haematophilum | GGQ61_RS02505 | 500790..504050 | 98513 | DUF499-domain-containing-protein-[Phenylobacterium-haematophilum] |
| WP_226745381.1 | DUF499_1.RM.like.systems | <-XIS-HTH<-SF1-SF2-helicase+RdgC+Beta-barrel+REase\|\|HTH+WYL->TUDOR+ SF1-SF2-Helicase+iREase+CTD+REase->MORC+DUF3684->METHYLASE->HEPN+DUF499*->MODE-HTH+PBPII-><-ParG_RHH | 1098 | Bacteria>Pseudomonadota>Alphaproteobacteria>Caulobacterales>Caulobacteraceae | GCF_020535245.1 | Asticcacaulis_sp._AND118 | LH365_RS06640 | 1378803..1382099 | 2840468 | DUF499-domain-containing-protein-[Asticcacaulis-sp.-AND118] |
| MBN8552584.1 | DUF499_1.RM.like.systems | ParG_RHH-><-MODE-HTH+PBPII<-HEPN+DUF499*<-METHYLASE<-MORC+DUF3684<-TUDOR+ SF1-SF2-Helicase+iREase+CTD+REase<-HTH+WYL\|\|SF1-SF2-Helicase+REase->SF1-SF2-Helicase+REase-> | 1098 | Bacteria>Pseudomonadota>Alphaproteobacteria>Caulobacterales>unclassified_Caulobacterales | GCA_017302105.1 | Caulobacterales_bacterium | JOL52_06785 | 4618..7914 | 2282150 | DUF499-domain-containing-protein-[Caulobacterales-bacterium] |
| WP_066690799.1 | DUF499_1.RM.like.systems | <-HicB\|\|HTH+WYL->TUDOR+ SF1-SF2-Helicase+iREase+CTD+REase->ASCH->METHYLASE->HEPN+DUF499*->MODE-HTH+PBPII-><-ParG_RHH | 1097 | Bacteria>Pseudomonadota>Alphaproteobacteria | GCF_001619955.1 | Sphingomonadales | BZF57_RS16495 | 60768..64061 | 204457 | MULTISPECIES:-DUF499-domain-containing-protein-[Sphingomonadales] |
| WP_106754608.1 | DUF499_1.RM.like.systems | ParG_RHH-><-TetR-HTH\|\|MODE-HTH+PBPII-><-HEPN+DUF499*<-METHYLASE<-MORC+DUF3684<-TUDOR+ SF1-SF2-Helicase+iREase+CTD+REase<-HTH+WYL\|\|SF1-SF2-Helicase+REase->SF1-SF2-Helicase+REase-> | 1098 | Bacteria>Pseudomonadota>Alphaproteobacteria | GCF_003012935.1 | Hyphomicrobiales | C8348_RS17400 | 71198..74494 | 356 | MULTISPECIES:-DUF499-domain-containing-protein-[Hyphomicrobiales] |
| WP_166015916.1 | DUF499_1.RM.like.systems | <-SF1-SF2-Helicase+REase\|\|HTH+WYL->TUDOR+ SF1-SF2-Helicase+iREase+CTD+REase->YtH->MORC+DUF3684->METHYLASE->HEPN+DUF499*->?->?->GDResolvase+Recombinase+ZnR-> | 1098 | Bacteria>Pseudomonadota>Alphaproteobacteria | GCF_015326295.1 | Hyphomicrobiales | GNP00_RS08405 | 1742058..1745354 | 356 | MULTISPECIES:-DUF499-domain-containing-protein-[Hyphomicrobiales] |
| MCL6609753.1 | DUF499_1.RM.like.systems | METHYLASE-p->HEPN+DUF499*-> | 1105 | Bacteria>Pseudomonadota>Alphaproteobacteria>Geminicoccales>Geminicoccaceae | GCA_023512415.1 | Geminicoccaceae_bacterium | K6T74_16850 | 1426..4743 | 2448052 | DUF499-domain-containing-protein-[Geminicoccaceae-bacterium] |
| MAS03479.1 | DUF499_1.RM.like.systems | Phage_integrase+Phage_integrase->Phage_integrase+Phage_integrase->TetR-HTH->XIS-HTH->TUDOR+ SF1-SF2-Helicase+iREase+CTD+REase->METHYLASE->HEPN+DUF499*->?->?-><-GNTR-HTH\|\|?->FtsK->XIS-HTH-> | 1123 | Bacteria>Pseudomonadota>Alphaproteobacteria>Hyphomicrobiales>Ahrensiaceae | GCA_002698425.1 | Ahrensia_sp. | CL534_02085 | 25586..28957 | 1872419 | AAA-family-ATPase-[Ahrensia-sp.] |
| MAS04609.1 | DUF499_1.RM.like.systems | <-TIR<-TIR<-?<-?\|\|TUDOR+ SF1-SF2-Helicase+iREase+CTD+REase-p->METHYLASE->HEPN+DUF499*->HTH_58-><-ParB+ParBC<-?<-?<-?<-HTH_36 | 1125 | Bacteria>Pseudomonadota>Alphaproteobacteria>Hyphomicrobiales>Ahrensiaceae | GCA_002698425.1 | Ahrensia_sp. | CL534_07935 | 36468..39845 | 1872419 | AAA-family-ATPase-[Ahrensia-sp.] |
| RAH96071.1 | DUF499_1.RM.like.systems | <-ArdCN+MPTase\|\|?-><-ParB+HTH<-PHP+SbcC<-ParA+HTH<-HEPN+DUF499*<-METHYLASE<-?<-TUDOR+ SF1-SF2-Helicase+iREase+CTD+REase<-Phage_int_SAM_1+Phage_integrase | 1116 | Bacteria>Pseudomonadota>Alphaproteobacteria>Hyphomicrobiales>Amorphaceae | GCA_003258595.1 | Acuticoccus_sediminis | DLJ53_33525 | 27428..30778 | 2184697 | ATPase-[Acuticoccus-sediminis] |
| WP_202913508.1 | DUF499_1.RM.like.systems | <-ArdCN+MPTase\|\|?-><-ParB+HTH\|\|?-><-PHP+SbcC<-ParA+HTH<-HEPN+DUF499*<-METHYLASE<-?<-TUDOR+ SF1-SF2-Helicase+iREase+CTD+REase<-Phage_int_SAM_1+Phage_integrase | 1115 | Bacteria>Pseudomonadota>Alphaproteobacteria>Hyphomicrobiales>Amorphaceae | GCF_003258595.1 | Acuticoccus_sediminis | DLJ53_RS33595 | 27428..30775 | 2184697 | DUF499-domain-containing-protein-[Acuticoccus-sediminis] |
| WP_235707141.1 | DUF499_1.RM.like.systems | HPT+HISKIN+REC->?->?->tRNA-><-HEPN+DUF499*<-METHYLASE<-?<-TUDOR+ SF1-SF2-Helicase+iREase+CTD+REase<-HTH+WYL<-HTH<-GDResolvase+Recombinase+ZnR | 1098 | Bacteria>Pseudomonadota>Alphaproteobacteria>Hyphomicrobiales>Amorphaceae | GCF_021556395.1 | Acuticoccus_kalidii | L1787_RS03610 | 494431..497727 | 2910977 | DUF499-domain-containing-protein-[Acuticoccus-kalidii] |
| WP_019994997.1 | DUF499_1.RM.like.systems | REase->ArdCN+MPTase-><-HEPN+DUF499*<-METHYLASE<-?<-TUDOR+ SF1-SF2-Helicase+iREase+CTD+REase<-GDResolvase<-SF1-SF2-Helicase<-REase | 1114 | Bacteria>Pseudomonadota>Alphaproteobacteria>Hyphomicrobiales>Aurantimonadaceae | GCF_000382705.1 | Aureimonas_ureilytica | B146_RS0103320 | 700135..703479 | 401562 | DUF499-domain-containing-protein-[Aureimonas-ureilytica] |
| WP_150971178.1 | DUF499_1.RM.like.systems | TUDOR+ SF1-SF2-Helicase+iREase+CTD+REase->METHYLASE->HEPN+DUF499*-><-SF1-SF2-Helicase+DUF1998+HKD-DNase+HKD-DNase<-?\|\|ParB+HNH-><-PIN<-HTH+MPTase<-PAPS_reduct | 1117 | Bacteria>Pseudomonadota>Alphaproteobacteria>Hyphomicrobiales>Aurantimonadaceae | GCF_008802405.1 | Aureimonas_leprariae | F6X38_RS15550 | 20910..24263 | 2615207 | DUF499-domain-containing-protein-[Aureimonas-leprariae] |
| WP_078709662.1 | DUF499_1.RM.like.systems | MODE-HTH+PBPII->?->ABC-ATPase+SbcC-><-METHYLASE-p<-?\|\|<-HEPN+DUF499*<-METHYLASE<-SIR2<-ParB+HNH<-ASCH<-TUDOR+ SF1-SF2-Helicase+iREase+CTD+REase\|\|REase-> | 1102 | Bacteria>Pseudomonadota>Alphaproteobacteria>Hyphomicrobiales>Aurantimonadaceae | GCF_900167365.1 | Consotaella_salsifontis | B5D38_RS17130 | 40507..43815 | 1365950 | DUF499-domain-containing-protein-[Consotaella-salsifontis] |
| TAJ30270.1 | DUF499_1.RM.like.systems | ParG_RHH-><-MODE-HTH+PBPII<-HIN-HTH<-HIN-HTH<-?<-SIGMA-HTH<-HEPN+DUF499*<-METHYLASE-p | 1098 | Bacteria>Pseudomonadota>Alphaproteobacteria>Hyphomicrobiales>Boseaceae | GCA_004297045.1 | Bosea_sp._(in:_a-proteobacteria) | EPO59_12315 | 16657..19953 | 1871050 | DUF499-domain-containing-protein-[Bosea-sp.] |
| WP_069053085.1 | DUF499_1.RM.like.systems | <-Response_reg+LuxR-HTH<-?\|\|?-><-?<-?<-?<-HEPN+DUF499*<-METHYLASE-p\|\|?->?->?->LON-AAA-> | 1097 | Bacteria>Pseudomonadota>Alphaproteobacteria>Hyphomicrobiales>Boseaceae | GCF_001713455.1 | Bosea_sp._RAC05 | BSY19_RS04405 | 191351..194644 | 1842539 | DUF499-domain-containing-protein-[Bosea-sp.-RAC05] |
| WP_069053609.1 | DUF499_1.RM.like.systems | Arm-DNA-bind_3+Phage_int_SAM_3+Phage_integrase->TUDOR+SF1-SF2-Helicase+iREase+CTD->?->TUDOR+ SF1-SF2-Helicase+iREase+CTD+REase->ASCH->METHYLASE->HEPN+DUF499*->?->XIS-HTH->Phage_int_SAM_4+Phage_integrase-><-ABC-ATPase+SbcC | 1113 | Bacteria>Pseudomonadota>Alphaproteobacteria>Hyphomicrobiales>Boseaceae | GCF_001713455.1 | Bosea_sp._RAC05 | BSY19_RS07510 | 857537..860878 | 1842539 | DUF499-domain-containing-protein-[Bosea-sp.-RAC05] |
| MCC0038267.1 | DUF499_1.RM.like.systems | <-TUDOR+SF1-SF2-Helicase+iREase+CTD<-?<-HHH_4+SF1-SF2-Helicase+REase<-HEPN+DUF499*<-METHYLASE | 1108 | Bacteria>Pseudomonadota>Alphaproteobacteria>Hyphomicrobiales>Brucellaceae | GCA_020637695.1 | Brucellaceae_bacterium | H6893_04600 | 209797..213123 | 2021378 | ATP-binding-protein-[Brucellaceae-bacterium] |
| WP_140022621.1 | DUF499_1.RM.like.systems | <-STYKinase-like<-HTH<-PIWI<-TIR+PNTD2<-HEPN+DUF499*<-METHYLASE<-MORC+DUF3684<-YtH<-ASCH<-TUDOR+ SF1-SF2-Helicase+iREase+CTD+REase<-HTH+WYL | 1098 | Bacteria>Pseudomonadota>Alphaproteobacteria>Hyphomicrobiales>Brucellaceae | GCF_006376675.1 | Brucella_pecoris | FIB18_RS21290 | 91464..94760 | 867683 | DUF499-domain-containing-protein-[Brucella-pecoris] |
| WP_210296176.1 | DUF499_1.RM.like.systems | HTH+WYL->TUDOR+ SF1-SF2-Helicase+iREase+CTD+REase->ASCH->YtH->MORC+DUF3684->METHYLASE->HEPN+DUF499*->TIR+PNTD2->PIWI->HTH->STYKinase-like-> | 1086 | Bacteria>Pseudomonadota>Alphaproteobacteria>Hyphomicrobiales>Brucellaceae | GCF_014197065.1 | Brucella_pecoris | GGQ79_RS19845 | 17581..20841 | 867683 | DUF499-domain-containing-protein-[Brucella-pecoris] |
| WP_258125110.1 | DUF499_1.RM.like.systems | <-SF1-SF2-Helicase+REase<-SF1-SF2-Helicase+REase\|\|HTH+WYL->TUDOR+ SF1-SF2-Helicase+iREase+CTD+REase->?->METHYLASE->HEPN+DUF499*->?->?->GDResolvase+Recombinase+ZnR-> | 1098 | Bacteria>Pseudomonadota>Alphaproteobacteria>Hyphomicrobiales>Brucellaceae | GCF_024720755.1 | Brucella_anthropi | NOF64_RS22210 | 68035..71331 | 529 | DUF499-domain-containing-protein-[Brucella-anthropi] |
| WP_259030292.1 | DUF499_1.RM.like.systems | <-SMODS\|\|HTH+WYL->TUDOR+ SF1-SF2-Helicase+iREase+CTD+REase->ParB+HNH->?->METHYLASE->HEPN+DUF499*->GDResolvase+HIN-HTH->HEPN->?->?-><-DDE_3<-PAIREDC-HTH | 1098 | Bacteria>Pseudomonadota>Alphaproteobacteria>Hyphomicrobiales>Brucellaceae | GCF_024799785.1 | Brucella_anthropi | NW321_RS01585 | 345006..348302 | 529 | DUF499-domain-containing-protein-[Brucella-anthropi] |
| WP_071923269.1 | DUF499_1.RM.like.systems | GDResolvase+Recombinase+ZnR->CopG/RHH->TUDOR+ SF1-SF2-Helicase+iREase+CTD+REase->METHYLASE->HEPN+DUF499*->OLD-ABC+TOPRIM->SF1-SF2-Helicase-><-HTH_17<-Phage_integrase<-HIN-HTH | 1097 | Bacteria>Pseudomonadota>Alphaproteobacteria>Hyphomicrobiales>Chelatococcaceae | GCF_001887265.1 | Chelatococcus_daeguensis | BOQ54_RS03320 | 711833..715126 | 444444 | DUF499-domain-containing-protein-[Chelatococcus-daeguensis] |
| WP_188611839.1 | DUF499_1.RM.like.systems | <-SF1-SF2-helicase+RdgC+Beta-barrel+REase\|\|HTH+WYL->TUDOR+ SF1-SF2-Helicase+iREase+CTD+REase->ASCH->METHYLASE->HEPN+DUF499*-><-?\|\|?-><-?\|\|MarR-HTH-> | 1097 | Bacteria>Pseudomonadota>Alphaproteobacteria>Hyphomicrobiales>Chelatococcaceae | GCF_014640075.1 | Chelatococcus_reniformis | IEW34_RS24475 | 395088..398381 | 1494448 | DUF499-domain-containing-protein-[Chelatococcus-reniformis] |
| MBN9362843.1 | DUF499_1.RM.like.systems | <-HicB\|\|HTH+WYL->TUDOR+ SF1-SF2-Helicase+iREase+CTD+REase->MORC+DUF3684->HEPN->METHYLASE->HEPN+DUF499*-><-?<-N6_MTase<-?<-Arm-DNA-bind_3+Phage_int_SAM_3+Phage_integrase<-GNTR-HTH | 1098 | Bacteria>Pseudomonadota>Alphaproteobacteria>Hyphomicrobiales>Devosiaceae | GCA_017307535.1 | Devosia_sp. | JOJ09_15200 | 310565..313861 | 1871048 | DUF499-domain-containing-protein-[Devosia-sp.] |
| WP_116654657.1 | DUF499_1.RM.like.systems | <-JAB<-UbE1+JAB<-SMODS\|\|HTH+WYL->TUDOR+ SF1-SF2-Helicase+iREase+CTD+REase->METHYLASE->HEPN+DUF499*->?->?-><-MODE-HTH+PBPII\|\|TetR-HTH-><-ParG_RHH | 1099 | Bacteria>Pseudomonadota>Alphaproteobacteria>Hyphomicrobiales>Devosiaceae | GCF_003390885.1 | Pelagibacterium_sediminicola | DQG18_RS12765 | 56546..59845 | 2248761 | DUF499-domain-containing-protein-[Pelagibacterium-sediminicola] |
| WP_152505130.1 | DUF499_1.RM.like.systems | Arm-DNA-bind_3+Phage_integrase->CopG/RHH->ACET-><-HEPN+DUF499*<-METHYLASE<-REase<-MORC+DUF3684<-YtH<-TUDOR+ SF1-SF2-Helicase+iREase+CTD+REase<-HTH+WYL | 1098 | Bacteria>Pseudomonadota>Alphaproteobacteria>Hyphomicrobiales | GCF_009363315.1 | Stappiaceae | FIV06_RS06815 | 1469673..1472969 | 2821832 | DUF499-domain-containing-protein-[Roseibium-sp.-Sym1] |
| MBA4172963.1 | DUF499_1.RM.like.systems | <-UbE1+JAB<-?\|\|HTH+WYL-><-HEPN+DUF499*<-METHYLASE<-TUDOR+ SF1-SF2-Helicase+iREase+CTD+REase<-HTH_37 | 1110 | Bacteria>Pseudomonadota>Alphaproteobacteria>Hyphomicrobiales>Hyphomicrobiaceae | GCA_013822135.1 | Hyphomicrobium_sp. | C0511_10000 | 5338..8670 | 82 | AAA+-family-ATPase-[Hyphomicrobium-sp.] |
| MBR2536595.1 | DUF499_1.RM.like.systems | ParB+N6_N4_MTase->?->?-><-REase<-REase\|\|HTH_3-><-HEPN+DUF499*<-METHYLASE<-HEPN<-TUDOR+ SF1-SF2-Helicase+iREase+CTD+REase<-HTH+WYL\|\|SMODS->SLATT-> | 1092 | Bacteria>Pseudomonadota>Alphaproteobacteria>Hyphomicrobiales>Hyphomicrobiaceae | GCA_017512425.1 | Hyphomicrobium_sp. | IKE66_11035 | 173240..176518 | 82 | DUF499-domain-containing-protein-[Hyphomicrobium-sp.] |
| WP_129609562.1 | DUF499_1.RM.like.systems | Phage_int_SAM_3+Phage_integrase->?->XIS-HTH->ZnR+TOPRIM->TOPRIM+SF1-SF2-Helicase+HTH-><-HEPN+DUF499*<-METHYLASE<-ASCH<-sGTP<-TUDOR+ SF1-SF2-Helicase+iREase+CTD+REase | 1121 | Bacteria>Pseudomonadota>Alphaproteobacteria>Hyphomicrobiales>Hyphomicrobiaceae | GCF_900604295.1 | Rhodoplanes_serenus | RHODPL_RS14920 | 10306..13671 | 200615 | Swt1-family-HEPN-domain-containing-protein-[Rhodoplanes-piscinae] |
| UYN95212.1 | DUF499_1.RM.like.systems | TUDOR+ SF1-SF2-Helicase+iREase+CTD+REase->?->ParB+HNH->METHYLASE->HEPN+DUF499*->AbiEii-><-?<-?<-SPOUT | 1104 | Bacteria>Pseudomonadota>Alphaproteobacteria>Hyphomicrobiales>Hyphomicrobiales_incertae_sedis | GCA_025808875.1 | Enhydrobacter_sp. | KIT25_24915 | 5028707..5032021 | 1894999 | MAG:-DUF499-domain-containing-protein-[Enhydrobacter-sp.] |
| WP_266282109.1 | DUF499_1.RM.like.systems | GNTR-HTH->?-><-?\|\|SIGMA-HTH-><-HEPN+DUF499*<-METHYLASE<-YtH<-MORC+DUF3684<-TUDOR+ SF1-SF2-Helicase+iREase+CTD+REase<-HTH+WYL | 1098 | Bacteria>Pseudomonadota>Alphaproteobacteria>Hyphomicrobiales>Kaistiaceae | GCF_026344055.1 | Kaistia_geumhonensis | OSH08_RS14490 | 3056555..3059851 | 410839 | DUF499-domain-containing-protein-[Kaistia-geumhonensis] |
| QGY06229.1 | DUF499_1.RM.like.systems | HTH+WYL-><-ParB+HNH<-ParB+HNH<-SF1-SF2-Helicase+DUF1998+HKD-DNase+HKD-DNase<-?<-HEPN+DUF499* | 1100 | Bacteria>Pseudomonadota>Alphaproteobacteria>Hyphomicrobiales>Methylobacteriaceae | GCA_000364445.2 | Methylobacterium_mesophilicum_SR1.6/6 | MMSR116_17605 | 3672852..3676154 | 908290 | ATP-binding-protein-[Methylobacterium-mesophilicum-SR1.6/6] |
| KQQ15548.1 | DUF499_1.RM.like.systems | PAIREDC-HTH->?->TUDOR+ SF1-SF2-Helicase+iREase+CTD+REase->ASCH->METHYLASE->HEPN+DUF499*->?->SF1-SF2-Helicase+DUF1998+HKD-DNase+HKD-DNase->Radical-SAM->?-><-Phage_integrase<-tRNA<-tRNA<-SIGMA-HTH+SIGMA-HTH | 1115 | Bacteria>Pseudomonadota>Alphaproteobacteria>Hyphomicrobiales>Methylobacteriaceae | GCA_001423385.1 | Methylobacterium_sp._Leaf121 | ASF59_14580 | 38300..41647 | 1736262 | ATPase-[Methylobacterium-sp.-Leaf121] |
| OAH31268.1 | DUF499_1.RM.like.systems | <-RVT+HNH<-MODE-HTH+PBPII<-HEPN+DUF499*<-METHYLASE<-MORC+DUF3684<-YtH<-TUDOR+ SF1-SF2-Helicase+iREase+CTD+REase<-HTH+WYL\|\|SF1-SF2-Helicase+REase-> | 1098 | Bacteria>Pseudomonadota>Alphaproteobacteria>Hyphomicrobiales>Methylobacteriaceae | GCA_001643095.1 | Methylorubrum_populi | AX289_21395 | 87979..91275 | 223967 | AAA-family-ATPase-[Methylorubrum-populi] |
| MCA3671902.1 | DUF499_1.RM.like.systems | TraJ-RHH->?->?-><-HEPN<-VirD4-TraG+VirD4-FtsK\|\|GDResolvase+HIN-HTH-><-HEPN+DUF499*<-METHYLASE<-TUDOR+ SF1-SF2-Helicase+iREase+CTD+REase<-?<-CHTH | 1106 | Bacteria>Pseudomonadota>Alphaproteobacteria>Hyphomicrobiales>Methylobacteriaceae | GCA_020253325.1 | Methylobacterium_sp. | INF11_08205 | 10966..14286 | 409 | DUF499-domain-containing-protein-[Methylobacterium-sp.] |
| WP_083920313.1 | DUF499_1.RM.like.systems | HTH+WYL-><-ParB+HNH<-ParB+HNH<-SF1-SF2-Helicase+DUF1998+HKD-DNase+HKD-DNase<-?<-HEPN+DUF499* | 1053 | Bacteria>Pseudomonadota>Alphaproteobacteria>Hyphomicrobiales>Methylobacteriaceae | GCF_000364445.2 | Methylobacterium_mesophilicum | MMSR116_RS17410 | 3672852..3676013 | 39956 | DUF499-domain-containing-protein-[Methylobacterium-mesophilicum] |
| WP_103711433.1 | DUF499_1.RM.like.systems | <-HEPN+DUF499*<-METHYLASE<-MORC+DUF3684<-TUDOR+ SF1-SF2-Helicase+iREase+CTD+REase<-HTH+WYL | 1098 | Bacteria>Pseudomonadota>Alphaproteobacteria>Hyphomicrobiales>Methylobacteriaceae | GCF_002917135.1 | Methylobacterium_sp._V23 | CRT23_RS20075 | 34063..37359 | 2044878 | DUF499-domain-containing-protein-[Methylobacterium-sp.-V23] |
| WP_210322592.1 | DUF499_1.RM.like.systems | TUDOR+ SF1-SF2-Helicase+iREase+CTD+REase->MORC+DUF3684->METHYLASE->HEPN+DUF499*-><-GDResolvase+Recombinase+ZnR<-RAMA<-?<-Phage_integrase<-URI-DNase | 1086 | Bacteria>Pseudomonadota>Alphaproteobacteria>Hyphomicrobiales>Methylobacteriaceae | GCF_014845095.1 | Methylobacterium_bullatum | MBTS_RS00690 | 11637..14897 | 570505 | DUF499-domain-containing-protein-[Methylobacterium-bullatum] |
| WP_207183158.1 | DUF499_1.RM.like.systems | TUDOR+ SF1-SF2-Helicase+iREase+CTD+REase->ASCH->METHYLASE-p->METHYLASE-p->METHYLASE-p->HEPN+DUF499*-> | 1117 | Bacteria>Pseudomonadota>Alphaproteobacteria>Hyphomicrobiales>Methylobacteriaceae | GCF_017347565.1 | Methylobacterium_indicum | miv_RS13675 | 2915960..2919313 | 1775910 | DUF499-domain-containing-protein-[Methylobacterium-indicum] |
| WP_192214889.1 | DUF499_1.RM.like.systems | URI-DNase->Phage_integrase->?->RAMA->GDResolvase+Recombinase+ZnR-><-HEPN+DUF499*<-METHYLASE<-MORC+DUF3684<-TUDOR+ SF1-SF2-Helicase+iREase+CTD+REase<-HTH+WYL | 1098 | Bacteria>Pseudomonadota>Alphaproteobacteria>Hyphomicrobiales>Methylobacteriaceae | GCF_022179105.1 | Methylobacterium_bullatum | MFF03_RS22740 | 7262..10558 | 570505 | DUF499-domain-containing-protein-[Methylobacterium-bullatum] |
| WP_262268834.1 | DUF499_1.RM.like.systems | RVT-><-HEPN+DUF499*<-METHYLASE<-TUDOR+ SF1-SF2-Helicase+iREase+CTD+REase\|\|?->Phage_integrase-> | 1110 | Bacteria>Pseudomonadota>Alphaproteobacteria>Hyphomicrobiales>Methylobacteriaceae | GCF_025460875.1 | Microvirga_sp._HBU65207 | NF876_RS10955 | 4117..7449 | 2953740 | DUF499-domain-containing-protein-[Microvirga-sp.-HBU65207] |
| WP_182180493.1 | DUF499_1.RM.like.systems | Arm-DNA-bind_3+Phage_int_SAM_3+Phage_integrase->TUDOR+ SF1-SF2-Helicase+iREase+CTD+REase->ASCH->METHYLASE->HEPN+DUF499*-> | 1111 | Bacteria>Pseudomonadota>Alphaproteobacteria>Hyphomicrobiales>Methylobacteriaceae | GCF_903971015.1 | Methylobacterium_radiotolerans | H7B86_RS28270 | 13923..17258 | 31998 | DUF499-domain-containing-protein-[Methylobacterium-radiotolerans] |
| CAA2157197.1 | DUF499_1.RM.like.systems | SF1-SF2-Helicase-p->?-><-GDResolvase+HIN-HTH\|\|TUDOR+ SF1-SF2-Helicase+iREase+CTD+REase->METHYLASE->HEPN+DUF499*-> | 1109 | Bacteria>Pseudomonadota>Alphaproteobacteria>Hyphomicrobiales>Methylobacteriaceae | - | Methylobacterium_brachiatum | MBRA_02601 | 587960..591289 | 269660 | hypothetical-protein-MBRA_02601-[Methylobacterium-brachiatum] |
| MBI5313198.1 | DUF499_1.RM.like.systems | TUDOR+ SF1-SF2-Helicase+iREase+CTD+REase->Ntox18->Peptidase_M78->METHYLASE->OLD-ABC->PIN->HEPN+DUF499*->?-><-HTH | 1107 | Bacteria>Pseudomonadota>Alphaproteobacteria>Hyphomicrobiales>Methylocystaceae | GCA_016219665.1 | Methylocystis_sp. | HZB28_08775 | 48507..51830 | 1911079 | DUF499-domain-containing-protein-[Methylocystis-sp.] |
| WP_064034073.1 | DUF499_1.RM.like.systems | MORC+DUF3684+REase-><-?<-SMODS\|\|?->?->MPTase-><-HEPN+DUF499*<-METHYLASE<-?<-TUDOR+ SF1-SF2-Helicase+iREase+CTD+REase<-HTH+WYL | 1109 | Bacteria>Pseudomonadota>Alphaproteobacteria>Hyphomicrobiales>Methylocystaceae | GCF_001644125.1 | Methylosinus_sp._R-45379 | A1351_RS23390 | 14763..18092 | 980563 | Swt1-family-HEPN-domain-containing-protein-[Methylosinus-sp.-R-45379] |
| WP_166143172.1 | DUF499_1.RM.like.systems | <-HEPN+DUF499*<-METHYLASE<-TUDOR+ SF1-SF2-Helicase+iREase+CTD+REase<-HTH+WYL\|\|?->ParB+HTH-> | 1109 | Bacteria>Pseudomonadota>Alphaproteobacteria>Hyphomicrobiales>Methylocystaceae | GCF_011378635.1 | Methylosinus_sp._RM1 | FMZ75_RS12510 | 924..4253 | 2583817 | Swt1-family-HEPN-domain-containing-protein-[Methylosinus-sp.-RM1] |
| OJU24986.1 | DUF499_1.RM.like.systems | PIN+OST-HTH->SF1-SF2-Helicase-p->TUDOR+ SF1-SF2-Helicase+iREase+CTD+REase->HEPN+DUF499*->ArdCN+MPTase-> | 1106 | Bacteria>Pseudomonadota>Alphaproteobacteria>Hyphomicrobiales>Nitrobacteraceae | GCA_001896955.1 | Nitrobacter_sp._62-13 | BGN91_09350 | 17137..20457 | 1895797 | MAG:-ATPase-[Nitrobacter-sp.-62-13] |
| TAH66859.1 | DUF499_1.RM.like.systems | <-SF1-SF2-Helicase+REase<-SF1-SF2-Helicase+REase\|\|HTH+WYL->TUDOR+ SF1-SF2-Helicase+iREase+CTD+REase->MORC+DUF3684->METHYLASE->HEPN+DUF499*-><-MODE-HTH+PBPII\|\|TetR-HTH->?->DUF488-l-><-ParG_RHH | 1098 | Bacteria>Pseudomonadota>Alphaproteobacteria>Hyphomicrobiales>Nitrobacteraceae | GCA_004294885.1 | Rhodopseudomonas_palustris | EWM45_09695 | 32006..35302 | 1076 | MAG:-ATP-binding-protein-[Rhodopseudomonas-palustris] |
| MCV0386470.1 | DUF499_1.RM.like.systems | Phage_integrase->XIS-HTH->TUDOR+ SF1-SF2-Helicase+iREase+CTD+REase->METHYLASE->HEPN+DUF499*->SF1-SF2-Helicase-p-> | 1096 | Bacteria>Pseudomonadota>Alphaproteobacteria>Hyphomicrobiales>Nitrobacteraceae | GCA_025698885.1 | Nitrobacter_sp. | K5821_08550 | 83290..86580 | 29420 | MAG:-DUF499-domain-containing-protein-[Nitrobacter-sp.] |
| WP_008960924.1 | DUF499_1.RM.like.systems | HEPN+DUF499*->?-><-Arm-DNA-bind_3+Phage_integrase | 1107 | Bacteria>Pseudomonadota>Alphaproteobacteria>Hyphomicrobiales>Nitrobacteraceae | GCF_000239795.1 | Bradyrhizobium_sp._STM_3809 | BRAS3809_RS04535 | 1765..5088 | 551936 | DUF499-domain-containing-protein-[Bradyrhizobium-sp.-STM-3809] |
| WP_036012275.1 | DUF499_1.RM.like.systems | URI-DNase-><-HEPN+DUF499*<-METHYLASE<-ASCH<-TUDOR+ SF1-SF2-Helicase+iREase+CTD+REase\|\|GDResolvase-> | 1104 | Bacteria>Pseudomonadota>Alphaproteobacteria>Hyphomicrobiales>Nitrobacteraceae | GCF_000261825.1 | Bradyrhizobium_yuanmingense | IC46_RS27575 | 9376..12690 | 108015 | DUF499-domain-containing-protein-[Bradyrhizobium-yuanmingense] |
| WP_028141996.1 | DUF499_1.RM.like.systems | REase-><-?<-MACRODOMAIN<-HEPN+DUF499*<-METHYLASE<-ASCH<-TUDOR+ SF1-SF2-Helicase+iREase+CTD+REase<-Phage_int_SAM_1+Phage_integrase | 1103 | Bacteria>Pseudomonadota>Alphaproteobacteria>Hyphomicrobiales>Nitrobacteraceae | GCF_000472725.1 | Bradyrhizobium_ottawaense | A3AQ_RS0117025 | 231238..234549 | 931866 | DUF499-domain-containing-protein-[Bradyrhizobium-ottawaense] |
| WP_025032482.1 | DUF499_1.RM.like.systems | ZnR+TOPRIM->TOPRIM+SF1-SF2-Helicase+HTH->?->?->REase->REase-><-HEPN+DUF499*<-METHYLASE<-ASCH<-TUDOR+ SF1-SF2-Helicase+iREase+CTD+REase<-?<-MULE-TRANSPOSASE<-tRNA<-HISKIN+MORC | 1106 | Bacteria>Pseudomonadota>Alphaproteobacteria>Hyphomicrobiales>Nitrobacteraceae | GCF_000617845.2 | Bradyrhizobium_sp._DOA9 | BDOA9_RS03050 | 657043..660363 | 1126627 | DUF499-domain-containing-protein-[Bradyrhizobium-sp.-DOA9] |
| WP_145832126.1 | DUF499_1.RM.like.systems | <-HEPN+DUF499*<-METHYLASE<-DDE_3 | 1113 | Bacteria>Pseudomonadota>Alphaproteobacteria>Hyphomicrobiales>Nitrobacteraceae | GCF_007830635.1 | Bradyrhizobium_huanghuaihaiense | IQ16_RS39330 | 51033..54374 | 990078 | DUF499-domain-containing-protein-[Bradyrhizobium-huanghuaihaiense] |
| WP_151650193.1 | DUF499_1.RM.like.systems | HTH_3->TUDOR+ SF1-SF2-Helicase+iREase+CTD+REase->METHYLASE->HEPN+DUF499*-> | 1111 | Bacteria>Pseudomonadota>Alphaproteobacteria>Hyphomicrobiales>Nitrobacteraceae | GCF_008932115.1 | Bradyrhizobium_betae | F8237_RS32705 | 6799441..6802776 | 244734 | DUF499-domain-containing-protein-[Bradyrhizobium-betae] |
| WP_166354634.1 | DUF499_1.RM.like.systems | <-HEPN+DUF499*<-METHYLASE<-MORC+DUF3684<-YtH<-ASCH<-ASCH<-TUDOR+ SF1-SF2-Helicase+iREase+CTD+REase | 1097 | Bacteria>Pseudomonadota>Alphaproteobacteria>Hyphomicrobiales>Nitrobacteraceae | GCF_011068405.1 | Bradyrhizobium_barranii | G6321_RS03310 | 151200..154493 | 2992140 | DUF499-domain-containing-protein-[Bradyrhizobium-barranii] |
| WP_049832574.1 | DUF499_1.RM.like.systems | <-AbiEii<-AbiEi_4\|\|CHTH->?-><-?\|\|REase-><-HEPN+DUF499*\|\|<-TUDOR+ SF1-SF2-Helicase+iREase+CTD+REase-p<-Recombinase<-REase<-tRNA<-HISKIN+MORC\|\|?->RNase-> | 1051 | Bacteria>Pseudomonadota>Alphaproteobacteria>Hyphomicrobiales>Nitrobacteraceae | GCF_011604625.1 | Bradyrhizobium_diazoefficiens | HAU57_RS27175 | 5742766..5745921 | 1355477 | DUF499-domain-containing-protein-[Bradyrhizobium-diazoefficiens] |
| WP_182868871.1 | DUF499_1.RM.like.systems | Classical-AAA->Peptidase_S8->?->?->?-><-HEPN+DUF499*<-METHYLASE<-sGTP<-TUDOR+ SF1-SF2-Helicase+iREase+CTD+REase<-?<-XIS-HTH | 1117 | Bacteria>Pseudomonadota>Alphaproteobacteria>Hyphomicrobiales>Nitrobacteraceae | GCF_014163475.1 | Bradyrhizobium_diazoefficiens | H7K76_RS29690 | 6385393..6388746 | 1355477 | Swt1-family-HEPN-domain-containing-protein-[Bradyrhizobium-diazoefficiens] |
| WP_011157774.1 | DUF499_1.RM.like.systems | <-SF1-SF2-Helicase+REase<-?\|\|HTH+WYL->TUDOR+ SF1-SF2-Helicase+iREase+CTD+REase->MORC+DUF3684->METHYLASE->HEPN+DUF499*->MODE-HTH+PBPII-><-ParG_RHH | 1099 | Bacteria>Pseudomonadota>Alphaproteobacteria>Hyphomicrobiales>Nitrobacteraceae | GCF_016584445.1 | Rhodopseudomonas_palustris | TX73_RS11465 | 2517133..2520432 | 1076 | DUF499-domain-containing-protein-[Rhodopseudomonas-palustris] |
| WP_211433350.1 | DUF499_1.RM.like.systems | <-DUF488-l<-REase\|\|TUDOR+ SF1-SF2-Helicase+iREase+CTD+REase->METHYLASE-p->METHYLASE-p->HEPN+DUF499*-> | 1097 | Bacteria>Pseudomonadota>Alphaproteobacteria>Hyphomicrobiales>Nitrobacteraceae | GCF_018130635.1 | Bradyrhizobium_diazoefficiens | JQ640_RS02510 | 177861..181154 | 1355477 | DUF499-domain-containing-protein-[Bradyrhizobium-diazoefficiens] |
| WP_212326836.1 | DUF499_1.RM.like.systems | Phage_integrase->?->XIS-HTH->?->DnaB->HIN-HTH-><-HEPN+DUF499*<-METHYLASE<-HNH<-OLD-ABC<-TUDOR+ SF1-SF2-Helicase+iREase+CTD+REase | 1108 | Bacteria>Pseudomonadota>Alphaproteobacteria>Hyphomicrobiales>Nitrobacteraceae | GCF_018131105.1 | Bradyrhizobium_manausense | JQ631_RS13545 | 2838930..2842256 | 989370 | Swt1-family-HEPN-domain-containing-protein-[Bradyrhizobium-manausense] |
| WP_225188922.1 | DUF499_1.RM.like.systems | <-REC<-REase+DUF4268\|\|?->HEPN->?->?->?-><-HEPN+DUF499*<-METHYLASE<-TUDOR+ SF1-SF2-Helicase+iREase+CTD+REase<-HTH_3\|\|?-><-XIS-HTH | 1094 | Bacteria>Pseudomonadota>Alphaproteobacteria>Hyphomicrobiales>Nitrobacteraceae | GCF_020193585.1 | Bradyrhizobium_sp._IC3195 | I6F09_RS20850 | 200244..203528 | 2793804 | DUF499-domain-containing-protein-[Bradyrhizobium-sp.-IC3195] |
| WP_225147255.1 | DUF499_1.RM.like.systems | REase->Recombinase->TUDOR+ SF1-SF2-Helicase+iREase+CTD+REase->ASCH->METHYLASE->HEPN+DUF499*-><-REase\|\|?-><-?<-CHTH\|\|AbiEi_4->AbiEii-> | 1113 | Bacteria>Pseudomonadota>Alphaproteobacteria>Hyphomicrobiales>Nitrobacteraceae | GCF_020193875.1 | Bradyrhizobium_sp._NBAIM16 | I6F29_RS15715 | 113590..116931 | 2793813 | DUF499-domain-containing-protein-[Bradyrhizobium-sp.-NBAIM16] |
| WP_225175154.1 | DUF499_1.RM.like.systems | REase->Recombinase->TUDOR+ SF1-SF2-Helicase+iREase+CTD+REase->ASCH->METHYLASE->HEPN+DUF499*-><-REase\|\|?-><-?<-CHTH\|\|AbiEi_4->AbiEii-> | 1113 | Bacteria>Pseudomonadota>Alphaproteobacteria>Hyphomicrobiales>Nitrobacteraceae | GCF_020193975.1 | Bradyrhizobium_sp._NBAIM02 | I6F23_RS23650 | 113567..116908 | 2793817 | DUF499-domain-containing-protein-[Bradyrhizobium-sp.-NBAIM02] |
| WP_231327536.1 | DUF499_1.RM.like.systems | <-Phage_int_SAM_1+Phage_integrase\|\|GNAT->?-><-?<-?<-HEPN+DUF499*<-METHYLASE<-TUDOR+ SF1-SF2-Helicase+iREase+CTD+REase | 1098 | Bacteria>Pseudomonadota>Alphaproteobacteria>Hyphomicrobiales>Nitrobacteraceae | GCF_021088345.1 | Bradyrhizobium_sp._A19 | LQG66_RS18125 | 4079105..4082401 | 2898149 | DUF499-domain-containing-protein-[Bradyrhizobium-sp.-A19] |
| WP_247344343.1 | DUF499_1.RM.like.systems | HTH+WYL->TUDOR+ SF1-SF2-Helicase+iREase+CTD+REase->ParB+HNH->?->METHYLASE->HEPN+DUF499*->HEPN-> | 1097 | Bacteria>Pseudomonadota>Alphaproteobacteria>Hyphomicrobiales>Nitrobacteraceae | GCF_023073075.1 | Bradyrhizobium_sp._61 | IVB46_RS33345 | 520098..523391 | 2782679 | DUF499-domain-containing-protein-[Bradyrhizobium-sp.-61] |
| WP_247407267.1 | DUF499_1.RM.like.systems | <-AbiEii<-AbiEi_4\|\|CHTH->?-><-?\|\|REase-><-HEPN+DUF499*<-METHYLASE<-ASCH<-TUDOR+ SF1-SF2-Helicase+iREase+CTD+REase<-Recombinase<-REase<-tRNA<-HISKIN+MORC | 1113 | Bacteria>Pseudomonadota>Alphaproteobacteria>Hyphomicrobiales>Nitrobacteraceae | GCF_023073575.1 | unclassified_Bradyrhizobium | IVB10_RS07520 | 491503..494844 | 2631580 | MULTISPECIES:-DUF499-domain-containing-protein-[unclassified-Bradyrhizobium] |
| WP_247531825.1 | DUF499_1.RM.like.systems | <-GDResolvase\|\|TUDOR+ SF1-SF2-Helicase+iREase+CTD+REase->ASCH->METHYLASE->HEPN+DUF499*-><-?<-PHP+SbcC | 1101 | Bacteria>Pseudomonadota>Alphaproteobacteria>Hyphomicrobiales>Nitrobacteraceae | GCF_023073775.1 | Bradyrhizobium_sp._162 | IVA99_RS22470 | 14137..17442 | 2782635 | DUF499-domain-containing-protein-[Bradyrhizobium-sp.-162] |
| WP_247438142.1 | DUF499_1.RM.like.systems | <-SF1-SF2-Helicase+REase-p\|\|?->TUDOR+ SF1-SF2-Helicase+iREase+CTD+REase->?->METHYLASE->HEPN+DUF499*-><-Peptidase\|\|?->VirD4-TraG+VirD4-FtsK-> | 1113 | Bacteria>Pseudomonadota>Alphaproteobacteria>Hyphomicrobiales>Nitrobacteraceae | GCF_023074175.1 | Bradyrhizobium_sp._139 | IVA80_RS29435 | 88222..91563 | 2782616 | DUF499-domain-containing-protein-[Bradyrhizobium-sp.-139] |
| WP_253593718.1 | DUF499_1.RM.like.systems | LZ_Tnp_IS66+zf-IS66+DDE_Tnp_IS66+DDE_Tnp_IS66_C-><-Phage_integrase\|\|HEPN+REase->TUDOR+ SF1-SF2-Helicase+iREase+CTD+REase->ASCH->METHYLASE->HEPN+DUF499*->?-><-HTH_3\|\|?-><-GDResolvase+Recombinase+ZnR | 1113 | Bacteria>Pseudomonadota>Alphaproteobacteria>Hyphomicrobiales>Nitrobacteraceae | GCF_024171365.1 | Bradyrhizobium_elkanii | M2199_RS04195 | 899505..902846 | 29448 | DUF499-domain-containing-protein-[Bradyrhizobium-elkanii] |
| WP_259172384.1 | DUF499_1.RM.like.systems | <-HEPN+DUF499*<-METHYLASE<-TUDOR+ SF1-SF2-Helicase+iREase+CTD+REase<-HTH_3 | 1098 | Bacteria>Pseudomonadota>Alphaproteobacteria>Hyphomicrobiales>Nitrobacteraceae | GCF_024806875.1 | Bradyrhizobium_betae | FHR88_RS05915 | 1186153..1189449 | 244734 | DUF499-domain-containing-protein-[Bradyrhizobium-betae] |
| WP_092683743.1 | DUF499_1.RM.like.systems | TUDOR+ SF1-SF2-Helicase+iREase+CTD+REase->MORC+DUF3684->METHYLASE->HEPN+DUF499*-><-?<-?<-Radical-SAM | 1116 | Bacteria>Pseudomonadota>Alphaproteobacteria>Hyphomicrobiales>Nitrobacteraceae | GCF_900110435.1 | Rhodopseudomonas_pseudopalustris | BMY71_RS11075 | 11290..14640 | 1513892 | Swt1-family-HEPN-domain-containing-protein-[Rhodopseudomonas-pseudopalustris] |
| WP_092298105.1 | DUF499_1.RM.like.systems | ZnR->?-><-SF1-SF2-Helicase-p<-SF1-SF2-Helicase<-Exonuc_V_gamma+REase<-HEPN+DUF499*<-METHYLASE<-MORC+DUF3684<-YtH<-TUDOR+ SF1-SF2-Helicase+iREase+CTD+REase<-HTH+WYL | 1094 | Bacteria>Pseudomonadota>Alphaproteobacteria>Hyphomicrobiales>Nitrobacteraceae | GCF_900115265.1 | Bradyrhizobium_sp._Ghvi | BM374_RS24865 | 13858..17142 | 1855319 | DUF499-domain-containing-protein-[Bradyrhizobium-sp.-Ghvi] |
| WP_079537876.1 | DUF499_1.RM.like.systems | <-ACYC\|\|?->OLD-ABC->?-><-HKD-DNase<-ParB+HNH<-HEPN+DUF499*<-METHYLASE<-ASCH<-TUDOR+ SF1-SF2-Helicase+iREase+CTD+REase\|\|?-><-GDResolvase+HIN-HTH | 1108 | Bacteria>Pseudomonadota>Alphaproteobacteria>Hyphomicrobiales>Nitrobacteraceae | GCF_900141755.1 | Bradyrhizobium_lablabi | B5526_RS08940 | 1910931..1914257 | 722472 | DUF499-domain-containing-protein-[Bradyrhizobium-lablabi] |
| WP_172236133.1 | DUF499_1.RM.like.systems | XIS-HTH->?->?->DnaB->HIN-HTH-><-HEPN+DUF499*<-METHYLASE<-HNH<-OLD-ABC<-TUDOR+ SF1-SF2-Helicase+iREase+CTD+REase | 1108 | Bacteria>Pseudomonadota>Alphaproteobacteria>Hyphomicrobiales>Nitrobacteraceae | GCF_902482845.1 | Bradyrhizobium | JQ619_RS32195 | 7694..11020 | 374 | MULTISPECIES:-Swt1-family-HEPN-domain-containing-protein-[Bradyrhizobium] |
| MAB12295.1 | DUF499_1.RM.like.systems | <-HEPN+DUF499*<-METHYLASE<-REase<-MORC+DUF3684<-YtH<-TUDOR+ SF1-SF2-Helicase+iREase+CTD+REase<-HTH+WYL | 1098 | Bacteria>Pseudomonadota>Alphaproteobacteria>Hyphomicrobiales>Parvibaculaceae | GCA_002684405.1 | Parvibaculum_sp. | CMI59_00100 | 13399..16695 | 2024848 | AAA-family-ATPase-[Parvibaculum-sp.] |
| MAM94938.1 | DUF499_1.RM.like.systems | <-SF1-SF2-Helicase+REase<-SF1-SF2-Helicase+REase\|\|HTH+WYL->TUDOR+ SF1-SF2-Helicase+iREase+CTD+REase->?->METHYLASE->HEPN+DUF499*-><-MODE-HTH+PBPII | 1098 | Bacteria>Pseudomonadota>Alphaproteobacteria>Hyphomicrobiales>Parvibaculaceae | GCA_002694985.1 | Parvibaculum_sp. | CMI61_09980 | 298093..301389 | 2024848 | AAA-family-ATPase-[Parvibaculum-sp.] |
| RWD48635.1 | DUF499_1.RM.like.systems | Peptidase_S11->tRNA-><-HEPN+DUF499*<-METHYLASE<-MORC+DUF3684<-YtH<-ASCH<-TUDOR+ SF1-SF2-Helicase+iREase+CTD+REase<-HTH+WYL | 1098 | Bacteria>Pseudomonadota>Alphaproteobacteria>Hyphomicrobiales>Phyllobacteriaceae | GCA_004016945.1 | Mesorhizobium_sp. | EOS59_16740 | 2840..6136 | 1871066 | ATP-binding-protein-[Mesorhizobium-sp.] |
| RWE54183.1 | DUF499_1.RM.like.systems | HTH+WYL->TUDOR+ SF1-SF2-Helicase+iREase+CTD+REase->ASCH->YtH->METHYLASE->HEPN+DUF499*->TUDOR+SF1-SF2-Helicase+iREase+CTD->?->HHH_4+SF1-SF2-Helicase+REase->?-><-tRNA<-Peptidase_S11 | 1100 | Bacteria>Pseudomonadota>Alphaproteobacteria>Hyphomicrobiales>Phyllobacteriaceae | GCA_004018175.1 | Mesorhizobium_sp. | EOS67_26215 | 294790..298092 | 1871066 | ATP-binding-protein-[Mesorhizobium-sp.] |
| RWJ00134.1 | DUF499_1.RM.like.systems | <-ZnR\|\|?-><-?<-?<-?\|\|<-HEPN+DUF499*<-METHYLASE<-MORC+DUF3684<-TUDOR+ SF1-SF2-Helicase+iREase+CTD+REase<-HTH+WYL\|\|SMODS->UbE1+JAB-> | 1110 | Bacteria>Pseudomonadota>Alphaproteobacteria>Hyphomicrobiales>Phyllobacteriaceae | GCA_004018865.1 | Mesorhizobium_sp. | EOR23_30945 | 21275..24607 | 1871066 | ATP-binding-protein-[Mesorhizobium-sp.] |
| RWL15103.1 | DUF499_1.RM.like.systems | HhH-RADC+JAB->?->\|\|Phage_integrase->?->METHYLASE->PIN-><-HEPN+DUF499*<-METHYLASE<-HEPN<-MORC+DUF3684<-TUDOR+ SF1-SF2-Helicase+iREase+CTD+REase<-HTH+WYL\|\|HicB-> | 1098 | Bacteria>Pseudomonadota>Alphaproteobacteria>Hyphomicrobiales>Phyllobacteriaceae | GCA_004019595.1 | Mesorhizobium_sp. | EOR57_31095 | 10515..13811 | 1871066 | DUF499-domain-containing-protein-[Mesorhizobium-sp.] |
| RWN27570.1 | DUF499_1.RM.like.systems | <-LD-peptidase<-?<-HEPN+DUF499*<-METHYLASE<-TUDOR+ SF1-SF2-Helicase+iREase+CTD+REase<-RVT<-RVT<-35exo | 1109 | Bacteria>Pseudomonadota>Alphaproteobacteria>Hyphomicrobiales>Phyllobacteriaceae | GCA_004020785.1 | Mesorhizobium_sp. | EOR95_24225 | 12019..15348 | 1871066 | ATP-binding-protein-[Mesorhizobium-sp.] |
| RWO89814.1 | DUF499_1.RM.like.systems | MarR-HTH->?-><-?\|\|TUDOR+ SF1-SF2-Helicase+iREase+CTD+REase->METHYLASE->HEPN+DUF499*-><-?\|\|REase-><-?<-?<-?<-tRNA<-Peptidase_S11 | 1105 | Bacteria>Pseudomonadota>Alphaproteobacteria>Hyphomicrobiales>Phyllobacteriaceae | GCA_004021485.1 | Mesorhizobium_sp. | EOQ95_16830 | 131837..135154 | 1871066 | ATP-binding-protein-[Mesorhizobium-sp.] |
| RWQ65486.1 | DUF499_1.RM.like.systems | <-ABC-ATPase+SbcC<-?<-?\|\|MarR-HTH->?-><-?<-HEPN+DUF499*<-METHYLASE<-TUDOR+ SF1-SF2-Helicase+iREase+CTD+REase<-tRNA<-Peptidase_S11 | 1103 | Bacteria>Pseudomonadota>Alphaproteobacteria>Hyphomicrobiales>Phyllobacteriaceae | GCA_004021745.1 | Mesorhizobium_sp. | EOS86_15645 | 123505..126816 | 1871066 | ATP-binding-protein-[Mesorhizobium-sp.] |
| TIS54880.1 | DUF499_1.RM.like.systems | <-HEPN+DUF499*<-METHYLASE<-Peptidase_M78<-Ntox18<-TUDOR+ SF1-SF2-Helicase+iREase+CTD+REase | 1115 | Bacteria>Pseudomonadota>Alphaproteobacteria>Hyphomicrobiales>Phyllobacteriaceae | GCA_004962755.1 | Mesorhizobium_sp. | E5W91_25095 | 74683..78030 | 1871066 | ATP-binding-protein-[Mesorhizobium-sp.] |
| TIX43843.1 | DUF499_1.RM.like.systems | METHYLASE->HEPN+DUF499*-><-XIS-HTH | 1132 | Bacteria>Pseudomonadota>Alphaproteobacteria>Hyphomicrobiales>Phyllobacteriaceae | GCA_004964855.1 | Mesorhizobium_sp. | E5V40_01625 | 4247..7645 | 1871066 | ATP-binding-protein-[Mesorhizobium-sp.] |
| TJW08146.1 | DUF499_1.RM.like.systems | Resolvase->SIR2+STALD->?->TUDOR+ SF1-SF2-Helicase+iREase+CTD+REase->METHYLASE->HEPN+DUF499*-><-XIS-HTH<-?\|\|?-><-tRNA<-?\|\|?->SAM-methylase-> | 1132 | Bacteria>Pseudomonadota>Alphaproteobacteria>Hyphomicrobiales>Phyllobacteriaceae | GCA_005047185.1 | Mesorhizobium_sp. | E5W97_02655 | 563673..567071 | 1871066 | ATP-binding-protein-[Mesorhizobium-sp.] |
| TPI12226.1 | DUF499_1.RM.like.systems | <-HicB\|\|HTH+WYL->TUDOR+ SF1-SF2-Helicase+iREase+CTD+REase->MORC+DUF3684->REase->METHYLASE->HEPN+DUF499*->TIR+PNTD2->PIWI-><-Arm-DNA-bind_3+Phage_int_SAM_3+Phage_integrase<-?<-?<-REase | 1098 | Bacteria>Pseudomonadota>Alphaproteobacteria>Hyphomicrobiales>Phyllobacteriaceae | GCA_006439355.1 | Mesorhizobium_sp._B4-1-3 | FJW06_17660 | 56845..60141 | 2589889 | ATP-binding-protein-[Mesorhizobium-sp.-B4-1-3] |
| MBP0439959.1 | DUF499_1.RM.like.systems | <-SF1-SF2-Helicase+REase\|\|HTH+WYL->TUDOR+ SF1-SF2-Helicase+iREase+CTD+REase->ASCH->YtH->METHYLASE->HEPN+DUF499*-><-ABC-ATPase+SbcC<-ABC-ATPase+SbcC | 1096 | Bacteria>Pseudomonadota>Alphaproteobacteria>Hyphomicrobiales>Phyllobacteriaceae | GCA_017813335.1 | Tianweitania_sediminis | J5Y06_14970 | 36806..40096 | 1502156 | DUF499-domain-containing-protein-[Tianweitania-sediminis] |
| MBZ9864227.1 | DUF499_1.RM.like.systems | <-BirA-HTH<-BirA-HTH<-BirA-HTH\|\|?->McrC-NTD+REase-><-REase<-HEPN+DUF499*<-?<-METHYLASE<-TUDOR+ SF1-SF2-Helicase+iREase+CTD+REase\|\|?->?-><-MPTase | 1112 | Bacteria>Pseudomonadota>Alphaproteobacteria>Hyphomicrobiales>Phyllobacteriaceae | GCA_020165215.1 | Mesorhizobium_sp._CA15 | LB515_02450 | 495752..499090 | 2876641 | DUF499-domain-containing-protein-[Mesorhizobium-sp.-CA15] |
| MBZ9939464.1 | DUF499_1.RM.like.systems | <-HEPN+DUF499*<-METHYLASE<-Peptidase_M78<-Ntox18<-TUDOR+ SF1-SF2-Helicase+iREase+CTD+REase<-Phage_integrase<-XIS-HTH | 1103 | Bacteria>Pseudomonadota>Alphaproteobacteria>Hyphomicrobiales>Phyllobacteriaceae | GCA_020165495.1 | Mesorhizobium_sp._BR1-1-16 | LB518_24480 | 798..4109 | 2876653 | DUF499-domain-containing-protein-[Mesorhizobium-sp.-BR1-1-16] |
| GLQ79184.1 | DUF499_1.RM.like.systems | HTH+WYL->TUDOR+ SF1-SF2-Helicase+iREase+CTD+REase->ASCH->YtH->METHYLASE->HEPN+DUF499*->TUDOR+SF1-SF2-Helicase+iREase+CTD->?->HHH_4+SF1-SF2-Helicase+REase-> | 1088 | Bacteria>Pseudomonadota>Alphaproteobacteria>Hyphomicrobiales>Phyllobacteriaceae | GCA_030160235.1 | Mesorhizobium_huakuii | GCM10007881_27020 | 53314..56580 | 28104 | hypothetical-protein-GCM10007881_27020-[Mesorhizobium-huakuii] |
| CAI2932244.1 | DUF499_1.RM.like.systems | ParG_RHH-><-MODE-HTH+PBPII<-HEPN+DUF499*<-METHYLASE<-ParB+HNH<-TUDOR+ SF1-SF2-Helicase+iREase+CTD+REase<-HTH+WYL\|\|SF1-SF2-Helicase+REase->SF1-SF2-Helicase+REase-> | 1098 | Bacteria>Pseudomonadota>Alphaproteobacteria>Hyphomicrobiales>Phyllobacteriaceae | GCA_946995915.1 | Aminobacter_niigataensis | AMD1_0962 | 943205..946501 | 83265 | ATPase-AAA-[Aminobacter-niigataensis] |
| WP_023805907.1 | DUF499_1.RM.like.systems | <-HEPN+DUF499*<-METHYLASE\|\|?->?->REase-> | 1110 | Bacteria>Pseudomonadota>Alphaproteobacteria>Hyphomicrobiales>Phyllobacteriaceae | GCF_000502355.1 | unclassified_Mesorhizobium | NLY44_RS22040 | 4450400..4453732 | 325217 | DUF499-domain-containing-protein-[Mesorhizobium-sp.-L2C089B000] |
| WP_023748119.1 | DUF499_1.RM.like.systems | <-HEPN+DUF499*<-?<-METHYLASE<-?<-TUDOR+ SF1-SF2-Helicase+iREase+CTD+REase<-RelE-ParE<-OLD-ABC | 1108 | Bacteria>Pseudomonadota>Alphaproteobacteria>Hyphomicrobiales>Phyllobacteriaceae | GCF_000502535.1 | Mesorhizobium_sp._LNJC391B00 | X750_RS144145 | 1284..4610 | 1287273 | Swt1-family-HEPN-domain-containing-protein-[Mesorhizobium-sp.-LNJC391B00] |
| WP_023715932.1 | DUF499_1.RM.like.systems | <-ParB<-?<-REase<-HEPN+DUF499*<-PIN<-OLD-ABC<-METHYLASE<-TUDOR+ SF1-SF2-Helicase+iREase+CTD+REase | 1115 | Bacteria>Pseudomonadota>Alphaproteobacteria>Hyphomicrobiales>Phyllobacteriaceae | GCF_000503075.1 | Mesorhizobium_sp._LSHC422A00 | X761_RS140575 | 209394..212741 | 1287294 | DUF499-domain-containing-protein-[Mesorhizobium-sp.-LSHC422A00] |
| WP_035530026.1 | DUF499_1.RM.like.systems | Phage_integrase-><-HEPN+DUF499*<-?<-METHYLASE<-?<-SIR2<-ASCH<-TUDOR+ SF1-SF2-Helicase+iREase+CTD+REase | 1103 | Bacteria>Pseudomonadota>Alphaproteobacteria>Hyphomicrobiales>Phyllobacteriaceae | GCF_000759435.1 | Hoeflea_sp._BAL378 | LL06_RS18050 | 3002..6313 | 1547437 | DUF499-domain-containing-protein-[Hoeflea-sp.-BAL378] |
| WP_056240747.1 | DUF499_1.RM.like.systems | <-SF1-SF2-Helicase<-PIWI<-TIR+PNTD2<-HicB\|\|HTH+WYL->HEPN+DUF499*->?-><-?\|\|PADR-HTH-> | 1103 | Bacteria>Pseudomonadota>Alphaproteobacteria>Hyphomicrobiales>Phyllobacteriaceae | GCF_001427285.1 | Mesorhizobium_sp._Root552 | ASD50_RS11320 | 108386..111697 | 1736555 | DUF499-domain-containing-protein-[Mesorhizobium-sp.-Root552] |
| WP_095831457.1 | DUF499_1.RM.like.systems | SF1-SF2-Helicase+REase->Resolvase->REase->TUDOR+ SF1-SF2-Helicase+iREase+CTD+REase->METHYLASE->HEPN+DUF499*-><-XIS-HTH\|\|?->?-><-tRNA<-?\|\|?->SAM-methylase-> | 1135 | Bacteria>Pseudomonadota>Alphaproteobacteria>Hyphomicrobiales>Phyllobacteriaceae | GCF_002294955.1 | Mesorhizobium_sp._WSM3864 | CK215_RS15785 | 236448..239855 | 2029404 | DUF499-domain-containing-protein-[Mesorhizobium-sp.-WSM3864] |
| WP_107649676.1 | DUF499_1.RM.like.systems | STYKinase-like->?-><-HEPN+DUF499*<-METHYLASE<-TUDOR+ SF1-SF2-Helicase+iREase+CTD+REase<-GDResolvase+HTH\|\|?-><-?\|\|HTH-> | 1105 | Bacteria>Pseudomonadota>Alphaproteobacteria>Hyphomicrobiales>Phyllobacteriaceae | GCF_003034915.1 | Mesorhizobium_helmanticense | C9427_RS13550 | 15712..19029 | 1776423 | DUF499-domain-containing-protein-[Mesorhizobium-helmanticense] |
| WP_120017243.1 | DUF499_1.RM.like.systems | TRPR-HTH+HTH->?->?->?->?->?-><-HEPN+DUF499*<-METHYLASE<-MORC+DUF3684<-TUDOR+ SF1-SF2-Helicase+iREase+CTD+REase<-?<-MORC+HISKIN<-METHYLASE-p | 1110 | Bacteria>Pseudomonadota>Alphaproteobacteria>Hyphomicrobiales>Phyllobacteriaceae | GCF_003601975.1 | Mesorhizobium_waimense | D3227_RS26610 | 4630..7962 | 1300307 | Swt1-family-HEPN-domain-containing-protein-[Mesorhizobium-waimense] |
| WP_125006738.1 | DUF499_1.RM.like.systems | <-SMODS\|\|HTH+WYL->?->TUDOR+ SF1-SF2-Helicase+iREase+CTD+REase->MORC+DUF3684->METHYLASE->HEPN+DUF499*-><-AbiEii<-AbiEi_4 | 1110 | Bacteria>Pseudomonadota>Alphaproteobacteria>Hyphomicrobiales>Phyllobacteriaceae | GCF_003863365.1 | Mesorhizobium_tamadayense | EH240_RS33970 | 18463..21795 | 425306 | Swt1-family-HEPN-domain-containing-protein-[Mesorhizobium-tamadayense] |
| WP_127367955.1 | DUF499_1.RM.like.systems | HTH_3-><-wHTH+SAD+HNH\|\|?->TUDOR+ SF1-SF2-Helicase+iREase+CTD+REase->wHTH+REase->METHYLASE->HEPN+DUF499*-> | 1118 | Bacteria>Pseudomonadota>Alphaproteobacteria>Hyphomicrobiales>Phyllobacteriaceae | GCF_003997975.1 | unclassified_Mesorhizobium | EN990_RS21670 | 38386..41742 | 325217 | MULTISPECIES:-DUF499-domain-containing-protein-[unclassified-Mesorhizobium] |
| WP_127864269.1 | DUF499_1.RM.like.systems | SF1-SF2-Helicase->TUDOR+ SF1-SF2-Helicase+iREase+CTD+REase->ASCH->METHYLASE->HEPN+DUF499*-><-tRNA<-Peptidase<-?\|\|MPTase-> | 1114 | Bacteria>Pseudomonadota>Alphaproteobacteria>Hyphomicrobiales>Phyllobacteriaceae | GCF_004010405.2 | unclassified_Mesorhizobium | EN797_RS08215 | 1680820..1684164 | 325217 | MULTISPECIES:-DUF499-domain-containing-protein-[unclassified-Mesorhizobium] |
| WP_164894099.1 | DUF499_1.RM.like.systems | <-HEPN+DUF499*<-METHYLASE<-wHTH+REase<-TUDOR+ SF1-SF2-Helicase+iREase+CTD+REase<-?\|\|wHTH+SAD+HNH-><-HTH_3 | 528 | Bacteria>Pseudomonadota>Alphaproteobacteria>Hyphomicrobiales>Phyllobacteriaceae | GCF_004022915.1 | Mesorhizobium_sp._M7A.F.Ca.US.006.04.2.1 | EN925_RS00840 | 1..1586 | 2496696 | Swt1-family-HEPN-domain-containing-protein,-partial-[Mesorhizobium-sp.-M7A.F.Ca.US.006.04.2.1] |
| WP_135867691.1 | DUF499_1.RM.like.systems | PAPS_reduct->HTH+MPTase->PIN-><-HEPN+DUF499*<-METHYLASE<-MORC+DUF3684<-TUDOR+ SF1-SF2-Helicase+iREase+CTD+REase<-REase<-?<-MAD-NTD+HNH | 1097 | Bacteria>Pseudomonadota>Alphaproteobacteria>Hyphomicrobiales>Phyllobacteriaceae | GCF_004791755.1 | unclassified_Mesorhizobium | EN861_RS33190 | 64871..68164 | 325217 | MULTISPECIES:-DUF499-domain-containing-protein-[unclassified-Mesorhizobium] |

<table>
<tr><th>Protein-ID</th><th>System</th><th>Gene-Neighbourhoods</th><th>Length</th><th>Taxonomy</th><th>Assembly-ID</th><th>Strain</th><th>Locus-Tag</th><th>Genomic-coordinates</th><th>TaxID</th><th>Fasta-defline</th></tr>
<tr><td>WP_135861107.1</td><td>DUF499_1.RM.like.systems</td><td>HTH+WYL->TUDOR+SF1-SF2-Helicase+iRease+CTD+REase->ASCH->YtH->MORC+DUF3684->METHYLASE->HEPN+DUF499*-><-TRPR-HTH+HTH<-?<-?<-?||TraJ-RHH-></td><td>1082</td><td>Bacteria>Pseudomonadota>Alphaproteobacteria>Hyphomicrobiales>Phyllobacteriaceae</td><td>GCF_004791825.1</td><td>unclassified_Mesorhizobium</td><td>EN833_RS30760</td><td>107531..110779</td><td>325217</td><td>MULTISPECIES:-DUF499-domain-containing-protein-[unclassified-Mesorhizobium]</td></tr>
<tr><td>WP_135929916.1</td><td>DUF499_1.RM.like.systems</td><td>TUDOR+SF1-SF2-Helicase+iRease+CTD+REase->METHYLASE->OLD-ABC->PIN->HEPN+DUF499*->?-><-?||Methylase-><-Radical-SAM<-Arm-DNA-bind_3+Phage_integrase</td><td>1110</td><td>Bacteria>Pseudomonadota>Alphaproteobacteria>Hyphomicrobiales>Phyllobacteriaceae</td><td>GCF_004791875.1</td><td>unclassified_Mesorhizobium</td><td>EN810_RS20200</td><td>141593..144925</td><td>325217</td><td>MULTISPECIES:-DUF499-domain-containing-protein-[unclassified-Mesorhizobium]</td></tr>
<tr><td>WP_136661243.1</td><td>DUF499_1.RM.like.systems</td><td>TUDOR+SF1-SF2-Helicase+iRease+CTD+REase->?->METHYLASE->HEPN+DUF499*->HIN-HTH-></td><td>1097</td><td>Bacteria>Pseudomonadota>Alphaproteobacteria>Hyphomicrobiales>Phyllobacteriaceae</td><td>GCF_004959855.1</td><td>Nitratireductor_sp._XY-223</td><td>E4M07_RS25460</td><td>21573..24866</td><td>2561926</td><td>DUF499-domain-containing-protein-[Nitratireductor-sp.-XY-223]</td></tr>
<tr><td>WP_246674663.1</td><td>DUF499_1.RM.like.systems</td><td><-HicB||HTH+WYL->TUDOR+SF1-SF2-Helicase+iRease+CTD+REase->MORC+DUF3684->REase->METHYLASE->HEPN+DUF499*->TIR+PNTD2->PIWI-><-Arm-DNA-bind_3+Phage_int_SAM_3+Phage_integrase<-?<-?<-REase</td><td>1086</td><td>Bacteria>Pseudomonadota>Alphaproteobacteria>Hyphomicrobiales>Phyllobacteriaceae</td><td>GCF_006439355.1</td><td>Mesorhizobium_sp._B4-1-3</td><td>FJW06_RS17660</td><td>56881..60141</td><td>2589889</td><td>DUF499-domain-containing-protein-[Mesorhizobium-sp.-B4-1-3]</td></tr>
<tr><td>WP_140871483.1</td><td>DUF499_1.RM.like.systems</td><td><-HEPN+DUF499*<-METHYLASE<-TUDOR+SF1-SF2-Helicase+iRease+CTD+REase<-?<-SIR2+STALD<-Resolvase</td><td>1134</td><td>Bacteria>Pseudomonadota>Alphaproteobacteria>Hyphomicrobiales>Phyllobacteriaceae</td><td>GCF_006440815.1</td><td>Mesorhizobium_sp._B2-8-3</td><td>FJ418_RS28720</td><td>4402..7806</td><td>2589905</td><td>DUF499-domain-containing-protein-[Mesorhizobium-sp.-B2-8-3]</td></tr>
<tr><td>WP_165021935.1</td><td>DUF499_1.RM.like.systems</td><td>TUDOR+SF1-SF2-Helicase+iRease+CTD+REase->METHYLASE->HEPN+DUF499*-><-?<-STYKinase-like<-?<-?||?-><-Phage_integrase</td><td>1105</td><td>Bacteria>Pseudomonadota>Alphaproteobacteria>Hyphomicrobiales>Phyllobacteriaceae</td><td>GCF_011045125.1</td><td>Mesorhizobium_camelthorni</td><td>G6N73_RS00955</td><td>189890..193207</td><td>475069</td><td>DUF499-domain-containing-protein-[Mesorhizobium-camelthorni]</td></tr>
<tr><td>WP_280515578.1</td><td>DUF499_1.RM.like.systems</td><td>TUDOR+SF1-SF2-Helicase+iRease+CTD+REase-p-><-HTH_Tnp_1+HTH_21+rve+rve_3||?->?->METHYLASE-p->METHYLASE->HEPN+DUF499*->DUF499->ParB+HTH->HTH_3->ParB+HNH-p->ParB+HNH-p->URI-DNase-></td><td>427</td><td>Bacteria>Pseudomonadota>Alphaproteobacteria>Hyphomicrobiales>Phyllobacteriaceae</td><td>GCF_013240295.1</td><td>Mesorhizobium_sediminum</td><td>HK436_RS32250</td><td>843839..845122</td><td>1849104</td><td>Swt1-family-HEPN-domain-containing-protein-[Mesorhizobium-sediminum]</td></tr>
<tr><td>WP_192357697.1</td><td>DUF499_1.RM.like.systems</td><td>REase->?->MACRODOMAIN->STYKinase-like->?-><-HEPN+DUF499*<-METHYLASE<-TUDOR+SF1-SF2-Helicase+iRease+CTD+REase</td><td>1105</td><td>Bacteria>Pseudomonadota>Alphaproteobacteria>Hyphomicrobiales>Phyllobacteriaceae</td><td>GCF_014843885.1</td><td>Mesorhizobium_mediterraneum</td><td>AK98_RS11975</td><td>23140..26457</td><td>43617</td><td>DUF499-domain-containing-protein-[Mesorhizobium-mediterraneum]</td></tr>
<tr><td>WP_202379562.1</td><td>DUF499_1.RM.like.systems</td><td>GDResolvase+Recombinase+ZnR-><-?||?->?-><-?||?-><-HEPN+DUF499*<-METHYLASE<-TUDOR+SF1-SF2-Helicase+iRease+CTD+REase||MerR-HTH->?-><-GDResolvase+Recombinase+ZnR</td><td>1114</td><td>Bacteria>Pseudomonadota>Alphaproteobacteria>Hyphomicrobiales>Phyllobacteriaceae</td><td>GCF_016756555.1</td><td>Mesorhizobium_sp._131-2-5</td><td>JG740_RS04010</td><td>865607..868951</td><td>2744519</td><td>DUF499-domain-containing-protein-[Mesorhizobium-sp.-131-2-5]</td></tr>
<tr><td>WP_245247928.1</td><td>DUF499_1.RM.like.systems</td><td><-SF1-SF2-Helicase+REase||HTH+WYL->TUDOR+SF1-SF2-Helicase+iRease+CTD+REase->ASCH->YtH->METHYLASE->HEPN+DUF499*-><-ABC-ATPase+SbcC<-ABC-ATPase+SbcC</td><td>1084</td><td>Bacteria>Pseudomonadota>Alphaproteobacteria>Hyphomicrobiales>Phyllobacteriaceae</td><td>GCF_017813335.1</td><td>Tianweitania_sediminis</td><td>J5Y06_RS14970</td><td>36842..40096</td><td>1502156</td><td>DUF499-domain-containing-protein-[Tianweitania-sediminis]</td></tr>
<tr><td>WP_224706639.1</td><td>DUF499_1.RM.like.systems</td><td><-BirA-HTH<-BirA-HTH<-BirA-HTH||?->McrC-NTD+REase-><-REase<-HEPN+DUF499*<-?<-METHYLASE<-TUDOR+SF1-SF2-Helicase+iRease+CTD+REase||?->?-><-MPTase</td><td>1112</td><td>Bacteria>Pseudomonadota>Alphaproteobacteria>Hyphomicrobiales>Phyllobacteriaceae</td><td>GCF_020165215.1</td><td>Mesorhizobium_sp._CA15</td><td>LB515_02450</td><td>495752..499090</td><td>2876641</td><td>DUF499-domain-containing-protein-[Mesorhizobium-sp.-CA15]</td></tr>
<tr><td>WP_224580510.1</td><td>DUF499_1.RM.like.systems</td><td><-HEPN+DUF499*<-METHYLASE<-Peptidase_M78<-Ntox18<-TUDOR+SF1-SF2-Helicase+iRease+CTD+REase<-Phage_integrase<-XIS-HTH</td><td>1103</td><td>Bacteria>Pseudomonadota>Alphaproteobacteria>Hyphomicrobiales>Phyllobacteriaceae</td><td>GCF_020165495.1</td><td>Mesorhizobium_sp._BR1-1-16</td><td>LB518_RS24470</td><td>798..4109</td><td>2876653</td><td>DUF499-domain-containing-protein-[Mesorhizobium-sp.-BR1-1-16]</td></tr>
<tr><td>WP_284273142.1</td><td>DUF499_1.RM.like.systems</td><td>HTH+WYL->TUDOR+SF1-SF2-Helicase+iRease+CTD+REase->ASCH->YtH->METHYLASE->HEPN+DUF499*->TUDOR+SF1-SF2-Helicase+iRease+CTD->?->HHH_4+SF1-SF2-Helicase+REase-><-?||HTH-></td><td>1100</td><td>Bacteria>Pseudomonadota>Alphaproteobacteria>Hyphomicrobiales>Phyllobacteriaceae</td><td>GCF_030160235.1</td><td>Mesorhizobium_huakuii</td><td>QQK95_RS13625</td><td>53278..56580</td><td>28104</td><td>DUF499-domain-containing-protein-[Mesorhizobium-huakuii]</td></tr>
<tr><td>WP_280515628.1</td><td>DUF499_1.RM.like.systems</td><td><-HTH_Tnp_1+HTH_21+rve+rve_3||?->?->METHYLASE-p->METHYLASE->HEPN+DUF499->DUF499*->ParB+HTH->HTH_3->ParB+HNH-p->ParB+HNH-p->URI-DNase-></td><td>678</td><td>Bacteria>Pseudomonadota>Alphaproteobacteria>Hyphomicrobiales>Phyllobacteriaceae</td><td>-</td><td>Mesorhizobium_sediminum</td><td>HK436_RS32255</td><td>845119..847155</td><td>1849104</td><td>DUF499-domain-containing-protein-[Mesorhizobium-sediminum]</td></tr>
<tr><td>WP_263490164.1</td><td>DUF499_1.RM.like.systems</td><td><-BirA-HTH<-BirA-HTH<-BirA-HTH||?->McrC-NTD+REase-><-REase<-HEPN+DUF499*<-?<-METHYLASE<-TUDOR+SF1-SF2-Helicase+iRease+CTD+REase||?->?-><-MPTase</td><td>1095</td><td>Bacteria>Pseudomonadota>Alphaproteobacteria>Hyphomicrobiales>Phyllobacteriaceae</td><td>-</td><td>Mesorhizobium_sp._CA15</td><td>LB515_RS02455</td><td>495752..499039</td><td>2876641</td><td>DUF499-domain-containing-protein-[Mesorhizobium-sp.-CA15]</td></tr>
<tr><td>MBY0324746.1</td><td>DUF499_1.RM.like.systems</td><td><-ABC-ATPase+SbcC<-ABC-ATPase+SbcC<-ABC-ATPase+SbcC<-?||?-><-?<-HEPN+DUF499*<-HEPN<-METHYLASE-p<-DDE_3<-METHYLASE<-YtH</td><td>1111</td><td>Bacteria>Pseudomonadota>Alphaproteobacteria>Hyphomicrobiales>Reyranellaceae</td><td>GCA_019750135.1</td><td>Reyranella_sp.</td><td>K2X72_38955</td><td>16276..19611</td><td>1929291</td><td>DUF499-domain-containing-protein-[Reyranella-sp.]</td></tr>
<tr><td>WP_174037242.1</td><td>DUF499_1.RM.like.systems</td><td>Phage_terminase->?-><-?<-HEPN+DUF499*<-METHYLASE<-MORC+DUF3684<-ASCH<-TUDOR+SF1-SF2-Helicase+iRease+CTD+REase<-HTH+WYL||HicB-></td><td>1098</td><td>Bacteria>Pseudomonadota>Alphaproteobacteria>Hyphomicrobiales>Rhizobiaceae</td><td>-</td><td>Agrobacterium_tumefaciens</td><td>G6M03_RS01055</td><td>214165..217461</td><td>358</td><td>DUF499-domain-containing-protein-[Agrobacterium-tumefaciens]</td></tr>
<tr><td>TBD87036.1</td><td>DUF499_1.RM.like.systems</td><td><-HEPN<-HEPN+DUF499*<-METHYLASE<-ASCH<-MORC+DUF3684<-YtH<-ASCH<-TUDOR+SF1-SF2-Helicase+iRease+CTD+REase</td><td>1097</td><td>Bacteria>Pseudomonadota>Alphaproteobacteria>Hyphomicrobiales>Rhizobiaceae</td><td>GCA_004306165.1</td><td>Rhizobium_ruizarguesonis</td><td>ELH13_20430</td><td>4103083..4106376</td><td>2081791</td><td>ATP-binding-protein-[Rhizobium-leguminosarum]</td></tr>
<tr><td>TCS04590.1</td><td>DUF499_1.RM.like.systems</td><td>XIS-HTH->?-><-?<-?<-?<-?<-HEPN+DUF499*<-METHYLASE<-MORC+DUF3684<-TUDOR+SF1-SF2-Helicase+iRease+CTD+REase<-?<-?||HTH_3-></td><td>1096</td><td>Bacteria>Pseudomonadota>Alphaproteobacteria>Hyphomicrobiales>Rhizobiaceae</td><td>GCA_004341105.1</td><td>Rhizobium_sp._BK418</td><td>EV281_103265</td><td>265866..269156</td><td>2512120</td><td>hypothetical-protein-EV281_103265-[Rhizobium-sp.-BK418]</td></tr>
<tr><td>NEH32401.1</td><td>DUF499_1.RM.like.systems</td><td>HTH+WYL->TUDOR+SF1-SF2-Helicase+iRease+CTD+REase->ASCH->YtH->MORC+DUF3684->METHYLASE->HEPN+DUF499*->HEPN-></td><td>1097</td><td>Bacteria>Pseudomonadota>Alphaproteobacteria>Hyphomicrobiales>Rhizobiaceae</td><td>GCA_010668085.1</td><td>Rhizobium_ruizarguesonis</td><td>GR203_31950</td><td>43158..46451</td><td>2081791</td><td>DUF499-domain-containing-protein-[Rhizobium-leguminosarum]</td></tr>
<tr><td>MBX4863812.1</td><td>DUF499_1.RM.like.systems</td><td><-HEPN+DUF499*<-METHYLASE<-YtH<-TUDOR+SF1-SF2-Helicase+iRease+CTD+REase<-HTH+WYL||?->UbE1+JAB-></td><td>1098</td><td>Bacteria>Pseudomonadota>Alphaproteobacteria>Hyphomicrobiales>Rhizobiaceae</td><td>GCA_019684005.1</td><td>Rhizobium_bangladeshense</td><td>HJA86_28440</td><td>35410..38706</td><td>1138189</td><td>ATP-binding-protein-[Rhizobium-bangladeshense]</td></tr>
<tr><td>MBY3260302.1</td><td>DUF499_1.RM.like.systems</td><td>HTH+WYL->TUDOR+SF1-SF2-Helicase+iRease+CTD+REase->YtH->MORC+DUF3684->METHYLASE->HEPN+DUF499*-><-wHTH+REase<-Arm-DNA-bind_3+Phage_integrase<-Peptidase_M24||TetR-HTH-></td><td>1098</td><td>Bacteria>Pseudomonadota>Alphaproteobacteria>Hyphomicrobiales>Rhizobiaceae</td><td>GCA_019779025.1</td><td>Rhizobium_laguerreae</td><td>HFO15_01275</td><td>256972..260268</td><td>1076926</td><td>ATP-binding-protein-[Rhizobium-laguerreae]</td></tr>
<tr><td>MBY5371752.1</td><td>DUF499_1.RM.like.systems</td><td><-PIN+OST-HTH||TUDOR+SF1-SF2-Helicase+iRease+CTD+REase->?->METHYLASE->?->HEPN+DUF499*-><-HTH_3</td><td>1103</td><td>Bacteria>Pseudomonadota>Alphaproteobacteria>Hyphomicrobiales>Rhizobiaceae</td><td>GCA_019798485.1</td><td>Rhizobium_leguminosarum</td><td>HFO92_34835</td><td>18961..22272</td><td>384</td><td>ATP-binding-protein-[Rhizobium-leguminosarum]</td></tr>
<tr><td>MCB1465833.1</td><td>DUF499_1.RM.like.systems</td><td>TUDOR+SF1-SF2-Helicase+iRease+CTD+REase-p->?->METHYLASE->HEPN+DUF499*-></td><td>428</td><td>Bacteria>Pseudomonadota>Alphaproteobacteria>Hyphomicrobiales>Rhizobiaceae</td><td>GCA_020441945.1</td><td>Rhizobiaceae_bacterium</td><td>KDK08_01465</td><td>5124..6408</td><td>1913961</td><td>DUF499-domain-containing-protein-[Rhizobiaceae-bacterium]</td></tr>
<tr><td>MCE6075329.1</td><td>DUF499_1.RM.like.systems</td><td><-HicB||HTH+WYL->TUDOR+SF1-SF2-Helicase+iRease+CTD+REase->ASCH->MORC+DUF3684->METHYLASE->HEPN+DUF499*->?-><-?<-?<-Phage_terminase<-TetR-HTH</td><td>1098</td><td>Bacteria>Pseudomonadota>Alphaproteobacteria>Hyphomicrobiales>Rhizobiaceae</td><td>GCA_021387325.1</td><td>Agrobacterium_vitis</td><td>FS799_10735</td><td>1060385..1063681</td><td>373</td><td>DUF499-domain-containing-protein-[Agrobacterium-vitis]</td></tr>
<tr><td>MCO5066228.1</td><td>DUF499_1.RM.like.systems</td><td>RelE-ParE->ParA+HTH->?->OLD-ABC+TOPRIM->SF1-SF2-Helicase-><-TIR<-HEPN+DUF499*<-METHYLASE<-?<-TUDOR+SF1-SF2-Helicase+iRease+CTD+REase<-HTH+WYL||HicB-></td><td>1098</td><td>Bacteria>Pseudomonadota>Alphaproteobacteria>Hyphomicrobiales>Rhizobiaceae</td><td>GCA_023953835.1</td><td>Rhizobiaceae_bacterium</td><td>M9924_17685</td><td>3745551..3748847</td><td>1913961</td><td>DUF499-domain-containing-protein-[Rhizobiaceae-bacterium]</td></tr>
<tr><td>SSC67554.1</td><td>DUF499_1.RM.like.systems</td><td>ParG_RHH-><-TetR-HTH||MODE-HTH+PBPII-><-HEPN+DUF499*<-METHYLASE<-MORC+DUF3684<-TUDOR+SF1-SF2-Helicase+iRease+CTD+REase<-HTH+WYL||SF1-SF2-Helicase+REase->SF1-SF2-Helicase+REase-></td><td>1086</td><td>Bacteria>Pseudomonadota>Alphaproteobacteria>Hyphomicrobiales>Rhizobiaceae</td><td>GCA_900492185.1</td><td>Ciceribacter_selenitireducens_ATCC_BAA-1503</td><td>RHIZ70_3262</td><td>225722..228982</td><td>1336235</td><td>unnamed-protein-product-[Ciceribacter-selenitireducens-ATCC-BAA-1503]</td></tr>
<tr><td>CAD6420434.1</td><td>DUF499_1.RM.like.systems</td><td><-LuxR-HTH||PIN+OST-HTH-><-SF1-SF2-Helicase<-HEPN+DUF499*<-?<-METHYLASE<-?<-TUDOR+SF1-SF2-Helicase+iRease+CTD+REase</td><td>1104</td><td>Bacteria>Pseudomonadota>Alphaproteobacteria>Hyphomicrobiales>Rhizobiaceae</td><td>GCA_902153285.2</td><td>Rhizobium_sp._Q54</td><td>REQ54_02098</td><td>2069964..2073278</td><td>1335060</td><td>hypothetical-protein-REQ54_02098-[Rhizobium-sp.-Q54]</td></tr>
<tr><td>WP_040958264.1</td><td>DUF499_1.RM.like.systems</td><td>Tox-RelA->TUDOR+SF1-SF2-Helicase+iRease+CTD+REase->METHYLASE->HEPN+DUF499*-><-SLATT<-SMODS<-RNase<-HTH_17<-DnaB+CITB-HTH<-ZnR+TOPRIM</td><td>1107</td><td>Bacteria>Pseudomonadota>Alphaproteobacteria>Hyphomicrobiales>Rhizobiaceae</td><td>GCF_000219415.3</td><td>Sinorhizobium_fredii</td><td>SFGR64A_RS01795</td><td>20467..23790</td><td>380</td><td>DUF499-domain-containing-protein-[Sinorhizobium-fredii]</td></tr>
<tr><td>WP_158499670.1</td><td>DUF499_1.RM.like.systems</td><td>Phage_integrase->XIS-HTH->TUDOR+SF1-SF2-Helicase+iRease+CTD+REase->METHYLASE->HEPN+DUF499*->?->SF1-SF2-Helicase-><-MORC||ArdCN+MPTase->ParB+HTH-></td><td>1096</td><td>Bacteria>Pseudomonadota>Alphaproteobacteria>Hyphomicrobiales>Rhizobiaceae</td><td>GCF_000219415.3</td><td>Sinorhizobium_fredii</td><td>SFGR64A_RS31000</td><td>11079..14369</td><td>380</td><td>DUF499-domain-containing-protein-[Sinorhizobium-fredii]</td></tr>
<tr><td>WP_028734362.1</td><td>DUF499_1.RM.like.systems</td><td><-HicB||HTH+WYL->TUDOR+SF1-SF2-Helicase+iRease+CTD+REase->ASCH->MORC+DUF3684->METHYLASE->HEPN+DUF499*->?-><-Arm-DNA-bind_3+Phage_integrase<-?<-?<-?<-HNH</td><td>1098</td><td>Bacteria>Pseudomonadota>Alphaproteobacteria>Hyphomicrobiales>Rhizobiaceae</td><td>GCF_000427705.1</td><td>Rhizobium_leguminosarum</td><td>YSW_RS0132760</td><td>58681..61977</td><td>384</td><td>DUF499-domain-containing-protein-[Rhizobium-leguminosarum]</td></tr>
<tr><td>WP_064332302.1</td><td>DUF499_1.RM.like.systems</td><td>ParG_RHH-><-MODE-HTH+PBPII<-HEPN+DUF499*<-METHYLASE<-ASCH<-?<-TUDOR+SF1-SF2-Helicase+iRease+CTD+REase<-HTH+WYL||SF1-SF2-Helicase+REase-></td><td>1098</td><td>Bacteria>Pseudomonadota>Alphaproteobacteria>Hyphomicrobiales>Rhizobiaceae</td><td>GCF_001652565.1</td><td>Shinella_sp._HZN7</td><td>shn_RS22015</td><td>4588511..4591807</td><td>879274</td><td>DUF499-domain-containing-protein-[Shinella-sp.-HZN7]</td></tr>
<tr><td>WP_064252405.1</td><td>DUF499_1.RM.like.systems</td><td>Terminase->?-><-HTH_3||?-><-?||<-HEPN+DUF499*<-METHYLASE<-TUDOR+SF1-SF2-Helicase+iRease+CTD+REase<-REase||35exo-></td><td>1109</td><td>Bacteria>Pseudomonadota>Alphaproteobacteria>Hyphomicrobiales>Rhizobiaceae</td><td>GCF_001889105.1</td><td>Sinorhizobium_americanum</td><td>SAMCFNEI73_RS12475</td><td>2595143..2598472</td><td>194963</td><td>DUF499-domain-containing-protein-[Sinorhizobium-americanum]</td></tr>
<tr><td>WP_077472796.1</td><td>DUF499_1.RM.like.systems</td><td><-REC<-?<-DOC+HTH<-?<-GDResolvase+HIN-HTH||<-HEPN+DUF499*<-METHYLASE<-ASCH<-TUDOR+SF1-SF2-Helicase+iRease+CTD+REase<-HTH+WYL<-McrC-NTD+REase</td><td>1095</td><td>Bacteria>Pseudomonadota>Alphaproteobacteria>Hyphomicrobiales>Rhizobiaceae</td><td>GCF_001999485.2</td><td>Rhizobium_sp._P44RR-XXIV</td><td>BSK43_RS00425</td><td>6721..10008</td><td>1921145</td><td>DUF499-domain-containing-protein-[Rhizobium-sp.-P44RR-XXIV]</td></tr>
<tr><td>WP_127665091.1</td><td>DUF499_1.RM.like.systems</td><td>Arm-DNA-bind_3+Phage_integrase->||HTH+PolB->?-><-?<-?<-ParB+HNH<-HEPN+DUF499*<-?<-METHYLASE<-?<-TUDOR+SF1-SF2-Helicase+iRease+CTD+REase<-SF1-SF2-Helicase+DUF1998+HKD-DNase+HKD-DNase</td><td>1104</td><td>Bacteria>Pseudomonadota>Alphaproteobacteria>Hyphomicrobiales>Rhizobiaceae</td><td>GCF_004001475.1</td><td>Sinorhizobium_meliloti</td><td>CN231_RS17685</td><td>56193..59507</td><td>382</td><td>DUF499-domain-containing-protein-[Sinorhizobium-meliloti]</td></tr>
<tr><td>WP_245582348.1</td><td>DUF499_1.RM.like.systems</td><td><-HEPN<-HEPN+DUF499*<-METHYLASE<-ASCH<-MORC+DUF3684<-YtH<-ASCH<-TUDOR+SF1-SF2-Helicase+iRease+CTD+REase</td><td>1085</td><td>Bacteria>Pseudomonadota>Alphaproteobacteria>Hyphomicrobiales>Rhizobiaceae</td><td>GCF_004306165.1</td><td>Rhizobium_ruizarguesonis</td><td>ELH13_RS20435</td><td>4103083..4106340</td><td>2081791</td><td>DUF499-domain-containing-protein-[Rhizobium-leguminosarum]</td></tr>
<tr><td>WP_132653484.1</td><td>DUF499_1.RM.like.systems</td><td>XIS-HTH->?-><-?<-?<-?<-?<-HEPN+DUF499*<-METHYLASE<-MORC+DUF3684<-TUDOR+SF1-SF2-Helicase+iRease+CTD+REase<-?||HTH_3->STYKinase-like-></td><td>1097</td><td>Bacteria>Pseudomonadota>Alphaproteobacteria>Hyphomicrobiales>Rhizobiaceae</td><td>GCF_004341105.1</td><td>Rhizobium_sp._BK418</td><td>EV281_RS12840</td><td>265866..269159</td><td>2512120</td><td>DUF499-domain-containing-protein-[Rhizobium-sp.-BK418]</td></tr>
<tr><td>WP_153314323.1</td><td>DUF499_1.RM.like.systems</td><td>TUDOR+SF1-SF2-Helicase+iRease+CTD+REase->METHYLASE->HEPN+DUF499*-><-GDResolvase+Recombinase+ZnR<-?<-?<-?||HTH_3-><-wHTH+SAD+HNH</td><td>1110</td><td>Bacteria>Pseudomonadota>Alphaproteobacteria>Hyphomicrobiales>Rhizobiaceae</td><td>GCF_009599625.1</td><td>Sinorhizobium_meliloti</td><td>GHK35_RS22105</td><td>11808..15140</td><td>382</td><td>DUF499-domain-containing-protein-[Sinorhizobium-meliloti]</td></tr>
<tr><td>WP_153320511.1</td><td>DUF499_1.RM.like.systems</td><td>wHTH+SAD+HNH-><-HTH_3||?->?->?->GDResolvase+Recombinase+ZnR-><-HEPN+DUF499*<-METHYLASE<-TUDOR+SF1-SF2-Helicase+iRease+CTD+REase</td><td>1110</td><td>Bacteria>Pseudomonadota>Alphaproteobacteria>Hyphomicrobiales>Rhizobiaceae</td><td>GCF_009599945.1</td><td>Sinorhizobium_meliloti</td><td>GHJ83_RS08885</td><td>35044..38376</td><td>382</td><td>DUF499-domain-containing-protein-[Sinorhizobium-meliloti]</td></tr>
<tr><td>WP_160871739.1</td><td>DUF499_1.RM.like.systems</td><td>Phage_integrase->?-><-ZnR||GDResolvase+HIN-HTH-><-?<-HEPN+DUF499*<-METHYLASE<-TUDOR+SF1-SF2-Helicase+iRease+CTD+REase||tRNA-><-?||?->?->ABC-ATPase+SbcC-></td><td>1118</td><td>Bacteria>Pseudomonadota>Alphaproteobacteria>Hyphomicrobiales>Rhizobiaceae</td><td>GCF_009826875.1</td><td>Shinella_sp._AETb1-6</td><td>GR158_RS15925</td><td>139338..142694</td><td>2692210</td><td>DUF499-domain-containing-protein-[Shinella-sp.-AETb1-6]</td></tr>
<tr><td>WP_168292930.1</td><td>DUF499_1.RM.like.systems</td><td><-OLD-ABC+TOPRIM||?-><-YtH<-HEPN+DUF499*<-METHYLASE<-TUDOR+SF1-SF2-Helicase+iRease+CTD+REase<-?<-STYKinase-like<-HTH_3<-TetR-HTH</td><td>1118</td><td>Bacteria>Pseudomonadota>Alphaproteobacteria>Hyphomicrobiales>Rhizobiaceae</td><td>GCF_012275615.1</td><td>Rhizobium_leguminosarum</td><td>GFL51_RS25220</td><td>66305..69661</td><td>384</td><td>DUF499-domain-containing-protein-[Rhizobium-leguminosarum]</td></tr>
<tr><td>WP_168332223.1</td><td>DUF499_1.RM.like.systems</td><td><-HEPN+DUF499*<-?<-METHYLASE<-?<-TUDOR+SF1-SF2-Helicase+iRease+CTD+REase</td><td>1103</td><td>Bacteria>Pseudomonadota>Alphaproteobacteria>Hyphomicrobiales>Rhizobiaceae</td><td>GCF_012276395.1</td><td>Rhizobium_leguminosarum</td><td>GFL80_RS35655</td><td>1353..4664</td><td>384</td><td>DUF499-domain-containing-protein-[Rhizobium-leguminosarum]</td></tr>
<tr><td>WP_173514149.1</td><td>DUF499_1.RM.like.systems</td><td><-UbE1+JAB<-?||HTH+WYL->TUDOR+SF1-SF2-Helicase+iRease+CTD+REase->YtH->METHYLASE->HEPN+DUF499*->?-><-?||?-><-?||REase-></td><td>1101</td><td>Bacteria>Pseudomonadota>Alphaproteobacteria>Hyphomicrobiales>Rhizobiaceae</td><td>GCF_013283645.1</td><td>Sinorhizobium_psoraleae</td><td>ILFOPFJJ_RS29980</td><td>72999..76304</td><td>520838</td><td>DUF499-domain-containing-protein-[Ensifer-psoraleae]</td></tr>
<tr><td>WP_174156950.1</td><td>DUF499_1.RM.like.systems</td><td><-XIS-HTH<-SF1-SF2-helicase+RdgC+Beta-barrel+REase||HTH+WYL->TUDOR+SF1-SF2-Helicase+iRease+CTD+REase->MORC+DUF3684->METHYLASE->HEPN+DUF499*->MODE-HTH+PBPII-><-ParG_RHH</td><td>1098</td><td>Bacteria>Pseudomonadota>Alphaproteobacteria>Hyphomicrobiales>Rhizobiaceae</td><td>GCF_013322055.1</td><td>Rhizobium_lusitanum</td><td>G6L40_RS07680</td><td>183867..187163</td><td>293958</td><td>DUF499-domain-containing-protein-[Rhizobium-lusitanum]</td></tr>
<tr><td>WP_209605924.1</td><td>DUF499_1.RM.like.systems</td><td><-Mbetalac<-?<-UbE1+JAB<-?||HTH+WYL-><-HEPN+DUF499*<-METHYLASE<-TUDOR+SF1-SF2-Helicase+iRease+CTD+REase<-SF1-SF2-Helicase+DUF1998+HKD-DNase+HKD-DNase<-?<-?<-REase</td><td>1109</td><td>Bacteria>Pseudomonadota>Alphaproteobacteria>Hyphomicrobiales>Rhizobiaceae</td><td>GCF_017874595.1</td><td>Sinorhizobium_kostiense</td><td>J2Z31_RS25385</td><td>30830..34159</td><td>76747</td><td>DUF499-domain-containing-protein-[Sinorhizobium-kostiense]</td></tr>
<tr><td>WP_220337433.1</td><td>DUF499_1.RM.like.systems</td><td>TUDOR+SF1-SF2-Helicase+iRease+CTD+REase->METHYLASE->HEPN+DUF499*->?-><-GDResolvase+HIN-HTH||ZnR-><-?||?-><-Phage_integrase</td><td>1118</td><td>Bacteria>Pseudomonadota>Alphaproteobacteria>Hyphomicrobiales>Rhizobiaceae</td><td>GCF_019511345.1</td><td>Rhizobium_mesosinicum</td><td>JNB85_RS26630</td><td>6855..10211</td><td>335017</td><td>DUF499-domain-containing-protein-[Rhizobium-mesosinicum]</td></tr>
<tr><td>WP_221096744.1</td><td>DUF499_1.RM.like.systems</td><td><-HicB||HTH+WYL->TUDOR+SF1-SF2-Helicase+iRease+CTD+REase->ASCH->MORC+DUF3684->METHYLASE->HEPN+DUF499*-></td><td>1098</td><td>Bacteria>Pseudomonadota>Alphaproteobacteria>Hyphomicrobiales>Rhizobiaceae</td><td>GCF_019684055.1</td><td>Rhizobium_bangladeshense</td><td>HJA85_RS24965</td><td>76913..80209</td><td>1138189</td><td>DUF499-domain-containing-protein-[Rhizobium-bangladeshense]</td></tr>
<tr><td>WP_222000518.1</td><td>DUF499_1.RM.like.systems</td><td>TetR-HTH->HTH_3->STYKinase-like->?-><-HEPN+DUF499*<-?<-METHYLASE<-?<-TUDOR+SF1-SF2-Helicase+iRease+CTD+REase</td><td>1103</td><td>Bacteria>Pseudomonadota>Alphaproteobacteria>Hyphomicrobiales>Rhizobiaceae</td><td>GCF_019779395.1</td><td>Rhizobium_laguerreae</td><td>HFN92_RS29725</td><td>56760..60071</td><td>1076926</td><td>DUF499-domain-containing-protein-[Rhizobium-laguerreae]</td></tr>
<tr><td>WP_261323154.1</td><td>DUF499_1.RM.like.systems</td><td><-PIN+OST-HTH||TUDOR+SF1-SF2-Helicase+iRease+CTD+REase->?->METHYLASE->?->HEPN+DUF499*-><-?<-STYKinase-like<-HTH_3</td><td>1091</td><td>Bacteria>Pseudomonadota>Alphaproteobacteria>Hyphomicrobiales>Rhizobiaceae</td><td>GCF_019798485.1</td><td>Rhizobium_leguminosarum</td><td>HFO91_RS33945</td><td>19091..22366</td><td>384</td><td>DUF499-domain-containing-protein-[Rhizobium-leguminosarum]</td></tr>
<tr><td>WP_222320084.1</td><td>DUF499_1.RM.like.systems</td><td><-OLD-ABC+TOPRIM<-?||?-><-YtH<-HEPN+DUF499*<-METHYLASE<-METHYLASE-p<-TUDOR+SF1-SF2-Helicase+iRease+CTD+REase</td><td>1108</td><td>Bacteria>Pseudomonadota>Alphaproteobacteria>Hyphomicrobiales>Rhizobiaceae</td><td>GCF_019798845.1</td><td>Rhizobium_leguminosarum</td><td>HFO61_RS22785</td><td>274165..277491</td><td>384</td><td>DUF499-domain-containing-protein-[Rhizobium-leguminosarum]</td></tr>
<tr><td>WP_222374442.1</td><td>DUF499_1.RM.like.systems</td><td>TetR-HTH->HTH_3->STYKinase-like->?->TUDOR+SF1-SF2-Helicase+iRease+CTD+REase->METHYLASE->HEPN+DUF499*-></td><td>1114</td><td>Bacteria>Pseudomonadota>Alphaproteobacteria>Hyphomicrobiales>Rhizobiaceae</td><td>GCF_019799295.1</td><td>Rhizobium_leguminosarum</td><td>HFO38_RS09705</td><td>496684..500028</td><td>384</td><td>DUF499-domain-containing-protein-[Rhizobium-leguminosarum]</td></tr>
<tr><td>WP_233741236.1</td><td>DUF499_1.RM.like.systems</td><td><-HicB||HTH+WYL->TUDOR+SF1-SF2-Helicase+iRease+CTD+REase->ASCH->MORC+DUF3684->METHYLASE->HEPN+DUF499*->?-><-?<-?<-Phage_terminase<-TetR-HTH</td><td>1098</td><td>Bacteria>Pseudomonadota>Alphaproteobacteria>Hyphomicrobiales>Rhizobiaceae</td><td>GCF_021387325.1</td><td>Agrobacterium_vitis</td><td>FS799_10735</td><td>1060385..1063681</td><td>373</td><td>DUF499-domain-containing-protein-[Agrobacterium-vitis]</td></tr>
<tr><td>WP_242219593.1</td><td>DUF499_1.RM.like.systems</td><td>ParG_RHH-><-MODE-HTH+PBPII<-HEPN+DUF499*<-METHYLASE<-?<-HIN-HTH||?->Phage_AlpA-><-Arm-DNA-bind_2+Phage_int_SAM_3+Phage_integrase</td><td>1098</td><td>Bacteria>Pseudomonadota>Alphaproteobacteria>Hyphomicrobiales>Rhizobiaceae</td><td>GCF_022682305.1</td><td>Shinella_zoogloeoides</td><td>MOE34_RS21255</td><td>4281309..4284605</td><td>352475</td><td>DUF499-domain-containing-protein-[Shinella-zoogloeoides]</td></tr>
<tr><td>WP_149746898.1</td><td>DUF499_1.RM.like.systems</td><td>ParG_RHH-><-MODE-HTH+PBPII<-HEPN+DUF499*<-METHYLASE<-REase<-TUDOR+SF1-SF2-Helicase+iRease+CTD+REase<-HTH+WYL||SF1-SF2-helicase+RdgC+Beta-barrel+REase->XIS-HTH-></td><td>1098</td><td>Bacteria>Pseudomonadota>Alphaproteobacteria>Hyphomicrobiales>Rhizobiaceae</td><td>GCF_900156005.1</td><td>Rhizobium_sp._RU35A</td><td>F3H95_RS17305</td><td>253422..256718</td><td>1907414</td><td>DUF499-domain-containing-protein-[Rhizobium-sp.-RU35A]</td></tr>
<tr><td>WP_097140486.1</td><td>DUF499_1.RM.like.systems</td><td>ParG_RHH-><-MODE-HTH+PBPII<-HEPN+DUF499*<-METHYLASE</td><td>1098</td><td>Bacteria>Pseudomonadota>Alphaproteobacteria>Hyphomicrobiales>Rhizobiaceae</td><td>GCF_900220975.1</td><td>Rhizobium_subbaraonis</td><td>CRO48_RS14615</td><td>291949..295245</td><td>908946</td><td>DUF499-domain-containing-protein-[Rhizobium-subbaraonis]</td></tr>
<tr><td>WP_142627962.1</td><td>DUF499_1.RM.like.systems</td><td>TPR-><-RNase+SF1-SF2-Helicase<-HEPN+DUF499*<-METHYLASE<-MORC+DUF3684<-YtH<-ASCH<-TUDOR+SF1-SF2-Helicase+iRease+CTD+REase<-HTH+WYL</td><td>1098</td><td>Bacteria>Pseudomonadota>Alphaproteobacteria>Hyphomicrobiales>Rhizobiaceae</td><td>GCF_902153225.2</td><td>Rhizobium_sp._P007</td><td>FN104_RS24455</td><td>262558..265854</td><td>285908</td><td>DUF499-domain-containing-protein-[Rhizobium-sp.-P007]</td></tr>
<tr><td>WP_246622248.1</td><td>DUF499_1.RM.like.systems</td><td>HTH+WYL->TUDOR+SF1-SF2-Helicase+iRease+CTD+REase->YtH->MORC+DUF3684->METHYLASE->HEPN+DUF499*-><-wHTH+REase<-Phage_integrase<-Peptidase_M24||TetR-HTH-></td><td>1086</td><td>Bacteria>Pseudomonadota>Alphaproteobacteria>Hyphomicrobiales>Rhizobiaceae</td><td>-</td><td>Rhizobium_laguerreae</td><td>HFO15_RS01270</td><td>257008..260268</td><td>1076926</td><td>DUF499-domain-containing-protein-[Rhizobium-laguerreae]</td></tr>
<tr><td>WP_222282260.1</td><td>DUF499_1.RM.like.systems</td><td><-PIN+OST-HTH||TUDOR+SF1-SF2-Helicase+iRease+CTD+REase->?->METHYLASE->?->HEPN+DUF499*-><-HTH_3</td><td>1103</td><td>Bacteria>Pseudomonadota>Alphaproteobacteria>Hyphomicrobiales>Rhizobiaceae</td><td>-</td><td>Rhizobium_leguminosarum</td><td>HFO92_34835</td><td>18961..22272</td><td>384</td><td>DUF499-domain-containing-protein-[Rhizobium-leguminosarum]</td></tr>
<tr><td>WP_166603209.1</td><td>DUF499_1.RM.like.systems</td><td>TUDOR+SF1-SF2-Helicase+iRease+CTD+REase-><-?<-?||METHYLASE->HEPN+DUF499*-></td><td>1119</td><td>Bacteria>Pseudomonadota>Alphaproteobacteria>Hyphomicrobiales>Rhizobiaceae</td><td>-</td><td>Rhizobium_sp._CRIBSB</td><td>GVN24_RS23375</td><td>29303..32662</td><td>2683264</td><td>DUF499-domain-containing-protein-[Rhizobium-sp.-CRIBSB]</td></tr>
<tr><td>WP_045535325.1</td><td>DUF499_1.RM.like.systems</td><td><-MORC+HisKin+REase<-HEPN+DUF499*<-?<-METHYLASE<-SIR2<-ASCH<-TUDOR+SF1-SF2-Helicase+iRease+CTD+REase<-HTH+WYL</td><td>1104</td><td>Bacteria>Pseudomonadota>Alphaproteobacteria>Hyphomicrobiales>Rhizobiaceae</td><td>-</td><td>Rhizobium_sp._LMB-1</td><td>VN14_RS23455</td><td>4128..7442</td><td>1605895</td><td>DUF499-domain-containing-protein-[Rhizobium-sp.-LMB-1]</td></tr>
<tr><td>WP_221153182.1</td><td>DUF499_1.RM.like.systems</td><td>TetR-HTH-><-tRNA<-Peptidase_M15<-?||TUDOR+SF1-SF2-Helicase+iRease+CTD+REase->METHYLASE->HEPN+DUF499*-><-GDResolvase+HIN-HTH||ZnR-></td><td>1114</td><td>Bacteria>Pseudomonadota>Alphaproteobacteria>Hyphomicrobiales>Rhizobiaceae</td><td>-</td><td>Rhizobium_sp._NLR10a</td><td>HJB90_RS14150</td><td>264416..267760</td><td>2731105</td><td>DUF499-domain-containing-protein-[Rhizobium-sp.-NLR10a]</td></tr>
<tr><td>MBC2861680.1</td><td>DUF499_1.RM.like.systems</td><td><-SF1-SF2-Helicase+REase<-SF1-SF2-Helicase+REase||HTH+WYL->TUDOR+SF1-SF2-Helicase+iRease+CTD+REase->?->METHYLASE->HEPN+DUF499*-><-MODE-HTH+PBPII||TetR-HTH-><-ParG_RHH</td><td>1098</td><td>Bacteria>Pseudomonadota>Alphaproteobacteria>Hyphomicrobiales>Stappiaceae</td><td>GCA_014252955.1</td><td>Stappia_sp._28M-7</td><td>H7H34_21795</td><td>99390..102686</td><td>2762596</td><td>ATP-binding-protein-[Stappia-sp.-28M-7]</td></tr>
<tr><td>MBO9421687.1</td><td>DUF499_1.RM.like.systems</td><td>HTH+WYL->TUDOR+SF1-SF2-Helicase+iRease+CTD+REase->YtH->MORC+DUF3684->REase->METHYLASE->HEPN+DUF499*-><-ACET<-CopG/RHH<-Arm-DNA-bind_3+Phage_integrase<-?<-MethyltransfD12</td><td>1098</td><td>Bacteria>Pseudomonadota>Alphaproteobacteria>Hyphomicrobiales>Stappiaceae</td><td>GCA_017743865.1</td><td>Labrenzia_sp._R4_2</td><td>J7481_19425</td><td>351032..354328</td><td>2821107</td><td>DUF499-domain-containing-protein-[Labrenzia-sp.-R4_2]</td></tr>
<tr><td>WP_055658296.1</td><td>DUF499_1.RM.like.systems</td><td>Arm-DNA-bind_3+Phage_integrase->?->Phage_AlpA->ZnR+TOPRIM->DnaB+CITB-HTH->?-><-HEPN+DUF499*<-?<-METHYLASE<-?<-ParB+HNH<-MORC+DUF3684<-TUDOR+SF1-SF2-Helicase+iRease+CTD+REase</td><td>1112</td><td>Bacteria>Pseudomonadota>Alphaproteobacteria>Hyphomicrobiales>Stappiaceae</td><td>GCF_001403915.1</td><td>Roseibium_aggregatum</td><td>LAL4801_RS18620</td><td>1338638..1341976</td><td>187304</td><td>DUF499-domain-containing-protein-[Roseibium-aggregatum]</td></tr>
<tr><td>WP_209006426.1</td><td>DUF499_1.RM.like.systems</td><td><-SF1-SF2-Helicase+REase<-SF1-SF2-Helicase+REase||HTH+WYL->TUDOR+SF1-SF2-Helicase+iRease+CTD+REase->?->METHYLASE->HEPN+DUF499*-><-MODE-HTH+PBPII||TetR-HTH-><-ParG_RHH</td><td>1086</td><td>Bacteria>Pseudomonadota>Alphaproteobacteria>Hyphomicrobiales>Stappiaceae</td><td>GCF_014252955.1</td><td>Stappia_sp._28M-7</td><td>H7H34_RS21795</td><td>99426..102686</td><td>2762596</td><td>DUF499-domain-containing-protein-[Stappia-sp.-28M-7]</td></tr>
<tr><td>WP_235988002.1</td><td>DUF499_1.RM.like.systems</td><td>ParG_RHH-><-TetR-HTH||MODE-HTH+PBPII-><-HEPN+DUF499*<-METHYLASE</td><td>1060</td><td>Bacteria>Pseudomonadota>Alphaproteobacteria>Hyphomicrobiales>Stappiaceae</td><td>GCF_014692675.1</td><td>Roseibium_aggregatum</td><td>HK439_RS18245</td><td>125935..129117</td><td>187304</td><td>DUF499-domain-containing-protein-[Roseibium-aggregatum]</td></tr>
<tr><td>WP_286190155.1</td><td>DUF499_1.RM.like.systems</td><td>HTH+WYL->TUDOR+SF1-SF2-Helicase+iRease+CTD+REase->YtH->MORC+DUF3684->REase->METHYLASE->HEPN+DUF499*-><-ACET<-CopG/RHH<-Arm-DNA-bind_3+Phage_integrase<-?<-MethyltransfD12</td><td>1086</td><td>Bacteria>Pseudomonadota>Alphaproteobacteria>Hyphomicrobiales>Stappiaceae</td><td>GCF_017743865.1</td><td>Labrenzia_sp._R4_2</td><td>J7481_RS19430</td><td>351068..354328</td><td>2821107</td><td>DUF499-domain-containing-protein-[Labrenzia-sp.-R4_2]</td></tr>
<tr><td>WP_186398691.1</td><td>DUF499_1.RM.like.systems</td><td><-AbiEii<-HEPN+DUF499*<-METHYLASE<-MORC+DUF3684<-TUDOR+SF1-SF2-Helicase+iRease+CTD+REase<-HTH+WYL<-XIS-HTH</td><td>1115</td><td>Bacteria>Pseudomonadota>Alphaproteobacteria>Hyphomicrobiales>Stappiaceae</td><td>GCF_900185735.1</td><td>Stappia_sp._P2PMeth1</td><td>P2PM1_RS07495</td><td>1079..4426</td><td>2003586</td><td>Swt1-family-HEPN-domain-containing-protein-[Stappia-sp.-P2PMeth1]</td></tr>
<tr><td>MBN8994771.1</td><td>DUF499_1.RM.like.systems</td><td><-SF1-SF2-Helicase+REase||HTH+WYL->TUDOR+SF1-SF2-Helicase+iRease+CTD+REase->ASCH->SIR2->METHYLASE->HEPN+DUF499*->MODE-HTH+PBPII-><-ParG_RHH</td><td>1100</td><td>Bacteria>Pseudomonadota>Alphaproteobacteria>Hyphomicrobiales>unclassified_Hyphomicrobiales</td><td>GCA_017305635.1</td><td>Hyphomicrobiales_bacterium</td><td>J0H94_06075</td><td>362415..365717</td><td>1909294</td><td>DUF499-domain-containing-protein-[Hyphomicrobiales-bacterium]</td></tr>
<tr><td>MBN9001777.1</td><td>DUF499_1.RM.like.systems</td><td>METHYLASE-p->HEPN+DUF499*-></td><td>379</td><td>Bacteria>Pseudomonadota>Alphaproteobacteria>Hyphomicrobiales>unclassified_Hyphomicrobiales</td><td>GCA_017305655.1</td><td>Hyphomicrobiales_bacterium</td><td>J0H75_06805</td><td>482..1619</td><td>1909294</td><td>DUF499-domain-containing-protein-[Hyphomicrobiales-bacterium]</td></tr>
<tr><td>MBN9010874.1</td><td>DUF499_1.RM.like.systems</td><td>METHYLASE-p->HEPN+DUF499*-></td><td>540</td><td>Bacteria>Pseudomonadota>Alphaproteobacteria>Hyphomicrobiales>unclassified_Hyphomicrobiales</td><td>GCA_017305735.1</td><td>Hyphomicrobiales_bacterium</td><td>J0H63_12355</td><td>981..2600</td><td>1909294</td><td>DUF499-domain-containing-protein-[Hyphomicrobiales-bacterium]</td></tr>
<tr><td>MBN9033564.1</td><td>DUF499_1.RM.like.systems</td><td>ABhydrolase->REase->Peptidase_M24->HhH-RADC+JAB->Arm-DNA-bind_3+Phage_integrase->?-><-HEPN+DUF499*<-METHYLASE</td><td>1098</td><td>Bacteria>Pseudomonadota>Alphaproteobacteria>Hyphomicrobiales>unclassified_Hyphomicrobiales</td><td>GCA_017305935.1</td><td>Hyphomicrobiales_bacterium</td><td>J0I23_27610</td><td>44902..48198</td><td>1909294</td><td>DUF499-domain-containing-protein-[Hyphomicrobiales-bacterium]</td></tr>
<tr><td>MBZ0262663.1</td><td>DUF499_1.RM.like.systems</td><td>SF1-SF2-Helicase+REase->SF1-SF2-Helicase+REase-><-?<-ACET<-HEPN+DUF499*<-METHYLASE<-TUDOR+SF1-SF2-Helicase+iRease+CTD+REase<-HTH+WYL||HicB->TIR+PNTD2->PIWI-></td><td>1104</td><td>Bacteria>Pseudomonadota>Alphaproteobacteria>Hyphomicrobiales>unclassified_Hyphomicrobiales</td><td>GCA_019912605.1</td><td>Hyphomicrobiales_bacterium</td><td>K8F90_18930</td><td>14586..17900</td><td>1909294</td><td>DUF499-domain-containing-protein-[Hyphomicrobiales-bacterium]</td></tr>
<tr><td>SDR39062.1</td><td>DUF499_1.RM.like.systems</td><td>TUDOR+SF1-SF2-Helicase+iRease+CTD+REase->YtH->METHYLASE->HEPN->HEPN+DUF499*->STAS+HisKin->?->?->?->?-><-ABhydrolase</td><td>1116</td><td>Bacteria>Pseudomonadota>Alphaproteobacteria>Hyphomicrobiales>unclassified_Hyphomicrobiales</td><td>GCA_900103895.1</td><td>Rhizobiales_bacterium_GAS113</td><td>SAMN05519103_02590</td><td>2751925..2755275</td><td>1884352</td><td>Predicted-ATPase,-AAA+-superfamily-[Rhizobiales-bacterium-GAS113]</td></tr>
<tr><td>ABS65938.1</td><td>DUF499_1.RM.like.systems</td><td>TUDOR+SF1-SF2-Helicase+iRease+CTD+REase->REase->METHYLASE-><-HTH_21+rve+rve_3<-HIN-HTH||METHYLASE-p->HEPN+DUF499*-><-MODE-HTH+PBPII||TetR-HTH-><-ParG_RHH</td><td>1098</td><td>Bacteria>Pseudomonadota>Alphaproteobacteria>Hyphomicrobiales>Xanthobacteraceae</td><td>GCA_000017645.1</td><td>Xanthobacter_autotrophicus_Py2</td><td>Xaut_0686</td><td>774507..777803</td><td>78245</td><td>ATPase-(AAA+-superfamily)-like-protein-[Xanthobacter-autotrophicus-Py2]</td></tr>
<tr><td>ABS68223.1</td><td>DUF499_1.RM.like.systems</td><td><-SF1-SF2-Helicase+REase||HTH+WYL->TUDOR+SF1-SF2-Helicase+iRease+CTD+REase->YtH->MORC+DUF3684->METHYLASE->HEPN+DUF499*->MODE-HTH+PBPII-></td><td>1098</td><td>Bacteria>Pseudomonadota>Alphaproteobacteria>Hyphomicrobiales>Xanthobacteraceae</td><td>GCA_000017645.1</td><td>Xanthobacter_autotrophicus_Py2</td><td>Xaut_2992</td><td>3328660..3331956</td><td>78245</td><td>conserved-hypothetical-protein-[Xanthobacter-autotrophicus-Py2]</td></tr>
<tr><td>MBX6329369.1</td><td>DUF499_1.RM.like.systems</td><td>TUDOR+SF1-SF2-Helicase+iRease+CTD+REase-p->?->?->PolB->HEPN->HEPN+DUF499*-></td><td>1114</td><td>Bacteria>Pseudomonadota>Alphaproteobacteria>Hyphomicrobiales>Xanthobacteraceae</td><td>GCA_019689975.1</td><td>Pseudolabrys_sp.</td><td>IRY89_12420</td><td>7103..10447</td><td>1960880</td><td>DUF499-domain-containing-protein-[Pseudolabrys-sp.]</td></tr>
<tr><td>MBX9829560.1</td><td>DUF499_1.RM.like.systems</td><td>TrwC+SF1-SF2-helicase->TUDOR+SF1-SF2-Helicase+iRease+CTD+REase->HEPN->METHYLASE-p->METHYLASE->?->HEPN+DUF499*->?->REase-></td><td>1101</td><td>Bacteria>Pseudomonadota>Alphaproteobacteria>Hyphomicrobiales>Xanthobacteraceae</td><td>GCA_019746355.1</td><td>Xanthobacteraceae_bacterium</td><td>K2Y27_31790</td><td>26343..29648</td><td>2021379</td><td>DUF499-domain-containing-protein-[Xanthobacteraceae-bacterium]</td></tr>
<tr><td>MBX9650469.1</td><td>DUF499_1.RM.like.systems</td><td><-SF1-SF2-Helicase+REase||HTH+WYL->TUDOR+SF1-SF2-Helicase+iRease+CTD+REase->MORC+DUF3684->METHYLASE->HEPN+DUF499*->MODE-HTH+PBPII-><-ParG_RHH</td><td>1098</td><td>Bacteria>Pseudomonadota>Alphaproteobacteria>Hyphomicrobiales>Xanthobacteraceae</td><td>GCA_019748335.1</td><td>Xanthobacteraceae_bacterium</td><td>K2X57_25835</td><td>14404..17700</td><td>2021379</td><td>DUF499-domain-containing-protein-[Xanthobacteraceae-bacterium]</td></tr>
<tr><td>MCW5685472.1</td><td>DUF499_1.RM.like.systems</td><td><-Peptidase_S11<-?||tRNA-><-?<-HEPN+DUF499*<-METHYLASE<-ParB+HNH<-TUDOR+SF1-SF2-Helicase+iRease+CTD+REase<-HTH+WYL||SF1-SF2-Helicase+REase-></td><td>1098</td><td>Bacteria>Pseudomonadota>Alphaproteobacteria>Hyphomicrobiales>Xanthobacteraceae</td><td>GCA_026125885.1</td><td>Pseudolabrys_sp.</td><td>KIT85_13790</td><td>135757..139053</td><td>1960880</td><td>MAG:-DUF499-domain-containing-protein-[Pseudolabrys-sp.]</td></tr>
<tr><td>MCZ7659369.1</td><td>DUF499_1.RM.like.systems</td><td>DEOR-HTH->TUDOR+SF1-SF2-Helicase+iRease+CTD+REase->TUDOR+SF1-SF2-Helicase+iRease+CTD+REase-p->?->REase->METHYLASE->HEPN+DUF499*->OLD-ABC+TOPRIM->SF1-SF2-Helicase-><-REase<-?<-N6_MTase<-DEOR-HTH</td><td>1120</td><td>Bacteria>Pseudomonadota>Alphaproteobacteria>Hyphomicrobiales>Xanthobacteraceae</td><td>GCA_027493405.1</td><td>Xanthobacteraceae_bacterium</td><td>M5U07_16605</td><td>9358..12720</td><td>2021379</td><td>MAG:-Swt1-family-HEPN-domain-containing-protein-[Xanthobacteraceae-bacterium]</td></tr>
<tr><td>WP_282888615.1</td><td>DUF499_1.RM.like.systems</td><td><-SF1-SF2-Helicase+REase<-?||HTH+WYL->TUDOR+SF1-SF2-Helicase+iRease+CTD+REase->MORC+DUF3684->METHYLASE->HEPN+DUF499*->TIR+PNTD2->PIWI->Arm-DNA-bind_3+Phage_integrase-><-XIS-HTH<-MobA_MobL+SF1-SF2-helicase||TraD-></td><td>1098</td><td>Bacteria>Pseudomonadota>Alphaproteobacteria>Hyphomicrobiales>Xanthobacteraceae</td><td>GCF_030015065.1</td><td>Xanthobacter_autotrophicus</td><td>K9U40_RS02195</td><td>72535..75831</td><td>280</td><td>DUF499-domain-containing-protein-[Xanthobacter-autotrophicus]</td></tr>
<tr><td>WP_159599724.1</td><td>DUF499_1.RM.like.systems</td><td>HTH+WYL->TUDOR+SF1-SF2-Helicase+iRease+CTD+REase->YtH->MORC+DUF3684->METHYLASE->HEPN+DUF499*-><-?<-Arm-DNA-bind_3+Phage_integrase<-HhH-RADC+JAB<-Peptidase_M24<-ACET||Aminotran_1_2-></td><td>1098</td><td>Bacteria>Pseudomonadota>Alphaproteobacteria>Hyphomicrobiales>Xanthobacteraceae</td><td>GCF_902703545.1</td><td>Starkeya_nomas</td><td>STARVERO_RS14985</td><td>3130835..3134131</td><td>2666134</td><td>DUF499-domain-containing-protein-[Starkeya-sp.-HF14-78462]</td></tr>
<tr><td>MCB9972622.1</td><td>DUF499_1.RM.like.systems</td><td>XIS-HTH->?->?-><-HEPN+DUF499*<-METHYLASE<-REase<-MORC+DUF3684<-TUDOR+SF1-SF2-Helicase+iRease+CTD+REase<-HTH+WYL||VirD4-TraG+VirD4-FtsK-></td><td>1097</td><td>Bacteria>Pseudomonadota>Alphaproteobacteria>Hyphomonadales>Hyphomonadaceae</td><td>GCA_020638455.1</td><td>Hyphomonas_sp.</td><td>H6847_14020</td><td>101452..104745</td><td>87</td><td>ATP-binding-protein-[Hyphomonas-sp.]</td></tr>
<tr><td>WP_188873657.1</td><td>DUF499_1.RM.like.systems</td><td>PAPS_reduct->HTH+MPTase->PIN-><-MalT<-HEPN<-HEPN<-HEPN+DUF499*<-TIR<-METHYLASE<-?<-TUDOR+SF1-SF2-Helicase+iRease+CTD+REase<-XIS-HTH||DOC+HTH-></td><td>1098</td><td>Bacteria>Pseudomonadota>Alphaproteobacteria>Iodidimonadales>Iodidimonadaceae</td><td>GCF_014647255.1</td><td>Iodidimonas_muriae</td><td>IEX96_RS06665</td><td>363502..366798</td><td>261467</td><td>DUF499-domain-containing-protein-[Iodidimonas-muriae]</td></tr>
<tr><td>GER06671.1</td><td>DUF499_1.RM.like.systems</td><td><-DOC+HTH||XIS-HTH->TUDOR+SF1-SF2-Helicase+iRease+CTD+REase->?->METHYLASE->TIR->HEPN+DUF499*->HEPN->HEPN->MalT-><-PIN<-HTH+MPTase<-PAPS_reduct</td><td>1074</td><td>Bacteria>Pseudomonadota>Alphaproteobacteria>Kordiimonadales>unclassified_Kordiimonadales</td><td>GCA_008579105.1</td><td>Kordiimonadales_bacterium_JCM_17843</td><td>JCM17843_09810</td><td>11822..15046</td><td>1236965</td><td>hypothetical-protein-JCM17843_09810-[Kordiimonadales-bacterium-JCM-17843]</td></tr>
<tr><td>MBF0108101.1</td><td>DUF499_1.RM.like.systems</td><td>PRTase->?->?->?-><-HEPN+DUF499*<-METHYLASE</td><td>1102</td><td>Bacteria>Pseudomonadota>Alphaproteobacteria>Magnetococcales>unclassified_Magnetococcales</td><td>GCA_015228815.1</td><td>Magnetococcales_bacterium</td><td>HQL76_02845</td><td>103029..106337</td><td>2026759</td><td>DUF499-domain-containing-protein-[Magnetococcales-bacterium]</td></tr>
<tr><td>MBF0262096.1</td><td>DUF499_1.RM.like.systems</td><td>XIS-HTH->TUDOR+SF1-SF2-Helicase+iRease+CTD+REase->SbcC->OLD-ABC->TOPRIM->METHYLASE->HEPN+DUF499*-></td><td>1048</td><td>Bacteria>Pseudomonadota>Alphaproteobacteria>Magnetococcales>unclassified_Magnetococcales</td><td>GCA_015231775.1</td><td>Magnetococcales_bacterium</td><td>HQL97_09720</td><td>11242..14384</td><td>2026759</td><td>DUF499-domain-containing-protein-[Magnetococcales-bacterium]</td></tr>
<tr><td>ADM09268.1</td><td>DUF499_1.RM.like.systems</td><td><-HEPN+DUF499*<-METHYLASE<-MORC+DUF3684<-TUDOR+SF1-SF2-Helicase+iRease+CTD+REase<-HTH+WYL||REase+PNTD1+PNTD2->PIWI-></td><td>1090</td><td>Bacteria>Pseudomonadota>Alphaproteobacteria>Parvularculales>Parvularculaceae</td><td>GCA_000152825.2</td><td>Parvularcula_bermudensis_HTCC2503</td><td>PB2503_05987</td><td>1154188..1157460</td><td>314260</td><td>hypothetical-protein-PB2503_05987-[Parvularcula-bermudensis-HTCC2503]</td></tr>
<tr><td>GGY37139.1</td><td>DUF499_1.RM.like.systems</td><td><-CopG/RHH<-VirD4-TraG+VirD4-FtsK||HTH+WYL->TUDOR+SF1-SF2-Helicase+iRease+CTD+REase->METHYLASE->HEPN+DUF499*-><-?||?-><-XIS-HTH<-?<-?<-ParB+HTH</td><td>1097</td><td>Bacteria>Pseudomonadota>Alphaproteobacteria>Parvularculales>Parvularculaceae</td><td>GCA_014651915.1</td><td>Parvularcula_lutaonensis</td><td>GCM10007148_01670</td><td>175474..178767</td><td>491923</td><td>hypothetical-protein-GCM10007148_01670-[Parvularcula-lutaonensis]</td></tr>
<tr><td>WP_041534905.1</td><td>DUF499_1.RM.like.systems</td><td><-HEPN+DUF499*<-METHYLASE<-MORC+DUF3684<-TUDOR+SF1-SF2-Helicase+iRease+CTD+REase<-HTH+WYL||REase+PNTD1+PNTD2->PIWI-></td><td>1102</td><td>Bacteria>Pseudomonadota>Alphaproteobacteria>Parvularculales>Parvularculaceae</td><td>GCF_000152825.2</td><td>Parvularcula_bermudensis</td><td>PB2503_RS05500</td><td>1154188..1157496</td><td>208216</td><td>DUF499-domain-containing-protein-[Parvularcula-bermudensis]</td></tr>
<tr><td>WP_229786002.1</td><td>DUF499_1.RM.like.systems</td><td><-CopG/RHH<-VirD4-TraG+VirD4-FtsK||HTH+WYL->TUDOR+SF1-SF2-Helicase+iRease+CTD+REase->METHYLASE->HEPN+DUF499*-><-?<-?<-XIS-HTH<-?<-?<-ParB+HTH</td><td>1085</td><td>Bacteria>Pseudomonadota>Alphaproteobacteria>Parvularculales>Parvularculaceae</td><td>GCF_014651915.1</td><td>Parvularcula_lutaonensis</td><td>IE053_RS00820</td><td>175510..178767</td><td>491923</td><td>DUF499-domain-containing-protein-[Parvularcula-lutaonensis]</td></tr>
<tr><td>KPP88834.1</td><td>DUF499_1.RM.like.systems</td><td>VirD4-TraG+VirD4-FtsK->CopG/RHH->TUDOR+SF1-SF2-Helicase+iRease+CTD+REase->PadR-HTH->METHYLASE->HEPN+DUF499*-><-PIN<-RHH<-?<-CHTH||VirB11+VirB4-FtsK->VirB2-></td><td>1100</td><td>Bacteria>Pseudomonadota>Alphaproteobacteria>Rhodobacterales>Paracoccaceae</td><td>GCA_001314655.1</td><td>Rhodobacteraceae_bacterium_HLUCCA08</td><td>HLUCCA08_13590</td><td>19280..22582</td><td>1666913</td><td>putative-ATPase-(AAA+-superfamily)-[Rhodobacteraceae-bacterium-HLUCCA08]</td></tr>
<tr><td>MBC6437846.1</td><td>DUF499_1.RM.like.systems</td><td><-ABC-ATPase+SbcC||?->ncRNA->?-><-HEPN+DUF499*<-METHYLASE-p</td><td>1099</td><td>Bacteria>Pseudomonadota>Alphaproteobacteria>Rhodobacterales>Paracoccaceae</td><td>GCA_014323785.1</td><td>Paracoccaceae_bacterium</td><td>GDA52_06875</td><td>42024..45323</td><td>1904441</td><td>ATP-binding-protein-[Rhodobacteraceae-bacterium]</td></tr>
<tr><td>MBC6443273.1</td><td>DUF499_1.RM.like.systems</td><td>Glycos_transf_4->?->HEPN+DUF499*->?-><-ParB+HNH||?->?->?->ABC-ATPase+SbcC-></td><td>1096</td><td>Bacteria>Pseudomonadota>Alphaproteobacteria>Rhodobacterales>Paracoccaceae</td><td>GCA_014323795.1</td><td>Paracoccaceae_bacterium</td><td>GDA53_09225</td><td>19361..22651</td><td>1904441</td><td>ATP-binding-protein-[Rhodobacteraceae-bacterium]</td></tr>
<tr><td>MBE2259559.1</td><td>DUF499_1.RM.like.systems</td><td><-HEPN+DUF499*<-HEPN<-PolB<-METHYLASE-p</td><td>485</td><td>Bacteria>Pseudomonadota>Alphaproteobacteria>Rhodobacterales>Paracoccaceae</td><td>GCA_014879755.1</td><td>Paracoccaceae_bacterium</td><td>IAE88_11910</td><td>1..1455</td><td>1904441</td><td>ATP-binding-protein-[Paracoccaceae-bacterium]</td></tr>
<tr><td>MCC5984903.1</td><td>DUF499_1.RM.like.systems</td><td><-HEPN+DUF499*<-METHYLASE<-REase<-TUDOR+SF1-SF2-Helicase+iRease+CTD+REase<-CopG/RHH<-VirD4-TraG+VirD4-FtsK||HNH-></td><td>351</td><td>Bacteria>Pseudomonadota>Alphaproteobacteria>Rhodobacterales>Paracoccaceae</td><td>GCA_020832705.1</td><td>Paracoccaceae_bacterium</td><td>JJU42_11110</td><td>1..1055</td><td>1904441</td><td>AAA-family-ATPase-[Paracoccaceae-bacterium]</td></tr>
<tr><td>MCQ0971011.1</td><td>DUF499_1.RM.like.systems</td><td>MODE-HTH+PBPII->||?->GDResolvase+Recombinase+ZnR->||DDE_Tnp_IS66->MODE-HTH+PBPII-><-HEPN+DUF499*<-METHYLASE<-ParB+HNH<-TUDOR+SF1-SF2-Helicase+iRease+CTD+REase<-HTH+WYL||SF1-SF2-Helicase+REase->SF1-SF2-Helicase+REase-></td><td>1097</td><td>Bacteria>Pseudomonadota>Alphaproteobacteria>Rhodobacterales>Paracoccaceae</td><td>GCA_024359525.1</td><td>Paracoccus_albicereus</td><td>MLD63_11305</td><td>50725..54018</td><td>2922394</td><td>DUF499-domain-containing-protein-[Paracoccus-sp.-TK19116]</td></tr>
<tr><td>GIX14656.1</td><td>DUF499_1.RM.like.systems</td><td>TUDOR+SF1-SF2-Helicase+iRease+CTD+REase->TUDOR+SF1-SF2-Helicase+iRease+CTD+REase-p->METHYLASE-p->METHYLASE-p->METHYLASE-p->HEPN+DUF499*-><-?<-?<-?||MULE-TRANSPOSASE-></td><td>1104</td><td>Bacteria>Pseudomonadota>Alphaproteobacteria>Rhodobacterales>Paracoccaceae</td><td>GCA_026004915.1</td><td>Paracoccaceae_bacterium</td><td>KatS3mg118_2615</td><td>175796..179110</td><td>1904441</td><td>MAG:-hypothetical-protein-KatS3mg118_2615-[Paracoccaceae-bacterium]</td></tr>
<tr><td>MCY4445744.1</td><td>DUF499_1.RM.like.systems</td><td><-Arm-DNA-bind_3+Phage_int_SAM_3+Phage_integrase<-Phage_int_SAM_3+Phage_integrase<-TUDOR+SF1-SF2-Helicase+iRease+CTD+REase<-HEPN+DUF499*||METHYLASE-p-></td><td>1118</td><td>Bacteria>Pseudomonadota>Alphaproteobacteria>Rhodobacterales>Paracoccaceae</td><td>GCA_026712645.1</td><td>Paracoccaceae_bacterium</td><td>OXC02_04685</td><td>7553..10909</td><td>1904441</td><td>MAG:-DUF499-domain-containing-protein-[Rhodobacteraceae-bacterium]</td></tr>
<tr><td>MCY4460275.1</td><td>DUF499_1.RM.like.systems</td><td><-HEPN+DUF499*<-Schlafen+HATPase+HTH<-METHYLASE<-TUDOR+SF1-SF2-Helicase+iRease+CTD+REase</td><td>1099</td><td>Bacteria>Pseudomonadota>Alphaproteobacteria>Rhodobacterales>Paracoccaceae</td><td>GCA_026712665.1</td><td>Albidovulum_sp.</td><td>OXC26_07750</td><td>691..3990</td><td>1872424</td><td>MAG:-DUF499-domain-containing-protein-[Albidovulum-sp.]</td></tr>
<tr><td>MDB4017580.1</td><td>DUF499_1.RM.like.systems</td><td>XIS-HTH->?->TUDOR+SF1-SF2-Helicase+iRease+CTD+REase->METHYLASE->METHYLASE->HEPN+DUF499*-><-URI-DNase<-RNase+35exo<-Schlafen||?->?->HTH-></td><td>1097</td><td>Bacteria>Pseudomonadota>Alphaproteobacteria>Rhodobacterales>Paracoccaceae</td><td>GCA_028244795.1</td><td>Amylibacter_sp.</td><td>N9493_06125</td><td>26402..29695</td><td>1872504</td><td>DUF499-domain-containing-protein-[Amylibacter-sp.]</td></tr>
<tr><td>MDC0095869.1</td><td>DUF499_1.RM.like.systems</td><td>XIS-HTH->?->TUDOR+SF1-SF2-Helicase+iRease+CTD+REase->METHYLASE->METHYLASE->HEPN+DUF499*-><-URI-DNase<-RNase+35exo<-RNase+SF1-SF2-Helicase||?->?->HTH-></td><td>1097</td><td>Bacteria>Pseudomonadota>Alphaproteobacteria>Rhodobacterales>Paracoccaceae</td><td>GCA_028342995.1</td><td>Amylibacter_sp.</td><td>OAI58_06255</td><td>84173..87466</td><td>1872504</td><td>DUF499-domain-containing-protein-[Amylibacter-sp.]</td></tr>
<tr><td>MDE0346579.1</td><td>DUF499_1.RM.like.systems</td><td><-HEPN+DUF499*<-PIN<-DUF499-p<-METHYLASE<-TUDOR+SF1-SF2-Helicase+iRease+CTD+REase-p</td><td>1102</td><td>Bacteria>Pseudomonadota>Alphaproteobacteria>Rhodobacterales>Paracoccaceae</td><td>GCA_028821095.1</td><td>Boseongicola_sp.</td><td>OXI66_12500</td><td>695..4003</td><td>1886600</td><td>MAG:-DUF499-domain-containing-protein-[Boseongicola-sp.]</td></tr>
<tr><td>WP_017929317.1</td><td>DUF499_1.RM.like.systems</td><td><-HEPN+DUF499*<-METHYLASE<-ASCH<-TUDOR+SF1-SF2-Helicase+iRease+CTD+REase<-?<-?<-SF1-SF2-Helicase-p</td><td>1110</td><td>Bacteria>Pseudomonadota>Alphaproteobacteria>Rhodobacterales>Paracoccaceae</td><td>GCF_000365005.1</td><td>Limimaricola_hongkongensis</td><td>A3EY_RS0112135</td><td>165688..169020</td><td>278132</td><td>DUF499-domain-containing-protein-[Limimaricola-hongkongensis]</td></tr>
<tr><td>WP_035746761.1</td><td>DUF499_1.RM.like.systems</td><td><-DDE_Tnp_1_5+DDE_Tnp_1<-HTH_Tnp_1<-HTH_Tnp_1<-HEPN+DUF499*<-METHYLASE<-ParB+HNH<-TUDOR+SF1-SF2-Helicase+iRease+CTD+REase<-HTH+WYL||SF1-SF2-Helicase+REase->SF1-SF2-Helicase+REase-></td><td>1098</td><td>Bacteria>Pseudomonadota>Alphaproteobacteria>Rhodobacterales>Paracoccaceae</td><td>GCF_000740775.1</td><td>Haematobacter_missouriensis</td><td>CG51_RS16225</td><td>8779..12075</td><td>366616</td><td>DUF499-domain-containing-protein-[Haematobacter-missouriensis]</td></tr>
<tr><td>WP_036708698.1</td><td>DUF499_1.RM.like.systems</td><td>XIS-HTH->?->?->MarR-HTH+REase->?->Enterotoxin_a-><-HEPN+DUF499*<-METHYLASE<-REase<-TUDOR+SF1-SF2-Helicase+iRease+CTD+REase||ParB+N6_N4_MTase->ParB+N6_N4_MTase-><-HNH</td><td>1100</td><td>Bacteria>Pseudomonadota>Alphaproteobacteria>Rhodobacterales>Paracoccaceae</td><td>GCF_000763845.1</td><td>Paracoccus_sanguinis</td><td>IX56_RS07055</td><td>10378..13680</td><td>1545044</td><td>DUF499-domain-containing-protein-[Paracoccus-sanguinis]</td></tr>
<tr><td>WP_058096461.1</td><td>DUF499_1.RM.like.systems</td><td><-SF1-SF2-Helicase+REase<-SF1-SF2-Helicase+REase||HTH+WYL->TUDOR+SF1-SF2-Helicase+iRease+CTD+REase->MORC+DUF3684->METHYLASE->HEPN+DUF499*->MODE-HTH+PBPII-><-ParG_RHH</td><td>1098</td><td>Bacteria>Pseudomonadota>Alphaproteobacteria>Rhodobacterales>Paracoccaceae</td><td>GCF_001447385.1</td><td>Paracoccus_sp._MKU1</td><td>AQY21_RS03730</td><td>27830..31126</td><td>1745182</td><td>DUF499-domain-containing-protein-[Paracoccus-sp.-MKU1]</td></tr>
<tr><td>WP_105291560.1</td><td>DUF499_1.RM.like.systems</td><td><-SbcC+SbcC<-Calcineurin||HTH+WYL->TUDOR+SF1-SF2-Helicase+iRease+CTD+REase->?->METHYLASE->HEPN+DUF499*->SbcC->?->RelE-ParE-><-?<-DUF4096+DDE_Tnp_1+DDE_Tnp<-DEDD_Tnp_IS110+Transposase_20</td><td>1101</td><td>Bacteria>Pseudomonadota>Alphaproteobacteria>Rhodobacterales>Paracoccaceae</td><td>GCF_002073635.2</td><td>Paracoccus_yeei</td><td>A6J80_RS16220</td><td>2818181..2821486</td><td>147645</td><td>DUF499-domain-containing-protein-[Paracoccus-yeei]</td></tr>
</table>

| Protein-ID | System | Gene-Neighbourhoods | Length | Taxonomy | Assembly-ID | Strain | Locus-Tag | Genomic-coordinates | TaxID | Fasta-defline |
|---|---|---|---|---|---|---|---|---|---|---|
| WP_096787622.1 | DUF499_1.RM.like.systems | METHYLASE-p-><-?\|\|?-><-SF1-SF2-Helicase+DUF1998+HKD-DNase+HKD-DNase<-?<-HEPN+DUF499*<-METHYLASE<-MACRODOMAIN<-TUDOR+SF1-SF2-Helicase+iREase+CTD+REase\|\|?-><-HTH+WYL | 1100 | Bacteria>Pseudomonadota>Alphaproteobacteria>Rhodobacterales>Paracoccaceae | GCF_002407205.1 | Rhodobacter_sp._CZR27 | CK951_RS18110 | 425347..428649 | 2033869 | DUF499-domain-containing-protein-[Rhodobacter-sp.-CZR27] |
| WP_099275178.1 | DUF499_1.RM.like.systems | rve_3+Mu-transpos_C->?-><-?<-HEPN+DUF499*<-METHYLASE<-ASCH<-TUDOR+SF1-SF2-Helicase+iREase+CTD+REase\|\|HIN-HTH->?->SF1-SF2-Helicase+DUF1998+HKD-DNase+HKD-DNase-> | 1110 | Bacteria>Pseudomonadota>Alphaproteobacteria>Rhodobacterales>Paracoccaceae | GCF_002723615.1 | Limimaricola_cinnabarinus | CJ301_RS05690 | 226587..229919 | 1125964 | DUF499-domain-containing-protein-[Limimaricola-cinnabarinus] |
| WP_108127680.1 | DUF499_1.RM.like.systems | CopG/RHH->TUDOR+SF1-SF2-Helicase+iREase+CTD+REase->PadR-HTH->METHYLASE->HEPN+DUF499*->DUF4011+SF1-SF2-Helicase+REase-><-?\|\|?->?->NTP_transf-> | 1112 | Bacteria>Pseudomonadota>Alphaproteobacteria>Rhodobacterales>Paracoccaceae | GCF_003054195.1 | Gemmobacter_caeni | C8N34_RS03245 | 88881..92219 | 589035 | DUF499-domain-containing-protein-[Gemmobacter-caeni] |
| WP_108644491.1 | DUF499_1.RM.like.systems | <-HNH<-ParB+N6_N4_MTase\|\|TUDOR+SF1-SF2-Helicase+iREase+CTD+REase->HEPN->METHYLASE->HEPN+DUF499*-><-LZ_Tnp_IS66+zf-IS66+DDE_Tnp_IS66+DDE_Tnp_IS66_C<-?<-HIN-HTH<-?<-MarR-HTH+REase | 1103 | Bacteria>Pseudomonadota>Alphaproteobacteria>Rhodobacterales>Paracoccaceae | GCF_003071405.1 | Fuscovulum_blasticum | B6K69_RS15915 | 3260938..3264249 | 1075 | DUF499-domain-containing-protein-[Fuscovulum-blasticum] |
| WP_118137057.1 | DUF499_1.RM.like.systems | <-UbE1+JAB<-?\|\|HTH+WYL->TUDOR+SF1-SF2-Helicase+iREase+CTD+REase->?->METHYLASE->HEPN+DUF499*-><-?\|\|REase->SF1-SF2-Helicase+REase->HNH->?->DOC-> | 1110 | Bacteria>Pseudomonadota>Alphaproteobacteria>Rhodobacterales>Paracoccaceae | GCF_003448965.1 | Oceanicella_sp._SM1341 | D3P35_RS19735 | 60544..63876 | 1548889 | DUF499-domain-containing-protein-[Oceanicella-sp.-SM1341] |
| WP_136646565.1 | DUF499_1.RM.like.systems | DUF2163+Phage_BR0599->?->?->RNase-><-DUF4236+TPR<-HEPN+DUF499*<-METHYLASE<-PadR-HTH<-TUDOR+SF1-SF2-Helicase+iREase+CTD+REase | 1097 | Bacteria>Pseudomonadota>Alphaproteobacteria>Rhodobacterales>Paracoccaceae | GCF_004959785.1 | Tabrizicola_sp._YIM_78059 | EZM81_RS16965 | 12465..15758 | 2529861 | DUF499-domain-containing-protein-[Tabrizicola-sp.-YIM-78059] |
| WP_139597753.1 | DUF499_1.RM.like.systems | <-RNase\|\|TUDOR+SF1-SF2-Helicase+iREase+CTD+REase->?->METHYLASE->HEPN+DUF499*->REase-><-PIN | 1100 | Bacteria>Pseudomonadota>Alphaproteobacteria>Rhodobacterales>Paracoccaceae | GCF_006239215.1 | Paracoccus_haeundaensis | FHD67_RS02835 | 188317..191619 | 225362 | DUF499-domain-containing-protein-[Paracoccus-haeundaensis] |
| WP_149139641.1 | DUF499_1.RM.like.systems | HEPN-><-ParB+N6_N4_MTase<-ParB+N6_N4_MTase\|\|TUDOR+SF1-SF2-Helicase+iREase+CTD+REase->ASCH->METHYLASE->HEPN+DUF499*-><-Enterotoxin_a<-?<-MarR-HTH+REase<-?<-?<-XIS-HTH | 1100 | Bacteria>Pseudomonadota>Alphaproteobacteria>Rhodobacterales>Paracoccaceae | GCF_008271655.1 | Gemmobacter_caeruleus | FOA17_RS00925 | 40432..43734 | 2595004 | DUF499-domain-containing-protein-[Gemmobacter-caeruleus] |
| WP_152367850.1 | DUF499_1.RM.like.systems | <-JAB<-UbE1+JAB<-SMODS\|\|HTH+WYL->TUDOR+SF1-SF2-Helicase+iREase+CTD+REase->METHYLASE->HEPN+DUF499*->?-><-MODE-HTH+PBPII\|\|TetR-HTH-><-ParG_RHH | 1099 | Bacteria>Pseudomonadota>Alphaproteobacteria>Rhodobacterales>Paracoccaceae | GCF_009301415.1 | Paracoccus_kondratievae | F8A10_RS15730 | 901669..904968 | 135740 | DUF499-domain-containing-protein-[Paracoccus-kondratievae] |
| WP_154336328.1 | DUF499_1.RM.like.systems | <-ABC-ATPase+SbcC<-?<-HEPN+DUF499*<-METHYLASE<-?<-TUDOR+SF1-SF2-Helicase+iREase+CTD+REase | 1097 | Bacteria>Pseudomonadota>Alphaproteobacteria>Rhodobacterales>Paracoccaceae | GCF_009674885.1 | Paracoccus_sp._S-4012 | GI374_RS07045 | 35979..39272 | 2665648 | DUF499-domain-containing-protein-[Paracoccus-sp.-S-4012] |
| WP_188408468.1 | DUF499_1.RM.like.systems | HTH+WYL->TUDOR+SF1-SF2-Helicase+iREase+CTD+REase->ASCH->YtH->METHYLASE->HEPN+DUF499*->ParB+HTH->HTH_3->STYKinase-like-> | 1102 | Bacteria>Pseudomonadota>Alphaproteobacteria>Rhodobacterales>Paracoccaceae | GCF_014635245.1 | Agaricicola_taiwanensis | IEV58_RS04465 | 943958..947266 | 591372 | DUF499-domain-containing-protein-[Agaricicola-taiwanensis] |
| WP_189381997.1 | DUF499_1.RM.like.systems | <-ParB+N6_N4_MTase<-ParB+N6_N4_MTase\|\|TUDOR+SF1-SF2-Helicase+iREase+CTD+REase->MACRODOMAIN->METHYLASE->?->HEPN+DUF499*-><-Enterotoxin_a<-?<-MarR-HTH+REase<-?<-?<-XIS-HTH | 1099 | Bacteria>Pseudomonadota>Alphaproteobacteria>Rhodobacterales>Paracoccaceae | GCF_014652195.1 | Gemmobacter_nanjingensis | IE126_RS20360 | 40653..43952 | 488454 | DUF499-domain-containing-protein-[Gemmobacter-nanjingensis] |
| WP_189635417.1 | DUF499_1.RM.like.systems | <-HEPN+DUF499*<-?<-METHYLASE<-?<-TUDOR+SF1-SF2-Helicase+iREase+CTD+REase\|\|ParB+N6_N4_MTase-> | 1099 | Bacteria>Pseudomonadota>Alphaproteobacteria>Rhodobacterales>Paracoccaceae | GCF_014652355.1 | Gemmobacter_lanyuensis | IE101_RS18950 | 648..3947 | 1054497 | DUF499-domain-containing-protein-[Gemmobacter-lanyuensis] |
| WP_233253227.1 | DUF499_1.RM.like.systems | ParG_RHH-><-TetR-HTH\|\|MODE-HTH+PBPII-><-HEPN+DUF499*<-METHYLASE<-ASCH<-TUDOR+SF1-SF2-Helicase+iREase+CTD+REase<-HTH+WYL\|\|SF1-SF2-Helicase+REase->SF1-SF2-Helicase+REase-> | 1086 | Bacteria>Pseudomonadota>Alphaproteobacteria>Rhodobacterales>Paracoccaceae | GCF_016446475.1 | Paracoccus_binzhouensis | I9P22_RS04960 | 53766..57026 | 2796149 | DUF499-domain-containing-protein-[Paracoccus-binzhouensis] |
| WP_209427763.1 | DUF499_1.RM.like.systems | <-AbiEii<-HEPN+DUF499*<-METHYLASE<-MORC+DUF3684<-TUDOR+SF1-SF2-Helicase+iREase+CTD+REase<-HTH+WYL<-Phage_int_SAM_3+Phage_integrase | 1112 | Bacteria>Pseudomonadota>Alphaproteobacteria>Rhodobacterales>Paracoccaceae | GCF_017811755.1 | Pararhodobacter_sp._SW119 | IM708_RS16910 | 1952..5290 | 2780075 | Swt1-family-HEPN-domain-containing-protein-[Pararhodobacter-sp.-SW119] |
| WP_213544567.1 | DUF499_1.RM.like.systems | TUDOR+SF1-SF2-Helicase+iREase+CTD+REase->?->METHYLASE->HEPN+DUF499*-><-?<-?<-XIS-HTH<-?<-Arm-DNA-bind_3+Phage_integrase | 1098 | Bacteria>Pseudomonadota>Alphaproteobacteria>Rhodobacterales>Paracoccaceae | GCF_018424695.1 | Vannielia_litorea | DYI42_RS03660 | 12366..15662 | 1217970 | DUF499-domain-containing-protein-[Vannielia-litorea] |
| WP_234869545.1 | DUF499_1.RM.like.systems | <-SF1-SF2-Helicase+DUF1998<-?\|\|?-><-?<-HEPN+DUF499*<-METHYLASE<-wHTH+REase<-TUDOR+SF1-SF2-Helicase+iREase+CTD+REase | 1100 | Bacteria>Pseudomonadota>Alphaproteobacteria>Rhodobacterales>Paracoccaceae | GCF_021491355.1 | Tabrizicola_rongguiensis | LOV05_RS07760 | 379884..383186 | 2909234 | DUF499-domain-containing-protein-[Tabrizicola-rongguiensis] |
| WP_255330034.1 | DUF499_1.RM.like.systems | MODE-HTH+PBPII->\|\|?->GDResolvase+Recombinase+ZnR->\|\|DDE_Tnp_IS66->MODE-HTH+PBPII-><-HEPN+DUF499*<-METHYLASE<-ParB+HNH<-TUDOR+SF1-SF2-Helicase+iREase+CTD+REase<-HTH+WYL\|\|SF1-SF2-Helicase+REase->SF1-SF2-Helicase+REase-> | 1097 | Bacteria>Pseudomonadota>Alphaproteobacteria>Rhodobacterales>Paracoccaceae | GCF_024359525.1 | Paracoccus_albicereus | MLD63_RS11355 | 50725..54018 | 2922394 | DUF499-domain-containing-protein-[Paracoccus-sp.-TK19116] |
| WP_259776874.1 | DUF499_1.RM.like.systems | REase->?->HTH+WYL->TUDOR+SF1-SF2-Helicase+iREase+CTD+REase->?->METHYLASE->HEPN+DUF499*->ArdCN+MPTase->?->?->?->?->VirD4-TraG+VirD4-FtsK-> | 1112 | Bacteria>Pseudomonadota>Alphaproteobacteria>Rhodobacterales>Paracoccaceae | GCF_025138535.1 | Aliiroseovarius_crassostreae | K3X25_RS03910 | 801884..805222 | 154981 | DUF499-domain-containing-protein-[Aliiroseovarius-crassostreae] |
| WP_090617184.1 | DUF499_1.RM.like.systems | <-HEPN+DUF499*<-METHYLASE<-ASCH<-TUDOR+SF1-SF2-Helicase+iREase+CTD+REase<-HTH+WYL<-?<-TetR-HTH | 1110 | Bacteria>Pseudomonadota>Alphaproteobacteria>Rhodobacterales>Paracoccaceae | GCF_028553725.1 | Paracoccus_alcaliphilus | BMW58_RS19895 | 488..3820 | 34002 | DUF499-domain-containing-protein-[Paracoccus-alcaliphilus] |
| WP_119887324.1 | DUF499_1.RM.like.systems | TUDOR+SF1-SF2-Helicase+iREase+CTD+REase->?->METHYLASE->HEPN+DUF499*-> | 1100 | Bacteria>Pseudomonadota>Alphaproteobacteria>Rhodobacterales>Paracoccaceae | GCF_028553885.1 | Paracoccus_aestuarii | D3P06_RS15025 | 7971..11273 | 453842 | DUF499-domain-containing-protein-[Paracoccus-aestuarii] |
| WP_104009245.1 | DUF499_1.RM.like.systems | XIS-HTH->?->Primpol->TUDOR+SF1-SF2-Helicase+iREase+CTD+REase->ASCH->METHYLASE->HEPN+DUF499*-><-?<-Peptidase_M15<-?<-RNase | 1104 | Bacteria>Pseudomonadota>Alphaproteobacteria>Rhodobacterales>Paracoccaceae | GCF_900108275.1 | Jhaorihella_thermophila | C4S70_RS16725 | 13940..17254 | 488547 | DUF499-domain-containing-protein-[Jhaorihella-thermophila] |
| WP_072748400.1 | DUF499_1.RM.like.systems | <-HEPN+DUF499*<-METHYLASE-p<-METHYLASE<-ASCH<-TUDOR+SF1-SF2-Helicase+iREase+CTD+REase<-?<-tRNA<-HEPN | 1104 | Bacteria>Pseudomonadota>Alphaproteobacteria>Rhodobacterales>Paracoccaceae | GCF_900111545.1 | Oceanicella_actignis | BMW67_RS14465 | 1463..4777 | 1189325 | DUF499-domain-containing-protein-[Oceanicella-actignis] |
| WP_094020052.1 | DUF499_1.RM.like.systems | XIS-HTH->HTH+WYL->ParB+HNH->TUDOR+SF1-SF2-Helicase+iREase+CTD+REase->?->METHYLASE->HEPN+DUF499*-><-RNase<-?<-?<-DUF2163+Phage_BR0599 | 1098 | Bacteria>Pseudomonadota>Alphaproteobacteria>Rhodobacterales>Paracoccaceae | GCF_900184945.1 | Maliponia_aquimaris | MAA8898_RS05760 | 37634..40930 | 1673631 | DUF499-domain-containing-protein-[Maliponia-aquimaris] |
| WP_088233907.1 | DUF499_1.RM.like.systems | <-SF1-SF2-Helicase+REase<-SF1-SF2-Helicase+REase\|\|HTH+WYL->TUDOR+SF1-SF2-Helicase+iREase+CTD+REase->ASCH->METHYLASE->HEPN+DUF499*-><-MODE-HTH+PBPII\|\|TetR-HTH-><-ParG_RHH | 1098 | Bacteria>Pseudomonadota>Alphaproteobacteria>Rhodobacterales>Paracoccaceae | GCF_943915015.1 | Haematobacter_missouriensis | CDV52_RS16275 | 17208..20504 | 366616 | DUF499-domain-containing-protein-[Haematobacter-missouriensis] |
| WP_276946692.1 | DUF499_1.RM.like.systems | ParG_RHH-><-TetR-HTH\|\|MODE-HTH+PBPII-><-HEPN+DUF499*<-METHYLASE<-ASCH<-TUDOR+SF1-SF2-Helicase+iREase+CTD+REase<-HTH+WYL\|\|SMODS->UbE1+JAB-> | 1098 | Bacteria>Pseudomonadota>Alphaproteobacteria>Rhodobacterales>Paracoccaceae | GCF_946222465.1 | Haematobacter_massiliensis | P6966_RS06770 | 8482..11778 | 195105 | DUF499-domain-containing-protein-[Haematobacter-massiliensis] |
| WP_181746701.1 | DUF499_1.RM.like.systems | NAD_binding_2+6PGD-><-?\|\|TUDOR+SF1-SF2-Helicase+iREase+CTD+REase->HEPN->METHYLASE->?->HEPN+DUF499*-><-Enterotoxin_a<-?<-?<-?<-Peptidase_M15 | 1099 | Bacteria>Pseudomonadota>Alphaproteobacteria>Rhodobacterales>Paracoccaceae | GCF_947502345.1 | Paracoccus_sp._S1E-3 | Q0821_RS11150 | 38877..42176 | 2756130 | DUF499-domain-containing-protein-[Paracoccus-sp.-S1E-3] |
| MBA96874.1 | DUF499_1.RM.like.systems | METHYLASE-p->HEPN+DUF499*-><-Glyco_transf_20<-?<-Glyco_transf_20 | 1111 | Bacteria>Pseudomonadota>Alphaproteobacteria>Rhodobacterales>Roseobacteraceae | GCA_002708965.1 | Roseobacter_sp. | CMN14_05170 | 1105..4440 | 1907202 | AAA-family-ATPase-[Roseobacter-sp.] |
| MBV47914.1 | DUF499_1.RM.like.systems | METHYLASE-p->HEPN+DUF499*-><-Glyco_transf_20<-?<-Glyco_transf_20 | 1111 | Bacteria>Pseudomonadota>Alphaproteobacteria>Rhodobacterales>Roseobacteraceae | GCA_002731875.1 | Roseobacter_sp. | CMN10_05040 | 1113..4448 | 1907202 | AAA-family-ATPase-[Roseobacter-sp.] |
| PWJ13798.1 | DUF499_1.RM.like.systems | MODE-HTH+PBPII->tRNA->TUDOR+SF1-SF2-Helicase+iREase+CTD+REase->HEPN->METHYLASE-p->HEPN+DUF499*->?-><-?<-?<-?<-?<-XIS-HTH | 1109 | Bacteria>Pseudomonadota>Alphaproteobacteria>Rhodobacterales>Roseobacteraceae | GCA_003149265.1 | Jannaschia_seohaensis | BCF38_11329 | 37396..40725 | 475081 | hypothetical-protein-BCF38_11329-[Jannaschia-seohaensis] |
| HCP81323.1 | DUF499_1.RM.like.systems | TUDOR+SF1-SF2-Helicase+iREase+CTD+REase->HEPN->METHYLASE->HEPN+DUF499*-><-?<-?<-Peptidase_M15 | 1102 | Bacteria>Pseudomonadota>Alphaproteobacteria>Rhodobacterales>Roseobacteraceae | GCA_003533975.1 | Octadecabacter_sp. | DIT67_06940 | 7896..11204 | 1920174 | AAA-family-ATPase-[Octadecabacter-sp.] |
| MBR2572727.1 | DUF499_1.RM.like.systems | <-ParB+N6_N4_MTase<-ParB+N6_N4_MTase\|\|TUDOR+SF1-SF2-Helicase+iREase+CTD+REase->ASCH->METHYLASE->HEPN+DUF499*-><-Enterotoxin_a<-DNA_methylase<-MarR-HTH+REase<-?<-?<-XIS-HTH | 1096 | Bacteria>Pseudomonadota>Alphaproteobacteria>Rhodobacterales>Roseobacteraceae | GCA_017542965.1 | Loktanella_sp. | IKE14_00205 | 39854..43144 | 1870901 | DUF499-domain-containing-protein-[Loktanella-sp.] |
| MCI5043002.1 | DUF499_1.RM.like.systems | <-HEPN+DUF499*<-METHYLASE<-?<-TUDOR+SF1-SF2-Helicase+iREase+CTD+REase<-HTH+WYL | 1111 | Bacteria>Pseudomonadota>Alphaproteobacteria>Rhodobacterales>Roseobacteraceae | GCA_022763195.1 | Donghicola_eburneus | MRY81_25480 | 833..4168 | 393278 | DUF499-domain-containing-protein-[Donghicola-eburneus] |
| WP_007798461.1 | DUF499_1.RM.like.systems | <-Peptidase_S11<-?\|\|tRNA->TUDOR+SF1-SF2-Helicase+iREase+CTD+REase->TUDOR+SF1-SF2-Helicase+iREase+CTD+REase-p->?->METHYLASE->HEPN+DUF499*-><-DDE_Tnp_1<-SF1-SF2-Helicase+DUF1998+HKD-DNase+HKD-DNase<-?<-?\|\|?-><-MULE-TRANSPOSASE | 1110 | Bacteria>Pseudomonadota>Alphaproteobacteria>Rhodobacterales>Roseobacteraceae | GCF_000153725.1 | Salipiger_bermudensis | R2601_RS11735 | 2461039..2464371 | 344736 | DUF499-domain-containing-protein-[Salipiger-bermudensis] |
| WP_026758864.1 | DUF499_1.RM.like.systems | GNTR-HTH->?->?-><-HEPN+DUF499*<-METHYLASE<-METHYLASE-p<-PadR-HTH<-TUDOR+SF1-SF2-Helicase+iREase+CTD+REase\|\|MarR-HTH->ABhydrolase-> | 1111 | Bacteria>Pseudomonadota>Alphaproteobacteria>Rhodobacterales>Roseobacteraceae | GCF_000423645.1 | Sediminimonas_qiaohouensis | G568_RS0115420 | 9768..13103 | 552061 | DUF499-domain-containing-protein-[Sediminimonas-qiaohouensis] |
| WP_050673791.1 | DUF499_1.RM.like.systems | Peptidase_M15->?->HTH+WYL->TUDOR+SF1-SF2-Helicase+iREase+CTD+REase->?->METHYLASE->HEPN+DUF499*->ArdCN+MPTase->?->?->?->?->VirD4-TraG+VirD4-FtsK-> | 1112 | Bacteria>Pseudomonadota>Alphaproteobacteria>Rhodobacterales>Roseobacteraceae | GCF_001251095.1 | Phaeobacter_italicus | NIT7321_RS13650 | 220898..224236 | 481446 | DUF499-domain-containing-protein-[Phaeobacter-italicus] |
| WP_068316335.1 | DUF499_1.RM.like.systems | <-HIN-HTH<-TPR<-HEPN+SF1-SF2-Helicase<-HIN-HTH\|\|?-><-AbiEii<-HEPN+DUF499*<-METHYLASE<-ASCH<-TUDOR+SF1-SF2-Helicase+iREase+CTD+REase<-Arm-DNA-bind_3+Phage_int_SAM_3+Phage_integrase\|\|TPR-> | 1099 | Bacteria>Pseudomonadota>Alphaproteobacteria>Rhodobacterales>Roseobacteraceae | GCF_001558155.1 | Aliiruegeria_sabulilitoris | AX756_RS20370 | 52266..55565 | 1510458 | DUF499-domain-containing-protein-[Aliiruegeria-sabulilitoris] |
| WP_109663884.1 | DUF499_1.RM.like.systems | VirD4-TraG+VirD4-FtsK->CopG/RHH->TUDOR+SF1-SF2-Helicase+iREase+CTD+REase->HKD-DNase->METHYLASE->HEPN+DUF499*-><-PIN<-RHH<-?<-CHTH\|\|VirB11+VirB4-FtsK->VirB2-> | 1100 | Bacteria>Pseudomonadota>Alphaproteobacteria>Rhodobacterales>Roseobacteraceae | GCF_003148775.1 | Roseicyclus_mahoneyensis | C7455_RS00150 | 34589..37891 | 164332 | DUF499-domain-containing-protein-[Roseicyclus-mahoneyensis] |
| WP_245947652.1 | DUF499_1.RM.like.systems | MODE-HTH+PBPII->tRNA->TUDOR+SF1-SF2-Helicase+iREase+CTD+REase->HEPN->METHYLASE->HEPN+DUF499*-> | 1097 | Bacteria>Pseudomonadota>Alphaproteobacteria>Rhodobacterales>Roseobacteraceae | GCF_003149265.1 | Jannaschia_seohaensis | BCF38_RS18825 | 37432..40725 | 475081 | DUF499-domain-containing-protein-[Jannaschia-seohaensis] |
| WP_240672187.1 | DUF499_1.RM.like.systems | TUDOR+SF1-SF2-Helicase+iREase+CTD+REase->PadR-HTH->METHYLASE->HEPN+DUF499*-><-?<-?<-?<-?<-ArsR-HTH+YTCD-HTH | 1098 | Bacteria>Pseudomonadota>Alphaproteobacteria>Rhodobacterales>Roseobacteraceae | GCF_003667315.2 | Tropicibacter_alexandrii | D8Z78_RS18955 | 49995..53291 | 2267683 | DUF499-domain-containing-protein-[Tropicibacter-sp.-LMIT003] |
| WP_170364042.1 | DUF499_1.RM.like.systems | Primpol->?->?->?-><-RNase<-REase<-HEPN+DUF499*<-METHYLASE<-TUDOR+SF1-SF2-Helicase+iREase+CTD+REase | 1107 | Bacteria>Pseudomonadota>Alphaproteobacteria>Rhodobacterales>Roseobacteraceae | GCF_013030285.1 | Ruegeria_arenilitoris | GOX59_RS08670 | 247066..250389 | 1173585 | DUF499-domain-containing-protein-[Ruegeria-arenilitoris] |
| WP_170755622.1 | DUF499_1.RM.like.systems | <-REase<-DNA_methylase\|\|HNH-><-?\|\|TUDOR+SF1-SF2-Helicase+iREase+CTD+REase->METHYLASE->HEPN+DUF499*->?->?->?-><-?<-DnaB+CITB-HTH | 1101 | Bacteria>Pseudomonadota>Alphaproteobacteria>Rhodobacterales>Roseobacteraceae | GCF_013031285.1 | Ruegeria_lacuscaerulensis | GOX67_RS18140 | 83323..86628 | 55218 | DUF499-domain-containing-protein-[Ruegeria-lacuscaerulensis] |
| WP_170518600.1 | DUF499_1.RM.like.systems | <-HEPN+DUF499*<-METHYLASE<-ASCH<-TUDOR+SF1-SF2-Helicase+iREase+CTD+REase<-HEPN<-REase | 1103 | Bacteria>Pseudomonadota>Alphaproteobacteria>Rhodobacterales>Roseobacteraceae | GCF_013032825.1 | Ruegeria_atlantica | GGW75_RS18355 | 50430..53741 | 81569 | DUF499-domain-containing-protein-[Ruegeria-atlantica] |
| WP_171727684.1 | DUF499_1.RM.like.systems | HEPN-><-REase<-HEPN+DUF499*<-METHYLASE<-?<-TUDOR+SF1-SF2-Helicase+iREase+CTD+REase | 1108 | Bacteria>Pseudomonadota>Alphaproteobacteria>Rhodobacterales>Roseobacteraceae | GCF_013110755.1 | Ruegeria_sp._HKCCD7318 | GS627_RS09025 | 215066..218392 | 2683014 | DUF499-domain-containing-protein-[Ruegeria-sp.-HKCCD7318] |
| WP_189682671.1 | DUF499_1.RM.like.systems | <-HicB\|\|HTH+WYL->TUDOR+SF1-SF2-Helicase+iREase+CTD+REase->MORC+DUF3684->METHYLASE->HEPN+DUF499*-><-Phage_integrase<-?<-?\|\|?->XIS-HTH-><-Phage_integrase | 1103 | Bacteria>Pseudomonadota>Alphaproteobacteria>Rhodobacterales>Roseobacteraceae | GCF_014656415.1 | Seohaeicola_zhoushanensis | IE266_RS23965 | 15420..18731 | 1569283 | DUF499-domain-containing-protein-[Seohaeicola-zhoushanensis] |
| WP_216044688.1 | DUF499_1.RM.like.systems | HTH+WYL->TUDOR+SF1-SF2-Helicase+iREase+CTD+REase->PadR-HTH->METHYLASE->HEPN+DUF499*-> | 1110 | Bacteria>Pseudomonadota>Alphaproteobacteria>Rhodobacterales>Roseobacteraceae | GCF_018860305.1 | Celeribacter_halophilus | KO492_RS09190 | 8991..12323 | 576117 | DUF499-domain-containing-protein-[Celeribacter-halophilus] |
| WP_219436397.1 | DUF499_1.RM.like.systems | HEPN+DUF499*-><-MODE-HTH+PBPII\|\|TetR-HTH-><-ParG_RHH | 1098 | Bacteria>Pseudomonadota>Alphaproteobacteria>Rhodobacterales>Roseobacteraceae | GCF_019392295.1 | Roseovarius_mucosus | KZZ08_RS20795 | 1456..4752 | 215743 | DUF499-domain-containing-protein-[Roseovarius-mucosus] |
| WP_090219105.1 | DUF499_1.RM.like.systems | <-ArdCN+MPTase\|\|TUDOR+SF1-SF2-Helicase+iREase+CTD+REase->?->METHYLASE->HEPN+DUF499*-><-?<-STYKinase-like<-HTH | 1105 | Bacteria>Pseudomonadota>Alphaproteobacteria>Rhodobacterales>Roseobacteraceae | GCF_900111945.1 | Litoreibacter_janthinus | BM352_RS16595 | 3306785..3310102 | 670154 | DUF499-domain-containing-protein-[Litoreibacter-janthinus] |
| WP_093359600.1 | DUF499_1.RM.like.systems | XIS-HTH->HTH+WYL->ParB+HNH->TUDOR+SF1-SF2-Helicase+iREase+CTD+REase->ASCH->METHYLASE->HEPN+DUF499*->RNase+SF1-SF2-Helicase-><-RNase<-?<-?<-DUF2163+Phage_BR0599 | 1109 | Bacteria>Pseudomonadota>Alphaproteobacteria>Rhodobacterales>Roseobacteraceae | GCF_900112335.1 | Tropicimonas_isoalkanivorans | BM075_RS05220 | 205092..208421 | 441112 | DUF499-domain-containing-protein-[Tropicimonas-isoalkanivorans] |
| MBF9033364.1 | DUF499_1.RM.like.systems | XIS-HTH->HTH+WYL->ParB+HNH->TUDOR+SF1-SF2-Helicase+iREase+CTD+REase->YtH->METHYLASE->HEPN+DUF499*->AbiEii-><-?<-Peptidase_M15 | 1098 | Bacteria>Pseudomonadota>Alphaproteobacteria>Rhodobacterales>unclassified_Rhodobacterales | GCA_015689685.1 | Rhodobacterales_bacterium_HKCCE2091 | HKCCE2091_03860 | 312307..315603 | 2792619 | DUF499-domain-containing-protein-[Rhodobacterales-bacterium-HKCCE2091] |
| QPI86373.1 | DUF499_1.RM.like.systems | TUDOR+SF1-SF2-Helicase+iREase+CTD+REase->METHYLASE->HEPN+DUF499*-><-HIN-HTH\|\|?->?-><-OBFOLD-NUCLEASE<-Aminotran_1_2 | 1111 | Bacteria>Pseudomonadota>Alphaproteobacteria>Rhodobacterales>unclassified_Rhodobacterales | GCA_015693905.1 | Rhodobacterales_bacterium_HKCCA1288 | I3V23_05250 | 1074611..1077946 | 2899152 | DUF499-domain-containing-protein-[Rhodobacterales-bacterium-HKCCA1288] |
| OZB26691.1 | DUF499_1.RM.like.systems | TUDOR+SF1-SF2-Helicase+iREase+CTD+REase->?->PIN->Ntox18->?->METHYLASE->HEPN+DUF499*->\|\|FAD-NAD-dep-oxidoreductase-><-NACHT+REase\|\|?->DnaB-><-NACHT+REase | 1091 | Bacteria>Pseudomonadota>Alphaproteobacteria>Rhodospirillales>Acetobacteraceae | GCA_002282635.1 | Acidiphilium_sp._34-64-41 | B7X49_12125 | 19315..22590 | 1970297 | MAG:-ATPase-[Acidiphilium-sp.-34-64-41] |
| NHO42826.1 | DUF499_1.RM.like.systems | HTH+WYL->TUDOR+SF1-SF2-Helicase+iREase+CTD+REase->?->METHYLASE->HEPN+DUF499*->VirB11->GTPase-AIG-> | 1097 | Bacteria>Pseudomonadota>Alphaproteobacteria>Rhodospirillales>Acetobacteraceae | GCA_011516925.1 | Acetobacter_fabarum | GOB83_11655 | 8419..11712 | 483199 | DUF499-domain-containing-protein-[Acetobacter-fabarum] |
| MBB3175065.1 | DUF499_1.RM.like.systems | rve_3->TUDOR+SF1-SF2-Helicase+iREase+CTD+REase->METHYLASE->HEPN+DUF499*->HTH_Tnp_1->?->?-><-HIN-HTH<-HIN-HTH<-HIN-HTH | 1110 | Bacteria>Pseudomonadota>Alphaproteobacteria>Rhodospirillales>Acetobacteraceae | GCA_014192375.1 | Endobacter_medicaginis | FHR90_002912 | 7080..10412 | 1181271 | hypothetical-protein-[Endobacter-medicaginis] |
| MBX6741130.1 | DUF499_1.RM.like.systems | METHYLASE-p->HEPN+DUF499*->?->?->?->?-><-SF1-SF2-Helicase+REase | 1108 | Bacteria>Pseudomonadota>Alphaproteobacteria>Rhodospirillales>Acetobacteraceae | GCA_019690105.1 | Acetobacteraceae_bacterium | IRY87_03665 | 460..3786 | 1909293 | DUF499-domain-containing-protein-[Acetobacteraceae-bacterium] |
| MBX6741668.1 | DUF499_1.RM.like.systems | TUDOR+SF1-SF2-Helicase+iREase+CTD+REase-p->ASCH->METHYLASE->HEPN->HEPN+DUF499*-> | 1110 | Bacteria>Pseudomonadota>Alphaproteobacteria>Rhodospirillales>Acetobacteraceae | GCA_019690105.1 | Acetobacteraceae_bacterium | IRY87_06425 | 5783..9115 | 1909293 | DUF499-domain-containing-protein-[Acetobacteraceae-bacterium] |
| MBX6743929.1 | DUF499_1.RM.like.systems | <-HEPN+DUF499*<-HEPN<-METHYLASE-p | 1111 | Bacteria>Pseudomonadota>Alphaproteobacteria>Rhodospirillales>Acetobacteraceae | GCA_019690105.1 | Acetobacteraceae_bacterium | IRY87_17990 | 402..3737 | 1909293 | DUF499-domain-containing-protein-[Acetobacteraceae-bacterium] |
| MBX9595850.1 | DUF499_1.RM.like.systems | MerR-HTH->TUDOR+SF1-SF2-Helicase+iREase+CTD+REase->HKD-DNase->METHYLASE->HEPN+DUF499*->MACRODOMAIN-><-HTH<-GDResolvase+Recombinase+ZnR | 1110 | Bacteria>Pseudomonadota>Alphaproteobacteria>Rhodospirillales>Acetobacteraceae | GCA_019748015.1 | Roseomonas_sp. | K2X46_15915 | 11392..14724 | 1869190 | DUF499-domain-containing-protein-[Roseomonas-sp.] |
| MCA3363599.1 | DUF499_1.RM.like.systems | METHYLASE->OLD-ABC->PIN->HEPN+DUF499*-> | 483 | Bacteria>Pseudomonadota>Alphaproteobacteria>Rhodospirillales>Acetobacteraceae | GCA_020249535.1 | Roseomonas_sp. | INF74_16815 | 3977..5424 | 1869190 | DUF499-domain-containing-protein-[Roseomonas-sp.] |
| MCA7120529.1 | DUF499_1.RM.like.systems | TUDOR+SF1-SF2-Helicase+iREase+CTD+REase->?->PIN->Ntox18->Peptidase_M78->METHYLASE->HEPN+DUF499*->?->?->HHH_5+SF1-SF2-helicase-><-RVT+HNH\|\|<-Phage_int_SAM_1+Phage_integrase<-Phage_integrase | 1113 | Bacteria>Pseudomonadota>Alphaproteobacteria>Rhodospirillales>Acetobacteraceae | GCA_020418435.1 | Acidibrevibacterium_fodinaquatile | LGL72_14265 | 2856594..2859935 | 1969806 | DUF499-domain-containing-protein-[Acidibrevibacterium-sp.] |
| MCH4022379.1 | DUF499_1.RM.like.systems | <-REase+CTD6\|\|HTH+WYL->TUDOR+SF1-SF2-Helicase+iREase+CTD+REase->?->METHYLASE->HEPN+DUF499*->?-><-?<-LexA-protease | 1097 | Bacteria>Pseudomonadota>Alphaproteobacteria>Rhodospirillales>Acetobacteraceae | GCA_022483365.1 | Acetobacter_sp. | LKE62_03305 | 718063..721356 | 440 | DUF499-domain-containing-protein-[Acetobacter-sp.] |
| WP_012553516.1 | DUF499_1.RM.like.systems | <-VirB2<-VirB11+VirB4-FtsK\|\|?->REase->?-><-HEPN+DUF499*<-METHYLASE<-TUDOR+SF1-SF2-Helicase+iREase+CTD+REase<-PIN<-?<-CopG/RHH<-VirD4-TraG+VirD4-FtsK | 1108 | Bacteria>Pseudomonadota>Alphaproteobacteria>Rhodospirillales>Acetobacteraceae | GCF_000021325.1 | Gluconacetobacter_diazotrophicus | GDIA_RS05030 | 1143198..1146524 | 33996 | DUF499-domain-containing-protein-[Gluconacetobacter-diazotrophicus] |
| WP_010666859.1 | DUF499_1.RM.like.systems | VirD4-TraG+VirD4-FtsK->CopG/RHH->?->PIN->TUDOR+SF1-SF2-Helicase+iREase+CTD+REase->METHYLASE->HEPN+DUF499*-><-?<-REase\|\|VirB11+VirB4-FtsK->VirB2->?->FtsK+VirB4-FtsK-> | 1108 | Bacteria>Pseudomonadota>Alphaproteobacteria>Rhodospirillales>Acetobacteraceae | GCF_000193495.2 | Acetobacter_aceti | EDC15_RS12660 | 41147..44473 | 435 | DUF499-domain-containing-protein-[Acetobacter-aceti] |
| WP_199897436.1 | DUF499_1.RM.like.systems | <-DUF499-p<-HEPN+DUF499*<-METHYLASE<-ASCH<-?<-TUDOR+SF1-SF2-Helicase+iREase+CTD+REase<-HTH+WYL<-METHYLASE-p | 1018 | Bacteria>Pseudomonadota>Alphaproteobacteria>Rhodospirillales>Acetobacteraceae | GCF_000613905.1 | Acetobacter_persici | JCM25330_RS21105 | 22243..25299 | 1076596 | DUF499-domain-containing-protein-[Acetobacter-persici] |
| WP_062273923.1 | DUF499_1.RM.like.systems | PIN->TUDOR+SF1-SF2-Helicase+iREase+CTD+REase->METHYLASE->HEPN+DUF499*-><-?<-?<-Radical-SAM<-PIN<-Phd-YefM | 1108 | Bacteria>Pseudomonadota>Alphaproteobacteria>Rhodospirillales>Acetobacteraceae | GCF_001581105.1 | Acetobacter_cerevisiae | AD954_RS12075 | 12356..15682 | 178900 | DUF499-domain-containing-protein-[Acetobacter-cerevisiae] |
| WP_114912917.1 | DUF499_1.RM.like.systems | Ntox18->Peptidase_M78->METHYLASE-><-HTH_Tnp_1+HTH_21+rve+rve_3\|\|MULE-TRANSPOSASE->METHYLASE-p->HEPN+DUF499*->TPR->?->?->HHH_5+SF1-SF2-helicase->?->PAIREDC-HTH-> | 1113 | Bacteria>Pseudomonadota>Alphaproteobacteria>Rhodospirillales>Acetobacteraceae | GCF_003352165.1 | Acidibrevibacterium_fodinaquatile | DEF76_RS14375 | 3023042..3026383 | 1969806 | DUF499-domain-containing-protein-[Acidibrevibacterium-fodinaquatile] |
| WP_132287513.1 | DUF499_1.RM.like.systems | TUDOR+SF1-SF2-Helicase+iREase+CTD+REase->?->METHYLASE->HEPN+DUF499*-><-?\|\|REase-><-HTH<-GDResolvase+Recombinase+ZnR<-RAMA | 1114 | Bacteria>Pseudomonadota>Alphaproteobacteria>Rhodospirillales>Acetobacteraceae | GCF_004343615.1 | Roseicella_aquatilis | EXY23_RS09345 | 90993..94337 | 2527868 | Swt1-family-HEPN-domain-containing-protein-[Paracraurococcus-sp.-NE82] |
| WP_135468376.1 | DUF499_1.RM.like.systems | <-LRP-HTH<-DnaJ<-?\|\|tRNA->TUDOR+SF1-SF2-Helicase+iREase+CTD+REase->?->METHYLASE->HEPN+DUF499*->ArdCN+MPTase-><-HTH<-GDResolvase+Recombinase+ZnR<-RAMA | 1112 | Bacteria>Pseudomonadota>Alphaproteobacteria>Rhodospirillales>Acetobacteraceae | GCF_004761865.1 | Crenalkalicoccus_roseus | ET526_RS14250 | 21738..25076 | 1485588 | DUF499-domain-containing-protein-[Crenalkalicoccus-roseus] |
| WP_153467943.1 | DUF499_1.RM.like.systems | <-GTPase-AIG<-VirB11\|\|<-VirB11<-DUF499-p<-HIN-HTH<-DUF4096+DDE_Tnp_1+DDE_Tnp_1_2<-HEPN+DUF499*<-METHYLASE<-DUF4096+DDE_Tnp_1+DDE_Tnp_1_2<-METHYLASE-p<-TUDOR+SF1-SF2-Helicase+iREase+CTD+REase<-HTH+WYL | 1047 | Bacteria>Pseudomonadota>Alphaproteobacteria>Rhodospirillales>Acetobacteraceae | GCF_009268135.1 | Komagataeibacter_medellinensis | D3W54_RS02225 | 408174..411317 | 1177712 | DUF499-domain-containing-protein-[Komagataeibacter-medellinensis] |
| WP_152832805.1 | DUF499_1.RM.like.systems | PIN->TUDOR+SF1-SF2-Helicase+iREase+CTD+REase->METHYLASE-p->METHYLASE-p->HEPN+DUF499*-><-PhdYeFM_antitox<-PIN<-Phd-YefM<-GDResolvase+Recombinase+ZnR<-XIS-HTH | 1108 | Bacteria>Pseudomonadota>Alphaproteobacteria>Rhodospirillales>Acetobacteraceae | GCF_009295745.1 | Acetobacter_senegalensis | GC248_RS02655 | 30596..33922 | 446692 | DUF499-domain-containing-protein-[Acetobacter-senegalensis] |
| WP_194301389.1 | DUF499_1.RM.like.systems | HTH+WYL->TUDOR+SF1-SF2-Helicase+iREase+CTD+REase->?->METHYLASE->HEPN+DUF499*->VirB11->GTPase-AIG-> | 1085 | Bacteria>Pseudomonadota>Alphaproteobacteria>Rhodospirillales>Acetobacteraceae | GCF_011516925.1 | Acetobacter_fabarum | GOB83_RS11695 | 8455..11712 | 483199 | DUF499-domain-containing-protein-[Acetobacter-fabarum] |
| WP_171617018.1 | DUF499_1.RM.like.systems | Phage_integrase-><-?\|\|TUDOR+SF1-SF2-Helicase+iREase+CTD+REase->?->METHYLASE->HEPN+DUF499*->HIN-HTH->?->LZ_Tnp_IS66+zf-IS66+DDE_Tnp_IS66+DDE_Tnp_IS66_C-><-DUF4158+DDE_Tnp_Tn3\|\|GDResolvase+HIN-HTH-> | 1113 | Bacteria>Pseudomonadota>Alphaproteobacteria>Rhodospirillales>Acetobacteraceae | GCF_013112485.1 | Roseicella_sp._DB1501 | HJG45_RS27180 | 18137..21478 | 2730925 | DUF499-domain-containing-protein-[Roseicella-sp.-DB1501] |
| WP_246330294.1 | DUF499_1.RM.like.systems | HIN-HTH-><-?<-HTH_Tnp_1<-HEPN+DUF499*<-METHYLASE<-TUDOR+SF1-SF2-Helicase+iREase+CTD+REase | 1093 | Bacteria>Pseudomonadota>Alphaproteobacteria>Rhodospirillales>Acetobacteraceae | GCF_014192375.1 | Endobacter_medicaginis | OR615_RS11695 | 24590..27871 | 1181271 | DUF499-domain-containing-protein-[Endobacter-medicaginis] |
| WP_211861077.1 | DUF499_1.RM.like.systems | TUDOR+SF1-SF2-Helicase+iREase+CTD+REase->HKD-DNase->?->METHYLASE->?->HEPN+DUF499*-><-HEPN+REase<-HTH<-GDResolvase+Recombinase+ZnR<-RAMA<-?<-HTH | 1110 | Bacteria>Pseudomonadota>Alphaproteobacteria>Rhodospirillales>Acetobacteraceae | GCF_018129065.1 | Neoroseomonas_soli | GXW76_RS05955 | 22990..26322 | 1081025 | DUF499-domain-containing-protein-[Roseomonas-soli] |
| WP_216840234.1 | DUF499_1.RM.like.systems | TUDOR+SF1-SF2-Helicase+iREase+CTD+REase->HKD-DNase->METHYLASE->HEPN+DUF499*-><-HTH<-GDResolvase+Recombinase+ZnR<-RAMA | 1109 | Bacteria>Pseudomonadota>Alphaproteobacteria>Rhodospirillales>Acetobacteraceae | GCF_019024325.1 | Falsiroseomonas_tokyonensis | J3S20_RS28040 | 39158..42487 | 430521 | DUF499-domain-containing-protein-[Falsiroseomonas-tokyonensis] |
| WP_274966108.1 | DUF499_1.RM.like.systems | TUDOR+SF1-SF2-Helicase+iREase+CTD+REase->?->PIN->Ntox18->Peptidase_M78->METHYLASE->HEPN+DUF499*->?->HHH_5+SF1-SF2-helicase-><-RVT+HNH\|\|<-Phage_int_SAM_1+Phage_integrase<-Phage_integrase<-Phage_integrase | 1113 | Bacteria>Pseudomonadota>Alphaproteobacteria>Rhodospirillales>Acetobacteraceae | GCF_020418435.1 | Acidibrevibacterium_fodinaquatile | LGL72_RS14205 | 2856594..2859935 | 1969806 | DUF499-domain-containing-protein-[Acidibrevibacterium-fodinaquatile] |
| WP_253545427.1 | DUF499_1.RM.like.systems | <-TraJ-RHH<-Phage_integrase\|\|?->PIN->TUDOR+SF1-SF2-Helicase+iREase+CTD+REase->METHYLASE->HEPN+DUF499*->?-><-?<-?<-?\|\|TrwC+SF1-SF2-helicase+TOPRIM-> | 1108 | Bacteria>Pseudomonadota>Alphaproteobacteria>Rhodospirillales>Acetobacteraceae | GCF_024158445.1 | Acetobacter_lovaniensis | NKW44_RS07985 | 22199..25525 | 104100 | DUF499-domain-containing-protein-[Acetobacter-lovaniensis] |
| WP_018981014.1 | DUF499_1.RM.like.systems | <-SF1-SF2-Helicase<-?<-?<-ABhydrolase<-?<-HEPN+DUF499*<-METHYLASE<-TUDOR+SF1-SF2-Helicase+iREase+CTD+REase | 1099 | Bacteria>Pseudomonadota>Alphaproteobacteria>Rhodospirillales>Acetobacteraceae | GCF_025994295.1 | Saccharibacter_floricola | F457_RS0110055 | 41499..44798 | 231053 | DUF499-domain-containing-protein-[Saccharibacter-floricola] |
| WP_045542211.1 | DUF499_1.RM.like.systems | METHYLASE-p->HEPN+DUF499*-><-?<-?<-Radical-SAM<-PIN<-Phd-YefM\|\|Phage_integrase-> | 1108 | Bacteria>Pseudomonadota>Alphaproteobacteria>Rhodospirillales>Acetobacteraceae | GCF_025995375.1 | Acetobacter_tropicalis | AD944_RS02385 | 1113..4439 | 104102 | DUF499-domain-containing-protein-[Acetobacter-tropicalis] |
| WP_165926191.1 | DUF499_1.RM.like.systems | <-GTPase-AIG<-VirB11<-HEPN+DUF499*<-METHYLASE<-?<-TUDOR+SF1-SF2-Helicase+iREase+CTD+REase-p | 1097 | Bacteria>Pseudomonadota>Alphaproteobacteria>Rhodospirillales>Acetobacteraceae | GCF_025995875.1 | Acetobacter_fabarum | AA19596_RS12715 | 13783..17076 | 483199 | DUF499-domain-containing-protein-[Acetobacter-fabarum] |
| WP_174502310.1 | DUF499_1.RM.like.systems | NUDIX->ABhydrolase->?->Arm-DNA-bind_3+Phage_integrase-><-Phage_AlpA<-HEPN+DUF499*<-METHYLASE<-MORC+DUF3684<-TUDOR+SF1-SF2-Helicase+iREase+CTD+REase<-HTH+WYL\|\|MODE-HTH+PBPII-><-ParG_RHH | 1110 | Bacteria>Pseudomonadota>Alphaproteobacteria>Rhodospirillales>Acetobacteraceae | GCF_902712915.1 | Acidiphilium_sp._C61 | HUW10_RS03970 | 224221..227553 | 1671485 | Swt1-family-HEPN-domain-containing-protein-[Acidiphilium-sp.-C61] |
| TWB14201.1 | DUF499_1.RM.like.systems | <-SF1-SF2-Helicase+REase\|\|HTH+WYL->TUDOR+SF1-SF2-Helicase+iREase+CTD+REase->YtH->MORC+DUF3684->METHYLASE-p->HEPN+DUF499*->MODE-HTH+PBPII-><-ParG_RHH | 1098 | Bacteria>Pseudomonadota>Alphaproteobacteria>Rhodospirillales>Azospirillaceae | GCA_007828045.1 | Nitrospirillum_amazonense | FBZ89_11767 | 103300..106596 | 28077 | hypothetical-protein-FBZ89_11767-[Nitrospirillum-amazonense] |
| WP_014199726.1 | DUF499_1.RM.like.systems | MODE-HTH+PBPII-><-Arm-DNA-bind_3+Phage_int_SAM_3+Phage_integrase<-?<-HEPN+DUF499*<-HEPN<-PolB<-METHYLASE<-HNH<-OLD-ABC<-TUDOR+SF1-SF2-Helicase+iREase+CTD+REase | 1115 | Bacteria>Pseudomonadota>Alphaproteobacteria>Rhodospirillales>Azospirillaceae | GCF_000237365.1 | Azospirillum | GAY33_RS05480 | 11187..14534 | 191 | MULTISPECIES:-Swt1-family-HEPN-domain-containing-protein-[Azospirillum] |
| WP_045580668.1 | DUF499_1.RM.like.systems | <-SF1-SF2-Helicase+REase\|\|TUDOR+SF1-SF2-Helicase+iREase+CTD+REase->METHYLASE->?->HEPN+DUF499*-><-?<-?<-LZ_Tnp_IS66+zf-IS66+DDE_Tnp_IS66+DDE_Tnp_IS66_C<-?<-HIN-HTH | 1118 | Bacteria>Pseudomonadota>Alphaproteobacteria>Rhodospirillales>Azospirillaceae | GCF_001305595.1 | Azospirillum_thiophilum | VY88_RS04045 | 887405..890761 | 528244 | Swt1-family-HEPN-domain-containing-protein-[Azospirillum-thiophilum] |
| WP_063633927.1 | DUF499_1.RM.like.systems | <-GDResolvase+HIN-HTH\|\|?-><-HEPN+DUF499*<-METHYLASE<-ASCH<-Cluster1090_5clades<-TUDOR+SF1-SF2-Helicase+iREase+CTD+REase<-HTH+WYL | 1115 | Bacteria>Pseudomonadota>Alphaproteobacteria>Rhodospirillales>Azospirillaceae | GCF_001639105.2 | Azospirillum_humicireducens | A6A40_RS02305 | 493921..497268 | 1226968 | Swt1-family-HEPN-domain-containing-protein-[Azospirillum-humicireducens] |
| WP_109152948.1 | DUF499_1.RM.like.systems | <-VirB4-FtsK+HerA<-?\|\|TUDOR+SF1-SF2-Helicase+iREase+CTD+REase->Cluster1090_5clades->ASCH->METHYLASE->HEPN+DUF499*->Lon_C-> | 1116 | Bacteria>Pseudomonadota>Alphaproteobacteria>Rhodospirillales>Azospirillaceae | GCF_003116065.1 | Azospirillum_sp._TSO5 | TSO5_RS18610 | 11435..14785 | 716760 | Swt1-family-HEPN-domain-containing-protein-[Azospirillum-sp.-TSO5] |
| WP_145610986.1 | DUF499_1.RM.like.systems | TUDOR+SF1-SF2-Helicase+iREase+CTD+REase->ParB+HNH->METHYLASE->PolB->HEPN->HEPN+DUF499*->?->PIN-> | 1115 | Bacteria>Pseudomonadota>Alphaproteobacteria>Rhodospirillales>Azospirillaceae | GCF_007828035.1 | Nitrospirillum_amazonense | FBZ87_RS14570 | 656187..659534 | 28077 | Swt1-family-HEPN-domain-containing-protein-[Nitrospirillum-amazonense] |
| WP_246172460.1 | DUF499_1.RM.like.systems | <-SF1-SF2-Helicase+REase\|\|HTH+WYL->TUDOR+SF1-SF2-Helicase+iREase+CTD+REase->YtH->MORC+DUF3684->METHYLASE->HEPN+DUF499*->MODE-HTH+PBPII-><-ParG_RHH | 1086 | Bacteria>Pseudomonadota>Alphaproteobacteria>Rhodospirillales>Azospirillaceae | GCF_007828045.1 | Nitrospirillum_amazonense | FBZ89_RS22360 | 103336..106596 | 28077 | DUF499-domain-containing-protein-[Nitrospirillum-amazonense] |
| WP_147041135.1 | DUF499_1.RM.like.systems | TPR->?->TUDOR+SF1-SF2-Helicase+iREase+CTD+REase->METHYLASE->HEPN+DUF499*->OLD-ABC+TOPRIM-><-RelE-ParE<-MetJarc-HTH | 1097 | Bacteria>Pseudomonadota>Alphaproteobacteria>Rhodospirillales>Azospirillaceae | GCF_007991775.1 | Skermanella_aerolata | SAE02_RS33460 | 15952..19245 | 393310 | DUF499-domain-containing-protein-[Skermanella-aerolata] |
| WP_246149069.1 | DUF499_1.RM.like.systems | REase->TUDOR+SF1-SF2-Helicase+iREase+CTD+REase->ASCH->ASCH->METHYLASE->METHYLASE-p->HEPN+DUF499*->DUF499-p-><-?<-DDE_Tnp_1\|\|REase-> | 926 | Bacteria>Pseudomonadota>Alphaproteobacteria>Rhodospirillales>Azospirillaceae | GCF_008843145.1 | Skermanella_pratensis | DPR14_RS09205 | 2041466..2044246 | 2233999 | DUF499-domain-containing-protein-[Skermanella-pratensis] |
| WP_180283596.1 | DUF499_1.RM.like.systems | <-PfkB\|\|DeoR-HTH+ISOCOT->Transposase_31+Yae1_N-><-HEPN+DUF499*<-PIN<-OLD-ABC<-METHYLASE<-TUDOR+SF1-SF2-Helicase+iREase+CTD+REase\|\|HTH->GDResolvase+PPP1-HTH-> | 1135 | Bacteria>Pseudomonadota>Alphaproteobacteria>Rhodospirillales>Azospirillaceae | GCF_013423495.1 | Azospirillum_oleiclasticum | HND93_RS19070 | 2984..6391 | 2735135 | DUF499-domain-containing-protein-[Azospirillum-oleiclasticum] |
| WP_211111227.1 | DUF499_1.RM.like.systems | <-GDResolvase+Recombinase+ZnR<-?\|\|?->?->Phage_integrase-><-HEPN+DUF499*<-METHYLASE<-TUDOR+SF1-SF2-Helicase+iREase+CTD+REase<-HTH+WYL\|\|HicB->TIR->REase-> | 1089 | Bacteria>Pseudomonadota>Alphaproteobacteria>Rhodospirillales>Azospirillaceae | GCF_016806835.2 | Skermanella_rosea | JL101_RS04580 | 983168..986437 | 1817965 | DUF499-domain-containing-protein-[Skermanella-rosea] |
| WP_209991153.1 | DUF499_1.RM.like.systems | TUDOR+SF1-SF2-Helicase+iREase+CTD+REase->OLD-ABC->HNH->METHYLASE->?->HEPN+DUF499*->?->Lon_C-> | 1114 | Bacteria>Pseudomonadota>Alphaproteobacteria>Rhodospirillales>Azospirillaceae | GCF_017876115.1 | Azospirillum_picis | J2850_RS30835 | 10080..13424 | 488438 | Swt1-family-HEPN-domain-containing-protein-[Azospirillum-picis] |
| WP_209875397.1 | DUF499_1.RM.like.systems | HIN-HTH->GDResolvase+Recombinase+ZnR->?->HIN-HTH->?->LZ_Tnp_IS66+zf-IS66+DDE_Tnp_IS66+DDE_Tnp_IS66_C-><-HEPN+DUF499*<-METHYLASE<-MORC+DUF3684<-TUDOR+SF1-SF2-Helicase+iREase+CTD+REase<-HTH+WYL | 1098 | Bacteria>Pseudomonadota>Alphaproteobacteria>Rhodospirillales>Azospirillaceae | GCF_017876165.1 | Azospirillum_soli | J2852_RS11360 | 44068..47364 | 1304799 | DUF499-domain-containing-protein-[Azospirillum-soli] |
| WP_238518868.1 | DUF499_1.RM.like.systems | <-SF1-SF2-Helicase+DUF1998+HKD-DNase+HKD-DNase<-?<-HEPN+DUF499*<-PIN<-OLD-ABC<-METHYLASE<-?<-Schlafen+HATPase+HTH<-TUDOR+SF1-SF2-Helicase+iREase+CTD+REase | 1154 | Bacteria>Pseudomonadota>Alphaproteobacteria>Rhodospirillales>Azospirillaceae | GCF_022214805.1 | Azospirillum_doebereinerae | MCW82_RS00020 | 10412..13876 | 92933 | DUF499-domain-containing-protein-[Azospirillum-doebereinerae] |
| WP_218285167.1 | DUF499_1.RM.like.systems | Peptidase->Peptidase_M24->TUDOR+SF1-SF2-Helicase+iREase+CTD+REase->METHYLASE->OLD-ABC->TOPRIM->HEPN+DUF499*->?-><-PAIREDC-HTH<-DUF4096+DDE_Tnp_1+DDE_Tnp_1_2\|\|DUF4096+DDE_Tnp_1+DDE_Tnp_1_2->PAIREDC-HTH-> | 1111 | Bacteria>Pseudomonadota>Alphaproteobacteria>Rhodospirillales>Elioraeaceae | GCF_019203965.1 | Elioraea_tepida | KO353_RS13040 | 2714596..2717931 | 2843330 | Swt1-family-HEPN-domain-containing-protein-[Elioraea-tepida] |
| MBC8269068.1 | DUF499_1.RM.like.systems | Y1_Tnp-><-?<-?\|\|HTH+WYL->TUDOR+SF1-SF2-Helicase+iREase+CTD+REase->METHYLASE->HEPN+DUF499*-> | 565 | Bacteria>Pseudomonadota>Alphaproteobacteria>Rhodospirillales>Rhodospirillaceae | GCA_014382375.1 | Rhodospirillaceae_bacterium | H8E36_09995 | 85324..87019 | 1898112 | ATP-binding-protein-[Rhodospirillaceae-bacterium] |
| MBT5356878.1 | DUF499_1.RM.like.systems | XIS-HTH->?->DnaB->?->Terminase-><-HEPN+DUF499*<-?<-?<-METHYLASE<-HEPN | 1096 | Bacteria>Pseudomonadota>Alphaproteobacteria>Rhodospirillales>Rhodospirillaceae | GCA_018663095.1 | Rhodospirillaceae_bacterium | HOL07_00895 | 173068..176358 | 1898112 | ATP-binding-protein-[Rhodospirillaceae-bacterium] |
| MBT6311358.1 | DUF499_1.RM.like.systems | XIS-HTH->?->DnaB->?->Terminase-><-HEPN+DUF499*<-?<-?<-METHYLASE<-HEPN | 1096 | Bacteria>Pseudomonadota>Alphaproteobacteria>Rhodospirillales>Rhodospirillaceae | GCA_018701575.1 | Rhodospirillaceae_bacterium | HOJ20_16125 | 146396..149686 | 1898112 | ATP-binding-protein-[Rhodospirillaceae-bacterium] |
| MCY4239105.1 | DUF499_1.RM.like.systems | METHYLASE->?->?->HEPN+DUF499*->SbcC-> | 1100 | Bacteria>Pseudomonadota>Alphaproteobacteria>Rhodospirillales>Rhodospirillaceae | GCA_026709535.1 | Rhodospirillaceae_bacterium | OXC93_12260 | 2987..6289 | 1898112 | MAG:-DUF499-domain-containing-protein-[Rhodospirillaceae-bacterium] |
| MCY4311819.1 | DUF499_1.RM.like.systems | <-SbcC<-HEPN+DUF499*<-?<-METHYLASE-p | 1100 | Bacteria>Pseudomonadota>Alphaproteobacteria>Rhodospirillales>Rhodospirillaceae | GCA_026710125.1 | Rhodospirillaceae_bacterium | OXC54_11025 | 2036..5338 | 1898112 | MAG:-DUF499-domain-containing-protein-[Rhodospirillaceae-bacterium] |
| MDE0176181.1 | DUF499_1.RM.like.systems | METHYLASE-p->METHYLASE->?->?->HEPN+DUF499*->HEPN->?->HEPN-> | 1096 | Bacteria>Pseudomonadota>Alphaproteobacteria>Rhodospirillales>Rhodospirillaceae | GCA_028819635.1 | Defluviicoccus_sp. | OXM60_22325 | 5691..8981 | 2682137 | MAG:-DUF499-domain-containing-protein-[Defluviicoccus-sp.] |
| MDE0333394.1 | DUF499_1.RM.like.systems | TUDOR+SF1-SF2-Helicase+iREase+CTD+REase->?->?->?->METHYLASE->HEPN->HEPN+DUF499*->REase-> | 1096 | Bacteria>Pseudomonadota>Alphaproteobacteria>Rhodospirillales>Rhodospirillaceae | GCA_028820945.1 | Defluviicoccus_sp. | OXI64_00425 | 9805..13095 | 2682137 | MAG:-DUF499-domain-containing-protein-[Defluviicoccus-sp.] |
| WP_190260570.1 | DUF499_1.RM.like.systems | <-SF1-SF2-Helicase+REase<-SF1-SF2-Helicase+REase\|\|HTH+WYL->TUDOR+SF1-SF2-Helicase+iREase+CTD+REase->YtH->METHYLASE->HEPN+DUF499*->MODE-HTH+PBPII-><-ParG_RHH | 1098 | Bacteria>Pseudomonadota>Alphaproteobacteria>Rhodospirillales>Rhodospirillaceae | GCF_014672695.1 | Defluviicoccus_vanus | HQ394_RS12700 | 2827067..2830363 | 111831 | DUF499-domain-containing-protein-[Defluviicoccus-vanus] |
| WP_173982139.1 | DUF499_1.RM.like.systems | HTH+WYL->TUDOR+SF1-SF2-Helicase+iREase+CTD+REase->METHYLASE->HEPN+DUF499*->TPR+TPR+PIN+PIN->TPR+TPR+PIN+PIN->REase+DUF4268-><-HTH<-GDResolvase+Recombinase+ZnR<-RAMA | 1097 | Bacteria>Pseudomonadota>Alphaproteobacteria>Rhodospirillales>Rhodospirillaceae | GCF_902729425.1 | Magnetospirillum_sp._SS-4 | MTBSS4_RS04180 | 57981..61274 | 2681465 | DUF499-domain-containing-protein-[Magnetospirillum-sp.-SS-4] |
| TVR79626.1 | DUF499_1.RM.like.systems | HDRnase-><-HEPN+DUF499*<-?<-PIN<-ParD<-METHYLASE<-OLD-ABC+TOPRIM<-TUDOR+SF1-SF2-Helicase+iREase+CTD+REase | 1117 | Bacteria>Pseudomonadota>Alphaproteobacteria>Rhodospirillales>unclassified_Rhodospirillales | GCA_007695045.1 | Rhodospirillales_bacterium | EA405_12420 | 709..4062 | 2026786 | ATP-binding-protein-[Rhodospirillales-bacterium] |
| MBC6439887.1 | DUF499_1.RM.like.systems | GATase->METHYLASE->HEPN+DUF499*-><-?<-?<-?<-ABC-ATPase+SbcC | 1142 | Bacteria>Pseudomonadota>Alphaproteobacteria>Rhodospirillales>unclassified_Rhodospirillales | GCA_014323865.1 | Rhodospirillales_bacterium | GDA49_05640 | 274906..278334 | 2026786 | ATP-binding-protein-[Rhodospirillales-bacterium] |
| MBK8177039.1 | DUF499_1.RM.like.systems | TrwC+SF1-SF2-helicase+TOPRIM->?->RHH->RelE-ParE-><-N6_N4_MTase<-?<-HEPN+DUF499*<-METHYLASE<-TUDOR+SF1-SF2-Helicase+iREase+CTD+REase<-?<-?<-ArdCN+MPTase\|\|DOC+HTH-> | 1106 | Bacteria>Pseudomonadota>Alphaproteobacteria>Rhodospirillales>unclassified_Rhodospirillales | GCA_016712595.1 | Rhodospirillales_bacterium | IPK66_17795 | 28556..31876 | 2026786 | DUF499-domain-containing-protein-[Rhodospirillales-bacterium] |
| MBL6940543.1 | DUF499_1.RM.like.systems | Y1_Tnp-><-?<-?\|\|HTH+WYL->TUDOR+SF1-SF2-Helicase+iREase+CTD+REase->METHYLASE->HEPN+DUF499*->HAD-> | 572 | Bacteria>Pseudomonadota>Alphaproteobacteria>Rhodospirillales>unclassified_Rhodospirillales | GCA_016776585.1 | Rhodospirillales_bacterium | ISR53_00145 | 38024..39742 | 2026786 | DUF499-domain-containing-protein-[Rhodospirillales-bacterium] |
| SEP49898.1 | DUF499_1.RM.like.systems | <-HEPN+DUF499*<-PIN<-OLD-ABC<-METHYLASE<-TUDOR+SF1-SF2-Helicase+iREase+CTD+REase<-HNH | 1099 | Bacteria>Pseudomonadota>Alphaproteobacteria>Rhodospirillales>unclassified_Rhodospirillales | GCA_900110395.1 | Rhodospirillales_bacterium_URHD0017 | SAMN02990966_07447 | 5640..8939 | 1380357 | hypothetical-protein-SAMN02990966_07447-[Rhodospirillales-bacterium-URHD0017] |

| Protein-ID | System | Gene-Neighbourhoods | Length | Taxonomy | Assembly-ID | Strain | Locus-Tag | Genomic-coordinates | TaxID | Fasta-define |
|---|---|---|---|---|---|---|---|---|---|---|
| MBA4046061.1 | DUF499_1.RM.like.systems | TUDOR+SF1-SF2-Helicase+iREase+CTD+REase->MORC+DUF3684->METHYLASE->HEPN+DUF499*->?-><-?<-TraH<-Thioredoxin | 1099 | Bacteria>Pseudomonadota>Alphaproteobacteria>Sphingomonadales>Erythrobacteraceae | GCA_013823025.1 | Erythrobacter_sp. | C0471_16820 | 9896..13195 | 1042 | AAA-family-ATPase-[Erythrobacter-sp.] |
| MCV0384725.1 | DUF499_1.RM.like.systems | <-HEPN+DUF499*<-METHYLASE<-HNH<-TUDOR+SF1-SF2-Helicase+iREase+CTD+REase-p | 421 | Bacteria>Pseudomonadota>Alphaproteobacteria>Sphingomonadales>Erythrobacteraceae | GCA_025698865.1 | Erythrobacter_sp. | K5799_14985 | 1..1263 | 1042 | MAG:-DUF499-domain-containing-protein,-partial-[Erythrobacter-sp.] |
| WP_017665279.1 | DUF499_1.RM.like.systems | <-HTH<-GDResolvase+Recombinase+ZnR<-RAMA<-?\|\|Peptidase_S74-><-HEPN+DUF499*<-METHYLASE<-TUDOR+SF1-SF2-Helicase+iREase+CTD+REase<-XIS-HTH | 1099 | Bacteria>Pseudomonadota>Alphaproteobacteria>Sphingomonadales>Erythrobacteraceae | GCF_000331285.1 | Porphyrobacter_sp._AAP82 | D620_RS0108005 | 32206..35505 | 1248917 | DUF499-domain-containing-protein-[Porphyrobacter-sp.-AAP82] |
| WP_034960305.1 | DUF499_1.RM.like.systems | <-TIR<-HicB\|\|HTH+WYL->TUDOR+SF1-SF2-Helicase+iREase+CTD+REase->MORC+DUF3684->METHYLASE->HEPN+DUF499*-><-?\|\|LexA-protease-><-wHTH+SAD+HNH<-TrwC+SF1-SF2-helicase<-VirD4-TraG+VirD4-FtsK<-CopG/RHH | 1100 | Bacteria>Pseudomonadota>Alphaproteobacteria>Sphingomonadales>Erythrobacteraceae | GCF_000715015.1 | Erythrobacter_longus | EH31_RS10535 | 101708..105010 | 1044 | DUF499-domain-containing-protein-[Erythrobacter-longus] |
| WP_073974519.1 | DUF499_1.RM.like.systems | <-HicB\|\|HTH+WYL->TUDOR+SF1-SF2-Helicase+iREase+CTD+REase->?->METHYLASE->HEPN+DUF499*->MODE-HTH+PBPII-><-ParG_RHH | 1097 | Bacteria>Pseudomonadota>Alphaproteobacteria>Sphingomonadales>Erythrobacteraceae | GCF_001556555.1 | Erythrobacter_donghaensis | AX698_RS16845 | 33517..36810 | 267135 | DUF499-domain-containing-protein-[Erythrobacter-donghaensis] |
| WP_120325010.1 | DUF499_1.RM.like.systems | CopG/RHH->VirD4-TraG+VirD4-FtsK->TrwC+SF1-SF2-helicase->wHTH+SAD+HNH-><-LexA-protease\|\|?-><-HEPN+DUF499*<-METHYLASE<-MORC+DUF3684<-TUDOR+SF1-SF2-Helicase+iREase+CTD+REase<-HTH+WYL\|\|HicB->TIR-> | 1100 | Bacteria>Pseudomonadota>Alphaproteobacteria>Sphingomonadales>Erythrobacteraceae | GCF_003610805.1 | Altericroceibacterium_spongiae | D6851_RS11415 | 253315..256617 | 2320269 | DUF499-domain-containing-protein-[Altericroceibacterium-spongiae] |
| WP_159791246.1 | DUF499_1.RM.like.systems | ZDNA-HTH->TUDOR+SF1-SF2-Helicase+iREase+CTD+REase->MarR-HTH->?->?->METHYLASE->HEPN+DUF499*-> | 1097 | Bacteria>Pseudomonadota>Alphaproteobacteria>Sphingomonadales>Erythrobacteraceae | GCF_009827435.1 | Croceibacterium_salegens | GRI89_RS00095 | 15268..18561 | 1737568 | DUF499-domain-containing-protein-[Croceibacterium-salegens] |
| WP_221424189.1 | DUF499_1.RM.like.systems | CopG/RHH->VirD4-TraG+VirD4-FtsK->TrwC+SF1-SF2-helicase->wHTH+SAD+HNH-><-LexA-protease\|\|?-><-HEPN+DUF499*<-METHYLASE<-MORC+DUF3684<-TUDOR+SF1-SF2-Helicase+iREase+CTD+REase<-HTH+WYL\|\|HicB->TIR-> | 1100 | Bacteria>Pseudomonadota>Alphaproteobacteria>Sphingomonadales>Erythrobacteraceae | GCF_019711375.1 | Qipengyuania_aurantiaca | K3148_RS07230 | 1481189..1484491 | 2867233 | DUF499-domain-containing-protein-[Qipengyuania-aurantiaca] |
| ABQ66570.1 | DUF499_1.RM.like.systems | <-HicB\|\|HTH+WYL->TUDOR+SF1-SF2-Helicase+iREase+CTD+REase->MORC+DUF3684->METHYLASE->HEPN+DUF499*->TIR+PNTD2->PIWI-><-tRNA<-?<-HHH+ENDO3+NUDIX | 1098 | Bacteria>Pseudomonadota>Alphaproteobacteria>Sphingomonadales>Sphingomonadaceae | GCA_000016765.1 | Rhizorhabdus_wittichii_RW1 | Swit_0199 | 198092..201388 | 392499 | ATPase-(AAA+-superfamily)-like-protein-[Rhizorhabdus-wittichii-RW1] |
| EPR17235.1 | DUF499_1.RM.like.systems | ParG_RHH-><-TetR-HTH\|\|MODE-HTH+PBPII-><-HEPN+DUF499*<-METHYLASE<-MORC+DUF3684<-TUDOR+SF1-SF2-Helicase+iREase+CTD+REase<-HTH+WYL\|\|SF1-SF2-Helicase+REase->SF1-SF2-Helicase+REase-> | 1086 | Bacteria>Pseudomonadota>Alphaproteobacteria>Sphingomonadales>Sphingomonadaceae | GCA_000421925.1 | Sphingobium_indicum_IP26 | M527_17965 | 110739..113999 | 1346790 | ATPase-AAA-[Sphingobium-chinhatense-IP26] |
| MAF61658.1 | DUF499_1.RM.like.systems | ParG_RHH-><-MODE-HTH+PBPII<-HEPN+DUF499*<-METHYLASE<-?<-TUDOR+SF1-SF2-Helicase+iREase+CTD+REase<-HTH+WYL\|\|HicB-> | 1097 | Bacteria>Pseudomonadota>Alphaproteobacteria>Sphingomonadales>Sphingomonadaceae | GCA_002687555.1 | Blastomonas_sp. | CL680_09370 | 351225..354518 | 1909299 | DUF499-domain-containing-protein-[Blastomonas-sp.] |
| PZU56403.1 | DUF499_1.RM.like.systems | METHYLASE->HEPN+DUF499*-><-?<-?<-?<-LON-AAA | 1100 | Bacteria>Pseudomonadota>Alphaproteobacteria>Sphingomonadales>Sphingomonadaceae | GCA_003248935.1 | Sphingobium_sp. | DI547_16590 | 2951..6253 | 1912891 | ATPase-[Sphingobium-sp.] |
| MBB3880227.1 | DUF499_1.RM.like.systems | <-SF1-SF2-Helicase+REase<-SF1-SF2-Helicase+REase\|\|HTH+WYL->TUDOR+SF1-SF2-Helicase+iREase+CTD+REase->REase->METHYLASE->HEPN+DUF499*->MODE-HTH+PBPII-><-ParG_RHH | 1098 | Bacteria>Pseudomonadota>Alphaproteobacteria>Sphingomonadales>Sphingomonadaceae | GCA_014196255.1 | Sphingomonas_pseudosanguinis | GGR48_002670 | 32767..36063 | 413712 | hypothetical-protein-[Sphingomonas-pseudosanguinis] |
| MBF7011871.1 | DUF499_1.RM.like.systems | Thioredoxin->TraH->?-><-?<-HEPN+DUF499*<-METHYLASE<-MORC+DUF3684<-TUDOR+SF1-SF2-Helicase+iREase+CTD+REase<-HTH+WYL\|\|HicB-> | 1097 | Bacteria>Pseudomonadota>Alphaproteobacteria>Sphingomonadales>Sphingomonadaceae | GCA_015594425.1 | Novosphingobium_sp._HR1a | HWN72_09985 | 40208..43501 | 1395637 | ATP-binding-protein-[Novosphingobium-sp.-HR1a] |
| MBP8234957.1 | DUF499_1.RM.like.systems | ParG_RHH-><-MODE-HTH+PBPII<-HEPN+DUF499*<-METHYLASE-p | 1098 | Bacteria>Pseudomonadota>Alphaproteobacteria>Sphingomonadales>Sphingomonadaceae | GCA_018006725.1 | Rhizorhabdus_sp. | KAY22_21925 | 3665..6961 | 1968843 | DUF499-domain-containing-protein-[Rhizorhabdus-sp.] |
| MBX9665921.1 | DUF499_1.RM.like.systems | <-TOPRIM+SF1-SF2-Helicase+HTH<-ZnR+TOPRIM<-?<-XIS-HTH<-?<-HEPN+DUF499*<-METHYLASE<-?<-TUDOR+SF1-SF2-Helicase+iREase+CTD+REase | 1100 | Bacteria>Pseudomonadota>Alphaproteobacteria>Sphingomonadales>Sphingomonadaceae | GCA_019748365.1 | Novosphingobium_sp. | K5Q19_21680 | 3115..6417 | 1874826 | DUF499-domain-containing-protein-[Novosphingobium-sp.] |
| MCB4860948.1 | DUF499_1.RM.like.systems | <-SF1-SF2-helicase+RdgC+Beta-barrel+REase<-HicB\|\|HTH+WYL->TUDOR+SF1-SF2-Helicase+iREase+CTD+REase->MORC+DUF3684->METHYLASE->HEPN+DUF499*-> | 1097 | Bacteria>Pseudomonadota>Alphaproteobacteria>Sphingomonadales>Sphingomonadaceae | GCA_019873065.1 | Sphingobium_sp._PNB | K7W03_15250 | 38941..42234 | 863934 | DUF499-domain-containing-protein-[Sphingobium-sp.-PNB] |
| MCL6741777.1 | DUF499_1.RM.like.systems | wHTH+REase->TUDOR+SF1-SF2-Helicase+iREase+CTD+REase->METHYLASE->OLD-ABC->PIN->HEPN+DUF499*-><-?<-BetaPropeller+ABhydrolase\|\|?-><-ABhydrolase<-MODE-HTH+PBPII | 1094 | Bacteria>Pseudomonadota>Alphaproteobacteria>Sphingomonadales>Sphingomonadaceae | GCA_023516505.1 | Sphingomonas_brevis | LZ518_11630 | 2249137..2252421 | 2908206 | DUF499-domain-containing-protein-[Sphingomonas-sp.-RB56-2] |
| MCX7283721.1 | DUF499_1.RM.like.systems | <-HEPN+DUF499*<-METHYLASE-p | 625 | Bacteria>Pseudomonadota>Alphaproteobacteria>Sphingomonadales>Sphingomonadaceae | GCA_026400375.1 | Novosphingobium_sp. | NTX28_06715 | 1..1874 | 1874826 | MAG:-DUF499-domain-containing-protein,-partial-[Novosphingobium-sp.] |
| WP_010188165.1 | DUF499_1.RM.like.systems | HTH+WYL->TUDOR+SF1-SF2-Helicase+iREase+CTD+REase->YtH->METHYLASE->HEPN+DUF499*->?->HIN-HTH->?->?->DOC->HTH_3-> | 1098 | Bacteria>Pseudomonadota>Alphaproteobacteria>Sphingomonadales>Sphingomonadaceae | GCF_000241485.1 | Sphingomonas_sp._PAMC_26605 | RH90_RS19210 | 226826..230122 | 1112214 | DUF499-domain-containing-protein-[Sphingomonas-sp.-PAMC-26605] |
| WP_017183457.1 | DUF499_1.RM.like.systems | <-HicB\|\|HTH+WYL->TUDOR+SF1-SF2-Helicase+iREase+CTD+REase->ParB+HNH->ASCH->METHYLASE->HEPN+DUF499*->MODE-HTH+PBPII-><-ParG_RHH | 1097 | Bacteria>Pseudomonadota>Alphaproteobacteria>Sphingomonadales>Sphingomonadaceae | GCF_000277525.1 | Sphingobium_xenophagum | QYYP_RS0113510 | 14048..17341 | 121428 | DUF499-domain-containing-protein-[Sphingobium-xenophagum] |
| WP_028055914.1 | DUF499_1.RM.like.systems | <-TIR<-HicB\|\|HTH+WYL->TUDOR+SF1-SF2-Helicase+iREase+CTD+REase->MORC+DUF3684->METHYLASE->HEPN+DUF499*-><-XIS-HTH | 1100 | Bacteria>Pseudomonadota>Alphaproteobacteria>Sphingomonadales>Sphingomonadaceae | GCF_000412635.1 | Sphingobium_bisphenolivorans | L858_RS0107580 | 162523..165825 | 1335760 | DUF499-domain-containing-protein-[Sphingobium-bisphenolivorans] |
| WP_025161153.1 | DUF499_1.RM.like.systems | ParG_RHH-><-TetR-HTH\|\|MODE-HTH+PBPII-><-HEPN+DUF499*<-METHYLASE<-MORC+DUF3684<-TUDOR+SF1-SF2-Helicase+iREase+CTD+REase<-HTH+WYL\|\|SF1-SF2-Helicase+REase->SF1-SF2-Helicase+REase-> | 1098 | Bacteria>Pseudomonadota>Alphaproteobacteria>Sphingomonadales>Sphingomonadaceae | GCF_000445085.1 | Sphingobium | L286_RS41375 | 45892..49188 | 165695 | MULTISPECIES:-DUF499-domain-containing-protein-[Sphingobium] |
| WP_037445586.1 | DUF499_1.RM.like.systems | Thioredoxin->TraH->?-><-?<-HEPN+DUF499*<-METHYLASE<-MORC+DUF3684<-TUDOR+SF1-SF2-Helicase+iREase+CTD+REase<-HTH+WYL\|\|HicB->SF1-SF2-helicase+RdgC+Beta-barrel+REase-> | 1101 | Bacteria>Pseudomonadota>Alphaproteobacteria>Sphingomonadales>Sphingomonadaceae | GCF_000633575.2 | Sphingobium_indicum | AL00_RS20465 | 14679..17984 | 332055 | DUF499-domain-containing-protein-[Sphingobium-indicum] |
| WP_066601098.1 | DUF499_1.RM.like.systems | ABhydrolase->tRNA->TUDOR+SF1-SF2-Helicase+iREase+CTD+REase->?->METHYLASE->HEPN+DUF499*-><-?<-MobA_MobL+SF1-SF2-helicase\|\|TraD->?-><-HEPN | 1100 | Bacteria>Pseudomonadota>Alphaproteobacteria>Sphingomonadales>Sphingomonadaceae | GCF_001046645.1 | Sphingobium_cupriresistens | V473_RS04750 | 974632..977934 | 1132417 | DUF499-domain-containing-protein-[Sphingobium-cupriresistens] |
| WP_053553316.1 | DUF499_1.RM.like.systems | <-SF1-SF2-Helicase+REase\|\|HTH+WYL->TUDOR+SF1-SF2-Helicase+iREase+CTD+REase->ASCH->SIR2->METHYLASE->HEPN+DUF499*->MODE-HTH+PBPII-><-ParG_RHH | 1100 | Bacteria>Pseudomonadota>Alphaproteobacteria>Sphingomonadales>Sphingomonadaceae | GCF_001278035.1 | Sphingopyxis_sp._113P3 | LH20_RS05260 | 1100261..1103563 | 292913 | DUF499-domain-containing-protein-[Sphingopyxis-sp.-113P3] |
| WP_053554356.1 | DUF499_1.RM.like.systems | HIN-HTH->?-><-?\|\|\|\|HIN-HTH->?->\|\|<-DEDD_Tnp_IS110+Transposase_20<-HEPN+DUF499*<-METHYLASE<-MORC+DUF3684<-TUDOR+SF1-SF2-Helicase+iREase+CTD+REase<-HTH+WYL<-XIS-HTH | 1115 | Bacteria>Pseudomonadota>Alphaproteobacteria>Sphingomonadales>Sphingomonadaceae | GCF_001278035.1 | Sphingopyxis_sp._113P3 | LH20_RS11780 | 2402847..2406194 | 292913 | Swt1-family-HEPN-domain-containing-protein-[Sphingopyxis-sp.-113P3] |
| WP_054108969.1 | DUF499_1.RM.like.systems | HIN-HTH->?-><-HEPN+DUF499*<-METHYLASE<-TUDOR+SF1-SF2-Helicase+iREase+CTD+REase<-HTH+WYL<-?\|\|HTH_3-> | 1099 | Bacteria>Pseudomonadota>Alphaproteobacteria>Sphingomonadales>Sphingomonadaceae | GCF_001295795.1 | Novosphingobium_sp._AAP83 | IP81_RS15755 | 15292..18591 | 1523425 | DUF499-domain-containing-protein-[Novosphingobium-sp.-AAP83] |
| WP_067912291.1 | DUF499_1.RM.like.systems | <-HicB\|\|HTH+WYL->TUDOR+SF1-SF2-Helicase+iREase+CTD+REase->MORC+DUF3684->METHYLASE->HEPN+DUF499*->?-><-?<-TraH<-Thioredoxin | 1099 | Bacteria>Pseudomonadota>Alphaproteobacteria>Sphingomonadales>Sphingomonadaceae | GCF_001519075.1 | Novosphingobium_fuchskuhlense | AQZ52_RS14075 | 23149..26448 | 1117702 | DUF499-domain-containing-protein-[Novosphingobium-fuchskuhlense] |
| WP_066778028.1 | DUF499_1.RM.like.systems | <-SF1-SF2-Helicase+REase\|\|HTH+WYL->TUDOR+SF1-SF2-Helicase+iREase+CTD+REase->YtH->MORC+DUF3684->METHYLASE->HEPN+DUF499*->MODE-HTH+PBPII-><-ParG_RHH | 1100 | Bacteria>Pseudomonadota>Alphaproteobacteria>Sphingomonadales>Sphingomonadaceae | GCF_001557395.1 | Sphingomonas_sp._CCH5-D11 | AX702_RS07275 | 80145..83447 | 1768786 | DUF499-domain-containing-protein-[Sphingomonas-sp.-CCH5-D11] |
| WP_069321300.1 | DUF499_1.RM.like.systems | ParG_RHH-><-MODE-HTH+PBPII<-HEPN+DUF499*<-METHYLASE<-MORC+DUF3684<-TUDOR+SF1-SF2-Helicase+iREase+CTD+REase<-HTH+WYL | 1099 | Bacteria>Pseudomonadota>Alphaproteobacteria>Sphingomonadales>Sphingomonadaceae | GCF_001721295.1 | Sphingomonas_turrisvirgatae | BFL28_RS14750 | 27774..31073 | 1888892 | DUF499-domain-containing-protein-[Sphingomonas-turrisvirgatae] |
| WP_066967630.1 | DUF499_1.RM.like.systems | ParG_RHH-><-MODE-HTH+PBPII<-HEPN+DUF499*<-METHYLASE<-MORC+DUF3684\|\|Arm-DNA-bind_3+Phage_integrase-><-XIS-HTH<-ParG_RHH | 1098 | Bacteria>Pseudomonadota>Alphaproteobacteria>Sphingomonadales>Sphingomonadaceae | GCF_002355275.1 | Rhizorhabdus_dicambivorans | CMV14_RS12985 | 2738842..2742138 | 1850238 | DUF499-domain-containing-protein-[Rhizorhabdus-dicambivorans] |
| WP_066520604.1 | DUF499_1.RM.like.systems | TUDOR+SF1-SF2-Helicase+iREase+CTD+REase-p->ASCH->SIR2->METHYLASE->HEPN+DUF499*->MODE-HTH+PBPII-><-ParG_RHH | 1098 | Bacteria>Pseudomonadota>Alphaproteobacteria>Sphingomonadales>Sphingomonadaceae | GCF_002355855.1 | Sphingobium_cloacae | SC4_RS16230 | 6611..9907 | 120107 | DUF499-domain-containing-protein-[Sphingobium-cloacae] |
| WP_100237044.1 | DUF499_1.RM.like.systems | ParG_RHH-><-TetR-HTH\|\|MODE-HTH+PBPII-><-HEPN+DUF499*<-METHYLASE<-MORC+DUF3684<-TUDOR+SF1-SF2-Helicase+iREase+CTD+REase<-HTH+WYL\|\|SF1-SF2-Helicase+REase->SF1-SF2-Helicase+REase-> | 1098 | Bacteria>Pseudomonadota>Alphaproteobacteria>Sphingomonadales>Sphingomonadaceae | GCF_002795205.1 | Sphingobium_sp._LB126 | CAF53_RS01415 | 292890..296186 | 1983755 | DUF499-domain-containing-protein-[Sphingobium-sp.-LB126] |
| WP_120216254.1 | DUF499_1.RM.like.systems | ParG_RHH-><-MODE-HTH+PBPII<-HEPN+DUF499*<-METHYLASE<-MORC+DUF3684<-YtH<-ASCH<-TUDOR+SF1-SF2-Helicase+iREase+CTD+REase<-HTH+WYL | 1098 | Bacteria>Pseudomonadota>Alphaproteobacteria>Sphingomonadales>Sphingomonadaceae | GCF_003609815.1 | Sphingopyxis_sp._EG6 | SPYCW_RS07385 | 1487775..1491071 | 1874061 | DUF499-domain-containing-protein-[Sphingopyxis-sp.-EG6] |
| WP_156842742.1 | DUF499_1.RM.like.systems | ParG_RHH-><-MODE-HTH+PBPII<-HEPN+DUF499*<-METHYLASE<-MORC+DUF3684<-YtH<-TUDOR+SF1-SF2-Helicase+iREase+CTD+REase<-HTH+WYL | 1098 | Bacteria>Pseudomonadota>Alphaproteobacteria>Sphingomonadales>Sphingomonadaceae | GCF_009746585.1 | Novosphingobium_aquimarinum | GQ094_RS17725 | 28376..31672 | 2682494 | DUF499-domain-containing-protein-[Novosphingobium-aquimarinum] |
| WP_182926476.1 | DUF499_1.RM.like.systems | <-SF1-SF2-Helicase+REase<-SF1-SF2-Helicase+REase\|\|HTH+WYL->TUDOR+SF1-SF2-Helicase+iREase+CTD+REase->?->METHYLASE->HEPN+DUF499*->MODE-HTH+PBPII-><-ParG_RHH | 1098 | Bacteria>Pseudomonadota>Alphaproteobacteria>Sphingomonadales>Sphingomonadaceae | GCF_014169955.1 | Sphingomonas_paucimobilis | SPKIRA_RS16090 | 3478703..3481999 | 13689 | DUF499-domain-containing-protein-[Sphingomonas-paucimobilis] |
| WP_221211510.1 | DUF499_1.RM.like.systems | <-SF1-SF2-Helicase+REase<-SF1-SF2-Helicase+REase\|\|HTH+WYL->TUDOR+SF1-SF2-Helicase+iREase+CTD+REase->REase->METHYLASE->HEPN+DUF499*->MODE-HTH+PBPII-><-ParG_RHH | 1086 | Bacteria>Pseudomonadota>Alphaproteobacteria>Sphingomonadales>Sphingomonadaceae | GCF_014196255.1 | Sphingomonas_pseudosanguinis | GGR48_RS13090 | 32803..36063 | 413712 | DUF499-domain-containing-protein-[Sphingomonas-pseudosanguinis] |
| WP_184477099.1 | DUF499_1.RM.like.systems | N6_MTase->REase-><-?<-?<-URI-DNase<-ParB+HTH<-HEPN+DUF499*<-METHYLASE<-ASCH<-HTH+WYL | 1098 | Bacteria>Pseudomonadota>Alphaproteobacteria>Sphingomonadales>Sphingomonadaceae | GCF_014200045.1 | Rhizorhapis_suberifaciens | HNQ99_RS14965 | 12710..16006 | 13656 | DUF499-domain-containing-protein-[Rhizorhapis-suberifaciens] |
| WP_145073130.1 | DUF499_1.RM.like.systems | TetR-HTH->XIS-HTH->TUDOR+SF1-SF2-Helicase+iREase+CTD+REase->ASCH->METHYLASE->HEPN+DUF499*-> | 1099 | Bacteria>Pseudomonadota>Alphaproteobacteria>Sphingomonadales>Sphingomonadaceae | GCF_014205735.1 | Sphingobium_wenxiniae | IQ35_RS10205 | 192114..195413 | 595605 | DUF499-domain-containing-protein-[Sphingobium-wenxiniae] |
| WP_194920702.1 | DUF499_1.RM.like.systems | METHYLASE-><-DDE_Tnp_1_4\|\|METHYLASE-p->OLD-ABC->?->HIN-HTH->HEPN+DUF499*->HIN-HTH->?-><-XIS-HTH | 1041 | Bacteria>Pseudomonadota>Alphaproteobacteria>Sphingomonadales>Sphingomonadaceae | GCF_015390225.1 | Novosphingobium_sp._NBM11 | F1640_RS18815 | 3882459..3885584 | 2596914 | DUF499-domain-containing-protein-[Novosphingobium-sp.-NBM11] |
| WP_205396700.1 | DUF499_1.RM.like.systems | ParG_RHH-><-MODE-HTH+PBPII<-HEPN+DUF499*<-METHYLASE<-?<-TUDOR+SF1-SF2-Helicase+iREase+CTD+REase<-HTH+WYL\|\|SF1-SF2-Helicase+REase->SF1-SF2-Helicase+REase-> | 1098 | Bacteria>Pseudomonadota>Alphaproteobacteria>Sphingomonadales>Sphingomonadaceae | GCF_016919545.1 | Sphingomonas_paucimobilis | JT366_RS09685 | 1990566..1993862 | 13689 | DUF499-domain-containing-protein-[Sphingomonas-paucimobilis] |
| WP_226632678.1 | DUF499_1.RM.like.systems | HTH+WYL->TUDOR+SF1-SF2-Helicase+iREase+CTD+REase->METHYLASE->HEPN+DUF499*-><-ACET<-?\|\|<-?<-STYKinase | 1110 | Bacteria>Pseudomonadota>Alphaproteobacteria>Sphingomonadales>Sphingomonadaceae | GCF_020524485.1 | Novosphingobium_profundi | LC060_RS03305 | 760563..763895 | 1774954 | DUF499-domain-containing-protein-[Novosphingobium-profundi] |
| WP_283938333.1 | DUF499_1.RM.like.systems | wHTH+REase->TUDOR+SF1-SF2-Helicase+iREase+CTD+REase->METHYLASE->OLD-ABC->PIN->HEPN+DUF499*-><-?<-BetaPropeller+ABhydrolase\|\|?-><-ABhydrolase<-MODE-HTH+PBPII | 1103 | Bacteria>Pseudomonadota>Alphaproteobacteria>Sphingomonadales>Sphingomonadaceae | GCF_023516505.1 | Sphingomonas_brevis | LZ518_RS11630 | 2249110..2252421 | 2908206 | DUF499-domain-containing-protein-[Sphingomonas-brevis] |
| WP_079648123.1 | DUF499_1.RM.like.systems | ParG_RHH-><-MODE-HTH+PBPII<-HEPN+DUF499*<-METHYLASE<-MORC+DUF3684<-YtH<-TUDOR+SF1-SF2-Helicase+iREase+CTD+REase<-HTH+WYL\|\|SF1-SF2-Helicase+REase-> | 1099 | Bacteria>Pseudomonadota>Alphaproteobacteria>Sphingomonadales>Sphingomonadaceae | GCF_900167915.1 | Rhizorhabdus_histidinilytica | B5X82_RS10580 | 218430..221729 | 439228 | DUF499-domain-containing-protein-[Rhizorhabdus-histidinilytica] |
| WP_201515748.1 | DUF499_1.RM.like.systems | <-TraJ-RHH<-?<-?<-?\|\|?->XIS-HTH-><-HEPN+DUF499*<-METHYLASE<-REase<-TUDOR+SF1-SF2-Helicase+iREase+CTD+REase<-HTH+WYL\|\|HicB-> | 1100 | Bacteria>Pseudomonadota>Alphaproteobacteria>Sphingomonadales>Sphingomonadaceae | GCF_904830875.1 | Sphingobium_sp._S6 | SPHS8_RS12190 | 306093..309395 | 2758386 | DUF499-domain-containing-protein-[Sphingobium-sp.-S6] |
| WP_248005921.1 | DUF499_1.RM.like.systems | Thioredoxin->TraH->?-><-?<-HEPN+DUF499*<-METHYLASE<-MORC+DUF3684<-TUDOR+SF1-SF2-Helicase+iREase+CTD+REase<-HTH+WYL\|\|HicB-> | 1085 | Bacteria>Pseudomonadota>Alphaproteobacteria>Sphingomonadales>Sphingomonadaceae | - | Novosphingobium_sp._HR1a | HWN72_RS09985 | 40208..43465 | 1395637 | DUF499-domain-containing-protein-[Novosphingobium-sp.-HR1a] |
| WP_226870194.1 | DUF499_1.RM.like.systems | <-SF1-SF2-helicase+RdgC+Beta-barrel+REase<-HicB\|\|HTH+WYL->TUDOR+SF1-SF2-Helicase+iREase+CTD+REase->MORC+DUF3684->METHYLASE->HEPN+DUF499*-> | 1085 | Bacteria>Pseudomonadota>Alphaproteobacteria>Sphingomonadales>Sphingomonadaceae | - | Sphingobium_sp._PNB | K7W03_RS15190 | 38977..42234 | 863934 | DUF499-domain-containing-protein-[Sphingobium-sp.-PNB] |
| WP_249916143.1 | DUF499_1.RM.like.systems | wHTH+REase->TUDOR+SF1-SF2-Helicase+iREase+CTD+REase->METHYLASE->OLD-ABC->PIN->HEPN+DUF499*-><-?<-BetaPropeller+ABhydrolase\|\|?-><-ABhydrolase<-MODE-HTH+PBPII | 1094 | Bacteria>Pseudomonadota>Alphaproteobacteria>Sphingomonadales>Sphingomonadaceae | - | Sphingomonas_brevis | LZ518_11630 | 2249137..2252421 | 2908206 | DUF499-domain-containing-protein-[Sphingomonas-sp.-RB56-2] |
| MBS3961271.1 | DUF499_1.RM.like.systems | <-TIR<-HEPN+DUF499*<-METHYLASE | 1098 | Bacteria>Pseudomonadota>Alphaproteobacteria>Sphingomonadales>Sphingosinicellaceae | GCA_018333935.1 | Sandarakinorhabdus_sp. | KGZ61_05485 | 446..3742 | 1916663 | DUF499-domain-containing-protein-[Sandarakinorhabdus-sp.] |
| WP_017667308.1 | DUF499_1.RM.like.systems | HEPN->Phage_integrase->TUDOR+SF1-SF2-Helicase+iREase+CTD+REase->METHYLASE->HEPN+DUF499*->PIN-> | 1100 | Bacteria>Pseudomonadota>Alphaproteobacteria>Sphingomonadales>Sphingosinicellaceae | GCF_000331225.1 | Sandarakinorhabdus_sp._AAP62 | D619_RS0104150 | 158738..162040 | 1248916 | DUF499-domain-containing-protein-[Sandarakinorhabdus-sp.-AAP62] |
| WP_094472303.1 | DUF499_1.RM.like.systems | <-HEPN+DUF499*<-?<-METHYLASE-p | 1098 | Bacteria>Pseudomonadota>Alphaproteobacteria>Sphingomonadales>Sphingosinicellaceae | GCF_002251755.1 | Sandarakinorhabdus_cyanobacteriorum | CHU93_RS00700 | 1379..4675 | 1981098 | DUF499-domain-containing-protein-[Sandarakinorhabdus-cyanobacteriorum] |
| WP_147041999.1 | DUF499_1.RM.like.systems | <-REase<-?\|\|HTH_3-><-?<-PIWI<-TIR+PNTD2<-HEPN+DUF499*<-METHYLASE<-?<-TUDOR+SF1-SF2-Helicase+iREase+CTD+REase<-HTH+WYL\|\|HicB-> | 1098 | Bacteria>Pseudomonadota>Alphaproteobacteria>Sphingomonadales>Sphingosinicellaceae | GCF_007995055.1 | Sphingosinicella_ginsenosidimutans | FRZ32_RS02405 | 490620..493916 | 1176539 | DUF499-domain-containing-protein-[Sphingosinicella-ginsenosidimutans] |
| TXG85100.1 | DUF499_1.RM.like.systems | ParG_RHH-><-MODE-HTH+PBPII<-HEPN+DUF499*<-METHYLASE<-MORC+DUF3684<-YtH<-TUDOR+SF1-SF2-Helicase+iREase+CTD+REase<-HTH+WYL\|\|SF1-SF2-Helicase+REase-> | 1098 | Bacteria>Pseudomonadota>Alphaproteobacteria>Sphingomonadales>unclassified_Sphingomonadales | GCA_008012005.1 | Sphingomonadales_bacterium | E6R12_02055 | 22905..26201 | 1978525 | MAG:-ATP-binding-protein-[Sphingomonadales-bacterium] |
| NCP12883.1 | DUF499_1.RM.like.systems | <-TIR<-HicB\|\|HTH+WYL->TUDOR+SF1-SF2-Helicase+iREase+CTD+REase->MORC+DUF3684->METHYLASE->HEPN+DUF499*-> | 1100 | Bacteria>Pseudomonadota>Alphaproteobacteria>Sphingomonadales>unclassified_Sphingomonadales | GCA_009993565.1 | Sphingomonadales_bacterium | GW858_01815 | 17021..20323 | 1978525 | ATP-binding-protein-[Sphingomonadales-bacterium] |
| KUO66709.1 | DUF499_1.RM.like.systems | ZnR+TOPRIM+D5ATPase-><-?\|\|?-><-HEPN+DUF499*<-METHYLASE<-TUDOR+SF1-SF2-Helicase+iREase+CTD+REase | 1101 | Bacteria>Pseudomonadota>Alphaproteobacteria>unclassified_Alphaproteobacteria | GCA_001516065.1 | Alphaproteobacteria_bacterium_BRH_c36 | APF80_06945 | 3407..6712 | 1734406 | hypothetical-protein-APF80_06945-[Alphaproteobacteria-bacterium-BRH_c36] |
| PKP67098.1 | DUF499_1.RM.like.systems | <-HEPN+DUF499*<-METHYLASE-p | 366 | Bacteria>Pseudomonadota>Alphaproteobacteria>unclassified_Alphaproteobacteria | GCA_002842795.1 | Alphaproteobacteria_bacterium_HGW-Alphaproteobacteria-8 | CVT86_02370 | 1..1098 | 2013671 | AAA-family-ATPase,-partial-[Alphaproteobacteria-bacterium-HGW-Alphaproteobacteria-8] |
| RMH48112.1 | DUF499_1.RM.like.systems | METHYLASE-p->PolB->HEPN->HEPN+DUF499*-> | 1114 | Bacteria>Pseudomonadota>Alphaproteobacteria>unclassified_Alphaproteobacteria | GCA_003696705.1 | Alphaproteobacteria_bacterium | D6688_02640 | 1365..4709 | 1913988 | ATP-binding-protein-[Alphaproteobacteria-bacterium] |
| MCL2716067.1 | DUF499_1.RM.like.systems | TUDOR+SF1-SF2-Helicase+iREase+CTD+REase-><-?\|\|METHYLASE->HEPN+DUF499*->REase+PNTD1+PNTD2->PIWI-> | 1073 | Bacteria>Pseudomonadota>Alphaproteobacteria>unclassified_Alphaproteobacteria | GCA_009781045.1 | Alphaproteobacteria_bacterium | FWD68_16235 | 9484..12705 | 1913988 | DUF499-domain-containing-protein-[Alphaproteobacteria-bacterium] |
| NBC34425.1 | DUF499_1.RM.like.systems | <-DUF29<-?\|\|?-><-HEPN+DUF499*<-METHYLASE<-TUDOR+SF1-SF2-Helicase+iREase+CTD+REase<-tRNA<-ZnR+TOPRIM+DnaB<-?\|\|?->Polbeta-> | 1148 | Bacteria>Pseudomonadota>Alphaproteobacteria>unclassified_Alphaproteobacteria | GCA_009909005.1 | Alphaproteobacteria_bacterium | GVY13_17240 | 19753..23199 | 1913988 | DUF499-domain-containing-protein-[Alphaproteobacteria-bacterium] |
| HHY48691.1 | DUF499_1.RM.like.systems | <-HEPN+DUF499*<-METHYLASE-p | 365 | Bacteria>Pseudomonadota>Alphaproteobacteria>unclassified_Alphaproteobacteria | GCA_012842825.1 | Alphaproteobacteria_bacterium | GYA10_02965 | 1..1094 | 1913988 | ATP-binding-protein-[Alphaproteobacteria-bacterium] |
| MBF0333240.1 | DUF499_1.RM.like.systems | TUDOR+SF1-SF2-Helicase+iREase+CTD+REase->HEPN->METHYLASE->PolB->HEPN->HEPN+DUF499*-> | 1111 | Bacteria>Pseudomonadota>Alphaproteobacteria>unclassified_Alphaproteobacteria | GCA_015232055.1 | Alphaproteobacteria_bacterium | HQL40_06270 | 8520..11855 | 1913988 | DUF499-domain-containing-protein-[Alphaproteobacteria-bacterium] |
| MBI1213755.1 | DUF499_1.RM.like.systems | DEOR-HTH->TUDOR+SF1-SF2-Helicase+iREase+CTD+REase->?->?->TPR->METHYLASE->HEPN+DUF499*->HEPN->DOC+HTH-><-?\|\|Phage_integrase-><-AbiEi_4\|\|AbiEi_4-> | 1117 | Bacteria>Pseudomonadota>Alphaproteobacteria>unclassified_Alphaproteobacteria | GCA_016124635.1 | Alphaproteobacteria_bacterium | GC190_20030 | 81379..84732 | 1913988 | DUF499-domain-containing-protein-[Alphaproteobacteria-bacterium] |
| MBM3508286.1 | DUF499_1.RM.like.systems | <-HEPN+DUF499*<-HEPN<-PolB<-METHYLASE | 566 | Bacteria>Pseudomonadota>Alphaproteobacteria>unclassified_Alphaproteobacteria | GCA_016869235.1 | Alphaproteobacteria_bacterium | FJX64_11435 | 1..1697 | 1913988 | ATP-binding-protein-[Alphaproteobacteria-bacterium] |
| MCB9797660.1 | DUF499_1.RM.like.systems | TUDOR+SF1-SF2-Helicase+iREase+CTD+REase->METHYLASE->OLD-ABC->TOPRIM->HEPN+DUF499*-> | 1109 | Bacteria>Pseudomonadota>Alphaproteobacteria>unclassified_Alphaproteobacteria | GCA_020632765.1 | Alphaproteobacteria_bacterium | H6741_33640 | 16618..19947 | 1913988 | ATP-binding-protein-[Alphaproteobacteria-bacterium] |
| MCB9746493.1 | DUF499_1.RM.like.systems | <-HEPN+DUF499*<-TOPRIM<-OLD-ABC<-METHYLASE<-TUDOR+SF1-SF2-Helicase+iREase+CTD+REase | 1109 | Bacteria>Pseudomonadota>Alphaproteobacteria>unclassified_Alphaproteobacteria | GCA_020632815.1 | Alphaproteobacteria_bacterium | H6740_28215 | 5203..8532 | 1913988 | ATP-binding-protein-[Alphaproteobacteria-bacterium] |
| MCR9130420.1 | DUF499_1.RM.like.systems | <-NTP_transferase<-?<-REase+REase<-Mbetalac<-REase<-?<-HEPN+DUF499*<-METHYLASE<-MarR-HTH<-TUDOR+SF1-SF2-Helicase+iREase+CTD+REase<-?\|\|XIS-HTH-> | 1090 | Bacteria>Pseudomonadota>Alphaproteobacteria>unclassified_Alphaproteobacteria | GCA_024742735.1 | Alphaproteobacteria_bacterium | NXI12_12945 | 97854..101126 | 1913988 | DUF499-domain-containing-protein-[Alphaproteobacteria-bacterium] |
| MDA8675490.1 | DUF499_1.RM.like.systems | RNase->?->tRNA->Phage_integrase->?->TUDOR+SF1-SF2-Helicase+iREase+CTD+REase->METHYLASE->HEPN+DUF499*-> | 1091 | Bacteria>Pseudomonadota>Alphaproteobacteria>unclassified_Alphaproteobacteria | GCA_028099405.1 | Alphaproteobacteria_bacterium | N9M53_02250 | 53918..57193 | 1913988 | DUF499-domain-containing-protein-[Alphaproteobacteria-bacterium] |
| MDA8776763.1 | DUF499_1.RM.like.systems | Arm-DNA-bind_3+Phage_int_SAM_3+Phage_integrase->XIS-HTH->DnaB-><-?<-REase<-HEPN+DUF499*<-METHYLASE | 1094 | Bacteria>Pseudomonadota>Alphaproteobacteria>unclassified_Alphaproteobacteria | GCA_028101585.1 | Alphaproteobacteria_bacterium | N9N58_04660 | 21038..24322 | 1913988 | DUF499-domain-containing-protein-[Alphaproteobacteria-bacterium] |
| EXI84547.1 | DUF499_1.RM.like.systems | TPR+TPR+TPR+TPR+TPR+TPR+TPR+TPR-><-HerA/FtsK<-DDE_Tnp_1<-HerA/FtsK<-?\|\|HEPN-><-HEPN+DUF499*<-METHYLASE | 1114 | Bacteria>Pseudomonadota>Betaproteobacteria>Betaproteobacteria_incertae_sedis>Candidatus_Accumulibacter | GCA_000585075.1 | Candidatus_Accumulibacter_regalis | AW11_03844 | 11867..15211 | 522306 | MAG:-hypothetical-protein-AW11_03844-[Candidatus-Accumulibacter-regalis] |
| HCZ17096.1 | DUF499_1.RM.like.systems | <-HEPN+DUF499*<-HEPN<-PolB<-METHYLASE-p | 1112 | Bacteria>Pseudomonadota>Betaproteobacteria>Betaproteobacteria_incertae_sedis>Candidatus_Accumulibacter | GCA_003538495.1 | Accumulibacter_sp. | DHV85_21500 | 171..3509 | 2053492 | AAA+-family-ATPase-[Candidatus-Accumulibacter-sp.] |
| QLH49283.1 | DUF499_1.RM.like.systems | TUDOR+SF1-SF2-Helicase+iREase+CTD+REase->OLD-ABC->TOPRIM->OLD-ABC->TOPRIM->METHYLASE->HEPN+DUF499*-><-tRNA<-?<-GNAT<-?<-?<-ABhydrolase | 1112 | Bacteria>Pseudomonadota>Betaproteobacteria>Betaproteobacteria_incertae_sedis>Candidatus_Accumulibacter | GCA_013414765.1 | Candidatus_Accumulibacter_cognatus | HWD57_05410 | 1172035..1175373 | 2954383 | MAG:-ATP-binding-protein-[Candidatus-Accumulibacter-cognatus] |
| MBK7674932.1 | DUF499_1.RM.like.systems | PIN->PolB->HEPN->PolB->HEPN->HEPN+DUF499*->?->Response_reg+LuxR-HTH-> | 1114 | Bacteria>Pseudomonadota>Betaproteobacteria>Betaproteobacteria_incertae_sedis>Candidatus_Accumulibacter | GCA_016709675.1 | Candidatus_Accumulibacter_proximus | IPJ27_09305 | 116820..120164 | 2954385 | DUF499-domain-containing-protein-[Candidatus-Accumulibacter-sp.] |
| MBK8115022.1 | DUF499_1.RM.like.systems | TRD+TRD->Schlafen+HATPase+HTH-><-?<-?<-TPR+TPR<-?<-HEPN+DUF499*<-HEPN<-PolB<-HEPN<-PolB<-METHYLASE<-TOPRIM | 1109 | Bacteria>Pseudomonadota>Betaproteobacteria>Betaproteobacteria_incertae_sedis>Candidatus_Accumulibacter | GCA_016712935.1 | Accumulibacter_sp. | IPK44_10990 | 647374..650703 | 2053492 | DUF499-domain-containing-protein-[Candidatus-Accumulibacter-sp.] |
| MBN8517040.1 | DUF499_1.RM.like.systems | TOPRIM->OLD-ABC->TOPRIM->METHYLASE->PolB->HEPN->HEPN+DUF499*-><-tRNA<-?<-GNAT<-?<-?<-ABhydrolase | 1112 | Bacteria>Pseudomonadota>Betaproteobacteria>Betaproteobacteria_incertae_sedis>Candidatus_Accumulibacter | GCA_017302385.1 | Accumulibacter_sp. | JOL71_04035 | 55735..59073 | 2053492 | DUF499-domain-containing-protein-[Candidatus-Accumulibacter-sp.] |
| MBN8437943.1 | DUF499_1.RM.like.systems | PIN->METHYLASE->PolB->HEPN->PolB->HEPN->HEPN+DUF499*->?->TPR+TPR-> | 1109 | Bacteria>Pseudomonadota>Betaproteobacteria>Betaproteobacteria_incertae_sedis>Candidatus_Accumulibacter | GCA_017302555.1 | Accumulibacter_sp. | JOL95_07850 | 90759..94088 | 2053492 | DUF499-domain-containing-protein-[Candidatus-Accumulibacter-sp.] |
| MCM8599234.1 | DUF499_1.RM.like.systems | MerR-HTH->wHTH+REase->METHYLASE->?->DUF488-l->?->HEPN+DUF499*-> | 512 | Bacteria>Pseudomonadota>Betaproteobacteria>Betaproteobacteria_incertae_sedis>Candidatus_Accumulibacter | GCA_023806585.1 | Accumulibacter_sp. | NFW04_11345 | 16520..18057 | 2053492 | Swt1-family-HEPN-domain-containing-protein-[Candidatus-Accumulibacter-sp.] |
| MCM8594943.1 | DUF499_1.RM.like.systems | <-NUDIX\|\|?->?-><-HEPN+DUF499*<-STAS<-HEPN<-PolB | 1108 | Bacteria>Pseudomonadota>Betaproteobacteria>Betaproteobacteria_incertae_sedis>Candidatus_Accumulibacter | GCA_023806595.1 | Accumulibacter_sp. | NFV71_06200 | 31893..35219 | 2053492 | Swt1-family-HEPN-domain-containing-protein-[Candidatus-Accumulibacter-sp.] |
| MCQ1549843.1 | DUF499_1.RM.like.systems | <-XIS-HTH<-?<-?<-?<-Arm-DNA-bind_3+Phage_integrase<-HEPN+DUF499*<-METHYLASE-p | 1112 | Bacteria>Pseudomonadota>Betaproteobacteria>Betaproteobacteria_incertae_sedis>Candidatus_Accumulibacter | GCA_024380085.1 | Candidatus_Accumulibacter_phosphatis | NOF05_13730 | 62282..65620 | 327160 | Swt1-family-HEPN-domain-containing-protein-[Candidatus-Accumulibacter-phosphatis] |
| WP_138677185.1 | DUF499_1.RM.like.systems | ABhydrolase->?->?->GNAT->?->tRNA-><-HEPN+DUF499*<-HEPN<-PolB<-METHYLASE<-TOPRIM<-OLD-ABC<-METHYLASE-p | 1112 | Bacteria>Pseudomonadota>Betaproteobacteria>Betaproteobacteria_incertae_sedis>Candidatus_Accumulibacter | GCF_005889575.1 | Candidatus_Accumulibacter_phosphatis | ACCUM_RS00275 | 63215..66553 | 327160 | Swt1-family-HEPN-domain-containing-protein-[Candidatus-Accumulibacter-phosphatis] |
| WP_169069333.1 | DUF499_1.RM.like.systems | <-HEPN+DUF499*<-HEPN<-PolB<-METHYLASE<-HNH<-OLD-ABC<-PemK_toxin | 1112 | Bacteria>Pseudomonadota>Betaproteobacteria>Betaproteobacteria_incertae_sedis>Candidatus_Accumulibacter | GCF_012940005.1 | Candidatus_Accumulibacter_contiguus | E4Q08_RS03235 | 1844..5182 | 2954381 | Swt1-family-HEPN-domain-containing-protein-[Candidatus-Accumulibacter-contiguus] |
| WP_169068632.1 | DUF499_1.RM.like.systems | TUDOR+SF1-SF2-Helicase+iREase+CTD+REase->MORC+DUF3684->wHTH+REase->METHYLASE->PolB->HEPN->HEPN+DUF499*->MarR-HTH->tRNA->?-><-GDResolvase+HIN-HTH\|\|MetJarc-HTH->RelE-ParE-> | 1113 | Bacteria>Pseudomonadota>Betaproteobacteria>Betaproteobacteria_incertae_sedis | GCF_907163155.1 | Candidatus_Accumulibacter | QZ648_RS13960 | 40235..43576 | 327159 | Swt1-family-HEPN-domain-containing-protein-[Candidatus-Accumulibacter-phosphatis] |
| OYY21358.1 | DUF499_1.RM.like.systems | <-ParB+HNH<-REase+35exo<-HTH+WYL<-Peptidase<-HEPN+DUF499*<-METHYLASE<-TUDOR+SF1-SF2-Helicase+iREase+CTD+REase\|\|MerR-HTH->?-><-REase | 1109 | Bacteria>Pseudomonadota>Betaproteobacteria>Burkholderiales>Burkholderiaceae | GCA_002281105.1 | Polynucleobacter_sp._35-46-11 | B7Y67_02075 | 7019..10348 | 1970425 | MAG:-AAA+-family-ATPase-[Polynucleobacter-sp.-35-46-11] |
| MBT8525405.1 | DUF499_1.RM.like.systems | Phage_integrase-><-?\|\|?->?->?-><-HEPN+DUF499*<-METHYLASE<-TUDOR+SF1-SF2-Helicase+iREase+CTD+REase | 1103 | Bacteria>Pseudomonadota>Betaproteobacteria>Burkholderiales>Burkholderiaceae | GCA_018705035.1 | Polynucleobacter_paneuropaeus | G6724_05515 | 1135363..1138674 | 2527775 | ATP-binding-protein-[Polynucleobacter-paneuropaeus] |
| MBT8573895.1 | DUF499_1.RM.like.systems | Phage_integrase-><-?\|\|?->?->?-><-HEPN+DUF499*<-METHYLASE<-TUDOR+SF1-SF2-Helicase+iREase+CTD+REase | 1100 | Bacteria>Pseudomonadota>Betaproteobacteria>Burkholderiales>Burkholderiaceae | GCA_018705385.1 | Polynucleobacter_paneuropaeus | G6701_00110 | 16919..20221 | 2527775 | ATP-binding-protein-[Polynucleobacter-paneuropaeus] |
| MCA3183095.1 | DUF499_1.RM.like.systems | METHYLASE-p->HEPN+DUF499*-> | 993 | Bacteria>Pseudomonadota>Betaproteobacteria>Burkholderiales>Burkholderiaceae | GCA_020248745.1 | Cupriavidus_sp. | INH13_05675 | 475..3455 | 1873897 | DUF499-domain-containing-protein-[Cupriavidus-sp.] |
| WP_062175042.1 | DUF499_1.RM.like.systems | <-PIN<-SF1-SF2-Helicase<-OLD-ABC+TOPRIM<-HEPN+DUF499*<-METHYLASE<-MORC+DUF3684<-DUF2326<-?<-?<-HTH_37 | 1108 | Bacteria>Pseudomonadota>Betaproteobacteria>Burkholderiales>Burkholderiaceae | GCF_001557535.2 | Burkholderia_sp._PAMC_26561 | AXG89_RS34140 | 126894..130220 | 1795043 | Swt1-family-HEPN-domain-containing-protein-[Burkholderia-sp.-PAMC-26561] |
| WP_205989436.1 | DUF499_1.RM.like.systems | RelB->RelE-ParE->TUDOR+SF1-SF2-Helicase+iREase+CTD+REase->METHYLASE->HEPN+DUF499*-><-HTH+WYL\|\|?->?->HTH+UbE1-> | 1110 | Bacteria>Pseudomonadota>Betaproteobacteria>Burkholderiales>Burkholderiaceae | GCF_017105025.1 | Paraburkholderia_sp._Ac-20336 | GXB81_RS13960 | 20393..23725 | 2703886 | Swt1-family-HEPN-domain-containing-protein-[Paraburkholderia-sp.-Ac-20336] |
| WP_035924723.1 | DUF499_1.RM.like.systems | Phage_integrase-><-HEPN+DUF499*<-METHYLASE<-TUDOR+SF1-SF2-Helicase+iREase+CTD+REase<-DUF2326<-?<-?<-HTH_37 | 1108 | Bacteria>Pseudomonadota>Betaproteobacteria>Burkholderiales>Burkholderiaceae | GCF_902833485.1 | Caballeronia_glathei | BG61_RS34410 | 1980..5306 | 60547 | Swt1-family-HEPN-domain-containing-protein-[Caballeronia-glathei] |
| MCP5281003.1 | DUF499_1.RM.like.systems | <-HEPN+DUF499*<-?<-METHYLASE<-DUF932<-ParB+HNH<-?<-TUDOR+SF1-SF2-Helicase+iREase+CTD+REase | 1107 | Bacteria>Pseudomonadota>Betaproteobacteria>Burkholderiales>Comamonadaceae | GCA_024235935.1 | Rhodoferax_sp. | H6930_05335 | 220759..224082 | 50421 | ATP-binding-protein-[Rhodoferax-sp.] |
| MCC6854228.1 | DUF499_1.RM.like.systems | <-HEPN+DUF499*<-HEPN<-PolB<-METHYLASE<-PemK_toxin<-PrlF_antitoxin<-TOPRIM | 1117 | Bacteria>Pseudomonadota>Betaproteobacteria>Burkholderiales>Sphaerotilaceae | GCA_020852865.1 | Rubrivivax_sp. | IT502_18195 | 74611..77964 | 50259 | DUF499-domain-containing-protein-[Rubrivivax-sp.] |
| WP_119292278.1 | DUF499_1.RM.like.systems | <-SbcC<-?<-?<-REase<-?<-HEPN+DUF499*<-HEPN<-PolB<-SbcC<-METHYLASE<-sGTP<-TUDOR+SF1-SF2-Helicase+iREase+CTD+REase | 1114 | Bacteria>Pseudomonadota>Betaproteobacteria>Burkholderiales>Sphaerotilaceae | GCF_003574675.1 | Azohydromonas_sediminis | DRV96_RS14230 | 10149..13493 | 2259674 | Swt1-family-HEPN-domain-containing-protein-[Calidifontimicrobium-sediminis] |
| RYG38846.1 | DUF499_1.RM.like.systems | METHYLASE->HEPN+DUF499*->?-><-HTH+WYL\|\|?->?->?->STYKIN-> | 1103 | Bacteria>Pseudomonadota>Betaproteobacteria>Burkholderiales>unclassified_Burkholderiales | GCA_004145455.1 | Burkholderiales_bacterium | EON93_01200 | 2210..5521 | 1891238 | ATP-binding-protein-[Burkholderiales-bacterium] |
| NMA26876.1 | DUF499_1.RM.like.systems | METHYLASE-p->OLD-ABC->TOPRIM->HEPN+DUF499*-> | 448 | Bacteria>Pseudomonadota>Betaproteobacteria>Burkholderiales>unclassified_Burkholderiales | GCA_012799975.1 | Burkholderiales_bacterium | GX934_03860 | 2623..3966 | 1891238 | ATP-binding-protein-[Burkholderiales-bacterium] |
| MBE0627115.1 | DUF499_1.RM.like.systems | TUDOR+SF1-SF2-Helicase+iREase+CTD+REase->OLD-ABC->TOPRIM->HEPN->METHYLASE->HEPN+DUF499*->?->?->HEPN->REase-> | 1113 | Bacteria>Pseudomonadota>Betaproteobacteria>Burkholderiales>unclassified_Burkholderiales | GCA_014860355.1 | Burkholderiales_bacterium | IH606_20135 | 9089..12430 | 1891238 | DUF499-domain-containing-protein-[Burkholderiales-bacterium] |
| MBE7523982.1 | DUF499_1.RM.like.systems | S1+RNase+2C->ABhydrolase->Glyoxalase-><-?<-HTH+PolB<-HEPN+DUF499*<-STAS<-?<-?<-METHYLASE<-?<-Schlafen+HATPase+HTH | 1119 | Bacteria>Pseudomonadota>Betaproteobacteria>Burkholderiales>unclassified_Burkholderiales | GCA_015075645.1 | Burkholderiales_bacterium | HS109_16570 | 209552..212911 | 1891238 | ATP-binding-protein-[Burkholderiales-bacterium] |
| MBI5907809.1 | DUF499_1.RM.like.systems | ABC-ATPase+SbcC->ABC-ATPase+SbcC->?-><-HEPN+DUF499*<-?<-METHYLASE | 1107 | Bacteria>Pseudomonadota>Betaproteobacteria>Burkholderiales>unclassified_Burkholderiales | GCA_016234645.1 | Burkholderiales_bacterium | HY853_04415 | 2095..5418 | 1891238 | ATP-binding-protein-[Burkholderiales-bacterium] |
| MBL8529542.1 | DUF499_1.RM.like.systems | Schlafen+HATPase+HTH->?->METHYLASE-p->HTH+Hiskin+STAS->PolB->HEPN+DUF499*-> | 1108 | Bacteria>Pseudomonadota>Betaproteobacteria>Burkholderiales>unclassified_Burkholderiales | GCA_016791285.1 | Burkholderiales_bacterium | JNL68_17825 | 6694..10020 | 1891238 | DUF499-domain-containing-protein-[Burkholderiales-bacterium] |
| MBN8477381.1 | DUF499_1.RM.like.systems | TUDOR+SF1-SF2-Helicase+iREase+CTD+REase->?->METHYLASE->?->?->HEPN+DUF499*->?->HTH->NTP_transf-> | 1117 | Bacteria>Pseudomonadota>Betaproteobacteria>Burkholderiales>unclassified_Burkholderiales | GCA_017302475.1 | Burkholderiales_bacterium | JOL91_00275 | 13639..16992 | 1891238 | DUF499-domain-containing-protein-[Burkholderiales-bacterium] |
| MBX9629971.1 | DUF499_1.RM.like.systems | RelE-ParE->?->TUDOR+SF1-SF2-Helicase+iREase+CTD+REase->?->?->METHYLASE->HEPN+DUF499*-><-?<-AbiEii<-AbiEi_4 | 1116 | Bacteria>Pseudomonadota>Betaproteobacteria>Burkholderiales>unclassified_Burkholderiales | GCA_019747905.1 | Burkholderiales_bacterium | K2X67_05560 | 12511..15861 | 1891238 | DUF499-domain-containing-protein-[Burkholderiales-bacterium] |
| MCA3160290.1 | DUF499_1.RM.like.systems | HTH_Tnp_1+HTH_21+rve+rve_3->METHYLASE-p->METHYLASE-p->HEPN+DUF499*->?-><-SAM-methylase | 1098 | Bacteria>Pseudomonadota>Betaproteobacteria>Burkholderiales>unclassified_Burkholderiales | GCA_020248485.1 | Burkholderiales_bacterium | ING31_11990 | 9686..12982 | 1891238 | DUF499-domain-containing-protein-[Burkholderiales-bacterium] |
| MCA3160409.1 | DUF499_1.RM.like.systems | <-HEPN+DUF499*<-METHYLASE | 815 | Bacteria>Pseudomonadota>Betaproteobacteria>Burkholderiales>unclassified_Burkholderiales | GCA_020248485.1 | Burkholderiales_bacterium | ING31_12605 | 1..2446 | 1891238 | DUF499-domain-containing-protein-[Burkholderiales-bacterium] |
| MCC6474792.1 | DUF499_1.RM.like.systems | <-HEPN+DUF499*<-HEPN<-PolB<-METHYLASE<-HEPN<-ParB+HNH\|\|tRNA-><-PIN | 1120 | Bacteria>Pseudomonadota>Betaproteobacteria>Burkholderiales>unclassified_Burkholderiales | GCA_020846575.1 | Burkholderiales_bacterium | IT514_13720 | 1159..4521 | 1891238 | DUF499-domain-containing-protein-[Burkholderiales-bacterium] |
| MCC6611669.1 | DUF499_1.RM.like.systems | VirD4-TraG+VirD4-FtsK->TrwC+SF1-SF2-helicase+TOPRIM-><-PIWI<-TIR+PNTD2<-HEPN+DUF499*<-METHYLASE<-PIN<-?<-STAS+HisKin<-TUDOR+SF1-SF2-Helicase+iREase+CTD+REase<-PIN | 1116 | Bacteria>Pseudomonadota>Betaproteobacteria>Burkholderiales>unclassified_Burkholderiales | GCA_020848075.1 | Burkholderiales_bacterium | IT515_18605 | 34273..37623 | 1891238 | DUF499-domain-containing-protein-[Burkholderiales-bacterium] |
| MCC7486352.1 | DUF499_1.RM.like.systems | ORC-CDC6+REase->HEPN+DUF499*->?->?-><-?<-ABhydrolase | 1107 | Bacteria>Pseudomonadota>Betaproteobacteria>Burkholderiales>unclassified_Burkholderiales | GCA_020854035.1 | Burkholderiales_bacterium | IT529_15375 | 2503..5826 | 1891238 | DUF499-domain-containing-protein-[Burkholderiales-bacterium] |
| MCI4428467.1 | DUF499_1.RM.like.systems | <-HEPN+DUF499*<-HEPN<-HTH+Hiskin+STAS<-HEPN<-PolB<-METHYLASE-p | 342 | Bacteria>Pseudomonadota>Betaproteobacteria>Burkholderiales>unclassified_Burkholderiales | GCA_022758925.1 | Burkholderiales_bacterium | JHC40_04765 | 1..1025 | 1891238 | ATP-binding-protein-[Burkholderiales-bacterium] |
| MCL4799144.1 | DUF499_1.RM.like.systems | <-HEPN+DUF499*<-HEPN<-PolB<-METHYLASE<-?<-TUDOR+SF1-SF2-Helicase+iREase+CTD+REase<-TIR | 1113 | Bacteria>Pseudomonadota>Betaproteobacteria>Burkholderiales>unclassified_Burkholderiales | GCA_023383795.1 | Burkholderiales_bacterium | KJ025_06120 | 926..4267 | 1891238 | DUF499-domain-containing-protein-[Burkholderiales-bacterium] |
| MCL6557122.1 | DUF499_1.RM.like.systems | <-HEPN+DUF499*<-PolB<-?<-ORC-CDC6+REase<-TUDOR+SF1-SF2-Helicase+iREase+CTD+REase-p | 1098 | Bacteria>Pseudomonadota>Betaproteobacteria>Burkholderiales>unclassified_Burkholderiales | GCA_023511995.1 | Burkholderiales_bacterium | K6T56_12270 | 1861..5157 | 1891238 | DUF499-domain-containing-protein-[Burkholderiales-bacterium] |
| GIX27884.1 | DUF499_1.RM.like.systems | METHYLASE-p-><-?\|\|METHYLASE-p->ParD->PIN->HEPN+DUF499*->DUF499-p->?->?->ORC-CDC6+REase-> | 389 | Bacteria>Pseudomonadota>Betaproteobacteria>Burkholderiales>unclassified_Burkholderiales | GCA_026005015.1 | Burkholderiales_bacterium | KatS3mg123_1765 | 1711823..1712992 | 1891238 | MAG:-hypothetical-protein-KatS3mg123_1765-[Burkholderiales-bacterium] |
| MCZ2419561.1 | DUF499_1.RM.like.systems | TPR+TPR-><-?<-HTH+UbE1<-?\|\|HTH->MPTase-><-HEPN+DUF499*<-?<-DUF488-l<-?<-METHYLASE<-TOPRIM<-OLD-ABC | 1108 | Bacteria>Pseudomonadota>Betaproteobacteria>Burkholderiales>unclassified_Burkholderiales | GCA_026931465.1 | Burkholderiales_bacterium | LC123_06960 | 80916..84242 | 1891238 | MAG:-DUF499-domain-containing-protein-[Burkholderiales-bacterium] |
| NDU85854.1 | DUF499_1.RM.like.systems | <-HTH+UbE1<-?\|\|?->?->MPTase-><-HEPN+DUF499*<-METHYLASE<-TOPRIM<-OLD-ABC<-TUDOR+SF1-SF2-Helicase+iREase+CTD+REase<-SF1-SF2-Helicase<-OLD-ABC+TOPRIM | 1110 | Bacteria>Pseudomonadota>Betaproteobacteria>Ferrovales>Ferrovaceae | GCA_010436015.1 | Ferrovum_sp. | G3H99_04485 | 6034..9366 | 2609467 | ATP-binding-protein-[Ferrovum-sp.] |
| QKE41374.1 | DUF499_1.RM.like.systems | HTH+UbE1->?->?->?->Phage_integrase-><-HEPN+DUF499*<-ORC-CDC6<-ParA-Soj-PloopNTPase<-HEPN<-METHYLASE<-TOPRIM<-OLD-ABC | 1109 | Bacteria>Pseudomonadota>Betaproteobacteria>Ferrovales>Ferrovaceae | GCA_013181415.1 | Ferrovum_myxofaciens | HO274_08630 | 1781795..1785124 | 416213 | ATP-binding-protein-[Ferrovum-myxofaciens] |
| WP_031597360.1 | DUF499_1.RM.like.systems | TUDOR+SF1-SF2-Helicase+iREase+CTD+REase->TOPRIM->METHYLASE->HEPN->ParA-Soj-PloopNTPase->ORC-CDC6->HEPN+DUF499*-><-Phage_integrase<-?<-?\|\|<-?<-HTH+UbE1 | 1109 | Bacteria>Pseudomonadota>Betaproteobacteria>Ferrovales>Ferrovaceae | GCF_000735045.1 | Ferrovum_myxofaciens | IV91_RS0107710 | 11022..14351 | 416213 | Swt1-family-HEPN-domain-containing-protein-[Ferrovum-myxofaciens] |
| WP_273118797.1 | DUF499_1.RM.like.systems | HTH+UbE1->?->\|\|?->?->Phage_integrase-><-HEPN+DUF499*<-ORC-CDC6<-ParA-Soj-PloopNTPase<-HEPN<-METHYLASE<-TOPRIM<-OLD-ABC | 1109 | Bacteria>Pseudomonadota>Betaproteobacteria>Ferrovales>Ferrovaceae | GCF_013181415.1 | Ferrovum_myxofaciens | HO274_RS08660 | 1781795..1785124 | 416213 | Swt1-family-HEPN-domain-containing-protein-[Ferrovum-myxofaciens] |
| CAG1003423.1 | DUF499_1.RM.like.systems | TUDOR+SF1-SF2-Helicase+iREase+CTD+REase->sGTP->ParB+HNH->METHYLASE->HEPN+DUF499*-> | 1107 | Bacteria>Pseudomonadota>Betaproteobacteria>Nitrosomonadales>Methylophilaceae | GCA_905339275.1 | Methylophilaceae_bacterium | MTYP_03091 | 15685..19008 | 2030816 | hypothetical-protein-MTYP_03091-[Methylophilaceae-bacterium] |
| MBP0133287.1 | DUF499_1.RM.like.systems | XIS-HTH->?->TUDOR+SF1-SF2-Helicase+iREase+CTD+REase->Cluster1090_5clades->METHYLASE->HEPN+DUF499*-><-HTH+WYL | 1105 | Bacteria>Pseudomonadota>Betaproteobacteria>Nitrosomonadales>Nitrosomonadaceae | GCA_016106425.1 | Nitrosospira_sp. | ITD31_07875 | 10694..14011 | 38080 | DUF499-domain-containing-protein-[Nitrosospira-sp.] |
| WP_107785979.1 | DUF499_1.RM.like.systems | <-HIN-HTH\|\|TUDOR+SF1-SF2-Helicase+iREase+CTD+REase->ParB+HNH->METHYLASE->HEPN+DUF499*-><-tRNA\|\|FURR-HTH-> | 1112 | Bacteria>Pseudomonadota>Betaproteobacteria>Nitrosomonadales>Nitrosomonadaceae | GCF_003051045.1 | Nitrosomonas_ureae | C8R28_RS01190 | 8316..11654 | 44577 | Swt1-family-HEPN-domain-containing-protein-[Nitrosomonas-ureae] |
| MBK7234801.1 | DUF499_1.RM.like.systems | <-SIGMA-HTH+SIGMA-HTH\|\|DDE_Tnp_1->DDE_Tnp_1_assoc+DDE_Tnp_1->DDE_Tnp_1_assoc+DDE_Tnp_1-><-35exo+SF1-SF2-Helicase<-HEPN+DUF499*<-HEPN<-PolB<-wHTH+REase | 1112 | Bacteria>Pseudomonadota>Betaproteobacteria>Nitrosomonadales>Sterolibacteriaceae | GCA_016707455.1 | Sterolibacteriaceae_bacterium | IPI02_03880 | 820379..823717 | 2682146 | DUF499-domain-containing-protein-[Sterolibacteriaceae-bacterium] |

| Protein-ID | System | Gene-Neighbourhoods | Length | Taxonomy | Assembly-ID | Strain | Locus-Tag | Genomic-coordinates | TaxID | Fasta-defline |
|---|---|---|---|---|---|---|---|---|---|---|
| MBK6974203.1 | DUF499_1.RM.like.systems | <-SIGMA-HTH+SIGMA-HTH\|\|DDE_Tnp_1_assoc+DDE_Tnp_1-><-35exo+SF1-SF2-Helicase<-HEPN+DUF499*<-HEPN<-PolB<-wHTH+REase<-TUDOR+SF1-SF2-Helicase+iREase+CTD+REase | 1112 | Bacteria>Pseudomonadota>Betaproteobacteria>Nitrosomonadales>Sterolibacteriaceae | GCA_016708905.1 | Candidatus_Methylophosphatis_roskildensis | IPH26_15090 | 451210..454548 | 2899263 | DUF499-domain-containing-protein-[Sterolibacteriaceae-bacterium] |
| MBK7665767.1 | DUF499_1.RM.like.systems | TUDOR+SF1-SF2-Helicase+iREase+CTD+REase->Schlafen+HATPase+HTH->?->PolB->HEPN->HEPN+DUF499*->35exo+SF1-SF2-Helicase->SIGMA-HTH+SIGMA-HTH-> | 1112 | Bacteria>Pseudomonadota>Betaproteobacteria>Nitrosomonadales>Sterolibacteriaceae | GCA_016709795.1 | Sterolibacteriaceae_bacterium | IPJ21_19955 | 4305507..4308845 | 2682146 | DUF499-domain-containing-protein-[Sterolibacteriaceae-bacterium] |
| ODU50157.1 | DUF499_1.RM.like.systems | <-HEPN+DUF499*<-?<-DUF488-l<-?<-METHYLASE<-sGTP<-TUDOR+SF1-SF2-Helicase+iREase+CTD+REase | 1110 | Bacteria>Pseudomonadota>Betaproteobacteria>Nitrosomonadales>Thiobacillaceae | GCA_001725135.1 | Thiobacillus_sp._SCN_63-374 | ABS92_03900 | 1302..4634 | 1660149 | MAG:-AAA+-family-ATPase-[Thiobacillus-sp.-SCN-63-374] |
| MBI1887192.1 | DUF499_1.RM.like.systems | <-Glycos_trans_3N+Glycos_transf_3<-?<-Schlafen+HATPase+HTH<-?<-?<-HEPN+DUF499*<-METHYLASE<-HEPN<-TOPRIM<-OLD-ABC<-TUDOR+SF1-SF2-Helicase+iREase+CTD+REase<-ParB | 1112 | Bacteria>Pseudomonadota>Betaproteobacteria>Nitrosomonadales>unclassified_Nitrosomonadales | GCA_016177795.1 | Nitrosomonadales_bacterium | HYS19_02300 | 86364..89702 | 2026761 | ATP-binding-protein-[Nitrosomonadales-bacterium] |
| MBI3902214.1 | DUF499_1.RM.like.systems | <-Glycos_trans_3N+Glycos_transf_3<-?<-?<-Schlafen+HATPase+HTH<-?<-HEPN+DUF499*<-METHYLASE<-?<-?<-TOPRIM<-OLD-ABC<-TUDOR+SF1-SF2-Helicase+iREase+CTD+REase | 1112 | Bacteria>Pseudomonadota>Betaproteobacteria>Nitrosomonadales>unclassified_Nitrosomonadales | GCA_016199835.1 | Nitrosomonadales_bacterium | HY306_04630 | 118339..121677 | 2026761 | ATP-binding-protein-[Nitrosomonadales-bacterium] |
| MBI5658243.1 | DUF499_1.RM.like.systems | TUDOR+SF1-SF2-Helicase+iREase+CTD+REase->OLD-ABC->TOPRIM->METHYLASE->?->?->HEPN+DUF499*-> | 1125 | Bacteria>Pseudomonadota>Betaproteobacteria>Nitrosomonadales>unclassified_Nitrosomonadales | GCA_016223025.1 | Nitrosomonadales_bacterium | HZC43_01545 | 64771..68148 | 2026761 | ATP-binding-protein-[Nitrosomonadales-bacterium] |
| MBN8462739.1 | DUF499_1.RM.like.systems | PrlF_antitoxin->PemK_toxin->REase->METHYLASE->PolB->HEPN->HEPN+DUF499*->?-><-?<-?\|\|?->sGTP-> | 1110 | Bacteria>Pseudomonadota>Betaproteobacteria>Rhodocyclales>Azonexaceae | GCA_017302355.1 | Dechloromonas_sp. | JOM01_07975 | 12361..15693 | 1917218 | DUF499-domain-containing-protein-[Dechloromonas-sp.] |
| KAB2934118.1 | DUF499_1.RM.like.systems | <-PAPS_reduct<-?<-KAP-NTPase<-HEPN+DUF499*<-PIN<-OLD-ABC<-METHYLASE<-?<-TUDOR+SF1-SF2-Helicase+iREase+CTD+REase | 1109 | Bacteria>Pseudomonadota>Betaproteobacteria>Rhodocyclales>Rhodocyclaceae | GCA_008933825.1 | Rhodocyclaceae_bacterium | F9K21_11715 | 6014..9343 | 1898103 | ATP-binding-protein-[Rhodocyclaceae-bacterium] |
| KAB2933410.1 | DUF499_1.RM.like.systems | OLD-ABC->TOPRIM->?->?->METHYLASE->REase->HEPN+DUF499*->?-><-?\|\|HTH->MarR-HTH-> | 1096 | Bacteria>Pseudomonadota>Betaproteobacteria>Rhodocyclales>Rhodocyclaceae | GCA_008933825.1 | Rhodocyclaceae_bacterium | F9K21_12245 | 12685..15975 | 1898103 | ATP-binding-protein-[Rhodocyclaceae-bacterium] |
| MBP6422202.1 | DUF499_1.RM.like.systems | <-CASPASE<-TIR<-HEPN+DUF499*<-METHYLASE-p | 1110 | Bacteria>Pseudomonadota>Betaproteobacteria>Rhodocyclales>Rhodocyclaceae | GCA_017988215.1 | Propionivibrio_sp. | KA271_04855 | 1433..4765 | 2212460 | DUF499-domain-containing-protein-[Propionivibrio-sp.] |
| MCA3140850.1 | DUF499_1.RM.like.systems | HTH_3->TUDOR+SF1-SF2-Helicase+iREase+CTD+REase->Schlafen+HATPase+HTH->?->METHYLASE->HTH+Hiskin+STAS->HEPN+DUF499*->?->XIS-HTH->?->TOPRIM->MetJarc-HTH->RelE-ParE-> | 1113 | Bacteria>Pseudomonadota>Betaproteobacteria>Rhodocyclales>Rhodocyclaceae | GCA_020248385.1 | Rhodocyclaceae_bacterium | ING80_01400 | 55201..58542 | 1898103 | DUF499-domain-containing-protein-[Rhodocyclaceae-bacterium] |
| WP_018989132.1 | DUF499_1.RM.like.systems | <-PIN+PhoH<-Radical-SAM+TRAM\|\|tRNA->?-><-HEPN+DUF499*<-METHYLASE<-HNH<-OLD-ABC<-PIN<-?<-STAS+HisKin | 1121 | Bacteria>Pseudomonadota>Betaproteobacteria>Rhodocyclales>Rhodocyclaceae | GCF_000378245.1 | Aromatoleum_toluclasticum | F464_RS0107275 | 366039..369404 | 92003 | Swt1-family-HEPN-domain-containing-protein-[Aromatoleum-toluclasticum] |
| WP_153167206.1 | DUF499_1.RM.like.systems | MORC+DNA_gyraseB+TOPRIM-><-?<-?<-HEPN+DUF499*<-?<-METHYLASE<-?<-MORC+DUF3684<-TUDOR+SF1-SF2-Helicase+iREase+CTD+REase | 1120 | Bacteria>Pseudomonadota>Betaproteobacteria>Rhodocyclales>Zoogloeaceae | GCF_009469595.2 | Thauera_sp._2A1 | GH664_RS06300 | 29943..33305 | 2570191 | Swt1-family-HEPN-domain-containing-protein-[Thauera-sp.-2A1] |
| AGG33842.1 | DUF499_1.RM.like.systems | Peptidase_M20->PrmC_N+Methyltransf_25-><-SMS-KaiC<-?<-?\|\|tRNA-><-HEPN+DUF499*<-METHYLASE<-TUDOR+SF1-SF2-Helicase+iREase+CTD+REase\|\|MerR-HTH-> | 1110 | Bacteria>Pseudomonadota>Betaproteobacteria>unclassified_Betaproteobacteria | GCA_000342265.1 | beta_proteobacterium_CB | D521_1274 | 1257498..1260830 | 543913 | ATPase-(AAA+-superfamily)-like-protein-[beta-proteobacterium-CB] |
| OGA65437.1 | DUF499_1.RM.like.systems | <-HEPN+DUF499*<-PIN<-MPTase<-?<-?<-METHYLASE<-ParB+HNH | 1109 | Bacteria>Pseudomonadota>Betaproteobacteria>unclassified_Betaproteobacteria | GCA_001770585.1 | Betaproteobacteria_bacterium_RIFCSPLOWO2_12_FULL_65_14 | A3G81_12440 | 218..3547 | 1797501 | AAA+-family-ATPase-[Betaproteobacteria-bacterium-RIFCSPLOWO2_12_FULL_65_14] |
| MBI4756762.1 | DUF499_1.RM.like.systems | METHYLASE->?->?->?->PolB->HEPN->HEPN+DUF499*-> | 1123 | Bacteria>Pseudomonadota>Betaproteobacteria>unclassified_Betaproteobacteria | GCA_016208415.1 | Betaproteobacteria_bacterium | HY778_15365 | 56697..60068 | 1891241 | ATP-binding-protein-[Betaproteobacteria-bacterium] |
| MBL8540233.1 | DUF499_1.RM.like.systems | TUDOR+SF1-SF2-Helicase+iREase+CTD+REase->METHYLASE->HTH+Hiskin+STAS->PolB->PolB->HEPN->HEPN+DUF499*-><-?<-?<-?<-Response_reg+LuxR-HTH<-STYKIN | 1105 | Bacteria>Pseudomonadota>Betaproteobacteria>unclassified_Betaproteobacteria | GCA_016791345.1 | Betaproteobacteria_bacterium | JNK68_07660 | 10175..13492 | 1891241 | DUF499-domain-containing-protein-[Betaproteobacteria-bacterium] |
| MDA8257273.1 | DUF499_1.RM.like.systems | <-EVE<-HEPN+DUF499*<-HEPN<-PolB<-ORC-CDC6<-ParA-Soj-PloopNTPase<-HEPN<-METHYLASE | 1114 | Bacteria>Pseudomonadota>Betaproteobacteria>unclassified_Betaproteobacteria | GCA_028065725.1 | Betaproteobacteria_bacterium | MOZ99_16875 | 140..3484 | 1891241 | MAG:-Swt1-family-HEPN-domain-containing-protein-[Betaproteobacteria-bacterium] |
| MDE2342677.1 | DUF499_1.RM.like.systems | <-HEPN+DUF499*<-METHYLASE<-TUDOR+SF1-SF2-Helicase+iREase+CTD+REase | 1094 | Bacteria>Pseudomonadota>Betaproteobacteria>unclassified_Betaproteobacteria | GCA_028870015.1 | Betaproteobacteria_bacterium | KGL63_04675 | 257..3541 | 1891241 | MAG:-DUF499-domain-containing-protein-[Betaproteobacteria-bacterium] |
| WP_096360574.1 | DUF499_1.RM.like.systems | <-CHTH<-HTH_3<-RelE-ParE<-UPF0061<-?<-?<-HEPN+DUF499*<-METHYLASE<-?<-TUDOR+SF1-SF2-Helicase+iREase+CTD+REase<-?<-BirA-HTH<-HKD-DNase | 1113 | Bacteria>Pseudomonadota>Gammaproteobacteria>Acidiferrobacterales>Acidiferrobacteraceae | GCF_002355735.1 | Sulfuricaulis_limicola | SCL_RS07110 | 1437860..1441201 | 1620215 | Swt1-family-HEPN-domain-containing-protein-[Sulfuricaulis-limicola] |
| MBK8637657.1 | DUF499_1.RM.like.systems | TOPRIM->METHYLASE->PolB->HEPN->?->Y1_Tnp->HEPN+DUF499*->RADICAL-SAM-> | 1120 | Bacteria>Pseudomonadota>Gammaproteobacteria>Chromatiales>Chromatiaceae | GCA_016714645.1 | Chromatiaceae_bacterium | IPN92_04990 | 1074519..1077881 | 2049432 | DUF499-domain-containing-protein-[Chromatiaceae-bacterium] |
| MBP6807674.1 | DUF499_1.RM.like.systems | <-HEPN+DUF499*<-?<-?<-PIN<-?<-METHYLASE | 368 | Bacteria>Pseudomonadota>Gammaproteobacteria>Chromatiales>Chromatiaceae | GCA_017991215.1 | Chromatiaceae_bacterium | KA125_05205 | 1..1104 | 2049432 | DUF499-domain-containing-protein-[Chromatiaceae-bacterium] |
| QVL49217.1 | DUF499_1.RM.like.systems | OLD-ABC->TOPRIM->METHYLASE->?->PolB->HEPN->HEPN+DUF499*-><-?<-?\|\|HicB-><-sGTP\|\|?-><-TPR+TPR | 1114 | Bacteria>Pseudomonadota>Gammaproteobacteria>Chromatiales>Chromatiaceae | GCA_018399035.1 | Thiocapsa_sp. | KFB96_01415 | 301904..305248 | 2024551 | MAG:-DUF499-domain-containing-protein-[Thiocapsa-sp.] |
| MCC7277404.1 | DUF499_1.RM.like.systems | CHTH-><-HEPN+DUF499*<-HEPN<-PolB<-HEPN<-PolB | 1122 | Bacteria>Pseudomonadota>Gammaproteobacteria>Chromatiales>Chromatiaceae | GCA_020851955.1 | Chromatiaceae_bacterium | IT487_03685 | 4457..7825 | 2049432 | DUF499-domain-containing-protein-[Chromatiaceae-bacterium] |
| MCU0834934.1 | DUF499_1.RM.like.systems | <-DUF499-p<-HEPN+DUF499*<-HTH+Hiskin+STAS<-METHYLASE-p<-METHYLASE<-PemK_toxin<-PrlF_antitoxin<-TUDOR+SF1-SF2-Helicase+iREase+CTD+REase-p | 755 | Bacteria>Pseudomonadota>Gammaproteobacteria>Chromatiales>Chromatiaceae | GCA_025458165.1 | Chromatiaceae_bacterium | MUC77_10980 | 31032..33299 | 2049432 | Swt1-family-HEPN-domain-containing-protein-[Chromatiaceae-bacterium] |
| WP_020507743.1 | DUF499_1.RM.like.systems | OLD-ABC->TOPRIM->METHYLASE->PolB->PolB->HEPN->HEPN+DUF499*-><-?<-?<-?<-REC<-HPT+HISKIN+REC | 1124 | Bacteria>Pseudomonadota>Gammaproteobacteria>Chromatiales>Chromatiaceae | GCF_000379525.1 | Lamprocystis_purpurea | A39O_RS0126800 | 79657..83031 | 61598 | Swt1-family-HEPN-domain-containing-protein-[Lamprocystis-purpurea] |
| WP_093189479.1 | DUF499_1.RM.like.systems | METHYLASE->PolB->HEPN->HEPN+DUF499*-><-?\|\|?->?->?-><-35exo | 1113 | Bacteria>Pseudomonadota>Gammaproteobacteria>Chromatiales>Chromatiaceae | GCF_001499735.1 | Thiocapsa_sp._KS1 | THIOKS1_RS17085 | 3720907..3724248 | 610332 | Swt1-family-HEPN-domain-containing-protein-[Thiocapsa-sp.-KS1] |
| WP_100917910.1 | DUF499_1.RM.like.systems | TUDOR+SF1-SF2-Helicase+iREase+CTD+REase->OLD-ABC->TOPRIM->METHYLASE->PolB->HEPN+DUF499*->?-><-NUDIX<-?<-?\|\|?-><-PAIREDC-HTH | 1111 | Bacteria>Pseudomonadota>Gammaproteobacteria>Chromatiales>Chromatiaceae | GCF_002813775.1 | Candidatus_Thiodictyon_syntrophicum | THSYN_RS03450 | 792207..795542 | 1166950 | Swt1-family-HEPN-domain-containing-protein-[Candidatus-Thiodictyon-syntrophicum] |
| WP_120796610.1 | DUF499_1.RM.like.systems | PrlF_antitoxin->RHH_6->PIN->METHYLASE-p->?->PolB->HEPN+DUF499*-><-?\|\|?->RVT+HNH->RVT+HNH-> | 1114 | Bacteria>Pseudomonadota>Gammaproteobacteria>Chromatiales>Chromatiaceae | GCF_003634315.1 | Thiocapsa_rosea | BDD21_RS07415 | 1614812..1618156 | 69360 | Swt1-family-HEPN-domain-containing-protein-[Thiocapsa-rosea] |
| WP_153974954.1 | DUF499_1.RM.like.systems | <-ABhydrolase<-?<-HEPN+DUF499*<-HEPN<-PolB<-METHYLASE<-?<-TUDOR+SF1-SF2-Helicase+iREase+CTD+REase | 1103 | Bacteria>Pseudomonadota>Gammaproteobacteria>Chromatiales>Chromatiaceae | GCF_009664085.1 | Thermochromatium_tepidum | E6P07_RS07035 | 1550502..1553813 | 1050 | Swt1-family-HEPN-domain-containing-protein-[Thermochromatium-tepidum] |
| WP_091642911.1 | DUF499_1.RM.like.systems | <-AbiEi_1<-GDResolvase+HIN-HTH\|\|HIN-HTH->?->LZ_Tnp_IS66+zf-IS66+DDE_Tnp_IS66+DDE_Tnp_IS66_C->?-><-HEPN+DUF499*<-METHYLASE<-TUDOR+SF1-SF2-Helicase+iREase+CTD+REase\|\|HTH_Tnp_1+HTH_21+rve-> | 1097 | Bacteria>Pseudomonadota>Gammaproteobacteria>Chromatiales>Ectothiorhodospiraceae | GCF_900110585.1 | Aquisalimonas_asiatica | BMZ02_RS09865 | 435856..439149 | 406100 | Swt1-family-HEPN-domain-containing-protein-[Aquisalimonas-asiatica] |
| MCB1764984.1 | DUF499_1.RM.like.systems | TUDOR+SF1-SF2-Helicase+iREase+CTD+REase->?->Tox-RelE->HTH->METHYLASE->HEPN+DUF499*->?->REase->Radical-SAM-> | 1099 | Bacteria>Pseudomonadota>Gammaproteobacteria>Gammaproteobacteria_incertae_sedis>Candidatus_Competibacteraceae | GCA_020444895.1 | Candidatus_Competibacteraceae_bacterium | KDJ22_02835 | 9090..12389 | 2053538 | DUF499-domain-containing-protein-[Candidatus-Competibacteraceae-bacterium] |
| WP_048676961.1 | DUF499_1.RM.like.systems | HEPN+DUF499*->MORC-><-?<-?<-?<-Phage_integrase\|\|DUF4158+DDE_Tnp_Tn3-> | 1099 | Bacteria>Pseudomonadota>Gammaproteobacteria>Gammaproteobacteria_incertae_sedis>Candidatus_Competibacteraceae | GCF_001051235.1 | Candidatus_Competibacter_denitrificans | BN873_RS17980 | 1195..4494 | 1400862 | DUF499-domain-containing-protein-[Candidatus-Competibacter-denitrificans] |
| MBC7162924.1 | DUF499_1.RM.like.systems | PIN->HTH+Hiskin+STAS->HEPN+DUF499*-><-ZDNA-HTH<-?<-?<-?<-HTH_3<-RelE-ParE | 1109 | Bacteria>Pseudomonadota>Gammaproteobacteria>Immundisolibacterales>Immundisolibacteraceae | GCA_014359565.1 | Immundisolibacter_sp. | H5U26_12570 | 5259..8588 | 1934948 | ATP-binding-protein-[Immundisolibacter-sp.] |
| NMZ24831.1 | DUF499_1.RM.like.systems | Arm-DNA-bind_3+Phage_integrase->TOPRIM->?->Integrase_1->?->HTH+MPTase-><-HEPN+DUF499*<-METHYLASE<-TUDOR+SF1-SF2-Helicase+iREase+CTD+REase<-HTH+WYL | 1098 | Bacteria>Pseudomonadota>Gammaproteobacteria>Pseudomonadales>Pseudomonadaceae | GCA_012986045.1 | Pseudomonas_proteolytica | HBO02_20655 | 84255..87551 | 219574 | ATP-binding-protein-[Pseudomonas-proteolytica] |
| WP_262499199.1 | DUF499_1.RM.like.systems | TOPRIM->?->Integrase_1->Phage_int_SAM->?->HTH+MPTase-><-HEPN+DUF499*<-METHYLASE<-TUDOR+SF1-SF2-Helicase+iREase+CTD+REase<-HTH+WYL | 1083 | Bacteria>Pseudomonadota>Gammaproteobacteria>Pseudomonadales>Pseudomonadaceae | GCF_012986045.1 | Pseudomonas_proteolytica | HBO02_RS20625 | 84255..87506 | 219574 | DUF499-domain-containing-protein-[Pseudomonas-proteolytica] |
| WP_169910036.1 | DUF499_1.RM.like.systems | Arm-DNA-bind_3+Phage_integrase->TOPRIM->?->Integrase_1->?->HTH+MPTase-><-HEPN+DUF499*<-METHYLASE<-TUDOR+SF1-SF2-Helicase+iREase+CTD+REase<-HTH+WYL | 1098 | Bacteria>Pseudomonadota>Gammaproteobacteria>Pseudomonadales>Pseudomonadaceae | - | Pseudomonas_proteolytica | HBO02_20655 | 84255..87551 | 219574 | DUF499-domain-containing-protein-[Pseudomonas-proteolytica] |
| MCG3168781.1 | DUF499_1.RM.like.systems | ASCH->HKD-DNase->METHYLASE->HEPN+DUF499*-> | 1110 | Bacteria>Pseudomonadota>Gammaproteobacteria>Pseudomonadales>unclassified_Pseudomonadales | GCA_022072145.1 | Pseudomonadales_bacterium | CALGDGBN_00288 | 5040..8372 | 1891229 | hypothetical-protein-[Pseudomonadales-bacterium] |
| MCP5148712.1 | DUF499_1.RM.like.systems | Radical-SAM-><-?<-?<-?<-HEPN+DUF499*<-METHYLASE<-?<-Cluster1090_5clades<-TUDOR+SF1-SF2-Helicase+iREase+CTD+REase\|\|GDResolvase+HIN-HTH-> | 1108 | Bacteria>Pseudomonadota>Gammaproteobacteria>Pseudomonadales>unclassified_Pseudomonadales | GCA_024234795.1 | Pseudomonadales_bacterium | H6986_11380 | 65855..69181 | 1891229 | ATP-binding-protein-[Pseudomonadales-bacterium] |
| MCP5129192.1 | DUF499_1.RM.like.systems | TraG_N-><-?<-SbcC<-?<-?<-HEPN+DUF499*<-METHYLASE<-MORC+DUF3684<-TUDOR+SF1-SF2-Helicase+iREase+CTD+REase<-XIS-HTH<-tRNA\|\|RNB+RNase->SPOUT-> | 1098 | Bacteria>Pseudomonadota>Gammaproteobacteria>Pseudomonadales>unclassified_Pseudomonadales | GCA_024234815.1 | Pseudomonadales_bacterium | H6985_06385 | 1449819..1453115 | 1891229 | ATP-binding-protein-[Pseudomonadales-bacterium] |
| MBC53307.1 | DUF499_1.RM.like.systems | MerR-HTH-><-?<-?<-Calcineurin<-?<-JAB<-HEPN+DUF499*<-METHYLASE<-MORC+DUF3684<-TUDOR+SF1-SF2-Helicase+iREase+CTD+REase<-HTH+WYL\|\|SF1-SF2-helicase+RdgC+Beta-barrel+REase-> | 1098 | Bacteria>Pseudomonadota>Gammaproteobacteria>unclassified_Gammaproteobacteria | GCA_002711845.1 | Gammaproteobacteria_bacterium | CMQ34_05660 | 7518..10814 | 1913989 | AAA-family-ATPase-[Gammaproteobacteria-bacterium] |
| RMG33561.1 | DUF499_1.RM.like.systems | <-HEPN+DUF499*<-METHYLASE | 1107 | Bacteria>Pseudomonadota>Gammaproteobacteria>unclassified_Gammaproteobacteria | GCA_003696905.1 | Gammaproteobacteria_bacterium | D6720_11165 | 157..3480 | 1913989 | ATP-binding-protein-[Gammaproteobacteria-bacterium] |
| TAK54308.1 | DUF499_1.RM.like.systems | TUDOR+SF1-SF2-Helicase+iREase+CTD+REase->Schlafen+HATPase+HTH->METHYLASE->OLD-ABC->TOPRIM->HEPN+DUF499*-> | 1105 | Bacteria>Pseudomonadota>Gammaproteobacteria>unclassified_Gammaproteobacteria | GCA_004297335.1 | Gammaproteobacteria_bacterium | EPO25_07530 | 13135..16452 | 1913989 | ATP-binding-protein-[Gammaproteobacteria-bacterium] |
| TVS19395.1 | DUF499_1.RM.like.systems | <-HEPN+DUF499*<-METHYLASE-p | 1100 | Bacteria>Pseudomonadota>Gammaproteobacteria>unclassified_Gammaproteobacteria | GCA_007694905.1 | Gammaproteobacteria_bacterium | EA417_00670 | 96..3398 | 1913989 | ATP-binding-protein-[Gammaproteobacteria-bacterium] |
| MXW44722.1 | DUF499_1.RM.like.systems | MORC+DUF3684->?->?->HEPN+DUF499*->?->MazE_antitoxin-> | 959 | Bacteria>Pseudomonadota>Gammaproteobacteria>unclassified_Gammaproteobacteria | GCA_009836205.1 | Gammaproteobacteria_bacterium | F4Z63_01800 | 11560..14439 | 1913989 | ATP-binding-protein-[Gammaproteobacteria-bacterium] |
| MXY90447.1 | DUF499_1.RM.like.systems | <-PrmC_N+Methyltransf_25\|\|?-><-?<-?<-HEPN+DUF499*<-TOPRIM<-OLD-ABC | 1116 | Bacteria>Pseudomonadota>Gammaproteobacteria>unclassified_Gammaproteobacteria | GCA_009837675.1 | Gammaproteobacteria_bacterium | F4Y89_07905 | 34085..37435 | 1913989 | ATP-binding-protein-[Gammaproteobacteria-bacterium] |
| MXY57169.1 | DUF499_1.RM.like.systems | METHYLASE-p->HEPN+DUF499*-> | 413 | Bacteria>Pseudomonadota>Gammaproteobacteria>unclassified_Gammaproteobacteria | GCA_009838585.1 | Gammaproteobacteria_bacterium | F4Y41_12365 | 1564..2802 | 1913989 | ATP-binding-protein-[Gammaproteobacteria-bacterium] |
| MYB35869.1 | DUF499_1.RM.like.systems | TPR+TPR+PIN+PIN->TUDOR+SF1-SF2-Helicase+iREase+CTD+REase->METHYLASE->HEPN+DUF499*-> | 1114 | Bacteria>Pseudomonadota>Gammaproteobacteria>unclassified_Gammaproteobacteria | GCA_009838905.1 | Gammaproteobacteria_bacterium | F4Y26_00435 | 10511..13855 | 1913989 | ATP-binding-protein-[Gammaproteobacteria-bacterium] |
| MYE82983.1 | DUF499_1.RM.like.systems | <-Arm-DNA-bind_3+Phage_int_SAM_3+Phage_integrase<-Phage_int_SAM_3+Phage_integrase\|\|HEPN+DUF499*-> | 1107 | Bacteria>Pseudomonadota>Gammaproteobacteria>unclassified_Gammaproteobacteria | GCA_009840715.1 | Gammaproteobacteria_bacterium | F4X36_14330 | 1398..4721 | 1913989 | ATP-binding-protein-[Gammaproteobacteria-bacterium] |
| MYH50428.1 | DUF499_1.RM.like.systems | METHYLASE-p->HEPN+DUF499*-><-HicB_lk_antitox<-?<-?<-LexA-protease\|\|XIS-HTH->VirB2-> | 1107 | Bacteria>Pseudomonadota>Gammaproteobacteria>unclassified_Gammaproteobacteria | GCA_009843625.1 | Gammaproteobacteria_bacterium | F4151_13115 | 723..4046 | 1913989 | ATP-binding-protein-[Gammaproteobacteria-bacterium] |
| MYH70086.1 | DUF499_1.RM.like.systems | METHYLASE->MazE_antitoxin->PemK_toxin->HEPN+DUF499*-><-TPR+TPR | 1110 | Bacteria>Pseudomonadota>Gammaproteobacteria>unclassified_Gammaproteobacteria | GCA_009843675.1 | Gammaproteobacteria_bacterium | F4147_08315 | 2722..6054 | 1913989 | ATP-binding-protein-[Gammaproteobacteria-bacterium] |
| NKC12747.1 | DUF499_1.RM.like.systems | <-HEPN+DUF499*<-METHYLASE<-?<-TUDOR+SF1-SF2-Helicase+iREase+CTD+REase\|\|?->?->Phage_integrase-> | 1098 | Bacteria>Pseudomonadota>Gammaproteobacteria>unclassified_Gammaproteobacteria | GCA_012103455.1 | Gammaproteobacteria_bacterium | GKR94_11270 | 2223400..2226696 | 1913989 | DUF499-domain-containing-protein-[Gammaproteobacteria-bacterium] |
| NKC12883.1 | DUF499_1.RM.like.systems | <-HEPN+DUF499*\|\|Phage_int_SAM_3+Phage_integrase->Phage_integrase->METHYLASE-p-><-HicA_toxin<-HicB_lk_antitox<-METHYLASE | 1107 | Bacteria>Pseudomonadota>Gammaproteobacteria>unclassified_Gammaproteobacteria | GCA_012103455.1 | Gammaproteobacteria_bacterium | GKR94_11995 | 2363179..2366502 | 1913989 | DUF499-domain-containing-protein-[Gammaproteobacteria-bacterium] |
| MBA2592967.1 | DUF499_1.RM.like.systems | ABC-ATPase->?->TUDOR+SF1-SF2-Helicase+iREase+CTD+REase->METHYLASE->HEPN+DUF499*-><-?<-?<-ParB<-TOPRIM<-ZnR<-XIS-HTH | 1115 | Bacteria>Pseudomonadota>Gammaproteobacteria>unclassified_Gammaproteobacteria | GCA_013697045.1 | Gammaproteobacteria_bacterium | HOU97_12235 | 29064..32411 | 1913989 | ATP-binding-protein-[Gammaproteobacteria-bacterium] |
| MBI2800244.1 | DUF499_1.RM.like.systems | DUF932-><-Phage_int_SAM_2+Integrase_1\|\|TUDOR+SF1-SF2-Helicase+iREase+CTD+REase->MORC+DUF3684->METHYLASE->?->HEPN+DUF499*-><-DUF772+DDE_Tnp_1<-Arm-DNA-bind_3+Phage_integrase\|\|REase+N6_MTase-><-?\|\|SPOUT-> | 1108 | Bacteria>Pseudomonadota>Gammaproteobacteria>unclassified_Gammaproteobacteria | GCA_016191285.1 | Gammaproteobacteria_bacterium | HYX63_08145 | 89797..93123 | 1913989 | ATP-binding-protein-[Gammaproteobacteria-bacterium] |
| MBM4223246.1 | DUF499_1.RM.like.systems | TUDOR+SF1-SF2-Helicase+iREase+CTD+REase->sGTP->METHYLASE->HEPN+DUF499*->REase-> | 1092 | Bacteria>Pseudomonadota>Gammaproteobacteria>unclassified_Gammaproteobacteria | GCA_016874585.1 | Gammaproteobacteria_bacterium | FJ167_00320 | 8980..12258 | 1913989 | ATP-binding-protein-[Gammaproteobacteria-bacterium] |
| MBT4891857.1 | DUF499_1.RM.like.systems | METHYLASE-p->HEPN+DUF499*->TIR-> | 1099 | Bacteria>Pseudomonadota>Gammaproteobacteria>unclassified_Gammaproteobacteria | GCA_018664395.1 | Gammaproteobacteria_bacterium | HON60_02830 | 645..3944 | 1913989 | ATP-binding-protein-[Gammaproteobacteria-bacterium] |
| MBU1235382.1 | DUF499_1.RM.like.systems | <-VirD4-TraG+VirD4-FtsK\|\|TUDOR+SF1-SF2-Helicase+iREase+CTD+REase->STAS+HisKin->?->PIN->METHYLASE->HEPN+DUF499*-><-?<-?<-?<-ParB | 1113 | Bacteria>Pseudomonadota>Gammaproteobacteria>unclassified_Gammaproteobacteria | GCA_018819515.1 | Gammaproteobacteria_bacterium | KJ788_01195 | 188697..192038 | 1913989 | DUF499-domain-containing-protein-[Gammaproteobacteria-bacterium] |
| MCG3202826.1 | DUF499_1.RM.like.systems | N6_N4_MTase-><-?<-AbiEii<-AbiEi_4<-REase<-HEPN+DUF499*<-METHYLASE-p | 1115 | Bacteria>Pseudomonadota>Gammaproteobacteria>unclassified_Gammaproteobacteria | GCA_022071965.1 | Gammaproteobacteria_bacterium | NFCOHLIN_02712 | 302948..306295 | 1913989 | hypothetical-protein-[Gammaproteobacteria-bacterium] |
| GIX22526.1 | DUF499_1.RM.like.systems | Response_reg+LuxR-HTH->URI-DNase+HhH-RADC->?->tRNA->tRNA->tRNA-><-HEPN+DUF499*<-METHYLASE<-REase<-TUDOR+SF1-SF2-Helicase+iREase+CTD+REase-p | 1122 | Bacteria>Pseudomonadota>Gammaproteobacteria>unclassified_Gammaproteobacteria | GCA_026004975.1 | Gammaproteobacteria_bacterium | KatS3mg121_1309 | 525909..529277 | 1913989 | MAG:-hypothetical-protein-KatS3mg121_1309-[Gammaproteobacteria-bacterium] |
| MCY3837525.1 | DUF499_1.RM.like.systems | METHYLASE->CR-ATPase8+REase->HEPN+DUF499*->Tox-RelE->HTH+MPTase-><-VapB_antitoxin | 1112 | Bacteria>Pseudomonadota>Gammaproteobacteria>unclassified_Gammaproteobacteria | GCA_026706425.1 | Gammaproteobacteria_bacterium | OXH09_02545 | 4546..7884 | 1913989 | MAG:-Swt1-family-HEPN-domain-containing-protein-[Gammaproteobacteria-bacterium] |
| MCY3821975.1 | DUF499_1.RM.like.systems | <-sGTP\|\|?->?-><-?<-HEPN+DUF499*\|\|Phage_int_SAM_3+Phage_integrase->Phage_integrase->METHYLASE-p-> | 1101 | Bacteria>Pseudomonadota>Gammaproteobacteria>unclassified_Gammaproteobacteria | GCA_026706665.1 | Gammaproteobacteria_bacterium | OXH52_21920 | 46820..50125 | 1913989 | MAG:-Swt1-family-HEPN-domain-containing-protein-[Gammaproteobacteria-bacterium] |
| MCY3939524.1 | DUF499_1.RM.like.systems | MORC+DUF3684->TUDOR+SF1-SF2-Helicase+iREase+CTD+REase->METHYLASE->HEPN+DUF499*->MazE_antitoxin->DUF433->PIN-> | 1097 | Bacteria>Pseudomonadota>Gammaproteobacteria>unclassified_Gammaproteobacteria | GCA_026707445.1 | Gammaproteobacteria_bacterium | OXG29_00400 | 16758..20051 | 1913989 | MAG:-DUF499-domain-containing-protein-[Gammaproteobacteria-bacterium] |
| MCY3988741.1 | DUF499_1.RM.like.systems | METHYLASE-p->HEPN+DUF499*->MazE_antitoxin->DUF433-> | 1097 | Bacteria>Pseudomonadota>Gammaproteobacteria>unclassified_Gammaproteobacteria | GCA_026708205.1 | Gammaproteobacteria_bacterium | OXF94_09155 | 914..4207 | 1913989 | MAG:-DUF499-domain-containing-protein-[Gammaproteobacteria-bacterium] |
| MCY4181966.1 | DUF499_1.RM.like.systems | <-HEPN+DUF499*<-METHYLASE | 1105 | Bacteria>Pseudomonadota>Gammaproteobacteria>unclassified_Gammaproteobacteria | GCA_026709135.1 | Gammaproteobacteria_bacterium | OXF43_03630 | 102..3419 | 1913989 | MAG:-DUF499-domain-containing-protein-[Gammaproteobacteria-bacterium] |
| MCY4211280.1 | DUF499_1.RM.like.systems | SAM-methylase->TPR+TPR->DOC+DOC-><-DUF499-p<-HEPN+DUF499* | 757 | Bacteria>Pseudomonadota>Gammaproteobacteria>unclassified_Gammaproteobacteria | GCA_026709385.1 | Gammaproteobacteria_bacterium | OXE97_09605 | 24101..26373 | 1913989 | MAG:-Swt1-family-HEPN-domain-containing-protein,-partial-[Gammaproteobacteria-bacterium] |
| MCY4265599.1 | DUF499_1.RM.like.systems | TUDOR+SF1-SF2-Helicase+iREase+CTD+REase->HEPN+DUF499*-><-MNS | 1088 | Bacteria>Pseudomonadota>Gammaproteobacteria>unclassified_Gammaproteobacteria | GCA_026709755.1 | Gammaproteobacteria_bacterium | OXE78_12210 | 4990..8256 | 1913989 | MAG:-DUF499-domain-containing-protein-[Gammaproteobacteria-bacterium] |
| MCY4282353.1 | DUF499_1.RM.like.systems | SAM-methylase->TPR+TPR->DOC+DOC-><-HEPN+DUF499*\|\|?->ParB+HNH->OLD-ABC-> | 941 | Bacteria>Pseudomonadota>Gammaproteobacteria>unclassified_Gammaproteobacteria | GCA_026709915.1 | Gammaproteobacteria_bacterium | OXE03_05465 | 14307..17132 | 1913989 | MAG:-Swt1-family-HEPN-domain-containing-protein-[Gammaproteobacteria-bacterium] |
| MCY4343961.1 | DUF499_1.RM.like.systems | METHYLASE->OLD-ABC->Schlafen+HATPase+HTH->?->PIN->HEPN+DUF499*-><-MNS+ZDNA-HTH<-PRTase | 1115 | Bacteria>Pseudomonadota>Gammaproteobacteria>unclassified_Gammaproteobacteria | GCA_026710215.1 | Gammaproteobacteria_bacterium | OXE83_10365 | 11310..14657 | 1913989 | MAG:-Swt1-family-HEPN-domain-containing-protein-[Gammaproteobacteria-bacterium] |
| MCZ0943260.1 | DUF499_1.RM.like.systems | DUF433-><-HEPN+DUF499*<-Schlafen+HATPase+HTH<-TOPRIM<-OLD-ABC<-METHYLASE-p | 1115 | Bacteria>Pseudomonadota>Gammaproteobacteria>unclassified_Gammaproteobacteria | GCA_026914145.1 | Gammaproteobacteria_bacterium | OXJ53_09405 | 2539..5886 | 1913989 | MAG:-Swt1-family-HEPN-domain-containing-protein-[Gammaproteobacteria-bacterium] |
| MDE0036818.1 | DUF499_1.RM.like.systems | METHYLASE-p->?-><-PIN<-OLD-ABC<-?<-HEPN+DUF499*<-METHYLASE<-OLD-ABC<-TUDOR+SF1-SF2-Helicase+iREase+CTD+REase<-?<-?\|\|SF1-SF2-Helicase-> | 1118 | Bacteria>Pseudomonadota>Gammaproteobacteria>unclassified_Gammaproteobacteria | GCA_028818795.1 | Gammaproteobacteria_bacterium | OXU77_04540 | 7629..10985 | 1913989 | MAG:-Swt1-family-HEPN-domain-containing-protein-[Gammaproteobacteria-bacterium] |
| MDE0073871.1 | DUF499_1.RM.like.systems | METHYLASE->OLD-ABC->TOPRIM->HEPN+DUF499*-><-HicB_lk_antitox<-HicA_toxin\|\|XIS-HTH->VirB2-> | 1107 | Bacteria>Pseudomonadota>Gammaproteobacteria>unclassified_Gammaproteobacteria | GCA_028819075.1 | Gammaproteobacteria_bacterium | OXU32_07805 | 3301..6624 | 1913989 | MAG:-Swt1-family-HEPN-domain-containing-protein-[Gammaproteobacteria-bacterium] |
| MDE0156012.1 | DUF499_1.RM.like.systems | SAM-methylase->TPR+TPR-><-HEPN+DUF499*<-PemK_toxin<-MazE_antitoxin<-HEPN<-CopG_antitoxin<-BrnT_toxin\|\|HEPN-> | 1110 | Bacteria>Pseudomonadota>Gammaproteobacteria>unclassified_Gammaproteobacteria | GCA_028819495.1 | Gammaproteobacteria_bacterium | OXS28_10440 | 75078..78410 | 1913989 | MAG:-Swt1-family-HEPN-domain-containing-protein-[Gammaproteobacteria-bacterium] |
| MDE0284673.1 | DUF499_1.RM.like.systems | <-HEPN\|\|BrnT_toxin->CopG_antitoxin->HEPN->MazE_antitoxin->PemK_toxin->HEPN+DUF499*-> | 949 | Bacteria>Pseudomonadota>Gammaproteobacteria>unclassified_Gammaproteobacteria | GCA_028820555.1 | Gammaproteobacteria_bacterium | OXN26_09020 | 3552..6397 | 1913989 | MAG:-Swt1-family-HEPN-domain-containing-protein,-partial-[Gammaproteobacteria-bacterium] |
| MDE0441971.1 | DUF499_1.RM.like.systems | <-HEPN+DUF499*<-METHYLASE | 570 | Bacteria>Pseudomonadota>Gammaproteobacteria>unclassified_Gammaproteobacteria | GCA_028821355.1 | Gammaproteobacteria_bacterium | OXL38_07620 | 38..1750 | 1913989 | MAG:-Swt1-family-HEPN-domain-containing-protein-[Gammaproteobacteria-bacterium] |
| MDE0510268.1 | DUF499_1.RM.like.systems | SAM-methylase->TPR+TPR-><-HEPN+DUF499*<-PemK_toxin<-MazE_antitoxin<-HEPN<-CopG_antitoxin<-BrnT_toxin\|\|HEPN-> | 1110 | Bacteria>Pseudomonadota>Gammaproteobacteria>unclassified_Gammaproteobacteria | GCA_028823865.1 | Gammaproteobacteria_bacterium | OXI88_00540 | 10766..14098 | 1913989 | MAG:-Swt1-family-HEPN-domain-containing-protein-[Gammaproteobacteria-bacterium] |
| MDE0678898.1 | DUF499_1.RM.like.systems | METHYLASE-p->HEPN+DUF499*->SF1-SF2-Helicase+HRDC-> | 1109 | Bacteria>Pseudomonadota>Gammaproteobacteria>unclassified_Gammaproteobacteria | GCA_028824615.1 | Gammaproteobacteria_bacterium | OXI11_01560 | 1155..4484 | 1913989 | MAG:-Swt1-family-HEPN-domain-containing-protein-[Gammaproteobacteria-bacterium] |
| MDE0680618.1 | DUF499_1.RM.like.systems | MORC+DUF3684->TUDOR+SF1-SF2-Helicase+iREase+CTD+REase->METHYLASE->?->?->HEPN+DUF499*->MazE_antitoxin-> | 1100 | Bacteria>Pseudomonadota>Gammaproteobacteria>unclassified_Gammaproteobacteria | GCA_028824615.1 | Gammaproteobacteria_bacterium | OXI11_10470 | 19020..22322 | 1913989 | MAG:-DUF499-domain-containing-protein-[Gammaproteobacteria-bacterium] |
| MDE0479832.1 | DUF499_1.RM.like.systems | <-HEPN+DUF499*<-METHYLASE<-?<-RNase<-TUDOR+SF1-SF2-Helicase+iREase+CTD+REase<-HTH+UbE1 | 1108 | Bacteria>Pseudomonadota>Gammaproteobacteria>unclassified_Gammaproteobacteria | GCA_028824825.1 | Gammaproteobacteria_bacterium | OXI13_09405 | 28778..32104 | 1913989 | MAG:-DUF499-domain-containing-protein-[Gammaproteobacteria-bacterium] |
| GIX32370.1 | DUF499_1.RM.like.systems | REase->METHYLASE-p->METHYLASE-p->METHYLASE->ParD->PIN->HEPN+DUF499*-><-tRNA<-SIGMA-HTH+SIGMA-HTH | 1108 | Bacteria>Pseudomonadota>Gammaproteobacteria>Xanthomonadales>unclassified_Xanthomonadales | GCA_026005055.1 | Xanthomonadales_bacterium | KatS3mg125_0326 | 361925..365251 | 2006849 | MAG:-hypothetical-protein-KatS3mg125_0326-[Xanthomonadales-bacterium] |
| WP_035746468.1 | DUF499_1.RM.like.systems | ParG_RHH-><-TetR-HTH\|\|MODE-HTH+PBPII-><-?<-HEPN+DUF499*<-METHYLASE<-TUDOR+SF1-SF2-Helicase+iREase+CTD+REase<-HTH+WYL\|\|HicB->KAP-NTPase-> | 1099 | Bacteria>Pseudomonadota | GCF_000740775.1 | Alphaproteobacteria | CG51_RS15425 | 33987..37286 | 28211 | MULTISPECIES:-DUF499-domain-containing-protein-[Alphaproteobacteria] |
| WP_102114747.1 | DUF499_1.RM.like.systems | METHYLASE->HEPN+DUF499*->MODE-HTH+PBPII-><-ParG_RHH | 1098 | Bacteria>Pseudomonadota | GCF_014640215.1 | Alphaproteobacteria | KF818_RS31845 | 5542..8838 | 28211 | DUF499-domain-containing-protein-[Niveispirillum-cyanobacteriorum] |
| WP_054362466.1 | DUF499_1.RM.like.systems | ParG_RHH-><-MODE-HTH+PBPII<-HEPN+DUF499*<-METHYLASE<-REase | 1098 | Bacteria>Pseudomonadota | GCF_963522535.1 | Alphaproteobacteria | ABB55_RS28500 | 24847..28143 | 28211 | MULTISPECIES:-DUF499-domain-containing-protein-[Alphaproteobacteria] |
| OQW45528.1 | DUF499_1.RM.like.systems | <-HicB\|\|HTH+WYL->TUDOR+SF1-SF2-Helicase+iREase+CTD+REase-><-?\|\|METHYLASE->HEPN+DUF499*->MODE-HTH+PBPII-><-ParG_RHH | 1097 | Bacteria>Pseudomonadota>unclassified_Pseudomonadota | GCA_002083435.1 | Proteobacteria_bacterium_SG_bin6 | A4S16_12875 | 8645..11938 | 1827383 | AAA-family-ATPase-[Proteobacteria-bacterium-SG_bin6] |
| PWT88877.1 | DUF499_1.RM.like.systems | TUDOR+SF1-SF2-Helicase+iREase+CTD+REase-p->HEPN+DUF499*-> | 1096 | Bacteria>Pseudomonadota>unclassified_Pseudomonadota | GCA_003175845.1 | Pseudomonadota_bacterium | C5B56_08055 | 3674..6960 | 1977087 | AAA+-family-ATPase,-partial-[Proteobacteria-bacterium] |
| RIK94108.1 | DUF499_1.RM.like.systems | DUF488->ORC-CDC6+REase->?-><-PIWI<-TIR+PNTD2<-HEPN+DUF499*<-METHYLASE<-MORC+DUF3684<-TUDOR+SF1-SF2-Helicase+iREase+CTD+REase<-HTH+WYL\|\|SF1-SF2-Helicase+REase->SF1-SF2-Helicase+REase-> | 1098 | Bacteria>Pseudomonadota>unclassified_Pseudomonadota | GCA_003577275.1 | Pseudomonadota_bacterium | DCC73_05940 | 240099..243395 | 1977087 | AAA-family-ATPase-[Proteobacteria-bacterium] |
| NBQ33337.1 | DUF499_1.RM.like.systems | <-HEPN+DUF499*<-METHYLASE-p | 1140 | Bacteria>Pseudomonadota>unclassified_Pseudomonadota | GCA_009920185.1 | Pseudomonadota_bacterium | EBU21_15535 | 254..3676 | 1977087 | ATP-binding-protein-[Proteobacteria-bacterium] |
| NBT04715.1 | DUF499_1.RM.like.systems | <-REase<-HEPN+DUF499*<-METHYLASE-p | 1140 | Bacteria>Pseudomonadota>unclassified_Pseudomonadota | GCA_009922565.1 | Pseudomonadota_bacterium | EBS94_15100 | 2192..5614 | 1977087 | ATP-binding-protein-[Proteobacteria-bacterium] |
| MBE9569155.1 | DUF499_1.RM.like.systems | PIN->METHYLASE->HEPN+DUF499*-> | 1063 | Bacteria>Pseudomonadota>unclassified_Pseudomonadota | GCA_015222845.1 | Pseudomonadota_bacterium | IMF11_00880 | 3390..6577 | 1977087 | DUF499-domain-containing-protein-[Proteobacteria-bacterium] |
| MBI4030489.1 | DUF499_1.RM.like.systems | Peptidase_M15_4->?->?->?->?-><-HEPN+DUF499*<-METHYLASE | 1115 | Bacteria>Pseudomonadota>unclassified_Pseudomonadota | GCA_016198545.1 | Pseudomonadota_bacterium | HY370_02330 | 8251..11598 | 1977087 | ATP-binding-protein-[Proteobacteria-bacterium] |
| MBQ9395962.1 | DUF499_1.RM.like.systems | METHYLASE-p-><-PIN<-?\|\|HEPN+DUF499*-> | 729 | Bacteria>Pseudomonadota>unclassified_Pseudomonadota | GCA_017455165.1 | Pseudomonadota_bacterium | IJU23_10695 | 3126..5311 | 1977087 | DUF499-domain-containing-protein-[Proteobacteria-bacterium] |
| MBS0482554.1 | DUF499_1.RM.like.systems | Thioredoxin->TraH->?-><-?<-HEPN+DUF499*<-METHYLASE-p | 1098 | Bacteria>Pseudomonadota>unclassified_Pseudomonadota | GCA_018240895.1 | Pseudomonadota_bacterium | JSR96_10460 | 15112..18408 | 1977087 | DUF499-domain-containing-protein-[Proteobacteria-bacterium] |
| MBS0471294.1 | DUF499_1.RM.like.systems | <-AbiEii<-AbiEi_3_N+AbiEi_3\|\|XIS-HTH->TUDOR+SF1-SF2-Helicase+iREase+CTD+REase->MORC+DUF3684->METHYLASE->HEPN+DUF499*-><-XIS-HTH<-?<-ParG_RHH | 1118 | Bacteria>Pseudomonadota>unclassified_Pseudomonadota | GCA_018241025.1 | Pseudomonadota_bacterium | JSR60_09500 | 167872..171228 | 1977087 | DUF499-domain-containing-protein-[Proteobacteria-bacterium] |
| MBU0466040.1 | DUF499_1.RM.like.systems | <-HEPN+DUF499*<-METHYLASE-p | 1114 | Bacteria>Pseudomonadota>unclassified_Pseudomonadota | GCA_018812385.1 | Pseudomonadota_bacterium | KKE12_21020 | 197..3541 | 1977087 | DUF499-domain-containing-protein-[Proteobacteria-bacterium] |
| MBU1208887.1 | DUF499_1.RM.like.systems | TUDOR+SF1-SF2-Helicase+iREase+CTD+REase-p->DrHyd+BirA-HTH->METHYLASE->HEPN+DUF499*->REase-p->?->?->Mbetalac-> | 1107 | Bacteria>Pseudomonadota>unclassified_Pseudomonadota | GCA_018819135.1 | Pseudomonadota_bacterium | KKH04_18500 | 6409..9732 | 1977087 | DUF499-domain-containing-protein-[Proteobacteria-bacterium] |
| MBU1342385.1 | DUF499_1.RM.like.systems | MORC+DUF3684->TUDOR+SF1-SF2-Helicase+iREase+CTD+REase->HEPN->METHYLASE->HEPN+DUF499*-> | 1114 | Bacteria>Pseudomonadota>unclassified_Pseudomonadota | GCA_018820265.1 | Pseudomonadota_bacterium | KKD66_12250 | 12988..16332 | 1977087 | DUF499-domain-containing-protein-[Proteobacteria-bacterium] |
| MBU2547024.1 | DUF499_1.RM.like.systems | <-HEPN+DUF499*<-METHYLASE-p | 1093 | Bacteria>Pseudomonadota>unclassified_Pseudomonadota | GCA_018830135.1 | Pseudomonadota_bacterium | KKB20_01315 | 29847..33128 | 1977087 | DUF499-domain-containing-protein-[Proteobacteria-bacterium] |
| MBU4581761.1 | DUF499_1.RM.like.systems | DDE_5+DDE_Tnp_1->?-><-HEPN+DUF499*<-METHYLASE<-?<-?<-SF1-SF2-Helicase<-DrHyd+BirA-HTH<-MORC+DUF3684 | 1124 | Bacteria>Pseudomonadota>unclassified_Pseudomonadota | GCA_018902925.1 | Pseudomonadota_bacterium | KKH02_05005 | 2369..5743 | 1977087 | DUF499-domain-containing-protein-[Proteobacteria-bacterium] |
| MCA0247253.1 | DUF499_1.RM.like.systems | <-SF1-SF2-Helicase+REase<-SF1-SF2-Helicase+REase\|\|HTH+WYL->TUDOR+SF1-SF2-Helicase+iREase+CTD+REase->MORC+DUF3684->METHYLASE->HEPN+DUF499*->MODE-HTH+PBPII-><-ParG_RHH | 1098 | Bacteria>Pseudomonadota>unclassified_Pseudomonadota | GCA_020160995.1 | Pseudomonadota_bacterium | LCH93_11605 | 30891..34187 | 1977087 | DUF499-domain-containing-protein-[Proteobacteria-bacterium] |
| MCA0405125.1 | DUF499_1.RM.like.systems | ASCH->METHYLASE->HEPN+DUF499*->MODE-HTH+PBPII-><-ParG_RHH | 1097 | Bacteria>Pseudomonadota>unclassified_Pseudomonadota | GCA_020161475.1 | Pseudomonadota_bacterium | LCH39_03110 | 5235..8528 | 1977087 | DUF499-domain-containing-protein-[Proteobacteria-bacterium] |
| MCA0401434.1 | DUF499_1.RM.like.systems | ParG_RHH-><-DUF488-l<-?<-TetR-HTH\|\|MODE-HTH+PBPII-><-HEPN+DUF499*<-METHYLASE<-HEPN<-TUDOR+SF1-SF2-Helicase+iREase+CTD+REase<-HTH+WYL\|\|SF1-SF2-Helicase+REase->SF1-SF2-Helicase+REase-> | 1098 | Bacteria>Pseudomonadota>unclassified_Pseudomonadota | GCA_020161575.1 | Pseudomonadota_bacterium | LCH38_11530 | 127907..131203 | 1977087 | DUF499-domain-containing-protein-[Proteobacteria-bacterium] |
| MCG8554136.1 | DUF499_1.RM.like.systems | Cas_Cmr3->?->Cas_Cmr5->?->CARF+HEPN->HEPN+DUF499*-> | 1114 | Bacteria>Pseudomonadota>unclassified_Pseudomonadota | GCA_022361155.1 | Pseudomonadota_bacterium | MJD61_02430 | 12967..16311 | 1977087 | Swt1-family-HEPN-domain-containing-protein-[Proteobacteria-bacterium] |
| MCX7197671.1 | DUF499_1.RM.like.systems | TUDOR+SF1-SF2-Helicase+iREase+CTD+REase-p->METHYLASE->HTH+Hiskin+STAS->HEPN+DUF499*->?->HNH->SIR2->DnaB-> | 1113 | Bacteria>Pseudomonadota>unclassified_Pseudomonadota | GCA_026399745.1 | Pseudomonadota_bacterium | NTW37_06695 | 4467..7808 | 1977087 | MAG:-Swt1-family-HEPN-domain-containing-protein-[Proteobacteria-bacterium] |
| MDA1076682.1 | DUF499_1.RM.like.systems | <-HEPN+DUF499*<-METHYLASE | 1005 | Bacteria>Pseudomonadota>unclassified_Pseudomonadota | GCA_027622495.1 | Pseudomonadota_bacterium | O3A63_18310 | 1..3015 | 1977087 | MAG:-Swt1-family-HEPN-domain-containing-protein,-partial-[Proteobacteria-bacterium] |
| MBU2496199.1 | DUF499_1.RM.like.systems | METHYLASE->HEPN+DUF499*-> | 365 | Bacteria>PVC_group>Candidatus_Omnitrophota>unclassified_Candidatus_Omnitrophota | GCA_018829455.1 | Candidatus_Omnitrophota_bacterium | KJ935_06870 | 1920..3013 | 2035772 | DUF499-domain-containing-protein-[Candidatus-Omnitrophota-bacterium] |
| MBN1673169.1 | DUF499_1.RM.like.systems | <-DrHyd+BirA-HTH<-TUDOR+SF1-SF2-Helicase+iREase+CTD+REase-p<-?<-?\|\|?-><-HEPN+DUF499*<-Schlafen+HATPase+HTH<-METHYLASE-p\|\|BirA-HTH->HhH-RADC+JAB-> | 1118 | Bacteria>PVC_group>Kiritimatiellota>Kiritimatiellia>unclassified_Kiritimatiellia | GCA_016929905.1 | Kiritimatiellia_bacterium | JXR37_19145 | 44287..47643 | 2212473 | DUF499-domain-containing-protein-[Kiritimatiellae-bacterium] |
| MDD4116915.1 | DUF499_1.RM.like.systems | SAM-methylase-><-?<-?<-?\|\|tRNA-><-HEPN+DUF499*<-Peptidase_C47<-?<-METHYLASE<-ParB+HNH<-?<-TUDOR+SF1-SF2-Helicase+iREase+CTD+REase | 1113 | Bacteria>PVC_group>Kiritimatiellota>Kiritimatiellia>unclassified_Kiritimatiellia | GCA_028701395.1 | Kiritimatiellia_bacterium | PHI39_01725 | 13604..16945 | 2212473 | MAG:-Swt1-family-HEPN-domain-containing-protein-[Kiritimatiellae-bacterium] |
| MCL1920537.1 | DUF499_1.RM.like.systems | <-HEPN+DUF499*<-METHYLASE-p | 1094 | Bacteria>PVC_group>Kiritimatiellota>unclassified_Kiritimatiellota | GCA_009778475.1 | Kiritimatiellota_bacterium | FWG50_05580 | 29293..32577 | 2599938 | DUF499-domain-containing-protein-[Kiritimatiellaeota-bacterium] |
| HIV08682.1 | DUF499_1.RM.like.systems | TUDOR+SF1-SF2-Helicase+iREase+CTD+REase->HEPN+DUF499*->ParB+HNH-p-> | 1108 | Bacteria>PVC_group>Lentisphaerota>Lentisphaeria>Lentisphaerales | GCA_018713405.1 | Candidatus_Spyradenecus_faecavium | IAC79_01020 | 6980..10306 | 2840947 | DUF499-domain-containing-protein-[Candidatus-Spyradenecus-faecavium] |
| MBN2450533.1 | DUF499_1.RM.like.systems | REase->?->PIN->HEPN+DUF499*-> | 1121 | Bacteria>PVC_group>Lentisphaerota>Lentisphaeria>unclassified_Lentisphaeria | GCA_016935045.1 | Lentisphaeria_bacterium | JXR77_09085 | 1530..4895 | 2053569 | DUF499-domain-containing-protein-[Lentisphaeria-bacterium] |
| MCE9615077.1 | DUF499_1.RM.like.systems | <-XIS-HTH<-?<-?<-?<-HicB<-?<-HEPN+DUF499*<-METHYLASE<-DOC+HTH<-HTH_3<-sGTP<-TUDOR+SF1-SF2-Helicase+iREase+CTD+REase | 1128 | Bacteria>PVC_group>Lentisphaerota>unclassified_Lentisphaerota | GCA_021413965.1 | Lentisphaerota_bacterium | K8T26_12435 | 13053..16439 | 1932692 | DUF499-domain-containing-protein-[Lentisphaerae-bacterium] |
| MBC8553109.1 | DUF499_1.RM.like.systems | XIS-HTH->TUDOR+SF1-SF2-Helicase+iREase+CTD+REase->METHYLASE->HEPN+DUF499*->HNH-> | 1106 | Bacteria>PVC_group>Planctomycetota>Candidatus_Brocadiia>Candidatus_Brocadiales | GCA_014384865.1 | Candidatus_Brocadiales_bacterium | H8D23_26105 | 6653..9973 | 2760054 | ATP-binding-protein-[Candidatus-Brocadiales-bacterium] |
| MBK9119771.1 | DUF499_1.RM.like.systems | DEOR-HTH->TUDOR+SF1-SF2-Helicase+iREase+CTD+REase->Schlafen+HATPase+HTH->?->?->METHYLASE->HEPN+DUF499*->MORC->REase-><-Trypsin\|\|HEPN->HTH->MPTase-> | 1121 | Bacteria>PVC_group>Planctomycetota>Phycisphaerae>Phycisphaerales | GCA_016716475.1 | Phycisphaerales_bacterium | IPM18_09260 | 379276..382641 | 2052180 | DUF499-domain-containing-protein-[Phycisphaerales-bacterium] |
| MCA3006076.1 | DUF499_1.RM.like.systems | METHYLASE->?->HEPN+DUF499*->?->XIS-HTH->?->?->?-><-tRNA\|\|ABC-ATPase+SbcC-> | 1114 | Bacteria>PVC_group>Planctomycetota>Phycisphaerae>Phycisphaerales | GCA_020247235.1 | Phycisphaerales_bacterium | IOD15_12050 | 3800..7144 | 2052180 | DUF499-domain-containing-protein-[Phycisphaerales-bacterium] |
| MCC6908120.1 | DUF499_1.RM.like.systems | <-Hydrolase_3<-HEPN+DUF499*<-METHYLASE<-?<-?<-Schlafen+HATPase+HTH<-TUDOR+SF1-SF2-Helicase+iREase+CTD+REase<-DEOR-HTH | 1120 | Bacteria>PVC_group>Planctomycetota>Phycisphaerae>Phycisphaerales | GCA_020852515.1 | Phycisphaerales_bacterium | IT430_09285 | 88345..91707 | 2052180 | DUF499-domain-containing-protein-[Phycisphaerales-bacterium] |
| MCL4740947.1 | DUF499_1.RM.like.systems | <-XIS-HTH<-?<-MPTase<-HTH_3\|\|?-><-HEPN+DUF499*<-METHYLASE<-sGTP<-TUDOR+SF1-SF2-Helicase+iREase+CTD+REase<-Phage_integrase<-ParB+HTH<-HTH_3 | 1114 | Bacteria>PVC_group>Planctomycetota>Phycisphaerae>Phycisphaerales | GCA_023384195.1 | Phycisphaerales_bacterium | KJZ54_01975 | 143211..146555 | 2052180 | DUF499-domain-containing-protein-[Phycisphaerales-bacterium] |
| MBN2560619.1 | DUF499_1.RM.like.systems | <-HEPN+DUF499*<-?<-METHYLASE<-wHTH+REase<-TUDOR+SF1-SF2-Helicase+iREase+CTD+REase<-?<-Cluster182_14clades | 1112 | Bacteria>PVC_group>Planctomycetota>Phycisphaerae>unclassified_Phycisphaerae | GCA_016935885.1 | Phycisphaerae_bacterium | JXQ75_06790 | 1428..4766 | 2026778 | DUF499-domain-containing-protein-[Phycisphaerae-bacterium] |
| MBW7904855.1 | DUF499_1.RM.like.systems | <-HTH_3\|\|DEOR-HTH->TUDOR+SF1-SF2-Helicase+iREase+CTD+REase->Schlafen+HATPase+HTH->REase->METHYLASE->HEPN+DUF499*->Hydrolase_3-> | 1120 | Bacteria>PVC_group>Planctomycetota>Phycisphaerae>unclassified_Phycisphaerae | GCA_019454965.1 | Phycisphaerae_bacterium | H3C42_04715 | 502745..506107 | 2026778 | ATP-binding-protein-[Phycisphaerae-bacterium] |
| MCK6484528.1 | DUF499_1.RM.like.systems | ArdCN+MPTase->DEOR-HTH->TUDOR+SF1-SF2-Helicase+iREase+CTD+REase->wHTH+REase->REase->METHYLASE->HEPN+DUF499*->DNA_methylase->MORC+HISKIN->?->REase->CR-Kinase+ORC-CDC6-like+CARF-> | 1119 | Bacteria>PVC_group>Planctomycetota>Phycisphaerae>unclassified_Phycisphaerae | GCA_023150595.1 | Phycisphaerae_bacterium | L6R00_10395 | 56365..59724 | 2026778 | DUF499-domain-containing-protein-[Phycisphaerae-bacterium] |
| GIW77359.1 | DUF499_1.RM.like.systems | TUDOR+SF1-SF2-Helicase+iREase+CTD+REase-p->TUDOR+SF1-SF2-Helicase+iREase+CTD+REase->METHYLASE->METHYLASE-p->HEPN+DUF499*->DUF499-p->DUF499-p->?-><-tRNA\|\|?->METHYLASE-> | 529 | Bacteria>PVC_group>Planctomycetota>Phycisphaerae>unclassified_Phycisphaerae | GCA_026004655.1 | Phycisphaerae_bacterium | KatS3mg104_2422 | 7750..9339 | 2026778 | MAG:-hypothetical-protein-KatS3mg104_2422-[Phycisphaerae-bacterium] |
| NLX95047.1 | DUF499_1.RM.like.systems | TPR+TPR->?->?-><-?<-PHP+SbcC<-HEPN+DUF499*<-METHYLASE<-HNH<-OLD-ABC<-TUDOR+SF1-SF2-Helicase+iREase+CTD+REase-p | 1104 | Bacteria>PVC_group>Planctomycetota>Planctomycetia>Pirellulales | GCA_012728855.1 | Rhodopirellula_sp. | GXY83_02605 | 12933..16247 | 2024855 | ATP-binding-protein-[Rhodopirellula-sp.] |
| MBL7036971.1 | DUF499_1.RM.like.systems | <-HEPN+DUF499*<-?<-METHYLASE<-?<-TUDOR+SF1-SF2-Helicase+iREase+CTD+REase\|\|?->METHYLASE-> | 498 | Bacteria>PVC_group>Planctomycetota>Planctomycetia>Pirellulales | GCA_016784735.1 | Pirellulaceae_bacterium | ISR77_00005 | 1..1496 | 2699754 | DUF499-domain-containing-protein-[Pirellulaceae-bacterium] |
| MBQ6108779.1 | DUF499_1.RM.like.systems | <-HEPN+DUF499*<-METHYLASE-p<-TUDOR+SF1-SF2-Helicase+iREase+CTD+REase-p | 927 | Bacteria>PVC_group>Planctomycetota>Planctomycetia>Pirellulales | GCA_017427905.1 | Thermoguttaceae_bacterium | IJK97_11270 | 67..2850 | 2691551 | DUF499-domain-containing-protein-[Thermoguttaceae-bacterium] |
| MBR2005706.1 | DUF499_1.RM.like.systems | METHYLASE-p->HEPN+DUF499*->DUF499-p-> | 960 | Bacteria>PVC_group>Planctomycetota>Planctomycetia>Pirellulales | GCA_017506065.1 | Thermoguttaceae_bacterium | IJ991_16180 | 1159..4041 | 2691551 | DUF499-domain-containing-protein-[Thermoguttaceae-bacterium] |

| Protein-ID | System | Gene-Neighbourhoods | Length | Taxonomy | Assembly-ID | Strain | Locus-Tag | Genomic-coordinates | TaxID | Fasta-defline |
|---|---|---|---|---|---|---|---|---|---|---|
| MBR2694935.1 | DUF499_1.RM.like.systems | <-HEPN+DUF499*<-METHYLASE<-TUDOR+ SF1-SF2-Helicase+iREase+CTD+REase | 1133 | Bacteria>PVC_group>Planctomycetota>Planctomycetia>Pirellulales | GCA_017515305.1 | Thermoguttaceae_bacterium | IKE69_12060 | 24593..27994 | 2691551 | DUF499-domain-containing-protein-[Thermoguttaceae-bacterium] |
| MBR4105130.1 | DUF499_1.RM.like.systems | <-HEPN+DUF499*<-METHYLASE-p | 819 | Bacteria>PVC_group>Planctomycetota>Planctomycetia>Pirellulales | GCA_017521545.1 | Thermoguttaceae_bacterium | IKK39_13850 | 1..2456 | 2691551 | DUF499-domain-containing-protein-[Thermoguttaceae-bacterium] |
| MBR4752420.1 | DUF499_1.RM.like.systems | TUDOR+ SF1-SF2-Helicase+iREase+CTD+REase->?->METHYLASE->HEPN+DUF499*-> | 1123 | Bacteria>PVC_group>Planctomycetota>Planctomycetia>Pirellulales | GCA_017550965.1 | Thermoguttaceae_bacterium | IK077_11730 | 50107..53478 | 2691551 | DUF499-domain-containing-protein-[Thermoguttaceae-bacterium] |
| GIW99318.1 | DUF499_1.RM.like.systems | METHYLASE-p->?->MACRODOMAIN->METHYLASE->PHP+SbcC->Schlafen+HATPase+HTH->HEPN+DUF499*-><-URI-DNase<-?<-REase<-DDE_Tnp_IS66+DDE_Tnp_IS66_C | 1115 | Bacteria>PVC_group>Planctomycetota>Planctomycetia>Pirellulales | GCA_026004795.1 | Pirellulaceae_bacterium | KatS3mg111_2651 | 704885..708232 | 2699754 | MAG:-hypothetical-protein-KatS3mg111_2651-[Pirellulaceae-bacterium] |
| MDB4650128.1 | DUF499_1.RM.like.systems | METHYLASE->HEPN+DUF499*->Synaptojanin->?-><-?<-?\|\|?->SF1-SF2-Helicase-> | 1109 | Bacteria>PVC_group>Planctomycetota>Planctomycetia>Pirellulales | GCA_028266775.1 | Pirellulaceae_bacterium | OAE37_00010 | 2073..5402 | 2699754 | DUF499-domain-containing-protein-[Pirellulaceae-bacterium] |
| MDD3586122.1 | DUF499_1.RM.like.systems | <-HEPN+DUF499*<-METHYLASE<-METHYLASE-p | 1130 | Bacteria>PVC_group>Planctomycetota>Planctomycetia>Pirellulales | GCA_028695865.1 | Thermoguttaceae_bacterium | PHQ75_02970 | 4102..7494 | 2691551 | MAG:-DUF499-domain-containing-protein-[Thermoguttaceae-bacterium] |
| WP_008662002.1 | DUF499_1.RM.like.systems | <-XIS-HTH\|\|URI-DNase-><-HEPN<-HEPN+DUF499*<-HEPN<-METHYLASE<-?<-TUDOR+ SF1-SF2-Helicase+iREase+CTD+REase\|\|HKD-DNase+SF1-SF2-Helicase+TRD->URI-DNase-> | 1113 | Bacteria>PVC_group>Planctomycetota>Planctomycetia>Pirellulales | GCF_000338295.1 | Rhodopirellula_europaea | RE6C_RS27575 | 3450..6791 | 1263866 | DUF499-domain-containing-protein-[Rhodopirellula-europaea] |
| WP_283433169.1 | DUF499_1.RM.like.systems | <-ParB+HNH<-?<-HTH\|\|MORC+DUF3684->HKD-DNase-><-HEPN+DUF499*<-PIN<-OLD-ABC<-METHYLASE<-REase\|\|?-><-TUDOR+ SF1-SF2-Helicase+iREase+CTD+REase | 1109 | Bacteria>PVC_group>Planctomycetota>Planctomycetia>Pirellulales | GCF_900182915.1 | Neorhodopirellula_lusitana | QOL80_RS14710 | 91339..94668 | 445327 | DUF499-domain-containing-protein-[Rhodopirellula-lusitana] |
| MBV8676907.1 | DUF499_1.RM.like.systems | METHYLASE-p->REase->HEPN+DUF499*->Y1_Tnp-> | 1104 | Bacteria>PVC_group>Planctomycetota>Planctomycetia>Planctomycetales | GCA_019232645.1 | Planctomycetaceae_bacterium | JO355_07025 | 1198..4512 | 2026779 | DUF499-domain-containing-protein-[Planctomycetaceae-bacterium] |
| MBV8313321.1 | DUF499_1.RM.like.systems | <-HEPN+DUF499*<-REase<-METHYLASE<-HicB+ZnR<-TUDOR+ SF1-SF2-Helicase+iREase+CTD+REase-p | 939 | Bacteria>PVC_group>Planctomycetota>Planctomycetia>Planctomycetales | GCA_019236145.1 | Planctomycetaceae_bacterium | JOZ53_00025 | 1..2817 | 2026779 | DUF499-domain-containing-protein-[Planctomycetaceae-bacterium] |
| MBV8269357.1 | DUF499_1.RM.like.systems | <-HEPN+DUF499*<-REase<-METHYLASE-p | 1104 | Bacteria>PVC_group>Planctomycetota>Planctomycetia>Planctomycetales | GCA_019236385.1 | Planctomycetaceae_bacterium | JO252_23790 | 243..3557 | 2026779 | DUF499-domain-containing-protein-[Planctomycetaceae-bacterium] |
| MBV8232626.1 | DUF499_1.RM.like.systems | TUDOR+ SF1-SF2-Helicase+iREase+CTD+REase-p->HicB+ZnR->METHYLASE->REase->HEPN+DUF499*->Y1_Tnp-> | 1104 | Bacteria>PVC_group>Planctomycetota>Planctomycetia>Planctomycetales | GCA_019236625.1 | Planctomycetaceae_bacterium | JO329_21805 | 5478..8792 | 2026779 | DUF499-domain-containing-protein-[Planctomycetaceae-bacterium] |
| MCA9076582.1 | DUF499_1.RM.like.systems | sGTP->METHYLASE-p->REase->METHYLASE->OLD-ABC->PIN->HEPN+DUF499*-><-PAIREDC-HTH<-?<-?<-SIGMA-HTH | 1095 | Bacteria>PVC_group>Planctomycetota>Planctomycetia>Planctomycetales | GCA_020426505.1 | Planctomycetaceae_bacterium | KDA93_16275 | 17776..21063 | 2026779 | DUF499-domain-containing-protein-[Planctomycetaceae-bacterium] |
| CAG0999165.1 | DUF499_1.RM.like.systems | HKD-DNase->?->HEPN+DUF499*->NERD+SF1-SF2-Helicase-> | 1113 | Bacteria>PVC_group>Planctomycetota>Planctomycetia>Planctomycetales | GCA_905339245.1 | Planctomycetaceae_bacterium | PLCT2_02959 | 2861..6202 | 2026779 | hypothetical-protein-PLCT2_02959-[Planctomycetaceae-bacterium] |
| WP_145451931.1 | DUF499_1.RM.like.systems | HTH_3-><-?<-?<-HKD-DNase<-?<-?<-HEPN+DUF499*<-METHYLASE<-Schlafen+HATPase+HTH<-MORC+DUF3684<-TUDOR+ SF1-SF2-Helicase+iREase+CTD+REase<-XIS-HTH | 1094 | Bacteria>PVC_group>Planctomycetota>Planctomycetia>Planctomycetales | GCF_007748155.1 | Gimesia_panareensis | Enr10x_RS26145 | 6953187..6956471 | 2527978 | Swt1-family-HEPN-domain-containing-protein-[Gimesia-panareensis] |
| MBE7458022.1 | DUF499_1.RM.like.systems | DEOR-HTH->TUDOR+ SF1-SF2-Helicase+iREase+CTD+REase->Schlafen+HATPase+HTH->?->?->METHYLASE->HEPN+DUF499*->MORC-><-?<-HTH_3<-RelE-ParE<-?<-DDE_Tnp_1_4 | 1120 | Bacteria>PVC_group>Planctomycetota>Planctomycetia>unclassified_Planctomycetia | GCA_015075705.1 | Planctomycetia_bacterium | HS102_15595 | 57764..61126 | 2052181 | ATP-binding-protein-[Planctomycetia-bacterium] |
| MAW61410.1 | DUF499_1.RM.like.systems | HNH->Phage_int_SAM_4+Phage_integrase-><-XIS-HTH<-Phage_int_SAM_1+Phage_integrase<-?<-?<-HEPN+DUF499*<-METHYLASE<-TUDOR+ SF1-SF2-Helicase+iREase+CTD+REase | 1080 | Bacteria>PVC_group>Planctomycetota>unclassified_Planctomycetota | GCA_002705055.1 | Planctomycetota_bacterium | CMJ94_11320 | 35772..39014 | 2026780 | AAA+-family-ATPase-[Planctomycetota-bacterium] |
| PIE22549.1 | DUF499_1.RM.like.systems | SMODS->SLATT->TUDOR+ SF1-SF2-Helicase+iREase+CTD+REase->METHYLASE->PHP+SbcC->Schlafen+HATPase+HTH->HEPN+DUF499*-><-tRNA\|\|?->?->?->?->SF1-SF2-Helicase-> | 1122 | Bacteria>PVC_group>Planctomycetota>unclassified_Planctomycetota | GCA_002748355.1 | Planctomycetota_bacterium | CSA62_11865 | 35983..39351 | 2026780 | MAG:-AAA+-family-ATPase-[Planctomycetota-bacterium] |
| RLS88266.1 | DUF499_1.RM.like.systems | METHYLASE->HEPN+DUF499*->DUF499-p-> | 912 | Bacteria>PVC_group>Planctomycetota>unclassified_Planctomycetota | GCA_003671065.1 | Planctomycetota_bacterium | DWI10_09315 | 3242..5980 | 2026780 | MAG:-ATP-binding-protein-[Planctomycetota-bacterium] |
| RMD62695.1 | DUF499_1.RM.like.systems | <-HEPN+DUF499*<-METHYLASE-p | 397 | Bacteria>PVC_group>Planctomycetota>unclassified_Planctomycetota | GCA_003693915.1 | Planctomycetota_bacterium | D6824_06430 | 1..1192 | 2026780 | MAG:-ATP-binding-protein,-partial-[Planctomycetota-bacterium] |
| RMH31880.1 | DUF499_1.RM.like.systems | METHYLASE-p->HEPN+DUF499*->HIN-HTH-> | 1102 | Bacteria>PVC_group>Planctomycetota>unclassified_Planctomycetota | GCA_003696675.1 | Planctomycetota_bacterium | D6692_00155 | 1472..4780 | 2026780 | MAG:-ATP-binding-protein-[Planctomycetota-bacterium] |
| TAH38052.1 | DUF499_1.RM.like.systems | <-wHTH+REase\|\|AbiEi_4->AbiEii-><-PIN<-ParB+HNH\|\|HKD-DNase-><-HEPN+DUF499*<-METHYLASE<-HNH<-OLD-ABC<-TUDOR+ SF1-SF2-Helicase+iREase+CTD+REase<-HTH | 1098 | Bacteria>PVC_group>Planctomycetota>unclassified_Planctomycetota | GCA_004295085.1 | Planctomycetota_bacterium | EYC70_05360 | 66593..69889 | 2026780 | MAG:-ATP-binding-protein-[Planctomycetota-bacterium] |
| QDU83134.1 | DUF499_1.RM.like.systems | TUDOR+ SF1-SF2-Helicase+iREase+CTD+REase->Schlafen+HATPase+HTH->?->METHYLASE->OLD-ABC->TOPRIM->HEPN+DUF499*->?-><-REase+SF1-SF2-Helicase<-DUF2357+REase<-Preader7+McrB-NTD+McrB-AAA | 1111 | Bacteria>PVC_group>Planctomycetota>unclassified_Planctomycetota | GCA_007750655.1 | Planctomycetes_bacterium_Pla163 | Pla163_02310 | 293863..297198 | 2528008 | hypothetical-protein-Pla163_02310-[Planctomycetes-bacterium-Pla163] |
| KAA3609103.1 | DUF499_1.RM.like.systems | TUDOR+ SF1-SF2-Helicase+iREase+CTD+REase->Schlafen+HATPase+HTH->METHYLASE->PHP+SbcC->Schlafen+HATPase+HTH->HEPN+DUF499*->?->?-><-?<-CHTH\|\|?-><-HhH-RADC+JAB | 1120 | Bacteria>PVC_group>Planctomycetota>unclassified_Planctomycetota | GCA_008501825.1 | Planctomycetota_bacterium | DWQ01_09790 | 14904..18266 | 2026780 | MAG:-DUF499-domain-containing-protein-[Planctomycetota-bacterium] |
| MCL2001508.1 | DUF499_1.RM.like.systems | <-HEPN+DUF499*<-METHYLASE-p<-METHYLASE-p | 524 | Bacteria>PVC_group>Planctomycetota>unclassified_Planctomycetota | GCA_009778045.1 | Planctomycetota_bacterium | FWG74_08750 | 1..1573 | 2026780 | Swt1-family-HEPN-domain-containing-protein-[Planctomycetota-bacterium] |
| MBI3462535.1 | DUF499_1.RM.like.systems | PIWI-><-ParB+HNH<-HEPN+DUF499*<-METHYLASE<-wHTH+REase<-TUDOR+ SF1-SF2-Helicase+iREase+CTD+REase-p | 1111 | Bacteria>PVC_group>Planctomycetota>unclassified_Planctomycetota | GCA_016196255.1 | Planctomycetota_bacterium | HY000_05665 | 2780..6115 | 2026780 | ATP-binding-protein-[Planctomycetota-bacterium] |
| MBI3462766.1 | DUF499_1.RM.like.systems | <-HEPN+DUF499*<-METHYLASE | 1074 | Bacteria>PVC_group>Planctomycetota>unclassified_Planctomycetota | GCA_016196255.1 | Planctomycetota_bacterium | HY000_06860 | 1..3221 | 2026780 | ATP-binding-protein-[Planctomycetota-bacterium] |
| MBI4617608.1 | DUF499_1.RM.like.systems | Radical-SAM->?->RelE-ParE->?->?->?-><-HEPN+DUF499*<-ORC-CDC6+REase | 1115 | Bacteria>PVC_group>Planctomycetota>unclassified_Planctomycetota | GCA_016209595.1 | Planctomycetota_bacterium | HY720_28610 | 4559..7906 | 2026780 | ATP-binding-protein-[Planctomycetota-bacterium] |
| MBK8205305.1 | DUF499_1.RM.like.systems | HEPN->METHYLASE-p->HKD-DNase->?-><-?<-?\|\|HEPN+DUF499*-><-?<-?<-HTH_36<-XIS-HTH\|\|Phage_integrase-> | 1097 | Bacteria>PVC_group>Planctomycetota>unclassified_Planctomycetota | GCA_016712285.1 | Planctomycetota_bacterium | IPK87_00800 | 221312..224605 | 2026780 | DUF499-domain-containing-protein-[Planctomycetota-bacterium] |
| MBX3463236.1 | DUF499_1.RM.like.systems | <-UbE1+JAB<-?\|\|HTH_3->MPTase->TUDOR+ SF1-SF2-Helicase+iREase+CTD+REase->METHYLASE->HEPN+DUF499*->HD-> | 1101 | Bacteria>PVC_group>Planctomycetota>unclassified_Planctomycetota | GCA_019636245.1 | Planctomycetota_bacterium | KF830_08695 | 93755..97060 | 2026780 | DUF499-domain-containing-protein-[Planctomycetota-bacterium] |
| MCC7173369.1 | DUF499_1.RM.like.systems | METHYLASE-p->HEPN+DUF499*->?->?-><-PIN\|\|?->SIGMA-HTH-> | 1113 | Bacteria>PVC_group>Planctomycetota>unclassified_Planctomycetota | GCA_020851675.1 | Planctomycetota_bacterium | IT459_23190 | 1090..4431 | 2026780 | DUF499-domain-containing-protein-[Planctomycetota-bacterium] |
| MCG3135902.1 | DUF499_1.RM.like.systems | TUDOR+ SF1-SF2-Helicase+iREase+CTD+REase->OLD-ABC->TOPRIM->METHYLASE->HEPN+DUF499*->KAP-NTPase->?->PAPS_reduct->?-><-TPR+TPR | 1110 | Bacteria>PVC_group>Planctomycetota>unclassified_Planctomycetota | GCA_022072415.1 | Planctomycetota_bacterium | HMLKMBBP_03678 | 15324..18656 | 2026780 | hypothetical-protein-[Planctomycetota-bacterium] |
| MCK5940853.1 | DUF499_1.RM.like.systems | <-HEPN+DUF499*<-METHYLASE | 1099 | Bacteria>PVC_group>Planctomycetota>unclassified_Planctomycetota | GCA_023146405.1 | Planctomycetota_bacterium | KAI24_02690 | 255..3554 | 2026780 | DUF499-domain-containing-protein-[Planctomycetota-bacterium] |
| MCK6472712.1 | DUF499_1.RM.like.systems | TUDOR+ SF1-SF2-Helicase+iREase+CTD+REase->?->REase-><-?\|\|?->HEPN+DUF499*->SbcC->?->?->METHYLASE-><-?<-XIS-HTH | 1124 | Bacteria>PVC_group>Planctomycetota>unclassified_Planctomycetota | GCA_023150555.1 | Planctomycetota_bacterium | L6R28_13300 | 20467..23841 | 2026780 | DUF499-domain-containing-protein-[Planctomycetota-bacterium] |
| MCZ7645664.1 | DUF499_1.RM.like.systems | HTH+MPTase->PemK_toxin->HNH+SAVED->SMODS->?-><-PIN<-HEPN+DUF499*<-METHYLASE<-?<-ParB+HNH<-TUDOR+ SF1-SF2-Helicase+iREase+CTD+REase | 1122 | Bacteria>PVC_group>Planctomycetota>unclassified_Planctomycetota | GCA_027493525.1 | Planctomycetota_bacterium | M5U26_10325 | 2469927..2473295 | 2026780 | MAG:-Swt1-family-HEPN-domain-containing-protein-[Planctomycetota-bacterium] |
| MDA1195884.1 | DUF499_1.RM.like.systems | <-HEPN+DUF499*<-?<-?<-METHYLASE | 886 | Bacteria>PVC_group>Planctomycetota>unclassified_Planctomycetota | GCA_027625315.1 | Planctomycetota_bacterium | O2894_11960 | 1..2659 | 2026780 | MAG:-Swt1-family-HEPN-domain-containing-protein,-partial-[Planctomycetota-bacterium] |
| MBW7893701.1 | DUF499_1.RM.like.systems | TUDOR+ SF1-SF2-Helicase+iREase+CTD+REase->MORC+DUF3684->HKD-DNase->?->METHYLASE->HEPN+DUF499*-><-?\|\|HIN-HTH-> | 1122 | Bacteria>PVC_group>Verrucomicrobiota>Opitutae>Opitutales | GCA_019455245.1 | Opitutaceae_bacterium | H3C27_01205 | 267980..271348 | 2006848 | ATP-binding-protein-[Opitutaceae-bacterium] |
| MBX3751087.1 | DUF499_1.RM.like.systems | TUDOR+ SF1-SF2-Helicase+iREase+CTD+REase->?->METHYLASE->HEPN+DUF499*-><-S1+RNase+2C | 1122 | Bacteria>PVC_group>Verrucomicrobiota>Opitutae>Opitutales | GCA_019634615.1 | Opitutaceae_bacterium | KF897_13455 | 8367..11735 | 2006848 | DUF499-domain-containing-protein-[Opitutaceae-bacterium] |
| MCC5835099.1 | DUF499_1.RM.like.systems | RVT->METHYLASE->?->CHTH->HEPN+DUF499*-><-?\|\|wHTH+SAD+HNH-><-?\|\|REase+35exo+Beta-Barrel+SF1-SF2-Helicase-><-AbiEi_4 | 1100 | Bacteria>PVC_group>Verrucomicrobiota>Opitutae>Opitutales | GCA_020831485.1 | Opitutales_bacterium | JJU20_10230 | 15950..19252 | 2026772 | DUF499-domain-containing-protein-[Opitutales-bacterium] |
| MCC5805073.1 | DUF499_1.RM.like.systems | OLD-ABC->HNH->METHYLASE->CHTH->RHH->PIN->HEPN+DUF499*->?->PhdYeFM_antitox->PIN-> | 1107 | Bacteria>PVC_group>Verrucomicrobiota>Opitutae>Opitutales | GCA_020831525.1 | Opitutales_bacterium | JJU00_01980 | 48883..52206 | 2026772 | DUF499-domain-containing-protein-[Opitutales-bacterium] |
| WP_189517054.1 | DUF499_1.RM.like.systems | <-ParB+HNH<-HEPN+DUF499*<-?<-HTH<-?<-KAP-NTPase<-METHYLASE | 1101 | Bacteria>PVC_group>Verrucomicrobiota>Opitutae>Puniceicoccales | GCF_014651635.1 | Cerasicoccus_arenae | JIN80_RS05850 | 25614..28919 | 424488 | DUF499-domain-containing-protein-[Cerasicoccus-arenae] |
| PWU21781.1 | DUF499_1.RM.like.systems | SF1-SF2-Helicase->?->HKD-DNase->?->?->?-><-HEPN+DUF499*<-?<-PIWI<-REase+PNTD1+PNTD2 | 1128 | Bacteria>PVC_group>Verrucomicrobiota>unclassified_Verrucomicrobiota | GCA_003176045.1 | Verrucomicrobiota_bacterium | C5B50_01090 | 37039..40425 | 2026799 | AAA+-family-ATPase-[Verrucomicrobia-bacterium] |
| RFC50837.1 | DUF499_1.RM.like.systems | <-UPF0061\|\|TUDOR+ SF1-SF2-Helicase+iREase+CTD+REase->STAS+HisKin->?->PIN->METHYLASE->HEPN+DUF499*-><-?<-?<-?<-SIGMA-HTH | 1074 | Bacteria>PVC_group>Verrucomicrobiota>unclassified_Verrucomicrobiota | GCA_003402705.1 | Verrucomicrobiota_bacterium | DVB22_001798 | 10693..13917 | 2026799 | putative-ATPase,-AAA+-superfamily-[Verrucomicrobia-bacterium] |
| RME69044.1 | DUF499_1.RM.like.systems | <-HEPN+DUF499*<-METHYLASE-p | 1114 | Bacteria>PVC_group>Verrucomicrobiota>unclassified_Verrucomicrobiota | GCA_003694825.1 | Verrucomicrobiota_bacterium | D6781_09510 | 385..3729 | 2026799 | ATP-binding-protein-[Verrucomicrobia-bacterium] |
| MBI3869845.1 | DUF499_1.RM.like.systems | Phage_integrase-><-?\|\|HTH-><-?<-HEPN+DUF499*<-PIWI<-TIR+PNTD2<-METHYLASE<-TIR<-SIR2+STALD<-TIR | 1119 | Bacteria>PVC_group>Verrucomicrobiota>unclassified_Verrucomicrobiota | GCA_016199985.1 | Verrucomicrobiota_bacterium | HY299_15085 | 262919..266278 | 2026799 | ATP-binding-protein-[Verrucomicrobia-bacterium] |
| MBI5394224.1 | DUF499_1.RM.like.systems | <-PfkB<-ABC-ATPase+SbcC<-?<-HEPN+DUF499*<-HEPN<-HTH+PolB<-TUDOR+ SF1-SF2-Helicase+iREase+CTD+REase | 1121 | Bacteria>PVC_group>Verrucomicrobiota>unclassified_Verrucomicrobiota | GCA_016219905.1 | Verrucomicrobiota_bacterium | HZA91_02900 | 10748..14113 | 2026799 | DUF499-domain-containing-protein-[Verrucomicrobia-bacterium] |
| MBM3880418.1 | DUF499_1.RM.like.systems | <-HEPN+DUF499*<-ParB+HNH<-METHYLASE<-TOPRIM<-OLD-ABC<-TUDOR+ SF1-SF2-Helicase+iREase+CTD+REase | 1127 | Bacteria>PVC_group>Verrucomicrobiota>unclassified_Verrucomicrobiota | GCA_016871715.1 | Verrucomicrobiota_bacterium | FJ387_11990 | 332..3715 | 2026799 | ATP-binding-protein-[Verrucomicrobia-bacterium] |
| MBS0646999.1 | DUF499_1.RM.like.systems | <-HEPN+DUF499*<-METHYLASE<-ParB+HNH<-AbiEii | 1093 | Bacteria>PVC_group>Verrucomicrobiota>unclassified_Verrucomicrobiota | GCA_018239885.1 | Verrucomicrobiota_bacterium | JSR97_10490 | 874..4155 | 2026799 | DUF499-domain-containing-protein-[Verrucomicrobia-bacterium] |
| MBS0647016.1 | DUF499_1.RM.like.systems | <-HEPN+DUF499*<-METHYLASE<-SF1-SF2-Helicase-p | 1093 | Bacteria>PVC_group>Verrucomicrobiota>unclassified_Verrucomicrobiota | GCA_018239885.1 | Verrucomicrobiota_bacterium | JSR97_10575 | 874..4155 | 2026799 | DUF499-domain-containing-protein-[Verrucomicrobia-bacterium] |
| MCC5848502.1 | DUF499_1.RM.like.systems | <-HEPN+DUF499*<-?<-SF1-SF2-Helicase+wHTH+PRTase<-METHYLASE<-TOPRIM<-OLD-ABC<-HTH_3 | 1113 | Bacteria>PVC_group>Verrucomicrobiota>unclassified_Verrucomicrobiota | GCA_020831635.1 | Verrucomicrobiota_bacterium | JJU29_10430 | 13119..16460 | 2026799 | DUF499-domain-containing-protein-[Verrucomicrobia-bacterium] |
| MCD6339150.1 | DUF499_1.RM.like.systems | <-HEPN+DUF499*<-METHYLASE-p | 1113 | Bacteria>PVC_group>Verrucomicrobiota>unclassified_Verrucomicrobiota | GCA_021160185.1 | Verrucomicrobiota_bacterium | J7M29_07215 | 9794..13135 | 2026799 | DUF499-domain-containing-protein-[Verrucomicrobia-bacterium] |
| MCF7668203.1 | DUF499_1.RM.like.systems | TUDOR+ SF1-SF2-Helicase+iREase+CTD+REase->METHYLASE->OLD-ABC->TOPRIM->HEPN+DUF499*->?->?-><-PfkB | 1108 | Bacteria>PVC_group>Verrucomicrobiota>unclassified_Verrucomicrobiota | GCA_021733785.1 | Verrucomicrobiota_bacterium | K9N48_00295 | 29599..32925 | 2026799 | DUF499-domain-containing-protein-[Verrucomicrobia-bacterium] |
| MCP5487625.1 | DUF499_1.RM.like.systems | <-HEPN+DUF499*<-METHYLASE<-?<-SMODS<-?<-HTH_3 | 1116 | Bacteria>PVC_group>Verrucomicrobiota>unclassified_Verrucomicrobiota | GCA_024233345.1 | Verrucomicrobiota_bacterium | H7A43_03165 | 141762..145112 | 2026799 | ATP-binding-protein-[Verrucomicrobia-bacterium] |
| MCZ7639656.1 | DUF499_1.RM.like.systems | TIR->TIR->PIWI->AbiEi_4->AbiEii->HEPN+DUF499*->HTH->HTH->NTP_transf-> | 1127 | Bacteria>PVC_group>Verrucomicrobiota>unclassified_Verrucomicrobiota | GCA_027493615.1 | Verrucomicrobiota_bacterium | M5U12_28570 | 388044..391427 | 2026799 | MAG:-Swt1-family-HEPN-domain-containing-protein-[Verrucomicrobia-bacterium] |
| MDE3100243.1 | DUF499_1.RM.like.systems | <-Response_reg+LuxR-HTH<-?\|\|?-><-REase<-?<-HEPN+DUF499*<-AbiEii<-AbiEi_4<-REase<-PIWI | 1114 | Bacteria>PVC_group>Verrucomicrobiota>unclassified_Verrucomicrobiota | GCA_028872915.1 | Verrucomicrobiota_bacterium | KGJ88_12300 | 8441..11785 | 2026799 | MAG:-DUF499-domain-containing-protein-[Verrucomicrobiota-bacterium] |
| MCS1417413.1 | DUF499_1.RM.like.systems | <-HEPN+DUF499*<-?<-DUF488-1<-PIN<-?<-METHYLASE-p | 1121 | Bacteria>PVC_group>Verrucomicrobiota>Verrucomicrobiae>Limisphaerales | GCA_024761205.1 | Limisphaerales_bacterium | M2R46_05167 | 18516..21881 | 2052186 | hypothetical-protein-[Verrucomicrobia-subdivision-3-bacterium] |
| MCW5552136.1 | DUF499_1.RM.like.systems | GDResolvase+PPP1-HTH->ParB->ParB-><-MarR-HTH\|\|?->ACET-><-HEPN+DUF499*<-METHYLASE<-PIN<-?<-STAS+HisKin<-TUDOR+ SF1-SF2-Helicase+iREase+CTD+REase | 1120 | Bacteria>PVC_group>Verrucomicrobiota>Verrucomicrobiae>unclassified_Verrucomicrobiae | GCA_026125285.1 | Verrucomicrobiae_bacterium | KIS67_08205 | 85951..89313 | 2499141 | MAG:-DUF499-domain-containing-protein-[Verrucomicrobiae-bacterium] |
| HCN27197.1 | DUF499_1.RM.like.systems | METHYLASE->ParD->PIN->HEPN+DUF499*-> | 1113 | Bacteria>PVC_group>Verrucomicrobiota>Verrucomicrobiae>Verrucomicrobiales | GCA_003545395.1 | Verrucomicrobiales_bacterium | DIT64_00065 | 2852..6193 | 2026801 | AAA+-family-ATPase-[Verrucomicrobiales-bacterium] |
| MBE2287190.1 | DUF499_1.RM.like.systems | TIR+PNTD2->PIWI->DUF932->TUDOR+ SF1-SF2-Helicase+iREase+CTD+REase->NACHT->METHYLASE->HEPN+DUF499*->?->SPOUT-> | 1114 | Bacteria>PVC_group>Verrucomicrobiota>Verrucomicrobiae>Verrucomicrobiales | GCA_014879935.1 | Prosthecobacter_sp. | IAE77_27290 | 17760..21104 | 1965333 | ATP-binding-protein-[Prosthecobacter-sp.] |
| MBL9159968.1 | DUF499_1.RM.like.systems | <-HISKIN<-ArdCN+MPTase<-HNH<-HEPN+DUF499*<-?<-Mbetalac<-?<-?<-METHYLASE | 1114 | Bacteria>PVC_group>Verrucomicrobiota>Verrucomicrobiae>Verrucomicrobiales | GCA_016795385.1 | Verrucomicrobiales_bacterium | JNJ70_20975 | 159944..163288 | 2026801 | DUF499-domain-containing-protein-[Verrucomicrobiales-bacterium] |
| QTN34153.1 | DUF499_1.RM.like.systems | MqsA_antitoxin+HTH_3->?->METHYLASE->HEPN+DUF499*->HEPN+SF1-SF2-Helicase->HEPN+TPR+RhoN->ParB+HNH-> | 1101 | Bacteria>PVC_group>Verrucomicrobiota>Verrucomicrobiae>Verrucomicrobiales | GCA_017798145.1 | Akkermansiaceae_bacterium | HZ994_18120 | 4237936..4241241 | 2562705 | ATP-binding-protein-[Akkermansiaceae-bacterium] |
| MBP7951461.1 | DUF499_1.RM.like.systems | ACET-><-tRNA<-?<-HEPN+DUF499*<-METHYLASE<-PIN<-?<-STAS+HisKin<-TUDOR+ SF1-SF2-Helicase+iREase+CTD+REase<-DDE_Tnp_1 | 1112 | Bacteria>PVC_group>Verrucomicrobiota>Verrucomicrobiae>Verrucomicrobiales | GCA_018003315.1 | Verrucomicrobiales_bacterium | KA004_17600 | 4745..8083 | 2026801 | DUF499-domain-containing-protein-[Verrucomicrobiales-bacterium] |
| AEJ18699.1 | DUF499_1.RM.like.systems | PhdYeFM_antitox->Phd-YefM->RelE-ParE->?->?->ParB+HNH->HEPN+DUF499*->ABC-ATPase+SbcC-> | 1058 | Bacteria>Spirochaetota>Spirochaetia>Spirochaetales>Breznakiellaceae | GCA_000219725.1 | Gracilinema_caldarium_DSM_7334 | Spica_0539 | 580385..583561 | 744872 | ATPase-(AAA+-superfamily)-like-protein-[Gracilinema-caldarium-DSM-7334] |
| WP_215904902.1 | DUF499_1.RM.like.systems | PIN->Phd-YefM->RelE-ParE->?->ParB+HNH-><-?\|\|HEPN+DUF499*->ABC-ATPase+SbcC-> | 1077 | Bacteria>Spirochaetota>Spirochaetia>Spirochaetales>Breznakiellaceae | GCF_000219725.1 | Gracilinema_caldarium | SPICA_RS02735 | 580328..583561 | 215591 | DUF499-domain-containing-protein-[Gracilinema-caldarium] |
| MBI9094232.1 | DUF499_1.RM.like.systems | <-zf-IS66+DDE_Tnp_IS66<-?\|\|tRNA-><-?<-?<-HEPN+DUF499*<-METHYLASE<-TUDOR+ SF1-SF2-Helicase+iREase+CTD+REase\|\|URI-DNase->HEPN->?-><-HTH_3 | 1134 | Bacteria>Spirochaetota>Spirochaetia>Spirochaetales>Sphaerochaetaceae | GCA_016342655.1 | Sphaerochaeta_sp. | JEY71_05065 | 29138..32542 | 1972642 | DUF499-domain-containing-protein-[Sphaerochaeta-sp.] |
| MCK9286308.1 | DUF499_1.RM.like.systems | <-sGTP<-SF1-SF2-Helicase<-OLD-ABC+TOPRIM<-HEPN+DUF499*<-METHYLASE<-TUDOR+ SF1-SF2-Helicase+iREase+CTD+REase<-HTH_26<-SbcC+SbcC<-?<-REase+CTD6 | 1135 | Bacteria>Spirochaetota>Spirochaetia>Spirochaetales>Sphaerochaetaceae | GCA_023230445.1 | Sphaerochaetaceae_bacterium | MOP29_01920 | 121891..125298 | 2791017 | DUF499-domain-containing-protein-[Sphaerochaetaceae-bacterium] |
| HAL93442.1 | DUF499_1.RM.like.systems | SF1-SF2-Helicase+DUF1998-><-?<-PIN<-?\|\|?-><-HEPN+DUF499*<-PIN<-PhdYeFM_antitox<-?<-HEPN-p | 1142 | Bacteria>Spirochaetota>Spirochaetia>Spirochaetales>Spirochaetaceae | GCA_003452735.1 | Spirochaetaceae_bacterium | DCP56_01610 | 10292..13720 | 1898206 | AAA+-family-ATPase-[Spirochaetaceae-bacterium] |
| HDQ14361.1 | DUF499_1.RM.like.systems | METHYLASE-p->METHYLASE-p->HEPN+DUF499*->?->ORC-CDC6+REase-> | 1109 | Bacteria>Spirochaetota>Spirochaetia>Spirochaetales>Spirochaetaceae | GCA_011046055.1 | Sediminispirochaeta_sp. | ENN41_06065 | 1978..5307 | 1911557 | ATP-binding-protein-[Sediminispirochaeta-sp.] |
| MCX7774711.1 | DUF499_1.RM.like.systems | ACET->TUDOR+ SF1-SF2-Helicase+iREase+CTD+REase->ORC-CDC6+REase->METHYLASE->AbiEi_4->AbiEii->HEPN+DUF499*-> | 1103 | Bacteria>Spirochaetota>Spirochaetia>Spirochaetales>Spirochaetaceae | GCA_026418115.1 | Spirochaetaceae_bacterium | N2067_01230 | 29258..32569 | 1898206 | MAG:-DUF499-domain-containing-protein-[Spirochaetaceae-bacterium] |
| MCY4372401.1 | DUF499_1.RM.like.systems | MPTase->HEPN+DUF499*->?-><-?\|\|Glyco_hydro->DUF2961-> | 1110 | Bacteria>Spirochaetota>Spirochaetia>Spirochaetales>Spirochaetaceae | GCA_026710565.1 | Spirochaetaceae_bacterium | OXC31_01430 | 992..4324 | 1898206 | MAG:-Swt1-family-HEPN-domain-containing-protein-[Spirochaetaceae-bacterium] |
| MDD9984792.1 | DUF499_1.RM.like.systems | <-Glyco_hydro\|\|?-><-NUDIX<-DOC+HTH\|\|?-><-?<-HEPN+DUF499*\|\|DDE_Tnp_1-> | 1110 | Bacteria>Spirochaetota>Spirochaetia>Spirochaetales>Spirochaetaceae | GCA_028818385.1 | Spirochaetaceae_bacterium | OXQ31_00820 | 20813..24145 | 1898206 | MAG:-Swt1-family-HEPN-domain-containing-protein-[Spirochaetaceae-bacterium] |
| MDE0219724.1 | DUF499_1.RM.like.systems | SbcC-><-OLD-ABC\|\|TUDOR+ SF1-SF2-Helicase+iREase+CTD+REase-><-?\|\|HTH+MPTase->PIN->HEPN+DUF499*->DOC+HTH->?->XIS-HTH+RVT->Glyco_hydro->DUF2961->HNH-> | 1110 | Bacteria>Spirochaetota>Spirochaetia>Spirochaetales>Spirochaetaceae | GCA_028820215.1 | Spirochaetaceae_bacterium | OXJ90_10675 | 22427..25759 | 1898206 | MAG:-Swt1-family-HEPN-domain-containing-protein-[Spirochaetaceae-bacterium] |
| WP_083048603.1 | DUF499_1.RM.like.systems | <-PfkB<-?<-?<-?<-HEPN+DUF499*<-METHYLASE<-TUDOR+ SF1-SF2-Helicase+iREase+CTD+REase<-?<-ParB+HNH<-?<-SF1-SF2-Helicase+REase | 1101 | Bacteria>Spirochaetota>Spirochaetia>Spirochaetales>Spirochaetaceae | GCF_002087085.1 | Marispirochaeta_aestuarii | B4097_RS04115 | 245818..249123 | 1963862 | DUF499-domain-containing-protein-[Marispirochaeta-aestuarii] |
| OQB96942.1 | DUF499_1.RM.like.systems | SIGMA-HTH->Radical-SAM-><-?<-?\|\|?->?-><-HEPN+DUF499*<-ParB+HNH<-METHYLASE<-TOPRIM<-OLD-ABC<-HEPN<-TUDOR+ SF1-SF2-Helicase+iREase+CTD+REase | 1176 | Bacteria>Spirochaetota>unclassified_Spirochaetota | GCA_002070755.1 | Spirochaetes_bacterium_ADurb.Bin110 | BWX81_01211 | 5339..8869 | 1852908 | hypothetical-protein-BWX81_01211-[Spirochaetes-bacterium-ADurb.Bin110] |
| MCL1994394.1 | DUF499_1.RM.like.systems | <-HEPN+DUF499*<-METHYLASE<-?<-OLD-ABC<-ParB+HNH<-TUDOR+ SF1-SF2-Helicase+iREase+CTD+REase | 1116 | Bacteria>Spirochaetota>unclassified_Spirochaetota | GCA_009778155.1 | Spirochaetota_bacterium | FWG66_15730 | 17581..20931 | 2202144 | DUF499-domain-containing-protein-[Spirochaetes-bacterium] |
| WP_026368871.1 | DUF499_1.RM.like.systems | <-HTH_3\|\|TUDOR+ SF1-SF2-Helicase+iREase+CTD+REase->METHYLASE->HEPN+DUF499*-><-ABC-ATPase+SbcC<-ABC-ATPase+SbcC\|\|<-?\|\|?->ABhydrolase-> | 1123 | Bacteria>Synergistota>Synergistia>Synergistales>Aminiphilaceae | GCF_000526375.1 | Aminiphilus_circumscriptus | K349_RS0105690 | 1079637..1083008 | 290732 | Swt1-family-HEPN-domain-containing-protein-[Aminiphilus-circumscriptus] |
| MBR1604261.1 | DUF499_1.RM.like.systems | TUDOR+ SF1-SF2-Helicase+iREase+CTD+REase->METHYLASE->HEPN+DUF499*-><-?\|\|?->REase-> | 1094 | Bacteria>Synergistota>Synergistia>Synergistales>Synergistaceae | GCA_017470785.1 | Synergistaceae_bacterium | IJ667_12605 | 8017..11301 | 2053624 | DUF499-domain-containing-protein-[Synergistaceae-bacterium] |
| HIR18062.1 | DUF499_1.RM.like.systems | <-MNS+REase<-DOC+HTH<-HEPN+DUF499*<-METHYLASE<-TUDOR+ SF1-SF2-Helicase+iREase+CTD+REase | 1115 | Bacteria>Synergistota>Synergistia>Synergistales>Synergistaceae | GCA_018712745.1 | Candidatus_Caccocola_faecigallinarum | IAD35_08445 | 169850..173197 | 2840712 | DUF499-domain-containing-protein-[Candidatus-Caccocola-faecigallinarum] |
| GHS93871.1 | DUF499_1.RM.like.systems | TUDOR+ SF1-SF2-Helicase+iREase+CTD+REase->HEPN+REase->METHYLASE->OLD-ABC->NERD->?->HEPN+DUF499*->?->?->METHYLASE->?-><-DDE_Tnp_1 | 1116 | Bacteria>Synergistota>Synergistia>Synergistales>unclassified_Synergistales | GCA_025997955.1 | Synergistales_bacterium | AGMMS50276_05310 | 53205..56555 | 2268201 | hypothetical-protein-AGMMS50276_05310-[Synergistales-bacterium] |
| GHV35275.1 | DUF499_1.RM.like.systems | HicA_toxin->OLD-ABC->NERD->?->OLD-ABC->?->HEPN+DUF499*-> | 1117 | Bacteria>Synergistota>Synergistia>Synergistales>unclassified_Synergistales | GCA_026002715.1 | Synergistales_bacterium | FACS1894187_07640 | 12375..15728 | 2268201 | hypothetical-protein-FACS1894187_07640-[Synergistales-bacterium] |
| MXW61875.1 | DUF499_1.RM.like.systems | RelE-ParE->ParA+HTH-><-HEPN+DUF499*\|\|RNase->HTH_58-><-METHYLASE\|\|KAP-NTPase->HEPN-> | 1084 | Bacteria>Terrabacteria_group>Actinomycetota>Acidimicrobiia>Acidimicrobiales | GCA_009835385.1 | Acidimicrobiaceae_bacterium | F4003_08870 | 881..4135 | 2024894 | ATP-binding-protein-[Acidimicrobiaceae-bacterium] |
| MXW97638.1 | DUF499_1.RM.like.systems | <-HEPN+DUF499*<-?<-METHYLASE\|\|HEPN-><-PemK_toxin<-?\|\|RHH_1-> | 1085 | Bacteria>Terrabacteria_group>Actinomycetota>Acidimicrobiia>Acidimicrobiales | GCA_009835735.1 | Acidimicrobiaceae_bacterium | F4Z89_03360 | 789..4046 | 2024894 | ATP-binding-protein-[Acidimicrobiaceae-bacterium] |
| MXW95443.1 | DUF499_1.RM.like.systems | <-SbcC+SbcC<-HEPN+DUF499*\|\|TraJ-RHH+MarR-HTH->AbiEii-><-METHYLASE | 1096 | Bacteria>Terrabacteria_group>Actinomycetota>Acidimicrobiia>Acidimicrobiales | GCA_009836165.1 | Acidimicrobiaceae_bacterium | F4Z64_08035 | 3972..7262 | 2024894 | ATP-binding-protein-[Acidimicrobiaceae-bacterium] |
| MXW76241.1 | DUF499_1.RM.like.systems | <-KAP-NTPase\|\|METHYLASE-><-HTH_58<-RNase\|\|HEPN+DUF499*-><-ParA+HTH<-RelE-ParE<-Creatinase_N+Peptidase_M24 | 1084 | Bacteria>Terrabacteria_group>Actinomycetota>Acidimicrobiia>Acidimicrobiales | GCA_009836285.1 | Acidimicrobiaceae_bacterium | F4Z58_09425 | 4280..7534 | 2024894 | ATP-binding-protein-[Acidimicrobiaceae-bacterium] |
| MXZ99353.1 | DUF499_1.RM.like.systems | <-AbiEii<-TraJ-RHH+MarR-HTH\|\|HEPN+DUF499*->SbcC+SbcC-> | 1096 | Bacteria>Terrabacteria_group>Actinomycetota>Acidimicrobiia>Acidimicrobiales | GCA_009836995.1 | Acidimicrobiaceae_bacterium | F4Z23_09980 | 3513..6803 | 2024894 | ATP-binding-protein-[Acidimicrobiaceae-bacterium] |
| MXY09521.1 | DUF499_1.RM.like.systems | TUDOR+ SF1-SF2-Helicase+iREase+CTD+REase->SIR2+SIR2->PIWI->METHYLASE->MerR-HTH->PIN->HEPN+DUF499*->ASCH->HAD-><-?<-?<-ABhydrolase | 1086 | Bacteria>Terrabacteria_group>Actinomycetota>Acidimicrobiia>Acidimicrobiales | GCA_009838345.1 | Acidimicrobiaceae_bacterium | F4Y56_03045 | 76366..79626 | 2024894 | ATP-binding-protein-[Acidimicrobiaceae-bacterium] |
| MYJ29885.1 | DUF499_1.RM.like.systems | ABhydrolase->?->?-><-HAD<-ASCH<-HEPN+DUF499*<-PIN<-MerR-HTH<-METHYLASE-p | 1086 | Bacteria>Terrabacteria_group>Actinomycetota>Acidimicrobiia>Acidimicrobiales | GCA_009845185.1 | Acidimicrobiaceae_bacterium | F4072_07945 | 6572..9832 | 2024894 | ATP-binding-protein-[Acidimicrobiaceae-bacterium] |
| NNN07581.1 | DUF499_1.RM.like.systems | <-HEPN+DUF499*<-METHYLASE<-TUDOR+ SF1-SF2-Helicase+iREase+CTD+REase | 435 | Bacteria>Terrabacteria_group>Actinomycetota>Acidimicrobiia>Acidimicrobiales | GCA_013044265.1 | Acidimicrobiaceae_bacterium | HKL85_00090 | 1..1304 | 2024894 | ATP-binding-protein-[Acidimicrobiaceae-bacterium] |
| NNN09184.1 | DUF499_1.RM.like.systems | METHYLASE->HEPN+DUF499*-><-?<-?<-GDResolvase<-?<-XIS-HTH | 1108 | Bacteria>Terrabacteria_group>Actinomycetota>Acidimicrobiia>Acidimicrobiales | GCA_013044265.1 | Acidimicrobiaceae_bacterium | HKL85_08335 | 1610..4936 | 2024894 | ATP-binding-protein-[Acidimicrobiaceae-bacterium] |
| MCI2976467.1 | DUF499_1.RM.like.systems | TUDOR+ SF1-SF2-Helicase+iREase+CTD+REase->METHYLASE->HEPN+DUF499*-> | 1103 | Bacteria>Terrabacteria_group>Actinomycetota>Acidimicrobiia>Acidimicrobiales | GCA_022690815.1 | Ferrimicrobium_sp. | MP439_10410 | 9887..13198 | 2926050 | DUF499-domain-containing-protein-[Ferrimicrobium-sp.] |
| MCY3849326.1 | DUF499_1.RM.like.systems | <-HEPN+DUF499*<-?<-METHYLASE-p\|\|?-><-?<-MORC+HISKIN<-DNA_methylase | 1088 | Bacteria>Terrabacteria_group>Actinomycetota>Acidimicrobiia>Acidimicrobiales | GCA_026706815.1 | Acidimicrobiaceae_bacterium | OXF75_00810 | 86121..89387 | 2024894 | MAG:-DUF499-domain-containing-protein-[Acidimicrobiaceae-bacterium] |
| MCY3892500.1 | DUF499_1.RM.like.systems | METHYLASE-p->OLD-ABC->TOPRIM->HEPN+DUF499*->?->ABhydrolase-> | 1131 | Bacteria>Terrabacteria_group>Actinomycetota>Acidimicrobiia>Acidimicrobiales | GCA_026707105.1 | Acidimicrobiaceae_bacterium | OXF65_04295 | 3174..6569 | 2024894 | MAG:-Swt1-family-HEPN-domain-containing-protein-[Acidimicrobiaceae-bacterium] |
| MCY3894210.1 | DUF499_1.RM.like.systems | <-ORC-CDC6+REase<-?<-HEPN+DUF499*<-?<-CHTH<-?<-METHYLASE<-TUDOR+ SF1-SF2-Helicase+iREase+CTD+REase<-MORC+REase | 1154 | Bacteria>Terrabacteria_group>Actinomycetota>Acidimicrobiia>Acidimicrobiales | GCA_026707105.1 | Acidimicrobiaceae_bacterium | OXF65_13050 | 72169..75633 | 2024894 | MAG:-DUF499-domain-containing-protein-[Acidimicrobiaceae-bacterium] |
| MCY4517135.1 | DUF499_1.RM.like.systems | METHYLASE-p->?->?->?->HEPN+DUF499*-> | 1090 | Bacteria>Terrabacteria_group>Actinomycetota>Acidimicrobiia>Acidimicrobiales | GCA_026713105.1 | Acidimicrobiaceae_bacterium | OXB99_07860 | 4033..7305 | 2024894 | MAG:-DUF499-domain-containing-protein-[Acidimicrobiaceae-bacterium] |
| MDE0065239.1 | DUF499_1.RM.like.systems | METHYLASE-p->?->?->HEPN+DUF499*-><-?<-?<-?<-?<-Aminotran_1_2 | 1145 | Bacteria>Terrabacteria_group>Actinomycetota>Acidimicrobiia>Acidimicrobiales | GCA_028819005.1 | Acidimicrobiaceae_bacterium | OXN44_00020 | 1449..4886 | 2024894 | MAG:-DUF499-domain-containing-protein-[Acidimicrobiaceae-bacterium] |
| MDE0067328.1 | DUF499_1.RM.like.systems | METHYLASE-p->?-><-PIN<-?<-HTH_58\|\|HEPN+DUF499*-><-ParA+HTH<-RelE-ParE<-Creatinase_N+Peptidase_M24 | 1084 | Bacteria>Terrabacteria_group>Actinomycetota>Acidimicrobiia>Acidimicrobiales | GCA_028819005.1 | Acidimicrobiaceae_bacterium | OXN44_10730 | 3279..6533 | 2024894 | MAG:-DUF499-domain-containing-protein-[Acidimicrobiaceae-bacterium] |
| MDE0136614.1 | DUF499_1.RM.like.systems | MORC+REase->TUDOR+ SF1-SF2-Helicase+iREase+CTD+REase->METHYLASE->?->CHTH->?->HEPN+DUF499*-><-?\|\|?->?-><-?<-?\|\|DOC+HTH-> | 1152 | Bacteria>Terrabacteria_group>Actinomycetota>Acidimicrobiia>Acidimicrobiales | GCA_028819405.1 | Acidimicrobiaceae_bacterium | OXM54_17470 | 118886..122344 | 2024894 | MAG:-DUF499-domain-containing-protein-[Acidimicrobiaceae-bacterium] |
| MDE0163189.1 | DUF499_1.RM.like.systems | <-HEPN+DUF499*<-?<-?<-METHYLASE<-TUDOR+ SF1-SF2-Helicase+iREase+CTD+REase<-HicB | 1126 | Bacteria>Terrabacteria_group>Actinomycetota>Acidimicrobiia>Acidimicrobiales | GCA_028819505.1 | Acidimicrobiaceae_bacterium | OXL98_15820 | 25839..29219 | 2024894 | MAG:-DUF499-domain-containing-protein-[Acidimicrobiaceae-bacterium] |
| MDE0268154.1 | DUF499_1.RM.like.systems | METHYLASE-p->HEPN+DUF499*->?-><-?\|\|?->DDE_Tnp_1->AbiEi_1-><-HTH_3 | 1111 | Bacteria>Terrabacteria_group>Actinomycetota>Acidimicrobiia>Acidimicrobiales | GCA_028820455.1 | Acidimicrobiaceae_bacterium | OXI96_03835 | 408..3743 | 2024894 | MAG:-DUF499-domain-containing-protein-[Acidimicrobiaceae-bacterium] |
| MDE0320770.1 | DUF499_1.RM.like.systems | PIN->TUDOR+ SF1-SF2-Helicase+iREase+CTD+REase->METHYLASE->?->CHTH->?->HEPN+DUF499*-><-?\|\|?->?-><-?<-?\|\|DOC+HTH-> | 1152 | Bacteria>Terrabacteria_group>Actinomycetota>Acidimicrobiia>Acidimicrobiales | GCA_028820815.1 | Acidimicrobiaceae_bacterium | OXI97_12920 | 73823..77281 | 2024894 | MAG:-DUF499-domain-containing-protein-[Acidimicrobiaceae-bacterium] |
| MDE0803902.1 | DUF499_1.RM.like.systems | <-HTH+SF1-SF2-Helicase+REase-p<-REase<-MORC<-HEPN+DUF499*<-METHYLASE | 1136 | Bacteria>Terrabacteria_group>Actinomycetota>Acidimicrobiia>Acidimicrobiales | GCA_028822535.1 | Acidimicrobiales_bacterium | OSA99_11315 | 4439..7849 | 2201156 | MAG:-Swt1-family-HEPN-domain-containing-protein-[Acidimicrobiales-bacterium] |
| MDE0497130.1 | DUF499_1.RM.like.systems | DNA_methylase->MORC+HISKIN->?->?->?->HEPN+DUF499*-> | 1089 | Bacteria>Terrabacteria_group>Actinomycetota>Acidimicrobiia>Acidimicrobiales | GCA_028823965.1 | Acidimicrobiaceae_bacterium | OXH86_07240 | 49527..52796 | 2024894 | MAG:-DUF499-domain-containing-protein-[Acidimicrobiaceae-bacterium] |
| MDE0498271.1 | DUF499_1.RM.like.systems | <-DOC+HTH\|\|?->?-><-?<-?\|\|?-><-HEPN+DUF499*<-?<-CHTH<-?<-METHYLASE<-TUDOR+ SF1-SF2-Helicase+iREase+CTD+REase<-MORC+REase | 1152 | Bacteria>Terrabacteria_group>Actinomycetota>Acidimicrobiia>Acidimicrobiales | GCA_028823965.1 | Acidimicrobiaceae_bacterium | OXH86_13045 | 54884..58342 | 2024894 | MAG:-DUF499-domain-containing-protein-[Acidimicrobiaceae-bacterium] |
| MDE0606846.1 | DUF499_1.RM.like.systems | HEPN+DUF499*-><-ParA+HTH<-RelE-ParE<-Creatinase_N+Peptidase_M24 | 1081 | Bacteria>Terrabacteria_group>Actinomycetota>Acidimicrobiia>Acidimicrobiales | GCA_028824095.1 | Acidimicrobiaceae_bacterium | OXH78_06815 | 1..3246 | 2024894 | MAG:-DUF499-domain-containing-protein,-partial-[Acidimicrobiaceae-bacterium] |
| MDE0699638.1 | DUF499_1.RM.like.systems | <-HTH_58\|\|?->DNA_methylase->MORC+HISKIN->?-><-wHTH+SAD+HNH\|\|HEPN+DUF499*-><-ParA+HTH<-RelE-ParE<-Creatinase_N+Peptidase_M24 | 1084 | Bacteria>Terrabacteria_group>Actinomycetota>Acidimicrobiia>Acidimicrobiales | GCA_028824665.1 | Acidimicrobiaceae_bacterium | OXH61_02820 | 85956..89210 | 2024894 | MAG:-DUF499-domain-containing-protein-[Acidimicrobiaceae-bacterium] |
| MDE0495499.1 | DUF499_1.RM.like.systems | METHYLASE-p->MerR-HTH->PIN->HEPN+DUF499*->ASCH->HAD-><-?<-?<-ABhydrolase | 1084 | Bacteria>Terrabacteria_group>Actinomycetota>Acidimicrobiia>Acidimicrobiales | GCA_028824895.1 | Acidimicrobiaceae_bacterium | OXH54_16270 | 41170..44424 | 2024894 | MAG:-DUF499-domain-containing-protein-[Acidimicrobiaceae-bacterium] |
| WP_219944160.1 | DUF499_1.RM.like.systems | TUDOR+ SF1-SF2-Helicase+iREase+CTD+REase->METHYLASE->METHYLASE->?->HEPN+DUF499*->?->?-><-Phage_int_SAM_1+Phage_integrase<-?\|\|MerR-HTH-> | 1096 | Bacteria>Terrabacteria_group>Actinomycetota>Acidimicrobiia>Acidimicrobiales | GCF_019443745.1 | Iamia_sp._SCSIO_61187 | HC251_RS04740 | 976382..979672 | 2722752 | Swt1-family-HEPN-domain-containing-protein-[Iamia-sp.-SCSIO-61187] |
| WP_262322266.1 | DUF499_1.RM.like.systems | MPTase->?->?-><-?\|\|?-><-?<-HEPN+DUF499*<-HEPN<-PolB<-METHYLASE<-?<-TUDOR+ SF1-SF2-Helicase+iREase+CTD+REase | 1118 | Bacteria>Terrabacteria_group>Actinomycetota>Acidimicrobiia>Acidimicrobiales | GCF_025499425.1 | Acidiferrimicrobium_sp._IK | K6U06_RS12505 | 271531..274887 | 2871700 | Swt1-family-HEPN-domain-containing-protein-[Acidiferrimicrobium-sp.-IK] |
| MXV89812.1 | DUF499_1.RM.like.systems | ABhydrolase-><-REase<-?<-?<-SF1-SF2-Helicase\|\|ABhydrolase-><-HEPN+DUF499*<-METHYLASE<-SF1-SF2-Helicase<-TUDOR+ SF1-SF2-Helicase+iREase+CTD+REase<-?\|\|?->SF1-SF2-Helicase-> | 1083 | Bacteria>Terrabacteria_group>Actinomycetota>Acidimicrobiia>unclassified_Acidimicrobiia | GCA_009835395.1 | Acidimicrobiia_bacterium | F4004_03955 | 27589..30840 | 2080302 | ATP-binding-protein-[Acidimicrobiia-bacterium] |
| MXW42977.1 | DUF499_1.RM.like.systems | <-METHYLASE<-PIN<-?<-HEPN+DUF499*<-METHYLASE<-?\|\|?-><-PIN<-DUF433 | 1079 | Bacteria>Terrabacteria_group>Actinomycetota>Acidimicrobiia>unclassified_Acidimicrobiia | GCA_009835645.1 | Acidimicrobiia_bacterium | F4Z91_07815 | 3856..7095 | 2080302 | ATP-binding-protein-[Acidimicrobiia-bacterium] |
| MXX00359.1 | DUF499_1.RM.like.systems | <-HEPN+DUF499*<-TUDOR+ SF1-SF2-Helicase+iREase+CTD+REase<-CHTH<-?<-CHTH | 1005 | Bacteria>Terrabacteria_group>Actinomycetota>Acidimicrobiia>unclassified_Acidimicrobiia | GCA_009835875.1 | Acidimicrobiia_bacterium | F4Z79_01885 | 1..3016 | 2080302 | ATP-binding-protein-[Acidimicrobiia-bacterium] |
| MXW58366.1 | DUF499_1.RM.like.systems | ABC-ATPase+SbcC->?->?->?->?-><-HEPN+DUF499*<-?<-?<-?<-Pkinase<-METHYLASE | 1114 | Bacteria>Terrabacteria_group>Actinomycetota>Acidimicrobiia>unclassified_Acidimicrobiia | GCA_009836065.1 | Acidimicrobiia_bacterium | F4Z70_07245 | 33723..37067 | 2080302 | ATP-binding-protein-[Acidimicrobiia-bacterium] |
| MXW68944.1 | DUF499_1.RM.like.systems | METHYLASE-p->HEPN+DUF499*-> | 1100 | Bacteria>Terrabacteria_group>Actinomycetota>Acidimicrobiia>unclassified_Acidimicrobiia | GCA_009836245.1 | Acidimicrobiia_bacterium | F4Z61_03125 | 1254..4555 | 2080302 | ATP-binding-protein-[Acidimicrobiia-bacterium] |
| MXX46239.1 | DUF499_1.RM.like.systems | <-GNTR-HTH<-?\|\|ORC-CDC6+REase-><-?<-?<-RNase<-HEPN+DUF499*<-METHYLASE-p<-METHYLASE-p<-METHYLASE-p<-TUDOR+ SF1-SF2-Helicase+iREase+CTD+REase<-CHTH | 1090 | Bacteria>Terrabacteria_group>Actinomycetota>Acidimicrobiia>unclassified_Acidimicrobiia | GCA_009836655.1 | Acidimicrobiia_bacterium | F4Z41_08550 | 57400..60672 | 2080302 | ATP-binding-protein-[Acidimicrobiia-bacterium] |
| MXZ30232.1 | DUF499_1.RM.like.systems | <-ParD_antitoxin<-ParD_antitoxin\|\|HEPN+DUF499*-><-ABhydrolase | 1083 | Bacteria>Terrabacteria_group>Actinomycetota>Acidimicrobiia>unclassified_Acidimicrobiia | GCA_009837005.1 | Acidimicrobiia_bacterium | F4Z22_05205 | 787..4038 | 2080302 | ATP-binding-protein-[Acidimicrobiia-bacterium] |
| MXZ78951.1 | DUF499_1.RM.like.systems | HTH_31->RNase->HTH_58-><-?<-HEPN+DUF499*\|\|?->PhdYeFM_antitox->PIN-><-METHYLASE<-?\|\|CHTH-> | 1079 | Bacteria>Terrabacteria_group>Actinomycetota>Acidimicrobiia>unclassified_Acidimicrobiia | GCA_009837345.1 | Acidimicrobiia_bacterium | F4Z06_11870 | 8705..11944 | 2080302 | ATP-binding-protein-[Acidimicrobiia-bacterium] |
| MXZ86859.1 | DUF499_1.RM.like.systems | ABC-ATPase+SbcC->?->?->?->?-><-HEPN+DUF499*<-?\|\|?-><-METHYLASE\|\|ParB+HNH->HEPN-><-TUDOR+ SF1-SF2-Helicase+iREase+CTD+REase | 1114 | Bacteria>Terrabacteria_group>Actinomycetota>Acidimicrobiia>unclassified_Acidimicrobiia | GCA_009837425.1 | Acidimicrobiia_bacterium | F4Z02_14550 | 33484..36828 | 2080302 | ATP-binding-protein-[Acidimicrobiia-bacterium] |
| MXY73348.1 | DUF499_1.RM.like.systems | METHYLASE-p->HEPN+DUF499*->HTH_3->NTP_transf-><-?<-PIN | 1107 | Bacteria>Terrabacteria_group>Actinomycetota>Acidimicrobiia>unclassified_Acidimicrobiia | GCA_009837815.1 | Acidimicrobiia_bacterium | F4Y83_00240 | 420..3743 | 2080302 | ATP-binding-protein-[Acidimicrobiia-bacterium] |
| MXY74717.1 | DUF499_1.RM.like.systems | <-HEPN+DUF499*<-METHYLASE-p | 1044 | Bacteria>Terrabacteria_group>Actinomycetota>Acidimicrobiia>unclassified_Acidimicrobiia | GCA_009837815.1 | Acidimicrobiia_bacterium | F4Y83_07345 | 1..3131 | 2080302 | ATP-binding-protein-[Acidimicrobiia-bacterium] |
| MYA38651.1 | DUF499_1.RM.like.systems | METHYLASE-p->HEPN+DUF499*->HTH_3->NTP_transf-><-?<-PIN | 1107 | Bacteria>Terrabacteria_group>Actinomycetota>Acidimicrobiia>unclassified_Acidimicrobiia | GCA_009838695.1 | Acidimicrobiia_bacterium | F4Y36_03440 | 1539..4862 | 2080302 | ATP-binding-protein-[Acidimicrobiia-bacterium] |
| MYB44039.1 | DUF499_1.RM.like.systems | METHYLASE-><-PIN<-?\|\|SIR2+STALD->?->PRTase+PRTase->HEPN+DUF499*-><-?\|\|PIN-><-ABC-ATPase+SbcC<-ABC-ATPase+SbcC<-Peptidase_M20 | 1097 | Bacteria>Terrabacteria_group>Actinomycetota>Acidimicrobiia>unclassified_Acidimicrobiia | GCA_009839965.1 | Acidimicrobiia_bacterium | F4X74_03760 | 19073..22366 | 2080302 | ATP-binding-protein-[Acidimicrobiia-bacterium] |
| MYB25632.1 | DUF499_1.RM.like.systems | <-Classical-AAA\|\|ORC-CDC6+REase-><-PIN<-HTH+MPTase<-PIN\|\|METHYLASE->HEPN+DUF499*-> | 1108 | Bacteria>Terrabacteria_group>Actinomycetota>Acidimicrobiia>unclassified_Acidimicrobiia | GCA_009840735.1 | Acidimicrobiia_bacterium | F4X37_11130 | 50648..53974 | 2080302 | ATP-binding-protein-[Acidimicrobiia-bacterium] |
| MYD40244.1 | DUF499_1.RM.like.systems | <-RNase<-HEPN+DUF499*<-METHYLASE | 1090 | Bacteria>Terrabacteria_group>Actinomycetota>Acidimicrobiia>unclassified_Acidimicrobiia | GCA_009841565.1 | Acidimicrobiia_bacterium | F4W94_00700 | 1349..4621 | 2080302 | ATP-binding-protein-[Acidimicrobiia-bacterium] |
| MYG93975.1 | DUF499_1.RM.like.systems | METHYLASE->?->?-><-?\|\|?->HEPN+DUF499*->?->PIN->METHYLASE-> | 1079 | Bacteria>Terrabacteria_group>Actinomycetota>Acidimicrobiia>unclassified_Acidimicrobiia | GCA_009843865.1 | Acidimicrobiia_bacterium | F4138_03135 | 9114..12353 | 2080302 | ATP-binding-protein-[Acidimicrobiia-bacterium] |
| MYJ15813.1 | DUF499_1.RM.like.systems | <-HTH_3<-HEPN+DUF499*<-METHYLASE-p | 1107 | Bacteria>Terrabacteria_group>Actinomycetota>Acidimicrobiia>unclassified_Acidimicrobiia | GCA_009844925.1 | Acidimicrobiia_bacterium | F4085_04615 | 343..3666 | 2080302 | ATP-binding-protein-[Acidimicrobiia-bacterium] |
| MBA2283088.1 | DUF499_1.RM.like.systems | Phage_int_SAM_1+Phage_integrase-><-NTD+SF1-SF2-Helicase\|\|TUDOR+ SF1-SF2-Helicase+iREase+CTD+REase->?->METHYLASE->?->HEPN+DUF499*->REase->?-><-?<-XIS-HTH\|\|XIS-HTH-> | 1110 | Bacteria>Terrabacteria_group>Actinomycetota>Acidimicrobiia>unclassified_Acidimicrobiia | GCA_013694495.1 | Acidimicrobiia_bacterium | HOW25_17905 | 18303..21635 | 2080302 | ATP-binding-protein-[Acidimicrobiia-bacterium] |
| MBP8181356.1 | DUF499_1.RM.like.systems | <-HEPN+DUF499*<-METHYLASE | 1096 | Bacteria>Terrabacteria_group>Actinomycetota>Acidimicrobiia>unclassified_Acidimicrobiia | GCA_018006325.1 | Acidimicrobiia_bacterium | KAZ88_10225 | 2758..6048 | 2080302 | DUF499-domain-containing-protein-[Acidimicrobiia-bacterium] |
| MCB9390232.1 | DUF499_1.RM.like.systems | <-HEPN<-XIS-HTH+RVT<-?<-DUF6361<-?<-REase<-HEPN+DUF499*<-METHYLASE\|\|?-><-?<-?<-DUF772+DDE_Tnp_1+DDE_Tnp_1_6 | 1107 | Bacteria>Terrabacteria_group>Actinomycetota>Acidimicrobiia>unclassified_Acidimicrobiia | GCA_020637365.1 | Acidimicrobiia_bacterium | H6512_11920 | 836965..840288 | 2080302 | ATP-binding-protein-[Acidimicrobiia-bacterium] |

| Protein-ID | System | Gene-Neighbourhoods | Length | Taxonomy | Assembly-ID | Strain | Locus-Tag | Genomic-coordinates | TaxID | Fasta-defline |
|---|---|---|---|---|---|---|---|---|---|---|
| MCC7078886.1 | DUF499_1.RM.like.systems | <-Schlafen+HATPase+HTH<-?<-HEPN+DUF499*<-METHYLASE<-Schlafen+HATPase+HTH<-TUDOR+SF1-SF2-Helicase+iREase+CTD+REase | 1112 | Bacteria>Terrabacteria_group>Actinomycetota>Acidimicrobiia>unclassified_Acidimicrobiia | GCA_020850945.1 | Acidimicrobiia_bacterium | IT198_17355 | 29484..32822 | 2080302 | DUF499-domain-containing-protein-[Acidimicrobiia-bacterium] |
| MCQ3809088.1 | DUF499_1.RM.like.systems | Radical-SAM->?<-?<-?<-HEPN+DUF499*\|\|?-><-?\|\|?-><-METHYLASE | 1114 | Bacteria>Terrabacteria_group>Actinomycetota>Acidimicrobiia>unclassified_Acidimicrobiia | GCA_024421205.1 | Acidimicrobiia_bacterium | KTV68_00880 | 53316..56660 | 2080302 | DUF499-domain-containing-protein-[Acidimicrobiia-bacterium] |
| ERH33797.1 | DUF499_1.RM.like.systems | <-HEPN+DUF499*<-METHYLASE<-TUDOR+SF1-SF2-Helicase+iREase+CTD+REase<-TraJ-RHH<-?<-?\|\|TetR-HTH-> | 1139 | Bacteria>Terrabacteria_group>Actinomycetota>Actinomycetes>Actinomycetales | GCA_000466305.1 | Actinomyces_sp._oral_taxon_877_str._F0543 | HMPREF1550_00139 | 24107..27526 | 1227264 | hypothetical-protein-HMPREF1550_00139-[Actinomyces-sp.-oral-taxon-877-str.-F0543] |
| NLI19748.1 | DUF499_1.RM.like.systems | <-HTH_31<-?\|\|UDG-conflict-><-HEPN+DUF499*<-METHYLASE<-TUDOR+SF1-SF2-Helicase+iREase+CTD+REase<-?<-TrwC+SF1-SF2-Helicase<-TrwC+SF1-SF2-Helicase | 1114 | Bacteria>Terrabacteria_group>Actinomycetota>Actinomycetes>Actinomycetales | GCA_012517595.1 | Actinomycetales_bacterium | GX427_13235 | 2536..5880 | 1911520 | ATP-binding-protein-[Actinomycetales-bacterium] |
| MBE6482584.1 | DUF499_1.RM.like.systems | <-Schlafen+HATPase+HTH<-REase<-HTH+SF1-SF2-Helicase+REase<-REase<-MORC<-HEPN+DUF499*<-METHYLASE\|\|DNA_methylase->DNA_methylase->MORC+HISKIN-> | 1137 | Bacteria>Terrabacteria_group>Actinomycetota>Actinomycetes>Actinomycetales | GCA_015062785.1 | Actinomyces_ruminicola | E7Z94_09510 | 10121..13534 | 332524 | ATP-binding-protein-[Actinomyces-ruminicola] |
| MBF0963210.1 | DUF499_1.RM.like.systems | <-HEPN+DUF499*<-TUDOR+SF1-SF2-Helicase+iREase+CTD+REase<-TraJ-RHH | 1136 | Bacteria>Terrabacteria_group>Actinomycetota>Actinomycetes>Actinomycetales | GCA_015259385.1 | Actinomyces_sp. | HXK13_05250 | 38..3448 | 29317 | ATP-binding-protein-[Actinomyces-sp.] |
| QQO76952.1 | DUF499_1.RM.like.systems | GATase-><-?<-Schlafen+HATPase+HTH<-?<-DNA_methylase\|\|METHYLASE->HEPN+DUF499*->?->?->Peptidase_M50-> | 1129 | Bacteria>Terrabacteria_group>Actinomycetota>Actinomycetes>Actinomycetales | GCA_016626125.1 | Actinomyces_sp._HMT897 | JJJ15_07540 | 1917676..1921065 | 2789424 | DUF499-domain-containing-protein-[Actinomyces-sp.-HMT897] |
| MBK8733830.1 | DUF499_1.RM.like.systems | <-DUF499-p<-?<-DUF499-p<-?<-HEPN+DUF499*<-METHYLASE<-TUDOR+SF1-SF2-Helicase+iREase+CTD+REase\|\|?->Phage_integrase+Phage_integrase->PIN-> | 347 | Bacteria>Terrabacteria_group>Actinomycetota>Actinomycetes>Actinomycetales | GCA_016710645.1 | Actinomycetales_bacterium | IPL93_13505 | 220122..221165 | 1911520 | hypothetical-protein-[Actinomycetales-bacterium] |
| MCI6556423.1 | DUF499_1.RM.like.systems | TUDOR+SF1-SF2-Helicase+iREase+CTD+REase->METHYLASE->HEPN+DUF499*->HEPN->RelB->?->tRNA-><-?<-?<-DNA_methylase | 1149 | Bacteria>Terrabacteria_group>Actinomycetota>Actinomycetes>Actinomycetales | GCA_022780845.1 | Schaalia_hyovaginalis | MR422_02105 | 263803..267252 | 29316 | DUF499-domain-containing-protein-[Schaalia-hyovaginalis] |
| BDF97829.1 | DUF499_1.RM.like.systems | ACET+ACET-><-?<-?<-DNA_methylase\|\|METHYLASE->HEPN+DUF499*->MORC->REase->HTH+SF1-SF2-Helicase+REase->REase-><-?\|\|METHYLASE-p-> | 1139 | Bacteria>Terrabacteria_group>Actinomycetota>Actinomycetes>Actinomycetales | GCA_023169925.1 | Actinomyces_oris_K20 | AoKodu_01300 | 147963..151382 | 871541 | hypothetical-protein-AoKodu_01300-[Actinomyces-oris-K20] |
| WP_003788306.1 | DUF499_1.RM.like.systems | <-METHYLASE-p\|\|?-><-REase<-HTH+SF1-SF2-Helicase+REase<-REase<-MORC<-HEPN+DUF499*<-METHYLASE\|\|DNA_methylase->?->?-><-?<-HTH_3+PolB | 1139 | Bacteria>Terrabacteria_group>Actinomycetota>Actinomycetes>Actinomycetales | GCF_000175315.3 | Actinomyces_viscosus | HMPREF0059_RS11125 | 118878..122297 | 1656 | DUF499-domain-containing-protein-[Actinomyces-viscosus] |
| WP_005985559.1 | DUF499_1.RM.like.systems | <-ABC-ATPase+SbcC<-?\|\|TUDOR+SF1-SF2-Helicase+iREase+CTD+REase->METHYLASE->HEPN+DUF499*-><-?<-?<-?<-?<-ABC-ATPase+SbcC | 1128 | Bacteria>Terrabacteria_group>Actinomycetota>Actinomycetes>Actinomycetales | GCF_000239695.1 | Actinomyces_graevenitzii | HMPREF0045_RS03480 | 792644..796030 | 55565 | DUF499-domain-containing-protein-[Actinomyces-graevenitzii] |
| WP_003781297.1 | DUF499_1.RM.like.systems | <-HEPN+DUF499*<-METHYLASE\|\|DNA_methylase-> | 1040 | Bacteria>Terrabacteria_group>Actinomycetota>Actinomycetes>Actinomycetales | GCF_000285995.1 | Actinomyces_naeslundii | HMPREF1129_RS04410 | 1..3119 | 1655 | DUF499-domain-containing-protein,-partial-[Actinomyces-naeslundii] |
| WP_017177796.1 | DUF499_1.RM.like.systems | Phage_holin_4_2->?-><-REase<-HTH+SF1-SF2-Helicase+REase<-REase<-MORC<-HEPN+DUF499*<-METHYLASE\|\|DNA_methylase->?->?-><-?<-HTH_3+PolB | 1137 | Bacteria>Terrabacteria_group>Actinomycetota>Actinomycetes>Actinomycetales | GCF_000295095.1 | Actinomyces_timonensis | A1QA_RS0104850 | 1087937..1091350 | 1288391 | DUF499-domain-containing-protein-[Actinomyces-timonensis] |
| WP_034502149.1 | DUF499_1.RM.like.systems | <-HEPN+DUF499*<-METHYLASE<-TUDOR+SF1-SF2-Helicase+iREase+CTD+REase | 1128 | Bacteria>Terrabacteria_group>Actinomycetota>Actinomycetes>Actinomycetales | GCF_000758755.1 | Actinomyces_sp._S6-Spd3 | HMPREF1627_RS06325 | 1474..4860 | 1284680 | DUF499-domain-containing-protein-[Actinomyces-sp.-S6-Spd3] |
| WP_053587850.1 | DUF499_1.RM.like.systems | <-REase<-HTH+SF1-SF2-Helicase+REase<-REase<-MORC<-HEPN+DUF499*<-METHYLASE\|\|METHYLASE->BirA-HTH->?-><-TPR<-MULE-TRANSPOSASE | 1165 | Bacteria>Terrabacteria_group>Actinomycetota>Actinomycetes>Actinomycetales | GCF_001278845.1 | Actinomyces_sp._oral_taxon_414 | AM609_RS14685 | 3673757..3677254 | 712122 | DUF499-domain-containing-protein-[Actinomyces-sp.-oral-taxon-414] |
| WP_070510861.1 | DUF499_1.RM.like.systems | <-REase<-HTH+SF1-SF2-Helicase+REase<-REase<-MORC\|\|?-><-HEPN+DUF499*<-METHYLASE\|\|DNA_methylase->?->?-><-?<-HTH_3+PolB | 1137 | Bacteria>Terrabacteria_group>Actinomycetota>Actinomycetes>Actinomycetales | GCF_001807465.1 | Actinomyces_sp._HMSC08A09 | HMPREF3165_RS03500 | 45763..49176 | 1581133 | DUF499-domain-containing-protein-[Actinomyces-sp.-HMSC08A09] |
| WP_073824655.1 | DUF499_1.RM.like.systems | TUDOR+SF1-SF2-Helicase+iREase+CTD+REase-p->TUDOR+SF1-SF2-Helicase+iREase+CTD+REase->METHYLASE->HEPN+DUF499*-> | 1140 | Bacteria>Terrabacteria_group>Actinomycetota>Actinomycetes>Actinomycetales | GCF_001907235.1 | Buchananella_hordeovulneris | BSZ40_RS07210 | 166478..169900 | 52770 | DUF499-domain-containing-protein-[Buchananella-hordeovulneris] |
| WP_075412471.1 | DUF499_1.RM.like.systems | <-METHYLASE-p<-REase<-HTH+SF1-SF2-Helicase+REase<-REase<-MORC<-HEPN+DUF499*<-METHYLASE\|\|DNA_methylase->?-><-HTH_3+PolB<-ACET+ACET | 1137 | Bacteria>Terrabacteria_group>Actinomycetota>Actinomycetes>Actinomycetales | GCF_001937365.1 | Actinomyces_oris | BKH31_RS11955 | 29944..33357 | 544580 | DUF499-domain-containing-protein-[Actinomyces-oris] |
| WP_075413329.1 | DUF499_1.RM.like.systems | <-METHYLASE-p<-REase<-HTH+SF1-SF2-Helicase+REase<-REase<-MORC<-HEPN+DUF499*<-METHYLASE\|\|DNA_methylase->?->?-><-ACET+ACET | 1137 | Bacteria>Terrabacteria_group>Actinomycetota>Actinomycetes>Actinomycetales | GCF_001937475.1 | Actinomyces_naeslundii | BKH08_RS04545 | 28951..32364 | 1655 | DUF499-domain-containing-protein-[Actinomyces-naeslundii] |
| WP_075403442.1 | DUF499_1.RM.like.systems | ACET+ACET->HTH_3+PolB-><-?<-?<-DNA_methylase\|\|METHYLASE->HEPN+DUF499*->MORC->REase->HTH+SF1-SF2-Helicase+REase->REase-><-?\|\|METHYLASE-p-> | 1139 | Bacteria>Terrabacteria_group>Actinomycetota>Actinomycetes>Actinomycetales | GCF_001937505.1 | Actinomyces_oris | BKH21_RS10525 | 35903..39322 | 544580 | DUF499-domain-containing-protein-[Actinomyces-oris] |
| WP_075409839.1 | DUF499_1.RM.like.systems | ACET+ACET-><-?<-?<-DNA_methylase\|\|METHYLASE->HEPN+DUF499*-> | 826 | Bacteria>Terrabacteria_group>Actinomycetota>Actinomycetes>Actinomycetales | GCF_001937735.1 | Actinomyces_naeslundii | BKH13_RS04850 | 82823..85302 | 1655 | DUF499-domain-containing-protein,-partial-[Actinomyces-naeslundii] |
| WP_076255389.1 | DUF499_1.RM.like.systems | ACET+ACET-><-?<-?<-DNA_methylase\|\|METHYLASE->HEPN+DUF499*->MORC->REase->HTH+SF1-SF2-Helicase+REase->REase->METHYLASE-p-> | 1137 | Bacteria>Terrabacteria_group>Actinomycetota>Actinomycetes>Actinomycetales | GCF_001956555.1 | Actinomyces_naeslundii | BKH25_RS11260 | 118842..122255 | 1655 | DUF499-domain-containing-protein-[Actinomyces-naeslundii] |
| WP_077240589.1 | DUF499_1.RM.like.systems | <-REase<-HTH+SF1-SF2-Helicase+REase<-REase<-MORC<-HEPN+DUF499*<-METHYLASE\|\|DNA_methylase->MORC+HEPN->\|\|?-><-?<-Peptidase_M50 | 1137 | Bacteria>Terrabacteria_group>Actinomycetota>Actinomycetes>Actinomycetales | GCF_002356195.1 | Actinomyces | DY404_RS06925 | 1656821..1660234 | 1654 | MULTISPECIES:-DUF499-domain-containing-protein-[Actinomyces] |
| WP_106110118.1 | DUF499_1.RM.like.systems | <-REase\|\|GATase-><-?<-HTH_28+HTH_33+DDE_3\|\|METHYLASE->HEPN+DUF499*->?->?->Peptidase_M50->?->?->VirB11+SF1-SF2-Helicase-> | 1130 | Bacteria>Terrabacteria_group>Actinomycetota>Actinomycetes>Actinomycetales | GCF_002999235.1 | Actinomyces_sp._oral_taxon_897 | C3V41_RS09940 | 2509136..2512528 | 2081702 | DUF499-domain-containing-protein-[Actinomyces-sp.-oral-taxon-897] |
| WP_124852157.1 | DUF499_1.RM.like.systems | <-HEPN+DUF499*<-METHYLASE<-TUDOR+SF1-SF2-Helicase+iREase+CTD+REase<-TUDOR+SF1-SF2-Helicase+iREase+CTD+REase-p | 1140 | Bacteria>Terrabacteria_group>Actinomycetota>Actinomycetes>Actinomycetales | GCF_003858495.1 | Buchananella_hordeovulneris | EII12_RS05710 | 157..3579 | 52770 | DUF499-domain-containing-protein-[Buchananella-hordeovulneris] |
| WP_124913173.1 | DUF499_1.RM.like.systems | TUDOR+SF1-SF2-Helicase+iREase+CTD+REase->METHYLASE->HEPN+DUF499*-> | 1140 | Bacteria>Terrabacteria_group>Actinomycetota>Actinomycetes>Actinomycetales | GCF_003858505.1 | Buchananella_hordeovulneris | EII13_RS09935 | 35169..38591 | 52770 | DUF499-domain-containing-protein-[Buchananella-hordeovulneris] |
| WP_103061758.1 | DUF499_1.RM.like.systems | <-REase<-HTH+SF1-SF2-Helicase+REase<-REase<-MORC<-HEPN+DUF499*<-METHYLASE\|\|DNA_methylase->MORC+HisKin->HTH_Tnp_4+DDE_Tnp_4->MORC+HEPN-> | 1138 | Bacteria>Terrabacteria_group>Actinomycetota>Actinomycetes>Actinomycetales | GCF_004102025.1 | Actinomyces_qiguomingii | CWT10_RS16110 | 3847675..3851091 | 2057800 | DUF499-domain-containing-protein-[Actinomyces-qiguomingii] |
| WP_128773202.1 | DUF499_1.RM.like.systems | SF1-SF2-Helicase+REase-><-?<-TrwC+SF1-SF2-Helicase\|\|DNA_methylase-><-HEPN+REase<-REase<-HEPN+DUF499*<-wHTH+REase<-METHYLASE<-TUDOR+SF1-SF2-Helicase+iREase+CTD+REase<-XIS-HTH<-ABhydrolase | 1132 | Bacteria>Terrabacteria_group>Actinomycetota>Actinomycetes>Actinomycetales | GCF_004104015.1 | Actinomyces_oricola | EUC46_RS05555 | 19810..23208 | 206043 | DUF499-domain-containing-protein-[Actinomyces-oricola] |
| WP_136193072.1 | DUF499_1.RM.like.systems | HNH->?->ABhydrolase-><-HAD<-?<-HEPN+DUF499*<-METHYLASE<-METHYLASE-p<-TUDOR+SF1-SF2-Helicase+iREase+CTD+REase<-TetR-HTH | 1138 | Bacteria>Terrabacteria_group>Actinomycetota>Actinomycetes>Actinomycetales | GCF_004796555.1 | Actinomyces_procaprae | FAQ94_RS06890 | 215265..218681 | 2560010 | DUF499-domain-containing-protein-[Actinomyces-procaprae] |
| WP_161924011.1 | DUF499_1.RM.like.systems | TetR-HTH->TUDOR+SF1-SF2-Helicase+iREase+CTD+REase->METHYLASE-p->METHYLASE->HEPN+DUF499*-><-HTH_Tnp_4+DDE_Tnp_4\|\|?->HAD-><-?<-HNH | 1139 | Bacteria>Terrabacteria_group>Actinomycetota>Actinomycetes>Actinomycetales | GCF_009930875.1 | Actinomyces_sp._432 | CWT12_RS05510 | 1339354..1342773 | 2057798 | DUF499-domain-containing-protein-[Actinomyces-sp.-432] |
| WP_162678504.1 | DUF499_1.RM.like.systems | TetR-HTH->TUDOR+SF1-SF2-Helicase+iREase+CTD+REase->METHYLASE-p->METHYLASE->HEPN+DUF499*->?->HAD-><-HNH | 1139 | Bacteria>Terrabacteria_group>Actinomycetota>Actinomycetes>Actinomycetales | GCF_010365615.1 | Actinomyces_sp._565 | GZ997_RS01005 | 32665..36084 | 2057794 | DUF499-domain-containing-protein-[Actinomyces-sp.-565] |
| WP_165316371.1 | DUF499_1.RM.like.systems | ABC-ATPase+SbcC-><-RecR+TOPRIM<-?\|\|TUDOR+SF1-SF2-Helicase+iREase+CTD+REase->?->METHYLASE->HEPN+DUF499*-><-PIN\|\|?->?->DDE_Tnp_1->DDE_Tnp_1-><-RNase | 1143 | Bacteria>Terrabacteria_group>Actinomycetota>Actinomycetes>Actinomycetales | GCF_011038885.2 | Schaalia_sp._ZJ405 | G7Y41_RS09385 | 2197397..2200828 | 2709403 | DUF499-domain-containing-protein-[Schaalia-sp.-ZJ405] |
| WP_169762497.1 | DUF499_1.RM.like.systems | <-HTH_3<-RelE-ParE<-TraJ-RHH<-RNase\|\|DUF433->PIN-><-HEPN+DUF499*<-METHYLASE<-TUDOR+SF1-SF2-Helicase+iREase+CTD+REase<-ParA+HTH<-TOPRIM<-PIN | 1131 | Bacteria>Terrabacteria_group>Actinomycetota>Actinomycetes>Actinomycetales | GCF_012976645.1 | Mobiluncus_mulieris | HHJ77_RS04685 | 61231..64626 | 2052 | DUF499-domain-containing-protein-[Mobiluncus-mulieris] |
| WP_169773626.1 | DUF499_1.RM.like.systems | PIN->TOPRIM->ParA+HTH->TUDOR+SF1-SF2-Helicase+iREase+CTD+REase->METHYLASE-p->HEPN+DUF499*-><-PIN<-DUF433<-?<-HEPN\|\|RNase->TraJ-RHH-> | 1131 | Bacteria>Terrabacteria_group>Actinomycetota>Actinomycetes>Actinomycetales | GCF_012976895.1 | Mobiluncus_mulieris | HHJ79_RS06650 | 39416..42811 | 2052 | DUF499-domain-containing-protein-[Mobiluncus-mulieris] |
| WP_182050741.1 | DUF499_1.RM.like.systems | SF1-SF2-Helicase->TUDOR+SF1-SF2-Helicase+iREase+CTD+REase->METHYLASE->REase->HEPN->HEPN+DUF499*->SbcC-> | 1134 | Bacteria>Terrabacteria_group>Actinomycetota>Actinomycetes>Actinomycetales | GCF_014041995.1 | Changpingibacter_yushuensis | H2017_RS05160 | 1185005..1188409 | 2758440 | Swt1-family-HEPN-domain-containing-protein-[Changpingibacter-yushuensis] |
| WP_184451288.1 | DUF499_1.RM.like.systems | TUDOR+SF1-SF2-Helicase+iREase+CTD+REase-p-><-?\|\|TUDOR+SF1-SF2-Helicase+iREase+CTD+REase->METHYLASE->HEPN+DUF499*->HEPN->RelB->tRNA-><-Glyoxalase | 1150 | Bacteria>Terrabacteria_group>Actinomycetota>Actinomycetes>Actinomycetales | GCF_014208035.1 | Schaalia_hyovaginalis | HD592_RS00720 | 169664..173116 | 29316 | DUF499-domain-containing-protein-[Schaalia-hyovaginalis] |
| WP_076142650.1 | DUF499_1.RM.like.systems | <-METHYLASE-p<-REase<-HTH+SF1-SF2-Helicase+REase<-REase<-MORC<-HEPN+DUF499*<-METHYLASE\|\|DNA_methylase->?->?-><-ACET+ACET | 1139 | Bacteria>Terrabacteria_group>Actinomycetota>Actinomycetes>Actinomycetales | GCF_016127855.1 | Actinomyces_naeslundii | BKH33_RS06345 | 173705..177124 | 1655 | DUF499-domain-containing-protein-[Actinomyces-naeslundii] |
| WP_198776534.1 | DUF499_1.RM.like.systems | sGTP-><-?<-DNA_methylase\|\|METHYLASE->HEPN+DUF499*->MORC->REase->HTH+SF1-SF2-Helicase+REase->REase-> | 1127 | Bacteria>Terrabacteria_group>Actinomycetota>Actinomycetes>Actinomycetales | GCF_016405145.1 | Schaalia_cardiffensis | JFX23_RS08485 | 21486..24869 | 181487 | DUF499-domain-containing-protein-[Schaalia-cardiffensis] |
| WP_200510328.1 | DUF499_1.RM.like.systems | GATase-><-?<-Schlafen+HATPase+HTH<-?<-DNA_methylase\|\|METHYLASE->HEPN+DUF499*->?->?->Peptidase_M50->?->?->VirB11+SF1-SF2-Helicase-> | 1129 | Bacteria>Terrabacteria_group>Actinomycetota>Actinomycetes>Actinomycetales | GCF_016626125.1 | Actinomyces_sp._HMT897 | JJJ15_RS13470 | 1917676..1921065 | 2789424 | DUF499-domain-containing-protein-[Actinomyces-sp.-HMT897] |
| WP_204423781.1 | DUF499_1.RM.like.systems | <-HEPN+DUF499*<-METHYLASE<-TUDOR+SF1-SF2-Helicase+iREase+CTD+REase | 1130 | Bacteria>Terrabacteria_group>Actinomycetota>Actinomycetes>Actinomycetales | GCF_016904675.1 | Arcanobacterium_phocisimile | JTE88_RS06650 | 1475854..1479246 | 1302235 | DUF499-domain-containing-protein-[Arcanobacterium-phocisimile] |
| WP_219110028.1 | DUF499_1.RM.like.systems | HNH-><-HAD<-?<-HEPN+DUF499*<-METHYLASE<-METHYLASE-p<-TUDOR+SF1-SF2-Helicase+iREase+CTD+REase<-TetR-HTH | 1139 | Bacteria>Terrabacteria_group>Actinomycetota>Actinomycetes>Actinomycetales | GCF_019325145.1 | Actinomyces_sp._594 | GZ998_RS12335 | 131266..134685 | 2057793 | DUF499-domain-containing-protein-[Actinomyces-sp.-594] |
| WP_282568806.1 | DUF499_1.RM.like.systems | ACET+ACET-><-?<-?<-DNA_methylase\|\|METHYLASE->HEPN+DUF499*->MORC->REase->HTH+SF1-SF2-Helicase+REase->REase-><-?\|\|METHYLASE-p-> | 1139 | Bacteria>Terrabacteria_group>Actinomycetota>Actinomycetes>Actinomycetales | GCF_023169925.1 | Actinomyces_oris | AoKodu_RS00635 | 147963..151382 | 544580 | DUF499-domain-containing-protein-[Actinomyces-oris] |
| WP_251775558.1 | DUF499_1.RM.like.systems | HKD-DNase+HKD-DNase-><-?\|\|?->?-><-?\|\|?-><-HEPN+DUF499*<-METHYLASE<-TUDOR+SF1-SF2-Helicase+iREase+CTD+REase | 1129 | Bacteria>Terrabacteria_group>Actinomycetota>Actinomycetes>Actinomycetales | GCF_023716965.1 | Schaalia_meyeri | ND910_RS08155 | 14607..17996 | 52773 | DUF499-domain-containing-protein-[Schaalia-meyeri] |
| WP_268916840.1 | DUF499_1.RM.like.systems | <-REase<-HTH+SF1-SF2-Helicase+REase<-REase<-MORC<-HEPN+DUF499*<-METHYLASE\|\|DNA_methylase->MORC+HEPN->SLATT->SMODS-> | 1131 | Bacteria>Terrabacteria_group>Actinomycetota>Actinomycetes>Actinomycetales | GCF_026891875.1 | Actinomyces_israelii | OHJ16_RS04000 | 49558..52953 | 1659 | DUF499-domain-containing-protein-[Actinomyces-israelii] |
| WP_271288667.1 | DUF499_1.RM.like.systems | ACET+ACET->HTH_3+PolB-><-?<-?<-DNA_methylase\|\|METHYLASE->HEPN+DUF499*->MORC->REase->HTH+SF1-SF2-Helicase+REase->REase-><-?\|\|METHYLASE-p-> | 1132 | Bacteria>Terrabacteria_group>Actinomycetota>Actinomycetes>Actinomycetales | GCF_027945475.1 | Actinomyces_oris | PGE45_RS00580 | 136591..139989 | 544580 | DUF499-domain-containing-protein-[Actinomyces-oris] |
| WP_073332337.1 | DUF499_1.RM.like.systems | <-ABC-ATPase+SbcC<-?<-?<-?<-?<-?<-HEPN+DUF499*<-METHYLASE<-TUDOR+SF1-SF2-Helicase+iREase+CTD+REase<-TetR-HTH | 1138 | Bacteria>Terrabacteria_group>Actinomycetota>Actinomycetes>Actinomycetales | GCF_900098745.1 | Actinomyces_glycerinitolerans | ACGLYG10_RS12005 | 81593..85009 | 1892869 | DUF499-domain-containing-protein-[Actinomyces-glycerinitolerans] |
| WP_108833962.1 | DUF499_1.RM.like.systems | <-METHYLASE-p<-REase<-HTH+SF1-SF2-Helicase+REase<-REase<-MORC<-HEPN+DUF499*<-METHYLASE\|\|DNA_methylase->?-><-ACET+ACET | 1138 | Bacteria>Terrabacteria_group>Actinomycetota>Actinomycetes>Actinomycetales | GCF_900323545.1 | Actinomyces_sp._Marseille-P3109 | BQ8008_RS10580 | 2440018..2443434 | 2083009 | DUF499-domain-containing-protein-[Actinomyces-sp.-Marseille-P3109] |
| WP_114990037.1 | DUF499_1.RM.like.systems | TOPRIM->RelE-ParE->ParA+HTH->TUDOR+SF1-SF2-Helicase+iREase+CTD+REase->METHYLASE->HEPN+DUF499*-><-PIN<-DUF433\|\|RNase->TraJ-RHH->RelE-ParE->HTH_3-> | 1131 | Bacteria>Terrabacteria_group>Actinomycetota>Actinomycetes>Actinomycetales | GCF_900453275.1 | Mobiluncus_mulieris | DX043_RS05220 | 1134969..1138364 | 2052 | DUF499-domain-containing-protein-[Mobiluncus-mulieris] |
| WP_278601539.1 | DUF499_1.RM.like.systems | ACET+ACET-><-?<-?<-DNA_methylase\|\|METHYLASE->HEPN+DUF499*->MORC->REase->HTH+SF1-SF2-Helicase+REase->REase->METHYLASE-p-> | 1137 | Bacteria>Terrabacteria_group>Actinomycetota>Actinomycetes>Actinomycetales | GCF_905372775.1 | Actinomyces_naeslundii | QDH38_RS03495 | 39335..42748 | 1655 | DUF499-domain-containing-protein-[Actinomyces-naeslundii] |
| WP_278972061.1 | DUF499_1.RM.like.systems | <-HTH_3<-RelE-ParE\|\|DUF433->PIN-><-HEPN+DUF499*<-METHYLASE-p<-TUDOR+SF1-SF2-Helicase+iREase+CTD+REase<-ParA+HTH<-RelE-ParE<-TOPRIM | 1131 | Bacteria>Terrabacteria_group>Actinomycetota>Actinomycetes>Actinomycetales | GCF_946891785.1 | Mobiluncus_mulieris | QDN45_RS08450 | 23006..26401 | 2052 | DUF499-domain-containing-protein-[Mobiluncus-mulieris] |
| WP_070662487.1 | DUF499_1.RM.like.systems | HTH_3+PolB->?-><-?<-?<-DNA_methylase\|\|METHYLASE->HEPN+DUF499*->MORC->REase->HTH+SF1-SF2-Helicase+REase->REase-><-?\|\|METHYLASE-p-> | 1137 | Bacteria>Terrabacteria_group>Actinomycetota>Actinomycetes>Actinomycetales | GCF_963519085.1 | Actinomyces | BKH20_RS04780 | 16618..20031 | 1654 | MULTISPECIES:-DUF499-domain-containing-protein-[Actinomyces] |
| WP_154475624.1 | DUF499_1.RM.like.systems | <-HEPN<-?\|\|?-><-?<-tRNA<-RelB<-HEPN<-HEPN+DUF499*<-METHYLASE<-TUDOR+SF1-SF2-Helicase+iREase+CTD+REase<-TUDOR+SF1-SF2-Helicase+iREase+CTD+REase-p | 1151 | Bacteria>Terrabacteria_group>Actinomycetota>Actinomycetes>Actinomycetales | - | Schaalia_hyovaginalis | FYJ77_RS02065 | 517238..520693 | 29316 | DUF499-domain-containing-protein-[Schaalia-hyovaginalis] |
| WP_129881947.1 | DUF499_1.RM.like.systems | <-REase<-ABC-ATPase+SbcC<-tRNA\|\|ACET-><-?<-HEPN+DUF499*<-METHYLASE<-TUDOR+SF1-SF2-Helicase+iREase+CTD+REase\|\|?-><-?<-ABC-ATPase+SbcC<-GNTR-HTH-p | 1127 | Bacteria>Terrabacteria_group>Actinomycetota>Actinomycetes>Bifidobacteriales | - | Bifidobacterium_animalis | PG1528B_RS04215 | 581191..584574 | 28025 | DUF499-domain-containing-protein-[Bifidobacterium-animalis] |
| WP_107040639.1 | DUF499_1.RM.like.systems | <-ABC-ATPase+SbcC<-?<-?<-SF1-SF2-Helicase\|\|PRTase->?-><-HEPN+DUF499*<-METHYLASE<-METHYLASE-p<-TUDOR+SF1-SF2-Helicase+iREase+CTD+REase\|\|HTH-><-TraJ-RHH | 1158 | Bacteria>Terrabacteria_group>Actinomycetota>Actinomycetes>Bifidobacteriales | - | Bifidobacterium_callitrichos | COO72_RS01265 | 282673..286149 | 762209 | Swt1-family-HEPN-domain-containing-protein-[Bifidobacterium-callitrichos] |
| WP_118284360.1 | DUF499_1.RM.like.systems | <-Schlafen+HATPase+HTH<-?\|\|?-><-?\|\|MULE-TRANSPOSASE-><-?<-HEPN+DUF499*<-METHYLASE<-TUDOR+SF1-SF2-Helicase+iREase+CTD+REase<-TraJ-RHH\|\|HTH_Tnp-> | 1129 | Bacteria>Terrabacteria_group>Actinomycetota>Actinomycetes>Bifidobacteriales | - | Bifidobacterium_pseudocatenulatum | DW149_RS06110 | 233918..237307 | 28026 | DUF499-domain-containing-protein-[Bifidobacterium-pseudocatenulatum] |
| OZG52261.1 | DUF499_1.RM.like.systems | <-JetD<-HTH\|\|?->?->?-><-HEPN+DUF499*<-METHYLASE<-?<-TUDOR+SF1-SF2-Helicase+iREase+CTD+REase<-?<-ABC-ATPase+SbcC<-ABC-ATPase+SbcC | 1139 | Bacteria>Terrabacteria_group>Actinomycetota>Actinomycetes>Bifidobacteriales | GCA_002259605.1 | Pseudoscardovia_radai | PSRA_0450 | 23959..27378 | 987066 | ATPase-AAA-[Pseudoscardovia-radai] |
| MBO5627607.1 | DUF499_1.RM.like.systems | <-HEPN+DUF499*<-METHYLASE-p | 467 | Bacteria>Terrabacteria_group>Actinomycetota>Actinomycetes>Bifidobacteriales | GCA_017627815.1 | Aeriscardovia_sp. | J5965_00830 | 1..1400 | 1872397 | DUF499-domain-containing-protein-[Aeriscardovia-sp.] |
| GDZ11402.1 | DUF499_1.RM.like.systems | MULE-TRANSPOSASE-><-?<-HEPN+DUF499*<-METHYLASE-p<-METHYLASE-p<-TUDOR+SF1-SF2-Helicase+iREase+CTD+REase<-TraJ-RHH\|\|HTH_Tnp-> | 1129 | Bacteria>Terrabacteria_group>Actinomycetota>Actinomycetes>Bifidobacteriales | GCA_018326745.1 | Bifidobacteriaceae_bacterium_MCC01993 | MCC01993_15150 | 22460..25849 | 2569496 | hypothetical-protein-MCC01993_15150-[Bifidobacteriaceae-bacterium-MCC01993] |
| MCI2174819.1 | DUF499_1.RM.like.systems | <-ABC-ATPase+SbcC\|\|Peptidase-><-?<-TetR-HTH<-HEPN+DUF499*<-METHYLASE<-?<-TUDOR+SF1-SF2-Helicase+iREase+CTD+REase | 1134 | Bacteria>Terrabacteria_group>Actinomycetota>Actinomycetes>Bifidobacteriales | GCA_022655115.1 | Bifidobacterium_thermacidophilum | LKK47_03040 | 24047..27451 | 246618 | DUF499-domain-containing-protein-[Bifidobacterium-thermacidophilum] |
| MCI2185070.1 | DUF499_1.RM.like.systems | HTH+MPTase->?->TUDOR+SF1-SF2-Helicase+iREase+CTD+REase->METHYLASE->HEPN+DUF499*->SbcC+ABC-ATPase->MethyltransfD12->?->URI-DNase+RAMA-> | 1149 | Bacteria>Terrabacteria_group>Actinomycetota>Actinomycetes>Bifidobacteriales | GCA_022655455.1 | Bifidobacterium_tibiigranuli | LKK60_02355 | 563467..566916 | 2172043 | DUF499-domain-containing-protein-[Bifidobacterium-tibiigranuli] |
| MDE5641417.1 | DUF499_1.RM.like.systems | METHYLASE-p->METHYLASE-p->HEPN+DUF499*-> | 583 | Bacteria>Terrabacteria_group>Actinomycetota>Actinomycetes>Bifidobacteriales | GCA_029051405.1 | Bifidobacterium_castoris | K2I40_06975 | 1749..3499 | 2306972 | MAG:-DUF499-domain-containing-protein,-partial-[Bifidobacterium-castoris] |
| MDD6374029.1 | DUF499_1.RM.like.systems | METHYLASE->HEPN+DUF499*-> | 1120 | Bacteria>Terrabacteria_group>Actinomycetota>Actinomycetes>Bifidobacteriales | GCA_029338855.1 | Bifidobacteriaceae_bacterium | PUF97_06115 | 2208..5567 | 2291324 | MAG:-DUF499-domain-containing-protein,-partial-[Bifidobacteriaceae-bacterium] |
| WP_003839950.1 | DUF499_1.RM.like.systems | Aldedh->?->?->PRTase->tRNA->TUDOR+SF1-SF2-Helicase+iREase+CTD+REase->METHYLASE->HEPN+DUF499*->?->HNH->?->?-><-Peptidase | 1138 | Bacteria>Terrabacteria_group>Actinomycetota>Actinomycetes>Bifidobacteriales | GCF_000024445.1 | Bifidobacterium_dentium | BIFDEN_RS08405 | 2078532..2081948 | 1689 | DUF499-domain-containing-protein-[Bifidobacterium-dentium] |
| WP_024464663.1 | DUF499_1.RM.like.systems | Cluster2439_3clades-><-HEPN\|\|TUDOR+SF1-SF2-Helicase+iREase+CTD+REase->METHYLASE-p->METHYLASE->HEPN+DUF499*->\|\|PAIREDC-HTH-> | 1163 | Bacteria>Terrabacteria_group>Actinomycetota>Actinomycetes>Bifidobacteriales | GCF_000426405.1 | Bifidobacterium_subtile | OUO0_RS10725 | 10278..13769 | 77635 | DUF499-domain-containing-protein-[Bifidobacterium-subtile] |
| WP_245834842.1 | DUF499_1.RM.like.systems | <-JetD<-HTH\|\|?->?-><-HEPN+DUF499*<-METHYLASE<-?<-TUDOR+SF1-SF2-Helicase+iREase+CTD+REase<-ABC-ATPase+SbcC<-ABC-ATPase+SbcC<-MarR-HTH | 1130 | Bacteria>Terrabacteria_group>Actinomycetota>Actinomycetes>Bifidobacteriales | GCF_002259605.1 | Pseudoscardovia_radai | PSRA_RS02315 | 23959..27351 | 987066 | DUF499-domain-containing-protein-[Pseudoscardovia-radai] |
| WP_100496559.1 | DUF499_1.RM.like.systems | Phage_integrase->?->PadR-HTH->SF1-SF2-Helicase-><-HEPN+DUF499*<-METHYLASE<-TUDOR+SF1-SF2-Helicase+iREase+CTD+REase<-Cluster2498_1clades<-?\|\|BirA-HTH-> | 1148 | Bacteria>Terrabacteria_group>Actinomycetota>Actinomycetes>Bifidobacteriales | GCF_002802865.1 | Bifidobacterium_scaligerum | CUU80_RS06820 | 234586..238032 | 2052656 | DUF499-domain-containing-protein-[Bifidobacterium-scaligerum] |
| WP_100513170.1 | DUF499_1.RM.like.systems | TUDOR+SF1-SF2-Helicase+iREase+CTD+REase->?->METHYLASE->HEPN+DUF499*-><-?<-Phage_holin_4_2\|\|?->?->JetB-> | 1147 | Bacteria>Terrabacteria_group>Actinomycetota>Actinomycetes>Bifidobacteriales | GCF_002802905.1 | Bifidobacterium_simiarum | CSQ87_RS07025 | 60477..63920 | 2045441 | Swt1-family-HEPN-domain-containing-protein-[Bifidobacterium-simiarum] |
| WP_126031245.1 | DUF499_1.RM.like.systems | <-ABC-ATPase+SbcC<-tRNA\|\|ACET-><-HEPN+DUF499*<-METHYLASE<-METHYLASE-p<-TUDOR+SF1-SF2-Helicase+iREase+CTD+REase | 1127 | Bacteria>Terrabacteria_group>Actinomycetota>Actinomycetes>Bifidobacteriales | GCF_003952025.1 | Bifidobacterium_castoris | D2E22_RS00925 | 206849..210232 | 2306972 | DUF499-domain-containing-protein-[Bifidobacterium-castoris] |
| WP_130013693.1 | DUF499_1.RM.like.systems | <-GDResolvase+HIN-HTH\|\|Mbetalac->TUDOR+SF1-SF2-Helicase+iREase+CTD+REase->METHYLASE->HEPN+DUF499*->?-><-URI-DNase | 1133 | Bacteria>Terrabacteria_group>Actinomycetota>Actinomycetes>Bifidobacteriales | GCF_004155015.1 | Bifidobacterium_pseudolongum | PG2054B_RS08290 | 18590..21991 | 1694 | DUF499-domain-containing-protein-[Bifidobacterium-pseudolongum] |
| WP_150353887.1 | DUF499_1.RM.like.systems | REase-p->TUDOR+SF1-SF2-Helicase+iREase+CTD+REase->METHYLASE-p->METHYLASE->HEPN+DUF499*->Primpol+D5N+D5ATPase-><-?<-Cysteine_desulfurase<-?<-ABC-ATPase+SbcC | 1134 | Bacteria>Terrabacteria_group>Actinomycetota>Actinomycetes>Bifidobacteriales | GCF_008698145.1 | Bifidobacterium_vespertilionis | EM090_RS02025 | 149240..152644 | 2562524 | Swt1-family-HEPN-domain-containing-protein-[Bifidobacterium-vespertilionis] |
| WP_150380554.1 | DUF499_1.RM.like.systems | <-ABhydrolase<-HEPN+DUF499*<-METHYLASE<-TUDOR+SF1-SF2-Helicase+iREase+CTD+REase | 1156 | Bacteria>Terrabacteria_group>Actinomycetota>Actinomycetes>Bifidobacteriales | GCF_008698165.1 | Bifidobacterium_tissieri | EM089_RS01160 | 325418..328888 | 1630162 | DUF499-domain-containing-protein-[Bifidobacterium-tissieri] |
| WP_150395254.1 | DUF499_1.RM.like.systems | TUDOR+SF1-SF2-Helicase+iREase+CTD+REase->METHYLASE->HEPN+DUF499*->ABhydrolase-> | 1156 | Bacteria>Terrabacteria_group>Actinomycetota>Actinomycetes>Bifidobacteriales | GCF_008698195.1 | Bifidobacterium_tissieri | EM849_RS05800 | 47872..51342 | 1630162 | DUF499-domain-containing-protein-[Bifidobacterium-tissieri] |
| WP_152234686.1 | DUF499_1.RM.like.systems | TUDOR+SF1-SF2-Helicase+iREase+CTD+REase->?->TraJ-RHH->METHYLASE-p->METHYLASE->HEPN+DUF499*-> | 1149 | Bacteria>Terrabacteria_group>Actinomycetota>Actinomycetes>Bifidobacteriales | GCF_009193355.1 | Bifidobacterium_leontopitheci | F7D09_RS07000 | 42381..45830 | 2650774 | Swt1-family-HEPN-domain-containing-protein-[Bifidobacterium-leontopitheci] |
| WP_195552455.1 | DUF499_1.RM.like.systems | <-Phage_int_SAM_1+Phage_integrase\|\|?->?->?->?-><-HEPN+DUF499*<-METHYLASE<-TUDOR+SF1-SF2-Helicase+iREase+CTD+REase\|\|?->HEPN-> | 1129 | Bacteria>Terrabacteria_group>Actinomycetota>Actinomycetes>Bifidobacteriales | GCF_015559175.1 | Bifidobacterium_pseudocatenulatum | I2G66_RS01750 | 59559..62948 | 28026 | DUF499-domain-containing-protein-[Bifidobacterium-pseudocatenulatum] |
| WP_198229240.1 | DUF499_1.RM.like.systems | NUDIX->TetR-HTH->ABC-ATPase+SbcC->?->?-><-HEPN+DUF499*<-METHYLASE<-TUDOR+SF1-SF2-Helicase+iREase+CTD+REase | 1150 | Bacteria>Terrabacteria_group>Actinomycetota>Actinomycetes>Bifidobacteriales | GCF_016101585.1 | Bifidobacterium_polysaccharolyticum | H3T90_RS00470 | 119489..122941 | 2750967 | DUF499-domain-containing-protein-[Bifidobacterium-polysaccharolyticum] |
| WP_214309728.1 | DUF499_1.RM.like.systems | Phage_integrase->PadR-HTH->SF1-SF2-Helicase-><-?<-?<-HEPN+DUF499*<-METHYLASE<-TUDOR+SF1-SF2-Helicase+iREase+CTD+REase<-Cluster2498_1clades<-PIN<-XIS-HTH<-Schlafen+HATPase+HTH | 1147 | Bacteria>Terrabacteria_group>Actinomycetota>Actinomycetes>Bifidobacteriales | GCF_018555415.1 | Bifidobacterium_simiarum | JS539_RS02125 | 175290..178733 | 2045441 | Swt1-family-HEPN-domain-containing-protein-[Bifidobacterium-simiarum] |
| WP_214310646.1 | DUF499_1.RM.like.systems | <-JetB<-?<-?\|\|Phage_holin_4_2->?-><-HEPN+DUF499*<-METHYLASE<-METHYLASE-p<-?<-?<-TUDOR+SF1-SF2-Helicase+iREase+CTD+REase | 1147 | Bacteria>Terrabacteria_group>Actinomycetota>Actinomycetes>Bifidobacteriales | GCF_018555415.1 | Bifidobacterium_simiarum | JS539_RS08840 | 19593..23036 | 2045441 | Swt1-family-HEPN-domain-containing-protein-[Bifidobacterium-simiarum] |
| WP_214375421.1 | DUF499_1.RM.like.systems | Phage_integrase-><-?\|\|PadR-HTH->SF1-SF2-Helicase-><-HEPN+DUF499*<-METHYLASE<-METHYLASE-p<-TUDOR+SF1-SF2-Helicase+iREase+CTD+REase<-Cluster2498_1clades | 1147 | Bacteria>Terrabacteria_group>Actinomycetota>Actinomycetes>Bifidobacteriales | GCF_018555455.1 | Bifidobacterium_colobi | JS530_RS01265 | 287140..290583 | 2809026 | Swt1-family-HEPN-domain-containing-protein-[Bifidobacterium-colobi] |
| WP_214376699.1 | DUF499_1.RM.like.systems | <-HTH_3<-?<-?\|\|Calcineurin->?-><-HEPN+DUF499*<-METHYLASE<-METHYLASE-p<-TUDOR+SF1-SF2-Helicase+iREase+CTD+REase<-FtsK\|\|DOC-> | 1148 | Bacteria>Terrabacteria_group>Actinomycetota>Actinomycetes>Bifidobacteriales | GCF_018555455.1 | Bifidobacterium_colobi | JS530_RS08230 | 76001..79447 | 2809026 | Swt1-family-HEPN-domain-containing-protein-[Bifidobacterium-colobi] |
| WP_214383368.1 | DUF499_1.RM.like.systems | ABC-ATPase+SbcC->?->Cysteine_desulfurase->?-><-Primpol+D5N+D5ATPase<-HEPN+DUF499*<-METHYLASE<-METHYLASE-p<-TUDOR+SF1-SF2-Helicase+iREase+CTD+REase<-REase-p | 1134 | Bacteria>Terrabacteria_group>Actinomycetota>Actinomycetes>Bifidobacteriales | GCF_018555535.1 | Bifidobacterium_vespertilionis | JS538_RS07550 | 34017..37421 | 2562524 | Swt1-family-HEPN-domain-containing-protein-[Bifidobacterium-vespertilionis] |
| WP_214305620.1 | DUF499_1.RM.like.systems | TUDOR+SF1-SF2-Helicase+iREase+CTD+REase->METHYLASE-p->METHYLASE->HEPN+DUF499*->ABhydrolase-> | 1154 | Bacteria>Terrabacteria_group>Actinomycetota>Actinomycetes>Bifidobacteriales | GCF_018555655.1 | Bifidobacterium_callimiconis | JS532_RS00845 | 238975..242439 | 2306973 | DUF499-domain-containing-protein-[Bifidobacterium-callimiconis] |
| WP_226595328.1 | DUF499_1.RM.like.systems | <-HTH_Tnp\|\|TraJ-RHH->TUDOR+SF1-SF2-Helicase+iREase+CTD+REase->METHYLASE->HEPN+DUF499*->?->\|\|?-><-?\|\|?->Schlafen+HATPase+HTH-><-Phage_integrase | 1129 | Bacteria>Terrabacteria_group>Actinomycetota>Actinomycetes>Bifidobacteriales | GCF_020526365.1 | Bifidobacterium_pseudocatenulatum | KZP21_RS03935 | 291458..294847 | 28026 | DUF499-domain-containing-protein-[Bifidobacterium-pseudocatenulatum] |
| WP_226678304.1 | DUF499_1.RM.like.systems | <-HTH_Tnp\|\|TraJ-RHH->TUDOR+SF1-SF2-Helicase+iREase+CTD+REase->METHYLASE->HEPN+DUF499*->?->\|\|?-><-?\|\|?->Schlafen+HATPase+HTH-><-Phage_integrase | 1129 | Bacteria>Terrabacteria_group>Actinomycetota>Actinomycetes>Bifidobacteriales | GCF_020526445.1 | Bifidobacterium_pseudocatenulatum | KZO94_RS04060 | 291413..294802 | 28026 | DUF499-domain-containing-protein-[Bifidobacterium-pseudocatenulatum] |
| WP_118262158.1 | DUF499_1.RM.like.systems | Phage_integrase-><-Schlafen+HATPase+HTH<-?\|\|?-><-?\|\|<-?<-HEPN+DUF499*<-METHYLASE<-TUDOR+SF1-SF2-Helicase+iREase+CTD+REase<-TraJ-RHH\|\|HTH_Tnp-><-?\|\|SAM-methylase-> | 1129 | Bacteria>Terrabacteria_group>Actinomycetota>Actinomycetes>Bifidobacteriales | GCF_020541925.1 | Bifidobacterium_pseudocatenulatum | PGQ74_RS03980 | 281260..284649 | 28026 | DUF499-domain-containing-protein-[Bifidobacterium-pseudocatenulatum] |
| WP_278801769.1 | DUF499_1.RM.like.systems | <-SF1-SF2-Helicase<-?<-WYL<-?<-HEPN+DUF499*<-METHYLASE<-TUDOR+SF1-SF2-Helicase+iREase+CTD+REase<-?\|\|RelB-><-tRNA<-sGTP | 1135 | Bacteria>Terrabacteria_group>Actinomycetota>Actinomycetes>Bifidobacteriales | GCF_944322225.1 | Bifidobacterium_pullorum | QDJ94_RS03760 | 68735..72142 | 78448 | DUF499-domain-containing-protein-[Bifidobacterium-pullorum] |
| WP_194891200.1 | DUF499_1.RM.like.systems | <-PAIREDC-HTH\|\|PAIREDC-HTH-><-?\|\|?->PAIREDC-HTH->HEPN-><-HEPN+DUF499*<-METHYLASE<-TUDOR+SF1-SF2-Helicase+iREase+CTD+REase\|\|?-><-?<-?\|\|PAIREDC-HTH-> | 1134 | Bacteria>Terrabacteria_group>Actinomycetota>Actinomycetes>Catenulisporales | GCF_015356215.1 | Catenulispora_pinisilvae | IZR00_RS01870 | 88900..92304 | 2705253 | DUF499-domain-containing-protein-[Catenulispora-pinisilvae] |
| TCJ34611.1 | DUF499_1.RM.like.systems | <-ParA-Soj-PloopNTPase<-REase\|\|TUDOR+SF1-SF2-Helicase+iREase+CTD+REase->METHYLASE->HEPN+DUF499*-><-HTH+UbE1<-?<-SbcC\|\|HTH_3->MPTase-><-REase | 1116 | Bacteria>Terrabacteria_group>Actinomycetota>Actinomycetes>Frankiales | GCA_004337455.1 | Parafrankia_sp._BMG5.11 | E0504_31915 | 85407..88757 | 222540 | ATP-binding-protein-[Frankia-sp.-BMG5.11] |
| WP_009738155.1 | DUF499_1.RM.like.systems | TUDOR+SF1-SF2-Helicase+iREase+CTD+REase->UDG-conflict->?->SF1-SF2-Helicase+REase->METHYLASE->HEPN+DUF499*-><-ParA+HTH<-RelE-ParE\|\|MULE-TRANSPOSASE->MULE-TRANSPOSASE->?-><-TPR+TPR+TPR+TPR+TPR+TPR+TPR+TPR | 1119 | Bacteria>Terrabacteria_group>Actinomycetota>Actinomycetes>Frankiales | GCF_000262465.1 | Frankia_sp._QA3 | FRAQA3DRAFT_RS05925 | 1493291..1496650 | 710111 | Swt1-family-HEPN-domain-containing-protein-[Frankia-sp.-QA3] |
| WP_203013794.1 | DUF499_1.RM.like.systems | <-NTD+SF1-SF2-Helicase\|\|TUDOR+SF1-SF2-Helicase+iREase+CTD+REase->UDG-conflict->METHYLASE->HEPN+DUF499*-><-?<-?\|\|?->?-><-?<-TetR-HTH | 1128 | Bacteria>Terrabacteria_group>Actinomycetota>Actinomycetes>Frankiales | GCF_016786345.1 | unclassified_Frankia | I6A60_RS06180 | 96819..100205 | 2632575 | MULTISPECIES:-Swt1-family-HEPN-domain-containing-protein-[unclassified-Frankia] |
| WP_248817712.1 | DUF499_1.RM.like.systems | TUDOR+SF1-SF2-Helicase+iREase+CTD+REase->UDG-conflict->METHYLASE->HEPN+DUF499*-><-ncRNA\|\|?->Pkinase-> | 1126 | Bacteria>Terrabacteria_group>Actinomycetota>Actinomycetes>Frankiales | GCF_023243045.1 | Frankia_sp._AgB32 | MXD60_RS04685 | 52343..55723 | 631119 | Swt1-family-HEPN-domain-containing-protein-[Frankia-sp.-AgB32] |
| WP_248814552.1 | DUF499_1.RM.like.systems | METHYLASE-p->HEPN+DUF499*->?-><-TPR+TPR+TPR+TPR+TPR+TPR+TPR+TPR | 1122 | Bacteria>Terrabacteria_group>Actinomycetota>Actinomycetes>Frankiales | GCF_023243055.1 | Frankia_sp._AgPm24 | MXD61_RS17500 | 1626..4994 | 631128 | Swt1-family-HEPN-domain-containing-protein-[Frankia-sp.-AgPm24] |
| WP_251744665.1 | DUF499_1.RM.like.systems | TUDOR+SF1-SF2-Helicase+iREase+CTD+REase->METHYLASE->HEPN+DUF499*-><-?<-ParA+HTH<-RelE-ParE | 1122 | Bacteria>Terrabacteria_group>Actinomycetota>Actinomycetes>Frankiales | GCF_023716945.1 | Frankia_sp._AiPs1 | ND748_RS06005 | 10061..13429 | 573493 | Swt1-family-HEPN-domain-containing-protein-[Frankia-sp.-AiPs1] |
| WP_261570743.1 | DUF499_1.RM.like.systems | METHYLASE->HEPN+DUF499*-><-?\|\|DDE_5+DDE_Tnp_1->?->?-><-TPR+TPR+TPR+TPR+TPR+TPR+TPR+TPR | 1120 | Bacteria>Terrabacteria_group>Actinomycetota>Actinomycetes>Frankiales | GCF_025403355.1 | Frankia_gtarii | NBZ84_RS30015 | 1969..5331 | 2950102 | Swt1-family-HEPN-domain-containing-protein-[Frankia-gtarii] |
| WP_226436803.1 | DUF499_1.RM.like.systems | <-DUF222+HNH<-HEPN+DUF499*<-METHYLASE\|\|CHTH-><-TUDOR+SF1-SF2-Helicase+iREase+CTD+REase | 1141 | Bacteria>Terrabacteria_group>Actinomycetota>Actinomycetes | GCF_012515855.1 | Mycobacteriales | GX543_RS09675 | 1506..4931 | 85007 | Swt1-family-HEPN-domain-containing-protein-[Rhodococcus-yananensis] |
| RBY97952.1 | DUF499_1.RM.like.systems | <-NUDIX\|\|?-><-ABhydrolase<-DNA_methylase\|\|REase->METHYLASE->HEPN+DUF499*->MORC->REase->HTH+SF1-SF2-Helicase+REase-> | 1128 | Bacteria>Terrabacteria_group>Actinomycetota>Actinomycetes>Geodermatophilales | GCA_003319175.1 | Blastococcus_sp._TF02-8 | DQ237_03405 | 728169..731555 | 2250574 | AAA-family-ATPase-[Blastococcus-sp.-TF02-8] |
| SNR82321.1 | DUF499_1.RM.like.systems | <-DNA-glycosylase\|\|?-><-METHYLASE\|\|?->TUDOR+SF1-SF2-Helicase+iREase+CTD+REase->METHYLASE->HEPN+DUF499*->tRNA-><-Arm-DNA-bind_4+Phage_int_SAM_3+Phage_integrase<-ZDNA-HTH\|\|HTH_3->?->?->DnaB-> | 1132 | Bacteria>Terrabacteria_group>Actinomycetota>Actinomycetes>Geodermatophilales | GCA_900188205.1 | Geodermatophilus_saharensis | SAMN04488107_0059 | 63726..67124 | 1137994 | Predicted-ATPase,-AAA+-superfamily-[Geodermatophilus-saharensis] |
| WP_113812542.1 | DUF499_1.RM.like.systems | <-ABhydrolase\|\|TUDOR+SF1-SF2-Helicase+iREase+CTD+REase->Schlafen+HATPase+HTH->METHYLASE->ParB+HNH->?->HEPN+DUF499*->NTD+SF1-SF2-Helicase->wHTH(SBD)+HNH-><-?<-?\|\|?-><-REase+35exo+Beta-Barrel+SF1-SF2-Helicase | 1114 | Bacteria>Terrabacteria_group>Actinomycetota>Actinomycetes>Geodermatophilales | GCF_003319095.1 | Blastococcus_sp._TBT05-19 | DQ244_RS16250 | 131306..134650 | 2250581 | Swt1-family-HEPN-domain-containing-protein-[Blastococcus-sp.-TBT05-19] |
| WP_205730045.1 | DUF499_1.RM.like.systems | <-ABhydrolase<-DNA_methylase\|\|REase->METHYLASE->HEPN+DUF499*->MORC->REase->HTH+SF1-SF2-Helicase+REase-> | 1113 | Bacteria>Terrabacteria_group>Actinomycetota>Actinomycetes>Geodermatophilales | GCF_003319175.1 | Blastococcus_sp._TF02-8 | DQ237_RS03405 | 728214..731555 | 2250574 | Swt1-family-HEPN-domain-containing-protein-[Blastococcus-sp.-TF02-8] |
| WP_236827009.1 | DUF499_1.RM.like.systems | HKD-DNase+SF1-SF2-Helicase+TRD-><-SF1-SF2-Helicase<-?\|\|TUDOR+SF1-SF2-Helicase+iREase+CTD+REase->ArdCN+MPTase->METHYLASE->HEPN+DUF499*->ParB+HNH-> | 1109 | Bacteria>Terrabacteria_group>Actinomycetota>Actinomycetes>Geodermatophilales | GCF_021698995.1 | unclassified_Blastococcus | E9565_RS02685 | 57900..61229 | 2619396 | MULTISPECIES:-Swt1-family-HEPN-domain-containing-protein-[unclassified-Blastococcus] |
| WP_284493306.1 | DUF499_1.RM.like.systems | <-HTH_19\|\|TUDOR+SF1-SF2-Helicase+iREase+CTD+REase-><-?<-?<-?\|\|?->HEPN+DUF499*->DOC+HTH-> | 1121 | Bacteria>Terrabacteria_group>Actinomycetota>Actinomycetes>Geodermatophilales | GCF_030165005.1 | Blastococcus_capsensis | QOZ86_RS17755 | 63627..66992 | 1564163 | Swt1-family-HEPN-domain-containing-protein-[Blastococcus-capsensis] |
| WP_176449778.1 | DUF499_1.RM.like.systems | DDE_Tnp_1_4-><-DNA-glycosylase<-METHYLASE\|\|?->TUDOR+SF1-SF2-Helicase+iREase+CTD+REase->METHYLASE->HEPN+DUF499*->tRNA->?-><-ZDNA-HTH\|\|HTH_3->?->DnaB-> | 1120 | Bacteria>Terrabacteria_group>Actinomycetota>Actinomycetes>Geodermatophilales | GCF_900188205.1 | Geodermatophilus_saharensis | CHC04_RS00285 | 63762..67124 | 1137994 | DUF499-domain-containing-protein-[Geodermatophilus-saharensis] |
| WP_013016869.1 | DUF499_1.RM.like.systems | MoxR-AAA->?->?->?-><-REase\|\|METHYLASE->HEPN+DUF499*-><-TUDOR+SF1-SF2-Helicase+iREase+CTD+REase | 1109 | Bacteria>Terrabacteria_group>Actinomycetota>Actinomycetes>Glycomycetales | GCF_000024545.1 | Stackebrandtia_nassauensis | SNAS_RS07855 | 1675315..1678644 | 283811 | DUF499-domain-containing-protein-[Stackebrandtia-nassauensis] |
| MBK6873560.1 | DUF499_1.RM.like.systems | UDG-conflict-><-TUDOR+SF1-SF2-Helicase+iREase+CTD+REase\|\|METHYLASE->HEPN+DUF499*->HTH_Tnp_1+HTH_21+rve+rve_2-><-?<-?<-Phage_integrase | 1150 | Bacteria>Terrabacteria_group>Actinomycetota>Actinomycetes>Kineosporiales | GCA_016702615.1 | Kineosporiaceae_bacterium | IPG94_19890 | 267597..271049 | 2487580 | DUF499-domain-containing-protein-[Kineosporiaceae-bacterium] |
| MBK7622561.1 | DUF499_1.RM.like.systems | Phage_int_SAM_1+Phage_integrase->?-><-HTH_Tnp_1+HTH_21+rve+rve_2<-HEPN+DUF499*<-METHYLASE\|\|TUDOR+SF1-SF2-Helicase+iREase+CTD+REase-><-UDG-conflict | 1150 | Bacteria>Terrabacteria_group>Actinomycetota>Actinomycetes>Kineosporiales | GCA_016709945.1 | Kineosporiaceae_bacterium | IPJ14_07830 | 115869..119321 | 2487580 | DUF499-domain-containing-protein-[Kineosporiaceae-bacterium] |
| KFG04509.1 | DUF499_1.RM.like.systems | <-DUF6731<-HEPN+DUF499*<-METHYLASE<-TUDOR+SF1-SF2-Helicase+iREase+CTD+REase\|\|HTH+DUF5753->ST-Hiskin->SIGMA-HTH-> | 1123 | Bacteria>Terrabacteria_group>Actinomycetota>Actinomycetes>Kitasatosporales | GCA_000738715.1 | Streptomyces_scabiei | IQ61_35115 | 15067..18438 | 1930 | hypothetical-protein-IQ61_35115-[Streptomyces-scabiei] |
| KOG77665.1 | DUF499_1.RM.like.systems | ACET-><-?<-?\|\|?->?-><-HEPN+DUF499*<-METHYLASE<-TUDOR+SF1-SF2-Helicase+iREase+CTD+REase<-ZDNA-HTH\|\|?->MORC->Phage_terminase-> | 1126 | Bacteria>Terrabacteria_group>Actinomycetota>Actinomycetes>Kitasatosporales | GCA_001270675.1 | Streptomyces_griseus_subsp._rhodochrous | ADK33_30195 | 165445..168825 | 285575 | hypothetical-protein-ADK33_30195-[Streptomyces-griseus-subsp.-rhodochrous] |
| OKI62803.1 | DUF499_1.RM.like.systems | <-MORC\|\|ZDNA-HTH->TUDOR+SF1-SF2-Helicase+iREase+CTD+REase->METHYLASE->HEPN+DUF499*-> | 1126 | Bacteria>Terrabacteria_group>Actinomycetota>Actinomycetes>Kitasatosporales | GCA_001905405.1 | Streptomyces_sp._TSRI0445 | AMK14_33000 | 173099..176479 | 1703925 | hypothetical-protein-AMK14_33000-[Streptomyces-sp.-TSRI0445] |
| TJZ52107.1 | DUF499_1.RM.like.systems | HAMP+HISKIN-><-Phage_holin<-DUF6731<-HEPN+DUF499*<-METHYLASE<-TUDOR+SF1-SF2-Helicase+iREase+CTD+REase<-tRNA\|\|METHYLASE-p-><-?\|\|TetR-HTH-> | 1122 | Bacteria>Terrabacteria_group>Actinomycetota>Actinomycetes>Kitasatosporales | GCA_005048155.1 | Streptomyces_piniterrae | FCH28_19955 | 322483..325851 | 2571125 | ATP-binding-protein-[Streptomyces-piniterrae] |
| TRZ87911.1 | DUF499_1.RM.like.systems | REase->METHYLASE->HEPN+DUF499*-><-Phage_int_SAM_1+Phage_integrase | 1108 | Bacteria>Terrabacteria_group>Actinomycetota>Actinomycetes>Kitasatosporales | GCA_007280125.1 | Streptomycetaceae_bacterium | D4R83_01365 | 3739..7065 | 2021369 | ATP-binding-protein-[Streptomycetaceae-bacterium] |
| GGS96436.1 | DUF499_1.RM.like.systems | <-ST-Hiskin\|\|HTH+DUF5753->HTH+DUF5753->DUF397-><-HEPN+DUF499*<-METHYLASE<-TUDOR+SF1-SF2-Helicase+iREase+CTD+REase<-SF1-SF2-Helicase<-?<-REase | 1137 | Bacteria>Terrabacteria_group>Actinomycetota>Actinomycetes>Kitasatosporales | GCA_014649155.1 | Streptomyces_griseoviridis | GCM10010240_32240 | 242124..245537 | 45398 | hypothetical-protein-GCM10010240_32240-[Streptomyces-griseoviridis] |
| BBB01161.1 | DUF499_1.RM.like.systems | <-DDE_Tnp<-?<-?\|\|?->TUDOR+SF1-SF2-Helicase+iREase+CTD+REase->METHYLASE->HEPN+DUF499*->DUF6731-><-?<-HTH+DUF5753\|\|HTH+DUF5753->METHYLASE-p-> | 1128 | Bacteria>Terrabacteria_group>Actinomycetota>Actinomycetes>Kitasatosporales | GCA_016756395.1 | Streptomyces_sp._SN-593 | RVR_8434 | 7009798..7013184 | 659352 | hypothetical-protein-RVR_8434-[Streptomyces-sp.-SN-593] |
| MBX7473484.1 | DUF499_1.RM.like.systems | HTH_3->REase->?->TUDOR+SF1-SF2-Helicase+iREase+CTD+REase->METHYLASE->HEPN+DUF499*-><-UDG-conflict\|\|?->?->Phage_int_SAM_1->DDE_Tnp_1_4->DDE_Tnp_1_4-> | 1146 | Bacteria>Terrabacteria_group>Actinomycetota>Actinomycetes>Kitasatosporales | GCA_019711445.1 | Streptomyces_sp._MAG02 | K1Y80_46530 | 10650..14090 | 2866213 | DUF499-domain-containing-protein-[Streptomyces-sp.-MAG02] |
| MCP9962608.1 | DUF499_1.RM.like.systems | ZDNA-HTH->TUDOR+SF1-SF2-Helicase+iREase+CTD+REase->METHYLASE->HEPN+DUF499*-><-?\|\|?->?-><-?\|\|ESAT-> | 1115 | Bacteria>Terrabacteria_group>Actinomycetota>Actinomycetes>Kitasatosporales | GCA_024349105.1 | Streptomyces_somaliensis | LUX05_16750 | 3621706..3625053 | 78355 | DUF499-domain-containing-protein-[Streptomyces-somaliensis] |
| BBC36956.1 | DUF499_1.RM.like.systems | PIN-><-?<-?<-ST-Hiskin\|\|HTH+DUF5753->DUF397-><-HEPN+DUF499*<-METHYLASE<-TUDOR+SF1-SF2-Helicase+iREase+CTD+REase<-SF1-SF2-Helicase<-?<-REase | 1135 | Bacteria>Terrabacteria_group>Actinomycetota>Actinomycetes>Kitasatosporales | GCA_030294945.1 | Streptomyces_graminofaciens | SGFS_082500 | 9169123..9172530 | 68212 | hypothetical-protein-SGFS_082500-[Streptomyces-graminofaciens] |
| WP_019888360.1 | DUF499_1.RM.like.systems | Radical-SAM->RADICAL-SAM->?-><-?\|\|TUDOR+SF1-SF2-Helicase+iREase+CTD+REase->METHYLASE->HEPN+DUF499*->DUF6731-><-?\|\|ACET-><-?<-?<-LuxR-HTH | 1154 | Bacteria>Terrabacteria_group>Actinomycetota>Actinomycetes>Kitasatosporales | GCF_000384175.1 | Streptomyces_purpureus | STRPU_RS0118820 | 4066802..4070266 | 1951 | DUF499-domain-containing-protein-[Streptomyces-purpureus] |

| Protein-ID | System | Gene-Neighbourhoods | Length | Taxonomy | Assembly-ID | Strain | Locus-Tag | Genomic-coordinates | TaxID | Fasta-defline |
|---|---|---|---|---|---|---|---|---|---|---|
| WP_043673297.1 | DUF499_1.RM.like.systems | <-NTP_transferase<-?<-ABC-ATPase+SbcC\|\|tRNA->?->Phage_cap_E->UDG-conflict-><-HEPN+DUF499*<-METHYLASE<-TUDOR+ SF1-SF2-Helicase+iREase+CTD+REase<-?<-REase<-SLATT | 1146 | Bacteria>Terrabacteria_group>Actinomycetota>Actinomycetes>Kitasatosporales | GCF_000716435.1 | Streptomyces_xylophagus | OO70_RS20695 | 140518..143958 | 285514 | Swt1-family-HEPN-domain-containing-protein-[Streptomyces-xylophagus] |
| WP_050490834.1 | DUF499_1.RM.like.systems | ACET-><-?\|\|<-?<-?\|\|?-><-DUF4096+DDE_Tnp_1+DDE_Tnp_1_2<-HEPN+DUF499*<-METHYLASE<-TUDOR+ SF1-SF2-Helicase+iREase+CTD+REase<-ZDNA-HTH\|\|MORC->Phage_terminase-><-TRAM+Methyltransf_31 | 1115 | Bacteria>Terrabacteria_group>Actinomycetota>Actinomycetes>Kitasatosporales | GCF_000718915.1 | Streptomyces | ADK33_RS30095 | 165445..168792 | 1883 | MULTISPECIES:-DUF499-domain-containing-protein-[Streptomyces] |
| WP_031508838.1 | DUF499_1.RM.like.systems | HTH+DUF5753->DUF397-><-HEPN+DUF499*<-METHYLASE<-TUDOR+ SF1-SF2-Helicase+iREase+CTD+REase<-?<-?<-?<-ABC-ATPase+SbcC | 1145 | Bacteria>Terrabacteria_group>Actinomycetota>Actinomycetes>Kitasatosporales | GCF_000718985.1 | Streptomyces_megasporus | IF47_RS0112740 | 147736..151173 | 44060 | Swt1-family-HEPN-domain-containing-protein-[Streptomyces-megasporus] |
| WP_030969976.1 | DUF499_1.RM.like.systems | REase->?->TUDOR+ SF1-SF2-Helicase+iREase+CTD+REase->METHYLASE->HEPN+DUF499*-><-UDG-conflict\|\|HTH_31-> | 1145 | Bacteria>Terrabacteria_group>Actinomycetota>Actinomycetes>Kitasatosporales | GCF_000719575.1 | Streptomyces_sp._NRRL_S-1824 | IG01_RS0105040 | 662425..665862 | 1463889 | Swt1-family-HEPN-domain-containing-protein-[Streptomyces-sp.-NRRL-S-1824] |
| WP_030708645.1 | DUF499_1.RM.like.systems | <-Aminotran_3\|\|?->?->?-><-HEPN+DUF499*<-METHYLASE<-TUDOR+ SF1-SF2-Helicase+iREase+CTD+REase\|\|?->?->?->Peptidase_M50+PDZ-> | 1136 | Bacteria>Terrabacteria_group>Actinomycetota>Actinomycetes>Kitasatosporales | GCF_000720115.1 | Streptomyces_sp._NRRL_F-2580 | IF88_RS0107850 | 389452..392862 | 1463841 | Swt1-family-HEPN-domain-containing-protein-[Streptomyces-sp.-NRRL-F-2580] |
| WP_067071154.1 | DUF499_1.RM.like.systems | TUDOR+ SF1-SF2-Helicase+iREase+CTD+REase-p->METHYLASE->HEPN+DUF499*->ParB+HNH-><-?<-?<-HTH+DUF5753 | 1122 | Bacteria>Terrabacteria_group>Actinomycetota>Actinomycetes>Kitasatosporales | GCF_001543895.1 | Carbonactinospora_thermoautotrophica | TR74_RS19795 | 3234..6602 | 1469144 | Swt1-family-HEPN-domain-containing-protein-[Carbonactinospora-thermoautotrophica] |
| WP_064533938.1 | DUF499_1.RM.like.systems | <-SF1-SF2-Helicase<-?<-?\|\|?->tRNA->tRNA->tRNA->TUDOR+ SF1-SF2-Helicase+iREase+CTD+REase->METHYLASE->HEPN+DUF499*->DUF6731->Phage_holin->tRNA->tRNA->?->?->SF1-SF2-Helicase-> | 1131 | Bacteria>Terrabacteria_group>Actinomycetota>Actinomycetes>Kitasatosporales | GCF_001654495.1 | Streptomyces_sp._SAT1 | A8713_RS14845 | 3432444..3435839 | 1849967 | DUF499-domain-containing-protein-[Streptomyces-sp.-SAT1] |
| WP_073228437.1 | DUF499_1.RM.like.systems | <-Phage_holin<-DUF6731<-HEPN+DUF499*<-METHYLASE<-TUDOR+ SF1-SF2-Helicase+iREase+CTD+REase | 1131 | Bacteria>Terrabacteria_group>Actinomycetota>Actinomycetes>Kitasatosporales | GCF_001895105.1 | Streptomyces_sp._NBRC_110465 | BHQ08_RS31730 | 47724..51119 | 1897621 | DUF499-domain-containing-protein-[Streptomyces-sp.-NBRC-110465] |
| WP_198941888.1 | DUF499_1.RM.like.systems | <-MORC\|\|ZDNA-HTH->TUDOR+ SF1-SF2-Helicase+iREase+CTD+REase->METHYLASE->HEPN+DUF499*-> | 1115 | Bacteria>Terrabacteria_group>Actinomycetota>Actinomycetes>Kitasatosporales | GCF_001905405.1 | Streptomyces_sp._TSRI0445 | AMK14_RS33650 | 173132..176479 | 1703925 | DUF499-domain-containing-protein-[Streptomyces-sp.-TSRI0445] |
| WP_075268487.1 | DUF499_1.RM.like.systems | HTH_3->?->TUDOR+ SF1-SF2-Helicase+iREase+CTD+REase->METHYLASE->HEPN+DUF499*->DUF6731->Phage_holin->\|\|?-><-?\|\|?-><-Peptidase_M6+Peptidase_M7 | 1126 | Bacteria>Terrabacteria_group>Actinomycetota>Actinomycetes>Kitasatosporales | GCF_001931635.1 | Streptomyces_sp._Tue_6075 | TK78_RS20735 | 4682915..4686295 | 1661694 | DUF499-domain-containing-protein-[Streptomyces-sp.-Tue-6075] |
| WP_208640870.1 | DUF499_1.RM.like.systems | HTH_3-><-NUDIX\|\|SLATT+SLATT->TIR-><-?\|\|?-><-HEPN+DUF499*<-METHYLASE<-TUDOR+ SF1-SF2-Helicase+iREase+CTD+REase | 1148 | Bacteria>Terrabacteria_group>Actinomycetota>Actinomycetes>Kitasatosporales | GCF_002154505.1 | Streptomyces_carpinensis | BZY54_RS35300 | 9342..12788 | 66369 | DUF499-domain-containing-protein-[Streptomyces-carpinensis] |
| WP_103510051.1 | DUF499_1.RM.like.systems | TUDOR+ SF1-SF2-Helicase+iREase+CTD+REase->METHYLASE->HEPN+DUF499*-><-?<-?<-?<-VirB4-FtsK | 1135 | Bacteria>Terrabacteria_group>Actinomycetota>Actinomycetes>Kitasatosporales | GCF_002910875.1 | Streptomyces_sp._SM13 | B9S60_RS19800 | 69389..72796 | 1983803 | Swt1-family-HEPN-domain-containing-protein-[Streptomyces-sp.-SM13] |
| WP_109294977.1 | DUF499_1.RM.like.systems | Methyltransf_25-><-?<-?\|\|TUDOR+ SF1-SF2-Helicase+iREase+CTD+REase->METHYLASE->HEPN+DUF499*-><-?<-?<-?<-XIS-HTH<-?\|\|HTH-> | 1137 | Bacteria>Terrabacteria_group>Actinomycetota>Actinomycetes>Kitasatosporales | GCF_003122365.1 | Streptomyces_spongiicola | DDQ41_RS15240 | 3467115..3470528 | 1690221 | Swt1-family-HEPN-domain-containing-protein-[Streptomyces-spongiicola] |
| WP_120051255.1 | DUF499_1.RM.like.systems | <-Phage_holin<-DUF6731<-HEPN+DUF499*<-METHYLASE<-TUDOR+ SF1-SF2-Helicase+iREase+CTD+REase\|\|RHH_1->PIN-> | 1131 | Bacteria>Terrabacteria_group>Actinomycetota>Actinomycetes>Kitasatosporales | GCF_003595235.1 | Streptomyces_griseorubiginosus | DWG14_RS17550 | 3948403..3951798 | 67304 | DUF499-domain-containing-protein-[Streptomyces-griseorubiginosus] |
| WP_122182258.1 | DUF499_1.RM.like.systems | ABhydrolase->?->tRNA->REase->?->TUDOR+ SF1-SF2-Helicase+iREase+CTD+REase->METHYLASE->HEPN+DUF499*-><-REase<-?<-?<-?<-?<-Pkinase | 1151 | Bacteria>Terrabacteria_group>Actinomycetota>Actinomycetes>Kitasatosporales | GCF_003696235.1 | Streptomyces_triticirhizae | EBN88_RS03330 | 36535..39990 | 2483353 | Swt1-family-HEPN-domain-containing-protein-[Streptomyces-triticirhizae] |
| WP_122182269.1 | DUF499_1.RM.like.systems | <-AdoMet_Synthase<-Pkinase<-?<-HTH_31\|\|tRNA-><-DUF6731<-HEPN+DUF499*<-METHYLASE<-TUDOR+ SF1-SF2-Helicase+iREase+CTD+REase | 1129 | Bacteria>Terrabacteria_group>Actinomycetota>Actinomycetes>Kitasatosporales | GCF_003696235.1 | Streptomyces_triticirhizae | EBN88_RS03395 | 53009..56398 | 2483353 | DUF499-domain-containing-protein-[Streptomyces-triticirhizae] |
| WP_123099817.1 | DUF499_1.RM.like.systems | REase->?->SF1-SF2-Helicase->TUDOR+ SF1-SF2-Helicase+iREase+CTD+REase->METHYLASE->HEPN+DUF499*-><-ST-Hiskin\|\|HTH+DUF5753->?-><-SLATT | 1137 | Bacteria>Terrabacteria_group>Actinomycetota>Actinomycetes>Kitasatosporales | GCF_003721215.1 | Streptomyces_botrytidirepellens | EEJ42_RS11295 | 50582..53995 | 2486417 | Swt1-family-HEPN-domain-containing-protein-[Streptomyces-botrytidirepellens] |
| WP_124289019.1 | DUF499_1.RM.like.systems | <-Pkinase<-AdoMet_Synthase<-Pkinase\|\|HTH+TPR+TPR->?-><-HEPN+DUF499*<-METHYLASE<-TUDOR+ SF1-SF2-Helicase+iREase+CTD+REase<-?<-REase<-SF1-SF2-Helicase | 1148 | Bacteria>Terrabacteria_group>Actinomycetota>Actinomycetes>Kitasatosporales | GCF_003846165.1 | Streptomyces_sp._ADI91-18 | EES37_RS07515 | 1680083..1683529 | 1522755 | DUF499-domain-containing-protein-[Streptomyces-sp.-ADI91-18] |
| WP_136222929.1 | DUF499_1.RM.like.systems | <-Radical-SAM<-?<-?\|\|HTH_3-><-?\|\|?-><-HEPN+DUF499*<-METHYLASE<-TUDOR+ SF1-SF2-Helicase+iREase+CTD+REase\|\|?->XIS-HTH->Phage_int_SAM_1+Phage_integrase-><-DUF4011+SF1-SF2-Helicase+REase | 1135 | Bacteria>Terrabacteria_group>Actinomycetota>Actinomycetes>Kitasatosporales | GCF_004794975.1 | Streptomyces_sp._A1547 | E6W17_RS04020 | 256183..259590 | 2563105 | Swt1-family-HEPN-domain-containing-protein-[Streptomyces-sp.-A1547] |
| WP_136741406.1 | DUF499_1.RM.like.systems | HAMP+HISKIN-><-SF1-SF2-Helicase+HRDC<-tRNA<-Phage_holin<-DUF6731<-HEPN+DUF499*<-METHYLASE<-TUDOR+ SF1-SF2-Helicase+iREase+CTD+REase<-?<-?<-?<-tRNA\|\|METHYLASE-p-> | 1124 | Bacteria>Terrabacteria_group>Actinomycetota>Actinomycetes>Kitasatosporales | GCF_005048155.1 | Streptomyces_piniterrae | FCH28_RS19960 | 322483..325857 | 2571125 | DUF499-domain-containing-protein-[Streptomyces-piniterrae] |
| WP_222946975.1 | DUF499_1.RM.like.systems | Phage_cap_E->?->?-><-wHTH(SBD)+HNH\|\|TUDOR+ SF1-SF2-Helicase+iREase+CTD+REase->METHYLASE->HEPN+DUF499*->DUF6731->Phage_holin-> | 1153 | Bacteria>Terrabacteria_group>Actinomycetota>Actinomycetes>Kitasatosporales | GCF_008271805.1 | Streptomyces_sp._TRM49041 | FVS66_RS24245 | 260388..263849 | 2603216 | DUF499-domain-containing-protein-[Streptomyces-sp.-TRM49041] |
| WP_161236363.1 | DUF499_1.RM.like.systems | <-HEPN\|\|<-wHTH(SBD)+HNH\|\|TUDOR+ SF1-SF2-Helicase+iREase+CTD+REase->METHYLASE->HEPN+DUF499*->DUF6731->Phage_holin-><-MNS+HTH+REase | 1161 | Bacteria>Terrabacteria_group>Actinomycetota>Actinomycetes>Kitasatosporales | GCF_010548075.1 | unclassified_Streptomyces | G3I23_RS06870 | 11181..14666 | 2593676 | MULTISPECIES:-DUF499-domain-containing-protein-[unclassified-Streptomyces] |
| WP_189323067.1 | DUF499_1.RM.like.systems | OLD-ABC->?-><-DUF397<-HTH+DUF5753\|\|?->METHYLASE->HEPN+DUF499*->SF1-SF2-Helicase-><-?\|\|SF1-SF2-Helicase-> | 1137 | Bacteria>Terrabacteria_group>Actinomycetota>Actinomycetes>Kitasatosporales | GCF_014647675.1 | Streptomyces_flaveus | IEZ21_RS18795 | 178873..182286 | 66370 | DUF499-domain-containing-protein-[Streptomyces-flaveus] |
| WP_189838087.1 | DUF499_1.RM.like.systems | <-FtsK<-?\|\|HTH_3-><-NUDIX\|\|TUDOR+ SF1-SF2-Helicase+iREase+CTD+REase->METHYLASE->HEPN+DUF499*->DUF6731-><-?\|\|GNTR-HTH->HTH_Tnp_4+DDE_Tnp_4->Methyltransf-> | 1150 | Bacteria>Terrabacteria_group>Actinomycetota>Actinomycetes>Kitasatosporales | GCF_014650195.1 | Streptomyces_umbrinus | IE284_RS09050 | 2097992..2101444 | 67370 | DUF499-domain-containing-protein-[Streptomyces-umbrinus] |
| WP_199584383.1 | DUF499_1.RM.like.systems | DNA-glycosylase-><-HNH<-Phage_holin<-DUF6731<-HEPN+DUF499*<-METHYLASE<-TUDOR+ SF1-SF2-Helicase+iREase+CTD+REase<-IclR-HTH\|\|?-><-HTH_3\|\|Phage_integrase+Phage_integrase-> | 1117 | Bacteria>Terrabacteria_group>Actinomycetota>Actinomycetes>Kitasatosporales | GCF_016482685.1 | Streptomyces_sp._DSM_110735 | IF655_RS11870 | 37375..40728 | 2775031 | DUF499-domain-containing-protein-[Streptomyces-sp.-DSM-110735] |
| WP_202237096.1 | DUF499_1.RM.like.systems | <-DDE_Tnp<-?\|\|?->TUDOR+ SF1-SF2-Helicase+iREase+CTD+REase->METHYLASE->HEPN+DUF499*->DUF6731->Phage_holin-><-?<-HTH+DUF5753\|\|HTH+DUF5753->METHYLASE-p-> | 1126 | Bacteria>Terrabacteria_group>Actinomycetota>Actinomycetes>Kitasatosporales | GCF_016756395.1 | Streptomyces_sp._SN-593 | RVR_RS30315 | 7009804..7013184 | 659352 | DUF499-domain-containing-protein-[Streptomyces-sp.-SN-593] |
| WP_210966257.1 | DUF499_1.RM.like.systems | HTH_Tnp_4+DDE_Tnp_4-><-?<-?<-?<-Phage_holin<-DUF6731<-HEPN+DUF499*<-METHYLASE<-TUDOR+ SF1-SF2-Helicase+iREase+CTD+REase<-ST-Hiskin\|\|HTH+DUF5753->DUF397-><-REase | 1129 | Bacteria>Terrabacteria_group>Actinomycetota>Actinomycetes>Kitasatosporales | GCF_018070605.1 | Streptomyces_sp._B15 | KBY19_RS19140 | 34073..37462 | 1537797 | DUF499-domain-containing-protein-[Streptomyces-sp.-B15] |
| WP_217467331.1 | DUF499_1.RM.like.systems | HTH+HKD-DNase->NUDIX-><-HDRnase\|\|TUDOR+ SF1-SF2-Helicase+iREase+CTD+REase->METHYLASE->HEPN+DUF499*-><-?\|\|?->?-><-MerR-HTH\|\|ABhydrolase-> | 1135 | Bacteria>Terrabacteria_group>Actinomycetota>Actinomycetes>Kitasatosporales | GCF_019090125.1 | Streptomyces_sp._BV333 | KUG12_RS25645 | 32046..35453 | 2849673 | Swt1-family-HEPN-domain-containing-protein-[Streptomyces-sp.-BV333] |
| WP_224221478.1 | DUF499_1.RM.like.systems | MORC+HisKin+REase-><-Phage_holin<-DUF6731<-HEPN+DUF499*<-METHYLASE<-TUDOR+ SF1-SF2-Helicase+iREase+CTD+REase<-DUF397\|\|ST-Hiskin-> | 1127 | Bacteria>Terrabacteria_group>Actinomycetota>Actinomycetes>Kitasatosporales | GCF_020092375.1 | Streptomyces_olivaceus | KVH28_RS23645 | 380730..384113 | 47716 | DUF499-domain-containing-protein-[Streptomyces-olivaceus] |
| WP_225014234.1 | DUF499_1.RM.like.systems | <-HEPN+DUF499*<-METHYLASE<-TUDOR+ SF1-SF2-Helicase+iREase+CTD+REase | 753 | Bacteria>Terrabacteria_group>Actinomycetota>Actinomycetes>Kitasatosporales | GCF_020176795.1 | Streptomyces_sp._8L | LCE31_RS12240 | 76..2337 | 2877242 | DUF499-domain-containing-protein-[Streptomyces-sp.-8L] |
| WP_235049964.1 | DUF499_1.RM.like.systems | <-PspA+ParA-Soj-PloopNTPase<-HEPN+DUF499*<-METHYLASE<-TUDOR+ SF1-SF2-Helicase+iREase+CTD+REase\|\|REase-> | 1137 | Bacteria>Terrabacteria_group>Actinomycetota>Actinomycetes>Kitasatosporales | GCF_021502675.1 | Yinghuangia_soli | LZ495_RS01505 | 359747..363160 | 2908204 | Swt1-family-HEPN-domain-containing-protein-[Yinghuangia-sp.-KLBMP8922] |
| WP_257563884.1 | DUF499_1.RM.like.systems | REase->?->SF1-SF2-Helicase->TUDOR+ SF1-SF2-Helicase+iREase+CTD+REase->METHYLASE->HEPN+DUF499*-><-DUF397<-HTH+DUF5753\|\|ST-Hiskin->?-><-PIN<-TraJ-RHH | 1137 | Bacteria>Terrabacteria_group>Actinomycetota>Actinomycetes>Kitasatosporales | GCF_024612035.1 | Streptomyces_sp._DSM_40750 | JIX55_RS15360 | 3457896..3461309 | 2801030 | Swt1-family-HEPN-domain-containing-protein-[Streptomyces-sp.-DSM-40750] |
| WP_260825327.1 | DUF499_1.RM.like.systems | REase->?->SF1-SF2-Helicase->TUDOR+ SF1-SF2-Helicase+iREase+CTD+REase->METHYLASE->HEPN+DUF499*-><-DUF397<-HTH+DUF5753\|\|ST-Hiskin->?->?-><-PIN | 1137 | Bacteria>Terrabacteria_group>Actinomycetota>Actinomycetes>Kitasatosporales | GCF_025290915.1 | Streptomyces_sp._15-116A | N4P33_RS06645 | 23393..26806 | 2259035 | Swt1-family-HEPN-domain-containing-protein-[Streptomyces-sp.-15-116A] |
| WP_266631931.1 | DUF499_1.RM.like.systems | METHYLASE->?->?->SLATT->TUDOR+ SF1-SF2-Helicase+iREase+CTD+REase->METHYLASE->HEPN+DUF499*->DUF6731->Phage_holin-> | 1159 | Bacteria>Terrabacteria_group>Actinomycetota>Actinomycetes>Kitasatosporales | GCF_026339655.1 | unclassified_Streptomyces | OOK58_RS43255 | 9431725..9435204 | 2593676 | MULTISPECIES:-DUF499-domain-containing-protein-[unclassified-Streptomyces] |
| WP_275780742.1 | DUF499_1.RM.like.systems | <-MORC+DNA_gyraseB+TOPRIM\|\|?-><-?\|\|HEPN-><-Phage_holin<-DUF6731<-HEPN+DUF499*<-METHYLASE<-TUDOR+ SF1-SF2-Helicase+iREase+CTD+REase | 1150 | Bacteria>Terrabacteria_group>Actinomycetota>Actinomycetes>Kitasatosporales | GCF_029223485.1 | Streptomyces_coacervatus | P2Q00_RS45645 | 83477..86929 | 647381 | DUF499-domain-containing-protein-[Streptomyces-coacervatus] |
| WP_158953066.1 | DUF499_1.RM.like.systems | <-SIGMA-HTH<-ST-Hiskin<-HTH+DUF5753\|\|TUDOR+ SF1-SF2-Helicase+iREase+CTD+REase->METHYLASE->HEPN+DUF499*->DUF6731-><-Phage_integrase<-XIS-HTH | 1150 | Bacteria>Terrabacteria_group>Actinomycetota>Actinomycetes>Kitasatosporales | GCF_033792825.1 | Streptomyces_scabiei | AA092_RS33625 | 12095..15547 | 1930 | DUF499-domain-containing-protein-[Streptomyces-scabiei] |
| WP_093789469.1 | DUF499_1.RM.like.systems | MORC+HisKin+REase-><-Phage_holin<-DUF6731<-HEPN+DUF499*<-METHYLASE<-TUDOR+ SF1-SF2-Helicase+iREase+CTD+REase\|\|VapB_antitoxin->PIN-><-DUF397<-HTH+DUF5753 | 1139 | Bacteria>Terrabacteria_group>Actinomycetota>Actinomycetes>Kitasatosporales | GCF_900114955.1 | Streptomyces_sp._cf124 | BM134_RS11480 | 95921..99340 | 1761903 | DUF499-domain-containing-protein-[Streptomyces-sp.-cf124] |
| WP_250287638.1 | DUF499_1.RM.like.systems | VirB4-FtsK->?->?->?-><-HEPN+DUF499*<-METHYLASE<-TUDOR+ SF1-SF2-Helicase+iREase+CTD+REase\|\|?->DnaB+DnaB-> | 1135 | Bacteria>Terrabacteria_group>Actinomycetota>Actinomycetes>Kitasatosporales | GCF_928722305.1 | Streptomyces_atroolivaceus | NBY68_RS04240 | 83190..86597 | 66869 | Swt1-family-HEPN-domain-containing-protein-[Streptomyces-atroolivaceus] |
| WP_052687173.1 | DUF499_1.RM.like.systems | <-PIN<-Phage_integrase+Phage_integrase\|\|HTH_3-><-?<-DUF6731<-HEPN+DUF499*<-METHYLASE<-TUDOR+ SF1-SF2-Helicase+iREase+CTD+REase<-?\|\|REase-> | 1168 | Bacteria>Terrabacteria_group>Actinomycetota>Actinomycetes>Kitasatosporales | - | Streptomyces_lydicus | T261_RS23850 | 5194255..5197761 | 47763 | DUF499-domain-containing-protein-[Streptomyces-lydicus] |
| WP_167559767.1 | DUF499_1.RM.like.systems | <-HEPN+REase<-REase<-HEPN+DUF499*<-wHTH+REase<-METHYLASE<-TUDOR+ SF1-SF2-Helicase+iREase+CTD+REase<-XIS-HTH\|\|MORC+HisKin+REase-><-TrwC+SF1-SF2-helicase | 1132 | Bacteria>Terrabacteria_group>Actinomycetota>Actinomycetes>Micrococcales | - | Brevibacterium_sp._LS14 | EZE58_RS02710 | 16654..20052 | 2528962 | DUF499-domain-containing-protein-[Brevibacterium-sp.-LS14] |
| AFR31400.1 | DUF499_1.RM.like.systems | <-NTD+SF1-SF2-Helicase\|\|TUDOR+ SF1-SF2-Helicase+iREase+CTD+REase->METHYLASE->HEPN+DUF499*-><-HEPN\|\|Phage_int_SAM_4->HEPN-> | 1120 | Bacteria>Terrabacteria_group>Actinomycetota>Actinomycetes>Micrococcales | GCA_000294695.2 | Arthrobacter_sp._Rue61a | ARUE_232p01920 | 197474..200836 | 1118963 | hypothetical-protein-ARUE_232p01920-[Arthrobacter-sp.-Rue61a] |
| KQY96844.1 | DUF499_1.RM.like.systems | Phage_holin->METHYLASE->?->ACET->HEPN+DUF499*->DUF4096->DDE_Tnp_1_2-><-UDG-conflict<-?<-REase<-MobL | 1102 | Bacteria>Terrabacteria_group>Actinomycetota>Actinomycetes>Micrococcales | GCA_001426925.1 | Microbacterium_sp._Root53 | ASD19_09760 | 99084..102392 | 1736553 | AAA-family-ATPase-[Microbacterium-sp.-Root53] |
| OFE15977.1 | DUF499_1.RM.like.systems | CopG/RHH->?->tRNA-><-SF1-SF2-Helicase<-?<-HEPN+DUF499*<-ParB+HNH<-METHYLASE<-TUDOR+ SF1-SF2-Helicase+iREase+CTD+REase<-?<-wHTH(SBD)+HNH<-REase+SF1-SF2-Helicase | 1100 | Bacteria>Terrabacteria_group>Actinomycetota>Actinomycetes>Micrococcales | GCA_001758225.1 | Humibacillus_sp._DSM_29435 | BA895_05295 | 164736..168038 | 1869167 | AAA-family-ATPase-[Humibacillus-sp.-DSM-29435] |
| MAT19458.1 | DUF499_1.RM.like.systems | <-ABhydrolase<-?\|\|?->?-><-SLATT+SLATT<-TIR<-HEPN+DUF499*<-METHYLASE<-TUDOR+ SF1-SF2-Helicase+iREase+CTD+REase<-XIS-HTH | 1130 | Bacteria>Terrabacteria_group>Actinomycetota>Actinomycetes>Micrococcales | GCA_002702785.1 | Leifsonia_sp. | CMF56_13050 | 20496..23888 | 1870902 | AAA-family-ATPase-[Leifsonia-sp.] |
| PPH45732.1 | DUF499_1.RM.like.systems | MACRODOMAIN+SIR2->TUDOR+ SF1-SF2-Helicase+iREase+CTD+REase->METHYLASE->HEPN+DUF499*-><-?<-?\|\|?->?->Peptidase->Peptidase_M73-> | 1120 | Bacteria>Terrabacteria_group>Actinomycetota>Actinomycetes>Micrococcales | GCA_002932315.1 | Rathayibacter_sp._AY1C9 | C5D09_10000 | 51639..55001 | 2080541 | AAA-family-ATPase-[Rathayibacter-sp.-AY1C9] |
| PZU36627.1 | DUF499_1.RM.like.systems | <-MODE-HTH+PBPII\|\|?->?->?->DUF4096+DDE_Tnp_1+DDE_Tnp_1_2-><-HEPN+DUF499*<-METHYLASE-p | 1129 | Bacteria>Terrabacteria_group>Actinomycetota>Actinomycetes>Micrococcales | GCA_003248605.1 | Microbacterium_sp. | DI573_13315 | 12880..16269 | 51671 | AAA-family-ATPase-[Microbacterium-sp.] |
| PZU44600.1 | DUF499_1.RM.like.systems | DnaB+SMS-KaiC->?->?-><-SLATT+SLATT<-TIR<-HEPN+DUF499*<-METHYLASE<-TUDOR+ SF1-SF2-Helicase+iREase+CTD+REase<-XIS-HTH | 1131 | Bacteria>Terrabacteria_group>Actinomycetota>Actinomycetes>Micrococcales | GCA_003248655.1 | Microbacterium_sp. | DI566_12010 | 17678..21073 | 51671 | AAA-family-ATPase-[Microbacterium-sp.] |
| PZU44169.1 | DUF499_1.RM.like.systems | TetR-HTH-><-?<-?\|\|?->\|\|HEPN+DUF499*->OLD-ABC+TOPRIM-><-?<-HIN-HTH\|\|HTH_19->Phd-YefM->PIN-> | 1129 | Bacteria>Terrabacteria_group>Actinomycetota>Actinomycetes>Micrococcales | GCA_003248655.1 | Microbacterium_sp. | DI566_12710 | 6913..10302 | 51671 | AAA-family-ATPase-[Microbacterium-sp.] |
| HBS75978.1 | DUF499_1.RM.like.systems | <-TrwC+SF1-SF2-Helicase<-wHTH+REase<-?\|\|NTD+SF1-SF2-Helicase-><-HEPN+DUF499* | 1125 | Bacteria>Terrabacteria_group>Actinomycetota>Actinomycetes>Micrococcales | GCA_003510265.1 | Microbacterium_sp. | DEB55_16590 | 13006..16383 | 51671 | AAA-family-ATPase-[Microbacterium-sp.] |
| QAY71606.1 | DUF499_1.RM.like.systems | <-AbiEi_4<-PIN<-XIS-HTH<-HEPN<-HEPN+DUF499*<-ParB+HNH<-METHYLASE<-?<-TUDOR+ SF1-SF2-Helicase+iREase+CTD+REase\|\|NTD+SF1-SF2-Helicase-> | 1123 | Bacteria>Terrabacteria_group>Actinomycetota>Actinomycetes>Micrococcales | GCA_004135385.1 | Xylanimonas_protaetiae | ET471_17505 | 3776288..3779659 | 2509457 | ATP-binding-protein-[Xylanimonas-protaetiae] |
| TFC67798.1 | DUF499_1.RM.like.systems | UDG-conflict-><-HTH+SF1-SF2-Helicase+REase<-REase<-MORC<-HEPN+DUF499*<-METHYLASE\|\|MORC+SIGMA-HTH->ASCH->?->DDE_Tnp_1->XIS-HTH-> | 1124 | Bacteria>Terrabacteria_group>Actinomycetota>Actinomycetes>Micrococcales | GCA_004402575.1 | Cryobacterium_sp._TMT2-4 | E3054_08285 | 76881..80255 | 1259254 | ATP-binding-protein-[Cryobacterium-sp.-TMT2-4] |
| TLM86009.1 | DUF499_1.RM.like.systems | HKD-DNase+SF1-SF2-Helicase+TRD->?->?->?->?-><-HEPN+DUF499*<-METHYLASE<-TUDOR+ SF1-SF2-Helicase+iREase+CTD+REase<-XIS-HTH<-SIR2+SIR2<-?\|\|XIS-HTH-> | 1120 | Bacteria>Terrabacteria_group>Actinomycetota>Actinomycetes>Micrococcales | GCA_005796225.1 | Pseudarthrobacter_sp._NamE5 | FDW84_06950 | 47724..51086 | 2576839 | ATP-binding-protein-[Pseudarthrobacter-sp.-NamE5] |
| QEO13112.1 | DUF499_1.RM.like.systems | VirB4-FtsK->?->VirD4-TraG+VirD4-FtsK->?->?->?-><-HEPN+DUF499*<-METHYLASE<-TUDOR+ SF1-SF2-Helicase+iREase+CTD+REase<-Phage_int_SAM_1+Phage_integrase<-XIS-HTH | 1128 | Bacteria>Terrabacteria_group>Actinomycetota>Actinomycetes>Micrococcales | GCA_008365295.1 | Agromyces_intestinalis | FLP10_00780 | 160366..163752 | 2592652 | ATP-binding-protein-[Agromyces-intestinalis] |
| MCL1800684.1 | DUF499_1.RM.like.systems | REase->PhdYeFM_antitox->RelE-ParE->METHYLASE->HEPN+DUF499*-> | 635 | Bacteria>Terrabacteria_group>Actinomycetota>Actinomycetes>Micrococcales | GCA_009782595.1 | Promicromonosporaceae_bacterium | FWG25_04890 | 4477..6383 | 2304674 | DUF499-domain-containing-protein-[Promicromonosporaceae-bacterium] |
| NDC48896.1 | DUF499_1.RM.like.systems | TUDOR+ SF1-SF2-Helicase+iREase+CTD+REase->METHYLASE->HEPN+DUF499*-> | 1108 | Bacteria>Terrabacteria_group>Actinomycetota>Actinomycetes>Micrococcales | GCA_010027415.1 | Micrococcales_bacterium | EBZ61_07440 | 7302..10628 | 2026762 | ATP-binding-protein-[Micrococcales-bacterium] |
| MBD1536843.1 | DUF499_1.RM.like.systems | HNH-><-Glyoxalase<-?\|\|ACET->TUDOR+ SF1-SF2-Helicase+iREase+CTD+REase->METHYLASE->HEPN+DUF499*->NTD+SF1-SF2-Helicase-> | 1126 | Bacteria>Terrabacteria_group>Actinomycetota>Actinomycetes>Micrococcales | GCA_014694415.1 | Arthrobacter_sp._S13_S34 | HC749_01495 | 347639..351019 | 2720227 | ATP-binding-protein-[Arthrobacter-sp.-S13_S34] |
| MBD3753304.1 | DUF499_1.RM.like.systems | <-HEPN+DUF499*<-METHYLASE-p | 452 | Bacteria>Terrabacteria_group>Actinomycetota>Actinomycetes>Micrococcales | GCA_014763555.1 | Micrococcales_bacterium | IE935_13015 | 1..1358 | 2026762 | ATP-binding-protein-[Micrococcales-bacterium] |
| MBK6441773.1 | DUF499_1.RM.like.systems | PrmC_N+Methyltransf_25-><-?<-PadR-HTH\|\|TUDOR+ SF1-SF2-Helicase+iREase+CTD+REase->METHYLASE->ParB+HNH->HEPN+DUF499*-> | 922 | Bacteria>Terrabacteria_group>Actinomycetota>Actinomycetes>Micrococcales | GCA_016704565.1 | Candidatus_Phosphoribacter_baldrii | IPF90_06370 | 1387205..1389971 | 2953744 | DUF499-domain-containing-protein-[Actinomycetales-bacterium] |
| MBK6299925.1 | DUF499_1.RM.like.systems | <-ABC-ATPase+SbcC<-ABC-ATPase+SbcC<-TetR-HTH<-NTD+SF1-SF2-Helicase<-HEPN+DUF499*<-Schlafen+HATPase+HTH<-METHYLASE<-TUDOR+ SF1-SF2-Helicase+iREase+CTD+REase\|\|PadR-HTH->?-><-PrmC_N+Methyltransf_25 | 1117 | Bacteria>Terrabacteria_group>Actinomycetota>Actinomycetes>Micrococcales | GCA_016705565.1 | Candidatus_Phosphoribacter_hodrii | IPF40_02340 | 519882..523235 | 2953743 | DUF499-domain-containing-protein-[Actinomycetales-bacterium] |
| MBK6954412.1 | DUF499_1.RM.like.systems | ArsR-HTH->PrmC_N+Methyltransf_25-><-?<-PadR-HTH\|\|TUDOR+ SF1-SF2-Helicase+iREase+CTD+REase->ParB+HNH->HEPN+DUF499*->NTD+SF1-SF2-Helicase->TetR-HTH->ABC-ATPase+SbcC->ABC-ATPase+SbcC-> | 1118 | Bacteria>Terrabacteria_group>Actinomycetota>Actinomycetes>Micrococcales | GCA_016708855.1 | Candidatus_Phosphoribacter_baldrii | IPH27_02570 | 528671..532027 | 2953744 | DUF499-domain-containing-protein-[Actinomycetales-bacterium] |
| MBK8435770.1 | DUF499_1.RM.like.systems | Phage_int_SAM_1+Phage_integrase->Phage_integrase->?->DDE_Tnp_1-><-HEPN+REase<-REase<-HEPN+DUF499*<-wHTH+REase<-METHYLASE<-XIS-HTH<-ABhydrolase<-MPTase<-HTH | 1139 | Bacteria>Terrabacteria_group>Actinomycetota>Actinomycetes>Micrococcales | GCA_016711265.1 | Austwickia_sp. | IPL37_04405 | 982795..986214 | 1872664 | DUF499-domain-containing-protein-[Austwickia-sp.] |
| MBN2177583.1 | DUF499_1.RM.like.systems | <-AbiEii<-AbiEi_4<-?<-HEPN+DUF499*<-METHYLASE\|\|?->HTH_31->?-><-TUDOR+ SF1-SF2-Helicase+iREase+CTD+REase | 1113 | Bacteria>Terrabacteria_group>Actinomycetota>Actinomycetes>Micrococcales | GCA_016933155.1 | Demequinaceae_bacterium | JW722_08015 | 69321..72662 | 2683256 | DUF499-domain-containing-protein-[Demequinaceae-bacterium] |
| MBN9213516.1 | DUF499_1.RM.like.systems | <-HTH+SF1-SF2-Helicase+REase<-REase<-MORC<-HEPN+DUF499*<-METHYLASE\|\|MORC+SIGMA-HTH->DNA_methylase->?->Phage_int_SAM_1+Phage_integrase-><-tRNA\|\|DUF222+HNH-> | 1119 | Bacteria>Terrabacteria_group>Actinomycetota>Actinomycetes>Micrococcales | GCA_017306385.1 | Microbacterium_sp. | JOJ04_01635 | 353134..356493 | 51671 | DUF499-domain-containing-protein-[Microbacterium-sp.] |
| MBS3179176.1 | DUF499_1.RM.like.systems | GDResolvase+HIN-HTH-><-?<-HNH\|\|TUDOR+ SF1-SF2-Helicase+iREase+CTD+REase->METHYLASE->HEPN+DUF499*->?-><-UDG-conflict\|\|?-><-?<-EVE | 1142 | Bacteria>Terrabacteria_group>Actinomycetota>Actinomycetes>Micrococcales | GCA_018304855.1 | Pseudoclavibacter_sp._Marseille-Q4354 | JSQ82_10750 | 821869..825297 | 2810515 | DUF499-domain-containing-protein-[Pseudoclavibacter-sp.-Marseille-Q4354] |
| MBW8172946.1 | DUF499_1.RM.like.systems | GNTR-HTH-><-?<-?<-DNA_methylase\|\|SIGMA-HTH->METHYLASE->HEPN+DUF499*->MORC->REase->HTH+SF1-SF2-Helicase+REase->NERD+SF1-SF2-Helicase-><-?<-ABC-ATPase+SbcC | 1128 | Bacteria>Terrabacteria_group>Actinomycetota>Actinomycetes>Micrococcales | GCA_019457945.1 | Ornithinimicrobium_laminariae | KO651_07785 | 644333..647719 | 2864873 | DUF499-domain-containing-protein-[Ornithinimicrobium-laminariae] |
| MCE1178998.1 | DUF499_1.RM.like.systems | ABhydrolase->XIS-HTH->TUDOR+ SF1-SF2-Helicase+iREase+CTD+REase->METHYLASE->wHTH+REase->HEPN+DUF499*->REase->?->?-><-DNA_methylase\|\|MULE-TRANSPOSASE-> | 1133 | Bacteria>Terrabacteria_group>Actinomycetota>Actinomycetes>Micrococcales | GCA_021287075.1 | Micrococcales_bacterium | LWW86_08210 | 123178..126579 | 2026762 | DUF499-domain-containing-protein-[Micrococcales-bacterium] |
| MCE7482742.1 | DUF499_1.RM.like.systems | REase+SF1-SF2-Helicase-><-SbcC<-HEPN+DUF499*<-METHYLASE-p | 1108 | Bacteria>Terrabacteria_group>Actinomycetota>Actinomycetes>Micrococcales | GCA_021395155.1 | Microbacterium_profundi | LZG07_12530 | 143258..146584 | 450380 | DUF499-domain-containing-protein-[Microbacterium-profundi] |
| MCI1262624.1 | DUF499_1.RM.like.systems | <-wHTH(SBD)+HNH<-Glyoxalase\|\|?-><-?<-HEPN<-PolB<-HEPN+DUF499*<-ParB+HNH<-METHYLASE<-TUDOR+ SF1-SF2-Helicase+iREase+CTD+REase\|\|NTD+SF1-SF2-Helicase-> | 1116 | Bacteria>Terrabacteria_group>Actinomycetota>Actinomycetes>Micrococcales | GCA_022641475.1 | Tetrasphaera_jenkinsii | LKG20_10180 | 13344..16694 | 330834 | Swt1-family-HEPN-domain-containing-protein-[Tetrasphaera-jenkinsii] |
| MCV7624012.1 | DUF499_1.RM.like.systems | <-REase<-?<-?<-REase\|\|DNA_methylase->METHYLASE->HEPN+DUF499*->MORC->REase->HTH+SF1-SF2-Helicase+REase-><-Peptidase_S66+Peptidase_S66C | 1157 | Bacteria>Terrabacteria_group>Actinomycetota>Actinomycetes>Micrococcales | GCA_025145605.2 | Micrococcus_luteus | M3B28_003235 | 32627..36100 | 1270 | Swt1-family-HEPN-domain-containing-protein-[Micrococcus-luteus] |
| MCV7527924.1 | DUF499_1.RM.like.systems | <-REase\|\|DNA_methylase->METHYLASE->HEPN+DUF499*->MORC->REase-> | 1157 | Bacteria>Terrabacteria_group>Actinomycetota>Actinomycetes>Micrococcales | GCA_025149725.2 | Micrococcus_luteus | M3D91_010335 | 6032..9505 | 1270 | Swt1-family-HEPN-domain-containing-protein-[Micrococcus-luteus] |
| MCV7506294.1 | DUF499_1.RM.like.systems | <-HTH+SF1-SF2-Helicase+REase<-REase<-MORC<-HEPN+DUF499*<-METHYLASE<-DNA_methylase\|\|REase-> | 1157 | Bacteria>Terrabacteria_group>Actinomycetota>Actinomycetes>Micrococcales | GCA_025151425.2 | Micrococcus_luteus | M3D82_001030 | 202363..205836 | 1270 | Swt1-family-HEPN-domain-containing-protein-[Micrococcus-luteus] |
| MCU1509501.1 | DUF499_1.RM.like.systems | <-SbcC<-DUF1679<-SIR2<-REase<-URI-DNase<-?<-HEPN+DUF499*<-METHYLASE<-STYKinase-like<-HTH_3\|\|TUDOR+ SF1-SF2-Helicase+iREase+CTD+REase-><-NTD+SF1-SF2-Helicase | 1121 | Bacteria>Terrabacteria_group>Actinomycetota>Actinomycetes>Micrococcales | GCA_025464095.1 | Glaciihabitans_sp. | JWQ12_1766 | 110555..113920 | 1965328 | hypothetical-protein-[Glaciihabitans-sp.] |
| CEA09674.1 | DUF499_1.RM.like.systems | <-McrC-NTD+REase<-?\|\|TUDOR+ SF1-SF2-Helicase+iREase+CTD+REase-><-?<-?\|\|METHYLASE-p->HEPN+DUF499*-><-?<-HKD-DNase+SF1-SF2-Helicase+TRD\|\|URI-DNase-><-SF1-SF2-Helicase\|\|NERD+SF1-SF2-Helicase-> | 1141 | Bacteria>Terrabacteria_group>Actinomycetota>Actinomycetes>Micrococcales | GCA_900016675.1 | Arthrobacter_saudimassiliensis | BN1051_03046 | 420821..424246 | 1461584 | hypothetical-protein-BN1051_03046-[Arthrobacter-saudimassiliensis] |
| SDF11628.1 | DUF499_1.RM.like.systems | <-wHTH(SBD)+HNH<-HTH_3<-PIN<-MPTase\|\|METHYLASE->ParB+HNH->HEPN+DUF499*-> | 1151 | Bacteria>Terrabacteria_group>Actinomycetota>Actinomycetes>Micrococcales | GCA_900102445.1 | Cellulosimicrobium_cellulans | SAMN04487781_0340 | 373492..376947 | 1710 | Predicted-ATPase,-AAA+-superfamily-[Cellulosimicrobium-cellulans] |
| SNS99685.1 | DUF499_1.RM.like.systems | <-HTH_19\|\|TUDOR+ SF1-SF2-Helicase+iREase+CTD+REase->Phd-YefM->RelE-ParE->METHYLASE->HEPN+DUF499*->?-><-?\|\|?->?->?->HNH-> | 1175 | Bacteria>Terrabacteria_group>Actinomycetota>Actinomycetes>Micrococcales | GCA_900188465.1 | Micrococcales_bacterium_KH10 | SAMN06309944_1702 | 505184..508711 | 1945885 | Predicted-ATPase,-AAA+-superfamily-[Micrococcales-bacterium-KH10] |
| WP_014923538.1 | DUF499_1.RM.like.systems | <-NTD+SF1-SF2-Helicase\|\|TUDOR+ SF1-SF2-Helicase+iREase+CTD+REase->METHYLASE->HEPN+DUF499*-><-HEPN\|\|Phage_int_SAM_4-> | 1120 | Bacteria>Terrabacteria_group>Actinomycetota>Actinomycetes>Micrococcales | GCF_000294695.2 | Arthrobacter_sp._Rue61a | ARUE_RS22630 | 197474..200836 | 1118963 | Swt1-family-HEPN-domain-containing-protein-[Arthrobacter-sp.-Rue61a] |
| WP_009381273.1 | DUF499_1.RM.like.systems | <-HEPN+REase<-REase<-HEPN+DUF499*<-wHTH+REase<-METHYLASE | 1132 | Bacteria>Terrabacteria_group>Actinomycetota>Actinomycetes>Micrococcales | GCF_000314575.1 | Brevibacterium_casei | C272_RS16125 | 18996..22394 | 33889 | DUF499-domain-containing-protein-[Brevibacterium-casei] |
| WP_026937806.1 | DUF499_1.RM.like.systems | <-HEPN+DUF499*<-METHYLASE\|\|DNA_methylase-> | 1131 | Bacteria>Terrabacteria_group>Actinomycetota>Actinomycetes>Micrococcales | GCF_000425685.1 | Gulosibacter_molinativorax | C7K25_RS15310 | 538..3933 | 256821 | Swt1-family-HEPN-domain-containing-protein-[Gulosibacter-molinativorax] |
| WP_029212622.1 | DUF499_1.RM.like.systems | SF1-SF2-Helicase->tRNA->TUDOR+ SF1-SF2-Helicase+iREase+CTD+REase->METHYLASE->ParB+HNH->HEPN+DUF499*-> | 1119 | Bacteria>Terrabacteria_group>Actinomycetota>Actinomycetes>Micrococcales | GCF_000426385.1 | Arsenicicoccus_bolidensis | K313_RS0116910 | 19529..22888 | 229480 | Swt1-family-HEPN-domain-containing-protein-[Arsenicicoccus-bolidensis] |
| WP_034802772.1 | DUF499_1.RM.like.systems | REase+SF1-SF2-Helicase->wHTH(SBD)+HNH->DndB->TUDOR+ SF1-SF2-Helicase+iREase+CTD+REase->METHYLASE->ParB+HNH->HEPN+DUF499*->?->SF1-SF2-Helicase-> | 1107 | Bacteria>Terrabacteria_group>Actinomycetota>Actinomycetes>Micrococcales | GCF_000576595.1 | Intrasporangium_oryzae | N865_RS05960 | 110230..113553 | 412687 | DUF499-domain-containing-protein-[Intrasporangium-oryzae] |
| WP_036294304.1 | DUF499_1.RM.like.systems | DnaB+SMS-KaiC->?-><-SLATT+SLATT<-TIR<-HEPN+DUF499*<-METHYLASE<-TUDOR+ SF1-SF2-Helicase+iREase+CTD+REase<-XIS-HTH | 1131 | Bacteria>Terrabacteria_group>Actinomycetota>Actinomycetes>Micrococcales | GCF_000582705.2 | Microbacterium | MIC448_RS00375 | 9420..12815 | 33882 | MULTISPECIES:-DUF499-domain-containing-protein-[Microbacterium] |
| WP_034226753.1 | DUF499_1.RM.like.systems | <-HTH+SF1-SF2-Helicase+REase<-REase<-MORC<-HEPN+DUF499*<-METHYLASE | 1144 | Bacteria>Terrabacteria_group>Actinomycetota>Actinomycetes>Micrococcales | GCF_000603945.1 | Actinotalea_ferrariae | N866_RS12275 | 5788..9222 | 1386098 | Swt1-family-HEPN-domain-containing-protein-[Actinotalea-ferrariae] |
| WP_029263985.1 | DUF499_1.RM.like.systems | <-Radical-SAM\|\|?-><-SF1-SF2-Helicase<-REase<-HEPN+DUF499*<-METHYLASE<-TUDOR+ SF1-SF2-Helicase+iREase+CTD+REase\|\|?-><-MerR-HTH | 1128 | Bacteria>Terrabacteria_group>Actinomycetota>Actinomycetes>Micrococcales | GCF_000708325.1 | unclassified_Microbacterium | DW303_RS18520 | 62853..66239 | 2609290 | MULTISPECIES:-DUF499-domain-containing-protein-[unclassified-Microbacterium] |
| WP_047687850.1 | DUF499_1.RM.like.systems | DNA_methylase-><-HEPN+REase<-REase<-HEPN+DUF499*<-wHTH+REase<-METHYLASE<-TUDOR+ SF1-SF2-Helicase+iREase+CTD+REase | 1133 | Bacteria>Terrabacteria_group>Actinomycetota>Actinomycetes>Micrococcales | GCF_000800955.1 | Kocuria_sp._ZOR0020 | L979_RS00020 | 3003..6404 | 1339234 | DUF499-domain-containing-protein-[Kocuria-sp.-ZOR0020] |
| WP_047691765.1 | DUF499_1.RM.like.systems | <-DNA_methylase\|\|REase->METHYLASE->HEPN+DUF499*-> | 452 | Bacteria>Terrabacteria_group>Actinomycetota>Actinomycetes>Micrococcales | GCF_000800955.1 | Kocuria_sp._ZOR0020 | L979_RS08865 | 18602..19957 | 1339234 | Swt1-family-HEPN-domain-containing-protein,-partial-[Kocuria-sp.-ZOR0020] |
| WP_043123141.1 | DUF499_1.RM.like.systems | <-NTD+SF1-SF2-Helicase\|\|TUDOR+ SF1-SF2-Helicase+iREase+CTD+REase->METHYLASE->HEPN+DUF499*-><-TrwC+SF1-SF2-helicase\|\|?->?->?-><-?<-ParB+HNH | 1115 | Bacteria>Terrabacteria_group>Actinomycetota>Actinomycetes>Micrococcales | GCF_000802235.1 | Sinomonas_humi | LK10_RS10225 | 159632..162979 | 1338436 | Swt1-family-HEPN-domain-containing-protein-[Sinomonas-humi] |
| WP_044658741.1 | DUF499_1.RM.like.systems | Peptidase_S66+Peptidase_S66C-><-HTH+SF1-SF2-Helicase+REase<-REase<-MORC<-HEPN+DUF499*<-METHYLASE<-DNA_methylase\|\|REase-> | 1157 | Bacteria>Terrabacteria_group>Actinomycetota>Actinomycetes>Micrococcales | GCF_000946635.1 | Micrococcus | B8W89_RS05705 | 32626..36099 | 1269 | MULTISPECIES:-Swt1-family-HEPN-domain-containing-protein-[Micrococcus] |
| WP_062529103.1 | DUF499_1.RM.like.systems | <-REase+SF1-SF2-Helicase\|\|MazG-><-ParB+HNH<-HTH+SF1-SF2-Helicase+REase<-REase<-MORC<-HEPN+DUF499*<-METHYLASE\|\|DNA_methylase-> | 1118 | Bacteria>Terrabacteria_group>Actinomycetota>Actinomycetes>Micrococcales | GCF_000971455.1 | Demequina_rhizosphaerae | AOM12_RS04465 | 350500..353856 | 1638985 | DUF499-domain-containing-protein-[Demequina-rhizosphaerae] |
| WP_046529475.1 | DUF499_1.RM.like.systems | XIS-HTH->PIN->HNH-><-HTH+SF1-SF2-Helicase+REase<-REase<-MORC<-HEPN+DUF499*<-METHYLASE<-REase\|\|DNA_methylase-><-OLD-ABC+TOPRIM<-DNA-glycosylase | 1124 | Bacteria>Terrabacteria_group>Actinomycetota>Actinomycetes>Micrococcales | GCF_000986825.1 | Cellulomonas_sp._FA1 | GY42_RS11845 | 2630948..2634322 | 1346710 | Swt1-family-HEPN-domain-containing-protein-[Cellulomonas-sp.-FA1] |
| WP_048693914.1 | DUF499_1.RM.like.systems | <-Radical-SAM+TRAM\|\|?->?-><-DNA_methylase\|\|PIN->METHYLASE->HEPN+DUF499*->MORC->REase->HTH+SF1-SF2-Helicase+REase->?->CR-Kinase+ORC-CDC6-like+CARF-> | 1134 | Bacteria>Terrabacteria_group>Actinomycetota>Actinomycetes>Micrococcales | GCF_001050535.1 | Tetrasphaera_australiensis | BN11_RS02235 | 472611..476015 | 99480 | DUF499-domain-containing-protein-[Tetrasphaera-australiensis] |
| WP_054682987.1 | DUF499_1.RM.like.systems | XIS-HTH->TUDOR+ SF1-SF2-Helicase+iREase+CTD+REase->METHYLASE->HIN-HTH->METHYLASE-p->HEPN+DUF499*->OLD-ABC+TOPRIM->HNH-><-HIN-HTH | 1129 | Bacteria>Terrabacteria_group>Actinomycetota>Actinomycetes>Micrococcales | GCF_001314225.1 | Microbacterium_sp._No._7 | AOA12_RS21690 | 98036..101425 | 1714373 | DUF499-domain-containing-protein-[Microbacterium-sp.-No.-7] |
| WP_055898767.1 | DUF499_1.RM.like.systems | <-wHTH+SAD+HNH\|\|NERD+SF1-SF2-Helicase-><-REase\|\|?->METHYLASE->HEPN+DUF499*->MORC->REase->HTH+SF1-SF2-Helicase+REase-><-?<-SF1-SF2-Helicase+REase<-Exonuc_V_gamma+REase | 1109 | Bacteria>Terrabacteria_group>Actinomycetota>Actinomycetes>Micrococcales | GCF_001425665.1 | Yonghaparkia_sp._Root332 | ASC54_RS09975 | 479255..482584 | 1736516 | DUF499-domain-containing-protein-[Yonghaparkia-sp.-Root332] |
| WP_200941118.1 | DUF499_1.RM.like.systems | Phage_holin->METHYLASE->?->ACET->HEPN+DUF499*->DUF4096+DDE_Tnp_1+DDE_Tnp_1_2-><-?<-REase<-MobL | 1112 | Bacteria>Terrabacteria_group>Actinomycetota>Actinomycetes>Micrococcales | GCF_001426925.1 | Microbacterium_sp._Root53 | ASD19_RS13260 | 99054..102392 | 1736553 | Swt1-family-HEPN-domain-containing-protein-[Microbacterium-sp.-Root53] |
| WP_197462106.1 | DUF499_1.RM.like.systems | HEPN+DUF499*->DUF499-p->OLD-ABC+TOPRIM-p->OLD-ABC+TOPRIM-p-> | 968 | Bacteria>Terrabacteria_group>Actinomycetota>Actinomycetes>Micrococcales | GCF_001595495.1 | Microbacterium_sp._CH1 | AUV07_RS03260 | 609..3515 | 1770208 | DUF499-domain-containing-protein-[Microbacterium-sp.-CH1] |
| WP_062860586.1 | DUF499_1.RM.like.systems | <-OLD-ABC+TOPRIM<-DNA_methylase\|\|DDE_Tnp_1_4->METHYLASE->HEPN+DUF499*->MORC->REase->HTH+SF1-SF2-Helicase+REase->SF1-SF2-Helicase+REase-><-?\|\|ZnR-> | 1132 | Bacteria>Terrabacteria_group>Actinomycetota>Actinomycetes>Micrococcales | GCF_001606005.1 | Brevibacterium_linens | A2T55_RS01955 | 443137..446535 | 1703 | DUF499-domain-containing-protein-[Brevibacterium-linens] |
| WP_063250773.1 | DUF499_1.RM.like.systems | <-HEPN+REase<-REase<-HEPN+DUF499*<-wHTH+REase<-METHYLASE<-TUDOR+ SF1-SF2-Helicase+iREase+CTD+REase<-XIS-HTH\|\|MORC+HisKin+REase-><-TrwC+SF1-SF2-helicase | 1132 | Bacteria>Terrabacteria_group>Actinomycetota>Actinomycetes>Micrococcales | GCF_001619685.1 | Brevibacterium_casei | AVW13_RS15020 | 17875..21273 | 33889 | DUF499-domain-containing-protein-[Brevibacterium-casei] |
| WP_069599267.1 | DUF499_1.RM.like.systems | <-SF1-SF2-Helicase+REase<-?<-HTH+SF1-SF2-Helicase+REase<-REase<-MORC<-HEPN+DUF499*<-METHYLASE<-REase<-MORC+HEPN\|\|DNA_methylase-><-?<-SF1-SF2-Helicase | 1131 | Bacteria>Terrabacteria_group>Actinomycetota>Actinomycetes>Micrococcales | GCF_001729525.1 | Brevibacterium_aurantiacum | CIK60_RS17585 | 12727..16122 | 273384 | DUF499-domain-containing-protein-[Brevibacterium-aurantiacum] |
| WP_139142668.1 | DUF499_1.RM.like.systems | CopG/RHH->?->CopG/RHH->tRNA-><-SF1-SF2-Helicase<-?<-HEPN+DUF499*<-ParB+HNH<-METHYLASE<-TUDOR+ SF1-SF2-Helicase+iREase+CTD+REase<-?<-wHTH(SBD)+HNH<-REase+SF1-SF2-Helicase | 1111 | Bacteria>Terrabacteria_group>Actinomycetota>Actinomycetes>Micrococcales | GCF_001758225.1 | Humibacillus_sp._DSM_29435 | BA895_RS14645 | 164736..168071 | 1869167 | Swt1-family-HEPN-domain-containing-protein-[Humibacillus-sp.-DSM-29435] |
| WP_076703521.1 | DUF499_1.RM.like.systems | <-HEPN+DUF499*<-wHTH+REase | 1131 | Bacteria>Terrabacteria_group>Actinomycetota>Actinomycetes>Micrococcales | GCF_001968835.1 | Tersicoccus_phoenicis | BKD30_RS07045 | 193..3588 | 554083 | Swt1-family-HEPN-domain-containing-protein-[Tersicoccus-phoenicis] |
| WP_232711167.1 | DUF499_1.RM.like.systems | Phage_holin->METHYLASE->?->?->HEPN+DUF499*->?->?->wHTH+REase->TrwC+SF1-SF2-Helicase-> | 1114 | Bacteria>Terrabacteria_group>Actinomycetota>Actinomycetes>Micrococcales | GCF_002812725.1 | unclassified_Microbacterium | CGU35_RS05955 | 370107..373451 | 2609290 | MULTISPECIES:-DUF499-domain-containing-protein-[unclassified-Microbacterium] |
| WP_101853235.1 | DUF499_1.RM.like.systems | <-HTH+SF1-SF2-Helicase+REase-p<-PAIREDC-HTH<-HTH+SF1-SF2-Helicase+REase-p<-REase<-MORC<-HEPN+DUF499*<-METHYLASE\|\|?->HTH+MPTase->PIN-> | 1144 | Bacteria>Terrabacteria_group>Actinomycetota>Actinomycetes>Micrococcales | GCF_002849405.1 | Kocuria_flava | AUQ48_RS15985 | 10331..13765 | 446860 | DUF499-domain-containing-protein-[Kocuria-flava] |
| WP_102209830.1 | DUF499_1.RM.like.systems | <-HEPN+DUF499*<-?<-METHYLASE<-Phage_holin<-?<-?<-TUDOR+ SF1-SF2-Helicase+iREase+CTD+REase | 1123 | Bacteria>Terrabacteria_group>Actinomycetota>Actinomycetes>Micrococcales | GCF_002871415.1 | Microbacterium_sp._UMB0228 | CJ226_RS16275 | 2567..5938 | 2029109 | DUF499-domain-containing-protein-[Microbacterium-sp.-UMB0228] |
| WP_104088073.1 | DUF499_1.RM.like.systems | <-HEPN+DUF499*<-METHYLASE<-TUDOR+ SF1-SF2-Helicase+iREase+CTD+REase<-XIS-HTH\|\|GDResolvase+HIN-HTH-> | 1133 | Bacteria>Terrabacteria_group>Actinomycetota>Actinomycetes>Micrococcales | GCF_002929595.1 | Cryobacterium_sp._N19 | CXY78_RS03325 | 298..3699 | 2048288 | DUF499-domain-containing-protein-[Cryobacterium-sp.-N19] |
| WP_104267247.1 | DUF499_1.RM.like.systems | <-GDResolvase+Recombinase+ZnR<-?\|\|ABhydrolase->XIS-HTH->TUDOR+ SF1-SF2-Helicase+iREase+CTD+REase->METHYLASE->HEPN+DUF499*->?->?->HerA/FtsK-> | 1135 | Bacteria>Terrabacteria_group>Actinomycetota>Actinomycetes>Micrococcales | GCF_002932235.1 | Rathayibacter_sp._AY1E3 | C5C53_RS07820 | 81195..84602 | 2080551 | Swt1-family-HEPN-domain-containing-protein-[Rathayibacter-sp.-AY1E3] |
| WP_258066970.1 | DUF499_1.RM.like.systems | MACRODOMAIN+SIR2->TUDOR+ SF1-SF2-Helicase+iREase+CTD+REase->METHYLASE->HEPN+DUF499*-><-?<-REase<-?<-?\|\|?->Peptidase_M73-> | 1112 | Bacteria>Terrabacteria_group>Actinomycetota>Actinomycetes>Micrococcales | GCF_002932315.1 | Rathayibacter_sp._AY1C9 | C5D09_RS10010 | 51663..55001 | 2080541 | Swt1-family-HEPN-domain-containing-protein-[Rathayibacter-sp.-AY1C9] |
| WP_105034124.1 | DUF499_1.RM.like.systems | TUDOR+ SF1-SF2-Helicase+iREase+CTD+REase->HTH_3->STYKinase-like->METHYLASE->HEPN+DUF499*->?-><-ABhydrolase<-?<-MPTase<-HTH\|\|Primpol+PriCT1-> | 1121 | Bacteria>Terrabacteria_group>Actinomycetota>Actinomycetes>Micrococcales | GCF_002954245.1 | Cryobacterium_aureum | CJ028_RS04835 | 402642..406007 | 995037 | Swt1-family-HEPN-domain-containing-protein-[Cryobacterium-aureum] |
| WP_106298858.1 | DUF499_1.RM.like.systems | MazG-><-?\|\|NTD+SF1-SF2-Helicase->TUDOR+ SF1-SF2-Helicase+iREase+CTD+REase->METHYLASE->ParB+HNH->HEPN+DUF499*->ACET->TraJ-RHH-> | 1112 | Bacteria>Terrabacteria_group>Actinomycetota>Actinomycetes>Micrococcales | GCF_003002895.1 | Knoellia_remsis | BCF74_RS15510 | 21779..25117 | 407159 | Swt1-family-HEPN-domain-containing-protein-[Knoellia-remsis] |
| WP_159096312.1 | DUF499_1.RM.like.systems | <-DDE_Tnp_1<-HEPN+DUF499*<-METHYLASE<-TUDOR+ SF1-SF2-Helicase+iREase+CTD+REase\|\|?-><-?<-?<-REase | 1126 | Bacteria>Terrabacteria_group>Actinomycetota>Actinomycetes>Micrococcales | GCF_003074035.1 | Miniimonas_sp._S16 | DCS86_RS00720 | 144486..147866 | 2171623 | DUF499-domain-containing-protein-[Miniimonas-sp.-S16] |
| WP_108718589.1 | DUF499_1.RM.like.systems | <-JetB<-JetA\|\|?->PIN->?-><-?<-HEPN+DUF499*<-ParB+HNH<-HEPN+HEPN<-METHYLASE\|\|HTH->Schlafen+HATPase+HTH-><-TUDOR+ SF1-SF2-Helicase+iREase+CTD+REase | 1134 | Bacteria>Terrabacteria_group>Actinomycetota>Actinomycetes>Micrococcales | GCF_003074035.1 | Miniimonas_sp._S16 | DCS86_RS08145 | 1885887..1889291 | 2171623 | Swt1-family-HEPN-domain-containing-protein-[Miniimonas-sp.-S16] |
| WP_109686041.1 | DUF499_1.RM.like.systems | PAIREDC-HTH->?->?->PAIREDC-HTH-><-?<-HEPN+DUF499*<-?<-METHYLASE<-TUDOR+ SF1-SF2-Helicase+iREase+CTD+REase\|\|NTD+SF1-SF2-Helicase->?-><-MazG | 1124 | Bacteria>Terrabacteria_group>Actinomycetota>Actinomycetes>Micrococcales | GCF_003149195.1 | Branchiibius_hedensis | ATK17_RS11720 | 2422832..2426206 | 672460 | Swt1-family-HEPN-domain-containing-protein-[Branchiibius-hedensis] |
| WP_112235018.1 | DUF499_1.RM.like.systems | <-MORC+HisKin+REase\|\|XIS-HTH->TUDOR+ SF1-SF2-Helicase+iREase+CTD+REase->METHYLASE->wHTH+REase->HEPN+DUF499*->OLD-ABC+TOPRIM->SF1-SF2-Helicase-> | 1132 | Bacteria>Terrabacteria_group>Actinomycetota>Actinomycetes>Micrococcales | GCF_003290245.1 | Kocuria_sp._BT304 | DNL27_RS06220 | 1382048..1385446 | 1702043 | DUF499-domain-containing-protein-[Kocuria-sp.-BT304] |
| WP_126874170.1 | DUF499_1.RM.like.systems | <-DNA_methylase\|\|REase->METHYLASE->HEPN+DUF499*->MORC->REase->HTH+SF1-SF2-Helicase+REase-><-?\|\|MarR-HTH-> | 1141 | Bacteria>Terrabacteria_group>Actinomycetota>Actinomycetes>Micrococcales | GCF_003989235.1 | unclassified_Kocuria | D8M38_RS00615 | 20820..24245 | 2649579 | MULTISPECIES:-DUF499-domain-containing-protein-[unclassified-Kocuria] |
| WP_128365158.1 | DUF499_1.RM.like.systems | Calcineurin->tRNA-><-Phage_integrase<-?<-HTH+SF1-SF2-Helicase+REase<-REase<-MORC<-HEPN+DUF499*<-METHYLASE<-REase\|\|DNA_methylase-><-?\|\|Peptidase_M48-> | 1135 | Bacteria>Terrabacteria_group>Actinomycetota>Actinomycetes>Micrococcales | GCF_004025965.2 | Phycicoccus_flavus | EPD83_RS11930 | 11514..14921 | 2502783 | DUF499-domain-containing-protein-[Phycicoccus-flavus] |
| WP_129190457.1 | DUF499_1.RM.like.systems | <-AbiEi_4<-PIN<-XIS-HTH<-HEPN<-HEPN+DUF499*<-ParB+HNH<-METHYLASE<-?<-TUDOR+ SF1-SF2-Helicase+iREase+CTD+REase\|\|NTD+SF1-SF2-Helicase-> | 1123 | Bacteria>Terrabacteria_group>Actinomycetota>Actinomycetes>Micrococcales | GCF_004135385.1 | Xylanimonas_protaetiae | ET471_RS17505 | 3776288..3779659 | 2509457 | Swt1-family-HEPN-domain-containing-protein-[Xylanimonas-protaetiae] |
| WP_208008530.1 | DUF499_1.RM.like.systems | <-MULE-TRANSPOSASE\|\|NTD+SF1-SF2-Helicase-><-HEPN+DUF499*<-REase<-METHYLASE<-Phage_holin<-?<-?<-TUDOR+ SF1-SF2-Helicase+iREase+CTD+REase | 1101 | Bacteria>Terrabacteria_group>Actinomycetota>Actinomycetes>Micrococcales | GCF_004135405.1 | Agromyces_protaetiae | ET445_RS03710 | 798698..802003 | 2509455 | Swt1-family-HEPN-domain-containing-protein-[Agromyces-protaetiae] |
| WP_131132843.1 | DUF499_1.RM.like.systems | <-HTH+SF1-SF2-Helicase+REase<-REase<-MORC<-HEPN+DUF499*<-METHYLASE\|\|DNA_methylase-><-SbcC<-DUF1679<-SIR2 | 1130 | Bacteria>Terrabacteria_group>Actinomycetota>Actinomycetes>Micrococcales | GCF_004320525.1 | Arthrobacter_sp._S39 | EYS21_RS09020 | 558026..561418 | 2509720 | Swt1-family-HEPN-domain-containing-protein-[Arthrobacter-sp.-S39] |

| Protein-ID | System | Gene-Neighbourhoods | Length | Taxonomy | Assembly-ID | Strain | Locus-Tag | Genomic-coordinates | TaxID | Fasta-define |
|---|---|---|---|---|---|---|---|---|---|---|
| WP_133351310.1 | DUF499_1.RM.like.systems | <-XIS-HTH<-NTD+SF1-SF2-Helicase\|\|TUDOR+SF1-SF2-Helicase+iREase+CTD+REase->METHYLASE->HEPN+DUF499*-><-?<-HIN-HTH | 1123 | Bacteria>Terrabacteria_group>Actinomycetota>Actinomycetes>Micrococcales | GCF_004358765.1 | Arthrobacter_nitrophenolicus | E2R57_RS17850 | 242800..246171 | 683150 | Swt1-family-HEPN-domain-containing-protein-[Arthrobacter-nitrophenolicus] |
| WP_134401373.1 | DUF499_1.RM.like.systems | UDG-conflict-><-HTH+SF1-SF2-Helicase+REase<-REase<-MORC<-HEPN+DUF499*<-METHYLASE\|\|MORC+SIGMA-HTH->ASCH->?->DDE_Tnp_1->XIS-HTH-> | 1124 | Bacteria>Terrabacteria_group>Actinomycetota>Actinomycetes>Micrococcales | GCF_004402575.1 | Cryobacterium_sp._TMT2-4 | E3054_RS08280 | 76881..80255 | 1259254 | Swt1-family-HEPN-domain-containing-protein-[Cryobacterium-sp.-TMT2-4] |
| WP_137123200.1 | DUF499_1.RM.like.systems | ABhydrolase->XIS-HTH->TUDOR+SF1-SF2-Helicase+iREase+CTD+REase->METHYLASE->wHTH+REase->HEPN+DUF499*->REase->HEPN+REase-><-DNA_methylase\|\|TrwC+SF1-SF2-Helicase->?->SMODS-> | 1132 | Bacteria>Terrabacteria_group>Actinomycetota>Actinomycetes>Micrococcales | GCF_005222685.1 | Ornithinicoccus_soli | FCN22_RS19310 | 12208..15606 | 2574732 | DUF499-domain-containing-protein-[Ornithinicoccus-soli] |
| WP_137320273.1 | DUF499_1.RM.like.systems | <-HEPN+DUF499*<-wHTH+REase<-METHYLASE<-TUDOR+SF1-SF2-Helicase+iREase+CTD+REase<-XIS-HTH<-ABhydrolase | 1132 | Bacteria>Terrabacteria_group>Actinomycetota>Actinomycetes>Micrococcales | GCF_005280295.1 | Brevibacterium_sp._CS2 | FDF13_RS00005 | 3239176..2834 | 2575923 | DUF499-domain-containing-protein-[Brevibacterium-sp.-CS2] |
| WP_137369040.1 | DUF499_1.RM.like.systems | <-NTD+SF1-SF2-Helicase\|\|TUDOR+SF1-SF2-Helicase+iREase+CTD+REase->METHYLASE->HEPN+DUF499*->REase+CTD6->?->HNH-><-URI-DNase\|\|URI-DNase-><-JetD | 1129 | Bacteria>Terrabacteria_group>Actinomycetota>Actinomycetes>Micrococcales | GCF_005281365.1 | Arthrobacter_sp._NamB2 | FDK12_RS07770 | 280727..284116 | 2576035 | Swt1-family-HEPN-domain-containing-protein-[Arthrobacter-sp.-NamB2] |
| WP_138608335.1 | DUF499_1.RM.like.systems | HKD-DNase+SF1-SF2-Helicase+TRD->?->?->?->?-><-HEPN+DUF499*<-METHYLASE<-TUDOR+SF1-SF2-Helicase+iREase+CTD+REase<-XIS-HTH<-SIR2+SIR2<-?\|\|XIS-HTH-> | 1120 | Bacteria>Terrabacteria_group>Actinomycetota>Actinomycetes>Micrococcales | GCF_005796225.1 | Pseudarthrobacter_sp._NamE5 | FDW84_RS08080 | 47724..51086 | 2576839 | DUF499-domain-containing-protein-[Pseudarthrobacter-sp.-NamE5] |
| WP_141391091.1 | DUF499_1.RM.like.systems | <-HEPN+DUF499*<-METHYLASE<-TUDOR+SF1-SF2-Helicase+iREase+CTD+REase<-tRNA<-Aminotran_4 | 1129 | Bacteria>Terrabacteria_group>Actinomycetota>Actinomycetes>Micrococcales | GCF_006539945.1 | Cellulosimicrobium_cellulans | CCE02nite_RS18570 | 1298..4687 | 1710 | DUF499-domain-containing-protein-[Cellulosimicrobium-cellulans] |
| WP_141842659.1 | DUF499_1.RM.like.systems | <-HIN-HTH<-SF1-SF2-Helicase<-OLD-ABC+TOPRIM<-HEPN+DUF499*<-wHTH+REase<-METHYLASE<-TUDOR+SF1-SF2-Helicase+iREase+CTD+REase<-XIS-HTH<-PAIREDC-HTH<-HIN-HTH | 1132 | Bacteria>Terrabacteria_group>Actinomycetota>Actinomycetes>Micrococcales | GCF_006704135.1 | Micrococcus_sp._R8502A1 | FLM69_RS00645 | 3870..7268 | 2583239 | Swt1-family-HEPN-domain-containing-protein-[Micrococcus-sp.-R8502A1] |
| WP_141784692.1 | DUF499_1.RM.like.systems | <-MACRODOMAIN+SIR2\|\|REase-><-DDE_Tnp_1_4\|\|METHYLASE->?->ParB+HNH->HEPN+DUF499*->?-><-Phage_int_SAM_1+Phage_integrase<-?\|\|HTH-> | 1119 | Bacteria>Terrabacteria_group>Actinomycetota>Actinomycetes>Micrococcales | GCF_006716185.1 | Ornithinicoccus_hortensis | FB467_RS08335 | 1805419..1808778 | 82346 | DUF499-domain-containing-protein-[Ornithinicoccus-hortensis] |
| WP_144831472.1 | DUF499_1.RM.like.systems | ACET-><-?<-SF1-SF2-Helicase<-OLD-ABC+TOPRIM<-HEPN+DUF499*<-wHTH+REase<-METHYLASE<-TUDOR+SF1-SF2-Helicase+iREase+CTD+REase<-XIS-HTH\|\|MORC+HisKin+REase-><-TrwC+SF1-SF2-helicase | 1133 | Bacteria>Terrabacteria_group>Actinomycetota>Actinomycetes>Micrococcales | GCF_007677595.1 | Kocuria_rhizophila | FS577_RS10180 | 96606..100007 | 72000 | DUF499-domain-containing-protein-[Kocuria-rhizophila] |
| WP_144794196.1 | DUF499_1.RM.like.systems | TetR-HTH->XIS-HTH->TUDOR+SF1-SF2-Helicase+iREase+CTD+REase->METHYLASE->HEPN+DUF499*->ParB+HNH->?-><-N6_MTase-p<-McrC-NTD+REase | 1132 | Bacteria>Terrabacteria_group>Actinomycetota>Actinomycetes>Micrococcales | GCF_007679925.1 | Microbacterium_paludicola | FS695_RS03635 | 729234..732632 | 300019 | DUF499-domain-containing-protein-[Microbacterium-paludicola] |
| WP_145227579.1 | DUF499_1.RM.like.systems | <-NERD<-?<-?<-?<-HEPN+DUF499*<-METHYLASE<-NERD<-TUDOR+SF1-SF2-Helicase+iREase+CTD+REase\|\|NTD+SF1-SF2-Helicase->SF1-SF2-Helicase+HNH-> | 1121 | Bacteria>Terrabacteria_group>Actinomycetota>Actinomycetes>Micrococcales | GCF_007829035.1 | Rudaeicoccus_suwonensis | BKA23_RS09875 | 2146112..2149477 | 657409 | DUF499-domain-containing-protein-[Rudaeicoccus-suwonensis] |
| WP_187368407.1 | DUF499_1.RM.like.systems | TUDOR+SF1-SF2-Helicase+iREase+CTD+REase-><-PIN<-DUF433<-wHTH(SBD)+HNH\|\|METHYLASE->ParB+HNH->HEPN+DUF499*-><-?<-?\|\|ACET+ACET-><-Lsr2 | 1133 | Bacteria>Terrabacteria_group>Actinomycetota>Actinomycetes>Micrococcales | GCF_008000995.1 | Ruania_zhangjianzhongii | FU260_RS03480 | 750431..753832 | 2603206 | DUF499-domain-containing-protein-[Ruania-zhangjianzhongii] |
| WP_147826310.1 | DUF499_1.RM.like.systems | <-SAM-methylase\|\|OLD-ABC+TOPRIM->URI-DNase->XIS-HTH->TUDOR+SF1-SF2-Helicase+iREase+CTD+REase->METHYLASE->HEPN+DUF499*->HTH_19->?-><-?<-?\|\|?->ParB+HNH-> | 1122 | Bacteria>Terrabacteria_group>Actinomycetota>Actinomycetes>Micrococcales | GCF_008017445.1 | Microbacterium_mitrae | FVP60_RS10835 | 35691..39059 | 664640 | DUF499-domain-containing-protein-[Microbacterium-mitrae] |
| WP_210418449.1 | DUF499_1.RM.like.systems | VirB4-FtsK->?->VirD4-TraG+VirD4-FtsK->?->?->?-><-HEPN+DUF499*<-METHYLASE<-TUDOR+SF1-SF2-Helicase+iREase+CTD+REase<-Phage_int_SAM_1+Phage_integrase<-XIS-HTH | 1116 | Bacteria>Terrabacteria_group>Actinomycetota>Actinomycetes>Micrococcales | GCF_008365295.1 | Agromyces_intestinalis | FLP10_RS00780 | 160366..163716 | 2592652 | DUF499-domain-containing-protein-[Agromyces-intestinalis] |
| WP_156242696.1 | DUF499_1.RM.like.systems | <-SbcC\|\|?->REase-><-HTH+SF1-SF2-Helicase+REase<-REase<-MORC<-HEPN+DUF499*<-METHYLASE\|\|MORC+SIGMA-HTH->DNA_methylase->?-><-DeoR-HTH+ISOCOT | 1129 | Bacteria>Terrabacteria_group>Actinomycetota>Actinomycetes>Micrococcales | GCF_009735645.1 | Microbacterium_oryzae | D7D94_RS11265 | 2403131..2406520 | 743009 | Swt1-family-HEPN-domain-containing-protein-[Microbacterium-oryzae] |
| WP_246200745.1 | DUF499_1.RM.like.systems | <-REase<-HTH+SF1-SF2-Helicase+REase<-REase<-MORC<-HEPN+DUF499*<-METHYLASE<-MORC+HEPN\|\|MORC+SIGMA-HTH->DNA_methylase->REase->HEPN+SF1-SF2-Helicase-> | 1131 | Bacteria>Terrabacteria_group>Actinomycetota>Actinomycetes>Micrococcales | GCF_009749385.1 | Agromyces_allii | GQR16_RS11060 | 282353..285748 | 393607 | Swt1-family-HEPN-domain-containing-protein-[Agromyces-allii] |
| WP_157002068.1 | DUF499_1.RM.like.systems | <-DNA_methylase<-SIGMA-HTH+SIGMA-HTH\|\|?->?->METHYLASE->HEPN+DUF499*->MORC->REase-><-?\|\|HTH+SF1-SF2-Helicase+REase->REase-> | 1143 | Bacteria>Terrabacteria_group>Actinomycetota>Actinomycetes>Micrococcales | GCF_009749475.1 | Agromyces_laixinhei | GQQ90_RS09115 | 49199..52630 | 2585717 | DUF499-domain-containing-protein-[Agromyces-laixinhei] |
| WP_163540895.1 | DUF499_1.RM.like.systems | PemK_toxin-><-HEPN+DUF499*<-METHYLASE-p<-METHYLASE-p\|\|DUF433-><-?<-TUDOR+SF1-SF2-Helicase+iREase+CTD+REase\|\|HTH_3-> | 1130 | Bacteria>Terrabacteria_group>Actinomycetota>Actinomycetes>Micrococcales | GCF_010994285.1 | Occultella_kanbiaonis | G7B41_RS01575 | 244678..248070 | 2675754 | Swt1-family-HEPN-domain-containing-protein-[Occultella-kanbiaonis] |
| WP_166844731.1 | DUF499_1.RM.like.systems | <-MORC+HEPN<-HTH+SF1-SF2-Helicase+REase<-REase<-MORC<-HEPN+DUF499*<-METHYLASE\|\|DNA_methylase->?-><-?<-?<-HTH_3+PolB | 1156 | Bacteria>Terrabacteria_group>Actinomycetota>Actinomycetes>Micrococcales | GCF_011634835.1 | Isoptericola_sp._BMS4 | EYB28_RS03030 | 693115..696585 | 2527875 | Swt1-family-HEPN-domain-containing-protein-[Isoptericola-sp.-BMS4] |
| WP_168429935.1 | DUF499_1.RM.like.systems | <-MerR-HTH<-?\|\|MORC+DUF3684+REase-><-?<-HEPN+DUF499*<-METHYLASE<-TUDOR+SF1-SF2-Helicase+iREase+CTD+REase | 1113 | Bacteria>Terrabacteria_group>Actinomycetota>Actinomycetes>Micrococcales | GCF_012396205.1 | Cellulomonas_hominis | HGA09_RS00165 | 40962..44303 | 156981 | DUF499-domain-containing-protein-[Cellulomonas-hominis] |
| WP_168888196.1 | DUF499_1.RM.like.systems | CopG_antitoxin-><-?<-HEPN+DUF499*<-wHTH+REase<-METHYLASE<-TUDOR+SF1-SF2-Helicase+iREase+CTD+REase | 1127 | Bacteria>Terrabacteria_group>Actinomycetota>Actinomycetes>Micrococcales | GCF_012641515.1 | Nesterenkonia_sedimenti | HGQ17_RS12070 | 43839..47222 | 1463632 | DUF499-domain-containing-protein-[Nesterenkonia-sedimenti] |
| WP_169323296.1 | DUF499_1.RM.like.systems | <-NTD+SF1-SF2-Helicase<-TUDOR+SF1-SF2-Helicase+iREase+CTD+REase\|\|Schlafen+HATPase+HTH->METHYLASE->HEPN+DUF499*->HD->TrwC+SF1-SF2-Helicase->?->?-><-DNA-glycosylase<-ArdCN+MPTase | 1120 | Bacteria>Terrabacteria_group>Actinomycetota>Actinomycetes>Micrococcales | GCF_012952065.1 | Cellulomonas_fimi | HIR71_RS02970 | 36548..39910 | 1708 | DUF499-domain-containing-protein-[Cellulomonas-fimi] |
| WP_176337038.1 | DUF499_1.RM.like.systems | URI-DNase-><-?\|\|XIS-HTH->TUDOR+SF1-SF2-Helicase+iREase+CTD+REase->METHYLASE->HEPN+DUF499*-><-?\|\|?->Mbetalac-><-BirA-HTH\|\|Aminotran_1_2-> | 1149 | Bacteria>Terrabacteria_group>Actinomycetota>Actinomycetes>Micrococcales | GCF_013368965.1 | Kocuria_salina | E7Z53_RS01815 | 401527..404976 | 1929416 | DUF499-domain-containing-protein-[Kocuria-salina] |
| WP_188034675.1 | DUF499_1.RM.like.systems | MPTase->?->ABhydrolase->XIS-HTH->TUDOR+SF1-SF2-Helicase+iREase+CTD+REase->METHYLASE->HEPN+DUF499*->REase+SAVED->?->TrwC+SF1-SF2-Helicase->?->?-><-HTH+MPTase | 1143 | Bacteria>Terrabacteria_group>Actinomycetota>Actinomycetes>Micrococcales | GCF_014530035.1 | unclassified_Leucobacter | KOE72_RS09585 | 56127..59558 | 2621730 | MULTISPECIES:-DUF499-domain-containing-protein-[unclassified-Leucobacter] |
| WP_188427984.1 | DUF499_1.RM.like.systems | <-MazG<-wHTH(SBD)+HNH<-REase+SF1-SF2-Helicase\|\|?->?-><-?<-HEPN+DUF499*<-ParB+HNH<-METHYLASE<-TUDOR+SF1-SF2-Helicase+iREase+CTD+REase<-?\|\|SF1-SF2-Helicase-> | 1123 | Bacteria>Terrabacteria_group>Actinomycetota>Actinomycetes>Micrococcales | GCF_014636495.1 | Ornithinimicrobium_tianjinense | IEV65_RS02295 | 470126..473497 | 1195761 | Swt1-family-HEPN-domain-containing-protein-[Ornithinimicrobium-tianjinense] |
| WP_192910841.1 | DUF499_1.RM.like.systems | HTH_Tnp_1+HTH_21+rve+rve_3->ACET->MULE-TRANSPOSASE-><-?<-HEPN<-HEPN+DUF499*<-METHYLASE<-TUDOR+SF1-SF2-Helicase+iREase+CTD+REase\|\|?->LPD36+METHYLASE+SF1-SF2-Helicase->?->HTH_Tnp_1+HTH_21+rve+rve_2-> | 1123 | Bacteria>Terrabacteria_group>Actinomycetota>Actinomycetes>Micrococcales | GCF_014883915.1 | Janibacter_indicus | IGS73_RS14740 | 2999354..3002725 | 857417 | DUF499-domain-containing-protein-[Janibacter-indicus] |
| WP_194396698.1 | DUF499_1.RM.like.systems | METHYLASE-><-HNH<-OLD-ABC+TOPRIM<-HEPN+DUF499*<-METHYLASE<-TUDOR+SF1-SF2-Helicase+iREase+CTD+REase<-XIS-HTH | 1130 | Bacteria>Terrabacteria_group>Actinomycetota>Actinomycetes>Micrococcales | GCF_015277815.1 | Microbacterium_atlanticum | IR212_RS15230 | 3329381..3332773 | 2782168 | DUF499-domain-containing-protein-[Microbacterium-atlanticum] |
| WP_194413436.1 | DUF499_1.RM.like.systems | <-HTH<-?<-?<-TrwC+SF1-SF2-helicase+TOPRIM\|\|NTD+SF1-SF2-Helicase-><-HEPN+CBS<-HEPN+DUF499*<-METHYLASE<-?<-TUDOR+SF1-SF2-Helicase+iREase+CTD+REase<-?<-ParB+HNH<-ParB+HNH | 1119 | Bacteria>Terrabacteria_group>Actinomycetota>Actinomycetes>Micrococcales | GCF_015278355.1 | Microbacterium_yannicii | NCG93_RS19890 | 89985..93344 | 671622 | Swt1-family-HEPN-domain-containing-protein-[Microbacterium-yannicii] |
| WP_195692263.1 | DUF499_1.RM.like.systems | <-Phage_int_SAM_1+Phage_integrase\|\|REase-><-NUDIX<-SbcC<-HEPN+DUF499*<-METHYLASE<-TUDOR+SF1-SF2-Helicase+iREase+CTD+REase\|\|GDResolvase+Recombinase+ZnR-><-PAIREDC-HTH | 1129 | Bacteria>Terrabacteria_group>Actinomycetota>Actinomycetes>Micrococcales | GCF_015565955.1 | Microbacterium | JOJ05_RS14720 | 85515..88904 | 33882 | DUF499-domain-containing-protein-[Microbacterium-schleiferi] |
| WP_197156507.1 | DUF499_1.RM.like.systems | <-HD<-?\|\|NUDIX-><-HTH+SF1-SF2-Helicase+REase<-REase<-MORC<-HEPN+DUF499*<-METHYLASE<-?<-?<-URI-DNase+SF1-SF2-Helicase<-MazG<-REase+35exo+Beta-Barrel+SF1-SF2-Helicase | 1110 | Bacteria>Terrabacteria_group>Actinomycetota>Actinomycetes>Micrococcales | GCF_015864945.1 | Salinibacterium_sp._NG253 | I6E52_RS09085 | 1888777..1892109 | 2792039 | DUF499-domain-containing-protein-[Salinibacterium-sp.-NG253] |
| WP_200569160.1 | DUF499_1.RM.like.systems | <-DNA_methylase\|\|REase->METHYLASE->HEPN+DUF499*->MORC->REase->HTH+SF1-SF2-Helicase+REase-> | 1144 | Bacteria>Terrabacteria_group>Actinomycetota>Actinomycetes>Micrococcales | GCF_016620465.1 | Kocuria_rhizophila | JHD97_RS04255 | 38629..42063 | 72000 | DUF499-domain-containing-protein-[Kocuria-rhizophila] |
| WP_201671845.1 | DUF499_1.RM.like.systems | <-HEPN+REase<-REase<-HEPN+DUF499*<-wHTH+REase<-METHYLASE-p<-METHYLASE-p<-TUDOR+SF1-SF2-Helicase+iREase+CTD+REase<-XIS-HTH\|\|MORC+HisKin+REase-> | 1132 | Bacteria>Terrabacteria_group>Actinomycetota>Actinomycetes>Micrococcales | GCF_016726245.1 | Brevibacterium_casei | I6I57_RS03275 | 754914..758312 | 33889 | DUF499-domain-containing-protein-[Brevibacterium-casei] |
| WP_204409507.1 | DUF499_1.RM.like.systems | <-HIN-HTH<-OLD-ABC+TOPRIM<-HEPN+DUF499*<-wHTH+REase<-METHYLASE<-TUDOR+SF1-SF2-Helicase+iREase+CTD+REase<-XIS-HTH | 1132 | Bacteria>Terrabacteria_group>Actinomycetota>Actinomycetes>Micrococcales | GCF_016907455.1 | Brachybacterium_muris | JOD52_RS08675 | 1871612..1875010 | 219301 | DUF499-domain-containing-protein-[Brachybacterium-muris] |
| WP_210233975.1 | DUF499_1.RM.like.systems | TUDOR+SF1-SF2-Helicase+iREase+CTD+REase->METHYLASE->REase->HEPN+DUF499*-><-REase<-HKD-DNase+SF1-SF2-Helicase+TRD\|\|DndB-> | 1116 | Bacteria>Terrabacteria_group>Actinomycetota>Actinomycetes>Micrococcales | GCF_017898105.1 | Arthrobacter_jiangjiafuii | J2M53_RS14475 | 604648..607998 | 2817475 | Swt1-family-HEPN-domain-containing-protein-[Arthrobacter-jiangjiafuii] |
| WP_249312439.1 | DUF499_1.RM.like.systems | GDResolvase+HIN-HTH-><-?<-HNH\|\|TUDOR+SF1-SF2-Helicase+iREase+CTD+REase->METHYLASE->HEPN+DUF499*->?-><-UDG-conflict\|\|?-><-?<-EVE | 1140 | Bacteria>Terrabacteria_group>Actinomycetota>Actinomycetes>Micrococcales | GCF_018304855.1 | Pseudoclavibacter_sp._Marseille-Q4354 | JSQ82_RS10750 | 821875..825297 | 2810515 | DUF499-domain-containing-protein-[Pseudoclavibacter-sp.-Marseille-Q4354] |
| WP_216886275.1 | DUF499_1.RM.like.systems | Peptidase_S66+Peptidase_S66C-><-HTH+SF1-SF2-Helicase+REase<-REase<-MORC<-HEPN+DUF499*<-METHYLASE<-DNA_methylase\|\|REase-> | 1157 | Bacteria>Terrabacteria_group>Actinomycetota>Actinomycetes>Micrococcales | GCF_019037785.1 | Micrococcus_luteus | KM867_RS03170 | 29276..32749 | 1270 | Swt1-family-HEPN-domain-containing-protein-[Micrococcus-luteus] |
| WP_217316614.1 | DUF499_1.RM.like.systems | N6_MTase-p->?->?-><-zf-ISL3+DDE_Tnp_ISL3\|\|<-REase+ParB<-?<-HEPN+DUF499*<-METHYLASE<-METHYLASE-p<-TUDOR+SF1-SF2-Helicase+iREase+CTD+REase<-XIS-HTH | 1128 | Bacteria>Terrabacteria_group>Actinomycetota>Actinomycetes>Micrococcales | GCF_019056515.1 | Microbacterium_paraoxydans | IZR02_RS16970 | 42865..46251 | 199592 | DUF499-domain-containing-protein-[Microbacterium-paraoxydans] |
| WP_223984660.1 | DUF499_1.RM.like.systems | REase-><-?\|\|HKD-DNase+SF1-SF2-Helicase+TRD->?-><-?<-HEPN+DUF499*<-METHYLASE<-TUDOR+SF1-SF2-Helicase+iREase+CTD+REase | 1139 | Bacteria>Terrabacteria_group>Actinomycetota>Actinomycetes>Micrococcales | GCF_019977355.1 | Arthrobacter_sp._NicSoilB8 | LDO15_RS05065 | 1116619..1120038 | 2830998 | Swt1-family-HEPN-domain-containing-protein-[Arthrobacter-sp.-NicSoilB8] |
| WP_224781058.1 | DUF499_1.RM.like.systems | <-MORC+HEPN<-DNA_methylase\|\|?->METHYLASE->HEPN+DUF499*->MORC->REase->HTH+SF1-SF2-Helicase+REase->REase-> | 1130 | Bacteria>Terrabacteria_group>Actinomycetota>Actinomycetes>Micrococcales | GCF_020096995.1 | Leucobacter_sp._Psy1 | K8P10_RS06885 | 1465839..1469231 | 2875729 | Swt1-family-HEPN-domain-containing-protein-[Leucobacter-sp.-Psy1] |
| WP_229577768.1 | DUF499_1.RM.like.systems | <-HTH+SF1-SF2-Helicase+REase<-REase<-MORC<-HEPN+DUF499*<-METHYLASE-p<-METHYLASE-p<-REase\|\|DNA_methylase-> | 1141 | Bacteria>Terrabacteria_group>Actinomycetota>Actinomycetes>Micrococcales | GCF_020829765.1 | Kocuria_rhizophila | JNE43_RS06540 | 5673..9098 | 72000 | DUF499-domain-containing-protein-[Kocuria-rhizophila] |
| WP_231831079.1 | DUF499_1.RM.like.systems | MORC+SIGMA-HTH-><-REase<-HTH+SF1-SF2-Helicase+REase<-REase<-MORC<-HEPN+DUF499*<-METHYLASE\|\|DNA_methylase->?->wHTH+SAD+HNH-> | 1128 | Bacteria>Terrabacteria_group>Actinomycetota>Actinomycetes>Micrococcales | GCF_021168475.1 | Pseudoclavibacter_sp._13-3 | LRX77_RS01510 | 334724..338110 | 2901228 | Swt1-family-HEPN-domain-containing-protein-[Pseudoclavibacter-sp.-13-3] |
| WP_276315051.1 | DUF499_1.RM.like.systems | REase+SF1-SF2-Helicase-><-SbcC<-HEPN+DUF499*<-METHYLASE-p | 1127 | Bacteria>Terrabacteria_group>Actinomycetota>Actinomycetes>Micrococcales | GCF_021395155.1 | Microbacterium_profundi | LZG07_RS12530 | 143258..146641 | 450380 | DUF499-domain-containing-protein-[Microbacterium-profundi] |
| WP_236803031.1 | DUF499_1.RM.like.systems | <-ABC-ATPase+SbcC<-?<-HEPN+DUF499*<-METHYLASE<-TUDOR+SF1-SF2-Helicase+iREase+CTD+REase<-XIS-HTH\|\|REase-> | 1127 | Bacteria>Terrabacteria_group>Actinomycetota>Actinomycetes>Micrococcales | GCF_021801725.1 | Arthrobacter_sp._FW306-05-C | LFT44_RS20985 | 51567..54950 | 2879620 | Swt1-family-HEPN-domain-containing-protein-[Arthrobacter-sp.-FW306-05-C] |
| WP_241710057.1 | DUF499_1.RM.like.systems | HIN-HTH->?-><-?<-HTH+SF1-SF2-Helicase+REase<-REase<-MORC<-HEPN+DUF499*<-METHYLASE\|\|DNA_methylase-><-MORC+HEPN\|\|?-><-SF1-SF2-Helicase\|\|wHTH+SAD+HNH-> | 1123 | Bacteria>Terrabacteria_group>Actinomycetota>Actinomycetes>Micrococcales | GCF_022606295.1 | Arthrobacter_bambusae | KNN17_RS16570 | 21950..25321 | 1338426 | DUF499-domain-containing-protein-[Arthrobacter-bambusae] |
| WP_134416003.1 | DUF499_1.RM.like.systems | Peptidase_S66+Peptidase_S66C-><-HTH+SF1-SF2-Helicase+REase<-REase<-MORC<-HEPN+DUF499*<-METHYLASE<-DNA_methylase\|\|REase-> | 1157 | Bacteria>Terrabacteria_group>Actinomycetota>Actinomycetes>Micrococcales | GCF_022664405.1 | Micrococcus_aloeverae | E2F92_RS02100 | 40430..43903 | 1391911 | Swt1-family-HEPN-domain-containing-protein-[Micrococcus-aloeverae] |
| WP_243591605.1 | DUF499_1.RM.like.systems | ACET-><-?<-REase<-HTH+SF1-SF2-Helicase+REase<-REase<-MORC<-HEPN+DUF499*<-METHYLASE\|\|DNA_methylase->MORC+HisKin-> | 1123 | Bacteria>Terrabacteria_group>Actinomycetota>Actinomycetes>Micrococcales | GCF_022815585.1 | Rathayibacter_caricis | MT349_RS09900 | 122292..125663 | 110936 | Swt1-family-HEPN-domain-containing-protein-[Rathayibacter-caricis] |
| WP_247619184.1 | DUF499_1.RM.like.systems | <-OLD-ABC+TOPRIM<-HEPN+DUF499*<-wHTH+REase<-METHYLASE<-TUDOR+SF1-SF2-Helicase+iREase+CTD+REase<-XIS-HTH | 1135 | Bacteria>Terrabacteria_group>Actinomycetota>Actinomycetes>Micrococcales | GCF_023072605.1 | Brevibacterium_sp._R8603A2 | MTQ12_RS14500 | 1781..5188 | 2929779 | Swt1-family-HEPN-domain-containing-protein-[Brevibacterium-sp.-R8603A2] |
| WP_247963105.1 | DUF499_1.RM.like.systems | DDE_Tnp_1->METHYLASE-p->HEPN+DUF499*->ParB+HNH->?-><-N6_MTase-p<-McrC-NTD+REase | 1132 | Bacteria>Terrabacteria_group>Actinomycetota>Actinomycetes>Micrococcales | GCF_023108875.1 | Microbacterium_aerolatum | MZK47_RS13285 | 1211..4609 | 153731 | DUF499-domain-containing-protein-[Microbacterium-aerolatum] |
| WP_247963551.1 | DUF499_1.RM.like.systems | DDE_Tnp_1->HIN-HTH->HTH_21+rve+rve_3->METHYLASE-p->HEPN+DUF499*->REase+SAVED-> | 1143 | Bacteria>Terrabacteria_group>Actinomycetota>Actinomycetes>Micrococcales | GCF_023108875.1 | Microbacterium_aerolatum | MZK47_RS16315 | 3209..6640 | 153731 | DUF499-domain-containing-protein-[Microbacterium-aerolatum] |
| WP_282449258.1 | DUF499_1.RM.like.systems | XIS-HTH->TUDOR+SF1-SF2-Helicase+iREase+CTD+REase->METHYLASE-p->METHYLASE->HEPN+DUF499*->HEPN-><-Glyoxalase | 1126 | Bacteria>Terrabacteria_group>Actinomycetota>Actinomycetes>Micrococcales | GCF_023155745.1 | Microbacterium_kunmingense | MA875_RS17035 | 50795..54175 | 2915939 | DUF499-domain-containing-protein-[Microbacterium-kunmingensis] |
| WP_249521258.1 | DUF499_1.RM.like.systems | <-HNH+SAVED<-HTH_3\|\|?->?->TUDOR+SF1-SF2-Helicase+iREase+CTD+REase->METHYLASE->HEPN+DUF499*->SbcC-><-N6_MTase-p\|\|?-><-MerR-HTH\|\|?->ACET-> | 1131 | Bacteria>Terrabacteria_group>Actinomycetota>Actinomycetes>Micrococcales | GCF_023383675.1 | Microbacterium_sp._ER1 | BV453_RS07415 | 50443..53838 | 1932846 | DUF499-domain-containing-protein-[Microbacterium-sp.-ER1] |
| WP_252619626.1 | DUF499_1.RM.like.systems | <-VirB11+VirB4-FtsK\|\|GDResolvase+Recombinase+ZnR-><-?<-HEPN+DUF499*<-METHYLASE<-TUDOR+SF1-SF2-Helicase+iREase+CTD+REase\|\|Lsr2-> | 1118 | Bacteria>Terrabacteria_group>Actinomycetota>Actinomycetes>Micrococcales | GCF_023923205.1 | Ornithinimicrobium_cryptoxanthini | NF557_RS11930 | 2576031..2579387 | 2934161 | DUF499-domain-containing-protein-[Ornithinimicrobium-sp.-JY.X270] |
| WP_253867645.1 | DUF499_1.RM.like.systems | <-HTH+SF1-SF2-Helicase+REase<-REase<-MORC<-HEPN+DUF499*<-METHYLASE\|\|HTH_3->wHTH+SAD+HNH->HTH-> | 1150 | Bacteria>Terrabacteria_group>Actinomycetota>Actinomycetes>Micrococcales | GCF_024171995.1 | Promicromonospora_umidemergens | APR04_RS02600 | 547565..551017 | 629679 | Swt1-family-HEPN-domain-containing-protein-[Promicromonospora-umidemergens] |
| WP_255162094.1 | DUF499_1.RM.like.systems | DDE_Tnp_IS240-><-Classical-AAA<-TrwC+SF1-SF2-Helicase<-OLD-ABC+TOPRIM<-?<-HEPN+DUF499*<-METHYLASE<-TUDOR+SF1-SF2-Helicase+iREase+CTD+REase<-XIS-HTH<-ABhydrolase<-?<-MPTase | 1143 | Bacteria>Terrabacteria_group>Actinomycetota>Actinomycetes>Micrococcales | GCF_024362265.1 | Microbacterium_liquefaciens | NMQ05_RS13065 | 2679727..2683158 | 33918 | DUF499-domain-containing-protein-[Microbacterium-liquefaciens] |
| WP_255770908.1 | DUF499_1.RM.like.systems | <-DNA_methylase\|\|METHYLASE->HEPN+DUF499*->MORC->REase->HTH+SF1-SF2-Helicase+REase->ParB+HNH-><-?<-NUDIX | 1122 | Bacteria>Terrabacteria_group>Actinomycetota>Actinomycetes>Micrococcales | GCF_024391045.1 | Pseudarthrobacter_sulfonivorans | NQJ30_RS15540 | 5007..8375 | 121292 | DUF499-domain-containing-protein-[Pseudarthrobacter-sulfonivorans] |
| WP_257744324.1 | DUF499_1.RM.like.systems | <-DNA_methylase\|\|REase->METHYLASE->HEPN+DUF499*->MORC->REase->HTH+SF1-SF2-Helicase+REase-><-?\|\|MarR-HTH-> | 1137 | Bacteria>Terrabacteria_group>Actinomycetota>Actinomycetes>Micrococcales | GCF_024655785.1 | Kocuria_rhizophila | NUV30_RS04805 | 31091..34504 | 72000 | DUF499-domain-containing-protein-[Kocuria-rhizophila] |
| WP_260073607.1 | DUF499_1.RM.like.systems | <-HIN-HTH\|\|?->TUDOR+SF1-SF2-Helicase+iREase+CTD+REase->METHYLASE->HEPN->HEPN+DUF499*-> | 1129 | Bacteria>Terrabacteria_group>Actinomycetota>Actinomycetes>Micrococcales | GCF_025146225.1 | Nesterenkonia_massiliensis | M3B43_RS10510 | 14559..17948 | 1232429 | DUF499-domain-containing-protein-[Nesterenkonia-massiliensis] |
| WP_259878499.1 | DUF499_1.RM.like.systems | <-Peptidase<-?<-?<-?<-HEPN+DUF499*<-wHTH+REase<-METHYLASE<-TUDOR+SF1-SF2-Helicase+iREase+CTD+REase<-XIS-HTH<-ABhydrolase\|\|DEDD_Tnp_IS110+Transposase_20-> | 1132 | Bacteria>Terrabacteria_group>Actinomycetota>Actinomycetes>Micrococcales | GCF_025148185.1 | Brachybacterium_muris | M3C58_RS02745 | 15014..18412 | 219301 | DUF499-domain-containing-protein-[Brachybacterium-muris] |
| WP_274500272.1 | DUF499_1.RM.like.systems | <-Mbetalac\|\|MarR-HTH->?->?->NERD-><-OLD-ABC+TOPRIM<-HEPN+DUF499*<-METHYLASE<-TUDOR+SF1-SF2-Helicase+iREase+CTD+REase<-UDG-conflict\|\|?-><-REase\|\|HEPN+SF1-SF2-Helicase-> | 1131 | Bacteria>Terrabacteria_group>Actinomycetota>Actinomycetes>Micrococcales | GCF_028765365.1 | Plantibacter_flavus | PUY80_RS13440 | 281410..284805 | 150123 | DUF499-domain-containing-protein-[Plantibacter-flavus] |
| WP_282859602.1 | DUF499_1.RM.like.systems | <-SbcC<-HEPN+DUF499*<-METHYLASE<-REase | 1126 | Bacteria>Terrabacteria_group>Actinomycetota>Actinomycetes>Micrococcales | GCF_030011915.1 | Pseudoclavibacter_helvolus | QM024_RS14940 | 2044..5424 | 255205 | DUF499-domain-containing-protein-[Pseudoclavibacter-helvolus] |
| WP_285329939.1 | DUF499_1.RM.like.systems | <-RelA<-?\|\|35exo->tRNA->TUDOR+SF1-SF2-Helicase+iREase+CTD+REase->METHYLASE->HEPN+DUF499*-> | 1139 | Bacteria>Terrabacteria_group>Actinomycetota>Actinomycetes>Micrococcales | GCF_030216795.1 | Pseudoglutamicibacter_cumminsii | QP411_RS02105 | 150205..153624 | 156979 | DUF499-domain-containing-protein-[Pseudoglutamicibacter-cumminsii] |
| WP_285140308.1 | DUF499_1.RM.like.systems | <-HTH_Tnp_1+HTH_21+rve+rve_3<-HIN-HTH\|\|XIS-HTH->TUDOR+SF1-SF2-Helicase+iREase+CTD+REase->METHYLASE->wHTH+REase->HEPN+DUF499*->OLD-ABC+TOPRIM->SF1-SF2-Helicase-><-DNA_methylase | 1132 | Bacteria>Terrabacteria_group>Actinomycetota>Actinomycetes>Micrococcales | GCF_030226745.1 | Micrococcus_luteus | QP726_RS10470 | 9707..13105 | 1270 | Swt1-family-HEPN-domain-containing-protein-[Micrococcus-luteus] |
| WP_068492369.1 | DUF499_1.RM.like.systems | <-HTH+SF1-SF2-Helicase+REase<-REase<-MORC<-HEPN+DUF499*<-METHYLASE\|\|DNA_methylase-> | 1129 | Bacteria>Terrabacteria_group>Actinomycetota>Actinomycetes>Micrococcales | GCF_900014995.1 | Pseudoclavibacter_helvolus | BN2259_RS00510 | 5683..9072 | 255205 | Swt1-family-HEPN-domain-containing-protein-[Pseudoclavibacter-helvolus] |
| WP_074694547.1 | DUF499_1.RM.like.systems | HEPN+SF1-SF2-Helicase->35exo-><-?<-?<-?<-HEPN+REase<-HEPN+DUF499*<-METHYLASE<-Mbetalac<-TUDOR+SF1-SF2-Helicase+iREase+CTD+REase<-HTH | 1131 | Bacteria>Terrabacteria_group>Actinomycetota>Actinomycetes>Micrococcales | GCF_900104345.1 | Microbacterium_testaceum | BLS25_RS04240 | 198218..201613 | 2033 | DUF499-domain-containing-protein-[Microbacterium-testaceum] |
| WP_076404375.1 | DUF499_1.RM.like.systems | REase-><-?<-?<-?<-HEPN+DUF499*<-ParB+HNH<-METHYLASE\|\|MPTase->PIN->HTH_3->wHTH(SBD)+HNH-> | 1151 | Bacteria>Terrabacteria_group>Actinomycetota>Actinomycetes>Micrococcales | GCF_900155925.1 | Cellulosimicrobium_aquatile | BXA05_RS06490 | 508556..512011 | 1612203 | DUF499-domain-containing-protein-[Cellulosimicrobium-aquatile] |
| WP_101639485.1 | DUF499_1.RM.like.systems | <-OLD-ABC+TOPRIM<-DNA_methylase\|\|MORC+REase->MORC+HEPN->REase->METHYLASE->HEPN+DUF499*->MORC->REase->HTH+SF1-SF2-Helicase+REase-> | 1135 | Bacteria>Terrabacteria_group>Actinomycetota>Actinomycetes>Micrococcales | GCF_900169095.1 | Brevibacterium_aurantiacum | BAUR920_RS16220 | 10995..14402 | 273384 | DUF499-domain-containing-protein-[Brevibacterium-aurantiacum] |
| WP_126499814.1 | DUF499_1.RM.like.systems | Response_reg+LuxR-HTH-><-?<-TetR-HTH\|\|ABhydrolase->?->METHYLASE->HEPN+DUF499*-> | 1125 | Bacteria>Terrabacteria_group>Actinomycetota>Actinomycetes>Micrococcales | GCF_900637985.1 | Rothia_aeria | EL250_RS03685 | 841376..844753 | 172042 | DUF499-domain-containing-protein-[Rothia-aeria] |
| WP_191562882.1 | DUF499_1.RM.like.systems | NTD+SF1-SF2-Helicase->TUDOR+SF1-SF2-Helicase+iREase+CTD+REase-><-?\|\|METHYLASE->ParB+HNH->HEPN+DUF499*->?->SF1-SF2-Helicase->?-><-URI-DNase | 1117 | Bacteria>Terrabacteria_group>Actinomycetota>Actinomycetes>Micrococcales | GCF_904423795.1 | Janibacter_melonis | IHQ27_RS00715 | 71564..74917 | 262209 | Swt1-family-HEPN-domain-containing-protein-[Janibacter-melonis] |
| WP_218115672.1 | DUF499_1.RM.like.systems | <-URI-DNase+SF1-SF2-Helicase<-MazG\|\|?->RVT-><-DNA_methylase\|\|METHYLASE->HEPN+DUF499*->MORC->REase->HTH+SF1-SF2-Helicase+REase->REase-><-MORC+SIGMA-HTH | 1133 | Bacteria>Terrabacteria_group>Actinomycetota>Actinomycetes>Micrococcales | GCF_910593785.1 | Leucobacter_soli | KXF26_RS09170 | 42150..45551 | 2812850 | Swt1-family-HEPN-domain-containing-protein-[Leucobacter-soli] |
| WP_225755648.1 | DUF499_1.RM.like.systems | REase->MORC->?->?->METHYLASE-p->METHYLASE-p->HEPN+DUF499*->MORC->REase->HTH+SF1-SF2-Helicase+REase->TUDOR+SF1-SF2-Helicase+iREase+CTD->SF1-SF2-Helicase-><-HNH | 1137 | Bacteria>Terrabacteria_group>Actinomycetota>Actinomycetes>Micrococcales | GCF_916618755.1 | Actinotalea_sp._Marseille-Q4924 | LG379_RS16320 | 268678..272091 | 2866571 | DUF499-domain-containing-protein-[Actinotalea-sp.-Marseille-Q4924] |
| WP_251355553.1 | DUF499_1.RM.like.systems | <-HNH<-?\|\|TUDOR+SF1-SF2-Helicase+iREase+CTD+REase->METHYLASE->HEPN+DUF499*->?-><-MULE-TRANSPOSASE\|\|?->?->zf-ISL3+DDE_Tnp_ISL3-> | 1104 | Bacteria>Terrabacteria_group>Actinomycetota>Actinomycetes>Micrococcales | - | Kocuria_rosea | M3684_RS03240 | 225275..228589 | 1275 | DUF499-domain-containing-protein-[Kocuria-rosea] |
| WP_234001366.1 | DUF499_1.RM.like.systems | REase+SF1-SF2-Helicase-><-SbcC<-HEPN+DUF499*<-METHYLASE-p | 1108 | Bacteria>Terrabacteria_group>Actinomycetota>Actinomycetes>Micrococcales | - | Microbacterium_profundi | LZG07_12530 | 143258..146584 | 450380 | DUF499-domain-containing-protein-[Microbacterium-profundi] |
| WP_220122165.1 | DUF499_1.RM.like.systems | GNTR-HTH-><-?<-?<-DNA_methylase\|\|SIGMA-HTH->METHYLASE->HEPN+DUF499*->MORC->REase->HTH+SF1-SF2-Helicase+REase->NERD+SF1-SF2-Helicase-><-?<-ABC-ATPase+SbcC | 1128 | Bacteria>Terrabacteria_group>Actinomycetota>Actinomycetes>Micrococcales | - | Ornithinimicrobium_laminariae | K0651_07785 | 644333..647719 | 2864873 | Swt1-family-HEPN-domain-containing-protein-[Ornithinimicrobium-laminariae] |
| WP_104269222.1 | DUF499_1.RM.like.systems | MACRODOMAIN+SIR2->TUDOR+SF1-SF2-Helicase+iREase+CTD+REase->METHYLASE->HEPN+DUF499*-><-?<-?\|\|?->?->Peptidase->Peptidase_M73-> | 1120 | Bacteria>Terrabacteria_group>Actinomycetota>Actinomycetes>Micrococcales | - | Rathayibacter_sp._AY1C9 | C5D09_10000 | 51639..55001 | 2080541 | Swt1-family-HEPN-domain-containing-protein-[Rathayibacter-sp.-AY1C9] |
| NMO53435.1 | DUF499_1.RM.like.systems | Pkinase->REase-><-?<-?<-HEPN+DUF499*<-TUDOR+SF1-SF2-Helicase+iREase+CTD+REase\|\|?->AAA_21-><-HEPN<-Primpol+D5N+D5ATPase | 1100 | Bacteria>Terrabacteria_group>Actinomycetota>Actinomycetes>Micromonosporales | GCA_012933445.1 | Actinoplanes_sp._TBRC_11911 | HH310_19870 | 150452..153754 | 2729386 | ATP-binding-protein-[Actinoplanes-sp.-TBRC-11911] |
| MBE1487514.1 | DUF499_1.RM.like.systems | HTH_17->Phage_int_SAM_1+Phage_integrase-><-?\|\|?->TUDOR+SF1-SF2-Helicase+iREase+CTD+REase->METHYLASE->HEPN+DUF499*-><-DDE_Tnp_1_2\|\|?->DNA-glycosylase-><-?<-DUF6361<-RNase | 1124 | Bacteria>Terrabacteria_group>Actinomycetota>Actinomycetes>Micromonosporales | GCA_014874095.1 | Plantactinospora_soyae | H4W31_003152 | 3518204..3521578 | 1544732 | putative-AAA+-superfamily-ATPase-[Plantactinospora-soyae] |
| MBN1173560.1 | DUF499_1.RM.like.systems | <-Exonuc_V_gamma+REase\|\|TUDOR+SF1-SF2-Helicase+iREase+CTD+REase->METHYLASE->HEPN+DUF499*->REase-><-?\|\|?->Cas_Cmr3-> | 1155 | Bacteria>Terrabacteria_group>Actinomycetota>Actinomycetes>Micromonosporales | GCA_016926535.1 | Micromonosporaceae_bacterium | JXA67_15425 | 10595..14062 | 1873464 | DUF499-domain-containing-protein-[Micromonosporaceae-bacterium] |
| BCJ64829.1 | DUF499_1.RM.like.systems | <-DDE_Tnp_1<-?<-?\|\|TUDOR+SF1-SF2-Helicase+iREase+CTD+REase->METHYLASE->HEPN+DUF499*-><-?<-?\|\|?->?->?->HIN-HTH-> | 1124 | Bacteria>Terrabacteria_group>Actinomycetota>Actinomycetes>Micromonosporales | GCA_018324255.1 | Polymorphospora_rubra | Prubr_18500 | 1986978..1990352 | 338584 | hypothetical-protein-Prubr_18500-[Polymorphospora-rubra] |
| WP_085067314.1 | DUF499_1.RM.like.systems | <-ABhydrolase\|\|SIGMA-HTH-><-?\|\|TUDOR+SF1-SF2-Helicase+iREase+CTD+REase->METHYLASE->HEPN+DUF499*-><-?<-?<-ABhydrolase\|\|HTH+WYL+WYL-C-> | 1111 | Bacteria>Terrabacteria_group>Actinomycetota>Actinomycetes>Micromonosporales | GCF_000717135.1 | Catenuloplanes_japonicus | OP04_RS45935 | 31135..34470 | 33876 | Swt1-family-HEPN-domain-containing-protein-[Catenuloplanes-japonicus] |
| WP_030271467.1 | DUF499_1.RM.like.systems | Cysteine_desulfurase->?->?->Phage_integrase-><-HEPN+DUF499*<-ParB+HNH<-METHYLASE<-TUDOR+SF1-SF2-Helicase+iREase+CTD+REase\|\|NTD+SF1-SF2-Helicase-><-?<-JetD | 1125 | Bacteria>Terrabacteria_group>Actinomycetota>Actinomycetes>Micromonosporales | GCF_000717585.1 | Micromonospora_globosa | IF06_RS0114120 | 15248..18625 | 47863 | Swt1-family-HEPN-domain-containing-protein-[Micromonospora-globosa] |
| WP_197494866.1 | DUF499_1.RM.like.systems | <-pPIWI+pPIWI<-REase<-NTD+SF1-SF2-Helicase\|\|TUDOR+SF1-SF2-Helicase+iREase+CTD+REase->METHYLASE->ParB+HNH->HEPN+DUF499*->?->DEDD_Tnp_IS110+Transposase_20->zf-ISL3+DDE_Tnp_ISL3-><-?<-HTH_3 | 1124 | Bacteria>Terrabacteria_group>Actinomycetota>Actinomycetes>Micromonosporales | GCF_001652995.1 | Micromonospora_sp._NBRC_110037 | A4R25_RS09725 | 2384437..2387811 | 1621261 | Swt1-family-HEPN-domain-containing-protein-[Micromonospora-sp.-NBRC-110037] |
| WP_107269540.1 | DUF499_1.RM.like.systems | <-ABhydrolase<-TetR-HTH\|\|?->HAMP+HISKIN->?-><-ParB+HNH<-HEPN+DUF499*\|\|?->?-><-?<-?\|\|HTH_3->Pkinase-> | 1110 | Bacteria>Terrabacteria_group>Actinomycetota>Actinomycetes>Micromonosporales | GCF_003030345.1 | Plantactinospora_sp._BC1 | C6361_RS28025 | 6382634..6385966 | 2108470 | Swt1-family-HEPN-domain-containing-protein-[Plantactinospora-sp.-BC1] |
| WP_107261307.1 | DUF499_1.RM.like.systems | <-ABhydrolase<-TetR-HTH\|\|?->HAMP+HISKIN->?-><-ParB+HNH<-HEPN+DUF499*\|\|?->?-><-?<-?\|\|HTH_3-> | 1110 | Bacteria>Terrabacteria_group>Actinomycetota>Actinomycetes>Micromonosporales | GCF_003030385.1 | Plantactinospora_sp._BB1 | C6W10_RS25965 | 5945851..5949183 | 2071627 | Swt1-family-HEPN-domain-containing-protein-[Plantactinospora-sp.-BB1] |
| WP_192902263.1 | DUF499_1.RM.like.systems | HTH+TPR+TPR->Pkinase-><-HEPN+DUF499*<-ParB+HNH<-METHYLASE<-TUDOR+SF1-SF2-Helicase+iREase+CTD+REase\|\|NTD+SF1-SF2-Helicase->REase->pPIWI+pPIWI-> | 1129 | Bacteria>Terrabacteria_group>Actinomycetota>Actinomycetes>Micromonosporales | GCF_003610785.1 | Micromonospora_globbae | D7I43_RS25240 | 101471..104860 | 1894969 | Swt1-family-HEPN-domain-containing-protein-[Micromonospora-globbae] |
| WP_211192376.1 | DUF499_1.RM.like.systems | Pkinase->REase-><-?<-?<-HEPN+DUF499*<-METHYLASE<-METHYLASE-p<-TUDOR+SF1-SF2-Helicase+iREase+CTD+REase\|\|?->AAA_21-><-HEPN | 1111 | Bacteria>Terrabacteria_group>Actinomycetota>Actinomycetes>Micromonosporales | GCF_012933445.1 | Actinoplanes_sp._TBRC_11911 | HH310_RS19870 | 150452..153787 | 2729386 | Swt1-family-HEPN-domain-containing-protein-[Actinoplanes-sp.-TBRC-11911] |
| WP_187690605.1 | DUF499_1.RM.like.systems | <-ParB+HNH+RAMA<-?<-HEPN+DUF499*<-METHYLASE<-TUDOR+SF1-SF2-Helicase+iREase+CTD+REase | 1128 | Bacteria>Terrabacteria_group>Actinomycetota>Actinomycetes>Micromonosporales | GCF_014472385.1 | Micromonospora_aurantiaca | H9X94_RS13515 | 9839..13225 | 47850 | Swt1-family-HEPN-domain-containing-protein-[Micromonospora-aurantiaca] |
| WP_187693213.1 | DUF499_1.RM.like.systems | SF1-SF2-helicase+RdgC+Beta-barrel+REase->\|\|HTH_Tnp_4+DDE_Tnp_4->\|\|DUF4096+DDE_Tnp_1+DDE_Tnp_1_2->TUDOR+SF1-SF2-Helicase+iREase+CTD+REase->METHYLASE->ParB+HNH->HEPN+DUF499*-><-Phage_integrase<-?<-?<-?<-Cysteine_desulfurase | 1129 | Bacteria>Terrabacteria_group>Actinomycetota>Actinomycetes>Micromonosporales | GCF_014472405.1 | Micromonospora_chalcea | H9X95_RS12030 | 19161..22550 | 1874 | Swt1-family-HEPN-domain-containing-protein-[Micromonospora-chalcea] |
| WP_190252034.1 | DUF499_1.RM.like.systems | <-ABC-ATPase+SbcC\|\|tRNA->?-><-?\|\|TUDOR+SF1-SF2-Helicase+iREase+CTD+REase->METHYLASE->HEPN+DUF499*->ParB+HNH->?-><-Phage_int_SAM_1+Phage_integrase<-HTH_17\|\|?-><-BirA-HTH | 1122 | Bacteria>Terrabacteria_group>Actinomycetota>Actinomycetes>Micromonosporales | GCF_014647515.1 | Dactylosporangium_sucinum | IEZ31_RS23190 | 135414..138782 | 1424081 | Swt1-family-HEPN-domain-containing-protein-[Dactylosporangium-sucinum] |
| WP_203769001.1 | DUF499_1.RM.like.systems | DNA_methylase-><-REase<-HTH+SF1-SF2-Helicase+REase<-REase<-MORC<-HEPN+DUF499*<-METHYLASE\|\|\|\|?->REase-><-?\|\|REase->MarR-HTH-> | 1138 | Bacteria>Terrabacteria_group>Actinomycetota>Actinomycetes>Micromonosporales | GCF_016862275.1 | Actinoplanes_nipponensis | Ani05nite_RS15610 | 113186..116602 | 135950 | Swt1-family-HEPN-domain-containing-protein-[Actinoplanes-nipponensis] |
| WP_184787881.1 | DUF499_1.RM.like.systems | TUDOR+SF1-SF2-Helicase+iREase+CTD+REase->METHYLASE->HEPN+DUF499*-><-TetR-HTH\|\|?-><-?\|\|ACET->?-><-HTH | 1112 | Bacteria>Terrabacteria_group>Actinomycetota>Actinomycetes>Micromonosporales | GCF_016862855.1 | Phytomonospora_endophytica | Pen01_RS22635 | 44949..48287 | 714109 | DUF499-domain-containing-protein-[Phytomonospora-endophytica] |
| WP_204020362.1 | DUF499_1.RM.like.systems | NUDIX->?-><-?<-REase<-ParB+HNH<-HEPN+DUF499*<-METHYLASE<-TUDOR+SF1-SF2-Helicase+iREase+CTD+REase\|\|RelE-ParE->ParA+HTH-><-PIN<-Phd-YefM | 1129 | Bacteria>Terrabacteria_group>Actinomycetota>Actinomycetes>Micromonosporales | GCF_016863595.1 | Micromonospora_andamanensis | Vwe01_RS03465 | 15617..19006 | 1287068 | Swt1-family-HEPN-domain-containing-protein-[Micromonospora-trujilloniae] |
| WP_208576856.1 | DUF499_1.RM.like.systems | HTH+TPR+TPR->Pkinase-><-HEPN+DUF499*<-ParB+HNH<-METHYLASE<-TUDOR+SF1-SF2-Helicase+iREase+CTD+REase\|\|NTD+SF1-SF2-Helicase->HEPN-> | 1126 | Bacteria>Terrabacteria_group>Actinomycetota>Actinomycetes>Micromonosporales | GCF_017599015.1 | Micromonospora_tulbaghiae | J5U46_RS08620 | 81650..85030 | 479978 | Swt1-family-HEPN-domain-containing-protein-[Micromonospora-tulbaghiae] |
| WP_212823736.1 | DUF499_1.RM.like.systems | Phage_int_SAM_1+Phage_integrase->?-><-HTH_21+rve+rve_3<-HIN-HTH\|\|TUDOR+SF1-SF2-Helicase+iREase+CTD+REase->METHYLASE->HEPN+DUF499*-><-?<-?\|\|?->HIN-HTH-> | 1124 | Bacteria>Terrabacteria_group>Actinomycetota>Actinomycetes>Micromonosporales | GCF_018324255.1 | Polymorphospora_rubra | Prubr_RS09020 | 1986978..1990352 | 338584 | DUF499-domain-containing-protein-[Polymorphospora-rubra] |
| WP_221380246.1 | DUF499_1.RM.like.systems | sGTP->tRNA->Phage_int_SAM_4+Phage_integrase->METHYLASE-><-CARF+HEPN<-?<-ParB+HNH<-HEPN+DUF499*<-METHYLASE<-TUDOR+SF1-SF2-Helicase+iREase+CTD+REase<-SF1-SF2-Helicase<-?<-REase | 1128 | Bacteria>Terrabacteria_group>Actinomycetota>Actinomycetes>Micromonosporales | GCF_019710575.1 | Actinoplanes_polyasparticus | KXC43_RS33640 | 36558..39944 | 2856853 | Swt1-family-HEPN-domain-containing-protein-[Actinoplanes-sp.-TRM66264-DLM] |
| WP_260034146.1 | DUF499_1.RM.like.systems | <-SF1-SF2-Helicase<-?\|\|METHYLASE->HEPN+DUF499*->MORC->REase->HTH+SF1-SF2-Helicase+REase-> | 1133 | Bacteria>Terrabacteria_group>Actinomycetota>Actinomycetes>Micromonosporales | GCF_025152325.1 | Micromonospora_chalcea | M3G91_RS14945 | 8947..12348 | 1874 | Swt1-family-HEPN-domain-containing-protein-[Micromonospora-chalcea] |
| WP_167666591.1 | DUF499_1.RM.like.systems | HTH+TPR+TPR->Pkinase-><-HEPN+DUF499*<-ParB+HNH<-METHYLASE<-TUDOR+SF1-SF2-Helicase+iREase+CTD+REase\|\|NTD+SF1-SF2-Helicase->\|\|HNH-> | 1124 | Bacteria>Terrabacteria_group>Actinomycetota>Actinomycetes>Micromonosporales | GCF_900089605.1 | Micromonospora_narathiwatensis | GA0070621_RS04900 | 1072112..1075486 | 299146 | Swt1-family-HEPN-domain-containing-protein-[Micromonospora-narathiwatensis] |
| WP_030502747.1 | DUF499_1.RM.like.systems | HTH_3->?-><-DEDD_Tnp_IS110+Transposase_20<-zf-ISL3+DDE_Tnp_ISL3\|\|MULE-TRANSPOSASE-><-HEPN+DUF499*<-ParB+HNH<-METHYLASE<-TUDOR+SF1-SF2-Helicase+iREase+CTD+REase\|\|NTD+SF1-SF2-Helicase->REase->pPIWI+pPIWI-> | 1124 | Bacteria>Terrabacteria_group>Actinomycetota>Actinomycetes>Micromonosporales | - | Micromonospora_purpureochromogenes | IH31_RS0121315 | 105944..109318 | 47872 | Swt1-family-HEPN-domain-containing-protein-[Micromonospora-purpureochromogenes] |
| WP_225945547.1 | DUF499_1.RM.like.systems | HTH_17->Phage_int_SAM_1+Phage_integrase->?->TUDOR+SF1-SF2-Helicase+iREase+CTD+REase->METHYLASE->HEPN+DUF499*->\|\|?->DNA-glycosylase-><-?<-DUF6361<-RNase<-REase | 1132 | Bacteria>Terrabacteria_group>Actinomycetota>Actinomycetes>Micromonosporales | - | Plantactinospora_soyae | H4W31_RS15750 | 3518180..3521578 | 1544732 | DUF499-domain-containing-protein-[Plantactinospora-soyae] |
| WP_259918390.1 | DUF499_1.RM.like.systems | <-ABhydrolase<-?<-Aminotran_1_2<-ncRNA\|\|tRNA->?->SF1-SF2-helicase+DUF1998-><-ORC-CDC6+REase<-HEPN+DUF499*<-METHYLASE<-TUDOR+SF1-SF2-Helicase+iREase+CTD+REase\|\|ABhydrolase-> | 1108 | Bacteria>Terrabacteria_group>Actinomycetota>Actinomycetes>Mycobacteriales | - | Corynebacterium_sanguinis | M3A76_RS07775 | 66134..69460 | 2594913 | DUF499-domain-containing-protein-[Corynebacterium-sanguinis] |
| ADK68968.1 | DUF499_1.RM.like.systems | TUDOR+SF1-SF2-Helicase+iREase+CTD+REase->NERD->?->METHYLASE->HEPN+DUF499*->REase+TPR->?->Peptidase_M48-> | 1120 | Bacteria>Terrabacteria_group>Actinomycetota>Actinomycetes>Mycobacteriales | GCA_000143885.2 | Gordonia_sp._KTR9 | KTR9_4887 | 112287..115649 | 337191 | Predicted-ATPase-(AAA+-superfamily)-[Gordonia-sp.-KTR9] |
| EHB57971.1 | DUF499_1.RM.like.systems | <-GNTR-HTH\|\|NUDIX-><-?<-HEPN<-Schlafen+HATPase+HTH<-HEPN+DUF499*<-REase<-METHYLASE<-TUDOR+SF1-SF2-Helicase+iREase+CTD+REase<-PIN<-XIS-HTH | 1108 | Bacteria>Terrabacteria_group>Actinomycetota>Actinomycetes>Mycobacteriales | GCA_000230935.2 | Mycolicibacterium_rhodesiae_JS60 | MycrhDRAFT_0407 | 390606..393932 | 931627 | hypothetical-protein-MycrhDRAFT_0407-[Mycolicibacterium-rhodesiae-JS60] |
| EHI39101.1 | DUF499_1.RM.like.systems | <-GDResolvase+HIN-HTH\|\|HEPN-><-?<-HEPN+DUF499*<-METHYLASE<-TUDOR+SF1-SF2-Helicase+iREase+CTD+REase\|\|DNA-glycosylase-> | 1127 | Bacteria>Terrabacteria_group>Actinomycetota>Actinomycetes>Mycobacteriales | GCA_000234335.1 | Rhodococcus_opacus_PD630 | OPAG_08353 | 61698..65081 | 543736 | hypothetical-protein-OPAG_08353-[Rhodococcus-opacus-PD630] |
| GAB08850.1 | DUF499_1.RM.like.systems | Amidohydro_3+Amidohydro_1-><-TetR-HTH<-?\|\|HTH_19-><-?<-HEPN+DUF499*<-METHYLASE<-TUDOR+SF1-SF2-Helicase+iREase+CTD+REase\|\|DOC+HTH-><-?<-?<-HNH | 1140 | Bacteria>Terrabacteria_group>Actinomycetota>Actinomycetes>Mycobacteriales | GCA_000241265.2 | Gordonia_araii_NBRC_100433 | GOARA_021_00870 | 82752..86174 | 1073574 | hypothetical-protein-GOARA_021_00870-[Gordonia-araii-NBRC-100433] |
| GAB20465.1 | DUF499_1.RM.like.systems | <-ABC-ATPase+SbcC\|\|HTH_3-><-?<-?<-?<-HEPN+DUF499*<-METHYLASE\|\|HTH_19-> | 1118 | Bacteria>Terrabacteria_group>Actinomycetota>Actinomycetes>Mycobacteriales | GCA_000241305.2 | Gordonia_effusa_NBRC_100432 | GOEFS_115_01050 | 108493..111849 | 1077974 | hypothetical-protein-GOEFS_115_01050-[Gordonia-effusa-NBRC-100432] |
| AFC52219.1 | DUF499_1.RM.like.systems | <-GNTR-HTH\|\|NUDIX-><-?<-?<-HEPN<-Schlafen+HATPase+HTH<-HEPN+DUF499*<-METHYLASE\|\|?-><-TUDOR+SF1-SF2-Helicase+iREase+CTD+REase<-PIN<-XIS-HTH | 1043 | Bacteria>Terrabacteria_group>Actinomycetota>Actinomycetes>Mycobacteriales | GCA_000276825.1 | Mycobacterium_paraintracellulare | OCQ_07060 | 694865..697996 | 1138383 | hypothetical-protein-OCQ_07060-[Mycobacterium-paraintracellulare] |
| EKF23780.1 | DUF499_1.RM.like.systems | ABC-ATPase->Cysteine_desulfurase->?->?->?-><-ABhydrolase<-HEPN+DUF499*<-METHYLASE<-TUDOR+SF1-SF2-Helicase+iREase+CTD+REase | 1105 | Bacteria>Terrabacteria_group>Actinomycetota>Actinomycetes>Mycobacteriales | GCA_000300375.1 | Mycolicibacterium_hassiacum_DSM_44199 | C731_2148 | 37750..41067 | 1122247 | hypothetical-protein-C731_2148-[Mycolicibacterium-hassiacum-DSM-44199] |
| AHK35548.1 | DUF499_1.RM.like.systems | <-DUF499-p\|\|?-><-?<-DUF499-p<-HEPN+DUF499*<-METHYLASE<-METHYLASE-p<-TUDOR+SF1-SF2-Helicase+iREase+CTD+REase<-TUDOR+SF1-SF2-Helicase+iREase+CTD+REase-p\|\|HEPN-p->DNA-glycosylase-> | 649 | Bacteria>Terrabacteria_group>Actinomycetota>Actinomycetes>Mycobacteriales | GCA_000599545.1 | Rhodococcus_opacus_PD630 | Pd630_LPD10098 | 71803..73752 | 543736 | hypothetical-protein-Pd630_LPD10098-[Rhodococcus-opacus-PD630] |
| KUI22690.1 | DUF499_1.RM.like.systems | <-TetR-HTH\|\|DapB_N-><-?<-NTD+SF1-SF2-Helicase\|\|TUDOR+SF1-SF2-Helicase+iREase+CTD+REase->METHYLASE->HEPN+DUF499*-><-?\|\|?->?-><-Cluster182_14clades<-MACRODOMAIN<-MACRODOMAIN | 1104 | Bacteria>Terrabacteria_group>Actinomycetota>Actinomycetes>Mycobacteriales | GCA_001499905.1 | Mycobacterium_sp._GA-1285 | AU193_20905 | 131956..135270 | 1772282 | AAA-family-ATPase-[Mycobacterium-sp.-GA-1285] |
| KUI38022.1 | DUF499_1.RM.like.systems | MACRODOMAIN->Cluster182_14clades-><-?<-?<-HEPN+DUF499*<-METHYLASE<-TUDOR+SF1-SF2-Helicase+iREase+CTD+REase\|\|NTD+SF1-SF2-Helicase-><-DapB_N\|\|TetR-HTH-> | 1104 | Bacteria>Terrabacteria_group>Actinomycetota>Actinomycetes>Mycobacteriales | GCA_001500025.1 | Mycobacterium_sp._IS-1590 | AU197_09575 | 351187..354501 | 1772286 | AAA-family-ATPase-[Mycobacterium-sp.-IS-1590] |
| KUI43016.1 | DUF499_1.RM.like.systems | MACRODOMAIN->Cluster182_14clades-><-?<-?<-HEPN+DUF499*<-METHYLASE<-TUDOR+SF1-SF2-Helicase+iREase+CTD+REase\|\|<-?\|\|REase->Mu-transpos_C-> | 1101 | Bacteria>Terrabacteria_group>Actinomycetota>Actinomycetes>Mycobacteriales | GCA_001500045.1 | Mycobacterium_sp._GA-1199 | AU198_07655 | 256994..260299 | 1772287 | AAA-family-ATPase-[Mycobacterium-sp.-GA-1199] |
| OAH45661.1 | DUF499_1.RM.like.systems | TUDOR+SF1-SF2-Helicase+iREase+CTD+REase-><-?\|\|METHYLASE->ParB+HNH->HEPN+DUF499*->ANK+ANK+RAMA-> | 1134 | Bacteria>Terrabacteria_group>Actinomycetota>Actinomycetes>Mycobacteriales | GCA_001643175.1 | Dietzia_cinnamea | AYJ66_16790 | 32668..36072 | 321318 | AAA-family-ATPase-[Dietzia-cinnamea] |
| OBA79911.1 | DUF499_1.RM.like.systems | <-LIP+Peptidase_S9\|\|?-><-?<-HTH_3\|\|TUDOR+SF1-SF2-Helicase+iREase+CTD+REase->METHYLASE->HEPN+DUF499*->VapB_antitoxin->PIN-><-JetD<-?<-JetB<-JetA | 1098 | Bacteria>Terrabacteria_group>Actinomycetota>Actinomycetes>Mycobacteriales | GCA_001665375.1 | Mycolicibacterium_mucogenicum | A5642_03430 | 86031..89327 | 56689 | AAA-family-ATPase-[Mycolicibacterium-mucogenicum] |
| OBB45420.1 | DUF499_1.RM.like.systems | ParB+HNH->?-><-HTH+SF1-SF2-Helicase+REase<-REase<-MORC<-HEPN+DUF499*<-METHYLASE\|\|DNA_methylase->OLD-ABC-><-REase<-SF1-SF2-Helicase+REase<-SF1-SF2-Helicase+REase-p | 1135 | Bacteria>Terrabacteria_group>Actinomycetota>Actinomycetes>Mycobacteriales | GCA_001665535.1 | Mycobacterium_sp._852002-51961_SCH5331710 | A5752_02020 | 17252..20659 | 1834105 | hypothetical-protein-A5752_02020-[Mycobacterium-sp.-852002-51961_SCH5331710] |
| OBB72728.1 | DUF499_1.RM.like.systems | HTH_Tnp->?->?->?->\|\|TUDOR+SF1-SF2-Helicase+iREase+CTD+REase->METHYLASE->HEPN+DUF499*->ParB+HNH->HIN-HTH->HTH_21+rve+rve_3->HEPN-> | 1110 | Bacteria>Terrabacteria_group>Actinomycetota>Actinomycetes>Mycobacteriales | GCA_001665615.1 | Mycobacterium_sp._852014-52144_SCH5372336 | A5759_18055 | 92771..96103 | 1834115 | AAA-family-ATPase-[Mycobacterium-sp.-852014-52144_SCH5372336] |
| OBK03541.1 | DUF499_1.RM.like.systems | <-NTD+SF1-SF2-Helicase\|\|TUDOR+SF1-SF2-Helicase+iREase+CTD+REase->METHYLASE->ParB+HNH->HEPN+DUF499*->?->?->RnaseIII+DSRBD+RnaseIII+DSRBD->REC+HEPN-> | 1103 | Bacteria>Terrabacteria_group>Actinomycetota>Actinomycetes>Mycobacteriales | GCA_001673265.1 | Mycolicibacterium_conceptionense | A5639_22740 | 13192..16503 | 451644 | AAA-family-ATPase-[Mycolicibacterium-conceptionense] |

| Protein-ID | System | Gene-Neighbourhoods | Length | Taxonomy | Assembly-ID | Strain | Locus-Tag | Genomic-coordinates | TaxID | Fasta-defline |
|---|---|---|---|---|---|---|---|---|---|---|
| OCB34496.1 | DUF499_1.RM.like.systems | <-HEPN+DUF499*<-METHYLASE<-TUDOR+SF1-SF2-Helicase+iREase+CTD+REase-p | 1101 | Bacteria>Terrabacteria_group>Actinomycetota>Actinomycetes>Mycobacteriales | GCA_001686725.1 | Mycobacterium_malmoense | A9X02_22585 | 3529..6834 | 1780 | AAA-family-ATPase-[Mycobacterium-malmoense] |
| OCC17284.1 | DUF499_1.RM.like.systems | FAD-NAD-dep-oxidoreductase-><-HEPN+DUF499*<-METHYLASE<-TUDOR+SF1-SF2-Helicase+iREase+CTD+REase | 1134 | Bacteria>Terrabacteria_group>Actinomycetota>Actinomycetes>Mycobacteriales | GCA_001687425.1 | Prescottella_equi | AS590_01070 | 160611..164015 | 43767 | AAA-family-ATPase-[Prescottella-equi] |
| OCW84481.1 | DUF499_1.RM.like.systems | <-SbcC<-SIR2<-HEPN+DUF499*<-METHYLASE<-?<-MULE-TRANSPOSASE<-?<-HTH_28+rve+Mu-transpos_C | 1113 | Bacteria>Terrabacteria_group>Actinomycetota>Actinomycetes>Mycobacteriales | GCA_001704135.1 | Nocardia_farcinica | A8M60_10225 | 5506..8847 | 37329 | AAA-family-ATPase-[Nocardia-farcinica] |
| OFJ52823.1 | DUF499_1.RM.like.systems | TUDOR+SF1-SF2-Helicase+iREase+CTD+REase->METHYLASE->ParB+HNH->HEPN+DUF499*->?->?-><-Cluster182_14clades<-MACRODOMAIN<-?<-IclR-HTH | 1109 | Bacteria>Terrabacteria_group>Actinomycetota>Actinomycetes>Mycobacteriales | GCA_001766635.1 | Mycobacterium_grossiae | BEL07_15685 | 42542..45871 | 1552759 | AAA-family-ATPase-[Mycobacterium-grossiae] |
| OHT48546.1 | DUF499_1.RM.like.systems | PAIREDC-HTH+HIN-HTH-><-?\|\|TUDOR+SF1-SF2-Helicase+iREase+CTD+REase->METHYLASE->ParB+HNH->HEPN+DUF499*->?->?->?-><-?<-METHYLASE-p | 1109 | Bacteria>Terrabacteria_group>Actinomycetota>Actinomycetes>Mycobacteriales | GCA_001853545.1 | Mycobacteroides_chelonae | BKG62_21080 | 210158..213487 | 1774 | AAA-family-ATPase-[Mycobacteroides-chelonae] |
| OHV00237.1 | DUF499_1.RM.like.systems | TUDOR+SF1-SF2-Helicase+iREase+CTD+REase->METHYLASE->ParB+HNH->HEPN+DUF499*->SIR2-><-?\|\|SbcC-> | 1095 | Bacteria>Terrabacteria_group>Actinomycetota>Actinomycetes>Mycobacteriales | GCA_001854525.1 | Mycobacterium_talmoniae | BKN37_18410 | 8707..11994 | 1858794 | AAA-family-ATPase-[Mycobacterium-talmoniae] |
| OKH62617.1 | DUF499_1.RM.like.systems | TUDOR+SF1-SF2-Helicase+iREase+CTD+REase->METHYLASE->ParB+HNH->HEPN+DUF499*->OLD-ABC-><-?<-McrC-NTD+REase<-MAD-NTD+EVE+McrB-AAA | 1117 | Bacteria>Terrabacteria_group>Actinomycetota>Actinomycetes>Mycobacteriales | GCA_001905305.1 | Mycobacterium_sp._SWH-M5 | EB74_16030 | 8207..11560 | 1490483 | ATPase-AAA-[Mycobacterium-sp.-SWH-M5] |
| OKH84136.1 | DUF499_1.RM.like.systems | <-LIP+Peptidase_S9\|\|?-><-HTH_31\|\|TUDOR+SF1-SF2-Helicase+iREase+CTD+REase->METHYLASE->HEPN+DUF499*->VapB_antitoxin->PIN-><-JetD<-?<-JetB<-JetA | 1109 | Bacteria>Terrabacteria_group>Actinomycetota>Actinomycetes>Mycobacteriales | GCA_001905655.1 | Mycobacterium_sp._ST-F2 | EB75_06540 | 184865..188194 | 1490484 | ATPase-AAA-[Mycobacterium-sp.-ST-F2] |
| OMC31131.1 | DUF499_1.RM.like.systems | <-ParB+HNH<-NTD+SF1-SF2-Helicase\|\|TUDOR+SF1-SF2-Helicase+iREase+CTD+REase->METHYLASE->ParB+HNH->HEPN+DUF499*-><-Glyoxalase\|\|?-><-TetR-HTH\|\|?->?-><-Pkinase | 1100 | Bacteria>Terrabacteria_group>Actinomycetota>Actinomycetes>Mycobacteriales | GCA_001954115.1 | Mycobacterium_colombiense | A5739_14025 | 105145..108447 | 339268 | AAA-family-ATPase-[Mycobacterium-colombiense] |
| OMB68868.1 | DUF499_1.RM.like.systems | <-HEPN+DUF499*<-ParB+HNH<-METHYLASE<-TUDOR+SF1-SF2-Helicase+iREase+CTD+REase\|\|NTD+SF1-SF2-Helicase-> | 1103 | Bacteria>Terrabacteria_group>Actinomycetota>Actinomycetes>Mycobacteriales | GCA_001954125.1 | Mycolicibacterium_conceptionense | A5741_09375 | 1839..5150 | 451644 | AAA-family-ATPase-[Mycolicibacterium-conceptionense] |
| OMC39072.1 | DUF499_1.RM.like.systems | McrC-NTD+REase-><-NTD+SF1-SF2-Helicase\|\|TUDOR+SF1-SF2-Helicase+iREase+CTD+REase->METHYLASE->HEPN+DUF499*->?->?->?->?-><-Primpol+D5N+D5ATPase<-REase | 1103 | Bacteria>Terrabacteria_group>Actinomycetota>Actinomycetes>Mycobacteriales | GCA_001954135.1 | Mycobacterium_sp._GA-1841 | A5740_02790 | 288168..291479 | 1834154 | AAA-family-ATPase-[Mycobacterium-sp.-GA-1841] |
| OOK63774.1 | DUF499_1.RM.like.systems | <-CNMP+TIR<-Schlafen+HATPase+HTH<-HEPN+DUF499*<-METHYLASE<-METHYLASE-p<-TUDOR+SF1-SF2-Helicase+iREase+CTD+REase-p<-?<-SF1-SF2-Helicase-p | 1104 | Bacteria>Terrabacteria_group>Actinomycetota>Actinomycetes>Mycobacteriales | GCA_002003585.1 | Mycobacterium_kansasii | BZL29_8433 | 3439..6753 | 1768 | hypothetical-protein-BZL29_8433-[Mycobacterium-kansasii] |
| ORA83573.1 | DUF499_1.RM.like.systems | REase->?->?->?->\|\|TUDOR+SF1-SF2-Helicase+iREase+CTD+REase->METHYLASE->HEPN+DUF499*->ParB+HNH->HEPN->?-><-SF1-SF2-Helicase | 1110 | Bacteria>Terrabacteria_group>Actinomycetota>Actinomycetes>Mycobacteriales | GCA_002086285.1 | Mycolicibacter_kumamotonensis | BST28_00055 | 16546..19878 | 354243 | AAA-family-ATPase-[Mycolicibacter-kumamotonensis] |
| ORB28066.1 | DUF499_1.RM.like.systems | HKD-DNase+SF1-SF2-Helicase+TRD-><-?<-?<-?<-HTH_Tnp_1+HTH_21+rve+rve_2<-HIN-HTH<-HEPN+DUF499*<-ParB+HNH<-METHYLASE<-TUDOR+SF1-SF2-Helicase+iREase+CTD+REase\|\|NTD+SF1-SF2-Helicase-> | 1100 | Bacteria>Terrabacteria_group>Actinomycetota>Actinomycetes>Mycobacteriales | GCA_002086815.1 | Mycolicibacterium_parafortuitum | BST38_21950 | 32526..35828 | 39692 | AAA-family-ATPase-[Mycolicibacterium-parafortuitum] |
| ORV96500.1 | DUF499_1.RM.like.systems | TUDOR+SF1-SF2-Helicase+iREase+CTD+REase->METHYLASE->HEPN+DUF499*->?->?->HTH_17->GDResolvase+Recombinase->Phage_int_SAM_1+Phage_integrase-> | 1110 | Bacteria>Terrabacteria_group>Actinomycetota>Actinomycetes>Mycobacteriales | GCA_002101745.1 | Mycobacterium_kubicae | AWC13_18590 | 6697..10029 | 120959 | AAA-family-ATPase-[Mycobacterium-kubicae] |
| ORV36019.1 | DUF499_1.RM.like.systems | HTH_Tnp->?->?->?->\|\|TUDOR+SF1-SF2-Helicase+iREase+CTD+REase->ParB+HNH->HEPN+DUF499*-> | 1111 | Bacteria>Terrabacteria_group>Actinomycetota>Actinomycetes>Mycobacteriales | GCA_002102115.1 | Mycolicibacterium_doricum | AWC01_00650 | 40820..44155 | 126673 | AAA-family-ATPase-[Mycobacterium-doricum] |
| OSC29643.1 | DUF499_1.RM.like.systems | <-GNTR-HTH\|\|NUDIX-><-?<-?<-Schlafen+HATPase+HTH<-HEPN+DUF499*<-METHYLASE\|\|?-><-TUDOR+SF1-SF2-Helicase+iREase+CTD+REase<-PIN<-XIS-HTH\|\|GNTR-HTH-p-> | 1097 | Bacteria>Terrabacteria_group>Actinomycetota>Actinomycetes>Mycobacteriales | GCA_002104735.1 | Mycobacterium_paraintracellulare | B8W68_04660 | 128474..131767 | 1138383 | AAA-family-ATPase-[Mycobacterium-paraintracellulare] |
| OYN79669.1 | DUF499_1.RM.like.systems | Abhydrolase_3-><-?<-?<-HEPN+DUF499*<-METHYLASE<-TUDOR+SF1-SF2-Helicase+iREase+CTD+REase\|\|NTD+SF1-SF2-Helicase-> | 1109 | Bacteria>Terrabacteria_group>Actinomycetota>Actinomycetes>Mycobacteriales | GCA_002250655.1 | Mycolicibacterium_sphagni | CG716_11635 | 96200..99529 | 1786 | AAA-family-ATPase-[Mycolicibacterium-sphagni] |
| ASW98156.1 | DUF499_1.RM.like.systems | Pkinase-><-?\|\|TetR-HTH-><-?\|\|Glyoxalase->?-><-HEPN+DUF499*<-METHYLASE<-TUDOR+SF1-SF2-Helicase+iREase+CTD+REase\|\|NTD+SF1-SF2-Helicase-><-SAD+HNH | 1096 | Bacteria>Terrabacteria_group>Actinomycetota>Actinomycetes>Mycobacteriales | GCA_002285675.1 | Mycobacterium_intracellulare | CKJ67_10690 | 2227205..2230495 | 1767 | AAA-family-ATPase-[Mycobacterium-intracellulare] |
| ATO62480.2 | DUF499_1.RM.like.systems | Glyoxalase->HIN-HTH-><-DNA_methylase\|\|REase->METHYLASE->HEPN+DUF499*-><-HIN-HTH | 1138 | Bacteria>Terrabacteria_group>Actinomycetota>Actinomycetes>Mycobacteriales | GCA_002716905.4 | Mycobacterium_avium_subsp._hominissuis | BEP52_09350 | 1958340..1961756 | 439334 | DUF499-domain-containing-protein-[Mycobacterium-avium-subsp.-hominissuis] |
| PND54237.1 | DUF499_1.RM.like.systems | METHYLASE->HEPN+DUF499*-><-HIN-HTH\|\|?-><-HTH_Tnp_1+HTH_21+rve+rve_2\|\|DUF499-p-> | 654 | Bacteria>Terrabacteria_group>Actinomycetota>Actinomycetes>Mycobacteriales | GCA_002887815.1 | Mycobacterium_sp._ENV421 | CRM90_29025 | 2865..4829 | 1213407 | hypothetical-protein-CRM90_29025-[Mycobacterium-sp.-ENV421] |
| PQP40021.1 | DUF499_1.RM.like.systems | <-GspA-ATPase<-HIN-HTH\|\|Cluster182_14clades-><-?<-?<-?<-HEPN+DUF499*<-ParB+HNH<-METHYLASE<-TUDOR+SF1-SF2-Helicase+iREase+CTD+REase\|\|NTD+SF1-SF2-Helicase-> | 1103 | Bacteria>Terrabacteria_group>Actinomycetota>Actinomycetes>Mycobacteriales | GCA_002968275.1 | Mycolicibacterium_austroafricanum | C6A88_31860 | 14193..17504 | 39687 | AAA-family-ATPase-[Mycolicibacterium-austroafricanum] |
| PZT93668.1 | DUF499_1.RM.like.systems | ABhydrolase->XIS-HTH->TUDOR+SF1-SF2-Helicase+iREase+CTD+REase->METHYLASE->wHTH+REase->HEPN+DUF499*->OLD-ABC+TOPRIM->SF1-SF2-Helicase-> | 1133 | Bacteria>Terrabacteria_group>Actinomycetota>Actinomycetes>Mycobacteriales | GCA_003248405.1 | Gordonia_sp._(in:_high_G+C_Gram-positive_bacteria) | DI630_26685 | 9643..13044 | 84139 | AAA-family-ATPase-[Gordonia-sp.] |
| AYJ07722.1 | DUF499_1.RM.like.systems | TIR->?->?->TUDOR+SF1-SF2-Helicase+iREase+CTD+REase-><-?\|\|METHYLASE->HEPN+DUF499*->Schlafen+HATPase+HTH->HEPN-><-NUDIX\|\|GNTR-HTH->?->FtsK-> | 1097 | Bacteria>Terrabacteria_group>Actinomycetota>Actinomycetes>Mycobacteriales | GCA_003640565.1 | Mycobacterium_avium | DBO90_17285 | 3694672..3697965 | 1764 | DUF499-domain-containing-protein-[Mycobacterium-avium] |
| RUP39290.1 | DUF499_1.RM.like.systems | TUDOR+SF1-SF2-Helicase+iREase+CTD+REase->METHYLASE->HEPN+DUF499*-><-HTH_3\|\|?->TetR-HTH-><-Amidohydro_3+Amidohydro_1 | 1112 | Bacteria>Terrabacteria_group>Actinomycetota>Actinomycetes>Mycobacteriales | GCA_003987705.1 | Gordonia_sp._(in:_high_G+C_Gram-positive_bacteria) | EKK60_07255 | 9352..12690 | 84139 | ATP-binding-protein-[Gordonia-sp.] |
| RUP02204.1 | DUF499_1.RM.like.systems | TUDOR+SF1-SF2-Helicase+iREase+CTD+REase->METHYLASE->HEPN+DUF499*->?->?-><-Phage_int_SAM_1+Phage_integrase\|\|ZnR->?-><-ABhydrolase | 1097 | Bacteria>Terrabacteria_group>Actinomycetota>Actinomycetes>Mycobacteriales | GCA_003987775.1 | Mycobacterium_sp. | EKK34_25285 | 13112..16405 | 1785 | MAG:-ATP-binding-protein-[Mycobacterium-sp.] |
| RUP31532.1 | DUF499_1.RM.like.systems | <-SF1-SF2-Helicase<-TRD+TRD<-?<-SMODS<-HEPN+DUF499*<-METHYLASE<-TUDOR+SF1-SF2-Helicase+iREase+CTD+REase<-XIS-HTH<-TetR-HTH<-Phage_integrase+Phage_integrase<-Phage_integrase+Phage_integrase | 1121 | Bacteria>Terrabacteria_group>Actinomycetota>Actinomycetes>Mycobacteriales | GCA_003987785.1 | Mycolicibacterium_sp. | EKK51_14275 | 206389..209754 | 2320850 | MAG:-ATP-binding-protein-[Mycolicibacterium-sp.] |
| TDK92004.1 | DUF499_1.RM.like.systems | <-PAIREDC-HTH<-HIN-HTH\|\|JetD-><-PIN<-VapB_antitoxin\|\|<-HEPN+DUF499*<-METHYLASE<-TUDOR+SF1-SF2-Helicase+iREase+CTD+REase\|\|HTH_31->?-><-?\|\|LIP+Peptidase_S9-> | 1099 | Bacteria>Terrabacteria_group>Actinomycetota>Actinomycetes>Mycobacteriales | GCA_004359045.1 | Mycolicibacterium_mucogenicum | EUA03_05700 | 2711..6010 | 56689 | ATP-binding-protein-[Mycolicibacterium-mucogenicum] |
| TXH28009.1 | DUF499_1.RM.like.systems | <-LIP+Peptidase_S9\|\|?-><-?<-HTH_31\|\|TUDOR+SF1-SF2-Helicase+iREase+CTD+REase->METHYLASE->HEPN+DUF499*->\|\|VapB_antitoxin->PIN-><-JetD<-?<-JetB<-JetA | 1098 | Bacteria>Terrabacteria_group>Actinomycetota>Actinomycetes>Mycobacteriales | GCA_008012155.1 | Mycobacterium_sp. | E6R06_02260 | 39291..42587 | 1785 | MAG:-ATP-binding-protein-[Mycobacterium-sp.] |
| TXH27607.1 | DUF499_1.RM.like.systems | TUDOR+SF1-SF2-Helicase+iREase+CTD+REase->METHYLASE->HEPN+DUF499*-> | 848 | Bacteria>Terrabacteria_group>Actinomycetota>Actinomycetes>Mycobacteriales | GCA_008012155.1 | Mycobacterium_sp. | E6R06_03465 | 7027..9570 | 1785 | MAG:-ATP-binding-protein,-partial-[Mycobacterium-sp.] |
| TXI55381.1 | DUF499_1.RM.like.systems | <-FtsK<-?<-GNTR-HTH\|\|NUDIX-><-HEPN<-Schlafen+HATPase+HTH<-HEPN+DUF499*<-ParB+HNH<-METHYLASE<-TUDOR+SF1-SF2-Helicase+iREase+CTD+REase<-?<-HEPN | 1103 | Bacteria>Terrabacteria_group>Actinomycetota>Actinomycetes>Mycobacteriales | GCA_008015745.1 | Mycolicibacter_arupensis | E6Q54_12720 | 7339..10650 | 342002 | ATP-binding-protein-[Mycolicibacter-arupensis] |
| TXI48376.1 | DUF499_1.RM.like.systems | <-HIN-HTH<-NTD+SF1-SF2-Helicase\|\|TUDOR+SF1-SF2-Helicase+iREase+CTD+REase-><-?\|\|METHYLASE->ParB+HNH->HEPN+DUF499*-> | 1102 | Bacteria>Terrabacteria_group>Actinomycetota>Actinomycetes>Mycobacteriales | GCA_008015745.1 | Mycolicibacter_arupensis | E6Q54_23710 | 12171..15479 | 342002 | ATP-binding-protein-[Mycolicibacter-arupensis] |
| GES35195.1 | DUF499_1.RM.like.systems | <-GDResolvase+HIN-HTH\|\|?->RNase-><-HEPN+DUF499*<-METHYLASE | 1137 | Bacteria>Terrabacteria_group>Actinomycetota>Actinomycetes>Mycobacteriales | GCA_009176285.1 | Rhodococcus_aetherivorans | RAJCM14343_0442 | 3451..6864 | 191292 | hypothetical-protein-RAJCM14343_0442-[Rhodococcus-aetherivorans] |
| NKZ77519.1 | DUF499_1.RM.like.systems | MarR-HTH->tRNA-><-Phage_integrase<-HTH\|\|XIS-HTH->?-><-HEPN+DUF499*<-METHYLASE<-TUDOR+SF1-SF2-Helicase+iREase+CTD+REase\|\|<-GDResolvase+HIN-HTH\|\|?-><-?\|\|ABhydrolase-> | 1133 | Bacteria>Terrabacteria_group>Actinomycetota>Actinomycetes>Mycobacteriales | GCA_012443725.1 | Prescottella_equi | GTA28_11640 | 113900..117301 | 43767 | DUF499-domain-containing-protein-[Prescottella-equi] |
| NLG46720.1 | DUF499_1.RM.like.systems | <-DUF222+HNH<-HEPN+DUF499*<-METHYLASE\|\|CHTH-><-TUDOR+SF1-SF2-Helicase+iREase+CTD+REase\|\|HTH-> | 1141 | Bacteria>Terrabacteria_group>Actinomycetota>Actinomycetes>Mycobacteriales | GCA_012515855.1 | Gordonia_sp._(in:_high_G+C_Gram-positive_bacteria) | GX543_09665 | 1506..4931 | 84139 | ATP-binding-protein-[Gordonia-sp.-(in:-high-G+C-Gram-positive-bacteria)] |
| NNG97058.1 | DUF499_1.RM.like.systems | Amidohydro_3+Amidohydro_1-><-TetR-HTH<-?\|\|HTH_19-><-?<-HEPN+DUF499*<-METHYLASE<-TUDOR+SF1-SF2-Helicase+iREase+CTD+REase\|\|DOC+HTH-><-?<-?<-HNH | 1118 | Bacteria>Terrabacteria_group>Actinomycetota>Actinomycetes>Mycobacteriales | GCA_013004535.1 | Gordonia_araii_NBRC_100433 | HLA97_07325 | 102425..105781 | 1073574 | ATP-binding-protein-[Gordonia-araii-NBRC-100433] |
| MBB4136224.1 | DUF499_1.RM.like.systems | <-ABhydrolase<-?\|\|?-><-ACET<-HEPN+DUF499*<-METHYLASE\|\|DUF222+HNH-><-TUDOR+SF1-SF2-Helicase+iREase+CTD+REase | 1130 | Bacteria>Terrabacteria_group>Actinomycetota>Actinomycetes>Mycobacteriales | GCA_014197435.1 | Gordonia_humi | BKA16_002776 | 3005759..3009151 | 686429 | hypothetical-protein-[Gordonia-humi] |
| MBF6311832.1 | DUF499_1.RM.like.systems | <-ACET<-?\|\|?-><-STAS<-?<-HEPN+DUF499*<-METHYLASE<-METHYLASE-p<-TUDOR+SF1-SF2-Helicase+iREase+CTD+REase<-ZDNA-HTH | 1125 | Bacteria>Terrabacteria_group>Actinomycetota>Actinomycetes>Mycobacteriales | GCA_015477685.1 | Nocardia_farcinica | IU455_12400 | 1148688..1152065 | 37329 | DUF499-domain-containing-protein-[Nocardia-farcinica] |
| MBF6378463.1 | DUF499_1.RM.like.systems | <-ACET<-?\|\|?-><-STAS<-?<-HEPN+DUF499*<-METHYLASE<-TUDOR+SF1-SF2-Helicase+iREase+CTD+REase<-ZDNA-HTH<-?<-?<-Response_reg+LuxR-HTH | 1125 | Bacteria>Terrabacteria_group>Actinomycetota>Actinomycetes>Mycobacteriales | GCA_015477815.1 | Nocardia_farcinica | IU446_07990 | 515758..519135 | 37329 | DUF499-domain-containing-protein-[Nocardia-farcinica] |
| MBM4528840.1 | DUF499_1.RM.like.systems | wHTH(SBD)+HNH-><-ParB<-HEPN+DUF499*<-ParB+HNH-p<-METHYLASE<-TUDOR+SF1-SF2-Helicase+iREase+CTD+REase\|\|NTD+SF1-SF2-Helicase-><-SF1-SF2-Helicase<-HKD-DNase | 1121 | Bacteria>Terrabacteria_group>Actinomycetota>Actinomycetes>Mycobacteriales | GCA_016879355.1 | Prescottella_equi | GS563_07715 | 1641397..1644762 | 43767 | DUF499-domain-containing-protein-[Prescottella-equi] |
| MBU8833314.1 | DUF499_1.RM.like.systems | McrC-NTD+REase-><-?<-HTH_Tnp_1+HTH_21+rve+rve_3\|\|HTH_Tnp_1+HTH_21+rve+rve_3->?-><-REase<-HEPN+DUF499*<-ParB+HNH<-METHYLASE<-TUDOR+SF1-SF2-Helicase+iREase+CTD+REase\|\|DDE_Tnp_1_4->?->HEPN+SF1-SF2-Helicase-> | 1119 | Bacteria>Terrabacteria_group>Actinomycetota>Actinomycetes>Mycobacteriales | GCA_018524295.1 | Mycolicibacterium_goodii | KL858_28165 | 17594..20953 | 134601 | DUF499-domain-containing-protein-[Mycobacterium-goodii] |
| HJC59998.1 | DUF499_1.RM.like.systems | <-PIN<-?\|\|?-><-?<-CHTH<-?<-HEPN+DUF499*<-ParB+HNH<-METHYLASE<-TUDOR+SF1-SF2-Helicase+iREase+CTD+REase\|\|HTH_19-><-Synaptojanin | 1131 | Bacteria>Terrabacteria_group>Actinomycetota>Actinomycetes>Mycobacteriales | GCA_019119875.1 | Candidatus_Dietzia_intestinigallinarum | H9755_06660 | 12112..15507 | 2838537 | DUF499-domain-containing-protein-[Candidatus-Dietzia-intestinigallinarum] |
| MBX7452224.1 | DUF499_1.RM.like.systems | <-REase\|\|DNA_methylase-><-NTD+SF1-SF2-Helicase\|\|TUDOR+SF1-SF2-Helicase+iREase+CTD+REase->METHYLASE->ParB+HNH->HEPN+DUF499*->MORC->REase->HTH+SF1-SF2-Helicase+REase->\|\|MORC+SF1-SF2-Helicase->NERD+SF1-SF2-Helicase->SF1-SF2-Helicase-> | 1122 | Bacteria>Terrabacteria_group>Actinomycetota>Actinomycetes>Mycobacteriales | GCA_019710595.1 | Mycolicibacterium_aurantiacum | GR927_29910 | 284651..288019 | 2692455 | DUF499-domain-containing-protein-[Mycolicibacterium-aurantiacum] |
| MBZ4519978.1 | DUF499_1.RM.like.systems | <-FtsK<-?<-GNTR-HTH\|\|NUDIX-><-SIR2<-HEPN+DUF499*<-METHYLASE<-TUDOR+SF1-SF2-Helicase+iREase+CTD+REase<-RNase | 1102 | Bacteria>Terrabacteria_group>Actinomycetota>Actinomycetes>Mycobacteriales | GCA_020055125.1 | Mycobacterium_avium_subsp._hominissuis | GBP86_08110 | 354734..358042 | 439334 | ATP-binding-protein-[Mycobacterium-avium-subsp.-hominissuis] |
| MBZ4613710.1 | DUF499_1.RM.like.systems | <-HEPN+DUF499*<-ParB+HNH<-METHYLASE<-TUDOR+SF1-SF2-Helicase+iREase+CTD+REase | 1103 | Bacteria>Terrabacteria_group>Actinomycetota>Actinomycetes>Mycobacteriales | GCA_020055445.1 | Mycobacterium_avium_subsp._hominissuis | GBP66_23845 | 83..3394 | 439334 | ATP-binding-protein-[Mycobacterium-avium-subsp.-hominissuis] |
| MCB0937449.1 | DUF499_1.RM.like.systems | ParB+ParBC-><-?<-HTH_58<-HEPN+DUF499*<-METHYLASE<-TUDOR+SF1-SF2-Helicase+iREase+CTD+REase-p | 1125 | Bacteria>Terrabacteria_group>Actinomycetota>Actinomycetes>Mycobacteriales | GCA_020438785.1 | Mycobacterium_sp. | KDB55_15800 | 12247..15624 | 1785 | DUF499-domain-containing-protein-[Mycobacterium-sp.] |
| MCB0932438.1 | DUF499_1.RM.like.systems | <-NTD+SF1-SF2-Helicase\|\|TUDOR+SF1-SF2-Helicase+iREase+CTD+REase->METHYLASE->ParB+HNH->HEPN+DUF499*->?->?->?-><-Cluster182_14clades<-MACRODOMAIN\|\|PIN-> | 1120 | Bacteria>Terrabacteria_group>Actinomycetota>Actinomycetes>Mycobacteriales | GCA_020438845.1 | Mycobacterium_sp. | KDB71_11130 | 13103..16465 | 1785 | DUF499-domain-containing-protein-[Mycobacterium-sp.] |
| MCB0924960.1 | DUF499_1.RM.like.systems | <-GDResolvase+Recombinase+ZnR<-?<-?<-HEPN+DUF499*<-METHYLASE-p | 1126 | Bacteria>Terrabacteria_group>Actinomycetota>Actinomycetes>Mycobacteriales | GCA_020438895.1 | Mycobacterium_sp. | KDB50_10535 | 13047..16427 | 1785 | DUF499-domain-containing-protein-[Mycobacterium-sp.] |
| MCB1297052.1 | DUF499_1.RM.like.systems | Peptidase_M20-><-?\|\|?->?->?-><-REase<-HEPN+DUF499*\|\|MazE_antitoxin->PIN-><-METHYLASE | 1119 | Bacteria>Terrabacteria_group>Actinomycetota>Actinomycetes>Mycobacteriales | GCA_020438995.1 | Gordonia_sp._(in:_high_G+C_Gram-positive_bacteria) | KDB67_20615 | 20423..23782 | 84139 | DUF499-domain-containing-protein-[Gordonia-sp.] |
| UHJ57678.1 | DUF499_1.RM.like.systems | REase->DNA-glycosylase->?-><-HTH+SF1-SF2-Helicase+REase<-REase<-MORC<-HEPN+DUF499*<-METHYLASE<-REase<-PIN\|\|DNA_methylase-><-?<-Phage_integrase+Phage_integrase | 1138 | Bacteria>Terrabacteria_group>Actinomycetota>Actinomycetes>Mycobacteriales | GCA_021249325.1 | Mycolicibacterium_fortuitum | LT337_13390 | 2620949..2624365 | 1766 | DUF499-domain-containing-protein-[Mycolicibacterium-fortuitum] |
| MCF8588060.1 | DUF499_1.RM.like.systems | TUDOR+SF1-SF2-Helicase+iREase+CTD+REase-><-DUF222+HNH\|\|METHYLASE->HEPN+DUF499*->ACET-><-?\|\|?->ABhydrolase-> | 1125 | Bacteria>Terrabacteria_group>Actinomycetota>Actinomycetes>Mycobacteriales | GCA_021738805.1 | Gordonia_liuliyuniae | L5G33_06210 | 131250..134627 | 2911517 | DUF499-domain-containing-protein-[Gordonia-liuliyuniae] |
| MCF8569704.1 | DUF499_1.RM.like.systems | <-ABhydrolase<-?\|\|?-><-ACET<-HEPN+DUF499*<-METHYLASE\|\|DUF222+HNH-><-TUDOR+SF1-SF2-Helicase+iREase+CTD+REase | 1115 | Bacteria>Terrabacteria_group>Actinomycetota>Actinomycetes>Mycobacteriales | GCA_021738965.1 | Gordonia_zhenghanii | L5G32_05410 | 142683..146030 | 2911516 | DUF499-domain-containing-protein-[Gordonia-zhenghanii] |
| MCF8610758.1 | DUF499_1.RM.like.systems | TUDOR+SF1-SF2-Helicase+iREase+CTD+REase-><-DUF222+HNH\|\|METHYLASE->HEPN+DUF499*->ACET-><-?\|\|?->ABhydrolase-> | 1125 | Bacteria>Terrabacteria_group>Actinomycetota>Actinomycetes>Mycobacteriales | GCA_021739025.1 | Gordonia_liuliyuniae | L5G28_11435 | 131354..134731 | 2911517 | DUF499-domain-containing-protein-[Gordonia-liuliyuniae] |
| MCI1256747.1 | DUF499_1.RM.like.systems | MPTase-><-HTH_3\|\|TUDOR+SF1-SF2-Helicase+iREase+CTD+REase->METHYLASE->HEPN+DUF499*-><-?\|\|HEPN+HEPN-> | 1166 | Bacteria>Terrabacteria_group>Actinomycetota>Actinomycetes>Mycobacteriales | GCA_022641305.1 | Corynebacterium_provencense | LKG15_08685 | 46466..49966 | 1737425 | DUF499-domain-containing-protein-[Corynebacterium-provencense] |
| MCQ4152489.1 | DUF499_1.RM.like.systems | HEPN-><-NTD+SF1-SF2-Helicase\|\|TUDOR+SF1-SF2-Helicase+iREase+CTD+REase->METHYLASE->ParB+HNH->HEPN+DUF499*-> | 1120 | Bacteria>Terrabacteria_group>Actinomycetota>Actinomycetes>Mycobacteriales | GCA_024438075.1 | Rhodococcus_qingshengii | NOS52_32470 | 22838..26200 | 334542 | Swt1-family-HEPN-domain-containing-protein-[Rhodococcus-qingshengii] |
| MCQ4364655.1 | DUF499_1.RM.like.systems | <-NTD+SF1-SF2-Helicase\|\|TUDOR+SF1-SF2-Helicase+iREase+CTD+REase->METHYLASE->ParB+HNH->HEPN+DUF499*->ParB+HNH->HEPN-> | 1118 | Bacteria>Terrabacteria_group>Actinomycetota>Actinomycetes>Mycobacteriales | GCA_024451565.1 | Mycobacterium_gordonae | KQR54_26610 | 18169..21525 | 1778 | Swt1-family-HEPN-domain-containing-protein-[Mycobacterium-gordonae] |
| MCT1882934.1 | DUF499_1.RM.like.systems | <-ABhydrolase<-?<-Aminotran_1_2<-ncRNA\|\|tRNA->?->SF1-SF2-helicase+DUF1998-><-ORC-CDC6+REase<-HEPN+DUF499*<-METHYLASE<-TUDOR+SF1-SF2-Helicase+iREase+CTD+REase\|\|ABhydrolase-> | 1081 | Bacteria>Terrabacteria_group>Actinomycetota>Actinomycetes>Mycobacteriales | GCA_025144605.1 | Corynebacterium_sanguinis | M3A76_07760 | 66134..69379 | 2594913 | DUF499-domain-containing-protein-[Corynebacterium-sanguinis] |
| QHN29206.1 | DUF499_1.RM.like.systems | <-MULE-TRANSPOSASE\|\|ParA-Soj-PloopNTPase->ORC-CDC6-><-HTH+SF1-SF2-Helicase+REase<-REase<-MORC<-HEPN+DUF499*<-METHYLASE\|\|XIS-HTH+RVT-><-HNH\|\|?-><-REase\|\|DNA_methylase-> | 1138 | Bacteria>Terrabacteria_group>Actinomycetota>Actinomycetes>Mycobacteriales | GCA_025268675.1 | Gordonia_amarae | GII32_01220 | 287584..291000 | 36821 | DUF499-domain-containing-protein-[Gordonia-amarae] |
| MCT7366341.1 | DUF499_1.RM.like.systems | JetA->JetB->?->JetD-><-?\|\|?-><-HEPN+DUF499*\|\|HIN-HTH->HIN-HTH-><-ParB+HNH<-HTH_21+rve+rve_3<-HIN-HTH<-TUDOR+SF1-SF2-Helicase+iREase+CTD+REase | 1099 | Bacteria>Terrabacteria_group>Actinomycetota>Actinomycetes>Mycobacteriales | GCA_025331195.1 | Mycolicibacterium_llatzerense | A7G45_26120 | 438962..442261 | 280871 | AAA-family-ATPase-[Mycolicibacterium-llatzerense] |
| SHV22416.1 | DUF499_1.RM.like.systems | <-NTD+SF1-SF2-Helicase\|\|TUDOR+SF1-SF2-Helicase+iREase+CTD+REase-><-?\|\|METHYLASE->ParB+HNH->HEPN+DUF499*->?->?-><-MACRODOMAIN | 1122 | Bacteria>Terrabacteria_group>Actinomycetota>Actinomycetes>Mycobacteriales | GCA_900132885.1 | Mycobacteroides_abscessus_subsp._abscessus | SAMEA2259605_05819 | 13396..16764 | 1185650 | Conserved-protein-of-uncharacterised-function-(part1)-[Mycobacteroides-abscessus-subsp.-abscessus] |
| WP_011210977.1 | DUF499_1.RM.like.systems | <-HTH_21+rve+rve_3\|\|?->HIN-HTH->?->rve+rve_3-><-HEPN+DUF499*<-ParB+HNH<-METHYLASE<-TUDOR+SF1-SF2-Helicase+iREase+CTD+REase\|\|GDResolvase+Recombinase+ZnR-> | 1127 | Bacteria>Terrabacteria_group>Actinomycetota>Actinomycetes>Mycobacteriales | GCF_000009805.1 | Nocardia_farcinica | NFA_RS22030 | 4605548..4608931 | 37329 | Swt1-family-HEPN-domain-containing-protein-[Nocardia-farcinica] |
| WP_044505526.1 | DUF499_1.RM.like.systems | TUDOR+SF1-SF2-Helicase+iREase+CTD+REase->NERD->METHYLASE->HEPN+DUF499*->REase+TPR->?->Peptidase_M48-> | 1115 | Bacteria>Terrabacteria_group>Actinomycetota>Actinomycetes>Mycobacteriales | GCF_000143885.2 | Gordonia | KTR9_RS00485 | 112302..115649 | 2053 | MULTISPECIES:-DUF499-domain-containing-protein-[Gordonia] |
| WP_005520661.1 | DUF499_1.RM.like.systems | TUDOR+SF1-SF2-Helicase+iREase+CTD+REase->METHYLASE->HEPN+DUF499*->HEPN-><-?<-?<-JetB<-JetA | 1186 | Bacteria>Terrabacteria_group>Actinomycetota>Actinomycetes>Mycobacteriales | GCF_000158635.1 | Corynebacterium_matruchotii | CORMATOL_RS04405 | 70263..73823 | 43768 | DUF499-domain-containing-protein-[Corynebacterium-matruchotii] |
| WP_009974960.1 | DUF499_1.RM.like.systems | TIR->?->?->TUDOR+SF1-SF2-Helicase+iREase+CTD+REase-><-?\|\|METHYLASE->HEPN+DUF499*->Schlafen+HATPase+HTH->HEPN-><-?<-NUDIX\|\|GNTR-HTH-> | 1108 | Bacteria>Terrabacteria_group>Actinomycetota>Actinomycetes>Mycobacteriales | GCF_000174035.1 | Mycobacterium_avium | MAA44156_RS17120 | 3691227..3694553 | 1764 | Swt1-family-HEPN-domain-containing-protein-[Mycobacterium-avium] |
| WP_005526420.1 | DUF499_1.RM.like.systems | TUDOR+SF1-SF2-Helicase+iREase+CTD+REase->METHYLASE->HEPN+DUF499*->HEPN->Classical-AAA->?->Classical-AAA-> | 1181 | Bacteria>Terrabacteria_group>Actinomycetota>Actinomycetes>Mycobacteriales | GCF_000175375.1 | Corynebacterium_matruchotii | HBA49_RS02175 | 486574..490119 | 43768 | DUF499-domain-containing-protein-[Corynebacterium-matruchotii] |
| WP_013470723.1 | DUF499_1.RM.like.systems | <-MORC<-HTH+SF1-SF2-Helicase+REase<-DDE_Tnp_1_4<-REase<-MORC<-HEPN+DUF499*<-ParB+HNH<-MULE-TRANSPOSASE<-TUDOR+SF1-SF2-Helicase+iREase+CTD+REase<-HIN-HTH | 1122 | Bacteria>Terrabacteria_group>Actinomycetota>Actinomycetes>Mycobacteriales | GCF_000184435.1 | Mycolicibacterium_gilvum | MSPYR1_RS04230 | 878356..881724 | 1804 | Swt1-family-HEPN-domain-containing-protein-[Mycolicibacterium-gilvum] |
| WP_009680352.1 | DUF499_1.RM.like.systems | TUDOR+SF1-SF2-Helicase+iREase+CTD+REase-><-DUF222+HNH\|\|METHYLASE->HEPN+DUF499*-><-?\|\|ACET-><-Peptidase_S51<-?\|\|?->ABhydrolase-> | 1137 | Bacteria>Terrabacteria_group>Actinomycetota>Actinomycetes>Mycobacteriales | GCF_000192435.1 | Gordonia_neofelifaecis | SCNU_RS15560 | 9488..12901 | 945692 | DUF499-domain-containing-protein-[Gordonia-neofelifaecis] |
| WP_010121755.1 | DUF499_1.RM.like.systems | <-HTH_3<-?\|\|TUDOR+SF1-SF2-Helicase+iREase+CTD+REase->METHYLASE->HEPN+DUF499*->\|\|zf-ISL3+DDE_Tnp_ISL3->HEPN+HEPN-> | 1182 | Bacteria>Terrabacteria_group>Actinomycetota>Actinomycetes>Mycobacteriales | GCF_000213935.1 | Corynebacterium_nuruki | FSW06_RS05470 | 1228839..1232387 | 1032851 | DUF499-domain-containing-protein-[Corynebacterium-nuruki] |
| WP_013827697.1 | DUF499_1.RM.like.systems | GDResolvase+HIN-HTH-><-REase<-HEPN+DUF499*<-METHYLASE<-TUDOR+SF1-SF2-Helicase+iREase+CTD+REase<-XIS-HTH\|\|ParA+HTH->HTH_58-> | 1138 | Bacteria>Terrabacteria_group>Actinomycetota>Actinomycetes>Mycobacteriales | GCF_000214155.1 | Mycolicibacter_sinensis | JDM601_RS03755 | 802521..805937 | 875328 | DUF499-domain-containing-protein-[Mycolicibacter-sinensis] |
| WP_006361012.1 | DUF499_1.RM.like.systems | <-ACYC<-ACYC<-HEPN+DUF499*<-METHYLASE\|\|?-><-TUDOR+SF1-SF2-Helicase+iREase+CTD+REase | 1113 | Bacteria>Terrabacteria_group>Actinomycetota>Actinomycetes>Mycobacteriales | GCF_000225505.1 | Gordonia_alkanivorans | GOALK_RS21795 | 2511..5852 | 84096 | DUF499-domain-containing-protein-[Gordonia-alkanivorans] |
| WP_026061790.1 | DUF499_1.RM.like.systems | <-DNA_methylase\|\|REase-><-HEPN+DUF499* | 1135 | Bacteria>Terrabacteria_group>Actinomycetota>Actinomycetes>Mycobacteriales | GCF_000239135.1 | Rhodococcus_rhodochrous | RHODODRAFT_RS0223985 | 1954..5361 | 1829 | Swt1-family-HEPN-domain-containing-protein-[Rhodococcus-rhodochrous] |
| WP_245548345.1 | DUF499_1.RM.like.systems | Amidohydro_3+Amidohydro_1-><-TetR-HTH<-?\|\|HTH_19-><-?<-HEPN+DUF499*<-METHYLASE<-TUDOR+SF1-SF2-Helicase+iREase+CTD+REase\|\|DOC+HTH-><-?<-?<-HNH | 1116 | Bacteria>Terrabacteria_group>Actinomycetota>Actinomycetes>Mycobacteriales | GCF_000241265.1 | Gordonia_araii | GOARA_RS03580 | 82752..86102 | 263909 | Swt1-family-HEPN-domain-containing-protein-[Gordonia-araii] |
| WP_005189162.1 | DUF499_1.RM.like.systems | <-MULE-TRANSPOSASE\|\|ParA-Soj-PloopNTPase->ORC-CDC6-><-HTH+SF1-SF2-Helicase+REase<-REase<-MORC<-HEPN+DUF499*<-METHYLASE\|\|XIS-HTH+RVT-><-HNH\|\|?-><-REase\|\|DNA_methylase-> | 1138 | Bacteria>Terrabacteria_group>Actinomycetota>Actinomycetes>Mycobacteriales | GCF_000241345.1 | Gordonia_amarae | GII35_RS01230 | 287739..291155 | 36821 | DUF499-domain-containing-protein-[Gordonia-amarae] |
| WP_014358285.1 | DUF499_1.RM.like.systems | MACRODOMAIN+SIR2->SF1-SF2-Helicase-><-HTH+SF1-SF2-Helicase+REase<-REase<-MORC<-HEPN+DUF499*<-METHYLASE<-REase\|\|DNA_methylase->SAM-methylase-><-CR-Kinase+ORC-CDC6-like+CARF | 1127 | Bacteria>Terrabacteria_group>Actinomycetota>Actinomycetes>Mycobacteriales | GCF_000247715.1 | Gordonia_polyisoprenivorans | GPOL_RS00855 | 197189..200572 | 84595 | DUF499-domain-containing-protein-[Gordonia-polyisoprenivorans] |
| WP_041784793.1 | DUF499_1.RM.like.systems | <-GNTR-HTH\|\|NUDIX-><-?<-?<-HEPN<-Schlafen+HATPase+HTH<-HEPN+DUF499*<-METHYLASE\|\|?-><-TUDOR+SF1-SF2-Helicase+iREase+CTD+REase<-PIN<-XIS-HTH\|\|GNTR-HTH-p-> | 1108 | Bacteria>Terrabacteria_group>Actinomycetota>Actinomycetes>Mycobacteriales | GCF_000276825.1 | Mycobacterium_paraintracellulare | B8W68_RS04660 | 128474..131800 | 1138383 | Swt1-family-HEPN-domain-containing-protein-[Mycobacterium-paraintracellulare] |
| WP_014941573.1 | DUF499_1.RM.like.systems | XIS-HTH->?-><-?\|\|?->TUDOR+SF1-SF2-Helicase+iREase+CTD+REase->METHYLASE->HEPN+DUF499*->HEPN->GDResolvase+HIN-HTH-><-?<-?<-?<-HIN-HTH | 1124 | Bacteria>Terrabacteria_group>Actinomycetota>Actinomycetes>Mycobacteriales | GCF_000298095.1 | Mycobacterium_avium_complex_(MAC) | MIP_RS08165 | 1729452..1732826 | 120793 | MULTISPECIES:-DUF499-domain-containing-protein-[Mycobacterium-avium-complex-(MAC)] |
| WP_026213067.1 | DUF499_1.RM.like.systems | ABC-ATPase->Cysteine_desulfurase->?->?->?-><-ABhydrolase<-HEPN+DUF499*<-METHYLASE<-TUDOR+SF1-SF2-Helicase+iREase+CTD+REase | 1116 | Bacteria>Terrabacteria_group>Actinomycetota>Actinomycetes>Mycobacteriales | GCF_000300375.1 | Mycolicibacterium_hassiacum | A3G5_RS0100810 | 158802..162152 | 46351 | Swt1-family-HEPN-domain-containing-protein-[Mycolicibacterium-hassiacum] |
| WP_015289521.1 | DUF499_1.RM.like.systems | Peptidase_S8->?->?->REase-><-DDE_Tnp_1<-HEPN<-HEPN+DUF499*\|\|RHH_1->PIN-><-METHYLASE<-TUDOR+SF1-SF2-Helicase+iREase+CTD+REase | 1115 | Bacteria>Terrabacteria_group>Actinomycetota>Actinomycetes>Mycobacteriales | GCF_000328785.1 | Mycobacterium_canettii | BN42_RS05705 | 1219765..1223112 | 78331 | Swt1-family-HEPN-domain-containing-protein-[Mycobacterium-canettii] |
| WP_005174963.1 | DUF499_1.RM.like.systems | GDSL-><-HTH_19\|\|TUDOR+SF1-SF2-Helicase+iREase+CTD+REase->METHYLASE-><-PIN<-MazE_antitoxin\|\|HEPN+DUF499*->?->ABhydrolase+ABhydrolase-> | 1134 | Bacteria>Terrabacteria_group>Actinomycetota>Actinomycetes>Mycobacteriales | GCF_000332975.1 | Gordonia_aichiensis | GOACH_RS12130 | 81836..85240 | 36820 | Swt1-family-HEPN-domain-containing-protein-[Gordonia-aichiensis] |
| WP_006896364.1 | DUF499_1.RM.like.systems | <-ABhydrolase<-?\|\|?->Peptidase_S51-><-ACET\|\|?-><-HEPN+DUF499*<-METHYLASE\|\|DUF222+HNH-><-TUDOR+SF1-SF2-Helicase+iREase+CTD+REase<-?<-tRNA<-tRNA<-tRNA<-?<-tRNA<-tRNA<-tRNA\|\|tRNA->ABC-ATPase+SbcC-> | 1124 | Bacteria>Terrabacteria_group>Actinomycetota>Actinomycetes>Mycobacteriales | GCF_000333035.1 | Gordonia | GSI01S_RS09635 | 144602..147976 | 2053 | MULTISPECIES:-DUF499-domain-containing-protein-[Gordonia] |
| WP_003939259.1 | DUF499_1.RM.like.systems | <-PRTase+PRTase<-?<-SIR2+STALD<-?\|\|TUDOR+SF1-SF2-Helicase+iREase+CTD+REase->METHYLASE->HEPN+DUF499*-><-RNase<-?\|\|?-><-GDResolvase+HIN-HTH<-?<-ArsR-HTH | 1138 | Bacteria>Terrabacteria_group>Actinomycetota>Actinomycetes>Mycobacteriales | GCF_000341965.1 | Rhodococcus_ruber | G352_RS25680 | 27793..31209 | 1830 | Swt1-family-HEPN-domain-containing-protein-[Rhodococcus-ruber] |
| WP_008379160.1 | DUF499_1.RM.like.systems | METHYLASE->HEPN+DUF499*->ACET-><-?\|\|?->ABhydrolase-> | 1125 | Bacteria>Terrabacteria_group>Actinomycetota>Actinomycetes>Mycobacteriales | GCF_000344135.1 | Gordonia_malaquae | GM1_RS11930 | 2383..5760 | 410332 | DUF499-domain-containing-protein-[Gordonia-malaquae] |
| WP_053777653.1 | DUF499_1.RM.like.systems | <-ABC-ATPase+SbcC<-?\|\|?-><-ABhydrolase+ABhydrolase<-?<-HEPN+DUF499*<-METHYLASE<-TUDOR+SF1-SF2-Helicase+iREase+CTD+REase\|\|HTH_19-> | 1120 | Bacteria>Terrabacteria_group>Actinomycetota>Actinomycetes>Mycobacteriales | GCF_000347295.2 | Gordonia_sp._NB41Y | ISGA_RS14495 | 38931..42293 | 875808 | DUF499-domain-containing-protein-[Gordonia-sp.-NB41Y] |
| WP_018177518.1 | DUF499_1.RM.like.systems | HTH_Tnp_1+HTH_21+rve+rve_2->HEPN+DUF499*->?-><-ACET | 1113 | Bacteria>Terrabacteria_group>Actinomycetota>Actinomycetes>Mycobacteriales | GCF_000380485.1 | Jongsikchunia_kroppenstedtii | G341_RS0102455 | 2656..5997 | 1121721 | Swt1-family-HEPN-domain-containing-protein-[Jongsikchunia-kroppenstedtii] |
| WP_006942562.1 | DUF499_1.RM.like.systems | <-HEPN+DUF499*<-ParB+HNH<-METHYLASE<-TUDOR+SF1-SF2-Helicase+iREase+CTD+REase\|\|NTD+SF1-SF2-Helicase-> | 1119 | Bacteria>Terrabacteria_group>Actinomycetota>Actinomycetes>Mycobacteriales | GCF_000382865.1 | Rhodococcus_aetherivorans | EBESD8_RS20690 | 433..3792 | 191292 | Swt1-family-HEPN-domain-containing-protein-[Rhodococcus-aetherivorans] |
| WP_005939379.1 | DUF499_1.RM.like.systems | ABhydrolase+ABhydrolase-><-?<-DUF222+HNH\|\|TUDOR+SF1-SF2-Helicase+iREase+CTD+REase->METHYLASE->HEPN+DUF499*->?->ACET-><-?\|\|?->ABhydrolase-> | 1129 | Bacteria>Terrabacteria_group>Actinomycetota>Actinomycetes>Mycobacteriales | GCF_000420685.1 | Gordonia_hirsuta | GOHSU_RS07845 | 157749..161138 | 53427 | Swt1-family-HEPN-domain-containing-protein-[Gordonia-hirsuta] |
| WP_193376452.1 | DUF499_1.RM.like.systems | <-Synaptojanin\|\|wHTH+SAD+HNH-><-NTD+SF1-SF2-Helicase\|\|TUDOR+SF1-SF2-Helicase+iREase+CTD+REase->METHYLASE->HEPN+DUF499*-><-SIR2+STALD<-?<-Glyoxalase | 1107 | Bacteria>Terrabacteria_group>Actinomycetota>Actinomycetes>Mycobacteriales | GCF_000455205.1 | Mycobacterium_sp._UM_CSW | N415_RS14335 | 81564..84887 | 1370119 | Swt1-family-HEPN-domain-containing-protein-[Mycobacterium-sp.-UM_CSW] |
| WP_024794486.1 | DUF499_1.RM.like.systems | <-TetR-HTH\|\|?->CHTH->Glyoxalase-><-?<-HEPN+DUF499*<-METHYLASE\|\|?->?-><-TUDOR+SF1-SF2-Helicase+iREase+CTD+REase\|\|HTH_3-> | 1134 | Bacteria>Terrabacteria_group>Actinomycetota>Actinomycetes>Mycobacteriales | GCF_000524475.1 | Tomitella_biformata | JJ684_RS09895 | 229365..232769 | 630403 | DUF499-domain-containing-protein-[Tomitella-biformata] |
| WP_036372468.1 | DUF499_1.RM.like.systems | <-SF1-SF2-Helicase<-TRD+TRD<-?<-HTH_Tnp_1+HTH_21+rve+rve_3<-?<-HEPN+DUF499*<-SIR2+SIR2<-BrxL<-PglZ<-METHYLASE-PglX | 1123 | Bacteria>Terrabacteria_group>Actinomycetota>Actinomycetes>Mycobacteriales | GCF_000612725.1 | Mycolicibacterium_austroafricanum | BN976_RS17325 | 44630..48001 | 39687 | DUF499-domain-containing-protein-[Mycolicibacterium-austroafricanum] |
| WP_032392811.1 | DUF499_1.RM.like.systems | HIN-HTH->?-><-DNA-glycosylase\|\|TUDOR+SF1-SF2-Helicase+iREase+CTD+REase-p->HEPN+DUF499*-><-?\|\|?->HNH-> | 1123 | Bacteria>Terrabacteria_group>Actinomycetota>Actinomycetes>Mycobacteriales | GCF_000760775.1 | Rhodococcus_fascians | BH98_RS0123030 | 109500..112871 | 1828 | Swt1-family-HEPN-domain-containing-protein-[Rhodococcus-fascians] |
| WP_032373157.1 | DUF499_1.RM.like.systems | <-HEPN+DUF499*<-METHYLASE<-TUDOR+SF1-SF2-Helicase+iREase+CTD+REase | 1122 | Bacteria>Terrabacteria_group>Actinomycetota>Actinomycetes>Mycobacteriales | GCF_000760815.1 | Rhodococcus_fascians | BI01_RS0106730 | 27469..30837 | 1828 | Swt1-family-HEPN-domain-containing-protein-[Rhodococcus-fascians] |
| WP_039584625.1 | DUF499_1.RM.like.systems | DNA-glycosylase-><-ZnR+ZnR<-TIR\|\|TUDOR+SF1-SF2-Helicase+iREase+CTD+REase->METHYLASE->HEPN+DUF499*-><-RNase | 1138 | Bacteria>Terrabacteria_group>Actinomycetota>Actinomycetes>Mycobacteriales | GCF_000801125.1 | Rhodococcus_sp._Chr-9 | QR64_RS06710 | 8521..11937 | 713612 | Swt1-family-HEPN-domain-containing-protein-[Rhodococcus-sp.-Chr-9] |
| WP_232512159.1 | DUF499_1.RM.like.systems | <-HEPN+DUF499*<-METHYLASE<-REase\|\|DNA_methylase-> | 1128 | Bacteria>Terrabacteria_group>Actinomycetota>Actinomycetes>Mycobacteriales | GCF_000829075.1 | Mycobacterium_avium | B9C26_RS25090 | 191..3577 | 1764 | DUF499-domain-containing-protein-[Mycobacterium-avium] |
| WP_045537629.1 | DUF499_1.RM.like.systems | <-ABhydrolase<-?\|\|?->Peptidase_S51-><-ACET\|\|?-><-HEPN+DUF499*<-METHYLASE\|\|DUF222+HNH-><-TUDOR+SF1-SF2-Helicase+iREase+CTD+REase<-?<-tRNA<-tRNA<-tRNA<-?<-tRNA<-tRNA<-tRNA\|\|tRNA->ABC-ATPase+SbcC-> | 1124 | Bacteria>Terrabacteria_group>Actinomycetota>Actinomycetes>Mycobacteriales | GCF_000960465.1 | Gordonia_sihwensis | UG54_RS06085 | 5143..8517 | 173559 | DUF499-domain-containing-protein-[Gordonia-sihwensis] |
| WP_047262804.1 | DUF499_1.RM.like.systems | ABC-ATPase+SbcC-><-tRNA<-tRNA<-?\|\|?-><-HEPN+DUF499*<-TUDOR+SF1-SF2-Helicase+iREase+CTD+REase\|\|XIS-HTH->SbcC->TetR-HTH-> | 1140 | Bacteria>Terrabacteria_group>Actinomycetota>Actinomycetes>Mycobacteriales | GCF_001020985.1 | Corynebacterium_mustelae | CMUST_RS12605 | 2821253..2824675 | 571915 | Swt1-family-HEPN-domain-containing-protein-[Corynebacterium-mustelae] |
| WP_048320042.1 | DUF499_1.RM.like.systems | <-DNA_methylase\|\|?->METHYLASE->HEPN+DUF499*->MORC->REase->HTH+SF1-SF2-Helicase+REase-><-wHTH+REase<-SF1-SF2-Helicase<-Primpol+D5N+D5ATPase | 1137 | Bacteria>Terrabacteria_group>Actinomycetota>Actinomycetes>Mycobacteriales | GCF_001037935.1 | Rhodococcus_fascians | ACG96_RS20770 | 139814..143227 | 1828 | DUF499-domain-containing-protein-[Rhodococcus-fascians] |
| WP_050067339.1 | DUF499_1.RM.like.systems | MarR-HTH->tRNA-><-Arm-DNA-bind_4+Phage_int_SAM_3+Phage_integrase<-HTH\|\|XIS-HTH->?-><-HEPN+DUF499*<-METHYLASE<-TUDOR+SF1-SF2-Helicase+iREase+CTD+REase<-?<-GDResolvase+HIN-HTH | 1134 | Bacteria>Terrabacteria_group>Actinomycetota>Actinomycetes>Mycobacteriales | GCF_001040705.1 | Rhodococcus_sp._RD6.2 | RHCRD62_RS24305 | 110292..113696 | 260936 | Swt1-family-HEPN-domain-containing-protein-[Rhodococcus-sp.-RD6.2] |
| WP_059384669.1 | DUF499_1.RM.like.systems | ParB+HNH-p->?-><-REase\|\|TUDOR+SF1-SF2-Helicase+iREase+CTD+REase->METHYLASE->HEPN+DUF499*-><-REase\|\|DNA_methylase-><-SF1-SF2-Helicase | 1131 | Bacteria>Terrabacteria_group>Actinomycetota>Actinomycetes>Mycobacteriales | GCF_001047055.1 | Rhodococcus_rhodochrous | RR2_RS22685 | 9470..12865 | 1829 | Swt1-family-HEPN-domain-containing-protein-[Rhodococcus-rhodochrous] |
| WP_174485135.1 | DUF499_1.RM.like.systems | <-NTD+SF1-SF2-Helicase\|\|TUDOR+SF1-SF2-Helicase+iREase+CTD+REase-><-?\|\|METHYLASE->ParB+HNH->HEPN+DUF499*->?->?-><-?<-REase<-?\|\|ABhydrolase-> | 1113 | Bacteria>Terrabacteria_group>Actinomycetota>Actinomycetes>Mycobacteriales | GCF_001214605.1 | Mycobacteroides_abscessus | IUT00_RS08255 | 1657384..1660725 | 36809 | Swt1-family-HEPN-domain-containing-protein-[Mycobacteroides-abscessus] |
| WP_172798208.1 | DUF499_1.RM.like.systems | DapB_N-><-?\|\|?-><-NTD+SF1-SF2-Helicase\|\|TUDOR+SF1-SF2-Helicase+iREase+CTD+REase->METHYLASE->HEPN+DUF499*-><-?\|\|?->?-><-Cluster182_14clades<-MACRODOMAIN<-MACRODOMAIN | 1115 | Bacteria>Terrabacteria_group>Actinomycetota>Actinomycetes>Mycobacteriales | GCF_001499905.1 | Mycobacterium_sp._GA-1285 | AU193_RS04065 | 131923..135270 | 1772282 | Swt1-family-HEPN-domain-containing-protein-[Mycobacterium-sp.-GA-1285] |
| WP_197415592.1 | DUF499_1.RM.like.systems | Cluster182_14clades-><-?<-?<-HEPN+DUF499*<-METHYLASE<-TUDOR+SF1-SF2-Helicase+iREase+CTD+REase\|\|NTD+SF1-SF2-Helicase-><-DapB_N\|\|TetR-HTH-><-PemK_toxin | 1115 | Bacteria>Terrabacteria_group>Actinomycetota>Actinomycetes>Mycobacteriales | GCF_001500025.1 | Mycobacterium_sp._IS-1590 | AU197_RS19315 | 351187..354534 | 1772286 | Swt1-family-HEPN-domain-containing-protein-[Mycobacterium-sp.-IS-1590] |
| WP_197416164.1 | DUF499_1.RM.like.systems | Cluster182_14clades-><-?<-?<-HEPN+DUF499*<-METHYLASE<-TUDOR+SF1-SF2-Helicase+iREase+CTD+REase\|\|NTD+SF1-SF2-Helicase-> | 1112 | Bacteria>Terrabacteria_group>Actinomycetota>Actinomycetes>Mycobacteriales | GCF_001500045.1 | Mycobacterium_sp._GA-1199 | AU198_RS19260 | 256994..260332 | 1772287 | Swt1-family-HEPN-domain-containing-protein-[Mycobacterium-sp.-GA-1199] |
| WP_066467289.1 | DUF499_1.RM.like.systems | TetR-HTH-><-RADICAL-SAM<-?<-?<-?<-HEPN+DUF499*<-METHYLASE\|\|DUF222+HNH-><-TUDOR+SF1-SF2-Helicase+iREase+CTD+REase\|\|HTH_19-> | 1118 | Bacteria>Terrabacteria_group>Actinomycetota>Actinomycetes>Mycobacteriales | GCF_001552855.1 | Skermania_piniformis | SP4_RS02850 | 6456..9812 | 39122 | DUF499-domain-containing-protein-[Skermania-piniformis] |
| WP_204161112.1 | DUF499_1.RM.like.systems | <-LIP+Peptidase_S9\|\|?-><-?<-HTH_31\|\|TUDOR+SF1-SF2-Helicase+iREase+CTD+REase->METHYLASE->HEPN+DUF499*->VapB_antitoxin->PIN-><-JetD<-?<-JetB<-JetA | 1109 | Bacteria>Terrabacteria_group>Actinomycetota>Actinomycetes>Mycobacteriales | GCF_001557045.1 | Mycolicibacterium_mucogenicum | AX749_RS05580 | 20286..23615 | 56689 | Swt1-family-HEPN-domain-containing-protein-[Mycolicibacterium-mucogenicum] |
| WP_062368091.1 | DUF499_1.RM.like.systems | <-DUF222+HNH\|\|METHYLASE->HEPN+DUF499*-><-?\|\|ACET-><-Peptidase_S51<-?\|\|?->ABhydrolase-> | 1124 | Bacteria>Terrabacteria_group>Actinomycetota>Actinomycetes>Mycobacteriales | GCF_001578675.1 | Gordonia_sp._QH-12 | Y710_RS17975 | 9303..12677 | 1437876 | DUF499-domain-containing-protein-[Gordonia-sp.-QH-12] |
| WP_066167603.1 | DUF499_1.RM.like.systems | ACET->TUDOR+SF1-SF2-Helicase+iREase+CTD+REase-><-DUF222+HNH\|\|METHYLASE->HEPN+DUF499*-><-Peptidase_S51<-?\|\|?->ABhydrolase-> | 1139 | Bacteria>Terrabacteria_group>Actinomycetota>Actinomycetes>Mycobacteriales | GCF_001592365.1 | Gordonia_hydrophobica | GH2_RS11425 | 77242..80661 | 40516 | DUF499-domain-containing-protein-[Gordonia-hydrophobica] |
| WP_062879806.1 | DUF499_1.RM.like.systems | <-GNTR-HTH\|\|NUDIX-><-?<-HEPN<-Schlafen+HATPase+HTH<-HEPN+DUF499*<-METHYLASE\|\|?-><-TUDOR+SF1-SF2-Helicase+iREase+CTD+REase<-PIN<-XIS-HTH | 1108 | Bacteria>Terrabacteria_group>Actinomycetota>Actinomycetes>Mycobacteriales | GCF_001606235.1 | Mycobacteroides_abscessus | B5431_RS12990 | 716837..720163 | 36809 | Swt1-family-HEPN-domain-containing-protein-[Mycobacteroides-abscessus] |
| WP_067473443.1 | DUF499_1.RM.like.systems | DDE_Tnp_1-><-?\|\|TUDOR+SF1-SF2-Helicase+iREase+CTD+REase->METHYLASE->HEPN+DUF499*-><-?<-Peptidase<-?<-?\|\|HEPN-p-> | 1116 | Bacteria>Terrabacteria_group>Actinomycetota>Actinomycetes>Mycobacteriales | GCF_001612745.1 | Nocardia_amamiensis | NA4_RS22590 | 30808..34158 | 404578 | DUF499-domain-containing-protein-[Nocardia-amamiensis] |
| WP_067488116.1 | DUF499_1.RM.like.systems | <-REase\|\|?->?->TUDOR+SF1-SF2-Helicase+iREase+CTD+REase->ParB-p->HEPN+DUF499*->NTD+SF1-SF2-Helicase-><-RnaseIII+DSRBD+RnaseIII+DSRBD<-?<-ABhydrolase | 1123 | Bacteria>Terrabacteria_group>Actinomycetota>Actinomycetes>Mycobacteriales | GCF_001612905.1 | Nocardia_ignorata | DFR75_RS26965 | 96694..100065 | 145285 | Swt1-family-HEPN-domain-containing-protein-[Nocardia-ignorata] |
| WP_063013730.1 | DUF499_1.RM.like.systems | UbE1+JAB->?->Lsr2->TUDOR+SF1-SF2-Helicase+iREase+CTD+REase->METHYLASE->HEPN+DUF499*-><-?\|\|ParB+HNH-><-?\|\|HTH_3-><-DEDD_Tnp_IS110+Transposase_20 | 1126 | Bacteria>Terrabacteria_group>Actinomycetota>Actinomycetes>Mycobacteriales | GCF_001613005.1 | Nocardia | NN2_RS13920 | 30731..34111 | 1817 | MULTISPECIES:-Swt1-family-HEPN-domain-containing-protein-[Nocardia] |
| WP_068276445.1 | DUF499_1.RM.like.systems | METHYLASE->ParB+HNH->HEPN+DUF499*-> | 1122 | Bacteria>Terrabacteria_group>Actinomycetota>Actinomycetes>Mycobacteriales | GCF_001646865.1 | Aldersonia_kunmingensis | A8G08_RS16790 | 4064..7432 | 408066 | Swt1-family-HEPN-domain-containing-protein-[Aldersonia-kunmingensis] |

<table>
<tr><th>Protein-ID</th><th>System</th><th>Gene-Neighbourhoods</th><th>Length</th><th>Taxonomy</th><th>Assembly-ID</th><th>Strain</th><th>Locus-Tag</th><th>Genomic-coordinates</th><th>TaxID</th><th>Fasta-defline</th></tr>
<tr><td>WP_156666271.1</td><td>DUF499_1.RM.like.systems</td><td><-SbcC||Tox-RelE->HTH+MPTase-> TUDOR+ SF1-SF2-Helicase+iREase+CTD+REase -> METHYLASE -> HEPN + DUF499 *-></td><td>536</td><td>Bacteria>Terrabacteria_group>Actinomycetota>Actinomycetes>Mycobacteriales</td><td>GCF_001651055.1</td><td>Rhodococcus_sp._HS-D2</td><td>A1E04_RS24870</td><td>10916..12523</td><td>1384636</td><td>Swt1-family-HEPN-domain-containing-protein,-partial-[Rhodococcus-sp.-HS-D2]</td></tr>
<tr><td>WP_082921219.1</td><td>DUF499_1.RM.like.systems</td><td>METHYLASE -p-> HEPN + DUF499 *->MORC->REase-></td><td>1121</td><td>Bacteria>Terrabacteria_group>Actinomycetota>Actinomycetes>Mycobacteriales</td><td>GCF_001651055.1</td><td>Rhodococcus_sp._HS-D2</td><td>A1E04_RS27825</td><td>597..3962</td><td>1384636</td><td>Swt1-family-HEPN-domain-containing-protein-[Rhodococcus-sp.-HS-D2]</td></tr>
<tr><td>WP_067083703.1</td><td>DUF499_1.RM.like.systems</td><td>ParB+HNH->?->?-><-HTH+SF1-SF2-Helicase+REase<-REase<-MORC<- HEPN + DUF499 *<- METHYLASE <-REase||DNA_methylase->OLD-ABC+TOPRIM-><-REase<-SF1-SF2-Helicase+REase</td><td>1135</td><td>Bacteria>Terrabacteria_group>Actinomycetota>Actinomycetes>Mycobacteriales</td><td>GCF_001665535.1</td><td>Mycobacterium_sp._852002-51961_SCH5331710</td><td>A5752_RS07195</td><td>17252..20659</td><td>1834105</td><td>Swt1-family-HEPN-domain-containing-protein-[Mycobacterium-sp.-852002-51961_SCH5331710]</td></tr>
<tr><td>WP_197506097.1</td><td>DUF499_1.RM.like.systems</td><td>HTH_Tnp->?->?-><-NTD+SF1-SF2-Helicase|| TUDOR+ SF1-SF2-Helicase+iREase+CTD+REase -> METHYLASE -> HEPN + DUF499 *->ParB+HNH->HTH_Tnp_1+HTH_21+rve+rve_2-> HEPN -></td><td>1121</td><td>Bacteria>Terrabacteria_group>Actinomycetota>Actinomycetes>Mycobacteriales</td><td>GCF_001665615.1</td><td>Mycobacterium_sp._852014-52144_SCH5372336</td><td>A5759_RS17450</td><td>92738..96103</td><td>1834115</td><td>Swt1-family-HEPN-domain-containing-protein-[Mycobacterium-sp.-852014-52144_SCH5372336]</td></tr>
<tr><td>WP_207544692.1</td><td>DUF499_1.RM.like.systems</td><td><-NTD+SF1-SF2-Helicase|| TUDOR+ SF1-SF2-Helicase+iREase+CTD+REase -> METHYLASE ->ParB+HNH-> HEPN + DUF499 *->?->?->RnaseIII+DSRBD+RnaseIII+DSRBD->REC+ HEPN -></td><td>1114</td><td>Bacteria>Terrabacteria_group>Actinomycetota>Actinomycetes>Mycobacteriales</td><td>GCF_001673265.1</td><td>Mycolicibacterium_conceptionense</td><td>A5639_RS11935</td><td>13159..16503</td><td>451644</td><td>Swt1-family-HEPN-domain-containing-protein-[Mycolicibacterium-conceptionense]</td></tr>
<tr><td>WP_065288965.1</td><td>DUF499_1.RM.like.systems</td><td><-HTH_58<-ParA+HTH||BirA-HTH+CRP-HTH-> TUDOR+ SF1-SF2-Helicase+iREase+CTD+REase -> METHYLASE -> HEPN + DUF499 *->REase-><-GDResolvase+HIN-HTH<-?<-?||HTH_3-></td><td>1141</td><td>Bacteria>Terrabacteria_group>Actinomycetota>Actinomycetes>Mycobacteriales</td><td>GCF_001679965.1</td><td>Mycolicibacter_kumamotonensis</td><td>ACT18_RS17285</td><td>28680..32105</td><td>354243</td><td>DUF499-domain-containing-protein-[Mycolicibacter-kumamotonensis]</td></tr>
<tr><td>WP_065923600.1</td><td>DUF499_1.RM.like.systems</td><td><-RVT+HNH||?-> HEPN -><- HEPN + DUF499 *<- METHYLASE <- TUDOR+ SF1-SF2-Helicase+iREase+CTD+REase <-?<-?<-?||HNH-></td><td>1138</td><td>Bacteria>Terrabacteria_group>Actinomycetota>Actinomycetes>Mycobacteriales</td><td>GCF_001700945.1</td><td>Rhodococcus_sp._WB1</td><td>A4U64_RS26630</td><td>5329..8745</td><td>1033922</td><td>Swt1-family-HEPN-domain-containing-protein-[Rhodococcus-sp.-WB1]</td></tr>
<tr><td>WP_207546421.1</td><td>DUF499_1.RM.like.systems</td><td><-NTD+SF1-SF2-Helicase||PAIREDC-HTH+HIN-HTH-><-?|| TUDOR+ SF1-SF2-Helicase+iREase+CTD+REase -> METHYLASE ->ParB+HNH-> HEPN + DUF499 *->?->?->?-><-?<- METHYLASE -p</td><td>1120</td><td>Bacteria>Terrabacteria_group>Actinomycetota>Actinomycetes>Mycobacteriales</td><td>GCF_001853545.1</td><td>Mycobacteroides_chelonae</td><td>BKG62_RS21080</td><td>210125..213487</td><td>1774</td><td>Swt1-family-HEPN-domain-containing-protein-[Mycobacteroides-chelonae]</td></tr>
<tr><td>WP_207547258.1</td><td>DUF499_1.RM.like.systems</td><td>TUDOR+ SF1-SF2-Helicase+iREase+CTD+REase -> METHYLASE ->ParB+HNH-> HEPN + DUF499 *->SIR2->?->SbcC-></td><td>1106</td><td>Bacteria>Terrabacteria_group>Actinomycetota>Actinomycetes>Mycobacteriales</td><td>GCF_001854525.1</td><td>Mycobacterium_talmoniae</td><td>BKN37_RS18415</td><td>8674..11994</td><td>1858794</td><td>DUF499-domain-containing-protein-[Mycobacterium-talmoniae]</td></tr>
<tr><td>WP_071935222.1</td><td>DUF499_1.RM.like.systems</td><td><-URI-DNase<-?<-?<-?||PAIREDC-HTH+HIN-HTH-><- HEPN + DUF499 *||HTH_Tnp_1+HTH_21+rve+rve_3-><-?||DDE_Tnp_1_4-><-GDResolvase+Recombinase+ZnR<-HTH_Tnp_1+HTH_21+rve+rve_3<-HTH_Tnp_1+HTH_21+rve+rve_3</td><td>1131</td><td>Bacteria>Terrabacteria_group>Actinomycetota>Actinomycetes>Mycobacteriales</td><td>GCF_001886355.1</td><td>Rhodococcus_sp._2G</td><td>BO226_RS11040</td><td>2363059..2366454</td><td>1570939</td><td>DUF499-domain-containing-protein-[Rhodococcus-sp.-2G]</td></tr>
<tr><td>WP_072804385.1</td><td>DUF499_1.RM.like.systems</td><td>TUDOR+ SF1-SF2-Helicase+iREase+CTD+REase ->?->AbiEii-> METHYLASE -><-PIN<-MazE_antitoxin|| HEPN + DUF499 *->?->?-><-?<-Phd-YefM</td><td>1125</td><td>Bacteria>Terrabacteria_group>Actinomycetota>Actinomycetes>Mycobacteriales</td><td>GCF_001895005.1</td><td>Rhodococcus_yunnanensis</td><td>RY1_RS14860</td><td>83069..86446</td><td>278209</td><td>Swt1-family-HEPN-domain-containing-protein-[Rhodococcus-yunnanensis]</td></tr>
<tr><td>WP_174557649.1</td><td>DUF499_1.RM.like.systems</td><td><-HTH_31|| TUDOR+ SF1-SF2-Helicase+iREase+CTD+REase -> METHYLASE -> HEPN + DUF499 *->VapB_antitoxin->PIN-><-JetD<-?<-JetB<-JetA</td><td>1109</td><td>Bacteria>Terrabacteria_group>Actinomycetota>Actinomycetes>Mycobacteriales</td><td>GCF_001905655.1</td><td>Mycobacterium_sp._ST-F2</td><td>EB75_RS10110</td><td>184865..188194</td><td>1490484</td><td>Swt1-family-HEPN-domain-containing-protein-[Mycobacterium-sp.-ST-F2]</td></tr>
<tr><td>WP_074506051.1</td><td>DUF499_1.RM.like.systems</td><td><-ABhydrolase||?-><- HEPN + DUF499 *<- METHYLASE <- TUDOR+ SF1-SF2-Helicase+iREase+CTD+REase ||HTH_3->MODE-HTH+PBPII-></td><td>1150</td><td>Bacteria>Terrabacteria_group>Actinomycetota>Actinomycetes>Mycobacteriales</td><td>GCF_001912755.2</td><td>Corynebacterium_glutamicum</td><td>AUO95_RS02900</td><td>202510..205962</td><td>1718</td><td>DUF499-domain-containing-protein-[Corynebacterium-glutamicum]</td></tr>
<tr><td>WP_075835625.1</td><td>DUF499_1.RM.like.systems</td><td>NUDIX->?-> TUDOR+ SF1-SF2-Helicase+iREase+CTD+REase -> METHYLASE -> HEPN + DUF499 *->?-><- HEPN ||PAIREDC-HTH->?->GDResolvase+Recombinase+ZnR->Phage_int_SAM_4+Phage_integrase-></td><td>1134</td><td>Bacteria>Terrabacteria_group>Actinomycetota>Actinomycetes>Mycobacteriales</td><td>GCF_001942025.1</td><td>unclassified_Rhodococcus_(in:_high_G+C_Gram-positive_bacteria)</td><td>BH682_RS29985</td><td>71816..75220</td><td>192944</td><td>MULTISPECIES:-Swt1-family-HEPN-domain-containing-protein-[unclassified-Rhodococcus]</td></tr>
<tr><td>WP_170880036.1</td><td>DUF499_1.RM.like.systems</td><td><-ParB+HNH<-NTD+SF1-SF2-Helicase|| TUDOR+ SF1-SF2-Helicase+iREase+CTD+REase -> METHYLASE ->ParB+HNH-> HEPN + DUF499 *-><-Glyoxalase||?-><-TetR-HTH||?->?-><-Pkinase</td><td>1111</td><td>Bacteria>Terrabacteria_group>Actinomycetota>Actinomycetes>Mycobacteriales</td><td>GCF_001954115.1</td><td>Mycobacterium_colombiense</td><td>A5739_RS13730</td><td>105112..108447</td><td>339268</td><td>Swt1-family-HEPN-domain-containing-protein-[Mycobacterium-colombiense]</td></tr>
<tr><td>WP_207553229.1</td><td>DUF499_1.RM.like.systems</td><td><- HEPN + DUF499 *<-ParB+HNH<- METHYLASE <- TUDOR+ SF1-SF2-Helicase+iREase+CTD+REase ||NTD+SF1-SF2-Helicase-></td><td>1114</td><td>Bacteria>Terrabacteria_group>Actinomycetota>Actinomycetes>Mycobacteriales</td><td>GCF_001954125.1</td><td>Mycolicibacterium_conceptionense</td><td>A5741_RS31960</td><td>1839..5183</td><td>451644</td><td>Swt1-family-HEPN-domain-containing-protein-[Mycolicibacterium-conceptionense]</td></tr>
<tr><td>WP_175616187.1</td><td>DUF499_1.RM.like.systems</td><td>McrC-NTD+REase-><-NTD+SF1-SF2-Helicase|| TUDOR+ SF1-SF2-Helicase+iREase+CTD+REase -> METHYLASE -> HEPN + DUF499 *->?->?->?->?-><-Primpol+D5N+D5ATPase<-REase</td><td>1114</td><td>Bacteria>Terrabacteria_group>Actinomycetota>Actinomycetes>Mycobacteriales</td><td>GCF_001954135.1</td><td>Mycobacterium_sp._GA-1841</td><td>A5740_RS07670</td><td>288135..291479</td><td>1834154</td><td>Swt1-family-HEPN-domain-containing-protein-[Mycobacterium-sp.-GA-1841]</td></tr>
<tr><td>WP_174682959.1</td><td>DUF499_1.RM.like.systems</td><td><- HEPN + DUF499 *<-ParB+HNH<- METHYLASE <-rve_3</td><td>1090</td><td>Bacteria>Terrabacteria_group>Actinomycetota>Actinomycetes>Mycobacteriales</td><td>GCF_001954235.1</td><td>Mycolicibacterium_conceptionense</td><td>A5746_RS13475</td><td>1..3270</td><td>451644</td><td>Swt1-family-HEPN-domain-containing-protein,-partial-[Mycolicibacterium-conceptionense]</td></tr>
<tr><td>WP_079929183.1</td><td>DUF499_1.RM.like.systems</td><td>TUDOR+ SF1-SF2-Helicase+iREase+CTD+REase -p-><-STYKinase-like<-HTH|| METHYLASE -> HEPN + DUF499 *-></td><td>762</td><td>Bacteria>Terrabacteria_group>Actinomycetota>Actinomycetes>Mycobacteriales</td><td>GCF_002043085.1</td><td>Gordonia_sp._i37</td><td>B1964_RS05750</td><td>6455..8741</td><td>1961707</td><td>Swt1-family-HEPN-domain-containing-protein,-partial-[Gordonia-sp.-i37]</td></tr>
<tr><td>WP_207569995.1</td><td>DUF499_1.RM.like.systems</td><td><-NTD+SF1-SF2-Helicase|| TUDOR+ SF1-SF2-Helicase+iREase+CTD+REase -> METHYLASE -> HEPN + DUF499 *->ParB+HNH->?-><-SF1-SF2-Helicase</td><td>1121</td><td>Bacteria>Terrabacteria_group>Actinomycetota>Actinomycetes>Mycobacteriales</td><td>GCF_002086285.1</td><td>Mycolicibacter_kumamotonensis</td><td>BST28_RS00055</td><td>16513..19878</td><td>354243</td><td>Swt1-family-HEPN-domain-containing-protein-[Mycolicibacter-kumamotonensis]</td></tr>
<tr><td>WP_181792003.1</td><td>DUF499_1.RM.like.systems</td><td><-35exo+SF1-SF2-Helicase||HKD-DNase+SF1-SF2-Helicase+TRD-><-?<-?<-?<-HTH_Tnp_1+HTH_21+rve+rve_2<- HEPN + DUF499 *<-ParB+HNH<- METHYLASE <- TUDOR+ SF1-SF2-Helicase+iREase+CTD+REase ||NTD+SF1-SF2-Helicase-></td><td>1111</td><td>Bacteria>Terrabacteria_group>Actinomycetota>Actinomycetes>Mycobacteriales</td><td>GCF_002086815.1</td><td>Mycolicibacterium_parafortuitum</td><td>MPP7335_RS09255</td><td>1901353..1904688</td><td>39692</td><td>Swt1-family-HEPN-domain-containing-protein-[Mycolicibacterium-parafortuitum]</td></tr>
<tr><td>WP_176213652.1</td><td>DUF499_1.RM.like.systems</td><td><- HEPN + DUF499 *<- METHYLASE <- TUDOR+ SF1-SF2-Helicase+iREase+CTD+REase <-RNase</td><td>1113</td><td>Bacteria>Terrabacteria_group>Actinomycetota>Actinomycetes>Mycobacteriales</td><td>GCF_002089315.1</td><td>Mycobacterium_avium_complex_(MAC)</td><td>BST46_RS22320</td><td>85..3426</td><td>120793</td><td>MULTISPECIES:-Swt1-family-HEPN-domain-containing-protein-[Mycobacterium-avium-complex-(MAC)]</td></tr>
<tr><td>WP_176217705.1</td><td>DUF499_1.RM.like.systems</td><td><-SIR2<- HEPN + DUF499 *<- METHYLASE <- TUDOR+ SF1-SF2-Helicase+iREase+CTD+REase <-RNase</td><td>1113</td><td>Bacteria>Terrabacteria_group>Actinomycetota>Actinomycetes>Mycobacteriales</td><td>GCF_002089755.1</td><td>Mycobacterium_avium</td><td>B9C05_RS24160</td><td>1185..4526</td><td>1764</td><td>Swt1-family-HEPN-domain-containing-protein-[Mycobacterium-avium]</td></tr>
<tr><td>WP_085176531.1</td><td>DUF499_1.RM.like.systems</td><td>Phage_integrase+Phage_integrase->?->HTH_Tnp_1+HTH_21+rve+rve_3-> TUDOR+ SF1-SF2-Helicase+iREase+CTD+REase -> METHYLASE -> HEPN + DUF499 *-></td><td>806</td><td>Bacteria>Terrabacteria_group>Actinomycetota>Actinomycetes>Mycobacteriales</td><td>GCF_002101705.1</td><td>Mycolicibacterium_iranicum</td><td>AWC12_RS19975</td><td>111693..114110</td><td>912594</td><td>DUF499-domain-containing-protein,-partial-[Mycolicibacterium-iranicum]</td></tr>
<tr><td>WP_085185294.1</td><td>DUF499_1.RM.like.systems</td><td><-PemK_toxin<-?<-?||Glyoxalase-><-?<-OLD-ABC<- HEPN + DUF499 *<- METHYLASE <- TUDOR+ SF1-SF2-Helicase+iREase+CTD+REase ||NTD+SF1-SF2-Helicase-><-wHTH+SAD+HNH||Synaptojanin-></td><td>1114</td><td>Bacteria>Terrabacteria_group>Actinomycetota>Actinomycetes>Mycobacteriales</td><td>GCF_002101935.1</td><td>Mycobacterium_sp._IEC1808</td><td>AWB92_RS10440</td><td>126977..130321</td><td>1743230</td><td>Swt1-family-HEPN-domain-containing-protein-[Mycobacterium-sp.-IEC1808]</td></tr>
<tr><td>WP_201263267.1</td><td>DUF499_1.RM.like.systems</td><td><-NTD+SF1-SF2-Helicase|| TUDOR+ SF1-SF2-Helicase+iREase+CTD+REase -> METHYLASE -> HEPN + DUF499 *->REase->SF1-SF2-Helicase-p->HTH_Tnp_1+HTH_21+rve+rve_2->SF1-SF2-Helicase-p-></td><td>1109</td><td>Bacteria>Terrabacteria_group>Actinomycetota>Actinomycetes>Mycobacteriales</td><td>GCF_002201735.1</td><td>Mycobacterium_avium</td><td>CEV14_RS17015</td><td>150524..153853</td><td>1764</td><td>Swt1-family-HEPN-domain-containing-protein-[Mycobacterium-avium]</td></tr>
<tr><td>WP_202931651.1</td><td>DUF499_1.RM.like.systems</td><td><- HEPN + DUF499 *<-ParB-p<- METHYLASE ||?-><- TUDOR+ SF1-SF2-Helicase+iREase+CTD+REase</td><td>1109</td><td>Bacteria>Terrabacteria_group>Actinomycetota>Actinomycetes>Mycobacteriales</td><td>GCF_002245615.1</td><td>Mycobacterium_neumannii</td><td>CEX88_RS25285</td><td>2442..5771</td><td>2048551</td><td>Swt1-family-HEPN-domain-containing-protein-[Mycobacterium-neumannii]</td></tr>
<tr><td>WP_207760645.1</td><td>DUF499_1.RM.like.systems</td><td>Abhydrolase_3-><-?<-?<- HEPN + DUF499 *<- METHYLASE <- TUDOR+ SF1-SF2-Helicase+iREase+CTD+REase ||NTD+SF1-SF2-Helicase-></td><td>1120</td><td>Bacteria>Terrabacteria_group>Actinomycetota>Actinomycetes>Mycobacteriales</td><td>GCF_002250655.1</td><td>Mycolicibacterium_sphagni</td><td>CG716_RS11640</td><td>96200..99562</td><td>1786</td><td>Swt1-family-HEPN-domain-containing-protein-[Mycolicibacterium-sphagni]</td></tr>
<tr><td>WP_094652354.1</td><td>DUF499_1.RM.like.systems</td><td><-ABhydrolase+ABhydrolase<-REase<-ParB+HNH+RAMA||Primpol+D5N+D5ATPase->SF1-SF2-Helicase-><- HEPN + DUF499 *<-ParB+HNH-p<- METHYLASE <- TUDOR+ SF1-SF2-Helicase+iREase+CTD+REase ||?->Exonuc_V_gamma+REase->SF1-SF2-Helicase+REase-></td><td>1120</td><td>Bacteria>Terrabacteria_group>Actinomycetota>Actinomycetes>Mycobacteriales</td><td>GCF_002259465.1</td><td>Rhodococcus_fascians</td><td>CH297_RS18200</td><td>13414..16776</td><td>1828</td><td>Swt1-family-HEPN-domain-containing-protein-[Rhodococcus-fascians]</td></tr>
<tr><td>WP_179300635.1</td><td>DUF499_1.RM.like.systems</td><td>Pkinase-><-?||TetR-HTH-><-?||Glyoxalase->?-><- HEPN + DUF499 *<- METHYLASE <- TUDOR+ SF1-SF2-Helicase+iREase+CTD+REase ||NTD+SF1-SF2-Helicase->||?->?->TetR-HTH-></td><td>1107</td><td>Bacteria>Terrabacteria_group>Actinomycetota>Actinomycetes>Mycobacteriales</td><td>GCF_002285675.1</td><td>Mycobacterium_intracellulare</td><td>CKJ67_RS10690</td><td>2227205..2230528</td><td>1767</td><td>Swt1-family-HEPN-domain-containing-protein-[Mycobacterium-intracellulare]</td></tr>
<tr><td>WP_099384398.1</td><td>DUF499_1.RM.like.systems</td><td>TUDOR+ SF1-SF2-Helicase+iREase+CTD+REase -> METHYLASE ->ParB+HNH-> HEPN + DUF499 *->NTD+SF1-SF2-Helicase-><-?<-?||REase-></td><td>1115</td><td>Bacteria>Terrabacteria_group>Actinomycetota>Actinomycetes>Mycobacteriales</td><td>GCF_002735905.1</td><td>Williamsia_marianensis</td><td>CSW57_RS21175</td><td>1638758..1642105</td><td>322609</td><td>Swt1-family-HEPN-domain-containing-protein-[Williamsia-marianensis]</td></tr>
<tr><td>WP_102999581.1</td><td>DUF499_1.RM.like.systems</td><td>TUDOR+ SF1-SF2-Helicase+iREase+CTD+REase -> METHYLASE ->ParB+HNH-> HEPN + DUF499 *->zf-ISL3+DDE_Tnp_ISL3->?-><-DNA_methylase<- HEPN <-ParB+HNH<-GDResolvase+HIN-HTH</td><td>1121</td><td>Bacteria>Terrabacteria_group>Actinomycetota>Actinomycetes>Mycobacteriales</td><td>GCF_002893965.1</td><td>Rhodococcus_qingshengii</td><td>C1M55_RS31235</td><td>10127..13492</td><td>334542</td><td>Swt1-family-HEPN-domain-containing-protein-[Rhodococcus-qingshengii]</td></tr>
<tr><td>WP_104363764.1</td><td>DUF499_1.RM.like.systems</td><td>HIN-HTH->HTH_21+rve+rve_3-><-?<-?||ParB+HNH-> HEPN + DUF499 *-></td><td>1121</td><td>Bacteria>Terrabacteria_group>Actinomycetota>Actinomycetes>Mycobacteriales</td><td>GCF_002933495.1</td><td>Nocardia_nova</td><td>C5F51_RS20340</td><td>202110..205475</td><td>37330</td><td>Swt1-family-HEPN-domain-containing-protein-[Nocardia-nova]</td></tr>
<tr><td>WP_205765478.1</td><td>DUF499_1.RM.like.systems</td><td><-HIN-HTH<-?||Cluster182_14clades-><-?<-?<-?<- HEPN + DUF499 *<-ParB+HNH<- METHYLASE <- TUDOR+ SF1-SF2-Helicase+iREase+CTD+REase ||NTD+SF1-SF2-Helicase-></td><td>1114</td><td>Bacteria>Terrabacteria_group>Actinomycetota>Actinomycetes>Mycobacteriales</td><td>GCF_002968275.1</td><td>Mycolicibacterium_austroafricanum</td><td>C6A88_RS31870</td><td>14193..17537</td><td>39687</td><td>Swt1-family-HEPN-domain-containing-protein-[Mycolicibacterium-austroafricanum]</td></tr>
<tr><td>WP_105942269.1</td><td>DUF499_1.RM.like.systems</td><td><-ABhydrolase<-?||?-><-ACET<-?||?-><- HEPN + DUF499 *<- METHYLASE <- TUDOR+ SF1-SF2-Helicase+iREase+CTD+REase ||DUF222+HNH-><-ABhydrolase+ABhydrolase</td><td>1121</td><td>Bacteria>Terrabacteria_group>Actinomycetota>Actinomycetes>Mycobacteriales</td><td>GCF_002993285.1</td><td>Gordonia_iterans</td><td>C6V83_RS09885</td><td>2154114..2157479</td><td>1004901</td><td>DUF499-domain-containing-protein-[Gordonia-iterans]</td></tr>
<tr><td>WP_106851264.1</td><td>DUF499_1.RM.like.systems</td><td>SIGMA-HTH+SIGMA-HTH-><-wHTH(SBD)+HNH<-REase+SF1-SF2-Helicase|| TUDOR+ SF1-SF2-Helicase+iREase+CTD+REase -> METHYLASE ->ParB+HNH-> HEPN + DUF499 *->NTD+SF1-SF2-Helicase-></td><td>1123</td><td>Bacteria>Terrabacteria_group>Actinomycetota>Actinomycetes>Mycobacteriales</td><td>GCF_003013675.1</td><td>Prescottella_equi</td><td>C7H75_RS09240</td><td>1958101..1961472</td><td>43767</td><td>Swt1-family-HEPN-domain-containing-protein-[Prescottella-equi]</td></tr>
<tr><td>WP_108404986.1</td><td>DUF499_1.RM.like.systems</td><td>GDSL->Schlafen-><-?|| TUDOR+ SF1-SF2-Helicase+iREase+CTD+REase -> METHYLASE -> HEPN + DUF499 *->REase-p-><-tRNA||ncRNA->Aminotran_1_2->?->GDSL-></td><td>1112</td><td>Bacteria>Terrabacteria_group>Actinomycetota>Actinomycetes>Mycobacteriales</td><td>GCF_003070865.1</td><td>Corynebacterium_liangguodongii</td><td>C3E79_RS00685</td><td>144858..148196</td><td>2079535</td><td>DUF499-domain-containing-protein-[Corynebacterium-liangguodongii]</td></tr>
<tr><td>WP_182858863.1</td><td>DUF499_1.RM.like.systems</td><td>TUDOR+ SF1-SF2-Helicase+iREase+CTD+REase ->AbiGi-> METHYLASE -> HEPN + DUF499 *->SbcC-><-NUDIX||GNTR-HTH->?->FtsK-></td><td>1108</td><td>Bacteria>Terrabacteria_group>Actinomycetota>Actinomycetes>Mycobacteriales</td><td>GCF_003581765.1</td><td>Mycobacteroides_abscessus</td><td>D2E28_RS06470</td><td>1322672..1325998</td><td>36809</td><td>Swt1-family-HEPN-domain-containing-protein-[Mycobacteroides-abscessus]</td></tr>
<tr><td>WP_119699599.1</td><td>DUF499_1.RM.like.systems</td><td><-JetA<-?<-?<-?<-?<-?<- HEPN + DUF499 *<-ParB+HNH<- METHYLASE <- TUDOR+ SF1-SF2-Helicase+iREase+CTD+REase ||NTD+SF1-SF2-Helicase-></td><td>1118</td><td>Bacteria>Terrabacteria_group>Actinomycetota>Actinomycetes>Mycobacteriales</td><td>GCF_003586525.1</td><td>Rhodococcus_ruber</td><td>TN91_RS02590</td><td>62027..65383</td><td>1830</td><td>Swt1-family-HEPN-domain-containing-protein-[Rhodococcus-ruber]</td></tr>
<tr><td>WP_124259269.1</td><td>DUF499_1.RM.like.systems</td><td><-GDResolvase+HIN-HTH||XIS-HTH->PIN-><- HEPN + DUF499 *<- METHYLASE <- TUDOR+ SF1-SF2-Helicase+iREase+CTD+REase <-DNA-glycosylase</td><td>1120</td><td>Bacteria>Terrabacteria_group>Actinomycetota>Actinomycetes>Mycobacteriales</td><td>GCF_003844175.1</td><td>Rhodococcus_ruber</td><td>TN91_RS21990</td><td>3527..6889</td><td>1830</td><td>DUF499-domain-containing-protein-[Rhodococcus-ruber]</td></tr>
<tr><td>WP_206613802.1</td><td>DUF499_1.RM.like.systems</td><td>BirA-HTH+CRP-HTH-><- HEPN + DUF499 *<-ParB+HNH<- METHYLASE <- TUDOR+ SF1-SF2-Helicase+iREase+CTD+REase</td><td>1113</td><td>Bacteria>Terrabacteria_group>Actinomycetota>Actinomycetes>Mycobacteriales</td><td>GCF_004014805.1</td><td>Mycolicibacterium_elephantis</td><td>H7I95_RS08850</td><td>1305..4646</td><td>81858</td><td>Swt1-family-HEPN-domain-containing-protein-[Mycolicibacterium-elephantis]</td></tr>
<tr><td>WP_129978122.1</td><td>DUF499_1.RM.like.systems</td><td><-DUF4192||?-><-GDResolvase+HIN-HTH||?->RNase-><- HEPN + DUF499 *<- METHYLASE <- TUDOR+ SF1-SF2-Helicase+iREase+CTD+REase ||TIR->ZnR+ZnR-><-DNA-glycosylase</td><td>1138</td><td>Bacteria>Terrabacteria_group>Actinomycetota>Actinomycetes>Mycobacteriales</td><td>GCF_004153645.1</td><td>Rhodococcus_sp._Q1</td><td>EQ874_RS21580</td><td>43740..47156</td><td>2508718</td><td>Swt1-family-HEPN-domain-containing-protein-[Rhodococcus-sp.-Q1]</td></tr>
<tr><td>WP_134205957.1</td><td>DUF499_1.RM.like.systems</td><td><-DUF222+HNH<-?<-?<-HTH+SF1-SF2-Helicase+REase<-REase<-MORC<- HEPN + DUF499 *<- METHYLASE -p<-HerA/FtsK<-?<-BrxL<-PglZ<- METHYLASE -PglX</td><td>1137</td><td>Bacteria>Terrabacteria_group>Actinomycetota>Actinomycetes>Mycobacteriales</td><td>GCF_004366875.1</td><td>Mycobacteroides_salmoniphilum</td><td>CCUG62472_RS09110</td><td>402187..405600</td><td>404941</td><td>DUF499-domain-containing-protein-[Mycobacteroides-salmoniphilum]</td></tr>
<tr><td>WP_208722091.1</td><td>DUF499_1.RM.like.systems</td><td><-TetR-HTH<-GDResolvase+HIN-HTH||Lsr2->?-><-?<- HEPN + DUF499 *<-ParB+HNH<- METHYLASE <-UDG-conflict<- TUDOR+ SF1-SF2-Helicase+iREase+CTD+REase ||NTD+SF1-SF2-Helicase-></td><td>1122</td><td>Bacteria>Terrabacteria_group>Actinomycetota>Actinomycetes>Mycobacteriales</td><td>GCF_006384225.1</td><td>Rhodococcus_qingshengii</td><td>IGTS8_RS29980</td><td>39886..43254</td><td>334542</td><td>Swt1-family-HEPN-domain-containing-protein-[Rhodococcus-qingshengii]</td></tr>
<tr><td>WP_141142680.1</td><td>DUF499_1.RM.like.systems</td><td>SF1-SF2-Helicase->?-><-REase||DNA_methylase-> METHYLASE -> HEPN + DUF499 *->MORC->REase->HTH+SF1-SF2-Helicase+REase->wHTH+REase-></td><td>1137</td><td>Bacteria>Terrabacteria_group>Actinomycetota>Actinomycetes>Mycobacteriales</td><td>GCF_006517415.1</td><td>Mycolicibacterium_fortuitum</td><td>FKW78_RS12965</td><td>17898..21311</td><td>1766</td><td>DUF499-domain-containing-protein-[Mycolicibacterium-fortuitum]</td></tr>
<tr><td>WP_143907483.1</td><td>DUF499_1.RM.like.systems</td><td><-PIN<-?<-?<- HEPN + DUF499 *<-Glyoxalase||ArsR-HTH->?->DNA_methylase-><-?<- METHYLASE</td><td>1132</td><td>Bacteria>Terrabacteria_group>Actinomycetota>Actinomycetes>Mycobacteriales</td><td>GCF_007559025.1</td><td>Tomitella_fengzijianii</td><td>M2W19_RS17095</td><td>32946..36344</td><td>2597660</td><td>DUF499-domain-containing-protein-[Tomitella-fengzijianii]</td></tr>
<tr><td>WP_144317543.1</td><td>DUF499_1.RM.like.systems</td><td><-ABhydrolase<-?<-Aminotran_1_2<-ncRNA||tRNA-><-ORC-CDC6+REase<- HEPN + DUF499 *<- METHYLASE <- TUDOR+ SF1-SF2-Helicase+iREase+CTD+REase ||ABhydrolase-></td><td>1111</td><td>Bacteria>Terrabacteria_group>Actinomycetota>Actinomycetes>Mycobacteriales</td><td>GCF_007641235.1</td><td>Corynebacterium_sanguinis</td><td>E3227_RS03440</td><td>690306..693641</td><td>2594913</td><td>DUF499-domain-containing-protein-[Corynebacterium-sanguinis]</td></tr>
<tr><td>WP_145941802.1</td><td>DUF499_1.RM.like.systems</td><td>ORC-CDC6+REase-><- HEPN + DUF499 *<- METHYLASE <- TUDOR+ SF1-SF2-Helicase+iREase+CTD+REase ||HTH_3-></td><td>1182</td><td>Bacteria>Terrabacteria_group>Actinomycetota>Actinomycetes>Mycobacteriales</td><td>GCF_007673095.1</td><td>Corynebacterium_glyciniphilum</td><td>FS507_RS05565</td><td>165529..169077</td><td>1404244</td><td>DUF499-domain-containing-protein-[Corynebacterium-glyciniphilum]</td></tr>
<tr><td>WP_276761078.1</td><td>DUF499_1.RM.like.systems</td><td><-FtsK<-?<-GNTR-HTH||NUDIX-><- HEPN <-Schlafen+HATPase+HTH<- HEPN + DUF499 *<-ParB+HNH<- METHYLASE <- TUDOR+ SF1-SF2-Helicase+iREase+CTD+REase <-?<- HEPN <- HEPN</td><td>1114</td><td>Bacteria>Terrabacteria_group>Actinomycetota>Actinomycetes>Mycobacteriales</td><td>GCF_008015745.1</td><td>Mycolicibacter_arupensis</td><td>E6Q54_RS12710</td><td>7339..10683</td><td>342002</td><td>Swt1-family-HEPN-domain-containing-protein-[Mycolicibacter-arupensis]</td></tr>
<tr><td>WP_276763638.1</td><td>DUF499_1.RM.like.systems</td><td><-NTD+SF1-SF2-Helicase|| TUDOR+ SF1-SF2-Helicase+iREase+CTD+REase -><-?|| METHYLASE ->ParB+HNH-> HEPN + DUF499 *-></td><td>1113</td><td>Bacteria>Terrabacteria_group>Actinomycetota>Actinomycetes>Mycobacteriales</td><td>GCF_008015745.1</td><td>Mycolicibacter_arupensis</td><td>E6Q54_RS23755</td><td>12138..15479</td><td>342002</td><td>Swt1-family-HEPN-domain-containing-protein-[Mycolicibacter-arupensis]</td></tr>
<tr><td>WP_149357518.1</td><td>DUF499_1.RM.like.systems</td><td>Lsr2-><-?<-?|| TUDOR+ SF1-SF2-Helicase+iREase+CTD+REase -> METHYLASE -> HEPN + DUF499 *->REase-><-?<-?||GDResolvase+HIN-HTH->URI-DNase-></td><td>1133</td><td>Bacteria>Terrabacteria_group>Actinomycetota>Actinomycetes>Mycobacteriales</td><td>GCF_008306195.1</td><td>Rhodococcus_qingshengii</td><td>D6M20_RS01760</td><td>367041..370442</td><td>334542</td><td>DUF499-domain-containing-protein-[Rhodococcus-qingshengii]</td></tr>
<tr><td>WP_176749585.1</td><td>DUF499_1.RM.like.systems</td><td><-NTD+SF1-SF2-Helicase|| TUDOR+ SF1-SF2-Helicase+iREase+CTD+REase -> METHYLASE -p->DDE_Tnp_1_4->ParB+HNH-> HEPN + DUF499 *->?->?->?-><-Cluster182_14clades<-?<-IclR-HTH</td><td>1120</td><td>Bacteria>Terrabacteria_group>Actinomycetota>Actinomycetes>Mycobacteriales</td><td>GCF_008329645.1</td><td>Mycobacterium_grossiae</td><td>FZ046_RS21205</td><td>4404866..4408228</td><td>1552759</td><td>Swt1-family-HEPN-domain-containing-protein-[Mycobacterium-grossiae]</td></tr>
<tr><td>WP_149772783.1</td><td>DUF499_1.RM.like.systems</td><td><-HTH_58<-ParA+HTH||XIS-HTH-> TUDOR+ SF1-SF2-Helicase+iREase+CTD+REase -> METHYLASE -> HEPN + DUF499 *->REase-><-GDResolvase+HIN-HTH</td><td>1138</td><td>Bacteria>Terrabacteria_group>Actinomycetota>Actinomycetes>Mycobacteriales</td><td>GCF_008373105.1</td><td>Mycolicibacter_arupensis</td><td>F0402_RS09270</td><td>8758..12174</td><td>342002</td><td>DUF499-domain-containing-protein-[Mycolicibacter-arupensis]</td></tr>
<tr><td>WP_043798089.1</td><td>DUF499_1.RM.like.systems</td><td><-GDResolvase+HIN-HTH||?->RNase-><- HEPN + DUF499 *</td><td>1139</td><td>Bacteria>Terrabacteria_group>Actinomycetota>Actinomycetes>Mycobacteriales</td><td>GCF_009176285.1</td><td>Rhodococcus</td><td>GDN71_RS02225</td><td>3451..6870</td><td>1827</td><td>MULTISPECIES:-Swt1-family-HEPN-domain-containing-protein-[Rhodococcus]</td></tr>
<tr><td>WP_153212242.1</td><td>DUF499_1.RM.like.systems</td><td><-NTD+SF1-SF2-Helicase|| TUDOR+ SF1-SF2-Helicase+iREase+CTD+REase ->UDG-conflict-> METHYLASE ->ParB+HNH-> HEPN + DUF499 *-><-NUDIX||?->REase->?-><-HAD<-ABC-ATPase+SbcC</td><td>1121</td><td>Bacteria>Terrabacteria_group>Actinomycetota>Actinomycetes>Mycobacteriales</td><td>GCF_009495805.1</td><td>Rhodococcus_erythropolis</td><td>GEO20_RS16000</td><td>38887..42252</td><td>1833</td><td>Swt1-family-HEPN-domain-containing-protein-[Rhodococcus-erythropolis]</td></tr>
<tr><td>WP_159371683.1</td><td>DUF499_1.RM.like.systems</td><td>REase+SPF-cons-><-?<-CHTH<-?<- HEPN + DUF499 *<-?<- METHYLASE <-?<- TUDOR+ SF1-SF2-Helicase+iREase+CTD+REase</td><td>1117</td><td>Bacteria>Terrabacteria_group>Actinomycetota>Actinomycetes>Mycobacteriales</td><td>GCF_009856645.1</td><td>Gordonia_sp._JH63</td><td>GR168_RS22070</td><td>4982408..4985761</td><td>2698900</td><td>DUF499-domain-containing-protein-[Gordonia-sp.-JH63]</td></tr>
<tr><td>WP_213249149.1</td><td>DUF499_1.RM.like.systems</td><td><-HTH+SF1-SF2-Helicase+REase<-REase<-MORC<- HEPN + DUF499 *<-SF1-SF2-Helicase<-OLD-ABC+TOPRIM||DNA_methylase-><-?||HNH-></td><td>1138</td><td>Bacteria>Terrabacteria_group>Actinomycetota>Actinomycetes>Mycobacteriales</td><td>GCF_009914535.1</td><td>Gordonia</td><td>IDH24_RS11690</td><td>142027..145443</td><td>2053</td><td>DUF499-domain-containing-protein-[Gordonia-pseudamarae]</td></tr>
<tr><td>WP_161926536.1</td><td>DUF499_1.RM.like.systems</td><td><-TetR-HTH<-?||HTH_19-><-?<- HEPN + DUF499 *<-DUF222+HNH<- METHYLASE <- TUDOR+ SF1-SF2-Helicase+iREase+CTD+REase ||DOC+HTH-></td><td>1124</td><td>Bacteria>Terrabacteria_group>Actinomycetota>Actinomycetes>Mycobacteriales</td><td>GCF_009932435.1</td><td>Gordonia_crocea</td><td>nbrcactino_RS06035</td><td>1265301..1268675</td><td>589162</td><td>DUF499-domain-containing-protein-[Gordonia-crocea]</td></tr>
<tr><td>WP_161896301.1</td><td>DUF499_1.RM.like.systems</td><td><-ABhydrolase<-?||?-><-ACET<- HEPN + DUF499 *<-AbiEii<-DDE_Tnp_1_4<-?<- METHYLASE ||DUF222+HNH-><- TUDOR+ SF1-SF2-Helicase+iREase+CTD+REase</td><td>1123</td><td>Bacteria>Terrabacteria_group>Actinomycetota>Actinomycetes>Mycobacteriales</td><td>GCF_009932475.1</td><td>Gordonia_spumicola</td><td>GYB83_RS15365</td><td>872250..875621</td><td>589161</td><td>DUF499-domain-containing-protein-[Gordonia-spumicola]</td></tr>
<tr><td>WP_174814236.1</td><td>DUF499_1.RM.like.systems</td><td>PAIREDC-HTH->HIN-HTH->?-> TUDOR+ SF1-SF2-Helicase+iREase+CTD+REase -> METHYLASE -> HEPN + DUF499 *->?->HTH_17->GDResolvase+Recombinase->Phage_int_SAM_1+Phage_integrase-></td><td>1121</td><td>Bacteria>Terrabacteria_group>Actinomycetota>Actinomycetes>Mycobacteriales</td><td>GCF_010723135.1</td><td>Mycobacterium_kubicae</td><td>G6N40_RS26550</td><td>2561020..2564385</td><td>120959</td><td>Swt1-family-HEPN-domain-containing-protein-[Mycobacterium-kubicae]</td></tr>
<tr><td>WP_179959972.1</td><td>DUF499_1.RM.like.systems</td><td><-NTD+SF1-SF2-Helicase|| TUDOR+ SF1-SF2-Helicase+iREase+CTD+REase ->ParB+HNH-> HEPN + DUF499 *->?->?->?-><-Cluster182_14clades</td><td>1122</td><td>Bacteria>Terrabacteria_group>Actinomycetota>Actinomycetes>Mycobacteriales</td><td>GCF_010728155.1</td><td>Mycolicibacterium_doricum</td><td>H7I39_RS15930</td><td>40769..44137</td><td>126673</td><td>Swt1-family-HEPN-domain-containing-protein-[Mycobacterium-doricum]</td></tr>
<tr><td>WP_083088684.1</td><td>DUF499_1.RM.like.systems</td><td>Recombinase->?->?-><- HEPN + DUF499 *<- METHYLASE <- TUDOR+ SF1-SF2-Helicase+iREase+CTD+REase ||NTD+SF1-SF2-Helicase->?-><-wHTH(SBD)+HNH</td><td>1108</td><td>Bacteria>Terrabacteria_group>Actinomycetota>Actinomycetes>Mycobacteriales</td><td>GCF_010731635.1</td><td>Mycobacterium_noviomagense</td><td>BST37_RS15635</td><td>32792..36118</td><td>459858</td><td>DUF499-domain-containing-protein-[Mycobacterium-noviomagense]</td></tr>
<tr><td>WP_179973625.1</td><td>DUF499_1.RM.like.systems</td><td><-PIN+ACET+EVE||<-REase<-NTD+SF1-SF2-Helicase|| TUDOR+ SF1-SF2-Helicase+iREase+CTD+REase -> METHYLASE ->ParB+HNH-> HEPN + DUF499 *-></td><td>1121</td><td>Bacteria>Terrabacteria_group>Actinomycetota>Actinomycetes>Mycobacteriales</td><td>GCF_010731815.2</td><td>Mycolicibacterium_arabiense</td><td>H5P33_RS16145</td><td>691085..694450</td><td>1286181</td><td>Swt1-family-HEPN-domain-containing-protein-[Mycolicibacterium-arabiense]</td></tr>
<tr><td>WP_166810353.1</td><td>DUF499_1.RM.like.systems</td><td>HEPN -><-NTD+SF1-SF2-Helicase|| TUDOR+ SF1-SF2-Helicase+iREase+CTD+REase -> HEPN + DUF499 *-><-?||?-><-?||<-GNTR-HTH</td><td>1121</td><td>Bacteria>Terrabacteria_group>Actinomycetota>Actinomycetes>Mycobacteriales</td><td>GCF_011602025.1</td><td>Rhodococcus_sp._IC4_135</td><td>G8767_RS12755</td><td>177145..180510</td><td>2715537</td><td>Swt1-family-HEPN-domain-containing-protein-[Rhodococcus-sp.-IC4_135]</td></tr>
<tr><td>WP_167106438.1</td><td>DUF499_1.RM.like.systems</td><td><-DNA_methylase||REase-> METHYLASE -> HEPN + DUF499 *->MORC->REase->HTH+SF1-SF2-Helicase+REase-><-?<-SF1-SF2-helicase+DUF1998</td><td>1137</td><td>Bacteria>Terrabacteria_group>Actinomycetota>Actinomycetes>Mycobacteriales</td><td>GCF_011694515.1</td><td>Mycobacterium_sp._DL592</td><td>HBE64_RS20620</td><td>4288954..4292367</td><td>2675524</td><td>DUF499-domain-containing-protein-[Mycobacterium-sp.-DL592]</td></tr>
<tr><td>WP_170194849.1</td><td>DUF499_1.RM.like.systems</td><td>HTH_3-><-?<-?||?-><-ABhydrolase+ABhydrolase<- HEPN + DUF499 *||DUF222+HNH-><-AbiEii<-?||?-><- METHYLASE ||HTH_19-></td><td>1113</td><td>Bacteria>Terrabacteria_group>Actinomycetota>Actinomycetes>Mycobacteriales</td><td>GCF_012933505.1</td><td>Gordonia_asplenii</td><td>HH308_RS14085</td><td>95388..98729</td><td>2725283</td><td>DUF499-domain-containing-protein-[Gordonia-asplenii]</td></tr>
<tr><td>WP_172936837.1</td><td>DUF499_1.RM.like.systems</td><td>REase+SF1-SF2-Helicase->wHTH+SAD+HNH-><-HTH+SF1-SF2-Helicase+REase<-REase<-MORC<- HEPN + DUF499 *<- METHYLASE ||DNA_methylase-><-?<-?<-?<-MORC</td><td>1125</td><td>Bacteria>Terrabacteria_group>Actinomycetota>Actinomycetes>Mycobacteriales</td><td>GCF_013277795.1</td><td>Rhodococcus_sp._MS16</td><td>FEZ60_RS20030</td><td>139457..142834</td><td>2579941</td><td>Swt1-family-HEPN-domain-containing-protein-[Rhodococcus-sp.-MS16]</td></tr>
<tr><td>WP_213934645.1</td><td>DUF499_1.RM.like.systems</td><td>McrC-NTD+REase-><-?<-?||CHTH->?-><- HEPN + DUF499 *||<- TUDOR+ SF1-SF2-Helicase+iREase+CTD+REase ||?->?-><-ACET</td><td>1138</td><td>Bacteria>Terrabacteria_group>Actinomycetota>Actinomycetes>Mycobacteriales</td><td>GCF_013602415.1</td><td>Rhodococcus_sp._B50</td><td>GDN09_RS24520</td><td>12924..16340</td><td>2682847</td><td>Swt1-family-HEPN-domain-containing-protein-[Rhodococcus-sp.-B50]</td></tr>
<tr><td>WP_181729591.1</td><td>DUF499_1.RM.like.systems</td><td><-ABhydrolase|| TUDOR+ SF1-SF2-Helicase+iREase+CTD+REase -> METHYLASE -> HEPN + DUF499 *->ORC-CDC6+REase-></td><td>1112</td><td>Bacteria>Terrabacteria_group>Actinomycetota>Actinomycetes>Mycobacteriales</td><td>GCF_013868425.1</td><td>Corynebacterium_sanguinis</td><td>H0H28_RS04340</td><td>15252..18590</td><td>2594913</td><td>DUF499-domain-containing-protein-[Corynebacterium-sanguinis]</td></tr>
<tr><td>WP_219849138.1</td><td>DUF499_1.RM.like.systems</td><td><-HTH+SF1-SF2-Helicase+REase<-REase<-MORC<- HEPN + DUF499 *<- METHYLASE <-REase||DNA_methylase-><- METHYLASE -p</td><td>1137</td><td>Bacteria>Terrabacteria_group>Actinomycetota>Actinomycetes>Mycobacteriales</td><td>GCF_014041935.1</td><td>Gordonia_jinghuaiqii</td><td>H1R19_RS12155</td><td>2732590..2736003</td><td>2758710</td><td>Swt1-family-HEPN-domain-containing-protein-[Gordonia-jinghuaiqii]</td></tr>
<tr><td>WP_281378415.1</td><td>DUF499_1.RM.like.systems</td><td><-ABhydrolase<-?||?-><- HEPN + DUF499 *<- METHYLASE ||DUF222+HNH-><- TUDOR+ SF1-SF2-Helicase+iREase+CTD+REase <-?<-?<-tRNA<-tRNA<-tRNA<-tRNA<-tRNA<-tRNA||tRNA->DDE_Tnp_1_4-></td><td>1142</td><td>Bacteria>Terrabacteria_group>Actinomycetota>Actinomycetes>Mycobacteriales</td><td>GCF_014197435.1</td><td>Gordonia_humi</td><td>BKA16_RS13810</td><td>3005759..3009187</td><td>686429</td><td>DUF499-domain-containing-protein-[Gordonia-humi]</td></tr>
<tr><td>WP_185712603.1</td><td>DUF499_1.RM.like.systems</td><td>Phage_integrase->?-><- HEPN ||REase-><- HEPN + DUF499 *<- TUDOR+ SF1-SF2-Helicase+iREase+CTD+REase <-HTH+MPTase<-Tox-RelE||SbcC->?->HTH+UbE1-></td><td>1138</td><td>Bacteria>Terrabacteria_group>Actinomycetota>Actinomycetes>Mycobacteriales</td><td>GCF_014230025.1</td><td>Rhodococcus_aetherivorans</td><td>H0B42_RS00215</td><td>49328..52744</td><td>191292</td><td>Swt1-family-HEPN-domain-containing-protein-[Rhodococcus-aetherivorans]</td></tr>
<tr><td>WP_193603884.1</td><td>DUF499_1.RM.like.systems</td><td>HTH_28+rve+Mu-transpos_C->?->?->?-> TUDOR+ SF1-SF2-Helicase+iREase+CTD+REase -> METHYLASE -> HEPN + DUF499 *-><-HAD||Classical-AAA-></td><td>1120</td><td>Bacteria>Terrabacteria_group>Actinomycetota>Actinomycetes>Mycobacteriales</td><td>GCF_015023815.1</td><td>Mycobacteroides_abscessus</td><td>E3G55_RS12995</td><td>2677503..2680865</td><td>36809</td><td>DUF499-domain-containing-protein-[Mycobacteroides-abscessus]</td></tr>
<tr><td>WP_193904012.1</td><td>DUF499_1.RM.like.systems</td><td>MODE-HTH+PBPII-><-IclR-HTH||?->RNase-><- HEPN + DUF499 *<- METHYLASE -p<-zf-ISL3+DDE_Tnp_ISL3<- METHYLASE <- TUDOR+ SF1-SF2-Helicase+iREase+CTD+REase ||DDE_Tnp_1_4-><-HTH+MPTase</td><td>1142</td><td>Bacteria>Terrabacteria_group>Actinomycetota>Actinomycetes>Mycobacteriales</td><td>GCF_015169995.1</td><td>Rhodococcus_pyridinivorans</td><td>INP59_RS25070</td><td>140646..144074</td><td>103816</td><td>Swt1-family-HEPN-domain-containing-protein-[Rhodococcus-pyridinivorans]</td></tr>
<tr><td>WP_194806632.1</td><td>DUF499_1.RM.like.systems</td><td>Phage_int_SAM_1+Phage_integrase->XIS-HTH->?->?->?-><- HEPN + DUF499 *<-ParB+HNH<- METHYLASE <-UDG-conflict<- TUDOR+ SF1-SF2-Helicase+iREase+CTD+REase</td><td>1121</td><td>Bacteria>Terrabacteria_group>Actinomycetota>Actinomycetes>Mycobacteriales</td><td>GCF_015354405.1</td><td>Nocardia_sp._BSTN01</td><td>IRT45_RS15950</td><td>106038..109403</td><td>2783665</td><td>Swt1-family-HEPN-domain-containing-protein-[Nocardia-sp.-BSTN01]</td></tr>
<tr><td>WP_228819156.1</td><td>DUF499_1.RM.like.systems</td><td><-STAS<-?<-Phage_integrase<-?<- HEPN + DUF499 *<- METHYLASE <- TUDOR+ SF1-SF2-Helicase+iREase+CTD+REase <-ZDNA-HTH<-?<-?<-Response_reg+LuxR-HTH</td><td>1114</td><td>Bacteria>Terrabacteria_group>Actinomycetota>Actinomycetes>Mycobacteriales</td><td>GCF_015477815.1</td><td>Nocardia_farcinica</td><td>QH106_RS06910</td><td>129402..132746</td><td>37329</td><td>DUF499-domain-containing-protein-[Nocardia-farcinica]</td></tr>
<tr><td>WP_197940422.1</td><td>DUF499_1.RM.like.systems</td><td><-HIN-HTH|| HEPN -><-NTD+SF1-SF2-Helicase|| TUDOR+ SF1-SF2-Helicase+iREase+CTD+REase -> METHYLASE ->ParB+HNH-> HEPN + DUF499 *->PAIREDC-HTH-></td><td>1121</td><td>Bacteria>Terrabacteria_group>Actinomycetota>Actinomycetes>Mycobacteriales</td><td>GCF_016019085.1</td><td>Rhodococcus_erythropolis</td><td>I3517_RS00940</td><td>13889..17254</td><td>1833</td><td>Swt1-family-HEPN-domain-containing-protein-[Rhodococcus-erythropolis]</td></tr>
<tr><td>WP_048892803.1</td><td>DUF499_1.RM.like.systems</td><td>DEDD_Tnp_IS110->DEDD_Tnp_IS110+Transposase_20-> TUDOR+ SF1-SF2-Helicase+iREase+CTD+REase -> METHYLASE -><-?<-?|| HEPN + DUF499 *-><-HKD-DNase+SF1-SF2-Helicase+TRD<-?<-PIN<-PhdYeFM_antitox<- HEPN <- Polbeta</td><td>1117</td><td>Bacteria>Terrabacteria_group>Actinomycetota>Actinomycetes>Mycobacteriales</td><td>GCF_016592155.1</td><td>Mycobacterium_heckeshornense</td><td>BMW24_RS01365</td><td>271817..275170</td><td>110505</td><td>DUF499-domain-containing-protein-[Mycobacterium-heckeshornense]</td></tr>
<tr><td>WP_200172468.1</td><td>DUF499_1.RM.like.systems</td><td><-CHTH|| METHYLASE ->?->HTH+MPTase-> HEPN + DUF499 *->?-><-?||?->?-><-?<-ABC-ATPase+SbcC</td><td>1132</td><td>Bacteria>Terrabacteria_group>Actinomycetota>Actinomycetes>Mycobacteriales</td><td>GCF_016599145.1</td><td>Tomitella_cavernea</td><td>JJ683_RS06825</td><td>52491..55889</td><td>1387982</td><td>DUF499-domain-containing-protein-[Tomitella-cavernea]</td></tr>
<tr><td>WP_203801307.1</td><td>DUF499_1.RM.like.systems</td><td><-SF1-SF2-helicase+DUF1998<-?|| TUDOR+ SF1-SF2-Helicase+iREase+CTD+REase -> METHYLASE -><-?<-?|| HEPN + DUF499 *-><-HKD-DNase+SF1-SF2-Helicase+TRD<-?<-PIN<-PhdYeFM_antitox<- HEPN <- Polbeta</td><td>1117</td><td>Bacteria>Terrabacteria_group>Actinomycetota>Actinomycetes>Mycobacteriales</td><td>GCF_016861545.1</td><td>Mycobacterium_heckeshornense</td><td>JUH53_RS19960</td><td>4159291..4162644</td><td>110505</td><td>DUF499-domain-containing-protein-[Mycobacterium-heckeshornense]</td></tr>
<tr><td>WP_205317328.1</td><td>DUF499_1.RM.like.systems</td><td>GDResolvase+HTH->?-><-?|| TUDOR+ SF1-SF2-Helicase+iREase+CTD+REase -> METHYLASE -> HEPN + DUF499 *-><-?||HTH-><-tRNA<-MarR-HTH</td><td>1133</td><td>Bacteria>Terrabacteria_group>Actinomycetota>Actinomycetes>Mycobacteriales</td><td>GCF_016921075.1</td><td>Prescottella_equi</td><td>J0861_RS22230</td><td>9979..13380</td><td>43767</td><td>Swt1-family-HEPN-domain-containing-protein-[Prescottella-equi]</td></tr>
<tr><td>WP_205874823.1</td><td>DUF499_1.RM.like.systems</td><td><-Cluster2439_3clades||?->XIS-HTH->PIN-> TUDOR+ SF1-SF2-Helicase+iREase+CTD+REase -> METHYLASE -> HEPN + DUF499 *-><-?<-HISKIN<-SbcC<-NUDIX||GNTR-HTH-></td><td>1108</td><td>Bacteria>Terrabacteria_group>Actinomycetota>Actinomycetes>Mycobacteriales</td><td>GCF_017086405.1</td><td>Mycobacterium_gordonae</td><td>JX552_RS26920</td><td>6279372..6282698</td><td>1778</td><td>Swt1-family-HEPN-domain-containing-protein-[Mycobacterium-gordonae]</td></tr>
<tr><td>WP_094602238.1</td><td>DUF499_1.RM.like.systems</td><td>GDSL-><-HTH_19|| TUDOR+ SF1-SF2-Helicase+iREase+CTD+REase -><-STYKinase-like<-HTH|| METHYLASE -> HEPN + DUF499 *->?->ABhydrolase+ABhydrolase-><-?||JetA->JetB-></td><td>1127</td><td>Bacteria>Terrabacteria_group>Actinomycetota>Actinomycetes>Mycobacteriales</td><td>GCF_018135605.1</td><td>Gordonia_polyisoprenivorans</td><td>J8M97_RS16335</td><td>3622924..3626307</td><td>84595</td><td>Swt1-family-HEPN-domain-containing-protein-[Gordonia-polyisoprenivorans]</td></tr>
<tr><td>WP_211919214.1</td><td>DUF499_1.RM.like.systems</td><td>HEPN -><-NTD+SF1-SF2-Helicase|| TUDOR+ SF1-SF2-Helicase+iREase+CTD+REase ->ParB+HNH-> HEPN + DUF499 *-><-NUDIX||?->REase->?->?->GDResolvase+HIN-HTH-></td><td>1121</td><td>Bacteria>Terrabacteria_group>Actinomycetota>Actinomycetes>Mycobacteriales</td><td>GCF_018283555.1</td><td>Rhodococcus_erythropolis</td><td>AC1659_RS28475</td><td>46414..49779</td><td>1833</td><td>Swt1-family-HEPN-domain-containing-protein-[Rhodococcus-erythropolis]</td></tr>
<tr><td>WP_205848809.1</td><td>DUF499_1.RM.like.systems</td><td><-PAIREDC-HTH<-HIN-HTH||JetD-><-PIN<-VapB_antitoxin<- HEPN + DUF499 *<- METHYLASE <- TUDOR+ SF1-SF2-Helicase+iREase+CTD+REase ||HTH_31->?-><-?||LIP+Peptidase_S9-></td><td>1110</td><td>Bacteria>Terrabacteria_group>Actinomycetota>Actinomycetes>Mycobacteriales</td><td>GCF_018326145.1</td><td>Mycolicibacterium</td><td>EUA03_RS05700</td><td>2711..6043</td><td>1866885</td><td>MULTISPECIES:-Swt1-family-HEPN-domain-containing-protein-[Mycolicibacterium]</td></tr>
<tr><td>WP_213184733.1</td><td>DUF499_1.RM.like.systems</td><td>TUDOR+ SF1-SF2-Helicase+iREase+CTD+REase -><-?|| METHYLASE ->ParB+HNH-> HEPN + DUF499 *->ANK+ANK+RAMA-></td><td>1134</td><td>Bacteria>Terrabacteria_group>Actinomycetota>Actinomycetes>Mycobacteriales</td><td>GCF_018390595.1</td><td>Dietzia_massiliensis</td><td>KHA85_RS02755</td><td>11826..15230</td><td>2697499</td><td>Swt1-family-HEPN-domain-containing-protein-[Dietzia-massiliensis]</td></tr>
<tr><td>WP_062887256.1</td><td>DUF499_1.RM.like.systems</td><td><-FtsK<-?||NUDIX-><- HEPN <-Schlafen+HATPase+HTH<- HEPN + DUF499 *<- METHYLASE ||?-><- TUDOR+ SF1-SF2-Helicase+iREase+CTD+REase</td><td>1107</td><td>Bacteria>Terrabacteria_group>Actinomycetota>Actinomycetes>Mycobacteriales</td><td>GCF_018524295.1</td><td>Mycobacteriaceae</td><td>KL858_RS23375</td><td>11898..15221</td><td>1762</td><td>Swt1-family-HEPN-domain-containing-protein-[Mycobacterium-avium]</td></tr>
<tr><td>WP_214389765.1</td><td>DUF499_1.RM.like.systems</td><td>EVE+McrB-AAA->McrC-NTD+REase-><-?<-HTH_Tnp_1+HTH_21+rve+rve_3||HTH_Tnp_1+HTH_21+rve+rve_3->?-><- HEPN + DUF499 *<-ParB+HNH<- METHYLASE <- TUDOR+ SF1-SF2-Helicase+iREase+CTD+REase ||NTD+SF1-SF2-Helicase->DDE_Tnp_1_4-></td><td>1119</td><td>Bacteria>Terrabacteria_group>Actinomycetota>Actinomycetes>Mycobacteriales</td><td>GCF_018524295.1</td><td>Mycolicibacterium_goodii</td><td>KL858_RS28165</td><td>17594..20953</td><td>134601</td><td>Swt1-family-HEPN-domain-containing-protein-[Mycolicibacterium-goodii]</td></tr>
<tr><td>WP_099696878.1</td><td>DUF499_1.RM.like.systems</td><td>FAD-NAD-dep-oxidoreductase->?-><- HEPN + DUF499 *<- METHYLASE <- TUDOR+ SF1-SF2-Helicase+iREase+CTD+REase</td><td>1134</td><td>Bacteria>Terrabacteria_group>Actinomycetota>Actinomycetes>Mycobacteriales</td><td>GCF_018588185.1</td><td>Rhodococcus</td><td>G4D79_RS05030</td><td>1105468..1108872</td><td>1827</td><td>MULTISPECIES:-Swt1-family-HEPN-domain-containing-protein-[Rhodococcus]</td></tr>
<tr><td>WP_214515005.1</td><td>DUF499_1.RM.like.systems</td><td>ABC-ATPase+SbcC->HAD-><-?||?-> TUDOR+ SF1-SF2-Helicase+iREase+CTD+REase -> METHYLASE -> HEPN + DUF499 *->?->||?-><-DDE_Tnp_IS240</td><td>1134</td><td>Bacteria>Terrabacteria_group>Actinomycetota>Actinomycetes>Mycobacteriales</td><td>GCF_018588185.1</td><td>Rhodococcus_sp._WY5</td><td>G4D79_RS33795</td><td>40006..43410</td><td>2708349</td><td>Swt1-family-HEPN-domain-containing-protein-[Rhodococcus-sp.-WY5]</td></tr>
<tr><td>WP_218681519.1</td><td>DUF499_1.RM.like.systems</td><td>NUDIX->?-> TUDOR+ SF1-SF2-Helicase+iREase+CTD+REase -> METHYLASE -> HEPN + DUF499 *-></td><td>1134</td><td>Bacteria>Terrabacteria_group>Actinomycetota>Actinomycetes>Mycobacteriales</td><td>GCF_019279115.1</td><td>Rhodococcus_qingshengii</td><td>KZA18_RS32415</td><td>9085..12489</td><td>334542</td><td>Swt1-family-HEPN-domain-containing-protein-[Rhodococcus-qingshengii]</td></tr>
<tr><td>WP_174714145.1</td><td>DUF499_1.RM.like.systems</td><td>XIS-HTH->PIN-> TUDOR+ SF1-SF2-Helicase+iREase+CTD+REase -><-?|| METHYLASE ->ParB+HNH-> HEPN + DUF499 *->?->?-><-NUDIX||GNTR-HTH->?->FtsK-></td><td>1108</td><td>Bacteria>Terrabacteria_group>Actinomycetota>Actinomycetes>Mycobacteriales</td><td>GCF_019456675.1</td><td>Mycolicibacterium_pallens</td><td>K0O64_RS24915</td><td>5121618..5124944</td><td>370524</td><td>MULTISPECIES:-Swt1-family-HEPN-domain-containing-protein-[Mycobacterium]</td></tr>
<tr><td>WP_220667114.1</td><td>DUF499_1.RM.like.systems</td><td><-REase<-PRTase+PRTase<-?<-SIR2+STALD|| TUDOR+ SF1-SF2-Helicase+iREase+CTD+REase -> HEPN + DUF499 *-><-RNase</td><td>1142</td><td>Bacteria>Terrabacteria_group>Actinomycetota>Actinomycetes>Mycobacteriales</td><td>GCF_019641445.1</td><td>Rhodococcus_sp._DMU2021</td><td>K3M35_RS24125</td><td>14618..18046</td><td>2866997</td><td>Swt1-family-HEPN-domain-containing-protein-[Rhodococcus-sp.-DMU2021]</td></tr>
<tr><td>WP_225506778.1</td><td>DUF499_1.RM.like.systems</td><td>XIS-HTH->PIN-> TUDOR+ SF1-SF2-Helicase+iREase+CTD+REase -> METHYLASE -> HEPN + DUF499 *->Schlafen+HATPase+HTH->||DDE_Tnp_1_4-> HEPN ->?-><-NUDIX||GNTR-HTH-></td><td>1108</td><td>Bacteria>Terrabacteria_group>Actinomycetota>Actinomycetes>Mycobacteriales</td><td>GCF_020227835.1</td><td>Mycolicibacterium_fortuitum</td><td>H8Z59_RS27040</td><td>5490569..5493895</td><td>1766</td><td>Swt1-family-HEPN-domain-containing-protein-[Mycolicibacterium-fortuitum]</td></tr>
<tr><td>WP_207546857.1</td><td>DUF499_1.RM.like.systems</td><td>JetA->JetB->?->JetD-><-PIN<-VapB_antitoxin<- HEPN + DUF499 *<- METHYLASE <- TUDOR+ SF1-SF2-Helicase+iREase+CTD+REase ||HTH_31->?-><-?||LIP+Peptidase_S9-></td><td>1109</td><td>Bacteria>Terrabacteria_group>Actinomycetota>Actinomycetes>Mycobacteriales</td><td>GCF_020520165.1</td><td>Mycolicibacterium</td><td>EKK51_RS16805</td><td>41858..45187</td><td>1866885</td><td>MULTISPECIES:-Swt1-family-HEPN-domain-containing-protein-[Mycolicibacterium]</td></tr>
<tr><td>WP_037183436.1</td><td>DUF499_1.RM.like.systems</td><td><-GDResolvase+HIN-HTH|| HEPN -><-?<- HEPN + DUF499 *<- METHYLASE <- TUDOR+ SF1-SF2-Helicase+iREase+CTD+REase ||DNA-glycosylase-></td><td>1136</td><td>Bacteria>Terrabacteria_group>Actinomycetota>Actinomycetes>Mycobacteriales</td><td>GCF_020542785.1</td><td>Rhodococcus_opacus</td><td>OPAG_RS41845</td><td>61698..65108</td><td>37919</td><td>Swt1-family-HEPN-domain-containing-protein-[Rhodococcus-opacus]</td></tr>
<tr><td>WP_227974732.1</td><td>DUF499_1.RM.like.systems</td><td>TIR->?->?-> TUDOR+ SF1-SF2-Helicase+iREase+CTD+REase -><-?|| METHYLASE -> HEPN + DUF499 *->Schlafen+HATPase+HTH-> HEPN -><-?<-NUDIX||GNTR-HTH-></td><td>1108</td><td>Bacteria>Terrabacteria_group>Actinomycetota>Actinomycetes>Mycobacteriales</td><td>GCF_020735405.1</td><td>Mycobacterium_avium</td><td>LT348_RS02565</td><td>526704..530030</td><td>1764</td><td>Swt1-family-HEPN-domain-containing-protein-[Mycobacterium-avium]</td></tr>
<tr><td>WP_228232863.1</td><td>DUF499_1.RM.like.systems</td><td><-GDResolvase+HIN-HTH<-?||ABhydrolase-><-GDResolvase+HIN-HTH||zf-ISL3+DDE_Tnp_ISL3-><-?<- HEPN + DUF499 *<-ParB+HNH<- METHYLASE <-UDG-conflict<- TUDOR+ SF1-SF2-Helicase+iREase+CTD+REase ||NTD+SF1-SF2-Helicase-><- HEPN</td><td>1121</td><td>Bacteria>Terrabacteria_group>Actinomycetota>Actinomycetes>Mycobacteriales</td><td>GCF_020782825.1</td><td>Rhodococcus_sp._3-2</td><td>LL008_RS29695</td><td>195028..198393</td><td>2890836</td><td>Swt1-family-HEPN-domain-containing-protein-[Rhodococcus-sp.-3-2]</td></tr>
<tr><td>WP_228833824.1</td><td>DUF499_1.RM.like.systems</td><td>ZDNA-HTH-> TUDOR+ SF1-SF2-Helicase+iREase+CTD+REase -> METHYLASE -p-> METHYLASE -> HEPN + DUF499 *->?->Phage_integrase->?->STAS-></td><td>1123</td><td>Bacteria>Terrabacteria_group>Actinomycetota>Actinomycetes>Mycobacteriales</td><td>GCF_020883195.1</td><td>Nocardia</td><td>LMJ57_RS17620</td><td>3762816..3766187</td><td>1817</td><td>MULTISPECIES:-DUF499-domain-containing-protein-[Nocardia]</td></tr>
<tr><td>WP_229481506.1</td><td>DUF499_1.RM.like.systems</td><td><-NTD+SF1-SF2-Helicase|| TUDOR+ SF1-SF2-Helicase+iREase+CTD+REase -> METHYLASE ->ParB+HNH-> HEPN + DUF499 *->?->?-><-SF1-SF2-Helicase<-OLD-ABC+TOPRIM<-?<-Primpol+D5N+D5ATPase</td><td>1114</td><td>Bacteria>Terrabacteria_group>Actinomycetota>Actinomycetes>Mycobacteriales</td><td>GCF_020884715.1</td><td>Mycolicibacterium_mageritense</td><td>MTY414_RS35935</td><td>11397..14741</td><td>53462</td><td>Swt1-family-HEPN-domain-containing-protein-[Mycolicibacterium-mageritense]</td></tr>
<tr><td>WP_231782199.1</td><td>DUF499_1.RM.like.systems</td><td><-MULE-TRANSPOSASE<-ZnR+ZnR<-TIR|| TUDOR+ SF1-SF2-Helicase+iREase+CTD+REase -> METHYLASE -> HEPN + DUF499 *-><-RNase<-?||GDResolvase+Recombinase+ZnR-><-?||DUF4192-></td><td>1139</td><td>Bacteria>Terrabacteria_group>Actinomycetota>Actinomycetes>Mycobacteriales</td><td>GCF_021165995.1</td><td>Rhodococcus_pyridinivorans</td><td>LSF60_RS23505</td><td>9142..12561</td><td>103816</td><td>Swt1-family-HEPN-domain-containing-protein-[Rhodococcus-pyridinivorans]</td></tr>
<tr><td>WP_233059597.1</td><td>DUF499_1.RM.like.systems</td><td>TIR->?->?-> TUDOR+ SF1-SF2-Helicase+iREase+CTD+REase -><-?|| METHYLASE -> HEPN + DUF499 *->Schlafen+HATPase+HTH-> HEPN -><-?<-NUDIX||GNTR-HTH-></td><td>1108</td><td>Bacteria>Terrabacteria_group>Actinomycetota>Actinomycetes>Mycobacteriales</td><td>GCF_021183805.1</td><td>Mycobacterium_avium</td><td>LTQ57_RS17660</td><td>3790685..3794011</td><td>1764</td><td>Swt1-family-HEPN-domain-containing-protein-[Mycobacterium-avium]</td></tr>
<tr><td>WP_236997292.1</td><td>DUF499_1.RM.like.systems</td><td>TUDOR+ SF1-SF2-Helicase+iREase+CTD+REase -><-DUF222+HNH|| METHYLASE -> HEPN + DUF499 *->ACET-><-?||?->ABhydrolase-></td><td>1125</td><td>Bacteria>Terrabacteria_group>Actinomycetota>Actinomycetes>Mycobacteriales</td><td>GCF_021738805.1</td><td>Gordonia_liuliyuniae</td><td>L5G33_06210</td><td>131250..134627</td><td>2911517</td><td>DUF499-domain-containing-protein-[Gordonia-sp.-HY366]</td></tr>
<tr><td>WP_237010003.1</td><td>DUF499_1.RM.like.systems</td><td><-ABhydrolase<-?||?-><-ACET<- HEPN + DUF499 *<- METHYLASE ||DUF222+HNH-><- TUDOR+ SF1-SF2-Helicase+iREase+CTD+REase</td><td>1115</td><td>Bacteria>Terrabacteria_group>Actinomycetota>Actinomycetes>Mycobacteriales</td><td>GCF_021738965.1</td><td>Gordonia_zhenghanii</td><td>L5G32_05410</td><td>142683..146030</td><td>2911516</td><td>MULTISPECIES:-DUF499-domain-containing-protein-[unclassified-Gordonia]</td></tr>
<tr><td>WP_237001659.1</td><td>DUF499_1.RM.like.systems</td><td>TUDOR+ SF1-SF2-Helicase+iREase+CTD+REase -><-DUF222+HNH|| METHYLASE -> HEPN + DUF499 *->ACET-><-?||?->ABhydrolase-></td><td>1125</td><td>Bacteria>Terrabacteria_group>Actinomycetota>Actinomycetes>Mycobacteriales</td><td>GCF_021739025.1</td><td>Gordonia_liuliyuniae</td><td>L5G28_11435</td><td>131354..134731</td><td>2911517</td><td>DUF499-domain-containing-protein-[Gordonia-sp.-HY285]</td></tr>
<tr><td>WP_237211675.1</td><td>DUF499_1.RM.like.systems</td><td><-REase<-PRTase+PRTase<-?<-SIR2+STALD|| TUDOR+ SF1-SF2-Helicase+iREase+CTD+REase -> METHYLASE -> HEPN + DUF499 *-><-RNase<-?||GDResolvase+HIN-HTH-></td><td>1138</td><td>Bacteria>Terrabacteria_group>Actinomycetota>Actinomycetes>Mycobacteriales</td><td>GCF_021852985.1</td><td>Rhodococcus_ruber</td><td>K7Z00_RS28700</td><td>67558..70974</td><td>1830</td><td>Swt1-family-HEPN-domain-containing-protein-[Rhodococcus-ruber]</td></tr>
<tr><td>WP_238054205.1</td><td>DUF499_1.RM.like.systems</td><td><-REase||?->DOC->GDResolvase+HIN-HTH-><- HEPN + DUF499 *<- METHYLASE <- TUDOR+ SF1-SF2-Helicase+iREase+CTD+REase <-?||?-><-?<-REase</td><td>1134</td><td>Bacteria>Terrabacteria_group>Actinomycetota>Actinomycetes>Mycobacteriales</td><td>GCF_022059865.1</td><td>Rhodococcus_erythropolis</td><td>ITJ47_RS31985</td><td>3376..6780</td><td>1833</td><td>Swt1-family-HEPN-domain-containing-protein-[Rhodococcus-erythropolis]</td></tr>
<tr><td>WP_238886618.1</td><td>DUF499_1.RM.like.systems</td><td>HIN-HTH->?->Cluster182_14clades-><-?<-?<-?<- HEPN + DUF499 *<-ParB+HNH<- METHYLASE <- TUDOR+ SF1-SF2-Helicase+iREase+CTD+REase ||NTD+SF1-SF2-Helicase-></td><td>1120</td><td>Bacteria>Terrabacteria_group>Actinomycetota>Actinomycetes>Mycobacteriales</td><td>GCF_022317005.1</td><td>Mycobacterium_sp._IDR2000157661</td><td>K3G64_RS19485</td><td>3782160..3785522</td><td>2867005</td><td>Swt1-family-HEPN-domain-containing-protein-[Mycobacterium-sp.-IDR2000157661]</td></tr>
<tr><td>WP_239342823.1</td><td>DUF499_1.RM.like.systems</td><td><-NTD+SF1-SF2-Helicase|| TUDOR+ SF1-SF2-Helicase+iREase+CTD+REase -> METHYLASE ->ParB+HNH-> HEPN + DUF499 *->REase->SF1-SF2-Helicase-></td><td>1109</td><td>Bacteria>Terrabacteria_group>Actinomycetota>Actinomycetes>Mycobacteriales</td><td>GCF_022348005.1</td><td>Mycolicibacterium_sp._OfavD-34-C</td><td>MHN82_RS23245</td><td>18852..22181</td><td>2917746</td><td>Swt1-family-HEPN-domain-containing-protein-[Mycolicibacterium-sp.-OfavD-34-C]</td></tr>
<tr><td>WP_244804062.1</td><td>DUF499_1.RM.like.systems</td><td>ABhydrolase->?-><-?<-?||?->ORC-CDC6+REase-><- HEPN + DUF499 *<- TUDOR+ SF1-SF2-Helicase+iREase+CTD+REase ||HTH_3->?->?->?-><-GNAT</td><td>1194</td><td>Bacteria>Terrabacteria_group>Actinomycetota>Actinomycetes>Mycobacteriales</td><td>GCF_022869005.1</td><td>Corynebacterium_kalidii</td><td>MUN33_RS06320</td><td>104934..108518</td><td>2931982</td><td>DUF499-domain-containing-protein-[Corynebacterium-sp.-LD5P10]</td></tr>
</table>

| Protein-ID | System | Gene-Neighbourhoods | Length | Taxonomy | Assembly-ID | Strain | Locus-Tag | Genomic-coordinates | TaxID | Fasta-defline |
|---|---|---|---|---|---|---|---|---|---|---|
| WP_247450629.1 | DUF499_1.RM.like.systems | HNH-><-REase+TPR<-HEPN+DUF499*<-METHYLASE<-TUDOR+ SF1-SF2-Helicase+iREase+CTD+REase | 1115 | Bacteria>Terrabacteria_group>Actinomycetota>Actinomycetes>Mycobacteriales | GCF_023078355.1 | Gordonia_hongkongensis | MVF96_RS23000 | 5106874..5110221 | 1701090 | DUF499-domain-containing-protein-[Gordonia-hongkongensis] |
| WP_248488617.1 | DUF499_1.RM.like.systems | SbcC->?-><-HTH_19<-HTH_3\|\|<-HTH\|\|METHYLASE->HEPN+DUF499*->?->?-><-?<-?\|\|?-><-RVT | 1120 | Bacteria>Terrabacteria_group>Actinomycetota>Actinomycetes>Mycobacteriales | GCF_023168325.1 | Tsukamurella_sp._PLM1 | MTP03_RS12765 | 2471846..2475208 | 2929795 | DUF499-domain-containing-protein-[Tsukamurella-sp.-PLM1] |
| WP_248689418.1 | DUF499_1.RM.like.systems | MULE-TRANSPOSASE->?->RNase-><-HEPN+DUF499*<-METHYLASE<-TUDOR+ SF1-SF2-Helicase+iREase+CTD+REase\|\|SF1-SF2-Helicase-><-DNA-glycosylase<-DNA-glycosylase | 1138 | Bacteria>Terrabacteria_group>Actinomycetota>Actinomycetes>Mycobacteriales | GCF_023238485.1 | Rhodococcus_pyridinivorans | M1C57_RS14915 | 3265410..3268826 | 103816 | Swt1-family-HEPN-domain-containing-protein-[Rhodococcus-pyridinivorans] |
| WP_249617808.1 | DUF499_1.RM.like.systems | TUDOR+ SF1-SF2-Helicase+iREase+CTD+REase->METHYLASE->HEPN+DUF499*-><-?<-ABhydrolase+ABhydrolase<-?<-?<-?<-ABhydrolase | 1124 | Bacteria>Terrabacteria_group>Actinomycetota>Actinomycetes>Mycobacteriales | GCF_023509275.1 | Corynebacterium_pseudodiphtheriticum | L9H27_RS09685 | 2151202..2154576 | 37637 | DUF499-domain-containing-protein-[Corynebacterium-pseudodiphtheriticum] |
| WP_249619951.1 | DUF499_1.RM.like.systems | TUDOR+ SF1-SF2-Helicase+iREase+CTD+REase->METHYLASE->HEPN+DUF499*-><-?<-ABhydrolase+ABhydrolase<-?<-?<-?<-ABhydrolase | 1124 | Bacteria>Terrabacteria_group>Actinomycetota>Actinomycetes>Mycobacteriales | GCF_023509295.1 | Corynebacterium_pseudodiphtheriticum | L9H78_RS09595 | 2140657..2144031 | 37637 | DUF499-domain-containing-protein-[Corynebacterium-pseudodiphtheriticum] |
| WP_256083185.1 | DUF499_1.RM.like.systems | HEPN-><-NTD+SF1-SF2-Helicase\|\|TUDOR+ SF1-SF2-Helicase+iREase+CTD+REase->METHYLASE->ParB+HNH->HEPN+DUF499*-> | 1120 | Bacteria>Terrabacteria_group>Actinomycetota>Actinomycetes>Mycobacteriales | GCF_024438075.1 | Rhodococcus_qingshengii | NOS52_RS32465 | 22838..26200 | 334542 | Swt1-family-HEPN-domain-containing-protein-[Rhodococcus-qingshengii] |
| WP_256101683.1 | DUF499_1.RM.like.systems | <-NTD+SF1-SF2-Helicase\|\|TUDOR+ SF1-SF2-Helicase+iREase+CTD+REase->METHYLASE->ParB+HNH->HEPN+DUF499*->ParB+HNH->HEPN-> | 1118 | Bacteria>Terrabacteria_group>Actinomycetota>Actinomycetes>Mycobacteriales | GCF_024451565.1 | Mycobacterium_gordonae | KQR54_RS26625 | 18169..21525 | 1778 | Swt1-family-HEPN-domain-containing-protein-[Mycobacterium-gordonae] |
| WP_160903143.1 | DUF499_1.RM.like.systems | RelE-ParE-><-?<-HEPN+DUF499*<-METHYLASE\|\|?-><-TUDOR+ SF1-SF2-Helicase+iREase+CTD+REase\|\|HTH_3-> | 1126 | Bacteria>Terrabacteria_group>Actinomycetota>Actinomycetes>Mycobacteriales | GCF_024734075.1 | Gordonia_mangrovi | GIY30_RS16680 | 372248..375628 | 2665643 | Swt1-family-HEPN-domain-containing-protein-[Gordonia-mangrovi] |
| WP_160904533.1 | DUF499_1.RM.like.systems | <-Exonuc_V_gamma+REase<-?<-NTD+SF1-SF2-Helicase\|\|TUDOR+ SF1-SF2-Helicase+iREase+CTD+REase->METHYLASE->?->HEPN+DUF499*->?->?-><-HKD-DNase+HKD-DNase<-?<-wHTH+REase<-wHTH+REase | 1121 | Bacteria>Terrabacteria_group>Actinomycetota>Actinomycetes>Mycobacteriales | GCF_024734075.1 | Gordonia_mangrovi | GIY30_RS23685 | 18846..22211 | 2665643 | Swt1-family-HEPN-domain-containing-protein-[Gordonia-mangrovi] |
| WP_258913985.1 | DUF499_1.RM.like.systems | <-MULE-TRANSPOSASE\|\|<-?<-ZnR+ZnR<-TIR\|\|TUDOR+ SF1-SF2-Helicase+iREase+CTD+REase->METHYLASE->HEPN+DUF499*-><-RNase<-?<-?<-?\|\|OLD-ABC-> | 1138 | Bacteria>Terrabacteria_group>Actinomycetota>Actinomycetes>Mycobacteriales | GCF_024760325.1 | Rhodococcus_pyridinivorans | NXT08_RS24750 | 68689..72105 | 103816 | Swt1-family-HEPN-domain-containing-protein-[Rhodococcus-pyridinivorans] |
| WP_259916185.1 | DUF499_1.RM.like.systems | <-ABhydrolase\|\|TUDOR+ SF1-SF2-Helicase+iREase+CTD+REase->METHYLASE->HEPN+DUF499*->ORC-CDC6+REase-><-tRNA\|\|ncRNA->Aminotran_1_2->?->ABhydrolase-> | 1111 | Bacteria>Terrabacteria_group>Actinomycetota>Actinomycetes>Mycobacteriales | GCF_025144095.1 | Corynebacterium_sanguinis | M3G11_RS01890 | 80632..83967 | 2594913 | DUF499-domain-containing-protein-[Corynebacterium-sanguinis] |
| WP_259795206.1 | DUF499_1.RM.like.systems | <-ABhydrolase\|\|TUDOR+ SF1-SF2-Helicase+iREase+CTD+REase->METHYLASE->HEPN+DUF499*->ORC-CDC6+REase-> | 1112 | Bacteria>Terrabacteria_group>Actinomycetota>Actinomycetes>Mycobacteriales | GCF_025144435.1 | Corynebacterium_sanguinis | M3G32_RS10715 | 21189..24527 | 2594913 | DUF499-domain-containing-protein-[Corynebacterium-sanguinis] |
| WP_259896744.1 | DUF499_1.RM.like.systems | <-ABhydrolase\|\|TUDOR+ SF1-SF2-Helicase+iREase+CTD+REase->METHYLASE->HEPN+DUF499*->ORC-CDC6+REase-> | 1112 | Bacteria>Terrabacteria_group>Actinomycetota>Actinomycetes>Mycobacteriales | GCF_025144475.1 | Corynebacterium_sanguinis | M3G45_RS07405 | 84077..87415 | 2594913 | DUF499-domain-containing-protein-[Corynebacterium-sanguinis] |
| WP_144823668.1 | DUF499_1.RM.like.systems | <-ORC-CDC6+REase<-HEPN+DUF499*<-METHYLASE<-TUDOR+ SF1-SF2-Helicase+iREase+CTD+REase-p<-TUDOR+ SF1-SF2-Helicase+iREase+CTD+REase-p\|\|ABhydrolase-> | 1112 | Bacteria>Terrabacteria_group>Actinomycetota>Actinomycetes>Mycobacteriales | GCF_025145725.1 | Corynebacterium_sanguinis | EKI50_RS10380 | 1436..4774 | 2594913 | DUF499-domain-containing-protein-[Corynebacterium-sanguinis] |
| WP_259825954.1 | DUF499_1.RM.like.systems | <-ABhydrolase<-?<-Aminotran_1_2<-ncRNA\|\|tRNA->?->SF1-SF2-helicase+DUF1998-><-ORC-CDC6+REase<-HEPN+DUF499*<-METHYLASE<-TUDOR+ SF1-SF2-Helicase+iREase+CTD+REase\|\|ABhydrolase-> | 1112 | Bacteria>Terrabacteria_group>Actinomycetota>Actinomycetes>Mycobacteriales | GCF_025146125.1 | Corynebacterium_sanguinis | QON61_RS06385 | 69194..72532 | 2594913 | DUF499-domain-containing-protein-[Corynebacterium-sanguinis] |
| WP_260836619.1 | DUF499_1.RM.like.systems | <-MULE-TRANSPOSASE\|\|ParA-Soj-PloopNTPase->ORC-CDC6-><-HTH+SF1-SF2-Helicase+REase<-REase<-MORC<-HEPN+DUF499*<-METHYLASE\|\|XIS-HTH+RVT-><-HNH\|\|?-><-REase\|\|DNA_methylase-> | 1138 | Bacteria>Terrabacteria_group>Actinomycetota>Actinomycetes>Mycobacteriales | GCF_025268675.1 | Gordonia_amarae | GII32_RS01225 | 287584..291000 | 36821 | DUF499-domain-containing-protein-[Gordonia-amarae] |
| WP_260941407.1 | DUF499_1.RM.like.systems | JetA->JetB->?->JetD-><-?\|\|?-><-HEPN+DUF499*<-ParB+HNH\|\|HIN-HTH-><-ParB+HNH<-HIN-HTH\|\|?-><-TUDOR+ SF1-SF2-Helicase+iREase+CTD+REase | 1110 | Bacteria>Terrabacteria_group>Actinomycetota>Actinomycetes>Mycobacteriales | GCF_025331195.1 | Mycolicibacterium_llatzerense | A7G45_RS26855 | 438962..442294 | 280871 | Swt1-family-HEPN-domain-containing-protein-[Mycolicibacterium-llatzerense] |
| WP_211284609.1 | DUF499_1.RM.like.systems | <-FtsK<-?<-GNTR-HTH\|\|NUDIX-><-HEPN<-Schlafen+HATPase+HTH<-HEPN+DUF499*<-METHYLASE<-TUDOR+ SF1-SF2-Helicase+iREase+CTD+REase<-?<-RNaseH | 1113 | Bacteria>Terrabacteria_group>Actinomycetota>Actinomycetes>Mycobacteriales | GCF_025823205.1 | Mycobacterium_alsense | BST11_RS03260 | 100678..104019 | 324058 | Swt1-family-HEPN-domain-containing-protein-[Mycobacterium-alsense] |
| WP_264721883.1 | DUF499_1.RM.like.systems | TUDOR+ SF1-SF2-Helicase+iREase+CTD+REase->METHYLASE->HEPN+DUF499*-><-RNase<-?\|\|?-><-?<-Phage_integrase | 1139 | Bacteria>Terrabacteria_group>Actinomycetota>Actinomycetes>Mycobacteriales | GCF_025989245.1 | Rhodococcus_pyridinivorans | OMK68_RS15380 | 34680..38099 | 103816 | Swt1-family-HEPN-domain-containing-protein-[Rhodococcus-pyridinivorans] |
| WP_267626547.1 | DUF499_1.RM.like.systems | XIS-HTH->?->?->?->TUDOR+ SF1-SF2-Helicase+iREase+CTD+REase->METHYLASE->HEPN+DUF499*-> | 1110 | Bacteria>Terrabacteria_group>Actinomycetota>Actinomycetes>Mycobacteriales | GCF_026616935.1 | Gordonia_sputi | NZL84_RS17670 | 45237..48569 | 36823 | DUF499-domain-containing-protein-[Gordonia-sputi] |
| WP_267616414.1 | DUF499_1.RM.like.systems | <-JetB<-JetA\|\|?->DUF222+HNH->?-><-?<-HEPN+DUF499*\|\|MazE_antitoxin->PIN-><-METHYLASE<-TUDOR+ SF1-SF2-Helicase+iREase+CTD+REase\|\|?->AbiEii-> | 1127 | Bacteria>Terrabacteria_group>Actinomycetota>Actinomycetes>Mycobacteriales | GCF_026616955.1 | Gordonia_bronchialis | NOB45_RS12880 | 105537..108920 | 2054 | Swt1-family-HEPN-domain-containing-protein-[Gordonia-bronchialis] |
| WP_268656219.1 | DUF499_1.RM.like.systems | DUF4158+DDE_Tnp_Tn3-><-TetR-HTH\|\|?->?->GDResolvase+HIN-HTH->\|\|RNase-><-HEPN+DUF499*<-METHYLASE<-TUDOR+ SF1-SF2-Helicase+iREase+CTD+REase<-HTH+MPTase<-Tox-RelE\|\|SbcC-> | 1138 | Bacteria>Terrabacteria_group>Actinomycetota>Actinomycetes>Mycobacteriales | GCF_026802115.1 | Rhodococcus_pyridinivorans | OQN32_RS28100 | 23296..26712 | 103816 | Swt1-family-HEPN-domain-containing-protein-[Rhodococcus-pyridinivorans] |
| WP_269614895.1 | DUF499_1.RM.like.systems | HEPN-><-NTD+SF1-SF2-Helicase\|\|TUDOR+ SF1-SF2-Helicase+iREase+CTD+REase->METHYLASE->ParB+HNH->HEPN+DUF499*-><-NUDIX\|\|?->REase->?->?->GDResolvase+HIN-HTH-> | 1121 | Bacteria>Terrabacteria_group>Actinomycetota>Actinomycetes>Mycobacteriales | GCF_027270755.1 | Rhodococcus_qingshengii | O4271_RS30885 | 26346..29711 | 334542 | Swt1-family-HEPN-domain-containing-protein-[Rhodococcus-qingshengii] |
| WP_270484774.1 | DUF499_1.RM.like.systems | <-HEPN+DUF499*<-METHYLASE<-TUDOR+ SF1-SF2-Helicase+iREase+CTD+REase<-?<-?<-?<-XIS-HTH | 1110 | Bacteria>Terrabacteria_group>Actinomycetota>Actinomycetes>Mycobacteriales | GCF_027659065.1 | Gordonia_jacobaea | PGQ40_RS12000 | 183639..186971 | 122202 | DUF499-domain-containing-protein-[Gordonia-jacobaea] |
| WP_276127103.1 | DUF499_1.RM.like.systems | <-TetR-HTH\|\|?->?-><-HTH+SF1-SF2-Helicase+REase<-REase<-MORC<-HEPN+DUF499*<-METHYLASE<-REase\|\|DNA_methylase-><-CR-Kinase+ORC-CDC6-like+CARF | 1137 | Bacteria>Terrabacteria_group>Actinomycetota>Actinomycetes>Mycobacteriales | GCF_029270025.1 | Mycolicibacterium_septicum | P3H80_RS10735 | 384555..387968 | 98668 | DUF499-domain-containing-protein-[Mycolicibacterium-septicum] |
| WP_277242420.1 | DUF499_1.RM.like.systems | DnaB+SMS-KaiC-><-?<-?<-?<-?<-?<-HEPN+DUF499*<-ParB+HNH<-METHYLASE<-TUDOR+ SF1-SF2-Helicase+iREase+CTD+REase<-?\|\|NTD+SF1-SF2-Helicase-> | 1119 | Bacteria>Terrabacteria_group>Actinomycetota>Actinomycetes>Mycobacteriales | GCF_029427115.1 | Gordonia_hongkongensis | L2299_RS00560 | 120321..123680 | 1701090 | Swt1-family-HEPN-domain-containing-protein-[Gordonia-hongkongensis] |
| WP_277371905.1 | DUF499_1.RM.like.systems | MODE-HTH+PBPII-><-IclR-HTH\|\|DDE_Tnp_1_4->?->RNase-><-HEPN+DUF499*<-METHYLASE<-TUDOR+ SF1-SF2-Helicase+iREase+CTD+REase\|\|SF1-SF2-Helicase-><-DNA-glycosylase | 1138 | Bacteria>Terrabacteria_group>Actinomycetota>Actinomycetes>Mycobacteriales | GCF_029457455.1 | Rhodococcus_rhodochrous | PVE95_RS25585 | 81370..84786 | 1829 | Swt1-family-HEPN-domain-containing-protein-[Rhodococcus-rhodochrous] |
| WP_278127257.1 | DUF499_1.RM.like.systems | <-ABhydrolase<-?\|\|?->Peptidase_S51-><-ACET\|\|?-><-HEPN+DUF499*<-METHYLASE\|\|DUF222+HNH-><-TUDOR+ SF1-SF2-Helicase+iREase+CTD+REase<-?<-tRNA<-tRNA<-tRNA<-?<-tRNA<-tRNA<-tRNA\|\|tRNA->ABC-ATPase+SbcC-> | 1124 | Bacteria>Terrabacteria_group>Actinomycetota>Actinomycetes>Mycobacteriales | GCF_029632925.1 | Gordonia_sihwensis | P5P27_RS08530 | 1762026..1765400 | 173559 | DUF499-domain-containing-protein-[Gordonia-sihwensis] |
| WP_278334519.1 | DUF499_1.RM.like.systems | SF1-SF2-Helicase-><-DNA_methylase\|\|REase-><-HEPN+DUF499*<-METHYLASE<-TUDOR+ SF1-SF2-Helicase+iREase+CTD+REase\|\|REase-> | 1138 | Bacteria>Terrabacteria_group>Actinomycetota>Actinomycetes>Mycobacteriales | GCF_029691645.1 | Rhodococcus_aetherivorans | P9K37_RS00335 | 73561..76977 | 191292 | Swt1-family-HEPN-domain-containing-protein-[Rhodococcus-aetherivorans] |
| WP_283569686.1 | DUF499_1.RM.like.systems | HEPN-><-NTD+SF1-SF2-Helicase\|\|TUDOR+ SF1-SF2-Helicase+iREase+CTD+REase->METHYLASE->ParB+HNH->HEPN+DUF499*-> | 1120 | Bacteria>Terrabacteria_group>Actinomycetota>Actinomycetes>Mycobacteriales | GCF_030063715.1 | Rhodococcus_qingshengii | QNA24_RS33665 | 11874..15236 | 334542 | Swt1-family-HEPN-domain-containing-protein-[Rhodococcus-qingshengii] |
| WP_284588418.1 | DUF499_1.RM.like.systems | TUDOR+ SF1-SF2-Helicase+iREase+CTD+REase->METHYLASE->HEPN+DUF499*-><-?<-ABhydrolase+ABhydrolase<-?<-?<-?<-ABhydrolase | 1124 | Bacteria>Terrabacteria_group>Actinomycetota>Actinomycetes>Mycobacteriales | GCF_030176355.1 | Corynebacterium_pseudodiphtheriticum | QPX18_RS08860 | 15682..19056 | 37637 | DUF499-domain-containing-protein-[Corynebacterium-pseudodiphtheriticum] |
| WP_284860121.1 | DUF499_1.RM.like.systems | ABhydrolase->?->?->?->ABhydrolase+ABhydrolase->?-><-HEPN+DUF499*<-METHYLASE<-TUDOR+ SF1-SF2-Helicase+iREase+CTD+REase | 1124 | Bacteria>Terrabacteria_group>Actinomycetota>Actinomycetes>Mycobacteriales | GCF_030229165.1 | Corynebacterium_pseudodiphtheriticum | QP898_RS02760 | 58935..62309 | 37637 | DUF499-domain-containing-protein-[Corynebacterium-pseudodiphtheriticum] |
| WP_074345357.1 | DUF499_1.RM.like.systems | <-MPTase<-HTH<-?<-?<-?<-?<-HEPN+DUF499*<-METHYLASE<-TUDOR+ SF1-SF2-Helicase+iREase+CTD+REase<-HTH+ZnR\|\|HTH_Tnp_1+HTH_21+rve+rve_2-> | 1115 | Bacteria>Terrabacteria_group>Actinomycetota>Actinomycetes>Mycobacteriales | GCF_900130695.1 | Mycobacteroides_abscessus | B5Z17_RS24110 | 72774..76121 | 36809 | DUF499-domain-containing-protein-[Mycobacteroides-abscessus] |
| WP_207554210.1 | DUF499_1.RM.like.systems | <-NTD+SF1-SF2-Helicase\|\|TUDOR+ SF1-SF2-Helicase+iREase+CTD+REase-><-?\|\|METHYLASE->ParB-p->ParB+HNH->HEPN+DUF499*->DUF499-p->?->?->\|\|MULE-TRANSPOSASE-> | 605 | Bacteria>Terrabacteria_group>Actinomycetota>Actinomycetes>Mycobacteriales | GCF_900141575.1 | Mycobacteroides_abscessus | B5Y25_RS21325 | 33267..35081 | 36809 | Swt1-family-HEPN-domain-containing-protein,-partial-[Mycobacteroides-abscessus] |
| WP_207554378.1 | DUF499_1.RM.like.systems | <-NTD+SF1-SF2-Helicase\|\|TUDOR+ SF1-SF2-Helicase+iREase+CTD+REase-><-?\|\|METHYLASE->ParB+HNH->HEPN+DUF499*->?->?->\|\|MULE-TRANSPOSASE-> | 1113 | Bacteria>Terrabacteria_group>Actinomycetota>Actinomycetes>Mycobacteriales | GCF_900141585.1 | Mycobacteroides_abscessus | B5Z43_RS22925 | 52451..55792 | 36809 | Swt1-family-HEPN-domain-containing-protein-[Mycobacteroides-abscessus] |
| WP_085470756.1 | DUF499_1.RM.like.systems | DNA-glycosylase->?-><-NTD+SF1-SF2-Helicase<-HEPN+DUF499*<-ParB+HNH<-METHYLASE<-TUDOR+ SF1-SF2-Helicase+iREase+CTD+REase\|\|?-><-PIN<-XIS-HTH | 1125 | Bacteria>Terrabacteria_group>Actinomycetota>Actinomycetes>Mycobacteriales | GCF_900177695.1 | Rhodococcus_rhodochrous | M502_RS25910 | 14095..17472 | 1829 | Swt1-family-HEPN-domain-containing-protein-[Rhodococcus-rhodochrous] |
| WP_089250630.1 | DUF499_1.RM.like.systems | <-DUF433<-?<-HEPN+DUF499*\|\|MazE_antitoxin->PIN-><-METHYLASE<-AbiEii<-?<-TUDOR+ SF1-SF2-Helicase+iREase+CTD+REase | 1122 | Bacteria>Terrabacteria_group>Actinomycetota>Actinomycetes>Mycobacteriales | GCF_900188125.1 | Rhodococcus_kyotonensis | CHB96_RS22960 | 65313..68681 | 398843 | Swt1-family-HEPN-domain-containing-protein-[Rhodococcus-kyotonensis] |
| WP_183155213.1 | DUF499_1.RM.like.systems | <-HTH+ZnR<-?\|\|HEPN->TUDOR+ SF1-SF2-Helicase+iREase+CTD+REase->METHYLASE->HEPN+DUF499*->Pkinase+SF1-SF2-Helicase->HTH+HKD-DNase-> | 1114 | Bacteria>Terrabacteria_group>Actinomycetota>Actinomycetes>Mycobacteriales | GCF_900565985.1 | Mycobacterium_kansasii | EET11_RS25330 | 51078..54422 | 1768 | DUF499-domain-containing-protein-[Mycobacterium-kansasii] |
| WP_125079869.1 | DUF499_1.RM.like.systems | ACET->GDResolvase+HIN-HTH-><-REase<-HEPN+DUF499* | 1124 | Bacteria>Terrabacteria_group>Actinomycetota>Actinomycetes>Mycobacteriales | GCF_900604305.1 | Mycobacterium_sp._P7213 | EJK91_RS20970 | 3382..6756 | 2478465 | DUF499-domain-containing-protein-[Mycobacterium-sp.-P7213] |
| WP_126299909.1 | DUF499_1.RM.like.systems | <-HEPN\|\|?-><-Classical-AAA<-?<-?<-Classical-AAA<-HEPN+DUF499*<-METHYLASE<-TUDOR+ SF1-SF2-Helicase+iREase+CTD+REase | 1141 | Bacteria>Terrabacteria_group>Actinomycetota>Actinomycetes>Mycobacteriales | GCF_900638255.1 | Corynebacterium_matruchotii | EL293_RS12755 | 2814253..2817678 | 43768 | DUF499-domain-containing-protein-[Corynebacterium-matruchotii] |
| WP_210731572.1 | DUF499_1.RM.like.systems | METHYLASE->ParB+HNH->HEPN+DUF499*-><-?\|\|?->?->SbcC->SbcC-> | 1116 | Bacteria>Terrabacteria_group>Actinomycetota>Actinomycetes>Mycobacteriales | GCF_900960235.1 | Mycobacterium_intracellulare | FAX65_RS26670 | 4893..8243 | 1767 | Swt1-family-HEPN-domain-containing-protein-[Mycobacterium-intracellulare] |
| WP_204801726.1 | DUF499_1.RM.like.systems | <-GNTR-HTH\|\|NUDIX-><-?<-HEPN<-Schlafen+HATPase+HTH<-HEPN+DUF499*<-METHYLASE\|\|?-><-TUDOR+ SF1-SF2-Helicase+iREase+CTD+REase<-HEPN+REase<-NTP_transf | 1109 | Bacteria>Terrabacteria_group>Actinomycetota>Actinomycetes>Mycobacteriales | GCF_905219495.1 | Mycobacterium_riyadhense | JX354_RS03520 | 789692..793021 | 486698 | Swt1-family-HEPN-domain-containing-protein-[Mycobacterium-riyadhense] |
| WP_278720226.1 | DUF499_1.RM.like.systems | <-Classical-AAA<-?<-?<-Classical-AAA<-HEPN<-HEPN+DUF499*<-METHYLASE<-TUDOR+ SF1-SF2-Helicase+iREase+CTD+REase\|\|?->?->?->?-><-HTH_28+HTH_33+DDE_3 | 1184 | Bacteria>Terrabacteria_group>Actinomycetota>Actinomycetes>Mycobacteriales | GCF_905372825.1 | Corynebacterium_matruchotii | QDH69_RS07065 | 79539..83093 | 43768 | DUF499-domain-containing-protein-[Corynebacterium-matruchotii] |
| WP_219068412.1 | DUF499_1.RM.like.systems | PIN-><-HEPN+DUF499*<-METHYLASE<-TUDOR+ SF1-SF2-Helicase+iREase+CTD+REase\|\|NTD+SF1-SF2-Helicase->?-><-MORC+DUF3684+REase | 1113 | Bacteria>Terrabacteria_group>Actinomycetota>Actinomycetes>Mycobacteriales | GCF_910591625.1 | Candidatus_Mycobacterium_methanotrophicum | M5I08_RS02450 | 450660..454001 | 2943498 | MULTISPECIES:-Swt1-family-HEPN-domain-containing-protein-[unclassified-Mycobacterium] |
| WP_230074521.1 | DUF499_1.RM.like.systems | <-MNS+HTH\|\|?->HNH->TIR->ZnR+ZnR-><-HEPN+DUF499*<-METHYLASE<-TUDOR+ SF1-SF2-Helicase+iREase+CTD+REase\|\|RnaseIII+DSRBD+RnaseIII+DSRBD-> | 1127 | Bacteria>Terrabacteria_group>Actinomycetota>Actinomycetes>Mycobacteriales | GCF_918697885.1 | Rhodococcus_fascians | LQV52_RS22465 | 73515..76898 | 1828 | DUF499-domain-containing-protein-[Rhodococcus-fascians] |
| WP_040525708.1 | DUF499_1.RM.like.systems | <-ABC-ATPase+SbcC\|\|HTH_3-><-?<-?<-?<-HEPN+DUF499*<-METHYLASE\|\|HTH_19-> | 1111 | Bacteria>Terrabacteria_group>Actinomycetota>Actinomycetes>Mycobacteriales | - | Gordonia_effusa | GOEFS_RS20135 | 108493..111828 | 263908 | DUF499-domain-containing-protein-[Gordonia-effusa] |
| WP_239588858.1 | DUF499_1.RM.like.systems | <-ABhydrolase<-?\|\|?->Peptidase_S51-><-HEPN+DUF499*<-METHYLASE\|\|DUF222+HNH-><-TUDOR+ SF1-SF2-Helicase+iREase+CTD+REase<-ACET | 1125 | Bacteria>Terrabacteria_group>Actinomycetota>Actinomycetes>Mycobacteriales | - | Gordonia_hydrophobica | JOC45_RS12260 | 227782..231159 | 40516 | DUF499-domain-containing-protein-[Gordonia-hydrophobica] |
| WP_260868497.1 | DUF499_1.RM.like.systems | TUDOR+ SF1-SF2-Helicase+iREase+CTD+REase-><-DUF222+HNH\|\|METHYLASE->HEPN+DUF499*-><-?\|\|ACET-><-Peptidase_S51<-?\|\|?->ABhydrolase-> | 1110 | Bacteria>Terrabacteria_group>Actinomycetota>Actinomycetes>Mycobacteriales | - | Gordonia_sihwensis | ACN95_RS06460 | 9637..12969 | 173559 | DUF499-domain-containing-protein-[Gordonia-sihwensis] |
| WP_023870228.1 | DUF499_1.RM.like.systems | Glyoxalase->HIN-HTH-><-DNA_methylase\|\|REase->METHYLASE->HEPN+DUF499*-><-HIN-HTH | 1138 | Bacteria>Terrabacteria_group>Actinomycetota>Actinomycetes>Mycobacteriales | - | Mycobacterium_avium | BEP52_09350 | 1958340..1961756 | 1764 | DUF499-domain-containing-protein-[Mycobacterium-avium] |
| WP_085075211.1 | DUF499_1.RM.like.systems | <-Phage_int_SAM_1+Phage_integrase<-GDResolvase+Recombinase<-HTH_17<-?<-HEPN+DUF499*<-METHYLASE<-TUDOR+ SF1-SF2-Helicase+iREase+CTD+REase | 1110 | Bacteria>Terrabacteria_group>Actinomycetota>Actinomycetes>Mycobacteriales | - | Mycobacterium_kubicae | H7H99_RS20905 | 74999..78331 | 120959 | Swt1-family-HEPN-domain-containing-protein-[Mycobacterium-kubicae] |
| WP_170826146.1 | DUF499_1.RM.like.systems | <-HEPN+DUF499*<-METHYLASE<-TUDOR+ SF1-SF2-Helicase+iREase+CTD+REase-p | 1112 | Bacteria>Terrabacteria_group>Actinomycetota>Actinomycetes>Mycobacteriales | - | Mycobacterium_malmoense | A9X02_RS19855 | 3529..6867 | 1780 | Swt1-family-HEPN-domain-containing-protein-[Mycobacterium-malmoense] |
| WP_176126766.1 | DUF499_1.RM.like.systems | <-Phage_integrase\|\|Cluster182_14clades->?-><-?<-?<-?<-HEPN+DUF499*<-ParB+HNH<-METHYLASE<-TUDOR+ SF1-SF2-Helicase+iREase+CTD+REase\|\|NTD+SF1-SF2-Helicase-> | 1120 | Bacteria>Terrabacteria_group>Actinomycetota>Actinomycetes>Mycobacteriales | - | Mycobacterium_sp._D16Q20 | B4386_RS22320 | 7597..10959 | 1855660 | Swt1-family-HEPN-domain-containing-protein-[Mycobacterium-sp.-D16Q20] |
| WP_005139269.1 | DUF499_1.RM.like.systems | <-GNTR-HTH\|\|NUDIX-><-?<-HEPN<-Schlafen+HATPase+HTH<-HEPN+DUF499*<-REase<-METHYLASE<-TUDOR+ SF1-SF2-Helicase+iREase+CTD+REase<-PIN<-XIS-HTH | 1108 | Bacteria>Terrabacteria_group>Actinomycetota>Actinomycetes>Mycobacteriales | - | Mycolicibacterium_rhodesiae | MycrhDRAFT_0407 | 390606..393932 | 36814 | Swt1-family-HEPN-domain-containing-protein-[Mycolicibacterium-rhodesiae] |
| WP_094024119.1 | DUF499_1.RM.like.systems | Peptidase_S66+Peptidase_S66C->SAM-methylase->?-><-ParB+HNH\|\|?-><-HEPN+DUF499*<-METHYLASE<-TUDOR+ SF1-SF2-Helicase+iREase+CTD+REase<-Lsr2<-?<-UbE1+JAB | 1126 | Bacteria>Terrabacteria_group>Actinomycetota>Actinomycetes>Mycobacteriales | - | Nocardia_cerradoensis | B7C42_RS01205 | 260073..263453 | 85688 | DUF499-domain-containing-protein-[Nocardia-cerradoensis] |
| WP_065456397.1 | DUF499_1.RM.like.systems | FAD-NAD-dep-oxidoreductase-><-HEPN+DUF499*<-METHYLASE<-TUDOR+ SF1-SF2-Helicase+iREase+CTD+REase | 1134 | Bacteria>Terrabacteria_group>Actinomycetota>Actinomycetes>Mycobacteriales | - | Prescottella_equi | AS590_01070 | 160611..164015 | 43767 | Swt1-family-HEPN-domain-containing-protein-[Prescottella-equi] |
| WP_116057355.1 | DUF499_1.RM.like.systems | <-URI-DNase<-DNA-glycosylase\|\|?->TUDOR+ SF1-SF2-Helicase+iREase+CTD+REase->METHYLASE->HEPN+DUF499*->?-><-HNH | 1122 | Bacteria>Terrabacteria_group>Actinomycetota>Actinomycetes>Mycobacteriales | - | Rhodococcus_erythropolis | DVG80_RS18290 | 3917488..3920856 | 1833 | Swt1-family-HEPN-domain-containing-protein-[Rhodococcus-erythropolis] |
| WP_116058404.1 | DUF499_1.RM.like.systems | HEPN-><-NTD+SF1-SF2-Helicase\|\|TUDOR+ SF1-SF2-Helicase+iREase+CTD+REase->METHYLASE->ParB+HNH->HEPN+DUF499*->?-><-Lsr2\|\|Phage_integrase->?->PAIREDC-HTH-> | 1121 | Bacteria>Terrabacteria_group>Actinomycetota>Actinomycetes>Mycobacteriales | - | Rhodococcus_erythropolis | DVG80_RS32980 | 11800..15165 | 1833 | Swt1-family-HEPN-domain-containing-protein-[Rhodococcus-erythropolis] |
| WP_194699134.1 | DUF499_1.RM.like.systems | SF1-SF2-Helicase->\|\|REase-><-HEPN+DUF499*<-TUDOR+ SF1-SF2-Helicase+iREase+CTD+REase\|\|PAIREDC-HTH-> | 1135 | Bacteria>Terrabacteria_group>Actinomycetota>Actinomycetes>Mycobacteriales | - | Rhodococcus_rhodochrous | IRZ34_RS06155 | 243205..246612 | 1829 | Swt1-family-HEPN-domain-containing-protein-[Rhodococcus-rhodochrous] |
| WP_264184695.1 | DUF499_1.RM.like.systems | <-HTH\|\|TUDOR+ SF1-SF2-Helicase+iREase+CTD+REase-><-CHTH\|\|METHYLASE->HEPN+DUF499*->DUF222+HNH-><-?<-?<-Peptidase_A8 | 1139 | Bacteria>Terrabacteria_group>Actinomycetota>Actinomycetes>Mycobacteriales | - | Rhodococcus_yananensis | LDQ05_RS09980 | 11697..15116 | 2879464 | Swt1-family-HEPN-domain-containing-protein-[Rhodococcus-yananensis] |
| MAO81794.1 | DUF499_1.RM.like.systems | <-DUF1679<-SIR2<-NERD<-?<-?<-?<-HEPN+DUF499*<-METHYLASE<-TUDOR+ SF1-SF2-Helicase+iREase+CTD+REase\|\|NTD+SF1-SF2-Helicase-><-?<-?<-TrwC+SF1-SF2-Helicase | 1117 | Bacteria>Terrabacteria_group>Actinomycetota>Actinomycetes>Propionibacteriales | GCA_002693615.1 | Nocardioides_sp. | CMH82_14275 | 37872..41225 | 35761 | AAA-family-ATPase-[Nocardioides-sp.] |
| MAS55988.1 | DUF499_1.RM.like.systems | XIS-HTH->METHYLASE-p-><-?<-MORC+HISKIN<-HEPN+DUF499*<-ParB+HNH<-METHYLASE<-UDG-conflict<-TUDOR+ SF1-SF2-Helicase+iREase+CTD+REase-p | 1116 | Bacteria>Terrabacteria_group>Actinomycetota>Actinomycetes>Propionibacteriales | GCA_002698575.1 | Pimelobacter_sp. | CMJ44_15460 | 17699..21049 | 51662 | AAA-family-ATPase-[Pimelobacter-sp.] |
| MCL2489469.1 | DUF499_1.RM.like.systems | <-TrwC+SF1-SF2-Helicase<-?<-PAPS_reduct<-?<-KAP-NTPase<-HEPN+DUF499*<-HEPN<-METHYLASE | 1120 | Bacteria>Terrabacteria_group>Actinomycetota>Actinomycetes>Propionibacteriales | GCA_009778875.1 | Propionibacteriaceae_bacterium | FWF36_01890 | 13226..16588 | 2021380 | DUF499-domain-containing-protein-[Propionibacteriaceae-bacterium] |
| NNG19379.1 | DUF499_1.RM.like.systems | LexA-HTH+LexA-protease-><-?<-35exo+SF1-SF2-Helicase<-?<-HEPN+DUF499*<-METHYLASE\|\|DUF222+HNH-><-TUDOR+ SF1-SF2-Helicase+iREase+CTD+REase\|\|HTH_3-> | 1110 | Bacteria>Terrabacteria_group>Actinomycetota>Actinomycetes>Propionibacteriales | GCA_013002685.1 | Naumannella_sp._ID2617S | HJ590_07255 | 1496426..1499758 | 1656893 | ATP-binding-protein-[Naumannella-sp.-ID2617S] |
| MBE6477658.1 | DUF499_1.RM.like.systems | TUDOR+ SF1-SF2-Helicase+iREase+CTD+REase->METHYLASE->HEPN->wHTH+REase->?->PemK_toxin->HEPN+DUF499*->?->TetR-HTH->?-><-?\|\|ArsR-HTH-><-ABhydrolase | 1088 | Bacteria>Terrabacteria_group>Actinomycetota>Actinomycetes>Propionibacteriales | GCA_015062705.1 | Propionibacteriaceae_bacterium | E7Z97_06285 | 37437..40699 | 2021380 | ATP-binding-protein-[Propionibacteriaceae-bacterium] |
| MBK8462833.1 | DUF499_1.RM.like.systems | <-REase<-?<-HEPN\|\|TUDOR+ SF1-SF2-Helicase+iREase+CTD+REase->METHYLASE->ParB+HNH->HEPN+DUF499*->?-><-Peptidase_M48<-?<-?\|\|TPR-> | 1117 | Bacteria>Terrabacteria_group>Actinomycetota>Actinomycetes>Propionibacteriales | GCA_016711325.1 | Nigerium_sp. | IPL36_07335 | 626569..629922 | 2042655 | DUF499-domain-containing-protein-[Nigerium-sp.] |
| MBM0123458.1 | DUF499_1.RM.like.systems | <-XIS-HTH<-XIS-HTH<-TrwC+SF1-SF2-helicase\|\|HEPN+Mrr_N-><-HEPN+DUF499*<-ParB+HNH<-METHYLASE<-TUDOR+ SF1-SF2-Helicase+iREase+CTD+REase\|\|NTD+SF1-SF2-Helicase-> | 1119 | Bacteria>Terrabacteria_group>Actinomycetota>Actinomycetes>Propionibacteriales | GCA_016803235.1 | Pimelobacter_simplex | IMY96_01775 | 51221..54580 | 2045 | DUF499-domain-containing-protein-[Pimelobacter-simplex] |
| MBN9103777.1 | DUF499_1.RM.like.systems | TUDOR+ SF1-SF2-Helicase+iREase+CTD+REase->UDG-conflict->METHYLASE->HEPN+DUF499*->ParB+HNH->?->SF1-SF2-Helicase-><-REase+PNTD1+PNTD2\|\|ACET-><-Phage_integrase | 1117 | Bacteria>Terrabacteria_group>Actinomycetota>Actinomycetes>Propionibacteriales | GCA_017306015.1 | Propionibacteriaceae_bacterium | JOI14_02120 | 371284..374637 | 2021380 | DUF499-domain-containing-protein-[Propionibacteriaceae-bacterium] |
| WP_013863116.1 | DUF499_1.RM.like.systems | PglZ-><-?\|\|?-><-DUF222+HNH\|\|?-><-HEPN+DUF499*<-ParB+HNH<-?<-METHYLASE\|\|wHTH(SBD)+HNH->HTH_19-> | 1117 | Bacteria>Terrabacteria_group>Actinomycetota>Actinomycetes>Propionibacteriales | GCF_000270245.1 | Microlunatus_phosphovorus | MLP_RS10775 | 2379873..2383226 | 29405 | Swt1-family-HEPN-domain-containing-protein-[Microlunatus-phosphovorus] |
| WP_014846821.1 | DUF499_1.RM.like.systems | REase->TUDOR+ SF1-SF2-Helicase+iREase+CTD+REase->METHYLASE->wHTH+REase->HEPN+DUF499*->?->ACET-><-?\|\|?-><-TetR-HTH | 1129 | Bacteria>Terrabacteria_group>Actinomycetota>Actinomycetes>Propionibacteriales | GCF_000277715.1 | Arachnia_propionica | J5A53_RS06805 | 1506504..1509893 | 1750 | DUF499-domain-containing-protein-[Arachnia-propionica] |
| WP_020385147.1 | DUF499_1.RM.like.systems | NUDIX->tRNA-><-?<-?\|\|TUDOR+ SF1-SF2-Helicase+iREase+CTD+REase->METHYLASE->ParB+HNH->HEPN+DUF499*-><-?<-?<-?<-METHYLASE-p | 1116 | Bacteria>Terrabacteria_group>Actinomycetota>Actinomycetes>Propionibacteriales | GCF_000372465.1 | Kribbella_catacumbae | A3ES_RS0102270 | 28038..31388 | 460086 | Swt1-family-HEPN-domain-containing-protein-[Kribbella-catacumbae] |
| WP_028701426.1 | DUF499_1.RM.like.systems | <-SF1-SF2-Helicase<-MORC+SF1-SF2-Helicase\|\|DNA_methylase-><-DNA_methylase\|\|METHYLASE->HEPN+DUF499*->MORC->REase->HTH+SF1-SF2-Helicase+REase->REase-><-ParB+HNH-p<-ParB+HNH-p | 1129 | Bacteria>Terrabacteria_group>Actinomycetota>Actinomycetes>Propionibacteriales | GCF_000427845.1 | Acidipropionibacterium_acidipropionici | ASQ49_RS03445 | 772026..775415 | 1748 | DUF499-domain-containing-protein-[Acidipropionibacterium-acidipropionici] |
| WP_021798346.1 | DUF499_1.RM.like.systems | TUDOR+ SF1-SF2-Helicase+iREase+CTD+REase->METHYLASE->wHTH+REase->HEPN+DUF499*->SF1-SF2-Helicase+REase-p->SF1-SF2-Helicase+REase-p->HTH_28+HTH_33+DDE_3->SF1-SF2-Helicase-p-><-Synaptojanin | 1128 | Bacteria>Terrabacteria_group>Actinomycetota>Actinomycetes>Propionibacteriales | GCF_000478805.1 | Propionibacterium_acidifaciens | HMPREF0682_RS10330 | 20512..23898 | 556499 | Swt1-family-HEPN-domain-containing-protein-[Propionibacterium-acidifaciens] |
| WP_061786876.1 | DUF499_1.RM.like.systems | REase->TUDOR+ SF1-SF2-Helicase+iREase+CTD+REase->METHYLASE->wHTH+REase->HEPN+DUF499*->KAP-NTPase->ACET-><-?\|\|?-><-TetR-HTH | 1129 | Bacteria>Terrabacteria_group>Actinomycetota>Actinomycetes>Propionibacteriales | GCF_001592325.1 | Arachnia_propionica | PP13_RS01005 | 243859..247248 | 1750 | DUF499-domain-containing-protein-[Arachnia-propionica] |
| WP_078920357.1 | DUF499_1.RM.like.systems | <-TrwC+SF1-SF2-helicase+TOPRIM<-DNA_methylase\|\|REase->METHYLASE->HEPN+DUF499*->MORC->REase->HTH+SF1-SF2-Helicase+REase-> | 1132 | Bacteria>Terrabacteria_group>Actinomycetota>Actinomycetes>Propionibacteriales | GCF_002022225.1 | Tessaracoccus | BKM78_RS14510 | 3153779..3157177 | 72763 | MULTISPECIES:-Swt1-family-HEPN-domain-containing-protein-[Tessaracoccus] |
| WP_094451589.1 | DUF499_1.RM.like.systems | <-SbcC<-HEPN+DUF499*<-METHYLASE<-TUDOR+ SF1-SF2-Helicase+iREase+CTD+REase<-?\|\|HTH-> | 1135 | Bacteria>Terrabacteria_group>Actinomycetota>Actinomycetes>Propionibacteriales | GCF_002250475.1 | Parenemella_sanctibonifatiensis | CGZ92_RS11810 | 2653..6060 | 2016505 | DUF499-domain-containing-protein-[Parenemella-sanctibonifatiensis] |
| WP_094363616.1 | DUF499_1.RM.like.systems | <-rve_3<-HIN-HTH\|\|?-><-?\|\|TUDOR+ SF1-SF2-Helicase+iREase+CTD+REase->METHYLASE->HEPN+DUF499*->SbcC-><-?<-?\|\|?->HEPN->DOC+HTH-> | 1125 | Bacteria>Terrabacteria_group>Actinomycetota>Actinomycetes>Propionibacteriales | GCF_002250625.1 | Enemella_dayhoffiae | CGZ93_RS07965 | 9213..12590 | 2016507 | DUF499-domain-containing-protein-[Enemella-dayhoffiae] |
| WP_115850048.1 | DUF499_1.RM.like.systems | <-Pkinase<-NTP_transf_5\|\|Pkinase->?->?-><-SF1-SF2-Helicase<-HEPN+DUF499*<-METHYLASE<-TUDOR+ SF1-SF2-Helicase+iREase+CTD+REase | 1124 | Bacteria>Terrabacteria_group>Actinomycetota>Actinomycetes>Propionibacteriales | GCF_003386795.1 | Thermasporomyces_composti | DFJ64_RS08950 | 2047152..2050526 | 696763 | Swt1-family-HEPN-domain-containing-protein-[Thermasporomyces-composti] |
| WP_123224702.1 | DUF499_1.RM.like.systems | <-NERD\|\|REase+SAVED->?->?-><-TIR<-HEPN+DUF499*<-METHYLASE\|\|?-><-TUDOR+ SF1-SF2-Helicase+iREase+CTD+REase\|\|NTD+SF1-SF2-Helicase-> | 1120 | Bacteria>Terrabacteria_group>Actinomycetota>Actinomycetes>Propionibacteriales | GCF_003725535.1 | Nocardioides_pocheonensis | EFL26_RS20130 | 110530..113892 | 661485 | DUF499-domain-containing-protein-[Marmoricola-pocheonensis] |
| WP_124237129.1 | DUF499_1.RM.like.systems | <-REase\|\|NTP_transf_2->tRNA-><-NTD+SF1-SF2-Helicase\|\|TUDOR+ SF1-SF2-Helicase+iREase+CTD+REase->METHYLASE->ParB+HNH->HEPN+DUF499*-> | 1124 | Bacteria>Terrabacteria_group>Actinomycetota>Actinomycetes>Propionibacteriales | GCF_003850045.1 | Aeromicrobium_camelliae | EHW97_RS10560 | 83306..86680 | 1538144 | Swt1-family-HEPN-domain-containing-protein-[Aeromicrobium-camelliae] |
| WP_125568181.1 | DUF499_1.RM.like.systems | HTH_3->?->?-><-HEPN+DUF499*<-METHYLASE<-TUDOR+ SF1-SF2-Helicase+iREase+CTD+REase\|\|NTD+SF1-SF2-Helicase-> | 1121 | Bacteria>Terrabacteria_group>Actinomycetota>Actinomycetes>Propionibacteriales | GCF_003945325.1 | Nocardioides_baekrokdamisoli | KCTC_RS07355 | 1515846..1519211 | 1804624 | Swt1-family-HEPN-domain-containing-protein-[Nocardioides-baekrokdamisoli] |
| WP_154690370.1 | DUF499_1.RM.like.systems | PAIREDC-HTH-><-?<-HTH+SF1-SF2-Helicase+REase<-REase<-MORC<-HEPN+DUF499*<-METHYLASE-p<-METHYLASE<-REase<-PIN\|\|DNA_methylase-> | 1132 | Bacteria>Terrabacteria_group>Actinomycetota>Actinomycetes>Propionibacteriales | GCF_009707365.1 | Microlunatus_sp._Gsoil_973 | GJV80_RS18445 | 3917369..3920767 | 2672569 | DUF499-domain-containing-protein-[Microlunatus-sp.-Gsoil-973] |
| WP_166139838.1 | DUF499_1.RM.like.systems | <-SLATT\|\|?-><-NTD+SF1-SF2-Helicase\|\|TUDOR+ SF1-SF2-Helicase+iREase+CTD+REase->METHYLASE->ParB+HNH->HEPN+DUF499*-><-?<-HNH\|\|?->?->?->REase+35exo+Beta-Barrel+SF1-SF2-Helicase-> | 1120 | Bacteria>Terrabacteria_group>Actinomycetota>Actinomycetes>Propionibacteriales | GCF_011420305.2 | Nocardioides_ochotonae | HBO46_RS08105 | 1673933..1677295 | 2685869 | Swt1-family-HEPN-domain-containing-protein-[Nocardioides-ochotonae] |
| WP_167165953.1 | DUF499_1.RM.like.systems | <-Pkinase<-?<-?\|\|TUDOR+ SF1-SF2-Helicase+iREase+CTD+REase->METHYLASE->HEPN+DUF499*->?->PIN->?->?-><-Aminotran_1_2 | 1131 | Bacteria>Terrabacteria_group>Actinomycetota>Actinomycetes>Propionibacteriales | GCF_011758645.1 | Brooklawnia_cerclae | FB473_RS07080 | 1539391..1542786 | 349934 | Swt1-family-HEPN-domain-containing-protein-[Brooklawnia-cerclae] |
| WP_246322323.1 | DUF499_1.RM.like.systems | <-REase<-?\|\|?->SIGMA-HTH-><-HEPN+DUF499*<-ParB+HNH<-METHYLASE<-TUDOR+ SF1-SF2-Helicase+iREase+CTD+REase\|\|REase-> | 1096 | Bacteria>Terrabacteria_group>Actinomycetota>Actinomycetes>Propionibacteriales | GCF_013407955.1 | Microlunatus_parietis | BKA15_RS11400 | 2395766..2399056 | 682979 | DUF499-domain-containing-protein-[Microlunatus-parietis] |
| WP_182784909.1 | DUF499_1.RM.like.systems | <-SF1-SF2-Helicase<-OLD-ABC+TOPRIM<-HEPN+DUF499*<-METHYLASE<-TUDOR+ SF1-SF2-Helicase+iREase+CTD+REase<-SF1-SF2-Helicase-p\|\|Glyoxalase-> | 1133 | Bacteria>Terrabacteria_group>Actinomycetota>Actinomycetes>Propionibacteriales | GCF_014164725.1 | Tessaracoccus_sp._MC1627 | H5399_RS17220 | 4312..7713 | 2760312 | DUF499-domain-containing-protein-[Tessaracoccus-sp.-MC1627] |
| WP_182766932.1 | DUF499_1.RM.like.systems | <-HTH+SF1-SF2-Helicase+REase<-REase<-HEPN+DUF499*<-METHYLASE<-REase\|\|DNA_methylase-> | 1129 | Bacteria>Terrabacteria_group>Actinomycetota>Actinomycetes>Propionibacteriales | GCF_014164735.1 | Tessaracoccus_sp._MC1756 | H5393_RS12480 | 209092..212481 | 2760311 | Swt1-family-HEPN-domain-containing-protein-[Tessaracoccus-sp.-MC1756] |
| WP_187768939.1 | DUF499_1.RM.like.systems | <-PIN+PhoH<-?\|\|?-><-?<-?<-HEPN+DUF499*<-METHYLASE<-TUDOR+ SF1-SF2-Helicase+iREase+CTD+REase\|\|NTD+SF1-SF2-Helicase-> | 1119 | Bacteria>Terrabacteria_group>Actinomycetota>Actinomycetes>Propionibacteriales | GCF_014490085.1 | Aeromicrobium | GL325_RS06400 | 1283810..1287169 | 2040 | MULTISPECIES:-Swt1-family-HEPN-domain-containing-protein-[unclassified-Aeromicrobium] |
| WP_194694759.1 | DUF499_1.RM.like.systems | <-DOC<-?<-tRNA<-DDE_Tnp_1_4<-?\|\|TUDOR+ SF1-SF2-Helicase+iREase+CTD+REase->METHYLASE->HEPN+DUF499*-><-?<-?\|\|?->ParB+HNH->?->Phage_int_SAM_1+Phage_integrase-> | 1113 | Bacteria>Terrabacteria_group>Actinomycetota>Actinomycetes>Propionibacteriales | GCF_015352445.1 | Nocardioides_agariphilus | ISU10_RS02340 | 86950..90291 | 433664 | DUF499-domain-containing-protein-[Nocardioides-agariphilus] |
| WP_207473688.1 | DUF499_1.RM.like.systems | ABhydrolase->XIS-HTH->TUDOR+ SF1-SF2-Helicase+iREase+CTD+REase->METHYLASE->wHTH+REase->HEPN+DUF499*->REase->HEPN+REase-><-DNA_methylase\|\|TrwC+SF1-SF2-Helicase-> | 1131 | Bacteria>Terrabacteria_group>Actinomycetota>Actinomycetes>Propionibacteriales | GCF_017355265.1 | Tessaracoccus_sp._SD287 | IPV09_RS03380 | 69383..72778 | 2782008 | DUF499-domain-containing-protein-[Tessaracoccus-sp.-SD287] |
| WP_211798072.1 | DUF499_1.RM.like.systems | TetR-HTH-><-?\|\|?-><-ACET<-?<-HEPN+DUF499*<-wHTH+REase<-METHYLASE<-TUDOR+ SF1-SF2-Helicase+iREase+CTD+REase | 1129 | Bacteria>Terrabacteria_group>Actinomycetota>Actinomycetes>Propionibacteriales | GCF_018128345.1 | Arachnia_propionica | J5A61_RS08580 | 1914777..1918166 | 1750 | DUF499-domain-containing-protein-[Arachnia-propionica] |
| WP_219081771.1 | DUF499_1.RM.like.systems | ABC-ATPase+SbcC->ABC-ATPase+SbcC-><-NTD+SF1-SF2-Helicase<-NACHT\|\|TUDOR+ SF1-SF2-Helicase+iREase+CTD+REase->METHYLASE->HEPN+DUF499*->?-><-ABhydrolase<-TetR-HTH | 1121 | Bacteria>Terrabacteria_group>Actinomycetota>Actinomycetes>Propionibacteriales | GCF_019316905.1 | Tessaracoccus_palaemonis | KDB89_RS13130 | 2884932..2888297 | 2829499 | Swt1-family-HEPN-domain-containing-protein-[Tessaracoccus-sp.-J1M15] |
| WP_227735039.1 | DUF499_1.RM.like.systems | TUDOR+ SF1-SF2-Helicase+iREase+CTD+REase-><-PIN<-DUF433\|\|METHYLASE-><-DOC+HTH\|\|HEPN+DUF499*-> | 1104 | Bacteria>Terrabacteria_group>Actinomycetota>Actinomycetes>Propionibacteriales | GCF_020687685.1 | Tessaracoccus_sp._OS52 | LKO27_RS00980 | 227595..230909 | 2886691 | DUF499-domain-containing-protein-[Tessaracoccus-sp.-OS52] |
| WP_240236717.1 | DUF499_1.RM.like.systems | <-wHTH(SBD)+HNH<-NTD+SF1-SF2-Helicase\|\|TUDOR+ SF1-SF2-Helicase+iREase+CTD+REase->METHYLASE->ParB+HNH->HEPN+DUF499*-><-Peptidase\|\|?->?->?->METHYLASE-> | 1117 | Bacteria>Terrabacteria_group>Actinomycetota>Actinomycetes>Propionibacteriales | GCF_022436705.1 | Nocardioides_sp._CFH_31398 | MJ876_RS12955 | 81599..84952 | 2919579 | Swt1-family-HEPN-domain-containing-protein-[Nocardioides-sp.-CFH-31398] |
| WP_255892000.1 | DUF499_1.RM.like.systems | <-NTD+SF1-SF2-Helicase\|\|TUDOR+ SF1-SF2-Helicase+iREase+CTD+REase->METHYLASE->HEPN+DUF499*->MORC+HISKIN->?-><-METHYLASE-p | 1104 | Bacteria>Terrabacteria_group>Actinomycetota>Actinomycetes>Propionibacteriales | GCF_024436375.1 | Nocardioides_astragali | J3P44_RS18825 | 368600..371914 | 1776736 | Swt1-family-HEPN-domain-containing-protein-[Nocardioides-astragali] |
| WP_232416936.1 | DUF499_1.RM.like.systems | <-HTH+SF1-SF2-Helicase+REase<-REase<-MORC<-HEPN+DUF499*<-METHYLASE<-REase\|\|DNA_methylase-><-?\|\|DUF222+HNH-> | 1136 | Bacteria>Terrabacteria_group>Actinomycetota>Actinomycetes>Propionibacteriales | GCF_024508295.1 | Aeromicrobium_duanguangcaii | NP095_RS05665 | 1110875..1114285 | 2968086 | MULTISPECIES:-Swt1-family-HEPN-domain-containing-protein-[unclassified-Aeromicrobium] |
| WP_257750656.1 | DUF499_1.RM.like.systems | TrwC+SF1-SF2-Helicase->?-><-NTD+SF1-SF2-Helicase\|\|TUDOR+ SF1-SF2-Helicase+iREase+CTD+REase->METHYLASE->HEPN+DUF499*->KAP-NTPase->?->PAPS_reduct->?->NERD-> | 1119 | Bacteria>Terrabacteria_group>Actinomycetota>Actinomycetes>Propionibacteriales | GCF_024662235.1 | Pimelobacter_simplex | MOM48_RS07490 | 1563266..1566625 | 2045 | Swt1-family-HEPN-domain-containing-protein-[Pimelobacter-simplex] |
| WP_131171419.1 | DUF499_1.RM.like.systems | <-MACRODOMAIN+SIR2<-HTH+SF1-SF2-Helicase+REase<-REase<-MORC<-HEPN+DUF499*<-METHYLASE<-METHYLASE-p\|\|DNA_methylase-><-SbcC<-SbcC\|\|TrwC+SF1-SF2-helicase+TOPRIM-> | 1146 | Bacteria>Terrabacteria_group>Actinomycetota>Actinomycetes>Propionibacteriales | GCF_900182665.1 | Propioniciclava_tarda | FN821_RS04755 | 6296..9736 | 433330 | Swt1-family-HEPN-domain-containing-protein-[Propioniciclava-tarda] |
| WP_119715651.1 | DUF499_1.RM.like.systems | TUDOR+ SF1-SF2-Helicase+iREase+CTD+REase->METHYLASE->HEPN->wHTH+REase->?->PemK_toxin->HEPN+DUF499*->?->?-><-?\|\|ArsR-HTH-><-ABhydrolase<-TetR-HTH | 1133 | Bacteria>Terrabacteria_group>Actinomycetota>Actinomycetes>Propionibacteriales | GCF_900289195.1 | Propionibacterium_ruminifibrarum | PROPJV5_RS07045 | 37471..40872 | 1962131 | Swt1-family-HEPN-domain-containing-protein-[Propionibacterium-ruminifibrarum] |
| WP_130874284.1 | DUF499_1.RM.like.systems | TetR-HTH-><-?\|\|?-><-ACET<-KAP-NTPase<-HEPN+DUF499*<-wHTH+REase<-METHYLASE<-TUDOR+ SF1-SF2-Helicase+iREase+CTD+REase<-MORC<-METHYLASE-p | 1129 | Bacteria>Terrabacteria_group>Actinomycetota>Actinomycetes>Propionibacteriales | GCF_900607225.1 | [Pseudopropionibacterium]_massiliense | EOE39_RS08095 | 1820154..1823543 | 2220000 | DUF499-domain-containing-protein-[[Pseudopropionibacterium] |
| WP_119162165.1 | DUF499_1.RM.like.systems | <-TetR-HTH\|\|?-><-SF1-SF2-Helicase<-OLD-ABC+TOPRIM<-HEPN+DUF499*<-PemK_toxin<-?<-wHTH+REase<-METHYLASE<-TUDOR+ SF1-SF2-Helicase+iREase+CTD+REase | 1133 | Bacteria>Terrabacteria_group>Actinomycetota>Actinomycetes>Propionibacteriales | GCF_900637915.1 | Propionibacterium_australiense | EL282_RS10960 | 2474046..2477447 | 119981 | DUF499-domain-containing-protein-[Propionibacterium-australiense] |
| WP_126378241.1 | DUF499_1.RM.like.systems | TetR-HTH-><-?\|\|?-><-ACET<-?<-HEPN+DUF499*<-wHTH+REase<-METHYLASE-p<-METHYLASE-p<-TUDOR+ SF1-SF2-Helicase+iREase+CTD+REase | 1129 | Bacteria>Terrabacteria_group>Actinomycetota>Actinomycetes>Propionibacteriales | GCF_900638645.1 | Arachnia_propionica | EL396_RS08525 | 1937379..1940768 | 1750 | DUF499-domain-containing-protein-[Arachnia-propionica] |
| WP_040158482.1 | DUF499_1.RM.like.systems | HTH+MPTase->?->?->HTH_Tnp_1+HTH_21+rve+rve_2-><-HEPN+DUF499*<-wHTH+REase<-METHYLASE<-TUDOR+ SF1-SF2-Helicase+iREase+CTD+REase<-XIS-HTH<-tRNA<-?<-ACET | 1132 | Bacteria>Terrabacteria_group>Actinomycetota>Actinomycetes>Propionibacteriales | GCF_902375435.1 | Nigerium_massiliense | BN1187_RS02660 | 259497..262895 | 1522317 | DUF499-domain-containing-protein-[Nigerium-massiliense] |
| WP_278693137.1 | DUF499_1.RM.like.systems | REase->TUDOR+ SF1-SF2-Helicase+iREase+CTD+REase->METHYLASE->wHTH+REase->HEPN+DUF499*-> | 1129 | Bacteria>Terrabacteria_group>Actinomycetota>Actinomycetes>Propionibacteriales | GCF_915063725.1 | Arachnia_propionica | QDI34_RS08815 | 15722..19111 | 1750 | DUF499-domain-containing-protein-[Arachnia-propionica] |
| WP_277066086.1 | DUF499_1.RM.like.systems | Synaptojanin-><-SF1-SF2-Helicase+REase<-HEPN+DUF499*<-wHTH+REase<-METHYLASE<-TUDOR+ SF1-SF2-Helicase+iREase+CTD+REase\|\|?-><-?\|\|Peptidase-> | 1128 | Bacteria>Terrabacteria_group>Actinomycetota>Actinomycetes>Propionibacteriales | GCF_938034675.1 | Propionibacterium_acidifaciens | P7A72_RS09150 | 66970..70356 | 556499 | Swt1-family-HEPN-domain-containing-protein-[Propionibacterium-acidifaciens] |
| WP_203043111.1 | DUF499_1.RM.like.systems | <-XIS-HTH<-XIS-HTH<-TrwC+SF1-SF2-helicase<-HEPN+DUF499*<-ParB+HNH<-METHYLASE<-TUDOR+ SF1-SF2-Helicase+iREase+CTD+REase\|\|NTD+SF1-SF2-Helicase-> | 1119 | Bacteria>Terrabacteria_group>Actinomycetota>Actinomycetes>Propionibacteriales | - | Pimelobacter_simplex | IMY96_RS01775 | 51221..54580 | 2045 | Swt1-family-HEPN-domain-containing-protein-[Pimelobacter-simplex] |
| PSL51415.1 | DUF499_1.RM.like.systems | HTH+DUF5753->ZnR->?->TPR+TPR+TPR-><-?<-HEPN+DUF499*<-METHYLASE<-TUDOR+ SF1-SF2-Helicase+iREase+CTD+REase\|\|XIS-HTH->Phage_int_SAM_1+Phage_integrase-> | 1099 | Bacteria>Terrabacteria_group>Actinomycetota>Actinomycetes>Pseudonocardiales | GCA_003014735.1 | Saccharothrix_carnea | BOI31_12144 | 38628..41927 | 1280637 | putative-AAA+-superfamily-ATPase-[Saccharothrix-carnea] |
| RAS61288.1 | DUF499_1.RM.like.systems | TUDOR+ SF1-SF2-Helicase+iREase+CTD+REase->METHYLASE-p->METHYLASE-p->HEPN+DUF499*-><-?<-HTH+DUF5753\|\|?->?->?->HTH-> | 1111 | Bacteria>Terrabacteria_group>Actinomycetota>Actinomycetes>Pseudonocardiales | GCA_003269295.1 | Lentzea_atacamensis | C8D87_110236 | 241893..245228 | 531938 | putative-AAA+-superfamily-ATPase-[Lentzea-atacamensis] |
| MBV9142989.1 | DUF499_1.RM.like.systems | <-HEPN+DUF499*<-METHYLASE-p<-TUDOR+ SF1-SF2-Helicase+iREase+CTD+REase<-Schlafen+HATPase+HTH<-Methyltransf\|\|METHYLASE-p-><-PIN | 1073 | Bacteria>Terrabacteria_group>Actinomycetota>Actinomycetes>Pseudonocardiales | GCA_019247305.1 | Pseudonocardiales_bacterium | JO115_19100 | 30..3251 | 2201155 | DUF499-domain-containing-protein-[Pseudonocardiales-bacterium] |
| WP_211261518.1 | DUF499_1.RM.like.systems | <-ArsR-HTH<-?<-?\|\|?->TUDOR+ SF1-SF2-Helicase+iREase+CTD+REase->METHYLASE->HEPN+DUF499*->?-><-?<-HTH+DUF5753<-TetR-HTH | 1100 | Bacteria>Terrabacteria_group>Actinomycetota>Actinomycetes>Pseudonocardiales | GCF_000620785.1 | Pseudonocardia_acaciae | N912_RS0136125 | 176490..179792 | 551276 | DUF499-domain-containing-protein-[Pseudonocardia-acaciae] |
| WP_106619913.1 | DUF499_1.RM.like.systems | HTH+DUF5753->ZnR->?->TPR+TPR+TPR-><-?<-HEPN+DUF499*<-METHYLASE<-TUDOR+ SF1-SF2-Helicase+iREase+CTD+REase\|\|XIS-HTH->Phage_int_SAM_1+Phage_integrase-> | 1114 | Bacteria>Terrabacteria_group>Actinomycetota>Actinomycetes>Pseudonocardiales | GCF_003014735.1 | Saccharothrix_carnea | BOI31_RS36295 | 38628..41972 | 1280637 | DUF499-domain-containing-protein-[Saccharothrix-carnea] |

| Protein-ID | System | Gene-Neighbourhoods | Length | Taxonomy | Assembly-ID | Strain | Locus-Tag | Genomic-coordinates | TaxID | Fasta-defline |
|---|---|---|---|---|---|---|---|---|---|---|
| WP_215732591.1 | DUF499_1.RM.like.systems | TUDOR+SF1-SF2-Helicase+iREase+CTD+REase->METHYLASE-p->METHYLASE-p->HEPN+DUF499*->?-><-?<-HTH+DUF5753 | 1100 | Bacteria>Terrabacteria_group>Actinomycetota>Actinomycetes>Pseudonocardiales | GCF_003269295.1 | Lentzea_atacamensis | C8D87_RS36395 | 241926..245228 | 531938 | DUF499-domain-containing-protein-[Lentzea-atacamensis] |
| WP_085916424.1 | DUF499_1.RM.like.systems | DNA-glycosylase->TUDOR+SF1-SF2-Helicase+iREase+CTD+REase->METHYLASE->ParB+HNH->HEPN+DUF499*->?-><-TrwC+SF1-SF2-helicase+TOPRIM | 1135 | Bacteria>Terrabacteria_group>Actinomycetota>Actinomycetes>Pseudonocardiales | GCF_003945385.1 | Pseudonocardia | PSA01_RS26720 | 8547..11954 | 1847 | MULTISPECIES:-Swt1-family-HEPN-domain-containing-protein-[Pseudonocardia] |
| WP_139094500.1 | DUF499_1.RM.like.systems | <-HIN-HTH+HIN-HTH\|\|?->?->?->?-><-HEPN+DUF499*<-METHYLASE<-TUDOR+SF1-SF2-Helicase+iREase+CTD+REase<-?<-HIN-HTH+HIN-HTH\|\|ABC-ATPase+SbcC->IclR-HTH-> | 1111 | Bacteria>Terrabacteria_group>Actinomycetota>Actinomycetes>Pseudonocardiales | GCF_006152065.1 | Amycolatopsis_alkalitolerans | FG385_RS00135 | 35621..38956 | 2547244 | DUF499-domain-containing-protein-[Amycolatopsis-alkalitolerans] |
| WP_184862258.1 | DUF499_1.RM.like.systems | HIN-HTH+HIN-HTH->?->TUDOR+SF1-SF2-Helicase+iREase+CTD+REase->UDG-conflict->METHYLASE->HEPN+DUF499*-><-?<-?<-PAIREDC-HTH<-?<-HTH<-SIGMA-HTH | 1111 | Bacteria>Terrabacteria_group>Actinomycetota>Actinomycetes>Pseudonocardiales | GCF_014203355.1 | Kutzneria_kofuensis | BJ998_RS15140 | 3332051..3335386 | 103725 | DUF499-domain-containing-protein-[Kutzneria-kofuensis] |
| WP_218590226.1 | DUF499_1.RM.like.systems | TUDOR+SF1-SF2-Helicase+iREase+CTD+REase->METHYLASE->ParB+HNH->HEPN+DUF499*->?-><-TrwC+SF1-SF2-helicase+TOPRIM<-?<-VirD4-TraG+VirD4-FtsK | 1126 | Bacteria>Terrabacteria_group>Actinomycetota>Actinomycetes>Pseudonocardiales | GCF_019263625.1 | Pseudonocardia_oceani | I4I79_RS18830 | 8746..12126 | 2792013 | Swt1-family-HEPN-domain-containing-protein-[Pseudonocardia-oceani] |
| WP_061989441.1 | DUF499_1.RM.like.systems | HIN-HTH+HIN-HTH->?->TUDOR+SF1-SF2-Helicase+iREase+CTD+REase->UDG-conflict->METHYLASE->HEPN+DUF499*-><-?\|\|Phage_integrase->?->?-><-tRNA<-?<-ACET | 1112 | Bacteria>Terrabacteria_group>Actinomycetota>Actinomycetes>Pseudonocardiales | GCF_900113515.1 | Amycolatopsis_regifaucium | AVL48_RS05675 | 397234..400572 | 546365 | DUF499-domain-containing-protein-[Amycolatopsis-regifaucium] |
| WP_146174093.1 | DUF499_1.RM.like.systems | HTH+DUF5753->ZnR->?->TPR+TPR+TPR-><-?<-HEPN+DUF499*<-METHYLASE<-TUDOR+SF1-SF2-Helicase+iREase+CTD+REase\|\|XIS-HTH->Phage_int_SAM_1+Phage_integrase-> | 1099 | Bacteria>Terrabacteria_group>Actinomycetota>Actinomycetes>Pseudonocardiales | - | Saccharothrix_carnea | B0I31_12144 | 38628..41927 | 1280637 | DUF499-domain-containing-protein-[Saccharothrix-carnea] |
| WP_019873098.1 | DUF499_1.RM.like.systems | <-ArdCN+MPTase<-?<-HEPN+DUF499*<-METHYLASE<-TUDOR+SF1-SF2-Helicase+iREase+CTD+REase-p<-DnaB+DnaB<-VirB4-FtsK<-?<-LysM+TPR | 1113 | Bacteria>Terrabacteria_group>Actinomycetota>Actinomycetes>Sporichthyales | GCF_000384115.1 | Sporichthya_polymorpha | SPOPO_RS0101950 | 384127..387468 | 35751 | DUF499-domain-containing-protein-[Sporichthya-polymorpha] |
| GIH50617.1 | DUF499_1.RM.like.systems | <-Primpol+D5N+D5ATPase<-DUF4011+SF1-SF2-Helicase+REase<-SF1-SF2-helicase+DUF1998<-?<-?<-HEPN+DUF499*<-METHYLASE<-TUDOR+SF1-SF2-Helicase+iREase+CTD+REase<-?<-REase+N6_MTase<-Pkinase | 1125 | Bacteria>Terrabacteria_group>Actinomycetota>Actinomycetes>Streptosporangiales | GCA_016863055.1 | Microbispora_rosea_subsp._rosea | Mro03_57960 | 50675..54052 | 58118 | hypothetical-protein-Mro03_57960-[Microbispora-rosea-subsp.-rosea] |
| GII94898.1 | DUF499_1.RM.like.systems | <-HEPN<-ParB+HNH<-HEPN+DUF499*<-METHYLASE<-TUDOR+SF1-SF2-Helicase+iREase+CTD+REase | 1137 | Bacteria>Terrabacteria_group>Actinomycetota>Actinomycetes>Streptosporangiales | GCA_016863435.1 | Sinosporangium_siamense | Ssi02_51290 | 167948..171361 | 1367973 | hypothetical-protein-Ssi02_51290-[Sinosporangium-siamense] |
| GHJ54460.1 | DUF499_1.RM.like.systems | <-REase<-?<-NTD+SF1-SF2-Helicase\|\|TUDOR+SF1-SF2-Helicase+iREase+CTD+REase->METHYLASE->ParB+HNH->HEPN+DUF499*->?->?->?-><-?<-SbcC\|\|HIN-HTH-> | 1122 | Bacteria>Terrabacteria_group>Actinomycetota>Actinomycetes>Streptosporangiales | GCA_018327725.1 | Nonomuraea_sp._TT08I-71 | Nm8I071_37670 | 3194026..3197394 | 2733864 | hypothetical-protein-Nm8I071_37670-[Nonomuraea-sp.-TT08I-71] |
| GHJ56382.1 | DUF499_1.RM.like.systems | <-HEPN+DUF499*<-METHYLASE<-TUDOR+SF1-SF2-Helicase+iREase+CTD+REase<-?\|\|SAD+HNH-><-Peptidase | 1124 | Bacteria>Terrabacteria_group>Actinomycetota>Actinomycetes>Streptosporangiales | GCA_018327725.1 | Nonomuraea_sp._TT08I-71 | Nm8I071_56890 | 5264667..5268041 | 2733864 | hypothetical-protein-Nm8I071_56890-[Nonomuraea-sp.-TT08I-71] |
| MCF6470725.1 | DUF499_1.RM.like.systems | GNTR-HTH-p->NUDIX->TUDOR+SF1-SF2-Helicase+iREase+CTD+REase->METHYLASE->HEPN+DUF499*->ParB+HNH->?->REase-><-?\|\|SIGMA-HTH-> | 1139 | Bacteria>Terrabacteria_group>Actinomycetota>Actinomycetes>Streptosporangiales | GCA_021699195.1 | Nonomuraea_sp._MG754425 | FAF44_20335 | 17638..21057 | 2570319 | ATP-binding-protein-[Nonomuraea-sp.-MG754425] |
| WP_017595710.1 | DUF499_1.RM.like.systems | <-JetB<-JetA<-HEPN+DUF499*<-METHYLASE<-TUDOR+SF1-SF2-Helicase+iREase+CTD+REase<-PIN<-XIS-HTH<-tRNA<-SIGMA-HTH | 1136 | Bacteria>Terrabacteria_group>Actinomycetota>Actinomycetes>Streptosporangiales | GCF_000341105.1 | Nocardiopsis_potens | D459_RS0124835 | 11224..14634 | 1246458 | Swt1-family-HEPN-domain-containing-protein-[Nocardiopsis-potens] |
| WP_030913330.1 | DUF499_1.RM.like.systems | <-REase+SF1-SF2-Helicase<-MazG\|\|UDG-conflict->?->TUDOR+SF1-SF2-Helicase+iREase+CTD+REase->METHYLASE->HEPN+DUF499*->?->?->?-><-DUF397<-HTH+DUF5753\|\|ST-Hiskin-> | 1133 | Bacteria>Terrabacteria_group>Actinomycetota>Actinomycetes>Streptosporangiales | GCF_000721075.1 | Streptosporangium_amethystogenes | IG98_RS0118225 | 190587..193988 | 2002 | Swt1-family-HEPN-domain-containing-protein-[Streptosporangium-amethystogenes] |
| WP_061082585.1 | DUF499_1.RM.like.systems | REase->?->McrC-NTD+REase->TUDOR+SF1-SF2-Helicase+iREase+CTD+REase->METHYLASE->HEPN+DUF499*-><-?<-ST-Hiskin\|\|HTH+DUF5753-> | 1135 | Bacteria>Terrabacteria_group>Actinomycetota>Actinomycetes>Streptosporangiales | GCF_001552695.1 | Nocardiopsis_alborubida | ND1_RS24360 | 24019..27426 | 146802 | Swt1-family-HEPN-domain-containing-protein-[Nocardiopsis-alborubida] |
| WP_131964568.1 | DUF499_1.RM.like.systems | <-Schlafen+HATPase+HTH<-Schlafen+HATPase+HTH<-ParB+HNH\|\|TUDOR+SF1-SF2-Helicase+iREase+CTD+REase->METHYLASE->HEPN+DUF499*->Phage_int_SAM_1+Phage_integrase->Phage_integrase-><-NUDIX<-SAM-methylase | 1135 | Bacteria>Terrabacteria_group>Actinomycetota>Actinomycetes>Streptosporangiales | GCF_004348735.1 | Actinomadura_sp._KC06 | E1287_RS09895 | 20684..24091 | 2530369 | DUF499-domain-containing-protein-[Actinomadura-sp.-KC06] |
| WP_132634803.1 | DUF499_1.RM.like.systems | <-REase+N6_MTase<-TRD+TRD<-REase+SF1-SF2-Helicase<-REase+SF1-SF2-Helicase\|\|TUDOR+SF1-SF2-Helicase+iREase+CTD+REase->METHYLASE->HEPN+DUF499*-><-DndB | 1126 | Bacteria>Terrabacteria_group>Actinomycetota>Actinomycetes>Streptosporangiales | GCF_004352805.1 | Nonomuraea_mesophila | E1295_RS29355 | 30406..33786 | 2530382 | Swt1-family-HEPN-domain-containing-protein-[Nonomuraea-mesophila] |
| WP_155999755.1 | DUF499_1.RM.like.systems | <-HTH+DUF5753\|\|?-><-?<-?<-METHYLASE<-?\|\|HEPN+DUF499*->ParB+HNH-><-?<-Glycos_trans_3N+Glycos_transf_3 | 1141 | Bacteria>Terrabacteria_group>Actinomycetota>Actinomycetes>Streptosporangiales | GCF_009728995.1 | Streptomonospora_sp._PA3 | FZ103_RS07070 | 271290..274715 | 2607326 | Swt1-family-HEPN-domain-containing-protein-[Streptomonospora-sp.-PA3] |
| WP_179824532.1 | DUF499_1.RM.like.systems | <-ST-Hiskin\|\|HTH+DUF5753->DUF397-><-?<-ParB+HNH<-HEPN+DUF499*<-METHYLASE<-TUDOR+SF1-SF2-Helicase+iREase+CTD+REase<-?<-UDG-conflict | 1132 | Bacteria>Terrabacteria_group>Actinomycetota>Actinomycetes>Streptosporangiales | GCF_013409935.1 | Streptosporangium_sandarakinum | HDA43_RS21565 | 651273..654671 | 1260955 | Swt1-family-HEPN-domain-containing-protein-[Streptosporangium-sandarakinum] |
| WP_185051091.1 | DUF499_1.RM.like.systems | <-HDRnase<-GNTR-HTH<-?\|\|HTH+HKD-DNase->TUDOR+SF1-SF2-Helicase+iREase+CTD+REase->METHYLASE->HEPN+DUF499*->tRNA-><-?\|\|MODE-HTH+PBPII->ABhydrolase-> | 1126 | Bacteria>Terrabacteria_group>Actinomycetota>Actinomycetes>Streptosporangiales | GCF_014202315.1 | Thermocatellispora_tengchongensis | HNP84_RS19220 | 438143..441523 | 1073253 | DUF499-domain-containing-protein-[Thermocatellispora-tengchongensis] |
| WP_184983644.1 | DUF499_1.RM.like.systems | 35exo-><-wHTH(SBD)+HNH<-?<-SF1-SF2-helicase+DUF1998<-?<-ParB+HNH<-HEPN+DUF499*<-METHYLASE<-TUDOR+SF1-SF2-Helicase+iREase+CTD+REase<-?<-REase+SF1-SF2-Helicase<-HTH_3<-ABhydrolase | 1124 | Bacteria>Terrabacteria_group>Actinomycetota>Actinomycetes>Streptosporangiales | GCF_014207705.1 | Sphaerisporangium_rubeum | BJ992_RS21150 | 4926161..4929535 | 321317 | Swt1-family-HEPN-domain-containing-protein-[Sphaerisporangium-rubeum] |
| WP_189237444.1 | DUF499_1.RM.like.systems | METHYLASE-p->HEPN+DUF499*-><-DUF397<-HTH+DUF5753\|\|ST-Hiskin->SIGMA-HTH->ParB+HNH-> | 1133 | Bacteria>Terrabacteria_group>Actinomycetota>Actinomycetes>Streptosporangiales | GCF_014647955.1 | Planomonospora_parontospora | Ppa05_RS15805 | 840..4241 | 58119 | Swt1-family-HEPN-domain-containing-protein-[Planomonospora-parontospora] |
| WP_191055573.1 | DUF499_1.RM.like.systems | HTH+HKD-DNase->?->?->TUDOR+SF1-SF2-Helicase+iREase+CTD+REase->METHYLASE->HEPN+DUF499*->ParB+HNH->HEPN-> | 1138 | Bacteria>Terrabacteria_group>Actinomycetota>Actinomycetes>Streptosporangiales | GCF_014712745.1 | Microbispora_camponoti | IEQ31_RS35675 | 16717..20133 | 1677852 | DUF499-domain-containing-protein-[Microbispora-camponoti] |
| WP_203980270.1 | DUF499_1.RM.like.systems | <-SF1-SF2-helicase+DUF1998<-?<-ParB+HNH<-?<-HEPN+DUF499*<-METHYLASE<-TUDOR+SF1-SF2-Helicase+iREase+CTD+REase<-?<-REase+N6_MTase | 1129 | Bacteria>Terrabacteria_group>Actinomycetota>Actinomycetes>Streptosporangiales | GCF_016863315.1 | Planotetraspora_silvatica | Psi02_RS35595 | 24804..28193 | 234614 | Swt1-family-HEPN-domain-containing-protein-[Planotetraspora-silvatica] |
| WP_203946296.1 | DUF499_1.RM.like.systems | REase+N6_MTase->?->TUDOR+SF1-SF2-Helicase+iREase+CTD+REase->METHYLASE->HEPN+DUF499*->?->ParB+HNH->?->SF1-SF2-helicase+DUF1998-><-?\|\|OLD-ABC+TOPRIM-> | 1129 | Bacteria>Terrabacteria_group>Actinomycetota>Actinomycetes>Streptosporangiales | GCF_016863335.1 | Planotetraspora_thailandica | Pth03_RS22295 | 43771..47160 | 487172 | Swt1-family-HEPN-domain-containing-protein-[Planotetraspora-thailandica] |
| WP_204029818.1 | DUF499_1.RM.like.systems | <-GNTR-HTH-p<-?<-?<-HEPN<-ParB+HNH<-HEPN+DUF499*<-METHYLASE<-TUDOR+SF1-SF2-Helicase+iREase+CTD+REase | 1127 | Bacteria>Terrabacteria_group>Actinomycetota>Actinomycetes>Streptosporangiales | GCF_016863435.1 | Sinosporangium_siamense | Ssi02_RS24940 | 167948..171331 | 1367973 | DUF499-domain-containing-protein-[Sinosporangium-siamense] |
| WP_237104850.1 | DUF499_1.RM.like.systems | GNTR-HTH-p->NUDIX->TUDOR+SF1-SF2-Helicase+iREase+CTD+REase->METHYLASE->HEPN+DUF499*->ParB+HNH->?->REase-><-?<-?\|\|SIGMA-HTH-> | 1137 | Bacteria>Terrabacteria_group>Actinomycetota>Actinomycetes>Streptosporangiales | GCF_021699195.1 | Nonomuraea_sp._MG754425 | FAF44_RS20630 | 17644..21057 | 2570319 | DUF499-domain-containing-protein-[Nonomuraea-sp.-MG754425] |
| WP_076440368.1 | DUF499_1.RM.like.systems | <-Primpol+D5N+D5ATPase<-DUF4011+SF1-SF2-Helicase+REase<-SF1-SF2-helicase+DUF1998<-?<-?<-HEPN+DUF499*<-METHYLASE<-TUDOR+SF1-SF2-Helicase+iREase+CTD+REase<-?<-REase+N6_MTase<-Pkinase | 1125 | Bacteria>Terrabacteria_group>Actinomycetota>Actinomycetes>Streptosporangiales | GCF_900156315.1 | Microbispora_rosea | BXA11_RS31445 | 30499..33876 | 58117 | Swt1-family-HEPN-domain-containing-protein-[Microbispora-rosea] |
| EAR26285.1 | DUF499_1.RM.like.systems | REase-><-REase<-DNA_methylase<-MORC+SIGMA-HTH\|\|METHYLASE->HEPN+DUF499*->MORC->REase->HTH+SF1-SF2-Helicase+REase->?-><-?<-ABC-ATPase+SbcC | 1127 | Bacteria>Terrabacteria_group>Actinomycetota>Actinomycetes>unclassified_Actinomycetes | GCA_000153145.1 | marine_actinobacterium_PHSC20C1 | A20C1_10399 | 2131040..2134423 | 312284 | hypothetical-protein-A20C1_10399-[marine-actinobacterium-PHSC20C1] |
| KLR61150.1 | DUF499_1.RM.like.systems | <-ParB+Schlafen\|\|TUDOR+SF1-SF2-Helicase+iREase+CTD+REase->METHYLASE-p->METHYLASE-p->HEPN+DUF499*->tRNA-><-?\|\|RelE-ParE->HTH-> | 1095 | Bacteria>Terrabacteria_group>Actinomycetota>Actinomycetes>unclassified_Actinomycetes | GCA_001025035.1 | Actinobacteria_bacterium_IMCC26207 | IMCC26207_109668 | 707438..710725 | 1641811 | putative-ATPase-(AAA+-superfamily)-[Actinobacteria-bacterium-IMCC26207] |
| MCL2317098.1 | DUF499_1.RM.like.systems | <-HEPN+DUF499*<-PIN<-PhdYeFM_antitox<-METHYLASE-p | 623 | Bacteria>Terrabacteria_group>Actinomycetota>Actinomycetes>unclassified_Actinomycetes | GCA_009785885.1 | Actinomycetes_bacterium | FWC46_08465 | 1..1871 | 1883427 | Swt1-family-HEPN-domain-containing-protein-[Actinomycetia-bacterium] |
| MCH9729512.1 | DUF499_1.RM.like.systems | PIN->TUDOR+SF1-SF2-Helicase+iREase+CTD+REase->METHYLASE->ParB+HNH->HEPN+DUF499*->SIR2-><-DndB\|\|?->Recombinase-><-NUDIX\|\|GNTR-HTH-> | 1114 | Bacteria>Terrabacteria_group>Actinomycetota>Actinomycetes>unclassified_Actinomycetes | GCA_022600605.1 | Actinomycetes_bacterium | K0U84_07525 | 8492..11836 | 1883427 | DUF499-domain-containing-protein-[Actinomycetia-bacterium] |
| PKQ27862.1 | DUF499_1.RM.like.systems | REase->TUDOR+SF1-SF2-Helicase+iREase+CTD+REase->Schlafen+HATPase+HTH->METHYLASE->HEPN+DUF499*->?->TIR-> | 1105 | Bacteria>Terrabacteria_group>Actinomycetota>Candidatus_Geothermincolia>Candidatus_Geothermincolales | GCA_002843005.1 | Candidatus_Anoxymicrobium_japonicum | CVT63_05765 | 8955..12272 | 2013648 | AAA+-family-ATPase-[Candidatus-Anoxymicrobium-japonicum] |
| MCL4078096.1 | DUF499_1.RM.like.systems | <-Peptidase_C39_2<-?<-?\|\|PspA+PIN->?-><-HEPN+DUF499*<-?<-MqsA_antitoxin+HTH_3<-?<-METHYLASE<-ParB+HNH<-TUDOR+SF1-SF2-Helicase+iREase+CTD+REase | 1106 | Bacteria>Terrabacteria_group>Actinomycetota>Coriobacteriia>Anaerosomatales | GCA_023371205.1 | Parvivirga_hydrogeniphila | MX659_00500 | 98516..101836 | 2939460 | Swt1-family-HEPN-domain-containing-protein-[Coriobacteriia-bacterium-Es71-Z0120] |
| WP_267192478.1 | DUF499_1.RM.like.systems | <-Peptidase_C39_2<-?<-?\|\|PspA+PIN->?-><-HEPN+DUF499*<-?<-MqsA_antitoxin+HTH_3<-?<-METHYLASE<-ParB+HNH<-TUDOR+SF1-SF2-Helicase+iREase+CTD+REase | 1106 | Bacteria>Terrabacteria_group>Actinomycetota>Coriobacteriia>Anaerosomatales | - | Parvivirga_hydrogeniphila | MX659_RS00500 | 98516..101836 | 2939460 | Swt1-family-HEPN-domain-containing-protein-[Parvivirga-hydrogeniphila] |
| TGY60542.1 | DUF499_1.RM.like.systems | MazG->URI-DNase+SF1-SF2-Helicase->HEPN->HEPN->TUDOR+SF1-SF2-Helicase+iREase+CTD+REase->METHYLASE->HEPN+DUF499*->ORC-CDC6+REase->?-><-?\|\|?->?->REase-> | 1126 | Bacteria>Terrabacteria_group>Actinomycetota>Coriobacteriia>Coriobacteriales | GCA_004793665.1 | Coriobacteriaceae_bacterium | E5335_00400 | 90399..93779 | 2011094 | ATP-binding-protein-[Coriobacteriaceae-bacterium] |
| MBQ6523654.1 | DUF499_1.RM.like.systems | <-HEPN+DUF499*<-METHYLASE-p | 1129 | Bacteria>Terrabacteria_group>Actinomycetota>Coriobacteriia>Coriobacteriales | GCA_017433555.1 | Atopobiaceae_bacterium | IJJ12_03205 | 1663..5052 | 2485926 | DUF499-domain-containing-protein-[Atopobiaceae-bacterium] |
| MBR3315601.1 | DUF499_1.RM.like.systems | ASCH-><-PIN+ACET+EVE<-HEPN+DUF499*<-METHYLASE<-TUDOR+SF1-SF2-Helicase+iREase+CTD+REase | 1137 | Bacteria>Terrabacteria_group>Actinomycetota>Coriobacteriia>Coriobacteriales | GCA_017510745.1 | Atopobiaceae_bacterium | IKG18_15850 | 6540..9953 | 2485926 | DUF499-domain-containing-protein-[Atopobiaceae-bacterium] |
| MBS6858281.1 | DUF499_1.RM.like.systems | MNS+HTH+REase->35exo->TUDOR+SF1-SF2-Helicase+iREase+CTD+REase->METHYLASE->HEPN+DUF499*-><-?<-?<-?\|\|?->Peptidase_M20-> | 1137 | Bacteria>Terrabacteria_group>Actinomycetota>Coriobacteriia>Coriobacteriales | GCA_018380015.1 | Collinsella_sp. | KH187_03945 | 112102..115515 | 1965294 | DUF499-domain-containing-protein-[Collinsella-sp.] |
| MCI1646461.1 | DUF499_1.RM.like.systems | Phage-portal-><-?<-REase<-?\|\|TUDOR+SF1-SF2-Helicase+iREase+CTD+REase->METHYLASE->HEPN+DUF499*-> | 1135 | Bacteria>Terrabacteria_group>Actinomycetota>Coriobacteriia>Coriobacteriales | GCA_022648325.1 | Olsenella_sp. | LKI40_09700 | 33450..36857 | 2042683 | Swt1-family-HEPN-domain-containing-protein-[Olsenella-sp.] |
| GJM55560.1 | DUF499_1.RM.like.systems | <-REase<-?<-?\|\|?-><-?<-ORC-CDC6+REase<-HEPN+DUF499*<-METHYLASE<-TUDOR+SF1-SF2-Helicase+iREase+CTD+REase<-HEPN<-URI-DNase+SF1-SF2-Helicase<-MazG | 1126 | Bacteria>Terrabacteria_group>Actinomycetota>Coriobacteriia>Coriobacteriales | GCA_022834715.1 | Granulimonas_faecalis | ATOP_12150 | 1358904..1362284 | 2894155 | hypothetical-protein-ATOP_12150-[Granulimonas-faecalis] |
| WP_059053651.1 | DUF499_1.RM.like.systems | <-HEPN+DUF499*<-METHYLASE<-TUDOR+SF1-SF2-Helicase+iREase+CTD+REase\|\|REase->?-><-Phage-portal | 1133 | Bacteria>Terrabacteria_group>Actinomycetota>Coriobacteriia>Coriobacteriales | GCF_001494635.1 | Tractidigestivibacter_scatoligenes | AUL39_RS03515 | 793561..796962 | 1299998 | DUF499-domain-containing-protein-[Tractidigestivibacter-scatoligenes] |
| WP_117794170.1 | DUF499_1.RM.like.systems | <-MNS+REase\|\|?-><-?<-HEPN+DUF499*<-METHYLASE<-TUDOR+SF1-SF2-Helicase+iREase+CTD+REase\|\|?-><-HEPN<-REase-p<-REase-p | 1137 | Bacteria>Terrabacteria_group>Actinomycetota>Coriobacteriia>Coriobacteriales | GCF_003466445.1 | Collinsella_sp._TM05-38 | DXD49_RS03005 | 54584..57997 | 2292341 | DUF499-domain-containing-protein-[Collinsella-sp.-TM05-38] |
| WP_218957899.1 | DUF499_1.RM.like.systems | <-Peptidase<-?<-?<-?<-?<-DUF499-p<-HEPN+DUF499*<-METHYLASE<-TUDOR+SF1-SF2-Helicase+iREase+CTD+REase\|\|OLD-ABC+TOPRIM->SF1-SF2-Helicase-> | 864 | Bacteria>Terrabacteria_group>Actinomycetota>Coriobacteriia>Coriobacteriales | GCF_007655325.1 | Enorma_shizhengliae | GJE22_RS07705 | 20738..23332 | 2606615 | DUF499-domain-containing-protein-[Enorma-shizhengliae] |
| WP_154324032.1 | DUF499_1.RM.like.systems | Classical-AAA->Peptidase_S8-><-HEPN+DUF499*<-METHYLASE<-TUDOR+SF1-SF2-Helicase+iREase+CTD+REase<-?<-Phage-portal | 1146 | Bacteria>Terrabacteria_group>Actinomycetota>Coriobacteriia>Coriobacteriales | GCF_009676995.1 | Enorma_shizhengliae | GJE22_RS00805 | 201758..205198 | 2606615 | DUF499-domain-containing-protein-[Enorma-shizhengliae] |
| WP_195644120.1 | DUF499_1.RM.like.systems | <-MNS+REase\|\|?-><-?<-HEPN+DUF499*<-METHYLASE<-TUDOR+SF1-SF2-Helicase+iREase+CTD+REase<-?<-35exo<-MNS+HTH+REase | 1137 | Bacteria>Terrabacteria_group>Actinomycetota>Coriobacteriia>Coriobacteriales | GCF_015561195.1 | Collinsella_aerofaciens | PMX13_RS01910 | 103698..107111 | 74426 | DUF499-domain-containing-protein-[Collinsella-aerofaciens] |
| WP_204556581.1 | DUF499_1.RM.like.systems | HTH_3->?->?-><-?\|\|TUDOR+SF1-SF2-Helicase+iREase+CTD+REase->METHYLASE->HEPN+DUF499*->?->XIS-HTH-><-SF1-SF2-Helicase\|\|?->?->MNS+HTH-> | 1118 | Bacteria>Terrabacteria_group>Actinomycetota>Coriobacteriia>Coriobacteriales | GCF_016901585.1 | Collinsella_intestinalis | H7U33_RS06515 | 78334..81690 | 147207 | DUF499-domain-containing-protein-[Collinsella-intestinalis] |
| WP_265590834.1 | DUF499_1.RM.like.systems | <-REase<-?<-?\|\|?-><-?<-ORC-CDC6+REase<-HEPN+DUF499*<-METHYLASE<-TUDOR+SF1-SF2-Helicase+iREase+CTD+REase<-HEPN<-URI-DNase+SF1-SF2-Helicase<-MazG | 1126 | Bacteria>Terrabacteria_group>Actinomycetota>Coriobacteriia>Coriobacteriales | GCF_022834715.1 | Granulimonas_faecalis | OR600_RS06205 | 1358904..1362284 | 2894155 | DUF499-domain-containing-protein-[Granulimonas-faecalis] |
| WP_270472295.1 | DUF499_1.RM.like.systems | HISKIN-><-?<-ORC-CDC6+REase\|\|TUDOR+SF1-SF2-Helicase+iREase+CTD+REase->METHYLASE->HEPN+DUF499*->?->?->?->?->ABC-ATPase+SbcC-> | 1129 | Bacteria>Terrabacteria_group>Actinomycetota>Coriobacteriia>Coriobacteriales | GCF_027683425.1 | Collinsella_intestinalis | PGX73_RS07325 | 61994..65383 | 147207 | DUF499-domain-containing-protein-[Collinsella-intestinalis] |
| WP_072371784.1 | DUF499_1.RM.like.systems | <-ABC-ATPase+SbcC<-MarR-HTH<-?\|\|ORC-CDC6+REase-><-ORC-CDC6+REase<-?<-HEPN+DUF499*<-METHYLASE<-TUDOR+SF1-SF2-Helicase+iREase+CTD+REase\|\|HTH_3->HAD-> | 1133 | Bacteria>Terrabacteria_group>Actinomycetota>Coriobacteriia>Coriobacteriales | GCF_900119385.1 | Thermophilibacter_mediterraneus | BQ7106_RS04355 | 522048..525449 | 1871031 | DUF499-domain-containing-protein-[Thermophilibacter-mediterraneus] |
| WP_251158236.1 | DUF499_1.RM.like.systems | <-ORC-CDC6+REase<-HEPN+DUF499*<-METHYLASE<-TUDOR+SF1-SF2-Helicase+iREase+CTD+REase<-HEPN<-URI-DNase+SF1-SF2-Helicase<-MazG | 1126 | Bacteria>Terrabacteria_group>Actinomycetota>Coriobacteriia>Coriobacteriales | GCF_943193035.1 | Caniella_muris | NFR49_RS03270 | 716481..719861 | 2941502 | DUF499-domain-containing-protein-[Caniella-sp.-DSM-110323] |
| WP_273395403.1 | DUF499_1.RM.like.systems | <-ABC-ATPase+SbcC<-MarR-HTH<-?\|\|ORC-CDC6+REase-><-ORC-CDC6+REase<-?<-HEPN+DUF499*<-METHYLASE<-TUDOR+SF1-SF2-Helicase+iREase+CTD+REase\|\|HTH_3->HAD-> | 1133 | Bacteria>Terrabacteria_group>Actinomycetota>Coriobacteriia>Coriobacteriales | GCF_944322105.1 | Thermophilibacter_mediterraneus | PUQ24_RS02970 | 23710..27111 | 1871031 | DUF499-domain-containing-protein-[Thermophilibacter-mediterraneus] |
| MBQ9000666.1 | DUF499_1.RM.like.systems | TUDOR+SF1-SF2-Helicase+iREase+CTD+REase->METHYLASE->HEPN+DUF499*->ParB-> | 1133 | Bacteria>Terrabacteria_group>Actinomycetota>Coriobacteriia>Eggerthellales | GCA_017460365.1 | Eggerthellaceae_bacterium | IJ087_02285 | 5494..8895 | 1972561 | DUF499-domain-containing-protein-[Eggerthellaceae-bacterium] |
| MCD8317268.1 | DUF499_1.RM.like.systems | TUDOR+SF1-SF2-Helicase+iREase+CTD+REase->METHYLASE-p-><-Phage_int_SAM_1+Phage_integrase<-Phage_integrase<-Phage_integrase\|\|METHYLASE-p->HEPN+DUF499*-> | 1113 | Bacteria>Terrabacteria_group>Actinomycetota>Coriobacteriia>Eggerthellales | GCA_021204985.1 | Eggerthellaceae_bacterium | LUB61_07720 | 22349..25690 | 1972561 | DUF499-domain-containing-protein-[Eggerthellaceae-bacterium] |
| MDD5805721.1 | DUF499_1.RM.like.systems | METHYLASE->HEPN+DUF499*-> | 557 | Bacteria>Terrabacteria_group>Actinomycetota>Coriobacteriia>Eggerthellales | GCA_029101185.1 | Eggerthellales_bacterium | PUD02_00035 | 2228..3899 | 2137879 | MAG:-Swt1-family-HEPN-domain-containing-protein,-partial-[Eggerthellales-bacterium] |
| WP_206214419.1 | DUF499_1.RM.like.systems | <-wHTH+REase<-HTH_3<-Tox-RelE\|\|Classical-AAA->Peptidase_S8-><-HEPN+DUF499*<-METHYLASE<-TUDOR+SF1-SF2-Helicase+iREase+CTD+REase<-?\|\|?-><-?<-ABC-ATPase | 1145 | Bacteria>Terrabacteria_group>Actinomycetota>Coriobacteriia>Eggerthellales | GCF_011038965.1 | Adlercreutzia_sp._ZJ141 | G7Y44_RS06490 | 70626..74063 | 2709406 | DUF499-domain-containing-protein-[Adlercreutzia-sp.-ZJ141] |
| WP_165173327.1 | DUF499_1.RM.like.systems | TUDOR+SF1-SF2-Helicase+iREase+CTD+REase->METHYLASE->HEPN+DUF499*->?->UDG-> | 1132 | Bacteria>Terrabacteria_group>Actinomycetota>Coriobacteriia>Eggerthellales | GCF_011038995.1 | Adlercreutzia_sp._ZJ242 | G7Y49_RS09770 | 7632..11030 | 2709409 | DUF499-domain-containing-protein-[Adlercreutzia-sp.-ZJ242] |
| MBC7265381.1 | DUF499_1.RM.like.systems | <-Peptidase_C39_2<-?\|\|PspA+PIN->?-><-HEPN+DUF499*<-?<-MqsA_antitoxin+HTH_3<-?<-METHYLASE<-ParB+HNH<-TUDOR+SF1-SF2-Helicase+iREase+CTD+REase | 1106 | Bacteria>Terrabacteria_group>Actinomycetota>Coriobacteriia>unclassified_Coriobacteriia | GCA_014360605.1 | Coriobacteriia_bacterium | H5T75_00145 | 31010..34330 | 2052159 | ATP-binding-protein-[Coriobacteriia-bacterium] |
| MBE0418219.1 | DUF499_1.RM.like.systems | METHYLASE-p->HEPN+DUF499*-> | 630 | Bacteria>Terrabacteria_group>Actinomycetota>Coriobacteriia>unclassified_Coriobacteriia | GCA_014859265.1 | Coriobacteriia_bacterium | IBX63_10685 | 617..2507 | 2052159 | ATP-binding-protein-[Coriobacteriia-bacterium] |
| MCP9484920.1 | DUF499_1.RM.like.systems | METHYLASE-p->?->METHYLASE-p->SbcC->PIN->?->HEPN+DUF499*->HIN-HTH->?->\|\|?-><-Peptidase_M20 | 1087 | Bacteria>Terrabacteria_group>Actinomycetota>Rubrobacteria>Gaiellales | GCA_024330215.1 | Candidatus_Gaiellasilicea_maunaloa | MSC30_03590 | 46535..49798 | 2928748 | DUF499-domain-containing-protein-[Candidatus-Gaiellasilicea-maunaloa] |
| GIU94958.1 | DUF499_1.RM.like.systems | Schlafen+HATPase+HTH->?->OLD-ABC->TOPRIM->METHYLASE->METHYLASE-p->HEPN+DUF499*->?->TPR->DUF222+HNH->REase+35exo+Beta-Barrel+SF1-SF2-Helicase-> | 1082 | Bacteria>Terrabacteria_group>Actinomycetota>Rubrobacteria>Gaiellales | GCA_026002995.1 | Gaiellaceae_bacterium | KatS3mg012_1415 | 1404220..1407468 | 2838783 | MAG:-hypothetical-protein-KatS3mg012_1415-[Gaiellaceae-bacterium] |
| MBX6764706.1 | DUF499_1.RM.like.systems | METHYLASE-p->MazE_antitoxin->PIN->HEPN+DUF499*->REase-> | 1106 | Bacteria>Terrabacteria_group>Actinomycetota>Rubrobacteria>Rubrobacterales | GCA_019690035.1 | Rubrobacteraceae_bacterium | IRY88_13660 | 1711..5031 | 2740537 | DUF499-domain-containing-protein-[Rubrobacteraceae-bacterium] |
| WP_219975492.1 | DUF499_1.RM.like.systems | <-SIGMA-HTH<-?<-MULE-TRANSPOSASE\|\|?->?-><-REase<-HEPN+DUF499*<-PIN<-MazE_antitoxin<-METHYLASE<-PIN<-TUDOR+SF1-SF2-Helicase+iREase+CTD+REase\|\|DUF4096-> | 1106 | Bacteria>Terrabacteria_group>Actinomycetota>Rubrobacteria>Rubrobacterales | GCF_019448335.1 | Rubrobacter_xylanophilus | Rxycam_RS04975 | 969402..972722 | 49319 | Swt1-family-HEPN-domain-containing-protein-[Rubrobacter-xylanophilus] |
| WP_217923107.1 | DUF499_1.RM.like.systems | <-HEPN+DUF499*<-METHYLASE<-TUDOR+SF1-SF2-Helicase+iREase+CTD+REase | 1108 | Bacteria>Terrabacteria_group>Actinomycetota>Thermoleophilia>Miltoncostaeales | GCF_018141545.1 | Miltoncostaea_oceani | IU369_RS03095 | 622243..625569 | 2843216 | Swt1-family-HEPN-domain-containing-protein-[Miltoncostaea-oceani] |
| MBI2692601.1 | DUF499_1.RM.like.systems | <-HTH+UbE1<-?\|\|HTH_31->MPTase->SF1-SF2-Helicase+REase-p->?-><-HEPN+DUF499*<-ParB+HNH<-METHYLASE<-TUDOR+SF1-SF2-Helicase+iREase+CTD+REase<-HTH_3\|\|?-><-HTH | 1109 | Bacteria>Terrabacteria_group>Actinomycetota>Thermoleophilia>Solirubrobacterales | GCA_016185345.1 | Solirubrobacterales_bacterium | HYX29_11735 | 159016..162345 | 1909295 | ATP-binding-protein-[Solirubrobacterales-bacterium] |
| MCW3047079.1 | DUF499_1.RM.like.systems | TUDOR+SF1-SF2-Helicase+iREase+CTD+REase->METHYLASE->HEPN+DUF499*->?->?-><-SF1-SF2-helicase<-SF1-SF2-Helicase+REase | 1109 | Bacteria>Terrabacteria_group>Actinomycetota>Thermoleophilia>Solirubrobacterales | GCA_025963905.1 | Solirubrobacterales_bacterium | JW074_1363 | 11829..15158 | 1909295 | MAG:-hypothetical-protein-JW074_1363-[Solirubrobacterales-bacterium] |
| MCZ7663785.1 | DUF499_1.RM.like.systems | <-NTP_transf<-HEPN<-PolB<-?<-TPR<-HEPN+DUF499*<-RelE-ParE<-?<-PIN<-?<-PIN | 1114 | Bacteria>Terrabacteria_group>Actinomycetota>Thermoleophilia>unclassified_Thermoleophilia | GCA_027493445.1 | Thermoleophilia_bacterium | M5U22_13030 | 2549788..2553132 | 2026888 | MAG:-Swt1-family-HEPN-domain-containing-protein-[Thermoleophilia-bacterium] |
| TAK69428.1 | DUF499_1.RM.like.systems | ParB+HNH->HEPN-><-TUDOR+SF1-SF2-Helicase+iREase+CTD+REase\|\|?->?->HEPN+DUF499*->SF1-SF2-Helicase->?-><-?<-REase+35exo+Beta-Barrel+SF1-SF2-Helicase | 1114 | Bacteria>Terrabacteria_group>Actinomycetota>unclassified_Actinomycetota | GCA_004297305.1 | Actinomycetota_bacterium | EPO13_05945 | 61030..64374 | 2900548 | ATP-binding-protein-[Actinobacteria-bacterium] |
| TAK68533.1 | DUF499_1.RM.like.systems | ABC-ATPase+SbcC->?-><-ACET+ACET\|\|?->?-><-HEPN+DUF499*<-METHYLASE<-TUDOR+SF1-SF2-Helicase+iREase+CTD+REase<-?\|\|SIR2+SIR2->PIWI-><-SF1-SF2-Helicase | 1113 | Bacteria>Terrabacteria_group>Actinomycetota>unclassified_Actinomycetota | GCA_004297305.1 | Actinomycetota_bacterium | EPO13_10510 | 141712..145053 | 2900548 | ATP-binding-protein-[Actinobacteria-bacterium] |
| TXH45529.1 | DUF499_1.RM.like.systems | Mbetalac-><-?<-?\|\|Peptidase-><-HEPN+DUF499*\|\|Phd-YefM->PIN-><-?<-METHYLASE\|\|?->PIN-> | 1130 | Bacteria>Terrabacteria_group>Actinomycetota>unclassified_Actinomycetota | GCA_008012465.1 | Actinomycetota_bacterium | E6Q90_00220 | 20268..23660 | 2900548 | ATP-binding-protein-[Actinobacteria-bacterium] |
| MSW27210.1 | DUF499_1.RM.like.systems | Phage_int_SAM_1+Phage_integrase->BirA-HTH-><-REase+DUF4268\|\|TUDOR+SF1-SF2-Helicase+iREase+CTD+REase->METHYLASE->METHYLASE->HEPN+DUF499*->RNase+SF1-SF2-Helicase->HEPN+SF1-SF2-Helicase->?->REase-> | 1092 | Bacteria>Terrabacteria_group>Actinomycetota>unclassified_Actinomycetota | GCA_009698865.1 | Actinomycetota_bacterium | F2849_00700 | 14320..17598 | 2900548 | DUF499-domain-containing-protein-[Actinomycetia-bacterium] |
| MSW31301.1 | DUF499_1.RM.like.systems | PadR-HTH->?->?-><-?<-RNase<-REase+DUF4268<-HEPN+DUF499*<-METHYLASE<-TUDOR+SF1-SF2-Helicase+iREase+CTD+REase<-?\|\|?->?->Phage_int_SAM_1-> | 1098 | Bacteria>Terrabacteria_group>Actinomycetota>unclassified_Actinomycetota | GCA_009698915.1 | Actinomycetota_bacterium | F2847_00635 | 27684..30980 | 2900548 | DUF499-domain-containing-protein-[Actinomycetia-bacterium] |
| MSW80815.1 | DUF499_1.RM.like.systems | <-REase<-?\|\|TUDOR+SF1-SF2-Helicase+iREase+CTD+REase->METHYLASE->HEPN+DUF499*-> | 1104 | Bacteria>Terrabacteria_group>Actinomycetota>unclassified_Actinomycetota | GCA_009699015.1 | Actinomycetota_bacterium | F2840_06650 | 17897..21211 | 2900548 | DUF499-domain-containing-protein-[Actinomycetia-bacterium] |
| NCD16052.1 | DUF499_1.RM.like.systems | <-MORC+REase<-HTH+SF1-SF2-Helicase+REase<-REase<-MORC<-HEPN+DUF499*<-METHYLASE<-REase\|\|DNA_methylase-><-?<-?<-TrwC+SF1-SF2-Helicase | 1131 | Bacteria>Terrabacteria_group>Actinomycetota>unclassified_Actinomycetota | GCA_009930385.1 | Actinomycetota_bacterium | EOL91_01830 | 7837..11232 | 2900548 | ATP-binding-protein-[Actinomycetia-bacterium] |
| NCD16598.1 | DUF499_1.RM.like.systems | <-HEPN+REase<-REase<-HEPN+DUF499*<-wHTH+REase<-METHYLASE | 1133 | Bacteria>Terrabacteria_group>Actinomycetota>unclassified_Actinomycetota | GCA_009930385.1 | Actinomycetota_bacterium | EOL91_04665 | 3104..6505 | 2900548 | ATP-binding-protein-[Actinomycetia-bacterium] |
| NLE73128.1 | DUF499_1.RM.like.systems | AbiEi_1->AbiEii->HIN-HTH-><-?<-Schlafen<-HEPN+DUF499*<-DOC+DOC<-HTH_3<-Tox-RelE<-RelE-ParE<-?<-METHYLASE-p | 1130 | Bacteria>Terrabacteria_group>Actinomycetota>unclassified_Actinomycetota | GCA_012514635.1 | Actinomycetota_bacterium | GX604_00625 | 12665..16057 | 2900548 | ATP-binding-protein-[Actinomycetia-bacterium] |
| NLO26762.1 | DUF499_1.RM.like.systems | PIN->METHYLASE->PIN->?->?->?->HEPN+DUF499*-> | 401 | Bacteria>Terrabacteria_group>Actinomycetota>unclassified_Actinomycetota | GCA_012523095.1 | Actinomycetota_bacterium | GX113_01060 | 16345..17548 | 2900548 | ATP-binding-protein-[Actinomycetia-bacterium] |
| MBC7252723.1 | DUF499_1.RM.like.systems | HIN-HTH->?-><-?<-Schlafen+HATPase+HTH<-ParB+HNH+Schlafen<-HEPN+DUF499*<-METHYLASE<-PIN<-TUDOR+SF1-SF2-Helicase+iREase+CTD+REase | 1115 | Bacteria>Terrabacteria_group>Actinomycetota>unclassified_Actinomycetota | GCA_014360685.1 | Actinomycetota_bacterium | H5T72_01965 | 75820..79167 | 2900548 | ATP-binding-protein-[Actinomycetia-bacterium] |
| QQS24220.1 | DUF499_1.RM.like.systems | TUDOR+SF1-SF2-Helicase+iREase+CTD+REase->METHYLASE->HEPN+DUF499*->NTD+SF1-SF2-Helicase->REase-><-ACET\|\|METHYLASE-p-><-PIN<-Phd-YefM | 1101 | Bacteria>Terrabacteria_group>Actinomycetota>unclassified_Actinomycetota | GCA_016700055.1 | Actinomycetota_bacterium | IPM43_12505 | 2586473..2589778 | 2900548 | DUF499-domain-containing-protein-[Actinobacteria-bacterium] |
| MBS1906106.1 | DUF499_1.RM.like.systems | N6_MTase-p->?->HTH+MPTase-><-SF1-SF2-Helicase<-REase<-HEPN+DUF499*<-METHYLASE<-TUDOR+SF1-SF2-Helicase+iREase+CTD+REase<-XIS-HTH<-?<-?<-HTH_3 | 1133 | Bacteria>Terrabacteria_group>Actinomycetota>unclassified_Actinomycetota | GCA_018268515.1 | Actinomycetota_bacterium | JST33_05955 | 17244..20645 | 2900548 | DUF499-domain-containing-protein-[Actinomycetia-bacterium] |
| MBU2074163.1 | DUF499_1.RM.like.systems | <-HTH+SF1-SF2-Helicase+REase<-REase<-MORC<-HEPN+DUF499*<-METHYLASE<-REase\|\|DNA_methylase->MORC+HEPN-><-?<-TrwC+SF1-SF2-Helicase | 1135 | Bacteria>Terrabacteria_group>Actinomycetota>unclassified_Actinomycetota | GCA_018826065.1 | Actinomycetota_bacterium | KJ938_06990 | 8952..12359 | 2900548 | DUF499-domain-containing-protein-[Actinomycetia-bacterium] |
| MBU2601803.1 | DUF499_1.RM.like.systems | <-Mbetalac<-?<-?<-Radical-SAM<-HEPN+DUF499*<-METHYLASE<-?<-?<-TUDOR+SF1-SF2-Helicase+iREase+CTD+REase<-AbiEii | 1115 | Bacteria>Terrabacteria_group>Actinomycetota>unclassified_Actinomycetota | GCA_018830645.1 | Actinomycetota_bacterium | KKA32_06500 | 57657..61004 | 2900548 | DUF499-domain-containing-protein-[Actinomycetia-bacterium] |
| MBU2602478.1 | DUF499_1.RM.like.systems | <-MNS+HTH\|\|?->HerA/FtsK-><-?<-HEPN+DUF499*<-?<-NTP_transf<-HTH | 1112 | Bacteria>Terrabacteria_group>Actinomycetota>unclassified_Actinomycetota | GCA_018830645.1 | Actinomycetota_bacterium | KKA32_09965 | 120264..123602 | 2900548 | DUF499-domain-containing-protein-[Actinomycetia-bacterium] |
| MBW3642892.1 | DUF499_1.RM.like.systems | <-HEPN+DUF499*<-?<-PIN<-SbcC<-METHYLASE | 1106 | Bacteria>Terrabacteria_group>Actinomycetota>unclassified_Actinomycetota | GCA_019348235.1 | Actinomycetota_bacterium | KY447_08275 | 1669..4989 | 2900548 | DUF499-domain-containing-protein-[Actinomycetia-bacterium] |
| MCA0180524.1 | DUF499_1.RM.like.systems | <-SMODS<-?<-TrwC+SF1-SF2-Helicase\|\|DNA_methylase-><-SF1-SF2-Helicase<-OLD-ABC+TOPRIM<-HEPN+DUF499*<-wHTH+REase<-METHYLASE<-TUDOR+SF1-SF2-Helicase+iREase+CTD+REase<-XIS-HTH<-?\|\|GDResolvase+Recombinase+ZnR-> | 1132 | Bacteria>Terrabacteria_group>Actinomycetota>unclassified_Actinomycetota | GCA_020160805.1 | Actinomycetota_bacterium | LCH77_13240 | 54204..57602 | 2900548 | DUF499-domain-containing-protein-[Actinomycetia-bacterium] |
| MCA0180887.1 | DUF499_1.RM.like.systems | <-VirD4-TraG+VirD4-FtsK<-?\|\|XIS-HTH->TUDOR+SF1-SF2-Helicase+iREase+CTD+REase->METHYLASE->wHTH+REase->HEPN+DUF499*->REase->HEPN+REase-> | 1132 | Bacteria>Terrabacteria_group>Actinomycetota>unclassified_Actinomycetota | GCA_020160805.1 | Actinomycetota_bacterium | LCH77_15145 | 46365..49763 | 2900548 | DUF499-domain-containing-protein-[Actinomycetia-bacterium] |
| MCA0294159.1 | DUF499_1.RM.like.systems | <-NTD+SF1-SF2-Helicase\|\|TUDOR+SF1-SF2-Helicase+iREase+CTD+REase->METHYLASE->HEPN+DUF499*->ParB+HNH->?->SF1-SF2-Helicase-><-REase+PNTD1+PNTD2\|\|MULE-TRANSPOSASE->ACET-> | 1117 | Bacteria>Terrabacteria_group>Actinomycetota>unclassified_Actinomycetota | GCA_020161755.1 | Actinomycetota_bacterium | LCH96_02385 | 200193..203546 | 2900548 | DUF499-domain-containing-protein-[Actinomycetia-bacterium] |
| MCL4554262.1 | DUF499_1.RM.like.systems | TUDOR+SF1-SF2-Helicase+iREase+CTD+REase-p->OLD-ABC+TOPRIM->METHYLASE->HEPN->ParA-Soj-PloopNTPase->ORC-CDC6->HEPN+DUF499*-> | 468 | Bacteria>Terrabacteria_group>Actinomycetota>unclassified_Actinomycetota | GCA_023382015.1 | Actinomycetota_bacterium | M1565_03890 | 9031..10434 | 2900548 | Swt1-family-HEPN-domain-containing-protein-[Actinomycetota-bacterium] |
| MCL5734830.1 | DUF499_1.RM.like.systems | TUDOR+SF1-SF2-Helicase+iREase+CTD+REase-p->MORC+DUF3684->METHYLASE->REase->SF1-SF2-Helicase->HEPN+DUF499*-><-HTH+DSBH | 1116 | Bacteria>Terrabacteria_group>Actinomycetota>unclassified_Actinomycetota | GCA_023483915.1 | Actinomycetota_bacterium | M1274_04450 | 22427..25777 | 2900548 | Swt1-family-HEPN-domain-containing-protein-[Actinobacteria-bacterium] |
| MCL5447091.1 | DUF499_1.RM.like.systems | <-HEPN+DUF499*<-METHYLASE<-?<-TUDOR+SF1-SF2-Helicase+iREase+CTD+REase<-REase<-?<-rve+rve_3 | 1178 | Bacteria>Terrabacteria_group>Actinomycetota>unclassified_Actinomycetota | GCA_023486005.1 | Actinomycetota_bacterium | M1350_06795 | 8812..12348 | 2900548 | DUF499-domain-containing-protein-[Actinobacteria-bacterium] |
| MCX6403861.1 | DUF499_1.RM.like.systems | REase->METHYLASE->HEPN+DUF499*-> | 1088 | Bacteria>Terrabacteria_group>Actinomycetota>unclassified_Actinomycetota | GCA_026392535.1 | Actinomycetota_bacterium | NT032_08225 | 3485..6751 | 2900548 | MAG:-DUF499-domain-containing-protein-[Actinobacteria-bacterium] |
| MDA8063428.1 | DUF499_1.RM.like.systems | <-HEPN+DUF499*<-HEPN<-PolB<-METHYLASE-p | 641 | Bacteria>Terrabacteria_group>Actinomycetota>unclassified_Actinomycetota | GCA_028064035.1 | Actinomycetota_bacterium | M0T80_13560 | 1..1922 | 2900548 | MAG:-Swt1-family-HEPN-domain-containing-protein,-partial-[Actinomycetota-bacterium] |
| MDA8117679.1 | DUF499_1.RM.like.systems | <-MerR-HTH<-?<-HEPN+DUF499*<-TOPRIM<-OLD-ABC<-METHYLASE<-HEPN<-ParB+HNH<-HicB+ZnR | 1122 | Bacteria>Terrabacteria_group>Actinomycetota>unclassified_Actinomycetota | GCA_028064745.1 | Actinomycetota_bacterium | M0000_09960 | 1809..5177 | 2900548 | MAG:-Swt1-family-HEPN-domain-containing-protein-[Actinomycetota-bacterium] |
| MDA8210090.1 | DUF499_1.RM.like.systems | METHYLASE->HEPN+DUF499*-> | 900 | Bacteria>Terrabacteria_group>Actinomycetota>unclassified_Actinomycetota | GCA_028065575.1 | Actinomycetota_bacterium | M0Z92_13975 | 2687..5388 | 2900548 | MAG:-Swt1-family-HEPN-domain-containing-protein,-partial-[Actinomycetota-bacterium] |
| MDA8316124.1 | DUF499_1.RM.like.systems | <-HEPN+DUF499*<-METHYLASE<-TUDOR+SF1-SF2-Helicase+iREase+CTD+REase-p | 1027 | Bacteria>Terrabacteria_group>Actinomycetota>unclassified_Actinomycetota | GCA_028067765.1 | Actinomycetota_bacterium | M0010_13295 | 109..3192 | 2900548 | MAG:-Swt1-family-HEPN-domain-containing-protein-[Actinomycetota-bacterium] |
| MDA8316555.1 | DUF499_1.RM.like.systems | TUDOR+SF1-SF2-Helicase+iREase+CTD+REase->METHYLASE->PolB->HEPN->PolB->HEPN+DUF499*-><-RNase<-XIS-HTH | 1117 | Bacteria>Terrabacteria_group>Actinomycetota>unclassified_Actinomycetota | GCA_028067765.1 | Actinomycetota_bacterium | M0010_15505 | 14614..17967 | 2900548 | MAG:-Swt1-family-HEPN-domain-containing-protein-[Actinomycetota-bacterium] |
| MDA8317108.1 | DUF499_1.RM.like.systems | <-HEPN+DUF499*<-METHYLASE\|\|?-><-TUDOR+SF1-SF2-Helicase+iREase+CTD+REase<-HTH+MPTase | 493 | Bacteria>Terrabacteria_group>Actinomycetota>unclassified_Actinomycetota | GCA_028067765.1 | Actinomycetota_bacterium | M0010_18355 | 6..1487 | 2900548 | MAG:-Swt1-family-HEPN-domain-containing-protein-[Actinomycetota-bacterium] |
| MDA8312150.1 | DUF499_1.RM.like.systems | Phage_integrase-><-HEPN+DUF499*<-METHYLASE<-TUDOR+SF1-SF2-Helicase+iREase+CTD+REase<-SIR2 | 1097 | Bacteria>Terrabacteria_group>Actinomycetota>unclassified_Actinomycetota | GCA_028067795.1 | Actinomycetota_bacterium | M0Z46_16395 | 628..3921 | 2900548 | MAG:-Swt1-family-HEPN-domain-containing-protein-[Actinomycetota-bacterium] |
| MDA8358565.1 | DUF499_1.RM.like.systems | TUDOR+SF1-SF2-Helicase+iREase+CTD+REase->METHYLASE->HEPN+DUF499*->DNA-glycosylase-><-PIWI<-SIR2+SIR2\|\|SbcC->SF1-SF2-Helicase-> | 1122 | Bacteria>Terrabacteria_group>Actinomycetota>unclassified_Actinomycetota | GCA_028067995.1 | Actinomycetota_bacterium | M0Z95_20250 | 17381..20749 | 2900548 | MAG:-DUF499-domain-containing-protein-[Actinomycetota-bacterium] |
| NLH98010.1 | DUF499_1.RM.like.systems | TUDOR+SF1-SF2-Helicase+iREase+CTD+REase->?->METHYLASE->OLD-ABC->TOPRIM->HEPN+DUF499*-> | 1111 | Bacteria>Terrabacteria_group>Armatimonadota>Chthonomonadetes>Chthonomonadales | GCA_012517255.1 | Chthonomonadales_bacterium | GX446_00800 | 20611..23946 | 2282151 | ATP-binding-protein-[Chthonomonadales-bacterium] |
| MBL8066214.1 | DUF499_1.RM.like.systems | <-HEPN+DUF499*<-HEPN<-METHYLASE-p | 1119 | Bacteria>Terrabacteria_group>Armatimonadota>Chthonomonadetes>Chthonomonadales | GCA_016788345.1 | Chthonomonadaceae_bacterium | JNM34_10200 | 1590..4949 | 2739509 | DUF499-domain-containing-protein-[Chthonomonadaceae-bacterium] |
| MBW7926628.1 | DUF499_1.RM.like.systems | TUDOR+SF1-SF2-Helicase+iREase+CTD+REase->?->SMODS->METHYLASE->HEPN+DUF499*->?->?->JAB-><-D5N+viral-AAA | 1119 | Bacteria>Terrabacteria_group>Armatimonadota>Fimbriimonadia>Fimbriimonadales | GCA_019454665.1 | Fimbriimonadaceae_bacterium | H3C58_00785 | 10553..13912 | 2682144 | ATP-binding-protein-[Fimbriimonadaceae-bacterium] |
| MBX7131086.1 | DUF499_1.RM.like.systems | Phage_int_SAM_1+Phage_integrase-><-Peptidase<-HEPN+DUF499*<-?<-HTH<-HKD-DNase<-METHYLASE<-ParB+HNH<-TUDOR+SF1-SF2-Helicase+iREase+CTD+REase | 1104 | Bacteria>Terrabacteria_group>Armatimonadota>Fimbriimonadia>Fimbriimonadales | GCA_019694835.1 | Fimbriimonadaceae_bacterium | K1X67_00255 | 55329..58643 | 2682144 | DUF499-domain-containing-protein-[Fimbriimonadaceae-bacterium] |
| MCC7230625.1 | DUF499_1.RM.like.systems | TIR->?->?->TUDOR+SF1-SF2-Helicase+iREase+CTD+REase->Schlafen+HATPase+HTH->METHYLASE->HEPN+DUF499*-><-JAB\|\|?->ArdCN+MPTase->?->N6_N4_MTase-><-PIN | 1116 | Bacteria>Terrabacteria_group>Armatimonadota>Fimbriimonadia>Fimbriimonadales | GCA_020851885.1 | Fimbriimonadaceae_bacterium | IT203_09555 | 16596..19946 | 2682144 | DUF499-domain-containing-protein-[Fimbriimonadaceae-bacterium] |
| GIV04239.1 | DUF499_1.RM.like.systems | METHYLASE-p->?->RAMPs->RAMPs->RAMPs->HEPN+HEPN->HEPN+DUF499*->MORC+DUF3684-> | 891 | Bacteria>Terrabacteria_group>Armatimonadota>Fimbriimonadia>Fimbriimonadales | GCA_026003055.1 | Fimbriimonadales_bacterium | KatS3mg015_3069 | 13495..16170 | 2682145 | MAG:-hypothetical-protein-KatS3mg015_3069-[Fimbriimonadales-bacterium] |
| MCX6342680.1 | DUF499_1.RM.like.systems | <-URI-DNase\|\|MazG->URI-DNase+SF1-SF2-Helicase-><-?<-REase<-HEPN+DUF499*<-METHYLASE<-TUDOR+SF1-SF2-Helicase+iREase+CTD+REase | 1119 | Bacteria>Terrabacteria_group>Armatimonadota>Fimbriimonadia>Fimbriimonadales | GCA_026392145.1 | Fimbriimonadales_bacterium | NTU72_09890 | 5893..9252 | 2682145 | MAG:-Swt1-family-HEPN-domain-containing-protein-[Fimbriimonadales-bacterium] |
| HCD99868.1 | DUF499_1.RM.like.systems | METHYLASE-p->HEPN+DUF499*->TUDOR+SF1-SF2-Helicase+iREase+CTD+REase-p-> | 1116 | Bacteria>Terrabacteria_group>Armatimonadota>unclassified_Armatimonadota | GCA_003516955.1 | Armatimonadota_bacterium | DER07_02370 | 679..4029 | 2033014 | AAA+-family-ATPase-[Armatimonadetes-bacterium] |
| MBI3912159.1 | DUF499_1.RM.like.systems | METHYLASE-p->DUF433->PIN->HEPN+DUF499*->?-><-NTP_transf<-Peptidase<-?<-Radical-SAM | 1099 | Bacteria>Terrabacteria_group>Armatimonadota>unclassified_Armatimonadota | GCA_016199525.1 | Armatimonadota_bacterium | HY320_14665 | 1429..4728 | 2033014 | ATP-binding-protein-[Armatimonadetes-bacterium] |
| MBM3457180.1 | DUF499_1.RM.like.systems | METHYLASE-p->HEPN+DUF499*-> | 1128 | Bacteria>Terrabacteria_group>Armatimonadota>unclassified_Armatimonadota | GCA_016868995.1 | Armatimonadota_bacterium | FJX77_01400 | 1230..4616 | 2033014 | DUF499-domain-containing-protein-[Armatimonadetes-bacterium] |
| MBS1724660.1 | DUF499_1.RM.like.systems | TUDOR+SF1-SF2-Helicase+iREase+CTD+REase->METHYLASE->HEPN+DUF499*->MORC-><-?\|\|N6_N4_MTase-><-Phage_integrase<-MAD-NTD+HNH | 1107 | Bacteria>Terrabacteria_group>Armatimonadota>unclassified_Armatimonadota | GCA_018268375.1 | Armatimonadota_bacterium | JSS66_17085 | 550844..554167 | 2033014 | DUF499-domain-containing-protein-[Armatimonadetes-bacterium] |
| MCX6379947.1 | DUF499_1.RM.like.systems | REase->HEPN+DUF499*-><-?<-?<-?<-?<-?\|\|Phage_int_SAM_1+Phage_integrase-> | 1119 | Bacteria>Terrabacteria_group>Armatimonadota>unclassified_Armatimonadota | GCA_026392135.1 | Armatimonadota_bacterium | NT023_10805 | 3514..6873 | 2033014 | MAG:-Swt1-family-HEPN-domain-containing-protein-[Armatimonadetes-bacterium] |
| AKM18157.1 | DUF499_1.RM.like.systems | <-GspA-ATPase<-HTH_Tnp<-ZnR\|\|?->TUDOR+SF1-SF2-Helicase+iREase+CTD+REase->METHYLASE->HEPN+DUF499*-><-?<-DDE_5+DDE_Tnp_2\|\|REase-><-Arm-DNA-bind_4+Phage_int_SAM_3+Phage_integrase<-DDE_5+DDE_Tnp_2 | 1126 | Bacteria>Terrabacteria_group>Bacillota>Bacilli>Bacillales | GCA_001028085.1 | Geobacillus_sp._12AMOR1 | GARCT_00860 | 838569..841949 | 1629723 | hypothetical-protein-GARCT_00860-[Geobacillus-sp.-12AMOR1] |
| PTQ56106.1 | DUF499_1.RM.like.systems | TUDOR+SF1-SF2-Helicase+iREase+CTD+REase->METHYLASE->HEPN+DUF499*-><-?\|\|?-><-?<-?<-Primpol+D5N+D5ATPase | 1116 | Bacteria>Terrabacteria_group>Bacillota>Bacilli>Bacillales | GCA_003050645.1 | Candidatus_Carbobacillus_altaicus | BSOLF_0824 | 6920..10270 | 2163959 | hypothetical-protein-BSOLF_0824-[Candidatus-Carbobacillus-altaicus] |

| Protein-ID | System | Gene-Neighbourhoods | Length | Taxonomy | Assembly-ID | Strain | Locus-Tag | Genomic-coordinates | TaxID | Fasta-define |
|---|---|---|---|---|---|---|---|---|---|---|
| RYL92593.1 | DUF499_1.RM.like.systems | REase+N6_MTase->?->TUDOR+SF1-SF2-Helicase+iREase+CTD+REase->METHYLASE->HEPN+DUF499*->ParB+HNH-><-DDE_Tnp_1<-?<-?\|\|DDE_Tnp_1-> | 1123 | Bacteria>Terrabacteria_group>Bacillota>Bacilli>Bacillales | GCA_004153195.1 | Sporolactobacillus_sp._THM19-2 | EWH91_06945 | 33831..37202 | 2511171 | ATP-binding-protein-[Sporolactobacillus-sp.-THM19-2] |
| MBO2534075.1 | DUF499_1.RM.like.systems | <-MqsA_antitoxin+HTH_3<-?<-HEPN+DUF499*<-METHYLASE-p | 1117 | Bacteria>Terrabacteria_group>Bacillota>Bacilli>Bacillales | GCA_017577425.1 | Thermoactinomycetaceae_bacterium | CW342_14595 | 2341..5694 | 2056641 | AAA+-family-ATPase-[Thermoactinomycetaceae-bacterium] |
| MCG0314785.1 | DUF499_1.RM.like.systems | <-HEPN+DUF499*<-METHYLASE-p | 616 | Bacteria>Terrabacteria_group>Bacillota>Bacilli>Bacillales | GCA_021890815.1 | Calditerricola_sp. | L5657_09065 | 1..1847 | 2885161 | DUF499-domain-containing-protein-[Calditerricola-sp.] |
| STO36143.1 | DUF499_1.RM.like.systems | <-REase\|\|DDE_5+DDE_Tnp_2->HIN-HTH->?->?-><-HEPN+DUF499*<-METHYLASE<-DUF4277+DDE_Tnp_1<-METHYLASE-p<-TUDOR+SF1-SF2-Helicase+iREase+CTD+REase<-?\|\|ZnR-> | 1026 | Bacteria>Terrabacteria_group>Bacillota>Bacilli>Bacillales | GCA_900450595.1 | [Flavobacterium]_thermophilum | NCTC11526_03101 | 455797..458877 | 1414643 | Protein-of-uncharacterised-function-(DUF499)-[[Flavobacterium] |
| WP_054669274.1 | DUF499_1.RM.like.systems | <-DUF499-p<-HEPN+DUF499*<-METHYLASE-p<-METHYLASE-p<-METHYLASE-p<-TUDOR+SF1-SF2-Helicase+iREase+CTD+REase-p | 833 | Bacteria>Terrabacteria_group>Bacillota>Bacilli>Bacillales | GCF_001311905.1 | Calditerricola_satsumensis | JCM14719_RS01550 | 8456..10957 | 373054 | Swt1-family-HEPN-domain-containing-protein-[Calditerricola-satsumensis] |
| WP_124727907.1 | DUF499_1.RM.like.systems | Mbetalac->?-><-?<-?<-ABhydrolase<-HEPN+DUF499*<-METHYLASE<-TUDOR+SF1-SF2-Helicase+iREase+CTD+REase\|\|?->ACET-><-CHTH | 1107 | Bacteria>Terrabacteria_group>Bacillota>Bacilli>Bacillales | GCF_003856495.1 | Staphylospora_marina | EG886_RS09450 | 1890646..1893969 | 2490858 | Swt1-family-HEPN-domain-containing-protein-[Staphylospora-marina] |
| WP_188816795.1 | DUF499_1.RM.like.systems | <-Radical-SAM+TRAM<-?<-?<-?<-?<-?<-HEPN+DUF499*<-METHYLASE<-TUDOR+SF1-SF2-Helicase+iREase+CTD+REase<-Peptidase | 1124 | Bacteria>Terrabacteria_group>Bacillota>Bacilli>Bacillales | GCF_014646935.1 | Calditerricola_satsumensis | IEX61_RS03410 | 40382..43756 | 373054 | DUF499-domain-containing-protein-[Calditerricola-satsumensis] |
| WP_220678456.1 | DUF499_1.RM.like.systems | TUDOR+SF1-SF2-Helicase+iREase+CTD+REase->METHYLASE-p-><-MULE-TRANSPOSASE\|\|METHYLASE->HEPN+DUF499*->?->DUF772+DDE_Tnp_1+DDE_Tnp_1_6->?->zf-ISL3+DDE_Tnp_ISL3->?->Glycos_trans_3N+Glycos_transf_3-> | 1124 | Bacteria>Terrabacteria_group>Bacillota>Bacilli>Bacillales | GCF_019645935.1 | Hydrogenibacillus_sp._N12 | K2M58_RS04470 | 965696..969070 | 2866627 | Swt1-family-HEPN-domain-containing-protein-[Hydrogenibacillus-sp.-N12] |
| WP_236408338.1 | DUF499_1.RM.like.systems | XIS-HTH->TUDOR+SF1-SF2-Helicase+iREase+CTD+REase->METHYLASE->HEPN+DUF499*-> | 1134 | Bacteria>Terrabacteria_group>Bacillota>Bacilli>Bacillales | GCF_021608135.1 | Microaerobacter_geothermalis | L1765_RS15200 | 6759..10163 | 674972 | Swt1-family-HEPN-domain-containing-protein-[Microaerobacter-geothermalis] |
| WP_244410622.1 | DUF499_1.RM.like.systems | <-HEPN+DUF499*<-METHYLASE<-TUDOR+SF1-SF2-Helicase+iREase+CTD+REase<-?\|\|?-><-MULE-TRANSPOSASE | 1126 | Bacteria>Terrabacteria_group>Bacillota>Bacilli>Bacillales | GCF_022846475.1 | Parageobacillus_thermoglucosidasius | MWM14_RS16385 | 3230827..3234207 | 1426 | Swt1-family-HEPN-domain-containing-protein-[Parageobacillus-thermoglucosidasius] |
| WP_258111975.1 | DUF499_1.RM.like.systems | <-HicB<-HicA_toxin<-?<-PIN<-?\|\|METHYLASE-p->HEPN+DUF499*->RVT-> | 1128 | Bacteria>Terrabacteria_group>Bacillota>Bacilli>Bacillales | GCF_024706985.1 | Alicyclobacillus_sp._SP_1 | M4I36_RS11945 | 3650..7036 | 2942475 | DUF499-domain-containing-protein-[Alicyclobacillus-sp.-SP_1] |
| WP_129929317.1 | DUF499_1.RM.like.systems | REase+N6_MTase->?->TUDOR+SF1-SF2-Helicase+iREase+CTD+REase->METHYLASE->HEPN+DUF499*->ParB+HNH-><-DDE_Tnp_1<-?<-?\|\|DDE_Tnp_1-> | 1123 | Bacteria>Terrabacteria_group>Bacillota>Bacilli>Bacillales | - | Sporolactobacillus_sp._THM19-2 | EWH91_06945 | 33831..37202 | 2511171 | DUF499-domain-containing-protein-[Sporolactobacillus-sp.-THM19-2] |
| WP_181339998.1 | DUF499_1.RM.like.systems | DeoR-HTH+ISOCOT->?->?->?->?-><-HEPN+DUF499*<-METHYLASE<-TUDOR+SF1-SF2-Helicase+iREase+CTD+REase\|\|OLD-ABC+TOPRIM-> | 1104 | Bacteria>Terrabacteria_group>Bacillota>Bacillota_sensu_stricto_incertae_sedis>Capillibacterium | GCF_013664685.1 | Capillibacterium_thermochitinicola | G5B42_RS08290 | 130685..133999 | 2699427 | Swt1-family-HEPN-domain-containing-protein-[Capillibacterium-thermochitinicola] |
| HIR07141.1 | DUF499_1.RM.like.systems | TUDOR+SF1-SF2-Helicase+iREase+CTD+REase->METHYLASE->HEPN+DUF499*-> | 987 | Bacteria>Terrabacteria_group>Bacillota>Clostridia>Clostridia_incertae_sedis | GCA_018712885.1 | Candidatus_Pullichristensenella_stercoripullorum | IAA71_02960 | 11277..14237 | 2840910 | DUF499-domain-containing-protein-[Candidatus-Pullichristensenella-stercoripullorum] |
| WP_073238382.1 | DUF499_1.RM.like.systems | <-TPR<-?<-Radical-SAM<-?<-tRNA<-?<-HEPN+DUF499*<-METHYLASE<-TUDOR+SF1-SF2-Helicase+iREase+CTD+REase\|\|tRNA->RNaseH->ZnR->PAIREDC-HTH-> | 1123 | Bacteria>Terrabacteria_group>Bacillota>Clostridia>Eubacteriales | - | Desulforamulus_putei | BUB67_RS07930 | 94636..98007 | 74701 | Swt1-family-HEPN-domain-containing-protein-[Desulforamulus-putei] |
| KLU38984.1 | DUF499_1.RM.like.systems | TUDOR+SF1-SF2-Helicase+iREase+CTD+REase->METHYLASE->METHYLASE->METHYLASE->HEPN+DUF499*->?->?->METHYLASE-p->?-><-?\|\|CHTH-> | 1130 | Bacteria>Terrabacteria_group>Bacillota>Clostridia>Eubacteriales | GCA_001028815.1 | Peptococcaceae_bacterium_1109 | AA931_11700 | 75012..78404 | 1655638 | hypothetical-protein-AA931_11700-[Peptococcaceae-bacterium-1109] |
| HCW03906.1 | DUF499_1.RM.like.systems | <-HEPN+DUF499*<-?<-?<-?<-ABC-ATPase+SbcC<-ABC-ATPase+SbcC | 1132 | Bacteria>Terrabacteria_group>Bacillota>Clostridia>Eubacteriales | GCA_003541575.1 | Clostridium_sp. | DGK91_04815 | 19..3417 | 1506 | hypothetical-protein-[Clostridium-sp.] |
| MCL1790896.1 | DUF499_1.RM.like.systems | <-Peptidase\|\|TUDOR+SF1-SF2-Helicase+iREase+CTD+REase->METHYLASE->Schlafen+HATPase+HTH->HEPN+DUF499*-> | 1122 | Bacteria>Terrabacteria_group>Bacillota>Clostridia>Eubacteriales | GCA_009778595.1 | Peptococcaceae_bacterium | FWG40_05990 | 9595..12963 | 2052179 | Swt1-family-HEPN-domain-containing-protein-[Peptococcaceae-bacterium] |
| MCL1794475.1 | DUF499_1.RM.like.systems | <-RelE-ParE<-MazE_antitoxin<-HEPN+DUF499*\|\|?->GDResolvase->?-><-DUF772+DDE_Tnp_1 | 1135 | Bacteria>Terrabacteria_group>Bacillota>Clostridia>Eubacteriales | GCA_009782405.1 | Oscillospiraceae_bacterium | FWG34_11475 | 15838..19245 | 2485925 | DUF499-domain-containing-protein-[Oscillospiraceae-bacterium] |
| MCL2699418.1 | DUF499_1.RM.like.systems | <-HEPN+DUF499*<-METHYLASE | 645 | Bacteria>Terrabacteria_group>Bacillota>Clostridia>Eubacteriales | GCA_009783765.1 | Defluviitaleaceae_bacterium | FWE68_03815 | 1..1937 | 2660712 | Swt1-family-HEPN-domain-containing-protein-[Defluviitaleaceae-bacterium] |
| NLI53236.1 | DUF499_1.RM.like.systems | WYL->?->TUDOR+SF1-SF2-Helicase+iREase+CTD+REase->METHYLASE->HEPN+DUF499*->OLD-ABC+TOPRIM->?->?-><-tRNA<-SMS-KaiC | 1124 | Bacteria>Terrabacteria_group>Bacillota>Clostridia>Eubacteriales | GCA_012517905.1 | Clostridiales_bacterium | GX417_02810 | 9381..12755 | 1898207 | ATP-binding-protein-[Clostridiales-bacterium] |
| NLI59701.1 | DUF499_1.RM.like.systems | METHYLASE-p->HEPN+DUF499*-> | 985 | Bacteria>Terrabacteria_group>Bacillota>Clostridia>Eubacteriales | GCA_012517995.1 | Clostridium_sp. | GX387_14560 | 352..3307 | 1506 | ATP-binding-protein-[Clostridium-sp.] |
| NLJ76878.1 | DUF499_1.RM.like.systems | <-ParB+HNH<-TPR<-?<-HEPN+DUF499*<-METHYLASE-p | 1133 | Bacteria>Terrabacteria_group>Bacillota>Clostridia>Eubacteriales | GCA_012518875.1 | Peptococcaceae_bacterium | GX325_06435 | 3363..6764 | 2052179 | ATP-binding-protein-[Peptococcaceae-bacterium] |
| NLW38082.1 | DUF499_1.RM.like.systems | <-MethyltransfD12\|\|?->PIN-><-HEPN+DUF499*<-METHYLASE<-TPR+TPR<-MetJarc-HTH<-SF1-SF2-Helicase<-EVE | 1120 | Bacteria>Terrabacteria_group>Bacillota>Clostridia>Eubacteriales | GCA_012728715.1 | Peptococcaceae_bacterium | GXY90_05855 | 3552..6914 | 2052179 | ATP-binding-protein-[Peptococcaceae-bacterium] |
| NLV15838.1 | DUF499_1.RM.like.systems | PRTase-><-?\|\|TUDOR+SF1-SF2-Helicase+iREase+CTD+REase->METHYLASE->HEPN+DUF499*->REase-> | 1154 | Bacteria>Terrabacteria_group>Bacillota>Clostridia>Eubacteriales | GCA_012729505.1 | Syntrophomonadaceae_bacterium | GXY50_01325 | 75931..79395 | 2093811 | ATP-binding-protein-[Syntrophomonadaceae-bacterium] |
| HHW47352.1 | DUF499_1.RM.like.systems | XIS-HTH->TUDOR+SF1-SF2-Helicase+iREase+CTD+REase->METHYLASE->HEPN+DUF499*->?->ParB+HNH-><-Aminotran_1_2\|\|?->ABC-ATPase+SbcC-> | 1135 | Bacteria>Terrabacteria_group>Bacillota>Clostridia>Eubacteriales | GCA_012840535.1 | Clostridiaceae_bacterium | GXX14_01895 | 11508..14915 | 1898204 | ATP-binding-protein-[Clostridiaceae-bacterium] |
| NQS74830.1 | DUF499_1.RM.like.systems | Peptidase_M24->?->ZnR->?->TUDOR+SF1-SF2-Helicase+iREase+CTD+REase->METHYLASE->HEPN+DUF499*->?->XIS-HTH-><-HTH+MPTase | 1135 | Bacteria>Terrabacteria_group>Bacillota>Clostridia>Eubacteriales | GCA_013201395.1 | Candidatus_Syntrophopropionicum_ammoniitolerans | HQP61_00110 | 30439..33846 | 2807340 | ATP-binding-protein-[Peptococcaceae-bacterium] |
| QNK42634.1 | DUF499_1.RM.like.systems | HTH_36-><-?<-?<-?<-JetA<-HEPN+DUF499*<-TUDOR+SF1-SF2-Helicase+iREase+CTD+REase<-CHTH | 1113 | Bacteria>Terrabacteria_group>Bacillota>Clostridia>Eubacteriales | GCA_014303955.1 | Caproicibacter_fermentans | HCR03_14035 | 2844426..2847767 | 2576756 | ATP-binding-protein-[Caproicibacter-fermentans] |
| MBC7075381.1 | DUF499_1.RM.like.systems | MazE_antitoxin->RelE-ParE->\|\|?->?->TUDOR+SF1-SF2-Helicase+iREase+CTD+REase->METHYLASE->HEPN+DUF499*->ParB+HNH-> | 1122 | Bacteria>Terrabacteria_group>Bacillota>Clostridia>Eubacteriales | GCA_014360155.1 | Syntrophomonadaceae_bacterium | H5T98_04805 | 11477..14845 | 2093811 | ATP-binding-protein-[Syntrophomonadaceae-bacterium] |
| MBD9257204.1 | DUF499_1.RM.like.systems | TRAM+Methyltransf_31->AbiGi->GDResolvase+Recombinase+ZnR->TUDOR+SF1-SF2-Helicase+iREase+CTD+REase->?->METHYLASE->HEPN+DUF499*-> | 1129 | Bacteria>Terrabacteria_group>Bacillota>Clostridia>Eubacteriales | GCA_014848725.1 | Faecalibacterium_sp. | EGR99_01350 | 18059..21448 | 1971605 | ATP-binding-protein-[Faecalibacterium-sp.] |
| MBQ3841838.1 | DUF499_1.RM.like.systems | <-XIS-HTH<-HNH+HNH\|\|?-><-HTH_3\|\|TUDOR+SF1-SF2-Helicase+iREase+CTD+REase->METHYLASE->HEPN+DUF499*->?->?->HTH_3-><-?<-HTH_3 | 1114 | Bacteria>Terrabacteria_group>Bacillota>Clostridia>Eubacteriales | GCA_017395555.1 | Ruminiclostridium_sp. | II820_04000 | 63719..67063 | 2053608 | DUF499-domain-containing-protein-[Ruminiclostridium-sp.] |
| MBQ9662099.1 | DUF499_1.RM.like.systems | TUDOR+SF1-SF2-Helicase+iREase+CTD+REase->METHYLASE->HEPN+DUF499*-><-?<-?<-?<-Glycos_transf_4 | 1119 | Bacteria>Terrabacteria_group>Bacillota>Clostridia>Eubacteriales | GCA_017458065.1 | Oscillospiraceae_bacterium | IJV40_02965 | 18565..21924 | 2485925 | DUF499-domain-containing-protein-[Oscillospiraceae-bacterium] |
| MBR3354573.1 | DUF499_1.RM.like.systems | HTH->tRNA->tRNA->tRNA->tRNA->tRNA->tRNA->tRNA->?->TUDOR+SF1-SF2-Helicase+iREase+CTD+REase->METHYLASE->METHYLASE->HEPN+DUF499*->REase-> | 1111 | Bacteria>Terrabacteria_group>Bacillota>Clostridia>Eubacteriales | GCA_017510465.1 | Oscillospiraceae_bacterium | IKG47_04355 | 36923..40258 | 2485925 | DUF499-domain-containing-protein-[Oscillospiraceae-bacterium] |
| MBR3354625.1 | DUF499_1.RM.like.systems | <-HTH\|\|HTH_26->TUDOR+SF1-SF2-Helicase+iREase+CTD+REase->METHYLASE->METHYLASE->METHYLASE->HEPN+DUF499*-> | 1125 | Bacteria>Terrabacteria_group>Bacillota>Clostridia>Eubacteriales | GCA_017510465.1 | Oscillospiraceae_bacterium | IKG47_04625 | 20932..24309 | 2485925 | DUF499-domain-containing-protein-[Oscillospiraceae-bacterium] |
| MBR3240816.1 | DUF499_1.RM.like.systems | <-MORC<-HEPN+DUF499*<-METHYLASE<-TUDOR+SF1-SF2-Helicase+iREase+CTD+REase<-?\|\|HTH->HTH_31->MPTase-> | 1133 | Bacteria>Terrabacteria_group>Bacillota>Clostridia>Eubacteriales | GCA_017511105.1 | Oscillospiraceae_bacterium | IKF99_20575 | 19684..23085 | 2485925 | DUF499-domain-containing-protein-[Oscillospiraceae-bacterium] |
| MBO6231158.1 | DUF499_1.RM.like.systems | <-HNH+HNH\|\|?-><-HTH_3\|\|TUDOR+SF1-SF2-Helicase+iREase+CTD+REase->METHYLASE->HEPN+DUF499*->?->Classical-AAA->REase->SMODS->HTH_3-><-Phd-YefM | 1114 | Bacteria>Terrabacteria_group>Bacillota>Clostridia>Eubacteriales | GCA_017635095.1 | Ruminiclostridium_sp. | J6050_11400 | 23257..26601 | 2053608 | DUF499-domain-containing-protein-[Ruminiclostridium-sp.] |
| MBS3975902.1 | DUF499_1.RM.like.systems | <-Phage_terminase<-NACHT<-HEPN+DUF499*<-METHYLASE<-TUDOR+SF1-SF2-Helicase+iREase+CTD+REase-p | 1133 | Bacteria>Terrabacteria_group>Bacillota>Clostridia>Eubacteriales | GCA_018333865.1 | Syntrophomonadaceae_bacterium | KGZ75_04135 | 3808..7209 | 2093811 | DUF499-domain-containing-protein-[Syntrophomonadaceae-bacterium] |
| MBS4922099.1 | DUF499_1.RM.like.systems | AbiGi->GDResolvase+Recombinase+ZnR->TUDOR+SF1-SF2-Helicase+iREase+CTD+REase->MNS->METHYLASE->HEPN+DUF499*->ParB->CRP-HTH+GNTR-HTH->?->ZDNA-HTH->GDResolvase+Recombinase+ZnR-> | 1129 | Bacteria>Terrabacteria_group>Bacillota>Clostridia>Eubacteriales | GCA_018364195.1 | Faecalibacterium_prausnitzii | KHZ84_09455 | 17444..20833 | 853 | DUF499-domain-containing-protein-[Faecalibacterium-prausnitzii] |
| MBU5249610.1 | DUF499_1.RM.like.systems | DUF772+DDE_Tnp_1_6->Schlafen+HATPase+HTH->TUDOR+SF1-SF2-Helicase+iREase+CTD+REase->METHYLASE->METHYLASE->HEPN+DUF499*->?->CRP-HTH+GNTR-HTH->?->?->?->REase-> | 1128 | Bacteria>Terrabacteria_group>Bacillota>Clostridia>Eubacteriales | GCA_018916865.1 | Coprococcus_comes | KQH93_07505 | 29424..32810 | 410072 | DUF499-domain-containing-protein-[Coprococcus-comes] |
| HJB61498.1 | DUF499_1.RM.like.systems | TRAM+Methyltransf_31-><-HEPN<-HEPN+DUF499*<-METHYLASE-p | 1134 | Bacteria>Terrabacteria_group>Bacillota>Clostridia>Eubacteriales | GCA_019119145.1 | Candidatus_Ruminococcus_gallistercoris | IAA38_07560 | 7302..10706 | 2838749 | DUF499-domain-containing-protein-[Candidatus-Ruminococcus-gallistercoris] |
| HJB96184.1 | DUF499_1.RM.like.systems | METHYLASE-p->HEPN+DUF499*-><-?<-PemK_toxin\|\|NACHT-> | 1128 | Bacteria>Terrabacteria_group>Bacillota>Clostridia>Eubacteriales | GCA_019119265.1 | Candidatus_Mediterraneibacter_intestinigallinarum | H9765_11315 | 720..4106 | 2838674 | DUF499-domain-containing-protein-[Candidatus-Mediterraneibacter-intestinigallinarum] |
| MBZ4669808.1 | DUF499_1.RM.like.systems | ABhydrolase+ABhydrolase->?->?->TUDOR+SF1-SF2-Helicase+iREase+CTD+REase->METHYLASE->HEPN+DUF499*->?-><-PIN<-PhdYeFM_antitox | 1133 | Bacteria>Terrabacteria_group>Bacillota>Clostridia>Eubacteriales | GCA_020054675.1 | Oscillospiraceae_bacterium | JG769_112 | 107395..110796 | 2485925 | hypothetical-protein-[Oscillospiraceae-bacterium] |
| MCC8151343.1 | DUF499_1.RM.like.systems | METHYLASE->HEPN+DUF499*->?->?->HTH+MPTase->PIN->SIGMA-HTH-> | 1132 | Bacteria>Terrabacteria_group>Bacillota>Clostridia>Eubacteriales | GCA_020860745.1 | Lachnospiraceae_bacterium | LIO96_07800 | 2635..6033 | 1898203 | DUF499-domain-containing-protein-[Lachnospiraceae-bacterium] |
| MCD7717395.1 | DUF499_1.RM.like.systems | REase->?->?-><-NACHT\|\|?-><-REase+ParB<-HEPN+DUF499*<-METHYLASE<-Schlafen+HATPase+HTH<-METHYLASE<-TUDOR+SF1-SF2-Helicase+iREase+CTD+REase<-XIS-HTH | 1132 | Bacteria>Terrabacteria_group>Bacillota>Clostridia>Eubacteriales | GCA_021199235.1 | Lachnospiraceae_bacterium | LUI39_13305 | 18966..22364 | 1898203 | DUF499-domain-containing-protein-[Lachnospiraceae-bacterium] |
| MCD7750767.1 | DUF499_1.RM.like.systems | <-NERD<-?<-?<-HEPN+DUF499*<-METHYLASE<-TUDOR+SF1-SF2-Helicase+iREase+CTD+REase<-XIS-HTH | 1127 | Bacteria>Terrabacteria_group>Bacillota>Clostridia>Eubacteriales | GCA_021199495.1 | Lachnospiraceae_bacterium | LUI10_03350 | 11139..14522 | 1898203 | DUF499-domain-containing-protein-[Lachnospiraceae-bacterium] |
| MCD7762788.1 | DUF499_1.RM.like.systems | <-NACHT\|\|?-><-MORC<-HEPN+DUF499*<-METHYLASE<-Schlafen+HATPase+HTH<-METHYLASE | 1132 | Bacteria>Terrabacteria_group>Bacillota>Clostridia>Eubacteriales | GCA_021199655.1 | Lachnospiraceae_bacterium | LUI14_06240 | 6036..9434 | 1898203 | DUF499-domain-containing-protein-[Lachnospiraceae-bacterium] |
| MCD7843035.1 | DUF499_1.RM.like.systems | METHYLASE->RelB->RelE-ParE->HEPN+DUF499*-> | 1126 | Bacteria>Terrabacteria_group>Bacillota>Clostridia>Eubacteriales | GCA_021200795.1 | Clostridiales_bacterium | LUG17_00205 | 3743..7123 | 1898207 | DUF499-domain-containing-protein-[Clostridiales-bacterium] |
| MCD8011085.1 | DUF499_1.RM.like.systems | <-HEPN+DUF499*<-RelE-ParE<-RelB<-METHYLASE-p<-METHYLASE-p<-GDResolvase<-GDResolvase+Recombinase+ZnR | 819 | Bacteria>Terrabacteria_group>Bacillota>Clostridia>Eubacteriales | GCA_021202235.1 | Lachnospiraceae_bacterium | LUF34_10055 | 34..2493 | 1898203 | DUF499-domain-containing-protein-[Lachnospiraceae-bacterium] |
| MCD8117856.1 | DUF499_1.RM.like.systems | <-HEPN+DUF499*<-ORC-CDC6+REase<-METHYLASE<-?<-?<-TUDOR+SF1-SF2-Helicase+iREase+CTD+REase | 1126 | Bacteria>Terrabacteria_group>Bacillota>Clostridia>Eubacteriales | GCA_021203045.1 | Oscillospiraceae_bacterium | LUE21_12275 | 415..3795 | 2485925 | DUF499-domain-containing-protein-[Oscillospiraceae-bacterium] |
| MCD8379771.1 | DUF499_1.RM.like.systems | <-HEPN+DUF499*<-METHYLASE-p | 474 | Bacteria>Terrabacteria_group>Bacillota>Clostridia>Eubacteriales | GCA_021205565.1 | Lachnospiraceae_bacterium | LUC95_05545 | 1..1422 | 1898203 | DUF499-domain-containing-protein-[Lachnospiraceae-bacterium] |
| MCH3999398.1 | DUF499_1.RM.like.systems | SbcC->HTH_26->TUDOR+SF1-SF2-Helicase+iREase+CTD+REase->METHYLASE->METHYLASE->HEPN+DUF499*->?->REase->HNH->?-><-?\|\|MarR-HTH-> | 1138 | Bacteria>Terrabacteria_group>Bacillota>Clostridia>Eubacteriales | GCA_022483165.1 | Lachnospiraceae_bacterium | LKE66_07755 | 1772421..1775837 | 1898203 | DUF499-domain-containing-protein-[Lachnospiraceae-bacterium] |
| MCI6268195.1 | DUF499_1.RM.like.systems | <-MerR-HTH<-?\|\|?->tRNA->HTH_26->TUDOR+SF1-SF2-Helicase+iREase+CTD+REase->METHYLASE->HEPN+DUF499*-> | 1130 | Bacteria>Terrabacteria_group>Bacillota>Clostridia>Eubacteriales | GCA_022772795.1 | Clostridiales_bacterium | MR637_06175 | 165246..168638 | 1898207 | DUF499-domain-containing-protein-[Clostridiales-bacterium] |
| MCI7107135.1 | DUF499_1.RM.like.systems | REase->?->?->?-><-ParB+HNH<-HEPN+DUF499*<-METHYLASE<-METHYLASE<-TUDOR+SF1-SF2-Helicase+iREase+CTD+REase<-HTH_26<-SbcC+SbcC | 1129 | Bacteria>Terrabacteria_group>Bacillota>Clostridia>Eubacteriales | GCA_022775045.1 | Agathobacter_sp. | MR945_04560 | 37097..40486 | 2021311 | DUF499-domain-containing-protein-[Agathobacter-sp.] |
| MCI7280004.1 | DUF499_1.RM.like.systems | METHYLASE->HEPN+DUF499*-><-?\|\|ZDNA-HTH->GDResolvase+Recombinase+ZnR-><-tRNA<-Peptidase_U32+Peptidase_U32_C<-?<-GNAT | 1129 | Bacteria>Terrabacteria_group>Bacillota>Clostridia>Eubacteriales | GCA_022783405.1 | Faecalibacterium_prausnitzii | MR538_06465 | 2347..5736 | 853 | DUF499-domain-containing-protein-[Faecalibacterium-prausnitzii] |
| MCI8809285.1 | DUF499_1.RM.like.systems | <-ABC-ATPase+SbcC<-?<-?\|\|TUDOR+SF1-SF2-Helicase+iREase+CTD+REase->METHYLASE->HEPN+DUF499*->Resolvase-> | 1137 | Bacteria>Terrabacteria_group>Bacillota>Clostridia>Eubacteriales | GCA_022795445.1 | Oscillibacter_sp. | HFF84_04010 | 14987..18400 | 1945593 | ATP-binding-protein-[Oscillibacter-sp.] |
| MCI9350231.1 | DUF499_1.RM.like.systems | TUDOR+SF1-SF2-Helicase+iREase+CTD+REase-p->?->METHYLASE->HEPN+DUF499*-> | 1134 | Bacteria>Terrabacteria_group>Bacillota>Clostridia>Eubacteriales | GCA_022799735.1 | Oscillibacter_sp. | HFF83_13605 | 3993..7397 | 1945593 | ATP-binding-protein-[Oscillibacter-sp.] |
| MCI9425971.1 | DUF499_1.RM.like.systems | METHYLASE->HEPN+DUF499*-> | 444 | Bacteria>Terrabacteria_group>Bacillota>Clostridia>Eubacteriales | GCA_022800375.1 | Flavonifractor_sp. | HFF30_10520 | 3535..4866 | 2049025 | ATP-binding-protein-[Flavonifractor-sp.] |
| MCM1579586.1 | DUF499_1.RM.like.systems | METHYLASE-p->HEPN+DUF499*-> | 410 | Bacteria>Terrabacteria_group>Bacillota>Clostridia>Eubacteriales | GCA_023667745.1 | Ruminococcus_sp. | NC078_12395 | 2153..3383 | 41978 | Swt1-family-HEPN-domain-containing-protein-[Ruminococcus-sp.] |
| MCR5263712.1 | DUF499_1.RM.like.systems | <-HTH\|\|?->?->TUDOR+SF1-SF2-Helicase+iREase+CTD+REase->METHYLASE->HEPN+DUF499*-> | 1118 | Bacteria>Terrabacteria_group>Bacillota>Clostridia>Eubacteriales | GCA_024695945.1 | Clostridiales_bacterium | K6D94_07550 | 8144..11500 | 1898207 | DUF499-domain-containing-protein-[Clostridiales-bacterium] |
| MCR5122540.1 | DUF499_1.RM.like.systems | METHYLASE-p->HEPN+DUF499*-> | 458 | Bacteria>Terrabacteria_group>Bacillota>Clostridia>Eubacteriales | GCA_024697705.1 | Ruminococcus_sp. | K6B74_09000 | 1756..3129 | 41978 | DUF499-domain-containing-protein-[Ruminococcus-sp.] |
| MCR5825960.1 | DUF499_1.RM.like.systems | <-HEPN+DUF499*<-METHYLASE | 853 | Bacteria>Terrabacteria_group>Bacillota>Clostridia>Eubacteriales | GCA_024698225.1 | Oscillospiraceae_bacterium | K6G54_05305 | 1..2559 | 2485925 | DUF499-domain-containing-protein-[Oscillospiraceae-bacterium] |
| MCR5807676.1 | DUF499_1.RM.like.systems | BirA-HTH->METHYLASE->HEPN+DUF499*-> | 732 | Bacteria>Terrabacteria_group>Bacillota>Clostridia>Eubacteriales | GCA_024698785.1 | Oscillospiraceae_bacterium | K6G68_11685 | 3761..5956 | 2485925 | DUF499-domain-containing-protein-[Oscillospiraceae-bacterium] |
| UYJ02009.1 | DUF499_1.RM.like.systems | TRAM+Methyltransf_31->AbiGi->TUDOR+SF1-SF2-Helicase+iREase+CTD+REase->MNS->METHYLASE->HEPN+DUF499*->ParB->CRP-HTH+GNTR-HTH->CRP-HTH+GNTR-HTH->?->?->ZDNA-HTH-> | 1130 | Bacteria>Terrabacteria_group>Bacillota>Clostridia>Eubacteriales | GCA_025757765.1 | Faecalibacterium_prausnitzii | OGM65_02940 | 601933..605325 | 853 | MAG:-Swt1-family-HEPN-domain-containing-protein-[Faecalibacterium-prausnitzii] |
| MDA8201143.1 | DUF499_1.RM.like.systems | <-RelE-ParE<-PhdYeFM_antitox\|\|DOC+HTH-><-?<-HEPN+DUF499*<-METHYLASE | 1093 | Bacteria>Terrabacteria_group>Bacillota>Clostridia>Eubacteriales | GCA_028065455.1 | Thermaerobacter_sp. | MOZ54_17205 | 4106..7387 | 1937527 | MAG:-Swt1-family-HEPN-domain-containing-protein-[Thermaerobacter-sp.] |
| WP_015942906.1 | DUF499_1.RM.like.systems | SF1-SF2-Helicase+HKD-DNase->MORC+SF1-SF2-Helicase->?->HTH_26->TUDOR+SF1-SF2-Helicase+iREase+CTD+REase->METHYLASE->HEPN+DUF499*->OLD-ABC+TOPRIM->SF1-SF2-Helicase->METHYLASE-p->ParB+HNH->HEPN->pPIWI+pPIWI-> | 1137 | Bacteria>Terrabacteria_group>Bacillota>Clostridia>Eubacteriales | GCF_000021925.1 | Desulfitobacterium_hafniense | DPCES_RS04010 | 854740..858153 | 49338 | DUF499-domain-containing-protein-[Desulfitobacterium-hafniense] |
| WP_006903458.1 | DUF499_1.RM.like.systems | TUDOR+SF1-SF2-Helicase+iREase+CTD+REase->METHYLASE->HEPN+DUF499*-> | 1128 | Bacteria>Terrabacteria_group>Bacillota>Clostridia>Eubacteriales | GCF_000183545.2 | Thermaerobacter_subterraneus | THESUDRAFT_RS04080 | 969973..973359 | 175696 | Swt1-family-HEPN-domain-containing-protein-[Thermaerobacter-subterraneus] |
| WP_008413292.1 | DUF499_1.RM.like.systems | <-HIN-HTH<-?<-TraJ-RHH<-HEPN+DUF499*<-METHYLASE<-TUDOR+SF1-SF2-Helicase+iREase+CTD+REase<-XIS-HTH<-?<-?<-HTH | 1135 | Bacteria>Terrabacteria_group>Bacillota>Clostridia>Eubacteriales | GCF_000315365.1 | Desulforamulus_hydrothermalis | BUA68_RS05960 | 1191..4598 | 412895 | Swt1-family-HEPN-domain-containing-protein-[Desulforamulus-hydrothermalis] |
| WP_018214270.1 | DUF499_1.RM.like.systems | <-pPIWI+pPIWI<-HEPN<-ParB+HNH<-METHYLASE-p<-Schlafen<-MORC<-HEPN+DUF499*<-METHYLASE<-TUDOR+SF1-SF2-Helicase+iREase+CTD+REase<-HTH_26<-?<-MORC+SF1-SF2-Helicase<-SF1-SF2-Helicase+HKD-DNase | 1137 | Bacteria>Terrabacteria_group>Bacillota>Clostridia>Eubacteriales | GCF_000378805.1 | Desulfitobacterium_hafniense | DESHAF_RS0122600 | 1962937..1966350 | 49338 | DUF499-domain-containing-protein-[Desulfitobacterium-hafniense] |
| WP_033117941.1 | DUF499_1.RM.like.systems | <-HTH\|\|?-><-?<-Response_reg+LuxR-HTH<-HISKIN+MORC<-?<-HEPN+DUF499*<-METHYLASE<-TUDOR+SF1-SF2-Helicase+iREase+CTD+REase<-XIS-HTH<-?\|\|?->GDResolvase-> | 1116 | Bacteria>Terrabacteria_group>Bacillota>Clostridia>Eubacteriales | GCF_000765135.1 | Intestinimonas | EOE41_RS12870 | 2605213..2608563 | 1392389 | MULTISPECIES:-DUF499-domain-containing-protein-[Intestinimonas] |
| WP_066668116.1 | DUF499_1.RM.like.systems | <-HD<-?\|\|?->?->?-><-HEPN+DUF499*<-METHYLASE<-PIN<-MetJarc-HTH<-TUDOR+SF1-SF2-Helicase+iREase+CTD+REase | 1130 | Bacteria>Terrabacteria_group>Bacillota>Clostridia>Eubacteriales | GCF_001655685.1 | Desulfotomaculum_copahuensis | A6M21_RS09370 | 52458..55850 | 1838280 | Swt1-family-HEPN-domain-containing-protein-[Desulfotomaculum-copahuensis] |
| WP_087179373.1 | DUF499_1.RM.like.systems | <-NACHT\|\|PemK_toxin->?-><-HEPN+DUF499*<-METHYLASE<-METHYLASE<-TUDOR+SF1-SF2-Helicase+iREase+CTD+REase<-XIS-HTH | 1128 | Bacteria>Terrabacteria_group>Bacillota>Clostridia>Eubacteriales | GCF_002161355.1 | Lachnoclostridium_sp._An76 | B5G27_RS13410 | 49376..52762 | 1965654 | DUF499-domain-containing-protein-[Lachnoclostridium-sp.-An76] |
| WP_135225812.1 | DUF499_1.RM.like.systems | <-HEPN+DUF499*<-METHYLASE<-TUDOR+SF1-SF2-Helicase+iREase+CTD+REase\|\|MarR-HTH->ABC-ATPase+SbcC-> | 1129 | Bacteria>Terrabacteria_group>Bacillota>Clostridia>Eubacteriales | GCF_004634385.1 | Thermaerobacter_sp._FW80 | E1B22_RS11885 | 2873077..2876466 | 2546351 | Swt1-family-HEPN-domain-containing-protein-[Thermaerobacter-sp.-FW80] |
| WP_166510781.1 | DUF499_1.RM.like.systems | <-HISKIN<-?<-?<-MPTase<-HTH<-?<-HEPN+DUF499*<-METHYLASE-p<-TUDOR+SF1-SF2-Helicase+iREase+CTD+REase-p | 1122 | Bacteria>Terrabacteria_group>Bacillota>Clostridia>Eubacteriales | GCF_008124625.1 | Desulfallas_thermosapovorans | LX24_RS03620 | 91122..94490 | 58137 | Swt1-family-HEPN-domain-containing-protein-[Desulfallas-thermosapovorans] |
| WP_166512732.1 | DUF499_1.RM.like.systems | <-SF1-SF2-Helicase+DUF1998<-HEPN+Mrr_N<-HDRnase<-?<-Primpol+D5N+D5ATPase<-?<-HEPN+DUF499*<-METHYLASE<-TUDOR+SF1-SF2-Helicase+iREase+CTD+REase<-Peptidase_M15 | 1110 | Bacteria>Terrabacteria_group>Bacillota>Clostridia>Eubacteriales | GCF_008124625.1 | Desulfallas_thermosapovorans | LX24_RS13825 | 12605..15937 | 58137 | Swt1-family-HEPN-domain-containing-protein-[Desulfallas-thermosapovorans] |
| WP_161846233.1 | DUF499_1.RM.like.systems | TUDOR+SF1-SF2-Helicase+iREase+CTD+REase->?->METHYLASE->HEPN+DUF499*->REase-><-GDResolvase | 1134 | Bacteria>Terrabacteria_group>Bacillota>Clostridia>Eubacteriales | GCF_009935845.1 | Pseudoflavonifractor_sp._524-17 | D1159_RS13725 | 7029..10433 | 2304577 | DUF499-domain-containing-protein-[Pseudoflavonifractor-sp.-524-17] |
| WP_168124741.1 | DUF499_1.RM.like.systems | <-HEPN+DUF499*<-METHYLASE<-METHYLASE<-Schlafen+HATPase+HTH<-TUDOR+SF1-SF2-Helicase+iREase+CTD+REase<-HTH_26<-SIGMA-HTH | 1132 | Bacteria>Terrabacteria_group>Bacillota>Clostridia>Eubacteriales | GCF_012027765.1 | Pseudoflavonifractor_sp._SW1122 | E0L15_RS04955 | 37276..40674 | 2530044 | DUF499-domain-containing-protein-[Pseudoflavonifractor-sp.-SW1122] |
| WP_228724972.1 | DUF499_1.RM.like.systems | HTH_36-><-?<-?<-?<-JetA<-HEPN+DUF499*<-METHYLASE<-METHYLASE-p<-TUDOR+SF1-SF2-Helicase+iREase+CTD+REase<-CHTH | 1117 | Bacteria>Terrabacteria_group>Bacillota>Clostridia>Eubacteriales | GCF_014303955.1 | Caproicibacter_fermentans | HCR03_RS14035 | 2844426..2847779 | 2576756 | DUF499-domain-containing-protein-[Caproicibacter-fermentans] |
| WP_195975511.1 | DUF499_1.RM.like.systems | HTH_26->TUDOR+SF1-SF2-Helicase+iREase+CTD+REase->METHYLASE->METHYLASE->HEPN+DUF499*-><-?<-GDResolvase+Recombinase+ZnR<-CHTH | 1134 | Bacteria>Terrabacteria_group>Bacillota>Clostridia>Eubacteriales | GCF_015670165.1 | Hydrogeniiclostridium_mannosilyticum | I4V57_RS07180 | 144125..147529 | 2764322 | DUF499-domain-containing-protein-[Hydrogeniiclostridium-mannosilyticum] |
| WP_204872058.1 | DUF499_1.RM.like.systems | SIGMA-HTH->HTH_26->TUDOR+SF1-SF2-Helicase+iREase+CTD+REase->Schlafen+HATPase+HTH->METHYLASE->METHYLASE->HEPN+DUF499*-> | 1132 | Bacteria>Terrabacteria_group>Bacillota>Clostridia>Eubacteriales | GCF_016899615.1 | Pseudoflavonifractor_capillosus | H9X87_RS10190 | 14789..18187 | 106588 | DUF499-domain-containing-protein-[Pseudoflavonifractor-capillosus] |
| WP_204889591.1 | DUF499_1.RM.like.systems | HTH_26->TUDOR+SF1-SF2-Helicase+iREase+CTD+REase->METHYLASE->Schlafen+HATPase+HTH->HEPN+DUF499*-><-?<-HIN-HTH\|\|DUF772+DDE_Tnp_1+DDE_Tnp_1_6-> | 1123 | Bacteria>Terrabacteria_group>Bacillota>Clostridia>Eubacteriales | GCF_016902175.1 | Pseudoflavonifractor_phocaeensis | H6B27_RS11850 | 9172..12543 | 1870988 | Swt1-family-HEPN-domain-containing-protein-[Pseudoflavonifractor-phocaeensis] |
| WP_207684444.1 | DUF499_1.RM.like.systems | GDResolvase->AbiGi->GDResolvase+Recombinase+ZnR->TUDOR+SF1-SF2-Helicase+iREase+CTD+REase->?->METHYLASE->HEPN+DUF499*->ParB->CRP-HTH+GNTR-HTH->?->?->ZDNA-HTH->GDResolvase+Recombinase+ZnR-> | 1129 | Bacteria>Terrabacteria_group>Bacillota>Clostridia>Eubacteriales | GCF_017377535.1 | Faecalibacterium_sp._Marseille-Q4137 | J3353_RS06385 | 45904..49293 | 2817021 | DUF499-domain-containing-protein-[Faecalibacterium-sp.-Marseille-Q4137] |
| WP_211802648.1 | DUF499_1.RM.like.systems | REase+CTD6->?->SbcC+SbcC->HTH_26->TUDOR+SF1-SF2-Helicase+iREase+CTD+REase->METHYLASE->HEPN+DUF499*->SF1-SF2-Helicase-p-> | 1134 | Bacteria>Terrabacteria_group>Bacillota>Clostridia>Eubacteriales | GCF_018128025.1 | Proteiniclasticum_sediminis | KCG48_RS13075 | 34929..38333 | 2804028 | DUF499-domain-containing-protein-[Proteiniclasticum-sediminis] |
| WP_156058230.1 | DUF499_1.RM.like.systems | AbiGi->GDResolvase+Recombinase+ZnR->TUDOR+SF1-SF2-Helicase+iREase+CTD+REase->TUDOR+SF1-SF2-Helicase+iREase+CTD+REase-p->MNS->METHYLASE->HEPN+DUF499*->?->ParB->CRP-HTH+GNTR-HTH->?->CHTH->GDResolvase+Recombinase+ZnR-> | 1130 | Bacteria>Terrabacteria_group>Bacillota>Clostridia>Eubacteriales | GCF_019969385.1 | Faecalibacterium_prausnitzii | GD822_RS02960 | 605262..608654 | 853 | Swt1-family-HEPN-domain-containing-protein-[Faecalibacterium-prausnitzii] |
| WP_092487583.1 | DUF499_1.RM.like.systems | HIN-HTH-><-DDE_Tnp_1<-HEPN+DUF499*<-PIN<-SbcC<-METHYLASE<-TUDOR+SF1-SF2-Helicase+iREase+CTD+REase | 888 | Bacteria>Terrabacteria_group>Bacillota>Clostridia>Eubacteriales | GCF_900115975.1 | Desulfoscipio_geothermicus | BM299_RS17670 | 2600..5266 | 39060 | Swt1-family-HEPN-domain-containing-protein-[Desulfoscipio-geothermicus] |
| WP_076780282.1 | DUF499_1.RM.like.systems | HDRnase->TRAM+Methyltransf_31-><-?<-?<-HEPN+DUF499*<-METHYLASE<-METHYLASE<-TUDOR+SF1-SF2-Helicase+iREase+CTD+REase<-HTH_26<-SIGMA-HTH+SIGMA-HTH<-HTH | 1134 | Bacteria>Terrabacteria_group>Bacillota>Clostridia>Eubacteriales | GCF_900120345.1 | Lachnoclostridium_phocaeense | BQ7370_RS15745 | 3233710..3237114 | 1871021 | DUF499-domain-containing-protein-[Lachnoclostridium-phocaeense] |
| WP_087282257.1 | DUF499_1.RM.like.systems | XIS-HTH->TUDOR+SF1-SF2-Helicase+iREase+CTD+REase->METHYLASE->METHYLASE->HEPN+DUF499*->?->SbcC->SF1-SF2-Helicase->WYL->?->LexA-HTH+LexA-protease-> | 1134 | Bacteria>Terrabacteria_group>Bacillota>Clostridia | GCF_016902345.1 | Eubacteriales | H6B11_RS01180 | 245745..249149 | 186802 | MULTISPECIES:-DUF499-domain-containing-protein-[Eubacteriales] |
| WP_071522349.1 | DUF499_1.RM.like.systems | PolB->HEPN->\|\|?->?->TUDOR+SF1-SF2-Helicase+iREase+CTD+REase->METHYLASE->HEPN+DUF499*->NERD-> | 1122 | Bacteria>Terrabacteria_group>Bacillota>Clostridia>Moorellales | GCF_001874085.1 | Moorella_thermoacetica | MOOTH_RS08230 | 71675..75043 | 1525 | Swt1-family-HEPN-domain-containing-protein-[Moorella-thermoacetica] |
| WP_156275504.1 | DUF499_1.RM.like.systems | <-CopG_antitoxin\|\|NTP_transf_2-><-?<-?<-Schlafen<-HEPN+DUF499*<-METHYLASE<-TUDOR+SF1-SF2-Helicase+iREase+CTD+REase\|\|?-><-?<-Phage_terminase | 1122 | Bacteria>Terrabacteria_group>Bacillota>Clostridia>Moorellales | GCF_009735625.1 | Moorella_glycerini | MGLY_RS15905 | 3178613..3181981 | 55779 | DUF499-domain-containing-protein-[Moorella-glycerini] |
| WP_013782413.1 | DUF499_1.RM.like.systems | TUDOR+SF1-SF2-Helicase+iREase+CTD+REase->METHYLASE->HEPN+DUF499*->?->XIS-HTH->BirA-HTH->?->Primpol+PriCT1->ZnR+TOPRIM+D5ATPase-> | 1118 | Bacteria>Terrabacteria_group>Bacillota>Clostridia>Thermoanaerobacterales | GCF_000213255.1 | Mahella_australiensis | MAHAU_RS14260 | 3047947..3051303 | 252966 | Swt1-family-HEPN-domain-containing-protein-[Mahella-australiensis] |
| WP_221039750.1 | DUF499_1.RM.like.systems | <-REase<-HEPN+DUF499*<-METHYLASE<-TUDOR+SF1-SF2-Helicase+iREase+CTD+REase<-McrC-NTD+REase | 1133 | Bacteria>Terrabacteria_group>Bacillota>Clostridia>Thermoanaerobacterales | GCF_019668485.1 | Gelria_sp._Kuro-4 | K5554_RS03460 | 654715..658116 | 2796927 | Swt1-family-HEPN-domain-containing-protein-[Gelria-sp.-Kuro-4] |
| HHY05947.1 | DUF499_1.RM.like.systems | <-TRAM+Methyltransf_31<-?<-HEPN+DUF499*<-METHYLASE<-TUDOR+SF1-SF2-Helicase+iREase+CTD+REase | 1125 | Bacteria>Terrabacteria_group>Bacillota>Clostridia>unclassified_Clostridia | GCA_012842035.1 | Clostridia_bacterium | GX532_03075 | 2394..5771 | 2044939 | ATP-binding-protein-[Clostridia-bacterium] |
| MBQ6961345.1 | DUF499_1.RM.like.systems | <-SF1-SF2-Helicase-p<-OLD-ABC+TOPRIM<-HEPN+DUF499*<-HEPN<-PolB<-METHYLASE<-MNS<-TUDOR+SF1-SF2-Helicase+iREase+CTD+REase<-HTH | 1126 | Bacteria>Terrabacteria_group>Bacillota>Clostridia>unclassified_Clostridia | GCA_017448975.1 | Clostridia_bacterium | IJP78_10255 | 13529..16909 | 2044939 | DUF499-domain-containing-protein-[Clostridia-bacterium] |
| GHU66463.1 | DUF499_1.RM.like.systems | ABhydrolase->TUDOR+SF1-SF2-Helicase+iREase+CTD+REase->NTP_transf->HEPN->METHYLASE->HEPN+DUF499*->?->?->?-><-PADR-HTH | 1148 | Bacteria>Terrabacteria_group>Bacillota>Clostridia>unclassified_Clostridia | GCA_026000995.1 | Clostridia_bacterium | FACS1894184_04510 | 37803..41249 | 2044939 | hypothetical-protein-FACS1894184_04510-[Clostridia-bacterium] |
| MBS3947521.1 | DUF499_1.RM.like.systems | <-HEPN+DUF499*<-HEPN<-Polbeta<-METHYLASE-p | 1118 | Bacteria>Terrabacteria_group>Bacillota>Dethiobacteria>Dethiobacterales | GCA_018333395.1 | Dethiobacter_sp. | KGZ57_04310 | 478..3834 | 2093370 | DUF499-domain-containing-protein-[Dethiobacter-sp.] |
| WP_281790372.1 | DUF499_1.RM.like.systems | <-HEPN+DUF499*<-METHYLASE | 1106 | Bacteria>Terrabacteria_group>Bacillota>Erysipelotrichia>Erysipelotrichales | GCF_937916425.1 | Faecalibaculum_rodentium | QU443_RS10445 | 250..3570 | 1702221 | DUF499-domain-containing-protein-[Faecalibaculum-rodentium] |
| WP_285944649.1 | DUF499_1.RM.like.systems | <-HEPN+DUF499*<-METHYLASE | 1106 | Bacteria>Terrabacteria_group>Bacillota>Erysipelotrichia>Erysipelotrichales | GCF_949383055.1 | Faecalibaculum_rodentium | QTM32_RS09735 | 250..3570 | 1702221 | DUF499-domain-containing-protein-[Faecalibaculum-rodentium] |
| MBE3584147.1 | DUF499_1.RM.like.systems | <-METHYLASE-p\|\|?->TUDOR+SF1-SF2-Helicase+iREase+CTD+REase->OLD-ABC+TOPRIM->METHYLASE->HEPN+DUF499*-><-?\|\|ABC-ATPase+SbcC->ABC-ATPase+SbcC-><-MazE_antitoxin | 1113 | Bacteria>Terrabacteria_group>Bacillota>Limnochordia>Limnochordales | GCA_014896305.1 | Limnochordaceae_bacterium | IMX01_08555 | 27896..31237 | 2773236 | DUF499-domain-containing-protein-[Limnochordaceae-bacterium] |
| MCK9524538.1 | DUF499_1.RM.like.systems | ArdCN+MPTase->?-><-?<-REase<-?<-REase<-HEPN+DUF499*<-METHYLASE<-TUDOR+SF1-SF2-Helicase+iREase+CTD+REase-p | 1113 | Bacteria>Terrabacteria_group>Bacillota>Limnochordia>unclassified_Limnochordia | GCA_023228555.1 | Limnochordia_bacterium | MOR49_01235 | 6495..9836 | 2497621 | Swt1-family-HEPN-domain-containing-protein-[Limnochordia-bacterium] |
| PWM79769.1 | DUF499_1.RM.like.systems | Peptidase_S8->?-><-?<-?<-STYKinase<-HEPN+DUF499*<-METHYLASE<-METHYLASE<-Schlafen+HATPase+HTH<-TUDOR+SF1-SF2-Helicase+iREase+CTD+REase<-HTH_26 | 1130 | Bacteria>Terrabacteria_group>Bacillota>Negativicutes>Acidaminococcales | GCA_003150755.1 | Phascolarctobacterium_sp. | DBY32_02635 | 546389..549781 | 2049039 | MAG:-hypothetical-protein-DBY32_02635-[Phascolarctobacterium-sp.] |
| MBR6861287.1 | DUF499_1.RM.like.systems | RADICAL-SAM-><-HEPN+DUF499*<-METHYLASE<-METHYLASE<-TUDOR+SF1-SF2-Helicase+iREase+CTD+REase<-?<-?<-REase+35exo | 1123 | Bacteria>Terrabacteria_group>Bacillota>Negativicutes>Acidaminococcales | GCA_017531245.1 | Acidaminococcaceae_bacterium | IKM73_08200 | 27143..30514 | 2049052 | DUF499-domain-containing-protein-[Acidaminococcaceae-bacterium] |
| WP_232709101.1 | DUF499_1.RM.like.systems | NUDIX->TUDOR+SF1-SF2-Helicase+iREase+CTD+REase->METHYLASE->METHYLASE-p->HIN-HTH->METHYLASE->HEPN+DUF499*->HTH_3->?->SIGMA-HTH-> | 1130 | Bacteria>Terrabacteria_group>Bacillota>Negativicutes>Acidaminococcales | GCF_902376135.1 | Acidaminococcus_massiliensis | BQ2657_RS09005 | 1875868..1879260 | 1852375 | DUF499-domain-containing-protein-[Acidaminococcus-massiliensis] |
| WP_279002420.1 | DUF499_1.RM.like.systems | NUDIX->?->TUDOR+SF1-SF2-Helicase+iREase+CTD+REase->METHYLASE->METHYLASE->HEPN+DUF499*->HTH_3-> | 1145 | Bacteria>Terrabacteria_group>Bacillota>Negativicutes>Acidaminococcales | GCF_945915305.1 | Acidaminococcus_fermentans | QDM86_RS09090 | 15622..19059 | 905 | DUF499-domain-containing-protein-[Acidaminococcus-fermentans] |
| MBF1662768.1 | DUF499_1.RM.like.systems | Classical-AAA->Peptidase_S8-><-?<-?<-?<-HEPN+DUF499*<-METHYLASE<-TUDOR+SF1-SF2-Helicase+iREase+CTD+REase<-?<-PfkB | 1145 | Bacteria>Terrabacteria_group>Bacillota>Negativicutes>Selenomonadales | GCA_015259755.1 | Selenomonas_noxia | HXO76_05010 | 11846..15283 | 135083 | ATP-binding-protein-[Selenomonas-noxia] |
| MBF1688626.1 | DUF499_1.RM.like.systems | TUDOR+SF1-SF2-Helicase+iREase+CTD+REase-p->METHYLASE->HEPN+DUF499*-> | 887 | Bacteria>Terrabacteria_group>Bacillota>Negativicutes>Selenomonadales | GCA_015265195.1 | Selenomonas_sp. | HXO83_10375 | 4201..6861 | 2053611 | ATP-binding-protein-[Selenomonas-sp.] |
| MBF1693417.1 | DUF499_1.RM.like.systems | TUDOR+SF1-SF2-Helicase+iREase+CTD+REase-p->HEPN+DUF499*-><-ASCH<-PIN+ACET+EVE<-?<-Classical-AAA | 1142 | Bacteria>Terrabacteria_group>Bacillota>Negativicutes>Selenomonadales | GCA_015265235.1 | Selenomonas_sp. | HXO82_03045 | 3480..6908 | 2053611 | ATP-binding-protein-[Selenomonas-sp.] |
| MBQ6005594.1 | DUF499_1.RM.like.systems | TUDOR+SF1-SF2-Helicase+iREase+CTD+REase->METHYLASE->HEPN+DUF499*-> | 1081 | Bacteria>Terrabacteria_group>Bacillota>Negativicutes>Selenomonadales | GCA_017427545.1 | Selenomonadaceae_bacterium | IJL14_05025 | 8141..11386 | 2652289 | DUF499-domain-containing-protein-[Selenomonadaceae-bacterium] |
| MBR0260489.1 | DUF499_1.RM.like.systems | METHYLASE->HEPN+DUF499*->sGTP->SIGMA-HTH+SIGMA-HTH->SIGMA-HTH+SIGMA-HTH-> | 1077 | Bacteria>Terrabacteria_group>Bacillota>Negativicutes>Selenomonadales | GCA_017446845.1 | Selenomonadaceae_bacterium | IJQ85_01715 | 15691..18924 | 2652289 | DUF499-domain-containing-protein-[Selenomonadaceae-bacterium] |
| MBR0290429.1 | DUF499_1.RM.like.systems | TUDOR+SF1-SF2-Helicase+iREase+CTD+REase->METHYLASE->HEPN+DUF499*->DUF499-p-> | 334 | Bacteria>Terrabacteria_group>Bacillota>Negativicutes>Selenomonadales | GCA_017446965.1 | Selenomonadaceae_bacterium | IJQ82_15765 | 5468..6472 | 2652289 | hypothetical-protein-[Selenomonadaceae-bacterium] |
| MBR0060910.1 | DUF499_1.RM.like.systems | METHYLASE->HEPN+DUF499*-> | 1084 | Bacteria>Terrabacteria_group>Bacillota>Negativicutes>Selenomonadales | GCA_017449105.1 | Selenomonadaceae_bacterium | IJP68_05465 | 1758..5012 | 2652289 | DUF499-domain-containing-protein-[Selenomonadaceae-bacterium] |
| MBQ9480498.1 | DUF499_1.RM.like.systems | <-HEPN+DUF499*<-?<-BrnT_toxin<-METHYLASE-p | 465 | Bacteria>Terrabacteria_group>Bacillota>Negativicutes>Selenomonadales | GCA_017454185.1 | Selenomonadaceae_bacterium | IJU71_13220 | 1..1395 | 2652289 | DUF499-domain-containing-protein-[Selenomonadaceae-bacterium] |
| MBQ7497714.1 | DUF499_1.RM.like.systems | <-DOC+HTH\|\|DeoR-HTH+ISOCOT-><-HEPN+DUF499*<-METHYLASE | 1125 | Bacteria>Terrabacteria_group>Bacillota>Negativicutes>Selenomonadales | GCA_017455465.1 | Selenomonas_sp. | IJU00_07780 | 3332..6709 | 2053611 | DUF499-domain-containing-protein-[Selenomonas-sp.] |
| MBR1580270.1 | DUF499_1.RM.like.systems | <-Peptidase_M30<-HEPN+DUF499*<-?<-NTP_transf<-METHYLASE | 1098 | Bacteria>Terrabacteria_group>Bacillota>Negativicutes>Selenomonadales | GCA_017477125.1 | Selenomonadaceae_bacterium | IJ668_07210 | 12971..16267 | 2652289 | hypothetical-protein-[Selenomonadaceae-bacterium] |
| MBR1807330.1 | DUF499_1.RM.like.systems | <-HEPN+DUF499*<-METHYLASE<-METHYLASE-p | 1085 | Bacteria>Terrabacteria_group>Bacillota>Negativicutes>Selenomonadales | GCA_017479925.1 | Selenomonadaceae_bacterium | IJ774_13210 | 2314..5571 | 2652289 | DUF499-domain-containing-protein-[Selenomonadaceae-bacterium] |
| MBR3051424.1 | DUF499_1.RM.like.systems | ABC-ATPase+SbcC-><-HEPN+DUF499*<-METHYLASE-p | 1077 | Bacteria>Terrabacteria_group>Bacillota>Negativicutes>Selenomonadales | GCA_017510185.1 | Selenomonadaceae_bacterium | IKG61_08245 | 624..3857 | 2652289 | DUF499-domain-containing-protein-[Selenomonadaceae-bacterium] |
| MBR7025847.1 | DUF499_1.RM.like.systems | ABC-ATPase+SbcC-><-?\|\|TUDOR+SF1-SF2-Helicase+iREase+CTD+REase->METHYLASE->HEPN+DUF499*-><-ABC-ATPase+SbcC<-ABC-ATPase+SbcC | 1077 | Bacteria>Terrabacteria_group>Bacillota>Negativicutes>Selenomonadales | GCA_017518905.1 | Selenomonadaceae_bacterium | IKI08_07585 | 26267..29500 | 2652289 | DUF499-domain-containing-protein-[Selenomonadaceae-bacterium] |
| MBR3623059.1 | DUF499_1.RM.like.systems | <-HEPN+DUF499*<-NERD<-OLD-ABC<-METHYLASE-p | 509 | Bacteria>Terrabacteria_group>Bacillota>Negativicutes>Selenomonadales | GCA_017529085.1 | Selenomonadaceae_bacterium | IKN43_06910 | 1..1529 | 2652289 | DUF499-domain-containing-protein-[Selenomonadaceae-bacterium] |

| Protein-ID | System | Gene-Neighbourhoods | Length | Taxonomy | Assembly-ID | Strain | Locus-Tag | Genomic-coordinates | TaxID | Fasta-defline |
|---|---|---|---|---|---|---|---|---|---|---|
| MBR3746746.1 | DUF499_1.RM.like.systems | <-HEPN+DUF499*<-METHYLASE-p | 1077 | Bacteria>Terrabacteria_group>Bacillota>Negativicutes>Selenomonadales | GCA_017529545.1 | Selenomonadaceae_bacterium | IKN27_07285 | 344..3577 | 2652289 | DUF499-domain-containing-protein-[Selenomonadaceae-bacterium] |
| MBR3723747.1 | DUF499_1.RM.like.systems | <-HEPN+DUF499*<-NERD<-OLD-ABC<-METHYLASE | 503 | Bacteria>Terrabacteria_group>Bacillota>Negativicutes>Selenomonadales | GCA_017530025.1 | Selenomonadaceae_bacterium | IKN12_13455 | 1..1509 | 2652289 | DUF499-domain-containing-protein-[Selenomonadaceae-bacterium] |
| MBR4642051.1 | DUF499_1.RM.like.systems | SF1-SF2-Helicase-p->REase->?->?->METHYLASE-p->METHYLASE->HEPN+DUF499*-> | 1118 | Bacteria>Terrabacteria_group>Bacillota>Negativicutes>Selenomonadales | GCA_017535925.1 | Selenomonadaceae_bacterium | IKO74_04930 | 6896..10252 | 2652289 | hypothetical-protein-[Selenomonadaceae-bacterium] |
| MCI6099016.1 | DUF499_1.RM.like.systems | TUDOR+SF1-SF2-Helicase+iREase+CTD+REase->Transposase_31->METHYLASE->HEPN+DUF499*-> | 1147 | Bacteria>Terrabacteria_group>Bacillota>Negativicutes>Selenomonadales | GCA_022770905.1 | Selenomonas_sp. | MR698_00025 | 8556..11999 | 2053611 | DUF499-domain-containing-protein-[Selenomonas-sp.] |
| MCI6231067.1 | DUF499_1.RM.like.systems | TUDOR+SF1-SF2-Helicase+iREase+CTD+REase->Transposase_31->METHYLASE->HEPN+DUF499*-> | 1147 | Bacteria>Terrabacteria_group>Bacillota>Negativicutes>Selenomonadales | GCA_022773585.1 | Selenomonas_sp. | MR630_00380 | 8523..11966 | 2053611 | DUF499-domain-containing-protein-[Selenomonas-sp.] |
| WP_007288290.1 | DUF499_1.RM.like.systems | Calcineurin->SbcC+SbcC-><-?<-?<-HEPN+DUF499*<-METHYLASE\|\|DDE_Tnp-><-?<-HTH_21+rve+rve_3<-HIN-HTH<-RNase | 1126 | Bacteria>Terrabacteria_group>Bacillota>Negativicutes>Selenomonadales | GCF_000169155.1 | Thermosinus_carboxydivorans | TCARDRAFT_RS01740 | 31494..34874 | 261685 | Swt1-family-HEPN-domain-containing-protein-[Thermosinus-carboxydivorans] |
| WP_009350157.1 | DUF499_1.RM.like.systems | XIS-HTH->HIN-HTH->?-><-CHTH<-HEPN+DUF499*<-METHYLASE<-TUDOR+SF1-SF2-Helicase+iREase+CTD+REase<-?\|\|DOC+HTH-><-MACRODOMAIN | 1142 | Bacteria>Terrabacteria_group>Bacillota>Negativicutes>Selenomonadales | GCF_000187125.1 | Selenomonas_artemidis | HMPREF9555_RS07020 | 6182..9610 | 671224 | DUF499-domain-containing-protein-[Selenomonas-artemidis] |
| WP_031587665.1 | DUF499_1.RM.like.systems | <-HEPN+DUF499*<-METHYLASE<-?<-TUDOR+SF1-SF2-Helicase+iREase+CTD+REase | 1132 | Bacteria>Terrabacteria_group>Bacillota>Negativicutes>Selenomonadales | GCF_000701885.1 | Selenomonas_bovis | T485_RS0110635 | 1077..4475 | 416586 | DUF499-domain-containing-protein-[Selenomonas-bovis] |
| WP_061945835.1 | DUF499_1.RM.like.systems | METHYLASE-p->?->METHYLASE-p->METHYLASE-p->METHYLASE-p->HEPN+DUF499*->tRNA-><-REase<-Phage_integrase\|\|?-><-?\|\|DUF772+DDE_Tnp_1_6-> | 377 | Bacteria>Terrabacteria_group>Bacillota>Negativicutes>Selenomonadales | GCF_001554015.1 | Selenomonas_sp._oral_taxon_136 | AXE86_RS06860 | 1418306..1419439 | 713030 | Swt1-family-HEPN-domain-containing-protein-[Selenomonas-sp.-oral-taxon-136] |
| WP_264918710.1 | DUF499_1.RM.like.systems | <-Mbetalac\|\|GDResolvase+HIN-HTH->TUDOR+SF1-SF2-Helicase+iREase+CTD+REase->METHYLASE->HEPN+DUF499*->?->Peptidase->?->sGTP->?->Aldedh-> | 1127 | Bacteria>Terrabacteria_group>Bacillota>Negativicutes>Selenomonadales | GCF_026013845.1 | Selenomonas_sputigena | OL240_RS12405 | 2695256..2698639 | 69823 | DUF499-domain-containing-protein-[Selenomonas-sputigena] |
| WP_270837292.1 | DUF499_1.RM.like.systems | <-MULE-TRANSPOSASE\|\|HTH_OrfB_IS605+OrfB_IS605+OrfB_Zn_ribbon->?->TUDOR+SF1-SF2-Helicase+iREase+CTD+REase->METHYLASE->METHYLASE->HEPN+DUF499*-><-DDE_Tnp_IS66+DDE_Tnp_IS66_C<-?<-HIN-HTH | 1134 | Bacteria>Terrabacteria_group>Bacillota>Negativicutes>Selenomonadales | GCF_027670005.1 | Mitsuokella_multacida | PHA28_RS02250 | 15218..18622 | 52226 | DUF499-domain-containing-protein-[Mitsuokella-multacida] |
| WP_270528574.1 | DUF499_1.RM.like.systems | <-35exo\|\|?-><-?\|\|?->?-><-HEPN+DUF499*<-METHYLASE<-TUDOR+SF1-SF2-Helicase+iREase+CTD+REase<-?<-?<-?<-Methylase | 1161 | Bacteria>Terrabacteria_group>Bacillota>Negativicutes>Selenomonadales | GCF_027681165.1 | Mitsuokella_jalaludinii | PGX61_RS07200 | 41318..44803 | 187979 | DUF499-domain-containing-protein-[Mitsuokella-jalaludinii] |
| WP_278503476.1 | DUF499_1.RM.like.systems | Pkinase-><-?<-HEPN+DUF499*<-METHYLASE<-METHYLASE<-TUDOR+SF1-SF2-Helicase+iREase+CTD+REase<-?<-HTH_OrfB_IS605+OrfB_IS605+OrfB_Zn_ribbon<-PhdYeFM_antitox | 1134 | Bacteria>Terrabacteria_group>Bacillota>Negativicutes>Selenomonadales | GCF_905193285.1 | Mitsuokella_multacida | QMF84_RS08000 | 50365..53769 | 52226 | DUF499-domain-containing-protein-[Mitsuokella-multacida] |
| WP_204967821.1 | DUF499_1.RM.like.systems | TRD+TRD->GDResolvase+Recombinase+ZnR->HTH_26->TUDOR+SF1-SF2-Helicase+iREase+CTD+REase->METHYLASE->METHYLASE->HEPN+DUF499*->?->?->?->TPR+TPR-> | 1132 | Bacteria>Terrabacteria_group>Bacillota>Negativicutes>Veillonellales | GCF_016900675.1 | Megasphaera_stantonii | H6G85_RS06205 | 65633..69031 | 2144175 | DUF499-domain-containing-protein-[Megasphaera-stantonii] |
| WP_008860390.1 | DUF499_1.RM.like.systems | Phage_integrase->Phage_integrase->?->TUDOR+SF1-SF2-Helicase+iREase+CTD+REase->METHYLASE->METHYLASE->HEPN+DUF499*->?-><-REase+SF1-SF2-Helicase<-REase+SF1-SF2-Helicase\|\|MazG-> | 1135 | Bacteria>Terrabacteria_group>Bacillota>Negativicutes>Veillonellales | GCF_902374635.1 | Dialister_succinatiphilus | HMPREF9453_RS09730 | 179520..182927 | 487173 | DUF499-domain-containing-protein-[Dialister-succinatiphilus] |
| OQB47592.1 | DUF499_1.RM.like.systems | DSBH-><-?\|\|tRNA->REase-><-HEPN+DUF499*<-METHYLASE-p<-TUDOR+SF1-SF2-Helicase+iREase+CTD+REase | 1103 | Bacteria>Terrabacteria_group>Bacillota>unclassified_Bacillota | GCA_002070415.1 | Firmicutes_bacterium_ADurb.Bin153 | BWY00_01229 | 26495..29806 | 1852883 | hypothetical-protein-BWY00_01229-[Firmicutes-bacterium-ADurb.Bin153] |
| HBL50377.1 | DUF499_1.RM.like.systems | <-SF1-SF2-Helicase<-OLD-ABC+TOPRIM<-HEPN+DUF499*<-METHYLASE<-TUDOR+SF1-SF2-Helicase+iREase+CTD+REase<-?<-MORC+DUF3684+HNH<-SF1-SF2-Helicase-p<-HEPN+SF1-SF2-Helicase | 1129 | Bacteria>Terrabacteria_group>Bacillota>unclassified_Bacillota | GCA_003504715.1 | Bacillota_bacterium | DDZ65_09075 | 4304..7693 | 1879010 | hypothetical-protein-[Firmicutes-bacterium] |
| MTI83942.1 | DUF499_1.RM.like.systems | <-SF1-SF2-Helicase+REase<-PIWI<-HEPN+DUF499*<-METHYLASE-p | 1111 | Bacteria>Terrabacteria_group>Bacillota>unclassified_Bacillota | GCA_009711705.1 | Bacillota_bacterium | FH756_08540 | 83034..86369 | 1879010 | ATP-binding-protein-[Firmicutes-bacterium] |
| NLG83533.1 | DUF499_1.RM.like.systems | REase-><-?<-HEPN+DUF499*<-METHYLASE<-Schlafen+HATPase+HTH<-TUDOR+SF1-SF2-Helicase+iREase+CTD+REase | 1127 | Bacteria>Terrabacteria_group>Bacillota>unclassified_Bacillota | GCA_012516115.1 | Bacillota_bacterium | GX493_02745 | 877..4260 | 1879010 | ATP-binding-protein-[Firmicutes-bacterium] |
| NLG85967.1 | DUF499_1.RM.like.systems | <-HHH_5+SF1-SF2-helicase<-?<-?<-?<-Cysteine_desulfurase<-HEPN+DUF499*<-METHYLASE-p | 1132 | Bacteria>Terrabacteria_group>Bacillota>unclassified_Bacillota | GCA_012516165.1 | Bacillota_bacterium | GX489_01905 | 9843..13241 | 1879010 | ATP-binding-protein-[Firmicutes-bacterium] |
| NLN07229.1 | DUF499_1.RM.like.systems | <-HEPN+DUF499*<-METHYLASE<-TUDOR+SF1-SF2-Helicase+iREase+CTD+REase<-REase | 1130 | Bacteria>Terrabacteria_group>Bacillota>unclassified_Bacillota | GCA_012522075.1 | Bacillota_bacterium | GX167_06390 | 552..3944 | 1879010 | ATP-binding-protein-[Firmicutes-bacterium] |
| HHU30916.1 | DUF499_1.RM.like.systems | METHYLASE->HEPN+DUF499*-> | 1077 | Bacteria>Terrabacteria_group>Bacillota>unclassified_Bacillota | GCA_012838605.1 | Bacillota_bacterium | GXZ53_09580 | 2024..5256 | 1879010 | ATP-binding-protein-[Firmicutes-bacterium] |
| HHU75981.1 | DUF499_1.RM.like.systems | <-HEPN+DUF499*<-METHYLASE-p | 1124 | Bacteria>Terrabacteria_group>Bacillota>unclassified_Bacillota | GCA_012839215.1 | Bacillota_bacterium | GXZ24_03690 | 12668..16042 | 1879010 | ATP-binding-protein-[Firmicutes-bacterium] |
| HHV07782.1 | DUF499_1.RM.like.systems | TUDOR+SF1-SF2-Helicase+iREase+CTD+REase-p->?->METHYLASE->HEPN+DUF499*-> | 441 | Bacteria>Terrabacteria_group>Bacillota>unclassified_Bacillota | GCA_012839445.1 | Bacillota_bacterium | GXX69_06175 | 4070..5392 | 1879010 | ATP-binding-protein-[Bacillota-bacterium] |
| HHV56346.1 | DUF499_1.RM.like.systems | <-DUF772+DDE_Tnp_1+DDE_Tnp_1_6\|\|METHYLASE-p->HEPN+DUF499*->REase-> | 1131 | Bacteria>Terrabacteria_group>Bacillota>unclassified_Bacillota | GCA_012839755.1 | Bacillota_bacterium | GXX50_01080 | 3217..6612 | 1879010 | ATP-binding-protein-[Firmicutes-bacterium] |
| HHW13082.1 | DUF499_1.RM.like.systems | TUDOR+SF1-SF2-Helicase+iREase+CTD+REase->METHYLASE->HEPN+DUF499*-><-?<-?<-?<-DeoR-HTH+ISOCOT | 1104 | Bacteria>Terrabacteria_group>Bacillota>unclassified_Bacillota | GCA_012840155.1 | Bacillota_bacterium | GXX33_08800 | 5884..9198 | 1879010 | ATP-binding-protein-[Firmicutes-bacterium] |
| NPV89870.1 | DUF499_1.RM.like.systems | GDResolvase->?->GDResolvase->TUDOR+SF1-SF2-Helicase+iREase+CTD+REase->AbiGi->METHYLASE->HEPN+DUF499*->OLD-ABC+TOPRIM->SF1-SF2-Helicase->?->Peptidase_S8-><-HTH_3 | 1137 | Bacteria>Terrabacteria_group>Bacillota>unclassified_Bacillota | GCA_013178275.1 | Bacillota_bacterium | HPY50_03730 | 179885..183298 | 1879010 | ATP-binding-protein-[Firmicutes-bacterium] |
| NPV80543.1 | DUF499_1.RM.like.systems | REase->?-><-DOC\|\|TUDOR+SF1-SF2-Helicase+iREase+CTD+REase->Schlafen+HATPase+HTH->METHYLASE->HEPN+DUF499*->?-><-HEPN-p | 1115 | Bacteria>Terrabacteria_group>Bacillota>unclassified_Bacillota | GCA_013178415.1 | Bacillota_bacterium | HPY52_09760 | 58381..61728 | 1879010 | ATP-binding-protein-[Firmicutes-bacterium] |
| MBO2495132.1 | DUF499_1.RM.like.systems | METHYLASE-p->HEPN+DUF499*->?->MqsA_antitoxin+HTH_3->ABhydrolase-> | 1117 | Bacteria>Terrabacteria_group>Bacillota>unclassified_Bacillota | GCA_017577845.1 | Bacillota_bacterium | COP67_00010 | 2134..5487 | 1879010 | AAA+-family-ATPase-[Firmicutes-bacterium] |
| MCL5982191.1 | DUF499_1.RM.like.systems | <-HEPN+DUF499*<-METHYLASE-p | 1131 | Bacteria>Terrabacteria_group>Bacillota>unclassified_Bacillota | GCA_023488665.1 | Bacillota_bacterium | M1571_06745 | 19839..23234 | 1879010 | DUF499-domain-containing-protein-[Firmicutes-bacterium] |
| MCL6590158.1 | DUF499_1.RM.like.systems | HicB_lk_antitox->TUDOR+SF1-SF2-Helicase+iREase+CTD+REase->?->MACRODOMAIN->METHYLASE->Y1_Tnp->HEPN+DUF499*->REase->SF1-SF2-Helicase->JetD-> | 1116 | Bacteria>Terrabacteria_group>Bacillota>unclassified_Bacillota | GCA_023512195.1 | Bacillota_bacterium | K6U80_09420 | 20552..23902 | 1879010 | DUF499-domain-containing-protein-[Firmicutes-bacterium] |
| MBJ7608107.1 | DUF499_1.RM.like.systems | Phd-YefM->PIN->?->?->?-><-HEPN+DUF499*<-METHYLASE<-TUDOR+SF1-SF2-Helicase+iREase+CTD+REase<-MPTase<-HTH\|\|?->SMODS-> | 1110 | Bacteria>Terrabacteria_group>Candidatus_Dormibacteraeota>unclassified_Candidatus_Dormibacteraeota | GCA_016464795.1 | Candidatus_Dormibacteraeota_bacterium | JF887_01575 | 11366..14698 | 2052315 | DUF499-domain-containing-protein-[Candidatus-Dormibacteraeota-bacterium] |
| MDA8331677.1 | DUF499_1.RM.like.systems | TUDOR+SF1-SF2-Helicase+iREase+CTD+REase->?->METHYLASE->HEPN+DUF499*-> | 1075 | Bacteria>Terrabacteria_group>Candidatus_Dormibacteraeota>unclassified_Candidatus_Dormibacteraeota | GCA_028067975.1 | Candidatus_Dormibacteraeota_bacterium | M0027_10845 | 5705..8930 | 2052315 | MAG:-DUF499-domain-containing-protein,-partial-[Candidatus-Dormibacteraeota-bacterium] |
| MDA8332521.1 | DUF499_1.RM.like.systems | <-HEPN+DUF499*<-METHYLASE<-?<-TUDOR+SF1-SF2-Helicase+iREase+CTD+REase<-TIR | 485 | Bacteria>Terrabacteria_group>Candidatus_Dormibacteraeota>unclassified_Candidatus_Dormibacteraeota | GCA_028067975.1 | Candidatus_Dormibacteraeota_bacterium | M0027_15220 | 1..1455 | 2052315 | MAG:-Swt1-family-HEPN-domain-containing-protein,-partial-[Candidatus-Dormibacteraeota-bacterium] |
| MAS36305.1 | DUF499_1.RM.like.systems | Glyoxalase->?-><-PIN<-?<-HEPN+DUF499*<-REase<-?<-METHYLASE<-Schlafen+HATPase+HTH | 1094 | Bacteria>Terrabacteria_group>Chloroflexota>Anaerolineae>Anaerolineales | GCA_002702065.1 | Anaerolineaceae_bacterium | CL610_20035 | 35853..39137 | 2024896 | AAA+-family-ATPase-[Anaerolineaceae-bacterium] |
| NLF02984.1 | DUF499_1.RM.like.systems | <-HEPN+DUF499*<-METHYLASE<-?<-MORC<-TUDOR+SF1-SF2-Helicase+iREase+CTD+REase | 1091 | Bacteria>Terrabacteria_group>Chloroflexota>Anaerolineae>Anaerolineales | GCA_012514695.1 | Anaerolineales_bacterium | GX601_18630 | 3316..6591 | 2073117 | ATP-binding-protein-[Anaerolineales-bacterium] |
| MBE2198717.1 | DUF499_1.RM.like.systems | METHYLASE-p->?->HEPN+DUF499*-> | 1099 | Bacteria>Terrabacteria_group>Chloroflexota>Anaerolineae>Anaerolineales | GCA_014879875.1 | Anaerolinea_sp. | IAE79_08915 | 3861..7160 | 1872519 | ATP-binding-protein-[Anaerolinea-sp.] |
| MBE7511263.1 | DUF499_1.RM.like.systems | Peptidase_M1->?->TUDOR+SF1-SF2-Helicase+iREase+CTD+REase->?->PIN->METHYLASE-p->HEPN+DUF499*-><-tRNA<-?<-ABhydrolase\|\|TPR+TPR->ACET+ACET->ACET-> | 1098 | Bacteria>Terrabacteria_group>Chloroflexota>Anaerolineae>Anaerolineales | GCA_015075625.1 | Anaerolineales_bacterium | HS103_00415 | 94825..98121 | 2073117 | ATP-binding-protein-[Anaerolineales-bacterium] |
| MBL6983853.1 | DUF499_1.RM.like.systems | <-REase<-HEPN+DUF499*<-SF1-SF2-Helicase-p | 1095 | Bacteria>Terrabacteria_group>Chloroflexota>Anaerolineae>Anaerolineales | GCA_016785015.1 | Anaerolineales_bacterium | ISR58_21930 | 2169..5456 | 2073117 | DUF499-domain-containing-protein-[Anaerolineales-bacterium] |
| MBQ6503986.1 | DUF499_1.RM.like.systems | METHYLASE-p->METHYLASE->HEPN+DUF499*->KAP-NTPase-> | 1120 | Bacteria>Terrabacteria_group>Chloroflexota>Anaerolineae>Anaerolineales | GCA_017432685.1 | Flexilinea_sp. | IJI57_08765 | 4499..7861 | 2017390 | DUF499-domain-containing-protein-[Flexilinea-sp.] |
| QYK51438.1 | DUF499_1.RM.like.systems | DOC+HTH-><-HEPN+DUF499*<-METHYLASE<-SIR2<-TUDOR+SF1-SF2-Helicase+iREase+CTD+REase<-?<-TIR\|\|PRTase-> | 1117 | Bacteria>Terrabacteria_group>Chloroflexota>Anaerolineae>Anaerolineales | GCA_019454165.1 | Anaerolineales_bacterium | KF701_02830 | 580231..583584 | 2073117 | MAG:-DUF499-domain-containing-protein-[Anaerolineales-bacterium] |
| MCB0010245.1 | DUF499_1.RM.like.systems | METHYLASE-p->TIR->TPR->?->?-><-?\|\|HEPN+DUF499*->?->HKD-DNase+HKD-DNase-><-nSTAND1<-?<-SIR2+CASPASE<-SIR2+nSTAND1+FGS | 1088 | Bacteria>Terrabacteria_group>Chloroflexota>Anaerolineae>Anaerolineales | GCA_020433005.1 | Anaerolineales_bacterium | KDE34_00040 | 5582..8848 | 2073117 | DUF499-domain-containing-protein-[Anaerolineales-bacterium] |
| MCB9452414.1 | DUF499_1.RM.like.systems | <-MetJarc-HTH\|\|?->TPR+TPR+TPR+TPR+TPR+TPR+TPR+TPR-><-HEPN+DUF499*<-ParB+HNH<-METHYLASE<-Schlafen+HATPase+HTH<-PIN<-?<-TUDOR+SF1-SF2-Helicase+iREase+CTD+REase | 1106 | Bacteria>Terrabacteria_group>Chloroflexota>Anaerolineae>Anaerolineales | GCA_020634115.1 | Anaerolineaceae_bacterium | H6672_13335 | 112307..115627 | 2024896 | ATP-binding-protein-[Anaerolineaceae-bacterium] |
| MCB9078004.1 | DUF499_1.RM.like.systems | <-HEPN+DUF499*<-REase<-?<-METHYLASE-p<-METHYLASE<-PIN<-DUF433 | 385 | Bacteria>Terrabacteria_group>Chloroflexota>Anaerolineae>Anaerolineales | GCA_020634955.1 | Anaerolineaceae_bacterium | H6631_10465 | 1..1157 | 2024896 | ATP-binding-protein-[Anaerolineaceae-bacterium] |
| MCC6958025.1 | DUF499_1.RM.like.systems | <-MarR-HTH<-HEPN+DUF499*<-?<-METHYLASE<-?<-PIN<-REase | 1114 | Bacteria>Terrabacteria_group>Chloroflexota>Anaerolineae>Anaerolineales | GCA_020853835.1 | Anaerolineales_bacterium | IT316_14590 | 1878..5222 | 2073117 | DUF499-domain-containing-protein-[Anaerolineales-bacterium] |
| MCW5886897.1 | DUF499_1.RM.like.systems | <-PRTase\|\|TIR->?->TUDOR+SF1-SF2-Helicase+iREase+CTD+REase->METHYLASE->HEPN+DUF499*-><-DOC+HTH | 1117 | Bacteria>Terrabacteria_group>Chloroflexota>Anaerolineae>Anaerolineales | GCA_026129325.1 | Anaerolineales_bacterium | KIT07_02110 | 8706..12059 | 2073117 | MAG:-DUF499-domain-containing-protein-[Anaerolineales-bacterium] |
| MCZ7550961.1 | DUF499_1.RM.like.systems | <-NACHT\|\|<-?<-?<-ParB+HNH<-?<-HEPN+DUF499*<-METHYLASE<-MORC+DUF3684<-MORC+DUF3684<-TUDOR+SF1-SF2-Helicase+iREase+CTD+REase-p | 1112 | Bacteria>Terrabacteria_group>Chloroflexota>Anaerolineae>Anaerolineales | GCA_027494025.1 | Anaerolineales_bacterium | M5U11_17685 | 3656371..3659709 | 2073117 | MAG:-Swt1-family-HEPN-domain-containing-protein-[Anaerolineales-bacterium] |
| NPV86424.1 | DUF499_1.RM.like.systems | <-REC<-tRNA\|\|TUDOR+SF1-SF2-Helicase+iREase+CTD+REase->PIN->METHYLASE->HEPN+DUF499*-> | 1115 | Bacteria>Terrabacteria_group>Chloroflexota>Anaerolineae>unclassified_Anaerolineae | GCA_013178295.1 | Anaerolineae_bacterium | HPY45_10505 | 184598..187945 | 2052143 | ATP-binding-protein-[Anaerolineae-bacterium] |
| MBK6429951.1 | DUF499_1.RM.like.systems | OLD-ABC->TOPRIM->Schlafen+HATPase+HTH->METHYLASE-p->Schlafen+HATPase+HTH->REase->HEPN+DUF499*->?->?->PemK_toxin->ABhydrolase->?->ABhydrolase+ABhydrolase-> | 1113 | Bacteria>Terrabacteria_group>Chloroflexota>Anaerolineae>unclassified_Anaerolineae | GCA_016704685.1 | Anaerolineae_bacterium | IPF85_04125 | 226121..229462 | 2052143 | DUF499-domain-containing-protein-[Anaerolineae-bacterium] |
| MBK9093182.1 | DUF499_1.RM.like.systems | <-ABhydrolase+ABhydrolase<-?<-ABhydrolase<-?<-?<-HEPN+DUF499*<-Schlafen+HATPase+HTH<-METHYLASE-p<-Schlafen+HATPase+HTH<-TUDOR+SF1-SF2-Helicase+iREase+CTD+REase<-TOPRIM<-OLD-ABC | 1133 | Bacteria>Terrabacteria_group>Chloroflexota>Anaerolineae>unclassified_Anaerolineae | GCA_016719785.1 | Anaerolineae_bacterium | IPM84_10425 | 127540..130941 | 2052143 | DUF499-domain-containing-protein-[Anaerolineae-bacterium] |
| MBN1136865.1 | DUF499_1.RM.like.systems | APATPase+iSTAND3->METHYLASE->?->?->HEPN+DUF499*-> | 684 | Bacteria>Terrabacteria_group>Chloroflexota>Anaerolineae>unclassified_Anaerolineae | GCA_016926595.1 | Anaerolineae_bacterium | JXM73_09775 | 6509..8561 | 2052143 | DUF499-domain-containing-protein-[Anaerolineae-bacterium] |
| MBN1429664.1 | DUF499_1.RM.like.systems | <-HEPN+DUF499*<-?<-?<-?<-?<-METHYLASE | 352 | Bacteria>Terrabacteria_group>Chloroflexota>Anaerolineae>unclassified_Anaerolineae | GCA_016928235.1 | Anaerolineae_bacterium | JXB07_14940 | 1..1058 | 2052143 | AAA-family-ATPase-[Anaerolineae-bacterium] |
| MBN2002049.1 | DUF499_1.RM.like.systems | <-HEPN+DUF499*<-Y1_Tnp<-METHYLASE<-MACRODOMAIN<-?<-?<-TUDOR+SF1-SF2-Helicase+iREase+CTD+REase | 1119 | Bacteria>Terrabacteria_group>Chloroflexota>Anaerolineae>unclassified_Anaerolineae | GCA_016931835.1 | Anaerolineae_bacterium | JXA21_01725 | 49309..52668 | 2052143 | DUF499-domain-containing-protein-[Anaerolineae-bacterium] |
| MCB0250740.1 | DUF499_1.RM.like.systems | <-HEPN+DUF499*<-Y1_Tnp<-METHYLASE-p | 1124 | Bacteria>Terrabacteria_group>Chloroflexota>Anaerolineae>unclassified_Anaerolineae | GCA_020434015.1 | Anaerolineae_bacterium | KDI07_19355 | 427..3801 | 2052143 | DUF499-domain-containing-protein-[Anaerolineae-bacterium] |
| MCC6801844.1 | DUF499_1.RM.like.systems | <-HNH<-CopG/RHH<-DUF2726+ZnR+TOPRIM<-HEPN+DUF499*<-?<-METHYLASE | 1095 | Bacteria>Terrabacteria_group>Chloroflexota>Anaerolineae>unclassified_Anaerolineae | GCA_020848855.1 | Anaerolineae_bacterium | IT319_03080 | 2285..5572 | 2052143 | DUF499-domain-containing-protein-[Anaerolineae-bacterium] |
| MCC6973771.1 | DUF499_1.RM.like.systems | PIN->?->METHYLASE->?->HEPN+DUF499*->HEPN-> | 1096 | Bacteria>Terrabacteria_group>Chloroflexota>Anaerolineae>unclassified_Anaerolineae | GCA_020853695.1 | Anaerolineae_bacterium | IT322_07165 | 5474..8764 | 2052143 | DUF499-domain-containing-protein-[Anaerolineae-bacterium] |
| MCK6580687.1 | DUF499_1.RM.like.systems | <-GDResolvase<-?<-PIN<-HEPN+DUF499*<-PIN<-DUF433<-?<-METHYLASE<-?<-TUDOR+SF1-SF2-Helicase+iREase+CTD+REase | 1104 | Bacteria>Terrabacteria_group>Chloroflexota>Anaerolineae>unclassified_Anaerolineae | GCA_023151025.1 | Anaerolineae_bacterium | L6Q98_21555 | 21655..24969 | 2052143 | DUF499-domain-containing-protein-[Anaerolineae-bacterium] |
| MCL4299450.1 | DUF499_1.RM.like.systems | TPR->?->?->METHYLASE->URI-DNase->HEPN+DUF499*->DUF2726+ZnR+TOPRIM-> | 1096 | Bacteria>Terrabacteria_group>Chloroflexota>Anaerolineae>unclassified_Anaerolineae | GCA_023382535.1 | Anaerolineae_bacterium | KJ077_27185 | 26645..29935 | 2052143 | DUF499-domain-containing-protein-[Anaerolineae-bacterium] |
| MCL4302340.1 | DUF499_1.RM.like.systems | <-TraJ-RHH\|\|TUDOR+SF1-SF2-Helicase+iREase+CTD+REase->?->?->METHYLASE->REase->HEPN+DUF499*-><-?\|\|?->GDResolvase+Recombinase+ZnR-> | 1117 | Bacteria>Terrabacteria_group>Chloroflexota>Anaerolineae>unclassified_Anaerolineae | GCA_023382535.1 | Anaerolineae_bacterium | KJ077_42015 | 34544..37897 | 2052143 | DUF499-domain-containing-protein-[Anaerolineae-bacterium] |
| MCL4237388.1 | DUF499_1.RM.like.systems | MULE-TRANSPOSASE->REase->SF1-SF2-Helicase-p->MULE-TRANSPOSASE-><-HEPN+DUF499*<-METHYLASE<-PIN<-?<-TUDOR+SF1-SF2-Helicase+iREase+CTD+REase | 1111 | Bacteria>Terrabacteria_group>Chloroflexota>Anaerolineae>unclassified_Anaerolineae | GCA_023382715.1 | Anaerolineae_bacterium | KJ047_03985 | 66352..69687 | 2052143 | DUF499-domain-containing-protein-[Anaerolineae-bacterium] |
| MCL4805199.1 | DUF499_1.RM.like.systems | <-HEPN+DUF499*<-METHYLASE<-?<-TUDOR+SF1-SF2-Helicase+iREase+CTD+REase<-McrC-NTD+REase | 1098 | Bacteria>Terrabacteria_group>Chloroflexota>Anaerolineae>unclassified_Anaerolineae | GCA_023384355.1 | Anaerolineae_bacterium | KJ046_12935 | 9107..12403 | 2052143 | DUF499-domain-containing-protein-[Anaerolineae-bacterium] |
| MCL4871680.1 | DUF499_1.RM.like.systems | <-AbiEii<-AbiEii<-AbiEi_4\|\|TUDOR+SF1-SF2-Helicase+iREase+CTD+REase->REase->METHYLASE->HEPN+DUF499*-><-?<-?<-SIGMA-HTH<-RNB+RNase\|\|?->PadR-HTH-> | 1115 | Bacteria>Terrabacteria_group>Chloroflexota>Anaerolineae>unclassified_Anaerolineae | GCA_023384415.1 | Anaerolineae_bacterium | KJ063_22200 | 16187..19534 | 2052143 | DUF499-domain-containing-protein-[Anaerolineae-bacterium] |
| MCS6835397.1 | DUF499_1.RM.like.systems | METHYLASE-p->OLD-ABC+TOPRIM-p->HEPN+DUF499*-> | 1089 | Bacteria>Terrabacteria_group>Chloroflexota>Anaerolineae>unclassified_Anaerolineae | GCA_025055175.1 | Anaerolineae_bacterium | NZ750_05190 | 1306..4575 | 2052143 | DUF499-domain-containing-protein-[Anaerolineae-bacterium] |
| MCU0500065.1 | DUF499_1.RM.like.systems | <-Glyco_transf_28<-TetR-HTH\|\|?-><-HEPN+DUF499*<-METHYLASE-p | 1097 | Bacteria>Terrabacteria_group>Chloroflexota>Anaerolineae>unclassified_Anaerolineae | GCA_025454525.1 | Anaerolineae_bacterium | MUF87_22175 | 4596..7889 | 2052143 | DUF499-domain-containing-protein-[Anaerolineae-bacterium] |
| MCU0476247.1 | DUF499_1.RM.like.systems | <-HEPN+DUF499*<-METHYLASE-p | 1076 | Bacteria>Terrabacteria_group>Chloroflexota>Anaerolineae>unclassified_Anaerolineae | GCA_025454715.1 | Anaerolineae_bacterium | MUC99_09085 | 257..3487 | 2052143 | DUF499-domain-containing-protein-[Anaerolineae-bacterium] |
| MCW5847519.1 | DUF499_1.RM.like.systems | METHYLASE-p->REase->HEPN+DUF499*-><-?<-?<-?<-?<-ABC-ATPase+SbcC | 1097 | Bacteria>Terrabacteria_group>Chloroflexota>Anaerolineae>unclassified_Anaerolineae | GCA_026129205.1 | Anaerolineae_bacterium | KIS95_09830 | 2201..5494 | 2052143 | MAG:-DUF499-domain-containing-protein-[Anaerolineae-bacterium] |
| MCZ2096700.1 | DUF499_1.RM.like.systems | SF1-SF2-Helicase->wHTH+SAD+HNH->ORC-CDC6+REase->HEPN+DUF499*-> | 1112 | Bacteria>Terrabacteria_group>Chloroflexota>Anaerolineae>unclassified_Anaerolineae | GCA_026954235.1 | Anaerolineae_bacterium | LC121_10630 | 39414..42752 | 2052143 | MAG:-DUF499-domain-containing-protein-[Anaerolineae-bacterium] |
| MCZ2114537.1 | DUF499_1.RM.like.systems | <-HTH+UbE1<-?\|\|HTH->MPTase-><-HEPN+DUF499*<-PIN<-OLD-ABC<-METHYLASE<-MORC+DUF3684<-TUDOR+SF1-SF2-Helicase+iREase+CTD+REase | 1108 | Bacteria>Terrabacteria_group>Chloroflexota>Anaerolineae>unclassified_Anaerolineae | GCA_026954395.1 | Anaerolineae_bacterium | LC131_12000 | 4338..7664 | 2052143 | MAG:-DUF499-domain-containing-protein-[Anaerolineae-bacterium] |
| MDD3828881.1 | DUF499_1.RM.like.systems | TUDOR+SF1-SF2-Helicase+iREase+CTD+REase->METHYLASE->HEPN+DUF499*-> | 350 | Bacteria>Terrabacteria_group>Chloroflexota>Anaerolineae>unclassified_Anaerolineae | GCA_028698515.1 | Anaerolineae_bacterium | PHY79_23180 | 6005..7056 | 2052143 | MAG:-Swt1-family-HEPN-domain-containing-protein,-partial-[Anaerolineae-bacterium] |
| MCB9423196.1 | DUF499_1.RM.like.systems | TUDOR+SF1-SF2-Helicase+iREase+CTD+REase->?->SF1-SF2-Helicase+HRDC->?->METHYLASE->TIR->HEPN+DUF499*->XIS-HTH-> | 1096 | Bacteria>Terrabacteria_group>Chloroflexota>Ardenticatenia>Ardenticatenales | GCA_020634245.1 | Ardenticatenaceae_bacterium | H6667_25585 | 35052..38342 | 2614247 | ATP-binding-protein-[Ardenticatenaceae-bacterium] |
| MCB8944346.1 | DUF499_1.RM.like.systems | GDResolvase+HIN-HTH-><-?\|\|?-><-?\|\|<-?<-?<-HEPN+DUF499*<-?<-METHYLASE<-ParB+HNH<-TUDOR+SF1-SF2-Helicase+iREase+CTD+REase | 1098 | Bacteria>Terrabacteria_group>Chloroflexota>Ardenticatenia>Ardenticatenales | GCA_020634415.1 | Ardenticatenaceae_bacterium | H6658_11405 | 5286..8582 | 2614247 | ATP-binding-protein-[Ardenticatenaceae-bacterium] |
| MCZ7570815.1 | DUF499_1.RM.like.systems | DEDD_Tnp_IS110+Transposase_20->?->TUDOR+SF1-SF2-Helicase+iREase+CTD+REase->METHYLASE->REase->HEPN+DUF499*-><-PIN<-PhdYeFM_antitox<-?<-HAD | 1117 | Bacteria>Terrabacteria_group>Chloroflexota>Ardenticatenia>Ardenticatenales | GCA_027493995.1 | Ardenticatenaceae_bacterium | M5U01_19825 | 4779576..4782929 | 2614247 | MAG:-Swt1-family-HEPN-domain-containing-protein-[Ardenticatenaceae-bacterium] |
| MXX28115.1 | DUF499_1.RM.like.systems | <-HEPN+DUF499*<-TOPRIM<-OLD-ABC<-HEPN<-PolB<-METHYLASE | 1121 | Bacteria>Terrabacteria_group>Chloroflexota>Caldilineae>Caldilineales | GCA_009835815.1 | Caldilineaceae_bacterium_SB0668_bin_21 | F4Z82_22075 | 67..3432 | 2605264 | ATP-binding-protein-[Caldilineaceae-bacterium-SB0668_bin_21] |
| MYA05496.1 | DUF499_1.RM.like.systems | <-METHYLASE<-TUDOR+SF1-SF2-Helicase+iREase+CTD+REase<-HEPN+DUF499* | 1079 | Bacteria>Terrabacteria_group>Chloroflexota>Caldilineae>Caldilineales | GCA_009838685.1 | Caldilineaceae_bacterium_SB0664_bin_22 | F4Y37_12945 | 5308..8547 | 2605259 | ATP-binding-protein-[Caldilineaceae-bacterium-SB0664_bin_22] |
| MBP7961438.1 | DUF499_1.RM.like.systems | TUDOR+SF1-SF2-Helicase+iREase+CTD+REase->?->?->METHYLASE->OLD-ABC->PIN->HEPN+DUF499*->?->SF1-SF2-Helicase->SF1-SF2-Helicase+DUF1998-> | 1102 | Bacteria>Terrabacteria_group>Chloroflexota>Caldilineae>Caldilineales | GCA_018003515.1 | Caldilineaceae_bacterium | KA003_05670 | 15367..18675 | 2614250 | DUF499-domain-containing-protein-[Caldilineaceae-bacterium] |
| MCC6169408.1 | DUF499_1.RM.like.systems | PIN->METHYLASE->HEPN+DUF499*-> | 1104 | Bacteria>Terrabacteria_group>Chloroflexota>Caldilineae>Caldilineales | GCA_020845455.1 | Caldilineaceae_bacterium | IT329_19460 | 3543..6857 | 2614250 | DUF499-domain-containing-protein-[Caldilineaceae-bacterium] |
| MCE7987213.1 | DUF499_1.RM.like.systems | <-HAMP+HISKIN<-?<-HEPN+DUF499*<-Y1_Tnp<-METHYLASE<-Schlafen+HATPase+HTH | 1167 | Bacteria>Terrabacteria_group>Chloroflexota>Caldilineae>Caldilineales | GCA_021405325.1 | Caldilinea_sp._CFX5 | DYG89_39080 | 104570..108073 | 2293554 | ATP-binding-protein-[Caldilinea-sp.-CFX5] |
| MCY3902338.1 | DUF499_1.RM.like.systems | TIR->TUDOR+SF1-SF2-Helicase+iREase+CTD+REase-p->PIN->METHYLASE->PolB->HEPN->HEPN+DUF499*->?->?-><-?<-DSBH | 1125 | Bacteria>Terrabacteria_group>Chloroflexota>Caldilineae>Caldilineales | GCA_026707125.1 | Caldilineaceae_bacterium | OXF76_04160 | 61061..64438 | 2614250 | MAG:-Swt1-family-HEPN-domain-containing-protein-[Caldilineaceae-bacterium] |
| MCY4079348.1 | DUF499_1.RM.like.systems | METHYLASE->PolB->?->?->OLD-ABC->TOPRIM->HEPN+DUF499*->?->?-><-?\|\|MazE_antitoxin-><-DSBH | 1121 | Bacteria>Terrabacteria_group>Chloroflexota>Caldilineae>Caldilineales | GCA_026707715.1 | Caldilineaceae_bacterium | OXF54_03840 | 16452..19817 | 2614250 | MAG:-Swt1-family-HEPN-domain-containing-protein-[Caldilineaceae-bacterium] |
| MCY4117637.1 | DUF499_1.RM.like.systems | TUDOR+SF1-SF2-Helicase+iREase+CTD+REase->METHYLASE->PolB->HEPN->OLD-ABC->TOPRIM->HEPN+DUF499*->?->?->?-><-DSBH | 1124 | Bacteria>Terrabacteria_group>Chloroflexota>Caldilineae>Caldilineales | GCA_026708105.1 | Caldilineaceae_bacterium | OXF55_12190 | 34492..37866 | 2614250 | MAG:-Swt1-family-HEPN-domain-containing-protein-[Caldilineaceae-bacterium] |
| MCZ0940408.1 | DUF499_1.RM.like.systems | REase+NACHT-><-?\|\|?->?-><-MAD-NTD+HNH<-HEPN+DUF499*<-METHYLASE-p | 1114 | Bacteria>Terrabacteria_group>Chloroflexota>Caldilineae>Caldilineales | GCA_026914085.1 | Caldilineaceae_bacterium | OXJ55_17325 | 15852..19196 | 2614250 | MAG:-DUF499-domain-containing-protein-[Caldilineaceae-bacterium] |
| MDE0069281.1 | DUF499_1.RM.like.systems | DSBH->?-><-?<-?<-HEPN+DUF499*<-?<-?<-METHYLASE-p | 1125 | Bacteria>Terrabacteria_group>Chloroflexota>Caldilineae>Caldilineales | GCA_028819015.1 | Caldilineaceae_bacterium | OX048_06170 | 52227..55604 | 2614250 | MAG:-Swt1-family-HEPN-domain-containing-protein-[Caldilineaceae-bacterium] |
| MDE0183518.1 | DUF499_1.RM.like.systems | METHYLASE->?->?->HEPN+DUF499*->?->?-><-?<-DSBH | 1122 | Bacteria>Terrabacteria_group>Chloroflexota>Caldilineae>Caldilineales | GCA_028819655.1 | Caldilineaceae_bacterium | OXL39_19355 | 3803..7171 | 2614250 | MAG:-Swt1-family-HEPN-domain-containing-protein-[Caldilineaceae-bacterium] |
| MDE0200338.1 | DUF499_1.RM.like.systems | METHYLASE->PolB->HEPN->OLD-ABC->TOPRIM->HEPN+DUF499*-> | 1112 | Bacteria>Terrabacteria_group>Chloroflexota>Caldilineae>Caldilineales | GCA_028819675.1 | Caldilineaceae_bacterium | OXK78_19185 | 5066..8404 | 2614250 | MAG:-Swt1-family-HEPN-domain-containing-protein-[Caldilineaceae-bacterium] |
| MDE0338616.1 | DUF499_1.RM.like.systems | <-HAD+HAD\|\|?-><-?\|\|TUDOR+SF1-SF2-Helicase+iREase+CTD+REase->METHYLASE->PolB->HEPN+DUF499*->?->?->MazE_antitoxin-><-DSBH | 1121 | Bacteria>Terrabacteria_group>Chloroflexota>Caldilineae>Caldilineales | GCA_028820895.1 | Caldilineaceae_bacterium | OXI80_13170 | 100490..103855 | 2614250 | MAG:-Swt1-family-HEPN-domain-containing-protein-[Caldilineaceae-bacterium] |
| MDE0431259.1 | DUF499_1.RM.like.systems | DSBH->?-><-?<-?<-HEPN+DUF499*<-TOPRIM<-OLD-ABC<-?<-HEPN<-PolB<-METHYLASE | 1123 | Bacteria>Terrabacteria_group>Chloroflexota>Caldilineae>Caldilineales | GCA_028821555.1 | Caldilineaceae_bacterium | OXH98_15965 | 76934..80305 | 2614250 | MAG:-Swt1-family-HEPN-domain-containing-protein-[Caldilineaceae-bacterium] |
| WP_014433685.1 | DUF499_1.RM.like.systems | <-Peptidase\|\|TUDOR+SF1-SF2-Helicase+iREase+CTD+REase->METHYLASE->METHYLASE->HEPN+DUF499*-> | 1108 | Bacteria>Terrabacteria_group>Chloroflexota>Caldilineae>Caldilineales | GCF_000281175.1 | Caldilinea | QVE71_RS01780 | 444218..447544 | 233191 | Swt1-family-HEPN-domain-containing-protein-[Caldilinea-aerophila] |
| MBK9709775.1 | DUF499_1.RM.like.systems | TUDOR+SF1-SF2-Helicase+iREase+CTD+REase->REase->METHYLASE->URI-DNase->HEPN+DUF499*->?->?-><-tRNA<-?<-?<-RnaseIII+DSRBD | 1115 | Bacteria>Terrabacteria_group>Chloroflexota>Chloroflexia>Chloroflexales | GCA_016717335.1 | Kouleothrix_sp. | IPO81_00330 | 71152..74499 | 2779161 | DUF499-domain-containing-protein-[Kouleothrix-sp.] |
| MBO9374240.1 | DUF499_1.RM.like.systems | <-TUDOR+SF1-SF2-Helicase+iREase+CTD+REase<-HEPN+DUF499* | 1112 | Bacteria>Terrabacteria_group>Chloroflexota>Chloroflexia>Chloroflexales | GCA_017743265.1 | Chloroflexus_sp. | J7459_12885 | 2503..5841 | 1904827 | DUF499-domain-containing-protein-[Chloroflexus-sp.] |
| MBO9390433.1 | DUF499_1.RM.like.systems | METHYLASE-p->?->RAMPs->RAMPs->RAMPs->HEPN+HEPN->HEPN+DUF499*-> | 1118 | Bacteria>Terrabacteria_group>Chloroflexota>Chloroflexia>Chloroflexales | GCA_017743475.1 | Roseiflexus_sp. | J7482_16995 | 17677..21033 | 2562120 | DUF499-domain-containing-protein-[Roseiflexus-sp.] |
| MBO9321562.1 | DUF499_1.RM.like.systems | <-TUDOR+SF1-SF2-Helicase+iREase+CTD+REase<-HEPN+DUF499* | 1112 | Bacteria>Terrabacteria_group>Chloroflexota>Chloroflexia>Chloroflexales | GCA_017743635.1 | Roseiflexus_sp. | J7457_05465 | 3598..6936 | 2562120 | DUF499-domain-containing-protein-[Roseiflexus-sp.] |
| MCS6937885.1 | DUF499_1.RM.like.systems | METHYLASE-p->HEPN+DUF499*-> | 531 | Bacteria>Terrabacteria_group>Chloroflexota>Chloroflexia>Chloroflexales | GCA_025055265.1 | Roseiflexus_sp. | NZM94_01360 | 1244..2838 | 2562120 | Swt1-family-HEPN-domain-containing-protein-[Roseiflexus-sp.] |
| MCS6883598.1 | DUF499_1.RM.like.systems | PolB->HEPN->?->METHYLASE->METHYLASE-p->REase->HEPN+DUF499*-> | 1130 | Bacteria>Terrabacteria_group>Chloroflexota>Chloroflexia>Chloroflexales | GCA_025057155.1 | Chloroflexaceae_bacterium | NZU74_19900 | 13790..17182 | 2562117 | Swt1-family-HEPN-domain-containing-protein-[Chloroflexaceae-bacterium] |
| MCS6889332.1 | DUF499_1.RM.like.systems | <-HEPN+DUF499*<-METHYLASE-p | 1115 | Bacteria>Terrabacteria_group>Chloroflexota>Chloroflexia>Chloroflexales | GCA_025057175.1 | Chloroflexus_sp. | NZQ09_14085 | 968..4315 | 1904827 | Swt1-family-HEPN-domain-containing-protein-[Chloroflexus-sp.] |
| MCX7860988.1 | DUF499_1.RM.like.systems | <-HEPN+DUF499*<-METHYLASE-p | 1115 | Bacteria>Terrabacteria_group>Chloroflexota>Chloroflexia>Chloroflexales | GCA_026417255.1 | Chloroflexus_sp. | N2385_12910 | 223..3570 | 1904827 | MAG:-Swt1-family-HEPN-domain-containing-protein-[Chloroflexus-sp.] |
| WP_011958262.1 | DUF499_1.RM.like.systems | HEPN->?->?->DDE_Tnp_1->DDE_Tnp_1->METHYLASE-p->HEPN+DUF499*->ABC-ATPase+SbcC-> | 1122 | Bacteria>Terrabacteria_group>Chloroflexota>Chloroflexia>Chloroflexales | GCF_000016665.1 | Roseiflexus_sp._RS-1 | ROSERS_RS18370 | 4458924..4462292 | 357808 | Swt1-family-HEPN-domain-containing-protein-[Roseiflexus-sp.-RS-1] |
| WP_255346773.1 | DUF499_1.RM.like.systems | HEPN+DUF499*-><-?<-MODE-HTH+PBPII\|\|?->?->ABC-ATPase+SbcC-> | 501 | Bacteria>Terrabacteria_group>Chloroflexota>Chloroflexia>Chloroflexales | GCF_000516515.1 | Chloroflexus_sp._Y-396-1 | CHY396_RS19730 | 597760..599265 | 867845 | Swt1-family-HEPN-domain-containing-protein-[Chloroflexus-sp.-Y-396-1] |
| WP_031460600.1 | DUF499_1.RM.like.systems | <-STAS\|\|?-><-?<-?<-?\|\|HEPN+DUF499*->TUDOR+SF1-SF2-Helicase+iREase+CTD+REase->METHYLASE-> | 1112 | Bacteria>Terrabacteria_group>Chloroflexota>Chloroflexia>Chloroflexales | GCF_000735195.1 | Chloroflexus_sp._MS-G | JE47_RS0115345 | 6861..10199 | 1521187 | Swt1-family-HEPN-domain-containing-protein-[Chloroflexus-sp.-MS-G] |
| WP_066790944.1 | DUF499_1.RM.like.systems | <-HEPN+DUF499*<-METHYLASE<-METHYLASE<-REase<-TUDOR+SF1-SF2-Helicase+iREase+CTD+REase | 1104 | Bacteria>Terrabacteria_group>Chloroflexota>Chloroflexia>Chloroflexales | GCF_001650695.1 | Chloroflexus_islandicus | A6A03_RS17620 | 125816..129130 | 1707952 | Swt1-family-HEPN-domain-containing-protein-[Chloroflexus-islandicus] |
| WP_097642123.1 | DUF499_1.RM.like.systems | <-DUF488-l<-?<-?<-Phage_integrase<-HKD-DNase+HKD-DNase<-?<-HEPN+DUF499*<-Polbeta<-PolB-NTase<-METHYLASE<-HEPN<-?<-TUDOR+SF1-SF2-Helicase+iREase+CTD+REase | 1103 | Bacteria>Terrabacteria_group>Chloroflexota>Chloroflexia>Chloroflexales | GCF_002532535.1 | Candidatus_Viridilinea_mediisalina | CJ255_RS00480 | 82670..85981 | 2024553 | Swt1-family-HEPN-domain-containing-protein-[Candidatus-Viridilinea-mediisalina] |
| WP_097643072.1 | DUF499_1.RM.like.systems | <-PIN\|\|TUDOR+SF1-SF2-Helicase+iREase+CTD+REase-><-?\|\|?->METHYLASE->DUF433->HEPN+DUF499*->?-><-MarR-HTH\|\|?-><-?<-ST-Hiskin<-NTP_transferase | 1128 | Bacteria>Terrabacteria_group>Chloroflexota>Chloroflexia>Chloroflexales | GCF_002532535.1 | Candidatus_Viridilinea_mediisalina | CJ255_RS05405 | 56277..59663 | 2024553 | Swt1-family-HEPN-domain-containing-protein-[Candidatus-Viridilinea-mediisalina] |
| WP_205703984.1 | DUF499_1.RM.like.systems | RAMPs->?->?->?->HEPN+DUF499*->?->?->Cas_Cas1->CRISPR_Cas2-> | 1169 | Bacteria>Terrabacteria_group>Chloroflexota>Chloroflexia>Chloroflexales | GCF_004138165.1 | Candidatus_Chloroploca_sp._Khr17 | EKD05_RS19000 | 28531..32040 | 2496869 | Swt1-family-HEPN-domain-containing-protein-[Candidatus-Chloroploca-sp.-Khr17] |
| WP_026369943.1 | DUF499_1.RM.like.systems | ABC-ATPase+SbcC->ABC-ATPase+SbcC->?->tRNA-><-HEPN+DUF499*\|\|PAIREDC-HTH->?->?->?->?-><-METHYLASE | 1108 | Bacteria>Terrabacteria_group>Chloroflexota>Chloroflexia>Kallotenuales | GCF_000526415.1 | Kallotenue_papyrolyticum | K361_RS0107050 | 1033569..1036895 | 1325125 | Swt1-family-HEPN-domain-containing-protein-[Kallotenue-papyrolyticum] |
| KAB8142895.1 | DUF499_1.RM.like.systems | PAPS_reduct->?->TUDOR+SF1-SF2-Helicase+iREase+CTD+REase->?->METHYLASE->HEPN+DUF499*-><-?<-HNH\|\|?->Phage_int_SAM_4+Phage_integrase-> | 1122 | Bacteria>Terrabacteria_group>Chloroflexota>Chloroflexia>unclassified_Chloroflexia | GCA_009268125.1 | Chloroflexia_bacterium_SDU3-3 | F8S13_11620 | 110122..113490 | 2651839 | ATP-binding-protein-[Chloroflexia-bacterium-SDU3-3] |
| KAB8139622.1 | DUF499_1.RM.like.systems | <-HEPN+DUF499*<-METHYLASE<-TUDOR+SF1-SF2-Helicase+iREase+CTD+REase | 1122 | Bacteria>Terrabacteria_group>Chloroflexota>Chloroflexia>unclassified_Chloroflexia | GCA_009268125.1 | Chloroflexia_bacterium_SDU3-3 | F8S13_27385 | 758..4126 | 2651839 | ATP-binding-protein-[Chloroflexia-bacterium-SDU3-3] |
| WP_012876078.1 | DUF499_1.RM.like.systems | <-GNTR-HTH\|\|?->TUDOR+SF1-SF2-Helicase+iREase+CTD+REase->METHYLASE->HEPN+DUF499*-> | 1099 | Bacteria>Terrabacteria_group>Chloroflexota>Chloroflexota_incertae_sedis>Thermobaculum | GCF_000025005.1 | Thermobaculum_terrenum | TTER_RS10895 | 291668..294967 | 166501 | Swt1-family-HEPN-domain-containing-protein-[Thermobaculum-terrenum] |
| MBN4064100.1 | DUF499_1.RM.like.systems | METHYLASE-p->HEPN+DUF499*-><-REase\|\|XIS-HTH-> | 1109 | Bacteria>Terrabacteria_group>Chloroflexota>Dehalococcoidia>Dehalococcoidales | GCA_017114025.1 | Dehalococcoides_mccartyi | JYU04_00010 | 654..3983 | 61435 | DUF499-domain-containing-protein-[Dehalococcoides-mccartyi] |
| WP_046961399.1 | DUF499_1.RM.like.systems | <-XIS-HTH<-HEPN+DUF499*<-METHYLASE<-MORC+DUF3684<-TUDOR+SF1-SF2-Helicase+iREase+CTD+REase<-HTH<-REase+SF1-SF2-Helicase<-HTH+WYL | 1114 | Bacteria>Terrabacteria_group>Chloroflexota>Dehalococcoidia>Dehalococcoidales | GCF_001010485.1 | Dehalococcoides | CWV2_RS06675 | 1251842..1255186 | 61434 | MULTISPECIES:-Swt1-family-HEPN-domain-containing-protein-[Dehalococcoides] |
| WP_072555801.1 | DUF499_1.RM.like.systems | <-XIS-HTH<-HEPN+DUF499*<-METHYLASE<-MORC+DUF3684<-TUDOR+SF1-SF2-Helicase+iREase+CTD+REase<-HTH<-REase+SF1-SF2-Helicase<-HTH+WYL | 1114 | Bacteria>Terrabacteria_group>Chloroflexota>Dehalococcoidia>Dehalococcoidales | GCF_001889305.1 | Dehalococcoides_mccartyi | ASJ33_RS07850 | 1416330..1419674 | 61435 | Swt1-family-HEPN-domain-containing-protein-[Dehalococcoides-mccartyi] |
| PWB45748.1 | DUF499_1.RM.like.systems | LysM+TPR-><-HEPN\|\|TUDOR+SF1-SF2-Helicase+iREase+CTD+REase->MORC+DUF3684->HEPN+DUF499*-><-?<-HISKIN<-SbcC<-sGTP | 1122 | Bacteria>Terrabacteria_group>Chloroflexota>Dehalococcoidia>unclassified_Dehalococcoidia | GCA_003104995.1 | Dehalococcoidia_bacterium | C3F10_05490 | 126089..129457 | 2026734 | AAA+-family-ATPase-[Dehalococcoidia-bacterium] |
| MPZ22055.1 | DUF499_1.RM.like.systems | <-CR-Kinase<-?<-HEPN+DUF499*\|\|<-TUDOR+SF1-SF2-Helicase+iREase+CTD+REase\|\|Peptidase_M20->?-><-DNA-glycosylase<-HAD | 1112 | Bacteria>Terrabacteria_group>Chloroflexota>Dehalococcoidia>unclassified_Dehalococcoidia | GCA_009377665.1 | Dehalococcoidia_bacterium | GEU28_00570 | 3174..6512 | 2026734 | DUF499-domain-containing-protein-[Dehalococcoidia-bacterium] |
| MXW25933.1 | DUF499_1.RM.like.systems | ABhydrolase-><-?<-URI-DNase\|\|tRNA->\|\|?-><-PadR-HTH\|\|?-><-HEPN+DUF499*<-HEPN<-PolB<-METHYLASE-p | 1120 | Bacteria>Terrabacteria_group>Chloroflexota>Dehalococcoidia>unclassified_Dehalococcoidia | GCA_009835935.1 | Dehalococcoidia_bacterium | F4Z77_06550 | 10102..13464 | 2026734 | ATP-binding-protein-[Dehalococcoidia-bacterium] |
| MXX18045.1 | DUF499_1.RM.like.systems | TUDOR+SF1-SF2-Helicase+iREase+CTD+REase->Trypsin->METHYLASE->PolB->HEPN->HEPN+DUF499*->?-><-ABhydrolase\|\|STYKinase-> | 1125 | Bacteria>Terrabacteria_group>Chloroflexota>Dehalococcoidia>unclassified_Dehalococcoidia | GCA_009836455.1 | Dehalococcoidia_bacterium | F4Z48_00945 | 10112..13489 | 2026734 | ATP-binding-protein-[Dehalococcoidia-bacterium] |
| MXY35173.1 | DUF499_1.RM.like.systems | <-HEPN+DUF499*<-HEPN<-PolB<-METHYLASE<-METHYLASE<-HEPN<-TUDOR+SF1-SF2-Helicase+iREase+CTD+REase | 1125 | Bacteria>Terrabacteria_group>Chloroflexota>Dehalococcoidia>unclassified_Dehalococcoidia | GCA_009838385.1 | Dehalococcoidia_bacterium | F4Y54_00090 | 1779..5156 | 2026734 | ATP-binding-protein-[Dehalococcoidia-bacterium] |
| MCA9831553.1 | DUF499_1.RM.like.systems | TUDOR+SF1-SF2-Helicase+iREase+CTD+REase->REase->?->?->?->METHYLASE->HEPN+DUF499*->?->HerA/FtsK->PIN-> | 1134 | Bacteria>Terrabacteria_group>Chloroflexota>Dehalococcoidia>unclassified_Dehalococcoidia | GCA_020432045.1 | Dehalococcoidia_bacterium | KC495_13860 | 27821..31225 | 2026734 | ATP-binding-protein-[Dehalococcoidia-bacterium] |
| MCL4230333.1 | DUF499_1.RM.like.systems | <-HTH_17\|\|HTH->Peptidase-><-HEPN+DUF499*<-METHYLASE<-HEPN<-TUDOR+SF1-SF2-Helicase+iREase+CTD+REase | 1139 | Bacteria>Terrabacteria_group>Chloroflexota>Dehalococcoidia>unclassified_Dehalococcoidia | GCA_023382735.1 | Dehalococcoidia_bacterium | KJ053_02015 | 402494..405913 | 2026734 | DUF499-domain-containing-protein-[Dehalococcoidia-bacterium] |
| MCL4239733.1 | DUF499_1.RM.like.systems | <-SF1-SF2-Helicase+REase<-?<-SIGMA-HTH<-?<-?<-HEPN+DUF499*<-METHYLASE<-?<-MORC+DUF3684<-TUDOR+SF1-SF2-Helicase+iREase+CTD+REase\|\|HEPN-><-LysM+TPR | 1122 | Bacteria>Terrabacteria_group>Chloroflexota>Dehalococcoidia>unclassified_Dehalococcoidia | GCA_023382755.1 | Dehalococcoidia_bacterium | KJ048_00120 | 28707..32075 | 2026734 | DUF499-domain-containing-protein-[Dehalococcoidia-bacterium] |
| MCL4240066.1 | DUF499_1.RM.like.systems | <-HEPN+DUF499*<-?<-HTH<-?<-METHYLASE<-ParB+HNH | 1138 | Bacteria>Terrabacteria_group>Chloroflexota>Dehalococcoidia>unclassified_Dehalococcoidia | GCA_023382755.1 | Dehalococcoidia_bacterium | KJ048_01835 | 17815..21231 | 2026734 | DUF499-domain-containing-protein-[Dehalococcoidia-bacterium] |
| MCS6803437.1 | DUF499_1.RM.like.systems | METHYLASE-><-?\|\|HEPN+DUF499*-> | 1109 | Bacteria>Terrabacteria_group>Chloroflexota>Dehalococcoidia>unclassified_Dehalococcoidia | GCA_025054935.1 | Dehalococcoidia_bacterium | NZ773_16035 | 2535..5864 | 2026734 | Swt1-family-HEPN-domain-containing-protein-[Dehalococcoidia-bacterium] |
| WP_126627899.1 | DUF499_1.RM.like.systems | ParB->TUDOR+SF1-SF2-Helicase+iREase+CTD+REase->METHYLASE-p->DDE_Tnp_1->DDE_Tnp_1+DDE_Tnp_1_2->METHYLASE->HEPN+DUF499*->?->?-><-?<-?<-NERD+Pkinase+Pkinase+nSTAND3 | 1124 | Bacteria>Terrabacteria_group>Chloroflexota>Ktedonobacteria>Ktedonobacterales | GCF_003967575.1 | Dictyobacter_alpinus | EL633_RS14700 | 3690225..3693599 | 2014873 | DUF499-domain-containing-protein-[Dictyobacter-alpinus] |
| WP_129891798.1 | DUF499_1.RM.like.systems | MPTase-><-METHYLASE-p<-SF1-SF2-helicase+DUF1998<-?\|\|TUDOR+SF1-SF2-Helicase+iREase+CTD+REase->METHYLASE->HEPN+DUF499*-><-?<-?<-?<-?\|\|?->ABhydrolase-> | 1138 | Bacteria>Terrabacteria_group>Chloroflexota>Ktedonobacteria>Ktedonobacterales | GCF_004208415.1 | Ktedonosporobacter_rubrisoli | EPA93_RS34115 | 7851540..7854956 | 2509675 | Swt1-family-HEPN-domain-containing-protein-[Ktedonosporobacter-rubrisoli] |

| Protein-ID | System | Gene-Neighbourhoods | Length | Taxonomy | Assembly-ID | Strain | Locus-Tag | Genomic-coordinates | TaxID | Fasta-defline |
|---|---|---|---|---|---|---|---|---|---|---|
| WP_111324027.1 | DUF499_1.RM.like.systems | TUDOR+ SF1-SF2-Helicase+iREase+CTD+REase ->METHYLASE->HEPN+DUF499*-> | 1126 | Bacteria>Terrabacteria_group>Chloroflexota>Ktedonobacteria>Ktedonobacterales | GCF_005402645.1 | Thermosporothrix_hazakensis | EI42_RS18025 | 56767..60147 | 644383 | Swt1-family-HEPN-domain-containing-protein-[Thermosporothrix-hazakensis] |
| WP_201387653.1 | DUF499_1.RM.like.systems | TUDOR+ SF1-SF2-Helicase+iREase+CTD+REase ->METHYLASE-p-><-DDE_3<-PAIREDC-HTH<-PAIREDC-HTH\|\|METHYLASE->HEPN+DUF499*->ParB+HNH-><-?<-?<-?\|\|HISKIN->REC-> | 1128 | Bacteria>Terrabacteria_group>Chloroflexota>Ktedonobacteria>Ktedonobacterales | GCF_016587395.1 | Ktedonobacter_sp._SOSP1-52 | JMJ16_RS30300 | 1126374..1129760 | 2778366 | DUF499-domain-containing-protein-[Ktedonobacter-sp.-SOSP1-52] |
| WP_201359963.1 | DUF499_1.RM.like.systems | NERD+Pkinase+Pkinase+nSTAND3->?->?->APATPase+iSTAND3-><-HEPN+DUF499*<-METHYLASE<-TUDOR+ SF1-SF2-Helicase+iREase+CTD+REase \|\|?->?->?->STYKIN-> | 1127 | Bacteria>Terrabacteria_group>Chloroflexota>Ktedonobacteria>Ktedonobacterales | GCF_016587435.1 | Dictyobacter_formicarum | JMI88_RS01395 | 354947..358330 | 2778368 | DUF499-domain-containing-protein-[Dictyobacter-formicarum] |
| WP_201365551.1 | DUF499_1.RM.like.systems | <-NUDIX<-?<-?<-HEPN+DUF499*<-METHYLASE<-TUDOR+ SF1-SF2-Helicase+iREase+CTD+REase | 1131 | Bacteria>Terrabacteria_group>Chloroflexota>Ktedonobacteria>Ktedonobacterales | GCF_016587435.1 | Dictyobacter_formicarum | JMI88_RS29565 | 30848..34243 | 2778368 | Swt1-family-HEPN-domain-containing-protein-[Dictyobacter-formicarum] |
| BBH87966.1 | DUF499_1.RM.like.systems | ACET->TUDOR+ SF1-SF2-Helicase+iREase+CTD+REase ->METHYLASE->HEPN+DUF499*-><-PADR-HTH | 1126 | Bacteria>Terrabacteria_group>Chloroflexota>Ktedonobacteria>Ktedonobacterales | - | Thermosporothrix_sp._COM3 | KTC_27170 | 3109376..3112756 | 2490863 | hypothetical-protein-KTC_27170-[Thermosporothrix-sp.-COM3] |
| MBX5459182.1 | DUF499_1.RM.like.systems | LuxR-HTH+LuxR-HTH->?-><-HEPN+DUF499*<-METHYLASE-p | 1136 | Bacteria>Terrabacteria_group>Chloroflexota>Ktedonobacteria>Thermogemmatisporales | GCA_019683635.1 | Thermogemmatispora_sp. | IRZ31_20000 | 1031..4441 | 1968838 | DUF499-domain-containing-protein-[Thermogemmatispora-sp.] |
| MBX5450396.1 | DUF499_1.RM.like.systems | TUDOR+ SF1-SF2-Helicase+iREase+CTD+REase ->METHYLASE->HEPN+DUF499*-><-?<-LuxR-HTH+LuxR-HTH | 1139 | Bacteria>Terrabacteria_group>Chloroflexota>Ktedonobacteria>Thermogemmatisporales | GCA_019683755.1 | Thermogemmatispora_sp. | IRZ24_10040 | 7541..10960 | 1968838 | DUF499-domain-containing-protein-[Thermogemmatispora-sp.] |
| WP_069804546.1 | DUF499_1.RM.like.systems | TUDOR+ SF1-SF2-Helicase+iREase+CTD+REase ->METHYLASE->HEPN+DUF499*-><-?<-LuxR-HTH+LuxR-HTH<-?\|\|XIS-HTH->?->HTH+TPR+TPR-> | 1138 | Bacteria>Terrabacteria_group>Chloroflexota>Ktedonobacteria>Thermogemmatisporales | GCF_001748285.1 | Thermogemmatispora_onikobensis | BGC09_RS13650 | 65405..68821 | 732234 | Swt1-family-HEPN-domain-containing-protein-[Thermogemmatispora-onikobensis] |
| WP_151729028.1 | DUF499_1.RM.like.systems | <-HTH+TPR+TPR\|\|?->LuxR-HTH+LuxR-HTH->?-><-HEPN+DUF499*<-METHYLASE<-TUDOR+ SF1-SF2-Helicase+iREase+CTD+REase | 1139 | Bacteria>Terrabacteria_group>Chloroflexota>Ktedonobacteria>Thermogemmatisporales | GCF_008974285.1 | Thermogemmatispora_aurantia | KTAU_RS14380 | 459463..462882 | 2045279 | Swt1-family-HEPN-domain-containing-protein-[Thermogemmatispora-aurantia] |
| BBH92994.1 | DUF499_1.RM.like.systems | TUDOR+ SF1-SF2-Helicase+iREase+CTD+REase ->METHYLASE->HEPN+DUF499*-><-?<-LuxR-HTH+LuxR-HTH<-?\|\|HTH+TPR+TPR-> | 1130 | Bacteria>Terrabacteria_group>Chloroflexota>Ktedonobacteria>Thermogemmatisporales | - | Thermogemmatispora_argillosa | KTA_11930 | 1512698..1516090 | 2045280 | hypothetical-protein-KTA_11930-[Thermogemmatispora-argillosa] |
| PFG73567.1 | DUF499_1.RM.like.systems | PolB->HEPN->?->TUDOR+ SF1-SF2-Helicase+iREase+CTD+REase ->OLD-ABC+TOPRIM->METHYLASE->HEPN+DUF499*->?-><-HAD | 1111 | Bacteria>Terrabacteria_group>Chloroflexota>Tepidiformia>Tepidiformales | GCA_002563855.1 | Tepidiforma_thermophila | A9A59_0767 | 812331..815666 | 2761530 | putative-AAA+-superfamily-ATPase-[Chloroflexi-bacterium-G233] |
| WP_098503018.1 | DUF499_1.RM.like.systems | PolB->HEPN->?->TUDOR+ SF1-SF2-Helicase+iREase+CTD+REase ->OLD-ABC+TOPRIM->METHYLASE->HEPN+DUF499*->?-><-HAD | 1111 | Bacteria>Terrabacteria_group>Chloroflexota>Tepidiformia>Tepidiformales | GCA_002563855.1 | Tepidiforma_thermophila | A9A59_RS03825 | 812331..815666 | 2761530 | Swt1-family-HEPN-domain-containing-protein-[Tepidiforma-thermophila] |
| WP_158067647.1 | DUF499_1.RM.like.systems | RNase+SF1-SF2-Helicase->CRISPR_Cas2->TUDOR+ SF1-SF2-Helicase+iREase+CTD+REase ->?->METHYLASE-p->METHYLASE->HEPN+DUF499*-><-?<-?\|\|?-><-?\|\|?-><-ZnR | 1109 | Bacteria>Terrabacteria_group>Chloroflexota>Tepidiformia>Tepidiformales | GCF_008838325.1 | Tepidiforma_bonchosmolovskayae | Tbon_RS10440 | 2086716..2090045 | 2601677 | Swt1-family-HEPN-domain-containing-protein-[Tepidiforma-bonchosmolovskayae] |
| MBE7520719.1 | DUF499_1.RM.like.systems | sGTP-><-PAIREDC-HTH<-?<-PAPS_reduct<-?<-KAP-NTPase<-HEPN+DUF499*<-METHYLASE<-?<-?<-TUDOR+ SF1-SF2-Helicase+iREase+CTD+REase <-LysM+TPR | 1132 | Bacteria>Terrabacteria_group>Chloroflexota>Thermoflexia>Thermoflexales | GCA_015075555.1 | Thermoflexaceae_bacterium | HS107_15925 | 106294..109692 | 2739507 | DUF499-domain-containing-protein-[Thermoflexaceae-bacterium] |
| MBO9362157.1 | DUF499_1.RM.like.systems | <-HEPN+DUF499*<-HEPN<-PolB<-METHYLASE | 1119 | Bacteria>Terrabacteria_group>Chloroflexota>Thermoflexia>Thermoflexales | GCA_017743605.1 | Thermoflexus_sp. | J7452_08145 | 638..3997 | 1969742 | DUF499-domain-containing-protein-[Thermoflexus-sp.] |
| QWK10277.1 | DUF499_1.RM.like.systems | <-OrfB_IS605+OrfB_Zn_ribbon<-?\|\|ABC-ATPase+SbcC-><-?<-?<-?<-HEPN+DUF499*<-HEPN<-PolB<-METHYLASE<-REase<-OLD-ABC+TOPRIM<-TUDOR+ SF1-SF2-Helicase+iREase+CTD+REase | 1119 | Bacteria>Terrabacteria_group>Chloroflexota>Thermoflexia>Thermoflexales | GCA_018771565.1 | Thermoflexus_hugenholtzii | KNN16_13035 | 2858619..2861978 | 1495650 | MAG:-DUF499-domain-containing-protein-[Thermoflexus-hugenholtzii] |
| MCE7937169.1 | DUF499_1.RM.like.systems | <-HEPN+DUF499*<-SF1-SF2-Helicase+HNH<-?<-?<-ORC-CDC6+REase | 1145 | Bacteria>Terrabacteria_group>Chloroflexota>unclassified_Chloroflexota | - | Chloroflexi_bacterium_CFX6 | DYG90_00940 | 4..3441 | 2293710 | ATP-binding-protein-[Chloroflexi-bacterium-CFX6] |
| MBM2811009.1 | DUF499_1.RM.like.systems | METHYLASE-p-><-?\|\|?->HicB_lk_antitox->HicA_toxin->HEPN+DUF499->DUF499*-> | 696 | Bacteria>Terrabacteria_group>Chloroflexota>unclassified_Chloroflexota | - | Chloroflexota_bacterium | HW416_1768 | 4823..6913 | 2026724 | ATPase-superfamily-[Chloroflexota-bacterium] |
| MBG54827.1 | DUF499_1.RM.like.systems | TUDOR+ SF1-SF2-Helicase+iREase+CTD+REase ->METHYLASE->HEPN+DUF499*-><-tRNA<-?<-?<-?<-?<-?<-RNase | 1093 | Bacteria>Terrabacteria_group>Chloroflexota>unclassified_Chloroflexota | GCA_002715705.1 | Chloroflexota_bacterium | CL795_03415 | 7874..11155 | 2026724 | AAA+-family-ATPase-[Chloroflexi-bacterium] |
| HAF11542.1 | DUF499_1.RM.like.systems | <-GDResolvase+Recombinase+ZnR<-?<-HEPN+DUF499*<-METHYLASE<-TUDOR+ SF1-SF2-Helicase+iREase+CTD+REase <-Schlafen+HATPase+HTH | 1109 | Bacteria>Terrabacteria_group>Chloroflexota>unclassified_Chloroflexota | GCA_003446795.1 | Chloroflexota_bacterium | DCK98_15880 | 2135..5464 | 2026724 | AAA+-family-ATPase-[Chloroflexi-bacterium] |
| MXW23202.1 | DUF499_1.RM.like.systems | TUDOR+ SF1-SF2-Helicase+iREase+CTD+REase -p->PolB->HEPN->HEPN+DUF499*-> | 1125 | Bacteria>Terrabacteria_group>Chloroflexota>unclassified_Chloroflexota | GCA_009835505.1 | Chloroflexota_bacterium | F4Z96_00705 | 5471..8848 | 2026724 | ATP-binding-protein-[Chloroflexi-bacterium] |
| MXX31451.1 | DUF499_1.RM.like.systems | <-REase\|\|TUDOR+ SF1-SF2-Helicase+iREase+CTD+REase ->HEPN->METHYLASE->Polbeta->HEPN->HEPN+DUF499*-> | 1112 | Bacteria>Terrabacteria_group>Chloroflexota>unclassified_Chloroflexota | GCA_009836445.1 | Chloroflexota_bacterium | F4Z51_04295 | 8660..11998 | 2026724 | ATP-binding-protein-[Chloroflexi-bacterium] |
| MXX47637.1 | DUF499_1.RM.like.systems | VirD4-TraG+HerA->PadR-HTH->RADICAL-SAM->?->?-><-HEPN+DUF499*<-HEPN<-Polbeta<-METHYLASE<-Trypsin<-TUDOR+ SF1-SF2-Helicase+iREase+CTD+REase \|\|REase-> | 1112 | Bacteria>Terrabacteria_group>Chloroflexota>unclassified_Chloroflexota | GCA_009836695.1 | Chloroflexota_bacterium | F4Z38_04900 | 11069..14407 | 2026724 | ATP-binding-protein-[Chloroflexi-bacterium] |
| MXZ46491.1 | DUF499_1.RM.like.systems | <-GDResolvase+Recombinase+ZnR<-?\|\|?->?-><-BirA-HTH+CRP-HTH\|\|?-><-HEPN+DUF499*<-HEPN<-PolB<-METHYLASE<-TUDOR+ SF1-SF2-Helicase+iREase+CTD+REase \|\|PIN+ACET+EVE->ASCH-> | 1125 | Bacteria>Terrabacteria_group>Chloroflexota>unclassified_Chloroflexota | GCA_009837285.1 | Chloroflexota_bacterium | F4Z08_05790 | 8633..12010 | 2026724 | ATP-binding-protein-[Chloroflexi-bacterium] |
| MXY00663.1 | DUF499_1.RM.like.systems | <-REase\|\|TUDOR+ SF1-SF2-Helicase+iREase+CTD+REase ->METHYLASE->METHYLASE-p->HEPN+DUF499*-><-?\|\|?-><-ABhydrolase | 1058 | Bacteria>Terrabacteria_group>Chloroflexota>unclassified_Chloroflexota | GCA_009838085.1 | Chloroflexota_bacterium | F4Y67_07570 | 133498..136674 | 2026724 | ATP-binding-protein-[Chloroflexi-bacterium] |
| MXY13704.1 | DUF499_1.RM.like.systems | METHYLASE-p->HEPN+DUF499*-> | 1057 | Bacteria>Terrabacteria_group>Chloroflexota>unclassified_Chloroflexota | GCA_009838555.1 | Chloroflexota_bacterium | F4Y46_06845 | 861..4034 | 2026724 | ATP-binding-protein-[Chloroflexi-bacterium] |
| MYD93547.1 | DUF499_1.RM.like.systems | DUF429->DOC-><-HEPN+DUF499*<-METHYLASE<-TUDOR+ SF1-SF2-Helicase+iREase+CTD+REase \|\|MORC+DUF3684-> | 1098 | Bacteria>Terrabacteria_group>Chloroflexota>unclassified_Chloroflexota | GCA_009839425.1 | Chloroflexota_bacterium | F4Y02_07625 | 2005..5301 | 2026724 | ATP-binding-protein-[Chloroflexi-bacterium] |
| MYD10935.1 | DUF499_1.RM.like.systems | HEPN+DUF499*-><-DUF2961<-ABC-ATPase+SbcC<-ABC-ATPase+SbcC | 1106 | Bacteria>Terrabacteria_group>Chloroflexota>unclassified_Chloroflexota | GCA_009841405.1 | Chloroflexota_bacterium | F4X02_12950 | 92..3412 | 2026724 | ATP-binding-protein-[Chloroflexi-bacterium] |
| MYJ01661.1 | DUF499_1.RM.like.systems | METHYLASE-p->Polbeta->HEPN->HEPN+DUF499*-><-PIN | 1112 | Bacteria>Terrabacteria_group>Chloroflexota>unclassified_Chloroflexota | GCA_009844585.1 | Chloroflexota_bacterium | F4102_05150 | 3394..6732 | 2026724 | ATP-binding-protein-[Chloroflexi-bacterium] |
| NJD65682.1 | DUF499_1.RM.like.systems | sGTP->SbcC->HISKIN->?-><-HEPN+DUF499*<-MORC+DUF3684<-TUDOR+ SF1-SF2-Helicase+iREase+CTD+REase \|\|HEPN-><-LysM+TPR | 1122 | Bacteria>Terrabacteria_group>Chloroflexota>unclassified_Chloroflexota | GCA_012027015.1 | Chloroflexota_bacterium | FIB00_10630 | 11912..15280 | 2026724 | ATP-binding-protein-[Chloroflexi-bacterium] |
| NJD66806.1 | DUF499_1.RM.like.systems | sGTP-><-SF1-SF2-Helicase<-OLD-ABC+TOPRIM<-HEPN+DUF499*<-METHYLASE<-DUF4326<-?<-DUF488-1<-TUDOR+ SF1-SF2-Helicase+iREase+CTD+REase <-LysM+TPR | 1122 | Bacteria>Terrabacteria_group>Chloroflexota>unclassified_Chloroflexota | GCA_012027015.1 | Chloroflexota_bacterium | FIB00_16460 | 5569..8937 | 2026724 | ATP-binding-protein-[Chloroflexi-bacterium] |
| NLE83803.1 | DUF499_1.RM.like.systems | TUDOR+ SF1-SF2-Helicase+iREase+CTD+REase -p->METHYLASE->HEPN+DUF499*-> | 944 | Bacteria>Terrabacteria_group>Chloroflexota>unclassified_Chloroflexota | GCA_012514645.1 | Chloroflexota_bacterium | GX603_04735 | 3343..6175 | 2026724 | ATP-binding-protein-[Chloroflexi-bacterium] |
| NLG97290.1 | DUF499_1.RM.like.systems | METHYLASE->Peptidase->HEPN+DUF499*-><-MULE-TRANSPOSASE<-MULE-TRANSPOSASE<-?<-tRNA<-REase<-Abi<-Glyoxalase | 1131 | Bacteria>Terrabacteria_group>Chloroflexota>unclassified_Chloroflexota | GCA_012516155.1 | Chloroflexota_bacterium | GX491_08015 | 33163..36558 | 2026724 | ATP-binding-protein-[Chloroflexi-bacterium] |
| NLH06920.1 | DUF499_1.RM.like.systems | METHYLASE-p->?->HEPN+DUF499*-> | 445 | Bacteria>Terrabacteria_group>Chloroflexota>unclassified_Chloroflexota | GCA_012516245.1 | Chloroflexota_bacterium | GX484_04875 | 1862..3195 | 2026724 | ATP-binding-protein-[Chloroflexota-bacterium] |
| NLS78478.1 | DUF499_1.RM.like.systems | <-HEPN+DUF499*<-?<-METHYLASE-p | 1075 | Bacteria>Terrabacteria_group>Chloroflexota>unclassified_Chloroflexota | GCA_012689325.1 | Chloroflexota_bacterium | GXY76_14575 | 1..3224 | 2026724 | ATP-binding-protein-[Chloroflexi-bacterium] |
| NOK58906.1 | DUF499_1.RM.like.systems | STAS->SAM-methylase->?-><-HEPN+DUF499*<-?<-METHYLASE<-TUDOR+ SF1-SF2-Helicase+iREase+CTD+REase | 1106 | Bacteria>Terrabacteria_group>Chloroflexota>unclassified_Chloroflexota | GCA_013122435.1 | Chloroflexi_bacterium_AL-N1 | GFH23_1086616n269 | 267583..270903 | 2686367 | ATP-binding-protein-[Chloroflexi-bacterium-AL-N1] |
| NWF79207.1 | DUF499_1.RM.like.systems | TUDOR+ SF1-SF2-Helicase+iREase+CTD+REase -><-?\|\|METHYLASE-><-PIN<-DUF433\|\|HEPN+DUF499*-> | 825 | Bacteria>Terrabacteria_group>Chloroflexota>unclassified_Chloroflexota | GCA_013388805.1 | Chloroflexota_bacterium | HXY37_04055 | 14161..16634 | 2026724 | ATP-binding-protein-[Chloroflexi-bacterium] |
| NWJ47051.1 | DUF499_1.RM.like.systems | <-Peptidase<-?<-SIGMA-HTH<-?<-?<-HEPN+DUF499*<-Y1_Tnp<-?<-TUDOR+ SF1-SF2-Helicase+iREase+CTD+REase <-?<-CRISPR_Cas2<-Cas_Cas1 | 1122 | Bacteria>Terrabacteria_group>Chloroflexota>unclassified_Chloroflexota | GCA_013390565.1 | Chloroflexota_bacterium | HXX08_14415 | 53237..56605 | 2026724 | ATP-binding-protein-[Chloroflexi-bacterium] |
| MBC8453263.1 | DUF499_1.RM.like.systems | METHYLASE-p->METHYLASE-p->HEPN+DUF499*-> | 404 | Bacteria>Terrabacteria_group>Chloroflexota>unclassified_Chloroflexota | GCA_014383785.1 | Chloroflexota_bacterium | H8D69_02180 | 1481..2692 | 2026724 | ATP-binding-protein-[Chloroflexota-bacterium] |
| MBF6606506.1 | DUF499_1.RM.like.systems | TUDOR+ SF1-SF2-Helicase+iREase+CTD+REase ->SIGMA-HTH+CR-ATPase9+Schlafen->METHYLASE->HEPN+DUF499*->?->?-><-Schlafen+HATPase+HTH | 1085 | Bacteria>Terrabacteria_group>Chloroflexota>unclassified_Chloroflexota | GCA_015478725.1 | Chloroflexota_bacterium | IVW53_13110 | 14708..17965 | 2026724 | DUF499-domain-containing-protein-[Chloroflexi-bacterium] |
| MBI1882242.1 | DUF499_1.RM.like.systems | <-HEPN+DUF499*<-REase<-METHYLASE-p<-METHYLASE-p | 401 | Bacteria>Terrabacteria_group>Chloroflexota>unclassified_Chloroflexota | GCA_016178315.1 | Chloroflexota_bacterium | HYR94_29060 | 1..1204 | 2026724 | ATP-binding-protein-[Chloroflexota-bacterium] |
| MBI3978251.1 | DUF499_1.RM.like.systems | <-RNase<-?\|\|?-><-?<-HEPN+DUF499*<-METHYLASE | 1129 | Bacteria>Terrabacteria_group>Chloroflexota>unclassified_Chloroflexota | GCA_016199325.1 | Chloroflexota_bacterium | HY331_08705 | 13275..16664 | 2026724 | ATP-binding-protein-[Chloroflexi-bacterium] |
| MBI3979239.1 | DUF499_1.RM.like.systems | <-PemK_toxin<-?\|\|Peptidase_M20-><-?<-?<-HEPN+DUF499*<-REase | 1136 | Bacteria>Terrabacteria_group>Chloroflexota>unclassified_Chloroflexota | GCA_016199325.1 | Chloroflexota_bacterium | HY331_13735 | 11922..15332 | 2026724 | ATP-binding-protein-[Chloroflexi-bacterium] |
| MBI3961042.1 | DUF499_1.RM.like.systems | METHYLASE-p->OLD-ABC->PIN->HEPN+DUF499*-> | 1102 | Bacteria>Terrabacteria_group>Chloroflexota>unclassified_Chloroflexota | GCA_016199425.1 | Chloroflexota_bacterium | HY328_19695 | 3483..6791 | 2026724 | ATP-binding-protein-[Chloroflexi-bacterium] |
| MBI4673984.1 | DUF499_1.RM.like.systems | TPR+TPR+TPR+TPR-><-HEPN+DUF499*<-NTP_transf<-?<-DOC+HTH<-TUDOR+ SF1-SF2-Helicase+iREase+CTD+REase | 1130 | Bacteria>Terrabacteria_group>Chloroflexota>unclassified_Chloroflexota | GCA_016209205.1 | Chloroflexota_bacterium | HY741_20245 | 1845..5237 | 2026724 | ATP-binding-protein-[Chloroflexi-bacterium] |
| MBI4332978.1 | DUF499_1.RM.like.systems | <-PIN<-MazE_antitoxin<-HicA_toxin<-HEPN+DUF499*<-METHYLASE<-Schlafen+HATPase+HTH<-TUDOR+ SF1-SF2-Helicase+iREase+CTD+REase | 1127 | Bacteria>Terrabacteria_group>Chloroflexota>unclassified_Chloroflexota | GCA_016210565.1 | Chloroflexota_bacterium | HY673_17055 | 9786..13169 | 2026724 | ATP-binding-protein-[Chloroflexi-bacterium] |
| MBI5949977.1 | DUF499_1.RM.like.systems | SIGMA-HTH-><-Pkinase<-?<-Phage_integrase<-Phage_integrase<-HEPN+DUF499*<-?<-METHYLASE<-?<-?<-?<-YtH | 1133 | Bacteria>Terrabacteria_group>Chloroflexota>unclassified_Chloroflexota | GCA_016234445.1 | Chloroflexota_bacterium | HY875_17715 | 143137..146538 | 2026724 | ATP-binding-protein-[Chloroflexi-bacterium] |
| MBK7919048.1 | DUF499_1.RM.like.systems | DDE_5-><-DDE_Tnp_IS1\|\|METHYLASE-p->?->?->HEPN-p->HEPN+DUF499*->DUF499-><-DDE_5<-DDE_5\|\|?-><-DDE_5+DDE_Tnp_2 | 582 | Bacteria>Terrabacteria_group>Chloroflexota>unclassified_Chloroflexota | GCA_016713795.1 | Chloroflexota_bacterium | IPJ94_22860 | 29678..31426 | 2026724 | DUF499-domain-containing-protein-[Chloroflexota-bacterium] |
| MBK7919049.1 | DUF499_1.RM.like.systems | <-DDE_Tnp_IS1\|\|METHYLASE-p->?->?->HEPN-p->HEPN+DUF499->DUF499*-><-DDE_5<-DDE_5\|\|?-><-DDE_5+DDE_Tnp_2 | 452 | Bacteria>Terrabacteria_group>Chloroflexota>unclassified_Chloroflexota | GCA_016713795.1 | Chloroflexota_bacterium | IPJ94_22865 | 31366..32724 | 2026724 | hypothetical-protein-[Chloroflexota-bacterium] |
| MBK8988277.1 | DUF499_1.RM.like.systems | TUDOR+ SF1-SF2-Helicase+iREase+CTD+REase ->?->PIN->CopG/RHH->PIN->METHYLASE-p->HEPN+DUF499*-><-AbiEii<-?\|\|?->?->?->ABC-ATPase+SbcC-> | 1097 | Bacteria>Terrabacteria_group>Chloroflexota>unclassified_Chloroflexota | GCA_016716065.1 | Chloroflexota_bacterium | IPM39_19790 | 148627..151920 | 2026724 | DUF499-domain-containing-protein-[Chloroflexi-bacterium] |
| MBM2811008.1 | DUF499_1.RM.like.systems | METHYLASE-p->METHYLASE-p-><-?\|\|?->HicB_lk_antitox->HicA_toxin->HEPN+DUF499*->DUF499-> | 384 | Bacteria>Terrabacteria_group>Chloroflexota>unclassified_Chloroflexota | GCA_016844585.1 | Chloroflexota_bacterium | HW416_1767 | 3649..4803 | 2026724 | ATPase-of-the-superfamily-[Chloroflexota-bacterium] |
| MBN9390798.1 | DUF499_1.RM.like.systems | <-HEPN+DUF499*<-METHYLASE<-YtH<-?<-TUDOR+ SF1-SF2-Helicase+iREase+CTD+REase <-HTH_17<-ArdCN+MPTase | 1111 | Bacteria>Terrabacteria_group>Chloroflexota>unclassified_Chloroflexota | GCA_017307685.1 | Chloroflexota_bacterium | J0I20_22395 | 2393..5728 | 2026724 | DUF499-domain-containing-protein-[Chloroflexi-bacterium] |
| MBP8000880.1 | DUF499_1.RM.like.systems | <-XIS-HTH<-?\|\|SF1-SF2-helicase+RdgC+Beta-barrel+REase-><-HEPN+DUF499*<-?<-?<-METHYLASE<-ParB+HNH<-TUDOR+ SF1-SF2-Helicase+iREase+CTD+REase | 1097 | Bacteria>Terrabacteria_group>Chloroflexota>unclassified_Chloroflexota | GCA_018004095.1 | Chloroflexota_bacterium | KA338_16630 | 13590..16883 | 2026724 | DUF499-domain-containing-protein-[Chloroflexi-bacterium] |
| MBU1746748.1 | DUF499_1.RM.like.systems | <-HEPN+DUF499*<-METHYLASE<-?<-ParB+HNH<-TUDOR+ SF1-SF2-Helicase+iREase+CTD+REase | 551 | Bacteria>Terrabacteria_group>Chloroflexota>unclassified_Chloroflexota | GCA_018823415.1 | Chloroflexota_bacterium | KKA73_03595 | 1..1652 | 2026724 | DUF499-domain-containing-protein-[Chloroflexota-bacterium] |
| MBU6150404.1 | DUF499_1.RM.like.systems | <-HEPN+DUF499*<-METHYLASE | 1130 | Bacteria>Terrabacteria_group>Chloroflexota>unclassified_Chloroflexota | GCA_018969665.1 | Chloroflexota_bacterium | KGR25_03080 | 672..4064 | 2026724 | DUF499-domain-containing-protein-[Chloroflexi-bacterium] |
| MBV9281297.1 | DUF499_1.RM.like.systems | METHYLASE-p->Glyoxalase->HEPN+DUF499*->REase-><-SIGMA-HTH+SIGMA-HTH<-?<-Glyoxalase | 1102 | Bacteria>Terrabacteria_group>Chloroflexota>unclassified_Chloroflexota | GCA_019245855.1 | Chloroflexota_bacterium | JOZ41_14550 | 2324..5632 | 2026724 | DUF499-domain-containing-protein-[Chloroflexi-bacterium] |
| MBX3030766.1 | DUF499_1.RM.like.systems | <-NUDIX<-?<-HEPN+DUF499*<-?\|\|HTH_31-><-TUDOR+ SF1-SF2-Helicase+iREase+CTD+REase <-?<-DOC+HTH\|\|ABC-ATPase+SbcC-> | 1095 | Bacteria>Terrabacteria_group>Chloroflexota>unclassified_Chloroflexota | GCA_019637815.1 | Chloroflexota_bacterium | KF809_11435 | 95688..98975 | 2026724 | DUF499-domain-containing-protein-[Chloroflexi-bacterium] |
| MBX6773122.1 | DUF499_1.RM.like.systems | <-HEPN+DUF499*<-METHYLASE | 328 | Bacteria>Terrabacteria_group>Chloroflexota>unclassified_Chloroflexota | GCA_019690095.1 | Chloroflexota_bacterium | IRY83_15455 | 1..985 | 2026724 | AAA+-family-ATPase-[Chloroflexota-bacterium] |
| MCA0453442.1 | DUF499_1.RM.like.systems | HTH-><-?<-?<-?<-?<-HEPN+DUF499*<-METHYLASE<-HEPN<-TUDOR+ SF1-SF2-Helicase+iREase+CTD+REase | 1099 | Bacteria>Terrabacteria_group>Chloroflexota>unclassified_Chloroflexota | GCA_020161935.1 | Chloroflexota_bacterium | LCI00_05665 | 1284524..1287823 | 2026724 | DUF499-domain-containing-protein-[Chloroflexi-bacterium] |
| MCA0453853.1 | DUF499_1.RM.like.systems | <-GDResolvase<-?<-zf-ISL3+DDE_Tnp_ISL3<-?<-?<-?<-HEPN+DUF499*<-PIN<-DUF433<-METHYLASE<-PIN<-?<-TUDOR+ SF1-SF2-Helicase+iREase+CTD+REase | 1103 | Bacteria>Terrabacteria_group>Chloroflexota>unclassified_Chloroflexota | GCA_020161935.1 | Chloroflexota_bacterium | LCI00_07765 | 53613..56924 | 2026724 | DUF499-domain-containing-protein-[Chloroflexi-bacterium] |
| MCC6174343.1 | DUF499_1.RM.like.systems | TUDOR+ SF1-SF2-Helicase+iREase+CTD+REase ->Schlafen->BirA-HTH->METHYLASE->ParD->PIN->HEPN+DUF499*-><-?\|\|?->Aldedh-> | 1160 | Bacteria>Terrabacteria_group>Chloroflexota>unclassified_Chloroflexota | GCA_020845495.1 | Chloroflexota_bacterium | IT305_03475 | 199239..202721 | 2026724 | DUF499-domain-containing-protein-[Chloroflexi-bacterium] |
| MCC6175191.1 | DUF499_1.RM.like.systems | <-METHYLASE-p<-HEPN+DUF499*<-METHYLASE<-TUDOR+ SF1-SF2-Helicase+iREase+CTD+REase | 1129 | Bacteria>Terrabacteria_group>Chloroflexota>unclassified_Chloroflexota | GCA_020845495.1 | Chloroflexota_bacterium | IT305_07815 | 409..3798 | 2026724 | DUF499-domain-containing-protein-[Chloroflexi-bacterium] |
| MCC6176203.1 | DUF499_1.RM.like.systems | TUDOR+ SF1-SF2-Helicase+iREase+CTD+REase ->Schlafen+HATPase+HTH->Cluster1090_5clades->METHYLASE->HEPN+DUF499*-><-REase | 1129 | Bacteria>Terrabacteria_group>Chloroflexota>unclassified_Chloroflexota | GCA_020845495.1 | Chloroflexota_bacterium | IT305_12930 | 12450..15839 | 2026724 | DUF499-domain-containing-protein-[Chloroflexi-bacterium] |
| MCC6178177.1 | DUF499_1.RM.like.systems | <-HTH_3<-Tox-RelE<-SF1-SF2-Helicase<-OLD-ABC+TOPRIM<-REase<-HEPN+DUF499*<-METHYLASE<-wHTH+REase<-TUDOR+ SF1-SF2-Helicase+iREase+CTD+REase \|\|?->Peptidase_S41-> | 1117 | Bacteria>Terrabacteria_group>Chloroflexota>unclassified_Chloroflexota | GCA_020845495.1 | Chloroflexota_bacterium | IT305_22975 | 6439..9792 | 2026724 | DUF499-domain-containing-protein-[Chloroflexi-bacterium] |
| MCC6567532.1 | DUF499_1.RM.like.systems | <-SF1-SF2-Helicase<-?<-?<-HEPN+DUF499*<-REase<-?<-METHYLASE<-?<-?<-TUDOR+ SF1-SF2-Helicase+iREase+CTD+REase | 1096 | Bacteria>Terrabacteria_group>Chloroflexota>unclassified_Chloroflexota | GCA_020848335.1 | Chloroflexota_bacterium | IT298_17380 | 16185..19475 | 2026724 | DUF499-domain-containing-protein-[Chloroflexi-bacterium] |
| MCC6627902.1 | DUF499_1.RM.like.systems | METHYLASE->?->?->HEPN+DUF499*-> | 524 | Bacteria>Terrabacteria_group>Chloroflexota>unclassified_Chloroflexota | GCA_020849725.1 | Chloroflexota_bacterium | IT340_10930 | 2805..4377 | 2026724 | DUF499-domain-containing-protein-[Chloroflexota-bacterium] |
| MCE7858272.1 | DUF499_1.RM.like.systems | HTH+UbE1-><-Schlafen<-?<-NACHT<-ParB+HNH<-HEPN+DUF499*<-METHYLASE<-MORC+DUF3684<-TUDOR+ SF1-SF2-Helicase+iREase+CTD+REase <-?<-?<-HTH+WYL | 1112 | Bacteria>Terrabacteria_group>Chloroflexota>unclassified_Chloroflexota | GCA_021404705.1 | Chloroflexi_bacterium_CFX2 | DYG86_00620 | 111078..114416 | 2293708 | ATP-binding-protein-[Chloroflexi-bacterium-CFX2] |
| MCQ3930198.1 | DUF499_1.RM.like.systems | ParB->?-><-?<-?<-STYKinase<-TIR<-HEPN+DUF499*<-HEPN<-PolB<-METHYLASE<-Pkinase<-TUDOR+ SF1-SF2-Helicase+iREase+CTD+REase | 1098 | Bacteria>Terrabacteria_group>Chloroflexota>unclassified_Chloroflexota | GCA_024434115.1 | Chloroflexota_bacterium | DPW16_07035 | 13947..17243 | 2026724 | AAA+-family-ATPase-[Chloroflexi-bacterium] |
| MCQ3935712.1 | DUF499_1.RM.like.systems | SF1-SF2-Helicase-><-PIWI<-REase+PNTD1+PNTD2<-HEPN+DUF499*<-METHYLASE<-SF1-SF2-Helicase<-?<-TUDOR+ SF1-SF2-Helicase+iREase+CTD+REase \|\|OLD-ABC+TOPRIM->SF1-SF2-Helicase-> | 1108 | Bacteria>Terrabacteria_group>Chloroflexota>unclassified_Chloroflexota | GCA_024434165.1 | Chloroflexota_bacterium | DPW18_01570 | 11480..14806 | 2026724 | AAA+-family-ATPase-[Chloroflexi-bacterium] |
| MCX6028868.1 | DUF499_1.RM.like.systems | TPR+TPR->?->HEPN-p->RNaseH->?->HEPN+DUF499*-> | 1119 | Bacteria>Terrabacteria_group>Chloroflexota>unclassified_Chloroflexota | GCA_026389845.1 | Chloroflexota_bacterium | NT169_06150 | 82672..86031 | 2026724 | MAG:-Swt1-family-HEPN-domain-containing-protein-[Chloroflexi-bacterium] |
| MCX6066946.1 | DUF499_1.RM.like.systems | <-REase<-?<-?<-?<-DUF499-p<-HEPN+DUF499*<-PIN<-MazE_antitoxin | 972 | Bacteria>Terrabacteria_group>Chloroflexota>unclassified_Chloroflexota | GCA_026389995.1 | Chloroflexota_bacterium | NT121_14520 | 4682..7598 | 2026724 | MAG:-Swt1-family-HEPN-domain-containing-protein,-partial-[Chloroflexi-bacterium] |
| MCY3559222.1 | DUF499_1.RM.like.systems | TUDOR+ SF1-SF2-Helicase+iREase+CTD+REase -p->Trypsin->METHYLASE->Polbeta->HEPN->HEPN+DUF499*->MORC+REase-><-REase<-PadR-HTH | 1111 | Bacteria>Terrabacteria_group>Chloroflexota>unclassified_Chloroflexota | GCA_026702785.1 | Chloroflexota_bacterium | OXH13_11370 | 6173..9508 | 2026724 | MAG:-Swt1-family-HEPN-domain-containing-protein-[Chloroflexi-bacterium] |
| MCY3570526.1 | DUF499_1.RM.like.systems | TUDOR+ SF1-SF2-Helicase+iREase+CTD+REase -p->HEPN->METHYLASE->Polbeta->HEPN->HEPN+DUF499*-><-?<-METHYLASE-p<-?<-ParB+HNH<-wHTH+REase<-PadR-HTH | 1110 | Bacteria>Terrabacteria_group>Chloroflexota>unclassified_Chloroflexota | GCA_026705025.1 | Chloroflexota_bacterium | OXH19_04200 | 6390..9722 | 2026724 | MAG:-Swt1-family-HEPN-domain-containing-protein-[Chloroflexi-bacterium] |
| MCY3747797.1 | DUF499_1.RM.like.systems | PolB->METHYLASE->PolB->?->?->HEPN+DUF499*->?->PADR-HTH->?-><-REase<-?<-HTH+UbE1 | 1113 | Bacteria>Terrabacteria_group>Chloroflexota>unclassified_Chloroflexota | GCA_026706265.1 | Chloroflexota_bacterium | OXG64_00755 | 49727..53068 | 2026724 | MAG:-Swt1-family-HEPN-domain-containing-protein-[Chloroflexi-bacterium] |
| MCY3801423.1 | DUF499_1.RM.like.systems | HEPN+DUF499*->METHYLASE->URI-DNase-><-?\|\|METHYLASE-p-><-ABhydrolase\|\|NUDIX-> | 1121 | Bacteria>Terrabacteria_group>Chloroflexota>unclassified_Chloroflexota | GCA_026706785.1 | Chloroflexota_bacterium | OXG46_07570 | 255383..258748 | 2026724 | MAG:-DUF499-domain-containing-protein-[Chloroflexi-bacterium] |
| MCY3923477.1 | DUF499_1.RM.like.systems | SF1-SF2-Helicase+REase-><-HEPN+DUF499*<-HEPN<-PolB<-METHYLASE<-HEPN<-TUDOR+ SF1-SF2-Helicase+iREase+CTD+REase | 1114 | Bacteria>Terrabacteria_group>Chloroflexota>unclassified_Chloroflexota | GCA_026707265.1 | Chloroflexota_bacterium | OXG27_13945 | 4231..7575 | 2026724 | MAG:-Swt1-family-HEPN-domain-containing-protein-[Chloroflexi-bacterium] |
| MCY3938336.1 | DUF499_1.RM.like.systems | HEPN+DUF499*->METHYLASE->URI-DNase-><-?\|\|METHYLASE-p-><-ABhydrolase\|\|NUDIX-> | 1121 | Bacteria>Terrabacteria_group>Chloroflexota>unclassified_Chloroflexota | GCA_026707405.1 | Chloroflexota_bacterium | OXG22_08290 | 17127..20492 | 2026724 | MAG:-DUF499-domain-containing-protein-[Chloroflexi-bacterium] |
| MCY3957107.1 | DUF499_1.RM.like.systems | TUDOR+ SF1-SF2-Helicase+iREase+CTD+REase ->METHYLASE->HEPN+DUF499*-> | 1113 | Bacteria>Terrabacteria_group>Chloroflexota>unclassified_Chloroflexota | GCA_026707585.1 | Chloroflexota_bacterium | OXG65_02325 | 8124..11465 | 2026724 | MAG:-DUF499-domain-containing-protein-[Chloroflexi-bacterium] |
| MCY4393097.1 | DUF499_1.RM.like.systems | SF1-SF2-Helicase-p-><-REase<-N6_MTase<-HEPN+DUF499*<-HEPN<-?<-METHYLASE<-HEPN<-TUDOR+ SF1-SF2-Helicase+iREase+CTD+REase | 1125 | Bacteria>Terrabacteria_group>Chloroflexota>unclassified_Chloroflexota | GCA_026710655.1 | Chloroflexota_bacterium | OXE43_13775 | 6581..9958 | 2026724 | MAG:-Swt1-family-HEPN-domain-containing-protein-[Chloroflexi-bacterium] |
| MCY4474618.1 | DUF499_1.RM.like.systems | <-METHYLASE<-PIN<-?<-?<-HEPN<-HEPN<-HEPN+DUF499* | 1144 | Bacteria>Terrabacteria_group>Chloroflexota>unclassified_Chloroflexota | GCA_026712855.1 | Chloroflexota_bacterium | OXC83_04195 | 178388..181822 | 2026724 | MAG:-DUF499-domain-containing-protein-[Chloroflexi-bacterium] |
| MCY4557669.1 | DUF499_1.RM.like.systems | <-NADAR<-?<-HIN-HTH<-HEPN+DUF499*<-METHYLASE<-OLD-ABC<-TUDOR+ SF1-SF2-Helicase+iREase+CTD+REase <-?<-CHTH | 1117 | Bacteria>Terrabacteria_group>Chloroflexota>unclassified_Chloroflexota | GCA_026713265.1 | Chloroflexota_bacterium | OXF79_15050 | 3333..6686 | 2026724 | MAG:-Swt1-family-HEPN-domain-containing-protein-[Chloroflexi-bacterium] |
| MCY4583299.1 | DUF499_1.RM.like.systems | METHYLASE->HEPN+DUF499*->?->?-><-ABC-ATPase+SbcC | 1128 | Bacteria>Terrabacteria_group>Chloroflexota>unclassified_Chloroflexota | GCA_026713505.1 | Chloroflexota_bacterium | OXE50_10970 | 2507..5893 | 2026724 | MAG:-Swt1-family-HEPN-domain-containing-protein-[Chloroflexi-bacterium] |
| MCY4618411.1 | DUF499_1.RM.like.systems | <-SF1-SF2-Helicase-p<-SF1-SF2-Helicase<-OLD-ABC+TOPRIM<-?<-?<-?<-HEPN+DUF499*<-METHYLASE<-TUDOR+ SF1-SF2-Helicase+iREase+CTD+REase <-?<-CHTH | 1114 | Bacteria>Terrabacteria_group>Chloroflexota>unclassified_Chloroflexota | GCA_026713765.1 | Chloroflexota_bacterium | OXD50_07650 | 54962..58306 | 2026724 | MAG:-DUF499-domain-containing-protein-[Chloroflexi-bacterium] |
| MDC0253131.1 | DUF499_1.RM.like.systems | REase->METHYLASE->HEPN+DUF499*-><-Mbetalac\|\|?-><-?<-?<-RNase | 1088 | Bacteria>Terrabacteria_group>Chloroflexota>unclassified_Chloroflexota | GCA_028346115.1 | Chloroflexota_bacterium | OAK52_03100 | 3358..6624 | 2026724 | DUF499-domain-containing-protein-[Chloroflexi-bacterium] |
| MDE2935729.1 | DUF499_1.RM.like.systems | <-HNH<-?<-OrfB_IS605+OrfB_Zn_ribbon\|\|?-><-?<-HEPN+DUF499*<-METHYLASE-p<-METHYLASE-p<-TUDOR+ SF1-SF2-Helicase+iREase+CTD+REase | 1054 | Bacteria>Terrabacteria_group>Chloroflexota>unclassified_Chloroflexota | GCA_028874095.1 | Chloroflexota_bacterium | OXP37_02700 | 49007..52171 | 2026724 | MAG:-DUF499-domain-containing-protein-[Chloroflexota-bacterium] |
| MDE2892396.1 | DUF499_1.RM.like.systems | <-REase\|\|TUDOR+ SF1-SF2-Helicase+iREase+CTD+REase ->Trypsin->METHYLASE->Polbeta->HEPN->HEPN+DUF499*-><-?<-?<-McrC-NTD+REase<-?<-wHTH+REase<-PadR-HTH | 1110 | Bacteria>Terrabacteria_group>Chloroflexota>unclassified_Chloroflexota | GCA_028874365.1 | Chloroflexota_bacterium | OXN86_07855 | 154158..157490 | 2026724 | MAG:-Swt1-family-HEPN-domain-containing-protein-[Chloroflexota-bacterium] |
| MDE2824105.1 | DUF499_1.RM.like.systems | METHYLASE->HEPN+DUF499*->SIR2-> | 1105 | Bacteria>Terrabacteria_group>Chloroflexota>unclassified_Chloroflexota | GCA_028874805.1 | Chloroflexota_bacterium | OXK79_11455 | 2958..6275 | 2026724 | MAG:-DUF499-domain-containing-protein-[Chloroflexota-bacterium] |
| MDE2668025.1 | DUF499_1.RM.like.systems | <-PIN<-AbrB<-HEPN+DUF499*<-METHYLASE<-TUDOR+ SF1-SF2-Helicase+iREase+CTD+REase <-REase<-MORC+REase<-DNA_methylase\|\|AbiEi_4-> | 1121 | Bacteria>Terrabacteria_group>Chloroflexota>unclassified_Chloroflexota | GCA_028875515.1 | Chloroflexota_bacterium | OXI51_00025 | 2709..6074 | 2026724 | MAG:-DUF499-domain-containing-protein-[Chloroflexota-bacterium] |
| WP_006616387.1 | DUF499_1.RM.like.systems | <-ABhydrolase<-?\|\|TUDOR+ SF1-SF2-Helicase+iREase+CTD+REase ->METHYLASE->?-><-?\|\|HEPN+DUF499*-> | 1135 | Bacteria>Terrabacteria_group>Cyanobacteriota/Melainabacteria_group>Cyanobacteriota>Cyanophyceae | - | Arthrospira_platensis | AP285_RS21915 | 4862811..4866218 | 118562 | Swt1-family-HEPN-domain-containing-protein-[Arthrospira-platensis] |
| WP_006624320.1 | DUF499_1.RM.like.systems | <-ABhydrolase\|\|TUDOR+ SF1-SF2-Helicase+iREase+CTD+REase ->METHYLASE-p->METHYLASE->METHYLASE-p-><-?\|\|HEPN+DUF499*-><-?\|\|?->?->?->SPOUT-> | 1123 | Bacteria>Terrabacteria_group>Cyanobacteriota/Melainabacteria_group>Cyanobacteriota>Cyanophyceae | - | Arthrospira_platensis | APLC1_RS09865 | 2007550..2010921 | 118562 | Swt1-family-HEPN-domain-containing-protein-[Arthrospira-platensis] |
| WP_096720868.1 | DUF499_1.RM.like.systems | DUF29-><-?<-?<-?\|\|?-><-HEPN+DUF499*<-PIN<-?<-METHYLASE\|\|REase-><-TUDOR+ SF1-SF2-Helicase+iREase+CTD+REase | 1125 | Bacteria>Terrabacteria_group>Cyanobacteriota/Melainabacteria_group>Cyanobacteriota>Cyanophyceae | - | Chondrocystis_sp._NIES-4102 | CDC35_RS11850 | 2748341..2751718 | 2005460 | Swt1-family-HEPN-domain-containing-protein-[Chondrocystis-sp.-NIES-4102] |
| EKD07858.1 | DUF499_1.RM.like.systems | <-ABhydrolase<-ABhydrolase\|\|TUDOR+ SF1-SF2-Helicase+iREase+CTD+REase -><-?\|\|METHYLASE-><-?\|\|HEPN+DUF499*-><-?\|\|?->?->?->?->SPOUT-> | 1123 | Bacteria>Terrabacteria_group>Cyanobacteriota/Melainabacteria_group>Cyanobacteriota>Cyanophyceae | GCA_000307915.1 | Arthrospira_platensis_C1 | SPLC1_S361100 | 3984646..3988017 | 459495 | hypothetical-protein-SPLC1_S361100-[Arthrospira-platensis-C1] |
| EKQ69986.1 | DUF499_1.RM.like.systems | TUDOR+ SF1-SF2-Helicase+iREase+CTD+REase ->REase->PolB->HEPN-><-?\|\|REase->HEPN+DUF499*-> | 1113 | Bacteria>Terrabacteria_group>Cyanobacteriota/Melainabacteria_group>Cyanobacteriota>Cyanophyceae | GCA_000309945.1 | Leptolyngbyaceae_cyanobacterium_JSC-12 | OsccyDRAFT_0242 | 268586..271927 | 864702 | putative-ATPase-(AAA+-superfamily)-[Leptolyngbyaceae-cyanobacterium-JSC-12] |
| KAB8314079.1 | DUF499_1.RM.like.systems | TUDOR+ SF1-SF2-Helicase+iREase+CTD+REase -p->REase-><-?\|\|METHYLASE->HEPN+DUF499*-><-?<-?\|\|FAD-NAD-dep-oxidoreductase-> | 1113 | Bacteria>Terrabacteria_group>Cyanobacteriota/Melainabacteria_group>Cyanobacteriota>Cyanophyceae | GCA_000828075.3 | Tolypothrix_campylonemoides_VB511288 | SD81_039415 | 98698..102039 | 1245935 | ATP-binding-protein-[Tolypothrix-campylonemoides-VB511288] |
| KPQ40173.1 | DUF499_1.RM.like.systems | <-CARF+HEPN+CARF<-REase<-HEPN+DUF499*<-?<-?<-?<-METHYLASE\|\|METHYLASE-><-TUDOR+ SF1-SF2-Helicase+iREase+CTD+REase | 1133 | Bacteria>Terrabacteria_group>Cyanobacteriota/Melainabacteria_group>Cyanobacteriota>Cyanophyceae | GCA_001314905.1 | Phormidium_sp._OSCR | HLUCCO16_04190 | 45328..48729 | 1666905 | putative-ATPase-(AAA+-superfamily)-[Phormidium-sp.-OSCR] |
| AMW31767.1 | DUF499_1.RM.like.systems | <-THUMP+Methylase<-ABhydrolase<-?\|\|TUDOR+ SF1-SF2-Helicase+iREase+CTD+REase ->METHYLASE->HEPN+DUF499*-> | 1119 | Bacteria>Terrabacteria_group>Cyanobacteriota/Melainabacteria_group>Cyanobacteriota>Cyanophyceae | GCA_001611905.1 | Arthrospira_platensis_YZ | AP285_21895 | 4862859..4866218 | 1738638 | AAA+-family-ATPase-[Arthrospira-platensis-YZ] |
| RFP57752.1 | DUF499_1.RM.like.systems | HISKIN+MORC->?-><-?\|\|TUDOR+ SF1-SF2-Helicase+iREase+CTD+REase ->METHYLASE->HEPN+DUF499*-> | 1119 | Bacteria>Terrabacteria_group>Cyanobacteriota/Melainabacteria_group>Cyanobacteriota>Cyanophyceae | GCA_001913845.2 | Limnothrix_sp._CACIAM_69d | BJG00_013040 | 63508..66867 | 1890733 | MAG:-ATP-binding-protein-[Limnothrix-sp.-CACIAM-69d] |
| ARI79893.1 | DUF499_1.RM.like.systems | TUDOR+ SF1-SF2-Helicase+iREase+CTD+REase ->REase->REase->METHYLASE->PAIREDC-HTH->PIN->HEPN+DUF499*-><-Cluster164_18clades<-?\|\|?->?->?->DUF29-> | 1155 | Bacteria>Terrabacteria_group>Cyanobacteriota/Melainabacteria_group>Cyanobacteriota>Cyanophyceae | GCA_002095975.1 | Microcystis_aeruginosa_PCC_7806SL | BH695_0612 | 541107..544574 | 1903187 | hypothetical-protein-BH695_0612-[Microcystis-aeruginosa-PCC-7806SL] |
| ARV61678.1 | DUF499_1.RM.like.systems | <-Trypsin\|\|?-><-?\|\|DDE_Tnp_1_3->TUDOR+ SF1-SF2-Helicase+iREase+CTD+REase ->METHYLASE->HEPN+DUF499*-><-tRNA\|\|?->LexA-HTH+LexA-protease->SF1-SF2-Helicase->SbcC->REase-> | 1115 | Bacteria>Terrabacteria_group>Cyanobacteriota/Melainabacteria_group>Cyanobacteriota>Cyanophyceae | GCA_002163975.1 | Nostocales_cyanobacterium_HT-58-2 | BZZ01_26385 | 6380631..6383978 | 1940762 | AAA+-family-ATPase-[Nostocales-cyanobacterium-HT-58-2] |
| BAZ89419.1 | DUF499_1.RM.like.systems | <-PolB\|\|TUDOR+ SF1-SF2-Helicase+iREase+CTD+REase -p->?-><-HTH+MPTase<-Tox-RelE\|\|TUDOR+ SF1-SF2-Helicase+iREase+CTD+REase ->HEPN+DUF499*-><-CR-ATPase9+REase\|\|ABhydrolase->?->?->Cysteine_desulfurase-> | 1015 | Bacteria>Terrabacteria_group>Cyanobacteriota/Melainabacteria_group>Cyanobacteriota>Cyanophyceae | GCA_002368135.1 | Raphidiopsis_curvata_NIES-932 | NIES932_08970 | 961455..964502 | 1973482 | hypothetical-protein-NIES932_08970-[Raphidiopsis-curvata-NIES-932] |
| GBE76047.1 | DUF499_1.RM.like.systems | REase->AbrB->PIN->METHYLASE->RHH_6->PIN->HEPN+DUF499*-><-Cluster164_18clades<-?\|\|?->?->?-><-TPR+TPR+TPR+TPR+TPR+TPR+TPR+TPR | 1111 | Bacteria>Terrabacteria_group>Cyanobacteriota/Melainabacteria_group>Cyanobacteriota>Cyanophyceae | GCA_002933835.1 | Microcystis_aeruginosa_NIES-87 | myaer87_32740 | 119164..122499 | 449440 | hypothetical-protein-myaer87_32740-[Microcystis-aeruginosa-NIES-87] |
| PSB44266.1 | DUF499_1.RM.like.systems | <-HEPN+DUF499*<-?<-METHYLASE<-REase<-PIN<-AbrB<-TUDOR+ SF1-SF2-Helicase+iREase+CTD+REase | 1113 | Bacteria>Terrabacteria_group>Cyanobacteriota/Melainabacteria_group>Cyanobacteriota>Cyanophyceae | GCA_003004015.1 | Cyanosarcina_cf._burmensis_CCALA_770 | C7B80_21165 | 280..3621 | 2107697 | AAA+-family-ATPase-[Cyanosarcina-cf.-burmensis-CCALA-770] |
| REJ55686.1 | DUF499_1.RM.like.systems | TUDOR+ SF1-SF2-Helicase+iREase+CTD+REase ->\|\|AbrB->PIN->METHYLASE->RHH_6->PIN->HEPN+DUF499*->?-><-Cluster164_18clades<-?\|\|?->?->DUF29-> | 1111 | Bacteria>Terrabacteria_group>Cyanobacteriota/Melainabacteria_group>Cyanobacteriota>Cyanophyceae | GCA_003388675.1 | Microcystis_aeruginosa_DA14 | DWQ56_15555 | 244128..247463 | 1987506 | DUF499-domain-containing-protein-[Microcystis-aeruginosa-DA14] |
| HBH74016.1 | DUF499_1.RM.like.systems | TUDOR+ SF1-SF2-Helicase+iREase+CTD+REase -><-PIN<-MazE_antitoxin\|\|?->wHTH(SBD)+HNH->HEPN+DUF499*-> | 1110 | Bacteria>Terrabacteria_group>Cyanobacteriota/Melainabacteria_group>Cyanobacteriota>Cyanophyceae | GCA_003508015.1 | Synechococcales_bacterium_UBA10510 | DDY43_11405 | 28893..32225 | 2055795 | AAA+-family-ATPase-[Synechococcales-bacterium-UBA10510] |
| HBP53069.1 | DUF499_1.RM.like.systems | METHYLASE->HEPN+DUF499*-> | 1057 | Bacteria>Terrabacteria_group>Cyanobacteriota/Melainabacteria_group>Cyanobacteriota>Cyanophyceae | GCA_003523965.1 | Synechococcus_sp._UBA8638 | DD643_01360 | 4940..8110 | 2055762 | AAA-family-ATPase-[Synechococcus-sp.-UBA8638] |
| TAD76280.1 | DUF499_1.RM.like.systems | METHYLASE-p->HEPN+DUF499*-> | 1114 | Bacteria>Terrabacteria_group>Cyanobacteriota/Melainabacteria_group>Cyanobacteriota>Cyanophyceae | GCA_004292515.1 | Oscillatoriales_cyanobacterium | EA001_12365 | 1816..5160 | 1151 | ATP-binding-protein-[Oscillatoriales-cyanobacterium] |
| TAE60284.1 | DUF499_1.RM.like.systems | <-HEPN+DUF499*\|\|<-METHYLASE | 1113 | Bacteria>Terrabacteria_group>Cyanobacteriota/Melainabacteria_group>Cyanobacteriota>Cyanophyceae | GCA_004292695.1 | Nostocales_cyanobacterium | EAZ87_06610 | 35276..38617 | 2447902 | ATP-binding-protein-[Nostocales-cyanobacterium] |
| TGH19526.1 | DUF499_1.RM.like.systems | TUDOR+ SF1-SF2-Helicase+iREase+CTD+REase -p->METHYLASE-p->METHYLASE-p->HEPN+DUF499*-> | 1094 | Bacteria>Terrabacteria_group>Cyanobacteriota/Melainabacteria_group>Cyanobacteriota>Cyanophyceae | GCA_004768385.2 | Aphanocapsa_feldmannii_277cI | ERJ68_08375 | 4147..7431 | 2507554 | ATP-binding-protein-[Aphanocapsa-feldmannii-277cI] |
| TGG91622.1 | DUF499_1.RM.like.systems | REase+NACHT-><-?<-?<-HEPN+DUF499*<-METHYLASE<-REase<-?<-?<-ParB+HNH | 1109 | Bacteria>Terrabacteria_group>Cyanobacteriota/Melainabacteria_group>Cyanobacteriota>Cyanophyceae | GCA_004768415.1 | Aphanocapsa_feldmannii_277cV | ERJ67_07705 | 17520..20849 | 2507553 | ATP-binding-protein-[Aphanocapsa-feldmannii-277cV] |
| GCL39965.1 | DUF499_1.RM.like.systems | <-METHYLASE-p\|\|HEPN+DUF499*-> | 931 | Bacteria>Terrabacteria_group>Cyanobacteriota/Melainabacteria_group>Cyanobacteriota>Cyanophyceae | GCA_005402885.1 | Sphaerospermopsis_reniformis | SR1949_50960 | 663..3458 | 531300 | hypothetical-protein-SR1949_50960-[Sphaerospermopsis-reniformis] |
| TRT82535.1 | DUF499_1.RM.like.systems | Cluster164_18clades-><-?<-HEPN+DUF499*<-ACET<-?<-METHYLASE<-PIN<-AbrB<-REase | 1130 | Bacteria>Terrabacteria_group>Cyanobacteriota/Melainabacteria_group>Cyanobacteriota>Cyanophyceae | GCA_007095595.1 | Microcystis_aeruginosa_Ma_AC_P_19900807_S299 | EWV82_11170 | 1786..5178 | 2486264 | ATP-binding-protein-[Microcystis-aeruginosa-Ma_AC_P_19900807_S299] |
| TRU11470.1 | DUF499_1.RM.like.systems | <-DUF29<-?<-?\|\|?->Cluster164_18clades-><-?<-HEPN+DUF499*<-ACET<-?<-METHYLASE<-AbrB<-PIN<-AbrB | 1117 | Bacteria>Terrabacteria_group>Cyanobacteriota/Melainabacteria_group>Cyanobacteriota>Cyanophyceae | GCA_007095705.1 | Microcystis_sp._Msp_OC_L_20101000_S702 | EWV60_07295 | 28722..32075 | 2486218 | MAG:-ATP-binding-protein-[Microcystis-sp.-Msp_OC_L_20101000_S702] |
| TRU24939.1 | DUF499_1.RM.like.systems | ABhydrolase->TUDOR+ SF1-SF2-Helicase+iREase+CTD+REase ->REase->METHYLASE->?->PIN->HEPN+DUF499*-><-Cluster164_18clades<-?\|\|?->?->DUF29->PolB-> | 1130 | Bacteria>Terrabacteria_group>Cyanobacteriota/Melainabacteria_group>Cyanobacteriota>Cyanophyceae | GCA_007095795.1 | Microcystis_aeruginosa_Ma_MB_S_20031200_S102D | EWV79_09040 | 24486..27878 | 2486255 | ATP-binding-protein-[Microcystis-aeruginosa-Ma_MB_S_20031200_S102D] |
| TVR05251.1 | DUF499_1.RM.like.systems | METHYLASE-p->?->?->?->HEPN+DUF499*-> | 1026 | Bacteria>Terrabacteria_group>Cyanobacteriota/Melainabacteria_group>Cyanobacteriota>Cyanophyceae | GCA_007693465.1 | Phormidium_sp._GEM2.Bin31 | EA395_15935 | 1874..4951 | 2480216 | ATP-binding-protein,-partial-[Phormidium-sp.-GEM2.Bin31] |
| MXW41265.1 | DUF499_1.RM.like.systems | HEPN->METHYLASE->HEPN+DUF499*-><-?<-N6_N4_MTase | 1110 | Bacteria>Terrabacteria_group>Cyanobacteriota/Melainabacteria_group>Cyanobacteriota>Cyanophyceae | GCA_009835965.1 | Synechococcus_sp._SB0668_bin_15 | F4Z75_09040 | 35350..38682 | 2604860 | ATP-binding-protein-[Synechococcus-sp.-SB0668_bin_15] |
| NCR40139.1 | DUF499_1.RM.like.systems | ABhydrolase->TUDOR+ SF1-SF2-Helicase+iREase+CTD+REase ->REase->METHYLASE->?->PIN->HEPN+DUF499*-><-Cluster164_18clades | 1130 | Bacteria>Terrabacteria_group>Cyanobacteriota/Melainabacteria_group>Cyanobacteriota>Cyanophyceae | GCA_009995705.1 | Microcystis_aeruginosa_W13-11 | GPJ21_09840 | 22820..26212 | 2685991 | ATP-binding-protein-[Microcystis-aeruginosa-W13-11] |
| NJM67133.1 | DUF499_1.RM.like.systems | METHYLASE->HEPN+DUF499*-> | 1106 | Bacteria>Terrabacteria_group>Cyanobacteriota/Melainabacteria_group>Cyanobacteriota>Cyanophyceae | GCA_012032145.1 | Acaryochloris_sp._RU_4_1 | HC851_16415 | 2448..5768 | 2720442 | ATP-binding-protein-[Acaryochloris-sp.-RU_4_1] |
| NJO72425.1 | DUF499_1.RM.like.systems | CHTH->TUDOR+ SF1-SF2-Helicase+iREase+CTD+REase ->REase->METHYLASE->?->PIN->HEPN+DUF499*->?-><-?<-?<-REase\|\|Phage_int_SAM_4+Phage_integrase-> | 1125 | Bacteria>Terrabacteria_group>Cyanobacteriota/Melainabacteria_group>Cyanobacteriota>Cyanophyceae | GCA_012033355.1 | Leptolyngbyaceae_cyanobacterium_RM1_406_9 | HC833_00760 | 33344..36721 | 2720434 | ATP-binding-protein-[Leptolyngbyaceae-cyanobacterium-RM1_406_9] |
| NJO76740.1 | DUF499_1.RM.like.systems | METHYLASE-p->REase->HEPN+DUF499*-> | 1125 | Bacteria>Terrabacteria_group>Cyanobacteriota/Melainabacteria_group>Cyanobacteriota>Cyanophyceae | GCA_012033355.1 | Leptolyngbyaceae_cyanobacterium_RM1_406_9 | HC833_25190 | 1227..4604 | 2720434 | ATP-binding-protein-[Leptolyngbyaceae-cyanobacterium-RM1_406_9] |

| Protein-ID | System | Gene-Neighbourhoods | Length | Taxonomy | Assembly-ID | Strain | Locus-Tag | Genomic-coordinates | TaxID | Fasta-defline |
|---|---|---|---|---|---|---|---|---|---|---|
| QJB25544.1 | DUF499_1.RM.like.systems | SF1-SF2-Helicase-><-THUMP+Methylase<-ABhydrolase\|\|TUDOR+SF1-SF2-Helicase+iREase+CTD+REase->METHYLASE-><-?\|\|HEPN+DUF499*-><-?\|\|?->?->?->?->SPOUT-> | 1123 | Bacteria>Terrabacteria_group>Cyanobacteriota/Melainabacteria_group>Cyanobacteriota>Cyanophyceae | GCA_012516315.1 | Limnospira_fusiformis_SAG_85.79 | HFV01_06765 | 1374121..1377492 | 1521533 | ATP-binding-protein-[Limnospira-fusiformis-SAG-85.79] |
| NMG59376.1 | DUF499_1.RM.like.systems | TUDOR+SF1-SF2-Helicase+iREase+CTD+REase->SbcC->?->METHYLASE->?->PIN->HEPN+DUF499*-> | 1130 | Bacteria>Terrabacteria_group>Cyanobacteriota/Melainabacteria_group>Cyanobacteriota>Cyanophyceae | GCA_012911965.1 | Geitlerinema_sp._P-1104 | E1H12_12830 | 9854..13246 | 2546230 | ATP-binding-protein-[Geitlerinema-sp.-P-1104] |
| MBD2550060.1 | DUF499_1.RM.like.systems | LRP-HTH->METHYLASE->HEPN+DUF499*->?->?->?->?->Peptidase_M48-><-HNH | 1126 | Bacteria>Terrabacteria_group>Cyanobacteriota/Melainabacteria_group>Cyanobacteriota>Cyanophyceae | GCA_014698445.1 | Microcystis_elabens_FACHB-917 | H6G65_10800 | 7559..10939 | 2692898 | ATP-binding-protein-[Microcystis-elabens-FACHB-917] |
| MBF2000219.1 | DUF499_1.RM.like.systems | <-HEPN+DUF499*<-REase<-REase | 1113 | Bacteria>Terrabacteria_group>Cyanobacteriota/Melainabacteria_group>Cyanobacteriota>Cyanophyceae | GCA_015272295.1 | Synechococcales_cyanobacterium_M58_A2018_015 | IGS38_05830 | 38711..42052 | 2774348 | DUF499-domain-containing-protein-[Synechococcales-cyanobacterium-M58_A2018_015] |
| MBF2075191.1 | DUF499_1.RM.like.systems | AbrB->HicB_lk_antitox->REase->REase->REase->REase->HEPN+DUF499*-> | 1114 | Bacteria>Terrabacteria_group>Cyanobacteriota/Melainabacteria_group>Cyanobacteriota>Cyanophyceae | GCA_015272325.1 | Synechococcales_cyanobacterium_C42_A2020_086 | IGS50_15730 | 92250..95594 | 2774344 | DUF499-domain-containing-protein-[Synechococcales-cyanobacterium-C42_A2020_086] |
| MBR8832157.1 | DUF499_1.RM.like.systems | <-PIN<-RHH<-HEPN+DUF499*<-METHYLASE<-?<-?<-?<-?<-TUDOR+SF1-SF2-Helicase+iREase+CTD+REase | 1101 | Bacteria>Terrabacteria_group>Cyanobacteriota/Melainabacteria_group>Cyanobacteriota>Cyanophyceae | GCA_016903695.1 | Chlorogloea_purpurea_SAG_13.99 | Cpurp_13245 | 34415..37720 | 1521637 | DUF499-domain-containing-protein-[Chlorogloea-purpurea-SAG-13.99] |
| MBV6627478.1 | DUF499_1.RM.like.systems | <-HEPN+DUF499*<-REase<-METHYLASE-p\|\|wHTH+REase-><-TUDOR+SF1-SF2-Helicase+iREase+CTD+REase-p<-REase | 1116 | Bacteria>Terrabacteria_group>Cyanobacteriota/Melainabacteria_group>Cyanobacteriota>Cyanophyceae | GCA_019192545.1 | Rivularia_sp._MS3 | KI793_31915 | 3971..7321 | 3012223 | DUF499-domain-containing-protein-[Rivularia-sp.-(in:-Bacteria)] |
| MBW4532455.1 | DUF499_1.RM.like.systems | <-HTH_OrfB_IS605+OrfB_IS605+OrfB_Zn_ribbon\|\|<-?<-?<-?<-HEPN+DUF499*<-CopG/RHH\|\|DDE_Tnp_1-><-PIN<-?<-?<-METHYLASE | 1104 | Bacteria>Terrabacteria_group>Cyanobacteriota/Melainabacteria_group>Cyanobacteriota>Cyanophyceae | GCA_019359095.1 | Pleurocapsa_minor_HA4230-MV1 | KME09_00805 | 42655..45969 | 2839664 | DUF499-domain-containing-protein-[Pleurocapsa-minor-HA4230-MV1] |
| MBW4455562.1 | DUF499_1.RM.like.systems | HEPN-><-?\|\|TUDOR+SF1-SF2-Helicase+iREase+CTD+REase->HicB+ZnR->METHYLASE-p->REase->HEPN+DUF499*-><-Phage_int_SAM_1+Phage_integrase\|\|?->?-><-Response_reg+LuxR-HTH<-HerA/FtsK | 1112 | Bacteria>Terrabacteria_group>Cyanobacteriota/Melainabacteria_group>Cyanobacteriota>Cyanophyceae | GCA_019359215.1 | Nostoc_indistinguendum_CM1-VF10 | KME55_24210 | 27093..30431 | 287863 | DUF499-domain-containing-protein-[Nostoc-indistinguendum-CM1-VF10] |
| MCA2627579.1 | DUF499_1.RM.like.systems | REase->METHYLASE->RHH_6->PIN->OLD-ABC->PIN->HEPN+DUF499*-><-Cluster164_18clades | 1130 | Bacteria>Terrabacteria_group>Cyanobacteriota/Melainabacteria_group>Cyanobacteriota>Cyanophyceae | GCA_020245485.1 | Microcystis_sp._M091S2 | IM505_05380 | 45571..48963 | 2771177 | DUF499-domain-containing-protein-[Microcystis-sp.-M091S2] |
| MCA2653775.1 | DUF499_1.RM.like.systems | Cluster164_18clades-><-HEPN+DUF499*<-PIN<-RHH_6<-METHYLASE<-AbrB<-PIN<-AbrB | 1130 | Bacteria>Terrabacteria_group>Cyanobacteriota/Melainabacteria_group>Cyanobacteriota>Cyanophyceae | GCA_020245605.1 | Microcystis_sp._M061S2 | IM475_05875 | 1597..4989 | 2771171 | DUF499-domain-containing-protein-[Microcystis-sp.-M061S2] |
| MCA2660027.1 | DUF499_1.RM.like.systems | Cluster164_18clades-><-HEPN+DUF499*<-ACET<-?<-METHYLASE | 1130 | Bacteria>Terrabacteria_group>Cyanobacteriota/Melainabacteria_group>Cyanobacteriota>Cyanophyceae | GCA_020245625.1 | Microcystis_sp._M049S2 | IM496_16500 | 8003..11395 | 2771169 | DUF499-domain-containing-protein-[Microcystis-sp.-M049S2] |
| MCA2716396.1 | DUF499_1.RM.like.systems | REase->AbrB->PIN->METHYLASE->RHH_6->HEPN+DUF499*-><-Cluster164_18clades<-?\|\|?->?->?-><-TPR+TPR+TPR+TPR+TPR+TPR+TPR+TPR | 1130 | Bacteria>Terrabacteria_group>Cyanobacteriota/Melainabacteria_group>Cyanobacteriota>Cyanophyceae | GCA_020245865.1 | Microcystis_sp._M169S2 | IM485_01040 | 5502..8894 | 2771157 | DUF499-domain-containing-protein-[Microcystis-sp.-M169S2] |
| MCA6503991.1 | DUF499_1.RM.like.systems | METHYLASE-p->?->PIN->REase->REase->REase->HEPN+DUF499*-><-Thioredoxin\|\|tRNA-><-PIN<-Phd-YefM\|\|?->?->ABhydrolase-> | 1118 | Bacteria>Terrabacteria_group>Cyanobacteriota/Melainabacteria_group>Cyanobacteriota>Cyanophyceae | GCA_020403835.1 | Pseudanabaena_sp._M090S1SP2A07QC | IM580_15730 | 4170..7526 | 2771254 | DUF499-domain-containing-protein-[Pseudanabaena-sp.-M090S1SP2A07QC] |
| MCA6581759.1 | DUF499_1.RM.like.systems | <-HEPN+DUF499*<-REase<-REase<-REase<-PIN | 1118 | Bacteria>Terrabacteria_group>Cyanobacteriota/Melainabacteria_group>Cyanobacteriota>Cyanophyceae | GCA_020404925.1 | Pseudanabaena_sp._M34BS1SP1A06MG | IM536_06930 | 164..3520 | 2771240 | DUF499-domain-containing-protein-[Pseudanabaena-sp.-M34BS1SP1A06MG] |
| MCA6600378.1 | DUF499_1.RM.like.systems | Phd-YefM->PIN-><-tRNA\|\|Thioredoxin-><-HEPN+DUF499*<-REase<-REase<-REase<-PIN<-?<-METHYLASE-p | 1118 | Bacteria>Terrabacteria_group>Cyanobacteriota/Melainabacteria_group>Cyanobacteriota>Cyanophyceae | GCA_020404965.1 | Pseudanabaena_sp._M57BS1SP1A06MG | IM537_09260 | 3869..7225 | 2771237 | DUF499-domain-containing-protein-[Pseudanabaena-sp.-M57BS1SP1A06MG] |
| MCA6597113.1 | DUF499_1.RM.like.systems | Phd-YefM->PIN-><-tRNA\|\|Thioredoxin-><-HEPN+DUF499*<-REase<-REase<-REase<-?<-REase<-REase | 1118 | Bacteria>Terrabacteria_group>Cyanobacteriota/Melainabacteria_group>Cyanobacteriota>Cyanophyceae | GCA_020405005.1 | Pseudanabaena_sp._M046S1SP1A06QC | IM539_12615 | 33014..36370 | 2771236 | DUF499-domain-containing-protein-[Pseudanabaena-sp.-M046S1SP1A06QC] |
| MCA6605401.1 | DUF499_1.RM.like.systems | PIN->REase->REase->REase->HEPN+DUF499*-> | 895 | Bacteria>Terrabacteria_group>Cyanobacteriota/Melainabacteria_group>Cyanobacteriota>Cyanophyceae | GCA_020405035.1 | Pseudanabaena_sp._M007S1SP1A06QC | IM533_13245 | 2810..5496 | 2771232 | DUF499-domain-containing-protein-[Pseudanabaena-sp.-M007S1SP1A06QC] |
| MCC5647010.1 | DUF499_1.RM.like.systems | <-SIGMA-HTH\|\|?-><-?<-PIWI\|\|TraJ-RHH-><-HEPN+DUF499*<-METHYLASE-p<-METHYLASE-p<-DOC<-?<-TUDOR+SF1-SF2-Helicase+iREase+CTD+REase | 1108 | Bacteria>Terrabacteria_group>Cyanobacteriota/Melainabacteria_group>Cyanobacteriota>Cyanophyceae | GCA_020829665.1 | Nostoc_sp._CHAB_5824 | LC607_29645 | 82754..86080 | 2780402 | DUF499-domain-containing-protein-[Nostoc-sp.-CHAB-5824] |
| MCD8490337.1 | DUF499_1.RM.like.systems | pPIWI+pPIWI->pPIWI->pPIWI+MID+RNaseH->?->?->?-><-HEPN+DUF499*<-METHYLASE<-PIN<-AbrB<-TUDOR+SF1-SF2-Helicase+iREase+CTD+REase<-TOPRIM+SF1-SF2-Helicase | 1118 | Bacteria>Terrabacteria_group>Cyanobacteriota/Melainabacteria_group>Cyanobacteriota>Cyanophyceae | GCA_021208125.1 | Desertifilum_sp. | LRZ84_27325 | 10272..13628 | 2497424 | DUF499-domain-containing-protein-[Desertifilum-sp.] |
| MCE2977896.1 | DUF499_1.RM.like.systems | METHYLASE-p->?->PIN->REase->REase->REase->HEPN+DUF499*-><-Thioredoxin\|\|tRNA-><-PIN | 1118 | Bacteria>Terrabacteria_group>Cyanobacteriota/Melainabacteria_group>Cyanobacteriota>Cyanophyceae | GCA_021298775.1 | Pseudanabaena_sp._CoA8_M7 | LW851_16625 | 3735..7091 | 2904992 | DUF499-domain-containing-protein-[Pseudanabaena-sp.-CoA8_M7] |
| MCE2669063.1 | DUF499_1.RM.like.systems | Cluster164_18clades-><-HEPN+DUF499*<-PIN<-OLD-ABC<-PIN<-RHH_6<-METHYLASE<-REase | 1130 | Bacteria>Terrabacteria_group>Cyanobacteriota/Melainabacteria_group>Cyanobacteriota>Cyanophyceae | GCA_021300285.1 | Microcystis_sp._49638_E5 | LW728_07560 | 10523..13915 | 2904986 | DUF499-domain-containing-protein-[Microcystis-sp.-49638_E5] |
| MCL6436540.1 | DUF499_1.RM.like.systems | <-HEPN+DUF499*<-METHYLASE-p | 1113 | Bacteria>Terrabacteria_group>Cyanobacteriota/Melainabacteria_group>Cyanobacteriota>Cyanophyceae | GCA_023511615.1 | Leptolyngbyaceae_cyanobacterium_HOT.MB2.61 | K6T90_20505 | 755..4096 | 2855467 | DUF499-domain-containing-protein-[Leptolyngbyaceae-cyanobacterium-HOT.MB2.61] |
| WJI23189.1 | DUF499_1.RM.like.systems | <-HEPN+DUF499*<-REase<-METHYLASE<-TUDOR+SF1-SF2-Helicase+iREase+CTD+REase\|\|REase-><-DDE_Tnp_1 | 1098 | Bacteria>Terrabacteria_group>Cyanobacteriota/Melainabacteria_group>Cyanobacteriota>Cyanophyceae | GCA_030322965.1 | Thermosynechococcus_sp._BO | MZ909_08100 | 1635834..1639130 | 2937284 | Swt1-family-HEPN-domain-containing-protein-[Thermosynechococcus-sp.-BO] |
| WP_006668432.1 | DUF499_1.RM.like.systems | SF1-SF2-Helicase-><-THUMP+Methylase\|\|TUDOR+SF1-SF2-Helicase+iREase+CTD+REase-><-?\|\|METHYLASE-><-?\|\|HEPN+DUF499*-><-?\|\|?->?->?->?->SPOUT-> | 1123 | Bacteria>Terrabacteria_group>Cyanobacteriota/Melainabacteria_group>Cyanobacteriota>Cyanophyceae | GCF_000173555.1 | Limnospira_maxima | AMAXDRAFT_RS04610 | 145363..148734 | 129910 | Swt1-family-HEPN-domain-containing-protein-[Limnospira-maxima] |
| WP_011318979.1 | DUF499_1.RM.like.systems | HTH_28+rve+Mu-transpos_C->?->?->TUDOR+SF1-SF2-Helicase+iREase+CTD+REase->METHYLASE->METHYLASE-p->HEPN+DUF499*-><-TraJ-RHH\|\|PIWI-><-HTH_3\|\|?-><-tRNA\|\|?->LexA-HTH+LexA-protease-> | 1109 | Bacteria>Terrabacteria_group>Cyanobacteriota/Melainabacteria_group>Cyanobacteriota>Cyanophyceae | GCF_000204075.1 | Nostocaceae | GSQ19_RS16260 | 3995693..3999022 | 1162 | MULTISPECIES:-Swt1-family-HEPN-domain-containing-protein-[Nostocaceae] |
| WP_008200990.1 | DUF499_1.RM.like.systems | REase->METHYLASE->?->ACET->HEPN+DUF499*->?-><-Cluster164_18clades | 1130 | Bacteria>Terrabacteria_group>Cyanobacteriota/Melainabacteria_group>Cyanobacteriota>Cyanophyceae | GCF_000297435.1 | Microcystis_sp._T1-4 | MICAI_RS09635 | 4446..7838 | 1160279 | Swt1-family-HEPN-domain-containing-protein-[Microcystis-sp.-T1-4] |
| WP_002754187.1 | DUF499_1.RM.like.systems | AbrB->PemK_toxin->METHYLASE->RHH_6->PIN->HEPN+DUF499*-><-Cluster164_18clades<-?\|\|?->?->DUF29->sGTP-> | 1130 | Bacteria>Terrabacteria_group>Cyanobacteriota/Melainabacteria_group>Cyanobacteriota>Cyanophyceae | GCF_000307995.1 | Microcystis_aeruginosa | MICCA_RS15375 | 7336..10728 | 1126 | Swt1-family-HEPN-domain-containing-protein-[Microcystis-aeruginosa] |
| WP_002777134.1 | DUF499_1.RM.like.systems | METHYLASE->RHH_6->PIN->OLD-ABC->PIN->HEPN+DUF499*-><-Cluster164_18clades<-?\|\|?->?->DUF29->sGTP-> | 1130 | Bacteria>Terrabacteria_group>Cyanobacteriota/Melainabacteria_group>Cyanobacteriota>Cyanophyceae | GCF_000312205.1 | Microcystis_aeruginosa | MICAD_RS14460 | 204713..208105 | 1126 | Swt1-family-HEPN-domain-containing-protein-[Microcystis-aeruginosa] |
| WP_002801405.1 | DUF499_1.RM.like.systems | REase->METHYLASE->RHH_6->PIN->OLD-ABC->PIN->HEPN+DUF499*-><-Cluster164_18clades | 1117 | Bacteria>Terrabacteria_group>Cyanobacteriota/Melainabacteria_group>Cyanobacteriota>Cyanophyceae | GCF_000312285.1 | Microcystis_aeruginosa | MICAK_RS07420 | 6579..9932 | 1126 | Swt1-family-HEPN-domain-containing-protein-[Microcystis-aeruginosa] |
| WP_015155777.1 | DUF499_1.RM.like.systems | Phage_integrase-><-Phage_integrase\|\|?-><-HEPN+DUF499*<-?<-METHYLASE<-REase<-PIN<-AbrB<-TUDOR+SF1-SF2-Helicase+iREase+CTD+REase | 1113 | Bacteria>Terrabacteria_group>Cyanobacteriota/Melainabacteria_group>Cyanobacteriota>Cyanophyceae | GCF_000317125.1 | Chroococcidiopsis_thermalis | CHRO_RS18600 | 4302197..4305538 | 54299 | Swt1-family-HEPN-domain-containing-protein-[Chroococcidiopsis-thermalis] |
| WP_015211462.1 | DUF499_1.RM.like.systems | DUF29->AbrB->OLD-ABC->?->METHYLASE->?->HEPN+DUF499*-><-?<-MobL+ZnR+TOPRIM | 1117 | Bacteria>Terrabacteria_group>Cyanobacteriota/Melainabacteria_group>Cyanobacteriota>Cyanophyceae | GCF_000317555.1 | Gloeocapsa_sp._PCC_7428 | GLO7428_RS25580 | 12645..15998 | 1173026 | Swt1-family-HEPN-domain-containing-protein-[Gloeocapsa-sp.-PCC-7428] |
| WP_015227928.1 | DUF499_1.RM.like.systems | <-HEPN+DUF499*<-HEPN<-PolB<-METHYLASE<-?<-HicB_lk_antitox\|\|BrnA_antitoxin-> | 1113 | Bacteria>Terrabacteria_group>Cyanobacteriota/Melainabacteria_group>Cyanobacteriota>Cyanophyceae | GCF_000317615.1 | Dactylococcopsis_salina | DACSA_RS00655 | 98261..101602 | 292566 | Swt1-family-HEPN-domain-containing-protein-[Dactylococcopsis-salina] |
| WP_006527613.1 | DUF499_1.RM.like.systems | <-METHYLASE-p<-HEPN+DUF499*<-?<-ABhydrolase\|\|sGTP-> | 1093 | Bacteria>Terrabacteria_group>Cyanobacteriota/Melainabacteria_group>Cyanobacteriota>Cyanophyceae | GCF_000332035.1 | Gloeocapsa_sp._PCC_73106 | GLO73106_RS03445 | 648..3929 | 102232 | Swt1-family-HEPN-domain-containing-protein-[Gloeocapsa-sp.-PCC-73106] |
| WP_019503187.1 | DUF499_1.RM.like.systems | <-DDE_3<-PAIREDC-HTH\|\|?->\|\|<-?\|\|PIWI-><-HEPN+DUF499*\|\|?-><-PIN<-?<-METHYLASE | 1127 | Bacteria>Terrabacteria_group>Cyanobacteriota/Melainabacteria_group>Cyanobacteriota>Cyanophyceae | GCF_000332195.1 | Pleurocapsa_sp._PCC_7319 | PLEUR7319_RS0100190 | 37513..40896 | 118161 | Swt1-family-HEPN-domain-containing-protein-[Pleurocapsa-sp.-PCC-7319] |
| WP_002748938.1 | DUF499_1.RM.like.systems | PAIREDC-HTH->PIN->HEPN+DUF499*-><-Cluster164_18clades<-?\|\|?->?->DUF29-> | 1111 | Bacteria>Terrabacteria_group>Cyanobacteriota/Melainabacteria_group>Cyanobacteriota>Cyanophyceae | GCF_000332585.1 | Microcystis_aeruginosa | C789_RS21375 | 1161..4496 | 1126 | Swt1-family-HEPN-domain-containing-protein-[Microcystis-aeruginosa] |
| WP_017744970.1 | DUF499_1.RM.like.systems | OrfB_IS605+OrfB_Zn_ribbon-><-HEPN+DUF499*\|\|?-><-MqsA_antitoxin+HTH_3<-Polbeta<-?<-METHYLASE | 1113 | Bacteria>Terrabacteria_group>Cyanobacteriota/Melainabacteria_group>Cyanobacteriota>Cyanophyceae | GCF_000346485.2 | Scytonema_hofmannii | WA1_RS46765 | 11110546..11113887 | 34078 | Swt1-family-HEPN-domain-containing-protein-[Scytonema-hofmannii] |
| WP_016515479.1 | DUF499_1.RM.like.systems | Cluster164_18clades-><-HEPN+DUF499*<-PIN<-OLD-ABC<-PIN<-RHH_6<-METHYLASE<-AbrB | 1130 | Bacteria>Terrabacteria_group>Cyanobacteriota/Melainabacteria_group>Cyanobacteriota>Cyanophyceae | GCF_000412595.1 | Microcystis_aeruginosa | MAESPC_RS19915 | 50545..53937 | 1126 | Swt1-family-HEPN-domain-containing-protein-[Microcystis-aeruginosa] |
| WP_026733571.1 | DUF499_1.RM.like.systems | TUDOR+SF1-SF2-Helicase+iREase+CTD+REase->AbrB->PIN->METHYLASE->REase->HEPN+DUF499*->?->TraJ-RHH-> | 1122 | Bacteria>Terrabacteria_group>Cyanobacteriota/Melainabacteria_group>Cyanobacteriota>Cyanophyceae | GCF_000517105.1 | Fischerella_sp._PCC_9605 | FIS9605_RS0116420 | 337165..340533 | 1173024 | Swt1-family-HEPN-domain-containing-protein-[Fischerella-sp.-PCC-9605] |
| WP_024971076.1 | DUF499_1.RM.like.systems | REase->METHYLASE->RHH_6->PIN->OLD-ABC->PIN->HEPN+DUF499*-> | 1130 | Bacteria>Terrabacteria_group>Cyanobacteriota/Melainabacteria_group>Cyanobacteriota>Cyanophyceae | GCF_000599945.1 | Microcystis_aeruginosa | MIC7005_RS0124495 | 6350..9742 | 1126 | Swt1-family-HEPN-domain-containing-protein-[Microcystis-aeruginosa] |
| WP_008052582.1 | DUF499_1.RM.like.systems | <-SPOUT<-?<-?<-?<-?\|\|?-><-HEPN+DUF499*<-METHYLASE<-TUDOR+SF1-SF2-Helicase+iREase+CTD+REase\|\|ABhydrolase->THUMP+Methylase-><-SF1-SF2-Helicase | 1120 | Bacteria>Terrabacteria_group>Cyanobacteriota/Melainabacteria_group>Cyanobacteriota>Cyanophyceae | GCF_000973065.1 | Limnospira_indica | XZ46_RS18610 | 4259944..4263306 | 147322 | Swt1-family-HEPN-domain-containing-protein-[Limnospira-indica] |
| WP_046663117.1 | DUF499_1.RM.like.systems | PP_kinase+HKD-DNase-><-?<-?\|\|TUDOR+SF1-SF2-Helicase+iREase+CTD+REase->REase->METHYLASE->HEPN+DUF499*-><-?<-?\|\|?->?-><-THUMP+Methylase | 1130 | Bacteria>Terrabacteria_group>Cyanobacteriota/Melainabacteria_group>Cyanobacteriota>Cyanophyceae | GCF_000981785.2 | Microcystis_aeruginosa | myaer_RS18245 | 3707749..3711141 | 1126 | Swt1-family-HEPN-domain-containing-protein-[Microcystis-aeruginosa] |
| WP_057177165.1 | DUF499_1.RM.like.systems | HKD-DNase+SF1-SF2-Helicase-p-><-HTH+MPTase<-?<-?\|\|TUDOR+SF1-SF2-Helicase+iREase+CTD+REase->METHYLASE->HEPN+DUF499*->REase->?->Peptidase_C14-> | 1103 | Bacteria>Terrabacteria_group>Cyanobacteriota/Melainabacteria_group>Cyanobacteriota>Cyanophyceae | GCF_001432185.1 | Cylindrospermopsis_sp._CR12 | ASL19_RS02560 | 94062..97373 | 1747196 | Swt1-family-HEPN-domain-containing-protein-[Cylindrospermopsis-sp.-CR12] |
| WP_059001129.1 | DUF499_1.RM.like.systems | PIN->?-><-HEPN+DUF499*<-METHYLASE\|\|REase+HTH-><-TUDOR+SF1-SF2-Helicase+iREase+CTD+REase<-TrwC+SF1-SF2-helicase+TOPRIM\|\|?->ParB+HNH-> | 1124 | Bacteria>Terrabacteria_group>Cyanobacteriota/Melainabacteria_group>Cyanobacteriota>Cyanophyceae | GCF_001485215.1 | Leptolyngbya_sp._NIES-2104 | NIES2104_RS25560 | 5320050..5323424 | 1552121 | Swt1-family-HEPN-domain-containing-protein-[Leptolyngbya-sp.-NIES-2104] |
| WP_066030118.1 | DUF499_1.RM.like.systems | PP_kinase+HKD-DNase-><-?<-?\|\|TUDOR+SF1-SF2-Helicase+iREase+CTD+REase->REase->METHYLASE->HEPN+DUF499*-><-?<-?\|\|?->?-><-THUMP+Methylase | 1130 | Bacteria>Terrabacteria_group>Cyanobacteriota/Melainabacteria_group>Cyanobacteriota>Cyanophyceae | GCF_001704955.2 | Microcystis_aeruginosa | amyaer_RS18255 | 3708666..3712058 | 1126 | Swt1-family-HEPN-domain-containing-protein-[Microcystis-aeruginosa] |
| WP_073632965.1 | DUF499_1.RM.like.systems | TUDOR+SF1-SF2-Helicase+iREase+CTD+REase->METHYLASE->REase->?->HEPN+DUF499*-><-HTH\|\|PIWI-> | 1113 | Bacteria>Terrabacteria_group>Cyanobacteriota/Melainabacteria_group>Cyanobacteriota>Cyanophyceae | GCF_001904675.1 | Scytonema_sp._HK-05 | NIES2130_RS19405 | 80454..83795 | 1137095 | Swt1-family-HEPN-domain-containing-protein-[Scytonema-sp.-HK-05] |
| WP_075601490.1 | DUF499_1.RM.like.systems | <-HEPN+DUF499*<-METHYLASE<-?<-SbcC<-TUDOR+SF1-SF2-Helicase+iREase+CTD+REase<-MPTase<-HTH | 834 | Bacteria>Terrabacteria_group>Cyanobacteriota/Melainabacteria_group>Cyanobacteriota>Cyanophyceae | GCF_001939115.1 | Leptolyngbya_sp._'hensonii' | BST81_RS26530 | 1..2502 | 1922337 | Swt1-family-HEPN-domain-containing-protein,-partial-[Leptolyngbya-sp.-'hensonii'] |
| WP_231828504.1 | DUF499_1.RM.like.systems | TUDOR+SF1-SF2-Helicase+iREase+CTD+REase->REase->REase->METHYLASE->PAIREDC-HTH->PIN->HEPN+DUF499*-><-Cluster164_18clades<-?\|\|?->?->DUF29-><-Y1_Tnp | 1129 | Bacteria>Terrabacteria_group>Cyanobacteriota/Melainabacteria_group>Cyanobacteriota>Cyanophyceae | GCF_002095975.1 | Microcystis_aeruginosa | BH695_RS02870 | 541185..544574 | 1126 | Swt1-family-HEPN-domain-containing-protein-[Microcystis-aeruginosa] |
| WP_223278246.1 | DUF499_1.RM.like.systems | TUDOR+SF1-SF2-Helicase+iREase+CTD+REase->Phd-YefM->PIN-><-HTH_3\|\|METHYLASE->HEPN+DUF499*-><-TraJ-RHH\|\|PIWI-><-HTH_3 | 1023 | Bacteria>Terrabacteria_group>Cyanobacteriota/Melainabacteria_group>Cyanobacteriota>Cyanophyceae | GCF_002245985.1 | Nostoc_sp._'Peltigera_membranacea_cyanobiont'_232 | CDG79_RS21465 | 10493..13564 | 2014531 | DUF499-domain-containing-protein-[Nostoc-sp.-'Peltigera-membranacea-cyanobiont'-232] |
| WP_094555433.1 | DUF499_1.RM.like.systems | GDResolvase-><-D5N+D5ATPase\|\|?->ArdCN+MPTase-><-?<-HEPN+DUF499*<-METHYLASE<-?<-PIN<-RHH<-TUDOR+SF1-SF2-Helicase+iREase+CTD+REase\|\|DUF4236-> | 1103 | Bacteria>Terrabacteria_group>Cyanobacteriota/Melainabacteria_group>Cyanobacteriota>Cyanophyceae | GCF_002252625.1 | Synechococcus_sp._1G10 | CJZ82_RS08935 | 55076..58387 | 2025605 | DUF499-domain-containing-protein-[Synechococcus-sp.-1G10] |
| WP_094561768.1 | DUF499_1.RM.like.systems | METHYLASE->HEPN+DUF499*-> | 902 | Bacteria>Terrabacteria_group>Cyanobacteriota/Melainabacteria_group>Cyanobacteriota>Cyanophyceae | GCF_002252665.1 | Synechococcus_sp._8F6 | CJZ78_RS13255 | 2049..4755 | 2025606 | DUF499-domain-containing-protein,-partial-[Synechococcus-sp.-8F6] |
| WP_100904446.1 | DUF499_1.RM.like.systems | <-DDE_Tnp_1_3<-HEPN+DUF499*<-REase<-TUDOR+SF1-SF2-Helicase+iREase+CTD+REase<-?<-?\|\|DndB-> | 1112 | Bacteria>Terrabacteria_group>Cyanobacteriota/Melainabacteria_group>Cyanobacteriota>Cyanophyceae | GCF_002813575.1 | Nostoc_flagelliforme | COO91_RS47960 | 745879..749217 | 1306274 | Swt1-family-HEPN-domain-containing-protein-[Nostoc-flagelliforme] |
| WP_103113079.1 | DUF499_1.RM.like.systems | PemK_toxin->METHYLASE->RHH_6->PIN->OLD-ABC->PIN->HEPN+DUF499*-><-Cluster164_18clades<-?\|\|?->?->DUF29->sGTP-> | 1111 | Bacteria>Terrabacteria_group>Cyanobacteriota/Melainabacteria_group>Cyanobacteriota>Cyanophyceae | GCF_002897275.1 | Microcystis_aeruginosa | D3800_RS23535 | 4968568..4971903 | 1126 | Swt1-family-HEPN-domain-containing-protein-[Microcystis-aeruginosa] |
| WP_104908185.1 | DUF499_1.RM.like.systems | TUDOR+SF1-SF2-Helicase+iREase+CTD+REase->Phd-YefM->PIN-><-HTH_3\|\|METHYLASE->REase->HEPN+DUF499*-> | 1109 | Bacteria>Terrabacteria_group>Cyanobacteriota/Melainabacteria_group>Cyanobacteriota>Cyanophyceae | GCF_002949795.1 | Nostoc_sp._'Lobaria_pulmonaria_(5183)_cyanobiont' | NLP_RS21725 | 4933532..4936861 | 1618022 | Swt1-family-HEPN-domain-containing-protein-[Nostoc-sp.-'Lobaria-pulmonaria-(5183)-cyanobiont'] |
| WP_106908219.1 | DUF499_1.RM.like.systems | Cluster164_18clades-><-?<-HEPN+DUF499*<-ACET<-?<-METHYLASE<-REase\|\|?-><-TUDOR+SF1-SF2-Helicase+iREase+CTD+REase | 1130 | Bacteria>Terrabacteria_group>Cyanobacteriota/Melainabacteria_group>Cyanobacteriota>Cyanophyceae | GCF_003019735.1 | Microcystis_sp._MC19 | B5D77_RS02535 | 488054..491446 | 1967666 | Swt1-family-HEPN-domain-containing-protein-[Microcystis-sp.-MC19] |
| WP_110545657.1 | DUF499_1.RM.like.systems | TUDOR+SF1-SF2-Helicase+iREase+CTD+REase->REase->METHYLASE->METHYLASE-p->?->ACET->HEPN+DUF499*-><-Cluster164_18clades | 1111 | Bacteria>Terrabacteria_group>Cyanobacteriota/Melainabacteria_group>Cyanobacteriota>Cyanophyceae | GCF_003206625.1 | Microcystis_aeruginosa | MTo_RS18515 | 13505..16840 | 1126 | Swt1-family-HEPN-domain-containing-protein-[Microcystis-aeruginosa] |
| WP_115538087.1 | DUF499_1.RM.like.systems | <-Peptidase_C14<-?<-REase<-DUF499-p\|\|DDE_Tnp_IS1-><-HEPN+DUF499*<-METHYLASE<-HTH_OrfB_IS605+OrfB_IS605+OrfB_Zn_ribbon\|\|<-TUDOR+SF1-SF2-Helicase+iREase+CTD+REase\|\|HTH+MPTase-><-?<-TUDOR+SF1-SF2-Helicase+iREase+CTD+REase-p | 1013 | Bacteria>Terrabacteria_group>Cyanobacteriota/Melainabacteria_group>Cyanobacteriota>Cyanophyceae | GCF_003367075.2 | Cylindrospermopsis_raciborskii | C6N34_RS14575 | 3182311..3185352 | 77022 | Swt1-family-HEPN-domain-containing-protein-[Cylindrospermopsis-raciborskii] |
| WP_130128840.1 | DUF499_1.RM.like.systems | <-MORC+DUF3684<-HEPN+DUF499*<-METHYLASE<-METHYLASE<-PIN<-RHH\|\|?->RelE-ParE-> | 1113 | Bacteria>Terrabacteria_group>Cyanobacteriota/Melainabacteria_group>Cyanobacteriota>Cyanophyceae | GCF_004209775.1 | Synechococcus_sp._WH_8101 | SynWH8101_RS05090 | 1007085..1010426 | 59932 | DUF499-domain-containing-protein-[Synechococcus-sp.-WH-8101] |
| WP_130756579.1 | DUF499_1.RM.like.systems | REase-p->AbrB->PIN->METHYLASE->RHH_6->PIN->HEPN+DUF499*-><-Cluster164_18clades<-?\|\|?->?->DUF29->sGTP-> | 1111 | Bacteria>Terrabacteria_group>Cyanobacteriota/Melainabacteria_group>Cyanobacteriota>Cyanophyceae | GCF_004305995.1 | Microcystis_aeruginosa | MiAbB_RS04285 | 58496..61831 | 1126 | Swt1-family-HEPN-domain-containing-protein-[Microcystis-aeruginosa] |
| WP_137669391.1 | DUF499_1.RM.like.systems | <-METHYLASE-p\|\|HEPN+DUF499*-> | 965 | Bacteria>Terrabacteria_group>Cyanobacteriota/Melainabacteria_group>Cyanobacteriota>Cyanophyceae | GCF_005402885.1 | Sphaerospermopsis_reniformis | FFR48_RS26320 | 561..3458 | 531300 | Swt1-family-HEPN-domain-containing-protein-[Sphaerospermopsis-reniformis] |
| WP_149107756.1 | DUF499_1.RM.like.systems | <-sGTP<-DUF29<-?\|\|?->Cluster164_18clades-><-HEPN+DUF499*<-PIN<-RHH_6<-METHYLASE<-PemK_toxin<-AbrB<-PIN | 1130 | Bacteria>Terrabacteria_group>Cyanobacteriota/Melainabacteria_group>Cyanobacteriota>Cyanophyceae | GCF_008257445.1 | Microcystis_aeruginosa | FXO09_RS20205 | 21434..24826 | 1126 | Swt1-family-HEPN-domain-containing-protein-[Microcystis-aeruginosa] |
| WP_149979248.1 | DUF499_1.RM.like.systems | Cluster164_18clades-><-HEPN+DUF499*<-ACET<-?<-METHYLASE<-REase\|\|?-><-TUDOR+SF1-SF2-Helicase+iREase+CTD+REase | 1130 | Bacteria>Terrabacteria_group>Cyanobacteriota/Melainabacteria_group>Cyanobacteriota>Cyanophyceae | GCF_008579225.1 | Microcystis_aeruginosa | MiYa_RS12425 | 10420..13812 | 1126 | Swt1-family-HEPN-domain-containing-protein-[Microcystis-aeruginosa] |
| WP_149976793.1 | DUF499_1.RM.like.systems | <-HEPN+DUF499*<-METHYLASE<-REase<-TUDOR+SF1-SF2-Helicase+iREase+CTD+REase | 1130 | Bacteria>Terrabacteria_group>Cyanobacteriota/Melainabacteria_group>Cyanobacteriota>Cyanophyceae | GCF_008579445.1 | Microcystis_aeruginosa | MiTs_RS18945 | 1418..4810 | 1126 | Swt1-family-HEPN-domain-containing-protein-[Microcystis-aeruginosa] |
| WP_149989607.1 | DUF499_1.RM.like.systems | Cluster164_18clades-><-?<-HEPN+DUF499*<-ACET<-?<-METHYLASE<-REase\|\|?-><-TUDOR+SF1-SF2-Helicase+iREase+CTD+REase | 1117 | Bacteria>Terrabacteria_group>Cyanobacteriota/Melainabacteria_group>Cyanobacteriota>Cyanophyceae | GCF_008579565.1 | Microcystis_aeruginosa | MiHa_RS15955 | 12499..15852 | 1126 | Swt1-family-HEPN-domain-containing-protein-[Microcystis-aeruginosa] |
| WP_149977735.1 | DUF499_1.RM.like.systems | Cluster164_18clades-><-?<-?<-HEPN+DUF499*<-ACET<-?<-METHYLASE<-REase<-TUDOR+SF1-SF2-Helicase+iREase+CTD+REase<-ABhydrolase | 1130 | Bacteria>Terrabacteria_group>Cyanobacteriota/Melainabacteria_group>Cyanobacteriota>Cyanophyceae | GCF_008579685.1 | Microcystis_aeruginosa | MiTa_RS10465 | 30953..34345 | 1126 | Swt1-family-HEPN-domain-containing-protein-[Microcystis-aeruginosa] |
| WP_150976007.1 | DUF499_1.RM.like.systems | REase->AbrB->PIN->METHYLASE->RHH_6->PIN->HEPN+DUF499*-><-DDE_3 | 1111 | Bacteria>Terrabacteria_group>Cyanobacteriota/Melainabacteria_group>Cyanobacteriota>Cyanophyceae | GCF_008757435.1 | Microcystis_aeruginosa | EZJ55_RS08120 | 1414057..1417392 | 1126 | Swt1-family-HEPN-domain-containing-protein-[Microcystis-aeruginosa] |
| WP_151696808.1 | DUF499_1.RM.like.systems | <-PRTase<-sGTP<-?\|\|?->?->?-><-HEPN+DUF499*<-PIN<-?<-METHYLASE-p | 1130 | Bacteria>Terrabacteria_group>Cyanobacteriota/Melainabacteria_group>Cyanobacteriota>Cyanophyceae | GCF_008974145.1 | Microcystis_aeruginosa | MiAbW_RS15325 | 39137..42529 | 1126 | Swt1-family-HEPN-domain-containing-protein-[Microcystis-aeruginosa] |
| WP_014276704.1 | DUF499_1.RM.like.systems | METHYLASE->HEPN+DUF499*-> | 1119 | Bacteria>Terrabacteria_group>Cyanobacteriota/Melainabacteria_group>Cyanobacteriota>Cyanophyceae | GCF_009176225.1 | Arthrospira_platensis | H6G70_RS20095 | 3941..7300 | 118562 | Swt1-family-HEPN-domain-containing-protein-[Arthrospira-platensis] |
| WP_152590042.1 | DUF499_1.RM.like.systems | TUDOR+SF1-SF2-Helicase+iREase+CTD+REase-><-HTH_3<-RelE-ParE\|\|METHYLASE->HEPN+DUF499*-><-TraJ-RHH\|\|PIWI-><-HTH_3\|\|?-><-tRNA<-?<-SPOUT | 1108 | Bacteria>Terrabacteria_group>Cyanobacteriota/Melainabacteria_group>Cyanobacteriota>Cyanophyceae | GCF_009372195.1 | Nostoc_sphaeroides | GXM_RS22730 | 5029149..5032475 | 446679 | Swt1-family-HEPN-domain-containing-protein-[Nostoc-sphaeroides] |
| WP_159253859.1 | DUF499_1.RM.like.systems | TUDOR+SF1-SF2-Helicase+iREase+CTD+REase->REase->METHYLASE->OLD-ABC->PIN->HEPN+DUF499*-><-?<-?\|\|?->?-><-THUMP+Methylase | 1130 | Bacteria>Terrabacteria_group>Cyanobacteriota/Melainabacteria_group>Cyanobacteriota>Cyanophyceae | GCF_009811855.1 | Microcystis_aeruginosa | GVO78_RS16540 | 16290..19682 | 1126 | Swt1-family-HEPN-domain-containing-protein-[Microcystis-aeruginosa] |
| WP_159297746.1 | DUF499_1.RM.like.systems | TUDOR+SF1-SF2-Helicase+iREase+CTD+REase->REase->METHYLASE->OLD-ABC->PIN->HEPN+DUF499*-><-?<-?\|\|?->?-><-THUMP+Methylase | 1130 | Bacteria>Terrabacteria_group>Cyanobacteriota/Melainabacteria_group>Cyanobacteriota>Cyanophyceae | GCF_009811875.1 | Microcystis_aeruginosa | GVO69_RS15755 | 16338..19730 | 1126 | Swt1-family-HEPN-domain-containing-protein-[Microcystis-aeruginosa] |
| WP_159816100.1 | DUF499_1.RM.like.systems | <-DUF772+DDE_Tnp_1<-?<-HIN-HTH<-MORC+DUF3684<-HEPN+DUF499*<-METHYLASE-p<-DUF772+DDE_Tnp_1+DDE_Tnp_1_6<-METHYLASE<-METHYLASE-p<-PIN | 1104 | Bacteria>Terrabacteria_group>Cyanobacteriota/Melainabacteria_group>Cyanobacteriota>Cyanophyceae | GCF_009834675.1 | Cyanobium_sp._Copco_Reservoir_LC18 | L107_RS01345 | 7355..10669 | 1328305 | DUF499-domain-containing-protein-[Cyanobium-sp.-Copco_Reservoir_LC18] |
| WP_163928210.1 | DUF499_1.RM.like.systems | <-TPR+TPR\|\|TUDOR+SF1-SF2-Helicase+iREase+CTD+REase->Phd-YefM->PIN-><-HTH_3\|\|METHYLASE->HEPN+DUF499*-><-TraJ-RHH\|\|PIWI-><-?<-?\|\|?->HTH_OrfB_IS605+OrfB_IS605+OrfB_Zn_ribbon-> | 1108 | Bacteria>Terrabacteria_group>Cyanobacteriota/Melainabacteria_group>Cyanobacteriota>Cyanophyceae | GCF_010747425.1 | Nostoc_sp._UIC_10630 | G1O98_RS07815 | 12096..15422 | 2100146 | Swt1-family-HEPN-domain-containing-protein-[Nostoc-sp.-UIC-10630] |
| WP_170190015.1 | DUF499_1.RM.like.systems | TUDOR+SF1-SF2-Helicase+iREase+CTD+REase->SbcC->?->METHYLASE->?->PIN->HEPN+DUF499*-> | 1130 | Bacteria>Terrabacteria_group>Cyanobacteriota/Melainabacteria_group>Cyanobacteriota>Cyanophyceae | GCF_012911965.1 | Geitlerinema_sp._P-1104 | E1H12_RS13025 | 9854..13246 | 2546230 | Swt1-family-HEPN-domain-containing-protein-[Geitlerinema-sp.-P-1104] |
| WP_172355891.1 | DUF499_1.RM.like.systems | REase->REase->AbrB->PIN->METHYLASE->?->HEPN+DUF499*->?-><-REase\|\|?->CR-Trypsin1->?-><-REase | 1105 | Bacteria>Terrabacteria_group>Cyanobacteriota/Melainabacteria_group>Cyanobacteriota>Cyanophyceae | GCF_013177315.1 | Thermoleptolyngbya_sichuanensis | HPC62_RS11805 | 2814598..2817915 | 2885951 | Swt1-family-HEPN-domain-containing-protein-[Thermoleptolyngbya-sichuanensis] |
| WP_273247370.1 | DUF499_1.RM.like.systems | <-SPOUT<-?<-?<-?<-?\|\|?-><-HEPN+DUF499*<-?<-METHYLASE<-TUDOR+SF1-SF2-Helicase+iREase+CTD+REase<-?\|\|ABhydrolase->THUMP+Methylase-> | 1056 | Bacteria>Terrabacteria_group>Cyanobacteriota/Melainabacteria_group>Cyanobacteriota>Cyanophyceae | GCF_014250495.1 | Limnospira_indica | H2674_RS19805 | 4259944..4263114 | 147322 | DUF499-domain-containing-protein,-partial-[Limnospira-indica] |
| WP_187757730.1 | DUF499_1.RM.like.systems | SF1-SF2-Helicase-><-THUMP+Methylase<-ABhydrolase\|\|TUDOR+SF1-SF2-Helicase+iREase+CTD+REase->METHYLASE->HEPN+DUF499*-><-?\|\|?->?->?->?->SPOUT-> | 1123 | Bacteria>Terrabacteria_group>Cyanobacteriota/Melainabacteria_group>Cyanobacteriota>Cyanophyceae | GCF_014489865.2 | Limnospira | H4N54_RS05785 | 1241805..1245176 | 2596745 | Swt1-family-HEPN-domain-containing-protein-[Limnospira-fusiformis] |
| WP_190676125.1 | DUF499_1.RM.like.systems | <-pPIWI+pPIWI\|\|ParB-><-DDE_3<-PAIREDC-HTH\|\|ParB-><-TUDOR+SF1-SF2-Helicase+iREase+CTD+REase<-HEPN+DUF499*<-PIN<-OLD-ABC<-METHYLASE<-REase\|\|?-><-TUDOR+SF1-SF2-Helicase+iREase+CTD+REase | 1112 | Bacteria>Terrabacteria_group>Cyanobacteriota/Melainabacteria_group>Cyanobacteriota>Cyanophyceae | GCF_014695435.1 | Coleofasciculus_sp._FACHB-129 | H6F60_RS14220 | 70669..74007 | 2692785 | Swt1-family-HEPN-domain-containing-protein-[Coleofasciculus-sp.-FACHB-129] |
| WP_190449861.1 | DUF499_1.RM.like.systems | CHTH->TUDOR+SF1-SF2-Helicase+iREase+CTD+REase->HEPN->METHYLASE->OLD-ABC->PIN->HEPN+DUF499*->ParB+HNH->?->CopG/RHH->HNH-><-RHH | 1121 | Bacteria>Terrabacteria_group>Cyanobacteriota/Melainabacteria_group>Cyanobacteriota>Cyanophyceae | GCF_014695795.1 | Phormidium_sp._FACHB-592 | H6F86_RS16680 | 58473..61838 | 2692850 | Swt1-family-HEPN-domain-containing-protein-[Phormidium-sp.-FACHB-592] |
| WP_190576377.1 | DUF499_1.RM.like.systems | RelE-ParE-><-?<-URI-DNase<-?<-?<-HEPN+DUF499*<-METHYLASE<-REase<-AbrB<-TUDOR+SF1-SF2-Helicase+iREase+CTD+REase | 1117 | Bacteria>Terrabacteria_group>Cyanobacteriota/Melainabacteria_group>Cyanobacteriota>Cyanophyceae | GCF_014697025.1 | Alkalinema_sp._FACHB-956 | H6G21_RS22810 | 35052..38405 | 2692768 | Swt1-family-HEPN-domain-containing-protein-[Alkalinema-sp.-FACHB-956] |
| WP_190513827.1 | DUF499_1.RM.like.systems | CHTH->TUDOR+SF1-SF2-Helicase+iREase+CTD+REase->?->METHYLASE->RHH_6->PIN->HEPN+DUF499*->URI-DNase-><-RelE-ParE<-?\|\|pPIWI+pPIWI->pPIWI->pPIWI+MID+RNaseH-> | 1117 | Bacteria>Terrabacteria_group>Cyanobacteriota/Melainabacteria_group>Cyanobacteriota>Cyanophyceae | GCF_014697545.1 | Oscillatoria_sp._FACHB-1407 | H6G89_RS30695 | 49800..53153 | 2692847 | Swt1-family-HEPN-domain-containing-protein-[Oscillatoria-sp.-FACHB-1407] |
| WP_194014089.1 | DUF499_1.RM.like.systems | AbrB->PIN->AbrB->METHYLASE->?->ACET->HEPN+DUF499*-><-?\|\|?->?->?-><-TPR+TPR+TPR+TPR+TPR+TPR+TPR+TPR<-ORC-CDC6 | 1130 | Bacteria>Terrabacteria_group>Cyanobacteriota/Melainabacteria_group>Cyanobacteriota>Cyanophyceae | GCF_015207115.1 | Microcystis_aeruginosa | IQ232_RS04140 | 10022..13414 | 1126 | Swt1-family-HEPN-domain-containing-protein-[Microcystis-aeruginosa] |
| WP_193965432.1 | DUF499_1.RM.like.systems | Phage_integrase-><-?\|\|?-><-HTH\|\|?-><-HEPN+DUF499*\|\|?-><-ParB+HNH<-?<-PIN<-?<-METHYLASE | 1114 | Bacteria>Terrabacteria_group>Cyanobacteriota/Melainabacteria_group>Cyanobacteriota>Cyanophyceae | GCF_015207395.1 | Nodosilinea_sp._LEGE_07088 | IQ254_RS06885 | 5859..9203 | 2777968 | Swt1-family-HEPN-domain-containing-protein-[Nodosilinea-sp.-LEGE-07088] |
| WP_200988810.1 | DUF499_1.RM.like.systems | PRTase-><-Y1_Tnp\|\|?->?->?-><-HEPN+DUF499*<-METHYLASE\|\|REase+HTH->REase+HTH-><-TUDOR+SF1-SF2-Helicase+iREase+CTD+REase | 1108 | Bacteria>Terrabacteria_group>Cyanobacteriota/Melainabacteria_group>Cyanobacteriota>Cyanophyceae | GCF_015245355.1 | Anabaenopsis_elenkinii | IM676_RS02385 | 515316..518642 | 156213 | Swt1-family-HEPN-domain-containing-protein-[Anabaenopsis-elenkinii] |
| WP_196514030.1 | DUF499_1.RM.like.systems | HTH+HKD-DNase->SPOUT->?->tRNA-><-?\|\|<-PIWI\|\|TraJ-RHH-><-HEPN+DUF499*<-METHYLASE\|\|HTH_3-><-TUDOR+SF1-SF2-Helicase+iREase+CTD+REase<-?<-?<-HTH_28+rve+Mu-transpos_C | 1107 | Bacteria>Terrabacteria_group>Cyanobacteriota/Melainabacteria_group>Cyanobacteriota>Cyanophyceae | GCF_015714705.1 | Nostoc_sp._NZL | F8R90_RS31675 | 50056..53379 | 2650612 | Swt1-family-HEPN-domain-containing-protein-[Nostoc-sp.-NZL] |
| WP_211941813.1 | DUF499_1.RM.like.systems | HKD-DNase+SF1-SF2-Helicase-p-><-HTH+MPTase<-?<-?\|\|TUDOR+SF1-SF2-Helicase+iREase+CTD+REase->METHYLASE->HEPN+DUF499*->REase->?->Peptidase_C14-> | 1103 | Bacteria>Terrabacteria_group>Cyanobacteriota/Melainabacteria_group>Cyanobacteriota>Cyanophyceae | GCF_018139025.1 | Cylindrospermopsis_raciborskii | J9B92_RS12735 | 2844714..2848025 | 77022 | Swt1-family-HEPN-domain-containing-protein-[Cylindrospermopsis-raciborskii] |
| WP_216916461.1 | DUF499_1.RM.like.systems | <-ArdCN+MPTase\|\|HNH-><-Peptidase_M48<-?<-?<-?<-HEPN+DUF499*<-METHYLASE<-TUDOR+SF1-SF2-Helicase+iREase+CTD+REase\|\|DUF4236->?->?-><-ABhydrolase | 1099 | Bacteria>Terrabacteria_group>Cyanobacteriota/Melainabacteria_group>Cyanobacteriota>Cyanophyceae | GCF_019038475.1 | unclassified_Synechococcus | KUG39_RS15775 | 99576..102875 | 2626047 | MULTISPECIES:-DUF499-domain-containing-protein-[unclassified-Synechococcus] |
| WP_230840692.1 | DUF499_1.RM.like.systems | TUDOR+SF1-SF2-Helicase+iREase+CTD+REase->REase->METHYLASE->DDE_Tnp_1->METHYLASE-p->?->HEPN+DUF499*-><-?<-Peptidase_U32+Peptidase_U32_C | 1101 | Bacteria>Terrabacteria_group>Cyanobacteriota/Melainabacteria_group>Cyanobacteriota>Cyanophyceae | GCF_021018745.1 | Gloeobacter_morelensis | ISF26_RS17965 | 3744120..3747425 | 2907343 | Swt1-family-HEPN-domain-containing-protein-[Gloeobacter-morelensis] |
| WP_233748345.1 | DUF499_1.RM.like.systems | <-Peptidase<-HEPN+DUF499*<-CopG/RHH<-PemK_toxin<-?<-REase<-REase<-METHYLASE | 1125 | Bacteria>Terrabacteria_group>Cyanobacteriota/Melainabacteria_group>Cyanobacteriota>Cyanophyceae | GCF_021379005.1 | Leptodesmis_sichuanensis | KIKO2_RS09445 | 2037020..2040397 | 2906798 | Swt1-family-HEPN-domain-containing-protein-[Leptodesmis-sichuanensis] |
| WP_235278697.1 | DUF499_1.RM.like.systems | METHYLASE->?->PemK_toxin->REase->REase->CopG/RHH->HEPN+DUF499*->?->?->MORC+DNA_gyraseB+TOPRIM-> | 1109 | Bacteria>Terrabacteria_group>Cyanobacteriota/Melainabacteria_group>Cyanobacteriota>Cyanophyceae | GCF_021533835.1 | Synechococcus_sp._Nb3U1 | L1047_RS09935 | 2086840..2090169 | 1914529 | Swt1-family-HEPN-domain-containing-protein-[Synechococcus-sp.-Nb3U1] |
| WP_240867511.1 | DUF499_1.RM.like.systems | <-PolB<-?<-DDE_Tnp_1<-?<-?\|\|TUDOR+SF1-SF2-Helicase+iREase+CTD+REase->HEPN+DUF499*->?->?->?-><-?\|\|Peptidase_C14-> | 1102 | Bacteria>Terrabacteria_group>Cyanobacteriota/Melainabacteria_group>Cyanobacteriota>Cyanophyceae | GCF_022496815.1 | Cylindrospermopsis_raciborskii | GSN00_RS07900 | 80955..84263 | 77022 | Swt1-family-HEPN-domain-containing-protein-[Cylindrospermopsis-raciborskii] |
| WP_244349767.1 | DUF499_1.RM.like.systems | TUDOR+SF1-SF2-Helicase+iREase+CTD+REase->METHYLASE->?->PemK_toxin->REase->REase->HEPN+DUF499*->?->ABC-ATPase+SbcC->?->?-><-ABC-ATPase+SbcC<-ABC-ATPase+SbcC | 1107 | Bacteria>Terrabacteria_group>Cyanobacteriota/Melainabacteria_group>Cyanobacteriota>Cyanophyceae | GCF_022848905.1 | Thermostichus_vulcanus | JX360_RS06145 | 28078..31401 | 32053 | Swt1-family-HEPN-domain-containing-protein-[Thermostichus-vulcanus] |
| WP_247214939.1 | DUF499_1.RM.like.systems | DDE_Tnp_1-><-Peptidase_M24\|\|?->tRNA-><-HEPN+DUF499*<-REase<-REase<-REase<-METHYLASE<-PIN<-AbrB | 1107 | Bacteria>Terrabacteria_group>Cyanobacteriota/Melainabacteria_group>Cyanobacteriota>Cyanophyceae | GCF_022984075.1 | Synechococcus_sp._C9 | MLD66_RS00195 | 39889..43212 | 102119 | Swt1-family-HEPN-domain-containing-protein-[Synechococcus-sp.-C9] |
| WP_250012765.1 | DUF499_1.RM.like.systems | AbrB->PIN->REase->METHYLASE-p->METHYLASE->?->HEPN+DUF499*->?-><-?\|\|Phage_integrase-><-Phage_integrase | 1113 | Bacteria>Terrabacteria_group>Cyanobacteriota/Melainabacteria_group>Cyanobacteriota>Cyanophyceae | GCF_023546805.1 | Chroococcidiopsis_sp._CCNUC1 | M5J74_RS19090 | 4105692..4109033 | 2653189 | Swt1-family-HEPN-domain-containing-protein-[Chroococcidiopsis-sp.-CCNUC1] |
| WP_250682698.1 | DUF499_1.RM.like.systems | <-HEPN+DUF499*<-METHYLASE<-?<-GATase<-?\|\|?->RNB+RNase-> | 1053 | Bacteria>Terrabacteria_group>Cyanobacteriota/Melainabacteria_group>Cyanobacteriota>Cyanophyceae | GCF_023650955.1 | Thermosynechococcus_sp._HN-54 | NBE99_RS01185 | 246244..249405 | 2933959 | DUF499-domain-containing-protein-[Thermosynechococcus-sp.-HN-54] |
| WP_251956292.1 | DUF499_1.RM.like.systems | <-HEPN+DUF499*<-METHYLASE-p<-METHYLASE-p\|\|HTH_3-><-PIN<-Phd-YefM<-TUDOR+SF1-SF2-Helicase+iREase+CTD+REase-p | 1108 | Bacteria>Terrabacteria_group>Cyanobacteriota/Melainabacteria_group>Cyanobacteriota>Cyanophyceae | GCF_023734775.1 | Nostoc_commune | ANSO36C_RS22370 | 4938684..4942010 | 1178 | Swt1-family-HEPN-domain-containing-protein-[Nostoc-commune] |
| WP_254935737.1 | DUF499_1.RM.like.systems | <-HEPN+DUF499*<-METHYLASE<-SF1-SF2-Helicase<-TUDOR+SF1-SF2-Helicase+iREase+CTD+REase<-LRP-HTH\|\|WYL-> | 1105 | Bacteria>Terrabacteria_group>Cyanobacteriota/Melainabacteria_group>Cyanobacteriota>Cyanophyceae | GCF_024345395.1 | unclassified_Cyanobium | KBZ17_RS00995 | 40704..44021 | 2627006 | MULTISPECIES:-DUF499-domain-containing-protein-[unclassified-Cyanobium] |
| WP_106502064.1 | DUF499_1.RM.like.systems | <-WYL\|\|LRP-HTH->TUDOR+SF1-SF2-Helicase+iREase+CTD+REase->METHYLASE->HEPN+DUF499*->?->?->?->?->Peptidase_M48-><-HNH | 1113 | Bacteria>Terrabacteria_group>Cyanobacteriota/Melainabacteria_group>Cyanobacteriota>Cyanophyceae | GCF_024345435.1 | Cyanobium | C7K55_RS03685 | 97796..101137 | 167375 | MULTISPECIES:-DUF499-domain-containing-protein-[Cyanobium] |
| WP_254940710.1 | DUF499_1.RM.like.systems | <-DUF4236\|\|TUDOR+SF1-SF2-Helicase+iREase+CTD+REase->?->PIN->METHYLASE->HEPN+DUF499*->?->?-><-ArdCN+MPTase | 1103 | Bacteria>Terrabacteria_group>Cyanobacteriota/Melainabacteria_group>Cyanobacteriota>Cyanophyceae | GCF_024345575.1 | Cyanobium_sp._Morenito_9A2 | KBY88_RS14795 | 41582..44893 | 2823718 | DUF499-domain-containing-protein-[Cyanobium-sp.-Morenito-9A2] |
| WP_255090875.1 | DUF499_1.RM.like.systems | <-ACET<-CopG/RHH<-WYL\|\|LRP-HTH->TUDOR+SF1-SF2-Helicase+iREase+CTD+REase->METHYLASE->HEPN+DUF499*->Peptidase_M48-><-HNH\|\|ArdCN+MPTase-> | 1122 | Bacteria>Terrabacteria_group>Cyanobacteriota/Melainabacteria_group>Cyanobacteriota>Cyanophyceae | GCF_024346175.1 | Vulcanococcus_limneticus | KBZ20_RS12915 | 25528..28896 | 2170428 | DUF499-domain-containing-protein-[Vulcanococcus-limneticus] |
| WP_045875055.1 | DUF499_1.RM.like.systems | <-HTH_37<-GDResolvase+Recombinase<-TraJ-RHH\|\|?->Phage_int_SAM_1+Phage_integrase-><-?<-HEPN+DUF499*<-REase<-?<-REase<-METHYLASE<-HicB+ZnR<-TUDOR+SF1-SF2-Helicase+iREase+CTD+REase | 1112 | Bacteria>Terrabacteria_group>Cyanobacteriota/Melainabacteria_group>Cyanobacteriota>Cyanophyceae | GCF_025860405.1 | unclassified_Tolypothrix | FDUTEX481_RS39715 | 28585..31923 | 2649714 | MULTISPECIES:-Swt1-family-HEPN-domain-containing-protein-[unclassified-Tolypothrix] |
| WP_265264586.1 | DUF499_1.RM.like.systems | NTP_transferase-><-HEPN+DUF499*<-PIN<-AbrB<-TUDOR+SF1-SF2-Helicase+iREase+CTD+REase | 1112 | Bacteria>Terrabacteria_group>Cyanobacteriota/Melainabacteria_group>Cyanobacteriota>Cyanophyceae | GCF_026130165.1 | Spirulina_subsalsa | K4A83_RS11005 | 13314..16652 | 54311 | Swt1-family-HEPN-domain-containing-protein-[Spirulina-subsalsa] |
| WP_190359815.1 | DUF499_1.RM.like.systems | REase-><-DUF29<-?<-?\|\|?->Cluster164_18clades-><-HEPN+DUF499*<-PIN<-RHH_6<-METHYLASE<-REase<-TUDOR+SF1-SF2-Helicase+iREase+CTD+REase<-ABhydrolase | 1117 | Bacteria>Terrabacteria_group>Cyanobacteriota/Melainabacteria_group>Cyanobacteriota>Cyanophyceae | GCF_026222535.1 | Microcystis | H6F92_RS24355 | 16318..19671 | 1125 | Swt1-family-HEPN-domain-containing-protein-[Microcystis-wesenbergii] |
| WP_265900465.1 | DUF499_1.RM.like.systems | REase->REase->?->?-><-sGTP<-HEPN+DUF499*<-PIN<-?<-REase<-REase<-REase | 1128 | Bacteria>Terrabacteria_group>Cyanobacteriota/Melainabacteria_group>Cyanobacteriota>Cyanophyceae | GCF_026240635.1 | Trichothermofontia_sichuanensis | OOK60_RS10505 | 2477245..2480631 | 3045816 | DUF499-domain-containing-protein-[Leptolyngbya-sp.-PKUAC-SCTB231] |
| WP_271935711.1 | DUF499_1.RM.like.systems | AbrB->METHYLASE->RHH_6->PIN->OLD-ABC->PIN->HEPN+DUF499*-><-Cluster164_18clades<-?\|\|?->?->DUF29->sGTP-> | 1111 | Bacteria>Terrabacteria_group>Cyanobacteriota/Melainabacteria_group>Cyanobacteriota>Cyanophyceae | GCF_028329095.1 | Microcystis | PN470_RS21190 | 25157..28492 | 1125 | MULTISPECIES:-Swt1-family-HEPN-domain-containing-protein-[Microcystis] |
| WP_272094882.1 | DUF499_1.RM.like.systems | <-HEPN+DUF499*<-PIN<-OLD-ABC<-PIN<-RHH_6<-METHYLASE<-REase | 1065 | Bacteria>Terrabacteria_group>Cyanobacteriota/Melainabacteria_group>Cyanobacteriota>Cyanophyceae | GCF_028330705.1 | Microcystis_aeruginosa | PN479_RS02395 | 1..3194 | 1126 | Swt1-family-HEPN-domain-containing-protein,-partial-[Microcystis-aeruginosa] |
| WP_272014891.1 | DUF499_1.RM.like.systems | METHYLASE-p->RHH_6->PIN->HEPN+DUF499*-><-Cluster164_18clades<-?\|\|?->?->DUF29->sGTP-> | 1111 | Bacteria>Terrabacteria_group>Cyanobacteriota/Melainabacteria_group>Cyanobacteriota>Cyanophyceae | GCF_028330745.1 | Microcystis_aeruginosa | PN482_RS11750 | 1666..5001 | 1126 | Swt1-family-HEPN-domain-containing-protein-[Microcystis-aeruginosa] |
| WP_272078194.1 | DUF499_1.RM.like.systems | DDE_3-><-HEPN+DUF499*<-PIN<-PIN<-RHH_6<-METHYLASE<-REase<-TUDOR+SF1-SF2-Helicase+iREase+CTD+REase | 1111 | Bacteria>Terrabacteria_group>Cyanobacteriota/Melainabacteria_group>Cyanobacteriota>Cyanophyceae | GCF_028331105.1 | Microcystis_aeruginosa | PN487_RS19700 | 7822..11157 | 1126 | Swt1-family-HEPN-domain-containing-protein-[Microcystis-aeruginosa] |
| WP_271953564.1 | DUF499_1.RM.like.systems | PIN->HEPN+DUF499*-><-Cluster164_18clades | 1130 | Bacteria>Terrabacteria_group>Cyanobacteriota/Melainabacteria_group>Cyanobacteriota>Cyanophyceae | GCF_028331205.1 | Microcystis_aeruginosa | PN437_RS04350 | 1001..4393 | 1126 | Swt1-family-HEPN-domain-containing-protein-[Microcystis-aeruginosa] |
| WP_280653420.1 | DUF499_1.RM.like.systems | DNA_methylase->?->?-><-?<-HEPN+DUF499* | 1111 | Bacteria>Terrabacteria_group>Cyanobacteriota/Melainabacteria_group>Cyanobacteriota>Cyanophyceae | GCF_029891735.1 | Chrysosporum_bergii | NWP17_RS02960 | 23805..27140 | 105352 | Swt1-family-HEPN-domain-containing-protein-[Chrysosporum-bergii] |
| WP_281006617.1 | DUF499_1.RM.like.systems | METHYLASE->?->PIN->REase->REase->REase->HEPN+DUF499*-> | 1118 | Bacteria>Terrabacteria_group>Cyanobacteriota/Melainabacteria_group>Cyanobacteriota>Cyanophyceae | GCF_029910235.1 | Pseudanabaena_galeata | OA858_RS18405 | 4059827..4063183 | 1112103 | Swt1-family-HEPN-domain-containing-protein-[Pseudanabaena-galeata] |
| WP_285965028.1 | DUF499_1.RM.like.systems | XIS-HTH-><-?<-Phage_integrase\|\|tRNA-><-?\|\|REase-><-DUF499-p<-HEPN+DUF499*<-AbiEii | 1005 | Bacteria>Terrabacteria_group>Cyanobacteriota/Melainabacteria_group>Cyanobacteriota>Cyanophyceae | GCF_030285465.1 | Oscillatoria_laete-virens | QQ056_RS17150 | 57885..60902 | 1565316 | Swt1-family-HEPN-domain-containing-protein-[Oscillatoria-laete-virens] |
| WP_087068561.1 | DUF499_1.RM.like.systems | TUDOR+SF1-SF2-Helicase+iREase+CTD+REase->?->PIN->METHYLASE-p->DUF772+DDE_Tnp_1_2->METHYLASE->HEPN+DUF499*->SF1-SF2-Helicase+REase->SF1-SF2-Helicase-p-><-?<-DUF772+DDE_Tnp_1_2 | 1101 | Bacteria>Terrabacteria_group>Cyanobacteriota/Melainabacteria_group>Cyanobacteriota>Cyanophyceae | GCF_900088535.1 | Cyanobium_sp._NIES-981 | CBM981_RS11760 | 2335245..2338550 | 1851505 | DUF499-domain-containing-protein-[Cyanobium-sp.-NIES-981] |
| WP_104397640.1 | DUF499_1.RM.like.systems | REase->AbrB->PIN->METHYLASE->RHH_6->PIN->HEPN+DUF499*-><-Cluster164_18clades<-?\|\|?->?->?-><-TPR+TPR+TPR+TPR+TPR+TPR+TPR+TPR | 1111 | Bacteria>Terrabacteria_group>Cyanobacteriota/Melainabacteria_group>Cyanobacteriota>Cyanophyceae | - | Microcystis_aeruginosa | myaer87_32740 | 119164..122499 | 1126 | Swt1-family-HEPN-domain-containing-protein-[Microcystis-aeruginosa] |
| WP_190546091.1 | DUF499_1.RM.like.systems | LRP-HTH->METHYLASE->HEPN+DUF499*->?->?->?->?->Peptidase_M48-><-HNH | 1126 | Bacteria>Terrabacteria_group>Cyanobacteriota/Melainabacteria_group>Cyanobacteriota>Cyanophyceae | - | Microcystis_elabens | H6G65_10800 | 7559..10939 | 44824 | DUF499-domain-containing-protein-[Microcystis-elabens] |
| WP_229465572.1 | DUF499_1.RM.like.systems | <-SIGMA-HTH\|\|?-><-?<-PIWI\|\|TraJ-RHH-><-HEPN+DUF499*<-METHYLASE-p<-METHYLASE-p<-DOC<-?<-TUDOR+SF1-SF2-Helicase+iREase+CTD+REase | 1108 | Bacteria>Terrabacteria_group>Cyanobacteriota/Melainabacteria_group>Cyanobacteriota>Cyanophyceae | - | Nostoc_sp._CHAB_5824 | LC607_29645 | 82754..86080 | 2780402 | Swt1-family-HEPN-domain-containing-protein-[Nostoc-sp.-CHAB-5824] |
| WP_255548761.1 | DUF499_1.RM.like.systems | REase-><-?<-CR-Trypsin1<-?\|\|REase-><-?<-HEPN+DUF499*<-?<-METHYLASE<-PIN<-AbrB<-REase<-REase | 1078 | Bacteria>Terrabacteria_group>Cyanobacteriota/Melainabacteria_group>Cyanobacteriota>Cyanophyceae | - | Thermoleptolyngbya_sp._PKUAC-SCTB121 | JTM80_RS13275 | 3238289..3241525 | 2811482 | DUF499-domain-containing-protein-[Thermoleptolyngbya-sp.-PKUAC-SCTB121] |
| OWY70620.1 | DUF499_1.RM.like.systems | TUDOR+SF1-SF2-Helicase+iREase+CTD+REase->?->METHYLASE->?->DUF488-1->?->HEPN+DUF499*->?-><-?\|\|ZnR+TOPRIM+D5ATPase-> | 1113 | Bacteria>Terrabacteria_group>Cyanobacteriota/Melainabacteria_group>Cyanobacteriota>unclassified_Cyanobacteriota | GCA_002213405.1 | cyanobacterium_TDX16 | B7486_13460 | 127054..130395 | 1503470 | AAA+-family-ATPase-[cyanobacterium-TDX16] |
| OWY65325.1 | DUF499_1.RM.like.systems | Phage_integrase-><-Phage_integrase<-HEPN+DUF499*<-DUF29<-REase<-PIN<-AbrB<-TUDOR+SF1-SF2-Helicase+iREase+CTD+REase\|\|Classical-AAA-> | 1119 | Bacteria>Terrabacteria_group>Cyanobacteriota/Melainabacteria_group>Cyanobacteriota>unclassified_Cyanobacteriota | GCA_002213405.1 | cyanobacterium_TDX16 | B7486_42500 | 6012..9371 | 1503470 | AAA+-family-ATPase-[cyanobacterium-TDX16] |
| PSO73463.1 | DUF499_1.RM.like.systems | <-HEPN+DUF499*<-TUDOR+SF1-SF2-Helicase+iREase+CTD+REase<-?\|\|?-><-BrnA_antitoxin\|\|?-><-DDE_Tnp_ISL3 | 1112 | Bacteria>Terrabacteria_group>Cyanobacteriota/Melainabacteria_group>Cyanobacteriota>unclassified_Cyanobacteriota | GCA_003021535.1 | Cyanobacteria_bacterium_QH_3_48_40 | BRC37_09455 | 1052..4390 | 1919122 | AAA+-family-ATPase-[Cyanobacteria-bacterium-QH_3_48_40] |
| PSO86698.1 | DUF499_1.RM.like.systems | REase->?-><-?\|\|HEPN+DUF499*-> | 1112 | Bacteria>Terrabacteria_group>Cyanobacteriota/Melainabacteria_group>Cyanobacteriota>unclassified_Cyanobacteriota | GCA_003021925.1 | Cyanobacteria_bacterium_QH_9_48_43 | BRC41_06380 | 3561..6899 | 1919126 | AAA+-family-ATPase-[Cyanobacteria-bacterium-QH_9_48_43] |
| PSP10430.1 | DUF499_1.RM.like.systems | DDE_Tnp_ISL3->BrnA_antitoxin-><-?\|\|?->HEPN+DUF499*-> | 1112 | Bacteria>Terrabacteria_group>Cyanobacteriota/Melainabacteria_group>Cyanobacteriota>unclassified_Cyanobacteriota | GCA_003022175.1 | Cyanobacteria_bacterium_SW_10_48_33 | BRC49_10060 | 15348..18686 | 1919136 | AAA+-family-ATPase-[Cyanobacteria-bacterium-SW_10_48_33] |
| PSO64590.1 | DUF499_1.RM.like.systems | <-HEPN+DUF499*<-TUDOR+SF1-SF2-Helicase+iREase+CTD+REase<-?\|\|?-><-BrnA_antitoxin\|\|?-><-DDE_Tnp_ISL3 | 1112 | Bacteria>Terrabacteria_group>Cyanobacteriota/Melainabacteria_group>Cyanobacteriota>unclassified_Cyanobacteriota | GCA_003022265.1 | Cyanobacteria_bacterium_QH_6_48_35 | BRC38_11110 | 999..4337 | 1919123 | AAA+-family-ATPase-[Cyanobacteria-bacterium-QH_6_48_35] |

| Protein-ID | System | Gene-Neighbourhoods | Length | Taxonomy | Assembly-ID | Strain | Locus-Tag | Genomic-coordinates | TaxID | Fasta-defline |
|---|---|---|---|---|---|---|---|---|---|---|
| MBJ7900527.1 | DUF499_1.RM.like.systems | <-Glyco_hydro_57+DUF3536<-?\|\|?->HAD-><-HEPN+DUF499*<-REase<-PIN<-OLD-ABC<-CopG/RHH<-?<-TUDOR+SF1-SF2-Helicase+iREase+CTD+REase | 1111 | Bacteria>Terrabacteria_group>Cyanobacteriota/Melainabacteria_group>Cyanobacteriota>unclassified_Cyanobacteriota | GCA_016471375.1 | Cyanobacteria_bacterium_RI_101 | GC158_11540 | 10083..13418 | 2775578 | DUF499-domain-containing-protein-[Cyanobacteria-bacterium-RI_101] |
| MBM5817894.1 | DUF499_1.RM.like.systems | HicB_lk_antitox->HicA_toxin->HEPN+DUF499*-> | 1124 | Bacteria>Terrabacteria_group>Cyanobacteriota/Melainabacteria_group>Cyanobacteriota>unclassified_Cyanobacteriota | GCA_016882305.1 | Cyanobacteria_bacterium_K_Offshore_surface_m2_239 | FJ083_15335 | 1681..5055 | 2775555 | ATP-binding-protein-[Cyanobacteria-bacterium-K_Offshore_surface_m2_239] |
| MBU6346401.1 | DUF499_1.RM.like.systems | <-Peptidase_C14<-?<-REase<-HEPN+DUF499*<-METHYLASE | 1103 | Bacteria>Terrabacteria_group>Cyanobacteriota/Melainabacteria_group>Cyanobacteriota>unclassified_Cyanobacteriota | GCA_018969285.1 | Cyanobacteria_bacterium_REEB494 | KGR70_14045 | 15811..19122 | 2800107 | DUF499-domain-containing-protein-[Cyanobacteria-bacterium-REEB494] |
| MBY0405416.1 | DUF499_1.RM.like.systems | METHYLASE-p->HEPN+DUF499*-> | 1058 | Bacteria>Terrabacteria_group>Cyanobacteriota/Melainabacteria_group>Cyanobacteriota>unclassified_Cyanobacteriota | GCA_019751245.1 | Cyanobacteriota_bacterium | K2X66_16060 | 763..3936 | 2847862 | DUF499-domain-containing-protein-[Cyanobacteria-bacterium] |
| MCY4055150.1 | DUF499_1.RM.like.systems | ABC-ATPase+SbcC-><-HEPN+DUF499*<-METHYLASE<-SIR2 | 1113 | Bacteria>Terrabacteria_group>Cyanobacteriota/Melainabacteria_group>Cyanobacteriota>unclassified_Cyanobacteriota | GCA_026708625.1 | Cyanobacteria_bacterium_MAG_CAR4_bin_6 | OXF67_04645 | 13192..16533 | 2775592 | MAG:-DUF499-domain-containing-protein-[Cyanobacteria-bacterium-MAG-CAR4_bin_6] |
| MCY4173696.1 | DUF499_1.RM.like.systems | ABC-ATPase+SbcC-><-HEPN+DUF499*<-METHYLASE<-SIR2<-?\|\|?->?-><-TUDOR+SF1-SF2-Helicase+iREase+CTD+REase | 1112 | Bacteria>Terrabacteria_group>Cyanobacteriota/Melainabacteria_group>Cyanobacteriota>unclassified_Cyanobacteriota | GCA_026709085.1 | Cyanobacteria_bacterium_MAG_CAR3_bin_5 | OXF25_06475 | 22273..25611 | 2775591 | MAG:-DUF499-domain-containing-protein-[Cyanobacteria-bacterium-MAG-CAR3_bin_5] |
| MCY4234708.1 | DUF499_1.RM.like.systems | HEPN->METHYLASE->HEPN+DUF499*-> | 347 | Bacteria>Terrabacteria_group>Cyanobacteriota/Melainabacteria_group>Cyanobacteriota>unclassified_Cyanobacteriota | GCA_026709585.1 | Cyanobacteria_bacterium_MAG_CAR2_bin_4 | OXE74_00055 | 3576..4616 | 2775590 | MAG:-Swt1-family-HEPN-domain-containing-protein,-partial-[Cyanobacteria-bacterium-MAG-CAR2_bin_4] |
| MCY4235602.1 | DUF499_1.RM.like.systems | SIR2->METHYLASE->HEPN+DUF499*-> | 348 | Bacteria>Terrabacteria_group>Cyanobacteriota/Melainabacteria_group>Cyanobacteriota>unclassified_Cyanobacteriota | GCA_026709585.1 | Cyanobacteria_bacterium_MAG_CAR2_bin_4 | OXE74_04750 | 4969..6012 | 2775590 | MAG:-Swt1-family-HEPN-domain-containing-protein,-partial-[Cyanobacteria-bacterium-MAG-CAR2_bin_4] |
| MCY4332863.1 | DUF499_1.RM.like.systems | ABC-ATPase+SbcC-><-HEPN+DUF499*<-METHYLASE<-wHTH+REase | 1113 | Bacteria>Terrabacteria_group>Cyanobacteriota/Melainabacteria_group>Cyanobacteriota>unclassified_Cyanobacteriota | GCA_026710375.1 | Cyanobacteria_bacterium_MAG_CAR1_bin_15 | OXC96_10300 | 26378..29719 | 2775589 | MAG:-DUF499-domain-containing-protein-[Cyanobacteria-bacterium-MAG-CAR1_bin_15] |
| OLV20266.1 | DUF499_1.RM.like.systems | TUDOR+SF1-SF2-Helicase+iREase+CTD+REase-><-?<-?\|\|METHYLASE->?->HEPN+DUF499*->?-><-NERD+PspA | 1092 | Bacteria>Terrabacteria_group>Deinococcota>Deinococci>Deinococcales | GCA_001949125.1 | Deinococcus_marmoris | BOO71_0000262 | 38538..41816 | 249408 | hypothetical-protein-BOO71_0000262-[Deinococcus-marmoris] |
| WP_075830034.1 | DUF499_1.RM.like.systems | TUDOR+SF1-SF2-Helicase+iREase+CTD+REase-><-?<-?\|\|METHYLASE->?->HEPN+DUF499*->?->?-><-NERD+PspA | 1100 | Bacteria>Terrabacteria_group>Deinococcota>Deinococci>Deinococcales | GCF_001949125.1 | Deinococcus_marmoris | BOO71_RS00480 | 38514..41816 | 249408 | DUF499-domain-containing-protein-[Deinococcus-marmoris] |
| WP_103130569.1 | DUF499_1.RM.like.systems | ABhydrolase->NTP_transferase->?->IclR-HTH-><-TraJ-RHH<-HEPN+DUF499*\|\|?-><-METHYLASE\|\|?->?-><-TUDOR+SF1-SF2-Helicase+iREase+CTD+REase | 1099 | Bacteria>Terrabacteria_group>Deinococcota>Deinococci>Deinococcales | GCF_002897375.1 | Deinococcus_aerius | DAERI_RS16670 | 55118..58417 | 200253 | DUF499-domain-containing-protein-[Deinococcus-aerius] |
| WP_107138393.1 | DUF499_1.RM.like.systems | TUDOR+SF1-SF2-Helicase+iREase+CTD+REase->HKD-DNase->Schlafen+HATPase+HTH->?->METHYLASE->HEPN+DUF499*->?->HIN-HTH-><-Phage_int_SAM_4+Phage_integrase | 1079 | Bacteria>Terrabacteria_group>Deinococcota>Deinococci>Deinococcales | GCF_003028415.1 | Deinococcus_arcticus | C8263_RS12115 | 79347..82586 | 2136176 | DUF499-domain-containing-protein-[Deinococcus-arcticus] |
| WP_135230540.1 | DUF499_1.RM.like.systems | Phage_integrase-><-?<-GDResolvase+HIN-HTH<-HEPN+DUF499*<-METHYLASE\|\|?-><-Schlafen+HATPase+HTH<-HKD-DNase<-Phage_integrase | 1098 | Bacteria>Terrabacteria_group>Deinococcota>Deinococci>Deinococcales | GCF_004634215.1 | Deinococcus_fonticola | E5Z01_RS17500 | 5706..9002 | 2528713 | DUF499-domain-containing-protein-[Deinococcus-fonticola] |
| WP_184025303.1 | DUF499_1.RM.like.systems | Phage_int_SAM_1+Phage_integrase->TUDOR+SF1-SF2-Helicase+iREase+CTD+REase->DUF433->PIN->METHYLASE->?->HEPN+DUF499*-><-?<-TOPRIM+primpol<-OrfB_Zn_ribbon<-XIS-HTH\|\|HTH->Phage_integrase-> | 1097 | Bacteria>Terrabacteria_group>Deinococcota>Deinococci>Deinococcales | GCF_014201885.1 | Deinococcus_budaensis | HNQ09_RS02950 | 29745..33038 | 1665626 | DUF499-domain-containing-protein-[Deinococcus-budaensis] |
| WP_189066307.1 | DUF499_1.RM.like.systems | Phage_int_SAM_4+Phage_integrase->?-><-HIN-HTH<-?<-HEPN+DUF499*<-METHYLASE-p<-METHYLASE<-?<-Schlafen+HATPase+HTH<-HKD-DNase<-TUDOR+SF1-SF2-Helicase+iREase+CTD+REase | 1082 | Bacteria>Terrabacteria_group>Deinococcota>Deinococci>Deinococcales | GCF_014648115.1 | Deinococcus_seoulensis | IEY70_RS17415 | 18426..21674 | 1837379 | DUF499-domain-containing-protein-[Deinococcus-seoulensis] |
| WP_239048765.1 | DUF499_1.RM.like.systems | <-HIN-HTH<-?<-?<-?<-McrC-NTD+REase<-DUF499-p<-HEPN+DUF499*<-HEPN-p<-METHYLASE\|\|?-><-Schlafen+HATPase+HTH<-HKD-DNase<-TUDOR+SF1-SF2-Helicase+iREase+CTD+REase | 559 | Bacteria>Terrabacteria_group>Deinococcota>Deinococci>Deinococcales | GCF_022344005.1 | Deinococcus_sp._KNUC1210 | MF271_RS10550 | 1554668..1556347 | 2917691 | DUF499-domain-containing-protein-[Deinococcus-sp.-KNUC1210] |
| WP_253581497.1 | DUF499_1.RM.like.systems | TUDOR+SF1-SF2-Helicase+iREase+CTD+REase->HKD-DNase->Schlafen+HATPase+HTH-><-?\|\|METHYLASE->HEPN+DUF499*-> | 1096 | Bacteria>Terrabacteria_group>Deinococcota>Deinococci>Deinococcales | GCF_024171495.1 | Deinococcus_sp._HSC-46F16 | L1280_RS07655 | 450556..453846 | 2910968 | DUF499-domain-containing-protein-[Deinococcus-sp.-HSC-46F16] |
| WP_129117970.1 | DUF499_1.RM.like.systems | <-HEPN+DUF499*<-METHYLASE-p<-METHYLASE-p<-?<-Schlafen+HATPase+HTH<-?<-HKD-DNase | 1076 | Bacteria>Terrabacteria_group>Deinococcota>Deinococci>Deinococcales | GCF_030161515.1 | Deinococcus_metallilatus | QQK08_RS18410 | 2112..5342 | 1211322 | DUF499-domain-containing-protein-[Deinococcus-metallilatus] |
| AEG34458.1 | DUF499_1.RM.like.systems | METHYLASE->HicB_lk_antitox->?->HigB/RHH->HTH_3->HIN-HTH+AT-HTH-HYD2-2+SecB->HEPN+DUF499*->?->?->?->?->Exonuc_V_gamma+REase->SF1-SF2-Helicase+REase-> | 1084 | Bacteria>Terrabacteria_group>Deinococcota>Deinococci>Thermales | GCA_000214845.1 | Thermus_thermophilus_SG0.5JP17-16 | Tthe16_2072 | 92847..96101 | 762633 | ATPase-(AAA+-superfamily)-like-protein-[Thermus-thermophilus-SG0.5JP17-16] |
| AEG34698.1 | DUF499_1.RM.like.systems | DDE_Tnp_1->HTH_3->HIN-HTH+SecB->HEPN+DUF499*->HTH+DSBH->?->?->?->?->ABC-ATPase+SbcC-> | 1084 | Bacteria>Terrabacteria_group>Deinococcota>Deinococci>Thermales | GCA_000214845.1 | Thermus_thermophilus_SG0.5JP17-16 | Tthe16_2339 | 352760..356014 | 762633 | ATPase-(AAA+-superfamily)-like-protein-[Thermus-thermophilus-SG0.5JP17-16] |
| AEV15179.1 | DUF499_1.RM.like.systems | <-DUF499-p\|\|MULE-TRANSPOSASE-><-HEPN+DUF499*<-HicB_lk_antitox<-HicA_toxin<-METHYLASE<-HKD-DNase+SF1-SF2-Helicase-p | 355 | Bacteria>Terrabacteria_group>Deinococcota>Deinococci>Thermales | GCA_000236585.1 | Thermus_sp._CCB_US3_UF1 | TCCBUS3UF1_1290 | 146394..147461 | 1111069 | hypothetical-protein-TCCBUS3UF1_1290-[Thermus-sp.-CCB_US3_UF1] |
| MBF6594441.1 | DUF499_1.RM.like.systems | TUDOR+SF1-SF2-Helicase+iREase+CTD+REase->Schlafen+HATPase+HTH->METHYLASE-><-PIN<-MazE_antitoxin\|\|HEPN+DUF499*-><-HTH_Tnp_4\|\|<-?<-?\|\|METHYLASE-p-> | 1117 | Bacteria>Terrabacteria_group>Deinococcota>Deinococci>Thermales | GCA_015478585.1 | Thermaceae_bacterium | IVW51_08335 | 21188..24541 | 2614257 | DUF499-domain-containing-protein-[Thermaceae-bacterium] |
| BCP99115.1 | DUF499_1.RM.like.systems | <-ABC-ATPase+SbcC<-?\|\|?-><-DDE_Tnp_1<-HEPN+DUF499*<-HIN-HTH+AT-HTH-HYD2-2+SecB<-HTH_3<-HicB_lk_antitox<-HicA_toxin<-METHYLASE<-DOC+HTH | 1083 | Bacteria>Terrabacteria_group>Deinococcota>Deinococci>Thermales | GCA_016865275.1 | Thermus_thermophilus | TthHB5002_b22180 | 17958..21209 | 274 | hypothetical-protein-TthHB5002_b22180-[Thermus-thermophilus] |
| BCZ90733.1 | DUF499_1.RM.like.systems | <-MULE-TRANSPOSASE\|\|HIN-HTH->HTH_21+rve+rve_3-><-?\|\|PAIREDC-HTH+HIN-HTH->DDE_3-><-HEPN+DUF499*<-HIN-HTH+AT-HTH-HYD2-2+SecB<-HTH_3<-HicB_lk_antitox<-HicA_toxin<-METHYLASE<-DOC+HTH | 1084 | Bacteria>Terrabacteria_group>Deinococcota>Deinococci>Thermales | GCA_019974355.1 | Thermus_thermophilus | TthAA22_25380 | 104954..108208 | 274 | hypothetical-protein-TthAA22_25380-[Thermus-thermophilus] |
| BDG20195.1 | DUF499_1.RM.like.systems | DDE_Tnp_1_3->DDE_Tnp_1->?->DDE_Tnp_1_3->DDE_Tnp_1-><-HEPN+DUF499*<-HIN-HTH+AT-HTH-HYD2-2+SecB<-HTH_3<-HigB/RHH<-HicB_lk_antitox<-METHYLASE | 1084 | Bacteria>Terrabacteria_group>Deinococcota>Deinococci>Thermales | GCA_022846275.1 | Thermus_thermophilus | TthSNM11_23980 | 20817..24071 | 274 | hypothetical-protein-TthSNM11_23980-[Thermus-thermophilus] |
| BDG27741.1 | DUF499_1.RM.like.systems | DDE_Tnp_1+DDE_Tnp_1_2->PAIREDC-HTH+HIN-HTH->DDE_3-><-HicA_toxin<-HicB<-DDE_Tnp_1<-HEPN+DUF499*<-HIN-HTH+AT-HTH-HYD2-2+SecB<-HTH_3<-HigB/RHH<-HicB_lk_antitox<-METHYLASE | 1084 | Bacteria>Terrabacteria_group>Deinococcota>Deinococci>Thermales | GCA_022846315.1 | Thermus_thermophilus | TthSNM66_23770 | 50871..54125 | 274 | hypothetical-protein-TthSNM66_23770-[Thermus-thermophilus] |
| BDG22724.1 | DUF499_1.RM.like.systems | METHYLASE->HicA_toxin->HicB_lk_antitox->HTH_3->HIN-HTH+AT-HTH-HYD2-2+SecB->HEPN+DUF499*-><-DDE_Tnp_1_3\|\|HTH_3->HIN-HTH+AT-HTH-HYD2-2+SecB->?->ABC-ATPase+SbcC-> | 1084 | Bacteria>Terrabacteria_group>Deinococcota>Deinococci>Thermales | GCA_022846355.1 | Thermus_thermophilus | TthSNM17_23860 | 105744..108998 | 274 | hypothetical-protein-TthSNM17_23860-[Thermus-thermophilus] |
| GIW28837.1 | DUF499_1.RM.like.systems | TPR-><-ABhydrolase<-HDRnase<-?\|\|HEPN+DUF499*->TUDOR+SF1-SF2-Helicase+iREase+CTD+REase->SF1-SF2-Helicase+wHTH+PRTase->?->Schlafen+HATPase+HTH->METHYLASE-><-TPR | 1063 | Bacteria>Terrabacteria_group>Deinococcota>Deinococci>Thermales | GCA_026004075.1 | Meiothermus_sp. | KatS3mg070_2200 | 2304208..2307399 | 1955249 | MAG:-hypothetical-protein-KatS3mg070_2200-[Meiothermus-sp.] |
| GIW32506.1 | DUF499_1.RM.like.systems | MerR-HTH->?->TUDOR+SF1-SF2-Helicase+iREase+CTD+REase->METHYLASE->HEPN+DUF499*-> | 1105 | Bacteria>Terrabacteria_group>Deinococcota>Deinococci>Thermales | GCA_026004095.1 | Meiothermus_sp. | KatS3mg071_2680 | 19805..23122 | 1955249 | MAG:-hypothetical-protein-KatS3mg071_2680-[Meiothermus-sp.] |
| GIW33264.1 | DUF499_1.RM.like.systems | Pkinase->DUF4388-><-?<-?\|\|?-><-HEPN+DUF499*<-HEPN<-PolB<-METHYLASE<-ORC-CDC6+REase<-TUDOR+SF1-SF2-Helicase+iREase+CTD+REase | 1104 | Bacteria>Terrabacteria_group>Deinococcota>Deinococci>Thermales | GCA_026004115.1 | Meiothermus_sp. | KatS3mg072_0597 | 655659..658973 | 1955249 | MAG:-hypothetical-protein-KatS3mg072_0597-[Meiothermus-sp.] |
| WP_013458840.1 | DUF499_1.RM.like.systems | TUDOR+SF1-SF2-Helicase+iREase+CTD+REase->HicA_toxin->HicB_lk_antitox->PIN->METHYLASE->HEPN+DUF499*->?-><-?<-?<-?\|\|?->METHYLASE-p-> | 1120 | Bacteria>Terrabacteria_group>Deinococcota>Deinococci>Thermales | GCF_000183745.1 | Oceanithermus_profundus | OCEPR_RS11190 | 2275579..2278941 | 187137 | DUF499-domain-containing-protein-[Oceanithermus-profundus] |
| WP_015716390.1 | DUF499_1.RM.like.systems | <-Phage_integrase\|\|?->?-><-?\|\|OLD-ABC+TOPRIM-><-HEPN+DUF499*<-METHYLASE<-MACRODOMAIN<-?<-TUDOR+SF1-SF2-Helicase+iREase+CTD+REase<-?<-ArsR-HTH | 1116 | Bacteria>Terrabacteria_group>Deinococcota>Deinococci>Thermales | GCF_000187005.1 | Thermus_scotoductus | TSC_RS02330 | 419073..422423 | 37636 | Swt1-family-HEPN-domain-containing-protein-[Thermus-scotoductus] |
| WP_014511304.1 | DUF499_1.RM.like.systems | DDE_Tnp_1->HTH_3->HIN-HTH+SecB->HEPN+DUF499*->\|\|HTH+DSBH->DSBH-> | 1084 | Bacteria>Terrabacteria_group>Deinococcota>Deinococci>Thermales | GCF_000214845.1 | Thermus_thermophilus | THTHE16_RS11840 | 352760..356014 | 274 | Swt1-family-HEPN-domain-containing-protein-[Thermus-thermophilus] |
| WP_014511084.1 | DUF499_1.RM.like.systems | METHYLASE->HicB_lk_antitox->?->HigB/RHH->HTH_3->HIN-HTH+AT-HTH-HYD2-2+SecB->HEPN+DUF499*->?->?->?->?->Exonuc_V_gamma+REase->SF1-SF2-Helicase+REase-> | 1084 | Bacteria>Terrabacteria_group>Deinococcota>Deinococci>Thermales | GCF_000214845.1 | Thermus_thermophilus | THTHE16_RS13550 | 92847..96101 | 274 | Swt1-family-HEPN-domain-containing-protein-[Thermus-thermophilus] |
| WP_014514547.1 | DUF499_1.RM.like.systems | 35exo+SF1-SF2-Helicase->?->?->?-><-?<-DUF499*\|\|MULE-TRANSPOSASE-><-HEPN+DUF499<-HicB_lk_antitox<-HicA_toxin<-METHYLASE<-HKD-DNase+SF1-SF2-Helicase-p | 742 | Bacteria>Terrabacteria_group>Deinococcota>Deinococci>Thermales | GCF_000236585.1 | Thermus_sp._CCB_US3_UF1 | TCCBUS3UF1_RS00625 | 142056..144284 | 1111069 | DUF499-domain-containing-protein-[Thermus-sp.-CCB_US3_UF1] |
| WP_155983233.1 | DUF499_1.RM.like.systems | <-DUF499\|\|MULE-TRANSPOSASE-><-HEPN+DUF499*<-HicB_lk_antitox<-HicA_toxin<-METHYLASE<-HKD-DNase+SF1-SF2-Helicase-p\|\|?->PIN-> | 341 | Bacteria>Terrabacteria_group>Deinococcota>Deinococci>Thermales | GCF_000236585.1 | Thermus_sp._CCB_US3_UF1 | TCCBUS3UF1_RS00635 | 146394..147419 | 1111069 | Swt1-family-HEPN-domain-containing-protein-[Thermus-sp.-CCB_US3_UF1] |
| WP_014632301.1 | DUF499_1.RM.like.systems | <-SF1-SF2-Helicase+REase<-Exonuc_V_gamma+REase<-?<-?<-?<-?<-HEPN+DUF499*<-HIN-HTH+AT-HTH-HYD2-2+SecB<-HTH_3<-HigB/RHH<-HicB_lk_antitox<-METHYLASE<-DOC+HTH | 1084 | Bacteria>Terrabacteria_group>Deinococcota>Deinococci>Thermales | GCF_000258245.1 | Thermus_thermophilus | TTJL18_RS12200 | 95863..99117 | 274 | Swt1-family-HEPN-domain-containing-protein-[Thermus-thermophilus] |
| WP_015065389.1 | DUF499_1.RM.like.systems | ParB->TUDOR+SF1-SF2-Helicase+iREase+CTD+REase->PIN->DDE_Tnp_1->HEPN+DUF499*->SF1-SF2-Helicase->IclR-HTH-> | 1099 | Bacteria>Terrabacteria_group>Deinococcota>Deinococci>Thermales | GCF_000309885.1 | Thermus_oshimai | THEOS_RS11985 | 210648..213947 | 56957 | Swt1-family-HEPN-domain-containing-protein-[Thermus-oshimai] |
| WP_018461866.1 | DUF499_1.RM.like.systems | ParB->TUDOR+SF1-SF2-Helicase+iREase+CTD+REase->PIN->METHYLASE->HEPN+DUF499*->SF1-SF2-Helicase->?-><-?<-Peptidase_C26 | 1097 | Bacteria>Terrabacteria_group>Deinococcota>Deinococci>Thermales | GCF_000373145.1 | Thermus_oshimai | B043_RS0109800 | 87375..90668 | 56957 | Swt1-family-HEPN-domain-containing-protein-[Thermus-oshimai] |
| WP_018110668.1 | DUF499_1.RM.like.systems | <-MULE-TRANSPOSASE\|\|METHYLASE->HigB/RHH->HTH_3->HIN-HTH+SecB->HEPN+DUF499*->?-><-?<-XIS-HTH\|\|Primpol->TOPRIM+primpol-> | 1095 | Bacteria>Terrabacteria_group>Deinococcota>Deinococci>Thermales | GCF_000376265.1 | Thermus_igniterrae | B128_RS0102275 | 11501..14788 | 88189 | Swt1-family-HEPN-domain-containing-protein-[Thermus-igniterrae] |
| WP_083922075.1 | DUF499_1.RM.like.systems | Hpr-Kinase-C->?->?->?->HEPN+DUF499*->TUDOR+SF1-SF2-Helicase+iREase+CTD+REase->METHYLASE-><-?\|\|?->TraJ-RHH->PIN-> | 1058 | Bacteria>Terrabacteria_group>Deinococcota>Deinococci>Thermales | GCF_000376265.1 | Thermus_igniterrae | B128_RS0103820 | 7701..10877 | 88189 | DUF499-domain-containing-protein-[Thermus-igniterrae] |
| WP_036270380.1 | DUF499_1.RM.like.systems | <-METHYLASE<-MACRODOMAIN<-?<-TUDOR+SF1-SF2-Helicase+iREase+CTD+REase<-HEPN+DUF499*<-?<-BirA-HTH | 1062 | Bacteria>Terrabacteria_group>Deinococcota>Deinococci>Thermales | GCF_000423425.1 | Meiothermus_rufus | G552_RS0106115 | 16057..19245 | 604332 | DUF499-domain-containing-protein-[Meiothermus-rufus] |
| WP_028494585.1 | DUF499_1.RM.like.systems | <-HEPN+DUF499*<-HIN-HTH+SecB<-HTH_3<-HigB/RHH | 1095 | Bacteria>Terrabacteria_group>Deinococcota>Deinococci>Thermales | GCF_000423905.1 | Thermus | CSW24_RS01285 | 852..4139 | 270 | MULTISPECIES:-Swt1-family-HEPN-domain-containing-protein-[Thermus] |
| WP_245587643.1 | DUF499_1.RM.like.systems | <-REase<-?\|\|HNH-><-?\|\|HEPN+DUF499*->TUDOR+SF1-SF2-Helicase+iREase+CTD+REase->?->MACRODOMAIN->METHYLASE-> | 1065 | Bacteria>Terrabacteria_group>Deinococcota>Deinococci>Thermales | GCF_000430045.1 | Calidithermus_chliarophilus | MEICH_RS0111510 | 2368714..2371911 | 52023 | DUF499-domain-containing-protein-[Calidithermus-chliarophilus] |
| WP_252829645.1 | DUF499_1.RM.like.systems | <-DDE_Tnp_1<-?<-?\|\|?->?->\|\|<-HTH+DSBH\|\|<-DUF499*<-HEPN+DUF499<-HIN-HTH+AT-HTH-HYD2-2+SecB<-HTH_3<-HigB/RHH<-HicB_lk_antitox<-METHYLASE-p | 779 | Bacteria>Terrabacteria_group>Deinococcota>Deinococci>Thermales | GCF_000744155.1 | Thermus_scotoductus | N686_RS12530 | 46025..48364 | 37636 | DUF499-domain-containing-protein-[Thermus-scotoductus] |
| WP_252829575.1 | DUF499_1.RM.like.systems | <-HTH+DSBH\|\|<-DUF499<-HEPN+DUF499*<-HIN-HTH+AT-HTH-HYD2-2+SecB<-HTH_3<-HigB/RHH<-HicB_lk_antitox<-METHYLASE-p<-METHYLASE | 335 | Bacteria>Terrabacteria_group>Deinococcota>Deinococci>Thermales | GCF_000744155.1 | Thermus_scotoductus | N686_RS14355 | 48271..49278 | 37636 | Swt1-family-HEPN-domain-containing-protein-[Thermus-scotoductus] |
| WP_236622573.1 | DUF499_1.RM.like.systems | HicB_lk_antitox->HigB/RHH->HTH_3->HIN-HTH+AT-HTH-HYD2-2+SecB->?->HEPN+DUF499->DUF499*->DUF499->\|\|DDE_Tnp_1_3->RADICAL-SAM-> | 235 | Bacteria>Terrabacteria_group>Deinococcota>Deinococci>Thermales | GCF_000744175.1 | Thermus_tengchongensis | BS82_RS15080 | 76525..77232 | 1214928 | DUF499-domain-containing-protein-[Thermus-tengchongensis] |
| WP_269077413.1 | DUF499_1.RM.like.systems | METHYLASE-p->HicB_lk_antitox->HigB/RHH->HTH_3->HIN-HTH+AT-HTH-HYD2-2+SecB->?->HEPN+DUF499*->DUF499->DUF499->\|\|DDE_Tnp_1_3->RADICAL-SAM-> | 335 | Bacteria>Terrabacteria_group>Deinococcota>Deinococci>Thermales | GCF_000744175.1 | Thermus_tengchongensis | BS82_RS15525 | 75605..76612 | 1214928 | Swt1-family-HEPN-domain-containing-protein-[Thermus-tengchongensis] |
| WP_269077414.1 | DUF499_1.RM.like.systems | HigB/RHH->HTH_3->HIN-HTH+AT-HTH-HYD2-2+SecB->?->HEPN+DUF499->DUF499->DUF499*->\|\|DDE_Tnp_1_3->RADICAL-SAM->?->?->?-><-DUF4277+DDE_Tnp_1 | 488 | Bacteria>Terrabacteria_group>Deinococcota>Deinococci>Thermales | GCF_000744175.1 | Thermus_tengchongensis | BS82_RS15530 | 77388..78854 | 1214928 | hypothetical-protein-[Thermus-tengchongensis] |
| WP_038055761.1 | DUF499_1.RM.like.systems | <-Radical-SAM<-tRNA<-rRNA<-rRNA\|\|DDE_Tnp_1_5->tRNA-><-HEPN+DUF499*<-HIN-HTH+SecB<-HTH_3<-HigB/RHH<-METHYLASE<-TUDOR+SF1-SF2-Helicase+iREase+CTD+REase-p<-TUDOR+SF1-SF2-Helicase+iREase+CTD+REase | 1086 | Bacteria>Terrabacteria_group>Deinococcota>Deinococci>Thermales | GCF_000744885.1 | Thermus_amyloliquefaciens | BS74_RS12920 | 400983..404243 | 1449080 | Swt1-family-HEPN-domain-containing-protein-[Thermus-amyloliquefaciens] |
| WP_223299558.1 | DUF499_1.RM.like.systems | <-HEPN+DUF499*<-METHYLASE<-TUDOR+SF1-SF2-Helicase+iREase+CTD+REase-p | 451 | Bacteria>Terrabacteria_group>Deinococcota>Deinococci>Thermales | GCF_001623175.1 | Meiothermus_taiwanensis | A3962_RS01225 | 1..1353 | 172827 | Swt1-family-HEPN-domain-containing-protein,-partial-[Meiothermus-taiwanensis] |
| WP_278359523.1 | DUF499_1.RM.like.systems | <-HEPN+DUF499*<-HIN-HTH+AT-HTH-HYD2-2+SecB<-HTH_3<-HicB_lk_antitox<-HicA_toxin<-METHYLASE<-DOC+HTH | 601 | Bacteria>Terrabacteria_group>Deinococcota>Deinococci>Thermales | GCF_002378965.1 | Thermus_thermophilus | B3N72_RS06335 | 2095..3898 | 274 | Swt1-family-HEPN-domain-containing-protein,-partial-[Thermus-thermophilus] |
| WP_114799026.1 | DUF499_1.RM.like.systems | ABC-ATPase+SbcC-><-?\|\|?->?->?-><-?\|\|tRNA->tRNA->HEPN+DUF499*->TUDOR+SF1-SF2-Helicase+iREase+CTD+REase->METHYLASE-><-TPR | 1057 | Bacteria>Terrabacteria_group>Deinococcota>Deinococci>Thermales | GCF_003351145.1 | Meiothermus_sp._QL-1 | DV704_RS07840 | 44133..47306 | 2058095 | DUF499-domain-containing-protein-[Meiothermus-sp.-QL-1] |
| WP_027888723.1 | DUF499_1.RM.like.systems | <-PAIREDC-HTH<-BrxL<-?<-?<-?<-ORC-CDC6+REase<-HEPN+DUF499*<-METHYLASE<-TUDOR+SF1-SF2-Helicase+iREase+CTD+REase\|\|DDE_Tnp_IS1-><-?\|\|?->TPR+TPR-> | 1105 | Bacteria>Terrabacteria_group>Deinococcota>Deinococci>Thermales | GCF_003568585.1 | Meiothermus_taiwanensis | MTAI_RS14270 | 92295..95612 | 172827 | Swt1-family-HEPN-domain-containing-protein-[Meiothermus-taiwanensis] |
| WP_119361752.1 | DUF499_1.RM.like.systems | TUDOR+SF1-SF2-Helicase+iREase+CTD+REase->METHYLASE->HEPN+DUF499*->ORC-CDC6+REase-> | 1108 | Bacteria>Terrabacteria_group>Deinococcota>Deinococci>Thermales | GCF_003574045.1 | Meiothermus_taiwanensis | Mcate_RS11630 | 6951..10277 | 172827 | Swt1-family-HEPN-domain-containing-protein-[Meiothermus-taiwanensis] |
| WP_119360281.1 | DUF499_1.RM.like.systems | ACET->?-><-?<-ABhydrolase<-HEPN+DUF499*<-?<-HTH_3<-METHYLASE<-?<-TUDOR+SF1-SF2-Helicase+iREase+CTD+REase | 1091 | Bacteria>Terrabacteria_group>Deinococcota>Deinococci>Thermales | GCF_003574085.1 | Meiothermus_luteus | Mlute_RS08485 | 5782..9057 | 2026184 | Swt1-family-HEPN-domain-containing-protein-[Meiothermus-luteus] |
| WP_119313343.1 | DUF499_1.RM.like.systems | TUDOR+SF1-SF2-Helicase+iREase+CTD+REase->METHYLASE->HEPN+DUF499*->ZnR+TOPRIM+primpol+HTH-> | 1117 | Bacteria>Terrabacteria_group>Deinococcota>Deinococci>Thermales | GCF_003574345.1 | Calidithermus_terrae | Mterra_RS00325 | 80239..83592 | 1408545 | Swt1-family-HEPN-domain-containing-protein-[Calidithermus-terrae] |
| WP_126164480.1 | DUF499_1.RM.like.systems | METHYLASE->HigB/RHH->HTH_3->HIN-HTH+SecB->HEPN+DUF499*-> | 412 | Bacteria>Terrabacteria_group>Deinococcota>Deinococci>Thermales | GCF_003958975.1 | Thermus_scotoductus | CSW48_RS02070 | 4415..5652 | 37636 | Swt1-family-HEPN-domain-containing-protein,-partial-[Thermus-scotoductus] |
| WP_126193807.1 | DUF499_1.RM.like.systems | METHYLASE->HigB/RHH->HTH_3->HIN-HTH+SecB->HEPN+DUF499*-> | 416 | Bacteria>Terrabacteria_group>Deinococcota>Deinococci>Thermales | GCF_003959075.1 | Thermus_scotoductus | CSW46_RS01260 | 4967..6215 | 37636 | Swt1-family-HEPN-domain-containing-protein,-partial-[Thermus-scotoductus] |
| WP_126199710.1 | DUF499_1.RM.like.systems | HigB/RHH->HTH_3->HIN-HTH+SecB->HEPN+DUF499*-> | 425 | Bacteria>Terrabacteria_group>Deinococcota>Deinococci>Thermales | GCF_003959105.1 | Thermus_scotoductus | CSW44_RS03290 | 2683..3958 | 37636 | Swt1-family-HEPN-domain-containing-protein,-partial-[Thermus-scotoductus] |
| WP_253724412.1 | DUF499_1.RM.like.systems | <-HEPN+DUF499*<-HIN-HTH+SecB<-HTH_3<-HigB/RHH<-METHYLASE | 424 | Bacteria>Terrabacteria_group>Deinococcota>Deinococci>Thermales | GCF_003959115.1 | Thermus_scotoductus | CSW43_RS02865 | 82..1356 | 37636 | Swt1-family-HEPN-domain-containing-protein-[Thermus-scotoductus] |
| WP_126209517.1 | DUF499_1.RM.like.systems | METHYLASE->HicA_toxin->HicB_lk_antitox->HIN-HTH+AT-HTH-HYD2-2+SecB->HEPN+DUF499*->DDE_Tnp_1_3-><-MULE-TRANSPOSASE | 1085 | Bacteria>Terrabacteria_group>Deinococcota>Deinococci>Thermales | GCF_003959225.1 | Thermus_scotoductus | CSW37_RS13630 | 3217..6474 | 37636 | DUF499-domain-containing-protein-[Thermus-scotoductus] |
| WP_126212451.1 | DUF499_1.RM.like.systems | HigB/RHH->HTH_3->HIN-HTH+SecB->HEPN+DUF499*-> | 419 | Bacteria>Terrabacteria_group>Deinococcota>Deinococci>Thermales | GCF_003959265.1 | Thermus_scotoductus | CSW34_RS03430 | 2738..3995 | 37636 | Swt1-family-HEPN-domain-containing-protein,-partial-[Thermus-scotoductus] |
| WP_253724062.1 | DUF499_1.RM.like.systems | <-HEPN+DUF499*<-HIN-HTH+SecB<-HTH_3<-HigB/RHH | 446 | Bacteria>Terrabacteria_group>Deinococcota>Deinococci>Thermales | GCF_003959305.1 | Thermus_scotoductus | CSW32_RS03250 | 10..1350 | 37636 | Swt1-family-HEPN-domain-containing-protein-[Thermus-scotoductus] |
| WP_126210662.1 | DUF499_1.RM.like.systems | <-HEPN+DUF499*<-HIN-HTH+SecB<-HTH_3<-HigB/RHH | 424 | Bacteria>Terrabacteria_group>Deinococcota>Deinococci>Thermales | GCF_003959395.1 | Thermus_scotoductus | CSW26_RS03555 | 1..1273 | 37636 | Swt1-family-HEPN-domain-containing-protein,-partial-[Thermus-scotoductus] |
| WP_253730420.1 | DUF499_1.RM.like.systems | METHYLASE-p->HigB/RHH->HTH_3->HIN-HTH+SecB->HEPN+DUF499*-> | 1004 | Bacteria>Terrabacteria_group>Deinococcota>Deinococci>Thermales | GCF_003959405.1 | Thermus_scotoductus | CSW25_RS02145 | 3152..6166 | 37636 | Swt1-family-HEPN-domain-containing-protein-[Thermus-scotoductus] |
| WP_153185240.1 | DUF499_1.RM.like.systems | <-HEPN+DUF499*<-HIN-HTH+SecB<-HTH_3<-HigB/RHH<-METHYLASE | 1011 | Bacteria>Terrabacteria_group>Deinococcota>Deinococci>Thermales | GCF_003959445.1 | Thermus_scotoductus | CSW20_RS00300 | 1..3032 | 37636 | Swt1-family-HEPN-domain-containing-protein,-partial-[Thermus-scotoductus] |
| WP_126212240.1 | DUF499_1.RM.like.systems | METHYLASE->HigB/RHH->HTH_3->HIN-HTH+SecB->HEPN+DUF499*-> | 418 | Bacteria>Terrabacteria_group>Deinococcota>Deinococci>Thermales | GCF_003959525.1 | Thermus_scotoductus | CSW17_RS01375 | 4891..6146 | 37636 | Swt1-family-HEPN-domain-containing-protein,-partial-[Thermus-scotoductus] |
| WP_253665106.1 | DUF499_1.RM.like.systems | HigB/RHH->HTH_3->HIN-HTH+SecB->HEPN+DUF499*-> | 1022 | Bacteria>Terrabacteria_group>Deinococcota>Deinococci>Thermales | GCF_003959545.1 | Thermus_scotoductus | CSW16_RS01280 | 2718..5783 | 37636 | Swt1-family-HEPN-domain-containing-protein,-partial-[Thermus-scotoductus] |
| WP_253665784.1 | DUF499_1.RM.like.systems | <-HEPN+DUF499*<-HIN-HTH+SecB<-HTH_3<-HigB/RHH<-METHYLASE | 429 | Bacteria>Terrabacteria_group>Deinococcota>Deinococci>Thermales | GCF_003959565.1 | Thermus_scotoductus | CSW39_RS03125 | 1..1288 | 37636 | Swt1-family-HEPN-domain-containing-protein,-partial-[Thermus-scotoductus] |
| WP_126216336.1 | DUF499_1.RM.like.systems | <-HEPN+DUF499*<-HIN-HTH+SecB<-HTH_3<-HigB/RHH | 413 | Bacteria>Terrabacteria_group>Deinococcota>Deinococci>Thermales | GCF_003959575.1 | Thermus_scotoductus | CSW36_RS03285 | 1..1238 | 37636 | Swt1-family-HEPN-domain-containing-protein,-partial-[Thermus-scotoductus] |
| WP_126252475.1 | DUF499_1.RM.like.systems | HigB/RHH->HTH_3->HIN-HTH+SecB->HEPN+DUF499*-> | 415 | Bacteria>Terrabacteria_group>Deinococcota>Deinococci>Thermales | GCF_003959675.1 | Thermus_scotoductus | CSW15_RS03250 | 2694..3937 | 37636 | Swt1-family-HEPN-domain-containing-protein,-partial-[Thermus-scotoductus] |
| WP_130842394.1 | DUF499_1.RM.like.systems | TUDOR+SF1-SF2-Helicase+iREase+CTD+REase->DOC+HTH->METHYLASE->HigB/RHH->HTH_3->HIN-HTH+SecB->HEPN+DUF499*-> | 1088 | Bacteria>Terrabacteria_group>Deinococcota>Deinococci>Thermales | GCF_004307015.1 | Thermus_thermamylovorans | ETP66_RS09490 | 48126..51392 | 2509362 | Swt1-family-HEPN-domain-containing-protein-[Thermus-thermamylovorans] |
| WP_135343460.1 | DUF499_1.RM.like.systems | <-HEPN+DUF499*<-METHYLASE<-PIN<-TUDOR+SF1-SF2-Helicase+iREase+CTD+REase<-ParB | 1099 | Bacteria>Terrabacteria_group>Deinococcota>Deinococci>Thermales | GCF_004685085.1 | Thermus_tengchongensis | E0489_RS07460 | 252..3551 | 1214928 | Swt1-family-HEPN-domain-containing-protein-[Thermus-tengchongensis] |
| WP_119341253.1 | DUF499_1.RM.like.systems | Pkinase->DUF4388-><-?<-?\|\|?-><-RHH<-HEPN+DUF499*<-METHYLASE<-TUDOR+SF1-SF2-Helicase+iREase+CTD+REase\|\|?-><-?<-PfkB | 1099 | Bacteria>Terrabacteria_group>Deinococcota>Deinococci>Thermales | GCF_007990975.1 | Meiothermus_hypogaeus | MHY01S_RS03040 | 15508..18807 | 884155 | Swt1-family-HEPN-domain-containing-protein-[Meiothermus-hypogaeus] |
| WP_240637265.1 | DUF499_1.RM.like.systems | DDE_Tnp_1->HEPN+DUF499*->TUDOR+SF1-SF2-Helicase+iREase+CTD+REase->SF1-SF2-Helicase+wHTH+PRTase->?->Schlafen+HATPase+HTH->METHYLASE-> | 1066 | Bacteria>Terrabacteria_group>Deinococcota>Deinococci>Thermales | GCF_007990975.1 | Meiothermus_hypogaeus | MHY01S_RS11115 | 3652..6852 | 884155 | DUF499-domain-containing-protein-[Meiothermus-hypogaeus] |
| WP_119356451.1 | DUF499_1.RM.like.systems | PIN->TUDOR+SF1-SF2-Helicase+iREase+CTD+REase->XIS-HTH->METHYLASE-><-PIN<-MazE_antitoxin\|\|HEPN+DUF499*-> | 1110 | Bacteria>Terrabacteria_group>Deinococcota>Deinococci>Thermales | GCF_007990995.1 | Meiothermus_granaticius | MGR01S_RS15620 | 8907..12239 | 863370 | Swt1-family-HEPN-domain-containing-protein-[Meiothermus-granaticius] |
| WP_202819508.1 | DUF499_1.RM.like.systems | <-ABC-ATPase+SbcC<-?\|\|?-><-DDE_Tnp_1<-HEPN+DUF499*<-HIN-HTH+AT-HTH-HYD2-2+SecB<-HTH_3<-HicB_lk_antitox<-HicA_toxin<-METHYLASE<-DOC+HTH | 1083 | Bacteria>Terrabacteria_group>Deinococcota>Deinococci>Thermales | GCF_016865275.1 | Thermus_thermophilus | TthHB5002_b22180 | 17958..21209 | 274 | Swt1-family-HEPN-domain-containing-protein-[Thermus-thermophilus] |
| WP_219760305.1 | DUF499_1.RM.like.systems | SLATT->METHYLASE->HEPN+DUF499*->?->?->DDE_3-> | 1116 | Bacteria>Terrabacteria_group>Deinococcota>Deinococci>Thermales | GCF_019430925.1 | Thermus_brevis | KZX47_RS11605 | 3607..6957 | 2862456 | Swt1-family-HEPN-domain-containing-protein-[Thermus-brevis] |
| WP_267461447.1 | DUF499_1.RM.like.systems | HIN-HTH-><-?\|\|PAIREDC-HTH+HIN-HTH->DDE_3-><-HEPN+DUF499*<-HIN-HTH+AT-HTH-HYD2-2+SecB<-HTH_3<-HicA_toxin<-METHYLASE<-DOC+HTH\|\|TUDOR+SF1-SF2-Helicase+iREase+CTD+REase-> | 1084 | Bacteria>Terrabacteria_group>Deinococcota>Deinococci>Thermales | GCF_019974355.1 | Thermus_thermophilus | LDD10_RS13400 | 104954..108208 | 274 | Swt1-family-HEPN-domain-containing-protein-[Thermus-thermophilus] |
| WP_224065603.1 | DUF499_1.RM.like.systems | PP_kinase+HKD-DNase-><-HEPN+DUF499*<-HIN-HTH+AT-HTH-HYD2-2+SecB<-HTH_3<-HicB_lk_antitox<-METHYLASE<-?\|\|TUDOR+SF1-SF2-Helicase+iREase+CTD+REase-p-> | 1086 | Bacteria>Terrabacteria_group>Deinococcota>Deinococci>Thermales | GCF_019974375.1 | Thermus_thermophilus | LDD80_RS12010 | 2426..5686 | 274 | Swt1-family-HEPN-domain-containing-protein-[Thermus-thermophilus] |
| WP_224065568.1 | DUF499_1.RM.like.systems | <-TUDOR+SF1-SF2-Helicase+iREase+CTD+REase\|\|DOC+HTH->METHYLASE->HicA_toxin->HTH_3->HIN-HTH+AT-HTH-HYD2-2+SecB->HEPN+DUF499*->DDE_Tnp_1+DDE_Tnp_1_2->?-><-HIN-HTH\|\|?-><-?\|\|DDE_Tnp_1_3-> | 1084 | Bacteria>Terrabacteria_group>Deinococcota>Deinococci>Thermales | GCF_019974375.1 | Thermus_thermophilus | LDD80_RS12530 | 90011..93265 | 274 | Swt1-family-HEPN-domain-containing-protein-[Thermus-thermophilus] |
| WP_234507999.1 | DUF499_1.RM.like.systems | DOC+HTH->METHYLASE->HigB/RHH->HTH_3->HIN-HTH+SecB->HEPN+DUF499*-><-Hpr-Kinase-C | 1090 | Bacteria>Terrabacteria_group>Deinococcota>Deinococci>Thermales | GCF_021462505.1 | Thermus_brockianus | LXN00_RS10430 | 5041..8313 | 56956 | DUF499-domain-containing-protein-[Thermus-brockianus] |
| WP_243028145.1 | DUF499_1.RM.like.systems | RNB+RNase-><-?\|\|?->?->tRNA-><-HEPN+DUF499*<-HicB_lk_antitox<-METHYLASE<-TUDOR+SF1-SF2-Helicase+iREase+CTD+REase\|\|Radical-SAM+TRAM-> | 1076 | Bacteria>Terrabacteria_group>Deinococcota>Deinococci>Thermales | GCF_022760855.1 | Thermus_albus | LOD18_RS06900 | 53799..57029 | 2908146 | DUF499-domain-containing-protein-[Thermus-albus] |
| WP_243092374.1 | DUF499_1.RM.like.systems | Hpr-Kinase-C-><-HEPN+DUF499*<-HIN-HTH+SecB<-HTH_3<-HigB/RHH<-METHYLASE<-DOC+HTH<-TUDOR+SF1-SF2-Helicase+iREase+CTD+REase | 1092 | Bacteria>Terrabacteria_group>Deinococcota>Deinococci>Thermales | GCF_022760925.1 | Thermus_hydrothermalis | LOC60_RS00500 | 88186..91464 | 2908148 | DUF499-domain-containing-protein-[Thermus-hydrothermalis] |
| WP_275973822.1 | DUF499_1.RM.like.systems | <-HTH_3<-?\|\|DDE_Tnp_1_3->DDE_Tnp_1->DDE_Tnp_1_3->DDE_Tnp_1-><-HEPN+DUF499*<-HIN-HTH+AT-HTH-HYD2-2+SecB<-HTH_3<-HigB/RHH<-HicB_lk_antitox<-METHYLASE | 1084 | Bacteria>Terrabacteria_group>Deinococcota>Deinococci>Thermales | GCF_022846275.1 | Thermus_thermophilus | MWM06_RS12915 | 20817..24071 | 274 | Swt1-family-HEPN-domain-containing-protein-[Thermus-thermophilus] |
| WP_244348266.1 | DUF499_1.RM.like.systems | DOC+HTH->METHYLASE->HicB_lk_antitox->HigB/RHH->HTH_3->HIN-HTH+AT-HTH-HYD2-2+SecB->HEPN+DUF499*->HNH->Peptidase_M48-> | 1084 | Bacteria>Terrabacteria_group>Deinococcota>Deinococci>Thermales | GCF_022846295.1 | Thermus_thermophilus | MWM13_RS12110 | 24920..28174 | 274 | Swt1-family-HEPN-domain-containing-protein-[Thermus-thermophilus] |
| WP_275974790.1 | DUF499_1.RM.like.systems | DUF4096+DDE_Tnp_1+DDE_Tnp->PAIREDC-HTH+HIN-HTH->DDE_3-><-HicA_toxin<-HicB<-DDE_Tnp_1<-HEPN+DUF499*<-HIN-HTH+AT-HTH-HYD2-2+SecB<-HTH_3<-HigB/RHH<-HicB_lk_antitox<-METHYLASE | 1084 | Bacteria>Terrabacteria_group>Deinococcota>Deinococci>Thermales | GCF_022846315.1 | Thermus_thermophilus | SNM_RS12085 | 50871..54125 | 274 | Swt1-family-HEPN-domain-containing-protein-[Thermus-thermophilus] |
| WP_275974710.1 | DUF499_1.RM.like.systems | <-TUDOR+SF1-SF2-Helicase+iREase+CTD+REase\|\|?->METHYLASE->HicA_toxin->HTH_3->HIN-HTH+AT-HTH-HYD2-2+SecB->HEPN+DUF499*-><-DDE_Tnp_1_3\|\|HTH_3->HIN-HTH+AT-HTH-HYD2-2+SecB->?->ABC-ATPase+SbcC-> | 1084 | Bacteria>Terrabacteria_group>Deinococcota>Deinococci>Thermales | GCF_022846355.1 | Thermus_thermophilus | MWM12_RS12790 | 105744..108998 | 274 | Swt1-family-HEPN-domain-containing-protein-[Thermus-thermophilus] |
| WP_071678256.1 | DUF499_1.RM.like.systems | TUDOR+SF1-SF2-Helicase+iREase+CTD+REase->DOC+HTH->METHYLASE->HigB/RHH->HTH_3->HIN-HTH+SecB->HEPN+DUF499*-><-Hpr-Kinase-C | 1092 | Bacteria>Terrabacteria_group>Deinococcota>Deinococci>Thermales | GCF_022846375.1 | Thermus_brockianus | MWM05_RS11465 | 93356..96634 | 56956 | DUF499-domain-containing-protein-[Thermus-brockianus] |
| WP_267262111.1 | DUF499_1.RM.like.systems | <-HEPN<-DDE_3<-?<-?<-HEPN+DUF499*<-METHYLASE<-OLD-ABC+TOPRIM<-TUDOR+SF1-SF2-Helicase+iREase+CTD+REase<-?<-ArsR-HTH | 1116 | Bacteria>Terrabacteria_group>Deinococcota>Deinococci>Thermales | GCF_026427635.1 | Thermus_sp._PS18 | KQ693_RS05000 | 868393..871743 | 2849039 | Swt1-family-HEPN-domain-containing-protein-[Thermus-sp.-PS18] |
| WP_014677588.1 | DUF499_1.RM.like.systems | <-HEPN+DUF499*<-HIN-HTH+AT-HTH-HYD2-2+SecB<-HTH_3<-HicB_lk_antitox<-HicA_toxin<-METHYLASE<-DOC+HTH | 1084 | Bacteria>Terrabacteria_group>Deinococcota>Deinococci>Thermales | - | Thermus | DCM56_RS11460 | 979..4233 | 270 | Swt1-family-HEPN-domain-containing-protein-[Thermus-thermophilus] |
| WP_126211477.1 | DUF499_1.RM.like.systems | <-HEPN+DUF499*<-HIN-HTH+SecB<-HTH_3<-HigB/RHH | 421 | Bacteria>Terrabacteria_group>Deinococcota>Deinococci>Thermales | - | Thermus_scotoductus | CSW19_RS03320 | 1..1262 | 37636 | Swt1-family-HEPN-domain-containing-protein,-partial-[Thermus-scotoductus] |
| MCY4354897.1 | DUF499_1.RM.like.systems | HicA_toxin->HicB->TUDOR+SF1-SF2-Helicase+iREase+CTD+REase->METHYLASE->PHP+SbcC->Schlafen+HATPase+HTH->HEPN+DUF499*->?->PIN->?->?->PrmC_N+Methyltransf_25-><-Peptidase_A8 | 1107 | Bacteria>Terrabacteria_group>Deinococcota>Deinococci>Trueperales | GCA_026710445.1 | Truepera_sp. | OXC09_08995 | 25870..29193 | 2478917 | MAG:-Swt1-family-HEPN-domain-containing-protein-[Truepera-sp.] |
| WP_013177687.1 | DUF499_1.RM.like.systems | PUA+METHYLASE->?->?->?-><-sGTP<-Aminotran_3<-HEPN+DUF499*<-METHYLASE-p\|\|Y1_Tnp-><-METHYLASE<-PIN | 1108 | Bacteria>Terrabacteria_group>Deinococcota>Deinococci>Trueperales | GCF_000092425.1 | Truepera_radiovictrix | TRAD_RS05920 | 1278601..1281927 | 332249 | Swt1-family-HEPN-domain-containing-protein-[Truepera-radiovictrix] |
| WP_013179180.1 | DUF499_1.RM.like.systems | TUDOR+SF1-SF2-Helicase+iREase+CTD+REase->Phd-YefM->PIN->METHYLASE->?->PIN->HEPN+DUF499*-><-Peptidase_M20 | 1069 | Bacteria>Terrabacteria_group>Deinococcota>Deinococci>Trueperales | GCF_000092425.1 | Truepera_radiovictrix | TRAD_RS13560 | 2957587..2960796 | 332249 | DUF499-domain-containing-protein-[Truepera-radiovictrix] |
| RMH56280.1 | DUF499_1.RM.like.systems | <-HAD<-?<-?\|\|TUDOR+SF1-SF2-Helicase+iREase+CTD+REase->METHYLASE->HTH_3->HEPN+DUF499*->ABhydrolase->?-><-?<-ACET | 1091 | Bacteria>Terrabacteria_group>Deinococcota>unclassified_Deinococcota | GCA_003695925.1 | Deinococcota_bacterium | D6684_05775 | 12315..15590 | 1913990 | ATP-binding-protein-[Deinococcus-Thermus-bacterium] |
| MCA9833507.1 | DUF499_1.RM.like.systems | Peptidase-><-?<-?\|\|TUDOR+SF1-SF2-Helicase+iREase+CTD+REase->HEPN->METHYLASE->HEPN+DUF499*->DDE_3->?-><-tRNA<-?\|\|?-><-?<-NACHT | 1122 | Bacteria>Terrabacteria_group>Thermomicrobiota>Thermomicrobia>Thermomicrobiales | GCA_020432165.1 | Thermomicrobiales_bacterium | KC435_06155 | 74622..77990 | 2482790 | DUF499-domain-containing-protein-[Thermomicrobiales-bacterium] |
| MCC6945442.1 | DUF499_1.RM.like.systems | METHYLASE-p->HEPN+DUF499*-> | 1060 | Bacteria>Terrabacteria_group>Thermomicrobiota>Thermomicrobia>Thermomicrobiales | GCA_020853875.1 | Thermomicrobiales_bacterium | IT335_12740 | 291..3470 | 2482790 | DUF499-domain-containing-protein-[Thermomicrobiales-bacterium] |
| MCC6945870.1 | DUF499_1.RM.like.systems | METHYLASE-p->HEPN+DUF499*-> | 1058 | Bacteria>Terrabacteria_group>Thermomicrobiota>Thermomicrobia>Thermomicrobiales | GCA_020853875.1 | Thermomicrobiales_bacterium | IT335_14920 | 865..4040 | 2482790 | DUF499-domain-containing-protein-[Thermomicrobiales-bacterium] |
| MBI5524652.1 | DUF499_1.RM.like.systems | <-Peptidase\|\|tRNA->TUDOR+SF1-SF2-Helicase+iREase+CTD+REase->DOC+HTH->METHYLASE->HEPN+DUF499*-> | 1110 | Bacteria>Thermodesulfobacteriota>Desulfarculia>Desulfarculales>Desulfarculaceae | GCA_016218705.1 | Desulfarculus_sp. | HY910_18660 | 18095..21427 | 2093367 | ATP-binding-protein-[Desulfarculus-sp.] |
| VBB42850.1 | DUF499_1.RM.like.systems | <-Phage_integrase<-?\|\|TUDOR+SF1-SF2-Helicase+iREase+CTD+REase->ParB+HNH->METHYLASE->HEPN+DUF499*->?->?->?-><-DDE_Tnp_1\|\|HNH-> | 1117 | Bacteria>Thermodesulfobacteriota>Desulfobacteria>Desulfatiglandales>Desulfatiglandaceae | GCA_900498135.1 | uncultured_Desulfatiglans_sp. | TRIP_B220098 | 96105..99458 | 1748965 | conserved-hypothetical-protein-[uncultured-Desulfatiglans-sp.] |
| WP_028322621.1 | DUF499_1.RM.like.systems | METHYLASE->HEPN+DUF499*->?->?->Primpol+D5N+D5ATPase-> | 1114 | Bacteria>Thermodesulfobacteriota>Desulfobacteria>Desulfatiglandales>Desulfatiglandaceae | GCF_000422285.1 | Desulfatiglans_anilini | H567_RS0119135 | 2085..5429 | 90728 | Swt1-family-HEPN-domain-containing-protein-[Desulfatiglans-anilini] |
| OPX33848.1 | DUF499_1.RM.like.systems | <-HicB_lk_antitox<-?<-HEPN+DUF499*<-METHYLASE-p | 957 | Bacteria>Thermodesulfobacteriota>Desulfobacteria>Desulfobacterales>Desulfobacteraceae | GCA_002049905.1 | Desulfobacteraceae_bacterium_4484_190.1 | B1H11_11870 | 1782..4655 | 1956167 | hypothetical-protein-B1H11_11870-[Desulfobacteraceae-bacterium-4484_190.1] |
| HHO48401.1 | DUF499_1.RM.like.systems | <-HEPN+DUF499*<-METHYLASE<-TUDOR+SF1-SF2-Helicase+iREase+CTD+REase | 1112 | Bacteria>Thermodesulfobacteriota>Desulfobacteria>Desulfobacterales>Desulfobacteraceae | GCA_011372855.1 | Desulfobacteraceae_bacterium | ENN06_08115 | 325..3663 | 2049433 | ATP-binding-protein-[Desulfobacteraceae-bacterium] |
| MBA3017256.1 | DUF499_1.RM.like.systems | <-N6_N4_MTase<-Peptidase_C47<-?\|\|TUDOR+SF1-SF2-Helicase+iREase+CTD+REase->MORC+HisKin+DUF3684->HEPN+DUF499*->RNase->ParB+HNH->HEPN-> | 1092 | Bacteria>Thermodesulfobacteriota>Desulfobacteria>Desulfobacterales>Desulfobacteraceae | GCA_013792055.1 | Desulfobacteraceae_bacterium | FP811_03735 | 27065..30343 | 2049433 | ATP-binding-protein-[Desulfobacteraceae-bacterium] |
| MBL7176677.1 | DUF499_1.RM.like.systems | TUDOR+SF1-SF2-Helicase+iREase+CTD+REase-p->OLD-ABC->PIN->?->HEPN+DUF499*->?->OLD-ABC->SF1-SF2-Helicase->?->MNS+HTH-> | 1094 | Bacteria>Thermodesulfobacteriota>Desulfobacteria>Desulfobacterales>Desulfobacteraceae | GCA_016782925.1 | Desulfobacteraceae_bacterium | ISS66_12705 | 13240..16524 | 2049433 | DUF499-domain-containing-protein-[Desulfobacteraceae-bacterium] |
| WP_004073680.1 | DUF499_1.RM.like.systems | SPOUT->PRTase->TUDOR+SF1-SF2-Helicase+iREase+CTD+REase->HEPN->METHYLASE->HEPN+DUF499*-> | 1113 | Bacteria>Thermodesulfobacteriota>Desulfobacteria>Desulfobacterales>Desulfobacteraceae | GCF_000233695.2 | Desulfobacter_postgatei | DESPODRAFT_RS11525 | 2528550..2531891 | 2293 | Swt1-family-HEPN-domain-containing-protein-[Desulfobacter-postgatei] |
| CBX30683.1 | DUF499_1.RM.like.systems | <-SbcC\|\|TUDOR+SF1-SF2-Helicase+iREase+CTD+REase->MqsR_toxin->MqsA_antitoxin+HTH_3->?->METHYLASE->HEPN+DUF499*-> | 1111 | Bacteria>Thermodesulfobacteriota>Desulfobacteria>Desulfobacterales>Desulfobacteriaceae | - | uncultured_Desulfobacterium_sp. | N47_E41950 | 104821..108156 | 201089 | hypothetical-protein-N47_E41950-[uncultured-Desulfobacterium-sp.] |
| MBC2710211.1 | DUF499_1.RM.like.systems | REase-><-Phage_integrase<-?\|\|?->?->?-><-HEPN+DUF499*<-METHYLASE<-TOPRIM<-OLD-ABC<-MORC+DUF3684<-TUDOR+SF1-SF2-Helicase+iREase+CTD+REase\|\|DDE_Tnp_1-> | 1102 | Bacteria>Thermodesulfobacteriota>Desulfobacteria>Desulfobacterales>Desulfosarcinaceae | GCA_014237405.1 | Desulfosarcina_sp. | HGJ94_04170 | 9166..12474 | 2027861 | ATP-binding-protein-[Desulfosarcina-sp.] |
| MBC2711122.1 | DUF499_1.RM.like.systems | TUDOR+SF1-SF2-Helicase+iREase+CTD+REase->?->OLD-ABC->HNH->METHYLASE->HEPN+DUF499*->?->GNTR-HTH->?->?->ABC-ATPase+SbcC->ABC-ATPase+SbcC-> | 1122 | Bacteria>Thermodesulfobacteriota>Desulfobacteria>Desulfobacterales>Desulfosarcinaceae | GCA_014237405.1 | Desulfosarcina_sp. | HGJ94_09035 | 43660..47028 | 2027861 | ATP-binding-protein-[Desulfosarcina-sp.] |
| PIU50309.1 | DUF499_1.RM.like.systems | METHYLASE->?->HEPN+DUF499*->?-><-?\|\|SF1-SF2-Helicase-> | 1109 | Bacteria>Thermodesulfobacteriota>Desulfobacteria>Desulfobacterales>unclassified_Desulfobacterales | GCA_002779465.1 | Desulfobacterales_bacterium_CG07_land_8_20_14_0_80_52_14 | COS92_01840 | 3954..7283 | 1973982 | AAA+-family-ATPase-[Desulfobacterales-bacterium-CG07_land_8_20_14_0_80_52_14] |
| MCU0558984.1 | DUF499_1.RM.like.systems | <-HEPN+DUF499*<-TOPRIM<-OLD-ABC<-METHYLASE-p | 1112 | Bacteria>Thermodesulfobacteriota>Desulfobacteria>Desulfobacterales>unclassified_Desulfobacterales | GCA_025454255.1 | Desulfobacterales_bacterium | MUD16_02165 | 66226..69564 | 2044940 | Swt1-family-HEPN-domain-containing-protein-[Desulfobacterales-bacterium] |
| WP_028319484.1 | DUF499_1.RM.like.systems | DDE_Tnp_1_2-><-HEPN+DUF499*<-METHYLASE<-TOPRIM<-OLD-ABC<-MORC+DUF3684<-TUDOR+SF1-SF2-Helicase+iREase+CTD+REase<-XIS-HTH | 1112 | Bacteria>Thermodesulfobacteriota>Desulfobulbia>Desulfobulbales>Desulfobulbaceae | GCF_000621145.1 | Desulfobulbus_elongatus | Q362_RS0116010 | 475..3813 | 53332 | Swt1-family-HEPN-domain-containing-protein-[Desulfobulbus-elongatus] |
| MBD5607241.1 | DUF499_1.RM.like.systems | TUDOR+SF1-SF2-Helicase+iREase+CTD+REase->METHYLASE->METHYLASE->HEPN+DUF499*->?->?->SF1-SF2-Helicase+REase-p-> | 1123 | Bacteria>Thermodesulfobacteriota>Desulfovibrionia>Desulfovibrionales>Desulfovibrionaceae | GCA_014804865.1 | Desulfovibrio_sp. | HDQ93_00095 | 9665..13036 | 885 | ATP-binding-protein-[Desulfovibrio-sp.] |
| MBR3880305.1 | DUF499_1.RM.like.systems | METHYLASE->HEPN+DUF499*-> | 1086 | Bacteria>Thermodesulfobacteriota>Desulfovibrionia>Desulfovibrionales>Desulfovibrionaceae | GCA_017522745.1 | Mailhella_sp. | IKJ34_01740 | 2844..6102 | 1981029 | DUF499-domain-containing-protein-[Mailhella-sp.] |
| HJB66433.1 | DUF499_1.RM.like.systems | TraJ-RHH->TraI-VirD2-RCRE->?->?->?-><-HEPN<-HEPN+DUF499*<-METHYLASE<-TOPRIM | 1112 | Bacteria>Thermodesulfobacteriota>Desulfovibrionia>Desulfovibrionales>Desulfovibrionaceae | GCA_019119045.1 | Candidatus_Mailhella_merdavium | H9768_09155 | 39301..42639 | 2838657 | DUF499-domain-containing-protein-[Candidatus-Mailhella-merdavium] |
| HJH15746.1 | DUF499_1.RM.like.systems | TOPRIM+SF1-SF2-Helicase+HTH->?->?->GDResolvase+Recombinase+ZnR->TUDOR+SF1-SF2-Helicase+iREase+CTD+REase->METHYLASE->HEPN+DUF499*-> | 865 | Bacteria>Thermodesulfobacteriota>Desulfovibrionia>Desulfovibrionales>Desulfovibrionaceae | GCA_020743955.1 | Bilophila_wadsworthia | K8U76_10820 | 19353..21949 | 35833 | DUF499-domain-containing-protein-[Bilophila-wadsworthia] |
| MDD6180986.1 | DUF499_1.RM.like.systems | CR-ATPase8+REase-><-HEPN+DUF499*<-METHYLASE<-TUDOR+SF1-SF2-Helicase+iREase+CTD+REase<-GDResolvase+Recombinase+ZnR<-Recombinase+ZnR<-GDResolvase+Recombinase+ZnR<-RAMA | 1147 | Bacteria>Thermodesulfobacteriota>Desulfovibrionia>Desulfovibrionales>Desulfovibrionaceae | GCA_029008115.1 | Desulfovibrionaceae_bacterium | PUB01_02780 | 4125..7568 | 2049043 | MAG:-Swt1-family-HEPN-domain-containing-protein-[Desulfovibrionaceae-bacterium] |
| WP_165067758.1 | DUF499_1.RM.like.systems | GDResolvase+Recombinase+ZnR->TUDOR+SF1-SF2-Helicase+iREase+CTD+REase-><-?\|\|METHYLASE->HEPN+DUF499*-><-ACET<-?<-?<-?<-?\|\|Arm-DNA-bind_3+Phage_integrase-> | 1129 | Bacteria>Thermodesulfobacteriota>Desulfovibrionia>Desulfovibrionales>Desulfovibrionaceae | GCF_011039095.1 | Desulfovibrio_sp._ZJ200 | G7Y52_RS05935 | 42436..45825 | 2709792 | Swt1-family-HEPN-domain-containing-protein-[Desulfovibrio-sp.-ZJ200] |
| WP_168936425.1 | DUF499_1.RM.like.systems | <-HEPN+DUF499*<-METHYLASE<-TUDOR+SF1-SF2-Helicase+iREase+CTD+REase-p<-TUDOR+SF1-SF2-Helicase+iREase+CTD+REase | 1121 | Bacteria>Thermodesulfobacteriota>Desulfovibrionia>Desulfovibrionales>Desulfovibrionaceae | GCF_012843875.1 | Desulfovibrio_piger | HF854_RS11535 | 110..3475 | 901 | Swt1-family-HEPN-domain-containing-protein-[Desulfovibrio-piger] |
| WP_278732740.1 | DUF499_1.RM.like.systems | TOPRIM+SF1-SF2-Helicase+HTH->?->?->GDResolvase+Recombinase+ZnR->TUDOR+SF1-SF2-Helicase+iREase+CTD+REase->METHYLASE->HEPN+DUF499*-> | 865 | Bacteria>Thermodesulfobacteriota>Desulfovibrionia>Desulfovibrionales>Desulfovibrionaceae | GCF_020743955.1 | Bilophila_wadsworthia | K8U76_RS10810 | 19353..21949 | 35833 | Swt1-family-HEPN-domain-containing-protein,-partial-[Bilophila-wadsworthia] |
| WP_279060950.1 | DUF499_1.RM.like.systems | GDResolvase+Recombinase+ZnR->TUDOR+SF1-SF2-Helicase+iREase+CTD+REase->METHYLASE->METHYLASE-p->?->HEPN+DUF499*-> | 1135 | Bacteria>Thermodesulfobacteriota>Desulfovibrionia>Desulfovibrionales>Desulfovibrionaceae | GCF_025543515.1 | Bilophila_wadsworthia | QDD56_RS08660 | 91751..95158 | 35833 | DUF499-domain-containing-protein-[Bilophila-wadsworthia] |

| Protein-ID | System | Gene-Neighbourhoods | Length | Taxonomy | Assembly-ID | Strain | Locus-Tag | Genomic-coordinates | TaxID | Fasta-defline |
|---|---|---|---|---|---|---|---|---|---|---|
| WP_273524160.1 | DUF499_1.RM.like.systems | DOC+HTH->REase+SAVED->?->?-><-?<-HEPN+DUF499*<-METHYLASE<-TUDOR+SF1-SF2-Helicase+iREase+CTD+REase<-GDResolvase+Recombinase+ZnR<-?<-?<-Phage_AlpA | 1120 | Bacteria>Thermodesulfobacteriota>Desulfovibrionia>Desulfovibrionales>Desulfovibrionaceae | GCF_944324605.1 | Mailhella_massiliensis | PUQ52_RS06600 | 138638..142000 | 1903261 | Swt1-family-HEPN-domain-containing-protein-[Mailhella-massiliensis] |
| WP_077072828.1 | DUF499_1.RM.like.systems | DOC+HTH->REase+SAVED->?->?-><-?<-HEPN+DUF499*<-METHYLASE<-TUDOR+SF1-SF2-Helicase+iREase+CTD+REase<-GDResolvase+Recombinase+ZnR<-?<-?<-Phage_AlpA | 1120 | Bacteria>Thermodesulfobacteriota>Desulfovibrionia>Desulfovibrionales>Desulfovibrionaceae | GCF_944324615.1 | Mailhella_massiliensis | QOK07_RS04875 | 39206..42568 | 1903261 | Swt1-family-HEPN-domain-containing-protein-[Mailhella-massiliensis] |
| TVR02140.1 | DUF499_1.RM.like.systems | <-HEPN+DUF499*<-?<-METHYLASE<-PemK_toxin<-?<-Y1_Tnp<-TUDOR+SF1-SF2-Helicase+iREase+CTD+REase | 1114 | Bacteria>Thermodesulfobacteriota>Desulfovibrionia>Desulfovibrionales>unclassified_Desulfovibrionales | GCA_007693505.1 | Desulfovibrionales_bacterium | EA399_00610 | 1087..4431 | 2448050 | ATP-binding-protein-[Desulfovibrionales-bacterium] |
| WP_067619295.1 | DUF499_1.RM.like.systems | LexA-HTH+LexA-protease->TUDOR+SF1-SF2-Helicase+iREase+CTD+REase->Schlafen+HATPase+HTH->?->METHYLASE->DOC+HTH->HEPN+DUF499*->?->Peptidase_C47->?->PIN-> | 1099 | Bacteria>Thermodesulfobacteriota>Dissulfuribacteria>Dissulfuribacterales>Dissulfuribacteraceae | GCF_001687335.1 | Dissulfuribacter_thermophilus | DBT_RS08725 | 80186..83485 | 1156395 | Swt1-family-HEPN-domain-containing-protein-[Dissulfuribacter-thermophilus] |
| NLA40358.1 | DUF499_1.RM.like.systems | <-HEPN+DUF499*<-METHYLASE | 1123 | Bacteria>Thermodesulfobacteriota>Syntrophia>Syntrophales>Smithellaceae | GCA_012510495.1 | Smithella_sp. | GX874_02955 | 108..3479 | 2053614 | ATP-binding-protein-[Smithella-sp.] |
| NLD79955.1 | DUF499_1.RM.like.systems | <-HEPN+DUF499*<-METHYLASE-p | 1123 | Bacteria>Thermodesulfobacteriota>Syntrophia>Syntrophales>Smithellaceae | GCA_012513865.1 | Smithella_sp. | GX642_02135 | 555..3926 | 2053614 | ATP-binding-protein-[Smithella-sp.] |
| ABC75909.1 | DUF499_1.RM.like.systems | <-sGTP\|\|?-><-DrHyd+BirA-HTH<-DUF499-p<-HEPN+DUF499*<-TOPRIM<-OLD-ABC<-METHYLASE-p<-METHYLASE-p<-HKD-DNase | 700 | Bacteria>Thermodesulfobacteriota>Syntrophia>Syntrophales>Syntrophaceae | GCA_000013405.1 | Syntrophus_aciditrophicus_SB | SYN_00008 | 32188..34290 | 56780 | ATPase-of-the-AAA+-superfamily-[Syntrophus-aciditrophicus-SB] |
| HBL52555.1 | DUF499_1.RM.like.systems | TUDOR+SF1-SF2-Helicase+iREase+CTD+REase-p->ORC-CDC6+REase->MqsA_antitoxin+HTH_3->HEPN+DUF499*-> | 1127 | Bacteria>Thermodesulfobacteriota>Syntrophia>Syntrophales>Syntrophaceae | GCA_003506745.1 | Syntrophaceae_bacterium | DDZ34_00420 | 4849..8232 | 2052184 | AAA+-family-ATPase-[Syntrophaceae-bacterium] |
| MBN1365905.1 | DUF499_1.RM.like.systems | HKD-DNase->METHYLASE->HEPN+DUF499*->?->?->REase-><-?\|\|HNH-><-REase | 1119 | Bacteria>Thermodesulfobacteriota>Syntrophia>Syntrophales>Syntrophaceae | GCA_016927935.1 | Syntrophaceae_bacterium | JW976_13940 | 4360..7719 | 2052184 | DUF499-domain-containing-protein-[Syntrophaceae-bacterium] |
| MCG2738715.1 | DUF499_1.RM.like.systems | MORC+DUF3684->DrHyd+BirA-HTH->SF1-SF2-Helicase->?->?->METHYLASE->HEPN+DUF499*-> | 1124 | Bacteria>Thermodesulfobacteriota>Syntrophia>Syntrophales>Syntrophaceae | GCA_022013455.1 | Syntrophaceae_bacterium | L6300_00555 | 19371..22745 | 2052184 | DUF499-domain-containing-protein-[Syntrophaceae-bacterium] |
| MCG2740517.1 | DUF499_1.RM.like.systems | MerR-HTH->METHYLASE->PIN->?-><-HEPN+DUF499*<-METHYLASE<-MerR-HTH<-TUDOR+SF1-SF2-Helicase+iREase+CTD+REase | 1109 | Bacteria>Thermodesulfobacteriota>Syntrophia>Syntrophales>Syntrophaceae | GCA_022013455.1 | Syntrophaceae_bacterium | L6300_09825 | 5770..9099 | 2052184 | DUF499-domain-containing-protein-[Syntrophaceae-bacterium] |
| WP_011415944.1 | DUF499_1.RM.like.systems | <-sGTP<-DrHyd+BirA-HTH<-DUF499-p<-HEPN+DUF499*<-TOPRIM<-OLD-ABC<-METHYLASE-p<-METHYLASE-p<-HKD-DNase | 700 | Bacteria>Thermodesulfobacteriota>Syntrophia>Syntrophales>Syntrophaceae | GCF_000013405.1 | Syntrophus_aciditrophicus | SYN_RS16675 | 32188..34290 | 316277 | Swt1-family-HEPN-domain-containing-protein-[Syntrophus-aciditrophicus] |
| MCK9528171.1 | DUF499_1.RM.like.systems | TUDOR+SF1-SF2-Helicase+iREase+CTD+REase->PHP+SbcC->Schlafen+HATPase+HTH->METHYLASE->OLD-ABC->TOPRIM->HEPN+DUF499*->ABhydrolase-> | 1108 | Bacteria>Thermodesulfobacteriota>Syntrophia>Syntrophales>unclassified_Syntrophales | GCA_023228425.1 | Syntrophales_bacterium | MOQ23_05915 | 66900..70226 | 2928017 | Swt1-family-HEPN-domain-containing-protein-[Syntrophales-bacterium] |
| MCK9230422.1 | DUF499_1.RM.like.systems | <-HEPN+DUF499*<-TOPRIM<-OLD-ABC<-METHYLASE<-Schlafen+HATPase+HTH<-PHP+SbcC<-TUDOR+SF1-SF2-Helicase+iREase+CTD+REase | 638 | Bacteria>Thermodesulfobacteriota>Syntrophia>Syntrophales>unclassified_Syntrophales | GCA_023230845.1 | Syntrophales_bacterium | MOP18_07070 | 1..1913 | 2928017 | Swt1-family-HEPN-domain-containing-protein-[Syntrophales-bacterium] |
| SPF49808.1 | DUF499_1.RM.like.systems | ParB+HNH->DrHyd+BirA-HTH->STAS+HisKin->METHYLASE->HEPN+DUF499*-> | 1116 | Bacteria>Thermodesulfobacteriota>Syntrophobacteria>Syntrophobacterales>Syntrophobacteraceae | GCA_900290365.1 | Syntrophobacter_sp._SbD1 | SBDP1_830032 | 43636..46986 | 2043167 | conserved-hypothetical-protein-[Syntrophobacter-sp.-SbD1] |
| OPY85236.1 | DUF499_1.RM.like.systems | METHYLASE->HEPN+DUF499*-> | 1175 | Bacteria>Thermodesulfobacteriota>Syntrophorhabdia>Syntrophorhabdales>Syntrophorhabdaceae | GCA_002067475.1 | Syntrophorhabdus_sp._PtaU1.Bin153 | A4E65_00163 | 2092..5619 | 1811705 | MAG:-hypothetical-protein-A4E65_00163-[Syntrophorhabdus-sp.-PtaU1.Bin153] |
| MDD3846877.1 | DUF499_1.RM.like.systems | <-HEPN<-ParB+HNH<-HEPN+DUF499*<-METHYLASE<-MORC+DUF3684<-TUDOR+SF1-SF2-Helicase+iREase+CTD+REase<-DUF932 | 1114 | Bacteria>Thermodesulfobacteriota>Syntrophorhabdia>Syntrophorhabdales>Syntrophorhabdaceae | GCA_028698615.1 | Syntrophorhabdaceae_bacterium | PHC90_11025 | 3963..7307 | 2660714 | MAG:-Swt1-family-HEPN-domain-containing-protein-[Syntrophorhabdaceae-bacterium] |
| WP_028893190.1 | DUF499_1.RM.like.systems | TUDOR+SF1-SF2-Helicase+iREase+CTD+REase->MORC+DUF3684->SbcC->OLD-ABC->TOPRIM->METHYLASE->HEPN+DUF499*->DUF4338->NurA->HerA/FtsK-><-?<-AbiEii<-AbiEi_1 | 1108 | Bacteria>Thermodesulfobacteriota>Syntrophorhabdia>Syntrophorhabdales>Syntrophorhabdaceae | GCF_000512235.1 | Syntrophorhabdus_aromaticivorans | SYNAR_RS0100650 | 131724..135050 | 328301 | Swt1-family-HEPN-domain-containing-protein-[Syntrophorhabdus-aromaticivorans] |
| OQA20075.1 | DUF499_1.RM.like.systems | DUF4236+TPR-><-Peptidase_M20<-?<-HEPN+DUF499*<-?<-DUF433<-METHYLASE<-TUDOR+SF1-SF2-Helicase+iREase+CTD+REase | 1128 | Bacteria>unclassified_Bacteria | GCA_002069435.1 | bacterium_ADurb.Bin363 | BWY64_00439 | 13366..16752 | 1866927 | hypothetical-protein-BWY64_00439-[bacterium-ADurb.Bin363] |
| PZN89122.1 | DUF499_1.RM.like.systems | SF1-SF2-Helicase+REase-p->SF1-SF2-Helicase+REase-><-?\|\|?-><-?<-?<-HEPN+DUF499*<-RNaseH<-METHYLASE-p | 1114 | Bacteria>unclassified_Bacteria | GCA_003242735.1 | bacterium | DIU52_15115 | 17855..21199 | 1869227 | AAA+-family-ATPase-[bacterium] |
| TRZ47091.1 | DUF499_1.RM.like.systems | <-HEPN+DUF499*<-?<-METHYLASE-p | 1109 | Bacteria>unclassified_Bacteria | GCA_007279595.1 | bacterium | D4S03_11515 | 927..4256 | 1869227 | ATP-binding-protein-[bacterium] |
| KAF0250487.1 | DUF499_1.RM.like.systems | Phd-YefM->PIN->METHYLASE->REase->OLD-ABC->TOPRIM->HEPN+DUF499*->?-><-Phage_integrase\|\|?->MqsA_antitoxin+HTH_3-> | 1130 | Bacteria>unclassified_Bacteria | GCA_009773835.1 | bacterium | FD167_113 | 11053..14445 | 1869227 | hypothetical-protein-FD167_113-[bacterium] |
| NJL57104.1 | DUF499_1.RM.like.systems | METHYLASE-p->DUF433->PIN->HEPN+DUF499*->DUF499-> | 411 | Bacteria>unclassified_Bacteria | GCA_012031555.1 | bacterium | HC928_19635 | 992..2227 | 1869227 | ATP-binding-protein-[bacterium] |
| NJL57105.1 | DUF499_1.RM.like.systems | METHYLASE-p->DUF433->PIN->HEPN+DUF499->DUF499*-> | 478 | Bacteria>unclassified_Bacteria | GCA_012031555.1 | bacterium | HC928_19640 | 2241..3676 | 1869227 | ATP-binding-protein-[bacterium] |
| MBK8128933.1 | DUF499_1.RM.like.systems | DDE_3->?-><-?\|\|?-><-?<-?<-HEPN+DUF499*<-METHYLASE-p<-PIN<-CopG/RHH<-PIN<-?<-TUDOR+SF1-SF2-Helicase+iREase+CTD+REase-p | 1097 | Bacteria>unclassified_Bacteria | GCA_016712825.1 | bacterium | IPK53_08290 | 27062..30355 | 1869227 | DUF499-domain-containing-protein-[bacterium] |
| MBQ7502581.1 | DUF499_1.RM.like.systems | <-HEPN+DUF499*\|\|REase->?->?-><-METHYLASE<-TUDOR+SF1-SF2-Helicase+iREase+CTD+REase\|\|REase-> | 1127 | Bacteria>unclassified_Bacteria | GCA_017458675.1 | bacterium | IJT93_07695 | 1961..5344 | 1869227 | DUF499-domain-containing-protein-[bacterium] |
| MBU2648361.1 | DUF499_1.RM.like.systems | TUDOR+SF1-SF2-Helicase+iREase+CTD+REase->wHTH+REase->METHYLASE->HEPN+DUF499*->?->Phage_integrase-><-Arm-DNA-bind_3+Phage_int_SAM_3+Phage_integrase | 1091 | Bacteria>unclassified_Bacteria | GCA_018830885.1 | bacterium | KKI24_26885 | 8390..11665 | 1869227 | DUF499-domain-containing-protein-[bacterium] |
| MCG3151370.1 | DUF499_1.RM.like.systems | TUDOR+SF1-SF2-Helicase+iREase+CTD+REase->SIR2->METHYLASE->HEPN+DUF499*->HTH->MPTase->?->?->Primpol+D5N+D5ATPase->HIN-HTH-> | 1134 | Bacteria>unclassified_Bacteria | GCA_022072165.1 | bacterium | GEEBNDBF_00641 | 748382..751786 | 1869227 | hypothetical-protein-[bacterium] |
| MCG3156467.1 | DUF499_1.RM.like.systems | TUDOR+SF1-SF2-Helicase+iREase+CTD+REase->Schlafen+HATPase+HTH->DOC+HTH->METHYLASE-><-PIN<-?\|\|HEPN+DUF499*->TIR-> | 1113 | Bacteria>unclassified_Bacteria | GCA_022072305.1 | bacterium | DKINENOH_03091 | 110830..114171 | 1869227 | hypothetical-protein-[bacterium] |
| MCI0414205.1 | DUF499_1.RM.like.systems | <-HEPN<-HEPN+DUF499*<-METHYLASE | 1107 | Bacteria>unclassified_Bacteria | GCA_022615055.1 | bacterium | L0222_15590 | 1218..4541 | 1869227 | DUF499-domain-containing-protein-[bacterium] |
| MCP4152612.1 | DUF499_1.RM.like.systems | Radical-SAM->?-><-HEPN+DUF499*<-METHYLASE-p | 1101 | Bacteria>unclassified_Bacteria | GCA_024224895.1 | bacterium | GY757_33055 | 6237..9542 | 1869227 | ATP-binding-protein-[bacterium] |
| MCP3995426.1 | DUF499_1.RM.like.systems | zf-ISL3+HTH_Tnp_ISL3+DDE_Tnp_ISL3->METHYLASE-p->HEPN+DUF499*-> | 479 | Bacteria>unclassified_Bacteria | GCA_024226295.1 | bacterium | GY722_10220 | 2321..3757 | 1869227 | ATP-binding-protein-[bacterium] |
| MCP4656470.1 | DUF499_1.RM.like.systems | <-HEPN+DUF499*<-?<-METHYLASE-p | 1058 | Bacteria>unclassified_Bacteria | GCA_024231315.1 | bacterium | GY856_13735 | 1..3174 | 1869227 | ATP-binding-protein-[bacterium] |
| MCR5845017.1 | DUF499_1.RM.like.systems | REase-><-?\|\|?-><-?<-HEPN+DUF499*<-METHYLASE | 1130 | Bacteria>unclassified_Bacteria | GCA_024698685.1 | bacterium | K6G78_00395 | 19668..23060 | 1869227 | DUF499-domain-containing-protein-[bacterium] |
| MCY3562591.1 | DUF499_1.RM.like.systems | METHYLASE->HEPN+DUF499*->HTH_3->NTP_transf-><-?<-PIN | 1107 | Bacteria>unclassified_Bacteria | GCA_026702805.1 | bacterium | OXH26_01515 | 1911..5234 | 1869227 | MAG:-DUF499-domain-containing-protein-[bacterium] |
| MCY3563828.1 | DUF499_1.RM.like.systems | <-HEPN+DUF499*<-METHYLASE-p | 754 | Bacteria>unclassified_Bacteria | GCA_026702805.1 | bacterium | OXH26_07915 | 1..2262 | 1869227 | MAG:-DUF499-domain-containing-protein,-partial-[bacterium] |
| MCY3560578.1 | DUF499_1.RM.like.systems | METHYLASE-p->HEPN+DUF499*-> | 1108 | Bacteria>unclassified_Bacteria | GCA_026702815.1 | bacterium | OXH20_05350 | 698..4024 | 1869227 | MAG:-Swt1-family-HEPN-domain-containing-protein-[bacterium] |
| MCY3562206.1 | DUF499_1.RM.like.systems | <-ABhydrolase\|\|?->?-><-HEPN+DUF499*\|\|ParD_antitoxin-><-?<-METHYLASE<-PIN<-DUF433 | 1084 | Bacteria>unclassified_Bacteria | GCA_026702815.1 | bacterium | OXH20_13745 | 3210..6464 | 1869227 | MAG:-DUF499-domain-containing-protein-[bacterium] |
| MCY3637183.1 | DUF499_1.RM.like.systems | <-HEPN+DUF499*<-METHYLASE<-OLD-ABC+TOPRIM<-?<-TOPRIM<-OLD-ABC | 1079 | Bacteria>unclassified_Bacteria | GCA_026705625.1 | bacterium | OXH23_16400 | 13524..16763 | 1869227 | MAG:-DUF499-domain-containing-protein-[bacterium] |
| MCY3785681.1 | DUF499_1.RM.like.systems | <-CHTH<-?\|\|BirA-HTH->CopG/RHH->METHYLASE->HEPN+DUF499*-> | 1126 | Bacteria>unclassified_Bacteria | GCA_026706565.1 | bacterium | OXG47_03025 | 15510..18890 | 1869227 | MAG:-Swt1-family-HEPN-domain-containing-protein-[bacterium] |
| MCY3806661.1 | DUF499_1.RM.like.systems | METHYLASE->MetJarc-HTH->PemK_toxin->HEPN+DUF499*->DUF499-><-?\|\|ABhydrolase-> | 355 | Bacteria>unclassified_Bacteria | GCA_026706585.1 | bacterium | OXG91_09240 | 4057..5121 | 1869227 | MAG:-Swt1-family-HEPN-domain-containing-protein,-partial-[bacterium] |
| MCY3806662.1 | DUF499_1.RM.like.systems | METHYLASE->MetJarc-HTH->PemK_toxin->HEPN+DUF499->DUF499*-><-?\|\|ABhydrolase-> | 710 | Bacteria>unclassified_Bacteria | GCA_026706585.1 | bacterium | OXG91_09245 | 5222..7356 | 1869227 | MAG:-AAA+-family-ATPase,-partial-[bacterium] |
| MCY3862385.1 | DUF499_1.RM.like.systems | <-PIN<-PhdYeFM_antitox<-?\|\|DUF222+HNH->?->PIN->HEPN+DUF499*-><-?<-DOC+DOC<-HTH_31 | 1079 | Bacteria>unclassified_Bacteria | GCA_026706925.1 | bacterium | OXG67_10535 | 19896..23135 | 1869227 | MAG:-DUF499-domain-containing-protein-[bacterium] |
| MCY3889489.1 | DUF499_1.RM.like.systems | <-PIN<-PhdYeFM_antitox<-?\|\|DUF222+HNH->?->PIN->HEPN+DUF499*-><-?<-DOC+DOC<-HTH_31 | 1079 | Bacteria>unclassified_Bacteria | GCA_026707085.1 | bacterium | OXG50_06285 | 19658..22897 | 1869227 | MAG:-DUF499-domain-containing-protein-[bacterium] |
| MCY3923936.1 | DUF499_1.RM.like.systems | <-CHTH\|\|DOC->CopG/RHH->METHYLASE->HEPN+DUF499*-> | 1131 | Bacteria>unclassified_Bacteria | GCA_026707305.1 | bacterium | OXG52_00270 | 5305..8700 | 1869227 | MAG:-Swt1-family-HEPN-domain-containing-protein-[bacterium] |
| MCY3924850.1 | DUF499_1.RM.like.systems | PIN->DUF433->PIN->METHYLASE->MetJarc-HTH->PemK_toxin->HEPN+DUF499*-><-?\|\|ABhydrolase->?-><-?<-Glyoxalase | 1099 | Bacteria>unclassified_Bacteria | GCA_026707305.1 | bacterium | OXG52_04985 | 27802..31101 | 1869227 | MAG:-DUF499-domain-containing-protein-[bacterium] |
| MCY3963126.1 | DUF499_1.RM.like.systems | <-PIN<-PhdYeFM_antitox<-?\|\|DUF222+HNH->?->PIN->HEPN+DUF499*-><-?<-HTH_31 | 1079 | Bacteria>unclassified_Bacteria | GCA_026707505.1 | bacterium | OXG34_15910 | 28430..31669 | 1869227 | MAG:-DUF499-domain-containing-protein-[bacterium] |
| MCY3951956.1 | DUF499_1.RM.like.systems | <-NUDIX<-?<-?<-HEPN+DUF499*<-METHYLASE\|\|CopG/RHH->PIN-><-REase-p<-PemK_toxin<-CopG/RHH | 1108 | Bacteria>unclassified_Bacteria | GCA_026707535.1 | bacterium | OXG69_05350 | 3578..6904 | 1869227 | MAG:-Swt1-family-HEPN-domain-containing-protein-[bacterium] |
| MCY3953257.1 | DUF499_1.RM.like.systems | DUF433->PIN->METHYLASE->MetJarc-HTH->PemK_toxin->HEPN+DUF499*-><-?\|\|ABhydrolase-> | 1099 | Bacteria>unclassified_Bacteria | GCA_026707535.1 | bacterium | OXG69_12060 | 23481..26780 | 1869227 | MAG:-DUF499-domain-containing-protein-[bacterium] |
| MCY4135752.1 | DUF499_1.RM.like.systems | <-HEPN+DUF499*\|\|?-><-METHYLASE\|\|DOC-><-HEPN+REase | 1079 | Bacteria>unclassified_Bacteria | GCA_026708015.1 | bacterium | OXG30_12705 | 27919..31158 | 1869227 | MAG:-DUF499-domain-containing-protein-[bacterium] |
| MCY4102170.1 | DUF499_1.RM.like.systems | CopG/RHH->PemK_toxin->REase-p-><-PIN<-CopG/RHH\|\|METHYLASE->HEPN+DUF499*->?->?->NUDIX-><-?<-?<-ABC-ATPase+SbcC | 1108 | Bacteria>unclassified_Bacteria | GCA_026708055.1 | bacterium | OXG55_02720 | 179181..182507 | 1869227 | MAG:-Swt1-family-HEPN-domain-containing-protein-[bacterium] |
| MCY4103460.1 | DUF499_1.RM.like.systems | DUF433->PIN->METHYLASE->MetJarc-HTH->PemK_toxin->HEPN+DUF499*-><-?\|\|ABhydrolase-><-?\|\|?-><-?\|\|REase-> | 1099 | Bacteria>unclassified_Bacteria | GCA_026708055.1 | bacterium | OXG55_09400 | 103234..106533 | 1869227 | MAG:-DUF499-domain-containing-protein-[bacterium] |
| MCY4163400.1 | DUF499_1.RM.like.systems | DUF433->PIN->METHYLASE->METHYLASE-p-><-?\|\|?->HEPN+DUF499*-><-?\|\|?->?->?->ABC-ATPase+SbcC->PfkB-> | 1091 | Bacteria>unclassified_Bacteria | GCA_026709025.1 | bacterium | OXE93_04200 | 10413..13688 | 1869227 | MAG:-DUF499-domain-containing-protein-[bacterium] |
| MCY4258449.1 | DUF499_1.RM.like.systems | <-HEPN+DUF499*<-?<-METHYLASE<-?<-TUDOR+SF1-SF2-Helicase+iREase+CTD+REase | 1116 | Bacteria>unclassified_Bacteria | GCA_026709785.1 | bacterium | OXE04_09265 | 10514..13864 | 1869227 | MAG:-Swt1-family-HEPN-domain-containing-protein-[bacterium] |
| MCY4258519.1 | DUF499_1.RM.like.systems | <-DUF222+HNH<-PIN<-DUF433\|\|?-><-HEPN+DUF499*<-?\|\|?-><-METHYLASE<-SbcC<-SbcC<-TUDOR+SF1-SF2-Helicase+iREase+CTD+REase | 1091 | Bacteria>unclassified_Bacteria | GCA_026709785.1 | bacterium | OXE04_09620 | 60604..63879 | 1869227 | MAG:-DUF499-domain-containing-protein-[bacterium] |
| MCY4370194.1 | DUF499_1.RM.like.systems | <-HEPN+DUF499*<-METHYLASE\|\|HEPN->DDE_Tnp_1-><-?<-HerA/FtsK | 1095 | Bacteria>unclassified_Bacteria | GCA_026710435.1 | bacterium | OXF41_12460 | 874..4161 | 1869227 | MAG:-DUF499-domain-containing-protein-[bacterium] |
| MCY4630164.1 | DUF499_1.RM.like.systems | <-HEPN+DUF499*<-METHYLASE | 332 | Bacteria>unclassified_Bacteria | GCA_026713745.1 | bacterium | OXE75_04665 | 1..996 | 1869227 | MAG:-Swt1-family-HEPN-domain-containing-protein,-partial-[bacterium] |
| MCY4632748.1 | DUF499_1.RM.like.systems | <-HEPN+DUF499*<-METHYLASE | 487 | Bacteria>unclassified_Bacteria | GCA_026713745.1 | bacterium | OXE75_17955 | 1..1460 | 1869227 | MAG:-Swt1-family-HEPN-domain-containing-protein,-partial-[bacterium] |
| MDC0992039.1 | DUF499_1.RM.like.systems | <-OLD-ABC<-HEPN+DUF499*<-METHYLASE | 1135 | Bacteria>unclassified_Bacteria | GCA_028375025.1 | bacterium | OAR33_00540 | 2470..5877 | 1869227 | Swt1-family-HEPN-domain-containing-protein-[bacterium] |
| MDD2328800.1 | DUF499_1.RM.like.systems | METHYLASE-p->HEPN+DUF499*-> | 1132 | Bacteria>unclassified_Bacteria | GCA_028683535.1 | bacterium | PHZ13_10910 | 283..3681 | 1869227 | MAG:-DUF499-domain-containing-protein-[bacterium] |
| MDD3925571.1 | DUF499_1.RM.like.systems | <-SF1-SF2-Helicase<-HEPN+DUF499*<-METHYLASE<-Schlafen+HATPase+HTH<-TUDOR+SF1-SF2-Helicase+iREase+CTD+REase | 1106 | Bacteria>unclassified_Bacteria | GCA_028699365.1 | bacterium | PHT33_02815 | 1591..4911 | 1869227 | MAG:-Swt1-family-HEPN-domain-containing-protein-[bacterium] |
| MDE0115147.1 | DUF499_1.RM.like.systems | <-HEPN+DUF499*<-DUF222+HNH\|\|?->PhdYeFM_antitox->PIN->?-><-METHYLASE | 1079 | Bacteria>unclassified_Bacteria | GCA_028819295.1 | bacterium | OXT07_00780 | 18092..21331 | 1869227 | MAG:-DUF499-domain-containing-protein-[bacterium] |
| MDE0170777.1 | DUF499_1.RM.like.systems | <-HEPN+DUF499*<-METHYLASE<-XIS-HTH+RVT<-?<-?<-REase | 1096 | Bacteria>unclassified_Bacteria | GCA_028819595.1 | bacterium | OXS29_14930 | 550..3840 | 1869227 | MAG:-DUF499-domain-containing-protein-[bacterium] |
| MDE0193912.1 | DUF499_1.RM.like.systems | METHYLASE->HEPN+DUF499*->?->?->NUDIX-> | 1108 | Bacteria>unclassified_Bacteria | GCA_028819945.1 | bacterium | OXP08_00010 | 2300..5626 | 1869227 | MAG:-Swt1-family-HEPN-domain-containing-protein-[bacterium] |
| MDE0377403.1 | DUF499_1.RM.like.systems | <-Glyoxalase<-?\|\|?->?-><-?<-HEPN+DUF499*<-METHYLASE | 1095 | Bacteria>unclassified_Bacteria | GCA_028821235.1 | bacterium | OXK16_15785 | 33048..36335 | 1869227 | MAG:-DUF499-domain-containing-protein-[bacterium] |
| MDE0368588.1 | DUF499_1.RM.like.systems | <-FAD-NAD-dep-oxidoreductase<-?<-?<-?\|\|?-><-HEPN+DUF499*<-METHYLASE\|\|?-><-MORC+HisKin+REase<-MNS+HTH<-REase\|\|HEPN-> | 1095 | Bacteria>unclassified_Bacteria | GCA_028821265.1 | bacterium | OXI84_00435 | 13176..16463 | 1869227 | MAG:-DUF499-domain-containing-protein-[bacterium] |
| MDE0668308.1 | DUF499_1.RM.like.systems | CopG/RHH-><-PIN<-MPTase<-PIN<-CopG/RHH\|\|METHYLASE->HEPN+DUF499*-> | 1108 | Bacteria>unclassified_Bacteria | GCA_028824385.1 | bacterium | OXI48_04695 | 5107..8433 | 1869227 | MAG:-Swt1-family-HEPN-domain-containing-protein-[bacterium] |
| MDE0578824.1 | DUF499_1.RM.like.systems | <-HEPN+DUF499*\|\|?-><-METHYLASE\|\|HTH_58-> | 1079 | Bacteria>unclassified_Bacteria | GCA_028824815.1 | bacterium | OXI29_03025 | 42794..46033 | 1869227 | MAG:-DUF499-domain-containing-protein-[bacterium] |
| MDE2480824.1 | DUF499_1.RM.like.systems | <-SF1-SF2-Helicase<-OLD-ABC+TOPRIM<-HEPN+DUF499*<-METHYLASE<-TUDOR+SF1-SF2-Helicase+iREase+CTD+REase\|\|PIN->AbiEi_3_N+AbiEi_3-><-SF1-SF2-helicase | 1110 | Bacteria>unclassified_Bacteria | GCA_028869385.1 | bacterium | KGN02_01370 | 63070..66402 | 1869227 | MAG:-DUF499-domain-containing-protein-[bacterium] |
| WP_248539435.1 | DUF499_1.RM.like.systems | MerR-HTH-><-?\|\|N6_MTase-p-><-OLD-ABC+TOPRIM<-HEPN+DUF499*<-METHYLASE<-TUDOR+SF1-SF2-Helicase+iREase+CTD+REase<-XIS-HTH<-?<-HTH_Tnp_1+HTH_21+rve+rve_2 | 1132 | - | GCF_023195655.1 | Not | M0721_RS02800 | 592569..595967 | 2851643 | DUF499-domain-containing-protein-[Microbacterium-sp.-SSW1-7] |
| -- | | | | | | | | | | |
| --HEPN(PolB->HEPN->) | | | | | | | | | | |
| -- | | | | | | | | | | |
| RLF61780.1 | DUF499_1.RM.like.systems | METHYLASE-p->PolB->HEPN->DUF499*-> | 863 | Archaea>Candidatus_Thermoplasmatota>Thermoplasmata>unclassified_Thermoplasmata | GCA_003650005.1 | Thermoplasmata_archaeon | DRN16_03190 | 1440..4028 | 1906666 | MAG:-hypothetical-protein-DRN16_03190,-partial-[Thermoplasmata-archaeon] |
| WP_015733362.1 | DUF499_1.RM.like.systems | METHYLASE->MNS->DUF499*->Cluster35_66clades->TUDOR+SF1-SF2-Helicase+iREase+CTD+REase->HEPN->RAMPs->HDRnase-> | 1104 | Archaea>Euryarchaeota>Methanomada_group>Methanococci>Methanococcales | GCF_000024625.1 | Methanocaldococcus_vulcanius | METVU_RS06315 | 1284462..1287776 | 73913 | DUF499-domain-containing-protein-[Methanocaldococcus-vulcanius] |
| RLG35314.1 | DUF499_1.RM.like.systems | METHYLASE-p->HEPN->DUF499*-> | 886 | Archaea>Euryarchaeota>Stenosarchaea_group>Methanomicrobia>Methanosarcinales | GCA_003661135.1 | Methanosarcinales_archaeon | DRN98_01510 | 3349..6009 | 2250255 | hypothetical-protein-DRN98_01510-[Methanosarcinales-archaeon] |
| MQY54544.1 | DUF499_1.RM.like.systems | PolB->HEPN->DUF499*->Cysteine_desulfurase-> | 840 | Archaea>Euryarchaeota>Stenosarchaea_group>Methanomicrobia>Methanosarcinales | GCA_009618475.1 | Methanosarcinales_archaeon | GH149_04580 | 2436..4958 | 2250255 | AAA-family-ATPase-[Methanosarcinales-archaeon] |
| OYT64646.1 | DUF499_1.RM.like.systems | <-SbcC<-DUF499*<-PIN<-SbcC<-METHYLASE-p<-HEPN<-PolB | 851 | Archaea>TACK_group>Candidatus_Bathyarchaeota>unclassified_Candidatus_Bathyarchaeota | GCA_002255045.1 | Candidatus_Bathyarchaeota_archaeon_ex4484_205 | B6U74_05075 | 1011..3566 | 2012510 | AAA-family-ATPase-[Candidatus-Bathyarchaeota-archaeon-ex4484_205] |
| MBO3798316.1 | DUF499_1.RM.like.systems | <-EVE<-HEPN<-DUF499*<-METHYLASE<-TUDOR+SF1-SF2-Helicase+iREase+CTD+REase<-URI-DNase+RAMA\|\|?->?->URI-DNase-> | 887 | Archaea>TACK_group>Candidatus_Brockarchaeota>unclassified_Candidatus_Brockarchaeota | GCA_017601225.1 | Candidatus_Brockarchaeota_archaeon | JTT13_05580 | 80557..83220 | 2815303 | ATP-binding-protein-[Candidatus-Brockarchaeota-archaeon] |
| KEQ56705.1 | DUF499_1.RM.like.systems | TUDOR+SF1-SF2-Helicase+iREase+CTD+REase->METHYLASE->DUF499*->HEPN-> | 855 | Archaea>TACK_group>Nitrososphaerota>Nitrososphaerota_incertae_sedis>Marine_Group_I | GCA_000722915.1 | Marine_Group_I_thaumarchaeote_SCGC_AAA799-N04 | AAA799N04_00839 | 23778..26345 | 1502293 | ATPase-AAA-protein-[Marine-Group-I-thaumarchaeote-SCGC-AAA799-N04] |
| KFM20182.1 | DUF499_1.RM.like.systems | <-HEPN<-DUF499*<-METHYLASE<-TUDOR+SF1-SF2-Helicase+iREase+CTD+REase<-?\|\|?-><-REase+DUF5655 | 852 | Archaea>TACK_group>Nitrososphaerota>Nitrososphaerota_incertae_sedis>Marine_Group_I | GCA_000746685.1 | Marine_Group_I_thaumarchaeote_SCGC_AAA799-P11 | AAA799P11_00258 | 97012..99570 | 1502295 | ATPase-AAA-protein-[Marine-Group-I-thaumarchaeote-SCGC-AAA799-P11] |
| MBT6647711.1 | DUF499_1.RM.like.systems | <-HEPN<-DUF499*<-METHYLASE-p\|\|METHYLASE-p-> | 834 | Archaea>TACK_group>Nitrososphaerota>unclassified_Nitrososphaerota | GCA_018697615.1 | Nitrososphaerota_archaeon | HOB51_09415 | 855..3359 | 2026795 | ATP-binding-protein-[Thaumarchaeota-archaeon] |
| MCH7560423.1 | DUF499_1.RM.like.systems | <-HEPN<-DUF499*<-METHYLASE | 852 | Archaea>TACK_group>Nitrososphaerota>unclassified_Nitrososphaerota | GCA_022561135.1 | Nitrososphaerota_archaeon | IIC67_03480 | 947..3505 | 2026795 | ATP-binding-protein-[Thaumarchaeota-archaeon] |
| PMB78266.1 | DUF499_1.RM.like.systems | TUDOR+SF1-SF2-Helicase+iREase+CTD+REase->METHYLASE-p->HEPN->NTP_transf_2->PolB->HEPN->DUF499*-> | 859 | Archaea>TACK_group>Thermoproteota>Thermoprotei>Fervidicoccales | GCA_002871295.1 | Fervidicoccus_fontis | C0177_00595 | 9295..11874 | 683846 | AAA-family-ATPase-[Fervidicoccus-fontis] |
| MBP1626439.1 | DUF499_1.RM.like.systems | HhH-RADC+JAB-><-PHP+SbcC<-DUF499*<-METHYLASE<-HEPN<-TUDOR+SF1-SF2-Helicase+iREase+CTD+REase<-HTH\|\|XIS-HTH-> | 952 | Bacteria>Acidobacteriota>Holophagae>Holophagales>Holophagaceae | GCA_017859955.1 | Holophagaceae_bacterium | H6Q00_914 | 10441..13299 | 2052165 | hypothetical-protein-[Holophagaceae-bacterium] |
| MBZ5523276.1 | DUF499_1.RM.like.systems | <-ABhydrolase<-?\|\|ParB+HNH->METHYLASE-><-HEPN\|\|DUF499*-><-PIN<-?\|\|MazE_antitoxin->PemK_toxin->?->PIN-> | 924 | Bacteria>Acidobacteriota>Terriglobia>unclassified_Terriglobia | GCA_020072595.1 | Terriglobia_bacterium | LAP21_13650 | 15770..18544 | 2081523 | DUF499-domain-containing-protein-[Acidobacteriia-bacterium] |
| MCL6564987.1 | DUF499_1.RM.like.systems | TUDOR+SF1-SF2-Helicase+iREase+CTD+REase->?->METHYLASE->HEPN->DUF499*-><-?\|\|?-><-?<-HTH_17 | 920 | Bacteria>Acidobacteriota>Terriglobia>unclassified_Terriglobia | GCA_023512135.1 | Terriglobia_bacterium | K6U09_01035 | 242502..245264 | 2081523 | DUF499-domain-containing-protein-[Acidobacteriia-bacterium] |
| MXX86607.1 | DUF499_1.RM.like.systems | TUDOR+SF1-SF2-Helicase+iREase+CTD+REase->PolB->HEPN->METHYLASE->DUF499*-><-?<-?<-?<-?\|\|PP_kinase+HKD-DNase-> | 922 | Bacteria>Acidobacteriota>unclassified_Acidobacteriota | GCA_009838025.1 | Acidobacteriota_bacterium | F4Y71_09150 | 13575..16343 | 1978231 | ATP-binding-protein-[Acidobacteria-bacterium] |
| MXY24404.1 | DUF499_1.RM.like.systems | TUDOR+SF1-SF2-Helicase+iREase+CTD+REase->SIR2+SIR2->METHYLASE->DUF499*-><-HEPN\|\|ParB+HNH-> | 934 | Bacteria>Acidobacteriota>unclassified_Acidobacteriota | GCA_009838525.1 | Acidobacteriota_bacterium | F4Y45_07765 | 419511..422315 | 1978231 | ATP-binding-protein-[Acidobacteria-bacterium] |
| MYN68213.1 | DUF499_1.RM.like.systems | TUDOR+SF1-SF2-Helicase+iREase+CTD+REase->Schlafen+HATPase+HTH->HEPN->METHYLASE->OLD-ABC->TOPRIM->DUF499*-><-?<-SAM-methylase<-ACET+ACET<-GspA-ATPase | 1101 | Bacteria>Acidobacteriota>unclassified_Acidobacteriota | GCA_009861545.1 | Acidobacteriota_bacterium | F4X11_24880 | 56553..59858 | 1978231 | ATP-binding-protein-[Acidobacteria-bacterium] |
| MBI4730026.1 | DUF499_1.RM.like.systems | TUDOR+SF1-SF2-Helicase+iREase+CTD+REase->?->HEPN+REase->METHYLASE->?->HEPN->DUF499*->?->?-><-?<-sGTP<-?<-SAM-methylase | 918 | Bacteria>Acidobacteriota>unclassified_Acidobacteriota | GCA_016208485.1 | Acidobacteriota_bacterium | HY775_11110 | 79906..82662 | 1978231 | DUF499-domain-containing-protein-[Acidobacteria-bacterium] |
| MCY3965602.1 | DUF499_1.RM.like.systems | PIN-><-?<-?\|\|OLD-ABC->TOPRIM-><-DUF499*\|\|HEPN-><-TOPRIM<-OLD-ABC<-METHYLASE\|\|?->PAPS_reduct-> | 1095 | Bacteria>Acidobacteriota>unclassified_Acidobacteriota | GCA_026707545.1 | Acidobacteriota_bacterium | OXG83_11235 | 2822742..2826029 | 1978231 | MAG:-DUF499-domain-containing-protein-[Acidobacteria-bacterium] |
| MDA2929041.1 | DUF499_1.RM.like.systems | JAB->GDResolvase+HTH->HicB+ZnR->?->TUDOR+SF1-SF2-Helicase+iREase+CTD+REase->METHYLASE->DUF499*->HEPN-> | 981 | Bacteria>Acidobacteriota>unclassified_Acidobacteriota | GCA_027717865.1 | Acidobacteria_bacterium_AH-259-006 | MYX84_03670 | 12694..15639 | 2935772 | DUF499-domain-containing-protein-[Acidobacteria-bacterium-AH-259-006] |
| MDE2979244.1 | DUF499_1.RM.like.systems | <-HicB_lk_antitox<-HicA_toxin\|\|METHYLASE-><-HEPN<-REase-p\|\|DUF499*-> | 1124 | Bacteria>Acidobacteriota>unclassified_Acidobacteriota | GCA_028873735.1 | Acidobacteriota_bacterium | OXU63_17235 | 11921..15295 | 1978231 | MAG:-DUF499-domain-containing-protein-[Acidobacteriota-bacterium] |
| PKP59483.1 | DUF499_1.RM.like.systems | METHYLASE->DUF499*->HEPN->RNase-> | 876 | Bacteria>Atribacterota>unclassified_Atribacterota | GCA_002841055.1 | Candidatus_Atribacteria_bacterium_HGW-Atribacteria-1 | CVT91_06625 | 3132..5762 | 2013676 | AAA-family-ATPase-[Candidatus-Atribacteria-bacterium-HGW-Atribacteria-1] |
| MBI4572390.1 | DUF499_1.RM.like.systems | <-ABC-ATPase+SbcC<-?<-?\|\|Polbeta->HEPN-><-DUF499*<-AAA-ATPase+REase | 1106 | Bacteria>Bacteria_incertae_sedis>Bacteria_candidate_phyla>candidate_division_NC10>unclassified_candidate_division_NC10 | GCA_016209725.1 | candidate_division_NC10_bacterium | HY713_03775 | 5635..8955 | 2072417 | ATP-binding-protein-[candidate-division-NC10-bacterium] |
| MCD6257887.1 | DUF499_1.RM.like.systems | METHYLASE-p->PolB->HEPN->DUF499*-> | 846 | Bacteria>Bacteria_incertae_sedis>Bacteria_candidate_phyla>Candidatus_Aerophobetes>unclassified_Candidatus_Aerophobetes | GCA_021163905.1 | Candidatus_Aerophobetes_bacterium | J7J45_07510 | 2315..4855 | 2030807 | ATP-binding-protein-[Candidatus-Aerophobetes-bacterium] |
| KUK26300.1 | DUF499_1.RM.like.systems | <-DUF499*<-HEPN<-PolB<-METHYLASE<-TUDOR+SF1-SF2-Helicase+iREase+CTD+REase | 889 | Bacteria>Bacteria_incertae_sedis>Bacteria_candidate_phyla>Candidatus_Bipolaricaulota>unclassified_Candidatus_Bipolaricaulota | GCA_001508155.1 | Acetothermia_bacterium_64_32 | XD60_1536 | 19316..21985 | 1635286 | Uncharacterized-protein-XD60_1536-[Acetothermia-bacterium-64_32] |
| HAZ27921.1 | DUF499_1.RM.like.systems | METHYLASE-p->PolB->HEPN->DUF499*-> | 742 | Bacteria>Bacteria_incertae_sedis>Bacteria_candidate_phyla>Candidatus_Bipolaricaulota>unclassified_Candidatus_Bipolaricaulota | GCA_003485525.1 | Candidatus_Acetothermia_bacterium | DCY67_04830 | 1017..3242 | 2053493 | AAA-family-ATPase-[Candidatus-Acetothermia-bacterium] |
| RLE33859.1 | DUF499_1.RM.like.systems | <-DUF499*<-HEPN<-PolB<-METHYLASE-p | 930 | Bacteria>Bacteria_incertae_sedis>Bacteria_candidate_phyla>Candidatus_Bipolaricaulota>unclassified_Candidatus_Bipolaricaulota | GCA_003650505.1 | Candidatus_Acetothermia_bacterium | DRJ24_04700 | 120..2912 | 2053493 | hypothetical-protein-DRJ24_04700-[Candidatus-Acetothermia-bacterium] |
| MXY29380.1 | DUF499_1.RM.like.systems | HEPN->DUF499*->REase-> | 1115 | Bacteria>Bacteria_incertae_sedis>Bacteria_candidate_phyla>Candidatus_Poribacteria>unclassified_Candidatus_Poribacteria | GCA_009838315.1 | Candidatus_Poribacteria_bacterium | F4Y59_14620 | 1265..4612 | 2026781 | ATP-binding-protein-[Candidatus-Poribacteria-bacterium] |
| MCG9130919.1 | DUF499_1.RM.like.systems | TUDOR+SF1-SF2-Helicase+iREase+CTD+REase->OLD-ABC->HNH->METHYLASE->PolB->HEPN->DUF499*->HEAT_2->SAM-methylase->SIGMA-HTH->ZnR->ABC-ATPase+SbcC->ABC-ATPase+SbcC-> | 1114 | Bacteria>Bacteria_incertae_sedis>Bacteria_candidate_phyla>Candidatus_Poribacteria>unclassified_Candidatus_Poribacteria | GCA_022394775.1 | Candidatus_Poribacteria_bacterium | J5I95_04465 | 27738..31082 | 2026781 | ATP-binding-protein-[Candidatus-Poribacteria-bacterium] |
| MDE0014698.1 | DUF499_1.RM.like.systems | <-ABC-ATPase+SbcC<-ABC-ATPase+SbcC<-ZnR<-SIGMA-HTH<-SAM-methylase<-HEAT_2<-DUF499*<-HEPN<-?<-METHYLASE<-HEPN+REase<-?<-SbcC | 1116 | Bacteria>Bacteria_incertae_sedis>Bacteria_candidate_phyla>Candidatus_Poribacteria>unclassified_Candidatus_Poribacteria | GCA_028816655.1 | Candidatus_Poribacteria_bacterium | OXU51_00820 | 15793..19143 | 2026781 | MAG:-DUF499-domain-containing-protein-[Candidatus-Poribacteria-bacterium] |
| MDE0324945.1 | DUF499_1.RM.like.systems | <-HEAT_2<-DUF499*<-HEPN<-?<-METHYLASE<-?<-HNH<-OLD-ABC | 1118 | Bacteria>Bacteria_incertae_sedis>Bacteria_candidate_phyla>Candidatus_Poribacteria>unclassified_Candidatus_Poribacteria | GCA_028820875.1 | Candidatus_Poribacteria_bacterium | OXN27_13610 | 421..3777 | 2026781 | MAG:-DUF499-domain-containing-protein-[Candidatus-Poribacteria-bacterium] |
| MDE0688672.1 | DUF499_1.RM.like.systems | <-PspA+ParA-Soj-PloopNTPase<-?<-?<-DUF499*<-METHYLASE<-?<-?<-HEPN<-TUDOR+SF1-SF2-Helicase+iREase+CTD+REase\|\|OrfB_IS605+OrfB_Zn_ribbon-> | 937 | Bacteria>Bacteria_incertae_sedis>Bacteria_candidate_phyla>Candidatus_Poribacteria>unclassified_Candidatus_Poribacteria | GCA_028824525.1 | Candidatus_Poribacteria_bacterium | OXI61_10940 | 39625..42438 | 2026781 | MAG:-DUF499-domain-containing-protein-[Candidatus-Poribacteria-bacterium] |
| MBI2567876.1 | DUF499_1.RM.like.systems | TUDOR+SF1-SF2-Helicase+iREase+CTD+REase->PolB->HEPN->?->DUF499*-> | 808 | Bacteria>Bacteria_incertae_sedis>Bacteria_candidate_phyla>Candidatus_Schekmanbacteria>unclassified_Candidatus_Schekmanbacteria | GCA_016188575.1 | Candidatus_Schekmanbacteria_bacterium | HYV63_12665 | 36519..38945 | 2419984 | DUF499-domain-containing-protein-[Candidatus-Schekmanbacteria-bacterium] |
| MCC6545645.1 | DUF499_1.RM.like.systems | <-DUF499*<-?<-PIWI<-METHYLASE<-?<-HEPN<-METHYLASE-p | 1108 | Bacteria>Bacteria_incertae_sedis>Bacteria_candidate_phyla>Candidatus_Sumerlaeota>unclassified_Candidatus_Sumerlaeota | GCA_020848155.1 | Candidatus_Sumerlaeota_bacterium | IT570_00645 | 2114..5440 | 2107588 | ATP-binding-protein-[Candidatus-Sumerlaeota-bacterium] |
| MBM3272924.1 | DUF499_1.RM.like.systems | HIN-HTH->TUDOR+SF1-SF2-Helicase+iREase+CTD+REase->HEPN->METHYLASE->DUF499*-> | 220 | Bacteria>Bacteria_incertae_sedis>Bacteria_candidate_phyla>Patescibacteria_group>Parcubacteria_group | GCA_016867345.1 | Candidatus_Kaiserbacteria_bacterium | FJY94_06720 | 8221..8881 | 2030810 | ATP-binding-protein-[Candidatus-Kaiserbacteria-bacterium] |
| PIV20448.1 | DUF499_1.RM.like.systems | MazE_antitoxin->PIN->?->METHYLASE->PolB->HEPN->DUF499*-> | 921 | Bacteria>delta/epsilon_subdivisions>Deltaproteobacteria>unclassified_Deltaproteobacteria | GCA_002780715.1 | Deltaproteobacteria_bacterium_CG03_land_8_20_14_0_80_45_14 | CDS40_12540 | 8362..11127 | 1973969 | AAA-family-ATPase-[Deltaproteobacteria-bacterium-CG03_land_8_20_14_0_80_45_14] |
| PKN69055.1 | DUF499_1.RM.like.systems | <-HKD-DNase<-?<-?<-?<-?<-?<-DUF499*<-METHYLASE<-TUDOR+SF1-SF2-Helicase+iREase+CTD+REase<-HEPN<-REase | 913 | Bacteria>delta/epsilon_subdivisions>Deltaproteobacteria>unclassified_Deltaproteobacteria | GCA_002840565.1 | Deltaproteobacteria_bacterium_HGW-Deltaproteobacteria-12 | CVU54_12145 | 60436..63177 | 2013739 | hypothetical-protein-CVU54_12145-[Deltaproteobacteria-bacterium-HGW-Deltaproteobacteria-12] |
| MBI5244845.1 | DUF499_1.RM.like.systems | <-DUF499*<-HEPN<-PolB<-METHYLASE | 944 | Bacteria>Elusimicrobiota>unclassified_Elusimicrobiota | GCA_016212885.1 | Elusimicrobiota_bacterium | HY922_14340 | 21906..24740 | 2030800 | ATP-binding-protein-[Elusimicrobia-bacterium] |
| MCS2505372.1 | DUF499_1.RM.like.systems | TUDOR+SF1-SF2-Helicase+iREase+CTD+REase->TUDOR+SF1-SF2-Helicase+iREase+CTD+REase-p->METHYLASE-p->METHYLASE-p->METHYLASE-p->HEPN-p->HEPN+DUF499*->DUF499-><-?<-HicB<-TOPRIM<-SbcC\|\|BirA-HTH-> | 434 | Bacteria>FCB_group>Bacteroidota/Chlorobiota_group>Bacteroidota>Bacteroidia | GCA_024788865.1 | Phocaeicola_vulgatus | NXW48_05305 | 1159689..1160993 | 821 | DUF499-domain-containing-protein-[Phocaeicola-vulgatus] |
| MCS2505373.1 | DUF499_1.RM.like.systems | TUDOR+SF1-SF2-Helicase+iREase+CTD+REase-p->METHYLASE-p->METHYLASE-p->METHYLASE-p->HEPN-p->HEPN+DUF499->DUF499*-><-?<-HicB<-TOPRIM<-SbcC\|\|BirA-HTH-><-Phage_int_SAM_5+Phage_integrase | 640 | Bacteria>FCB_group>Bacteroidota/Chlorobiota_group>Bacteroidota>Bacteroidia | GCA_024788865.1 | Phocaeicola_vulgatus | NXW48_05310 | 1160902..1162824 | 821 | ATP-binding-protein-[Phocaeicola-vulgatus] |
| MCS2524521.1 | DUF499_1.RM.like.systems | Peptidase-><-?<-?<-?<-DUF499-p<-DUF499*<-HEPN-p<-METHYLASE-p<-METHYLASE<-TUDOR+SF1-SF2-Helicase+iREase+CTD+REase<-TUDOR+SF1-SF2-Helicase+iREase+CTD+REase-p | 704 | Bacteria>FCB_group>Bacteroidota/Chlorobiota_group>Bacteroidota>Bacteroidia | GCA_024789005.1 | Bacteroides_ovatus | NXV02_14125 | 3438684..3440798 | 28116 | DUF499-domain-containing-protein-[Bacteroides-ovatus] |
| MBK9661552.1 | DUF499_1.RM.like.systems | <-Integrase_1+Phage_integrase\|\|?->DUF488-1-><-?<-DUF499*<-wHTH+REase<-METHYLASE<-HEPN<-TUDOR+SF1-SF2-Helicase+iREase+CTD+REase<-HTH+WYL\|\|RnaseHII-> | 943 | Bacteria>FCB_group>Bacteroidota/Chlorobiota_group>Bacteroidota>Chitinophagia | GCA_016717645.1 | Chitinophagaceae_bacterium | IPO68_16915 | 5414..8245 | 1869212 | ATP-binding-protein-[Chitinophagaceae-bacterium] |
| MCR4416749.1 | DUF499_1.RM.like.systems | Cas_Cmr3->?->?->?->?->?-><-DUF499*<-HEPN<-PolB<-METHYLASE<-TUDOR+SF1-SF2-Helicase+iREase+CTD+REase<-tRNA\|\|?->Peptidase_M16+Peptidase_M16_C-> | 847 | Bacteria>FCB_group>Bacteroidota/Chlorobiota_group>Ignavibacteriota>Ignavibacteria | GCA_024653255.1 | Ignavibacteria_bacterium | NUV92_03270 | 797868..800411 | 2053306 | DUF499-domain-containing-protein-[Ignavibacteria-bacterium] |
| MCW8961935.1 | DUF499_1.RM.like.systems | METHYLASE-p->HEPN->DUF499*-> | 543 | Bacteria>FCB_group>Bacteroidota/Chlorobiota_group>Ignavibacteriota>Ignavibacteria | GCA_026196095.1 | Ignavibacteriaceae_bacterium | OQK29_10260 | 1268..2896 | 2651163 | MAG:-DUF499-domain-containing-protein,-partial-[Ignavibacteriaceae-bacterium] |
| NLH93351.1 | DUF499_1.RM.like.systems | HEPN->TUDOR+SF1-SF2-Helicase+iREase+CTD+REase->ZDNA-HTH->METHYLASE->DUF499*->?->SF1-SF2-Helicase-><-DUF772+DDE_Tnp_1 | 945 | Bacteria>FCB_group>Candidatus_Cloacimonadota>unclassified_Candidatus_Cloacimonadota | GCA_012516855.1 | Candidatus_Cloacimonadota_bacterium | GX466_03915 | 18260..21097 | 2030808 | ATP-binding-protein-[Candidatus-Cloacimonetes-bacterium] |
| MBW7864504.1 | DUF499_1.RM.like.systems | PolB->HEPN->HEPN->METHYLASE->?->?->DUF499*->?->?->XIS-HTH->?->Primpol+PriCT1->HTH-> | 944 | Bacteria>FCB_group>Candidatus_Hydrogenedentes>unclassified_Candidatus_Hydrogenedentes | GCA_019455225.1 | Candidatus_Hydrogenedentes_bacterium | H3C30_08835 | 18999..21833 | 2030809 | ATP-binding-protein-[Candidatus-Hydrogenedentes-bacterium] |
| MBR2089170.1 | DUF499_1.RM.like.systems | <-DUF4417<-PIN<-HTH+MPTase<-?<-?<-HEPN<-DUF499*<-METHYLASE<-TUDOR+SF1-SF2-Helicase+iREase+CTD+REase-p | 933 | Bacteria>FCB_group>Fibrobacterota>Fibrobacteria>Fibrobacterales | - | Fibrobacter_sp. | IJ905_02095 | 11314..14115 | 35828 | ATP-binding-protein-[Fibrobacter-sp.] |
| MXW55543.1 | DUF499_1.RM.like.systems | <-DUF499*<-HEPN-p<-METHYLASE-p | 977 | Bacteria>FCB_group>Gemmatimonadota>Gemmatimonadetes>Gemmatimonadales | GCA_009836275.1 | Gemmatimonadales_bacterium | F4Z59_02325 | 190..3123 | 2448054 | ATP-binding-protein-[Gemmatimonadales-bacterium] |
| MYA34704.1 | DUF499_1.RM.like.systems | HEPN->TUDOR+SF1-SF2-Helicase+iREase+CTD+REase->METHYLASE->OLD-ABC->TOPRIM->ParB+HNH+Schlafen->DUF499*-><-?\|\|HTH->MPTase->?->SMODS-><-ABhydrolase | 928 | Bacteria>FCB_group>Gemmatimonadota>Gemmatimonadetes>Gemmatimonadales | GCA_009838765.1 | Gemmatimonadales_bacterium | F4Y33_13800 | 70561..73347 | 2448054 | ATP-binding-protein-[Gemmatimonadales-bacterium] |
| MBA4158336.1 | DUF499_1.RM.like.systems | HEPN->DUF499*->SF1-SF2-Helicase+wHTH+PRTase->?->?->XIS-HTH-> | 907 | Bacteria>FCB_group>Gemmatimonadota>unclassified_Gemmatimonadota | GCA_013820915.1 | Gemmatimonadota_bacterium | HOX65_12770 | 579..3302 | 2026742 | ATP-binding-protein-[Gemmatimonadetes-bacterium] |
| RLC09317.1 | DUF499_1.RM.like.systems | <-DUF499*<-REase<-HEPN<-METHYLASE-p | 517 | Bacteria>Myxococcota>Myxococcia>unclassified_Myxococcia | GCA_003647435.1 | Deltaproteobacteria_bacterium | DRH43_08475 | 1..1550 | 2026735 | AAA-family-ATPase,-partial-[Deltaproteobacteria-bacterium] |
| MBC8458333.1 | DUF499_1.RM.like.systems | METHYLASE->PolB->HEPN->DUF499*-> | 802 | Bacteria>Myxococcota>Myxococcia>unclassified_Myxococcia | GCA_014383825.1 | Deltaproteobacteria_bacterium | H8D67_10095 | 2315..4719 | 2026735 | ATP-binding-protein-[Deltaproteobacteria-bacterium] |
| MBC8460283.1 | DUF499_1.RM.like.systems | HEPN+REase->METHYLASE->DUF499*-> | 926 | Bacteria>Myxococcota>Myxococcia>unclassified_Myxococcia | GCA_014383825.1 | Deltaproteobacteria_bacterium | H8D67_20045 | 4060..6840 | 2026735 | ATP-binding-protein-[Deltaproteobacteria-bacterium] |
| MBW2135777.1 | DUF499_1.RM.like.systems | METHYLASE-p->PolB->HEPN->DUF499*-> | 916 | Bacteria>Myxococcota>Myxococcia>unclassified_Myxococcia | GCA_019309545.1 | Deltaproteobacteria_bacterium | JRG72_11235 | 2815..5565 | 2026735 | ATP-binding-protein-[Deltaproteobacteria-bacterium] |
| MBW2309256.1 | DUF499_1.RM.like.systems | METHYLASE-p->HEPN->DUF499*-> | 920 | Bacteria>Myxococcota>Myxococcia>unclassified_Myxococcia | GCA_019310535.1 | Deltaproteobacteria_bacterium | JRG73_20230 | 2438..5200 | 2026735 | ATP-binding-protein-[Deltaproteobacteria-bacterium] |
| GIW47700.1 | DUF499_1.RM.like.systems | <-HEPN<-PolB\|\|?-><-DUF499*<-HEPN<-PolB<-METHYLASE<-Schlafen+HATPase+HTH<-TUDOR+SF1-SF2-Helicase+iREase+CTD+REase | 857 | Bacteria>Myxococcota>Myxococcia>unclassified_Myxococcia | GCA_026004235.1 | Deltaproteobacteria_bacterium | KatS3mg078_1577 | 289556..292129 | 2026735 | MAG:-hypothetical-protein-KatS3mg078_1577-[Deltaproteobacteria-bacterium] |
| MDD9873633.1 | DUF499_1.RM.like.systems | TUDOR+SF1-SF2-Helicase+iREase+CTD+REase->HEPN->METHYLASE->OLD-ABC->TOPRIM->DUF499*->?->?->?->?->ABC-ATPase+SbcC-> | 1106 | Bacteria>Myxococcota>Myxococcia>unclassified_Myxococcia | GCA_028817295.1 | Deltaproteobacteria_bacterium | OXU65_09090 | 9437..12757 | 2026735 | MAG:-DUF499-domain-containing-protein-[Deltaproteobacteria-bacterium] |

| Protein-ID | System | Gene-Neighbourhoods | Length | Taxonomy | Assembly-ID | Strain | Locus-Tag | Genomic-coordinates | TaxID | Fasta-defline |
|---|---|---|---|---|---|---|---|---|---|---|
| NGZ60735.1 | DUF499_1.RM.like.systems | <-DUF499*<-HEPN<-PolB<-METHYLASE-p | 916 | Bacteria>Nitrospirota>Nitrospiria>Nitrospirales>Nitrospiraceae | GCA_011090395.1 | Nitrospira_sp._LK265 | CV081_09575 | 239..2989 | 2039810 | AAA-family-ATPase-[Nitrospira-sp.-LK265] |
| MBS0175039.1 | DUF499_1.RM.like.systems | HTH+WYL->TUDOR+ SF1-SF2-Helicase+iREase+CTD+REase->HEPN->METHYLASE->DUF499*->?->?->?-><-Phage_lysozyme | 941 | Bacteria>Nitrospirota>Nitrospiria>Nitrospirales>Nitrospiraceae | GCA_018242785.1 | Nitrospira_sp. | JSR64_13460 | 8684..11509 | 70125 | ATP-binding-protein-[Nitrospira-sp.] |
| MCP9448661.1 | DUF499_1.RM.like.systems | sGTP->?->?->?->?->?-><-DUF499*<-HEPN<-ORC-CDC6+REase<-METHYLASE-p\|\|?-><-Radical-SAM | 987 | Bacteria>Nitrospirota>Nitrospiria>Nitrospirales>Nitrospiraceae | GCA_024331085.1 | Nitrospira_sp. | NNA21_01185 | 6072..9035 | 70125 | DUF499-domain-containing-protein-[Nitrospira-sp.] |
| MCP9469842.1 | DUF499_1.RM.like.systems | <-Glyoxalase\|\|?->HTH->?-><-DUF499*<-HEPN<-PolB<-METHYLASE<-?<-TUDOR+ SF1-SF2-Helicase+iREase+CTD+REase | 919 | Bacteria>Nitrospirota>Nitrospiria>Nitrospirales>Nitrospiraceae | GCA_024331125.1 | Nitrospira_sp. | NNA31_07565 | 179179..181938 | 70125 | DUF499-domain-containing-protein-[Nitrospira-sp.] |
| WP_059175576.1 | DUF499_1.RM.like.systems | REase->TUDOR+ SF1-SF2-Helicase+iREase+CTD+REase->METHYLASE->PolB->HEPN->METHYLASE-p->DUF499*-><-?<-?\|\|SAM-methylase-> | 842 | Bacteria>Nitrospirota>Thermodesulfovibrionia>Thermodesulfovibrionales>Thermodesulfovibrionaceae | GCF_001514535.1 | Thermodesulfovibrio_aggregans | TAGGR_RS01355 | 271390..273918 | 86166 | DUF499-domain-containing-protein-[Thermodesulfovibrio-aggregans] |
| MBF0565077.1 | DUF499_1.RM.like.systems | <-DUF499*<-NTP_transf_2<-HEPN<-METHYLASE | 1116 | Bacteria>Nitrospirota>unclassified_Nitrospirota | GCA_015233615.1 | Nitrospirota_bacterium | HQK89_07535 | 20523..23873 | 2026887 | ATP-binding-protein-[Nitrospirae-bacterium] |
| MBI5634975.1 | DUF499_1.RM.like.systems | TUDOR+ SF1-SF2-Helicase+iREase+CTD+REase->?->METHYLASE->HEPN->DUF499*->HTH+MPTase->?->?-><-Schlafen+HATPase+HTH<-MqsA_antitoxin+HTH_3\|\|REase-> | 927 | Bacteria>Nitrospirota>unclassified_Nitrospirota | GCA_016217675.1 | Nitrospirota_bacterium | HZA15_16020 | 27832..30615 | 2026887 | ATP-binding-protein-[Nitrospirae-bacterium] |
| MBI5740568.1 | DUF499_1.RM.like.systems | SbcC->METHYLASE->PolB->HEPN->DUF499*->?->?->?-><-Peptidase_U32+Peptidase_U32_C | 919 | Bacteria>Nitrospirota>unclassified_Nitrospirota | GCA_016234115.1 | Nitrospirota_bacterium | HZA16_07595 | 59463..62222 | 2026887 | ATP-binding-protein-[Nitrospirae-bacterium] |
| WP_244532343.1 | DUF499_1.RM.like.systems | METHYLASE->HEPN->DUF499*-><-?\|\|MODE-HTH+PBPII->\|\|?->?->HTH_24->ZnR-> | 752 | Bacteria>Pseudomonadota>Alphaproteobacteria>Hyphomicrobiales>Beijerinckiaceae | GCF_900114285.1 | Methylocapsa_palsarum | BM177_RS13845 | 3654..5912 | 1612308 | DUF499-domain-containing-protein-[Methylocapsa-palsarum] |
| MCW5716290.1 | DUF499_1.RM.like.systems | Peptidase_M15->?-><-HEPN<-DUF499*<-METHYLASE<-SF1-SF2-Helicase<-TUDOR+ SF1-SF2-Helicase+iREase+CTD+REase<-XIS-HTH<-GDResolvase+Recombinase | 1108 | Bacteria>Pseudomonadota>Alphaproteobacteria>Hyphomicrobiales>Kaistiaceae | GCA_026126085.1 | Bauldia_sp. | KIS68_00515 | 97641..100967 | 2575872 | MAG:-ATP-binding-protein-[Bauldia-sp.] |
| WP_280515578.1 | DUF499_1.RM.like.systems | TUDOR+ SF1-SF2-Helicase+iREase+CTD+REase-p-><-HTH_Tnp_1+HTH_21+rve+rve_3\|\|?->?->METHYLASE-p->METHYLASE->HEPN+DUF499*->DUF499->ParB+HTH->HTH_3->ParB+HNH-p->ParB+HNH-p->URI-DNase-> | 427 | Bacteria>Pseudomonadota>Alphaproteobacteria>Hyphomicrobiales>Phyllobacteriaceae | GCF_013240295.1 | Mesorhizobium_sediminum | HK436_RS32250 | 843839..845122 | 1849104 | Swt1-family-HEPN-domain-containing-protein-[Mesorhizobium-sediminum] |
| WP_280515628.1 | DUF499_1.RM.like.systems | <-HTH_Tnp_1+HTH_21+rve+rve_3\|\|?->?->METHYLASE-p->METHYLASE->HEPN+DUF499->DUF499*->ParB+HTH->HTH_3->ParB+HNH-p->ParB+HNH-p->URI-DNase-> | 678 | Bacteria>Pseudomonadota>Alphaproteobacteria>Hyphomicrobiales>Phyllobacteriaceae | - | Mesorhizobium_sediminum | HK436_RS32255 | 845119..847155 | 1849104 | DUF499-domain-containing-protein-[Mesorhizobium-sediminum] |
| HCX68724.1 | DUF499_1.RM.like.systems | TUDOR+ SF1-SF2-Helicase+iREase+CTD+REase->?->METHYLASE->HEPN->DUF499*-> | 525 | Bacteria>Pseudomonadota>Alphaproteobacteria>Hyphomicrobiales>Rhodobiaceae | GCA_003538155.1 | Rhodobiaceae_bacterium | DHK64_14660 | 8558..10133 | 2026785 | AAA-family-ATPase-[Rhodobiaceae-bacterium] |
| MBF0185782.1 | DUF499_1.RM.like.systems | REase+35exo+Beta-Barrel+SF1-SF2-Helicase->TUDOR+ SF1-SF2-Helicase+iREase+CTD+REase->HEPN->METHYLASE->?->PIN->DUF499*-> | 943 | Bacteria>Pseudomonadota>Alphaproteobacteria>Magnetococcales>unclassified_Magnetococcales | GCA_015231215.1 | Magnetococcales_bacterium | HQM06_15545 | 19234..22065 | 2026759 | ATP-binding-protein-[Magnetococcales-bacterium] |
| MBF0340191.1 | DUF499_1.RM.like.systems | <-DUF499*<-PIN<-?<-HEPN<-TUDOR+ SF1-SF2-Helicase+iREase+CTD+REase<-HTH+WYL\|\|REase+SAVED-> | 705 | Bacteria>Pseudomonadota>Alphaproteobacteria>Magnetococcales>unclassified_Magnetococcales | GCA_015232245.1 | Magnetococcales_bacterium | HQL95_04415 | 72..2189 | 2026759 | ATP-binding-protein-[Magnetococcales-bacterium] |
| WP_013654352.1 | DUF499_1.RM.like.systems | <-PIN<-XIS-HTH\|\|?-><-ArdCN+MPTase<-HEPN<-DUF499*<-METHYLASE<-HNH<-OLD-ABC<-TUDOR+ SF1-SF2-Helicase+iREase+CTD+REase<-HTH_3<-HigB | 944 | Bacteria>Pseudomonadota>Alphaproteobacteria>Rhodobacterales>Paracoccaceae | GCF_000192745.1 | Polymorphum_gilvum | SL003B_RS18265 | 3920112..3922946 | 991904 | DUF499-domain-containing-protein-[Polymorphum-gilvum] |
| MBT3398604.1 | DUF499_1.RM.like.systems | REase->METHYLASE->DUF499*->HEPN->Methylase-><-Radical-SAM<-?<-Peptidase_M15_4 | 922 | Bacteria>Pseudomonadota>Alphaproteobacteria>Rhodospirillales>Rhodospirillaceae | GCA_018648085.1 | Rhodospirillaceae_bacterium | HN420_01275 | 4129..6897 | 1898112 | ATP-binding-protein-[Rhodospirillaceae-bacterium] |
| NWG45252.1 | DUF499_1.RM.like.systems | Peptidase_M15->?->?-><-?<-HEPN<-DUF499*<-METHYLASE-p | 1108 | Bacteria>Pseudomonadota>Alphaproteobacteria>unclassified_Alphaproteobacteria | GCA_013389075.1 | Alphaproteobacteria_bacterium | HXY25_01725 | 2157..5483 | 1913988 | ATP-binding-protein-[Alphaproteobacteria-bacterium] |
| QKS30678.1 | DUF499_1.RM.like.systems | <-MerR-HTH\|\|TUDOR+ SF1-SF2-Helicase+iREase+CTD+REase->?->?->METHYLASE->DUF499*->HEPN+SF1-SF2-Helicase->MORC+DUF3684+HNH->?-><-?<-TraH<-Thioredoxin | 1092 | Bacteria>Pseudomonadota>Betaproteobacteria>Betaproteobacteria_incertae_sedis>Candidatus_Accumulibacter | GCA_013347225.1 | Candidatus_Accumulibacter_similis | HT579_18190 | 4129453..4132731 | 2954388 | MAG:-ATP-binding-protein-[Candidatus-Accumulibacter-similis] |
| MBL8396550.1 | DUF499_1.RM.like.systems | HTH+WYL->TUDOR+ SF1-SF2-Helicase+iREase+CTD+REase->HEPN->METHYLASE->wHTH+REase->DUF499*->DDE_5+DDE_Tnp_1->?-><-MORC+DNA_gyraseB+TOPRIM | 939 | Bacteria>Pseudomonadota>Betaproteobacteria>Betaproteobacteria_incertae_sedis>Candidatus_Accumulibacter | GCA_016789495.1 | Accumulibacter_sp. | JNL84_00165 | 43696..46515 | 2053492 | ATP-binding-protein-[Candidatus-Accumulibacter-sp.] |
| MCM8600113.1 | DUF499_1.RM.like.systems | <-SbcC<-DUF499*<-Schlafen+HATPase+HTH<-METHYLASE<-METHYLASE<-HEPN<-TUDOR+ SF1-SF2-Helicase+iREase+CTD+REase<-HTH_Tnp | 949 | Bacteria>Pseudomonadota>Betaproteobacteria>Betaproteobacteria_incertae_sedis>Candidatus_Accumulibacter | GCA_023806585.1 | Accumulibacter_sp. | NFW04_15895 | 29772..32621 | 2053492 | DUF499-domain-containing-protein-[Candidatus-Accumulibacter-sp.] |
| WP_033446622.1 | DUF499_1.RM.like.systems | TetR-HTH->?->MarR-HTH->Phage_int_SAM_2+Integrase_1-><-DUF499*<-Schlafen+HATPase+HTH<-METHYLASE<-METHYLASE<-HEPN<-TUDOR+ SF1-SF2-Helicase+iREase+CTD+REase<-HTH+WYL | 947 | Bacteria>Pseudomonadota>Betaproteobacteria>Burkholderiales>Alcaligenaceae | GCF_000690475.1 | Bordetella_bronchiseptica | EL288_RS19650 | 4193914..4196757 | 518 | DUF499-domain-containing-protein-[Bordetella-bronchiseptica] |
| TAL53973.1 | DUF499_1.RM.like.systems | HTH->STYKinase-like-><-DUF2357+REase<-Preader7+McrB-NTD+McrB-AAA\|\|HTH->STYKinase-like-><-DUF499*<-METHYLASE<-HEPN<-HEPN<-TUDOR+ SF1-SF2-Helicase+iREase+CTD+REase | 966 | Bacteria>Pseudomonadota>Betaproteobacteria>Burkholderiales>Burkholderiaceae | GCA_004297875.1 | Pandoraea_sp. | EPN80_13305 | 57479..60379 | 1883445 | MAG:-ATP-binding-protein-[Pandoraea-sp.] |
| MBX9834804.1 | DUF499_1.RM.like.systems | METHYLASE->HEPN->DUF499*->?->?-><-SAM-methylase | 945 | Bacteria>Pseudomonadota>Betaproteobacteria>Burkholderiales>Burkholderiaceae | GCA_019746655.1 | Burkholderiaceae_bacterium | K2X65_01340 | 3559..6396 | 2030806 | DUF499-domain-containing-protein-[Burkholderiaceae-bacterium] |
| WP_028211840.1 | DUF499_1.RM.like.systems | <-DUF499*<-METHYLASE<-TUDOR+ SF1-SF2-Helicase+iREase+CTD+REase\|\|HEPN-p-> | 939 | Bacteria>Pseudomonadota>Betaproteobacteria>Burkholderiales>Burkholderiaceae | GCF_000472845.1 | Paraburkholderia_mimosarum | A19S_RS0115645 | 1157..3976 | 312026 | DUF499-domain-containing-protein-[Paraburkholderia-mimosarum] |
| WP_065497253.1 | DUF499_1.RM.like.systems | TUDOR+ SF1-SF2-Helicase+iREase+CTD+REase-p->TUDOR+ SF1-SF2-Helicase+iREase+CTD+REase->PIN->METHYLASE->METHYLASE->Schlafen+HATPase+HTH->DUF499*->HEPN-> | 955 | Bacteria>Pseudomonadota>Betaproteobacteria>Burkholderiales>Burkholderiaceae | GCF_001685525.1 | Burkholderia_sp._CCA53 | BCR55_RS29355 | 26270..29137 | 1776288 | DUF499-domain-containing-protein-[Burkholderia-sp.-CCA53] |
| WP_226267509.1 | DUF499_1.RM.like.systems | <-MAD-NTD+HNH<-?\|\|HTH_3-><-Peptidase_C70<-?<-DUF499*<-METHYLASE<-HEPN<-TUDOR+ SF1-SF2-Helicase+iREase+CTD+REase<-HTH+WYL\|\|REase->Resolvase-> | 943 | Bacteria>Pseudomonadota>Betaproteobacteria>Burkholderiales>Burkholderiaceae | GCF_020419455.1 | Burkholderia_arboris | LGM95_RS31475 | 204448..207279 | 488730 | DUF499-domain-containing-protein-[Burkholderia-arboris] |
| WP_150698381.1 | DUF499_1.RM.like.systems | HTH->GDResolvase+HIN-HTH->URI-DNase-><-DUF499*<-METHYLASE<-HEPN<-TUDOR+ SF1-SF2-Helicase+iREase+CTD+REase | 944 | Bacteria>Pseudomonadota>Betaproteobacteria>Burkholderiales>Burkholderiaceae | GCF_902459695.1 | Pandoraea_terrae | PTE30175_RS17675 | 118123..120957 | 1537710 | DUF499-domain-containing-protein-[Pandoraea-terrae] |
| WP_186144236.1 | DUF499_1.RM.like.systems | HTH+WYL->TIR+PNTD2->PIWI->TUDOR+ SF1-SF2-Helicase+iREase+CTD+REase->HEPN->METHYLASE->DUF499*->REase-><-tRNA\|\|?->?-><-?\|\|MODE-HTH+PBPII-><-TOPRIM+ZnR | 955 | Bacteria>Pseudomonadota>Betaproteobacteria>Burkholderiales>Burkholderiaceae | GCF_902831945.1 | Burkholderia_gladioli | H9E34_RS32225 | 51067..53934 | 28095 | DUF499-domain-containing-protein-[Burkholderia-gladioli] |
| WP_102632169.1 | DUF499_1.RM.like.systems | HTH-><-Peptidase_C70<-?<-DUF499*<-METHYLASE<-HEPN<-TUDOR+ SF1-SF2-Helicase+iREase+CTD+REase<-HTH+WYL\|\|Resolvase-><-Peptidase_S8 | 943 | Bacteria>Pseudomonadota>Betaproteobacteria>Burkholderiales>Burkholderiaceae | GCF_902859775.1 | Paraburkholderia_rhynchosiae | HV734_RS08115 | 141591..144422 | 487049 | DUF499-domain-containing-protein-[Paraburkholderia-rhynchosiae] |
| WP_211950277.1 | DUF499_1.RM.like.systems | <-STYKinase-like\|\|HTH+WYL->TUDOR+ SF1-SF2-Helicase+iREase+CTD+REase->HEPN->METHYLASE->DUF499*->?->HEPN+CBS->?-><-?<-?<-Phage_int_SAM_1+Phage_integrase | 966 | Bacteria>Pseudomonadota>Betaproteobacteria>Burkholderiales>Burkholderiaceae | GCF_905397275.1 | Cupriavidus_yeoncheonensis | KHO22_RS27195 | 69520..72420 | 1462994 | DUF499-domain-containing-protein-[Cupriavidus-yeoncheonensis] |
| WP_143969665.1 | DUF499_1.RM.like.systems | HTH+WYL->TUDOR+ SF1-SF2-Helicase+iREase+CTD+REase->PIN->METHYLASE->OLD-ABC->TOPRIM->DUF499*->HEPN-> | 952 | Bacteria>Pseudomonadota>Betaproteobacteria>Burkholderiales>Burkholderiales_genera_incertae_sedis | GCF_007556755.1 | Tepidimonas_fonticaldi | Tfont_RS12360 | 12901..15759 | 1101373 | DUF499-domain-containing-protein-[Tepidimonas-fonticaldi] |
| NMM11921.1 | DUF499_1.RM.like.systems | <-HEPN<-REase<-?<-?<-DUF499*<-METHYLASE<-HEPN<-TUDOR+ SF1-SF2-Helicase+iREase+CTD+REase<-HTH+WYL\|\|STYKinase-like-> | 961 | Bacteria>Pseudomonadota>Betaproteobacteria>Burkholderiales>Comamonadaceae | GCA_012927445.1 | Rhodoferax_sp. | HHJ17_00055 | 10474..13359 | 50421 | ATP-binding-protein-[Rhodoferax-sp.] |
| NMM07407.1 | DUF499_1.RM.like.systems | TUDOR+ SF1-SF2-Helicase+iREase+CTD+REase->HEPN->METHYLASE->DUF499*-> | 948 | Bacteria>Pseudomonadota>Betaproteobacteria>Burkholderiales>Comamonadaceae | GCA_012927455.1 | Polaromonas_sp. | HHJ18_14460 | 504043..506889 | 1869339 | ATP-binding-protein-[Polaromonas-sp.] |
| MBX3657558.1 | DUF499_1.RM.like.systems | HTH+WYL->TUDOR+ SF1-SF2-Helicase+iREase+CTD+REase->HEPN->METHYLASE->?->?->DUF499*->SbcC->HIN-HTH->?-><-?<-Phage_lysozyme<-Phage_holin | 951 | Bacteria>Pseudomonadota>Betaproteobacteria>Burkholderiales>Comamonadaceae | GCA_019635415.1 | Ramlibacter_sp. | KF740_03900 | 804820..807675 | 1917967 | ATP-binding-protein-[Ramlibacter-sp.] |
| WP_012839779.1 | DUF499_1.RM.like.systems | <-MORC+HisKin<-REase<-?<-DUF499*<-METHYLASE<-HEPN<-TUDOR+ SF1-SF2-Helicase+iREase+CTD+REase<-HTH+WYL | 955 | Bacteria>Pseudomonadota>Betaproteobacteria>Burkholderiales>Comamonadaceae | GCF_000093145.3 | Comamonas_thiooxydans | HNB49_RS10950 | 10966..13833 | 363952 | DUF499-domain-containing-protein-[Comamonas-thiooxydans] |
| WP_042427102.1 | DUF499_1.RM.like.systems | Phage_lysozyme->?->HTH_3->Radical-SAM-><-?<-TIR<-DUF499*<-?<-?<-METHYLASE<-HEPN<-PIN<-TUDOR+ SF1-SF2-Helicase+iREase+CTD+REase | 951 | Bacteria>Pseudomonadota>Betaproteobacteria>Burkholderiales>Comamonadaceae | GCF_000739995.1 | Comamonas_granuli | CGR01S_RS11705 | 67732..70587 | 290309 | DUF499-domain-containing-protein-[Comamonas-granuli] |
| WP_045536717.1 | DUF499_1.RM.like.systems | TUDOR+ SF1-SF2-Helicase+iREase+CTD+REase->PIN->HEPN->METHYLASE-p->METHYLASE->ParB+HNH->DUF499*-> | 950 | Bacteria>Pseudomonadota>Betaproteobacteria>Burkholderiales>Comamonadaceae | GCF_000828915.1 | Serpentinimonas_maccroryi | SMCB_RS09950 | 2162226..2165078 | 1458426 | DUF499-domain-containing-protein-[Serpentinimonas-maccroryi] |
| WP_095541484.1 | DUF499_1.RM.like.systems | METHYLASE-p->?->HAD->Arm-DNA-bind_3->?-><-DUF499*<-ParB+HNH<-METHYLASE<-HEPN<-TUDOR+ SF1-SF2-Helicase+iREase+CTD+REase<-HTH_Tnp<-HTH+WYL | 948 | Bacteria>Pseudomonadota>Betaproteobacteria>Burkholderiales>Comamonadaceae | GCF_002285255.1 | Vandammella_animalimorsus | CLI93_RS07370 | 83443..86289 | 2029117 | DUF499-domain-containing-protein-[Vandammella-animalimorsus] |
| WP_127807271.1 | DUF499_1.RM.like.systems | TUDOR+ SF1-SF2-Helicase+iREase+CTD+REase->PIN->HEPN->METHYLASE->?->?->DUF499*->TIR->?-><-Radical-SAM<-HTH_3<-?<-Phage_lysozyme | 950 | Bacteria>Pseudomonadota>Betaproteobacteria>Burkholderiales>Comamonadaceae | GCF_004006415.1 | Hydrogenophaga_sp._NH-16 | DD542_RS03485 | 729405..732257 | 2184519 | DUF499-domain-containing-protein-[Hydrogenophaga-sp.-NH-16] |
| WP_210930504.1 | DUF499_1.RM.like.systems | HTH->?->Phage_integrase->STYKinase-like-><-?\|\|DDE_Tnp_1_5+DDE_Tnp_1-><-DUF499*<-REase<-OLD-ABC<-HEPN<-METHYLASE<-HNH<-OLD-ABC | 945 | Bacteria>Pseudomonadota>Betaproteobacteria>Burkholderiales>Comamonadaceae | GCF_018064205.1 | Comamonas_aquatica | KAQ61_RS03430 | 737208..740045 | 225991 | DUF499-domain-containing-protein-[Comamonas-aquatica] |
| WP_265282594.1 | DUF499_1.RM.like.systems | TUDOR+ SF1-SF2-Helicase+iREase+CTD+REase->HEPN->METHYLASE->DUF499*->OLD-ABC+TOPRIM->?->?-><-?<-?<-Peptidase_C26 | 951 | Bacteria>Pseudomonadota>Betaproteobacteria>Burkholderiales>Comamonadaceae | GCF_026127365.1 | Verminephrobacter_aporrectodeae | D5039_RS14590 | 1238377..1241232 | 1110389 | DUF499-domain-containing-protein-[Verminephrobacter-aporrectodeae] |
| WP_265654212.1 | DUF499_1.RM.like.systems | Peptidase_C26->?->?-><-?<-OLD-ABC+TOPRIM<-DUF499*<-METHYLASE<-HEPN<-TUDOR+ SF1-SF2-Helicase+iREase+CTD+REase | 951 | Bacteria>Pseudomonadota>Betaproteobacteria>Burkholderiales>Comamonadaceae | GCF_026192715.1 | Verminephrobacter_aporrectodeae | D8B27_RS10360 | 21500..24355 | 1110389 | DUF499-domain-containing-protein-[Verminephrobacter-aporrectodeae] |
| WP_088958124.1 | DUF499_1.RM.like.systems | ParB->HTH+WYL->HTH_Tnp->TUDOR+ SF1-SF2-Helicase+iREase+CTD+REase->HEPN->METHYLASE->DUF499*->TIR->SLATT->SMODS->Methylase-><-Radical-SAM<-HTH_3 | 951 | Bacteria>Pseudomonadota>Betaproteobacteria>Burkholderiales>Comamonadaceae | GCF_900090195.1 | Variovorax_sp._HW608 | VAR608DRAFT_RS34345 | 7257646..7260501 | 1034889 | DUF499-domain-containing-protein-[Variovorax-sp.-HW608] |
| WP_206111008.1 | DUF499_1.RM.like.systems | HTH+WYL->HTH_Tnp->TUDOR+ SF1-SF2-Helicase+iREase+CTD+REase->PIN->HEPN->METHYLASE->DUF499*->TIR->SLATT->SMODS->Methylase-><-Radical-SAM<-Phage_lysozyme | 955 | Bacteria>Pseudomonadota>Betaproteobacteria>Burkholderiales | GCF_017150735.1 | Burkholderiaceae | KTD10_RS22775 | 57014..59881 | 119060 | MULTISPECIES:-DUF499-domain-containing-protein-[Burkholderiaceae] |
| UUZ52278.1 | DUF499_1.RM.like.systems | SF1-SF2-Helicase-p->?-><-REase<-DUF499-p<-DUF499-p<-DUF499*<-METHYLASE-p<-METHYLASE-p<-HEPN<-TUDOR+ SF1-SF2-Helicase+iREase+CTD+REase-p<-TUDOR+ SF1-SF2-Helicase+iREase+CTD+REase<-TUDOR+ SF1-SF2-Helicase+iREase+CTD+REase-p | 325 | Bacteria>Pseudomonadota>Betaproteobacteria>Burkholderiales>Oxalobacteraceae | GCA_024703915.1 | Massilia_sp._H-1 | LP419_21985 | 3096112..3097089 | 2895566 | DUF499-domain-containing-protein-[Massilia-sp.-H-1] |
| UUZ46680.1 | DUF499_1.RM.like.systems | SF1-SF2-Helicase-p->?->?-><-REase<-DUF499-p<-DUF499-p<-DUF499*<-METHYLASE-p<-METHYLASE-p<-METHYLASE-p<-HEPN\|\|?->RNaseP-> | 339 | Bacteria>Pseudomonadota>Betaproteobacteria>Burkholderiales>Oxalobacteraceae | GCA_024703935.1 | Massilia_sp._B-10 | LP420_22520 | 3094423..3095442 | 2895567 | DUF499-domain-containing-protein-[Massilia-sp.-B-10] |
| WP_170204580.1 | DUF499_1.RM.like.systems | <-RVT\|\|HTH_Tnp_1+HTH_21+rve+rve_3-><-?<-?<-DUF499*<-METHYLASE<-HEPN<-TUDOR+ SF1-SF2-Helicase+iREase+CTD+REase\|\|STYKinase-like-> | 956 | Bacteria>Pseudomonadota>Betaproteobacteria>Burkholderiales>Oxalobacteraceae | GCF_012849555.1 | Massilia_forsythiae | HH212_RS22790 | 5438600..5441470 | 2728020 | DUF499-domain-containing-protein-[Massilia-forsythiae] |
| WP_072453399.1 | DUF499_1.RM.like.systems | <-STYKinase-like\|\|HTH+WYL->TUDOR+ SF1-SF2-Helicase+iREase+CTD+REase->HEPN->METHYLASE->DUF499*->REase-> | 961 | Bacteria>Pseudomonadota>Betaproteobacteria>Burkholderiales>Oxalobacteraceae | GCF_900119665.1 | Janthinobacterium_lividum | BT188_RS05025 | 1132446..1135331 | 29581 | DUF499-domain-containing-protein-[Janthinobacterium-lividum] |
| RZL03828.1 | DUF499_1.RM.like.systems | HTH+WYL->TIR+PNTD2->PIWI->TUDOR+ SF1-SF2-Helicase+iREase+CTD+REase->HEPN->METHYLASE->DUF499*->?->KAP-NTPase->?->PAPS_reduct-> | 956 | Bacteria>Pseudomonadota>Betaproteobacteria>Burkholderiales>Sphaerotilaceae | GCA_004211355.1 | Rubrivivax_sp. | EOP36_03025 | 28587..31457 | 50259 | MAG:-ATP-binding-protein-[Rubrivivax-sp.] |
| WP_281915166.1 | DUF499_1.RM.like.systems | Phage_lysozyme-><-HEPN<-DUF499*<-Schlafen+HATPase+HTH<-METHYLASE<-METHYLASE | 955 | Bacteria>Pseudomonadota>Betaproteobacteria>Burkholderiales>Sphaerotilaceae | GCF_937957785.1 | Caldimonas_thermodepolymerans | QMG53_RS04175 | 1459..4326 | 215580 | DUF499-domain-containing-protein-[Caldimonas-thermodepolymerans] |
| MBX9611601.1 | DUF499_1.RM.like.systems | TUDOR+ SF1-SF2-Helicase+iREase+CTD+REase->HEPN->METHYLASE->DUF499*-><-?<-?<-SF1-SF2-Helicase+HRDC | 953 | Bacteria>Pseudomonadota>Betaproteobacteria>Burkholderiales>unclassified_Burkholderiales | GCA_019747995.1 | Burkholderiales_bacterium | K2X51_08260 | 22656..25517 | 1891238 | DUF499-domain-containing-protein-[Burkholderiales-bacterium] |
| MCA3215581.1 | DUF499_1.RM.like.systems | <-MetJarc-HTH\|\|HTH+WYL->HTH_Tnp->TUDOR+ SF1-SF2-Helicase+iREase+CTD+REase->HEPN->METHYLASE->DUF499*->?->Methylase-><-Radical-SAM<-HTH_3 | 948 | Bacteria>Pseudomonadota>Betaproteobacteria>Burkholderiales>unclassified_Burkholderiales | GCA_020248845.1 | Burkholderiales_bacterium | ING39_09185 | 33455..36301 | 1891238 | ATP-binding-protein-[Burkholderiales-bacterium] |
| MCZ2134305.1 | DUF499_1.RM.like.systems | <-RAMA<-?<-?<-HTH_21+rve+rve_3<-HIN-HTH<-?<-DUF499*<-wHTH+REase<-METHYLASE<-HEPN<-PIN<-TUDOR+ SF1-SF2-Helicase+iREase+CTD+REase\|\|HIN-HTH-> | 956 | Bacteria>Pseudomonadota>Betaproteobacteria>Burkholderiales>unclassified_Burkholderiales | GCA_026932525.1 | Burkholderiales_bacterium | LC098_02580 | 42253..45123 | 1891238 | MAG:-DUF499-domain-containing-protein-[Burkholderiales-bacterium] |
| MBP6189859.1 | DUF499_1.RM.like.systems | DUF488-1-><-?<-DUF499*<-wHTH+REase<-METHYLASE<-HEPN | 943 | Bacteria>Pseudomonadota>Betaproteobacteria>Rhodocyclales>Azonexaceae | GCA_017985755.1 | Azonexus_sp. | KA440_12990 | 2046..4877 | 1872668 | ATP-binding-protein-[Azonexus-sp.] |
| WP_169202394.1 | DUF499_1.RM.like.systems | TUDOR+ SF1-SF2-Helicase+iREase+CTD+REase->?-><-PAIREDC-HTH+HIN-HTH\|\|METHYLASE->HEPN->wHTH+REase->DUF499*-><-?<-?<-NADAR<-?<-NUDIX | 941 | Bacteria>Pseudomonadota>Betaproteobacteria>Rhodocyclales>Rhodocyclaceae | GCF_017894365.1 | Aromatoleum_bremense | GPA24_RS09420 | 37549..40374 | 76115 | DUF499-domain-containing-protein-[Aromatoleum-bremense] |
| WP_191658157.1 | DUF499_1.RM.like.systems | <-DUF499*<-wHTH+REase<-HEPN<-METHYLASE<-?<-ParB+HNH<-TUDOR+ SF1-SF2-Helicase+iREase+CTD+REase | 944 | Bacteria>Pseudomonadota>Betaproteobacteria>Rhodocyclales>Zoogloeaceae | - | Azoarcus_sp._Aa7 | AZOA_12030 | 1309..4143 | 2303328 | DUF499-domain-containing-protein-[Azoarcus-sp.-Aa7] |
| MBD5801779.1 | DUF499_1.RM.like.systems | <-DUF499*<-wHTH+REase<-HEPN<-METHYLASE<-?<-ParB+HNH<-TUDOR+ SF1-SF2-Helicase+iREase+CTD+REase | 944 | Bacteria>Pseudomonadota>Betaproteobacteria>Rhodocyclales>Zoogloeaceae | GCA_014822185.1 | Azoarcus_sp._Aa7 | AZOA_12030 | 1309..4143 | 2303328 | hypothetical-protein-[Azoarcus-sp.-Aa7] |
| WP_002936068.1 | DUF499_1.RM.like.systems | <-RNaseP<-?\|\|?->?->MORC+DNA_gyraseB+TOPRIM->?-><-DUF499*<-wHTH+REase<-METHYLASE<-HEPN<-ParB+HNH<-TUDOR+ SF1-SF2-Helicase+iREase+CTD+REase | 941 | Bacteria>Pseudomonadota>Betaproteobacteria>Rhodocyclales>Zoogloeaceae | GCF_000310125.2 | Thauera_sp._27 | B447_RS02400 | 104287..107112 | 305700 | DUF499-domain-containing-protein-[Thauera-sp.-27] |
| WP_153168444.1 | DUF499_1.RM.like.systems | <-Radical-SAM+TRAM<-?\|\|tRNA->?->?->?-><-?<-DUF499*<-wHTH+REase<-REase<-OLD-ABC<-METHYLASE<-HEPN<-TUDOR+ SF1-SF2-Helicase+iREase+CTD+REase | 941 | Bacteria>Pseudomonadota>Betaproteobacteria>Rhodocyclales>Zoogloeaceae | GCF_009469595.2 | Thauera_sp._2A1 | GH664_RS12700 | 85060..87885 | 2570191 | DUF499-domain-containing-protein-[Thauera-sp.-2A1] |
| OGA43321.1 | DUF499_1.RM.like.systems | METHYLASE->PolB->HEPN->REase->DUF499*-> | 927 | Bacteria>Pseudomonadota>Betaproteobacteria>unclassified_Betaproteobacteria | GCA_001771985.1 | Betaproteobacteria_bacterium_RIFCSPLOWO2_12_FULL_62_13 | A3G24_09290 | 4491..7274 | 1797495 | AAA-family-ATPase-[Betaproteobacteria-bacterium-RIFCSPLOWO2_12_FULL_62_13] |
| WP_047727733.1 | DUF499_1.RM.like.systems | Arm-DNA-bind_3+Phage_int_SAM_3+Phage_integrase-><-?<-?<-DUF499*<-METHYLASE<-HEPN<-TUDOR+ SF1-SF2-Helicase+iREase+CTD+REase<-HTH+WYL | 942 | Bacteria>Pseudomonadota>Gammaproteobacteria>Enterobacterales>Yersiniaceae | GCF_001022215.1 | Serratia | DKC05_RS04355 | 483774..486602 | 613 | MULTISPECIES:-DUF499-domain-containing-protein-[Serratia] |
| WP_064023787.1 | DUF499_1.RM.like.systems | PIN-><-?<-RelE-ParE<-TraJ-RHH\|\|Phage_int_SAM_1+Phage_integrase-><-HEPN<-DUF499*<-METHYLASE<-TUDOR+ SF1-SF2-Helicase+iREase+CTD+REase<-RHH<-HIN-HTH | 936 | Bacteria>Pseudomonadota>Gammaproteobacteria>Methylococcales>Methylococcaceae | GCF_001644685.1 | Methylomonas | A1356_RS16225 | 62920..65730 | 416 | MULTISPECIES:-DUF499-domain-containing-protein-[Methylomonas] |
| MDB9958356.1 | DUF499_1.RM.like.systems | <-DUF499*<-METHYLASE<-HEPN<-TUDOR+ SF1-SF2-Helicase+iREase+CTD+REase | 942 | Bacteria>Pseudomonadota>Gammaproteobacteria>Oceanospirillales>Oceanospirillaceae | GCA_028340495.1 | Oceanospirillaceae_bacterium | OAD42_04655 | 239..3067 | 1899355 | DUF499-domain-containing-protein-[Oceanospirillaceae-bacterium] |
| EPL60615.1 | DUF499_1.RM.like.systems | <-RelE-ParE<-RelB<-PIN+OST-HTH<-?\|\|?->RlpA-><-DUF499*<-HEPN<-METHYLASE<-PIN<-TUDOR+ SF1-SF2-Helicase+iREase+CTD+REase<-MORC+DNA_gyraseB+TOPRIM<-SbcC | 946 | Bacteria>Pseudomonadota>Gammaproteobacteria>Pseudomonadales>Pseudomonadaceae | GCA_000416345.1 | Stutzerimonas_stutzeri_B1SMN1 | B382_19940 | 6557..9397 | 1212549 | hypothetical-protein-B382_19940-[Stutzerimonas-stutzeri-B1SMN1] |
| MBG7398186.1 | DUF499_1.RM.like.systems | SF1-SF2-Helicase-><-SF1-SF2-helicase+RdgC+Beta-barrel+REase<-35exo+SF1-SF2-Helicase\|\|ABhydrolase->?->RlpA-><-DUF499*<-REase<-OLD-ABC<-METHYLASE<-HEPN<-TUDOR+ SF1-SF2-Helicase+iREase+CTD+REase | 942 | Bacteria>Pseudomonadota>Gammaproteobacteria>Pseudomonadales>Pseudomonadaceae | GCA_015766545.1 | Pseudomonas_aeruginosa | GHU66_27420 | 38429..41257 | 287 | ATP-binding-protein-[Pseudomonas-aeruginosa] |
| HBO5114778.1 | DUF499_1.RM.like.systems | TUDOR+ SF1-SF2-Helicase+iREase+CTD+REase->HEPN->METHYLASE->OLD-ABC->REase->DUF499*-><-RlpA<-?<-ABhydrolase | 946 | Bacteria>Pseudomonadota>Gammaproteobacteria>Pseudomonadales>Pseudomonadaceae | GCA_021662155.1 | Pseudomonas_aeruginosa | L4W37_005387 | 12730..15570 | 287 | ATP-binding-protein-[Pseudomonas-aeruginosa] |
| MCO2926705.1 | DUF499_1.RM.like.systems | TUDOR+ SF1-SF2-Helicase+iREase+CTD+REase->Schlafen+HATPase+HTH->?->METHYLASE->HEPN->DUF499*-><-RlpA<-?<-?\|\|HTH_3-><-ZnR+ZnR | 943 | Bacteria>Pseudomonadota>Gammaproteobacteria>Pseudomonadales>Pseudomonadaceae | GCA_023909145.1 | Pseudomonas_aeruginosa | FA260_26715 | 18266..21097 | 287 | ATP-binding-protein-[Pseudomonas-aeruginosa] |
| HCF1493245.1 | DUF499_1.RM.like.systems | <-MORC+HisKin<-RVT\|\|?->?->RlpA-><-?<-DUF499*<-HEPN<-METHYLASE-p | 948 | Bacteria>Pseudomonadota>Gammaproteobacteria>Pseudomonadales>Pseudomonadaceae | GCA_023941105.1 | Pseudomonas_aeruginosa | NH848_005990 | 4619..7465 | 287 | ATP-binding-protein-[Pseudomonas-aeruginosa] |
| HCI3160560.1 | DUF499_1.RM.like.systems | ABhydrolase->?->RlpA-><-DUF499*<-REase<-OLD-ABC<-METHYLASE<-HEPN<-TUDOR+ SF1-SF2-Helicase+iREase+CTD+REase | 942 | Bacteria>Pseudomonadota>Gammaproteobacteria>Pseudomonadales>Pseudomonadaceae | GCA_024356045.1 | Pseudomonas_aeruginosa | NOV63_004506 | 69244..72072 | 287 | ATP-binding-protein-[Pseudomonas-aeruginosa] |
| MCR7692359.1 | DUF499_1.RM.like.systems | TUDOR+ SF1-SF2-Helicase+iREase+CTD+REase->HEPN->METHYLASE->OLD-ABC->REase->DUF499*-> | 417 | Bacteria>Pseudomonadota>Gammaproteobacteria>Pseudomonadales>Pseudomonadaceae | GCA_024717035.1 | Pseudomonas_aeruginosa | NTA33_29240 | 17686..18936 | 287 | DUF499-domain-containing-protein-[Pseudomonas-aeruginosa] |
| HCK4430924.1 | DUF499_1.RM.like.systems | TUDOR+ SF1-SF2-Helicase+iREase+CTD+REase->HEPN->METHYLASE->OLD-ABC->TOPRIM->DUF499*-><-RlpA<-?<-JAB<-ParB<-?\|\|MazE_antitoxin-> | 942 | Bacteria>Pseudomonadota>Gammaproteobacteria>Pseudomonadales>Pseudomonadaceae | GCA_024830635.1 | Pseudomonas_aeruginosa | NOG72_005469 | 46146..48974 | 287 | ATP-binding-protein-[Pseudomonas-aeruginosa] |
| WP_023835622.1 | DUF499_1.RM.like.systems | TUDOR+ SF1-SF2-Helicase+iREase+CTD+REase->HEPN->METHYLASE->OLD-ABC->REase->DUF499*-><-RlpA<-?<-JAB<-ParB<-?\|\|HIN-HTH-> | 944 | Bacteria>Pseudomonadota>Gammaproteobacteria>Pseudomonadales>Pseudomonadaceae | GCF_000503175.1 | Pseudomonas_aeruginosa | CF344_RS23300 | 12270..15104 | 287 | DUF499-domain-containing-protein-[Pseudomonas-aeruginosa] |
| WP_031643596.1 | DUF499_1.RM.like.systems | TUDOR+ SF1-SF2-Helicase+iREase+CTD+REase->HEPN->METHYLASE->OLD-ABC->REase->DUF499*-><-RlpA<-?<-ABhydrolase<-?<-?\|\|HTH_3-> | 950 | Bacteria>Pseudomonadota>Gammaproteobacteria>Pseudomonadales>Pseudomonadaceae | GCF_000629305.1 | Pseudomonas_aeruginosa | AJ61_RS31490 | 12829..15681 | 287 | DUF499-domain-containing-protein-[Pseudomonas-aeruginosa] |
| WP_049308056.1 | DUF499_1.RM.like.systems | TUDOR+ SF1-SF2-Helicase+iREase+CTD+REase->HEPN->METHYLASE->OLD-ABC->REase->DUF499*-><-RlpA<-?<-?\|\|SF1-SF2-helicase+RdgC+Beta-barrel+REase-><-SF1-SF2-Helicase<-HKD-DNase | 942 | Bacteria>Pseudomonadota>Gammaproteobacteria>Pseudomonadales>Pseudomonadaceae | GCF_001061475.1 | Pseudomonas_aeruginosa | ADM76_RS18680 | 17855..20683 | 287 | DUF499-domain-containing-protein-[Pseudomonas-aeruginosa] |
| WP_054902927.1 | DUF499_1.RM.like.systems | DUF932->?->ParB->JAB->?->RlpA-><-DUF499*<-REase<-OLD-ABC<-METHYLASE<-HEPN<-TUDOR+ SF1-SF2-Helicase+iREase+CTD+REase\|\|HTH+WYL-> | 944 | Bacteria>Pseudomonadota>Gammaproteobacteria>Pseudomonadales>Pseudomonadaceae | GCF_001320525.1 | Pseudomonas_sp._NBRC_111131 | APH34_RS22145 | 20096..22930 | 1661046 | DUF499-domain-containing-protein-[Pseudomonas-sp.-NBRC-111131] |
| WP_074197337.1 | DUF499_1.RM.like.systems | DUF932->?->ParB->JAB->?->RlpA-><-DUF499*<-REase<-OLD-ABC<-METHYLASE<-HEPN<-TUDOR+ SF1-SF2-Helicase+iREase+CTD+REase | 946 | Bacteria>Pseudomonadota>Gammaproteobacteria>Pseudomonadales>Pseudomonadaceae | GCF_001909445.1 | Pseudomonas_aeruginosa | BH596_RS15220 | 11473..14313 | 287 | DUF499-domain-containing-protein-[Pseudomonas-aeruginosa] |
| WP_128472827.1 | DUF499_1.RM.like.systems | RlpA-><-DUF499*<-TOPRIM<-OLD-ABC<-METHYLASE<-HEPN<-TUDOR+ SF1-SF2-Helicase+iREase+CTD+REase | 943 | Bacteria>Pseudomonadota>Gammaproteobacteria>Pseudomonadales>Pseudomonadaceae | GCF_004053955.1 | Pseudomonas_aeruginosa | EQH74_RS32310 | 490..3321 | 287 | DUF499-domain-containing-protein-[Pseudomonas-aeruginosa] |
| WP_154355052.1 | DUF499_1.RM.like.systems | TUDOR+ SF1-SF2-Helicase+iREase+CTD+REase->HEPN->METHYLASE->OLD-ABC->REase->DUF499*-><-RlpA<-?<-JAB<-ParB<-?<-DUF932 | 942 | Bacteria>Pseudomonadota>Gammaproteobacteria>Pseudomonadales>Pseudomonadaceae | GCF_009676785.1 | Pseudomonas_aeruginosa | GJV02_RS03270 | 700187..703015 | 287 | DUF499-domain-containing-protein-[Pseudomonas-aeruginosa] |
| WP_192907661.1 | DUF499_1.RM.like.systems | TUDOR+ SF1-SF2-Helicase+iREase+CTD+REase->OLD-ABC->HNH->METHYLASE->HEPN->?->DUF499*-><-RlpA\|\|SF1-SF2-helicase+RdgC+Beta-barrel+REase-><-REase<-Phage_int_SAM<-Integrase_1<-Arm-DNA-bind_3+Phage_integrase | 941 | Bacteria>Pseudomonadota>Gammaproteobacteria>Pseudomonadales>Pseudomonadaceae | GCF_014883875.1 | Pseudomonas_taiwanensis | ICN73_RS10045 | 2229206..2232031 | 470150 | DUF499-domain-containing-protein-[Pseudomonas-taiwanensis] |
| WP_198580799.1 | DUF499_1.RM.like.systems | TUDOR+ SF1-SF2-Helicase+iREase+CTD+REase->HEPN->METHYLASE->OLD-ABC->REase->DUF499*-><-RlpA<-?<-JAB<-ParB<-?<-DUF932 | 944 | Bacteria>Pseudomonadota>Gammaproteobacteria>Pseudomonadales>Pseudomonadaceae | GCF_016134915.1 | Pseudomonas_aeruginosa | QMY34_RS12045 | 381175..384009 | 287 | DUF499-domain-containing-protein-[Pseudomonas-aeruginosa] |
| WP_200582036.1 | DUF499_1.RM.like.systems | TUDOR+ SF1-SF2-Helicase+iREase+CTD+REase->HEPN->METHYLASE->OLD-ABC->REase->DUF499*-><-RlpA<-?<-SymE_toxin<-?<-ZNR+Pkinase+SF1-SF2-Helicase+REase | 943 | Bacteria>Pseudomonadota>Gammaproteobacteria>Pseudomonadales>Pseudomonadaceae | GCF_016622185.1 | Pseudomonas_aeruginosa | GFM12_RS08425 | 1738931..1741762 | 287 | DUF499-domain-containing-protein-[Pseudomonas-aeruginosa] |
| WP_213918359.1 | DUF499_1.RM.like.systems | <-rve_3+Mu-transpos_C<-REase\|\|ParB->RlpA-><-DUF499*<-HEPN<-METHYLASE<-PIN<-TUDOR+ SF1-SF2-Helicase+iREase+CTD+REase<-MORC+DNA_gyraseB+TOPRIM<-SbcC | 946 | Bacteria>Pseudomonadota>Gammaproteobacteria>Pseudomonadales>Pseudomonadaceae | GCF_018457005.1 | Stutzerimonas_stutzeri | JR047_RS05385 | 4622..7462 | 316 | DUF499-domain-containing-protein-[Stutzerimonas-stutzeri] |
| WP_219806875.1 | DUF499_1.RM.like.systems | <-MazE_antitoxin\|\|?->ParB->JAB->?->RlpA-><-DUF499*<-REase<-OLD-ABC<-METHYLASE<-HEPN<-TUDOR+ SF1-SF2-Helicase+iREase+CTD+REase<-Peptidase | 943 | Bacteria>Pseudomonadota>Gammaproteobacteria>Pseudomonadales>Pseudomonadaceae | GCF_019429735.1 | Pseudomonas_aeruginosa | KZ843_RS17265 | 5657..8488 | 287 | DUF499-domain-containing-protein-[Pseudomonas-aeruginosa] |
| WP_223119133.1 | DUF499_1.RM.like.systems | ZnR+ZnR-><-HTH_3\|\|?->?->RlpA-><-DUF499*<-HEPN<-METHYLASE<-?<-Schlafen+HATPase+HTH<-TUDOR+ SF1-SF2-Helicase+iREase+CTD+REase | 942 | Bacteria>Pseudomonadota>Gammaproteobacteria>Pseudomonadales>Pseudomonadaceae | GCF_020036175.1 | Pseudomonas_aeruginosa | NFY03_RS10210 | 2202477..2205305 | 287 | DUF499-domain-containing-protein-[Pseudomonas-aeruginosa] |
| WP_033975677.1 | DUF499_1.RM.like.systems | TUDOR+ SF1-SF2-Helicase+iREase+CTD+REase->HEPN->METHYLASE->OLD-ABC->REase->DUF499*-><-RlpA<-?<-SymE_toxin<-?<-ZNR+Pkinase+SF1-SF2-Helicase+REase | 943 | Bacteria>Pseudomonadota>Gammaproteobacteria>Pseudomonadales>Pseudomonadaceae | GCF_022569915.1 | Pseudomonas_aeruginosa | CGU39_RS16575 | 614189..617020 | 287 | DUF499-domain-containing-protein-[Pseudomonas-aeruginosa] |
| WP_150035379.1 | DUF499_1.RM.like.systems | TUDOR+ SF1-SF2-Helicase+iREase+CTD+REase->Schlafen+HATPase+HTH->?->METHYLASE->HEPN->DUF499*-><-RlpA<-?<-?\|\|HTH_3-><-ZnR+ZnR | 942 | Bacteria>Pseudomonadota>Gammaproteobacteria>Pseudomonadales>Pseudomonadaceae | GCF_022627285.1 | Pseudomonas_aeruginosa | JNB05_RS00615 | 132976..135804 | 287 | DUF499-domain-containing-protein-[Pseudomonas-aeruginosa] |
| WP_266022655.1 | DUF499_1.RM.like.systems | TUDOR+ SF1-SF2-Helicase+iREase+CTD+REase->HEPN->METHYLASE->OLD-ABC->REase->DUF499*-><-RlpA<-?<-JAB<-ParB<-?<-DUF932 | 948 | Bacteria>Pseudomonadota>Gammaproteobacteria>Pseudomonadales>Pseudomonadaceae | GCF_026241065.1 | unclassified_Pseudomonas | OO409_RS22205 | 32872..35718 | 196821 | MULTISPECIES:-DUF499-domain-containing-protein-[unclassified-Pseudomonas] |
| WP_270067442.1 | DUF499_1.RM.like.systems | TUDOR+ SF1-SF2-Helicase+iREase+CTD+REase->HEPN->METHYLASE->SF1-SF2-Helicase+wHTH+PRTase->?->DUF499*-><-RlpA<-?<-JAB<-ParB<-?<-DUF932 | 944 | Bacteria>Pseudomonadota>Gammaproteobacteria>Pseudomonadales>Pseudomonadaceae | GCF_027594645.1 | Stutzerimonas_stutzeri | N7388_RS04725 | 1052728..1055562 | 316 | DUF499-domain-containing-protein-[Stutzerimonas-stutzeri] |
| WP_272438616.1 | DUF499_1.RM.like.systems | HKD-DNase->SF1-SF2-Helicase-><-SF1-SF2-helicase+RdgC+Beta-barrel+REase\|\|?->?->RlpA-><-DUF499*<-REase<-OLD-ABC<-METHYLASE<-HEPN<-TUDOR+ SF1-SF2-Helicase+iREase+CTD+REase | 942 | Bacteria>Pseudomonadota>Gammaproteobacteria>Pseudomonadales>Pseudomonadaceae | GCF_028435625.1 | Pseudomonas_aeruginosa | KGZ09_RS30315 | 38429..41257 | 287 | DUF499-domain-containing-protein-[Pseudomonas-aeruginosa] |
| WP_213870614.1 | DUF499_1.RM.like.systems | JAB->?-><-HTH_3\|\|?->?->RlpA-><-DUF499*<-HEPN<-METHYLASE<-HNH<-OLD-ABC<-TUDOR+ SF1-SF2-Helicase+iREase+CTD+REase | 952 | Bacteria>Pseudomonadota>Gammaproteobacteria>Pseudomonadales>Pseudomonadaceae | GCF_029201245.1 | Pseudomonas_aeruginosa | KKT79_RS19565 | 32215..35073 | 287 | DUF499-domain-containing-protein-[Pseudomonas-aeruginosa] |
| WP_279947325.1 | DUF499_1.RM.like.systems | DUF932->?->ParB->JAB->?->RlpA-><-DUF499*<-?<-SF1-SF2-Helicase+wHTH+PRTase<-METHYLASE<-HEPN<-TUDOR+ SF1-SF2-Helicase+iREase+CTD+REase | 944 | Bacteria>Pseudomonadota>Gammaproteobacteria>Pseudomonadales>Pseudomonadaceae | GCF_029843075.1 | Stutzerimonas_stutzeri | N5E83_RS18215 | 56624..59458 | 316 | DUF499-domain-containing-protein-[Stutzerimonas-stutzeri] |
| WP_033982644.1 | DUF499_1.RM.like.systems | ABhydrolase->?->RlpA-><-DUF499*<-REase<-OLD-ABC<-METHYLASE<-HEPN<-TUDOR+ SF1-SF2-Helicase+iREase+CTD+REase | 942 | Bacteria>Pseudomonadota>Gammaproteobacteria>Pseudomonadales>Pseudomonadaceae | GCF_900707645.1 | Pseudomonas_aeruginosa | BUH55_RS18155 | 7471..10299 | 287 | DUF499-domain-containing-protein-[Pseudomonas-aeruginosa] |
| WP_280091279.1 | DUF499_1.RM.like.systems | Arm-DNA-bind_3+Phage_integrase->Phage_int_SAM-><-HNH+SAVED<-SF1-SF2-helicase+RdgC+Beta-barrel+REase\|\|RlpA-><-DUF499*<-HEPN<-METHYLASE<-HNH<-OLD-ABC<-TUDOR+ SF1-SF2-Helicase+iREase+CTD+REase | 941 | Bacteria>Pseudomonadota>Gammaproteobacteria>Pseudomonadales>Pseudomonadaceae | GCF_947311875.1 | Pseudomonas | QME24_RS13335 | 66050..68875 | 286 | DUF499-domain-containing-protein-[Pseudomonas-juntendi] |
| WP_033979009.1 | DUF499_1.RM.like.systems | ZnR+ZnR-><-HTH_3\|\|?->?->RlpA-><-DUF499*<-HEPN<-METHYLASE<-?<-Schlafen+HATPase+HTH<-TUDOR+ SF1-SF2-Helicase+iREase+CTD+REase | 943 | Bacteria>Pseudomonadota>Gammaproteobacteria>Pseudomonadales>Pseudomonadaceae | - | Pseudomonas_aeruginosa | CJT50_RS28650 | 19614..22445 | 287 | DUF499-domain-containing-protein-[Pseudomonas-aeruginosa] |
| WP_132674023.1 | DUF499_1.RM.like.systems | DUF932->?->ParB->JAB->?->RlpA-><-DUF499*<-REase<-OLD-ABC<-METHYLASE<-HEPN<-TUDOR+ SF1-SF2-Helicase+iREase+CTD+REase | 944 | Bacteria>Pseudomonadota>Gammaproteobacteria>Pseudomonadales>Pseudomonadaceae | - | Pseudomonas_aeruginosa | EQ623_RS00180 | 30601..33435 | 287 | DUF499-domain-containing-protein-[Pseudomonas-aeruginosa] |
| WP_020308343.1 | DUF499_1.RM.like.systems | <-RelE-ParE<-RelB<-PIN+OST-HTH<-?\|\|?->RlpA-><-DUF499*<-HEPN<-METHYLASE<-PIN<-TUDOR+ SF1-SF2-Helicase+iREase+CTD+REase<-MORC+DNA_gyraseB+TOPRIM<-SbcC | 946 | Bacteria>Pseudomonadota>Gammaproteobacteria>Pseudomonadales>Pseudomonadaceae | - | Stutzerimonas_stutzeri | B382_19940 | 6557..9397 | 316 | DUF499-domain-containing-protein-[Pseudomonas-stutzeri] |
| MXY51727.1 | DUF499_1.RM.like.systems | <-DUF499*<-METHYLASE<-METHYLASE-p<-HEPN<-DNA_methylase<-MORC+REase<-TUDOR+ SF1-SF2-Helicase+iREase+CTD+REase | 951 | Bacteria>Pseudomonadota>Gammaproteobacteria>unclassified_Gammaproteobacteria | GCA_009837725.1 | Gammaproteobacteria_bacterium | F4Y86_04245 | 1..2853 | 1913989 | ATP-binding-protein-[Gammaproteobacteria-bacterium] |
| WP_182517583.1 | DUF499_1.RM.like.systems | XIS-HTH->TUDOR+ SF1-SF2-Helicase+iREase+CTD+REase->HEPN->METHYLASE->OLD-ABC->TOPRIM->DUF499*-> | 930 | Bacteria>Pseudomonadota>Gammaproteobacteria>Xanthomonadales>Rhodanobacteraceae | GCF_014138295.1 | Dokdonella_fugitiva | FHW13_RS05515 | 304049..306841 | 328517 | DUF499-domain-containing-protein-[Dokdonella-fugitiva] |
| WP_091239202.1 | DUF499_1.RM.like.systems | Phage_holin->Phage_lysozyme->?->HTH_3->Radical-SAM-><-?<-DUF499*<-ParB+HNH<-METHYLASE<-HEPN<-TUDOR+ SF1-SF2-Helicase+iREase+CTD+REase<-HTH_Tnp<-HTH+WYL | 949 | Bacteria>Pseudomonadota>Gammaproteobacteria>Xanthomonadales>Rhodanobacteraceae | GCF_900101825.1 | Aquimonas_voraii | BLP83_RS04255 | 981108..983957 | 265719 | DUF499-domain-containing-protein-[Aquimonas-voraii] |
| MCK9487878.1 | DUF499_1.RM.like.systems | <-SIGMA-HTH<-OLD-ABC<-?<-?<-?<-DUF499*<-REase<-OLD-ABC<-METHYLASE<-HEPN<-PIN<-PemK_toxin | 944 | Bacteria>Pseudomonadota>Gammaproteobacteria>Xanthomonadales>unclassified_Xanthomonadales | GCA_023228845.1 | Xanthomonadales_bacterium | MOQ42_00470 | 105333..108167 | 2006849 | DUF499-domain-containing-protein-[Xanthomonadales-bacterium] |
| MBH2008931.1 | DUF499_1.RM.like.systems | <-SF1-SF2-Helicase<-OLD-ABC+TOPRIM<-DUF499*<-METHYLASE<-HEPN<-TUDOR+ SF1-SF2-Helicase+iREase+CTD+REase<-?\|\|DDE_Tnp_1_2-> | 969 | Bacteria>Pseudomonadota>Gammaproteobacteria>Xanthomonadales>Xanthomonadaceae | GCA_015999665.1 | Xanthomonadaceae_bacterium | I8H71_04430 | 4236..7145 | 1926873 | ATP-binding-protein-[Xanthomonadaceae-bacterium] |
| MBP7655321.1 | DUF499_1.RM.like.systems | HEPN->METHYLASE->?->?->DUF499*->TIR->Methylase-><-Radical-SAM<-HTH_3<-?<-Phage_lysozyme | 950 | Bacteria>Pseudomonadota>Gammaproteobacteria>Xanthomonadales>Xanthomonadaceae | GCA_017999045.1 | Pseudoxanthomonas_sp. | KA742_02095 | 7019..9871 | 1871049 | ATP-binding-protein-[Pseudoxanthomonas-sp.] |
| MBU6247856.1 | DUF499_1.RM.like.systems | <-DUF499*<-ParB+HNH<-?<-HEPN+CBS<-METHYLASE<-PIN | 934 | Bacteria>Pseudomonadota>Gammaproteobacteria>Xanthomonadales>Xanthomonadaceae | GCA_018970895.1 | Xanthomonadaceae_bacterium | KGN77_08870 | 9374..12178 | 1926873 | DUF499-domain-containing-protein-[Xanthomonadaceae-bacterium] |
| WP_106991537.1 | DUF499_1.RM.like.systems | <-DUF499*<-?<-METHYLASE<-HEPN<-PIN<-TUDOR+ SF1-SF2-Helicase+iREase+CTD+REase<-HTH_26 | 930 | Bacteria>Pseudomonadota>Gammaproteobacteria>Xanthomonadales>Xanthomonadaceae | GCF_003024235.1 | Arenimonas_caeni | C6N40_RS13415 | 668..3460 | 2058085 | DUF499-domain-containing-protein-[Arenimonas-caeni] |
| WP_161614005.1 | DUF499_1.RM.like.systems | TUDOR+ SF1-SF2-Helicase+iREase+CTD+REase->HEPN->METHYLASE->?->?->?->DUF499*->?->?->?->?->?->D5N+D5ATPase-> | 929 | Bacteria>Pseudomonadota>Gammaproteobacteria>Xanthomonadales>Xanthomonadaceae | GCF_009996725.1 | Stenotrophomonas_acidaminiphila | GXB84_RS06675 | 54604..57393 | 128780 | DUF499-domain-containing-protein-[Stenotrophomonas-acidaminiphila] |
| WP_028873821.1 | DUF499_1.RM.like.systems | DDE_Tnp_1_assoc+DDE_Tnp_1->PIN->HEPN->METHYLASE->?->DUF499*->?->PIN->?-><-?<-RuvC | 947 | Bacteria>Pseudomonadota>Hydrogenophilia>Hydrogenophilales>Hydrogenophilaceae | GCF_000425565.1 | Tepidiphilus_margaritifer | H607_RS0100570 | 5909..8752 | 203471 | DUF499-domain-containing-protein-[Tepidiphilus-margaritifer] |
| MBS0576259.1 | DUF499_1.RM.like.systems | RNase->?->?->Phage_lysozyme-><-SbcC<-DUF499*<-METHYLASE<-HEPN<-PIN<-TUDOR+ SF1-SF2-Helicase+iREase+CTD+REase<-HTH_Tnp<-HTH+WYL | 955 | Bacteria>Pseudomonadota>unclassified_Pseudomonadota | GCA_018240435.1 | Pseudomonadota_bacterium | JSS45_07535 | 37839..40706 | 1977087 | ATP-binding-protein-[Proteobacteria-bacterium] |
| MBU2056000.1 | DUF499_1.RM.like.systems | wHTH+REase->SF1-SF2-Helicase->METHYLASE->REase->PolB->HEPN->DUF499*-><-?<-?<-HTH | 950 | Bacteria>Pseudomonadota>unclassified_Pseudomonadota | GCA_018825885.1 | Pseudomonadota_bacterium | KKC25_13705 | 11854..14706 | 1977087 | DUF499-domain-containing-protein-[Proteobacteria-bacterium] |
| MBI3319953.1 | DUF499_1.RM.like.systems | <-ORC-CDC6+REase<-?<-?<-MqsA_antitoxin+HTH_3<-?<-DUF499*<-HEPN<-PolB<-METHYLASE | 925 | Bacteria>PVC_group>Candidatus_Omnitrophota>unclassified_Candidatus_Omnitrophota | GCA_016196435.1 | Candidatus_Omnitrophota_bacterium | HYZ89_05150 | 6165..8942 | 2035772 | ATP-binding-protein-[Candidatus-Omnitrophica-bacterium] |
| OGV65464.1 | DUF499_1.RM.like.systems | BrnT_toxin->CopG_antitoxin->?-><-?<-?<-?<-DUF499*<-?<-METHYLASE<-?<-HEPN<-TUDOR+ SF1-SF2-Helicase+iREase+CTD+REase<-HEPN | 939 | Bacteria>PVC_group>Lentisphaerota>unclassified_Lentisphaerota | GCA_001803235.1 | Lentisphaerae_bacterium_RIFOXYA12_64_32 | A3K18_32130 | 5290..8109 | 1798577 | MAG:-hypothetical-protein-A3K18_32130-[Lentisphaerae-bacterium-RIFOXYA12_64_32] |
| MCG3177990.1 | DUF499_1.RM.like.systems | Radical-SAM-><-DOC+HTH\|\|HTH->STYKinase-like-><-SF1-SF2-Helicase<-DUF499*<-METHYLASE<-HEPN<-TUDOR+ SF1-SF2-Helicase+iREase+CTD+REase | 1112 | Bacteria>PVC_group>Planctomycetota>Phycisphaerae>unclassified_Phycisphaerae | GCA_022072085.1 | Phycisphaerae_bacterium | BIFFINMI_00313 | 58540..61878 | 2026778 | hypothetical-protein-[Phycisphaerae-bacterium] |

| Protein-ID | System | Gene-Neighbourhoods | Length | Taxonomy | Assembly-ID | Strain | Locus-Tag | Genomic-coordinates | TaxID | Fasta-defline |
|---|---|---|---|---|---|---|---|---|---|---|
| GJQ28342.1 | DUF499_1.RM.like.systems | Phage_integrase-><-?<-?<-?<-N6_N4_MTase<-DUF499*<-METHYLASE<-HEPN<-TUDOR+SF1-SF2-Helicase+iREase+CTD+REase | 1114 | Bacteria>PVC_group>Planctomycetota>Phycisphaerae>unclassified_Phycisphaerae | GCA_022180725.1 | Phycisphaerae_bacterium | HBSAPP03_02260 | 258751..262095 | 2026778 | hypothetical-protein-HBSAPP03_02260-[Phycisphaerae-bacterium] |
| NLX57996.1 | DUF499_1.RM.like.systems | TUDOR+SF1-SF2-Helicase+iREase+CTD+REase->HEPN->?->METHYLASE->DUF499*-> | 975 | Bacteria>PVC_group>Planctomycetota>Phycisphaerae>unclassified_Phycisphaerae | - | Phycisphaerae_bacterium | GXY74_02785 | 9938..12865 | 2026778 | ATP-binding-protein-[Phycisphaerae-bacterium] |
| WP_146679086.1 | DUF499_1.RM.like.systems | XIS-HTH->TUDOR+SF1-SF2-Helicase+iREase+CTD+REase->HEPN->METHYLASE->METHYLASE->REase->DUF499*->REase+ParB->RVT->?->RAMA->GDResolvase+Recombinase+ZnR->GDResolvase+Recombinase+ZnR-> | 981 | Bacteria>PVC_group>Planctomycetota>Planctomycetia>Pirellulales | GCF_001610875.1 | Pirellula_sp._SH-Sr6A | VN12_RS23070 | 5951575..5954520 | 1632865 | DUF499-domain-containing-protein-[Pirellula-sp.-SH-Sr6A] |
| WP_262025600.1 | DUF499_1.RM.like.systems | TUDOR+SF1-SF2-Helicase+iREase+CTD+REase->OLD-ABC->?->HEPN->METHYLASE->?->DUF499*->?->?->?->?->HKD-DNase->SF1-SF2-Helicase-> | 962 | Bacteria>PVC_group>Planctomycetota>Planctomycetia>Pirellulales | GCF_024584705.1 | Blastopirellula_sp._J2-11 | M4951_RS06120 | 1475319..1478207 | 2943192 | DUF499-domain-containing-protein-[Blastopirellula-sp.-J2-11] |
| MDA8378650.1 | DUF499_1.RM.like.systems | <-35exo\|\|?->TUDOR+SF1-SF2-Helicase+iREase+CTD+REase->HEPN->METHYLASE->wHTH+REase->DUF499*-> | 945 | Bacteria>PVC_group>Planctomycetota>Planctomycetia>unclassified_Planctomycetia | GCA_028068345.1 | Planctomycetia_bacterium | MOZ50_16785 | 11833..14670 | 2052181 | MAG:-DUF499-domain-containing-protein-[Planctomycetia-bacterium] |
| MBA3683762.1 | DUF499_1.RM.like.systems | <-DUF499*<-METHYLASE<-PolB<-HEPN<-TUDOR+SF1-SF2-Helicase+iREase+CTD+REase<-tRNA<-tRNA\|\|PRTase-> | 945 | Bacteria>PVC_group>Planctomycetota>unclassified_Planctomycetota | GCA_013816645.1 | Planctomycetota_bacterium | HOW72_00725 | 4185..7022 | 2026780 | ATP-binding-protein-[Planctomycetota-bacterium] |
| MBI2191311.1 | DUF499_1.RM.like.systems | <-DUF499*<-REase<-METHYLASE<-HEPN<-HEPN<-Schlafen+HATPase+HTH\|\|DEDD_Tnp_IS110+Transposase_20-> | 926 | Bacteria>PVC_group>Planctomycetota>unclassified_Planctomycetota | GCA_016179685.1 | Planctomycetota_bacterium | HYU36_04935 | 72260..75040 | 2026780 | ATP-binding-protein-[Planctomycetota-bacterium] |
| MBI2806086.1 | DUF499_1.RM.like.systems | <-ParB+HNH<-?<-PIWI<-?<-DUF499*<-HTH<-METHYLASE<-REase<-HEPN<-TUDOR+SF1-SF2-Helicase+iREase+CTD+REase\|\|HTH-> | 936 | Bacteria>PVC_group>Planctomycetota>unclassified_Planctomycetota | GCA_016191165.1 | Planctomycetota_bacterium | HYX68_13985 | 123843..126653 | 2026780 | ATP-binding-protein-[Planctomycetota-bacterium] |
| MBI3271453.1 | DUF499_1.RM.like.systems | <-Phage_int_SAM_1+Phage_integrase<-?<-?<-ParB+HNH<-?<-DUF499*<-PIN<-?<-?<-METHYLASE<-HEPN<-TUDOR+SF1-SF2-Helicase+iREase+CTD+REase | 953 | Bacteria>PVC_group>Planctomycetota>unclassified_Planctomycetota | GCA_016197205.1 | Planctomycetota_bacterium | HYZ53_20855 | 61698..64559 | 2026780 | ATP-binding-protein-[Planctomycetota-bacterium] |
| MCX5669700.1 | DUF499_1.RM.like.systems | TUDOR+SF1-SF2-Helicase+iREase+CTD+REase->OLD-ABC+TOPRIM->METHYLASE->PolB->HEPN->DUF499*-> | 924 | Bacteria>PVC_group>Planctomycetota>unclassified_Planctomycetota | GCA_026387055.1 | Planctomycetota_bacterium | NTU94_00045 | 8049..10823 | 2026780 | MAG:-DUF499-domain-containing-protein-[Planctomycetota-bacterium] |
| OQC32217.1 | DUF499_1.RM.like.systems | DDE_Tnp_IS66+DDE_Tnp_IS66_C-><-?<-Radical-SAM<-?<-DUF499*<-?<-?<-METHYLASE<-HEPN<-ParB+HNH<-TUDOR+SF1-SF2-Helicase+iREase+CTD+REase | 945 | Bacteria>PVC_group>Verrucomicrobiota>unclassified_Verrucomicrobiota | GCA_002071025.1 | Verrucomicrobia_bacterium_ADurb.Bin070 | BWX70_00665 | 3737..6574 | 1852927 | hypothetical-protein-BWX70_00665-[Verrucomicrobia-bacterium-ADurb.Bin070] |
| MBI3850446.1 | DUF499_1.RM.like.systems | <-ACET+PIN+ACET\|\|TUDOR+SF1-SF2-Helicase+iREase+CTD+REase->?->SF1-SF2-Helicase->?->METHYLASE->DUF499*->HEPN->?->REase+HTH->NTP_transf->?-><-35exo | 961 | Bacteria>PVC_group>Verrucomicrobiota>unclassified_Verrucomicrobiota | GCA_016200005.1 | Verrucomicrobiota_bacterium | HY298_09175 | 22264..25149 | 2026799 | DUF499-domain-containing-protein-[Verrucomicrobia-bacterium] |
| MBM3880845.1 | DUF499_1.RM.like.systems | TUDOR+SF1-SF2-Helicase+iREase+CTD+REase->HEPN->ParA-Soj-PloopNTPase->ORC-CDC6->REase->DUF499*-> | 1105 | Bacteria>PVC_group>Verrucomicrobiota>unclassified_Verrucomicrobiota | GCA_016871715.1 | Verrucomicrobiota_bacterium | FJ387_14200 | 23630..26947 | 2026799 | DUF499-domain-containing-protein-[Verrucomicrobiota-bacterium] |
| MCX7997938.1 | DUF499_1.RM.like.systems | <-DUF499*<-HEPN<-PolB<-METHYLASE-p | 838 | Bacteria>Spirochaetota>Spirochaetia>Leptospirales>Leptospiraceae | GCA_026415905.1 | Leptospiraceae_bacterium | N3A69_03160 | 811..3327 | 2026757 | MAG:-hypothetical-protein-N3A69_03160-[Leptospiraceae-bacterium] |
| MDD4082191.1 | DUF499_1.RM.like.systems | <-SF1-SF2-Helicase-p<-HEPN<-DUF499*<-METHYLASE-p<-METHYLASE<-TUDOR+SF1-SF2-Helicase+iREase+CTD+REase | 924 | Bacteria>Spirochaetota>Spirochaetia>Spirochaetales>Sphaerochaetaceae | GCA_028701295.1 | Sphaerochaetaceae_bacterium | PHD05_02270 | 2146..4920 | 2791017 | MAG:-DUF499-domain-containing-protein-[Sphaerochaetaceae-bacterium] |
| MBV5322918.1 | DUF499_1.RM.like.systems | HEPN->OLD-ABC->HNH->METHYLASE->METHYLASE->HEPN->DUF499*-> | 872 | Bacteria>Terrabacteria_group>Actinomycetota>Acidimicrobiia>Acidimicrobiales | GCA_019162935.1 | Ilumatobacteraceae_bacterium | JZU60_03690 | 12690..15310 | 2497619 | ATP-binding-protein-[Ilumatobacteraceae-bacterium] |
| WP_252863026.1 | DUF499_1.RM.like.systems | <-METHYLASE-p<-REase<-HTH+SF1-SF2-Helicase+REase<-REase<-MORC<-DUF499*<-HEPN<-METHYLASE\|\|DNA_methylase-> | 912 | Bacteria>Terrabacteria_group>Actinomycetota>Actinomycetes>Actinomycetales | - | Actinomyces_oris | BKH30_RS08650 | 21386..24124 | 544580 | DUF499-domain-containing-protein-[Actinomyces-oris] |
| NUQ96612.1 | DUF499_1.RM.like.systems | <-HNH<-ABC-ATPase<-HEPN<-DUF499*<-METHYLASE<-TUDOR+SF1-SF2-Helicase+iREase+CTD+REase\|\|?->?-><-REase | 910 | Bacteria>Terrabacteria_group>Actinomycetota>Actinomycetes>Kitasatosporales | GCA_013361825.1 | Streptomyces_sp. | HOY79_08625 | 38052..40784 | 1931 | ATP-binding-protein-[Streptomyces-sp.] |
| WP_037620150.1 | DUF499_1.RM.like.systems | REase->Peptidase-S8+Subtilisin->?->?->TUDOR+SF1-SF2-Helicase+iREase+CTD+REase->METHYLASE->DUF499*->HEPN->ABC-ATPase->HNH-> | 910 | Bacteria>Terrabacteria_group>Actinomycetota>Actinomycetes>Kitasatosporales | GCF_000725495.1 | Streptomyces_aureus | IO36_RS15000 | 183930..186662 | 193461 | DUF499-domain-containing-protein-[Streptomyces-aureus] |
| WP_190197152.1 | DUF499_1.RM.like.systems | ABhydrolase+ABhydrolase+ABhydrolase->Peptidase->?-><-HNH<-ABC-ATPase<-HEPN<-DUF499*<-METHYLASE<-TUDOR+SF1-SF2-Helicase+iREase+CTD+REase\|\|HIN-HTH-> | 910 | Bacteria>Terrabacteria_group>Actinomycetota>Actinomycetes>Kitasatosporales | GCF_014651075.1 | Streptomyces_djakartensis | IE212_RS08750 | 190992..193724 | 68193 | DUF499-domain-containing-protein-[Streptomyces-djakartensis] |
| MDD0856750.1 | DUF499_1.RM.like.systems | ABC-ATPase+SbcC-><-?<-TrwC+SF1-SF2-Helicase<-?<-?\|\|<-DUF499-p<-DUF499*<-DUF499-p<-HEPN<-METHYLASE<-?<-METHYLASE-p | 458 | Bacteria>Terrabacteria_group>Actinomycetota>Actinomycetes>Micrococcales | GCA_024137885.2 | Arthrobacter_alpinus | NHF46_00925 | 169502..170878 | 656366 | DUF499-domain-containing-protein-[Arthrobacter-alpinus] |
| WBM79848.1 | DUF499_1.RM.like.systems | XIS-HTH->TUDOR+SF1-SF2-Helicase+iREase+CTD+REase-p->TUDOR+SF1-SF2-Helicase+iREase+CTD+REase->METHYLASE->ParB+HNH->HEPN->DUF499*->DUF499-p->DUF499-p-><-?\|\|REase->REase-> | 285 | Bacteria>Terrabacteria_group>Actinomycetota>Actinomycetes>Micrococcales | GCA_027594785.1 | Cryobacterium_breve | KIV56_17020 | 3683119..3683976 | 1259258 | DUF499-domain-containing-protein-[Cryobacterium-breve] |
| WP_262391126.1 | DUF499_1.RM.like.systems | Peptidase_S66+Peptidase_S66C-><-HTH+SF1-SF2-Helicase+REase<-REase<-MORC<-MORC<-DUF499*<-HEPN<-METHYLASE<-DNA_methylase\|\|REase-> | 679 | Bacteria>Terrabacteria_group>Actinomycetota>Actinomycetes>Micrococcales | GCF_003071525.2 | Micrococcus_luteus | CO205_RS12620 | 704510..706549 | 1270 | DUF499-domain-containing-protein-[Micrococcus-luteus] |
| WP_281534461.1 | DUF499_1.RM.like.systems | TUDOR+SF1-SF2-Helicase+iREase+CTD+REase-p->TUDOR+SF1-SF2-Helicase+iREase+CTD+REase->METHYLASE->ParB+HNH->HEPN->DUF499-p->DUF499*->DUF499-p-><-?\|\|REase->?->?-><-RelA | 430 | Bacteria>Terrabacteria_group>Actinomycetota>Actinomycetes>Micrococcales | GCF_027594785.1 | Cryobacterium_breve | KIV56_RS17895 | 3683885..3685177 | 1259258 | DUF499-domain-containing-protein-[Cryobacterium-breve] |
| CCK55004.1 | DUF499_1.RM.like.systems | STYKinase->HIN-HTH->?-><-?<-REase<-DUF499*<-HEPN<-?<-?<-HIN-HTH<-HIN-HTH<-METHYLASE-p | 792 | Bacteria>Terrabacteria_group>Actinomycetota>Actinomycetes>Mycobacteriales | GCA_000328845.1 | Mycobacterium_canettii_CIPT_140070008 | BN43_30077 | 1164706..1167084 | 1205675 | Conserved-protein-of-unknown-function-[Mycobacterium-canettii-CIPT-140070008] |
| WP_267324306.1 | DUF499_1.RM.like.systems | <-DUF499*<-zf-ISL3+DDE_Tnp_ISL3<-HEPN<-METHYLASE<-TUDOR+SF1-SF2-Helicase+iREase+CTD+REase-p | 810 | Bacteria>Terrabacteria_group>Actinomycetota>Actinomycetes>Mycobacteriales | GCF_026501155.1 | Mycolicibacterium_mucogenicum | OS121_RS10005 | 2088715..2091147 | 56689 | DUF499-domain-containing-protein-[Mycolicibacterium-mucogenicum] |
| WP_142566095.1 | DUF499_1.RM.like.systems | PAIREDC-HTH-><-tRNA\|\|TUDOR+SF1-SF2-Helicase+iREase+CTD+REase->METHYLASE->DUF499*->HEPN-><-HTH+HKD-DNase\|\|?->GNTR-HTH->HDRnase-> | 976 | Bacteria>Terrabacteria_group>Actinomycetota>Actinomycetes>Streptosporangiales | GCF_006874435.1 | Microbispora_sp._KK1-11 | FLW16_RS39395 | 9114..12044 | 2053005 | DUF499-domain-containing-protein-[Microbispora-sp.-KK1-11] |
| WP_147945511.1 | DUF499_1.RM.like.systems | <-HEPN<-DUF499*<-METHYLASE | 976 | Bacteria>Terrabacteria_group>Actinomycetota>Actinomycetes>Streptosporangiales | GCF_008033685.1 | Microbispora_sp._CSR-4 | FLV34_RS38505 | 1003..3933 | 2592813 | DUF499-domain-containing-protein-[Microbispora-sp.-CSR-4] |
| WP_238498630.1 | DUF499_1.RM.like.systems | <-TetR-HTH<-tRNA<-?<-DUF499*<-METHYLASE<-TUDOR+SF1-SF2-Helicase+iREase+CTD+REase<-HEPN<-XIS-HTH+RVT<-?<-HTH+DUF5753 | 939 | Bacteria>Terrabacteria_group>Actinomycetota>Actinomycetes>Streptosporangiales | - | Streptosporangium_sp._KLBMP_9127 | MBA17_23095 | 277278..280097 | 2916433 | DUF499-domain-containing-protein-[Streptosporangium-sp.-KLBMP-9127] |
| GFP32491.1 | DUF499_1.RM.like.systems | <-DUF499*<-HEPN<-PolB<-METHYLASE | 577 | Bacteria>Terrabacteria_group>Actinomycetota>Actinomycetota_incertae_sedis>Candidatus_Hakubellales | GCA_013281995.1 | Candidatus_Hakubella_thermalkaliphila | HKBW3S42_00795 | 50..1783 | 2754717 | hypothetical-protein-HKBW3S42_00795-[Candidatus-Hakubanella-thermoalkaliphilus] |
| WP_176233286.1 | DUF499_1.RM.like.systems | METHYLASE->PolB->HEPN->DUF499*-><-tRNA\|\|?->NTP_transferase->?->?-><-ACET | 876 | Bacteria>Terrabacteria_group>Actinomycetota>Actinomycetota_incertae_sedis>Candidatus_Hakubellales | GCF_013281955.1 | Candidatus_Hakubella_thermalkaliphila | HW440_RS04150 | 3204..5834 | 2754717 | DUF499-domain-containing-protein-[Candidatus-Hakubanella-thermoalkaliphilus] |
| MCS7301717.1 | DUF499_1.RM.like.systems | <-TUDOR+SF1-SF2-Helicase+iREase+CTD+REase-p<-DUF499*<-HEPN<-PolB<-HEPN<-PolB<-METHYLASE-p | 950 | Bacteria>Terrabacteria_group>Armatimonadota>Fimbriimonadia>Fimbriimonadales | GCA_025062205.1 | Fimbriimonadales_bacterium | NZ556_09225 | 1592..4444 | 2682145 | DUF499-domain-containing-protein-[Fimbriimonadales-bacterium] |
| GIV02394.1 | DUF499_1.RM.like.systems | <-URI-DNase+RAMA<-?<-HEPN+REase<-?<-REase<-Schlafen+HATPase+HTH<-DUF499*<-REase<-?<-METHYLASE-p\|\|?-><-?<-METHYLASE | 920 | Bacteria>Terrabacteria_group>Armatimonadota>Fimbriimonadia>Fimbriimonadales | GCA_026003055.1 | Fimbriimonadales_bacterium | KatS3mg015_1224 | 1249734..1252496 | 2682145 | MAG:-hypothetical-protein-KatS3mg015_1224-[Fimbriimonadales-bacterium] |
| PIU67978.1 | DUF499_1.RM.like.systems | HEPN-p->METHYLASE->DUF499*->?-><-?<-?<-SF1-SF2-Helicase<-Schlafen+HATPase+HTH | 944 | Bacteria>Terrabacteria_group>Armatimonadota>unclassified_Armatimonadota | GCA_002779575.1 | Armatimonadetes_bacterium_CG07_land_8_20_14_0_80_40_9 | COS84_02435 | 3211..6045 | 1973916 | AAA-family-ATPase-[Armatimonadetes-bacterium-CG07_land_8_20_14_0_80_40_9] |
| NLB94469.1 | DUF499_1.RM.like.systems | REase->HEPN->METHYLASE->DUF499*->DNA-glycosylase-> | 991 | Bacteria>Terrabacteria_group>Armatimonadota>unclassified_Armatimonadota | GCA_012512265.1 | Armatimonadota_bacterium | GX785_02050 | 5194..8169 | 2033014 | DUF499-domain-containing-protein-[Armatimonadetes-bacterium] |
| MCA1947578.1 | DUF499_1.RM.like.systems | URI-DNase-><-DUF499*<-HEPN<-PolB<-METHYLASE | 919 | Bacteria>Terrabacteria_group>Armatimonadota>unclassified_Armatimonadota | GCA_020217215.1 | Armatimonadota_bacterium | LDL55_05595 | 1010..3769 | 2033014 | DUF499-domain-containing-protein-[Armatimonadetes-bacterium] |
| BAF59044.1 | DUF499_1.RM.like.systems | TUDOR+SF1-SF2-Helicase+iREase+CTD+REase->?->METHYLASE->HEPN->HEPN+Mrr_N->DUF499*-><-?\|\|?->?->DNA_pol3_delta+DNA_pol3_gamma3-> | 885 | Bacteria>Terrabacteria_group>Bacillota>Clostridia>Eubacteriales | GCA_000010565.1 | Pelotomaculum_thermopropionicum_SI | PTH_0863 | 877158..879815 | 370438 | predicted-ATPase-[Pelotomaculum-thermopropionicum-SI] |
| WP_166349185.1 | DUF499_1.RM.like.systems | ArsR-HTH->BirA-HTH->?-><-DUF499*<-N6_N4_MTase<-?<-HEPN<-METHYLASE<-TUDOR+SF1-SF2-Helicase+iREase+CTD+REase | 1104 | Bacteria>Terrabacteria_group>Bacillota>Clostridia>Eubacteriales | GCF_011393055.1 | Desulfofundulus_sp._TPOSR | G7K71_RS17495 | 347545..350859 | 2714340 | DUF499-domain-containing-protein-[Desulfofundulus-sp.-TPOSR] |
| MCL5054662.1 | DUF499_1.RM.like.systems | METHYLASE->DUF499*-><-HEPN<-Polbeta | 1100 | Bacteria>Terrabacteria_group>Bacillota>unclassified_Bacillota | GCA_023475635.1 | Bacillota_bacterium | M1169_01785 | 1736..5038 | 1879010 | DUF499-domain-containing-protein-[Firmicutes-bacterium] |
| MBE7469119.1 | DUF499_1.RM.like.systems | METHYLASE-p->?->?->?-><-DUF499*<-REase<-METHYLASE<-HEPN<-PolB<-HEPN<-Polbeta | 906 | Bacteria>Terrabacteria_group>Chloroflexota>Anaerolineae>Anaerolineales | GCA_015075405.1 | Anaerolineales_bacterium | HS114_08270 | 1909271..1911991 | 2073117 | ATP-binding-protein-[Anaerolineales-bacterium] |
| MBN8655123.1 | DUF499_1.RM.like.systems | PIN->REase+NACHT->TUDOR+SF1-SF2-Helicase+iREase+CTD+REase->METHYLASE->HEPN->?->DUF499*-><-SF1-SF2-Helicase | 853 | Bacteria>Terrabacteria_group>Chloroflexota>Anaerolineae>unclassified_Anaerolineae | GCA_017303735.1 | Anaerolineae_bacterium | JOM11_05235 | 15902..18463 | 2052143 | DUF499-domain-containing-protein-[Anaerolineae-bacterium] |
| MCQ3973997.1 | DUF499_1.RM.like.systems | PIN->CHTH->METHYLASE-p->?->METHYLASE->CR-ATPase9->DUF499*->?->HEPN-><-?<-Phage_integrase | 1225 | Bacteria>Terrabacteria_group>Chloroflexota>Anaerolineae>unclassified_Anaerolineae | GCA_024434315.1 | Anaerolineae_bacterium | DPW09_11170 | 46079..49756 | 2052143 | hypothetical-protein-[Anaerolineae-bacterium] |
| CAG1014975.1 | DUF499_1.RM.like.systems | PHP+SbcC->Schlafen+HATPase+HTH->METHYLASE-p->?->METHYLASE->HEPN->DUF499*-> | 923 | Bacteria>Terrabacteria_group>Chloroflexota>Anaerolineae>unclassified_Anaerolineae | GCA_905339325.1 | Anaerolineae_bacterium | ANRL4_05364 | 17229..20000 | 2052143 | hypothetical-protein-ANRL4_05364-[Anaerolineae-bacterium] |
| MDE0464749.1 | DUF499_1.RM.like.systems | METHYLASE->PolB->HEPN->OLD-ABC->TOPRIM->HEPN->DUF499*->?->?-><-?\|\|MazE_antitoxin-><-DSBH | 775 | Bacteria>Terrabacteria_group>Chloroflexota>Caldilineae>Caldilineales | GCA_028821465.1 | Caldilineaceae_bacterium | OXH93_20190 | 7230..9557 | 2614250 | MAG:-DUF499-domain-containing-protein-[Caldilineaceae-bacterium] |
| MYD51869.1 | DUF499_1.RM.like.systems | TUDOR+SF1-SF2-Helicase+iREase+CTD+REase->PolB->HEPN->METHYLASE->DUF433->PIN->DUF499*-> | 928 | Bacteria>Terrabacteria_group>Chloroflexota>Dehalococcoidia>unclassified_Dehalococcoidia | GCA_009841605.1 | Dehalococcoidia_bacterium | F4W93_10360 | 50124..52910 | 2026734 | ATP-binding-protein-[Dehalococcoidia-bacterium] |
| MBI4295304.1 | DUF499_1.RM.like.systems | TUDOR+SF1-SF2-Helicase+iREase+CTD+REase->HEPN->Pkinase->METHYLASE->DUF433->PIN->DUF499*-> | 770 | Bacteria>Terrabacteria_group>Chloroflexota>unclassified_Chloroflexota | GCA_016210605.1 | Chloroflexota_bacterium | HY669_03965 | 9180..11489 | 2026724 | ATP-binding-protein-[Chloroflexi-bacterium] |
| MBK7919048.1 | DUF499_1.RM.like.systems | DDE_5-><-DDE_Tnp_IS1\|\|METHYLASE-p->?->?->HEPN-p->HEPN+DUF499*->DUF499-><-DDE_5<-DDE_5\|\|?-><-DDE_5+DDE_Tnp_2 | 582 | Bacteria>Terrabacteria_group>Chloroflexota>unclassified_Chloroflexota | GCA_016713795.1 | Chloroflexota_bacterium | IPJ94_22860 | 29678..31426 | 2026724 | DUF499-domain-containing-protein-[Chloroflexota-bacterium] |
| MBK7919049.1 | DUF499_1.RM.like.systems | <-DDE_Tnp_IS1\|\|METHYLASE-p->?->?->HEPN-p->HEPN+DUF499->DUF499*-><-DDE_5<-DDE_5\|\|?-><-DDE_5+DDE_Tnp_2 | 452 | Bacteria>Terrabacteria_group>Chloroflexota>unclassified_Chloroflexota | GCA_016713795.1 | Chloroflexota_bacterium | IPJ94_22865 | 31366..32724 | 2026724 | hypothetical-protein-[Chloroflexota-bacterium] |
| MBM3143888.1 | DUF499_1.RM.like.systems | TUDOR+SF1-SF2-Helicase+iREase+CTD+REase->HEPN->METHYLASE->DUF499*->?->PIN-><-?\|\|?->Peptidase_M24-> | 927 | Bacteria>Terrabacteria_group>Chloroflexota>unclassified_Chloroflexota | GCA_016865865.1 | Chloroflexota_bacterium | FJ010_02740 | 17191..19974 | 2026724 | ATP-binding-protein-[Chloroflexi-bacterium] |
| MCL4543926.1 | DUF499_1.RM.like.systems | TUDOR+SF1-SF2-Helicase+iREase+CTD+REase->METHYLASE->PolB->HEPN->DUF499*->TraJ-RHH->PIN->DDE_Tnp_1_3-> | 860 | Bacteria>Terrabacteria_group>Chloroflexota>unclassified_Chloroflexota | GCA_023381895.1 | Chloroflexota_bacterium | M1118_04890 | 7735..10317 | 2026724 | DUF499-domain-containing-protein-[Chloroflexi-bacterium] |
| MCL5107378.1 | DUF499_1.RM.like.systems | TUDOR+SF1-SF2-Helicase+iREase+CTD+REase->?->HEPN->METHYLASE->DOC+HTH->DUF499*->Schlafen+HATPase+HTH-> | 1102 | Bacteria>Terrabacteria_group>Chloroflexota>unclassified_Chloroflexota | GCA_023473795.1 | Chloroflexota_bacterium | M1401_00595 | 20015..23323 | 2026724 | DUF499-domain-containing-protein-[Chloroflexi-bacterium] |
| MCZ7670577.1 | DUF499_1.RM.like.systems | Peptidase_C26->?->tRNA-><-?<-?<-DUF499-p<-DUF499*<-HEPN<-?<-METHYLASE-p\|\|?-><-METHYLASE-p<-METHYLASE-p | 431 | Bacteria>Terrabacteria_group>Chloroflexota>unclassified_Chloroflexota | GCA_027493515.1 | Chloroflexota_bacterium | M5U34_27215 | 48065..49360 | 2026724 | MAG:-DUF499-domain-containing-protein-[Chloroflexi-bacterium] |
| KKZ10364.1 | DUF499_1.RM.like.systems | METHYLASE->HEPN->DUF499*-> | 965 | Bacteria>Terrabacteria_group>Cyanobacteriota/Melainabacteria_group>Cyanobacteriota>Cyanophyceae | GCA_001007625.1 | Candidatus_Synechococcus_spongiarum_142 | TH68_11020 | 5098..7996 | 1608213 | ATPase-AAA,-partial-[Candidatus-Synechococcus-spongiarum-142] |
| WP_143780302.1 | DUF499_1.RM.like.systems | <-HHH+ENDO3+NUDIX<-Peptidase_M50+PDZ\|\|?-><-METHYLASE-p<-DUF499*<-HEPN<-?<-?<-REase<-METHYLASE-p | 735 | Bacteria>Terrabacteria_group>Cyanobacteriota/Melainabacteria_group>Cyanobacteriota>Cyanophyceae | GCF_001939115.1 | Leptolyngbya_sp._'hensonii' | BST81_RS10560 | 80963..83170 | 1922337 | DUF499-domain-containing-protein-[Leptolyngbya-sp.-'hensonii'] |
| WP_106370048.1 | DUF499_1.RM.like.systems | RNase-><-?\|\|?->TIR-><-HKD-DNase+SF1-SF2-Helicase<-HEPN<-DUF499*<-METHYLASE<-PIN | 1004 | Bacteria>Terrabacteria_group>Cyanobacteriota/Melainabacteria_group>Cyanobacteriota>Cyanophyceae | GCF_003003885.1 | Chlorogloea_sp._CCALA_695 | C7B70_RS16210 | 39604..42618 | 2107693 | DUF499-domain-containing-protein-[Chlorogloea-sp.-CCALA-695] |
| WP_250683155.1 | DUF499_1.RM.like.systems | <-DUF499*<-HEPN<-REase<-METHYLASE<-PemK_toxin<-AbrB<-PolB | 788 | Bacteria>Terrabacteria_group>Cyanobacteriota/Melainabacteria_group>Cyanobacteriota>Cyanophyceae | - | Thermosynechococcus_sp._HN-54 | NBE99_RS03710 | 771476..773842 | 2933959 | DUF499-domain-containing-protein-[Thermosynechococcus-sp.-HN-54] |
| WP_252829645.1 | DUF499_1.RM.like.systems | <-DDE_Tnp_1<-?<-?\|\|?->?->\|\|<-HTH+DSBH\|\|<-DUF499*<-HEPN+DUF499<-HIN-HTH+AT-HTH-HYD2-2+SecB<-HTH_3<-HigB/RHH<-HicB_lk_antitox<-METHYLASE-p | 779 | Bacteria>Terrabacteria_group>Deinococcota>Deinococci>Thermales | GCF_000744155.1 | Thermus_scotoductus | N686_RS12530 | 46025..48364 | 37636 | DUF499-domain-containing-protein-[Thermus-scotoductus] |
| WP_252829575.1 | DUF499_1.RM.like.systems | <-HTH+DSBH\|\|<-DUF499<-HEPN+DUF499*<-HIN-HTH+AT-HTH-HYD2-2+SecB<-HTH_3<-HigB/RHH<-HicB_lk_antitox<-METHYLASE-p<-METHYLASE | 335 | Bacteria>Terrabacteria_group>Deinococcota>Deinococci>Thermales | GCF_000744155.1 | Thermus_scotoductus | N686_RS14355 | 48271..49278 | 37636 | Swt1-family-HEPN-domain-containing-protein-[Thermus-scotoductus] |
| MDD3580510.1 | DUF499_1.RM.like.systems | BrnT_toxin->BrnA_antitoxin->TUDOR+SF1-SF2-Helicase+iREase+CTD+REase->METHYLASE->PolB->HEPN->DUF499*-> | 920 | Bacteria>Thermodesulfobacteriota>Desulfobaccia>Desulfobaccales>Desulfobaccaceae | GCA_028695935.1 | Desulfobacca_sp. | PHW74_05770 | 25341..28103 | 2067990 | MAG:-DUF499-domain-containing-protein-[Desulfobacca-sp.] |
| MBL7204915.1 | DUF499_1.RM.like.systems | Schlafen->METHYLASE->PolB->HEPN->DUF499*->HEPN+Mrr_N->Schlafen+HATPase+HTH-> | 945 | Bacteria>Thermodesulfobacteriota>Desulfobacteria>Desulfobacterales>Desulfobacteraceae | GCA_016783005.1 | Desulfobacteraceae_bacterium | ISS63_11385 | 7746..10583 | 2049433 | ATP-binding-protein-[Desulfobacteraceae-bacterium] |
| MDA8161932.1 | DUF499_1.RM.like.systems | <-DUF499*<-HEPN<-PolB<-METHYLASE-p | 922 | Bacteria>Thermodesulfobacteriota>Desulfobacteria>Desulfobacterales>Desulfobacteraceae | GCA_028065225.1 | Desulfobacteraceae_bacterium | MOO22_03365 | 1133..3901 | 2049433 | MAG:-DUF499-domain-containing-protein-[Desulfobacteraceae-bacterium] |
| VEN75475.1 | DUF499_1.RM.like.systems | 35exo->?->Pterin_bind->?->tRNA-><-URI-DNase<-DUF499*<-METHYLASE<-HEPN<-TUDOR+SF1-SF2-Helicase+iREase+CTD+REase<-wHTH+REase<-?<-METHYLASE | 988 | Bacteria>Thermodesulfobacteriota>Desulfobacteria>Desulfobacterales>Desulfobacteraceae | GCA_900659855.1 | uncultured_Desulfobacteraceae_bacterium | EPICR_90073 | 73527..76493 | 218296 | conserved-hypothetical-protein-[uncultured-Desulfobacteraceae-bacterium] |
| MBA3013967.1 | DUF499_1.RM.like.systems | DUF499*->REase->HEPN->OLD-ABC+TOPRIM->SF1-SF2-Helicase-><-METHYLASE+SF1-SF2-Helicase | 944 | Bacteria>Thermodesulfobacteriota>Desulfobulbia>Desulfobulbales>Desulfobulbaceae | GCA_013792005.1 | Desulfobulbaceae_bacterium | FP815_03320 | 13..2847 | 2053307 | ATP-binding-protein-[Desulfobulbaceae-bacterium] |
| WP_205241970.1 | DUF499_1.RM.like.systems | TUDOR+SF1-SF2-Helicase+iREase+CTD+REase->HD->DUF4065->METHYLASE->OLD-ABC->TOPRIM->DUF499*->HEPN+SF1-SF2-Helicase->MORC+DUF3684+HNH->?->?-><-?<-HEPN | 985 | Bacteria>Thermodesulfobacteriota>Desulfobulbia>Desulfobulbales>Desulfobulbaceae | GCF_016918545.1 | Desulfobulbus_alkaliphilus | JWG43_RS06300 | 125818..128775 | 869814 | DUF499-domain-containing-protein-[Desulfobulbus-alkaliphilus] |
| HCF05840.1 | DUF499_1.RM.like.systems | TUDOR+SF1-SF2-Helicase+iREase+CTD+REase->PIN->HEPN->METHYLASE->OLD-ABC->TOPRIM->DUF499*-> | 934 | Bacteria>Thermodesulfobacteriota>Desulfovibrionia>Desulfovibrionales>Desulfomicrobiaceae | GCA_003530935.1 | Desulfomicrobiaceae_bacterium | DEU72_06355 | 12752..15556 | 2053547 | hypothetical-protein-[Desulfomicrobiaceae-bacterium] |
| MBZ4685271.1 | DUF499_1.RM.like.systems | FAD-NAD-dep-oxidoreductase-><-?<-?<-?\|\|?-><-DUF499*<-TOPRIM<-OLD-ABC<-METHYLASE<-HEPN<-PIN | 934 | Bacteria>Thermodesulfobacteriota>Desulfovibrionia>Desulfovibrionales>Desulfomicrobiaceae | GCA_020063575.1 | Desulfomicrobiaceae_bacterium | JG774_1016 | 59895..62699 | 2053547 | putative-ATPase-[Desulfomicrobiaceae-bacterium] |
| WP_160963929.1 | DUF499_1.RM.like.systems | TUDOR+SF1-SF2-Helicase+iREase+CTD+REase->HEPN->METHYLASE->DUF499*-> | 956 | Bacteria>Thermodesulfobacteriota>Desulfovibrionia>Desulfovibrionales>Desulfovibrionaceae | GCF_009856865.1 | Solidesulfovibrio_aerotolerans | GTA51_RS18990 | 10316..13186 | 295255 | DUF499-domain-containing-protein-[Solidesulfovibrio-aerotolerans] |
| WP_179981475.1 | DUF499_1.RM.like.systems | ABC-ATPase+SbcC->?->?->GDResolvase+HIN-HTH->DUF4158+DDE_Tnp_Tn3-><-OLD-ABC<-DUF499*<-NERD<-OLD-ABC<-METHYLASE<-HEPN<-TUDOR+SF1-SF2-Helicase+iREase+CTD+REase<-SF1-SF2-Helicase-p | 944 | Bacteria>Thermodesulfobacteriota>Desulfovibrionia>Desulfovibrionales>Desulfovibrionaceae | GCF_902702915.1 | Desulfovibrio_sp._86 | QZL82_RS00255 | 53858..56692 | 2666132 | DUF499-domain-containing-protein-[Desulfovibrio-sp.-86] |
| MBN1472033.1 | DUF499_1.RM.like.systems | METHYLASE-p->PolB->HEPN->?->DUF499*-> | 791 | Bacteria>Thermodesulfobacteriota>Syntrophia>Syntrophales>Syntrophaceae | GCA_016928665.1 | Syntrophaceae_bacterium | JW925_09645 | 2090..4465 | 2052184 | ATP-binding-protein-[Syntrophaceae-bacterium] |
| NSW87838.1 | DUF499_1.RM.like.systems | <-DUF499*<-METHYLASE-p<-HEPN<-wHTH+REase<-REase | 698 | Bacteria>Thermodesulfobacteriota>Syntrophobacteria>Syntrophobacterales>Syntrophobacteraceae | GCA_013314815.1 | Syntrophobacteraceae_bacterium | HPY84_16135 | 1106..3202 | 2052185 | DUF499-domain-containing-protein-[Syntrophobacteraceae-bacterium] |
| MBZ4682110.1 | DUF499_1.RM.like.systems | PRTase->?->Calcineurin->Radical-SAM-><-?<-?<-DUF499*<-METHYLASE-p<-HEPN<-PolB<-PolB<-METHYLASE<-TUDOR+SF1-SF2-Helicase+iREase+CTD+REase | 856 | Bacteria>Thermodesulfobacteriota>Thermodesulfobacteria>Thermodesulfobacteriales>Thermodesulfobacteriaceae | GCA_020055045.1 | Thermodesulfobacterium_sp. | JG778_1185 | 26969..29539 | 1965289 | hypothetical-protein-[Thermodesulfobacterium-sp.] |
| MCC6048503.1 | DUF499_1.RM.like.systems | <-DUF499*<-METHYLASE-p<-HEPN<-PolB<-METHYLASE-p | 454 | Bacteria>Thermodesulfobacteriota>Thermodesulfobacteria>Thermodesulfobacteriales>Thermodesulfobacteriaceae | GCA_020833455.1 | Thermodesulfobacterium_sp. | LM579_03100 | 1..1362 | 1965289 | DUF499-domain-containing-protein-[Thermodesulfobacterium-sp.] |
| WP_038060516.1 | DUF499_1.RM.like.systems | TUDOR+SF1-SF2-Helicase+iREase+CTD+REase->METHYLASE->PolB->PolB->HEPN->METHYLASE-p->DUF499*->?->?-><-Radical-SAM<-Calcineurin<-?<-PRTase | 856 | Bacteria>Thermodesulfobacteriota>Thermodesulfobacteria>Thermodesulfobacteriales>Thermodesulfobacteriaceae | GCF_000734015.1 | Thermodesulfobacterium_commune | CC99_RS02005 | 41791..44361 | 1741 | ATP-binding-protein-[Thermodesulfobacterium-commune] |
| KUK23759.1 | DUF499_1.RM.like.systems | REase-><-?<-?<-?<-DUF499-p<-DUF499*<-METHYLASE-p<-HEPN<-PolB<-METHYLASE<-TUDOR+SF1-SF2-Helicase+iREase+CTD+REase | 604 | Bacteria>Thermotogota>Thermotogae>Thermotogales>Thermotogaceae | GCA_001508125.1 | Thermotoga_petrophila | XD57_0150 | 12939..14753 | 93929 | MAG:-ATPase-(AAA+-superfamily)-like-protein-[Thermotoga-petrophila] |
| MBC7121832.1 | DUF499_1.RM.like.systems | REase-><-?<-?<-?<-DUF499*<-HEPN<-PolB<-METHYLASE-p | 847 | Bacteria>Thermotogota>Thermotogae>Thermotogales>Thermotogaceae | GCA_014360265.1 | Pseudothermotoga_sp. | H5T94_01065 | 9865..12408 | 2033661 | ATP-binding-protein-[Pseudothermotoga-sp.] |
| WP_031503324.1 | DUF499_1.RM.like.systems | TUDOR+SF1-SF2-Helicase+iREase+CTD+REase->METHYLASE-p->PolB->HEPN->METHYLASE-p->DUF499*->?->?->?-><-REase | 847 | Bacteria>Thermotogota>Thermotogae>Thermotogales>Thermotogaceae | GCF_000504105.1 | Pseudothermotoga_hypogea | THYO1S_RS07620 | 1580734..1583277 | 57487 | ATP-binding-protein-[Pseudothermotoga-hypogea] |
| GBD39155.1 | DUF499_1.RM.like.systems | <-RelE-ParE\|\|TUDOR+SF1-SF2-Helicase+iREase+CTD+REase->Schlafen->METHYLASE->PolB->HEPN->DUF499*->?-><-?\|\|?->NAD_binding_2+6PGD-> | 872 | Bacteria>unclassified_Bacteria | GCA_002898375.1 | bacterium_HR37 | HRbin37_01422 | 79215..81833 | 2035432 | hypothetical-protein-HRbin37_01422-[bacterium-HR37] |
| NJL57104.1 | DUF499_1.RM.like.systems | METHYLASE-p->DUF433->PIN->HEPN+DUF499*->DUF499-> | 411 | Bacteria>unclassified_Bacteria | GCA_012031555.1 | bacterium | HC928_19635 | 992..2227 | 1869227 | ATP-binding-protein-[bacterium] |
| NJL57105.1 | DUF499_1.RM.like.systems | METHYLASE-p->DUF433->PIN->HEPN+DUF499->DUF499*-> | 478 | Bacteria>unclassified_Bacteria | GCA_012031555.1 | bacterium | HC928_19640 | 2241..3676 | 1869227 | ATP-binding-protein-[bacterium] |
| MBU1261643.1 | DUF499_1.RM.like.systems | METHYLASE-p->PolB->HEPN->DUF499*-> | 515 | Bacteria>unclassified_Bacteria | GCA_018819855.1 | bacterium | KJ640_01640 | 1911..3456 | 1869227 | DUF499-domain-containing-protein-[bacterium] |
| MBU1261806.1 | DUF499_1.RM.like.systems | <-DUF499*<-TOPRIM<-OLD-ABC<-HEPN<-PolB<-METHYLASE-p | 494 | Bacteria>unclassified_Bacteria | GCA_018819855.1 | bacterium | KJ640_02495 | 1..1484 | 1869227 | DUF499-domain-containing-protein-[bacterium] |
| MBU1598556.1 | DUF499_1.RM.like.systems | METHYLASE-p->PolB->HEPN->OLD-ABC->TOPRIM->DUF499*-> | 494 | Bacteria>unclassified_Bacteria | GCA_018822205.1 | bacterium | KKG61_00350 | 3347..4830 | 1869227 | DUF499-domain-containing-protein-[bacterium] |
| MBU2461874.1 | DUF499_1.RM.like.systems | METHYLASE-p->PolB->HEPN->DUF499*-> | 513 | Bacteria>unclassified_Bacteria | GCA_018828965.1 | bacterium | KKH65_03235 | 1911..3449 | 1869227 | DUF499-domain-containing-protein-[bacterium] |
| MBU2462380.1 | DUF499_1.RM.like.systems | METHYLASE-p->PolB->HEPN->OLD-ABC->TOPRIM->DUF499*-> | 441 | Bacteria>unclassified_Bacteria | GCA_018828965.1 | bacterium | KKH65_05855 | 2757..4081 | 1869227 | DUF499-domain-containing-protein-[bacterium] |
| MCY3806661.1 | DUF499_1.RM.like.systems | METHYLASE->MetJarc-HTH->PemK_toxin->HEPN+DUF499*->DUF499-><-?\|\|ABhydrolase-> | 355 | Bacteria>unclassified_Bacteria | GCA_026706585.1 | bacterium | OXG91_09240 | 4057..5121 | 1869227 | MAG:-Swt1-family-HEPN-domain-containing-protein,-partial-[bacterium] |
| MCY3806662.1 | DUF499_1.RM.like.systems | METHYLASE->MetJarc-HTH->PemK_toxin->HEPN+DUF499->DUF499*-><-?\|\|ABhydrolase-> | 710 | Bacteria>unclassified_Bacteria | GCA_026706585.1 | bacterium | OXG91_09245 | 5222..7356 | 1869227 | MAG:-AAA+-family-ATPase,-partial-[bacterium] |
| MCY4104683.1 | DUF499_1.RM.like.systems | TUDOR+SF1-SF2-Helicase+iREase+CTD+REase->DUF4527->?->?->METHYLASE->DUF499*->HEPN-><-?<-?<-35exo<-?\|\|N6_N4_MTase-> | 985 | Bacteria>unclassified_Bacteria | GCA_026708055.1 | bacterium | OXG55_15720 | 11834..14791 | 1869227 | MAG:-DUF499-domain-containing-protein-[bacterium] |
| MCY4369121.1 | DUF499_1.RM.like.systems | <-HEPN<-DUF499*<-METHYLASE-p | 984 | Bacteria>unclassified_Bacteria | GCA_026710435.1 | bacterium | OXF41_06785 | 3276..6230 | 1869227 | MAG:-DUF499-domain-containing-protein-[bacterium] |

| Protein-ID | System | Gene-Neighbourhoods | Length | Taxonomy | Assembly-ID | Strain | Locus-Tag | Genomic-coordinates | TaxID | Fasta-defline |
|---|---|---|---|---|---|---|---|---|---|---|
| -- | | | | | | | | | | |
| -- ParB+HNH-(GmrSD) | | | | | | | | | | |
| -- | | | | | | | | | | |
| WP_146418283.1 | DUF499_1.RM.like.systems | <-RNase<-METHYLASE\|\|ParB+HNH-><-DUF499*\|\|SF1-SF2-Helicase->?-><-ORC/CDC6+HTH | 1068 | Archaea>Euryarchaeota>Stenosarchaea_group>Halobacteria>Halobacteriales | GCF_902209435.1 | Haloarcula_hispanica | FUT35_RS18485 | 5617..8823 | 51589 | DUF499-domain-containing-protein-[Haloarcula-hispanica] |
| WP_004062079.1 | DUF499_1.RM.like.systems | ORC/CDC6+HTH->?-><-?<-SF1-SF2-Helicase\|\|DUF499*-><-?\|\|?->?->METHYLASE->DUF429->ParB+HNH-> | 1039 | Archaea>Euryarchaeota>Stenosarchaea_group>Halobacteria>Haloferacales | GCF_000336795.1 | Haloferax_lucentense | C456_RS02980 | 68527..71646 | 2254 | DUF499-domain-containing-protein-[Haloferax-lucentense] |
| WP_224338217.1 | DUF499_1.RM.like.systems | <-PIWI<-SF1-SF2-Helicase+REase<-Exonuc_V_gamma+REase<-?<-?<-SF1-SF2-Helicase\|\|DUF499*->?-><-ParB+HNH<-ParB+HNH-p<-?\|\|METHYLASE->RNase-> | 1069 | Archaea>Euryarchaeota>Stenosarchaea_group>Halobacteria>Haloferacales | GCF_020097835.1 | Haloprofundus_halobius | LAQ74_RS18925 | 18862..22071 | 2876194 | DUF499-domain-containing-protein-[Haloprofundus-halobius] |
| WP_159663625.1 | DUF499_1.RM.like.systems | <-PIN<-METHYLASE<-?\|\|ParB+HNH->?-><-DUF499*\|\|SF1-SF2-Helicase-> | 1071 | Archaea>Euryarchaeota>Stenosarchaea_group>Halobacteria>Haloferacales | - | Halobaculum_saliterrae | GRX01_RS03890 | 204032..207247 | 2073113 | DUF499-domain-containing-protein-[Halobaculum-saliterrae] |
| CCJ36896.1 | DUF499_1.RM.like.systems | TUDOR+ SF1-SF2-Helicase+iREase+CTD+REase->METHYLASE-p->wHTH+REase->METHYLASE->ParB+HNH->DUF499*->?->PolB->HEPN-> | 944 | Archaea>Euryarchaeota>Stenosarchaea_group>Methanomicrobia>Methanomicrobiales | GCA_000304355.2 | Methanoculleus_bourgensis_MS2 | BN140_1973 | 2039579..2042413 | 1201294 | ATPase-[Methanoculleus-bourgensis-MS2] |
| RXE56548.1 | DUF499_1.RM.like.systems | ORC/CDC6+HTH-><-DUF499*<-ParB+HNH<-METHYLASE<-TUDOR+ SF1-SF2-Helicase+iREase+CTD+REase\|\|REase-> | 935 | Archaea>Euryarchaeota>Stenosarchaea_group>Methanomicrobia>Methanomicrobiales | GCA_004102725.1 | Methanoculleus_taiwanensis | ABH15_07530 | 552550..555357 | 1550565 | hypothetical-protein-ABH15_07530-[Methanoculleus-taiwanensis] |
| NLM31070.1 | DUF499_1.RM.like.systems | METHYLASE->ParB+HNH->DUF499*-> | 942 | Archaea>Euryarchaeota>Stenosarchaea_group>Methanomicrobia>Methanomicrobiales | GCA_012521035.1 | Methanomicrobiales_archaeon | GX216_11755 | 4804..7632 | 2184052 | ATP-binding-protein-[Methanomicrobiales-archaeon] |
| MCE5337191.1 | DUF499_1.RM.like.systems | ORC/CDC6+HTH->DOC+HTH-><-?<-PolB<-?<-PolB<-DUF499*<-ParB+HNH<-METHYLASE<-?<-wHTH+REase<-METHYLASE-p<-TUDOR+ SF1-SF2-Helicase+iREase+CTD+REase | 944 | Archaea>Euryarchaeota>Stenosarchaea_group>Methanomicrobia>Methanomicrobiales | GCA_021371835.1 | Methanomicrobiaceae_archaeon | LLF90_00675 | 34057..36891 | 2716262 | DUF499-domain-containing-protein-[Methanomicrobiaceae-archaeon] |
| WP_156147608.1 | DUF499_1.RM.like.systems | TUDOR+ SF1-SF2-Helicase+iREase+CTD+REase->METHYLASE-p->wHTH+REase->?->METHYLASE->ParB+HNH->DUF499*->PolB->HEPN-><-?<-Ntox49 | 942 | Archaea>Euryarchaeota>Stenosarchaea_group>Methanomicrobia>Methanomicrobiales | GCF_000304355.2 | Methanoculleus_bourgensis | BN140_RS09865 | 2039585..2042413 | 83986 | DUF499-domain-containing-protein-[Methanoculleus-bourgensis] |
| WP_235808115.1 | DUF499_1.RM.like.systems | <-SF1-SF2-Helicase<-?<-?\|\|TUDOR+ SF1-SF2-Helicase+iREase+CTD+REase->METHYLASE->ParB+HNH->DUF499*-><-?<-?<-?<-?<-?<-XIS-HTH | 945 | Archaea>Euryarchaeota>Stenosarchaea_group>Methanomicrobia>Methanomicrobiales | GCF_001602375.1 | Methanoculleus_horonobensis | MCUHO_RS01275 | 28363..31200 | 528314 | DUF499-domain-containing-protein-[Methanoculleus-horonobensis] |
| WP_206633427.1 | DUF499_1.RM.like.systems | ORC/CDC6+HTH-><-DUF499*<-ParB+HNH<-METHYLASE<-TUDOR+ SF1-SF2-Helicase+iREase+CTD+REase\|\|REase-> | 937 | Archaea>Euryarchaeota>Stenosarchaea_group>Methanomicrobia>Methanomicrobiales | GCF_004102725.1 | Methanoculleus_taiwanensis | ABH15_RS07615 | 552550..555363 | 1550565 | DUF499-domain-containing-protein-[Methanoculleus-taiwanensis] |
| WP_253456790.1 | DUF499_1.RM.like.systems | TUDOR+ SF1-SF2-Helicase+iREase+CTD+REase->METHYLASE->ParB+HNH->DUF499*-><-?<-tRNA<-?\|\|Radical-SAM->?->tRNA-><-METHYLASE | 947 | Archaea>Euryarchaeota>Stenosarchaea_group>Methanomicrobia>Methanomicrobiales | GCF_024170265.1 | Methanocalculus_sp._AMF5 | J2T61_RS02245 | 78750..81593 | 1198257 | DUF499-domain-containing-protein-[Methanocalculus-sp.-AMF5] |
| MBZ5523276.1 | DUF499_1.RM.like.systems | <-ABhydrolase<-?\|\|ParB+HNH->METHYLASE-><-HEPN\|\|DUF499*-><-PIN<-?\|\|MazE_antitoxin->PemK_toxin->?->PIN-> | 924 | Bacteria>Acidobacteriota>Terriglobia>unclassified_Terriglobia | GCA_020072595.1 | Terriglobia_bacterium | LAP21_13650 | 15770..18544 | 2081523 | DUF499-domain-containing-protein-[Acidobacteriia-bacterium] |
| KAB2964430.1 | DUF499_1.RM.like.systems | REase->TUDOR+ SF1-SF2-Helicase+iREase+CTD+REase->ParB+HNH->HEPN->METHYLASE->?->HEPN+ DUF499*->HNH-> | 1109 | Bacteria>Acidobacteriota>Thermoanaerobaculia>unclassified_Thermoanaerobaculia | GCA_009026165.1 | Thermoanaerobaculia_bacterium | F9K16_04350 | 12106..15435 | 2651171 | MAG:-ATP-binding-protein-[Thermoanaerobaculia-bacterium] |
| MBA4124439.1 | DUF499_1.RM.like.systems | DDE_Tnp_1_assoc+DDE_Tnp_1->CHTH->?->TUDOR+ SF1-SF2-Helicase+iREase+CTD+REase->ParB+HNH->METHYLASE->HEPN+ DUF499*->?->?->BirA-HTH-><-35exo<-Phage_integrase | 1091 | Bacteria>Acidobacteriota>unclassified_Acidobacteriota | GCA_013820745.1 | Acidobacteriota_bacterium | HOX72_18505 | 11663..14938 | 1978231 | DUF499-domain-containing-protein-[Acidobacteria-bacterium] |
| MBI4902187.1 | DUF499_1.RM.like.systems | TUDOR+ SF1-SF2-Helicase+iREase+CTD+REase->ParB+HNH->METHYLASE->OLD-ABC->TOPRIM->HEPN+ DUF499*->Cluster2725_2clades->Terminase-> | 1115 | Bacteria>Acidobacteriota>unclassified_Acidobacteriota | GCA_016213775.1 | Acidobacteriota_bacterium | HY820_01040 | 17603..20950 | 1978231 | ATP-binding-protein-[Acidobacteria-bacterium] |
| MBP7777875.1 | DUF499_1.RM.like.systems | TUDOR+ SF1-SF2-Helicase+iREase+CTD+REase->DUF932->METHYLASE->HEPN+ DUF499*->?->Pkinase->ParB+HNH->REase->?->HicB_lk_antitox-> | 1114 | Bacteria>Acidobacteriota>unclassified_Acidobacteriota | GCA_018000455.1 | Acidobacteriota_bacterium | KA371_12150 | 120064..123408 | 1978231 | DUF499-domain-containing-protein-[Acidobacteria-bacterium] |
| MCY3932061.1 | DUF499_1.RM.like.systems | TUDOR+ SF1-SF2-Helicase+iREase+CTD+REase->ParB+HNH->Schlafen->?->METHYLASE-p->METHYLASE->DUF499*-><-?<-?\|\|Peptidase_M28-> | 937 | Bacteria>Acidobacteriota>unclassified_Acidobacteriota | GCA_026707325.1 | Acidobacteriota_bacterium | OXH70_10095 | 67644..70457 | 1978231 | MAG:-DUF499-domain-containing-protein-[Acidobacteria-bacterium] |
| MCY4659240.1 | DUF499_1.RM.like.systems | <-ORC-CDC6+REase<-DUF499*<-PIN<-OLD-ABC<-METHYLASE<-ParB+HNH+Schlafen<-TOPRIM<-OLD-ABC | 941 | Bacteria>Acidobacteriota>unclassified_Acidobacteriota | GCA_026713985.1 | Acidobacteriota_bacterium | OXF93_05425 | 1121..3946 | 1978231 | MAG:-DUF499-domain-containing-protein-[Acidobacteria-bacterium] |
| MBE0461501.1 | DUF499_1.RM.like.systems | <-HEPN<-ParB+HNH<-DUF499*<-METHYLASE-p | 878 | Bacteria>Bacteria_incertae_sedis>Bacteria_candidate_phyla>Candidatus_Aminicenantes>unclassified_Candidatus_Aminicenantes | GCA_014859315.1 | Candidatus_Aminicenantes_bacterium | IBX60_07695 | 3318..5954 | 2052149 | ATP-binding-protein-[Candidatus-Aminicenantes-bacterium] |
| MDD5453217.1 | DUF499_1.RM.like.systems | SF1-SF2-Helicase-p->PIN->ParB+HNH+Schlafen->METHYLASE->PolB->HEPN->DUF499*-><-ABC-ATPase | 932 | Bacteria>Bacteria_incertae_sedis>Bacteria_candidate_phyla>Candidatus_Bipolaricaulota>Bipolaricaulis | GCA_028715685.1 | Candidatus_Bipolaricaulis_sp. | PHZ21_02925 | 36651..39449 | 2814139 | MAG:-DUF499-domain-containing-protein-[Candidatus-Bipolaricaulis-sp.] |
| MCF7889851.1 | DUF499_1.RM.like.systems | TUDOR+ SF1-SF2-Helicase+iREase+CTD+REase->METHYLASE->DUF499*->ParB+HNH->XIS-HTH-> | 911 | Bacteria>Bacteria_incertae_sedis>Bacteria_candidate_phyla>Candidatus_Bipolaricaulota>unclassified_Candidatus_Bipolaricaulota | GCA_021735245.1 | Candidatus_Bipolaricaulota_bacterium | K9M78_01380 | 69042..71777 | 2306012 | DUF499-domain-containing-protein-[Candidatus-Bipolaricaulota-bacterium] |
| NPV88406.1 | DUF499_1.RM.like.systems | TUDOR+ SF1-SF2-Helicase+iREase+CTD+REase->METHYLASE->DUF499*->ParB+HNH->HEPN+Mrr_N->HEPN+Mrr_N-> | 865 | Bacteria>Coprothermobacterota>Coprothermobacteria>Coprothermobacterales>Coprothermobacteraceae | GCA_013178255.1 | Coprothermobacteraceae_bacterium | HPY42_02620 | 513240..515837 | 2736298 | ATP-binding-protein-[Coprothermobacteraceae-bacterium] |
| MBP6872709.1 | DUF499_1.RM.like.systems | TUDOR+ SF1-SF2-Helicase+iREase+CTD+REase->AbiEi_1->AbiEii->ParB+HNH->METHYLASE->HEPN+ DUF499*->?->SbcC->?-><-?<-REase+SF1-SF2-Helicase<-MazG | 1092 | Bacteria>FCB_group>Bacteroidota/Chlorobiota_group>Bacteroidota>Bacteroidia | GCA_017991835.1 | Bacteroidales_bacterium | KBC43_11940 | 58567..61845 | 2030927 | ATP-binding-protein-[Bacteroidales-bacterium] |
| NCU04700.1 | DUF499_1.RM.like.systems | wHTH+AbiEi->AbiEii->METHYLASE->ParB+HNH->OLD-ABC->?->DUF499*-> | 237 | Bacteria>FCB_group>Bacteroidota/Chlorobiota_group>Bacteroidota>Chitinophagia | GCA_009996555.1 | Chitinophagaceae_bacterium | GXC73_12005 | 16910..17621 | 1869212 | ATP-binding-protein-[Chitinophagaceae-bacterium] |
| MBN8676639.1 | DUF499_1.RM.like.systems | REase+35exo+Beta-Barrel+SF1-SF2-Helicase->?->?->TUDOR+ SF1-SF2-Helicase+iREase+CTD+REase->ParB+HNH->METHYLASE->HEPN+ DUF499*->REase-> | 1087 | Bacteria>FCB_group>Bacteroidota/Chlorobiota_group>Bacteroidota>Chitinophagia | GCA_017303475.1 | Chitinophagales_bacterium | JOM29_00350 | 102288..105551 | 2448779 | DUF499-domain-containing-protein-[Chitinophagales-bacterium] |
| MBP8114174.1 | DUF499_1.RM.like.systems | HTH->TUDOR+ SF1-SF2-Helicase+iREase+CTD+REase->METHYLASE->ParB+HNH->OLD-ABC->?->DUF499*-> | 925 | Bacteria>FCB_group>Bacteroidota/Chlorobiota_group>Bacteroidota>Chitinophagia | GCA_018005575.1 | Chitinophagaceae_bacterium | KAY50_02395 | 14241..17018 | 1869212 | ATP-binding-protein-[Chitinophagaceae-bacterium] |
| MBX7108627.1 | DUF499_1.RM.like.systems | HTH->TUDOR+ SF1-SF2-Helicase+iREase+CTD+REase->METHYLASE->ParB+HNH->OLD-ABC->?->DUF499*->?->PIWI-> | 925 | Bacteria>FCB_group>Bacteroidota/Chlorobiota_group>Bacteroidota>Chitinophagia | GCA_019694975.1 | Chitinophagales_bacterium | K1X61_08290 | 480306..483083 | 2448779 | DUF499-domain-containing-protein-[Chitinophagales-bacterium] |
| WP_086597021.1 | DUF499_1.RM.like.systems | TUDOR+ SF1-SF2-Helicase+iREase+CTD+REase->ParB+HNH->METHYLASE->DUF499*-><-?\|\|?->Peptidase_S74-> | 972 | Bacteria>FCB_group>Bacteroidota/Chlorobiota_group>Bacteroidota>Cytophagia | GCF_002154225.1 | Hymenobacter_crusticola | BXP70_RS25835 | 30420..33338 | 1770526 | DUF499-domain-containing-protein-[Hymenobacter-crusticola] |
| WP_114065477.1 | DUF499_1.RM.like.systems | <-TUDOR+SF1-SF2-Helicase+iREase+CTD<-HEPN+ DUF499*<-?<-OLD-ABC<-ParB+HNH<-METHYLASE<-REase<-TUDOR+ SF1-SF2-Helicase+iREase+CTD+REase | 1094 | Bacteria>FCB_group>Bacteroidota/Chlorobiota_group>Bacteroidota>Cytophagia | GCF_003325355.1 | Runella_rosea | DR864_RS02565 | 579082..582366 | 2259595 | DUF499-domain-containing-protein-[Runella-rosea] |
| WP_209329595.1 | DUF499_1.RM.like.systems | <-BirA-HTH<-?<-?<-HEPN+ DUF499*<-AAA-ATPase+REase<-METHYLASE<-ParB+HNH<-TUDOR+ SF1-SF2-Helicase+iREase+CTD+REase | 1097 | Bacteria>FCB_group>Bacteroidota/Chlorobiota_group>Bacteroidota>Cytophagia | GCF_017811155.1 | Lunatimonas_salinarum | IAQ59_RS04505 | 160980..164273 | 1774590 | DUF499-domain-containing-protein-[Lunatimonas-salinarum] |
| MBN1449251.1 | DUF499_1.RM.like.systems | <-DUF499*<-ParB+HNH<-METHYLASE | 939 | Bacteria>FCB_group>Bacteroidota/Chlorobiota_group>Bacteroidota>unclassified_Bacteroidota | GCA_016928515.1 | Bacteroidota_bacterium | JXA28_15095 | 364..3183 | 1898104 | ATP-binding-protein-[Bacteroidetes-bacterium] |
| MBS1776511.1 | DUF499_1.RM.like.systems | XIS-HTH->?->HTH->TUDOR+ SF1-SF2-Helicase+iREase+CTD+REase->METHYLASE->DUF499*->PIWI->ParB+HNH-> | 925 | Bacteria>FCB_group>Bacteroidota/Chlorobiota_group>Bacteroidota>unclassified_Bacteroidota | GCA_018267925.1 | Bacteroidota_bacterium | JSS64_09560 | 68237..71014 | 1898104 | ATP-binding-protein-[Bacteroidetes-bacterium] |
| GIV55103.1 | DUF499_1.RM.like.systems | TUDOR+ SF1-SF2-Helicase+iREase+CTD+REase->Schlafen->BirA-HTH->METHYLASE-p->METHYLASE-p->DUF499*->ParB+HNH-> | 875 | Bacteria>FCB_group>Bacteroidota/Chlorobiota_group>Candidatus_Kapabacteria>unclassified_Candidatus_Kapabacteria | GCA_026003475.1 | Candidatus_Kapabacteria_bacterium | KatS3mg039_1621 | 1981096..1983723 | 2447898 | MAG:-hypothetical-protein-KatS3mg039_1621-[Candidatus-Kapabacteria-bacterium] |
| MYA34704.1 | DUF499_1.RM.like.systems | HEPN->TUDOR+ SF1-SF2-Helicase+iREase+CTD+REase->METHYLASE->OLD-ABC->TOPRIM->ParB+HNH+Schlafen->DUF499*-><-?\|\|HTH->MPTase->?->SMODS-><-ABhydrolase | 928 | Bacteria>FCB_group>Gemmatimonadota>Gemmatimonadetes>Gemmatimonadales | GCA_009838765.1 | Gemmatimonadales_bacterium | F4Y33_13800 | 70561..73347 | 2448054 | ATP-binding-protein-[Gemmatimonadales-bacterium] |
| MXZ76449.1 | DUF499_1.RM.like.systems | Synaptojanin->TUDOR+ SF1-SF2-Helicase+iREase+CTD+REase->OLD-ABC->METHYLASE->HEPN+ DUF499*->ParB+HNH-><-?<-MazE_antitoxin | 1113 | Bacteria>FCB_group>Gemmatimonadota>unclassified_Gemmatimonadota | GCA_009837395.1 | Gemmatimonadota_bacterium | F4Z03_15245 | 64974..68315 | 2026742 | ATP-binding-protein-[Gemmatimonadetes-bacterium] |
| MYH20375.1 | DUF499_1.RM.like.systems | TUDOR+ SF1-SF2-Helicase+iREase+CTD+REase->ParB+HNH->METHYLASE->HEPN+ DUF499*->?->TPR-><-Peptidase-S8<-Classical-AAA<-Peptidase_M1 | 1108 | Bacteria>FCB_group>Gemmatimonadota>unclassified_Gemmatimonadota | GCA_009844015.1 | Gemmatimonadota_bacterium | F4132_14865 | 91560..94886 | 2026742 | ATP-binding-protein-[Gemmatimonadetes-bacterium] |
| MYL00076.1 | DUF499_1.RM.like.systems | TUDOR+ SF1-SF2-Helicase+iREase+CTD+REase->ParB+HNH-p->METHYLASE->HEPN+ DUF499*->?->TPR-><-Peptidase-S8 | 1108 | Bacteria>FCB_group>Gemmatimonadota>unclassified_Gemmatimonadota | GCA_009846305.1 | Gemmatimonadota_bacterium | F4014_15170 | 26347..29673 | 2026742 | ATP-binding-protein-[Gemmatimonadetes-bacterium] |
| MBI5504530.1 | DUF499_1.RM.like.systems | MazE_antitoxin->DOC-><-HNH<-?<-?<-HEPN+ DUF499*<-METHYLASE<-?<-ParB+HNH<-TUDOR+ SF1-SF2-Helicase+iREase+CTD+REase<-?\|\|HIN-HTH-> | 1116 | Bacteria>Myxococcota>Myxococcia>unclassified_Myxococcia | GCA_016218925.1 | Deltaproteobacteria_bacterium | HY899_06990 | 27440..30790 | 2026735 | ATP-binding-protein-[Deltaproteobacteria-bacterium] |
| NGZ08749.1 | DUF499_1.RM.like.systems | Glycos_transf_4->TUDOR+ SF1-SF2-Helicase+iREase+CTD+REase->ParB+HNH->METHYLASE->HEPN+ DUF499*-> | 1121 | Bacteria>Nitrospirota>Nitrospiria>Nitrospirales>Nitrospiraceae | GCA_011090425.1 | Nitrospira_sp._LX70 | CV088_05085 | 57956..61321 | 2039809 | ATP-binding-protein-[Nitrospira-sp.-LX70] |
| MBP6261842.1 | DUF499_1.RM.like.systems | <-XIS-HTH\|\|TUDOR+ SF1-SF2-Helicase+iREase+CTD+REase->AbiGi->ParB+HNH->METHYLASE->HEPN+ DUF499*->HEPN+CBS->?-><-?\|\|HIN-HTH-> | 1121 | Bacteria>Nitrospirota>Nitrospiria>Nitrospirales>Nitrospiraceae | GCA_017986555.1 | Nitrospira_sp. | KA426_01230 | 11559..14924 | 70125 | DUF499-domain-containing-protein-[Nitrospira-sp.] |
| MBS0184610.1 | DUF499_1.RM.like.systems | TUDOR+ SF1-SF2-Helicase+iREase+CTD+REase-p->TUDOR+ SF1-SF2-Helicase+iREase+CTD+REase-p->ParB+HNH->METHYLASE->METHYLASE-p->HEPN->HEPN+ DUF499*-> | 1121 | Bacteria>Nitrospirota>Nitrospiria>Nitrospirales>Nitrospiraceae | GCA_018242685.1 | Nitrospira_sp. | JSS39_19650 | 30690..34055 | 70125 | DUF499-domain-containing-protein-[Nitrospira-sp.] |
| MCS6288012.1 | DUF499_1.RM.like.systems | <-XIS-HTH\|\|TUDOR+ SF1-SF2-Helicase+iREase+CTD+REase->ParB+HNH->METHYLASE->HEPN+ DUF499*->RelE-ParE->HTH->SF1-SF2-Helicase-> | 1121 | Bacteria>Nitrospirota>Nitrospiria>Nitrospirales>Nitrospiraceae | GCA_024998625.1 | Nitrospira_sp. | H8K10_03430 | 260842..264207 | 70125 | ATP-binding-protein-[Nitrospira-sp.] |
| KAF0185697.1 | DUF499_1.RM.like.systems | MazE_antitoxin->PIN->TUDOR+ SF1-SF2-Helicase+iREase+CTD+REase->METHYLASE->HEPN+ DUF499*-><-ParB+HNH\|\|?->?->?->Phage_AlpA-><-Arm-DNA-bind_3+Phage_integrase | 1077 | Bacteria>Pseudomonadota>Alphaproteobacteria>Caulobacterales>Caulobacteraceae | GCA_009773335.1 | Caulobacteraceae_bacterium | FD160_506 | 42091..45324 | 1978230 | ATPase--like-protein-[Caulobacteraceae-bacterium] |
| WP_078709662.1 | DUF499_1.RM.like.systems | MODE-HTH+PBPII->?->ABC-ATPase+SbcC-><-METHYLASE-p<-?\|\|<-HEPN+ DUF499*<-METHYLASE<-SIR2<-ParB+HNH<-ASCH<-TUDOR+ SF1-SF2-Helicase+iREase+CTD+REase\|\|REase-> | 1102 | Bacteria>Pseudomonadota>Alphaproteobacteria>Hyphomicrobiales>Aurantimonadaceae | GCF_900167365.1 | Consotaella_salsifontis | B5D38_RS17130 | 40507..43815 | 1365950 | DUF499-domain-containing-protein-[Consotaella-salsifontis] |
| WP_259030292.1 | DUF499_1.RM.like.systems | <-SMODS\|\|HTH+WYL->TUDOR+ SF1-SF2-Helicase+iREase+CTD+REase->ParB+HNH->?->METHYLASE->HEPN+ DUF499*->GDResolvase+HIN-HTH->HEPN->?->?-><-DDE_3<-PAIREDC-HTH | 1098 | Bacteria>Pseudomonadota>Alphaproteobacteria>Hyphomicrobiales>Brucellaceae | GCF_024799785.1 | Brucella_anthropi | NW321_RS01585 | 345006..348302 | 529 | DUF499-domain-containing-protein-[Brucella-anthropi] |
| UYN95212.1 | DUF499_1.RM.like.systems | TUDOR+ SF1-SF2-Helicase+iREase+CTD+REase->?->ParB+HNH->METHYLASE->HEPN+ DUF499*->AbiEii-><-?<-?<-SPOUT | 1104 | Bacteria>Pseudomonadota>Alphaproteobacteria>Hyphomicrobiales>Hyphomicrobiales_incertae_sedis | GCA_025808875.1 | Enhydrobacter_sp. | KIT25_24915 | 5028707..5032021 | 1894999 | MAG:-DUF499-domain-containing-protein-[Enhydrobacter-sp.] |
| WP_244479882.1 | DUF499_1.RM.like.systems | TUDOR+ SF1-SF2-Helicase+iREase+CTD+REase->OLD-ABC->HNH->METHYLASE->DUF499*->GDResolvase+HTH->ParB+HNH-><-?\|\|HTH_3-><-?\|\|REase-> | 950 | Bacteria>Pseudomonadota>Alphaproteobacteria>Hyphomicrobiales>Methylobacteriaceae | GCF_001426045.1 | Methylobacterium_sp._Leaf94 | ASG63_RS11585 | 13925..16777 | 1736250 | ATP-binding-protein-[Methylobacterium-sp.-Leaf94] |
| WP_247344343.1 | DUF499_1.RM.like.systems | HTH+WYL->TUDOR+ SF1-SF2-Helicase+iREase+CTD+REase->ParB+HNH->?->METHYLASE->HEPN+ DUF499*->HEPN-> | 1097 | Bacteria>Pseudomonadota>Alphaproteobacteria>Hyphomicrobiales>Nitrobacteraceae | GCF_023073075.1 | Bradyrhizobium_sp._61 | IVB46_RS33345 | 520098..523391 | 2782679 | DUF499-domain-containing-protein-[Bradyrhizobium-sp.-61] |
| WP_079537876.1 | DUF499_1.RM.like.systems | <-ACYC\|\|?->OLD-ABC->?-><-HKD-DNase<-ParB+HNH<-HEPN+ DUF499*<-METHYLASE<-ASCH<-TUDOR+ SF1-SF2-Helicase+iREase+CTD+REase\|\|?-><-GDResolvase+HIN-HTH | 1108 | Bacteria>Pseudomonadota>Alphaproteobacteria>Hyphomicrobiales>Nitrobacteraceae | GCF_900141755.1 | Bradyrhizobium_lablabi | B5526_RS08940 | 1910931..1914257 | 722472 | DUF499-domain-containing-protein-[Bradyrhizobium-lablabi] |
| CAI2932244.1 | DUF499_1.RM.like.systems | ParG_RHH-><-MODE-HTH+PBPII<-HEPN+ DUF499*<-METHYLASE<-ParB+HNH<-TUDOR+ SF1-SF2-Helicase+iREase+CTD+REase<-HTH+WYL\|\|SF1-SF2-Helicase+REase->SF1-SF2-Helicase+REase-> | 1098 | Bacteria>Pseudomonadota>Alphaproteobacteria>Hyphomicrobiales>Phyllobacteriaceae | GCA_946995915.1 | Aminobacter_niigataensis | AMD1_0962 | 943205..946501 | 83265 | ATPase-AAA-[Aminobacter-niigataensis] |
| WP_127665091.1 | DUF499_1.RM.like.systems | Arm-DNA-bind_3+Phage_integrase->\|\|HTH+PolB->?-><-?<-?<-ParB+HNH<-HEPN+ DUF499*<-?<-METHYLASE<-?<-TUDOR+ SF1-SF2-Helicase+iREase+CTD+REase<-SF1-SF2-Helicase+DUF1998+HKD-DNase+HKD-DNase | 1104 | Bacteria>Pseudomonadota>Alphaproteobacteria>Hyphomicrobiales>Rhizobiaceae | GCF_004001475.1 | Sinorhizobium_meliloti | CN231_RS17685 | 56193..59507 | 382 | DUF499-domain-containing-protein-[Sinorhizobium-meliloti] |
| WP_055658296.1 | DUF499_1.RM.like.systems | Arm-DNA-bind_3+Phage_integrase->?->Phage_AlpA->ZnR+TOPRIM->DnaB+CITB-HTH->?-><-HEPN+ DUF499*<-?<-METHYLASE<-?<-ParB+HNH<-MORC+DUF3684<-TUDOR+ SF1-SF2-Helicase+iREase+CTD+REase | 1112 | Bacteria>Pseudomonadota>Alphaproteobacteria>Hyphomicrobiales>Stappiaceae | GCF_001403915.1 | Roseibium_aggregatum | LAL4801_RS18620 | 1338638..1341976 | 187304 | DUF499-domain-containing-protein-[Roseibium-aggregatum] |
| MCW5685472.1 | DUF499_1.RM.like.systems | <-Peptidase_S11<-?\|\|tRNA-><-?<-HEPN+ DUF499*<-METHYLASE<-ParB+HNH<-TUDOR+ SF1-SF2-Helicase+iREase+CTD+REase<-HTH+WYL\|\|SF1-SF2-Helicase+REase-> | 1098 | Bacteria>Pseudomonadota>Alphaproteobacteria>Hyphomicrobiales>Xanthobacteraceae | GCA_026125885.1 | Pseudolabrys_sp. | KIT85_13790 | 135757..139053 | 1960880 | MAG:-DUF499-domain-containing-protein-[Pseudolabrys-sp.] |
| MCQ0971011.1 | DUF499_1.RM.like.systems | MODE-HTH+PBPII->\|\|?->GDResolvase+Recombinase+ZnR->\|\|DDE_Tnp_IS66->MODE-HTH+PBPII-><-HEPN+ DUF499*<-METHYLASE<-ParB+HNH<-TUDOR+ SF1-SF2-Helicase+iREase+CTD+REase<-HTH+WYL\|\|SF1-SF2-Helicase+REase->SF1-SF2-Helicase+REase-> | 1097 | Bacteria>Pseudomonadota>Alphaproteobacteria>Rhodobacterales>Paracoccaceae | GCA_024359525.1 | Paracoccus_albicereus | MLD63_11305 | 50725..54018 | 2922394 | DUF499-domain-containing-protein-[Paracoccus-sp.-TK19116] |
| WP_035746761.1 | DUF499_1.RM.like.systems | <-DDE_Tnp_1_5+DDE_Tnp_1<-HTH_Tnp_1<-HTH_Tnp_1<-HEPN+ DUF499*<-METHYLASE<-ParB+HNH<-TUDOR+ SF1-SF2-Helicase+iREase+CTD+REase<-HTH+WYL\|\|SF1-SF2-Helicase+REase->SF1-SF2-Helicase+REase-> | 1098 | Bacteria>Pseudomonadota>Alphaproteobacteria>Rhodobacterales>Paracoccaceae | GCF_000740775.1 | Haematobacter_missouriensis | CG51_RS16225 | 8779..12075 | 366616 | DUF499-domain-containing-protein-[Haematobacter-missouriensis] |
| WP_255330034.1 | DUF499_1.RM.like.systems | MODE-HTH+PBPII->\|\|?->GDResolvase+Recombinase+ZnR->\|\|DDE_Tnp_IS66->MODE-HTH+PBPII-><-HEPN+ DUF499*<-METHYLASE<-ParB+HNH<-TUDOR+ SF1-SF2-Helicase+iREase+CTD+REase<-HTH+WYL\|\|SF1-SF2-Helicase+REase->SF1-SF2-Helicase+REase-> | 1097 | Bacteria>Pseudomonadota>Alphaproteobacteria>Rhodobacterales>Paracoccaceae | GCF_024359525.1 | Paracoccus_albicereus | MLD63_RS11355 | 50725..54018 | 2922394 | DUF499-domain-containing-protein-[Paracoccus-sp.-TK19116] |
| WP_094020052.1 | DUF499_1.RM.like.systems | XIS-HTH->HTH+WYL->ParB+HNH->TUDOR+ SF1-SF2-Helicase+iREase+CTD+REase->?->METHYLASE->HEPN+ DUF499*-><-RNase<-?<-?<-DUF2163+Phage_BR0599 | 1098 | Bacteria>Pseudomonadota>Alphaproteobacteria>Rhodobacterales>Paracoccaceae | GCF_900184945.1 | Maliponia_aquimaris | MAA8898_RS05760 | 37634..40930 | 1673631 | DUF499-domain-containing-protein-[Maliponia-aquimaris] |
| WP_093359600.1 | DUF499_1.RM.like.systems | XIS-HTH->HTH+WYL->ParB+HNH->TUDOR+ SF1-SF2-Helicase+iREase+CTD+REase->ASCH->METHYLASE->HEPN+ DUF499*->RNase+SF1-SF2-Helicase-><-RNase<-?<-?<-DUF2163+Phage_BR0599 | 1109 | Bacteria>Pseudomonadota>Alphaproteobacteria>Rhodobacterales>Roseobacteraceae | GCF_900112335.1 | Tropicimonas_isoalkanivorans | BM075_RS05220 | 205092..208421 | 441112 | DUF499-domain-containing-protein-[Tropicimonas-isoalkanivorans] |
| MBF9033364.1 | DUF499_1.RM.like.systems | XIS-HTH->HTH+WYL->ParB+HNH->TUDOR+ SF1-SF2-Helicase+iREase+CTD+REase->YtH->METHYLASE->HEPN+ DUF499*->AbiEii-><-?<-Peptidase_M15 | 1098 | Bacteria>Pseudomonadota>Alphaproteobacteria>Rhodobacterales>unclassified_Rhodobacterales | GCA_015689685.1 | Rhodobacterales_bacterium_HKCCE2091 | HKCCE2091_03860 | 312307..315603 | 2792619 | DUF499-domain-containing-protein-[Rhodobacterales-bacterium-HKCCE2091] |
| WP_145610986.1 | DUF499_1.RM.like.systems | TUDOR+ SF1-SF2-Helicase+iREase+CTD+REase->ParB+HNH->METHYLASE->PolB->HEPN->HEPN+ DUF499*->?->PIN-> | 1115 | Bacteria>Pseudomonadota>Alphaproteobacteria>Rhodospirillales>Azospirillaceae | GCF_007828035.1 | Nitrospirillum_amazonense | FBZ87_RS14570 | 656187..659534 | 28077 | Swt1-family-HEPN-domain-containing-protein-[Nitrospirillum-amazonense] |
| WP_017183457.1 | DUF499_1.RM.like.systems | <-HicB\|\|HTH+WYL->TUDOR+ SF1-SF2-Helicase+iREase+CTD+REase->ParB+HNH->ASCH->METHYLASE->HEPN+ DUF499*->MODE-HTH+PBPII-><-ParG_RHH | 1097 | Bacteria>Pseudomonadota>Alphaproteobacteria>Sphingomonadales>Sphingomonadaceae | GCF_000277525.1 | Sphingobium_xenophagum | QYYP_RS0113510 | 14048..17341 | 121428 | DUF499-domain-containing-protein-[Sphingobium-xenophagum] |
| WP_166837252.1 | DUF499_1.RM.like.systems | TUDOR+ SF1-SF2-Helicase+iREase+CTD+REase->ParB+HNH->PIN->METHYLASE->DUF499*-> | 936 | Bacteria>Pseudomonadota>Betaproteobacteria>Burkholderiales>Burkholderiales_genera_incertae_sedis | GCF_011742065.1 | Aquabacterium_sp._A08 | HA630_RS20340 | 28723..31533 | 2718532 | DUF499-domain-containing-protein-[Aquabacterium-sp.-A08] |
| MCP5281003.1 | DUF499_1.RM.like.systems | <-HEPN+ DUF499*<-?<-METHYLASE<-DUF932<-ParB+HNH<-?<-TUDOR+ SF1-SF2-Helicase+iREase+CTD+REase | 1107 | Bacteria>Pseudomonadota>Betaproteobacteria>Burkholderiales>Comamonadaceae | GCA_024235935.1 | Rhodoferax_sp. | H6930_05335 | 220759..224082 | 50421 | ATP-binding-protein-[Rhodoferax-sp.] |
| WP_045536717.1 | DUF499_1.RM.like.systems | TUDOR+ SF1-SF2-Helicase+iREase+CTD+REase->PIN->HEPN->METHYLASE-p->METHYLASE->ParB+HNH->DUF499*-> | 950 | Bacteria>Pseudomonadota>Betaproteobacteria>Burkholderiales>Comamonadaceae | GCF_000828915.1 | Serpentinimonas_maccroryi | SMCB_RS09950 | 2162226..2165078 | 1458426 | DUF499-domain-containing-protein-[Serpentinimonas-maccroryi] |
| WP_095541484.1 | DUF499_1.RM.like.systems | METHYLASE-p->?->HAD->Arm-DNA-bind_3->?-><-DUF499*<-ParB+HNH<-METHYLASE<-HEPN<-TUDOR+ SF1-SF2-Helicase+iREase+CTD+REase<-HTH_Tnp<-HTH+WYL | 948 | Bacteria>Pseudomonadota>Betaproteobacteria>Burkholderiales>Comamonadaceae | GCF_002285255.1 | Vandammella_animalimorsus | CLI93_RS07370 | 83443..86289 | 2029117 | DUF499-domain-containing-protein-[Vandammella-animalimorsus] |
| WP_157271680.1 | DUF499_1.RM.like.systems | <-MACRODOMAIN<-DUF499*<-ParB+HNH<-METHYLASE<-TUDOR+ SF1-SF2-Helicase+iREase+CTD+REase | 943 | Bacteria>Pseudomonadota>Betaproteobacteria>Burkholderiales>Sphaerotilaceae | GCF_009760915.1 | Azohydromonas_aeria | FJ694_RS32510 | 1271..4102 | 2590212 | DUF499-domain-containing-protein-[Azohydromonas-aeria] |
| MCC6474792.1 | DUF499_1.RM.like.systems | <-HEPN+ DUF499*<-HEPN<-PolB<-METHYLASE<-HEPN<-ParB+HNH\|\|tRNA-><-PIN | 1120 | Bacteria>Pseudomonadota>Betaproteobacteria>Burkholderiales>unclassified_Burkholderiales | GCA_020846575.1 | Burkholderiales_bacterium | IT514_13720 | 1159..4521 | 1891238 | DUF499-domain-containing-protein-[Burkholderiales-bacterium] |
| CAG1003423.1 | DUF499_1.RM.like.systems | TUDOR+ SF1-SF2-Helicase+iREase+CTD+REase->sGTP->ParB+HNH->METHYLASE->HEPN+ DUF499*-> | 1107 | Bacteria>Pseudomonadota>Betaproteobacteria>Nitrosomonadales>Methylophilaceae | GCA_905339275.1 | Methylophilaceae_bacterium | MTYP_03091 | 15685..19008 | 2030816 | hypothetical-protein-MTYP_03091-[Methylophilaceae-bacterium] |
| WP_107785979.1 | DUF499_1.RM.like.systems | <-HIN-HTH\|\|TUDOR+ SF1-SF2-Helicase+iREase+CTD+REase->ParB+HNH->METHYLASE->HEPN+ DUF499*-><-tRNA\|\|FURR-HTH-> | 1112 | Bacteria>Pseudomonadota>Betaproteobacteria>Nitrosomonadales>Nitrosomonadaceae | GCF_003051045.1 | Nitrosomonas_ureae | C8R28_RS01190 | 8316..11654 | 44577 | Swt1-family-HEPN-domain-containing-protein-[Nitrosomonas-ureae] |
| MBQ7625733.1 | DUF499_1.RM.like.systems | METHYLASE-p->ParB+HNH->DUF499*-><-REase\|\|TP_methylase->?->?->ncRNA->?->Methyltransf_5-> | 945 | Bacteria>Pseudomonadota>Betaproteobacteria>Rhodocyclales>Rhodocyclaceae | GCA_017444505.1 | Rhodocyclaceae_bacterium | IJS87_01550 | 2639..5476 | 1898103 | ATP-binding-protein-[Rhodocyclaceae-bacterium] |
| WP_191658157.1 | DUF499_1.RM.like.systems | <-DUF499*<-wHTH+REase<-HEPN<-METHYLASE<-?<-ParB+HNH<-TUDOR+ SF1-SF2-Helicase+iREase+CTD+REase | 944 | Bacteria>Pseudomonadota>Betaproteobacteria>Rhodocyclales>Zoogloeaceae | - | Azoarcus_sp._Aa7 | AZOA_12030 | 1309..4143 | 2303328 | DUF499-domain-containing-protein-[Azoarcus-sp.-Aa7] |
| MBD5801779.1 | DUF499_1.RM.like.systems | <-DUF499*<-wHTH+REase<-HEPN<-METHYLASE<-?<-ParB+HNH<-TUDOR+ SF1-SF2-Helicase+iREase+CTD+REase | 944 | Bacteria>Pseudomonadota>Betaproteobacteria>Rhodocyclales>Zoogloeaceae | GCA_014822185.1 | Azoarcus_sp._Aa7 | AZOA_12030 | 1309..4143 | 2303328 | hypothetical-protein-[Azoarcus-sp.-Aa7] |
| WP_002936068.1 | DUF499_1.RM.like.systems | <-RNaseP<-?\|\|?->?->MORC+DNA_gyraseB+TOPRIM->?-><-DUF499*<-wHTH+REase<-METHYLASE<-HEPN<-ParB+HNH<-TUDOR+ SF1-SF2-Helicase+iREase+CTD+REase | 941 | Bacteria>Pseudomonadota>Betaproteobacteria>Rhodocyclales>Zoogloeaceae | GCF_000310125.2 | Thauera_sp._27 | B447_RS02400 | 104287..107112 | 305700 | DUF499-domain-containing-protein-[Thauera-sp.-27] |
| WP_168986546.1 | DUF499_1.RM.like.systems | Phage_integrase-><-DUF499*<-?<-?<-ParB+HNH<-METHYLASE<-PIN<-TUDOR+ SF1-SF2-Helicase+iREase+CTD+REase | 944 | Bacteria>Pseudomonadota>Betaproteobacteria>Rhodocyclales>Zoogloeaceae | GCF_012910885.1 | Azoarcus_taiwanensis | GPA21_RS02025 | 735..3569 | 666964 | DUF499-domain-containing-protein-[Azoarcus-taiwanensis] |
| TQE98724.1 | DUF499_1.RM.like.systems | ParB+HNH->DUF499*-><-?<-REase+35exo<-HTH+WYL | 939 | Bacteria>Pseudomonadota>Gammaproteobacteria>Chromatiales>Ectothiorhodospiraceae | GCA_006569195.1 | Spiribacter_salinus | FKY71_12295 | 713..3532 | 1335746 | ATP-binding-protein-[Spiribacter-salinus] |
| WP_220756270.1 | DUF499_1.RM.like.systems | TUDOR+ SF1-SF2-Helicase+iREase+CTD+REase->OLD-ABC->HNH->METHYLASE->DUF499*-><-RlpA<-RVT+HNH<-?<-ParB+HNH<-ParB+HNH | 947 | Bacteria>Pseudomonadota>Gammaproteobacteria>Enterobacterales | GCF_019655755.1 | Enterobacteriaceae | K5R72_RS10955 | 43358..46201 | 543 | DUF499-domain-containing-protein-[Escherichia-coli] |
| RUQ36603.1 | DUF499_1.RM.like.systems | ParB+HNH->DUF499*-><-ABhydrolase+ABhydrolase<-?<-tRNA<-?<-HHH+ENDO3+NUDIX | 937 | Bacteria>Pseudomonadota>Gammaproteobacteria>Gammaproteobacteria_incertae_sedis>Candidatus_Competibacteraceae | GCA_003989065.1 | Candidatus_Competibacteraceae_bacterium | EKK71_07765 | 2719..5532 | 2053538 | ATP-binding-protein-[Candidatus-Competibacteraceae-bacterium] |
| WP_107866351.1 | DUF499_1.RM.like.systems | METHYLASE->OLD-ABC->REase->DUF499*-><-ParB+HNH<-?<-ParB | 941 | Bacteria>Pseudomonadota>Gammaproteobacteria>Moraxellales>Moraxellaceae | GCF_003051055.1 | Agitococcus_lubricus | C8N29_RS12225 | 4668..7493 | 1077255 | DUF499-domain-containing-protein-[Agitococcus-lubricus] |
| WP_194364110.1 | DUF499_1.RM.like.systems | ParB+HNH->ParB+HNH->TUDOR+ SF1-SF2-Helicase+iREase+CTD+REase->METHYLASE->DUF499*-><-?<-?<-Arm-DNA-bind_2+Phage_int_SAM_3+Phage_integrase<-?\|\|HTH_26-> | 949 | Bacteria>Pseudomonadota>Gammaproteobacteria>Moraxellales>Moraxellaceae | GCF_015277735.1 | Psychrobacter_sp._NG254 | IP510_RS07350 | 481008..483857 | 2782003 | DUF499-domain-containing-protein-[Psychrobacter-sp.-NG254] |
| WP_231663150.1 | DUF499_1.RM.like.systems | HTH+WYL->TUDOR+ SF1-SF2-Helicase+iREase+CTD+REase->METHYLASE->OLD-ABC->REase->ParB+HNH->DUF499*->Primpol+D5N+D5ATPase-><-Arm-DNA-bind_3+Phage_integrase\|\|?->FAD-NAD-dep-oxidoreductase-> | 957 | Bacteria>Pseudomonadota>Gammaproteobacteria>Oceanospirillales>Halomonadaceae | GCF_021151495.1 | Halomonas_sp._IOP_31 | K7I12_RS04335 | 15378..18251 | 2876584 | DUF499-domain-containing-protein-[Halomonas-sp.-IOP_31] |
| WP_239285448.1 | DUF499_1.RM.like.systems | TUDOR+ SF1-SF2-Helicase+iREase+CTD+REase->OLD-ABC->HNH->METHYLASE->ParB+HNH->DUF499*->HTH_Tnp_1+HTH_21+rve+rve_3->?->?->AbiEi-><-CHTH<-RelE | 962 | Bacteria>Pseudomonadota>Gammaproteobacteria>Oceanospirillales>Halomonadaceae | GCF_022348135.1 | unclassified_Halomonas | MHM85_RS05495 | 207037..209925 | 2609666 | MULTISPECIES:-DUF499-domain-containing-protein-[unclassified-Halomonas] |
| MBE0508133.1 | DUF499_1.RM.like.systems | <-DUF499*<-ParB+HNH<-METHYLASE | 954 | Bacteria>Pseudomonadota>Gammaproteobacteria>Oceanospirillales>Oceanospirillaceae | GCA_014859565.1 | Marinospirillum_sp. | IBX50_15700 | 354..3218 | 2183934 | ATP-binding-protein-[Marinospirillum-sp.] |
| WP_098424145.1 | DUF499_1.RM.like.systems | XIS-HTH->TUDOR+ SF1-SF2-Helicase+iREase+CTD+REase->OLD-ABC->HNH->METHYLASE->ParB+HNH->DUF499*->GDResolvase+HIN-HTH-><-?<-REase<-HNH | 946 | Bacteria>Pseudomonadota>Gammaproteobacteria>Pseudomonadales>Marinobacteraceae | GCF_002563815.1 | Marinobacter_sp._LV10R520-4 | ATG98_RS19590 | 4036541..4039381 | 1761796 | DUF499-domain-containing-protein-[Marinobacter-sp.-LV10R520-4] |
| WP_098418633.1 | DUF499_1.RM.like.systems | XIS-HTH->TUDOR+ SF1-SF2-Helicase+iREase+CTD+REase->OLD-ABC->HNH->METHYLASE->ParB+HNH->DUF499*->GDResolvase+HIN-HTH-><-?<-REase<-HNH<-?\|\|HIN-HTH-> | 946 | Bacteria>Pseudomonadota>Gammaproteobacteria>Pseudomonadales>Marinobacteraceae | GCF_002563885.1 | Marinobacter_sp._LV10MA510-1 | ATI45_RS05630 | 1155568..1158408 | 1415567 | DUF499-domain-containing-protein-[Marinobacter-sp.-LV10MA510-1] |
| WP_199457594.1 | DUF499_1.RM.like.systems | <-DUF499*<-ParB+HNH<-METHYLASE | 953 | Bacteria>Pseudomonadota>Gammaproteobacteria>Pseudomonadales>Marinobacteraceae | GCF_016463955.1 | unclassified_Marinobacter | JIF54_RS10850 | 749..3610 | 83889 | MULTISPECIES:-DUF499-domain-containing-protein-[unclassified-Marinobacter] |
| MTI93108.1 | DUF499_1.RM.like.systems | ParB->ParB->Classical-AAA->Peptidase_S8->?->RlpA-><-DUF499*<-ParB+HNH<-METHYLASE<-PIN<-TUDOR+ SF1-SF2-Helicase+iREase+CTD+REase<-MORC+DNA_gyraseB+TOPRIM<-SbcC | 946 | Bacteria>Pseudomonadota>Gammaproteobacteria>Pseudomonadales>Pseudomonadaceae | GCA_009711815.1 | Stutzerimonas_stutzeri | FH750_17245 | 14123..16963 | 316 | ATP-binding-protein-[Stutzerimonas-stutzeri] |
| HIQ43647.1 | DUF499_1.RM.like.systems | ParB->ParB-><-MORC+HisKin\|\|<-RVT\|\|?->RlpA-><-DUF499*<-ParB+HNH<-METHYLASE<-PIN<-TUDOR+ SF1-SF2-Helicase+iREase+CTD+REase<-MORC+DNA_gyraseB+TOPRIM<-SbcC | 946 | Bacteria>Pseudomonadota>Gammaproteobacteria>Pseudomonadales>Pseudomonadaceae | GCA_015661735.1 | Pseudomonas_oleovorans | EYH47_12960 | 31988..34828 | 301 | ATP-binding-protein-[Pseudomonas-oleovorans] |
| MBG3917698.1 | DUF499_1.RM.like.systems | SbcC->MORC+DNA_gyraseB+TOPRIM->TUDOR+ SF1-SF2-Helicase+iREase+CTD+REase->PIN->METHYLASE->ParB+HNH->DUF499*-><-RlpA<-?<-Peptidase_S8<-Classical-AAA<-ParB<-ParB | 946 | Bacteria>Pseudomonadota>Gammaproteobacteria>Pseudomonadales>Pseudomonadaceae | GCA_015732935.1 | Pseudomonas_aeruginosa | I4931_21710 | 92367..95207 | 287 | ATP-binding-protein-[Pseudomonas-aeruginosa] |
| MCF6804930.1 | DUF499_1.RM.like.systems | ParB->Classical-AAA->Peptidase_S8->?->?->RlpA-><-DUF499*<-ParB+HNH<-METHYLASE<-PIN<-TUDOR+ SF1-SF2-Helicase+iREase+CTD+REase<-MORC+DNA_gyraseB+TOPRIM<-SbcC | 889 | Bacteria>Pseudomonadota>Gammaproteobacteria>Pseudomonadales>Pseudomonadaceae | GCA_021713635.1 | Stutzerimonas_stutzeri | L3X13_09150 | 18126..20795 | 316 | DUF499-domain-containing-protein-[Stutzerimonas-stutzeri] |
| MCF6781034.1 | DUF499_1.RM.like.systems | ParB->Classical-AAA->Peptidase_S8->?->?->RlpA-><-DUF499*<-ParB+HNH<-METHYLASE<-PIN<-TUDOR+ SF1-SF2-Helicase+iREase+CTD+REase<-MORC+DNA_gyraseB+TOPRIM<-SbcC | 946 | Bacteria>Pseudomonadota>Gammaproteobacteria>Pseudomonadales>Pseudomonadaceae | GCA_021713665.1 | Stutzerimonas_stutzeri | L3X17_04510 | 18122..20962 | 316 | DUF499-domain-containing-protein-[Stutzerimonas-stutzeri] |
| WP_079328502.1 | DUF499_1.RM.like.systems | ParB+HNH->DUF499*-><-RlpA<-?<-?\|\|RVT->RelB-><-ParB | 946 | Bacteria>Pseudomonadota>Gammaproteobacteria>Pseudomonadales>Pseudomonadaceae | GCF_002027175.1 | Stutzerimonas_stutzeri | B2J73_RS08375 | 2891..5731 | 316 | DUF499-domain-containing-protein-[Stutzerimonas-stutzeri] |
| WP_125852814.1 | DUF499_1.RM.like.systems | SbcC->MORC+DNA_gyraseB+TOPRIM->TUDOR+ SF1-SF2-Helicase+iREase+CTD+REase->PIN->METHYLASE->ParB+HNH->DUF499*-><-RlpA<-ParB\|\|REase-> | 946 | Bacteria>Pseudomonadota>Gammaproteobacteria>Pseudomonadales>Pseudomonadaceae | GCF_003936145.1 | Stutzerimonas_stutzeri | EGJ42_RS05720 | 52381..55221 | 316 | DUF499-domain-containing-protein-[Stutzerimonas-stutzeri] |
| WP_143508093.1 | DUF499_1.RM.like.systems | SbcC->MORC+DNA_gyraseB+TOPRIM->TUDOR+ SF1-SF2-Helicase+iREase+CTD+REase->PIN->METHYLASE->ParB+HNH->DUF499*-><-RlpA<-?<-Peptidase_S8<-Classical-AAA<-ParB<-ParB | 946 | Bacteria>Pseudomonadota>Gammaproteobacteria>Pseudomonadales>Pseudomonadaceae | GCF_007049885.1 | Pseudomonas_mendocina | EQ828_RS12690 | 220581..223421 | 300 | DUF499-domain-containing-protein-[Pseudomonas-mendocina] |
| WP_165892256.1 | DUF499_1.RM.like.systems | TPR->RlpA-><-DUF499*<-ParB+HNH<-METHYLASE<-HNH<-OLD-ABC<-TUDOR+ SF1-SF2-Helicase+iREase+CTD+REase | 933 | Bacteria>Pseudomonadota>Gammaproteobacteria>Pseudomonadales>Pseudomonadaceae | GCF_011470035.1 | Azotobacter_chroococcum | HA520_RS07840 | 19913..22714 | 353 | DUF499-domain-containing-protein-[Azotobacter-chroococcum] |
| WP_262384438.1 | DUF499_1.RM.like.systems | PIN->METHYLASE->METHYLASE-p->ParB+HNH->DUF499*-><-?\|\|HTH+MPTase->PIN->HAD->HAD-><-ParB | 415 | Bacteria>Pseudomonadota>Gammaproteobacteria>Pseudomonadales>Pseudomonadaceae | GCF_012986835.1 | Pseudomonas_fragi | HBN88_RS02020 | 44440..45687 | 296 | DUF499-domain-containing-protein-[Pseudomonas-fragi] |
| WP_213582008.1 | DUF499_1.RM.like.systems | ParB-><-HAD\|\|Classical-AAA->Peptidase_S8-><-?\|\|RlpA-><-DUF499*<-ParB+HNH<-METHYLASE<-TUDOR+ SF1-SF2-Helicase+iREase+CTD+REase\|\|?-><-MORC+DNA_gyraseB+TOPRIM<-SbcC | 946 | Bacteria>Pseudomonadota>Gammaproteobacteria>Pseudomonadales>Pseudomonadaceae | GCF_018383465.1 | Pseudomonas_syringae | KID98_RS09080 | 2027689..2030529 | 317 | DUF499-domain-containing-protein-[Pseudomonas-syringae] |
| WP_226045701.1 | DUF499_1.RM.like.systems | SbcC->MORC+DNA_gyraseB+TOPRIM->TUDOR+ SF1-SF2-Helicase+iREase+CTD+REase->PIN->METHYLASE->ParB+HNH->DUF499*-><-RlpA<-?<-Peptidase_S8<-Classical-AAA<-ParB<-ParB | 946 | Bacteria>Pseudomonadota>Gammaproteobacteria>Pseudomonadales>Pseudomonadaceae | GCF_020401845.1 | Pseudomonas_sp._HS-18 | LDJ84_RS00045 | 14958..17798 | 2879114 | DUF499-domain-containing-protein-[Pseudomonas-sp.-HS-18] |
| WP_129484085.1 | DUF499_1.RM.like.systems | SbcC->MORC+DNA_gyraseB+TOPRIM->TUDOR+ SF1-SF2-Helicase+iREase+CTD+REase->PIN->METHYLASE->ParB+HNH->DUF499*-><-RlpA<-?<-Peptidase_S8<-Classical-AAA<-ParB<-ParB | 946 | Bacteria>Pseudomonadota>Gammaproteobacteria>Pseudomonadales>Pseudomonadaceae | GCF_021495345.1 | Pseudomonas_hydrolytica | L1F06_RS00045 | 14914..17754 | 2493633 | DUF499-domain-containing-protein-[Pseudomonas-hydrolytica] |
| WP_279641602.1 | DUF499_1.RM.like.systems | SbcC->MORC+DNA_gyraseB+TOPRIM->TUDOR+ SF1-SF2-Helicase+iREase+CTD+REase->PIN->METHYLASE->ParB+HNH->DUF499*->?-><-RlpA<-?\|\|RVT-><-ParB<-ParB | 946 | Bacteria>Pseudomonadota>Gammaproteobacteria>Pseudomonadales>Pseudomonadaceae | GCF_029839225.1 | Stutzerimonas_stutzeri | N5C32_RS13925 | 184183..187023 | 316 | DUF499-domain-containing-protein-[Stutzerimonas-stutzeri] |
| WP_283258799.1 | DUF499_1.RM.like.systems | HIN-HTH->?->RlpA-><-DUF499*<-ParB+HNH<-METHYLASE<-PIN<-TUDOR+ SF1-SF2-Helicase+iREase+CTD+REase<-MORC+DNA_gyraseB+TOPRIM<-SbcC | 946 | Bacteria>Pseudomonadota>Gammaproteobacteria>Pseudomonadales>Pseudomonadaceae | GCF_030056015.1 | Stutzerimonas_stutzeri | QM290_RS03915 | 3700..6540 | 316 | DUF499-domain-containing-protein-[Stutzerimonas-stutzeri] |
| WP_046835095.1 | DUF499_1.RM.like.systems | TUDOR+ SF1-SF2-Helicase+iREase+CTD+REase->PIN->METHYLASE-p->DUF772+DDE_Tnp_1->METHYLASE->ParB+HNH->DUF499*-><-RlpA<-?\|\|CHTH-><-LZ_Tnp_IS66+zf-IS66+DDE_Tnp_IS66+DDE_Tnp_IS66_C | 944 | Bacteria>Pseudomonadota>Gammaproteobacteria>Pseudomonadales>Pseudomonadaceae | GCF_030252675.1 | Pseudomonas_amygdali | QO021_RS27770 | 6030244..6033078 | 47877 | DUF499-domain-containing-protein-[Pseudomonas-amygdali] |
| WP_110599711.1 | DUF499_1.RM.like.systems | ParB-><-PIN<-HTH+MPTase\|\|?->?->RlpA-><-DUF499*<-ParB+HNH<-METHYLASE<-PIN<-TUDOR+ SF1-SF2-Helicase+iREase+CTD+REase<-MORC+DNA_gyraseB+TOPRIM<-SbcC | 945 | Bacteria>Pseudomonadota>Gammaproteobacteria>Pseudomonadales>Pseudomonadaceae | - | Pseudomonas_koreensis | DMX04_RS04460 | 134660..137497 | 198620 | DUF499-domain-containing-protein-[Pseudomonas-koreensis] |
| WP_003246716.1 | DUF499_1.RM.like.systems | SbcC->MORC+DNA_gyraseB+TOPRIM->TUDOR+ SF1-SF2-Helicase+iREase+CTD+REase->PIN->METHYLASE->ParB+HNH->DUF499*-><-RlpA<-?<-?\|\|RVT-> | 946 | Bacteria>Pseudomonadota>Gammaproteobacteria>Pseudomonadales>Pseudomonadaceae | - | Pseudomonas_mendocina | A471_RS19815 | 19321..22161 | 300 | DUF499-domain-containing-protein-[Pseudomonas-mendocina] |
| WP_236481327.1 | DUF499_1.RM.like.systems | <-NERD+SF1-SF2-Helicase\|\|RlpA-><-DUF499*<-ParB+HNH<-METHYLASE<-HNH<-OLD-ABC<-TUDOR+ SF1-SF2-Helicase+iREase+CTD+REase<-MORC+DNA_gyraseB+TOPRIM | 946 | Bacteria>Pseudomonadota>Gammaproteobacteria>Pseudomonadales>Pseudomonadaceae | - | Pseudomonas_syringae | GIW69_RS06575 | 2105..4945 | 317 | DUF499-domain-containing-protein-[Pseudomonas-syringae] |
| WP_122622773.1 | DUF499_1.RM.like.systems | <-ParB+HNH\|\|HTH+WYL->TUDOR+ SF1-SF2-Helicase+iREase+CTD+REase->OLD-ABC->HNH->METHYLASE->DUF499*-><-RlpA<-Phage_int_SAM<-?<-TOPRIM<-Mbetalac<-Arm-DNA-bind_3+Phage_integrase | 942 | Bacteria>Pseudomonadota>Gammaproteobacteria>Pseudomonadales>Pseudomonadaceae | - | Pseudomonas_viridiflava | EEX91_RS04620 | 101897..104725 | 33069 | DUF499-domain-containing-protein-[Pseudomonas-viridiflava] |
| WP_125867554.1 | DUF499_1.RM.like.systems | SbcC->MORC+DNA_gyraseB+TOPRIM->TUDOR+ SF1-SF2-Helicase+iREase+CTD+REase->PIN->METHYLASE->ParB+HNH->DUF499*->HIN-HTH->?->LZ_Tnp_IS66+zf-IS66+DDE_Tnp_IS66+DDE_Tnp_IS66_C-><-?<-?<-?<-Peptidase_S8 | 946 | Bacteria>Pseudomonadota>Gammaproteobacteria>Pseudomonadales>Pseudomonadaceae | - | Stutzerimonas_stutzeri | EGI99_RS04500 | 994408..997248 | 316 | DUF499-domain-containing-protein-[Stutzerimonas-stutzeri] |
| WP_161612279.1 | DUF499_1.RM.like.systems | METHYLASE->ParB+HNH->DUF499*->?-><-RlpA<-?\|\|RVT-><-ParB<-ParB | 946 | Bacteria>Pseudomonadota>Gammaproteobacteria>Pseudomonadales>Pseudomonadaceae | - | Stutzerimonas_stutzeri | GXB82_RS15495 | 4443..7283 | 316 | DUF499-domain-containing-protein-[Stutzerimonas-stutzeri] |
| MCB1656848.1 | DUF499_1.RM.like.systems | <-HIN-HTH\|\|ParB-><-DUF499*<-ParB+HNH<-METHYLASE | 939 | Bacteria>Pseudomonadota>Gammaproteobacteria>Pseudomonadales>unclassified_Pseudomonadales | GCA_020443205.1 | Pseudomonadales_bacterium | KDI39_01315 | 4825..7644 | 1891229 | ATP-binding-protein-[Pseudomonadales-bacterium] |
| MCY4286579.1 | DUF499_1.RM.like.systems | PolB->ParB+HNH->METHYLASE->?->OLD-ABC->TOPRIM->DUF499*->?->?-><-?<-?<-ABC-ATPase+SbcC | 1107 | Bacteria>Pseudomonadota>Gammaproteobacteria>Thiotrichales>unclassified_Thiotrichales | GCA_026709935.1 | Thiotrichales_bacterium | OXC65_14725 | 7747..11070 | 2026796 | MAG:-DUF499-domain-containing-protein-[Thiotrichales-bacterium] |

| Protein-ID | System | Gene-Neighbourhoods | Length | Taxonomy | Assembly-ID | Strain | Locus-Tag | Genomic-coordinates | TaxID | Fasta-defline |
|---|---|---|---|---|---|---|---|---|---|---|
| MCP5421179.1 | DUF499_1.RM.like.systems | GDResolvase+HIN-HTH->?->TUDOR+SF1-SF2-Helicase+iREase+CTD+REase->METHYLASE->ParB+HNH->DUF499*->wHTH+REase->SbcC-><-?\|\|BrnT_toxin->BrnA_antitoxin-> | 941 | Bacteria>Pseudomonadota>Gammaproteobacteria>unclassified_Gammaproteobacteria | GCA_024235155.1 | Gammaproteobacteria_bacterium | H6969_11925 | 8981..11806 | 1913989 | ATP-binding-protein-[Gammaproteobacteria-bacterium] |
| WP_091239202.1 | DUF499_1.RM.like.systems | Phage_holin->Phage_lysozyme->?->HTH_3->Radical-SAM-><-?<-DUF499*<-ParB+HNH<-METHYLASE<-HEPN<-TUDOR+SF1-SF2-Helicase+iREase+CTD+REase<-HTH_Tnp<-HTH+WYL | 949 | Bacteria>Pseudomonadota>Gammaproteobacteria>Xanthomonadales>Rhodanobacteraceae | GCF_900101825.1 | Aquimonas_voraii | BLP83_RS04255 | 981108..983957 | 265719 | DUF499-domain-containing-protein-[Aquimonas-voraii] |
| MCU0753612.1 | DUF499_1.RM.like.systems | <-ArsR-HTH\|\|?->?->?->?->?-><-DUF499*<-Schlafen+HATPase+HTH<-ParB+HNH<-REase<-OLD-ABC<-METHYLASE<-HNH | 955 | Bacteria>Pseudomonadota>Gammaproteobacteria>Xanthomonadales>unclassified_Xanthomonadales | GCA_025458445.1 | Xanthomonadales_bacterium | MUE46_00615 | 13163..16030 | 2006849 | DUF499-domain-containing-protein-[Xanthomonadales-bacterium] |
| PJK14600.1 | DUF499_1.RM.like.systems | <-DUF499*<-ParB+HNH<-METHYLASE<-HNH<-OLD-ABC<-TUDOR+SF1-SF2-Helicase+iREase+CTD+REase | 945 | Bacteria>Pseudomonadota>Gammaproteobacteria>Xanthomonadales>Xanthomonadaceae | GCA_002798295.1 | Xanthomonadaceae_bacterium_NML07-0707 | CO613_05135 | 84524..87361 | 2032574 | hypothetical-protein-CO613_05135-[Xanthomonadaceae-bacterium-NML07-0707] |
| MBU6247856.1 | DUF499_1.RM.like.systems | <-DUF499*<-ParB+HNH<-?<-HEPN+CBS<-METHYLASE<-PIN | 934 | Bacteria>Pseudomonadota>Gammaproteobacteria>Xanthomonadales>Xanthomonadaceae | GCA_018970895.1 | Xanthomonadaceae_bacterium | KGN77_08870 | 9374..12178 | 1926873 | DUF499-domain-containing-protein-[Xanthomonadaceae-bacterium] |
| WP_049462969.1 | DUF499_1.RM.like.systems | HTH_26->TUDOR+SF1-SF2-Helicase+iREase+CTD+REase->OLD-ABC->HNH->METHYLASE->ParB+HNH->DUF499*->?->?->?->?->TPR-> | 930 | Bacteria>Pseudomonadota>Gammaproteobacteria>Xanthomonadales>Xanthomonadaceae | GCF_001075485.1 | Stenotrophomonas_maltophilia | ADM82_RS13495 | 40138..42930 | 40324 | DUF499-domain-containing-protein-[Stenotrophomonas-maltophilia] |
| WP_055771167.1 | DUF499_1.RM.like.systems | <-D5N+D5ATPase<-?<-?<-?<-?<-?<-DUF499*<-ParB+HNH<-?<-METHYLASE<-PIN<-TUDOR+SF1-SF2-Helicase+iREase+CTD+REase<-Phage_integrase | 929 | Bacteria>Pseudomonadota>Gammaproteobacteria>Xanthomonadales>Xanthomonadaceae | GCF_001422025.1 | Stenotrophomonas_sp._Leaf70 | ASF01_RS13960 | 54995..57784 | 1736233 | DUF499-domain-containing-protein-[Stenotrophomonas-sp.-Leaf70] |
| WP_144815145.1 | DUF499_1.RM.like.systems | METHYLASE-p->?->ParB+HNH->DUF499*->?->?->?->?->?->D5N+D5ATPase-> | 930 | Bacteria>Pseudomonadota>Gammaproteobacteria>Xanthomonadales>Xanthomonadaceae | GCF_007830115.1 | Lysobacter_ruishenii | IP93_RS09870 | 5117..7909 | 686800 | DUF499-domain-containing-protein-[Lysobacter-ruishenii] |
| WP_053505566.1 | DUF499_1.RM.like.systems | HEPN+TPR+RhoN->HicA_toxin->METHYLASE->?->?->ParB+HNH->DUF499*->?->NACHT-> | 930 | Bacteria>Pseudomonadota>Gammaproteobacteria>Xanthomonadales>Xanthomonadaceae | - | Stenotrophomonas_maltophilia | ABW45_RS08210 | 26513..29305 | 40324 | DUF499-domain-containing-protein-[Stenotrophomonas-maltophilia] |
| MDD4116915.1 | DUF499_1.RM.like.systems | SAM-methylase-><-?<-?<-?\|\|tRNA-><-HEPN+DUF499*<-Peptidase_C47<-?<-METHYLASE<-ParB+HNH<-?<-TUDOR+SF1-SF2-Helicase+iREase+CTD+REase | 1113 | Bacteria>PVC_group>Kiritimatiellota>Kiritimatiellia>unclassified_Kiritimatiellia | GCA_028701395.1 | Kiritimatiellia_bacterium | PHI39_01725 | 13604..16945 | 2212473 | MAG:-Swt1-family-HEPN-domain-containing-protein-[Kiritimatiellae-bacterium] |
| HIV08682.1 | DUF499_1.RM.like.systems | TUDOR+SF1-SF2-Helicase+iREase+CTD+REase->HEPN+DUF499*->ParB+HNH-p-> | 1108 | Bacteria>PVC_group>Lentisphaerota>Lentisphaeria>Lentisphaerales | GCA_018713405.1 | Candidatus_Spyradenecus_faecavium | IAC79_01020 | 6980..10306 | 2840947 | DUF499-domain-containing-protein-[Candidatus-Spyradenecus-faecavium] |
| MCG3126049.1 | DUF499_1.RM.like.systems | HEPN+REase->METHYLASE->ParB+HNH->DUF499*->?->SF1-SF2-Helicase->?->?->CASPASE+APATPase+TPR+TPR+TPR-> | 1107 | Bacteria>PVC_group>Planctomycetota>Phycisphaerae>unclassified_Phycisphaerae | GCA_022072445.1 | Phycisphaerae_bacterium | CHACPFDD_00877 | 44646..47969 | 2026778 | hypothetical-protein-[Phycisphaerae-bacterium] |
| NLX98932.1 | DUF499_1.RM.like.systems | TUDOR+SF1-SF2-Helicase+iREase+CTD+REase->ParB+HNH->?->METHYLASE->REase->DUF499*->REase->MORC+REase-><-DNA_methylase<-HTH | 983 | Bacteria>PVC_group>Planctomycetota>Planctomycetia>Pirellulales | GCA_012728855.1 | Rhodopirellula_sp. | GXY83_22530 | 8787..11738 | 2024855 | ATP-binding-protein-[Rhodopirellula-sp.] |
| MCE2752488.1 | DUF499_1.RM.like.systems | Methyltransf_21->TUDOR+SF1-SF2-Helicase+iREase+CTD+REase->?->?->METHYLASE->ParB+HNH->DUF499*->REase->ParB+HNH->?->URI-DNase-> | 957 | Bacteria>PVC_group>Planctomycetota>Planctomycetia>Pirellulales | GCA_021299695.1 | Pirellula_sp. | LW720_11440 | 12221..15094 | 117 | ATP-binding-protein-[Pirellula-sp.] |
| QDT36019.1 | DUF499_1.RM.like.systems | <-DUF499*<-?<-METHYLASE<-ParB+HNH<-TUDOR+SF1-SF2-Helicase+iREase+CTD+REase<-GDResolvase+Recombinase+ZnR | 961 | Bacteria>PVC_group>Planctomycetota>Planctomycetia>Planctomycetales | GCA_007744515.1 | Stratiformator_vulcanicus | Pan189_03740 | 483899..486784 | 2527980 | hypothetical-protein-Pan189_03740-[Planctomycetes-bacterium-Pan189] |
| MCA9232427.1 | DUF499_1.RM.like.systems | <-ParB+HNH<-DUF499*<-METHYLASE<-TOPRIM<-OLD-ABC<-TUDOR+SF1-SF2-Helicase+iREase+CTD+REase-p | 953 | Bacteria>PVC_group>Planctomycetota>Planctomycetia>Planctomycetales | GCA_020427155.1 | Planctomycetales_bacterium | KDA57_17390 | 2105..4966 | 2053591 | ATP-binding-protein-[Planctomycetales-bacterium] |
| TAH38052.1 | DUF499_1.RM.like.systems | <-wHTH+REase\|\|AbiEi_4->AbiEii-><-PIN<-ParB+HNH\|\|HKD-DNase-><-HEPN+DUF499*<-METHYLASE<-HNH<-OLD-ABC<-TUDOR+SF1-SF2-Helicase+iREase+CTD+REase<-HTH | 1098 | Bacteria>PVC_group>Planctomycetota>unclassified_Planctomycetota | GCA_004295085.1 | Planctomycetota_bacterium | EYC70_05360 | 66593..69889 | 2026780 | MAG:-ATP-binding-protein-[Planctomycetota-bacterium] |
| MBI3462535.1 | DUF499_1.RM.like.systems | PIWI-><-ParB+HNH<-HEPN+DUF499*<-METHYLASE<-wHTH+REase<-TUDOR+SF1-SF2-Helicase+iREase+CTD+REase-p | 1111 | Bacteria>PVC_group>Planctomycetota>unclassified_Planctomycetota | GCA_016196255.1 | Planctomycetota_bacterium | HY000_05665 | 2780..6115 | 2026780 | ATP-binding-protein-[Planctomycetota-bacterium] |
| MBI3271453.1 | DUF499_1.RM.like.systems | <-Phage_int_SAM_1+Phage_integrase<-?<-?<-ParB+HNH<-?<-DUF499*<-PIN<-?<-?<-METHYLASE<-HEPN<-TUDOR+SF1-SF2-Helicase+iREase+CTD+REase | 953 | Bacteria>PVC_group>Planctomycetota>unclassified_Planctomycetota | GCA_016197205.1 | Planctomycetota_bacterium | HYZ53_20855 | 61698..64559 | 2026780 | ATP-binding-protein-[Planctomycetota-bacterium] |
| MCE9567305.1 | DUF499_1.RM.like.systems | <-SF1-SF2-Helicase+REase<-?<-SIGMA-HTH\|\|ParB+HNH->METHYLASE->REase->DUF499*->ParB+HNH->ParB+HNH-> | 986 | Bacteria>PVC_group>Planctomycetota>unclassified_Planctomycetota | GCA_021413735.1 | Planctomycetota_bacterium | K8U57_35335 | 37179..40139 | 2026780 | DUF499-domain-containing-protein-[Planctomycetota-bacterium] |
| MCZ7645664.1 | DUF499_1.RM.like.systems | HTH+MPTase->PemK_toxin->HNH+SAVED->SMODS->?-><-PIN<-HEPN+DUF499*<-METHYLASE<-?<-ParB+HNH<-TUDOR+SF1-SF2-Helicase+iREase+CTD+REase | 1122 | Bacteria>PVC_group>Planctomycetota>unclassified_Planctomycetota | GCA_027493525.1 | Planctomycetota_bacterium | M5U26_10325 | 2469927..2473295 | 2026780 | MAG:-Swt1-family-HEPN-domain-containing-protein-[Planctomycetota-bacterium] |
| WP_189517054.1 | DUF499_1.RM.like.systems | <-ParB+HNH<-HEPN+DUF499*<-?<-HTH<-?<-KAP-NTPase<-METHYLASE | 1101 | Bacteria>PVC_group>Verrucomicrobiota>Opitutae>Puniceicoccales | GCF_014651635.1 | Cerasicoccus_arenae | JIN80_RS05850 | 25614..28919 | 424488 | DUF499-domain-containing-protein-[Cerasicoccus-arenae] |
| OQC32217.1 | DUF499_1.RM.like.systems | DDE_Tnp_IS66+DDE_Tnp_IS66_C-><-?<-Radical-SAM<-?<-DUF499*<-?<-?<-METHYLASE<-HEPN<-ParB+HNH<-TUDOR+SF1-SF2-Helicase+iREase+CTD+REase | 945 | Bacteria>PVC_group>Verrucomicrobiota>unclassified_Verrucomicrobiota | GCA_002071025.1 | Verrucomicrobia_bacterium_ADurb.Bin070 | BWX70_00665 | 3737..6574 | 1852927 | hypothetical-protein-BWX70_00665-[Verrucomicrobia-bacterium-ADurb.Bin070] |
| MBM3880418.1 | DUF499_1.RM.like.systems | <-HEPN+DUF499*<-ParB+HNH<-METHYLASE<-TOPRIM<-OLD-ABC<-TUDOR+SF1-SF2-Helicase+iREase+CTD+REase | 1127 | Bacteria>PVC_group>Verrucomicrobiota>unclassified_Verrucomicrobiota | GCA_016871715.1 | Verrucomicrobiota_bacterium | FJ387_11990 | 332..3715 | 2026799 | ATP-binding-protein-[Verrucomicrobia-bacterium] |
| MBS0646999.1 | DUF499_1.RM.like.systems | <-HEPN+DUF499*<-METHYLASE<-ParB+HNH<-AbiEii | 1093 | Bacteria>PVC_group>Verrucomicrobiota>unclassified_Verrucomicrobiota | GCA_018239885.1 | Verrucomicrobiota_bacterium | JSR97_10490 | 874..4155 | 2026799 | DUF499-domain-containing-protein-[Verrucomicrobia-bacterium] |
| MBE7497142.1 | DUF499_1.RM.like.systems | HKD-DNase->ParB+HNH->METHYLASE->?->nSTAND1+sGTP->REase->DUF499*->tRNA-><-?\|\|?-><-NTP_transf | 920 | Bacteria>PVC_group>Verrucomicrobiota>Verrucomicrobiae>Verrucomicrobiales | GCA_015075365.1 | Verrucomicrobiaceae_bacterium | HS117_19545 | 529489..532251 | 2026800 | ATP-binding-protein-[Verrucomicrobiaceae-bacterium] |
| MBK9498689.1 | DUF499_1.RM.like.systems | METHYLASE->ParB+HNH->OLD-ABC->?->?->DUF499*-><-?\|\|?-><-STYKIN<-HKD-DNase | 972 | Bacteria>Spirochaetota>Spirochaetia>Leptospirales>Leptospiraceae | GCA_016718635.1 | Leptospiraceae_bacterium | IPO06_04755 | 70687..73605 | 2026757 | ATP-binding-protein-[Leptospiraceae-bacterium] |
| MBP6738374.1 | DUF499_1.RM.like.systems | <-Phage_integrase<-DUF499*<-?<-OLD-ABC<-ParB+HNH<-METHYLASE | 972 | Bacteria>Spirochaetota>Spirochaetia>Leptospirales>Leptospiraceae | GCA_017990775.1 | Leptospiraceae_bacterium | KA146_00215 | 1347..4265 | 2026757 | ATP-binding-protein-[Leptospiraceae-bacterium] |
| MCB1144525.1 | DUF499_1.RM.like.systems | <-DUF499*<-?<-?<-OLD-ABC<-ParB+HNH<-METHYLASE | 972 | Bacteria>Spirochaetota>Spirochaetia>Leptospirales>Leptospiraceae | GCA_020440865.1 | Leptospiraceae_bacterium | KDK54_19880 | 628..3546 | 2026757 | ATP-binding-protein-[Leptospiraceae-bacterium] |
| AEJ18699.1 | DUF499_1.RM.like.systems | PhdYeFM_antitox->Phd-YefM->RelE-ParE->?->?->ParB+HNH->HEPN+DUF499*->ABC-ATPase+SbcC-> | 1058 | Bacteria>Spirochaetota>Spirochaetia>Spirochaetales>Breznakiellaceae | GCA_000219725.1 | Gracilinema_caldarium_DSM_7334 | Spica_0539 | 580385..583561 | 744872 | ATPase-(AAA+-superfamily)-like-protein-[Gracilinema-caldarium-DSM-7334] |
| WP_083048603.1 | DUF499_1.RM.like.systems | <-PfkB<-?<-?<-?<-HEPN+DUF499*<-METHYLASE<-TUDOR+SF1-SF2-Helicase+iREase+CTD+REase<-?<-ParB+HNH<-?<-SF1-SF2-Helicase+REase | 1101 | Bacteria>Spirochaetota>Spirochaetia>Spirochaetales>Spirochaetaceae | GCF_002087085.1 | Marispirochaeta_aestuarii | B4097_RS04115 | 245818..249123 | 1963862 | DUF499-domain-containing-protein-[Marispirochaeta-aestuarii] |
| OQB96942.1 | DUF499_1.RM.like.systems | SIGMA-HTH->Radical-SAM-><-?<-?\|\|?->?-><-HEPN+DUF499*<-ParB+HNH<-METHYLASE<-TOPRIM<-OLD-ABC<-HEPN<-TUDOR+SF1-SF2-Helicase+iREase+CTD+REase | 1176 | Bacteria>Spirochaetota>unclassified_Spirochaetota | GCA_002070755.1 | Spirochaetes_bacterium_ADurb.Bin110 | BWX81_01211 | 5339..8869 | 1852908 | hypothetical-protein-BWX81_01211-[Spirochaetes-bacterium-ADurb.Bin110] |
| MCL1994394.1 | DUF499_1.RM.like.systems | <-HEPN+DUF499*<-METHYLASE<-?<-OLD-ABC<-ParB+HNH<-TUDOR+SF1-SF2-Helicase+iREase+CTD+REase | 1116 | Bacteria>Spirochaetota>unclassified_Spirochaetota | GCA_009778155.1 | Spirochaetota_bacterium | FWG66_15730 | 17581..20931 | 2202144 | DUF499-domain-containing-protein-[Spirochaetes-bacterium] |
| MXZ86859.1 | DUF499_1.RM.like.systems | ABC-ATPase+SbcC->?->?->?->?-><-HEPN+DUF499*<-?\|\|?-><-METHYLASE\|\|ParB+HNH->HEPN-><-TUDOR+SF1-SF2-Helicase+iREase+CTD+REase | 1114 | Bacteria>Terrabacteria_group>Actinomycetota>Acidimicrobiia>unclassified_Acidimicrobiia | GCA_009837425.1 | Acidimicrobiia_bacterium | F4Z02_14550 | 33484..36828 | 2080302 | ATP-binding-protein-[Acidimicrobiia-bacterium] |
| WP_284940342.1 | DUF499_1.RM.like.systems | HEPN-><-HEPN<-ParB+HNH<-DUF499*<-TraJ-RHH | 797 | Bacteria>Terrabacteria_group>Actinomycetota>Actinomycetes>Bifidobacteriales | GCF_030250405.1 | Bifidobacterium_catenulatum | K1513_RS02315 | 519682..522075 | 1686 | SNF2-related-protein-[Bifidobacterium-catenulatum] |
| WP_113812542.1 | DUF499_1.RM.like.systems | <-ABhydrolase\|\|TUDOR+SF1-SF2-Helicase+iREase+CTD+REase->Schlafen+HATPase+HTH->METHYLASE->ParB+HNH->?->HEPN+DUF499*->NTD+SF1-SF2-Helicase->wHTH(SBD)+HNH-><-?<-?\|\|?-><-REase+35exo+Beta-Barrel+SF1-SF2-Helicase | 1114 | Bacteria>Terrabacteria_group>Actinomycetota>Actinomycetes>Geodermatophilales | GCF_003319095.1 | Blastococcus_sp._TBT05-19 | DQ244_RS16250 | 131306..134650 | 2250581 | Swt1-family-HEPN-domain-containing-protein-[Blastococcus-sp.-TBT05-19] |
| WP_236827009.1 | DUF499_1.RM.like.systems | HKD-DNase+SF1-SF2-Helicase+TRD-><-SF1-SF2-Helicase<-?\|\|TUDOR+SF1-SF2-Helicase+iREase+CTD+REase->ArdCN+MPTase->METHYLASE->HEPN+DUF499*->ParB+HNH-> | 1109 | Bacteria>Terrabacteria_group>Actinomycetota>Actinomycetes>Geodermatophilales | GCF_021698995.1 | unclassified_Blastococcus | E9565_RS02685 | 57900..61229 | 2619396 | MULTISPECIES:-Swt1-family-HEPN-domain-containing-protein-[unclassified-Blastococcus] |
| WP_067071154.1 | DUF499_1.RM.like.systems | TUDOR+SF1-SF2-Helicase+iREase+CTD+REase-p->METHYLASE->HEPN+DUF499*->ParB+HNH-><-?<-?<-HTH+DUF5753 | 1122 | Bacteria>Terrabacteria_group>Actinomycetota>Actinomycetes>Kitasatosporales | GCF_001543895.1 | Carbonactinospora_thermoautotrophica | TR74_RS19795 | 3234..6602 | 1469144 | Swt1-family-HEPN-domain-containing-protein-[Carbonactinospora-thermoautotrophica] |
| OFE15977.1 | DUF499_1.RM.like.systems | CopG/RHH->?->tRNA-><-SF1-SF2-Helicase<-?<-HEPN+DUF499*<-ParB+HNH<-METHYLASE<-TUDOR+SF1-SF2-Helicase+iREase+CTD+REase<-?<-wHTH(SBD)+HNH<-REase+SF1-SF2-Helicase | 1100 | Bacteria>Terrabacteria_group>Actinomycetota>Actinomycetes>Micrococcales | GCA_001758225.1 | Humibacillus_sp._DSM_29435 | BA895_05295 | 164736..168038 | 1869167 | AAA-family-ATPase-[Humibacillus-sp.-DSM-29435] |
| QAY71606.1 | DUF499_1.RM.like.systems | <-AbiEi_4<-PIN<-XIS-HTH<-HEPN<-HEPN+DUF499*<-ParB+HNH<-METHYLASE<-?<-TUDOR+SF1-SF2-Helicase+iREase+CTD+REase\|\|NTD+SF1-SF2-Helicase-> | 1123 | Bacteria>Terrabacteria_group>Actinomycetota>Actinomycetes>Micrococcales | GCA_004135385.1 | Xylanimonas_protaetiae | ET471_17505 | 3776288..3779659 | 2509457 | ATP-binding-protein-[Xylanimonas-protaetiae] |
| MBK6441773.1 | DUF499_1.RM.like.systems | PrmC_N+Methyltransf_25-><-?<-PadR-HTH\|\|TUDOR+SF1-SF2-Helicase+iREase+CTD+REase->METHYLASE->ParB+HNH->HEPN+DUF499*-> | 922 | Bacteria>Terrabacteria_group>Actinomycetota>Actinomycetes>Micrococcales | GCA_016704565.1 | Candidatus_Phosphoribacter_baldrii | IPF90_06370 | 1387205..1389971 | 2953744 | DUF499-domain-containing-protein-[Actinomycetales-bacterium] |
| MBK6954412.1 | DUF499_1.RM.like.systems | ArsR-HTH->PrmC_N+Methyltransf_25-><-?<-PadR-HTH\|\|TUDOR+SF1-SF2-Helicase+iREase+CTD+REase->ParB+HNH->HEPN+DUF499*->NTD+SF1-SF2-Helicase->TetR-HTH->ABC-ATPase+SbcC->ABC-ATPase+SbcC-> | 1118 | Bacteria>Terrabacteria_group>Actinomycetota>Actinomycetes>Micrococcales | GCA_016708855.1 | Candidatus_Phosphoribacter_baldrii | IPH27_02570 | 528671..532027 | 2953744 | DUF499-domain-containing-protein-[Actinomycetales-bacterium] |
| MCI1262624.1 | DUF499_1.RM.like.systems | <-wHTH(SBD)+HNH<-Glyoxalase\|\|?-><-?<-HEPN<-PolB<-HEPN+DUF499*<-ParB+HNH<-METHYLASE<-TUDOR+SF1-SF2-Helicase+iREase+CTD+REase\|\|NTD+SF1-SF2-Helicase-> | 1116 | Bacteria>Terrabacteria_group>Actinomycetota>Actinomycetes>Micrococcales | GCA_022641475.1 | Tetrasphaera_jenkinsii | LKG20_10180 | 13344..16694 | 330834 | Swt1-family-HEPN-domain-containing-protein-[Tetrasphaera-jenkinsii] |
| WBM79848.1 | DUF499_1.RM.like.systems | XIS-HTH->TUDOR+SF1-SF2-Helicase+iREase+CTD+REase-p->TUDOR+SF1-SF2-Helicase+iREase+CTD+REase->METHYLASE->ParB+HNH->HEPN->DUF499*->DUF499-p->DUF499-p-><-?\|\|REase->REase-> | 285 | Bacteria>Terrabacteria_group>Actinomycetota>Actinomycetes>Micrococcales | GCA_027594785.1 | Cryobacterium_breve | KIV56_17020 | 3683119..3683976 | 1259258 | DUF499-domain-containing-protein-[Cryobacterium-breve] |
| SDF11628.1 | DUF499_1.RM.like.systems | <-wHTH(SBD)+HNH<-HTH_3<-PIN<-MPTase\|\|METHYLASE->ParB+HNH->HEPN+DUF499*-> | 1151 | Bacteria>Terrabacteria_group>Actinomycetota>Actinomycetes>Micrococcales | GCA_900102445.1 | Cellulosimicrobium_cellulans | SAMN04487781_0340 | 373492..376947 | 1710 | Predicted-ATPase,-AAA+-superfamily-[Cellulosimicrobium-cellulans] |
| WP_029212622.1 | DUF499_1.RM.like.systems | SF1-SF2-Helicase->tRNA->TUDOR+SF1-SF2-Helicase+iREase+CTD+REase->METHYLASE->ParB+HNH->HEPN+DUF499*-> | 1119 | Bacteria>Terrabacteria_group>Actinomycetota>Actinomycetes>Micrococcales | GCF_000426385.1 | Arsenicicoccus_bolidensis | K313_RS0116910 | 19529..22888 | 229480 | Swt1-family-HEPN-domain-containing-protein-[Arsenicicoccus-bolidensis] |
| WP_034802772.1 | DUF499_1.RM.like.systems | REase+SF1-SF2-Helicase->wHTH(SBD)+HNH->DndB->TUDOR+SF1-SF2-Helicase+iREase+CTD+REase->METHYLASE->ParB+HNH->HEPN+DUF499*->?->SF1-SF2-Helicase-> | 1107 | Bacteria>Terrabacteria_group>Actinomycetota>Actinomycetes>Micrococcales | GCF_000576595.1 | Intrasporangium_oryzae | N865_RS05960 | 110230..113553 | 412687 | DUF499-domain-containing-protein-[Intrasporangium-oryzae] |
| WP_139142668.1 | DUF499_1.RM.like.systems | CopG/RHH->?->CopG/RHH->tRNA-><-SF1-SF2-Helicase<-?<-HEPN+DUF499*<-ParB+HNH<-METHYLASE<-TUDOR+SF1-SF2-Helicase+iREase+CTD+REase<-?<-wHTH(SBD)+HNH<-REase+SF1-SF2-Helicase | 1111 | Bacteria>Terrabacteria_group>Actinomycetota>Actinomycetes>Micrococcales | GCF_001758225.1 | Humibacillus_sp._DSM_29435 | BA895_RS14645 | 164736..168071 | 1869167 | Swt1-family-HEPN-domain-containing-protein-[Humibacillus-sp.-DSM-29435] |
| WP_106298858.1 | DUF499_1.RM.like.systems | MazG-><-?\|\|NTD+SF1-SF2-Helicase->TUDOR+SF1-SF2-Helicase+iREase+CTD+REase->METHYLASE->ParB+HNH->HEPN+DUF499*->ACET->TraJ-RHH-> | 1112 | Bacteria>Terrabacteria_group>Actinomycetota>Actinomycetes>Micrococcales | GCF_003002895.1 | Knoellia_remsis | BCF74_RS15510 | 21779..25117 | 407159 | Swt1-family-HEPN-domain-containing-protein-[Knoellia-remsis] |
| WP_108718589.1 | DUF499_1.RM.like.systems | <-JetB<-JetA\|\|?->PIN->?-><-?<-HEPN+DUF499*<-ParB+HNH<-HEPN+HEPN<-METHYLASE\|\|HTH->Schlafen+HATPase+HTH-><-TUDOR+SF1-SF2-Helicase+iREase+CTD+REase | 1134 | Bacteria>Terrabacteria_group>Actinomycetota>Actinomycetes>Micrococcales | GCF_003074035.1 | Miniimonas_sp._S16 | DCS86_RS08145 | 1885887..1889291 | 2171623 | Swt1-family-HEPN-domain-containing-protein-[Miniimonas-sp.-S16] |
| WP_129190457.1 | DUF499_1.RM.like.systems | <-AbiEi_4<-PIN<-XIS-HTH<-HEPN<-HEPN+DUF499*<-ParB+HNH<-METHYLASE<-?<-TUDOR+SF1-SF2-Helicase+iREase+CTD+REase\|\|NTD+SF1-SF2-Helicase-> | 1123 | Bacteria>Terrabacteria_group>Actinomycetota>Actinomycetes>Micrococcales | GCF_004135385.1 | Xylanimonas_protaetiae | ET471_RS17505 | 3776288..3779659 | 2509457 | Swt1-family-HEPN-domain-containing-protein-[Xylanimonas-protaetiae] |
| WP_141784692.1 | DUF499_1.RM.like.systems | <-MACRODOMAIN+SIR2\|\|REase-><-DDE_Tnp_1_4\|\|METHYLASE->?->ParB+HNH->HEPN+DUF499*->?-><-Phage_int_SAM_1+Phage_integrase<-?\|\|HTH-> | 1119 | Bacteria>Terrabacteria_group>Actinomycetota>Actinomycetes>Micrococcales | GCF_006716185.1 | Ornithinicoccus_hortensis | FB467_RS08335 | 1805419..1808778 | 82346 | DUF499-domain-containing-protein-[Ornithinicoccus-hortensis] |
| WP_144794196.1 | DUF499_1.RM.like.systems | TetR-HTH->XIS-HTH->TUDOR+SF1-SF2-Helicase+iREase+CTD+REase->METHYLASE->HEPN+DUF499*->ParB+HNH->?-><-N6_MTase-p<-McrC-NTD+REase | 1132 | Bacteria>Terrabacteria_group>Actinomycetota>Actinomycetes>Micrococcales | GCF_007679925.1 | Microbacterium_paludicola | FS695_RS03635 | 729234..732632 | 300019 | DUF499-domain-containing-protein-[Microbacterium-paludicola] |
| WP_187368407.1 | DUF499_1.RM.like.systems | TUDOR+SF1-SF2-Helicase+iREase+CTD+REase-><-PIN<-DUF433<-wHTH(SBD)+HNH\|\|METHYLASE->ParB+HNH->HEPN+DUF499*-><-?<-?\|\|ACET+ACET-><-Lsr2 | 1133 | Bacteria>Terrabacteria_group>Actinomycetota>Actinomycetes>Micrococcales | GCF_008000995.1 | Ruania_zhangjianzhongii | FU260_RS03480 | 750431..753832 | 2603206 | DUF499-domain-containing-protein-[Ruania-zhangjianzhongii] |
| WP_188427984.1 | DUF499_1.RM.like.systems | <-MazG<-wHTH(SBD)+HNH<-REase+SF1-SF2-Helicase\|\|?->?-><-?<-HEPN+DUF499*<-ParB+HNH<-METHYLASE<-TUDOR+SF1-SF2-Helicase+iREase+CTD+REase<-?\|\|SF1-SF2-Helicase-> | 1123 | Bacteria>Terrabacteria_group>Actinomycetota>Actinomycetes>Micrococcales | GCF_014636495.1 | Ornithinimicrobium_tianjinense | IEV65_RS02295 | 470126..473497 | 1195761 | Swt1-family-HEPN-domain-containing-protein-[Ornithinimicrobium-tianjinense] |
| WP_247963105.1 | DUF499_1.RM.like.systems | DDE_Tnp_1->METHYLASE-p->HEPN+DUF499*->ParB+HNH->?-><-N6_MTase-p<-McrC-NTD+REase | 1132 | Bacteria>Terrabacteria_group>Actinomycetota>Actinomycetes>Micrococcales | GCF_023108875.1 | Microbacterium_aerolatum | MZK47_RS13285 | 1211..4609 | 153731 | DUF499-domain-containing-protein-[Microbacterium-aerolatum] |
| WP_281534461.1 | DUF499_1.RM.like.systems | TUDOR+SF1-SF2-Helicase+iREase+CTD+REase-p->TUDOR+SF1-SF2-Helicase+iREase+CTD+REase->METHYLASE->ParB+HNH->HEPN->DUF499-p->DUF499*->DUF499-p-><-?\|\|REase->?->?-><-RelA | 430 | Bacteria>Terrabacteria_group>Actinomycetota>Actinomycetes>Micrococcales | GCF_027594785.1 | Cryobacterium_breve | KIV56_RS17895 | 3683885..3685177 | 1259258 | DUF499-domain-containing-protein-[Cryobacterium-breve] |
| WP_218122143.1 | DUF499_1.RM.like.systems | HNH->\|\|?-><-?<-?<-MPTase<-DUF499*<-METHYLASE<-TUDOR+SF1-SF2-Helicase+iREase+CTD+REase<-ParB+HNH\|\|?-><-?\|\|METHYLASE-> | 977 | Bacteria>Terrabacteria_group>Actinomycetota>Actinomycetes>Micrococcales | GCF_900102895.1 | Arthrobacter_sp._ov407 | BLS42_RS19705 | 16403..19336 | 1761748 | DUF499-domain-containing-protein-[Arthrobacter-sp.-ov407] |
| WP_076404375.1 | DUF499_1.RM.like.systems | REase-><-?<-?<-?<-HEPN+DUF499*<-ParB+HNH<-METHYLASE\|\|MPTase->PIN->HTH_3->wHTH(SBD)+HNH-> | 1151 | Bacteria>Terrabacteria_group>Actinomycetota>Actinomycetes>Micrococcales | GCF_900155925.1 | Cellulosimicrobium_aquatile | BXA05_RS06490 | 508556..512011 | 1612203 | DUF499-domain-containing-protein-[Cellulosimicrobium-aquatile] |
| WP_191562882.1 | DUF499_1.RM.like.systems | NTD+SF1-SF2-Helicase->TUDOR+SF1-SF2-Helicase+iREase+CTD+REase-><-?\|\|METHYLASE->ParB+HNH->HEPN+DUF499*->?->SF1-SF2-Helicase->?-><-URI-DNase | 1117 | Bacteria>Terrabacteria_group>Actinomycetota>Actinomycetes>Micrococcales | GCF_904423795.1 | Janibacter_melonis | IHQ27_RS00715 | 71564..74917 | 262209 | Swt1-family-HEPN-domain-containing-protein-[Janibacter-melonis] |
| WP_030271467.1 | DUF499_1.RM.like.systems | Cysteine_desulfurase->?->?->Phage_integrase-><-HEPN+DUF499*<-ParB+HNH<-METHYLASE<-TUDOR+SF1-SF2-Helicase+iREase+CTD+REase\|\|NTD+SF1-SF2-Helicase-><-?<-JetD | 1125 | Bacteria>Terrabacteria_group>Actinomycetota>Actinomycetes>Micromonosporales | GCF_000717585.1 | Micromonospora_globosa | IF06_RS0114120 | 15248..18625 | 47863 | Swt1-family-HEPN-domain-containing-protein-[Micromonospora-globosa] |
| WP_197494866.1 | DUF499_1.RM.like.systems | <-pPIWI+pPIWI<-REase<-NTD+SF1-SF2-Helicase\|\|TUDOR+SF1-SF2-Helicase+iREase+CTD+REase->METHYLASE->ParB+HNH->HEPN+DUF499*->?->DEDD_Tnp_IS110+Transposase_20->zf-ISL3+DDE_Tnp_ISL3-><-?<-HTH_3 | 1124 | Bacteria>Terrabacteria_group>Actinomycetota>Actinomycetes>Micromonosporales | GCF_001652995.1 | Micromonospora_sp._NBRC_110037 | A4R25_RS09725 | 2384437..2387811 | 1621261 | Swt1-family-HEPN-domain-containing-protein-[Micromonospora-sp.-NBRC-110037] |
| WP_107269540.1 | DUF499_1.RM.like.systems | <-ABhydrolase<-TetR-HTH\|\|?->HAMP+HISKIN->?-><-ParB+HNH<-HEPN+DUF499*\|\|?->?-><-?<-?\|\|HTH_3->Pkinase-> | 1110 | Bacteria>Terrabacteria_group>Actinomycetota>Actinomycetes>Micromonosporales | GCF_003030345.1 | Plantactinospora_sp._BC1 | C6361_RS28025 | 6382634..6385966 | 2108470 | Swt1-family-HEPN-domain-containing-protein-[Plantactinospora-sp.-BC1] |
| WP_107261307.1 | DUF499_1.RM.like.systems | <-ABhydrolase<-TetR-HTH\|\|?->HAMP+HISKIN->?-><-ParB+HNH<-HEPN+DUF499*\|\|?->?-><-?<-?\|\|HTH_3-> | 1110 | Bacteria>Terrabacteria_group>Actinomycetota>Actinomycetes>Micromonosporales | GCF_003030385.1 | Plantactinospora_sp._BB1 | C6W10_RS25965 | 5945851..5949183 | 2071627 | Swt1-family-HEPN-domain-containing-protein-[Plantactinospora-sp.-BB1] |
| WP_192902263.1 | DUF499_1.RM.like.systems | HTH+TPR+TPR->Pkinase-><-HEPN+DUF499*<-ParB+HNH<-METHYLASE<-TUDOR+SF1-SF2-Helicase+iREase+CTD+REase\|\|NTD+SF1-SF2-Helicase->REase->pPIWI+pPIWI-> | 1129 | Bacteria>Terrabacteria_group>Actinomycetota>Actinomycetes>Micromonosporales | GCF_003610785.1 | Micromonospora_globbae | D7I43_RS25240 | 101471..104860 | 1894969 | Swt1-family-HEPN-domain-containing-protein-[Micromonospora-globbae] |
| WP_187693213.1 | DUF499_1.RM.like.systems | SF1-SF2-helicase+RdgC+Beta-barrel+REase->\|\|HTH_Tnp_4+DDE_Tnp_4->\|\|DUF4096+DDE_Tnp_1+DDE_Tnp_1_2->TUDOR+SF1-SF2-Helicase+iREase+CTD+REase->METHYLASE->ParB+HNH->HEPN+DUF499*-><-Phage_integrase<-?<-?<-?<-Cysteine_desulfurase | 1129 | Bacteria>Terrabacteria_group>Actinomycetota>Actinomycetes>Micromonosporales | GCF_014472405.1 | Micromonospora_chalcea | H9X95_RS12030 | 19161..22550 | 1874 | Swt1-family-HEPN-domain-containing-protein-[Micromonospora-chalcea] |
| WP_190252034.1 | DUF499_1.RM.like.systems | <-ABC-ATPase+SbcC\|\|tRNA->?-><-?\|\|TUDOR+SF1-SF2-Helicase+iREase+CTD+REase->METHYLASE->HEPN+DUF499*->ParB+HNH->?-><-Phage_int_SAM_1+Phage_integrase<-HTH_17\|\|?-><-BirA-HTH | 1122 | Bacteria>Terrabacteria_group>Actinomycetota>Actinomycetes>Micromonosporales | GCF_014647515.1 | Dactylosporangium_sucinum | IEZ31_RS23190 | 135414..138782 | 1424081 | Swt1-family-HEPN-domain-containing-protein-[Dactylosporangium-sucinum] |
| WP_204020362.1 | DUF499_1.RM.like.systems | NUDIX->?-><-?<-REase<-ParB+HNH<-HEPN+DUF499*<-METHYLASE<-TUDOR+SF1-SF2-Helicase+iREase+CTD+REase\|\|RelE-ParE->ParA+HTH-><-PIN<-Phd-YefM | 1129 | Bacteria>Terrabacteria_group>Actinomycetota>Actinomycetes>Micromonosporales | GCF_016863595.1 | Micromonospora_andamanensis | Vwe01_RS03465 | 15617..19006 | 1287068 | Swt1-family-HEPN-domain-containing-protein-[Micromonospora-trujilloniae] |
| WP_208576856.1 | DUF499_1.RM.like.systems | HTH+TPR+TPR->Pkinase-><-HEPN+DUF499*<-ParB+HNH<-METHYLASE<-TUDOR+SF1-SF2-Helicase+iREase+CTD+REase\|\|NTD+SF1-SF2-Helicase->HEPN-> | 1126 | Bacteria>Terrabacteria_group>Actinomycetota>Actinomycetes>Micromonosporales | GCF_017599015.1 | Micromonospora_tulbaghiae | J5U46_RS08620 | 81650..85030 | 479978 | Swt1-family-HEPN-domain-containing-protein-[Micromonospora-tulbaghiae] |
| WP_221380246.1 | DUF499_1.RM.like.systems | sGTP->tRNA->Phage_int_SAM_4+Phage_integrase->METHYLASE-><-CARF+HEPN<-?<-ParB+HNH<-HEPN+DUF499*<-METHYLASE<-TUDOR+SF1-SF2-Helicase+iREase+CTD+REase<-SF1-SF2-Helicase<-?<-REase | 1128 | Bacteria>Terrabacteria_group>Actinomycetota>Actinomycetes>Micromonosporales | GCF_019710575.1 | Actinoplanes_polyasparticus | KXC43_RS33640 | 36558..39944 | 2856853 | Swt1-family-HEPN-domain-containing-protein-[Actinoplanes-sp.-TRM66264-DLM] |
| WP_167666591.1 | DUF499_1.RM.like.systems | HTH+TPR+TPR->Pkinase-><-HEPN+DUF499*<-ParB+HNH<-METHYLASE<-TUDOR+SF1-SF2-Helicase+iREase+CTD+REase\|\|NTD+SF1-SF2-Helicase->\|\|HNH-> | 1124 | Bacteria>Terrabacteria_group>Actinomycetota>Actinomycetes>Micromonosporales | GCF_900089605.1 | Micromonospora_narathiwatensis | GA0070621_RS04900 | 1072112..1075486 | 299146 | Swt1-family-HEPN-domain-containing-protein-[Micromonospora-narathiwatensis] |
| WP_030502747.1 | DUF499_1.RM.like.systems | HTH_3->?-><-DEDD_Tnp_IS110+Transposase_20<-zf-ISL3+DDE_Tnp_ISL3\|\|MULE-TRANSPOSASE-><-HEPN+DUF499*<-ParB+HNH<-METHYLASE<-TUDOR+SF1-SF2-Helicase+iREase+CTD+REase\|\|NTD+SF1-SF2-Helicase->REase->pPIWI+pPIWI-> | 1124 | Bacteria>Terrabacteria_group>Actinomycetota>Actinomycetes>Micromonosporales | - | Micromonospora_purpureochromogenes | IH31_RS0121315 | 105944..109318 | 47872 | Swt1-family-HEPN-domain-containing-protein-[Micromonospora-purpureochromogenes] |
| OAH45661.1 | DUF499_1.RM.like.systems | TUDOR+SF1-SF2-Helicase+iREase+CTD+REase-><-?\|\|METHYLASE->ParB+HNH->HEPN+DUF499*->ANK+ANK+RAMA-> | 1134 | Bacteria>Terrabacteria_group>Actinomycetota>Actinomycetes>Mycobacteriales | GCA_001643175.1 | Dietzia_cinnamea | AYJ66_16790 | 32668..36072 | 321318 | AAA-family-ATPase-[Dietzia-cinnamea] |
| OBB72728.1 | DUF499_1.RM.like.systems | HTH_Tnp->?->?->?->\|\|TUDOR+SF1-SF2-Helicase+iREase+CTD+REase->METHYLASE->HEPN+DUF499*->ParB+HNH->HIN-HTH->HTH_21+rve+rve_3->HEPN-> | 1110 | Bacteria>Terrabacteria_group>Actinomycetota>Actinomycetes>Mycobacteriales | GCA_001665615.1 | Mycobacterium_sp._852014-52144_SCH5372336 | A5759_18055 | 92771..96103 | 1834115 | AAA-family-ATPase-[Mycobacterium-sp.-852014-52144_SCH5372336] |
| OBK03541.1 | DUF499_1.RM.like.systems | <-NTD+SF1-SF2-Helicase\|\|TUDOR+SF1-SF2-Helicase+iREase+CTD+REase->METHYLASE->ParB+HNH->HEPN+DUF499*->?->?->RnaseIII+DSRBD+RnaseIII+DSRBD->REC+HEPN-> | 1103 | Bacteria>Terrabacteria_group>Actinomycetota>Actinomycetes>Mycobacteriales | GCA_001673265.1 | Mycolicibacterium_conceptionense | A5639_22740 | 13192..16503 | 451644 | AAA-family-ATPase-[Mycolicibacterium-conceptionense] |
| OFJ52823.1 | DUF499_1.RM.like.systems | TUDOR+SF1-SF2-Helicase+iREase+CTD+REase->METHYLASE->ParB+HNH->HEPN+DUF499*->?->?-><-Cluster182_14clades<-MACRODOMAIN<-?<-IclR-HTH | 1109 | Bacteria>Terrabacteria_group>Actinomycetota>Actinomycetes>Mycobacteriales | GCA_001766635.1 | Mycobacterium_grossiae | BEL07_15685 | 42542..45871 | 1552759 | AAA-family-ATPase-[Mycobacterium-grossiae] |
| OHT48546.1 | DUF499_1.RM.like.systems | PAIREDC-HTH+HIN-HTH-><-?\|\|TUDOR+SF1-SF2-Helicase+iREase+CTD+REase->METHYLASE->ParB+HNH->HEPN+DUF499*->?->?->?-><-?<-METHYLASE-p | 1109 | Bacteria>Terrabacteria_group>Actinomycetota>Actinomycetes>Mycobacteriales | GCA_001853545.1 | Mycobacteroides_chelonae | BKG62_21080 | 210158..213487 | 1774 | AAA-family-ATPase-[Mycobacteroides-chelonae] |
| OHV00237.1 | DUF499_1.RM.like.systems | TUDOR+SF1-SF2-Helicase+iREase+CTD+REase->METHYLASE->ParB+HNH->HEPN+DUF499*->SIR2-><-?\|\|SbcC-> | 1095 | Bacteria>Terrabacteria_group>Actinomycetota>Actinomycetes>Mycobacteriales | GCA_001854525.1 | Mycobacterium_talmoniae | BKN37_18410 | 8707..11994 | 1858794 | AAA-family-ATPase-[Mycobacterium-talmoniae] |
| OKH62617.1 | DUF499_1.RM.like.systems | TUDOR+SF1-SF2-Helicase+iREase+CTD+REase->METHYLASE->ParB+HNH->HEPN+DUF499*->OLD-ABC-><-?<-McrC-NTD+REase<-MAD-NTD+EVE+McrB-AAA | 1117 | Bacteria>Terrabacteria_group>Actinomycetota>Actinomycetes>Mycobacteriales | GCA_001905305.1 | Mycobacterium_sp._SWH-M5 | EB74_16030 | 8207..11560 | 1490483 | ATPase-AAA-[Mycobacterium-sp.-SWH-M5] |
| OMC31131.1 | DUF499_1.RM.like.systems | <-ParB+HNH<-NTD+SF1-SF2-Helicase\|\|TUDOR+SF1-SF2-Helicase+iREase+CTD+REase->METHYLASE->ParB+HNH->HEPN+DUF499*-><-Glyoxalase\|\|?-><-TetR-HTH\|\|?->?-><-Pkinase | 1100 | Bacteria>Terrabacteria_group>Actinomycetota>Actinomycetes>Mycobacteriales | GCA_001954115.1 | Mycobacterium_colombiense | A5739_14025 | 105145..108447 | 339268 | AAA-family-ATPase-[Mycobacterium-colombiense] |
| OMB68868.1 | DUF499_1.RM.like.systems | <-HEPN+DUF499*<-ParB+HNH<-METHYLASE<-TUDOR+SF1-SF2-Helicase+iREase+CTD+REase\|\|NTD+SF1-SF2-Helicase-> | 1103 | Bacteria>Terrabacteria_group>Actinomycetota>Actinomycetes>Mycobacteriales | GCA_001954125.1 | Mycolicibacterium_conceptionense | A5741_09375 | 1839..5150 | 451644 | AAA-family-ATPase-[Mycolicibacterium-conceptionense] |
| ORA83573.1 | DUF499_1.RM.like.systems | REase->?->?->?->\|\|TUDOR+SF1-SF2-Helicase+iREase+CTD+REase->METHYLASE->HEPN+DUF499*->ParB+HNH->HEPN->?-><-SF1-SF2-Helicase | 1110 | Bacteria>Terrabacteria_group>Actinomycetota>Actinomycetes>Mycobacteriales | GCA_002086285.1 | Mycolicibacter_kumamotonensis | BST28_00055 | 16546..19878 | 354243 | AAA-family-ATPase-[Mycolicibacter-kumamotonensis] |
| ORB28066.1 | DUF499_1.RM.like.systems | HKD-DNase+SF1-SF2-Helicase+TRD-><-?<-?<-?<-HTH_Tnp_1+HTH_21+rve+rve_2<-HIN-HTH<-HEPN+DUF499*<-ParB+HNH<-METHYLASE<-TUDOR+SF1-SF2-Helicase+iREase+CTD+REase\|\|NTD+SF1-SF2-Helicase-> | 1100 | Bacteria>Terrabacteria_group>Actinomycetota>Actinomycetes>Mycobacteriales | GCA_002086815.1 | Mycolicibacterium_parafortuitum | BST38_21950 | 32526..35828 | 39692 | AAA-family-ATPase-[Mycolicibacterium-parafortuitum] |
| ORV36019.1 | DUF499_1.RM.like.systems | HTH_Tnp->?->?->?->\|\|TUDOR+SF1-SF2-Helicase+iREase+CTD+REase->ParB+HNH->HEPN+DUF499*-> | 1111 | Bacteria>Terrabacteria_group>Actinomycetota>Actinomycetes>Mycobacteriales | GCA_002102115.1 | Mycolicibacterium_doricum | AWC01_00650 | 40820..44155 | 126673 | AAA-family-ATPase-[Mycobacterium-doricum] |
| PQP40021.1 | DUF499_1.RM.like.systems | <-GspA-ATPase<-HIN-HTH\|\|Cluster182_14clades-><-?<-?<-?<-HEPN+DUF499*<-ParB+HNH<-METHYLASE<-TUDOR+SF1-SF2-Helicase+iREase+CTD+REase\|\|NTD+SF1-SF2-Helicase-> | 1103 | Bacteria>Terrabacteria_group>Actinomycetota>Actinomycetes>Mycobacteriales | GCA_002968275.1 | Mycolicibacterium_austroafricanum | C6A88_31860 | 14193..17504 | 39687 | AAA-family-ATPase-[Mycolicibacterium-austroafricanum] |
| TXI55381.1 | DUF499_1.RM.like.systems | <-FtsK<-?<-GNTR-HTH\|\|NUDIX-><-HEPN<-Schlafen+HATPase+HTH<-HEPN+DUF499*<-ParB+HNH<-METHYLASE<-TUDOR+SF1-SF2-Helicase+iREase+CTD+REase<-?<-HEPN | 1103 | Bacteria>Terrabacteria_group>Actinomycetota>Actinomycetes>Mycobacteriales | GCA_008015745.1 | Mycolicibacter_arupensis | E6Q54_12720 | 7339..10650 | 342002 | ATP-binding-protein-[Mycolicibacter-arupensis] |
| TXI48376.1 | DUF499_1.RM.like.systems | <-HIN-HTH<-NTD+SF1-SF2-Helicase\|\|TUDOR+SF1-SF2-Helicase+iREase+CTD+REase-><-?\|\|METHYLASE->ParB+HNH->HEPN+DUF499*-> | 1102 | Bacteria>Terrabacteria_group>Actinomycetota>Actinomycetes>Mycobacteriales | GCA_008015745.1 | Mycolicibacter_arupensis | E6Q54_23710 | 12171..15479 | 342002 | ATP-binding-protein-[Mycolicibacter-arupensis] |
| MBM4528840.1 | DUF499_1.RM.like.systems | wHTH(SBD)+HNH-><-ParB<-HEPN+DUF499*<-ParB+HNH-p<-METHYLASE<-TUDOR+SF1-SF2-Helicase+iREase+CTD+REase\|\|NTD+SF1-SF2-Helicase-><-SF1-SF2-Helicase<-HKD-DNase | 1121 | Bacteria>Terrabacteria_group>Actinomycetota>Actinomycetes>Mycobacteriales | GCA_016879355.1 | Prescottella_equi | GS563_07715 | 1641397..1644762 | 43767 | DUF499-domain-containing-protein-[Prescottella-equi] |
| MBU8833314.1 | DUF499_1.RM.like.systems | McrC-NTD+REase-><-?<-HTH_Tnp_1+HTH_21+rve+rve_3\|\|HTH_Tnp_1+HTH_21+rve+rve_3->?-><-REase<-HEPN+DUF499*<-ParB+HNH<-METHYLASE<-TUDOR+SF1-SF2-Helicase+iREase+CTD+REase\|\|DDE_Tnp_1_4->?->HEPN+SF1-SF2-Helicase-> | 1119 | Bacteria>Terrabacteria_group>Actinomycetota>Actinomycetes>Mycobacteriales | GCA_018524295.1 | Mycolicibacterium_goodii | KL858_28165 | 17594..20953 | 134601 | DUF499-domain-containing-protein-[Mycobacterium-goodii] |
| HJC59998.1 | DUF499_1.RM.like.systems | <-PIN<-?\|\|?-><-?<-CHTH<-?<-HEPN+DUF499*<-ParB+HNH<-METHYLASE<-TUDOR+SF1-SF2-Helicase+iREase+CTD+REase\|\|HTH_19-><-Synaptojanin | 1131 | Bacteria>Terrabacteria_group>Actinomycetota>Actinomycetes>Mycobacteriales | GCA_019119875.1 | Candidatus_Dietzia_intestinigallinarum | H9755_06660 | 12112..15507 | 2838537 | DUF499-domain-containing-protein-[Candidatus-Dietzia-intestinigallinarum] |
| MBX7452224.1 | DUF499_1.RM.like.systems | <-REase\|\|DNA_methylase-><-NTD+SF1-SF2-Helicase\|\|TUDOR+SF1-SF2-Helicase+iREase+CTD+REase->METHYLASE->ParB+HNH->HEPN+DUF499*->MORC->REase->HTH+SF1-SF2-Helicase+REase->\|\|MORC+SF1-SF2-Helicase->NERD+SF1-SF2-Helicase->SF1-SF2-Helicase-> | 1122 | Bacteria>Terrabacteria_group>Actinomycetota>Actinomycetes>Mycobacteriales | GCA_019710595.1 | Mycolicibacterium_aurantiacum | GR927_29910 | 284651..288019 | 2692455 | DUF499-domain-containing-protein-[Mycolicibacterium-aurantiacum] |
| MBZ4613710.1 | DUF499_1.RM.like.systems | <-HEPN+DUF499*<-ParB+HNH<-METHYLASE<-TUDOR+SF1-SF2-Helicase+iREase+CTD+REase | 1103 | Bacteria>Terrabacteria_group>Actinomycetota>Actinomycetes>Mycobacteriales | GCA_020055445.1 | Mycobacterium_avium_subsp._hominissuis | GBP66_23845 | 83..3394 | 439334 | ATP-binding-protein-[Mycobacterium-avium-subsp.-hominissuis] |
| MCB0932438.1 | DUF499_1.RM.like.systems | <-NTD+SF1-SF2-Helicase\|\|TUDOR+SF1-SF2-Helicase+iREase+CTD+REase->METHYLASE->ParB+HNH->HEPN+DUF499*->?->?->?-><-Cluster182_14clades<-MACRODOMAIN\|\|PIN-> | 1120 | Bacteria>Terrabacteria_group>Actinomycetota>Actinomycetes>Mycobacteriales | GCA_020438845.1 | Mycobacterium_sp. | KDB71_11130 | 13103..16465 | 1785 | DUF499-domain-containing-protein-[Mycobacterium-sp.] |
| MCQ4152489.1 | DUF499_1.RM.like.systems | HEPN-><-NTD+SF1-SF2-Helicase\|\|TUDOR+SF1-SF2-Helicase+iREase+CTD+REase->METHYLASE->ParB+HNH->HEPN+DUF499*-> | 1120 | Bacteria>Terrabacteria_group>Actinomycetota>Actinomycetes>Mycobacteriales | GCA_024438075.1 | Rhodococcus_qingshengii | NOS52_32470 | 22838..26200 | 334542 | Swt1-family-HEPN-domain-containing-protein-[Rhodococcus-qingshengii] |
| MCQ4364655.1 | DUF499_1.RM.like.systems | <-NTD+SF1-SF2-Helicase\|\|TUDOR+SF1-SF2-Helicase+iREase+CTD+REase->METHYLASE->ParB+HNH->HEPN+DUF499*->ParB+HNH->HEPN-> | 1118 | Bacteria>Terrabacteria_group>Actinomycetota>Actinomycetes>Mycobacteriales | GCA_024451565.1 | Mycobacterium_gordonae | KQR54_26610 | 18169..21525 | 1778 | Swt1-family-HEPN-domain-containing-protein-[Mycobacterium-gordonae] |
| MCT7366341.1 | DUF499_1.RM.like.systems | JetA->JetB->?->JetD-><-?\|\|?-><-HEPN+DUF499*\|\|HIN-HTH->HIN-HTH-><-ParB+HNH<-HTH_21+rve+rve_3<-HIN-HTH<-TUDOR+SF1-SF2-Helicase+iREase+CTD+REase | 1099 | Bacteria>Terrabacteria_group>Actinomycetota>Actinomycetes>Mycobacteriales | GCA_025331195.1 | Mycolicibacterium_llatzerense | A7G45_26120 | 438962..442261 | 280871 | AAA-family-ATPase-[Mycolicibacterium-llatzerense] |
| SHV22416.1 | DUF499_1.RM.like.systems | <-NTD+SF1-SF2-Helicase\|\|TUDOR+SF1-SF2-Helicase+iREase+CTD+REase-><-?\|\|METHYLASE->ParB+HNH->HEPN+DUF499*->?->?-><-MACRODOMAIN | 1122 | Bacteria>Terrabacteria_group>Actinomycetota>Actinomycetes>Mycobacteriales | GCA_900132885.1 | Mycobacteroides_abscessus_subsp._abscessus | SAMEA2259605_05819 | 13396..16764 | 1185650 | Conserved-protein-of-uncharacterised-function-(part1)-[Mycobacteroides-abscessus-subsp.-abscessus] |
| WP_011210977.1 | DUF499_1.RM.like.systems | <-HTH_21+rve+rve_3\|\|?->HIN-HTH->?->rve+rve_3-><-HEPN+DUF499*<-ParB+HNH<-METHYLASE<-TUDOR+SF1-SF2-Helicase+iREase+CTD+REase\|\|GDResolvase+Recombinase+ZnR-> | 1127 | Bacteria>Terrabacteria_group>Actinomycetota>Actinomycetes>Mycobacteriales | GCF_000009805.1 | Nocardia_farcinica | NFA_RS22030 | 4605548..4608931 | 37329 | Swt1-family-HEPN-domain-containing-protein-[Nocardia-farcinica] |
| WP_013470723.1 | DUF499_1.RM.like.systems | <-MORC<-HTH+SF1-SF2-Helicase+REase<-DDE_Tnp_1_4<-REase<-MORC<-HEPN+DUF499*<-ParB+HNH<-MULE-TRANSPOSASE<-TUDOR+SF1-SF2-Helicase+iREase+CTD+REase<-HIN-HTH | 1122 | Bacteria>Terrabacteria_group>Actinomycetota>Actinomycetes>Mycobacteriales | GCF_000184435.1 | Mycolicibacterium_gilvum | MSPYR1_RS04230 | 878356..881724 | 1804 | Swt1-family-HEPN-domain-containing-protein-[Mycolicibacterium-gilvum] |
| WP_006942562.1 | DUF499_1.RM.like.systems | <-HEPN+DUF499*<-ParB+HNH<-METHYLASE<-TUDOR+SF1-SF2-Helicase+iREase+CTD+REase\|\|NTD+SF1-SF2-Helicase-> | 1119 | Bacteria>Terrabacteria_group>Actinomycetota>Actinomycetes>Mycobacteriales | GCF_000382865.1 | Rhodococcus_aetherivorans | EBESD8_RS20690 | 433..3792 | 191292 | Swt1-family-HEPN-domain-containing-protein-[Rhodococcus-aetherivorans] |
| WP_174485135.1 | DUF499_1.RM.like.systems | <-NTD+SF1-SF2-Helicase\|\|TUDOR+SF1-SF2-Helicase+iREase+CTD+REase-><-?\|\|METHYLASE->ParB+HNH->HEPN+DUF499*->?->?-><-?<-REase<-?\|\|ABhydrolase-> | 1113 | Bacteria>Terrabacteria_group>Actinomycetota>Actinomycetes>Mycobacteriales | GCF_001214605.1 | Mycobacteroides_abscessus | IUT00_RS08255 | 1657384..1660725 | 36809 | Swt1-family-HEPN-domain-containing-protein-[Mycobacteroides-abscessus] |
| WP_068276445.1 | DUF499_1.RM.like.systems | METHYLASE->ParB+HNH->HEPN+DUF499*-> | 1122 | Bacteria>Terrabacteria_group>Actinomycetota>Actinomycetes>Mycobacteriales | GCF_001646865.1 | Aldersonia_kunmingensis | A8G08_RS16790 | 4064..7432 | 408066 | Swt1-family-HEPN-domain-containing-protein-[Aldersonia-kunmingensis] |
| WP_197506097.1 | DUF499_1.RM.like.systems | HTH_Tnp->?->?-><-NTD+SF1-SF2-Helicase\|\|TUDOR+SF1-SF2-Helicase+iREase+CTD+REase->METHYLASE->HEPN+DUF499*->ParB+HNH->HTH_Tnp_1+HTH_21+rve+rve_2->HEPN-> | 1121 | Bacteria>Terrabacteria_group>Actinomycetota>Actinomycetes>Mycobacteriales | GCF_001665615.1 | Mycobacterium_sp._852014-52144_SCH5372336 | A5759_RS17450 | 92738..96103 | 1834115 | Swt1-family-HEPN-domain-containing-protein-[Mycobacterium-sp.-852014-52144_SCH5372336] |
| WP_207544692.1 | DUF499_1.RM.like.systems | <-NTD+SF1-SF2-Helicase\|\|TUDOR+SF1-SF2-Helicase+iREase+CTD+REase->METHYLASE->ParB+HNH->HEPN+DUF499*->?->?->RnaseIII+DSRBD+RnaseIII+DSRBD->REC+HEPN-> | 1114 | Bacteria>Terrabacteria_group>Actinomycetota>Actinomycetes>Mycobacteriales | GCF_001673265.1 | Mycolicibacterium_conceptionense | A5639_RS11935 | 13159..16503 | 451644 | Swt1-family-HEPN-domain-containing-protein-[Mycolicibacterium-conceptionense] |
| WP_207546421.1 | DUF499_1.RM.like.systems | <-NTD+SF1-SF2-Helicase\|\|PAIREDC-HTH+HIN-HTH-><-?\|\|TUDOR+SF1-SF2-Helicase+iREase+CTD+REase->METHYLASE->ParB+HNH->HEPN+DUF499*->?->?->?-><-?<-METHYLASE-p | 1120 | Bacteria>Terrabacteria_group>Actinomycetota>Actinomycetes>Mycobacteriales | GCF_001853545.1 | Mycobacteroides_chelonae | BKG62_RS21080 | 210125..213487 | 1774 | Swt1-family-HEPN-domain-containing-protein-[Mycobacteroides-chelonae] |
| WP_207547258.1 | DUF499_1.RM.like.systems | TUDOR+SF1-SF2-Helicase+iREase+CTD+REase->METHYLASE->ParB+HNH->HEPN+DUF499*->SIR2->?->SbcC-> | 1106 | Bacteria>Terrabacteria_group>Actinomycetota>Actinomycetes>Mycobacteriales | GCF_001854525.1 | Mycobacterium_talmoniae | BKN37_RS18415 | 8674..11994 | 1858794 | DUF499-domain-containing-protein-[Mycobacterium-talmoniae] |
| WP_170880036.1 | DUF499_1.RM.like.systems | <-ParB+HNH<-NTD+SF1-SF2-Helicase\|\|TUDOR+SF1-SF2-Helicase+iREase+CTD+REase->METHYLASE->ParB+HNH->HEPN+DUF499*-><-Glyoxalase\|\|?-><-TetR-HTH\|\|?->?-><-Pkinase | 1111 | Bacteria>Terrabacteria_group>Actinomycetota>Actinomycetes>Mycobacteriales | GCF_001954115.1 | Mycobacterium_colombiense | A5739_RS13730 | 105112..108447 | 339268 | Swt1-family-HEPN-domain-containing-protein-[Mycobacterium-colombiense] |
| WP_207553229.1 | DUF499_1.RM.like.systems | <-HEPN+DUF499*<-ParB+HNH<-METHYLASE<-TUDOR+SF1-SF2-Helicase+iREase+CTD+REase\|\|NTD+SF1-SF2-Helicase-> | 1114 | Bacteria>Terrabacteria_group>Actinomycetota>Actinomycetes>Mycobacteriales | GCF_001954125.1 | Mycolicibacterium_conceptionense | A5741_RS31960 | 1839..5183 | 451644 | Swt1-family-HEPN-domain-containing-protein-[Mycolicibacterium-conceptionense] |
| WP_174682959.1 | DUF499_1.RM.like.systems | <-HEPN+DUF499*<-ParB+HNH<-METHYLASE<-rve_3 | 1090 | Bacteria>Terrabacteria_group>Actinomycetota>Actinomycetes>Mycobacteriales | GCF_001954235.1 | Mycolicibacterium_conceptionense | A5746_RS13475 | 1..3270 | 451644 | Swt1-family-HEPN-domain-containing-protein,-partial-[Mycolicibacterium-conceptionense] |

| Protein-ID | System | Gene-Neighbourhoods | Length | Taxonomy | Assembly-ID | Strain | Locus-Tag | Genomic-coordinates | TaxID | Fasta-defline |
|---|---|---|---|---|---|---|---|---|---|---|
| WP_207569995.1 | DUF499_1.RM.like.systems | <-NTD+SF1-SF2-Helicase\|\|TUDOR+ SF1-SF2-Helicase+iREase+CTD+REase->METHYLASE->HEPN+ DUF499 *->ParB+HNH->?-><-SF1-SF2-Helicase | 1121 | Bacteria>Terrabacteria_group>Actinomycetota>Actinomycetes>Mycobacteriales | GCF_002086285.1 | Mycolicibacter_kumamotonensis | BST28_RS00055 | 16513..19878 | 354243 | Swt1-family-HEPN-domain-containing-protein-[Mycolicibacter-kumamotonensis] |
| WP_181792003.1 | DUF499_1.RM.like.systems | <-35exo+SF1-SF2-Helicase\|\|HKD-DNase+SF1-SF2-Helicase+TRD-><-?<-?<-?<-HTH_Tnp_1+HTH_21+rve+rve_2<-HEPN+ DUF499 *<-ParB+HNH<-METHYLASE<-TUDOR+ SF1-SF2-Helicase+iREase+CTD+REase\|\|NTD+SF1-SF2-Helicase-> | 1111 | Bacteria>Terrabacteria_group>Actinomycetota>Actinomycetes>Mycobacteriales | GCF_002086815.1 | Mycolicibacterium_parafortuitum | MPP7335_RS09255 | 1901353..1904688 | 39692 | Swt1-family-HEPN-domain-containing-protein-[Mycolicibacterium-parafortuitum] |
| WP_094652354.1 | DUF499_1.RM.like.systems | <-ABhydrolase+ABhydrolase<-REase<-ParB+HNH+RAMA\|\|Primpol+D5N+D5ATPase->SF1-SF2-Helicase-><-HEPN+ DUF499 *<-ParB+HNH-p<-METHYLASE<-TUDOR+ SF1-SF2-Helicase+iREase+CTD+REase\|\|?->Exonuc_V_gamma+REase->SF1-SF2-Helicase+REase-> | 1120 | Bacteria>Terrabacteria_group>Actinomycetota>Actinomycetes>Mycobacteriales | GCF_002259465.1 | Rhodococcus_fascians | CH297_RS18200 | 13414..16776 | 1828 | Swt1-family-HEPN-domain-containing-protein-[Rhodococcus-fascians] |
| WP_099384398.1 | DUF499_1.RM.like.systems | TUDOR+ SF1-SF2-Helicase+iREase+CTD+REase->METHYLASE->ParB+HNH->HEPN+ DUF499 *->NTD+SF1-SF2-Helicase-><-?<-?\|\|REase-> | 1115 | Bacteria>Terrabacteria_group>Actinomycetota>Actinomycetes>Mycobacteriales | GCF_002735905.1 | Williamsia_marianensis | CSW57_RS21175 | 1638758..1642105 | 322609 | Swt1-family-HEPN-domain-containing-protein-[Williamsia-marianensis] |
| WP_102999581.1 | DUF499_1.RM.like.systems | TUDOR+ SF1-SF2-Helicase+iREase+CTD+REase->METHYLASE->ParB+HNH->HEPN+ DUF499 *->zf-ISL3+DDE_Tnp_ISL3->?-><-DNA_methylase<-HEPN<-ParB+HNH<-GDResolvase+HIN-HTH | 1121 | Bacteria>Terrabacteria_group>Actinomycetota>Actinomycetes>Mycobacteriales | GCF_002893965.1 | Rhodococcus_qingshengii | C1M55_RS31235 | 10127..13492 | 334542 | Swt1-family-HEPN-domain-containing-protein-[Rhodococcus-qingshengii] |
| WP_104363764.1 | DUF499_1.RM.like.systems | HIN-HTH->HTH_21+rve+rve_3-><-?<-?\|\|ParB+HNH->HEPN+ DUF499 *-> | 1121 | Bacteria>Terrabacteria_group>Actinomycetota>Actinomycetes>Mycobacteriales | GCF_002933495.1 | Nocardia_nova | C5F51_RS20340 | 202110..205475 | 37330 | Swt1-family-HEPN-domain-containing-protein-[Nocardia-nova] |
| WP_205765478.1 | DUF499_1.RM.like.systems | <-HIN-HTH<-?\|\|Cluster182_14clades-><-?<-?<-?<-HEPN+ DUF499 *<-ParB+HNH<-METHYLASE<-TUDOR+ SF1-SF2-Helicase+iREase+CTD+REase\|\|NTD+SF1-SF2-Helicase-> | 1114 | Bacteria>Terrabacteria_group>Actinomycetota>Actinomycetes>Mycobacteriales | GCF_002968275.1 | Mycolicibacterium_austroafricanum | C6A88_RS31870 | 14193..17537 | 39687 | Swt1-family-HEPN-domain-containing-protein-[Mycolicibacterium-austroafricanum] |
| WP_106851264.1 | DUF499_1.RM.like.systems | SIGMA-HTH+SIGMA-HTH-><-wHTH(SBD)+HNH<-REase+SF1-SF2-Helicase\|\|TUDOR+ SF1-SF2-Helicase+iREase+CTD+REase->METHYLASE->ParB+HNH->HEPN+ DUF499 *->NTD+SF1-SF2-Helicase-> | 1123 | Bacteria>Terrabacteria_group>Actinomycetota>Actinomycetes>Mycobacteriales | GCF_003013675.1 | Prescottella_equi | C7H75_RS09240 | 1958101..1961472 | 43767 | Swt1-family-HEPN-domain-containing-protein-[Prescottella-equi] |
| WP_119699599.1 | DUF499_1.RM.like.systems | <-JetA<-?<-?<-?<-?<-?<-HEPN+ DUF499 *<-ParB+HNH<-METHYLASE<-TUDOR+ SF1-SF2-Helicase+iREase+CTD+REase\|\|NTD+SF1-SF2-Helicase-> | 1118 | Bacteria>Terrabacteria_group>Actinomycetota>Actinomycetes>Mycobacteriales | GCF_003586525.1 | Rhodococcus_ruber | TN91_RS02590 | 62027..65383 | 1830 | Swt1-family-HEPN-domain-containing-protein-[Rhodococcus-ruber] |
| WP_206613802.1 | DUF499_1.RM.like.systems | BirA-HTH+CRP-HTH-><-HEPN+ DUF499 *<-ParB+HNH<-METHYLASE<-TUDOR+ SF1-SF2-Helicase+iREase+CTD+REase | 1113 | Bacteria>Terrabacteria_group>Actinomycetota>Actinomycetes>Mycobacteriales | GCF_004014805.1 | Mycolicibacterium_elephantis | H7I95_RS08850 | 1305..4646 | 81858 | Swt1-family-HEPN-domain-containing-protein-[Mycolicibacterium-elephantis] |
| WP_208722091.1 | DUF499_1.RM.like.systems | <-TetR-HTH<-GDResolvase+HIN-HTH\|\|Lsr2->?-><-?<-HEPN+ DUF499 *<-ParB+HNH<-METHYLASE<-UDG-conflict<-TUDOR+ SF1-SF2-Helicase+iREase+CTD+REase\|\|NTD+SF1-SF2-Helicase-> | 1122 | Bacteria>Terrabacteria_group>Actinomycetota>Actinomycetes>Mycobacteriales | GCF_006384225.1 | Rhodococcus_qingshengii | IGTS8_RS29980 | 39886..43254 | 334542 | Swt1-family-HEPN-domain-containing-protein-[Rhodococcus-qingshengii] |
| WP_276761078.1 | DUF499_1.RM.like.systems | <-FtsK<-?<-GNTR-HTH\|\|NUDIX-><-HEPN<-Schlafen+HATPase+HTH<-HEPN+ DUF499 *<-ParB+HNH<-METHYLASE<-TUDOR+ SF1-SF2-Helicase+iREase+CTD+REase<-?<-HEPN<-HEPN | 1114 | Bacteria>Terrabacteria_group>Actinomycetota>Actinomycetes>Mycobacteriales | GCF_008015745.1 | Mycolicibacter_arupensis | E6Q54_RS12710 | 7339..10683 | 342002 | Swt1-family-HEPN-domain-containing-protein-[Mycolicibacter-arupensis] |
| WP_276763638.1 | DUF499_1.RM.like.systems | <-NTD+SF1-SF2-Helicase\|\|TUDOR+ SF1-SF2-Helicase+iREase+CTD+REase-><-?\|\|METHYLASE->ParB+HNH->HEPN+ DUF499 *-> | 1113 | Bacteria>Terrabacteria_group>Actinomycetota>Actinomycetes>Mycobacteriales | GCF_008015745.1 | Mycolicibacter_arupensis | E6Q54_RS23755 | 12138..15479 | 342002 | Swt1-family-HEPN-domain-containing-protein-[Mycolicibacter-arupensis] |
| WP_176749585.1 | DUF499_1.RM.like.systems | <-NTD+SF1-SF2-Helicase\|\|TUDOR+ SF1-SF2-Helicase+iREase+CTD+REase->METHYLASE-p->DDE_Tnp_1_4->ParB+HNH->HEPN+ DUF499 *->?->?->?-><-Cluster182_14clades<-?<-IclR-HTH | 1120 | Bacteria>Terrabacteria_group>Actinomycetota>Actinomycetes>Mycobacteriales | GCF_008329645.1 | Mycobacterium_grossiae | FZ046_RS21205 | 4404866..4408228 | 1552759 | Swt1-family-HEPN-domain-containing-protein-[Mycobacterium-grossiae] |
| WP_153212242.1 | DUF499_1.RM.like.systems | <-NTD+SF1-SF2-Helicase\|\|TUDOR+ SF1-SF2-Helicase+iREase+CTD+REase->UDG-conflict->METHYLASE->ParB+HNH->HEPN+ DUF499 *-><-NUDIX\|\|?->REase->?-><-HAD<-ABC-ATPase+SbcC | 1121 | Bacteria>Terrabacteria_group>Actinomycetota>Actinomycetes>Mycobacteriales | GCF_009495805.1 | Rhodococcus_erythropolis | GEO20_RS16000 | 38887..42252 | 1833 | Swt1-family-HEPN-domain-containing-protein-[Rhodococcus-erythropolis] |
| WP_179959972.1 | DUF499_1.RM.like.systems | <-NTD+SF1-SF2-Helicase\|\|TUDOR+ SF1-SF2-Helicase+iREase+CTD+REase->ParB+HNH->HEPN+ DUF499 *->?->?->?-><-Cluster182_14clades | 1122 | Bacteria>Terrabacteria_group>Actinomycetota>Actinomycetes>Mycobacteriales | GCF_010728155.1 | Mycolicibacterium_doricum | H7I39_RS15930 | 40769..44137 | 126673 | Swt1-family-HEPN-domain-containing-protein-[Mycobacterium-doricum] |
| WP_179973625.1 | DUF499_1.RM.like.systems | <-PIN+ACET+EVE\|\|<-REase<-NTD+SF1-SF2-Helicase\|\|TUDOR+ SF1-SF2-Helicase+iREase+CTD+REase->METHYLASE->ParB+HNH->HEPN+ DUF499 *-> | 1121 | Bacteria>Terrabacteria_group>Actinomycetota>Actinomycetes>Mycobacteriales | GCF_010731815.2 | Mycolicibacterium_arabiense | H5P33_RS16145 | 691085..694450 | 1286181 | Swt1-family-HEPN-domain-containing-protein-[Mycolicibacterium-arabiense] |
| WP_194806632.1 | DUF499_1.RM.like.systems | Phage_int_SAM_1+Phage_integrase->XIS-HTH->?->?->?-><-HEPN+ DUF499 *<-ParB+HNH<-METHYLASE<-UDG-conflict<-TUDOR+ SF1-SF2-Helicase+iREase+CTD+REase | 1121 | Bacteria>Terrabacteria_group>Actinomycetota>Actinomycetes>Mycobacteriales | GCF_015354405.1 | Nocardia_sp._BSTN01 | IRT45_RS15950 | 106038..109403 | 2783665 | Swt1-family-HEPN-domain-containing-protein-[Nocardia-sp.-BSTN01] |
| WP_197940422.1 | DUF499_1.RM.like.systems | <-HIN-HTH\|\|HEPN-><-NTD+SF1-SF2-Helicase\|\|TUDOR+ SF1-SF2-Helicase+iREase+CTD+REase->METHYLASE->ParB+HNH->HEPN+ DUF499 *->PAIREDC-HTH-> | 1121 | Bacteria>Terrabacteria_group>Actinomycetota>Actinomycetes>Mycobacteriales | GCF_016019085.1 | Rhodococcus_erythropolis | I3517_RS00940 | 13889..17254 | 1833 | Swt1-family-HEPN-domain-containing-protein-[Rhodococcus-erythropolis] |
| WP_211919214.1 | DUF499_1.RM.like.systems | HEPN-><-NTD+SF1-SF2-Helicase\|\|TUDOR+ SF1-SF2-Helicase+iREase+CTD+REase->ParB+HNH->HEPN+ DUF499 *-><-NUDIX\|\|?->REase->?->?->GDResolvase+HIN-HTH-> | 1121 | Bacteria>Terrabacteria_group>Actinomycetota>Actinomycetes>Mycobacteriales | GCF_018283555.1 | Rhodococcus_erythropolis | AC1659_RS28475 | 46414..49779 | 1833 | Swt1-family-HEPN-domain-containing-protein-[Rhodococcus-erythropolis] |
| WP_213184733.1 | DUF499_1.RM.like.systems | TUDOR+ SF1-SF2-Helicase+iREase+CTD+REase-><-?\|\|METHYLASE->ParB+HNH->HEPN+ DUF499 *->ANK+ANK+RAMA-> | 1134 | Bacteria>Terrabacteria_group>Actinomycetota>Actinomycetes>Mycobacteriales | GCF_018390595.1 | Dietzia_massiliensis | KHA85_RS02755 | 11826..15230 | 2697499 | Swt1-family-HEPN-domain-containing-protein-[Dietzia-massiliensis] |
| WP_214389765.1 | DUF499_1.RM.like.systems | EVE+McrB-AAA->McrC-NTD+REase-><-?<-HTH_Tnp_1+HTH_21+rve+rve_3\|\|HTH_Tnp_1+HTH_21+rve+rve_3->?-><-HEPN+ DUF499 *<-ParB+HNH<-METHYLASE<-TUDOR+ SF1-SF2-Helicase+iREase+CTD+REase\|\|NTD+SF1-SF2-Helicase->DDE_Tnp_1_4-> | 1119 | Bacteria>Terrabacteria_group>Actinomycetota>Actinomycetes>Mycobacteriales | GCF_018524295.1 | Mycolicibacterium_goodii | KL858_RS28165 | 17594..20953 | 134601 | Swt1-family-HEPN-domain-containing-protein-[Mycolicibacterium-goodii] |
| WP_174714145.1 | DUF499_1.RM.like.systems | XIS-HTH->PIN->TUDOR+ SF1-SF2-Helicase+iREase+CTD+REase-><-?\|\|METHYLASE->ParB+HNH->HEPN+ DUF499 *->?->?-><-NUDIX\|\|GNTR-HTH->?->FtsK-> | 1108 | Bacteria>Terrabacteria_group>Actinomycetota>Actinomycetes>Mycobacteriales | GCF_019456675.1 | Mycolicibacterium_pallens | KO064_RS24915 | 5121618..5124944 | 370524 | MULTISPECIES:-Swt1-family-HEPN-domain-containing-protein-[Mycobacterium] |
| WP_228232863.1 | DUF499_1.RM.like.systems | <-GDResolvase+HIN-HTH<-?\|\|ABhydrolase-><-GDResolvase+HIN-HTH\|\|zf-ISL3+DDE_Tnp_ISL3-><-?<-HEPN+ DUF499 *<-ParB+HNH<-METHYLASE<-UDG-conflict<-TUDOR+ SF1-SF2-Helicase+iREase+CTD+REase\|\|NTD+SF1-SF2-Helicase-><-HEPN | 1121 | Bacteria>Terrabacteria_group>Actinomycetota>Actinomycetes>Mycobacteriales | GCF_020782825.1 | Rhodococcus_sp._3-2 | LL008_RS29695 | 195028..198393 | 2890836 | Swt1-family-HEPN-domain-containing-protein-[Rhodococcus-sp.-3-2] |
| WP_229481506.1 | DUF499_1.RM.like.systems | <-NTD+SF1-SF2-Helicase\|\|TUDOR+ SF1-SF2-Helicase+iREase+CTD+REase->METHYLASE->ParB+HNH->HEPN+ DUF499 *->?->?-><-SF1-SF2-Helicase<-OLD-ABC+TOPRIM<-?<-Primpol+D5N+D5ATPase | 1114 | Bacteria>Terrabacteria_group>Actinomycetota>Actinomycetes>Mycobacteriales | GCF_020884715.1 | Mycolicibacterium_mageritense | MTY414_RS35935 | 11397..14741 | 53462 | Swt1-family-HEPN-domain-containing-protein-[Mycolicibacterium-mageritense] |
| WP_238886618.1 | DUF499_1.RM.like.systems | HIN-HTH->?->Cluster182_14clades-><-?<-?<-?<-HEPN+ DUF499 *<-ParB+HNH<-METHYLASE<-TUDOR+ SF1-SF2-Helicase+iREase+CTD+REase\|\|NTD+SF1-SF2-Helicase-> | 1120 | Bacteria>Terrabacteria_group>Actinomycetota>Actinomycetes>Mycobacteriales | GCF_022317005.1 | Mycobacterium_sp._IDR2000157661 | K3G64_RS19485 | 3782160..3785522 | 2867005 | Swt1-family-HEPN-domain-containing-protein-[Mycobacterium-sp.-IDR2000157661] |
| WP_239342823.1 | DUF499_1.RM.like.systems | <-NTD+SF1-SF2-Helicase\|\|TUDOR+ SF1-SF2-Helicase+iREase+CTD+REase->METHYLASE->ParB+HNH->HEPN+ DUF499 *->REase->SF1-SF2-Helicase-> | 1109 | Bacteria>Terrabacteria_group>Actinomycetota>Actinomycetes>Mycobacteriales | GCF_022348005.1 | Mycolicibacterium_sp._OfavD-34-C | MHN82_RS23245 | 18852..22181 | 2917746 | Swt1-family-HEPN-domain-containing-protein-[Mycolicibacterium-sp.-OfavD-34-C] |
| WP_256083185.1 | DUF499_1.RM.like.systems | HEPN-><-NTD+SF1-SF2-Helicase\|\|TUDOR+ SF1-SF2-Helicase+iREase+CTD+REase->METHYLASE->ParB+HNH->HEPN+ DUF499 *-> | 1120 | Bacteria>Terrabacteria_group>Actinomycetota>Actinomycetes>Mycobacteriales | GCF_024438075.1 | Rhodococcus_qingshengii | NOS52_RS32465 | 22838..26200 | 334542 | Swt1-family-HEPN-domain-containing-protein-[Rhodococcus-qingshengii] |
| WP_256101683.1 | DUF499_1.RM.like.systems | <-NTD+SF1-SF2-Helicase\|\|TUDOR+ SF1-SF2-Helicase+iREase+CTD+REase->METHYLASE->ParB+HNH->HEPN+ DUF499 *->ParB+HNH->HEPN-> | 1118 | Bacteria>Terrabacteria_group>Actinomycetota>Actinomycetes>Mycobacteriales | GCF_024451565.1 | Mycobacterium_gordonae | KQR54_RS26625 | 18169..21525 | 1778 | Swt1-family-HEPN-domain-containing-protein-[Mycobacterium-gordonae] |
| WP_260941407.1 | DUF499_1.RM.like.systems | JetA->JetB->?->JetD-><-?\|\|?-><-HEPN+ DUF499 *<-ParB+HNH\|\|HIN-HTH-><-ParB+HNH<-HIN-HTH\|\|?-><-TUDOR+ SF1-SF2-Helicase+iREase+CTD+REase | 1110 | Bacteria>Terrabacteria_group>Actinomycetota>Actinomycetes>Mycobacteriales | GCF_025331195.1 | Mycolicibacterium_llatzerense | A7G45_RS26855 | 438962..442294 | 280871 | Swt1-family-HEPN-domain-containing-protein-[Mycolicibacterium-llatzerense] |
| WP_269614895.1 | DUF499_1.RM.like.systems | HEPN-><-NTD+SF1-SF2-Helicase\|\|TUDOR+ SF1-SF2-Helicase+iREase+CTD+REase->METHYLASE->ParB+HNH->HEPN+ DUF499 *-><-NUDIX\|\|?->REase->?->?->GDResolvase+HIN-HTH-> | 1121 | Bacteria>Terrabacteria_group>Actinomycetota>Actinomycetes>Mycobacteriales | GCF_027270755.1 | Rhodococcus_qingshengii | O4271_RS30885 | 26346..29711 | 334542 | Swt1-family-HEPN-domain-containing-protein-[Rhodococcus-qingshengii] |
| WP_277242420.1 | DUF499_1.RM.like.systems | DnaB+SMS-KaiC-><-?<-?<-?<-?<-HEPN+ DUF499 *<-ParB+HNH<-METHYLASE<-TUDOR+ SF1-SF2-Helicase+iREase+CTD+REase<-?\|\|NTD+SF1-SF2-Helicase-> | 1119 | Bacteria>Terrabacteria_group>Actinomycetota>Actinomycetes>Mycobacteriales | GCF_029427115.1 | Gordonia_hongkongensis | L2299_RS00560 | 120321..123680 | 1701090 | Swt1-family-HEPN-domain-containing-protein-[Gordonia-hongkongensis] |
| WP_283569686.1 | DUF499_1.RM.like.systems | HEPN-><-NTD+SF1-SF2-Helicase\|\|TUDOR+ SF1-SF2-Helicase+iREase+CTD+REase->METHYLASE->ParB+HNH->HEPN+ DUF499 *-> | 1120 | Bacteria>Terrabacteria_group>Actinomycetota>Actinomycetes>Mycobacteriales | GCF_030063715.1 | Rhodococcus_qingshengii | QNA24_RS33665 | 11874..15236 | 334542 | Swt1-family-HEPN-domain-containing-protein-[Rhodococcus-qingshengii] |
| WP_207554210.1 | DUF499_1.RM.like.systems | <-NTD+SF1-SF2-Helicase\|\|TUDOR+ SF1-SF2-Helicase+iREase+CTD+REase-><-?\|\|METHYLASE->ParB-p->ParB+HNH->HEPN+ DUF499 *->DUF499-p->?->?->\|\|MULE-TRANSPOSASE-> | 605 | Bacteria>Terrabacteria_group>Actinomycetota>Actinomycetes>Mycobacteriales | GCF_900141575.1 | Mycobacteroides_abscessus | B5Y25_RS21325 | 33267..35081 | 36809 | Swt1-family-HEPN-domain-containing-protein,-partial-[Mycobacteroides-abscessus] |
| WP_207554378.1 | DUF499_1.RM.like.systems | <-NTD+SF1-SF2-Helicase\|\|TUDOR+ SF1-SF2-Helicase+iREase+CTD+REase-><-?\|\|METHYLASE->ParB+HNH->HEPN+ DUF499 *->?->?->\|\|MULE-TRANSPOSASE-> | 1113 | Bacteria>Terrabacteria_group>Actinomycetota>Actinomycetes>Mycobacteriales | GCF_900141585.1 | Mycobacteroides_abscessus | B5Z43_RS22925 | 52451..55792 | 36809 | Swt1-family-HEPN-domain-containing-protein-[Mycobacteroides-abscessus] |
| WP_085470756.1 | DUF499_1.RM.like.systems | DNA-glycosylase->?-><-NTD+SF1-SF2-Helicase<-HEPN+ DUF499 *<-ParB+HNH<-METHYLASE<-TUDOR+ SF1-SF2-Helicase+iREase+CTD+REase\|\|?-><-PIN<-XIS-HTH | 1125 | Bacteria>Terrabacteria_group>Actinomycetota>Actinomycetes>Mycobacteriales | GCF_900177695.1 | Rhodococcus_rhodochrous | M502_RS25910 | 14095..17472 | 1829 | Swt1-family-HEPN-domain-containing-protein-[Rhodococcus-rhodochrous] |
| WP_210731572.1 | DUF499_1.RM.like.systems | METHYLASE->ParB+HNH->HEPN+ DUF499 *-><-?\|\|?->?->SbcC->SbcC-> | 1116 | Bacteria>Terrabacteria_group>Actinomycetota>Actinomycetes>Mycobacteriales | GCF_900960235.1 | Mycobacterium_intracellulare | FAX65_RS26670 | 4893..8243 | 1767 | Swt1-family-HEPN-domain-containing-protein-[Mycobacterium-intracellulare] |
| WP_176126766.1 | DUF499_1.RM.like.systems | <-Phage_integrase\|\|Cluster182_14clades->?-><-?<-?<-?<-HEPN+ DUF499 *<-ParB+HNH<-METHYLASE<-TUDOR+ SF1-SF2-Helicase+iREase+CTD+REase\|\|NTD+SF1-SF2-Helicase-> | 1120 | Bacteria>Terrabacteria_group>Actinomycetota>Actinomycetes>Mycobacteriales | - | Mycobacterium_sp._D16Q20 | B4386_RS22320 | 7597..10959 | 1855660 | Swt1-family-HEPN-domain-containing-protein-[Mycobacterium-sp.-D16Q20] |
| WP_116058404.1 | DUF499_1.RM.like.systems | HEPN-><-NTD+SF1-SF2-Helicase\|\|TUDOR+ SF1-SF2-Helicase+iREase+CTD+REase->METHYLASE->ParB+HNH->HEPN+ DUF499 *->?-><-Lsr2\|\|Phage_integrase->?->PAIREDC-HTH-> | 1121 | Bacteria>Terrabacteria_group>Actinomycetota>Actinomycetes>Mycobacteriales | - | Rhodococcus_erythropolis | DVG80_RS32980 | 11800..15165 | 1833 | Swt1-family-HEPN-domain-containing-protein-[Rhodococcus-erythropolis] |
| MAS55988.1 | DUF499_1.RM.like.systems | XIS-HTH->METHYLASE-p-><-?<-MORC+HISKIN<-HEPN+ DUF499 *<-ParB+HNH<-METHYLASE<-UDG-conflict<-TUDOR+ SF1-SF2-Helicase+iREase+CTD+REase-p | 1116 | Bacteria>Terrabacteria_group>Actinomycetota>Actinomycetes>Propionibacteriales | GCA_002698575.1 | Pimelobacter_sp. | CMJ44_15460 | 17699..21049 | 51662 | AAA-family-ATPase-[Pimelobacter-sp.] |
| MBK8462833.1 | DUF499_1.RM.like.systems | <-REase<-?<-HEPN\|\|TUDOR+ SF1-SF2-Helicase+iREase+CTD+REase->METHYLASE->ParB+HNH->HEPN+ DUF499 *->?-><-Peptidase_M48<-?<-?\|\|TPR-> | 1117 | Bacteria>Terrabacteria_group>Actinomycetota>Actinomycetes>Propionibacteriales | GCA_016711325.1 | Nigerium_sp. | IPL36_07335 | 626569..629922 | 2042655 | DUF499-domain-containing-protein-[Nigerium-sp.] |
| MBM0123458.1 | DUF499_1.RM.like.systems | <-XIS-HTH<-XIS-HTH<-TrwC+SF1-SF2-helicase\|\|HEPN+Mrr_N-><-HEPN+ DUF499 *<-ParB+HNH<-METHYLASE<-TUDOR+ SF1-SF2-Helicase+iREase+CTD+REase\|\|NTD+SF1-SF2-Helicase-> | 1119 | Bacteria>Terrabacteria_group>Actinomycetota>Actinomycetes>Propionibacteriales | GCA_016803235.1 | Pimelobacter_simplex | IMY96_01775 | 51221..54580 | 2045 | DUF499-domain-containing-protein-[Pimelobacter-simplex] |
| MBN9103777.1 | DUF499_1.RM.like.systems | TUDOR+ SF1-SF2-Helicase+iREase+CTD+REase->UDG-conflict->METHYLASE->HEPN+ DUF499 *->ParB+HNH->?->SF1-SF2-Helicase-><-REase+PNTD1+PNTD2\|\|ACET-><-Phage_integrase | 1117 | Bacteria>Terrabacteria_group>Actinomycetota>Actinomycetes>Propionibacteriales | GCA_017306015.1 | Propionibacteriaceae_bacterium | JOI14_02120 | 371284..374637 | 2021380 | DUF499-domain-containing-protein-[Propionibacteriaceae-bacterium] |
| WP_013863116.1 | DUF499_1.RM.like.systems | PglZ-><-?\|\|?-><-DUF222+HNH\|\|?-><-HEPN+ DUF499 *<-ParB+HNH<-?<-METHYLASE\|\|wHTH(SBD)+HNH->HTH_19-> | 1117 | Bacteria>Terrabacteria_group>Actinomycetota>Actinomycetes>Propionibacteriales | GCF_000270245.1 | Microlunatus_phosphovorus | MLP_RS10775 | 2379873..2383226 | 29405 | Swt1-family-HEPN-domain-containing-protein-[Microlunatus-phosphovorus] |
| WP_020385147.1 | DUF499_1.RM.like.systems | NUDIX->tRNA-><-?<-?\|\|TUDOR+ SF1-SF2-Helicase+iREase+CTD+REase->METHYLASE->ParB+HNH->HEPN+ DUF499 *-><-?<-?<-?<-METHYLASE-p | 1116 | Bacteria>Terrabacteria_group>Actinomycetota>Actinomycetes>Propionibacteriales | GCF_000372465.1 | Kribbella_catacumbae | A3ES_RS0102270 | 28038..31388 | 460086 | Swt1-family-HEPN-domain-containing-protein-[Kribbella-catacumbae] |
| WP_124237129.1 | DUF499_1.RM.like.systems | <-REase\|\|NTP_transf_2->tRNA-><-NTD+SF1-SF2-Helicase\|\|TUDOR+ SF1-SF2-Helicase+iREase+CTD+REase->METHYLASE->ParB+HNH->HEPN+ DUF499 *-> | 1124 | Bacteria>Terrabacteria_group>Actinomycetota>Actinomycetes>Propionibacteriales | GCF_003850045.1 | Aeromicrobium_camelliae | EHW97_RS10560 | 83306..86680 | 1538144 | Swt1-family-HEPN-domain-containing-protein-[Aeromicrobium-camelliae] |
| WP_166139838.1 | DUF499_1.RM.like.systems | <-SLATT\|\|?-><-NTD+SF1-SF2-Helicase\|\|TUDOR+ SF1-SF2-Helicase+iREase+CTD+REase->METHYLASE->ParB+HNH->HEPN+ DUF499 *-><-?<-HNH\|\|?->?->?->REase+35exo+Beta-Barrel+SF1-SF2-Helicase-> | 1120 | Bacteria>Terrabacteria_group>Actinomycetota>Actinomycetes>Propionibacteriales | GCF_011420305.2 | Nocardioides_ochotonae | HB046_RS08105 | 1673933..1677295 | 2685869 | Swt1-family-HEPN-domain-containing-protein-[Nocardioides-ochotonae] |
| WP_246322323.1 | DUF499_1.RM.like.systems | <-REase<-?\|\|?->SIGMA-HTH-><-HEPN+ DUF499 *<-ParB+HNH<-METHYLASE<-TUDOR+ SF1-SF2-Helicase+iREase+CTD+REase\|\|REase-> | 1096 | Bacteria>Terrabacteria_group>Actinomycetota>Actinomycetes>Propionibacteriales | GCF_013407955.1 | Microlunatus_parietis | BKA15_RS11400 | 2395766..2399056 | 682979 | DUF499-domain-containing-protein-[Microlunatus-parietis] |
| WP_240236717.1 | DUF499_1.RM.like.systems | <-wHTH(SBD)+HNH<-NTD+SF1-SF2-Helicase\|\|TUDOR+ SF1-SF2-Helicase+iREase+CTD+REase->METHYLASE->ParB+HNH->HEPN+ DUF499 *-><-Peptidase\|\|?->?->?->METHYLASE-> | 1117 | Bacteria>Terrabacteria_group>Actinomycetota>Actinomycetes>Propionibacteriales | GCF_022436705.1 | Nocardioides_sp._CFH_31398 | MJ876_RS12955 | 81599..84952 | 2919579 | Swt1-family-HEPN-domain-containing-protein-[Nocardioides-sp.-CFH-31398] |
| WP_203043111.1 | DUF499_1.RM.like.systems | <-XIS-HTH<-XIS-HTH<-TrwC+SF1-SF2-helicase<-HEPN+ DUF499 *<-ParB+HNH<-METHYLASE<-TUDOR+ SF1-SF2-Helicase+iREase+CTD+REase\|\|NTD+SF1-SF2-Helicase-> | 1119 | Bacteria>Terrabacteria_group>Actinomycetota>Actinomycetes>Propionibacteriales | - | Pimelobacter_simplex | IMY96_RS01775 | 51221..54580 | 2045 | Swt1-family-HEPN-domain-containing-protein-[Pimelobacter-simplex] |
| WP_085916424.1 | DUF499_1.RM.like.systems | DNA-glycosylase->TUDOR+ SF1-SF2-Helicase+iREase+CTD+REase->METHYLASE->ParB+HNH->HEPN+ DUF499 *->?-><-TrwC+SF1-SF2-helicase+TOPRIM | 1135 | Bacteria>Terrabacteria_group>Actinomycetota>Actinomycetes>Pseudonocardiales | GCF_003945385.1 | Pseudonocardia | PSA01_RS26720 | 8547..11954 | 1847 | MULTISPECIES:-Swt1-family-HEPN-domain-containing-protein-[Pseudonocardia] |
| WP_218590226.1 | DUF499_1.RM.like.systems | TUDOR+ SF1-SF2-Helicase+iREase+CTD+REase->METHYLASE->ParB+HNH->HEPN+ DUF499 *->?-><-TrwC+SF1-SF2-helicase+TOPRIM<-?<-VirD4-TraG+VirD4-FtsK | 1126 | Bacteria>Terrabacteria_group>Actinomycetota>Actinomycetes>Pseudonocardiales | GCF_019263625.1 | Pseudonocardia_oceani | I4I79_RS18830 | 8746..12126 | 2792013 | Swt1-family-HEPN-domain-containing-protein-[Pseudonocardia-oceani] |
| GII94898.1 | DUF499_1.RM.like.systems | <-HEPN<-ParB+HNH<-HEPN+ DUF499 *<-METHYLASE<-TUDOR+ SF1-SF2-Helicase+iREase+CTD+REase | 1137 | Bacteria>Terrabacteria_group>Actinomycetota>Actinomycetes>Streptosporangiales | GCA_016863435.1 | Sinosporangium_siamense | Ssi02_51290 | 167948..171361 | 1367973 | hypothetical-protein-Ssi02_51290-[Sinosporangium-siamense] |
| GHJ54460.1 | DUF499_1.RM.like.systems | <-REase<-?<-NTD+SF1-SF2-Helicase\|\|TUDOR+ SF1-SF2-Helicase+iREase+CTD+REase->METHYLASE->ParB+HNH->HEPN+ DUF499 *->?->?->?-><-?<-SbcC\|\|HIN-HTH-> | 1122 | Bacteria>Terrabacteria_group>Actinomycetota>Actinomycetes>Streptosporangiales | GCA_018327725.1 | Nonomuraea_sp._TT08I-71 | Nm8I071_37670 | 3194026..3197394 | 2733864 | hypothetical-protein-Nm8I071_37670-[Nonomuraea-sp.-TT08I-71] |
| MCF6470725.1 | DUF499_1.RM.like.systems | GNTR-HTH-p->NUDIX->TUDOR+ SF1-SF2-Helicase+iREase+CTD+REase->METHYLASE->HEPN+ DUF499 *->ParB+HNH->?->REase-><-?\|\|SIGMA-HTH-> | 1139 | Bacteria>Terrabacteria_group>Actinomycetota>Actinomycetes>Streptosporangiales | GCA_021699195.1 | Nonomuraea_sp._MG754425 | FAF44_20335 | 17638..21057 | 2570319 | ATP-binding-protein-[Nonomuraea-sp.-MG754425] |
| WP_155999755.1 | DUF499_1.RM.like.systems | <-HTH+DUF5753\|\|?-><-?<-?<-METHYLASE<-?\|\|HEPN+ DUF499 *->ParB+HNH-><-?<-Glycos_trans_3N+Glycos_transf_3 | 1141 | Bacteria>Terrabacteria_group>Actinomycetota>Actinomycetes>Streptosporangiales | GCF_009728995.1 | Streptomonospora_sp._PA3 | FZ103_RS07070 | 271290..274715 | 2607326 | Swt1-family-HEPN-domain-containing-protein-[Streptomonospora-sp.-PA3] |
| WP_179824532.1 | DUF499_1.RM.like.systems | <-ST-Hiskin\|\|HTH+DUF5753->DUF397-><-?<-ParB+HNH<-HEPN+ DUF499 *<-METHYLASE<-TUDOR+ SF1-SF2-Helicase+iREase+CTD+REase<-?<-UDG-conflict | 1132 | Bacteria>Terrabacteria_group>Actinomycetota>Actinomycetes>Streptosporangiales | GCF_013409935.1 | Streptosporangium_sandarakinum | HDA43_RS21565 | 651273..654671 | 1260955 | Swt1-family-HEPN-domain-containing-protein-[Streptosporangium-sandarakinum] |
| WP_184983644.1 | DUF499_1.RM.like.systems | 35exo-><-wHTH(SBD)+HNH<-?<-SF1-SF2-helicase+DUF1998<-?<-ParB+HNH<-HEPN+ DUF499 *<-METHYLASE<-TUDOR+ SF1-SF2-Helicase+iREase+CTD+REase<-?<-REase+SF1-SF2-Helicase<-HTH_3<-ABhydrolase | 1124 | Bacteria>Terrabacteria_group>Actinomycetota>Actinomycetes>Streptosporangiales | GCF_014207705.1 | Sphaerisporangium_rubeum | BJ992_RS21150 | 4926161..4929535 | 321317 | Swt1-family-HEPN-domain-containing-protein-[Sphaerisporangium-rubeum] |
| WP_191055573.1 | DUF499_1.RM.like.systems | HTH+HKD-DNase->?->?->TUDOR+ SF1-SF2-Helicase+iREase+CTD+REase->METHYLASE->HEPN+ DUF499 *->ParB+HNH->HEPN-> | 1138 | Bacteria>Terrabacteria_group>Actinomycetota>Actinomycetes>Streptosporangiales | GCF_014712745.1 | Microbispora_camponoti | IEQ31_RS35675 | 16717..20133 | 1677852 | DUF499-domain-containing-protein-[Microbispora-camponoti] |
| WP_203980270.1 | DUF499_1.RM.like.systems | <-SF1-SF2-helicase+DUF1998<-?<-ParB+HNH<-?<-HEPN+ DUF499 *<-METHYLASE<-TUDOR+ SF1-SF2-Helicase+iREase+CTD+REase<-?<-REase+N6_MTase | 1129 | Bacteria>Terrabacteria_group>Actinomycetota>Actinomycetes>Streptosporangiales | GCF_016863315.1 | Planotetraspora_silvatica | Psi02_RS35595 | 24804..28193 | 234614 | Swt1-family-HEPN-domain-containing-protein-[Planotetraspora-silvatica] |
| WP_203946296.1 | DUF499_1.RM.like.systems | REase+N6_MTase->?->TUDOR+ SF1-SF2-Helicase+iREase+CTD+REase->METHYLASE->HEPN+ DUF499 *->?->ParB+HNH->?->SF1-SF2-helicase+DUF1998-><-?\|\|OLD-ABC+TOPRIM-> | 1129 | Bacteria>Terrabacteria_group>Actinomycetota>Actinomycetes>Streptosporangiales | GCF_016863335.1 | Planotetraspora_thailandica | Pth03_RS22295 | 43771..47160 | 487172 | Swt1-family-HEPN-domain-containing-protein-[Planotetraspora-thailandica] |
| WP_204029818.1 | DUF499_1.RM.like.systems | <-GNTR-HTH-p<-?<-?<-?<-HEPN<-ParB+HNH<-HEPN+ DUF499 *<-METHYLASE<-TUDOR+ SF1-SF2-Helicase+iREase+CTD+REase | 1127 | Bacteria>Terrabacteria_group>Actinomycetota>Actinomycetes>Streptosporangiales | GCF_016863435.1 | Sinosporangium_siamense | Ssi02_RS24940 | 167948..171331 | 1367973 | DUF499-domain-containing-protein-[Sinosporangium-siamense] |
| WP_237104850.1 | DUF499_1.RM.like.systems | GNTR-HTH-p->NUDIX->TUDOR+ SF1-SF2-Helicase+iREase+CTD+REase->METHYLASE->HEPN+ DUF499 *->ParB+HNH->?->REase-><-?<-?\|\|SIGMA-HTH-> | 1137 | Bacteria>Terrabacteria_group>Actinomycetota>Actinomycetes>Streptosporangiales | GCF_021699195.1 | Nonomuraea_sp._MG754425 | FAF44_RS20630 | 17644..21057 | 2570319 | DUF499-domain-containing-protein-[Nonomuraea-sp.-MG754425] |
| MCH9729512.1 | DUF499_1.RM.like.systems | PIN->TUDOR+ SF1-SF2-Helicase+iREase+CTD+REase->METHYLASE->ParB+HNH->HEPN+ DUF499 *->SIR2-><-DndB\|\|?->Recombinase-><-NUDIX\|\|GNTR-HTH-> | 1114 | Bacteria>Terrabacteria_group>Actinomycetota>Actinomycetes>unclassified_Actinomycetes | GCA_022600605.1 | Actinomycetes_bacterium | KOU84_07525 | 8492..11836 | 1883427 | DUF499-domain-containing-protein-[Actinomycetia-bacterium] |
| MCL4078096.1 | DUF499_1.RM.like.systems | <-Peptidase_C39_2<-?<-?\|\|PspA+PIN->?-><-HEPN+ DUF499 *<-?<-MqsA_antitoxin+HTH_3<-?<-METHYLASE<-ParB+HNH<-TUDOR+ SF1-SF2-Helicase+iREase+CTD+REase | 1106 | Bacteria>Terrabacteria_group>Actinomycetota>Coriobacteriia>Anaerosomatales | GCA_023371205.1 | Parvivirga_hydrogeniphila | MX659_00500 | 98516..101836 | 2939460 | Swt1-family-HEPN-domain-containing-protein-[Coriobacteriia-bacterium-Es71-Z0120] |
| WP_267192478.1 | DUF499_1.RM.like.systems | <-Peptidase_C39_2<-?<-?\|\|PspA+PIN->?-><-HEPN+ DUF499 *<-?<-MqsA_antitoxin+HTH_3<-?<-METHYLASE<-ParB+HNH<-TUDOR+ SF1-SF2-Helicase+iREase+CTD+REase | 1106 | Bacteria>Terrabacteria_group>Actinomycetota>Coriobacteriia>Anaerosomatales | - | Parvivirga_hydrogeniphila | MX659_RS00500 | 98516..101836 | 2939460 | Swt1-family-HEPN-domain-containing-protein-[Parvivirga-hydrogeniphila] |
| GAV31939.1 | DUF499_1.RM.like.systems | PspA+PIN->?-><-DUF499 *<-?<-MqsA_antitoxin+HTH_3<-METHYLASE<-ParB+HNH<-TUDOR+ SF1-SF2-Helicase+iREase+CTD+REase | 950 | Bacteria>Terrabacteria_group>Actinomycetota>Coriobacteriia>Coriobacteriales | GCA_001950395.1 | Coriobacteriaceae_bacterium_EMTCatB1 | emb_1c0561 | 1370215..1373067 | 1927122 | predicted-ATPase-[Coriobacteriaceae-bacterium-EMTCatB1] |
| MBC7265381.1 | DUF499_1.RM.like.systems | <-Peptidase_C39_2<-?\|\|PspA+PIN->?-><-HEPN+ DUF499 *<-?<-MqsA_antitoxin+HTH_3<-?<-METHYLASE<-ParB+HNH<-TUDOR+ SF1-SF2-Helicase+iREase+CTD+REase | 1106 | Bacteria>Terrabacteria_group>Actinomycetota>Coriobacteriia>unclassified_Coriobacteriia | GCA_014360605.1 | Coriobacteriia_bacterium | H5T75_00145 | 31010..34330 | 2052159 | ATP-binding-protein-[Coriobacteriia-bacterium] |
| MBI2692601.1 | DUF499_1.RM.like.systems | <-HTH+UbE1<-?\|\|HTH_31->MPTase->SF1-SF2-Helicase+REase-p->?-><-HEPN+ DUF499 *<-ParB+HNH<-METHYLASE<-TUDOR+ SF1-SF2-Helicase+iREase+CTD+REase<-HTH_3\|\|?-><-HTH | 1109 | Bacteria>Terrabacteria_group>Actinomycetota>Thermoleophilia>Solirubrobacterales | GCA_016185345.1 | Solirubrobacterales_bacterium | HYX29_11735 | 159016..162345 | 1909295 | ATP-binding-protein-[Solirubrobacterales-bacterium] |
| TAK69428.1 | DUF499_1.RM.like.systems | ParB+HNH->HEPN-><-TUDOR+ SF1-SF2-Helicase+iREase+CTD+REase\|\|?->?->HEPN+ DUF499 *->SF1-SF2-Helicase->?-><-?<-REase+35exo+Beta-Barrel+SF1-SF2-Helicase | 1114 | Bacteria>Terrabacteria_group>Actinomycetota>unclassified_Actinomycetota | GCA_004297305.1 | Actinomycetota_bacterium | EP013_05945 | 61030..64374 | 2900548 | ATP-binding-protein-[Actinobacteria-bacterium] |
| MBC7252723.1 | DUF499_1.RM.like.systems | HIN-HTH->?-><-?<-Schlafen+HATPase+HTH<-ParB+HNH+Schlafen<-HEPN+ DUF499 *<-METHYLASE<-PIN<-TUDOR+ SF1-SF2-Helicase+iREase+CTD+REase | 1115 | Bacteria>Terrabacteria_group>Actinomycetota>unclassified_Actinomycetota | GCA_014360685.1 | Actinomycetota_bacterium | H5T72_01965 | 75820..79167 | 2900548 | ATP-binding-protein-[Actinomycetia-bacterium] |
| MCA0294159.1 | DUF499_1.RM.like.systems | <-NTD+SF1-SF2-Helicase\|\|TUDOR+ SF1-SF2-Helicase+iREase+CTD+REase->METHYLASE->HEPN+ DUF499 *->ParB+HNH->?->SF1-SF2-Helicase-><-REase+PNTD1+PNTD2\|\|MULE-TRANSPOSASE->ACET-> | 1117 | Bacteria>Terrabacteria_group>Actinomycetota>unclassified_Actinomycetota | GCA_020161755.1 | Actinomycetota_bacterium | LCH96_02385 | 200193..203546 | 2900548 | DUF499-domain-containing-protein-[Actinomycetia-bacterium] |
| MDA8117679.1 | DUF499_1.RM.like.systems | <-MerR-HTH<-?<-HEPN+ DUF499 *<-TOPRIM<-OLD-ABC<-METHYLASE<-HEPN<-ParB+HNH<-HicB+ZnR | 1122 | Bacteria>Terrabacteria_group>Actinomycetota>unclassified_Actinomycetota | GCA_028064745.1 | Actinomycetota_bacterium | MO000_09960 | 1809..5177 | 2900548 | MAG:-Swt1-family-HEPN-domain-containing-protein-[Actinomycetota-bacterium] |
| MDA8281345.1 | DUF499_1.RM.like.systems | Phage_integrase->PIN-><-XIS-HTH\|\|TUDOR+ SF1-SF2-Helicase+iREase+CTD+REase->ParB+HNH->METHYLASE->DUF499 *->DUF4527->?->?->HTH-> | 939 | Bacteria>Terrabacteria_group>Actinomycetota>unclassified_Actinomycetota | GCA_028067595.1 | Actinomycetota_bacterium | MOZ63_13180 | 23767..26586 | 2900548 | MAG:-DUF499-domain-containing-protein-[Actinomycetota-bacterium] |
| MBX7131086.1 | DUF499_1.RM.like.systems | Phage_int_SAM_1+Phage_integrase-><-Peptidase<-HEPN+ DUF499 *<-?<-HTH<-HKD-DNase<-METHYLASE<-ParB+HNH<-TUDOR+ SF1-SF2-Helicase+iREase+CTD+REase | 1104 | Bacteria>Terrabacteria_group>Armatimonadota>Fimbriimonadia>Fimbriimonadales | GCA_019694835.1 | Fimbriimonadaceae_bacterium | K1X67_00255 | 55329..58643 | 2682144 | DUF499-domain-containing-protein-[Fimbriimonadaceae-bacterium] |
| RYL92593.1 | DUF499_1.RM.like.systems | REase+N6_MTase->?->TUDOR+ SF1-SF2-Helicase+iREase+CTD+REase->METHYLASE->HEPN+ DUF499 *->ParB+HNH-><-DDE_Tnp_1<-?<-?\|\|DDE_Tnp_1-> | 1123 | Bacteria>Terrabacteria_group>Bacillota>Bacilli>Bacillales | GCA_004153195.1 | Sporolactobacillus_sp._THM19-2 | EWH91_06945 | 33831..37202 | 2511171 | ATP-binding-protein-[Sporolactobacillus-sp.-THM19-2] |
| WP_129929317.1 | DUF499_1.RM.like.systems | REase+N6_MTase->?->TUDOR+ SF1-SF2-Helicase+iREase+CTD+REase->METHYLASE->HEPN+ DUF499 *->ParB+HNH-><-DDE_Tnp_1<-?<-?\|\|DDE_Tnp_1-> | 1123 | Bacteria>Terrabacteria_group>Bacillota>Bacilli>Bacillales | - | Sporolactobacillus_sp._THM19-2 | EWH91_06945 | 33831..37202 | 2511171 | DUF499-domain-containing-protein-[Sporolactobacillus-sp.-THM19-2] |
| WP_013432067.1 | DUF499_1.RM.like.systems | MazE_antitoxin->?->\|\|REase+NACHT->REase+NACHT->TUDOR+ SF1-SF2-Helicase+iREase+CTD+REase->METHYLASE->DUF499 *->ParB+HNH-p->DDE_Tnp_1->ParB+HNH-p-> | 913 | Bacteria>Terrabacteria_group>Bacillota>Clostridia>Caldicellulosiruptorales | GCF_000166695.1 | Caldicellulosiruptor_acetigenus | CALKR_RS03530 | 770817..773558 | 301953 | DUF499-domain-containing-protein-[Caldicellulosiruptor-acetigenus] |
| NLJ76878.1 | DUF499_1.RM.like.systems | <-ParB+HNH<-TPR<-?<-HEPN+ DUF499 *<-METHYLASE-p | 1133 | Bacteria>Terrabacteria_group>Bacillota>Clostridia>Eubacteriales | GCA_012518875.1 | Peptococcaceae_bacterium | GX325_06435 | 3363..6764 | 2052179 | ATP-binding-protein-[Peptococcaceae-bacterium] |
| HHW47352.1 | DUF499_1.RM.like.systems | XIS-HTH->TUDOR+ SF1-SF2-Helicase+iREase+CTD+REase->METHYLASE->HEPN+ DUF499 *->?->ParB+HNH-><-Aminotran_1_2\|\|?->ABC-ATPase+SbcC-> | 1135 | Bacteria>Terrabacteria_group>Bacillota>Clostridia>Eubacteriales | GCA_012840535.1 | Clostridiaceae_bacterium | GXX14_01895 | 11508..14915 | 1898204 | ATP-binding-protein-[Clostridiaceae-bacterium] |
| MBC7075381.1 | DUF499_1.RM.like.systems | MazE_antitoxin->RelE-ParE->\|\|?->?->TUDOR+ SF1-SF2-Helicase+iREase+CTD+REase->METHYLASE->HEPN+ DUF499 *->ParB+HNH-> | 1122 | Bacteria>Terrabacteria_group>Bacillota>Clostridia>Eubacteriales | GCA_014360155.1 | Syntrophomonadaceae_bacterium | H5T98_04805 | 11477..14845 | 2093811 | ATP-binding-protein-[Syntrophomonadaceae-bacterium] |
| MBO8129694.1 | DUF499_1.RM.like.systems | TUDOR+ SF1-SF2-Helicase+iREase+CTD+REase->ParB+HNH->METHYLASE->DUF499 *-> | 984 | Bacteria>Terrabacteria_group>Bacillota>Clostridia>Eubacteriales | GCA_017656185.1 | Peptococcaceae_bacterium | HOZ39_10965 | 8170..11124 | 2052179 | DUF499-domain-containing-protein-[Peptococcaceae-bacterium] |
| MCI7107135.1 | DUF499_1.RM.like.systems | REase->?->?->?-><-ParB+HNH<-HEPN+ DUF499 *<-METHYLASE<-METHYLASE<-TUDOR+ SF1-SF2-Helicase+iREase+CTD+REase<-HTH_26<-SbcC+SbcC | 1129 | Bacteria>Terrabacteria_group>Bacillota>Clostridia>Eubacteriales | GCA_022775045.1 | Agathobacter_sp. | MR945_04560 | 37097..40486 | 2021311 | DUF499-domain-containing-protein-[Agathobacter-sp.] |
| WP_018214270.1 | DUF499_1.RM.like.systems | <-pPIWI+pPIWI<-HEPN<-ParB+HNH<-METHYLASE-p<-Schlafen<-MORC<-HEPN+ DUF499 *<-METHYLASE<-TUDOR+ SF1-SF2-Helicase+iREase+CTD+REase<-HTH_26<-?<-MORC+SF1-SF2-Helicase<-SF1-SF2-Helicase+HKD-DNase | 1137 | Bacteria>Terrabacteria_group>Bacillota>Clostridia>Eubacteriales | GCF_000378805.1 | Desulfitobacterium_hafniense | DESHAF_RS0122600 | 1962937..1966350 | 49338 | DUF499-domain-containing-protein-[Desulfitobacterium-hafniense] |
| WP_088552559.1 | DUF499_1.RM.like.systems | HKD-DNase+SF1-SF2-Helicase+TRD->?->?->TUDOR+ SF1-SF2-Helicase+iREase+CTD+REase->METHYLASE->DUF499 *->ParB+HNH->?->PIN-><-35exo | 850 | Bacteria>Terrabacteria_group>Bacillota>Clostridia>Moorellales | GCF_002207765.1 | Calderihabitans_maritimus | KKC1_RS00515 | 36542..39094 | 1246530 | DUF499-domain-containing-protein-[Calderihabitans-maritimus] |
| WP_054937991.1 | DUF499_1.RM.like.systems | DEDD_Tnp_IS110+Transposase_20->TUDOR+ SF1-SF2-Helicase+iREase+CTD+REase->?->HEPN->METHYLASE->DUF499 *->ParB+HNH->?->?->?-><-?<-ArsR-HTH | 1103 | Bacteria>Terrabacteria_group>Bacillota>Clostridia>Moorellales | - | Moorella | MOST_RS05225 | 66804..70115 | 44260 | MULTISPECIES:-DUF499-domain-containing-protein-[Moorella] |
| MCB9452414.1 | DUF499_1.RM.like.systems | <-MetJarc-HTH\|\|?->TPR+TPR+TPR+TPR+TPR+TPR+TPR+TPR-><-HEPN+ DUF499 *<-ParB+HNH<-METHYLASE<-Schlafen+HATPase+HTH<-PIN<-?<-TUDOR+ SF1-SF2-Helicase+iREase+CTD+REase | 1106 | Bacteria>Terrabacteria_group>Chloroflexota>Anaerolineae>Anaerolineales | GCA_020634115.1 | Anaerolineaceae_bacterium | H6672_13335 | 112307..115627 | 2024896 | ATP-binding-protein-[Anaerolineaceae-bacterium] |
| MCZ7550961.1 | DUF499_1.RM.like.systems | <-NACHT\|\|<-?<-?<-ParB+HNH<-?<-HEPN+ DUF499 *<-METHYLASE<-MORC+DUF3684<-MORC+DUF3684<-TUDOR+ SF1-SF2-Helicase+iREase+CTD+REase-p | 1112 | Bacteria>Terrabacteria_group>Chloroflexota>Anaerolineae>Anaerolineales | GCA_027494025.1 | Anaerolineales_bacterium | M5U11_17685 | 3656371..3659709 | 2073117 | MAG:-Swt1-family-HEPN-domain-containing-protein-[Anaerolineales-bacterium] |
| MBK9093501.1 | DUF499_1.RM.like.systems | TUDOR+ SF1-SF2-Helicase+iREase+CTD+REase->OLD-ABC->HNH->METHYLASE->OLD-ABC->REase->DUF499 *->?->ParB+HNH->?->REase-p-> | 943 | Bacteria>Terrabacteria_group>Chloroflexota>Anaerolineae>unclassified_Anaerolineae | GCA_016719785.1 | Anaerolineae_bacterium | IPM84_12145 | 210722..213553 | 2052143 | ATP-binding-protein-[Anaerolineae-bacterium] |
| MBN1139287.1 | DUF499_1.RM.like.systems | MazE_antitoxin->PIN-><-?<-DUF433<-DUF499 *\|\|?->PIN-><-ParB+HNH<-METHYLASE | 1237 | Bacteria>Terrabacteria_group>Chloroflexota>Anaerolineae>unclassified_Anaerolineae | GCA_016926595.1 | Anaerolineae_bacterium | JXM73_22110 | 53334..57047 | 2052143 | ATP-binding-protein-[Anaerolineae-bacterium] |
| MCB8944346.1 | DUF499_1.RM.like.systems | GDResolvase+HIN-HTH-><-?\|\|?-><-?\|\|<-?<-?<-HEPN+ DUF499 *<-?<-METHYLASE<-ParB+HNH<-TUDOR+ SF1-SF2-Helicase+iREase+CTD+REase | 1098 | Bacteria>Terrabacteria_group>Chloroflexota>Ardenticatenia>Ardenticatenales | GCA_020634415.1 | Ardenticatenaceae_bacterium | H6658_11405 | 5286..8582 | 2614247 | ATP-binding-protein-[Ardenticatenaceae-bacterium] |
| MDE0078893.1 | DUF499_1.RM.like.systems | ParB+HNH+SF1-SF2-Helicase+HDRC->?->?->HEPN-><-?<-ParB+HNH<-DUF499 *<-OLD-ABC<-?<-?<-?<-HEPN<-METHYLASE | 1109 | Bacteria>Terrabacteria_group>Chloroflexota>Caldilineae>Caldilineales | GCA_028819025.1 | Caldilineaceae_bacterium | OXO50_15340 | 14318..17647 | 2614250 | MAG:-DUF499-domain-containing-protein-[Caldilineaceae-bacterium] |
| MCL4240066.1 | DUF499_1.RM.like.systems | <-HEPN+ DUF499 *<-?<-HTH<-?<-METHYLASE<-ParB+HNH | 1138 | Bacteria>Terrabacteria_group>Chloroflexota>Dehalococcoidia>unclassified_Dehalococcoidia | GCA_023382755.1 | Dehalococcoidia_bacterium | KJ048_01835 | 17815..21231 | 2026734 | DUF499-domain-containing-protein-[Dehalococcoidia-bacterium] |
| WP_201387653.1 | DUF499_1.RM.like.systems | TUDOR+ SF1-SF2-Helicase+iREase+CTD+REase->METHYLASE-p-><-DDE_3<-PAIREDC-HTH<-PAIREDC-HTH\|\|METHYLASE->HEPN+ DUF499 *->ParB+HNH-><-?<-?<-?\|\|HISKIN->REC-> | 1128 | Bacteria>Terrabacteria_group>Chloroflexota>Ktedonobacteria>Ktedonobacterales | GCF_016587395.1 | Ktedonobacter_sp._SOSP1-52 | JMJ16_RS30300 | 1126374..1129760 | 2778366 | DUF499-domain-containing-protein-[Ktedonobacter-sp.-SOSP1-52] |
| MBP8000880.1 | DUF499_1.RM.like.systems | <-XIS-HTH<-?\|\|SF1-SF2-helicase+RdgC+Beta-barrel+REase-><-HEPN+ DUF499 *<-?<-?<-METHYLASE<-ParB+HNH<-TUDOR+ SF1-SF2-Helicase+iREase+CTD+REase | 1097 | Bacteria>Terrabacteria_group>Chloroflexota>unclassified_Chloroflexota | GCA_018004095.1 | Chloroflexota_bacterium | KA338_16630 | 13590..16883 | 2026724 | DUF499-domain-containing-protein-[Chloroflexi-bacterium] |
| MBU1746748.1 | DUF499_1.RM.like.systems | <-HEPN+ DUF499 *<-METHYLASE<-?<-ParB+HNH<-TUDOR+ SF1-SF2-Helicase+iREase+CTD+REase | 551 | Bacteria>Terrabacteria_group>Chloroflexota>unclassified_Chloroflexota | GCA_018823415.1 | Chloroflexota_bacterium | KKA73_03595 | 1..1652 | 2026724 | DUF499-domain-containing-protein-[Chloroflexota-bacterium] |
| MCE7858272.1 | DUF499_1.RM.like.systems | HTH+UbE1-><-Schlafen<-?<-NACHT<-ParB+HNH<-HEPN+ DUF499 *<-METHYLASE<-MORC+DUF3684<-TUDOR+ SF1-SF2-Helicase+iREase+CTD+REase<-?<-?<-HTH+WYL | 1112 | Bacteria>Terrabacteria_group>Chloroflexota>unclassified_Chloroflexota | GCA_021404705.1 | Chloroflexi_bacterium_CFX2 | DYG86_00620 | 111078..114416 | 2293708 | ATP-binding-protein-[Chloroflexi-bacterium-CFX2] |
| MCX6033329.1 | DUF499_1.RM.like.systems | REase->?->METHYLASE->ParB+HNH->APATPase+iSTAND3->DUF499 *->STYKIN-> | 993 | Bacteria>Terrabacteria_group>Chloroflexota>unclassified_Chloroflexota | GCA_026389845.1 | Chloroflexota_bacterium | NT169_29110 | 10350..13331 | 2026724 | MAG:-DUF499-domain-containing-protein-[Chloroflexi-bacterium] |
| MCY3570526.1 | DUF499_1.RM.like.systems | TUDOR+ SF1-SF2-Helicase+iREase+CTD+REase-p->HEPN->METHYLASE->Polbeta->HEPN->HEPN+ DUF499 *-><-?<-METHYLASE-p<-?<-ParB+HNH<-wHTH+REase<-PadR-HTH | 1110 | Bacteria>Terrabacteria_group>Chloroflexota>unclassified_Chloroflexota | GCA_026705025.1 | Chloroflexota_bacterium | OXH19_04200 | 6390..9722 | 2026724 | MAG:-Swt1-family-HEPN-domain-containing-protein-[Chloroflexi-bacterium] |
| MCY3783979.1 | DUF499_1.RM.like.systems | MORC->?->?->?-><-?<-DUF499 *<-METHYLASE<-ParB+HNH+Schlafen<-SF1-SF2-Helicase-p<-VirB4-FtsK+HerA<-?<-TUDOR+ SF1-SF2-Helicase+iREase+CTD+REase | 920 | Bacteria>Terrabacteria_group>Chloroflexota>unclassified_Chloroflexota | GCA_026706485.1 | Chloroflexota_bacterium | OXG79_09360 | 384253..387015 | 2026724 | MAG:-DUF499-domain-containing-protein-[Chloroflexi-bacterium] |
| WP_190449861.1 | DUF499_1.RM.like.systems | CHTH->TUDOR+ SF1-SF2-Helicase+iREase+CTD+REase->HEPN->METHYLASE->OLD-ABC->PIN->HEPN+ DUF499 *->ParB+HNH->?->CopG/RHH->HNH-><-RHH | 1121 | Bacteria>Terrabacteria_group>Cyanobacteriota/Melainabacteria_group>Cyanobacteriota>Cyanophyceae | GCF_014695795.1 | Phormidium_sp._FACHB-592 | H6F86_RS16680 | 58473..61838 | 2692850 | Swt1-family-HEPN-domain-containing-protein-[Phormidium-sp.-FACHB-592] |
| WP_193965432.1 | DUF499_1.RM.like.systems | Phage_integrase-><-?\|\|?->?-><-HTH\|\|?-><-HEPN+ DUF499 *\|\|?-><-ParB+HNH<-?<-PIN<-?<-METHYLASE | 1114 | Bacteria>Terrabacteria_group>Cyanobacteriota/Melainabacteria_group>Cyanobacteriota>Cyanophyceae | GCF_015207395.1 | Nodosilinea_sp._LEGE_07088 | IQ254_RS06885 | 5859..9203 | 2777968 | Swt1-family-HEPN-domain-containing-protein-[Nodosilinea-sp.-LEGE-07088] |
| VBB42850.1 | DUF499_1.RM.like.systems | <-Phage_integrase<-?\|\|TUDOR+ SF1-SF2-Helicase+iREase+CTD+REase->ParB+HNH->METHYLASE->HEPN+ DUF499 *->?->?->?-><-DDE_Tnp_1\|\|HNH-> | 1117 | Bacteria>Thermodesulfobacteriota>Desulfobacteria>Desulfatiglandales>Desulfatiglandaceae | GCA_900498135.1 | uncultured_Desulfatiglans_sp. | TRIP_B220098 | 96105..99458 | 1748965 | conserved-hypothetical-protein-[uncultured-Desulfatiglans-sp.] |
| MBA3017256.1 | DUF499_1.RM.like.systems | <-N6_N4_MTase<-Peptidase_C47<-?\|\|TUDOR+ SF1-SF2-Helicase+iREase+CTD+REase->MORC+HisKin+DUF3684->HEPN+ DUF499 *->RNase->ParB+HNH->HEPN-> | 1092 | Bacteria>Thermodesulfobacteriota>Desulfobacteria>Desulfobacterales>Desulfobacteraceae | GCA_013792055.1 | Desulfobacteraceae_bacterium | FP811_03735 | 27065..30343 | 2049433 | ATP-binding-protein-[Desulfobacteraceae-bacterium] |
| MDD5204467.1 | DUF499_1.RM.like.systems | <-DUF499 *<-METHYLASE<-ParB+HNH | 968 | Bacteria>Thermodesulfobacteriota>Desulfobacteria>Desulfobacterales>unclassified_Desulfobacterales | GCA_028713085.1 | Desulfobacterales_bacterium | PHS17_03550 | 258..3164 | 2044940 | MAG:-DUF499-domain-containing-protein-[Desulfobacterales-bacterium] |
| RII28688.1 | DUF499_1.RM.like.systems | <-REase<-RelE-ParE<-RHH_5<-Cysteine_desulfurase\|\|DUF4007->DUF499 *->PAPS_reduct-><-ParB+HNH<-?<-DUF2726+ZnR+TOPRIM<-?\|\|XIS-HTH-> | 1131 | Bacteria>Thermodesulfobacteriota>Desulfuromonadia>Geobacterales>Geobacteraceae | GCA_003574885.1 | Geobacter_sp. | CXR31_02060 | 447432..450827 | 46610 | MAG:-ATP-binding-protein-[Geobacter-sp.] |
| MDD3846877.1 | DUF499_1.RM.like.systems | <-HEPN<-ParB+HNH<-HEPN+ DUF499 *<-METHYLASE<-MORC+DUF3684<-TUDOR+ SF1-SF2-Helicase+iREase+CTD+REase<-DUF932 | 1114 | Bacteria>Thermodesulfobacteriota>Syntrophorhabdia>Syntrophorhabdales>Syntrophorhabdaceae | GCA_028698615.1 | Syntrophorhabdaceae_bacterium | PHC90_11025 | 3963..7307 | 2660714 | MAG:-Swt1-family-HEPN-domain-containing-protein-[Syntrophorhabdaceae-bacterium] |

| Protein-ID | System | Gene-Neighbourhoods | Length | Taxonomy | Assembly-ID | Strain | Locus-Tag | Genomic-coordinates | TaxID | Fasta-defline |
|---|---|---|---|---|---|---|---|---|---|---|
| ACM23765.1 | DUF499_1.RM.like.systems | ParB+HNH-><-TUDOR+SF1-SF2-Helicase+iREase+CTD+REase<-Cluster35_66clades<-DUF499*<-METHYLASE<-?<-?<-Peptidase_M9 | 1123 | Bacteria>Thermotogota>Thermotogae>Thermotogales>Thermotogaceae | GCA_000018945.1 | Thermotoga_neapolitana_DSM_4359 | CTN_1589 | 1557052..1560423 | 309803 | Putative-uncharacterized-protein-[Thermotoga-neapolitana-DSM-4359] |
| KUK23102.1 | DUF499_1.RM.like.systems | METHYLASE->DUF499*->Cluster35_66clades->TUDOR+SF1-SF2-Helicase+iREase+CTD+REase-><-ParB+HNH | 1124 | Bacteria>Thermotogota>Thermotogae>Thermotogales>Thermotogaceae | GCA_001508125.1 | Thermotoga_petrophila | XD57_0797 | 2049..5423 | 93929 | MAG:-Uncharacterized-protein-XD57_0797-[Thermotoga-petrophila] |
| MBZ4661022.1 | DUF499_1.RM.like.systems | ParB+HNH-><-TUDOR+SF1-SF2-Helicase+iREase+CTD+REase<-Cluster35_66clades<-DUF499*<-DUF499<-METHYLASE<-?<-Peptidase_M9 | 347 | Bacteria>Thermotogota>Thermotogae>Thermotogales>Thermotogaceae | GCA_020054995.1 | Thermotoga_sp. | JG779_176 | 11603..12646 | 28240 | hypothetical-protein-[Thermotoga-sp.] |
| MBZ4661023.1 | DUF499_1.RM.like.systems | ParB+HNH-><-TUDOR+SF1-SF2-Helicase+iREase+CTD+REase<-Cluster35_66clades<-DUF499<-DUF499*<-METHYLASE<-?<-Peptidase_M9 | 704 | Bacteria>Thermotogota>Thermotogae>Thermotogales>Thermotogaceae | GCA_020054995.1 | Thermotoga_sp. | JG779_177 | 12744..14858 | 28240 | hypothetical-protein-[Thermotoga-sp.] |
| WP_004080572.1 | DUF499_1.RM.like.systems | Peptidase_M9->?->METHYLASE->DUF499*->Cluster35_66clades->TUDOR+SF1-SF2-Helicase+iREase+CTD+REase-><-ParB+HNH | 1123 | Bacteria>Thermotogota>Thermotogae>Thermotogales>Thermotogaceae | GCF_000008545.1 | Thermotoga_maritima | TM_RS05035 | 1002898..1006269 | 2336 | ATP-binding-protein-[Thermotoga-maritima] |
| WP_012311354.1 | DUF499_1.RM.like.systems | ParB+HNH-><-TUDOR+SF1-SF2-Helicase+iREase+CTD+REase<-Cluster35_66clades<-DUF499*<-METHYLASE<-?<-Peptidase_M9 | 1122 | Bacteria>Thermotogota>Thermotogae>Thermotogales>Thermotogaceae | GCF_000019625.1 | Thermotoga | TRQ2_RS09320 | 1826657..1830025 | 2335 | MULTISPECIES:-ATP-binding-protein-[Thermotoga] |
| WP_255347884.1 | DUF499_1.RM.like.systems | ParB+HNH->ParB+HNH-><-REase<-TUDOR+SF1-SF2-Helicase+iREase+CTD+REase<-Cluster35_66clades<-DUF499<-DUF499*<-DUF499<-METHYLASE<-?<-Peptidase_M9 | 391 | Bacteria>Thermotogota>Thermotogae>Thermotogales>Thermotogaceae | GCF_000832145.1 | Thermotoga_sp._RQ7 | TRQ7_RS09840 | 1801298..1802473 | 126738 | DUF499-domain-containing-protein-[Thermotoga-sp.-RQ7] |
| WP_236681711.1 | DUF499_1.RM.like.systems | ParB+HNH-><-REase<-TUDOR+SF1-SF2-Helicase+iREase+CTD+REase<-Cluster35_66clades<-DUF499<-DUF499<-DUF499*<-METHYLASE<-?<-Peptidase_M9 | 807 | Bacteria>Thermotogota>Thermotogae>Thermotogales>Thermotogaceae | GCF_000832145.1 | Thermotoga_sp._RQ7 | TRQ7_RS09855 | 1802757..1805180 | 126738 | ATP-binding-protein-[Thermotoga-sp.-RQ7] |
| WP_255347883.1 | DUF499_1.RM.like.systems | ParB+HNH->ParB+HNH-><-REase<-TUDOR+SF1-SF2-Helicase+iREase+CTD+REase<-Cluster35_66clades<-DUF499*<-DUF499<-DUF499<-METHYLASE<-?<-Peptidase_M9 | 78 | Bacteria>Thermotogota>Thermotogae>Thermotogales>Thermotogaceae | GCF_000832145.1 | Thermotoga_sp._RQ7 | TRQ7_RS09895 | 1801093..1801329 | 126738 | DUF499-domain-containing-protein-[Thermotoga-sp.-RQ7] |
| WP_165304882.1 | DUF499_1.RM.like.systems | Peptidase_M9->?-><-DUF499*<-METHYLASE<-TUDOR+SF1-SF2-Helicase+iREase+CTD+REase<-TUDOR+SF1-SF2-Helicase+iREase+CTD+REase-p<-ParB+HNH<-ParB+HNH | 842 | Bacteria>Thermotogota>Thermotogae>Thermotogales>Thermotogaceae | GCF_011057665.1 | Thermotoga_sp._38H-to | AS158_RS08895 | 65896..68424 | 1755812 | ATP-binding-protein-[Thermotoga-sp.-38H-to] |
| MBC7327984.1 | DUF499_1.RM.like.systems | ParB+HNH->TUDOR+SF1-SF2-Helicase+iREase+CTD+REase->METHYLASE->DUF499*-><-?<-ABC-ATPase+SbcC | 922 | Bacteria>unclassified_Bacteria | GCA_014360465.1 | bacterium | H5T87_07705 | 9838..12606 | 1869227 | ATP-binding-protein-[bacterium] |
| MCP3963401.1 | DUF499_1.RM.like.systems | TUDOR+SF1-SF2-Helicase+iREase+CTD+REase-p->ParB+HNH+Schlafen->METHYLASE->DUF499*-><-?\|\|?->?-><-?<-?\|\|JAB-> | 946 | Bacteria>unclassified_Bacteria | GCA_024226315.1 | bacterium | GY719_36660 | 6844..9684 | 1869227 | ATP-binding-protein-[bacterium] |
| -- | | | | | | | | | | |
| --wHTH(SBD)+HNH | | | | | | | | | | |
| -- | | | | | | | | | | |
| WP_099185493.1 | DUF499_1.RM.like.systems | HTH+WYL->CopG/RHH->TUDOR+SF1-SF2-Helicase+iREase+CTD+REase->OLD-ABC->HNH->METHYLASE->DUF499*-><-?\|\|LexA-protease-><-wHTH(SBD)+HNH<-TrwC+SF1-SF2-helicase<-VirD4-TraG+VirD4-FtsK<-CopG/RHH | 940 | Bacteria>Pseudomonadota>Alphaproteobacteria>Sphingomonadales>Sphingomonadaceae | GCF_002723855.1 | Sphingobium | SFOMI_RS05000 | 866506..869328 | 165695 | ATP-binding-protein-[Sphingobium-fuliginis] |
| HAV2024006.1 | DUF499_1.RM.like.systems | <-ParB+BAM<-XIS-HTH<-?\|\|wHTH(SBD)+HNH->TUDOR+SF1-SF2-Helicase+iREase+CTD+REase->METHYLASE->DUF499*-><-RlpA | 941 | Bacteria>Pseudomonadota>Gammaproteobacteria>Enterobacterales>Enterobacteriaceae | GCA_016444345.1 | Citrobacter_koseri | JHJ49_004194 | 16997..19822 | 545 | ATP-binding-protein-[Citrobacter-koseri] |
| WP_048233987.1 | DUF499_1.RM.like.systems | SymE_toxin->wHTH(SBD)+HNH-><-AbiEii<-AbiEi_4\|\|?->RlpA-><-DUF499*<-METHYLASE<-TUDOR+SF1-SF2-Helicase+iREase+CTD+REase<-wHTH(SBD)+HNH\|\|?->XIS-HTH->ParB+BAM-> | 941 | Bacteria>Pseudomonadota>Gammaproteobacteria>Enterobacterales>Enterobacteriaceae | GCF_001034485.1 | Citrobacter_sp._MGH104 | SK34_RS19955 | 343965..346790 | 1686379 | DUF499-domain-containing-protein-[Citrobacter-sp.-MGH104] |
| WP_121585419.1 | DUF499_1.RM.like.systems | <-ParB+BAM<-XIS-HTH<-?\|\|wHTH(SBD)+HNH->TUDOR+SF1-SF2-Helicase+iREase+CTD+REase->METHYLASE->DUF499*-><-RlpA<-?\|\|AbiEi_4->AbiEii-><-wHTH(SBD)+HNH\|\|MAD-NTD+HNH-> | 941 | Bacteria>Pseudomonadota>Gammaproteobacteria>Enterobacterales>Enterobacteriaceae | GCF_003665575.1 | Citrobacter | CUC49_RS22185 | 4547789..4550614 | 544 | MULTISPECIES:-DUF499-domain-containing-protein-[Citrobacter] |
| WP_061423982.1 | DUF499_1.RM.like.systems | wHTH(SBD)+HNH->TUDOR+SF1-SF2-Helicase+iREase+CTD+REase->METHYLASE->DUF499*-><-RlpA<-?\|\|AbiEi_4->AbiEii-> | 941 | Bacteria>Pseudomonadota>Gammaproteobacteria>Enterobacterales>Enterobacteriaceae | GCF_004194635.3 | Salmonella_enterica | DBZ85_RS25390 | 353324..356149 | 28901 | DUF499-domain-containing-protein-[Salmonella-enterica] |
| WP_155022959.1 | DUF499_1.RM.like.systems | <-ParB+BAM<-XIS-HTH<-?\|\|wHTH(SBD)+HNH->TUDOR+SF1-SF2-Helicase+iREase+CTD+REase->METHYLASE->DUF499*-><-RlpA<-?\|\|AbiEi_4->AbiEii-><-wHTH(SBD)+HNH\|\|SymE_toxin-> | 941 | Bacteria>Pseudomonadota>Gammaproteobacteria>Enterobacterales>Enterobacteriaceae | GCF_015704165.1 | Enterobacter_hormaechei | I4P33_RS13240 | 224794..227619 | 158836 | MULTISPECIES:-DUF499-domain-containing-protein-[Enterobacter-cloacae-complex] |
| WP_251853646.1 | DUF499_1.RM.like.systems | <-SymE_toxin\|\|wHTH(SBD)+HNH-><-AbiEii<-AbiEi_4\|\|?->RlpA-><-DUF499*<-METHYLASE<-TUDOR+SF1-SF2-Helicase+iREase+CTD+REase<-wHTH(SBD)+HNH\|\|?->XIS-HTH->ParB+BAM-> | 941 | Bacteria>Pseudomonadota>Gammaproteobacteria>Enterobacterales>Enterobacteriaceae | GCF_903935915.1 | Enterobacter_cloacae | NGL52_RS07590 | 54377..57202 | 550 | DUF499-domain-containing-protein-[Enterobacter-cloacae] |
| MBX3448931.1 | DUF499_1.RM.like.systems | TUDOR+SF1-SF2-Helicase+iREase+CTD+REase->OLD-ABC->HNH->METHYLASE->OLD-ABC->REase->DUF499*->?->MORC->?->?-><-wHTH(SBD)+HNH<-DNA_methylase | 948 | Bacteria>PVC_group>Planctomycetota>Planctomycetia>Planctomycetales | GCA_019636285.1 | Planctomycetaceae_bacterium | KF777_05185 | 10482..13328 | 2026779 | ATP-binding-protein-[Planctomycetaceae-bacterium] |
| WP_222946975.1 | DUF499_1.RM.like.systems | Phage_cap_E->?->?-><-wHTH(SBD)+HNH\|\|TUDOR+SF1-SF2-Helicase+iREase+CTD+REase->METHYLASE->HEPN+DUF499*->DUF6731->Phage_holin-> | 1153 | Bacteria>Terrabacteria_group>Actinomycetota>Actinomycetes>Kitasatosporales | GCF_008271805.1 | Streptomyces_sp._TRM49041 | FVS66_RS24245 | 260388..263849 | 2603216 | DUF499-domain-containing-protein-[Streptomyces-sp.-TRM49041] |
| WP_161236363.1 | DUF499_1.RM.like.systems | <-HEPN\|\|<-wHTH(SBD)+HNH\|\|TUDOR+SF1-SF2-Helicase+iREase+CTD+REase->METHYLASE->HEPN+DUF499*->DUF6731->Phage_holin-><-MNS+HTH+REase | 1161 | Bacteria>Terrabacteria_group>Actinomycetota>Actinomycetes>Kitasatosporales | GCF_010548075.1 | unclassified_Streptomyces | G3I23_RS06870 | 11181..14666 | 2593676 | MULTISPECIES:-DUF499-domain-containing-protein-[unclassified-Streptomyces] |
| WP_257017806.1 | DUF499_1.RM.like.systems | SF1-SF2-Helicase+REase->SF1-SF2-helicase-><-?<-?\|\|TUDOR+SF1-SF2-Helicase+iREase+CTD+REase->METHYLASE->DUF499*-><-wHTH(SBD)+HNH<-REase+SF1-SF2-Helicase<-MazG | 1022 | Bacteria>Terrabacteria_group>Actinomycetota>Actinomycetes>Mycobacteriales | GCF_900215125.1 | unclassified_Rhodococcus_(in:_high_G+C_Gram-positive_bacteria) | C8K38_RS25080 | 92076..95144 | 192944 | MULTISPECIES:-DUF499-domain-containing-protein-[unclassified-Rhodococcus-(in:-high-G+C-Gram-positive-bacteria)] |
| HBH74016.1 | DUF499_1.RM.like.systems | TUDOR+SF1-SF2-Helicase+iREase+CTD+REase-><-PIN<-MazE_antitoxin\|\|?->wHTH(SBD)+HNH->HEPN+DUF499*-> | 1110 | Bacteria>Terrabacteria_group>Cyanobacteriota/Melainabacteria_group>Cyanobacteriota>Cyanophyceae | GCA_003508015.1 | Synechococcales_bacterium_UBA10510 | DDY43_11405 | 28893..32225 | 2055795 | AAA+-family-ATPase-[Synechococcales-bacterium-UBA10510] |
| MCP9859835.1 | DUF499_1.RM.like.systems | <-SIGMA-HTH<-?<-?<-DUF499*<-wHTH(SBD)+HNH<-METHYLASE-p<-HNH<-OLD-ABC<-TUDOR+SF1-SF2-Helicase+iREase+CTD+REase<-LRP-HTH | 819 | Bacteria>Terrabacteria_group>Cyanobacteriota/Melainabacteria_group>Cyanobacteriota>Cyanophyceae | GCA_024345955.1 | Cyanobium_sp._Cruz-8H5 | KBZ34_12405 | 60686..63145 | 2823712 | ATP-binding-protein-[Cyanobium-sp.-Cruz-8H5] |
| WP_254953294.1 | DUF499_1.RM.like.systems | <-SIGMA-HTH<-?<-?<-DUF499*<-wHTH(SBD)+HNH<-METHYLASE-p<-HNH<-OLD-ABC<-TUDOR+SF1-SF2-Helicase+iREase+CTD+REase<-LRP-HTH | 819 | Bacteria>Terrabacteria_group>Cyanobacteriota/Melainabacteria_group>Cyanobacteriota>Cyanophyceae | GCF_024345955.1 | unclassified_Cyanobium | KBZ34_12405 | 60686..63145 | 2627006 | MULTISPECIES:-DUF499-domain-containing-protein-[unclassified-Cyanobium] |
| WP_261362403.1 | DUF499_1.RM.like.systems | <-SIGMA-HTH<-?<-?<-DUF499*<-wHTH(SBD)+HNH<-METHYLASE-p<-HNH<-OLD-ABC<-TUDOR+SF1-SF2-Helicase+iREase+CTD+REase<-LRP-HTH | 820 | Bacteria>Terrabacteria_group>Cyanobacteriota/Melainabacteria_group>Cyanobacteriota>Cyanophyceae | - | unclassified_Cyanobium | KBZ34_RS12460 | 60686..63148 | 2627006 | MULTISPECIES:-DUF499-domain-containing-protein-[unclassified-Cyanobium] |
| -- | | | | | | | | | | |
| --HNH(OLD->HNH->) | | | | | | | | | | |
| -- | | | | | | | | | | |
| MCL4551940.1 | DUF499_1.RM.like.systems | XIS-HTH->OLD-ABC->HNH->METHYLASE->DUF499*->DUF499-p->SF1-SF2-Helicase+REase-> | 336 | Archaea>TACK_group>Candidatus_Marsarchaeota>unclassified_Candidatus_Marsarchaeota | GCA_023381795.1 | Candidatus_Marsarchaeota_archaeon | M1305_00065 | 10946..11954 | 2823144 | DUF499-domain-containing-protein-[Candidatus-Marsarchaeota-archaeon] |
| WP_194449891.1 | DUF499_1.RM.like.systems | <-XIS-HTH<-?<-?\|\|TUDOR+SF1-SF2-Helicase+iREase+CTD+REase->?->METHYLASE->DUF499*->HNH->?->?-><-ParB<-GDResolvase+Recombinase+ZnR<-RAMA | 1110 | Bacteria>Acidobacteriota>Terriglobia>Bryobacterales>Bryobacteraceae | GCF_015277775.1 | Paludibaculum_fermentans | IRI77_RS36750 | 9264025..9267357 | 1473598 | DUF499-domain-containing-protein-[Paludibaculum-fermentans] |
| MYC75109.1 | DUF499_1.RM.like.systems | <-ABC-ATPase+SbcC<-ABC-ATPase+SbcC<-ZnR<-SIGMA-HTH<-HEAT_2<-DUF499*<-?<-METHYLASE<-?<-HNH<-OLD-ABC<-TUDOR+SF1-SF2-Helicase+iREase+CTD+REase | 1114 | Bacteria>Bacteria_incertae_sedis>Bacteria_candidate_phyla>Candidatus_Poribacteria>unclassified_Candidatus_Poribacteria | GCA_009841255.1 | Candidatus_Poribacteria_bacterium | F4X10_04955 | 70263..73607 | 2026781 | ATP-binding-protein-[Candidatus-Poribacteria-bacterium] |
| MYK19441.1 | DUF499_1.RM.like.systems | <-DUF499*<-HNH<-?<-METHYLASE | 934 | Bacteria>Bacteria_incertae_sedis>Bacteria_candidate_phyla>Candidatus_Poribacteria>unclassified_Candidatus_Poribacteria | GCA_009845505.1 | Candidatus_Poribacteria_bacterium | F4055_14930 | 451..3255 | 2026781 | ATP-binding-protein-[Candidatus-Poribacteria-bacterium] |
| MCG9130919.1 | DUF499_1.RM.like.systems | TUDOR+SF1-SF2-Helicase+iREase+CTD+REase->OLD-ABC->HNH->METHYLASE->PolB->HEPN->DUF499*->HEAT_2->SAM-methylase->SIGMA-HTH->ZnR->ABC-ATPase+SbcC->ABC-ATPase+SbcC-> | 1114 | Bacteria>Bacteria_incertae_sedis>Bacteria_candidate_phyla>Candidatus_Poribacteria>unclassified_Candidatus_Poribacteria | GCA_022394775.1 | Candidatus_Poribacteria_bacterium | J5I95_04465 | 27738..31082 | 2026781 | ATP-binding-protein-[Candidatus-Poribacteria-bacterium] |
| MCY3550002.1 | DUF499_1.RM.like.systems | <-Polbeta<-ABC-ATPase+SbcC<-ABC-ATPase+SbcC<-ZnR<-SIGMA-HTH<-HEAT_2<-DUF499*<-PolB<-METHYLASE<-?<-HNH<-OLD-ABC<-TUDOR+SF1-SF2-Helicase+iREase+CTD+REase | 1122 | Bacteria>Bacteria_incertae_sedis>Bacteria_candidate_phyla>Candidatus_Poribacteria>unclassified_Candidatus_Poribacteria | GCA_026702755.1 | Candidatus_Poribacteria_bacterium | OXH39_06030 | 100519..103887 | 2026781 | MAG:-DUF499-domain-containing-protein-[Candidatus-Poribacteria-bacterium] |
| MDE0324945.1 | DUF499_1.RM.like.systems | <-HEAT_2<-DUF499*<-HEPN<-?<-METHYLASE<-?<-HNH<-OLD-ABC | 1118 | Bacteria>Bacteria_incertae_sedis>Bacteria_candidate_phyla>Candidatus_Poribacteria>unclassified_Candidatus_Poribacteria | GCA_028820875.1 | Candidatus_Poribacteria_bacterium | OXN27_13610 | 421..3777 | 2026781 | MAG:-DUF499-domain-containing-protein-[Candidatus-Poribacteria-bacterium] |
| MDE0634775.1 | DUF499_1.RM.like.systems | TUDOR+SF1-SF2-Helicase+iREase+CTD+REase->SMODS->HNH+SAVED->HEPN+REase->?->METHYLASE->DUF499*->?->TPR+TPR->SbcC-> | 935 | Bacteria>Bacteria_incertae_sedis>Bacteria_candidate_phyla>Candidatus_Poribacteria>unclassified_Candidatus_Poribacteria | GCA_028824125.1 | Candidatus_Poribacteria_bacterium | OXI43_02830 | 28592..31399 | 2026781 | MAG:-DUF499-domain-containing-protein-[Candidatus-Poribacteria-bacterium] |
| MDE0555374.1 | DUF499_1.RM.like.systems | <-DUF499*<-HNH<-?<-METHYLASE | 546 | Bacteria>Bacteria_incertae_sedis>Bacteria_candidate_phyla>Candidatus_Poribacteria>unclassified_Candidatus_Poribacteria | GCA_028824855.1 | Candidatus_Poribacteria_bacterium | OXI24_14220 | 20..1660 | 2026781 | MAG:-DUF499-domain-containing-protein-[Candidatus-Poribacteria-bacterium] |
| MBI4801911.1 | DUF499_1.RM.like.systems | STAS+HisKin->?->PIN->OLD-ABC->HNH->METHYLASE->HEPN+DUF499*->Glycos_trans_3N+Glycos_transf_3->Radical-SAM->Radical-SAM->?->?->RADICAL-SAM-> | 1132 | Bacteria>Elusimicrobiota>unclassified_Elusimicrobiota | GCA_016208085.1 | Elusimicrobiota_bacterium | HY796_05240 | 17514..20912 | 2030800 | ATP-binding-protein-[Elusimicrobia-bacterium] |
| WP_070725327.1 | DUF499_1.RM.like.systems | <-SF1-SF2-Helicase+REase<-?\|\|TUDOR+SF1-SF2-Helicase+iREase+CTD+REase->OLD-ABC->HNH->METHYLASE->HEPN+DUF499*-><-HTH | 1110 | Bacteria>FCB_group>Bacteroidota/Chlorobiota_group>Bacteroidota>Cytophagia | GCF_001816145.1 | Hymenobacter_lapidarius | BEN47_RS03945 | 71126..74458 | 1908237 | DUF499-domain-containing-protein-[Hymenobacter-lapidarius] |
| MCB0808336.1 | DUF499_1.RM.like.systems | <-DUF499*<-METHYLASE<-HNH<-OLD-ABC<-TUDOR+SF1-SF2-Helicase+iREase+CTD+REase | 759 | Bacteria>FCB_group>Bacteroidota/Chlorobiota_group>Bacteroidota>Flavobacteriia | GCA_020438005.1 | Flavobacteriales_bacterium | KDB96_03570 | 1..2279 | 2021391 | ATP-binding-protein-[Flavobacteriales-bacterium] |
| NUQ24578.1 | DUF499_1.RM.like.systems | TUDOR+SF1-SF2-Helicase+iREase+CTD+REase->OLD-ABC->HNH->METHYLASE->AAA-ATPase+REase->HEPN+DUF499*->?->PolB->HEPN-><-ORC-CDC6+REase | 1095 | Bacteria>FCB_group>Bacteroidota/Chlorobiota_group>Bacteroidota>Saprospiria | GCA_013360405.1 | Saprospiraceae_bacterium | HUU34_11550 | 44219..47506 | 2202734 | DUF499-domain-containing-protein-[Saprospiraceae-bacterium] |
| MBK6932297.1 | DUF499_1.RM.like.systems | <-Glyco_hydro_1\|\|?-><-DUF499-p<-HEPN+DUF499*<-URI-DNase<-METHYLASE-p<-?<-METHYLASE-p<-HNH | 951 | Bacteria>FCB_group>Bacteroidota/Chlorobiota_group>Bacteroidota>Saprospiria | GCA_016709195.1 | Saprospirales_bacterium | IPH12_16140 | 92714..95569 | 2026790 | DUF499-domain-containing-protein-[Saprospirales-bacterium] |
| MBI2502153.1 | DUF499_1.RM.like.systems | MACRODOMAIN->METHYLASE->?->DUF488-1->?->HEPN+DUF499*->HNH->?->DDE_Tnp_1_4-> | 1125 | Bacteria>FCB_group>Candidatus_Latescibacteria>unclassified_Candidatus_Latescibacteria | GCA_016187725.1 | Candidatus_Latescibacteria_bacterium | HYW07_02835 | 5709..9086 | 2053570 | ATP-binding-protein-[Candidatus-Latescibacteria-bacterium] |
| WP_171805995.1 | DUF499_1.RM.like.systems | TUDOR+SF1-SF2-Helicase+iREase+CTD+REase->OLD-ABC->HNH->METHYLASE->DUF499*->DNA-glycosylase->CHTH->?->LZ_Tnp_IS66+zf-IS66+DDE_Tnp_IS66+DDE_Tnp_IS66_C->DUF4096+DDE_Tnp_1+DDE_Tnp_1_2-> | 942 | Bacteria>Myxococcota>Myxococcia>Myxococcales>Cystobacterineae | GCF_013155195.1 | Corallococcus_exiguus | HPP06_RS00265 | 63781..66609 | 83462 | DUF499-domain-containing-protein-[Corallococcus-exiguus] |
| WP_174489000.1 | DUF499_1.RM.like.systems | <-METHYLASE+DNA_methylase\|\|tRNA-><-DNA-glycosylase<-?<-Pkinase<-REase<-DUF499*<-METHYLASE<-HNH<-OLD-ABC<-TUDOR+SF1-SF2-Helicase+iREase+CTD+REase<-XIS-HTH<-PAIREDC-HTH | 943 | Bacteria>Myxococcota>Myxococcia>Myxococcales>Cystobacterineae | GCF_013336725.1 | Myxococcus_sp._CA040A | HUA75_RS36310 | 164891..167722 | 2741738 | DUF499-domain-containing-protein-[Myxococcus-sp.-CA040A] |
| MDE0344143.1 | DUF499_1.RM.like.systems | <-Arm-DNA-bind_3+Phage_integrase<-?<-HNH<-HEPN+DUF499*<-?\|\|Phage_integrase->METHYLASE-p-> | 1111 | Bacteria>Myxococcota>Myxococcia>unclassified_Myxococcia | GCA_028821055.1 | Deltaproteobacteria_bacterium | OXK82_13420 | 9792..13127 | 2026735 | MAG:-Swt1-family-HEPN-domain-containing-protein-[Deltaproteobacteria-bacterium] |
| KYF87240.1 | DUF499_1.RM.like.systems | <-DUF499*<-REase<-OLD-ABC<-METHYLASE<-HNH<-OLD-ABC<-TUDOR+SF1-SF2-Helicase+iREase+CTD+REase | 954 | Bacteria>Myxococcota>Polyangia>Polyangiales>Polyangiaceae | GCA_001589185.1 | Sorangium_cellulosum | BE17_46125 | 4063..6927 | 56 | hypothetical-protein-BE17_46125-[Sorangium-cellulosum] |
| MBZ0154723.1 | DUF499_1.RM.like.systems | PIN->OLD-ABC->HNH->METHYLASE->?->RelE-ParE->HEPN+DUF499*-> | 1114 | Bacteria>Nitrospirota>Nitrospiria>Nitrospirales>Nitrospiraceae | - | Candidatus_Nitrobium_versatile | K8I29_00740 | 55978..59322 | 2884831 | DUF499-domain-containing-protein-[Alphaproteobacteria-bacterium] |
| MBU2768209.1 | DUF499_1.RM.like.systems | <-DUF499*<-REase<-OLD-ABC<-wHTH+REase<-METHYLASE<-HNH<-OLD-ABC | 542 | Bacteria>Pseudomonadota>Acidithiobacillia>Acidithiobacillales>Acidithiobacillaceae | GCA_018853935.1 | Acidithiobacillus_ferrivorans | HAP94_19055 | 1..1626 | 160808 | ATP-binding-protein-[Acidithiobacillus-ferrivorans] |
| WP_166097056.1 | DUF499_1.RM.like.systems | MerR-HTH->tRNA->TUDOR+SF1-SF2-Helicase+iREase+CTD+REase->OLD-ABC->HNH->METHYLASE->DUF499*->DUF433->PIN-><-Phage_integrase | 945 | Bacteria>Pseudomonadota>Alphaproteobacteria>Hyphomicrobiales>Nitrobacteraceae | GCF_011516635.5 | Bradyrhizobium_barranii | HAP41_RS22275 | 4684908..4687745 | 2992140 | ATP-binding-protein-[Bradyrhizobium-barranii] |
| WP_212326836.1 | DUF499_1.RM.like.systems | Phage_integrase->?->XIS-HTH->?->DnaB->HIN-HTH-><-HEPN+DUF499*<-METHYLASE<-HNH<-OLD-ABC<-TUDOR+SF1-SF2-Helicase+iREase+CTD+REase | 1108 | Bacteria>Pseudomonadota>Alphaproteobacteria>Hyphomicrobiales>Nitrobacteraceae | GCF_018131105.1 | Bradyrhizobium_manausense | JQ631_RS13545 | 2838930..2842256 | 989370 | Swt1-family-HEPN-domain-containing-protein-[Bradyrhizobium-manausense] |
| WP_172236133.1 | DUF499_1.RM.like.systems | XIS-HTH->?->?->DnaB->HIN-HTH-><-HEPN+DUF499*<-METHYLASE<-HNH<-OLD-ABC<-TUDOR+SF1-SF2-Helicase+iREase+CTD+REase | 1108 | Bacteria>Pseudomonadota>Alphaproteobacteria>Hyphomicrobiales>Nitrobacteraceae | GCF_902482845.1 | Bradyrhizobium | JQ619_RS32195 | 7694..11020 | 374 | MULTISPECIES:-Swt1-family-HEPN-domain-containing-protein-[Bradyrhizobium] |
| WP_116622075.1 | DUF499_1.RM.like.systems | HTH_3->TUDOR+SF1-SF2-Helicase+iREase+CTD+REase->OLD-ABC->HNH->ASCH->METHYLASE->DUF499*-><-?<-TraD\|\|MobA_MobL+SF1-SF2-helicase->XIS-HTH->?->PhdYeFM_antitox-> | 945 | Bacteria>Pseudomonadota>Alphaproteobacteria>Hyphomicrobiales>Phyllobacteriaceae | GCF_003403035.1 | Mesorhizobium_denitrificans | DY251_RS01610 | 349492..352329 | 2294114 | DUF499-domain-containing-protein-[Mesorhizobium-denitrificans] |
| WP_140734688.1 | DUF499_1.RM.like.systems | Phage_terminase->TUDOR+SF1-SF2-Helicase+iREase+CTD+REase->OLD-ABC->HNH->METHYLASE->DUF499*->?->\|\|?-><-?<-?<-?\|\|Phage_integrase-> | 942 | Bacteria>Pseudomonadota>Alphaproteobacteria>Hyphomicrobiales>Phyllobacteriaceae | GCF_006442905.1 | Mesorhizobium_sp._B2-4-9 | FJ945_RS19675 | 32509..35337 | 2589940 | ATP-binding-protein-[Mesorhizobium-sp.-B2-4-9] |
| NEJ60793.1 | DUF499_1.RM.like.systems | XIS-HTH->TUDOR+SF1-SF2-Helicase+iREase+CTD+REase->OLD-ABC->HNH->METHYLASE->DUF499*-><-?<-Radical-SAM | 943 | Bacteria>Pseudomonadota>Alphaproteobacteria>Hyphomicrobiales>Rhizobiaceae | GCA_010668765.1 | Rhizobium_ruizarguesonis | GUK21_31970 | 69157..71988 | 2081791 | DUF499-domain-containing-protein-[Rhizobium-leguminosarum] |
| MCJ8508804.1 | DUF499_1.RM.like.systems | <-XIS-HTH\|\|HTH_3->TUDOR+SF1-SF2-Helicase+iREase+CTD+REase->OLD-ABC->HNH->METHYLASE->DUF499*->MODE-HTH+PBPII-><-ParG_RHH | 937 | Bacteria>Pseudomonadota>Alphaproteobacteria>Hyphomicrobiales>Rhizobiaceae | GCA_022968395.1 | Rhizobium_lemnae | MUU53_12875 | 44346..47159 | 1214924 | ATP-binding-protein-[Rhizobium-lemnae] |
| WP_010968623.1 | DUF499_1.RM.like.systems | HTH_3->TUDOR+SF1-SF2-Helicase+iREase+CTD+REase->OLD-ABC->HNH->METHYLASE->DUF499*-><-?<-REase<-HTH_3 | 942 | Bacteria>Pseudomonadota>Alphaproteobacteria>Hyphomicrobiales>Rhizobiaceae | GCF_000006965.1 | Sinorhizobium_meliloti | J3S94_RS16455 | 3478625..3481453 | 382 | ATP-binding-protein-[Sinorhizobium-meliloti] |
| WP_012755925.1 | DUF499_1.RM.like.systems | Phage_integrase->?->?->OLD-ABC+TOPRIM->?-><-DUF499*<-METHYLASE<-HNH<-OLD-ABC<-TUDOR+SF1-SF2-Helicase+iREase+CTD+REase<-HerA/FtsK | 945 | Bacteria>Pseudomonadota>Alphaproteobacteria>Hyphomicrobiales>Rhizobiaceae | GCF_000023185.1 | Rhizobium_leguminosarum | RLEG_RS01005 | 199729..202566 | 384 | ATP-binding-protein-[Rhizobium-leguminosarum] |
| WP_069615760.1 | DUF499_1.RM.like.systems | HTH_3->TUDOR+SF1-SF2-Helicase+iREase+CTD+REase->OLD-ABC->HNH->METHYLASE->DUF499*-><-?<-N6_MTase<-HTH<-?\|\|?->McrC-NTD+REase-> | 945 | Bacteria>Pseudomonadota>Alphaproteobacteria>Hyphomicrobiales>Rhizobiaceae | GCF_001729315.1 | Rhizobium_sp._YK2 | A9Z06_RS30975 | 24492..27329 | 1860096 | ATP-binding-protein-[Rhizobium-sp.-YK2] |
| WP_130838769.1 | DUF499_1.RM.like.systems | <-REase\|\|TUDOR+SF1-SF2-Helicase+iREase+CTD+REase->OLD-ABC->HNH->METHYLASE->DUF499*-> | 944 | Bacteria>Pseudomonadota>Alphaproteobacteria>Hyphomicrobiales>Rhizobiaceae | GCF_004305895.1 | Rhizobium_ruizarguesonis | HA464_RS06765 | 1375987..1378821 | 2081791 | ATP-binding-protein-[Rhizobium-leguminosarum] |
| WP_164568970.1 | DUF499_1.RM.like.systems | XIS-HTH->TUDOR+SF1-SF2-Helicase+iREase+CTD+REase->OLD-ABC->HNH->METHYLASE->DUF499*-><-?<-Radical-SAM | 943 | Bacteria>Pseudomonadota>Alphaproteobacteria>Hyphomicrobiales>Rhizobiaceae | GCF_010668765.1 | Rhizobium_ruizarguesonis | GUK21_31970 | 69157..71988 | 2081791 | ATP-binding-protein-[Rhizobium-leguminosarum] |
| WP_164056349.1 | DUF499_1.RM.like.systems | Arm-DNA-bind_3+Phage_integrase-><-XIS-HTH<-MobA_MobL+SF1-SF2-helicase\|\|TraD->?-><-DUF499*<-METHYLASE<-HNH<-OLD-ABC<-TUDOR+SF1-SF2-Helicase+iREase+CTD+REase<-HTH_3<-HigB | 944 | Bacteria>Pseudomonadota>Alphaproteobacteria>Hyphomicrobiales>Rhizobiaceae | GCF_010669145.1 | Rhizobium_oryzihabitans | G3A56_RS10085 | 1920588..1923422 | 2267833 | ATP-binding-protein-[Rhizobium-oryzihabitans] |
| WP_242223016.1 | DUF499_1.RM.like.systems | HigB->HTH_3->TUDOR+SF1-SF2-Helicase+iREase+CTD+REase->OLD-ABC->HNH->METHYLASE->DUF499*-><-?<-TraD\|\|MobA_MobL+SF1-SF2-helicase-><-?\|\|XIS-HTH-> | 946 | Bacteria>Pseudomonadota>Alphaproteobacteria>Hyphomicrobiales>Rhizobiaceae | GCF_022682305.1 | Shinella_zoogloeoides | MOE34_RS09030 | 1765134..1767974 | 352475 | ATP-binding-protein-[Shinella-zoogloeoides] |
| WP_281435123.1 | DUF499_1.RM.like.systems | <-XIS-HTH\|\|HTH_3->TUDOR+SF1-SF2-Helicase+iREase+CTD+REase->OLD-ABC->HNH->METHYLASE->DUF499*->MODE-HTH+PBPII-><-ParG_RHH | 943 | Bacteria>Pseudomonadota>Alphaproteobacteria>Hyphomicrobiales>Rhizobiaceae | GCF_022968395.1 | Rhizobium_lemnae | MUU53_RS12880 | 44328..47159 | 1214924 | ATP-binding-protein-[Rhizobium-lemnae] |
| WP_248132970.1 | DUF499_1.RM.like.systems | <-DUF499*<-METHYLASE<-HNH<-OLD-ABC<-TUDOR+SF1-SF2-Helicase+iREase+CTD+REase<-HTH_3 | 506 | Bacteria>Pseudomonadota>Alphaproteobacteria>Hyphomicrobiales>Rhizobiaceae | GCF_023119065.1 | Sinorhizobium_meliloti | J3S92_RS00005 | 1..1518 | 382 | DUF499-domain-containing-protein,-partial-[Sinorhizobium-meliloti] |
| WP_247261244.1 | DUF499_1.RM.like.systems | <-XIS-HTH\|\|HTH_3->TUDOR+SF1-SF2-Helicase+iREase+CTD+REase->OLD-ABC->HNH->METHYLASE->DUF499*->MODE-HTH+PBPII-><-ParG_RHH | 937 | Bacteria>Pseudomonadota>Alphaproteobacteria>Hyphomicrobiales>Rhizobiaceae | - | Rhizobium_lemnae | MUU53_12875 | 44346..47159 | 1214924 | ATP-binding-protein-[Rhizobium-lemnae] |
| WP_200985559.1 | DUF499_1.RM.like.systems | <-VirD4-TraG+VirD4-FtsK<-?<-Classical-AAA<-PIN+OST-HTH\|\|MobA_MobL+SF1-SF2-helicase-><-DUF499*<-METHYLASE<-HNH<-OLD-ABC<-TUDOR+SF1-SF2-Helicase+iREase+CTD+REase<-HTH_3 | 944 | Bacteria>Pseudomonadota>Alphaproteobacteria>Hyphomicrobiales>Rhizobiaceae | - | Rhizobium_rhizogenes | JL018_RS00080 | 14669..17503 | 359 | ATP-binding-protein-[Agrobacterium-rhizogenes] |
| SDR63348.1 | DUF499_1.RM.like.systems | Phage_int_SAM_1+Phage_integrase->REase-><-?<-REase+35exo+Beta-Barrel+SF1-SF2-Helicase<-DUF499*<-METHYLASE<-HNH<-OLD-ABC<-TUDOR+SF1-SF2-Helicase+iREase+CTD+REase | 942 | Bacteria>Pseudomonadota>Alphaproteobacteria>Hyphomicrobiales>unclassified_Hyphomicrobiales | GCA_900103895.1 | Rhizobiales_bacterium_GAS113 | SAMN05519103_08517 | 30774..33602 | 1884352 | hypothetical-protein-SAMN05519103_08517-[Rhizobiales-bacterium-GAS113] |
| WP_013654352.1 | DUF499_1.RM.like.systems | <-PIN<-XIS-HTH\|\|?-><-ArdCN+MPTase<-HEPN<-DUF499*<-METHYLASE<-HNH<-OLD-ABC<-TUDOR+SF1-SF2-Helicase+iREase+CTD+REase<-HTH_3<-HigB | 944 | Bacteria>Pseudomonadota>Alphaproteobacteria>Rhodobacterales>Paracoccaceae | GCF_000192745.1 | Polymorphum_gilvum | SL003B_RS18265 | 3920112..3922946 | 991904 | DUF499-domain-containing-protein-[Polymorphum-gilvum] |
| WP_014199726.1 | DUF499_1.RM.like.systems | MODE-HTH+PBPII-><-Arm-DNA-bind_3+Phage_int_SAM_3+Phage_integrase<-?<-HEPN+DUF499*<-HEPN<-PolB<-METHYLASE<-HNH<-OLD-ABC<-TUDOR+SF1-SF2-Helicase+iREase+CTD+REase | 1115 | Bacteria>Pseudomonadota>Alphaproteobacteria>Rhodospirillales>Azospirillaceae | GCF_000237365.1 | Azospirillum | GAY33_RS05480 | 11187..14534 | 191 | MULTISPECIES:-Swt1-family-HEPN-domain-containing-protein-[Azospirillum] |
| WP_209991153.1 | DUF499_1.RM.like.systems | TUDOR+SF1-SF2-Helicase+iREase+CTD+REase->OLD-ABC->HNH->METHYLASE->?->HEPN+DUF499*->?->Lon_C-> | 1114 | Bacteria>Pseudomonadota>Alphaproteobacteria>Rhodospirillales>Azospirillaceae | GCF_017876115.1 | Azospirillum_picis | J2850_RS30835 | 10080..13424 | 488438 | Swt1-family-HEPN-domain-containing-protein-[Azospirillum-picis] |
| TXH33435.1 | DUF499_1.RM.like.systems | HTH_3->TUDOR+SF1-SF2-Helicase+iREase+CTD+REase->OLD-ABC->HNH->ASCH->METHYLASE->DUF499*-><-?<-TraD\|\|MobA_MobL+SF1-SF2-helicase-> | 945 | Bacteria>Pseudomonadota>Alphaproteobacteria>Rhodospirillales>Rhodospirillaceae | GCA_008012315.1 | Rhodospirillaceae_bacterium | E6Q98_22210 | 66565..69402 | 1898112 | DUF499-domain-containing-protein-[Rhodospirillaceae-bacterium] |
| SEP49898.1 | DUF499_1.RM.like.systems | <-HEPN+DUF499*<-PIN<-OLD-ABC<-METHYLASE<-TUDOR+SF1-SF2-Helicase+iREase+CTD+REase<-HNH | 1099 | Bacteria>Pseudomonadota>Alphaproteobacteria>Rhodospirillales>unclassified_Rhodospirillales | GCA_900110395.1 | Rhodospirillales_bacterium_URHD0017 | SAMN02990966_07447 | 5640..8939 | 1380357 | hypothetical-protein-SAMN02990966_07447-[Rhodospirillales-bacterium-URHD0017] |
| MBL8657365.1 | DUF499_1.RM.like.systems | <-DUF499*<-METHYLASE<-ASCH<-HNH<-OLD-ABC<-TUDOR+SF1-SF2-Helicase+iREase+CTD+REase<-HTH_3 | 944 | Bacteria>Pseudomonadota>Alphaproteobacteria>Sphingomonadales>Erythrobacteraceae | GCA_016793865.1 | Altererythrobacter_sp. | JNJ92_08435 | 1147..3981 | 1872480 | DUF499-domain-containing-protein-[Altererythrobacter-sp.] |
| MCV0384725.1 | DUF499_1.RM.like.systems | <-HEPN+DUF499*<-METHYLASE<-HNH<-TUDOR+SF1-SF2-Helicase+iREase+CTD+REase-p | 421 | Bacteria>Pseudomonadota>Alphaproteobacteria>Sphingomonadales>Erythrobacteraceae | GCA_025698865.1 | Erythrobacter_sp. | K5799_14985 | 1..1263 | 1042 | MAG:-DUF499-domain-containing-protein,-partial-[Erythrobacter-sp.] |
| RXZ34416.1 | DUF499_1.RM.like.systems | TUDOR+SF1-SF2-Helicase+iREase+CTD+REase->OLD-ABC->HNH->METHYLASE->DUF499*-> | 940 | Bacteria>Pseudomonadota>Alphaproteobacteria>Sphingomonadales>Sphingomonadaceae | GCA_004135605.1 | Sphingomonas_desiccabilis | EO081_01620 | 357116..359938 | 429134 | DUF499-domain-containing-protein-[Sphingomonas-desiccabilis] |
| WP_130751563.1 | DUF499_1.RM.like.systems | CopG/RHH->VirD4-TraG+VirD4-FtsK->TrwC+SF1-SF2-helicase->wHTH(SBD)+HNH-><-LexA-protease\|\|?-><-DUF499*<-METHYLASE<-HNH<-OLD-ABC<-TUDOR+SF1-SF2-Helicase+iREase+CTD+REase<-CopG/RHH<-HTH+WYL | 940 | Bacteria>Pseudomonadota>Alphaproteobacteria>Sphingomonadales>Sphingomonadaceae | GCF_004305355.1 | Sphingobium_xenophagum | MBESOW_RS00075 | 12724..15546 | 121428 | ATP-binding-protein-[Sphingobium-xenophagum] |
| WP_159367773.1 | DUF499_1.RM.like.systems | <-DDE_Tnp_IS240<-?<-LZ_Tnp_IS66+zf-IS66+DDE_Tnp_IS66+DDE_Tnp_IS66_C<-?<-HIN-HTH<-PAIREDC-HTH+HIN-HTH<-DUF499*<-METHYLASE<-HNH<-OLD-ABC<-TUDOR+SF1-SF2-Helicase+iREase+CTD+REase<-CopG/RHH<-HTH+WYL | 940 | Bacteria>Pseudomonadota>Alphaproteobacteria>Sphingomonadales>Sphingomonadaceae | GCF_009856545.1 | Sphingobium_yanoikuyae | GS397_RS23435 | 4919345..4922167 | 13690 | ATP-binding-protein-[Sphingobium-yanoikuyae] |
| WP_168853122.1 | DUF499_1.RM.like.systems | CopG/RHH->VirD4-TraG+VirD4-FtsK->TrwC+SF1-SF2-helicase->wHTH(SBD)+HNH-><-LexA-protease\|\|?-><-DUF499*<-METHYLASE<-HNH<-OLD-ABC<-TUDOR+SF1-SF2-Helicase+iREase+CTD+REase<-CopG/RHH<-HTH+WYL | 940 | Bacteria>Pseudomonadota>Alphaproteobacteria>Sphingomonadales>Sphingomonadaceae | GCF_012641335.1 | Novosphingobium_sp._ERN07 | HGI47_RS12585 | 95856..98678 | 2726187 | ATP-binding-protein-[Novosphingobium-sp.-ERN07] |
| WP_206364336.1 | DUF499_1.RM.like.systems | TUDOR+SF1-SF2-Helicase+iREase+CTD+REase->OLD-ABC->HNH->METHYLASE->DUF499*-> | 934 | Bacteria>Pseudomonadota>Alphaproteobacteria>Sphingomonadales>Sphingomonadaceae | GCF_014196135.1 | Sphingomonas_desiccabilis | EO081_RS01620 | 357134..359938 | 429134 | ATP-binding-protein-[Sphingomonas-desiccabilis] |
| WP_260153764.1 | DUF499_1.RM.like.systems | TUDOR+SF1-SF2-Helicase+iREase+CTD+REase->OLD-ABC->HNH->METHYLASE->DUF499*-> | 822 | Bacteria>Pseudomonadota>Alphaproteobacteria>Sphingomonadales>Sphingomonadaceae | - | Sphingomonas_desiccabilis | GGQ81_RS01635 | 357438..359906 | 429134 | DUF499-domain-containing-protein-[Sphingomonas-desiccabilis] |
| WP_248595613.1 | DUF499_1.RM.like.systems | RelE-ParE->Phage_int_SAM_4+Phage_integrase->?->OLD-ABC->HNH->METHYLASE->DUF499*->DUF499-p->Phage_int_SAM_4+Phage_integrase-> | 644 | Bacteria>Pseudomonadota>Betaproteobacteria>Betaproteobacteria_incertae_sedis>Candidatus_Accumulibacter | GCF_012939955.1 | Candidatus_Accumulibacter_phosphatis | E4Q23_RS20285 | 20006..21940 | 327160 | DUF499-domain-containing-protein-[Candidatus-Accumulibacter-phosphatis] |
| WP_169069333.1 | DUF499_1.RM.like.systems | <-HEPN+DUF499*<-HEPN<-PolB<-METHYLASE<-HNH<-OLD-ABC<-PemK_toxin | 1112 | Bacteria>Pseudomonadota>Betaproteobacteria>Betaproteobacteria_incertae_sedis>Candidatus_Accumulibacter | GCF_012940005.1 | Candidatus_Accumulibacter_contiguus | E4Q08_RS03235 | 1844..5182 | 2954381 | Swt1-family-HEPN-domain-containing-protein-[Candidatus-Accumulibacter-contiguus] |
| WP_217096864.1 | DUF499_1.RM.like.systems | <-ACET\|\|?->?->MarR-HTH->Phage_int_SAM_2+Integrase_1-><-DUF499*<-Schlafen+HATPase+HTH<-METHYLASE<-METHYLASE<-HNH<-OLD-ABC<-TUDOR+SF1-SF2-Helicase+iREase+CTD+REase | 947 | Bacteria>Pseudomonadota>Betaproteobacteria>Burkholderiales>Burkholderiaceae | GCF_019041095.1 | Burkholderia_multivorans | KTE23_RS28415 | 22880..25723 | 87883 | DUF499-domain-containing-protein-[Burkholderia-multivorans] |
| MCW5668677.1 | DUF499_1.RM.like.systems | TUDOR+SF1-SF2-Helicase+iREase+CTD+REase->OLD-ABC->HNH->METHYLASE->?->?->DUF499*-><-HTH_3<-?<-HD\|\|NUDIX-> | 937 | Bacteria>Pseudomonadota>Betaproteobacteria>Burkholderiales>Comamonadaceae | GCA_026125825.1 | Hydrogenophaga_sp. | KIT86_03380 | 51119..53932 | 1904254 | MAG:-ATP-binding-protein-[Hydrogenophaga-sp.] |
| MDD2610199.1 | DUF499_1.RM.like.systems | TUDOR+SF1-SF2-Helicase+iREase+CTD+REase->OLD-ABC->HNH->METHYLASE-p->METHYLASE->?->DUF499*->?-><-?\|\|<-JAB<-UbE1+JAB<-SMODS | 951 | Bacteria>Pseudomonadota>Betaproteobacteria>Burkholderiales>Comamonadaceae | GCA_028686475.1 | Giesbergeria_sp. | PHX60_11025 | 19718..22573 | 2818473 | MAG:-DUF499-domain-containing-protein-[Giesbergeria-sp.] |
| WP_087746901.1 | DUF499_1.RM.like.systems | <-SPOUT\|\|?-><-?<-?<-?<-DUF499*<-REase<-OLD-ABC<-METHYLASE<-HNH<-OLD-ABC<-TUDOR+SF1-SF2-Helicase+iREase+CTD+REase | 945 | Bacteria>Pseudomonadota>Betaproteobacteria>Burkholderiales>Comamonadaceae | GCF_002176815.1 | unclassified_Acidovorax | CCX87_RS13165 | 2797139..2799976 | 2684926 | MULTISPECIES:-DUF499-domain-containing-protein-[unclassified-Acidovorax] |
| WP_055397337.1 | DUF499_1.RM.like.systems | <-ArdCN+MPTase<-?<-?<-MobA_MobL+SF1-SF2-helicase\|\|?-><-?<-DUF499*<-REase<-OLD-ABC<-METHYLASE<-HNH<-OLD-ABC<-TUDOR+SF1-SF2-Helicase+iREase+CTD+REase | 945 | Bacteria>Pseudomonadota>Betaproteobacteria>Burkholderiales>Comamonadaceae | GCF_017355385.1 | Acidovorax | AE621_RS12465 | 7782..10619 | 12916 | MULTISPECIES:-DUF499-domain-containing-protein-[Acidovorax] |
| WP_210568820.1 | DUF499_1.RM.like.systems | TUDOR+SF1-SF2-Helicase+iREase+CTD+REase->OLD-ABC->HNH->METHYLASE->OLD-ABC->REase->DUF499*->?->?->?-><-?\|\|SPOUT-> | 945 | Bacteria>Pseudomonadota>Betaproteobacteria>Burkholderiales>Comamonadaceae | GCF_017939685.1 | Acidovorax_sp._JG5 | J8G26_RS10065 | 96032..98869 | 2822718 | DUF499-domain-containing-protein-[Acidovorax-sp.-JG5] |
| WP_210930504.1 | DUF499_1.RM.like.systems | HTH->?->Phage_integrase->STYKinase-like-><-?\|\|DDE_Tnp_1_5+DDE_Tnp_1-><-DUF499*<-REase<-OLD-ABC<-HEPN<-METHYLASE<-HNH<-OLD-ABC | 945 | Bacteria>Pseudomonadota>Betaproteobacteria>Burkholderiales>Comamonadaceae | GCF_018064205.1 | Comamonas_aquatica | KAQ61_RS03430 | 737208..740045 | 225991 | DUF499-domain-containing-protein-[Comamonas-aquatica] |
| MCM1512954.1 | DUF499_1.RM.like.systems | HTH_3->?->?->?->TUDOR+SF1-SF2-Helicase+iREase+CTD+REase->METHYLASE->DUF499*->OLD-ABC->HNH-><-?\|\|MazE_antitoxin-> | 937 | Bacteria>Pseudomonadota>Betaproteobacteria>Burkholderiales>Oxalobacteraceae | GCA_023667225.1 | Oxalobacter_formigenes | NC112_07605 | 9124..11937 | 847 | DUF499-domain-containing-protein-[Oxalobacter-formigenes] |
| WP_182994400.1 | DUF499_1.RM.like.systems | HTH+WYL->TUDOR+SF1-SF2-Helicase+iREase+CTD+REase->OLD-ABC->HNH->METHYLASE->DUF499*-><-?\|\|?-><-?<-Arm-DNA-bind_3+Phage_integrase | 941 | Bacteria>Pseudomonadota>Betaproteobacteria>Burkholderiales>Oxalobacteraceae | GCF_014171515.1 | Herbaspirillum_frisingense | G5S34_RS08630 | 1936607..1939432 | 92645 | DUF499-domain-containing-protein-[Herbaspirillum-frisingense] |
| NRT58384.1 | DUF499_1.RM.like.systems | HAD->?-><-rve_3<-ParB+HTH<-?\|\|?-><-DUF499*<-TOPRIM<-OLD-ABC<-METHYLASE<-HNH<-OLD-ABC<-TUDOR+SF1-SF2-Helicase+iREase+CTD+REase | 953 | Bacteria>Pseudomonadota>Betaproteobacteria>Burkholderiales>Sphaerotilaceae | GCA_013294065.1 | Leptothrix_sp._C29 | HNQ01_004152 | 18472..21333 | 2735897 | hypothetical-protein-[Leptothrix-sp.-C29] |
| WP_140628337.1 | DUF499_1.RM.like.systems | PIN+OST-HTH-><-?\|\|HTH_3-><-?<-?<-?<-DUF499*<-?<-?<-METHYLASE<-HNH<-OLD-ABC<-TUDOR+SF1-SF2-Helicase+iREase+CTD+REase | 936 | Bacteria>Pseudomonadota>Betaproteobacteria>Burkholderiales>Sphaerotilaceae | GCF_006386545.1 | Methylibium_rhizosphaerae | E8D38_RS09385 | 117386..120196 | 2570323 | DUF499-domain-containing-protein-[Pseudorivibacter-rhizosphaerae] |
| WP_173807403.1 | DUF499_1.RM.like.systems | HAD->?-><-ParB+HTH<-?\|\|?-><-DUF499*<-TOPRIM<-OLD-ABC<-METHYLASE<-HNH<-OLD-ABC<-TUDOR+SF1-SF2-Helicase+iREase+CTD+REase | 953 | Bacteria>Pseudomonadota>Betaproteobacteria>Burkholderiales>Sphaerotilaceae | GCF_013294065.1 | Leptothrix_sp._C29 | HNQ01_RS20655 | 18472..21333 | 2735897 | DUF499-domain-containing-protein-[Leptothrix-sp.-C29] |
| WP_110402335.1 | DUF499_1.RM.like.systems | <-ABhydrolase\|\|?->?-><-?<-DUF499*<-REase<-OLD-ABC<-METHYLASE<-HNH<-OLD-ABC<-TUDOR+SF1-SF2-Helicase+iREase+CTD+REase | 958 | Bacteria>Pseudomonadota>Betaproteobacteria>Burkholderiales>Sphaerotilaceae | - | Sphaerotilus_hippei | C7444_RS18460 | 22920..25796 | 744406 | DUF499-domain-containing-protein-[Sphaerotilus-hippei] |
| QKE41939.1 | DUF499_1.RM.like.systems | DDE_Tnp_1_assoc+DDE_Tnp_1->HTH+WYL->TUDOR+SF1-SF2-Helicase+iREase+CTD+REase->OLD-ABC->HNH->wHTH+REase->DUF499*->DDE_Tnp_ISAZ013-><-?<-Hydrolase_3 | 942 | Bacteria>Pseudomonadota>Betaproteobacteria>Ferrovales>Ferrovaceae | GCA_013181415.1 | Ferrovum_myxofaciens | HO274_11940 | 2423876..2426704 | 416213 | ATP-binding-protein-[Ferrovum-myxofaciens] |
| WP_031597390.1 | DUF499_1.RM.like.systems | HTH+WYL->TUDOR+SF1-SF2-Helicase+iREase+CTD+REase->OLD-ABC->HNH->METHYLASE->DUF499*->?->?->?-><-?<-?<-DUF2163+Phage_BR0599 | 944 | Bacteria>Pseudomonadota>Betaproteobacteria>Ferrovales>Ferrovaceae | GCF_000735045.1 | Ferrovum_myxofaciens | IV91_RS0107820 | 14389..17223 | 416213 | DUF499-domain-containing-protein-[Ferrovum-myxofaciens] |
| WP_273119931.1 | DUF499_1.RM.like.systems | TUDOR+SF1-SF2-Helicase+iREase+CTD+REase->OLD-ABC->HNH->METHYLASE-p->METHYLASE-p->wHTH+REase->DUF499*->DDE_Tnp_ISAZ013-><-?<-?<-Hydrolase_3 | 942 | Bacteria>Pseudomonadota>Betaproteobacteria>Ferrovales>Ferrovaceae | GCF_013181415.1 | Ferrovum_myxofaciens | HO274_RS11980 | 2423876..2426704 | 416213 | DUF499-domain-containing-protein-[Ferrovum-myxofaciens] |
| MBX3631056.1 | DUF499_1.RM.like.systems | <-DUF499*<-METHYLASE<-HNH<-OLD-ABC<-TUDOR+SF1-SF2-Helicase+iREase+CTD+REase<-REase+RNase | 949 | Bacteria>Pseudomonadota>Betaproteobacteria>Nitrosomonadales>Nitrosomonadaceae | GCA_019635215.1 | Nitrosomonas_sp. | KF908_14345 | 13134..15983 | 42353 | ATP-binding-protein-[Nitrosomonas-sp.] |
| MCW5608748.1 | DUF499_1.RM.like.systems | HNH->METHYLASE->DUF499*-> | 949 | Bacteria>Pseudomonadota>Betaproteobacteria>Nitrosomonadales>Nitrosomonadaceae | GCA_026125505.1 | Nitrosomonas_sp. | KIT26_11955 | 3990..6839 | 42353 | MAG:-ATP-binding-protein-[Nitrosomonas-sp.] |
| MBX3668836.1 | DUF499_1.RM.like.systems | HTH_Tnp->TUDOR+SF1-SF2-Helicase+iREase+CTD+REase->OLD-ABC->HNH->METHYLASE->DUF499*->?-><-Phage_lysozyme<-PIN<-?\|\|?->HTH-> | 945 | Bacteria>Pseudomonadota>Betaproteobacteria>Rhodocyclales>Rhodocyclaceae | GCA_019635235.1 | Rhodocyclaceae_bacterium | KF778_10575 | 185865..188702 | 1898103 | ATP-binding-protein-[Rhodocyclaceae-bacterium] |
| WP_018989132.1 | DUF499_1.RM.like.systems | <-PIN+PhoH<-Radical-SAM+TRAM\|\|tRNA->?-><-HEPN+DUF499*<-METHYLASE<-HNH<-OLD-ABC<-PIN<-?<-STAS+HisKin | 1121 | Bacteria>Pseudomonadota>Betaproteobacteria>Rhodocyclales>Rhodocyclaceae | GCF_000378245.1 | Aromatoleum_toluclasticum | F464_RS0107275 | 366039..369404 | 92003 | Swt1-family-HEPN-domain-containing-protein-[Aromatoleum-toluclasticum] |

| Protein-ID | System | Gene-Neighbourhoods | Length | Taxonomy | Assembly-ID | Strain | Locus-Tag | Genomic-coordinates | TaxID | Fasta-defline |
|---|---|---|---|---|---|---|---|---|---|---|
| WP_169202998.1 | DUF499 _1.RM.like.systems | <- DUF499 *<-REase<-OLD-ABC<- METHYLASE <- HNH <-OLD-ABC <- TUDOR+ SF1-SF2-Helicase+iREase+CTD+REase | 951 | Bacteria>Pseudomonadota>Betaproteobacteria>Rhodocyclales>Rhodocyclaceae | GCF_017894365.1 | Aromatoleum_bremense | GPA24_RS12870 | 12346..15201 | 76115 | DUF499 -domain-containing-protein-[Aromatoleum-bremense] |
| TMW75521.1 | DUF499 _1.RM.like.systems | <- DUF499 *<- METHYLASE <- HNH <-OLD-ABC <-DOC+HTH<- TUDOR+ SF1-SF2-Helicase+iREase+CTD+REase \|\|HTH_31-> | 622 | Bacteria>Pseudomonadota>Betaproteobacteria>Rhodocyclales>Zoogloeaceae | GCA_006007845.1 | Thauera_sp._UPWRP | FG147_05165 | 1..1865 | 2488794 | ATP-binding-protein,-partial-[Thauera-sp.-UPWRP] |
| WP_159691715.1 | DUF499 _1.RM.like.systems | MODE-HTH+PBPII-><-TPR<- DUF499 *<-REase<-OLD-ABC<- METHYLASE <- HNH <-OLD-ABC <- TUDOR+ SF1-SF2-Helicase+iREase+CTD+REase | 943 | Bacteria>Pseudomonadota>Betaproteobacteria>Rhodocyclales>Zoogloeaceae | GCF_009800965.1 | Cognatazoarcus_halotolerans | GQB03_RS16405 | 71425..74256 | 2686016 | DUF499 -domain-containing-protein-[Cognatazoarcus-halotolerans] |
| WP_212351537.1 | DUF499 _1.RM.like.systems | <-Phage_integrase\|\|HTH+WYL-> TUDOR+ SF1-SF2-Helicase+iREase+CTD+REase -> OLD-ABC-> HNH -> METHYLASE -> DUF499 *->?-><-?<-RelA | 954 | Bacteria>Pseudomonadota>Betaproteobacteria>Rhodocyclales>Zoogloeaceae | GCF_018122735.1 | Azoarcus_sp._L1K30 | J5J83_RS19190 | 4181146..4184010 | 2820277 | DUF499 -domain-containing-protein-[Azoarcus-sp.-L1K30] |
| MDA8417189.1 | DUF499 _1.RM.like.systems | TUDOR+ SF1-SF2-Helicase+iREase+CTD+REase ->RVT-> OLD-ABC-> HNH -> METHYLASE ->wHTH+REase-> DUF499 *-><-?\|\|?->?->?-><-Response_reg+LuxR-HTH | 946 | Bacteria>Pseudomonadota>Betaproteobacteria>unclassified_Betaproteobacteria | GCA_028069095.1 | Betaproteobacteria_bacterium | M0Z78_09075 | 11618..14458 | 1891241 | MAG:- DUF499 -domain-containing-protein-[Betaproteobacteria-bacterium] |
| QWL60397.1 | DUF499 _1.RM.like.systems | URI-DNase-><-PAIREDC-HTH\|\|?-> OLD-ABC-> HNH -> METHYLASE -> DUF499 *-><-?\|\|HTH-><-RlpA<-JAB | 954 | Bacteria>Pseudomonadota>Gammaproteobacteria>Aeromonadales>Aeromonadaceae | GCA_018802265.1 | Aeromonas_jandaei | HQ400_01350 | 291564..294428 | 650 | ATP-binding-protein-[Aeromonas-jandaei] |
| MCF3097928.1 | DUF499 _1.RM.like.systems | TUDOR+ SF1-SF2-Helicase+iREase+CTD+REase -> HNH -> METHYLASE ->OLD-ABC->TOPRIM-> DUF499 *-> | 954 | Bacteria>Pseudomonadota>Gammaproteobacteria>Aeromonadales>Aeromonadaceae | GCA_021538905.1 | Aeromonas_australiensis | EHZ86_11640 | 15807..18671 | 1114880 | ATP-binding-protein-[Aeromonas-australiensis] |
| WP_064338617.1 | DUF499 _1.RM.like.systems | JAB->RlpA-><-HTH\|\|?-><- DUF499 *<-TOPRIM<-OLD-ABC<-DDE_Tnp_1_assoc+DDE_Tnp_1<-OLD-ABC-p<- METHYLASE <- HNH | 954 | Bacteria>Pseudomonadota>Gammaproteobacteria>Aeromonadales>Aeromonadaceae | GCF_001634325.1 | Aeromonas_veronii | WM43_RS07825 | 1773937..1776801 | 654 | DUF499 -domain-containing-protein-[Aeromonas-veronii] |
| WP_103858986.1 | DUF499 _1.RM.like.systems | JAB->CbeA_antitoxin->RlpA-><- DUF499 *<-TOPRIM<-OLD-ABC<- METHYLASE <- HNH <-OLD-ABC <- TUDOR+ SF1-SF2-Helicase+iREase+CTD+REase | 948 | Bacteria>Pseudomonadota>Gammaproteobacteria>Aeromonadales>Aeromonadaceae | GCF_002919855.1 | Aeromonas_sp._ASNIH8 | C3418_RS17745 | 3201..6047 | 1920113 | DUF499 -domain-containing-protein-[Aeromonas-sp.-ASNIH8] |
| WP_064341063.1 | DUF499 _1.RM.like.systems | <-HTH\|\|?-><-HIN-HTH\|\|?->?->DDE_Tnp_1-><- DUF499 *<-TOPRIM<-OLD-ABC<-OLD-ABC-p<- METHYLASE <- METHYLASE -p<- HNH | 954 | Bacteria>Pseudomonadota>Gammaproteobacteria>Aeromonadales>Aeromonadaceae | GCF_014169835.1 | Aeromonas_veronii | QBS04_RS15305 | 3271066..3273930 | 654 | DUF499 -domain-containing-protein-[Aeromonas-veronii] |
| WP_201871420.1 | DUF499 _1.RM.like.systems | JAB->CbeA_antitoxin->RlpA-><- DUF499 *<-TOPRIM<-OLD-ABC<- METHYLASE <- HNH <-OLD-ABC <- TUDOR+ SF1-SF2-Helicase+iREase+CTD+REase | 948 | Bacteria>Pseudomonadota>Gammaproteobacteria>Aeromonadales>Aeromonadaceae | GCF_016728905.1 | Aeromonas_caviae | JD551_RS01070 | 83459..86305 | 648 | DUF499 -domain-containing-protein-[Aeromonas-caviae] |
| WP_216991887.1 | DUF499 _1.RM.like.systems | TUDOR+ SF1-SF2-Helicase+iREase+CTD+REase -> OLD-ABC-> HNH -> METHYLASE ->OLD-ABC->TOPRIM-> DUF499 *-><-RlpA<-CbeA_antitoxin<-JAB | 948 | Bacteria>Pseudomonadota>Gammaproteobacteria>Aeromonadales>Aeromonadaceae | GCF_019048725.1 | Aeromonas_sp._FDAARGOS_1407 | I6L37_RS11490 | 2395827..2398673 | 2778056 | DUF499 -domain-containing-protein-[Aeromonas-sp.-FDAARGOS-1407] |
| WP_219296379.1 | DUF499 _1.RM.like.systems | JAB->RlpA-><-HTH\|\|?-><- DUF499 *<-TOPRIM<-OLD-ABC<- METHYLASE <- HNH <-OLD-ABC <- TUDOR+ SF1-SF2-Helicase+iREase+CTD+REase | 952 | Bacteria>Pseudomonadota>Gammaproteobacteria>Aeromonadales>Aeromonadaceae | GCF_019348625.1 | Aeromonas_veronii | GL270_RS12655 | 107539..110397 | 654 | DUF499 -domain-containing-protein-[Aeromonas-veronii] |
| WP_223918005.1 | DUF499 _1.RM.like.systems | TUDOR+ SF1-SF2-Helicase+iREase+CTD+REase -> OLD-ABC-> HNH -> METHYLASE ->OLD-ABC->TOPRIM-> DUF499 *-><-RlpA<-CbeA_antitoxin<-JAB | 948 | Bacteria>Pseudomonadota>Gammaproteobacteria>Aeromonadales>Aeromonadaceae | GCF_019971655.1 | Aeromonas_caviae | LDN21_RS02455 | 71531..74377 | 648 | DUF499 -domain-containing-protein-[Aeromonas-caviae] |
| WP_223954701.1 | DUF499 _1.RM.like.systems | JAB->CbeA_antitoxin->RlpA-><- DUF499 *<-REase<-OLD-ABC<- METHYLASE <- HNH <-OLD-ABC <- TUDOR+ SF1-SF2-Helicase+iREase+CTD+REase | 946 | Bacteria>Pseudomonadota>Gammaproteobacteria>Aeromonadales>Aeromonadaceae | GCF_019972775.1 | Aeromonas_caviae | LDN68_RS17105 | 3642842..3645682 | 648 | DUF499 -domain-containing-protein-[Aeromonas-caviae] |
| WP_235462717.1 | DUF499 _1.RM.like.systems | TUDOR+ SF1-SF2-Helicase+iREase+CTD+REase -> OLD-ABC-> HNH -> METHYLASE ->OLD-ABC->TOPRIM-> DUF499 *-> | 897 | Bacteria>Pseudomonadota>Gammaproteobacteria>Aeromonadales>Aeromonadaceae | GCF_021538905.1 | Aeromonas_australiensis | EHZ86_RS11610 | 15978..18671 | 1114880 | DUF499 -domain-containing-protein-[Aeromonas-australiensis] |
| WP_257709589.1 | DUF499 _1.RM.like.systems | TUDOR+ SF1-SF2-Helicase+iREase+CTD+REase -> OLD-ABC-> HNH -> METHYLASE ->OLD-ABC->TOPRIM-> DUF499 *-><-RlpA<-CbeA_antitoxin<-JAB<-?\|\|<-Arm-DNA-bind_3+Phage_integrase | 948 | Bacteria>Pseudomonadota>Gammaproteobacteria>Aeromonadales>Aeromonadaceae | GCF_024649925.1 | Aeromonas_caviae | NUK49_RS00060 | 12274..15120 | 648 | DUF499 -domain-containing-protein-[Aeromonas-caviae] |
| WP_270668501.1 | DUF499 _1.RM.like.systems | <- DUF499 *<- METHYLASE <- HNH <-OLD-ABC <-?<- TUDOR+ SF1-SF2-Helicase+iREase+CTD+REase | 954 | Bacteria>Pseudomonadota>Gammaproteobacteria>Aeromonadales>Aeromonadaceae | GCF_027705205.1 | unclassified_Aeromonas | OKB91_RS16400 | 106..2970 | 257493 | MULTISPECIES:- DUF499 -domain-containing-protein-[unclassified-Aeromonas] |
| WP_101201433.1 | DUF499 _1.RM.like.systems | HTH+WYL-> TUDOR+ SF1-SF2-Helicase+iREase+CTD+REase -> OLD-ABC-> HNH -> METHYLASE -> DUF499 *-><-RlpA<-?\|\|?->Phage_integrase-> | 944 | Bacteria>Pseudomonadota>Gammaproteobacteria>Alteromonadales>Alteromonadaceae | GCF_002835935.1 | Paraglaciecola_sp._MB-3u-78 | CXF95_RS26560 | 32531..35365 | 2058332 | DUF499 -domain-containing-protein-[Paraglaciecola-sp.-MB-3u-78] |
| WP_250655185.1 | DUF499 _1.RM.like.systems | HTH+WYL-> TUDOR+ SF1-SF2-Helicase+iREase+CTD+REase -> OLD-ABC-> HNH -> TUDOR+ SF1-SF2-Helicase+iREase+CTD+REase -p-> METHYLASE -> DUF499 *->SMODS->?-><-RlpA<-Phage_lysozyme | 945 | Bacteria>Pseudomonadota>Gammaproteobacteria>Alteromonadales>Alteromonadaceae | GCF_023650515.1 | Alkalimarinus_coralli | MY525_RS07060 | 1557790..1560627 | 2935863 | MULTISPECIES:- DUF499 -domain-containing-protein-[unclassified-Alkalimarinus] |
| BCV35016.1 | DUF499 _1.RM.like.systems | <-PRTase<-RNase\|\|?->Arm-DNA-bind_3+Phage_integrase->RlpA-><- DUF499 *<- METHYLASE <- HNH <-OLD-ABC <- TUDOR+ SF1-SF2-Helicase+iREase+CTD+REase <-HTH_3 | 943 | Bacteria>Pseudomonadota>Gammaproteobacteria>Alteromonadales>Shewanellaceae | GCA_019670565.1 | Shewanella_chilikensis | TUM17377_03440 | 374214..377045 | 558541 | hypothetical-protein-TUM17377_03440-[Shewanella-chilikensis] |
| WP_186265255.1 | DUF499 _1.RM.like.systems | HTH_3-> TUDOR+ SF1-SF2-Helicase+iREase+CTD+REase -> OLD-ABC-> HNH -> METHYLASE -> DUF499 *-><-RlpA<-Arm-DNA-bind_3+Phage_integrase<-?\|\|RNase->PRTase-> | 943 | Bacteria>Pseudomonadota>Gammaproteobacteria>Alteromonadales>Shewanellaceae | GCF_014263185.1 | Shewanella_algae | HU689_RS02010 | 444450..447281 | 38313 | DUF499 -domain-containing-protein-[Shewanella-algae] |
| WP_115016378.1 | DUF499 _1.RM.like.systems | <-JAB\|\|XIS-HTH->REase-p->REase->RlpA-><- DUF499 *<- METHYLASE <- HNH <-OLD-ABC <- TUDOR+ SF1-SF2-Helicase+iREase+CTD+REase \|\|?->DDE_Tnp_1_3-> | 943 | Bacteria>Pseudomonadota>Gammaproteobacteria>Alteromonadales>Shewanellaceae | GCF_019266125.1 | Shewanella | DX079_RS11245 | 2521097..2523928 | 22 | MULTISPECIES:- DUF499 -domain-containing-protein-[Shewanella] |
| WP_219013226.1 | DUF499 _1.RM.like.systems | HTH_3-> TUDOR+ SF1-SF2-Helicase+iREase+CTD+REase -> OLD-ABC-> HNH -> METHYLASE -> DUF499 *-><-RlpA<-Arm-DNA-bind_3+Phage_integrase<-?\|\|RNase->PRTase-> | 943 | Bacteria>Pseudomonadota>Gammaproteobacteria>Alteromonadales>Shewanellaceae | GCF_019303275.1 | Shewanella_algae | KOE54_RS14875 | 19508..22339 | 38313 | DUF499 -domain-containing-protein-[Shewanella-algae] |
| WP_079541822.1 | DUF499 _1.RM.like.systems | TUDOR+ SF1-SF2-Helicase+iREase+CTD+REase -> OLD-ABC-> HNH -> METHYLASE ->OLD-ABC->TOPRIM-> DUF499 *-> | 937 | Bacteria>Pseudomonadota>Gammaproteobacteria>Cardiobacteriales>Cardiobacteriaceae | GCF_900079085.1 | Cardiobacterium_hominis | CHUV0807_RS10775 | 37520..40333 | 2718 | DUF499 -domain-containing-protein-[Cardiobacterium-hominis] |
| MTW22277.1 | DUF499 _1.RM.like.systems | TUDOR+ SF1-SF2-Helicase+iREase+CTD+REase -> OLD-ABC-> HNH ->OLD-ABC->TOPRIM-> DUF499 *-> | 885 | Bacteria>Pseudomonadota>Gammaproteobacteria>Chromatiales>Chromatiaceae | GCA_009720725.1 | Allochromatium_palmeri | GJ668_14455 | 38860..41517 | 231048 | DUF499 -domain-containing-protein-[Allochromatium-palmeri] |
| WP_155450846.1 | DUF499 _1.RM.like.systems | OLD-ABC-> HNH -> METHYLASE -p-> METHYLASE -p->OLD-ABC->TOPRIM-> DUF499 *-> | 883 | Bacteria>Pseudomonadota>Gammaproteobacteria>Chromatiales>Chromatiaceae | GCF_009720725.1 | Allochromatium_palmeri | GJ668_RS14460 | 38866..41517 | 231048 | DUF499 -domain-containing-protein-[Allochromatium-palmeri] |
| EFO1856148.1 | DUF499 _1.RM.like.systems | TUDOR+ SF1-SF2-Helicase+iREase+CTD+REase -> OLD-ABC-> HNH -> METHYLASE -> DUF499 *-><-HTH_Tnp_1+HTH_21+rve+rve_2<-HTH_Tnp_1+HTH_21+rve+rve_2\|\|<-?<-?\|\|XIS-HTH->DUF3987-> | 947 | Bacteria>Pseudomonadota>Gammaproteobacteria>Enterobacterales>Enterobacteriaceae | GCA_013065125.1 | Escherichia_coli | DUZ37_17320 | 27712..30555 | 562 | DUF499 -domain-containing-protein-[Escherichia-coli] |
| EGQ7342643.1 | DUF499 _1.RM.like.systems | RlpA-><- DUF499 *<- METHYLASE <- HNH <-OLD-ABC <- TUDOR+ SF1-SF2-Helicase+iREase+CTD+REase <-?\|\| ParB -> | 947 | Bacteria>Pseudomonadota>Gammaproteobacteria>Enterobacterales>Enterobacteriaceae | GCA_015730265.1 | Enterobacter_cloacae | I6C89_001957 | 175814..178657 | 550 | ATP-binding-protein-[Enterobacter-cloacae] |
| HAU4285776.1 | DUF499 _1.RM.like.systems | TUDOR+ SF1-SF2-Helicase+iREase+CTD+REase -> OLD-ABC-> HNH -> METHYLASE -> DUF499 *-><-RlpA | 947 | Bacteria>Pseudomonadota>Gammaproteobacteria>Enterobacterales>Enterobacteriaceae | GCA_016065535.1 | Citrobacter_freundii | F6W46_24070 | 10411..13254 | 546 | ATP-binding-protein-[Citrobacter-freundii] |
| HAU5791339.1 | DUF499 _1.RM.like.systems | Phage_ASH->?->?->TOPRIM+DNAG+D5N+D5ATPase+P4PRIMASE-HTH->?->RlpA-><- DUF499 *<- METHYLASE <- HNH <-OLD-ABC <- TUDOR+ SF1-SF2-Helicase+iREase+CTD+REase | 947 | Bacteria>Pseudomonadota>Gammaproteobacteria>Enterobacterales>Enterobacteriaceae | GCA_016076255.1 | Citrobacter_freundii | JD367_15405 | 174239..177082 | 546 | ATP-binding-protein-[Citrobacter-freundii] |
| HAW1058251.1 | DUF499 _1.RM.like.systems | <-nSTAND1+sGTP\|\|?-> TUDOR+ SF1-SF2-Helicase+iREase+CTD+REase -> OLD-ABC-> HNH -> METHYLASE -> DUF499 *-><-DEDD_Tnp_IS110+Transposase_20 | 947 | Bacteria>Pseudomonadota>Gammaproteobacteria>Enterobacterales>Enterobacteriaceae | GCA_016672425.1 | Escherichia_coli | JLK83_004317 | 23199..26042 | 562 | ATP-binding-protein-[Escherichia-coli] |
| HBC9812127.1 | DUF499 _1.RM.like.systems | TUDOR+ SF1-SF2-Helicase+iREase+CTD+REase -p-> OLD-ABC-> HNH -> METHYLASE -> DUF499 *-><-RlpA | 951 | Bacteria>Pseudomonadota>Gammaproteobacteria>Enterobacterales>Enterobacteriaceae | GCA_018195115.1 | Escherichia_coli | KFP54_004194 | 5317..8172 | 562 | ATP-binding-protein-[Escherichia-coli] |
| HBW9592181.1 | DUF499 _1.RM.like.systems | TUDOR+ SF1-SF2-Helicase+iREase+CTD+REase -> OLD-ABC-> HNH -> METHYLASE -> DUF499 *-><-RlpA | 953 | Bacteria>Pseudomonadota>Gammaproteobacteria>Enterobacterales>Enterobacteriaceae | GCA_022207645.1 | Klebsiella_pneumoniae | MFS54_26490 | 9564..12425 | 573 | DUF499 -domain-containing-protein-[Klebsiella-pneumoniae] |
| HCI7559692.1 | DUF499 _1.RM.like.systems | <-HIN-HTH\|\|?-> TUDOR+ SF1-SF2-Helicase+iREase+CTD+REase -> OLD-ABC-> HNH -> METHYLASE -> DUF499 *-><-RlpA<-?<-Arm-DNA-bind_3+Phage_integrase<-tRNA\|\|?->?->ADH_N+ADH_zinc_N-> | 947 | Bacteria>Pseudomonadota>Gammaproteobacteria>Enterobacterales>Enterobacteriaceae | GCA_024446695.1 | Klebsiella_pneumoniae | NPW42_004570 | 42168..45011 | 573 | ATP-binding-protein-[Klebsiella-pneumoniae] |
| MCQ4415112.1 | DUF499 _1.RM.like.systems | Arm-DNA-bind_3+Phage_integrase->RlpA-><- DUF499 *<- METHYLASE <- HNH <-OLD-ABC <- TUDOR+ SF1-SF2-Helicase+iREase+CTD+REase | 947 | Bacteria>Pseudomonadota>Gammaproteobacteria>Enterobacterales>Enterobacteriaceae | GCA_024451305.1 | Enterobacter_kobei | NOX26_15660 | 3292884..3295727 | 208224 | DUF499 -domain-containing-protein-[Enterobacter-kobei] |
| WP_045284468.1 | DUF499 _1.RM.like.systems | RlpA-><- DUF499 *<- METHYLASE <- HNH <-OLD-ABC <- TUDOR+ SF1-SF2-Helicase+iREase+CTD+REase | 947 | Bacteria>Pseudomonadota>Gammaproteobacteria>Enterobacterales>Enterobacteriaceae | GCF_000958085.1 | Enterobacter_sichuanensis | SS37_RS01295 | 119849..122692 | 2071710 | DUF499 -domain-containing-protein-[Enterobacter-sichuanensis] |
| WP_097584906.1 | DUF499 _1.RM.like.systems | <-REC+LuxR-HTH<-SAM-methylase<- wHTH(SBD)+ HNH \|\|tRNA->RlpA-><- DUF499 *<- METHYLASE <-SbcC<- TUDOR+ SF1-SF2-Helicase+iREase+CTD+REase | 947 | Bacteria>Pseudomonadota>Gammaproteobacteria>Enterobacterales>Enterobacteriaceae | GCF_002532355.1 | Cronobacter_sakazakii | FC600_RS12565 | 210517..213360 | 28141 | DUF499 -domain-containing-protein-[Cronobacter-sakazakii] |
| WP_086660584.1 | DUF499 _1.RM.like.systems | <-nSTAND1+sGTP\|\|?-> TUDOR+ SF1-SF2-Helicase+iREase+CTD+REase -> OLD-ABC-> HNH -> METHYLASE -> DUF499 *->\|\|?->CbeA_antitoxin->?-><-tRNA<-Aldedh | 947 | Bacteria>Pseudomonadota>Gammaproteobacteria>Enterobacterales>Enterobacteriaceae | GCF_014216435.1 | Escherichia_coli | GVI19_RS16185 | 90398..93241 | 562 | DUF499 -domain-containing-protein-[Escherichia-coli] |
| WP_196359062.1 | DUF499 _1.RM.like.systems | RlpA-><- DUF499 *<- METHYLASE <- HNH <-OLD-ABC <- TUDOR+ SF1-SF2-Helicase+iREase+CTD+REase <-?\|\| ParB -> | 947 | Bacteria>Pseudomonadota>Gammaproteobacteria>Enterobacterales>Enterobacteriaceae | GCF_015701305.1 | Enterobacter_hormaechei | I4P56_RS16415 | 165030..167873 | 158836 | DUF499 -domain-containing-protein-[Enterobacter-hormaechei] |
| WP_214577408.1 | DUF499 _1.RM.like.systems | RlpA-><- DUF499 *<- METHYLASE <- HNH <-OLD-ABC <- TUDOR+ SF1-SF2-Helicase+iREase+CTD+REase | 947 | Bacteria>Pseudomonadota>Gammaproteobacteria>Enterobacterales>Enterobacteriaceae | GCF_018597265.1 | Enterobacter_dykesii | KK116_RS19460 | 158324..161167 | 2797506 | DUF499 -domain-containing-protein-[Enterobacter-dykesii] |
| WP_222563661.1 | DUF499 _1.RM.like.systems | Arm-DNA-bind_3+Phage_integrase->RlpA-><- DUF499 *<- METHYLASE <- HNH <-OLD-ABC <- TUDOR+ SF1-SF2-Helicase+iREase+CTD+REase | 947 | Bacteria>Pseudomonadota>Gammaproteobacteria>Enterobacterales>Enterobacteriaceae | GCF_019803045.1 | Citrobacter_farmeri | Cf24236_RS16360 | 3446361..3449204 | 67824 | MULTISPECIES:- DUF499 -domain-containing-protein-[Enterobacteriaceae] |
| WP_229532552.1 | DUF499 _1.RM.like.systems | ACET-><-Phage_integrase\|\|RlpA-><- DUF499 *<- METHYLASE <- HNH <-?<-OLD-ABC<- TUDOR+ SF1-SF2-Helicase+iREase+CTD+REase | 947 | Bacteria>Pseudomonadota>Gammaproteobacteria>Enterobacterales>Enterobacteriaceae | GCF_020577415.1 | Klebsiella_variicola | KAG03_RS12050 | 331978..334821 | 244366 | DUF499 -domain-containing-protein-[Klebsiella-variicola] |
| WP_048997346.1 | DUF499 _1.RM.like.systems | RlpA-><- DUF499 *<- METHYLASE <- HNH <-OLD-ABC <- TUDOR+ SF1-SF2-Helicase+iREase+CTD+REase | 947 | Bacteria>Pseudomonadota>Gammaproteobacteria>Enterobacterales>Enterobacteriaceae | GCF_021869685.1 | Enterobacter | L3068_RS02450 | 520532..523375 | 547 | MULTISPECIES:- DUF499 -domain-containing-protein-[Enterobacter] |
| WP_256043565.1 | DUF499 _1.RM.like.systems | Arm-DNA-bind_3+Phage_integrase->RlpA-><- DUF499 *<- METHYLASE <- HNH <-OLD-ABC <- TUDOR+ SF1-SF2-Helicase+iREase+CTD+REase | 947 | Bacteria>Pseudomonadota>Gammaproteobacteria>Enterobacterales>Enterobacteriaceae | GCF_024451305.1 | Enterobacter_kobei | NOX26_RS15660 | 3292884..3295727 | 208224 | DUF499 -domain-containing-protein-[Enterobacter-kobei] |
| WP_279936012.1 | DUF499 _1.RM.like.systems | <- DUF499 *<- METHYLASE <- HNH | 951 | Bacteria>Pseudomonadota>Gammaproteobacteria>Enterobacterales>Enterobacteriaceae | GCF_029837515.1 | Klebsiella_oxytoca | N7642_RS28020 | 124..2979 | 571 | DUF499 -domain-containing-protein-[Klebsiella-oxytoca] |
| WP_193132801.1 | DUF499 _1.RM.like.systems | HTH->?->HEPN->RlpA-><- DUF499 *<- METHYLASE <- HNH <-OLD-ABC <- TUDOR+ SF1-SF2-Helicase+iREase+CTD+REase <-?\|\| ParB -> | 942 | Bacteria>Pseudomonadota>Gammaproteobacteria>Enterobacterales>Enterobacteriaceae | GCF_032369155.1 | Enterobacter_hormaechei | CXM24_RS20100 | 4287724..4290552 | 158836 | DUF499 -domain-containing-protein-[Enterobacter-hormaechei] |
| WP_086872901.1 | DUF499 _1.RM.like.systems | TUDOR+ SF1-SF2-Helicase+iREase+CTD+REase -> OLD-ABC-> HNH -> METHYLASE -> DUF499 *-><-RlpA<-Arm-DNA-bind_3+Phage_integrase<-tRNA\|\|REC+LuxR-HTH-> | 953 | Bacteria>Pseudomonadota>Gammaproteobacteria>Enterobacterales>Enterobacteriaceae | GCF_900184035.1 | Kosakonia_pseudosacchari | BN2591_RS15655 | 26880..29741 | 1646340 | DUF499 -domain-containing-protein-[Kosakonia-pseudosacchari] |
| WP_008325059.1 | DUF499 _1.RM.like.systems | TUDOR+ SF1-SF2-Helicase+iREase+CTD+REase -> OLD-ABC-> HNH -> METHYLASE -> DUF499 *-><-RlpA | 947 | Bacteria>Pseudomonadota>Gammaproteobacteria>Enterobacterales | GCF_000277565.1 | Enterobacteriaceae | B397_RS01195 | 10694..13537 | 543 | MULTISPECIES:- DUF499 -domain-containing-protein-[Enterobacteriaceae] |
| WP_077999168.1 | DUF499 _1.RM.like.systems | JAB->RlpA-><- DUF499 *<-REase<-OLD-ABC<- METHYLASE <- HNH <-OLD-ABC <- TUDOR+ SF1-SF2-Helicase+iREase+CTD+REase | 949 | Bacteria>Pseudomonadota>Gammaproteobacteria>Enterobacterales>Hafniaceae | GCF_002006945.1 | Edwardsiella_tarda | BTN58_RS02535 | 130022..132871 | 636 | DUF499 -domain-containing-protein-[Edwardsiella-tarda] |
| WP_205545050.1 | DUF499 _1.RM.like.systems | <-PAIREDC-HTH+HIN-HTH\|\|?-> TUDOR+ SF1-SF2-Helicase+iREase+CTD+REase -> OLD-ABC-> HNH -> METHYLASE -> DUF499 *-><-?<- HNH <-?<-DHTH+WYL\|\|nSTAND1+sGTP->HTH+WYL-> | 951 | Bacteria>Pseudomonadota>Gammaproteobacteria>Enterobacterales>Pectobacteriaceae | GCF_016949795.1 | Pectobacterium_brasiliense | H4F47_RS10595 | 47611..50466 | 180957 | DUF499 -domain-containing-protein-[Pectobacterium-brasiliense] |
| SUI50248.1 | DUF499 _1.RM.like.systems | GDResolvase+SIGMA-HTH->?->SF1-SF2-Helicase->RlpA-><- DUF499 *<- METHYLASE <- HNH <-OLD-ABC <- TUDOR+ SF1-SF2-Helicase+iREase+CTD+REase | 975 | Bacteria>Pseudomonadota>Gammaproteobacteria>Enterobacterales>Yersiniaceae | GCA_900457075.1 | Serratia_quinivorans | NCTC11544_01113 | 1025353..1028280 | 137545 | Protein-of-uncharacterised-function-( DUF499 )-[Serratia-quinivorans] |
| CAI1753142.1 | DUF499 _1.RM.like.systems | TUDOR+ SF1-SF2-Helicase+iREase+CTD+REase -> OLD-ABC-> HNH -> METHYLASE -> DUF499 *-><-RlpA<-SF1-SF2-Helicase<-?<-GDResolvase+SIGMA-HTH<-?<-XIS-HTH | 975 | Bacteria>Pseudomonadota>Gammaproteobacteria>Enterobacterales>Yersiniaceae | GCA_946228205.1 | Serratia_quinivorans | SAMEA4857275_02169 | 318450..321377 | 137545 | Protein-of-uncharacterised-function-( DUF499 )-[Serratia-quinivorans] |
| WP_115183101.1 | DUF499 _1.RM.like.systems | <-nSTAND1+sGTP\|\|?-> TUDOR+ SF1-SF2-Helicase+iREase+CTD+REase -> OLD-ABC-> HNH -> METHYLASE -> DUF499 *-><-RlpA<-SF1-SF2-Helicase<-?<-GDResolvase+SIGMA-HTH<-?<-XIS-HTH | 951 | Bacteria>Pseudomonadota>Gammaproteobacteria>Enterobacterales>Yersiniaceae | GCF_946228205.1 | Serratia | BSR03_RS13425 | 143107..145962 | 613 | MULTISPECIES:- DUF499 -domain-containing-protein-[Serratia] |
| PZN71299.1 | DUF499 _1.RM.like.systems | <- DUF499 *<-PIN<- METHYLASE <- HNH <-OLD-ABC <-PemK_toxin<-PrlF_antitoxin | 955 | Bacteria>Pseudomonadota>Gammaproteobacteria>Methylococcales>Methylococcales_incertae_sedis | GCA_003242955.1 | Candidatus_Methylumidiphilus_alinenensis | DM484_26720 | 13367..16234 | 2202197 | hypothetical-protein-DM484_26720-[Candidatus-Methylumidiphilus-alinenensis] |
| WP_142947823.1 | DUF499 _1.RM.like.systems | Arm-DNA-bind_3+Phage_integrase-><-Phage_AlpA<-ZnR+TOPRIM+primpol+HTH<-XIS-HTH\|\|MobI-><- DUF499 *<-Schlafen+HATPase+HTH<- METHYLASE <- METHYLASE <- HNH <-OLD-ABC <- TUDOR+ SF1-SF2-Helicase+iREase+CTD+REase | 953 | Bacteria>Pseudomonadota>Gammaproteobacteria>Oceanospirillales>Alcanivoracaceae | - | Alcanivorax_sp._DSM_26295 | FN816_RS03075 | 107194..110055 | 1938738 | DUF499 -domain-containing-protein-[Alcanivorax-sp.-DSM-26295] |
| MCP5162750.1 | DUF499 _1.RM.like.systems | <-ABC-ATPase+SbcC<-?<- DUF499 *<- METHYLASE <- HNH <-OLD-ABC <- TUDOR+ SF1-SF2-Helicase+iREase+CTD+REase <-HTH+WYL | 944 | Bacteria>Pseudomonadota>Gammaproteobacteria>Oceanospirillales>Hahellaceae | GCA_024234505.1 | Hahellaceae_bacterium | H7A00_13970 | 74140..76974 | 2048971 | ATP-binding-protein-[Hahellaceae-bacterium] |
| MCD6438846.1 | DUF499 _1.RM.like.systems | HNH -> METHYLASE ->OLD-ABC->REase->EVE+McrB-AAA->McrC-NTD+REase-> DUF499 *-> | 282 | Bacteria>Pseudomonadota>Gammaproteobacteria>Oceanospirillales>Halomonadaceae | GCA_021161325.1 | Halomonas_sp. | J7L56_11360 | 10325..11171 | 1486246 | ATP-binding-protein-[Halomonas-sp.] |
| WP_253017818.1 | DUF499 _1.RM.like.systems | HNH -> METHYLASE ->OLD-ABC->REase->EVE+McrB-AAA->McrC-NTD+REase-> DUF499 *->?->AbiEi-><-CHTH<-RelE | 961 | Bacteria>Pseudomonadota>Gammaproteobacteria>Oceanospirillales>Halomonadaceae | GCF_024124605.1 | Halomonas_sp._CnH100-B | NH398_RS08385 | 365976..368861 | 2954490 | DUF499 -domain-containing-protein-[Halomonas-sp.-CnH100-B] |
| WP_137094585.1 | DUF499 _1.RM.like.systems | TUDOR+ SF1-SF2-Helicase+iREase+CTD+REase -> OLD-ABC-> HNH -> METHYLASE ->OLD-ABC->REase-> DUF499 *-><-MobI\|\|XIS-HTH->ZnR+TOPRIM+primpol+HTH->Phage_AlpA-><-Arm-DNA-bind_3+Phage_integrase | 953 | Bacteria>Pseudomonadota>Gammaproteobacteria>Oceanospirillales>Halomonadaceae | GCF_028580285.1 | Halomonas | E8Q34_RS13770 | 13695..16556 | 2745 | DUF499 -domain-containing-protein-[Halomonas-sp.-15WGF] |
| WP_284453091.1 | DUF499 _1.RM.like.systems | Arm-DNA-bind_3+Phage_integrase-><-Phage_AlpA<-ZnR+TOPRIM+primpol+HTH<-XIS-HTH\|\|MobI-><- DUF499 *<-REase<-OLD-ABC<- METHYLASE <- HNH <-OLD-ABC <- TUDOR+ SF1-SF2-Helicase+iREase+CTD+REase | 953 | Bacteria>Pseudomonadota>Gammaproteobacteria>Oceanospirillales>Halomonadaceae | GCF_030145725.1 | Halomonas_meridiana | KYX60_RS01235 | 106598..109459 | 29570 | DUF499 -domain-containing-protein-[Halomonas-meridiana] |
| WP_072841218.1 | DUF499 _1.RM.like.systems | TUDOR+ SF1-SF2-Helicase+iREase+CTD+REase -> OLD-ABC-> HNH -> METHYLASE ->OLD-ABC->REase-> DUF499 *->SMODS->?-><-RlpA\|\|?-><-HTH_26 | 942 | Bacteria>Pseudomonadota>Gammaproteobacteria>Oceanospirillales>Oceanospirillaceae | GCF_900129155.1 | Marinomonas_polaris | BUA59_RS18830 | 76459..79287 | 293552 | DUF499 -domain-containing-protein-[Marinomonas-polaris] |
| WP_116685595.1 | DUF499 _1.RM.like.systems | HTH+WYL-> TUDOR+ SF1-SF2-Helicase+iREase+CTD+REase -> OLD-ABC-> HNH -> METHYLASE -> DUF499 *-><-?<-?\|\|?->TIR-> | 935 | Bacteria>Pseudomonadota>Gammaproteobacteria>Oceanospirillales>Oceanospirillales_incertae_sedis | GCF_003097315.1 | Pelagibaculum_spongiae | DC094_RS03025 | 690947..693754 | 2080658 | DUF499 -domain-containing-protein-[Pelagibaculum-spongiae] |
| AKV97706.1 | DUF499 _1.RM.like.systems | ParB ->?->REase->?->DDE_Tnp_1_4->VirB4-FtsK+HerA-><- DUF499 *<- METHYLASE -p\|\|?-><- HNH <-OLD-ABC <- TUDOR+ SF1-SF2-Helicase+iREase+CTD+REase <-XIS-HTH | 951 | Bacteria>Pseudomonadota>Gammaproteobacteria>Pseudomonadales>Marinobacteraceae | GCA_001266795.1 | Marinobacter_sp._CP1 | ACP86_16945 | 3679225..3682080 | 1671721 | hypothetical-protein-ACP86_16945-[Marinobacter-sp.-CP1] |
| WP_200905073.1 | DUF499 _1.RM.like.systems | REase->?->DDE_Tnp_1_4->?->VirB4-FtsK+HerA-><- DUF499 *<- METHYLASE -p\|\|?-><- METHYLASE <- HNH <-OLD-ABC <- TUDOR+ SF1-SF2-Helicase+iREase+CTD+REase | 935 | Bacteria>Pseudomonadota>Gammaproteobacteria>Pseudomonadales>Marinobacteraceae | GCF_001266795.1 | Marinobacter_sp._CP1 | ACP86_RS16950 | 3679225..3682032 | 1671721 | DUF499 -domain-containing-protein-[Marinobacter-sp.-CP1] |
| WP_091985110.1 | DUF499 _1.RM.like.systems | TUDOR+ SF1-SF2-Helicase+iREase+CTD+REase -> OLD-ABC-> HNH -> METHYLASE ->MAD-NTD+EVE+McrB-AAA->McrC-NTD+REase-> DUF499 *->SAM-methylase->HTH+ HNH -><-?<-CHTH | 947 | Bacteria>Pseudomonadota>Gammaproteobacteria>Pseudomonadales>Marinobacteraceae | GCF_900115175.1 | Marinobacter_gudaonensis | BM344_RS01255 | 264147..266990 | 375760 | DUF499 -domain-containing-protein-[Marinobacter-gudaonensis] |
| MBI8725677.1 | DUF499 _1.RM.like.systems | TUDOR+ SF1-SF2-Helicase+iREase+CTD+REase -> OLD-ABC-> HNH -> METHYLASE ->SF1-SF2-Helicase+wHTH+PRTase->?-> DUF499 *-><-RlpA<-?<-?\|\|HTH_3-><-ZnR+ZnR | 952 | Bacteria>Pseudomonadota>Gammaproteobacteria>Pseudomonadales>Pseudomonadaceae | GCA_016342885.1 | Pseudomonas_aeruginosa | JEZ69_03815 | 204270..207128 | 287 | ATP-binding-protein-[Pseudomonas-aeruginosa] |
| HCL3757805.1 | DUF499 _1.RM.like.systems | TUDOR+ SF1-SF2-Helicase+iREase+CTD+REase -> OLD-ABC-> HNH -> METHYLASE ->OLD-ABC->TOPRIM-> DUF499 *-> | 667 | Bacteria>Pseudomonadota>Gammaproteobacteria>Pseudomonadales>Pseudomonadaceae | GCA_025003865.1 | Pseudomonas_aeruginosa | N2B17_005733 | 10859..12860 | 287 | ATP-binding-protein-[Pseudomonas-aeruginosa] |
| WCY08688.1 | DUF499 _1.RM.like.systems | TUDOR+ SF1-SF2-Helicase+iREase+CTD+REase -> OLD-ABC-> HNH ->SF1-SF2-Helicase+wHTH+PRTase->?-> DUF499 *-><-RlpA<-?<-JAB<- ParB <-?\|\|MazE_antitoxin-> | 943 | Bacteria>Pseudomonadota>Gammaproteobacteria>Pseudomonadales>Pseudomonadaceae | GCA_028609905.1 | Pseudomonas_aeruginosa | KK229_11705 | 2446793..2449624 | 287 | DUF499 -domain-containing-protein-[Pseudomonas-aeruginosa] |
| WP_015271883.1 | DUF499 _1.RM.like.systems | DUF932->?-> ParB ->JAB-><-?\|\|RlpA-><- DUF499 *<-?<-SF1-SF2-Helicase+wHTH+PRTase<- METHYLASE <- HNH <-OLD-ABC <- TUDOR+ SF1-SF2-Helicase+iREase+CTD+REase | 943 | Bacteria>Pseudomonadota>Gammaproteobacteria>Pseudomonadales>Pseudomonadaceae | GCF_000325725.1 | Pseudomonas | LU639_RS23750 | 5103653..5106484 | 286 | MULTISPECIES:- DUF499 -domain-containing-protein-[Pseudomonas] |
| WP_033888172.1 | DUF499 _1.RM.like.systems | TUDOR+ SF1-SF2-Helicase+iREase+CTD+REase -> OLD-ABC-> HNH -> METHYLASE ->SF1-SF2-Helicase+wHTH+PRTase->?-> DUF499 *-><-RlpA<-SymE_toxin<-?<-ZNR+Pkinase+SF1-SF2-Helicase+REase | 942 | Bacteria>Pseudomonadota>Gammaproteobacteria>Pseudomonadales>Pseudomonadaceae | GCF_000506025.1 | Pseudomonas_aeruginosa | HB728_RS10975 | 2312198..2315026 | 287 | DUF499 -domain-containing-protein-[Pseudomonas-aeruginosa] |
| WP_031691594.1 | DUF499 _1.RM.like.systems | ZnR+ZnR-><-HTH_3\|\|?->?->RlpA-><- DUF499 *<-?<-SF1-SF2-Helicase+wHTH+PRTase<- METHYLASE <- HNH <-OLD-ABC <- TUDOR+ SF1-SF2-Helicase+iREase+CTD+REase | 954 | Bacteria>Pseudomonadota>Gammaproteobacteria>Pseudomonadales>Pseudomonadaceae | GCF_000629125.1 | Pseudomonas_aeruginosa | AJ70_RS18015 | 30417..33281 | 287 | DUF499 -domain-containing-protein-[Pseudomonas-aeruginosa] |
| WP_033983017.1 | DUF499 _1.RM.like.systems | TUDOR+ SF1-SF2-Helicase+iREase+CTD+REase -> OLD-ABC-> HNH -> METHYLASE ->SF1-SF2-Helicase+wHTH+PRTase->?-> DUF499 *-><-RlpA<-?<-JAB<- ParB <-?\|\|MazE_antitoxin-> | 943 | Bacteria>Pseudomonadota>Gammaproteobacteria>Pseudomonadales>Pseudomonadaceae | GCF_000789805.1 | Pseudomonas_aeruginosa | NR66_RS02500 | 146772..149603 | 287 | DUF499 -domain-containing-protein-[Pseudomonas-aeruginosa] |
| WP_064616121.1 | DUF499 _1.RM.like.systems | HTH+WYL-> TUDOR+ SF1-SF2-Helicase+iREase+CTD+REase -> OLD-ABC-> HNH -> METHYLASE -> DUF499 *-><-RlpA<-Phage_int_SAM<-Integrase_1<-?<-TOPRIM | 931 | Bacteria>Pseudomonadota>Gammaproteobacteria>Pseudomonadales>Pseudomonadaceae | GCF_001655615.1 | Pseudomonas_sp._GR_6-02 | PGR6_RS03540 | 827904..830699 | 1659194 | DUF499 -domain-containing-protein-[Pseudomonas-sp.-GR-6-02] |
| WP_104882974.1 | DUF499 _1.RM.like.systems | DUF932->?-> ParB ->JAB->RlpA-><- DUF499 *<-?<-SF1-SF2-Helicase+wHTH+PRTase<- METHYLASE <- HNH <-OLD-ABC <- TUDOR+ SF1-SF2-Helicase+iREase+CTD+REase | 943 | Bacteria>Pseudomonadota>Gammaproteobacteria>Pseudomonadales>Pseudomonadaceae | GCF_002947915.1 | Pseudomonas_sp._SWI44 | C4Q26_RS08410 | 1754987..1757818 | 2083053 | DUF499 -domain-containing-protein-[Pseudomonas-sp.-SWI44] |
| WP_126634673.1 | DUF499 _1.RM.like.systems | <-DDE_Tnp_1\|\|?-><-LZ_Tnp_IS66+zf-IS66+DDE_Tnp_IS66+DDE_Tnp_IS66_C<-?<-HIN-HTH\|\|RlpA-><- DUF499 *<-?<-SF1-SF2-Helicase+wHTH+PRTase<- METHYLASE <- HNH <-OLD-ABC <- TUDOR+ SF1-SF2-Helicase+iREase+CTD+REase | 943 | Bacteria>Pseudomonadota>Gammaproteobacteria>Pseudomonadales>Pseudomonadaceae | GCF_003970625.1 | Pseudomonas_aeruginosa | E5D53_RS31320 | 11116..13947 | 287 | DUF499 -domain-containing-protein-[Pseudomonas-aeruginosa] |
| WP_131349395.1 | DUF499 _1.RM.like.systems | TUDOR+ SF1-SF2-Helicase+iREase+CTD+REase -> OLD-ABC-> HNH -> METHYLASE ->OLD-ABC->REase-> DUF499 *-><-RlpA<-TPR<-?<-?\|\|?-><-Mbetalac | 929 | Bacteria>Pseudomonadota>Gammaproteobacteria>Pseudomonadales>Pseudomonadaceae | GCF_004327885.1 | Azotobacter_chroococcum | E0E52_RS08250 | 59995..62784 | 353 | DUF499 -domain-containing-protein-[Azotobacter-chroococcum] |
| WP_135265733.1 | DUF499 _1.RM.like.systems | TUDOR+ SF1-SF2-Helicase+iREase+CTD+REase -> OLD-ABC-> HNH -> METHYLASE ->SF1-SF2-Helicase+wHTH+PRTase->?-> DUF499 *-><-RlpA | 943 | Bacteria>Pseudomonadota>Gammaproteobacteria>Pseudomonadales>Pseudomonadaceae | GCF_004683755.1 | Stutzerimonas_stutzeri | AK6_RS01615 | 334422..337253 | 316 | DUF499 -domain-containing-protein-[Stutzerimonas-stutzeri] |
| WP_146538006.1 | DUF499 _1.RM.like.systems | TUDOR+ SF1-SF2-Helicase+iREase+CTD+REase -> OLD-ABC-> HNH -> METHYLASE ->SF1-SF2-Helicase+wHTH+PRTase->?-> DUF499 *-><-RlpA | 942 | Bacteria>Pseudomonadota>Gammaproteobacteria>Pseudomonadales>Pseudomonadaceae | GCF_007896855.1 | Pseudomonas_aeruginosa | FQA91_RS32330 | 37996..40824 | 287 | DUF499 -domain-containing-protein-[Pseudomonas-aeruginosa] |
| WP_151883645.1 | DUF499 _1.RM.like.systems | TUDOR+ SF1-SF2-Helicase+iREase+CTD+REase -> OLD-ABC-> HNH -> METHYLASE -> DUF499 *-><-RlpA\|\|?-><-JAB<- ParB <-?<-DUF932 | 943 | Bacteria>Pseudomonadota>Gammaproteobacteria>Pseudomonadales>Pseudomonadaceae | GCF_009012945.1 | Pseudomonas | GB470_RS16305 | 7805..10636 | 286 | MULTISPECIES:- DUF499 -domain-containing-protein-[Pseudomonas] |
| WP_103518020.1 | DUF499 _1.RM.like.systems | <-HTH+WYL\|\|?-> TUDOR+ SF1-SF2-Helicase+iREase+CTD+REase -> OLD-ABC-> HNH -> METHYLASE -> DUF499 *-><-RlpA\|\|?-><-JAB<- ParB <-?<-DUF932 | 942 | Bacteria>Pseudomonadota>Gammaproteobacteria>Pseudomonadales>Pseudomonadaceae | GCF_013305665.1 | Pseudomonas | CW309_RS13950 | 90223..93051 | 286 | MULTISPECIES:- DUF499 -domain-containing-protein-[Pseudomonas] |
| WP_093986953.1 | DUF499 _1.RM.like.systems | TUDOR+ SF1-SF2-Helicase+iREase+CTD+REase -> OLD-ABC-> HNH -> METHYLASE ->SF1-SF2-Helicase+wHTH+PRTase->?-> DUF499 *-><-RlpA<-JAB<- ParB | 943 | Bacteria>Pseudomonadota>Gammaproteobacteria>Pseudomonadales>Pseudomonadaceae | GCF_014635625.1 | Pseudomonas_fluvialis | CFY91_RS13770 | 27997..30828 | 1793966 | DUF499 -domain-containing-protein-[Pseudomonas-fluvialis] |
| WP_192907661.1 | DUF499 _1.RM.like.systems | TUDOR+ SF1-SF2-Helicase+iREase+CTD+REase -> OLD-ABC-> HNH -> METHYLASE ->HEPN->?-> DUF499 *-><-RlpA\|\|SF1-SF2-helicase+RdgC+Beta-barrel+REase-><-REase<-Phage_int_SAM<-Integrase_1<-Arm-DNA-bind_3+Phage_integrase | 941 | Bacteria>Pseudomonadota>Gammaproteobacteria>Pseudomonadales>Pseudomonadaceae | GCF_014883875.1 | Pseudomonas_taiwanensis | ICN73_RS10045 | 2229206..2232031 | 470150 | DUF499 -domain-containing-protein-[Pseudomonas-taiwanensis] |
| WP_232392569.1 | DUF499 _1.RM.like.systems | DUF932->?-> ParB ->JAB->RlpA-><- DUF499 *<- METHYLASE <- HNH <-OLD-ABC <- TUDOR+ SF1-SF2-Helicase+iREase+CTD+REase -p<- TUDOR+ SF1-SF2-Helicase+iREase+CTD+REase <-HTH+WYL | 921 | Bacteria>Pseudomonadota>Gammaproteobacteria>Pseudomonadales>Pseudomonadaceae | GCF_021228675.1 | Pseudomonas_sp._NIBR-H-19 | LS633_RS06540 | 1376164..1378929 | 2901380 | DUF499 -domain-containing-protein-[Pseudomonas-sp.-NIBR-H-19] |
| WP_232885556.1 | DUF499 _1.RM.like.systems | <-HTH+WYL\|\|?-> TUDOR+ SF1-SF2-Helicase+iREase+CTD+REase -> OLD-ABC-> HNH -> METHYLASE -> DUF499 *-><-RlpA\|\|?-><-JAB<- ParB <-?<-DUF932 | 943 | Bacteria>Pseudomonadota>Gammaproteobacteria>Pseudomonadales>Pseudomonadaceae | GCF_021282765.1 | Pseudomonas | QK328_RS05450 | 308623..311454 | 286 | DUF499 -domain-containing-protein-[Pseudomonas-sp.-NMI795_08] |
| WP_232896897.1 | DUF499 _1.RM.like.systems | DUF932->?-> ParB ->JAB-><-?\|\|RlpA-><- DUF499 *<-?<-SF1-SF2-Helicase+wHTH+PRTase<- METHYLASE <- HNH <-OLD-ABC <- TUDOR+ SF1-SF2-Helicase+iREase+CTD+REase | 928 | Bacteria>Pseudomonadota>Gammaproteobacteria>Pseudomonadales>Pseudomonadaceae | GCF_021283465.2 | Pseudomonas | LU687_RS25005 | 5299214..5302000 | 286 | MULTISPECIES:- DUF499 -domain-containing-protein-[Pseudomonas] |
| WP_213870614.1 | DUF499 _1.RM.like.systems | JAB->?-><-HTH_3\|\|?->?->RlpA-><- DUF499 *<-HEPN<- METHYLASE <- HNH <-OLD-ABC <- TUDOR+ SF1-SF2-Helicase+iREase+CTD+REase | 952 | Bacteria>Pseudomonadota>Gammaproteobacteria>Pseudomonadales>Pseudomonadaceae | GCF_029201245.1 | Pseudomonas_aeruginosa | KKT79_RS19565 | 32215..35073 | 287 | DUF499 -domain-containing-protein-[Pseudomonas-aeruginosa] |
| WP_276226492.1 | DUF499 _1.RM.like.systems | TUDOR+ SF1-SF2-Helicase+iREase+CTD+REase -> OLD-ABC-> HNH -> METHYLASE ->SF1-SF2-Helicase+wHTH+PRTase->?-> DUF499 *-><-RlpA\|\|<- METHYLASE \|\|HTH->SbcC-> | 939 | Bacteria>Pseudomonadota>Gammaproteobacteria>Pseudomonadales>Pseudomonadaceae | GCF_029335315.1 | Pseudomonas_citronellolis | P3W75_RS06225 | 49283..52102 | 53408 | DUF499 -domain-containing-protein-[Pseudomonas-citronellolis] |
| WP_128722906.1 | DUF499 _1.RM.like.systems | TUDOR+ SF1-SF2-Helicase+iREase+CTD+REase -> OLD-ABC-> HNH -> METHYLASE ->SF1-SF2-Helicase+wHTH+PRTase->?-> DUF499 *-><-RlpA<-SymE_toxin<-?<-ZNR+Pkinase+SF1-SF2-Helicase+REase | 942 | Bacteria>Pseudomonadota>Gammaproteobacteria>Pseudomonadales>Pseudomonadaceae | GCF_029954295.1 | Pseudomonas_aeruginosa | OLM70_RS05770 | 1186505..1189333 | 287 | DUF499 -domain-containing-protein-[Pseudomonas-aeruginosa] |
| WP_285534180.1 | DUF499 _1.RM.like.systems | ZnR+ZnR-><-HTH_3\|\|?->?->RlpA-><- DUF499 *<-?<-SF1-SF2-Helicase+wHTH+PRTase<- METHYLASE <- HNH <-OLD-ABC <- TUDOR+ SF1-SF2-Helicase+iREase+CTD+REase | 942 | Bacteria>Pseudomonadota>Gammaproteobacteria>Pseudomonadales>Pseudomonadaceae | GCF_030269585.1 | Stutzerimonas_stutzeri | QSJ82_RS21895 | 7754..10582 | 316 | DUF499 -domain-containing-protein-[Stutzerimonas-stutzeri] |
| WP_124078188.1 | DUF499 _1.RM.like.systems | ZnR+ZnR-><-HTH_3\|\|?->?->RlpA-><- DUF499 *<-TOPRIM<-OLD-ABC<- METHYLASE <- HNH <-OLD-ABC <- TUDOR+ SF1-SF2-Helicase+iREase+CTD+REase | 943 | Bacteria>Pseudomonadota>Gammaproteobacteria>Pseudomonadales>Pseudomonadaceae | GCF_033140865.1 | Pseudomonas_aeruginosa | IPC1239_RS13835 | 644387..647218 | 287 | DUF499 -domain-containing-protein-[Pseudomonas-aeruginosa] |
| WP_090941615.1 | DUF499 _1.RM.like.systems | TUDOR+ SF1-SF2-Helicase+iREase+CTD+REase -> OLD-ABC-> HNH -> METHYLASE ->OLD-ABC->REase-> DUF499 *-><-RlpA<-?<-?<-ZnR+TOPRIM+D5ATPase | 929 | Bacteria>Pseudomonadota>Gammaproteobacteria>Pseudomonadales>Pseudomonadaceae | GCF_900114395.1 | Azotobacter_beijerinckii | BM240_RS15650 | 17867..20656 | 170623 | DUF499 -domain-containing-protein-[Azotobacter-beijerinckii] |
| WP_280091279.1 | DUF499 _1.RM.like.systems | Arm-DNA-bind_3+Phage_integrase->Phage_int_SAM-><- HNH +SAVED<-SF1-SF2-helicase+RdgC+Beta-barrel+REase\|\|RlpA-><- DUF499 *<-HEPN<- METHYLASE <- HNH <-OLD-ABC <- TUDOR+ SF1-SF2-Helicase+iREase+CTD+REase | 941 | Bacteria>Pseudomonadota>Gammaproteobacteria>Pseudomonadales>Pseudomonadaceae | GCF_947311875.1 | Pseudomonas | QME24_RS13335 | 66050..68875 | 286 | DUF499 -domain-containing-protein-[Pseudomonas-juntendi] |
| WP_029406945.1 | DUF499 _1.RM.like.systems | Phage_integrase-><-GDResolvase+HIN-HTH<- DUF499 *<- METHYLASE <- HNH <-OLD-ABC <- TUDOR+ SF1-SF2-Helicase+iREase+CTD+REase <-?<-HTH | 941 | Bacteria>Pseudomonadota>Gammaproteobacteria>Thiotrichales>Piscirickettsiaceae | GCF_000702325.1 | Thiomicrorhabdus_sp._Milos-T2 | BS34_RS0102315 | 537290..540115 | 90814 | DUF499 -domain-containing-protein-[Thiomicrorhabdus-sp.-Milos-T2] |
| WP_020558458.1 | DUF499 _1.RM.like.systems | 35exo->?->?->?-><- DUF499 *<- METHYLASE <- HNH <-OLD-ABC <- TUDOR+ SF1-SF2-Helicase+iREase+CTD+REase <-SF1-SF2-Helicase | 942 | Bacteria>Pseudomonadota>Gammaproteobacteria>Thiotrichales>Thiofilaceae | GCF_000380185.1 | Thiofilum_flexile | A3IK_RS0102855 | 95361..98189 | 125627 | DUF499 -domain-containing-protein-[Thiofilum-flexile] |
| SEH06532.1 | DUF499 _1.RM.like.systems | TUDOR+ SF1-SF2-Helicase+iREase+CTD+REase -> OLD-ABC-> HNH -> METHYLASE -> DUF499 *->?->?->?->ParA-Soj-PloopNTPase-> | 934 | Bacteria>Pseudomonadota>Gammaproteobacteria>Thiotrichales>Thiotrichaceae | GCA_900099115.2 | Candidatus_Venteria_ishoeyi | MBHS_02395 | 35251..38055 | 1899563 | Uncharacterised-protein-[Thiotrichales-bacterium-HS_08] |
| WP_109837975.1 | DUF499 _1.RM.like.systems | <-ORC-CDC6+REase\|\| TUDOR+ SF1-SF2-Helicase+iREase+CTD+REase -> OLD-ABC-> HNH -> METHYLASE -> DUF499 *-> | 944 | Bacteria>Pseudomonadota>Gammaproteobacteria>Thiotrichales>Thiotrichaceae | GCF_003172875.1 | Leucothrix_pacifica | DKW60_RS12435 | 56874..59708 | 1247513 | DUF499 -domain-containing-protein-[Leucothrix-pacifica] |
| WP_103920297.1 | DUF499 _1.RM.like.systems | TUDOR+ SF1-SF2-Helicase+iREase+CTD+REase -> OLD-ABC-> HNH -> METHYLASE -> DUF499 *->?->?->ParA-Soj-PloopNTPase-> | 934 | Bacteria>Pseudomonadota>Gammaproteobacteria>Thiotrichales>Thiotrichaceae | GCF_900099115.1 | Candidatus_Venteria_ishoeyi | MBHS_RS13590 | 35251..38055 | 1899563 | DUF499 -domain-containing-protein-[Candidatus-Venteria-ishoeyi] |
| WP_078924154.1 | DUF499 _1.RM.like.systems | TUDOR+ SF1-SF2-Helicase+iREase+CTD+REase -> OLD-ABC-> HNH -> METHYLASE -> DUF499 *-> | 937 | Bacteria>Pseudomonadota>Gammaproteobacteria>Thiotrichales>Thiotrichaceae | GCF_900167255.1 | Thiothrix_eikelboomii | B5D50_RS18550 | 28151..30964 | 92487 | DUF499 -domain-containing-protein-[Thiothrix-eikelboomii] |
| MCL4555400.1 | DUF499 _1.RM.like.systems | <- DUF499 *<-?<- METHYLASE <- HNH | 513 | Bacteria>Pseudomonadota>Gammaproteobacteria>unclassified_Gammaproteobacteria | GCA_023382035.1 | Gammaproteobacteria_bacterium | M1572_00725 | 1..1540 | 1913989 | DUF499 -domain-containing-protein-[Gammaproteobacteria-bacterium] |
| TBT35889.1 | DUF499 _1.RM.like.systems | Arm-DNA-bind_3+Phage_integrase->RlpA-><- DUF499 *<- TUDOR+ SF1-SF2-Helicase+iREase+CTD+REase -p<- HNH <-OLD-ABC <-HTH+WYL<-XIS-HTH\|\|JAB-> | 940 | Bacteria>Pseudomonadota>Gammaproteobacteria>Vibrionales>Vibrionaceae | GCA_004324405.1 | Vibrio_parahaemolyticus | D5E85_08495 | 430725..433547 | 670 | ATP-binding-protein-[Vibrio-parahaemolyticus] |
| HCH6462476.1 | DUF499 _1.RM.like.systems | XIS-HTH->HTH+WYL-> TUDOR+ SF1-SF2-Helicase+iREase+CTD+REase -> OLD-ABC-> HNH -> METHYLASE -> DUF499 *-><-RlpA<-Arm-DNA-bind_3+Phage_integrase | 940 | Bacteria>Pseudomonadota>Gammaproteobacteria>Vibrionales>Vibrionaceae | GCA_024102875.1 | Vibrio_parahaemolyticus | NK401_003629 | 41421..44243 | 670 | ATP-binding-protein-[Vibrio-parahaemolyticus] |
| EJL6982897.1 | DUF499 _1.RM.like.systems | XIS-HTH->HTH+WYL-> TUDOR+ SF1-SF2-Helicase+iREase+CTD+REase -> OLD-ABC-> HNH -> METHYLASE -> DUF499 *-><-RlpA<-Arm-DNA-bind_3+Phage_integrase | 915 | Bacteria>Pseudomonadota>Gammaproteobacteria>Vibrionales>Vibrionaceae | GCA_024219855.1 | Vibrio_cholerae | NMT40_002449 | 120290..123037 | 666 | ATP-binding-protein-[Vibrio-cholerae] |
| EJR0961152.1 | DUF499 _1.RM.like.systems | Arm-DNA-bind_3+Phage_integrase->RlpA-><- DUF499 *<- METHYLASE <- HNH <-OLD-ABC <- TUDOR+ SF1-SF2-Helicase+iREase+CTD+REase <-HTH+WYL<-XIS-HTH | 940 | Bacteria>Pseudomonadota>Gammaproteobacteria>Vibrionales>Vibrionaceae | GCA_024870505.1 | Vibrio_parahaemolyticus | NYK10_002560 | 278932..281754 | 670 | ATP-binding-protein-[Vibrio-parahaemolyticus] |
| WP_213964958.1 | DUF499 _1.RM.like.systems | XIS-HTH->HTH+WYL-> TUDOR+ SF1-SF2-Helicase+iREase+CTD+REase -> OLD-ABC-> HNH -> METHYLASE -> DUF499 *-><-RlpA<-Arm-DNA-bind_3+Phage_integrase | 940 | Bacteria>Pseudomonadota>Gammaproteobacteria>Vibrionales>Vibrionaceae | GCF_018457925.1 | Vibrio_alginolyticus | J4H38_RS12950 | 94691..97513 | 663 | DUF499 -domain-containing-protein-[Vibrio-alginolyticus] |
| WP_062660547.1 | DUF499 _1.RM.like.systems | HTH+WYL-> TUDOR+ SF1-SF2-Helicase+iREase+CTD+REase -> OLD-ABC-> HNH -> TUDOR+ SF1-SF2-Helicase+iREase+CTD+REase -p-> METHYLASE -> DUF499 *-><-RlpA<-?\|\|?->?-><- HNH <-OBFOLD-NUCLEASE | 947 | Bacteria>Pseudomonadota>Gammaproteobacteria>Vibrionales>Vibrionaceae | GCF_900055185.1 | Grimontia_celer | GCE9029_RS00270 | 67941..70784 | 1796497 | DUF499 -domain-containing-protein-[Grimontia-celer] |
| WP_254620468.1 | DUF499 _1.RM.like.systems | Arm-DNA-bind_3+Phage_integrase->RlpA-><- DUF499 *<- METHYLASE <- HNH <-OLD-ABC <- TUDOR+ SF1-SF2-Helicase+iREase+CTD+REase <-HTH+WYL<-XIS-HTH | 856 | Bacteria>Pseudomonadota>Gammaproteobacteria>Vibrionales>Vibrionaceae | - | Vibrio_cholerae | KVS75_RS02420 | 551801..554371 | 666 | DUF499 -domain-containing-protein-[Vibrio-cholerae] |

<table>
<tr><th>Protein-ID</th><th>System</th><th>Gene-Neighbourhoods</th><th>Length</th><th>Taxonomy</th><th>Assembly-ID</th><th>Strain</th><th>Locus-Tag</th><th>Genomic-coordinates</th><th>TaxID</th><th>Fasta-defline</th></tr>
<tr><td>MCC7256721.1</td><td>DUF499_1.RM.like.systems</td><td><-KAP-NTPase<-?<- DUF499 *<-KAP-NTPase<- METHYLASE <- HNH <-OLD-ABC <- TUDOR+ SF1-SF2-Helicase+iREase+CTD+REase <-PIWI</td><td>930</td><td>Bacteria>Pseudomonadota>Gammaproteobacteria>Xanthomonadales>Rhodanobacteraceae</td><td>GCA_020852005.1</td><td>Dokdonella_sp.</td><td>IT467_12405</td><td>2246..5038</td><td>2291710</td><td>ATP-binding-protein-[Dokdonella-sp.]</td></tr>
<tr><td>MCX7197671.1</td><td>DUF499_1.RM.like.systems</td><td>TUDOR+ SF1-SF2-Helicase+iREase+CTD+REase -p-> METHYLASE ->HTH+Hiskin+STAS-> HEPN+ DUF499 *->?-> HNH ->SIR2->DnaB-></td><td>1113</td><td>Bacteria>Pseudomonadota>unclassified_Pseudomonadota</td><td>GCA_026399745.1</td><td>Pseudomonadota_bacterium</td><td>NTW37_06695</td><td>4467..7808</td><td>1977087</td><td>MAG:-Swt1-family-HEPN-domain-containing-protein-[Proteobacteria-bacterium]</td></tr>
<tr><td>MBQ6473657.1</td><td>DUF499_1.RM.like.systems</td><td>Schlafen+HATPase+HTH-> OLD-ABC-> HNH -> METHYLASE ->OLD-ABC->NERD-> DUF499 *-></td><td>944</td><td>Bacteria>PVC_group>Lentisphaerota>Lentisphaeria>Victivallales</td><td>GCA_017431185.1</td><td>Victivallales_bacterium</td><td>IJJ33_16845</td><td>18968..21802</td><td>2212475</td><td>ATP-binding-protein-[Victivallales-bacterium]</td></tr>
<tr><td>WP_168963703.1</td><td>DUF499_1.RM.like.systems</td><td>XIS-HTH-><-?<-Phage_integrase||ORC-CDC6+REase->?-><-?<- DUF499 *<-NERD<-OLD-ABC<- METHYLASE <- HNH <-OLD-ABC <- TUDOR+ SF1-SF2-Helicase+iREase+CTD+REase</td><td>935</td><td>Bacteria>PVC_group>Lentisphaerota>Lentisphaeria>Victivallales</td><td>GCF_012843305.1</td><td>Victivallis_vadensis</td><td>HF882_RS18870</td><td>10950..13757</td><td>172901</td><td>DUF499-domain-containing-protein-[Victivallis-vadensis]</td></tr>
<tr><td>MBE6357833.1</td><td>DUF499_1.RM.like.systems</td><td>ParB -HTH->?-><-?<-?<- DUF499 *<-REase<-OLD-ABC<- METHYLASE <- HNH <-OLD-ABC <- TUDOR+ SF1-SF2-Helicase+iREase+CTD+REase</td><td>938</td><td>Bacteria>PVC_group>Lentisphaerota>unclassified_Lentisphaerota</td><td>GCA_015061525.1</td><td>Lentisphaerota_bacterium</td><td>E7057_01125</td><td>3177..5993</td><td>1932692</td><td>ATP-binding-protein-[Lentisphaerae-bacterium]</td></tr>
<tr><td>MBC8553109.1</td><td>DUF499_1.RM.like.systems</td><td>XIS-HTH-> TUDOR+ SF1-SF2-Helicase+iREase+CTD+REase -> METHYLASE -> HEPN+ DUF499 *-> HNH -></td><td>1106</td><td>Bacteria>PVC_group>Planctomycetota>Candidatus_Brocadiia>Candidatus_Brocadiales</td><td>GCA_014384865.1</td><td>Candidatus_Brocadiales_bacterium</td><td>H8D23_26105</td><td>6653..9973</td><td>2760054</td><td>ATP-binding-protein-[Candidatus-Brocadiales-bacterium]</td></tr>
<tr><td>MBX3408154.1</td><td>DUF499_1.RM.like.systems</td><td>RAMA->GDResolvase+Recombinase+ZnR-> ParB -><-?<-Radical-SAM<- DUF499 *<- METHYLASE <- HNH <-OLD-ABC <- TUDOR+ SF1-SF2-Helicase+iREase+CTD+REase</td><td>858</td><td>Bacteria>PVC_group>Planctomycetota>Phycisphaerae>Phycisphaerales</td><td>GCA_019636415.1</td><td>Phycisphaeraceae_bacterium</td><td>KF869_15485</td><td>6654..9230</td><td>2026777</td><td>ATP-binding-protein-[Phycisphaeraceae-bacterium]</td></tr>
<tr><td>MBX3409892.1</td><td>DUF499_1.RM.like.systems</td><td>RAMA->GDResolvase+Recombinase+ZnR-> ParB -><-?<-Radical-SAM<- DUF499 *<- METHYLASE <- HNH <-OLD-ABC <- TUDOR+ SF1-SF2-Helicase+iREase+CTD+REase ||HTH->MPTase-></td><td>947</td><td>Bacteria>PVC_group>Planctomycetota>Phycisphaerae>Phycisphaerales</td><td>GCA_019636575.1</td><td>Phycisphaeraceae_bacterium</td><td>KF859_08400</td><td>137437..140280</td><td>2026777</td><td>ATP-binding-protein-[Phycisphaeraceae-bacterium]</td></tr>
<tr><td>OYP37234.1</td><td>DUF499_1.RM.like.systems</td><td><- wHTH(SBD)+ HNH <-URI-DNase+RAMA<-REase<-EVE<-REase<- DUF499 *<- HNH <-OLD-ABC <- METHYLASE <-Peptidase_S8<-Classical-AAA<- TUDOR+ SF1-SF2-Helicase+iREase+CTD+REase</td><td>948</td><td>Bacteria>PVC_group>Planctomycetota>Planctomycetia>Pirellulales</td><td>GCA_002251265.1</td><td>Rhodopirellula_sp._MGV</td><td>CGZ80_05855</td><td>9695..12541</td><td>2023130</td><td>hypothetical-protein-CGZ80_05855-[Rhodopirellula-sp.-MGV]</td></tr>
<tr><td>NLX95047.1</td><td>DUF499_1.RM.like.systems</td><td>TPR+TPR->?->?-><-?<-PHP+SbcC<- HEPN+ DUF499 *<- METHYLASE <- HNH <-OLD-ABC <- TUDOR+ SF1-SF2-Helicase+iREase+CTD+REase -p</td><td>1104</td><td>Bacteria>PVC_group>Planctomycetota>Planctomycetia>Pirellulales</td><td>GCA_012728855.1</td><td>Rhodopirellula_sp.</td><td>GXY83_02605</td><td>12933..16247</td><td>2024855</td><td>ATP-binding-protein-[Rhodopirellula-sp.]</td></tr>
<tr><td>WP_198953633.1</td><td>DUF499_1.RM.like.systems</td><td><- wHTH(SBD)+ HNH <-URI-DNase+RAMA<-REase<-EVE<-REase<- DUF499 *<- HNH <-OLD-ABC <- METHYLASE <-Peptidase_S8<-Classical-AAA<- TUDOR+ SF1-SF2-Helicase+iREase+CTD+REase</td><td>951</td><td>Bacteria>PVC_group>Planctomycetota>Planctomycetia>Pirellulales</td><td>GCF_002251265.1</td><td>Rhodopirellula_sp._MGV</td><td>CGZ80_RS05865</td><td>9695..12550</td><td>2023130</td><td>DUF499-domain-containing-protein-[Rhodopirellula-sp.-MGV]</td></tr>
<tr><td>MDB5334583.1</td><td>DUF499_1.RM.like.systems</td><td>TUDOR+ SF1-SF2-Helicase+iREase+CTD+REase -> OLD-ABC-> HNH -> METHYLASE -> DUF499 *-></td><td>942</td><td>Bacteria>PVC_group>Planctomycetota>Planctomycetia>Planctomycetales</td><td>GCA_028291605.1</td><td>Planctomycetaceae_bacterium</td><td>JWN70_202</td><td>14767..17595</td><td>2026779</td><td>MAG:-hypothetical-protein-JWN70_202-[Planctomycetaceae-bacterium]</td></tr>
<tr><td>MCC5805073.1</td><td>DUF499_1.RM.like.systems</td><td>OLD-ABC-> HNH -> METHYLASE ->CHTH->RHH->PIN-> HEPN+ DUF499 *->?->PhdYeFM_antitox->PIN-></td><td>1107</td><td>Bacteria>PVC_group>Verrucomicrobiota>Opitutae>Opitutales</td><td>GCA_020831525.1</td><td>Opitutales_bacterium</td><td>JJU00_01980</td><td>48883..52206</td><td>2026772</td><td>DUF499-domain-containing-protein-[Opitutales-bacterium]</td></tr>
<tr><td>TAE90777.1</td><td>DUF499_1.RM.like.systems</td><td><- DUF499 *<-REase<- METHYLASE <- HNH <-OLD-ABC <-TOPRIM<-OLD-ABC</td><td>939</td><td>Bacteria>PVC_group>Verrucomicrobiota>unclassified_Verrucomicrobiota</td><td>GCA_004293285.1</td><td>Verrucomicrobiota_bacterium</td><td>EAZ81_08635</td><td>280..3099</td><td>2026799</td><td>ATP-binding-protein-[Verrucomicrobia-bacterium]</td></tr>
<tr><td>MBI5767835.1</td><td>DUF499_1.RM.like.systems</td><td>HTH-> TUDOR+ SF1-SF2-Helicase+iREase+CTD+REase -> OLD-ABC-> HNH -> METHYLASE -> DUF499 *->Radical-SAM-></td><td>944</td><td>Bacteria>PVC_group>Verrucomicrobiota>unclassified_Verrucomicrobiota</td><td>GCA_016235555.1</td><td>Verrucomicrobiota_bacterium</td><td>HZA93_08560</td><td>101523..104357</td><td>2026799</td><td>ATP-binding-protein-[Verrucomicrobia-bacterium]</td></tr>
<tr><td>MCX6879289.1</td><td>DUF499_1.RM.like.systems</td><td><-HISKIN<- DUF499 *<-REase<-OLD-ABC<- METHYLASE <- HNH <-OLD-ABC <- TUDOR+ SF1-SF2-Helicase+iREase+CTD+REase</td><td>941</td><td>Bacteria>PVC_group>Verrucomicrobiota>unclassified_Verrucomicrobiota</td><td>GCA_026397725.1</td><td>Verrucomicrobiota_bacterium</td><td>NTW21_36620</td><td>9837..12662</td><td>2026799</td><td>MAG:-DUF499-domain-containing-protein-[Verrucomicrobia-bacterium]</td></tr>
<tr><td>MCX6953048.1</td><td>DUF499_1.RM.like.systems</td><td>TUDOR+ SF1-SF2-Helicase+iREase+CTD+REase -> OLD-ABC-> HNH -> METHYLASE ->REase-> DUF499 *->?->?->?->ABhydrolase-></td><td>941</td><td>Bacteria>PVC_group>Verrucomicrobiota>unclassified_Verrucomicrobiota</td><td>GCA_026398065.1</td><td>Verrucomicrobiota_bacterium</td><td>NTV51_12915</td><td>8536..11361</td><td>2026799</td><td>MAG:-DUF499-domain-containing-protein-[Verrucomicrobia-bacterium]</td></tr>
<tr><td>HAO79056.1</td><td>DUF499_1.RM.like.systems</td><td>METHYLASE -p->RNaseH->?-> DUF499 *-><- HNH ||TPR+TPR-></td><td>924</td><td>Bacteria>PVC_group>Verrucomicrobiota>Verrucomicrobiae>Limisphaerales</td><td>GCA_003455565.1</td><td>Limisphaerales_bacterium</td><td>DCQ92_08790</td><td>3206..5980</td><td>2052186</td><td>AAA-family-ATPase-[Verrucomicrobia-subdivision-3-bacterium]</td></tr>
<tr><td>MBL9159968.1</td><td>DUF499_1.RM.like.systems</td><td><-HISKIN<-ArdCN+MPTase<- HNH <- HEPN+ DUF499 *<-?<-Mbetalac<-?<-?<- METHYLASE</td><td>1114</td><td>Bacteria>PVC_group>Verrucomicrobiota>Verrucomicrobiae>Verrucomicrobiales</td><td>GCA_016795385.1</td><td>Verrucomicrobiales_bacterium</td><td>JNJ70_20975</td><td>159944..163288</td><td>2026801</td><td>DUF499-domain-containing-protein-[Verrucomicrobiales-bacterium]</td></tr>
<tr><td>MYH43296.1</td><td>DUF499_1.RM.like.systems</td><td>HNH ->?->?-> TUDOR+ SF1-SF2-Helicase+iREase+CTD+REase -> METHYLASE -> DUF499 *-><-?<-HEPN<-?||?->Aminotran_1_2->ACET+PIN+ACET-></td><td>923</td><td>Bacteria>Terrabacteria_group>Actinomycetota>Acidimicrobiia>Acidimicrobiales</td><td>GCA_009843995.1</td><td>Acidimicrobiaceae_bacterium</td><td>F4131_06280</td><td>9069..11840</td><td>2024894</td><td>ATP-binding-protein-[Acidimicrobiaceae-bacterium]</td></tr>
<tr><td>MBV5322918.1</td><td>DUF499_1.RM.like.systems</td><td>HEPN-> OLD-ABC-> HNH -> METHYLASE -> METHYLASE ->HEPN-> DUF499 *-></td><td>872</td><td>Bacteria>Terrabacteria_group>Actinomycetota>Acidimicrobiia>Acidimicrobiales</td><td>GCA_019162935.1</td><td>Ilumatobacteraceae_bacterium</td><td>JZU60_03690</td><td>12690..15310</td><td>2497619</td><td>ATP-binding-protein-[Ilumatobacteraceae-bacterium]</td></tr>
<tr><td>WP_003839950.1</td><td>DUF499_1.RM.like.systems</td><td>Aldedh->?->?->PRTase->tRNA-> TUDOR+ SF1-SF2-Helicase+iREase+CTD+REase -> METHYLASE -> HEPN+ DUF499 *->?-> HNH ->?->?-><-Peptidase</td><td>1138</td><td>Bacteria>Terrabacteria_group>Actinomycetota>Actinomycetes>Bifidobacteriales</td><td>GCF_000024445.1</td><td>Bifidobacterium_dentium</td><td>BIFDEN_RS08405</td><td>2078532..2081948</td><td>1689</td><td>DUF499-domain-containing-protein-[Bifidobacterium-dentium]</td></tr>
<tr><td>WP_226436803.1</td><td>DUF499_1.RM.like.systems</td><td><-DUF222+ HNH <- HEPN+ DUF499 *<- METHYLASE ||CHTH-><- TUDOR+ SF1-SF2-Helicase+iREase+CTD+REase</td><td>1141</td><td>Bacteria>Terrabacteria_group>Actinomycetota>Actinomycetes</td><td>GCF_012515855.1</td><td>Mycobacteriales</td><td>GX543_RS09675</td><td>1506..4931</td><td>85007</td><td>Swt1-family-HEPN-domain-containing-protein-[Rhodococcus-yananensis]</td></tr>
<tr><td>WP_242328934.1</td><td>DUF499_1.RM.like.systems</td><td>Phage_terminase-> TUDOR+ SF1-SF2-Helicase+iREase+CTD+REase -> METHYLASE -> DUF499 *->?-> HNH ->?->?->REase-></td><td>999</td><td>Bacteria>Terrabacteria_group>Actinomycetota>Actinomycetes>Kitasatosporales</td><td>GCF_022647665.1</td><td>Streptomyces_formicae</td><td>J4032_RS01965</td><td>417762..420761</td><td>1616117</td><td>DUF499-domain-containing-protein-[Streptomyces-formicae]</td></tr>
<tr><td>MBS3179176.1</td><td>DUF499_1.RM.like.systems</td><td>GDResolvase+HIN-HTH-><-?<- HNH || TUDOR+ SF1-SF2-Helicase+iREase+CTD+REase -> METHYLASE -> HEPN+ DUF499 *->?-><-UDG-conflict||?-><-?<-EVE</td><td>1142</td><td>Bacteria>Terrabacteria_group>Actinomycetota>Actinomycetes>Micrococcales</td><td>GCA_018304855.1</td><td>Pseudoclavibacter_sp._Marseille-Q4354</td><td>JSQ82_10750</td><td>821869..825297</td><td>2810515</td><td>DUF499-domain-containing-protein-[Pseudoclavibacter-sp.-Marseille-Q4354]</td></tr>
<tr><td>WP_054682987.1</td><td>DUF499_1.RM.like.systems</td><td>XIS-HTH-> TUDOR+ SF1-SF2-Helicase+iREase+CTD+REase -> METHYLASE ->HIN-HTH-> METHYLASE -p-> HEPN+ DUF499 *->OLD-ABC+TOPRIM-> HNH -><-HIN-HTH</td><td>1129</td><td>Bacteria>Terrabacteria_group>Actinomycetota>Actinomycetes>Micrococcales</td><td>GCF_001314225.1</td><td>Microbacterium_sp._No._7</td><td>AOA12_RS21690</td><td>98036..101425</td><td>1714373</td><td>DUF499-domain-containing-protein-[Microbacterium-sp.-No.-7]</td></tr>
<tr><td>WP_194396698.1</td><td>DUF499_1.RM.like.systems</td><td>METHYLASE -><- HNH <-OLD-ABC +TOPRIM<- HEPN+ DUF499 *<- METHYLASE <- TUDOR+ SF1-SF2-Helicase+iREase+CTD+REase <-XIS-HTH</td><td>1130</td><td>Bacteria>Terrabacteria_group>Actinomycetota>Actinomycetes>Micrococcales</td><td>GCF_015277815.1</td><td>Microbacterium_atlanticum</td><td>IR212_RS15230</td><td>3329381..3332773</td><td>2782168</td><td>DUF499-domain-containing-protein-[Microbacterium-atlanticum]</td></tr>
<tr><td>WP_249312439.1</td><td>DUF499_1.RM.like.systems</td><td>GDResolvase+HIN-HTH-><-?<- HNH || TUDOR+ SF1-SF2-Helicase+iREase+CTD+REase -> METHYLASE -> HEPN+ DUF499 *->?-><-UDG-conflict||?-><-?<-EVE</td><td>1140</td><td>Bacteria>Terrabacteria_group>Actinomycetota>Actinomycetes>Micrococcales</td><td>GCF_018304855.1</td><td>Pseudoclavibacter_sp._Marseille-Q4354</td><td>JSQ82_RS10750</td><td>821875..825297</td><td>2810515</td><td>DUF499-domain-containing-protein-[Pseudoclavibacter-sp.-Marseille-Q4354]</td></tr>
<tr><td>NLG46720.1</td><td>DUF499_1.RM.like.systems</td><td><-DUF222+ HNH <- HEPN+ DUF499 *<- METHYLASE ||CHTH-><- TUDOR+ SF1-SF2-Helicase+iREase+CTD+REase ||HTH-></td><td>1141</td><td>Bacteria>Terrabacteria_group>Actinomycetota>Actinomycetes>Mycobacteriales</td><td>GCA_012515855.1</td><td>Gordonia_sp._(in:_high_G+C_Gram-positive_bacteria)</td><td>GX543_09665</td><td>1506..4931</td><td>84139</td><td>ATP-binding-protein-[Gordonia-sp.-(in:-high-G+C-Gram-positive-bacteria)]</td></tr>
<tr><td>MBB4136224.1</td><td>DUF499_1.RM.like.systems</td><td><-ABhydrolase<-?||?-><-ACET<- HEPN+ DUF499 *<- METHYLASE ||DUF222+ HNH -><- TUDOR+ SF1-SF2-Helicase+iREase+CTD+REase</td><td>1130</td><td>Bacteria>Terrabacteria_group>Actinomycetota>Actinomycetes>Mycobacteriales</td><td>GCA_014197435.1</td><td>Gordonia_humi</td><td>BKA16_002776</td><td>3005759..3009151</td><td>686429</td><td>hypothetical-protein-[Gordonia-humi]</td></tr>
<tr><td>MCF8588060.1</td><td>DUF499_1.RM.like.systems</td><td>TUDOR+ SF1-SF2-Helicase+iREase+CTD+REase -><-DUF222+ HNH || METHYLASE -> HEPN+ DUF499 *->ACET-><-?||?->ABhydrolase-></td><td>1125</td><td>Bacteria>Terrabacteria_group>Actinomycetota>Actinomycetes>Mycobacteriales</td><td>GCA_021738805.1</td><td>Gordonia_liuliyuniae</td><td>L5G33_06210</td><td>131250..134627</td><td>2911517</td><td>DUF499-domain-containing-protein-[Gordonia-liuliyuniae]</td></tr>
<tr><td>MCF8569704.1</td><td>DUF499_1.RM.like.systems</td><td><-ABhydrolase<-?||?-><-ACET<- HEPN+ DUF499 *<- METHYLASE ||DUF222+ HNH -><- TUDOR+ SF1-SF2-Helicase+iREase+CTD+REase</td><td>1115</td><td>Bacteria>Terrabacteria_group>Actinomycetota>Actinomycetes>Mycobacteriales</td><td>GCA_021738965.1</td><td>Gordonia_zhenghanii</td><td>L5G32_05410</td><td>142683..146030</td><td>2911516</td><td>DUF499-domain-containing-protein-[Gordonia-zhenghanii]</td></tr>
<tr><td>MCF8610758.1</td><td>DUF499_1.RM.like.systems</td><td>TUDOR+ SF1-SF2-Helicase+iREase+CTD+REase -><-DUF222+ HNH || METHYLASE -> HEPN+ DUF499 *->ACET-><-?||?->ABhydrolase-></td><td>1125</td><td>Bacteria>Terrabacteria_group>Actinomycetota>Actinomycetes>Mycobacteriales</td><td>GCA_021739025.1</td><td>Gordonia_liuliyuniae</td><td>L5G28_11435</td><td>131354..134731</td><td>2911517</td><td>DUF499-domain-containing-protein-[Gordonia-liuliyuniae]</td></tr>
<tr><td>WP_009680352.1</td><td>DUF499_1.RM.like.systems</td><td>TUDOR+ SF1-SF2-Helicase+iREase+CTD+REase -><-DUF222+ HNH || METHYLASE -> HEPN+ DUF499 *-><-?||ACET-><-Peptidase_S51<-?||?->ABhydrolase-></td><td>1137</td><td>Bacteria>Terrabacteria_group>Actinomycetota>Actinomycetes>Mycobacteriales</td><td>GCF_000192435.1</td><td>Gordonia_neofelifaecis</td><td>SCNU_RS15560</td><td>9488..12901</td><td>945692</td><td>DUF499-domain-containing-protein-[Gordonia-neofelifaecis]</td></tr>
<tr><td>WP_006896364.1</td><td>DUF499_1.RM.like.systems</td><td><-ABhydrolase<-?||?->Peptidase_S51-><-ACET||?-><- HEPN+ DUF499 *<- METHYLASE ||DUF222+ HNH -><- TUDOR+ SF1-SF2-Helicase+iREase+CTD+REase <-?<-tRNA<-tRNA<-tRNA<-?<-tRNA<-tRNA<-tRNA||tRNA->ABC-ATPase+SbcC-></td><td>1124</td><td>Bacteria>Terrabacteria_group>Actinomycetota>Actinomycetes>Mycobacteriales</td><td>GCF_000333035.1</td><td>Gordonia</td><td>GSI01S_RS09635</td><td>144602..147976</td><td>2053</td><td>MULTISPECIES:-DUF499-domain-containing-protein-[Gordonia]</td></tr>
<tr><td>WP_045537629.1</td><td>DUF499_1.RM.like.systems</td><td><-ABhydrolase<-?||?->Peptidase_S51-><-ACET||?-><- HEPN+ DUF499 *<- METHYLASE ||DUF222+ HNH -><- TUDOR+ SF1-SF2-Helicase+iREase+CTD+REase <-?<-tRNA<-tRNA<-tRNA<-?<-tRNA<-tRNA<-tRNA||tRNA->ABC-ATPase+SbcC-></td><td>1124</td><td>Bacteria>Terrabacteria_group>Actinomycetota>Actinomycetes>Mycobacteriales</td><td>GCF_000960465.1</td><td>Gordonia_sihwensis</td><td>UG54_RS06085</td><td>5143..8517</td><td>173559</td><td>DUF499-domain-containing-protein-[Gordonia-sihwensis]</td></tr>
<tr><td>WP_066467289.1</td><td>DUF499_1.RM.like.systems</td><td>TetR-HTH-><-RADICAL-SAM<-?<-?<-?<- HEPN+ DUF499 *<- METHYLASE ||DUF222+ HNH -><- TUDOR+ SF1-SF2-Helicase+iREase+CTD+REase ||HTH_19-></td><td>1118</td><td>Bacteria>Terrabacteria_group>Actinomycetota>Actinomycetes>Mycobacteriales</td><td>GCF_001552855.1</td><td>Skermania_piniformis</td><td>SP4_RS02850</td><td>6456..9812</td><td>39122</td><td>DUF499-domain-containing-protein-[Skermania-piniformis]</td></tr>
<tr><td>WP_066167603.1</td><td>DUF499_1.RM.like.systems</td><td>ACET-> TUDOR+ SF1-SF2-Helicase+iREase+CTD+REase -><-DUF222+ HNH || METHYLASE -> HEPN+ DUF499 *-><-Peptidase_S51<-?||?->ABhydrolase-></td><td>1139</td><td>Bacteria>Terrabacteria_group>Actinomycetota>Actinomycetes>Mycobacteriales</td><td>GCF_001592365.1</td><td>Gordonia_hydrophobica</td><td>GH2_RS11425</td><td>77242..80661</td><td>40516</td><td>DUF499-domain-containing-protein-[Gordonia-hydrophobica]</td></tr>
<tr><td>WP_161926536.1</td><td>DUF499_1.RM.like.systems</td><td><-TetR-HTH<-?||HTH_19-><-?<- HEPN+ DUF499 *<-DUF222+ HNH <- METHYLASE <- TUDOR+ SF1-SF2-Helicase+iREase+CTD+REase ||DOC+HTH-></td><td>1124</td><td>Bacteria>Terrabacteria_group>Actinomycetota>Actinomycetes>Mycobacteriales</td><td>GCF_009932435.1</td><td>Gordonia_crocea</td><td>nbrcactino_RS06035</td><td>1265301..1268675</td><td>589162</td><td>DUF499-domain-containing-protein-[Gordonia-crocea]</td></tr>
<tr><td>WP_083088684.1</td><td>DUF499_1.RM.like.systems</td><td>Recombinase->?->?-><- HEPN+ DUF499 *<- METHYLASE <- TUDOR+ SF1-SF2-Helicase+iREase+CTD+REase ||NTD+SF1-SF2-Helicase->?-><- wHTH(SBD)+ HNH</td><td>1108</td><td>Bacteria>Terrabacteria_group>Actinomycetota>Actinomycetes>Mycobacteriales</td><td>GCF_010731635.1</td><td>Mycobacterium_noviomagense</td><td>BST37_RS15635</td><td>32792..36118</td><td>459858</td><td>DUF499-domain-containing-protein-[Mycobacterium-noviomagense]</td></tr>
<tr><td>WP_170194849.1</td><td>DUF499_1.RM.like.systems</td><td>HTH_3-><-?<-?||?-><-ABhydrolase+ABhydrolase<- HEPN+ DUF499 *||DUF222+ HNH -><-AbiEii<-?||?-><- METHYLASE ||HTH_19-></td><td>1113</td><td>Bacteria>Terrabacteria_group>Actinomycetota>Actinomycetes>Mycobacteriales</td><td>GCF_012933505.1</td><td>Gordonia_asplenii</td><td>HH308_RS14085</td><td>95388..98729</td><td>2725283</td><td>DUF499-domain-containing-protein-[Gordonia-asplenii]</td></tr>
<tr><td>WP_281378415.1</td><td>DUF499_1.RM.like.systems</td><td><-ABhydrolase<-?||?-><- HEPN+ DUF499 *<- METHYLASE ||DUF222+ HNH -><- TUDOR+ SF1-SF2-Helicase+iREase+CTD+REase <-?<-?<-tRNA<-tRNA<-tRNA<-tRNA<-tRNA||tRNA->DDE_Tnp_1_4-></td><td>1142</td><td>Bacteria>Terrabacteria_group>Actinomycetota>Actinomycetes>Mycobacteriales</td><td>GCF_014197435.1</td><td>Gordonia_humi</td><td>BKA16_RS13810</td><td>3005759..3009187</td><td>686429</td><td>DUF499-domain-containing-protein-[Gordonia-humi]</td></tr>
<tr><td>WP_236997292.1</td><td>DUF499_1.RM.like.systems</td><td>TUDOR+ SF1-SF2-Helicase+iREase+CTD+REase -><-DUF222+ HNH || METHYLASE -> HEPN+ DUF499 *->ACET-><-?||?->ABhydrolase-></td><td>1125</td><td>Bacteria>Terrabacteria_group>Actinomycetota>Actinomycetes>Mycobacteriales</td><td>GCF_021738805.1</td><td>Gordonia_liuliyuniae</td><td>L5G33_06210</td><td>131250..134627</td><td>2911517</td><td>DUF499-domain-containing-protein-[Gordonia-sp.-HY366]</td></tr>
<tr><td>WP_237010003.1</td><td>DUF499_1.RM.like.systems</td><td><-ABhydrolase<-?||?-><-ACET<- HEPN+ DUF499 *<- METHYLASE ||DUF222+ HNH -><- TUDOR+ SF1-SF2-Helicase+iREase+CTD+REase</td><td>1115</td><td>Bacteria>Terrabacteria_group>Actinomycetota>Actinomycetes>Mycobacteriales</td><td>GCF_021738965.1</td><td>Gordonia_zhenghanii</td><td>L5G32_05410</td><td>142683..146030</td><td>2911516</td><td>MULTISPECIES:-DUF499-domain-containing-protein-[unclassified-Gordonia]</td></tr>
<tr><td>WP_237001659.1</td><td>DUF499_1.RM.like.systems</td><td>TUDOR+ SF1-SF2-Helicase+iREase+CTD+REase -><-DUF222+ HNH || METHYLASE -> HEPN+ DUF499 *->ACET-><-?||?->ABhydrolase-></td><td>1125</td><td>Bacteria>Terrabacteria_group>Actinomycetota>Actinomycetes>Mycobacteriales</td><td>GCF_021739025.1</td><td>Gordonia_liuliyuniae</td><td>L5G28_11435</td><td>131354..134731</td><td>2911517</td><td>DUF499-domain-containing-protein-[Gordonia-sp.-HY285]</td></tr>
<tr><td>WP_278127257.1</td><td>DUF499_1.RM.like.systems</td><td><-ABhydrolase<-?||?->Peptidase_S51-><-ACET||?-><- HEPN+ DUF499 *<- METHYLASE ||DUF222+ HNH -><- TUDOR+ SF1-SF2-Helicase+iREase+CTD+REase <-?<-tRNA<-tRNA<-tRNA<-?<-tRNA<-tRNA<-tRNA||tRNA->ABC-ATPase+SbcC-></td><td>1124</td><td>Bacteria>Terrabacteria_group>Actinomycetota>Actinomycetes>Mycobacteriales</td><td>GCF_029632925.1</td><td>Gordonia_sihwensis</td><td>P5P27_RS08530</td><td>1762026..1765400</td><td>173559</td><td>DUF499-domain-containing-protein-[Gordonia-sihwensis]</td></tr>
<tr><td>WP_239588858.1</td><td>DUF499_1.RM.like.systems</td><td><-ABhydrolase<-?||?->Peptidase_S51-><- HEPN+ DUF499 *<- METHYLASE ||DUF222+ HNH -><- TUDOR+ SF1-SF2-Helicase+iREase+CTD+REase <-ACET</td><td>1125</td><td>Bacteria>Terrabacteria_group>Actinomycetota>Actinomycetes>Mycobacteriales</td><td>-</td><td>Gordonia_hydrophobica</td><td>JOC45_RS12260</td><td>227782..231159</td><td>40516</td><td>DUF499-domain-containing-protein-[Gordonia-hydrophobica]</td></tr>
<tr><td>WP_260868497.1</td><td>DUF499_1.RM.like.systems</td><td>TUDOR+ SF1-SF2-Helicase+iREase+CTD+REase -><-DUF222+ HNH || METHYLASE -> HEPN+ DUF499 *-><-?||ACET-><-Peptidase_S51<-?||?->ABhydrolase-></td><td>1110</td><td>Bacteria>Terrabacteria_group>Actinomycetota>Actinomycetes>Mycobacteriales</td><td>-</td><td>Gordonia_sihwensis</td><td>ACN95_RS06460</td><td>9637..12969</td><td>173559</td><td>DUF499-domain-containing-protein-[Gordonia-sihwensis]</td></tr>
<tr><td>WP_116057355.1</td><td>DUF499_1.RM.like.systems</td><td><-URI-DNase<-DNA-glycosylase||?-> TUDOR+ SF1-SF2-Helicase+iREase+CTD+REase -> METHYLASE -> HEPN+ DUF499 *->?-><- HNH</td><td>1122</td><td>Bacteria>Terrabacteria_group>Actinomycetota>Actinomycetes>Mycobacteriales</td><td>-</td><td>Rhodococcus_erythropolis</td><td>DVG80_RS18290</td><td>3917488..3920856</td><td>1833</td><td>Swt1-family-HEPN-domain-containing-protein-[Rhodococcus-erythropolis]</td></tr>
<tr><td>WP_264184695.1</td><td>DUF499_1.RM.like.systems</td><td><-HTH|| TUDOR+ SF1-SF2-Helicase+iREase+CTD+REase -><-CHTH|| METHYLASE -> HEPN+ DUF499 *->DUF222+ HNH -><-?<-?<-Peptidase_A8</td><td>1139</td><td>Bacteria>Terrabacteria_group>Actinomycetota>Actinomycetes>Mycobacteriales</td><td>-</td><td>Rhodococcus_yananensis</td><td>LDQ05_RS09980</td><td>11697..15116</td><td>2879464</td><td>Swt1-family-HEPN-domain-containing-protein-[Rhodococcus-yananensis]</td></tr>
<tr><td>NNG19379.1</td><td>DUF499_1.RM.like.systems</td><td>LexA-HTH+LexA-protease-><-?<-35exo+SF1-SF2-Helicase<-?<- HEPN+ DUF499 *<- METHYLASE ||DUF222+ HNH -><- TUDOR+ SF1-SF2-Helicase+iREase+CTD+REase ||HTH_3-></td><td>1110</td><td>Bacteria>Terrabacteria_group>Actinomycetota>Actinomycetes>Propionibacteriales</td><td>GCA_013002685.1</td><td>Naumannella_sp._ID2617S</td><td>HJ590_07255</td><td>1496426..1499758</td><td>1656893</td><td>ATP-binding-protein-[Naumannella-sp.-ID2617S]</td></tr>
<tr><td>WP_232416936.1</td><td>DUF499_1.RM.like.systems</td><td><-HTH+SF1-SF2-Helicase+REase<-REase<-MORC<- HEPN+ DUF499 *<- METHYLASE <-REase||DNA_methylase-><-?||DUF222+ HNH -></td><td>1136</td><td>Bacteria>Terrabacteria_group>Actinomycetota>Actinomycetes>Propionibacteriales</td><td>GCF_024508295.1</td><td>Aeromicrobium_duanguangcaii</td><td>NP095_RS05665</td><td>1110875..1114285</td><td>2968086</td><td>MULTISPECIES:-Swt1-family-HEPN-domain-containing-protein-[unclassified-Aeromicrobium]</td></tr>
<tr><td>WP_016481846.1</td><td>DUF499_1.RM.like.systems</td><td>TUDOR+ SF1-SF2-Helicase+iREase+CTD+REase ->SMS-KaiC-> HNH -> METHYLASE -> DUF499 *-><-?||?->?-><-HAD||?->REase-></td><td>937</td><td>Bacteria>Terrabacteria_group>Armatimonadota>Chthonomonadetes>Chthonomonadales</td><td>GCF_000427095.1</td><td>Chthonomonas_calidirosea</td><td>CCALI_RS02245</td><td>511927..514740</td><td>454171</td><td>DUF499-domain-containing-protein-[Chthonomonas-calidirosea]</td></tr>
<tr><td>WP_075162425.1</td><td>DUF499_1.RM.like.systems</td><td><-REase<-?||HAD-><-?<-?||?-><- DUF499 *<- METHYLASE -p<- METHYLASE -p<- HNH <-SMS-KaiC<- TUDOR+ SF1-SF2-Helicase+iREase+CTD+REase</td><td>937</td><td>Bacteria>Terrabacteria_group>Armatimonadota>Chthonomonadetes>Chthonomonadales</td><td>GCF_001457975.1</td><td>Chthonomonas_calidirosea</td><td>CP488_RS03500</td><td>839037..841850</td><td>454171</td><td>DUF499-domain-containing-protein-[Chthonomonas-calidirosea]</td></tr>
<tr><td>HBL50377.1</td><td>DUF499_1.RM.like.systems</td><td><-SF1-SF2-Helicase<-OLD-ABC+TOPRIM<- HEPN+ DUF499 *<- METHYLASE <- TUDOR+ SF1-SF2-Helicase+iREase+CTD+REase <-?<-MORC+DUF3684+ HNH <-SF1-SF2-Helicase-p<-HEPN+SF1-SF2-Helicase</td><td>1129</td><td>Bacteria>Terrabacteria_group>Bacillota>unclassified_Bacillota</td><td>GCA_003504715.1</td><td>Bacillota_bacterium</td><td>DDZ65_09075</td><td>4304..7693</td><td>1879010</td><td>hypothetical-protein-[Firmicutes-bacterium]</td></tr>
<tr><td>MBK6433234.1</td><td>DUF499_1.RM.like.systems</td><td><- DUF499 *<-REase<-OLD-ABC<- HNH</td><td>943</td><td>Bacteria>Terrabacteria_group>Chloroflexota>Anaerolineae>unclassified_Anaerolineae</td><td>GCA_016704685.1</td><td>Anaerolineae_bacterium</td><td>IPF85_21385</td><td>32175..35006</td><td>2052143</td><td>ATP-binding-protein-[Anaerolineae-bacterium]</td></tr>
<tr><td>WP_244348266.1</td><td>DUF499_1.RM.like.systems</td><td>DOC+HTH-> METHYLASE ->HicB_lk_antitox->HigB/RHH->HTH_3->HIN-HTH+AT-HTH-HYD2-2+SecB-> HEPN+ DUF499 *-> HNH ->Peptidase_M48-></td><td>1084</td><td>Bacteria>Terrabacteria_group>Deinococcota>Deinococci>Thermales</td><td>GCF_022846295.1</td><td>Thermus_thermophilus</td><td>MWM13_RS12110</td><td>24920..28174</td><td>274</td><td>Swt1-family-HEPN-domain-containing-protein-[Thermus-thermophilus]</td></tr>
<tr><td>MBC2711122.1</td><td>DUF499_1.RM.like.systems</td><td>TUDOR+ SF1-SF2-Helicase+iREase+CTD+REase ->?-> OLD-ABC-> HNH -> METHYLASE -> HEPN+ DUF499 *->?->GNTR-HTH->?->?->ABC-ATPase+SbcC->ABC-ATPase+SbcC-></td><td>1122</td><td>Bacteria>Thermodesulfobacteriota>Desulfobacteria>Desulfobacterales>Desulfosarcinaceae</td><td>GCA_014237405.1</td><td>Desulfosarcina_sp.</td><td>HGJ94_09035</td><td>43660..47028</td><td>2027861</td><td>ATP-binding-protein-[Desulfosarcina-sp.]</td></tr>
<tr><td>MBF0121377.1</td><td>DUF499_1.RM.like.systems</td><td>HNH -> TUDOR+ SF1-SF2-Helicase+iREase+CTD+REase -> METHYLASE ->?->RelE-> DUF499 *-></td><td>942</td><td>Bacteria>Thermodesulfobacteriota>Desulfobacteria>Desulfobacterales>unclassified_Desulfobacterales</td><td>GCA_015228685.1</td><td>Desulfobacterales_bacterium</td><td>HQK79_21310</td><td>12197..15025</td><td>2044940</td><td>ATP-binding-protein-[Desulfobacterales-bacterium]</td></tr>
<tr><td>MBF0121417.1</td><td>DUF499_1.RM.like.systems</td><td><- HNH <- DUF499 *<-PIN<-?<- METHYLASE <- TUDOR+ SF1-SF2-Helicase+iREase+CTD+REase</td><td>973</td><td>Bacteria>Thermodesulfobacteriota>Desulfobacteria>Desulfobacterales>unclassified_Desulfobacterales</td><td>GCA_015228685.1</td><td>Desulfobacterales_bacterium</td><td>HQK79_21515</td><td>1603..4524</td><td>2044940</td><td>ATP-binding-protein-[Desulfobacterales-bacterium]</td></tr>
<tr><td>WP_168935605.1</td><td>DUF499_1.RM.like.systems</td><td><-HTH_3||?-><-?||?->?-><- DUF499 *<-REase<-OLD-ABC<- METHYLASE <- HNH <-OLD-ABC <- TUDOR+ SF1-SF2-Helicase+iREase+CTD+REase</td><td>933</td><td>Bacteria>Thermodesulfobacteriota>Desulfovibrionia>Desulfovibrionales>Desulfovibrionaceae</td><td>GCF_012843875.1</td><td>Desulfovibrio_piger</td><td>HF854_RS06725</td><td>17522..20323</td><td>901</td><td>DUF499-domain-containing-protein-[Desulfovibrio-piger]</td></tr>
<tr><td>MCI0602497.1</td><td>DUF499_1.RM.like.systems</td><td><-XIS-HTH||?-><-?<- HNH <- DUF499 *<-REase<- METHYLASE <-HEPN+REase</td><td>922</td><td>Bacteria>unclassified_Bacteria</td><td>GCA_022616075.1</td><td>bacterium</td><td>L0156_05740</td><td>5054..7822</td><td>1869227</td><td>DUF499-domain-containing-protein-[bacterium]</td></tr>
</table>

| Protein-ID | System | Gene-Neighbourhoods | Length | Taxonomy | Assembly-ID | Strain | Locus-Tag | Genomic-coordinates | TaxID | Fasta-defline |
|---|---|---|---|---|---|---|---|---|---|---|
| -- | | | | | | | | | | |
| --PIN-EndoRNase | | | | | | | | | | |
| --OLD-ABC->PIN-> | | | | | | | | | | |
| -- | | | | | | | | | | |
| AGB03431.1 | DUF499_1.RM.like.systems | TUDOR+ SF1-SF2-Helicase+iREase+CTD+REase->METHYLASE->?->PADR-HTH->PIN->DUF499*-> | 981 | Archaea>Euryarchaeota>Stenosarchaea_group>Methanomicrobia>Methanomicrobiales | GCA_000327485.1 | Methanoregula_formicica_SMSP | Metfor_2431 | 2388276..2391221 | 593750 | putative-ATPase-(AAA+-superfamily)-[Methanoregula-formicica-SMSP] |
| WP_233504506.1 | DUF499_1.RM.like.systems | TUDOR+ SF1-SF2-Helicase+iREase+CTD+REase->METHYLASE->?->PADR-HTH->PIN->DUF499*-> | 867 | Archaea>Euryarchaeota>Stenosarchaea_group>Methanomicrobia>Methanomicrobiales | GCF_000327485.1 | Methanoregula_formicica | METFOR_RS11875 | 2388618..2391221 | 882104 | DUF499-domain-containing-protein-[Methanoregula-formicica] |
| WP_010477366.1 | DUF499_1.RM.like.systems | <-Phage_int_SAM_1+Phage_integrase<-TOPRIM\|\|tRNA-><-DUF499*\|\|?->PIN-><-METHYLASE<-TUDOR+ SF1-SF2-Helicase+iREase+CTD+REase | 887 | Archaea>Euryarchaeota>Thermococci>Thermococcales>Thermococcaceae | GCF_000258515.1 | Thermococcus_zilligii | TZI_RS0101400 | 259596..262259 | 54076 | DUF499-domain-containing-protein-[Thermococcus-zilligii] |
| WP_139681370.1 | DUF499_1.RM.like.systems | <-GDResolvase+HIN-HTH\|\|TUDOR+ SF1-SF2-Helicase+iREase+CTD+REase->METHYLASE-><-PIN<-?\|\|?->DUF499*->?->?->SF1-SF2-Helicase-><-tRNA\|\|?->REase-> | 883 | Archaea>Euryarchaeota>Thermococci>Thermococcales>Thermococcaceae | GCF_006274605.1 | Thermococcus_indicus | FH039_RS11140 | 2049351..2052002 | 2586643 | DUF499-domain-containing-protein-[Thermococcus-indicus] |
| WP_167893361.1 | DUF499_1.RM.like.systems | <-REase<-?\|\|tRNA-><-SF1-SF2-Helicase<-?<-?<-DUF499*<-?\|\|?->PIN-><-METHYLASE<-?<-TUDOR+ SF1-SF2-Helicase+iREase+CTD+REase | 880 | Archaea>Euryarchaeota>Thermococci>Thermococcales>Thermococcaceae | GCF_012027305.1 | Thermococcus_sp._MAR1 | E3E25_RS11215 | 186255..188897 | 1638263 | DUF499-domain-containing-protein-[Thermococcus-sp.-MAR1] |
| WP_167910221.1 | DUF499_1.RM.like.systems | <-PadR-HTH\|\|?->Phage_integrase->?->?->?-><-DUF499*<-?\|\|?->PIN-><-METHYLASE<-?<-TUDOR+ SF1-SF2-Helicase+iREase+CTD+REase | 885 | Archaea>Euryarchaeota>Thermococci>Thermococcales>Thermococcaceae | GCF_012027595.1 | Thermococcus_sp._Bubb.Bath | E3E29_RS06900 | 296042..298699 | 1638242 | DUF499-domain-containing-protein-[Thermococcus-sp.-Bubb.Bath] |
| OYT64646.1 | DUF499_1.RM.like.systems | <-SbcC<-DUF499*<-PIN<-SbcC<-METHYLASE-p<-PolB+HEPN<-PolB | 851 | Archaea>TACK_group>Candidatus_Bathyarchaeota>unclassified_Candidatus_Bathyarchaeota | GCA_002255045.1 | Candidatus_Bathyarchaeota_archaeon_ex4484_205 | B6U74_05075 | 1011..3566 | 2012510 | AAA-family-ATPase-[Candidatus-Bathyarchaeota-archaeon-ex4484_205] |
| MXY61028.1 | DUF499_1.RM.like.systems | PIN->DUF499*->METHYLASE->SF1-SF2-Helicase+iREase+CTD+REase-> | 820 | Archaea>TACK_group>Nitrososphaerota>Cenarchaeales>Cenarchaeaceae | GCA_009837825.1 | Cenarchaeum_sp._SB0665_bin_23 | F4Y82_02795 | 8011..10473 | 2604876 | ATP-binding-protein-[Cenarchaeum-sp.-SB0665_bin_23] |
| MYG32491.1 | DUF499_1.RM.like.systems | BirA-HTH->?->PIN->DUF499*->METHYLASE->SF1-SF2-Helicase+iREase+CTD+REase-> | 806 | Archaea>TACK_group>Nitrososphaerota>Cenarchaeales>Cenarchaeaceae | GCA_009842575.1 | Cenarchaeum_sp._SB0677_bin_16 | F4202_00565 | 1815..4235 | 2604883 | ATP-binding-protein-[Cenarchaeum-sp.-SB0677_bin_16] |
| MYG32966.1 | DUF499_1.RM.like.systems | PIN->DUF499*->METHYLASE->SF1-SF2-Helicase+iREase+CTD+REase-> | 820 | Archaea>TACK_group>Nitrososphaerota>Cenarchaeales>Cenarchaeaceae | GCA_009842575.1 | Cenarchaeum_sp._SB0677_bin_16 | F4202_03070 | 8355..10817 | 2604883 | ATP-binding-protein-[Cenarchaeum-sp.-SB0677_bin_16] |
| WP_009070048.1 | DUF499_1.RM.like.systems | <-ORC-CDC6+REase\|\|?->?->?->PIN-><-DUF499*<-METHYLASE<-?<-REase+SF1-SF2-Helicase\|\|?->MethyltransfD12-><-HKD-DNase | 137 | Archaea>TACK_group>Thermoproteota>Thermoprotei>Sulfolobales | GCF_000243315.1 | Metallosphaera_yellowstonensis | METMK1DRAFT_RS18680 | 133361..133774 | 1111107 | DUF499-domain-containing-protein-[Metallosphaera-yellowstonensis] |
| MBA3712812.1 | DUF499_1.RM.like.systems | Schlafen+HATPase+HTH->?->METHYLASE->DUF433->PIN->HEPN+ DUF499*->TIR+PNTD2->PIWI->?->Phage_integrase-> | 1118 | Bacteria>Acidobacteriota>Blastocatellia>Blastocatellales>Pyrinomonadaceae | GCA_013816505.1 | Pyrinomonadaceae_bacterium | H0W76_10240 | 5538..8894 | 2283092 | ATP-binding-protein-[Pyrinomonadaceae-bacterium] |
| WP_211425996.1 | DUF499_1.RM.like.systems | SAM-methylase-><-?<-?\|\|TUDOR+ SF1-SF2-Helicase+iREase+CTD+REase->PIN->DUF499*->sGTP-> | 933 | Bacteria>Acidobacteriota>Blastocatellia>Chloracidobacterium | GCF_018304805.1 | Chloracidobacterium_aggregatum | J8C02_RS10960 | 2617767..2620568 | 2851959 | DUF499-domain-containing-protein-[Chloracidobacterium-aggregatum] |
| MDE0626645.1 | DUF499_1.RM.like.systems | METHYLASE-p->?->PIN->DUF499*->REase-> | 1105 | Bacteria>Acidobacteriota>Terriglobia>Bryobacterales>unclassified_Bryobacterales | GCA_028824065.1 | Bryobacterales_bacterium | OXH99_09625 | 1430..4747 | 2026791 | MAG:-DUF499-domain-containing-protein-[Bryobacterales-bacterium] |
| MBP7799276.1 | DUF499_1.RM.like.systems | <-HEPN+ DUF499*<-PIN<-OLD-ABC<-METHYLASE<-TOPRIM<-OLD-ABC | 1114 | Bacteria>Acidobacteriota>Thermoanaerobaculia>Thermoanaerobaculales>Thermoanaerobaculaceae | GCA_018000955.1 | Thermoanaerobaculaceae_bacterium | KA072_11695 | 22806..26150 | 2699758 | DUF499-domain-containing-protein-[Thermoanaerobaculaceae-bacterium] |
| RMH20600.1 | DUF499_1.RM.like.systems | METHYLASE-p->DUF433->PIN->HEPN+ DUF499*-> | 1096 | Bacteria>Acidobacteriota>unclassified_Acidobacteriota | GCA_003697015.1 | Acidobacteriota_bacterium | D6696_07825 | 1571..4858 | 1978231 | ATP-binding-protein,-partial-[Acidobacteria-bacterium] |
| MXW39171.1 | DUF499_1.RM.like.systems | <-DUF499*<-PIN<-DUF433<-DUF433<-METHYLASE<-HEPN | 922 | Bacteria>Acidobacteriota>unclassified_Acidobacteriota | GCA_009836105.1 | Acidobacteriota_bacterium | F4Z65_13025 | 3012..5780 | 1978231 | DUF499-domain-containing-protein-[Acidobacteria-bacterium] |
| MYN64243.1 | DUF499_1.RM.like.systems | OLD-ABC->PIN->OLD-ABC->TOPRIM->MetJarc-HTH->PIN->DUF499*-><-?\|\|?->?->PIN-> | 935 | Bacteria>Acidobacteriota>unclassified_Acidobacteriota | GCA_009861545.1 | Acidobacteriota_bacterium | F4X11_04340 | 45133..47940 | 1978231 | ATP-binding-protein-[Acidobacteria-bacterium] |
| QQS46399.1 | DUF499_1.RM.like.systems | TUDOR+ SF1-SF2-Helicase+iREase+CTD+REase->Phd-YefM->PIN->METHYLASE->?->DUF499*->Phd-YefM-> | 931 | Bacteria>Acidobacteriota>unclassified_Acidobacteriota | GCA_016700195.1 | Acidobacteriota_bacterium | IPM66_21230 | 5343713..5346508 | 1978231 | ATP-binding-protein-[Acidobacteria-bacterium] |
| MCI0623777.1 | DUF499_1.RM.like.systems | HTH-><-HEPN+ DUF499*<-METHYLASE<-PIN<-?<-STAS+HisKin<-TUDOR+ SF1-SF2-Helicase+iREase+CTD+REase<-XIS-HTH | 1110 | Bacteria>Acidobacteriota>unclassified_Acidobacteriota | GCA_022615935.1 | Acidobacteriota_bacterium | L0387_19325 | 508..3840 | 1978231 | DUF499-domain-containing-protein-[Acidobacteria-bacterium] |
| MCY3931995.1 | DUF499_1.RM.like.systems | <-HicB_lk_antitox<-HicA_toxin\|\|METHYLASE->Phd-YefM->PIN->TPR+TPR->DUF499*-> | 1099 | Bacteria>Acidobacteriota>unclassified_Acidobacteriota | GCA_026707325.1 | Acidobacteriota_bacterium | OXH70_09765 | 12744..16043 | 1978231 | MAG:-DUF499-domain-containing-protein-[Acidobacteria-bacterium] |
| MCY4659240.1 | DUF499_1.RM.like.systems | <-ORC-CDC6+REase<-DUF499*<-PIN<-OLD-ABC<-METHYLASE<-ParB+HNH+Schlafen<-TOPRIM<-OLD-ABC | 941 | Bacteria>Acidobacteriota>unclassified_Acidobacteriota | GCA_026713985.1 | Acidobacteriota_bacterium | OXF93_05425 | 1121..3946 | 1978231 | MAG:-DUF499-domain-containing-protein-[Acidobacteria-bacterium] |
| MCL4821144.1 | DUF499_1.RM.like.systems | STYKinase-><-HEPN+ DUF499*<-?<-ORC-CDC6<-ParA-Soj-PloopNTPase<-HEPN<-METHYLASE<-PIN | 1116 | Bacteria>Acidobacteriota>Vicinamibacteria>unclassified_Vicinamibacteria | GCA_023384615.1 | Vicinamibacteria_bacterium | KJ067_18565 | 2540..5890 | 2793154 | DUF499-domain-containing-protein-[Vicinamibacteria-bacterium] |
| MCL4821456.1 | DUF499_1.RM.like.systems | TUDOR+ SF1-SF2-Helicase+iREase+CTD+REase-p->XIS-HTH+RVT->METHYLASE->OLD-ABC->PIN->HEPN+ DUF499*->SF1-SF2-Helicase-><-?<-?\|\|Pkinase-> | 1115 | Bacteria>Acidobacteriota>Vicinamibacteria>unclassified_Vicinamibacteria | GCA_023384615.1 | Vicinamibacteria_bacterium | KJ067_20160 | 8492..11839 | 2793154 | DUF499-domain-containing-protein-[Vicinamibacteria-bacterium] |
| MPZ20368.1 | DUF499_1.RM.like.systems | TUDOR+ SF1-SF2-Helicase+iREase+CTD+REase->PIN->?->SIR2->SIGMA-HTH->METHYLASE->DUF499*->?->HEPN-> | 1103 | Bacteria>Acidobacteriota>Vicinamibacteria>Vicinamibacterales>Vicinamibacteraceae | GCA_009377605.1 | Luteitalea_sp. | GEV06_21000 | 17307..20618 | 2004800 | DUF499-domain-containing-protein-[Luteitalea-sp.] |
| RUM61916.1 | DUF499_1.RM.like.systems | METHYLASE-p->MazE_antitoxin->PIN->DUF499*-> | 858 | Bacteria>Aquificota>Aquificae>Aquificales>Hydrogenothermaceae | GCA_003978555.1 | Persephonella_sp. | DSY59_01055 | 1949..4525 | 2060922 | MAG:-AAA-family-ATPase-[Persephonella-sp.] |
| KAA0227462.1 | DUF499_1.RM.like.systems | TUDOR+ SF1-SF2-Helicase+iREase+CTD+REase->ArdCN+MPTase-><-PIN<-?\|\|HEPN+ DUF499*-><-?<-STYKIN<-?<-?\|\|?->HAD-> | 1113 | Bacteria>Bacteria_incertae_sedis>Bacteria_candidate_phyla>candidate_division_KSB1>unclassified_candidate_division_KSB1 | GCA_008363195.1 | candidate_division_KSB1_bacterium | EDS67_15790 | 88921..92262 | 2172550 | ATP-binding-protein-[candidate-division-KSB1-bacterium] |
| MBZ0160824.1 | DUF499_1.RM.like.systems | <-DUF499*<-PIN<-DUF433<-REase<-HTH<-BrnT_toxin<-METHYLASE | 920 | Bacteria>Bacteria_incertae_sedis>Bacteria_candidate_phyla>candidate_division_NC10>Methylomirabilis | GCA_019912425.1 | Candidatus_Methylomirabilis_sp. | K8G79_11930 | 646..3408 | 2032687 | DUF499-domain-containing-protein-[bacterium] |
| NIM17954.1 | DUF499_1.RM.like.systems | PIN->?->?->RADICAL-SAM-><-MNS+REase<-DUF499*<-PIN<-MazE_antitoxin<-METHYLASE-p | 846 | Bacteria>Bacteria_incertae_sedis>Bacteria_candidate_phyla>Candidatus_Aminicenantes>unclassified_Candidatus_Aminicenantes | GCA_011771225.1 | Candidatus_Aminicenantes_bacterium | GT081_38750 | 19109..21649 | 2052149 | DUF499-domain-containing-protein-[Candidatus-Aminicenantes-bacterium] |
| MBF6571348.1 | DUF499_1.RM.like.systems | METHYLASE->?->XIS-HTH->?->PIN->DUF499*-> | 924 | Bacteria>Bacteria_incertae_sedis>Bacteria_candidate_phyla>Candidatus_Binatota>Candidatus_Binatia | GCA_015478625.1 | Candidatus_Binataceae_bacterium | IVW54_21020 | 19322..22096 | 2785449 | ATP-binding-protein-[Candidatus-Binataceae-bacterium] |
| MBF6571679.1 | DUF499_1.RM.like.systems | <-DUF499*<-DUF499-p<-PIN<-DUF433<-?<-METHYLASE | 905 | Bacteria>Bacteria_incertae_sedis>Bacteria_candidate_phyla>Candidatus_Binatota>Candidatus_Binatia | GCA_015478625.1 | Candidatus_Binataceae_bacterium | IVW54_22790 | 1..2714 | 2785449 | ATP-binding-protein-[Candidatus-Binataceae-bacterium] |
| WP_122031531.1 | DUF499_1.RM.like.systems | TUDOR+ SF1-SF2-Helicase+iREase+CTD+REase->ORC-CDC6+REase->METHYLASE->OLD-ABC->PIN->HEPN+ DUF499*->Radical-SAM+TRAM->PfkB->?->35exo+SF1-SF2-Helicase->?-><-NTP_transf_2 | 1110 | Bacteria>Bacteria_incertae_sedis>Bacteria_candidate_phyla>Candidatus_Bipolaricaulota>Bipolaricaulis | GCF_900465355.1 | Candidatus_Bipolaricaulis_anaerobius | BARAN1_RS05370 | 1116813..1120145 | 2026885 | Swt1-family-HEPN-domain-containing-protein-[Candidatus-Bipolaricaulis-anaerobius] |
| KUK26942.1 | DUF499_1.RM.like.systems | <-DUF499-p<-HEPN+ DUF499*<-PolB-NTase<-METHYLASE-p<-PolB+HEPN<-PolB<-METHYLASE-p<-PIN | 493 | Bacteria>Bacteria_incertae_sedis>Bacteria_candidate_phyla>Candidatus_Bipolaricaulota>unclassified_Candidatus_Bipolaricaulota | GCA_001508155.1 | Acetothermia_bacterium_64_32 | XD60_0831 | 38456..39937 | 1635286 | Uncharacterized-protein-XD60_0831-[Acetothermia-bacterium-64_32] |
| HAF70679.1 | DUF499_1.RM.like.systems | <-DUF499-p<-HEPN+ DUF499*<-PolB-NTase<-METHYLASE-p<-PolB+HEPN<-PolB<-METHYLASE-p<-PIN | 515 | Bacteria>Bacteria_incertae_sedis>Bacteria_candidate_phyla>Candidatus_Bipolaricaulota>unclassified_Candidatus_Bipolaricaulota | GCA_003445465.1 | Candidatus_Acetothermia_bacterium | DCL37_04935 | 22341..23888 | 2053493 | hypothetical-protein-[Candidatus-Acetothermia-bacterium] |
| MXV83700.1 | DUF499_1.RM.like.systems | <-URI-DNase<-?<-METHYLASE<-REase<-REase<-REase<-HEPN+ DUF499*<-PIN | 1088 | Bacteria>Bacteria_incertae_sedis>Bacteria_candidate_phyla>Candidatus_Poribacteria>unclassified_Candidatus_Poribacteria | GCA_009835485.1 | Candidatus_Poribacteria_bacterium | F4Z98_10065 | 15224..18490 | 2026781 | ATP-binding-protein-[Candidatus-Poribacteria-bacterium] |
| MYC78293.1 | DUF499_1.RM.like.systems | <-HEPN<-URI-DNase<-?<-REase<-REase<-REase<-HEPN+ DUF499*<-PIN<-HTH+MPTase<-?<-?<-?<-SF1-SF2-Helicase+REase | 1093 | Bacteria>Bacteria_incertae_sedis>Bacteria_candidate_phyla>Candidatus_Poribacteria>unclassified_Candidatus_Poribacteria | GCA_009841255.1 | Candidatus_Poribacteria_bacterium | F4X10_21230 | 47302..50583 | 2026781 | ATP-binding-protein-[Candidatus-Poribacteria-bacterium] |
| MBI5115316.1 | DUF499_1.RM.like.systems | <-SF1-SF2-Helicase<-?<-HEPN+REase<-?<-PIN<-HTH+MPTase<-HEPN+ DUF499*<-METHYLASE-p | 1118 | Bacteria>Bacteria_incertae_sedis>Bacteria_candidate_phyla>Candidatus_Poribacteria>unclassified_Candidatus_Poribacteria | GCA_016215705.1 | Candidatus_Poribacteria_bacterium | HZA56_02480 | 5349..8705 | 2026781 | ATP-binding-protein-[Candidatus-Poribacteria-bacterium] |
| MBM3210709.1 | DUF499_1.RM.like.systems | <-XIS-HTH<-DUF499*<-PIN<-?<-METHYLASE<-TUDOR+ SF1-SF2-Helicase+iREase+CTD+REase-p | 903 | Bacteria>Bacteria_incertae_sedis>Bacteria_candidate_phyla>Candidatus_Poribacteria>unclassified_Candidatus_Poribacteria | GCA_016866835.1 | Candidatus_Poribacteria_bacterium | FJZ33_00700 | 511..3222 | 2026781 | DUF499-domain-containing-protein-[Candidatus-Poribacteria-bacterium] |
| MCY3724179.1 | DUF499_1.RM.like.systems | PIN->HEPN+ DUF499*->REase+HTH->REase->METHYLASE->?->URI-DNase-> | 1093 | Bacteria>Bacteria_incertae_sedis>Bacteria_candidate_phyla>Candidatus_Poribacteria>unclassified_Candidatus_Poribacteria | GCA_026705955.1 | Candidatus_Poribacteria_bacterium | OXG97_18330 | 699..3980 | 2026781 | MAG:-DUF499-domain-containing-protein-[Candidatus-Poribacteria-bacterium] |
| MDE0014209.1 | DUF499_1.RM.like.systems | HTH+MPTase->PIN->HEPN+ DUF499*->REase+HTH->REase->METHYLASE->URI-DNase-> | 1095 | Bacteria>Bacteria_incertae_sedis>Bacteria_candidate_phyla>Candidatus_Poribacteria>unclassified_Candidatus_Poribacteria | GCA_028818705.1 | Candidatus_Poribacteria_bacterium | OXU36_23900 | 11254..14541 | 2026781 | MAG:-DUF499-domain-containing-protein-[Candidatus-Poribacteria-bacterium] |
| MDE0681304.1 | DUF499_1.RM.like.systems | HTH+MPTase->PIN->HEPN+ DUF499*->METHYLASE-> | 1094 | Bacteria>Bacteria_incertae_sedis>Bacteria_candidate_phyla>Candidatus_Poribacteria>unclassified_Candidatus_Poribacteria | GCA_028824575.1 | Candidatus_Poribacteria_bacterium | OXI63_00195 | 3088..6372 | 2026781 | MAG:-DUF499-domain-containing-protein-[Candidatus-Poribacteria-bacterium] |
| GAK56408.1 | DUF499_1.RM.like.systems | TUDOR+ SF1-SF2-Helicase+iREase+CTD+REase->?->PIN->METHYLASE->DUF499*->?->?-><-?<-?<-REase | 1005 | Bacteria>Candidatus_Moduliflexota>Candidatus_Vecturitrichia>Candidatus_Vecturitrichales>Candidatus_Vecturitrichaceae | GCA_000739535.1 | Candidatus_Vecturithrix_granuli | U27_03370 | 807271..810288 | 1499967 | ATPase-(AAA+-superfamily)-like-protein-[Candidatus-Vecturithrix-granuli] |
| MBI5243147.1 | DUF499_1.RM.like.systems | <-ORC-CDC6+REase\|\|?->PIN-><-TPR\|\|?-><-HEPN+ DUF499*<-METHYLASE<-PIN<-?<-STAS+HisKin<-TUDOR+ SF1-SF2-Helicase+iREase+CTD+REase | 1133 | Bacteria>Elusimicrobiota>unclassified_Elusimicrobiota | GCA_016212885.1 | Elusimicrobiota_bacterium | HY922_05595 | 9033..12434 | 2030800 | ATP-binding-protein-[Elusimicrobia-bacterium] |
| MCX5796464.1 | DUF499_1.RM.like.systems | TUDOR+ SF1-SF2-Helicase+iREase+CTD+REase->STAS+HisKin->?->PIN->METHYLASE->OLD-ABC+TOPRIM->HEPN+ DUF499*-> | 1133 | Bacteria>Elusimicrobiota>unclassified_Elusimicrobiota | GCA_026388095.1 | Elusimicrobiota_bacterium | NTY77_13305 | 62095..65496 | 2030800 | MAG:-Swt1-family-HEPN-domain-containing-protein-[Elusimicrobia-bacterium] |
| MBR1880072.1 | DUF499_1.RM.like.systems | TUDOR+ SF1-SF2-Helicase+iREase+CTD+REase->METHYLASE-><-PIN<-?\|\|HEPN+ DUF499*-> | 593 | Bacteria>FCB_group>Bacteroidota/Chlorobiota_group>Bacteroidota>Bacteroidia | GCA_017478995.1 | Prevotella_sp. | IJ804_04915 | 6915..8693 | 59823 | DUF499-domain-containing-protein-[Prevotella-sp.] |
| MCZ2341133.1 | DUF499_1.RM.like.systems | TUDOR+ SF1-SF2-Helicase+iREase+CTD+REase->?->METHYLASE->Phd-YefM->PIN->DUF499*-> | 944 | Bacteria>FCB_group>Bacteroidota/Chlorobiota_group>Bacteroidota>Bacteroidia | GCA_026931725.1 | Bacteroidales_bacterium | LC104_04965 | 19853..22687 | 2030927 | MAG:-DUF499-domain-containing-protein-[Bacteroidales-bacterium] |
| HCE56000.1 | DUF499_1.RM.like.systems | <-PIN<-HEPN+ DUF499*<-METHYLASE<-AbiEii<-AbiEi_1<-TUDOR+ SF1-SF2-Helicase+iREase+CTD+REase<-AbiEii<-AbiEi_4 | 1089 | Bacteria>FCB_group>Bacteroidota/Chlorobiota_group>Bacteroidota>Flavobacteriia | GCA_003516285.1 | Lutibacter_sp. | DER05_13810 | 23309..26578 | 1925666 | hypothetical-protein-[Lutibacter-sp.] |
| MBK7697113.1 | DUF499_1.RM.like.systems | PIN->PIN->METHYLASE-p->METHYLASE-p->?->METHYLASE-p->DUF499*->DUF499-p->?-><-?<-?\|\|?->HTH_3-> | 749 | Bacteria>FCB_group>Bacteroidota/Chlorobiota_group>Bacteroidota>Saprospiria | GCA_016706925.1 | Candidatus_Vicinibacter_affinis | IPI30_23260 | 68066..70315 | 2982026 | ATP-binding-protein-[Saprospiraceae-bacterium] |
| MBK8644365.1 | DUF499_1.RM.like.systems | <-HTH_3<-?\|\|?->?-><-?<-DUF499*<-METHYLASE<-METHYLASE-p<-PIN<-TUDOR+ SF1-SF2-Helicase+iREase+CTD+REase<-HTH<-SIR2 | 923 | Bacteria>FCB_group>Bacteroidota/Chlorobiota_group>Bacteroidota>Saprospiria | GCA_016715705.1 | Candidatus_Vicinibacter_affinis | IPN15_19785 | 428646..431417 | 2982026 | ATP-binding-protein-[Saprospiraceae-bacterium] |
| MCY3594931.1 | DUF499_1.RM.like.systems | Peptidase_S41-><-?<-DUF499*<-REase<-PIN<-OLD-ABC<-METHYLASE<-TUDOR+ SF1-SF2-Helicase+iREase+CTD+REase | 913 | Bacteria>FCB_group>Bacteroidota/Chlorobiota_group>Bacteroidota>unclassified_Bacteroidota | GCA_026705205.1 | Bacteroidota_bacterium | OXH01_06810 | 22578..25319 | 1898104 | MAG:-DUF499-domain-containing-protein-[Bacteroidetes-bacterium] |
| MCY3614378.1 | DUF499_1.RM.like.systems | SWACOS-><-DUF499*<-PIN<-OLD-ABC<-METHYLASE-p | 924 | Bacteria>FCB_group>Bacteroidota/Chlorobiota_group>Bacteroidota>unclassified_Bacteroidota | GCA_026705365.1 | Bacteroidota_bacterium | OXH03_04690 | 2086..4860 | 1898104 | MAG:-DUF499-domain-containing-protein-[Bacteroidetes-bacterium] |
| MCY3630037.1 | DUF499_1.RM.like.systems | <-DUF499*<-PIN<-OLD-ABC<-METHYLASE<-wHTH+REase<-TUDOR+ SF1-SF2-Helicase+iREase+CTD+REase | 884 | Bacteria>FCB_group>Bacteroidota/Chlorobiota_group>Bacteroidota>unclassified_Bacteroidota | GCA_026705375.1 | Bacteroidota_bacterium | OXG94_08505 | 1..2653 | 1898104 | MAG:-DUF499-domain-containing-protein,-partial-[Bacteroidetes-bacterium] |
| MCY4001546.1 | DUF499_1.RM.like.systems | wHTH+REase->METHYLASE->OLD-ABC->PIN->DUF499*-> | 903 | Bacteria>FCB_group>Bacteroidota/Chlorobiota_group>Bacteroidota>unclassified_Bacteroidota | GCA_026708685.1 | Bacteroidota_bacterium | OXF84_12180 | 5800..8511 | 1898104 | MAG:-DUF499-domain-containing-protein-[Bacteroidetes-bacterium] |
| MDE2646784.1 | DUF499_1.RM.like.systems | ABhydrolase-><-DUF499*<-PIN<-OLD-ABC<-METHYLASE<-METHYLASE-p<-wHTH+REase<-TUDOR+ SF1-SF2-Helicase+iREase+CTD+REase | 924 | Bacteria>FCB_group>Bacteroidota/Chlorobiota_group>Bacteroidota>unclassified_Bacteroidota | GCA_028875465.1 | Bacteroidota_bacterium | OXI05_13240 | 2968..5742 | 1898104 | MAG:-DUF499-domain-containing-protein-[Bacteroidota-bacterium] |
| MXW15222.1 | DUF499_1.RM.like.systems | TUDOR+ SF1-SF2-Helicase+iREase+CTD+REase->wHTH+REase->OLD-ABC->PIN->DUF499*-><-ABhydrolase | 924 | Bacteria>FCB_group>Bacteroidota/Chlorobiota_group>Rhodothermota>Rhodothermia | GCA_009835915.1 | Rhodothermaceae_bacterium | F4Z76_09955 | 11969..14743 | 2026787 | ATP-binding-protein-[Rhodothermaceae-bacterium] |
| MXZ58767.1 | DUF499_1.RM.like.systems | ABhydrolase-><-DUF499*<-PIN<-OLD-ABC<-METHYLASE<-wHTH+REase<-TUDOR+ SF1-SF2-Helicase+iREase+CTD+REase | 924 | Bacteria>FCB_group>Bacteroidota/Chlorobiota_group>Rhodothermota>Rhodothermia | GCA_009837125.1 | Rhodothermaceae_bacterium | F4Z16_09865 | 2884..5658 | 2026787 | ATP-binding-protein-[Rhodothermaceae-bacterium] |
| MXX97465.1 | DUF499_1.RM.like.systems | TUDOR+ SF1-SF2-Helicase+iREase+CTD+REase->wHTH+REase->METHYLASE->OLD-ABC->PIN->DUF499*-><-ABhydrolase | 907 | Bacteria>FCB_group>Bacteroidota/Chlorobiota_group>Rhodothermota>Rhodothermia | GCA_009838195.1 | Rhodothermaceae_bacterium | F4Y64_07620 | 10395..13118 | 2026787 | ATP-binding-protein-[Rhodothermaceae-bacterium] |
| MYC03829.1 | DUF499_1.RM.like.systems | ABhydrolase-><-DUF499*<-PIN<-OLD-ABC<-wHTH+REase<-TUDOR+ SF1-SF2-Helicase+iREase+CTD+REase | 907 | Bacteria>FCB_group>Bacteroidota/Chlorobiota_group>Rhodothermota>Rhodothermia | GCA_009840265.1 | Rhodothermaceae_bacterium | F4X61_04285 | 3864..6587 | 2026787 | ATP-binding-protein-[Rhodothermaceae-bacterium] |
| MYH11281.1 | DUF499_1.RM.like.systems | TUDOR+ SF1-SF2-Helicase+iREase+CTD+REase->wHTH+REase->METHYLASE->OLD-ABC->PIN->DUF499*-><-ABhydrolase | 924 | Bacteria>FCB_group>Bacteroidota/Chlorobiota_group>Rhodothermota>Rhodothermia | GCA_009842805.1 | Rhodothermaceae_bacterium | F4191_00350 | 10395..13169 | 2026787 | ATP-binding-protein-[Rhodothermaceae-bacterium] |
| MBX3180306.1 | DUF499_1.RM.like.systems | <-AbiEii<-AbiEi_4\|\|?-><-ArdCN+MPTase<-DUF499*<-PIN<-DUF433<-METHYLASE<-?<-?<-SF1-SF2-Helicase-p | 1107 | Bacteria>FCB_group>Candidatus_Hydrogenedentes>unclassified_Candidatus_Hydrogenedentes | GCA_019637335.1 | Candidatus_Hydrogenedentes_bacterium | KF886_23390 | 5509..8832 | 2030809 | ATP-binding-protein-[Candidatus-Hydrogenedentes-bacterium] |
| MBO8151135.1 | DUF499_1.RM.like.systems | TUDOR+ SF1-SF2-Helicase+iREase+CTD+REase->?->PIN->DUF499*-> | 854 | Bacteria>FCB_group>Candidatus_Marinimicrobia>unclassified_Candidatus_Marinimicrobia | GCA_017656455.1 | Candidatus_Marinimicrobia_bacterium | HOZ30_01400 | 332624..335188 | 2026760 | ATP-binding-protein-[Candidatus-Marinimicrobia-bacterium] |
| MCY4604211.1 | DUF499_1.RM.like.systems | METHYLASE->METHYLASE-p->DUF433->PIN->METHYLASE-p->REase->DUF499*-> | 922 | Bacteria>FCB_group>Gemmatimonadota>unclassified_Gemmatimonadota | GCA_026713655.1 | Gemmatimonadota_bacterium | OXE49_08275 | 9321..12089 | 2026742 | MAG:-DUF499-domain-containing-protein-[Gemmatimonadetes-bacterium] |
| MDE2653631.1 | DUF499_1.RM.like.systems | TUDOR+ SF1-SF2-Helicase+iREase+CTD+REase->METHYLASE->METHYLASE-p->?->PIN->DUF499*-> | 1103 | Bacteria>FCB_group>Gemmatimonadota>unclassified_Gemmatimonadota | GCA_028875795.1 | Gemmatimonadota_bacterium | OXI71_07420 | 11724..15035 | 2026742 | MAG:-DUF499-domain-containing-protein-[Gemmatimonadota-bacterium] |
| NMB74978.1 | DUF499_1.RM.like.systems | XIS-HTH->?->?-><-?<-?<-?<-DUF499*<-METHYLASE<-PIN<-?<-TUDOR+ SF1-SF2-Helicase+iREase+CTD+REase<-XIS-HTH | 1126 | Bacteria>Myxococcota>Myxococcia>Myxococcales>unclassified_Myxococcales | GCA_012798435.1 | Myxococcales_bacterium | GYA21_07595 | 16528..19908 | 2026763 | ATP-binding-protein-[Myxococcales-bacterium] |
| MCG5052821.1 | DUF499_1.RM.like.systems | METHYLASE-p->DUF499-p->TUDOR+ SF1-SF2-Helicase+iREase+CTD+REase->PIN->METHYLASE->METHYLASE-p->DUF499*->?-><-DDE_Tnp_ISAZ013<-?\|\|?->SIGMA-HTH-> | 941 | Bacteria>Myxococcota>Myxococcia>Myxococcales>unclassified_Myxococcales | GCA_022184915.1 | Myxococcales_bacterium | KA712_07655 | 10615..13440 | 2026763 | DUF499-domain-containing-protein-[Myxococcales-bacterium] |
| MBI2179305.1 | DUF499_1.RM.like.systems | METHYLASE-p->DUF433->PIN->PIN->DUF499*->?-><-?\|\|?-><-Glyoxalase | 937 | Bacteria>Myxococcota>Myxococcia>unclassified_Myxococcia | GCA_016179825.1 | Deltaproteobacteria_bacterium | HYU31_00605 | 1817..4630 | 2026735 | ATP-binding-protein-[Deltaproteobacteria-bacterium] |
| MBI3073658.1 | DUF499_1.RM.like.systems | <-HTH_3\|\|TUDOR+ SF1-SF2-Helicase+iREase+CTD+REase->?->METHYLASE-><-PIN<-MetJarc-HTH\|\|HEPN+ DUF499*->sGTP->?->Phage_integrase-> | 1105 | Bacteria>Myxococcota>Myxococcia>unclassified_Myxococcia | GCA_016192705.1 | Deltaproteobacteria_bacterium | HYY84_16220 | 33822..37139 | 2026735 | ATP-binding-protein-[Deltaproteobacteria-bacterium] |
| MBI3799269.1 | DUF499_1.RM.like.systems | METHYLASE->DUF1828+REase->METHYLASE-p->DUF433->DUF433->PIN->DUF499*-> | 939 | Bacteria>Myxococcota>Myxococcia>unclassified_Myxococcia | GCA_016200595.1 | Deltaproteobacteria_bacterium | HY268_20180 | 14299..17118 | 2026735 | ATP-binding-protein-[Deltaproteobacteria-bacterium] |
| MBI4814845.1 | DUF499_1.RM.like.systems | <-DNA-glycosylase\|\|TUDOR+ SF1-SF2-Helicase+iREase+CTD+REase->METHYLASE->?->PIN->METHYLASE->DUF499*->REase+NACHT-> | 920 | Bacteria>Myxococcota>Myxococcia>unclassified_Myxococcia | GCA_016208175.1 | Deltaproteobacteria_bacterium | HY791_01225 | 36261..39023 | 2026735 | ATP-binding-protein-[Deltaproteobacteria-bacterium] |
| MBI4527351.1 | DUF499_1.RM.like.systems | Schlafen+HATPase+HTH->METHYLASE->?->PIN->HicB_lk_antitox->?->DUF499*-> | 920 | Bacteria>Myxococcota>Myxococcia>unclassified_Myxococcia | GCA_016210115.1 | Deltaproteobacteria_bacterium | HY695_26450 | 25107..27869 | 2026735 | ATP-binding-protein-[Deltaproteobacteria-bacterium] |
| MBN2361667.1 | DUF499_1.RM.like.systems | TUDOR+ SF1-SF2-Helicase+iREase+CTD+REase->PIN->METHYLASE->DUF499*->Schlafen->EVE-> | 935 | Bacteria>Myxococcota>Myxococcia>unclassified_Myxococcia | GCA_016934455.1 | Deltaproteobacteria_bacterium | JXR83_19595 | 9461..12268 | 2026735 | ATP-binding-protein-[Deltaproteobacteria-bacterium] |
| MBS3919360.1 | DUF499_1.RM.like.systems | SF1-SF2-Helicase->METHYLASE->DUF433->PIN->DUF499*-> | 921 | Bacteria>Myxococcota>Myxococcia>unclassified_Myxococcia | GCA_018333535.1 | Deltaproteobacteria_bacterium | KG012_10765 | 28833..31598 | 2026735 | ATP-binding-protein-[Deltaproteobacteria-bacterium] |
| MBW1911012.1 | DUF499_1.RM.like.systems | Radical-SAM-><-DUF499*<-PIN<-?<-DUF433<-METHYLASE<-TOPRIM<-OLD-ABC-p | 941 | Bacteria>Myxococcota>Myxococcia>unclassified_Myxococcia | GCA_019307765.1 | Deltaproteobacteria_bacterium | JRJ11_15995 | 905..3730 | 2026735 | ATP-binding-protein-[Deltaproteobacteria-bacterium] |
| MBW2149119.1 | DUF499_1.RM.like.systems | METHYLASE-p->METHYLASE-p->DUF433->PIN->DUF499*-> | 939 | Bacteria>Myxococcota>Myxococcia>unclassified_Myxococcia | GCA_019309515.1 | Deltaproteobacteria_bacterium | JRI22_19100 | 2839..5658 | 2026735 | ATP-binding-protein-[Deltaproteobacteria-bacterium] |
| MBW2309273.1 | DUF499_1.RM.like.systems | <-DUF499*<-PIN<-DUF433<-METHYLASE-p<-METHYLASE | 908 | Bacteria>Myxococcota>Myxococcia>unclassified_Myxococcia | GCA_019310535.1 | Deltaproteobacteria_bacterium | JRG73_20315 | 275..3001 | 2026735 | ATP-binding-protein-[Deltaproteobacteria-bacterium] |
| MDD9852708.1 | DUF499_1.RM.like.systems | TUDOR+ SF1-SF2-Helicase+iREase+CTD+REase->?->?->METHYLASE->OLD-ABC->PIN->HEPN+ DUF499*-> | 632 | Bacteria>Myxococcota>Myxococcia>unclassified_Myxococcia | GCA_028817165.1 | Deltaproteobacteria_bacterium | OXU78_02000 | 21524..23422 | 2026735 | MAG:-Swt1-family-HEPN-domain-containing-protein-[Deltaproteobacteria-bacterium] |
| MBX3269805.1 | DUF499_1.RM.like.systems | PIN->TUDOR+ SF1-SF2-Helicase+iREase+CTD+REase->?->HEPN->METHYLASE->METHYLASE->HEPN+ DUF499*->?-><-HIN-HTH | 1090 | Bacteria>Myxococcota>Polyangia>Polyangiales>Sandaracinaceae | GCA_019637095.1 | Sandaracinaceae_bacterium | KF729_06070 | 123033..126305 | 2510337 | DUF499-domain-containing-protein-[Sandaracinaceae-bacterium] |
| MBU0551512.1 | DUF499_1.RM.like.systems | TUDOR+ SF1-SF2-Helicase+iREase+CTD+REase-p->HTH+MPTase->PIN->METHYLASE->OLD-ABC->REase->DUF499*-><-?<-?<-RecR+TOPRIM | 942 | Bacteria>Myxococcota>unclassified_Myxococcota | GCA_018813065.1 | Myxococcota_bacterium | KJ940_08450 | 6900..9728 | 2818507 | DUF499-domain-containing-protein-[Myxococcota-bacterium] |

| Protein-ID | System | Gene-Neighbourhoods | Length | Taxonomy | Assembly-ID | Strain | Locus-Tag | Genomic-coordinates | TaxID | Fasta-defline |
|---|---|---|---|---|---|---|---|---|---|---|
| MBS0166207.1 | DUF499_1.RM.like.systems | <-TrwC+SF1-SF2-helicase+TOPRIM<-VirD4-TraG+VirD4-FtsK<-?<-?<-?<-?<- DUF499 *<- PIN <-?<- METHYLASE <-HicA_toxin<-HicB_lk_antitox<- TUDOR+ SF1-SF2-Helicase+iREase+CTD+REase | 920 | Bacteria>Nitrospirota>Nitrospiria>Nitrospirales>Nitrospiraceae | GCA_018242765.1 | Nitrospira_sp. | JSR29_09005 | 53219..55981 | 70125 | ATP-binding-protein-[Nitrospira-sp.] |
| MCW5796793.1 | DUF499_1.RM.like.systems | DUF4096+DDE_Tnp_1+DDE_Tnp->?-><- DUF499 *<- PIN <-?<-?<- METHYLASE | 852 | Bacteria>Nitrospirota>Nitrospiria>Nitrospirales>Nitrospiraceae | GCA_026128925.1 | Nitrospira_sp. | KIS97_20860 | 1483..4041 | 70125 | MAG:-DUF499-domain-containing-protein-[Nitrospira-sp.] |
| MDC8448550.1 | DUF499_1.RM.like.systems | rve+rve_3->?-><-?<- DUF499 *<- PIN <-?<- METHYLASE -p<- METHYLASE -p<-HicA_toxin<-HicB_lk_antitox | 922 | Bacteria>Nitrospirota>Nitrospiria>Nitrospirales>Nitrospiraceae | GCA_028580895.1 | Nitrospira_sp. | LV473_09340 | 1848..4616 | 70125 | MAG:-DUF499-domain-containing-protein-[Nitrospira-sp.] |
| MDD9860639.1 | DUF499_1.RM.like.systems | SPOUT-><-LuxR-HTH\|\|?->?->?->?-><- DUF499 *<- PIN <-OLD-ABC <- METHYLASE -p | 939 | Bacteria>Nitrospirota>Nitrospiria>Nitrospirales>Nitrospiraceae | GCA_028817215.1 | Nitrospira_sp. | OXU40_09370 | 15116..17935 | 70125 | MAG:-DUF499-domain-containing-protein-[Nitrospira-sp.] |
| MDD9819374.1 | DUF499_1.RM.like.systems | <- DUF499 *<- PIN <-OLD-ABC <- METHYLASE -p | 939 | Bacteria>Nitrospirota>Nitrospiria>Nitrospirales>Nitrospiraceae | GCA_028817545.1 | Nitrospira_sp. | OXR07_00640 | 2590..5409 | 70125 | MAG:-DUF499-domain-containing-protein-[Nitrospira-sp.] |
| WP_090899592.1 | DUF499_1.RM.like.systems | PIN -> TUDOR+ SF1-SF2-Helicase+iREase+CTD+REase ->STAS+HisKin->?-> PIN -> METHYLASE -> HEPN+ DUF499 *->?->KAP-NTPase->?->PAPS_reduct-> | 1117 | Bacteria>Nitrospirota>Nitrospiria>Nitrospirales>Nitrospiraceae | GCF_001458775.1 | Candidatus_Nitrospira_nitrificans | COMA2_RS14290 | 32189..35542 | 1742973 | Swt1-family-HEPN-domain-containing-protein-[Candidatus-Nitrospira-nitrificans] |
| WP_281755187.1 | DUF499_1.RM.like.systems | <-Phage_integrase<-XIS-HTH<-?\|\|?-><- DUF499 *<- METHYLASE <- PIN <-?<- TUDOR+ SF1-SF2-Helicase+iREase+CTD+REase | 846 | Bacteria>Nitrospirota>Thermodesulfovibrionia>Thermodesulfovibrionales>Thermodesulfovibrionaceae | GCF_027925485.1 | Thermodesulfovibrio_islandicus | QMF67_RS07335 | 1419604..1422144 | 53423 | DUF499-domain-containing-protein-[Thermodesulfovibrio-islandicus] |
| RMH36427.1 | DUF499_1.RM.like.systems | <- DUF499 *<- PIN <-DUF433<-REase<-HTH | 933 | Bacteria>Nitrospirota>unclassified_Nitrospirota | GCA_003695915.1 | Nitrospirota_bacterium | D6690_05850 | 2045..4846 | 2026887 | ATP-binding-protein-[Nitrospirae-bacterium] |
| WP_215870431.1 | DUF499_1.RM.like.systems | <- DUF499 *<- METHYLASE <- PIN <- TUDOR+ SF1-SF2-Helicase+iREase+CTD+REase | 949 | Bacteria>Pseudomonadota>Acidithiobacillia>Acidithiobacillales>Acidithiobacillaceae | GCF_018853695.1 | Acidithiobacillus_caldus | HAQ02_RS11835 | 505..3354 | 33059 | DUF499-domain-containing-protein-[Acidithiobacillus-caldus] |
| KAF0185697.1 | DUF499_1.RM.like.systems | MazE_antitoxin-> PIN -> TUDOR+ SF1-SF2-Helicase+iREase+CTD+REase -> METHYLASE -> HEPN+ DUF499 *-><-ParB+HNH\|\|?->?->?->Phage_AlpA-><-Arm-DNA-bind_3+Phage_integrase | 1077 | Bacteria>Pseudomonadota>Alphaproteobacteria>Caulobacterales>Caulobacteraceae | GCA_009773335.1 | Caulobacteraceae_bacterium | FD160_506 | 42091..45324 | 1978230 | ATPase-like-protein-[Caulobacteraceae-bacterium] |
| MBI5313198.1 | DUF499_1.RM.like.systems | TUDOR+ SF1-SF2-Helicase+iREase+CTD+REase ->Ntox18->Peptidase_M78-> METHYLASE -> OLD-ABC-> PIN -> HEPN+ DUF499 *->?-><-HTH | 1107 | Bacteria>Pseudomonadota>Alphaproteobacteria>Hyphomicrobiales>Methylocystaceae | GCA_016219665.1 | Methylocystis_sp. | HZB28_08775 | 48507..51830 | 1911079 | DUF499-domain-containing-protein-[Methylocystis-sp.] |
| WP_166097056.1 | DUF499_1.RM.like.systems | MerR-HTH->tRNA-> TUDOR+ SF1-SF2-Helicase+iREase+CTD+REase ->OLD-ABC->HNH-> METHYLASE -> DUF499 *->DUF433-> PIN -><-Phage_integrase | 945 | Bacteria>Pseudomonadota>Alphaproteobacteria>Hyphomicrobiales>Nitrobacteraceae | GCF_011516635.5 | Bradyrhizobium_barranii | HAP41_RS22275 | 4684908..4687745 | 2992140 | ATP-binding-protein-[Bradyrhizobium-barranii] |
| RWL15103.1 | DUF499_1.RM.like.systems | HhH-RADC+JAB->?->\|\|Phage_integrase->?-> METHYLASE -> PIN -><- HEPN+ DUF499 *<- METHYLASE <-HEPN<-MORC+DUF3684<- TUDOR+ SF1-SF2-Helicase+iREase+CTD+REase <-HTH+WYL\|\|HicB-> | 1098 | Bacteria>Pseudomonadota>Alphaproteobacteria>Hyphomicrobiales>Phyllobacteriaceae | GCA_004019595.1 | Mesorhizobium_sp. | EOR57_31095 | 10515..13811 | 1871066 | DUF499-domain-containing-protein-[Mesorhizobium-sp.] |
| WP_023715932.1 | DUF499_1.RM.like.systems | <-ParB<-?<-REase<- HEPN+ DUF499 *<- PIN <-OLD-ABC <- METHYLASE <- TUDOR+ SF1-SF2-Helicase+iREase+CTD+REase | 1115 | Bacteria>Pseudomonadota>Alphaproteobacteria>Hyphomicrobiales>Phyllobacteriaceae | GCF_000503075.1 | Mesorhizobium_sp._LSHC422A00 | X761_RS140575 | 209394..212741 | 1287294 | DUF499-domain-containing-protein-[Mesorhizobium-sp.-LSHC422A00] |
| WP_135929916.1 | DUF499_1.RM.like.systems | TUDOR+ SF1-SF2-Helicase+iREase+CTD+REase -> METHYLASE -> OLD-ABC-> PIN -> HEPN+ DUF499 *->?-><-?\|\|Methylase-><-Radical-SAM<-Arm-DNA-bind_3+Phage_integrase | 1110 | Bacteria>Pseudomonadota>Alphaproteobacteria>Hyphomicrobiales>Phyllobacteriaceae | GCF_004791875.1 | unclassified_Mesorhizobium | EN810_RS20200 | 141593..144925 | 325217 | MULTISPECIES:-DUF499-domain-containing-protein-[unclassified-Mesorhizobium] |
| MBF0143013.1 | DUF499_1.RM.like.systems | TUDOR+ SF1-SF2-Helicase+iREase+CTD+REase -> PIN -> METHYLASE ->?-> PIN -> DUF499 *-> | 945 | Bacteria>Pseudomonadota>Alphaproteobacteria>Magnetococcales>unclassified_Magnetococcales | GCA_015229005.1 | Magnetococcales_bacterium | HQL59_06090 | 11314..14151 | 2026759 | ATP-binding-protein-[Magnetococcales-bacterium] |
| MBF0185782.1 | DUF499_1.RM.like.systems | REase+35exo+Beta-Barrel+SF1-SF2-Helicase-> TUDOR+ SF1-SF2-Helicase+iREase+CTD+REase ->HEPN-> METHYLASE ->?-> PIN -> DUF499 *-> | 943 | Bacteria>Pseudomonadota>Alphaproteobacteria>Magnetococcales>unclassified_Magnetococcales | GCA_015231215.1 | Magnetococcales_bacterium | HQM06_15545 | 19234..22065 | 2026759 | ATP-binding-protein-[Magnetococcales-bacterium] |
| MBF0214027.1 | DUF499_1.RM.like.systems | TUDOR+ SF1-SF2-Helicase+iREase+CTD+REase -> PIN -> METHYLASE ->?-> PIN -> DUF499 *-> | 946 | Bacteria>Pseudomonadota>Alphaproteobacteria>Magnetococcales>unclassified_Magnetococcales | GCA_015231295.1 | Magnetococcales_bacterium | HQM00_10780 | 8228..11068 | 2026759 | ATP-binding-protein-[Magnetococcales-bacterium] |
| MBF0340191.1 | DUF499_1.RM.like.systems | <- DUF499 *<- PIN <-?<-HEPN<- TUDOR+ SF1-SF2-Helicase+iREase+CTD+REase <-HTH+WYL\|\|REase+SAVED-> | 705 | Bacteria>Pseudomonadota>Alphaproteobacteria>Magnetococcales>unclassified_Magnetococcales | GCA_015232245.1 | Magnetococcales_bacterium | HQL95_04415 | 72..2189 | 2026759 | ATP-binding-protein-[Magnetococcales-bacterium] |
| MYA89893.1 | DUF499_1.RM.like.systems | Arm-DNA-bind_3+Phage_int_SAM_3+Phage_integrase->Arm-DNA-bind_3+Phage_int_SAM_3+Phage_integrase-><-?<- DUF499 *<- PIN <-DUF433<- METHYLASE -p | 926 | Bacteria>Pseudomonadota>Alphaproteobacteria>Rhodobacterales>Paracoccaceae | GCA_009839535.1 | Boseongicola_sp._SB0662_bin_57 | F4X97_15820 | 7656..10436 | 2604843 | ATP-binding-protein-[Boseongicola-sp.-SB0662_bin_57] |
| MDE0346579.1 | DUF499_1.RM.like.systems | <- HEPN+ DUF499 *<- PIN <- DUF499 -p<- METHYLASE <- TUDOR+ SF1-SF2-Helicase+iREase+CTD+REase -p | 1102 | Bacteria>Pseudomonadota>Alphaproteobacteria>Rhodobacterales>Paracoccaceae | GCA_028821095.1 | Boseongicola_sp. | OXI66_12500 | 695..4003 | 1886600 | MAG:-DUF499-domain-containing-protein-[Boseongicola-sp.] |
| OZB26691.1 | DUF499_1.RM.like.systems | TUDOR+ SF1-SF2-Helicase+iREase+CTD+REase ->?-> PIN ->Ntox18->?-> METHYLASE -> HEPN+ DUF499 *->\|\|FAD-NAD-dep-oxidoreductase-><-NACHT+REase\|\|?->DnaB-><-NACHT+REase | 1091 | Bacteria>Pseudomonadota>Alphaproteobacteria>Rhodospirillales>Acetobacteraceae | GCA_002282635.1 | Acidiphilium_sp._34-64-41 | B7X49_12125 | 19315..22590 | 1970297 | MAG:-ATPase-[Acidiphilium-sp.-34-64-41] |
| MCA3363599.1 | DUF499_1.RM.like.systems | METHYLASE -> OLD-ABC-> PIN -> HEPN+ DUF499 *-> | 483 | Bacteria>Pseudomonadota>Alphaproteobacteria>Rhodospirillales>Acetobacteraceae | GCA_020249535.1 | Roseomonas_sp. | INF74_16815 | 3977..5424 | 1869190 | DUF499-domain-containing-protein-[Roseomonas-sp.] |
| MCA7120529.1 | DUF499_1.RM.like.systems | TUDOR+ SF1-SF2-Helicase+iREase+CTD+REase ->?-> PIN ->Ntox18->Peptidase_M78-> METHYLASE -> HEPN+ DUF499 *->?->?->HHH_5+SF1-SF2-helicase-><-RVT+HNH\|\|<-Phage_int_SAM_1+Phage_integrase<-Phage_integrase | 1113 | Bacteria>Pseudomonadota>Alphaproteobacteria>Rhodospirillales>Acetobacteraceae | GCA_020418435.1 | Acidibrevibacterium_fodinaquatile | LGL72_14265 | 2856594..2859935 | 1969806 | DUF499-domain-containing-protein-[Acidibrevibacterium-sp.] |
| WP_012553516.1 | DUF499_1.RM.like.systems | <-VirB2<-VirB11+VirB4-FtsK\|\|?->REase->?-><- HEPN+ DUF499 *<- METHYLASE <- TUDOR+ SF1-SF2-Helicase+iREase+CTD+REase <- PIN <-?<-CopG/RHH<-VirD4-TraG+VirD4-FtsK | 1108 | Bacteria>Pseudomonadota>Alphaproteobacteria>Rhodospirillales>Acetobacteraceae | GCF_000021325.1 | Gluconacetobacter_diazotrophicus | GDIA_RS05030 | 1143198..1146524 | 33996 | DUF499-domain-containing-protein-[Gluconacetobacter-diazotrophicus] |
| WP_010666859.1 | DUF499_1.RM.like.systems | VirD4-TraG+VirD4-FtsK->CopG/RHH->?-> PIN -> TUDOR+ SF1-SF2-Helicase+iREase+CTD+REase -> METHYLASE -> HEPN+ DUF499 *-><-?<-REase\|\|VirB11+VirB4-FtsK->VirB2->?->FtsK+VirB4-FtsK-> | 1108 | Bacteria>Pseudomonadota>Alphaproteobacteria>Rhodospirillales>Acetobacteraceae | GCF_000193495.2 | Acetobacter_aceti | EDC15_RS12660 | 41147..44473 | 435 | DUF499-domain-containing-protein-[Acetobacter-aceti] |
| WP_062273923.1 | DUF499_1.RM.like.systems | PIN -> TUDOR+ SF1-SF2-Helicase+iREase+CTD+REase -> METHYLASE -> HEPN+ DUF499 *-><-?<-?<-Radical-SAM<- PIN <-Phd-YefM | 1108 | Bacteria>Pseudomonadota>Alphaproteobacteria>Rhodospirillales>Acetobacteraceae | GCF_001581105.1 | Acetobacter_cerevisiae | AD954_RS12075 | 12356..15682 | 178900 | DUF499-domain-containing-protein-[Acetobacter-cerevisiae] |
| WP_274966108.1 | DUF499_1.RM.like.systems | TUDOR+ SF1-SF2-Helicase+iREase+CTD+REase ->?-> PIN ->Ntox18->Peptidase_M78-> METHYLASE -> HEPN+ DUF499 *->?->HHH_5+SF1-SF2-helicase-><-RVT+HNH\|\|<-Phage_int_SAM_1+Phage_integrase<-Phage_integrase<-Phage_integrase | 1113 | Bacteria>Pseudomonadota>Alphaproteobacteria>Rhodospirillales>Acetobacteraceae | GCF_020418435.1 | Acidibrevibacterium_fodinaquatile | LGL72_RS14205 | 2856594..2859935 | 1969806 | DUF499-domain-containing-protein-[Acidibrevibacterium-fodinaquatile] |
| WP_253545427.1 | DUF499_1.RM.like.systems | <-TraJ-RHH<-Phage_integrase\|\|?-> PIN -> TUDOR+ SF1-SF2-Helicase+iREase+CTD+REase -> METHYLASE -> HEPN+ DUF499 *->?-><-?<-?<-?\|\|TrwC+SF1-SF2-helicase+TOPRIM-> | 1108 | Bacteria>Pseudomonadota>Alphaproteobacteria>Rhodospirillales>Acetobacteraceae | GCF_024158445.1 | Acetobacter_lovaniensis | NKW44_RS07985 | 22199..25525 | 104100 | DUF499-domain-containing-protein-[Acetobacter-lovaniensis] |
| WP_180283596.1 | DUF499_1.RM.like.systems | <-PfkB\|\|DeoR-HTH+ISOCOT->Transposase_31+Yae1_N-><- HEPN+ DUF499 *<- PIN <-OLD-ABC <- METHYLASE <- TUDOR+ SF1-SF2-Helicase+iREase+CTD+REase \|\|HTH->GDResolvase+PPP1-HTH-> | 1135 | Bacteria>Pseudomonadota>Alphaproteobacteria>Rhodospirillales>Azospirillaceae | GCF_013423495.1 | Azospirillum_oleiclasticum | HND93_RS19070 | 2984..6391 | 2735135 | DUF499-domain-containing-protein-[Azospirillum-oleiclasticum] |
| WP_238518868.1 | DUF499_1.RM.like.systems | <-SF1-SF2-Helicase+DUF1998+HKD-DNase+HKD-DNase<-?<- HEPN+ DUF499 *<- PIN <-OLD-ABC <- METHYLASE <-?<-Schlafen+HATPase+HTH<- TUDOR+ SF1-SF2-Helicase+iREase+CTD+REase | 1154 | Bacteria>Pseudomonadota>Alphaproteobacteria>Rhodospirillales>Azospirillaceae | GCF_022214805.1 | Azospirillum_doebereinerae | MCW82_RS00020 | 10412..13876 | 92933 | DUF499-domain-containing-protein-[Azospirillum-doebereinerae] |
| TVR79626.1 | DUF499_1.RM.like.systems | HDRnase-><- HEPN+ DUF499 *<-?<- PIN <-ParD<- METHYLASE <-OLD-ABC+TOPRIM<- TUDOR+ SF1-SF2-Helicase+iREase+CTD+REase | 1117 | Bacteria>Pseudomonadota>Alphaproteobacteria>Rhodospirillales>unclassified_Rhodospirillales | GCA_007695045.1 | Rhodospirillales_bacterium | EA405_12420 | 709..4062 | 2026786 | ATP-binding-protein-[Rhodospirillales-bacterium] |
| SEP49898.1 | DUF499_1.RM.like.systems | <- HEPN+ DUF499 *<- PIN <-OLD-ABC <- METHYLASE <- TUDOR+ SF1-SF2-Helicase+iREase+CTD+REase <-HNH | 1099 | Bacteria>Pseudomonadota>Alphaproteobacteria>Rhodospirillales>unclassified_Rhodospirillales | GCA_900110395.1 | Rhodospirillales_bacterium_URHD0017 | SAMN02990966_07447 | 5640..8939 | 1380357 | hypothetical-protein-SAMN02990966_07447-[Rhodospirillales-bacterium-URHD0017] |
| MCL6741777.1 | DUF499_1.RM.like.systems | wHTH+REase-> TUDOR+ SF1-SF2-Helicase+iREase+CTD+REase -> METHYLASE -> OLD-ABC-> PIN -> HEPN+ DUF499 *-><-?<-BetaPropeller+ABhydrolase\|\|?-><-ABhydrolase<-MODE-HTH+PBPII | 1094 | Bacteria>Pseudomonadota>Alphaproteobacteria>Sphingomonadales>Sphingomonadaceae | GCA_023516505.1 | Sphingomonas_brevis | LZ518_11630 | 2249137..2252421 | 2908206 | DUF499-domain-containing-protein-[Sphingomonas-sp.-RB56-2] |
| WP_283938333.1 | DUF499_1.RM.like.systems | wHTH+REase-> TUDOR+ SF1-SF2-Helicase+iREase+CTD+REase -> METHYLASE -> OLD-ABC-> PIN -> HEPN+ DUF499 *-><-?<-BetaPropeller+ABhydrolase\|\|?-><-ABhydrolase<-MODE-HTH+PBPII | 1103 | Bacteria>Pseudomonadota>Alphaproteobacteria>Sphingomonadales>Sphingomonadaceae | GCF_023516505.1 | Sphingomonas_brevis | LZ518_RS11630 | 2249110..2252421 | 2908206 | DUF499-domain-containing-protein-[Sphingomonas-brevis] |
| WP_249916143.1 | DUF499_1.RM.like.systems | wHTH+REase-> TUDOR+ SF1-SF2-Helicase+iREase+CTD+REase -> METHYLASE -> OLD-ABC-> PIN -> HEPN+ DUF499 *-><-?<-BetaPropeller+ABhydrolase\|\|?-><-ABhydrolase<-MODE-HTH+PBPII | 1094 | Bacteria>Pseudomonadota>Alphaproteobacteria>Sphingomonadales>Sphingomonadaceae | - | Sphingomonas_brevis | LZ518_11630 | 2249137..2252421 | 2908206 | DUF499-domain-containing-protein-[Sphingomonas-sp.-RB56-2] |
| WP_017667308.1 | DUF499_1.RM.like.systems | HEPN->Phage_integrase-> TUDOR+ SF1-SF2-Helicase+iREase+CTD+REase -> METHYLASE -> HEPN+ DUF499 *-> PIN -> | 1100 | Bacteria>Pseudomonadota>Alphaproteobacteria>Sphingomonadales>Sphingosinicellaceae | GCF_000331225.1 | Sandarakinorhabdus_sp._AAP62 | D619_RS0104150 | 158738..162040 | 1248916 | DUF499-domain-containing-protein-[Sandarakinorhabdus-sp.-AAP62] |
| MDD3182285.1 | DUF499_1.RM.like.systems | <-ACET\|\|Peptidase_M15->?-><-SF1-SF2-Helicase<-OLD-ABC+TOPRIM<- DUF499 *<- METHYLASE <- PIN <- TUDOR+ SF1-SF2-Helicase+iREase+CTD+REase <-HTH+WYL | 940 | Bacteria>Pseudomonadota>Alphaproteobacteria>unclassified_Alphaproteobacteria | GCA_028691835.1 | Alphaproteobacteria_bacterium | PHD48_05740 | 6780..9602 | 1913988 | MAG:-DUF499-domain-containing-protein-[Alphaproteobacteria-bacterium] |
| MDD3183268.1 | DUF499_1.RM.like.systems | <- DUF499 *<- METHYLASE <- PIN <- TUDOR+ SF1-SF2-Helicase+iREase+CTD+REase <-HTH+WYL | 795 | Bacteria>Pseudomonadota>Alphaproteobacteria>unclassified_Alphaproteobacteria | GCA_028691835.1 | Alphaproteobacteria_bacterium | PHD48_10775 | 1..2386 | 1913988 | MAG:-DUF499-domain-containing-protein,-partial-[Alphaproteobacteria-bacterium] |
| KFB77813.1 | DUF499_1.RM.like.systems | Hpr-Kinase-C-><-JetD<-?<-JetB<-JetA<- DUF499 *<-Schlafen+HATPase+HTH<- METHYLASE <- PIN \|\|?-><- TUDOR+ SF1-SF2-Helicase+iREase+CTD+REase <-HAMP+HISKIN | 1093 | Bacteria>Pseudomonadota>Betaproteobacteria>Betaproteobacteria_incertae_sedis>Candidatus_Accumulibacter | GCA_000584975.2 | Candidatus_Accumulibacter_cognatus | AW06_000996 | 10153..13434 | 2954383 | MAG:-hypothetical-protein-AW06_000996-[Candidatus-Accumulibacter-cognatus] |
| EXI65857.1 | DUF499_1.RM.like.systems | TUDOR+ SF1-SF2-Helicase+iREase+CTD+REase -> PIN ->?-> METHYLASE -> DUF499 *-> | 1091 | Bacteria>Pseudomonadota>Betaproteobacteria>Betaproteobacteria_incertae_sedis>Candidatus_Accumulibacter | GCA_000585015.1 | Candidatus_Accumulibacter_adjunctus | AW08_02882 | 60181..63456 | 1454001 | MAG:-hypothetical-protein-AW08_02882-[Candidatus-Accumulibacter-adjunctus] |
| MBN8437943.1 | DUF499_1.RM.like.systems | PIN -> METHYLASE ->PolB->PolB+HEPN->PolB->HEPN-> HEPN+ DUF499 *->?->TPR+TPR-> | 1109 | Bacteria>Pseudomonadota>Betaproteobacteria>Betaproteobacteria_incertae_sedis>Candidatus_Accumulibacter | GCA_017302555.1 | Accumulibacter_sp. | JOL95_07850 | 90759..94088 | 2053492 | DUF499-domain-containing-protein-[Candidatus-Accumulibacter-sp.] |
| MCM8638973.1 | DUF499_1.RM.like.systems | METHYLASE -p-> PIN -> DUF499 *-> | 937 | Bacteria>Pseudomonadota>Betaproteobacteria>Betaproteobacteria_incertae_sedis>Candidatus_Accumulibacter | GCA_023806635.1 | Accumulibacter_sp. | NFW30_04525 | 1720..4533 | 2053492 | DUF499-domain-containing-protein-[Candidatus-Accumulibacter-sp.] |
| MCQ1548298.1 | DUF499_1.RM.like.systems | HAMP+HISKIN-> TUDOR+ SF1-SF2-Helicase+iREase+CTD+REase -> PIN -> METHYLASE ->Schlafen+HATPase+HTH-> DUF499 *-> | 1093 | Bacteria>Pseudomonadota>Betaproteobacteria>Betaproteobacteria_incertae_sedis>Candidatus_Accumulibacter | GCA_024380085.1 | Candidatus_Accumulibacter_phosphatis | NOF05_05620 | 18582..21863 | 327160 | DUF499-domain-containing-protein-[Candidatus-Accumulibacter-phosphatis] |
| WP_034945897.1 | DUF499_1.RM.like.systems | Hpr-Kinase-C-><-JetD<-?<-JetB<-JetA<- DUF499 *<-Schlafen+HATPase+HTH<- METHYLASE <- PIN <- TUDOR+ SF1-SF2-Helicase+iREase+CTD+REase <-HAMP+HISKIN | 1093 | Bacteria>Pseudomonadota>Betaproteobacteria>Betaproteobacteria_incertae_sedis>Candidatus_Accumulibacter | GCF_000584975.1 | Candidatus_Accumulibacter_cognatus | AW06_RS03920 | 10153..13434 | 2954383 | DUF499-domain-containing-protein-[Candidatus-Accumulibacter-cognatus] |
| MCX8099543.1 | DUF499_1.RM.like.systems | <-SF1-SF2-Helicase+DUF1998<-ORC-CDC6+REase<-?<- DUF499 *<- PIN <-HTH<- METHYLASE <-TOPRIM<-OLD-ABC<- TUDOR+ SF1-SF2-Helicase+iREase+CTD+REase | 918 | Bacteria>Pseudomonadota>Betaproteobacteria>Betaproteobacteria_incertae_sedis>Casimicrobiaceae | GCA_026414825.1 | Casimicrobiaceae_bacterium | N3F11_11190 | 5518..8274 | 2906524 | MAG:-DUF499-domain-containing-protein-[Casimicrobiaceae-bacterium] |
| WP_065497253.1 | DUF499_1.RM.like.systems | TUDOR+ SF1-SF2-Helicase+iREase+CTD+REase -p-> TUDOR+ SF1-SF2-Helicase+iREase+CTD+REase -> PIN -> METHYLASE -> METHYLASE ->Schlafen+HATPase+HTH-> DUF499 *->HEPN-> | 955 | Bacteria>Pseudomonadota>Betaproteobacteria>Burkholderiales>Burkholderiaceae | GCF_001685525.1 | Burkholderia_sp._CCA53 | BCR55_RS29355 | 26270..29137 | 1776288 | DUF499-domain-containing-protein-[Burkholderia-sp.-CCA53] |
| WP_254661892.1 | DUF499_1.RM.like.systems | PIN -> METHYLASE ->AAA_21->Schlafen+HATPase+HTH->STYKIN-><-?\|\| DUF499 *-> | 314 | Bacteria>Pseudomonadota>Betaproteobacteria>Burkholderiales>Burkholderiaceae | GCF_001685525.1 | Burkholderia_sp._CCA53 | BCR55_RS29865 | 130717..131658 | 1776288 | DUF499-domain-containing-protein,-partial-[Burkholderia-sp.-CCA53] |
| WP_132962623.1 | DUF499_1.RM.like.systems | GST_N+GST_C->tRNA->tRNA-><- DUF499 *<-REase<-OLD-ABC<- METHYLASE <- PIN <- TUDOR+ SF1-SF2-Helicase+iREase+CTD+REase \|\|Peptidase_M24-> | 949 | Bacteria>Pseudomonadota>Betaproteobacteria>Burkholderiales>Burkholderiales_genera_incertae_sedis | GCF_004342625.1 | Tepidimonas_ignava | EDC36_RS07390 | 30173..33022 | 114249 | DUF499-domain-containing-protein-[Tepidimonas-ignava] |
| WP_143969665.1 | DUF499_1.RM.like.systems | HTH+WYL-> TUDOR+ SF1-SF2-Helicase+iREase+CTD+REase -> PIN -> METHYLASE ->OLD-ABC->TOPRIM-> DUF499 *->HEPN-> | 952 | Bacteria>Pseudomonadota>Betaproteobacteria>Burkholderiales>Burkholderiales_genera_incertae_sedis | GCF_007556755.1 | Tepidimonas_fonticaldi | Tfont_RS12360 | 12901..15759 | 1101373 | DUF499-domain-containing-protein-[Tepidimonas-fonticaldi] |
| WP_166837252.1 | DUF499_1.RM.like.systems | TUDOR+ SF1-SF2-Helicase+iREase+CTD+REase ->ParB+HNH-> PIN -> METHYLASE -> DUF499 *-> | 936 | Bacteria>Pseudomonadota>Betaproteobacteria>Burkholderiales>Burkholderiales_genera_incertae_sedis | GCF_011742065.1 | Aquabacterium_sp._A08 | HA630_RS20340 | 28723..31533 | 2718532 | DUF499-domain-containing-protein-[Aquabacterium-sp.-A08] |
| WP_043699941.1 | DUF499_1.RM.like.systems | PIN -> METHYLASE ->OLD-ABC->TOPRIM-> METHYLASE ->Schlafen+HATPase+HTH-> DUF499 *-><-?<-?<-MORC<-?<-VirB11+MoxR-AAA | 945 | Bacteria>Pseudomonadota>Betaproteobacteria>Burkholderiales>Burkholderiales_genera_incertae_sedis | GCF_020162115.1 | Tepidimonas_taiwanensis | PL21_RS04270 | 89059..91896 | 307486 | DUF499-domain-containing-protein-[Tepidimonas-taiwanensis] |
| TFI45392.1 | DUF499_1.RM.like.systems | <-MobA_MobL+SF1-SF2-helicase\|\|?-><-?<- DUF499 *<-REase<-OLD-ABC<- METHYLASE <- PIN <- TUDOR+ SF1-SF2-Helicase+iREase+CTD+REase | 945 | Bacteria>Pseudomonadota>Betaproteobacteria>Burkholderiales>Comamonadaceae | - | Diaphorobacter_sp._DS2 | E4093_19980 | 3252..6089 | 1302548 | ATP-binding-protein-[Diaphorobacter-sp.-DS2] |
| WP_042427102.1 | DUF499_1.RM.like.systems | Phage_lysozyme->?->HTH_3->Radical-SAM-><-?<-TIR<- DUF499 *<-?<-?<- METHYLASE <-HEPN<- PIN <- TUDOR+ SF1-SF2-Helicase+iREase+CTD+REase | 951 | Bacteria>Pseudomonadota>Betaproteobacteria>Burkholderiales>Comamonadaceae | GCF_000739995.1 | Comamonas_granuli | CGR01S_RS11705 | 67732..70587 | 290309 | DUF499-domain-containing-protein-[Comamonas-granuli] |
| WP_045536717.1 | DUF499_1.RM.like.systems | TUDOR+ SF1-SF2-Helicase+iREase+CTD+REase -> PIN ->HEPN-> METHYLASE -p-> METHYLASE ->ParB+HNH-> DUF499 *-> | 950 | Bacteria>Pseudomonadota>Betaproteobacteria>Burkholderiales>Comamonadaceae | GCF_000828915.1 | Serpentinimonas_maccroryi | SMCB_RS09950 | 2162226..2165078 | 1458426 | DUF499-domain-containing-protein-[Serpentinimonas-maccroryi] |
| WP_127807271.1 | DUF499_1.RM.like.systems | TUDOR+ SF1-SF2-Helicase+iREase+CTD+REase -> PIN ->HEPN-> METHYLASE ->?->?-> DUF499 *->TIR->?-><-Radical-SAM<-HTH_3<-?<-Phage_lysozyme | 950 | Bacteria>Pseudomonadota>Betaproteobacteria>Burkholderiales>Comamonadaceae | GCF_004006415.1 | Hydrogenophaga_sp._NH-16 | DD542_RS03485 | 729405..732257 | 2184519 | DUF499-domain-containing-protein-[Hydrogenophaga-sp.-NH-16] |
| WP_265283911.1 | DUF499_1.RM.like.systems | TUDOR+ SF1-SF2-Helicase+iREase+CTD+REase -> PIN -> METHYLASE -> DUF499 *->Methylase-><-Radical-SAM<-THUMP+Methylase<-?<-ABC-ATPase+SbcC | 1094 | Bacteria>Pseudomonadota>Betaproteobacteria>Burkholderiales>Comamonadaceae | GCF_026127385.1 | Verminephrobacter_aporrectodeae | D5041_RS03845 | 914052..917336 | 1110389 | DUF499-domain-containing-protein-[Verminephrobacter-aporrectodeae] |
| WP_265663937.1 | DUF499_1.RM.like.systems | ABC-ATPase+SbcC->?->THUMP+Methylase->Radical-SAM-><-Methylase<- DUF499 *<- METHYLASE <- PIN <- TUDOR+ SF1-SF2-Helicase+iREase+CTD+REase | 1094 | Bacteria>Pseudomonadota>Betaproteobacteria>Burkholderiales>Comamonadaceae | GCF_026192565.1 | Verminephrobacter_aporrectodeae | D8B23_RS01230 | 11909..15193 | 1110389 | DUF499-domain-containing-protein-[Verminephrobacter-aporrectodeae] |
| WP_265653305.1 | DUF499_1.RM.like.systems | TUDOR+ SF1-SF2-Helicase+iREase+CTD+REase -> PIN -> METHYLASE -> DUF499 *->\|\|?->HerA/FtsK-><-THUMP+Methylase<-?<-ABC-ATPase+SbcC | 1094 | Bacteria>Pseudomonadota>Betaproteobacteria>Burkholderiales>Comamonadaceae | GCF_026192715.1 | Verminephrobacter_aporrectodeae | D8B27_RS01360 | 119835..123119 | 1110389 | DUF499-domain-containing-protein-[Verminephrobacter-aporrectodeae] |
| WP_206111008.1 | DUF499_1.RM.like.systems | HTH+WYL->HTH_Tnp-> TUDOR+ SF1-SF2-Helicase+iREase+CTD+REase -> PIN ->HEPN-> METHYLASE -> DUF499 *->TIR->SLATT->SMODS->Methylase-><-Radical-SAM<-Phage_lysozyme | 955 | Bacteria>Pseudomonadota>Betaproteobacteria>Burkholderiales | GCF_017150735.1 | Burkholderiaceae | KTD10_RS22775 | 57014..59881 | 119060 | MULTISPECIES:-DUF499-domain-containing-protein-[Burkholderiaceae] |
| MCK6404267.1 | DUF499_1.RM.like.systems | <-zf-IS66+DDE_Tnp_IS66<-DDE_Tnp_1_assoc+DDE_Tnp_1<-ParB+HTH<-?\|\|?-><- DUF499 *<-TOPRIM<-OLD-ABC<- METHYLASE <-?<- PIN <-OLD-ABC | 955 | Bacteria>Pseudomonadota>Betaproteobacteria>Burkholderiales>Sphaerotilaceae | GCA_023150255.1 | Sphaerotilus_sulfidivorans | L6Q74_20550 | 5829..8696 | 639200 | DUF499-domain-containing-protein-[Sphaerotilus-sulfidivorans] |
| WP_245558353.1 | DUF499_1.RM.like.systems | RNase->?->?->Phage_lysozyme-><-TIR<- DUF499 *<-?<-?<- METHYLASE <- PIN <- TUDOR+ SF1-SF2-Helicase+iREase+CTD+REase <-HTH_Tnp | 934 | Bacteria>Pseudomonadota>Betaproteobacteria>Burkholderiales>Sphaerotilaceae | GCF_000381125.1 | Caldimonas_manganoxidans | F615_RS0111630 | 4122..6926 | 196015 | DUF499-domain-containing-protein-[Caldimonas-manganoxidans] |
| WP_273473498.1 | DUF499_1.RM.like.systems | <-zf-IS66+DDE_Tnp_IS66<-DDE_Tnp_1_assoc+DDE_Tnp_1<-ParB+HTH<-?\|\|?-><- DUF499 *<-TOPRIM<-OLD-ABC<- METHYLASE <-?<- PIN <-OLD-ABC | 955 | Bacteria>Pseudomonadota>Betaproteobacteria>Burkholderiales>Sphaerotilaceae | GCF_023150255.1 | Sphaerotilus_sulfidivorans | L6Q74_RS20510 | 5829..8696 | 639200 | DUF499-domain-containing-protein-[Sphaerotilus-sulfidivorans] |
| MCC6611669.1 | DUF499_1.RM.like.systems | VirD4-TraG+VirD4-FtsK->TrwC+SF1-SF2-helicase+TOPRIM-><-PIWI<-TIR+PNTD2<- HEPN+ DUF499 *<- METHYLASE <- PIN <-?<-STAS+HisKin<- TUDOR+ SF1-SF2-Helicase+iREase+CTD+REase <- PIN | 1116 | Bacteria>Pseudomonadota>Betaproteobacteria>Burkholderiales>unclassified_Burkholderiales | GCA_020848075.1 | Burkholderiales_bacterium | IT515_18605 | 34273..37623 | 1891238 | DUF499-domain-containing-protein-[Burkholderiales-bacterium] |
| GIX27884.1 | DUF499_1.RM.like.systems | METHYLASE -p-><-?\|\| METHYLASE -p->ParD-> PIN -> HEPN+ DUF499 *-> DUF499 -p->?->?->ORC-CDC6+REase-> | 389 | Bacteria>Pseudomonadota>Betaproteobacteria>Burkholderiales>unclassified_Burkholderiales | GCA_026005015.1 | Burkholderiales_bacterium | KatS3mg123_1765 | 1711823..1712992 | 1891238 | MAG:-hypothetical-protein-KatS3mg123_1765-[Burkholderiales-bacterium] |
| MCZ2134305.1 | DUF499_1.RM.like.systems | <-RAMA<-?<-?<-HTH_21+rve+rve_3<-HIN-HTH<-?<- DUF499 *<-wHTH+REase<- METHYLASE <-HEPN<- PIN <- TUDOR+ SF1-SF2-Helicase+iREase+CTD+REase \|\|HIN-HTH-> | 956 | Bacteria>Pseudomonadota>Betaproteobacteria>Burkholderiales>unclassified_Burkholderiales | GCA_026932525.1 | Burkholderiales_bacterium | LC098_02580 | 42253..45123 | 1891238 | MAG:-DUF499-domain-containing-protein-[Burkholderiales-bacterium] |
| KAB2934118.1 | DUF499_1.RM.like.systems | <-PAPS_reduct<-?<-KAP-NTPase<- HEPN+ DUF499 *<- PIN <-OLD-ABC <- METHYLASE <-?<- TUDOR+ SF1-SF2-Helicase+iREase+CTD+REase | 1109 | Bacteria>Pseudomonadota>Betaproteobacteria>Rhodocyclales>Rhodocyclaceae | GCA_008933825.1 | Rhodocyclaceae_bacterium | F9K21_11715 | 6014..9343 | 1898103 | ATP-binding-protein-[Rhodocyclaceae-bacterium] |
| WP_168986546.1 | DUF499_1.RM.like.systems | Phage_integrase-><- DUF499 *<-?<-?<-ParB+HNH<- METHYLASE <- PIN <- TUDOR+ SF1-SF2-Helicase+iREase+CTD+REase | 944 | Bacteria>Pseudomonadota>Betaproteobacteria>Rhodocyclales>Zoogloeaceae | GCF_012910885.1 | Azoarcus_taiwanensis | GPA21_RS02025 | 735..3569 | 666964 | DUF499-domain-containing-protein-[Azoarcus-taiwanensis] |
| OGA65437.1 | DUF499_1.RM.like.systems | <- HEPN+ DUF499 *<- PIN <-MPTase<-?<-?<- METHYLASE <-ParB+HNH | 1109 | Bacteria>Pseudomonadota>Betaproteobacteria>unclassified_Betaproteobacteria | GCA_001770585.1 | Betaproteobacteria_bacterium_RIFCSPLOWO2_12_FULL_65_14 | A3G81_12440 | 218..3547 | 1797501 | AAA+-family-ATPase-[Betaproteobacteria-bacterium-RIFCSPLOWO2_12_FULL_65_14] |
| OGA54123.1 | DUF499_1.RM.like.systems | <- DUF499 *<- METHYLASE <- PIN <- TUDOR+ SF1-SF2-Helicase+iREase+CTD+REase \|\|?->?->MqsA_antitoxin+HTH_3-> | 1091 | Bacteria>Pseudomonadota>Betaproteobacteria>unclassified_Betaproteobacteria | GCA_001772005.1 | Betaproteobacteria_bacterium_RIFCSPLOWO2_12_FULL_62_58 | A3F74_24210 | 1834..5109 | 1797497 | hypothetical-protein-A3F74_24210-[Betaproteobacteria-bacterium-RIFCSPLOWO2_12_FULL_62_58] |
| MSQ72376.1 | DUF499_1.RM.like.systems | ABC-ATPase+SbcC->?->?->?->Phd-YefM-> PIN -> DUF499 *-><-DOC+HTH<-HTH<-RelE-ParE\|\|?-><-BirA-HTH | 104 | Bacteria>Pseudomonadota>Betaproteobacteria>unclassified_Betaproteobacteria | GCA_009693435.1 | Betaproteobacteria_bacterium | EXR27_13930 | 25540..25854 | 1891241 | hypothetical-protein-[Betaproteobacteria-bacterium] |
| GHU44749.1 | DUF499_1.RM.like.systems | <-Aminotran_1_2<- DUF499 *<-Schlafen+HATPase+HTH<- METHYLASE <- METHYLASE <- PIN <- TUDOR+ SF1-SF2-Helicase+iREase+CTD+REase \|\|Aminotran_3-> | 945 | Bacteria>Pseudomonadota>Betaproteobacteria>unclassified_Betaproteobacteria | GCA_025997095.1 | Betaproteobacteria_bacterium | AGMMS50289_13810 | 2255..5092 | 1891241 | hypothetical-protein-AGMMS50289_13810-[Betaproteobacteria-bacterium] |
| WP_273009210.1 | DUF499_1.RM.like.systems | <- DUF499 *<- METHYLASE <- PIN <- TUDOR+ SF1-SF2-Helicase+iREase+CTD+REase | 943 | Bacteria>Pseudomonadota>Gammaproteobacteria>Cellvibrionales>Spongiibacteraceae | GCF_002435105.1 | Spongiibacter_tropicus | B3U69_RS04085 | 395..3226 | 454602 | DUF499-domain-containing-protein-[Spongiibacter-tropicus] |
| MBP6807674.1 | DUF499_1.RM.like.systems | <- HEPN+ DUF499 *<-?<-?<- PIN <-?<- METHYLASE | 368 | Bacteria>Pseudomonadota>Gammaproteobacteria>Chromatiales>Chromatiaceae | GCA_017991215.1 | Chromatiaceae_bacterium | KA125_05205 | 1..1104 | 2049432 | DUF499-domain-containing-protein-[Chromatiaceae-bacterium] |
| WP_120796610.1 | DUF499_1.RM.like.systems | PrlF_antitoxin->RHH_6-> PIN -> METHYLASE -p->?->PolB-> HEPN+ DUF499 *-><-?\|\|?->RVT+HNH->RVT+HNH-> | 1114 | Bacteria>Pseudomonadota>Gammaproteobacteria>Chromatiales>Chromatiaceae | GCF_003634315.1 | Thiocapsa_rosea | BDD21_RS07415 | 1614812..1618156 | 69360 | Swt1-family-HEPN-domain-containing-protein-[Thiocapsa-rosea] |
| WP_019627826.1 | DUF499_1.RM.like.systems | Arm-DNA-bind_2+Phage_int_SAM_3+Phage_integrase->HDRnase->ZnR-> TUDOR+ SF1-SF2-Helicase+iREase+CTD+REase -> PIN -> METHYLASE -> DUF499 *->?->SF1-SF2-Helicase->?-><-?<-Phage_integrase | 951 | Bacteria>Pseudomonadota>Gammaproteobacteria>Chromatiales>Ectothiorhodospiraceae | GCF_000381845.1 | Thioalkalivibrio_sp._AKL10 | G333_RS0104940 | 1030421..1033276 | 1158158 | DUF499-domain-containing-protein-[Thioalkalivibrio-sp.-AKL10] |
| TVR61882.1 | DUF499_1.RM.like.systems | <- DUF499 *<- PIN <-DUF433<- METHYLASE -p | 1105 | Bacteria>Pseudomonadota>Gammaproteobacteria>Gammaproteobacteria_incertae_sedis>Candidatus_Competibacteraceae | GCA_007695245.1 | Candidatus_Competibacteraceae_bacterium | EA420_10930 | 75..3392 | 2053538 | ATP-binding-protein-[Candidatus-Competibacteraceae-bacterium] |

| Protein-ID | System | Gene-Neighbourhoods | Length | Taxonomy | Assembly-ID | Strain | Locus-Tag | Genomic-coordinates | TaxID | Fasta-defline |
|---|---|---|---|---|---|---|---|---|---|---|
| MBK8754773.1 | DUF499_1.RM.like.systems | METHYLASE ->OLD-ABC->TOPRIM->?->PIN->DUF499*->?-><-?<-CASPASE<-TIR<-?<-ADH_N+ADH_zinc_N | 946 | Bacteria>Pseudomonadota>Gammaproteobacteria>Gammaproteobacteria_incertae_sedis>Candidatus_Competibacteraceae | GCA_016710535.1 | Candidatus_Competibacteraceae_bacterium | IPL99_25425 | 33622..36462 | 2053538 | ATP-binding-protein-[Candidatus-Competibacteraceae-bacterium] |
| MBL8253032.1 | DUF499_1.RM.like.systems | <-DUF499*<-PIN<-DUF433<-METHYLASE<-TUDOR+ SF1-SF2-Helicase+iREase+CTD+REase | 937 | Bacteria>Pseudomonadota>Gammaproteobacteria>Gammaproteobacteria_incertae_sedis>Candidatus_Competibacteraceae | GCA_016789525.1 | Candidatus_Competibacter_sp. | JNJ76_05450 | 1..2812 | 2800415 | ATP-binding-protein-[Candidatus-Competibacter-sp.] |
| MCY4450795.1 | DUF499_1.RM.like.systems | <-STYKinase-like<-HTH<-DUF499*<-PIN<-?<-METHYLASE \|\|HicB_lk_antitox-><-Schlafen+HATPase+HTH<-HEPN | 1074 | Bacteria>Pseudomonadota>Gammaproteobacteria>Immundisolibacterales>unclassified_Immundisolibacterales | - | Immundisolibacterales_bacterium | OXC01_02485 | 119191..122415 | 2764111 | MAG:-DUF499-domain-containing-protein-[Immundisolibacterales-bacterium] |
| WP_218800673.1 | DUF499_1.RM.like.systems | <-TIR<-DUF499*<-REase<-OLD-ABC<-METHYLASE<-PIN<-TUDOR+ SF1-SF2-Helicase+iREase+CTD+REase <-HTH_Tnp | 942 | Bacteria>Pseudomonadota>Gammaproteobacteria>Methylococcales>Methylococcaceae | GCF_019285535.1 | Methylococcus_capsulatus | M3M30_RS10970 | 2380003..2382831 | 414 | DUF499-domain-containing-protein-[Methylococcus-capsulatus] |
| PZN71299.1 | DUF499_1.RM.like.systems | <-DUF499*<-PIN<-METHYLASE<-HNH<-OLD-ABC<-PemK_toxin<-PrlF_antitoxin | 955 | Bacteria>Pseudomonadota>Gammaproteobacteria>Methylococcales>Methylococcales_incertae_sedis | GCA_003242955.1 | Candidatus_Methylumidiphilus_alinenensis | DM484_26720 | 13367..16234 | 2202197 | hypothetical-protein-DM484_26720-[Candidatus-Methylumidiphilus-alinenensis] |
| MCK0538078.1 | DUF499_1.RM.like.systems | HTH+WYL->TUDOR+ SF1-SF2-Helicase+iREase+CTD+REase ->PIN->METHYLASE->DUF499*->SIR2-><-Phage_lysozyme<-?<-SF1-SF2-Helicase<-Schlafen+HATPase+HTH<-TRD+TRD | 947 | Bacteria>Pseudomonadota>Gammaproteobacteria>Oceanospirillales>Alcanivoracaceae | GCA_023051175.1 | Alcanivorax_quisquiliarum | MU846_10190 | 134529..137372 | 2933565 | DUF499-domain-containing-protein-[Alcanivorax-sp.-CY1518] |
| WP_246952339.1 | DUF499_1.RM.like.systems | HTH+WYL->TUDOR+ SF1-SF2-Helicase+iREase+CTD+REase ->PIN->METHYLASE->DUF499*->SIR2-><-Phage_lysozyme<-?<-SF1-SF2-Helicase<-Schlafen+HATPase+HTH<-TRD+TRD | 947 | Bacteria>Pseudomonadota>Gammaproteobacteria>Oceanospirillales>Alcanivoracaceae | GCF_023051175.1 | Alcanivorax_quisquiliarum | MU846_10190 | 134529..137372 | 2933565 | DUF499-domain-containing-protein-[Alcanivorax-sp.-CY1518] |
| EPL60615.1 | DUF499_1.RM.like.systems | <-RelE-ParE<-RelB<-PIN+OST-HTH<-?\|\|?->RlpA-><-DUF499*<-HEPN<-METHYLASE<-PIN<-TUDOR+ SF1-SF2-Helicase+iREase+CTD+REase <-MORC+DNA_gyraseB+TOPRIM<-SbcC | 946 | Bacteria>Pseudomonadota>Gammaproteobacteria>Pseudomonadales>Pseudomonadaceae | GCA_000416345.1 | Stutzerimonas_stutzeri_B1SMN1 | B382_19940 | 6557..9397 | 1212549 | hypothetical-protein-B382_19940-[Stutzerimonas-stutzeri-B1SMN1] |
| MTI93108.1 | DUF499_1.RM.like.systems | ParB->ParB->Classical-AAA->Peptidase_S8->?->RlpA-><-DUF499*<-ParB+HNH<-METHYLASE<-PIN<-TUDOR+ SF1-SF2-Helicase+iREase+CTD+REase <-MORC+DNA_gyraseB+TOPRIM<-SbcC | 946 | Bacteria>Pseudomonadota>Gammaproteobacteria>Pseudomonadales>Pseudomonadaceae | GCA_009711815.1 | Stutzerimonas_stutzeri | FH750_17245 | 14123..16963 | 316 | ATP-binding-protein-[Stutzerimonas-stutzeri] |
| HIQ43647.1 | DUF499_1.RM.like.systems | ParB->ParB-><-MORC+HisKin\|\|<-RVT\|\|?->RlpA-><-DUF499*<-ParB+HNH<-METHYLASE<-PIN<-TUDOR+ SF1-SF2-Helicase+iREase+CTD+REase <-MORC+DNA_gyraseB+TOPRIM<-SbcC | 946 | Bacteria>Pseudomonadota>Gammaproteobacteria>Pseudomonadales>Pseudomonadaceae | GCA_015661735.1 | Pseudomonas_oleovorans | EYH47_12960 | 31988..34828 | 301 | ATP-binding-protein-[Pseudomonas-oleovorans] |
| MBG3917698.1 | DUF499_1.RM.like.systems | SbcC->MORC+DNA_gyraseB+TOPRIM->TUDOR+ SF1-SF2-Helicase+iREase+CTD+REase ->PIN->METHYLASE->ParB+HNH->DUF499*-><-RlpA<-?<-Peptidase_S8<-Classical-AAA<-ParB<-ParB | 946 | Bacteria>Pseudomonadota>Gammaproteobacteria>Pseudomonadales>Pseudomonadaceae | GCA_015732935.1 | Pseudomonas_aeruginosa | I4931_21710 | 92367..95207 | 287 | ATP-binding-protein-[Pseudomonas-aeruginosa] |
| MCF6804930.1 | DUF499_1.RM.like.systems | ParB->Classical-AAA->Peptidase_S8->?->?->RlpA-><-DUF499*<-ParB+HNH<-METHYLASE<-PIN<-TUDOR+ SF1-SF2-Helicase+iREase+CTD+REase <-MORC+DNA_gyraseB+TOPRIM<-SbcC | 889 | Bacteria>Pseudomonadota>Gammaproteobacteria>Pseudomonadales>Pseudomonadaceae | GCA_021713635.1 | Stutzerimonas_stutzeri | L3X13_09150 | 18126..20795 | 316 | DUF499-domain-containing-protein-[Stutzerimonas-stutzeri] |
| MCF6781034.1 | DUF499_1.RM.like.systems | ParB->Classical-AAA->Peptidase_S8->?->?->RlpA-><-DUF499*<-ParB+HNH<-METHYLASE<-PIN<-TUDOR+ SF1-SF2-Helicase+iREase+CTD+REase <-MORC+DNA_gyraseB+TOPRIM<-SbcC | 946 | Bacteria>Pseudomonadota>Gammaproteobacteria>Pseudomonadales>Pseudomonadaceae | GCA_021713665.1 | Stutzerimonas_stutzeri | L3X17_04510 | 18122..20962 | 316 | DUF499-domain-containing-protein-[Stutzerimonas-stutzeri] |
| WP_125852814.1 | DUF499_1.RM.like.systems | SbcC->MORC+DNA_gyraseB+TOPRIM->TUDOR+ SF1-SF2-Helicase+iREase+CTD+REase ->PIN->METHYLASE->ParB+HNH->DUF499*-><-RlpA<-ParB\|\|REase-> | 946 | Bacteria>Pseudomonadota>Gammaproteobacteria>Pseudomonadales>Pseudomonadaceae | GCF_003936145.1 | Stutzerimonas_stutzeri | EGJ42_RS05720 | 52381..55221 | 316 | DUF499-domain-containing-protein-[Stutzerimonas-stutzeri] |
| WP_143508093.1 | DUF499_1.RM.like.systems | SbcC->MORC+DNA_gyraseB+TOPRIM->TUDOR+ SF1-SF2-Helicase+iREase+CTD+REase ->PIN->METHYLASE->ParB+HNH->DUF499*-><-RlpA<-?<-Peptidase_S8<-Classical-AAA<-ParB<-ParB | 946 | Bacteria>Pseudomonadota>Gammaproteobacteria>Pseudomonadales>Pseudomonadaceae | GCF_007049885.1 | Pseudomonas_mendocina | EQ828_RS12690 | 220581..223421 | 300 | DUF499-domain-containing-protein-[Pseudomonas-mendocina] |
| WP_213918359.1 | DUF499_1.RM.like.systems | <-rve_3+Mu-transpos_C<-REase\|\|ParB->RlpA-><-DUF499*<-HEPN<-METHYLASE<-PIN<-TUDOR+ SF1-SF2-Helicase+iREase+CTD+REase <-MORC+DNA_gyraseB+TOPRIM<-SbcC | 946 | Bacteria>Pseudomonadota>Gammaproteobacteria>Pseudomonadales>Pseudomonadaceae | GCF_018457005.1 | Stutzerimonas_stutzeri | JR047_RS05385 | 4622..7462 | 316 | DUF499-domain-containing-protein-[Stutzerimonas-stutzeri] |
| WP_226045701.1 | DUF499_1.RM.like.systems | SbcC->MORC+DNA_gyraseB+TOPRIM->TUDOR+ SF1-SF2-Helicase+iREase+CTD+REase ->PIN->METHYLASE->ParB+HNH->DUF499*-><-RlpA<-?<-Peptidase_S8<-Classical-AAA<-ParB<-ParB | 946 | Bacteria>Pseudomonadota>Gammaproteobacteria>Pseudomonadales>Pseudomonadaceae | GCF_020401845.1 | Pseudomonas_sp._HS-18 | LDJ84_RS00045 | 14958..17798 | 2879114 | DUF499-domain-containing-protein-[Pseudomonas-sp.-HS-18] |
| WP_129484085.1 | DUF499_1.RM.like.systems | SbcC->MORC+DNA_gyraseB+TOPRIM->TUDOR+ SF1-SF2-Helicase+iREase+CTD+REase ->PIN->METHYLASE->ParB+HNH->DUF499*-><-RlpA<-?<-Peptidase_S8<-Classical-AAA<-ParB<-ParB | 946 | Bacteria>Pseudomonadota>Gammaproteobacteria>Pseudomonadales>Pseudomonadaceae | GCF_021495345.1 | Pseudomonas_hydrolytica | L1F06_RS00045 | 14914..17754 | 2493633 | DUF499-domain-containing-protein-[Pseudomonas-hydrolytica] |
| WP_279641602.1 | DUF499_1.RM.like.systems | SbcC->MORC+DNA_gyraseB+TOPRIM->TUDOR+ SF1-SF2-Helicase+iREase+CTD+REase ->PIN->METHYLASE->ParB+HNH->DUF499*->?-><-RlpA<-?\|\|RVT-><-ParB<-ParB | 946 | Bacteria>Pseudomonadota>Gammaproteobacteria>Pseudomonadales>Pseudomonadaceae | GCF_029839225.1 | Stutzerimonas_stutzeri | N5C32_RS13925 | 184183..187023 | 316 | DUF499-domain-containing-protein-[Stutzerimonas-stutzeri] |
| WP_283258799.1 | DUF499_1.RM.like.systems | HIN-HTH->?->RlpA-><-DUF499*<-ParB+HNH<-METHYLASE<-PIN<-TUDOR+ SF1-SF2-Helicase+iREase+CTD+REase <-MORC+DNA_gyraseB+TOPRIM<-SbcC | 946 | Bacteria>Pseudomonadota>Gammaproteobacteria>Pseudomonadales>Pseudomonadaceae | GCF_030056015.1 | Stutzerimonas_stutzeri | QM290_RS03915 | 3700..6540 | 316 | DUF499-domain-containing-protein-[Stutzerimonas-stutzeri] |
| WP_046835095.1 | DUF499_1.RM.like.systems | TUDOR+ SF1-SF2-Helicase+iREase+CTD+REase ->PIN->METHYLASE-p->DUF772+DDE_Tnp_1->METHYLASE->ParB+HNH->DUF499*-><-RlpA<-?\|\|CHTH-><-LZ_Tnp_IS66+zf-IS66+DDE_Tnp_IS66+DDE_Tnp_IS66_C | 944 | Bacteria>Pseudomonadota>Gammaproteobacteria>Pseudomonadales>Pseudomonadaceae | GCF_030252675.1 | Pseudomonas_amygdali | QO021_RS27770 | 6030244..6033078 | 47877 | DUF499-domain-containing-protein-[Pseudomonas-amygdali] |
| WP_110599711.1 | DUF499_1.RM.like.systems | ParB-><-PIN<-HTH+MPTase\|\|?->?->RlpA-><-DUF499*<-ParB+HNH<-METHYLASE<-PIN<-TUDOR+ SF1-SF2-Helicase+iREase+CTD+REase <-MORC+DNA_gyraseB+TOPRIM<-SbcC | 945 | Bacteria>Pseudomonadota>Gammaproteobacteria>Pseudomonadales>Pseudomonadaceae | - | Pseudomonas_koreensis | DMX04_RS04460 | 134660..137497 | 198620 | DUF499-domain-containing-protein-[Pseudomonas-koreensis] |
| WP_003246716.1 | DUF499_1.RM.like.systems | SbcC->MORC+DNA_gyraseB+TOPRIM->TUDOR+ SF1-SF2-Helicase+iREase+CTD+REase ->PIN->METHYLASE->ParB+HNH->DUF499*-><-RlpA<-?<-?\|\|RVT-> | 946 | Bacteria>Pseudomonadota>Gammaproteobacteria>Pseudomonadales>Pseudomonadaceae | - | Pseudomonas_mendocina | A471_RS19815 | 19321..22161 | 300 | DUF499-domain-containing-protein-[Pseudomonas-mendocina] |
| WP_020308343.1 | DUF499_1.RM.like.systems | <-RelE-ParE<-RelB<-PIN+OST-HTH<-?\|\|?->RlpA-><-DUF499*<-HEPN<-METHYLASE<-PIN<-TUDOR+ SF1-SF2-Helicase+iREase+CTD+REase <-MORC+DNA_gyraseB+TOPRIM<-SbcC | 946 | Bacteria>Pseudomonadota>Gammaproteobacteria>Pseudomonadales>Pseudomonadaceae | - | Stutzerimonas_stutzeri | B382_19940 | 6557..9397 | 316 | DUF499-domain-containing-protein-[Stutzerimonas-stutzeri] |
| WP_125867554.1 | DUF499_1.RM.like.systems | SbcC->MORC+DNA_gyraseB+TOPRIM->TUDOR+ SF1-SF2-Helicase+iREase+CTD+REase ->PIN->METHYLASE->ParB+HNH->DUF499*->HIN-HTH->?->LZ_Tnp_IS66+zf-IS66+DDE_Tnp_IS66+DDE_Tnp_IS66_C-><-?<-?<-Peptidase_S8 | 946 | Bacteria>Pseudomonadota>Gammaproteobacteria>Pseudomonadales>Pseudomonadaceae | - | Stutzerimonas_stutzeri | EGI99_RS04500 | 994408..997248 | 316 | DUF499-domain-containing-protein-[Stutzerimonas-stutzeri] |
| MAS10374.1 | DUF499_1.RM.like.systems | <-HhH-RADC+JAB\|\|?->?->?-><-DUF499*<-TOPRIM<-OLD-ABC<-METHYLASE<-PIN<-TUDOR+ SF1-SF2-Helicase+iREase+CTD+REase <-JetD | 942 | Bacteria>Pseudomonadota>Gammaproteobacteria>Salinisphaerales>Salinisphaeraceae | GCA_002702145.1 | Salinisphaera_sp. | CMN26_09985 | 5272..8100 | 1914330 | hypothetical-protein-[Salinisphaera-sp.] |
| MBU1235382.1 | DUF499_1.RM.like.systems | <-VirD4-TraG+VirD4-FtsK\|\|TUDOR+ SF1-SF2-Helicase+iREase+CTD+REase ->STAS+HisKin->?->PIN->METHYLASE->HEPN+ DUF499 *-><-?<-?<-?<-ParB | 1113 | Bacteria>Pseudomonadota>Gammaproteobacteria>unclassified_Gammaproteobacteria | GCA_018819515.1 | Gammaproteobacteria_bacterium | KJ788_01195 | 188697..192038 | 1913989 | DUF499-domain-containing-protein-[Gammaproteobacteria-bacterium] |
| MCY4157213.1 | DUF499_1.RM.like.systems | TUDOR+ SF1-SF2-Helicase+iREase+CTD+REase ->?->?->METHYLASE->DUF433->PIN->DUF499*-><-LD-peptidase | 954 | Bacteria>Pseudomonadota>Gammaproteobacteria>unclassified_Gammaproteobacteria | GCA_026708965.1 | Gammaproteobacteria_bacterium | OXF66_07255 | 51365..54229 | 1913989 | MAG:-DUF499-domain-containing-protein-[Gammaproteobacteria-bacterium] |
| MCY4164747.1 | DUF499_1.RM.like.systems | TUDOR+ SF1-SF2-Helicase+iREase+CTD+REase ->?->?->METHYLASE->DUF433->PIN->DUF499*-><-LD-peptidase | 953 | Bacteria>Pseudomonadota>Gammaproteobacteria>unclassified_Gammaproteobacteria | GCA_026709045.1 | Gammaproteobacteria_bacterium | OXF03_01150 | 156657..159518 | 1913989 | MAG:-DUF499-domain-containing-protein-[Gammaproteobacteria-bacterium] |
| MCY4199355.1 | DUF499_1.RM.like.systems | AbiEi_1->?->?-><-HEPN<-?<-TPR+TPR+PIN+PIN<-DUF499*<-PIN<-DUF433<-METHYLASE<-?<-TUDOR+ SF1-SF2-Helicase+iREase+CTD+REase \|\|RVT-> | 924 | Bacteria>Pseudomonadota>Gammaproteobacteria>unclassified_Gammaproteobacteria | GCA_026709225.1 | Gammaproteobacteria_bacterium | OXF31_05975 | 33360..36134 | 1913989 | MAG:-DUF499-domain-containing-protein-[Gammaproteobacteria-bacterium] |
| MCY4200247.1 | DUF499_1.RM.like.systems | AbiEii->?->?-><-DUF499*<-METHYLASE<-?<-PIN<-TUDOR+ SF1-SF2-Helicase+iREase+CTD+REase \|\|?->PIN-> | 1089 | Bacteria>Pseudomonadota>Gammaproteobacteria>unclassified_Gammaproteobacteria | GCA_026709225.1 | Gammaproteobacteria_bacterium | OXF31_10650 | 19603..22872 | 1913989 | MAG:-hypothetical-protein-OXF31_10650-[Gammaproteobacteria-bacterium] |
| MCY4255936.1 | DUF499_1.RM.like.systems | TUDOR+ SF1-SF2-Helicase+iREase+CTD+REase ->?->?->METHYLASE->DUF433->PIN->DUF499*-><-LD-peptidase | 953 | Bacteria>Pseudomonadota>Gammaproteobacteria>unclassified_Gammaproteobacteria | GCA_026709685.1 | Gammaproteobacteria_bacterium | OXE51_07470 | 8826..11687 | 1913989 | MAG:-DUF499-domain-containing-protein-[Gammaproteobacteria-bacterium] |
| MCY4278451.1 | DUF499_1.RM.like.systems | <-PIN<-?\|\|TUDOR+ SF1-SF2-Helicase+iREase+CTD+REase ->PIN->?->METHYLASE->DUF499*-><-?<-?<-AbiEii | 1093 | Bacteria>Pseudomonadota>Gammaproteobacteria>unclassified_Gammaproteobacteria | GCA_026709965.1 | Gammaproteobacteria_bacterium | OXE81_11575 | 12290..15571 | 1913989 | MAG:-hypothetical-protein-OXE81_11575-[Gammaproteobacteria-bacterium] |
| MCY4342454.1 | DUF499_1.RM.like.systems | ORC-CDC6+REase-><-DUF499*<-PIN<-DUF433<-METHYLASE-p<-METHYLASE<-Peptidase_S8<-Classical-AAA | 924 | Bacteria>Pseudomonadota>Gammaproteobacteria>unclassified_Gammaproteobacteria | GCA_026710215.1 | Gammaproteobacteria_bacterium | OXE83_02620 | 2601..5375 | 1913989 | MAG:-DUF499-domain-containing-protein-[Gammaproteobacteria-bacterium] |
| MCY4343961.1 | DUF499_1.RM.like.systems | METHYLASE->OLD-ABC->Schlafen+HATPase+HTH->?->PIN->HEPN+ DUF499 *-><-MNS+ZDNA-HTH<-PRTase | 1115 | Bacteria>Pseudomonadota>Gammaproteobacteria>unclassified_Gammaproteobacteria | GCA_026710215.1 | Gammaproteobacteria_bacterium | OXE83_10365 | 11310..14657 | 1913989 | MAG:-Swt1-family-HEPN-domain-containing-protein-[Gammaproteobacteria-bacterium] |
| MCY4340924.1 | DUF499_1.RM.like.systems | LD-peptidase-><-DUF499*<-PIN<-DUF433<-METHYLASE<-?<-?<-TUDOR+ SF1-SF2-Helicase+iREase+CTD+REase | 954 | Bacteria>Pseudomonadota>Gammaproteobacteria>unclassified_Gammaproteobacteria | GCA_026710305.1 | Gammaproteobacteria_bacterium | OXE48_06110 | 63766..66630 | 1913989 | MAG:-DUF499-domain-containing-protein-[Gammaproteobacteria-bacterium] |
| MCY4533098.1 | DUF499_1.RM.like.systems | HTH+WYL->TUDOR+ SF1-SF2-Helicase+iREase+CTD+REase ->PIN->METHYLASE->DUF499*->?-><-Peptidase_S8<-Classical-AAA | 931 | Bacteria>Pseudomonadota>Gammaproteobacteria>unclassified_Gammaproteobacteria | GCA_026713185.1 | Gammaproteobacteria_bacterium | OXC84_12365 | 15998..18793 | 1913989 | MAG:-DUF499-domain-containing-protein-[Gammaproteobacteria-bacterium] |
| MDD9824620.1 | DUF499_1.RM.like.systems | <-DUF499*<-PIN<-OLD-ABC<-OLD-ABC+TOPRIM<-METHYLASE<-TPR+TPR | 928 | Bacteria>Pseudomonadota>Gammaproteobacteria>unclassified_Gammaproteobacteria | GCA_028817255.1 | Gammaproteobacteria_bacterium | OXU43_05565 | 147..2933 | 1913989 | MAG:-DUF499-domain-containing-protein-[Gammaproteobacteria-bacterium] |
| MDE0036818.1 | DUF499_1.RM.like.systems | METHYLASE-p->?-><-PIN<-OLD-ABC<-?<-HEPN+ DUF499 *<-METHYLASE<-OLD-ABC<-TUDOR+ SF1-SF2-Helicase+iREase+CTD+REase <-?<-?\|\|SF1-SF2-Helicase-> | 1118 | Bacteria>Pseudomonadota>Gammaproteobacteria>unclassified_Gammaproteobacteria | GCA_028818795.1 | Gammaproteobacteria_bacterium | OXU77_04540 | 7629..10985 | 1913989 | MAG:-Swt1-family-HEPN-domain-containing-protein-[Gammaproteobacteria-bacterium] |
| MDE0258535.1 | DUF499_1.RM.like.systems | <-TraI-VirD2-RCRE<-TraJ-RHH<-Phage_integrase<-DUF499*<-PIN<-?<-METHYLASE<-TUDOR+ SF1-SF2-Helicase+iREase+CTD+REase | 1103 | Bacteria>Pseudomonadota>Gammaproteobacteria>unclassified_Gammaproteobacteria | GCA_028820325.1 | Gammaproteobacteria_bacterium | OXR82_09155 | 1794..5105 | 1913989 | MAG:-DUF499-domain-containing-protein-[Gammaproteobacteria-bacterium] |
| WP_008437332.1 | DUF499_1.RM.like.systems | <-AbiEi_4\|\|?->?->XIS-HTH->XIS-HTH->Phage_int_SAM_1+Phage_integrase-><-DUF499*<-?<-METHYLASE<-?<-PIN<-TUDOR+ SF1-SF2-Helicase+iREase+CTD+REase | 933 | Bacteria>Pseudomonadota>Gammaproteobacteria>Xanthomonadales>Rhodanobacteraceae | GCF_000264375.1 | Rhodanobacter_thiooxydans | RHOFW104T7_RS11960 | 24308..27109 | 416169 | DUF499-domain-containing-protein-[Rhodanobacter-thiooxydans] |
| WP_243041017.1 | DUF499_1.RM.like.systems | HTH+WYL->HTH_Tnp->TUDOR+ SF1-SF2-Helicase+iREase+CTD+REase ->PIN->METHYLASE->DUF499*->SbcC-><-Phage_lysozyme<-?<-?<-RNase | 955 | Bacteria>Pseudomonadota>Gammaproteobacteria>Xanthomonadales>Rhodanobacteraceae | GCF_022760995.1 | Dyella_sedimenti | MJ605_RS08555 | 206646..209513 | 2919947 | DUF499-domain-containing-protein-[Dyella-sp.-NS14] |
| MCK9487878.1 | DUF499_1.RM.like.systems | <-SIGMA-HTH<-OLD-ABC<-?<-?<-?<-DUF499*<-REase<-OLD-ABC<-METHYLASE<-HEPN<-PIN<-PemK_toxin | 944 | Bacteria>Pseudomonadota>Gammaproteobacteria>Xanthomonadales>unclassified_Xanthomonadales | GCA_023228845.1 | Xanthomonadales_bacterium | MOQ42_00470 | 105333..108167 | 2006849 | DUF499-domain-containing-protein-[Xanthomonadales-bacterium] |
| GIX32370.1 | DUF499_1.RM.like.systems | REase->METHYLASE-p->METHYLASE-p->METHYLASE->ParD->PIN->HEPN+ DUF499 *-><-tRNA<-SIGMA-HTH+SIGMA-HTH | 1108 | Bacteria>Pseudomonadota>Gammaproteobacteria>Xanthomonadales>unclassified_Xanthomonadales | GCA_026005055.1 | Xanthomonadales_bacterium | KatS3mg125_0326 | 361925..365251 | 2006849 | MAG:-hypothetical-protein-KatS3mg125_0326-[Xanthomonadales-bacterium] |
| PZO64463.1 | DUF499_1.RM.like.systems | <-DUF499*<-TOPRIM<-OLD-ABC<-METHYLASE<-PIN<-TUDOR+ SF1-SF2-Helicase+iREase+CTD+REase | 932 | Bacteria>Pseudomonadota>Gammaproteobacteria>Xanthomonadales>Xanthomonadaceae | GCA_003240695.1 | Pseudoxanthomonas_suwonensis | DI635_05295 | 731..3529 | 314722 | hypothetical-protein-DI635_05295-[Pseudoxanthomonas-suwonensis] |
| KAA2283481.1 | DUF499_1.RM.like.systems | HTH+WYL->HTH_Tnp->TUDOR+ SF1-SF2-Helicase+iREase+CTD+REase ->PIN->METHYLASE->DUF499*->SIR2+STALD->TIR->Methylase-><-Radical-SAM<-HTH_3<-Phage_lysozyme | 955 | Bacteria>Pseudomonadota>Gammaproteobacteria>Xanthomonadales>Xanthomonadaceae | GCA_008386465.1 | Arenimonas_fontis | F0415_12220 | 12040..14907 | 2608255 | ATP-binding-protein-[Arenimonas-fontis] |
| MBU6247856.1 | DUF499_1.RM.like.systems | <-DUF499*<-ParB+HNH<-?<-HEPN+CBS<-METHYLASE<-PIN | 934 | Bacteria>Pseudomonadota>Gammaproteobacteria>Xanthomonadales>Xanthomonadaceae | GCA_018970895.1 | Xanthomonadaceae_bacterium | KGN77_08870 | 9374..12178 | 1926873 | DUF499-domain-containing-protein-[Xanthomonadaceae-bacterium] |
| WP_034214097.1 | DUF499_1.RM.like.systems | <-DUF499*<-?<-METHYLASE<-PIN<-DOC+HTH<-TUDOR+ SF1-SF2-Helicase+iREase+CTD+REase | 933 | Bacteria>Pseudomonadota>Gammaproteobacteria>Xanthomonadales>Xanthomonadaceae | GCF_000747155.1 | Arenimonas_metalli | N787_RS11310 | 49441..52242 | 948077 | DUF499-domain-containing-protein-[Arenimonas-metalli] |
| WP_055771167.1 | DUF499_1.RM.like.systems | <-D5N+D5ATPase<-?<-?<-?<-?<-?<-DUF499*<-ParB+HNH<-?<-METHYLASE<-PIN<-TUDOR+ SF1-SF2-Helicase+iREase+CTD+REase <-Phage_integrase | 929 | Bacteria>Pseudomonadota>Gammaproteobacteria>Xanthomonadales>Xanthomonadaceae | GCF_001422025.1 | Stenotrophomonas_sp._Leaf70 | ASF01_RS13960 | 54995..57784 | 1736233 | DUF499-domain-containing-protein-[Stenotrophomonas-sp.-Leaf70] |
| WP_106991537.1 | DUF499_1.RM.like.systems | <-DUF499*<-?<-METHYLASE<-HEPN<-PIN<-TUDOR+ SF1-SF2-Helicase+iREase+CTD+REase <-HTH_26 | 930 | Bacteria>Pseudomonadota>Gammaproteobacteria>Xanthomonadales>Xanthomonadaceae | GCF_003024235.1 | Arenimonas_caeni | C6N40_RS13415 | 668..3460 | 2058085 | DUF499-domain-containing-protein-[Arenimonas-caeni] |
| WP_246149960.1 | DUF499_1.RM.like.systems | HTH+WYL->HTH_Tnp->TUDOR+ SF1-SF2-Helicase+iREase+CTD+REase ->PIN->METHYLASE->DUF499*->SIR2+STALD->TIR->Methylase-><-Radical-SAM<-HTH_3<-Phage_lysozyme | 959 | Bacteria>Pseudomonadota>Gammaproteobacteria>Xanthomonadales>Xanthomonadaceae | GCF_008386465.1 | Arenimonas_fontis | F0415_RS12225 | 12028..14907 | 2608255 | DUF499-domain-containing-protein-[Arenimonas-fontis] |
| WP_028873821.1 | DUF499_1.RM.like.systems | DDE_Tnp_1_assoc+DDE_Tnp_1->PIN->HEPN->METHYLASE->?->DUF499*->?->PIN->?-><-?<-RuvC | 947 | Bacteria>Pseudomonadota>Hydrogenophilia>Hydrogenophilales>Hydrogenophilaceae | GCF_000425565.1 | Tepidiphilus_margaritifer | H607_RS0100570 | 5909..8752 | 203471 | DUF499-domain-containing-protein-[Tepidiphilus-margaritifer] |
| WP_169116186.1 | DUF499_1.RM.like.systems | HEPN->PolB->TUDOR+ SF1-SF2-Helicase+iREase+CTD+REase ->PIN->METHYLASE->DUF499*->?-><-?<-RuvC<-?\|\|SF1-SF2-Helicase-> | 948 | Bacteria>Pseudomonadota>Hydrogenophilia>Hydrogenophilales>Hydrogenophilaceae | GCF_012911495.1 | Tepidiphilus_baoligensis | GV368_RS08285 | 50966..53812 | 2698687 | DUF499-domain-containing-protein-[Tepidiphilus-baoligensis] |
| WP_219972353.1 | DUF499_1.RM.like.systems | MODE-HTH+PBPII->?-><-tRNA\|\|?->?-><-?<-DUF499*<-?<-METHYLASE<-PIN<-TUDOR+ SF1-SF2-Helicase+iREase+CTD+REase | 955 | Bacteria>Pseudomonadota>Hydrogenophilia>Hydrogenophilales>Hydrogenophilaceae | GCF_019448605.1 | Hydrogenophilus_thermoluteolus | K1I42_RS04670 | 179917..182784 | 297 | DUF499-domain-containing-protein-[Hydrogenophilus-thermoluteolus] |
| MCL1861026.1 | DUF499_1.RM.like.systems | PIN->METHYLASE->?->DUF499*-><-?<-?<-ZnR+TOPRIM+D5ATPase<-?<-HTH_17 | 943 | Bacteria>Pseudomonadota>unclassified_Pseudomonadota | GCA_009778425.1 | Pseudomonadota_bacterium | FWG52_05760 | 4102..6933 | 1977087 | DUF499-domain-containing-protein-[Proteobacteria-bacterium] |
| MBE9569155.1 | DUF499_1.RM.like.systems | PIN->METHYLASE->HEPN+ DUF499 *-> | 1063 | Bacteria>Pseudomonadota>unclassified_Pseudomonadota | GCA_015222845.1 | Pseudomonadota_bacterium | IMF11_00880 | 3390..6577 | 1977087 | DUF499-domain-containing-protein-[Proteobacteria-bacterium] |
| MBQ9395962.1 | DUF499_1.RM.like.systems | METHYLASE-p-><-PIN<-?\|\|HEPN+ DUF499 *-> | 729 | Bacteria>Pseudomonadota>unclassified_Pseudomonadota | GCA_017455165.1 | Pseudomonadota_bacterium | IJU23_10695 | 3126..5311 | 1977087 | DUF499-domain-containing-protein-[Proteobacteria-bacterium] |
| MBS0576259.1 | DUF499_1.RM.like.systems | RNase->?->?->Phage_lysozyme-><-SbcC<-DUF499*<-METHYLASE<-HEPN<-PIN<-TUDOR+ SF1-SF2-Helicase+iREase+CTD+REase <-HTH_Tnp<-HTH+WYL | 955 | Bacteria>Pseudomonadota>unclassified_Pseudomonadota | GCA_018240435.1 | Pseudomonadota_bacterium | JSS45_07535 | 37839..40706 | 1977087 | ATP-binding-protein-[Proteobacteria-bacterium] |
| MBS0225815.1 | DUF499_1.RM.like.systems | ACET->?-><-?\|\|HEPN-><-DUF499*<-?<-?<-PIN<-TUDOR+ SF1-SF2-Helicase+iREase+CTD+REase | 929 | Bacteria>Pseudomonadota>unclassified_Pseudomonadota | GCA_018242465.1 | Pseudomonadota_bacterium | JSS25_05685 | 37623..40412 | 1977087 | ATP-binding-protein-[Proteobacteria-bacterium] |
| MBU0673024.1 | DUF499_1.RM.like.systems | TUDOR+ SF1-SF2-Helicase+iREase+CTD+REase ->PIN->?->METHYLASE->OLD-ABC->REase->DUF499*-> | 943 | Bacteria>Pseudomonadota>unclassified_Pseudomonadota | GCA_018814305.1 | Pseudomonadota_bacterium | KJ950_00090 | 19515..22346 | 1977087 | DUF499-domain-containing-protein-[Proteobacteria-bacterium] |
| MDE3269952.1 | DUF499_1.RM.like.systems | REase-><-TUDOR+ SF1-SF2-Helicase+iREase+CTD+REase <-?<-DUF499*<-PIN<-MPTase<-HTH<-?<-METHYLASE | 917 | Bacteria>Pseudomonadota>unclassified_Pseudomonadota | GCA_028877195.1 | Pseudomonadota_bacterium | OYH77_06680 | 5373..8126 | 1977087 | MAG:-hypothetical-protein-OYH77_06680-[Pseudomonadota-bacterium] |
| MBP7275753.1 | DUF499_1.RM.like.systems | <-DUF499*<-?<-METHYLASE<-PIN<-?<-REase | 1001 | Bacteria>PVC_group>Kiritimatiellota>Kiritimatiellia>unclassified_Kiritimatiellia | GCA_017996315.1 | Kiritimatiellia_bacterium | KBA51_06085 | 1..3004 | 2212473 | ATP-binding-protein-[Kiritimatiellae-bacterium] |
| MBN2450533.1 | DUF499_1.RM.like.systems | REase->?->PIN->HEPN+ DUF499 *-> | 1121 | Bacteria>PVC_group>Lentisphaerota>Lentisphaeria>unclassified_Lentisphaeria | GCA_016935045.1 | Lentisphaeria_bacterium | JXR77_09085 | 1530..4895 | 2053569 | DUF499-domain-containing-protein-[Lentisphaeria-bacterium] |
| MBT5607699.1 | DUF499_1.RM.like.systems | PIN->METHYLASE->DUF499*->?->DUF6361->HKD-DNase->SF1-SF2-Helicase->ORC-CDC6+HTH-> | 1109 | Bacteria>PVC_group>Lentisphaerota>unclassified_Lentisphaerota | GCA_018666605.1 | Lentisphaerota_bacterium | HOJ57_17295 | 4653..7982 | 1932692 | ATP-binding-protein-[Lentisphaerae-bacterium] |
| KXK32838.1 | DUF499_1.RM.like.systems | <-DUF1264\|\|?-><-?<-?<-DUF499*<-DUF433<-METHYLASE-p<-METHYLASE<-PIN | 925 | Bacteria>PVC_group>Planctomycetota>Candidatus_Brocadiia>Candidatus_Brocadiales | GCA_001567345.1 | Candidatus_Brocadia_sinica | UZ01_00260 | 122708..125485 | 795830 | hypothetical-protein-UZ01_00260-[Candidatus-Brocadia-sinica] |
| WP_052562727.1 | DUF499_1.RM.like.systems | TUDOR+ SF1-SF2-Helicase+iREase+CTD+REase ->?->?->PIN->METHYLASE->DUF433->DUF499*->?->?-><-?\|\|DUF1264-> | 925 | Bacteria>PVC_group>Planctomycetota>Candidatus_Brocadiia>Candidatus_Brocadiales | GCF_000949635.1 | Brocadia | FBQ80_RS01160 | 33537..36314 | 380240 | DUF499-domain-containing-protein-[Candidatus-Brocadia-sinica] |
| MBX3391865.1 | DUF499_1.RM.like.systems | <-ParA-Soj-PloopNTPase\|\|?->?-><-?<-N6_N4_MTase<-DUF499*<-PIN<-METHYLASE<-?<-TUDOR+ SF1-SF2-Helicase+iREase+CTD+REase | 1114 | Bacteria>PVC_group>Planctomycetota>Phycisphaerae>Phycisphaerales | GCA_019636515.1 | Phycisphaeraceae_bacterium | KF787_04385 | 1038912..1042256 | 2026777 | ATP-binding-protein-[Phycisphaeraceae-bacterium] |
| MCK4341360.1 | DUF499_1.RM.like.systems | HTH_3-><-GDResolvase+Recombinase+ZnR<-?<-DUF499*<-PIN<-?<-?<-METHYLASE-p | 947 | Bacteria>PVC_group>Planctomycetota>Phycisphaerae>unclassified_Phycisphaerae | GCA_023128695.1 | Phycisphaerae_bacterium | KAY37_06525 | 2745..5588 | 2026778 | ATP-binding-protein-[Phycisphaerae-bacterium] |
| MSR55922.1 | DUF499_1.RM.like.systems | <-DUF499*<-REase<-PIN<-DUF433<-METHYLASE-p | 984 | Bacteria>PVC_group>Planctomycetota>Planctomycetia>Gemmatales | GCA_009691845.1 | Gemmataceae_bacterium | EXS09_22015 | 769..3723 | 2052164 | ATP-binding-protein-[Gemmataceae-bacterium] |
| GIW84013.1 | DUF499_1.RM.like.systems | GDResolvase+Recombinase+ZnR->ParB->TUDOR+ SF1-SF2-Helicase+iREase+CTD+REase ->METHYLASE->DUF433->PIN->DUF499*->HTH->MPTase-><-DUF4096+DDE_Tnp_1+DDE_Tnp | 924 | Bacteria>PVC_group>Planctomycetota>Planctomycetia>Gemmatales | GCA_026004695.1 | Gemmataceae_bacterium | KatS3mg106_526 | 15876..18650 | 2052164 | MAG:-hypothetical-protein-KatS3mg106_526-[Gemmataceae-bacterium] |
| MDB5306879.1 | DUF499_1.RM.like.systems | <-DUF499*<-PIN<-Phd-YefM<-METHYLASE<-TUDOR+ SF1-SF2-Helicase+iREase+CTD+REase -p<-TUDOR+ SF1-SF2-Helicase+iREase+CTD+REase | 207 | Bacteria>PVC_group>Planctomycetota>Planctomycetia>Gemmatales | GCA_028288765.1 | Gemmataceae_bacterium | JWO38_1081 | 91..714 | 2052164 | MAG:-hypothetical-protein-JWO38_1081-[Gemmataceae-bacterium] |
| MBV8611623.1 | DUF499_1.RM.like.systems | METHYLASE-p->DUF433->PIN->HTH->DUF499*-> | 454 | Bacteria>PVC_group>Planctomycetota>Planctomycetia>Isosphaerales | GCA_019233025.1 | Singulisphaera_sp. | JO034_29790 | 1731..3093 | 1906241 | ATP-binding-protein-[Singulisphaera-sp.] |
| WP_013566219.1 | DUF499_1.RM.like.systems | <-NurA<-DUF4338<-?\|\|?-><-REase<-?<-DUF499*<-PIN<-DUF433<-METHYLASE<-TUDOR+ SF1-SF2-Helicase+iREase+CTD+REase <-SF1-SF2-Helicase-p<-METHYLASE | 959 | Bacteria>PVC_group>Planctomycetota>Planctomycetia>Isosphaerales | GCF_000186345.1 | Isosphaera_pallida | ISOP_RS17995 | 4934004..4936883 | 128 | ATP-binding-protein-[Isosphaera-pallida] |
| NLX99940.1 | DUF499_1.RM.like.systems | <-DUF499*<-PIN<-?<-REase<-?<-METHYLASE<-REase | 981 | Bacteria>PVC_group>Planctomycetota>Planctomycetia>Pirellulales | GCA_012728855.1 | Rhodopirellula_sp. | GXY83_27900 | 2887..5832 | 2024855 | ATP-binding-protein-[Rhodopirellula-sp.] |
| NMC20499.1 | DUF499_1.RM.like.systems | <-DUF499*<-PIN<-DUF433<-METHYLASE-p | 919 | Bacteria>PVC_group>Planctomycetota>Planctomycetia>Pirellulales | GCA_012798155.1 | Thermogutta_sp. | GYA33_08775 | 1111..3870 | 1962930 | ATP-binding-protein-[Thermogutta-sp.] |
| WP_283433169.1 | DUF499_1.RM.like.systems | <-ParB+HNH<-?<-HTH\|\|MORC+DUF3684->HKD-DNase-><-HEPN+ DUF499 *<-PIN<-OLD-ABC<-METHYLASE<-REase\|\|?-><-TUDOR+ SF1-SF2-Helicase+iREase+CTD+REase | 1109 | Bacteria>PVC_group>Planctomycetota>Planctomycetia>Pirellulales | GCF_900182915.1 | Neorhodopirellula_lusitana | QOL80_RS14710 | 91339..94668 | 445327 | DUF499-domain-containing-protein-[Rhodopirellula-lusitana] |
| MCA9013093.1 | DUF499_1.RM.like.systems | <-DUF499*<-PIN<-?<-METHYLASE<-METHYLASE-p<-TUDOR+ SF1-SF2-Helicase+iREase+CTD+REase | 944 | Bacteria>PVC_group>Planctomycetota>Planctomycetia>Planctomycetales | GCA_020426105.1 | Planctomycetaceae_bacterium | KDB01_25255 | 1526..4360 | 2026779 | ATP-binding-protein-[Planctomycetaceae-bacterium] |
| MCA9076582.1 | DUF499_1.RM.like.systems | sGTP->METHYLASE-p->REase->METHYLASE->OLD-ABC->PIN->HEPN+ DUF499 *-><-PAIREDC-HTH<-?<-?<-SIGMA-HTH | 1095 | Bacteria>PVC_group>Planctomycetota>Planctomycetia>Planctomycetales | GCA_020426505.1 | Planctomycetaceae_bacterium | KDA93_16275 | 17776..21063 | 2026779 | DUF499-domain-containing-protein-[Planctomycetaceae-bacterium] |
| MDA8378780.1 | DUF499_1.RM.like.systems | TUDOR+ SF1-SF2-Helicase+iREase+CTD+REase ->RNase->?->METHYLASE->PIN->DUF499*-><-AAA_21 | 1112 | Bacteria>PVC_group>Planctomycetota>Planctomycetia>unclassified_Planctomycetia | GCA_028068345.1 | Planctomycetia_bacterium | MOZ50_17440 | 9415..12753 | 2052181 | MAG:-DUF499-domain-containing-protein-[Planctomycetia-bacterium] |

| Protein-ID | System | Gene-Neighbourhoods | Length | Taxonomy | Assembly-ID | Strain | Locus-Tag | Genomic-coordinates | TaxID | Fasta-defline |
|---|---|---|---|---|---|---|---|---|---|---|
| MBI3271453.1 | DUF499_1.RM.like.systems | <-Phage_int_SAM_1+Phage_integrase<-?<-?<-ParB+HNH<-?<-DUF499*<-PIN<-?<-?<-METHYLASE<-HEPN<-TUDOR+ SF1-SF2-Helicase+iREase+CTD+REase | 953 | Bacteria>PVC_group>Planctomycetota>unclassified_Planctomycetota | GCA_016197205.1 | Planctomycetota_bacterium | HYZ53_20855 | 61698..64559 | 2026780 | ATP-binding-protein-[Planctomycetota-bacterium] |
| MBM4022804.1 | DUF499_1.RM.like.systems | <-DUF499*<-PIN<-METHYLASE-p | 620 | Bacteria>PVC_group>Planctomycetota>unclassified_Planctomycetota | GCA_016872955.1 | Planctomycetota_bacterium | FJ284_11300 | 1..1859 | 2026780 | ATP-binding-protein-[Planctomycetota-bacterium] |
| MCH7685161.1 | DUF499_1.RM.like.systems | METHYLASE-p->?->DUF433->PIN->HTH->DUF499*-> | 617 | Bacteria>PVC_group>Planctomycetota>unclassified_Planctomycetota | GCA_022562015.1 | Planctomycetota_bacterium | IH899_00515 | 3772..5625 | 2026780 | ATP-binding-protein-[Planctomycetota-bacterium] |
| MCY2991648.1 | DUF499_1.RM.like.systems | <-DUF499*<-PIN<-DUF433<-DUF433<-METHYLASE<-REase<-TUDOR+ SF1-SF2-Helicase+iREase+CTD+REase | 983 | Bacteria>PVC_group>Planctomycetota>unclassified_Planctomycetota | GCA_026692885.1 | Planctomycetota_bacterium | NTY19_27810 | 2128..5079 | 2026780 | MAG:-DUF499-domain-containing-protein-[Planctomycetota-bacterium] |
| MCZ7645664.1 | DUF499_1.RM.like.systems | HTH+MPTase->PemK_toxin->HNH+SAVED->SMODS->?-><-PIN<-HEPN+ DUF499*<-METHYLASE<-?<-ParB+HNH<-TUDOR+ SF1-SF2-Helicase+iREase+CTD+REase | 1122 | Bacteria>PVC_group>Planctomycetota>unclassified_Planctomycetota | GCA_027493525.1 | Planctomycetota_bacterium | M5U26_10325 | 2469927..2473295 | 2026780 | MAG:-Swt1-family-HEPN-domain-containing-protein-[Planctomycetota-bacterium] |
| MCC5805073.1 | DUF499_1.RM.like.systems | OLD-ABC->HNH->METHYLASE->CHTH->RHH->PIN->HEPN+ DUF499*->?->PhdYeFM_antitox->PIN-> | 1107 | Bacteria>PVC_group>Verrucomicrobiota>Opitutae>Opitutales | GCA_020831525.1 | Opitutales_bacterium | JJU00_01980 | 48883..52206 | 2026772 | DUF499-domain-containing-protein-[Opitutales-bacterium] |
| RFC50837.1 | DUF499_1.RM.like.systems | <-UPF0061\|\|TUDOR+ SF1-SF2-Helicase+iREase+CTD+REase->STAS+HisKin->?->PIN->METHYLASE->HEPN+ DUF499*-><-?<-?<-?<-SIGMA-HTH | 1074 | Bacteria>PVC_group>Verrucomicrobiota>unclassified_Verrucomicrobiota | GCA_003402705.1 | Verrucomicrobiota_bacterium | DVB22_001798 | 10693..13917 | 2026799 | putative-ATPase,-AAA+-superfamily-[Verrucomicrobia-bacterium] |
| MBI2947997.1 | DUF499_1.RM.like.systems | METHYLASE-p->SF1-SF2-Helicase+REase->?->PIN->HicB->DUF499*-> | 264 | Bacteria>PVC_group>Verrucomicrobiota>unclassified_Verrucomicrobiota | GCA_016190065.1 | Verrucomicrobiota_bacterium | HYY23_10135 | 3501..4294 | 2026799 | ATP-binding-protein-[Verrucomicrobiota-bacterium] |
| MCL4785388.1 | DUF499_1.RM.like.systems | <-URI-DNase<-?<-?\|\|ABhydrolase+ABhydrolase->HHH_5+SF1-SF2-helicase-><-ASCH<-DUF499*<-?<-METHYLASE<-?<-TUDOR+ SF1-SF2-Helicase+iREase+CTD+REase<-PIN<-HTH+MPTase | 915 | Bacteria>PVC_group>Verrucomicrobiota>unclassified_Verrucomicrobiota | GCA_023383755.1 | Verrucomicrobiota_bacterium | KJ070_01140 | 19812..22559 | 2026799 | DUF499-domain-containing-protein-[Verrucomicrobia-bacterium] |
| MCS1417413.1 | DUF499_1.RM.like.systems | <-HEPN+ DUF499*<-?<-DUF488-l<-PIN<-?<-METHYLASE-p | 1121 | Bacteria>PVC_group>Verrucomicrobiota>Verrucomicrobiae>Limisphaerales | GCA_024761205.1 | Limisphaerales_bacterium | M2R46_05167 | 18516..21881 | 2052186 | hypothetical-protein-[Verrucomicrobia-subdivision-3-bacterium] |
| MCW5552136.1 | DUF499_1.RM.like.systems | GDResolvase+PPP1-HTH->ParB->ParB-><-MarR-HTH\|\|?->ACET-><-HEPN+ DUF499*<-METHYLASE<-PIN<-?<-STAS+HisKin<-TUDOR+ SF1-SF2-Helicase+iREase+CTD+REase | 1120 | Bacteria>PVC_group>Verrucomicrobiota>Verrucomicrobiae>unclassified_Verrucomicrobiae | GCA_026125285.1 | Verrucomicrobiae_bacterium | KIS67_08205 | 85951..89313 | 2499141 | MAG:-DUF499-domain-containing-protein-[Verrucomicrobiae-bacterium] |
| HCN27197.1 | DUF499_1.RM.like.systems | METHYLASE->ParD->PIN->HEPN+ DUF499*-> | 1113 | Bacteria>PVC_group>Verrucomicrobiota>Verrucomicrobiae>Verrucomicrobiales | GCA_003545395.1 | Verrucomicrobiales_bacterium | DIT64_00065 | 2852..6193 | 2026801 | AAA+-family-ATPase-[Verrucomicrobiales-bacterium] |
| MBP7951461.1 | DUF499_1.RM.like.systems | ACET-><-tRNA<-?<-HEPN+ DUF499*<-METHYLASE<-PIN<-?<-STAS+HisKin<-TUDOR+ SF1-SF2-Helicase+iREase+CTD+REase<-DDE_Tnp_1 | 1112 | Bacteria>PVC_group>Verrucomicrobiota>Verrucomicrobiae>Verrucomicrobiales | GCA_018003315.1 | Verrucomicrobiales_bacterium | KA004_17600 | 4745..8083 | 2026801 | DUF499-domain-containing-protein-[Verrucomicrobiales-bacterium] |
| HAL93442.1 | DUF499_1.RM.like.systems | SF1-SF2-Helicase+DUF1998-><-?<-PIN<-?\|\|?-><-HEPN+ DUF499*<-PIN<-PhdYeFM_antitox<-?<-HEPN-p | 1142 | Bacteria>Spirochaetota>Spirochaetia>Spirochaetales>Spirochaetaceae | GCA_003452735.1 | Spirochaetaceae_bacterium | DCP56_01610 | 10292..13720 | 1898206 | AAA+-family-ATPase-[Spirochaetaceae-bacterium] |
| MDE0219724.1 | DUF499_1.RM.like.systems | SbcC-><-OLD-ABC\|\|TUDOR+ SF1-SF2-Helicase+iREase+CTD+REase-><-?\|\|HTH+MPTase->PIN->HEPN+ DUF499*->DOC+HTH->?->XIS-HTH+RVT->Glyco_hydro->DUF2961->HNH-> | 1110 | Bacteria>Spirochaetota>Spirochaetia>Spirochaetales>Spirochaetaceae | GCA_028820215.1 | Spirochaetaceae_bacterium | OXJ90_10675 | 22427..25759 | 1898206 | MAG:-Swt1-family-HEPN-domain-containing-protein-[Spirochaetaceae-bacterium] |
| MCL2137974.1 | DUF499_1.RM.like.systems | TUDOR+ SF1-SF2-Helicase+iREase+CTD+REase->SbcC->?->METHYLASE->MetJarc-HTH->PIN->DUF499*-> | 821 | Bacteria>Spirochaetota>Spirochaetia>Spirochaetales>Treponemataceae | GCA_009776715.1 | Treponema_sp. | FWH41_00405 | 14879..17342 | 166 | DUF499-domain-containing-protein-[Treponema-sp.] |
| MXY09521.1 | DUF499_1.RM.like.systems | TUDOR+ SF1-SF2-Helicase+iREase+CTD+REase->SIR2+SIR2->PIWI->METHYLASE->MerR-HTH->PIN->HEPN+ DUF499*->ASCH->HAD-><-?<-?<-ABhydrolase | 1086 | Bacteria>Terrabacteria_group>Actinomycetota>Acidimicrobiia>Acidimicrobiales | GCA_009838345.1 | Acidimicrobiaceae_bacterium | F4Y56_03045 | 76366..79626 | 2024894 | ATP-binding-protein-[Acidimicrobiaceae-bacterium] |
| MYJ29885.1 | DUF499_1.RM.like.systems | ABhydrolase->?->?-><-HAD<-ASCH<-HEPN+ DUF499*<-PIN<-MerR-HTH<-METHYLASE-p | 1086 | Bacteria>Terrabacteria_group>Actinomycetota>Acidimicrobiia>Acidimicrobiales | GCA_009845185.1 | Acidimicrobiaceae_bacterium | F4072_07945 | 6572..9832 | 2024894 | ATP-binding-protein-[Acidimicrobiaceae-bacterium] |
| MDE0067328.1 | DUF499_1.RM.like.systems | METHYLASE-p->?-><-PIN<-?<-HTH_58\|\|HEPN+ DUF499*-><-ParA+HTH<-RelE-ParE<-Creatinase_N+Peptidase_M24 | 1084 | Bacteria>Terrabacteria_group>Actinomycetota>Acidimicrobiia>Acidimicrobiales | GCA_028819005.1 | Acidimicrobiaceae_bacterium | OXN44_10730 | 3279..6533 | 2024894 | MAG:-DUF499-domain-containing-protein-[Acidimicrobiaceae-bacterium] |
| MDE0160529.1 | DUF499_1.RM.like.systems | TUDOR+ SF1-SF2-Helicase+iREase+CTD+REase->DUF433->PIN->METHYLASE->DUF433->PIN->DUF499*->?-><-tRNA\|\|?->?->FAD-NAD-dep-oxidoreductase-><-HEPN-p | 923 | Bacteria>Terrabacteria_group>Actinomycetota>Acidimicrobiia>Acidimicrobiales | GCA_028819505.1 | Acidimicrobiaceae_bacterium | OXL98_02215 | 9144..11915 | 2024894 | MAG:-DUF499-domain-containing-protein-[Acidimicrobiaceae-bacterium] |
| MDE0320770.1 | DUF499_1.RM.like.systems | PIN->TUDOR+ SF1-SF2-Helicase+iREase+CTD+REase->METHYLASE->?->CHTH->?->HEPN+ DUF499*-><-?\|\|?->?-><-?<-?\|\|DOC+HTH-> | 1152 | Bacteria>Terrabacteria_group>Actinomycetota>Acidimicrobiia>Acidimicrobiales | GCA_028820815.1 | Acidimicrobiaceae_bacterium | OXI97_12920 | 73823..77281 | 2024894 | MAG:-DUF499-domain-containing-protein-[Acidimicrobiaceae-bacterium] |
| MDE0495499.1 | DUF499_1.RM.like.systems | METHYLASE-p->MerR-HTH->PIN->HEPN+ DUF499*->ASCH->HAD-><-?<-?<-ABhydrolase | 1084 | Bacteria>Terrabacteria_group>Actinomycetota>Acidimicrobiia>Acidimicrobiales | GCA_028824895.1 | Acidimicrobiaceae_bacterium | OXH54_16270 | 41170..44424 | 2024894 | MAG:-DUF499-domain-containing-protein-[Acidimicrobiaceae-bacterium] |
| MXW42977.1 | DUF499_1.RM.like.systems | <-METHYLASE<-PIN<-?<-HEPN+ DUF499*<-METHYLASE<-?\|\|?-><-PIN<-DUF433 | 1079 | Bacteria>Terrabacteria_group>Actinomycetota>Acidimicrobiia>unclassified_Acidimicrobiia | GCA_009835645.1 | Acidimicrobiia_bacterium | F4Z91_07815 | 3856..7095 | 2080302 | ATP-binding-protein-[Acidimicrobiia-bacterium] |
| MXZ78951.1 | DUF499_1.RM.like.systems | HTH_31->RNase->HTH_58-><-?<-HEPN+ DUF499*\|\|?->PhdYeFM_antitox->PIN-><-METHYLASE<-?\|\|CHTH-> | 1079 | Bacteria>Terrabacteria_group>Actinomycetota>Acidimicrobiia>unclassified_Acidimicrobiia | GCA_009837345.1 | Acidimicrobiia_bacterium | F4Z06_11870 | 8705..11944 | 2080302 | ATP-binding-protein-[Acidimicrobiia-bacterium] |
| MYB44039.1 | DUF499_1.RM.like.systems | METHYLASE-><-PIN<-?\|\|SIR2+STALD->?->PRTase+PRTase->HEPN+ DUF499*-><-?\|\|PIN-><-ABC-ATPase+SbcC<-ABC-ATPase+SbcC<-Peptidase_M20 | 1097 | Bacteria>Terrabacteria_group>Actinomycetota>Acidimicrobiia>unclassified_Acidimicrobiia | GCA_009839965.1 | Acidimicrobiia_bacterium | F4X74_03760 | 19073..22366 | 2080302 | ATP-binding-protein-[Acidimicrobiia-bacterium] |
| MYG93975.1 | DUF499_1.RM.like.systems | METHYLASE->?->?-><-?\|\|?->HEPN+ DUF499*->?->PIN->METHYLASE-> | 1079 | Bacteria>Terrabacteria_group>Actinomycetota>Acidimicrobiia>unclassified_Acidimicrobiia | GCA_009843865.1 | Acidimicrobiia_bacterium | F4138_03135 | 9114..12353 | 2080302 | ATP-binding-protein-[Acidimicrobiia-bacterium] |
| WP_165316371.1 | DUF499_1.RM.like.systems | ABC-ATPase+SbcC-><-RecR+TOPRIM<-?\|\|TUDOR+ SF1-SF2-Helicase+iREase+CTD+REase->?->METHYLASE->HEPN+ DUF499*-><-PIN\|\|?->?->DDE_Tnp_1->DDE_Tnp_1-><-RNase | 1143 | Bacteria>Terrabacteria_group>Actinomycetota>Actinomycetes>Actinomycetales | GCF_011038885.2 | Schaalia_sp._ZJ405 | G7Y41_RS09385 | 2197397..2200828 | 2709403 | DUF499-domain-containing-protein-[Schaalia-sp.-ZJ405] |
| WP_169762497.1 | DUF499_1.RM.like.systems | <-HTH_3<-RelE-ParE<-TraJ-RHH<-RNase\|\|DUF433->PIN-><-HEPN+ DUF499*<-METHYLASE<-TUDOR+ SF1-SF2-Helicase+iREase+CTD+REase<-ParA+HTH<-TOPRIM<-PIN | 1131 | Bacteria>Terrabacteria_group>Actinomycetota>Actinomycetes>Actinomycetales | GCF_012976645.1 | Mobiluncus_mulieris | HHJ77_RS04685 | 61231..64626 | 2052 | DUF499-domain-containing-protein-[Mobiluncus-mulieris] |
| NEB00544.1 | DUF499_1.RM.like.systems | VapB_antitoxin->PIN-><-DUF499*<-METHYLASE<-TUDOR+ SF1-SF2-Helicase+iREase+CTD+REase\|\|Schlafen+HATPase+HTH-> | 931 | Bacteria>Terrabacteria_group>Actinomycetota>Actinomycetes>Kitasatosporales | GCA_010548375.1 | Streptomyces_sp._SID13726 | G3I78_15960 | 365366..368161 | 2706058 | ATP-binding-protein-[Streptomyces-sp.-SID13726] |
| WP_048693914.1 | DUF499_1.RM.like.systems | <-Radical-SAM+TRAM\|\|?->?-><-DNA_methylase\|\|PIN->METHYLASE->HEPN+ DUF499*->MORC->REase->HTH+SF1-SF2-Helicase+REase->?->CR-Kinase+ORC-CDC6-like+CARF-> | 1134 | Bacteria>Terrabacteria_group>Actinomycetota>Actinomycetes>Micrococcales | GCF_001050535.1 | Tetrasphaera_australiensis | BN11_RS02235 | 472611..476015 | 99480 | DUF499-domain-containing-protein-[Tetrasphaera-australiensis] |
| WP_187368407.1 | DUF499_1.RM.like.systems | TUDOR+ SF1-SF2-Helicase+iREase+CTD+REase-><-PIN<-DUF433<-wHTH+SAD+HNH\|\|METHYLASE->ParB+HNH->HEPN+ DUF499*-><-?<-?\|\|ACET+ACET-><-Lsr2 | 1133 | Bacteria>Terrabacteria_group>Actinomycetota>Actinomycetes>Micrococcales | GCF_008000995.1 | Ruania_zhangjianzhongii | FU260_RS03480 | 750431..753832 | 2603206 | DUF499-domain-containing-protein-[Ruania-zhangjianzhongii] |
| EHB57971.1 | DUF499_1.RM.like.systems | <-GNTR-HTH\|\|NUDIX-><-?<-HEPN<-Schlafen+HATPase+HTH<-HEPN+ DUF499*<-REase<-METHYLASE<-TUDOR+ SF1-SF2-Helicase+iREase+CTD+REase<-PIN<-XIS-HTH | 1108 | Bacteria>Terrabacteria_group>Actinomycetota>Actinomycetes>Mycobacteriales | GCA_000230935.2 | Mycolicibacterium_rhodesiae_JS60 | MycrhDRAFT_0407 | 390606..393932 | 931627 | hypothetical-protein-MycrhDRAFT_0407-[Mycolicibacterium-rhodesiae-JS60] |
| AFC52219.1 | DUF499_1.RM.like.systems | <-GNTR-HTH\|\|NUDIX-><-?<-?<-HEPN<-Schlafen+HATPase+HTH<-HEPN+ DUF499*<-METHYLASE\|\|?-><-TUDOR+ SF1-SF2-Helicase+iREase+CTD+REase<-PIN<-XIS-HTH | 1043 | Bacteria>Terrabacteria_group>Actinomycetota>Actinomycetes>Mycobacteriales | GCA_000276825.1 | Mycobacterium_paraintracellulare | OCQ_07060 | 694865..697996 | 1138383 | hypothetical-protein-OCQ_07060-[Mycobacterium-paraintracellulare] |
| OSC29643.1 | DUF499_1.RM.like.systems | <-GNTR-HTH\|\|NUDIX-><-?<-?<-Schlafen+HATPase+HTH<-HEPN+ DUF499*<-METHYLASE\|\|?-><-TUDOR+ SF1-SF2-Helicase+iREase+CTD+REase<-PIN<-XIS-HTH\|\|GNTR-HTH-p-> | 1097 | Bacteria>Terrabacteria_group>Actinomycetota>Actinomycetes>Mycobacteriales | GCA_002104735.1 | Mycobacterium_paraintracellulare | B8W68_04660 | 128474..131767 | 1138383 | AAA-family-ATPase-[Mycobacterium-paraintracellulare] |
| UHJ57678.1 | DUF499_1.RM.like.systems | REase->DNA-glycosylase->?-><-HTH+SF1-SF2-Helicase+REase<-REase<-MORC<-HEPN+ DUF499*<-METHYLASE<-REase<-PIN\|\|DNA_methylase-><-?<-Phage_integrase+Phage_integrase | 1138 | Bacteria>Terrabacteria_group>Actinomycetota>Actinomycetes>Mycobacteriales | GCA_021249325.1 | Mycolicibacterium_fortuitum | LT337_13390 | 2620949..2624365 | 1766 | DUF499-domain-containing-protein-[Mycolicibacterium-fortuitum] |
| WP_041784793.1 | DUF499_1.RM.like.systems | <-GNTR-HTH\|\|NUDIX-><-?<-?<-HEPN<-Schlafen+HATPase+HTH<-HEPN+ DUF499*<-METHYLASE\|\|?-><-TUDOR+ SF1-SF2-Helicase+iREase+CTD+REase<-PIN<-XIS-HTH\|\|GNTR-HTH-p-> | 1108 | Bacteria>Terrabacteria_group>Actinomycetota>Actinomycetes>Mycobacteriales | GCF_000276825.1 | Mycobacterium_paraintracellulare | B8W68_RS04660 | 128474..131800 | 1138383 | Swt1-family-HEPN-domain-containing-protein-[Mycobacterium-paraintracellulare] |
| WP_015289521.1 | DUF499_1.RM.like.systems | Peptidase_S8->?->?->REase-><-DDE_Tnp_1<-HEPN<-HEPN+ DUF499*\|\|RHH_1->PIN-><-METHYLASE<-TUDOR+ SF1-SF2-Helicase+iREase+CTD+REase | 1115 | Bacteria>Terrabacteria_group>Actinomycetota>Actinomycetes>Mycobacteriales | GCF_000328785.1 | Mycobacterium_canettii | BN42_RS05705 | 1219765..1223112 | 78331 | Swt1-family-HEPN-domain-containing-protein-[Mycobacterium-canettii] |
| WP_005174963.1 | DUF499_1.RM.like.systems | GDSL-><-HTH_19\|\|TUDOR+ SF1-SF2-Helicase+iREase+CTD+REase->METHYLASE-><-PIN<-MazE_antitoxin\|\|HEPN+ DUF499*->?->ABhydrolase+ABhydrolase-> | 1134 | Bacteria>Terrabacteria_group>Actinomycetota>Actinomycetes>Mycobacteriales | GCF_000332975.1 | Gordonia_aichiensis | GOACH_RS12130 | 81836..85240 | 36820 | Swt1-family-HEPN-domain-containing-protein-[Gordonia-aichiensis] |
| WP_062879806.1 | DUF499_1.RM.like.systems | <-GNTR-HTH\|\|NUDIX-><-?<-HEPN<-Schlafen+HATPase+HTH<-HEPN+ DUF499*<-METHYLASE\|\|?-><-TUDOR+ SF1-SF2-Helicase+iREase+CTD+REase<-PIN<-XIS-HTH | 1108 | Bacteria>Terrabacteria_group>Actinomycetota>Actinomycetes>Mycobacteriales | GCF_001606235.1 | Mycobacteroides_abscessus | B5431_RS12990 | 716837..720163 | 36809 | Swt1-family-HEPN-domain-containing-protein-[Mycobacteroides-abscessus] |
| WP_072804385.1 | DUF499_1.RM.like.systems | TUDOR+ SF1-SF2-Helicase+iREase+CTD+REase->?->AbiEii->METHYLASE-><-PIN<-MazE_antitoxin\|\|HEPN+ DUF499*->?->?-><-?<-Phd-YefM | 1125 | Bacteria>Terrabacteria_group>Actinomycetota>Actinomycetes>Mycobacteriales | GCF_001895005.1 | Rhodococcus_yunnanensis | RY1_RS14860 | 83069..86446 | 278209 | Swt1-family-HEPN-domain-containing-protein-[Rhodococcus-yunnanensis] |
| WP_124259269.1 | DUF499_1.RM.like.systems | <-GDResolvase+HIN-HTH\|\|XIS-HTH->PIN-><-HEPN+ DUF499*<-METHYLASE<-TUDOR+ SF1-SF2-Helicase+iREase+CTD+REase<-DNA-glycosylase | 1120 | Bacteria>Terrabacteria_group>Actinomycetota>Actinomycetes>Mycobacteriales | GCF_003844175.1 | Rhodococcus_ruber | TN91_RS21990 | 3527..6889 | 1830 | DUF499-domain-containing-protein-[Rhodococcus-ruber] |
| WP_205874823.1 | DUF499_1.RM.like.systems | <-Cluster2439_3clades\|\|?->XIS-HTH->PIN->TUDOR+ SF1-SF2-Helicase+iREase+CTD+REase->METHYLASE->HEPN+ DUF499*-><-?<-HISKIN<-SbcC<-NUDIX\|\|GNTR-HTH-> | 1108 | Bacteria>Terrabacteria_group>Actinomycetota>Actinomycetes>Mycobacteriales | GCF_017086405.1 | Mycobacterium_gordonae | JX552_RS26920 | 6279372..6282698 | 1778 | Swt1-family-HEPN-domain-containing-protein-[Mycobacterium-gordonae] |
| WP_174714145.1 | DUF499_1.RM.like.systems | XIS-HTH->PIN->TUDOR+ SF1-SF2-Helicase+iREase+CTD+REase-><-?\|\|METHYLASE->ParB+HNH->HEPN+ DUF499*->?->?-><-NUDIX\|\|GNTR-HTH->?->FtsK-> | 1108 | Bacteria>Terrabacteria_group>Actinomycetota>Actinomycetes>Mycobacteriales | GCF_019456675.1 | Mycolicibacterium_pallens | K0O64_RS24915 | 5121618..5124944 | 370524 | MULTISPECIES:-Swt1-family-HEPN-domain-containing-protein-[Mycobacterium] |
| WP_225506778.1 | DUF499_1.RM.like.systems | XIS-HTH->PIN->TUDOR+ SF1-SF2-Helicase+iREase+CTD+REase->METHYLASE->HEPN+ DUF499*->Schlafen+HATPase+HTH->\|\|DDE_Tnp_1_4->HEPN->?-><-NUDIX\|\|GNTR-HTH-> | 1108 | Bacteria>Terrabacteria_group>Actinomycetota>Actinomycetes>Mycobacteriales | GCF_020227835.1 | Mycolicibacterium_fortuitum | H8Z59_RS27040 | 5490569..5493895 | 1766 | Swt1-family-HEPN-domain-containing-protein-[Mycolicibacterium-fortuitum] |
| WP_267616414.1 | DUF499_1.RM.like.systems | <-JetB<-JetA\|\|?->DUF222+HNH->?-><-?<-HEPN+ DUF499*\|\|MazE_antitoxin->PIN-><-METHYLASE<-TUDOR+ SF1-SF2-Helicase+iREase+CTD+REase\|\|?->AbiEii-> | 1127 | Bacteria>Terrabacteria_group>Actinomycetota>Actinomycetes>Mycobacteriales | GCF_026616955.1 | Gordonia_bronchialis | NOB45_RS12880 | 105537..108920 | 2054 | Swt1-family-HEPN-domain-containing-protein-[Gordonia-bronchialis] |
| WP_089250630.1 | DUF499_1.RM.like.systems | <-DUF433<-?<-HEPN+ DUF499*\|\|MazE_antitoxin->PIN-><-METHYLASE<-AbiEii<-?<-TUDOR+ SF1-SF2-Helicase+iREase+CTD+REase | 1122 | Bacteria>Terrabacteria_group>Actinomycetota>Actinomycetes>Mycobacteriales | GCF_900188125.1 | Rhodococcus_kyotonensis | CHB96_RS22960 | 65313..68681 | 398843 | Swt1-family-HEPN-domain-containing-protein-[Rhodococcus-kyotonensis] |
| WP_005139269.1 | DUF499_1.RM.like.systems | <-GNTR-HTH\|\|NUDIX-><-?<-HEPN<-Schlafen+HATPase+HTH<-HEPN+ DUF499*<-REase<-METHYLASE<-TUDOR+ SF1-SF2-Helicase+iREase+CTD+REase<-PIN<-XIS-HTH | 1108 | Bacteria>Terrabacteria_group>Actinomycetota>Actinomycetes>Mycobacteriales | - | Mycolicibacterium_rhodesiae | MycrhDRAFT_0407 | 390606..393932 | 36814 | Swt1-family-HEPN-domain-containing-protein-[Mycolicibacterium-rhodesiae] |
| WP_227735039.1 | DUF499_1.RM.like.systems | TUDOR+ SF1-SF2-Helicase+iREase+CTD+REase-><-PIN<-DUF433\|\|METHYLASE-><-DOC+HTH\|\|HEPN+ DUF499*-> | 1104 | Bacteria>Terrabacteria_group>Actinomycetota>Actinomycetes>Propionibacteriales | GCF_020687685.1 | Tessaracoccus_sp._OS52 | LK027_RS00980 | 227595..230909 | 2886691 | DUF499-domain-containing-protein-[Tessaracoccus-sp.-OS52] |
| WP_017595710.1 | DUF499_1.RM.like.systems | <-JetB<-JetA<-HEPN+ DUF499*<-METHYLASE<-TUDOR+ SF1-SF2-Helicase+iREase+CTD+REase<-PIN<-XIS-HTH<-tRNA<-SIGMA-HTH | 1136 | Bacteria>Terrabacteria_group>Actinomycetota>Actinomycetes>Streptosporangiales | GCF_000341105.1 | Nocardiopsis_potens | D459_RS0124835 | 11224..14634 | 1246458 | Swt1-family-HEPN-domain-containing-protein-[Nocardiopsis-potens] |
| MCL2317098.1 | DUF499_1.RM.like.systems | <-HEPN+ DUF499*<-PIN<-PhdYeFM_antitox<-METHYLASE-p | 623 | Bacteria>Terrabacteria_group>Actinomycetota>Actinomycetes>unclassified_Actinomycetes | GCA_009785885.1 | Actinomycetes_bacterium | FWC46_08465 | 1..1871 | 1883427 | Swt1-family-HEPN-domain-containing-protein-[Actinomycetia-bacterium] |
| MCH9729512.1 | DUF499_1.RM.like.systems | PIN->TUDOR+ SF1-SF2-Helicase+iREase+CTD+REase->METHYLASE->ParB+HNH->HEPN+ DUF499*->SIR2-><-DndB\|\|?->Recombinase-><-NUDIX\|\|GNTR-HTH-> | 1114 | Bacteria>Terrabacteria_group>Actinomycetota>Actinomycetes>unclassified_Actinomycetes | GCA_022600605.1 | Actinomycetes_bacterium | KOU84_07525 | 8492..11836 | 1883427 | DUF499-domain-containing-protein-[Actinomycetia-bacterium] |
| MCP9484920.1 | DUF499_1.RM.like.systems | METHYLASE-p->?->METHYLASE-p->SbcC->PIN->?->HEPN+ DUF499*->HIN-HTH->?->\|\|?-><-Peptidase_M20 | 1087 | Bacteria>Terrabacteria_group>Actinomycetota>Rubrobacteria>Gaiellales | GCA_024330215.1 | Candidatus_Gaiellasilicea_maunaloa | MSC30_03590 | 46535..49798 | 2928748 | DUF499-domain-containing-protein-[Candidatus-Gaiellasilicea-maunaloa] |
| MBX6764706.1 | DUF499_1.RM.like.systems | METHYLASE-p->MazE_antitoxin->PIN->HEPN+ DUF499*->REase-> | 1106 | Bacteria>Terrabacteria_group>Actinomycetota>Rubrobacteria>Rubrobacterales | GCA_019690035.1 | Rubrobacteraceae_bacterium | IRY88_13660 | 1711..5031 | 2740537 | DUF499-domain-containing-protein-[Rubrobacteraceae-bacterium] |
| WP_219975492.1 | DUF499_1.RM.like.systems | <-SIGMA-HTH<-?<-MULE-TRANSPOSASE\|\|?->?-><-REase<-HEPN+ DUF499*<-PIN<-MazE_antitoxin<-METHYLASE<-PIN<-TUDOR+ SF1-SF2-Helicase+iREase+CTD+REase\|\|DUF4096-> | 1106 | Bacteria>Terrabacteria_group>Actinomycetota>Rubrobacteria>Rubrobacterales | GCF_019448335.1 | Rubrobacter_xylanophilus | Rxycam_RS04975 | 969402..972722 | 49319 | Swt1-family-HEPN-domain-containing-protein-[Rubrobacter-xylanophilus] |
| UGS34698.1 | DUF499_1.RM.like.systems | PIN->tRNA->TUDOR+ SF1-SF2-Helicase+iREase+CTD+REase->METHYLASE->DUF499*-><-?\|\|?->?->Phage_int_SAM_1+Phage_integrase-><-HTH_3 | 940 | Bacteria>Terrabacteria_group>Actinomycetota>Thermoleophilia>Solirubrobacterales | GCA_021172045.1 | Capillimicrobium_parvum | DSM104329_01080 | 1087018..1089840 | 2884022 | hypothetical-protein-DSM104329_01080-[Solirubrobacterales-bacterium-0166_1] |
| WP_259314362.1 | DUF499_1.RM.like.systems | PIN->tRNA->TUDOR+ SF1-SF2-Helicase+iREase+CTD+REase->METHYLASE->DUF499*->?-><-?\|\|XIS-HTH->?->?->Phage_int_SAM_1+Phage_integrase-> | 940 | Bacteria>Terrabacteria_group>Actinomycetota>Thermoleophilia>Solirubrobacterales | GCF_021172045.1 | Capillimicrobium_parvum | DSM104329_RS05360 | 1087018..1089840 | 2884022 | DUF499-domain-containing-protein-[Capillimicrobium-parvum] |
| TXH45529.1 | DUF499_1.RM.like.systems | Mbetalac-><-?<-?\|\|Peptidase-><-HEPN+ DUF499*\|\|Phd-YefM->PIN-><-?<-METHYLASE\|\|?->PIN-> | 1130 | Bacteria>Terrabacteria_group>Actinomycetota>unclassified_Actinomycetota | GCA_008012465.1 | Actinomycetota_bacterium | E6Q90_00220 | 20268..23660 | 2900548 | ATP-binding-protein-[Actinobacteria-bacterium] |
| NLO26762.1 | DUF499_1.RM.like.systems | PIN->METHYLASE->PIN->?->?->?->HEPN+ DUF499*-> | 401 | Bacteria>Terrabacteria_group>Actinomycetota>unclassified_Actinomycetota | GCA_012523095.1 | Actinomycetota_bacterium | GX113_01060 | 16345..17548 | 2900548 | ATP-binding-protein-[Actinomycetia-bacterium] |
| MBC7252723.1 | DUF499_1.RM.like.systems | HIN-HTH->?-><-?<-Schlafen+HATPase+HTH<-ParB+HNH+Schlafen<-HEPN+ DUF499*<-METHYLASE<-PIN<-TUDOR+ SF1-SF2-Helicase+iREase+CTD+REase | 1115 | Bacteria>Terrabacteria_group>Actinomycetota>unclassified_Actinomycetota | GCA_014360685.1 | Actinomycetota_bacterium | H5T72_01965 | 75820..79167 | 2900548 | ATP-binding-protein-[Actinomycetia-bacterium] |
| MBW3642892.1 | DUF499_1.RM.like.systems | <-HEPN+ DUF499*<-?<-PIN<-SbcC<-METHYLASE | 1106 | Bacteria>Terrabacteria_group>Actinomycetota>unclassified_Actinomycetota | GCA_019348235.1 | Actinomycetota_bacterium | KY447_08275 | 1669..4989 | 2900548 | DUF499-domain-containing-protein-[Actinomycetia-bacterium] |
| PIY44270.1 | DUF499_1.RM.like.systems | METHYLASE->DUF433->PIN->DUF499*-> | 318 | Bacteria>Terrabacteria_group>Armatimonadota>unclassified_Armatimonadota | GCA_002784925.1 | Armatimonadetes_bacterium_CG_4_10_14_3_um_filter_59_10 | COZ05_08660 | 3346..4299 | 1973909 | hypothetical-protein-COZ05_08660,-partial-[Armatimonadetes-bacterium-CG_4_10_14_3_um_filter_59_10] |
| NCO42947.1 | DUF499_1.RM.like.systems | METHYLASE-p->DUF433->PIN->DUF499*->?->Tox-RelE->HTH_3-><-?<-HicA_toxin<-HicB_lk_antitox | 1098 | Bacteria>Terrabacteria_group>Armatimonadota>unclassified_Armatimonadota | GCA_009992845.1 | Armatimonadota_bacterium | GW892_30120 | 1561..4857 | 2033014 | ATP-binding-protein-[Armatimonadetes-bacterium] |
| MBI3912159.1 | DUF499_1.RM.like.systems | METHYLASE-p->DUF433->PIN->HEPN+ DUF499*->?-><-NTP_transf<-Peptidase<-?<-Radical-SAM | 1099 | Bacteria>Terrabacteria_group>Armatimonadota>unclassified_Armatimonadota | GCA_016199525.1 | Armatimonadota_bacterium | HY320_14665 | 1429..4728 | 2033014 | ATP-binding-protein-[Armatimonadetes-bacterium] |
| HHW19260.1 | DUF499_1.RM.like.systems | TUDOR+ SF1-SF2-Helicase+iREase+CTD+REase->SbcC->METHYLASE->REase->DUF499*->DUF433->PIN-> | 932 | Bacteria>Terrabacteria_group>Bacillota>Candidatus_Fermentithermobacillia>Candidatus_Fermentithermobacillales | GCA_012840505.1 | Candidatus_Fermentithermobacillaceae_bacterium | GXX30_10280 | 57551..60349 | 3085339 | ATP-binding-protein-[Firmicutes-bacterium] |
| NLW38082.1 | DUF499_1.RM.like.systems | <-MethyltransfD12\|\|?->PIN-><-HEPN+ DUF499*<-METHYLASE<-TPR+TPR<-MetJarc-HTH<-SF1-SF2-Helicase<-EVE | 1120 | Bacteria>Terrabacteria_group>Bacillota>Clostridia>Eubacteriales | GCA_012728715.1 | Peptococcaceae_bacterium | GXY90_05855 | 3552..6914 | 2052179 | ATP-binding-protein-[Peptococcaceae-bacterium] |
| WP_027356905.1 | DUF499_1.RM.like.systems | TUDOR+ SF1-SF2-Helicase+iREase+CTD+REase->METHYLASE->DUF499*-><-Polbeta<-PIN<-PhdYeFM_antitox\|\|?->HIN-HTH-> | 896 | Bacteria>Terrabacteria_group>Bacillota>Clostridia>Eubacteriales | GCF_000686645.1 | Desulfofundulus_thermocisternus | BR83_RS0109605 | 50288..52978 | 42471 | DUF499-domain-containing-protein-[Desulfofundulus-thermocisternus] |
| WP_066668116.1 | DUF499_1.RM.like.systems | <-HD<-?\|\|?->?->?-><-HEPN+ DUF499*<-METHYLASE<-PIN<-MetJarc-HTH<-TUDOR+ SF1-SF2-Helicase+iREase+CTD+REase | 1130 | Bacteria>Terrabacteria_group>Bacillota>Clostridia>Eubacteriales | GCF_001655685.1 | Desulfotomaculum_copahuensis | A6M21_RS09370 | 52458..55850 | 1838280 | Swt1-family-HEPN-domain-containing-protein-[Desulfotomaculum-copahuensis] |
| WP_092487583.1 | DUF499_1.RM.like.systems | HIN-HTH-><-DDE_Tnp_1<-HEPN+ DUF499*<-PIN<-SbcC<-METHYLASE<-TUDOR+ SF1-SF2-Helicase+iREase+CTD+REase | 888 | Bacteria>Terrabacteria_group>Bacillota>Clostridia>Eubacteriales | GCF_900115975.1 | Desulfoscipio_geothermicus | BM299_RS17670 | 2600..5266 | 39060 | Swt1-family-HEPN-domain-containing-protein-[Desulfoscipio-geothermicus] |
| MCL6474638.1 | DUF499_1.RM.like.systems | PIN->DUF499*->TUDOR+ SF1-SF2-Helicase+iREase+CTD+REase->?-><-REase | 879 | Bacteria>Terrabacteria_group>Bacillota>unclassified_Bacillota | GCA_023511455.1 | Bacillota_bacterium | K6U75_06255 | 18500..21139 | 1879010 | DUF499-domain-containing-protein-[Firmicutes-bacterium] |
| MBE2269142.1 | DUF499_1.RM.like.systems | <-DUF499*<-METHYLASE<-PIN<-DUF433<-TUDOR+ SF1-SF2-Helicase+iREase+CTD+REase | 1106 | Bacteria>Terrabacteria_group>Chloroflexota>Anaerolineae>Anaerolineales | GCA_014879865.1 | Anaerolinea_sp. | IAE80_13000 | 195..3515 | 1872519 | ATP-binding-protein-[Anaerolinea-sp.] |
| MBE7511263.1 | DUF499_1.RM.like.systems | Peptidase_M1->?->TUDOR+ SF1-SF2-Helicase+iREase+CTD+REase->?->PIN->METHYLASE-p->HEPN+ DUF499*-><-tRNA<-?<-ABhydrolase\|\|TPR+TPR->ACET+ACET->ACET-> | 1098 | Bacteria>Terrabacteria_group>Chloroflexota>Anaerolineae>Anaerolineales | GCA_015075625.1 | Anaerolineales_bacterium | HS103_00415 | 94825..98121 | 2073117 | ATP-binding-protein-[Anaerolineales-bacterium] |
| MCB9452414.1 | DUF499_1.RM.like.systems | <-MetJarc-HTH\|\|?->TPR+TPR+TPR+TPR+TPR+TPR+TPR+TPR-><-HEPN+ DUF499*<-ParB+HNH<-METHYLASE<-Schlafen+HATPase+HTH<-PIN<-?<-TUDOR+ SF1-SF2-Helicase+iREase+CTD+REase | 1106 | Bacteria>Terrabacteria_group>Chloroflexota>Anaerolineae>Anaerolineales | GCA_020634115.1 | Anaerolineaceae_bacterium | H6672_13335 | 112307..115627 | 2024896 | ATP-binding-protein-[Anaerolineaceae-bacterium] |
| MCB9078004.1 | DUF499_1.RM.like.systems | <-HEPN+ DUF499*<-REase<-?<-METHYLASE-p<-METHYLASE<-PIN<-DUF433 | 385 | Bacteria>Terrabacteria_group>Chloroflexota>Anaerolineae>Anaerolineales | GCA_020634955.1 | Anaerolineaceae_bacterium | H6631_10465 | 1..1157 | 2024896 | ATP-binding-protein-[Anaerolineaceae-bacterium] |
| MCC6958025.1 | DUF499_1.RM.like.systems | <-MarR-HTH<-HEPN+ DUF499*<-?<-METHYLASE<-?<-PIN<-REase | 1114 | Bacteria>Terrabacteria_group>Chloroflexota>Anaerolineae>Anaerolineales | GCA_020853835.1 | Anaerolineales_bacterium | IT316_14590 | 1878..5222 | 2073117 | DUF499-domain-containing-protein-[Anaerolineales-bacterium] |
| NPV86424.1 | DUF499_1.RM.like.systems | <-REC<-tRNA\|\|TUDOR+ SF1-SF2-Helicase+iREase+CTD+REase->PIN->METHYLASE->HEPN+ DUF499*-> | 1115 | Bacteria>Terrabacteria_group>Chloroflexota>Anaerolineae>unclassified_Anaerolineae | GCA_013178295.1 | Anaerolineae_bacterium | HPY45_10505 | 184598..187945 | 2052143 | ATP-binding-protein-[Anaerolineae-bacterium] |

| Protein-ID | System | Gene-Neighbourhoods | Length | Taxonomy | Assembly-ID | Strain | Locus-Tag | Genomic-coordinates | TaxID | Fasta-defline |
|---|---|---|---|---|---|---|---|---|---|---|
| MBN1139681.1 | DUF499_1.RM.like.systems | <-DUF433<-DUF499 *<-METHYLASE <-PIN<-CopG/RHH<-TUDOR+ SF1-SF2-Helicase+iREase+CTD+REase \|\|?-><-Methyltransf | 1104 | Bacteria>Terrabacteria_group>Chloroflexota>Anaerolineae>unclassified_Anaerolineae | GCA_016926595.1 | Anaerolineae_bacterium | JXM73_24120 | 8352..11666 | 2052143 | ATP-binding-protein-[Anaerolineae-bacterium] |
| MCK6580687.1 | DUF499_1.RM.like.systems | <-GDResolvase<-?<-PIN <-HEPN+ DUF499 *<-PIN<-DUF433<-?<-METHYLASE <-?<-TUDOR+ SF1-SF2-Helicase+iREase+CTD+REase | 1104 | Bacteria>Terrabacteria_group>Chloroflexota>Anaerolineae>unclassified_Anaerolineae | GCA_023151025.1 | Anaerolineae_bacterium | L6Q98_21555 | 21655..24969 | 2052143 | DUF499-domain-containing-protein-[Anaerolineae-bacterium] |
| MCL4264803.1 | DUF499_1.RM.like.systems | TUDOR+ SF1-SF2-Helicase+iREase+CTD+REase ->?-><-DUF4096+DDE_Tnp_1+DDE_Tnp_1_2\|\|?->PIN-> METHYLASE -> DUF499 *->?->?->?->XIS-HTH-><-?\|\|HhH-RADC+JAB-> | 1222 | Bacteria>Terrabacteria_group>Chloroflexota>Anaerolineae>unclassified_Anaerolineae | GCA_023382655.1 | Anaerolineae_bacterium | KJ069_16390 | 143359..147027 | 2052143 | DUF499-domain-containing-protein-[Anaerolineae-bacterium] |
| MCL4237388.1 | DUF499_1.RM.like.systems | MULE-TRANSPOSASE->REase->SF1-SF2-Helicase-p->MULE-TRANSPOSASE-><-HEPN+ DUF499 *<-METHYLASE <-PIN<-?<-TUDOR+ SF1-SF2-Helicase+iREase+CTD+REase | 1111 | Bacteria>Terrabacteria_group>Chloroflexota>Anaerolineae>unclassified_Anaerolineae | GCA_023382715.1 | Anaerolineae_bacterium | KJ047_03985 | 66352..69687 | 2052143 | DUF499-domain-containing-protein-[Anaerolineae-bacterium] |
| MCR4407531.1 | DUF499_1.RM.like.systems | Phage_integrase-><-?\|\|<-DUF499 *<-TOPRIM-p<-SbcC<-METHYLASE <-PIN<-MazE_antitoxin<-TUDOR+ SF1-SF2-Helicase+iREase+CTD+REase | 1132 | Bacteria>Terrabacteria_group>Chloroflexota>Anaerolineae>unclassified_Anaerolineae | GCA_024653325.1 | Anaerolineae_bacterium | NUW24_11530 | 3238..6636 | 2052143 | DUF499-domain-containing-protein-[Anaerolineae-bacterium] |
| MCS7282892.1 | DUF499_1.RM.like.systems | METHYLASE -p->DUF499 *-><-PIN<-PhdYeFM_antitox | 1035 | Bacteria>Terrabacteria_group>Chloroflexota>Anaerolineae>unclassified_Anaerolineae | GCA_025062415.1 | Anaerolineae_bacterium | NZ769_06035 | 1264..4371 | 2052143 | DUF499-domain-containing-protein-[Anaerolineae-bacterium] |
| MCZ2114537.1 | DUF499_1.RM.like.systems | <-HTH+UbE1<-?\|\|HTH->MPTase-><-HEPN+ DUF499 *<-PIN<-OLD-ABC <-METHYLASE <-MORC+DUF3684<-TUDOR+ SF1-SF2-Helicase+iREase+CTD+REase | 1108 | Bacteria>Terrabacteria_group>Chloroflexota>Anaerolineae>unclassified_Anaerolineae | GCA_026954395.1 | Anaerolineae_bacterium | LC131_12000 | 4338..7664 | 2052143 | MAG:-DUF499-domain-containing-protein-[Anaerolineae-bacterium] |
| MCZ7570815.1 | DUF499_1.RM.like.systems | DEDD_Tnp_IS110+Transposase_20->?->TUDOR+ SF1-SF2-Helicase+iREase+CTD+REase -> METHYLASE ->REase->HEPN+ DUF499 *-><-PIN<-PhdYeFM_antitox<-?<-HAD | 1117 | Bacteria>Terrabacteria_group>Chloroflexota>Ardenticatenia>Ardenticatenales | GCA_027493995.1 | Ardenticatenaceae_bacterium | M5U01_19825 | 4779576..4782929 | 2614247 | MAG:-Swt1-family-HEPN-domain-containing-protein-[Ardenticatenaceae-bacterium] |
| MBP7961438.1 | DUF499_1.RM.like.systems | TUDOR+ SF1-SF2-Helicase+iREase+CTD+REase ->?->?->METHYLASE -> OLD-ABC->PIN -> HEPN+ DUF499 *->?->SF1-SF2-Helicase->SF1-SF2-Helicase+DUF1998-> | 1102 | Bacteria>Terrabacteria_group>Chloroflexota>Caldilineae>Caldilineales | GCA_018003515.1 | Caldilineaceae_bacterium | KA003_05670 | 15367..18675 | 2614250 | DUF499-domain-containing-protein-[Caldilineaceae-bacterium] |
| MCC6169408.1 | DUF499_1.RM.like.systems | PIN-> METHYLASE -> HEPN+ DUF499 *-> | 1104 | Bacteria>Terrabacteria_group>Chloroflexota>Caldilineae>Caldilineales | GCA_020845455.1 | Caldilineaceae_bacterium | IT329_19460 | 3543..6857 | 2614250 | DUF499-domain-containing-protein-[Caldilineaceae-bacterium] |
| MCY3902338.1 | DUF499_1.RM.like.systems | TIR->TUDOR+ SF1-SF2-Helicase+iREase+CTD+REase -p->PIN-> METHYLASE ->PolB->HEPN->HEPN+ DUF499 *->?->?-><-?<-DSBH | 1125 | Bacteria>Terrabacteria_group>Chloroflexota>Caldilineae>Caldilineales | GCA_026707125.1 | Caldilineaceae_bacterium | OXF76_04160 | 61061..64438 | 2614250 | MAG:-Swt1-family-HEPN-domain-containing-protein-[Caldilineaceae-bacterium] |
| MDE0079138.1 | DUF499_1.RM.like.systems | <-DUF499 *<-METHYLASE <-PIN<-TUDOR+ SF1-SF2-Helicase+iREase+CTD+REase | 1096 | Bacteria>Terrabacteria_group>Chloroflexota>Caldilineae>Caldilineales | GCA_028819025.1 | Caldilineaceae_bacterium | OX050_16575 | 525..3815 | 2614250 | MAG:-DUF499-domain-containing-protein-[Caldilineaceae-bacterium] |
| MYD51869.1 | DUF499_1.RM.like.systems | TUDOR+ SF1-SF2-Helicase+iREase+CTD+REase ->PolB->HEPN->METHYLASE ->DUF433->PIN-> DUF499 *-> | 928 | Bacteria>Terrabacteria_group>Chloroflexota>Dehalococcoidia>unclassified_Dehalococcoidia | GCA_009841605.1 | Dehalococcoidia_bacterium | F4W93_10360 | 50124..52910 | 2026734 | ATP-binding-protein-[Dehalococcoidia-bacterium] |
| NWF79207.1 | DUF499_1.RM.like.systems | TUDOR+ SF1-SF2-Helicase+iREase+CTD+REase -><-?\|\| METHYLASE -><-PIN<-DUF433\|\| HEPN+ DUF499 *-> | 825 | Bacteria>Terrabacteria_group>Chloroflexota>unclassified_Chloroflexota | GCA_013388805.1 | Chloroflexota_bacterium | HXY37_04055 | 14161..16634 | 2026724 | ATP-binding-protein-[Chloroflexi-bacterium] |
| MBI2958637.1 | DUF499_1.RM.like.systems | METHYLASE -p->DUF433->PIN-> DUF499 -p-> DUF499 *-> | 1006 | Bacteria>Terrabacteria_group>Chloroflexota>unclassified_Chloroflexota | GCA_016189825.1 | Chloroflexota_bacterium | HYY32_07340 | 1155..4175 | 2026724 | DUF499-domain-containing-protein-[Chloroflexi-bacterium] |
| MBI3961042.1 | DUF499_1.RM.like.systems | METHYLASE -p-> OLD-ABC->PIN -> HEPN+ DUF499 *-> | 1102 | Bacteria>Terrabacteria_group>Chloroflexota>unclassified_Chloroflexota | GCA_016199425.1 | Chloroflexota_bacterium | HY328_19695 | 3483..6791 | 2026724 | ATP-binding-protein-[Chloroflexi-bacterium] |
| MBI4295304.1 | DUF499_1.RM.like.systems | TUDOR+ SF1-SF2-Helicase+iREase+CTD+REase ->HEPN->Pkinase->METHYLASE ->DUF433->PIN-> DUF499 *-> | 770 | Bacteria>Terrabacteria_group>Chloroflexota>unclassified_Chloroflexota | GCA_016210605.1 | Chloroflexota_bacterium | HY669_03965 | 9180..11489 | 2026724 | ATP-binding-protein-[Chloroflexi-bacterium] |
| MBK8988277.1 | DUF499_1.RM.like.systems | TUDOR+ SF1-SF2-Helicase+iREase+CTD+REase ->?->PIN->CopG/RHH->PIN-> METHYLASE -p->HEPN+ DUF499 *-><-AbiEii<-?\|\|?->?->?->ABC-ATPase+SbcC-> | 1097 | Bacteria>Terrabacteria_group>Chloroflexota>unclassified_Chloroflexota | GCA_016716065.1 | Chloroflexota_bacterium | IPM39_19790 | 148627..151920 | 2026724 | DUF499-domain-containing-protein-[Chloroflexi-bacterium] |
| MBK9748864.1 | DUF499_1.RM.like.systems | <-VirD4-TraG+VirD4-FtsK<-?<-?<-MetJarc-HTH<-DUF499 *<-METHYLASE <-PIN<-DUF433<-TUDOR+ SF1-SF2-Helicase+iREase+CTD+REase \|\|?-><-PIWI | 1104 | Bacteria>Terrabacteria_group>Chloroflexota>unclassified_Chloroflexota | GCA_016717205.1 | Chloroflexota_bacterium | IPO91_19080 | 75969..79283 | 2026724 | ATP-binding-protein-[Chloroflexi-bacterium] |
| MBM3143888.1 | DUF499_1.RM.like.systems | TUDOR+ SF1-SF2-Helicase+iREase+CTD+REase ->HEPN->METHYLASE -> DUF499 *->?->PIN-><-?\|\|?->Peptidase_M24-> | 927 | Bacteria>Terrabacteria_group>Chloroflexota>unclassified_Chloroflexota | GCA_016865865.1 | Chloroflexota_bacterium | FJ010_02740 | 17191..19974 | 2026724 | ATP-binding-protein-[Chloroflexi-bacterium] |
| MCA0453853.1 | DUF499_1.RM.like.systems | <-GDResolvase<-?<-zf-ISL3+DDE_Tnp_ISL3<-?<-?<-?<-HEPN+ DUF499 *<-PIN<-DUF433<-METHYLASE <-PIN<-?<-TUDOR+ SF1-SF2-Helicase+iREase+CTD+REase | 1103 | Bacteria>Terrabacteria_group>Chloroflexota>unclassified_Chloroflexota | GCA_020161935.1 | Chloroflexota_bacterium | LCI00_07765 | 53613..56924 | 2026724 | DUF499-domain-containing-protein-[Chloroflexi-bacterium] |
| MCC6174343.1 | DUF499_1.RM.like.systems | TUDOR+ SF1-SF2-Helicase+iREase+CTD+REase ->Schlafen->BirA-HTH->METHYLASE ->ParD->PIN-> HEPN+ DUF499 *-><-?\|\|?->Aldedh-> | 1160 | Bacteria>Terrabacteria_group>Chloroflexota>unclassified_Chloroflexota | GCA_020845495.1 | Chloroflexota_bacterium | IT305_03475 | 199239..202721 | 2026724 | DUF499-domain-containing-protein-[Chloroflexi-bacterium] |
| GIW20263.1 | DUF499_1.RM.like.systems | TUDOR+ SF1-SF2-Helicase+iREase+CTD+REase -> PIN-> METHYLASE ->?-> DUF499 *-><-?\|\|?->?-><-tRNA<-Peptidase<-tRNA<-tRNA<-tRNA<-tRNA<-REase | 906 | Bacteria>Terrabacteria_group>Chloroflexota>unclassified_Chloroflexota | GCA_026004015.1 | Chloroflexota_bacterium | KatS3mg065_0559 | 209249..211969 | 2026724 | MAG:-hypothetical-protein-KatS3mg065_0559-[Chloroflexota-bacterium] |
| MCX6066946.1 | DUF499_1.RM.like.systems | <-REase<-?<-?<-?<-DUF499 -p<-HEPN+ DUF499 *<-PIN<-MazE_antitoxin | 972 | Bacteria>Terrabacteria_group>Chloroflexota>unclassified_Chloroflexota | GCA_026389995.1 | Chloroflexota_bacterium | NT121_14520 | 4682..7598 | 2026724 | MAG:-Swt1-family-HEPN-domain-containing-protein,-partial-[Chloroflexi-bacterium] |
| MCY3958689.1 | DUF499_1.RM.like.systems | SF1-SF2-Helicase-p->Peptidase->?-><-DUF499 *<-PIN<-DUF433<-METHYLASE <-?<-TUDOR+ SF1-SF2-Helicase+iREase+CTD+REase | 926 | Bacteria>Terrabacteria_group>Chloroflexota>unclassified_Chloroflexota | GCA_026707585.1 | Chloroflexota_bacterium | OXG65_10380 | 10889..13669 | 2026724 | MAG:-DUF499-domain-containing-protein-[Chloroflexi-bacterium] |
| MCY4474618.1 | DUF499_1.RM.like.systems | <-METHYLASE <-PIN<-?<-?<-HEPN<-HEPN<-HEPN+ DUF499 * | 1144 | Bacteria>Terrabacteria_group>Chloroflexota>unclassified_Chloroflexota | GCA_026712855.1 | Chloroflexota_bacterium | OXC83_04195 | 178388..181822 | 2026724 | MAG:-DUF499-domain-containing-protein-[Chloroflexi-bacterium] |
| MDE2669829.1 | DUF499_1.RM.like.systems | <-REase+NACHT\|\|TUDOR+ SF1-SF2-Helicase+iREase+CTD+REase -> METHYLASE ->DUF433->PIN-> DUF499 *-> | 918 | Bacteria>Terrabacteria_group>Chloroflexota>unclassified_Chloroflexota | GCA_028875515.1 | Chloroflexota_bacterium | OXI51_09280 | 52276..55032 | 2026724 | MAG:-DUF499-domain-containing-protein-[Chloroflexota-bacterium] |
| WP_096720868.1 | DUF499_1.RM.like.systems | DUF29-><-?<-?<-?\|\|?-><-HEPN+ DUF499 *<-PIN<-?<-METHYLASE \|\|REase-><-TUDOR+ SF1-SF2-Helicase+iREase+CTD+REase | 1125 | Bacteria>Terrabacteria_group>Cyanobacteriota/Melainabacteria_group>Cyanobacteriota>Cyanophyceae | - | Chondrocystis_sp._NIES-4102 | CDC35_RS11850 | 2748341..2751718 | 2005460 | Swt1-family-HEPN-domain-containing-protein-[Chondrocystis-sp.-NIES-4102] |
| BAQ63189.1 | DUF499_1.RM.like.systems | TUDOR+ SF1-SF2-Helicase+iREase+CTD+REase -><-ZnR\|\| METHYLASE ->?->PIN-> DUF499 *-><-Phage_integrase<-DUF4158+DDE_Tnp_Tn3<-Phage_integrase\|\|?->N6_MTase-p-> | 1005 | Bacteria>Terrabacteria_group>Cyanobacteriota/Melainabacteria_group>Cyanobacteriota>Cyanophyceae | GCA_001548095.1 | Geminocystis_sp._NIES-3708 | GM3708_3595 | 38002..41019 | 1615909 | predicted-ATPase-[Geminocystis-sp.-NIES-3708] |
| OBQ34507.1 | DUF499_1.RM.like.systems | <-DUF499 *<-PIN<-MetJarc-HTH<-?<-METHYLASE -p | 966 | Bacteria>Terrabacteria_group>Cyanobacteriota/Melainabacteria_group>Cyanobacteriota>Cyanophyceae | GCA_001672075.1 | Anabaena_sp._CRKS33 | AN487_17930 | 14..2914 | 1710888 | MAG:-AAA-family-ATPase-[Anabaena-sp.-CRKS33] |
| OBQ20276.1 | DUF499_1.RM.like.systems | <-DnaJ<-?<-NTP_transferase<-?<-DUF499 *<-PIN<-?<-REase+HTH<-TUDOR+ SF1-SF2-Helicase+iREase+CTD+REase <-CHTH | 916 | Bacteria>Terrabacteria_group>Cyanobacteriota/Melainabacteria_group>Cyanobacteriota>Cyanophyceae | GCA_001672085.1 | Anabaena_sp._AL93 | AN486_07300 | 48734..51484 | 1678133 | MAG:-ATPase-[Anabaena-sp.-AL93] |
| ARI79893.1 | DUF499_1.RM.like.systems | TUDOR+ SF1-SF2-Helicase+iREase+CTD+REase ->REase->REase->METHYLASE ->PAIREDC-HTH->PIN-> HEPN+ DUF499 *-><-Cluster164_18clades<-?\|\|?->?->?->DUF29-> | 1155 | Bacteria>Terrabacteria_group>Cyanobacteriota/Melainabacteria_group>Cyanobacteriota>Cyanophyceae | GCA_002095975.1 | Microcystis_aeruginosa_PCC_7806SL | BH695_0612 | 541107..544574 | 1903187 | hypothetical-protein-BH695_0612-[Microcystis-aeruginosa-PCC-7806SL] |
| GBE76047.1 | DUF499_1.RM.like.systems | REase->AbrB->PIN-> METHYLASE ->RHH_6->PIN-> HEPN+ DUF499 *-><-Cluster164_18clades<-?\|\|?->?->?-><-TPR+TPR+TPR+TPR+TPR+TPR+TPR+TPR | 1111 | Bacteria>Terrabacteria_group>Cyanobacteriota/Melainabacteria_group>Cyanobacteriota>Cyanophyceae | GCA_002933835.1 | Microcystis_aeruginosa_NIES-87 | myaer87_32740 | 119164..122499 | 449440 | hypothetical-protein-myaer87_32740-[Microcystis-aeruginosa-NIES-87] |
| PSB44266.1 | DUF499_1.RM.like.systems | <-HEPN+ DUF499 *<-?<-METHYLASE <-REase<-PIN<-AbrB<-TUDOR+ SF1-SF2-Helicase+iREase+CTD+REase | 1113 | Bacteria>Terrabacteria_group>Cyanobacteriota/Melainabacteria_group>Cyanobacteriota>Cyanophyceae | GCA_003004015.1 | Cyanosarcina_cf._burmensis_CCALA_770 | C7B80_21165 | 280..3621 | 2107697 | AAA+-family-ATPase-[Cyanosarcina-cf.-burmensis-CCALA-770] |
| REJ55686.1 | DUF499_1.RM.like.systems | TUDOR+ SF1-SF2-Helicase+iREase+CTD+REase ->\|\|AbrB->PIN-> METHYLASE ->RHH_6->PIN-> HEPN+ DUF499 *->?-><-Cluster164_18clades<-?\|\|?->?->DUF29-> | 1111 | Bacteria>Terrabacteria_group>Cyanobacteriota/Melainabacteria_group>Cyanobacteriota>Cyanophyceae | GCA_003388675.1 | Microcystis_aeruginosa_DA14 | DWQ56_15555 | 244128..247463 | 1987506 | DUF499-domain-containing-protein-[Microcystis-aeruginosa-DA14] |
| BBD57116.1 | DUF499_1.RM.like.systems | MethyltransfD12->MethyltransfD12-><-?<-HTH_3\|\|DDE_Tnp_1->?-><-DUF499 *<-REase<-REase<-REase<-METHYLASE <-PIN | 1002 | Bacteria>Terrabacteria_group>Cyanobacteriota/Melainabacteria_group>Cyanobacteriota>Cyanophyceae | GCA_003609755.1 | Planktothrix_agardhii_NIES-204 | NIES204_44520 | 39433..42441 | 282423 | AAA-family-ATPase-[Planktothrix-agardhii-NIES-204] |
| TRT82535.1 | DUF499_1.RM.like.systems | Cluster164_18clades-><-?<-HEPN+ DUF499 *<-ACET<-?<-METHYLASE <-PIN<-AbrB<-REase | 1130 | Bacteria>Terrabacteria_group>Cyanobacteriota/Melainabacteria_group>Cyanobacteriota>Cyanophyceae | GCA_007095595.1 | Microcystis_aeruginosa_Ma_AC_P_19900807_S299 | EWV82_11170 | 1786..5178 | 2486264 | ATP-binding-protein-[Microcystis-aeruginosa-Ma_AC_P_19900807_S299] |
| TRU24939.1 | DUF499_1.RM.like.systems | ABhydrolase->TUDOR+ SF1-SF2-Helicase+iREase+CTD+REase ->REase->METHYLASE ->?->PIN-> HEPN+ DUF499 *-><-Cluster164_18clades<-?\|\|?->?->DUF29->PolB-> | 1130 | Bacteria>Terrabacteria_group>Cyanobacteriota/Melainabacteria_group>Cyanobacteriota>Cyanophyceae | GCA_007095795.1 | Microcystis_aeruginosa_Ma_MB_S_20031200_S102D | EWV79_09040 | 24486..27878 | 2486255 | ATP-binding-protein-[Microcystis-aeruginosa-Ma_MB_S_20031200_S102D] |
| NCR40139.1 | DUF499_1.RM.like.systems | ABhydrolase->TUDOR+ SF1-SF2-Helicase+iREase+CTD+REase ->REase->METHYLASE ->?->PIN-> HEPN+ DUF499 *-><-Cluster164_18clades | 1130 | Bacteria>Terrabacteria_group>Cyanobacteriota/Melainabacteria_group>Cyanobacteriota>Cyanophyceae | GCA_009995705.1 | Microcystis_aeruginosa_W13-11 | GPJ21_09840 | 22820..26212 | 2685991 | ATP-binding-protein-[Microcystis-aeruginosa-W13-11] |
| NJO72425.1 | DUF499_1.RM.like.systems | CHTH->TUDOR+ SF1-SF2-Helicase+iREase+CTD+REase ->REase->METHYLASE ->?->PIN-> HEPN+ DUF499 *->?-><-?<-?<-REase\|\|Phage_int_SAM_4+Phage_integrase-> | 1125 | Bacteria>Terrabacteria_group>Cyanobacteriota/Melainabacteria_group>Cyanobacteriota>Cyanophyceae | GCA_012033355.1 | Leptolyngbyaceae_cyanobacterium_RM1_406_9 | HC833_00760 | 33344..36721 | 2720434 | ATP-binding-protein-[Leptolyngbyaceae-cyanobacterium-RM1_406_9] |
| NMG59376.1 | DUF499_1.RM.like.systems | TUDOR+ SF1-SF2-Helicase+iREase+CTD+REase ->SbcC->?->METHYLASE ->?->PIN-> HEPN+ DUF499 *-> | 1130 | Bacteria>Terrabacteria_group>Cyanobacteriota/Melainabacteria_group>Cyanobacteriota>Cyanophyceae | GCA_012911965.1 | Geitlerinema_sp._P-1104 | E1H12_12830 | 9854..13246 | 2546230 | ATP-binding-protein-[Geitlerinema-sp.-P-1104] |
| MBC6477472.1 | DUF499_1.RM.like.systems | REase->REase->?->?->?->PIN-><-DUF499 *<-METHYLASE <-TUDOR+ SF1-SF2-Helicase+iREase+CTD+REase \|\|ABC-ATPase+SbcC->OLD-ABC->TOPRIM-> | 976 | Bacteria>Terrabacteria_group>Cyanobacteriota/Melainabacteria_group>Cyanobacteriota>Cyanophyceae | GCA_014323675.1 | Hormoscilla_sp._GM7CHS1pb | GDA56_06445 | 1108802..1111732 | 2653677 | ATP-binding-protein-[Hormoscilla-sp.-GM7CHS1pb] |
| MBO1068320.1 | DUF499_1.RM.like.systems | <-NTP_transferase\|\|N6_N4_MTase-><-HTH_3<-?<-?<-?<-DUF499 *<-PIN<-?<-METHYLASE <-?<-REase+HTH<-TUDOR+ SF1-SF2-Helicase+iREase+CTD+REase | 911 | Bacteria>Terrabacteria_group>Cyanobacteriota/Melainabacteria_group>Cyanobacteriota>Cyanophyceae | GCA_017355655.1 | Dolichospermum_sp._DEX189 | HEQ13_02485 | 530022..532757 | 2717870 | ATP-binding-protein-[Dolichospermum-sp.-DEX189] |
| MBS9772311.1 | DUF499_1.RM.like.systems | <-DUF499 *<-REase<-PIN<-?<-METHYLASE <-TUDOR+ SF1-SF2-Helicase+iREase+CTD+REase | 1016 | Bacteria>Terrabacteria_group>Cyanobacteriota/Melainabacteria_group>Cyanobacteriota>Cyanophyceae | GCA_018457135.1 | Trichodesmium_erythraeum_GBRTRLIN201 | J7F05_18140 | 304..3354 | 1045867 | ATP-binding-protein-[Trichodesmium-erythraeum-GBRTRLIN201] |
| MCA2627579.1 | DUF499_1.RM.like.systems | REase->METHYLASE ->RHH_6->PIN-> OLD-ABC->PIN -> HEPN+ DUF499 *-><-Cluster164_18clades | 1130 | Bacteria>Terrabacteria_group>Cyanobacteriota/Melainabacteria_group>Cyanobacteriota>Cyanophyceae | GCA_020245485.1 | Microcystis_sp._M091S2 | IM505_05380 | 45571..48963 | 2771177 | DUF499-domain-containing-protein-[Microcystis-sp.-M091S2] |
| MCA2653775.1 | DUF499_1.RM.like.systems | Cluster164_18clades-><-HEPN+ DUF499 *<-PIN<-RHH_6<-METHYLASE <-AbrB<-PIN<-AbrB | 1130 | Bacteria>Terrabacteria_group>Cyanobacteriota/Melainabacteria_group>Cyanobacteriota>Cyanophyceae | GCA_020245605.1 | Microcystis_sp._M061S2 | IM475_05875 | 1597..4989 | 2771171 | DUF499-domain-containing-protein-[Microcystis-sp.-M061S2] |
| MCA2716396.1 | DUF499_1.RM.like.systems | REase->AbrB->PIN-> METHYLASE ->RHH_6->HEPN+ DUF499 *-><-Cluster164_18clades<-?\|\|?->?->?-><-TPR+TPR+TPR+TPR+TPR+TPR+TPR+TPR | 1130 | Bacteria>Terrabacteria_group>Cyanobacteriota/Melainabacteria_group>Cyanobacteriota>Cyanophyceae | GCA_020245865.1 | Microcystis_sp._M169S2 | IM485_01040 | 5502..8894 | 2771157 | DUF499-domain-containing-protein-[Microcystis-sp.-M169S2] |
| MCD8490337.1 | DUF499_1.RM.like.systems | pPIWI+pPIWI->pPIWI->pPIWI+MID+RNaseH->?->?->?-><-HEPN+ DUF499 *<-METHYLASE <-PIN<-AbrB<-TUDOR+ SF1-SF2-Helicase+iREase+CTD+REase <-TOPRIM+SF1-SF2-Helicase | 1118 | Bacteria>Terrabacteria_group>Cyanobacteriota/Melainabacteria_group>Cyanobacteriota>Cyanophyceae | GCA_021208125.1 | Desertifilum_sp. | LRZ84_27325 | 10272..13628 | 2497424 | DUF499-domain-containing-protein-[Desertifilum-sp.] |
| MCE2669063.1 | DUF499_1.RM.like.systems | Cluster164_18clades-><-HEPN+ DUF499 *<-PIN<-OLD-ABC <-PIN<-RHH_6<-METHYLASE <-REase | 1130 | Bacteria>Terrabacteria_group>Cyanobacteriota/Melainabacteria_group>Cyanobacteriota>Cyanophyceae | GCA_021300285.1 | Microcystis_sp._49638_E5 | LW728_07560 | 10523..13915 | 2904986 | DUF499-domain-containing-protein-[Microcystis-sp.-49638_E5] |
| WP_002754187.1 | DUF499_1.RM.like.systems | AbrB->PemK_toxin->METHYLASE ->RHH_6->PIN-> HEPN+ DUF499 *-><-Cluster164_18clades<-?\|\|?->?->DUF29->sGTP-> | 1130 | Bacteria>Terrabacteria_group>Cyanobacteriota/Melainabacteria_group>Cyanobacteriota>Cyanophyceae | GCF_000307995.1 | Microcystis_aeruginosa | MICCA_RS15375 | 7336..10728 | 1126 | Swt1-family-HEPN-domain-containing-protein-[Microcystis-aeruginosa] |
| WP_002777134.1 | DUF499_1.RM.like.systems | METHYLASE ->RHH_6->PIN-> OLD-ABC->PIN -> HEPN+ DUF499 *-><-Cluster164_18clades<-?\|\|?->?->DUF29->sGTP-> | 1130 | Bacteria>Terrabacteria_group>Cyanobacteriota/Melainabacteria_group>Cyanobacteriota>Cyanophyceae | GCF_000312205.1 | Microcystis_aeruginosa | MICAD_RS14460 | 204713..208105 | 1126 | Swt1-family-HEPN-domain-containing-protein-[Microcystis-aeruginosa] |
| WP_002801405.1 | DUF499_1.RM.like.systems | REase->METHYLASE ->RHH_6->PIN-> OLD-ABC->PIN -> HEPN+ DUF499 *-><-Cluster164_18clades | 1117 | Bacteria>Terrabacteria_group>Cyanobacteriota/Melainabacteria_group>Cyanobacteriota>Cyanophyceae | GCF_000312285.1 | Microcystis_aeruginosa | MICAK_RS07420 | 6579..9932 | 1126 | Swt1-family-HEPN-domain-containing-protein-[Microcystis-aeruginosa] |
| WP_015155777.1 | DUF499_1.RM.like.systems | Phage_integrase-><-Phage_integrase\|\|?-><-HEPN+ DUF499 *<-?<-METHYLASE <-REase<-PIN<-AbrB<-TUDOR+ SF1-SF2-Helicase+iREase+CTD+REase | 1113 | Bacteria>Terrabacteria_group>Cyanobacteriota/Melainabacteria_group>Cyanobacteriota>Cyanophyceae | GCF_000317125.1 | Chroococcidiopsis_thermalis | CHRO_RS18600 | 4302197..4305538 | 54299 | Swt1-family-HEPN-domain-containing-protein-[Chroococcidiopsis-thermalis] |
| WP_019503187.1 | DUF499_1.RM.like.systems | <-DDE_3<-PAIREDC-HTH\|\|?->\|\|<-?\|\|PIWI-><-HEPN+ DUF499 *\|\|?-><-PIN<-?<-METHYLASE | 1127 | Bacteria>Terrabacteria_group>Cyanobacteriota/Melainabacteria_group>Cyanobacteriota>Cyanophyceae | GCF_000332195.1 | Pleurocapsa_sp._PCC_7319 | PLEUR7319_RS0100190 | 37513..40896 | 118161 | Swt1-family-HEPN-domain-containing-protein-[Pleurocapsa-sp.-PCC-7319] |
| WP_002748938.1 | DUF499_1.RM.like.systems | PAIREDC-HTH->PIN-> HEPN+ DUF499 *-><-Cluster164_18clades<-?\|\|?->?->DUF29-> | 1111 | Bacteria>Terrabacteria_group>Cyanobacteriota/Melainabacteria_group>Cyanobacteriota>Cyanophyceae | GCF_000332585.1 | Microcystis_aeruginosa | C789_RS21375 | 1161..4496 | 1126 | Swt1-family-HEPN-domain-containing-protein-[Microcystis-aeruginosa] |
| WP_016515479.1 | DUF499_1.RM.like.systems | Cluster164_18clades-><-?<-HEPN+ DUF499 *<-PIN<-OLD-ABC <-PIN<-RHH_6<-METHYLASE <-AbrB | 1130 | Bacteria>Terrabacteria_group>Cyanobacteriota/Melainabacteria_group>Cyanobacteriota>Cyanophyceae | GCF_000412595.1 | Microcystis_aeruginosa | MAESPC_RS19915 | 50545..53937 | 1126 | Swt1-family-HEPN-domain-containing-protein-[Microcystis-aeruginosa] |
| WP_200865503.1 | DUF499_1.RM.like.systems | TUDOR+ SF1-SF2-Helicase+iREase+CTD+REase ->?->PIN-> METHYLASE ->DUF433->PIN-> DUF499 *-> DUF499 -p->tRNA->?->?-><-?\|\|NADAR-><-MORC | 732 | Bacteria>Terrabacteria_group>Cyanobacteriota/Melainabacteria_group>Cyanobacteriota>Cyanophyceae | GCF_000505665.1 | Thermosynechococcus_sp._NK55a | NK55_RS09315 | 1926061..1928259 | 1394889 | DUF499-domain-containing-protein-[Thermosynechococcus-sp.-NK55a] |
| WP_026733571.1 | DUF499_1.RM.like.systems | TUDOR+ SF1-SF2-Helicase+iREase+CTD+REase ->AbrB->PIN-> METHYLASE ->REase->HEPN+ DUF499 *->?->TraJ-RHH-> | 1122 | Bacteria>Terrabacteria_group>Cyanobacteriota/Melainabacteria_group>Cyanobacteriota>Cyanophyceae | GCF_000517105.1 | Fischerella_sp._PCC_9605 | FIS9605_RS0116420 | 337165..340533 | 1173024 | Swt1-family-HEPN-domain-containing-protein-[Fischerella-sp.-PCC-9605] |
| WP_024971076.1 | DUF499_1.RM.like.systems | REase->METHYLASE ->RHH_6->PIN-> OLD-ABC->PIN -> HEPN+ DUF499 *-> | 1130 | Bacteria>Terrabacteria_group>Cyanobacteriota/Melainabacteria_group>Cyanobacteriota>Cyanophyceae | GCF_000599945.1 | Microcystis_aeruginosa | MIC7005_RS0124495 | 6350..9742 | 1126 | Swt1-family-HEPN-domain-containing-protein-[Microcystis-aeruginosa] |
| WP_042158736.1 | DUF499_1.RM.like.systems | PIN-> METHYLASE ->REase->REase->REase->DUF499 *-><-?\|\|RelB_antitoxin->PemK_toxin-> | 1002 | Bacteria>Terrabacteria_group>Cyanobacteriota/Melainabacteria_group>Cyanobacteriota>Cyanophyceae | GCF_000710505.1 | Planktothrix_agardhii | NMK38_RS22725 | 84961..87969 | 1160 | DUF499-domain-containing-protein-[Planktothrix-agardhii] |
| WP_053539516.1 | DUF499_1.RM.like.systems | <-NTP_transferase\|\|N6_N4_MTase-><-HTH_3<-?<-?<-DUF499 *<-PIN<-?<-METHYLASE <-REase+HTH<-TUDOR+ SF1-SF2-Helicase+iREase+CTD+REase <-CHTH | 911 | Bacteria>Terrabacteria_group>Cyanobacteriota/Melainabacteria_group>Cyanobacteriota>Cyanophyceae | GCF_001277295.1 | Anabaena_sp._WA102 | AA650_RS14270 | 3307169..3309904 | 1647413 | DUF499-domain-containing-protein-[Anabaena-sp.-WA102] |
| WP_069791018.1 | DUF499_1.RM.like.systems | METHYLASE ->?->?->?->PIN->?-> DUF499 *-> DUF499 -p-><-GDResolvase+HIN-HTH\|\|DUF4158+DDE_Tnp_Tn3-> | 838 | Bacteria>Terrabacteria_group>Cyanobacteriota/Melainabacteria_group>Cyanobacteriota>Cyanophyceae | GCF_001747005.1 | Cyanobacterium_sp._IPPAS_B-1200 | A5482_RS09625 | 19329..21845 | 1562720 | DUF499-domain-containing-protein-[Cyanobacterium-sp.-IPPAS-B-1200] |
| WP_075888510.1 | DUF499_1.RM.like.systems | <-GDResolvase+HIN-HTH\|\|DUF4158+DDE_Tnp_Tn3-><-?<-?<-?<-?<-DUF499 *<-Polbeta<-PolB-NTase<-PIN<-?<-METHYLASE | 965 | Bacteria>Terrabacteria_group>Cyanobacteriota/Melainabacteria_group>Cyanobacteriota>Cyanophyceae | GCF_001904615.1 | [Limnothrix_rosea]_IAM_M-220 | NIES208_RS00240 | 58146..61043 | 454133 | DUF499-domain-containing-protein-[Limnothrix-rosea] |
| WP_231828504.1 | DUF499_1.RM.like.systems | TUDOR+ SF1-SF2-Helicase+iREase+CTD+REase ->REase->REase->METHYLASE ->PAIREDC-HTH->PIN-> HEPN+ DUF499 *-><-Cluster164_18clades<-?\|\|?->?->DUF29-><-Y1_Tnp | 1129 | Bacteria>Terrabacteria_group>Cyanobacteriota/Melainabacteria_group>Cyanobacteriota>Cyanophyceae | GCF_002095975.1 | Microcystis_aeruginosa | BH695_RS02870 | 541185..544574 | 1126 | Swt1-family-HEPN-domain-containing-protein-[Microcystis-aeruginosa] |
| WP_223278246.1 | DUF499_1.RM.like.systems | TUDOR+ SF1-SF2-Helicase+iREase+CTD+REase ->Phd-YefM->PIN-><-HTH_3\|\| METHYLASE -> HEPN+ DUF499 *-><-TraJ-RHH\|\|PIWI-><-HTH_3 | 1023 | Bacteria>Terrabacteria_group>Cyanobacteriota/Melainabacteria_group>Cyanobacteriota>Cyanophyceae | GCF_002245985.1 | Nostoc_sp._'Peltigera_membranacea_cyanobiont'_232 | CDG79_RS21465 | 10493..13564 | 2014531 | DUF499-domain-containing-protein-[Nostoc-sp.-'Peltigera-membranacea-cyanobiont'-232] |
| WP_094555433.1 | DUF499_1.RM.like.systems | GDResolvase-><-D5N+D5ATPase\|\|?->ArdCN+MPTase-><-?<-HEPN+ DUF499 *<-METHYLASE <-?<-PIN<-RHH<-TUDOR+ SF1-SF2-Helicase+iREase+CTD+REase \|\|DUF4236-> | 1103 | Bacteria>Terrabacteria_group>Cyanobacteriota/Melainabacteria_group>Cyanobacteriota>Cyanophyceae | GCF_002252625.1 | Synechococcus_sp._1G10 | CJZ82_RS08935 | 55076..58387 | 2025605 | DUF499-domain-containing-protein-[Synechococcus-sp.-1G10] |
| WP_096671658.1 | DUF499_1.RM.like.systems | <-DUF499 *<-PIN<-?<-METHYLASE <-METHYLASE -p<-REase+HTH<-TUDOR+ SF1-SF2-Helicase+iREase+CTD+REase | 912 | Bacteria>Terrabacteria_group>Cyanobacteriota/Melainabacteria_group>Cyanobacteriota>Cyanophyceae | GCF_002368115.1 | Dolichospermum_compactum | CA730_RS19465 | 4197332..4200070 | 136073 | DUF499-domain-containing-protein-[Dolichospermum-compactum] |
| WP_103113079.1 | DUF499_1.RM.like.systems | PemK_toxin->METHYLASE ->RHH_6->PIN-> OLD-ABC->PIN -> HEPN+ DUF499 *-><-Cluster164_18clades<-?\|\|?->?->DUF29->sGTP-> | 1111 | Bacteria>Terrabacteria_group>Cyanobacteriota/Melainabacteria_group>Cyanobacteriota>Cyanophyceae | GCF_002897275.1 | Microcystis_aeruginosa | D3800_RS23535 | 4968568..4971903 | 1126 | Swt1-family-HEPN-domain-containing-protein-[Microcystis-aeruginosa] |
| WP_104908185.1 | DUF499_1.RM.like.systems | TUDOR+ SF1-SF2-Helicase+iREase+CTD+REase ->Phd-YefM->PIN-><-HTH_3\|\| METHYLASE ->REase->HEPN+ DUF499 *-> | 1109 | Bacteria>Terrabacteria_group>Cyanobacteriota/Melainabacteria_group>Cyanobacteriota>Cyanophyceae | GCF_002949795.1 | Nostoc_sp._'Lobaria_pulmonaria_(5183)_cyanobiont' | NLP_RS21725 | 4933532..4936861 | 1618022 | Swt1-family-HEPN-domain-containing-protein-[Nostoc-sp.-'Lobaria-pulmonaria-(5183)-cyanobiont'] |
| WP_106370048.1 | DUF499_1.RM.like.systems | RNase-><-?\|\|?->TIR-><-HKD-DNase+SF1-SF2-Helicase<-HEPN<-DUF499 *<-METHYLASE <-PIN | 1004 | Bacteria>Terrabacteria_group>Cyanobacteriota/Melainabacteria_group>Cyanobacteriota>Cyanophyceae | GCF_003003885.1 | Chlorogloea_sp._CCALA_695 | C7B70_RS16210 | 39604..42618 | 2107693 | DUF499-domain-containing-protein-[Chlorogloea-sp.-CCALA-695] |
| WP_106371305.1 | DUF499_1.RM.like.systems | <-STYKIN\|\|?-><-SIR2+SIR2<-DUF499 *<-REase<-METHYLASE <-PIN | 994 | Bacteria>Terrabacteria_group>Cyanobacteriota/Melainabacteria_group>Cyanobacteriota>Cyanophyceae | GCF_003003885.1 | Chlorogloea_sp._CCALA_695 | C7B70_RS23190 | 7463..10447 | 2107693 | DUF499-domain-containing-protein-[Chlorogloea-sp.-CCALA-695] |
| WP_130128840.1 | DUF499_1.RM.like.systems | <-MORC+DUF3684<-HEPN+ DUF499 *<-METHYLASE <-METHYLASE <-PIN<-RHH\|\|?->RelE-ParE-> | 1113 | Bacteria>Terrabacteria_group>Cyanobacteriota/Melainabacteria_group>Cyanobacteriota>Cyanophyceae | GCF_004209775.1 | Synechococcus_sp._WH_8101 | SynWH8101_RS05090 | 1007085..1010426 | 59932 | DUF499-domain-containing-protein-[Synechococcus-sp.-WH-8101] |
| WP_130756579.1 | DUF499_1.RM.like.systems | REase-p->AbrB->PIN-> METHYLASE ->RHH_6->PIN-> HEPN+ DUF499 *-><-Cluster164_18clades<-?\|\|?->?->DUF29->sGTP-> | 1111 | Bacteria>Terrabacteria_group>Cyanobacteriota/Melainabacteria_group>Cyanobacteriota>Cyanophyceae | GCF_004305995.1 | Microcystis_aeruginosa | MiAbB_RS04285 | 58496..61831 | 1126 | Swt1-family-HEPN-domain-containing-protein-[Microcystis-aeruginosa] |
| WP_137908144.1 | DUF499_1.RM.like.systems | <-REase+HTH<-?\|\|?-><-?<-?<-DUF499 *<-PIN<-?<-MetJarc-HTH<-MetJarc-HTH<-OrfB_Zn_ribbon<-REase | 969 | Bacteria>Terrabacteria_group>Cyanobacteriota/Melainabacteria_group>Cyanobacteriota>Cyanophyceae | GCF_005402965.1 | Dolichospermum_planctonicum | FF006_RS11405 | 62234..65143 | 136072 | DUF499-domain-containing-protein-[Dolichospermum-planctonicum] |
| WP_149107756.1 | DUF499_1.RM.like.systems | <-sGTP<-DUF29<-?\|\|?->Cluster164_18clades-><-HEPN+ DUF499 *<-PIN<-RHH_6<-METHYLASE <-PemK_toxin<-AbrB<-PIN | 1130 | Bacteria>Terrabacteria_group>Cyanobacteriota/Melainabacteria_group>Cyanobacteriota>Cyanophyceae | GCF_008257445.1 | Microcystis_aeruginosa | FXO09_RS20205 | 21434..24826 | 1126 | Swt1-family-HEPN-domain-containing-protein-[Microcystis-aeruginosa] |
| WP_150976007.1 | DUF499_1.RM.like.systems | REase->AbrB->PIN-> METHYLASE ->RHH_6->PIN-> HEPN+ DUF499 *-><-DDE_3 | 1111 | Bacteria>Terrabacteria_group>Cyanobacteriota/Melainabacteria_group>Cyanobacteriota>Cyanophyceae | GCF_008757435.1 | Microcystis_aeruginosa | EZJ55_RS08120 | 1414057..1417392 | 1126 | Swt1-family-HEPN-domain-containing-protein-[Microcystis-aeruginosa] |
| WP_151696808.1 | DUF499_1.RM.like.systems | <-PRTase<-sGTP<-?\|\|?->?->?-><-HEPN+ DUF499 *<-PIN<-?<-METHYLASE -p | 1130 | Bacteria>Terrabacteria_group>Cyanobacteriota/Melainabacteria_group>Cyanobacteriota>Cyanophyceae | GCF_008974145.1 | Microcystis_aeruginosa | MiAbW_RS15325 | 39137..42529 | 1126 | Swt1-family-HEPN-domain-containing-protein-[Microcystis-aeruginosa] |

| Protein-ID | System | Gene-Neighbourhoods | Length | Taxonomy | Assembly-ID | Strain | Locus-Tag | Genomic-coordinates | TaxID | Fasta-defline |
|---|---|---|---|---|---|---|---|---|---|---|
| WP_168507185.1 | DUF499_1.RM.like.systems | <- DUF499 *<-REase<- METHYLASE <- PIN <- TUDOR+ SF1-SF2-Helicase+iREase+CTD+REase <-CHTH | 962 | Bacteria>Terrabacteria_group>Cyanobacteriota/Melainabacteria_group>Cyanobacteriota>Cyanophyceae | GCF_009712035.1 | Anabaena_sp._UHCC_0187 | FJR11_RS20795 | 2169..5057 | 2590018 | DUF499-domain-containing-protein-[Anabaena-sp.-UHCC-0187] |
| WP_254434496.1 | DUF499_1.RM.like.systems | SF1-SF2-Helicase-p-> METHYLASE -p->MetJarc-HTH-> PIN ->MetJarc-HTH-> DUF499 *-> DUF499 -p->?->HISKIN+MORC->Methyltransf_21->?->HTH_3-> | 386 | Bacteria>Terrabacteria_group>Cyanobacteriota/Melainabacteria_group>Cyanobacteriota>Cyanophyceae | GCF_009712065.1 | Aphanizomenon_sp._UHCC_0183 | FJR37_RS03355 | 2964..4124 | 2590028 | DUF499-domain-containing-protein-[Aphanizomenon-sp.-UHCC-0183] |
| WP_159253859.1 | DUF499_1.RM.like.systems | TUDOR+ SF1-SF2-Helicase+iREase+CTD+REase ->REase-> METHYLASE -> OLD-ABC-> PIN -> HEPN+ DUF499 *-><-?<-?\|\|?->?-><-THUMP+Methylase | 1130 | Bacteria>Terrabacteria_group>Cyanobacteriota/Melainabacteria_group>Cyanobacteriota>Cyanophyceae | GCF_009811855.1 | Microcystis_aeruginosa | GVO78_RS16540 | 16290..19682 | 1126 | Swt1-family-HEPN-domain-containing-protein-[Microcystis-aeruginosa] |
| WP_159297746.1 | DUF499_1.RM.like.systems | TUDOR+ SF1-SF2-Helicase+iREase+CTD+REase ->REase-> METHYLASE -> OLD-ABC-> PIN -> HEPN+ DUF499 *-><-?<-?\|\|?->?-><-THUMP+Methylase | 1130 | Bacteria>Terrabacteria_group>Cyanobacteriota/Melainabacteria_group>Cyanobacteriota>Cyanophyceae | GCF_009811875.1 | Microcystis_aeruginosa | GVO69_RS15755 | 16338..19730 | 1126 | Swt1-family-HEPN-domain-containing-protein-[Microcystis-aeruginosa] |
| WP_163928210.1 | DUF499_1.RM.like.systems | <-TPR+TPR\|\| TUDOR+ SF1-SF2-Helicase+iREase+CTD+REase ->Phd-YefM-> PIN -><-HTH_3\|\| METHYLASE -> HEPN+ DUF499 *-><-TraJ-RHH\|\|PIWI-><-?<-?\|\|?->HTH_OrfB_IS605+OrfB_IS605+OrfB_Zn_ribbon-> | 1108 | Bacteria>Terrabacteria_group>Cyanobacteriota/Melainabacteria_group>Cyanobacteriota>Cyanophyceae | GCF_010747425.1 | Nostoc_sp._UIC_10630 | G1098_RS07815 | 12096..15422 | 2100146 | Swt1-family-HEPN-domain-containing-protein-[Nostoc-sp.-UIC-10630] |
| WP_170190015.1 | DUF499_1.RM.like.systems | TUDOR+ SF1-SF2-Helicase+iREase+CTD+REase ->SbcC->?-> METHYLASE ->?-> PIN -> HEPN+ DUF499 *-> | 1130 | Bacteria>Terrabacteria_group>Cyanobacteriota/Melainabacteria_group>Cyanobacteriota>Cyanophyceae | GCF_012911965.1 | Geitlerinema_sp._P-1104 | E1H12_RS13025 | 9854..13246 | 2546230 | Swt1-family-HEPN-domain-containing-protein-[Geitlerinema-sp.-P-1104] |
| WP_172355891.1 | DUF499_1.RM.like.systems | REase->REase->AbrB-> PIN -> METHYLASE ->?-> HEPN+ DUF499 *->?-><-REase\|\|?->CR-Trypsin1->?-><-REase | 1105 | Bacteria>Terrabacteria_group>Cyanobacteriota/Melainabacteria_group>Cyanobacteriota>Cyanophyceae | GCF_013177315.1 | Thermoleptolyngbya_sichuanensis | HPC62_RS11805 | 2814598..2817915 | 2885951 | Swt1-family-HEPN-domain-containing-protein-[Thermoleptolyngbya-sichuanensis] |
| WP_222620775.1 | DUF499_1.RM.like.systems | <- DUF499 *<- PIN <-?<- METHYLASE -p\|\|?->?-><- TUDOR+ SF1-SF2-Helicase+iREase+CTD+REase | 616 | Bacteria>Terrabacteria_group>Cyanobacteriota/Melainabacteria_group>Cyanobacteriota>Cyanophyceae | GCF_014299995.1 | Sphaerospermopsis_sp._LEGE_00249 | H5968_RS22665 | 435..2285 | 1380707 | DUF499-domain-containing-protein-[Sphaerospermopsis-sp.-LEGE-00249] |
| WP_190676125.1 | DUF499_1.RM.like.systems | <-pPIWI+pPIWI\|\|ParB-><-DDE_3<-PAIREDC-HTH\|\|ParB-><- TUDOR+ SF1-SF2-Helicase+iREase+CTD+REase <- HEPN+ DUF499 *<- PIN <-OLD-ABC <- METHYLASE <-REase\|\|?-><- TUDOR+ SF1-SF2-Helicase+iREase+CTD+REase | 1112 | Bacteria>Terrabacteria_group>Cyanobacteriota/Melainabacteria_group>Cyanobacteriota>Cyanophyceae | GCF_014695435.1 | Coleofasciculus_sp._FACHB-129 | H6F60_RS14220 | 70669..74007 | 2692785 | Swt1-family-HEPN-domain-containing-protein-[Coleofasciculus-sp.-FACHB-129] |
| WP_190449861.1 | DUF499_1.RM.like.systems | CHTH-> TUDOR+ SF1-SF2-Helicase+iREase+CTD+REase ->HEPN-> METHYLASE -> OLD-ABC-> PIN -> HEPN+ DUF499 *->ParB+HNH->?->CopG/RHH->HNH-><-RHH | 1121 | Bacteria>Terrabacteria_group>Cyanobacteriota/Melainabacteria_group>Cyanobacteriota>Cyanophyceae | GCF_014695795.1 | Phormidium_sp._FACHB-592 | H6F86_RS16680 | 58473..61838 | 2692850 | Swt1-family-HEPN-domain-containing-protein-[Phormidium-sp.-FACHB-592] |
| WP_190368028.1 | DUF499_1.RM.like.systems | <- DUF499 *<- PIN <-MetJarc-HTH\|\|?->?-><- TUDOR+ SF1-SF2-Helicase+iREase+CTD+REase | 977 | Bacteria>Terrabacteria_group>Cyanobacteriota/Melainabacteria_group>Cyanobacteriota>Cyanophyceae | GCF_014696225.1 | Anabaena_sp._FACHB-1237 | H6F32_RS03895 | 63906..66839 | 2692769 | ATP-binding-protein-[Anabaena-sp.-FACHB-1237] |
| WP_190347798.1 | DUF499_1.RM.like.systems | ABC-ATPase+SbcC-><- DUF499 *<- PIN <-?<- METHYLASE \|\|?->?-><- TUDOR+ SF1-SF2-Helicase+iREase+CTD+REase | 963 | Bacteria>Terrabacteria_group>Cyanobacteriota/Melainabacteria_group>Cyanobacteriota>Cyanophyceae | GCF_014696235.1 | Sphaerospermopsis_sp._FACHB-1194 | H6F45_RS16760 | 52263..55154 | 2692862 | DUF499-domain-containing-protein-[Sphaerospermopsis-sp.-FACHB-1194] |
| WP_190383648.1 | DUF499_1.RM.like.systems | METHYLASE ->MetJarc-HTH-> PIN ->MetJarc-HTH-> DUF499 *->HISKIN+MORC->?->HTH_3->REase-><-N6_N4_MTase | 963 | Bacteria>Terrabacteria_group>Cyanobacteriota/Melainabacteria_group>Cyanobacteriota>Cyanophyceae | GCF_014696815.1 | Aphanizomenon_flos-aquae | H6F99_RS16810 | 3730..6621 | 1176 | DUF499-domain-containing-protein-[Aphanizomenon-flos-aquae] |
| WP_190513827.1 | DUF499_1.RM.like.systems | CHTH-> TUDOR+ SF1-SF2-Helicase+iREase+CTD+REase ->?-> METHYLASE ->RHH_6-> PIN -> HEPN+ DUF499 *->URI-DNase-><-RelE-ParE<-?\|\|pPIWI+pPIWI->pPIWI->pPIWI+MID+RNaseH-> | 1117 | Bacteria>Terrabacteria_group>Cyanobacteriota/Melainabacteria_group>Cyanobacteriota>Cyanophyceae | GCF_014697545.1 | Oscillatoria_sp._FACHB-1407 | H6G89_RS30695 | 49800..53153 | 2692847 | Swt1-family-HEPN-domain-containing-protein-[Oscillatoria-sp.-FACHB-1407] |
| WP_242054772.1 | DUF499_1.RM.like.systems | TUDOR+ SF1-SF2-Helicase+iREase+CTD+REase ->Phd-YefM-> PIN -><-HTH_3\|\| METHYLASE -p-> DUF499 *-><-TraJ-RHH\|\|PIWI->\|\|?-><-tRNA<-?<-SPOUT | 1065 | Bacteria>Terrabacteria_group>Cyanobacteriota/Melainabacteria_group>Cyanobacteriota>Cyanophyceae | GCF_014698065.1 | Nostoc_flagelliforme | H6G97_RS01005 | 252887..256084 | 1306274 | DUF499-domain-containing-protein-[Nostoc-flagelliforme] |
| WP_193965432.1 | DUF499_1.RM.like.systems | Phage_integrase-><-?\|\|?->?-><-HTH\|\|?-><- HEPN+ DUF499 *\|\|?-><-ParB+HNH<-?<- PIN <-?<- METHYLASE | 1114 | Bacteria>Terrabacteria_group>Cyanobacteriota/Melainabacteria_group>Cyanobacteriota>Cyanophyceae | GCF_015207395.1 | Nodosilinea_sp._LEGE_07088 | IQ254_RS06885 | 5859..9203 | 2777968 | Swt1-family-HEPN-domain-containing-protein-[Nodosilinea-sp.-LEGE-07088] |
| WP_200988278.1 | DUF499_1.RM.like.systems | <-REase<-?<-?<- DUF499 *<-CR-ATPase9+REase<- DUF499 <-CR-ATPase9+REase<- METHYLASE <-REase<- PIN | 950 | Bacteria>Terrabacteria_group>Cyanobacteriota/Melainabacteria_group>Cyanobacteriota>Cyanophyceae | GCF_015245355.1 | Anabaenopsis_elenkinii | IM676_RS18785 | 4299000..4301852 | 156213 | DUF499-domain-containing-protein-[Anabaenopsis-elenkinii] |
| WP_225880403.1 | DUF499_1.RM.like.systems | <-REase<-?<-?<- DUF499 <-CR-ATPase9+REase<- DUF499 *<-CR-ATPase9+REase<- METHYLASE <-REase<- PIN <-MetJarc-HTH | 821 | Bacteria>Terrabacteria_group>Cyanobacteriota/Melainabacteria_group>Cyanobacteriota>Cyanophyceae | GCF_015245355.1 | Anabaenopsis_elenkinii | IM676_RS18795 | 4304033..4306498 | 156213 | DUF499-domain-containing-protein-[Anabaenopsis-elenkinii] |
| WP_168636740.1 | DUF499_1.RM.like.systems | TUDOR+ SF1-SF2-Helicase+iREase+CTD+REase ->REase+HTH->?-> METHYLASE ->?-> PIN -> DUF499 *->?->?->HTH_3->REase-><-N6_N4_MTase\|\|NTP_transferase-> | 911 | Bacteria>Terrabacteria_group>Cyanobacteriota/Melainabacteria_group>Cyanobacteriota>Cyanophyceae | GCF_017355725.1 | Nostocales | HEQ36_RS21575 | 43162..45897 | 1161 | DUF499-domain-containing-protein-[Dolichospermum-flos-aquae] |
| WP_227382638.1 | DUF499_1.RM.like.systems | PIN -> METHYLASE ->REase->REase->REase-> DUF499 *-><-?\|\|RelB_antitoxin->PemK_toxin->?->RHH_1-><-TOPRIM+SF1-SF2-Helicase | 1002 | Bacteria>Terrabacteria_group>Cyanobacteriota/Melainabacteria_group>Cyanobacteriota>Cyanophyceae | GCF_020594575.1 | Planktothrix_agardhii | LJE16_RS21930 | 10143..13151 | 1160 | DUF499-domain-containing-protein-[Planktothrix-agardhii] |
| WP_227398372.1 | DUF499_1.RM.like.systems | ZnR->TOPRIM+SF1-SF2-Helicase-><-PemK_toxin<-RelB_antitoxin<-?\|\|?-><- DUF499 *<-REase<-REase<-REase<- METHYLASE <- PIN | 1002 | Bacteria>Terrabacteria_group>Cyanobacteriota/Melainabacteria_group>Cyanobacteriota>Cyanophyceae | GCF_020594695.1 | Planktothrix_agardhii | L2E74_RS21495 | 55743..58751 | 1160 | DUF499-domain-containing-protein-[Planktothrix-agardhii] |
| WP_235753326.1 | DUF499_1.RM.like.systems | PIN -> METHYLASE ->REase->REase->REase->REase-> DUF499 *-><-?\|\|RelB_antitoxin->HTH_OrfB_IS605+OrfB_IS605+OrfB_Zn_ribbon-><-PAIREDC-HTH | 1002 | Bacteria>Terrabacteria_group>Cyanobacteriota/Melainabacteria_group>Cyanobacteriota>Cyanophyceae | GCF_021554825.1 | Planktothrix_agardhii | L2E72_RS21865 | 12950..15958 | 1160 | DUF499-domain-containing-protein-[Planktothrix-agardhii] |
| WP_250012765.1 | DUF499_1.RM.like.systems | AbrB-> PIN ->REase-> METHYLASE -p-> METHYLASE ->?-> HEPN+ DUF499 *->?-><-?\|\|Phage_integrase-><-Phage_integrase | 1113 | Bacteria>Terrabacteria_group>Cyanobacteriota/Melainabacteria_group>Cyanobacteriota>Cyanophyceae | GCF_023546805.1 | Chroococcidiopsis_sp._CCNUC1 | M5J74_RS19090 | 4105692..4109033 | 2653189 | Swt1-family-HEPN-domain-containing-protein-[Chroococcidiopsis-sp.-CCNUC1] |
| WP_251956292.1 | DUF499_1.RM.like.systems | <- HEPN+ DUF499 *<- METHYLASE -p<- METHYLASE -p\|\|HTH_3-><- PIN <-Phd-YefM<- TUDOR+ SF1-SF2-Helicase+iREase+CTD+REase -p | 1108 | Bacteria>Terrabacteria_group>Cyanobacteriota/Melainabacteria_group>Cyanobacteriota>Cyanophyceae | GCF_023734775.1 | Nostoc_commune | ANS036C_RS22370 | 4938684..4942010 | 1178 | Swt1-family-HEPN-domain-containing-protein-[Nostoc-commune] |
| WP_254940710.1 | DUF499_1.RM.like.systems | <-DUF4236\|\| TUDOR+ SF1-SF2-Helicase+iREase+CTD+REase ->?-> PIN -> METHYLASE -> HEPN+ DUF499 *->?->?-><-ArdCN+MPTase | 1103 | Bacteria>Terrabacteria_group>Cyanobacteriota/Melainabacteria_group>Cyanobacteriota>Cyanophyceae | GCF_024345575.1 | Cyanobium_sp._Morenito_9A2 | KBY88_RS14795 | 41582..44893 | 2823718 | DUF499-domain-containing-protein-[Cyanobium-sp.-Morenito-9A2] |
| WP_265264586.1 | DUF499_1.RM.like.systems | NTP_transferase-><- HEPN+ DUF499 *<- PIN <-AbrB<- TUDOR+ SF1-SF2-Helicase+iREase+CTD+REase | 1112 | Bacteria>Terrabacteria_group>Cyanobacteriota/Melainabacteria_group>Cyanobacteriota>Cyanophyceae | GCF_026130165.1 | Spirulina_subsalsa | K4A83_RS11005 | 13314..16652 | 54311 | Swt1-family-HEPN-domain-containing-protein-[Spirulina-subsalsa] |
| WP_190359815.1 | DUF499_1.RM.like.systems | REase-><-DUF29<-?<-?\|\|?->Cluster164_18clades-><- HEPN+ DUF499 *<- PIN <-RHH_6<- METHYLASE <-REase<- TUDOR+ SF1-SF2-Helicase+iREase+CTD+REase <-ABhydrolase | 1117 | Bacteria>Terrabacteria_group>Cyanobacteriota/Melainabacteria_group>Cyanobacteriota>Cyanophyceae | GCF_026222535.1 | Microcystis | H6F92_RS24355 | 16318..19671 | 1125 | Swt1-family-HEPN-domain-containing-protein-[Microcystis-wesenbergii] |
| WP_265900465.1 | DUF499_1.RM.like.systems | REase->REase->?->?-><-sGTP<- HEPN+ DUF499 *<- PIN <-?<-REase<-REase<-REase | 1128 | Bacteria>Terrabacteria_group>Cyanobacteriota/Melainabacteria_group>Cyanobacteriota>Cyanophyceae | GCF_026240635.1 | Trichothermofontia_sichuanensis | OOK60_RS10505 | 2477245..2480631 | 3045816 | DUF499-domain-containing-protein-[Leptolyngbya-sp.-PKUAC-SCTB231] |
| WP_271935711.1 | DUF499_1.RM.like.systems | AbrB-> METHYLASE ->RHH_6-> PIN -> OLD-ABC-> PIN -> HEPN+ DUF499 *-><-Cluster164_18clades<-?\|\|?->?->DUF29->sGTP-> | 1111 | Bacteria>Terrabacteria_group>Cyanobacteriota/Melainabacteria_group>Cyanobacteriota>Cyanophyceae | GCF_028329095.1 | Microcystis | PN470_RS21190 | 25157..28492 | 1125 | MULTISPECIES:-Swt1-family-HEPN-domain-containing-protein-[Microcystis] |
| WP_272056151.1 | DUF499_1.RM.like.systems | <-SF1-SF2-Helicase-p<-?\|\|?-> PIN -> TUDOR+ SF1-SF2-Helicase+iREase+CTD+REase -> METHYLASE -> DUF499 *-><-Phage_integrase\|\|RelE-ParE->HTH_37->MORC-> | 1005 | Bacteria>Terrabacteria_group>Cyanobacteriota/Melainabacteria_group>Cyanobacteriota>Cyanophyceae | GCF_028330355.1 | Nodularia_spumigena | PN490_RS03030 | 15639..18656 | 70799 | DUF499-domain-containing-protein-[Nodularia-spumigena] |
| WP_272094882.1 | DUF499_1.RM.like.systems | <- HEPN+ DUF499 *<- PIN <-OLD-ABC <- PIN <-RHH_6<- METHYLASE <-REase | 1065 | Bacteria>Terrabacteria_group>Cyanobacteriota/Melainabacteria_group>Cyanobacteriota>Cyanophyceae | GCF_028330705.1 | Microcystis_aeruginosa | PN479_RS02395 | 1..3194 | 1126 | Swt1-family-HEPN-domain-containing-protein,-partial-[Microcystis-aeruginosa] |
| WP_272014891.1 | DUF499_1.RM.like.systems | METHYLASE -p->RHH_6-> PIN -> HEPN+ DUF499 *-><-Cluster164_18clades<-?\|\|?->?->DUF29->sGTP-> | 1111 | Bacteria>Terrabacteria_group>Cyanobacteriota/Melainabacteria_group>Cyanobacteriota>Cyanophyceae | GCF_028330745.1 | Microcystis_aeruginosa | PN482_RS11750 | 1666..5001 | 1126 | Swt1-family-HEPN-domain-containing-protein-[Microcystis-aeruginosa] |
| WP_272078194.1 | DUF499_1.RM.like.systems | DDE_3-><- HEPN+ DUF499 *<- PIN <- PIN <-RHH_6<- METHYLASE <-REase<- TUDOR+ SF1-SF2-Helicase+iREase+CTD+REase | 1111 | Bacteria>Terrabacteria_group>Cyanobacteriota/Melainabacteria_group>Cyanobacteriota>Cyanophyceae | GCF_028331105.1 | Microcystis_aeruginosa | PN487_RS19700 | 7822..11157 | 1126 | Swt1-family-HEPN-domain-containing-protein-[Microcystis-aeruginosa] |
| WP_271953564.1 | DUF499_1.RM.like.systems | PIN -> HEPN+ DUF499 *-><-Cluster164_18clades | 1130 | Bacteria>Terrabacteria_group>Cyanobacteriota/Melainabacteria_group>Cyanobacteriota>Cyanophyceae | GCF_028331205.1 | Microcystis_aeruginosa | PN437_RS04350 | 1001..4393 | 1126 | Swt1-family-HEPN-domain-containing-protein-[Microcystis-aeruginosa] |
| WP_281006617.1 | DUF499_1.RM.like.systems | METHYLASE ->?-> PIN ->REase->REase->REase-> HEPN+ DUF499 *-> | 1118 | Bacteria>Terrabacteria_group>Cyanobacteriota/Melainabacteria_group>Cyanobacteriota>Cyanophyceae | GCF_029910235.1 | Pseudanabaena_galeata | OA858_RS18405 | 4059827..4063183 | 1112103 | Swt1-family-HEPN-domain-containing-protein-[Pseudanabaena-galeata] |
| WP_087068561.1 | DUF499_1.RM.like.systems | TUDOR+ SF1-SF2-Helicase+iREase+CTD+REase ->?-> PIN -> METHYLASE -p->DUF772+DDE_Tnp_1_2-> METHYLASE -> HEPN+ DUF499 *->SF1-SF2-Helicase+REase->SF1-SF2-Helicase-p-><-?<-DUF772+DDE_Tnp_1_2 | 1101 | Bacteria>Terrabacteria_group>Cyanobacteriota/Melainabacteria_group>Cyanobacteriota>Cyanophyceae | GCF_900088535.1 | Cyanobium_sp._NIES-981 | CBM981_RS11760 | 2335245..2338550 | 1851505 | DUF499-domain-containing-protein-[Cyanobium-sp.-NIES-981] |
| WP_254171845.1 | DUF499_1.RM.like.systems | PIN -> METHYLASE ->REase->REase->REase->REase-> DUF499 *-><-?\|\|RelB_antitoxin->HTH_OrfB_IS605+OrfB_IS605+OrfB_Zn_ribbon->?->RHH_1-> | 1002 | Bacteria>Terrabacteria_group>Cyanobacteriota/Melainabacteria_group>Cyanobacteriota>Cyanophyceae | GCF_904830885.1 | Planktothrix_agardhii | NMK31_RS00300 | 70820..73828 | 1160 | DUF499-domain-containing-protein-[Planktothrix-agardhii] |
| WP_254035908.1 | DUF499_1.RM.like.systems | PIN -> PIN -> METHYLASE ->REase->REase->REase-> DUF499 *-><-?\|\|RelB_antitoxin->PemK_toxin->RHH_1-><-TOPRIM+SF1-SF2-Helicase | 999 | Bacteria>Terrabacteria_group>Cyanobacteriota/Melainabacteria_group>Cyanobacteriota>Cyanophyceae | GCF_904830935.1 | Planktothrix_agardhii | NMG88_RS22325 | 51359..54358 | 1160 | DUF499-domain-containing-protein-[Planktothrix-agardhii] |
| WP_066349645.1 | DUF499_1.RM.like.systems | TUDOR+ SF1-SF2-Helicase+iREase+CTD+REase -><-ZnR\|\| METHYLASE ->?-> PIN -> DUF499 *-><-DUF4158+DDE_Tnp_Tn3<-Phage_integrase\|\|?->N6_MTase-p-> | 1007 | Bacteria>Terrabacteria_group>Cyanobacteriota/Melainabacteria_group>Cyanobacteriota>Cyanophyceae | - | Geminocystis_sp._NIES-3708 | GM3708_RS17435 | 37996..41019 | 1615909 | DUF499-domain-containing-protein-[Geminocystis-sp.-NIES-3708] |
| WP_104397640.1 | DUF499_1.RM.like.systems | REase->AbrB-> PIN -> METHYLASE ->RHH_6-> PIN -> HEPN+ DUF499 *-><-Cluster164_18clades<-?\|\|?->?->?-><-TPR+TPR+TPR+TPR+TPR+TPR+TPR+TPR | 1111 | Bacteria>Terrabacteria_group>Cyanobacteriota/Melainabacteria_group>Cyanobacteriota>Cyanophyceae | - | Microcystis_aeruginosa | myaer87_32740 | 119164..122499 | 1126 | Swt1-family-HEPN-domain-containing-protein-[Microcystis-aeruginosa] |
| WP_235758599.1 | DUF499_1.RM.like.systems | TUDOR+ SF1-SF2-Helicase+iREase+CTD+REase -p-> PIN -> METHYLASE -p->REase->REase->REase-> DUF499 *-><-TOPRIM+SF1-SF2-Helicase<-ZnR<-?<-?<-URI-DNase | 1002 | Bacteria>Terrabacteria_group>Cyanobacteriota/Melainabacteria_group>Cyanobacteriota>Cyanophyceae | - | Planktothrix_agardhii | L2E81_RS19965 | 6975..9983 | 1160 | DUF499-domain-containing-protein-[Planktothrix-agardhii] |
| WP_255548761.1 | DUF499_1.RM.like.systems | REase-><-?<-CR-Trypsin1<-?\|\|REase-><-?<- HEPN+ DUF499 *<-?<- METHYLASE <- PIN <-AbrB<-REase<-REase | 1078 | Bacteria>Terrabacteria_group>Cyanobacteriota/Melainabacteria_group>Cyanobacteriota>Cyanophyceae | - | Thermoleptolyngbya_sp._PKUAC-SCTB121 | JTM80_RS13275 | 3238289..3241525 | 2811482 | DUF499-domain-containing-protein-[Thermoleptolyngbya-sp.-PKUAC-SCTB121] |
| MDA0267772.1 | DUF499_1.RM.like.systems | METHYLASE ->?-> PIN ->REase-> DUF499 *-> | 1000 | Bacteria>Terrabacteria_group>Cyanobacteriota/Melainabacteria_group>Cyanobacteriota>unclassified_Cyanobacteriota | GCA_027592485.1 | Cyanobacteriota_bacterium | O3A14_12630 | 3513..6515 | 2847862 | MAG:-DUF499-domain-containing-protein-[Cyanobacteria-bacterium] |
| WP_184025303.1 | DUF499_1.RM.like.systems | Phage_int_SAM_1+Phage_integrase-> TUDOR+ SF1-SF2-Helicase+iREase+CTD+REase ->DUF433-> PIN -> METHYLASE ->?-> HEPN+ DUF499 *-><-?<-TOPRIM+primpol<-OrfB_Zn_ribbon<-XIS-HTH\|\|HTH->Phage_integrase-> | 1097 | Bacteria>Terrabacteria_group>Deinococcota>Deinococci>Deinococcales | GCF_014201885.1 | Deinococcus_budaensis | HNQ09_RS02950 | 29745..33038 | 1665626 | DUF499-domain-containing-protein-[Deinococcus-budaensis] |
| MBF6594441.1 | DUF499_1.RM.like.systems | TUDOR+ SF1-SF2-Helicase+iREase+CTD+REase ->Schlafen+HATPase+HTH-> METHYLASE -><- PIN <-MazE_antitoxin\|\| HEPN+ DUF499 *-><-HTH_Tnp_4\|\|<-?<-?\|\| METHYLASE -p-> | 1117 | Bacteria>Terrabacteria_group>Deinococcota>Deinococci>Thermales | GCA_015478585.1 | Thermaceae_bacterium | IVW51_08335 | 21188..24541 | 2614257 | DUF499-domain-containing-protein-[Thermaceae-bacterium] |
| WP_013458840.1 | DUF499_1.RM.like.systems | TUDOR+ SF1-SF2-Helicase+iREase+CTD+REase ->HicA_toxin->HicB_lk_antitox-> PIN -> METHYLASE -> HEPN+ DUF499 *->?-><-?<-?<-?\|\|?-> METHYLASE -p-> | 1120 | Bacteria>Terrabacteria_group>Deinococcota>Deinococci>Thermales | GCF_000183745.1 | Oceanithermus_profundus | OCEPR_RS11190 | 2275579..2278941 | 187137 | DUF499-domain-containing-protein-[Oceanithermus-profundus] |
| WP_015065389.1 | DUF499_1.RM.like.systems | ParB-> TUDOR+ SF1-SF2-Helicase+iREase+CTD+REase -> PIN ->DDE_Tnp_1-> HEPN+ DUF499 *->SF1-SF2-Helicase->IclR-HTH-> | 1099 | Bacteria>Terrabacteria_group>Deinococcota>Deinococci>Thermales | GCF_000309885.1 | Thermus_oshimai | THEOS_RS11985 | 210648..213947 | 56957 | Swt1-family-HEPN-domain-containing-protein-[Thermus-oshimai] |
| WP_018461866.1 | DUF499_1.RM.like.systems | ParB-> TUDOR+ SF1-SF2-Helicase+iREase+CTD+REase -> PIN -> METHYLASE -> HEPN+ DUF499 *->SF1-SF2-Helicase->?-><-?<-Peptidase_C26 | 1097 | Bacteria>Terrabacteria_group>Deinococcota>Deinococci>Thermales | GCF_000373145.1 | Thermus_oshimai | B043_RS0109800 | 87375..90668 | 56957 | Swt1-family-HEPN-domain-containing-protein-[Thermus-oshimai] |
| WP_135343460.1 | DUF499_1.RM.like.systems | <- HEPN+ DUF499 *<- METHYLASE <- PIN <- TUDOR+ SF1-SF2-Helicase+iREase+CTD+REase <-ParB | 1099 | Bacteria>Terrabacteria_group>Deinococcota>Deinococci>Thermales | GCF_004685085.1 | Thermus_tengchongensis | E0489_RS07460 | 252..3551 | 1214928 | Swt1-family-HEPN-domain-containing-protein-[Thermus-tengchongensis] |
| WP_119356451.1 | DUF499_1.RM.like.systems | PIN -> TUDOR+ SF1-SF2-Helicase+iREase+CTD+REase ->XIS-HTH-> METHYLASE -><- PIN <-MazE_antitoxin\|\| HEPN+ DUF499 *-> | 1110 | Bacteria>Terrabacteria_group>Deinococcota>Deinococci>Thermales | GCF_007990995.1 | Meiothermus_granaticius | MGR01S_RS15620 | 8907..12239 | 863370 | Swt1-family-HEPN-domain-containing-protein-[Meiothermus-granaticius] |
| WP_013177687.1 | DUF499_1.RM.like.systems | PUA+ METHYLASE ->?->?->?-><-sGTP<-Aminotran_3<- HEPN+ DUF499 *<- METHYLASE -p\|\|Y1_Tnp-><- METHYLASE <- PIN | 1108 | Bacteria>Terrabacteria_group>Deinococcota>Deinococci>Trueperales | GCF_000092425.1 | Truepera_radiovictrix | TRAD_RS05920 | 1278601..1281927 | 332249 | Swt1-family-HEPN-domain-containing-protein-[Truepera-radiovictrix] |
| WP_013179180.1 | DUF499_1.RM.like.systems | TUDOR+ SF1-SF2-Helicase+iREase+CTD+REase ->Phd-YefM-> PIN -> METHYLASE ->?-> PIN -> HEPN+ DUF499 *-><-Peptidase_M20 | 1069 | Bacteria>Terrabacteria_group>Deinococcota>Deinococci>Trueperales | GCF_000092425.1 | Truepera_radiovictrix | TRAD_RS13560 | 2957587..2960796 | 332249 | DUF499-domain-containing-protein-[Truepera-radiovictrix] |
| OQX20475.1 | DUF499_1.RM.like.systems | METHYLASE -p->PhdYefM_antitox->RelE-ParE-><-?\|\|MazE_antitoxin-> PIN -> DUF499 *->?->?->?->Peptidase_M48-> | 913 | Bacteria>Thermodesulfobacteriota>Desulfobacteria>Desulfobacterales>Desulfobacteraceae | GCA_002083955.1 | Desulfobacteraceae_bacterium_IS3 | BWK80_35715 | 2623..5364 | 1934248 | hypothetical-protein-BWK80_35715-[Desulfobacteraceae-bacterium-IS3] |
| MBF0572396.1 | DUF499_1.RM.like.systems | <-CHASE2+ACYC<-?<-?<- DUF499 *<-REase<- PIN <-?<-?<- METHYLASE -p | 936 | Bacteria>Thermodesulfobacteriota>Desulfobacteria>Desulfobacterales>Desulfobacteraceae | GCA_015233755.1 | Desulfamplus_sp. | HQK69_01355 | 13339..16149 | 2006184 | ATP-binding-protein-[Desulfamplus-sp.] |
| MBL7176677.1 | DUF499_1.RM.like.systems | TUDOR+ SF1-SF2-Helicase+iREase+CTD+REase -p-> OLD-ABC-> PIN ->?-> HEPN+ DUF499 *->?->OLD-ABC->SF1-SF2-Helicase->?->MNS+HTH-> | 1094 | Bacteria>Thermodesulfobacteriota>Desulfobacteria>Desulfobacterales>Desulfobacteraceae | GCA_016782925.1 | Desulfobacteraceae_bacterium | ISS66_12705 | 13240..16524 | 2049433 | DUF499-domain-containing-protein-[Desulfobacteraceae-bacterium] |
| MBF0121417.1 | DUF499_1.RM.like.systems | <-HNH<- DUF499 *<- PIN <-?<- METHYLASE <- TUDOR+ SF1-SF2-Helicase+iREase+CTD+REase | 973 | Bacteria>Thermodesulfobacteriota>Desulfobacteria>Desulfobacterales>unclassified_Desulfobacterales | GCA_015228685.1 | Desulfobacterales_bacterium | HQK79_21515 | 1603..4524 | 2044940 | ATP-binding-protein-[Desulfobacterales-bacterium] |
| MCK4391114.1 | DUF499_1.RM.like.systems | MNS+HTH-><-?\|\| METHYLASE ->DUF433-> PIN ->HicB-> DUF499 *-> | 914 | Bacteria>Thermodesulfobacteriota>Desulfobacteria>Desulfobacterales>unclassified_Desulfobacterales | GCA_023129535.1 | Desulfobacterales_bacterium | KAV83_12860 | 5895..8639 | 2044940 | ATP-binding-protein-[Desulfobacterales-bacterium] |
| HCF05840.1 | DUF499_1.RM.like.systems | TUDOR+ SF1-SF2-Helicase+iREase+CTD+REase -> PIN ->HEPN-> METHYLASE ->OLD-ABC->TOPRIM-> DUF499 *-> | 934 | Bacteria>Thermodesulfobacteriota>Desulfovibrionia>Desulfovibrionales>Desulfomicrobiaceae | GCA_003530935.1 | Desulfomicrobiaceae_bacterium | DEU72_06355 | 12752..15556 | 2053547 | hypothetical-protein-[Desulfomicrobiaceae-bacterium] |
| MBZ4685271.1 | DUF499_1.RM.like.systems | FAD-NAD-dep-oxidoreductase-><-?<-?<-?\|\|?-><- DUF499 *<-TOPRIM<-OLD-ABC<- METHYLASE <-HEPN<- PIN | 934 | Bacteria>Thermodesulfobacteriota>Desulfovibrionia>Desulfovibrionales>Desulfomicrobiaceae | GCA_020063575.1 | Desulfomicrobiaceae_bacterium | JG774_1016 | 59895..62699 | 2053547 | putative-ATPase-[Desulfomicrobiaceae-bacterium] |
| MCG2740517.1 | DUF499_1.RM.like.systems | MerR-HTH-> METHYLASE -> PIN ->?-><- HEPN+ DUF499 *<- METHYLASE <-MerR-HTH<- TUDOR+ SF1-SF2-Helicase+iREase+CTD+REase | 1109 | Bacteria>Thermodesulfobacteriota>Syntrophia>Syntrophales>Syntrophaceae | GCA_022013455.1 | Syntrophaceae_bacterium | L6300_09825 | 5770..9099 | 2052184 | DUF499-domain-containing-protein-[Syntrophaceae-bacterium] |
| WP_073037944.1 | DUF499_1.RM.like.systems | TUDOR+ SF1-SF2-Helicase+iREase+CTD+REase ->TIR-> METHYLASE -> DUF499 *-> PIN -> | 928 | Bacteria>Thermodesulfobacteriota>Syntrophobacteria>Syntrophobacterales>Syntrophobacteraceae | GCF_900129305.1 | Desulfacinum_infernum | BUB04_RS06060 | 31912..34698 | 35837 | DUF499-domain-containing-protein-[Desulfacinum-infernum] |
| OQA20379.1 | DUF499_1.RM.like.systems | PIN ->HicB_lk_antitox->HicB_lk_antitox->?-> PIN ->?-> DUF499 *-><-NUDIX<-?<-?\|\|TPR+TPR-> | 993 | Bacteria>unclassified_Bacteria | GCA_002069435.1 | bacterium_ADurb.Bin363 | BWY64_00213 | 12863..15844 | 1866927 | hypothetical-protein-BWY64_00213-[bacterium-ADurb.Bin363] |
| KAF0250487.1 | DUF499_1.RM.like.systems | Phd-YefM-> PIN -> METHYLASE ->REase->OLD-ABC->TOPRIM-> HEPN+ DUF499 *->?-><-Phage_integrase\|\|?->MqsA_antitoxin+HTH_3-> | 1130 | Bacteria>unclassified_Bacteria | GCA_009773835.1 | bacterium | FD167_113 | 11053..14445 | 1869227 | hypothetical-protein-FD167_113-[bacterium] |
| NJL57104.1 | DUF499_1.RM.like.systems | METHYLASE -p->DUF433-> PIN -> HEPN+ DUF499 *-> DUF499 -> | 411 | Bacteria>unclassified_Bacteria | GCA_012031555.1 | bacterium | HC928_19635 | 992..2227 | 1869227 | ATP-binding-protein-[bacterium] |
| NJL57105.1 | DUF499_1.RM.like.systems | METHYLASE -p->DUF433-> PIN -> HEPN+ DUF499 -> DUF499 *-> | 478 | Bacteria>unclassified_Bacteria | GCA_012031555.1 | bacterium | HC928_19640 | 2241..3676 | 1869227 | ATP-binding-protein-[bacterium] |
| MBE3604313.1 | DUF499_1.RM.like.systems | <-HIN-HTH\|\|tRNA-><- DUF499 *<- PIN <-DUF433\|\|CopG_antitoxin->?-><-?<- METHYLASE | 1108 | Bacteria>unclassified_Bacteria | GCA_014896235.1 | bacterium | IMX07_11830 | 1248..4574 | 1869227 | ATP-binding-protein-[bacterium] |
| MBK8128933.1 | DUF499_1.RM.like.systems | DDE_3->?-><-?\|\|?-><-?<-?<- HEPN+ DUF499 *<- METHYLASE -p<- PIN <-CopG/RHH<- PIN <-?<- TUDOR+ SF1-SF2-Helicase+iREase+CTD+REase -p | 1097 | Bacteria>unclassified_Bacteria | GCA_016712825.1 | bacterium | IPK53_08290 | 27062..30355 | 1869227 | DUF499-domain-containing-protein-[bacterium] |
| MBN2381437.1 | DUF499_1.RM.like.systems | GDResolvase+Recombinase+ZnR->?-> TUDOR+ SF1-SF2-Helicase+iREase+CTD+REase ->?-> PIN -> METHYLASE -> DUF499 *-> | 970 | Bacteria>unclassified_Bacteria | GCA_016934675.1 | bacterium | JXQ70_01010 | 9963..12875 | 1869227 | ATP-binding-protein-[bacterium] |
| MCD6327181.1 | DUF499_1.RM.like.systems | wHTH+REase->SbcC->?-> METHYLASE ->?-> PIN -> DUF499 *-> | 928 | Bacteria>unclassified_Bacteria | GCA_021160245.1 | bacterium | J7M28_06475 | 15142..17928 | 1869227 | ATP-binding-protein-[bacterium] |
| MCG3156467.1 | DUF499_1.RM.like.systems | TUDOR+ SF1-SF2-Helicase+iREase+CTD+REase ->Schlafen+HATPase+HTH->DOC+HTH-> METHYLASE -><- PIN <-?\|\| HEPN+ DUF499 *->TIR-> | 1113 | Bacteria>unclassified_Bacteria | GCA_022072305.1 | bacterium | DKINENOH_03091 | 110830..114171 | 1869227 | hypothetical-protein-[bacterium] |
| MCY3924850.1 | DUF499_1.RM.like.systems | PIN ->DUF433-> PIN -> METHYLASE ->MetJarc-HTH->PemK_toxin-> HEPN+ DUF499 *-><-?\|\|ABhydrolase->?-><-?<-Glyoxalase | 1099 | Bacteria>unclassified_Bacteria | GCA_026707305.1 | bacterium | OXG52_04985 | 27802..31101 | 1869227 | MAG:-DUF499-domain-containing-protein-[bacterium] |
| MCY3953257.1 | DUF499_1.RM.like.systems | DUF433-> PIN -> METHYLASE ->MetJarc-HTH->PemK_toxin-> HEPN+ DUF499 *-><-?\|\|ABhydrolase-> | 1099 | Bacteria>unclassified_Bacteria | GCA_026707535.1 | bacterium | OXG69_12060 | 23481..26780 | 1869227 | MAG:-DUF499-domain-containing-protein-[bacterium] |
| MCY4103460.1 | DUF499_1.RM.like.systems | DUF433-> PIN -> METHYLASE ->MetJarc-HTH->PemK_toxin-> HEPN+ DUF499 *-><-?\|\|ABhydrolase-><-?\|\|?-><-?\|\|REase-> | 1099 | Bacteria>unclassified_Bacteria | GCA_026708055.1 | bacterium | OXG55_09400 | 103234..106533 | 1869227 | MAG:-DUF499-domain-containing-protein-[bacterium] |
| MCY4163400.1 | DUF499_1.RM.like.systems | DUF433-> PIN -> METHYLASE -> METHYLASE -p-><-?\|\|?-> HEPN+ DUF499 *-><-?\|\|?->?->?->ABC-ATPase+SbcC->PfkB-> | 1091 | Bacteria>unclassified_Bacteria | GCA_026709025.1 | bacterium | OXE93_04200 | 10413..13688 | 1869227 | MAG:-DUF499-domain-containing-protein-[bacterium] |

| Protein-ID | System | Gene-Neighbourhoods | Length | Taxonomy | Assembly-ID | Strain | Locus-Tag | Genomic-coordinates | TaxID | Fasta-defline |
|---|---|---|---|---|---|---|---|---|---|---|
| -- | | | | | | | | | | |
| -- TUDOR+SF1-SF2-Helicase+i REase +CTD+ REase | and | additional | REase ; | | | | | | | |
| -- OLD-ABC-> REase | | | | | | | | | | |
| -- | | | | | | | | | | |
| RLI54042.1 | DUF499_1.RM.like.systems | TUDOR+SF1-SF2-Helicase+i REase +CTD+ REase -> REase -> METHYLASE -> DUF499 *-> | 866 | Archaea>Asgard_group>Candidatus_Thorarchaeota>unclassified_Candidatus_Thorarchaeota | GCA_003662775.1 | Candidatus_Thorarchaeota_archaeon | DRP09_14185 | 7004..9603 | 2053491 | AAA-family-ATPase,-partial-[Candidatus-Thorarchaeota-archaeon] |
| CCJ36896.1 | DUF499_1.RM.like.systems | TUDOR+SF1-SF2-Helicase+i REase +CTD+ REase -> METHYLASE -p-> w HTH+ REase -> METHYLASE ->ParB+HNH-> DUF499 *->?->PolB->HEPN-> | 944 | Archaea>Euryarchaeota>Stenosarchaea_group>Methanomicrobia>Methanomicrobiales | GCA_000304355.2 | Methanoculleus_bourgensis_MS2 | BN140_1973 | 2039579..2042413 | 1201294 | ATPase-[Methanoculleus-bourgensis-MS2] |
| MCE5337191.1 | DUF499_1.RM.like.systems | ORC/CDC6+HTH->DOC+HTH-><-?<-PolB<-?<-PolB<- DUF499 *<-ParB+HNH<- METHYLASE <-?<- w HTH+ REase <- METHYLASE -p<- TUDOR+SF1-SF2-Helicase+i REase +CTD+ REase | 944 | Archaea>Euryarchaeota>Stenosarchaea_group>Methanomicrobia>Methanomicrobiales | GCA_021371835.1 | Methanomicrobiaceae_archaeon | LLF90_00675 | 34057..36891 | 2716262 | DUF499-domain-containing-protein-[Methanomicrobiaceae-archaeon] |
| WP_156147608.1 | DUF499_1.RM.like.systems | TUDOR+SF1-SF2-Helicase+i REase +CTD+ REase -> METHYLASE -p-> w HTH+ REase ->?-> METHYLASE ->ParB+HNH-> DUF499 *->PolB->HEPN-><-?<-Ntox49 | 942 | Archaea>Euryarchaeota>Stenosarchaea_group>Methanomicrobia>Methanomicrobiales | GCF_000304355.2 | Methanoculleus_bourgensis | BN140_RS09865 | 2039585..2042413 | 83986 | DUF499-domain-containing-protein-[Methanoculleus-bourgensis] |
| AKB47809.1 | DUF499_1.RM.like.systems | TUDOR+SF1-SF2-Helicase+i REase +CTD+ REase -> METHYLASE -> REase -> DUF499 *-><-Radical-SAM<-?<-Radical-SAM<-Radical-SAM<-?\|\|DDE_Tnp_1_2-> | 939 | Archaea>Euryarchaeota>Stenosarchaea_group>Methanomicrobia>Methanosarcinales | GCA_000969945.1 | Methanosarcina_sp._Kolksee | MSKOL_2032 | 2634692..2637511 | 1434099 | putative-ATPase-(AAA+-superfamily)-[Methanosarcina-sp.-Kolksee] |
| PXF61726.1 | DUF499_1.RM.like.systems | TUDOR+SF1-SF2-Helicase+i REase +CTD+ REase ->HD->DUF4065-> REase ->PolB->HEPN-> DUF499 *-><-Mbetalac<-?<-?<-?<-CR-Trypsin1 | 968 | Archaea>Euryarchaeota>Stenosarchaea_group>Methanomicrobia>Methanosarcinales | GCA_003194445.1 | ANME-2_cluster_archaeon | C4B59_02400 | 71391..74297 | 2056317 | AAA-family-ATPase-[ANME-2-cluster-archaeon] |
| WP_197078759.1 | DUF499_1.RM.like.systems | TUDOR+SF1-SF2-Helicase+i REase +CTD+ REase ->?-> METHYLASE -> REase -> DUF499 *-><-Radical-SAM<-?<-Radical-SAM<-Radical-SAM | 943 | Archaea>Euryarchaeota>Stenosarchaea_group>Methanomicrobia>Methanosarcinales | GCF_000969945.1 | Methanosarcina_sp._Kolksee | MSKOL_RS10880 | 2634680..2637511 | 1434099 | DUF499-domain-containing-protein-[Methanosarcina-sp.-Kolksee] |
| AET64040.1 | DUF499_1.RM.like.systems | TUDOR+SF1-SF2-Helicase+i REase +CTD+ REase ->Schlafen+HATPase+HTH-> METHYLASE -> REase -><-HicB_lk_antitox\|\| DUF499 *->HicB_lk_antitox-> | 1010 | Archaea>Euryarchaeota>Stenosarchaea_group>Methanomicrobia>Methanotrichales | GCA_000235565.1 | Methanothrix_harundinacea_6Ac | Mhar_0662 | 698704..701736 | 1110509 | ATPase-(AAA+-superfamily)-like-protein-[Methanosaeta-harundinacea-6Ac] |
| KUK43676.1 | DUF499_1.RM.like.systems | TUDOR+SF1-SF2-Helicase+i REase +CTD+ REase ->HD->DUF4065-> METHYLASE -> REase -> DUF499 *->RelB->?-><-HicB_lk_antitox | 987 | Archaea>Euryarchaeota>Stenosarchaea_group>Methanomicrobia>Methanotrichales | GCA_001509375.1 | Methanothrix_harundinacea | XD72_1943 | 10548..13511 | 301375 | ATPase-(AAA+-superfamily)-like-protein-[Methanothrix-harundinacea] |
| NLX39269.1 | DUF499_1.RM.like.systems | Y1_Tnp-><- DUF499 *<- REase <- METHYLASE <- TUDOR+SF1-SF2-Helicase+i REase +CTD+ REase | 1044 | Archaea>Euryarchaeota>Stenosarchaea_group>Methanomicrobia>Methanotrichales | GCA_012729025.1 | Methanothrix_sp. | GXY73_07140 | 1114..4248 | 90426 | ATP-binding-protein-[Methanothrix-sp.] |
| MBN1235613.1 | DUF499_1.RM.like.systems | <- DUF499 *<- REase <-ASCH<-PIN+ACET+EVE<- METHYLASE <-Schlafen+HATPase+HTH<- TUDOR+SF1-SF2-Helicase+i REase +CTD+ REase | 1051 | Archaea>Euryarchaeota>Stenosarchaea_group>Methanomicrobia>Methanotrichales | GCA_016927055.1 | Methanotrichaceae_archaeon | JW999_06125 | 575..3730 | 2595067 | DUF499-domain-containing-protein-[Methanotrichaceae-archaeon] |
| MDD3564060.1 | DUF499_1.RM.like.systems | DUF6361->HKD-DNase->SF1-SF2-Helicase-><- SF1-SF2-Helicase+ REase <- REase <-?<- DUF499 *<- REase <- METHYLASE <-DUF4065<-HD<-HD<- TUDOR+SF1-SF2-Helicase+i REase +CTD+ REase | 1047 | Archaea>Euryarchaeota>Stenosarchaea_group>Methanomicrobia>Methanotrichales | GCA_028695755.1 | Methanothrix_sp. | PHN90_00125 | 45442..48585 | 90426 | MAG:-DUF499-domain-containing-protein-[Methanothrix-sp.] |
| WP_048144800.1 | DUF499_1.RM.like.systems | TUDOR+SF1-SF2-Helicase+i REase +CTD+ REase ->Schlafen+HATPase+HTH-> METHYLASE -> REase -><-HicB_lk_antitox\|\| DUF499 *->RelB->HicB_lk_antitox-> | 987 | Archaea>Euryarchaeota>Stenosarchaea_group>Methanomicrobia>Methanotrichales | GCF_000235565.1 | Methanothrix_harundinacea | MHAR_RS03425 | 698773..701736 | 301375 | DUF499-domain-containing-protein-[Methanothrix-harundinacea] |
| MCD6241711.1 | DUF499_1.RM.like.systems | <-OLD-ABC+TOPRIM<-?<-PIWI<-?<-?\|\|?-><- DUF499 *<- METHYLASE <- REase <- TUDOR+SF1-SF2-Helicase+i REase +CTD+ REase | 896 | Archaea>TACK_group>Candidatus_Bathyarchaeota>unclassified_Candidatus_Bathyarchaeota | GCA_021158145.1 | Candidatus_Bathyarchaeota_archaeon | J7K27_09445 | 7902..10592 | 2026714 | ATP-binding-protein-[Candidatus-Bathyarchaeota-archaeon] |
| MBZ5490311.1 | DUF499_1.RM.like.systems | <- DUF499 *<- REase <- METHYLASE <-OLD-ABC<- TUDOR+SF1-SF2-Helicase+i REase +CTD+ REase | 921 | Bacteria>Acidobacteriota>Terriglobia>unclassified_Terriglobia | GCA_020072645.1 | Terriglobia_bacterium | LAO76_05190 | 365622..368387 | 2081523 | DUF499-domain-containing-protein-[Acidobacteriia-bacterium] |
| MBZ5536629.1 | DUF499_1.RM.like.systems | TUDOR+SF1-SF2-Helicase+i REase +CTD+ REase ->DOC+HTH->?-> METHYLASE -> REase -> DUF499 *-><-?<-?<-?<-?<-TPR+TPR<-NTP_transf_3 | 926 | Bacteria>Acidobacteriota>Terriglobia>unclassified_Terriglobia | GCA_020072965.1 | Terriglobia_bacterium | LAO31_11795 | 35949..38729 | 2081523 | DUF499-domain-containing-protein-[Acidobacteriia-bacterium] |
| KAB2964430.1 | DUF499_1.RM.like.systems | REase -> TUDOR+SF1-SF2-Helicase+i REase +CTD+ REase ->ParB+HNH->HEPN-> METHYLASE ->?-> HEPN+ DUF499 *->HNH-> | 1109 | Bacteria>Acidobacteriota>Thermoanaerobaculia>unclassified_Thermoanaerobaculia | GCA_009026165.1 | Thermoanaerobaculia_bacterium | F9K16_04350 | 12106..15435 | 2651171 | MAG:-ATP-binding-protein-[Thermoanaerobaculia-bacterium] |
| PYV70338.1 | DUF499_1.RM.like.systems | REase +35exo-><-?\|\| TUDOR+SF1-SF2-Helicase+i REase +CTD+ REase -p-> METHYLASE -> METHYLASE -p-> REase -> DUF499 *-> | 691 | Bacteria>Acidobacteriota>unclassified_Acidobacteriota | GCA_003225375.1 | Acidobacteriota_bacterium | DMG97_19575 | 8894..10968 | 1978231 | AAA-family-ATPase,-partial-[Acidobacteriota-bacterium] |
| HBL27432.1 | DUF499_1.RM.like.systems | <- DUF499 *<-?<- DUF499 -p<- REase <- REase <- METHYLASE <- TUDOR+SF1-SF2-Helicase+i REase +CTD+ REase | 720 | Bacteria>Acidobacteriota>unclassified_Acidobacteriota | GCA_003504885.1 | Acidobacteriota_bacterium | DD490_11410 | 1..2162 | 1978231 | AAA-family-ATPase-[Acidobacteria-bacterium] |
| MBI2841722.1 | DUF499_1.RM.like.systems | TUDOR+SF1-SF2-Helicase+i REase +CTD+ REase ->?->AAA-ATPase+ REase -> METHYLASE -> DUF499 *-> | 1100 | Bacteria>Acidobacteriota>unclassified_Acidobacteriota | GCA_016191015.1 | Acidobacteriota_bacterium | HYX75_25690 | 46699..50001 | 1978231 | ATP-binding-protein-[Acidobacteria-bacterium] |
| MBI4730026.1 | DUF499_1.RM.like.systems | TUDOR+SF1-SF2-Helicase+i REase +CTD+ REase ->?->HEPN+ REase -> METHYLASE ->?->HEPN-> DUF499 *->?->?-><-?<-sGTP<-?<-SAM-methylase | 918 | Bacteria>Acidobacteriota>unclassified_Acidobacteriota | GCA_016208485.1 | Acidobacteriota_bacterium | HY775_11110 | 79906..82662 | 1978231 | DUF499-domain-containing-protein-[Acidobacteria-bacterium] |
| MBN1568514.1 | DUF499_1.RM.like.systems | ABC-ATPase+SbcC->ABC-ATPase+SbcC->ABC-ATPase+SbcC-> TUDOR+SF1-SF2-Helicase+i REase +CTD+ REase -> METHYLASE -> REase -> DUF499 *->?->?->?->HEPN->OLD-ABC+TOPRIM->SF1-SF2-Helicase-> | 971 | Bacteria>Acidobacteriota>unclassified_Acidobacteriota | GCA_016929335.1 | Acidobacteriota_bacterium | JXA73_11765 | 13094..16009 | 1978231 | ATP-binding-protein-[Acidobacteria-bacterium] |
| MCL5262739.1 | DUF499_1.RM.like.systems | <- HEPN+ DUF499 *<- METHYLASE <-?<- TUDOR+SF1-SF2-Helicase+i REase +CTD+ REase -p<- REase <-HKD-DNase+SF1-SF2-Helicase+TRD | 1103 | Bacteria>Acidobacteriota>unclassified_Acidobacteriota | GCA_023480755.1 | Acidobacteriota_bacterium | M1568_00545 | 1..3309 | 1978231 | Swt1-family-HEPN-domain-containing-protein-[Acidobacteria-bacterium] |
| MDE2058883.1 | DUF499_1.RM.like.systems | <- HEPN+ DUF499 *<-?<- METHYLASE <-TOPRIM<-OLD-ABC<-ORC-CDC6+ REase <- TUDOR+SF1-SF2-Helicase+i REase +CTD+ REase | 1122 | Bacteria>Bacteria_incertae_sedis>Bacteria_candidate_phyla>candidate_division_NC10>unclassified_candidate_division_NC10 | GCA_028864815.1 | candidate_division_NC10_bacterium | KGJ27_04040 | 4506..7874 | 2072417 | MAG:-DUF499-domain-containing-protein-[candidate-division-NC10-bacterium] |
| MCX6842949.1 | DUF499_1.RM.like.systems | <-PIN<-?<- DUF499 *<- METHYLASE <-TOPRIM<-OLD-ABC<- REase <- TUDOR+SF1-SF2-Helicase+i REase +CTD+ REase | 912 | Bacteria>Bacteria_incertae_sedis>Bacteria_candidate_phyla>candidate_division_WOR-3 | GCA_026397595.1 | candidate_division_WOR-3_bacterium | NTX53_11780 | 2277..5015 | 2052148 | MAG:-DUF499-domain-containing-protein-[candidate-division-WOR-3-bacterium] |
| HDP95525.1 | DUF499_1.RM.like.systems | <- DUF499 *<- REase <-?<- METHYLASE <-DOC+HTH<- TUDOR+SF1-SF2-Helicase+i REase +CTD+ REase | 933 | Bacteria>Bacteria_incertae_sedis>Bacteria_candidate_phyla>Candidatus_Aminicenantes>unclassified_Candidatus_Aminicenantes | GCA_011049425.1 | Candidatus_Aminicenantes_bacterium | ENN40_09225 | 209..3010 | 2052149 | ATP-binding-protein-[Candidatus-Aminicenantes-bacterium] |
| NQT80771.1 | DUF499_1.RM.like.systems | <-URI-DNase<-HEPN<- REase <- McrC-NTD+ REase <-?<- REase <- DUF499 *<- METHYLASE <- TUDOR+SF1-SF2-Helicase+i REase +CTD+ REase | 981 | Bacteria>Bacteria_incertae_sedis>Bacteria_candidate_phyla>Candidatus_Aminicenantes>unclassified_Candidatus_Aminicenantes | GCA_013203055.1 | Candidatus_Aminicenantes_bacterium | HQ555_10335 | 19157..22102 | 2052149 | ATP-binding-protein-[Candidatus-Aminicenantes-bacterium] |
| QAA76831.1 | DUF499_1.RM.like.systems | METHYLASE -p-> TUDOR+SF1-SF2-Helicase+i REase +CTD+ REase ->?->SbcC-> METHYLASE -> REase -> DUF499 *->DEDD_Tnp_IS110+Transposase_20->Glyco_transf_20-><-ABC-ATPase+SbcC<-ABC-ATPase+SbcC | 920 | Bacteria>Bacteria_incertae_sedis>Bacteria_candidate_phyla>Candidatus_Bipolaricaulota>Bipolaricaulis | GCA_004102645.1 | Candidatus_Bipolaricaulis_sibiricus | BIP78_1065 | 1085101..1087863 | 2501609 | hypothetical-protein-BIP78_1065-[Candidatus-Bipolaricaulis-sibiricus] |
| WP_122031531.1 | DUF499_1.RM.like.systems | TUDOR+SF1-SF2-Helicase+i REase +CTD+ REase ->ORC-CDC6+ REase -> METHYLASE ->OLD-ABC->PIN-> HEPN+ DUF499 *->Radical-SAM+TRAM->PfkB->?->35exo+SF1-SF2-Helicase->?-><-NTP_transf_2 | 1110 | Bacteria>Bacteria_incertae_sedis>Bacteria_candidate_phyla>Candidatus_Bipolaricaulota>Bipolaricaulis | GCF_900465355.1 | Candidatus_Bipolaricaulis_anaerobius | BARAN1_RS05370 | 1116813..1120145 | 2026885 | Swt1-family-HEPN-domain-containing-protein-[Candidatus-Bipolaricaulis-anaerobius] |
| MCD6320860.1 | DUF499_1.RM.like.systems | <- DUF499 *<- METHYLASE -p<-HEPN<-PolB<- METHYLASE -p<- REase <- TUDOR+SF1-SF2-Helicase+i REase +CTD+ REase | 867 | Bacteria>Bacteria_incertae_sedis>Bacteria_candidate_phyla>Candidatus_Bipolaricaulota>unclassified_Candidatus_Bipolaricaulota | GCA_021160365.1 | Candidatus_Bipolaricaulota_bacterium | J7L84_01245 | 3909..6512 | 2306012 | DUF499-domain-containing-protein-[Candidatus-Bipolaricaulota-bacterium] |
| MDE0634775.1 | DUF499_1.RM.like.systems | TUDOR+SF1-SF2-Helicase+i REase +CTD+ REase ->SMODS->HNH+SAVED->HEPN+ REase ->?-> METHYLASE -> DUF499 *->?->TPR+TPR->SbcC-> | 935 | Bacteria>Bacteria_incertae_sedis>Bacteria_candidate_phyla>Candidatus_Poribacteria>unclassified_Candidatus_Poribacteria | GCA_028824125.1 | Candidatus_Poribacteria_bacterium | OXI43_02830 | 28592..31399 | 2026781 | MAG:-DUF499-domain-containing-protein-[Candidatus-Poribacteria-bacterium] |
| MCP6718634.1 | DUF499_1.RM.like.systems | TUDOR+SF1-SF2-Helicase+i REase +CTD+ REase -> METHYLASE -> DUF499 *-> w HTH+ REase -><-?<-?\|\|?-><-ZnR | 947 | Bacteria>Bacteria_incertae_sedis>Bacteria_candidate_phyla>Patescibacteria_group>unclassified_Patescibacteria_group | GCA_024239015.1 | Patescibacteria_group_bacterium | KJI71_00195 | 43190..46033 | 2052139 | hypothetical-protein-[Patescibacteria-group-bacterium] |
| MDE2100040.1 | DUF499_1.RM.like.systems | HTH_26->Phage_integrase-> TUDOR+SF1-SF2-Helicase+i REase +CTD+ REase -> METHYLASE -> REase -> DUF499 *-> | 970 | Bacteria>Bacteria_incertae_sedis>Bacteria_candidate_phyla>Patescibacteria_group>unclassified_Patescibacteria_group | GCA_028872085.1 | Patescibacteria_group_bacterium | KGL39_22490 | 13393..16305 | 2052139 | MAG:-ATP-binding-protein-[Patescibacteria-group-bacterium] |
| WP_142699382.1 | DUF499_1.RM.like.systems | HTH-><-?\|\|?-> TUDOR+SF1-SF2-Helicase+i REase +CTD+ REase -> METHYLASE -> HEPN+ DUF499 *-> REase -><-ASCH | 1099 | Bacteria>Bdellovibrionota>Bdellovibrionia>Bdellovibrionales>Pseudobdellovibrionaceae | GCF_006874645.1 | Bdellovibrio_sp._ZAP7 | DOM22_RS05315 | 1082808..1086107 | 2231053 | Swt1-family-HEPN-domain-containing-protein-[Bdellovibrio-sp.-ZAP7] |
| MCR4433941.1 | DUF499_1.RM.like.systems | TUDOR+SF1-SF2-Helicase+i REase +CTD+ REase ->?-> METHYLASE -> REase -> DUF499 *-> | 924 | Bacteria>Caldisericota/Cryosericota_group>Caldisericota>unclassified_Caldisericota | GCA_024655865.1 | Caldisericota_bacterium | NUV70_07825 | 12514..15288 | 2052147 | DUF499-domain-containing-protein-[Caldisericota-bacterium] |
| OGP05895.1 | DUF499_1.RM.like.systems | ParA-Soj-PloopNTPase->Peptidase-> TUDOR+SF1-SF2-Helicase+i REase +CTD+ REase -> w HTH+ REase -> METHYLASE -> DUF499 *-> | 918 | Bacteria>delta/epsilon_subdivisions>Deltaproteobacteria>unclassified_Deltaproteobacteria | GCA_001795495.1 | Deltaproteobacteria_bacterium_GWA2_42_85 | A2056_04335 | 11127..13883 | 1797803 | AAA-family-ATPase-[Deltaproteobacteria-bacterium-GWA2_42_85] |
| PKN69055.1 | DUF499_1.RM.like.systems | <-HKD-DNase<-?<-?<-?<-?<-?<- DUF499 *<- METHYLASE <- TUDOR+SF1-SF2-Helicase+i REase +CTD+ REase <-HEPN<- REase | 913 | Bacteria>delta/epsilon_subdivisions>Deltaproteobacteria>unclassified_Deltaproteobacteria | GCA_002840565.1 | Deltaproteobacteria_bacterium_HGW-Deltaproteobacteria-12 | CVU54_12145 | 60436..63177 | 2013739 | hypothetical-protein-CVU54_12145-[Deltaproteobacteria-bacterium-HGW-Deltaproteobacteria-12] |
| CDD51558.1 | DUF499_1.RM.like.systems | <-XIS-HTH<-?<-Phage_integrase\|\|TIR->TIR-><-?<- HEPN+ DUF499 *<- METHYLASE <- REase +ParB<- TUDOR+SF1-SF2-Helicase+i REase +CTD+ REase <-HTH_3<-DnaJ | 1098 | Bacteria>FCB_group>Bacteroidota/Chlorobiota_group>Bacteroidota>Bacteroidia | GCA_000435415.1 | Bacteroides_sp._CAG:875 | BN800_02603 | 61429..64725 | 1262752 | putative-uncharacterized-protein-[Bacteroides-sp.-CAG:875] |
| CDD33920.1 | DUF499_1.RM.like.systems | <-Peptidase_M3<-?\|\|?-><-?<- HEPN+ DUF499 *<- METHYLASE <- REase +ParB<- TUDOR+SF1-SF2-Helicase+i REase +CTD+ REase <-Radical-SAM | 1098 | Bacteria>FCB_group>Bacteroidota/Chlorobiota_group>Bacteroidota>Bacteroidia | GCA_000436055.1 | Bacteroides_sp._CAG:714 | BN762_01576 | 11019..14315 | 1262749 | putative-uncharacterized-protein-[Bacteroides-sp.-CAG:714] |
| NCB25654.1 | DUF499_1.RM.like.systems | TUDOR+SF1-SF2-Helicase+i REase +CTD+ REase -> METHYLASE -> HEPN+ DUF499 *-> REase ->Pkinase-> | 1134 | Bacteria>FCB_group>Bacteroidota/Chlorobiota_group>Bacteroidota>Bacteroidia | GCA_009929165.1 | Bacteroidia_bacterium | EOM62_09315 | 4747..8151 | 2044936 | ATP-binding-protein-[Bacteroidia-bacterium] |
| MBN1340543.1 | DUF499_1.RM.like.systems | TUDOR+SF1-SF2-Helicase+i REase +CTD+ REase ->BirA-HTH-> METHYLASE -> w HTH+ REase -> DUF499 *-><-?<-tRNA\|\|sGTP->sGTP-> | 974 | Bacteria>FCB_group>Bacteroidota/Chlorobiota_group>Bacteroidota>Bacteroidia | GCA_016927695.1 | Bacteroidales_bacterium | JXA03_14535 | 29128..32052 | 2030927 | ATP-binding-protein-[Bacteroidales-bacterium] |
| WP_018709397.1 | DUF499_1.RM.like.systems | REase +SF1-SF2-Helicase ->?->Radical-SAM-> TUDOR+SF1-SF2-Helicase+i REase +CTD+ REase -> REase +ParB-> METHYLASE -> HEPN+ DUF499 *->?-><-?\|\|?->Peptidase_M3-> | 1098 | Bacteria>FCB_group>Bacteroidota/Chlorobiota_group>Bacteroidota>Bacteroidia | GCF_000374585.1 | Phocaeicola_barnesiae | C510_RS0100610 | 147306..150602 | 376804 | DUF499-domain-containing-protein-[Phocaeicola-barnesiae] |
| WP_087396459.1 | DUF499_1.RM.like.systems | REase ->?-><-?<-DOC+HTH<- HEPN+ DUF499 *<- METHYLASE <- REase +ParB<- TUDOR+SF1-SF2-Helicase+i REase +CTD+ REase <-DOC<-HTH_3<-FAD-NAD-dep-oxidoreductase | 1097 | Bacteria>FCB_group>Bacteroidota/Chlorobiota_group>Bacteroidota>Bacteroidia | GCF_002161565.1 | Bacteroides_sp._An51A | B5G04_RS06890 | 141402..144695 | 1965640 | DUF499-domain-containing-protein-[Bacteroides-sp.-An51A] |
| WP_278365558.1 | DUF499_1.RM.like.systems | <-Peptidase_M3<-?\|\|?-><-?<- HEPN+ DUF499 *<- METHYLASE <- REase +ParB<- TUDOR+SF1-SF2-Helicase+i REase +CTD+ REase <-Radical-SAM<-?<-SF1-SF2-Helicase | 1098 | Bacteria>FCB_group>Bacteroidota/Chlorobiota_group>Bacteroidota>Bacteroidia | GCF_002359135.1 | Phocaeicola_barnesiae | B3R84_RS06870 | 398812..402108 | 376804 | DUF499-domain-containing-protein-[Phocaeicola-barnesiae] |
| WP_117747735.1 | DUF499_1.RM.like.systems | <- HEPN+ DUF499 *<- METHYLASE <- REase +ParB<- TUDOR+SF1-SF2-Helicase+i REase +CTD+ REase <-HTH_3<-DnaJ | 1098 | Bacteria>FCB_group>Bacteroidota/Chlorobiota_group>Bacteroidota>Bacteroidia | GCF_003438135.1 | Phocaeicola_plebeius | DXC17_RS06660 | 1179..4475 | 310297 | DUF499-domain-containing-protein-[Phocaeicola-plebeius] |
| WP_195204977.1 | DUF499_1.RM.like.systems | DnaJ->HTH_3-> TUDOR+SF1-SF2-Helicase+i REase +CTD+ REase -> REase +ParB-> METHYLASE -p-> METHYLASE -> HEPN+ DUF499 *-> | 1098 | Bacteria>FCB_group>Bacteroidota/Chlorobiota_group>Bacteroidota>Bacteroidia | GCF_015546785.1 | Phocaeicola_plebeius | I1Q75_RS16665 | 24617..27913 | 310297 | DUF499-domain-containing-protein-[Phocaeicola-plebeius] |
| WP_235301263.1 | DUF499_1.RM.like.systems | TUDOR+SF1-SF2-Helicase+i REase +CTD+ REase -> REase +ParB-> METHYLASE -> HEPN+ DUF499 *->?-><-?\|\|?->Peptidase_M3-> | 1098 | Bacteria>FCB_group>Bacteroidota/Chlorobiota_group>Bacteroidota>Bacteroidia | GCF_021531035.1 | Phocaeicola_barnesiae | I6E10_RS09100 | 8385..11681 | 376804 | DUF499-domain-containing-protein-[Phocaeicola-barnesiae] |
| WP_118026126.1 | DUF499_1.RM.like.systems | DnaJ->HTH_3-> TUDOR+SF1-SF2-Helicase+i REase +CTD+ REase -> REase +ParB-> METHYLASE -> HEPN+ DUF499 *-><-tRNA<-tRNA<-?<-?<-PRTase<-PRTase<-?<-PrmC_N+Methyltransf_25 | 1098 | Bacteria>FCB_group>Bacteroidota/Chlorobiota_group>Bacteroidota>Bacteroidia | GCF_027686295.1 | Phocaeicola_plebeius | PGZ43_RS01830 | 134398..137694 | 310297 | DUF499-domain-containing-protein-[Phocaeicola-plebeius] |
| WP_118432131.1 | DUF499_1.RM.like.systems | PrmC_N+Methyltransf_25->?->PRTase->PRTase->?->?->tRNA->tRNA-><- HEPN+ DUF499 *<- METHYLASE <- REase +ParB<- TUDOR+SF1-SF2-Helicase+i REase +CTD+ REase <-HTH_3<-DnaJ | 1098 | Bacteria>FCB_group>Bacteroidota/Chlorobiota_group>Bacteroidota>Bacteroidia | GCF_958447975.1 | Phocaeicola_plebeius | PGY45_RS13550 | 13557..16853 | 310297 | DUF499-domain-containing-protein-[Phocaeicola-plebeius] |
| WP_118338055.1 | DUF499_1.RM.like.systems | <- HEPN+ DUF499 *<- METHYLASE <- REase +ParB<- TUDOR+SF1-SF2-Helicase+i REase +CTD+ REase <-HTH_3<-DnaJ | 1098 | Bacteria>FCB_group>Bacteroidota/Chlorobiota_group>Bacteroidota>Bacteroidia | GCF_959025615.1 | Phocaeicola_plebeius | PG845_RS15420 | 1179..4475 | 310297 | DUF499-domain-containing-protein-[Phocaeicola-plebeius] |
| WP_284525582.1 | DUF499_1.RM.like.systems | <- DUF499 -p<- DUF499 -p<- HEPN+ DUF499 *<- METHYLASE -p<- METHYLASE <- REase +ParB<-?<- TUDOR+SF1-SF2-Helicase+i REase +CTD+ REase <-HTH_3 | 518 | Bacteria>FCB_group>Bacteroidota/Chlorobiota_group>Bacteroidota>Bacteroidia | - | Phocaeicola_plebeius | DW982_RS17975 | 3045..4601 | 310297 | DUF499-domain-containing-protein-[Phocaeicola-plebeius] |
| MBK9661552.1 | DUF499_1.RM.like.systems | <-Integrase_1+Phage_integrase\|\|?->DUF488-1-><-?<- DUF499 *<- w HTH+ REase <- METHYLASE <-HEPN<- TUDOR+SF1-SF2-Helicase+i REase +CTD+ REase <-HTH+WYL\|\|RnaseHII-> | 943 | Bacteria>FCB_group>Bacteroidota/Chlorobiota_group>Bacteroidota>Chitinophagia | GCA_016717645.1 | Chitinophagaceae_bacterium | IPO68_16915 | 5414..8245 | 1869212 | ATP-binding-protein-[Chitinophagaceae-bacterium] |
| WP_027303393.1 | DUF499_1.RM.like.systems | MNS+ REase ->CHTH->?->?-><-?\|\|MORC+HisKin+ REase -><- HEPN+ DUF499 *<- w HTH+ REase <- METHYLASE \|\|?-><- TUDOR+SF1-SF2-Helicase+i REase +CTD+ REase <-PAIREDC-HTH\|\|Phage_integrase-> | 1100 | Bacteria>FCB_group>Bacteroidota/Chlorobiota_group>Bacteroidota>Cytophagia | GCF_000383955.1 | Rudanella_lutea | RUDLU_RS0125935 | 41364..44666 | 451374 | DUF499-domain-containing-protein-[Rudanella-lutea] |
| WP_114065477.1 | DUF499_1.RM.like.systems | <-TUDOR+SF1-SF2-Helicase+i REase +CTD<- HEPN+ DUF499 *<-?<-OLD-ABC<-ParB+HNH<- METHYLASE <- REase <- TUDOR+SF1-SF2-Helicase+i REase +CTD+ REase | 1094 | Bacteria>FCB_group>Bacteroidota/Chlorobiota_group>Bacteroidota>Cytophagia | GCF_003325355.1 | Runella_rosea | DR864_RS02565 | 579082..582366 | 2259595 | DUF499-domain-containing-protein-[Runella-rosea] |
| WP_130024109.1 | DUF499_1.RM.like.systems | <-HTH<-XIS-HTH<-?<- HEPN+ DUF499 *<- METHYLASE <-AbiEii<-wHTH+AbiEi<- REase <- TUDOR+SF1-SF2-Helicase+i REase +CTD+ REase | 1096 | Bacteria>FCB_group>Bacteroidota/Chlorobiota_group>Bacteroidota>Cytophagia | GCF_004168285.1 | Emticicia_agri | EWM59_RS25780 | 4303..7593 | 2492393 | DUF499-domain-containing-protein-[Emticicia-agri] |
| WP_209329595.1 | DUF499_1.RM.like.systems | <-BirA-HTH<-?<-?<- HEPN+ DUF499 *<-AAA-ATPase+ REase <- METHYLASE <-ParB+HNH<- TUDOR+SF1-SF2-Helicase+i REase +CTD+ REase | 1097 | Bacteria>FCB_group>Bacteroidota/Chlorobiota_group>Bacteroidota>Cytophagia | GCF_017811155.1 | Lunatimonas_salinarum | IAQ59_RS04505 | 160980..164273 | 1774590 | DUF499-domain-containing-protein-[Lunatimonas-salinarum] |
| NUQ24578.1 | DUF499_1.RM.like.systems | TUDOR+SF1-SF2-Helicase+i REase +CTD+ REase ->OLD-ABC->HNH-> METHYLASE ->AAA-ATPase+ REase -> HEPN+ DUF499 *->?->PolB->HEPN-><-ORC-CDC6+ REase | 1095 | Bacteria>FCB_group>Bacteroidota/Chlorobiota_group>Bacteroidota>Saprospiria | GCA_013360405.1 | Saprospiraceae_bacterium | HUU34_11550 | 44219..47506 | 2202734 | DUF499-domain-containing-protein-[Saprospiraceae-bacterium] |
| MCY3594931.1 | DUF499_1.RM.like.systems | Peptidase_S41-><-?<- DUF499 *<- REase <-PIN<-OLD-ABC<- METHYLASE <- TUDOR+SF1-SF2-Helicase+i REase +CTD+ REase | 913 | Bacteria>FCB_group>Bacteroidota/Chlorobiota_group>Bacteroidota>unclassified_Bacteroidota | GCA_026705205.1 | Bacteroidota_bacterium | OXH01_06810 | 22578..25319 | 1898104 | MAG:-DUF499-domain-containing-protein-[Bacteroidetes-bacterium] |
| MCY3630037.1 | DUF499_1.RM.like.systems | <- DUF499 *<-PIN<-OLD-ABC<- METHYLASE <- w HTH+ REase <- TUDOR+SF1-SF2-Helicase+i REase +CTD+ REase | 884 | Bacteria>FCB_group>Bacteroidota/Chlorobiota_group>Bacteroidota>unclassified_Bacteroidota | GCA_026705375.1 | Bacteroidota_bacterium | OXG94_08505 | 1..2653 | 1898104 | MAG:-DUF499-domain-containing-protein,-partial-[Bacteroidetes-bacterium] |
| MCY4170340.1 | DUF499_1.RM.like.systems | OLD-ABC-><- DUF499 *<- REase <-?<- METHYLASE <- TUDOR+SF1-SF2-Helicase+i REase +CTD+ REase | 912 | Bacteria>FCB_group>Bacteroidota/Chlorobiota_group>Bacteroidota>unclassified_Bacteroidota | GCA_026709065.1 | Bacteroidota_bacterium | OXF08_01935 | 1316..4054 | 1898104 | MAG:-DUF499-domain-containing-protein-[Bacteroidetes-bacterium] |
| MDE2646784.1 | DUF499_1.RM.like.systems | ABhydrolase-><- DUF499 *<-PIN<-OLD-ABC<- METHYLASE <- METHYLASE -p<- w HTH+ REase <- TUDOR+SF1-SF2-Helicase+i REase +CTD+ REase | 924 | Bacteria>FCB_group>Bacteroidota/Chlorobiota_group>Bacteroidota>unclassified_Bacteroidota | GCA_028875465.1 | Bacteroidota_bacterium | OXI05_13240 | 2968..5742 | 1898104 | MAG:-DUF499-domain-containing-protein-[Bacteroidota-bacterium] |
| MBL6956529.1 | DUF499_1.RM.like.systems | <-HTH+MPTase<-Tox-RelE<-?<- HEPN+ DUF499 *<- METHYLASE <- w HTH+ REase <- TUDOR+SF1-SF2-Helicase+i REase +CTD+ REase | 1115 | Bacteria>FCB_group>Bacteroidota/Chlorobiota_group>Chlorobiota>Chlorobiia | GCA_016776555.1 | Chlorobium_phaeobacteroides | ISR54_06890 | 24108..27455 | 1096 | DUF499-domain-containing-protein-[Chlorobium-phaeobacteroides] |
| WP_059138717.1 | DUF499_1.RM.like.systems | RelE-ParE-> TUDOR+SF1-SF2-Helicase+i REase +CTD+ REase -> w HTH+ REase -> METHYLASE -> HEPN+ DUF499 *->?->?->?->TraJ-RHH-> | 1115 | Bacteria>FCB_group>Bacteroidota/Chlorobiota_group>Chlorobiota>Chlorobiia | GCF_001509575.1 | Chlorobium_limicola | ASB62_RS03910 | 18289..21636 | 1092 | Swt1-family-HEPN-domain-containing-protein-[Chlorobium-limicola] |
| KXB97080.1 | DUF499_1.RM.like.systems | TUDOR+SF1-SF2-Helicase+i REase +CTD+ REase -> REase ->?-> METHYLASE -> DUF499 *-> | 876 | Bacteria>FCB_group>Bacteroidota/Chlorobiota_group>Chlorobiota>unclassified_Chlorobiota | GCA_001302225.1 | Chlorobi_bacterium_NICIL-2 | AA908_08510 | 7880..10510 | 1662223 | hypothetical-protein-AA908_08510-[Chlorobi-bacterium-NICIL-2] |
| MCX6133013.1 | DUF499_1.RM.like.systems | <-DUF932<-?\|\| TUDOR+SF1-SF2-Helicase+i REase +CTD+ REase -> METHYLASE -> REase -> DUF499 *->?->DUF488-1-> | 972 | Bacteria>FCB_group>Bacteroidota/Chlorobiota_group>Ignavibacteriota>Ignavibacteria | GCA_026390595.1 | Ignavibacteriales_bacterium | NTU47_04275 | 82490..85408 | 2049428 | MAG:-DUF499-domain-containing-protein-[Ignavibacteriales-bacterium] |
| MXW15222.1 | DUF499_1.RM.like.systems | TUDOR+SF1-SF2-Helicase+i REase +CTD+ REase -> w HTH+ REase ->OLD-ABC->PIN-> DUF499 *-><-ABhydrolase | 924 | Bacteria>FCB_group>Bacteroidota/Chlorobiota_group>Rhodothermota>Rhodothermia | GCA_009835915.1 | Rhodothermaceae_bacterium | F4Z76_09955 | 11969..14743 | 2026787 | ATP-binding-protein-[Rhodothermaceae-bacterium] |
| MXZ58767.1 | DUF499_1.RM.like.systems | ABhydrolase-><- DUF499 *<-PIN<-OLD-ABC<- METHYLASE <- w HTH+ REase <- TUDOR+SF1-SF2-Helicase+i REase +CTD+ REase | 924 | Bacteria>FCB_group>Bacteroidota/Chlorobiota_group>Rhodothermota>Rhodothermia | GCA_009837125.1 | Rhodothermaceae_bacterium | F4Z16_09865 | 2884..5658 | 2026787 | ATP-binding-protein-[Rhodothermaceae-bacterium] |
| MXX97465.1 | DUF499_1.RM.like.systems | TUDOR+SF1-SF2-Helicase+i REase +CTD+ REase -> w HTH+ REase -> METHYLASE ->OLD-ABC->PIN-> DUF499 *-><-ABhydrolase | 907 | Bacteria>FCB_group>Bacteroidota/Chlorobiota_group>Rhodothermota>Rhodothermia | GCA_009838195.1 | Rhodothermaceae_bacterium | F4Y64_07620 | 10395..13118 | 2026787 | ATP-binding-protein-[Rhodothermaceae-bacterium] |
| MYC03829.1 | DUF499_1.RM.like.systems | ABhydrolase-><- DUF499 *<-PIN<-OLD-ABC<- w HTH+ REase <- TUDOR+SF1-SF2-Helicase+i REase +CTD+ REase | 907 | Bacteria>FCB_group>Bacteroidota/Chlorobiota_group>Rhodothermota>Rhodothermia | GCA_009840265.1 | Rhodothermaceae_bacterium | F4X61_04285 | 3864..6587 | 2026787 | ATP-binding-protein-[Rhodothermaceae-bacterium] |
| MYH11281.1 | DUF499_1.RM.like.systems | TUDOR+SF1-SF2-Helicase+i REase +CTD+ REase -> w HTH+ REase -> METHYLASE ->OLD-ABC->PIN-> DUF499 *-><-ABhydrolase | 924 | Bacteria>FCB_group>Bacteroidota/Chlorobiota_group>Rhodothermota>Rhodothermia | GCA_009842805.1 | Rhodothermaceae_bacterium | F4191_00350 | 10395..13169 | 2026787 | ATP-binding-protein-[Rhodothermaceae-bacterium] |
| MCC6694832.1 | DUF499_1.RM.like.systems | TUDOR+SF1-SF2-Helicase+i REase +CTD+ REase -> REase ->HEPN-> METHYLASE -p->?-> METHYLASE -> HEPN+ DUF499 *->?->?->ArdCN+MPTase-><-Arm-DNA-bind_3+Phage_integrase | 1102 | Bacteria>FCB_group>Candidatus_Hydrogenedentes>unclassified_Candidatus_Hydrogenedentes | GCA_020849535.1 | Candidatus_Hydrogenedentes_bacterium | IT365_04280 | 13299..16607 | 2030809 | DUF499-domain-containing-protein-[Candidatus-Hydrogenedentes-bacterium] |
| MBN1601227.1 | DUF499_1.RM.like.systems | <-PIN<-OLD-ABC<-?<-?<- DUF499 *<- w HTH+ REase <-?<- METHYLASE <- TUDOR+SF1-SF2-Helicase+i REase +CTD+ REase <- REase | 922 | Bacteria>FCB_group>Fibrobacterota>Chitinispirillia>Chitinispirillales | GCA_016929595.1 | Chitinispirillaceae_bacterium | JW915_06440 | 4163..6931 | 2813024 | ATP-binding-protein-[Chitinispirillaceae-bacterium] |
| MYC87147.1 | DUF499_1.RM.like.systems | TUDOR+SF1-SF2-Helicase+i REase +CTD+ REase ->OLD-ABC-p->OLD-ABC-p-> REase -> DUF499 *->?-><-?<-?<-SF1-SF2-Helicase<-HKD-DNase<-DUF6361 | 922 | Bacteria>FCB_group>Gemmatimonadota>Gemmatimonadetes>Candidatus_Palauibacterales | GCA_009841015.1 | Candidatus_Palauibacter_denitrificans | F4X22_02775 | 10288..13056 | 3056734 | ATP-binding-protein-[Gemmatimonadales-bacterium] |
| MBV6522081.1 | DUF499_1.RM.like.systems | TUDOR+SF1-SF2-Helicase+i REase +CTD+ REase ->ORC-CDC6+ REase -> METHYLASE ->ORC-CDC6+ REase -> HEPN+ DUF499 *-><-Pkinase | 1116 | Bacteria>FCB_group>Gemmatimonadota>Gemmatimonadetes>Gemmatimonadales | GCA_019186945.1 | Gemmatimonadaceae_bacterium | MNPFHGCM_02226 | 129597..132947 | 2026741 | hypothetical-protein-[Gemmatimonadaceae-bacterium] |
| MDE2973202.1 | DUF499_1.RM.like.systems | TUDOR+SF1-SF2-Helicase+i REase +CTD+ REase -> w HTH+ REase -> METHYLASE -> DUF499 *-> | 933 | Bacteria>FCB_group>Gemmatimonadota>unclassified_Gemmatimonadota | GCA_028873825.1 | Gemmatimonadota_bacterium | OXU64_00560 | 8847..11648 | 2026742 | MAG:-DUF499-domain-containing-protein-[Gemmatimonadota-bacterium] |
| MDE2733147.1 | DUF499_1.RM.like.systems | <- TUDOR+SF1-SF2-Helicase+i REase +CTD+ REase <- HEPN+ DUF499 *\|\| REase ->?-><-?<- METHYLASE <- REase <-OLD-ABC | 1112 | Bacteria>FCB_group>Gemmatimonadota>unclassified_Gemmatimonadota | GCA_028875335.1 | Gemmatimonadota_bacterium | OXI72_02050 | 10955..14293 | 2026742 | MAG:-Swt1-family-HEPN-domain-containing-protein-[Gemmatimonadota-bacterium] |
| WP_012098801.1 | DUF499_1.RM.like.systems | <-Phage_integrase<-XIS-HTH<-?<-wHTH+SAD+HNH<-HKD-DNase+SF1-SF2-Helicase+TRD<- DUF499 *<- REase <- METHYLASE <- METHYLASE <- TUDOR+SF1-SF2-Helicase+i REase +CTD+ REase \|\|DUF6361->HKD-DNase-> | 932 | Bacteria>Myxococcota>Myxococcia>Myxococcales>Cystobacterineae | GCF_000017505.1 | Anaeromyxobacter_sp._Fw109-5 | ANAE109_RS20460 | 4648491..4651289 | 404589 | DUF499-domain-containing-protein-[Anaeromyxobacter-sp.-Fw109-5] |
| WP_121761665.1 | DUF499_1.RM.like.systems | TUDOR+SF1-SF2-Helicase+i REase +CTD+ REase -> METHYLASE -> DUF499 *-> REase -> | 947 | Bacteria>Myxococcota>Myxococcia>Myxococcales>Cystobacterineae | GCF_003668955.1 | Corallococcus_sp._AB038B | D7Y04_RS31655 | 364596..367439 | 2316718 | DUF499-domain-containing-protein-[Corallococcus-sp.-AB038B] |
| WP_174489000.1 | DUF499_1.RM.like.systems | <- METHYLASE +DNA_methylase\|\|tRNA-><-DNA-glycosylase<-?<-Pkinase<- REase <- DUF499 *<- METHYLASE <-HNH<-OLD-ABC<- TUDOR+SF1-SF2-Helicase+i REase +CTD+ REase <-XIS-HTH<-PAIREDC-HTH | 943 | Bacteria>Myxococcota>Myxococcia>Myxococcales>Cystobacterineae | GCF_013336725.1 | Myxococcus_sp._CA040A | HUA75_RS36310 | 164891..167722 | 2741738 | DUF499-domain-containing-protein-[Myxococcus-sp.-CA040A] |
| MBI3184665.1 | DUF499_1.RM.like.systems | <-HAD<-?\|\|XIS-HTH-> TUDOR+SF1-SF2-Helicase+i REase +CTD+ REase -> METHYLASE -> REase -> DUF499 *->?->Radical-SAM->ParB->?->XIS-HTH-><- REase | 930 | Bacteria>Myxococcota>Myxococcia>Myxococcales>unclassified_Myxococcales | GCA_016191815.1 | Myxococcales_bacterium | HYZ28_21220 | 50523..53315 | 2026763 | ATP-binding-protein-[Myxococcales-bacterium] |
| MBK7583793.1 | DUF499_1.RM.like.systems | <-HTH\|\|?-> REase -> TUDOR+SF1-SF2-Helicase+i REase +CTD+ REase -> METHYLASE -> HEPN+ DUF499 *-> REase -> | 1105 | Bacteria>Myxococcota>Myxococcia>Myxococcales>unclassified_Myxococcales | GCA_016706225.1 | Myxococcales_bacterium | IPI67_26815 | 461421..464738 | 2026763 | DUF499-domain-containing-protein-[Myxococcales-bacterium] |
| MBL9006089.1 | DUF499_1.RM.like.systems | XIS-HTH-> TUDOR+SF1-SF2-Helicase+i REase +CTD+ REase ->OLD-ABC-> METHYLASE -> METHYLASE -> REase -> DUF499 *-> | 928 | Bacteria>Myxococcota>Myxococcia>Myxococcales>unclassified_Myxococcales | GCA_016794505.1 | Myxococcales_bacterium | JNJ46_17680 | 14814..17600 | 2026763 | ATP-binding-protein-[Myxococcales-bacterium] |
| HBA39938.1 | DUF499_1.RM.like.systems | TUDOR+SF1-SF2-Helicase+i REase +CTD+ REase ->AAA-ATPase+ REase -> REase -p-> METHYLASE -> REase -> DUF499 *-> | 1099 | Bacteria>Myxococcota>Myxococcia>unclassified_Myxococcia | GCA_003490035.1 | Deltaproteobacteria_bacterium | DCZ05_09410 | 9752..13051 | 2026735 | hypothetical-protein-[Deltaproteobacteria-bacterium] |
| MBC8466666.1 | DUF499_1.RM.like.systems | <- DUF499 *<- REase <- METHYLASE <-?<-DrHyd+BirA-HTH<- TUDOR+SF1-SF2-Helicase+i REase +CTD+ REase | 918 | Bacteria>Myxococcota>Myxococcia>unclassified_Myxococcia | GCA_014384065.1 | Deltaproteobacteria_bacterium | H8D55_02390 | 3412..6168 | 2026735 | ATP-binding-protein-[Deltaproteobacteria-bacterium] |
| MBI4514358.1 | DUF499_1.RM.like.systems | <-sGTP<-?<-BirA-HTH<-?<-?\|\|tRNA-><- DUF499 *<- REase <- METHYLASE <-AAA-ATPase+ REase <- TUDOR+SF1-SF2-Helicase+i REase +CTD+ REase \|\|?->\|\|tRNA->NTP_transferase+MannoseP_isomer-> | 1100 | Bacteria>Myxococcota>Myxococcia>unclassified_Myxococcia | GCA_016210005.1 | Deltaproteobacteria_bacterium | HY699_00880 | 115685..118987 | 2026735 | ATP-binding-protein-[Deltaproteobacteria-bacterium] |
| MBM4330413.1 | DUF499_1.RM.like.systems | TUDOR+SF1-SF2-Helicase+i REase +CTD+ REase -> w HTH+ REase ->?-> METHYLASE -> DUF499 *-><-?<-PIN | 1106 | Bacteria>Myxococcota>Myxococcia>unclassified_Myxococcia | GCA_016875165.1 | Deltaproteobacteria_bacterium | FJ117_04155 | 62692..66012 | 2026735 | ATP-binding-protein-[Deltaproteobacteria-bacterium] |
| MBW1705706.1 | DUF499_1.RM.like.systems | <-SMS-KaiC<-RelE-ParE<-MetJarc-HTH\|\| TUDOR+SF1-SF2-Helicase+i REase +CTD+ REase -> w HTH+ REase -> METHYLASE -> DUF499 *-> | 670 | Bacteria>Myxococcota>Myxococcia>unclassified_Myxococcia | GCA_019306165.1 | Deltaproteobacteria_bacterium | JRJ86_11145 | 11472..13481 | 2026735 | ATP-binding-protein-[Deltaproteobacteria-bacterium] |
| MBW2343658.1 | DUF499_1.RM.like.systems | <-RelE-ParE<-MetJarc-HTH\|\|?-> TUDOR+SF1-SF2-Helicase+i REase +CTD+ REase -> w HTH+ REase -> METHYLASE -> DUF499 *-> | 1106 | Bacteria>Myxococcota>Myxococcia>unclassified_Myxococcia | GCA_019310785.1 | Deltaproteobacteria_bacterium | JRF53_06530 | 24941..28261 | 2026735 | DUF499-domain-containing-protein-[Deltaproteobacteria-bacterium] |
| KYF87240.1 | DUF499_1.RM.like.systems | <- DUF499 *<- REase <-OLD-ABC <- METHYLASE <-HNH<-OLD-ABC<- TUDOR+SF1-SF2-Helicase+i REase +CTD+ REase | 954 | Bacteria>Myxococcota>Polyangia>Polyangiales>Polyangiaceae | GCA_001589185.1 | Sorangium_cellulosum | BE17_46125 | 4063..6927 | 56 | hypothetical-protein-BE17_46125-[Sorangium-cellulosum] |
| WP_050434136.1 | DUF499_1.RM.like.systems | <-SF1-SF2-Helicase<-PAIREDC-HTH<-DNA-glycosylase<- DUF499 *<- REase <- METHYLASE <-BirA-HTH<- METHYLASE <-OLD-ABC<- TUDOR+SF1-SF2-Helicase+i REase +CTD+ REase | 936 | Bacteria>Myxococcota>Polyangia>Polyangiales>Polyangiaceae | GCF_001189295.1 | Chondromyces_crocatus | CMC5_RS33060 | 9153103..9155913 | 52 | DUF499-domain-containing-protein-[Chondromyces-crocatus] |
| MBU0551512.1 | DUF499_1.RM.like.systems | TUDOR+SF1-SF2-Helicase+i REase +CTD+ REase -p->HTH+MPTase->PIN-> METHYLASE -> OLD-ABC-> REase -> DUF499 *-><-?<-?<-RecR+TOPRIM | 942 | Bacteria>Myxococcota>unclassified_Myxococcota | GCA_018813065.1 | Myxococcota_bacterium | KJ940_08450 | 6900..9728 | 2818507 | DUF499-domain-containing-protein-[Myxococcota-bacterium] |
| MBS0151038.1 | DUF499_1.RM.like.systems | TUDOR+SF1-SF2-Helicase+i REase +CTD+ REase -> REase -> METHYLASE ->?->DUF488-1->?-> HEPN+ DUF499 *->?-><-?\|\|?-><-?<-SMODS | 1121 | Bacteria>Nitrospirota>Nitrospiria>Nitrospirales>Nitrospiraceae | GCA_018242725.1 | Nitrospira_sp. | JSR31_08860 | 34905..38270 | 70125 | DUF499-domain-containing-protein-[Nitrospira-sp.] |
| MCP9446717.1 | DUF499_1.RM.like.systems | <- REase <-?<-?<-?<-N6_N4_MTase<-?<- DUF499 *<- REase <- METHYLASE <-Schlafen+HATPase+HTH<- TUDOR+SF1-SF2-Helicase+i REase +CTD+ REase | 941 | Bacteria>Nitrospirota>Nitrospiria>Nitrospirales>Nitrospiraceae | GCA_024330945.1 | Nitrospira_sp. | NNA22_03990 | 9048..11873 | 70125 | DUF499-domain-containing-protein-[Nitrospira-sp.] |
| MDA8090379.1 | DUF499_1.RM.like.systems | ACET-><-?<-?<- REase <- HEPN+ DUF499 *<- METHYLASE <-TIR+PNTD2<- TUDOR+SF1-SF2-Helicase+i REase +CTD+ REase <-?\|\|Tox-RelE->HTH_3-> | 1107 | Bacteria>Nitrospirota>Nitrospiria>Nitrospirales>Nitrospiraceae | GCA_028064195.1 | Nitrospiraceae_bacterium | MOZ61_09200 | 18029..21352 | 2026770 | MAG:-Swt1-family-HEPN-domain-containing-protein-[Nitrospiraceae-bacterium] |
| WP_059175576.1 | DUF499_1.RM.like.systems | REase -> TUDOR+SF1-SF2-Helicase+i REase +CTD+ REase -> METHYLASE ->PolB->HEPN-> METHYLASE -p-> DUF499 *-><-?<-?\|\|SAM-methylase-> | 842 | Bacteria>Nitrospirota>Thermodesulfovibrionia>Thermodesulfovibrionales>Thermodesulfovibrionaceae | GCF_001514535.1 | Thermodesulfovibrio_aggregans | TAGGR_RS01355 | 271390..273918 | 86166 | DUF499-domain-containing-protein-[Thermodesulfovibrio-aggregans] |

| Protein-ID | System | Gene-Neighbourhoods | Length | Taxonomy | Assembly-ID | Strain | Locus-Tag | Genomic-coordinates | TaxID | Fasta-define |
|---|---|---|---|---|---|---|---|---|---|---|
| MBB3889803.1 | DUF499_1.RM.like.systems | <-SF1-SF2-Helicase+REase\|\|HTH+WYL->TUDOR+SF1-SF2-Helicase+iREase+CTD+REase->REase->METHYLASE->HEPN+DUF499*->MODE-HTH+PBPII-><-ParG_RHH | 1098 | Bacteria>Pseudomonadota>Alphaproteobacteria>Caulobacterales>Caulobacteraceae | GCA_014196295.1 | Phenylobacterium_haematophilum | GGQ61_000500 | 500754..504050 | 98513 | hypothetical-protein-[Phenylobacterium-haematophilum] |
| WP_152505130.1 | DUF499_1.RM.like.systems | Arm-DNA-bind_3+Phage_integrase->CopG/RHH->ACET-><-HEPN+DUF499*<-METHYLASE<-REase<-MORC+DUF3684<-YtH<-TUDOR+SF1-SF2-Helicase+iREase+CTD+REase<-HTH+WYL | 1098 | Bacteria>Pseudomonadota>Alphaproteobacteria>Hyphomicrobiales | GCF_009363315.1 | Stappiaceae | FIV06_RS06815 | 1469673..1472969 | 2821832 | DUF499-domain-containing-protein-[Roseibium-sp.-Sym1] |
| WP_253593718.1 | DUF499_1.RM.like.systems | LZ_Tnp_IS66+zf-IS66+DDE_Tnp_IS66+DDE_Tnp_IS66_C-><-Phage_integrase\|\|HEPN+REase->TUDOR+SF1-SF2-Helicase+iREase+CTD+REase->ASCH->METHYLASE->HEPN+DUF499*->?-><-HTH_3\|\|?-><-GDResolvase+Recombinase+ZnR | 1113 | Bacteria>Pseudomonadota>Alphaproteobacteria>Hyphomicrobiales>Nitrobacteraceae | GCF_024171365.1 | Bradyrhizobium_elkanii | M2199_RS04195 | 899505..902846 | 29448 | DUF499-domain-containing-protein-[Bradyrhizobium-elkanii] |
| WP_092298105.1 | DUF499_1.RM.like.systems | ZnR->?-><-SF1-SF2-Helicase-p<-SF1-SF2-Helicase<-Exonuc_V_gamma+REase<-HEPN+DUF499*<-METHYLASE<-MORC+DUF3684<-YtH<-TUDOR+SF1-SF2-Helicase+iREase+CTD+REase<-HTH+WYL | 1094 | Bacteria>Pseudomonadota>Alphaproteobacteria>Hyphomicrobiales>Nitrobacteraceae | GCF_900115265.1 | Bradyrhizobium_sp._Ghvi | BM374_RS24865 | 13858..17142 | 1855319 | DUF499-domain-containing-protein-[Bradyrhizobium-sp.-Ghvi] |
| TPI12226.1 | DUF499_1.RM.like.systems | <-HicB\|\|HTH+WYL->TUDOR+SF1-SF2-Helicase+iREase+CTD+REase->MORC+DUF3684->REase->METHYLASE->HEPN+DUF499*->TIR+PNTD2->PIWI-><-Arm-DNA-bind_3+Phage_int_SAM_3+Phage_integrase<-?<-?<-REase | 1098 | Bacteria>Pseudomonadota>Alphaproteobacteria>Hyphomicrobiales>Phyllobacteriaceae | GCA_006439355.1 | Mesorhizobium_sp._B4-1-3 | FJW06_17660 | 56845..60141 | 2589889 | ATP-binding-protein-[Mesorhizobium-sp.-B4-1-3] |
| MBZ9864227.1 | DUF499_1.RM.like.systems | <-BirA-HTH<-BirA-HTH<-BirA-HTH\|\|?->McrC-NTD+REase-><-REase<-HEPN+DUF499*<-?<-METHYLASE<-TUDOR+SF1-SF2-Helicase+iREase+CTD+REase\|\|?->?-><-MPTase | 1112 | Bacteria>Pseudomonadota>Alphaproteobacteria>Hyphomicrobiales>Phyllobacteriaceae | GCA_020165215.1 | Mesorhizobium_sp._CA15 | LB515_02450 | 495752..499090 | 2876641 | DUF499-domain-containing-protein-[Mesorhizobium-sp.-CA15] |
| WP_023715932.1 | DUF499_1.RM.like.systems | <-ParB<-?<-REase<-HEPN+DUF499*<-PIN<-OLD-ABC<-METHYLASE<-TUDOR+SF1-SF2-Helicase+iREase+CTD+REase | 1115 | Bacteria>Pseudomonadota>Alphaproteobacteria>Hyphomicrobiales>Phyllobacteriaceae | GCF_000503075.1 | Mesorhizobium_sp._LSHC422A00 | X761_RS140575 | 209394..212741 | 1287294 | DUF499-domain-containing-protein-[Mesorhizobium-sp.-LSHC422A00] |
| WP_095831457.1 | DUF499_1.RM.like.systems | SF1-SF2-Helicase+REase->Resolvase->REase->TUDOR+SF1-SF2-Helicase+iREase+CTD+REase->METHYLASE->HEPN+DUF499*-><-XIS-HTH\|\|?->?-><-tRNA<-?\|\|?->SAM-methylase-> | 1135 | Bacteria>Pseudomonadota>Alphaproteobacteria>Hyphomicrobiales>Phyllobacteriaceae | GCF_002294955.1 | Mesorhizobium_sp._WSM3864 | CK215_RS15785 | 236448..239855 | 2029404 | DUF499-domain-containing-protein-[Mesorhizobium-sp.-WSM3864] |
| WP_127367955.1 | DUF499_1.RM.like.systems | HTH_3-><-wHTH+SAD+HNH\|\|?->TUDOR+SF1-SF2-Helicase+iREase+CTD+REase->wHTH+REase->METHYLASE->HEPN+DUF499*-> | 1118 | Bacteria>Pseudomonadota>Alphaproteobacteria>Hyphomicrobiales>Phyllobacteriaceae | GCF_003997975.1 | unclassified_Mesorhizobium | EN990_RS21670 | 38386..41742 | 325217 | MULTISPECIES:-DUF499-domain-containing-protein-[unclassified-Mesorhizobium] |
| WP_164894099.1 | DUF499_1.RM.like.systems | <-HEPN+DUF499*<-METHYLASE<-wHTH+REase<-TUDOR+SF1-SF2-Helicase+iREase+CTD+REase<-?\|\|wHTH+SAD+HNH-><-HTH_3 | 528 | Bacteria>Pseudomonadota>Alphaproteobacteria>Hyphomicrobiales>Phyllobacteriaceae | GCF_004022915.1 | Mesorhizobium_sp._M7A.F.Ca.US.006.04.2.1 | EN925_RS00840 | 1..1586 | 2496696 | Swt1-family-HEPN-domain-containing-protein,-partial-[Mesorhizobium-sp.-M7A.F.Ca.US.006.04.2.1] |
| WP_135867691.1 | DUF499_1.RM.like.systems | PAPS_reduct->HTH+MPTase->PIN-><-HEPN+DUF499*<-METHYLASE<-MORC+DUF3684<-TUDOR+SF1-SF2-Helicase+iREase+CTD+REase<-REase<-?<-MAD-NTD+HNH | 1097 | Bacteria>Pseudomonadota>Alphaproteobacteria>Hyphomicrobiales>Phyllobacteriaceae | GCF_004791755.1 | unclassified_Mesorhizobium | EN861_RS33190 | 64871..68164 | 325217 | MULTISPECIES:-DUF499-domain-containing-protein-[unclassified-Mesorhizobium] |
| WP_246674663.1 | DUF499_1.RM.like.systems | <-HicB\|\|HTH+WYL->TUDOR+SF1-SF2-Helicase+iREase+CTD+REase->MORC+DUF3684->REase->METHYLASE->HEPN+DUF499*->TIR+PNTD2->PIWI-><-Arm-DNA-bind_3+Phage_int_SAM_3+Phage_integrase<-?<-?<-REase | 1086 | Bacteria>Pseudomonadota>Alphaproteobacteria>Hyphomicrobiales>Phyllobacteriaceae | GCF_006439355.1 | Mesorhizobium_sp._B4-1-3 | FJW06_RS17660 | 56881..60141 | 2589889 | DUF499-domain-containing-protein-[Mesorhizobium-sp.-B4-1-3] |
| WP_224706639.1 | DUF499_1.RM.like.systems | <-BirA-HTH<-BirA-HTH<-BirA-HTH\|\|?->McrC-NTD+REase-><-REase<-HEPN+DUF499*<-?<-METHYLASE<-TUDOR+SF1-SF2-Helicase+iREase+CTD+REase\|\|?->?-><-MPTase | 1112 | Bacteria>Pseudomonadota>Alphaproteobacteria>Hyphomicrobiales>Phyllobacteriaceae | GCF_020165215.1 | Mesorhizobium_sp._CA15 | LB515_02450 | 495752..499090 | 2876641 | DUF499-domain-containing-protein-[Mesorhizobium-sp.-CA15] |
| WP_263490164.1 | DUF499_1.RM.like.systems | <-BirA-HTH<-BirA-HTH<-BirA-HTH\|\|?->McrC-NTD+REase-><-REase<-HEPN+DUF499*<-?<-METHYLASE<-TUDOR+SF1-SF2-Helicase+iREase+CTD+REase\|\|?->?-><-MPTase | 1095 | Bacteria>Pseudomonadota>Alphaproteobacteria>Hyphomicrobiales>Phyllobacteriaceae | - | Mesorhizobium_sp._CA15 | LB515_RS02455 | 495752..499039 | 2876641 | DUF499-domain-containing-protein-[Mesorhizobium-sp.-CA15] |
| WP_064252405.1 | DUF499_1.RM.like.systems | Terminase->?-><-HTH_3\|\|?-><-?\|\|<-HEPN+DUF499*<-METHYLASE<-TUDOR+SF1-SF2-Helicase+iREase+CTD+REase<-REase\|\|35exo-> | 1109 | Bacteria>Pseudomonadota>Alphaproteobacteria>Hyphomicrobiales>Rhizobiaceae | GCF_001889105.1 | Sinorhizobium_americanum | SAMCFNEI73_RS12475 | 2595143..2598472 | 194963 | DUF499-domain-containing-protein-[Sinorhizobium-americanum] |
| WP_149746898.1 | DUF499_1.RM.like.systems | ParG_RHH-><-MODE-HTH+PBPII<-HEPN+DUF499*<-METHYLASE<-REase<-TUDOR+SF1-SF2-Helicase+iREase+CTD+REase<-HTH+WYL\|\|SF1-SF2-helicase+RdgC+Beta-barrel+REase->XIS-HTH-> | 1098 | Bacteria>Pseudomonadota>Alphaproteobacteria>Hyphomicrobiales>Rhizobiaceae | GCF_900156005.1 | Rhizobium_sp._RU35A | F3H95_RS17305 | 253422..256718 | 1907414 | DUF499-domain-containing-protein-[Rhizobium-sp.-RU35A] |
| ABS65938.1 | DUF499_1.RM.like.systems | TUDOR+SF1-SF2-Helicase+iREase+CTD+REase->REase->METHYLASE-><-HTH_21+rve+rve_3<-HIN-HTH\|\|METHYLASE-p->HEPN+DUF499*-><-MODE-HTH+PBPII\|\|TetR-HTH-><-ParG_RHH | 1098 | Bacteria>Pseudomonadota>Alphaproteobacteria>Hyphomicrobiales>Xanthobacteraceae | GCA_000017645.1 | Xanthobacter_autotrophicus_Py2 | Xaut_0686 | 774507..777803 | 78245 | ATPase-(AAA+-superfamily)-like-protein-[Xanthobacter-autotrophicus-Py2] |
| MCZ7659369.1 | DUF499_1.RM.like.systems | DEOR-HTH->TUDOR+SF1-SF2-Helicase+iREase+CTD+REase->TUDOR+SF1-SF2-Helicase+iREase+CTD+REase-p->?->REase->METHYLASE->HEPN+DUF499*->OLD-ABC+TOPRIM->SF1-SF2-Helicase-><-REase<-?<-N6_MTase<-DEOR-HTH | 1120 | Bacteria>Pseudomonadota>Alphaproteobacteria>Hyphomicrobiales>Xanthobacteraceae | GCA_027493405.1 | Xanthobacteraceae_bacterium | M5U07_16605 | 9358..12720 | 2021379 | MAG:-Swt1-family-HEPN-domain-containing-protein-[Xanthobacteraceae-bacterium] |
| WP_011714306.1 | DUF499_1.RM.like.systems | <-HISKIN+MORC<-SbcC\|\|HTH+WYL->REase+35exo+Beta-Barrel+SF1-SF2-Helicase->TUDOR+SF1-SF2-Helicase+iREase+CTD+REase->METHYLASE->DUF499*->\|\|Y1_Tnp-><-DDE_5 | 948 | Bacteria>Pseudomonadota>Alphaproteobacteria>Magnetococcales>Magnetococcaceae | GCF_000014865.1 | Magnetococcus_marinus | MMC1_RS13905 | 3444637..3447483 | 1124597 | DUF499-domain-containing-protein-[Magnetococcus-marinus] |
| MCC5984903.1 | DUF499_1.RM.like.systems | <-HEPN+DUF499*<-METHYLASE<-REase<-TUDOR+SF1-SF2-Helicase+iREase+CTD+REase<-CopG/RHH<-VirD4-TraG+VirD4-FtsK\|\|HNH-> | 351 | Bacteria>Pseudomonadota>Alphaproteobacteria>Rhodobacterales>Paracoccaceae | GCA_020832705.1 | Paracoccaceae_bacterium | JJU42_11110 | 1..1055 | 1904441 | AAA-family-ATPase-[Paracoccaceae-bacterium] |
| WP_036708698.1 | DUF499_1.RM.like.systems | XIS-HTH->?->?->MarR-HTH+REase->?->Enterotoxin_a-><-HEPN+DUF499*<-METHYLASE<-REase<-TUDOR+SF1-SF2-Helicase+iREase+CTD+REase\|\|ParB+N6_N4_MTase->ParB+N6_N4_MTase-><-HNH | 1100 | Bacteria>Pseudomonadota>Alphaproteobacteria>Rhodobacterales>Paracoccaceae | GCF_000763845.1 | Paracoccus_sanguinis | IX56_RS07055 | 10378..13680 | 1545044 | DUF499-domain-containing-protein-[Paracoccus-sanguinis] |
| WP_139597753.1 | DUF499_1.RM.like.systems | <-RNase\|\|TUDOR+SF1-SF2-Helicase+iREase+CTD+REase->?->METHYLASE->HEPN+DUF499*->REase-><-PIN | 1100 | Bacteria>Pseudomonadota>Alphaproteobacteria>Rhodobacterales>Paracoccaceae | GCF_006239215.1 | Paracoccus_haeundaensis | FHD67_RS02835 | 188317..191619 | 225362 | DUF499-domain-containing-protein-[Paracoccus-haeundaensis] |
| WP_234869545.1 | DUF499_1.RM.like.systems | <-SF1-SF2-Helicase+DUF1998<-?\|\|?-><-?<-HEPN+DUF499*<-METHYLASE<-wHTH+REase<-TUDOR+SF1-SF2-Helicase+iREase+CTD+REase | 1100 | Bacteria>Pseudomonadota>Alphaproteobacteria>Rhodobacterales>Paracoccaceae | GCF_021491355.1 | Tabrizicola_rongguiensis | LOV05_RS07760 | 379884..383186 | 2909234 | DUF499-domain-containing-protein-[Tabrizicola-rongguiensis] |
| WP_170364042.1 | DUF499_1.RM.like.systems | Primpol->?->?->?-><-RNase<-REase<-HEPN+DUF499*<-METHYLASE<-TUDOR+SF1-SF2-Helicase+iREase+CTD+REase | 1107 | Bacteria>Pseudomonadota>Alphaproteobacteria>Rhodobacterales>Roseobacteraceae | GCF_013030285.1 | Ruegeria_arenilitoris | GOX59_RS08670 | 247066..250389 | 1173585 | DUF499-domain-containing-protein-[Ruegeria-arenilitoris] |
| WP_170518600.1 | DUF499_1.RM.like.systems | <-HEPN+DUF499*<-METHYLASE<-ASCH<-TUDOR+SF1-SF2-Helicase+iREase+CTD+REase<-HEPN<-REase | 1103 | Bacteria>Pseudomonadota>Alphaproteobacteria>Rhodobacterales>Roseobacteraceae | GCF_013032825.1 | Ruegeria_atlantica | GOW75_RS18355 | 50430..53741 | 81569 | DUF499-domain-containing-protein-[Ruegeria-atlantica] |
| WP_171727684.1 | DUF499_1.RM.like.systems | HEPN-><-REase<-HEPN+DUF499*<-METHYLASE<-?<-TUDOR+SF1-SF2-Helicase+iREase+CTD+REase | 1108 | Bacteria>Pseudomonadota>Alphaproteobacteria>Rhodobacterales>Roseobacteraceae | GCF_013110755.1 | Ruegeria_sp._HKCCD7318 | GS627_RS09025 | 215066..218392 | 2683014 | DUF499-domain-containing-protein-[Ruegeria-sp.-HKCCD7318] |
| MDE0333394.1 | DUF499_1.RM.like.systems | TUDOR+SF1-SF2-Helicase+iREase+CTD+REase->?->?->?->METHYLASE->HEPN->HEPN+DUF499*->REase-> | 1096 | Bacteria>Pseudomonadota>Alphaproteobacteria>Rhodospirillales>Rhodospirillaceae | GCA_028820945.1 | Defluviicoccus_sp. | OXI64_00425 | 9805..13095 | 2682137 | MAG:-DUF499-domain-containing-protein-[Defluviicoccus-sp.] |
| MBB3880227.1 | DUF499_1.RM.like.systems | <-SF1-SF2-Helicase+REase<-SF1-SF2-Helicase+REase\|\|HTH+WYL->TUDOR+SF1-SF2-Helicase+iREase+CTD+REase->REase->METHYLASE->HEPN+DUF499*->MODE-HTH+PBPII-><-ParG_RHH | 1098 | Bacteria>Pseudomonadota>Alphaproteobacteria>Sphingomonadales>Sphingomonadaceae | GCA_014196255.1 | Sphingomonas_pseudosanguinis | GGR48_002670 | 32767..36063 | 413712 | hypothetical-protein-[Sphingomonas-pseudosanguinis] |
| MCL6741777.1 | DUF499_1.RM.like.systems | wHTH+REase->TUDOR+SF1-SF2-Helicase+iREase+CTD+REase->METHYLASE->OLD-ABC->PIN->HEPN+DUF499*-><-?<-BetaPropeller+ABhydrolase\|\|?-><-ABhydrolase<-MODE-HTH+PBPII | 1094 | Bacteria>Pseudomonadota>Alphaproteobacteria>Sphingomonadales>Sphingomonadaceae | GCA_023516505.1 | Sphingomonas_brevis | LZ518_11630 | 2249137..2252421 | 2908206 | DUF499-domain-containing-protein-[Sphingomonas-sp.-RB56-2] |
| WP_221211510.1 | DUF499_1.RM.like.systems | <-SF1-SF2-Helicase+REase<-SF1-SF2-Helicase+REase\|\|HTH+WYL->TUDOR+SF1-SF2-Helicase+iREase+CTD+REase->REase->METHYLASE->HEPN+DUF499*->MODE-HTH+PBPII-><-ParG_RHH | 1086 | Bacteria>Pseudomonadota>Alphaproteobacteria>Sphingomonadales>Sphingomonadaceae | GCF_014196255.1 | Sphingomonas_pseudosanguinis | GGR48_RS13090 | 32803..36063 | 413712 | DUF499-domain-containing-protein-[Sphingomonas-pseudosanguinis] |
| WP_283938333.1 | DUF499_1.RM.like.systems | wHTH+REase->TUDOR+SF1-SF2-Helicase+iREase+CTD+REase->METHYLASE->OLD-ABC->PIN->HEPN+DUF499*-><-?<-BetaPropeller+ABhydrolase\|\|?-><-ABhydrolase<-MODE-HTH+PBPII | 1103 | Bacteria>Pseudomonadota>Alphaproteobacteria>Sphingomonadales>Sphingomonadaceae | GCF_023516505.1 | Sphingomonas_brevis | LZ518_RS11630 | 2249110..2252421 | 2908206 | DUF499-domain-containing-protein-[Sphingomonas-brevis] |
| WP_201515748.1 | DUF499_1.RM.like.systems | <-TraJ-RHH<-?<-?<-?\|\|?->XIS-HTH-><-HEPN+DUF499*<-METHYLASE<-REase<-TUDOR+SF1-SF2-Helicase+iREase+CTD+REase<-HTH+WYL\|\|HicB-> | 1100 | Bacteria>Pseudomonadota>Alphaproteobacteria>Sphingomonadales>Sphingomonadaceae | GCF_904830875.1 | Sphingobium_sp._S6 | SPHS8_RS12190 | 306093..309395 | 2758386 | DUF499-domain-containing-protein-[Sphingobium-sp.-S6] |
| WP_249916143.1 | DUF499_1.RM.like.systems | wHTH+REase->TUDOR+SF1-SF2-Helicase+iREase+CTD+REase->METHYLASE->OLD-ABC->PIN->HEPN+DUF499*-><-?<-BetaPropeller+ABhydrolase\|\|?-><-ABhydrolase<-MODE-HTH+PBPII | 1094 | Bacteria>Pseudomonadota>Alphaproteobacteria>Sphingomonadales>Sphingomonadaceae | - | Sphingomonas_brevis | LZ518_11630 | 2249137..2252421 | 2908206 | DUF499-domain-containing-protein-[Sphingomonas-sp.-RB56-2] |
| MCL2716067.1 | DUF499_1.RM.like.systems | TUDOR+SF1-SF2-Helicase+iREase+CTD+REase-><-?\|\|METHYLASE->HEPN+DUF499*->REase+PNTD1+PNTD2->PIWI-> | 1073 | Bacteria>Pseudomonadota>Alphaproteobacteria>unclassified_Alphaproteobacteria | GCA_009781045.1 | Alphaproteobacteria_bacterium | FWD68_16235 | 9484..12705 | 1913988 | DUF499-domain-containing-protein-[Alphaproteobacteria-bacterium] |
| MBL8396550.1 | DUF499_1.RM.like.systems | HTH+WYL->TUDOR+SF1-SF2-Helicase+iREase+CTD+REase->HEPN->METHYLASE->wHTH+REase->DUF499*->DDE_5+DDE_Tnp_1->?-><-MORC+DNA_gyraseB+TOPRIM | 939 | Bacteria>Pseudomonadota>Betaproteobacteria>Betaproteobacteria_incertae_sedis>Candidatus_Accumulibacter | GCA_016789495.1 | Accumulibacter_sp. | JNL84_00165 | 43696..46515 | 2053492 | ATP-binding-protein-[Candidatus-Accumulibacter-sp.] |
| WP_169068632.1 | DUF499_1.RM.like.systems | TUDOR+SF1-SF2-Helicase+iREase+CTD+REase->MORC+DUF3684->wHTH+REase->METHYLASE->PolB->HEPN->HEPN+DUF499*->MarR-HTH->tRNA->?-><-GDResolvase+HIN-HTH\|\|MetJarc-HTH->RelE-ParE-> | 1113 | Bacteria>Pseudomonadota>Betaproteobacteria>Betaproteobacteria_incertae_sedis | GCF_907163155.1 | Candidatus_Accumulibacter | QZ648_RS13960 | 40235..43576 | 327159 | Swt1-family-HEPN-domain-containing-protein-[Candidatus-Accumulibacter-phosphatis] |
| WP_116311964.1 | DUF499_1.RM.like.systems | <-RNaseP<-?\|\|?->?->MORC+DNA_gyraseB+TOPRIM-><-REase<-DUF499*<-METHYLASE<-TUDOR+SF1-SF2-Helicase+iREase+CTD+REase<-HTH+WYL\|\|STYKinase-like->?->DUF6361-> | 963 | Bacteria>Pseudomonadota>Betaproteobacteria>Burkholderiales>Burkholderiaceae | GCF_900249825.1 | Cupriavidus_taiwanensis | CBM2633_RS00535 | 131730..134621 | 164546 | DUF499-domain-containing-protein-[Cupriavidus-taiwanensis] |
| WP_186144236.1 | DUF499_1.RM.like.systems | HTH+WYL->TIR+PNTD2->PIWI->TUDOR+SF1-SF2-Helicase+iREase+CTD+REase->HEPN->METHYLASE->DUF499*->REase-><-tRNA\|\|?->?-><-?\|\|MODE-HTH+PBPII-><-TOPRIM+ZnR | 955 | Bacteria>Pseudomonadota>Betaproteobacteria>Burkholderiales>Burkholderiaceae | GCF_902831945.1 | Burkholderia_gladioli | H9E34_RS32225 | 51067..53934 | 28095 | DUF499-domain-containing-protein-[Burkholderia-gladioli] |
| WP_132962623.1 | DUF499_1.RM.like.systems | GST_N+GST_C->tRNA->tRNA-><-DUF499*<-REase<-OLD-ABC<-METHYLASE<-PIN<-TUDOR+SF1-SF2-Helicase+iREase+CTD+REase\|\|Peptidase_M24-> | 949 | Bacteria>Pseudomonadota>Betaproteobacteria>Burkholderiales>Burkholderiales_genera_incertae_sedis | GCF_004342625.1 | Tepidimonas_ignava | EDC36_RS07390 | 30173..33022 | 114249 | DUF499-domain-containing-protein-[Tepidimonas-ignava] |
| WP_143969782.1 | DUF499_1.RM.like.systems | <-DUF772+DDE_Tnp_1\|\|<-DUF499*<-REase<-METHYLASE<-TOPRIM<-OLD-ABC<-TUDOR+SF1-SF2-Helicase+iREase+CTD+REase<-XIS-HTH | 948 | Bacteria>Pseudomonadota>Betaproteobacteria>Burkholderiales>Burkholderiales_genera_incertae_sedis | GCF_007556755.1 | Tepidimonas_fonticaldi | Tfont_RS13255 | 1599..4445 | 1101373 | DUF499-domain-containing-protein-[Tepidimonas-fonticaldi] |
| WP_180684522.1 | DUF499_1.RM.like.systems | XIS-HTH->TUDOR+SF1-SF2-Helicase+iREase+CTD+REase->OLD-ABC->METHYLASE->REase->DUF499*-><-Phage_lysozyme | 939 | Bacteria>Pseudomonadota>Betaproteobacteria>Burkholderiales>Burkholderiales_genera_incertae_sedis | GCF_013432195.1 | Tepidicella_baoligensis | HZS63_RS13535 | 8909..11728 | 2707016 | DUF499-domain-containing-protein-[Tepidicella-baoligensis] |
| WP_252101816.1 | DUF499_1.RM.like.systems | TUDOR+SF1-SF2-Helicase+iREase+CTD+REase->Schlafen+HATPase+HTH->?->METHYLASE->OLD-ABC->REase->DUF499*-><-?<-?<-Phage_integrase+Phage_integrase<-?<-HIN-HTH<-Phage_integrase+Phage_integrase | 945 | Bacteria>Pseudomonadota>Betaproteobacteria>Burkholderiales>Comamonadaceae | - | [Acidovorax]_ebreus | MRB47_RS02565 | 530947..533784 | 721785 | DUF499-domain-containing-protein-[[Acidovorax] |
| TFI45392.1 | DUF499_1.RM.like.systems | <-MobA_MobL+SF1-SF2-helicase\|\|?-><-?<-DUF499*<-REase<-OLD-ABC<-METHYLASE<-PIN<-TUDOR+SF1-SF2-Helicase+iREase+CTD+REase | 945 | Bacteria>Pseudomonadota>Betaproteobacteria>Burkholderiales>Comamonadaceae | - | Diaphorobacter_sp._DS2 | E4093_19980 | 3252..6089 | 1302548 | ATP-binding-protein-[Diaphorobacter-sp.-DS2] |
| WP_087746901.1 | DUF499_1.RM.like.systems | <-SPOUT\|\|?-><-?<-?<-?<-DUF499*<-REase<-OLD-ABC<-METHYLASE<-HNH<-OLD-ABC<-TUDOR+SF1-SF2-Helicase+iREase+CTD+REase | 945 | Bacteria>Pseudomonadota>Betaproteobacteria>Burkholderiales>Comamonadaceae | GCF_002176815.1 | unclassified_Acidovorax | CCX87_RS13165 | 2797139..2799976 | 2684926 | MULTISPECIES:-DUF499-domain-containing-protein-[unclassified-Acidovorax] |
| WP_055397337.1 | DUF499_1.RM.like.systems | <-ArdCN+MPTase<-?<-?<-MobA_MobL+SF1-SF2-helicase\|\|?-><-?<-DUF499*<-REase<-OLD-ABC<-METHYLASE<-HNH<-OLD-ABC<-TUDOR+SF1-SF2-Helicase+iREase+CTD+REase | 945 | Bacteria>Pseudomonadota>Betaproteobacteria>Burkholderiales>Comamonadaceae | GCF_017355385.1 | Acidovorax | AE621_RS12465 | 7782..10619 | 12916 | MULTISPECIES:-DUF499-domain-containing-protein-[Acidovorax] |
| WP_210568820.1 | DUF499_1.RM.like.systems | TUDOR+SF1-SF2-Helicase+iREase+CTD+REase->OLD-ABC->HNH->METHYLASE->OLD-ABC->REase->DUF499*->?->?->?-><-?\|\|SPOUT-> | 945 | Bacteria>Pseudomonadota>Betaproteobacteria>Burkholderiales>Comamonadaceae | GCF_017939685.1 | Acidovorax_sp._JG5 | J8G26_RS10065 | 96032..98869 | 2822718 | DUF499-domain-containing-protein-[Acidovorax-sp.-JG5] |
| WP_265286858.1 | DUF499_1.RM.like.systems | ZnR+TOPRIM+D5ATPase->TUDOR+SF1-SF2-Helicase+iREase+CTD+REase->SMODS->?->wHTH+REase->METHYLASE->DUF499*->REase->REase->?-><-GNAT | 1092 | Bacteria>Pseudomonadota>Betaproteobacteria>Burkholderiales>Comamonadaceae | GCF_026127525.1 | Verminephrobacter_eiseniae | D5047_RS05820 | 1238736..1242014 | 364317 | DUF499-domain-containing-protein-[Verminephrobacter-eiseniae] |
| UUZ52278.1 | DUF499_1.RM.like.systems | SF1-SF2-Helicase-p->?-><-REase<-DUF499-p<-DUF499-p<-DUF499*<-METHYLASE-p<-METHYLASE-p<-HEPN<-TUDOR+SF1-SF2-Helicase+iREase+CTD+REase-p<-TUDOR+SF1-SF2-Helicase+iREase+CTD+REase<-TUDOR+SF1-SF2-Helicase+iREase+CTD+REase-p | 325 | Bacteria>Pseudomonadota>Betaproteobacteria>Burkholderiales>Oxalobacteraceae | GCA_024703915.1 | Massilia_sp._H-1 | LP419_21985 | 3096112..3097089 | 2895566 | DUF499-domain-containing-protein-[Massilia-sp.-H-1] |
| WP_072453399.1 | DUF499_1.RM.like.systems | <-STYKinase-like\|\|HTH+WYL->TUDOR+SF1-SF2-Helicase+iREase+CTD+REase->HEPN->METHYLASE->DUF499*->REase-> | 961 | Bacteria>Pseudomonadota>Betaproteobacteria>Burkholderiales>Oxalobacteraceae | GCF_900119665.1 | Janthinobacterium_lividum | BT188_RS05025 | 1132446..1135331 | 29581 | DUF499-domain-containing-protein-[Janthinobacterium-lividum] |
| WP_066335883.1 | DUF499_1.RM.like.systems | <-RNase+SF1-SF2-Helicase<-?<-?<-MACRODOMAIN<-DUF499*<-REase<-OLD-ABC<-METHYLASE<-?<-TUDOR+SF1-SF2-Helicase+iREase+CTD+REase | 939 | Bacteria>Pseudomonadota>Betaproteobacteria>Burkholderiales>Sphaerotilaceae | GCF_001571085.1 | Azohydromonas_lata | ALA01S_RS10810 | 44580..47399 | 45677 | DUF499-domain-containing-protein-[Azohydromonas-lata] |
| WP_110402335.1 | DUF499_1.RM.like.systems | <-ABhydrolase\|\|?->?-><-?<-DUF499*<-REase<-OLD-ABC<-METHYLASE<-HNH<-OLD-ABC<-TUDOR+SF1-SF2-Helicase+iREase+CTD+REase | 958 | Bacteria>Pseudomonadota>Betaproteobacteria>Burkholderiales>Sphaerotilaceae | - | Sphaerotilus_hippei | C7444_RS18460 | 22920..25796 | 744406 | DUF499-domain-containing-protein-[Sphaerotilus-hippei] |
| MCL6557122.1 | DUF499_1.RM.like.systems | <-HEPN+DUF499*<-PolB<-?<-ORC-CDC6+REase<-TUDOR+SF1-SF2-Helicase+iREase+CTD+REase-p | 1098 | Bacteria>Pseudomonadota>Betaproteobacteria>Burkholderiales>unclassified_Burkholderiales | GCA_023511995.1 | Burkholderiales_bacterium | K6T56_12270 | 1861..5157 | 1891238 | DUF499-domain-containing-protein-[Burkholderiales-bacterium] |
| MCZ2134305.1 | DUF499_1.RM.like.systems | <-RAMA<-?<-?<-HTH_21+rve+rve_3<-HIN-HTH<-?<-DUF499*<-wHTH+REase<-METHYLASE<-HEPN<-PIN<-TUDOR+SF1-SF2-Helicase+iREase+CTD+REase\|\|HIN-HTH-> | 956 | Bacteria>Pseudomonadota>Betaproteobacteria>Burkholderiales>unclassified_Burkholderiales | GCA_026932525.1 | Burkholderiales_bacterium | LC098_02580 | 42253..45123 | 1891238 | MAG:-DUF499-domain-containing-protein-[Burkholderiales-bacterium] |
| QKE41939.1 | DUF499_1.RM.like.systems | DDE_Tnp_1_assoc+DDE_Tnp_1->HTH+WYL->TUDOR+SF1-SF2-Helicase+iREase+CTD+REase->OLD-ABC->HNH->wHTH+REase->DUF499*->DDE_Tnp_ISAZ013-><-?<-Hydrolase_3 | 942 | Bacteria>Pseudomonadota>Betaproteobacteria>Ferrovales>Ferrovaceae | GCA_013181415.1 | Ferrovum_myxofaciens | HO274_11940 | 2423876..2426704 | 416213 | ATP-binding-protein-[Ferrovum-myxofaciens] |
| WP_273119931.1 | DUF499_1.RM.like.systems | TUDOR+SF1-SF2-Helicase+iREase+CTD+REase->OLD-ABC->HNH->METHYLASE-p->METHYLASE-p->wHTH+REase->DUF499*->DDE_Tnp_ISAZ013-><-?<-?<-Hydrolase_3 | 942 | Bacteria>Pseudomonadota>Betaproteobacteria>Ferrovales>Ferrovaceae | GCF_013181415.1 | Ferrovum_myxofaciens | HO274_RS11980 | 2423876..2426704 | 416213 | DUF499-domain-containing-protein-[Ferrovum-myxofaciens] |
| MBX3631056.1 | DUF499_1.RM.like.systems | <-DUF499*<-METHYLASE<-HNH<-OLD-ABC<-TUDOR+SF1-SF2-Helicase+iREase+CTD+REase<-REase+RNase | 949 | Bacteria>Pseudomonadota>Betaproteobacteria>Nitrosomonadales>Nitrosomonadaceae | GCA_019635215.1 | Nitrosomonas_sp. | KF908_14345 | 13134..15983 | 42353 | ATP-binding-protein-[Nitrosomonas-sp.] |
| MBK6974203.1 | DUF499_1.RM.like.systems | <-SIGMA-HTH+SIGMA-HTH\|\|DDE_Tnp_1_assoc+DDE_Tnp_1-><-35exo+SF1-SF2-Helicase<-HEPN+DUF499*<-HEPN<-PolB<-wHTH+REase<-TUDOR+SF1-SF2-Helicase+iREase+CTD+REase | 1112 | Bacteria>Pseudomonadota>Betaproteobacteria>Nitrosomonadales>Sterolibacteriaceae | GCA_016708905.1 | Candidatus_Methylophosphatis_roskildensis | IPH26_15090 | 451210..454548 | 2899263 | DUF499-domain-containing-protein-[Sterolibacteriaceae-bacterium] |
| WP_169202394.1 | DUF499_1.RM.like.systems | TUDOR+SF1-SF2-Helicase+iREase+CTD+REase->?-><-PAIREDC-HTH+HIN-HTH\|\|METHYLASE->HEPN->wHTH+REase->DUF499*-><-?<-?<-NADAR<-?<-NUDIX | 941 | Bacteria>Pseudomonadota>Betaproteobacteria>Rhodocyclales>Rhodocyclaceae | GCF_017894365.1 | Aromatoleum_bremense | GPA24_RS09420 | 37549..40374 | 76115 | DUF499-domain-containing-protein-[Aromatoleum-bremense] |
| WP_169202998.1 | DUF499_1.RM.like.systems | <-DUF499*<-REase<-OLD-ABC<-METHYLASE<-HNH<-OLD-ABC<-TUDOR+SF1-SF2-Helicase+iREase+CTD+REase | 951 | Bacteria>Pseudomonadota>Betaproteobacteria>Rhodocyclales>Rhodocyclaceae | GCF_017894365.1 | Aromatoleum_bremense | GPA24_RS12870 | 12346..15201 | 76115 | DUF499-domain-containing-protein-[Aromatoleum-bremense] |
| WP_191658157.1 | DUF499_1.RM.like.systems | <-DUF499*<-wHTH+REase<-HEPN<-METHYLASE<-?<-ParB+HNH<-TUDOR+SF1-SF2-Helicase+iREase+CTD+REase | 944 | Bacteria>Pseudomonadota>Betaproteobacteria>Rhodocyclales>Zoogloeaceae | - | Azoarcus_sp._Aa7 | AZOA_12030 | 1309..4143 | 2303328 | DUF499-domain-containing-protein-[Azoarcus-sp.-Aa7] |
| MBD5801779.1 | DUF499_1.RM.like.systems | <-DUF499*<-wHTH+REase<-HEPN<-METHYLASE<-?<-ParB+HNH<-TUDOR+SF1-SF2-Helicase+iREase+CTD+REase | 944 | Bacteria>Pseudomonadota>Betaproteobacteria>Rhodocyclales>Zoogloeaceae | GCA_014822185.1 | Azoarcus_sp._Aa7 | AZOA_12030 | 1309..4143 | 2303328 | hypothetical-protein-[Azoarcus-sp.-Aa7] |
| WP_002936068.1 | DUF499_1.RM.like.systems | <-RNaseP<-?\|\|?->?->MORC+DNA_gyraseB+TOPRIM->?-><-DUF499*<-wHTH+REase<-METHYLASE<-HEPN<-ParB+HNH<-TUDOR+SF1-SF2-Helicase+iREase+CTD+REase | 941 | Bacteria>Pseudomonadota>Betaproteobacteria>Rhodocyclales>Zoogloeaceae | GCF_000310125.2 | Thauera_sp._27 | B447_RS02400 | 104287..107112 | 305700 | DUF499-domain-containing-protein-[Thauera-sp.-27] |
| WP_153168444.1 | DUF499_1.RM.like.systems | <-Radical-SAM+TRAM<-?\|\|tRNA->?->?->?-><-?<-DUF499*<-wHTH+REase<-REase<-OLD-ABC<-METHYLASE<-HEPN<-TUDOR+SF1-SF2-Helicase+iREase+CTD+REase | 941 | Bacteria>Pseudomonadota>Betaproteobacteria>Rhodocyclales>Zoogloeaceae | GCF_009469595.2 | Thauera_sp._2A1 | GH664_RS12700 | 85060..87885 | 2570191 | DUF499-domain-containing-protein-[Thauera-sp.-2A1] |
| WP_159691715.1 | DUF499_1.RM.like.systems | MODE-HTH+PBPII-><-TPR<-DUF499*<-REase<-OLD-ABC<-METHYLASE<-HNH<-OLD-ABC<-TUDOR+SF1-SF2-Helicase+iREase+CTD+REase | 943 | Bacteria>Pseudomonadota>Betaproteobacteria>Rhodocyclales>Zoogloeaceae | GCF_009800965.1 | Cognatazoarcus_halotolerans | GQB03_RS16405 | 71425..74256 | 2686016 | DUF499-domain-containing-protein-[Cognatazoarcus-halotolerans] |
| OGA55362.1 | DUF499_1.RM.like.systems | <-DUF499*<-REase<-METHYLASE<-Schlafen+HATPase+HTH<-TUDOR+SF1-SF2-Helicase+iREase+CTD+REase | 941 | Bacteria>Pseudomonadota>Betaproteobacteria>unclassified_Betaproteobacteria | GCA_001772005.1 | Betaproteobacteria_bacterium_RIFCSPLOWO2_12_FULL_62_58 | A3F74_07155 | 338..3163 | 1797497 | AAA-family-ATPase-[Betaproteobacteria-bacterium-RIFCSPLOWO2_12_FULL_62_58] |
| MBI5911562.1 | DUF499_1.RM.like.systems | <-DOC+HTH\|\|TUDOR+SF1-SF2-Helicase+iREase+CTD+REase->AAA-ATPase+REase->?->?->METHYLASE->DUF499*-><-?<-?<-Aminotran_4 | 1090 | Bacteria>Pseudomonadota>Betaproteobacteria>unclassified_Betaproteobacteria | GCA_016234705.1 | Betaproteobacteria_bacterium | HY848_16630 | 24102..27374 | 1891241 | ATP-binding-protein-[Betaproteobacteria-bacterium] |
| MDA8417189.1 | DUF499_1.RM.like.systems | TUDOR+SF1-SF2-Helicase+iREase+CTD+REase->RVT->OLD-ABC->HNH->METHYLASE->wHTH+REase->DUF499*-><-?\|\|?->?->?-><-Response_reg+LuxR-HTH | 946 | Bacteria>Pseudomonadota>Betaproteobacteria>unclassified_Betaproteobacteria | GCA_028069095.1 | Betaproteobacteria_bacterium | MOZ78_09075 | 11618..14458 | 1891241 | MAG:-DUF499-domain-containing-protein-[Betaproteobacteria-bacterium] |
| WP_096459285.1 | DUF499_1.RM.like.systems | TUDOR+SF1-SF2-Helicase+iREase+CTD+REase->OLD-ABC->METHYLASE->REase->DUF499*->?->?->?->?->?-><-HTH_3 | 943 | Bacteria>Pseudomonadota>Gammaproteobacteria>Acidiferrobacterales>Acidiferrobacteraceae | GCF_002355415.1 | Sulfurifustis_variabilis | SVA_RS04285 | 892754..895585 | 1675686 | DUF499-domain-containing-protein-[Sulfurifustis-variabilis] |
| WP_223954701.1 | DUF499_1.RM.like.systems | JAB->CbeA_antitoxin->RlpA-><-DUF499*<-REase<-OLD-ABC<-METHYLASE<-HNH<-OLD-ABC<-TUDOR+SF1-SF2-Helicase+iREase+CTD+REase | 946 | Bacteria>Pseudomonadota>Gammaproteobacteria>Aeromonadales>Aeromonadaceae | GCF_019972775.1 | Aeromonas_caviae | LDN68_RS17105 | 3642842..3645682 | 648 | DUF499-domain-containing-protein-[Aeromonas-caviae] |
| WP_087504467.1 | DUF499_1.RM.like.systems | <-Phage_integrase\|\|HTH+WYL->TUDOR+SF1-SF2-Helicase+iREase+CTD+REase->METHYLASE->OLD-ABC->REase->DUF499*-><-?<-?<-?<-?\|\|REase-> | 940 | Bacteria>Pseudomonadota>Gammaproteobacteria>Alteromonadales>Echinimonadaceae | GCF_014636195.1 | Neiella_marina | CBE68_RS02320 | 109774..112596 | 508461 | DUF499-domain-containing-protein-[Neiella-marina] |
| KTG26834.1 | DUF499_1.RM.like.systems | Arm-DNA-bind_3+Phage_integrase-><-Phage_AlpA<-ZnR+TOPRIM+primpol+HTH<-XIS-HTH\|\|MobI-><-DUF499*<-REase<-OLD-ABC<-METHYLASE<-OLD-ABC<-TUDOR+SF1-SF2-Helicase+iREase+CTD+REase<-HTH+WYL | 953 | Bacteria>Pseudomonadota>Gammaproteobacteria>Alteromonadales>Idiomarinaceae | GCA_001469885.1 | Idiomarina_sp._H105 | AUR68_16335 | 105470..108331 | 1766622 | hypothetical-protein-AUR68_16335-[Idiomarina-sp.-H105] |
| WP_020507360.1 | DUF499_1.RM.like.systems | <-McrC-NTD+REase<-?<-DUF499*<-wHTH+REase<-METHYLASE<-TOPRIM<-OLD-ABC<-METHYLASE-p<-TUDOR+SF1-SF2-Helicase+iREase+CTD+REase | 961 | Bacteria>Pseudomonadota>Gammaproteobacteria>Chromatiales>Chromatiaceae | GCF_000379525.1 | Lamprocystis_purpurea | A390_RS0124805 | 120589..123474 | 61598 | DUF499-domain-containing-protein-[Lamprocystis-purpurea] |
| WP_213378539.1 | DUF499_1.RM.like.systems | MerR-HTH+GDResolvase->HTH_OrfB_IS605+OrfB_IS605+OrfB_Zn_ribbon->?-><-DUF499*<-REase<-OLD-ABC<-METHYLASE<-TUDOR+SF1-SF2-Helicase+iREase+CTD+REase<-HTH+WYL\|\|HTH_OrfB_IS605+OrfB_IS605+OrfB_Zn_ribbon-> | 943 | Bacteria>Pseudomonadota>Gammaproteobacteria>Chromatiales>Chromatiaceae | GCF_018409545.1 | Allochromatium_tepidum | Atep_RS10840 | 2243146..2245977 | 553982 | DUF499-domain-containing-protein-[Allochromatium-tepidum] |
| WP_025282662.1 | DUF499_1.RM.like.systems | TraH->?->TraG_N-><-?\|\|AbiEi_4-><-DUF499*<-REase<-OLD-ABC<-METHYLASE<-?<-TUDOR+SF1-SF2-Helicase+iREase+CTD+REase\|\|TPR+TPR-> | 956 | Bacteria>Pseudomonadota>Gammaproteobacteria>Chromatiales>Ectothiorhodospiraceae | GCF_000583815.1 | Ectothiorhodospira_haloalkaliphila | ECTHA_RS0102365 | 117439..120309 | 421628 | DUF499-domain-containing-protein-[Ectothiorhodospira-haloalkaliphila] |
| WP_186934038.1 | DUF499_1.RM.like.systems | <-DUF499*<-REase<-OLD-ABC<-METHYLASE<-SbcC<-TUDOR+SF1-SF2-Helicase+iREase+CTD+REase | 946 | Bacteria>Pseudomonadota>Gammaproteobacteria>Enterobacterales>Enterobacteriaceae | GCF_014285945.1 | Klebsiella | H8M57_RS27310 | 138..2978 | 570 | MULTISPECIES:-DUF499-domain-containing-protein-[Klebsiella] |
| WP_219173436.1 | DUF499_1.RM.like.systems | HIN-HTH->TUDOR+SF1-SF2-Helicase+iREase+CTD+REase->SbcC->METHYLASE->OLD-ABC->REase->DUF499*-><-RlpA<-Arm-DNA-bind_3+Phage_integrase\|\|HIN-HTH->\|\|?->?->ParB+HNH-> | 946 | Bacteria>Pseudomonadota>Gammaproteobacteria>Enterobacterales>Enterobacteriaceae | GCF_019334445.1 | Klebsiella/Raoultella_group | LVJ43_RS03725 | 746872..749712 | 2890311 | DUF499-domain-containing-protein-[Raoultella-ornithinolytica] |
| WP_224381981.1 | DUF499_1.RM.like.systems | TUDOR+SF1-SF2-Helicase+iREase+CTD+REase->SbcC->METHYLASE->OLD-ABC->REase->DUF499*-> | 350 | Bacteria>Pseudomonadota>Gammaproteobacteria>Enterobacterales>Enterobacteriaceae | GCF_020118495.1 | Klebsiella_michiganensis | FMK66_RS24360 | 60161..61211 | 1134687 | DUF499-domain-containing-protein,-partial-[Klebsiella-michiganensis] |
| WP_134906061.1 | DUF499_1.RM.like.systems | Arm-DNA-bind_3+Phage_integrase->RlpA-><-DUF499*<-REase<-OLD-ABC<-METHYLASE<-SbcC<-TUDOR+SF1-SF2-Helicase+iREase+CTD+REase<-HIN-HTH | 946 | Bacteria>Pseudomonadota>Gammaproteobacteria>Enterobacterales | GCF_004801595.1 | Enterobacteriaceae | I8Q62_RS26385 | 2639..5479 | 543 | MULTISPECIES:-DUF499-domain-containing-protein-[Enterobacteriaceae] |
| WP_077999168.1 | DUF499_1.RM.like.systems | JAB->RlpA-><-DUF499*<-REase<-OLD-ABC<-METHYLASE<-HNH<-OLD-ABC<-TUDOR+SF1-SF2-Helicase+iREase+CTD+REase | 949 | Bacteria>Pseudomonadota>Gammaproteobacteria>Enterobacterales>Hafniaceae | GCF_002006945.1 | Edwardsiella_tarda | BTN58_RS02535 | 130022..132871 | 636 | DUF499-domain-containing-protein-[Edwardsiella-tarda] |
| WP_156881498.1 | DUF499_1.RM.like.systems | <-SLATT\|\|XIS-HTH->TUDOR+SF1-SF2-Helicase+iREase+CTD+REase->OLD-ABC->METHYLASE->REase->DUF499*->?-><-Radical-SAM<-HTH_3<-Phage_lysozyme | 939 | Bacteria>Pseudomonadota>Gammaproteobacteria>Methylococcales>Methylococcaceae | GCF_017832375.1 | Methylocaldum_sp._RMAD-M | JOD69_RS15380 | 37425..40244 | 2806557 | DUF499-domain-containing-protein-[Methylocaldum-sp.-RMAD-M] |
| WP_218800673.1 | DUF499_1.RM.like.systems | <-TIR<-DUF499*<-REase<-OLD-ABC<-METHYLASE<-PIN<-TUDOR+SF1-SF2-Helicase+iREase+CTD+REase<-HTH_Tnp | 942 | Bacteria>Pseudomonadota>Gammaproteobacteria>Methylococcales>Methylococcaceae | GCF_019285535.1 | Methylococcus_capsulatus | M3M30_RS10970 | 2380003..2382831 | 414 | DUF499-domain-containing-protein-[Methylococcus-capsulatus] |
| WP_218796637.1 | DUF499_1.RM.like.systems | XIS-HTH->TUDOR+SF1-SF2-Helicase+iREase+CTD+REase->OLD-ABC->TOPRIM->METHYLASE->REase->DUF499*->MORC+REase->HEPN-><-Phage_lysozyme<-?<-?<-RNase | 939 | Bacteria>Pseudomonadota>Gammaproteobacteria>Methylococcales>Methylococcaceae | GCF_019285555.1 | Methylococcus_capsulatus | KW114_RS08340 | 1841753..1844572 | 414 | DUF499-domain-containing-protein-[Methylococcus-capsulatus] |
| WP_218797031.1 | DUF499_1.RM.like.systems | HHH_5+SF1-SF2-helicase->XIS-HTH->TUDOR+SF1-SF2-Helicase+iREase+CTD+REase->TIR->METHYLASE->REase->DUF499*->Methylase-><-Radical-SAM<-HTH_3 | 939 | Bacteria>Pseudomonadota>Gammaproteobacteria>Methylococcales>Methylococcaceae | GCF_019285555.1 | Methylococcus_capsulatus | KW114_RS10930 | 2381240..2384059 | 414 | DUF499-domain-containing-protein-[Methylococcus-capsulatus] |
| WP_282213184.1 | DUF499_1.RM.like.systems | HHH_5+SF1-SF2-helicase->XIS-HTH->TUDOR+SF1-SF2-Helicase+iREase+CTD+REase->REase->METHYLASE->REase->DUF499*->Methylase-><-Radical-SAM<-HTH_3 | 939 | Bacteria>Pseudomonadota>Gammaproteobacteria>Methylococcales>Methylococcaceae | GCF_949769275.1 | Methylococcus_capsulatus | QMX71_RS08995 | 1893622..1896441 | 414 | DUF499-domain-containing-protein-[Methylococcus-capsulatus] |
| WP_022949219.1 | DUF499_1.RM.like.systems | LON+AAA->TUDOR+SF1-SF2-Helicase+iREase+CTD+REase->METHYLASE->OLD-ABC->REase->DUF499*->?->RADICAL-SAM->Glyco_transf_8-> | 952 | Bacteria>Pseudomonadota>Gammaproteobacteria>Methylococcales>Methylothermaceae | GCF_000421465.1 | Methylohalobius_crimeensis | H035_RS0111990 | 2436080..2438938 | 244365 | DUF499-domain-containing-protein-[Methylohalobius-crimeensis] |
| MAE39903.1 | DUF499_1.RM.like.systems | HTH_26->TUDOR+SF1-SF2-Helicase+iREase+CTD+REase->METHYLASE->OLD-ABC->REase->DUF499*->?->?-><-Phage_integrase<-Phage_integrase | 941 | Bacteria>Pseudomonadota>Gammaproteobacteria>Moraxellales>Moraxellaceae | GCA_002686735.1 | Psychrobacter_sp. | CML07_03075 | 11656..14481 | 56811 | hypothetical-protein-[Psychrobacter-sp.] |
| WP_231663150.1 | DUF499_1.RM.like.systems | HTH+WYL->TUDOR+SF1-SF2-Helicase+iREase+CTD+REase->METHYLASE->OLD-ABC->REase->ParB+HNH->DUF499*->Primpol+D5N+D5ATPase-><-Arm-DNA-bind_3+Phage_integrase\|\|?->FAD-NAD-dep-oxidoreductase-> | 957 | Bacteria>Pseudomonadota>Gammaproteobacteria>Oceanospirillales>Halomonadaceae | GCF_021151495.1 | Halomonas_sp._IOP_31 | K7I12_RS04335 | 15378..18251 | 2876584 | DUF499-domain-containing-protein-[Halomonas-sp.-IOP_31] |
| WP_239295914.1 | DUF499_1.RM.like.systems | <-HTH_Tnp_1+HTH_21+rve+rve_2<-?<-DUF499*<-?<-REase<-OLD-ABC<-METHYLASE<-?<-TUDOR+SF1-SF2-Helicase+iREase+CTD+REase | 958 | Bacteria>Pseudomonadota>Gammaproteobacteria>Oceanospirillales>Halomonadaceae | GCF_022348215.1 | unclassified_Halomonas | MHM38_RS03720 | 8315..11191 | 2609666 | MULTISPECIES:-DUF499-domain-containing-protein-[unclassified-Halomonas] |
| WP_253027667.1 | DUF499_1.RM.like.systems | Phage_integrase-><-Phage_AlpA<-ZnR+TOPRIM+primpol+HTH<-XIS-HTH\|\|MobI-><-DUF499*<-REase<-OLD-ABC<-METHYLASE<-OLD-ABC<-TUDOR+SF1-SF2-Helicase+iREase+CTD+REase<-HTH+WYL | 953 | Bacteria>Pseudomonadota>Gammaproteobacteria>Oceanospirillales>Halomonadaceae | GCF_024124725.1 | Halomonas_sp._Ps84H-12 | NH344_RS06635 | 49336..52197 | 2954501 | DUF499-domain-containing-protein-[Halomonas-sp.-Ps84H-12] |
| WP_137094585.1 | DUF499_1.RM.like.systems | TUDOR+SF1-SF2-Helicase+iREase+CTD+REase->OLD-ABC->HNH->METHYLASE->OLD-ABC->REase->DUF499*-><-MobI\|\|XIS-HTH->ZnR+TOPRIM+primpol+HTH->Phage_AlpA-><-Arm-DNA-bind_3+Phage_integrase | 953 | Bacteria>Pseudomonadota>Gammaproteobacteria>Oceanospirillales>Halomonadaceae | GCF_028580285.1 | Halomonas | E8Q34_RS13770 | 13695..16556 | 2745 | DUF499-domain-containing-protein-[Halomonas-sp.-15WGF] |
| WP_284453091.1 | DUF499_1.RM.like.systems | Arm-DNA-bind_3+Phage_integrase-><-Phage_AlpA<-ZnR+TOPRIM+primpol+HTH<-XIS-HTH\|\|MobI-><-DUF499*<-REase<-OLD-ABC<-METHYLASE<-HNH<-OLD-ABC<-TUDOR+SF1-SF2-Helicase+iREase+CTD+REase | 953 | Bacteria>Pseudomonadota>Gammaproteobacteria>Oceanospirillales>Halomonadaceae | GCF_030145725.1 | Halomonas_meridiana | KYX60_RS01235 | 106598..109459 | 29570 | DUF499-domain-containing-protein-[Halomonas-meridiana] |
| WP_072841218.1 | DUF499_1.RM.like.systems | TUDOR+SF1-SF2-Helicase+iREase+CTD+REase->OLD-ABC->HNH->METHYLASE->OLD-ABC->REase->DUF499*->SMODS->?-><-RlpA\|\|?-><-HTH_26 | 942 | Bacteria>Pseudomonadota>Gammaproteobacteria>Oceanospirillales>Oceanospirillaceae | GCF_900129155.1 | Marinomonas_polaris | BUA59_RS18830 | 76459..79287 | 293552 | DUF499-domain-containing-protein-[Marinomonas-polaris] |
| WP_230713447.1 | DUF499_1.RM.like.systems | Phage_integrase-><-Phage_AlpA<-ZnR+TOPRIM+primpol+HTH<-XIS-HTH\|\|MobI-><-DUF499*<-REase<-OLD-ABC<-METHYLASE<-OLD-ABC<-TUDOR+SF1-SF2-Helicase+iREase+CTD+REase<-HTH+WYL | 953 | Bacteria>Pseudomonadota>Gammaproteobacteria>Pseudomonadales>Marinobacteraceae | GCF_021021515.1 | Marinobacter_shengliensis | K7I15_RS06080 | 103063..105924 | 1389223 | DUF499-domain-containing-protein-[Marinobacter-shengliensis] |
| WP_091985110.1 | DUF499_1.RM.like.systems | TUDOR+SF1-SF2-Helicase+iREase+CTD+REase->OLD-ABC->HNH->METHYLASE->MAD-NTD+EVE+McrB-AAA->McrC-NTD+REase->DUF499*->SAM-methylase->HTH+HNH-><-?<-CHTH | 947 | Bacteria>Pseudomonadota>Gammaproteobacteria>Pseudomonadales>Marinobacteraceae | GCF_900115175.1 | Marinobacter_gudaonensis | BM344_RS01255 | 264147..266990 | 375760 | DUF499-domain-containing-protein-[Marinobacter-gudaonensis] |
| MBG7398186.1 | DUF499_1.RM.like.systems | SF1-SF2-Helicase-><-SF1-SF2-helicase+RdgC+Beta-barrel+REase<-35exo+SF1-SF2-Helicase\|\|ABhydrolase->?->RlpA-><-DUF499*<-REase<-OLD-ABC<-METHYLASE<-HEPN<-TUDOR+SF1-SF2-Helicase+iREase+CTD+REase | 942 | Bacteria>Pseudomonadota>Gammaproteobacteria>Pseudomonadales>Pseudomonadaceae | GCA_015766545.1 | Pseudomonas_aeruginosa | GHU66_27420 | 38429..41257 | 287 | ATP-binding-protein-[Pseudomonas-aeruginosa] |
| HBO5114778.1 | DUF499_1.RM.like.systems | TUDOR+SF1-SF2-Helicase+iREase+CTD+REase->HEPN->METHYLASE->OLD-ABC->REase->DUF499*-><-RlpA<-?<-ABhydrolase | 946 | Bacteria>Pseudomonadota>Gammaproteobacteria>Pseudomonadales>Pseudomonadaceae | GCA_021662155.1 | Pseudomonas_aeruginosa | L4W37_005387 | 12730..15570 | 287 | ATP-binding-protein-[Pseudomonas-aeruginosa] |
| MCO2346212.1 | DUF499_1.RM.like.systems | TUDOR+SF1-SF2-Helicase+iREase+CTD+REase-><-?\|\|METHYLASE->OLD-ABC->REase->DUF499*-><-RlpA<-?\|\|35exo+SF1-SF2-Helicase-><-?<-HIN-HTH | 942 | Bacteria>Pseudomonadota>Gammaproteobacteria>Pseudomonadales>Pseudomonadaceae | GCA_023907105.1 | Pseudomonas_aeruginosa | FA362_14075 | 17744..20572 | 287 | ATP-binding-protein-[Pseudomonas-aeruginosa] |
| HCI3160560.1 | DUF499_1.RM.like.systems | ABhydrolase->?->RlpA-><-DUF499*<-REase<-OLD-ABC<-METHYLASE<-HEPN<-TUDOR+SF1-SF2-Helicase+iREase+CTD+REase | 942 | Bacteria>Pseudomonadota>Gammaproteobacteria>Pseudomonadales>Pseudomonadaceae | GCA_024356045.1 | Pseudomonas_aeruginosa | NOV63_004506 | 69244..72072 | 287 | ATP-binding-protein-[Pseudomonas-aeruginosa] |
| MCR7692359.1 | DUF499_1.RM.like.systems | TUDOR+SF1-SF2-Helicase+iREase+CTD+REase->HEPN->METHYLASE->OLD-ABC->REase->DUF499*-> | 417 | Bacteria>Pseudomonadota>Gammaproteobacteria>Pseudomonadales>Pseudomonadaceae | GCA_024717035.1 | Pseudomonas_aeruginosa | NTA33_29240 | 17686..18936 | 287 | DUF499-domain-containing-protein-[Pseudomonas-aeruginosa] |

| Protein-ID | System | Gene-Neighbourhoods | Length | Taxonomy | Assembly-ID | Strain | Locus-Tag | Genomic-coordinates | TaxID | Fasta-defline |
|---|---|---|---|---|---|---|---|---|---|---|
| WP_023835622.1 | DUF499_1.RM.like.systems | TUDOR+SF1-SF2-Helicase+iREase+CTD+REase->HEPN->METHYLASE->OLD-ABC->REase->DUF499*-><-RlpA<-?<-JAB<-ParB<-?\|\|HIN-HTH-> | 944 | Bacteria>Pseudomonadota>Gammaproteobacteria>Pseudomonadales>Pseudomonadaceae | GCF_000503175.1 | Pseudomonas_aeruginosa | CF344_RS23300 | 12270..15104 | 287 | DUF499-domain-containing-protein-[Pseudomonas-aeruginosa] |
| WP_031643596.1 | DUF499_1.RM.like.systems | TUDOR+SF1-SF2-Helicase+iREase+CTD+REase->HEPN->METHYLASE->OLD-ABC->REase->DUF499*-><-RlpA<-?<-ABhydrolase<-?<-?\|\|HTH_3-> | 950 | Bacteria>Pseudomonadota>Gammaproteobacteria>Pseudomonadales>Pseudomonadaceae | GCF_000629305.1 | Pseudomonas_aeruginosa | AJ61_RS31490 | 12829..15681 | 287 | DUF499-domain-containing-protein-[Pseudomonas-aeruginosa] |
| WP_049308056.1 | DUF499_1.RM.like.systems | TUDOR+SF1-SF2-Helicase+iREase+CTD+REase->HEPN->METHYLASE->OLD-ABC->REase->DUF499*-><-RlpA<-?<-?\|SF1-SF2-helicase+RdgC+Beta-barrel+REase-><-SF1-SF2-Helicase<-HKD-DNase | 942 | Bacteria>Pseudomonadota>Gammaproteobacteria>Pseudomonadales>Pseudomonadaceae | GCF_001061475.1 | Pseudomonas_aeruginosa | ADM76_RS18680 | 17855..20683 | 287 | DUF499-domain-containing-protein-[Pseudomonas-aeruginosa] |
| WP_054902927.1 | DUF499_1.RM.like.systems | DUF932->?->ParB->JAB->?->RlpA-><-DUF499*<-REase<-OLD-ABC<-METHYLASE<-HEPN<-TUDOR+SF1-SF2-Helicase+iREase+CTD+REase\|\|HTH+WYL-> | 944 | Bacteria>Pseudomonadota>Gammaproteobacteria>Pseudomonadales>Pseudomonadaceae | GCF_001320525.1 | Pseudomonas_sp._NBRC_111131 | APH34_RS22145 | 20096..22930 | 1661046 | DUF499-domain-containing-protein-[Pseudomonas-sp.-NBRC-111131] |
| WP_074197337.1 | DUF499_1.RM.like.systems | DUF932->?->ParB->JAB->?->RlpA-><-DUF499*<-REase<-OLD-ABC<-METHYLASE<-HEPN<-TUDOR+SF1-SF2-Helicase+iREase+CTD+REase | 946 | Bacteria>Pseudomonadota>Gammaproteobacteria>Pseudomonadales>Pseudomonadaceae | GCF_001909445.1 | Pseudomonas_aeruginosa | BH596_RS15220 | 11473..14313 | 287 | DUF499-domain-containing-protein-[Pseudomonas-aeruginosa] |
| WP_085592307.1 | DUF499_1.RM.like.systems | MerR-HTH->tRNA->TUDOR+SF1-SF2-Helicase+iREase+CTD+REase->METHYLASE->OLD-ABC->REase->DUF499*-><-RlpA<-REase+SF1-SF2-Helicase<-N6_N4_MTase\|\|MAD-NTD+HNH-> | 941 | Bacteria>Pseudomonadota>Gammaproteobacteria>Pseudomonadales>Pseudomonadaceae | GCF_002113255.1 | Pseudomonas_sp._B14(2017) | CAG34_RS21645 | 108391..111216 | 1981745 | DUF499-domain-containing-protein-[Pseudomonas-sp.-B14(2017)] |
| WP_131349395.1 | DUF499_1.RM.like.systems | TUDOR+SF1-SF2-Helicase+iREase+CTD+REase->OLD-ABC->HNH->METHYLASE->OLD-ABC->REase->DUF499*-><-RlpA<-TPR<-?<-?\|\|?-><-Mbetalac | 929 | Bacteria>Pseudomonadota>Gammaproteobacteria>Pseudomonadales>Pseudomonadaceae | GCF_004327885.1 | Azotobacter_chroococcum | EOE52_RS08250 | 59995..62784 | 353 | DUF499-domain-containing-protein-[Azotobacter-chroococcum] |
| WP_154355052.1 | DUF499_1.RM.like.systems | TUDOR+SF1-SF2-Helicase+iREase+CTD+REase->HEPN->METHYLASE->OLD-ABC->REase->DUF499*-><-RlpA<-?<-JAB<-ParB<-?<-DUF932 | 942 | Bacteria>Pseudomonadota>Gammaproteobacteria>Pseudomonadales>Pseudomonadaceae | GCF_009676785.1 | Pseudomonas_aeruginosa | GJV02_RS03270 | 700187..703015 | 287 | DUF499-domain-containing-protein-[Pseudomonas-aeruginosa] |
| WP_192069117.1 | DUF499_1.RM.like.systems | Integrase_1->Phage_int_SAM->N6_N4_MTase->REase+SF1-SF2-Helicase->RlpA-><-DUF499*<-REase<-OLD-ABC<-METHYLASE<-TUDOR+SF1-SF2-Helicase+iREase+CTD+REase\|\|?-><-HTH_3 | 941 | Bacteria>Pseudomonadota>Gammaproteobacteria>Pseudomonadales>Pseudomonadaceae | GCF_014842375.1 | Pseudomonas_coleopterorum | IFT40_RS20700 | 77168..79993 | 1605838 | DUF499-domain-containing-protein-[Pseudomonas-coleopterorum] |
| WP_198580799.1 | DUF499_1.RM.like.systems | TUDOR+SF1-SF2-Helicase+iREase+CTD+REase->HEPN->METHYLASE->OLD-ABC->REase->DUF499*-><-RlpA<-?<-JAB<-ParB<-?<-DUF932 | 944 | Bacteria>Pseudomonadota>Gammaproteobacteria>Pseudomonadales>Pseudomonadaceae | GCF_016134915.1 | Pseudomonas_aeruginosa | QMY34_RS12045 | 381175..384009 | 287 | DUF499-domain-containing-protein-[Pseudomonas-aeruginosa] |
| WP_200582036.1 | DUF499_1.RM.like.systems | TUDOR+SF1-SF2-Helicase+iREase+CTD+REase->HEPN->METHYLASE->OLD-ABC->REase->DUF499*-><-RlpA<-?<-SymE_toxin<-?<-ZNR+Pkinase+SF1-SF2-Helicase+REase | 943 | Bacteria>Pseudomonadota>Gammaproteobacteria>Pseudomonadales>Pseudomonadaceae | GCF_016622185.1 | Pseudomonas_aeruginosa | GFM12_RS08425 | 1738931..1741762 | 287 | DUF499-domain-containing-protein-[Pseudomonas-aeruginosa] |
| WP_219806875.1 | DUF499_1.RM.like.systems | <-MazE_antitoxin\|\|?->ParB->JAB->?->RlpA-><-DUF499*<-REase<-OLD-ABC<-METHYLASE<-HEPN<-TUDOR+SF1-SF2-Helicase+iREase+CTD+REase<-Peptidase | 943 | Bacteria>Pseudomonadota>Gammaproteobacteria>Pseudomonadales>Pseudomonadaceae | GCF_019429735.1 | Pseudomonas_aeruginosa | KZ843_RS17265 | 5657..8488 | 287 | DUF499-domain-containing-protein-[Pseudomonas-aeruginosa] |
| WP_033975677.1 | DUF499_1.RM.like.systems | TUDOR+SF1-SF2-Helicase+iREase+CTD+REase->HEPN->METHYLASE->OLD-ABC->REase->DUF499*-><-RlpA<-?<-SymE_toxin<-?<-ZNR+Pkinase+SF1-SF2-Helicase+REase | 943 | Bacteria>Pseudomonadota>Gammaproteobacteria>Pseudomonadales>Pseudomonadaceae | GCF_022569915.1 | Pseudomonas_aeruginosa | CGU39_RS16575 | 614189..617020 | 287 | DUF499-domain-containing-protein-[Pseudomonas-aeruginosa] |
| WP_266022655.1 | DUF499_1.RM.like.systems | TUDOR+SF1-SF2-Helicase+iREase+CTD+REase->HEPN->METHYLASE->OLD-ABC->REase->DUF499*-><-RlpA<-?<-JAB<-ParB<-?<-DUF932 | 948 | Bacteria>Pseudomonadota>Gammaproteobacteria>Pseudomonadales>Pseudomonadaceae | GCF_026241065.1 | unclassified_Pseudomonas | OO409_RS22205 | 32872..35718 | 196821 | MULTISPECIES:-DUF499-domain-containing-protein-[unclassified-Pseudomonas] |
| WP_272438616.1 | DUF499_1.RM.like.systems | HKD-DNase->SF1-SF2-Helicase-><-SF1-SF2-helicase+RdgC+Beta-barrel+REase\|\|?->?->RlpA-><-DUF499*<-REase<-OLD-ABC<-METHYLASE<-HEPN<-TUDOR+SF1-SF2-Helicase+iREase+CTD+REase | 942 | Bacteria>Pseudomonadota>Gammaproteobacteria>Pseudomonadales>Pseudomonadaceae | GCF_028435625.1 | Pseudomonas_aeruginosa | KGZ09_RS30315 | 38429..41257 | 287 | DUF499-domain-containing-protein-[Pseudomonas-aeruginosa] |
| WP_090734260.1 | DUF499_1.RM.like.systems | TUDOR+SF1-SF2-Helicase+iREase+CTD+REase->?->METHYLASE->OLD-ABC->REase->DUF499*-><-RlpA<-?\|\|PAIREDC-HTH-> | 929 | Bacteria>Pseudomonadota>Gammaproteobacteria>Pseudomonadales>Pseudomonadaceae | GCF_900108885.1 | Azotobacter_beijerinckii | BMW56_RS18070 | 16681..19470 | 170623 | DUF499-domain-containing-protein-[Azotobacter-beijerinckii] |
| WP_090941615.1 | DUF499_1.RM.like.systems | TUDOR+SF1-SF2-Helicase+iREase+CTD+REase->OLD-ABC->HNH->METHYLASE->OLD-ABC->REase->DUF499*-><-RlpA<-?<-?<-ZnR+TOPRIM+D5ATPase | 929 | Bacteria>Pseudomonadota>Gammaproteobacteria>Pseudomonadales>Pseudomonadaceae | GCF_900114395.1 | Azotobacter_beijerinckii | BM240_RS15650 | 17867..20656 | 170623 | DUF499-domain-containing-protein-[Azotobacter-beijerinckii] |
| WP_033982644.1 | DUF499_1.RM.like.systems | ABhydrolase->?->RlpA-><-DUF499*<-REase<-OLD-ABC<-METHYLASE<-HEPN<-TUDOR+SF1-SF2-Helicase+iREase+CTD+REase | 942 | Bacteria>Pseudomonadota>Gammaproteobacteria>Pseudomonadales>Pseudomonadaceae | GCF_900707645.1 | Pseudomonas_aeruginosa | BUH55_RS18155 | 7471..10299 | 287 | DUF499-domain-containing-protein-[Pseudomonas-aeruginosa] |
| WP_132674023.1 | DUF499_1.RM.like.systems | DUF932->?->ParB->JAB->?->RlpA-><-DUF499*<-REase<-OLD-ABC<-METHYLASE<-HEPN<-TUDOR+SF1-SF2-Helicase+iREase+CTD+REase | 944 | Bacteria>Pseudomonadota>Gammaproteobacteria>Pseudomonadales>Pseudomonadaceae | - | Pseudomonas_aeruginosa | EQ623_RS00180 | 30601..33435 | 287 | DUF499-domain-containing-protein-[Pseudomonas-aeruginosa] |
| WP_203168599.1 | DUF499_1.RM.like.systems | SF1-SF2-Helicase+DUF1998->SF1-SF2-Helicase+DUF1998->?-><-?<-DUF499*<-REase<-METHYLASE<-TUDOR+SF1-SF2-Helicase+iREase+CTD+REase<-XIS-HTH | 913 | Bacteria>Pseudomonadota>Gammaproteobacteria>Steroidobacterales>Steroidobacteraceae | GCF_016801985.1 | Steroidobacter_gossypii | JM946_RS17390 | 830297..833038 | 2805490 | DUF499-domain-containing-protein-[Steroidobacter-gossypii] |
| MBM4223246.1 | DUF499_1.RM.like.systems | TUDOR+SF1-SF2-Helicase+iREase+CTD+REase->sGTP->METHYLASE->HEPN+DUF499*->REase-> | 1092 | Bacteria>Pseudomonadota>Gammaproteobacteria>unclassified_Gammaproteobacteria | GCA_016874585.1 | Gammaproteobacteria_bacterium | FJ167_00320 | 8980..12258 | 1913989 | ATP-binding-protein-[Gammaproteobacteria-bacterium] |
| MCP5421179.1 | DUF499_1.RM.like.systems | GDResolvase+HIN-HTH->?->TUDOR+SF1-SF2-Helicase+iREase+CTD+REase->METHYLASE->ParB+HNH->DUF499*->w HTH+REase->SbcC-><-?\|\|BrnT_toxin->BrnA_antitoxin-> | 941 | Bacteria>Pseudomonadota>Gammaproteobacteria>unclassified_Gammaproteobacteria | GCA_024235155.1 | Gammaproteobacteria_bacterium | H6969_11925 | 8981..11806 | 1913989 | ATP-binding-protein-[Gammaproteobacteria-bacterium] |
| GIX22526.1 | DUF499_1.RM.like.systems | Response_reg+LuxR-HTH->URI-DNase+HhH-RADC->?->tRNA->tRNA->tRNA-><-HEPN+DUF499*<-METHYLASE<-REase<-TUDOR+SF1-SF2-Helicase+iREase+CTD+REase-p | 1122 | Bacteria>Pseudomonadota>Gammaproteobacteria>unclassified_Gammaproteobacteria | GCA_026004975.1 | Gammaproteobacteria_bacterium | KatS3mg121_1309 | 525909..529277 | 1913989 | MAG:-hypothetical-protein-KatS3mg121_1309-[Gammaproteobacteria-bacterium] |
| MDE0157838.1 | DUF499_1.RM.like.systems | TUDOR+SF1-SF2-Helicase+iREase+CTD+REase->N6_N4_MTase->?->?->METHYLASE->REase->DUF499*-> | 926 | Bacteria>Pseudomonadota>Gammaproteobacteria>unclassified_Gammaproteobacteria | GCA_028819495.1 | Gammaproteobacteria_bacterium | OXS28_19845 | 41740..44520 | 1913989 | MAG:-DUF499-domain-containing-protein-[Gammaproteobacteria-bacterium] |
| MBS0213168.1 | DUF499_1.RM.like.systems | ZnR+TOPRIM+D5ATPase->TUDOR+SF1-SF2-Helicase+iREase+CTD+REase->SMODS->?->w HTH+REase->METHYLASE->DUF499*->REase->?->ParB+HNH-p-> | 1093 | Bacteria>Pseudomonadota>unclassified_Pseudomonadota | GCA_018242445.1 | Pseudomonadota_bacterium | JSR26_08340 | 14587..17868 | 1977087 | ATP-binding-protein-[Proteobacteria-bacterium] |
| MBU0673024.1 | DUF499_1.RM.like.systems | TUDOR+SF1-SF2-Helicase+iREase+CTD+REase->PIN->?->METHYLASE->OLD-ABC->REase->DUF499*-> | 943 | Bacteria>Pseudomonadota>unclassified_Pseudomonadota | GCA_018814305.1 | Pseudomonadota_bacterium | KJ950_00090 | 19515..22346 | 1977087 | DUF499-domain-containing-protein-[Proteobacteria-bacterium] |
| MBU1208887.1 | DUF499_1.RM.like.systems | TUDOR+SF1-SF2-Helicase+iREase+CTD+REase-p->DrHyd+BirA-HTH->METHYLASE->HEPN+DUF499*->REase-p->?->?->Mbetalac-> | 1107 | Bacteria>Pseudomonadota>unclassified_Pseudomonadota | GCA_018819135.1 | Pseudomonadota_bacterium | KKH04_18500 | 6409..9732 | 1977087 | DUF499-domain-containing-protein-[Proteobacteria-bacterium] |
| MCG8418451.1 | DUF499_1.RM.like.systems | <-DUF499*<-REase<-METHYLASE<-METHYLASE<-TUDOR+SF1-SF2-Helicase+iREase+CTD+REase\|\|Phage_integrase-> | 934 | Bacteria>Pseudomonadota>unclassified_Pseudomonadota | GCA_022360315.1 | Pseudomonadota_bacterium | MJE77_10965 | 1001..3805 | 1977087 | DUF499-domain-containing-protein-[Proteobacteria-bacterium] |
| MCX7113477.1 | DUF499_1.RM.like.systems | <-DUF499*<-?<-METHYLASE<-?<-ORC-CDC6+REase<-TUDOR+SF1-SF2-Helicase+iREase+CTD+REase | 938 | Bacteria>Pseudomonadota>unclassified_Pseudomonadota | GCA_026399135.1 | Pseudomonadota_bacterium | NTX45_25930 | 1011..3827 | 1977087 | MAG:-DUF499-domain-containing-protein-[Proteobacteria-bacterium] |
| MDE3269952.1 | DUF499_1.RM.like.systems | REase-><-TUDOR+SF1-SF2-Helicase+iREase+CTD+REase<-?<-DUF499*<-PIN<-MPTase<-HTH<-?<-METHYLASE | 917 | Bacteria>Pseudomonadota>unclassified_Pseudomonadota | GCA_028877195.1 | Pseudomonadota_bacterium | OYH77_06680 | 5373..8126 | 1977087 | MAG:-hypothetical-protein-OYH77_06680-[Pseudomonadota-bacterium] |
| OGW72082.1 | DUF499_1.RM.like.systems | ParB+HNH->REase->TUDOR+SF1-SF2-Helicase+iREase+CTD+REase->?->METHYLASE->DUF499-p->DUF499*->REase-><-DNA_methylase\|\|MORC+HISKIN->REC-><-Phage_integrase+ZnR | 616 | Bacteria>PVC_group>Candidatus_Omnitrophota>unclassified_Candidatus_Omnitrophota | GCA_001803975.1 | Omnitrophica_bacterium_GWA2_41_15 | A2047_01995 | 16552..18402 | 1801814 | MAG:-hypothetical-protein-A2047_01995-[Omnitrophica-bacterium-GWA2_41_15] |
| MBE6357833.1 | DUF499_1.RM.like.systems | ParB-HTH->?-><-?<-?<-DUF499*<-REase<-OLD-ABC<-METHYLASE<-HNH<-OLD-ABC<-TUDOR+SF1-SF2-Helicase+iREase+CTD+REase | 938 | Bacteria>PVC_group>Lentisphaerota>unclassified_Lentisphaerota | GCA_015061525.1 | Lentisphaerota_bacterium | E7057_01125 | 3177..5993 | 1932692 | ATP-binding-protein-[Lentisphaerae-bacterium] |
| TVL98635.1 | DUF499_1.RM.like.systems | TUDOR+SF1-SF2-Helicase+iREase+CTD+REase->OLD-ABC->METHYLASE->REase->DUF499*-> | 981 | Bacteria>PVC_group>Planctomycetota>Candidatus_Brocadiia>Candidatus_Brocadiales | GCA_007618135.1 | Candidatus_Brocadia_sp._WS118 | CV087_21015 | 12418..15363 | 2040625 | MAG:-AAA-family-ATPase-[Candidatus-Brocadia-sp.-WS118] |
| NUO09865.1 | DUF499_1.RM.like.systems | <-DUF499*<-ORC-CDC6+REase<-METHYLASE<-TUDOR+SF1-SF2-Helicase+iREase+CTD+REase<-Aldedh | 1099 | Bacteria>PVC_group>Planctomycetota>Candidatus_Brocadiia>Candidatus_Brocadiales | GCA_013360945.1 | Candidatus_Brocadia_sp. | HUU08_14505 | 241..3540 | 2172549 | ATP-binding-protein-[Candidatus-Brocadia-sp.] |
| UJS18027.1 | DUF499_1.RM.like.systems | Aldedh->TUDOR+SF1-SF2-Helicase+iREase+CTD+REase->METHYLASE->ORC-CDC6+REase->DUF499*->DUF499-p->SLATT-><-?<-URI-DNase+HhH-RADC | 841 | Bacteria>PVC_group>Planctomycetota>Candidatus_Brocadiia>Candidatus_Brocadiales | GCA_021650895.1 | Candidatus_Jettenia_sp. | L3J17_02955 | 714724..717249 | 2170669 | MAG:-DUF499-domain-containing-protein-[Candidatus-Jettenia-sp.] |
| MBN1125969.1 | DUF499_1.RM.like.systems | Phage_integrase-><-?<-DUF499*<-REase<-METHYLASE<-SF1-SF2-Helicase<-TUDOR+SF1-SF2-Helicase+iREase+CTD+REase\|\|?->PAPS_reduct-> | 912 | Bacteria>PVC_group>Planctomycetota>Phycisphaerae>Sedimentisphaerales | GCA_016926155.1 | Sedimentisphaerales_bacterium | JXA82_13255 | 1576..4314 | 2813034 | ATP-binding-protein-[Sedimentisphaerales-bacterium] |
| MAO24327.1 | DUF499_1.RM.like.systems | TUDOR+SF1-SF2-Helicase+iREase+CTD+REase->TIR->METHYLASE->REase->DUF499*-><-GDResolvase+Recombinase+ZnR<-RAMA | 991 | Bacteria>PVC_group>Planctomycetota>Phycisphaerae>unclassified_Phycisphaerae | GCA_002696165.1 | Phycisphaerae_bacterium | CMJ25_26550 | 12971..15946 | 2026778 | AAA-family-ATPase-[Phycisphaerae-bacterium] |
| MBE0537368.1 | DUF499_1.RM.like.systems | TUDOR+SF1-SF2-Helicase+iREase+CTD+REase->?->REase->DUF499*-> | 1115 | Bacteria>PVC_group>Planctomycetota>Phycisphaerae>unclassified_Phycisphaerae | GCA_014859965.1 | Phycisphaerae_bacterium | IH624_17025 | 11714..15061 | 2026778 | ATP-binding-protein-[Phycisphaerae-bacterium] |
| MBN2560619.1 | DUF499_1.RM.like.systems | <-HEPN+DUF499*<-?<-METHYLASE<-w HTH+REase<-TUDOR+SF1-SF2-Helicase+iREase+CTD+REase<-?<-Cluster182_14clades | 1112 | Bacteria>PVC_group>Planctomycetota>Phycisphaerae>unclassified_Phycisphaerae | GCA_016935885.1 | Phycisphaerae_bacterium | JXQ75_06790 | 1428..4766 | 2026778 | DUF499-domain-containing-protein-[Phycisphaerae-bacterium] |
| MBW7904855.1 | DUF499_1.RM.like.systems | <-HTH_3\|\|DEOR-HTH->TUDOR+SF1-SF2-Helicase+iREase+CTD+REase->Schlafen+HATPase+HTH->REase->METHYLASE->HEPN+DUF499*->Hydrolase_3-> | 1120 | Bacteria>PVC_group>Planctomycetota>Phycisphaerae>unclassified_Phycisphaerae | GCA_019454965.1 | Phycisphaerae_bacterium | H3C42_04715 | 502745..506107 | 2026778 | ATP-binding-protein-[Phycisphaerae-bacterium] |
| MCK6484528.1 | DUF499_1.RM.like.systems | ArdCN+MPTase->DEOR-HTH->TUDOR+SF1-SF2-Helicase+iREase+CTD+REase->w HTH+REase->REase->METHYLASE->HEPN+DUF499*->DNA_methylase->MORC+HISKIN->?->REase->CR-Kinase+ORC-CDC6-like+CARF-> | 1119 | Bacteria>PVC_group>Planctomycetota>Phycisphaerae>unclassified_Phycisphaerae | GCA_023150595.1 | Phycisphaerae_bacterium | L6R00_10395 | 56365..59724 | 2026778 | DUF499-domain-containing-protein-[Phycisphaerae-bacterium] |
| MCI0639191.1 | DUF499_1.RM.like.systems | TUDOR+SF1-SF2-Helicase+iREase+CTD+REase-p->TUDOR+SF1-SF2-Helicase+iREase+CTD+REase->METHYLASE->DUF499*-> | 949 | Bacteria>PVC_group>Planctomycetota>Planctomycetia>Gemmatales | GCA_022616245.1 | Gemmataceae_bacterium | LOY70_08955 | 7374..10223 | 2052164 | DUF499-domain-containing-protein-[Gemmataceae-bacterium] |
| WP_210660196.1 | DUF499_1.RM.like.systems | <-PadR-HTH\|\|?->?-><-?<-DUF499*<-REase<-METHYLASE<-DUF4065<-HD<-TUDOR+SF1-SF2-Helicase+iREase+CTD+REase<-ZnR+TOPRIM+D5ATPase | 985 | Bacteria>PVC_group>Planctomycetota>Planctomycetia>Gemmatales | GCF_017939745.1 | Gemmata_palustris | J8F10_RS30110 | 7260785..7263742 | 2822762 | DUF499-domain-containing-protein-[Gemmata-palustris] |
| WP_076345234.1 | DUF499_1.RM.like.systems | Glyoxalase->?->?->?->?->tRNA-><-XIS-HTH<-DUF499*<-REase<-OLD-ABC<-METHYLASE<-TUDOR+SF1-SF2-Helicase+iREase+CTD+REase<-ParB | 941 | Bacteria>PVC_group>Planctomycetota>Planctomycetia>Isosphaerales | GCF_001956985.1 | Paludisphaera_borealis | BSF38_RS09985 | 2556227..2559052 | 1387353 | DUF499-domain-containing-protein-[Paludisphaera-borealis] |
| WP_250846938.1 | DUF499_1.RM.like.systems | <-DUF499*<-REase<-OLD-ABC<-METHYLASE<-TUDOR+SF1-SF2-Helicase+iREase+CTD+REase | 939 | Bacteria>PVC_group>Planctomycetota>Planctomycetia>Isosphaerales | GCF_011064615.1 | Aquisphaera_insulae | G6054_RS16920 | 61801..64620 | 2712864 | DUF499-domain-containing-protein-[Aquisphaera-sp.-JC669] |
| NLX98932.1 | DUF499_1.RM.like.systems | TUDOR+SF1-SF2-Helicase+iREase+CTD+REase->ParB+HNH->?->METHYLASE->REase->DUF499*->REase->MORC+REase-><-DNA_methylase<-HTH | 983 | Bacteria>PVC_group>Planctomycetota>Planctomycetia>Pirellulales | GCA_012728855.1 | Rhodopirellula_sp. | GXY83_22530 | 8787..11738 | 2024855 | ATP-binding-protein-[Rhodopirellula-sp.] |
| MBN1909337.1 | DUF499_1.RM.like.systems | TUDOR+SF1-SF2-Helicase+iREase+CTD+REase->REase->METHYLASE->?->?->HTH->DUF499*-><-PIN<-HTH+MPTase<-PAPS_reduct<-?\|\|HEPN->HTH-> | 936 | Bacteria>PVC_group>Planctomycetota>Planctomycetia>Pirellulales | GCA_016931075.1 | Pirellulales_bacterium | JW818_06335 | 16839..19649 | 2715253 | ATP-binding-protein-[Pirellulales-bacterium] |
| MBX9790027.1 | DUF499_1.RM.like.systems | Classical-AAA->Peptidase_S8->TUDOR+SF1-SF2-Helicase+iREase+CTD+REase->REase->METHYLASE->?->DUF499*-><-SF1-SF2-Helicase | 926 | Bacteria>PVC_group>Planctomycetota>Planctomycetia>Pirellulales | GCA_019745975.1 | Pirellulales_bacterium | K2Y37_14000 | 59028..61808 | 2715253 | DUF499-domain-containing-protein-[Pirellulales-bacterium] |
| MCE2752488.1 | DUF499_1.RM.like.systems | Methyltransf_21->TUDOR+SF1-SF2-Helicase+iREase+CTD+REase->?->?->METHYLASE->ParB+HNH->DUF499*->REase->ParB+HNH->?->URI-DNase-> | 957 | Bacteria>PVC_group>Planctomycetota>Planctomycetia>Pirellulales | GCA_021299695.1 | Pirellula_sp. | LW720_11440 | 12221..15094 | 117 | ATP-binding-protein-[Pirellula-sp.] |
| MCL4190101.1 | DUF499_1.RM.like.systems | TUDOR+SF1-SF2-Helicase+iREase+CTD+REase->?->METHYLASE->DOC+HTH->w HTH+REase->DUF499*->SbcC->?->?->LZ_Tnp_IS66+zf-IS66+DDE_Tnp_IS66+DDE_Tnp_IS66_C-> | 1115 | Bacteria>PVC_group>Planctomycetota>Planctomycetia>Pirellulales | GCA_023382995.1 | Thermoguttaceae_bacterium | KJZ87_00035 | 12107..15454 | 2691551 | DUF499-domain-containing-protein-[Thermoguttaceae-bacterium] |
| WP_146679086.1 | DUF499_1.RM.like.systems | XIS-HTH->TUDOR+SF1-SF2-Helicase+iREase+CTD+REase->HEPN->METHYLASE->METHYLASE->REase->DUF499*->REase+ParB->RVT->?->RAMA->GDResolvase+Recombinase+ZnR->GDResolvase+Recombinase+ZnR-> | 981 | Bacteria>PVC_group>Planctomycetota>Planctomycetia>Pirellulales | GCF_001610875.1 | Pirellula_sp._SH-Sr6A | VN12_RS23070 | 5951575..5954520 | 1632865 | DUF499-domain-containing-protein-[Pirellula-sp.-SH-Sr6A] |
| WP_146436433.1 | DUF499_1.RM.like.systems | ST-Hiskin->tRNA-><-DUF499*<-REase<-METHYLASE<-PRTase+PRTase<-TUDOR+SF1-SF2-Helicase+iREase+CTD+REase<-XIS-HTH<-ParB | 983 | Bacteria>PVC_group>Planctomycetota>Planctomycetia>Pirellulales | GCF_007859755.1 | Blastopirellula_retiformator | Enr8_RS23480 | 388216..391167 | 2527970 | DUF499-domain-containing-protein-[Blastopirellula-retiformator] |
| WP_230262563.1 | DUF499_1.RM.like.systems | <-RAMA<-URI-DNase+RAMA<-REase<-EVE<-REase<-DUF499*<-REase<-OLD-ABC<-METHYLASE<-Peptidase_S8<-Classical-AAA<-TUDOR+SF1-SF2-Helicase+iREase+CTD+REase | 950 | Bacteria>PVC_group>Planctomycetota>Planctomycetia>Pirellulales | GCF_020966635.1 | Stieleria_sp._JC731 | LOC67_RS10680 | 1300044..1302896 | 2894195 | DUF499-domain-containing-protein-[Stieleria-sp.-JC731] |
| WP_283433169.1 | DUF499_1.RM.like.systems | <-ParB+HNH<-?<-HTH\|\|MORC+DUF3684->HKD-DNase-><-HEPN+DUF499*<-PIN<-OLD-ABC<-METHYLASE<-REase\|\|?-><-TUDOR+SF1-SF2-Helicase+iREase+CTD+REase | 1109 | Bacteria>PVC_group>Planctomycetota>Planctomycetia>Pirellulales | GCF_900182915.1 | Neorhodopirellula_lusitana | QOL80_RS14710 | 91339..94668 | 445327 | DUF499-domain-containing-protein-[Rhodopirellula-lusitana] |
| MBV8313321.1 | DUF499_1.RM.like.systems | <-HEPN+DUF499*<-REase<-METHYLASE<-HicB+ZnR<-TUDOR+SF1-SF2-Helicase+iREase+CTD+REase-p | 939 | Bacteria>PVC_group>Planctomycetota>Planctomycetia>Planctomycetales | GCA_019236145.1 | Planctomycetaceae_bacterium | JOZ53_00025 | 1..2817 | 2026779 | DUF499-domain-containing-protein-[Planctomycetaceae-bacterium] |
| MBV8232626.1 | DUF499_1.RM.like.systems | TUDOR+SF1-SF2-Helicase+iREase+CTD+REase-p->HicB+ZnR->METHYLASE->REase->HEPN+DUF499*->Y1_Tnp-> | 1104 | Bacteria>PVC_group>Planctomycetota>Planctomycetia>Planctomycetales | GCA_019236625.1 | Planctomycetaceae_bacterium | JO329_21805 | 5478..8792 | 2026779 | DUF499-domain-containing-protein-[Planctomycetaceae-bacterium] |
| MBX3448931.1 | DUF499_1.RM.like.systems | TUDOR+SF1-SF2-Helicase+iREase+CTD+REase->OLD-ABC->HNH->METHYLASE->OLD-ABC->REase->DUF499*->?->MORC->?->?-><-wHTH(SBD)+HNH<-DNA_methylase | 948 | Bacteria>PVC_group>Planctomycetota>Planctomycetia>Planctomycetales | GCA_019636285.1 | Planctomycetaceae_bacterium | KF777_05185 | 10482..13328 | 2026779 | ATP-binding-protein-[Planctomycetaceae-bacterium] |
| MCB9950010.1 | DUF499_1.RM.like.systems | <-JAB+UbE1<-?<-?<-MPTase<-?<-SIGMA-HTH<-DUF499*<-REase<-?<-METHYLASE<-SAM-methylase<-TUDOR+SF1-SF2-Helicase+iREase+CTD+REase<-XIS-HTH | 984 | Bacteria>PVC_group>Planctomycetota>Planctomycetia>Planctomycetales | GCA_020638955.1 | Planctomycetaceae_bacterium | H6824_03425 | 868726..871680 | 2026779 | ATP-binding-protein-[Planctomycetaceae-bacterium] |
| WP_145201095.1 | DUF499_1.RM.like.systems | <-JAB+UbE1<-?<-?<-MPTase<-?<-SIGMA-HTH<-DUF499*<-REase<-?<-METHYLASE<-OLD-ABC<-TUDOR+SF1-SF2-Helicase+iREase+CTD+REase<-XIS-HTH | 984 | Bacteria>PVC_group>Planctomycetota>Planctomycetia>Planctomycetales | GCF_007744255.1 | Thalassoglobus_polymorphus | Mal48_RS14940 | 4184383..4187337 | 2527994 | DUF499-domain-containing-protein-[Thalassoglobus-polymorphus] |
| MDA8378650.1 | DUF499_1.RM.like.systems | <-35exo\|\|?->TUDOR+SF1-SF2-Helicase+iREase+CTD+REase->HEPN->METHYLASE->w HTH+REase->DUF499*-> | 945 | Bacteria>PVC_group>Planctomycetota>Planctomycetia>unclassified_Planctomycetia | GCA_028068345.1 | Planctomycetia_bacterium | MOZ50_16785 | 11833..14670 | 2052181 | MAG:-DUF499-domain-containing-protein-[Planctomycetia-bacterium] |
| RIK66096.1 | DUF499_1.RM.like.systems | TUDOR+SF1-SF2-Helicase+iREase+CTD+REase-><-?\|\|?->METHYLASE->MORC->REase->DUF499*-><-?\|\|HTH_3-><-?<-?\|\|HTH_3->MPTase-> | 974 | Bacteria>PVC_group>Planctomycetota>unclassified_Planctomycetota | GCA_003576905.1 | Planctomycetota_bacterium | DCC65_10760 | 38314..41238 | 2026780 | MAG:-AAA-family-ATPase-[Planctomycetota-bacterium] |
| MBE7491676.1 | DUF499_1.RM.like.systems | PAIREDC-HTH+HIN-HTH->?-><-DDE_Tnp_1+DDE_Tnp_1_2<-DUF4096<-?<-DUF499*<-REase<-METHYLASE<-OLD-ABC<-OLD-ABC-p<-TUDOR+SF1-SF2-Helicase+iREase+CTD+REase | 974 | Bacteria>PVC_group>Planctomycetota>unclassified_Planctomycetota | GCA_015075515.1 | Planctomycetota_bacterium | HS108_07990 | 570856..573780 | 2026780 | ATP-binding-protein-[Planctomycetota-bacterium] |
| MBI2930414.1 | DUF499_1.RM.like.systems | <-DUF499*<-METHYLASE<-REase<-TUDOR+SF1-SF2-Helicase+iREase+CTD+REase-p | 940 | Bacteria>PVC_group>Planctomycetota>unclassified_Planctomycetota | GCA_016190155.1 | Planctomycetota_bacterium | HYY16_02075 | 259..3081 | 2026780 | ATP-binding-protein-[Planctomycetota-bacterium] |
| MBI2806086.1 | DUF499_1.RM.like.systems | <-ParB+HNH<-?<-PIWI<-?<-DUF499*<-HTH<-METHYLASE<-REase<-HEPN<-TUDOR+SF1-SF2-Helicase+iREase+CTD+REase\|\|HTH-> | 936 | Bacteria>PVC_group>Planctomycetota>unclassified_Planctomycetota | GCA_016191165.1 | Planctomycetota_bacterium | HYX68_13985 | 123843..126653 | 2026780 | ATP-binding-protein-[Planctomycetota-bacterium] |
| MBI3462535.1 | DUF499_1.RM.like.systems | PIWI-><-ParB+HNH<-HEPN+DUF499*<-METHYLASE<-w HTH+REase<-TUDOR+SF1-SF2-Helicase+iREase+CTD+REase-p | 1111 | Bacteria>PVC_group>Planctomycetota>unclassified_Planctomycetota | GCA_016196255.1 | Planctomycetota_bacterium | HYO00_05665 | 2780..6115 | 2026780 | ATP-binding-protein-[Planctomycetota-bacterium] |
| MBI5364695.1 | DUF499_1.RM.like.systems | Phage_integrase->TUDOR+SF1-SF2-Helicase+iREase+CTD+REase->OLD-ABC->METHYLASE->REase->DUF499*->DNA-glycosylase-> | 936 | Bacteria>PVC_group>Planctomycetota>unclassified_Planctomycetota | GCA_016220035.1 | Planctomycetota_bacterium | HZA53_16080 | 12813..15623 | 2026780 | ATP-binding-protein-[Planctomycetota-bacterium] |
| MBI5763662.1 | DUF499_1.RM.like.systems | <-ParB+N6_N4_MTase<-?<-ZnR+TOPRIM+D5ATPase<-?<-?<-HIN-HTH<-DUF499*<-REase<-?<-METHYLASE<-?\|\|?-><-TUDOR+SF1-SF2-Helicase+iREase+CTD+REase | 921 | Bacteria>PVC_group>Planctomycetota>unclassified_Planctomycetota | GCA_016235815.1 | Planctomycetota_bacterium | HZA51_09085 | 13223..15988 | 2026780 | ATP-binding-protein-[Planctomycetota-bacterium] |
| MBN8597233.1 | DUF499_1.RM.like.systems | TUDOR+SF1-SF2-Helicase+iREase+CTD+REase->?->?->METHYLASE->?->REase->DUF499*-><-?<-?\|\|?->?->RAMA->GDResolvase+Recombinase+ZnR-> | 987 | Bacteria>PVC_group>Planctomycetota>unclassified_Planctomycetota | GCA_017303555.1 | Planctomycetota_bacterium | JOL78_06125 | 35545..38508 | 2026780 | ATP-binding-protein-[Planctomycetota-bacterium] |
| MCE9554597.1 | DUF499_1.RM.like.systems | SMS-KaiC->?->GDResolvase->?-><-SF1-SF2-Helicase-p<-SF1-SF2-Helicase<-DUF499*<-w HTH+REase<-METHYLASE<-DUF4065<-HD<-TUDOR+SF1-SF2-Helicase+iREase+CTD+REase | 950 | Bacteria>PVC_group>Planctomycetota>unclassified_Planctomycetota | GCA_021413705.1 | Planctomycetota_bacterium | K8T91_14665 | 30922..33774 | 2026780 | DUF499-domain-containing-protein-[Planctomycetota-bacterium] |
| MCH8343169.1 | DUF499_1.RM.like.systems | <-METHYLASE\|\|TUDOR+SF1-SF2-Helicase+iREase+CTD+REase->METHYLASE->DUF499*->REase->?->XIS-HTH->ZnR+TOPRIM+D5ATPase->XIS-HTH->HTH-> | 982 | Bacteria>PVC_group>Planctomycetota>unclassified_Planctomycetota | GCA_022568135.1 | Planctomycetota_bacterium | IH983_04190 | 73866..76814 | 2026780 | ATP-binding-protein-[Planctomycetota-bacterium] |
| MCK6472712.1 | DUF499_1.RM.like.systems | TUDOR+SF1-SF2-Helicase+iREase+CTD+REase->?->REase-><-?\|\|?->HEPN+DUF499*->SbcC->?->?->METHYLASE-><-?<-XIS-HTH | 1124 | Bacteria>PVC_group>Planctomycetota>unclassified_Planctomycetota | GCA_023150555.1 | Planctomycetota_bacterium | L6R28_13300 | 20467..23841 | 2026780 | DUF499-domain-containing-protein-[Planctomycetota-bacterium] |
| MCY2991648.1 | DUF499_1.RM.like.systems | <-DUF499*<-PIN<-DUF433<-DUF433<-METHYLASE<-REase<-TUDOR+SF1-SF2-Helicase+iREase+CTD+REase | 983 | Bacteria>PVC_group>Planctomycetota>unclassified_Planctomycetota | GCA_026692885.1 | Planctomycetota_bacterium | NTY19_27810 | 2128..5079 | 2026780 | MAG:-DUF499-domain-containing-protein-[Planctomycetota-bacterium] |
| MCY2978566.1 | DUF499_1.RM.like.systems | <-DUF499*<-REase<-OLD-ABC<-METHYLASE<-TUDOR+SF1-SF2-Helicase+iREase+CTD+REase<-XIS-HTH | 968 | Bacteria>PVC_group>Planctomycetota>unclassified_Planctomycetota | GCA_026692945.1 | Planctomycetota_bacterium | NTU79_07885 | 10024..12930 | 2026780 | MAG:-ATP-binding-protein-[Planctomycetota-bacterium] |
| MCY2928835.1 | DUF499_1.RM.like.systems | HTH_3->MPTase->?->?->?->?-><-DUF499*<-REase<-METHYLASE<-?<-?<-TUDOR+SF1-SF2-Helicase+iREase+CTD+REase\|\|ParB+HNH-> | 970 | Bacteria>PVC_group>Planctomycetota>unclassified_Planctomycetota | GCA_026693065.1 | Planctomycetota_bacterium | NTV86_04950 | 16719..19631 | 2026780 | MAG:-DUF499-domain-containing-protein-[Planctomycetota-bacterium] |
| MBE2203055.1 | DUF499_1.RM.like.systems | <-JAB\|\|?-><-REase<-?<-?<-MORC+HisKin+REase<-DUF499*<-METHYLASE<-TOPRIM<-OLD-ABC<-TUDOR+SF1-SF2-Helicase+iREase+CTD+REase | 938 | Bacteria>PVC_group>Verrucomicrobiota>Spartobacteria>Chthoniobacterales | GCA_014879575.1 | Chthoniobacterales_bacterium | IAE94_01770 | 19431..22247 | 2201154 | ATP-binding-protein-[Chthoniobacterales-bacterium] |
| PWU13198.1 | DUF499_1.RM.like.systems | TUDOR+SF1-SF2-Helicase+iREase+CTD+REase->METHYLASE->REase->OLD-ABC->TOPRIM->DUF499*->?->?->?->?->Cluster1029_5clades-> | 928 | Bacteria>PVC_group>Verrucomicrobiota>unclassified_Verrucomicrobiota | GCA_003176045.1 | Verrucomicrobiota_bacterium | C5B50_19760 | 19033..21819 | 2026799 | AAA-family-ATPase-[Verrucomicrobia-bacterium] |
| MBM3880845.1 | DUF499_1.RM.like.systems | TUDOR+SF1-SF2-Helicase+iREase+CTD+REase->HEPN->ParA-Soj-PloopNTPase->ORC-CDC6->REase->DUF499*-> | 1105 | Bacteria>PVC_group>Verrucomicrobiota>unclassified_Verrucomicrobiota | GCA_016871715.1 | Verrucomicrobiota_bacterium | FJ387_14200 | 23630..26947 | 2026799 | DUF499-domain-containing-protein-[Verrucomicrobiota-bacterium] |
| MBU4201307.1 | DUF499_1.RM.like.systems | <-DUF499*<-METHYLASE<-TOPRIM<-OLD-ABC<-?<-TUDOR+SF1-SF2-Helicase+iREase+CTD+REase<-ORC-CDC6+REase | 941 | Bacteria>PVC_group>Verrucomicrobiota>unclassified_Verrucomicrobiota | GCA_018898455.1 | Verrucomicrobiota_bacterium | KJ919_15475 | 11057..13882 | 2026799 | DUF499-domain-containing-protein-[Verrucomicrobia-bacterium] |
| MCX6879289.1 | DUF499_1.RM.like.systems | <-HISKIN<-DUF499*<-REase<-OLD-ABC<-METHYLASE<-HNH<-OLD-ABC<-TUDOR+SF1-SF2-Helicase+iREase+CTD+REase | 941 | Bacteria>PVC_group>Verrucomicrobiota>unclassified_Verrucomicrobiota | GCA_026397725.1 | Verrucomicrobiota_bacterium | NTW21_36620 | 9837..12662 | 2026799 | MAG:-DUF499-domain-containing-protein-[Verrucomicrobia-bacterium] |
| MCX6953048.1 | DUF499_1.RM.like.systems | TUDOR+SF1-SF2-Helicase+iREase+CTD+REase->OLD-ABC->HNH->METHYLASE->REase->DUF499*->?->?->?->ABhydrolase-> | 941 | Bacteria>PVC_group>Verrucomicrobiota>unclassified_Verrucomicrobiota | GCA_026398065.1 | Verrucomicrobiota_bacterium | NTV51_12915 | 8536..11361 | 2026799 | MAG:-DUF499-domain-containing-protein-[Verrucomicrobia-bacterium] |
| WP_165107642.1 | DUF499_1.RM.like.systems | Cluster1029_5clades-><-?\|\|?-><-SF1-SF2-Helicase\|\|?-><-DUF499*<-REase<-HTH<-REase<-METHYLASE<-?<-TUDOR+SF1-SF2-Helicase+iREase+CTD+REase | 925 | Bacteria>PVC_group>Verrucomicrobiota>Verrucomicrobiae>Limisphaerales | GCF_011044775.1 | Limisphaera_ngatamarikiensis | G4L39_RS09160 | 324178..326955 | 1324935 | DUF499-domain-containing-protein-[Limisphaera-ngatamarikiensis] |
| MCI0539271.1 | DUF499_1.RM.like.systems | <-SF1-SF2-Helicase<-OLD-ABC+TOPRIM\|\|TUDOR+SF1-SF2-Helicase+iREase+CTD+REase->w HTH+REase->METHYLASE->DUF499*->PIWI->REase-> | 1107 | Bacteria>PVC_group>Verrucomicrobiota>Verrucomicrobiae>Verrucomicrobiales | GCA_022615925.1 | Verrucomicrobiales_bacterium | LOZ50_29035 | 12351..15674 | 2026801 | DUF499-domain-containing-protein-[Verrucomicrobiales-bacterium] |
| MCB1193039.1 | DUF499_1.RM.like.systems | <-REase<-PIN<-?<-REase<-REase<-DUF499*<-REase<-METHYLASE<-REase<-TUDOR+SF1-SF2-Helicase+iREase+CTD+REase | 1067 | Bacteria>Spirochaetota>Spirochaetia>Leptospirales>Leptospiraceae | GCA_020440835.1 | Leptospiraceae_bacterium | KDK90_21525 | 1949..5152 | 2026757 | ATP-binding-protein-[Leptospiraceae-bacterium] |
| MCX7774711.1 | DUF499_1.RM.like.systems | ACET->TUDOR+SF1-SF2-Helicase+iREase+CTD+REase->ORC-CDC6+REase->METHYLASE->AbiEi_4->AbiEii->HEPN+DUF499*-> | 1103 | Bacteria>Spirochaetota>Spirochaetia>Spirochaetales>Spirochaetaceae | GCA_026418115.1 | Spirochaetaceae_bacterium | N2067_01230 | 29258..32569 | 1898206 | MAG:-DUF499-domain-containing-protein-[Spirochaetaceae-bacterium] |
| WP_010252910.1 | DUF499_1.RM.like.systems | <-w HTH+REase<-DUF499*<-METHYLASE-p<-METHYLASE<-TUDOR+SF1-SF2-Helicase+iREase+CTD+REase\|\|?->ORC-CDC6+REase->Phage_integrase-> | 929 | Bacteria>Spirochaetota>Spirochaetia>Spirochaetales>Treponemataceae | GCF_000297095.1 | Treponema_primitia | TPRIMZ1_RS0100035 | 3238..6027 | 88058 | DUF499-domain-containing-protein-[Treponema-primitia] |
| HIR18062.1 | DUF499_1.RM.like.systems | <-MNS+REase<-DOC+HTH<-HEPN+DUF499*<-METHYLASE<-TUDOR+SF1-SF2-Helicase+iREase+CTD+REase | 1115 | Bacteria>Synergistota>Synergistia>Synergistales>Synergistaceae | GCA_018712745.1 | Candidatus_Caccocola_faecigallinarum | IAD35_08445 | 169850..173197 | 2840712 | DUF499-domain-containing-protein-[Candidatus-Caccocola-faecigallinarum] |
| GHS93871.1 | DUF499_1.RM.like.systems | TUDOR+SF1-SF2-Helicase+iREase+CTD+REase->HEPN+REase->METHYLASE->OLD-ABC->NERD->?->HEPN+DUF499*->?->?->METHYLASE->?-><-DDE_Tnp_1 | 1116 | Bacteria>Synergistota>Synergistia>Synergistales>unclassified_Synergistales | GCA_025997955.1 | Synergistales_bacterium | AGMMS50276_05310 | 53205..56555 | 2268201 | hypothetical-protein-AGMMS50276_05310-[Synergistales-bacterium] |
| MXZ14251.1 | DUF499_1.RM.like.systems | NTP_transf-><-DUF499*<-METHYLASE<-REase<-TUDOR+SF1-SF2-Helicase+iREase+CTD+REase | 970 | Bacteria>Terrabacteria_group>Actinomycetota>Acidimicrobiia>Acidimicrobiales | GCA_009837945.1 | Acidimicrobiales_bacterium | F4Y76_01880 | 8322..11234 | 2201156 | ATP-binding-protein-[Acidimicrobiales-bacterium] |
| MYD34460.1 | DUF499_1.RM.like.systems | NTP_transf-><-DUF499*<-METHYLASE<-REase<-TUDOR+SF1-SF2-Helicase+iREase+CTD+REase\|\|?-><-Cysteine_desulfurase\|\|Aminotran_3-> | 970 | Bacteria>Terrabacteria_group>Actinomycetota>Acidimicrobiia>Acidimicrobiales | GCA_009841525.1 | Acidimicrobiales_bacterium | F4W98_11365 | 6251..9163 | 2201156 | ATP-binding-protein-[Acidimicrobiales-bacterium] |
| MCY4516058.1 | DUF499_1.RM.like.systems | <-Aminotran_3\|\|Cysteine_desulfurase-><-?\|\|TUDOR+SF1-SF2-Helicase+iREase+CTD+REase->REase->METHYLASE->DUF499*-><-NTP_transf | 966 | Bacteria>Terrabacteria_group>Actinomycetota>Acidimicrobiia>Acidimicrobiales | GCA_026713105.1 | Acidimicrobiaceae_bacterium | OXB99_02335 | 12101..15001 | 2024894 | MAG:-DUF499-domain-containing-protein-[Acidimicrobiaceae-bacterium] |
| MDE0133758.1 | DUF499_1.RM.like.systems | <-Aminotran_3\|\|Cysteine_desulfurase->?->TUDOR+SF1-SF2-Helicase+iREase+CTD+REase->REase->METHYLASE->DUF499*-><-NTP_transf | 970 | Bacteria>Terrabacteria_group>Actinomycetota>Acidimicrobiia>Acidimicrobiales | GCA_028819405.1 | Acidimicrobiaceae_bacterium | OXM54_02860 | 72167..75079 | 2024894 | MAG:-DUF499-domain-containing-protein-[Acidimicrobiaceae-bacterium] |
| MDE0606020.1 | DUF499_1.RM.like.systems | MAD-NTD+HNH-><-MORC+REase<-DNA_methylase<-DUF499*<-METHYLASE<-REase<-TUDOR+SF1-SF2-Helicase+iREase+CTD+REase\|\|DUF222+HNH-><-PIN | 983 | Bacteria>Terrabacteria_group>Actinomycetota>Acidimicrobiia>Acidimicrobiales | GCA_028824095.1 | Acidimicrobiaceae_bacterium | OXH78_02565 | 4211..7162 | 2024894 | MAG:-DUF499-domain-containing-protein-[Acidimicrobiaceae-bacterium] |
| MYB24421.1 | DUF499_1.RM.like.systems | TUDOR+SF1-SF2-Helicase+iREase+CTD+REase->REase->?->?->METHYLASE->DUF499*-> | 1025 | Bacteria>Terrabacteria_group>Actinomycetota>Acidimicrobiia>unclassified_Acidimicrobiia | GCA_009840735.1 | Acidimicrobiia_bacterium | F4X37_04860 | 55279..58356 | 2080302 | DUF499-domain-containing-protein-[Acidimicrobiia-bacterium] |
| MBA2283088.1 | DUF499_1.RM.like.systems | Phage_int_SAM_1+Phage_integrase-><-NTD+SF1-SF2-Helicase\|\|TUDOR+SF1-SF2-Helicase+iREase+CTD+REase->?->METHYLASE->?->HEPN+DUF499*->REase->?-><-?<-XIS-HTH\|\|XIS-HTH-> | 1110 | Bacteria>Terrabacteria_group>Actinomycetota>Acidimicrobiia>unclassified_Acidimicrobiia | GCA_013694495.1 | Acidimicrobiia_bacterium | HOW25_17905 | 18303..21635 | 2080302 | ATP-binding-protein-[Acidimicrobiia-bacterium] |
| WP_128773202.1 | DUF499_1.RM.like.systems | SF1-SF2-Helicase+REase-><-?<-TrwC+SF1-SF2-Helicase\|\|DNA_methylase-><-HEPN+REase<-REase<-HEPN+DUF499*<-w HTH+REase<-METHYLASE<-TUDOR+SF1-SF2-Helicase+iREase+CTD+REase<-XIS-HTH<-ABhydrolase | 1132 | Bacteria>Terrabacteria_group>Actinomycetota>Actinomycetes>Actinomycetales | GCF_004104015.1 | Actinomyces_oricola | EUC46_RS05555 | 19810..23208 | 206043 | DUF499-domain-containing-protein-[Actinomyces-oricola] |
| WP_182050741.1 | DUF499_1.RM.like.systems | SF1-SF2-Helicase->TUDOR+SF1-SF2-Helicase+iREase+CTD+REase->METHYLASE->REase->HEPN->HEPN+DUF499*->SbcC-> | 1134 | Bacteria>Terrabacteria_group>Actinomycetota>Actinomycetes>Actinomycetales | GCF_014041995.1 | Changpingibacter_yushuensis | H2017_RS05160 | 1185005..1188409 | 2758440 | Swt1-family-HEPN-domain-containing-protein-[Changpingibacter-yushuensis] |
| WP_150353887.1 | DUF499_1.RM.like.systems | REase-p->TUDOR+SF1-SF2-Helicase+iREase+CTD+REase->METHYLASE-p->METHYLASE->HEPN+DUF499*->Primpol+D5N+D5ATPase-><-?<-Cysteine_desulfurase<-?<-ABC-ATPase+SbcC | 1134 | Bacteria>Terrabacteria_group>Actinomycetota>Actinomycetes>Bifidobacteriales | GCF_008698145.1 | Bifidobacterium_vespertilionis | EMO90_RS02025 | 149240..152644 | 2562524 | Swt1-family-HEPN-domain-containing-protein-[Bifidobacterium-vespertilionis] |
| WP_214383368.1 | DUF499_1.RM.like.systems | ABC-ATPase+SbcC->?->Cysteine_desulfurase->?-><-Primpol+D5N+D5ATPase<-HEPN+DUF499*<-METHYLASE<-METHYLASE-p<-TUDOR+SF1-SF2-Helicase+iREase+CTD+REase<-REase-p | 1134 | Bacteria>Terrabacteria_group>Actinomycetota>Actinomycetes>Bifidobacteriales | GCF_018555535.1 | Bifidobacterium_vespertilionis | JS538_RS07550 | 34017..37421 | 2562524 | Swt1-family-HEPN-domain-containing-protein-[Bifidobacterium-vespertilionis] |
| WP_009738155.1 | DUF499_1.RM.like.systems | TUDOR+SF1-SF2-Helicase+iREase+CTD+REase->UDG-conflict->?->SF1-SF2-Helicase+REase->METHYLASE->HEPN+DUF499*-><-ParA+HTH<-RelE-ParE\|\|MULE-TRANSPOSASE->MULE-TRANSPOSASE->?-><-TPR+TPR+TPR+TPR+TPR+TPR+TPR+TPR | 1119 | Bacteria>Terrabacteria_group>Actinomycetota>Actinomycetes>Frankiales | GCF_000262465.1 | Frankia_sp._QA3 | FRAQA3DRAFT_RS05925 | 1493291..1496650 | 710111 | Swt1-family-HEPN-domain-containing-protein-[Frankia-sp.-QA3] |
| WP_189449183.1 | DUF499_1.RM.like.systems | VirD4-TraG+VirD4-FtsK->DNA_methylase-><-HNH<-ABC-ATPase<-DUF499*<-METHYLASE<-TUDOR+SF1-SF2-Helicase+iREase+CTD+REase<-REase+DUF5655<-TRAM+Methyltransf_31 | 922 | Bacteria>Terrabacteria_group>Actinomycetota>Actinomycetes>Kitasatosporales | GCF_014648775.1 | Streptomyces_abikoensis | IE096_RS06315 | 1448410..1451178 | 97398 | DUF499-domain-containing-protein-[Streptomyces-abikoensis] |
| WP_167559767.1 | DUF499_1.RM.like.systems | <-HEPN+REase<-REase<-HEPN+DUF499*<-w HTH+REase<-METHYLASE<-TUDOR+SF1-SF2-Helicase+iREase+CTD+REase<-XIS-HTH\|\|MORC+HisKin+REase-><-TrwC+SF1-SF2-helicase | 1132 | Bacteria>Terrabacteria_group>Actinomycetota>Actinomycetes>Micrococcales | - | Brevibacterium_sp._LS14 | EZE58_RS02710 | 16654..20052 | 2528962 | DUF499-domain-containing-protein-[Brevibacterium-sp.-LS14] |
| MCE1178998.1 | DUF499_1.RM.like.systems | ABhydrolase->XIS-HTH->TUDOR+SF1-SF2-Helicase+iREase+CTD+REase->METHYLASE->w HTH+REase->HEPN+DUF499*->REase->?->?-><-DNA_methylase\|\|MULE-TRANSPOSASE-> | 1133 | Bacteria>Terrabacteria_group>Actinomycetota>Actinomycetes>Micrococcales | GCA_021287075.1 | Micrococcales_bacterium | LWW86_08210 | 123178..126579 | 2026762 | DUF499-domain-containing-protein-[Micrococcales-bacterium] |

| Protein-ID | System | Gene-Neighbourhoods | Length | Taxonomy | Assembly-ID | Strain | Locus-Tag | Genomic-coordinates | TaxID | Fasta-define |
|---|---|---|---|---|---|---|---|---|---|---|
| WP_029263985.1 | DUF499_1.RM.like.systems | <-Radical-SAM\|\|?-><-SF1-SF2-Helicase<- REase <- HEPN+ DUF499 *<- METHYLASE <- TUDOR+SF1-SF2-Helicase+i REase +CTD+ REase \|\|?-><-MerR-HTH | 1128 | Bacteria>Terrabacteria_group>Actinomycetota>Actinomycetes>Micrococcales | GCF_000708325.1 | unclassified_Microbacterium | DW303_RS18520 | 62853..66239 | 2609290 | MULTISPECIES:-DUF499-domain-containing-protein-[unclassified-Microbacterium] |
| WP_047687850.1 | DUF499_1.RM.like.systems | DNA_methylase-><-HEPN+ REase <- REase <- HEPN+ DUF499 *<- w HTH+ REase <- METHYLASE <- TUDOR+SF1-SF2-Helicase+i REase +CTD+ REase | 1133 | Bacteria>Terrabacteria_group>Actinomycetota>Actinomycetes>Micrococcales | GCF_000800955.1 | Kocuria_sp._ZOR0020 | L979_RS00020 | 3003..6404 | 1339234 | DUF499-domain-containing-protein-[Kocuria-sp.-ZOR0020] |
| WP_063250773.1 | DUF499_1.RM.like.systems | <-HEPN+ REase <- REase <- HEPN+ DUF499 *<- w HTH+ REase <- METHYLASE <- TUDOR+SF1-SF2-Helicase+i REase +CTD+ REase <-XIS-HTH\|\|MORC+HisKin+ REase -><-TrwC+SF1-SF2-helicase | 1132 | Bacteria>Terrabacteria_group>Actinomycetota>Actinomycetes>Micrococcales | GCF_001619685.1 | Brevibacterium_casei | AVW13_RS15020 | 17875..21273 | 33889 | DUF499-domain-containing-protein-[Brevibacterium-casei] |
| WP_112235018.1 | DUF499_1.RM.like.systems | <-MORC+HisKin+ REase \|\|XIS-HTH-> TUDOR+SF1-SF2-Helicase+i REase +CTD+ REase -> METHYLASE -> w HTH+ REase -> HEPN+ DUF499 *->OLD-ABC+TOPRIM->SF1-SF2-Helicase-> | 1132 | Bacteria>Terrabacteria_group>Actinomycetota>Actinomycetes>Micrococcales | GCF_003290245.1 | Kocuria_sp._BT304 | DNL27_RS06220 | 1382048..1385446 | 1702043 | DUF499-domain-containing-protein-[Kocuria-sp.-BT304] |
| WP_208008530.1 | DUF499_1.RM.like.systems | <-MULE-TRANSPOSASE\|\|NTD+SF1-SF2-Helicase-><- HEPN+ DUF499 *<- REase <- METHYLASE <-Phage_holin<-?<-?<- TUDOR+SF1-SF2-Helicase+i REase +CTD+ REase | 1101 | Bacteria>Terrabacteria_group>Actinomycetota>Actinomycetes>Micrococcales | GCF_004135405.1 | Agromyces_protaetiae | ET445_RS03710 | 798698..802003 | 2509455 | Swt1-family-HEPN-domain-containing-protein-[Agromyces-protaetiae] |
| WP_137123200.1 | DUF499_1.RM.like.systems | ABhydrolase->XIS-HTH-> TUDOR+SF1-SF2-Helicase+i REase +CTD+ REase -> METHYLASE -> w HTH+ REase -> HEPN+ DUF499 *-> REase ->HEPN+ REase -><-DNA_methylase\|\|TrwC+SF1-SF2-Helicase->?->SMODS-> | 1132 | Bacteria>Terrabacteria_group>Actinomycetota>Actinomycetes>Micrococcales | GCF_005222685.1 | Ornithinicoccus_soli | FCN22_RS19310 | 12208..15606 | 2574732 | DUF499-domain-containing-protein-[Ornithinicoccus-soli] |
| WP_137320273.1 | DUF499_1.RM.like.systems | <- HEPN+ DUF499 *<- w HTH+ REase <- METHYLASE <- TUDOR+SF1-SF2-Helicase+i REase +CTD+ REase <-XIS-HTH<-ABhydrolase | 1132 | Bacteria>Terrabacteria_group>Actinomycetota>Actinomycetes>Micrococcales | GCF_005280295.1 | Brevibacterium_sp._CS2 | FDF13_RS00005 | 3239176..2834 | 2575923 | DUF499-domain-containing-protein-[Brevibacterium-sp.-CS2] |
| WP_137369040.1 | DUF499_1.RM.like.systems | <-NTD+SF1-SF2-Helicase\|\| TUDOR+SF1-SF2-Helicase+i REase +CTD+ REase -> METHYLASE -> HEPN+ DUF499 *-> REase +CTD6->?->HNH-><-URI-DNase\|\|URI-DNase-><-JetD | 1129 | Bacteria>Terrabacteria_group>Actinomycetota>Actinomycetes>Micrococcales | GCF_005281365.1 | Arthrobacter_sp._NamB2 | FDK12_RS07770 | 280727..284116 | 2576035 | Swt1-family-HEPN-domain-containing-protein-[Arthrobacter-sp.-NamB2] |
| WP_141842659.1 | DUF499_1.RM.like.systems | <-HIN-HTH<-SF1-SF2-Helicase<-OLD-ABC+TOPRIM<- HEPN+ DUF499 *<- w HTH+ REase <- METHYLASE <- TUDOR+SF1-SF2-Helicase+i REase +CTD+ REase <-XIS-HTH<-PAIREDC-HTH<-HIN-HTH | 1132 | Bacteria>Terrabacteria_group>Actinomycetota>Actinomycetes>Micrococcales | GCF_006704135.1 | Micrococcus_sp._R8502A1 | FLM69_RS00645 | 3870..7268 | 2583239 | Swt1-family-HEPN-domain-containing-protein-[Micrococcus-sp.-R8502A1] |
| WP_144831472.1 | DUF499_1.RM.like.systems | ACET-><-?<-SF1-SF2-Helicase<-OLD-ABC+TOPRIM<- HEPN+ DUF499 *<- w HTH+ REase <- METHYLASE <- TUDOR+SF1-SF2-Helicase+i REase +CTD+ REase <-XIS-HTH\|\|MORC+HisKin+ REase -><-TrwC+SF1-SF2-helicase | 1133 | Bacteria>Terrabacteria_group>Actinomycetota>Actinomycetes>Micrococcales | GCF_007677595.1 | Kocuria_rhizophila | FS577_RS10180 | 96606..100007 | 72000 | DUF499-domain-containing-protein-[Kocuria-rhizophila] |
| WP_168888196.1 | DUF499_1.RM.like.systems | CopG_antitoxin-><-?<- HEPN+ DUF499 *<- w HTH+ REase <- METHYLASE <- TUDOR+SF1-SF2-Helicase+i REase +CTD+ REase | 1127 | Bacteria>Terrabacteria_group>Actinomycetota>Actinomycetes>Micrococcales | GCF_012641515.1 | Nesterenkonia_sedimenti | HGQ17_RS12070 | 43839..47222 | 1463632 | DUF499-domain-containing-protein-[Nesterenkonia-sedimenti] |
| WP_188034675.1 | DUF499_1.RM.like.systems | MPTase->?->ABhydrolase->XIS-HTH-> TUDOR+SF1-SF2-Helicase+i REase +CTD+ REase -> METHYLASE -> HEPN+ DUF499 *-> REase +SAVED->?->TrwC+SF1-SF2-Helicase->?->?-><-HTH+MPTase | 1143 | Bacteria>Terrabacteria_group>Actinomycetota>Actinomycetes>Micrococcales | GCF_014530035.1 | unclassified_Leucobacter | KOE72_RS09585 | 56127..59558 | 2621730 | MULTISPECIES:-DUF499-domain-containing-protein-[unclassified-Leucobacter] |
| WP_201671845.1 | DUF499_1.RM.like.systems | <-HEPN+ REase <- REase <- HEPN+ DUF499 *<- w HTH+ REase <- METHYLASE -p<- METHYLASE -p<- TUDOR+SF1-SF2-Helicase+i REase +CTD+ REase <-XIS-HTH\|\|MORC+HisKin+ REase -> | 1132 | Bacteria>Terrabacteria_group>Actinomycetota>Actinomycetes>Micrococcales | GCF_016726245.1 | Brevibacterium_casei | I6I57_RS03275 | 754914..758312 | 33889 | DUF499-domain-containing-protein-[Brevibacterium-casei] |
| WP_204409507.1 | DUF499_1.RM.like.systems | <-HIN-HTH<-OLD-ABC+TOPRIM<- HEPN+ DUF499 *<- w HTH+ REase <- METHYLASE <- TUDOR+SF1-SF2-Helicase+i REase +CTD+ REase <-XIS-HTH | 1132 | Bacteria>Terrabacteria_group>Actinomycetota>Actinomycetes>Micrococcales | GCF_016907455.1 | Brachybacterium_muris | JOD52_RS08675 | 1871612..1875010 | 219301 | DUF499-domain-containing-protein-[Brachybacterium-muris] |
| WP_210233975.1 | DUF499_1.RM.like.systems | TUDOR+SF1-SF2-Helicase+i REase +CTD+ REase -> METHYLASE -> REase -> HEPN+ DUF499 *-><- REase <-HKD-DNase+SF1-SF2-Helicase+TRD\|\|DndB-> | 1116 | Bacteria>Terrabacteria_group>Actinomycetota>Actinomycetes>Micrococcales | GCF_017898105.1 | Arthrobacter_jiangjiafuii | J2M53_RS14475 | 604648..607998 | 2817475 | Swt1-family-HEPN-domain-containing-protein-[Arthrobacter-jiangjiafuii] |
| WP_236803031.1 | DUF499_1.RM.like.systems | <-ABC-ATPase+SbcC<-?<- HEPN+ DUF499 *<- METHYLASE <- TUDOR+SF1-SF2-Helicase+i REase +CTD+ REase <-XIS-HTH\|\| REase -> | 1127 | Bacteria>Terrabacteria_group>Actinomycetota>Actinomycetes>Micrococcales | GCF_021801725.1 | Arthrobacter_sp._FW306-05-C | LFT44_RS20985 | 51567..54950 | 2879620 | Swt1-family-HEPN-domain-containing-protein-[Arthrobacter-sp.-FW306-05-C] |
| WP_247619184.1 | DUF499_1.RM.like.systems | <-OLD-ABC+TOPRIM<- HEPN+ DUF499 *<- w HTH+ REase <- METHYLASE <- TUDOR+SF1-SF2-Helicase+i REase +CTD+ REase <-XIS-HTH | 1135 | Bacteria>Terrabacteria_group>Actinomycetota>Actinomycetes>Micrococcales | GCF_023072605.1 | Brevibacterium_sp._R8603A2 | MTQ12_RS14500 | 1781..5188 | 2929779 | Swt1-family-HEPN-domain-containing-protein-[Brevibacterium-sp.-R8603A2] |
| WP_259878499.1 | DUF499_1.RM.like.systems | <-Peptidase<-?<-?<-?<- HEPN+ DUF499 *<- w HTH+ REase <- METHYLASE <- TUDOR+SF1-SF2-Helicase+i REase +CTD+ REase <-XIS-HTH<-ABhydrolase\|\|DEDD_Tnp_IS110+Transposase_20-> | 1132 | Bacteria>Terrabacteria_group>Actinomycetota>Actinomycetes>Micrococcales | GCF_025148185.1 | Brachybacterium_muris | M3C58_RS02745 | 15014..18412 | 219301 | DUF499-domain-containing-protein-[Brachybacterium-muris] |
| WP_285140308.1 | DUF499_1.RM.like.systems | <-HTH_Tnp_1+HTH_21+rve+rve_3<-HIN-HTH\|\|XIS-HTH-> TUDOR+SF1-SF2-Helicase+i REase +CTD+ REase -> METHYLASE -> w HTH+ REase -> HEPN+ DUF499 *->OLD-ABC+TOPRIM->SF1-SF2-Helicase-><-DNA_methylase | 1132 | Bacteria>Terrabacteria_group>Actinomycetota>Actinomycetes>Micrococcales | GCF_030226745.1 | Micrococcus_luteus | QP726_RS10470 | 9707..13105 | 1270 | Swt1-family-HEPN-domain-containing-protein-[Micrococcus-luteus] |
| WP_074694547.1 | DUF499_1.RM.like.systems | HEPN+SF1-SF2-Helicase->35exo-><-?<-?<-?<-HEPN+ REase <- HEPN+ DUF499 *<- METHYLASE <-Mbetalac<- TUDOR+SF1-SF2-Helicase+i REase +CTD+ REase <-HTH | 1131 | Bacteria>Terrabacteria_group>Actinomycetota>Actinomycetes>Micrococcales | GCF_900104345.1 | Microbacterium_testaceum | BLS25_RS04240 | 198218..201613 | 2033 | DUF499-domain-containing-protein-[Microbacterium-testaceum] |
| PWU48923.1 | DUF499_1.RM.like.systems | <-HTH_31<-?<- DUF499 *<- METHYLASE <- TUDOR+SF1-SF2-Helicase+i REase +CTD+ REase <- REase <-?\|\| REase -> | 962 | Bacteria>Terrabacteria_group>Actinomycetota>Actinomycetes>Micromonosporales | GCA_003176735.1 | Micromonospora_globispora | DLJ46_10740 | 15153..18041 | 1450148 | AAA-family-ATPase-[Micromonospora-globispora] |
| MBN1173560.1 | DUF499_1.RM.like.systems | <-Exonuc_V_gamma+ REase \|\| TUDOR+SF1-SF2-Helicase+i REase +CTD+ REase -> METHYLASE -> HEPN+ DUF499 *-> REase -><-?\|\|?->Cas_Cmr3-> | 1155 | Bacteria>Terrabacteria_group>Actinomycetota>Actinomycetes>Micromonosporales | GCA_016926535.1 | Micromonosporaceae_bacterium | JXA67_15425 | 10595..14062 | 1873464 | DUF499-domain-containing-protein-[Micromonosporaceae-bacterium] |
| WP_204020362.1 | DUF499_1.RM.like.systems | NUDIX->?-><-?<- REase <-ParB+HNH<- HEPN+ DUF499 *<- METHYLASE <- TUDOR+SF1-SF2-Helicase+i REase +CTD+ REase \|\|RelE-ParE->ParA+HTH-><-PIN<-Phd-YefM | 1129 | Bacteria>Terrabacteria_group>Actinomycetota>Actinomycetes>Micromonosporales | GCF_016863595.1 | Micromonospora_andamanensis | Vwe01_RS03465 | 15617..19006 | 1287068 | Swt1-family-HEPN-domain-containing-protein-[Micromonospora-trujilloniae] |
| WP_259918390.1 | DUF499_1.RM.like.systems | <-ABhydrolase<-?<-Aminotran_1_2<-ncRNA\|\|tRNA->?->SF1-SF2-helicase+DUF1998-><-ORC-CDC6+ REase <- HEPN+ DUF499 *<- METHYLASE <- TUDOR+SF1-SF2-Helicase+i REase +CTD+ REase \|\|ABhydrolase-> | 1108 | Bacteria>Terrabacteria_group>Actinomycetota>Actinomycetes>Mycobacteriales | - | Corynebacterium_sanguinis | M3A76_RS07775 | 66134..69460 | 2594913 | DUF499-domain-containing-protein-[Corynebacterium-sanguinis] |
| ADK68968.1 | DUF499_1.RM.like.systems | TUDOR+SF1-SF2-Helicase+i REase +CTD+ REase ->NERD->?-> METHYLASE -> HEPN+ DUF499 *-> REase +TPR->?->Peptidase_M48-> | 1120 | Bacteria>Terrabacteria_group>Actinomycetota>Actinomycetes>Mycobacteriales | GCA_000143885.2 | Gordonia_sp._KTR9 | KTR9_4887 | 112287..115649 | 337191 | Predicted-ATPase-(AAA+-superfamily)-[Gordonia-sp.-KTR9] |
| EHB57971.1 | DUF499_1.RM.like.systems | <-GNTR-HTH\|\|NUDIX-><-?<-HEPN<-Schlafen+HATPase+HTH<- HEPN+ DUF499 *<- REase <- METHYLASE <- TUDOR+SF1-SF2-Helicase+i REase +CTD+ REase <-PIN<-XIS-HTH | 1108 | Bacteria>Terrabacteria_group>Actinomycetota>Actinomycetes>Mycobacteriales | GCA_000230935.2 | Mycolicibacterium_rhodesiae_JS60 | MycrhDRAFT_0407 | 390606..393932 | 931627 | hypothetical-protein-MycrhDRAFT_0407-[Mycolicibacterium-rhodesiae-JS60] |
| PZT93668.1 | DUF499_1.RM.like.systems | ABhydrolase->XIS-HTH-> TUDOR+SF1-SF2-Helicase+i REase +CTD+ REase -> METHYLASE -> w HTH+ REase -> HEPN+ DUF499 *->OLD-ABC+TOPRIM->SF1-SF2-Helicase-> | 1133 | Bacteria>Terrabacteria_group>Actinomycetota>Actinomycetes>Mycobacteriales | GCA_003248405.1 | Gordonia_sp._(in:_high_G+C_Gram-positive_bacteria) | DI630_26685 | 9643..13044 | 84139 | AAA-family-ATPase-[Gordonia-sp.] |
| MBU8833314.1 | DUF499_1.RM.like.systems | McrC-NTD+ REase -><-?<-HTH_Tnp_1+HTH_21+rve+rve_3\|\|HTH_Tnp_1+HTH_21+rve+rve_3->?-><- REase <- HEPN+ DUF499 *<-ParB+HNH<- METHYLASE <- TUDOR+SF1-SF2-Helicase+i REase +CTD+ REase \|\|DDE_Tnp_1_4->?->HEPN+SF1-SF2-Helicase-> | 1119 | Bacteria>Terrabacteria_group>Actinomycetota>Actinomycetes>Mycobacteriales | GCA_018524295.1 | Mycolicibacterium_goodii | KL858_28165 | 17594..20953 | 134601 | DUF499-domain-containing-protein-[Mycobacterium-goodii] |
| MCT1882934.1 | DUF499_1.RM.like.systems | <-ABhydrolase<-?<-Aminotran_1_2<-ncRNA\|\|tRNA->?->SF1-SF2-helicase+DUF1998-><-ORC-CDC6+ REase <- HEPN+ DUF499 *<- METHYLASE <- TUDOR+SF1-SF2-Helicase+i REase +CTD+ REase \|\|ABhydrolase-> | 1081 | Bacteria>Terrabacteria_group>Actinomycetota>Actinomycetes>Mycobacteriales | GCA_025144605.1 | Corynebacterium_sanguinis | M3A76_07760 | 66134..69379 | 2594913 | DUF499-domain-containing-protein-[Corynebacterium-sanguinis] |
| SBS79264.1 | DUF499_1.RM.like.systems | TUDOR+SF1-SF2-Helicase+i REase +CTD+ REase -> REase -> METHYLASE -> DUF499 *-> | 986 | Bacteria>Terrabacteria_group>Actinomycetota>Actinomycetes>Mycobacteriales | GCA_900078375.1 | uncultured_Mycobacterium_sp. | MHPYR_700004 | 5438..8398 | 171292 | AAA-family-ATPase-[uncultured-Mycobacterium-sp.] |
| WP_044505526.1 | DUF499_1.RM.like.systems | TUDOR+SF1-SF2-Helicase+i REase +CTD+ REase ->NERD-> METHYLASE -> HEPN+ DUF499 *-> REase +TPR->?->Peptidase_M48-> | 1115 | Bacteria>Terrabacteria_group>Actinomycetota>Actinomycetes>Mycobacteriales | GCF_000143885.2 | Gordonia | KTR9_RS00485 | 112302..115649 | 2053 | MULTISPECIES:-DUF499-domain-containing-protein-[Gordonia] |
| WP_013827697.1 | DUF499_1.RM.like.systems | GDResolvase+HIN-HTH-><- REase <- HEPN+ DUF499 *<- METHYLASE <- TUDOR+SF1-SF2-Helicase+i REase +CTD+ REase <-XIS-HTH\|\|ParA+HTH->HTH_58-> | 1138 | Bacteria>Terrabacteria_group>Actinomycetota>Actinomycetes>Mycobacteriales | GCF_000214155.1 | Mycolicibacter_sinensis | JDM601_RS03755 | 802521..805937 | 875328 | DUF499-domain-containing-protein-[Mycolicibacter-sinensis] |
| WP_065288965.1 | DUF499_1.RM.like.systems | <-HTH_58<-ParA+HTH\|\|BirA-HTH+CRP-HTH-> TUDOR+SF1-SF2-Helicase+i REase +CTD+ REase -> METHYLASE -> HEPN+ DUF499 *-> REase -><-GDResolvase+HIN-HTH<-?<-?\|\|HTH_3-> | 1141 | Bacteria>Terrabacteria_group>Actinomycetota>Actinomycetes>Mycobacteriales | GCF_001679965.1 | Mycolicibacter_kumamotonensis | ACT18_RS17285 | 28680..32105 | 354243 | DUF499-domain-containing-protein-[Mycolicibacter-kumamotonensis] |
| WP_201263267.1 | DUF499_1.RM.like.systems | <-NTD+SF1-SF2-Helicase\|\| TUDOR+SF1-SF2-Helicase+i REase +CTD+ REase -> METHYLASE -> HEPN+ DUF499 *-> REase ->SF1-SF2-Helicase-p->HTH_Tnp_1+HTH_21+rve+rve_2->SF1-SF2-Helicase-p-> | 1109 | Bacteria>Terrabacteria_group>Actinomycetota>Actinomycetes>Mycobacteriales | GCF_002201735.1 | Mycobacterium_avium | CEV14_RS17015 | 150524..153853 | 1764 | Swt1-family-HEPN-domain-containing-protein-[Mycobacterium-avium] |
| WP_108404986.1 | DUF499_1.RM.like.systems | GDSL->Schlafen-><-?\|\| TUDOR+SF1-SF2-Helicase+i REase +CTD+ REase -> METHYLASE -> HEPN+ DUF499 *-> REase -p-><-tRNA\|\|ncRNA->Aminotran_1_2->?->GDSL-> | 1112 | Bacteria>Terrabacteria_group>Actinomycetota>Actinomycetes>Mycobacteriales | GCF_003070865.1 | Corynebacterium_liangguodongii | C3E79_RS00685 | 144858..148196 | 2079535 | DUF499-domain-containing-protein-[Corynebacterium-liangguodongii] |
| WP_144317543.1 | DUF499_1.RM.like.systems | <-ABhydrolase<-?<-Aminotran_1_2<-ncRNA\|\|tRNA-><-ORC-CDC6+ REase <- HEPN+ DUF499 *<- METHYLASE <- TUDOR+SF1-SF2-Helicase+i REase +CTD+ REase \|\|ABhydrolase-> | 1111 | Bacteria>Terrabacteria_group>Actinomycetota>Actinomycetes>Mycobacteriales | GCF_007641235.1 | Corynebacterium_sanguinis | E3227_RS03440 | 690306..693641 | 2594913 | DUF499-domain-containing-protein-[Corynebacterium-sanguinis] |
| WP_149357518.1 | DUF499_1.RM.like.systems | Lsr2-><-?<-?\|\| TUDOR+SF1-SF2-Helicase+i REase +CTD+ REase -> METHYLASE -> HEPN+ DUF499 *-> REase -><-?<-?\|\|GDResolvase+HIN-HTH->URI-DNase-> | 1133 | Bacteria>Terrabacteria_group>Actinomycetota>Actinomycetes>Mycobacteriales | GCF_008306195.1 | Rhodococcus_qingshengii | D6M20_RS01760 | 367041..370442 | 334542 | DUF499-domain-containing-protein-[Rhodococcus-qingshengii] |
| WP_149772783.1 | DUF499_1.RM.like.systems | <-HTH_58<-ParA+HTH\|\|XIS-HTH-> TUDOR+SF1-SF2-Helicase+i REase +CTD+ REase -> METHYLASE -> HEPN+ DUF499 *-> REase -><-GDResolvase+HIN-HTH | 1138 | Bacteria>Terrabacteria_group>Actinomycetota>Actinomycetes>Mycobacteriales | GCF_008373105.1 | Mycolicibacter_arupensis | F0402_RS09270 | 8758..12174 | 342002 | DUF499-domain-containing-protein-[Mycolicibacter-arupensis] |
| WP_239342823.1 | DUF499_1.RM.like.systems | <-NTD+SF1-SF2-Helicase\|\| TUDOR+SF1-SF2-Helicase+i REase +CTD+ REase -> METHYLASE ->ParB+HNH-> HEPN+ DUF499 *-> REase ->SF1-SF2-Helicase-> | 1109 | Bacteria>Terrabacteria_group>Actinomycetota>Actinomycetes>Mycobacteriales | GCF_022348005.1 | Mycolicibacterium_sp._OfavD-34-C | MHN82_RS23245 | 18852..22181 | 2917746 | Swt1-family-HEPN-domain-containing-protein-[Mycolicibacterium-sp.-OfavD-34-C] |
| WP_259916185.1 | DUF499_1.RM.like.systems | <-ABhydrolase\|\| TUDOR+SF1-SF2-Helicase+i REase +CTD+ REase -> METHYLASE -> HEPN+ DUF499 *->ORC-CDC6+ REase -><-tRNA\|\|ncRNA->Aminotran_1_2->?->ABhydrolase-> | 1111 | Bacteria>Terrabacteria_group>Actinomycetota>Actinomycetes>Mycobacteriales | GCF_025144095.1 | Corynebacterium_sanguinis | M3G11_RS01890 | 80632..83967 | 2594913 | DUF499-domain-containing-protein-[Corynebacterium-sanguinis] |
| WP_259795206.1 | DUF499_1.RM.like.systems | <-ABhydrolase\|\| TUDOR+SF1-SF2-Helicase+i REase +CTD+ REase -> METHYLASE -> HEPN+ DUF499 *->ORC-CDC6+ REase -> | 1112 | Bacteria>Terrabacteria_group>Actinomycetota>Actinomycetes>Mycobacteriales | GCF_025144435.1 | Corynebacterium_sanguinis | M3G32_RS10715 | 21189..24527 | 2594913 | DUF499-domain-containing-protein-[Corynebacterium-sanguinis] |
| WP_259896744.1 | DUF499_1.RM.like.systems | <-ABhydrolase\|\| TUDOR+SF1-SF2-Helicase+i REase +CTD+ REase -> METHYLASE -> HEPN+ DUF499 *->ORC-CDC6+ REase -> | 1112 | Bacteria>Terrabacteria_group>Actinomycetota>Actinomycetes>Mycobacteriales | GCF_025144475.1 | Corynebacterium_sanguinis | M3G45_RS07405 | 84077..87415 | 2594913 | DUF499-domain-containing-protein-[Corynebacterium-sanguinis] |
| WP_259825954.1 | DUF499_1.RM.like.systems | <-ABhydrolase<-?<-Aminotran_1_2<-ncRNA\|\|tRNA->?->SF1-SF2-helicase+DUF1998-><-ORC-CDC6+ REase <- HEPN+ DUF499 *<- METHYLASE <- TUDOR+SF1-SF2-Helicase+i REase +CTD+ REase \|\|ABhydrolase-> | 1112 | Bacteria>Terrabacteria_group>Actinomycetota>Actinomycetes>Mycobacteriales | GCF_025146125.1 | Corynebacterium_sanguinis | QON61_RS06385 | 69194..72532 | 2594913 | DUF499-domain-containing-protein-[Corynebacterium-sanguinis] |
| WP_283569455.1 | DUF499_1.RM.like.systems | <-rve+rve_3<-HIN-HTH\|\|?->PemK_toxin-> TUDOR+SF1-SF2-Helicase+i REase +CTD+ REase -> METHYLASE -> DUF499 *-> REase +TPR-> | 982 | Bacteria>Terrabacteria_group>Actinomycetota>Actinomycetes>Mycobacteriales | GCF_030063715.1 | Rhodococcus_qingshengii | QNA24_RS28365 | 79281..82229 | 334542 | DUF499-domain-containing-protein-[Rhodococcus-qingshengii] |
| WP_005139269.1 | DUF499_1.RM.like.systems | <-GNTR-HTH\|\|NUDIX-><-?<-HEPN<-Schlafen+HATPase+HTH<- HEPN+ DUF499 *<- REase <- METHYLASE <- TUDOR+SF1-SF2-Helicase+i REase +CTD+ REase <-PIN<-XIS-HTH | 1108 | Bacteria>Terrabacteria_group>Actinomycetota>Actinomycetes>Mycobacteriales | - | Mycolicibacterium_rhodesiae | MycrhDRAFT_0407 | 390606..393932 | 36814 | Swt1-family-HEPN-domain-containing-protein-[Mycolicibacterium-rhodesiae] |
| MBE6477658.1 | DUF499_1.RM.like.systems | TUDOR+SF1-SF2-Helicase+i REase +CTD+ REase -> METHYLASE ->HEPN-> w HTH+ REase ->?->PemK_toxin-> HEPN+ DUF499 *->?->TetR-HTH->?-><-?\|\|ArsR-HTH-><-ABhydrolase | 1088 | Bacteria>Terrabacteria_group>Actinomycetota>Actinomycetes>Propionibacteriales | GCA_015062705.1 | Propionibacteriaceae_bacterium | E7Z97_06285 | 37437..40699 | 2021380 | ATP-binding-protein-[Propionibacteriaceae-bacterium] |
| MBK8462833.1 | DUF499_1.RM.like.systems | <- REase <-?<-HEPN\|\| TUDOR+SF1-SF2-Helicase+i REase +CTD+ REase -> METHYLASE ->ParB+HNH-> HEPN+ DUF499 *->?-><-Peptidase_M48<-?<-?\|\|TPR-> | 1117 | Bacteria>Terrabacteria_group>Actinomycetota>Actinomycetes>Propionibacteriales | GCA_016711325.1 | Nigerium_sp. | IPL36_07335 | 626569..629922 | 2042655 | DUF499-domain-containing-protein-[Nigerium-sp.] |
| WP_014846821.1 | DUF499_1.RM.like.systems | REase -> TUDOR+SF1-SF2-Helicase+i REase +CTD+ REase -> METHYLASE -> w HTH+ REase -> HEPN+ DUF499 *->?->ACET-><-?\|\|?-><-TetR-HTH | 1129 | Bacteria>Terrabacteria_group>Actinomycetota>Actinomycetes>Propionibacteriales | GCF_000277715.1 | Arachnia_propionica | J5A53_RS06805 | 1506504..1509893 | 1750 | DUF499-domain-containing-protein-[Arachnia-propionica] |
| WP_021798346.1 | DUF499_1.RM.like.systems | TUDOR+SF1-SF2-Helicase+i REase +CTD+ REase -> METHYLASE -> w HTH+ REase -> HEPN+ DUF499 *-> SF1-SF2-Helicase+ REase -p-> SF1-SF2-Helicase+ REase -p->HTH_28+HTH_33+DDE_3->SF1-SF2-Helicase-p-><-Synaptojanin | 1128 | Bacteria>Terrabacteria_group>Actinomycetota>Actinomycetes>Propionibacteriales | GCF_000478805.1 | Propionibacterium_acidifaciens | HMPREF0682_RS10330 | 20512..23898 | 556499 | Swt1-family-HEPN-domain-containing-protein-[Propionibacterium-acidifaciens] |
| WP_061786876.1 | DUF499_1.RM.like.systems | REase -> TUDOR+SF1-SF2-Helicase+i REase +CTD+ REase -> METHYLASE -> w HTH+ REase -> HEPN+ DUF499 *->KAP-NTPase->ACET-><-?\|\|?-><-TetR-HTH | 1129 | Bacteria>Terrabacteria_group>Actinomycetota>Actinomycetes>Propionibacteriales | GCF_001592325.1 | Arachnia_propionica | PP13_RS01005 | 243859..247248 | 1750 | DUF499-domain-containing-protein-[Arachnia-propionica] |
| WP_207473688.1 | DUF499_1.RM.like.systems | ABhydrolase->XIS-HTH-> TUDOR+SF1-SF2-Helicase+i REase +CTD+ REase -> METHYLASE -> w HTH+ REase -> HEPN+ DUF499 *-> REase ->HEPN+ REase -><-DNA_methylase\|\|TrwC+SF1-SF2-Helicase-> | 1131 | Bacteria>Terrabacteria_group>Actinomycetota>Actinomycetes>Propionibacteriales | GCF_017355265.1 | Tessaracoccus_sp._SD287 | IPV09_RS03380 | 69383..72778 | 2782008 | DUF499-domain-containing-protein-[Tessaracoccus-sp.-SD287] |
| WP_211798072.1 | DUF499_1.RM.like.systems | TetR-HTH-><-?\|\|?-><-ACET<-?<- HEPN+ DUF499 *<- w HTH+ REase <- METHYLASE <- TUDOR+SF1-SF2-Helicase+i REase +CTD+ REase | 1129 | Bacteria>Terrabacteria_group>Actinomycetota>Actinomycetes>Propionibacteriales | GCF_018128345.1 | Arachnia_propionica | J5A61_RS08580 | 1914777..1918166 | 1750 | DUF499-domain-containing-protein-[Arachnia-propionica] |
| WP_119715651.1 | DUF499_1.RM.like.systems | TUDOR+SF1-SF2-Helicase+i REase +CTD+ REase -> METHYLASE ->HEPN-> w HTH+ REase ->?->PemK_toxin-> HEPN+ DUF499 *->?->?-><-?\|\|ArsR-HTH-><-ABhydrolase<-TetR-HTH | 1133 | Bacteria>Terrabacteria_group>Actinomycetota>Actinomycetes>Propionibacteriales | GCF_900289195.1 | Propionibacterium_ruminifibrarum | PROPJV5_RS07045 | 37471..40872 | 1962131 | Swt1-family-HEPN-domain-containing-protein-[Propionibacterium-ruminifibrarum] |
| WP_130874284.1 | DUF499_1.RM.like.systems | TetR-HTH-><-?\|\|?-><-ACET<-KAP-NTPase<- HEPN+ DUF499 *<- w HTH+ REase <- METHYLASE <- TUDOR+SF1-SF2-Helicase+i REase +CTD+ REase <-MORC<- METHYLASE -p | 1129 | Bacteria>Terrabacteria_group>Actinomycetota>Actinomycetes>Propionibacteriales | GCF_900607225.1 | [Pseudopropionibacterium]_massiliense | EOE39_RS08095 | 1820154..1823543 | 2220000 | DUF499-domain-containing-protein-[[Pseudopropionibacterium] |
| WP_119162165.1 | DUF499_1.RM.like.systems | <-TetR-HTH\|\|?-><-SF1-SF2-Helicase<-OLD-ABC+TOPRIM<- HEPN+ DUF499 *<-PemK_toxin<-?<- w HTH+ REase <- METHYLASE <- TUDOR+SF1-SF2-Helicase+i REase +CTD+ REase | 1133 | Bacteria>Terrabacteria_group>Actinomycetota>Actinomycetes>Propionibacteriales | GCF_900637915.1 | Propionibacterium_australiense | EL282_RS10960 | 2474046..2477447 | 119981 | DUF499-domain-containing-protein-[Propionibacterium-australiense] |
| WP_126378241.1 | DUF499_1.RM.like.systems | TetR-HTH-><-?\|\|?-><-ACET<-?<- HEPN+ DUF499 *<- w HTH+ REase <- METHYLASE -p<- METHYLASE -p<- TUDOR+SF1-SF2-Helicase+i REase +CTD+ REase | 1129 | Bacteria>Terrabacteria_group>Actinomycetota>Actinomycetes>Propionibacteriales | GCF_900638645.1 | Arachnia_propionica | EL396_RS08525 | 1937379..1940768 | 1750 | DUF499-domain-containing-protein-[Arachnia-propionica] |
| WP_040158482.1 | DUF499_1.RM.like.systems | HTH+MPTase->?->?->HTH_Tnp_1+HTH_21+rve+rve_2-><- HEPN+ DUF499 *<- w HTH+ REase <- METHYLASE <- TUDOR+SF1-SF2-Helicase+i REase +CTD+ REase <-XIS-HTH<-tRNA<-?<-ACET | 1132 | Bacteria>Terrabacteria_group>Actinomycetota>Actinomycetes>Propionibacteriales | GCF_902375435.1 | Nigerium_massiliense | BN1187_RS02660 | 259497..262895 | 1522317 | DUF499-domain-containing-protein-[Nigerium-massiliense] |
| WP_278693137.1 | DUF499_1.RM.like.systems | REase -> TUDOR+SF1-SF2-Helicase+i REase +CTD+ REase -> METHYLASE -> w HTH+ REase -> HEPN+ DUF499 *-> | 1129 | Bacteria>Terrabacteria_group>Actinomycetota>Actinomycetes>Propionibacteriales | GCF_915063725.1 | Arachnia_propionica | QDI34_RS08815 | 15722..19111 | 1750 | DUF499-domain-containing-protein-[Arachnia-propionica] |
| WP_277066086.1 | DUF499_1.RM.like.systems | Synaptojanin-><- SF1-SF2-Helicase+ REase <- HEPN+ DUF499 *<- w HTH+ REase <- METHYLASE <- TUDOR+SF1-SF2-Helicase+i REase +CTD+ REase \|\|?-><-?\|\|Peptidase-> | 1128 | Bacteria>Terrabacteria_group>Actinomycetota>Actinomycetes>Propionibacteriales | GCF_938034675.1 | Propionibacterium_acidifaciens | P7A72_RS09150 | 66970..70356 | 556499 | Swt1-family-HEPN-domain-containing-protein-[Propionibacterium-acidifaciens] |
| MCY4088355.1 | DUF499_1.RM.like.systems | HTH+UbE1->?-><-?\|\| TUDOR+SF1-SF2-Helicase+i REase +CTD+ REase -> REase -> METHYLASE -> DUF499 *-><-SF1-SF2-Helicase<-ParB+HNH\|\|DUF4065-><-HD\|\|Peptidase-> | 929 | Bacteria>Terrabacteria_group>Actinomycetota>Actinomycetes>unclassified_Actinomycetes | GCA_026708005.1 | Actinomycetes_bacterium | OXG37_16080 | 16647..19436 | 1883427 | MAG:-DUF499-domain-containing-protein-[Actinomycetia-bacterium] |
| PKQ27862.1 | DUF499_1.RM.like.systems | REase -> TUDOR+SF1-SF2-Helicase+i REase +CTD+ REase ->Schlafen+HATPase+HTH-> METHYLASE -> HEPN+ DUF499 *->?->TIR-> | 1105 | Bacteria>Terrabacteria_group>Actinomycetota>Candidatus_Geothermincolia>Candidatus_Geothermincolales | GCA_002843005.1 | Candidatus_Anoxymicrobium_japonicum | CVT63_05765 | 8955..12272 | 2013648 | AAA+-family-ATPase-[Candidatus-Anoxymicrobium-japonicum] |
| TGY60542.1 | DUF499_1.RM.like.systems | MazG->URI-DNase+SF1-SF2-Helicase->HEPN->HEPN-> TUDOR+SF1-SF2-Helicase+i REase +CTD+ REase -> METHYLASE -> HEPN+ DUF499 *->ORC-CDC6+ REase ->?-><-?\|\|?->?-> REase -> | 1126 | Bacteria>Terrabacteria_group>Actinomycetota>Coriobacteriia>Coriobacteriales | GCA_004793665.1 | Coriobacteriaceae_bacterium | E5335_00400 | 90399..93779 | 2011094 | ATP-binding-protein-[Coriobacteriaceae-bacterium] |
| GJM55560.1 | DUF499_1.RM.like.systems | <- REase <-?<-?\|\|?-><-?<-ORC-CDC6+ REase <- HEPN+ DUF499 *<- METHYLASE <- TUDOR+SF1-SF2-Helicase+i REase +CTD+ REase <-HEPN<-URI-DNase+SF1-SF2-Helicase<-MazG | 1126 | Bacteria>Terrabacteria_group>Actinomycetota>Coriobacteriia>Coriobacteriales | GCA_022834715.1 | Granulimonas_faecalis | ATOP_12150 | 1358904..1362284 | 2894155 | hypothetical-protein-ATOP_12150-[Granulimonas-faecalis] |
| WP_265590834.1 | DUF499_1.RM.like.systems | <- REase <-?<-?\|\|?-><-?<-ORC-CDC6+ REase <- HEPN+ DUF499 *<- METHYLASE <- TUDOR+SF1-SF2-Helicase+i REase +CTD+ REase <-HEPN<-URI-DNase+SF1-SF2-Helicase<-MazG | 1126 | Bacteria>Terrabacteria_group>Actinomycetota>Coriobacteriia>Coriobacteriales | GCA_022834715.1 | Granulimonas_faecalis | OR600_RS06205 | 1358904..1362284 | 2894155 | DUF499-domain-containing-protein-[Granulimonas-faecalis] |
| WP_251158236.1 | DUF499_1.RM.like.systems | <-ORC-CDC6+ REase <- HEPN+ DUF499 *<- METHYLASE <- TUDOR+SF1-SF2-Helicase+i REase +CTD+ REase <-HEPN<-URI-DNase+SF1-SF2-Helicase<-MazG | 1126 | Bacteria>Terrabacteria_group>Actinomycetota>Coriobacteriia>Coriobacteriales | GCF_943193035.1 | Caniella_muris | NFR49_RS03270 | 716481..719861 | 2941502 | DUF499-domain-containing-protein-[Caniella-sp.-DSM-110323] |
| WP_219975492.1 | DUF499_1.RM.like.systems | <-SIGMA-HTH<-?<-MULE-TRANSPOSASE\|\|?->?-><- REase <- HEPN+ DUF499 *<-PIN<-MazE_antitoxin<- METHYLASE <-PIN<- TUDOR+SF1-SF2-Helicase+i REase +CTD+ REase \|\|DUF4096-> | 1106 | Bacteria>Terrabacteria_group>Actinomycetota>Rubrobacteria>Rubrobacterales | GCF_019448335.1 | Rubrobacter_xylanophilus | Rxycam_RS04975 | 969402..972722 | 49319 | Swt1-family-HEPN-domain-containing-protein-[Rubrobacter-xylanophilus] |
| MBS1906106.1 | DUF499_1.RM.like.systems | N6_MTase-p->?->HTH+MPTase-><-SF1-SF2-Helicase<- REase <- HEPN+ DUF499 *<- METHYLASE <- TUDOR+SF1-SF2-Helicase+i REase +CTD+ REase <-XIS-HTH<-?<-?<-HTH_3 | 1133 | Bacteria>Terrabacteria_group>Actinomycetota>unclassified_Actinomycetota | GCA_018268515.1 | Actinomycetota_bacterium | JST33_05955 | 17244..20645 | 2900548 | DUF499-domain-containing-protein-[Actinomycetia-bacterium] |
| MCA0180524.1 | DUF499_1.RM.like.systems | <-SMODS<-?<-TrwC+SF1-SF2-Helicase\|\|DNA_methylase-><-SF1-SF2-Helicase<-OLD-ABC+TOPRIM<- HEPN+ DUF499 *<- w HTH+ REase <- METHYLASE <- TUDOR+SF1-SF2-Helicase+i REase +CTD+ REase <-XIS-HTH<-?\|\|GDResolvase+Recombinase+ZnR-> | 1132 | Bacteria>Terrabacteria_group>Actinomycetota>unclassified_Actinomycetota | GCA_020160805.1 | Actinomycetota_bacterium | LCH77_13240 | 54204..57602 | 2900548 | DUF499-domain-containing-protein-[Actinomycetia-bacterium] |
| MCA0180887.1 | DUF499_1.RM.like.systems | <-VirD4-TraG+VirD4-FtsK<-?\|\|XIS-HTH-> TUDOR+SF1-SF2-Helicase+i REase +CTD+ REase -> METHYLASE -> w HTH+ REase -> HEPN+ DUF499 *-> REase ->HEPN+ REase -> | 1132 | Bacteria>Terrabacteria_group>Actinomycetota>unclassified_Actinomycetota | GCA_020160805.1 | Actinomycetota_bacterium | LCH77_15145 | 46365..49763 | 2900548 | DUF499-domain-containing-protein-[Actinomycetia-bacterium] |
| MCL5734830.1 | DUF499_1.RM.like.systems | TUDOR+SF1-SF2-Helicase+i REase +CTD+ REase -p->MORC+DUF3684-> METHYLASE -> REase ->SF1-SF2-Helicase-> HEPN+ DUF499 *-><-HTH+DSBH | 1116 | Bacteria>Terrabacteria_group>Actinomycetota>unclassified_Actinomycetota | GCA_023483915.1 | Actinomycetota_bacterium | M1274_04450 | 22427..25777 | 2900548 | Swt1-family-HEPN-domain-containing-protein-[Actinobacteria-bacterium] |
| MCL5447091.1 | DUF499_1.RM.like.systems | <- HEPN+ DUF499 *<- METHYLASE <-?<- TUDOR+SF1-SF2-Helicase+i REase +CTD+ REase <- REase <-?<-rve+rve_3 | 1178 | Bacteria>Terrabacteria_group>Actinomycetota>unclassified_Actinomycetota | GCA_023486005.1 | Actinomycetota_bacterium | M1350_06795 | 8812..12348 | 2900548 | DUF499-domain-containing-protein-[Actinobacteria-bacterium] |
| MCX6342680.1 | DUF499_1.RM.like.systems | <-URI-DNase\|\|MazG->URI-DNase+SF1-SF2-Helicase-><-?<- REase <- HEPN+ DUF499 *<- METHYLASE <- TUDOR+SF1-SF2-Helicase+i REase +CTD+ REase | 1119 | Bacteria>Terrabacteria_group>Armatimonadota>Fimbriimonadia>Fimbriimonadales | GCA_026392145.1 | Fimbriimonadales_bacterium | NTU72_09890 | 5893..9252 | 2682145 | MAG:-Swt1-family-HEPN-domain-containing-protein-[Fimbriimonadales-bacterium] |
| NIM68640.1 | DUF499_1.RM.like.systems | TUDOR+SF1-SF2-Helicase+i REase +CTD+ REase ->NTP_transf-> REase -> METHYLASE -> DUF499 *->DNA-glycosylase-> | 918 | Bacteria>Terrabacteria_group>Armatimonadota>unclassified_Armatimonadota | GCA_011771675.1 | Armatimonadota_bacterium | GT055_11015 | 8511..11267 | 2033014 | DUF499-domain-containing-protein-[Armatimonadetes-bacterium] |
| WP_119798229.1 | DUF499_1.RM.like.systems | <-Phage_integrase<- REase <- DUF499 *<- METHYLASE <- TUDOR+SF1-SF2-Helicase+i REase +CTD+ REase \|\|TraJ-RHH-><-?<-HerA/FtsK | 952 | Bacteria>Terrabacteria_group>Bacillota>Bacilli>Bacillales | GCF_003591485.1 | Paenibacillus_sp._1011MAR3C5 | D3P07_RS00960 | 194996..197854 | 1675787 | DUF499-domain-containing-protein-[Paenibacillus-sp.-1011MAR3C5] |
| WP_082451901.1 | DUF499_1.RM.like.systems | <- REase <- DUF499 *<- METHYLASE <- TUDOR+SF1-SF2-Helicase+i REase +CTD+ REase <-?<-?<-?\|\|HTH_3-> | 838 | Bacteria>Terrabacteria_group>Bacillota>Bacilli>Bacillales | GCF_902375745.1 | Paenibacillus_ihuae | FY521_RS18285 | 3939778..3942291 | 1232431 | DUF499-domain-containing-protein,-partial-[Paenibacillus-ihuae] |
| HHW19260.1 | DUF499_1.RM.like.systems | TUDOR+SF1-SF2-Helicase+i REase +CTD+ REase ->SbcC-> METHYLASE -> REase -> DUF499 *->DUF433->PIN-> | 932 | Bacteria>Terrabacteria_group>Bacillota>Candidatus_Fermentithermobacillia>Candidatus_Fermentithermobacillales | GCA_012840505.1 | Candidatus_Fermentithermobacillaceae_bacterium | GXX30_10280 | 57551..60349 | 3085339 | ATP-binding-protein-[Firmicutes-bacterium] |
| NLV15838.1 | DUF499_1.RM.like.systems | PRTase-><-?\|\| TUDOR+SF1-SF2-Helicase+i REase +CTD+ REase -> METHYLASE -> HEPN+ DUF499 *-> REase -> | 1154 | Bacteria>Terrabacteria_group>Bacillota>Clostridia>Eubacteriales | GCA_012729505.1 | Syntrophomonadaceae_bacterium | GXY50_01325 | 75931..79395 | 2093811 | ATP-binding-protein-[Syntrophomonadaceae-bacterium] |
| NSW83680.1 | DUF499_1.RM.like.systems | <-DDE_Tnp\|\|HIN-HTH-><-DDE_Tnp_1<-NACHT+ REase <- DUF499 *<- METHYLASE <- TUDOR+SF1-SF2-Helicase+i REase +CTD+ REase <-PrlF_antitoxin | 905 | Bacteria>Terrabacteria_group>Bacillota>Clostridia>Eubacteriales | GCA_013314835.1 | Syntrophothermus_sp. | HPY90_10510 | 22691..25408 | 2736299 | ATP-binding-protein-[Syntrophothermus-sp.] |
| MBR3354573.1 | DUF499_1.RM.like.systems | HTH->tRNA->tRNA->tRNA->tRNA->tRNA->tRNA->tRNA->tRNA->?-> TUDOR+SF1-SF2-Helicase+i REase +CTD+ REase -> METHYLASE -> METHYLASE -> HEPN+ DUF499 *-> REase -> | 1111 | Bacteria>Terrabacteria_group>Bacillota>Clostridia>Eubacteriales | GCA_017510465.1 | Oscillospiraceae_bacterium | IKG47_04355 | 36923..40258 | 2485925 | DUF499-domain-containing-protein-[Oscillospiraceae-bacterium] |
| MCR4442858.1 | DUF499_1.RM.like.systems | TUDOR+SF1-SF2-Helicase+i REase +CTD+ REase -> REase ->SF1-SF2-Helicase-> METHYLASE ->OLD-ABC-> DUF499 *->?->?->TPR+TPR->?->PolB-> | 1104 | Bacteria>Terrabacteria_group>Bacillota>Clostridia>Eubacteriales | GCA_024655825.1 | Peptococcaceae_bacterium | NUV48_11985 | 21314..24628 | 2052179 | DUF499-domain-containing-protein-[Peptococcaceae-bacterium] |
| WP_012301643.1 | DUF499_1.RM.like.systems | REase -> TUDOR+SF1-SF2-Helicase+i REase +CTD+ REase ->?->OLD-ABC+TOPRIM-> METHYLASE -> REase -> DUF499 *->RNase->?->?->?->HicB_lk_antitox-> | 924 | Bacteria>Terrabacteria_group>Bacillota>Clostridia>Eubacteriales | GCF_000018425.1 | Candidatus_Desulforudis_audaxviator | Daudx_RS02685 | 550555..553329 | 471827 | DUF499-domain-containing-protein-[Candidatus-Desulforudis-audaxviator] |
| WP_161846233.1 | DUF499_1.RM.like.systems | TUDOR+SF1-SF2-Helicase+i REase +CTD+ REase ->?-> METHYLASE -> HEPN+ DUF499 *-> REase -><-GDResolvase | 1134 | Bacteria>Terrabacteria_group>Bacillota>Clostridia>Eubacteriales | GCF_009935845.1 | Pseudoflavonifractor_sp._524-17 | D1159_RS13725 | 7029..10433 | 2304577 | DUF499-domain-containing-protein-[Pseudoflavonifractor-sp.-524-17] |
| WP_221039750.1 | DUF499_1.RM.like.systems | <- REase <- HEPN+ DUF499 *<- METHYLASE <- TUDOR+SF1-SF2-Helicase+i REase +CTD+ REase <- McrC-NTD+ REase | 1133 | Bacteria>Terrabacteria_group>Bacillota>Clostridia>Thermoanaerobacterales | GCF_019668485.1 | Gelria_sp._Kuro-4 | K5554_RS03460 | 654715..658116 | 2796927 | Swt1-family-HEPN-domain-containing-protein-[Gelria-sp.-Kuro-4] |
| MCK9524538.1 | DUF499_1.RM.like.systems | ArdCN+MPTase->?-><-?<- REase <-?<- REase <- HEPN+ DUF499 *<- METHYLASE <- TUDOR+SF1-SF2-Helicase+i REase +CTD+ REase -p | 1113 | Bacteria>Terrabacteria_group>Bacillota>Limnochordia>unclassified_Limnochordia | GCA_023228555.1 | Limnochordia_bacterium | MOR49_01235 | 6495..9836 | 2497621 | Swt1-family-HEPN-domain-containing-protein-[Limnochordia-bacterium] |
| NLG79149.1 | DUF499_1.RM.like.systems | <-XIS-HTH+RVT\|\|?->?-><- REase <- DUF499 *<- METHYLASE <- TUDOR+SF1-SF2-Helicase+i REase +CTD+ REase | 946 | Bacteria>Terrabacteria_group>Bacillota>unclassified_Bacillota | GCA_012516125.1 | Bacillota_bacterium | GX492_04905 | 11356..14196 | 1879010 | ATP-binding-protein-[Firmicutes-bacterium] |
| NLN07229.1 | DUF499_1.RM.like.systems | <- HEPN+ DUF499 *<- METHYLASE <- TUDOR+SF1-SF2-Helicase+i REase +CTD+ REase <- REase | 1130 | Bacteria>Terrabacteria_group>Bacillota>unclassified_Bacillota | GCA_012522075.1 | Bacillota_bacterium | GX167_06390 | 552..3944 | 1879010 | ATP-binding-protein-[Firmicutes-bacterium] |
| MBC7083917.1 | DUF499_1.RM.like.systems | <- DUF499 *<- REase <- METHYLASE <- REase <- TUDOR+SF1-SF2-Helicase+i REase +CTD+ REase | 907 | Bacteria>Terrabacteria_group>Bacillota>unclassified_Bacillota | GCA_014360195.1 | Bacillota_bacterium | H5T95_10620 | 51..2774 | 1879010 | ATP-binding-protein-[Firmicutes-bacterium] |
| WP_075063424.1 | DUF499_1.RM.like.systems | LuxR-HTH->?->SF1-SF2-Helicase+i REase +CTD+ REase -> METHYLASE -> DUF499 *-> REase +SF1-SF2-Helicase ->N6_N4_MTase-><-SF1-SF2-Helicase | 1097 | Bacteria>Terrabacteria_group>Chloroflexota>Anaerolineae>Anaerolineales | GCF_001306115.1 | Ornatilinea_apprima | ADN00_RS13000 | 57976..61269 | 1134406 | DUF499-domain-containing-protein-[Ornatilinea-apprima] |
| MBK9093501.1 | DUF499_1.RM.like.systems | TUDOR+SF1-SF2-Helicase+i REase +CTD+ REase ->OLD-ABC->HNH-> METHYLASE -> OLD-ABC-> REase -> DUF499 *->?->ParB+HNH->?-> REase -p-> | 943 | Bacteria>Terrabacteria_group>Chloroflexota>Anaerolineae>unclassified_Anaerolineae | GCA_016719785.1 | Anaerolineae_bacterium | IPM84_12145 | 210722..213553 | 2052143 | ATP-binding-protein-[Anaerolineae-bacterium] |
| MBN2002245.1 | DUF499_1.RM.like.systems | STYKIN-> REase ->Trypsin-> REase ->SF1-SF2-Helicase+i REase +CTD+ REase -> METHYLASE -> DUF499 *->?-><-URI-DNase | 1003 | Bacteria>Terrabacteria_group>Chloroflexota>Anaerolineae>unclassified_Anaerolineae | GCA_016931835.1 | Anaerolineae_bacterium | JXA21_02715 | 14285..17296 | 2052143 | ATP-binding-protein-[Anaerolineae-bacterium] |
| MBN2007579.1 | DUF499_1.RM.like.systems | TUDOR+SF1-SF2-Helicase+i REase +CTD+ REase -> REase -> METHYLASE ->OLD-ABC->TOPRIM-> DUF499 *-> | 737 | Bacteria>Terrabacteria_group>Chloroflexota>Anaerolineae>unclassified_Anaerolineae | GCA_016931835.1 | Anaerolineae_bacterium | JXA21_29810 | 7893..10102 | 2052143 | ATP-binding-protein-[Anaerolineae-bacterium] |
| MBZ0275669.1 | DUF499_1.RM.like.systems | SF1-SF2-Helicase+i REase +CTD+ REase -> METHYLASE -> DUF499 *-> REase -> | 1097 | Bacteria>Terrabacteria_group>Chloroflexota>Anaerolineae>unclassified_Anaerolineae | GCA_019912915.1 | Anaerolineae_bacterium | K8I60_05975 | 6820..10113 | 2052143 | DUF499-domain-containing-protein-[Anaerolineae-bacterium] |
| MCL4300573.1 | DUF499_1.RM.like.systems | TUDOR+SF1-SF2-Helicase+i REase +CTD+ REase ->?->?-> METHYLASE -> REase ->?-> DUF499 *-><-?\|\|?->?->?-><-HNH | 929 | Bacteria>Terrabacteria_group>Chloroflexota>Anaerolineae>unclassified_Anaerolineae | GCA_023382535.1 | Anaerolineae_bacterium | KJ077_32855 | 82827..85616 | 2052143 | DUF499-domain-containing-protein-[Anaerolineae-bacterium] |
| MCL4302340.1 | DUF499_1.RM.like.systems | <-TraJ-RHH\|\| TUDOR+SF1-SF2-Helicase+i REase +CTD+ REase ->?->?-> METHYLASE -> REase -> HEPN+ DUF499 *-><-?\|\|?->GDResolvase+Recombinase+ZnR-> | 1117 | Bacteria>Terrabacteria_group>Chloroflexota>Anaerolineae>unclassified_Anaerolineae | GCA_023382535.1 | Anaerolineae_bacterium | KJ077_42015 | 34544..37897 | 2052143 | DUF499-domain-containing-protein-[Anaerolineae-bacterium] |
| MCL4871680.1 | DUF499_1.RM.like.systems | <-AbiEii<-AbiEii<-AbiEi_4\|\| TUDOR+SF1-SF2-Helicase+i REase +CTD+ REase -> REase -> METHYLASE -> HEPN+ DUF499 *-><-?<-?<-SIGMA-HTH<-RNB+RNase\|\|?->PadR-HTH-> | 1115 | Bacteria>Terrabacteria_group>Chloroflexota>Anaerolineae>unclassified_Anaerolineae | GCA_023384415.1 | Anaerolineae_bacterium | KJ063_22200 | 16187..19534 | 2052143 | DUF499-domain-containing-protein-[Anaerolineae-bacterium] |
| MCZ7570815.1 | DUF499_1.RM.like.systems | DEDD_Tnp_IS110+Transposase_20->?-> TUDOR+SF1-SF2-Helicase+i REase +CTD+ REase -> METHYLASE -> REase -> HEPN+ DUF499 *-><-PIN<-PhdYefM_antitox<-?<-HAD | 1117 | Bacteria>Terrabacteria_group>Chloroflexota>Ardenticatenia>Ardenticatenales | GCA_027493995.1 | Ardenticatenaceae_bacterium | M5U01_19825 | 4779576..4782929 | 2614247 | MAG:-Swt1-family-HEPN-domain-containing-protein-[Ardenticatenaceae-bacterium] |
| MBK9709775.1 | DUF499_1.RM.like.systems | TUDOR+SF1-SF2-Helicase+i REase +CTD+ REase -> REase -> METHYLASE ->URI-DNase-> HEPN+ DUF499 *->?->?-><-tRNA<-?<-?<-RnaseIII+DSRBD | 1115 | Bacteria>Terrabacteria_group>Chloroflexota>Chloroflexia>Chloroflexales | GCA_016717335.1 | Kouleothrix_sp. | IPO81_00330 | 71152..74499 | 2779161 | DUF499-domain-containing-protein-[Kouleothrix-sp.] |

| Protein-ID | System | Gene-Neighbourhoods | Length | Taxonomy | Assembly-ID | Strain | Locus-Tag | Genomic-coordinates | TaxID | Fasta-defline |
|---|---|---|---|---|---|---|---|---|---|---|
| WP_011955032.1 | DUF499_1.RM.like.systems | MethyltransfD12-><-?<-?\|\|MODE-HTH+PBPII-><- DUF499 *<- REase <- METHYLASE <- TUDOR+SF1-SF2-Helicase+i REase +CTD+ REase <-DDE_Tnp_1<-DDE_Tnp_1 | 865 | Bacteria>Terrabacteria_group>Chloroflexota>Chloroflexia>Chloroflexales | GCF_000016665.1 | Roseiflexus_sp._RS-1 | ROSERS_RS01210 | 293633..296230 | 357808 | DUF499-domain-containing-protein-[Roseiflexus-sp.-RS-1] |
| WP_066790944.1 | DUF499_1.RM.like.systems | <- HEPN+ DUF499 *<- METHYLASE <- METHYLASE <- REase <- TUDOR+SF1-SF2-Helicase+i REase +CTD+ REase | 1104 | Bacteria>Terrabacteria_group>Chloroflexota>Chloroflexia>Chloroflexales | GCF_001650695.1 | Chloroflexus_islandicus | A6A03_RS17620 | 125816..129130 | 1707952 | Swt1-family-HEPN-domain-containing-protein-[Chloroflexus-islandicus] |
| MCA9831553.1 | DUF499_1.RM.like.systems | TUDOR+SF1-SF2-Helicase+i REase +CTD+ REase -> REase ->?->?->?-> METHYLASE -> HEPN+ DUF499 *->?->HerA/FtsK->PIN-> | 1134 | Bacteria>Terrabacteria_group>Chloroflexota>Dehalococcoidia>unclassified_Dehalococcoidia | GCA_020432045.1 | Dehalococcoidia_bacterium | KC495_13860 | 27821..31225 | 2026734 | ATP-binding-protein-[Dehalococcoidia-bacterium] |
| MCL0075718.1 | DUF499_1.RM.like.systems | SF1-SF2-Helicase+DUF1998->?->SLATT->HicB-> TUDOR+SF1-SF2-Helicase+i REase +CTD+ REase -> METHYLASE -> DUF499 *-> REase ->Phage_int_SAM_4+Phage_integrase->?->HEPN-> | 847 | Bacteria>Terrabacteria_group>Chloroflexota>Dehalococcoidia>unclassified_Dehalococcoidia | GCA_023271455.1 | Dehalococcoidia_bacterium | M1017_02415 | 8590..11133 | 2026734 | DUF499-domain-containing-protein-[Dehalococcoidia-bacterium] |
| QWK10277.1 | DUF499_1.RM.like.systems | <-OrfB_IS605+OrfB_Zn_ribbon<-?\|\|ABC-ATPase+SbcC-><-?<-?<-?<- HEPN+ DUF499 *<-HEPN<-PolB<- METHYLASE <- REase <-OLD-ABC +TOPRIM<- TUDOR+SF1-SF2-Helicase+i REase +CTD+ REase | 1119 | Bacteria>Terrabacteria_group>Chloroflexota>Thermoflexia>Thermoflexales | GCA_018771565.1 | Thermoflexus_hugenholtzii | KNN16_13035 | 2858619..2861978 | 1495650 | MAG:-DUF499-domain-containing-protein-[Thermoflexus-hugenholtzii] |
| MBI2863770.1 | DUF499_1.RM.like.systems | TUDOR+SF1-SF2-Helicase+i REase +CTD+ REase -p->HicB_lk_antitox->MqsA_antitoxin->?-> METHYLASE -> REase -> DUF499 *-> | 1112 | Bacteria>Terrabacteria_group>Chloroflexota>unclassified_Chloroflexota | GCA_016190625.1 | Chloroflexota_bacterium | HYX94_04330 | 24989..28327 | 2026724 | ATP-binding-protein-[Chloroflexi-bacterium] |
| MBI4322283.1 | DUF499_1.RM.like.systems | TUDOR+SF1-SF2-Helicase+i REase +CTD+ REase ->HD->DUF4065-> METHYLASE -p-> REase -> METHYLASE -> DUF499 *->?-><-?<-?<-?\|\| TUDOR+SF1-SF2-Helicase+i REase +CTD+ REase ->Schlafen+HATPase+HTH-> | 979 | Bacteria>Terrabacteria_group>Chloroflexota>unclassified_Chloroflexota | GCA_016210495.1 | Chloroflexota_bacterium | HY675_27635 | 34699..37638 | 2026724 | ATP-binding-protein-[Chloroflexi-bacterium] |
| MBI4322294.1 | DUF499_1.RM.like.systems | TUDOR+SF1-SF2-Helicase+i REase +CTD+ REase ->Schlafen+HATPase+HTH->?-> METHYLASE -> OLD-ABC-> REase -> DUF499 *-> | 938 | Bacteria>Terrabacteria_group>Chloroflexota>unclassified_Chloroflexota | GCA_016210495.1 | Chloroflexota_bacterium | HY675_27690 | 51598..54414 | 2026724 | ATP-binding-protein-[Chloroflexi-bacterium] |
| MCC6178177.1 | DUF499_1.RM.like.systems | <-HTH_3<-Tox-RelE<-SF1-SF2-Helicase<-OLD-ABC+TOPRIM<- REase <- HEPN+ DUF499 *<- METHYLASE <- w HTH+ REase <- TUDOR+SF1-SF2-Helicase+i REase +CTD+ REase \|\|?->Peptidase_S41-> | 1117 | Bacteria>Terrabacteria_group>Chloroflexota>unclassified_Chloroflexota | GCA_020845495.1 | Chloroflexota_bacterium | IT305_22975 | 6439..9792 | 2026724 | DUF499-domain-containing-protein-[Chloroflexi-bacterium] |
| MCC6567532.1 | DUF499_1.RM.like.systems | <-SF1-SF2-Helicase<-?<-?<- HEPN+ DUF499 *<- REase <-?<- METHYLASE <-?<-?<- TUDOR+SF1-SF2-Helicase+i REase +CTD+ REase | 1096 | Bacteria>Terrabacteria_group>Chloroflexota>unclassified_Chloroflexota | GCA_020848335.1 | Chloroflexota_bacterium | IT298_17380 | 16185..19475 | 2026724 | DUF499-domain-containing-protein-[Chloroflexi-bacterium] |
| MCL4370916.1 | DUF499_1.RM.like.systems | <- DUF499 *<- METHYLASE <- REase <-?<-?<-Schlafen+HATPase+HTH<- TUDOR+SF1-SF2-Helicase+i REase +CTD+ REase -p | 584 | Bacteria>Terrabacteria_group>Chloroflexota>unclassified_Chloroflexota | GCA_023380955.1 | Chloroflexota_bacterium | M1380_08425 | 1..1753 | 2026724 | DUF499-domain-containing-protein-[Chloroflexota-bacterium] |
| MCL5961065.1 | DUF499_1.RM.like.systems | TUDOR+SF1-SF2-Helicase+i REase +CTD+ REase ->Schlafen+HATPase+HTH-> METHYLASE -> OLD-ABC-> REase -> DUF499 *->MerR-HTH->?-> REase ->IclR-HTH-> | 939 | Bacteria>Terrabacteria_group>Chloroflexota>unclassified_Chloroflexota | GCA_023489135.1 | Chloroflexota_bacterium | M1358_17465 | 13442..16261 | 2026724 | ATP-binding-protein-[Chloroflexi-bacterium] |
| MCL5950399.1 | DUF499_1.RM.like.systems | <-ParB+HNH<- REase <- REase <-PIWI<- DUF499 *<- REase <- METHYLASE <-MarR-HTH<-HKD-DNase<-HKD-DNase<- TUDOR+SF1-SF2-Helicase+i REase +CTD+ REase | 935 | Bacteria>Terrabacteria_group>Chloroflexota>unclassified_Chloroflexota | GCA_023489485.1 | Chloroflexota_bacterium | M1132_01520 | 20661..23468 | 2026724 | DUF499-domain-containing-protein-[Chloroflexi-bacterium] |
| MCX6032091.1 | DUF499_1.RM.like.systems | <- SF1-SF2-Helicase+ REase <-Exonuc_V_gamma+ REase <-SF1-SF2-Helicase+DUF1998<-?<-?<- DUF499 *<-?<-?<- METHYLASE <- REase <- TUDOR+SF1-SF2-Helicase+i REase +CTD+ REase | 1118 | Bacteria>Terrabacteria_group>Chloroflexota>unclassified_Chloroflexota | GCA_026389845.1 | Chloroflexota_bacterium | NT169_22690 | 15055..18411 | 2026724 | MAG:-DUF499-domain-containing-protein-[Chloroflexi-bacterium] |
| MDE2668025.1 | DUF499_1.RM.like.systems | <-PIN<-AbrB<- HEPN+ DUF499 *<- METHYLASE <- TUDOR+SF1-SF2-Helicase+i REase +CTD+ REase <- REase <-MORC+ REase <-DNA_methylase\|\|AbiEi_4-> | 1121 | Bacteria>Terrabacteria_group>Chloroflexota>unclassified_Chloroflexota | GCA_028875515.1 | Chloroflexota_bacterium | OXI51_00025 | 2709..6074 | 2026724 | MAG:-DUF499-domain-containing-protein-[Chloroflexota-bacterium] |
| MBY0357952.1 | DUF499_1.RM.like.systems | <- SF1-SF2-Helicase+ REase -p\|\| TUDOR+SF1-SF2-Helicase+i REase +CTD+ REase -> METHYLASE -> DUF499 *-> REase ->?->SIGMA-HTH-> | 939 | Bacteria>Terrabacteria_group>Cyanobacteriota/Melainabacteria_group>Candidatus_Melainabacteria>Candidatus_Obscuribacterales | GCA_019750335.1 | Candidatus_Obscuribacterales_bacterium | K2W82_08125 | 417056..419875 | 2364082 | DUF499-domain-containing-protein-[Candidatus-Obscuribacterales-bacterium] |
| WP_096720868.1 | DUF499_1.RM.like.systems | DUF29-><-?<-?<-?\|\|?-><- HEPN+ DUF499 *<-PIN<-?<- METHYLASE \|\| REase -><- TUDOR+SF1-SF2-Helicase+i REase +CTD+ REase | 1125 | Bacteria>Terrabacteria_group>Cyanobacteriota/Melainabacteria_group>Cyanobacteriota>Cyanophyceae | - | Chondrocystis_sp._NIES-4102 | CDC35_RS11850 | 2748341..2751718 | 2005460 | Swt1-family-HEPN-domain-containing-protein-[Chondrocystis-sp.-NIES-4102] |
| WP_102148297.1 | DUF499_1.RM.like.systems | ABC-ATPase+SbcC->?->?-><-?\|\|?-><- DUF499 *<- REase <- METHYLASE <-OLD-ABC-p<- TUDOR+SF1-SF2-Helicase+i REase +CTD+ REase | 975 | Bacteria>Terrabacteria_group>Cyanobacteriota/Melainabacteria_group>Cyanobacteriota>Cyanophyceae | - | Fischerella_thermalis | CEN42_03245 | 8470..11397 | 372787 | DUF499-domain-containing-protein-[Fischerella-thermalis] |
| EGK85305.1 | DUF499_1.RM.like.systems | NTP_transferase->ST-Hiskin->?-> TUDOR+SF1-SF2-Helicase+i REase +CTD+ REase -> METHYLASE -> REase -> DUF499 *-> | 998 | Bacteria>Terrabacteria_group>Cyanobacteriota/Melainabacteria_group>Cyanobacteriota>Cyanophyceae | GCA_000214075.2 | Microcoleus_vaginatus_FGP-2 | MicvaDRAFT_4687 | 60148..63144 | 756067 | AAA-family-ATPase-[Microcoleus-vaginatus-FGP-2] |
| EHC19655.1 | DUF499_1.RM.like.systems | TUDOR+SF1-SF2-Helicase+i REase +CTD+ REase ->?->OLD-ABC-p-> METHYLASE -> REase -> DUF499 *->?-><-?<-?<-ABC-ATPase+SbcC | 975 | Bacteria>Terrabacteria_group>Cyanobacteriota/Melainabacteria_group>Cyanobacteriota>Cyanophyceae | GCA_000231365.2 | Fischerella_thermalis_JSC-11 | FJSC11DRAFT_0472 | 574051..576978 | 741277 | AAA-family-ATPase-[Fischerella-thermalis-JSC-11] |
| EKQ69986.1 | DUF499_1.RM.like.systems | TUDOR+SF1-SF2-Helicase+i REase +CTD+ REase -> REase ->PolB->HEPN-><-?\|\| REase -> HEPN+ DUF499 *-> | 1113 | Bacteria>Terrabacteria_group>Cyanobacteriota/Melainabacteria_group>Cyanobacteriota>Cyanophyceae | GCA_000309945.1 | Leptolyngbyaceae_cyanobacterium_JSC-12 | OsccyDRAFT_0242 | 268586..271927 | 864702 | putative-ATPase-(AAA+-superfamily)-[Leptolyngbyaceae-cyanobacterium-JSC-12] |
| KAB8314079.1 | DUF499_1.RM.like.systems | TUDOR+SF1-SF2-Helicase+i REase +CTD+ REase -p-> REase -><-?\|\| METHYLASE -> HEPN+ DUF499 *-><-?<-?\|\|FAD-NAD-dep-oxidoreductase-> | 1113 | Bacteria>Terrabacteria_group>Cyanobacteriota/Melainabacteria_group>Cyanobacteriota>Cyanophyceae | GCA_000828075.3 | Tolypothrix_campylonemoides_VB511288 | SD81_039415 | 98698..102039 | 1245935 | ATP-binding-protein-[Tolypothrix-campylonemoides-VB511288] |
| MBO1045985.1 | DUF499_1.RM.like.systems | Aldedh-><-?\|\|?->?->?-><- DUF499 *<-PemK_toxin<-MetJarc-HTH<- METHYLASE <-?<- REase <- TUDOR+SF1-SF2-Helicase+i REase +CTD+ REase | 965 | Bacteria>Terrabacteria_group>Cyanobacteriota/Melainabacteria_group>Cyanobacteriota>Cyanophyceae | GCA_001593825.2 | Aphanizomenon_flos-aquae_UKL13-PB | A2T80_019100 | 69616..72513 | 1803587 | DUF499-domain-containing-protein-[Aphanizomenon-flos-aquae-UKL13-PB] |
| OAB62772.1 | DUF499_1.RM.like.systems | TUDOR+SF1-SF2-Helicase+i REase +CTD+ REase ->PRTase+PRTase->?-> METHYLASE -> REase -> DUF499 *-> | 1013 | Bacteria>Terrabacteria_group>Cyanobacteriota/Melainabacteria_group>Cyanobacteriota>Cyanophyceae | GCA_001637395.1 | Leptolyngbya_valderiana_BDU_20041 | AY599_12440 | 19718..22759 | 322866 | AAA-family-ATPase-[Leptolyngbya-valderiana-BDU-20041] |
| OBQ20276.1 | DUF499_1.RM.like.systems | <-DnaJ<-?<-NTP_transferase<-?<- DUF499 *<-PIN<-?<- REase +HTH<- TUDOR+SF1-SF2-Helicase+i REase +CTD+ REase <-CHTH | 916 | Bacteria>Terrabacteria_group>Cyanobacteriota/Melainabacteria_group>Cyanobacteriota>Cyanophyceae | GCA_001672085.1 | Anabaena_sp._AL93 | AN486_07300 | 48734..51484 | 1678133 | MAG:-ATPase-[Anabaena-sp.-AL93] |
| ARI79893.1 | DUF499_1.RM.like.systems | TUDOR+SF1-SF2-Helicase+i REase +CTD+ REase -> REase -> REase -> METHYLASE ->PAIREDC-HTH->PIN-> HEPN+ DUF499 *-><-Cluster164_18clades<-?\|\|?->?->?->DUF29-> | 1155 | Bacteria>Terrabacteria_group>Cyanobacteriota/Melainabacteria_group>Cyanobacteriota>Cyanophyceae | GCA_002095975.1 | Microcystis_aeruginosa_PCC_7806SL | BH695_0612 | 541107..544574 | 1903187 | hypothetical-protein-BH695_0612-[Microcystis-aeruginosa-PCC-7806SL] |
| PMB36920.1 | DUF499_1.RM.like.systems | ABC-ATPase+SbcC->?->?-><-?\|\|?-><- DUF499 *<- REase <- METHYLASE <-OLD-ABC-p<- TUDOR+SF1-SF2-Helicase+i REase +CTD+ REase | 975 | Bacteria>Terrabacteria_group>Cyanobacteriota/Melainabacteria_group>Cyanobacteriota>Cyanophyceae | GCA_002870755.1 | Fischerella_thermalis_CCMEE_5208 | CEN42_03245 | 8470..11397 | 2019669 | AAA-family-ATPase-[Fischerella-thermalis-CCMEE-5208] |
| PSB44266.1 | DUF499_1.RM.like.systems | <- HEPN+ DUF499 *<-?<- METHYLASE <- REase <-PIN<-AbrB<- TUDOR+SF1-SF2-Helicase+i REase +CTD+ REase | 1113 | Bacteria>Terrabacteria_group>Cyanobacteriota/Melainabacteria_group>Cyanobacteriota>Cyanophyceae | GCA_003004015.1 | Cyanosarcina_cf._burmensis_CCALA_770 | C7B80_21165 | 280..3621 | 2107697 | AAA+-family-ATPase-[Cyanosarcina-cf.-burmensis-CCALA-770] |
| TRU24939.1 | DUF499_1.RM.like.systems | ABhydrolase-> TUDOR+SF1-SF2-Helicase+i REase +CTD+ REase -> REase -> METHYLASE ->?->PIN-> HEPN+ DUF499 *-><-Cluster164_18clades<-?\|\|?->?->DUF29->PolB-> | 1130 | Bacteria>Terrabacteria_group>Cyanobacteriota/Melainabacteria_group>Cyanobacteriota>Cyanophyceae | GCA_007095795.1 | Microcystis_aeruginosa_Ma_MB_S_20031200_S102D | EWV79_09040 | 24486..27878 | 2486255 | ATP-binding-protein-[Microcystis-aeruginosa-Ma_MB_S_20031200_S102D] |
| NCR40139.1 | DUF499_1.RM.like.systems | ABhydrolase-> TUDOR+SF1-SF2-Helicase+i REase +CTD+ REase -> REase -> METHYLASE ->?->PIN-> HEPN+ DUF499 *-><-Cluster164_18clades | 1130 | Bacteria>Terrabacteria_group>Cyanobacteriota/Melainabacteria_group>Cyanobacteriota>Cyanophyceae | GCA_009995705.1 | Microcystis_aeruginosa_W13-11 | GPJ21_09840 | 22820..26212 | 2685991 | ATP-binding-protein-[Microcystis-aeruginosa-W13-11] |
| MBD1215623.1 | DUF499_1.RM.like.systems | TPR+TPR-><-?\|\|?-><- DUF499 *<- METHYLASE <-?<- REase +HTH<- TUDOR+SF1-SF2-Helicase+i REase +CTD+ REase | 913 | Bacteria>Terrabacteria_group>Cyanobacteriota/Melainabacteria_group>Cyanobacteriota>Cyanophyceae | GCA_014654715.1 | Dolichospermum_circinale_Clear-D4 | H9534_23500 | 3273..6014 | 2766971 | ATP-binding-protein-[Dolichospermum-circinale-Clear-D4] |
| MBI1240693.1 | DUF499_1.RM.like.systems | Cysteine_desulfurase-> TUDOR+SF1-SF2-Helicase+i REase +CTD+ REase -> METHYLASE -> REase -> DUF499 *-><-Peptidase_S51<-?<-?<-?\|\|sGTP-> | 973 | Bacteria>Terrabacteria_group>Cyanobacteriota/Melainabacteria_group>Cyanobacteriota>Cyanophyceae | GCA_016124525.1 | Nostoc_sp._RI_552 | GC195_03655 | 36340..39261 | 2654185 | DUF499-domain-containing-protein-[Nostoc-sp.-RI_552] |
| MBO1068320.1 | DUF499_1.RM.like.systems | <-NTP_transferase\|\|N6_N4_MTase-><-HTH_3<-?<-?<-?<- DUF499 *<-PIN<-?<- METHYLASE <-?<- REase +HTH<- TUDOR+SF1-SF2-Helicase+i REase +CTD+ REase | 911 | Bacteria>Terrabacteria_group>Cyanobacteriota/Melainabacteria_group>Cyanobacteriota>Cyanophyceae | GCA_017355655.1 | Dolichospermum_sp._DEX189 | HEQ13_02485 | 530022..532757 | 2717870 | ATP-binding-protein-[Dolichospermum-sp.-DEX189] |
| MBS9772311.1 | DUF499_1.RM.like.systems | <- DUF499 *<- REase <-PIN<-?<- METHYLASE <- TUDOR+SF1-SF2-Helicase+i REase +CTD+ REase | 1016 | Bacteria>Terrabacteria_group>Cyanobacteriota/Melainabacteria_group>Cyanobacteriota>Cyanophyceae | GCA_018457135.1 | Trichodesmium_erythraeum_GBRTRLIN201 | J7F05_18140 | 304..3354 | 1045867 | ATP-binding-protein-[Trichodesmium-erythraeum-GBRTRLIN201] |
| MBV6627478.1 | DUF499_1.RM.like.systems | <- HEPN+ DUF499 *<- REase <- METHYLASE -p\|\| w HTH+ REase -><- TUDOR+SF1-SF2-Helicase+i REase +CTD+ REase -p<- REase | 1116 | Bacteria>Terrabacteria_group>Cyanobacteriota/Melainabacteria_group>Cyanobacteriota>Cyanophyceae | GCA_019192545.1 | Rivularia_sp._MS3 | KI793_31915 | 3971..7321 | 3012223 | DUF499-domain-containing-protein-[Rivularia-sp.-(in:-Bacteria)] |
| MBW4455562.1 | DUF499_1.RM.like.systems | HEPN-><-?\|\| TUDOR+SF1-SF2-Helicase+i REase +CTD+ REase ->HicB+ZnR-> METHYLASE -p-> REase -> HEPN+ DUF499 *-><-Phage_int_SAM_1+Phage_integrase\|\|?->?-><-Response_reg+LuxR-HTH<-HerA/FtsK | 1112 | Bacteria>Terrabacteria_group>Cyanobacteriota/Melainabacteria_group>Cyanobacteriota>Cyanophyceae | GCA_019359215.1 | Nostoc_indistinguendum_CM1-VF10 | KME55_24210 | 27093..30431 | 287863 | DUF499-domain-containing-protein-[Nostoc-indistinguendum-CM1-VF10] |
| MBW4429556.1 | DUF499_1.RM.like.systems | TUDOR+SF1-SF2-Helicase+i REase +CTD+ REase -> METHYLASE -> REase -> DUF499 *->?->PIN-> | 1005 | Bacteria>Terrabacteria_group>Cyanobacteriota/Melainabacteria_group>Cyanobacteriota>Cyanophyceae | GCA_019359245.1 | Nostoc_desertorum_CM1-VF14 | KME50_35640 | 5971..8988 | 287862 | DUF499-domain-containing-protein-[Nostoc-desertorum-CM1-VF14] |
| WJI23189.1 | DUF499_1.RM.like.systems | <- HEPN+ DUF499 *<- REase <- METHYLASE <- TUDOR+SF1-SF2-Helicase+i REase +CTD+ REase \|\| REase -><-DDE_Tnp_1 | 1098 | Bacteria>Terrabacteria_group>Cyanobacteriota/Melainabacteria_group>Cyanobacteriota>Cyanophyceae | GCA_030322965.1 | Thermosynechococcus_sp._B0 | MZ909_08100 | 1635834..1639130 | 2937284 | Swt1-family-HEPN-domain-containing-protein-[Thermosynechococcus-sp.-B0] |
| CAH2571600.1 | DUF499_1.RM.like.systems | TUDOR+SF1-SF2-Helicase+i REase +CTD+ REase -> REase -> METHYLASE -> REase -> REase -> REase -> DUF499 *->?->?->RelB_antitoxin-> | 999 | Bacteria>Terrabacteria_group>Cyanobacteriota/Melainabacteria_group>Cyanobacteriota>Cyanophyceae | GCA_937024135.1 | Planktothrix_rubescens | PRNO82_00998 | 1099392..1102391 | 59512 | AAA-family-ATPase-[Planktothrix-rubescens] |
| WP_039990643.1 | DUF499_1.RM.like.systems | TUDOR+SF1-SF2-Helicase+i REase +CTD+ REase ->OLD-ABC-p-> METHYLASE -> REase -> DUF499 *-> | 955 | Bacteria>Terrabacteria_group>Cyanobacteriota/Melainabacteria_group>Cyanobacteriota>Cyanophyceae | GCF_000231365.1 | Fischerella_thermalis | FJSC11DRAFT_RS02260 | 574111..576978 | 372787 | DUF499-domain-containing-protein-[Fischerella-thermalis] |
| WP_015155777.1 | DUF499_1.RM.like.systems | Phage_integrase-><-Phage_integrase\|\|?-><- HEPN+ DUF499 *<-?<- METHYLASE <- REase <-PIN<-AbrB<- TUDOR+SF1-SF2-Helicase+i REase +CTD+ REase | 1113 | Bacteria>Terrabacteria_group>Cyanobacteriota/Melainabacteria_group>Cyanobacteriota>Cyanophyceae | GCF_000317125.1 | Chroococcidiopsis_thermalis | CHRO_RS18600 | 4302197..4305538 | 54299 | Swt1-family-HEPN-domain-containing-protein-[Chroococcidiopsis-thermalis] |
| WP_026797173.1 | DUF499_1.RM.like.systems | <-RelB_antitoxin<-?<-?<- DUF499 *<- REase <- REase <- REase <- METHYLASE <- REase <- TUDOR+SF1-SF2-Helicase+i REase +CTD+ REase | 999 | Bacteria>Terrabacteria_group>Cyanobacteriota/Melainabacteria_group>Cyanobacteriota>Cyanophyceae | GCF_000464765.1 | Planktothrix | PLAN_RS23755 | 2806320..2809319 | 54304 | MULTISPECIES:-DUF499-domain-containing-protein-[Planktothrix] |
| WP_023065229.1 | DUF499_1.RM.like.systems | TUDOR+SF1-SF2-Helicase+i REase +CTD+ REase -> METHYLASE -> REase -> DUF499 *-><-?\|\|?->?-><-NTP_transf_9<-TPR+TPR+TPR+TPR+TPR+TPR+TPR+TPR | 968 | Bacteria>Terrabacteria_group>Cyanobacteriota/Melainabacteria_group>Cyanobacteriota>Cyanophyceae | GCF_000478195.2 | Lyngbya_aestuarii | M595_RS06070 | 76585..79491 | 118322 | DUF499-domain-containing-protein-[Lyngbya-aestuarii] |
| WP_026733571.1 | DUF499_1.RM.like.systems | TUDOR+SF1-SF2-Helicase+i REase +CTD+ REase ->AbrB->PIN-> METHYLASE -> REase -> HEPN+ DUF499 *->?->TraJ-RHH-> | 1122 | Bacteria>Terrabacteria_group>Cyanobacteriota/Melainabacteria_group>Cyanobacteriota>Cyanophyceae | GCF_000517105.1 | Fischerella_sp._PCC_9605 | FIS9605_RS0116420 | 337165..340533 | 1173024 | Swt1-family-HEPN-domain-containing-protein-[Fischerella-sp.-PCC-9605] |
| WP_200888917.1 | DUF499_1.RM.like.systems | TUDOR+SF1-SF2-Helicase+i REase +CTD+ REase -> REase +HTH->?-> METHYLASE -p-> METHYLASE -p-> METHYLASE -p-> DUF499 *->?->Methyltransf_21->?->HTH_3-> REase -><-N6_N4_MTase | 909 | Bacteria>Terrabacteria_group>Cyanobacteriota/Melainabacteria_group>Cyanobacteriota>Cyanophyceae | GCF_000789435.1 | Aphanizomenon_flos-aquae | OA07_RS10570 | 7797..10526 | 1176 | DUF499-domain-containing-protein-[Aphanizomenon-flos-aquae] |
| WP_046663117.1 | DUF499_1.RM.like.systems | PP_kinase+HKD-DNase-><-?<-?\|\| TUDOR+SF1-SF2-Helicase+i REase +CTD+ REase -> REase -> METHYLASE -> HEPN+ DUF499 *-><-?<-?\|\|?->?-><-THUMP+Methylase | 1130 | Bacteria>Terrabacteria_group>Cyanobacteriota/Melainabacteria_group>Cyanobacteriota>Cyanophyceae | GCF_000981785.2 | Microcystis_aeruginosa | myaer_RS18245 | 3707749..3711141 | 1126 | Swt1-family-HEPN-domain-containing-protein-[Microcystis-aeruginosa] |
| WP_053539516.1 | DUF499_1.RM.like.systems | <-NTP_transferase\|\|N6_N4_MTase-><-HTH_3<-?<-?<- DUF499 *<-PIN<-?<- METHYLASE <- REase +HTH<- TUDOR+SF1-SF2-Helicase+i REase +CTD+ REase <-CHTH | 911 | Bacteria>Terrabacteria_group>Cyanobacteriota/Melainabacteria_group>Cyanobacteriota>Cyanophyceae | GCF_001277295.1 | Anabaena_sp._WA102 | AA650_RS14270 | 3307169..3309904 | 1647413 | DUF499-domain-containing-protein-[Anabaena-sp.-WA102] |
| WP_057177165.1 | DUF499_1.RM.like.systems | HKD-DNase+SF1-SF2-Helicase-p-><-HTH+MPTase<-?<-?\|\| TUDOR+SF1-SF2-Helicase+i REase +CTD+ REase -> METHYLASE -> HEPN+ DUF499 *-> REase ->?->Peptidase_C14-> | 1103 | Bacteria>Terrabacteria_group>Cyanobacteriota/Melainabacteria_group>Cyanobacteriota>Cyanophyceae | GCF_001432185.1 | Cylindrospermopsis_sp._CR12 | ASL19_RS02560 | 94062..97373 | 1747196 | Swt1-family-HEPN-domain-containing-protein-[Cylindrospermopsis-sp.-CR12] |
| WP_059001129.1 | DUF499_1.RM.like.systems | PIN->?-><- HEPN+ DUF499 *<- METHYLASE \|\| REase +HTH-><- TUDOR+SF1-SF2-Helicase+i REase +CTD+ REase <-TrwC+SF1-SF2-helicase+TOPRIM\|\|?->ParB+HNH-> | 1124 | Bacteria>Terrabacteria_group>Cyanobacteriota/Melainabacteria_group>Cyanobacteriota>Cyanophyceae | GCF_001485215.1 | Leptolyngbya_sp._NIES-2104 | NIES2104_RS25560 | 5320050..5323424 | 1552121 | Swt1-family-HEPN-domain-containing-protein-[Leptolyngbya-sp.-NIES-2104] |
| WP_065711823.1 | DUF499_1.RM.like.systems | REase -> TUDOR+SF1-SF2-Helicase+i REase +CTD+ REase ->OLD-ABC+TOPRIM->PolB-> DUF499 *->?->?->NACHT->?-><-HIN-HTH\|\|PIN-> | 989 | Bacteria>Terrabacteria_group>Cyanobacteriota/Melainabacteria_group>Cyanobacteriota>Cyanophyceae | GCF_001693275.1 | Picosynechococcus_sp._PCC_7117 | AWQ22_RS15050 | 32294..35263 | 195498 | DUF499-domain-containing-protein-[Synechococcus-sp.-PCC-7117] |
| WP_066030118.1 | DUF499_1.RM.like.systems | PP_kinase+HKD-DNase-><-?<-?\|\| TUDOR+SF1-SF2-Helicase+i REase +CTD+ REase -> REase -> METHYLASE -> HEPN+ DUF499 *-><-?<-?\|\|?->?-><-THUMP+Methylase | 1130 | Bacteria>Terrabacteria_group>Cyanobacteriota/Melainabacteria_group>Cyanobacteriota>Cyanophyceae | GCF_001704955.2 | Microcystis_aeruginosa | amyaer_RS18255 | 3708666..3712058 | 1126 | Swt1-family-HEPN-domain-containing-protein-[Microcystis-aeruginosa] |
| WP_073632965.1 | DUF499_1.RM.like.systems | TUDOR+SF1-SF2-Helicase+i REase +CTD+ REase -> METHYLASE -> REase ->?-> HEPN+ DUF499 *-><-HTH\|\|PIWI-> | 1113 | Bacteria>Terrabacteria_group>Cyanobacteriota/Melainabacteria_group>Cyanobacteriota>Cyanophyceae | GCF_001904675.1 | Scytonema_sp._HK-05 | NIES2130_RS19405 | 80454..83795 | 1137095 | Swt1-family-HEPN-domain-containing-protein-[Scytonema-sp.-HK-05] |
| WP_231828504.1 | DUF499_1.RM.like.systems | TUDOR+SF1-SF2-Helicase+i REase +CTD+ REase -> REase -> REase -> METHYLASE ->PAIREDC-HTH->PIN-> HEPN+ DUF499 *-><-Cluster164_18clades<-?\|\|?->?->DUF29-><-Y1_Tnp | 1129 | Bacteria>Terrabacteria_group>Cyanobacteriota/Melainabacteria_group>Cyanobacteriota>Cyanophyceae | GCF_002095975.1 | Microcystis_aeruginosa | BH695_RS02870 | 541185..544574 | 1126 | Swt1-family-HEPN-domain-containing-protein-[Microcystis-aeruginosa] |
| WP_096671658.1 | DUF499_1.RM.like.systems | <- DUF499 *<-PIN<-?<- METHYLASE <- METHYLASE -p<- REase +HTH<- TUDOR+SF1-SF2-Helicase+i REase +CTD+ REase | 912 | Bacteria>Terrabacteria_group>Cyanobacteriota/Melainabacteria_group>Cyanobacteriota>Cyanophyceae | GCF_002368115.1 | Dolichospermum_compactum | CA730_RS19465 | 4197332..4200070 | 136073 | DUF499-domain-containing-protein-[Dolichospermum-compactum] |
| WP_100904446.1 | DUF499_1.RM.like.systems | <-DDE_Tnp_1_3<- HEPN+ DUF499 *<- REase <- TUDOR+SF1-SF2-Helicase+i REase +CTD+ REase <-?<-?\|\|DndB-> | 1112 | Bacteria>Terrabacteria_group>Cyanobacteriota/Melainabacteria_group>Cyanobacteriota>Cyanophyceae | GCF_002813575.1 | Nostoc_flagelliforme | COO91_RS47960 | 745879..749217 | 1306274 | Swt1-family-HEPN-domain-containing-protein-[Nostoc-flagelliforme] |
| WP_257998857.1 | DUF499_1.RM.like.systems | ABC-ATPase+SbcC->?->?-><-?\|\|?-><- DUF499 *<- REase <- METHYLASE <-OLD-ABC-p<- TUDOR+SF1-SF2-Helicase+i REase +CTD+ REase | 955 | Bacteria>Terrabacteria_group>Cyanobacteriota/Melainabacteria_group>Cyanobacteriota>Cyanophyceae | GCF_002870755.1 | Fischerella_thermalis | CEN42_RS03265 | 8470..11337 | 372787 | DUF499-domain-containing-protein-[Fischerella-thermalis] |
| WP_104908185.1 | DUF499_1.RM.like.systems | TUDOR+SF1-SF2-Helicase+i REase +CTD+ REase ->Phd-YefM->PIN-><-HTH_3\|\| METHYLASE -> REase -> HEPN+ DUF499 *-> | 1109 | Bacteria>Terrabacteria_group>Cyanobacteriota/Melainabacteria_group>Cyanobacteriota>Cyanophyceae | GCF_002949795.1 | Nostoc_sp._'Lobaria_pulmonaria_(5183)_cyanobiont' | NLP_RS21725 | 4933532..4936861 | 1618022 | Swt1-family-HEPN-domain-containing-protein-[Nostoc-sp.-'Lobaria-pulmonaria-(5183)-cyanobiont'] |
| WP_106289188.1 | DUF499_1.RM.like.systems | <- DUF499 *<- REase <- METHYLASE <-OLD-ABC<- TUDOR+SF1-SF2-Helicase+i REase +CTD+ REase <-CHTH\|\|OBFOLD-NUCLEASE-> | 994 | Bacteria>Terrabacteria_group>Cyanobacteriota/Melainabacteria_group>Cyanobacteriota>Cyanophyceae | GCF_003003775.1 | Merismopedia_glauca | C7B64_RS13505 | 1286..4270 | 292586 | DUF499-domain-containing-protein-[Merismopedia-glauca] |
| WP_106908219.1 | DUF499_1.RM.like.systems | Cluster164_18clades-><-?<- HEPN+ DUF499 *<-ACET<-?<- METHYLASE <- REase \|\|?-><- TUDOR+SF1-SF2-Helicase+i REase +CTD+ REase | 1130 | Bacteria>Terrabacteria_group>Cyanobacteriota/Melainabacteria_group>Cyanobacteriota>Cyanophyceae | GCF_003019735.1 | Microcystis_sp._MC19 | B5D77_RS02535 | 488054..491446 | 1967666 | Swt1-family-HEPN-domain-containing-protein-[Microcystis-sp.-MC19] |
| WP_110545657.1 | DUF499_1.RM.like.systems | TUDOR+SF1-SF2-Helicase+i REase +CTD+ REase -> REase -> METHYLASE -> METHYLASE -p->?->ACET-> HEPN+ DUF499 *-><-Cluster164_18clades | 1111 | Bacteria>Terrabacteria_group>Cyanobacteriota/Melainabacteria_group>Cyanobacteriota>Cyanophyceae | GCF_003206625.1 | Microcystis_aeruginosa | MTo_RS18515 | 13505..16840 | 1126 | Swt1-family-HEPN-domain-containing-protein-[Microcystis-aeruginosa] |
| WP_141294973.1 | DUF499_1.RM.like.systems | TUDOR+SF1-SF2-Helicase+i REase +CTD+ REase -> REase -> METHYLASE -> REase -> REase -> REase -> DUF499 *->?->?->?-><-Radical-SAM<-?<-AbrB-like | 999 | Bacteria>Terrabacteria_group>Cyanobacteriota/Melainabacteria_group>Cyanobacteriota>Cyanophyceae | GCF_005402765.1 | Planktothrix_agardhii | FLZ02_RS17820 | 16554..19553 | 1160 | DUF499-domain-containing-protein-[Planktothrix-agardhii] |
| WP_137909444.1 | DUF499_1.RM.like.systems | TUDOR+SF1-SF2-Helicase+i REase +CTD+ REase -> REase +HTH->?-> METHYLASE -> DUF499 *-><-?\|\|?-><-TPR+TPR | 913 | Bacteria>Terrabacteria_group>Cyanobacteriota/Melainabacteria_group>Cyanobacteriota>Cyanophyceae | GCF_005402965.1 | Dolichospermum_planctonicum | FFO06_RS19070 | 11296..14037 | 136072 | DUF499-domain-containing-protein-[Dolichospermum-planctonicum] |
| WP_189523574.1 | DUF499_1.RM.like.systems | <-ArdA\|\|MetJarc-HTH->?-><-?<-?<-?<- DUF499 *<- REase <- METHYLASE <-OLD-ABC<- TUDOR+SF1-SF2-Helicase+i REase +CTD+ REase \|\|?-><-HKD-DNase+SF1-SF2-Helicase | 962 | Bacteria>Terrabacteria_group>Cyanobacteriota/Melainabacteria_group>Cyanobacteriota>Cyanophyceae | GCF_005869855.1 | Nostoc_sp._PA-18-2419 | FCR06_RS25145 | 41137..44025 | 2575443 | DUF499-domain-containing-protein-[Nostoc-sp.-PA-18-2419] |
| WP_149979248.1 | DUF499_1.RM.like.systems | Cluster164_18clades-><- HEPN+ DUF499 *<-ACET<-?<- METHYLASE <- REase \|\|?-><- TUDOR+SF1-SF2-Helicase+i REase +CTD+ REase | 1130 | Bacteria>Terrabacteria_group>Cyanobacteriota/Melainabacteria_group>Cyanobacteriota>Cyanophyceae | GCF_008579225.1 | Microcystis_aeruginosa | MiYa_RS12425 | 10420..13812 | 1126 | Swt1-family-HEPN-domain-containing-protein-[Microcystis-aeruginosa] |
| WP_149976793.1 | DUF499_1.RM.like.systems | <- HEPN+ DUF499 *<- METHYLASE <- REase <- TUDOR+SF1-SF2-Helicase+i REase +CTD+ REase | 1130 | Bacteria>Terrabacteria_group>Cyanobacteriota/Melainabacteria_group>Cyanobacteriota>Cyanophyceae | GCF_008579445.1 | Microcystis_aeruginosa | MiTs_RS18945 | 1418..4810 | 1126 | Swt1-family-HEPN-domain-containing-protein-[Microcystis-aeruginosa] |
| WP_149989607.1 | DUF499_1.RM.like.systems | Cluster164_18clades-><-?<- HEPN+ DUF499 *<-ACET<-?<- METHYLASE <- REase \|\|?-><- TUDOR+SF1-SF2-Helicase+i REase +CTD+ REase | 1117 | Bacteria>Terrabacteria_group>Cyanobacteriota/Melainabacteria_group>Cyanobacteriota>Cyanophyceae | GCF_008579565.1 | Microcystis_aeruginosa | MiHa_RS15955 | 12499..15852 | 1126 | Swt1-family-HEPN-domain-containing-protein-[Microcystis-aeruginosa] |
| WP_149977735.1 | DUF499_1.RM.like.systems | Cluster164_18clades-><-?<-?<- HEPN+ DUF499 *<-ACET<-?<- METHYLASE <- REase <- TUDOR+SF1-SF2-Helicase+i REase +CTD+ REase <-ABhydrolase | 1130 | Bacteria>Terrabacteria_group>Cyanobacteriota/Melainabacteria_group>Cyanobacteriota>Cyanophyceae | GCF_008579685.1 | Microcystis_aeruginosa | MiTa_RS10465 | 30953..34345 | 1126 | Swt1-family-HEPN-domain-containing-protein-[Microcystis-aeruginosa] |
| WP_168507185.1 | DUF499_1.RM.like.systems | <- DUF499 *<- REase <- METHYLASE <-PIN<- TUDOR+SF1-SF2-Helicase+i REase +CTD+ REase <-CHTH | 962 | Bacteria>Terrabacteria_group>Cyanobacteriota/Melainabacteria_group>Cyanobacteriota>Cyanophyceae | GCF_009712035.1 | Anabaena_sp._UHCC_0187 | FJR11_RS20795 | 2169..5057 | 2590018 | DUF499-domain-containing-protein-[Anabaena-sp.-UHCC-0187] |
| WP_159253859.1 | DUF499_1.RM.like.systems | TUDOR+SF1-SF2-Helicase+i REase +CTD+ REase -> REase -> METHYLASE ->OLD-ABC->PIN-> HEPN+ DUF499 *-><-?<-?\|\|?->?-><-THUMP+Methylase | 1130 | Bacteria>Terrabacteria_group>Cyanobacteriota/Melainabacteria_group>Cyanobacteriota>Cyanophyceae | GCF_009811855.1 | Microcystis_aeruginosa | GVO78_RS16540 | 16290..19682 | 1126 | Swt1-family-HEPN-domain-containing-protein-[Microcystis-aeruginosa] |
| WP_159297746.1 | DUF499_1.RM.like.systems | TUDOR+SF1-SF2-Helicase+i REase +CTD+ REase -> REase -> METHYLASE ->OLD-ABC->PIN-> HEPN+ DUF499 *-><-?<-?\|\|?->?-><-THUMP+Methylase | 1130 | Bacteria>Terrabacteria_group>Cyanobacteriota/Melainabacteria_group>Cyanobacteriota>Cyanophyceae | GCF_009811875.1 | Microcystis_aeruginosa | GVO69_RS15755 | 16338..19730 | 1126 | Swt1-family-HEPN-domain-containing-protein-[Microcystis-aeruginosa] |
| WP_181929825.1 | DUF499_1.RM.like.systems | <-TetR-HTH<- DUF499 *<- METHYLASE <- METHYLASE -p<- REase <- TUDOR+SF1-SF2-Helicase+i REase +CTD+ REase \|\|CHTH-> | 966 | Bacteria>Terrabacteria_group>Cyanobacteriota/Melainabacteria_group>Cyanobacteriota>Cyanophyceae | GCF_014023275.1 | Nostoc_edaphicum | HUN01_RS33840 | 7488405..7491305 | 264686 | DUF499-domain-containing-protein-[Nostoc-edaphicum] |
| WP_190667193.1 | DUF499_1.RM.like.systems | CHTH-> TUDOR+SF1-SF2-Helicase+i REase +CTD+ REase -> METHYLASE -> REase -> DUF499 *-><- REase <-CHTH | 1004 | Bacteria>Terrabacteria_group>Cyanobacteriota/Melainabacteria_group>Cyanobacteriota>Cyanophyceae | GCF_014695285.1 | Coleofasciculus_sp._FACHB-501 | H6F54_RS11880 | 20325..23339 | 2692786 | DUF499-domain-containing-protein-[Coleofasciculus-sp.-FACHB-501] |
| WP_190676125.1 | DUF499_1.RM.like.systems | <-pPIWI+pPIWI\|\|ParB-><-DDE_3<-PAIREDC-HTH\|\|ParB-><- TUDOR+SF1-SF2-Helicase+i REase +CTD+ REase <- HEPN+ DUF499 *<-PIN<-OLD-ABC<- METHYLASE <- REase \|\|?-><- TUDOR+SF1-SF2-Helicase+i REase +CTD+ REase | 1112 | Bacteria>Terrabacteria_group>Cyanobacteriota/Melainabacteria_group>Cyanobacteriota>Cyanophyceae | GCF_014695435.1 | Coleofasciculus_sp._FACHB-129 | H6F60_RS14220 | 70669..74007 | 2692785 | Swt1-family-HEPN-domain-containing-protein-[Coleofasciculus-sp.-FACHB-129] |
| WP_190447191.1 | DUF499_1.RM.like.systems | Prim-Pol->CHTH-> TUDOR+SF1-SF2-Helicase+i REase +CTD+ REase ->RnaseIII+DSRBD-> METHYLASE -> REase -> DUF499 *->?->PAPS_reduct->?->ParB-><- REase | 968 | Bacteria>Terrabacteria_group>Cyanobacteriota/Melainabacteria_group>Cyanobacteriota>Cyanophyceae | GCF_014695795.1 | Phormidium_sp._FACHB-592 | H6F86_RS05945 | 30309..33215 | 2692850 | DUF499-domain-containing-protein-[Phormidium-sp.-FACHB-592] |
| WP_190397164.1 | DUF499_1.RM.like.systems | TPR+TPR-><-?\|\|?-><- DUF499 *<- METHYLASE <-?<- REase +HTH<- TUDOR+SF1-SF2-Helicase+i REase +CTD+ REase | 913 | Bacteria>Terrabacteria_group>Cyanobacteriota/Melainabacteria_group>Cyanobacteriota>Cyanophyceae | GCF_014696335.1 | Anabaena_sp._FACHB-1250 | H6F39_RS18020 | 6551..9292 | 2692770 | DUF499-domain-containing-protein-[Anabaena-sp.-FACHB-1250] |
| WP_190372718.1 | DUF499_1.RM.like.systems | TPR+TPR-><-?\|\|?-><- DUF499 *<- METHYLASE <-?<-?<- REase +HTH<- TUDOR+SF1-SF2-Helicase+i REase +CTD+ REase | 913 | Bacteria>Terrabacteria_group>Cyanobacteriota/Melainabacteria_group>Cyanobacteriota>Cyanophyceae | GCF_014696755.1 | Anabaena_sp._FACHB-1391 | H6F62_RS19000 | 6552..9293 | 2692771 | DUF499-domain-containing-protein-[Anabaena-sp.-FACHB-1391] |
| WP_190576377.1 | DUF499_1.RM.like.systems | RelE-ParE-><-?<-URI-DNase<-?<-?<- HEPN+ DUF499 *<- METHYLASE <- REase <-AbrB<- TUDOR+SF1-SF2-Helicase+i REase +CTD+ REase | 1117 | Bacteria>Terrabacteria_group>Cyanobacteriota/Melainabacteria_group>Cyanobacteriota>Cyanophyceae | GCF_014697025.1 | Alkalinema_sp._FACHB-956 | H6G21_RS22810 | 35052..38405 | 2692768 | Swt1-family-HEPN-domain-containing-protein-[Alkalinema-sp.-FACHB-956] |
| WP_193849072.1 | DUF499_1.RM.like.systems | TUDOR+SF1-SF2-Helicase+i REase +CTD+ REase -><-?\|\| METHYLASE -> REase -> DUF499 *->?-><-?\|\|?-><-?\|\| REase +HTH-> | 969 | Bacteria>Terrabacteria_group>Cyanobacteriota/Melainabacteria_group>Cyanobacteriota>Cyanophyceae | GCF_015207785.1 | Dolichospermum_flos-aquae | IQ222_RS12495 | 19378..22287 | 1166 | DUF499-domain-containing-protein-[Dolichospermum-flos-aquae] |
| WP_193849784.1 | DUF499_1.RM.like.systems | TUDOR+SF1-SF2-Helicase+i REase +CTD+ REase -> REase +HTH->?-> METHYLASE -><-?\|\| DUF499 *-> | 913 | Bacteria>Terrabacteria_group>Cyanobacteriota/Melainabacteria_group>Cyanobacteriota>Cyanophyceae | GCF_015207785.1 | Dolichospermum_flos-aquae | IQ222_RS15370 | 28947..31688 | 1166 | DUF499-domain-containing-protein-[Dolichospermum-flos-aquae] |
| WP_200988810.1 | DUF499_1.RM.like.systems | PRTase-><-Y1_Tnp\|\|?->?->?-><- HEPN+ DUF499 *<- METHYLASE \|\| REase +HTH-> REase +HTH-><- TUDOR+SF1-SF2-Helicase+i REase +CTD+ REase | 1108 | Bacteria>Terrabacteria_group>Cyanobacteriota/Melainabacteria_group>Cyanobacteriota>Cyanophyceae | GCF_015245355.1 | Anabaenopsis_elenkinii | IM676_RS02385 | 515316..518642 | 156213 | Swt1-family-HEPN-domain-containing-protein-[Anabaenopsis-elenkinii] |
| WP_168636740.1 | DUF499_1.RM.like.systems | TUDOR+SF1-SF2-Helicase+i REase +CTD+ REase -> REase +HTH->?-> METHYLASE ->?->PIN-> DUF499 *->?->?->HTH_3-> REase -><-N6_N4_MTase\|\|NTP_transferase-> | 911 | Bacteria>Terrabacteria_group>Cyanobacteriota/Melainabacteria_group>Cyanobacteriota>Cyanophyceae | GCF_017355725.1 | Nostocales | HEQ36_RS21575 | 43162..45897 | 1161 | DUF499-domain-containing-protein-[Dolichospermum-flos-aquae] |
| WP_211941813.1 | DUF499_1.RM.like.systems | HKD-DNase+SF1-SF2-Helicase-p-><-HTH+MPTase<-?<-?\|\| TUDOR+SF1-SF2-Helicase+i REase +CTD+ REase -> METHYLASE -> HEPN+ DUF499 *-> REase ->?->Peptidase_C14-> | 1103 | Bacteria>Terrabacteria_group>Cyanobacteriota/Melainabacteria_group>Cyanobacteriota>Cyanophyceae | GCF_018139025.1 | Cylindrospermopsis_raciborskii | J9B92_RS12735 | 2844714..2848025 | 77022 | Swt1-family-HEPN-domain-containing-protein-[Cylindrospermopsis-raciborskii] |
| WP_245395605.1 | DUF499_1.RM.like.systems | <-DDE_Tnp_1\|\| TUDOR+SF1-SF2-Helicase+i REase +CTD+ REase -><-?\|\| METHYLASE -> REase -> DUF499 *->?->GDResolvase+HIN-HTH-><- REase +HTH | 977 | Bacteria>Terrabacteria_group>Cyanobacteriota/Melainabacteria_group>Cyanobacteriota>Cyanophyceae | GCF_018389385.1 | Anthocerotibacter_panamensis | IL331_RS05745 | 1255834..1258767 | 2857077 | DUF499-domain-containing-protein-[Anthocerotibacter-panamensis] |
| WP_230840692.1 | DUF499_1.RM.like.systems | TUDOR+SF1-SF2-Helicase+i REase +CTD+ REase -> REase -> METHYLASE ->DDE_Tnp_1-> METHYLASE -p->?-> HEPN+ DUF499 *-><-?<-Peptidase_U32+Peptidase_U32_C | 1101 | Bacteria>Terrabacteria_group>Cyanobacteriota/Melainabacteria_group>Cyanobacteriota>Cyanophyceae | GCF_021018745.1 | Gloeobacter_morelensis | ISF26_RS17965 | 3744120..3747425 | 2907343 | Swt1-family-HEPN-domain-containing-protein-[Gloeobacter-morelensis] |
| WP_244349767.1 | DUF499_1.RM.like.systems | TUDOR+SF1-SF2-Helicase+i REase +CTD+ REase -> METHYLASE ->?->PemK_toxin-> REase -> REase -> HEPN+ DUF499 *->?->ABC-ATPase+SbcC->?->?-><-ABC-ATPase+SbcC<-ABC-ATPase+SbcC | 1107 | Bacteria>Terrabacteria_group>Cyanobacteriota/Melainabacteria_group>Cyanobacteriota>Cyanophyceae | GCF_022848905.1 | Thermostichus_vulcanus | JX360_RS06145 | 28078..31401 | 32053 | Swt1-family-HEPN-domain-containing-protein-[Thermostichus-vulcanus] |
| WP_045875055.1 | DUF499_1.RM.like.systems | <-HTH_37<-GDResolvase+Recombinase<-TraJ-RHH\|\|?->Phage_int_SAM_1+Phage_integrase-><-?<- HEPN+ DUF499 *<- REase <-?<- REase <- METHYLASE <-HicB+ZnR<- TUDOR+SF1-SF2-Helicase+i REase +CTD+ REase | 1112 | Bacteria>Terrabacteria_group>Cyanobacteriota/Melainabacteria_group>Cyanobacteriota>Cyanophyceae | GCF_025860405.1 | unclassified_Tolypothrix | FDUTEX481_RS39715 | 28585..31923 | 2649714 | MULTISPECIES:-Swt1-family-HEPN-domain-containing-protein-[unclassified-Tolypothrix] |
| WP_190359815.1 | DUF499_1.RM.like.systems | REase -><-DUF29<-?<-?\|\|?->Cluster164_18clades-><- HEPN+ DUF499 *<-PIN<-RHH_6<- METHYLASE <- REase <- TUDOR+SF1-SF2-Helicase+i REase +CTD+ REase <-ABhydrolase | 1117 | Bacteria>Terrabacteria_group>Cyanobacteriota/Melainabacteria_group>Cyanobacteriota>Cyanophyceae | GCF_026222535.1 | Microcystis | H6F92_RS24355 | 16318..19671 | 1125 | Swt1-family-HEPN-domain-containing-protein-[Microcystis-wesenbergii] |
| WP_272078194.1 | DUF499_1.RM.like.systems | DDE_3-><- HEPN+ DUF499 *<-PIN<-PIN<-RHH_6<- METHYLASE <- REase <- TUDOR+SF1-SF2-Helicase+i REase +CTD+ REase | 1111 | Bacteria>Terrabacteria_group>Cyanobacteriota/Melainabacteria_group>Cyanobacteriota>Cyanophyceae | GCF_028331105.1 | Microcystis_aeruginosa | PN487_RS19700 | 7822..11157 | 1126 | Swt1-family-HEPN-domain-containing-protein-[Microcystis-aeruginosa] |
| WP_254175247.1 | DUF499_1.RM.like.systems | <- DUF499 *<- REase <- METHYLASE <-?<-SbcC<- TUDOR+SF1-SF2-Helicase+i REase +CTD+ REase <-RHH_1 | 998 | Bacteria>Terrabacteria_group>Cyanobacteriota/Melainabacteria_group>Cyanobacteriota>Cyanophyceae | GCF_904830925.1 | Planktothrix_pseudagardhii | NMK36_RS28460 | 5796..8792 | 132604 | DUF499-domain-containing-protein-[Planktothrix-pseudagardhii] |
| WP_006634723.1 | DUF499_1.RM.like.systems | NTP_transferase->ST-Hiskin->?-> TUDOR+SF1-SF2-Helicase+i REase +CTD+ REase -> METHYLASE -> REase -> DUF499 *-> | 998 | Bacteria>Terrabacteria_group>Cyanobacteriota/Melainabacteria_group>Cyanobacteriota>Cyanophyceae | - | Microcoleus_vaginatus | MicvaDRAFT_4687 | 60148..63144 | 119532 | DUF499-domain-containing-protein-[Microcoleus-vaginatus] |
| WP_235758599.1 | DUF499_1.RM.like.systems | TUDOR+SF1-SF2-Helicase+i REase +CTD+ REase -p->PIN-> METHYLASE -p-> REase -> REase -> REase -> DUF499 *-><-TOPRIM+SF1-SF2-Helicase<-ZnR<-?<-?<-URI-DNase | 1002 | Bacteria>Terrabacteria_group>Cyanobacteriota/Melainabacteria_group>Cyanobacteriota>Cyanophyceae | - | Planktothrix_agardhii | L2E81_RS19965 | 6975..9983 | 1160 | DUF499-domain-containing-protein-[Planktothrix-agardhii] |
| OWY65325.1 | DUF499_1.RM.like.systems | Phage_integrase-><-Phage_integrase<- HEPN+ DUF499 *<-DUF29<- REase <-PIN<-AbrB<- TUDOR+SF1-SF2-Helicase+i REase +CTD+ REase \|\|Classical-AAA-> | 1119 | Bacteria>Terrabacteria_group>Cyanobacteriota/Melainabacteria_group>Cyanobacteriota>unclassified_Cyanobacteriota | GCA_002213405.1 | cyanobacterium_TDX16 | B7486_42500 | 6012..9371 | 1503470 | AAA+-family-ATPase-[cyanobacterium-TDX16] |
| HBB33200.1 | DUF499_1.RM.like.systems | TUDOR+SF1-SF2-Helicase+i REase +CTD+ REase -> REase -> METHYLASE -p-> METHYLASE ->Polbeta->MqsA_antitoxin+HTH_3-> DUF499 *-> | 1007 | Bacteria>Terrabacteria_group>Cyanobacteriota/Melainabacteria_group>Cyanobacteriota>unclassified_Cyanobacteriota | GCA_003488275.1 | Cyanobacteria_bacterium_UBA9273 | DC064_15720 | 10674..13697 | 2055791 | AAA-family-ATPase-[Cyanobacteria-bacterium-UBA9273] |
| MBJ7900527.1 | DUF499_1.RM.like.systems | <-Glyco_hydro_57+DUF3536<-?\|\|?->HAD-><- HEPN+ DUF499 *<- REase <-PIN<-OLD-ABC<-CopG/RHH<-?<- TUDOR+SF1-SF2-Helicase+i REase +CTD+ REase | 1111 | Bacteria>Terrabacteria_group>Cyanobacteriota/Melainabacteria_group>Cyanobacteriota>unclassified_Cyanobacteriota | GCA_016471375.1 | Cyanobacteria_bacterium_RI_101 | GC158_11540 | 10083..13418 | 2775578 | DUF499-domain-containing-protein-[Cyanobacteria-bacterium-RI_101] |
| WP_229782279.1 | DUF499_1.RM.like.systems | OrfB_Zn_ribbon->?->ParB-><- DUF499 *<- REase <-OLD-ABC <- METHYLASE <- TUDOR+SF1-SF2-Helicase+i REase +CTD+ REase | 959 | Bacteria>Terrabacteria_group>Deinococcota>Deinococci>Deinococcales | GCF_014647175.1 | Deinococcus_daejeonensis | IEY59_RS18160 | 23471..26350 | 1007098 | DUF499-domain-containing-protein-[Deinococcus-daejeonensis] |
| GIW33264.1 | DUF499_1.RM.like.systems | Pkinase->DUF4388-><-?<-?\|\|?-><- HEPN+ DUF499 *<-HEPN<-PolB<- METHYLASE <-ORC-CDC6+ REase <- TUDOR+SF1-SF2-Helicase+i REase +CTD+ REase | 1104 | Bacteria>Terrabacteria_group>Deinococcota>Deinococci>Thermales | GCA_026004115.1 | Meiothermus_sp. | KatS3mg072_0597 | 655659..658973 | 1955249 | MAG:-hypothetical-protein-KatS3mg072_0597-[Meiothermus-sp.] |
| WP_027888723.1 | DUF499_1.RM.like.systems | <-PAIREDC-HTH<-BrxL<-?<-?<-?<-ORC-CDC6+ REase <- HEPN+ DUF499 *<- METHYLASE <- TUDOR+SF1-SF2-Helicase+i REase +CTD+ REase \|\|DDE_Tnp_IS1-><-?\|\|?->TPR+TPR-> | 1105 | Bacteria>Terrabacteria_group>Deinococcota>Deinococci>Thermales | GCF_003568585.1 | Meiothermus_taiwanensis | MTAI_RS14270 | 92295..95612 | 172827 | Swt1-family-HEPN-domain-containing-protein-[Meiothermus-taiwanensis] |
| WP_119361752.1 | DUF499_1.RM.like.systems | TUDOR+SF1-SF2-Helicase+i REase +CTD+ REase -> METHYLASE -> HEPN+ DUF499 *->ORC-CDC6+ REase -> | 1108 | Bacteria>Terrabacteria_group>Deinococcota>Deinococci>Thermales | GCF_003574045.1 | Meiothermus_taiwanensis | Mcate_RS11630 | 6951..10277 | 172827 | Swt1-family-HEPN-domain-containing-protein-[Meiothermus-taiwanensis] |
| WP_126203060.1 | DUF499_1.RM.like.systems | TUDOR+SF1-SF2-Helicase+i REase +CTD+ REase ->Schlafen+HATPase+HTH-> METHYLASE -> REase -> DUF499 *-><-ParB | 924 | Bacteria>Terrabacteria_group>Deinococcota>Deinococci>Thermales | GCF_003959165.1 | Thermus_scotoductus | CSW41_RS02895 | 15669..18443 | 37636 | DUF499-domain-containing-protein-[Thermus-scotoductus] |
| KPA15639.1 | DUF499_1.RM.like.systems | TUDOR+SF1-SF2-Helicase+i REase +CTD+ REase -p->HIN-HTH-> TUDOR+SF1-SF2-Helicase+i REase +CTD+ REase -> REase +HTH-> METHYLASE -> DUF499 *-><- REase | 941 | Bacteria>Thermodesulfobacteriota>Desulfobacteria>Desulfobacterales>Candidatus_Magnetomoraceae | GCA_001292585.1 | Candidatus_Magnetomorum_sp._HK-1 | MHK_004158 | 9986..12811 | 1509431 | ATPase-AAA-[Candidatus-Magnetomorum-sp.-HK-1] |
| VEN75475.1 | DUF499_1.RM.like.systems | 35exo->?->Pterin_bind->?->tRNA-><-URI-DNase<- DUF499 *<- METHYLASE <-HEPN<- TUDOR+SF1-SF2-Helicase+i REase +CTD+ REase <- w HTH+ REase <-?<- METHYLASE | 988 | Bacteria>Thermodesulfobacteriota>Desulfobacteria>Desulfobacterales>Desulfobacteraceae | GCA_900659855.1 | uncultured_Desulfobacteraceae_bacterium | EPICR_90073 | 73527..76493 | 218296 | conserved-hypothetical-protein-[uncultured-Desulfobacteraceae-bacterium] |
| WP_178366469.1 | DUF499_1.RM.like.systems | <- DUF499 *<- REase <-OLD-ABC <- METHYLASE <-OLD-ABC<- TUDOR+SF1-SF2-Helicase+i REase +CTD+ REase <-Radical-SAM+TRAM | 952 | Bacteria>Thermodesulfobacteriota>Desulfobacteria>Desulfobacterales>Desulfobacteraceae | GCF_013389365.1 | Desulfobacter_latus | HXW94_RS08505 | 32689..35547 | 2292 | DUF499-domain-containing-protein-[Desulfobacter-latus] |
| MBC8360097.1 | DUF499_1.RM.like.systems | TUDOR+SF1-SF2-Helicase+i REase +CTD+ REase ->Schlafen+HATPase+HTH->?->PHP+SbcC-> METHYLASE -> REase -> DUF499 *-> | 920 | Bacteria>Thermodesulfobacteriota>Desulfobacteria>Desulfobacterales>Desulfobacterales_incertae_sedis | GCA_014382665.1 | Candidatus_Desulfatibia_profunda | H8E23_01700 | 12931..15693 | 2841695 | ATP-binding-protein-[Desulfobacterales-bacterium] |
| MBC8432173.1 | DUF499_1.RM.like.systems | <- DUF499 *<- REase <- METHYLASE <-?<- TUDOR+SF1-SF2-Helicase+i REase +CTD+ REase | 979 | Bacteria>Thermodesulfobacteriota>Desulfobacteria>Desulfobacterales>Desulfobacterales_incertae_sedis | GCA_014383215.1 | Candidatus_Desulfatibia_vada | H8D96_09645 | 2855..5794 | 2841696 | DUF499-domain-containing-protein-[Desulfobacterales-bacterium] |
| WP_168935605.1 | DUF499_1.RM.like.systems | <-HTH_3\|\|?-><-?\|\|?->?-><- DUF499 *<- REase <-OLD-ABC <- METHYLASE <-HNH<-OLD-ABC<- TUDOR+SF1-SF2-Helicase+i REase +CTD+ REase | 933 | Bacteria>Thermodesulfobacteriota>Desulfovibrionia>Desulfovibrionales>Desulfovibrionaceae | GCF_012843875.1 | Desulfovibrio_piger | HF854_RS06725 | 17522..20323 | 901 | DUF499-domain-containing-protein-[Desulfovibrio-piger] |

<table>
<tr><th>Protein-ID</th><th>System</th><th>Gene-Neighbourhoods</th><th>Length</th><th>Taxonomy</th><th>Assembly-ID</th><th>Strain</th><th>Locus-Tag</th><th>Genomic-coordinates</th><th>TaxID</th><th>Fasta-define</th></tr>
<tr><td>HBL52555.1</td><td>DUF499_1.RM.like.systems</td><td>TUDOR+SF1-SF2-Helicase+i REase +CTD+ REase -p->ORC-CDC6+ REase ->MqsA_antitoxin+HTH_3-> HEPN+ DUF499 *-></td><td>1127</td><td>Bacteria>Thermodesulfobacteriota>Syntrophia>Syntrophales>Syntrophaceae</td><td>GCA_003506745.1</td><td>Syntrophaceae_bacterium</td><td>DDZ34_00420</td><td>4849..8232</td><td>2052184</td><td>AAA+-family-ATPase-[Syntrophaceae-bacterium]</td></tr>
<tr><td>MBS3905937.1</td><td>DUF499_1.RM.like.systems</td><td>TOPRIM-> TUDOR+SF1-SF2-Helicase+i REase +CTD+ REase ->Schlafen+HATPase+HTH->?-> METHYLASE ->AAA-ATPase+ REase -> DUF499 *->SF1-SF2-Helicase->HTH-><-TPR</td><td>1115</td><td>Bacteria>Thermodesulfobacteriota>Syntrophia>Syntrophales>Syntrophaceae</td><td>GCA_018333715.1</td><td>Syntrophaceae_bacterium</td><td>KGZ49_02765</td><td>19111..22458</td><td>2052184</td><td>ATP-binding-protein-[Syntrophaceae-bacterium]</td></tr>
<tr><td>MCK8600135.1</td><td>DUF499_1.RM.like.systems</td><td>REase +NACHT-><-Polbeta<-?<-?<- DUF499 *<- REase <- METHYLASE <-?<-SF1-SF2-Helicase<-SbcC<- TUDOR+SF1-SF2-Helicase+i REase +CTD+ REase</td><td>921</td><td>Bacteria>Thermodesulfobacteriota>Syntrophobacteria>Syntrophobacterales>Syntrophobacteraceae</td><td>GCA_022603395.1</td><td>Desulfoferrobacter_suflitae</td><td>K1V27_00335</td><td>59530..62295</td><td>2865782</td><td>DUF499-domain-containing-protein-[Syntrophobacteraceae-bacterium-DRH4]</td></tr>
<tr><td>WP_258927534.1</td><td>DUF499_1.RM.like.systems</td><td>REase +NACHT-><-Polbeta<-?<-?<- DUF499 *<- REase <- METHYLASE <-?<-SF1-SF2-Helicase<-SbcC<- TUDOR+SF1-SF2-Helicase+i REase +CTD+ REase</td><td>921</td><td>Bacteria>Thermodesulfobacteriota>Syntrophobacteria>Syntrophobacterales>Syntrophobacteraceae</td><td>GCF_022603395.1</td><td>Desulfoferrobacter_suflitae</td><td>K1V27_RS00340</td><td>59530..62295</td><td>2865782</td><td>DUF499-domain-containing-protein-[Desulfoferrobacter-suflitae]</td></tr>
<tr><td>WP_028895668.1</td><td>DUF499_1.RM.like.systems</td><td>XIS-HTH-> TUDOR+SF1-SF2-Helicase+i REase +CTD+ REase -> w HTH+ REase ->?->VirB4-FtsK+HerA-> METHYLASE -> DUF499 *->Classical-AAA->?->Peptidase_S8-><-?||HTH_3-></td><td>965</td><td>Bacteria>Thermodesulfobacteriota>Syntrophorhabdia>Syntrophorhabdales>Syntrophorhabdaceae</td><td>GCF_000512235.1</td><td>Syntrophorhabdus_aromaticivorans</td><td>SYNAR_RS0117510</td><td>3034488..3037385</td><td>328301</td><td>DUF499-domain-containing-protein-[Syntrophorhabdus-aromaticivorans]</td></tr>
<tr><td>WP_255347884.1</td><td>DUF499_1.RM.like.systems</td><td>ParB+HNH->ParB+HNH-><- REase <- TUDOR+SF1-SF2-Helicase+i REase +CTD+ REase <-Cluster35_66clades<- DUF499 <- DUF499 *<- DUF499 <- METHYLASE <-?<-Peptidase_M9</td><td>391</td><td>Bacteria>Thermotogota>Thermotogae>Thermotogales>Thermotogaceae</td><td>GCF_000832145.1</td><td>Thermotoga_sp._RQ7</td><td>TRQ7_RS09840</td><td>1801298..1802473</td><td>126738</td><td>DUF499-domain-containing-protein-[Thermotoga-sp.-RQ7]</td></tr>
<tr><td>WP_236681711.1</td><td>DUF499_1.RM.like.systems</td><td>ParB+HNH-><- REase <- TUDOR+SF1-SF2-Helicase+i REase +CTD+ REase <-Cluster35_66clades<- DUF499 <- DUF499 <- DUF499 *<- METHYLASE <-?<-Peptidase_M9</td><td>807</td><td>Bacteria>Thermotogota>Thermotogae>Thermotogales>Thermotogaceae</td><td>GCF_000832145.1</td><td>Thermotoga_sp._RQ7</td><td>TRQ7_RS09855</td><td>1802757..1805180</td><td>126738</td><td>ATP-binding-protein-[Thermotoga-sp.-RQ7]</td></tr>
<tr><td>WP_255347883.1</td><td>DUF499_1.RM.like.systems</td><td>ParB+HNH->ParB+HNH-><- REase <- TUDOR+SF1-SF2-Helicase+i REase +CTD+ REase <-Cluster35_66clades<- DUF499 *<- DUF499 <- DUF499 <- METHYLASE <-?<-Peptidase_M9</td><td>78</td><td>Bacteria>Thermotogota>Thermotogae>Thermotogales>Thermotogaceae</td><td>GCF_000832145.1</td><td>Thermotoga_sp._RQ7</td><td>TRQ7_RS09895</td><td>1801093..1801329</td><td>126738</td><td>DUF499-domain-containing-protein-[Thermotoga-sp.-RQ7]</td></tr>
<tr><td>MBN2383967.1</td><td>DUF499_1.RM.like.systems</td><td>TUDOR+SF1-SF2-Helicase+i REase +CTD+ REase ->?->?->OLD-ABC+TOPRIM-> METHYLASE -> REase -> DUF499 *-><-?||?-><-?||Aminotran_3-></td><td>920</td><td>Bacteria>unclassified_Bacteria</td><td>GCA_016934675.1</td><td>bacterium</td><td>JXQ70_13910</td><td>28377..31139</td><td>1869227</td><td>ATP-binding-protein-[bacterium]</td></tr>
<tr><td>MBU2648361.1</td><td>DUF499_1.RM.like.systems</td><td>TUDOR+SF1-SF2-Helicase+i REase +CTD+ REase -> w HTH+ REase -> METHYLASE -> HEPN+ DUF499 *->?->Phage_integrase-><-Arm-DNA-bind_3+Phage_int_SAM_3+Phage_integrase</td><td>1091</td><td>Bacteria>unclassified_Bacteria</td><td>GCA_018830885.1</td><td>bacterium</td><td>KKI24_26885</td><td>8390..11665</td><td>1869227</td><td>DUF499-domain-containing-protein-[bacterium]</td></tr>
<tr><td>--</td><td></td><td></td><td></td><td></td><td></td><td></td><td></td><td></td><td></td><td></td></tr>
<tr><td>-- REase</td><td>without</td><td>TUDOR+SF1-SF2-Helicase+i REase +CTD+ REase ;</td><td></td><td></td><td></td><td></td><td></td><td></td><td></td><td></td></tr>
<tr><td>-- OLD-ABC-> REase</td><td></td><td></td><td></td><td></td><td></td><td></td><td></td><td></td><td></td><td></td></tr>
<tr><td>--</td><td></td><td></td><td></td><td></td><td></td><td></td><td></td><td></td><td></td><td></td></tr>
<tr><td>MBU1103405.1</td><td>DUF499_1.RM.like.systems</td><td><- REase +35exo<- DUF499 *<-?<- METHYLASE <- REase</td><td>918</td><td>Archaea>DPANN_group>Nanoarchaeota>unclassified_Nanoarchaeota</td><td>GCA_018817905.1</td><td>Nanoarchaeota_archaeon</td><td>KJ600_02510</td><td>1498..4254</td><td>2026764</td><td>hypothetical-protein-[Nanoarchaeota-archaeon]</td></tr>
<tr><td>WP_092695686.1</td><td>DUF499_1.RM.like.systems</td><td>DDE_Tnp_1-><-?<- REase <- METHYLASE || DUF499 *-></td><td>805</td><td>Archaea>Euryarchaeota>Stenosarchaea_group>Halobacteria>Halobacteriales</td><td>GCF_900102305.1</td><td>Halorientalis_regularis</td><td>BLQ72_RS20055</td><td>9401..11817</td><td>660518</td><td>DUF499-domain-containing-protein,-partial-[Halorientalis-regularis]</td></tr>
<tr><td>ABD41477.1</td><td>DUF499_1.RM.like.systems</td><td>OLD-ABC-> OLD-ABC-> REase -> METHYLASE ->OLD-ABC->TOPRIM-> HEPN+ DUF499 *-></td><td>1094</td><td>Archaea>Euryarchaeota>Stenosarchaea_group>Methanomicrobia>Methanomicrobiales</td><td>GCA_000013445.1</td><td>Methanospirillum_hungatei_JF-1</td><td>Mhun_1756</td><td>1984858..1988142</td><td>323259</td><td>conserved-hypothetical-protein-[Methanospirillum-hungatei-JF-1]</td></tr>
<tr><td>WP_204222940.1</td><td>DUF499_1.RM.like.systems</td><td>OLD-ABC-> OLD-ABC-> REase -> METHYLASE ->OLD-ABC->TOPRIM-> HEPN+ DUF499 *-></td><td>1082</td><td>Archaea>Euryarchaeota>Stenosarchaea_group>Methanomicrobia>Methanomicrobiales</td><td>GCF_000013445.1</td><td>Methanospirillum_hungatei</td><td>MHUN_RS09170</td><td>1984894..1988142</td><td>2203</td><td>DUF499-domain-containing-protein-[Methanospirillum-hungatei]</td></tr>
<tr><td>MCP1392941.1</td><td>DUF499_1.RM.like.systems</td><td><-RelB<- DUF499 *||HicB_lk_antitox-><- REase <- METHYLASE -p</td><td>1008</td><td>Archaea>Euryarchaeota>Stenosarchaea_group>Methanomicrobia>Methanotrichales</td><td>GCA_024166385.1</td><td>Methanothrix_harundinacea</td><td>J5I35_09320</td><td>253..3279</td><td>301375</td><td>ATP-binding-protein-[Methanothrix-harundinacea]</td></tr>
<tr><td>MCX6669299.1</td><td>DUF499_1.RM.like.systems</td><td>METHYLASE -p->PIN->PIN+ACET+EVE->ASCH-> REase -> DUF499 *-></td><td>1023</td><td>Archaea>Euryarchaeota>Stenosarchaea_group>Methanomicrobia>Methanotrichales</td><td>GCA_026394475.1</td><td>Methanothrix_sp.</td><td>NTV25_05770</td><td>3501..6572</td><td>90426</td><td>MAG:-DUF499-domain-containing-protein-[Methanothrix-sp.]</td></tr>
<tr><td>WP_013719923.1</td><td>DUF499_1.RM.like.systems</td><td>PADR-HTH->?->?-> METHYLASE ->?-> REase -> DUF499 *-><-?<-SbcC+SbcC||RADICAL-SAM->Radical-SAM-></td><td>1047</td><td>Archaea>Euryarchaeota>Stenosarchaea_group>Methanomicrobia>Methanotrichales</td><td>GCF_000204415.1</td><td>Methanothrix</td><td>QVD88_RS00980</td><td>24822..27965</td><td>2222</td><td>DUF499-domain-containing-protein-[Methanothrix-soehngenii]</td></tr>
<tr><td>MCL4551940.1</td><td>DUF499_1.RM.like.systems</td><td>XIS-HTH->OLD-ABC->HNH-> METHYLASE -> DUF499 *-> DUF499 -p-> SF1-SF2-Helicase+ REase -></td><td>336</td><td>Archaea>TACK_group>Candidatus_Marsarchaeota>unclassified_Candidatus_Marsarchaeota</td><td>GCA_023381795.1</td><td>Candidatus_Marsarchaeota_archaeon</td><td>M1305_00065</td><td>10946..11954</td><td>2823144</td><td>DUF499-domain-containing-protein-[Candidatus-Marsarchaeota-archaeon]</td></tr>
<tr><td>MCC6018921.1</td><td>DUF499_1.RM.like.systems</td><td>METHYLASE -> DUF499 *->Cluster35_66clades->TUDOR+SF1-SF2-Helicase+i REase +CTD->?-><- REase</td><td>1124</td><td>Archaea>TACK_group>Candidatus_Verstraetearchaeota>unclassified_Candidatus_Verstraetearchaeota</td><td>GCA_020833005.1</td><td>Candidatus_Verstraetearchaeota_archaeon</td><td>LM601_07815</td><td>5184..8558</td><td>2250257</td><td>ATP-binding-protein-[Candidatus-Verstraetearchaeota-archaeon]</td></tr>
<tr><td>MCC6023551.1</td><td>DUF499_1.RM.like.systems</td><td>EVE-><- SF1-SF2-Helicase+ REase <-Cluster35_66clades<- DUF499 *<- METHYLASE</td><td>1124</td><td>Archaea>TACK_group>Nitrososphaerota>unclassified_Nitrososphaerota</td><td>GCA_020833125.1</td><td>Nitrososphaerota_archaeon</td><td>LM600_02495</td><td>4852..8226</td><td>2026795</td><td>ATP-binding-protein-[Thaumarchaeota-archaeon]</td></tr>
<tr><td>AFK51621.1</td><td>DUF499_1.RM.like.systems</td><td>ABC-ATPase+SbcC->?-> METHYLASE -> DUF499 *->?-> SF1-SF2-Helicase+ REase ->HHH+ENDO3+NUDIX-></td><td>1233</td><td>Archaea>TACK_group>Thermoproteota>Thermoprotei>Desulfurococcales</td><td>GCA_000264495.1</td><td>Thermogladius_calderae_1633</td><td>TCELL_1198</td><td>1136308..1140009</td><td>1184251</td><td>hypothetical-protein-TCELL_1198-[Thermogladius-calderae-1633]</td></tr>
<tr><td>NAZ12665.1</td><td>DUF499_1.RM.like.systems</td><td><- SF1-SF2-Helicase+ REase <-?<- DUF499 *<- METHYLASE</td><td>1252</td><td>Archaea>TACK_group>Thermoproteota>Thermoprotei>Desulfurococcales</td><td>GCA_009903825.1</td><td>Desulfurococcales_archaeon</td><td>GU359_00740</td><td>3566..7324</td><td>2480821</td><td>DUF499-domain-containing-protein-[Desulfurococcales-archaeon]</td></tr>
<tr><td>MCE4605485.1</td><td>DUF499_1.RM.like.systems</td><td>REase -><- DUF499 *<- METHYLASE || REase ->SF1-SF2-Helicase-><-?<-?||N6_N4_MTase-></td><td>1031</td><td>Archaea>TACK_group>Thermoproteota>Thermoprotei>Desulfurococcales</td><td>GCA_015521665.1</td><td>Desulfurococcales_archaeon</td><td>F7C08_03020</td><td>104683..107778</td><td>2480821</td><td>DUF499-domain-containing-protein-[Desulfurococcales-archaeon]</td></tr>
<tr><td>MCE4602804.1</td><td>DUF499_1.RM.like.systems</td><td><-N6_N4_MTase||?->?-><-SF1-SF2-Helicase<- REase || METHYLASE -> DUF499 *-></td><td>996</td><td>Archaea>TACK_group>Thermoproteota>Thermoprotei>Desulfurococcales</td><td>GCA_015523445.1</td><td>Desulfurococcales_archaeon</td><td>F7B18_06450</td><td>10084..13072</td><td>2480821</td><td>DUF499-domain-containing-protein-[Desulfurococcales-archaeon]</td></tr>
<tr><td>MCC6010980.1</td><td>DUF499_1.RM.like.systems</td><td><- REase <- SF1-SF2-Helicase+ REase <-?<- DUF499 *<- METHYLASE -p</td><td>1248</td><td>Archaea>TACK_group>Thermoproteota>Thermoprotei>Desulfurococcales</td><td>GCA_020832865.1</td><td>Desulfurococcaceae_archaeon</td><td>LM556_01090</td><td>5254..9000</td><td>2184738</td><td>ATP-binding-protein-[Desulfurococcaceae-archaeon]</td></tr>
<tr><td>WP_013129635.1</td><td>DUF499_1.RM.like.systems</td><td><- SF1-SF2-Helicase+ REase <-?<- DUF499 *<- METHYLASE ||?->MerR-HTH+GDResolvase->ZnR->HNH-></td><td>1246</td><td>Archaea>TACK_group>Thermoproteota>Thermoprotei>Desulfurococcales</td><td>GCF_000092185.1</td><td>Thermosphaera_aggregans</td><td>TAGG_RS03820</td><td>727041..730781</td><td>54254</td><td>DUF499-domain-containing-protein-[Thermosphaera-aggregans]</td></tr>
<tr><td>WP_048163335.1</td><td>DUF499_1.RM.like.systems</td><td>ABC-ATPase+SbcC-> METHYLASE -> DUF499 *->?-> SF1-SF2-Helicase+ REase ->HHH+ENDO3+NUDIX-></td><td>1244</td><td>Archaea>TACK_group>Thermoproteota>Thermoprotei>Desulfurococcales</td><td>GCF_000264495.1</td><td>Thermogladius_calderae</td><td>TCELL_RS06135</td><td>1136275..1140009</td><td>1200300</td><td>DUF499-domain-containing-protein-[Thermogladius-calderae]</td></tr>
<tr><td>WP_055408547.1</td><td>DUF499_1.RM.like.systems</td><td><- METHYLASE -p<- REase <-SF1-SF2-Helicase<- REase || METHYLASE -> DUF499 *-><-?<-HNH</td><td>1047</td><td>Archaea>TACK_group>Thermoproteota>Thermoprotei>Desulfurococcales</td><td>GCF_001412615.1</td><td>Pyrodictium_delaneyi</td><td>Pyrde_RS04390</td><td>857202..860345</td><td>1273541</td><td>DUF499-domain-containing-protein-[Pyrodictium-delaneyi]</td></tr>
<tr><td>EHP69856.1</td><td>DUF499_1.RM.like.systems</td><td><-ORC-CDC6+ REase <-?||?->PIN->?->Transposase-><- DUF499 *<- DUF499 -p<- METHYLASE <-?<- REase +SF1-SF2-Helicase ||?->MethyltransfD12-></td><td>129</td><td>Archaea>TACK_group>Thermoproteota>Thermoprotei>Sulfolobales</td><td>GCA_000243315.1</td><td>Metallosphaera_yellowstonensis_MK1</td><td>MetMK1DRAFT_00003580</td><td>133014..133403</td><td>671065</td><td>putative-ATPase-(AAA+-superfamily)-[Metallosphaera-yellowstonensis-MK1]</td></tr>
<tr><td>WP_009070048.1</td><td>DUF499_1.RM.like.systems</td><td><-ORC-CDC6+ REase ||?->?->?->PIN-><- DUF499 *<- METHYLASE <-?<- REase +SF1-SF2-Helicase ||?->MethyltransfD12-><-HKD-DNase</td><td>137</td><td>Archaea>TACK_group>Thermoproteota>Thermoprotei>Sulfolobales</td><td>GCF_000243315.1</td><td>Metallosphaera_yellowstonensis</td><td>METMK1DRAFT_RS18680</td><td>133361..133774</td><td>1111107</td><td>DUF499-domain-containing-protein-[Metallosphaera-yellowstonensis]</td></tr>
<tr><td>KUO92820.1</td><td>DUF499_1.RM.like.systems</td><td><-ABC-ATPase+SbcC||?->Radical-SAM->tRNA->?->?-> METHYLASE -> DUF499 *-> SF1-SF2-Helicase+ REase -><-?<-ZnR<-HKD-DNase<-SF1-SF2-Helicase<- METHYLASE</td><td>1052</td><td>Archaea>TACK_group>Thermoproteota>Thermoprotei>Thermoproteales</td><td>GCA_001516585.1</td><td>Thermocladium_sp._ECH_B</td><td>AT710_02110</td><td>21625..24783</td><td>1714261</td><td>MAG:-hypothetical-protein-AT710_02110-[Thermocladium-sp.-ECH_B]</td></tr>
<tr><td>NAZ33307.1</td><td>DUF499_1.RM.like.systems</td><td>METHYLASE -> DUF499 *->?-> SF1-SF2-Helicase+ REase -></td><td>1024</td><td>Archaea>TACK_group>Thermoproteota>Thermoprotei>Thermoproteales</td><td>GCA_009903905.1</td><td>Pyrobaculum_sp.</td><td>GU356_03295</td><td>3707..6781</td><td>2004705</td><td>DUF499-domain-containing-protein-[Pyrobaculum-sp.]</td></tr>
<tr><td>WP_013680508.1</td><td>DUF499_1.RM.like.systems</td><td><-MORC+DUF3684+ REase ||?-><-APATPase+TPR||<-PIWI|| METHYLASE -> DUF499 *->?-> SF1-SF2-Helicase+ REase -><-ZnR<-?<-?||HEPN-></td><td>1023</td><td>Archaea>TACK_group>Thermoproteota>Thermoprotei>Thermoproteales</td><td>GCF_000193375.1</td><td>Thermoproteus_uzoniensis</td><td>TUZN_RS08265</td><td>1481435..1484506</td><td>184117</td><td>DUF499-domain-containing-protein-[Thermoproteus-uzoniensis]</td></tr>
<tr><td>WP_014290093.1</td><td>DUF499_1.RM.like.systems</td><td><-SF1-SF2-Helicase||ZnR-><- SF1-SF2-Helicase+ REase <-?<- DUF499 *<- METHYLASE ||PIWI->||?->APATPase+TPR-></td><td>1023</td><td>Archaea>TACK_group>Thermoproteota>Thermoprotei>Thermoproteales</td><td>GCF_000234805.1</td><td>Pyrobaculum_ferrireducens</td><td>P186_RS13545</td><td>2438033..2441104</td><td>1104324</td><td>DUF499-domain-containing-protein-[Pyrobaculum-ferrireducens]</td></tr>
<tr><td>NPA84494.1</td><td>DUF499_1.RM.like.systems</td><td><-SF1-SF2-Helicase<- REase || METHYLASE -> DUF499 *-></td><td>785</td><td>Archaea>TACK_group>Thermoproteota>unclassified_Thermoproteota</td><td>GCA_013154085.1</td><td>Thermoproteota_archaeon</td><td>GXO07_00630</td><td>7378..9733</td><td>2056631</td><td>DUF499-domain-containing-protein-[Crenarchaeota-archaeon]</td></tr>
<tr><td>HDZ17009.1</td><td>DUF499_1.RM.like.systems</td><td>SF1-SF2-Helicase+ REase -> METHYLASE -> DUF499 *-></td><td>949</td><td>Archaea>unclassified_Archaea</td><td>GCA_011050315.1</td><td>archaeon</td><td>ENH75_01880</td><td>4806..7655</td><td>1906665</td><td>ATP-binding-protein-[archaeon]</td></tr>
<tr><td>MBA2524268.1</td><td>DUF499_1.RM.like.systems</td><td>REase -> METHYLASE -> REase -> DUF499 *-><-?<-N6_N4_MTase</td><td>930</td><td>Bacteria>Acidobacteriota>Blastocatellia>Blastocatellales>Pyrinomonadaceae</td><td>GCA_013696625.1</td><td>Pyrinomonadaceae_bacterium</td><td>HOV18_00595</td><td>5309..8101</td><td>2283092</td><td>ATP-binding-protein-[Pyrinomonadaceae-bacterium]</td></tr>
<tr><td>MDE0626645.1</td><td>DUF499_1.RM.like.systems</td><td>METHYLASE -p->?->PIN-> DUF499 *-> REase -></td><td>1105</td><td>Bacteria>Acidobacteriota>Terriglobia>Bryobacterales>unclassified_Bryobacterales</td><td>GCA_028824065.1</td><td>Bryobacterales_bacterium</td><td>OXH99_09625</td><td>1430..4747</td><td>2026791</td><td>MAG:-DUF499-domain-containing-protein-[Bryobacterales-bacterium]</td></tr>
<tr><td>RMG44917.1</td><td>DUF499_1.RM.like.systems</td><td>METHYLASE -p-> REase -> DUF499 *-></td><td>649</td><td>Bacteria>Acidobacteriota>unclassified_Acidobacteriota</td><td>GCA_003696865.1</td><td>Acidobacteriota_bacterium</td><td>D6723_18895</td><td>1491..3439</td><td>1978231</td><td>ATP-binding-protein,-partial-[Acidobacteriota-bacterium]</td></tr>
<tr><td>MBI2833279.1</td><td>DUF499_1.RM.like.systems</td><td>TOPRIM->OLD-ABC->TOPRIM-> REase ->?->?-> DUF499 *->sGTP-></td><td>917</td><td>Bacteria>Acidobacteriota>unclassified_Acidobacteriota</td><td>GCA_016190995.1</td><td>Acidobacteriota_bacterium</td><td>HYX76_02505</td><td>4163..6916</td><td>1978231</td><td>ATP-binding-protein-[Acidobacteria-bacterium]</td></tr>
<tr><td>MCI0622286.1</td><td>DUF499_1.RM.like.systems</td><td><- REase <- HEPN+ DUF499 *<-?<-DUF488-1<-?<- METHYLASE <-HEPN<- METHYLASE -p</td><td>1117</td><td>Bacteria>Acidobacteriota>unclassified_Acidobacteriota</td><td>GCA_022615935.1</td><td>Acidobacteriota_bacterium</td><td>L0387_11575</td><td>924..4277</td><td>1978231</td><td>DUF499-domain-containing-protein-[Acidobacteria-bacterium]</td></tr>
<tr><td>MCY4659240.1</td><td>DUF499_1.RM.like.systems</td><td><-ORC-CDC6+ REase <- DUF499 *<-PIN<-OLD-ABC<- METHYLASE <-ParB+HNH+Schlafen<-TOPRIM<-OLD-ABC</td><td>941</td><td>Bacteria>Acidobacteriota>unclassified_Acidobacteriota</td><td>GCA_026713985.1</td><td>Acidobacteriota_bacterium</td><td>OXF93_05425</td><td>1121..3946</td><td>1978231</td><td>MAG:-DUF499-domain-containing-protein-[Acidobacteria-bacterium]</td></tr>
<tr><td>MDE2906585.1</td><td>DUF499_1.RM.like.systems</td><td><- DUF499 *<- METHYLASE <- REase</td><td>963</td><td>Bacteria>Acidobacteriota>unclassified_Acidobacteriota</td><td>GCA_028874255.1</td><td>Acidobacteriota_bacterium</td><td>OXQ28_10940</td><td>1..2891</td><td>1978231</td><td>MAG:-DUF499-domain-containing-protein,-partial-[Acidobacteriota-bacterium]</td></tr>
<tr><td>OGB88303.1</td><td>DUF499_1.RM.like.systems</td><td>METHYLASE -p->AAA-ATPase+ REase -> DUF499 *-><-?<-PIN<-?<-?<-?<-NUDIX</td><td>1106</td><td>Bacteria>Bacteria_incertae_sedis>Bacteria_candidate_phyla>candidate_division_NC10>unclassified_candidate_division_NC10</td><td>GCA_001771245.1</td><td>candidate_division_NC10_bacterium_RIFCSPLOWO2_02_FULL_66_22</td><td>A3H39_08635</td><td>5174..8494</td><td>1801658</td><td>hypothetical-protein-A3H39_08635-[candidate-division-NC10-bacterium-RIFCSPLOWO2_02_FULL_66_22]</td></tr>
<tr><td>MBI4572390.1</td><td>DUF499_1.RM.like.systems</td><td><-ABC-ATPase+SbcC<-?<-?||Polbeta->HEPN-><- DUF499 *<-AAA-ATPase+ REase</td><td>1106</td><td>Bacteria>Bacteria_incertae_sedis>Bacteria_candidate_phyla>candidate_division_NC10>unclassified_candidate_division_NC10</td><td>GCA_016209725.1</td><td>candidate_division_NC10_bacterium</td><td>HY713_03775</td><td>5635..8955</td><td>2072417</td><td>ATP-binding-protein-[candidate-division-NC10-bacterium]</td></tr>
<tr><td>MBI4536775.1</td><td>DUF499_1.RM.like.systems</td><td><- REase <-sGTP<- DUF499 *<-?<-TOPRIM<-OLD-ABC<- METHYLASE -p<-?<- METHYLASE</td><td>938</td><td>Bacteria>Bacteria_incertae_sedis>Bacteria_candidate_phyla>candidate_division_NC10>unclassified_candidate_division_NC10</td><td>GCA_016209735.1</td><td>candidate_division_NC10_bacterium</td><td>HY712_02325</td><td>61380..64196</td><td>2072417</td><td>ATP-binding-protein-[candidate-division-NC10-bacterium]</td></tr>
<tr><td>MDE2058207.1</td><td>DUF499_1.RM.like.systems</td><td>OLD-ABC->TOPRIM-> METHYLASE ->BrnT_toxin->HTH-> REase -> DUF499 *->?->HTH+PolB->HEPN-></td><td>931</td><td>Bacteria>Bacteria_incertae_sedis>Bacteria_candidate_phyla>candidate_division_NC10>unclassified_candidate_division_NC10</td><td>GCA_028864815.1</td><td>candidate_division_NC10_bacterium</td><td>KGJ27_00545</td><td>34245..37040</td><td>2072417</td><td>MAG:-ATP-binding-protein-[candidate-division-NC10-bacterium]</td></tr>
<tr><td>MCX7973587.1</td><td>DUF499_1.RM.like.systems</td><td>METHYLASE -p-> METHYLASE -p-> DUF499 *-><- REase</td><td>845</td><td>Bacteria>Bacteria_incertae_sedis>Bacteria_candidate_phyla>Candidatus_Aminicenantes>unclassified_Candidatus_Aminicenantes</td><td>GCA_026416195.1</td><td>Candidatus_Aminicenantes_bacterium</td><td>N3B16_03705</td><td>1453..3990</td><td>2052149</td><td>MAG:-DUF499-domain-containing-protein-[Candidatus-Aminicenantes-bacterium]</td></tr>
<tr><td>MBC7170403.1</td><td>DUF499_1.RM.like.systems</td><td>ABC-ATPase+SbcC->ABC-ATPase+SbcC-><-Glyco_transf_20<- DUF499 *<- REase <- METHYLASE</td><td>920</td><td>Bacteria>Bacteria_incertae_sedis>Bacteria_candidate_phyla>Candidatus_Bipolaricaulota>unclassified_Candidatus_Bipolaricaulota</td><td>GCA_014361015.1</td><td>Candidatus_Bipolaricaulota_bacterium</td><td>H5T54_05700</td><td>11895..14657</td><td>2306012</td><td>ATP-binding-protein-[Candidatus-Bipolaricaulota-bacterium]</td></tr>
<tr><td>MBP6875772.1</td><td>DUF499_1.RM.like.systems</td><td>HerA/FtsK->?-> METHYLASE -> REase -> DUF499 *-><-ParB</td><td>924</td><td>Bacteria>Bacteria_incertae_sedis>Bacteria_candidate_phyla>Candidatus_Eisenbacteria>unclassified_Candidatus_Eisenbacteria</td><td>GCA_017991665.1</td><td>Candidatus_Eisenbacteria_bacterium</td><td>KBD56_06850</td><td>148901..151675</td><td>2212470</td><td>DUF499-domain-containing-protein-[Candidatus-Eisenbacteria-bacterium]</td></tr>
<tr><td>MAE19516.1</td><td>DUF499_1.RM.like.systems</td><td>ABhydrolase-><-?<-?<-Aminotran_4<-ACET<- DUF499 *<- METHYLASE <- REase</td><td>922</td><td>Bacteria>Bacteria_incertae_sedis>Bacteria_candidate_phyla>Candidatus_Poribacteria>unclassified_Candidatus_Poribacteria</td><td>GCA_002686625.1</td><td>Candidatus_Poribacteria_bacterium</td><td>CMK12_11405</td><td>49414..52182</td><td>2026781</td><td>AAA-family-ATPase-[Candidatus-Poribacteria-bacterium]</td></tr>
<tr><td>MXV83700.1</td><td>DUF499_1.RM.like.systems</td><td><-URI-DNase<-?<- METHYLASE <- REase <- REase <- REase <- HEPN+ DUF499 *<-PIN</td><td>1088</td><td>Bacteria>Bacteria_incertae_sedis>Bacteria_candidate_phyla>Candidatus_Poribacteria>unclassified_Candidatus_Poribacteria</td><td>GCA_009835485.1</td><td>Candidatus_Poribacteria_bacterium</td><td>F4Z98_10065</td><td>15224..18490</td><td>2026781</td><td>ATP-binding-protein-[Candidatus-Poribacteria-bacterium]</td></tr>
<tr><td>MXY28219.1</td><td>DUF499_1.RM.like.systems</td><td>DUF499 *-> REase +HTH-> REase -> METHYLASE ->URI-DNase-></td><td>895</td><td>Bacteria>Bacteria_incertae_sedis>Bacteria_candidate_phyla>Candidatus_Poribacteria>unclassified_Candidatus_Poribacteria</td><td>GCA_009838315.1</td><td>Candidatus_Poribacteria_bacterium</td><td>F4Y59_08690</td><td>1..2690</td><td>2026781</td><td>ATP-binding-protein-[Candidatus-Poribacteria-bacterium]</td></tr>
<tr><td>MXY29380.1</td><td>DUF499_1.RM.like.systems</td><td>HEPN-> DUF499 *-> REase -></td><td>1115</td><td>Bacteria>Bacteria_incertae_sedis>Bacteria_candidate_phyla>Candidatus_Poribacteria>unclassified_Candidatus_Poribacteria</td><td>GCA_009838315.1</td><td>Candidatus_Poribacteria_bacterium</td><td>F4Y59_14620</td><td>1265..4612</td><td>2026781</td><td>ATP-binding-protein-[Candidatus-Poribacteria-bacterium]</td></tr>
<tr><td>MYB95133.1</td><td>DUF499_1.RM.like.systems</td><td><- METHYLASE <- REase +HTH<- REase +HTH<- HEPN+ DUF499 *<-?<-?<-?<- SF1-SF2-Helicase+ REase <-ABC-ATPase+SbcC</td><td>1108</td><td>Bacteria>Bacteria_incertae_sedis>Bacteria_candidate_phyla>Candidatus_Poribacteria>unclassified_Candidatus_Poribacteria</td><td>GCA_009838805.1</td><td>Candidatus_Poribacteria_bacterium</td><td>F4Y43_14870</td><td>2954..6280</td><td>2026781</td><td>ATP-binding-protein-[Candidatus-Poribacteria-bacterium]</td></tr>
<tr><td>MYC78293.1</td><td>DUF499_1.RM.like.systems</td><td><-HEPN<-URI-DNase<-?<- REase <- REase <- REase <- HEPN+ DUF499 *<-PIN<-HTH+MPTase<-?<-?<-?<- SF1-SF2-Helicase+ REase</td><td>1093</td><td>Bacteria>Bacteria_incertae_sedis>Bacteria_candidate_phyla>Candidatus_Poribacteria>unclassified_Candidatus_Poribacteria</td><td>GCA_009841255.1</td><td>Candidatus_Poribacteria_bacterium</td><td>F4X10_21230</td><td>47302..50583</td><td>2026781</td><td>ATP-binding-protein-[Candidatus-Poribacteria-bacterium]</td></tr>
<tr><td>MYK17049.1</td><td>DUF499_1.RM.like.systems</td><td>HEPN+ DUF499 *-> REase +HTH-> REase -> METHYLASE ->URI-DNase-></td><td>1090</td><td>Bacteria>Bacteria_incertae_sedis>Bacteria_candidate_phyla>Candidatus_Poribacteria>unclassified_Candidatus_Poribacteria</td><td>GCA_009845505.1</td><td>Candidatus_Poribacteria_bacterium</td><td>F4055_02625</td><td>1958..5230</td><td>2026781</td><td>ATP-binding-protein-[Candidatus-Poribacteria-bacterium]</td></tr>
<tr><td>MCY3724179.1</td><td>DUF499_1.RM.like.systems</td><td>PIN-> HEPN+ DUF499 *-> REase +HTH-> REase -> METHYLASE ->?->URI-DNase-></td><td>1093</td><td>Bacteria>Bacteria_incertae_sedis>Bacteria_candidate_phyla>Candidatus_Poribacteria>unclassified_Candidatus_Poribacteria</td><td>GCA_026705955.1</td><td>Candidatus_Poribacteria_bacterium</td><td>OXG97_18330</td><td>699..3980</td><td>2026781</td><td>MAG:-DUF499-domain-containing-protein-[Candidatus-Poribacteria-bacterium]</td></tr>
<tr><td>MDE0014209.1</td><td>DUF499_1.RM.like.systems</td><td>HTH+MPTase->PIN-> HEPN+ DUF499 *-> REase +HTH-> REase -> METHYLASE ->URI-DNase-></td><td>1095</td><td>Bacteria>Bacteria_incertae_sedis>Bacteria_candidate_phyla>Candidatus_Poribacteria>unclassified_Candidatus_Poribacteria</td><td>GCA_028818705.1</td><td>Candidatus_Poribacteria_bacterium</td><td>OXU36_23900</td><td>11254..14541</td><td>2026781</td><td>MAG:-DUF499-domain-containing-protein-[Candidatus-Poribacteria-bacterium]</td></tr>
<tr><td>MDE0423419.1</td><td>DUF499_1.RM.like.systems</td><td>SF1-SF2-Helicase+ REase ->?->?->?->?->?-> HEPN+ DUF499 *-> REase +HTH-> REase -> METHYLASE ->URI-DNase-></td><td>1090</td><td>Bacteria>Bacteria_incertae_sedis>Bacteria_candidate_phyla>Candidatus_Poribacteria>unclassified_Candidatus_Poribacteria</td><td>GCA_028820845.1</td><td>Candidatus_Poribacteria_bacterium</td><td>OXN25_00975</td><td>9456..12728</td><td>2026781</td><td>MAG:-DUF499-domain-containing-protein-[Candidatus-Poribacteria-bacterium]</td></tr>
<tr><td>OGK75197.1</td><td>DUF499_1.RM.like.systems</td><td>AAA-ATPase+ REase -> METHYLASE -> DUF499 *->Radical-SAM->?->?->?->?->ABC-ATPase+SbcC-></td><td>1096</td><td>Bacteria>Bacteria_incertae_sedis>Bacteria_candidate_phyla>Candidatus_Rokubacteria>unclassified_Candidatus_Rokubacteria</td><td>GCA_001788395.1</td><td>Candidatus_Rokubacteria_bacterium_GWA2_70_23</td><td>A2X53_06550</td><td>6264..9554</td><td>1802095</td><td>MAG:-hypothetical-protein-A2X53_06550-[Candidatus-Rokubacteria-bacterium-GWA2_70_23]</td></tr>
<tr><td>HAM55582.1</td><td>DUF499_1.RM.like.systems</td><td><- DUF499 -p<- DUF499 *<- METHYLASE <-AAA-ATPase+ REase</td><td>911</td><td>Bacteria>Bacteria_incertae_sedis>Bacteria_candidate_phyla>Candidatus_Rokubacteria>unclassified_Candidatus_Rokubacteria</td><td>GCA_003452995.1</td><td>Candidatus_Rokubacteria_bacterium</td><td>DCQ64_09305</td><td>854..3589</td><td>2053607</td><td>hypothetical-protein-[Candidatus-Rokubacteria-bacterium]</td></tr>
<tr><td>MBU2264520.1</td><td>DUF499_1.RM.like.systems</td><td>METHYLASE -p-> REase ->PolB->HEPN-> DUF499 *-></td><td>810</td><td>Bacteria>Bacteria_incertae_sedis>Bacteria_candidate_phyla>Patescibacteria_group>unclassified_Patescibacteria_group</td><td>GCA_018827685.1</td><td>Patescibacteria_group_bacterium</td><td>KJ797_04325</td><td>2908..5337</td><td>2052139</td><td>DUF499-domain-containing-protein-[Patescibacteria-group-bacterium]</td></tr>
<tr><td>MBL0059014.1</td><td>DUF499_1.RM.like.systems</td><td>SMODS->?-> METHYLASE -> w HTH+ REase -> DUF499 -p-> DUF499 *-><-rve_3<-HTH_21+rve+rve_3<-HIN-HTH</td><td>1036</td><td>Bacteria>Elusimicrobiota>unclassified_Elusimicrobiota</td><td>GCA_016721625.1</td><td>Elusimicrobiota_bacterium</td><td>IPP35_07880</td><td>1838166..1841276</td><td>2030800</td><td>ATP-binding-protein-[Elusimicrobia-bacterium]</td></tr>
<tr><td>HBF05518.1</td><td>DUF499_1.RM.like.systems</td><td><-ORC-CDC6+ REase <- HEPN+ DUF499 *<- METHYLASE -p</td><td>1108</td><td>Bacteria>FCB_group>Bacteroidota/Chlorobiota_group>Bacteroidota>Bacteroidia</td><td>GCA_003501465.1</td><td>Candidatus_Segatella_violae</td><td>DDW28_05315</td><td>5202..8528</td><td>1947177</td><td>hypothetical-protein-[Prevotella-sp.]</td></tr>
<tr><td>MBO5419352.1</td><td>DUF499_1.RM.like.systems</td><td>XIS-HTH->?->?-><-?<-?||ORC-CDC6+ REase -><- HEPN+ DUF499 *<- METHYLASE -p</td><td>1102</td><td>Bacteria>FCB_group>Bacteroidota/Chlorobiota_group>Bacteroidota>Bacteroidia</td><td>GCA_017625725.1</td><td>Bacteroidales_bacterium</td><td>J6A22_04620</td><td>7056..10364</td><td>2030927</td><td>DUF499-domain-containing-protein-[Bacteroidales-bacterium]</td></tr>
<tr><td>MBK9465916.1</td><td>DUF499_1.RM.like.systems</td><td>ABhydrolase-><-?<-Integrase_1+Phage_integrase||?->DUF488-1-><-?<- DUF499 *<- w HTH+ REase <-HEPN<-HTH+WYL||RnaseHII-></td><td>943</td><td>Bacteria>FCB_group>Bacteroidota/Chlorobiota_group>Bacteroidota>Chitinophagia</td><td>GCA_016719015.1</td><td>Chitinophagaceae_bacterium</td><td>IPN56_14270</td><td>8966..11797</td><td>1869212</td><td>ATP-binding-protein-[Chitinophagaceae-bacterium]</td></tr>
<tr><td>MBK7652070.1</td><td>DUF499_1.RM.like.systems</td><td>HTH-> REase -> METHYLASE -p-> HEPN+ DUF499 *-> DUF499 -p-> DUF499 -p-></td><td>498</td><td>Bacteria>FCB_group>Bacteroidota/Chlorobiota_group>Bacteroidota>Cytophagia</td><td>GCA_016709845.1</td><td>Flammeovirgaceae_bacterium</td><td>IPJ20_16990</td><td>23879..25375</td><td>2026740</td><td>DUF499-domain-containing-protein-[Flammeovirgaceae-bacterium]</td></tr>
<tr><td>MCX7650091.1</td><td>DUF499_1.RM.like.systems</td><td><- DUF499 *<-AAA-ATPase+ REase <- METHYLASE -p</td><td>926</td><td>Bacteria>FCB_group>Bacteroidota/Chlorobiota_group>Bacteroidota>Flavobacteriia</td><td>GCA_026419215.1</td><td>Flavobacteriales_bacterium</td><td>N2050_06005</td><td>2592..5372</td><td>2021391</td><td>MAG:-DUF499-domain-containing-protein-[Flavobacteriales-bacterium]</td></tr>
<tr><td>MCY4001546.1</td><td>DUF499_1.RM.like.systems</td><td>w HTH+ REase -> METHYLASE ->OLD-ABC->PIN-> DUF499 *-></td><td>903</td><td>Bacteria>FCB_group>Bacteroidota/Chlorobiota_group>Bacteroidota>unclassified_Bacteroidota</td><td>GCA_026708685.1</td><td>Bacteroidota_bacterium</td><td>OXF84_12180</td><td>5800..8511</td><td>1898104</td><td>MAG:-DUF499-domain-containing-protein-[Bacteroidetes-bacterium]</td></tr>
<tr><td>MBL7989353.1</td><td>DUF499_1.RM.like.systems</td><td><- REase <-?<-?<-?||ORC-CDC6+ REase -><- DUF499 *</td><td>945</td><td>Bacteria>FCB_group>Bacteroidota/Chlorobiota_group>Chlorobiota>unclassified_Chlorobiota</td><td>GCA_016788565.1</td><td>Chlorobiota_bacterium</td><td>JNJ94_14925</td><td>66381..69218</td><td>2268192</td><td>ATP-binding-protein-[Chlorobi-bacterium]</td></tr>
<tr><td>MCB5246762.1</td><td>DUF499_1.RM.like.systems</td><td>METHYLASE -> OLD-ABC-> REase -> DUF499 *-></td><td>957</td><td>Bacteria>FCB_group>Candidatus_Cloacimonadota>unclassified_Candidatus_Cloacimonadota</td><td>GCA_020532215.1</td><td>Candidatus_Cloacimonadota_bacterium</td><td>LHW57_01885</td><td>3263..6136</td><td>2030808</td><td>DUF499-domain-containing-protein-[Candidatus-Cloacimonetes-bacterium]</td></tr>
<tr><td>MCB5253275.1</td><td>DUF499_1.RM.like.systems</td><td>SF1-SF2-Helicase-><- REase <-NERD<- DUF499 *</td><td>944</td><td>Bacteria>FCB_group>Candidatus_Cloacimonadota>unclassified_Candidatus_Cloacimonadota</td><td>GCA_020532345.1</td><td>Candidatus_Cloacimonadota_bacterium</td><td>LHW51_09920</td><td>5656..8490</td><td>2030808</td><td>DUF499-domain-containing-protein-[Candidatus-Cloacimonetes-bacterium]</td></tr>
<tr><td>MBI5093542.1</td><td>DUF499_1.RM.like.systems</td><td>DUF4065->HEPN-> REase -> METHYLASE -p-> METHYLASE ->?-> DUF499 *->?->?->HISKIN+MORC->REC-></td><td>944</td><td>Bacteria>FCB_group>Candidatus_Hydrogenedentes>unclassified_Candidatus_Hydrogenedentes</td><td>GCA_016215105.1</td><td>Candidatus_Hydrogenedentes_bacterium</td><td>HZB26_14005</td><td>11856..14690</td><td>2030809</td><td>ATP-binding-protein-[Candidatus-Hydrogenedentes-bacterium]</td></tr>
<tr><td>MBI2505138.1</td><td>DUF499_1.RM.like.systems</td><td>Glycos_transf_4->?->?->?-><- DUF499 *<-?<-PRTase<-SF1-SF2-Helicase-p<- w HTH+ REase <- METHYLASE <- METHYLASE -p</td><td>1105</td><td>Bacteria>FCB_group>Candidatus_Latescibacteria>unclassified_Candidatus_Latescibacteria</td><td>GCA_016187725.1</td><td>Candidatus_Latescibacteria_bacterium</td><td>HYW07_18115</td><td>13371..16688</td><td>2053570</td><td>ATP-binding-protein-[Candidatus-Latescibacteria-bacterium]</td></tr>
<tr><td>MCD6099133.1</td><td>DUF499_1.RM.like.systems</td><td>REase -> DUF499 *->OBFOLD-NUCLEASE->SIR2+SIR2->PIWI-></td><td>932</td><td>Bacteria>FCB_group>Candidatus_Marinimicrobia>unclassified_Candidatus_Marinimicrobia</td><td>GCA_021158485.1</td><td>Candidatus_Marinimicrobia_bacterium</td><td>J7K33_00965</td><td>724..3522</td><td>2026760</td><td>ATP-binding-protein-[Candidatus-Marinimicrobia-bacterium]</td></tr>
<tr><td>MCZ0935996.1</td><td>DUF499_1.RM.like.systems</td><td>SIGMA-HTH->?->?->OLD-ABC+TOPRIM->SF1-SF2-Helicase-> REase -><- HEPN+ DUF499 *<- METHYLASE -p||Zn_Tnp_IS1595+DDE_Tnp_IS1595-><-?|| METHYLASE +HNH->?-><- METHYLASE</td><td>1098</td><td>Bacteria>FCB_group>Gemmatimonadota>Gemmatimonadetes>Candidatus_Palauibacterales</td><td>GCA_026914155.1</td><td>Candidatus_Palauibacter_rhopaloidicola</td><td>OXJ54_12525</td><td>22300..25596</td><td>3056738</td><td>MAG:-DUF499-domain-containing-protein-[Gemmatimonadetes-bacterium]</td></tr>
<tr><td>MDE2877868.1</td><td>DUF499_1.RM.like.systems</td><td>DNA_methylase-> REase ->DNA_methylase-> SF1-SF2-Helicase+ REase ->OLD-ABC-> METHYLASE -> HEPN+ DUF499 *->?->?->?->MULE-TRANSPOSASE-></td><td>1098</td><td>Bacteria>FCB_group>Gemmatimonadota>Gemmatimonadetes>Candidatus_Palauibacterales</td><td>GCA_028874535.1</td><td>Candidatus_Palauibacter_soopunensis</td><td>OXO55_03070</td><td>20960..24256</td><td>3056739</td><td>MAG:-DUF499-domain-containing-protein-[Gemmatimonadota-bacterium]</td></tr>
<tr><td>MCC7194161.1</td><td>DUF499_1.RM.like.systems</td><td>METHYLASE -p->ORC-CDC6+ REase -> HEPN+ DUF499 *-><-?||SIGMA-HTH-></td><td>1104</td><td>Bacteria>FCB_group>Gemmatimonadota>Gemmatimonadetes>Gemmatimonadales</td><td>GCA_020851455.1</td><td>Gemmatimonadaceae_bacterium</td><td>IT356_01265</td><td>1930..5244</td><td>2026741</td><td>DUF499-domain-containing-protein-[Gemmatimonadaceae-bacterium]</td></tr>
<tr><td>MYE95017.1</td><td>DUF499_1.RM.like.systems</td><td><- DUF499 *<- METHYLASE <- REase <-DNA_methylase<-HNH</td><td>358</td><td>Bacteria>FCB_group>Gemmatimonadota>unclassified_Gemmatimonadota</td><td>GCA_009841825.1</td><td>Gemmatimonadota_bacterium</td><td>F4238_16835</td><td>1..1075</td><td>2026742</td><td>ATP-binding-protein-[Gemmatimonadota-bacterium]</td></tr>
<tr><td>MYE70563.1</td><td>DUF499_1.RM.like.systems</td><td>METHYLASE -> METHYLASE -p-> REase -> DUF499 *-></td><td>400</td><td>Bacteria>FCB_group>Gemmatimonadota>unclassified_Gemmatimonadota</td><td>GCA_009841855.1</td><td>Gemmatimonadota_bacterium</td><td>F4237_11010</td><td>2710..3911</td><td>2026742</td><td>ATP-binding-protein-[Gemmatimonadota-bacterium]</td></tr>
<tr><td>MCY3788288.1</td><td>DUF499_1.RM.like.systems</td><td>REase -> w HTH+ REase -> METHYLASE -> REase -> DUF499 *-></td><td>918</td><td>Bacteria>FCB_group>Gemmatimonadota>unclassified_Gemmatimonadota</td><td>GCA_026706505.1</td><td>Gemmatimonadota_bacterium</td><td>OXH63_05815</td><td>5478..8234</td><td>2026742</td><td>MAG:-DUF499-domain-containing-protein-[Gemmatimonadetes-bacterium]</td></tr>
<tr><td>MCY4604211.1</td><td>DUF499_1.RM.like.systems</td><td>METHYLASE -> METHYLASE -p->DUF433->PIN-> METHYLASE -p-> REase -> DUF499 *-></td><td>922</td><td>Bacteria>FCB_group>Gemmatimonadota>unclassified_Gemmatimonadota</td><td>GCA_026713655.1</td><td>Gemmatimonadota_bacterium</td><td>OXE49_08275</td><td>9321..12089</td><td>2026742</td><td>MAG:-DUF499-domain-containing-protein-[Gemmatimonadetes-bacterium]</td></tr>
<tr><td>MDE2981419.1</td><td>DUF499_1.RM.like.systems</td><td>ZDNA-HTH->SF1-SF2-Helicase->Schlafen+HATPase+HTH-><-TPR|| METHYLASE -> REase -> DUF499 *->?->URI-DNase->HEPN-></td><td>922</td><td>Bacteria>FCB_group>Gemmatimonadota>unclassified_Gemmatimonadota</td><td>GCA_028873815.1</td><td>Gemmatimonadota_bacterium</td><td>OXU74_09510</td><td>42276..45044</td><td>2026742</td><td>MAG:-DUF499-domain-containing-protein-[Gemmatimonadota-bacterium]</td></tr>
<tr><td>MDE2763067.1</td><td>DUF499_1.RM.like.systems</td><td><-HTH<-?||?-><-?||<-ORC-CDC6+ REase <-?<- DUF499 *<-MNS+ZDNA-HTH<-DDE_Tnp_1</td><td>934</td><td>Bacteria>FCB_group>Gemmatimonadota>unclassified_Gemmatimonadota</td><td>GCA_028874985.1</td><td>Gemmatimonadota_bacterium</td><td>OXK74_09825</td><td>20104..22908</td><td>2026742</td><td>MAG:-DUF499-domain-containing-protein-[Gemmatimonadota-bacterium]</td></tr>
<tr><td>RLC09317.1</td><td>DUF499_1.RM.like.systems</td><td><- DUF499 *<- REase <-HEPN<- METHYLASE -p</td><td>517</td><td>Bacteria>Myxococcota>Myxococcia>unclassified_Myxococcia</td><td>GCA_003647435.1</td><td>Deltaproteobacteria_bacterium</td><td>DRH43_08475</td><td>1..1550</td><td>2026735</td><td>AAA-family-ATPase,-partial-[Deltaproteobacteria-bacterium]</td></tr>
<tr><td>MBC8392014.1</td><td>DUF499_1.RM.like.systems</td><td>METHYLASE -> REase -> DUF499 *-></td><td>427</td><td>Bacteria>Myxococcota>Myxococcia>unclassified_Myxococcia</td><td>GCA_014382765.1</td><td>Deltaproteobacteria_bacterium</td><td>H8E17_05550</td><td>4033..5315</td><td>2026735</td><td>ATP-binding-protein-[Deltaproteobacteria-bacterium]</td></tr>
<tr><td>MBC8460283.1</td><td>DUF499_1.RM.like.systems</td><td>HEPN+ REase -> METHYLASE -> DUF499 *-></td><td>926</td><td>Bacteria>Myxococcota>Myxococcia>unclassified_Myxococcia</td><td>GCA_014383825.1</td><td>Deltaproteobacteria_bacterium</td><td>H8D67_20045</td><td>4060..6840</td><td>2026735</td><td>ATP-binding-protein-[Deltaproteobacteria-bacterium]</td></tr>
<tr><td>MBI2988835.1</td><td>DUF499_1.RM.like.systems</td><td><-sGTP<-?<-?<-?<-?<- DUF499 *<- METHYLASE <-?<- REase</td><td>931</td><td>Bacteria>Myxococcota>Myxococcia>unclassified_Myxococcia</td><td>GCA_016193465.1</td><td>Deltaproteobacteria_bacterium</td><td>HYY45_18895</td><td>27398..30193</td><td>2026735</td><td>DUF499-domain-containing-protein-[Deltaproteobacteria-bacterium]</td></tr>
<tr><td>MBI3249564.1</td><td>DUF499_1.RM.like.systems</td><td>METHYLASE -> REase -> HEPN+ DUF499 *-><-HicB_lk_antitox||Phage_integrase-><-tRNA<-ABhydrolase</td><td>1110</td><td>Bacteria>Myxococcota>Myxococcia>unclassified_Myxococcia</td><td>GCA_016197285.1</td><td>Deltaproteobacteria_bacterium</td><td>HYZ50_23930</td><td>2669..6001</td><td>2026735</td><td>ATP-binding-protein-[Deltaproteobacteria-bacterium]</td></tr>
<tr><td>MBI4525152.1</td><td>DUF499_1.RM.like.systems</td><td>REase +SAVED->SMODS->UbE1+JAB->JAB-> METHYLASE -> REase -> DUF499 *-> REase +nSTAND1-></td><td>1099</td><td>Bacteria>Myxococcota>Myxococcia>unclassified_Myxococcia</td><td>GCA_016210115.1</td><td>Deltaproteobacteria_bacterium</td><td>HY695_15215</td><td>17806..21105</td><td>2026735</td><td>DUF499-domain-containing-protein-[Deltaproteobacteria-bacterium]</td></tr>
<tr><td>MBW1926153.1</td><td>DUF499_1.RM.like.systems</td><td><- DUF499 *<-DUF1828+ REase <-?<- METHYLASE</td><td>912</td><td>Bacteria>Myxococcota>Myxococcia>unclassified_Myxococcia</td><td>GCA_019307845.1</td><td>Deltaproteobacteria_bacterium</td><td>JRI35_02965</td><td>2089..4827</td><td>2026735</td><td>ATP-binding-protein-[Deltaproteobacteria-bacterium]</td></tr>
<tr><td>MCL5406608.1</td><td>DUF499_1.RM.like.systems</td><td>DUF499 *-> REase ->?->?-><-?<-?<-GDResolvase+Recombinase+ZnR</td><td>564</td><td>Bacteria>Myxococcota>Myxococcia>unclassified_Myxococcia</td><td>GCA_023484975.1</td><td>Deltaproteobacteria_bacterium</td><td>M1398_07815</td><td>1..1696</td><td>2026735</td><td>DUF499-domain-containing-protein-[Deltaproteobacteria-bacterium]</td></tr>
<tr><td>MCP9448661.1</td><td>DUF499_1.RM.like.systems</td><td>sGTP->?->?->?->?->?-><- DUF499 *<-HEPN<-ORC-CDC6+ REase <- METHYLASE -p||?-><-Radical-SAM</td><td>987</td><td>Bacteria>Nitrospirota>Nitrospiria>Nitrospirales>Nitrospiraceae</td><td>GCA_024331085.1</td><td>Nitrospira_sp.</td><td>NNA21_01185</td><td>6072..9035</td><td>70125</td><td>DUF499-domain-containing-protein-[Nitrospira-sp.]</td></tr>
<tr><td>MCP9450544.1</td><td>DUF499_1.RM.like.systems</td><td>METHYLASE ->?-> DUF499 *-> REase +N6_MTase-></td><td>918</td><td>Bacteria>Nitrospirota>Nitrospiria>Nitrospirales>Nitrospiraceae</td><td>GCA_024331085.1</td><td>Nitrospira_sp.</td><td>NNA21_10860</td><td>3927..6683</td><td>70125</td><td>DUF499-domain-containing-protein-[Nitrospira-sp.]</td></tr>
<tr><td>OIP61731.1</td><td>DUF499_1.RM.like.systems</td><td><- DUF499 *<- REase <- REase <-DUF1828+ REase <-?<- METHYLASE -p</td><td>918</td><td>Bacteria>Nitrospirota>unclassified_Nitrospirota</td><td>GCA_001873285.1</td><td>Nitrospirae_bacterium_CG2_30_53_67</td><td>AUK29_09775</td><td>1402..4158</td><td>1805278</td><td>AAA-family-ATPase-[Nitrospirae-bacterium-CG2_30_53_67]</td></tr>
<tr><td>PIV85847.1</td><td>DUF499_1.RM.like.systems</td><td><- DUF499 *<- REase <- REase <-DUF1828+ REase <-?<- METHYLASE -p</td><td>916</td><td>Bacteria>Nitrospirota>unclassified_Nitrospirota</td><td>GCA_002781445.1</td><td>Nitrospirae_bacterium_CG17_big_fil_post_rev_8_21_14_2_50_50_9</td><td>COW52_00060</td><td>1390..4140</td><td>1974070</td><td>AAA-family-ATPase,-partial-[Nitrospirae-bacterium-CG17_big_fil_post_rev_8_21_14_2_50_50_9]</td></tr>
<tr><td>RMH36427.1</td><td>DUF499_1.RM.like.systems</td><td><- DUF499 *<-PIN<-DUF433<- REase <-HTH</td><td>933</td><td>Bacteria>Nitrospirota>unclassified_Nitrospirota</td><td>GCA_003695915.1</td><td>Nitrospirota_bacterium</td><td>D6690_05850</td><td>2045..4846</td><td>2026887</td><td>ATP-binding-protein-[Nitrospirae-bacterium]</td></tr>
<tr><td>MBI2060378.1</td><td>DUF499_1.RM.like.systems</td><td>PHP+SbcC-> METHYLASE ->?->DUF1828+ REase ->?-> DUF499 *-><-PAIREDC-HTH<-?<-?<-ZnR+TOPRIM+D5ATPase<-?<-XIS-HTH</td><td>918</td><td>Bacteria>Nitrospirota>unclassified_Nitrospirota</td><td>GCA_016180645.1</td><td>Nitrospirota_bacterium</td><td>HYT87_11470</td><td>28661..31417</td><td>2026887</td><td>ATP-binding-protein-[Nitrospirae-bacterium]</td></tr>
<tr><td>MBU2768209.1</td><td>DUF499_1.RM.like.systems</td><td><- DUF499 *<- REase <-OLD-ABC <- w HTH+ REase <- METHYLASE <-HNH<-OLD-ABC</td><td>542</td><td>Bacteria>Pseudomonadota>Acidithiobacillia>Acidithiobacillales>Acidithiobacillaceae</td><td>GCA_018853935.1</td><td>Acidithiobacillus_ferrivorans</td><td>HAP94_19055</td><td>1..1626</td><td>160808</td><td>ATP-binding-protein-[Acidithiobacillus-ferrivorans]</td></tr>
<tr><td>WP_198926060.1</td><td>DUF499_1.RM.like.systems</td><td><- DUF499 -p<- DUF499 *<- w HTH+ REase</td><td>460</td><td>Bacteria>Pseudomonadota>Acidithiobacillia>Acidithiobacillales>Acidithiobacillaceae</td><td>GCF_001705725.1</td><td>Acidithiobacillus_thiooxidans</td><td>A6M27_RS22710</td><td>4222..5604</td><td>930</td><td>DUF499-domain-containing-protein-[Acidithiobacillus-thiooxidans]</td></tr>
<tr><td>MCC0038267.1</td><td>DUF499_1.RM.like.systems</td><td><-TUDOR+SF1-SF2-Helicase+i REase +CTD<-?<-HHH_4+ SF1-SF2-Helicase+ REase <- HEPN+ DUF499 *<- METHYLASE</td><td>1108</td><td>Bacteria>Pseudomonadota>Alphaproteobacteria>Hyphomicrobiales>Brucellaceae</td><td>GCA_020637695.1</td><td>Brucellaceae_bacterium</td><td>H6893_04600</td><td>209797..213123</td><td>2021378</td><td>ATP-binding-protein-[Brucellaceae-bacterium]</td></tr>
<tr><td>RWB26422.1</td><td>DUF499_1.RM.like.systems</td><td><-GDResolvase+Recombinase<-?||?->N6_MTase-> REase -><- DUF499 *<- METHYLASE -p</td><td>944</td><td>Bacteria>Pseudomonadota>Alphaproteobacteria>Hyphomicrobiales>Phyllobacteriaceae</td><td>GCA_004016585.1</td><td>Mesorhizobium_sp.</td><td>EOQ43_30895</td><td>29681..32515</td><td>1871066</td><td>ATP-binding-protein-[Mesorhizobium-sp.]</td></tr>
<tr><td>MBF0177023.1</td><td>DUF499_1.RM.like.systems</td><td>SAM-methylase-><-?<-?<-?<-SF1-SF2-Helicase+HRDC<- DUF499 *<- REase <-OLD-ABC <- METHYLASE</td><td>950</td><td>Bacteria>Pseudomonadota>Alphaproteobacteria>Magnetococcales>unclassified_Magnetococcales</td><td>GCA_015231175.1</td><td>Magnetococcales_bacterium</td><td>HQL63_09275</td><td>22537..25389</td><td>2026759</td><td>ATP-binding-protein-[Magnetococcales-bacterium]</td></tr>
<tr><td>MBF8274722.1</td><td>DUF499_1.RM.like.systems</td><td><- REase +HTH<- DUF499 *<- METHYLASE</td><td>935</td><td>Bacteria>Pseudomonadota>Alphaproteobacteria>Magnetococcales>unclassified_Magnetococcales</td><td>GCA_015660855.1</td><td>Magnetococcales_bacterium</td><td>HW380_3827</td><td>1830..4637</td><td>2026759</td><td>hypothetical-protein-[Magnetococcales-bacterium]</td></tr>
<tr><td>MBF0416113.1</td><td>DUF499_1.RM.like.systems</td><td><-TOPRIM<-?<- REase +HTH<- DUF499 *<- METHYLASE -p</td><td>927</td><td>Bacteria>Pseudomonadota>Alphaproteobacteria>Magnetococcales>unclassified_Magnetococcales</td><td>-</td><td>Magnetococcales_bacterium</td><td>HQL79_10150</td><td>22485..25268</td><td>2026759</td><td>ATP-binding-protein-[Magnetococcales-bacterium]</td></tr>
<tr><td>MBT3398604.1</td><td>DUF499_1.RM.like.systems</td><td>REase -> METHYLASE -> DUF499 *->HEPN->Methylase-><-Radical-SAM<-?<-Peptidase_M15_4</td><td>922</td><td>Bacteria>Pseudomonadota>Alphaproteobacteria>Rhodospirillales>Rhodospirillaceae</td><td>GCA_018648085.1</td><td>Rhodospirillaceae_bacterium</td><td>HN420_01275</td><td>4129..6897</td><td>1898112</td><td>ATP-binding-protein-[Rhodospirillaceae-bacterium]</td></tr>
<tr><td>MCZ0954488.1</td><td>DUF499_1.RM.like.systems</td><td><- DUF499 *<- REase <- METHYLASE</td><td>920</td><td>Bacteria>Pseudomonadota>Alphaproteobacteria>Rhodospirillales>Rhodospirillaceae</td><td>GCA_026914105.1</td><td>Rhodospirillaceae_bacterium</td><td>OXJ56_18105</td><td>195..2957</td><td>1898112</td><td>MAG:-DUF499-domain-containing-protein-[Rhodospirillaceae-bacterium]</td></tr>
<tr><td>MDA8001255.1</td><td>DUF499_1.RM.like.systems</td><td><- DUF499 *<-DUF1828+ REase <-?<- METHYLASE</td><td>799</td><td>Bacteria>Pseudomonadota>Alphaproteobacteria>unclassified_Alphaproteobacteria</td><td>GCA_028066975.1</td><td>Alphaproteobacteria_bacterium</td><td>MPL62_08195</td><td>1..2396</td><td>1913988</td><td>MAG:-DUF499-domain-containing-protein,-partial-[Alphaproteobacteria-bacterium]</td></tr>
<tr><td>MDA8776763.1</td><td>DUF499_1.RM.like.systems</td><td>Arm-DNA-bind_3+Phage_int_SAM_3+Phage_integrase->XIS-HTH->DnaB-><-?<- REase <- HEPN+ DUF499 *<- METHYLASE</td><td>1094</td><td>Bacteria>Pseudomonadota>Alphaproteobacteria>unclassified_Alphaproteobacteria</td><td>GCA_028101585.1</td><td>Alphaproteobacteria_bacterium</td><td>N9N58_04660</td><td>21038..24322</td><td>1913988</td><td>DUF499-domain-containing-protein-[Alphaproteobacteria-bacterium]</td></tr>
<tr><td>MCM8599234.1</td><td>DUF499_1.RM.like.systems</td><td>MerR-HTH-> w HTH+ REase -> METHYLASE ->?->DUF488-1->?-> HEPN+ DUF499 *-></td><td>512</td><td>Bacteria>Pseudomonadota>Betaproteobacteria>Betaproteobacteria_incertae_sedis>Candidatus_Accumulibacter</td><td>GCA_023806585.1</td><td>Accumulibacter_sp.</td><td>NFW04_11345</td><td>16520..18057</td><td>2053492</td><td>Swt1-family-HEPN-domain-containing-protein-[Candidatus-Accumulibacter-sp.]</td></tr>
<tr><td>SBT04363.1</td><td>DUF499_1.RM.like.systems</td><td>TP_methylase-><- DUF499 -p<- DUF499 *<- REase <-OLD-ABC <- METHYLASE -p</td><td>671</td><td>Bacteria>Pseudomonadota>Betaproteobacteria>Betaproteobacteria_incertae_sedis>Candidatus_Accumulibacter</td><td>GCA_900089955.1</td><td>Candidatus_Accumulibacter_aalborgensis</td><td>ACCAA_130242</td><td>226085..228100</td><td>1860102</td><td>conserved-hypothetical-protein-[Candidatus-Accumulibacter-aalborgensis]</td></tr>
<tr><td>MBP6559243.1</td><td>DUF499_1.RM.like.systems</td><td><- DUF499 *<- REase <-OLD-ABC <- METHYLASE -p</td><td>945</td><td>Bacteria>Pseudomonadota>Betaproteobacteria>Burkholderiales>Burkholderiaceae</td><td>GCA_017989415.1</td><td>Burkholderiaceae_bacterium</td><td>KA212_02995</td><td>512..3349</td><td>2030806</td><td>ATP-binding-protein-[Burkholderiaceae-bacterium]</td></tr>
<tr><td>MBK6748240.1</td><td>DUF499_1.RM.like.systems</td><td><-ABhydrolase||<-?<-HEPN|| w HTH+ REase -> DUF499 *-><-tRNA<-?<-?||?->MoxR-AAA-></td><td>942</td><td>Bacteria>Pseudomonadota>Betaproteobacteria>Burkholderiales>Comamonadaceae</td><td>GCA_016703145.1</td><td>Ottowia_sp.</td><td>IPG65_18050</td><td>1102212..1105040</td><td>1898956</td><td>ATP-binding-protein-[Ottowia-sp.]</td></tr>
<tr><td>MBP7545005.1</td><td>DUF499_1.RM.like.systems</td><td><- DUF499 *<- REase <-OLD-ABC <- METHYLASE -p</td><td>949</td><td>Bacteria>Pseudomonadota>Betaproteobacteria>Burkholderiales>Comamonadaceae</td><td>GCA_017998835.1</td><td>Acidovorax_sp.</td><td>KA749_05805</td><td>238..3087</td><td>1872122</td><td>ATP-binding-protein-[Acidovorax-sp.]</td></tr>
<tr><td>WP_210930504.1</td><td>DUF499_1.RM.like.systems</td><td>HTH->?->Phage_integrase->STYKinase-like-><-?||DDE_Tnp_1_5+DDE_Tnp_1-><- DUF499 *<- REase <-OLD-ABC <-HEPN<- METHYLASE <-HNH<-OLD-ABC</td><td>945</td><td>Bacteria>Pseudomonadota>Betaproteobacteria>Burkholderiales>Comamonadaceae</td><td>GCF_018064205.1</td><td>Comamonas_aquatica</td><td>KAQ61_RS03430</td><td>737208..740045</td><td>225991</td><td>DUF499-domain-containing-protein-[Comamonas-aquatica]</td></tr>
</table>

<table>
<tr><th>Protein-ID</th><th>System</th><th>Gene-Neighbourhoods</th><th>Length</th><th>Taxonomy</th><th>Assembly-ID</th><th>Strain</th><th>Locus-Tag</th><th>Genomic-coordinates</th><th>TaxID</th><th>Fasta-defline</th></tr>
<tr><td>WP_209423136.1</td><td>DUF499_1.RM.like.systems</td><td>SF1-SF2-Helicase+DUF1998->?-><-?<-?<-?||?-><-DUF499*<-REase<-OLD-ABC<-TOPRIM<-OLD-ABC<-METHYLASE-p<-METHYLASE</td><td>943</td><td>Bacteria>Pseudomonadota>Betaproteobacteria>Burkholderiales>Comamonadaceae</td><td>GCF_021249385.1</td><td>Melaminivora_jejuensis</td><td>LVC68_RS01445</td><td>284242..287073</td><td>1267217</td><td>DUF499-domain-containing-protein-[Melaminivora-jejuensis]</td></tr>
<tr><td>UUZ46680.1</td><td>DUF499_1.RM.like.systems</td><td>SF1-SF2-Helicase-p->?->?-><-REase<-DUF499-p<-DUF499-p<-DUF499*<-METHYLASE-p<-METHYLASE-p<-METHYLASE-p<-HEPN||?->RNaseP-></td><td>339</td><td>Bacteria>Pseudomonadota>Betaproteobacteria>Burkholderiales>Oxalobacteraceae</td><td>GCA_024703935.1</td><td>Massilia_sp._B-10</td><td>LP420_22520</td><td>3094423..3095442</td><td>2895567</td><td>DUF499-domain-containing-protein-[Massilia-sp.-B-10]</td></tr>
<tr><td>MBK6716517.1</td><td>DUF499_1.RM.like.systems</td><td>METHYLASE-p->OLD-ABC->REase->DUF499*-><-?<-?||METHYLASE-p-></td><td>948</td><td>Bacteria>Pseudomonadota>Betaproteobacteria>Burkholderiales>unclassified_Burkholderiales</td><td>GCA_016703325.1</td><td>Burkholderiales_bacterium</td><td>IPG57_15845</td><td>2519..5365</td><td>1891238</td><td>ATP-binding-protein-[Burkholderiales-bacterium]</td></tr>
<tr><td>MCA3175663.1</td><td>DUF499_1.RM.like.systems</td><td>METHYLASE-p->w HTH+REase->DUF499*-><-tRNA||?->?->Methyltransf_4-></td><td>941</td><td>Bacteria>Pseudomonadota>Betaproteobacteria>Burkholderiales>unclassified_Burkholderiales</td><td>GCA_020248585.1</td><td>Burkholderiales_bacterium</td><td>ING36_09020</td><td>1791..4616</td><td>1891238</td><td>ATP-binding-protein-[Burkholderiales-bacterium]</td></tr>
<tr><td>MCC7486352.1</td><td>DUF499_1.RM.like.systems</td><td>ORC-CDC6+REase->HEPN+DUF499*->?-><-?<-ABhydrolase</td><td>1107</td><td>Bacteria>Pseudomonadota>Betaproteobacteria>Burkholderiales>unclassified_Burkholderiales</td><td>GCA_020854035.1</td><td>Burkholderiales_bacterium</td><td>IT529_15375</td><td>2503..5826</td><td>1891238</td><td>DUF499-domain-containing-protein-[Burkholderiales-bacterium]</td></tr>
<tr><td>MCZ2407521.1</td><td>DUF499_1.RM.like.systems</td><td>Phage_integrase+Phage_integrase->?-><-DUF499*<-REase<-OLD-ABC<-METHYLASE</td><td>945</td><td>Bacteria>Pseudomonadota>Betaproteobacteria>Burkholderiales>unclassified_Burkholderiales</td><td>GCA_026931285.1</td><td>Burkholderiales_bacterium</td><td>LC097_12425</td><td>115108..117945</td><td>1891238</td><td>MAG:-DUF499-domain-containing-protein-[Burkholderiales-bacterium]</td></tr>
<tr><td>NNM68915.1</td><td>DUF499_1.RM.like.systems</td><td>METHYLASE-p->w HTH+REase->DUF499*-></td><td>522</td><td>Bacteria>Pseudomonadota>Betaproteobacteria>Nitrosomonadales>Gallionellaceae</td><td>GCA_013043965.1</td><td>Gallionella_sp.</td><td>HKM00_02900</td><td>1745..3312</td><td>2052161</td><td>ATP-binding-protein-[Gallionella-sp.]</td></tr>
<tr><td>MBN8462739.1</td><td>DUF499_1.RM.like.systems</td><td>PrlF_antitoxin->PemK_toxin->REase->METHYLASE->PolB->HEPN->HEPN+DUF499*->?-><-?<-?||?->sGTP-></td><td>1110</td><td>Bacteria>Pseudomonadota>Betaproteobacteria>Rhodocyclales>Azonexaceae</td><td>GCA_017302355.1</td><td>Dechloromonas_sp.</td><td>JOM01_07975</td><td>12361..15693</td><td>1917218</td><td>DUF499-domain-containing-protein-[Dechloromonas-sp.]</td></tr>
<tr><td>MBP6189859.1</td><td>DUF499_1.RM.like.systems</td><td>DUF488-l-><-?<-DUF499*<-w HTH+REase<-METHYLASE<-HEPN</td><td>943</td><td>Bacteria>Pseudomonadota>Betaproteobacteria>Rhodocyclales>Azonexaceae</td><td>GCA_017985755.1</td><td>Azonexus_sp.</td><td>KA440_12990</td><td>2046..4877</td><td>1872668</td><td>ATP-binding-protein-[Azonexus-sp.]</td></tr>
<tr><td>KAB2933410.1</td><td>DUF499_1.RM.like.systems</td><td>OLD-ABC->TOPRIM->?->?->METHYLASE->REase->HEPN+DUF499*->?-><-?||HTH->MarR-HTH-></td><td>1096</td><td>Bacteria>Pseudomonadota>Betaproteobacteria>Rhodocyclales>Rhodocyclaceae</td><td>GCA_008933825.1</td><td>Rhodocyclaceae_bacterium</td><td>F9K21_12245</td><td>12685..15975</td><td>1898103</td><td>ATP-binding-protein-[Rhodocyclaceae-bacterium]</td></tr>
<tr><td>MBQ7625733.1</td><td>DUF499_1.RM.like.systems</td><td>METHYLASE-p->ParB+HNH->DUF499*-><-REase||TP_methylase->?->?->ncRNA->?->Methyltransf_5-></td><td>945</td><td>Bacteria>Pseudomonadota>Betaproteobacteria>Rhodocyclales>Rhodocyclaceae</td><td>GCA_017444505.1</td><td>Rhodocyclaceae_bacterium</td><td>IJS87_01550</td><td>2639..5476</td><td>1898103</td><td>ATP-binding-protein-[Rhodocyclaceae-bacterium]</td></tr>
<tr><td>MDD3354816.1</td><td>DUF499_1.RM.like.systems</td><td>PIN-><-DUF499*<-REase<-OLD-ABC<-METHYLASE</td><td>948</td><td>Bacteria>Pseudomonadota>Betaproteobacteria>Rhodocyclales>Zoogloeaceae</td><td>GCA_028693495.1</td><td>Zoogloea_sp.</td><td>PHD22_14435</td><td>4878..7724</td><td>49181</td><td>MAG:-DUF499-domain-containing-protein-[Zoogloea-sp.]</td></tr>
<tr><td>OGA43321.1</td><td>DUF499_1.RM.like.systems</td><td>METHYLASE->PolB->HEPN->REase->DUF499*-></td><td>927</td><td>Bacteria>Pseudomonadota>Betaproteobacteria>unclassified_Betaproteobacteria</td><td>GCA_001771985.1</td><td>Betaproteobacteria_bacterium_RIFCSPLOWO2_12_FULL_62_13</td><td>A3G24_09290</td><td>4491..7274</td><td>1797495</td><td>AAA-family-ATPase-[Betaproteobacteria-bacterium-RIFCSPLOWO2_12_FULL_62_13]</td></tr>
<tr><td>MPZ45705.1</td><td>DUF499_1.RM.like.systems</td><td>AAA-ATPase+REase->METHYLASE->DUF499*-></td><td>1088</td><td>Bacteria>Pseudomonadota>Betaproteobacteria>unclassified_Betaproteobacteria</td><td>GCA_009377585.1</td><td>Betaproteobacteria_bacterium</td><td>GEV05_20400</td><td>5826..9092</td><td>1891241</td><td>DUF499-domain-containing-protein-[Betaproteobacteria-bacterium]</td></tr>
<tr><td>MBI5922415.1</td><td>DUF499_1.RM.like.systems</td><td><-DUF499*<-REase<-METHYLASE<-?<-Schlafen+HATPase+HTH||URI-DNase-></td><td>933</td><td>Bacteria>Pseudomonadota>Betaproteobacteria>unclassified_Betaproteobacteria</td><td>GCA_016234745.1</td><td>Betaproteobacteria_bacterium</td><td>HY847_12340</td><td>39471..42272</td><td>1891241</td><td>ATP-binding-protein-[Betaproteobacteria-bacterium]</td></tr>
<tr><td>WP_135445665.1</td><td>DUF499_1.RM.like.systems</td><td>METHYLASE->?->?->OLD-ABC->TOPRIM->DUF499*->REase->MORC+DUF3684+REase-><-SF1-SF2-Helicase+REase<-SF1-SF2-Helicase+REase||?->HKD-DNase+HKD-DNase-></td><td>947</td><td>Bacteria>Pseudomonadota>Gammaproteobacteria>Cellvibrionales>Halieaceae</td><td>GCF_004745945.1</td><td>Haliea_sp._SAOS-164</td><td>E4634_RS16035</td><td>159390..162233</td><td>2562682</td><td>DUF499-domain-containing-protein-[Haliea-sp.-SAOS-164]</td></tr>
<tr><td>MCK9506376.1</td><td>DUF499_1.RM.like.systems</td><td>OLD-ABC->REase->DUF499*->Classical-AAA-><-?<-DUF932</td><td>951</td><td>Bacteria>Pseudomonadota>Gammaproteobacteria>Cellvibrionales>Porticoccaceae</td><td>GCA_023228865.1</td><td>Porticoccaceae_bacterium</td><td>MOQ95_19625</td><td>878..3733</td><td>2026782</td><td>DUF499-domain-containing-protein-[Porticoccaceae-bacterium]</td></tr>
<tr><td>MBV5272245.1</td><td>DUF499_1.RM.like.systems</td><td>REase->w HTH+REase->DUF499*->?->McrC-NTD+REase-></td><td>961</td><td>Bacteria>Pseudomonadota>Gammaproteobacteria>Chromatiales>Chromatiaceae</td><td>GCA_019163055.1</td><td>Lamprocystis_purpurea</td><td>JZU52_00945</td><td>1014..3899</td><td>61598</td><td>ATP-binding-protein-[Lamprocystis-purpurea]</td></tr>
<tr><td>TQE99812.1</td><td>DUF499_1.RM.like.systems</td><td>HEPN->ParA-Soj-PloopNTPase->ORC-CDC6->METHYLASE->OLD-ABC->REase->DUF499*->?->UbE1+JAB->?->Schlafen-></td><td>955</td><td>Bacteria>Pseudomonadota>Gammaproteobacteria>Chromatiales>Ectothiorhodospiraceae</td><td>GCA_006569195.1</td><td>Spiribacter_salinus</td><td>FKY71_06670</td><td>12316..15183</td><td>1335746</td><td>ATP-binding-protein-[Spiribacter-salinus]</td></tr>
<tr><td>MCC6374735.1</td><td>DUF499_1.RM.like.systems</td><td><-DUF499*<-REase<-OLD-ABC<-METHYLASE-p</td><td>878</td><td>Bacteria>Pseudomonadota>Gammaproteobacteria>Moraxellales>Moraxellaceae</td><td>GCA_020846355.1</td><td>Moraxellaceae_bacterium</td><td>IT465_07990</td><td>1..2635</td><td>1889775</td><td>ATP-binding-protein-[Moraxellaceae-bacterium]</td></tr>
<tr><td>WP_107866351.1</td><td>DUF499_1.RM.like.systems</td><td>METHYLASE->OLD-ABC->REase->DUF499*-><-ParB+HNH<-?<-ParB</td><td>941</td><td>Bacteria>Pseudomonadota>Gammaproteobacteria>Moraxellales>Moraxellaceae</td><td>GCF_003051055.1</td><td>Agitococcus_lubricus</td><td>C8N29_RS12225</td><td>4668..7493</td><td>1077255</td><td>DUF499-domain-containing-protein-[Agitococcus-lubricus]</td></tr>
<tr><td>EWH01940.1</td><td>DUF499_1.RM.like.systems</td><td>REase->MAD-NTD+EVE+McrB-AAA->?->McrC-NTD+REase->?->DUF499-p->DUF499*->?->?-><-?||?->REase-></td><td>676</td><td>Bacteria>Pseudomonadota>Gammaproteobacteria>Oceanospirillales>Halomonadaceae</td><td>GCA_000568385.1</td><td>Halomonas_sp._BC04</td><td>Q427_11510</td><td>32984..35014</td><td>1403540</td><td>hypothetical-protein-Q427_11510-[Halomonas-sp.-BC04]</td></tr>
<tr><td>BBI71108.2</td><td>DUF499_1.RM.like.systems</td><td>Phage_integrase-><-Phage_AlpA<-TOPRIM+primpol<-?<-XIS-HTH<-DUF499-p<-DUF499*<-DUF499-p<-REase<-OLD-ABC<-?<-METHYLASE-p<-METHYLASE-p</td><td>386</td><td>Bacteria>Pseudomonadota>Gammaproteobacteria>Oceanospirillales>Halomonadaceae</td><td>GCA_007163885.2</td><td>Halomonas_axialensis</td><td>HAALTHF_10150n</td><td>665831..666991</td><td>115555</td><td>hypothetical-protein-HAALTHF_10150n-[Halomonas-axialensis]</td></tr>
<tr><td>MCD6438846.1</td><td>DUF499_1.RM.like.systems</td><td>HNH->METHYLASE->OLD-ABC->REase->EVE+McrB-AAA->McrC-NTD+REase->DUF499*-></td><td>282</td><td>Bacteria>Pseudomonadota>Gammaproteobacteria>Oceanospirillales>Halomonadaceae</td><td>GCA_021161325.1</td><td>Halomonas_sp.</td><td>J7L56_11360</td><td>10325..11171</td><td>1486246</td><td>ATP-binding-protein-[Halomonas-sp.]</td></tr>
<tr><td>WP_269321216.1</td><td>DUF499_1.RM.like.systems</td><td>REase->MAD-NTD+EVE+McrB-AAA->?->McrC-NTD+REase->?->DUF499->DUF499*->?-><-?||?->REase-></td><td>680</td><td>Bacteria>Pseudomonadota>Gammaproteobacteria>Oceanospirillales>Halomonadaceae</td><td>GCF_000568385.1</td><td>Halomonas_sp._BC04</td><td>Q427_RS09405</td><td>32972..35014</td><td>1403540</td><td>DUF499-domain-containing-protein-[Halomonas-sp.-BC04]</td></tr>
<tr><td>WP_197030572.1</td><td>DUF499_1.RM.like.systems</td><td>OLD-ABC->REase->MAD-NTD+EVE+McrB-AAA->?->McrC-NTD+REase->?->DUF499*->DUF499->?-><-?||?->REase-></td><td>264</td><td>Bacteria>Pseudomonadota>Gammaproteobacteria>Oceanospirillales>Halomonadaceae</td><td>GCF_000568385.1</td><td>Halomonas_sp._BC04</td><td>Q427_RS35225</td><td>32151..32945</td><td>1403540</td><td>DUF499-domain-containing-protein-[Halomonas-sp.-BC04]</td></tr>
<tr><td>WP_030073674.1</td><td>DUF499_1.RM.like.systems</td><td>RelE->CHTH-><-AbiEi||?-><-DUF499*<-McrC-NTD+REase<-MAD-NTD+EVE+McrB-AAA<-OLD-ABC<-?<-METHYLASE<-HNH</td><td>961</td><td>Bacteria>Pseudomonadota>Gammaproteobacteria>Oceanospirillales>Halomonadaceae</td><td>GCF_000712975.1</td><td>Halomonas_alkaliantarctica</td><td>CR46_RS0113290</td><td>253918..256803</td><td>232346</td><td>DUF499-domain-containing-protein-[Halomonas-alkaliantarctica]</td></tr>
<tr><td>WP_097789710.1</td><td>DUF499_1.RM.like.systems</td><td>Peptidase_M20->Arm-DNA-bind_3+Phage_integrase->TPR+TPR-><-?<-DUF499*<-McrC-NTD+REase<-MAD-NTD+EVE+McrB-AAA<-REase<-OLD-ABC<-HEPN+CBS<-METHYLASE</td><td>957</td><td>Bacteria>Pseudomonadota>Gammaproteobacteria>Oceanospirillales>Halomonadaceae</td><td>GCF_002549795.1</td><td>Halomonas_beimenensis</td><td>BEI_RS11820</td><td>2512898..2515771</td><td>475662</td><td>DUF499-domain-containing-protein-[Halomonas-beimenensis]</td></tr>
<tr><td>WP_044628625.1</td><td>DUF499_1.RM.like.systems</td><td>METHYLASE->OLD-ABC->REase->DUF499*-><-MobI||XIS-HTH->ZnR+TOPRIM+primpol+HTH->Phage_AlpA-><-Arm-DNA-bind_3+Phage_integrase</td><td>953</td><td>Bacteria>Pseudomonadota>Gammaproteobacteria>Oceanospirillales>Halomonadaceae</td><td>GCF_011398715.1</td><td>Halomonas</td><td>QWE69_RS13020</td><td>3280..6141</td><td>2745</td><td>MULTISPECIES:-DUF499-domain-containing-protein-[Halomonas]</td></tr>
<tr><td>WP_234246274.1</td><td>DUF499_1.RM.like.systems</td><td>METHYLASE->METHYLASE->Schlafen+HATPase+HTH->HEPN+CBS->OLD-ABC->REase->DUF499*-></td><td>957</td><td>Bacteria>Pseudomonadota>Gammaproteobacteria>Oceanospirillales>Halomonadaceae</td><td>GCF_021404345.1</td><td>Halomonas_desiderata</td><td>HOP57_RS15495</td><td>85349..88222</td><td>52021</td><td>DUF499-domain-containing-protein-[Halomonas-desiderata]</td></tr>
<tr><td>WP_253017818.1</td><td>DUF499_1.RM.like.systems</td><td>HNH->METHYLASE->OLD-ABC->REase->EVE+McrB-AAA->McrC-NTD+REase->DUF499*->?->AbiEi-><-CHTH<-RelE</td><td>961</td><td>Bacteria>Pseudomonadota>Gammaproteobacteria>Oceanospirillales>Halomonadaceae</td><td>GCF_024124605.1</td><td>Halomonas_sp._CnH100-B</td><td>NH398_RS08385</td><td>365976..368861</td><td>2954490</td><td>DUF499-domain-containing-protein-[Halomonas-sp.-CnH100-B]</td></tr>
<tr><td>WP_230706347.1</td><td>DUF499_1.RM.like.systems</td><td>Phage_integrase-><-Phage_AlpA<-ZnR+TOPRIM+primpol+HTH<-XIS-HTH||MobI-><-DUF499*<-REase<-OLD-ABC||||?-><-?<-?<-Resolvase</td><td>953</td><td>Bacteria>Pseudomonadota>Gammaproteobacteria>Pseudomonadales>Marinobacteraceae</td><td>GCF_021021495.1</td><td>Marinobacter_adhaerens</td><td>K7I10_RS06135</td><td>1373639..1376500</td><td>1033846</td><td>DUF499-domain-containing-protein-[Marinobacter-adhaerens]</td></tr>
<tr><td>WP_049271643.1</td><td>DUF499_1.RM.like.systems</td><td>ParB->JAB->?->RlpA-><-DUF499*<-REase<-OLD-ABC<-METHYLASE</td><td>943</td><td>Bacteria>Pseudomonadota>Gammaproteobacteria>Pseudomonadales>Pseudomonadaceae</td><td>GCF_001077135.1</td><td>Pseudomonas_aeruginosa</td><td>ADL79_RS26490</td><td>2020..4851</td><td>287</td><td>DUF499-domain-containing-protein-[Pseudomonas-aeruginosa]</td></tr>
<tr><td>RTZ75007.1</td><td>DUF499_1.RM.like.systems</td><td><-NERD+SF1-SF2-Helicase<-DUF499*<-REase<-OLD-ABC<-METHYLASE-p</td><td>943</td><td>Bacteria>Pseudomonadota>Gammaproteobacteria>unclassified_Gammaproteobacteria</td><td>GCA_003973075.1</td><td>Gammaproteobacteria_bacterium</td><td>DSZ00_02995</td><td>2492..5323</td><td>1913989</td><td>hypothetical-protein-DSZ00_02995-[Gammaproteobacteria-bacterium]</td></tr>
<tr><td>MYA16811.1</td><td>DUF499_1.RM.like.systems</td><td>OLD-ABC-p->OLD-ABC-p->REase->DUF499*->URI-DNase-></td><td>922</td><td>Bacteria>Pseudomonadota>Gammaproteobacteria>unclassified_Gammaproteobacteria</td><td>GCA_009836885.1</td><td>Gammaproteobacteria_bacterium</td><td>F4Z28_08615</td><td>8657..11425</td><td>1913989</td><td>ATP-binding-protein-[Gammaproteobacteria-bacterium]</td></tr>
<tr><td>HFD86238.1</td><td>DUF499_1.RM.like.systems</td><td><-DUF499*<-REase<-OLD-ABC<-METHYLASE-p</td><td>692</td><td>Bacteria>Pseudomonadota>Gammaproteobacteria>unclassified_Gammaproteobacteria</td><td>GCA_011322685.1</td><td>Gammaproteobacteria_bacterium</td><td>ENJ35_01010</td><td>1..2075</td><td>1913989</td><td>ATP-binding-protein-[Gammaproteobacteria-bacterium]</td></tr>
<tr><td>MCG3202826.1</td><td>DUF499_1.RM.like.systems</td><td>N6_N4_MTase-><-?<-AbiEii<-AbiEi_4<-REase<-HEPN+DUF499*<-METHYLASE-p</td><td>1115</td><td>Bacteria>Pseudomonadota>Gammaproteobacteria>unclassified_Gammaproteobacteria</td><td>GCA_022071965.1</td><td>Gammaproteobacteria_bacterium</td><td>NFCOHLIN_02712</td><td>302948..306295</td><td>1913989</td><td>hypothetical-protein-[Gammaproteobacteria-bacterium]</td></tr>
<tr><td>MCY3837525.1</td><td>DUF499_1.RM.like.systems</td><td>METHYLASE->CR-ATPase8+REase->HEPN+DUF499*->Tox-RelE->HTH+MPTase-><-VapB_antitoxin</td><td>1112</td><td>Bacteria>Pseudomonadota>Gammaproteobacteria>unclassified_Gammaproteobacteria</td><td>GCA_026706425.1</td><td>Gammaproteobacteria_bacterium</td><td>OXH09_02545</td><td>4546..7884</td><td>1913989</td><td>MAG:-Swt1-family-HEPN-domain-containing-protein-[Gammaproteobacteria-bacterium]</td></tr>
<tr><td>MCY4342454.1</td><td>DUF499_1.RM.like.systems</td><td>ORC-CDC6+REase-><-DUF499*<-PIN<-DUF433<-METHYLASE-p<-METHYLASE<-Peptidase_S8<-Classical-AAA</td><td>924</td><td>Bacteria>Pseudomonadota>Gammaproteobacteria>unclassified_Gammaproteobacteria</td><td>GCA_026710215.1</td><td>Gammaproteobacteria_bacterium</td><td>OXE83_02620</td><td>2601..5375</td><td>1913989</td><td>MAG:-DUF499-domain-containing-protein-[Gammaproteobacteria-bacterium]</td></tr>
<tr><td>MDE0285439.1</td><td>DUF499_1.RM.like.systems</td><td><-DUF499*<-REase<-METHYLASE-p</td><td>924</td><td>Bacteria>Pseudomonadota>Gammaproteobacteria>unclassified_Gammaproteobacteria</td><td>GCA_028820555.1</td><td>Gammaproteobacteria_bacterium</td><td>OXN26_12940</td><td>1577..4351</td><td>1913989</td><td>MAG:-DUF499-domain-containing-protein-[Gammaproteobacteria-bacterium]</td></tr>
<tr><td>MCK9487878.1</td><td>DUF499_1.RM.like.systems</td><td><-SIGMA-HTH<-OLD-ABC<-?<-?<-?<-DUF499*<-REase<-OLD-ABC<-METHYLASE<-HEPN<-PIN<-PemK_toxin</td><td>944</td><td>Bacteria>Pseudomonadota>Gammaproteobacteria>Xanthomonadales>unclassified_Xanthomonadales</td><td>GCA_023228845.1</td><td>Xanthomonadales_bacterium</td><td>MOQ42_00470</td><td>105333..108167</td><td>2006849</td><td>DUF499-domain-containing-protein-[Xanthomonadales-bacterium]</td></tr>
<tr><td>MCU0753612.1</td><td>DUF499_1.RM.like.systems</td><td><-ArsR-HTH||?->?->?->?->?-><-DUF499*<-Schlafen+HATPase+HTH<-ParB+HNH<-REase<-OLD-ABC<-METHYLASE<-HNH</td><td>955</td><td>Bacteria>Pseudomonadota>Gammaproteobacteria>Xanthomonadales>unclassified_Xanthomonadales</td><td>GCA_025458445.1</td><td>Xanthomonadales_bacterium</td><td>MUE46_00615</td><td>13163..16030</td><td>2006849</td><td>DUF499-domain-containing-protein-[Xanthomonadales-bacterium]</td></tr>
<tr><td>GIX32370.1</td><td>DUF499_1.RM.like.systems</td><td>REase->METHYLASE-p->METHYLASE-p->METHYLASE->ParD->PIN->HEPN+DUF499*-><-tRNA<-SIGMA-HTH+SIGMA-HTH</td><td>1108</td><td>Bacteria>Pseudomonadota>Gammaproteobacteria>Xanthomonadales>unclassified_Xanthomonadales</td><td>GCA_026005055.1</td><td>Xanthomonadales_bacterium</td><td>KatS3mg125_0326</td><td>361925..365251</td><td>2006849</td><td>MAG:-hypothetical-protein-KatS3mg125_0326-[Xanthomonadales-bacterium]</td></tr>
<tr><td>GIX36093.1</td><td>DUF499_1.RM.like.systems</td><td>METHYLASE-p->METHYLASE-p->OLD-ABC->REase->DUF499*-><-?<-?<-?<-?<-Radical-SAM</td><td>950</td><td>Bacteria>Pseudomonadota>Gammaproteobacteria>Xanthomonadales>Xanthomonadaceae</td><td>GCA_026005075.1</td><td>Xanthomonadaceae_bacterium</td><td>KatS3mg126_1872</td><td>4497..7349</td><td>1926873</td><td>MAG:-hypothetical-protein-KatS3mg126_1872-[Xanthomonadaceae-bacterium]</td></tr>
<tr><td>WP_054362466.1</td><td>DUF499_1.RM.like.systems</td><td>ParG_RHH-><-MODE-HTH+PBPII<-HEPN+DUF499*<-METHYLASE<-REase</td><td>1098</td><td>Bacteria>Pseudomonadota</td><td>GCF_963522535.1</td><td>Alphaproteobacteria</td><td>ABB55_RS28500</td><td>24847..28143</td><td>28211</td><td>MULTISPECIES:-DUF499-domain-containing-protein-[Alphaproteobacteria]</td></tr>
<tr><td>NBT04715.1</td><td>DUF499_1.RM.like.systems</td><td><-REase<-HEPN+DUF499*<-METHYLASE-p</td><td>1140</td><td>Bacteria>Pseudomonadota>unclassified_Pseudomonadota</td><td>GCA_009922565.1</td><td>Pseudomonadota_bacterium</td><td>EBS94_15100</td><td>2192..5614</td><td>1977087</td><td>ATP-binding-protein-[Proteobacteria-bacterium]</td></tr>
<tr><td>MBU0652986.1</td><td>DUF499_1.RM.like.systems</td><td>AAA-ATPase+REase->DUF499*-></td><td>325</td><td>Bacteria>Pseudomonadota>unclassified_Pseudomonadota</td><td>GCA_018814185.1</td><td>Pseudomonadota_bacterium</td><td>KKG96_08920</td><td>1165..2139</td><td>1977087</td><td>DUF499-domain-containing-protein-[Pseudomonadota-bacterium]</td></tr>
<tr><td>MBU2056000.1</td><td>DUF499_1.RM.like.systems</td><td>w HTH+REase->SF1-SF2-Helicase->METHYLASE->REase->PolB->HEPN->DUF499*-><-?<-?<-HTH</td><td>950</td><td>Bacteria>Pseudomonadota>unclassified_Pseudomonadota</td><td>GCA_018825885.1</td><td>Pseudomonadota_bacterium</td><td>KKC25_13705</td><td>11854..14706</td><td>1977087</td><td>DUF499-domain-containing-protein-[Proteobacteria-bacterium]</td></tr>
<tr><td>MBU4372115.1</td><td>DUF499_1.RM.like.systems</td><td>Peptidase_M24->?->Aminotran_1_2->?->?->DNA_pol3_delta+DNA_pol3_gamma3-><-DUF499*<-AAA-ATPase+REase<-METHYLASE-p</td><td>1098</td><td>Bacteria>Pseudomonadota>unclassified_Pseudomonadota</td><td>GCA_018896355.1</td><td>Pseudomonadota_bacterium</td><td>KKD68_09540</td><td>35625..38921</td><td>1977087</td><td>DUF499-domain-containing-protein-[Proteobacteria-bacterium]</td></tr>
<tr><td>MBU4582289.1</td><td>DUF499_1.RM.like.systems</td><td>AAA-ATPase+REase->DUF499*-><-DNA_pol3_delta+DNA_pol3_gamma3<-?<-?<-Aminotran_1_2<-?<-Peptidase_M24</td><td>1098</td><td>Bacteria>Pseudomonadota>unclassified_Pseudomonadota</td><td>GCA_018902925.1</td><td>Pseudomonadota_bacterium</td><td>KKH02_07770</td><td>1387..4683</td><td>1977087</td><td>DUF499-domain-containing-protein-[Proteobacteria-bacterium]</td></tr>
<tr><td>MDA8349445.1</td><td>DUF499_1.RM.like.systems</td><td>METHYLASE-p->DUF499*->REase+nSTAND1-></td><td>941</td><td>Bacteria>Pseudomonadota>unclassified_Pseudomonadota</td><td>GCA_028067895.1</td><td>Pseudomonadota_bacterium</td><td>MO038_11690</td><td>747..3572</td><td>1977087</td><td>MAG:-DUF499-domain-containing-protein-[Pseudomonadota-bacterium]</td></tr>
<tr><td>MBP7275753.1</td><td>DUF499_1.RM.like.systems</td><td><-DUF499*<-?<-METHYLASE<-PIN<-?<-REase</td><td>1001</td><td>Bacteria>PVC_group>Kiritimatiellota>Kiritimatiellia>unclassified_Kiritimatiellia</td><td>GCA_017996315.1</td><td>Kiritimatiellia_bacterium</td><td>KBA51_06085</td><td>1..3004</td><td>2212473</td><td>ATP-binding-protein-[Kiritimatiellae-bacterium]</td></tr>
<tr><td>MBN2450533.1</td><td>DUF499_1.RM.like.systems</td><td>REase->?->PIN->HEPN+DUF499*-></td><td>1121</td><td>Bacteria>PVC_group>Lentisphaerota>Lentisphaeria>unclassified_Lentisphaeria</td><td>GCA_016935045.1</td><td>Lentisphaeria_bacterium</td><td>JXR77_09085</td><td>1530..4895</td><td>2053569</td><td>DUF499-domain-containing-protein-[Lentisphaeria-bacterium]</td></tr>
<tr><td>NLF19752.1</td><td>DUF499_1.RM.like.systems</td><td><-DUF499*<-w HTH+REase<-METHYLASE<-?<-?<-?<-Schlafen+HATPase+HTH</td><td>905</td><td>Bacteria>PVC_group>Lentisphaerota>unclassified_Lentisphaerota</td><td>GCA_012514865.1</td><td>Lentisphaerota_bacterium</td><td>GX595_21175</td><td>1..2716</td><td>1932692</td><td>ATP-binding-protein-[Lentisphaerae-bacterium]</td></tr>
<tr><td>NLG01020.1</td><td>DUF499_1.RM.like.systems</td><td><-DOC+HTH<-?<-DUF499*<-?<-TIR<-TIR<-METHYLASE<-REase<-SF1-SF2-Helicase</td><td>925</td><td>Bacteria>PVC_group>Lentisphaerota>unclassified_Lentisphaerota</td><td>GCA_012515415.1</td><td>Lentisphaerota_bacterium</td><td>GX565_12840</td><td>2112..4889</td><td>1932692</td><td>ATP-binding-protein-[Lentisphaerae-bacterium]</td></tr>
<tr><td>NLN03021.1</td><td>DUF499_1.RM.like.systems</td><td><-DUF499*<-REase<-METHYLASE-p</td><td>437</td><td>Bacteria>PVC_group>Lentisphaerota>unclassified_Lentisphaerota</td><td>GCA_012521875.1</td><td>Lentisphaerota_bacterium</td><td>GX174_14140</td><td>1..1313</td><td>1932692</td><td>ATP-binding-protein-[Lentisphaerota-bacterium]</td></tr>
<tr><td>MBN1509225.1</td><td>DUF499_1.RM.like.systems</td><td><-PAPS_reduct<-?<-?<-KAP-NTPase<-DUF499*<-HTH<-REase<-METHYLASE</td><td>922</td><td>Bacteria>PVC_group>Planctomycetota>Phycisphaerae>Sedimentisphaerales</td><td>GCA_016928975.1</td><td>Sedimentisphaerales_bacterium</td><td>JW955_20435</td><td>4212..6980</td><td>2813034</td><td>ATP-binding-protein-[Sedimentisphaerales-bacterium]</td></tr>
<tr><td>MCG3126049.1</td><td>DUF499_1.RM.like.systems</td><td>HEPN+REase->METHYLASE->ParB+HNH->DUF499*->?->SF1-SF2-Helicase->?->?->CASPASE+APATPase+TPR+TPR+TPR-></td><td>1107</td><td>Bacteria>PVC_group>Planctomycetota>Phycisphaerae>unclassified_Phycisphaerae</td><td>GCA_022072445.1</td><td>Phycisphaerae_bacterium</td><td>CHACPFDD_00877</td><td>44646..47969</td><td>2026778</td><td>hypothetical-protein-[Phycisphaerae-bacterium]</td></tr>
<tr><td>MSR55922.1</td><td>DUF499_1.RM.like.systems</td><td><-DUF499*<-REase<-PIN<-DUF433<-METHYLASE-p</td><td>984</td><td>Bacteria>PVC_group>Planctomycetota>Planctomycetia>Gemmatales</td><td>GCA_009691845.1</td><td>Gemmataceae_bacterium</td><td>EXS09_22015</td><td>769..3723</td><td>2052164</td><td>ATP-binding-protein-[Gemmataceae-bacterium]</td></tr>
<tr><td>MCI0378738.1</td><td>DUF499_1.RM.like.systems</td><td><-DUF499*<-METHYLASE<-REase</td><td>207</td><td>Bacteria>PVC_group>Planctomycetota>Planctomycetia>Gemmatales</td><td>GCA_022614915.1</td><td>Gemmataceae_bacterium</td><td>LO215_14110</td><td>1..621</td><td>2052164</td><td>DUF499-domain-containing-protein-[Gemmataceae-bacterium]</td></tr>
<tr><td>NLF68873.1</td><td>DUF499_1.RM.like.systems</td><td>ParB->ParB-><-HEPN<-PIWI<-REase+PNTD1+PNTD2<-?<-DUF499*</td><td>982</td><td>Bacteria>PVC_group>Planctomycetota>Planctomycetia>Pirellulales</td><td>GCA_012515235.1</td><td>Candidatus_Anammoximicrobium_sp.</td><td>GX575_07440</td><td>18006..20954</td><td>2699742</td><td>ATP-binding-protein-[Candidatus-Anammoximicrobium-sp.]</td></tr>
<tr><td>NLX99940.1</td><td>DUF499_1.RM.like.systems</td><td><-DUF499*<-PIN<-?<-REase<-?<-METHYLASE<-REase</td><td>981</td><td>Bacteria>PVC_group>Planctomycetota>Planctomycetia>Pirellulales</td><td>GCA_012728855.1</td><td>Rhodopirellula_sp.</td><td>GXY83_27900</td><td>2887..5832</td><td>2024855</td><td>ATP-binding-protein-[Rhodopirellula-sp.]</td></tr>
<tr><td>MBN1912210.1</td><td>DUF499_1.RM.like.systems</td><td>METHYLASE->w HTH+REase->DUF499*-><-Radical-SAM</td><td>1103</td><td>Bacteria>PVC_group>Planctomycetota>Planctomycetia>Pirellulales</td><td>GCA_016931075.1</td><td>Pirellulales_bacterium</td><td>JW818_20990</td><td>4913..8224</td><td>2715253</td><td>ATP-binding-protein-[Pirellulales-bacterium]</td></tr>
<tr><td>GIW94553.1</td><td>DUF499_1.RM.like.systems</td><td><-DUF499*<-REase<-?<-TOPRIM<-OLD-ABC<-METHYLASE-p<-METHYLASE-p</td><td>938</td><td>Bacteria>PVC_group>Planctomycetota>Planctomycetia>Pirellulales</td><td>GCA_026004775.1</td><td>Pirellulaceae_bacterium</td><td>KatS3mg110_2594</td><td>12784..15600</td><td>2699754</td><td>MAG:-hypothetical-protein-KatS3mg110_2594-[Pirellulaceae-bacterium]</td></tr>
<tr><td>HBI42788.1</td><td>DUF499_1.RM.like.systems</td><td>SIGMA-HTH->?-><-DUF499*<-REase<-OLD-ABC<-METHYLASE</td><td>939</td><td>Bacteria>PVC_group>Planctomycetota>Planctomycetia>Planctomycetales</td><td>GCA_003508275.1</td><td>Planctomycetales_bacterium</td><td>DDY78_08010</td><td>2578..5397</td><td>2053591</td><td>hypothetical-protein-[Planctomycetales-bacterium]</td></tr>
<tr><td>MBV8676907.1</td><td>DUF499_1.RM.like.systems</td><td>METHYLASE-p->REase->HEPN+DUF499*->Y1_Tnp-></td><td>1104</td><td>Bacteria>PVC_group>Planctomycetota>Planctomycetia>Planctomycetales</td><td>GCA_019232645.1</td><td>Planctomycetaceae_bacterium</td><td>JO355_07025</td><td>1198..4512</td><td>2026779</td><td>DUF499-domain-containing-protein-[Planctomycetaceae-bacterium]</td></tr>
<tr><td>MBV8269357.1</td><td>DUF499_1.RM.like.systems</td><td><-HEPN+DUF499*<-REase<-METHYLASE-p</td><td>1104</td><td>Bacteria>PVC_group>Planctomycetota>Planctomycetia>Planctomycetales</td><td>GCA_019236385.1</td><td>Planctomycetaceae_bacterium</td><td>JO252_23790</td><td>243..3557</td><td>2026779</td><td>DUF499-domain-containing-protein-[Planctomycetaceae-bacterium]</td></tr>
<tr><td>MCA9067325.1</td><td>DUF499_1.RM.like.systems</td><td><-DUF499*<-?<-REase<-REase</td><td>982</td><td>Bacteria>PVC_group>Planctomycetota>Planctomycetia>Planctomycetales</td><td>GCA_020426385.1</td><td>Planctomycetaceae_bacterium</td><td>KDA84_00275</td><td>118..3066</td><td>2026779</td><td>ATP-binding-protein-[Planctomycetaceae-bacterium]</td></tr>
<tr><td>MCA9076582.1</td><td>DUF499_1.RM.like.systems</td><td>sGTP->METHYLASE-p->REase->METHYLASE->OLD-ABC->PIN->HEPN+DUF499*-><-PAIREDC-HTH<-?<-?<-SIGMA-HTH</td><td>1095</td><td>Bacteria>PVC_group>Planctomycetota>Planctomycetia>Planctomycetales</td><td>GCA_020426505.1</td><td>Planctomycetaceae_bacterium</td><td>KDA93_16275</td><td>17776..21063</td><td>2026779</td><td>DUF499-domain-containing-protein-[Planctomycetaceae-bacterium]</td></tr>
<tr><td>MBI2825945.1</td><td>DUF499_1.RM.like.systems</td><td><-REase<-?<-?<-?<-DNA_methylase<-DUF499*<-HTH<-REase<-RNaseH<-METHYLASE<-METHYLASE-p</td><td>930</td><td>Bacteria>PVC_group>Planctomycetota>Planctomycetia>unclassified_Planctomycetia</td><td>GCA_016191115.1</td><td>Planctomycetia_bacterium</td><td>HYX69_14775</td><td>236780..239572</td><td>2052181</td><td>ATP-binding-protein-[Planctomycetia-bacterium]</td></tr>
<tr><td>NMD35173.1</td><td>DUF499_1.RM.like.systems</td><td><-DUF499*<-REase<-METHYLASE<-OLD-ABC+TOPRIM</td><td>932</td><td>Bacteria>PVC_group>Planctomycetota>unclassified_Planctomycetota</td><td>GCA_012797405.1</td><td>Planctomycetota_bacterium</td><td>GYA73_04750</td><td>409..3207</td><td>2026780</td><td>ATP-binding-protein-[Planctomycetota-bacterium]</td></tr>
<tr><td>MBI1914948.1</td><td>DUF499_1.RM.like.systems</td><td>ABC-ATPase-><-DUF499*<-REase<-METHYLASE<-?<-DUF4065<-?<-HIN-HTH</td><td>974</td><td>Bacteria>PVC_group>Planctomycetota>unclassified_Planctomycetota</td><td>GCA_016177925.1</td><td>Planctomycetota_bacterium</td><td>HYS12_09465</td><td>5117..8041</td><td>2026780</td><td>DUF499-domain-containing-protein-[Planctomycetota-bacterium]</td></tr>
<tr><td>MBI2191311.1</td><td>DUF499_1.RM.like.systems</td><td><-DUF499*<-REase<-METHYLASE<-HEPN<-HEPN<-Schlafen+HATPase+HTH||DEDD_Tnp_IS110+Transposase_20-></td><td>926</td><td>Bacteria>PVC_group>Planctomycetota>unclassified_Planctomycetota</td><td>GCA_016179685.1</td><td>Planctomycetota_bacterium</td><td>HYU36_04935</td><td>72260..75040</td><td>2026780</td><td>ATP-binding-protein-[Planctomycetota-bacterium]</td></tr>
<tr><td>MBI3467145.1</td><td>DUF499_1.RM.like.systems</td><td><-DUF499*<-HTH<-TIR<-METHYLASE<-REase<-?<-REase</td><td>942</td><td>Bacteria>PVC_group>Planctomycetota>unclassified_Planctomycetota</td><td>GCA_016196255.1</td><td>Planctomycetota_bacterium</td><td>HY000_29375</td><td>4772..7600</td><td>2026780</td><td>ATP-binding-protein-[Planctomycetota-bacterium]</td></tr>
<tr><td>MBI4617608.1</td><td>DUF499_1.RM.like.systems</td><td>Radical-SAM->?->RelE-ParE->?->?->?-><-HEPN+DUF499*<-ORC-CDC6+REase</td><td>1115</td><td>Bacteria>PVC_group>Planctomycetota>unclassified_Planctomycetota</td><td>GCA_016209595.1</td><td>Planctomycetota_bacterium</td><td>HY720_28610</td><td>4559..7906</td><td>2026780</td><td>ATP-binding-protein-[Planctomycetota-bacterium]</td></tr>
<tr><td>MBI4584119.1</td><td>DUF499_1.RM.like.systems</td><td>Schlafen+HATPase+HTH->?->METHYLASE->REase->OLD-ABC->TOPRIM->DUF499*->HicB_lk_antitox->HicA_toxin-></td><td>920</td><td>Bacteria>PVC_group>Planctomycetota>unclassified_Planctomycetota</td><td>GCA_016209605.1</td><td>Planctomycetota_bacterium</td><td>HY717_08860</td><td>13681..16443</td><td>2026780</td><td>ATP-binding-protein-[Planctomycetota-bacterium]</td></tr>
<tr><td>MBN8644244.1</td><td>DUF499_1.RM.like.systems</td><td><-McrC-NTD+REase<-?<-DUF499*<-REase<-METHYLASE</td><td>978</td><td>Bacteria>PVC_group>Planctomycetota>unclassified_Planctomycetota</td><td>GCA_017303565.1</td><td>Planctomycetota_bacterium</td><td>JOL61_03265</td><td>4226..7162</td><td>2026780</td><td>ATP-binding-protein-[Planctomycetota-bacterium]</td></tr>
<tr><td>MBP7751892.1</td><td>DUF499_1.RM.like.systems</td><td><-MoxR-AAA<-?||?->?->?-><-DUF499*<-REase<-METHYLASE-p</td><td>932</td><td>Bacteria>PVC_group>Planctomycetota>unclassified_Planctomycetota</td><td>GCA_018000135.1</td><td>Planctomycetota_bacterium</td><td>KA323_14485</td><td>46655..49453</td><td>2026780</td><td>ATP-binding-protein-[Planctomycetota-bacterium]</td></tr>
<tr><td>MCA8917196.1</td><td>DUF499_1.RM.like.systems</td><td>XIS-HTH->?-><-DUF499*<-REase<-OLD-ABC<-METHYLASE</td><td>951</td><td>Bacteria>PVC_group>Planctomycetota>unclassified_Planctomycetota</td><td>GCA_020425605.1</td><td>Planctomycetota_bacterium</td><td>KDB90_17530</td><td>3869..6724</td><td>2026780</td><td>ATP-binding-protein-[Planctomycetota-bacterium]</td></tr>
<tr><td>MCE9567305.1</td><td>DUF499_1.RM.like.systems</td><td><-SF1-SF2-Helicase+REase<-?<-SIGMA-HTH||ParB+HNH->METHYLASE->REase->DUF499*->ParB+HNH->ParB+HNH-></td><td>986</td><td>Bacteria>PVC_group>Planctomycetota>unclassified_Planctomycetota</td><td>GCA_021413735.1</td><td>Planctomycetota_bacterium</td><td>K8U57_35335</td><td>37179..40139</td><td>2026780</td><td>DUF499-domain-containing-protein-[Planctomycetota-bacterium]</td></tr>
<tr><td>GIW70817.1</td><td>DUF499_1.RM.like.systems</td><td><-Glyco_hydro_57+DUF3536<-?<-DUF499*<-REase<-REase<-TOPRIM<-OLD-ABC<-METHYLASE-p<-METHYLASE-p</td><td>942</td><td>Bacteria>PVC_group>Planctomycetota>unclassified_Planctomycetota</td><td>GCA_026004615.1</td><td>Planctomycetota_bacterium</td><td>KatS3mg102_0359</td><td>449544..452372</td><td>2026780</td><td>MAG:-hypothetical-protein-KatS3mg102_0359-[Planctomycetota-bacterium]</td></tr>
<tr><td>TAE90777.1</td><td>DUF499_1.RM.like.systems</td><td><-DUF499*<-REase<-METHYLASE<-HNH<-OLD-ABC<-TOPRIM<-OLD-ABC</td><td>939</td><td>Bacteria>PVC_group>Verrucomicrobiota>unclassified_Verrucomicrobiota</td><td>GCA_004293285.1</td><td>Verrucomicrobiota_bacterium</td><td>EAZ81_08635</td><td>280..3099</td><td>2026799</td><td>ATP-binding-protein-[Verrucomicrobia-bacterium]</td></tr>
<tr><td>MBI2925504.1</td><td>DUF499_1.RM.like.systems</td><td>ZnR->?->?->?-><-DUF499*<-REase<-METHYLASE-p</td><td>941</td><td>Bacteria>PVC_group>Verrucomicrobiota>unclassified_Verrucomicrobiota</td><td>GCA_016190045.1</td><td>Verrucomicrobiota_bacterium</td><td>HYY24_07350</td><td>8775..11600</td><td>2026799</td><td>ATP-binding-protein-[Verrucomicrobia-bacterium]</td></tr>
<tr><td>MBI2947997.1</td><td>DUF499_1.RM.like.systems</td><td>METHYLASE-p->AAA-ATPase+REase->?->PIN->HicB->DUF499*-></td><td>264</td><td>Bacteria>PVC_group>Verrucomicrobiota>unclassified_Verrucomicrobiota</td><td>GCA_016190065.1</td><td>Verrucomicrobiota_bacterium</td><td>HYY23_10135</td><td>3501..4294</td><td>2026799</td><td>ATP-binding-protein-[Verrucomicrobiota-bacterium]</td></tr>
<tr><td>MBM3880410.1</td><td>DUF499_1.RM.like.systems</td><td>HEPN->?->?->METHYLASE->ZDNA-HTH->REase->DUF499*->?-><-PIN<-?<-?<-?<-PIN</td><td>967</td><td>Bacteria>PVC_group>Verrucomicrobiota>unclassified_Verrucomicrobiota</td><td>GCA_016871715.1</td><td>Verrucomicrobiota_bacterium</td><td>FJ387_11950</td><td>21225..24128</td><td>2026799</td><td>ATP-binding-protein-[Verrucomicrobia-bacterium]</td></tr>
<tr><td>MCU0784170.1</td><td>DUF499_1.RM.like.systems</td><td>REase->?->METHYLASE->ORC-CDC6+REase->DUF499*->HTH_31->Peptidase-></td><td>935</td><td>Bacteria>PVC_group>Verrucomicrobiota>unclassified_Verrucomicrobiota</td><td>GCA_025457745.1</td><td>Verrucomicrobiota_bacterium</td><td>MUF81_09035</td><td>5492..8299</td><td>2026799</td><td>DUF499-domain-containing-protein-[Verrucomicrobiota-bacterium]</td></tr>
<tr><td>MCX6880519.1</td><td>DUF499_1.RM.like.systems</td><td><-DUF499*<-REase<-SPOUT<-METHYLASE<-DrHyd+BirA-HTH<-REase</td><td>689</td><td>Bacteria>PVC_group>Verrucomicrobiota>unclassified_Verrucomicrobiota</td><td>GCA_026397725.1</td><td>Verrucomicrobiota_bacterium</td><td>NTW21_42945</td><td>1..2067</td><td>2026799</td><td>MAG:-DUF499-domain-containing-protein,-partial-[Verrucomicrobia-bacterium]</td></tr>
<tr><td>MBE7497142.1</td><td>DUF499_1.RM.like.systems</td><td>HKD-DNase->ParB+HNH->METHYLASE->?->nSTAND1+sGTP->REase->DUF499*->tRNA-><-?||?-><-NTP_transf</td><td>920</td><td>Bacteria>PVC_group>Verrucomicrobiota>Verrucomicrobiae>Verrucomicrobiales</td><td>GCA_015075365.1</td><td>Verrucomicrobiaceae_bacterium</td><td>HS117_19545</td><td>529489..532251</td><td>2026800</td><td>ATP-binding-protein-[Verrucomicrobiaceae-bacterium]</td></tr>
<tr><td>HDQ14361.1</td><td>DUF499_1.RM.like.systems</td><td>METHYLASE-p->METHYLASE-p->HEPN+DUF499*->?->ORC-CDC6+REase-></td><td>1109</td><td>Bacteria>Spirochaetota>Spirochaetia>Spirochaetales>Spirochaetaceae</td><td>GCA_011046055.1</td><td>Sediminispirochaeta_sp.</td><td>ENN41_06065</td><td>1978..5307</td><td>1911557</td><td>ATP-binding-protein-[Sediminispirochaeta-sp.]</td></tr>
<tr><td>MXV87683.1</td><td>DUF499_1.RM.like.systems</td><td>NTP_transf-><-DUF499*<-METHYLASE<-REase</td><td>970</td><td>Bacteria>Terrabacteria_group>Actinomycetota>Acidimicrobiia>Acidimicrobiales</td><td>GCA_009835365.1</td><td>Acidimicrobiales_bacterium</td><td>F4005_09195</td><td>34597..37509</td><td>2201156</td><td>ATP-binding-protein-[Acidimicrobiales-bacterium]</td></tr>
<tr><td>MXX43802.1</td><td>DUF499_1.RM.like.systems</td><td>NTP_transf-><-DUF499*<-METHYLASE<-REase</td><td>970</td><td>Bacteria>Terrabacteria_group>Actinomycetota>Acidimicrobiia>Acidimicrobiales</td><td>GCA_009836375.1</td><td>Acidimicrobiales_bacterium</td><td>F4Z53_12210</td><td>6244..9156</td><td>2201156</td><td>ATP-binding-protein-[Acidimicrobiales-bacterium]</td></tr>
<tr><td>MXY02283.1</td><td>DUF499_1.RM.like.systems</td><td>NTP_transf-><-DUF499*<-METHYLASE<-REase</td><td>970</td><td>Bacteria>Terrabacteria_group>Actinomycetota>Acidimicrobiia>Acidimicrobiales</td><td>GCA_009838155.1</td><td>Acidimicrobiales_bacterium</td><td>F4Y66_05500</td><td>5633..8545</td><td>2201156</td><td>DUF499-domain-containing-protein-[Acidimicrobiales-bacterium]</td></tr>
<tr><td>MCY3894574.1</td><td>DUF499_1.RM.like.systems</td><td>REase->METHYLASE->DUF499*-><-NTP_transf</td><td>970</td><td>Bacteria>Terrabacteria_group>Actinomycetota>Acidimicrobiia>Acidimicrobiales</td><td>GCA_026707105.1</td><td>Acidimicrobiaceae_bacterium</td><td>OXF65_14910</td><td>3656..6568</td><td>2024894</td><td>MAG:-DUF499-domain-containing-protein-[Acidimicrobiaceae-bacterium]</td></tr>
<tr><td>MDE0319984.1</td><td>DUF499_1.RM.like.systems</td><td>NTP_transf-><-DUF499*<-METHYLASE<-REase</td><td>970</td><td>Bacteria>Terrabacteria_group>Actinomycetota>Acidimicrobiia>Acidimicrobiales</td><td>GCA_028820815.1</td><td>Acidimicrobiaceae_bacterium</td><td>OXI97_08880</td><td>1201..4113</td><td>2024894</td><td>MAG:-DUF499-domain-containing-protein-[Acidimicrobiaceae-bacterium]</td></tr>
<tr><td>MDE0803902.1</td><td>DUF499_1.RM.like.systems</td><td><-HTH+SF1-SF2-Helicase+REase-p<-REase<-MORC<-HEPN+DUF499*<-METHYLASE</td><td>1136</td><td>Bacteria>Terrabacteria_group>Actinomycetota>Acidimicrobiia>Acidimicrobiales</td><td>GCA_028822535.1</td><td>Acidimicrobiales_bacterium</td><td>OSA99_11315</td><td>4439..7849</td><td>2201156</td><td>MAG:-Swt1-family-HEPN-domain-containing-protein-[Acidimicrobiales-bacterium]</td></tr>
<tr><td>MDE0498678.1</td><td>DUF499_1.RM.like.systems</td><td>NTP_transf-><-DUF499*<-METHYLASE<-REase</td><td>970</td><td>Bacteria>Terrabacteria_group>Actinomycetota>Acidimicrobiia>Acidimicrobiales</td><td>GCA_028823965.1</td><td>Acidimicrobiaceae_bacterium</td><td>OXH86_15120</td><td>56164..59076</td><td>2024894</td><td>MAG:-DUF499-domain-containing-protein-[Acidimicrobiaceae-bacterium]</td></tr>
<tr><td>MCB9390232.1</td><td>DUF499_1.RM.like.systems</td><td><-HEPN<-XIS-HTH+RVT<-?<-DUF6361<-?<-REase<-HEPN+DUF499*<-METHYLASE||?-><-?<-?<-DUF772+DDE_Tnp_1+DDE_Tnp_1_6</td><td>1107</td><td>Bacteria>Terrabacteria_group>Actinomycetota>Acidimicrobiia>unclassified_Acidimicrobiia</td><td>GCA_020637365.1</td><td>Acidimicrobiia_bacterium</td><td>H6512_11920</td><td>836965..840288</td><td>2080302</td><td>ATP-binding-protein-[Acidimicrobiia-bacterium]</td></tr>
<tr><td>MCZ7536061.1</td><td>DUF499_1.RM.like.systems</td><td><-DUF499*<-DUF499<-DUF499<-METHYLASE-p<-METHYLASE-p<-REase</td><td>300</td><td>Bacteria>Terrabacteria_group>Actinomycetota>Acidimicrobiia>unclassified_Acidimicrobiia</td><td>GCA_027494135.1</td><td>Acidimicrobiia_bacterium</td><td>M5T61_09135</td><td>11348..12250</td><td>2080302</td><td>MAG:-hypothetical-protein-M5T61_09135-[Acidimicrobiia-bacterium]</td></tr>
<tr><td>MCZ7536062.1</td><td>DUF499_1.RM.like.systems</td><td><-DUF499<-DUF499*<-DUF499<-METHYLASE-p<-METHYLASE-p<-REase<-?<-HKD-DNase+SF1-SF2-Helicase-p</td><td>184</td><td>Bacteria>Terrabacteria_group>Actinomycetota>Acidimicrobiia>unclassified_Acidimicrobiia</td><td>GCA_027494135.1</td><td>Acidimicrobiia_bacterium</td><td>M5T61_09140</td><td>12615..13169</td><td>2080302</td><td>MAG:-hypothetical-protein-M5T61_09140-[Acidimicrobiia-bacterium]</td></tr>
<tr><td>MCZ7536063.1</td><td>DUF499_1.RM.like.systems</td><td><-DUF499<-DUF499<-DUF499*<-METHYLASE-p<-METHYLASE-p<-REase<-?<-HKD-DNase+SF1-SF2-Helicase-p</td><td>207</td><td>Bacteria>Terrabacteria_group>Actinomycetota>Acidimicrobiia>unclassified_Acidimicrobiia</td><td>GCA_027494135.1</td><td>Acidimicrobiia_bacterium</td><td>M5T61_09145</td><td>13163..13786</td><td>2080302</td><td>MAG:-hypothetical-protein-M5T61_09145-[Acidimicrobiia-bacterium]</td></tr>
<tr><td>WP_252863026.1</td><td>DUF499_1.RM.like.systems</td><td><-METHYLASE-p<-REase<-HTH+SF1-SF2-Helicase+REase<-REase<-MORC<-DUF499*<-HEPN<-METHYLASE||DNA_methylase-></td><td>912</td><td>Bacteria>Terrabacteria_group>Actinomycetota>Actinomycetes>Actinomycetales</td><td>-</td><td>Actinomyces_oris</td><td>BKH30_RS08650</td><td>21386..24124</td><td>544580</td><td>DUF499-domain-containing-protein-[Actinomyces-oris]</td></tr>
<tr><td>MBE6482584.1</td><td>DUF499_1.RM.like.systems</td><td><-Schlafen+HATPase+HTH<-REase<-HTH+SF1-SF2-Helicase+REase<-REase<-MORC<-HEPN+DUF499*<-METHYLASE||DNA_methylase->DNA_methylase->MORC+HISKIN-></td><td>1137</td><td>Bacteria>Terrabacteria_group>Actinomycetota>Actinomycetes>Actinomycetales</td><td>GCA_015062785.1</td><td>Actinomyces_ruminicola</td><td>E7Z94_09510</td><td>10121..13534</td><td>332524</td><td>ATP-binding-protein-[Actinomyces-ruminicola]</td></tr>
<tr><td>BDF97829.1</td><td>DUF499_1.RM.like.systems</td><td>ACET+ACET-><-?<-?<-DNA_methylase||METHYLASE->HEPN+DUF499*->MORC->REase->HTH+SF1-SF2-Helicase+REase->REase-><-?||METHYLASE-p-></td><td>1139</td><td>Bacteria>Terrabacteria_group>Actinomycetota>Actinomycetes>Actinomycetales</td><td>GCA_023169925.1</td><td>Actinomyces_oris_K20</td><td>AoKodu_01300</td><td>147963..151382</td><td>871541</td><td>hypothetical-protein-AoKodu_01300-[Actinomyces-oris-K20]</td></tr>
<tr><td>WP_003788306.1</td><td>DUF499_1.RM.like.systems</td><td><-METHYLASE-p||?-><-REase<-HTH+SF1-SF2-Helicase+REase<-REase<-MORC<-HEPN+DUF499*<-METHYLASE||DNA_methylase->?->?-><-?<-HTH_3+PolB</td><td>1139</td><td>Bacteria>Terrabacteria_group>Actinomycetota>Actinomycetes>Actinomycetales</td><td>GCF_000175315.3</td><td>Actinomyces_viscosus</td><td>HMPREF0059_RS11125</td><td>118878..122297</td><td>1656</td><td>DUF499-domain-containing-protein-[Actinomyces-viscosus]</td></tr>
<tr><td>WP_017177796.1</td><td>DUF499_1.RM.like.systems</td><td>Phage_holin_4_2->?-><-REase<-HTH+SF1-SF2-Helicase+REase<-REase<-MORC<-HEPN+DUF499*<-METHYLASE||DNA_methylase->?->?-><-?<-HTH_3+PolB</td><td>1137</td><td>Bacteria>Terrabacteria_group>Actinomycetota>Actinomycetes>Actinomycetales</td><td>GCF_000295095.1</td><td>Actinomyces_timonensis</td><td>A1QA_RS0104850</td><td>1087937..1091350</td><td>1288391</td><td>DUF499-domain-containing-protein-[Actinomyces-timonensis]</td></tr>
<tr><td>WP_053587850.1</td><td>DUF499_1.RM.like.systems</td><td><-REase<-HTH+SF1-SF2-Helicase+REase<-REase<-MORC<-HEPN+DUF499*<-METHYLASE||METHYLASE->BirA-HTH->?-><-TPR<-MULE-TRANSPOSASE</td><td>1165</td><td>Bacteria>Terrabacteria_group>Actinomycetota>Actinomycetes>Actinomycetales</td><td>GCF_001278845.1</td><td>Actinomyces_sp._oral_taxon_414</td><td>AM609_RS14685</td><td>3673757..3677254</td><td>712122</td><td>DUF499-domain-containing-protein-[Actinomyces-sp.-oral-taxon-414]</td></tr>
<tr><td>WP_075412471.1</td><td>DUF499_1.RM.like.systems</td><td><-METHYLASE-p<-REase<-HTH+SF1-SF2-Helicase+REase<-REase<-MORC<-HEPN+DUF499*<-METHYLASE||DNA_methylase->?-><-HTH_3+PolB<-ACET+ACET</td><td>1137</td><td>Bacteria>Terrabacteria_group>Actinomycetota>Actinomycetes>Actinomycetales</td><td>GCF_001937365.1</td><td>Actinomyces_oris</td><td>BKH31_RS11955</td><td>29944..33357</td><td>544580</td><td>DUF499-domain-containing-protein-[Actinomyces-oris]</td></tr>
<tr><td>WP_075413329.1</td><td>DUF499_1.RM.like.systems</td><td><-METHYLASE-p<-REase<-HTH+SF1-SF2-Helicase+REase<-REase<-MORC<-HEPN+DUF499*<-METHYLASE||DNA_methylase->?->?-><-ACET+ACET</td><td>1137</td><td>Bacteria>Terrabacteria_group>Actinomycetota>Actinomycetes>Actinomycetales</td><td>GCF_001937475.1</td><td>Actinomyces_naeslundii</td><td>BKH08_RS04545</td><td>28951..32364</td><td>1655</td><td>DUF499-domain-containing-protein-[Actinomyces-naeslundii]</td></tr>
<tr><td>WP_075403442.1</td><td>DUF499_1.RM.like.systems</td><td>ACET+ACET->HTH_3+PolB-><-?<-?<-DNA_methylase||METHYLASE->HEPN+DUF499*->MORC->REase->HTH+SF1-SF2-Helicase+REase->REase-><-?||METHYLASE-p-></td><td>1139</td><td>Bacteria>Terrabacteria_group>Actinomycetota>Actinomycetes>Actinomycetales</td><td>GCF_001937505.1</td><td>Actinomyces_oris</td><td>BKH21_RS10525</td><td>35903..39322</td><td>544580</td><td>DUF499-domain-containing-protein-[Actinomyces-oris]</td></tr>
<tr><td>WP_076255389.1</td><td>DUF499_1.RM.like.systems</td><td>ACET+ACET-><-?<-?<-DNA_methylase||METHYLASE->HEPN+DUF499*->MORC->REase->HTH+SF1-SF2-Helicase+REase->REase->METHYLASE-p-></td><td>1137</td><td>Bacteria>Terrabacteria_group>Actinomycetota>Actinomycetes>Actinomycetales</td><td>GCF_001956555.1</td><td>Actinomyces_naeslundii</td><td>BKH25_RS11260</td><td>118842..122255</td><td>1655</td><td>DUF499-domain-containing-protein-[Actinomyces-naeslundii]</td></tr>
<tr><td>WP_077240589.1</td><td>DUF499_1.RM.like.systems</td><td><-REase<-HTH+SF1-SF2-Helicase+REase<-REase<-MORC<-HEPN+DUF499*<-METHYLASE||DNA_methylase->MORC+HEPN->||?-><-?<-?<-Peptidase_M50</td><td>1137</td><td>Bacteria>Terrabacteria_group>Actinomycetota>Actinomycetes>Actinomycetales</td><td>GCF_002356195.1</td><td>Actinomyces</td><td>DY404_RS06925</td><td>1656821..1660234</td><td>1654</td><td>MULTISPECIES:-DUF499-domain-containing-protein-[Actinomyces]</td></tr>
<tr><td>WP_103061758.1</td><td>DUF499_1.RM.like.systems</td><td><-REase<-HTH+SF1-SF2-Helicase+REase<-REase<-MORC<-HEPN+DUF499*<-METHYLASE||DNA_methylase->MORC+HisKin->HTH_Tnp_4+DDE_Tnp_4->MORC+HEPN-></td><td>1138</td><td>Bacteria>Terrabacteria_group>Actinomycetota>Actinomycetes>Actinomycetales</td><td>GCF_004102025.1</td><td>Actinomyces_qiguomingii</td><td>CWT10_RS16110</td><td>3847675..3851091</td><td>2057800</td><td>DUF499-domain-containing-protein-[Actinomyces-qiguomingii]</td></tr>
<tr><td>WP_076142650.1</td><td>DUF499_1.RM.like.systems</td><td><-METHYLASE-p<-REase<-HTH+SF1-SF2-Helicase+REase<-REase<-MORC<-HEPN+DUF499*<-METHYLASE||DNA_methylase->?->?-><-ACET+ACET</td><td>1139</td><td>Bacteria>Terrabacteria_group>Actinomycetota>Actinomycetes>Actinomycetales</td><td>GCF_016127855.1</td><td>Actinomyces_naeslundii</td><td>BKH33_RS06345</td><td>173705..177124</td><td>1655</td><td>DUF499-domain-containing-protein-[Actinomyces-naeslundii]</td></tr>
<tr><td>WP_198776534.1</td><td>DUF499_1.RM.like.systems</td><td>sGTP-><-?<-DNA_methylase||METHYLASE->HEPN+DUF499*->MORC->REase->HTH+SF1-SF2-Helicase+REase->REase-></td><td>1127</td><td>Bacteria>Terrabacteria_group>Actinomycetota>Actinomycetes>Actinomycetales</td><td>GCF_016405145.1</td><td>Schaalia_cardiffensis</td><td>JFX23_RS08485</td><td>21486..24869</td><td>181487</td><td>DUF499-domain-containing-protein-[Schaalia-cardiffensis]</td></tr>
<tr><td>WP_282568806.1</td><td>DUF499_1.RM.like.systems</td><td>ACET+ACET-><-?<-?<-DNA_methylase||METHYLASE->HEPN+DUF499*->MORC->REase->HTH+SF1-SF2-Helicase+REase->REase-><-?||METHYLASE-p-></td><td>1139</td><td>Bacteria>Terrabacteria_group>Actinomycetota>Actinomycetes>Actinomycetales</td><td>GCF_023169925.1</td><td>Actinomyces_oris</td><td>AoKodu_RS00635</td><td>147963..151382</td><td>544580</td><td>DUF499-domain-containing-protein-[Actinomyces-oris]</td></tr>
<tr><td>WP_268916840.1</td><td>DUF499_1.RM.like.systems</td><td><-REase<-HTH+SF1-SF2-Helicase+REase<-REase<-MORC<-HEPN+DUF499*<-METHYLASE||DNA_methylase->MORC+HEPN->SLATT->SMODS-></td><td>1131</td><td>Bacteria>Terrabacteria_group>Actinomycetota>Actinomycetes>Actinomycetales</td><td>GCF_026891875.1</td><td>Actinomyces_israelii</td><td>OHJ16_RS04000</td><td>49558..52953</td><td>1659</td><td>DUF499-domain-containing-protein-[Actinomyces-israelii]</td></tr>
<tr><td>WP_271288667.1</td><td>DUF499_1.RM.like.systems</td><td>ACET+ACET->HTH_3+PolB-><-?<-?<-DNA_methylase||METHYLASE->HEPN+DUF499*->MORC->REase->HTH+SF1-SF2-Helicase+REase->REase-><-?||METHYLASE-p-></td><td>1132</td><td>Bacteria>Terrabacteria_group>Actinomycetota>Actinomycetes>Actinomycetales</td><td>GCF_027945475.1</td><td>Actinomyces_oris</td><td>PGE45_RS00580</td><td>136591..139989</td><td>544580</td><td>DUF499-domain-containing-protein-[Actinomyces-oris]</td></tr>
<tr><td>WP_108833962.1</td><td>DUF499_1.RM.like.systems</td><td><-METHYLASE-p<-REase<-HTH+SF1-SF2-Helicase+REase<-REase<-MORC<-HEPN+DUF499*<-METHYLASE||DNA_methylase->?->?-><-ACET+ACET</td><td>1138</td><td>Bacteria>Terrabacteria_group>Actinomycetota>Actinomycetes>Actinomycetales</td><td>GCF_900323545.1</td><td>Actinomyces_sp._Marseille-P3109</td><td>BQ8008_RS10580</td><td>2440018..2443434</td><td>2083009</td><td>DUF499-domain-containing-protein-[Actinomyces-sp.-Marseille-P3109]</td></tr>
</table>

<table>
<tr><th>Protein-ID</th><th>System</th><th>Gene-Neighbourhoods</th><th>Length</th><th>Taxonomy</th><th>Assembly-ID</th><th>Strain</th><th>Locus-Tag</th><th>Genomic-coordinates</th><th>TaxID</th><th>Fasta-define</th></tr>
<tr><td>WP_278601539.1</td><td>DUF499_1.RM.like.systems</td><td>ACET+ACET-><-?<-?<-DNA_methylase||METHYLASE->HEPN+DUF499*->MORC->REase->HTH+SF1-SF2-Helicase+REase->REase->METHYLASE-p-></td><td>1137</td><td>Bacteria>Terrabacteria_group>Actinomycetota>Actinomycetes>Actinomycetales</td><td>GCF_905372775.1</td><td>Actinomyces_naeslundii</td><td>QDH38_RS03495</td><td>39335..42748</td><td>1655</td><td>DUF499-domain-containing-protein-[Actinomyces-naeslundii]</td></tr>
<tr><td>WP_070662487.1</td><td>DUF499_1.RM.like.systems</td><td>HTH_3+PolB->?-><-?<-?<-DNA_methylase||METHYLASE->HEPN+DUF499*->MORC->REase->HTH+SF1-SF2-Helicase+REase->REase-><-?||METHYLASE-p-></td><td>1137</td><td>Bacteria>Terrabacteria_group>Actinomycetota>Actinomycetes>Actinomycetales</td><td>GCF_963519085.1</td><td>Actinomyces</td><td>BKH20_RS04780</td><td>16618..20031</td><td>1654</td><td>MULTISPECIES:-DUF499-domain-containing-protein-[Actinomyces]</td></tr>
<tr><td>RBY97952.1</td><td>DUF499_1.RM.like.systems</td><td><-NUDIX||?-><-ABhydrolase<-DNA_methylase||REase->METHYLASE->HEPN+DUF499*->MORC->REase->HTH+SF1-SF2-Helicase+REase-></td><td>1128</td><td>Bacteria>Terrabacteria_group>Actinomycetota>Actinomycetes>Geodermatophilales</td><td>GCA_003319175.1</td><td>Blastococcus_sp._TF02-8</td><td>DQ237_03405</td><td>728169..731555</td><td>2250574</td><td>AAA-family-ATPase-[Blastococcus-sp.-TF02-8]</td></tr>
<tr><td>WP_205730045.1</td><td>DUF499_1.RM.like.systems</td><td><-ABhydrolase<-DNA_methylase||REase->METHYLASE->HEPN+DUF499*->MORC->REase->HTH+SF1-SF2-Helicase+REase-></td><td>1113</td><td>Bacteria>Terrabacteria_group>Actinomycetota>Actinomycetes>Geodermatophilales</td><td>GCF_003319175.1</td><td>Blastococcus_sp._TF02-8</td><td>DQ237_RS03405</td><td>728214..731555</td><td>2250574</td><td>Swt1-family-HEPN-domain-containing-protein-[Blastococcus-sp.-TF02-8]</td></tr>
<tr><td>WP_158716411.1</td><td>DUF499_1.RM.like.systems</td><td>DUF499*->MORC->REase->HTH+SF1-SF2-Helicase+REase->SIGMA-HTH->HTH_Tnp_1+HTH_21+rve+rve_3-></td><td>941</td><td>Bacteria>Terrabacteria_group>Actinomycetota>Actinomycetes>Geodermatophilales</td><td>GCF_900292035.1</td><td>Blastococcus_sp._Marseille-P5729</td><td>DAA40_RS12375</td><td>801548..804373</td><td>2086582</td><td>DUF499-domain-containing-protein-[Blastococcus-sp.-Marseille-P5729]</td></tr>
<tr><td>TRZ87911.1</td><td>DUF499_1.RM.like.systems</td><td>REase->METHYLASE->HEPN+DUF499*-><-Phage_int_SAM_1+Phage_integrase</td><td>1108</td><td>Bacteria>Terrabacteria_group>Actinomycetota>Actinomycetes>Kitasatosporales</td><td>GCA_007280125.1</td><td>Streptomycetaceae_bacterium</td><td>D4R83_01365</td><td>3739..7065</td><td>2021369</td><td>ATP-binding-protein-[Streptomycetaceae-bacterium]</td></tr>
<tr><td>HBS75978.1</td><td>DUF499_1.RM.like.systems</td><td><-TrwC+SF1-SF2-Helicase<-wHTH+REase<-?||NTD+SF1-SF2-Helicase-><-HEPN+DUF499*</td><td>1125</td><td>Bacteria>Terrabacteria_group>Actinomycetota>Actinomycetes>Micrococcales</td><td>GCA_003510265.1</td><td>Microbacterium_sp.</td><td>DEB55_16590</td><td>13006..16383</td><td>51671</td><td>AAA-family-ATPase-[Microbacterium-sp.]</td></tr>
<tr><td>TFC67798.1</td><td>DUF499_1.RM.like.systems</td><td>UDG-conflict-><-HTH+SF1-SF2-Helicase+REase<-REase<-MORC<-HEPN+DUF499*<-METHYLASE||MORC+SIGMA-HTH->ASCH->?->DDE_Tnp_1->XIS-HTH-></td><td>1124</td><td>Bacteria>Terrabacteria_group>Actinomycetota>Actinomycetes>Micrococcales</td><td>GCA_004402575.1</td><td>Cryobacterium_sp._TMT2-4</td><td>E3054_08285</td><td>76881..80255</td><td>1259254</td><td>ATP-binding-protein-[Cryobacterium-sp.-TMT2-4]</td></tr>
<tr><td>MBK8435770.1</td><td>DUF499_1.RM.like.systems</td><td>Phage_int_SAM_1+Phage_integrase->Phage_integrase->?->DDE_Tnp_1-><-HEPN+REase<-REase<-HEPN+DUF499*<-wHTH+REase<-METHYLASE<-XIS-HTH<-ABhydrolase<-MPTase<-HTH</td><td>1139</td><td>Bacteria>Terrabacteria_group>Actinomycetota>Actinomycetes>Micrococcales</td><td>GCA_016711265.1</td><td>Austwickia_sp.</td><td>IPL37_04405</td><td>982795..986214</td><td>1872664</td><td>DUF499-domain-containing-protein-[Austwickia-sp.]</td></tr>
<tr><td>MBN9213516.1</td><td>DUF499_1.RM.like.systems</td><td><-HTH+SF1-SF2-Helicase+REase<-REase<-MORC<-HEPN+DUF499*<-METHYLASE||MORC+SIGMA-HTH->DNA_methylase->?->Phage_int_SAM_1+Phage_integrase-><-tRNA||DUF222+HNH-></td><td>1119</td><td>Bacteria>Terrabacteria_group>Actinomycetota>Actinomycetes>Micrococcales</td><td>GCA_017306385.1</td><td>Microbacterium_sp.</td><td>JOJ04_01635</td><td>353134..356493</td><td>51671</td><td>DUF499-domain-containing-protein-[Microbacterium-sp.]</td></tr>
<tr><td>MBW8172946.1</td><td>DUF499_1.RM.like.systems</td><td>GNTR-HTH-><-?<-?<-DNA_methylase||SIGMA-HTH->METHYLASE->HEPN+DUF499*->MORC->REase->HTH+SF1-SF2-Helicase+REase->NERD+SF1-SF2-Helicase-><-?<-ABC-ATPase+SbcC</td><td>1128</td><td>Bacteria>Terrabacteria_group>Actinomycetota>Actinomycetes>Micrococcales</td><td>GCA_019457945.1</td><td>Ornithinimicrobium_laminariae</td><td>K0651_07785</td><td>644333..647719</td><td>2864873</td><td>DUF499-domain-containing-protein-[Ornithinimicrobium-laminariae]</td></tr>
<tr><td>MCE7482742.1</td><td>DUF499_1.RM.like.systems</td><td>REase+SF1-SF2-Helicase-><-SbcC<-HEPN+DUF499*<-METHYLASE-p</td><td>1108</td><td>Bacteria>Terrabacteria_group>Actinomycetota>Actinomycetes>Micrococcales</td><td>GCA_021395155.1</td><td>Microbacterium_profundi</td><td>LZG07_12530</td><td>143258..146584</td><td>450380</td><td>DUF499-domain-containing-protein-[Microbacterium-profundi]</td></tr>
<tr><td>MCV7624012.1</td><td>DUF499_1.RM.like.systems</td><td><-REase<-?<-?<-REase||DNA_methylase->METHYLASE->HEPN+DUF499*->MORC->REase->HTH+SF1-SF2-Helicase+REase-><-Peptidase_S66+Peptidase_S66C</td><td>1157</td><td>Bacteria>Terrabacteria_group>Actinomycetota>Actinomycetes>Micrococcales</td><td>GCA_025145605.2</td><td>Micrococcus_luteus</td><td>M3B28_003235</td><td>32627..36100</td><td>1270</td><td>Swt1-family-HEPN-domain-containing-protein-[Micrococcus-luteus]</td></tr>
<tr><td>MCV7527924.1</td><td>DUF499_1.RM.like.systems</td><td><-REase||DNA_methylase->METHYLASE->HEPN+DUF499*->MORC->REase-></td><td>1157</td><td>Bacteria>Terrabacteria_group>Actinomycetota>Actinomycetes>Micrococcales</td><td>GCA_025149725.2</td><td>Micrococcus_luteus</td><td>M3D91_010335</td><td>6032..9505</td><td>1270</td><td>Swt1-family-HEPN-domain-containing-protein-[Micrococcus-luteus]</td></tr>
<tr><td>MCV7506294.1</td><td>DUF499_1.RM.like.systems</td><td><-HTH+SF1-SF2-Helicase+REase<-REase<-MORC<-HEPN+DUF499*<-METHYLASE<-DNA_methylase||REase-></td><td>1157</td><td>Bacteria>Terrabacteria_group>Actinomycetota>Actinomycetes>Micrococcales</td><td>GCA_025151425.2</td><td>Micrococcus_luteus</td><td>M3D82_001030</td><td>202363..205836</td><td>1270</td><td>Swt1-family-HEPN-domain-containing-protein-[Micrococcus-luteus]</td></tr>
<tr><td>WP_009381273.1</td><td>DUF499_1.RM.like.systems</td><td><-HEPN+REase<-REase<-HEPN+DUF499*<-wHTH+REase<-METHYLASE</td><td>1132</td><td>Bacteria>Terrabacteria_group>Actinomycetota>Actinomycetes>Micrococcales</td><td>GCF_000314575.1</td><td>Brevibacterium_casei</td><td>C272_RS16125</td><td>18996..22394</td><td>33889</td><td>DUF499-domain-containing-protein-[Brevibacterium-casei]</td></tr>
<tr><td>WP_034226753.1</td><td>DUF499_1.RM.like.systems</td><td><-HTH+SF1-SF2-Helicase+REase<-REase<-MORC<-HEPN+DUF499*<-METHYLASE</td><td>1144</td><td>Bacteria>Terrabacteria_group>Actinomycetota>Actinomycetes>Micrococcales</td><td>GCF_000603945.1</td><td>Actinotalea_ferrariae</td><td>N866_RS12275</td><td>5788..9222</td><td>1386098</td><td>Swt1-family-HEPN-domain-containing-protein-[Actinotalea-ferrariae]</td></tr>
<tr><td>WP_047688630.1</td><td>DUF499_1.RM.like.systems</td><td>Glyoxalase-><-?<-HTH+SF1-SF2-Helicase+REase<-REase<-MORC<-DUF499*</td><td>699</td><td>Bacteria>Terrabacteria_group>Actinomycetota>Actinomycetes>Micrococcales</td><td>GCF_000800955.1</td><td>Kocuria_sp._ZOR0020</td><td>L979_RS02200</td><td>64471..66571</td><td>1339234</td><td>DUF499-domain-containing-protein,-partial-[Kocuria-sp.-ZOR0020]</td></tr>
<tr><td>WP_047691765.1</td><td>DUF499_1.RM.like.systems</td><td><-DNA_methylase||REase->METHYLASE->HEPN+DUF499*-></td><td>452</td><td>Bacteria>Terrabacteria_group>Actinomycetota>Actinomycetes>Micrococcales</td><td>GCF_000800955.1</td><td>Kocuria_sp._ZOR0020</td><td>L979_RS08865</td><td>18602..19957</td><td>1339234</td><td>Swt1-family-HEPN-domain-containing-protein,-partial-[Kocuria-sp.-ZOR0020]</td></tr>
<tr><td>WP_044658741.1</td><td>DUF499_1.RM.like.systems</td><td>Peptidase_S66+Peptidase_S66C-><-HTH+SF1-SF2-Helicase+REase<-REase<-MORC<-HEPN+DUF499*<-METHYLASE<-DNA_methylase||REase-></td><td>1157</td><td>Bacteria>Terrabacteria_group>Actinomycetota>Actinomycetes>Micrococcales</td><td>GCF_000946635.1</td><td>Micrococcus</td><td>B8W89_RS05705</td><td>32626..36099</td><td>1269</td><td>MULTISPECIES:-Swt1-family-HEPN-domain-containing-protein-[Micrococcus]</td></tr>
<tr><td>WP_062529103.1</td><td>DUF499_1.RM.like.systems</td><td><-REase+SF1-SF2-Helicase||MazG-><-ParB+HNH<-HTH+SF1-SF2-Helicase+REase<-REase<-MORC<-HEPN+DUF499*<-METHYLASE||DNA_methylase-></td><td>1118</td><td>Bacteria>Terrabacteria_group>Actinomycetota>Actinomycetes>Micrococcales</td><td>GCF_000971455.1</td><td>Demequina_rhizosphaerae</td><td>AOM12_RS04465</td><td>350500..353856</td><td>1638985</td><td>DUF499-domain-containing-protein-[Demequina-rhizosphaerae]</td></tr>
<tr><td>WP_046529475.1</td><td>DUF499_1.RM.like.systems</td><td>XIS-HTH->PIN->HNH-><-HTH+SF1-SF2-Helicase+REase<-REase<-MORC<-HEPN+DUF499*<-METHYLASE<-REase||DNA_methylase-><-OLD-ABC+TOPRIM<-DNA-glycosylase</td><td>1124</td><td>Bacteria>Terrabacteria_group>Actinomycetota>Actinomycetes>Micrococcales</td><td>GCF_000986825.1</td><td>Cellulomonas_sp._FA1</td><td>GY42_RS11845</td><td>2630948..2634322</td><td>1346710</td><td>Swt1-family-HEPN-domain-containing-protein-[Cellulomonas-sp.-FA1]</td></tr>
<tr><td>WP_048693914.1</td><td>DUF499_1.RM.like.systems</td><td><-Radical-SAM+TRAM||?->?-><-DNA_methylase||PIN->METHYLASE->HEPN+DUF499*->MORC->REase->HTH+SF1-SF2-Helicase+REase->?->CR-Kinase+ORC-CDC6-like+CARF-></td><td>1134</td><td>Bacteria>Terrabacteria_group>Actinomycetota>Actinomycetes>Micrococcales</td><td>GCF_001050535.1</td><td>Tetrasphaera_australiensis</td><td>BN11_RS02235</td><td>472611..476015</td><td>99480</td><td>DUF499-domain-containing-protein-[Tetrasphaera-australiensis]</td></tr>
<tr><td>WP_055898767.1</td><td>DUF499_1.RM.like.systems</td><td><-wHTH+SAD+HNH||NERD+SF1-SF2-Helicase-><-REase||?->METHYLASE->HEPN+DUF499*->MORC->REase->HTH+SF1-SF2-Helicase+REase-><-?<-SF1-SF2-Helicase+REase<-Exonuc_V_gamma+REase</td><td>1109</td><td>Bacteria>Terrabacteria_group>Actinomycetota>Actinomycetes>Micrococcales</td><td>GCF_001425665.1</td><td>Yonghaparkia_sp._Root332</td><td>ASC54_RS09975</td><td>479255..482584</td><td>1736516</td><td>DUF499-domain-containing-protein-[Yonghaparkia-sp.-Root332]</td></tr>
<tr><td>WP_200941118.1</td><td>DUF499_1.RM.like.systems</td><td>Phage_holin->METHYLASE->?->ACET->HEPN+DUF499*->DUF4096+DDE_Tnp_1+DDE_Tnp_1_2-><-?<-REase<-MobL</td><td>1112</td><td>Bacteria>Terrabacteria_group>Actinomycetota>Actinomycetes>Micrococcales</td><td>GCF_001426925.1</td><td>Microbacterium_sp._Root53</td><td>ASD19_RS13260</td><td>99054..102392</td><td>1736553</td><td>Swt1-family-HEPN-domain-containing-protein-[Microbacterium-sp.-Root53]</td></tr>
<tr><td>WP_062860586.1</td><td>DUF499_1.RM.like.systems</td><td><-OLD-ABC+TOPRIM<-DNA_methylase||DDE_Tnp_1_4->METHYLASE->HEPN+DUF499*->MORC->REase->HTH+SF1-SF2-Helicase+REase->SF1-SF2-Helicase+REase-><-?||ZnR-></td><td>1132</td><td>Bacteria>Terrabacteria_group>Actinomycetota>Actinomycetes>Micrococcales</td><td>GCF_001606005.1</td><td>Brevibacterium_linens</td><td>A2T55_RS01955</td><td>443137..446535</td><td>1703</td><td>DUF499-domain-containing-protein-[Brevibacterium-linens]</td></tr>
<tr><td>WP_069599267.1</td><td>DUF499_1.RM.like.systems</td><td><-SF1-SF2-Helicase+REase<-?<-HTH+SF1-SF2-Helicase+REase<-REase<-MORC<-HEPN+DUF499*<-METHYLASE<-REase<-MORC+HEPN||DNA_methylase-><-?<-SF1-SF2-Helicase</td><td>1131</td><td>Bacteria>Terrabacteria_group>Actinomycetota>Actinomycetes>Micrococcales</td><td>GCF_001729525.1</td><td>Brevibacterium_aurantiacum</td><td>CIK60_RS17585</td><td>12727..16122</td><td>273384</td><td>DUF499-domain-containing-protein-[Brevibacterium-aurantiacum]</td></tr>
<tr><td>WP_076703521.1</td><td>DUF499_1.RM.like.systems</td><td><-HEPN+DUF499*<-wHTH+REase</td><td>1131</td><td>Bacteria>Terrabacteria_group>Actinomycetota>Actinomycetes>Micrococcales</td><td>GCF_001968835.1</td><td>Tersicoccus_phoenicis</td><td>BKD30_RS07045</td><td>193..3588</td><td>554083</td><td>Swt1-family-HEPN-domain-containing-protein-[Tersicoccus-phoenicis]</td></tr>
<tr><td>WP_232711167.1</td><td>DUF499_1.RM.like.systems</td><td>Phage_holin->METHYLASE->?->?->HEPN+DUF499*->?->?->wHTH+REase->TrwC+SF1-SF2-Helicase-></td><td>1114</td><td>Bacteria>Terrabacteria_group>Actinomycetota>Actinomycetes>Micrococcales</td><td>GCF_002812725.1</td><td>unclassified_Microbacterium</td><td>CGU35_RS05955</td><td>370107..373451</td><td>2609290</td><td>MULTISPECIES:-DUF499-domain-containing-protein-[unclassified-Microbacterium]</td></tr>
<tr><td>WP_101853235.1</td><td>DUF499_1.RM.like.systems</td><td><-HTH+SF1-SF2-Helicase+REase-p<-PAIREDC-HTH<-HTH+SF1-SF2-Helicase+REase-p<-REase<-MORC<-HEPN+DUF499*<-METHYLASE||?->HTH+MPTase->PIN-></td><td>1144</td><td>Bacteria>Terrabacteria_group>Actinomycetota>Actinomycetes>Micrococcales</td><td>GCF_002849405.1</td><td>Kocuria_flava</td><td>AUQ48_RS15985</td><td>10331..13765</td><td>446860</td><td>DUF499-domain-containing-protein-[Kocuria-flava]</td></tr>
<tr><td>WP_262391126.1</td><td>DUF499_1.RM.like.systems</td><td>Peptidase_S66+Peptidase_S66C-><-HTH+SF1-SF2-Helicase+REase<-REase<-MORC<-MORC<-DUF499*<-HEPN<-METHYLASE<-DNA_methylase||REase-></td><td>679</td><td>Bacteria>Terrabacteria_group>Actinomycetota>Actinomycetes>Micrococcales</td><td>GCF_003071525.2</td><td>Micrococcus_luteus</td><td>C0205_RS12620</td><td>704510..706549</td><td>1270</td><td>DUF499-domain-containing-protein-[Micrococcus-luteus]</td></tr>
<tr><td>WP_126874170.1</td><td>DUF499_1.RM.like.systems</td><td><-DNA_methylase||REase->METHYLASE->HEPN+DUF499*->MORC->REase->HTH+SF1-SF2-Helicase+REase-><-?||MarR-HTH-></td><td>1141</td><td>Bacteria>Terrabacteria_group>Actinomycetota>Actinomycetes>Micrococcales</td><td>GCF_003989235.1</td><td>unclassified_Kocuria</td><td>D8M38_RS00615</td><td>20820..24245</td><td>2649579</td><td>MULTISPECIES:-DUF499-domain-containing-protein-[unclassified-Kocuria]</td></tr>
<tr><td>WP_128365158.1</td><td>DUF499_1.RM.like.systems</td><td>Calcineurin->tRNA-><-Phage_integrase<-?<-HTH+SF1-SF2-Helicase+REase<-REase<-MORC<-HEPN+DUF499*<-METHYLASE<-REase||DNA_methylase-><-?||Peptidase_M48-></td><td>1135</td><td>Bacteria>Terrabacteria_group>Actinomycetota>Actinomycetes>Micrococcales</td><td>GCF_004025965.2</td><td>Phycicoccus_flavus</td><td>EPD83_RS11930</td><td>11514..14921</td><td>2502783</td><td>DUF499-domain-containing-protein-[Phycicoccus-flavus]</td></tr>
<tr><td>WP_131132843.1</td><td>DUF499_1.RM.like.systems</td><td><-HTH+SF1-SF2-Helicase+REase<-REase<-MORC<-HEPN+DUF499*<-METHYLASE||DNA_methylase-><-SbcC<-DUF1679<-SIR2</td><td>1130</td><td>Bacteria>Terrabacteria_group>Actinomycetota>Actinomycetes>Micrococcales</td><td>GCF_004320525.1</td><td>Arthrobacter_sp._S39</td><td>EYS21_RS09020</td><td>558026..561418</td><td>2509720</td><td>Swt1-family-HEPN-domain-containing-protein-[Arthrobacter-sp.-S39]</td></tr>
<tr><td>WP_134401373.1</td><td>DUF499_1.RM.like.systems</td><td>UDG-conflict-><-HTH+SF1-SF2-Helicase+REase<-REase<-MORC<-HEPN+DUF499*<-METHYLASE||MORC+SIGMA-HTH->ASCH->?->DDE_Tnp_1->XIS-HTH-></td><td>1124</td><td>Bacteria>Terrabacteria_group>Actinomycetota>Actinomycetes>Micrococcales</td><td>GCF_004402575.1</td><td>Cryobacterium_sp._TMT2-4</td><td>E3054_RS08280</td><td>76881..80255</td><td>1259254</td><td>Swt1-family-HEPN-domain-containing-protein-[Cryobacterium-sp.-TMT2-4]</td></tr>
<tr><td>WP_156242696.1</td><td>DUF499_1.RM.like.systems</td><td><-SbcC||?->REase-><-HTH+SF1-SF2-Helicase+REase<-REase<-MORC<-HEPN+DUF499*<-METHYLASE||MORC+SIGMA-HTH->DNA_methylase->?-><-DeoR-HTH+ISOCOT</td><td>1129</td><td>Bacteria>Terrabacteria_group>Actinomycetota>Actinomycetes>Micrococcales</td><td>GCF_009735645.1</td><td>Microbacterium_oryzae</td><td>D7D94_RS11265</td><td>2403131..2406520</td><td>743009</td><td>Swt1-family-HEPN-domain-containing-protein-[Microbacterium-oryzae]</td></tr>
<tr><td>WP_246200745.1</td><td>DUF499_1.RM.like.systems</td><td><-REase<-HTH+SF1-SF2-Helicase+REase<-REase<-MORC<-HEPN+DUF499*<-METHYLASE<-MORC+HEPN||MORC+SIGMA-HTH->DNA_methylase->REase->HEPN+SF1-SF2-Helicase-></td><td>1131</td><td>Bacteria>Terrabacteria_group>Actinomycetota>Actinomycetes>Micrococcales</td><td>GCF_009749385.1</td><td>Agromyces_allii</td><td>GQR16_RS11060</td><td>282353..285748</td><td>393607</td><td>Swt1-family-HEPN-domain-containing-protein-[Agromyces-allii]</td></tr>
<tr><td>WP_157002068.1</td><td>DUF499_1.RM.like.systems</td><td><-DNA_methylase<-SIGMA-HTH+SIGMA-HTH||?->?->METHYLASE->HEPN+DUF499*->MORC->REase-><-?||HTH+SF1-SF2-Helicase+REase->REase-></td><td>1143</td><td>Bacteria>Terrabacteria_group>Actinomycetota>Actinomycetes>Micrococcales</td><td>GCF_009749475.1</td><td>Agromyces_laixinhei</td><td>GQQ90_RS09115</td><td>49199..52630</td><td>2585717</td><td>DUF499-domain-containing-protein-[Agromyces-laixinhei]</td></tr>
<tr><td>WP_166844731.1</td><td>DUF499_1.RM.like.systems</td><td><-MORC+HEPN<-HTH+SF1-SF2-Helicase+REase<-REase<-MORC<-HEPN+DUF499*<-METHYLASE||DNA_methylase->?-><-?<-?<-HTH_3+PolB</td><td>1156</td><td>Bacteria>Terrabacteria_group>Actinomycetota>Actinomycetes>Micrococcales</td><td>GCF_011634835.1</td><td>Isoptericola_sp._BMS4</td><td>EYB28_RS03030</td><td>693115..696585</td><td>2527875</td><td>Swt1-family-HEPN-domain-containing-protein-[Isoptericola-sp.-BMS4]</td></tr>
<tr><td>WP_197156507.1</td><td>DUF499_1.RM.like.systems</td><td><-HD<-?||NUDIX-><-HTH+SF1-SF2-Helicase+REase<-REase<-MORC<-HEPN+DUF499*<-METHYLASE<-?<-?<-URI-DNase+SF1-SF2-Helicase<-MazG<-REase+35exo+Beta-Barrel+SF1-SF2-Helicase</td><td>1110</td><td>Bacteria>Terrabacteria_group>Actinomycetota>Actinomycetes>Micrococcales</td><td>GCF_015864945.1</td><td>Salinibacterium_sp._NG253</td><td>I6E52_RS09085</td><td>1888777..1892109</td><td>2792039</td><td>DUF499-domain-containing-protein-[Salinibacterium-sp.-NG253]</td></tr>
<tr><td>WP_200569160.1</td><td>DUF499_1.RM.like.systems</td><td><-DNA_methylase||REase->METHYLASE->HEPN+DUF499*->MORC->REase->HTH+SF1-SF2-Helicase+REase-></td><td>1144</td><td>Bacteria>Terrabacteria_group>Actinomycetota>Actinomycetes>Micrococcales</td><td>GCF_016620465.1</td><td>Kocuria_rhizophila</td><td>JHD97_RS04255</td><td>38629..42063</td><td>72000</td><td>DUF499-domain-containing-protein-[Kocuria-rhizophila]</td></tr>
<tr><td>WP_216886275.1</td><td>DUF499_1.RM.like.systems</td><td>Peptidase_S66+Peptidase_S66C-><-HTH+SF1-SF2-Helicase+REase<-REase<-MORC<-HEPN+DUF499*<-METHYLASE<-DNA_methylase||REase-></td><td>1157</td><td>Bacteria>Terrabacteria_group>Actinomycetota>Actinomycetes>Micrococcales</td><td>GCF_019037785.1</td><td>Micrococcus_luteus</td><td>KM867_RS03170</td><td>29276..32749</td><td>1270</td><td>Swt1-family-HEPN-domain-containing-protein-[Micrococcus-luteus]</td></tr>
<tr><td>WP_224781058.1</td><td>DUF499_1.RM.like.systems</td><td><-MORC+HEPN<-DNA_methylase||?->METHYLASE->HEPN+DUF499*->MORC->REase->HTH+SF1-SF2-Helicase+REase->REase-></td><td>1130</td><td>Bacteria>Terrabacteria_group>Actinomycetota>Actinomycetes>Micrococcales</td><td>GCF_020096995.1</td><td>Leucobacter_sp._Psy1</td><td>K8P10_RS06885</td><td>1465839..1469231</td><td>2875729</td><td>Swt1-family-HEPN-domain-containing-protein-[Leucobacter-sp.-Psy1]</td></tr>
<tr><td>WP_229577768.1</td><td>DUF499_1.RM.like.systems</td><td><-HTH+SF1-SF2-Helicase+REase<-REase<-MORC<-HEPN+DUF499*<-METHYLASE-p<-METHYLASE-p<-REase||DNA_methylase-></td><td>1141</td><td>Bacteria>Terrabacteria_group>Actinomycetota>Actinomycetes>Micrococcales</td><td>GCF_020829765.1</td><td>Kocuria_rhizophila</td><td>JNE43_RS06540</td><td>5673..9098</td><td>72000</td><td>DUF499-domain-containing-protein-[Kocuria-rhizophila]</td></tr>
<tr><td>WP_231831079.1</td><td>DUF499_1.RM.like.systems</td><td>MORC+SIGMA-HTH-><-REase<-HTH+SF1-SF2-Helicase+REase<-REase<-MORC<-HEPN+DUF499*<-METHYLASE||DNA_methylase->?->wHTH+SAD+HNH-></td><td>1128</td><td>Bacteria>Terrabacteria_group>Actinomycetota>Actinomycetes>Micrococcales</td><td>GCF_021168475.1</td><td>Pseudoclavibacter_sp._13-3</td><td>LRX77_RS01510</td><td>334724..338110</td><td>2901228</td><td>Swt1-family-HEPN-domain-containing-protein-[Pseudoclavibacter-sp.-13-3]</td></tr>
<tr><td>WP_276315051.1</td><td>DUF499_1.RM.like.systems</td><td>REase+SF1-SF2-Helicase-><-SbcC<-HEPN+DUF499*<-METHYLASE-p</td><td>1127</td><td>Bacteria>Terrabacteria_group>Actinomycetota>Actinomycetes>Micrococcales</td><td>GCF_021395155.1</td><td>Microbacterium_profundi</td><td>LZG07_RS12530</td><td>143258..146641</td><td>450380</td><td>DUF499-domain-containing-protein-[Microbacterium-profundi]</td></tr>
<tr><td>WP_241710057.1</td><td>DUF499_1.RM.like.systems</td><td>HIN-HTH->?-><-?<-HTH+SF1-SF2-Helicase+REase<-REase<-MORC<-HEPN+DUF499*<-METHYLASE||DNA_methylase-><-MORC+HEPN||?-><-SF1-SF2-Helicase||wHTH+SAD+HNH-></td><td>1123</td><td>Bacteria>Terrabacteria_group>Actinomycetota>Actinomycetes>Micrococcales</td><td>GCF_022606295.1</td><td>Arthrobacter_bambusae</td><td>KNN17_RS16570</td><td>21950..25321</td><td>1338426</td><td>DUF499-domain-containing-protein-[Arthrobacter-bambusae]</td></tr>
<tr><td>WP_134416003.1</td><td>DUF499_1.RM.like.systems</td><td>Peptidase_S66+Peptidase_S66C-><-HTH+SF1-SF2-Helicase+REase<-REase<-MORC<-HEPN+DUF499*<-METHYLASE<-DNA_methylase||REase-></td><td>1157</td><td>Bacteria>Terrabacteria_group>Actinomycetota>Actinomycetes>Micrococcales</td><td>GCF_022664405.1</td><td>Micrococcus_aloeverae</td><td>E2F92_RS02100</td><td>40430..43903</td><td>1391911</td><td>Swt1-family-HEPN-domain-containing-protein-[Micrococcus-aloeverae]</td></tr>
<tr><td>WP_243591605.1</td><td>DUF499_1.RM.like.systems</td><td>ACET-><-?<-REase<-HTH+SF1-SF2-Helicase+REase<-REase<-MORC<-HEPN+DUF499*<-METHYLASE||DNA_methylase->MORC+HisKin-></td><td>1123</td><td>Bacteria>Terrabacteria_group>Actinomycetota>Actinomycetes>Micrococcales</td><td>GCF_022815585.1</td><td>Rathayibacter_caricis</td><td>MT349_RS09900</td><td>122292..125663</td><td>110936</td><td>Swt1-family-HEPN-domain-containing-protein-[Rathayibacter-caricis]</td></tr>
<tr><td>WP_247963551.1</td><td>DUF499_1.RM.like.systems</td><td>DDE_Tnp_1->HIN-HTH->HTH_21+rve+rve_3->METHYLASE-p->HEPN+DUF499*->REase+SAVED-></td><td>1143</td><td>Bacteria>Terrabacteria_group>Actinomycetota>Actinomycetes>Micrococcales</td><td>GCF_023108875.1</td><td>Microbacterium_aerolatum</td><td>MZK47_RS16315</td><td>3209..6640</td><td>153731</td><td>DUF499-domain-containing-protein-[Microbacterium-aerolatum]</td></tr>
<tr><td>WP_253867645.1</td><td>DUF499_1.RM.like.systems</td><td><-HTH+SF1-SF2-Helicase+REase<-REase<-MORC<-HEPN+DUF499*<-METHYLASE||HTH_3->wHTH+SAD+HNH->HTH-></td><td>1150</td><td>Bacteria>Terrabacteria_group>Actinomycetota>Actinomycetes>Micrococcales</td><td>GCF_024171995.1</td><td>Promicromonospora_umidemergens</td><td>APR04_RS02600</td><td>547565..551017</td><td>629679</td><td>Swt1-family-HEPN-domain-containing-protein-[Promicromonospora-umidemergens]</td></tr>
<tr><td>WP_255770908.1</td><td>DUF499_1.RM.like.systems</td><td><-DNA_methylase||METHYLASE->HEPN+DUF499*->MORC->REase->HTH+SF1-SF2-Helicase+REase->ParB+HNH-><-?<-NUDIX</td><td>1122</td><td>Bacteria>Terrabacteria_group>Actinomycetota>Actinomycetes>Micrococcales</td><td>GCF_024391045.1</td><td>Pseudarthrobacter_sulfonivorans</td><td>NQJ30_RS15540</td><td>5007..8375</td><td>121292</td><td>DUF499-domain-containing-protein-[Pseudarthrobacter-sulfonivorans]</td></tr>
<tr><td>WP_257744324.1</td><td>DUF499_1.RM.like.systems</td><td><-DNA_methylase||REase->METHYLASE->HEPN+DUF499*->MORC->REase->HTH+SF1-SF2-Helicase+REase-><-?||MarR-HTH-></td><td>1137</td><td>Bacteria>Terrabacteria_group>Actinomycetota>Actinomycetes>Micrococcales</td><td>GCF_024655785.1</td><td>Kocuria_rhizophila</td><td>NUV30_RS04805</td><td>31091..34504</td><td>72000</td><td>DUF499-domain-containing-protein-[Kocuria-rhizophila]</td></tr>
<tr><td>WP_282859602.1</td><td>DUF499_1.RM.like.systems</td><td><-SbcC<-HEPN+DUF499*<-METHYLASE<-REase</td><td>1126</td><td>Bacteria>Terrabacteria_group>Actinomycetota>Actinomycetes>Micrococcales</td><td>GCF_030011915.1</td><td>Pseudoclavibacter_helvolus</td><td>QM024_RS14940</td><td>2044..5424</td><td>255205</td><td>DUF499-domain-containing-protein-[Pseudoclavibacter-helvolus]</td></tr>
<tr><td>WP_068492369.1</td><td>DUF499_1.RM.like.systems</td><td><-HTH+SF1-SF2-Helicase+REase<-REase<-MORC<-HEPN+DUF499*<-METHYLASE||DNA_methylase-></td><td>1129</td><td>Bacteria>Terrabacteria_group>Actinomycetota>Actinomycetes>Micrococcales</td><td>GCF_900014995.1</td><td>Pseudoclavibacter_helvolus</td><td>BN2259_RS00510</td><td>5683..9072</td><td>255205</td><td>Swt1-family-HEPN-domain-containing-protein-[Pseudoclavibacter-helvolus]</td></tr>
<tr><td>WP_101639485.1</td><td>DUF499_1.RM.like.systems</td><td><-OLD-ABC+TOPRIM<-DNA_methylase||MORC+REase->MORC+HEPN->REase->METHYLASE->HEPN+DUF499*->MORC->REase->HTH+SF1-SF2-Helicase+REase-></td><td>1135</td><td>Bacteria>Terrabacteria_group>Actinomycetota>Actinomycetes>Micrococcales</td><td>GCF_900169095.1</td><td>Brevibacterium_aurantiacum</td><td>BAUR920_RS16220</td><td>10995..14402</td><td>273384</td><td>DUF499-domain-containing-protein-[Brevibacterium-aurantiacum]</td></tr>
<tr><td>WP_218115672.1</td><td>DUF499_1.RM.like.systems</td><td><-URI-DNase+SF1-SF2-Helicase<-MazG||?->RVT-><-DNA_methylase||METHYLASE->HEPN+DUF499*->MORC->REase->HTH+SF1-SF2-Helicase+REase->REase-><-MORC+SIGMA-HTH</td><td>1133</td><td>Bacteria>Terrabacteria_group>Actinomycetota>Actinomycetes>Micrococcales</td><td>GCF_910593785.1</td><td>Leucobacter_soli</td><td>KXF26_RS09170</td><td>42150..45551</td><td>2812850</td><td>Swt1-family-HEPN-domain-containing-protein-[Leucobacter-soli]</td></tr>
<tr><td>WP_225755648.1</td><td>DUF499_1.RM.like.systems</td><td>REase->MORC->?->?->METHYLASE-p->METHYLASE-p->HEPN+DUF499*->MORC->REase->HTH+SF1-SF2-Helicase+REase->TUDOR+SF1-SF2-Helicase+iREase+CTD->SF1-SF2-Helicase-><-HNH</td><td>1137</td><td>Bacteria>Terrabacteria_group>Actinomycetota>Actinomycetes>Micrococcales</td><td>GCF_916618755.1</td><td>Actinotalea_sp._Marseille-Q4924</td><td>LG379_RS16320</td><td>268678..272091</td><td>2866571</td><td>DUF499-domain-containing-protein-[Actinotalea-sp.-Marseille-Q4924]</td></tr>
<tr><td>WP_234001366.1</td><td>DUF499_1.RM.like.systems</td><td>REase+SF1-SF2-Helicase-><-SbcC<-HEPN+DUF499*<-METHYLASE-p</td><td>1108</td><td>Bacteria>Terrabacteria_group>Actinomycetota>Actinomycetes>Micrococcales</td><td>-</td><td>Microbacterium_profundi</td><td>LZG07_12530</td><td>143258..146584</td><td>450380</td><td>DUF499-domain-containing-protein-[Microbacterium-profundi]</td></tr>
<tr><td>WP_220122165.1</td><td>DUF499_1.RM.like.systems</td><td>GNTR-HTH-><-?<-?<-DNA_methylase||SIGMA-HTH->METHYLASE->HEPN+DUF499*->MORC->REase->HTH+SF1-SF2-Helicase+REase->NERD+SF1-SF2-Helicase-><-?<-ABC-ATPase+SbcC</td><td>1128</td><td>Bacteria>Terrabacteria_group>Actinomycetota>Actinomycetes>Micrococcales</td><td>-</td><td>Ornithinimicrobium_laminariae</td><td>K0651_07785</td><td>644333..647719</td><td>2864873</td><td>Swt1-family-HEPN-domain-containing-protein-[Ornithinimicrobium-laminariae]</td></tr>
<tr><td>WP_203769001.1</td><td>DUF499_1.RM.like.systems</td><td>DNA_methylase-><-REase<-HTH+SF1-SF2-Helicase+REase<-REase<-MORC<-HEPN+DUF499*<-METHYLASE||||?->REase-><-?||REase->MarR-HTH-></td><td>1138</td><td>Bacteria>Terrabacteria_group>Actinomycetota>Actinomycetes>Micromonosporales</td><td>GCF_016862275.1</td><td>Actinoplanes_nipponensis</td><td>Ani05nite_RS15610</td><td>113186..116602</td><td>135950</td><td>Swt1-family-HEPN-domain-containing-protein-[Actinoplanes-nipponensis]</td></tr>
<tr><td>WP_260034146.1</td><td>DUF499_1.RM.like.systems</td><td><-SF1-SF2-Helicase<-?||METHYLASE->HEPN+DUF499*->MORC->REase->HTH+SF1-SF2-Helicase+REase-></td><td>1133</td><td>Bacteria>Terrabacteria_group>Actinomycetota>Actinomycetes>Micromonosporales</td><td>GCF_025152325.1</td><td>Micromonospora_chalcea</td><td>M3G91_RS14945</td><td>8947..12348</td><td>1874</td><td>Swt1-family-HEPN-domain-containing-protein-[Micromonospora-chalcea]</td></tr>
<tr><td>CCK55004.1</td><td>DUF499_1.RM.like.systems</td><td>STYKinase->HIN-HTH->?-><-?<-REase<-DUF499*<-HEPN<-?<-?<-HIN-HTH<-HIN-HTH<-METHYLASE-p</td><td>792</td><td>Bacteria>Terrabacteria_group>Actinomycetota>Actinomycetes>Mycobacteriales</td><td>GCA_000328845.1</td><td>Mycobacterium_canettii_CIPT_140070008</td><td>BN43_30077</td><td>1164706..1167084</td><td>1205675</td><td>Conserved-protein-of-unknown-function-[Mycobacterium-canettii-CIPT-140070008]</td></tr>
<tr><td>OBB45420.1</td><td>DUF499_1.RM.like.systems</td><td>ParB+HNH->?-><-HTH+SF1-SF2-Helicase+REase<-REase<-MORC<-HEPN+DUF499*<-METHYLASE||DNA_methylase->OLD-ABC-><-REase<-SF1-SF2-Helicase+REase<-SF1-SF2-Helicase+REase-p</td><td>1135</td><td>Bacteria>Terrabacteria_group>Actinomycetota>Actinomycetes>Mycobacteriales</td><td>GCA_001665535.1</td><td>Mycobacterium_sp._852002-51961_SCH5331710</td><td>A5752_02020</td><td>17252..20659</td><td>1834105</td><td>hypothetical-protein-A5752_02020-[Mycobacterium-sp.-852002-51961_SCH5331710]</td></tr>
<tr><td>ATO62480.2</td><td>DUF499_1.RM.like.systems</td><td>Glyoxalase->HIN-HTH-><-DNA_methylase||REase->METHYLASE->HEPN+DUF499*-><-HIN-HTH</td><td>1138</td><td>Bacteria>Terrabacteria_group>Actinomycetota>Actinomycetes>Mycobacteriales</td><td>GCA_002716905.4</td><td>Mycobacterium_avium_subsp._hominissuis</td><td>BEP52_09350</td><td>1958340..1961756</td><td>439334</td><td>DUF499-domain-containing-protein-[Mycobacterium-avium-subsp.-hominissuis]</td></tr>
<tr><td>MCB1297052.1</td><td>DUF499_1.RM.like.systems</td><td>Peptidase_M20-><-?||?->?->?-><-REase<-HEPN+DUF499*||MazE_antitoxin->PIN-><-METHYLASE</td><td>1119</td><td>Bacteria>Terrabacteria_group>Actinomycetota>Actinomycetes>Mycobacteriales</td><td>GCA_020438995.1</td><td>Gordonia_sp._(in:_high_G+C_Gram-positive_bacteria)</td><td>KDB67_20615</td><td>20423..23782</td><td>84139</td><td>DUF499-domain-containing-protein-[Gordonia-sp.]</td></tr>
<tr><td>UHJ57678.1</td><td>DUF499_1.RM.like.systems</td><td>REase->DNA-glycosylase->?-><-HTH+SF1-SF2-Helicase+REase<-REase<-MORC<-HEPN+DUF499*<-METHYLASE<-REase<-PIN||DNA_methylase-><-?<-Phage_integrase+Phage_integrase</td><td>1138</td><td>Bacteria>Terrabacteria_group>Actinomycetota>Actinomycetes>Mycobacteriales</td><td>GCA_021249325.1</td><td>Mycolicibacterium_fortuitum</td><td>LT337_13390</td><td>2620949..2624365</td><td>1766</td><td>DUF499-domain-containing-protein-[Mycolicibacterium-fortuitum]</td></tr>
<tr><td>QHN29206.1</td><td>DUF499_1.RM.like.systems</td><td><-MULE-TRANSPOSASE||ParA-Soj-PloopNTPase->ORC-CDC6-><-HTH+SF1-SF2-Helicase+REase<-REase<-MORC<-HEPN+DUF499*<-METHYLASE||XIS-HTH+RVT-><-HNH||?-><-REase||DNA_methylase-></td><td>1138</td><td>Bacteria>Terrabacteria_group>Actinomycetota>Actinomycetes>Mycobacteriales</td><td>GCA_025268675.1</td><td>Gordonia_amarae</td><td>GII32_01220</td><td>287584..291000</td><td>36821</td><td>DUF499-domain-containing-protein-[Gordonia-amarae]</td></tr>
<tr><td>WP_026061790.1</td><td>DUF499_1.RM.like.systems</td><td><-DNA_methylase||REase-><-HEPN+DUF499*</td><td>1135</td><td>Bacteria>Terrabacteria_group>Actinomycetota>Actinomycetes>Mycobacteriales</td><td>GCF_000239135.1</td><td>Rhodococcus_rhodochrous</td><td>RHODODRAFT_RS0223985</td><td>1954..5361</td><td>1829</td><td>Swt1-family-HEPN-domain-containing-protein-[Rhodococcus-rhodochrous]</td></tr>
<tr><td>WP_005189162.1</td><td>DUF499_1.RM.like.systems</td><td><-MULE-TRANSPOSASE||ParA-Soj-PloopNTPase->ORC-CDC6-><-HTH+SF1-SF2-Helicase+REase<-REase<-MORC<-HEPN+DUF499*<-METHYLASE||XIS-HTH+RVT-><-HNH||?-><-REase||DNA_methylase-></td><td>1138</td><td>Bacteria>Terrabacteria_group>Actinomycetota>Actinomycetes>Mycobacteriales</td><td>GCF_000241345.1</td><td>Gordonia_amarae</td><td>GII35_RS01230</td><td>287739..291155</td><td>36821</td><td>DUF499-domain-containing-protein-[Gordonia-amarae]</td></tr>
<tr><td>WP_014358285.1</td><td>DUF499_1.RM.like.systems</td><td>MACRODOMAIN+SIR2->SF1-SF2-Helicase-><-HTH+SF1-SF2-Helicase+REase<-REase<-MORC<-HEPN+DUF499*<-METHYLASE<-REase||DNA_methylase->SAM-methylase-><-CR-Kinase+ORC-CDC6-like+CARF</td><td>1127</td><td>Bacteria>Terrabacteria_group>Actinomycetota>Actinomycetes>Mycobacteriales</td><td>GCF_000247715.1</td><td>Gordonia_polyisoprenivorans</td><td>GPOL_RS00855</td><td>197189..200572</td><td>84595</td><td>DUF499-domain-containing-protein-[Gordonia-polyisoprenivorans]</td></tr>
<tr><td>WP_232512159.1</td><td>DUF499_1.RM.like.systems</td><td><-HEPN+DUF499*<-METHYLASE<-REase||DNA_methylase-></td><td>1128</td><td>Bacteria>Terrabacteria_group>Actinomycetota>Actinomycetes>Mycobacteriales</td><td>GCF_000829075.1</td><td>Mycobacterium_avium</td><td>B9C26_RS25090</td><td>191..3577</td><td>1764</td><td>DUF499-domain-containing-protein-[Mycobacterium-avium]</td></tr>
<tr><td>WP_048320042.1</td><td>DUF499_1.RM.like.systems</td><td><-DNA_methylase||?->METHYLASE->HEPN+DUF499*->MORC->REase->HTH+SF1-SF2-Helicase+REase-><-wHTH+REase<-SF1-SF2-Helicase<-Primpol+D5N+D5ATPase</td><td>1137</td><td>Bacteria>Terrabacteria_group>Actinomycetota>Actinomycetes>Mycobacteriales</td><td>GCF_001037935.1</td><td>Rhodococcus_fascians</td><td>ACG96_RS20770</td><td>139814..143227</td><td>1828</td><td>DUF499-domain-containing-protein-[Rhodococcus-fascians]</td></tr>
<tr><td>WP_082921219.1</td><td>DUF499_1.RM.like.systems</td><td>METHYLASE-p->HEPN+DUF499*->MORC->REase-></td><td>1121</td><td>Bacteria>Terrabacteria_group>Actinomycetota>Actinomycetes>Mycobacteriales</td><td>GCF_001651055.1</td><td>Rhodococcus_sp._HS-D2</td><td>A1E04_RS27825</td><td>597..3962</td><td>1384636</td><td>Swt1-family-HEPN-domain-containing-protein-[Rhodococcus-sp.-HS-D2]</td></tr>
<tr><td>WP_067083703.1</td><td>DUF499_1.RM.like.systems</td><td>ParB+HNH->?-><-HTH+SF1-SF2-Helicase+REase<-REase<-MORC<-HEPN+DUF499*<-METHYLASE<-REase||DNA_methylase->OLD-ABC+TOPRIM-><-REase<-SF1-SF2-Helicase+REase</td><td>1135</td><td>Bacteria>Terrabacteria_group>Actinomycetota>Actinomycetes>Mycobacteriales</td><td>GCF_001665535.1</td><td>Mycobacterium_sp._852002-51961_SCH5331710</td><td>A5752_RS07195</td><td>17252..20659</td><td>1834105</td><td>Swt1-family-HEPN-domain-containing-protein-[Mycobacterium-sp.-852002-51961_SCH5331710]</td></tr>
<tr><td>WP_134205957.1</td><td>DUF499_1.RM.like.systems</td><td><-DUF222+HNH<-?<-?<-HTH+SF1-SF2-Helicase+REase<-REase<-MORC<-HEPN+DUF499*<-METHYLASE-p<-HerA/FtsK<-?<-BrxL<-PglZ<-METHYLASE-PglX</td><td>1137</td><td>Bacteria>Terrabacteria_group>Actinomycetota>Actinomycetes>Mycobacteriales</td><td>GCF_004366875.1</td><td>Mycobacteroides_salmoniphilum</td><td>CCUG62472_RS09110</td><td>402187..405600</td><td>404941</td><td>DUF499-domain-containing-protein-[Mycobacteroides-salmoniphilum]</td></tr>
<tr><td>WP_141142680.1</td><td>DUF499_1.RM.like.systems</td><td>SF1-SF2-Helicase->?-><-REase||DNA_methylase->METHYLASE->HEPN+DUF499*->MORC->REase->HTH+SF1-SF2-Helicase+REase->wHTH+REase-></td><td>1137</td><td>Bacteria>Terrabacteria_group>Actinomycetota>Actinomycetes>Mycobacteriales</td><td>GCF_006517415.1</td><td>Mycolicibacterium_fortuitum</td><td>FKW78_RS12965</td><td>17898..21311</td><td>1766</td><td>DUF499-domain-containing-protein-[Mycolicibacterium-fortuitum]</td></tr>
<tr><td>WP_213249149.1</td><td>DUF499_1.RM.like.systems</td><td><-HTH+SF1-SF2-Helicase+REase<-REase<-MORC<-HEPN+DUF499*<-SF1-SF2-Helicase<-OLD-ABC+TOPRIM||DNA_methylase-><-?||HNH-></td><td>1138</td><td>Bacteria>Terrabacteria_group>Actinomycetota>Actinomycetes>Mycobacteriales</td><td>GCF_009914535.1</td><td>Gordonia</td><td>IDH24_RS11690</td><td>142027..145443</td><td>2053</td><td>DUF499-domain-containing-protein-[Gordonia-pseudamarae]</td></tr>
<tr><td>WP_162113231.1</td><td>DUF499_1.RM.like.systems</td><td>DUF499*->REase-><-GDResolvase+HIN-HTH</td><td>883</td><td>Bacteria>Terrabacteria_group>Actinomycetota>Actinomycetes>Mycobacteriales</td><td>GCF_010093495.1</td><td>Mycolicibacter_kumamotonensis</td><td>GWR20_RS22070</td><td>1..2652</td><td>354243</td><td>DUF499-domain-containing-protein,-partial-[Mycolicibacter-kumamotonensis]</td></tr>
<tr><td>WP_167106438.1</td><td>DUF499_1.RM.like.systems</td><td><-DNA_methylase||REase->METHYLASE->HEPN+DUF499*->MORC->REase->HTH+SF1-SF2-Helicase+REase-><-?<-SF1-SF2-helicase+DUF1998</td><td>1137</td><td>Bacteria>Terrabacteria_group>Actinomycetota>Actinomycetes>Mycobacteriales</td><td>GCF_011694515.1</td><td>Mycobacterium_sp._DL592</td><td>HBE64_RS20620</td><td>4288954..4292367</td><td>2675524</td><td>DUF499-domain-containing-protein-[Mycobacterium-sp.-DL592]</td></tr>
<tr><td>WP_172936837.1</td><td>DUF499_1.RM.like.systems</td><td>REase+SF1-SF2-Helicase->wHTH+SAD+HNH-><-HTH+SF1-SF2-Helicase+REase<-REase<-MORC<-HEPN+DUF499*<-METHYLASE||DNA_methylase-><-?<-?<-?<-MORC</td><td>1125</td><td>Bacteria>Terrabacteria_group>Actinomycetota>Actinomycetes>Mycobacteriales</td><td>GCF_013277795.1</td><td>Rhodococcus_sp._MS16</td><td>FEZ60_RS20030</td><td>139457..142834</td><td>2579941</td><td>Swt1-family-HEPN-domain-containing-protein-[Rhodococcus-sp.-MS16]</td></tr>
<tr><td>WP_219849138.1</td><td>DUF499_1.RM.like.systems</td><td><-HTH+SF1-SF2-Helicase+REase<-REase<-MORC<-HEPN+DUF499*<-METHYLASE<-REase||DNA_methylase-><-METHYLASE-p</td><td>1137</td><td>Bacteria>Terrabacteria_group>Actinomycetota>Actinomycetes>Mycobacteriales</td><td>GCF_014041935.1</td><td>Gordonia_jinghuaiqii</td><td>H1R19_RS12155</td><td>2732590..2736003</td><td>2758710</td><td>Swt1-family-HEPN-domain-containing-protein-[Gordonia-jinghuaiqii]</td></tr>
<tr><td>WP_260836619.1</td><td>DUF499_1.RM.like.systems</td><td><-MULE-TRANSPOSASE||ParA-Soj-PloopNTPase->ORC-CDC6-><-HTH+SF1-SF2-Helicase+REase<-REase<-MORC<-HEPN+DUF499*<-METHYLASE||XIS-HTH+RVT-><-HNH||?-><-REase||DNA_methylase-></td><td>1138</td><td>Bacteria>Terrabacteria_group>Actinomycetota>Actinomycetes>Mycobacteriales</td><td>GCF_025268675.1</td><td>Gordonia_amarae</td><td>GII32_RS01225</td><td>287584..291000</td><td>36821</td><td>DUF499-domain-containing-protein-[Gordonia-amarae]</td></tr>
<tr><td>WP_276127103.1</td><td>DUF499_1.RM.like.systems</td><td><-TetR-HTH||?->?-><-HTH+SF1-SF2-Helicase+REase<-REase<-MORC<-HEPN+DUF499*<-METHYLASE<-REase||DNA_methylase-><-CR-Kinase+ORC-CDC6-like+CARF</td><td>1137</td><td>Bacteria>Terrabacteria_group>Actinomycetota>Actinomycetes>Mycobacteriales</td><td>GCF_029270025.1</td><td>Mycolicibacterium_septicum</td><td>P3H80_RS10735</td><td>384555..387968</td><td>98668</td><td>DUF499-domain-containing-protein-[Mycolicibacterium-septicum]</td></tr>
<tr><td>WP_125079869.1</td><td>DUF499_1.RM.like.systems</td><td>ACET->GDResolvase+HIN-HTH-><-REase<-HEPN+DUF499*</td><td>1124</td><td>Bacteria>Terrabacteria_group>Actinomycetota>Actinomycetes>Mycobacteriales</td><td>GCF_900604305.1</td><td>Mycobacterium_sp._P7213</td><td>EJK91_RS20970</td><td>3382..6756</td><td>2478465</td><td>DUF499-domain-containing-protein-[Mycobacterium-sp.-P7213]</td></tr>
<tr><td>WP_023870228.1</td><td>DUF499_1.RM.like.systems</td><td>Glyoxalase->HIN-HTH-><-DNA_methylase||REase->METHYLASE->HEPN+DUF499*-><-HIN-HTH</td><td>1138</td><td>Bacteria>Terrabacteria_group>Actinomycetota>Actinomycetes>Mycobacteriales</td><td>-</td><td>Mycobacterium_avium</td><td>BEP52_09350</td><td>1958340..1961756</td><td>1764</td><td>DUF499-domain-containing-protein-[Mycobacterium-avium]</td></tr>
<tr><td>WP_028701426.1</td><td>DUF499_1.RM.like.systems</td><td><-SF1-SF2-Helicase<-MORC+SF1-SF2-Helicase||DNA_methylase-><-DNA_methylase||METHYLASE->HEPN+DUF499*->MORC->REase->HTH+SF1-SF2-Helicase+REase->REase-><-ParB+HNH-p<-ParB+HNH-p</td><td>1129</td><td>Bacteria>Terrabacteria_group>Actinomycetota>Actinomycetes>Propionibacteriales</td><td>GCF_000427845.1</td><td>Acidipropionibacterium_acidipropionici</td><td>ASQ49_RS03445</td><td>772026..775415</td><td>1748</td><td>DUF499-domain-containing-protein-[Acidipropionibacterium-acidipropionici]</td></tr>
<tr><td>WP_078920357.1</td><td>DUF499_1.RM.like.systems</td><td><-TrwC+SF1-SF2-helicase+TOPRIM<-DNA_methylase||REase->METHYLASE->HEPN+DUF499*->MORC->REase->HTH+SF1-SF2-Helicase+REase-></td><td>1132</td><td>Bacteria>Terrabacteria_group>Actinomycetota>Actinomycetes>Propionibacteriales</td><td>GCF_002022225.1</td><td>Tessaracoccus</td><td>BKM78_RS14510</td><td>3153779..3157177</td><td>72763</td><td>MULTISPECIES:-Swt1-family-HEPN-domain-containing-protein-[Tessaracoccus]</td></tr>
<tr><td>WP_154690370.1</td><td>DUF499_1.RM.like.systems</td><td>PAIREDC-HTH-><-?<-HTH+SF1-SF2-Helicase+REase<-REase<-MORC<-HEPN+DUF499*<-METHYLASE-p<-METHYLASE<-REase<-PIN||DNA_methylase-></td><td>1132</td><td>Bacteria>Terrabacteria_group>Actinomycetota>Actinomycetes>Propionibacteriales</td><td>GCF_009707365.1</td><td>Microlunatus_sp._Gsoil_973</td><td>GJV80_RS18445</td><td>3917369..3920767</td><td>2672569</td><td>DUF499-domain-containing-protein-[Microlunatus-sp.-Gsoil-973]</td></tr>
<tr><td>WP_182766932.1</td><td>DUF499_1.RM.like.systems</td><td><-HTH+SF1-SF2-Helicase+REase<-REase<-HEPN+DUF499*<-METHYLASE<-REase||DNA_methylase-></td><td>1129</td><td>Bacteria>Terrabacteria_group>Actinomycetota>Actinomycetes>Propionibacteriales</td><td>GCF_014164735.1</td><td>Tessaracoccus_sp._MC1756</td><td>H5393_RS12480</td><td>209092..212481</td><td>2760311</td><td>Swt1-family-HEPN-domain-containing-protein-[Tessaracoccus-sp.-MC1756]</td></tr>
<tr><td>WP_232416936.1</td><td>DUF499_1.RM.like.systems</td><td><-HTH+SF1-SF2-Helicase+REase<-REase<-MORC<-HEPN+DUF499*<-METHYLASE<-REase||DNA_methylase-><-?||DUF222+HNH-></td><td>1136</td><td>Bacteria>Terrabacteria_group>Actinomycetota>Actinomycetes>Propionibacteriales</td><td>GCF_024508295.1</td><td>Aeromicrobium_duanguangcaii</td><td>NP095_RS05665</td><td>1110875..1114285</td><td>2968086</td><td>MULTISPECIES:-Swt1-family-HEPN-domain-containing-protein-[unclassified-Aeromicrobium]</td></tr>
<tr><td>WP_131171419.1</td><td>DUF499_1.RM.like.systems</td><td><-MACRODOMAIN+SIR2<-HTH+SF1-SF2-Helicase+REase<-REase<-MORC<-HEPN+DUF499*<-METHYLASE<-METHYLASE-p||DNA_methylase-><-SbcC<-SbcC||TrwC+SF1-SF2-helicase+TOPRIM-></td><td>1146</td><td>Bacteria>Terrabacteria_group>Actinomycetota>Actinomycetes>Propionibacteriales</td><td>GCF_900182665.1</td><td>Propioniciclava_tarda</td><td>FN821_RS04755</td><td>6296..9736</td><td>433330</td><td>Swt1-family-HEPN-domain-containing-protein-[Propioniciclava-tarda]</td></tr>
<tr><td>EAR26285.1</td><td>DUF499_1.RM.like.systems</td><td>REase-><-REase<-DNA_methylase<-MORC+SIGMA-HTH||METHYLASE->HEPN+DUF499*->MORC->REase->HTH+SF1-SF2-Helicase+REase->?-><-?<-ABC-ATPase+SbcC</td><td>1127</td><td>Bacteria>Terrabacteria_group>Actinomycetota>Actinomycetes>unclassified_Actinomycetes</td><td>GCA_000153145.1</td><td>marine_actinobacterium_PHSC20C1</td><td>A20C1_10399</td><td>2131040..2134423</td><td>312284</td><td>hypothetical-protein-A20C1_10399-[marine-actinobacterium-PHSC20C1]</td></tr>
<tr><td>MBX6764706.1</td><td>DUF499_1.RM.like.systems</td><td>METHYLASE-p->MazE_antitoxin->PIN->HEPN+DUF499*->REase-></td><td>1106</td><td>Bacteria>Terrabacteria_group>Actinomycetota>Rubrobacteria>Rubrobacterales</td><td>GCA_019690035.1</td><td>Rubrobacteraceae_bacterium</td><td>IRY88_13660</td><td>1711..5031</td><td>2740537</td><td>DUF499-domain-containing-protein-[Rubrobacteraceae-bacterium]</td></tr>
<tr><td>NCD16052.1</td><td>DUF499_1.RM.like.systems</td><td><-MORC+REase<-HTH+SF1-SF2-Helicase+REase<-REase<-MORC<-HEPN+DUF499*<-METHYLASE<-REase||DNA_methylase-><-?<-?<-TrwC+SF1-SF2-Helicase</td><td>1131</td><td>Bacteria>Terrabacteria_group>Actinomycetota>unclassified_Actinomycetota</td><td>GCA_009930385.1</td><td>Actinomycetota_bacterium</td><td>EOL91_01830</td><td>7837..11232</td><td>2900548</td><td>ATP-binding-protein-[Actinomycetia-bacterium]</td></tr>
<tr><td>NCD16598.1</td><td>DUF499_1.RM.like.systems</td><td><-HEPN+REase<-REase<-HEPN+DUF499*<-wHTH+REase<-METHYLASE</td><td>1133</td><td>Bacteria>Terrabacteria_group>Actinomycetota>unclassified_Actinomycetota</td><td>GCA_009930385.1</td><td>Actinomycetota_bacterium</td><td>EOL91_04665</td><td>3104..6505</td><td>2900548</td><td>ATP-binding-protein-[Actinomycetia-bacterium]</td></tr>
<tr><td>MBU2074163.1</td><td>DUF499_1.RM.like.systems</td><td><-HTH+SF1-SF2-Helicase+REase<-REase<-MORC<-HEPN+DUF499*<-METHYLASE<-REase||DNA_methylase->MORC+HEPN-><-?<-TrwC+SF1-SF2-Helicase</td><td>1135</td><td>Bacteria>Terrabacteria_group>Actinomycetota>unclassified_Actinomycetota</td><td>GCA_018826065.1</td><td>Actinomycetota_bacterium</td><td>KJ938_06990</td><td>8952..12359</td><td>2900548</td><td>DUF499-domain-containing-protein-[Actinomycetia-bacterium]</td></tr>
<tr><td>MBU2110960.1</td><td>DUF499_1.RM.like.systems</td><td>DUF499*->MORC->REase->HTH+SF1-SF2-Helicase+REase-></td><td>991</td><td>Bacteria>Terrabacteria_group>Actinomycetota>unclassified_Actinomycetota</td><td>GCA_018826625.1</td><td>Actinomycetota_bacterium</td><td>KKE65_04805</td><td>1..2976</td><td>2900548</td><td>DUF499-domain-containing-protein-[Actinomycetia-bacterium]</td></tr>
<tr><td>MCE7883902.1</td><td>DUF499_1.RM.like.systems</td><td><-REase+SF1-SF2-Helicase<-DUF499*<-DUF499-p||XIS-HTH-></td><td>594</td><td>Bacteria>Terrabacteria_group>Actinomycetota>unclassified_Actinomycetota</td><td>GCA_021404985.1</td><td>Actinobacteria_bacterium_ATB1</td><td>DYH08_08590</td><td>2755..4539</td><td>2293627</td><td>ATP-binding-protein-[Actinobacteria-bacterium-ATB1]</td></tr>
<tr><td>MCX6403861.1</td><td>DUF499_1.RM.like.systems</td><td>REase->METHYLASE->HEPN+DUF499*-></td><td>1088</td><td>Bacteria>Terrabacteria_group>Actinomycetota>unclassified_Actinomycetota</td><td>GCA_026392535.1</td><td>Actinomycetota_bacterium</td><td>NT032_08225</td><td>3485..6751</td><td>2900548</td><td>MAG:-DUF499-domain-containing-protein-[Actinobacteria-bacterium]</td></tr>
<tr><td>GIV02394.1</td><td>DUF499_1.RM.like.systems</td><td><-URI-DNase+RAMA<-?<-HEPN+REase<-?<-REase<-Schlafen+HATPase+HTH<-DUF499*<-REase<-?<-METHYLASE-p||?-><-?<-METHYLASE</td><td>920</td><td>Bacteria>Terrabacteria_group>Armatimonadota>Fimbriimonadia>Fimbriimonadales</td><td>GCA_026003055.1</td><td>Fimbriimonadales_bacterium</td><td>KatS3mg015_1224</td><td>1249734..1252496</td><td>2682145</td><td>MAG:-hypothetical-protein-KatS3mg015_1224-[Fimbriimonadales-bacterium]</td></tr>
<tr><td>NIM06025.1</td><td>DUF499_1.RM.like.systems</td><td><-DNA-glycosylase<-DUF499*<-REase<-METHYLASE-p</td><td>976</td><td>Bacteria>Terrabacteria_group>Armatimonadota>unclassified_Armatimonadota</td><td>GCA_011771275.1</td><td>Armatimonadota_bacterium</td><td>GTO78_10545</td><td>837..3767</td><td>2033014</td><td>DUF499-domain-containing-protein-[Armatimonadetes-bacterium]</td></tr>
<tr><td>NLB94469.1</td><td>DUF499_1.RM.like.systems</td><td>REase->HEPN->METHYLASE->DUF499*->DNA-glycosylase-></td><td>991</td><td>Bacteria>Terrabacteria_group>Armatimonadota>unclassified_Armatimonadota</td><td>GCA_012512265.1</td><td>Armatimonadota_bacterium</td><td>GX785_02050</td><td>5194..8169</td><td>2033014</td><td>DUF499-domain-containing-protein-[Armatimonadetes-bacterium]</td></tr>
<tr><td>MBM3494074.1</td><td>DUF499_1.RM.like.systems</td><td><-REase+DUF4268<-DUF499*</td><td>989</td><td>Bacteria>Terrabacteria_group>Armatimonadota>unclassified_Armatimonadota</td><td>GCA_016869115.1</td><td>Armatimonadota_bacterium</td><td>FJX72_07105</td><td>2132..5102</td><td>2033014</td><td>ATP-binding-protein-[Armatimonadetes-bacterium]</td></tr>
<tr><td>MCL6449134.1</td><td>DUF499_1.RM.like.systems</td><td>METHYLASE->?->DUF1828+REase->DUF499*->REase->Radical-SAM-><-PIN<-PhdYeFM_antitox||?-><-PIN</td><td>1104</td><td>Bacteria>Terrabacteria_group>Armatimonadota>unclassified_Armatimonadota</td><td>GCA_023511535.1</td><td>Armatimonadota_bacterium</td><td>K6U04_13475</td><td>3761..7075</td><td>2033014</td><td>DUF499-domain-containing-protein-[Armatimonadetes-bacterium]</td></tr>
</table>

| Protein-ID | System | Gene-Neighbourhoods | Length | Taxonomy | Assembly-ID | Strain | Locus-Tag | Genomic-coordinates | TaxID | Fasta-defline |
|---|---|---|---|---|---|---|---|---|---|---|
| MCX6379947.1 | DUF499_1.RM.like.systems | REase->HEPN+DUF499*-><-?<-?<-?<-?<-?\|\|Phage_int_SAM_1+Phage_integrase-> | 1119 | Bacteria>Terrabacteria_group>Armatimonadota>unclassified_Armatimonadota | GCA_026392135.1 | Armatimonadota_bacterium | NT023_10805 | 3514..6873 | 2033014 | MAG:-Swt1-family-HEPN-domain-containing-protein-[Armatimonadetes-bacterium] |
| MDE2206237.1 | DUF499_1.RM.like.systems | DUF499*->HEPN+REase->HTH->MPTase->?->HTH+UbE1->MarR-HTH-> | 600 | Bacteria>Terrabacteria_group>Armatimonadota>unclassified_Armatimonadota | GCA_028871005.1 | Armatimonadota_bacterium | KGK12_06315 | 1..1805 | 2033014 | MAG:-ATP-binding-protein,-partial-[Armatimonadetes-bacterium] |
| HHW04285.1 | DUF499_1.RM.like.systems | REase->METHYLASE->DUF499*-> | 910 | Bacteria>Terrabacteria_group>Bacillota>Clostridia>Thermoanaerobacterales | GCA_012840135.1 | Thermoanaerobacterales_bacterium | GXX35_16055 | 4069..6801 | 2304039 | ATP-binding-protein-[Thermoanaerobacterales-bacterium] |
| MTI83942.1 | DUF499_1.RM.like.systems | <-SF1-SF2-Helicase+REase<-PIWI<-HEPN+DUF499*<-METHYLASE-p | 1111 | Bacteria>Terrabacteria_group>Bacillota>unclassified_Bacillota | GCA_009711705.1 | Bacillota_bacterium | FH756_08540 | 83034..86369 | 1879010 | ATP-binding-protein-[Firmicutes-bacterium] |
| HHV56346.1 | DUF499_1.RM.like.systems | <-DUF772+DDE_Tnp_1+DDE_Tnp_1_6\|\|METHYLASE-p->HEPN+DUF499*->REase-> | 1131 | Bacteria>Terrabacteria_group>Bacillota>unclassified_Bacillota | GCA_012839755.1 | Bacillota_bacterium | GXX50_01080 | 3217..6612 | 1879010 | ATP-binding-protein-[Firmicutes-bacterium] |
| MAS36305.1 | DUF499_1.RM.like.systems | Glyoxalase->?-><-PIN<-?<-HEPN+DUF499*<-REase<-?<-METHYLASE<-Schlafen+HATPase+HTH | 1094 | Bacteria>Terrabacteria_group>Chloroflexota>Anaerolineae>Anaerolineales | GCA_002702065.1 | Anaerolineaceae_bacterium | CL610_20035 | 35853..39137 | 2024896 | AAA+-family-ATPase-[Anaerolineaceae-bacterium] |
| MBE7469119.1 | DUF499_1.RM.like.systems | METHYLASE-p->?->?->?-><-DUF499*<-REase<-METHYLASE<-HEPN<-PolB<-HEPN<-Polbeta | 906 | Bacteria>Terrabacteria_group>Chloroflexota>Anaerolineae>Anaerolineales | GCA_015075405.1 | Anaerolineales_bacterium | HS114_08270 | 1909271..1911991 | 2073117 | ATP-binding-protein-[Anaerolineales-bacterium] |
| MBK8904525.1 | DUF499_1.RM.like.systems | <-HhH-RADC+JAB<-?<-?<-DDE_5<-DUF499*\|\|?-><-?<-?<-REase<-METHYLASE-p<-HTH_Tnp_4 | 948 | Bacteria>Terrabacteria_group>Chloroflexota>Anaerolineae>Anaerolineales | GCA_016710325.1 | Anaerolineaceae_bacterium | IPM53_25325 | 3764417..3767263 | 2024896 | ATP-binding-protein-[Anaerolineaceae-bacterium] |
| MBL6983853.1 | DUF499_1.RM.like.systems | <-REase<-HEPN+DUF499*<-SF1-SF2-Helicase-p | 1095 | Bacteria>Terrabacteria_group>Chloroflexota>Anaerolineae>Anaerolineales | GCA_016785015.1 | Anaerolineales_bacterium | ISR58_21930 | 2169..5456 | 2073117 | DUF499-domain-containing-protein-[Anaerolineales-bacterium] |
| MCB9078004.1 | DUF499_1.RM.like.systems | <-HEPN+DUF499*<-REase<-?<-METHYLASE-p<-METHYLASE<-PIN<-DUF433 | 385 | Bacteria>Terrabacteria_group>Chloroflexota>Anaerolineae>Anaerolineales | GCA_020634955.1 | Anaerolineaceae_bacterium | H6631_10465 | 1..1157 | 2024896 | ATP-binding-protein-[Anaerolineaceae-bacterium] |
| MCC6958025.1 | DUF499_1.RM.like.systems | <-MarR-HTH<-HEPN+DUF499*<-?<-METHYLASE<-?<-PIN<-REase | 1114 | Bacteria>Terrabacteria_group>Chloroflexota>Anaerolineae>Anaerolineales | GCA_020853835.1 | Anaerolineales_bacterium | IT316_14590 | 1878..5222 | 2073117 | DUF499-domain-containing-protein-[Anaerolineales-bacterium] |
| PJH76238.1 | DUF499_1.RM.like.systems | DUF499*->?->ORC-CDC6+REase-> | 750 | Bacteria>Terrabacteria_group>Chloroflexota>Anaerolineae>unclassified_Anaerolineae | GCA_002796565.1 | Anaerolineae_bacterium_CG_4_9_14_0_8_um_filter_58_9 | C0064_02300 | 1..2255 | 1973903 | MAG:-AAA-family-ATPase,-partial-[Anaerolineae-bacterium-CG_4_9_14_0_8_um_filter_58_9] |
| MBK6429951.1 | DUF499_1.RM.like.systems | OLD-ABC->TOPRIM->Schlafen+HATPase+HTH->METHYLASE-p->Schlafen+HATPase+HTH->REase->HEPN+DUF499*->?->?->PemK_toxin->ABhydrolase->?->ABhydrolase+ABhydrolase-> | 1113 | Bacteria>Terrabacteria_group>Chloroflexota>Anaerolineae>unclassified_Anaerolineae | GCA_016704685.1 | Anaerolineae_bacterium | IPF85_04125 | 226121..229462 | 2052143 | DUF499-domain-containing-protein-[Anaerolineae-bacterium] |
| MBK6433234.1 | DUF499_1.RM.like.systems | <-DUF499*<-REase<-OLD-ABC<-HNH | 943 | Bacteria>Terrabacteria_group>Chloroflexota>Anaerolineae>unclassified_Anaerolineae | GCA_016704685.1 | Anaerolineae_bacterium | IPF85_21385 | 32175..35006 | 2052143 | ATP-binding-protein-[Anaerolineae-bacterium] |
| MBL7062619.1 | DUF499_1.RM.like.systems | MazE_antitoxin->PIN->?->AbiEii->METHYLASE->REase->DUF499*-><-?<-ABC-ATPase+SbcC | 858 | Bacteria>Terrabacteria_group>Chloroflexota>Anaerolineae>unclassified_Anaerolineae | GCA_016783285.1 | Anaerolineae_bacterium | ISS49_00240 | 51879..54455 | 2052143 | ATP-binding-protein-[Anaerolineae-bacterium] |
| MBN1814375.1 | DUF499_1.RM.like.systems | DDE_3-><-REase<-DUF499* | 1242 | Bacteria>Terrabacteria_group>Chloroflexota>Anaerolineae>unclassified_Anaerolineae | GCA_016930605.1 | Anaerolineae_bacterium | JXA14_21225 | 14526..18254 | 2052143 | ATP-binding-protein-[Anaerolineae-bacterium] |
| MBN1921823.1 | DUF499_1.RM.like.systems | REase->DUF499*->McrC-NTD+REase-> | 924 | Bacteria>Terrabacteria_group>Chloroflexota>Anaerolineae>unclassified_Anaerolineae | GCA_016931315.1 | Anaerolineae_bacterium | JW892_11295 | 1846..4620 | 2052143 | ATP-binding-protein-[Anaerolineae-bacterium] |
| MBN2393686.1 | DUF499_1.RM.like.systems | <-DUF499*<-REase<-?<-?<-METHYLASE-p | 955 | Bacteria>Terrabacteria_group>Chloroflexota>Anaerolineae>unclassified_Anaerolineae | GCA_016934645.1 | Anaerolineae_bacterium | JXR84_23340 | 10232..13099 | 2052143 | DUF499-domain-containing-protein-[Anaerolineae-bacterium] |
| MCD6553270.1 | DUF499_1.RM.like.systems | <-DUF499*<-REase<-TPR+TPR+TPR<-?<-?<-METHYLASE | 901 | Bacteria>Terrabacteria_group>Chloroflexota>Anaerolineae>unclassified_Anaerolineae | GCA_021162125.1 | Anaerolineae_bacterium | J7M16_04575 | 6228..8933 | 2052143 | ATP-binding-protein-[Anaerolineae-bacterium] |
| MCG3212549.1 | DUF499_1.RM.like.systems | SMODS->?->METHYLASE->?->AbiEii->?->DUF499*->REase->?->?->?->?->JetA-> | 1128 | Bacteria>Terrabacteria_group>Chloroflexota>Anaerolineae>unclassified_Anaerolineae | GCA_022071955.1 | Anaerolineae_bacterium | FOGNACKC_06219 | 22074..25460 | 2052143 | hypothetical-protein-[Anaerolineae-bacterium] |
| MCG3142365.1 | DUF499_1.RM.like.systems | METHYLASE->REase->DUF499*-><-?<-?\|\|?->Phage_int_SAM_1+Phage_integrase->Phage_integrase-> | 937 | Bacteria>Terrabacteria_group>Chloroflexota>Anaerolineae>unclassified_Anaerolineae | GCA_022072245.1 | Anaerolineae_bacterium | HDKAJFGB_03811 | 2691..5504 | 2052143 | hypothetical-protein-[Anaerolineae-bacterium] |
| MCW5847519.1 | DUF499_1.RM.like.systems | METHYLASE-p->REase->HEPN+DUF499*-><-?<-?<-?<-?<-ABC-ATPase+SbcC | 1097 | Bacteria>Terrabacteria_group>Chloroflexota>Anaerolineae>unclassified_Anaerolineae | GCA_026129205.1 | Anaerolineae_bacterium | KIS95_09830 | 2201..5494 | 2052143 | MAG:-DUF499-domain-containing-protein-[Anaerolineae-bacterium] |
| MCZ2096700.1 | DUF499_1.RM.like.systems | SF1-SF2-Helicase->wHTH+SAD+HNH->ORC-CDC6+REase->HEPN+DUF499*-> | 1112 | Bacteria>Terrabacteria_group>Chloroflexota>Anaerolineae>unclassified_Anaerolineae | GCA_026954235.1 | Anaerolineae_bacterium | LC121_10630 | 39414..42752 | 2052143 | MAG:-DUF499-domain-containing-protein-[Anaerolineae-bacterium] |
| MCZ7573949.1 | DUF499_1.RM.like.systems | <-Pkinase<-?<-?<-SIGMA-HTH<-?<-DUF499*<-HTH<-PolB<-METHYLASE<-?<-PolB<-REase | 933 | Bacteria>Terrabacteria_group>Chloroflexota>Ardenticatenia>Ardenticatenales | GCA_027493995.1 | Ardenticatenaceae_bacterium | M5U01_35810 | 8592301..8595102 | 2614247 | MAG:-DUF499-domain-containing-protein-[Ardenticatenaceae-bacterium] |
| MYC61956.1 | DUF499_1.RM.like.systems | <-DUF499*<-METHYLASE-p<-zf-IS66+DDE_Tnp_IS66<-METHYLASE<-REase | 218 | Bacteria>Terrabacteria_group>Chloroflexota>Caldilineae>Caldilineales | GCA_009841145.1 | Caldilineaceae_bacterium_SB0661_bin_34 | F4X16_03815 | 38..694 | 2605256 | hypothetical-protein-[Caldilineaceae-bacterium-SB0661_bin_34] |
| GIV69804.1 | DUF499_1.RM.like.systems | Schlafen+HATPase+HTH->METHYLASE->REase-><-?\|\|DUF499*-> | 854 | Bacteria>Terrabacteria_group>Chloroflexota>Caldilineae>Caldilineales | GCA_026003635.1 | Caldilinea_sp. | KatS3mg048_2666 | 137004..139568 | 2293560 | MAG:-hypothetical-protein-KatS3mg048_2666-[Caldilinea-sp.] |
| MDE0182657.1 | DUF499_1.RM.like.systems | PIWI->METHYLASE->OLD-ABC->TPR->?->DOC+HTH->DUF499*->?->HEPN->REase+35exo+Beta-Barrel+SF1-SF2-Helicase->?->TPR+TPR-> | 1109 | Bacteria>Terrabacteria_group>Chloroflexota>Caldilineae>Caldilineales | GCA_028819655.1 | Caldilineaceae_bacterium | OXL39_15010 | 68027..71356 | 2614250 | MAG:-DUF499-domain-containing-protein-[Caldilineaceae-bacterium] |
| MCS6883598.1 | DUF499_1.RM.like.systems | PolB->HEPN->?->METHYLASE->METHYLASE-p->REase->HEPN+DUF499*-> | 1130 | Bacteria>Terrabacteria_group>Chloroflexota>Chloroflexia>Chloroflexales | GCA_025057155.1 | Chloroflexaceae_bacterium | NZU74_19900 | 13790..17182 | 2562117 | Swt1-family-HEPN-domain-containing-protein-[Chloroflexaceae-bacterium] |
| MCS7288799.1 | DUF499_1.RM.like.systems | METHYLASE-p->REase->DUF499*-><-TPR+TPR<-?<-STYKinase<-ZnR | 857 | Bacteria>Terrabacteria_group>Chloroflexota>Chloroflexia>Chloroflexales | GCA_025062325.1 | Roseiflexus_sp. | NZ699_06675 | 1458..4031 | 2562120 | DUF499-domain-containing-protein-[Roseiflexus-sp.] |
| MBN4064100.1 | DUF499_1.RM.like.systems | METHYLASE-p->HEPN+DUF499*-><-REase\|\|XIS-HTH-> | 1109 | Bacteria>Terrabacteria_group>Chloroflexota>Dehalococcoidia>Dehalococcoidales | GCA_017114025.1 | Dehalococcoides_mccartyi | JYU04_00010 | 654..3983 | 61435 | DUF499-domain-containing-protein-[Dehalococcoides-mccartyi] |
| MBK6562039.1 | DUF499_1.RM.like.systems | <-REase<-REase+DUF4268<-DUF499-p<-?<-DUF499*<-DUF499-p<-METHYLASE-p | 582 | Bacteria>Terrabacteria_group>Chloroflexota>Dehalococcoidia>unclassified_Dehalococcoidia | GCA_016704105.1 | Dehalococcoidia_bacterium | IPG14_10980 | 2567..4315 | 2026734 | DUF499-domain-containing-protein-[Dehalococcoidia-bacterium] |
| MCL0064885.1 | DUF499_1.RM.like.systems | <-TPR+TPR+TPR+TPR+TPR+TPR+TPR+TPR<-?<-DUF499*<-REase<-REase<-OLD-ABC<-METHYLASE | 925 | Bacteria>Terrabacteria_group>Chloroflexota>Dehalococcoidia>unclassified_Dehalococcoidia | GCA_023271695.1 | Dehalococcoidia_bacterium | M1N84_03205 | 1953..4730 | 2026734 | DUF499-domain-containing-protein-[Dehalococcoidia-bacterium] |
| MBN1890000.1 | DUF499_1.RM.like.systems | REase->DUF499*-><-?\|\|?->ABhydrolase->DOC+HTH-> | 987 | Bacteria>Terrabacteria_group>Chloroflexota>Thermoflexia>Thermoflexales | GCA_016931085.1 | Thermoflexales_bacterium | JW850_18535 | 987..3950 | 2497639 | ATP-binding-protein-[Thermoflexales-bacterium] |
| MBC8450530.1 | DUF499_1.RM.like.systems | <-HAD<-?\|\|METHYLASE->REase->DUF499*->ABhydrolase-> | 946 | Bacteria>Terrabacteria_group>Chloroflexota>unclassified_Chloroflexota | GCA_014383605.1 | Chloroflexota_bacterium | H8D78_22580 | 11778..14618 | 2026724 | ATP-binding-protein-[Chloroflexi-bacterium] |
| MBC8450556.1 | DUF499_1.RM.like.systems | METHYLASE-p->REase->DUF499*-> | 874 | Bacteria>Terrabacteria_group>Chloroflexota>unclassified_Chloroflexota | GCA_014383605.1 | Chloroflexota_bacterium | H8D78_22710 | 1848..4469 | 2026724 | ATP-binding-protein-[Chloroflexi-bacterium] |
| MBI1882242.1 | DUF499_1.RM.like.systems | <-HEPN+DUF499*<-REase<-METHYLASE-p<-METHYLASE-p | 401 | Bacteria>Terrabacteria_group>Chloroflexota>unclassified_Chloroflexota | GCA_016178315.1 | Chloroflexota_bacterium | HYR94_29060 | 1..1204 | 2026724 | ATP-binding-protein-[Chloroflexota-bacterium] |
| MBI3979239.1 | DUF499_1.RM.like.systems | <-PemK_toxin<-?\|\|Peptidase_M20-><-?<-?<-HEPN+DUF499*<-REase | 1136 | Bacteria>Terrabacteria_group>Chloroflexota>unclassified_Chloroflexota | GCA_016199325.1 | Chloroflexota_bacterium | HY331_13735 | 11922..15332 | 2026724 | ATP-binding-protein-[Chloroflexi-bacterium] |
| MBM3127627.1 | DUF499_1.RM.like.systems | Pkinase->?->?->?-><-SF1-SF2-Helicase+DUF1998<-?<-DUF499*<-REase<-TIR<-METHYLASE<-?<-REase+ParB<-RNase | 922 | Bacteria>Terrabacteria_group>Chloroflexota>unclassified_Chloroflexota | GCA_016865935.1 | Chloroflexota_bacterium | FJ009_03225 | 26548..29316 | 2026724 | ATP-binding-protein-[Chloroflexi-bacterium] |
| MBU1747544.1 | DUF499_1.RM.like.systems | <-DUF499*<-REase<-?<-METHYLASE | 437 | Bacteria>Terrabacteria_group>Chloroflexota>unclassified_Chloroflexota | GCA_018823415.1 | Chloroflexota_bacterium | KKA73_07640 | 1..1312 | 2026724 | DUF499-domain-containing-protein-[Chloroflexota-bacterium] |
| MBV9281297.1 | DUF499_1.RM.like.systems | METHYLASE-p->Glyoxalase->HEPN+DUF499*->REase-><-SIGMA-HTH+SIGMA-HTH<-?<-Glyoxalase | 1102 | Bacteria>Terrabacteria_group>Chloroflexota>unclassified_Chloroflexota | GCA_019245855.1 | Chloroflexota_bacterium | JOZ41_14550 | 2324..5632 | 2026724 | DUF499-domain-containing-protein-[Chloroflexi-bacterium] |
| MCX6066946.1 | DUF499_1.RM.like.systems | <-REase<-?<-?<-?<-DUF499-p<-HEPN+DUF499*<-PIN<-MazE_antitoxin | 972 | Bacteria>Terrabacteria_group>Chloroflexota>unclassified_Chloroflexota | GCA_026389995.1 | Chloroflexota_bacterium | NT121_14520 | 4682..7598 | 2026724 | MAG:-Swt1-family-HEPN-domain-containing-protein,-partial-[Chloroflexi-bacterium] |
| MDC0253131.1 | DUF499_1.RM.like.systems | REase->METHYLASE->HEPN+DUF499*-><-Mbetalac\|\|?-><-?<-?<-RNase | 1088 | Bacteria>Terrabacteria_group>Chloroflexota>unclassified_Chloroflexota | GCA_028346115.1 | Chloroflexota_bacterium | OAK52_03100 | 3358..6624 | 2026724 | DUF499-domain-containing-protein-[Chloroflexi-bacterium] |
| KAB8333560.1 | DUF499_1.RM.like.systems | REase->REase->SF1-SF2-Helicase+REase->METHYLASE->METHYLASE->?->DUF499*->WYL-> | 553 | Bacteria>Terrabacteria_group>Cyanobacteriota/Melainabacteria_group>Cyanobacteriota>Cyanophyceae | GCA_000828085.3 | Scytonema_tolypothrichoides_VB-61278 | SD80_012900 | 39454..41115 | 1233231 | hypothetical-protein-SD80_012900-[Scytonema-tolypothrichoides-VB-61278] |
| BBD57116.1 | DUF499_1.RM.like.systems | MethyltransfD12->MethyltransfD12-><-?<-HTH_3\|\|DDE_Tnp_1->?-><-DUF499*<-REase<-REase<-REase<-METHYLASE<-PIN | 1002 | Bacteria>Terrabacteria_group>Cyanobacteriota/Melainabacteria_group>Cyanobacteriota>Cyanophyceae | GCA_003609755.1 | Planktothrix_agardhii_NIES-204 | NIES204_44520 | 39433..42441 | 282423 | AAA-family-ATPase-[Planktothrix-agardhii-NIES-204] |
| TGG91622.1 | DUF499_1.RM.like.systems | REase+NACHT-><-?<-?<-HEPN+DUF499*<-METHYLASE<-REase<-?<-?<-ParB+HNH | 1109 | Bacteria>Terrabacteria_group>Cyanobacteriota/Melainabacteria_group>Cyanobacteriota>Cyanophyceae | GCA_004768415.1 | Aphanocapsa_feldmannii_277cV | ERJ67_07705 | 17520..20849 | 2507553 | ATP-binding-protein-[Aphanocapsa-feldmannii-277cV] |
| NJO76740.1 | DUF499_1.RM.like.systems | METHYLASE-p->REase->HEPN+DUF499*-> | 1125 | Bacteria>Terrabacteria_group>Cyanobacteriota/Melainabacteria_group>Cyanobacteriota>Cyanophyceae | GCA_012033355.1 | Leptolyngbyaceae_cyanobacterium_RM1_406_9 | HC833_25190 | 1227..4604 | 2720434 | ATP-binding-protein-[Leptolyngbyaceae-cyanobacterium-RM1_406_9] |
| NJR56033.1 | DUF499_1.RM.like.systems | METHYLASE-p->REase->DUF499*->Phage_integrase->?->MazE_antitoxin->DOC->DUF29-> | 968 | Bacteria>Terrabacteria_group>Cyanobacteriota/Melainabacteria_group>Cyanobacteriota>Cyanophyceae | GCA_012034865.1 | Acaryochloris_sp._CRU_2_0 | HC768_16565 | 11406..14312 | 2720402 | ATP-binding-protein-[Acaryochloris-sp.-CRU_2_0] |
| MBF2000219.1 | DUF499_1.RM.like.systems | <-HEPN+DUF499*<-REase<-REase | 1113 | Bacteria>Terrabacteria_group>Cyanobacteriota/Melainabacteria_group>Cyanobacteriota>Cyanophyceae | GCA_015272295.1 | Synechococcales_cyanobacterium_M58_A2018_015 | IGS38_05830 | 38711..42052 | 2774348 | DUF499-domain-containing-protein-[Synechococcales-cyanobacterium-M58_A2018_015] |
| MBF2075191.1 | DUF499_1.RM.like.systems | AbrB->HicB_lk_antitox->REase->REase->REase->REase->HEPN+DUF499*-> | 1114 | Bacteria>Terrabacteria_group>Cyanobacteriota/Melainabacteria_group>Cyanobacteriota>Cyanophyceae | GCA_015272325.1 | Synechococcales_cyanobacterium_C42_A2020_086 | IGS50_15730 | 92250..95594 | 2774344 | DUF499-domain-containing-protein-[Synechococcales-cyanobacterium-C42_A2020_086] |
| MBW4559185.1 | DUF499_1.RM.like.systems | METHYLASE-p->REase->DUF499*-><-RVT+HNH\|\|?-><-?<-Phage_integrase<-REase | 990 | Bacteria>Terrabacteria_group>Cyanobacteriota/Melainabacteria_group>Cyanobacteriota>Cyanophyceae | GCA_019358915.1 | Trichormus_sp._ATA11-4-KO1 | KME59_25360 | 2404..5376 | 2835091 | DUF499-domain-containing-protein-[Trichormus-sp.-ATA11-4-KO1] |
| MCA2627579.1 | DUF499_1.RM.like.systems | REase->METHYLASE->RHH_6->PIN->OLD-ABC->PIN->HEPN+DUF499*-><-Cluster164_18clades | 1130 | Bacteria>Terrabacteria_group>Cyanobacteriota/Melainabacteria_group>Cyanobacteriota>Cyanophyceae | GCA_020245485.1 | Microcystis_sp._M091S2 | IM505_05380 | 45571..48963 | 2771177 | DUF499-domain-containing-protein-[Microcystis-sp.-M091S2] |
| MCA6503991.1 | DUF499_1.RM.like.systems | METHYLASE-p->?->PIN->REase->REase->REase->HEPN+DUF499*-><-Thioredoxin\|\|tRNA-><-PIN<-Phd-YefM\|\|?->?->ABhydrolase-> | 1118 | Bacteria>Terrabacteria_group>Cyanobacteriota/Melainabacteria_group>Cyanobacteriota>Cyanophyceae | GCA_020403835.1 | Pseudanabaena_sp._M090S1SP2A07QC | IM580_15730 | 4170..7526 | 2771254 | DUF499-domain-containing-protein-[Pseudanabaena-sp.-M090S1SP2A07QC] |
| MCA6581759.1 | DUF499_1.RM.like.systems | <-HEPN+DUF499*<-REase<-REase<-REase<-PIN | 1118 | Bacteria>Terrabacteria_group>Cyanobacteriota/Melainabacteria_group>Cyanobacteriota>Cyanophyceae | GCA_020404925.1 | Pseudanabaena_sp._M34BS1SP1A06MG | IM536_06930 | 164..3520 | 2771240 | DUF499-domain-containing-protein-[Pseudanabaena-sp.-M34BS1SP1A06MG] |
| MCA6600378.1 | DUF499_1.RM.like.systems | Phd-YefM->PIN-><-tRNA\|\|Thioredoxin-><-HEPN+DUF499*<-REase<-REase<-REase<-PIN<-?<-METHYLASE-p | 1118 | Bacteria>Terrabacteria_group>Cyanobacteriota/Melainabacteria_group>Cyanobacteriota>Cyanophyceae | GCA_020404965.1 | Pseudanabaena_sp._M57BS1SP1A06MG | IM537_09260 | 3869..7225 | 2771237 | DUF499-domain-containing-protein-[Pseudanabaena-sp.-M57BS1SP1A06MG] |
| MCA6597113.1 | DUF499_1.RM.like.systems | Phd-YefM->PIN-><-tRNA\|\|Thioredoxin-><-HEPN+DUF499*<-REase<-REase<-REase<-?<-REase<-REase | 1118 | Bacteria>Terrabacteria_group>Cyanobacteriota/Melainabacteria_group>Cyanobacteriota>Cyanophyceae | GCA_020405005.1 | Pseudanabaena_sp._M046S1SP1A06QC | IM539_12615 | 33014..36370 | 2771236 | DUF499-domain-containing-protein-[Pseudanabaena-sp.-M046S1SP1A06QC] |
| MCA6605401.1 | DUF499_1.RM.like.systems | PIN->REase->REase->REase->HEPN+DUF499*-> | 895 | Bacteria>Terrabacteria_group>Cyanobacteriota/Melainabacteria_group>Cyanobacteriota>Cyanophyceae | GCA_020405035.1 | Pseudanabaena_sp._M007S1SP1A06QC | IM533_13245 | 2810..5496 | 2771232 | DUF499-domain-containing-protein-[Pseudanabaena-sp.-M007S1SP1A06QC] |
| MCE2977896.1 | DUF499_1.RM.like.systems | METHYLASE-p->?->PIN->REase->REase->REase->HEPN+DUF499*-><-Thioredoxin\|\|tRNA-><-PIN | 1118 | Bacteria>Terrabacteria_group>Cyanobacteriota/Melainabacteria_group>Cyanobacteriota>Cyanophyceae | GCA_021298775.1 | Pseudanabaena_sp._CoA8_M7 | LW851_16625 | 3735..7091 | 2904992 | DUF499-domain-containing-protein-[Pseudanabaena-sp.-CoA8_M7] |
| MCE2669063.1 | DUF499_1.RM.like.systems | Cluster164_18clades-><-HEPN+DUF499*<-PIN<-OLD-ABC<-PIN<-RHH_6<-METHYLASE<-REase | 1130 | Bacteria>Terrabacteria_group>Cyanobacteriota/Melainabacteria_group>Cyanobacteriota>Cyanophyceae | GCA_021300285.1 | Microcystis_sp._49638_E5 | LW728_07560 | 10523..13915 | 2904986 | DUF499-domain-containing-protein-[Microcystis-sp.-49638_E5] |
| CAD5939228.1 | DUF499_1.RM.like.systems | AbrB-like->?->Radical-SAM-><-?<-?<-DUF499*<-REase<-REase<-REase<-REase<-METHYLASE<-REase | 1002 | Bacteria>Terrabacteria_group>Cyanobacteriota/Melainabacteria_group>Cyanobacteriota>Cyanophyceae | GCA_904830785.1 | Planktothrix_rubescens | NO108_02187 | 2314018..2317026 | 59512 | Putative-AAA-family-ATPase-[Planktothrix-rubescens] |
| WP_008200990.1 | DUF499_1.RM.like.systems | REase->METHYLASE->?->ACET->HEPN+DUF499*->?-><-Cluster164_18clades | 1130 | Bacteria>Terrabacteria_group>Cyanobacteriota/Melainabacteria_group>Cyanobacteriota>Cyanophyceae | GCF_000297435.1 | Microcystis_sp._T1-4 | MICAI_RS09635 | 4446..7838 | 1160279 | Swt1-family-HEPN-domain-containing-protein-[Microcystis-sp.-T1-4] |
| WP_027249039.1 | DUF499_1.RM.like.systems | METHYLASE-p->METHYLASE->REase->REase->REase->REase->DUF499*->?-><-?\|\|?->RelB_antitoxin-> | 999 | Bacteria>Terrabacteria_group>Cyanobacteriota/Melainabacteria_group>Cyanobacteriota>Cyanophyceae | GCF_000464665.1 | Planktothrix_agardhii | N318_RS0105300 | 1257351..1260350 | 1160 | DUF499-domain-containing-protein-[Planktothrix-agardhii] |
| WP_026794456.1 | DUF499_1.RM.like.systems | <-RelB_antitoxin<-?\|\|?-><-?<-DUF499*<-REase<-REase<-REase<-REase<-METHYLASE<-METHYLASE-p | 999 | Bacteria>Terrabacteria_group>Cyanobacteriota/Melainabacteria_group>Cyanobacteriota>Cyanophyceae | GCF_000464725.1 | Planktothrix | N317_RS0110130 | 2122440..2125439 | 54304 | MULTISPECIES:-DUF499-domain-containing-protein-[Planktothrix] |
| WP_026788986.1 | DUF499_1.RM.like.systems | REase->METHYLASE->REase->REase->REase->REase->DUF499*->?->?->?->?-><-Radical-SAM | 999 | Bacteria>Terrabacteria_group>Cyanobacteriota/Melainabacteria_group>Cyanobacteriota>Cyanophyceae | GCF_000464785.1 | Planktothrix_rubescens | N313_RS0123225 | 1913145..1916144 | 59512 | ATP-binding-protein-[Planktothrix-rubescens] |
| WP_027255058.1 | DUF499_1.RM.like.systems | <-DUF499*<-REase<-REase<-REase<-REase<-METHYLASE<-REase | 996 | Bacteria>Terrabacteria_group>Cyanobacteriota/Melainabacteria_group>Cyanobacteriota>Cyanophyceae | GCF_000464825.1 | Planktothrix_agardhii | N316_RS0113720 | 328554..331544 | 1160 | DUF499-domain-containing-protein-[Planktothrix-agardhii] |
| WP_024971076.1 | DUF499_1.RM.like.systems | REase->METHYLASE->RHH_6->PIN->OLD-ABC->PIN->HEPN+DUF499*-> | 1130 | Bacteria>Terrabacteria_group>Cyanobacteriota/Melainabacteria_group>Cyanobacteriota>Cyanophyceae | GCF_000599945.1 | Microcystis_aeruginosa | MIC7005_RS0124495 | 6350..9742 | 1126 | Swt1-family-HEPN-domain-containing-protein-[Microcystis-aeruginosa] |
| WP_042158736.1 | DUF499_1.RM.like.systems | PIN->METHYLASE->REase->REase->REase->DUF499*-><-?\|\|RelB_antitoxin->PemK_toxin-> | 1002 | Bacteria>Terrabacteria_group>Cyanobacteriota/Melainabacteria_group>Cyanobacteriota>Cyanophyceae | GCF_000710505.1 | Planktothrix_agardhii | NMK38_RS22725 | 84961..87969 | 1160 | DUF499-domain-containing-protein-[Planktothrix-agardhii] |
| WP_046277358.1 | DUF499_1.RM.like.systems | METHYLASE-p->REase->DUF499*-><-Pkinase\|\|?->?-><-NTP_transf_9<-TPR+TPR+TPR+TPR+TPR+TPR+TPR+TPR | 968 | Bacteria>Terrabacteria_group>Cyanobacteriota/Melainabacteria_group>Cyanobacteriota>Cyanophyceae | GCF_000972705.2 | Limnoraphis_robusta | WN50_RS27095 | 1686..4592 | 1118279 | DUF499-domain-containing-protein-[Limnoraphis-robusta] |
| WP_073075160.1 | DUF499_1.RM.like.systems | CR-Trypsin1->?-><-RVT+HNH<-DUF499*<-REase<-REase<-METHYLASE<-DUF4926+DUF2281 | 965 | Bacteria>Terrabacteria_group>Cyanobacteriota/Melainabacteria_group>Cyanobacteriota>Cyanophyceae | GCF_001895925.1 | Phormidesmis_priestleyi | C7B65_RS24715 | 5123..8020 | 268141 | DUF499-domain-containing-protein-[Phormidesmis-priestleyi] |
| WP_106371305.1 | DUF499_1.RM.like.systems | <-STYKIN\|\|?-><-SIR2+SIR2<-DUF499*<-REase<-METHYLASE<-PIN | 994 | Bacteria>Terrabacteria_group>Cyanobacteriota/Melainabacteria_group>Cyanobacteriota>Cyanophyceae | GCF_003003885.1 | Chlorogloea_sp._CCALA_695 | C7B70_RS23190 | 7463..10447 | 2107693 | DUF499-domain-containing-protein-[Chlorogloea-sp.-CCALA-695] |
| WP_168360489.1 | DUF499_1.RM.like.systems | <-REase+HTH\|\|?-><-?\|\|?-><-?<-?<-DUF499*<-?<-?<-MetJarc-HTH<-REase<-METHYLASE | 967 | Bacteria>Terrabacteria_group>Cyanobacteriota/Melainabacteria_group>Cyanobacteriota>Cyanophyceae | GCF_009712075.2 | Dolichospermum_planctonicum | FJR41_RS21600 | 23214..26117 | 136072 | DUF499-domain-containing-protein-[Dolichospermum-planctonicum] |
| WP_190397100.1 | DUF499_1.RM.like.systems | METHYLASE->REase->MetJarc-HTH->PemK_toxin->DUF499*->?->?-><-?\|\|?-><-?\|\|REase+HTH-> | 969 | Bacteria>Terrabacteria_group>Cyanobacteriota/Melainabacteria_group>Cyanobacteriota>Cyanophyceae | GCF_014696335.1 | Anabaena_sp._FACHB-1250 | H6F39_RS17030 | 5832..8741 | 2692770 | DUF499-domain-containing-protein-[Anabaena-sp.-FACHB-1250] |
| WP_190371458.1 | DUF499_1.RM.like.systems | <-REase+HTH<-?\|\|?-><-?<-?<-DUF499*<-REase<-METHYLASE | 969 | Bacteria>Terrabacteria_group>Cyanobacteriota/Melainabacteria_group>Cyanobacteriota>Cyanophyceae | GCF_014696755.1 | Anabaena_sp._FACHB-1391 | H6F62_RS04790 | 144117..147026 | 2692771 | DUF499-domain-containing-protein-[Anabaena-sp.-FACHB-1391] |
| WP_192216664.1 | DUF499_1.RM.like.systems | <-HTH\|\|?->?-><-?<-GDResolvase+Recombinase+ZnR<-?<-DUF499*<-REase<-REase<-METHYLASE-p | 969 | Bacteria>Terrabacteria_group>Cyanobacteriota/Melainabacteria_group>Cyanobacteriota>Cyanophyceae | GCF_014849525.1 | Microcoleus_sp._FACHB-SPT15 | H6F98_RS00230 | 47740..50649 | 2692830 | DUF499-domain-containing-protein-[Microcoleus-sp.-FACHB-SPT15] |
| WP_200988278.1 | DUF499_1.RM.like.systems | <-REase<-?<-?<-DUF499*<-CR-ATPase9+REase<-DUF499<-CR-ATPase9+REase<-METHYLASE<-REase<-PIN | 950 | Bacteria>Terrabacteria_group>Cyanobacteriota/Melainabacteria_group>Cyanobacteriota>Cyanophyceae | GCF_015245355.1 | Anabaenopsis_elenkinii | IM676_RS18785 | 4299000..4301852 | 156213 | DUF499-domain-containing-protein-[Anabaenopsis-elenkinii] |
| WP_225880403.1 | DUF499_1.RM.like.systems | <-REase<-?<-?<-DUF499<-CR-ATPase9+REase<-DUF499*<-CR-ATPase9+REase<-METHYLASE<-REase<-PIN<-MetJarc-HTH | 821 | Bacteria>Terrabacteria_group>Cyanobacteriota/Melainabacteria_group>Cyanobacteriota>Cyanophyceae | GCF_015245355.1 | Anabaenopsis_elenkinii | IM676_RS18795 | 4304033..4306498 | 156213 | DUF499-domain-containing-protein-[Anabaenopsis-elenkinii] |
| WP_227366555.1 | DUF499_1.RM.like.systems | GspA-ATPase->METHYLASE-p->REase->REase->REase->DUF499*-><-TOPRIM+SF1-SF2-Helicase\|\|Phage_int_SAM_1+Phage_integrase->?->TUDOR+SF1-SF2-Helicase+iREase+CTD->METHYLASE-p->REase-> | 1002 | Bacteria>Terrabacteria_group>Cyanobacteriota/Melainabacteria_group>Cyanobacteriota>Cyanophyceae | GCF_020594525.1 | Planktothrix_agardhii | LJE15_RS21495 | 6203..9211 | 1160 | DUF499-domain-containing-protein-[Planktothrix-agardhii] |
| WP_227382638.1 | DUF499_1.RM.like.systems | PIN->METHYLASE->REase->REase->REase->DUF499*-><-?\|\|RelB_antitoxin->PemK_toxin->?->RHH_1-><-TOPRIM+SF1-SF2-Helicase | 1002 | Bacteria>Terrabacteria_group>Cyanobacteriota/Melainabacteria_group>Cyanobacteriota>Cyanophyceae | GCF_020594575.1 | Planktothrix_agardhii | LJE16_RS21930 | 10143..13151 | 1160 | DUF499-domain-containing-protein-[Planktothrix-agardhii] |
| WP_227398372.1 | DUF499_1.RM.like.systems | ZnR->TOPRIM+SF1-SF2-Helicase-><-PemK_toxin<-RelB_antitoxin<-?\|\|?-><-DUF499*<-REase<-REase<-REase<-METHYLASE<-PIN | 1002 | Bacteria>Terrabacteria_group>Cyanobacteriota/Melainabacteria_group>Cyanobacteriota>Cyanophyceae | GCF_020594695.1 | Planktothrix_agardhii | L2E74_RS21495 | 55743..58751 | 1160 | DUF499-domain-containing-protein-[Planktothrix-agardhii] |
| WP_233748345.1 | DUF499_1.RM.like.systems | <-Peptidase<-HEPN+DUF499*<-CopG/RHH<-PemK_toxin<-?<-REase<-REase<-METHYLASE | 1125 | Bacteria>Terrabacteria_group>Cyanobacteriota/Melainabacteria_group>Cyanobacteriota>Cyanophyceae | GCF_021379005.1 | Leptodesmis_sichuanensis | KIK02_RS09445 | 2037020..2040397 | 2906798 | Swt1-family-HEPN-domain-containing-protein-[Leptodesmis-sichuanensis] |
| WP_235278697.1 | DUF499_1.RM.like.systems | METHYLASE->?->PemK_toxin->REase->REase->CopG/RHH->HEPN+DUF499*->?->?->MORC+DNA_gyraseB+TOPRIM-> | 1109 | Bacteria>Terrabacteria_group>Cyanobacteriota/Melainabacteria_group>Cyanobacteriota>Cyanophyceae | GCF_021533835.1 | Synechococcus_sp._Nb3U1 | L1047_RS09935 | 2086840..2090169 | 1914529 | Swt1-family-HEPN-domain-containing-protein-[Synechococcus-sp.-Nb3U1] |
| WP_235753326.1 | DUF499_1.RM.like.systems | PIN->METHYLASE->REase->REase->REase->DUF499*-><-?\|\|RelB_antitoxin->HTH_OrfB_IS605+OrfB_IS605+OrfB_Zn_ribbon-><-PAIREDC-HTH | 1002 | Bacteria>Terrabacteria_group>Cyanobacteriota/Melainabacteria_group>Cyanobacteriota>Cyanophyceae | GCF_021554825.1 | Planktothrix_agardhii | L2E72_RS21865 | 12950..15958 | 1160 | DUF499-domain-containing-protein-[Planktothrix-agardhii] |
| WP_247214939.1 | DUF499_1.RM.like.systems | DDE_Tnp_1-><-Peptidase_M24\|\|?->tRNA-><-HEPN+DUF499*<-REase<-REase<-REase<-METHYLASE<-PIN<-AbrB | 1107 | Bacteria>Terrabacteria_group>Cyanobacteriota/Melainabacteria_group>Cyanobacteriota>Cyanophyceae | GCF_022984075.1 | Synechococcus_sp._C9 | MLD66_RS00195 | 39889..43212 | 102119 | Swt1-family-HEPN-domain-containing-protein-[Synechococcus-sp.-C9] |
| WP_250012765.1 | DUF499_1.RM.like.systems | AbrB->PIN->REase->METHYLASE-p->METHYLASE->?->HEPN+DUF499*->?-><-?\|\|Phage_integrase-><-Phage_integrase | 1113 | Bacteria>Terrabacteria_group>Cyanobacteriota/Melainabacteria_group>Cyanobacteriota>Cyanophyceae | GCF_023546805.1 | Chroococcidiopsis_sp._CCNUC1 | M5J74_RS19090 | 4105692..4109033 | 2653189 | Swt1-family-HEPN-domain-containing-protein-[Chroococcidiopsis-sp.-CCNUC1] |
| WP_272094882.1 | DUF499_1.RM.like.systems | <-HEPN+DUF499*<-PIN<-OLD-ABC<-PIN<-RHH_6<-METHYLASE<-REase | 1065 | Bacteria>Terrabacteria_group>Cyanobacteriota/Melainabacteria_group>Cyanobacteriota>Cyanophyceae | GCF_028330705.1 | Microcystis_aeruginosa | PN479_RS02395 | 1..3194 | 1126 | Swt1-family-HEPN-domain-containing-protein,-partial-[Microcystis-aeruginosa] |
| WP_280650398.1 | DUF499_1.RM.like.systems | <-sGTP\|\|?->?->?->Peptidase_S51-><-DUF499*<-REase<-METHYLASE-p<-Cysteine_desulfurase | 953 | Bacteria>Terrabacteria_group>Cyanobacteriota/Melainabacteria_group>Cyanobacteriota>Cyanophyceae | GCF_029891655.1 | Umezakia_ovalisporum | NWP27_RS05680 | 14158..17019 | 75695 | DUF499-domain-containing-protein-[Umezakia-ovalisporum] |
| WP_280699380.1 | DUF499_1.RM.like.systems | <-sGTP\|\|?->?->?->Peptidase_S51-><-DUF499*<-REase<-METHYLASE-p<-Cysteine_desulfurase | 953 | Bacteria>Terrabacteria_group>Cyanobacteriota/Melainabacteria_group>Cyanobacteriota>Cyanophyceae | GCF_029891685.1 | Umezakia_ovalisporum | NWP25_RS10975 | 18984..21845 | 75695 | DUF499-domain-containing-protein-[Umezakia-ovalisporum] |
| WP_280700572.1 | DUF499_1.RM.like.systems | Cysteine_desulfurase->METHYLASE-p->REase->DUF499*-><-Peptidase_S51<-?<-?<-?\|\|sGTP-> | 953 | Bacteria>Terrabacteria_group>Cyanobacteriota/Melainabacteria_group>Cyanobacteriota>Cyanophyceae | GCF_029891755.1 | Umezakia_ovalisporum | NWP23_RS06515 | 33643..36504 | 75695 | DUF499-domain-containing-protein-[Umezakia-ovalisporum] |
| WP_280656504.1 | DUF499_1.RM.like.systems | <-sGTP\|\|?->?->?->Peptidase_S51-><-DUF499*<-REase<-METHYLASE-p<-Cysteine_desulfurase | 953 | Bacteria>Terrabacteria_group>Cyanobacteriota/Melainabacteria_group>Cyanobacteriota>Cyanophyceae | GCF_029891765.1 | Umezakia_ovalisporum | NWP26_RS00065 | 19111..21972 | 75695 | DUF499-domain-containing-protein-[Umezakia-ovalisporum] |
| WP_280687902.1 | DUF499_1.RM.like.systems | <-sGTP\|\|?->?->?->Peptidase_S51-><-DUF499*<-REase<-METHYLASE-p<-Cysteine_desulfurase | 953 | Bacteria>Terrabacteria_group>Cyanobacteriota/Melainabacteria_group>Cyanobacteriota>Cyanophyceae | GCF_029891835.1 | Umezakia_ovalisporum | NWP18_RS13190 | 14598..17459 | 75695 | DUF499-domain-containing-protein-[Umezakia-ovalisporum] |
| WP_280689836.1 | DUF499_1.RM.like.systems | <-sGTP\|\|?->?->?->Peptidase_S51-><-DUF499*<-REase<-METHYLASE-p<-Cysteine_desulfurase | 973 | Bacteria>Terrabacteria_group>Cyanobacteriota/Melainabacteria_group>Cyanobacteriota>Cyanophyceae | GCF_029891865.1 | Umezakia_ovalisporum | NWP15_RS06365 | 19189..22110 | 75695 | DUF499-domain-containing-protein-[Umezakia-ovalisporum] |
| WP_281006617.1 | DUF499_1.RM.like.systems | METHYLASE->?->PIN->REase->REase->REase->HEPN+DUF499*-> | 1118 | Bacteria>Terrabacteria_group>Cyanobacteriota/Melainabacteria_group>Cyanobacteriota>Cyanophyceae | GCF_029910235.1 | Pseudanabaena_galeata | OA858_RS18405 | 4059827..4063183 | 1112103 | Swt1-family-HEPN-domain-containing-protein-[Pseudanabaena-galeata] |
| WP_285965028.1 | DUF499_1.RM.like.systems | XIS-HTH-><-?<-Phage_integrase\|\|tRNA-><-?\|\|REase-><-DUF499-p<-HEPN+DUF499*<-AbiEii | 1005 | Bacteria>Terrabacteria_group>Cyanobacteriota/Melainabacteria_group>Cyanobacteriota>Cyanophyceae | GCF_030285465.1 | Oscillatoria_laete-virens | QQ056_RS17150 | 57885..60902 | 1565316 | Swt1-family-HEPN-domain-containing-protein-[Oscillatoria-laete-virens] |
| WP_214559734.1 | DUF499_1.RM.like.systems | <-RelB_antitoxin<-?<-?\|\|?-><-?<-DUF499*<-REase<-REase<-REase<-REase<-METHYLASE<-METHYLASE-p | 999 | Bacteria>Terrabacteria_group>Cyanobacteriota/Melainabacteria_group>Cyanobacteriota>Cyanophyceae | GCF_903969095.1 | Planktothrix_agardhii | KM073_RS08965 | 67038..70037 | 1160 | DUF499-domain-containing-protein-[Planktothrix-agardhii] |
| WP_254171845.1 | DUF499_1.RM.like.systems | PIN->METHYLASE->REase->REase->REase->REase->DUF499*-><-?\|\|RelB_antitoxin->HTH_OrfB_IS605+OrfB_IS605+OrfB_Zn_ribbon->?->RHH_1-> | 1002 | Bacteria>Terrabacteria_group>Cyanobacteriota/Melainabacteria_group>Cyanobacteriota>Cyanophyceae | GCF_904830885.1 | Planktothrix_agardhii | NMK31_RS00300 | 70820..73828 | 1160 | DUF499-domain-containing-protein-[Planktothrix-agardhii] |
| WP_254035908.1 | DUF499_1.RM.like.systems | PIN->PIN->METHYLASE->REase->REase->REase->DUF499*-><-?\|\|RelB_antitoxin->PemK_toxin->RHH_1-><-TOPRIM+SF1-SF2-Helicase | 999 | Bacteria>Terrabacteria_group>Cyanobacteriota/Melainabacteria_group>Cyanobacteriota>Cyanophyceae | GCF_904830935.1 | Planktothrix_agardhii | NMG88_RS22325 | 51359..54358 | 1160 | DUF499-domain-containing-protein-[Planktothrix-agardhii] |
| WP_254031531.1 | DUF499_1.RM.like.systems | <-RelB_antitoxin<-?\|\|?-><-?<-DUF499*<-REase<-REase<-REase<-REase<-METHYLASE<-DDE_Tnp_1_assoc+DDE_Tnp_1 | 999 | Bacteria>Terrabacteria_group>Cyanobacteriota/Melainabacteria_group>Cyanobacteriota>Cyanophyceae | GCF_904830945.1 | Planktothrix_agardhii | NMG75_RS16735 | 3649445..3652444 | 1160 | DUF499-domain-containing-protein-[Planktothrix-agardhii] |
| WP_250683155.1 | DUF499_1.RM.like.systems | <-DUF499*<-HEPN<-REase<-METHYLASE<-PemK_toxin<-AbrB<-PolB | 788 | Bacteria>Terrabacteria_group>Cyanobacteriota/Melainabacteria_group>Cyanobacteriota>Cyanophyceae | - | Thermosynechococcus_sp._HN-54 | NBE99_RS03710 | 771476..773842 | 2933959 | DUF499-domain-containing-protein-[Thermosynechococcus-sp.-HN-54] |
| OWY64929.1 | DUF499_1.RM.like.systems | METHYLASE->REase->DUF499*-><-?\|\|HTH_3-><-?<-?<-SbcC+SidE<-METHYLASE+SF1-SF2-Helicase | 921 | Bacteria>Terrabacteria_group>Cyanobacteriota/Melainabacteria_group>Cyanobacteriota>unclassified_Cyanobacteriota | GCA_002213405.1 | cyanobacterium_TDX16 | B7486_44590 | 3155..5920 | 1503470 | AAA-family-ATPase-[cyanobacterium-TDX16] |
| MBU6346401.1 | DUF499_1.RM.like.systems | <-Peptidase_C14<-?<-REase<-HEPN+DUF499*<-METHYLASE | 1103 | Bacteria>Terrabacteria_group>Cyanobacteriota/Melainabacteria_group>Cyanobacteriota>unclassified_Cyanobacteriota | GCA_018969285.1 | Cyanobacteria_bacterium_REEB494 | KGR70_14045 | 15811..19122 | 2800107 | DUF499-domain-containing-protein-[Cyanobacteria-bacterium-REEB494] |
| MCY4332863.1 | DUF499_1.RM.like.systems | ABC-ATPase+SbcC-><-HEPN+DUF499*<-METHYLASE<-wHTH+REase | 1113 | Bacteria>Terrabacteria_group>Cyanobacteriota/Melainabacteria_group>Cyanobacteriota>unclassified_Cyanobacteriota | GCA_026710375.1 | Cyanobacteria_bacterium_MAG_CAR1_bin_15 | OXC96_10300 | 26378..29719 | 2775589 | MAG:-DUF499-domain-containing-protein-[Cyanobacteria-bacterium-MAG-CAR1_bin_15] |
| MDA0267772.1 | DUF499_1.RM.like.systems | METHYLASE->?->PIN->REase->DUF499*-> | 1000 | Bacteria>Terrabacteria_group>Cyanobacteriota/Melainabacteria_group>Cyanobacteriota>unclassified_Cyanobacteriota | GCA_027592485.1 | Cyanobacteriota_bacterium | O3A14_12630 | 3513..6515 | 2847862 | MAG:-DUF499-domain-containing-protein-[Cyanobacteria-bacterium] |
| WP_150118021.1 | DUF499_1.RM.like.systems | DUF499*->ORC-CDC6+REase-> | 664 | Bacteria>Terrabacteria_group>Deinococcota>Deinococci>Thermales | GCF_001623175.1 | Meiothermus_taiwanensis | A3962_RS14295 | 1..1997 | 172827 | DUF499-domain-containing-protein,-partial-[Meiothermus-taiwanensis] |
| WP_126188140.1 | DUF499_1.RM.like.systems | ParB-><-DUF499*<-REase\|\|DDE_Tnp_1_assoc-> | 924 | Bacteria>Terrabacteria_group>Deinococcota>Deinococci>Thermales | GCF_003958985.1 | Thermus_scotoductus | CSW50_RS14435 | 4946..7720 | 37636 | DUF499-domain-containing-protein-[Thermus-scotoductus] |
| MBF0572396.1 | DUF499_1.RM.like.systems | <-CHASE2+ACYC<-?<-?<-DUF499*<-REase<-PIN<-?<-?<-METHYLASE-p | 936 | Bacteria>Thermodesulfobacteriota>Desulfobacteria>Desulfobacterales>Desulfobacteraceae | GCA_015233755.1 | Desulfamplus_sp. | HQK69_01355 | 13339..16149 | 2006184 | ATP-binding-protein-[Desulfamplus-sp.] |
| PIP36530.1 | DUF499_1.RM.like.systems | <-ACET<-SAM-methylase<-?<-DUF499-p<-DUF499-p<-DUF499*<-ORC-CDC6+REase<-?<-MqsA_antitoxin+HTH_3<-MqsR_toxin | 497 | Bacteria>Thermodesulfobacteriota>Desulfobacteria>Desulfobacterales>unclassified_Desulfobacterales | GCA_002771315.1 | Desulfobacterales_bacterium_CG23_combo_of_CG06-09_8_20_14_all_52_9 | COX20_05265 | 9529..11022 | 1973984 | hypothetical-protein-COX20_05265-[Desulfobacterales-bacterium-CG23_combo_of_CG06-09_8_20_14_all_52_9] |
| MCG2778998.1 | DUF499_1.RM.like.systems | <-PolB<-?<-?<-DUF499*<-REase<-METHYLASE<-VirB4-FtsK+HerA | 920 | Bacteria>Thermodesulfobacteriota>Desulfobacteria>Desulfobacterales>unclassified_Desulfobacterales | GCA_022013615.1 | Desulfobacterales_bacterium | L6406_25260 | 2158..4920 | 2044940 | DUF499-domain-containing-protein-[Desulfobacterales-bacterium] |
| HHL35500.1 | DUF499_1.RM.like.systems | <-DUF499*<-DUF1828+REase | 826 | Bacteria>Thermodesulfobacteriota>Desulfobulbia>Desulfobulbales>Desulfobulbaceae | GCA_011322715.1 | Desulfobulbaceae_bacterium | ENJ30_14165 | 1..2478 | 2053307 | ATP-binding-protein-[Desulfobulbaceae-bacterium] |
| MBA3013967.1 | DUF499_1.RM.like.systems | DUF499*->REase->HEPN->OLD-ABC+TOPRIM->SF1-SF2-Helicase-><-METHYLASE+SF1-SF2-Helicase | 944 | Bacteria>Thermodesulfobacteriota>Desulfobulbia>Desulfobulbales>Desulfobulbaceae | GCA_013792005.1 | Desulfobulbaceae_bacterium | FP815_03320 | 13..2847 | 2053307 | ATP-binding-protein-[Desulfobulbaceae-bacterium] |
| MBQ9405898.1 | DUF499_1.RM.like.systems | PfkB->?-><-DUF499*\|\|?-><-REase<-REase<-METHYLASE-p<-METHYLASE-p<-HicB | 1096 | Bacteria>Thermodesulfobacteriota>Desulfovibrionia>Desulfovibrionales>Desulfovibrionaceae | GCA_017454855.1 | Desulfovibrio_sp. | IJU37_04050 | 4592..7882 | 885 | ATP-binding-protein-[Desulfovibrio-sp.] |
| MCG2742124.1 | DUF499_1.RM.like.systems | Aminotran_1_2->?->?->?->DNA_pol3_delta+DNA_pol3_gamma3-><-DUF499*<-AAA-ATPase+REase<-METHYLASE<-?<-Schlafen+HATPase+HTH | 1098 | Bacteria>Thermodesulfobacteriota>Syntrophia>Syntrophales>Syntrophaceae | GCA_022013455.1 | Syntrophaceae_bacterium | L6300_18065 | 15618..18914 | 2052184 | DUF499-domain-containing-protein-[Syntrophaceae-bacterium] |
| NSW87838.1 | DUF499_1.RM.like.systems | <-DUF499*<-METHYLASE-p<-HEPN<-wHTH+REase<-REase | 698 | Bacteria>Thermodesulfobacteriota>Syntrophobacteria>Syntrophobacterales>Syntrophobacteraceae | GCA_013314815.1 | Syntrophobacteraceae_bacterium | HPY84_16135 | 1106..3202 | 2052185 | DUF499-domain-containing-protein-[Syntrophobacteraceae-bacterium] |
| MCD6238356.1 | DUF499_1.RM.like.systems | <-DUF499*<-REase<-METHYLASE<-?\|\|?->?-><-PIWI | 935 | Bacteria>Thermotogota>unclassified_Thermotogota | GCA_021158015.1 | Thermotogota_bacterium | J7K51_03355 | 264..3071 | 2053689 | ATP-binding-protein-[Thermotogae-bacterium] |
| KAF0250487.1 | DUF499_1.RM.like.systems | Phd-YefM->PIN->METHYLASE->REase->OLD-ABC->TOPRIM->HEPN+DUF499*->?-><-Phage_integrase\|\|?->MqsA_antitoxin+HTH_3-> | 1130 | Bacteria>unclassified_Bacteria | GCA_009773835.1 | bacterium | FD167_113 | 11053..14445 | 1869227 | hypothetical-protein-FD167_113-[bacterium] |
| MCI0602497.1 | DUF499_1.RM.like.systems | <-XIS-HTH\|\|?-><-?<-HNH<-DUF499*<-REase<-METHYLASE<-HEPN+REase | 922 | Bacteria>unclassified_Bacteria | GCA_022616075.1 | bacterium | L0156_05740 | 5054..7822 | 1869227 | DUF499-domain-containing-protein-[bacterium] |
| MCR9296473.1 | DUF499_1.RM.like.systems | <-Phage_integrase<-?<-DUF499*<-REase | 985 | Bacteria>unclassified_Bacteria | GCA_024743355.1 | bacterium | NXI32_27500 | 1811..4768 | 1869227 | DUF499-domain-containing-protein-[bacterium] |
| MCY4630094.1 | DUF499_1.RM.like.systems | REase->METHYLASE-p->METHYLASE-p->DUF499*-><-?\|\|CR-ATPase9+REase->?-><-?<-REase-p | 1004 | Bacteria>unclassified_Bacteria | GCA_026713745.1 | bacterium | OXE75_04305 | 3540..6554 | 1869227 | MAG:-DUF499-domain-containing-protein-[bacterium] |

| Protein-ID | System | Gene-Neighbourhoods | Length | Taxonomy | Assembly-ID | Strain | Locus-Tag | Genomic-coordinates | TaxID | Fasta-defline |
|---|---|---|---|---|---|---|---|---|---|---|
| -- | | | | | | | | | | |
| -- Schlafen+HATPase+wHTH | | | | | | | | | | |
| -- | | | | | | | | | | |
| CBH38392.1 | DUF499_1.RM.like.systems | <-HicB_lk_antitox\|\|?->METHYLASE-p->TUDOR+SF1-SF2-Helicase+iREase+CTD+REase->Schlafen->METHYLASE->DUF499*->?-><-?<-?<-?<-SPOUT | 900 | Archaea>environmental_samples | - | uncultured_archaeon | BSM_18690 | 1759679..1762381 | 115547 | conserved-hypothetical-protein-[uncultured-archaeon] |
| WP_167816002.1 | DUF499_1.RM.like.systems | Schlafen+HATPase+wHTH->?->Schlafen+HATPase+wHTH->METHYLASE->OLD-ABC->TOPRIM->HEPN+DUF499*-> | 756 | Archaea>Euryarchaeota>Stenosarchaea_group>Methanomicrobia>Methanomicrobiales | GCF_012062385.1 | Methanocorpusculum_sp._GPch4 | EYA77_RS04465 | 15957..18225 | 2527877 | Swt1-family-HEPN-domain-containing-protein,-partial-[Methanocorpusculum-sp.-GPch4] |
| MCG7853928.1 | DUF499_1.RM.like.systems | <-DUF499*<-METHYLASE<-?<-Schlafen+HATPase+wHTH<-TUDOR+SF1-SF2-Helicase+iREase+CTD+REase | 377 | Archaea>Euryarchaeota>Stenosarchaea_group>Methanomicrobia>Methanosarcinales | GCA_022353215.1 | Methanosarcinaceae_archaeon | MI092_15510 | 1..1132 | 1945595 | DUF499-domain-containing-protein-[Methanosarcinaceae-archaeon] |
| AET64040.1 | DUF499_1.RM.like.systems | TUDOR+SF1-SF2-Helicase+iREase+CTD+REase->Schlafen+HATPase+wHTH->METHYLASE->REase-><-HicB_lk_antitox\|\|DUF499*->HicB_lk_antitox-> | 1010 | Archaea>Euryarchaeota>Stenosarchaea_group>Methanomicrobia>Methanotrichales | GCA_000235565.1 | Methanothrix_harundinacea_6Ac | Mhar_0662 | 698704..701736 | 1110509 | ATPase-(AAA+-superfamily)-like-protein-[Methanosaeta-harundinacea-6Ac] |
| MBN1235613.1 | DUF499_1.RM.like.systems | <-DUF499*<-REase<-ASCH<-PIN+ACET+EVE<-METHYLASE<-Schlafen+HATPase+wHTH<-TUDOR+SF1-SF2-Helicase+iREase+CTD+REase | 1051 | Archaea>Euryarchaeota>Stenosarchaea_group>Methanomicrobia>Methanotrichales | GCA_016927055.1 | Methanotrichaceae_archaeon | JW999_06125 | 575..3730 | 2595067 | DUF499-domain-containing-protein-[Methanotrichaceae-archaeon] |
| WP_048144800.1 | DUF499_1.RM.like.systems | TUDOR+SF1-SF2-Helicase+iREase+CTD+REase->Schlafen+HATPase+wHTH->METHYLASE->REase-><-HicB_lk_antitox\|\|DUF499*->RelB->HicB_lk_antitox-> | 987 | Archaea>Euryarchaeota>Stenosarchaea_group>Methanomicrobia>Methanotrichales | GCF_000235565.1 | Methanothrix_harundinacea | MHAR_RS03425 | 698773..701736 | 301375 | DUF499-domain-containing-protein-[Methanothrix-harundinacea] |
| MXX20696.1 | DUF499_1.RM.like.systems | PIN->Schlafen+HATPase+wHTH->DUF499*->METHYLASE->SF1-SF2-Helicase+iREase+CTD+REase-> | 822 | Archaea>TACK_group>Nitrososphaerota>Cenarchaeales>Cenarchaeaceae | GCA_009836545.1 | Cenarchaeum_sp._SB0667_bin_13 | F4Z46_02780 | 8230..10698 | 2604878 | ATP-binding-protein-[Cenarchaeum-sp.-SB0667_bin_13] |
| MBA3712812.1 | DUF499_1.RM.like.systems | Schlafen+HATPase+wHTH->?->METHYLASE->DUF433->PIN->HEPN+DUF499*->TIR+PNTD2->PIWI->?->Phage_integrase-> | 1118 | Bacteria>Acidobacteriota>Blastocatellia>Blastocatellales>Pyrinomonadaceae | GCA_013816505.1 | Pyrinomonadaceae_bacterium | HOW76_10240 | 5538..8894 | 2283092 | ATP-binding-protein-[Pyrinomonadaceae-bacterium] |
| MBK9166737.1 | DUF499_1.RM.like.systems | Schlafen+HATPase+wHTH->METHYLASE-p-><-rve+rve_3<-HTH_Tnp_1\|\|METHYLASE-p->ZDNA-HTH->HEPN+DUF499*->?->?->ABhydrolase-> | 1121 | Bacteria>Acidobacteriota>Terriglobia>Bryobacterales>unclassified_Bryobacterales | GCA_016716385.1 | Bryobacterales_bacterium | IPM24_04655 | 396638..400003 | 2026791 | DUF499-domain-containing-protein-[Bryobacterales-bacterium] |
| MCC6263216.1 | DUF499_1.RM.like.systems | <-DUF499*<-Schlafen+HATPase+wHTH<-METHYLASE<-TUDOR+SF1-SF2-Helicase+iREase+CTD+REase | 926 | Bacteria>Acidobacteriota>Terriglobia>Bryobacterales>unclassified_Bryobacterales | GCA_020847315.1 | Bryobacterales_bacterium | IT169_06530 | 310..3090 | 2026791 | ATP-binding-protein-[Bryobacterales-bacterium] |
| MCG3192277.1 | DUF499_1.RM.like.systems | TUDOR+SF1-SF2-Helicase+iREase+CTD+REase->OLD-ABC->TOPRIM->METHYLASE->PHP+SbcC->Schlafen+HATPase+wHTH->HEPN+DUF499*->?->?->PAIREDC-HTH-> | 1114 | Bacteria>Acidobacteriota>Thermoanaerobaculia>unclassified_Thermoanaerobaculia | GCA_022071945.1 | Thermoanaerobaculia_bacterium | DIJKHBIC_01515 | 140917..144261 | 2651171 | hypothetical-protein-[Thermoanaerobaculia-bacterium] |
| ODS52398.1 | DUF499_1.RM.like.systems | <-MethyltransfD12<-ParB<-HEPN+DUF499*<-METHYLASE<-Schlafen+HATPase+wHTH<-TUDOR+SF1-SF2-Helicase+iREase+CTD+REase<-?\|\|?->Peptidase_S41-> | 1114 | Bacteria>Acidobacteriota>unclassified_Acidobacteriota | GCA_001724025.1 | Acidobacteria_bacterium_SCN_69-37 | ABS36_17605 | 152750..156094 | 1660160 | AAA+-family-ATPase-[Acidobacteria-bacterium-SCN-69-37] |
| RIK10616.1 | DUF499_1.RM.like.systems | DOC->?->MPTase->METHYLASE->HEPN+DUF499*-><-?<-Schlafen<-?<-tRNA<-tRNA<-?\|\|tRNA-><-METHYLASE-p | 1113 | Bacteria>Acidobacteriota>unclassified_Acidobacteriota | GCA_003577105.1 | Acidobacteriota_bacterium | DCC49_02885 | 59433..62774 | 1978231 | AAA+-family-ATPase-[Acidobacteria-bacterium] |
| MYN68213.1 | DUF499_1.RM.like.systems | TUDOR+SF1-SF2-Helicase+iREase+CTD+REase->Schlafen+HATPase+wHTH->HEPN->METHYLASE->OLD-ABC->TOPRIM->DUF499*-><-?<-SAM-methylase<-ACET+ACET<-GspA-ATPase | 1101 | Bacteria>Acidobacteriota>unclassified_Acidobacteriota | GCA_009861545.1 | Acidobacteriota_bacterium | F4X11_24880 | 56553..59858 | 1978231 | ATP-binding-protein-[Acidobacteria-bacterium] |
| MBI1792208.1 | DUF499_1.RM.like.systems | TUDOR+SF1-SF2-Helicase+iREase+CTD+REase->Schlafen+HATPase+wHTH->METHYLASE->PolB->HEPN+DUF499*->OLD-ABC+TOPRIM-> | 1093 | Bacteria>Acidobacteriota>unclassified_Acidobacteriota | GCA_016178945.1 | Acidobacteriota_bacterium | HYR60_32190 | 8610..11891 | 1978231 | ATP-binding-protein-[Acidobacteria-bacterium] |
| MBI3951052.1 | DUF499_1.RM.like.systems | Schlafen+HATPase+wHTH->MNS->HEPN->METHYLASE->?->HicA_toxin->DUF499*-> | 965 | Bacteria>Acidobacteriota>unclassified_Acidobacteriota | GCA_016199695.1 | Acidobacteriota_bacterium | HY314_11430 | 12465..15362 | 1978231 | ATP-binding-protein-[Acidobacteria-bacterium] |
| MBI4454518.1 | DUF499_1.RM.like.systems | METHYLASE-p-><-?\|\|RADICAL-SAM->?-><-DUF499*<-?<-Schlafen+HATPase+wHTH<-METHYLASE\|\|?->?-><-TUDOR+SF1-SF2-Helicase+iREase+CTD+REase | 1101 | Bacteria>Acidobacteriota>unclassified_Acidobacteriota | GCA_016211105.1 | Acidobacteriota_bacterium | HY644_01320 | 37960..41265 | 1978231 | ATP-binding-protein-[Acidobacteria-bacterium] |
| MBP7569108.1 | DUF499_1.RM.like.systems | Cas_Cmr3->?->Cas_Cmr5->?->CARF+HEPN->HEPN+DUF499*->Schlafen-> | 1117 | Bacteria>Acidobacteriota>unclassified_Acidobacteriota | GCA_017998655.1 | Acidobacteriota_bacterium | KBA95_03500 | 22125..25478 | 1978231 | DUF499-domain-containing-protein-[Acidobacteria-bacterium] |
| MCC6131982.1 | DUF499_1.RM.like.systems | <-HEPN+DUF499*<-Schlafen+HATPase+wHTH<-PHP+SbcC<-METHYLASE<-TOPRIM<-OLD-ABC<-TUDOR+SF1-SF2-Helicase+iREase+CTD+REase | 1114 | Bacteria>Acidobacteriota>unclassified_Acidobacteriota | GCA_020845555.1 | Acidobacteriota_bacterium | IT186_18830 | 2559..5903 | 1978231 | DUF499-domain-containing-protein-[Acidobacteria-bacterium] |
| MCY3932061.1 | DUF499_1.RM.like.systems | TUDOR+SF1-SF2-Helicase+iREase+CTD+REase->ParB+HNH->Schlafen->?->METHYLASE-p->METHYLASE->DUF499*-><-?<-?\|\|Peptidase_M28-> | 937 | Bacteria>Acidobacteriota>unclassified_Acidobacteriota | GCA_026707325.1 | Acidobacteriota_bacterium | OXH70_10095 | 67644..70457 | 1978231 | MAG:-DUF499-domain-containing-protein-[Acidobacteria-bacterium] |
| MCY4659240.1 | DUF499_1.RM.like.systems | <-ORC-CDC6+REase<-DUF499*<-PIN<-OLD-ABC<-METHYLASE<-ParB+HNH+Schlafen<-TOPRIM<-OLD-ABC | 941 | Bacteria>Acidobacteriota>unclassified_Acidobacteriota | GCA_026713985.1 | Acidobacteriota_bacterium | OXF93_05425 | 1121..3946 | 1978231 | MAG:-DUF499-domain-containing-protein-[Acidobacteria-bacterium] |
| MDE2979142.1 | DUF499_1.RM.like.systems | ParB+HNH+Schlafen->PolB->METHYLASE->?->OLD-ABC->TOPRIM->DUF499*->XIS-HTH+RVT-> | 923 | Bacteria>Acidobacteriota>unclassified_Acidobacteriota | GCA_028873735.1 | Acidobacteriota_bacterium | OXU63_16720 | 13221..15992 | 1978231 | MAG:-DUF499-domain-containing-protein-[Acidobacteriota-bacterium] |
| MBE3120494.1 | DUF499_1.RM.like.systems | <-KAP-NTPase<-HEPN+DUF499*<-METHYLASE<-Schlafen+HATPase+wHTH | 1116 | Bacteria>Atribacterota>Atribacteria>unclassified_Atribacteria | GCA_014894575.1 | Candidatus_Atribacteria_bacterium | IMZ50_17295 | 4085..7435 | 2053509 | DUF499-domain-containing-protein-[Candidatus-Atribacteria-bacterium] |
| MBC7217419.1 | DUF499_1.RM.like.systems | Schlafen+HATPase+wHTH->Schlafen->METHYLASE->DUF499*-> | 870 | Bacteria>Atribacterota>Atribacterota_incertae_sedis>Candidatus_Caldatribacterium>unclassified_Candidatus_Caldatribacterium | GCA_014359405.1 | Candidatus_Caldatribacterium_sp. | H5U36_04510 | 5319..7931 | 2282143 | ATP-binding-protein-[Candidatus-Caldatribacterium-sp.] |
| MBN2055102.1 | DUF499_1.RM.like.systems | <-DUF499*<-METHYLASE<-Schlafen+HATPase+wHTH<-TUDOR+SF1-SF2-Helicase+iREase+CTD+REase | 1105 | Bacteria>Bacteria_incertae_sedis>Bacteria_candidate_phyla>candidate_division_CSSED10-310 | GCA_016932295.1 | candidate_division_CSSED10-310_bacterium | JW905_09275 | 5385..8702 | 2855610 | ATP-binding-protein-[bacterium] |
| PWB47246.1 | DUF499_1.RM.like.systems | TUDOR+SF1-SF2-Helicase+iREase+CTD+REase->Schlafen+HATPase+wHTH->METHYLASE->HEPN+DUF499*->?->?-><-?<-REase<-?<-CopG_antitoxin | 1102 | Bacteria>Bacteria_incertae_sedis>Bacteria_candidate_phyla>candidate_division_NC10>unclassified_candidate_division_NC10 | GCA_003104975.1 | candidate_division_NC10_bacterium | C3F12_04520 | 184009..187317 | 2072417 | AAA+-family-ATPase-[candidate-division-NC10-bacterium] |
| MCX6842110.1 | DUF499_1.RM.like.systems | NTP_transf->TUDOR+SF1-SF2-Helicase+iREase+CTD+REase->Schlafen+HATPase+wHTH->?->METHYLASE-p->METHYLASE->HEPN+DUF499*-> | 1030 | Bacteria>Bacteria_incertae_sedis>Bacteria_candidate_phyla>candidate_division_WOR-3 | GCA_026397595.1 | candidate_division_WOR-3_bacterium | NTX53_07520 | 11691..14783 | 2052148 | MAG:-Swt1-family-HEPN-domain-containing-protein-[candidate-division-WOR-3-bacterium] |
| OQB57508.1 | DUF499_1.RM.like.systems | TUDOR+SF1-SF2-Helicase+iREase+CTD+REase->Schlafen+HATPase+wHTH->METHYLASE->HEPN+DUF499*-><-?<-?<-Cysteine_desulfurase\|\|Peptidase_M24-> | 1114 | Bacteria>Bacteria_incertae_sedis>Bacteria_candidate_phyla>Candidatus_Aminicenantes>unclassified_Candidatus_Aminicenantes | GCA_002070465.1 | Candidatus_Aminicenantes_bacterium_ADurb.Bin147 | BWX98_01441 | 17398..20742 | 1852831 | hypothetical-protein-BWX98_01441-[Candidatus-Aminicenantes-bacterium-ADurb.Bin147] |
| MCX6557429.1 | DUF499_1.RM.like.systems | Schlafen+HATPase+wHTH->METHYLASE->TIR->TIR->?->DUF499*-> | 896 | Bacteria>Bacteria_incertae_sedis>Bacteria_candidate_phyla>Candidatus_Aminicenantes>unclassified_Candidatus_Aminicenantes | GCA_026393795.1 | Candidatus_Aminicenantes_bacterium | NTW95_08395 | 10594..13284 | 2052149 | MAG:-DUF499-domain-containing-protein-[Candidatus-Aminicenantes-bacterium] |
| MDD5263468.1 | DUF499_1.RM.like.systems | <-HEPN+DUF499*\|\|?-><-OLD-ABC<-METHYLASE<-Schlafen+HATPase+wHTH<-TUDOR+SF1-SF2-Helicase+iREase+CTD+REase-p | 1114 | Bacteria>Bacteria_incertae_sedis>Bacteria_candidate_phyla>Candidatus_Bipolaricaulota>Bipolaricaulis | GCA_028713635.1 | Candidatus_Bipolaricaulis_sp. | PHU43_01355 | 30296..33640 | 2814139 | MAG:-Swt1-family-HEPN-domain-containing-protein-[Candidatus-Bipolaricaulis-sp.] |
| MDD5453217.1 | DUF499_1.RM.like.systems | SF1-SF2-Helicase-p->PIN->ParB+HNH+Schlafen->METHYLASE->PolB->HEPN->DUF499*-><-ABC-ATPase | 932 | Bacteria>Bacteria_incertae_sedis>Bacteria_candidate_phyla>Candidatus_Bipolaricaulota>Bipolaricaulis | GCA_028715685.1 | Candidatus_Bipolaricaulis_sp. | PHZ21_02925 | 36651..39449 | 2814139 | MAG:-DUF499-domain-containing-protein-[Candidatus-Bipolaricaulis-sp.] |
| WP_259099629.1 | DUF499_1.RM.like.systems | TUDOR+SF1-SF2-Helicase+iREase+CTD+REase->Schlafen+HATPase+wHTH->METHYLASE->DUF499*->?->?->PfkB-> | 880 | Bacteria>Bacteria_incertae_sedis>Bacteria_candidate_phyla>Candidatus_Fervidibacteria>Candidatus_Fervidibacter | GCF_030520105.1 | Candidatus_Fervidibacter_sacchari | M2350_RS13990 | 282876..285518 | 1448929 | DUF499-domain-containing-protein-[Candidatus-Fervidibacter-sacchari] |
| MCD6507091.1 | DUF499_1.RM.like.systems | TUDOR+SF1-SF2-Helicase+iREase+CTD+REase->Schlafen->BirA-HTH->METHYLASE->DUF499*-> | 864 | Bacteria>Bacteria_incertae_sedis>Bacteria_candidate_phyla>Candidatus_Poribacteria>unclassified_Candidatus_Poribacteria | GCA_021162805.1 | Candidatus_Poribacteria_bacterium | J7M22_10765 | 25392..27986 | 2026781 | ATP-binding-protein-[Candidatus-Poribacteria-bacterium] |
| MBI2216638.1 | DUF499_1.RM.like.systems | <-nSTAND3<-nSTAND3\|\|TUDOR+SF1-SF2-Helicase+iREase+CTD+REase->PHP+SbcC->Schlafen+HATPase+wHTH->DUF499*->?->?-><-?<-Radical-SAM | 1102 | Bacteria>Bacteria_incertae_sedis>Bacteria_candidate_phyla>Candidatus_Rokubacteria>unclassified_Candidatus_Rokubacteria | GCA_016184125.1 | Candidatus_Rokubacteria_bacterium | HYU51_05015 | 42973..46281 | 2053607 | ATP-binding-protein-[Candidatus-Rokubacteria-bacterium] |
| MCX7047269.1 | DUF499_1.RM.like.systems | DDE_Tnp_1-><-?<-?<-?<-tRNA\|\|?->Radical-SAM->tRNA-><-DUF499*<-DUF499<-Schlafen+HATPase+wHTH<-?<-METHYLASE<-Schlafen+HATPase+wHTH | 616 | Bacteria>Bacteria_incertae_sedis>Bacteria_candidate_phyla>Candidatus_Sumerlaeota>unclassified_Candidatus_Sumerlaeota | GCA_026398575.1 | Candidatus_Sumerlaeota_bacterium | NTX50_17505 | 5846..7696 | 2107588 | MAG:-DUF499-domain-containing-protein-[Candidatus-Sumerlaeota-bacterium] |
| MCX7047270.1 | DUF499_1.RM.like.systems | Radical-SAM->tRNA-><-DUF499<-DUF499*<-Schlafen+HATPase+wHTH<-?<-METHYLASE<-Schlafen+HATPase+wHTH<-?<-TUDOR+SF1-SF2-Helicase+iREase+CTD+REase | 330 | Bacteria>Bacteria_incertae_sedis>Bacteria_candidate_phyla>Candidatus_Sumerlaeota>unclassified_Candidatus_Sumerlaeota | GCA_026398575.1 | Candidatus_Sumerlaeota_bacterium | NTX50_17510 | 7797..8789 | 2107588 | MAG:-DUF499-domain-containing-protein-[Candidatus-Sumerlaeota-bacterium] |
| MBN1883904.1 | DUF499_1.RM.like.systems | Peptidase_M48->TUDOR+SF1-SF2-Helicase+iREase+CTD+REase->Schlafen+HATPase+wHTH->METHYLASE->OLD-ABC->TOPRIM->HEPN+DUF499*->PIN+OST-HTH-><-?<-?<-REase<-?\|\|DOC+HTH-> | 1113 | Bacteria>Candidatus_Krumholzibacteriota>unclassified_Candidatus_Krumholzibacteriota | GCA_016931295.1 | Candidatus_Krumholzibacteriota_bacterium | JW876_00100 | 28696..32037 | 2725313 | DUF499-domain-containing-protein-[Candidatus-Krumholzibacteriota-bacterium] |
| WP_237889222.1 | DUF499_1.RM.like.systems | TUDOR+SF1-SF2-Helicase+iREase+CTD+REase->Schlafen+HATPase+wHTH->METHYLASE->DUF499*->?->?->tRNA->ABC-ATPase+SbcC-> | 970 | Bacteria>delta/epsilon_subdivisions>Deltaproteobacteria>Deltaproteobacteria_incertae_sedis>Dissulfurimicrobium | GCF_022026155.1 | Dissulfurimicrobium_hydrothermale | LGS26_RS03280 | 705149..708061 | 1750598 | DUF499-domain-containing-protein-[Dissulfurimicrobium-hydrothermale] |
| OGR05786.1 | DUF499_1.RM.like.systems | METHYLASE->DUF499*->?->Schlafen+HATPase+wHTH-> | 944 | Bacteria>delta/epsilon_subdivisions>Deltaproteobacteria>unclassified_Deltaproteobacteria | GCA_001799225.1 | Deltaproteobacteria_bacterium_RIFOXYD12_FULL_50_9 | A2511_04565 | 2695..5529 | 1797902 | hypothetical-protein-A2511_04565-[Deltaproteobacteria-bacterium-RIFOXYD12_FULL_50_9] |
| PJB39408.1 | DUF499_1.RM.like.systems | <-DUF499-p<-?<-HEPN+DUF499*<-Schlafen+HATPase+wHTH<-PHP+SbcC<-METHYLASE-p | 1117 | Bacteria>delta/epsilon_subdivisions>Deltaproteobacteria>unclassified_Deltaproteobacteria | GCA_002789955.1 | Deltaproteobacteria_bacterium_CG_4_9_14_3_um_filter_63_12 | CO108_17305 | 2362..5715 | 1973967 | AAA+-family-ATPase-[Deltaproteobacteria-bacterium-CG_4_9_14_3_um_filter_63_12] |
| CAG1064741.1 | DUF499_1.RM.like.systems | <-DOC<-MazE_antitoxin<-?<-TUDOR+SF1-SF2-Helicase+iREase+CTD<-?<-HEPN+DUF499*<-METHYLASE<-ArdCN+MPTase<-Schlafen+HATPase+wHTH<-ArdCN+MPTase<-TUDOR+SF1-SF2-Helicase+iREase+CTD+REase<-HTH_26 | 1109 | Bacteria>environmental_samples | GCA_905339405.1 | uncultured_bacterium | BAC1_00306 | 123828..127157 | 77133 | hypothetical-protein-BAC1_00306-[uncultured-bacterium] |
| MCC8143016.1 | DUF499_1.RM.like.systems | <-AraC-HTH+SHS2+SHS2<-Peptidase_M28<-?<-BirA-HTH<-Schlafen<-HEPN+DUF499*<-METHYLASE | 1102 | Bacteria>FCB_group>Bacteroidota/Chlorobiota_group>Bacteroidota>Bacteroidia | GCA_020860665.1 | Tannerellaceae_bacterium | LIO97_03250 | 4810..8118 | 2785930 | DUF499-domain-containing-protein-[Tannerellaceae-bacterium] |
| MCD8261448.1 | DUF499_1.RM.like.systems | <-STYKIN<-Schlafen<-HEPN+DUF499*<-METHYLASE-p<-METHYLASE-p | 1102 | Bacteria>FCB_group>Bacteroidota/Chlorobiota_group>Bacteroidota>Bacteroidia | GCA_021204375.1 | Bacteroides_sp. | LUD15_08080 | 1305..4613 | 29523 | DUF499-domain-containing-protein-[Bacteroides-sp.] |
| GIV23748.1 | DUF499_1.RM.like.systems | TUDOR+SF1-SF2-Helicase+iREase+CTD+REase->Schlafen+HATPase+wHTH->METHYLASE->DUF499*->DUF499-p->?->?->TUDOR+SF1-SF2-Helicase+iREase+CTD->?->METHYLASE-p-> | 393 | Bacteria>FCB_group>Bacteroidota/Chlorobiota_group>Bacteroidota>Bacteroidia | GCA_026003215.1 | Bacteroidia_bacterium | KatS3mg025_1407 | 1616391..1617572 | 2044936 | MAG:-hypothetical-protein-KatS3mg025_1407-[Bacteroidia-bacterium] |
| WP_074635742.1 | DUF499_1.RM.like.systems | DEDD_Tnp_IS110->HTH_3->TUDOR+SF1-SF2-Helicase+iREase+CTD+REase->METHYLASE->HEPN+DUF499*->Schlafen+HATPase+wHTH->?->HTH_3->DNA_methylase->MORC-> | 1096 | Bacteria>FCB_group>Bacteroidota/Chlorobiota_group>Bacteroidota>Bacteroidia | GCF_900100465.1 | Bacteroides_ovatus | BLS71_RS01410 | 96540..99830 | 28116 | DUF499-domain-containing-protein-[Bacteroides-ovatus] |
| WP_074760634.1 | DUF499_1.RM.like.systems | Schlafen+HATPase+wHTH->TUDOR+SF1-SF2-Helicase+iREase+CTD+REase->METHYLASE->DUF499*-> | 930 | Bacteria>FCB_group>Bacteroidota/Chlorobiota_group>Bacteroidota>Bacteroidia | GCF_900107775.1 | Xylanibacter_ruminicola | BLU97_RS05645 | 384764..387556 | 839 | DUF499-domain-containing-protein-[Prevotella-ruminicola] |
| MQY79011.1 | DUF499_1.RM.like.systems | TUDOR+SF1-SF2-Helicase+iREase+CTD+REase->Schlafen+HATPase+wHTH->METHYLASE->?->DUF499*-> | 491 | Bacteria>FCB_group>Bacteroidota/Chlorobiota_group>Bacteroidota>unclassified_Bacteroidota | GCA_009619225.1 | Bacteroidota_bacterium | GH151_07430 | 8478..9950 | 1898104 | DUF499-domain-containing-protein-[Bacteroidota-bacterium] |
| BCX00053.1 | DUF499_1.RM.like.systems | <-35exo\|\|?-><-Methyltransf_4<-?<-ABhydrolase<-DUF499*<-METHYLASE<-Schlafen+HATPase+wHTH<-TUDOR+SF1-SF2-Helicase+iREase+CTD+REase<-?<-?<-Endonuclease_5 | 878 | Bacteria>FCB_group>Bacteroidota/Chlorobiota_group>Bacteroidota>unclassified_Bacteroidota | GCA_025998875.1 | Bacteroidota_bacterium | KatS3mg041_0099 | 106704..109340 | 1898104 | MAG:-hypothetical-protein-KatS3mg041_0099-[Bacteroidota-bacterium] |
| GIV55103.1 | DUF499_1.RM.like.systems | TUDOR+SF1-SF2-Helicase+iREase+CTD+REase->Schlafen->BirA-HTH->METHYLASE-p->METHYLASE-p->DUF499*->ParB+HNH-> | 875 | Bacteria>FCB_group>Bacteroidota/Chlorobiota_group>Candidatus_Kapabacteria>unclassified_Candidatus_Kapabacteria | GCA_026003475.1 | Candidatus_Kapabacteria_bacterium | KatS3mg039_1621 | 1981096..1983723 | 2447898 | MAG:-hypothetical-protein-KatS3mg039_1621-[Candidatus-Kapabacteria-bacterium] |
| MCB5288062.1 | DUF499_1.RM.like.systems | <-SbcC<-DUF499*<-Schlafen+HATPase+wHTH<-METHYLASE | 940 | Bacteria>FCB_group>Candidatus_Cloacimonadota>unclassified_Candidatus_Cloacimonadota | GCA_020532145.1 | Candidatus_Cloacimonadota_bacterium | LHW64_09695 | 2385..5207 | 2030808 | DUF499-domain-containing-protein-[Candidatus-Cloacimonetes-bacterium] |
| MCB5262479.1 | DUF499_1.RM.like.systems | <-DUF499*<-Schlafen+HATPase+wHTH<-METHYLASE-p | 826 | Bacteria>FCB_group>Candidatus_Cloacimonadota>unclassified_Candidatus_Cloacimonadota | GCA_020532425.1 | Candidatus_Cloacimonadota_bacterium | LHW53_07100 | 1..2478 | 2030808 | DUF499-domain-containing-protein-[Candidatus-Cloacimonetes-bacterium] |
| NLO34316.1 | DUF499_1.RM.like.systems | <-Schlafen+HATPase+wHTH<-?<-DUF499*<-METHYLASE-p | 1113 | Bacteria>FCB_group>Candidatus_Hydrogenedentes>unclassified_Candidatus_Hydrogenedentes | GCA_012523015.1 | Candidatus_Hydrogenedentes_bacterium | GX117_13355 | 3475..6816 | 2030809 | ATP-binding-protein-[Candidatus-Hydrogenedentes-bacterium] |
| MCF7823046.1 | DUF499_1.RM.like.systems | <-MazG<-?<-?<-MORC+HISKIN<-METHYLASE<-HEPN+DUF499*<-METHYLASE<-MORC+HisKin+DUF3684<-TUDOR+SF1-SF2-Helicase+iREase+CTD+REase<-XIS-HTH<-?<-Schlafen+HATPase+wHTH | 1111 | Bacteria>FCB_group>Candidatus_Marinimicrobia>unclassified_Candidatus_Marinimicrobia | GCA_021734325.1 | Candidatus_Marinimicrobia_bacterium | K9N35_02645 | 72966..76301 | 2026760 | DUF499-domain-containing-protein-[Candidatus-Marinimicrobia-bacterium] |
| ODT03617.1 | DUF499_1.RM.like.systems | TUDOR+SF1-SF2-Helicase+iREase+CTD+REase->Schlafen+HATPase+wHTH->METHYLASE->HEPN+DUF499*->Primpol+D5N+D5ATPase->PAIREDC-HTH->ORC-CDC6+REase-> | 1109 | Bacteria>FCB_group>Gemmatimonadota>unclassified_Gemmatimonadota | GCA_001724275.1 | Gemmatimonadetes_bacterium_SCN_70-22 | ABS52_08400 | 10589..13918 | 1660163 | AAA+-family-ATPase-[Gemmatimonadetes-bacterium-SCN-70-22] |
| MYG34810.1 | DUF499_1.RM.like.systems | <-XIS-HTH\|\|LexA-protease->TUDOR+SF1-SF2-Helicase+iREase+CTD+REase->METHYLASE->DUF499*->Schlafen+HATPase+wHTH-> | 963 | Bacteria>FCB_group>Gemmatimonadota>unclassified_Gemmatimonadota | GCA_009842515.1 | Gemmatimonadota_bacterium | F4205_04900 | 8108..10999 | 2026742 | ATP-binding-protein-[Gemmatimonadetes-bacterium] |
| MCB9507837.1 | DUF499_1.RM.like.systems | REase-><-Schlafen+HATPase+wHTH<-SF1-SF2-Helicase+HRDC<-?<-NERD+SF1-SF2-Helicase<-HEPN+DUF499*<-METHYLASE<-?<-TUDOR+SF1-SF2-Helicase+iREase+CTD+REase\|\|DNA-glycosylase->?->HTH-> | 1103 | Bacteria>Myxococcota>Myxococcia>Myxococcales>unclassified_Myxococcales | GCA_020633615.1 | Myxococcales_bacterium | H6697_09260 | 18345..21656 | 2026763 | ATP-binding-protein-[Myxococcales-bacterium] |
| MAG32379.1 | DUF499_1.RM.like.systems | METHYLASE-p->OLD-ABC->Schlafen+HATPase+wHTH->HEPN+DUF499*->?-><-METHYLASE-p<-TOPRIM<-OLD-ABC\|\|?->METHYLASE-p-> | 1115 | Bacteria>Myxococcota>Myxococcia>unclassified_Myxococcia | GCA_002688255.1 | Deltaproteobacteria_bacterium | CL908_15970 | 4485..7832 | 2026735 | AAA+-family-ATPase-[Deltaproteobacteria-bacterium] |
| PXF55798.1 | DUF499_1.RM.like.systems | TUDOR+SF1-SF2-Helicase+iREase+CTD+REase->Schlafen+HATPase+wHTH->?->METHYLASE-p-><-MULE-TRANSPOSASE\|\|METHYLASE-p->HEPN+DUF499*->MULE-TRANSPOSASE->?-><-LD-peptidase | 1108 | Bacteria>Myxococcota>Myxococcia>unclassified_Myxococcia | GCA_003194485.1 | Deltaproteobacteria_bacterium | C4B57_02005 | 126317..129643 | 2026735 | AAA+-family-ATPase-[Deltaproteobacteria-bacterium] |
| PXF54128.1 | DUF499_1.RM.like.systems | Phage_int_SAM_1+Phage_integrase-><-SbcC+SbcC<-Calcineurin<-?<-?<-DUF499*<-Schlafen+HATPase+wHTH<-METHYLASE-p<-TOPRIM<-OLD-ABC | 1011 | Bacteria>Myxococcota>Myxococcia>unclassified_Myxococcia | GCA_003194485.1 | Deltaproteobacteria_bacterium | C4B57_08240 | 17848..20883 | 2026735 | AAA-family-ATPase-[Deltaproteobacteria-bacterium] |
| PXF60387.1 | DUF499_1.RM.like.systems | PAIREDC-HTH->Schlafen+HATPase+wHTH->METHYLASE->METHYLASE-p->HEPN+DUF499*-> | 1108 | Bacteria>Myxococcota>Myxococcia>unclassified_Myxococcia | GCA_003194495.1 | Deltaproteobacteria_bacterium | C4B58_00710 | 172327..175653 | 2026735 | AAA+-family-ATPase-[Deltaproteobacteria-bacterium] |
| RLC12751.1 | DUF499_1.RM.like.systems | <-DUF499*<-Schlafen+HATPase+wHTH<-METHYLASE-p<-METHYLASE<-MNS<-MACRODOMAIN<-TUDOR+SF1-SF2-Helicase+iREase+CTD+REase | 645 | Bacteria>Myxococcota>Myxococcia>unclassified_Myxococcia | GCA_003647435.1 | Deltaproteobacteria_bacterium | DRH43_00745 | 1..1934 | 2026735 | AAA-family-ATPase,-partial-[Deltaproteobacteria-bacterium] |
| MBC8466732.1 | DUF499_1.RM.like.systems | Phage_integrase->TUDOR+SF1-SF2-Helicase+iREase+CTD+REase->Schlafen->METHYLASE->DUF499*-> | 1095 | Bacteria>Myxococcota>Myxococcia>unclassified_Myxococcia | GCA_014384065.1 | Deltaproteobacteria_bacterium | H8D55_02725 | 8487..11774 | 2026735 | ATP-binding-protein-[Deltaproteobacteria-bacterium] |
| MBF0104316.1 | DUF499_1.RM.like.systems | <-Radical-SAM\|\|METHYLASE->Methyltrans_RNA-><-TPR+TPR<-TPR+TPR<-?<-DUF499*<-Schlafen+HATPase+wHTH<-METHYLASE<-METHYLASE-p<-TUDOR+SF1-SF2-Helicase+iREase+CTD+REase-p<-TPR+TPR | 375 | Bacteria>Myxococcota>Myxococcia>unclassified_Myxococcia | GCA_015228875.1 | Deltaproteobacteria_bacterium | HQM16_03205 | 33942..35069 | 2026735 | ATP-binding-protein-[Deltaproteobacteria-bacterium] |
| MBI4527351.1 | DUF499_1.RM.like.systems | Schlafen+HATPase+wHTH->METHYLASE->?->PIN->HicB_lk_antitox->?->DUF499*-> | 920 | Bacteria>Myxococcota>Myxococcia>unclassified_Myxococcia | GCA_016210115.1 | Deltaproteobacteria_bacterium | HY695_26450 | 25107..27869 | 2026735 | ATP-binding-protein-[Deltaproteobacteria-bacterium] |
| MBM4341420.1 | DUF499_1.RM.like.systems | <-DUF499*<-Schlafen+HATPase+wHTH<-METHYLASE<-SIR2<-TUDOR+SF1-SF2-Helicase+iREase+CTD+REase | 454 | Bacteria>Myxococcota>Myxococcia>unclassified_Myxococcia | GCA_016875195.1 | Deltaproteobacteria_bacterium | FJ110_17985 | 1..1363 | 2026735 | ATP-binding-protein-[Deltaproteobacteria-bacterium] |
| MBN2361667.1 | DUF499_1.RM.like.systems | TUDOR+SF1-SF2-Helicase+iREase+CTD+REase->PIN->METHYLASE->DUF499*->Schlafen->EVE-> | 935 | Bacteria>Myxococcota>Myxococcia>unclassified_Myxococcia | GCA_016934455.1 | Deltaproteobacteria_bacterium | JXR83_19595 | 9461..12268 | 2026735 | ATP-binding-protein-[Deltaproteobacteria-bacterium] |
| MBW1922080.1 | DUF499_1.RM.like.systems | ORC-CDC6+REase->TUDOR+SF1-SF2-Helicase+iREase+CTD+REase-p->Schlafen+HATPase+wHTH->METHYLASE->PolB->HEPN->HEPN+DUF499*-> | 1062 | Bacteria>Myxococcota>Myxococcia>unclassified_Myxococcia | GCA_019307945.1 | Deltaproteobacteria_bacterium | JRJ35_01275 | 87250..90435 | 2026735 | DUF499-domain-containing-protein-[Deltaproteobacteria-bacterium] |
| MBW1949838.1 | DUF499_1.RM.like.systems | TUDOR+SF1-SF2-Helicase+iREase+CTD+REase-p->Schlafen+HATPase+wHTH->?->METHYLASE->PolB->HEPN->HEPN+DUF499*-> | 1092 | Bacteria>Myxococcota>Myxococcia>unclassified_Myxococcia | GCA_019308065.1 | Deltaproteobacteria_bacterium | JRJ18_09875 | 33770..37048 | 2026735 | DUF499-domain-containing-protein-[Deltaproteobacteria-bacterium] |
| MBW2077703.1 | DUF499_1.RM.like.systems | TUDOR+SF1-SF2-Helicase+iREase+CTD+REase-p->Schlafen+HATPase+wHTH->METHYLASE->URI-DNase->PolB->HEPN->HEPN+DUF499*-> | 1093 | Bacteria>Myxococcota>Myxococcia>unclassified_Myxococcia | GCA_019309065.1 | Deltaproteobacteria_bacterium | JRI71_09200 | 7259..10540 | 2026735 | DUF499-domain-containing-protein-[Deltaproteobacteria-bacterium] |
| GIW47700.1 | DUF499_1.RM.like.systems | <-PolB+HEPN<-PolB\|\|?-><-DUF499*<-PolB+HEPN<-PolB<-METHYLASE<-Schlafen+HATPase+wHTH<-TUDOR+SF1-SF2-Helicase+iREase+CTD+REase | 857 | Bacteria>Myxococcota>Myxococcia>unclassified_Myxococcia | GCA_026004235.1 | Deltaproteobacteria_bacterium | KatS3mg078_1577 | 289556..292129 | 2026735 | MAG:-hypothetical-protein-KatS3mg078_1577-[Deltaproteobacteria-bacterium] |
| MDD9854174.1 | DUF499_1.RM.like.systems | <-DUF499*<-OLD-ABC<-METHYLASE<-TPR+TPR<-Schlafen+HATPase+wHTH<-TUDOR+SF1-SF2-Helicase+iREase+CTD+REase | 946 | Bacteria>Myxococcota>Myxococcia>unclassified_Myxococcia | GCA_028817165.1 | Deltaproteobacteria_bacterium | OXU78_09525 | 86..2926 | 2026735 | MAG:-DUF499-domain-containing-protein-[Deltaproteobacteria-bacterium] |
| MBP7125459.1 | DUF499_1.RM.like.systems | TUDOR+SF1-SF2-Helicase+iREase+CTD+REase->Schlafen+HATPase+wHTH->METHYLASE->OLD-ABC->TOPRIM->HEPN+DUF499*-> | 1039 | Bacteria>Myxococcota>unclassified_Myxococcota | GCA_017994695.1 | Myxococcota_bacterium | KBD49_03725 | 64170..67286 | 2818507 | DUF499-domain-containing-protein-[Myxococcota-bacterium] |
| MDE0208007.1 | DUF499_1.RM.like.systems | <-HEPN+DUF499*<-Schlafen+HATPase+wHTH<-METHYLASE<-TUDOR+SF1-SF2-Helicase+iREase+CTD+REase<-BrnA_antitoxin | 1111 | Bacteria>Nitrospinae/Tectomicrobia_group>Candidatus_Tectomicrobia>unclassified_Candidatus_Tectomicrobia | GCA_028819935.1 | Candidatus_Tectomicrobia_bacterium | OXP66_18475 | 1406..4741 | 2528274 | MAG:-Swt1-family-HEPN-domain-containing-protein-[Candidatus-Tectomicrobia-bacterium] |

| Protein-ID | System | Gene-Neighbourhoods | Length | Taxonomy | Assembly-ID | Strain | Locus-Tag | Genomic-coordinates | TaxID | Fasta-defline |
|---|---|---|---|---|---|---|---|---|---|---|
| MBZ0154906.1 | DUF499 _1.RM.like.systems | TIR->PAIREDC-HTH+HIN-HTH->DDE_3-><-SF1-SF2-Helicase<-HKD-DNase<-?<- HEPN+ DUF499 *<- METHYLASE <-?<-?<- Schlafen +HATPase+wHTH \|\|REase-> | 1108 | Bacteria>Nitrospirota>Nitrospiria>Nitrospirales>Nitrospiraceae | GCA_019912415.1 | Candidatus_Nitrobium_versatile | K8I29_01665 | 56123..59449 | 2884831 | DUF499 -domain-containing-protein-[Alphaproteobacteria-bacterium] |
| MCP9446717.1 | DUF499 _1.RM.like.systems | <-REase<-?<-?<-?<-N6_N4_MTase<-?<- DUF499 *<-REase<- METHYLASE <- Schlafen +HATPase+wHTH <- TUDOR+ SF1-SF2-Helicase+iREase+CTD+REase | 941 | Bacteria>Nitrospirota>Nitrospiria>Nitrospirales>Nitrospiraceae | GCA_024330945.1 | Nitrospira_sp. | NNA22_03990 | 9048..11873 | 70125 | DUF499 -domain-containing-protein-[Nitrospira-sp.] |
| MCY4460275.1 | DUF499 _1.RM.like.systems | <- HEPN+ DUF499 *<- Schlafen +HATPase+wHTH <- METHYLASE <- TUDOR+ SF1-SF2-Helicase+iREase+CTD+REase | 1099 | Bacteria>Pseudomonadota>Alphaproteobacteria>Rhodobacterales>Paracoccaceae | GCA_026712665.1 | Albidovulum_sp. | OXC26_07750 | 691..3990 | 1872424 | MAG:- DUF499 -domain-containing-protein-[Albidovulum-sp.] |
| WP_238518868.1 | DUF499 _1.RM.like.systems | <-SF1-SF2-Helicase+DUF1998+HKD-DNase+HKD-DNase<-?<- HEPN+ DUF499 *<-PIN<-OLD-ABC<- METHYLASE <-?<- Schlafen +HATPase+wHTH <- TUDOR+ SF1-SF2-Helicase+iREase+CTD+REase | 1154 | Bacteria>Pseudomonadota>Alphaproteobacteria>Rhodospirillales>Azospirillaceae | GCF_022214805.1 | Azospirillum_doebereinerae | MCW82_RS00020 | 10412..13876 | 92933 | DUF499 -domain-containing-protein-[Azospirillum-doebereinerae] |
| KFB77813.1 | DUF499 _1.RM.like.systems | Hpr-Kinase-C-><-JetD<-?<-JetB<-JetA<- DUF499 *<- Schlafen +HATPase+wHTH <- METHYLASE <-PIN\|\|?-><- TUDOR+ SF1-SF2-Helicase+iREase+CTD+REase <-HAMP+HISKIN | 1093 | Bacteria>Pseudomonadota>Betaproteobacteria>Betaproteobacteria_incertae_sedis>Candidatus_Accumulibacter | GCA_000584975.2 | Candidatus_Accumulibacter_cognatus | AW06_000996 | 10153..13434 | 2954383 | MAG:-hypothetical-protein-AW06_000996-[Candidatus-Accumulibacter-cognatus] |
| KFB70630.1 | DUF499 _1.RM.like.systems | <- DUF499 -p<- DUF499 *<- Schlafen +HATPase+wHTH <- Schlafen <-AAA_21 | 484 | Bacteria>Pseudomonadota>Betaproteobacteria>Betaproteobacteria_incertae_sedis>Candidatus_Accumulibacter | GCA_000585035.2 | Candidatus_Accumulibacter_phosphatis | AW09_004270 | 1814..3268 | 327160 | MAG:-hypothetical-protein-AW09_004270-[Candidatus-Accumulibacter-phosphatis] |
| MCM8600113.1 | DUF499 _1.RM.like.systems | <-SbcC<- DUF499 *<- Schlafen +HATPase+wHTH <- METHYLASE <- METHYLASE <-HEPN<- TUDOR+ SF1-SF2-Helicase+iREase+CTD+REase <-HTH_Tnp | 949 | Bacteria>Pseudomonadota>Betaproteobacteria>Betaproteobacteria_incertae_sedis>Candidatus_Accumulibacter | GCA_023806585.1 | Accumulibacter_sp. | NFW04_15895 | 29772..32621 | 2053492 | DUF499 -domain-containing-protein-[Candidatus-Accumulibacter-sp.] |
| MCQ1548298.1 | DUF499 _1.RM.like.systems | HAMP+HISKIN-> TUDOR+ SF1-SF2-Helicase+iREase+CTD+REase ->PIN-> METHYLASE -> Schlafen +HATPase+wHTH -> DUF499 *-> | 1093 | Bacteria>Pseudomonadota>Betaproteobacteria>Betaproteobacteria_incertae_sedis>Candidatus_Accumulibacter | GCA_024380085.1 | Candidatus_Accumulibacter_phosphatis | NOF05_05620 | 18582..21863 | 327160 | DUF499 -domain-containing-protein-[Candidatus-Accumulibacter-phosphatis] |
| WP_034945897.1 | DUF499 _1.RM.like.systems | Hpr-Kinase-C-><-JetD<-?<-JetB<-JetA<- DUF499 *<- Schlafen +HATPase+wHTH <- METHYLASE <-PIN<- TUDOR+ SF1-SF2-Helicase+iREase+CTD+REase <-HAMP+HISKIN | 1093 | Bacteria>Pseudomonadota>Betaproteobacteria>Betaproteobacteria_incertae_sedis>Candidatus_Accumulibacter | GCF_000584975.1 | Candidatus_Accumulibacter_cognatus | AW06_RS03920 | 10153..13434 | 2954383 | DUF499 -domain-containing-protein-[Candidatus-Accumulibacter-cognatus] |
| WP_033446622.1 | DUF499 _1.RM.like.systems | TetR-HTH->?->MarR-HTH->Phage_int_SAM_2+Integrase_1-><- DUF499 *<- Schlafen +HATPase+wHTH <- METHYLASE <- METHYLASE <-HEPN<- TUDOR+ SF1-SF2-Helicase+iREase+CTD+REase <-HTH+WYL | 947 | Bacteria>Pseudomonadota>Betaproteobacteria>Burkholderiales>Alcaligenaceae | GCF_000690475.1 | Bordetella_bronchiseptica | EL288_RS19650 | 4193914..4196757 | 518 | DUF499 -domain-containing-protein-[Bordetella-bronchiseptica] |
| WP_065497253.1 | DUF499 _1.RM.like.systems | TUDOR+ SF1-SF2-Helicase+iREase+CTD+REase -p-> TUDOR+ SF1-SF2-Helicase+iREase+CTD+REase ->PIN-> METHYLASE -> METHYLASE -> Schlafen +HATPase+wHTH -> DUF499 *->HEPN-> | 955 | Bacteria>Pseudomonadota>Betaproteobacteria>Burkholderiales>Burkholderiaceae | GCF_001685525.1 | Burkholderia_sp._CCA53 | BCR55_RS29355 | 26270..29137 | 1776288 | DUF499 -domain-containing-protein-[Burkholderia-sp.-CCA53] |
| WP_217096864.1 | DUF499 _1.RM.like.systems | <-ACET\|\|?->?->MarR-HTH->Phage_int_SAM_2+Integrase_1-><- DUF499 *<- Schlafen +HATPase+wHTH <- METHYLASE <- METHYLASE <-HNH<-OLD-ABC<- TUDOR+ SF1-SF2-Helicase+iREase+CTD+REase | 947 | Bacteria>Pseudomonadota>Betaproteobacteria>Burkholderiales>Burkholderiaceae | GCF_019041095.1 | Burkholderia_multivorans | KTE23_RS28415 | 22880..25723 | 87883 | DUF499 -domain-containing-protein-[Burkholderia-multivorans] |
| WP_043699941.1 | DUF499 _1.RM.like.systems | PIN-> METHYLASE ->OLD-ABC->TOPRIM-> METHYLASE -> Schlafen +HATPase+wHTH -> DUF499 *-><-?<-?<-MORC<-?<-VirB11+MoxR-AAA | 945 | Bacteria>Pseudomonadota>Betaproteobacteria>Burkholderiales>Burkholderiales_genera_incertae_sedis | GCF_020162115.1 | Tepidimonas_taiwanensis | PL21_RS04270 | 89059..91896 | 307486 | DUF499 -domain-containing-protein-[Tepidimonas-taiwanensis] |
| WP_252101816.1 | DUF499 _1.RM.like.systems | TUDOR+ SF1-SF2-Helicase+iREase+CTD+REase -> Schlafen +HATPase+wHTH ->?-> METHYLASE ->OLD-ABC->REase-> DUF499 *-><-?<-?<-Phage_integrase+Phage_integrase<-?<-HIN-HTH<-Phage_integrase+Phage_integrase | 945 | Bacteria>Pseudomonadota>Betaproteobacteria>Burkholderiales>Comamonadaceae | - | [Acidovorax]_ebreus | MRB47_RS02565 | 530947..533784 | 721785 | DUF499 -domain-containing-protein-[[Acidovorax] |
| WP_011810371.1 | DUF499 _1.RM.like.systems | <-S1+RNase+2C<-?\|\|?-><-SPOUT_MTase<-?\|\|?-><- DUF499 *<- Schlafen +HATPase+wHTH <- METHYLASE <- METHYLASE <-DUF4096+DDE_Tnp_1+DDE_Tnp_1_2<-PAIREDC-HTH+HIN-HTH<-HAMP+HISKIN | 936 | Bacteria>Pseudomonadota>Betaproteobacteria>Burkholderiales>Comamonadaceae | GCF_000015565.1 | Verminephrobacter_eiseniae | VEIS_RS12785 | 2967373..2970183 | 364317 | DUF499 -domain-containing-protein-[Verminephrobacter-eiseniae] |
| WP_265259651.1 | DUF499 _1.RM.like.systems | METHYLASE -> METHYLASE -> Schlafen +HATPase+wHTH -> DUF499 *-><-?\|\|?->SPOUT_MTase-><-?\|\|?->S1+RNase+2C-> | 936 | Bacteria>Pseudomonadota>Betaproteobacteria>Burkholderiales>Comamonadaceae | GCF_026127445.1 | Verminephrobacter_eiseniae | D8B29_RS12175 | 5874..8684 | 364317 | DUF499 -domain-containing-protein-[Verminephrobacter-eiseniae] |
| WP_265289361.1 | DUF499 _1.RM.like.systems | METHYLASE -p-> METHYLASE -><-?\|\| METHYLASE -> Schlafen +HATPase+wHTH -> Schlafen +HATPase+wHTH -> DUF499 *-><-?\|\|?->SPOUT_MTase->?->S1+RNase+2C-> | 942 | Bacteria>Pseudomonadota>Betaproteobacteria>Burkholderiales>Comamonadaceae | GCF_026127525.1 | Verminephrobacter_eiseniae | D5047_RS20580 | 4396867..4399695 | 364317 | DUF499 -domain-containing-protein-[Verminephrobacter-eiseniae] |
| WP_281915166.1 | DUF499 _1.RM.like.systems | Phage_lysozyme-><-HEPN<- DUF499 *<- Schlafen +HATPase+wHTH <- METHYLASE <- METHYLASE | 955 | Bacteria>Pseudomonadota>Betaproteobacteria>Burkholderiales>Sphaerotilaceae | GCF_937957785.1 | Caldimonas_thermodepolymerans | QMG53_RS04175 | 1459..4326 | 215580 | DUF499 -domain-containing-protein-[Caldimonas-thermodepolymerans] |
| MBL8529542.1 | DUF499 _1.RM.like.systems | Schlafen +HATPase+wHTH ->?-> METHYLASE -p->HTH+Hiskin+STAS->PolB-> HEPN+ DUF499 *-> | 1108 | Bacteria>Pseudomonadota>Betaproteobacteria>Burkholderiales>unclassified_Burkholderiales | GCA_016791285.1 | Burkholderiales_bacterium | JNL68_17825 | 6694..10020 | 1891238 | DUF499 -domain-containing-protein-[Burkholderiales-bacterium] |
| MBK7665767.1 | DUF499 _1.RM.like.systems | TUDOR+ SF1-SF2-Helicase+iREase+CTD+REase -> Schlafen +HATPase+wHTH ->?->PolB->PolB+HEPN-> HEPN+ DUF499 *->35exo+SF1-SF2-Helicase->SIGMA-HTH+SIGMA-HTH-> | 1112 | Bacteria>Pseudomonadota>Betaproteobacteria>Nitrosomonadales>Sterolibacteriaceae | GCA_016709795.1 | Sterolibacteriaceae_bacterium | IPJ21_19955 | 4305507..4308845 | 2682146 | DUF499 -domain-containing-protein-[Sterolibacteriaceae-bacterium] |
| ODU50877.1 | DUF499 _1.RM.like.systems | <-HAD<-?\|\|?-><-?<-?\|\|?-><- DUF499 *<-?<- METHYLASE <-?<- Schlafen +HATPase+wHTH <- TUDOR+ SF1-SF2-Helicase+iREase+CTD+REase | 937 | Bacteria>Pseudomonadota>Betaproteobacteria>Nitrosomonadales>Thiobacillaceae | GCA_001725135.1 | Thiobacillus_sp._SCN_63-374 | ABS92_01640 | 11556..14369 | 1660149 | MAG:-hypothetical-protein-ABS92_01640-[Thiobacillus-sp.-SCN-63-374] |
| MCX7674148.1 | DUF499 _1.RM.like.systems | NTP_transf_2->HEPN->Mbetalac->tRNA->?-><- DUF499 *<- Schlafen +HATPase+wHTH <- METHYLASE <-?<-TOPRIM<-OLD-ABC<- METHYLASE | 1093 | Bacteria>Pseudomonadota>Betaproteobacteria>Nitrosomonadales>Thiobacillaceae | GCA_026419045.1 | Thiobacillaceae_bacterium | N2Z63_11245 | 10361..13642 | 2528409 | MAG:- DUF499 -domain-containing-protein-[Thiobacillaceae-bacterium] |
| MBI1887192.1 | DUF499 _1.RM.like.systems | <-Glycos_trans_3N+Glycos_transf_3<-?<- Schlafen +HATPase+wHTH <-?<-?<- HEPN+ DUF499 *<- METHYLASE <-HEPN<-TOPRIM<-OLD-ABC<- TUDOR+ SF1-SF2-Helicase+iREase+CTD+REase <-ParB | 1112 | Bacteria>Pseudomonadota>Betaproteobacteria>Nitrosomonadales>unclassified_Nitrosomonadales | GCA_016177795.1 | Nitrosomonadales_bacterium | HYS19_02300 | 86364..89702 | 2026761 | ATP-binding-protein-[Nitrosomonadales-bacterium] |
| MBI3902214.1 | DUF499 _1.RM.like.systems | <-Glycos_trans_3N+Glycos_transf_3<-?<-?<- Schlafen +HATPase+wHTH <-?<- HEPN+ DUF499 *<- METHYLASE <-?<-?<-TOPRIM<-OLD-ABC<- TUDOR+ SF1-SF2-Helicase+iREase+CTD+REase | 1112 | Bacteria>Pseudomonadota>Betaproteobacteria>Nitrosomonadales>unclassified_Nitrosomonadales | GCA_016199835.1 | Nitrosomonadales_bacterium | HY306_04630 | 118339..121677 | 2026761 | ATP-binding-protein-[Nitrosomonadales-bacterium] |
| MCA3140850.1 | DUF499 _1.RM.like.systems | HTH_3-> TUDOR+ SF1-SF2-Helicase+iREase+CTD+REase -> Schlafen +HATPase+wHTH ->?-> METHYLASE ->HTH+Hiskin+STAS-> HEPN+ DUF499 *->?->XIS-HTH->?->TOPRIM->MetJarc-HTH->RelE-ParE-> | 1113 | Bacteria>Pseudomonadota>Betaproteobacteria>Rhodocyclales>Rhodocyclaceae | GCA_020248385.1 | Rhodocyclaceae_bacterium | ING80_01400 | 55201..58542 | 1898103 | DUF499 -domain-containing-protein-[Rhodocyclaceae-bacterium] |
| MCX8086911.1 | DUF499 _1.RM.like.systems | <-Phage_integrase\|\|tRNA-><- DUF499 *<- Schlafen +HATPase+wHTH <- METHYLASE <-TOPRIM<-OLD-ABC<- METHYLASE | 943 | Bacteria>Pseudomonadota>Betaproteobacteria>Rhodocyclales>Rhodocyclaceae | GCA_026415155.1 | Rhodocyclaceae_bacterium | N3C63_08430 | 24540..27371 | 1898103 | MAG:- DUF499 -domain-containing-protein-[Rhodocyclaceae-bacterium] |
| OGA30960.1 | DUF499 _1.RM.like.systems | TUDOR+ SF1-SF2-Helicase+iREase+CTD+REase ->PHP+SbcC-> Schlafen +HATPase+wHTH -> METHYLASE ->DOC+HTH-> DUF499 *-><-?\|\|?->?-><-?<-?\|\|SPOUT-> | 1081 | Bacteria>Pseudomonadota>Betaproteobacteria>unclassified_Betaproteobacteria | GCA_001771975.1 | Betaproteobacteria_bacterium_RIFCSPLOWO2_02_FULL_68_150 | A3I65_01690 | 23801..27046 | 1797493 | hypothetical-protein-A3I65_01690-[Betaproteobacteria-bacterium-RIFCSPLOWO2_02_FULL_68_150] |
| OGA55362.1 | DUF499 _1.RM.like.systems | <- DUF499 *<-REase<- METHYLASE <- Schlafen +HATPase+wHTH <- TUDOR+ SF1-SF2-Helicase+iREase+CTD+REase | 941 | Bacteria>Pseudomonadota>Betaproteobacteria>unclassified_Betaproteobacteria | GCA_001772005.1 | Betaproteobacteria_bacterium_RIFCSPLOWO2_12_FULL_62_58 | A3F74_07155 | 338..3163 | 1797497 | AAA-family-ATPase-[Betaproteobacteria-bacterium-RIFCSPLOWO2_12_FULL_62_58] |
| MBI3917901.1 | DUF499 _1.RM.like.systems | OLD-ABC->PHP+SbcC-> Schlafen +HATPase+wHTH -> METHYLASE ->OLD-ABC->TOPRIM-> DUF499 *-> | 927 | Bacteria>Pseudomonadota>Betaproteobacteria>unclassified_Betaproteobacteria | GCA_016199545.1 | Betaproteobacteria_bacterium | HY322_12940 | 28133..30916 | 1891241 | ATP-binding-protein-[Betaproteobacteria-bacterium] |
| MBI5922415.1 | DUF499 _1.RM.like.systems | <- DUF499 *<-REase<- METHYLASE <-?<- Schlafen +HATPase+wHTH \|\|URI-DNase-> | 933 | Bacteria>Pseudomonadota>Betaproteobacteria>unclassified_Betaproteobacteria | GCA_016234745.1 | Betaproteobacteria_bacterium | HY847_12340 | 39471..42272 | 1891241 | ATP-binding-protein-[Betaproteobacteria-bacterium] |
| GHU44749.1 | DUF499 _1.RM.like.systems | <-Aminotran_1_2<- DUF499 *<- Schlafen +HATPase+wHTH <- METHYLASE <- METHYLASE <-PIN<- TUDOR+ SF1-SF2-Helicase+iREase+CTD+REase \|\|Aminotran_3-> | 945 | Bacteria>Pseudomonadota>Betaproteobacteria>unclassified_Betaproteobacteria | GCA_025997095.1 | Betaproteobacteria_bacterium | AGMMS50289_13810 | 2255..5092 | 1891241 | hypothetical-protein-AGMMS50289_13810-[Betaproteobacteria-bacterium] |
| WP_020742688.1 | DUF499 _1.RM.like.systems | <- METHYLASE <-Methyltransf_4<-ABhydrolase\|\|HHH+ENDO3+NUDIX->?->tRNA-><- Schlafen <- DUF499 *<- METHYLASE -p\|\|<-PAIREDC-HTH\|\|?->Resolvase->HTH_28+rve+Mu-transpos_C->GspA-ATPase-> | 939 | Bacteria>Pseudomonadota>Gammaproteobacteria>Alteromonadales>Alteromonadaceae | GCF_000439515.1 | Alteromonas_mediterranea | I876_RS01280 | 288971..291790 | 314275 | DUF499 -domain-containing-protein-[Alteromonas-mediterranea] |
| WP_182173095.1 | DUF499 _1.RM.like.systems | Phage_integrase-><-Phage_AlpA<-ZnR+TOPRIM+primpol+HTH<-XIS-HTH\|\|MobI-><- DUF499 *<- Schlafen +HATPase+wHTH <- METHYLASE <- METHYLASE <-SbcC+SbcC<- TUDOR+ SF1-SF2-Helicase+iREase+CTD+REase <-HTH+WYL | 951 | Bacteria>Pseudomonadota>Gammaproteobacteria>Cellvibrionales>Halieaceae | GCF_014077625.1 | Sediminihaliea_albiluteola | H2508_RS10780 | 844621..847476 | 2758564 | DUF499 -domain-containing-protein-[Sediminihaliea-albiluteola] |
| MAT52642.1 | DUF499 _1.RM.like.systems | <-KAP-NTPase\|\|HTH+WYL-> TUDOR+ SF1-SF2-Helicase+iREase+CTD+REase -> METHYLASE -> METHYLASE -> Schlafen +HATPase+wHTH -> DUF499 *-><-?<-TOPRIM<-XIS-HTH<-?<-Arm-DNA-bind_3+Phage_int_SAM_3+Phage_integrase | 949 | Bacteria>Pseudomonadota>Gammaproteobacteria>Cellvibrionales>Porticoccaceae | GCA_002702725.1 | Porticoccaceae_bacterium | CMK32_15795 | 17542..20391 | 2026782 | hypothetical-protein-[Porticoccaceae-bacterium] |
| WP_120796693.1 | DUF499 _1.RM.like.systems | TUDOR+ SF1-SF2-Helicase+iREase+CTD+REase -> METHYLASE -> Schlafen +HATPase+wHTH -> DUF499 *->?->?->?->CHTH-> | 1093 | Bacteria>Pseudomonadota>Gammaproteobacteria>Chromatiales>Chromatiaceae | GCF_003634315.1 | Thiocapsa_rosea | BDD21_RS07920 | 1738895..1742176 | 69360 | DUF499 -domain-containing-protein-[Thiocapsa-rosea] |
| WP_010962200.1 | DUF499 _1.RM.like.systems | TUDOR+ SF1-SF2-Helicase+iREase+CTD+REase ->PrlF_antitoxin->PemK_toxin-> METHYLASE -> METHYLASE -> Schlafen +HATPase+wHTH -> DUF499 *->TIR-> | 942 | Bacteria>Pseudomonadota>Gammaproteobacteria>Methylococcales>Methylococcaceae | GCF_000008325.1 | Methylococcus_capsulatus | MCA_RS14770 | 3194105..3196933 | 414 | DUF499 -domain-containing-protein-[Methylococcus-capsulatus] |
| WP_017365774.1 | DUF499 _1.RM.like.systems | XIS-HTH->?->?->?->?-><-TIR<- DUF499 *<- Schlafen +HATPase+wHTH <- METHYLASE <- METHYLASE <-PemK_toxin<-PrlF_antitoxin<- TUDOR+ SF1-SF2-Helicase+iREase+CTD+REase | 942 | Bacteria>Pseudomonadota>Gammaproteobacteria>Methylococcales>Methylococcaceae | GCF_000297615.1 | Methylococcus_capsulatus | QMX71_RS07545 | 1581839..1584667 | 414 | DUF499 -domain-containing-protein-[Methylococcus-capsulatus] |
| WP_142947823.1 | DUF499 _1.RM.like.systems | Arm-DNA-bind_3+Phage_integrase-><-Phage_AlpA<-ZnR+TOPRIM+primpol+HTH<-XIS-HTH\|\|MobI-><- DUF499 *<- Schlafen +HATPase+wHTH <- METHYLASE <- METHYLASE <-HNH<-OLD-ABC<- TUDOR+ SF1-SF2-Helicase+iREase+CTD+REase | 953 | Bacteria>Pseudomonadota>Gammaproteobacteria>Oceanospirillales>Alcanivoracaceae | - | Alcanivorax_sp._DSM_26295 | FN816_RS03075 | 107194..110055 | 1938738 | DUF499 -domain-containing-protein-[Alcanivorax-sp.-DSM-26295] |
| WP_234246274.1 | DUF499 _1.RM.like.systems | METHYLASE -> METHYLASE -> Schlafen +HATPase+wHTH ->HEPN+CBS->OLD-ABC->REase-> DUF499 *-> | 957 | Bacteria>Pseudomonadota>Gammaproteobacteria>Oceanospirillales>Halomonadaceae | GCF_021404345.1 | Halomonas_desiderata | HOP57_RS15495 | 85349..88222 | 52021 | DUF499 -domain-containing-protein-[Halomonas-desiderata] |
| WP_200203476.1 | DUF499 _1.RM.like.systems | Arm-DNA-bind_3+Phage_integrase-><-Phage_AlpA<-ZnR+TOPRIM+primpol+HTH<-?<-XIS-HTH\|\|<- DUF499 *<- Schlafen +HATPase+wHTH <- METHYLASE <- METHYLASE <-?<- Schlafen +HATPase+wHTH <- TUDOR+ SF1-SF2-Helicase+iREase+CTD+REase | 951 | Bacteria>Pseudomonadota>Gammaproteobacteria>Pseudomonadales>Marinobacteraceae | GCF_016597775.1 | Marinobacter_sp._1-3A | FE848_RS12575 | 132618..135473 | 2582920 | DUF499 -domain-containing-protein-[Marinobacter-sp.-1-3A] |
| MCO2926705.1 | DUF499 _1.RM.like.systems | TUDOR+ SF1-SF2-Helicase+iREase+CTD+REase -> Schlafen +HATPase+wHTH ->?-> METHYLASE ->HEPN-> DUF499 *-><-RlpA<-?<-?\|\|HTH_3-><-ZnR+ZnR | 943 | Bacteria>Pseudomonadota>Gammaproteobacteria>Pseudomonadales>Pseudomonadaceae | GCA_023909145.1 | Pseudomonas_aeruginosa | FA260_26715 | 18266..21097 | 287 | ATP-binding-protein-[Pseudomonas-aeruginosa] |
| WP_223119133.1 | DUF499 _1.RM.like.systems | ZnR+ZnR-><-HTH_3\|\|?->?->RlpA-><- DUF499 *<-HEPN<- METHYLASE <-?<- Schlafen +HATPase+wHTH <- TUDOR+ SF1-SF2-Helicase+iREase+CTD+REase | 942 | Bacteria>Pseudomonadota>Gammaproteobacteria>Pseudomonadales>Pseudomonadaceae | GCF_020036175.1 | Pseudomonas_aeruginosa | NFY03_RS10210 | 2202477..2205305 | 287 | DUF499 -domain-containing-protein-[Pseudomonas-aeruginosa] |
| WP_150035379.1 | DUF499 _1.RM.like.systems | TUDOR+ SF1-SF2-Helicase+iREase+CTD+REase -> Schlafen +HATPase+wHTH ->?-> METHYLASE ->HEPN-> DUF499 *-><-RlpA<-?<-?\|\|HTH_3-><-ZnR+ZnR | 942 | Bacteria>Pseudomonadota>Gammaproteobacteria>Pseudomonadales>Pseudomonadaceae | GCF_022627285.1 | Pseudomonas_aeruginosa | JNB05_RS00615 | 132976..135804 | 287 | DUF499 -domain-containing-protein-[Pseudomonas-aeruginosa] |
| WP_033979009.1 | DUF499 _1.RM.like.systems | ZnR+ZnR-><-HTH_3\|\|?->?->RlpA-><- DUF499 *<-HEPN<- METHYLASE <-?<- Schlafen +HATPase+wHTH <- TUDOR+ SF1-SF2-Helicase+iREase+CTD+REase | 943 | Bacteria>Pseudomonadota>Gammaproteobacteria>Pseudomonadales>Pseudomonadaceae | - | Pseudomonas_aeruginosa | CJT50_RS28650 | 19614..22445 | 287 | DUF499 -domain-containing-protein-[Pseudomonas-aeruginosa] |
| TAK54308.1 | DUF499 _1.RM.like.systems | TUDOR+ SF1-SF2-Helicase+iREase+CTD+REase -> Schlafen +HATPase+wHTH -> METHYLASE ->OLD-ABC->TOPRIM-> HEPN+ DUF499 *-> | 1105 | Bacteria>Pseudomonadota>Gammaproteobacteria>unclassified_Gammaproteobacteria | GCA_004297335.1 | Gammaproteobacteria_bacterium | EPO25_07530 | 13135..16452 | 1913989 | ATP-binding-protein-[Gammaproteobacteria-bacterium] |
| MCG3203220.1 | DUF499 _1.RM.like.systems | <-NTP_transf<-MORC+DUF3684+REase<- DUF499 *<- METHYLASE <- Schlafen +HATPase+wHTH <-PHP+SbcC<- TUDOR+ SF1-SF2-Helicase+iREase+CTD+REase \|\|MerR-HTH-> | 1092 | Bacteria>Pseudomonadota>Gammaproteobacteria>unclassified_Gammaproteobacteria | GCA_022071965.1 | Gammaproteobacteria_bacterium | NFCOHLIN_03153 | 6649..9927 | 1913989 | hypothetical-protein-[Gammaproteobacteria-bacterium] |
| MCP5127473.1 | DUF499 _1.RM.like.systems | 35exo->DOC+HTH-> TUDOR+ SF1-SF2-Helicase+iREase+CTD+REase -> Schlafen +HATPase+wHTH -> METHYLASE -> DUF499 *-> METHYLASE ->XIS-HTH+RVT-><-MobL+ZnR+TOPRIM<-DOC<-MazE_antitoxin<-Phage_int_SAM_4+Phage_integrase | 986 | Bacteria>Pseudomonadota>Gammaproteobacteria>unclassified_Gammaproteobacteria | GCA_024235095.1 | Gammaproteobacteria_bacterium | H6973_18120 | 278648..281608 | 1913989 | ATP-binding-protein-[Gammaproteobacteria-bacterium] |
| MCY4343961.1 | DUF499 _1.RM.like.systems | METHYLASE ->OLD-ABC-> Schlafen +HATPase+wHTH ->?->PIN-> HEPN+ DUF499 *-><-MNS+ZDNA-HTH<-PRTase | 1115 | Bacteria>Pseudomonadota>Gammaproteobacteria>unclassified_Gammaproteobacteria | GCA_026710215.1 | Gammaproteobacteria_bacterium | OXE83_10365 | 11310..14657 | 1913989 | MAG:-Swt1-family-HEPN-domain-containing-protein-[Gammaproteobacteria-bacterium] |
| MCZ0943260.1 | DUF499 _1.RM.like.systems | DUF433-><- HEPN+ DUF499 *<- Schlafen +HATPase+wHTH <-TOPRIM<-OLD-ABC<- METHYLASE -p | 1115 | Bacteria>Pseudomonadota>Gammaproteobacteria>unclassified_Gammaproteobacteria | GCA_026914145.1 | Gammaproteobacteria_bacterium | OXJ53_09405 | 2539..5886 | 1913989 | MAG:-Swt1-family-HEPN-domain-containing-protein-[Gammaproteobacteria-bacterium] |
| MDD9817274.1 | DUF499 _1.RM.like.systems | <- DUF499 *<-?<-TOPRIM<-OLD-ABC<- METHYLASE <- Schlafen +HATPase+wHTH <-RelE-ParE | 1113 | Bacteria>Pseudomonadota>Gammaproteobacteria>unclassified_Gammaproteobacteria | GCA_028817495.1 | Gammaproteobacteria_bacterium | OXU61_03925 | 191..3532 | 1913989 | MAG:-hypothetical-protein-OXU61_03925-[Gammaproteobacteria-bacterium] |
| MDE0129995.1 | DUF499 _1.RM.like.systems | <-SF1-SF2-Helicase<-OLD-ABC+TOPRIM<- Schlafen +HATPase+wHTH <- DUF499 *<- METHYLASE <- TUDOR+ SF1-SF2-Helicase+iREase+CTD+REase <-?<-OLD-ABC+TOPRIM | 968 | Bacteria>Pseudomonadota>Gammaproteobacteria>unclassified_Gammaproteobacteria | GCA_028819355.1 | Gammaproteobacteria_bacterium | OXQ86_10575 | 58066..60972 | 1913989 | MAG:- DUF499 -domain-containing-protein-[Gammaproteobacteria-bacterium] |
| MDE0453907.1 | DUF499 _1.RM.like.systems | <-HEPN+REase<- Schlafen +HATPase+wHTH <- Schlafen <- DUF499 *<- METHYLASE <-SF1-SF2-Helicase<-OLD-ABC+TOPRIM<- TUDOR+ SF1-SF2-Helicase+iREase+CTD+REase <-AbiEii<-AbiEi_4 | 967 | Bacteria>Pseudomonadota>Gammaproteobacteria>unclassified_Gammaproteobacteria | GCA_028821145.1 | Gammaproteobacteria_bacterium | OXJ63_01205 | 18382..21285 | 1913989 | MAG:- DUF499 -domain-containing-protein-[Gammaproteobacteria-bacterium] |
| TAN08689.1 | DUF499 _1.RM.like.systems | <-STYKinase-like<-HTH\|\|OLD-ABC+TOPRIM-> METHYLASE -> METHYLASE -> Schlafen +HATPase+wHTH -> DUF499 *-><-BirA-HTH<-?<-Phage_integrase<-?<-ZnR | 934 | Bacteria>Pseudomonadota>Gammaproteobacteria>Xanthomonadales>Rhodanobacteraceae | GCA_004299555.1 | Rhodanobacteraceae_bacterium | EPN38_01845 | 144698..147502 | 2052182 | ATP-binding-protein-[Rhodanobacteraceae-bacterium] |
| WP_043696850.1 | DUF499 _1.RM.like.systems | HTH+WYL-> TUDOR+ SF1-SF2-Helicase+iREase+CTD+REase ->SbcC+SbcC-> METHYLASE -> METHYLASE -> Schlafen +HATPase+wHTH -> DUF499 *-><-Phage_int_SAM_2+Integrase_1<-MarR-HTH\|\|?-><-?<-MODE-HTH+PBPII\|\|ADH_N+ADH_zinc_N-> | 947 | Bacteria>Pseudomonadota>Gammaproteobacteria>Xanthomonadales>Rhodanobacteraceae | GCF_000800155.1 | Luteibacter_sp._9133 | FA87_RS19045 | 4371799..4374642 | 1500891 | DUF499 -domain-containing-protein-[Luteibacter-sp.-9133] |
| MCU0753612.1 | DUF499 _1.RM.like.systems | <-ArsR-HTH\|\|?->?->?->?->?-><- DUF499 *<- Schlafen +HATPase+wHTH <-ParB+HNH<-REase<-OLD-ABC<- METHYLASE <-HNH | 955 | Bacteria>Pseudomonadota>Gammaproteobacteria>Xanthomonadales>unclassified_Xanthomonadales | GCA_025458445.1 | Xanthomonadales_bacterium | MUE46_00615 | 13163..16030 | 2006849 | DUF499 -domain-containing-protein-[Xanthomonadales-bacterium] |
| MCX5809780.1 | DUF499 _1.RM.like.systems | TUDOR+ SF1-SF2-Helicase+iREase+CTD+REase -> METHYLASE -> Schlafen +HATPase+wHTH -> DUF499 *->ParB-> | 1105 | Bacteria>Pseudomonadota>unclassified_Pseudomonadota | GCA_026388245.1 | Pseudomonadota_bacterium | NTX36_10495 | 9204..12521 | 1977087 | MAG:- DUF499 -domain-containing-protein-[Proteobacteria-bacterium] |
| MBN1673169.1 | DUF499 _1.RM.like.systems | <-DrHyd+BirA-HTH<- TUDOR+ SF1-SF2-Helicase+iREase+CTD+REase -p<-?<-?\|\|?->?-><- HEPN+ DUF499 *<- Schlafen +HATPase+wHTH <- METHYLASE -p\|\|BirA-HTH->HhH-RADC+JAB-> | 1118 | Bacteria>PVC_group>Kiritimatiellota>Kiritimatiellia>unclassified_Kiritimatiellia | GCA_016929905.1 | Kiritimatiellia_bacterium | JXR37_19145 | 44287..47643 | 2212473 | DUF499 -domain-containing-protein-[Kiritimatiellae-bacterium] |
| MBQ6473657.1 | DUF499 _1.RM.like.systems | Schlafen +HATPase+wHTH ->OLD-ABC->HNH-> METHYLASE ->OLD-ABC->NERD-> DUF499 *-> | 944 | Bacteria>PVC_group>Lentisphaerota>Lentisphaeria>Victivallales | GCA_017431185.1 | Victivallales_bacterium | IJJ33_16845 | 18968..21802 | 2212475 | ATP-binding-protein-[Victivallales-bacterium] |
| NLE53431.1 | DUF499 _1.RM.like.systems | METHYLASE -p-> Schlafen +HATPase+wHTH -> DUF499 *-> | 656 | Bacteria>PVC_group>Lentisphaerota>unclassified_Lentisphaerota | GCA_012514365.1 | Lentisphaerota_bacterium | GX617_00715 | 3231..5197 | 1932692 | ATP-binding-protein-[Lentisphaerota-bacterium] |
| NLF19752.1 | DUF499 _1.RM.like.systems | <- DUF499 *<-wHTH+REase<- METHYLASE <-?<-?<-?<- Schlafen +HATPase+wHTH | 905 | Bacteria>PVC_group>Lentisphaerota>unclassified_Lentisphaerota | GCA_012514865.1 | Lentisphaerota_bacterium | GX595_21175 | 1..2716 | 1932692 | ATP-binding-protein-[Lentisphaerae-bacterium] |
| MBK9119771.1 | DUF499 _1.RM.like.systems | DEOR-HTH-> TUDOR+ SF1-SF2-Helicase+iREase+CTD+REase -> Schlafen +HATPase+wHTH ->?->?-> METHYLASE -> HEPN+ DUF499 *->MORC->REase-><-Trypsin\|\|HEPN->HTH->MPTase-> | 1121 | Bacteria>PVC_group>Planctomycetota>Phycisphaerae>Phycisphaerales | GCA_016716475.1 | Phycisphaerales_bacterium | IPM18_09260 | 379276..382641 | 2052180 | DUF499 -domain-containing-protein-[Phycisphaerales-bacterium] |
| MCC6908120.1 | DUF499 _1.RM.like.systems | <-Hydrolase_3<- HEPN+ DUF499 *<- METHYLASE <-?<-?<- Schlafen +HATPase+wHTH <- TUDOR+ SF1-SF2-Helicase+iREase+CTD+REase <-DEOR-HTH | 1120 | Bacteria>PVC_group>Planctomycetota>Phycisphaerae>Phycisphaerales | GCA_020852515.1 | Phycisphaerales_bacterium | IT430_09285 | 88345..91707 | 2052180 | DUF499 -domain-containing-protein-[Phycisphaerales-bacterium] |
| MBW7904855.1 | DUF499 _1.RM.like.systems | <-HTH_3\|\|DEOR-HTH-> TUDOR+ SF1-SF2-Helicase+iREase+CTD+REase -> Schlafen +HATPase+wHTH ->REase-> METHYLASE -> HEPN+ DUF499 *->Hydrolase_3-> | 1120 | Bacteria>PVC_group>Planctomycetota>Phycisphaerae>unclassified_Phycisphaerae | GCA_019454965.1 | Phycisphaerae_bacterium | H3C42_04715 | 502745..506107 | 2026778 | ATP-binding-protein-[Phycisphaerae-bacterium] |
| GJQ27840.1 | DUF499 _1.RM.like.systems | <-N6_N4_MTase<-MPTase<-HTH<- DUF499 *<-?<- METHYLASE <- Schlafen +HATPase+wHTH \|\|?-><-?<- TUDOR+ SF1-SF2-Helicase+iREase+CTD+REase | 926 | Bacteria>PVC_group>Planctomycetota>Phycisphaerae>unclassified_Phycisphaerae | GCA_022180675.1 | Phycisphaerae_bacterium | HBSAPP02_28720 | 66529..69309 | 2026778 | hypothetical-protein-HBSAPP02_28720-[Phycisphaerae-bacterium] |
| NLF73683.1 | DUF499 _1.RM.like.systems | HD->Peptidase_M48-> TUDOR+ SF1-SF2-Helicase+iREase+CTD+REase -> Schlafen +HATPase+wHTH -> METHYLASE ->?-> DUF499 *->?->?->Primpol->Phage_integrase-> | 979 | Bacteria>PVC_group>Planctomycetota>Planctomycetia>Pirellulales | GCA_012515235.1 | Candidatus_Anammoximicrobium_sp. | GX575_31980 | 17079..20018 | 2699742 | ATP-binding-protein-[Candidatus-Anammoximicrobium-sp.] |
| MCL6502992.1 | DUF499 _1.RM.like.systems | <-OBFOLD-NUCLEASE<- DUF499 *<-?<- Schlafen +HATPase+wHTH <- METHYLASE -p | 973 | Bacteria>PVC_group>Planctomycetota>Planctomycetia>Pirellulales | GCA_023511675.1 | Pirellulales_bacterium | K6T86_09935 | 15292..18213 | 2715253 | DUF499 -domain-containing-protein-[Pirellulales-bacterium] |
| GIW99318.1 | DUF499 _1.RM.like.systems | METHYLASE -p->?->MACRODOMAIN-> METHYLASE ->PHP+SbcC-> Schlafen +HATPase+wHTH -> HEPN+ DUF499 *-><-URI-DNase<-?<-REase<-DDE_Tnp_IS66+DDE_Tnp_IS66_C | 1115 | Bacteria>PVC_group>Planctomycetota>Planctomycetia>Pirellulales | GCA_026004795.1 | Pirellulaceae_bacterium | KatS3mg111_2651 | 704885..708232 | 2699754 | MAG:-hypothetical-protein-KatS3mg111_2651-[Pirellulaceae-bacterium] |
| GHT10305.1 | DUF499 _1.RM.like.systems | METHYLASE -p-> METHYLASE -p->ZDNA-HTH-> DUF499 *->Peptidase-> Schlafen ->BirA-HTH-> | 1093 | Bacteria>PVC_group>Planctomycetota>Planctomycetia>Planctomycetales | GCA_026001785.1 | Planctomycetales_bacterium | FACS1894170_02190 | 13810..17091 | 2053591 | hypothetical-protein-FACS1894170_02190-[Planctomycetales-bacterium] |
| WP_145451931.1 | DUF499 _1.RM.like.systems | HTH_3-><-?<-?<-HKD-DNase<-?<-?<- HEPN+ DUF499 *<- METHYLASE <- Schlafen +HATPase+wHTH <-MORC+DUF3684<- TUDOR+ SF1-SF2-Helicase+iREase+CTD+REase <-XIS-HTH | 1094 | Bacteria>PVC_group>Planctomycetota>Planctomycetia>Planctomycetales | GCF_007748155.1 | Gimesia_panareensis | Enr10x_RS26145 | 6953187..6956471 | 2527978 | Swt1-family-HEPN-domain-containing-protein-[Gimesia-panareensis] |
| MBE7458022.1 | DUF499 _1.RM.like.systems | DEOR-HTH-> TUDOR+ SF1-SF2-Helicase+iREase+CTD+REase -> Schlafen +HATPase+wHTH ->?->?-> METHYLASE -> HEPN+ DUF499 *->MORC-><-?<-HTH_3<-RelE-ParE<-?<-DDE_Tnp_1_4 | 1120 | Bacteria>PVC_group>Planctomycetota>Planctomycetia>unclassified_Planctomycetia | GCA_015075705.1 | Planctomycetia_bacterium | HS102_15595 | 57764..61126 | 2052181 | ATP-binding-protein-[Planctomycetia-bacterium] |
| PIE22549.1 | DUF499 _1.RM.like.systems | SMODS->SLATT-> TUDOR+ SF1-SF2-Helicase+iREase+CTD+REase -> METHYLASE ->PHP+SbcC-> Schlafen +HATPase+wHTH -> HEPN+ DUF499 *-><-tRNA\|\|?->?->?->?->SF1-SF2-Helicase-> | 1122 | Bacteria>PVC_group>Planctomycetota>unclassified_Planctomycetota | GCA_002748355.1 | Planctomycetota_bacterium | CSA62_11865 | 35983..39351 | 2026780 | MAG:-AAA+-family-ATPase-[Planctomycetota-bacterium] |
| QDU83134.1 | DUF499 _1.RM.like.systems | TUDOR+ SF1-SF2-Helicase+iREase+CTD+REase -> Schlafen +HATPase+wHTH ->?-> METHYLASE ->OLD-ABC->TOPRIM-> HEPN+ DUF499 *->?-><-REase+SF1-SF2-Helicase<-DUF2357+REase<-Preader7+McrB-NTD+McrB-AAA | 1111 | Bacteria>PVC_group>Planctomycetota>unclassified_Planctomycetota | GCA_007750655.1 | Planctomycetes_bacterium_Pla163 | Pla163_02310 | 293863..297198 | 2528008 | hypothetical-protein-Pla163_02310-[Planctomycetes-bacterium-Pla163] |
| KAA3609103.1 | DUF499 _1.RM.like.systems | TUDOR+ SF1-SF2-Helicase+iREase+CTD+REase -> Schlafen +HATPase+wHTH -> METHYLASE ->PHP+SbcC-> Schlafen +HATPase+wHTH -> HEPN+ DUF499 *->?->?-><-?<-CHTH\|\|?-><-HhH-RADC+JAB | 1120 | Bacteria>PVC_group>Planctomycetota>unclassified_Planctomycetota | GCA_008501825.1 | Planctomycetota_bacterium | DWQ01_09790 | 14904..18266 | 2026780 | MAG:- DUF499 -domain-containing-protein-[Planctomycetota-bacterium] |
| MBI2191311.1 | DUF499 _1.RM.like.systems | <- DUF499 *<-REase<- METHYLASE <-HEPN<-HEPN<- Schlafen +HATPase+wHTH \|\|DEDD_Tnp_IS110+Transposase_20-> | 926 | Bacteria>PVC_group>Planctomycetota>unclassified_Planctomycetota | GCA_016179685.1 | Planctomycetota_bacterium | HYU36_04935 | 72260..75040 | 2026780 | ATP-binding-protein-[Planctomycetota-bacterium] |
| MBI2193014.1 | DUF499 _1.RM.like.systems | TUDOR+ SF1-SF2-Helicase+iREase+CTD+REase -> Schlafen +HATPase+wHTH -> METHYLASE -> METHYLASE ->Y1_Tnp->DUF433-> DUF499 *->ABhydrolase-><-TPR<-SIGMA-HTH | 946 | Bacteria>PVC_group>Planctomycetota>unclassified_Planctomycetota | GCA_016179685.1 | Planctomycetota_bacterium | HYU36_13630 | 54036..56876 | 2026780 | ATP-binding-protein-[Planctomycetota-bacterium] |

| Protein-ID | System | Gene-Neighbourhoods | Length | Taxonomy | Assembly-ID | Strain | Locus-Tag | Genomic-coordinates | TaxID | Fasta-defline |
|---|---|---|---|---|---|---|---|---|---|---|
| MBM4038866.1 | DUF499_1.RM.like.systems | <- DUF499 *<-?<- METHYLASE -p<- METHYLASE <- Schlafen +HATPase+wHTH <- TUDOR+ SF1-SF2-Helicase+iREase+CTD+REase | 938 | Bacteria>PVC_group>Planctomycetota>unclassified_Planctomycetota | GCA_016872855.1 | Planctomycetota_bacterium | FJ290_10155 | 3372..6188 | 2026780 | ATP-binding-protein-[Planctomycetota-bacterium] |
| MCF7707718.1 | DUF499_1.RM.like.systems | TUDOR+ SF1-SF2-Helicase+iREase+CTD+REase ->OLD-ABC->TOPRIM->?-> METHYLASE -> Schlafen +HATPase+wHTH -> DUF499 *-> | 1113 | Bacteria>PVC_group>Verrucomicrobiota>unclassified_Verrucomicrobiota | GCA_021733835.1 | Verrucomicrobiota_bacterium | K9N52_02315 | 37691..41032 | 2026799 | DUF499-domain-containing-protein-[Verrucomicrobia-bacterium] |
| WP_142661086.1 | DUF499_1.RM.like.systems | Schlafen +HATPase+wHTH -> METHYLASE -> DUF499 *-> | 945 | Bacteria>PVC_group>Verrucomicrobiota>Verrucomicrobiota_incertae_sedis>Methylacidimicrobium | GCF_902143375.2 | Methylacidimicrobium_tartarophylax | MAMT_RS11075 | 5444..8281 | 1041768 | DUF499-domain-containing-protein-[Methylacidimicrobium-tartarophylax] |
| MDD6486202.1 | DUF499_1.RM.like.systems | <-HTH\|\| TUDOR+ SF1-SF2-Helicase+iREase+CTD+REase -> METHYLASE ->APATPase+iSTAND3-> Schlafen +HATPase+wHTH -> DUF499 *->HKD-DNase+HKD-DNase->?->HicB_lk_antitox->tRNA->tRNA->?->Mbetalac-> | 948 | Bacteria>Spirochaetota>Spirochaetia>Spirochaetales>unclassified_Spirochaetales | GCA_029227075.1 | Spirochaetales_bacterium | PUF61_04585 | 321927..324773 | 2026792 | MAG:-DUF499-domain-containing-protein-[Spirochaetales-bacterium] |
| MCC7078886.1 | DUF499_1.RM.like.systems | <- Schlafen +HATPase+wHTH <-?<- HEPN+ DUF499 *<- METHYLASE <- Schlafen +HATPase+wHTH <- TUDOR+ SF1-SF2-Helicase+iREase+CTD+REase | 1112 | Bacteria>Terrabacteria_group>Actinomycetota>Acidimicrobiia>unclassified_Acidimicrobiia | GCA_020850945.1 | Acidimicrobiia_bacterium | IT198_17355 | 29484..32822 | 2080302 | DUF499-domain-containing-protein-[Acidimicrobiia-bacterium] |
| WP_113812542.1 | DUF499_1.RM.like.systems | <-ABhydrolase\|\| TUDOR+ SF1-SF2-Helicase+iREase+CTD+REase -> Schlafen +HATPase+wHTH -> METHYLASE ->ParB+HNH->?-> HEPN+ DUF499 *->NTD+SF1-SF2-Helicase->wHTH+SAD+HNH-><-?<-?\|\|?-><-REase+35exo+Beta-Barrel+SF1-SF2-Helicase | 1114 | Bacteria>Terrabacteria_group>Actinomycetota>Actinomycetes>Geodermatophilales | GCF_003319095.1 | Blastococcus_sp._TBT05-19 | DQ244_RS16250 | 131306..134650 | 2250581 | Swt1-family-HEPN-domain-containing-protein-[Blastococcus-sp.-TBT05-19] |
| NEB00544.1 | DUF499_1.RM.like.systems | VapB_antitoxin->PIN-><- DUF499 *<- METHYLASE <- TUDOR+ SF1-SF2-Helicase+iREase+CTD+REase \|\| Schlafen +HATPase+wHTH -> | 931 | Bacteria>Terrabacteria_group>Actinomycetota>Actinomycetes>Kitasatosporales | GCA_010548375.1 | Streptomyces_sp._SID13726 | G3I78_15960 | 365366..368161 | 2706058 | ATP-binding-protein-[Streptomyces-sp.-SID13726] |
| MBV7696079.1 | DUF499_1.RM.like.systems | <-REase<-?<- Schlafen +HATPase+wHTH \|\| TUDOR+ SF1-SF2-Helicase+iREase+CTD+REase -> METHYLASE -> DUF499 *-> | 931 | Bacteria>Terrabacteria_group>Actinomycetota>Actinomycetes>Kitasatosporales | GCA_019219825.1 | Streptomyces_sp._TRM70350 | KWI83_10510 | 185854..188649 | 2856165 | DUF499-domain-containing-protein-[Streptomyces-sp.-TRM70350] |
| MBX9397445.1 | DUF499_1.RM.like.systems | <- Schlafen +HATPase+wHTH \|\| TUDOR+ SF1-SF2-Helicase+iREase+CTD+REase -> METHYLASE -> DUF499 *-> | 932 | Bacteria>Terrabacteria_group>Actinomycetota>Actinomycetes>Kitasatosporales | GCA_019733235.1 | Streptomyces_sp._TRM72054 | K4749_28635 | 8261..11059 | 2870562 | DUF499-domain-containing-protein-[Streptomyces-sp.-TRM72054] |
| WP_030324200.1 | DUF499_1.RM.like.systems | wHTH+SAD+HNH->MetJarc-HTH->?-><- Schlafen +HATPase+wHTH \|\| TUDOR+ SF1-SF2-Helicase+iREase+CTD+REase -> METHYLASE -> DUF499 *-> | 931 | Bacteria>Terrabacteria_group>Actinomycetota>Actinomycetes>Kitasatosporales | GCF_000719505.1 | Streptomyces_sp._NRRL_B-3229 | IF54_RS0129580 | 225412..228207 | 1463836 | DUF499-domain-containing-protein-[Streptomyces-sp.-NRRL-B-3229] |
| WP_166682464.1 | DUF499_1.RM.like.systems | <- Schlafen +HATPase+wHTH \|\| TUDOR+ SF1-SF2-Helicase+iREase+CTD+REase -> METHYLASE -> DUF499 *-><-HAD\|\|HTH_31->?->?->RADICAL-SAM-> | 955 | Bacteria>Terrabacteria_group>Actinomycetota>Actinomycetes>Kitasatosporales | GCF_004365705.1 | Streptomyces_sp._846.5 | EDD99_RS20240 | 4433112..4435979 | 2485156 | DUF499-domain-containing-protein-[Streptomyces-sp.-846.5] |
| WP_239157864.1 | DUF499_1.RM.like.systems | VapB_antitoxin-><- DUF499 *<- METHYLASE <- TUDOR+ SF1-SF2-Helicase+iREase+CTD+REase \|\| Schlafen +HATPase+wHTH ->REase-> | 850 | Bacteria>Terrabacteria_group>Actinomycetota>Actinomycetes>Kitasatosporales | GCF_010548375.1 | Streptomyces_sp._SID13726 | G3I78_RS15960 | 365366..367918 | 2706058 | DUF499-domain-containing-protein-[Streptomyces-sp.-SID13726] |
| WP_218514040.1 | DUF499_1.RM.like.systems | <-REase<-?<- Schlafen +HATPase+wHTH \|\| TUDOR+ SF1-SF2-Helicase+iREase+CTD+REase -> METHYLASE -> DUF499 *-> | 931 | Bacteria>Terrabacteria_group>Actinomycetota>Actinomycetes>Kitasatosporales | GCF_019219825.1 | Streptomyces_sp._TRM70350 | KWI83_10510 | 185854..188649 | 2856165 | DUF499-domain-containing-protein-[Streptomyces-sp.-TRM70350] |
| WP_221752200.1 | DUF499_1.RM.like.systems | <- Schlafen +HATPase+wHTH \|\| TUDOR+ SF1-SF2-Helicase+iREase+CTD+REase -> METHYLASE -> DUF499 *-> | 932 | Bacteria>Terrabacteria_group>Actinomycetota>Actinomycetes>Kitasatosporales | GCF_019733235.1 | Streptomyces_sp._TRM72054 | K4749_28635 | 8261..11059 | 2870562 | DUF499-domain-containing-protein-[Streptomyces-sp.-TRM72054] |
| WP_254370730.1 | DUF499_1.RM.like.systems | Methyltransf->?-><-?\|\|?-><- DUF499 *<- METHYLASE <- TUDOR+ SF1-SF2-Helicase+iREase+CTD+REase \|\| Schlafen +HATPase+wHTH ->?-><-?\|\|ABhydrolase-> | 953 | Bacteria>Terrabacteria_group>Actinomycetota>Actinomycetes>Kitasatosporales | GCF_024213665.1 | Streptomyces_sp._A3M-1-3 | NLX86_07020 | 25922..28783 | 2962044 | DUF499-domain-containing-protein-[Streptomyces-sp.-A3M-1-3] |
| WP_263976853.1 | DUF499_1.RM.like.systems | <- DUF499 *<- TUDOR+ SF1-SF2-Helicase+iREase+CTD+REase \|\| Schlafen +HATPase+wHTH -><-?<-HTH_3\|\|ST-Hiskin-> | 851 | Bacteria>Terrabacteria_group>Actinomycetota>Actinomycetes>Kitasatosporales | - | Streptomyces_sp._NK08204 | K7G59_RS27400 | 90033..92588 | 2873260 | DUF499-domain-containing-protein-[Streptomyces-sp.-NK08204] |
| WP_256362104.1 | DUF499_1.RM.like.systems | HTH+DUF5753->?-><-REase<-?<- Schlafen +HATPase+wHTH \|\| TUDOR+ SF1-SF2-Helicase+iREase+CTD+REase -> DUF499 *-> | 811 | Bacteria>Terrabacteria_group>Actinomycetota>Actinomycetes>Kitasatosporales | - | Streptomyces_sp._TRM70350 | KWI83_RS10510 | 186214..188649 | 2856165 | DUF499-domain-containing-protein-[Streptomyces-sp.-TRM70350] |
| WP_260463288.1 | DUF499_1.RM.like.systems | <-REase<- Schlafen +HATPase+wHTH \|\| TUDOR+ SF1-SF2-Helicase+iREase+CTD+REase -> DUF499 *-> | 851 | Bacteria>Terrabacteria_group>Actinomycetota>Actinomycetes>Kitasatosporales | - | Streptomyces_sp._TRM72054 | K4749_RS28650 | 8504..11059 | 2870562 | DUF499-domain-containing-protein-[Streptomyces-sp.-TRM72054] |
| MBK6299925.1 | DUF499_1.RM.like.systems | <-ABC-ATPase+SbcC<-ABC-ATPase+SbcC<-TetR-HTH<-NTD+SF1-SF2-Helicase<- HEPN+ DUF499 *<- Schlafen +HATPase+wHTH <- METHYLASE <- TUDOR+ SF1-SF2-Helicase+iREase+CTD+REase \|\|PadR-HTH->?-><-PrmC_N+Methyltransf_25 | 1117 | Bacteria>Terrabacteria_group>Actinomycetota>Actinomycetes>Micrococcales | GCA_016705565.1 | Candidatus_Phosphoribacter_hodrii | IPF40_02340 | 519882..523235 | 2953743 | DUF499-domain-containing-protein-[Actinomycetales-bacterium] |
| WP_108718589.1 | DUF499_1.RM.like.systems | <-JetB<-JetA\|\|?->PIN->?-><-?<- HEPN+ DUF499 *<-ParB+HNH<-HEPN+HEPN<- METHYLASE \|\|HTH-> Schlafen +HATPase+wHTH -><- TUDOR+ SF1-SF2-Helicase+iREase+CTD+REase | 1134 | Bacteria>Terrabacteria_group>Actinomycetota>Actinomycetes>Micrococcales | GCF_003074035.1 | Miniimonas_sp._S16 | DCS86_RS08145 | 1885887..1889291 | 2171623 | Swt1-family-HEPN-domain-containing-protein-[Miniimonas-sp.-S16] |
| WP_169323296.1 | DUF499_1.RM.like.systems | <-NTD+SF1-SF2-Helicase<- TUDOR+ SF1-SF2-Helicase+iREase+CTD+REase \|\| Schlafen +HATPase+wHTH -> METHYLASE -> HEPN+ DUF499 *->HD->TrwC+SF1-SF2-Helicase->?->?-><-DNA-glycosylase<-ArdCN+MPTase | 1120 | Bacteria>Terrabacteria_group>Actinomycetota>Actinomycetes>Micrococcales | GCF_012952065.1 | Cellulomonas_fimi | HIR71_RS02970 | 36548..39910 | 1708 | DUF499-domain-containing-protein-[Cellulomonas-fimi] |
| WP_119663874.1 | DUF499_1.RM.like.systems | <- DUF499 *<- METHYLASE <- TUDOR+ SF1-SF2-Helicase+iREase+CTD+REase <-ZnR<-GDresolvase<-ParB+HTH\|\| Schlafen +HATPase+wHTH -> | 581 | Bacteria>Terrabacteria_group>Actinomycetota>Actinomycetes>Micromonosporales | GCF_003581605.1 | Micromonospora_endophytica | D3H59_RS20815 | 1..1744 | 515350 | DUF499-domain-containing-protein,-partial-[Micromonospora-endophytica] |
| EHB57971.1 | DUF499_1.RM.like.systems | <-GNTR-HTH\|\|NUDIX-><-?<-HEPN<- Schlafen +HATPase+wHTH <- HEPN+ DUF499 *<-REase<- METHYLASE <- TUDOR+ SF1-SF2-Helicase+iREase+CTD+REase <-PIN<-XIS-HTH | 1108 | Bacteria>Terrabacteria_group>Actinomycetota>Actinomycetes>Mycobacteriales | GCA_000230935.2 | Mycolicibacterium_rhodesiae_JS60 | MycrhDRAFT_0407 | 390606..393932 | 931627 | hypothetical-protein-MycrhDRAFT_0407-[Mycolicibacterium-rhodesiae-JS60] |
| AFC52219.1 | DUF499_1.RM.like.systems | <-GNTR-HTH\|\|NUDIX-><-?<-?<-HEPN<- Schlafen +HATPase+wHTH <- HEPN+ DUF499 *<- METHYLASE \|\|?-><- TUDOR+ SF1-SF2-Helicase+iREase+CTD+REase <-PIN<-XIS-HTH | 1043 | Bacteria>Terrabacteria_group>Actinomycetota>Actinomycetes>Mycobacteriales | GCA_000276825.1 | Mycobacterium_paraintracellulare | OCQ_07060 | 694865..697996 | 1138383 | hypothetical-protein-OCQ_07060-[Mycobacterium-paraintracellulare] |
| OOK63774.1 | DUF499_1.RM.like.systems | <-CNMP+TIR<- Schlafen +HATPase+wHTH <- HEPN+ DUF499 *<- METHYLASE <- METHYLASE -p<- TUDOR+ SF1-SF2-Helicase+iREase+CTD+REase -p<-?<-SF1-SF2-Helicase-p | 1104 | Bacteria>Terrabacteria_group>Actinomycetota>Actinomycetes>Mycobacteriales | GCA_002003585.1 | Mycobacterium_kansasii | BZL29_8433 | 3439..6753 | 1768 | hypothetical-protein-BZL29_8433-[Mycobacterium-kansasii] |
| OSC29643.1 | DUF499_1.RM.like.systems | <-GNTR-HTH\|\|NUDIX-><-?<-?<- Schlafen +HATPase+wHTH <- HEPN+ DUF499 *<- METHYLASE \|\|?-><- TUDOR+ SF1-SF2-Helicase+iREase+CTD+REase <-PIN<-XIS-HTH\|\|GNTR-HTH-p-> | 1097 | Bacteria>Terrabacteria_group>Actinomycetota>Actinomycetes>Mycobacteriales | GCA_002104735.1 | Mycobacterium_paraintracellulare | B8W68_04660 | 128474..131767 | 1138383 | AAA-family-ATPase-[Mycobacterium-paraintracellulare] |
| AYJ07722.1 | DUF499_1.RM.like.systems | TIR->?->?-> TUDOR+ SF1-SF2-Helicase+iREase+CTD+REase -><-?\|\| METHYLASE -> HEPN+ DUF499 *-> Schlafen +HATPase+wHTH ->HEPN-><-NUDIX\|\|GNTR-HTH->?->FtsK-> | 1097 | Bacteria>Terrabacteria_group>Actinomycetota>Actinomycetes>Mycobacteriales | GCA_003640565.1 | Mycobacterium_avium | DBO90_17285 | 3694672..3697965 | 1764 | DUF499-domain-containing-protein-[Mycobacterium-avium] |
| TXI55381.1 | DUF499_1.RM.like.systems | <-FtsK<-?<-GNTR-HTH\|\|NUDIX-><-HEPN<- Schlafen +HATPase+wHTH <- HEPN+ DUF499 *<-ParB+HNH<- METHYLASE <- TUDOR+ SF1-SF2-Helicase+iREase+CTD+REase <-?<-HEPN | 1103 | Bacteria>Terrabacteria_group>Actinomycetota>Actinomycetes>Mycobacteriales | GCA_008015745.1 | Mycolicibacter_arupensis | E6Q54_12720 | 7339..10650 | 342002 | ATP-binding-protein-[Mycolicibacter-arupensis] |
| WP_009974960.1 | DUF499_1.RM.like.systems | TIR->?->?-> TUDOR+ SF1-SF2-Helicase+iREase+CTD+REase -><-?\|\| METHYLASE -> HEPN+ DUF499 *-> Schlafen +HATPase+wHTH ->HEPN-><-?<-NUDIX\|\|GNTR-HTH-> | 1108 | Bacteria>Terrabacteria_group>Actinomycetota>Actinomycetes>Mycobacteriales | GCF_000174035.1 | Mycobacterium_avium | MAA44156_RS17120 | 3691227..3694553 | 1764 | Swt1-family-HEPN-domain-containing-protein-[Mycobacterium-avium] |
| WP_041784793.1 | DUF499_1.RM.like.systems | <-GNTR-HTH\|\|NUDIX-><-?<-?<-HEPN<- Schlafen +HATPase+wHTH <- HEPN+ DUF499 *<- METHYLASE \|\|?-><- TUDOR+ SF1-SF2-Helicase+iREase+CTD+REase <-PIN<-XIS-HTH\|\|GNTR-HTH-p-> | 1108 | Bacteria>Terrabacteria_group>Actinomycetota>Actinomycetes>Mycobacteriales | GCF_000276825.1 | Mycobacterium_paraintracellulare | B8W68_RS04660 | 128474..131800 | 1138383 | Swt1-family-HEPN-domain-containing-protein-[Mycobacterium-paraintracellulare] |
| WP_062879806.1 | DUF499_1.RM.like.systems | <-GNTR-HTH\|\|NUDIX-><-?<-HEPN<- Schlafen +HATPase+wHTH <- HEPN+ DUF499 *<- METHYLASE \|\|?-><- TUDOR+ SF1-SF2-Helicase+iREase+CTD+REase <-PIN<-XIS-HTH | 1108 | Bacteria>Terrabacteria_group>Actinomycetota>Actinomycetes>Mycobacteriales | GCF_001606235.1 | Mycobacteroides_abscessus | B5431_RS12990 | 716837..720163 | 36809 | Swt1-family-HEPN-domain-containing-protein-[Mycobacteroides-abscessus] |
| WP_157762129.1 | DUF499_1.RM.like.systems | BirA-HTH-><- Schlafen +HATPase+wHTH \|\| TUDOR+ SF1-SF2-Helicase+iREase+CTD+REase -> METHYLASE -> DUF499 *-><-?<-tRNA\|\|CR-Kinase-><-TetR-HTH | 911 | Bacteria>Terrabacteria_group>Actinomycetota>Actinomycetes>Mycobacteriales | GCF_001613325.1 | Nocardia_yamanashiensis | NY1_RS09400 | 81283..84018 | 209247 | DUF499-domain-containing-protein-[Nocardia-yamanashiensis] |
| WP_276761078.1 | DUF499_1.RM.like.systems | <-FtsK<-?<-GNTR-HTH\|\|NUDIX-><-HEPN<- Schlafen +HATPase+wHTH <- HEPN+ DUF499 *<-ParB+HNH<- METHYLASE <- TUDOR+ SF1-SF2-Helicase+iREase+CTD+REase <-?<-HEPN<-HEPN | 1114 | Bacteria>Terrabacteria_group>Actinomycetota>Actinomycetes>Mycobacteriales | GCF_008015745.1 | Mycolicibacter_arupensis | E6Q54_RS12710 | 7339..10683 | 342002 | Swt1-family-HEPN-domain-containing-protein-[Mycolicibacter-arupensis] |
| WP_062887256.1 | DUF499_1.RM.like.systems | <-FtsK<-?\|\|NUDIX-><-HEPN<- Schlafen +HATPase+wHTH <- HEPN+ DUF499 *<- METHYLASE \|\|?-><- TUDOR+ SF1-SF2-Helicase+iREase+CTD+REase | 1107 | Bacteria>Terrabacteria_group>Actinomycetota>Actinomycetes>Mycobacteriales | GCF_018524295.1 | Mycobacteriaceae | KL858_RS23375 | 11898..15221 | 1762 | Swt1-family-HEPN-domain-containing-protein-[Mycobacterium-avium] |
| WP_225506778.1 | DUF499_1.RM.like.systems | XIS-HTH->PIN-> TUDOR+ SF1-SF2-Helicase+iREase+CTD+REase -> METHYLASE -> HEPN+ DUF499 *-> Schlafen +HATPase+wHTH ->\|\|DDE_Tnp_1_4->HEPN->?-><-NUDIX\|\|GNTR-HTH-> | 1108 | Bacteria>Terrabacteria_group>Actinomycetota>Actinomycetes>Mycobacteriales | GCF_020227835.1 | Mycolicibacterium_fortuitum | H8Z59_RS27040 | 5490569..5493895 | 1766 | Swt1-family-HEPN-domain-containing-protein-[Mycolicibacterium-fortuitum] |
| WP_227974732.1 | DUF499_1.RM.like.systems | TIR->?->?-> TUDOR+ SF1-SF2-Helicase+iREase+CTD+REase -><-?\|\| METHYLASE -> HEPN+ DUF499 *-> Schlafen +HATPase+wHTH ->HEPN-><-?<-NUDIX\|\|GNTR-HTH-> | 1108 | Bacteria>Terrabacteria_group>Actinomycetota>Actinomycetes>Mycobacteriales | GCF_020735405.1 | Mycobacterium_avium | LT348_RS02565 | 526704..530030 | 1764 | Swt1-family-HEPN-domain-containing-protein-[Mycobacterium-avium] |
| WP_233059597.1 | DUF499_1.RM.like.systems | TIR->?->?-> TUDOR+ SF1-SF2-Helicase+iREase+CTD+REase -><-?\|\| METHYLASE -> HEPN+ DUF499 *-> Schlafen +HATPase+wHTH ->HEPN-><-?<-NUDIX\|\|GNTR-HTH-> | 1108 | Bacteria>Terrabacteria_group>Actinomycetota>Actinomycetes>Mycobacteriales | GCF_021183805.1 | Mycobacterium_avium | LTQ57_RS17660 | 3790685..3794011 | 1764 | Swt1-family-HEPN-domain-containing-protein-[Mycobacterium-avium] |
| WP_211284609.1 | DUF499_1.RM.like.systems | <-FtsK<-?<-GNTR-HTH\|\|NUDIX-><-HEPN<- Schlafen +HATPase+wHTH <- HEPN+ DUF499 *<- METHYLASE <- TUDOR+ SF1-SF2-Helicase+iREase+CTD+REase <-?<-RNaseH | 1113 | Bacteria>Terrabacteria_group>Actinomycetota>Actinomycetes>Mycobacteriales | GCF_025823205.1 | Mycobacterium_alsense | BST11_RS03260 | 100678..104019 | 324058 | Swt1-family-HEPN-domain-containing-protein-[Mycobacterium-alsense] |
| WP_204801726.1 | DUF499_1.RM.like.systems | <-GNTR-HTH\|\|NUDIX-><-?<-HEPN<- Schlafen +HATPase+wHTH <- HEPN+ DUF499 *<- METHYLASE \|\|?-><- TUDOR+ SF1-SF2-Helicase+iREase+CTD+REase <-HEPN+REase<-NTP_transf | 1109 | Bacteria>Terrabacteria_group>Actinomycetota>Actinomycetes>Mycobacteriales | GCF_905219495.1 | Mycobacterium_riyadhense | JX354_RS03520 | 789692..793021 | 486698 | Swt1-family-HEPN-domain-containing-protein-[Mycobacterium-riyadhense] |
| WP_005139269.1 | DUF499_1.RM.like.systems | <-GNTR-HTH\|\|NUDIX-><-?<-HEPN<- Schlafen +HATPase+wHTH <- HEPN+ DUF499 *<-REase<- METHYLASE <- TUDOR+ SF1-SF2-Helicase+iREase+CTD+REase <-PIN<-XIS-HTH | 1108 | Bacteria>Terrabacteria_group>Actinomycetota>Actinomycetes>Mycobacteriales | - | Mycolicibacterium_rhodesiae | MycrhDRAFT_0407 | 390606..393932 | 36814 | Swt1-family-HEPN-domain-containing-protein-[Mycolicibacterium-rhodesiae] |
| MBV9142989.1 | DUF499_1.RM.like.systems | <- HEPN+ DUF499 *<- METHYLASE -p<- TUDOR+ SF1-SF2-Helicase+iREase+CTD+REase <- Schlafen +HATPase+wHTH <-Methyltransf\|\| METHYLASE -p-><-PIN | 1073 | Bacteria>Terrabacteria_group>Actinomycetota>Actinomycetes>Pseudonocardiales | GCA_019247305.1 | Pseudonocardiales_bacterium | JO115_19100 | 30..3251 | 2201155 | DUF499-domain-containing-protein-[Pseudonocardiales-bacterium] |
| GFP42119.1 | DUF499_1.RM.like.systems | <- DUF499 *<- Schlafen +HATPase+wHTH <-PHP+SbcC<- METHYLASE <- METHYLASE -p<- TUDOR+ SF1-SF2-Helicase+iREase+CTD+REase -p | 603 | Bacteria>Terrabacteria_group>Actinomycetota>Actinomycetota_incertae_sedis>Candidatus_Hakubellales | GCA_013282075.1 | Candidatus_Hakubella_thermalkaliphila | HKBW3C_01245 | 1..1809 | 2754717 | hypothetical-protein-HKBW3C_01245,-partial-[Candidatus-Hakubanella-thermoalkaliphilus] |
| WP_176229285.1 | DUF499_1.RM.like.systems | Peptidase_M29-><-PemK_toxin<-?<- DUF499 *<- Schlafen +HATPase+wHTH <-PHP+SbcC<- METHYLASE <- TUDOR+ SF1-SF2-Helicase+iREase+CTD+REase | 867 | Bacteria>Terrabacteria_group>Actinomycetota>Actinomycetota_incertae_sedis>Candidatus_Hakubellales | GCA_013282015.1 | Candidatus_Hakubella_thermalkaliphila | HW437_RS01600 | 1284..3887 | 2754717 | DUF499-domain-containing-protein-[Candidatus-Hakubanella-thermoalkaliphilus] |
| PKQ27862.1 | DUF499_1.RM.like.systems | REase-> TUDOR+ SF1-SF2-Helicase+iREase+CTD+REase -> Schlafen +HATPase+wHTH -> METHYLASE -> HEPN+ DUF499 *->?->TIR-> | 1105 | Bacteria>Terrabacteria_group>Actinomycetota>Candidatus_Geothermincolia>Candidatus_Geothermincolales | GCA_002843005.1 | Candidatus_Anoxymicrobium_japonicum | CVT63_05765 | 8955..12272 | 2013648 | AAA+-family-ATPase-[Candidatus-Anoxymicrobium-japonicum] |
| NLE73128.1 | DUF499_1.RM.like.systems | AbiEi_1->AbiEii->HIN-HTH-><-?<- Schlafen <- HEPN+ DUF499 *<-DOC+DOC<-HTH_3<-Tox-RelE<-RelE-ParE<-?<- METHYLASE -p | 1130 | Bacteria>Terrabacteria_group>Actinomycetota>unclassified_Actinomycetota | GCA_012514635.1 | Actinomycetota_bacterium | GX604_00625 | 12665..16057 | 2900548 | ATP-binding-protein-[Actinomycetia-bacterium] |
| MCW3097907.1 | DUF499_1.RM.like.systems | TUDOR+ SF1-SF2-Helicase+iREase+CTD+REase ->?-> METHYLASE ->?->MACRODOMAIN-> DUF499 *-> Schlafen +TUDOR+PIWI->?-><-?<-ParB-HTH | 1081 | Bacteria>Terrabacteria_group>Armatimonadota>Chthonomonadetes>Chthonomonadales | GCA_025963795.1 | Chthonomonadaceae_bacterium | JWL77_3525 | 19451..22696 | 2739509 | MAG:-hypothetical-protein-JWL77_3525-[Chthonomonadaceae-bacterium] |
| MCC7230625.1 | DUF499_1.RM.like.systems | TIR->?->?-> TUDOR+ SF1-SF2-Helicase+iREase+CTD+REase -> Schlafen +HATPase+wHTH -> METHYLASE -> HEPN+ DUF499 *-><-JAB\|\|?->ArdCN+MPTase->?->N6_N4_MTase-><-PIN | 1116 | Bacteria>Terrabacteria_group>Armatimonadota>Fimbriimonadia>Fimbriimonadales | GCA_020851885.1 | Fimbriimonadaceae_bacterium | IT203_09555 | 16596..19946 | 2682144 | DUF499-domain-containing-protein-[Fimbriimonadaceae-bacterium] |
| GIV02394.1 | DUF499_1.RM.like.systems | <-URI-DNase+RAMA<-?<-HEPN+REase<-?<-REase<- Schlafen +HATPase+wHTH <- DUF499 *<-REase<-?<- METHYLASE -p\|\|?-><-?<- METHYLASE | 920 | Bacteria>Terrabacteria_group>Armatimonadota>Fimbriimonadia>Fimbriimonadales | GCA_026003055.1 | Fimbriimonadales_bacterium | KatS3mg015_1224 | 1249734..1252496 | 2682145 | MAG:-hypothetical-protein-KatS3mg015_1224-[Fimbriimonadales-bacterium] |
| GIV04311.1 | DUF499_1.RM.like.systems | <- DUF499 *<-HTH<- METHYLASE <-OLD-ABC+TOPRIM<-SF1-SF2-Helicase-p<- Schlafen +HATPase+wHTH <- TUDOR+ SF1-SF2-Helicase+iREase+CTD+REase | 944 | Bacteria>Terrabacteria_group>Armatimonadota>Fimbriimonadia>Fimbriimonadales | GCA_026003055.1 | Fimbriimonadales_bacterium | KatS3mg015_3141 | 322..3156 | 2682145 | MAG:-hypothetical-protein-KatS3mg015_3141-[Fimbriimonadales-bacterium] |
| MCX7689026.1 | DUF499_1.RM.like.systems | TUDOR+ SF1-SF2-Helicase+iREase+CTD+REase -p-> Schlafen +HATPase+wHTH -> METHYLASE -> METHYLASE -p-> DUF499 *-> | 872 | Bacteria>Terrabacteria_group>Armatimonadota>Fimbriimonadia>Fimbriimonadales | GCA_026418845.1 | Fimbriimonadales_bacterium | N2045_13770 | 4406..7024 | 2682145 | MAG:-DUF499-domain-containing-protein-[Fimbriimonadales-bacterium] |
| GIV15264.1 | DUF499_1.RM.like.systems | TUDOR+ SF1-SF2-Helicase+iREase+CTD+REase ->?-> Schlafen +HATPase+wHTH -> Schlafen +HATPase+wHTH -> METHYLASE -p-> METHYLASE -p-> DUF499 *-> | 873 | Bacteria>Terrabacteria_group>Armatimonadota>unclassified_Armatimonadota | GCA_026003175.1 | Armatimonadota_bacterium | KatS3mg022_0699 | 793472..796093 | 2033014 | MAG:-hypothetical-protein-KatS3mg022_0699-[Armatimonadota-bacterium] |
| MCX6379001.1 | DUF499_1.RM.like.systems | <- DUF499 *<- Schlafen +HATPase+wHTH <- METHYLASE <-MORC+DUF3684+REase<- TUDOR+ SF1-SF2-Helicase+iREase+CTD+REase <-?\|\|XIS-HTH-> | 625 | Bacteria>Terrabacteria_group>Armatimonadota>unclassified_Armatimonadota | GCA_026392135.1 | Armatimonadota_bacterium | NT023_05910 | 1..1875 | 2033014 | MAG:-DUF499-domain-containing-protein,-partial-[Armatimonadetes-bacterium] |
| MCL1790896.1 | DUF499_1.RM.like.systems | <-Peptidase\|\| TUDOR+ SF1-SF2-Helicase+iREase+CTD+REase -> METHYLASE -> Schlafen +HATPase+wHTH -> HEPN+ DUF499 *-> | 1122 | Bacteria>Terrabacteria_group>Bacillota>Clostridia>Eubacteriales | GCA_009778595.1 | Peptococcaceae_bacterium | FWG40_05990 | 9595..12963 | 2052179 | Swt1-family-HEPN-domain-containing-protein-[Peptococcaceae-bacterium] |
| MBU5249610.1 | DUF499_1.RM.like.systems | DUF772+DDE_Tnp_1_6-> Schlafen +HATPase+wHTH -> TUDOR+ SF1-SF2-Helicase+iREase+CTD+REase -> METHYLASE -> METHYLASE -> HEPN+ DUF499 *->?->CRP-HTH+GNTR-HTH->?->?->?->REase-> | 1128 | Bacteria>Terrabacteria_group>Bacillota>Clostridia>Eubacteriales | GCA_018916865.1 | Coprococcus_comes | KQH93_07505 | 29424..32810 | 410072 | DUF499-domain-containing-protein-[Coprococcus-comes] |
| MCD7717395.1 | DUF499_1.RM.like.systems | REase->?->?-><-NACHT\|\|?-><-REase+ParB<- HEPN+ DUF499 *<- METHYLASE <- Schlafen +HATPase+wHTH <- METHYLASE <- TUDOR+ SF1-SF2-Helicase+iREase+CTD+REase <-XIS-HTH | 1132 | Bacteria>Terrabacteria_group>Bacillota>Clostridia>Eubacteriales | GCA_021199235.1 | Lachnospiraceae_bacterium | LUI39_13305 | 18966..22364 | 1898203 | DUF499-domain-containing-protein-[Lachnospiraceae-bacterium] |
| MCD7762788.1 | DUF499_1.RM.like.systems | <-NACHT\|\|?-><-MORC<- HEPN+ DUF499 *<- METHYLASE <- Schlafen +HATPase+wHTH <- METHYLASE | 1132 | Bacteria>Terrabacteria_group>Bacillota>Clostridia>Eubacteriales | GCA_021199655.1 | Lachnospiraceae_bacterium | LUI14_06240 | 6036..9434 | 1898203 | DUF499-domain-containing-protein-[Lachnospiraceae-bacterium] |
| WP_018214270.1 | DUF499_1.RM.like.systems | <-pPIWI+pPIWI<-HEPN<-ParB+HNH<- METHYLASE -p<- Schlafen <-MORC<- HEPN+ DUF499 *<- METHYLASE <- TUDOR+ SF1-SF2-Helicase+iREase+CTD+REase <-HTH_26<-?<-MORC+SF1-SF2-Helicase<-SF1-SF2-Helicase+HKD-DNase | 1137 | Bacteria>Terrabacteria_group>Bacillota>Clostridia>Eubacteriales | GCF_000378805.1 | Desulfitobacterium_hafniense | DESHAF_RS0122600 | 1962937..1966350 | 49338 | DUF499-domain-containing-protein-[Desulfitobacterium-hafniense] |
| WP_168124741.1 | DUF499_1.RM.like.systems | <- HEPN+ DUF499 *<- METHYLASE <- METHYLASE <- Schlafen +HATPase+wHTH <- TUDOR+ SF1-SF2-Helicase+iREase+CTD+REase <-HTH_26<-SIGMA-HTH | 1132 | Bacteria>Terrabacteria_group>Bacillota>Clostridia>Eubacteriales | GCF_012027765.1 | Pseudoflavonifractor_sp._SW1122 | EOL15_RS04955 | 37276..40674 | 2530044 | DUF499-domain-containing-protein-[Pseudoflavonifractor-sp.-SW1122] |
| WP_204872058.1 | DUF499_1.RM.like.systems | SIGMA-HTH->HTH_26-> TUDOR+ SF1-SF2-Helicase+iREase+CTD+REase -> Schlafen +HATPase+wHTH -> METHYLASE -> METHYLASE -> HEPN+ DUF499 *-> | 1132 | Bacteria>Terrabacteria_group>Bacillota>Clostridia>Eubacteriales | GCF_016899615.1 | Pseudoflavonifractor_capillosus | H9X87_RS10190 | 14789..18187 | 106588 | DUF499-domain-containing-protein-[Pseudoflavonifractor-capillosus] |
| WP_204889591.1 | DUF499_1.RM.like.systems | HTH_26-> TUDOR+ SF1-SF2-Helicase+iREase+CTD+REase -> METHYLASE -> Schlafen +HATPase+wHTH -> HEPN+ DUF499 *-><-?<-HIN-HTH\|\|DUF772+DDE_Tnp_1+DDE_Tnp_1_6-> | 1123 | Bacteria>Terrabacteria_group>Bacillota>Clostridia>Eubacteriales | GCF_016902175.1 | Pseudoflavonifractor_phocaeensis | H6B27_RS11850 | 9172..12543 | 1870988 | Swt1-family-HEPN-domain-containing-protein-[Pseudoflavonifractor-phocaeensis] |
| WP_156275504.1 | DUF499_1.RM.like.systems | <-CopG_antitoxin\|\|NTP_transf_2-><-?<-?<- Schlafen <- HEPN+ DUF499 *<- METHYLASE <- TUDOR+ SF1-SF2-Helicase+iREase+CTD+REase \|\|?-><-?<-Phage_terminase | 1122 | Bacteria>Terrabacteria_group>Bacillota>Clostridia>Moorellales | GCF_009735625.1 | Moorella_glycerini | MGLY_RS15905 | 3178613..3181981 | 55779 | DUF499-domain-containing-protein-[Moorella-glycerini] |
| PWM79769.1 | DUF499_1.RM.like.systems | Peptidase_S8->?-><-?<-?<-STYKinase<- HEPN+ DUF499 *<- METHYLASE <- METHYLASE <- Schlafen +HATPase+wHTH <- TUDOR+ SF1-SF2-Helicase+iREase+CTD+REase <-HTH_26 | 1130 | Bacteria>Terrabacteria_group>Bacillota>Negativicutes>Acidaminococcales | GCA_003150755.1 | Phascolarctobacterium_sp. | DBY32_02635 | 546389..549781 | 2049039 | MAG:-hypothetical-protein-DBY32_02635-[Phascolarctobacterium-sp.] |
| NLG83533.1 | DUF499_1.RM.like.systems | REase-><-?<- HEPN+ DUF499 *<- METHYLASE <- Schlafen +HATPase+wHTH <- TUDOR+ SF1-SF2-Helicase+iREase+CTD+REase | 1127 | Bacteria>Terrabacteria_group>Bacillota>unclassified_Bacillota | GCA_012516115.1 | Bacillota_bacterium | GX493_02745 | 877..4260 | 1879010 | ATP-binding-protein-[Firmicutes-bacterium] |
| NPV80543.1 | DUF499_1.RM.like.systems | REase->?-><-DOC\|\| TUDOR+ SF1-SF2-Helicase+iREase+CTD+REase -> Schlafen +HATPase+wHTH -> METHYLASE -> HEPN+ DUF499 *->?-><-HEPN-p | 1115 | Bacteria>Terrabacteria_group>Bacillota>unclassified_Bacillota | GCA_013178415.1 | Bacillota_bacterium | HPY52_09760 | 58381..61728 | 1879010 | ATP-binding-protein-[Firmicutes-bacterium] |
| MAS36305.1 | DUF499_1.RM.like.systems | Glyoxalase->?-><-PIN<-?<- HEPN+ DUF499 *<-REase<-?<- METHYLASE <- Schlafen +HATPase+wHTH | 1094 | Bacteria>Terrabacteria_group>Chloroflexota>Anaerolineae>Anaerolineales | GCA_002702065.1 | Anaerolineaceae_bacterium | CL610_20035 | 35853..39137 | 2024896 | AAA+-family-ATPase-[Anaerolineaceae-bacterium] |
| MCB9452414.1 | DUF499_1.RM.like.systems | <-MetJarc-HTH\|\|?->TPR+TPR+TPR+TPR+TPR+TPR+TPR+TPR-><- HEPN+ DUF499 *<-ParB+HNH<- METHYLASE <- Schlafen +HATPase+wHTH <-PIN<-?<- TUDOR+ SF1-SF2-Helicase+iREase+CTD+REase | 1106 | Bacteria>Terrabacteria_group>Chloroflexota>Anaerolineae>Anaerolineales | GCA_020634115.1 | Anaerolineaceae_bacterium | H6672_13335 | 112307..115627 | 2024896 | ATP-binding-protein-[Anaerolineaceae-bacterium] |
| MBK6429951.1 | DUF499_1.RM.like.systems | OLD-ABC->TOPRIM-> Schlafen +HATPase+wHTH -> METHYLASE -p-> Schlafen +HATPase+wHTH ->REase-> HEPN+ DUF499 *->?->?->PemK_toxin->ABhydrolase->?->ABhydrolase+ABhydrolase-> | 1113 | Bacteria>Terrabacteria_group>Chloroflexota>Anaerolineae>unclassified_Anaerolineae | GCA_016704685.1 | Anaerolineae_bacterium | IPF85_04125 | 226121..229462 | 2052143 | DUF499-domain-containing-protein-[Anaerolineae-bacterium] |
| MBK9093182.1 | DUF499_1.RM.like.systems | <-ABhydrolase+ABhydrolase<-?<-ABhydrolase<-?<-?<- HEPN+ DUF499 *<- Schlafen +HATPase+wHTH <- METHYLASE -p<- Schlafen +HATPase+wHTH <- TUDOR+ SF1-SF2-Helicase+iREase+CTD+REase <-TOPRIM<-OLD-ABC | 1133 | Bacteria>Terrabacteria_group>Chloroflexota>Anaerolineae>unclassified_Anaerolineae | GCA_016719785.1 | Anaerolineae_bacterium | IPM84_10425 | 127540..130941 | 2052143 | DUF499-domain-containing-protein-[Anaerolineae-bacterium] |
| CAG1014975.1 | DUF499_1.RM.like.systems | PHP+SbcC-> Schlafen +HATPase+wHTH -> METHYLASE -p->?-> METHYLASE ->HEPN-> DUF499 *-> | 923 | Bacteria>Terrabacteria_group>Chloroflexota>Anaerolineae>unclassified_Anaerolineae | GCA_905339325.1 | Anaerolineae_bacterium | ANRL4_05364 | 17229..20000 | 2052143 | hypothetical-protein-ANRL4_05364-[Anaerolineae-bacterium] |
| MCE7987213.1 | DUF499_1.RM.like.systems | <-HAMP+HISKIN<-?<- HEPN+ DUF499 *<-Y1_Tnp<- METHYLASE <- Schlafen +HATPase+wHTH | 1167 | Bacteria>Terrabacteria_group>Chloroflexota>Caldilineae>Caldilineales | GCA_021405325.1 | Caldilinea_sp._CFX5 | DYG89_39080 | 104570..108073 | 2293554 | ATP-binding-protein-[Caldilinea-sp.-CFX5] |
| GIV69804.1 | DUF499_1.RM.like.systems | Schlafen +HATPase+wHTH -> METHYLASE ->REase-><-?\|\| DUF499 *-> | 854 | Bacteria>Terrabacteria_group>Chloroflexota>Caldilineae>Caldilineales | GCA_026003635.1 | Caldilinea_sp. | KatS3mg048_2666 | 137004..139568 | 2293560 | MAG:-hypothetical-protein-KatS3mg048_2666-[Caldilinea-sp.] |
| WP_012120125.1 | DUF499_1.RM.like.systems | <-Peptidase_C14<-Peptidase_C14<- DUF499 *<- Schlafen +HATPase+wHTH <-PHP+SbcC+PspA+SbcC<- METHYLASE <- TUDOR+ SF1-SF2-Helicase+iREase+CTD+REase <-AbiEii<-AbiEi_4 | 854 | Bacteria>Terrabacteria_group>Chloroflexota>Chloroflexia>Chloroflexales | GCF_000017805.1 | Roseiflexus_castenholzii | RCAS_RS08215 | 2067227..2069791 | 120962 | DUF499-domain-containing-protein-[Roseiflexus-castenholzii] |
| HAF11542.1 | DUF499_1.RM.like.systems | <-GDResolvase+Recombinase+ZnR<-?<- HEPN+ DUF499 *<- METHYLASE <- TUDOR+ SF1-SF2-Helicase+iREase+CTD+REase <- Schlafen +HATPase+wHTH | 1109 | Bacteria>Terrabacteria_group>Chloroflexota>unclassified_Chloroflexota | GCA_003446795.1 | Chloroflexota_bacterium | DCK98_15880 | 2135..5464 | 2026724 | AAA+-family-ATPase-[Chloroflexi-bacterium] |
| MBI4322283.1 | DUF499_1.RM.like.systems | TUDOR+ SF1-SF2-Helicase+iREase+CTD+REase ->HD->DUF4065-> METHYLASE -p->REase-> METHYLASE -> DUF499 *->?-><-?<-?<-?\|\| TUDOR+ SF1-SF2-Helicase+iREase+CTD+REase -> Schlafen +HATPase+wHTH -> | 979 | Bacteria>Terrabacteria_group>Chloroflexota>unclassified_Chloroflexota | GCA_016210495.1 | Chloroflexota_bacterium | HY675_27635 | 34699..37638 | 2026724 | ATP-binding-protein-[Chloroflexi-bacterium] |

--
-- OLD-ABC -> TOPRIM ->
-- OLD-ABC + TOPRIM ->SF1-SF2-Helicase
--

| Protein-ID | System | Gene-Neighbourhoods | Length | Taxonomy | Assembly-ID | Strain | Locus-Tag | Genomic-coordinates | TaxID | Fasta-defline |
|---|---|---|---|---|---|---|---|---|---|---|
| RLF71961.1 | DUF499_1.RM.like.systems | <-DUF499*<-TOPRIM<-OLD-ABC<-METHYLASE<-?<-SbcC<-TUDOR+SF1-SF2-Helicase+iREase+CTD+REase | 691 | Archaea>Candidatus_Thermoplasmatota>Thermoplasmata>unclassified_Thermoplasmata | GCA_003663625.1 | Thermoplasmata_archaeon | DRN40_01330 | 1..2074 | 1906666 | MAG:-AAA-family-ATPase,-partial-[Thermoplasmata-archaeon] |
| RLI77774.1 | DUF499_1.RM.like.systems | METHYLASE-p-><-TOPRIM<-OLD-ABC\|\|MazE_antitoxin->PIN->DUF499*-> | 227 | Archaea>Euryarchaeota>Archaeoglobi>Archaeoglobales>unclassified_Archaeoglobales | GCA_003662925.1 | Archaeoglobales_archaeon | DRP04_10895 | 3032..3711 | 2250258 | hypothetical-protein-DRP04_10895,-partial-[Archaeoglobales-archaeon] |
| WP_214400399.1 | DUF499_1.RM.like.systems | PolB+HEPN->METHYLASE->SbcC-><-TOPRIM<-TOPRIM<-OLD-ABC\|\|DUF499*-><-?<-?\|\|MarR-HTH->?-><-Peptidase_M42 | 878 | Archaea>Euryarchaeota>Methanomada_group>Methanococci>Methanococcales | GCF_902827225.1 | Methanocaldococcus_lauensis | KMP69_RS02590 | 459984..462620 | 2546128 | DUF499-domain-containing-protein-[Methanocaldococcus-sp.-SG7] |
| NLL10045.1 | DUF499_1.RM.like.systems | METHYLASE-p->OLD-ABC->TOPRIM->DUF499*-><-?\|\|Transposase_31+DUF4351-> | 970 | Archaea>Euryarchaeota>Stenosarchaea_group>Methanomicrobia>Methanomicrobiales | GCA_012520015.1 | Methanomicrobiales_archaeon | GX268_03955 | 2465..5377 | 2184052 | ATP-binding-protein-[Methanomicrobiales-archaeon] |
| WP_167816002.1 | DUF499_1.RM.like.systems | Schlafen+HATPase+HTH->?->Schlafen+HATPase+HTH->METHYLASE->OLD-ABC->TOPRIM->HEPN+DUF499*-> | 756 | Archaea>Euryarchaeota>Stenosarchaea_group>Methanomicrobia>Methanomicrobiales | GCF_012062385.1 | Methanocorpusculum_sp._GPch4 | EYA77_RS04465 | 15957..18225 | 2527877 | Swt1-family-HEPN-domain-containing-protein,-partial-[Methanocorpusculum-sp.-GPch4] |
| WP_204222940.1 | DUF499_1.RM.like.systems | OLD-ABC->OLD-ABC->REase->METHYLASE->OLD-ABC->TOPRIM->HEPN+DUF499*-> | 1082 | Archaea>Euryarchaeota>Stenosarchaea_group>Methanomicrobia>Methanomicrobiales | GCF_000013445.1 | Methanospirillum_hungatei | MHUN_RS09170 | 1984894..1988142 | 2203 | DUF499-domain-containing-protein-[Methanospirillum-hungatei] |
| ABD41477.1 | DUF499_1.RM.like.systems | OLD-ABC->OLD-ABC->REase->METHYLASE->OLD-ABC->TOPRIM->HEPN+DUF499*-> | 1094 | Archaea>Euryarchaeota>Stenosarchaea_group>Methanomicrobia>Methanomicrobiales | GCA_000013445.1 | Methanospirillum_hungatei_JF-1 | Mhun_1756 | 1984858..1988142 | 323259 | conserved-hypothetical-protein-[Methanospirillum-hungatei-JF-1] |
| MCW3134161.1 | DUF499_1.RM.like.systems | <-DUF499*<-TOPRIM<-OLD-ABC<-METHYLASE-p<-METHYLASE<-SF1-SF2-Helicase | 434 | Archaea>Euryarchaeota>Stenosarchaea_group>Methanomicrobia>Methanophagales | GCA_025966595.1 | Methanophagales_archaeon | N2V78_07540 | 1..1304 | 2056316 | MAG:-DUF499-domain-containing-protein,-partial-[Methanophagales-archaeon] |
| MCW3129356.1 | DUF499_1.RM.like.systems | METHYLASE->OLD-ABC->TOPRIM->DUF499*-><-?<-?<-Aminotran_1_2<-?<-?<-sGTP | 843 | Archaea>Euryarchaeota>Stenosarchaea_group>Methanomicrobia>Methanophagales | GCA_025966515.1 | Methanophagales_archaeon | N2V75_04565 | 3652..6183 | 2056316 | MAG:-hypothetical-protein-N2V75_04565-[Methanophagales-archaeon] |
| QNO47559.1 | DUF499_1.RM.like.systems | <-DDE_Tnp_1\|\|METHYLASE-p->METHYLASE-p->OLD-ABC->TOPRIM->DUF499*->?->?-><-?<-?<-?<-Phage_integrase | 854 | Archaea>Euryarchaeota>Stenosarchaea_group>Methanomicrobia>Methanosarcinales | - | Methanosarcinales_archaeon_ANME-2c_ERB4 | GGGHDLIA_00049 | 33141..35705 | 2759911 | hypothetical-protein-GGGHDLIA_00049-[Methanosarcinales-archaeon-ANME-2c-ERB4] |
| OFV65717.1 | DUF499_1.RM.like.systems | <-DUF499*<-TOPRIM<-OLD-ABC<-METHYLASE<-?\|\|PADR-HTH->PIN-> | 860 | Archaea>Euryarchaeota>Stenosarchaea_group>Methanomicrobia>Methanosarcinales | GCA_001766825.1 | Candidatus_Syntrophoarchaeum_butanivorans | SBU_001300 | 796..3378 | 1839936 | ATPase-AAA-[Candidatus-Syntrophoarchaeum-butanivorans] |
| WP_273278519.1 | DUF499_1.RM.like.systems | Primpol->?->ArdCN+MPTase->?->?-><-REase<-HEPN+DUF499*<-TOPRIM<-OLD-ABC<-PolB<-METHYLASE<-HEPN<-TUDOR+SF1-SF2-Helicase+iREase+CTD+REase | 1113 | Archaea>Euryarchaeota>Stenosarchaea_group>Methanomicrobia>Methanotrichales | GCF_913757535.1 | Methanothrix_soehngenii | NW278_RS08460 | 1661104..1664445 | 2223 | DUF499-domain-containing-protein-[Methanothrix-soehngenii] |
| MBK7385730.1 | DUF499_1.RM.like.systems | Methyltransf-><-HEPN+DUF499*<-TOPRIM<-OLD-ABC<-METHYLASE\|\|?->?-><-TUDOR+SF1-SF2-Helicase+iREase+CTD+REase | 1119 | Archaea>Euryarchaeota>Stenosarchaea_group>Methanomicrobia>Methanotrichales | GCA_016706325.1 | Methanothrix_sp. | IPI63_02955 | 568305..571664 | 90426 | DUF499-domain-containing-protein-[Methanothrix-sp.] |
| MBU4220744.1 | DUF499_1.RM.like.systems | <-PolB+HEPN<-PolB\|\|?-><-DUF499*<-TOPRIM<-OLD-ABC<-METHYLASE<-?<-SbcC<-METHYLASE-p | 846 | Archaea>Euryarchaeota>unclassified_Euryarchaeota | GCA_018898145.1 | Euryarchaeota_archaeon | KKA10_03835 | 1706..4246 | 2026739 | DUF499-domain-containing-protein-[Euryarchaeota-archaeon] |
| OQA55746.1 | DUF499_1.RM.like.systems | METHYLASE->?->OLD-ABC->?->TOPRIM->HEPN+DUF499*-> | 1082 | Archaea>Euryarchaeota>unclassified_Euryarchaeota | GCA_002069575.1 | Euryarchaeota_archaeon_ADurb.Bin294 | BWY45_02073 | 4487..7735 | 1852878 | hypothetical-protein-BWY45_02073-[Euryarchaeota-archaeon-ADurb.Bin294] |
| QQR73412.1 | DUF499_1.RM.like.systems | STAS+HisKin->?->?->METHYLASE->OLD-ABC->TOPRIM->HEPN+DUF499*-><-?<-Aminotran_1_2<-?<-?\|\|ACET-> | 1117 | Bacteria>Acidobacteriota>Holophagae>Holophagales>unclassified_Holophagales | GCA_016699405.1 | Holophagales_bacterium | IPJ17_18310 | 4435335..4438688 | 2478486 | DUF499-domain-containing-protein-[Holophagales-bacterium] |
| MBP7799241.1 | DUF499_1.RM.like.systems | OLD-ABC+TOPRIM->METHYLASE->OLD-ABC->TOPRIM->?->?->DUF499*-> | 919 | Bacteria>Acidobacteriota>Thermoanaerobaculia>Thermoanaerobaculales>Thermoanaerobaculaceae | GCA_018000955.1 | Thermoanaerobaculaceae_bacterium | KA072_11485 | 15680..18439 | 2699758 | ATP-binding-protein-[Thermoanaerobaculaceae-bacterium] |
| MCU0292531.1 | DUF499_1.RM.like.systems | <-PIN<-?\|\|?->?-><-DUF499-p<-DUF499*<-TOPRIM<-OLD-ABC<-METHYLASE | 869 | Bacteria>Acidobacteriota>Thermoanaerobaculia>Thermoanaerobaculales>Thermoanaerobaculaceae | GCA_025456225.1 | Thermoanaerobaculaceae_bacterium | MUF10_11190 | 5913..8522 | 2699758 | DUF499-domain-containing-protein-[Thermoanaerobaculaceae-bacterium] |
| MBP7799276.1 | DUF499_1.RM.like.systems | <-HEPN+DUF499*<-PIN<-OLD-ABC<-METHYLASE<-TOPRIM<-OLD-ABC | 1114 | Bacteria>Acidobacteriota>Thermoanaerobaculia>Thermoanaerobaculales>Thermoanaerobaculaceae | GCA_018000955.1 | Thermoanaerobaculaceae_bacterium | KA072_11695 | 22806..26150 | 2699758 | DUF499-domain-containing-protein-[Thermoanaerobaculaceae-bacterium] |
| MCG3190932.1 | DUF499_1.RM.like.systems | <-DUF499*<-TOPRIM<-OLD-ABC<-METHYLASE | 1102 | Bacteria>Acidobacteriota>Thermoanaerobaculia>unclassified_Thermoanaerobaculia | GCA_022071945.1 | Thermoanaerobaculia_bacterium | DIJKHBIC_00155 | 188742..192050 | 2651171 | hypothetical-protein-[Thermoanaerobaculia-bacterium] |
| MCG3192277.1 | DUF499_1.RM.like.systems | TUDOR+SF1-SF2-Helicase+iREase+CTD+REase->OLD-ABC->TOPRIM->METHYLASE->PHP+SbcC->Schlafen+HATPase+HTH->HEPN+DUF499*->?->?->PAIREDC-HTH-> | 1114 | Bacteria>Acidobacteriota>Thermoanaerobaculia>unclassified_Thermoanaerobaculia | GCA_022071945.1 | Thermoanaerobaculia_bacterium | DIJKHBIC_01515 | 140917..144261 | 2651171 | hypothetical-protein-[Thermoanaerobaculia-bacterium] |
| MCK6683888.1 | DUF499_1.RM.like.systems | TUDOR+SF1-SF2-Helicase+iREase+CTD+REase->?->?->MACRODOMAIN->METHYLASE->HEPN+DUF499*->?->OLD-ABC+TOPRIM->DUF932->?->ASCH+DUF3780-> | 1093 | Bacteria>Acidobacteriota>Thermoanaerobaculia>unclassified_Thermoanaerobaculia | GCA_023151375.1 | Thermoanaerobaculia_bacterium | L6R30_15920 | 12714..15995 | 2651171 | DUF499-domain-containing-protein-[Thermoanaerobaculia-bacterium] |
| MCY4659240.1 | DUF499_1.RM.like.systems | <-ORC-CDC6+REase<-DUF499*<-PIN<-OLD-ABC<-METHYLASE<-ParB+HNH+Schlafen<-TOPRIM<-OLD-ABC | 941 | Bacteria>Acidobacteriota>unclassified_Acidobacteriota | GCA_026713985.1 | Acidobacteriota_bacterium | OXF93_05425 | 1121..3946 | 1978231 | MAG:-DUF499-domain-containing-protein-[Acidobacteria-bacterium] |
| MDE2979142.1 | DUF499_1.RM.like.systems | ParB+HNH+Schlafen->PolB->METHYLASE->?->OLD-ABC->TOPRIM->DUF499*->XIS-HTH+RVT-> | 923 | Bacteria>Acidobacteriota>unclassified_Acidobacteriota | GCA_028873735.1 | Acidobacteriota_bacterium | OXU63_16720 | 13221..15992 | 1978231 | MAG:-DUF499-domain-containing-protein-[Acidobacteriota-bacterium] |
| MCY4077553.1 | DUF499_1.RM.like.systems | <-Radical-SAM<-?\|\|?->?-><-OLD-ABC+TOPRIM<-DUF499*<-METHYLASE-p | 876 | Bacteria>Acidobacteriota>unclassified_Acidobacteriota | GCA_026707615.1 | Acidobacteriota_bacterium | OXH04_19220 | 9433..12063 | 1978231 | MAG:-DUF499-domain-containing-protein-[Acidobacteria-bacterium] |
| MYN68213.1 | DUF499_1.RM.like.systems | TUDOR+SF1-SF2-Helicase+iREase+CTD+REase->Schlafen+HATPase+HTH->HEPN->METHYLASE->OLD-ABC->TOPRIM->DUF499*-><-?<-SAM-methylase<-ACET+ACET<-GspA-ATPase | 1101 | Bacteria>Acidobacteriota>unclassified_Acidobacteriota | GCA_009861545.1 | Acidobacteriota_bacterium | F4X11_24880 | 56553..59858 | 1978231 | ATP-binding-protein-[Acidobacteria-bacterium] |
| MCY3965602.1 | DUF499_1.RM.like.systems | PIN-><-?<-?\|\|OLD-ABC->TOPRIM-><-DUF499*\|\|HEPN-><-TOPRIM<-OLD-ABC<-METHYLASE\|\|?->PAPS_reduct-> | 1095 | Bacteria>Acidobacteriota>unclassified_Acidobacteriota | GCA_026707545.1 | Acidobacteriota_bacterium | OXG83_11235 | 2822742..2826029 | 1978231 | MAG:-DUF499-domain-containing-protein-[Acidobacteria-bacterium] |
| MYN67290.1 | DUF499_1.RM.like.systems | <-REase\|\|?-><-DUF499*<-TOPRIM<-OLD-ABC<-?<-TIR<-OLD-ABC<-SF1-SF2-Helicase | 1106 | Bacteria>Acidobacteriota>unclassified_Acidobacteriota | GCA_009861545.1 | Acidobacteriota_bacterium | F4X11_20045 | 66058..69378 | 1978231 | ATP-binding-protein-[Acidobacteria-bacterium] |
| MBI4902187.1 | DUF499_1.RM.like.systems | TUDOR+SF1-SF2-Helicase+iREase+CTD+REase->ParB+HNH->METHYLASE->OLD-ABC->TOPRIM->HEPN+DUF499*->Cluster2725_2clades->Terminase-> | 1115 | Bacteria>Acidobacteriota>unclassified_Acidobacteriota | GCA_016213775.1 | Acidobacteriota_bacterium | HY820_01040 | 17603..20950 | 1978231 | ATP-binding-protein-[Acidobacteria-bacterium] |
| MBS1816795.1 | DUF499_1.RM.like.systems | <-XIS-HTH<-?\|\|HTH+PolB->TUDOR+SF1-SF2-Helicase+iREase+CTD+REase->METHYLASE->?->HEPN+DUF499*->?-><-OLD-ABC+TOPRIM<-?<-?<-AbiEi_4\|\|REase-> | 1111 | Bacteria>Acidobacteriota>unclassified_Acidobacteriota | GCA_018268955.1 | Acidobacteriota_bacterium | JSU08_02600 | 186783..190118 | 1978231 | DUF499-domain-containing-protein-[Acidobacteria-bacterium] |
| MCC6131982.1 | DUF499_1.RM.like.systems | <-HEPN+DUF499*<-Schlafen+HATPase+HTH<-PHP+SbcC<-METHYLASE<-TOPRIM<-OLD-ABC<-TUDOR+SF1-SF2-Helicase+iREase+CTD+REase | 1114 | Bacteria>Acidobacteriota>unclassified_Acidobacteriota | GCA_020845555.1 | Acidobacteriota_bacterium | IT186_18830 | 2559..5903 | 1978231 | DUF499-domain-containing-protein-[Acidobacteria-bacterium] |
| MBI1792208.1 | DUF499_1.RM.like.systems | TUDOR+SF1-SF2-Helicase+iREase+CTD+REase->Schlafen+HATPase+HTH->METHYLASE->PolB->HEPN+DUF499*->OLD-ABC+TOPRIM-> | 1093 | Bacteria>Acidobacteriota>unclassified_Acidobacteriota | GCA_016178945.1 | Acidobacteriota_bacterium | HYR60_32190 | 8610..11891 | 1978231 | ATP-binding-protein-[Acidobacteria-bacterium] |
| MBS1819352.1 | DUF499_1.RM.like.systems | <-SF1-SF2-Helicase<-OLD-ABC+TOPRIM<-HEPN+DUF499*<-METHYLASE-p | 1115 | Bacteria>Acidobacteriota>unclassified_Acidobacteriota | GCA_018268955.1 | Acidobacteriota_bacterium | JSU08_15565 | 76293..79640 | 1978231 | DUF499-domain-containing-protein-[Acidobacteria-bacterium] |
| MDD5559009.1 | DUF499_1.RM.like.systems | HD->DUF4065->METHYLASE-p->OLD-ABC->TOPRIM->METHYLASE->DUF499*-><-?<-REase<-?<-?<-?<-CopG_antitoxin | 950 | Bacteria>Bacteria_incertae_sedis>Bacteria_candidate_phyla>candidate_division_NC10>Methylomirabilis | GCA_028716865.1 | Candidatus_Methylomirabilis_sp. | PHV01_05045 | 55831..58683 | 2032687 | MAG:-DUF499-domain-containing-protein-[Candidatus-Methylomirabilis-sp.] |
| MDE2058207.1 | DUF499_1.RM.like.systems | OLD-ABC->TOPRIM->METHYLASE->BrnT_toxin->HTH->REase->DUF499*->?->HTH+PolB->HEPN-> | 931 | Bacteria>Bacteria_incertae_sedis>Bacteria_candidate_phyla>candidate_division_NC10>unclassified_candidate_division_NC10 | GCA_028864815.1 | candidate_division_NC10_bacterium | KGJ27_00545 | 34245..37040 | 2072417 | MAG:-ATP-binding-protein-[candidate-division-NC10-bacterium] |
| MBI4536775.1 | DUF499_1.RM.like.systems | <-REase<-sGTP<-DUF499*<-?<-TOPRIM<-OLD-ABC<-METHYLASE-p<-?<-METHYLASE | 938 | Bacteria>Bacteria_incertae_sedis>Bacteria_candidate_phyla>candidate_division_NC10>unclassified_candidate_division_NC10 | GCA_016209735.1 | candidate_division_NC10_bacterium | HY712_02325 | 61380..64196 | 2072417 | ATP-binding-protein-[candidate-division-NC10-bacterium] |
| MDE2058883.1 | DUF499_1.RM.like.systems | <-HEPN+DUF499*<-?<-METHYLASE<-TOPRIM<-OLD-ABC<-ORC-CDC6+REase<-TUDOR+SF1-SF2-Helicase+iREase+CTD+REase | 1122 | Bacteria>Bacteria_incertae_sedis>Bacteria_candidate_phyla>candidate_division_NC10>unclassified_candidate_division_NC10 | GCA_028864815.1 | candidate_division_NC10_bacterium | KGJ27_04040 | 4506..7874 | 2072417 | MAG:-DUF499-domain-containing-protein-[candidate-division-NC10-bacterium] |
| MCX6842949.1 | DUF499_1.RM.like.systems | <-PIN<-?<-DUF499*<-METHYLASE<-TOPRIM<-OLD-ABC<-REase<-TUDOR+SF1-SF2-Helicase+iREase+CTD+REase | 912 | Bacteria>Bacteria_incertae_sedis>Bacteria_candidate_phyla>candidate_division_WOR-3 | GCA_026397595.1 | candidate_division_WOR-3_bacterium | NTX53_11780 | 2277..5015 | 2052148 | MAG:-DUF499-domain-containing-protein-[candidate-division-WOR-3-bacterium] |
| QAA75938.1 | DUF499_1.RM.like.systems | <-RNaseH<-?<-?<-?<-?<-?<-HEPN+DUF499*<-?<-METHYLASE<-TOPRIM<-OLD-ABC<-TUDOR+SF1-SF2-Helicase+iREase+CTD+REase | 1113 | Bacteria>Bacteria_incertae_sedis>Bacteria_candidate_phyla>Candidatus_Bipolaricaulota>Bipolaricaulis | GCA_004102645.1 | Candidatus_Bipolaricaulis_sibiricus | BIP78_0170 | 158925..162266 | 2501609 | hypothetical-protein-BIP78_0170-[Candidatus-Bipolaricaulis-sibiricus] |
| HDN19517.1 | DUF499_1.RM.like.systems | TUDOR+SF1-SF2-Helicase+iREase+CTD+REase->OLD-ABC+TOPRIM->METHYLASE->PolB->HEPN+DUF499*->MazE_antitoxin->PIN-> | 1111 | Bacteria>Bacteria_incertae_sedis>Bacteria_candidate_phyla>Candidatus_Bipolaricaulota>unclassified_Candidatus_Bipolaricaulota | GCA_011042375.1 | Candidatus_Acetothermia_bacterium | ENF46_00270 | 10022..13357 | 2053493 | ATP-binding-protein-[Candidatus-Acetothermia-bacterium] |
| MDE0183895.1 | DUF499_1.RM.like.systems | OLD-ABC->REase-><-Radical-SAM<-?<-?<-REase+35exo+Beta-Barrel+SF1-SF2-Helicase<-DUF499*<-Y1_Tnp<-TOPRIM<-OLD-ABC<-METHYLASE | 1100 | Bacteria>Bacteria_incertae_sedis>Bacteria_candidate_phyla>Candidatus_Poribacteria>unclassified_Candidatus_Poribacteria | GCA_028819845.1 | Candidatus_Poribacteria_bacterium | OXP71_00395 | 88990..92292 | 2026781 | MAG:-DUF499-domain-containing-protein-[Candidatus-Poribacteria-bacterium] |
| MBI2567004.1 | DUF499_1.RM.like.systems | <-REase\|\|XIS-HTH->HhH-RADC+JAB->TUDOR+SF1-SF2-Helicase+iREase+CTD+REase->METHYLASE-p->METHYLASE->DUF499*->OLD-ABC->TOPRIM->?->McrC-NTD+McrC-> | 962 | Bacteria>Bacteria_incertae_sedis>Bacteria_candidate_phyla>Candidatus_Schekmanbacteria>unclassified_Candidatus_Schekmanbacteria | GCA_016188575.1 | Candidatus_Schekmanbacteria_bacterium | HYV63_08250 | 57213..60101 | 2419984 | ATP-binding-protein-[Candidatus-Schekmanbacteria-bacterium] |
| MBN1883904.1 | DUF499_1.RM.like.systems | Peptidase_M48->TUDOR+SF1-SF2-Helicase+iREase+CTD+REase->Schlafen+HATPase+HTH->METHYLASE->OLD-ABC->TOPRIM->HEPN+DUF499*->PIN+OST-HTH-><-?<-?<-REase<-?\|\|DOC+HTH-> | 1113 | Bacteria>Candidatus_Krumholzibacteriota>unclassified_Candidatus_Krumholzibacteriota | GCA_016931295.1 | Candidatus_Krumholzibacteriota_bacterium | JW876_00100 | 28696..32037 | 2725313 | DUF499-domain-containing-protein-[Candidatus-Krumholzibacteriota-bacterium] |
| OGQ97428.1 | DUF499_1.RM.like.systems | REase->?->?->?-><-?<-SF1-SF2-Helicase<-HEPN+DUF499*<-METHYLASE<-?<-?<-ParB+HNH<-TOPRIM<-OLD-ABC | 1111 | Bacteria>delta/epsilon_subdivisions>Deltaproteobacteria>unclassified_Deltaproteobacteria | GCA_001799275.1 | Deltaproteobacteria_bacterium_RIFOXYD12_FULL_57_12 | A2521_08565 | 7962..11297 | 1797906 | AAA+-family-ATPase-[Deltaproteobacteria-bacterium-RIFOXYD12_FULL_57_12] |
| MBI4676018.1 | DUF499_1.RM.like.systems | <-Peptidase<-DUF499*<-TOPRIM<-OLD-ABC<-TOPRIM<-OLD-ABC<-METHYLASE<-ORC-CDC6 | 925 | Bacteria>Elusimicrobiota>unclassified_Elusimicrobiota | GCA_016209035.1 | Elusimicrobiota_bacterium | HY748_00360 | 72321..75098 | 2030800 | ATP-binding-protein-[Elusimicrobia-bacterium] |
| MBI5244011.1 | DUF499_1.RM.like.systems | OLD-ABC->TOPRIM->REase->SF1-SF2-Helicase+wHTH+PRTase->?->DUF499*->?-><-Methyltrans_RNA<-DnaJ | 1107 | Bacteria>Elusimicrobiota>unclassified_Elusimicrobiota | GCA_016212885.1 | Elusimicrobiota_bacterium | HY922_10100 | 5875..9198 | 2030800 | ATP-binding-protein-[Elusimicrobia-bacterium] |
| MCX5796464.1 | DUF499_1.RM.like.systems | TUDOR+SF1-SF2-Helicase+iREase+CTD+REase->STAS+HisKin->?->PIN->METHYLASE->OLD-ABC+TOPRIM->HEPN+DUF499*-> | 1133 | Bacteria>Elusimicrobiota>unclassified_Elusimicrobiota | GCA_026388095.1 | Elusimicrobiota_bacterium | NTY77_13305 | 62095..65496 | 2030800 | MAG:-Swt1-family-HEPN-domain-containing-protein-[Elusimicrobia-bacterium] |
| MCL2726873.1 | DUF499_1.RM.like.systems | <-ABhydrolase\|\|REase->Peptidase-><-Peptidase_M24<-Peptidase<-?<-HEPN+DUF499*<-TOPRIM<-OLD-ABC<-METHYLASE<-TUDOR+SF1-SF2-Helicase+iREase+CTD+REase<-LysE | 1124 | Bacteria>FCB_group>Bacteroidota/Chlorobiota_group>Bacteroidota>Bacteroidia | GCA_009781275.1 | Bacteroidales_bacterium | FWD56_00650 | 13721..17095 | 2030927 | DUF499-domain-containing-protein-[Bacteroidales-bacterium] |
| MBX7052699.1 | DUF499_1.RM.like.systems | MazG->REase+SF1-SF2-Helicase->TUDOR+SF1-SF2-Helicase+iREase+CTD+REase->OLD-ABC->TOPRIM->METHYLASE->HEPN+DUF499*-> | 1088 | Bacteria>FCB_group>Bacteroidota/Chlorobiota_group>Bacteroidota>Flavobacteriia | GCA_019694805.1 | Flavobacteriales_bacterium | K1X54_11735 | 22264..25530 | 2021391 | DUF499-domain-containing-protein-[Flavobacteriales-bacterium] |
| MBK8920115.1 | DUF499_1.RM.like.systems | <-Glyco_hydro_1<-?<-HEPN+DUF499*<-URI-DNase<-METHYLASE-p<-METHYLASE-p<-METHYLASE-p<-TOPRIM<-OLD-ABC | 1088 | Bacteria>FCB_group>Bacteroidota/Chlorobiota_group>Bacteroidota>Saprospiria | GCA_016719895.1 | Saprospirales_bacterium | IPM81_01165 | 280177..283443 | 2026790 | DUF499-domain-containing-protein-[Saprospirales-bacterium] |
| MBP7719545.1 | DUF499_1.RM.like.systems | TUDOR+SF1-SF2-Helicase+iREase+CTD+REase->METHYLASE->OLD-ABC->TOPRIM->HEPN+DUF499*-> | 1111 | Bacteria>FCB_group>Candidatus_Fermentibacteria>Candidatus_Fermentibacteria_(class)>Candidatus_Fermentibacterales | GCA_017999735.1 | Candidatus_Fermentibacter_sp. | KA108_02790 | 14441..17776 | 2779163 | DUF499-domain-containing-protein-[Candidatus-Fermentibacter-sp.] |
| MCF7837530.1 | DUF499_1.RM.like.systems | Phage_integrase-><-ZnR+TOPRIM+D5ATPase<-XIS-HTH\|\|HTH->XIS-HTH-><-HEPN+DUF499*<-OLD-ABC+TOPRIM<-METHYLASE-p | 1110 | Bacteria>FCB_group>Candidatus_Marinimicrobia>unclassified_Candidatus_Marinimicrobia | GCA_021734645.1 | Candidatus_Marinimicrobia_bacterium | K9N49_02780 | 12805..16137 | 2026760 | DUF499-domain-containing-protein-[Candidatus-Marinimicrobia-bacterium] |
| MYA34704.1 | DUF499_1.RM.like.systems | HEPN->TUDOR+SF1-SF2-Helicase+iREase+CTD+REase->METHYLASE->OLD-ABC->TOPRIM->ParB+HNH+Schlafen->DUF499*-><-?\|\|HTH->MPTase->?->SMODS-><-ABhydrolase | 928 | Bacteria>FCB_group>Gemmatimonadota>Gemmatimonadetes>Gemmatimonadales | GCA_009838765.1 | Gemmatimonadales_bacterium | F4Y33_13800 | 70561..73347 | 2448054 | ATP-binding-protein-[Gemmatimonadales-bacterium] |
| MXX77944.1 | DUF499_1.RM.like.systems | OLD-ABC->TOPRIM->HEPN+DUF499*-><-RNase<-?<-REase | 1102 | Bacteria>FCB_group>Gemmatimonadota>Gemmatimonadetes>Gemmatimonadales | GCA_009836785.1 | Gemmatimonadales_bacterium | F4Z33_02945 | 1211..4519 | 2448054 | ATP-binding-protein-[Gemmatimonadales-bacterium] |
| MYB67636.1 | DUF499_1.RM.like.systems | <-DUF499*<-TOPRIM<-OLD-ABC<-HEPN<-PolB<-?<-TraJ-RHH | 924 | Bacteria>FCB_group>Gemmatimonadota>unclassified_Gemmatimonadota | GCA_009840005.1 | Gemmatimonadota_bacterium | F4X75_03955 | 7160..9934 | 2026742 | ATP-binding-protein-[Gemmatimonadetes-bacterium] |
| MYA24522.1 | DUF499_1.RM.like.systems | <-DUF499*<-TOPRIM | 924 | Bacteria>FCB_group>Gemmatimonadota>unclassified_Gemmatimonadota | GCA_009836855.1 | Gemmatimonadota_bacterium | F4Z30_17850 | 5045..7819 | 2026742 | ATP-binding-protein-[Gemmatimonadetes-bacterium] |
| MDE2723058.1 | DUF499_1.RM.like.systems | <-DUF499*<-TOPRIM<-OLD-ABC<-HTH_3<-METHYLASE<-TOPRIM | 656 | Bacteria>FCB_group>Gemmatimonadota>unclassified_Gemmatimonadota | GCA_028875455.1 | Gemmatimonadota_bacterium | OXI59_06750 | 1..1968 | 2026742 | MAG:-DUF499-domain-containing-protein,-partial-[Gemmatimonadota-bacterium] |
| MYD59776.1 | DUF499_1.RM.like.systems | TUDOR+SF1-SF2-Helicase+iREase+CTD+REase->?->METHYLASE->HTH_3->OLD-ABC->TOPRIM->DUF499*->?->?->HigB_toxin->HTH->?->Peptidase-> | 1104 | Bacteria>FCB_group>Gemmatimonadota>unclassified_Gemmatimonadota | GCA_009841665.1 | Gemmatimonadota_bacterium | F4W91_01925 | 35019..38333 | 2026742 | ATP-binding-protein-[Gemmatimonadetes-bacterium] |
| MCY4603567.1 | DUF499_1.RM.like.systems | CR-ATPase9+REase->PhdYeFM_antitox->METHYLASE->OLD-ABC->TOPRIM->DUF499*->Glyco_hydro_42-> | 1097 | Bacteria>FCB_group>Gemmatimonadota>unclassified_Gemmatimonadota | GCA_026713655.1 | Gemmatimonadota_bacterium | OXE49_04985 | 6802..10095 | 2026742 | MAG:-DUF499-domain-containing-protein-[Gemmatimonadetes-bacterium] |
| MXX38675.1 | DUF499_1.RM.like.systems | OLD-ABC->TOPRIM->DUF499*->OLD-ABC->TOPRIM->Glyco_hydro_42->?-><-MqsA_antitoxin+HTH_3<-Polbeta | 1097 | Bacteria>FCB_group>Gemmatimonadota>unclassified_Gemmatimonadota | GCA_009835785.1 | Gemmatimonadota_bacterium | F4Z85_11545 | 1202..4495 | 2026742 | ATP-binding-protein-[Gemmatimonadetes-bacterium] |
| MDE3256982.1 | DUF499_1.RM.like.systems | <-SbcC<-OLD-ABC+TOPRIM<-?<-Phd-YefM<-DUF499*\|\|?-><-?<-?<-METHYLASE<-SbcC | 950 | Bacteria>FCB_group>Gemmatimonadota>unclassified_Gemmatimonadota | GCA_028877215.1 | Gemmatimonadota_bacterium | OYM47_04020 | 48493..51345 | 2026742 | MAG:-DUF499-domain-containing-protein-[Gemmatimonadota-bacterium] |
| MDE2654306.1 | DUF499_1.RM.like.systems | <-REC<-HEPN+DUF499*<-TOPRIM<-OLD-ABC<-METHYLASE-p | 1116 | Bacteria>FCB_group>Gemmatimonadota>unclassified_Gemmatimonadota | GCA_028875795.1 | Gemmatimonadota_bacterium | OXI71_10900 | 83236..86586 | 2026742 | MAG:-Swt1-family-HEPN-domain-containing-protein-[Gemmatimonadota-bacterium] |
| MBL9045676.1 | DUF499_1.RM.like.systems | TUDOR+SF1-SF2-Helicase+iREase+CTD+REase->METHYLASE->OLD-ABC->TOPRIM->HEPN+DUF499*-><-?<-HTH | 1115 | Bacteria>Myxococcota>Myxococcia>Myxococcales>unclassified_Myxococcales | GCA_016794465.1 | Myxococcales_bacterium | JNM83_28965 | 9575..12922 | 2026763 | DUF499-domain-containing-protein-[Myxococcales-bacterium] |
| NLN59460.1 | DUF499_1.RM.like.systems | <-SF1-SF2-helicase<-?<-PAIREDC-HTH+HIN-HTH<-Methyltransf_25<-?<-?<-DUF499*<-METHYLASE<-TOPRIM<-OLD-ABC<-DOC+HTH<-TUDOR+SF1-SF2-Helicase+iREase+CTD+REase<-XIS-HTH | 920 | Bacteria>Myxococcota>Myxococcia>unclassified_Myxococcia | GCA_012522415.1 | Deltaproteobacteria_bacterium | GX147_01890 | 29834..32596 | 2026735 | ATP-binding-protein-[Deltaproteobacteria-bacterium] |
| PXF54128.1 | DUF499_1.RM.like.systems | Phage_int_SAM_1+Phage_integrase-><-SbcC+SbcC<-Calcineurin<-?<-?<-DUF499*<-Schlafen+HATPase+HTH<-METHYLASE-p<-TOPRIM<-OLD-ABC | 1011 | Bacteria>Myxococcota>Myxococcia>unclassified_Myxococcia | GCA_003194485.1 | Deltaproteobacteria_bacterium | C4B57_08240 | 17848..20883 | 2026735 | AAA-family-ATPase-[Deltaproteobacteria-bacterium] |
| MCD6198158.1 | DUF499_1.RM.like.systems | <-DUF499*<-OLD-ABC+TOPRIM<-MACRODOMAIN<-?<-TUDOR+SF1-SF2-Helicase+iREase+CTD+REase<-XIS-HTH | 992 | Bacteria>Myxococcota>Myxococcia>unclassified_Myxococcia | GCA_021159445.1 | Deltaproteobacteria_bacterium | J7K15_06235 | 2443..5421 | 2026735 | ATP-binding-protein-[Deltaproteobacteria-bacterium] |
| MBW1911012.1 | DUF499_1.RM.like.systems | Radical-SAM-><-DUF499*<-PIN<-?<-DUF433<-METHYLASE<-TOPRIM<-OLD-ABC-p | 941 | Bacteria>Myxococcota>Myxococcia>unclassified_Myxococcia | GCA_019307765.1 | Deltaproteobacteria_bacterium | JRJ11_15995 | 905..3730 | 2026735 | ATP-binding-protein-[Deltaproteobacteria-bacterium] |
| MBI4706258.1 | DUF499_1.RM.like.systems | <-DUF499*<-?<-SbcC<-OLD-ABC+TOPRIM<-METHYLASE<-TOPRIM<-OLD-ABC | 925 | Bacteria>Myxococcota>Myxococcia>unclassified_Myxococcia | GCA_016209145.1 | Deltaproteobacteria_bacterium | HY744_34610 | 6205..8982 | 2026735 | ATP-binding-protein-[Deltaproteobacteria-bacterium] |
| MBW1922689.1 | DUF499_1.RM.like.systems | METHYLASE->OLD-ABC->TOPRIM->DUF499*->EVE-><-HIN-HTH | 965 | Bacteria>Myxococcota>Myxococcia>unclassified_Myxococcia | GCA_019307945.1 | Deltaproteobacteria_bacterium | JRJ35_04385 | 5613..8510 | 2026735 | DUF499-domain-containing-protein-[Deltaproteobacteria-bacterium] |
| RLC08398.1 | DUF499_1.RM.like.systems | <-DUF499*<-TOPRIM<-OLD-ABC<-METHYLASE-p | 353 | Bacteria>Myxococcota>Myxococcia>unclassified_Myxococcia | GCA_003647435.1 | Deltaproteobacteria_bacterium | DRH43_10115 | 1..1059 | 2026735 | AAA-family-ATPase,-partial-[Deltaproteobacteria-bacterium] |
| MCX5910053.1 | DUF499_1.RM.like.systems | OLD-ABC-p->TOPRIM->DUF499-p->DUF499*->DUF499-p->AbiEii-> | 258 | Bacteria>Myxococcota>Myxococcia>unclassified_Myxococcia | GCA_026388955.1 | Deltaproteobacteria_bacterium | NTY64_23470 | 1666..2438 | 2026735 | MAG:-DUF499-domain-containing-protein,-partial-[Deltaproteobacteria-bacterium] |
| MDD9873633.1 | DUF499_1.RM.like.systems | TUDOR+SF1-SF2-Helicase+iREase+CTD+REase->HEPN->METHYLASE->OLD-ABC->TOPRIM->DUF499*->?->?->?->?->ABC-ATPase+SbcC-> | 1106 | Bacteria>Myxococcota>Myxococcia>unclassified_Myxococcia | GCA_028817295.1 | Deltaproteobacteria_bacterium | OXU65_09090 | 9437..12757 | 2026735 | MAG:-DUF499-domain-containing-protein-[Deltaproteobacteria-bacterium] |
| MBM4396669.1 | DUF499_1.RM.like.systems | TUDOR+SF1-SF2-Helicase+iREase+CTD+REase-p->METHYLASE->OLD-ABC->TOPRIM->DUF499*-> | 1103 | Bacteria>Myxococcota>Myxococcia>unclassified_Myxococcia | GCA_016875475.1 | Deltaproteobacteria_bacterium | FJ087_13395 | 5583..8894 | 2026735 | ATP-binding-protein-[Deltaproteobacteria-bacterium] |
| MBI3795502.1 | DUF499_1.RM.like.systems | <-DUF499*<-TOPRIM<-OLD-ABC<-METHYLASE | 1108 | Bacteria>Myxococcota>Myxococcia>unclassified_Myxococcia | GCA_016200595.1 | Deltaproteobacteria_bacterium | HY268_00830 | 2141..5467 | 2026735 | ATP-binding-protein-[Deltaproteobacteria-bacterium] |
| MAG32379.1 | DUF499_1.RM.like.systems | METHYLASE-p->OLD-ABC->Schlafen+HATPase+HTH->HEPN+DUF499*->?-><-METHYLASE-p<-TOPRIM<-OLD-ABC\|\|?->METHYLASE-p-> | 1115 | Bacteria>Myxococcota>Myxococcia>unclassified_Myxococcia | GCA_002688255.1 | Deltaproteobacteria_bacterium | CL908_15970 | 4485..7832 | 2026735 | AAA+-family-ATPase-[Deltaproteobacteria-bacterium] |
| MDD3553583.1 | DUF499_1.RM.like.systems | XIS-HTH->TUDOR+SF1-SF2-Helicase+iREase+CTD+REase->METHYLASE->OLD-ABC->TOPRIM->HEPN+DUF499*-> | 1108 | Bacteria>Myxococcota>Myxococcia>unclassified_Myxococcia | GCA_028695735.1 | Deltaproteobacteria_bacterium | PHC35_03575 | 35937..39263 | 2026735 | MAG:-Swt1-family-HEPN-domain-containing-protein-[Deltaproteobacteria-bacterium] |
| TRZ75307.1 | DUF499_1.RM.like.systems | METHYLASE->OLD-ABC->TOPRIM->HEPN+DUF499*-> | 362 | Bacteria>Myxococcota>Myxococcia>unclassified_Myxococcia | GCA_007279985.1 | Deltaproteobacteria_bacterium | D4R93_05190 | 4209..5295 | 2026735 | ATP-binding-protein,-partial-[Deltaproteobacteria-bacterium] |
| MCS6797210.1 | DUF499_1.RM.like.systems | Peptidase->?-><-?<-?<-DUF499*<-METHYLASE-p<-METHYLASE<-?<-?<-TOPRIM<-OLD-ABC | 1007 | Bacteria>Myxococcota>unclassified_Myxococcota | GCA_025054695.1 | Myxococcota_bacterium | NZ898_01550 | 47913..50936 | 2818507 | DUF499-domain-containing-protein-[Myxococcota-bacterium] |
| MBP7125459.1 | DUF499_1.RM.like.systems | TUDOR+SF1-SF2-Helicase+iREase+CTD+REase->Schlafen+HATPase+HTH->METHYLASE->OLD-ABC->TOPRIM->HEPN+DUF499*-> | 1039 | Bacteria>Myxococcota>unclassified_Myxococcota | GCA_017994695.1 | Myxococcota_bacterium | KBD49_03725 | 64170..67286 | 2818507 | DUF499-domain-containing-protein-[Myxococcota-bacterium] |
| MDE0331736.1 | DUF499_1.RM.like.systems | <-OLD-ABC+TOPRIM<-DUF499*<-METHYLASE<-TUDOR+SF1-SF2-Helicase+iREase+CTD+REase | 967 | Bacteria>Nitrospinae/Tectomicrobia_group>Nitrospinota>unclassified_Nitrospinota | GCA_028820795.1 | Nitrospinota_bacterium | OXL41_07695 | 2738..5641 | 2026769 | MAG:-DUF499-domain-containing-protein-[Nitrospinae-bacterium] |
| PIX82647.1 | DUF499_1.RM.like.systems | METHYLASE->OLD-ABC->TOPRIM->HEPN+DUF499*-> | 561 | Bacteria>Nitrospirota>unclassified_Nitrospirota | GCA_002783945.1 | Nitrospirae_bacterium_CG_4_10_14_3_um_filter_70_108 | COZ33_09565 | 4155..5838 | 1974054 | AAA+-family-ATPase,-partial-[Nitrospirae-bacterium-CG_4_10_14_3_um_filter_70_108] |
| WP_071923269.1 | DUF499_1.RM.like.systems | GDResolvase+Recombinase+ZnR->CopG/RHH->TUDOR+SF1-SF2-Helicase+iREase+CTD+REase->METHYLASE->HEPN+DUF499*->OLD-ABC+TOPRIM->SF1-SF2-Helicase-><-HTH_17<-Phage_integrase<-HIN-HTH | 1097 | Bacteria>Pseudomonadota>Alphaproteobacteria>Hyphomicrobiales>Chelatococcaceae | GCF_001887265.1 | Chelatococcus_daeguensis | BOQ54_RS03320 | 711833..715126 | 444444 | DUF499-domain-containing-protein-[Chelatococcus-daeguensis] |
| WP_012755925.1 | DUF499_1.RM.like.systems | Phage_integrase->?->?->OLD-ABC+TOPRIM->?-><-DUF499*<-METHYLASE<-HNH<-OLD-ABC<-TUDOR+SF1-SF2-Helicase+iREase+CTD+REase<-HerA/FtsK | 945 | Bacteria>Pseudomonadota>Alphaproteobacteria>Hyphomicrobiales>Rhizobiaceae | GCF_000023185.1 | Rhizobium_leguminosarum | RLEG_RS01005 | 199729..202566 | 384 | ATP-binding-protein-[Rhizobium-leguminosarum] |
| MCZ7659369.1 | DUF499_1.RM.like.systems | DEOR-HTH->TUDOR+SF1-SF2-Helicase+iREase+CTD+REase->TUDOR+SF1-SF2-Helicase+iREase+CTD+REase-p->?->REase->METHYLASE->HEPN+DUF499*->OLD-ABC+TOPRIM->SF1-SF2-Helicase-><-REase<-?<-N6_MTase<-DEOR-HTH | 1120 | Bacteria>Pseudomonadota>Alphaproteobacteria>Hyphomicrobiales>Xanthobacteraceae | GCA_027493405.1 | Xanthobacteraceae_bacterium | M5U07_16605 | 9358..12720 | 2021379 | MAG:-Swt1-family-HEPN-domain-containing-protein-[Xanthobacteraceae-bacterium] |
| MBF0262096.1 | DUF499_1.RM.like.systems | XIS-HTH->TUDOR+SF1-SF2-Helicase+iREase+CTD+REase->SbcC->OLD-ABC->TOPRIM->METHYLASE->HEPN+DUF499*-> | 1048 | Bacteria>Pseudomonadota>Alphaproteobacteria>Magnetococcales>unclassified_Magnetococcales | GCA_015231775.1 | Magnetococcales_bacterium | HQL97_09720 | 11242..14384 | 2026759 | DUF499-domain-containing-protein-[Magnetococcales-bacterium] |
| WP_147041135.1 | DUF499_1.RM.like.systems | TPR->?->TUDOR+SF1-SF2-Helicase+iREase+CTD+REase->METHYLASE->HEPN+DUF499*->OLD-ABC+TOPRIM-><-RelE-ParE<-MetJarc-HTH | 1097 | Bacteria>Pseudomonadota>Alphaproteobacteria>Rhodospirillales>Azospirillaceae | GCF_007991775.1 | Skermanella_aerolata | SAE02_RS33460 | 15952..19245 | 393310 | DUF499-domain-containing-protein-[Skermanella-aerolata] |
| GIX11744.1 | DUF499_1.RM.like.systems | <-Peptidase_S8<-Classical-AAA\|\|TUDOR+SF1-SF2-Helicase+iREase+CTD+REase->RNase->METHYLASE->DUF499*->OLD-ABC+TOPRIM-><-?<-Peptidase_M15 | 1107 | Bacteria>Pseudomonadota>Alphaproteobacteria>Rhodospirillales>Elioraeaceae | GCA_026004895.1 | Elioraea_sp. | KatS3mg116_3454 | 526065..529388 | 2185103 | MAG:-hypothetical-protein-KatS3mg116_3454-[Elioraea-sp.] |
| WP_218285167.1 | DUF499_1.RM.like.systems | Peptidase->Peptidase_M24->TUDOR+SF1-SF2-Helicase+iREase+CTD+REase->METHYLASE->OLD-ABC->TOPRIM->HEPN+DUF499*->?-><-PAIREDC-HTH<-DUF4096+DDE_Tnp_1+DDE_Tnp_1_2\|\|DUF4096+DDE_Tnp_1+DDE_Tnp_1_2->PAIREDC-HTH-> | 1111 | Bacteria>Pseudomonadota>Alphaproteobacteria>Rhodospirillales>Elioraeaceae | GCF_019203965.1 | Elioraea_tepida | KO353_RS13040 | 2714596..2717931 | 2843330 | Swt1-family-HEPN-domain-containing-protein-[Elioraea-tepida] |
| MDE0384393.1 | DUF499_1.RM.like.systems | TUDOR+SF1-SF2-Helicase+iREase+CTD+REase->RNase->?->METHYLASE->OLD-ABC->TOPRIM->DUF499*-><-?<-SF1-SF2-Helicase+DUF1998<-?<-PADR-HTH | 1108 | Bacteria>Pseudomonadota>Alphaproteobacteria>Rhodospirillales>Rhodospirillaceae | GCA_028821395.1 | Defluviicoccus_sp. | OXI22_10955 | 42906..46232 | 2682137 | MAG:-DUF499-domain-containing-protein-[Defluviicoccus-sp.] |
| MDA8051502.1 | DUF499_1.RM.like.systems | TUDOR+SF1-SF2-Helicase+iREase+CTD+REase->OLD-ABC->TOPRIM->METHYLASE->DUF499*-><-?\|\|?->Arm-DNA-bind_3+Phage_int_SAM_3+Phage_integrase->?->ZnR+TOPRIM->D5N+D5ATPase-> | 933 | Bacteria>Pseudomonadota>Alphaproteobacteria>Rhodospirillales>unclassified_Rhodospirillales | GCA_028063655.1 | Rhodospirillales_bacterium | M0002_16125 | 8854..11655 | 2026786 | MAG:-DUF499-domain-containing-protein-[Rhodospirillales-bacterium] |

| Protein-ID | System | Gene-Neighbourhoods | Length | Taxonomy | Assembly-ID | Strain | Locus-Tag | Genomic-coordinates | TaxID | Fasta-define |
|---|---|---|---|---|---|---|---|---|---|---|
| TVR79626.1 | DUF499_1.RM.like.systems | HDRnase-><- HEPN+ DUF499 *<-?<-PIN<-ParD<- METHYLASE <- OLD-ABC + TOPRIM <- TUDOR+ SF1-SF2-Helicase+iREase+CTD +REase | 1117 | Bacteria>Pseudomonadota>Alphaproteobacteria>Rhodospirillales>unclassified_Rhodospirillales | GCA_007695045.1 | Rhodospirillales_bacterium | EA405_12420 | 709..4062 | 2026786 | ATP-binding-protein-[Rhodospirillales-bacterium] |
| MDD3182285.1 | DUF499_1.RM.like.systems | <-ACET\|\|Peptidase_M15->?-><- SF1-SF2-Helicase<- OLD-ABC + TOPRIM <- DUF499 *<- METHYLASE <-PIN<- TUDOR+ SF1-SF2-Helicase+iREase+CTD +REase <-HTH+WYL | 940 | Bacteria>Pseudomonadota>Alphaproteobacteria>unclassified_Alphaproteobacteria | GCA_028691835.1 | Alphaproteobacteria_bacterium | PHD48_05740 | 6780..9602 | 1913988 | MAG:-DUF499-domain-containing-protein-[Alphaproteobacteria-bacterium] |
| MCY3753479.1 | DUF499_1.RM.like.systems | <-HTH_3\|\| TUDOR+ SF1-SF2-Helicase+iREase+CTD +REase ->?-> METHYLASE -> OLD-ABC -> TOPRIM -> DUF499 *-><-MPTase<-HTH_3\|\|SMODS->?->?-><-Arm-DNA-bind_3+Phage_int_SAM_3+Phage_integrase | 1104 | Bacteria>Pseudomonadota>Alphaproteobacteria>unclassified_Alphaproteobacteria | GCA_026706245.1 | Alphaproteobacteria_bacterium | OXG99_05240 | 13033..16347 | 1913988 | MAG:-DUF499-domain-containing-protein-[Alphaproteobacteria-bacterium] |
| MCB9797660.1 | DUF499_1.RM.like.systems | TUDOR+ SF1-SF2-Helicase+iREase+CTD +REase -> METHYLASE -> OLD-ABC -> TOPRIM -> HEPN+ DUF499 *-> | 1109 | Bacteria>Pseudomonadota>Alphaproteobacteria>unclassified_Alphaproteobacteria | GCA_020632765.1 | Alphaproteobacteria_bacterium | H6741_33640 | 16618..19947 | 1913988 | ATP-binding-protein-[Alphaproteobacteria-bacterium] |
| MCB9746493.1 | DUF499_1.RM.like.systems | <- HEPN+ DUF499 *<- TOPRIM <- OLD-ABC <- METHYLASE <- TUDOR+ SF1-SF2-Helicase+iREase+CTD +REase | 1109 | Bacteria>Pseudomonadota>Alphaproteobacteria>unclassified_Alphaproteobacteria | GCA_020632815.1 | Alphaproteobacteria_bacterium | H6740_28215 | 5203..8532 | 1913988 | ATP-binding-protein-[Alphaproteobacteria-bacterium] |
| QLH49283.1 | DUF499_1.RM.like.systems | TUDOR+ SF1-SF2-Helicase+iREase+CTD +REase -> OLD-ABC -> TOPRIM -> OLD-ABC -> TOPRIM -> METHYLASE -> HEPN+ DUF499 *-><-tRNA<-?<-GNAT<-?<-?<-ABhydrolase | 1112 | Bacteria>Pseudomonadota>Betaproteobacteria>Betaproteobacteria_incertae_sedis>Candidatus_Accumulibacter | GCA_013414765.1 | Candidatus_Accumulibacter_cognatus | HWD57_05410 | 1172035..1175373 | 2954383 | MAG:-ATP-binding-protein-[Candidatus-Accumulibacter-cognatus] |
| MBK8115022.1 | DUF499_1.RM.like.systems | TRD+TRD->Schlafen+HATPase+HTH-><-?<-?<-TPR+TPR<-?<- HEPN+ DUF499 *<-HEPN<-PolB<-PolB+HEPN<-PolB<- METHYLASE <- TOPRIM | 1109 | Bacteria>Pseudomonadota>Betaproteobacteria>Betaproteobacteria_incertae_sedis>Candidatus_Accumulibacter | GCA_016712935.1 | Accumulibacter_sp. | IPK44_10990 | 647374..650703 | 2053492 | DUF499-domain-containing-protein-[Candidatus-Accumulibacter-sp.] |
| WP_138677185.1 | DUF499_1.RM.like.systems | ABhydrolase->?->?->GNAT->?->tRNA-><- HEPN+ DUF499 *<-PolB+HEPN<-PolB<- METHYLASE <- TOPRIM <- OLD-ABC <- METHYLASE -p | 1112 | Bacteria>Pseudomonadota>Betaproteobacteria>Betaproteobacteria_incertae_sedis>Candidatus_Accumulibacter | GCF_005889575.1 | Candidatus_Accumulibacter_phosphatis | ACCUM_RS00275 | 63215..66553 | 327160 | Swt1-family-HEPN-domain-containing-protein-[Candidatus-Accumulibacter-phosphatis] |
| MBN8517040.1 | DUF499_1.RM.like.systems | TOPRIM -> OLD-ABC -> TOPRIM -> METHYLASE ->PolB->PolB+HEPN-> HEPN+ DUF499 *-><-tRNA<-?<-GNAT<-?<-?<-ABhydrolase | 1112 | Bacteria>Pseudomonadota>Betaproteobacteria>Betaproteobacteria_incertae_sedis>Candidatus_Accumulibacter | GCA_017302385.1 | Accumulibacter_sp. | JOL71_04035 | 55735..59073 | 2053492 | DUF499-domain-containing-protein-[Candidatus-Accumulibacter-sp.] |
| MCX8099543.1 | DUF499_1.RM.like.systems | <-SF1-SF2-Helicase+DUF1998<-ORC-CDC6+REase<-?<- DUF499 *<-PIN<-HTH<- METHYLASE <- TOPRIM <- OLD-ABC <- TUDOR+ SF1-SF2-Helicase+iREase+CTD +REase | 918 | Bacteria>Pseudomonadota>Betaproteobacteria>Betaproteobacteria_incertae_sedis>Casimicrobiaceae | GCA_026414825.1 | Casimicrobiaceae_bacterium | N3F11_11190 | 5518..8274 | 2906524 | MAG:-DUF499-domain-containing-protein-[Casimicrobiaceae-bacterium] |
| WP_062175042.1 | DUF499_1.RM.like.systems | <-PIN<- SF1-SF2-Helicase<- OLD-ABC + TOPRIM <- HEPN+ DUF499 *<- METHYLASE <-MORC+DUF3684<-DUF2326<-?<-?<-HTH_37 | 1108 | Bacteria>Pseudomonadota>Betaproteobacteria>Burkholderiales>Burkholderiaceae | GCF_001557535.2 | Burkholderia_sp._PAMC_26561 | AXG89_RS34140 | 126894..130220 | 1795043 | Swt1-family-HEPN-domain-containing-protein-[Burkholderia-sp.-PAMC-26561] |
| WP_143969665.1 | DUF499_1.RM.like.systems | HTH+WYL-> TUDOR+ SF1-SF2-Helicase+iREase+CTD +REase ->PIN-> METHYLASE -> OLD-ABC -> TOPRIM -> DUF499 *->HEPN-> | 952 | Bacteria>Pseudomonadota>Betaproteobacteria>Burkholderiales>Burkholderiales_genera_incertae_sedis | GCF_007556755.1 | Tepidimonas_fonticaldi | Tfont_RS12360 | 12901..15759 | 1101373 | DUF499-domain-containing-protein-[Tepidimonas-fonticaldi] |
| WP_043699941.1 | DUF499_1.RM.like.systems | PIN-> METHYLASE -> OLD-ABC -> TOPRIM -> METHYLASE ->Schlafen+HATPase+HTH-> DUF499 *-><-?<-?<-MORC<-?<-VirB11+MoxR-AAA | 945 | Bacteria>Pseudomonadota>Betaproteobacteria>Burkholderiales>Burkholderiales_genera_incertae_sedis | GCF_020162115.1 | Tepidimonas_taiwanensis | PL21_RS04270 | 89059..91896 | 307486 | DUF499-domain-containing-protein-[Tepidimonas-taiwanensis] |
| WP_143969782.1 | DUF499_1.RM.like.systems | <-DUF772+DDE_Tnp_1\|\|<- DUF499 *<-REase<- METHYLASE <- TOPRIM <- OLD-ABC <- TUDOR+ SF1-SF2-Helicase+iREase+CTD +REase <-XIS-HTH | 948 | Bacteria>Pseudomonadota>Betaproteobacteria>Burkholderiales>Burkholderiales_genera_incertae_sedis | GCF_007556755.1 | Tepidimonas_fonticaldi | Tfont_RS13255 | 1599..4445 | 1101373 | DUF499-domain-containing-protein-[Tepidimonas-fonticaldi] |
| WP_265654212.1 | DUF499_1.RM.like.systems | Peptidase_C26->?->?-><-?<- OLD-ABC + TOPRIM <- DUF499 *<- METHYLASE <-HEPN<- TUDOR+ SF1-SF2-Helicase+iREase+CTD +REase | 951 | Bacteria>Pseudomonadota>Betaproteobacteria>Burkholderiales>Comamonadaceae | GCF_026192715.1 | Verminephrobacter_aporrectodeae | D8B27_RS10360 | 21500..24355 | 1110389 | DUF499-domain-containing-protein-[Verminephrobacter-aporrectodeae] |
| WP_265282594.1 | DUF499_1.RM.like.systems | TUDOR+ SF1-SF2-Helicase+iREase+CTD +REase ->HEPN-> METHYLASE -> DUF499 *-> OLD-ABC + TOPRIM ->?->?-><-?<-?<-Peptidase_C26 | 951 | Bacteria>Pseudomonadota>Betaproteobacteria>Burkholderiales>Comamonadaceae | GCF_026127365.1 | Verminephrobacter_aporrectodeae | D5039_RS14590 | 1238377..1241232 | 1110389 | DUF499-domain-containing-protein-[Verminephrobacter-aporrectodeae] |
| WP_209423136.1 | DUF499_1.RM.like.systems | SF1-SF2-Helicase+DUF1998->?-><-?<-?<-?\|\|?-><- DUF499 *<-REase<- OLD-ABC <- TOPRIM <- OLD-ABC <- METHYLASE -p<- METHYLASE | 943 | Bacteria>Pseudomonadota>Betaproteobacteria>Burkholderiales>Comamonadaceae | GCF_021249385.1 | Melaminivora_jejuensis | LVC68_RS01445 | 284242..287073 | 1267217 | DUF499-domain-containing-protein-[Melaminivora-jejuensis] |
| WP_086929120.1 | DUF499_1.RM.like.systems | TUDOR+ SF1-SF2-Helicase+iREase+CTD +REase -> METHYLASE -> DUF499 *->?-> OLD-ABC + TOPRIM -> | 970 | Bacteria>Pseudomonadota>Betaproteobacteria>Burkholderiales>Comamonadaceae | GCF_002157185.1 | Acidovorax_carolinensis | CBP35_RS20910 | 151033..153945 | 553814 | DUF499-domain-containing-protein-[Acidovorax-carolinensis] |
| NRT58384.1 | DUF499_1.RM.like.systems | HAD->?-><-rve_3<-ParB+HTH<-?\|\|?-><- DUF499 *<- TOPRIM <- OLD-ABC <- METHYLASE <-HNH<- OLD-ABC <- TUDOR+ SF1-SF2-Helicase+iREase+CTD +REase | 953 | Bacteria>Pseudomonadota>Betaproteobacteria>Burkholderiales>Sphaerotilaceae | GCA_013294065.1 | Leptothrix_sp._C29 | HNQ01_004152 | 18472..21333 | 2735897 | hypothetical-protein-[Leptothrix-sp.-C29] |
| WP_273473498.1 | DUF499_1.RM.like.systems | <-zf-IS66+DDE_Tnp_IS66<-DDE_Tnp_1_assoc+DDE_Tnp_1<-ParB+HTH<-?\|\|?-><- DUF499 *<- TOPRIM <- OLD-ABC <- METHYLASE <-?<-PIN<- OLD-ABC | 955 | Bacteria>Pseudomonadota>Betaproteobacteria>Burkholderiales>Sphaerotilaceae | GCF_023150255.1 | Sphaerotilus_sulfidivorans | L6Q74_RS20510 | 5829..8696 | 639200 | DUF499-domain-containing-protein-[Sphaerotilus-sulfidivorans] |
| MCK6404267.1 | DUF499_1.RM.like.systems | <-zf-IS66+DDE_Tnp_IS66<-DDE_Tnp_1_assoc+DDE_Tnp_1<-ParB+HTH<-?\|\|?-><- DUF499 *<- TOPRIM <- OLD-ABC <- METHYLASE <-?<-PIN<- OLD-ABC | 955 | Bacteria>Pseudomonadota>Betaproteobacteria>Burkholderiales>Sphaerotilaceae | GCA_023150255.1 | Sphaerotilus_sulfidivorans | L6Q74_20550 | 5829..8696 | 639200 | DUF499-domain-containing-protein-[Sphaerotilus-sulfidivorans] |
| WP_173807403.1 | DUF499_1.RM.like.systems | HAD->?-><-ParB+HTH<-?\|\|?-><- DUF499 *<- TOPRIM <- OLD-ABC <- METHYLASE <-HNH<- OLD-ABC <- TUDOR+ SF1-SF2-Helicase+iREase+CTD +REase | 953 | Bacteria>Pseudomonadota>Betaproteobacteria>Burkholderiales>Sphaerotilaceae | GCF_013294065.1 | Leptothrix_sp._C29 | HNQ01_RS20655 | 18472..21333 | 2735897 | DUF499-domain-containing-protein-[Leptothrix-sp.-C29] |
| NNM65793.1 | DUF499_1.RM.like.systems | HTH+WYL-> TUDOR+ SF1-SF2-Helicase+iREase+CTD +REase -> METHYLASE -> DUF499 *-> OLD-ABC + TOPRIM ->SF1-SF2-Helicase ->?-> OLD-ABC + TOPRIM -> | 971 | Bacteria>Pseudomonadota>Betaproteobacteria>Burkholderiales>unclassified_Burkholderiales | GCA_013044035.1 | Burkholderiales_bacterium | HKL99_14520 | 53445..56360 | 1891238 | ATP-binding-protein-[Burkholderiales-bacterium] |
| MBE0627115.1 | DUF499_1.RM.like.systems | TUDOR+ SF1-SF2-Helicase+iREase+CTD +REase -> OLD-ABC -> TOPRIM ->HEPN-> METHYLASE -> HEPN+ DUF499 *->?->?->HEPN->REase-> | 1113 | Bacteria>Pseudomonadota>Betaproteobacteria>Burkholderiales>unclassified_Burkholderiales | GCA_014860355.1 | Burkholderiales_bacterium | IH606_20135 | 9089..12430 | 1891238 | DUF499-domain-containing-protein-[Burkholderiales-bacterium] |
| MCZ2419561.1 | DUF499_1.RM.like.systems | TPR+TPR-><-?<-HTH+UbE1<-?\|\|HTH->MPTase-><- HEPN+ DUF499 *<-?<-DUF488-1<-?<- METHYLASE <- TOPRIM <- OLD-ABC | 1108 | Bacteria>Pseudomonadota>Betaproteobacteria>Burkholderiales>unclassified_Burkholderiales | GCA_026931465.1 | Burkholderiales_bacterium | LC123_06960 | 80916..84242 | 1891238 | MAG:-DUF499-domain-containing-protein-[Burkholderiales-bacterium] |
| NMA26876.1 | DUF499_1.RM.like.systems | METHYLASE -p-> OLD-ABC -> TOPRIM -> HEPN+ DUF499 *-> | 448 | Bacteria>Pseudomonadota>Betaproteobacteria>Burkholderiales>unclassified_Burkholderiales | GCA_012799975.1 | Burkholderiales_bacterium | GX934_03860 | 2623..3966 | 1891238 | ATP-binding-protein-[Burkholderiales-bacterium] |
| WP_031597360.1 | DUF499_1.RM.like.systems | TUDOR+ SF1-SF2-Helicase+iREase+CTD +REase -> TOPRIM -> METHYLASE ->HEPN->ParA-Soj-PloopNTPase->ORC-CDC6-> HEPN+ DUF499 *-><-Phage_integrase<-?<-?\|\|<-?<-HTH+UbE1 | 1109 | Bacteria>Pseudomonadota>Betaproteobacteria>Ferrovales>Ferrovaceae | GCF_000735045.1 | Ferrovum_myxofaciens | IV91_RS0107710 | 11022..14351 | 416213 | Swt1-family-HEPN-domain-containing-protein-[Ferrovum-myxofaciens] |
| NDU85854.1 | DUF499_1.RM.like.systems | <-HTH+UbE1<-?\|\|?->?->MPTase-><- HEPN+ DUF499 *<- METHYLASE <- TOPRIM <- OLD-ABC <- TUDOR+ SF1-SF2-Helicase+iREase+CTD +REase <- SF1-SF2-Helicase<- OLD-ABC + TOPRIM | 1110 | Bacteria>Pseudomonadota>Betaproteobacteria>Ferrovales>Ferrovaceae | GCA_010436015.1 | Ferrovum_sp. | G3H99_04485 | 6034..9366 | 2609467 | ATP-binding-protein-[Ferrovum-sp.] |
| WP_273118797.1 | DUF499_1.RM.like.systems | HTH+UbE1->?->\|\|?->?->Phage_integrase-><- HEPN+ DUF499 *<-ORC-CDC6<-ParA-Soj-PloopNTPase<-HEPN<- METHYLASE <- TOPRIM <- OLD-ABC | 1109 | Bacteria>Pseudomonadota>Betaproteobacteria>Ferrovales>Ferrovaceae | GCF_013181415.1 | Ferrovum_myxofaciens | HO274_RS08660 | 1781795..1785124 | 416213 | Swt1-family-HEPN-domain-containing-protein-[Ferrovum-myxofaciens] |
| QKE41374.1 | DUF499_1.RM.like.systems | HTH+UbE1->?->?->?->Phage_integrase-><- HEPN+ DUF499 *<-ORC-CDC6<-ParA-Soj-PloopNTPase<-HEPN<- METHYLASE <- TOPRIM <- OLD-ABC | 1109 | Bacteria>Pseudomonadota>Betaproteobacteria>Ferrovales>Ferrovaceae | GCA_013181415.1 | Ferrovum_myxofaciens | HO274_08630 | 1781795..1785124 | 416213 | ATP-binding-protein-[Ferrovum-myxofaciens] |
| MCX7674148.1 | DUF499_1.RM.like.systems | NTP_transf_2->HEPN->Mbetalac->tRNA->?-><- DUF499 *<-Schlafen+HATPase+HTH<- METHYLASE <-?<- TOPRIM <- OLD-ABC <- METHYLASE | 1093 | Bacteria>Pseudomonadota>Betaproteobacteria>Nitrosomonadales>Thiobacillaceae | GCA_026419045.1 | Thiobacillaceae_bacterium | N2Z63_11245 | 10361..13642 | 2528409 | MAG:-DUF499-domain-containing-protein-[Thiobacillaceae-bacterium] |
| MBI1887192.1 | DUF499_1.RM.like.systems | <-Glycos_trans_3N+Glycos_transf_3<-?<-Schlafen+HATPase+HTH<-?<-?<- HEPN+ DUF499 *<- METHYLASE <-HEPN<- TOPRIM <- OLD-ABC <- TUDOR+ SF1-SF2-Helicase+iREase+CTD +REase <-ParB | 1112 | Bacteria>Pseudomonadota>Betaproteobacteria>Nitrosomonadales>unclassified_Nitrosomonadales | GCA_016177795.1 | Nitrosomonadales_bacterium | HYS19_02300 | 86364..89702 | 2026761 | ATP-binding-protein-[Nitrosomonadales-bacterium] |
| MBI3902214.1 | DUF499_1.RM.like.systems | <-Glycos_trans_3N+Glycos_transf_3<-?<-?<-Schlafen+HATPase+HTH<-?<- HEPN+ DUF499 *<- METHYLASE <-?<-?<- TOPRIM <- OLD-ABC <- TUDOR+ SF1-SF2-Helicase+iREase+CTD +REase | 1112 | Bacteria>Pseudomonadota>Betaproteobacteria>Nitrosomonadales>unclassified_Nitrosomonadales | GCA_016199835.1 | Nitrosomonadales_bacterium | HY306_04630 | 118339..121677 | 2026761 | ATP-binding-protein-[Nitrosomonadales-bacterium] |
| MBI5658243.1 | DUF499_1.RM.like.systems | TUDOR+ SF1-SF2-Helicase+iREase+CTD +REase -> OLD-ABC -> TOPRIM -> METHYLASE ->?->?-> HEPN+ DUF499 *-> | 1125 | Bacteria>Pseudomonadota>Betaproteobacteria>Nitrosomonadales>unclassified_Nitrosomonadales | GCA_016223025.1 | Nitrosomonadales_bacterium | HZC43_01545 | 64771..68148 | 2026761 | ATP-binding-protein-[Nitrosomonadales-bacterium] |
| MCX8086911.1 | DUF499_1.RM.like.systems | <-Phage_integrase\|\|tRNA-><- DUF499 *<-Schlafen+HATPase+HTH<- METHYLASE <- TOPRIM <- OLD-ABC <- METHYLASE | 943 | Bacteria>Pseudomonadota>Betaproteobacteria>Rhodocyclales>Rhodocyclaceae | GCA_026415155.1 | Rhodocyclaceae_bacterium | N3C63_08430 | 24540..27371 | 1898103 | MAG:-DUF499-domain-containing-protein-[Rhodocyclaceae-bacterium] |
| MBI3917901.1 | DUF499_1.RM.like.systems | OLD-ABC ->PHP+SbcC->Schlafen+HATPase+HTH-> METHYLASE -> OLD-ABC -> TOPRIM -> DUF499 *-> | 927 | Bacteria>Pseudomonadota>Betaproteobacteria>unclassified_Betaproteobacteria | GCA_016199545.1 | Betaproteobacteria_bacterium | HY322_12940 | 28133..30916 | 1891241 | ATP-binding-protein-[Betaproteobacteria-bacterium] |
| MBI3370955.1 | DUF499_1.RM.like.systems | <- OLD-ABC + TOPRIM <-?<- DUF499 *<- METHYLASE | 1089 | Bacteria>Pseudomonadota>Betaproteobacteria>unclassified_Betaproteobacteria | GCA_016195915.1 | Betaproteobacteria_bacterium | HYO17_04235 | 112207..115476 | 1891241 | ATP-binding-protein-[Betaproteobacteria-bacterium] |
| WP_257709589.1 | DUF499_1.RM.like.systems | TUDOR+ SF1-SF2-Helicase+iREase+CTD +REase -> OLD-ABC ->HNH-> METHYLASE -> OLD-ABC -> TOPRIM -> DUF499 *-><-RlpA<-CbeA_antitoxin<-JAB<-?\|\|<-Arm-DNA-bind_3+Phage_integrase | 948 | Bacteria>Pseudomonadota>Gammaproteobacteria>Aeromonadales>Aeromonadaceae | GCF_024649925.1 | Aeromonas_caviae | NUK49_RS00060 | 12274..15120 | 648 | DUF499-domain-containing-protein-[Aeromonas-caviae] |
| WP_223918005.1 | DUF499_1.RM.like.systems | TUDOR+ SF1-SF2-Helicase+iREase+CTD +REase -> OLD-ABC ->HNH-> METHYLASE -> OLD-ABC -> TOPRIM -> DUF499 *-><-RlpA<-CbeA_antitoxin<-JAB | 948 | Bacteria>Pseudomonadota>Gammaproteobacteria>Aeromonadales>Aeromonadaceae | GCF_019971655.1 | Aeromonas_caviae | LDN21_RS02455 | 71531..74377 | 648 | DUF499-domain-containing-protein-[Aeromonas-caviae] |
| WP_216991887.1 | DUF499_1.RM.like.systems | TUDOR+ SF1-SF2-Helicase+iREase+CTD +REase -> OLD-ABC ->HNH-> METHYLASE -> OLD-ABC -> TOPRIM -> DUF499 *-><-RlpA<-CbeA_antitoxin<-JAB | 948 | Bacteria>Pseudomonadota>Gammaproteobacteria>Aeromonadales>Aeromonadaceae | GCF_019048725.1 | Aeromonas_sp._FDAARGOS_1407 | I6L37_RS11490 | 2395827..2398673 | 2778056 | DUF499-domain-containing-protein-[Aeromonas-sp.-FDAARGOS-1407] |
| WP_201871420.1 | DUF499_1.RM.like.systems | JAB->CbeA_antitoxin->RlpA-><- DUF499 *<- TOPRIM <- OLD-ABC <- METHYLASE <-HNH<- OLD-ABC <- TUDOR+ SF1-SF2-Helicase+iREase+CTD +REase | 948 | Bacteria>Pseudomonadota>Gammaproteobacteria>Aeromonadales>Aeromonadaceae | GCF_016728905.1 | Aeromonas_caviae | JD551_RS01070 | 83459..86305 | 648 | DUF499-domain-containing-protein-[Aeromonas-caviae] |
| WP_103858986.1 | DUF499_1.RM.like.systems | JAB->CbeA_antitoxin->RlpA-><- DUF499 *<- TOPRIM <- OLD-ABC <- METHYLASE <-HNH<- OLD-ABC <- TUDOR+ SF1-SF2-Helicase+iREase+CTD +REase | 948 | Bacteria>Pseudomonadota>Gammaproteobacteria>Aeromonadales>Aeromonadaceae | GCF_002919855.1 | Aeromonas_sp._ASNIH8 | C3418_RS17745 | 3201..6047 | 1920113 | DUF499-domain-containing-protein-[Aeromonas-sp.-ASNIH8] |
| WP_219296379.1 | DUF499_1.RM.like.systems | JAB->RlpA-><-HTH\|\|?-><- DUF499 *<- TOPRIM <- OLD-ABC <- METHYLASE <-HNH<- OLD-ABC <- TUDOR+ SF1-SF2-Helicase+iREase+CTD +REase | 952 | Bacteria>Pseudomonadota>Gammaproteobacteria>Aeromonadales>Aeromonadaceae | GCF_019348625.1 | Aeromonas_veronii | GL270_RS12655 | 107539..110397 | 654 | DUF499-domain-containing-protein-[Aeromonas-veronii] |
| WP_264787584.1 | DUF499_1.RM.like.systems | HTH+WYL-> TUDOR+ SF1-SF2-Helicase+iREase+CTD +REase ->?-> METHYLASE -> OLD-ABC -> TOPRIM -> DUF499 *-><-RlpA<-?<-JAB<-?<-?<-ParB | 945 | Bacteria>Pseudomonadota>Gammaproteobacteria>Aeromonadales>Aeromonadaceae | GCF_026000055.1 | Aeromonas_caviae | OPI30_RS08280 | 1646827..1649664 | 648 | DUF499-domain-containing-protein-[Aeromonas-caviae] |
| WP_244812704.1 | DUF499_1.RM.like.systems | METHYLASE -p-> OLD-ABC -> TOPRIM -> DUF499 *-><-RlpA<-CbeA_antitoxin<-JAB<-?\|\|<-Arm-DNA-bind_3+Phage_integrase | 948 | Bacteria>Pseudomonadota>Gammaproteobacteria>Aeromonadales>Aeromonadaceae | GCF_022869435.1 | Aeromonas | MUW49_RS10495 | 3086..5932 | 642 | DUF499-domain-containing-protein-[Aeromonas-sp.-LsrichE-8G] |
| WP_064338617.1 | DUF499_1.RM.like.systems | JAB->RlpA-><-HTH\|\|?-><- DUF499 *<- TOPRIM <- OLD-ABC <-DDE_Tnp_1_assoc+DDE_Tnp_1<- OLD-ABC -p<- METHYLASE <-HNH | 954 | Bacteria>Pseudomonadota>Gammaproteobacteria>Aeromonadales>Aeromonadaceae | GCF_001634325.1 | Aeromonas_veronii | WM43_RS07825 | 1773937..1776801 | 654 | DUF499-domain-containing-protein-[Aeromonas-veronii] |
| WP_064341063.1 | DUF499_1.RM.like.systems | <-HTH\|\|?-><-HIN-HTH\|\|?->?->DDE_Tnp_1-><- DUF499 *<- TOPRIM <- OLD-ABC <- OLD-ABC -p<- METHYLASE <- METHYLASE -p<-HNH | 954 | Bacteria>Pseudomonadota>Gammaproteobacteria>Aeromonadales>Aeromonadaceae | GCF_014169835.1 | Aeromonas_veronii | QBS04_RS15305 | 3271066..3273930 | 654 | DUF499-domain-containing-protein-[Aeromonas-veronii] |
| WP_270793588.1 | DUF499_1.RM.like.systems | TUDOR+ SF1-SF2-Helicase+iREase+CTD +REase ->?-> METHYLASE -> OLD-ABC -> TOPRIM -> DUF499 *-><-RlpA<-?<-JAB | 945 | Bacteria>Pseudomonadota>Gammaproteobacteria>Aeromonadales>Aeromonadaceae | GCF_027705385.1 | unclassified_Aeromonas | OKC33_RS16320 | 9044..11881 | 257493 | MULTISPECIES:-DUF499-domain-containing-protein-[unclassified-Aeromonas] |
| WP_235462717.1 | DUF499_1.RM.like.systems | TUDOR+ SF1-SF2-Helicase+iREase+CTD +REase -> OLD-ABC ->HNH-> METHYLASE -> OLD-ABC -> TOPRIM -> DUF499 *-> | 897 | Bacteria>Pseudomonadota>Gammaproteobacteria>Aeromonadales>Aeromonadaceae | GCF_021538905.1 | Aeromonas_australiensis | EHZ86_RS11610 | 15978..18671 | 1114880 | DUF499-domain-containing-protein-[Aeromonas-australiensis] |
| MCF3097928.1 | DUF499_1.RM.like.systems | TUDOR+ SF1-SF2-Helicase+iREase+CTD +REase ->HNH-> METHYLASE -> OLD-ABC -> TOPRIM -> DUF499 *-> | 954 | Bacteria>Pseudomonadota>Gammaproteobacteria>Aeromonadales>Aeromonadaceae | GCA_021538905.1 | Aeromonas_australiensis | EHZ86_11640 | 15807..18671 | 1114880 | ATP-binding-protein-[Aeromonas-australiensis] |
| WP_079541822.1 | DUF499_1.RM.like.systems | TUDOR+ SF1-SF2-Helicase+iREase+CTD +REase -> OLD-ABC ->HNH-> METHYLASE -> OLD-ABC -> TOPRIM -> DUF499 *-> | 937 | Bacteria>Pseudomonadota>Gammaproteobacteria>Cardiobacteriales>Cardiobacteriaceae | GCF_900079085.1 | Cardiobacterium_hominis | CHUV0807_RS10775 | 37520..40333 | 2718 | DUF499-domain-containing-protein-[Cardiobacterium-hominis] |
| WP_135445665.1 | DUF499_1.RM.like.systems | METHYLASE ->?->?-> OLD-ABC -> TOPRIM -> DUF499 *->REase->MORC+DUF3684+REase-><-SF1-SF2-Helicase+REase<-SF1-SF2-Helicase+REase\|\|?->HKD-DNase+HKD-DNase-> | 947 | Bacteria>Pseudomonadota>Gammaproteobacteria>Cellvibrionales>Halieaceae | GCF_004745945.1 | Haliea_sp._SAOS-164 | E4634_RS16035 | 159390..162233 | 2562682 | DUF499-domain-containing-protein-[Haliea-sp.-SAOS-164] |
| MTW22277.1 | DUF499_1.RM.like.systems | TUDOR+ SF1-SF2-Helicase+iREase+CTD +REase -> OLD-ABC ->HNH-> OLD-ABC -> TOPRIM -> DUF499 *-> | 885 | Bacteria>Pseudomonadota>Gammaproteobacteria>Chromatiales>Chromatiaceae | GCA_009720725.1 | Allochromatium_palmeri | GJ668_14455 | 38860..41517 | 231048 | DUF499-domain-containing-protein-[Allochromatium-palmeri] |
| WP_155450846.1 | DUF499_1.RM.like.systems | OLD-ABC ->HNH-> METHYLASE -p-> METHYLASE -p-> OLD-ABC -> TOPRIM -> DUF499 *-> | 883 | Bacteria>Pseudomonadota>Gammaproteobacteria>Chromatiales>Chromatiaceae | GCF_009720725.1 | Allochromatium_palmeri | GJ668_RS14460 | 38866..41517 | 231048 | DUF499-domain-containing-protein-[Allochromatium-palmeri] |
| WP_020507360.1 | DUF499_1.RM.like.systems | <-McrC-NTD+McrC<-?<- DUF499 *<-wHTH+REase<- METHYLASE <- TOPRIM <- OLD-ABC <- METHYLASE -p<- TUDOR+ SF1-SF2-Helicase+iREase+CTD +REase | 961 | Bacteria>Pseudomonadota>Gammaproteobacteria>Chromatiales>Chromatiaceae | GCF_000379525.1 | Lamprocystis_purpurea | A390_RS0124805 | 120589..123474 | 61598 | DUF499-domain-containing-protein-[Lamprocystis-purpurea] |
| WP_157817756.1 | DUF499_1.RM.like.systems | <-Abhydrolase_4\|\|DDE_Tnp_IS1-><-DUF29<- DUF499 *<- METHYLASE <- TOPRIM <- OLD-ABC <- TUDOR+ SF1-SF2-Helicase+iREase+CTD +REase | 1094 | Bacteria>Pseudomonadota>Gammaproteobacteria>Chromatiales>Chromatiaceae | GCF_002813775.1 | Candidatus_Thiodictyon_syntrophicum | THSYN_RS17695 | 4223024..4226308 | 1166950 | DUF499-domain-containing-protein-[Candidatus-Thiodictyon-syntrophicum] |
| AUB82572.1 | DUF499_1.RM.like.systems | <-Abhydrolase_4\|\|DDE_Tnp_IS1-><-DUF29<- DUF499 *<- METHYLASE <- TOPRIM <- OLD-ABC <- TUDOR+ SF1-SF2-Helicase+iREase+CTD +REase | 1083 | Bacteria>Pseudomonadota>Gammaproteobacteria>Chromatiales>Chromatiaceae | GCA_002813775.1 | Candidatus_Thiodictyon_syntrophicum | THSYN_17560 | 4223024..4226275 | 1166950 | hypothetical-protein-THSYN_17560-[Candidatus-Thiodictyon-syntrophicum] |
| WP_100917910.1 | DUF499_1.RM.like.systems | TUDOR+ SF1-SF2-Helicase+iREase+CTD +REase -> OLD-ABC -> TOPRIM -> METHYLASE ->PolB-> HEPN+ DUF499 *->?-><-NUDIX<-?<-?\|\|?-><-PAIREDC-HTH | 1111 | Bacteria>Pseudomonadota>Gammaproteobacteria>Chromatiales>Chromatiaceae | GCF_002813775.1 | Candidatus_Thiodictyon_syntrophicum | THSYN_RS03450 | 792207..795542 | 1166950 | Swt1-family-HEPN-domain-containing-protein-[Candidatus-Thiodictyon-syntrophicum] |
| WP_020507743.1 | DUF499_1.RM.like.systems | OLD-ABC -> TOPRIM -> METHYLASE ->PolB->PolB->PolB+HEPN-> HEPN+ DUF499 *-><-?<-?<-?<-REC<-HPT+HISKIN+REC | 1124 | Bacteria>Pseudomonadota>Gammaproteobacteria>Chromatiales>Chromatiaceae | GCF_000379525.1 | Lamprocystis_purpurea | A390_RS0126800 | 79657..83031 | 61598 | Swt1-family-HEPN-domain-containing-protein-[Lamprocystis-purpurea] |
| QVL49217.1 | DUF499_1.RM.like.systems | OLD-ABC -> TOPRIM -> METHYLASE ->?->PolB->PolB+HEPN-> HEPN+ DUF499 *-><-?<-?\|\|HicB-><-sGTP\|\|?-><-TPR+TPR | 1114 | Bacteria>Pseudomonadota>Gammaproteobacteria>Chromatiales>Chromatiaceae | GCA_018399035.1 | Thiocapsa_sp. | KFB96_01415 | 301904..305248 | 2024551 | MAG:-DUF499-domain-containing-protein-[Thiocapsa-sp.] |
| MBK8637657.1 | DUF499_1.RM.like.systems | TOPRIM -> METHYLASE ->PolB->HEPN->?->Y1_Tnp-> HEPN+ DUF499 *->RADICAL-SAM-> | 1120 | Bacteria>Pseudomonadota>Gammaproteobacteria>Chromatiales>Chromatiaceae | GCA_016714645.1 | Chromatiaceae_bacterium | IPN92_04990 | 1074519..1077881 | 2049432 | DUF499-domain-containing-protein-[Chromatiaceae-bacterium] |
| MBF2761740.1 | DUF499_1.RM.like.systems | <- TOPRIM <- OLD-ABC <-N6_N4_MTase<- DUF499 *<- METHYLASE <- METHYLASE -p<- TOPRIM <- OLD-ABC <-?<-N6_N4_MTase | 1092 | Bacteria>Pseudomonadota>Gammaproteobacteria>Chromatiales>Ectothiorhodospiraceae | GCA_015278165.1 | Ectothiorhodospiraceae_bacterium_AqS1 | ISN28_16015 | 2939..6217 | 1725326 | ATP-binding-protein-[Ectothiorhodospiraceae-bacterium-AqS1] |
| BAL53124.1 | DUF499_1.RM.like.systems | TUDOR+ SF1-SF2-Helicase+iREase+CTD +REase -> OLD-ABC -> TOPRIM -> METHYLASE ->?->?-> DUF499 *-><-?<-?<-tRNA\|\|?-><-?<-TRAM+Methyltransf_31 | 967 | Bacteria>Pseudomonadota>Gammaproteobacteria>environmental_samples | - | uncultured_Gammaproteobacteria_bacterium | HGMM_F04D09C21 | 20323..23226 | 86473 | AAA-family-ATPase-[uncultured-Gammaproteobacteria-bacterium] |
| MBK8754773.1 | DUF499_1.RM.like.systems | METHYLASE -> OLD-ABC -> TOPRIM ->?->PIN-> DUF499 *->?-><-?<-CASPASE<-TIR<-?<-ADH_N+ADH_zinc_N | 946 | Bacteria>Pseudomonadota>Gammaproteobacteria>Gammaproteobacteria_incertae_sedis>Candidatus_Competibacteraceae | GCA_016710535.1 | Candidatus_Competibacteraceae_bacterium | IPL99_25425 | 33622..36462 | 2053538 | ATP-binding-protein-[Candidatus-Competibacteraceae-bacterium] |
| NOT11404.1 | DUF499_1.RM.like.systems | PemK_toxin-> OLD-ABC -> TOPRIM -> METHYLASE -> OLD-ABC -> TOPRIM -> DUF499 *-> | 939 | Bacteria>Pseudomonadota>Gammaproteobacteria>Methylococcales>Methylococcaceae | GCA_013140705.1 | Methylococcaceae_bacterium | HOP23_06170 | 47500..50319 | 1933926 | ATP-binding-protein-[Methylococcaceae-bacterium] |
| WP_218796637.1 | DUF499_1.RM.like.systems | XIS-HTH-> TUDOR+ SF1-SF2-Helicase+iREase+CTD +REase -> OLD-ABC -> TOPRIM -> METHYLASE ->REase-> DUF499 *->MORC+REase->HEPN-><-Phage_lysozyme<-?<-?<-RNase | 939 | Bacteria>Pseudomonadota>Gammaproteobacteria>Methylococcales>Methylococcaceae | GCF_019285555.1 | Methylococcus_capsulatus | KW114_RS08340 | 1841753..1844572 | 414 | DUF499-domain-containing-protein-[Methylococcus-capsulatus] |
| HCK4430924.1 | DUF499_1.RM.like.systems | TUDOR+ SF1-SF2-Helicase+iREase+CTD +REase ->HEPN-> METHYLASE -> OLD-ABC -> TOPRIM -> DUF499 *-><-RlpA<-?<-JAB<-ParB<-?\|\|MazE_antitoxin-> | 942 | Bacteria>Pseudomonadota>Gammaproteobacteria>Pseudomonadales>Pseudomonadaceae | GCA_024830635.1 | Pseudomonas_aeruginosa | NOG72_005469 | 46146..48974 | 287 | ATP-binding-protein-[Pseudomonas-aeruginosa] |
| WP_124078188.1 | DUF499_1.RM.like.systems | ZnR+ZnR-><-HTH_3\|\|?->?->RlpA-><- DUF499 *<- TOPRIM <- OLD-ABC <- METHYLASE <-HNH<- OLD-ABC <- TUDOR+ SF1-SF2-Helicase+iREase+CTD +REase | 943 | Bacteria>Pseudomonadota>Gammaproteobacteria>Pseudomonadales>Pseudomonadaceae | GCF_033140865.1 | Pseudomonas_aeruginosa | IPC1239_RS13835 | 644387..647218 | 287 | DUF499-domain-containing-protein-[Pseudomonas-aeruginosa] |
| WP_128472827.1 | DUF499_1.RM.like.systems | RlpA-><- DUF499 *<- TOPRIM <- OLD-ABC <- METHYLASE <-HEPN<- TUDOR+ SF1-SF2-Helicase+iREase+CTD +REase | 943 | Bacteria>Pseudomonadota>Gammaproteobacteria>Pseudomonadales>Pseudomonadaceae | GCF_004053955.1 | Pseudomonas_aeruginosa | EQH74_RS32310 | 490..3321 | 287 | DUF499-domain-containing-protein-[Pseudomonas-aeruginosa] |
| HCL3757805.1 | DUF499_1.RM.like.systems | TUDOR+ SF1-SF2-Helicase+iREase+CTD +REase -> OLD-ABC ->HNH-> METHYLASE -> OLD-ABC -> TOPRIM -> DUF499 *-> | 667 | Bacteria>Pseudomonadota>Gammaproteobacteria>Pseudomonadales>Pseudomonadaceae | GCA_025003865.1 | Pseudomonas_aeruginosa | N2B17_005733 | 10859..12860 | 287 | ATP-binding-protein-[Pseudomonas-aeruginosa] |
| TXH86520.1 | DUF499_1.RM.like.systems | JAB->RlpA-><- DUF499 *<- TOPRIM <- OLD-ABC <- METHYLASE | 942 | Bacteria>Pseudomonadota>Gammaproteobacteria>Pseudomonadales>Pseudomonadaceae | GCA_008015405.1 | Pseudomonas_sp. | E6Q71_06445 | 770..3598 | 306 | ATP-binding-protein-[Pseudomonas-sp.] |
| MCG8614350.1 | DUF499_1.RM.like.systems | HISKIN-><-?\|\|?->Phage_int_SAM_4-><-?\|\|?-><- DUF499 *<- TOPRIM <- OLD-ABC <- METHYLASE <- TUDOR+ SF1-SF2-Helicase+iREase+CTD +REase -p | 947 | Bacteria>Pseudomonadota>Gammaproteobacteria>Pseudomonadales>unclassified_Pseudomonadales | GCA_022361325.1 | Pseudomonadales_bacterium | MI864_27885 | 9196..12039 | 1891229 | DUF499-domain-containing-protein-[Pseudomonadales-bacterium] |
| MAS10374.1 | DUF499_1.RM.like.systems | <-HhH-RADC+JAB\|\|?->?->?-><- DUF499 *<- TOPRIM <- OLD-ABC <- METHYLASE <-PIN<- TUDOR+ SF1-SF2-Helicase+iREase+CTD +REase <-JetD | 942 | Bacteria>Pseudomonadota>Gammaproteobacteria>Salinisphaerales>Salinisphaeraceae | GCA_002702145.1 | Salinisphaera_sp. | CMN26_09985 | 5272..8100 | 1914330 | hypothetical-protein-[Salinisphaera-sp.] |
| MCY4286579.1 | DUF499_1.RM.like.systems | PolB->ParB+HNH-> METHYLASE ->?-> OLD-ABC -> TOPRIM -> DUF499 *->?->?-><-?<-?<-ABC-ATPase+SbcC | 1107 | Bacteria>Pseudomonadota>Gammaproteobacteria>Thiotrichales>unclassified_Thiotrichales | GCA_026709935.1 | Thiotrichales_bacterium | OXC65_14725 | 7747..11070 | 2026796 | MAG:-DUF499-domain-containing-protein-[Thiotrichales-bacterium] |
| MDE0453907.1 | DUF499_1.RM.like.systems | <-HEPN+REase<-Schlafen+HATPase+HTH<-Schlafen<- DUF499 *<- METHYLASE <- SF1-SF2-Helicase<- OLD-ABC + TOPRIM <- TUDOR+ SF1-SF2-Helicase+iREase+CTD +REase <-AbiEii<-AbiEi_4 | 967 | Bacteria>Pseudomonadota>Gammaproteobacteria>unclassified_Gammaproteobacteria | GCA_028821145.1 | Gammaproteobacteria_bacterium | OXJ63_01205 | 18382..21285 | 1913989 | MAG:-DUF499-domain-containing-protein-[Gammaproteobacteria-bacterium] |
| MXY51432.1 | DUF499_1.RM.like.systems | <-PhdYeFM_antitox\|\| TUDOR+ SF1-SF2-Helicase+iREase+CTD +REase -> OLD-ABC + TOPRIM ->SF1-SF2-Helicase -> METHYLASE -p-> METHYLASE -> DUF499 *->TIR-> | 968 | Bacteria>Pseudomonadota>Gammaproteobacteria>unclassified_Gammaproteobacteria | GCA_009837725.1 | Gammaproteobacteria_bacterium | F4Y86_02730 | 44418..47324 | 1913989 | ATP-binding-protein-[Gammaproteobacteria-bacterium] |
| MDE0129995.1 | DUF499_1.RM.like.systems | <- SF1-SF2-Helicase<- OLD-ABC + TOPRIM <-Schlafen+HATPase+HTH<- DUF499 *<- METHYLASE <- TUDOR+ SF1-SF2-Helicase+iREase+CTD +REase <-?<- OLD-ABC + TOPRIM | 968 | Bacteria>Pseudomonadota>Gammaproteobacteria>unclassified_Gammaproteobacteria | GCA_028819355.1 | Gammaproteobacteria_bacterium | OXQ86_10575 | 58066..60972 | 1913989 | MAG:-DUF499-domain-containing-protein-[Gammaproteobacteria-bacterium] |
| MDD9824620.1 | DUF499_1.RM.like.systems | <- DUF499 *<-PIN<- OLD-ABC <- OLD-ABC + TOPRIM <- METHYLASE <-TPR+TPR | 928 | Bacteria>Pseudomonadota>Gammaproteobacteria>unclassified_Gammaproteobacteria | GCA_028817255.1 | Gammaproteobacteria_bacterium | OXU43_05565 | 147..2933 | 1913989 | MAG:-DUF499-domain-containing-protein-[Gammaproteobacteria-bacterium] |
| MDD9822476.1 | DUF499_1.RM.like.systems | METHYLASE -p-> OLD-ABC + TOPRIM -> DUF499 *-> | 928 | Bacteria>Pseudomonadota>Gammaproteobacteria>unclassified_Gammaproteobacteria | GCA_028816685.1 | Gammaproteobacteria_bacterium | OXU98_06290 | 2678..5464 | 1913989 | MAG:-DUF499-domain-containing-protein-[Gammaproteobacteria-bacterium] |
| MDE0157713.1 | DUF499_1.RM.like.systems | OLD-ABC -> TOPRIM -> METHYLASE -p->?->?-> DUF499 *-> | 952 | Bacteria>Pseudomonadota>Gammaproteobacteria>unclassified_Gammaproteobacteria | GCA_028819495.1 | Gammaproteobacteria_bacterium | OXS28_19190 | 7492..10350 | 1913989 | MAG:-DUF499-domain-containing-protein-[Gammaproteobacteria-bacterium] |
| MDE0512489.1 | DUF499_1.RM.like.systems | <- DUF499 *<-?<-?<- METHYLASE <- TOPRIM <- OLD-ABC | 952 | Bacteria>Pseudomonadota>Gammaproteobacteria>unclassified_Gammaproteobacteria | GCA_028823865.1 | Gammaproteobacteria_bacterium | OXI88_11955 | 1695..4553 | 1913989 | MAG:-DUF499-domain-containing-protein-[Gammaproteobacteria-bacterium] |
| MDE0038699.1 | DUF499_1.RM.like.systems | OLD-ABC -> TOPRIM -><-?<-TIR\|\|TIR-> METHYLASE -> DUF499 *-><-SF1-SF2-Helicase | 1089 | Bacteria>Pseudomonadota>Gammaproteobacteria>unclassified_Gammaproteobacteria | GCA_028818795.1 | Gammaproteobacteria_bacterium | OXU77_14270 | 23040..26309 | 1913989 | MAG:-hypothetical-protein-OXU77_14270-[Gammaproteobacteria-bacterium] |
| MDD9817274.1 | DUF499_1.RM.like.systems | <- DUF499 *<-?<- TOPRIM <- OLD-ABC <- METHYLASE <-Schlafen+HATPase+HTH<-RelE-ParE | 1113 | Bacteria>Pseudomonadota>Gammaproteobacteria>unclassified_Gammaproteobacteria | GCA_028817495.1 | Gammaproteobacteria_bacterium | OXU61_03925 | 191..3532 | 1913989 | MAG:-hypothetical-protein-OXU61_03925-[Gammaproteobacteria-bacterium] |
| MDD9864539.1 | DUF499_1.RM.like.systems | METHYLASE -> OLD-ABC -> TOPRIM -> DUF499 *-><-?<-?\|\|?->?->?->ABC-ATPase+SbcC-> | 1113 | Bacteria>Pseudomonadota>Gammaproteobacteria>unclassified_Gammaproteobacteria | GCA_028817305.1 | Gammaproteobacteria_bacterium | OXU54_08205 | 4537..7878 | 1913989 | MAG:-DUF499-domain-containing-protein-[Gammaproteobacteria-bacterium] |
| TAK54308.1 | DUF499_1.RM.like.systems | TUDOR+ SF1-SF2-Helicase+iREase+CTD +REase ->Schlafen+HATPase+HTH-> METHYLASE -> OLD-ABC -> TOPRIM -> HEPN+ DUF499 *-> | 1105 | Bacteria>Pseudomonadota>Gammaproteobacteria>unclassified_Gammaproteobacteria | GCA_004297335.1 | Gammaproteobacteria_bacterium | EPO25_07530 | 13135..16452 | 1913989 | ATP-binding-protein-[Gammaproteobacteria-bacterium] |
| MDE0073871.1 | DUF499_1.RM.like.systems | METHYLASE -> OLD-ABC -> TOPRIM -> HEPN+ DUF499 *-><-HicB_lk_antitox<-HicA_toxin\|\|XIS-HTH->VirB2-> | 1107 | Bacteria>Pseudomonadota>Gammaproteobacteria>unclassified_Gammaproteobacteria | GCA_028819075.1 | Gammaproteobacteria_bacterium | OXU32_07805 | 3301..6624 | 1913989 | MAG:-Swt1-family-HEPN-domain-containing-protein-[Gammaproteobacteria-bacterium] |
| MCZ0943260.1 | DUF499_1.RM.like.systems | DUF433-><- HEPN+ DUF499 *<-Schlafen+HATPase+HTH<- TOPRIM <- OLD-ABC <- METHYLASE -p | 1115 | Bacteria>Pseudomonadota>Gammaproteobacteria>unclassified_Gammaproteobacteria | GCA_026914145.1 | Gammaproteobacteria_bacterium | OXJ53_09405 | 2539..5886 | 1913989 | MAG:-Swt1-family-HEPN-domain-containing-protein-[Gammaproteobacteria-bacterium] |
| MXY90447.1 | DUF499_1.RM.like.systems | <-PrmC_N+Methyltransf_25\|\|?-><-?<-?<- HEPN+ DUF499 *<- TOPRIM <- OLD-ABC | 1116 | Bacteria>Pseudomonadota>Gammaproteobacteria>unclassified_Gammaproteobacteria | GCA_009837675.1 | Gammaproteobacteria_bacterium | F4Y89_07905 | 34085..37435 | 1913989 | ATP-binding-protein-[Gammaproteobacteria-bacterium] |
| TAN08689.1 | DUF499_1.RM.like.systems | <-STYKinase-like<-HTH\|\| OLD-ABC + TOPRIM -> METHYLASE -> METHYLASE ->Schlafen+HATPase+HTH-> DUF499 *-><-BirA-HTH<-?<-Phage_integrase<-?<-ZnR | 934 | Bacteria>Pseudomonadota>Gammaproteobacteria>Xanthomonadales>Rhodanobacteraceae | GCA_004299555.1 | Rhodanobacteraceae_bacterium | EPN38_01845 | 144698..147502 | 2052182 | ATP-binding-protein-[Rhodanobacteraceae-bacterium] |
| WP_182517583.1 | DUF499_1.RM.like.systems | XIS-HTH-> TUDOR+ SF1-SF2-Helicase+iREase+CTD +REase ->HEPN-> METHYLASE -> OLD-ABC -> TOPRIM -> DUF499 *-> | 930 | Bacteria>Pseudomonadota>Gammaproteobacteria>Xanthomonadales>Rhodanobacteraceae | GCF_014138295.1 | Dokdonella_fugitiva | FHW13_RS05515 | 304049..306841 | 328517 | DUF499-domain-containing-protein-[Dokdonella-fugitiva] |

| Protein-ID | System | Gene-Neighbourhoods | Length | Taxonomy | Assembly-ID | Strain | Locus-Tag | Genomic-coordinates | TaxID | Fasta-defline |
|---|---|---|---|---|---|---|---|---|---|---|
| MBH2008931.1 | DUF499 _1.RM.like.systems | <- SF1-SF2-Helicase<- OLD-ABC + TOPRIM <- DUF499 *<- METHYLASE <-HEPN<- TUDOR+ SF1-SF2-Helicase+iREase+CTD +REase <-?\|\|DDE_Tnp_1_2-> | 969 | Bacteria>Pseudomonadota>Gammaproteobacteria>Xanthomonadales>Xanthomonadaceae | GCA_015999665.1 | Xanthomonadaceae_bacterium | I8H71_04430 | 4236..7145 | 1926873 | ATP-binding-protein-[Xanthomonadaceae-bacterium] |
| MTI74440.1 | DUF499 _1.RM.like.systems | <-JAB<-?<-D5N+D5ATPase<-?<-?<-?<- DUF499 *<-?<-?<- TOPRIM <- OLD-ABC <- TUDOR+ SF1-SF2-Helicase+iREase+CTD +REase <-Phage_integrase | 931 | Bacteria>Pseudomonadota>Gammaproteobacteria>Xanthomonadales>Xanthomonadaceae | GCA_009711865.1 | Stenotrophomonas_sp. | FH747_12410 | 103914..106709 | 69392 | ATP-binding-protein-[Stenotrophomonas-sp.] |
| PZO64463.1 | DUF499 _1.RM.like.systems | <- DUF499 *<- TOPRIM <- OLD-ABC <- METHYLASE <-PIN<- TUDOR+ SF1-SF2-Helicase+iREase+CTD +REase | 932 | Bacteria>Pseudomonadota>Gammaproteobacteria>Xanthomonadales>Xanthomonadaceae | GCA_003240695.1 | Pseudoxanthomonas_suwonensis | DI635_05295 | 731..3529 | 314722 | hypothetical-protein-DI635_05295-[Pseudoxanthomonas-suwonensis] |
| OQC12437.1 | DUF499 _1.RM.like.systems | TUDOR+ SF1-SF2-Helicase+iREase+CTD +REase ->MqsA_antitoxin+HTH_3->?-> METHYLASE -> OLD-ABC -> TOPRIM -> DUF499 *-><-?<-?<-ZnR<-35exo+SF1-SF2-Helicase | 976 | Bacteria>PVC_group>Lentisphaerota>unclassified_Lentisphaerota | GCA_002070945.1 | Lentisphaerae_bacterium_ADurb.Bin082 | BWX73_02930 | 53685..56615 | 1852899 | MAG:-hypothetical-protein-BWX73_02930-[Lentisphaerae-bacterium-ADurb.Bin082] |
| MCK6483011.1 | DUF499 _1.RM.like.systems | N6_N4_MTase->?->?-><-ParB<-GDResolvase+Recombinase+ZnR<-RAMA<- DUF499 *<- TOPRIM <- OLD-ABC <- METHYLASE <-?<-?<- TUDOR+ SF1-SF2-Helicase+iREase+CTD +REase | 1108 | Bacteria>PVC_group>Planctomycetota>Phycisphaerae>unclassified_Phycisphaerae | GCA_023150595.1 | Phycisphaerae_bacterium | L6R00_02615 | 131087..134413 | 2026778 | DUF499 -domain-containing-protein-[Phycisphaerae-bacterium] |
| GIW78771.1 | DUF499 _1.RM.like.systems | <-Phage_int_SAM_4+Phage_integrase\|\|tRNA-><-?<-XIS-HTH<-?<-?<- DUF499 *<- TOPRIM <- OLD-ABC <- METHYLASE <-?<- TUDOR+ SF1-SF2-Helicase+iREase+CTD +REase <-SF1-SF2-Helicase-p | 1113 | Bacteria>PVC_group>Planctomycetota>Planctomycetia>Gemmatales | GCA_026004675.1 | Gemmatales_bacterium | KatS3mg105_0578 | 670726..674067 | 2838789 | MAG:-hypothetical-protein-KatS3mg105_0578-[Gemmatales-bacterium] |
| MCF7962597.1 | DUF499 _1.RM.like.systems | <-Pkinase\|\|Phage_int_SAM_4+Phage_integrase-><- OLD-ABC + TOPRIM <-?<- DUF499 *<- METHYLASE | 956 | Bacteria>PVC_group>Planctomycetota>Planctomycetia>Pirellulales | GCA_021736575.1 | Pirellula_sp. | K9M08_17805 | 7810..10680 | 117 | ATP-binding-protein-[Pirellula-sp.] |
| GIW94553.1 | DUF499 _1.RM.like.systems | <- DUF499 *<-REase<-?<- TOPRIM <- OLD-ABC <- METHYLASE -p<- METHYLASE -p | 938 | Bacteria>PVC_group>Planctomycetota>Planctomycetia>Pirellulales | GCA_026004775.1 | Pirellulaceae_bacterium | KatS3mg110_2594 | 12784..15600 | 2699754 | MAG:-hypothetical-protein-KatS3mg110_2594-[Pirellulaceae-bacterium] |
| NLY99303.1 | DUF499 _1.RM.like.systems | <- DUF499 *<-?<- TOPRIM <- OLD-ABC <- METHYLASE -p | 1112 | Bacteria>PVC_group>Planctomycetota>Planctomycetia>Pirellulales | GCA_012744395.1 | Pirellulaceae_bacterium | GXY25_02065 | 2321..5659 | 2699754 | ATP-binding-protein-[Pirellulaceae-bacterium] |
| MDD4270717.1 | DUF499 _1.RM.like.systems | METHYLASE -> OLD-ABC -> TOPRIM ->?-> DUF499 *-> | 1112 | Bacteria>PVC_group>Planctomycetota>Planctomycetia>Pirellulales | GCA_028703175.1 | Thermoguttaceae_bacterium | PHN77_21030 | 4016..7354 | 2691551 | MAG:- DUF499 -domain-containing-protein-[Thermoguttaceae-bacterium] |
| MCA9232427.1 | DUF499 _1.RM.like.systems | <-ParB+HNH<- DUF499 *<- METHYLASE <- TOPRIM <- OLD-ABC <- TUDOR+ SF1-SF2-Helicase+iREase+CTD +REase -p | 953 | Bacteria>PVC_group>Planctomycetota>Planctomycetia>Planctomycetales | GCA_020427155.1 | Planctomycetales_bacterium | KDA57_17390 | 2105..4966 | 2053591 | ATP-binding-protein-[Planctomycetales-bacterium] |
| MBL8860740.1 | DUF499 _1.RM.like.systems | TUDOR+ SF1-SF2-Helicase+iREase+CTD +REase -> OLD-ABC + TOPRIM -> METHYLASE -> DUF499 *->DNA-glycosylase->?->?-><-?<-TPR+TPR\|\|Phage_integrase-> | 888 | Bacteria>PVC_group>Planctomycetota>unclassified_Planctomycetota | GCA_016795205.1 | Planctomycetota_bacterium | JNK02_01915 | 8775..11441 | 2026780 | DUF499 -domain-containing-protein-[Planctomycetota-bacterium] |
| GIW70817.1 | DUF499 _1.RM.like.systems | <-Glyco_hydro_57+DUF3536<-?<- DUF499 *<-REase<-REase<- TOPRIM <- OLD-ABC <- METHYLASE -p<- METHYLASE -p | 942 | Bacteria>PVC_group>Planctomycetota>unclassified_Planctomycetota | GCA_026004615.1 | Planctomycetota_bacterium | KatS3mg102_0359 | 449544..452372 | 2026780 | MAG:-hypothetical-protein-KatS3mg102_0359-[Planctomycetota-bacterium] |
| MBI4584119.1 | DUF499 _1.RM.like.systems | Schlafen+HATPase+HTH->?-> METHYLASE ->REase-> OLD-ABC -> TOPRIM -> DUF499 *->HicB_lk_antitox->HicA_toxin-> | 920 | Bacteria>PVC_group>Planctomycetota>unclassified_Planctomycetota | GCA_016209605.1 | Planctomycetota_bacterium | HY717_08860 | 13681..16443 | 2026780 | ATP-binding-protein-[Planctomycetota-bacterium] |
| MBI1826653.1 | DUF499 _1.RM.like.systems | <- DUF499 *<- TOPRIM <- OLD-ABC <- METHYLASE <-PHP+SbcC<- TUDOR+ SF1-SF2-Helicase+iREase+CTD +REase <-ArdCN+MPTase | 948 | Bacteria>PVC_group>Planctomycetota>unclassified_Planctomycetota | GCA_016178855.1 | Planctomycetota_bacterium | HYR83_09750 | 14494..17340 | 2026780 | ATP-binding-protein-[Planctomycetota-bacterium] |
| MCX5669700.1 | DUF499 _1.RM.like.systems | TUDOR+ SF1-SF2-Helicase+iREase+CTD +REase -> OLD-ABC + TOPRIM -> METHYLASE ->PolB->HEPN-> DUF499 *-> | 924 | Bacteria>PVC_group>Planctomycetota>unclassified_Planctomycetota | GCA_026387055.1 | Planctomycetota_bacterium | NTU94_00045 | 8049..10823 | 2026780 | MAG:- DUF499 -domain-containing-protein-[Planctomycetota-bacterium] |
| NMD35173.1 | DUF499 _1.RM.like.systems | <- DUF499 *<-REase<- METHYLASE <- OLD-ABC + TOPRIM | 932 | Bacteria>PVC_group>Planctomycetota>unclassified_Planctomycetota | GCA_012797405.1 | Planctomycetota_bacterium | GYA73_04750 | 409..3207 | 2026780 | ATP-binding-protein-[Planctomycetota-bacterium] |
| NQW48683.1 | DUF499 _1.RM.like.systems | TUDOR+ SF1-SF2-Helicase+iREase+CTD +REase ->SF1-SF2-Helicase-> METHYLASE -> OLD-ABC -> TOPRIM -> DUF499 *-> | 332 | Bacteria>PVC_group>Planctomycetota>unclassified_Planctomycetota | GCA_013205375.1 | Planctomycetota_bacterium | HQ464_13060 | 11784..12781 | 2026780 | ATP-binding-protein-[Planctomycetota-bacterium] |
| MBM4083197.1 | DUF499 _1.RM.like.systems | <- DUF499 *<- TOPRIM <- OLD-ABC <- METHYLASE <-BirA-HTH | 878 | Bacteria>PVC_group>Planctomycetota>unclassified_Planctomycetota | GCA_016873055.1 | Planctomycetota_bacterium | FJ272_00230 | 1..2634 | 2026780 | ATP-binding-protein-[Planctomycetota-bacterium] |
| QDU83134.1 | DUF499 _1.RM.like.systems | TUDOR+ SF1-SF2-Helicase+iREase+CTD +REase ->Schlafen+HATPase+HTH->?-> METHYLASE -> OLD-ABC -> TOPRIM -> HEPN+ DUF499 *->?-><-REase+SF1-SF2-Helicase<-DUF2357+REase<-Preader7+McrB-NTD+McrB-AAA | 1111 | Bacteria>PVC_group>Planctomycetota>unclassified_Planctomycetota | GCA_007750655.1 | Planctomycetes_bacterium_Pla163 | Pla163_02310 | 293863..297198 | 2528008 | hypothetical-protein-Pla163_02310-[Planctomycetes-bacterium-Pla163] |
| MCG3135902.1 | DUF499 _1.RM.like.systems | TUDOR+ SF1-SF2-Helicase+iREase+CTD +REase -> OLD-ABC -> TOPRIM -> METHYLASE -> HEPN+ DUF499 *->KAP-NTPase->?->PAPS_reduct->?-><-TPR+TPR | 1110 | Bacteria>PVC_group>Planctomycetota>unclassified_Planctomycetota | GCA_022072415.1 | Planctomycetota_bacterium | HMLKMBBP_03678 | 15324..18656 | 2026780 | hypothetical-protein-[Planctomycetota-bacterium] |
| MBE2203055.1 | DUF499 _1.RM.like.systems | <-JAB\|\|?-><-REase<-?<-?<-MORC+HisKin+REase<- DUF499 *<- METHYLASE <- TOPRIM <- OLD-ABC <- TUDOR+ SF1-SF2-Helicase+iREase+CTD +REase | 938 | Bacteria>PVC_group>Verrucomicrobiota>Spartobacteria>Chthoniobacterales | GCA_014879575.1 | Chthoniobacterales_bacterium | IAE94_01770 | 19431..22247 | 2201154 | ATP-binding-protein-[Chthoniobacterales-bacterium] |
| MCX6966137.1 | DUF499 _1.RM.like.systems | TUDOR+ SF1-SF2-Helicase+iREase+CTD +REase -> OLD-ABC -> TOPRIM -> METHYLASE ->?-> DUF499 *->?->?-><-Peptidase | 941 | Bacteria>PVC_group>Verrucomicrobiota>unclassified_Verrucomicrobiota | GCA_026398255.1 | Verrucomicrobiota_bacterium | NTW41_12430 | 37940..40765 | 2026799 | MAG:- DUF499 -domain-containing-protein-[Verrucomicrobia-bacterium] |
| TAE90777.1 | DUF499 _1.RM.like.systems | <- DUF499 *<-REase<- METHYLASE <-HNH<- OLD-ABC <- TOPRIM <- OLD-ABC | 939 | Bacteria>PVC_group>Verrucomicrobiota>unclassified_Verrucomicrobiota | GCA_004293285.1 | Verrucomicrobiota_bacterium | EAZ81_08635 | 280..3099 | 2026799 | ATP-binding-protein-[Verrucomicrobia-bacterium] |
| PWU13198.1 | DUF499 _1.RM.like.systems | TUDOR+ SF1-SF2-Helicase+iREase+CTD +REase -> METHYLASE ->REase-> OLD-ABC -> TOPRIM -> DUF499 *->?->?->?->?->Cluster1029_5clades-> | 928 | Bacteria>PVC_group>Verrucomicrobiota>unclassified_Verrucomicrobiota | GCA_003176045.1 | Verrucomicrobiota_bacterium | C5B50_19760 | 19033..21819 | 2026799 | AAA-family-ATPase-[Verrucomicrobia-bacterium] |
| MBU4201307.1 | DUF499 _1.RM.like.systems | <- DUF499 *<- METHYLASE <- TOPRIM <- OLD-ABC <-?<- TUDOR+ SF1-SF2-Helicase+iREase+CTD +REase <-ORC-CDC6+REase | 941 | Bacteria>PVC_group>Verrucomicrobiota>unclassified_Verrucomicrobiota | GCA_018898455.1 | Verrucomicrobiota_bacterium | KJ919_15475 | 11057..13882 | 2026799 | DUF499 -domain-containing-protein-[Verrucomicrobia-bacterium] |
| MBU6399160.1 | DUF499 _1.RM.like.systems | METHYLASE -p-> OLD-ABC -> TOPRIM -> DUF499 *->MORC+HisKin-><-SIGMA-HTH | 908 | Bacteria>PVC_group>Verrucomicrobiota>unclassified_Verrucomicrobiota | GCA_018969165.1 | Verrucomicrobiota_bacterium | KGS61_02480 | 2710..5436 | 2026799 | DUF499 -domain-containing-protein-[Verrucomicrobia-bacterium] |
| RME92999.1 | DUF499 _1.RM.like.systems | DUF499 *->?-> TUDOR+ SF1-SF2-Helicase+iREase+CTD +REase -p->?-> OLD-ABC -> TOPRIM -> | 774 | Bacteria>PVC_group>Verrucomicrobiota>unclassified_Verrucomicrobiota | GCA_003695015.1 | Verrucomicrobiota_bacterium | D6766_09020 | 1..2326 | 2026799 | DUF499 -domain-containing-protein,-partial-[Verrucomicrobia-bacterium] |
| NLH73611.1 | DUF499 _1.RM.like.systems | TPR-> METHYLASE -p->?->?-><- DUF499 *<- TOPRIM <- OLD-ABC <- METHYLASE <- TUDOR+ SF1-SF2-Helicase+iREase+CTD +REase | 1104 | Bacteria>PVC_group>Verrucomicrobiota>unclassified_Verrucomicrobiota | GCA_012517065.1 | Verrucomicrobiota_bacterium | GX456_11205 | 15913..19227 | 2026799 | ATP-binding-protein-[Verrucomicrobia-bacterium] |
| MCF7707718.1 | DUF499 _1.RM.like.systems | TUDOR+ SF1-SF2-Helicase+iREase+CTD +REase -> OLD-ABC -> TOPRIM ->?-> METHYLASE ->Schlafen+HATPase+HTH-> DUF499 *-> | 1113 | Bacteria>PVC_group>Verrucomicrobiota>unclassified_Verrucomicrobiota | GCA_021733835.1 | Verrucomicrobiota_bacterium | K9N52_02315 | 37691..41032 | 2026799 | DUF499 -domain-containing-protein-[Verrucomicrobia-bacterium] |
| MBP7947571.1 | DUF499 _1.RM.like.systems | <- DUF499 *<- TOPRIM <- OLD-ABC <- METHYLASE <-?<- TUDOR+ SF1-SF2-Helicase+iREase+CTD +REase | 1105 | Bacteria>PVC_group>Verrucomicrobiota>unclassified_Verrucomicrobiota | GCA_018003215.1 | Verrucomicrobiota_bacterium | KA191_16095 | 1214..4531 | 2026799 | DUF499 -domain-containing-protein-[Verrucomicrobiota-bacterium] |
| NMD22053.1 | DUF499 _1.RM.like.systems | METHYLASE -p-> OLD-ABC -> TOPRIM -> OLD-ABC -> TOPRIM -> DUF499 *-> | 1068 | Bacteria>PVC_group>Verrucomicrobiota>unclassified_Verrucomicrobiota | GCA_012797305.1 | Verrucomicrobiota_bacterium | GYA76_17670 | 5686..8891 | 2026799 | ATP-binding-protein-[Verrucomicrobia-bacterium] |
| MBP7947738.1 | DUF499 _1.RM.like.systems | <- DUF499 *<- TOPRIM <- OLD-ABC <- TOPRIM <- OLD-ABC <- METHYLASE | 1104 | Bacteria>PVC_group>Verrucomicrobiota>unclassified_Verrucomicrobiota | GCA_018003215.1 | Verrucomicrobiota_bacterium | KA191_16985 | 4010..7324 | 2026799 | ATP-binding-protein-[Verrucomicrobia-bacterium] |
| MBM3880418.1 | DUF499 _1.RM.like.systems | <- HEPN+ DUF499 *<-ParB+HNH<- METHYLASE <- TOPRIM <- OLD-ABC <- TUDOR+ SF1-SF2-Helicase+iREase+CTD +REase | 1127 | Bacteria>PVC_group>Verrucomicrobiota>unclassified_Verrucomicrobiota | GCA_016871715.1 | Verrucomicrobiota_bacterium | FJ387_11990 | 332..3715 | 2026799 | ATP-binding-protein-[Verrucomicrobia-bacterium] |
| MCF7668203.1 | DUF499 _1.RM.like.systems | TUDOR+ SF1-SF2-Helicase+iREase+CTD +REase -> METHYLASE -> OLD-ABC -> TOPRIM -> HEPN+ DUF499 *->?->?-><-PfkB | 1108 | Bacteria>PVC_group>Verrucomicrobiota>unclassified_Verrucomicrobiota | GCA_021733785.1 | Verrucomicrobiota_bacterium | K9N48_00295 | 29599..32925 | 2026799 | DUF499 -domain-containing-protein-[Verrucomicrobia-bacterium] |
| MCC5848502.1 | DUF499 _1.RM.like.systems | <- HEPN+ DUF499 *<-?<-SF1-SF2-Helicase+wHTH+PRTase<- METHYLASE <- TOPRIM <- OLD-ABC <-HTH_3 | 1113 | Bacteria>PVC_group>Verrucomicrobiota>unclassified_Verrucomicrobiota | GCA_020831635.1 | Verrucomicrobiota_bacterium | JJU29_10430 | 13119..16460 | 2026799 | DUF499 -domain-containing-protein-[Verrucomicrobia-bacterium] |
| WP_165106894.1 | DUF499 _1.RM.like.systems | Cluster1029_5clades->?-><- DUF499 *<-DUF488-1<- METHYLASE <-MNS<- TOPRIM <- OLD-ABC <- TUDOR+ SF1-SF2-Helicase+iREase+CTD +REase | 963 | Bacteria>PVC_group>Verrucomicrobiota>Verrucomicrobiae>Limisphaerales | GCF_011044775.1 | Limisphaera_ngatamarikiensis | G4L39_RS06720 | 76974..79865 | 1324935 | DUF499 -domain-containing-protein-[Limisphaera-ngatamarikiensis] |
| MBE7503149.1 | DUF499 _1.RM.like.systems | wHTH+REase->?->?-> METHYLASE -> OLD-ABC -> TOPRIM -> DUF499 *-> DUF499 -> | 393 | Bacteria>PVC_group>Verrucomicrobiota>Verrucomicrobiae>Verrucomicrobiales | GCA_015075445.1 | Verrucomicrobiales_bacterium | HS113_23265 | 55757..56938 | 2026801 | ATP-binding-protein-[Verrucomicrobiales-bacterium] |
| MCI0539271.1 | DUF499 _1.RM.like.systems | <- SF1-SF2-Helicase<- OLD-ABC + TOPRIM \|\| TUDOR+ SF1-SF2-Helicase+iREase+CTD +REase ->wHTH+REase-> METHYLASE -> DUF499 *->PIWI->REase-> | 1107 | Bacteria>PVC_group>Verrucomicrobiota>Verrucomicrobiae>Verrucomicrobiales | GCA_022615925.1 | Verrucomicrobiales_bacterium | LOZ50_29035 | 12351..15674 | 2026801 | DUF499 -domain-containing-protein-[Verrucomicrobiales-bacterium] |
| MBE7503150.1 | DUF499 _1.RM.like.systems | METHYLASE -> OLD-ABC -> TOPRIM -> DUF499 -> DUF499 *-> | 747 | Bacteria>PVC_group>Verrucomicrobiota>Verrucomicrobiae>Verrucomicrobiales | GCA_015075445.1 | Verrucomicrobiales_bacterium | HS113_23270 | 56827..59070 | 2026801 | hypothetical-protein-[Verrucomicrobiales-bacterium] |
| MCK9286308.1 | DUF499 _1.RM.like.systems | <-sGTP<- SF1-SF2-Helicase<- OLD-ABC + TOPRIM <- HEPN+ DUF499 *<- METHYLASE <- TUDOR+ SF1-SF2-Helicase+iREase+CTD +REase <-HTH_26<-SbcC+SbcC<-?<-REase+CTD6 | 1135 | Bacteria>Spirochaetota>Spirochaetia>Spirochaetales>Sphaerochaetaceae | GCA_023230445.1 | Sphaerochaetaceae_bacterium | MOP29_01920 | 121891..125298 | 2791017 | DUF499 -domain-containing-protein-[Sphaerochaetaceae-bacterium] |
| MDE0447651.1 | DUF499 _1.RM.like.systems | Cluster182_14clades->TPR+TPR-><-?<- DUF499 *<- TOPRIM <- OLD-ABC <-HTH_3<- METHYLASE <-SbcC<- TUDOR+ SF1-SF2-Helicase+iREase+CTD +REase | 1107 | Bacteria>Spirochaetota>Spirochaetia>Spirochaetales>Spirochaetaceae | GCA_028821475.1 | Spirochaetaceae_bacterium | OXH96_13340 | 24817..28140 | 1898206 | MAG:- DUF499 -domain-containing-protein-[Spirochaetaceae-bacterium] |
| MDD9989001.1 | DUF499 _1.RM.like.systems | Cluster182_14clades-><-CR-ATPase8+REase<-?<- DUF499 *<- TOPRIM <- OLD-ABC <-HTH_3<- METHYLASE <-TIR | 1107 | Bacteria>Spirochaetota>Spirochaetia>Spirochaetales>Spirochaetaceae | GCA_028818385.1 | Spirochaetaceae_bacterium | OXQ31_22210 | 3073..6396 | 1898206 | MAG:- DUF499 -domain-containing-protein-[Spirochaetaceae-bacterium] |
| OQB96942.1 | DUF499 _1.RM.like.systems | SIGMA-HTH->Radical-SAM-><-?<-?\|\|?->?-><- HEPN+ DUF499 *<-ParB+HNH<- METHYLASE <- TOPRIM <- OLD-ABC <-HEPN<- TUDOR+ SF1-SF2-Helicase+iREase+CTD +REase | 1176 | Bacteria>Spirochaetota>unclassified_Spirochaetota | GCA_002070755.1 | Spirochaetes_bacterium_ADurb.Bin110 | BWX81_01211 | 5339..8869 | 1852908 | hypothetical-protein-BWX81_01211-[Spirochaetes-bacterium-ADurb.Bin110] |
| MXW97851.1 | DUF499 _1.RM.like.systems | TUDOR+ SF1-SF2-Helicase+iREase+CTD +REase ->RelE-ParE->HTH_Tnp+HTH_Tnp-> METHYLASE -> DUF499 *-> OLD-ABC -> TOPRIM -> | 977 | Bacteria>Terrabacteria_group>Actinomycetota>Acidimicrobiia>Acidimicrobiales | GCA_009835735.1 | Acidimicrobiaceae_bacterium | F4Z89_04460 | 7235..10168 | 2024894 | ATP-binding-protein-[Acidimicrobiaceae-bacterium] |
| MCY3892500.1 | DUF499 _1.RM.like.systems | METHYLASE -p-> OLD-ABC -> TOPRIM -> HEPN+ DUF499 *->?->ABhydrolase-> | 1131 | Bacteria>Terrabacteria_group>Actinomycetota>Acidimicrobiia>Acidimicrobiales | GCA_026707105.1 | Acidimicrobiaceae_bacterium | OXF65_04295 | 3174..6569 | 2024894 | MAG:-Swt1-family-HEPN-domain-containing-protein-[Acidimicrobiaceae-bacterium] |
| WP_285140308.1 | DUF499 _1.RM.like.systems | <-HTH_Tnp_1+HTH_21+rve+rve_3<-HIN-HTH\|\|XIS-HTH-> TUDOR+ SF1-SF2-Helicase+iREase+CTD +REase -> METHYLASE ->wHTH+REase-> HEPN+ DUF499 *-> OLD-ABC + TOPRIM ->SF1-SF2-Helicase -><-DNA_methylase | 1132 | Bacteria>Terrabacteria_group>Actinomycetota>Actinomycetes>Micrococcales | GCF_030226745.1 | Micrococcus_luteus | QP726_RS10470 | 9707..13105 | 1270 | Swt1-family-HEPN-domain-containing-protein-[Micrococcus-luteus] |
| WP_144831472.1 | DUF499 _1.RM.like.systems | ACET-><-?<- SF1-SF2-Helicase<- OLD-ABC + TOPRIM <- HEPN+ DUF499 *<-wHTH+REase<- METHYLASE <- TUDOR+ SF1-SF2-Helicase+iREase+CTD +REase <-XIS-HTH\|\|MORC+HisKin+REase-><-TrwC+SF1-SF2-helicase | 1133 | Bacteria>Terrabacteria_group>Actinomycetota>Actinomycetes>Micrococcales | GCF_007677595.1 | Kocuria_rhizophila | FS577_RS10180 | 96606..100007 | 72000 | DUF499 -domain-containing-protein-[Kocuria-rhizophila] |
| WP_274500272.1 | DUF499 _1.RM.like.systems | <-Mbetalac\|\|MarR-HTH->?->?->NERD-><- OLD-ABC + TOPRIM <- HEPN+ DUF499 *<- METHYLASE <- TUDOR+ SF1-SF2-Helicase+iREase+CTD +REase <-UDG-conflict\|\|?-><-REase\|\|HEPN+SF1-SF2-Helicase-> | 1131 | Bacteria>Terrabacteria_group>Actinomycetota>Actinomycetes>Micrococcales | GCF_028765365.1 | Plantibacter_flavus | PUY80_RS13440 | 281410..284805 | 150123 | DUF499 -domain-containing-protein-[Plantibacter-flavus] |
| WP_255162094.1 | DUF499 _1.RM.like.systems | DDE_Tnp_IS240-><-Classical-AAA<-TrwC+ SF1-SF2-Helicase<- OLD-ABC + TOPRIM <-?<- HEPN+ DUF499 *<- METHYLASE <- TUDOR+ SF1-SF2-Helicase+iREase+CTD +REase <-XIS-HTH<-ABhydrolase<-?<-MPTase | 1143 | Bacteria>Terrabacteria_group>Actinomycetota>Actinomycetes>Micrococcales | GCF_024362265.1 | Microbacterium_liquefaciens | NMQ05_RS13065 | 2679727..2683158 | 33918 | DUF499 -domain-containing-protein-[Microbacterium-liquefaciens] |
| WP_141842659.1 | DUF499 _1.RM.like.systems | <-HIN-HTH<- SF1-SF2-Helicase<- OLD-ABC + TOPRIM <- HEPN+ DUF499 *<-wHTH+REase<- METHYLASE <- TUDOR+ SF1-SF2-Helicase+iREase+CTD +REase <-XIS-HTH<-PAIREDC-HTH<-HIN-HTH | 1132 | Bacteria>Terrabacteria_group>Actinomycetota>Actinomycetes>Micrococcales | GCF_006704135.1 | Micrococcus_sp._R8502A1 | FLM69_RS00645 | 3870..7268 | 2583239 | Swt1-family-HEPN-domain-containing-protein-[Micrococcus-sp.-R8502A1] |
| WP_112235018.1 | DUF499 _1.RM.like.systems | <-MORC+HisKin+REase\|\|XIS-HTH-> TUDOR+ SF1-SF2-Helicase+iREase+CTD +REase -> METHYLASE ->wHTH+REase-> HEPN+ DUF499 *-> OLD-ABC + TOPRIM ->SF1-SF2-Helicase -> | 1132 | Bacteria>Terrabacteria_group>Actinomycetota>Actinomycetes>Micrococcales | GCF_003290245.1 | Kocuria_sp._BT304 | DNL27_RS06220 | 1382048..1385446 | 1702043 | DUF499 -domain-containing-protein-[Kocuria-sp.-BT304] |
| WP_046529475.1 | DUF499 _1.RM.like.systems | XIS-HTH->PIN->HNH-><-HTH+SF1-SF2-Helicase+REase<-REase<-MORC<- HEPN+ DUF499 *<- METHYLASE <-REase\|\|DNA_methylase-><- OLD-ABC + TOPRIM <-DNA-glycosylase | 1124 | Bacteria>Terrabacteria_group>Actinomycetota>Actinomycetes>Micrococcales | GCF_000986825.1 | Cellulomonas_sp._FA1 | GY42_RS11845 | 2630948..2634322 | 1346710 | Swt1-family-HEPN-domain-containing-protein-[Cellulomonas-sp.-FA1] |
| WP_054682987.1 | DUF499 _1.RM.like.systems | XIS-HTH-> TUDOR+ SF1-SF2-Helicase+iREase+CTD +REase -> METHYLASE ->HIN-HTH-> METHYLASE -p-> HEPN+ DUF499 *-> OLD-ABC + TOPRIM ->HNH-><-HIN-HTH | 1129 | Bacteria>Terrabacteria_group>Actinomycetota>Actinomycetes>Micrococcales | GCF_001314225.1 | Microbacterium_sp._No._7 | AOA12_RS21690 | 98036..101425 | 1714373 | DUF499 -domain-containing-protein-[Microbacterium-sp.-No.-7] |
| WP_204409507.1 | DUF499 _1.RM.like.systems | <-HIN-HTH<- OLD-ABC + TOPRIM <- HEPN+ DUF499 *<-wHTH+REase<- METHYLASE <- TUDOR+ SF1-SF2-Helicase+iREase+CTD +REase <-XIS-HTH | 1132 | Bacteria>Terrabacteria_group>Actinomycetota>Actinomycetes>Micrococcales | GCF_016907455.1 | Brachybacterium_muris | JOD52_RS08675 | 1871612..1875010 | 219301 | DUF499 -domain-containing-protein-[Brachybacterium-muris] |
| WP_247619184.1 | DUF499 _1.RM.like.systems | <- OLD-ABC + TOPRIM <- HEPN+ DUF499 *<-wHTH+REase<- METHYLASE <- TUDOR+ SF1-SF2-Helicase+iREase+CTD +REase <-XIS-HTH | 1135 | Bacteria>Terrabacteria_group>Actinomycetota>Actinomycetes>Micrococcales | GCF_023072605.1 | Brevibacterium_sp._R8603A2 | MTQ12_RS14500 | 1781..5188 | 2929779 | Swt1-family-HEPN-domain-containing-protein-[Brevibacterium-sp.-R8603A2] |
| WP_194396698.1 | DUF499 _1.RM.like.systems | METHYLASE -><-HNH<- OLD-ABC + TOPRIM <- HEPN+ DUF499 *<- METHYLASE <- TUDOR+ SF1-SF2-Helicase+iREase+CTD +REase <-XIS-HTH | 1130 | Bacteria>Terrabacteria_group>Actinomycetota>Actinomycetes>Micrococcales | GCF_015277815.1 | Microbacterium_atlanticum | IR212_RS15230 | 3329381..3332773 | 2782168 | DUF499 -domain-containing-protein-[Microbacterium-atlanticum] |
| PZU44169.1 | DUF499 _1.RM.like.systems | TetR-HTH-><-?<-?\|\|?->\|\| HEPN+ DUF499 *-> OLD-ABC + TOPRIM -><-?<-HIN-HTH\|\|HTH_19->Phd-YefM->PIN-> | 1129 | Bacteria>Terrabacteria_group>Actinomycetota>Actinomycetes>Micrococcales | GCA_003248655.1 | Microbacterium_sp. | DI566_12710 | 6913..10302 | 51671 | AAA-family-ATPase-[Microbacterium-sp.] |
| WP_197462106.1 | DUF499 _1.RM.like.systems | HEPN+ DUF499 *-> DUF499 -p-> OLD-ABC + TOPRIM -p-> OLD-ABC + TOPRIM -p-> | 968 | Bacteria>Terrabacteria_group>Actinomycetota>Actinomycetes>Micrococcales | GCF_001595495.1 | Microbacterium_sp._CH1 | AUV07_RS03260 | 609..3515 | 1770208 | DUF499 -domain-containing-protein-[Microbacterium-sp.-CH1] |
| PZT93668.1 | DUF499 _1.RM.like.systems | ABhydrolase->XIS-HTH-> TUDOR+ SF1-SF2-Helicase+iREase+CTD +REase -> METHYLASE ->wHTH+REase-> HEPN+ DUF499 *-> OLD-ABC + TOPRIM ->SF1-SF2-Helicase -> | 1133 | Bacteria>Terrabacteria_group>Actinomycetota>Actinomycetes>Mycobacteriales | GCA_003248405.1 | Gordonia_sp._(in:_high_G+C_Gram-positive_bacteria) | DI630_26685 | 9643..13044 | 84139 | AAA-family-ATPase-[Gordonia-sp.] |
| WP_213249149.1 | DUF499 _1.RM.like.systems | <-HTH+SF1-SF2-Helicase+REase<-REase<-MORC<- HEPN+ DUF499 *<- SF1-SF2-Helicase<- OLD-ABC + TOPRIM \|\|DNA_methylase-><-?\|\|HNH-> | 1138 | Bacteria>Terrabacteria_group>Actinomycetota>Actinomycetes>Mycobacteriales | GCF_009914535.1 | Gordonia | IDH24_RS11690 | 142027..145443 | 2053 | DUF499 -domain-containing-protein-[Gordonia-pseudamarae] |
| WP_119162165.1 | DUF499 _1.RM.like.systems | <-TetR-HTH\|\|?-><- SF1-SF2-Helicase<- OLD-ABC + TOPRIM <- HEPN+ DUF499 *<-PemK_toxin<-?<-wHTH+REase<- METHYLASE <- TUDOR+ SF1-SF2-Helicase+iREase+CTD +REase | 1133 | Bacteria>Terrabacteria_group>Actinomycetota>Actinomycetes>Propionibacteriales | GCF_900637915.1 | Propionibacterium_australiense | EL282_RS10960 | 2474046..2477447 | 119981 | DUF499 -domain-containing-protein-[Propionibacterium-australiense] |
| WP_182784909.1 | DUF499 _1.RM.like.systems | <- SF1-SF2-Helicase<- OLD-ABC + TOPRIM <- HEPN+ DUF499 *<- METHYLASE <- TUDOR+ SF1-SF2-Helicase+iREase+CTD +REase <-SF1-SF2-Helicase-p\|\|Glyoxalase-> | 1133 | Bacteria>Terrabacteria_group>Actinomycetota>Actinomycetes>Propionibacteriales | GCF_014164725.1 | Tessaracoccus_sp._MC1627 | H5399_RS17220 | 4312..7713 | 2760312 | DUF499 -domain-containing-protein-[Tessaracoccus-sp.-MC1627] |
| WP_176233419.1 | DUF499 _1.RM.like.systems | DDE_Tnp_1->Phage_int_SAM_1-><-?<-sGTP<-?<-?<- DUF499 *<- TOPRIM <- OLD-ABC <- METHYLASE -p | 683 | Bacteria>Terrabacteria_group>Actinomycetota>Actinomycetota_incertae_sedis>Candidatus_Hakubellales | GCF_013281955.1 | Candidatus_Hakubella_thermalkaliphila | HW440_RS05440 | 5834..7885 | 2754717 | DUF499 -domain-containing-protein-[Candidatus-Hakubanella-thermoalkaliphilus] |
| GFP42415.1 | DUF499 _1.RM.like.systems | DDE_Tnp_1-><-?\|\|Phage_int_SAM_1-><-?<-sGTP<-?<- DUF499 *<- TOPRIM | 683 | Bacteria>Terrabacteria_group>Actinomycetota>Actinomycetota_incertae_sedis>Candidatus_Hakubellales | GCA_013282075.1 | Candidatus_Hakubella_thermalkaliphila | HKBW3C_01541 | 5550..7601 | 2754717 | hypothetical-protein-HKBW3C_01541-[Candidatus-Hakubanella-thermoalkaliphilus] |
| WP_218957899.1 | DUF499 _1.RM.like.systems | <-Peptidase<-?<-?<-?<-?<- DUF499 -p<- HEPN+ DUF499 *<- METHYLASE <- TUDOR+ SF1-SF2-Helicase+iREase+CTD +REase \|\| OLD-ABC + TOPRIM ->SF1-SF2-Helicase -> | 864 | Bacteria>Terrabacteria_group>Actinomycetota>Coriobacteriia>Coriobacteriales | GCF_007655325.1 | Enorma_shizhengliae | GJE22_RS07705 | 20738..23332 | 2606615 | DUF499 -domain-containing-protein-[Enorma-shizhengliae] |
| GIU94958.1 | DUF499 _1.RM.like.systems | Schlafen+HATPase+HTH->?-> OLD-ABC -> TOPRIM -> METHYLASE -> METHYLASE -p-> HEPN+ DUF499 *->?->TPR->DUF222+HNH->REase+35exo+Beta-Barrel+SF1-SF2-Helicase-> | 1082 | Bacteria>Terrabacteria_group>Actinomycetota>Rubrobacteria>Gaiellales | GCA_026002995.1 | Gaiellaceae_bacterium | KatS3mg012_1415 | 1404220..1407468 | 2838783 | MAG:-hypothetical-protein-KatS3mg012_1415-[Gaiellaceae-bacterium] |
| MDA8047538.1 | DUF499 _1.RM.like.systems | TUDOR+ SF1-SF2-Helicase+iREase+CTD +REase -> METHYLASE -> DUF499 *-> OLD-ABC + TOPRIM -><-PIWI<-REase+PNTD1+PNTD2\|\|HicB-> | 941 | Bacteria>Terrabacteria_group>Actinomycetota>unclassified_Actinomycetota | GCA_028063615.1 | Actinomycetota_bacterium | MOZ30_20285 | 6559..9384 | 2900548 | MAG:- DUF499 -domain-containing-protein-[Actinomycetota-bacterium] |
| MCA0180524.1 | DUF499 _1.RM.like.systems | <-SMODS<-?<-TrwC+SF1-SF2-Helicase\|\|DNA_methylase-><- SF1-SF2-Helicase<- OLD-ABC + TOPRIM <- HEPN+ DUF499 *<-wHTH+REase<- METHYLASE <- TUDOR+ SF1-SF2-Helicase+iREase+CTD +REase <-XIS-HTH<-?\|\|GDResolvase+Recombinase+ZnR-> | 1132 | Bacteria>Terrabacteria_group>Actinomycetota>unclassified_Actinomycetota | GCA_020160805.1 | Actinomycetota_bacterium | LCH77_13240 | 54204..57602 | 2900548 | DUF499 -domain-containing-protein-[Actinomycetia-bacterium] |
| MCL4554262.1 | DUF499 _1.RM.like.systems | TUDOR+ SF1-SF2-Helicase+iREase+CTD +REase -p-> OLD-ABC + TOPRIM -> METHYLASE ->HEPN->ParA-Soj-PloopNTPase->ORC-CDC6-> HEPN+ DUF499 *-> | 468 | Bacteria>Terrabacteria_group>Actinomycetota>unclassified_Actinomycetota | GCA_023382015.1 | Actinomycetota_bacterium | M1565_03890 | 9031..10434 | 2900548 | Swt1-family-HEPN-domain-containing-protein-[Actinomycetota-bacterium] |
| MDA8117679.1 | DUF499 _1.RM.like.systems | <-MerR-HTH<-?<- HEPN+ DUF499 *<- TOPRIM <- OLD-ABC <- METHYLASE <-HEPN<-ParB+HNH<-HicB+ZnR | 1122 | Bacteria>Terrabacteria_group>Actinomycetota>unclassified_Actinomycetota | GCA_028064745.1 | Actinomycetota_bacterium | M0000_09960 | 1809..5177 | 2900548 | MAG:-Swt1-family-HEPN-domain-containing-protein-[Actinomycetota-bacterium] |
| NLH98010.1 | DUF499 _1.RM.like.systems | TUDOR+ SF1-SF2-Helicase+iREase+CTD +REase ->?-> METHYLASE -> OLD-ABC -> TOPRIM -> HEPN+ DUF499 *-> | 1111 | Bacteria>Terrabacteria_group>Armatimonadota>Chthonomonadetes>Chthonomonadales | GCA_012517255.1 | Chthonomonadales_bacterium | GX446_00800 | 20611..23946 | 2282151 | ATP-binding-protein-[Chthonomonadales-bacterium] |
| GIV04311.1 | DUF499 _1.RM.like.systems | <- DUF499 *<-HTH<- METHYLASE <- OLD-ABC + TOPRIM <-SF1-SF2-Helicase-p<-Schlafen+HATPase+HTH<- TUDOR+ SF1-SF2-Helicase+iREase+CTD +REase | 944 | Bacteria>Terrabacteria_group>Armatimonadota>Fimbriimonadia>Fimbriimonadales | GCA_026003055.1 | Fimbriimonadales_bacterium | KatS3mg015_3141 | 322..3156 | 2682145 | MAG:-hypothetical-protein-KatS3mg015_3141-[Fimbriimonadales-bacterium] |
| WP_181339998.1 | DUF499 _1.RM.like.systems | DeoR-HTH+ISOCOT->?->?->?->?-><- HEPN+ DUF499 *<- METHYLASE <- TUDOR+ SF1-SF2-Helicase+iREase+CTD +REase \|\| OLD-ABC + TOPRIM -> | 1104 | Bacteria>Terrabacteria_group>Bacillota>Bacillota_sensu_stricto_incertae_sedis>Capillibacterium | GCF_013664685.1 | Capillibacterium_thermochitinicola | G5B42_RS08290 | 130685..133999 | 2699427 | Swt1-family-HEPN-domain-containing-protein-[Capillibacterium-thermochitinicola] |
| WP_012301643.1 | DUF499 _1.RM.like.systems | REase-> TUDOR+ SF1-SF2-Helicase+iREase+CTD +REase ->?-> OLD-ABC + TOPRIM -> METHYLASE ->REase-> DUF499 *->RNase->?->?->?->HicB_lk_antitox-> | 924 | Bacteria>Terrabacteria_group>Bacillota>Clostridia>Eubacteriales | GCF_000018425.1 | Candidatus_Desulforudis_audaxviator | Daudx_RS02685 | 550555..553329 | 471827 | DUF499 -domain-containing-protein-[Candidatus-Desulforudis-audaxviator] |
| WP_015942906.1 | DUF499 _1.RM.like.systems | SF1-SF2-Helicase+HKD-DNase->MORC+SF1-SF2-Helicase->?->HTH_26-> TUDOR+ SF1-SF2-Helicase+iREase+CTD +REase -> METHYLASE -> HEPN+ DUF499 *-> OLD-ABC + TOPRIM ->SF1-SF2-Helicase -> METHYLASE -p->ParB+HNH->HEPN->pPIWI+pPIWI-> | 1137 | Bacteria>Terrabacteria_group>Bacillota>Clostridia>Eubacteriales | GCF_000021925.1 | Desulfitobacterium_hafniense | DPCES_RS04010 | 854740..858153 | 49338 | DUF499 -domain-containing-protein-[Desulfitobacterium-hafniense] |
| NLI53236.1 | DUF499 _1.RM.like.systems | WYL->?-> TUDOR+ SF1-SF2-Helicase+iREase+CTD +REase -> METHYLASE -> HEPN+ DUF499 *-> OLD-ABC + TOPRIM ->?->?-><-tRNA<-SMS-KaiC | 1124 | Bacteria>Terrabacteria_group>Bacillota>Clostridia>Eubacteriales | GCA_012517905.1 | Clostridiales_bacterium | GX417_02810 | 9381..12755 | 1898207 | ATP-binding-protein-[Clostridiales-bacterium] |
| MCR4419670.1 | DUF499 _1.RM.like.systems | <- DUF499 *<- TOPRIM <- OLD-ABC <-?<- METHYLASE <- TUDOR+ SF1-SF2-Helicase+iREase+CTD +REase | 1104 | Bacteria>Terrabacteria_group>Bacillota>Clostridia>unclassified_Clostridia | GCA_024653205.1 | Clostridia_bacterium | NUV99_05980 | 77810..81124 | 2044939 | DUF499 -domain-containing-protein-[Clostridia-bacterium] |
| MBQ6961345.1 | DUF499 _1.RM.like.systems | <-SF1-SF2-Helicase-p<- OLD-ABC + TOPRIM <- HEPN+ DUF499 *<-PolB+HEPN<-PolB<- METHYLASE <-MNS<- TUDOR+ SF1-SF2-Helicase+iREase+CTD +REase <-HTH | 1126 | Bacteria>Terrabacteria_group>Bacillota>Clostridia>unclassified_Clostridia | GCA_017448975.1 | Clostridia_bacterium | IJP78_10255 | 13529..16909 | 2044939 | DUF499 -domain-containing-protein-[Clostridia-bacterium] |
| MBE3584147.1 | DUF499 _1.RM.like.systems | <- METHYLASE -p\|\|?-> TUDOR+ SF1-SF2-Helicase+iREase+CTD +REase -> OLD-ABC + TOPRIM -> METHYLASE -> HEPN+ DUF499 *-><-?\|\|ABC-ATPase+SbcC->ABC-ATPase+SbcC-><-MazE_antitoxin | 1113 | Bacteria>Terrabacteria_group>Bacillota>Limnochordia>Limnochordales | GCA_014896305.1 | Limnochordaceae_bacterium | IMX01_08555 | 27896..31237 | 2773236 | DUF499 -domain-containing-protein-[Limnochordaceae-bacterium] |
| NCB76905.1 | DUF499 _1.RM.like.systems | Ntox49->?->Resolvase->Recombinase+ZnR->?->PIN+OST-HTH-><- DUF499 *<- TOPRIM <- OLD-ABC <- METHYLASE <- TUDOR+ SF1-SF2-Helicase+iREase+CTD +REase <-?\|\|NACHT-> | 964 | Bacteria>Terrabacteria_group>Bacillota>Negativicutes>unclassified_Negativicutes | GCA_009929485.1 | Negativicutes_bacterium | EOM42_07415 | 5128..8022 | 2499140 | ATP-binding-protein-[Negativicutes-bacterium] |
| HBL50377.1 | DUF499 _1.RM.like.systems | <- SF1-SF2-Helicase<- OLD-ABC + TOPRIM <- HEPN+ DUF499 *<- METHYLASE <- TUDOR+ SF1-SF2-Helicase+iREase+CTD +REase <-?<-MORC+DUF3684+HNH<-SF1-SF2-Helicase-p<-HEPN+SF1-SF2-Helicase | 1129 | Bacteria>Terrabacteria_group>Bacillota>unclassified_Bacillota | GCA_003504715.1 | Bacillota_bacterium | DDZ65_09075 | 4304..7693 | 1879010 | hypothetical-protein-[Firmicutes-bacterium] |
| NPV89870.1 | DUF499 _1.RM.like.systems | GDResolvase->?->GDResolvase-> TUDOR+ SF1-SF2-Helicase+iREase+CTD +REase ->AbiGi-> METHYLASE -> HEPN+ DUF499 *-> OLD-ABC + TOPRIM ->SF1-SF2-Helicase ->?->Peptidase_S8-><-HTH_3 | 1137 | Bacteria>Terrabacteria_group>Bacillota>unclassified_Bacillota | GCA_013178275.1 | Bacillota_bacterium | HPY50_03730 | 179885..183298 | 1879010 | ATP-binding-protein-[Firmicutes-bacterium] |
| MBT9164708.1 | DUF499 _1.RM.like.systems | <- DUF499 *<- TOPRIM <- OLD-ABC <- METHYLASE | 873 | Bacteria>Terrabacteria_group>Candidatus_Lithacetigenota>Candidatus_Lithacetigena | GCA_018725485.1 | Candidatus_Lithacetigena_glycinireducens | DDT23_00712 | 1258..3879 | 2986055 | hypothetical-protein-[candidate-division-WS2-bacterium] |
| MBN1317038.1 | DUF499 _1.RM.like.systems | METHYLASE -> OLD-ABC -> TOPRIM ->?->SWACOS-> DUF499 *-> | 449 | Bacteria>Terrabacteria_group>Chloroflexota>Anaerolineae>Anaerolineales | GCA_016927355.1 | Anaerolineales_bacterium | JXA42_16270 | 39202..40547 | 2073117 | ATP-binding-protein-[Anaerolineales-bacterium] |
| MBN1920387.1 | DUF499 _1.RM.like.systems | DUF433->?->?-> OLD-ABC + TOPRIM -> METHYLASE ->?-> DUF499 *-> | 909 | Bacteria>Terrabacteria_group>Chloroflexota>Anaerolineae>unclassified_Anaerolineae | GCA_016931315.1 | Anaerolineae_bacterium | JW892_04030 | 10544..13273 | 2052143 | ATP-binding-protein-[Anaerolineae-bacterium] |
| MBN2007579.1 | DUF499 _1.RM.like.systems | TUDOR+ SF1-SF2-Helicase+iREase+CTD +REase ->REase-> METHYLASE -> OLD-ABC -> TOPRIM -> DUF499 *-> | 737 | Bacteria>Terrabacteria_group>Chloroflexota>Anaerolineae>unclassified_Anaerolineae | GCA_016931835.1 | Anaerolineae_bacterium | JXA21_29810 | 7893..10102 | 2052143 | ATP-binding-protein-[Anaerolineae-bacterium] |
| MCR4407531.1 | DUF499 _1.RM.like.systems | Phage_integrase-><-?\|\|<- DUF499 *<- TOPRIM -p<-SbcC<- METHYLASE <-PIN<-MazE_antitoxin<- TUDOR+ SF1-SF2-Helicase+iREase+CTD +REase | 1132 | Bacteria>Terrabacteria_group>Chloroflexota>Anaerolineae>unclassified_Anaerolineae | GCA_024653325.1 | Anaerolineae_bacterium | NUW24_11530 | 3238..6636 | 2052143 | DUF499 -domain-containing-protein-[Anaerolineae-bacterium] |

| Protein-ID | System | Gene-Neighbourhoods | Length | Taxonomy | Assembly-ID | Strain | Locus-Tag | Genomic-coordinates | TaxID | Fasta-defline |
|---|---|---|---|---|---|---|---|---|---|---|
| -- | | | | | | | | | | |
| --HEPN-EndoRNase | | | | | | | | | | |
| -- | | | | | | | | | | |
| WP_228841862.1 | DUF499_2.RM.like.systems | <-HKD-DNase+SF1-SF2-Helicase+iREase+CTD<-HEPN<-METHYLASE<-DUF3780<-DUF499*<-AbrB-HTH | 1108 | Archaea>Euryarchaeota>Stenosarchaea_group>Halobacteria>Halobacteriales | GCF_001485575.1 | Haloarcula_sp._CBA1127 | AV059_RS20935 | 42218..45544 | 1765055 | DUF499-domain-containing-protein-[Haloarcula-sp.-CBA1127] |
| ELZ56264.1 | DUF499_2.RM.like.systems | DDE_Tnp_1-><-HKD-DNase+SF1-SF2-Helicase+iREase+CTD<-HEPN<-METHYLASE-p<-DUF3780<-DUF499*<-?\|\|Resolvase->?->PadR-HTH->?->BirA-HTH-> | 935 | Archaea>Euryarchaeota>Stenosarchaea_group>Halobacteria>Haloferacales | GCA_000337075.1 | Halorubrum_hochstenium_ATCC_700873 | C467_08795 | 110652..113459 | 1227481 | hypothetical-protein-C467_08795-[Halorubrum-hochstenium-ATCC-700873] |
| MBB6645927.1 | DUF499_2.RM.like.systems | Phage_integrase-><-HKD-DNase+SF1-SF2-Helicase+iREase+CTD<-HEPN<-METHYLASE<-DUF3780<-DUF499* | 935 | Archaea>Euryarchaeota>Stenosarchaea_group>Halobacteria>Haloferacales | GCA_014212355.1 | Halobellus_ruber | H5V44_06430 | 284116..286923 | 2761102 | DUF499-domain-containing-protein-[Halobellus-ruber] |
| WP_239638715.1 | DUF499_2.RM.like.systems | <-PadR-HTH\|\|?->?-><-Resolvase\|\|?->AbrB-HTH->DUF499*->DUF3780->METHYLASE->HEPN->HKD-DNase+SF1-SF2-Helicase+iREase+CTD->REase-><-DDE_Tnp_1 | 1054 | Archaea>Euryarchaeota>Stenosarchaea_group>Halobacteria>Haloferacales | GCF_000337075.1 | Halorubrum | NAF06_RS08930 | 1779031..1782195 | 56688 | MULTISPECIES:-DUF499-domain-containing-protein-[Halorubrum] |
| WP_246403805.1 | DUF499_2.RM.like.systems | Phage_integrase-><-HKD-DNase+SF1-SF2-Helicase+iREase+CTD<-HEPN<-METHYLASE<-DUF3780<-DUF499* | 1109 | Archaea>Euryarchaeota>Stenosarchaea_group>Halobacteria>Haloferacales | GCF_014212355.1 | Halobellus_ruber | H5V44_RS06435 | 284116..287445 | 2761102 | DUF499-domain-containing-protein-[Halobellus-ruber] |
| WP_256397181.1 | DUF499_2.RM.like.systems | <-DDE_Tnp_IS240\|\|AbrB-HTH->DUF499*->DUF3780->METHYLASE->HKD-DNase+SF1-SF2-Helicase+iREase+CTD->HEPN-><-?\|\|ORC/CDC6+HTH-> | 1045 | Archaea>Euryarchaeota>Stenosarchaea_group>Halobacteria>Haloferacales | GCF_024494645.1 | Halorubraceae | NO887_RS09745 | 233066..236203 | 1963271 | MULTISPECIES:-DUF499-domain-containing-protein-[Halorubraceae] |
| WP_267623061.1 | DUF499_2.RM.like.systems | <-REase<-ORC/CDC6+HTH\|\|CoiA-><-DOC\|\|AbrB-HTH->DUF499*->DUF3780->METHYLASE->HEPN->HKD-DNase+SF1-SF2-Helicase+iREase+CTD->Cluster197_21clades-> | 1103 | Archaea>Euryarchaeota>Stenosarchaea_group>Halobacteria>Haloferacales | GCF_026586625.1 | Halobium_salinum | N5095_RS11700 | 30435..33746 | 1364940 | DUF499-domain-containing-protein-[Halobium-salinum] |
| WP_211427121.1 | DUF499_2.RM.like.systems | SIGMA-HTH->?->?->DUF499*->DUF3780->?->?->PemK_toxin->METHYLASE->HEPN-> | 1041 | Bacteria>Acidobacteriota>Blastocatellia>Chloracidobacterium | GCF_018304805.1 | Chloracidobacterium_aggregatum | J8C02_RS13600 | 632551..635676 | 2851959 | DUF499-domain-containing-protein-[Chloracidobacterium-aggregatum] |
| MBV9397053.1 | DUF499_2.RM.like.systems | <-Cluster2725_2clades\|\|Terminase->PolB->HEPN->DUF499*->DUF3780->METHYLASE->HKD-DNase+SF1-SF2-Helicase+iREase+CTD-> | 1087 | Bacteria>Acidobacteriota>Terriglobia>Bryobacterales>unclassified_Bryobacterales | GCA_019261985.1 | Bryobacterales_bacterium | J0062_03680 | 122094..125357 | 2026791 | ATP-binding-protein-[Bryobacterales-bacterium] |
| ODS53422.1 | DUF499_2.RM.like.systems | RVT-><-HEPN<-HKD-DNase+SF1-SF2-Helicase+iREase+CTD<-METHYLASE<-DOC+HTH<-DUF3780<-DUF499* | 997 | Bacteria>Acidobacteriota>unclassified_Acidobacteriota | GCA_001724025.1 | Acidobacteria_bacterium_SCN_69-37 | ABS36_12275 | 12058..15051 | 1660160 | hypothetical-protein-ABS36_12275-[Acidobacteria-bacterium-SCN-69-37] |
| WP_109646585.1 | DUF499_2.RM.like.systems | ParB+HNH->DUF499*->DUF3780->METHYLASE->HKD-DNase+SF1-SF2-Helicase->HEPN+CBS-> | 1023 | Bacteria>FCB_group>Bacteroidota/Chlorobiota_group>Balneolota>Balneolia | GCF_003150675.1 | Rhodohalobacter_mucosus | DDZ15_RS08075 | 64787..67858 | 2079485 | DUF499-domain-containing-protein-[Rhodohalobacter-mucosus] |
| MDD3554381.1 | DUF499_2.RM.like.systems | <-ParA-Soj-PloopNTPase<-ParA-Soj-PloopNTPase<-?\|\|DUF499*->DUF3780->?->MACRODOMAIN->METHYLASE->HEPN->ParA-Soj-PloopNTPase-> | 1040 | Bacteria>Myxococcota>Myxococcia>unclassified_Myxococcia | GCA_028695735.1 | Deltaproteobacteria_bacterium | PHC35_07690 | 76771..79893 | 2026735 | MAG:-DUF499-domain-containing-protein-[Deltaproteobacteria-bacterium] |
| MDD5374995.1 | DUF499_2.RM.like.systems | <-ParA-Soj-PloopNTPase<-HEPN<-METHYLASE<-MACRODOMAIN<-?<-DUF3780<-DUF499*<-RHH_5 | 1041 | Bacteria>Pseudomonadota>Acidithiobacillia>Acidithiobacillales>Acidithiobacillaceae | GCA_028714835.1 | Acidithiobacillus_sp. | PHD38_05125 | 22024..25149 | 1872118 | MAG:-DUF499-domain-containing-protein-[Acidithiobacillus-sp.] |
| MCZ8170923.1 | DUF499_2.RM.like.systems | <-SMODS<-OBFOLD-NUCLEASE<-HEPN+REase<-HKD-DNase+SF1-SF2-Helicase<-METHYLASE<-DUF3780<-DUF499*<-CopG/RHH\|\|?-><-?<-HTH_3 | 1034 | Bacteria>Pseudomonadota>Alphaproteobacteria>Caulobacterales>Caulobacteraceae | GCA_027531485.1 | Brevundimonas_sp. | O9272_04190 | 53052..56156 | 1871086 | MAG:-DUF499-domain-containing-protein-[Brevundimonas-sp.] |
| WP_203443311.1 | DUF499_2.RM.like.systems | <-GDResolvase+HIN-HTH<-HEPN<-HKD-DNase+SF1-SF2-Helicase+iREase+CTD-p<-DUF932<-METHYLASE<-DUF3780<-DUF499*\|\|?->?->?->?-><-?\|\|MarR-HTH-> | 1021 | Bacteria>Pseudomonadota>Alphaproteobacteria>Hyphomicrobiales>Methylobacteriaceae | GCF_016864375.1 | Microvirga_sp._VF16 | J0965_RS08880 | 1887910..1890975 | 2807101 | DUF499-domain-containing-protein-[Microvirga-sp.-VF16] |
| WP_230534162.1 | DUF499_2.RM.like.systems | <-HEPN<-HKD-DNase+SF1-SF2-Helicase+iREase+CTD<-DUF932<-METHYLASE<-DUF3780<-DUF499* | 1021 | Bacteria>Pseudomonadota>Alphaproteobacteria>Hyphomicrobiales>Methylobacteriaceae | GCF_020866965.1 | Microvirga_roseola | LG661_RS22040 | 8580..11645 | 2883126 | DUF499-domain-containing-protein-[Microvirga-roseola] |
| RWG75421.1 | DUF499_2.RM.like.systems | UbE1+JAB->?-><-HEPN<-HKD-DNase+SF1-SF2-Helicase<-METHYLASE<-DUF3780<-DUF499*<-CopG/RHH<-HEPN | 830 | Bacteria>Pseudomonadota>Alphaproteobacteria>Hyphomicrobiales>Phyllobacteriaceae | GCA_004018415.1 | Mesorhizobium_sp. | EOQ66_00050 | 16547..19039 | 1871066 | DUF499-domain-containing-protein-[Mesorhizobium-sp.] |
| RWK94990.1 | DUF499_2.RM.like.systems | MULE-TRANSPOSASE-><-Resolvase<-?<-HKD-DNase+SF1-SF2-Helicase<-METHYLASE<-DUF3780<-DUF499*\|\|HEPN->?->HTH+MPTase-><-?<-TRPR-HTH+HTH | 1036 | Bacteria>Pseudomonadota>Alphaproteobacteria>Hyphomicrobiales>Phyllobacteriaceae | GCA_004020105.1 | Mesorhizobium_sp. | EOR53_16295 | 117341..120451 | 1871066 | ATP-binding-protein-[Mesorhizobium-sp.] |
| WP_127299913.1 | DUF499_2.RM.like.systems | <-HKD-DNase+SF1-SF2-Helicase+iREase+CTD<-METHYLASE<-DUF3780<-DUF499*<-?<-HEPN | 1017 | Bacteria>Pseudomonadota>Alphaproteobacteria>Hyphomicrobiales>Phyllobacteriaceae | GCF_003997355.1 | Mesorhizobium_sp._M1E.F.Ca.ET.041.01.1.1 | EOA38_RS18715 | 4307..7360 | 2496759 | DUF499-domain-containing-protein-[Mesorhizobium-sp.-M1E.F.Ca.ET.041.01.1.1] |
| WP_128294455.1 | DUF499_2.RM.like.systems | HEPN->CopG/RHH->DUF499*->DUF3780->METHYLASE->HKD-DNase+SF1-SF2-Helicase->HKD-DNase+SF1-SF2-Helicase-p->HEPN-> | 1042 | Bacteria>Pseudomonadota>Alphaproteobacteria>Hyphomicrobiales>Phyllobacteriaceae | GCF_004023225.1 | unclassified_Mesorhizobium | EOQ59_RS13085 | 2651..5779 | 325217 | DUF499-domain-containing-protein-[Mesorhizobium-sp.-M1A.F.Ca.IN.020.30.1.1] |
| QYK40247.1 | DUF499_2.RM.like.systems | <-ACET<-?<-HEPN+REase<-HKD-DNase+SF1-SF2-Helicase<-METHYLASE<-DUF3780<-DUF499*<-CopG/RHH\|\|?-><-tRNA\|\|?->?-><-tRNA\|\|?->Peptidase_M48-> | 1034 | Bacteria>Pseudomonadota>Alphaproteobacteria>Rhodobacterales>Paracoccaceae | GCA_019454225.1 | Paracoccaceae_bacterium | KF887_12470 | 2639699..2642803 | 1904441 | MAG:-DUF499-domain-containing-protein-[Rhodobacteraceae-bacterium] |
| WP_203075822.1 | DUF499_2.RM.like.systems | RAMA->GDResolvase+Recombinase+ZnR->HTH->?->DUF499*->DUF3780->METHYLASE->HKD-DNase+SF1-SF2-Helicase->HEPN-> | 1045 | Bacteria>Pseudomonadota>Alphaproteobacteria>Rhodospirillales>Acetobacteraceae | GCF_016805305.1 | Roseomonas_ponticola | JJI54_RS23740 | 272470..275607 | 2786951 | DUF499-domain-containing-protein-[Roseomonas-ponticola] |
| WP_144094495.1 | DUF499_2.RM.like.systems | <-LexA-protease\|\|HTH+WYL->DUF499*->DUF3780->METHYLASE->HEPN->HKD-DNase+SF1-SF2-Helicase->?->TUDOR+SF1-SF2-Helicase+iREase+CTD-> | 1034 | Bacteria>Pseudomonadota>Alphaproteobacteria>Sphingomonadales>Erythrobacteraceae | GCF_007570835.1 | Croceicoccus_sediminis | FAX90_RS00355 | 90290..93394 | 2571150 | DUF499-domain-containing-protein-[Croceicoccus-sediminis] |
| WP_168134876.1 | DUF499_2.RM.like.systems | PIWI-><-?<-HKD-DNase+SF1-SF2-Helicase+iREase+CTD<-?<-METHYLASE<-DUF3780<-DUF499*<-?<-HEPN\|\|AAA_21->?-><-TniQ | 1031 | Bacteria>Pseudomonadota>Alphaproteobacteria>Sphingomonadales>Sphingomonadaceae | GCF_012035195.1 | Sphingomonas_corticis | HBH26_RS12185 | 283937..287032 | 2722791 | DUF499-domain-containing-protein-[Sphingomonas-corticis] |
| TXH35005.1 | DUF499_2.RM.like.systems | <-ParA-Soj-PloopNTPase<-HEPN<-METHYLASE<-MACRODOMAIN<-?<-DUF3780<-DUF499* | 828 | Bacteria>Pseudomonadota>Betaproteobacteria>Burkholderiales>Burkholderiaceae | GCA_008012415.1 | Burkholderiaceae_bacterium | E6Q92_12395 | 11663..14151 | 2030806 | DUF499-domain-containing-protein,-partial-[Burkholderiaceae-bacterium] |
| WP_138938924.1 | DUF499_2.RM.like.systems | <-SIGMA-HTH<-Transposase_mut\|\|HTH_3->RHH_1->DUF499*->DUF3780->Schlafen+HATPase+HTH->METHYLASE->HEPN->ParA-Soj-PloopNTPase->ORC-CDC6-> | 1022 | Bacteria>Pseudomonadota>Betaproteobacteria>Burkholderiales>Sphaerotilaceae | GCF_011365705.1 | Rubrivivax_benzoatilyticus | G7087_RS13130 | 24930..27998 | 316997 | DUF499-domain-containing-protein-[Rubrivivax-benzoatilyticus] |
| MCA3223201.1 | DUF499_2.RM.like.systems | sGTP->tRNA->?->DUF499*->DUF3780->METHYLASE->HKD-DNase+SF1-SF2-Helicase->HEPN->?->URI-DNase+RAMA-> | 1027 | Bacteria>Pseudomonadota>Betaproteobacteria>Burkholderiales>unclassified_Burkholderiales | GCA_020248865.1 | Burkholderiales_bacterium | ING41_12560 | 90260..93343 | 1891238 | DUF499-domain-containing-protein-[Burkholderiales-bacterium] |
| MBI3370534.1 | DUF499_2.RM.like.systems | <-MqsA_antitoxin+HTH_3\|\|?->HEPN->?->DUF499*->DUF3780->METHYLASE->?->?->HKD-DNase+SF1-SF2-Helicase-> | 1026 | Bacteria>Pseudomonadota>Betaproteobacteria>unclassified_Betaproteobacteria | GCA_016195915.1 | Betaproteobacteria_bacterium | HY017_02090 | 33561..36641 | 1891241 | DUF499-domain-containing-protein-[Betaproteobacteria-bacterium] |
| GHU10669.1 | DUF499_2.RM.like.systems | <-ParA-Soj-PloopNTPase<-HEPN<-METHYLASE<-MACRODOMAIN<-?<-DUF3780<-DUF499*<-RHH_5 | 1041 | Bacteria>Pseudomonadota>Betaproteobacteria>unclassified_Betaproteobacteria | GCA_025997015.1 | Betaproteobacteria_bacterium | AGMMS50225_14630 | 22175..25300 | 1891241 | hypothetical-protein-AGMMS50225_14630-[Betaproteobacteria-bacterium] |
| ESQ16532.1 | DUF499_2.RM.like.systems | <-HKD-DNase+SF1-SF2-Helicase+iREase+CTD<-HEPN<-PolB<-METHYLASE<-DUF3780<-REase<-DUF499* | 1087 | Bacteria>Pseudomonadota>Gammaproteobacteria>Chromatiales>Chromatiaceae | GCA_000495615.1 | Thiohalocapsa_sp._PB-PSB1 | N838_05685 | 9740..13003 | 1385625 | MAG:-hypothetical-protein-N838_05685-[Thiohalocapsa-sp.-PB-PSB1] |
| QQ057135.1 | DUF499_2.RM.like.systems | DDE_Tnp_1->DUF499*->REase->DUF3780->METHYLASE->PolB->HEPN->HKD-DNase+SF1-SF2-Helicase+iREase+CTD-> | 1085 | Bacteria>Pseudomonadota>Gammaproteobacteria>Chromatiales>Chromatiaceae | GCA_016745215.1 | Thiohalocapsa_sp._PB-PSB1 | N838_31100 | 7373531..7376788 | 1385625 | MAG:-ATP-binding-protein-[Thiohalocapsa-sp.-PB-PSB1] |
| MCU0836365.1 | DUF499_2.RM.like.systems | <-METHYLASE-p<-HEPN<-PolB<-PIN<-?<-DUF3780<-DUF499*<-RHH\|\|REase-><-Cluster182_14clades\|\|?->?->ZnR+TOPRIM-> | 1037 | Bacteria>Pseudomonadota>Gammaproteobacteria>Chromatiales>Chromatiaceae | GCA_025458165.1 | Chromatiaceae_bacterium | MUC77_18335 | 8879..11992 | 2049432 | DUF499-domain-containing-protein-[Chromatiaceae-bacterium] |
| WP_201244800.1 | DUF499_2.RM.like.systems | PrmC_N+Methyltransf_25->?->?-><-?\|\|DUF499*->DUF3780->HicA_toxin->?->PolB->HEPN->METHYLASE-> | 1086 | Bacteria>Pseudomonadota>Gammaproteobacteria>Chromatiales>Chromatiaceae | GCF_016653295.1 | Halochromatium_salexigens | CCR82_RS07690 | 81879..85139 | 49447 | DUF499-domain-containing-protein-[Halochromatium-salexigens] |
| OZB36169.1 | DUF499_2.RM.like.systems | <-ParA-Soj-PloopNTPase<-HEPN<-METHYLASE<-MACRODOMAIN<-?<-DUF3780<-DUF499*<-RHH_5<-?<-?\|\|XIS-HTH->DnaB-><-HAMP+HISKIN | 1041 | Bacteria>Pseudomonadota>Gammaproteobacteria>Chromatiales>Halothiobacillaceae | GCA_002282715.1 | Halothiobacillus_sp._15-55-196 | B7X44_07150 | 24460..27585 | 1970382 | MAG:-AAA-family-ATPase-[Halothiobacillus-sp.-15-55-196] |
| WP_198323836.1 | DUF499_2.RM.like.systems | ADH_N+ADH_zinc_N-><-SF1-SF2-Helicase\|\|?->?->RHH->DUF499*->DUF3780->MACRODOMAIN->METHYLASE->HEPN->ParA-Soj-PloopNTPase->ORC-CDC6-> | 1041 | Bacteria>Pseudomonadota>Gammaproteobacteria>Methylococcales>Methylococcaceae | GCF_016106025.1 | Methylococcus | ELQ36_RS13470 | 36069..39194 | 413 | DUF499-domain-containing-protein-[Methylococcus-sp.-BF19-07] |
| WP_218806438.1 | DUF499_2.RM.like.systems | ADH_N+ADH_zinc_N-><-SF1-SF2-Helicase\|\|?->?->CopG_antitoxin->DUF499*->DUF3780->METHYLASE->Schlafen+HATPase+HTH->?->METHYLASE->HEPN-> | 1041 | Bacteria>Pseudomonadota>Gammaproteobacteria>Methylococcales>Methylococcaceae | GCF_019285515.1 | Methylococcus_sp._Mc7 | KW115_RS14805 | 3051261..3054386 | 2860258 | DUF499-domain-containing-protein-[Methylococcus-sp.-Mc7] |
| WP_092844114.1 | DUF499_2.RM.like.systems | CHTH-><-?\|\|?->XIS-HTH->DUF499*->DUF3780->METHYLASE->PolB->HEPN->HKD-DNase+SF1-SF2-Helicase+iREase+CTD->ParB+HNH-> | 1088 | Bacteria>Pseudomonadota>Gammaproteobacteria>Oceanospirillales>Halomonadaceae | GCF_900113605.1 | Halomonas_xianhensis | BM194_RS04450 | 210680..213946 | 442341 | DUF499-domain-containing-protein-[Halomonas-xianhensis] |
| HBP0460596.1 | DUF499_2.RM.like.systems | ParB->JAB-><-HEPN<-HKD-DNase+SF1-SF2-Helicase<-METHYLASE<-DUF3780<-DUF499*<-RHH\|\|?->Phage_integrase->Phage_integrase-> | 1040 | Bacteria>Pseudomonadota>Gammaproteobacteria>Pseudomonadales>Pseudomonadaceae | GCA_021685755.1 | Pseudomonas_aeruginosa | L5Q34_003641 | 43212..46334 | 287 | DUF499-domain-containing-protein-[Pseudomonas-aeruginosa] |
| WP_023443648.1 | DUF499_2.RM.like.systems | <-Phage_integrase<-?\|\|RHH->DUF499*->DUF3780->METHYLASE->HKD-DNase+SF1-SF2-Helicase->HEPN-><-?<-JAB | 1040 | Bacteria>Pseudomonadota>Gammaproteobacteria>Pseudomonadales>Pseudomonadaceae | GCF_000495915.1 | Stutzerimonas_stutzeri_group | F753_RS25405 | 1974..5096 | 136846 | MULTISPECIES:-DUF499-domain-containing-protein-[Stutzerimonas-stutzeri-group] |
| WP_058147188.1 | DUF499_2.RM.like.systems | <-MPTase<-?\|\|?-><-?<-Arm-DNA-bind_3+Phage_integrase\|\|RHH->DUF499*->DUF3780->METHYLASE->HKD-DNase+SF1-SF2-Helicase->HEPN-><-?<-Phage_lysozyme | 1040 | Bacteria>Pseudomonadota>Gammaproteobacteria>Pseudomonadales>Pseudomonadaceae | GCF_001449155.1 | Pseudomonas_aeruginosa | A0881_RS09155 | 11696..14818 | 287 | DUF499-domain-containing-protein-[Pseudomonas-aeruginosa] |
| WP_254873384.1 | DUF499_2.RM.like.systems | <-ParA-Soj-PloopNTPase<-HEPN<-METHYLASE<-MACRODOMAIN<-?<-DUF3780<-DUF499*<-RHH | 1036 | Bacteria>Pseudomonadota>Gammaproteobacteria>Pseudomonadales>Pseudomonadaceae | GCF_003411965.2 | Pseudomonas_aeruginosa | DZ913_RS12395 | 105397..108507 | 287 | DUF499-domain-containing-protein-[Pseudomonas-aeruginosa] |
| WP_073667316.1 | DUF499_2.RM.like.systems | RHH_5->DUF499*->DUF3780->?->MACRODOMAIN->METHYLASE->HEPN->ParA-Soj-PloopNTPase-> | 1041 | Bacteria>Pseudomonadota>Gammaproteobacteria>Pseudomonadales>Pseudomonadaceae | GCF_900708645.1 | Pseudomonas_aeruginosa | CHH44_RS18170 | 263427..266552 | 287 | DUF499-domain-containing-protein-[Pseudomonas-aeruginosa] |
| QFY90794.1 | DUF499_2.RM.like.systems | XIS-HTH->DUF499*->DUF3780->METHYLASE->HEPN->ParA-Soj-PloopNTPase->ORC-CDC6->HKD-DNase+SF1-SF2-Helicase-> | 1040 | Bacteria>Pseudomonadota>Gammaproteobacteria>unclassified_Gammaproteobacteria | GCA_009497875.2 | gamma_proteobacterium_SS-5 | D5125_15735 | 2296694..2299816 | 947516 | DUF499-domain-containing-protein-[gamma-proteobacterium-SS-5] |
| MDD9983184.1 | DUF499_2.RM.like.systems | ABhydrolase->DUF499*->DUF3780->METHYLASE-p->?->METHYLASE-p->ParB+HNH->HEPN-> | 1021 | Bacteria>Pseudomonadota>Gammaproteobacteria>unclassified_Gammaproteobacteria | GCA_028818395.1 | Gammaproteobacteria_bacterium | OXU81_17805 | 2420..5485 | 1913989 | MAG:-DUF499-domain-containing-protein-[Gammaproteobacteria-bacterium] |
| WP_032129329.1 | DUF499_2.RM.like.systems | Methyltransf_11->RNase->35exo-><-?\|\|tRNA->RHH_5->DUF499*->DUF3780->?->METHYLASE->HEPN->ParA-Soj-PloopNTPase->ORC-CDC6-> | 1044 | Bacteria>Pseudomonadota>Gammaproteobacteria>Xanthomonadales>Xanthomonadaceae | GCF_000758465.1 | Stenotrophomonas_maltophilia | LI87_RS0113700 | 18740..21874 | 40324 | DUF499-domain-containing-protein-[Stenotrophomonas-maltophilia] |
| WP_099589177.1 | DUF499_2.RM.like.systems | RHH_5->DUF499*->DUF3780->?->MACRODOMAIN->METHYLASE->HEPN->ParA-Soj-PloopNTPase-> | 1041 | Bacteria>Pseudomonadota>Gammaproteobacteria>Xanthomonadales>Xanthomonadaceae | GCF_002738905.1 | Stenotrophomonas_maltophilia | BRB41_RS09090 | 797..3922 | 40324 | DUF499-domain-containing-protein-[Stenotrophomonas-maltophilia] |
| WP_053462285.1 | DUF499_2.RM.like.systems | Methyltransf_11->RNase->35exo-><-?\|\|tRNA->RHH_5->DUF499*->DUF3780->?->MACRODOMAIN->METHYLASE->HEPN->ParA-Soj-PloopNTPase-> | 1041 | Bacteria>Pseudomonadota>Gammaproteobacteria>Xanthomonadales>Xanthomonadaceae | GCF_021554495.1 | Stenotrophomonas | N5C25_RS09560 | 28481..31606 | 40323 | DUF499-domain-containing-protein-[Stenotrophomonas-maltophilia] |
| WP_261997736.1 | DUF499_2.RM.like.systems | Methyltransf_11->RNase->35exo-><-?\|\|tRNA->RHH_5->DUF499*->DUF3780->?->MACRODOMAIN->METHYLASE->HEPN->ParA-Soj-PloopNTPase-> | 1041 | Bacteria>Pseudomonadota>Gammaproteobacteria>Xanthomonadales>Xanthomonadaceae | GCF_024519495.1 | Stenotrophomonas_sp._Ste96 | OC373_RS11855 | 536568..539693 | 2926029 | DUF499-domain-containing-protein-[Stenotrophomonas-sp.-Ste96] |
| MBI3099182.1 | DUF499_2.RM.like.systems | <-HKD-DNase+SF1-SF2-Helicase+iREase+CTD<-METHYLASE<-DUF3780<-DUF499*<-HEPN<-?<-Phage_integrase\|\|tRNA->TPR-> | 977 | Bacteria>PVC_group>Planctomycetota>unclassified_Planctomycetota | GCA_016192525.1 | Planctomycetota_bacterium | HYY93_13215 | 4536..7469 | 2026780 | ATP-binding-protein-[Planctomycetota-bacterium] |
| MBM3879657.1 | DUF499_2.RM.like.systems | <-ABC-ATPase+SbcC<-?<-?<-?<-?<-ABC-ATPase+SbcC\|\|DUF499*->DUF3780->Schlafen+HATPase+HTH->?->Schlafen+HATPase+HTH->METHYLASE->HEPN-> | 1019 | Bacteria>PVC_group>Verrucomicrobiota>unclassified_Verrucomicrobiota | GCA_016871715.1 | Verrucomicrobiota_bacterium | FJ387_08035 | 7884..10943 | 2026799 | ATP-binding-protein-[Verrucomicrobia-bacterium] |
| MCX8496484.1 | DUF499_2.RM.like.systems | <-REase<-?\|\|HTH->MPTase->?->?->DUF499*->DUF3780->METHYLASE->METHYLASE->HKD-DNase+SF1-SF2-Helicase->REase->HEPN-> | 1044 | Bacteria>PVC_group>Verrucomicrobiota>Verrucomicrobiae>Verrucomicrobiales | GCA_026461385.1 | Akkermansiaceae_bacterium | ORN51_09920 | 5504..8638 | 2562705 | MAG:-DUF499-domain-containing-protein-[Akkermansiaceae-bacterium] |
| MDD9986492.1 | DUF499_2.RM.like.systems | <-HEPN<-PolB<-?\|\|?->?->?->DUF499*->DUF3780->HEPN->?->?->?->METHYLASE-p-> | 1007 | Bacteria>Spirochaetota>Spirochaetia>Spirochaetales>Spirochaetaceae | GCA_028818385.1 | Spirochaetaceae_bacterium | OXQ31_09490 | 10176..13199 | 1898206 | MAG:-DUF499-domain-containing-protein-[Spirochaetaceae-bacterium] |
| MBN2352085.1 | DUF499_2.RM.like.systems | <-HKD-DNase+SF1-SF2-Helicase<-AbiEii<-BirA-HTH<-HEPN<-METHYLASE<-DUF3780<-DUF499*\|\|?->?-><-?<-?<-HNH | 1039 | Bacteria>Spirochaetota>Spirochaetia>Spirochaetales>unclassified_Spirochaetales | GCA_016934415.1 | Spirochaetales_bacterium | JXD23_05895 | 38302..41421 | 2026792 | ATP-binding-protein-[Spirochaetales-bacterium] |
| MBB2482470.1 | DUF499_2.RM.like.systems | REase+DUF4268->?->DUF499*->DUF3780->METHYLASE->HKD-DNase+SF1-SF2-Helicase->HEPN-> | 1031 | Bacteria>Terrabacteria_group>Bacillota>Bacilli>Bacillales | GCA_014174455.1 | Bacillus_sp._APMAM | H5P36_20065 | 1869..4964 | 2761103 | DUF499-domain-containing-protein-[Bacillus-sp.-APMAM] |
| WP_075310335.1 | DUF499_2.RM.like.systems | Preader7+McrB-NTD+McrB-AAA->McrC-NTD+REase->REase->HNH-><-HEPN<-HEPN-p\|\|DUF499*->DUF3780->METHYLASE->HKD-DNase+SF1-SF2-Helicase->?->?->MerR-HTH-> | 1022 | Bacteria>Terrabacteria_group>Bacillota>Bacilli>Bacillales | GCF_001932005.1 | Bacillus_wiedmannii | BUC33_RS14890 | 28645..31713 | 1890302 | DUF499-domain-containing-protein-[Bacillus-wiedmannii] |
| WP_230127616.1 | DUF499_2.RM.like.systems | SPOUT_MTase-><-HTH+MPTase<-?\|\|DUF499*->DUF3780->METHYLASE->HKD-DNase+SF1-SF2-Helicase->HEPN-> | 1022 | Bacteria>Terrabacteria_group>Bacillota>Bacilli>Bacillales | GCF_918741275.1 | Bacillus_sp._CECT_9360 | LQ216_RS05715 | 45654..48722 | 2845821 | DUF499-domain-containing-protein-[Bacillus-sp.-CECT-9360] |
| WP_045165444.1 | DUF499_2.RM.like.systems | <-REase<-MethyltransfD12<-?<-HKD-DNase+SF1-SF2-Helicase<-METHYLASE<-DUF3780<-DUF499*<-HEPN<-PolB | 1080 | Bacteria>Terrabacteria_group>Bacillota>Clostridia>Caldicellulosiruptorales | GCF_026914285.1 | Caldicellulosiruptor_naganoensis | N907_RS06310 | 1198788..1202030 | 29324 | DUF499-domain-containing-protein-[Caldicellulosiruptor-naganoensis] |
| WP_045175072.1 | DUF499_2.RM.like.systems | <-PIN<-MPTase<-SbcC<-HKD-DNase+SF1-SF2-Helicase<-METHYLASE<-DUF3780<-DUF499*<-?<-HEPN<-PolB | 1081 | Bacteria>Terrabacteria_group>Bacillota>Clostridia>Caldicellulosiruptorales | GCF_034343125.1 | Caldicellulosiruptor_danielii | N913_RS08680 | 1859027..1862272 | 1387557 | DUF499-domain-containing-protein-[Caldicellulosiruptor-danielii] |
| WP_161821648.1 | DUF499_2.RM.like.systems | <-wHTH+SAD+HNH<-REase<-HEPN<-HKD-DNase+SF1-SF2-Helicase+iREase+CTD<-METHYLASE<-DUF3780<-DUF499*\|\|?-><-?<-?<-Glycos_trans_3N+Glycos_transf_3\|\|?->HAMP+HISKIN-> | 1082 | Bacteria>Terrabacteria_group>Bacillota>Clostridia>Eubacteriales | GCF_009932395.1 | Sporotomaculum_syntrophicum | SPSYN_RS06335 | 287290..290538 | 182264 | DUF499-domain-containing-protein-[Sporotomaculum-syntrophicum] |
| WP_080657027.1 | DUF499_2.RM.like.systems | <-HKD-DNase+SF1-SF2-Helicase+iREase+CTD<-HEPN<-METHYLASE<-DUF3780<-DUF499*<-RelE-ParE<-Phd-YefM<-?<-?<-?<-ABC-ATPase+SbcC | 1126 | Bacteria>Terrabacteria_group>Bacillota>Negativicutes>Selenomonadales | GCF_000424065.1 | Selenomonas_ruminantium | G598_RS0111655 | 42037..45417 | 971 | DUF499-domain-containing-protein-[Selenomonas-ruminantium] |
| WP_074570871.1 | DUF499_2.RM.like.systems | <-HIN-HTH\|\|?->DUF499*->DUF3780->METHYLASE->HEPN->HKD-DNase+SF1-SF2-Helicase+iREase+CTD-><-DOC+HTH | 1085 | Bacteria>Terrabacteria_group>Bacillota>Negativicutes>Selenomonadales | GCF_900104055.1 | Selenomonas_ruminantium | BLQ59_RS00930 | 972..4229 | 971 | DUF499-domain-containing-protein-[Selenomonas-ruminantium] |
| MCX7789960.1 | DUF499_2.RM.like.systems | ZnR->?->PolB->DUF499*->DUF499-p->DUF3780->PolB->HEPN-> | 910 | Bacteria>Terrabacteria_group>Chloroflexota>Chloroflexia>Chloroflexales | GCA_026417815.1 | Chloroflexaceae_bacterium | N2378_04905 | 2392..5124 | 2562117 | MAG:-DUF499-domain-containing-protein-[Chloroflexaceae-bacterium] |
| OQA44952.1 | DUF499_2.RM.like.systems | <-METHYLASE<-HEPN<-PolB<-PhdYeFM_antitox<-RelE-ParE<-ASCH+DUF3780<-DUF499* | 1029 | Bacteria>Terrabacteria_group>Chloroflexota>unclassified_Chloroflexota | GCA_002068935.1 | Chloroflexi_bacterium_ADurb.Bin325 | BWY52_01458 | 8239..11328 | 1852859 | hypothetical-protein-BWY52_01458-[Chloroflexi-bacterium-ADurb.Bin325] |
| OQB28032.1 | DUF499_2.RM.like.systems | <-PolB\|\|?-><-HEPN+Mrr_N<-HKD-DNase+SF1-SF2-Helicase+iREase+CTD<-METHYLASE<-DUF3780<-DUF499*<-?<-?<-REase<-?<-HHH_4+SF1-SF2-Helicase+REase | 1020 | Bacteria>Terrabacteria_group>Chloroflexota>unclassified_Chloroflexota | GCA_002070215.1 | Chloroflexi_bacterium_ADurb.Bin180 | BWY10_00836 | 30087..33149 | 1852857 | hypothetical-protein-BWY10_00836-[Chloroflexi-bacterium-ADurb.Bin180] |
| KKZ13991.1 | DUF499_2.RM.like.systems | HEPN->DUF499*->DUF3780->REase->?->ParB+HNH->OLD-ABC-> | 1040 | Bacteria>Terrabacteria_group>Cyanobacteriota/Melainabacteria_group>Cyanobacteriota>Cyanophyceae | GCA_001007635.1 | Candidatus_Synechococcus_spongiarum_15L | TQ37_02560 | 1477..4599 | 1608419 | hypothetical-protein-TQ37_02560-[Candidatus-Synechococcus-spongiarum-15L] |
| BCX11788.1 | DUF499_2.RM.like.systems | SF1-SF2-Helicase-><-REase\|\|TUDOR+SF1-SF2-Helicase+iREase+CTD+REase-p->DUF499*->DUF3780->METHYLASE->HKD-DNase+SF1-SF2-Helicase->HEPN+Mrr_N->HEPN+REase-> | 1027 | Bacteria>Terrabacteria_group>Cyanobacteriota/Melainabacteria_group>Cyanobacteriota>Cyanophyceae | GCA_025999095.1 | Thermosynechococcus_sp. | KatS3mg067_0726 | 755896..758979 | 2814275 | MAG:-hypothetical-protein-KatS3mg067_0726-[Thermosynechococcus-sp.] |
| MCY4234851.1 | DUF499_2.RM.like.systems | DUF499*->DUF3780->?->METHYLASE-><-HEPN<-?\|\|HKD-DNase+SF1-SF2-Helicase+iREase+CTD-> | 1020 | Bacteria>Terrabacteria_group>Cyanobacteriota/Melainabacteria_group>Cyanobacteriota>unclassified_Cyanobacteriota | GCA_026709585.1 | Cyanobacteria_bacterium_MAG_CAR2_bin_4 | OXE74_00800 | 134..3196 | 2775590 | MAG:-DUF499-domain-containing-protein-[Cyanobacteria-bacterium-MAG-CAR2_bin_4] |
| MCM8748012.1 | DUF499_2.RM.like.systems | <-Phage_integrase\|\|DUF499*->DUF3780->HEPN+Mrr_N->?->?->REase->MazE_antitoxin-> | 1016 | Bacteria>Terrabacteria_group>Thermomicrobiota>Thermomicrobia>Thermomicrobiales | GCA_023806485.1 | Thermalbibacter_longus | NET02_02505 | 568134..571184 | 2951981 | DUF499-domain-containing-protein-[Thermomicrobiaceae-bacterium-CFH-74404] |
| WP_284055791.1 | DUF499_2.RM.like.systems | <-Phage_integrase\|\|DUF499*->DUF3780->HEPN+Mrr_N->?->?->REase->MazE_antitoxin-> | 1016 | Bacteria>Terrabacteria_group>Thermomicrobiota>Thermomicrobia>Thermomicrobiales | GCF_023806485.1 | Thermalbibacter_longus | NET02_RS02500 | 568134..571184 | 2951981 | DUF499-domain-containing-protein-[Thermalbibacter-longus] |
| MBC8359968.1 | DUF499_2.RM.like.systems | Schlafen+HATPase+HTH->BirA-HTH->?->PolB->HEPN->DUF499*-> | 637 | Bacteria>Thermodesulfobacteriota>Desulfobacteria>Desulfobacterales>Desulfobacterales_incertae_sedis | GCA_014382665.1 | Candidatus_Desulfatibia_profunda | H8E23_01035 | 3975..5887 | 2841695 | ATP-binding-protein-[Desulfobacterales-bacterium] |
| MBF0120192.1 | DUF499_2.RM.like.systems | <-NERD<-OLD-ABC<-METHYLASE<-HEPN<-PolB<-DUF3780<-DUF499*<-?<-?<-ABC-ATPase+SbcC | 1037 | Bacteria>Thermodesulfobacteriota>Desulfobacteria>Desulfobacterales>unclassified_Desulfobacterales | GCA_015228685.1 | Desulfobacterales_bacterium | HQK79_15260 | 23007..26120 | 2044940 | DUF499-domain-containing-protein-[Desulfobacterales-bacterium] |
| TVQ96846.1 | DUF499_2.RM.like.systems | DUF4236+TPR->DUF499*->DUF3780-><-HEPN<-PolB<-?\|\|METHYLASE-p->REase+N6_MTase-> | 438 | Bacteria>Thermodesulfobacteriota>Desulfovibrionia>Desulfovibrionales>unclassified_Desulfovibrionales | GCA_007693505.1 | Desulfovibrionales_bacterium | EA399_14900 | 820..2136 | 2448050 | hypothetical-protein-EA399_14900-[Desulfovibrionales-bacterium] |

| Protein-ID | System | Gene-Neighbourhoods | Length | Taxonomy | Assembly-ID | Strain | Locus-Tag | Genomic-coordinates | TaxID | Fasta-defline |
|---|---|---|---|---|---|---|---|---|---|---|
| -- | | | | | | | | | | |
| -- ParB + HNH | | | | | | | | | | |
| -- | | | | | | | | | | |
| AOW80904.1 | DUF499_2.RM.like.systems | <-REase\|\|ParB+HNH-p->?-><-METHYLASE<-DUF3780<-DUF499* | 1107 | Archaea>Euryarchaeota>Stenosarchaea_group>Halobacteria>Halobacteriales | GCA_001767315.1 | Halodesulfurarchaeum_formicicum | HTSR_1734 | 1660006..1663329 | 1873524 | putative-ATPase-(AAA+-superfamily)-[Halodesulfurarchaeum-formicicum] |
| WP_070365960.1 | DUF499_2.RM.like.systems | <-REase\|\|ParB+HNH-p->?-><-METHYLASE<-DUF3780<-DUF499* | 933 | Archaea>Euryarchaeota>Stenosarchaea_group>Halobacteria>Halobacteriales | GCF_001767315.1 | Halodesulfurarchaeum_formicicum | HTSR_RS08490 | 1660006..1662807 | 1873524 | DUF499-domain-containing-protein-[Halodesulfurarchaeum-formicicum] |
| WP_226011842.1 | DUF499_2.RM.like.systems | GDResolvase+Recombinase->?->DUF499*->DUF3780->METHYLASE-><-?\|\|ParB+HNH->HKD-DNase+SF1-SF2-Helicase+iREase+CTD-> | 1101 | Archaea>Euryarchaeota>Stenosarchaea_group>Halobacteria>Halobacteriales | GCF_020405185.1 | Halomicrobium_salinisoli | LE162_RS01580 | 297593..300898 | 2878391 | DUF499-domain-containing-protein-[Halomicrobium-salinisoli] |
| WP_254824893.1 | DUF499_2.RM.like.systems | ZnR+REase-><-HKD-DNase+SF1-SF2-Helicase+iREase+CTD\|\|ParB+HNH-><-METHYLASE<-DUF3780<-DUF499*\|\|ZnR+REase-><-ParA-Soj-PloopNTPase | 1107 | Archaea>Euryarchaeota>Stenosarchaea_group>Halobacteria>Halobacteriales | GCF_024298885.1 | Haloglomus_halophilum | NL114_RS21530 | 126665..129988 | 2962672 | DUF499-domain-containing-protein-[Haloglomus-sp.-ZY41] |
| MBC9987873.1 | DUF499_2.RM.like.systems | <-Peptidase_M48<-?<-SF1-SF2-Helicase+REase<-Exonuc_V_gamma+REase<-CBS+HEPN\|\|?->DUF499*->DUF3780->METHYLASE-><-ParB+HNH\|\|HKD-DNase+SF1-SF2-Helicase+iREase+CTD-><-ZnR+REase | 933 | Archaea>Euryarchaeota>Stenosarchaea_group>Halobacteria>Haloferacales | GCA_014530665.1 | Haloferax_sp._AS1 | E4P24_16065 | 306929..309730 | 2562277 | DUF499-domain-containing-protein-[Haloferax-sp.-AS1] |
| WP_188148783.1 | DUF499_2.RM.like.systems | <-Peptidase_M48<-?<-SF1-SF2-Helicase+REase<-Exonuc_V_gamma+REase<-CBS+HEPN\|\|?->DUF499*->DUF3780->METHYLASE-><-ParB+HNH\|\|HKD-DNase+SF1-SF2-Helicase+iREase+CTD-><-ZnR+REase | 1107 | Archaea>Euryarchaeota>Stenosarchaea_group>Halobacteria>Haloferacales | GCF_014530665.1 | Haloferax_sp._AS1 | E4P24_RS16200 | 306407..309730 | 2562277 | DUF499-domain-containing-protein-[Haloferax-sp.-AS1] |
| WP_271963542.1 | DUF499_2.RM.like.systems | <-Schlafen<-HKD-DNase+SF1-SF2-Helicase+iREase+CTD\|\|ParB+HNH-><-METHYLASE<-DUF3780<-DUF499*\|\|?->?->?-><-Phage_integrase\|\|PadR-HTH->PadR-HTH-> | 1107 | Archaea>Euryarchaeota>Stenosarchaea_group>Halobacteria>Haloferacales | GCF_028211285.1 | Halorubrum_ezzemoulense | PM025_RS15440 | 14437..17760 | 337243 | DUF499-domain-containing-protein-[Halorubrum-ezzemoulense] |
| CAG0952062.1 | DUF499_2.RM.like.systems | <-Primpol+D5N+D5ATPase\|\|DUF499*->DUF3780->PADR-HTH->HD->METHYLASE->SF1-SF2-Helicase+iREase+CTD+REase->ParB+HNH-> | 980 | Archaea>Euryarchaeota>Stenosarchaea_group>Methanomicrobia>Methanosarcinales | GCA_905339305.1 | Methanosarcinales_archaeon | METP3_00272 | 13443..16385 | 2250255 | hypothetical-protein-METP3_00272-[Methanosarcinales-archaeon] |
| WP_156161107.1 | DUF499_2.RM.like.systems | <-ParA-Soj-PloopNTPase\|\|?->HD-><-?\|\|ParB+HNH->?->DUF499*->DUF3780->METHYLASE->HKD-DNase+SF1-SF2-Helicase-> | 1034 | Archaea>Euryarchaeota>Stenosarchaea_group>Methanomicrobia>Methanosarcinales | GCF_000979415.1 | unclassified_Methanosarcina | EO97_RS16030 | 24899..28003 | 2644672 | MULTISPECIES:-DUF499-domain-containing-protein-[unclassified-Methanosarcina] |
| MBP7066715.1 | DUF499_2.RM.like.systems | TPR+TPR->?->?-><-DUF3780<-DUF499*\|\|METHYLASE-><-?<-?<-HKD-DNase+SF1-SF2-Helicase+iREase+CTD<-ParB+HNH<-ArdCN+MPTase | 1011 | Archaea>Euryarchaeota>Stenosarchaea_group>Methanomicrobia>Methanotrichales | GCA_017994095.1 | Methanothrix_sp. | KBB04_00320 | 64883..67918 | 90426 | ATP-binding-protein-[Methanothrix-sp.] |
| NHV98109.1 | DUF499_2.RM.like.systems | <-Radical-SAM\|\|Radical-SAM-><-EVE\|\|ParB+HNH-><-DUF3780<-DUF499*\|\|METHYLASE-><-?<-HKD-DNase+SF1-SF2-Helicase+iREase+CTD\|\|?-><-DNA-glycosylase | 982 | Archaea>TACK_group>Nitrososphaerota>unclassified_Nitrososphaerota | GCA_011605725.1 | Nitrososphaerota_archaeon | HA496_00520 | 58939..61887 | 2026795 | ATP-binding-protein-[Thaumarchaeota-archaeon] |
| MCY4585111.1 | DUF499_2.RM.like.systems | <-HKD-DNase+SF1-SF2-Helicase+iREase+CTD<-METHYLASE<-?<-OLD-ABC<-ParB+HNH<-DUF3780<-DUF499*\|\|Phage_integrase-> | 1018 | Bacteria>Acidobacteriota>Terriglobia>Bryobacterales>unclassified_Bryobacterales | GCA_026713545.1 | Bryobacterales_bacterium | OXB98_03645 | 8005..11061 | 2026791 | MAG:-DUF499-domain-containing-protein-[Bryobacterales-bacterium] |
| MCY3933638.1 | DUF499_2.RM.like.systems | <-HNH<-OLD-ABC<-?<-OLD-ABC<-ParB+HNH<-DUF3780<-DUF499*<-?<-?<-SMODS\|\|HTH+MPTase-><-?<-Phd-YefM | 1021 | Bacteria>Acidobacteriota>unclassified_Acidobacteriota | GCA_026707325.1 | Acidobacteriota_bacterium | OXH70_18150 | 6925..9990 | 1978231 | MAG:-DUF499-domain-containing-protein-[Acidobacteria-bacterium] |
| MDE2976517.1 | DUF499_2.RM.like.systems | MetJarc-HTH->?->?->DUF499*->DUF3780->ParB+HNH->OLD-ABC->?->OLD-ABC->HNH-> | 1021 | Bacteria>Acidobacteriota>unclassified_Acidobacteriota | GCA_028873735.1 | Acidobacteriota_bacterium | OXU63_03230 | 1513..4578 | 1978231 | MAG:-DUF499-domain-containing-protein-[Acidobacteriota-bacterium] |
| MCB9367519.1 | DUF499_2.RM.like.systems | DUF499*->DUF3780->METHYLASE->HKD-DNase+SF1-SF2-Helicase+iREase+CTD->ParB+HNH->OLD-ABC-> | 970 | Bacteria>Calditrichota>unclassified_Calditrichota | GCA_020637445.1 | Calditrichota_bacterium | H6507_00185 | 43427..46339 | 2212469 | ATP-binding-protein-[Calditrichaeota-bacterium] |
| NLT02438.1 | DUF499_2.RM.like.systems | <-ParB+HNH<-?<-HKD-DNase+SF1-SF2-Helicase<-Schlafen+HATPase+HTH<-METHYLASE<-DUF3780<-DUF499*<-tRNA\|\|?->?->NADAR->?->Aminotran_1_2-> | 1038 | Bacteria>FCB_group>Bacteroidota/Chlorobiota_group>Bacteroidota>Bacteroidia | GCA_012718945.1 | Bacteroidales_bacterium | GXY09_01085 | 15955..19071 | 2030927 | DUF499-domain-containing-protein-[Bacteroidales-bacterium] |
| TAK42618.1 | DUF499_2.RM.like.systems | BirA-HTH->?->wHTH+REase->METHYLASE-p->DUF499*->DUF3780->ParB+HNH->OLD-ABC->RelE-ParE->METHYLASE-p-> | 1024 | Bacteria>FCB_group>Bacteroidota/Chlorobiota_group>Bacteroidota>Saprospiria | GCA_004297645.1 | Saprospiraceae_bacterium | EPO28_07065 | 17499..20573 | 2202734 | DUF499-domain-containing-protein-[Saprospiraceae-bacterium] |
| NUQ24407.1 | DUF499_2.RM.like.systems | <-ABhydrolase\|\|?->?-><-?<-?\|\|?->DUF499*->DUF3780->ParB+HNH->OLD-ABC->RelE-ParE->METHYLASE->OLD-ABC-> | 1024 | Bacteria>FCB_group>Bacteroidota/Chlorobiota_group>Bacteroidota>Saprospiria | GCA_013360405.1 | Saprospiraceae_bacterium | HUU34_10665 | 13446..16520 | 2202734 | DUF499-domain-containing-protein-[Saprospiraceae-bacterium] |
| WP_109646585.1 | DUF499_2.RM.like.systems | ParB+HNH->DUF499*->DUF3780->METHYLASE->HKD-DNase+SF1-SF2-Helicase->HEPN+CBS-> | 1023 | Bacteria>FCB_group>Bacteroidota/Chlorobiota_group>Balneolota>Balneolia | GCF_003150675.1 | Rhodohalobacter_mucosus | DDZ15_RS08075 | 64787..67858 | 2079485 | DUF499-domain-containing-protein-[Rhodohalobacter-mucosus] |
| MBM2814383.1 | DUF499_2.RM.like.systems | <-ParB+HNH<-?<-HKD-DNase+SF1-SF2-Helicase<-METHYLASE<-DUF3780<-DUF499*<-Peptidase<-Peptidase | 1037 | Bacteria>FCB_group>Bacteroidota/Chlorobiota_group>Ignavibacteriota>Ignavibacteria | GCA_016844745.1 | Ignavibacteria_bacterium | HW421_1145 | 16386..19499 | 2053306 | hypothetical-protein-[Ignavibacteria-bacterium] |
| MBK8059521.1 | DUF499_2.RM.like.systems | GDResolvase->GDResolvase-><-HKD-DNase+SF1-SF2-Helicase+iREase+CTD<-ParB+HNH<-METHYLASE<-DUF3780<-DUF499*\|\|HTH_3-><-?<-PAPS_reduct<-?<-KAP-NTPase<-HTH | 1026 | Bacteria>FCB_group>Gemmatimonadota>unclassified_Gemmatimonadota | GCA_016713015.1 | Gemmatimonadota_bacterium | IPK33_17185 | 755230..758310 | 2026742 | ATP-binding-protein-[Gemmatimonadetes-bacterium] |
| MBN1335625.1 | DUF499_2.RM.like.systems | REase->SF1-SF2-Helicase-><-ParB+HNH<-HKD-DNase+SF1-SF2-Helicase<-METHYLASE<-DUF3780<-DUF499*<-XIS-HTH<-?<-?<-HEPN | 1020 | Bacteria>Myxococcota>Myxococcia>unclassified_Myxococcia | GCA_016927815.1 | Deltaproteobacteria_bacterium | JXB39_06665 | 17124..20186 | 2026735 | DUF499-domain-containing-protein-[Deltaproteobacteria-bacterium] |
| MDE0030313.1 | DUF499_2.RM.like.systems | <-HKD-DNase+SF1-SF2-Helicase+iREase+CTD<-METHYLASE<-?<-OLD-ABC<-ParB+HNH<-DUF3780<-DUF499*\|\|?-><-HEPN\|\|HEPN-><-DUF772+DDE_Tnp_1+DDE_Tnp_1_6 | 1005 | Bacteria>Myxococcota>Myxococcia>unclassified_Myxococcia | GCA_028818775.1 | Deltaproteobacteria_bacterium | OXU42_13040 | 19997..23014 | 2026735 | MAG:-DUF499-domain-containing-protein-[Deltaproteobacteria-bacterium] |
| MBT3366889.1 | DUF499_2.RM.like.systems | ParB+HNH+Schlafen->DUF499*->DUF3780->METHYLASE->HicB_lk_antitox->HicA_toxin->HKD-DNase+SF1-SF2-Helicase->HNH-> | 1035 | Bacteria>Nitrospinae/Tectomicrobia_group>Nitrospinota>Nitrospinia>Nitrospinales | GCA_018648125.1 | Nitrospina_sp. | HN416_07030 | 2974..6081 | 2024844 | DUF499-domain-containing-protein-[Nitrospina-sp.] |
| HBP88391.1 | DUF499_2.RM.like.systems | DUF499*->DUF3780->METHYLASE->HKD-DNase+SF1-SF2-Helicase->ParB+HNH+RAMA->HEPN-> | 1027 | Bacteria>Nitrospirota>Nitrospiria>Nitrospirales>Nitrospiraceae | GCA_003523945.1 | Nitrospiraceae_bacterium | DD706_11920 | 779..3862 | 2026770 | hypothetical-protein-[Nitrospiraceae-bacterium] |
| WP_136498122.1 | DUF499_2.RM.like.systems | Phd-YefM->PIN->Phage_integrase->CopG/RHH->DUF499*->DUF3780->METHYLASE->HKD-DNase+SF1-SF2-Helicase->ParB+HNH-p->ParB+HNH-p-> | 1045 | Bacteria>Pseudomonadota>Alphaproteobacteria>Hyphomicrobiales>Methylocystaceae | GCF_004802635.2 | Methylocystis_heyeri | H2LOC_RS20845 | 59150..62287 | 391905 | DUF499-domain-containing-protein-[Methylocystis-heyeri] |
| RWI67698.1 | DUF499_2.RM.like.systems | <-HKD-DNase+SF1-SF2-Helicase<-METHYLASE<-DUF3780<-DUF499*<-?<-ParB+HNH+RAMA\|\|?->Resolvase-> | 1033 | Bacteria>Pseudomonadota>Alphaproteobacteria>Hyphomicrobiales>Phyllobacteriaceae | GCA_004020235.1 | Mesorhizobium_sp. | EOR19_30450 | 12024..15125 | 1871066 | ATP-binding-protein-[Mesorhizobium-sp.] |
| TPG55477.1 | DUF499_2.RM.like.systems | <-HKD-DNase+SF1-SF2-Helicase<-MACRODOMAIN<-ParB+HNH<-METHYLASE<-DUF3780<-DUF499*<-35exo<-?<-?\|\|HEPN+REase-> | 1086 | Bacteria>Pseudomonadota>Alphaproteobacteria>Rhodospirillales>Acetobacteraceae | GCA_006438825.1 | Roseomonas_nepalensis | EAH89_14595 | 46471..49731 | 1854500 | DUF499-domain-containing-protein-[Roseomonas-nepalensis] |
| MBE7157121.1 | DUF499_2.RM.like.systems | HTH->ParB+HNH->ParB+HNH->DUF499*->DUF3780-> | 1031 | Bacteria>Pseudomonadota>Alphaproteobacteria>Rhodospirillales>Acetobacteraceae | GCA_015069705.1 | Acetobacter_sp. | INR62_01575 | 6375..9470 | 440 | DUF499-domain-containing-protein-[Rhodospirillales-bacterium] |
| MCA3288087.1 | DUF499_2.RM.like.systems | <-sGTP<-?<-?\|\|tRNA->DUF499*->DUF3780->METHYLASE->HKD-DNase+SF1-SF2-Helicase-><-ParB+HNH<-HTH<-GDResolvase+Recombinase+ZnR | 1045 | Bacteria>Pseudomonadota>Alphaproteobacteria>Rhodospirillales>Acetobacteraceae | GCA_020249245.1 | Roseomonas_sp. | ING09_16215 | 121046..124183 | 1869190 | ATP-binding-protein-[Roseomonas-sp.] |
| MCA4920336.1 | DUF499_2.RM.like.systems | DUF499*->DUF3780->METHYLASE->HKD-DNase+SF1-SF2-Helicase-><-ParB+HNH<-HTH<-GDResolvase+Recombinase+ZnR | 1045 | Bacteria>Pseudomonadota>Alphaproteobacteria>Rhodospirillales>Acetobacteraceae | GCA_020285315.1 | Roseomonas_sp. | ING82_12990 | 1209..4346 | 1869190 | ATP-binding-protein-[Roseomonas-sp.] |
| WP_075822782.1 | DUF499_2.RM.like.systems | <-DDE_Tnp_1<-?<-?<-?<-?\|\|DUF499*->DUF3780->METHYLASE->ParB+HNH->MACRODOMAIN->HKD-DNase+SF1-SF2-Helicase-> | 1066 | Bacteria>Pseudomonadota>Alphaproteobacteria>Rhodospirillales>Acetobacteraceae | GCF_001950415.1 | Roseomonas | MHL39_RS17975 | 5795..8995 | 125216 | MULTISPECIES:-DUF499-domain-containing-protein-[Roseomonas] |
| WP_161993601.1 | DUF499_2.RM.like.systems | <-HKD-DNase+SF1-SF2-Helicase<-MACRODOMAIN<-ParB+HNH<-METHYLASE<-DUF3780<-DUF499*<-35exo<-?<-?\|\|HEPN+REase-> | 1022 | Bacteria>Pseudomonadota>Alphaproteobacteria>Rhodospirillales>Acetobacteraceae | GCF_006438825.1 | Roseomonas_nepalensis | EAH89_RS14655 | 46471..49539 | 1854500 | DUF499-domain-containing-protein-[Roseomonas-nepalensis] |
| WP_256444398.1 | DUF499_2.RM.like.systems | DUF499*->DUF3780->METHYLASE->ParB+HNH->?->?->HKD-DNase+SF1-SF2-Helicase-p-> | 869 | Bacteria>Pseudomonadota>Alphaproteobacteria>Rhodospirillales>Acetobacteraceae | GCF_019084075.1 | Siccirubricoccus_sp._G192 | KSF81_RS27995 | 936..3545 | 2849651 | DUF499-domain-containing-protein-[Siccirubricoccus-sp.-G192] |
| WP_217382201.1 | DUF499_2.RM.like.systems | Phage_integrase-><-?<-?<-ParA-Soj-PloopNTPase\|\|SF1-SF2-Helicase->?->\|\|DUF499*->DUF3780->METHYLASE-p->METHYLASE->ParB+HNH->HKD-DNase+SF1-SF2-Helicase-><-HicB | 1072 | Bacteria>Pseudomonadota>Alphaproteobacteria>Rhodospirillales>Acetobacteraceae | GCF_019084075.1 | Siccirubricoccus_sp._G192 | KSF81_RS28660 | 165691..168909 | 2849651 | DUF499-domain-containing-protein-[Siccirubricoccus-sp.-G192] |
| SUS04809.1 | DUF499_2.RM.like.systems | ParB+HNH->DUF499*->DUF3780->RelE-ParE->ParA+HTH->METHYLASE->HKD-DNase+SF1-SF2-Helicase-> | 1024 | Bacteria>Pseudomonadota>Alphaproteobacteria>Rhodospirillales>Rhodospirillaceae | GCA_900498315.1 | uncultured_Defluviicoccus_sp. | DF3PB_150002 | 1531..4605 | 543062 | conserved-hypothetical-protein-[uncultured-Defluviicoccus-sp.] |
| WP_130302965.1 | DUF499_2.RM.like.systems | <-RelB\|\|?->XIS-HTH->DUF499*->DUF3780->METHYLASE->HKD-DNase+SF1-SF2-Helicase-><-ParB+HNH<-MORC+REase\|\|HEPN-> | 1022 | Bacteria>Pseudomonadota>Betaproteobacteria>Burkholderiales>Alcaligenaceae | GCF_004217095.1 | Advenella_incenata | EV681_RS00025 | 2520..5588 | 267800 | DUF499-domain-containing-protein-[Advenella-incenata] |
| WP_216254672.1 | DUF499_2.RM.like.systems | <-HTH+WYL<-?\|\|?->?-><-?<-MerR-HTH\|\|DUF499*->DUF3780->METHYLASE->HKD-DNase+SF1-SF2-Helicase->ParB+HNH+Schlafen-><-PIN | 1038 | Bacteria>Pseudomonadota>Betaproteobacteria>Burkholderiales>Burkholderiaceae | GCF_018882365.1 | Polynucleobacter_sp._71A-WALBACH | ICN42_RS10945 | 39602..42718 | 2689097 | DUF499-domain-containing-protein-[Polynucleobacter-sp.-71A-WALBACH] |
| MBK7616707.1 | DUF499_2.RM.like.systems | <-HTH+WYL\|\|HNH-><-?\|\|Arm-DNA-bind_3+Phage_integrase-><-ParB+HNH<-DUF3780<-DUF499*<-SF1-SF2-Helicase+DUF1998<-?<-?<-?<-?\|\|HNH-> | 1045 | Bacteria>Pseudomonadota>Betaproteobacteria>Burkholderiales>unclassified_Burkholderiales | GCA_016710065.1 | Burkholderiales_bacterium | IPJ08_20475 | 317100..320237 | 1891238 | ATP-binding-protein-[Burkholderiales-bacterium] |
| TMO42407.1 | DUF499_2.RM.like.systems | <-MNS+REase\|\|HTH->STYKinase-like->XIS-HTH->ZnR+TOPRIM+primpol+HTH->HTH+WYL->DUF499*->DUF3780->METHYLASE->ParB+HNH->HKD-DNase+SF1-SF2-Helicase+iREase+CTD->?->MqsA_antitoxin+HTH_3-> | 1081 | Bacteria>Pseudomonadota>Gammaproteobacteria>Alteromonadales>Pseudoalteromonadaceae | GCA_005886815.1 | Pseudoalteromonas_sp._S4389 | CWC25_15720 | 9911..13156 | 579556 | AAA-family-ATPase-[Pseudoalteromonas-sp.-S4389] |
| WP_171040448.1 | DUF499_2.RM.like.systems | <-MNS+REase\|\|HTH->STYKinase-like->XIS-HTH->ZnR+TOPRIM+primpol+HTH->HTH+WYL->DUF499*->DUF3780->METHYLASE->ParB+HNH->HKD-DNase+SF1-SF2-Helicase+iREase+CTD->?->MqsA_antitoxin+HTH_3-> | 1077 | Bacteria>Pseudomonadota>Gammaproteobacteria>Alteromonadales>Pseudoalteromonadaceae | GCF_005886815.1 | Pseudoalteromonas_sp._S4389 | CWC25_RS15645 | 9923..13156 | 579556 | DUF499-domain-containing-protein-[Pseudoalteromonas-sp.-S4389] |
| WP_199525347.1 | DUF499_2.RM.like.systems | <-MqsA_antitoxin+HTH_3<-?<-HKD-DNase+SF1-SF2-Helicase+iREase+CTD<-ParB+HNH<-METHYLASE<-DUF3780<-DUF499*<-HTH+WYL<-ZnR+TOPRIM+primpol+HTH<-XIS-HTH<-STYKinase-like<-HTH\|\|MNS+REase-> | 1077 | Bacteria>Pseudomonadota>Gammaproteobacteria>Alteromonadales>Pseudoalteromonadaceae | GCF_016464045.1 | Pseudoalteromonas_sp._bablab_jr011 | JIG41_RS20360 | 15866..19099 | 2755062 | DUF499-domain-containing-protein-[Pseudoalteromonas-sp.-bablab_jr011] |
| WP_076922918.1 | DUF499_2.RM.like.systems | <-MqsA_antitoxin+HTH_3<-?<-HKD-DNase+SF1-SF2-Helicase+iREase+CTD<-ParB+HNH<-METHYLASE<-DUF3780<-DUF499*<-HTH+WYL<-ZnR+TOPRIM+primpol+HTH<-XIS-HTH<-STYKinase-like<-HTH\|\|MNS+REase-> | 1077 | Bacteria>Pseudomonadota>Gammaproteobacteria>Alteromonadales>Pseudoalteromonadaceae | GCF_030407565.1 | unclassified_Pseudoalteromonas | BW140_RS15975 | 19057..22290 | 194690 | DUF499-domain-containing-protein-[Pseudoalteromonas-sp.-SK20] |
| MCP5417446.1 | DUF499_2.RM.like.systems | TraH->?->TraG_N-><-?\|\|XIS-HTH->ParB+HNH->DUF499*->DUF3780->METHYLASE->HKD-DNase+SF1-SF2-Helicase->?->Hydrolase_3-> | 1043 | Bacteria>Pseudomonadota>Gammaproteobacteria>Chromatiales>Chromatiaceae | GCA_024235215.1 | Chromatiaceae_bacterium | H6965_10165 | 390461..393592 | 2049432 | DUF499-domain-containing-protein-[Chromatiaceae-bacterium] |
| WP_007193607.1 | DUF499_2.RM.like.systems | <-HKD-DNase+SF1-SF2-Helicase+iREase+CTD<-ParB+HNH<-METHYLASE-p<-METHYLASE<-Schlafen+HATPase+HTH<-DUF3780<-DUF499*\|\|?->?->?->CTHH-> | 1041 | Bacteria>Pseudomonadota>Gammaproteobacteria>Chromatiales>Chromatiaceae | GCF_000223985.1 | Thiocapsa_marina | THIMADRAFT_RS13475 | 269670..272795 | 244573 | DUF499-domain-containing-protein-[Thiocapsa-marina] |
| WP_009148233.1 | DUF499_2.RM.like.systems | <-ACET<-?\|\|DUF499*->DUF3780->RHH_1->?->PIN->ParB+HNH->METHYLASE-> | 1040 | Bacteria>Pseudomonadota>Gammaproteobacteria>Chromatiales>Chromatiaceae | GCF_033954835.1 | Thiorhodovibrio_frisius | THI970DRAFT_RS08785 | 535191..538313 | 631362 | DUF499-domain-containing-protein-[Thiorhodovibrio-sp.-970] |
| WP_005369522.1 | DUF499_2.RM.like.systems | SbcC->?->MetJarc-HTH->RelE-ParE->DUF499*->DUF3780->METHYLASE->ParB+HNH->HKD-DNase+SF1-SF2-Helicase+iREase+CTD->OLD-ABC+TOPRIM->SF1-SF2-Helicase-> | 1077 | Bacteria>Pseudomonadota>Gammaproteobacteria>Methylococcales>Methylococcaceae | GCF_000214275.2 | Methylomicrobium_album | METAL_RS03085 | 732822..736055 | 39775 | DUF499-domain-containing-protein-[Methylomicrobium-album] |
| WP_249692546.1 | DUF499_2.RM.like.systems | HTH+WYL->DUF499*->DUF3780->METHYLASE->ParB+HNH->HKD-DNase+SF1-SF2-Helicase+iREase+CTD-><-Phage_lysozyme | 1077 | Bacteria>Pseudomonadota>Gammaproteobacteria>Oceanospirillales>Aestuariirhabdaceae | GCF_023509475.1 | Aestuariirhabdus_haliotis | MIB40_RS07390 | 56644..59877 | 2918751 | DUF499-domain-containing-protein-[Aestuariirhabdus-haliotis] |
| WP_153843409.1 | DUF499_2.RM.like.systems | NERD+SF1-SF2-Helicase->DUF499*->DUF3780->METHYLASE->ParB+HNH->REase+ParB->HKD-DNase+SF1-SF2-Helicase+iREase+CTD-> | 1106 | Bacteria>Pseudomonadota>Gammaproteobacteria>Oceanospirillales>Halomonadaceae | GCF_012427705.1 | Halomonas_piezotolerans | GYM47_RS02295 | 484423..487743 | 2609667 | DUF499-domain-containing-protein-[Halomonas-piezotolerans] |
| WP_247619860.1 | DUF499_2.RM.like.systems | DUF499*->DUF3780->METHYLASE->ParB+HNH->?->HKD-DNase+SF1-SF2-Helicase+iREase+CTD-> | 1088 | Bacteria>Pseudomonadota>Gammaproteobacteria>Oceanospirillales>Halomonadaceae | GCF_025393375.1 | Chromohalobacter | KZO79_RS04210 | 936140..939406 | 42054 | MULTISPECIES:-DUF499-domain-containing-protein-[Chromohalobacter] |
| WP_092844114.1 | DUF499_2.RM.like.systems | CTHH-><-?\|\|?->XIS-HTH->DUF499*->DUF3780->METHYLASE->PolB->HEPN->HKD-DNase+SF1-SF2-Helicase+iREase+CTD->ParB+HNH-> | 1088 | Bacteria>Pseudomonadota>Gammaproteobacteria>Oceanospirillales>Halomonadaceae | GCF_900113605.1 | Halomonas_xianhensis | BM194_RS04450 | 210680..213946 | 442341 | DUF499-domain-containing-protein-[Halomonas-xianhensis] |
| WP_200371316.1 | DUF499_2.RM.like.systems | <-DUF4236+TPR<-?\|\|?->?->?->?->DUF499*->DUF3780->METHYLASE->ParB+HNH->OLD-ABC->PIN-> | 1081 | Bacteria>Pseudomonadota>Gammaproteobacteria>Pseudomonadales>Marinobacteraceae | GCF_016597815.1 | Marinobacter_sp._DY40_1A1 | FFI61_RS14505 | 345780..349025 | 2583229 | DUF499-domain-containing-protein-[Marinobacter-sp.-DY40_1A1] |
| WP_058181793.1 | DUF499_2.RM.like.systems | <-SF1-SF2-Helicase<-STYKIN<-?<-?<-?\|\|RHH_5->DUF499*->DUF3780->?->MACRODOMAIN->METHYLASE->HKD-DNase+SF1-SF2-Helicase->ParB+HNH-> | 1041 | Bacteria>Pseudomonadota>Gammaproteobacteria>Pseudomonadales>Pseudomonadaceae | GCF_001453595.1 | Pseudomonas_aeruginosa | APA84_RS22695 | 17823..20948 | 287 | DUF499-domain-containing-protein-[Pseudomonas-aeruginosa] |
| WP_236167518.1 | DUF499_2.RM.like.systems | <-ParB+HNH<-ParB+HNH<-HKD-DNase+SF1-SF2-Helicase<-METHYLASE<-DUF3780<-DUF499*<-?<-Phage_int_SAM<-Integrase_1<-?<-TOPRIM<-Arm-DNA-bind_3+Phage_integrase | 1038 | Bacteria>Pseudomonadota>Gammaproteobacteria>Pseudomonadales>Pseudomonadaceae | GCF_021602545.1 | Pseudomonas_fulva | L6D33_RS20200 | 11642..14758 | 47880 | DUF499-domain-containing-protein-[Pseudomonas-fulva] |
| MBU0657159.1 | DUF499_2.RM.like.systems | Phage_integrase-><-SF1-SF2-Helicase<-?<-HKD-DNase+SF1-SF2-Helicase<-METHYLASE<-DUF3780<-DUF499*<-ParB+HNH+Schlafen\|\|N6_N4_MTase->?->Mbetalac->?->XIS-HTH-> | 1043 | Bacteria>Pseudomonadota>Gammaproteobacteria>unclassified_Gammaproteobacteria | GCA_018813855.1 | Gammaproteobacteria_bacterium | KJ914_18710 | 11621..14752 | 1913989 | DUF499-domain-containing-protein-[Gammaproteobacteria-bacterium] |
| MCB1859932.1 | DUF499_2.RM.like.systems | <-HKD-DNase+SF1-SF2-Helicase<-METHYLASE<-DUF3780<-DUF499*<-ParB+HNH<-XIS-HTH\|\|TIR-><-TraG_N | 1043 | Bacteria>Pseudomonadota>Gammaproteobacteria>unclassified_Gammaproteobacteria | GCA_020445415.1 | Gammaproteobacteria_bacterium | KDI63_16790 | 5935..9066 | 1913989 | DUF499-domain-containing-protein-[Gammaproteobacteria-bacterium] |
| MDD9983184.1 | DUF499_2.RM.like.systems | ABhydrolase->DUF499*->DUF3780->METHYLASE-p->?->METHYLASE-p->ParB+HNH->HEPN-> | 1021 | Bacteria>Pseudomonadota>Gammaproteobacteria>unclassified_Gammaproteobacteria | GCA_028818395.1 | Gammaproteobacteria_bacterium | OXU81_17805 | 2420..5485 | 1913989 | MAG:-DUF499-domain-containing-protein-[Gammaproteobacteria-bacterium] |
| WP_285401339.1 | DUF499_2.RM.like.systems | <-JAB\|\|XIS-HTH->?->XIS-HTH->DUF499*->DUF3780->METHYLASE->ParB+HNH->HKD-DNase+SF1-SF2-Helicase+iREase+CTD-><-RlpA<-Arm-DNA-bind_3+Phage_integrase | 1076 | Bacteria>Pseudomonadota>Gammaproteobacteria>Vibrionales>Vibrionaceae | GCF_030248535.1 | unclassified_Vibrio | KJB41_RS10780 | 100358..103588 | 2614977 | MULTISPECIES:-DUF499-domain-containing-protein-[unclassified-Vibrio] |
| MBK8067117.1 | DUF499_2.RM.like.systems | <-REase+35exo<-REase+35exo\|\|?->HTH+WYL->XIS-HTH->DUF499*->DUF3780->METHYLASE->HKD-DNase+SF1-SF2-Helicase->ParB+HNH-> | 1046 | Bacteria>Pseudomonadota>Gammaproteobacteria>Xanthomonadales>Rhodanobacteraceae | GCA_016713135.1 | Rhodanobacteraceae_bacterium | IPK27_05685 | 321286..324426 | 2052182 | ATP-binding-protein-[Rhodanobacteraceae-bacterium] |
| MCE9554575.1 | DUF499_2.RM.like.systems | DUF499*->DUF3780->ParB+HNH->METHYLASE->?->HKD-DNase+SF1-SF2-Helicase+iREase+CTD-> | 885 | Bacteria>PVC_group>Planctomycetota>unclassified_Planctomycetota | GCA_021413705.1 | Planctomycetota_bacterium | K8T91_14550 | 155..2812 | 2026780 | ATP-binding-protein-[Planctomycetota-bacterium] |
| MBK9497791.1 | DUF499_2.RM.like.systems | <-HNH<-OLD-ABC<-?<-OLD-ABC<-ParB+HNH<-DUF3780<-DUF499*<-DUF499\|\|?->?->?->Peptidase-> | 736 | Bacteria>Spirochaetota>Spirochaetia>Leptospirales>Leptospiraceae | GCA_016718635.1 | Leptospiraceae_bacterium | IPO06_00155 | 42788..44998 | 2026757 | DUF499-domain-containing-protein-[Leptospiraceae-bacterium] |
| MBK9497792.1 | DUF499_2.RM.like.systems | <-OLD-ABC<-?<-OLD-ABC<-ParB+HNH<-DUF3780<-DUF499<-DUF499*\|\|?->?->?->Peptidase-> | 294 | Bacteria>Spirochaetota>Spirochaetia>Leptospirales>Leptospiraceae | GCA_016718635.1 | Leptospiraceae_bacterium | IPO06_00160 | 44986..45870 | 2026757 | DUF499-domain-containing-protein-[Leptospiraceae-bacterium] |
| MBP6739048.1 | DUF499_2.RM.like.systems | DUF499*->DUF3780->ParB+HNH->OLD-ABC->?->Y1_Tnp->METHYLASE-> | 1072 | Bacteria>Spirochaetota>Spirochaetia>Leptospirales>Leptospiraceae | GCA_017990775.1 | Leptospiraceae_bacterium | KA146_03605 | 102..3320 | 2026757 | ATP-binding-protein-[Leptospiraceae-bacterium] |
| ORC33988.1 | DUF499_2.RM.like.systems | <-HKD-DNase+SF1-SF2-Helicase<-METHYLASE<-ParB+HNH<-DUF3780<-DUF499*<-?<-Phage_integrase<-tRNA\|\|?-><-NTP_transf_3 | 1091 | Bacteria>Spirochaetota>Spirochaetia>Spirochaetales>Spirochaetaceae | GCA_002087085.1 | Marispirochaeta_aestuarii | B4097_13955 | 8718..11993 | 1963862 | hypothetical-protein-B4097_13955-[Marispirochaeta-aestuarii] |
| WP_158084306.1 | DUF499_2.RM.like.systems | <-HKD-DNase+SF1-SF2-Helicase<-METHYLASE<-ParB+HNH<-DUF3780<-DUF499*<-?<-Phage_integrase<-tRNA\|\|?-><-NTP_transf_3 | 1039 | Bacteria>Spirochaetota>Spirochaetia>Spirochaetales>Spirochaetaceae | GCF_002087085.1 | Marispirochaeta_aestuarii | B4097_RS13955 | 8718..11837 | 1963862 | DUF499-domain-containing-protein-[Marispirochaeta-aestuarii] |
| KOX56111.1 | DUF499_2.RM.like.systems | <-ABC-ATPase+SbcC\|\|?->?->DUF499*->METHYLASE->ParB+HNH->HKD-DNase+SF1-SF2-Helicase-> | 1045 | Bacteria>Terrabacteria_group>Actinomycetota>Actinomycetes>Kitasatosporales | GCA_001280155.1 | Streptomyces_purpurogeneiscleroticus | ADL19_10425 | 17672..20809 | 68259 | hypothetical-protein-ADL19_10425-[Streptomyces-purpurogeneiscleroticus] |
| MCB0857325.1 | DUF499_2.RM.like.systems | <-TUDOR+SF1-SF2-Helicase+iREase+CTD<-?<-?\|\|ParB+HNH->DUF499*->DUF3780->METHYLASE->HKD-DNase+SF1-SF2-Helicase+iREase+CTD->?-><-REase | 976 | Bacteria>Terrabacteria_group>Actinomycetota>Thermoleophilia>Solirubrobacterales | GCA_020438355.1 | Solirubrobacterales_bacterium | KDB57_04290 | 63088..66018 | 1909295 | DUF499-domain-containing-protein-[Solirubrobacterales-bacterium] |
| WP_015908040.1 | DUF499_2.RM.like.systems | <-OLD-ABC+TOPRIM<-ParB+HNH<-HKD-DNase+SF1-SF2-Helicase<-METHYLASE<-DUF3780<-DUF499*<-SF1-SF2-Helicase<-OLD-ABC+TOPRIM<-NACHT<-?<-NERD | 1080 | Bacteria>Terrabacteria_group>Bacillota>Clostridia>Caldicellulosiruptorales | GCF_000022325.1 | Caldicellulosiruptor_bescii | ATHE_RS08025 | 1698593..1701835 | 31899 | DUF499-domain-containing-protein-[Caldicellulosiruptor-bescii] |
| WP_013403049.1 | DUF499_2.RM.like.systems | NERD->?->NACHT->OLD-ABC+TOPRIM->SF1-SF2-Helicase->DUF499*->DUF3780->METHYLASE->HKD-DNase+SF1-SF2-Helicase->ParB+HNH-> | 1080 | Bacteria>Terrabacteria_group>Bacillota>Clostridia>Caldicellulosiruptorales | GCF_000166355.1 | Caldicellulosiruptor_hydrothermalis | CALHY_RS05800 | 1166413..1169655 | 413888 | DUF499-domain-containing-protein-[Caldicellulosiruptor-hydrothermalis] |
| WP_014042422.1 | DUF499_2.RM.like.systems | <-MethyltransfD12<-?<-ParB+HNH<-HKD-DNase+SF1-SF2-Helicase<-METHYLASE<-DUF3780<-DUF499*<-SbcC<-SF1-SF2-Helicase<-NERD<-?<-?<-Peptidase_M56+Peptidase_M48 | 1080 | Bacteria>Terrabacteria_group>Bacillota>Clostridia>Caldicellulosiruptorales | GCF_000193435.2 | Caldicellulosiruptor_acetigenus | CALLA_RS05120 | 1088104..1091346 | 301953 | DUF499-domain-containing-protein-[Caldicellulosiruptor-acetigenus] |
| HHY76914.1 | DUF499_2.RM.like.systems | <-URI-DNase<-ParB+HNH<-SbcC<-HKD-DNase+SF1-SF2-Helicase<-METHYLASE<-DUF3780<-DUF499*<-TraJ-RHH | 1098 | Bacteria>Terrabacteria_group>Bacillota>Clostridia>Eubacteriales | GCA_012842725.1 | Clostridiales_bacterium | GX498_00205 | 10822..14118 | 1898207 | DUF499-domain-containing-protein-[Clostridiales-bacterium] |
| MBM7867397.1 | DUF499_2.RM.like.systems | REase->DUF499*->DUF3780->METHYLASE->HKD-DNase+SF1-SF2-Helicase->ParB+HNH-p->?->Classical-AAA-> | 1023 | Bacteria>Terrabacteria_group>Bacillota>Clostridia>Eubacteriales | GCA_016908025.1 | Heliomicrobium_gestii | JOC69_002303 | 75173..78244 | 2699 | hypothetical-protein-[Heliomicrobium-gestii] |
| MCF8011662.1 | DUF499_2.RM.like.systems | <-TPR<-?<-ParB+HNH<-HKD-DNase+SF1-SF2-Helicase+iREase+CTD<-METHYLASE<-DUF3780<-DUF499*<-PrlF_antitoxin | 1080 | Bacteria>Terrabacteria_group>Bacillota>Clostridia>Eubacteriales | GCA_021736925.1 | Clostridiales_bacterium | K9L56_10290 | 19311..22553 | 1898207 | DUF499-domain-containing-protein-[Clostridiales-bacterium] |
| WP_029232559.1 | DUF499_2.RM.like.systems | <-ParB+HNH<-ParB+HNH<-HKD-DNase+SF1-SF2-Helicase<-METHYLASE<-DUF3780<-DUF499*<-XIS-HTH<-?<-?<-ZnR | 1039 | Bacteria>Terrabacteria_group>Bacillota>Clostridia>Eubacteriales | GCF_000420945.1 | Butyrivibrio_sp._VCB2006 | G613_RS0110145 | 46570..49689 | 1280679 | DUF499-domain-containing-protein-[Butyrivibrio-sp.-VCB2006] |
| WP_235920429.1 | DUF499_2.RM.like.systems | REase->DUF499*->DUF3780->METHYLASE->HKD-DNase+SF1-SF2-Helicase->ParB+HNH-p->?->Classical-AAA-> | 942 | Bacteria>Terrabacteria_group>Bacillota>Clostridia>Eubacteriales | GCF_016908025.1 | Heliomicrobium_gestii | JOC69_RS11260 | 75416..78244 | 2699 | DUF499-domain-containing-protein-[Heliomicrobium-gestii] |
| WP_089655503.1 | DUF499_2.RM.like.systems | MazF_antitoxin-><-RAMA<-?<-HKD-DNase+SF1-SF2-Helicase<-METHYLASE<-DUF3780<-DUF499*<-ParB+HNH<-SF1-SF2-Helicase | 1030 | Bacteria>Terrabacteria_group>Bacillota>Clostridia>Halanaerobiales | GCF_900100575.1 | Halanaerobium_congolense | BLS10_RS04565 | 105480..108572 | 54121 | DUF499-domain-containing-protein-[Halanaerobium-congolense] |
| WP_014063447.1 | DUF499_2.RM.like.systems | <-Primpol+D5N+D5ATPase<-ParB+HNH<-HKD-DNase+SF1-SF2-Helicase<-METHYLASE<-DUF3780<-DUF499*<-TraJ-RHH\|\|DDE_Tnp_1-> | 1037 | Bacteria>Terrabacteria_group>Bacillota>Clostridia>Thermoanaerobacterales | GCF_000147695.2 | Thermoanaerobacter_wiegelii | THEWI_RS10775 | 2213989..2217102 | 46354 | DUF499-domain-containing-protein-[Thermoanaerobacter-wiegelii] |
| WP_216384534.1 | DUF499_2.RM.like.systems | HTH_26->DUF499*->DUF3780->METHYLASE->HKD-DNase+SF1-SF2-Helicase->ParB+HNH->?->Cysteine_desulfurase-> | 1038 | Bacteria>Terrabacteria_group>Bacillota>Tissierellia>Tissierellales | GCF_018917195.1 | Anaerosalibacter_bizertensis | KQH90_RS07250 | 53730..56846 | 932217 | DUF499-domain-containing-protein-[Anaerosalibacter-bizertensis] |
| MBE7435441.1 | DUF499_2.RM.like.systems | <-URI-DNase<-ParB+HNH<-DUF499-p<-ASCH+DUF3780<-?<-?<-DUF499*<-?<-METHYLASE<-TUDOR+SF1-SF2-Helicase+iREase+CTD+REase | 1032 | Bacteria>Terrabacteria_group>Chloroflexota>Anaerolineae>Anaerolineales | GCA_015075725.1 | Anaerolineales_bacterium | HS100_16115 | 3331332..3334430 | 2073117 | DUF499-domain-containing-protein-[Anaerolineales-bacterium] |
| MCA9900782.1 | DUF499_2.RM.like.systems | DUF499*->DUF3780->METHYLASE->METHYLASE-p->HKD-DNase+SF1-SF2-Helicase->ParB+HNH-p-> | 1027 | Bacteria>Terrabacteria_group>Chloroflexota>Anaerolineae>Anaerolineales | GCA_020432515.1 | Anaerolineales_bacterium | KC433_21490 | 560..3643 | 2073117 | DUF499-domain-containing-protein-[Anaerolineales-bacterium] |
| MCC6456909.1 | DUF499_2.RM.like.systems | <-HIN-HTH<-HIN-HTH<-?\|\|MazE_antitoxin->DUF499*->MAD-NTD+HNH->DUF3780->DUF3780->ParB+HNH->METHYLASE->Calcineurin-> | 1021 | Bacteria>Terrabacteria_group>Chloroflexota>Caldilineae>Caldilineales | GCA_020846635.1 | Caldilineaceae_bacterium | IT328_18275 | 1966..5031 | 2614250 | DUF499-domain-containing-protein-[Caldilineaceae-bacterium] |
| MCX5991391.1 | DUF499_2.RM.like.systems | <-HTH+PolB<-?<-ParB+HNH<-HKD-DNase+SF1-SF2-Helicase<-METHYLASE<-DUF3780<-DUF499*\|\|HD->PADR-HTH-><-PIN<-HTH+MPTase | 1031 | Bacteria>Terrabacteria_group>Chloroflexota>unclassified_Chloroflexota | GCA_026389585.1 | Chloroflexota_bacterium | NTZ04_03550 | 87097..90192 | 2026724 | MAG:-DUF499-domain-containing-protein-[Chloroflexi-bacterium] |
| WP_224174984.1 | DUF499_2.RM.like.systems | <-DDE_Tnp_1<-HKD-DNase+SF1-SF2-Helicase+iREase+CTD<-ParB+HNH<-METHYLASE<-DUF3780<-DUF499*<-CHTH\|\|STYKIN-> | 1013 | Bacteria>Terrabacteria_group>Cyanobacteriota/Melainabacteria_group>Cyanobacteriota>Cyanophyceae | - | Calothrix_sp._PCC_7716 | CAL7716_066380 | 7620880..7623921 | 2853227 | DUF499-domain-containing-protein-[Calothrix-sp.-PCC-7716] |
| MBD2438833.1 | DUF499_2.RM.like.systems | <-Phage_int_SAM_1+Phage_integrase\|\|?->?->?->DUF499*->DUF3780->METHYLASE->ParB+HNH->HKD-DNase+SF1-SF2-Helicase+iREase+CTD-> | 1019 | Bacteria>Terrabacteria_group>Cyanobacteriota/Melainabacteria_group>Cyanobacteriota>Cyanophyceae | GCA_014697425.1 | Nostoc_sp._FACHB-110 | H6G69_18770 | 61408..64467 | 2692834 | ATP-binding-protein-[Nostoc-sp.-FACHB-110] |
| MBD2467864.1 | DUF499_2.RM.like.systems | <-CHTH<-TPR\|\|<-?\|\|?->?->\|\|DUF499*->DUF3780->METHYLASE->ParB+HNH->HKD-DNase+SF1-SF2-Helicase+iREase+CTD-> | 1019 | Bacteria>Terrabacteria_group>Cyanobacteriota/Melainabacteria_group>Cyanobacteriota>Cyanophyceae | GCA_014697455.1 | Nostoc_sp._FACHB-145 | H6G90_09455 | 23220..26279 | 2692836 | ATP-binding-protein-[Nostoc-sp.-FACHB-145] |
| MBE9018013.1 | DUF499_2.RM.like.systems | <-HKD-DNase+SF1-SF2-Helicase+iREase+CTD<-ParB+HNH<-METHYLASE<-DUF3780<-DUF499* | 1026 | Bacteria>Terrabacteria_group>Cyanobacteriota/Melainabacteria_group>Cyanobacteriota>Cyanophyceae | GCA_015206905.1 | Chroococcidiopsidales_cyanobacterium_LEGE_13417 | IQ272_18060 | 6249..9329 | 1828829 | ATP-binding-protein-[Chroococcidiopsidales-cyanobacterium-LEGE-13417] |
| BDA72472.1 | DUF499_2.RM.like.systems | <-DDE_Tnp_1<-HKD-DNase+SF1-SF2-Helicase+iREase+CTD<-ParB+HNH<-METHYLASE<-DUF3780<-DUF499*<-CHTH\|\|STYKIN-> | 1013 | Bacteria>Terrabacteria_group>Cyanobacteriota/Melainabacteria_group>Cyanobacteriota>Cyanophyceae | GCA_019977735.1 | Calothrix_sp._PCC_7716 | CAL7716_066380 | 7620880..7623921 | 2853227 | hypothetical-protein-CAL7716_066380-[Calothrix-sp.-PCC-7716] |
| WP_199338599.1 | DUF499_2.RM.like.systems | <-CHTH<-TPR\|\|<-?\|\|?->?->\|\|DUF499*->DUF3780->METHYLASE->ParB+HNH->HKD-DNase+SF1-SF2-Helicase+iREase+CTD-> | 1045 | Bacteria>Terrabacteria_group>Cyanobacteriota/Melainabacteria_group>Cyanobacteriota>Cyanophyceae | GCF_014697455.1 | Nostoc_sp._FACHB-145 | H6G90_RS09455 | 23142..26279 | 2692836 | DUF499-domain-containing-protein-[Nostoc-sp.-FACHB-145] |
| WP_201323620.1 | DUF499_2.RM.like.systems | <-HKD-DNase+SF1-SF2-Helicase<-ParB+HNH<-METHYLASE<-REase<-DUF3780<-DUF499* | 1027 | Bacteria>Terrabacteria_group>Cyanobacteriota/Melainabacteria_group>Cyanobacteriota>Cyanophyceae | GCF_016584665.1 | Pseudanabaena_sp._lw0831 | APA_RS17915 | 65254..68337 | 1357935 | DUF499-domain-containing-protein-[Pseudanabaena-sp.-lw0831] |
| WP_255094986.1 | DUF499_2.RM.like.systems | AbrB-like->?->HD-><-?<-DUF4236\|\|DUF499*->DUF3780->ParB+HNH->METHYLASE-p-> | 1037 | Bacteria>Terrabacteria_group>Cyanobacteriota/Melainabacteria_group>Cyanobacteriota>Cyanophyceae | GCF_024345795.1 | Synechococcus_sp._J7-Johnson | KBY93_RS13695 | 20943..24056 | 2823737 | DUF499-domain-containing-protein-[Synechococcus-sp.-J7-Johnson] |
| WP_255001865.1 | DUF499_2.RM.like.systems | DUF499*->DUF3780->METHYLASE->METHYLASE-p->HKD-DNase+SF1-SF2-Helicase-><-ParB+HNH | 1047 | Bacteria>Terrabacteria_group>Cyanobacteriota/Melainabacteria_group>Cyanobacteriota>Cyanophyceae | GCF_024346435.1 | Cyanobium_sp._Alchichica_3B3-8F6 | KBY65_RS13005 | 1719..4862 | 2823696 | DUF499-domain-containing-protein-[Cyanobium-sp.-Alchichica-3B3-8F6] |
| WP_268243228.1 | DUF499_2.RM.like.systems | <-HKD-DNase+SF1-SF2-Helicase<-ParB+HNH<-METHYLASE<-DUF3780<-DUF499*\|\|?->SF1-SF2-Helicase-><-Phage_int_SAM_4+Phage_integrase | 1031 | Bacteria>Terrabacteria_group>Cyanobacteriota/Melainabacteria_group>Cyanobacteriota>Cyanophyceae | GCF_026735145.1 | Leptolyngbya_sp._GGD | OU814_RS30730 | 9328..12423 | 2997907 | DUF499-domain-containing-protein-[Leptolyngbya-sp.-GGD] |
| WP_242072110.1 | DUF499_2.RM.like.systems | <-Phage_int_SAM_1+Phage_integrase\|\|?->?->?->DUF499*->DUF3780->METHYLASE->ParB+HNH->HKD-DNase+SF1-SF2-Helicase+iREase+CTD-> | 920 | Bacteria>Terrabacteria_group>Cyanobacteriota/Melainabacteria_group>Cyanobacteriota>Cyanophyceae | - | Nostoc_sp._FACHB-110 | H6G69_RS18775 | 61705..64467 | 2692834 | DUF499-domain-containing-protein-[Nostoc-sp.-FACHB-110] |
| KPA09724.1 | DUF499_2.RM.like.systems | DUF499*->DUF3780->Schlafen+HATPase+HTH->METHYLASE->ParB+HNH->OLD-ABC-> | 1021 | Bacteria>Thermodesulfobacteriota>Desulfobacteria>Desulfobacterales>Candidatus_Magnetomoraceae | GCA_001292585.1 | Candidatus_Magnetomorum_sp._HK-1 | MHK_010064 | 367..3432 | 1509431 | ATPase-AAA-[Candidatus-Magnetomorum-sp.-HK-1] |
| OQY49645.1 | DUF499_2.RM.like.systems | <-OLD-ABC<-ParB+HNH<-DUF3780<-DUF499*<-HTH+WYL\|\|SMODS-> | 1021 | Bacteria>Thermodesulfobacteriota>Desulfobacteria>Desulfobacterales>Desulfobacteraceae | GCA_002085465.1 | Desulfobacteraceae_bacterium_4572_89 | B6230_07770 | 5853..8918 | 1972445 | hypothetical-protein-B6230_07770-[Desulfobacteraceae-bacterium-4572_89] |
| MCF8025627.1 | DUF499_2.RM.like.systems | DUF499*->DUF3780->METHYLASE->HKD-DNase+SF1-SF2-Helicase->ParB+HNH-> | 1031 | Bacteria>Thermodesulfobacteriota>Desulfobacteria>Desulfobacterales>Desulfobacteraceae | GCA_021737095.1 | Desulfobacteraceae_bacterium | K9K82_09080 | 113..3208 | 2049433 | DUF499-domain-containing-protein-[Desulfobacteraceae-bacterium] |
| WP_205241274.1 | DUF499_2.RM.like.systems | <-HKD-DNase+SF1-SF2-Helicase+iREase+CTD<-METHYLASE<-DUF3780<-DUF499*<-ParB+HNH-p<-?<-DOC+DOC<-HTH_31 | 1077 | Bacteria>Thermodesulfobacteriota>Desulfobulbia>Desulfobulbales>Desulfobulbaceae | GCF_016918545.1 | Desulfobulbus_alkaliphilus | JWG43_RS02730 | 133485..136718 | 869814 | DUF499-domain-containing-protein-[Desulfobulbus-alkaliphilus] |
| WP_243366428.1 | DUF499_2.RM.like.systems | <-METHYLASE<-HKD-DNase+SF1-SF2-Helicase<-METHYLASE<-DUF3780<-DUF499*<-ParB+HNH<-?<-?<-REase | 1022 | Bacteria>Thermodesulfobacteriota>Desulfovibrionia>Desulfovibrionales>Desulfovibrionaceae | GCF_022808695.1 | Fundidesulfovibrio_soli | MLE18_RS00810 | 167395..170463 | 2922716 | DUF499-domain-containing-protein-[Fundidesulfovibrio-sp.-SG60] |
| OPX97686.1 | DUF499_2.RM.like.systems | <-REase<-?\|\|?->HEPN+Mrr_N->?->DUF499-p->DUF499*->DUF3780->METHYLASE->HKD-DNase+SF1-SF2-Helicase->ParB+HNH-> | 861 | Bacteria>Thermodesulfobacteriota>Syntrophorhabdia>Syntrophorhabdales>Syntrophorhabdaceae | GCA_002067585.1 | Syntrophorhabdus_sp._PtaB.Bin027 | A4E59_00667 | 16354..18939 | 1811699 | MAG:-hypothetical-protein-A4E59_00667-[Syntrophorhabdus-sp.-PtaB.Bin027] |

| Protein-ID | System | Gene-Neighbourhoods | Length | Taxonomy | Assembly-ID | Strain | Locus-Tag | Genomic-coordinates | TaxID | Fasta-defline |
|---|---|---|---|---|---|---|---|---|---|---|
| -- | | | | | | | | | | |
| --PIN-EndoRNase | | | | | | | | | | |
| -- | | | | | | | | | | |
| WP_202318894.1 | DUF499_2.RM.like.systems | <-METHYLASE-p<-PIN<-?<-MazE_antitoxin\|\|?-><-DUF3780<-DUF499*<-?<-tRNA\|\|?->?->Cysteine_desulfurase-> | 773 | Archaea>Euryarchaeota>Archaeoglobi>Archaeoglobales>Archaeoglobaceae | GCF_016757965.1 | Archaeoglobus_neptunius | JFQ59_RS02795 | 87809..90130 | 2798580 | ATP-binding-protein-[Archaeoglobus-neptunius] |
| GIU72777.1 | DUF499_2.RM.like.systems | DUF488->DUF499*->DUF3780-><-PIN<-?<-?\|\|METHYLASE->SF1-SF2-Helicase+iREase+CTD+REase-> | 894 | Archaea>TACK_group>Nitrososphaerota>Nitrososphaeria>Candidatus_Nitrosocaldales | GCA_026002835.1 | Candidatus_Nitrosocaldaceae_archaeon | KatS3mg003_2256 | 14146..16830 | 2838776 | MAG:-hypothetical-protein-KatS3mg003_2256-[Candidatus-Nitrosocaldaceae-archaeon] |
| MCS6954437.1 | DUF499_2.RM.like.systems | <-HicA_toxin<-HicB_lk_antitox\|\|?->?->DUF499*->DUF3780->MPTase->PIN->METHYLASE-p-> | 1077 | Bacteria>Acidobacteriota>Terriglobia>Bryobacterales>Bryobacteraceae | GCA_025055455.1 | Bryobacteraceae_bacterium | NZM33_16450 | 10420..13653 | 2212468 | DUF499-domain-containing-protein-[Bryobacteraceae-bacterium] |
| GIU78629.1 | DUF499_2.RM.like.systems | Arm-DNA-bind_3+Phage_integrase->DUF772+DDE_Tnp_1+DDE_Tnp_1_6-><-?\|\|DUF499*->DUF3780->HTH+MPTase->PIN->METHYLASE->HKD-DNase+SF1-SF2-Helicase+iREase+CTD-> | 1076 | Bacteria>Acidobacteriota>Terriglobia>Bryobacterales>Bryobacteraceae | GCA_026002875.1 | Bryobacteraceae_bacterium | KatS3mg005_1867 | 2212823..2216053 | 2212468 | MAG:-hypothetical-protein-KatS3mg005_1867-[Bryobacteraceae-bacterium] |
| MCY4597199.1 | DUF499_2.RM.like.systems | <-HKD-DNase+SF1-SF2-Helicase+iREase+CTD<-PIN<-?<-METHYLASE<-TOPRIM<-OLD-ABC\|\|DUF499*->DUF3780-> | 1006 | Bacteria>Acidobacteriota>Terriglobia>Bryobacterales>unclassified_Bryobacterales | GCA_026713455.1 | Bryobacterales_bacterium | OXC19_20655 | 14043..17063 | 2026791 | MAG:-DUF499-domain-containing-protein-[Bryobacterales-bacterium] |
| MBI1799725.1 | DUF499_2.RM.like.systems | <-ParA-Soj-PloopNTPase\|\|Radical-SAM->?-><-HKD-DNase+SF1-SF2-Helicase+iREase+CTD<-METHYLASE<-PIN<-DUF499*<-?<-ABC-ATPase+SbcC<-?<-?<-PRTase | 972 | Bacteria>Bacteria_incertae_sedis>Bacteria_candidate_phyla>Candidatus_Eisenbacteria>unclassified_Candidatus_Eisenbacteria | GCA_016178685.1 | Candidatus_Eisenbacteria_bacterium | HYR73_08560 | 14195..17113 | 2212470 | ATP-binding-protein-[Candidatus-Eisenbacteria-bacterium] |
| WP_259060935.1 | DUF499_2.RM.like.systems | <-URI-DNase+RAMA<-HKD-DNase+SF1-SF2-Helicase<-METHYLASE<-PIN<-HTH+MPTase<-DUF3780<-DUF499*<-RNase\|\|URI-DNase->REase-> | 1021 | Bacteria>FCB_group>Bacteroidota/Chlorobiota_group>Rhodothermota>Rhodothermia | GCF_024806505.1 | Salinibacter_ruber | GGQ15_RS12220 | 47661..50726 | 146919 | DUF499-domain-containing-protein-[Salinibacter-ruber] |
| WP_259082418.1 | DUF499_2.RM.like.systems | <-HKD-DNase+SF1-SF2-Helicase-p<-METHYLASE<-METHYLASE<-PIN<-HTH+MPTase<-DUF3780<-DUF499*<-RNase<-?\|\|REase-> | 1021 | Bacteria>FCB_group>Bacteroidota/Chlorobiota_group>Rhodothermota>Rhodothermia | GCF_024806735.1 | Salinibacter_ruber | GGP83_RS14580 | 9847..12912 | 146919 | DUF499-domain-containing-protein-[Salinibacter-ruber] |
| WP_259170624.1 | DUF499_2.RM.like.systems | <-HKD-DNase+SF1-SF2-Helicase<-METHYLASE<-METHYLASE<-PIN<-HTH+MPTase<-DUF3780<-DUF499*<-REase+35exo<-?<-?\|\|DEDD_Tnp_IS110+Transposase_20->?->DUF4096+DDE_Tnp_1+DDE_Tnp-> | 1021 | Bacteria>FCB_group>Bacteroidota/Chlorobiota_group>Rhodothermota>Rhodothermia | GCF_024807095.1 | Salinibacter_ruber | GGP42_RS07650 | 508801..511866 | 146919 | DUF499-domain-containing-protein-[Salinibacter-ruber] |
| WP_259066518.1 | DUF499_2.RM.like.systems | <-HKD-DNase+SF1-SF2-Helicase<-METHYLASE<-PIN<-HTH+MPTase<-DUF3780<-DUF499*<-?\|\|URI-DNase->REase-> | 1021 | Bacteria>FCB_group>Bacteroidota/Chlorobiota_group>Rhodothermota>Rhodothermia | GCF_024809675.1 | Salinibacter_ruber | GGP46_RS15095 | 82348..85413 | 146919 | DUF499-domain-containing-protein-[Salinibacter-ruber] |
| WP_251930233.1 | DUF499_2.RM.like.systems | <-REase<-URI-DNase\|\|?->RNase->DUF499*->DUF3780->HTH+MPTase->PIN->METHYLASE->HKD-DNase+SF1-SF2-Helicase-> | 1021 | Bacteria>FCB_group>Bacteroidota/Chlorobiota_group>Rhodothermota>Rhodothermia | GCF_943908145.1 | Salinibacter_ruber | NGL32_RS11815 | 75461..78526 | 146919 | DUF499-domain-containing-protein-[Salinibacter-ruber] |
| MCD4708414.1 | DUF499_2.RM.like.systems | MqsA_antitoxin+HTH_3->HTH+MPTase->PIN->?->DUF499*->DUF499-p->DUF3780->METHYLASE->RADICAL-SAM-><-METHYLASE | 647 | Bacteria>FCB_group>Candidatus_Fermentibacteria>Sabulitectum>unclassified_Candidatus_Sabulitectum | GCA_021107435.1 | Candidatus_Sabulitectum_sp. | K8S62_11845 | 5005..6948 | 2822438 | DUF499-domain-containing-protein-[Candidatus-Sabulitectum-sp.] |
| MBI5442127.1 | DUF499_2.RM.like.systems | <-METHYLASE-p<-?<-PIN<-METHYLASE<-DUF3780<-DUF499* | 842 | Bacteria>Myxococcota>Myxococcia>unclassified_Myxococcia | GCA_016218905.1 | Deltaproteobacteria_bacterium | HY900_13060 | 8281..10809 | 2026735 | DUF499-domain-containing-protein-[Deltaproteobacteria-bacterium] |
| ETX04990.1 | DUF499_2.RM.like.systems | <-HKD-DNase+SF1-SF2-Helicase<-PIN<-?<-METHYLASE<-DUF3780<-DUF499-p<-DUF499* | 495 | Bacteria>Nitrospinae/Tectomicrobia_group>Candidatus_Tectomicrobia>Entotheonella | GCA_000522445.1 | Candidatus_Entotheonella_gemina | ETSY2_25620 | 8747..10231 | 1429439 | hypothetical-protein-ETSY2_25620,-partial-[Candidatus-Entotheonella-gemina] |
| GIW54142.1 | DUF499_2.RM.like.systems | REase+HTH->NTP_transf-><-?<-?\|\|AbiEii->?->DUF499*->DUF3780->PIN->METHYLASE-> | 972 | Bacteria>Nitrospirota>Nitrospiria>Nitrospirales>Nitrospiraceae | GCA_026004315.1 | Nitrospiraceae_bacterium | KatS3mg082_0546 | 99014..101932 | 2026770 | MAG:-hypothetical-protein-KatS3mg082_0546-[Nitrospiraceae-bacterium] |
| WP_215851339.1 | DUF499_2.RM.like.systems | Phage_int_SAM_4+Phage_integrase->RHH->DUF499*->DUF3780->?->?->PIN->METHYLASE->REase+ParB-> | 1041 | Bacteria>Pseudomonadota>Acidithiobacillia>Acidithiobacillales>Acidithiobacillaceae | GCF_031200535.1 | Acidithiobacillus_caldus | HFV02_RS10450 | 3946..7071 | 33059 | DUF499-domain-containing-protein-[Acidithiobacillus-caldus] |
| WP_152828752.1 | DUF499_2.RM.like.systems | Peptidase_S11-><-HKD-DNase+SF1-SF2-Helicase<-METHYLASE<-DUF3780<-DUF499*<-PIN<-RHH\|\|RNase-> | 1044 | Bacteria>Pseudomonadota>Alphaproteobacteria>Rhodobacterales>Paracoccaceae | GCF_009380135.2 | Fertoeibacter_niger | GEU84_RS20380 | 38813..41947 | 2656921 | DUF499-domain-containing-protein-[Fertoeibacter-niger] |
| WP_237719425.1 | DUF499_2.RM.like.systems | MazE_antitoxin->PIN-><-HKD-DNase+SF1-SF2-Helicase+iREase+CTD<-?<-METHYLASE<-DUF3780<-DUF499* | 411 | Bacteria>Pseudomonadota>Alphaproteobacteria>Rhodospirillales>Acetobacteraceae | GCF_000292225.1 | Roseomonas_sp._B5 | B555_RS0120340 | 9698..10935 | 1217720 | ATP-binding-protein,-partial-[Roseomonas-sp.-B5] |
| WP_019463153.1 | DUF499_2.RM.like.systems | DUF499*->DUF3780->METHYLASE->?->HKD-DNase+SF1-SF2-Helicase+iREase+CTD-><-PIN<-MazE_antitoxin | 399 | Bacteria>Pseudomonadota>Alphaproteobacteria>Rhodospirillales>Acetobacteraceae | GCF_006384895.1 | Roseomonas_mucosa | BT98_RS0119345 | 343..1542 | 207340 | hypothetical-protein-[Roseomonas-mucosa] |
| EXI76478.1 | DUF499_2.RM.like.systems | <-PIN<-?<-METHYLASE<-PIN<-MazE_antitoxin<-DUF3780<-DUF499*<-?<-?\|\|?->?-><-?<-Peptidase | 1044 | Bacteria>Pseudomonadota>Betaproteobacteria>Betaproteobacteria_incertae_sedis>Candidatus_Accumulibacter | GCA_000584995.1 | Candidatus_Accumulibacter_sp._SK-11 | AW07_00426 | 125131..128265 | 1454000 | MAG:-hypothetical-protein-AW07_00426-[Candidatus-Accumulibacter-sp.-SK-11] |
| MBK8387660.1 | DUF499_2.RM.like.systems | <-HKD-DNase+SF1-SF2-Helicase-p<-TOPRIM<-METHYLASE<-PIN<-MazE_antitoxin<-DUF3780<-DUF499*<-DSBH\|\|?->?-><-?<-Methyltransf_25 | 1044 | Bacteria>Pseudomonadota>Betaproteobacteria>Betaproteobacteria_incertae_sedis>Candidatus_Accumulibacter | GCA_016711725.1 | Accumulibacter_sp. | IPL11_19685 | 6141..9275 | 2053492 | DUF499-domain-containing-protein-[Candidatus-Accumulibacter-sp.] |
| MBK8117091.1 | DUF499_2.RM.like.systems | <-HKD-DNase+SF1-SF2-Helicase<-?<-METHYLASE<-PIN<-MazE_antitoxin<-DUF3780<-DUF499*<-DSBH\|\|?->?-><-?<-Methyltransf_25 | 1044 | Bacteria>Pseudomonadota>Betaproteobacteria>Betaproteobacteria_incertae_sedis>Candidatus_Accumulibacter | GCA_016712935.1 | Accumulibacter_sp. | IPK44_22565 | 276336..279470 | 2053492 | DUF499-domain-containing-protein-[Candidatus-Accumulibacter-sp.] |
| MBK8576832.1 | DUF499_2.RM.like.systems | <-HicB<-PIN<-METHYLASE<-PIN<-MazE_antitoxin<-DUF3780<-DUF499*<-DSBH\|\|?->?-><-?<-Methyltransf_25 | 1044 | Bacteria>Pseudomonadota>Betaproteobacteria>Betaproteobacteria_incertae_sedis>Candidatus_Accumulibacter | GCA_016714935.1 | Candidatus_Accumulibacter_propinquus | IPN78_00515 | 111101..114235 | 2954380 | DUF499-domain-containing-protein-[Candidatus-Accumulibacter-sp.] |
| MBN8450450.1 | DUF499_2.RM.like.systems | <-HKD-DNase+SF1-SF2-Helicase<-PIN<-?<-METHYLASE<-DUF3780<-DUF499* | 1034 | Bacteria>Pseudomonadota>Betaproteobacteria>Betaproteobacteria_incertae_sedis>Candidatus_Accumulibacter | GCA_017302435.1 | Candidatus_Accumulibacter_necessarius | JOM13_15880 | 43840..46944 | 2954386 | ATP-binding-protein-[Candidatus-Accumulibacter-sp.] |
| MBN8436880.1 | DUF499_2.RM.like.systems | Methyltransf_25->?-><-?<-?\|\|?->DSBH->DUF499*->DUF3780->MazE_antitoxin->PIN->METHYLASE->?->PIN-> | 1044 | Bacteria>Pseudomonadota>Betaproteobacteria>Betaproteobacteria_incertae_sedis>Candidatus_Accumulibacter | GCA_017302555.1 | Accumulibacter_sp. | JOL95_02360 | 92935..96069 | 2053492 | DUF499-domain-containing-protein-[Candidatus-Accumulibacter-sp.] |
| MCM8613692.1 | DUF499_2.RM.like.systems | <-METHYLASE<-PIN<-MazE_antitoxin<-DUF3780<-DUF499*\|\|?->?-><-?<-?<-Radical-SAM | 1044 | Bacteria>Pseudomonadota>Betaproteobacteria>Betaproteobacteria_incertae_sedis>Candidatus_Accumulibacter | GCA_023806505.1 | Accumulibacter_sp. | NFW17_16745 | 3715..6849 | 2053492 | DUF499-domain-containing-protein-[Candidatus-Accumulibacter-sp.] |
| WP_186409049.1 | DUF499_2.RM.like.systems | DnaB->?->?->?->DUF499*->DUF3780->MazE_antitoxin->PIN->METHYLASE->?->PIN-> | 1044 | Bacteria>Pseudomonadota>Betaproteobacteria>Betaproteobacteria_incertae_sedis>Candidatus_Accumulibacter | GCF_900089955.1 | Candidatus_Accumulibacter_aalborgensis | ACCAA_RS20210 | 12255..15389 | 1860102 | DUF499-domain-containing-protein-[Candidatus-Accumulibacter-aalborgensis] |
| ODS67619.1 | DUF499_2.RM.like.systems | <-HKD-DNase+SF1-SF2-Helicase<-METHYLASE<-PIN<-MazE_antitoxin<-DUF3780<-DUF499*\|\|OLD-ABC+TOPRIM->SF1-SF2-Helicase-><-HTH_21+rve+rve_3<-HIN-HTH | 1045 | Bacteria>Pseudomonadota>Betaproteobacteria>Burkholderiales>Comamonadaceae | GCA_001724795.1 | Acidovorax_sp._SCN_65-28 | ABS39_19765 | 13942..17079 | 1660086 | MAG:-hypothetical-protein-ABS39_19765-[Acidovorax-sp.-SCN-65-28] |
| MBN9628764.1 | DUF499_2.RM.like.systems | HTH_3-><-?<-?\|\|HTH_Tnp_1+HTH_21+rve+rve_2-><-SF1-SF2-Helicase<-OLD-ABC+TOPRIM\|\|DUF499*->DUF3780->MazE_antitoxin->PIN->METHYLASE->HKD-DNase+SF1-SF2-Helicase-> | 1024 | Bacteria>Pseudomonadota>Betaproteobacteria>Burkholderiales>Comamonadaceae | GCA_017307415.1 | Acidovorax_sp. | JOG38_21240 | 6814..9888 | 1872122 | DUF499-domain-containing-protein-[Acidovorax-sp.] |
| WP_022771796.1 | DUF499_2.RM.like.systems | <-PIN<-?<-METHYLASE<-PIN<-MazE_antitoxin<-DUF3780<-DUF499*<-RHH<-tRNA<-?<-RecR+TOPRIM | 1061 | Bacteria>Pseudomonadota>Betaproteobacteria>Burkholderiales>Comamonadaceae | GCF_000477435.1 | Candidatus_Symbiobacter_mobilis | CENROD_RS03870 | 937551..940736 | 1436290 | DUF499-domain-containing-protein-[Candidatus-Symbiobacter-mobilis] |
| WP_169741491.1 | DUF499_2.RM.like.systems | <-REase<-?\|\|?->DUF499*->DUF3780->MazE_antitoxin->PIN->METHYLASE->HKD-DNase+SF1-SF2-Helicase-> | 1024 | Bacteria>Pseudomonadota>Betaproteobacteria>Burkholderiales>Comamonadaceae | GCF_000739875.1 | Comamonas_aquatica | CA2_RS16400 | 2260..5334 | 225991 | DUF499-domain-containing-protein-[Comamonas-aquatica] |
| WP_210931298.1 | DUF499_2.RM.like.systems | XIS-HTH->?->?->?->DUF499*->DUF3780->MazE_antitoxin->PIN->METHYLASE->HKD-DNase+SF1-SF2-Helicase-><-STYKinase-like | 1024 | Bacteria>Pseudomonadota>Betaproteobacteria>Burkholderiales>Comamonadaceae | GCF_018064205.1 | Comamonas_aquatica | KAQ61_RS08175 | 1732991..1736065 | 225991 | DUF499-domain-containing-protein-[Comamonas-aquatica] |
| MDE2076666.1 | DUF499_2.RM.like.systems | <-HTH+WYL\|\|?->?->DUF499*->DUF3780->?->PIN->METHYLASE->HKD-DNase+SF1-SF2-Helicase-p-> | 1056 | Bacteria>Pseudomonadota>Betaproteobacteria>Burkholderiales>unclassified_Burkholderiales | GCA_028864885.1 | Burkholderiales_bacterium | KGI91_06275 | 36073..39243 | 1891238 | MAG:-DUF499-domain-containing-protein-[Burkholderiales-bacterium] |
| MPZ42202.1 | DUF499_2.RM.like.systems | <-HKD-DNase+SF1-SF2-Helicase<-?<-Ntox49<-DUF3780<-PIN<-?<-DUF499* | 1021 | Bacteria>Pseudomonadota>Betaproteobacteria>unclassified_Betaproteobacteria | GCA_009377585.1 | Betaproteobacteria_bacterium | GEV05_02130 | 145657..148722 | 1891241 | DUF499-domain-containing-protein-[Betaproteobacteria-bacterium] |
| MAL94418.1 | DUF499_2.RM.like.systems | <-HKD-DNase+SF1-SF2-Helicase+iREase+CTD<-?<-PIN<-OLD-ABC<-METHYLASE<-DUF3780<-DUF499*\|\|SF1-SF2-Helicase-> | 1088 | Bacteria>Pseudomonadota>Gammaproteobacteria>Cellvibrionales>Halieaceae | GCA_002692965.1 | Haliea_sp. | CME40_05000 | 41677..44943 | 1932666 | hypothetical-protein-[Haliea-sp.] |
| AUB85626.1 | DUF499_2.RM.like.systems | <-HKD-DNase+SF1-SF2-Helicase<-PIN<-?<-?<-METHYLASE<-DUF3780<-DUF499* | 1026 | Bacteria>Pseudomonadota>Gammaproteobacteria>Chromatiales>Chromatiaceae | GCA_002813775.1 | Candidatus_Thiodictyon_syntrophicum | THSYN_32535 | 358287..361367 | 1166950 | AAA-family-ATPase-[Candidatus-Thiodictyon-syntrophicum] |
| TVQ88665.1 | DUF499_2.RM.like.systems | ABhydrolase-><-?<-?\|\|?->DUF499*->MazE_antitoxin->PIN->DUF3780->?->PIN->PolB-> | 1030 | Bacteria>Pseudomonadota>Gammaproteobacteria>Chromatiales>Chromatiaceae | GCA_007693535.1 | Chromatiaceae_bacterium | EA400_08510 | 10879..13971 | 2049432 | ATP-binding-protein-[Chromatiaceae-bacterium] |
| MBK1620020.1 | DUF499_2.RM.like.systems | <-METHYLASE<-PIN<-?<-DUF3780<-PIN<-MazE_antitoxin<-DUF499* | 1039 | Bacteria>Pseudomonadota>Gammaproteobacteria>Chromatiales>Chromatiaceae | GCA_016583905.1 | Lamprobacter_modestohalophilus | CK042_16545 | 30403..33522 | 1064514 | AAA-family-ATPase-[Lamprobacter-modestohalophilus] |
| MBV5274819.1 | DUF499_2.RM.like.systems | <-SF1-SF2-Helicase\|\|?->NTP_transferase->DUF499*->?->DUF3780-><-PIN<-?\|\|METHYLASE->OLD-ABC-p-> | 898 | Bacteria>Pseudomonadota>Gammaproteobacteria>Chromatiales>Chromatiaceae | GCA_019163055.1 | Lamprocystis_purpurea | JZU52_14660 | 4546..7239 | 61598 | ATP-binding-protein-[Lamprocystis-purpurea] |
| MCF8003666.1 | DUF499_2.RM.like.systems | <-DUF29<-HKD-DNase+SF1-SF2-Helicase<-PIN<-?<-METHYLASE<-DUF3780<-DUF499*<-CR-ATPase9+REase | 1047 | Bacteria>Pseudomonadota>Gammaproteobacteria>Chromatiales>Chromatiaceae | GCA_021736845.1 | Chromatiaceae_bacterium | K9L32_05580 | 9009..12152 | 2049432 | DUF499-domain-containing-protein-[Chromatiaceae-bacterium] |
| MCF8005387.1 | DUF499_2.RM.like.systems | <-DUF3780<-PIN<-MazE_antitoxin<-DUF499* | 1026 | Bacteria>Pseudomonadota>Gammaproteobacteria>Chromatiales>Chromatiaceae | GCA_021736845.1 | Chromatiaceae_bacterium | K9L32_14520 | 1092..4172 | 2049432 | DUF499-domain-containing-protein-[Chromatiaceae-bacterium] |
| WP_020503208.1 | DUF499_2.RM.like.systems | <-NUDIX\|\|?-><-SF1-SF2-Helicase\|\|?->?->NTP_transferase->DUF499*->DUF3780-><-PIN<-?\|\|METHYLASE->OLD-ABC-p->OLD-ABC-> | 1081 | Bacteria>Pseudomonadota>Gammaproteobacteria>Chromatiales>Chromatiaceae | GCF_000379525.1 | Lamprocystis_purpurea | A390_RS0102830 | 110926..114171 | 61598 | DUF499-domain-containing-protein-[Lamprocystis-purpurea] |
| WP_236849074.1 | DUF499_2.RM.like.systems | <-HKD-DNase+SF1-SF2-Helicase<-PIN<-?<-?<-METHYLASE<-DUF3780<-DUF499* | 1039 | Bacteria>Pseudomonadota>Gammaproteobacteria>Chromatiales>Chromatiaceae | GCF_002813775.1 | Candidatus_Thiodictyon_syntrophicum | THSYN_RS32810 | 358287..361406 | 1166950 | DUF499-domain-containing-protein-[Candidatus-Thiodictyon-syntrophicum] |
| WP_245969509.1 | DUF499_2.RM.like.systems | <-ABhydrolase<-?\|\|ABhydrolase-><-?\|\|?->DUF499*->REase->DUF3780->?->PIN->METHYLASE->METHYLASE-p-> | 1016 | Bacteria>Pseudomonadota>Gammaproteobacteria>Chromatiales>Chromatiaceae | GCF_003634315.1 | Thiocapsa_rosea | BDD21_RS09910 | 2229310..2232360 | 69360 | DUF499-domain-containing-protein-[Thiocapsa-rosea] |
| WP_245970045.1 | DUF499_2.RM.like.systems | <-PIN<-?<-METHYLASE<-PIN<-?<-DUF3780<-DUF499*<-?<-REase+NACHT<-GDResolvase+HIN-HTH | 1018 | Bacteria>Pseudomonadota>Gammaproteobacteria>Chromatiales>Chromatiaceae | GCF_003634315.1 | Thiocapsa_rosea | BDD21_RS27140 | 182504..185560 | 69360 | DUF499-domain-containing-protein-[Thiocapsa-rosea] |
| WP_132976384.1 | DUF499_2.RM.like.systems | <-HAD<-?<-METHYLASE<-PIN<-MazE_antitoxin<-DUF3780<-DUF499*\|\|?-><-?<-?<-HISKIN<-REC | 1086 | Bacteria>Pseudomonadota>Gammaproteobacteria>Chromatiales>Chromatiaceae | GCF_004342175.1 | Thiobaca_trueperi | EDC35_RS06210 | 66133..69393 | 127458 | DUF499-domain-containing-protein-[Thiobaca-trueperi] |
| WP_153975105.1 | DUF499_2.RM.like.systems | <-HKD-DNase+SF1-SF2-Helicase<-METHYLASE<-PIN<-?<-MACRODOMAIN<-DUF3780<-DUF499* | 1040 | Bacteria>Pseudomonadota>Gammaproteobacteria>Chromatiales>Chromatiaceae | GCF_009664085.1 | Thermochromatium_tepidum | E6P07_RS07905 | 1743171..1746293 | 1050 | DUF499-domain-containing-protein-[Thermochromatium-tepidum] |
| WP_200375560.1 | DUF499_2.RM.like.systems | <-MerR-HTH<-?\|\|?->?->?->SMS-KaiC->DUF499*->DUF3780->?->PIN->OLD-ABC->TOPRIM->METHYLASE-> | 1027 | Bacteria>Pseudomonadota>Gammaproteobacteria>Chromatiales>Chromatiaceae | GCF_016583575.1 | Thiocystis_violacea | CK027_RS14205 | 21670..24753 | 13725 | DUF499-domain-containing-protein-[Thiocystis-violacea] |
| WP_207148756.1 | DUF499_2.RM.like.systems | <-METHYLASE<-PIN<-?<-DUF3780<-PIN<-MazE_antitoxin<-DUF499* | 1020 | Bacteria>Pseudomonadota>Gammaproteobacteria>Chromatiales>Chromatiaceae | GCF_016583905.1 | Lamprobacter_modestohalophilus | CK042_RS16530 | 30403..33465 | 1064514 | DUF499-domain-containing-protein-[Lamprobacter-modestohalophilus] |
| WP_242518638.1 | DUF499_2.RM.like.systems | DUF499*->DUF3780->PIN->METHYLASE->DDE_Tnp_1-><-GDResolvase+Recombinase+ZnR | 1021 | Bacteria>Pseudomonadota>Gammaproteobacteria>Chromatiales>Chromatiaceae | GCF_016653315.1 | Halochromatium_roseum | CCR96_RS05805 | 17129..20194 | 391920 | DUF499-domain-containing-protein-[Halochromatium-roseum] |
| WP_201219751.1 | DUF499_2.RM.like.systems | CR-ATPase9+REase->DUF499*->DUF3780->METHYLASE->?->PIN->HKD-DNase+SF1-SF2-Helicase->DUF29-> | 1047 | Bacteria>Pseudomonadota>Gammaproteobacteria>Chromatiales>Chromatiaceae | GCF_016653315.1 | Halochromatium_roseum | CCR96_RS08785 | 123675..126818 | 391920 | DUF499-domain-containing-protein-[Halochromatium-roseum] |
| WP_009148233.1 | DUF499_2.RM.like.systems | <-ACET<-?\|\|DUF499*->DUF3780->RHH_1->?->PIN->ParB+HNH->METHYLASE-> | 1040 | Bacteria>Pseudomonadota>Gammaproteobacteria>Chromatiales>Chromatiaceae | GCF_033954835.1 | Thiorhodovibrio_frisius | THI970DRAFT_RS08785 | 535191..538313 | 631362 | DUF499-domain-containing-protein-[Thiorhodovibrio-sp.-970] |
| RUQ39484.1 | DUF499_2.RM.like.systems | <-FURR-HTH\|\|?-><-?<-?\|\|?->RHH_5->DUF499*->DUF3780->?->PIN->METHYLASE-p-> | 1041 | Bacteria>Pseudomonadota>Gammaproteobacteria>Gammaproteobacteria_incertae_sedis>Candidatus_Competibacteraceae | GCA_003989075.1 | Candidatus_Competibacteraceae_bacterium | EKK69_09085 | 13765..16890 | 2053538 | DUF499-domain-containing-protein-[Candidatus-Competibacteraceae-bacterium] |
| MBS3952485.1 | DUF499_2.RM.like.systems | HDRnase->?->DUF499*->DUF3780->?->RelE-ParE-><-?\|\|?->PIN-> | 1039 | Bacteria>Pseudomonadota>Gammaproteobacteria>Methylococcales>Methylococcaceae | GCA_018333785.1 | Methylomicrobium_sp. | KGZ88_06015 | 4130..7249 | 2303549 | DUF499-domain-containing-protein-[Methylomicrobium-sp.] |
| WP_026223744.1 | DUF499_2.RM.like.systems | <-METHYLASE<-PIN<-VapB_antitoxin<-PIN<-?<-DUF3780<-DUF499*\|\|?->?->?-><-?<-?<-LON-AAA | 1042 | Bacteria>Pseudomonadota>Gammaproteobacteria>Methylococcales>Methylococcaceae | GCF_000372865.1 | Methylosarcina_fibrata | A3OW_RS0119500 | 4143557..4146685 | 105972 | DUF499-domain-containing-protein-[Methylosarcina-fibrata] |
| WP_017842534.1 | DUF499_2.RM.like.systems | HNH-><-?\|\|?->?->MetJarc-HTH->RelE-ParE->DUF499*->DUF3780->?->PIN->?->PIN->METHYLASE-> | 1021 | Bacteria>Pseudomonadota>Gammaproteobacteria>Methylococcales>Methylococcaceae | GCF_005931095.1 | Methylotuvimicrobium_buryatense | EQU24_RS16455 | 3692788..3695853 | 95641 | DUF499-domain-containing-protein-[Methylotuvimicrobium-buryatense] |
| MSP27870.1 | DUF499_2.RM.like.systems | HIN-HTH->DUF499*->DUF3780->?->PIN->METHYLASE->HKD-DNase+SF1-SF2-Helicase-> | 1021 | Bacteria>Pseudomonadota>Gammaproteobacteria>Methylococcales>unclassified_Methylococcales | GCA_009692395.1 | Methylococcales_bacterium | EXR80_05400 | 1625..4690 | 2304002 | DUF499-domain-containing-protein-[Methylococcales-bacterium] |
| MDD5268641.1 | DUF499_2.RM.like.systems | <-HKD-DNase+SF1-SF2-Helicase<-PIN<-VapB_antitoxin<-METHYLASE<-DUF3780<-DUF499*<-DOC+HTH | 1043 | Bacteria>Pseudomonadota>Gammaproteobacteria>Methylococcales>unclassified_Methylococcales | GCA_028712795.1 | Methylococcales_bacterium | PHO08_16185 | 15697..18828 | 2304002 | MAG:-DUF499-domain-containing-protein-[Methylococcales-bacterium] |
| MDD5320442.1 | DUF499_2.RM.like.systems | DUF499*->DUF3780->REase->?->PIN->REase->METHYLASE-> | 1041 | Bacteria>Pseudomonadota>Gammaproteobacteria>Methylococcales>unclassified_Methylococcales | GCA_028714175.1 | Methylococcales_bacterium | PHD43_07485 | 17817..20942 | 2304002 | MAG:-DUF499-domain-containing-protein-[Methylococcales-bacterium] |
| MDD5321594.1 | DUF499_2.RM.like.systems | <-HKD-DNase+SF1-SF2-Helicase<-?<-PIN<-?<-METHYLASE<-DUF3780<-DUF499*<-tRNA<-MerR-HTH | 1047 | Bacteria>Pseudomonadota>Gammaproteobacteria>Methylococcales>unclassified_Methylococcales | GCA_028714175.1 | Methylococcales_bacterium | PHD43_13460 | 8074..11217 | 2304002 | MAG:-DUF499-domain-containing-protein-[Methylococcales-bacterium] |
| WP_257264226.1 | DUF499_2.RM.like.systems | DUF499*->DUF3780->?->METHYLASE-p->METHYLASE->PIN->XIS-HTH-> | 740 | Bacteria>Pseudomonadota>Gammaproteobacteria>Oceanospirillales>Endozoicomonadaceae | GCF_024606365.1 | Endozoicomonas_sp._ONNA2 | KDK14_RS06570 | 43..2265 | 2828741 | FN3-associated-domain-containing-protein-[Endozoicomonas-sp.-ONNA2] |
| KAE8440079.1 | DUF499_2.RM.like.systems | <-DUF932<-?\|\|<-?\|\|SH3->DUF499*->DUF3780->METHYLASE->OLD-ABC->PIN->?->ParB+HNH-> | 1113 | Bacteria>Pseudomonadota>Gammaproteobacteria>Oceanospirillales>Halomonadaceae | GCA_009660035.1 | Halomonas_piezotolerans | F1978_02220 | 465970..469311 | 2609667 | ATP-binding-protein-[Halomonas-piezotolerans] |
| WP_086622422.1 | DUF499_2.RM.like.systems | DUF932->?->?->?->DUF499*->DUF3780->METHYLASE->OLD-ABC->PIN->?->HKD-DNase+SF1-SF2-Helicase+iREase+CTD-> | 1088 | Bacteria>Pseudomonadota>Gammaproteobacteria>Oceanospirillales>Halomonadaceae | GCF_002155145.1 | Kushneria_konosiri | B9G99_RS12330 | 2656346..2659612 | 698828 | DUF499-domain-containing-protein-[Kushneria-konosiri] |
| WP_110069957.1 | DUF499_2.RM.like.systems | TraH->?->TraG_N-><-?\|\|DUF4236+TPR->DUF499*->DUF3780->METHYLASE->OLD-ABC->PIN->?->ParB+HNH-> | 1088 | Bacteria>Pseudomonadota>Gammaproteobacteria>Oceanospirillales>Halomonadaceae | GCF_003182195.1 | Halomonas_sp._A11-A | DER72_RS10560 | 14381..17647 | 2183985 | DUF499-domain-containing-protein-[Halomonas-sp.-A11-A] |
| WP_147183730.1 | DUF499_2.RM.like.systems | <-PIN<-DUF433\|\|DUF4236+TPR->DUF499*->DUF3780->METHYLASE->OLD-ABC->PIN->?->ParB+HNH-> | 1088 | Bacteria>Pseudomonadota>Gammaproteobacteria>Oceanospirillales>Halomonadaceae | GCF_007954585.1 | Pistricoccus_aurantiacus | FGL86_RS05990 | 1230922..1234188 | 1883414 | DUF499-domain-containing-protein-[Pistricoccus-aurantiacus] |
| WP_163558801.1 | DUF499_2.RM.like.systems | <-ParB+HNH<-?<-PIN<-OLD-ABC<-METHYLASE<-DUF3780<-DUF499*<-SH3\|\|?-><-CHTH<-?\|\|?->DUF932-> | 1088 | Bacteria>Pseudomonadota>Gammaproteobacteria>Oceanospirillales>Halomonadaceae | GCF_010994375.1 | Halomonas_sp._NO4 | FEF15_RS08845 | 14465..17731 | 2484813 | DUF499-domain-containing-protein-[Halomonas-sp.-NO4] |
| WP_168444437.1 | DUF499_2.RM.like.systems | <-DUF932\|\|?->?-><-?\|\|SH3->DUF499*->DUF3780->METHYLASE->OLD-ABC->PIN->?->ParB+HNH-> | 1088 | Bacteria>Pseudomonadota>Gammaproteobacteria>Oceanospirillales>Halomonadaceae | GCF_012427705.1 | Halomonas_piezotolerans | GYM47_RS05600 | 1204301..1207567 | 2609667 | DUF499-domain-containing-protein-[Halomonas-piezotolerans] |
| WP_223136851.1 | DUF499_2.RM.like.systems | CHTH->?-><-?\|\|SH3->DUF499*->DUF3780->METHYLASE->OLD-ABC->PIN->?->ParB+HNH-> | 1088 | Bacteria>Pseudomonadota>Gammaproteobacteria>Oceanospirillales>Halomonadaceae | GCF_019915905.1 | Halomonas_sp._ANAO-440 | KZO25_RS10115 | 168620..171886 | 2861360 | DUF499-domain-containing-protein-[Halomonas-sp.-ANAO-440] |
| WP_234259726.1 | DUF499_2.RM.like.systems | <-ParB+HNH<-?<-PIN<-OLD-ABC<-METHYLASE<-DUF3780<-DUF499*<-SH3 | 1088 | Bacteria>Pseudomonadota>Gammaproteobacteria>Oceanospirillales>Halomonadaceae | GCF_021404355.1 | Halomonas_sp._MCCC_1A11062 | HOP58_RS04945 | 176236..179502 | 2733485 | DUF499-domain-containing-protein-[Halomonas-sp.-MCCC-1A11062] |
| WP_133483952.1 | DUF499_2.RM.like.systems | TraH->?->TraG_N-><-?<-CHTH\|\|DUF4236+TPR->DUF499*->DUF3780->METHYLASE->OLD-ABC->PIN->?->ParB+HNH-> | 1088 | Bacteria>Pseudomonadota>Gammaproteobacteria>Oceanospirillales>Halomonadaceae | GCF_024137965.1 | Halomonas | NHL60_RS01495 | 294251..297517 | 2745 | MULTISPECIES:-DUF499-domain-containing-protein-[Halomonas] |
| WP_200371316.1 | DUF499_2.RM.like.systems | <-DUF4236+TPR<-?\|\|?->?->?->?->DUF499*->DUF3780->METHYLASE->ParB+HNH->OLD-ABC->PIN-> | 1081 | Bacteria>Pseudomonadota>Gammaproteobacteria>Pseudomonadales>Marinobacteraceae | GCF_016597815.1 | Marinobacter_sp._DY40_1A1 | FFI61_RS14505 | 345780..349025 | 2583229 | DUF499-domain-containing-protein-[Marinobacter-sp.-DY40_1A1] |
| MXX05925.1 | DUF499_2.RM.like.systems | <-HEPN<-?\|\|MazE_antitoxin->PIN->DUF499*->DUF3780->?->?->?->METHYLASE->HKD-DNase+SF1-SF2-Helicase+iREase+CTD-> | 1000 | Bacteria>Pseudomonadota>Gammaproteobacteria>unclassified_Gammaproteobacteria | GCA_009836015.1 | Gammaproteobacteria_bacterium | F4Z71_04120 | 93593..96595 | 1913989 | DUF499-domain-containing-protein-[Gammaproteobacteria-bacterium] |
| MDE0038229.1 | DUF499_2.RM.like.systems | <-METHYLASE<-PIN<-MPTase<-DUF3780<-DUF499*\|\|?-><-HicA_toxin<-SF1-SF2-Helicase+REase<-SF1-SF2-Helicase+REase<-?<-PIN | 1015 | Bacteria>Pseudomonadota>Gammaproteobacteria>unclassified_Gammaproteobacteria | GCA_028818795.1 | Gammaproteobacteria_bacterium | OXU77_11860 | 4307..7354 | 1913989 | MAG:-DUF499-domain-containing-protein-[Gammaproteobacteria-bacterium] |
| MDE0444961.1 | DUF499_2.RM.like.systems | <-HKD-DNase+SF1-SF2-Helicase+iREase+CTD<-METHYLASE<-PIN<-MPTase<-DUF3780<-DUF499*\|\|?-><-HicA_toxin<-SF1-SF2-Helicase+REase<-SF1-SF2-Helicase+REase<-?<-PIN | 1015 | Bacteria>Pseudomonadota>Gammaproteobacteria>unclassified_Gammaproteobacteria | GCA_028821355.1 | Gammaproteobacteria_bacterium | OXL38_23000 | 9764..12811 | 1913989 | MAG:-DUF499-domain-containing-protein-[Gammaproteobacteria-bacterium] |
| MDE0478449.1 | DUF499_2.RM.like.systems | <-METHYLASE<-?<-?<-?<-DUF3780<-DUF499*<-PIN<-MazE_antitoxin\|\|?->HEPN-> | 1000 | Bacteria>Pseudomonadota>Gammaproteobacteria>unclassified_Gammaproteobacteria | GCA_028824825.1 | Gammaproteobacteria_bacterium | OXI13_02365 | 5014..8016 | 1913989 | MAG:-DUF499-domain-containing-protein-[Gammaproteobacteria-bacterium] |
| NCC04604.1 | DUF499_2.RM.like.systems | <-HKD-DNase+SF1-SF2-Helicase+iREase+CTD<-METHYLASE<-PIN<-DUF3780<-DUF499* | 1015 | Bacteria>Pseudomonadota>unclassified_Pseudomonadota | GCA_009929575.1 | Pseudomonadota_bacterium | EOM37_11320 | 14987..18034 | 1977087 | ATP-binding-protein-[Proteobacteria-bacterium] |
| MBS0593923.1 | DUF499_2.RM.like.systems | <-HKD-DNase+SF1-SF2-Helicase<-HicB<-PIN<-METHYLASE<-DUF3780<-DUF499*<-?\|\|?->ABhydrolase-><-MerR-HTH | 1044 | Bacteria>Pseudomonadota>unclassified_Pseudomonadota | GCA_018240325.1 | Pseudomonadota_bacterium | JSR84_10695 | 72059..75193 | 1977087 | DUF499-domain-containing-protein-[Proteobacteria-bacterium] |
| WP_194535999.1 | DUF499_2.RM.like.systems | <-HKD-DNase+SF1-SF2-Helicase+iREase+CTD<-PIN<-?<-?<-METHYLASE<-DUF3780<-DUF499*<-?\|\|?->?->PAIREDC-HTH-> | 1013 | Bacteria>PVC_group>Planctomycetota>Planctomycetia>Gemmatales | GCF_013694095.1 | Thermogemmata_fonticola | H0921_RS00060 | 23590..26631 | 2755323 | DUF499-domain-containing-protein-[Thermogemmata-fonticola] |
| MCK6447147.1 | DUF499_2.RM.like.systems | <-ABhydrolase\|\|Phage_integrase->?->?->XIS-HTH->DUF499*->DUF3780->METHYLASE->?->?->PIN->HKD-DNase+SF1-SF2-Helicase+iREase+CTD-> | 1029 | Bacteria>PVC_group>Planctomycetota>unclassified_Planctomycetota | GCA_023150475.1 | Planctomycetota_bacterium | L6Q99_12220 | 25859..28948 | 2026780 | ATP-binding-protein-[Planctomycetota-bacterium] |
| MCY2953678.1 | DUF499_2.RM.like.systems | DUF499*->DUF3780->PIN->METHYLASE-> | 977 | Bacteria>PVC_group>Planctomycetota>unclassified_Planctomycetota | GCA_026693085.1 | Planctomycetota_bacterium | NTU53_17115 | 20539..23472 | 2026780 | MAG:-DUF499-domain-containing-protein-[Planctomycetota-bacterium] |
| MCC5838834.1 | DUF499_2.RM.like.systems | <-Radical-SAM<-Classical-AAA<-PIN<-?<-DUF3780<-DUF499*<-?<-PIN<-RHH\|\|?-><-?<-XIS-HTH+RVT | 1043 | Bacteria>PVC_group>Verrucomicrobiota>Opitutae>Opitutales | GCA_020831585.1 | Opitutales_bacterium | JJT96_01820 | 59223..62354 | 2026772 | DUF499-domain-containing-protein-[Opitutales-bacterium] |
| MCH8539497.1 | DUF499_2.RM.like.systems | Radical-SAM->?->?->DUF499*->DUF3780->PIN->OLD-ABC+TOPRIM->METHYLASE->OLD-ABC->TOPRIM-> | 1021 | Bacteria>PVC_group>Verrucomicrobiota>Opitutae>Opitutales | GCA_022565675.1 | Opitutales_bacterium | LAT58_01865 | 10372..13437 | 2026772 | DUF499-domain-containing-protein-[Opitutales-bacterium] |
| AKC83089.1 | DUF499_2.RM.like.systems | SbcC+SbcC-><-?\|\|DUF499*->DUF3780->SMODS->SLATT->METHYLASE->HKD-DNase+SF1-SF2-Helicase+iREase+CTD->PIN-> | 988 | Bacteria>PVC_group>Verrucomicrobiota>unclassified_Verrucomicrobiota | GCA_000972765.1 | Verrucomicrobia_bacterium_IMCC26134 | IMCC26134_10435 | 2769967..2772933 | 1637999 | hypothetical-protein-IMCC26134_10435-[Verrucomicrobia-bacterium-IMCC26134] |
| TGM42334.1 | DUF499_2.RM.like.systems | DUF499*->DUF3780->MPTase->PIN->METHYLASE->HKD-DNase+SF1-SF2-Helicase-> | 1030 | Bacteria>Spirochaetota>Spirochaetia>Leptospirales>Leptospiraceae | GCA_004770805.1 | Leptospira_biflexa | EHQ92_16185 | 338..3430 | 172 | DUF499-domain-containing-protein-[Leptospira-biflexa] |
| MCB1144576.1 | DUF499_2.RM.like.systems | <-HKD-DNase+SF1-SF2-Helicase<-METHYLASE<-?<-PIN<-MPTase<-DUF3780<-DUF499*<-?<-PIN<-ParB+HNH<-rve+rve_3 | 1030 | Bacteria>Spirochaetota>Spirochaetia>Leptospirales>Leptospiraceae | GCA_020440865.1 | Leptospiraceae_bacterium | KDK54_20140 | 8602..11694 | 2026757 | ATP-binding-protein-[Leptospiraceae-bacterium] |
| WP_167881406.1 | DUF499_2.RM.like.systems | DUF499*->DUF3780->MPTase->PIN->METHYLASE->HKD-DNase+SF1-SF2-Helicase-> | 998 | Bacteria>Spirochaetota>Spirochaetia>Leptospirales>Leptospiraceae | GCF_004770325.1 | Leptospira_biflexa | EHQ92_RS16185 | 434..3430 | 172 | DUF499-domain-containing-protein-[Leptospira-biflexa] |
| WP_135740495.1 | DUF499_2.RM.like.systems | DUF499*->DUF3780->MPTase->PIN->METHYLASE->HKD-DNase+SF1-SF2-Helicase-> | 1030 | Bacteria>Spirochaetota>Spirochaetia>Leptospirales>Leptospiraceae | GCF_004770445.1 | Leptospira_bouyouniensis | EHQ99_RS04855 | 387..3479 | 2484911 | DUF499-domain-containing-protein-[Leptospira-bouyouniensis] |
| AEJ20205.1 | DUF499_2.RM.like.systems | <-HTH+UbE1\|\|DUF499*->DUF3780->OLD-ABC+TOPRIM->DUF772+DDE_Tnp_1_2->HKD-DNase+SF1-SF2-Helicase->HTH+MPTase->PIN-> | 1024 | Bacteria>Spirochaetota>Spirochaetia>Spirochaetales>Breznakiellaceae | GCA_000219725.1 | Gracilinema_caldarium_DSM_7334 | Spica_2078 | 2335967..2339041 | 744872 | hypothetical-protein-Spica_2078-[Gracilinema-caldarium-DSM-7334] |
| MDD3916530.1 | DUF499_2.RM.like.systems | DUF932->DUF499*->DUF3780->METHYLASE->HKD-DNase+SF1-SF2-Helicase+iREase+CTD->HKD-DNase+SF1-SF2-Helicase+iREase+CTD->VapB_antitoxin->PIN-> | 1079 | Bacteria>Synergistota>Synergistia>Synergistales>Synergistaceae | GCA_028699385.1 | Synergistaceae_bacterium | PHX00_05210 | 3629..6868 | 2053624 | MAG:-DUF499-domain-containing-protein-[Synergistaceae-bacterium] |
| MXX42195.1 | DUF499_2.RM.like.systems | MODE-HTH+PBPII-><-PIN<-?<-HKD-DNase+SF1-SF2-Helicase+iREase+CTD<-METHYLASE<-DUF3780<-DUF499*\|\|REase-> | 1088 | Bacteria>Terrabacteria_group>Actinomycetota>Acidimicrobiia>Acidimicrobiales | GCA_009836375.1 | Acidimicrobiales_bacterium | F4Z53_04000 | 47327..50593 | 2201156 | ATP-binding-protein-[Acidimicrobiales-bacterium] |
| MBE3519594.1 | DUF499_2.RM.like.systems | <-ABC-ATPase+SbcC<-PAIREDC-HTH\|\|tRNA-><-Phage_integrase\|\|MazE_antitoxin->PIN->DUF499*->DUF3780->METHYLASE->HKD-DNase+SF1-SF2-Helicase+iREase+CTD-> | 1008 | Bacteria>Terrabacteria_group>Bacillota>Candidatus_Fermentithermobacillia>Candidatus_Fermentithermobacillales | GCA_014896535.1 | Candidatus_Fermentithermobacillaceae_bacterium | IMW97_04750 | 107753..110779 | 3085339 | ATP-binding-protein-[Firmicutes-bacterium] |
| WP_207712783.1 | DUF499_2.RM.like.systems | PolB-NTase->Polbeta-><-Peptidase_S51<-HKD-DNase+SF1-SF2-Helicase+iREase+CTD<-METHYLASE<-DUF3780<-DUF499*<-PIN<-MetJarc-HTH<-REase | 1069 | Bacteria>Terrabacteria_group>Bacillota>Clostridia>Eubacteriales | - | Desulfofundulus_sp._TPOSR | G7K71_RS06320 | 1300314..1303523 | 2714340 | DUF499-domain-containing-protein-[Desulfofundulus-sp.-TPOSR] |

| Protein-ID | System | Gene-Neighbourhoods | Length | Taxonomy | Assembly-ID | Strain | Locus-Tag | Genomic-coordinates | TaxID | Fasta-defline |
|---|---|---|---|---|---|---|---|---|---|---|
| -- | | | | | | | | | | |
| -- SF1-SF2-Helicase+i REase +CTD+ REase | | | | | | | | | | |
| --(Type-1_BR-systems-like) | | | | | | | | | | |
| -- | | | | | | | | | | |
| PMP75322.1 | DUF499 _2.RM.like.systems | <- SF1-SF2-Helicase+i REase +CTD+ REase <- METHYLASE <- DUF3780 <- DUF499 * | 883 | Archaea>Candidatus_Thermoplasmatota>DHVE2_group>Aciduliprofundum>unclassified_Aciduliprofundum | GCA_002878135.1 | Aciduliprofundum_sp. | C0180_01660 | 6420..9071 | 2060325 | MAG:-hypothetical-protein-C0180_01660-[Aciduliprofundum-sp.] |
| MBD6956366.1 | DUF499 _2.RM.like.systems | <- SF1-SF2-Helicase+i REase +CTD+ REase <- METHYLASE <- DUF3780 <- DUF499 * | 890 | Archaea>Candidatus_Thermoplasmatota>Thermoplasmata>Thermoplasmatales>unclassified_Thermoplasmatales | GCA_014836265.1 | Thermoplasmatales_archaeon | F9Y92_07015 | 7804..10476 | 2268204 | DUF499 -domain-containing-protein-[Thermoplasmatales-archaeon] |
| PKL69081.1 | DUF499 _2.RM.like.systems | <- REase <-?<-EVE<- SF1-SF2-Helicase+i REase +CTD+ REase <- METHYLASE <- DUF3780 <- DUF499 *\|\| REase ->?->?-><- REase \|\|?->ACET-> | 1018 | Archaea>Euryarchaeota>Methanomada_group>Methanobacteria>Methanobacteriales | GCA_002839705.1 | Methanobacteriales_archaeon_HGW-Methanobacteriales-1 | CVV28_02915 | 553077..556133 | 2013815 | hypothetical-protein-CVV28_02915-[Methanobacteriales-archaeon-HGW-Methanobacteriales-1] |
| WP_211533304.1 | DUF499 _2.RM.like.systems | <- REase <-?<-EVE<- SF1-SF2-Helicase+i REase +CTD+ REase <- METHYLASE <- DUF3780 <- DUF499 *\|\| REase ->?->?->?-><-?<-TPR+TPR | 965 | Archaea>Euryarchaeota>Methanomada_group>Methanobacteria>Methanobacteriales | GCF_018141185.1 | Methanobacterium_alkalithermotolerans | HYG87_RS00570 | 112143..115040 | 2731220 | DUF499 -domain-containing-protein-[Methanobacterium-alkalithermotolerans] |
| WP_256532744.1 | DUF499 _2.RM.like.systems | REase -> SF1-SF2-Helicase+ REase -> DUF499 *-> DUF3780 -> METHYLASE ->SF1-SF2-Helicase->?-> HKD-DNase+SF1-SF2-Helicase +i REase +CTD->iSTAND-> | 986 | Archaea>Euryarchaeota>Stenosarchaea_group>Halobacteria>Natrialbales | GCF_024498195.1 | Halovivax_cerinus | NO366_RS02535 | 531191..534151 | 1487865 | DUF499 -domain-containing-protein-[Halovivax-cerinus] |
| WAH99918.1 | DUF499 _2.RM.like.systems | <- SF1-SF2-Helicase+i REase +CTD+ REase <- METHYLASE <-?<- DUF3780 <- DUF499 * | 978 | Archaea>Euryarchaeota>Stenosarchaea_group>Methanomicrobia>Methanosarcinales | - | Candidatus_Methanoperedens_sp. | OI716_00725 | 153031..155967 | 2032688 | MAG:- DUF499 -domain-containing-protein-[Candidatus-Methanoperedens-sp.] |
| MCZ7400325.1 | DUF499 _2.RM.like.systems | RelE-ParE->?->?-> DUF499 *-> DUF3780 ->PADR-HTH->HD-> METHYLASE ->NUDIX-> SF1-SF2-Helicase+i REase +CTD+ REase -> | 980 | Archaea>Euryarchaeota>Stenosarchaea_group>Methanomicrobia>Methanosarcinales | GCA_027460605.1 | Candidatus_Methanoperedens_sp. | O8C62_11750 | 3097..6039 | 2032688 | MAG:- DUF499 -domain-containing-protein-[Candidatus-Methanoperedens-sp.] |
| CAG0952062.1 | DUF499 _2.RM.like.systems | <-Primpol+D5N+D5ATPase\|\| DUF499 *-> DUF3780 ->PADR-HTH->HD-> METHYLASE -> SF1-SF2-Helicase+i REase +CTD+ REase ->ParB+HNH-> | 980 | Archaea>Euryarchaeota>Stenosarchaea_group>Methanomicrobia>Methanosarcinales | GCA_905339305.1 | Methanosarcinales_archaeon | METP3_00272 | 13443..16385 | 2250255 | hypothetical-protein-METP3_00272-[Methanosarcinales-archaeon] |
| WP_048043536.1 | DUF499 _2.RM.like.systems | <- SF1-SF2-Helicase+i REase +CTD+ REase <- METHYLASE <- DUF3780 <- DUF499 *<-?<-RelE-ParE | 983 | Archaea>Euryarchaeota>Stenosarchaea_group>Methanomicrobia>Methanosarcinales | GCF_000970185.1 | Methanosarcina_mazei | MSMAP_RS10730 | 2555385..2558336 | 2209 | DUF499 -domain-containing-protein-[Methanosarcina-mazei] |
| WP_221068977.1 | DUF499 _2.RM.like.systems | <-HTH\|\|?->?-><-?<-?\|\|Phage_integrase+ZnR-> DUF499 *-> DUF3780 -> METHYLASE -> SF1-SF2-Helicase+i REase +CTD+ REase -> | 983 | Archaea>Euryarchaeota>Stenosarchaea_group>Methanomicrobia>Methanosarcinales | GCF_019669945.1 | Methanosarcina_mazei | MmazTMA_RS08615 | 2012144..2015095 | 2209 | DUF499 -domain-containing-protein-[Methanosarcina-mazei] |
| WP_013749028.1 | DUF499 _2.RM.like.systems | <- SF1-SF2-Helicase+i REase +CTD+ REase <- METHYLASE <- DUF3780 <- DUF499 *<-ORC-CDC6+ REase <-?<-?<-?<-LRP-HTH<-Peptidase | 1059 | Archaea>Euryarchaeota>Thermococci>Thermococcales>Thermococcaceae | GCF_000211475.1 | Pyrococcus_sp._NA2 | PNA2_RS07895 | 1443844..1447023 | 342949 | ATP-binding-protein-[Pyrococcus-sp.-NA2] |
| MBN1683203.1 | DUF499 _2.RM.like.systems | DUF499 *-> DUF3780 -> METHYLASE -> SF1-SF2-Helicase+i REase +CTD+ REase -> | 678 | Archaea>TACK_group>Candidatus_Bathyarchaeota>unclassified_Candidatus_Bathyarchaeota | GCA_016930055.1 | Candidatus_Bathyarchaeota_archaeon | JW865_06595 | 1..2037 | 2026714 | hypothetical-protein-[Candidatus-Bathyarchaeota-archaeon] |
| MCW3988812.1 | DUF499 _2.RM.like.systems | <-URI-DNase\|\|?->35exo->?-><-SF1-SF2-Helicase<-?\|\| DUF499 *-> DUF3780 -> METHYLASE -> SF1-SF2-Helicase+i REase +CTD+ REase ->?->?->HIN-HTH-> | 885 | Archaea>TACK_group>Candidatus_Bathyarchaeota>unclassified_Candidatus_Bathyarchaeota | GCA_026014875.1 | Candidatus_Bathyarchaeota_archaeon | NWE88_01905 | 126678..129335 | 2026714 | MAG:- DUF499 -domain-containing-protein-[Candidatus-Bathyarchaeota-archaeon] |
| RWX74207.1 | DUF499 _2.RM.like.systems | PIN->?-><-?\|\| DUF499 *-> DUF3780 -> METHYLASE -> SF1-SF2-Helicase+i REase +CTD+ REase ->?->?-><-PIN | 892 | Archaea>TACK_group>Candidatus_Verstraetearchaeota>Candidatus_Methanomethylicia>Candidatus_Methanomethylicales | GCA_004028775.1 | Candidatus_Methanosuratincola_subterraneus | Metus_0232 | 214945..217623 | 2593994 | hypothetical-protein-Metus_0232-[Candidatus-Methanosuratincola-subterraneus] |
| GIU72777.1 | DUF499 _2.RM.like.systems | DUF488-> DUF499 *-> DUF3780 -><-PIN<-?<-?\|\| METHYLASE -> SF1-SF2-Helicase+i REase +CTD+ REase -> | 894 | Archaea>TACK_group>Nitrososphaerota>Nitrososphaeria>Candidatus_Nitrosocaldales | GCA_026002835.1 | Candidatus_Nitrosocaldaceae_archaeon | KatS3mg003_2256 | 14146..16830 | 2838776 | MAG:-hypothetical-protein-KatS3mg003_2256-[Candidatus-Nitrosocaldaceae-archaeon] |
| MDE0525348.1 | DUF499 _2.RM.like.systems | <-MarR-HTH\|\| DUF499 *-> DUF3780 -> METHYLASE -> SF1-SF2-Helicase+i REase +CTD+ REase -> | 885 | Archaea>TACK_group>Nitrososphaerota>unclassified_Nitrososphaerota | GCA_028823795.1 | Nitrososphaerota_archaeon | OXI27_01935 | 36993..39650 | 2026795 | MAG:- DUF499 -domain-containing-protein-[Thaumarchaeota-archaeon] |
| WP_054837388.1 | DUF499 _2.RM.like.systems | DUF499 *-> DUF3780 -> METHYLASE -p-> METHYLASE -p-> SF1-SF2-Helicase+i REase +CTD+ REase -p-> REase -> | 877 | Archaea>TACK_group>Thermoproteota>Thermoprotei>Sulfolobales | GCF_001315825.1 | Metallosphaera_hakonensis | JCM8857_RS10980 | 481..3114 | 79601 | hypothetical-protein-[Metallosphaera-hakonensis] |
| MBI4750004.1 | DUF499 _2.RM.like.systems | TUDOR+ SF1-SF2-Helicase+i REase +CTD+ REase ->HicB_lk_antitox-> REase -> METHYLASE ->HicB_lk_antitox->HicA_toxin-> DUF499 *->ABhydrolase-><-?<-?<-?<-BirA-HTH | 939 | Bacteria>Acidobacteriota>unclassified_Acidobacteriota | GCA_016208495.1 | Acidobacteriota_bacterium | HY774_16075 | 69401..72220 | 1978231 | ATP-binding-protein-[Acidobacteria-bacterium] |
| WP_088441309.1 | DUF499 _2.RM.like.systems | <- HKD-DNase+SF1-SF2-Helicase +i REase +CTD<- METHYLASE <- DUF3780 <- DUF499 *<-HHH_4+ SF1-SF2-Helicase+ REase <- REase +35exo+Beta-Barrel+SF1-SF2-Helicase<-Ntox49\|\|HTH+WYL-> | 1022 | Bacteria>Pseudomonadota>Alphaproteobacteria>Sphingomonadales>Sphingomonadaceae | GCF_014645635.1 | Sphingopyxis_bauzanensis | CDQ92_RS10605 | 849090..852158 | 651663 | DUF499 -domain-containing-protein-[Sphingopyxis-bauzanensis] |
| NLF73182.1 | DUF499 _2.RM.like.systems | TUDOR+ SF1-SF2-Helicase+i REase +CTD+ REase ->SF1-SF2-Helicase-> METHYLASE ->?-> DUF499 *->?->ASCH+ DUF3780 -><-?\|\|?->XIS-HTH-> | 975 | Bacteria>PVC_group>Planctomycetota>Planctomycetia>Pirellulales | GCA_012515235.1 | Candidatus_Anammoximicrobium_sp. | GX575_29450 | 8853..11780 | 2699742 | ATP-binding-protein-[Candidatus-Anammoximicrobium-sp.] |
| WP_025333200.1 | DUF499 _2.RM.like.systems | HTH+WYL->?->?->?->Exonuc_V_gamma+ REase -> SF1-SF2-Helicase+ REase -> DUF499 *-> DUF3780 -> METHYLASE -> HKD-DNase+SF1-SF2-Helicase ->?-> REase ->pPIWI+pPIWI-> | 1022 | Bacteria>Terrabacteria_group>Bacillota>Bacilli>Bacillales | GCF_000612505.1 | Paenibacillus_sabinae | PSAB_RS03435 | 744096..747164 | 365617 | DUF499 -domain-containing-protein-[Paenibacillus-sabinae] |
| WP_042210529.1 | DUF499 _2.RM.like.systems | TPR->?->?->HTH+WYL->Exonuc_V_gamma+ REase -> SF1-SF2-Helicase+ REase -> DUF499 *-> DUF3780 -> METHYLASE -> HKD-DNase+SF1-SF2-Helicase ->Calcineurin->SbcC+SbcC-> | 1022 | Bacteria>Terrabacteria_group>Bacillota>Bacilli>Bacillales | GCF_000758665.1 | Paenibacillus_borealis | PBOR_RS03715 | 849881..852949 | 160799 | DUF499 -domain-containing-protein-[Paenibacillus-borealis] |
| WP_274435612.1 | DUF499 _2.RM.like.systems | <-RVT+HNH<-DUF4277+DDE_Tnp_1<-RVT<- HKD-DNase+SF1-SF2-Helicase <- METHYLASE <- DUF3780 <- DUF499 *<-TUDOR+ SF1-SF2-Helicase+i REase +CTD+ REase -p | 1058 | Bacteria>Terrabacteria_group>Bacillota>Bacilli>Bacillales | GCF_028751195.1 | Alicyclobacillus_sp._ALC3 | JC200_RS07360 | 1554106..1557282 | 2796143 | DUF499 -domain-containing-protein-[Alicyclobacillus-sp.-ALC3] |
| MBE7435441.1 | DUF499 _2.RM.like.systems | <-URI-DNase<-ParB+HNH<- DUF499 -p<-ASCH+ DUF3780 <-?<-?<- DUF499 *<-?<- METHYLASE <-TUDOR+ SF1-SF2-Helicase+i REase +CTD+ REase | 1032 | Bacteria>Terrabacteria_group>Chloroflexota>Anaerolineae>Anaerolineales | GCA_015075725.1 | Anaerolineales_bacterium | HS100_16115 | 3331332..3334430 | 2073117 | DUF499 -domain-containing-protein-[Anaerolineales-bacterium] |
| BCX11788.1 | DUF499 _2.RM.like.systems | SF1-SF2-Helicase-><- REase \|\|TUDOR+ SF1-SF2-Helicase+i REase +CTD+ REase -p-> DUF499 *-> DUF3780 -> METHYLASE -> HKD-DNase+SF1-SF2-Helicase ->HEPN+Mrr_N->HEPN+ REase -> | 1027 | Bacteria>Terrabacteria_group>Cyanobacteriota/Melainabacteria_group>Cyanobacteriota>Cyanophyceae | GCA_025999095.1 | Thermosynechococcus_sp. | KatS3mg067_0726 | 755896..758979 | 2814275 | MAG:-hypothetical-protein-KatS3mg067_0726-[Thermosynechococcus-sp.] |
| WP_045216075.1 | DUF499 _2.RM.like.systems | TUDOR+ SF1-SF2-Helicase+i REase +CTD+ REase ->?->PIN->?->?-> DUF499 *-> DUF3780 -> METHYLASE ->PolB->HEPN-> | 1078 | Bacteria>Thermodesulfobacteriota>Desulfovibrionia>Desulfovibrionales>Desulfonatronovibrionaceae | GCF_000934755.1 | Desulfonatronovibrio_magnus | LZ23_RS16895 | 146926..150162 | 698827 | DUF499 -domain-containing-protein-[Desulfonatronovibrio-magnus] |
| -- | | | | | | | | | | |
| -- HTH+ REase | | | | | | | | | | |
| -- | | | | | | | | | | |
| TAK42618.1 | DUF499 _2.RM.like.systems | BirA-HTH->?-> w HTH+ REase -> METHYLASE -p-> DUF499 *-> DUF3780 ->ParB+HNH->OLD-ABC->RelE-ParE-> METHYLASE -p-> | 1024 | Bacteria>FCB_group>Bacteroidota/Chlorobiota_group>Bacteroidota>Saprospiria | GCA_004297645.1 | Saprospiraceae_bacterium | EPO28_07065 | 17499..20573 | 2202734 | DUF499 -domain-containing-protein-[Saprospiraceae-bacterium] |
| WP_233943241.1 | DUF499 _2.RM.like.systems | <- HKD-DNase+SF1-SF2-Helicase <- METHYLASE <- DUF499 -p<- DUF499 *\|\| w HTH+ REase ->\|\|?-><-?<-PIN+OST-HTH<-?<-Arm-DNA-bind_3+Phage_integrase | 526 | Bacteria>Pseudomonadota>Alphaproteobacteria>Hyphomicrobiales>Rhizobiaceae | GCF_021391595.1 | Rhizobium_leguminosarum | LZK78_RS32110 | 712264..713844 | 384 | DUF499 -domain-containing-protein-[Rhizobium-leguminosarum] |
| QTL37180.1 | DUF499 _2.RM.like.systems | <-VirD4-TraG+VirB4-FtsK<-HDRNase<- w HTH+ REase <- HKD-DNase+SF1-SF2-Helicase <- METHYLASE <- DUF3780 +RAMA<- DUF499 *<-RHH_1<-35exo\|\|?->?->Classical-AAA->Peptidase_S8-> | 1022 | Bacteria>Pseudomonadota>Gammaproteobacteria>Alteromonadales>Pseudoalteromonadaceae | GCA_017742995.1 | Pseudoalteromonas_viridis | J5X90_01860 | 434218..437286 | 339617 | DUF499 -domain-containing-protein-[Pseudoalteromonas-viridis] |
| WP_036957217.1 | DUF499 _2.RM.like.systems | <-HNH<-ABC-ATPase<- w HTH+ REase <- HKD-DNase+SF1-SF2-Helicase <- METHYLASE <- DUF3780 +RAMA<- DUF499 *<-RHH<-?<-XIS-HTH<-?<-XIS-HTH\|\|JAB-> | 1041 | Bacteria>Pseudomonadota>Gammaproteobacteria>Alteromonadales>Pseudoalteromonadaceae | GCF_913044265.1 | Pseudoalteromonas | QL994_RS18060 | 15258..18383 | 53246 | DUF499 -domain-containing-protein-[Pseudoalteromonas-sp.-A2] |
| WP_230608232.1 | DUF499 _2.RM.like.systems | 35exo->RHH_1-> DUF499 *-> DUF3780 +RAMA-> METHYLASE -> HKD-DNase+SF1-SF2-Helicase -> w HTH+ REase ->HDRNase->VirD4-TraG+VirB4-FtsK-> | 1041 | Bacteria>Pseudomonadota>Gammaproteobacteria>Enterobacterales>Budviciaceae | GCF_021010575.1 | Limnobaculum_eriocheiris | LPW36_RS04285 | 61896..65021 | 2897391 | DUF499 -domain-containing-protein-[Jinshanibacter-sp.-LJY008] |
| WP_200893021.1 | DUF499 _2.RM.like.systems | Arm-DNA-bind_3+Phage_integrase->RHH-> DUF499 *-> DUF3780 +RAMA-> METHYLASE -> HKD-DNase+SF1-SF2-Helicase -> w HTH+ REase -><-?\|\|HTH_Tnp_1+HTH_21+rve+rve_2-> | 1022 | Bacteria>Pseudomonadota>Gammaproteobacteria>Enterobacterales>Enterobacteriaceae | - | Enterobacter_hormaechei | I4A92_000735 | 150340..153408 | 158836 | DUF499 -domain-containing-protein-[Enterobacter-hormaechei] |
| WP_218968419.1 | DUF499 _2.RM.like.systems | <- w HTH+ REase <- HKD-DNase+SF1-SF2-Helicase <- METHYLASE <- DUF3780 +RAMA<- DUF499 *<-RHH<-Arm-DNA-bind_3+Phage_integrase | 1022 | Bacteria>Pseudomonadota>Gammaproteobacteria>Enterobacterales>Enterobacteriaceae | - | Enterobacter_hormaechei | SAMEA2053862_03698 | 27017..30085 | 158836 | DUF499 -domain-containing-protein-[Enterobacter-hormaechei] |
| KDL80228.1 | DUF499 _2.RM.like.systems | <-ADH_N+ADH_zinc_N<-?<-?\|\|tRNA->Arm-DNA-bind_3+Phage_integrase->RHH-> DUF499 *-> DUF3780 +RAMA-> METHYLASE -> HKD-DNase+SF1-SF2-Helicase -> w HTH+ REase -> | 1022 | Bacteria>Pseudomonadota>Gammaproteobacteria>Enterobacterales>Enterobacteriaceae | GCA_000694675.1 | Klebsiella_pneumoniae_MGH_72 | AD98_04564 | 185426..188494 | 1438795 | hypothetical-protein-AD98_04564-[Klebsiella-pneumoniae-MGH-72] |
| ARA42377.1 | DUF499 _2.RM.like.systems | SF1-SF2-Helicase->?-><- w HTH+ REase <- HKD-DNase+SF1-SF2-Helicase <- METHYLASE <- DUF3780 +RAMA<- DUF499 *<-DDE_Tnp_1<- DUF499 -p<-RHH<-Arm-DNA-bind_3+Phage_integrase | 984 | Bacteria>Pseudomonadota>Gammaproteobacteria>Enterobacterales>Enterobacteriaceae | GCA_002055765.1 | Klebsiella_pneumoniae | AM364_19165 | 3781580..3784534 | 573 | AAA-family-ATPase-[Klebsiella-pneumoniae] |
| PMD76486.1 | DUF499 _2.RM.like.systems | <-ADH_N+ADH_zinc_N<-?<-?\|\|tRNA->Arm-DNA-bind_3+Phage_integrase->RHH-> DUF499 *-> DUF3780 +RAMA-> METHYLASE -> HKD-DNase+SF1-SF2-Helicase -> w HTH+ REase -><-?\|\|<-XIS-HTH | 1041 | Bacteria>Pseudomonadota>Gammaproteobacteria>Enterobacterales>Enterobacteriaceae | GCA_002872205.1 | Klebsiella_pneumoniae | A8A10_21890 | 132525..135650 | 573 | AAA-family-ATPase-[Klebsiella-pneumoniae] |
| ECA3794685.1 | DUF499 _2.RM.like.systems | <-DnaB+SMS-KaiC<-?<-ParA-Soj-PloopNTPase\|\|Arm-DNA-bind_3+Phage_integrase->RHH-> DUF499 *-> DUF3780 +RAMA-> METHYLASE -> HKD-DNase+SF1-SF2-Helicase -> w HTH+ REase -> | 1041 | Bacteria>Pseudomonadota>Gammaproteobacteria>Enterobacterales>Enterobacteriaceae | GCA_007378915.1 | Salmonella_enterica_subsp._enterica_serovar_Aqua | EKG95_23275 | 13505..16630 | 1302615 | DUF499 -domain-containing-protein-[Salmonella-enterica-subsp.-enterica-serovar-Aqua] |
| EFA6413129.1 | DUF499 _2.RM.like.systems | <-HIN-HTH<- w HTH+ REase <- HKD-DNase+SF1-SF2-Helicase <- METHYLASE <- DUF3780 +RAMA<- DUF499 *<-RHH_5\|\|<-?<-Aminotran_1_2\|\|<-Arm-DNA-bind_3+Phage_integrase<-tRNA\|\|ADH_N+ADH_zinc_N->FtsK+HerA+A32-> | 1041 | Bacteria>Pseudomonadota>Gammaproteobacteria>Enterobacterales>Enterobacteriaceae | GCA_012256865.1 | Escherichia_coli | CIW06_000361 | 54690..57815 | 562 | DUF499 -domain-containing-protein-[Escherichia-coli] |
| MBK1516290.1 | DUF499 _2.RM.like.systems | <-ADH_N+ADH_zinc_N<-?<-?\|\|tRNA->Arm-DNA-bind_3+Phage_integrase->RHH-> DUF499 *-> DUF3780 +RAMA-> METHYLASE -> HKD-DNase+SF1-SF2-Helicase -> w HTH+ REase -><-HTH+MPTase | 1022 | Bacteria>Pseudomonadota>Gammaproteobacteria>Enterobacterales>Enterobacteriaceae | GCA_014235385.1 | Klebsiella_variicola | H8S48_22830 | 314255..317323 | 244366 | DUF499 -domain-containing-protein-[Klebsiella-variicola] |
| HAV2294932.1 | DUF499 _2.RM.like.systems | HTH+MPTase-><- w HTH+ REase <- HKD-DNase+SF1-SF2-Helicase <- METHYLASE <- DUF3780 +RAMA<- DUF499 *<-RHH<-Arm-DNA-bind_3+Phage_integrase<-tRNA\|\|ADH_N+ADH_zinc_N->FtsK+HerA+A32-> | 1041 | Bacteria>Pseudomonadota>Gammaproteobacteria>Enterobacterales>Enterobacteriaceae | GCA_016445565.1 | Citrobacter_koseri | JHK52_001290 | 177195..180320 | 545 | DUF499 -domain-containing-protein-[Citrobacter-koseri] |
| MBJ8391833.1 | DUF499 _2.RM.like.systems | <-HIN-HTH<- w HTH+ REase <- HKD-DNase+SF1-SF2-Helicase <- METHYLASE <- DUF3780 +RAMA<- DUF499 *<-RHH<-Arm-DNA-bind_3+Phage_integrase<-tRNA\|\|ADH_N+ADH_zinc_N-> | 1022 | Bacteria>Pseudomonadota>Gammaproteobacteria>Enterobacterales>Enterobacteriaceae | GCA_016501115.1 | Citrobacter_cronae | I6M80_16495 | 16784..19852 | 1748967 | DUF499 -domain-containing-protein-[Citrobacter-cronae] |
| MBK2656985.1 | DUF499 _2.RM.like.systems | HTH+MPTase-><- w HTH+ REase <- HKD-DNase+SF1-SF2-Helicase <- METHYLASE <- DUF3780 +RAMA<- DUF499 *<-RHH<-Arm-DNA-bind_3+Phage_integrase<-tRNA\|\|?->?->ADH_N+ADH_zinc_N-> | 1039 | Bacteria>Pseudomonadota>Gammaproteobacteria>Enterobacterales>Enterobacteriaceae | GCA_016614755.1 | Klebsiella_pneumoniae | FKL12_18780 | 12401..15520 | 573 | DUF499 -domain-containing-protein-[Klebsiella-pneumoniae] |
| MBK5711249.1 | DUF499 _2.RM.like.systems | HTH+MPTase-><- w HTH+ REase <- HKD-DNase+SF1-SF2-Helicase <- METHYLASE <- DUF3780 +RAMA<- DUF499 *<-RHH<-Arm-DNA-bind_3+Phage_integrase<-tRNA\|\|?->?->ADH_N+ADH_zinc_N-> | 1022 | Bacteria>Pseudomonadota>Gammaproteobacteria>Enterobacterales>Enterobacteriaceae | GCA_016651545.1 | Klebsiella_pneumoniae | JKA42_15120 | 66258..69326 | 573 | DUF499 -domain-containing-protein-[Klebsiella-pneumoniae] |
| HBB8152959.1 | DUF499 _2.RM.like.systems | DEDD_Tnp_IS110+Transposase_20->?->RHH_1-> DUF499 -p-> DUF499 *-> DUF3780 +RAMA-> METHYLASE -> HKD-DNase+SF1-SF2-Helicase -> w HTH+ REase ->HIN-HTH-> | 958 | Bacteria>Pseudomonadota>Gammaproteobacteria>Enterobacterales>Enterobacteriaceae | GCA_018015815.1 | Escherichia_coli | J8710_004813 | 1587..4463 | 562 | DUF499 -domain-containing-protein-[Escherichia-coli] |
| HBC5231502.1 | DUF499 _2.RM.like.systems | SF1-SF2-Helicase->?-><- w HTH+ REase <- HKD-DNase+SF1-SF2-Helicase <- METHYLASE <- DUF3780 +RAMA<- DUF499 *<- DUF499 -p<-RHH<-Arm-DNA-bind_3+Phage_integrase<-tRNA\|\|ADH_N+ADH_zinc_N-> | 959 | Bacteria>Pseudomonadota>Gammaproteobacteria>Enterobacterales>Enterobacteriaceae | GCA_018155305.1 | Klebsiella_oxytoca | KEU23_000991 | 23793..26672 | 571 | DUF499 -domain-containing-protein-[Klebsiella-oxytoca] |
| EHN8715928.1 | DUF499 _2.RM.like.systems | Arm-DNA-bind_3+Phage_integrase->RHH-> DUF499 *-> DUF3780 +RAMA-> METHYLASE -> HKD-DNase+SF1-SF2-Helicase -> w HTH+ REase -><-?\|\|HTH_Tnp_1+HTH_21+rve+rve_2-> | 1022 | Bacteria>Pseudomonadota>Gammaproteobacteria>Enterobacterales>Enterobacteriaceae | GCA_018454885.1 | Enterobacter_hormaechei | I4A92_000735 | 150340..153408 | 158836 | DUF499 -domain-containing-protein-[Enterobacter-hormaechei] |
| HBE2428354.1 | DUF499 _2.RM.like.systems | RHH_1-> DUF499 *-> DUF3780 +RAMA-> METHYLASE -> HKD-DNase+SF1-SF2-Helicase -> w HTH+ REase -> | 1022 | Bacteria>Pseudomonadota>Gammaproteobacteria>Enterobacterales>Enterobacteriaceae | GCA_018545885.1 | Escherichia_coli | KL339_004657 | 1620..4688 | 562 | DUF499 -domain-containing-protein-[Escherichia-coli] |
| MBV4586514.1 | DUF499 _2.RM.like.systems | ZnR->ZnR->HTH_Tnp_1+HTH_21+rve+rve_3->?->RHH_1-> DUF499 *-> DUF3780 +RAMA-> METHYLASE -> HKD-DNase+SF1-SF2-Helicase -> w HTH+ REase ->HIN-HTH-> | 1022 | Bacteria>Pseudomonadota>Gammaproteobacteria>Enterobacterales>Enterobacteriaceae | GCA_019145775.1 | Escherichia_coli | I2922_14300 | 166455..169523 | 562 | DUF499 -domain-containing-protein-[Escherichia-coli] |
| HBM8437234.1 | DUF499 _2.RM.like.systems | <-HIN-HTH<- w HTH+ REase <- HKD-DNase+SF1-SF2-Helicase <- METHYLASE <- DUF3780 +RAMA<- DUF499 *<-RHH_5\|\|<-?<-Aminotran_1_2 | 1022 | Bacteria>Pseudomonadota>Gammaproteobacteria>Enterobacterales>Enterobacteriaceae | GCA_021425605.1 | Escherichia_coli | LOV60_003801 | 18779..21847 | 562 | DUF499 -domain-containing-protein-[Escherichia-coli] |
| HBQ8501082.1 | DUF499 _2.RM.like.systems | SF1-SF2-Helicase->?-><- w HTH+ REase <- HKD-DNase+SF1-SF2-Helicase <- METHYLASE <- DUF3780 +RAMA<- DUF499 *<-RHH<-Arm-DNA-bind_3+Phage_integrase | 1022 | Bacteria>Pseudomonadota>Gammaproteobacteria>Enterobacterales>Enterobacteriaceae | GCA_021895715.1 | Klebsiella_pneumoniae | L9A44_004853 | 54041..57109 | 573 | DUF499 -domain-containing-protein-[Klebsiella-pneumoniae] |
| HBX9967586.1 | DUF499 _2.RM.like.systems | HTH+MPTase-><- w HTH+ REase <- HKD-DNase+SF1-SF2-Helicase <- METHYLASE <- DUF3780 +RAMA<- DUF499 *<-RHH<-Arm-DNA-bind_3+Phage_integrase<-tRNA\|\|?->?->ADH_N+ADH_zinc_N-> | 1041 | Bacteria>Pseudomonadota>Gammaproteobacteria>Enterobacterales>Enterobacteriaceae | GCA_022266445.1 | Klebsiella_variicola | MIM73_23195 | 35789..38914 | 244366 | DUF499 -domain-containing-protein-[Klebsiella-variicola] |
| HBY8019970.1 | DUF499 _2.RM.like.systems | SF1-SF2-Helicase->?-><- w HTH+ REase <- HKD-DNase+SF1-SF2-Helicase <- METHYLASE <- DUF3780 +RAMA<- DUF499 * | 982 | Bacteria>Pseudomonadota>Gammaproteobacteria>Enterobacterales>Enterobacteriaceae | GCA_022301155.1 | Klebsiella_pneumoniae | MI822_22070 | 54206..57154 | 573 | DUF499 -domain-containing-protein-[Klebsiella-pneumoniae] |
| EIY1898012.1 | DUF499 _2.RM.like.systems | SF1-SF2-Helicase->?-><- w HTH+ REase <- HKD-DNase+SF1-SF2-Helicase <- METHYLASE <- DUF3780 +RAMA<- DUF499 *<-RHH<-Arm-DNA-bind_3+Phage_integrase<-tRNA\|\|?->?->ADH_N+ADH_zinc_N-> | 1022 | Bacteria>Pseudomonadota>Gammaproteobacteria>Enterobacterales>Enterobacteriaceae | GCA_022464035.1 | Klebsiella_pneumoniae | MMH51_002463 | 54491..57559 | 573 | DUF499 -domain-containing-protein-[Klebsiella-pneumoniae] |
| HBZ7479225.1 | DUF499 _2.RM.like.systems | <-ADH_N+ADH_zinc_N<-?<-?\|\|tRNA->Arm-DNA-bind_3+Phage_integrase->RHH-> DUF499 *-> DUF3780 +RAMA-> METHYLASE -> HKD-DNase+SF1-SF2-Helicase -> w HTH+ REase -><-HTH+MPTase | 1022 | Bacteria>Pseudomonadota>Gammaproteobacteria>Enterobacterales>Enterobacteriaceae | GCA_022497945.1 | Klebsiella_pneumoniae | MM066_002839 | 198069..201137 | 573 | DUF499 -domain-containing-protein-[Klebsiella-pneumoniae] |
| MCM7420480.1 | DUF499 _2.RM.like.systems | SF1-SF2-Helicase->?-><- w HTH+ REase <- HKD-DNase+SF1-SF2-Helicase <- METHYLASE <- DUF3780 +RAMA<- DUF499 *<-RHH<-Arm-DNA-bind_3+Phage_integrase | 1041 | Bacteria>Pseudomonadota>Gammaproteobacteria>Enterobacterales>Enterobacteriaceae | GCA_023751025.1 | Enterobacter_hormaechei | M8R61_11745 | 350884..354009 | 158836 | DUF499 -domain-containing-protein-[Enterobacter-hormaechei] |
| HCD9026459.1 | DUF499 _2.RM.like.systems | HTH+MPTase-><-HNH+SAVED<- w HTH+ REase <- HKD-DNase+SF1-SF2-Helicase <- METHYLASE <- DUF3780 +RAMA<- DUF499 *<-RHH<-Arm-DNA-bind_3+Phage_integrase<-tRNA\|\|wHTH+HNH->SAM-methylase-> | 1022 | Bacteria>Pseudomonadota>Gammaproteobacteria>Enterobacterales>Enterobacteriaceae | GCA_023760145.1 | Klebsiella_oxytoca | NEF91_003870 | 43874..46942 | 571 | DUF499 -domain-containing-protein-[Klebsiella-oxytoca] |
| HCJ8634260.1 | DUF499 _2.RM.like.systems | <-ADH_N+ADH_zinc_N<-?<-?\|\|tRNA->RHH-> DUF499 *-> DUF3780 +RAMA-> METHYLASE -> HKD-DNase+SF1-SF2-Helicase -> w HTH+ REase -><-?<-SF1-SF2-Helicase | 1022 | Bacteria>Pseudomonadota>Gammaproteobacteria>Enterobacterales>Enterobacteriaceae | GCA_024674695.1 | Klebsiella_pneumoniae | NV748_003018 | 174251..177319 | 573 | DUF499 -domain-containing-protein-[Klebsiella-pneumoniae] |
| HCM9136403.1 | DUF499 _2.RM.like.systems | ZnR->ZnR-><- w HTH+ REase <- HKD-DNase+SF1-SF2-Helicase <- METHYLASE <- DUF3780 +RAMA<- DUF499 *<-RHH<-Arm-DNA-bind_3+Phage_integrase | 1022 | Bacteria>Pseudomonadota>Gammaproteobacteria>Enterobacterales>Enterobacteriaceae | GCA_025209595.1 | Enterobacter_cloacae_subsp._cloacae | N5L02_000891 | 375093..378161 | 336306 | DUF499 -domain-containing-protein-[Enterobacter-cloacae-subsp.-cloacae] |
| UZF63336.1 | DUF499 _2.RM.like.systems | <-HIN-HTH<- w HTH+ REase <- HKD-DNase+SF1-SF2-Helicase <- METHYLASE <- DUF3780 +RAMA<- DUF499 -p<- DUF499 *<- DUF499 -p<-RHH_5<-?<-DEDD_Tnp_IS110+Transposase_20<-HTH_Tnp_1+HTH_21+rve+rve_3\|\|<-Arm-DNA-bind_3+Phage_integrase | 355 | Bacteria>Pseudomonadota>Gammaproteobacteria>Enterobacterales>Enterobacteriaceae | GCA_026016605.1 | Escherichia_coli | OM434_11360 | 2373962..2375029 | 562 | ATP-binding-protein-[Escherichia-coli] |
| HCW2948669.1 | DUF499 _2.RM.like.systems | <-HIN-HTH<- w HTH+ REase <- HKD-DNase+SF1-SF2-Helicase <- METHYLASE <- DUF3780 +RAMA<- DUF499 *<-RHH_1 | 1022 | Bacteria>Pseudomonadota>Gammaproteobacteria>Enterobacterales>Enterobacteriaceae | GCA_026548625.1 | Escherichia_coli | OXB36_004615 | 17018..20086 | 562 | TPA:- DUF499 -domain-containing-protein-[Escherichia-coli] |
| HCW3139350.1 | DUF499 _2.RM.like.systems | <-FtsK+HerA+A32<-ADH_N+ADH_zinc_N\|\|tRNA->Arm-DNA-bind_3+Phage_integrase->RHH-> DUF499 *-> DUF3780 +RAMA-> METHYLASE -> HKD-DNase+SF1-SF2-Helicase -> w HTH+ REase -><-HTH+MPTase | 1022 | Bacteria>Pseudomonadota>Gammaproteobacteria>Enterobacterales>Enterobacteriaceae | GCA_026549635.1 | Citrobacter_freundii | OXC88_001869 | 152209..155277 | 546 | TPA:- DUF499 -domain-containing-protein-[Citrobacter-freundii] |
| EKK5919799.1 | DUF499 _2.RM.like.systems | Arm-DNA-bind_3+Phage_integrase->RHH-> DUF499 *-> DUF3780 +RAMA-> METHYLASE -> HKD-DNase+SF1-SF2-Helicase -> w HTH+ REase -><-?<-SF1-SF2-Helicase | 1022 | Bacteria>Pseudomonadota>Gammaproteobacteria>Enterobacterales>Enterobacteriaceae | GCA_028182685.1 | Enterobacter_hormaechei | POC94_000676 | 161243..164311 | 158836 | DUF499 -domain-containing-protein-[Enterobacter-hormaechei] |
| WIJ56059.1 | DUF499 _2.RM.like.systems | Arm-DNA-bind_3+Phage_integrase->RHH-> DUF499 -p-> DUF499 *-> DUF3780 +RAMA-> METHYLASE -> HKD-DNase+SF1-SF2-Helicase -> w HTH+ REase -><-?\|\|HTH_Tnp_1+HTH_21+rve+rve_2-> | 894 | Bacteria>Pseudomonadota>Gammaproteobacteria>Enterobacterales>Enterobacteriaceae | GCA_030179615.1 | Enterobacter_hormaechei | OI979_06745 | 1401626..1404310 | 158836 | DUF499 -domain-containing-protein-[Enterobacter-hormaechei] |
| STR56259.1 | DUF499 _2.RM.like.systems | SF1-SF2-Helicase->?-><- w HTH+ REase <- HKD-DNase+SF1-SF2-Helicase <- METHYLASE <- DUF3780 +RAMA<- DUF499 *<- DUF499 -p<-RHH<-Arm-DNA-bind_3+Phage_integrase<-tRNA\|\|ADH_N+ADH_zinc_N-> | 984 | Bacteria>Pseudomonadota>Gammaproteobacteria>Enterobacterales>Enterobacteriaceae | GCA_900451255.1 | Klebsiella_oxytoca | NCTC11696_05224 | 5162019..5164973 | 571 | Protein-of-uncharacterised-function-( DUF499 )-[Klebsiella-oxytoca] |

| Protein-ID | System | Gene-Neighbourhoods | Length | Taxonomy | Assembly-ID | Strain | Locus-Tag | Genomic-coordinates | TaxID | Fasta-defline |
|---|---|---|---|---|---|---|---|---|---|---|
| VAE22474.1 | DUF499_2.RM.like.systems | <-w HTH+REase<-HKD-DNase+SF1-SF2-Helicase<-METHYLASE<-DUF3780+RAMA<-DUF499*<-RHH<-Arm-DNA-bind_3+Phage_integrase | 1022 | Bacteria>Pseudomonadota>Gammaproteobacteria>Enterobacterales>Enterobacteriaceae | GCA_900558075.1 | Enterobacter_hormaechei | SAMEA2053862_03698 | 27017..30085 | 158836 | Protein-of-uncharacterised-function-(DUF499)-[Enterobacter-hormaechei] |
| VTN25635.1 | DUF499_2.RM.like.systems | HTH+MPTase-><-w HTH+REase<-HKD-DNase+SF1-SF2-Helicase<-METHYLASE<-DUF3780+RAMA<-DUF499-p<-DUF499*<-RHH<-Arm-DNA-bind_3+Phage_integrase<-tRNA\|\|?->?->ADH_N+ADH_zinc_N-> | 247 | Bacteria>Pseudomonadota>Gammaproteobacteria>Enterobacterales>Enterobacteriaceae | GCA_901420815.1 | Klebsiella_pneumoniae | NCTC9129_01967 | 1513305..1514048 | 573 | Uncharacterised-protein-[Klebsiella-pneumoniae] |
| WP_024008845.1 | DUF499_2.RM.like.systems | <-VirD4-TraG+VirB4-FtsK<-HDRNase<-w HTH+REase<-HKD-DNase+SF1-SF2-Helicase<-METHYLASE<-DUF3780+RAMA<-DUF499*<-RHH_1<-35exo\|\|?->Phage_integrase-> | 1041 | Bacteria>Pseudomonadota>Gammaproteobacteria>Enterobacterales>Enterobacteriaceae | GCF_000507645.1 | Escherichia_coli | G831_RS00450 | 65264..68389 | 562 | DUF499-domain-containing-protein-[Escherichia-coli] |
| WP_227518579.1 | DUF499_2.RM.like.systems | SF1-SF2-Helicase->?-><-w HTH+REase<-HKD-DNase+SF1-SF2-Helicase<-METHYLASE<-DUF3780+RAMA<-DUF499*<-DDE_Tnp_1<-DUF499-p<-RHH<-Arm-DNA-bind_3+Phage_integrase | 973 | Bacteria>Pseudomonadota>Gammaproteobacteria>Enterobacterales>Enterobacteriaceae | GCF_002055765.1 | Klebsiella_pneumoniae | AM364_RS18410 | 3781580..3784501 | 573 | DUF499-domain-containing-protein-[Klebsiella-pneumoniae] |
| WP_102217321.1 | DUF499_2.RM.like.systems | <-ADH_N+ADH_zinc_N<-?<-?\|\|tRNA->Arm-DNA-bind_3+Phage_integrase->RHH->DUF499*->DUF3780+RAMA->METHYLASE->HKD-DNase+SF1-SF2-Helicase->w HTH+REase-><-HTH+MPTase | 1041 | Bacteria>Pseudomonadota>Gammaproteobacteria>Enterobacterales>Enterobacteriaceae | GCF_002872205.1 | Klebsiella_pneumoniae | A8A10_RS01535 | 132525..135650 | 573 | DUF499-domain-containing-protein-[Klebsiella-pneumoniae] |
| WP_120159929.1 | DUF499_2.RM.like.systems | RHH_5->DUF499*->DUF3780+RAMA->METHYLASE->HKD-DNase+SF1-SF2-Helicase->w HTH+REase->HIN-HTH-> | 1041 | Bacteria>Pseudomonadota>Gammaproteobacteria>Enterobacterales>Enterobacteriaceae | GCF_003605695.1 | Escherichia_coli | D3Y67_RS09395 | 1782..4907 | 562 | DUF499-domain-containing-protein-[Escherichia-coli] |
| WP_131009058.1 | DUF499_2.RM.like.systems | HTH+MPTase-><-w HTH+REase<-HKD-DNase+SF1-SF2-Helicase<-METHYLASE<-DUF3780+RAMA<-DUF499*<-RHH<-Arm-DNA-bind_3+Phage_integrase<-tRNA\|\|?->?->ADH_N+ADH_zinc_N-> | 1041 | Bacteria>Pseudomonadota>Gammaproteobacteria>Enterobacterales>Enterobacteriaceae | GCF_004312385.1 | Klebsiella_variicola | HZ923_RS12940 | 35774..38899 | 244366 | DUF499-domain-containing-protein-[Klebsiella-variicola] |
| WP_000077518.1 | DUF499_2.RM.like.systems | DEDD_Tnp_IS110+Transposase_20->?->RHH_5->DUF499*->DUF3780+RAMA->METHYLASE->HKD-DNase+SF1-SF2-Helicase->w HTH+REase->HIN-HTH-> | 1022 | Bacteria>Pseudomonadota>Gammaproteobacteria>Enterobacterales>Enterobacteriaceae | GCF_005390065.1 | Escherichia_coli | EXY46_RS20920 | 4095..7163 | 562 | DUF499-domain-containing-protein-[Escherichia-coli] |
| WP_153674499.1 | DUF499_2.RM.like.systems | rve+rve_3-><-HIN-HTH<-w HTH+REase<-HKD-DNase+SF1-SF2-Helicase<-METHYLASE<-DUF3780+RAMA<-DUF499*<-RHH_5<-?\|\|<-?<-?<-HTH_Tnp_1+HTH_21+rve+rve_3<-ZnR | 1041 | Bacteria>Pseudomonadota>Gammaproteobacteria>Enterobacterales>Enterobacteriaceae | GCF_009648255.1 | Escherichia_coli | GIJ00_RS05185 | 9599..12724 | 562 | DUF499-domain-containing-protein-[Escherichia-coli] |
| WP_185774377.1 | DUF499_2.RM.like.systems | <-ADH_N+ADH_zinc_N<-?\|\|tRNA->Arm-DNA-bind_3+Phage_integrase->RHH->DUF499*->DUF3780+RAMA->METHYLASE->HKD-DNase+SF1-SF2-Helicase->w HTH+REase-><-HTH+MPTase | 1041 | Bacteria>Pseudomonadota>Gammaproteobacteria>Enterobacterales>Enterobacteriaceae | GCF_014235385.1 | Klebsiella_variicola | H8S48_RS22840 | 314198..317323 | 244366 | DUF499-domain-containing-protein-[Klebsiella-variicola] |
| WP_227628590.1 | DUF499_2.RM.like.systems | <-ADH_N+ADH_zinc_N\|\|tRNA->Arm-DNA-bind_3+Phage_integrase->RHH->DUF499*->DUF3780+RAMA->METHYLASE->HKD-DNase+SF1-SF2-Helicase->w HTH+REase-> | 1041 | Bacteria>Pseudomonadota>Gammaproteobacteria>Enterobacterales>Enterobacteriaceae | GCF_015721555.1 | Klebsiella_oxytoca | QP770_RS13995 | 142941..146066 | 571 | DUF499-domain-containing-protein-[Klebsiella-oxytoca] |
| WP_233593808.1 | DUF499_2.RM.like.systems | <-w HTH+REase<-HKD-DNase+SF1-SF2-Helicase<-METHYLASE<-DUF3780+RAMA<-DUF499*<-RHH<-Arm-DNA-bind_3+Phage_integrase<-tRNA\|\|ADH_N+ADH_zinc_N-> | 1041 | Bacteria>Pseudomonadota>Gammaproteobacteria>Enterobacterales>Enterobacteriaceae | GCF_016501115.1 | Citrobacter | OGW14_RS13170 | 9006..12131 | 544 | DUF499-domain-containing-protein-[Citrobacter-cronae] |
| WP_227639547.1 | DUF499_2.RM.like.systems | HTH+MPTase-><-w HTH+REase<-HKD-DNase+SF1-SF2-Helicase<-METHYLASE<-DUF3780+RAMA<-DUF499*<-RHH<-Arm-DNA-bind_3+Phage_integrase<-tRNA\|\|?->?->ADH_N+ADH_zinc_N-> | 1041 | Bacteria>Pseudomonadota>Gammaproteobacteria>Enterobacterales>Enterobacteriaceae | GCF_016651545.1 | Klebsiella_pneumoniae | JKA42_RS15120 | 66258..69383 | 573 | DUF499-domain-containing-protein-[Klebsiella-pneumoniae] |
| WP_089553231.1 | DUF499_2.RM.like.systems | <-HIN-HTH<-w HTH+REase<-HKD-DNase+SF1-SF2-Helicase<-METHYLASE<-DUF3780+RAMA<-DUF499*<-RHH_5\|\|<-?<-Aminotran_1_2\|\|<-Arm-DNA-bind_3+Phage_integrase<-tRNA\|\|ADH_N+ADH_zinc_N->FtsK+HerA+A32-> | 1041 | Bacteria>Pseudomonadota>Gammaproteobacteria>Enterobacterales>Enterobacteriaceae | GCF_019957165.1 | Escherichia_coli | BEC12_RS18680 | 17132..20257 | 562 | DUF499-domain-containing-protein-[Escherichia-coli] |
| WP_196557776.1 | DUF499_2.RM.like.systems | Arm-DNA-bind_3+Phage_integrase->RHH->DUF499*->DUF3780+RAMA->METHYLASE->HKD-DNase+SF1-SF2-Helicase->w HTH+REase-><-?<-SF1-SF2-Helicase | 1022 | Bacteria>Pseudomonadota>Gammaproteobacteria>Enterobacterales>Enterobacteriaceae | GCF_020116575.1 | Klebsiella_oxytoca | FML32_RS25495 | 2110..5178 | 571 | DUF499-domain-containing-protein-[Klebsiella-oxytoca] |
| WP_232904627.1 | DUF499_2.RM.like.systems | PIN->?-><-w HTH+REase<-HKD-DNase+SF1-SF2-Helicase<-METHYLASE<-DUF3780+RAMA<-DUF499*<-RHH<-Arm-DNA-bind_3+Phage_integrase | 1041 | Bacteria>Pseudomonadota>Gammaproteobacteria>Enterobacterales>Enterobacteriaceae | GCF_021282435.1 | Buttiauxella_sp._W03-F01 | LVQ77_RS12925 | 38707..41832 | 2904524 | DUF499-domain-containing-protein-[Buttiauxella-sp.-W03-F01] |
| WP_201792481.1 | DUF499_2.RM.like.systems | RHH_5->DUF499*->DUF3780+RAMA->METHYLASE->HKD-DNase+SF1-SF2-Helicase->w HTH+REase->HIN-HTH-> | 1022 | Bacteria>Pseudomonadota>Gammaproteobacteria>Enterobacterales>Enterobacteriaceae | GCF_023516095.1 | Escherichia_coli | IXW39_RS10870 | 2714..5782 | 562 | DUF499-domain-containing-protein-[Escherichia-coli] |
| WP_252076363.1 | DUF499_2.RM.like.systems | Arm-DNA-bind_3+Phage_integrase->RHH->DUF499*->DUF3780+RAMA->METHYLASE->HKD-DNase+SF1-SF2-Helicase->w HTH+REase-><-HTH+MPTase | 1041 | Bacteria>Pseudomonadota>Gammaproteobacteria>Enterobacterales>Enterobacteriaceae | GCF_023751425.1 | Enterobacter_kobei | M8R77_RS12255 | 159585..162710 | 208224 | DUF499-domain-containing-protein-[Enterobacter-kobei] |
| WP_258301444.1 | DUF499_2.RM.like.systems | <-HIN-HTH<-w HTH+REase<-HKD-DNase+SF1-SF2-Helicase<-METHYLASE<-DUF3780+RAMA<-DUF499*<-RHH_5\|\|<-?<-Aminotran_1_2\|\|<-Arm-DNA-bind_3+Phage_integrase<-tRNA\|\|ADH_N+ADH_zinc_N->FtsK+HerA+A32-> | 1041 | Bacteria>Pseudomonadota>Gammaproteobacteria>Enterobacterales>Enterobacteriaceae | GCF_024733045.1 | Escherichia_coli | NX108_RS16810 | 9461..12586 | 562 | DUF499-domain-containing-protein-[Escherichia-coli] |
| WP_258442601.1 | DUF499_2.RM.like.systems | <-ADH_N+ADH_zinc_N\|\|tRNA->Arm-DNA-bind_3+Phage_integrase->RHH->DUF499*->DUF3780+RAMA->METHYLASE->HKD-DNase+SF1-SF2-Helicase->w HTH+REase-><-?<-SF1-SF2-Helicase | 1041 | Bacteria>Pseudomonadota>Gammaproteobacteria>Enterobacterales>Enterobacteriaceae | GCF_024741195.1 | Citrobacter_amalonaticus | NXA99_RS21100 | 70776..73901 | 35703 | DUF499-domain-containing-protein-[Citrobacter-amalonaticus] |
| WP_101974273.1 | DUF499_2.RM.like.systems | HTH+MPTase-><-w HTH+REase<-HKD-DNase+SF1-SF2-Helicase<-METHYLASE<-DUF3780+RAMA<-DUF499*<-RHH<-Arm-DNA-bind_3+Phage_integrase<-tRNA\|\|?->?->ADH_N+ADH_zinc_N-> | 1041 | Bacteria>Pseudomonadota>Gammaproteobacteria>Enterobacterales>Enterobacteriaceae | GCF_028894585.1 | Klebsiella_pneumoniae | PXW64_RS18870 | 24255..27380 | 573 | DUF499-domain-containing-protein-[Klebsiella-pneumoniae] |
| WP_275399056.1 | DUF499_2.RM.like.systems | <-FtsK+HerA+A32<-ADH_N+ADH_zinc_N\|\|tRNA->Arm-DNA-bind_3+Phage_integrase->RHH->DUF499*->DUF3780+RAMA->METHYLASE->HKD-DNase+SF1-SF2-Helicase->w HTH+REase-><-SF1-SF2-Helicase<-OLD-ABC+TOPRIM | 1041 | Bacteria>Pseudomonadota>Gammaproteobacteria>Enterobacterales>Enterobacteriaceae | GCF_029104325.1 | Citrobacter_portucalensis | L2173_RS05340 | 153527..156652 | 1639133 | DUF499-domain-containing-protein-[Citrobacter-portucalensis] |
| WP_045337753.1 | DUF499_2.RM.like.systems | Arm-DNA-bind_3+Phage_integrase->RHH->DUF499*->DUF3780+RAMA->METHYLASE->HKD-DNase+SF1-SF2-Helicase->w HTH+REase-><-?\|\|HTH_Tnp_1+HTH_21+rve+rve_2-> | 1041 | Bacteria>Pseudomonadota>Gammaproteobacteria>Enterobacterales>Enterobacteriaceae | GCF_029207635.1 | Enterobacter_hormaechei | H0762_RS10605 | 150283..153408 | 158836 | DUF499-domain-containing-protein-[Enterobacter-hormaechei] |
| WP_284106250.1 | DUF499_2.RM.like.systems | HTH+MPTase-><-HNH+SAVED<-w HTH+REase<-HKD-DNase+SF1-SF2-Helicase<-METHYLASE<-DUF3780+RAMA<-DUF499*<-RHH<-Arm-DNA-bind_3+Phage_integrase<-tRNA\|\|ADH_N+ADH_zinc_N->FtsK+HerA+A32-> | 1041 | Bacteria>Pseudomonadota>Gammaproteobacteria>Enterobacterales>Enterobacteriaceae | GCF_030132145.1 | Citrobacter_portucalensis | QL289_RS16600 | 141668..144793 | 1639133 | DUF499-domain-containing-protein-[Citrobacter-portucalensis] |
| WP_285145128.1 | DUF499_2.RM.like.systems | XIS-HTH->Phage_ASH->?->?->TOPRIM+DNAG+D5N+D5ATPase+P4PRIMASE-HTH->RHH_5->DUF499*->DUF3780+RAMA->METHYLASE->HKD-DNase+SF1-SF2-Helicase->w HTH+REase-><-ParB+HNH<-SymE_toxin | 1041 | Bacteria>Pseudomonadota>Gammaproteobacteria>Enterobacterales>Enterobacteriaceae | GCF_030234145.1 | unclassified_Lelliottia | QQF24_RS22360 | 130751..133876 | 2642424 | MULTISPECIES:-DUF499-domain-containing-protein-[unclassified-Lelliottia] |
| WP_180245885.1 | DUF499_2.RM.like.systems | SF1-SF2-Helicase->?-><-w HTH+REase<-HKD-DNase+SF1-SF2-Helicase<-METHYLASE<-DUF3780+RAMA<-DUF499*<-RHH<-Arm-DNA-bind_3+Phage_integrase | 1041 | Bacteria>Pseudomonadota>Gammaproteobacteria>Enterobacterales>Enterobacteriaceae | GCF_900558075.1 | Enterobacter_hormaechei | H0785_RS18375 | 27017..30142 | 158836 | DUF499-domain-containing-protein-[Enterobacter-hormaechei] |
| WP_204372837.1 | DUF499_2.RM.like.systems | SF1-SF2-Helicase->?-><-w HTH+REase<-HKD-DNase+SF1-SF2-Helicase<-METHYLASE<-DUF3780+RAMA<-DUF499*<-RHH<-Arm-DNA-bind_3+Phage_integrase<-tRNA\|\|?->?->ADH_N+ADH_zinc_N-> | 1022 | Bacteria>Pseudomonadota>Gammaproteobacteria>Enterobacterales>Enterobacteriaceae | - | Klebsiella_pneumoniae | MMH51_002463 | 54491..57559 | 573 | DUF499-domain-containing-protein-[Klebsiella-pneumoniae] |
| WP_198740177.1 | DUF499_2.RM.like.systems | Arm-DNA-bind_3+Phage_integrase->RHH->DUF499*->DUF3780+RAMA->METHYLASE->HKD-DNase+SF1-SF2-Helicase->w HTH+REase-><-?<-SF1-SF2-Helicase | 1022 | Bacteria>Pseudomonadota>Gammaproteobacteria>Enterobacterales>Enterobacteriaceae | - | Klebsiella_pneumoniae | POC94_000676 | 161243..164311 | 573 | MULTISPECIES:-DUF499-domain-containing-protein-[Enterobacteriaceae] |
| WP_048997689.1 | DUF499_2.RM.like.systems | <-ADH_N+ADH_zinc_N<-?<-?\|\|tRNA->Arm-DNA-bind_3+Phage_integrase->RHH->DUF499*->DUF3780+RAMA->METHYLASE->HKD-DNase+SF1-SF2-Helicase->w HTH+REase-><-?<-SF1-SF2-Helicase | 1041 | Bacteria>Pseudomonadota>Gammaproteobacteria>Enterobacterales>Enterobacteriaceae | - | Klebsiella_pneumoniae | QVO21_RS15045 | 200942..204067 | 573 | DUF499-domain-containing-protein-[Klebsiella-pneumoniae] |
| WP_116197723.1 | DUF499_2.RM.like.systems | <-NACHT+SF1-SF2-Helicase+HKD-DNase\|\|ParB+HNH-><-w HTH+REase<-HKD-DNase+SF1-SF2-Helicase<-METHYLASE<-DUF3780+RAMA<-DUF499*<-RHH | 1041 | Bacteria>Pseudomonadota>Gammaproteobacteria>Enterobacterales>Erwiniaceae | GCF_017920215.1 | Pantoea_ananatis | HG975_RS10020 | 359371..362496 | 553 | DUF499-domain-containing-protein-[Pantoea-ananatis] |
| WP_266093834.1 | DUF499_2.RM.like.systems | DUF1176->tRNA->RHH->DUF499*->DUF3780+RAMA->METHYLASE->HKD-DNase+SF1-SF2-Helicase->w HTH+REase-> | 1041 | Bacteria>Pseudomonadota>Gammaproteobacteria>Enterobacterales>Erwiniaceae | GCF_026241965.1 | [Curtobacterium]_plantarum | ORG41_RS07860 | 286061..289186 | 221276 | DUF499-domain-containing-protein-[[Curtobacterium] |
| WP_040132773.1 | DUF499_2.RM.like.systems | 35exo->RHH_1->DUF499*->DUF3780+RAMA->METHYLASE->HKD-DNase+SF1-SF2-Helicase->w HTH+REase->HDRNase->VirD4-TraG+VirB4-FtsK-> | 1041 | Bacteria>Pseudomonadota>Gammaproteobacteria>Enterobacterales | GCF_000754345.1 | Morganellaceae | I6H92_RS05395 | 1203958..1207083 | 1903414 | MULTISPECIES:-DUF499-domain-containing-protein-[Morganellaceae] |
| WP_032437326.1 | DUF499_2.RM.like.systems | SF1-SF2-Helicase->?-><-w HTH+REase<-HKD-DNase+SF1-SF2-Helicase<-METHYLASE<-DUF3780+RAMA<-DUF499*<-RHH<-Arm-DNA-bind_3+Phage_integrase<-tRNA\|\|?->?->ADH_N+ADH_zinc_N-> | 1041 | Bacteria>Pseudomonadota>Gammaproteobacteria>Enterobacterales | GCF_009387945.1 | Enterobacteriaceae | OMD39_RS22365 | 4570598..4573723 | 543 | MULTISPECIES:-DUF499-domain-containing-protein-[Enterobacteriaceae] |
| MBK1419820.1 | DUF499_2.RM.like.systems | 35exo->RHH_1->DUF499*->DUF3780+RAMA->METHYLASE->HKD-DNase+SF1-SF2-Helicase->w HTH+REase->HDRNase->VirD4-TraG+VirB4-FtsK-> | 1022 | Bacteria>Pseudomonadota>Gammaproteobacteria>Enterobacterales>Morganellaceae | GCA_016623625.1 | Providencia_stuartii | I6H92_05395 | 1204015..1207083 | 588 | DUF499-domain-containing-protein-[Providencia-stuartii] |
| MCL8535002.1 | DUF499_2.RM.like.systems | DUF499*->DUF3780->METHYLASE->HKD-DNase+SF1-SF2-Helicase->w HTH+REase->HDRNase->VirD4-TraG+VirB4-FtsK-> | 850 | Bacteria>Pseudomonadota>Gammaproteobacteria>Enterobacterales>Morganellaceae | GCA_023604365.1 | Proteus_mirabilis | NAE57_17165 | 121..2673 | 584 | ATP-binding-protein-[Proteus-mirabilis] |
| WP_038193804.1 | DUF499_2.RM.like.systems | <-w HTH+REase<-HKD-DNase+SF1-SF2-Helicase<-METHYLASE<-DUF3780+RAMA<-DUF499*<-RHH<-tRNA\|\|Beta-lactamase-><-?<-?\|\|Peptidase-> | 1041 | Bacteria>Pseudomonadota>Gammaproteobacteria>Enterobacterales>Morganellaceae | GCF_000736695.1 | Xenorhabdus_bovienii | XBKB1_RS06355 | 8159..11284 | 40576 | DUF499-domain-containing-protein-[Xenorhabdus-bovienii] |
| WP_152110353.1 | DUF499_2.RM.like.systems | <-Peptidase_S8<-?\|\|35exo->RHH_1->DUF499*->DUF3780+RAMA->METHYLASE->HKD-DNase+SF1-SF2-Helicase->w HTH+REase->HDRNase->VirD4-TraG+VirB4-FtsK-> | 1041 | Bacteria>Pseudomonadota>Gammaproteobacteria>Enterobacterales>Morganellaceae | GCF_009183685.1 | Proteus_mirabilis | GBN12_RS03240 | 63661..66786 | 584 | DUF499-domain-containing-protein-[Proteus-mirabilis] |
| WP_159262627.1 | DUF499_2.RM.like.systems | <-VirD4-TraG+VirB4-FtsK<-HDRNase<-w HTH+REase<-HKD-DNase+SF1-SF2-Helicase<-METHYLASE<-DUF3780<-DUF499*<-XIS-HTH<-35exo | 1022 | Bacteria>Pseudomonadota>Gammaproteobacteria>Enterobacterales>Morganellaceae | GCF_010692865.1 | Proteus_mirabilis | QN307_RS16755 | 34681..37749 | 584 | DUF499-domain-containing-protein-[Proteus-mirabilis] |
| WP_275358359.1 | DUF499_2.RM.like.systems | <-Peptidase\|\|?->?-><-Beta-lactamase\|\|tRNA->RHH->DUF499*->DUF3780+RAMA->METHYLASE->HKD-DNase+SF1-SF2-Helicase->w HTH+REase-> | 1041 | Bacteria>Pseudomonadota>Gammaproteobacteria>Enterobacterales>Morganellaceae | GCF_029104585.1 | Xenorhabdus_bovienii | KKI95_RS09990 | 130588..133713 | 40576 | DUF499-domain-containing-protein-[Xenorhabdus-bovienii] |
| WP_196578769.1 | DUF499_2.RM.like.systems | 35exo->RHH_1->DUF499*->DUF3780+RAMA->METHYLASE->HKD-DNase+SF1-SF2-Helicase->w HTH+REase->HDRNase->VirD4-TraG+VirB4-FtsK-> | 1022 | Bacteria>Pseudomonadota>Gammaproteobacteria>Enterobacterales>Morganellaceae | - | Proteus_mirabilis | I6H92_05395 | 1204015..1207083 | 584 | DUF499-domain-containing-protein-[Proteus-mirabilis] |
| WP_250269692.1 | DUF499_2.RM.like.systems | DUF499*->DUF3780->METHYLASE->HKD-DNase+SF1-SF2-Helicase->w HTH+REase->HDRNase->VirD4-TraG+VirB4-FtsK-> | 850 | Bacteria>Pseudomonadota>Gammaproteobacteria>Enterobacterales>Morganellaceae | - | Proteus_mirabilis | NAE57_17165 | 121..2673 | 584 | DUF499-domain-containing-protein-[Proteus-mirabilis] |
| WP_226974938.1 | DUF499_2.RM.like.systems | <-Peptidase\|\|?->?->tRNA->Arm-DNA-bind_3+Phage_integrase->RHH->DUF499*->METHYLASE-p->HKD-DNase+SF1-SF2-Helicase->w HTH+REase-><-OLD-ABC+TOPRIM<-SbcC<-HTH+MPTase | 777 | Bacteria>Pseudomonadota>Gammaproteobacteria>Enterobacterales>Yersiniaceae | GCF_001319705.1 | Yersinia_pseudotuberculosis | EGX52_RS11275 | 2325348..2327681 | 633 | DUF499-domain-containing-protein-[Yersinia-pseudotuberculosis] |
| WP_095924507.1 | DUF499_2.RM.like.systems | <-Peptidase\|\|?->?-><-Beta-lactamase\|\|tRNA->Arm-DNA-bind_3+Phage_integrase->RHH->DUF499*->DUF3780+RAMA->METHYLASE->HKD-DNase+SF1-SF2-Helicase->w HTH+REase-><-HIN-HTH | 1041 | Bacteria>Pseudomonadota>Gammaproteobacteria>Enterobacterales>Yersiniaceae | GCF_002303785.1 | Rahnella_victoriana | C0635_RS19350 | 4088068..4091193 | 1510570 | DUF499-domain-containing-protein-[Rahnella-victoriana] |
| WP_024250705.1 | DUF499_2.RM.like.systems | <-HIN-HTH<-w HTH+REase<-HKD-DNase+SF1-SF2-Helicase<-METHYLASE<-DUF3780+RAMA<-DUF499*<-RHH_5<-?<-DEDD_Tnp_IS110+Transposase_20<-HTH_Tnp_1+HTH_21+rve+rve_3\|\|<-Arm-DNA-bind_3+Phage_integrase | 1041 | Bacteria>Pseudomonadota>Gammaproteobacteria | GCF_000212715.2 | Enterobacterales | KT011_RS04530 | 922314..925439 | 91347 | MULTISPECIES:-DUF499-domain-containing-protein-[Enterobacterales] |
| WP_235045586.1 | DUF499_2.RM.like.systems | Phage_integrase->Phage_integrase->?->?->LON-AAA->RHH->DUF499*->DUF499-p->DUF3780->METHYLASE-p->METHYLASE->HKD-DNase+SF1-SF2-Helicase->w HTH+REase-> | 776 | Bacteria>Pseudomonadota>Gammaproteobacteria>Thiotrichales>Piscirickettsiaceae | GCF_000349205.1 | Methylophaga_lonarensis | MPL1_RS12025 | 17805..20135 | 999151 | DUF499-domain-containing-protein-[Methylophaga-lonarensis] |
| PUB90401.1 | DUF499_2.RM.like.systems | DUF499*->DUF3780->METHYLASE->HKD-DNase+SF1-SF2-Helicase->w HTH+REase->HTH-> | 1012 | Bacteria>Pseudomonadota>Gammaproteobacteria>unclassified_Gammaproteobacteria | GCA_003058535.1 | gamma_proteobacterium_symbiont_of_Ctena_orbiculata | DBP01_07750 | 1..3039 | 1968598 | AAA-family-ATPase,-partial-[gamma-proteobacterium-symbiont-of-Ctena-orbiculata] |
| WP_087868382.1 | DUF499_2.RM.like.systems | <-w HTH+REase<-HKD-DNase+SF1-SF2-Helicase<-METHYLASE<-?<-DUF3780<-DUF499*<-RHH<-XIS-HTH | 1041 | Bacteria>Pseudomonadota>Gammaproteobacteria>unclassified_Gammaproteobacteria | GCF_002189065.1 | Bathymodiolus_platifrons_methanotrophic_gill_symbiont | CDZ78_RS01565 | 51590..54715 | 113268 | DUF499-domain-containing-protein-[Bathymodiolus-platifrons-methanotrophic-gill-symbiont] |
| MBE4228752.1 | DUF499_2.RM.like.systems | <-VirD4-TraG+VirB4-FtsK<-HDRNase<-w HTH+REase<-HKD-DNase+SF1-SF2-Helicase<-DUF3780+RAMA<-DUF499*<-RHH_1<-35exo | 1041 | Bacteria>Pseudomonadota>Gammaproteobacteria>Vibrionales>Vibrionaceae | GCA_014920775.1 | Vibrio_parahaemolyticus | HJ082_06195 | 253692..256817 | 670 | DUF499-domain-containing-protein-[Vibrio-parahaemolyticus] |
| MBY7816961.1 | DUF499_2.RM.like.systems | 35exo->XIS-HTH->DUF499*->DUF3780->METHYLASE->HKD-DNase+SF1-SF2-Helicase->w HTH+REase->HDRNase->VirD4-TraG+VirB4-FtsK-> | 1022 | Bacteria>Pseudomonadota>Gammaproteobacteria>Vibrionales>Vibrionaceae | GCA_019852895.1 | Vibrio_fluvialis | KW416_13045 | 310292..313360 | 676 | DUF499-domain-containing-protein-[Vibrio-fluvialis] |
| EKO3646393.1 | DUF499_2.RM.like.systems | <-VirD4-TraG+VirB4-FtsK<-HDRNase<-w HTH+REase<-HKD-DNase+SF1-SF2-Helicase<-METHYLASE<-DUF3780+RAMA<-DUF499*<-RHH_1<-35exo | 1041 | Bacteria>Pseudomonadota>Gammaproteobacteria>Vibrionales>Vibrionaceae | GCA_028989625.1 | Vibrio_metschnikovii | M3925_003006 | 63497..66622 | 28172 | DUF499-domain-containing-protein-[Vibrio-metschnikovii] |
| EKO3692967.1 | DUF499_2.RM.like.systems | 35exo->XIS-HTH->DUF499*->DUF3780->METHYLASE->HKD-DNase+SF1-SF2-Helicase->w HTH+REase->HDRNase->VirD4-TraG+VirB4-FtsK-> | 1022 | Bacteria>Pseudomonadota>Gammaproteobacteria>Vibrionales>Vibrionaceae | GCA_028990005.1 | Vibrio_metschnikovii | POF02_001010 | 192750..195818 | 28172 | ATP-binding-protein-[Vibrio-metschnikovii] |
| WP_203342017.1 | DUF499_2.RM.like.systems | <-VirD4-TraG+VirB4-FtsK<-HDRNase<-w HTH+REase<-HKD-DNase+SF1-SF2-Helicase<-METHYLASE<-DUF3780+RAMA<-DUF499*<-?<-DUF499-p<-RHH_1<-35exo | 782 | Bacteria>Pseudomonadota>Gammaproteobacteria>Vibrionales>Vibrionaceae | GCF_014525255.1 | Vibrio_metschnikovii | GO180_RS04985 | 107142..109490 | 28172 | DUF499-domain-containing-protein-[Vibrio-metschnikovii] |
| WP_238130455.1 | DUF499_2.RM.like.systems | <-VirD4-TraG+VirB4-FtsK<-HDRNase<-w HTH+REase<-HKD-DNase+SF1-SF2-Helicase<-METHYLASE<-DUF3780+RAMA<-DUF499*<-RHH_1<-35exo | 1041 | Bacteria>Pseudomonadota>Gammaproteobacteria>Vibrionales>Vibrionaceae | GCF_022114045.1 | Vibrio_cincinnatiensis | EXA22_RS10090 | 12757..15882 | 675 | DUF499-domain-containing-protein-[Vibrio-cincinnatiensis] |
| WP_108717267.1 | DUF499_2.RM.like.systems | <-VirD4-TraG+VirB4-FtsK<-HDRNase<-w HTH+REase<-HKD-DNase+SF1-SF2-Helicase<-METHYLASE<-DUF3780+RAMA<-DUF499*<-RHH_1<-35exo<-DDE_Tnp_1 | 1041 | Bacteria>Pseudomonadota | GCF_003073935.1 | Gammaproteobacteria | QQJ65_RS05090 | 258052..261177 | 1236 | MULTISPECIES:-DUF499-domain-containing-protein-[Gammaproteobacteria] |
| WP_222132889.1 | DUF499_2.RM.like.systems | DDE_Tnp_1-><-Phage_integrase<-?\|\|35exo->XIS-HTH->DUF499*->DUF3780->METHYLASE->HKD-DNase+SF1-SF2-Helicase->w HTH+REase->HDRNase->VirD4-TraG+VirB4-FtsK-> | 1022 | Bacteria>Pseudomonadota | GCF_030388465.1 | Gammaproteobacteria | FG482_RS02520 | 511750..514818 | 1236 | DUF499-domain-containing-protein-[Vibrio-parahaemolyticus] |
| MCF7762652.1 | DUF499_2.RM.like.systems | <-HKD-DNase+SF1-SF2-Helicase<-Schlafen+HATPase+HTH<-METHYLASE<-w HTH+REase<-DUF3780<-DUF499*<-XIS-HTH\|\|SF1-SF2-Helicase-><-Glyoxalase<-?<-REC | 1027 | Bacteria>PVC_group>Verrucomicrobiota>unclassified_Verrucomicrobiota | GCA_021733945.1 | Verrucomicrobiota_bacterium | K9N62_03175 | 25626..28709 | 2026799 | DUF499-domain-containing-protein-[Verrucomicrobia-bacterium] |
| WP_098986579.1 | DUF499_2.RM.like.systems | <-NERD+SF1-SF2-Helicase\|\|Preader7+McrB-NTD+McrB-AAA->McrC-NTD+REase->?-><-HEPN\|\|?->DUF499*->DUF3780->METHYLASE->HKD-DNase+SF1-SF2-Helicase->?->?->MerR-HTH-> | 1022 | Bacteria>Terrabacteria_group>Bacillota>Bacilli>Bacillales | - | Bacillus_cereus | COD90_RS22430 | 19622..22690 | 1396 | DUF499-domain-containing-protein-[Bacillus-cereus] |
| PGW11809.1 | DUF499_2.RM.like.systems | <-NERD+SF1-SF2-Helicase\|\|Preader7+McrB-NTD+McrB-AAA->McrC-NTD+REase->?-><-HEPN\|\|?->DUF499*->DUF3780->METHYLASE->HKD-DNase+SF1-SF2-Helicase->?->?->MerR-HTH-> | 1022 | Bacteria>Terrabacteria_group>Bacillota>Bacilli>Bacillales | GCA_002585495.1 | Bacillus_cereus | COD90_22430 | 19622..22690 | 1396 | hypothetical-protein-COD90_22430-[Bacillus-cereus] |
| WP_098128686.1 | DUF499_2.RM.like.systems | HTH+WYL->?->Preader7+McrB-NTD+McrB-AAA->McrC-NTD+REase->REase->?->DUF499*->DUF3780->METHYLASE->HKD-DNase+SF1-SF2-Helicase-> | 1033 | Bacteria>Terrabacteria_group>Bacillota>Bacilli>Bacillales | GCF_002556745.1 | Bacillus_wiedmannii | CN576_RS24165 | 43423..46524 | 1890302 | DUF499-domain-containing-protein-[Bacillus-wiedmannii] |
| WP_156271943.1 | DUF499_2.RM.like.systems | DSBH-><-HKD-DNase+SF1-SF2-Helicase+iREase+CTD<-METHYLASE<-DUF3780<-DUF499*<-MNS+HTH+REase<-RVT<-?<-?\|\|OLD-ABC-p->OLD-ABC-> | 1074 | Bacteria>Terrabacteria_group>Bacillota>Clostridia>Moorellales | GCF_009735625.1 | Moorella_glycerini | MGLY_RS03970 | 823454..826678 | 55779 | DUF499-domain-containing-protein-[Moorella-glycerini] |
| HHV75152.1 | DUF499_2.RM.like.systems | <-McrC-NTD+REase<-?<-HKD-DNase+SF1-SF2-Helicase<-METHYLASE<-DUF3780<-DUF499*<-SF1-SF2-Helicase<-SIGMA-HTH+SIGMA-HTH<-?<-ACET | 1037 | Bacteria>Terrabacteria_group>Bacillota>Clostridia>Thermoanaerobacterales | GCA_012839995.1 | Thermoanaerobacterium_sp. | GXX41_11045 | 45158..48271 | 40549 | DUF499-domain-containing-protein-[Thermoanaerobacterium-sp.] |
| MCE7987123.1 | DUF499_2.RM.like.systems | RNaseH-><-HKD-DNase+SF1-SF2-Helicase<-w HTH+REase<-METHYLASE<-DUF3780<-DUF499* | 1028 | Bacteria>Terrabacteria_group>Chloroflexota>Caldilineae>Caldilineales | GCA_021405325.1 | Caldilinea_sp._CFX5 | DYG89_38625 | 47315..50401 | 2293554 | DUF499-domain-containing-protein-[Caldilinea-sp.-CFX5] |
| WP_026149087.1 | DUF499_2.RM.like.systems | Classical-AAA->Peptidase-S8-><-?<-?\|\|?-><-DUF3780<-DUF499*<-w HTH+REase | 1071 | Bacteria>Terrabacteria_group>Cyanobacteriota/Melainabacteria_group>Cyanobacteriota>Cyanophyceae | GCF_002142475.1 | Leptolyngbya_boryana | MCP04_RS33160 | 388752..391967 | 1184 | DUF499-domain-containing-protein-[Leptolyngbya-boryana] |

| Protein-ID | System | Gene-Neighbourhoods | Length | Taxonomy | Assembly-ID | Strain | Locus-Tag | Genomic-coordinates | TaxID | Fasta-defline |
|---|---|---|---|---|---|---|---|---|---|---|
| WP_088889980.1 | DUF499 _2.RM.like.systems | DEDD_Tnp_IS110+Transposase_20-><-REase<-PIN<-?<-DUF3780 <-DUF499 *\|\| w HTH+REase -><-?<-URI-DNase<-Prim-Pol | 1072 | Bacteria>Terrabacteria_group>Cyanobacteriota/Melainabacteria_group>Cyanobacteriota>Cyanophyceae | GCF_002215035.1 | Leptolyngbya_ohadii | CDV24_RS06815 | 190985..194203 | 1962290 | DUF499 -domain-containing-protein-[Leptolyngbya-ohadii] |
| WP_017291799.1 | DUF499 _2.RM.like.systems | w HTH+REase -> DUF499 *-> DUF3780 -><-?\|\|?->?-><-Peptidase-S8<-Classical-AAA | 1073 | Bacteria>Terrabacteria_group>Cyanobacteriota/Melainabacteria_group>Cyanobacteriota>Cyanophyceae | GCF_014697225.1 | Leptolyngbya | H6G55_RS21500 | 113566..116787 | 47251 | MULTISPECIES:-DUF499 -domain-containing-protein-[Leptolyngbya] |
| WP_242049455.1 | DUF499 _2.RM.like.systems | Classical-AAA->Peptidase-S8-><-?<-?\|\|?-><-DUF3780 <-DUF499 *<- w HTH+REase | 1070 | Bacteria>Terrabacteria_group>Cyanobacteriota/Melainabacteria_group>Cyanobacteriota>Cyanophyceae | GCF_014697245.1 | unclassified_Leptolyngbya | H6G61_RS21985 | 21330..24542 | 2650499 | MULTISPECIES:-DUF499 -domain-containing-protein-[unclassified-Leptolyngbya] |
| KPA18631.1 | DUF499 _2.RM.like.systems | <-HKD-DNase+SF1-SF2-Helicase <- w HTH+REase <-METHYLASE <-DUF3780 <-DUF499 * | 1032 | Bacteria>Thermodesulfobacteriota>Desulfobacteria>Desulfobacterales>Candidatus_Magnetomoraceae | GCA_001292585.1 | Candidatus_Magnetomorum_sp._HK-1 | MHK_001159 | 10914..14012 | 1509431 | ATPase-AAA-[Candidatus-Magnetomorum-sp.-HK-1] |
| MBK7550235.1 | DUF499 _2.RM.like.systems | <-HKD-DNase+SF1-SF2-Helicase <-DUF3780 <-DUF499 *<- w HTH+REase <-PIN<-?<-CHTH | 1026 | Bacteria>Thermodesulfobacteriota>Syntrophia>Syntrophales>Syntrophaceae | - | Syntrophaceae_bacterium | IPI61_00440 | 88025..91105 | 2052184 | DUF499 -domain-containing-protein-[Syntrophaceae-bacterium] |
| -- | | | | | | | | | | |
| --REase | | | | | | | | | | |
| -- | | | | | | | | | | |
| WP_220589967.1 | DUF499 _2.RM.like.systems | SWACOS-><-?<-?<-HNH\|\|REase -> DUF499 *-> DUF3780 -> METHYLASE -> HKD-DNase+SF1-SF2-Helicase +i REase +CTD->Resolvase-><-MarR-HTH | 1098 | Archaea>Euryarchaeota>Stenosarchaea_group>Halobacteria>Halobacteriales | GCF_019599405.1 | Halomicroarcula_salinisoli | EGD98_RS19205 | 187933..191229 | 2487746 | DUF499 -domain-containing-protein-[Halomicroarcula-salinisoli] |
| WP_239638715.1 | DUF499 _2.RM.like.systems | <-PadR-HTH\|\|?->?-><-Resolvase\|\|?->AbrB-HTH-> DUF499 *-> DUF3780 -> METHYLASE ->HEPN-> HKD-DNase+SF1-SF2-Helicase +i REase +CTD-> REase -><-DDE_Tnp_1 | 1054 | Archaea>Euryarchaeota>Stenosarchaea_group>Halobacteria>Haloferacales | GCF_000337075.1 | Halorubrum | NAF06_RS08930 | 1779031..1782195 | 56688 | MULTISPECIES:-DUF499 -domain-containing-protein-[Halorubrum] |
| WP_195155664.1 | DUF499 _2.RM.like.systems | Zn_Tnp_IS1595+DDE_Tnp_IS1595->HEPN->HEPN-><-?\|\|?->AbrB-HTH-> DUF499 *-> DUF3780 -> METHYLASE -> HKD-DNase+SF1-SF2-Helicase +i REase +CTD-> REase -p->HEPN-> | 1099 | Archaea>Euryarchaeota>Stenosarchaea_group>Halobacteria>Haloferacales | GCF_000513435.1 | Halorubrum_sp._AJ67 | BN903_RS00695 | 145466..148765 | 1173487 | DUF499 -domain-containing-protein-[Halorubrum-sp.-AJ67] |
| WP_205742829.1 | DUF499 _2.RM.like.systems | <-ORC/CDC6+HTH<-ORC/CDC6+HTH<-?<-?<-?\|\|AbrB-HTH-> DUF499 *-> DUF3780 -> METHYLASE -> HKD-DNase+SF1-SF2-Helicase +i REase +CTD-> REase -> | 1099 | Archaea>Euryarchaeota>Stenosarchaea_group>Halobacteria>Haloferacales | GCF_004799685.1 | Halobellus_limi | DV707_RS18320 | 167196..170495 | 699433 | DUF499 -domain-containing-protein-[Halobellus-limi] |
| WP_234699407.1 | DUF499 _2.RM.like.systems | AbrB-HTH-> DUF499 *-> DUF3780 -> METHYLASE -> HKD-DNase+SF1-SF2-Helicase +i REase +CTD-> REase -><-?<-BirA-HTH | 1099 | Archaea>Euryarchaeota>Stenosarchaea_group>Halobacteria>Haloferacales | GCF_016635065.1 | Halorubrum_sp._LN27 | JIN95_RS16120 | 50683..53982 | 2801032 | DUF499 -domain-containing-protein-[Halorubrum-sp.-LN27] |
| OAQ51496.1 | DUF499 _2.RM.like.systems | ZetaToxin-><-HEPN-p\|\|?->?->AbrB-HTH-> DUF499 *-> DUF3780 -> METHYLASE -p-> HKD-DNase+SF1-SF2-Helicase +i REase +CTD-> REase -><-?\|\|CoiA-> | 1099 | Archaea>Euryarchaeota>Stenosarchaea_group>Halobacteria>Natrialbales | GCA_000690595.2 | Natrinema_mahii | HTG_19000 | 43326..46625 | 1416969 | ATPase-AAA-[Haloterrigena-mahii] |
| WP_160063634.1 | DUF499 _2.RM.like.systems | <-REase <-HKD-DNase+SF1-SF2-Helicase +i REase +CTD<-METHYLASE <-DUF3780 <-DUF499 *<-AbrB-HTH<-?<-?\|\|?->HNH-> | 926 | Archaea>Euryarchaeota>Stenosarchaea_group>Halobacteria>Natrialbales | GCF_009834785.1 | Natronorubrum_halalkaliphilum | GS429_RS05865 | 14261..17041 | 2691917 | DUF499 -domain-containing-protein-[Natronorubrum-halalkaliphilum] |
| MCK9404871.1 | DUF499 _2.RM.like.systems | DUF499 *-> DUF3780 -> METHYLASE -> HKD-DNase+SF1-SF2-Helicase -><-SF1-SF2-Helicase<-REase \|\|PIN-> | 1027 | Archaea>Euryarchaeota>Stenosarchaea_group>Methanomicrobia>Methanotrichales | GCA_023229265.1 | Methanothrix_sp. | MOQ47_00445 | 3470..6553 | 90426 | DUF499 -domain-containing-protein-[Methanothrix-sp.] |
| MDE3261994.1 | DUF499 _2.RM.like.systems | REase -> DUF499 *-> DUF3780 ->OLD-ABC->HNH-> METHYLASE -> HKD-DNase+SF1-SF2-Helicase +i REase +CTD-> | 1006 | Bacteria>Acidobacteriota>unclassified_Acidobacteriota | GCA_028877265.1 | Acidobacteriota_bacterium | OYL41_08435 | 3377..6397 | 1978231 | MAG:-DUF499 -domain-containing-protein-[Acidobacteriota-bacterium] |
| QQS19533.1 | DUF499 _2.RM.like.systems | DDE_Tnp_1->Peptidase_S58-><-?\|\|DEDD_Tnp_IS110+Transposase_20->FtsK->ORC-CDC6+REase -> DUF499 *-><-?\|\|?->?-><-?\|\|ABhydrolase-> | 766 | Bacteria>Bacteria_incertae_sedis>Bacteria_candidate_phyla>Candidatus_Saccharibacteria>unclassified_Candidatus_Saccharibacteria | GCA_016699955.1 | Candidatus_Saccharibacteria_bacterium | IPL85_04630 | 901666..903966 | 2026720 | ATP-binding-protein-[Candidatus-Saccharibacteria-bacterium] |
| MBI2571108.1 | DUF499 _2.RM.like.systems | <-DUF499 *\|\|REase ->AbiEi_1->AbiEi_1-><-?<-REase | 526 | Bacteria>Bacteria_incertae_sedis>Bacteria_candidate_phyla>Candidatus_Schekmanbacteria>unclassified_Candidatus_Schekmanbacteria | GCA_016188575.1 | Candidatus_Schekmanbacteria_bacterium | HYV63_29230 | 1..1578 | 2419984 | ATP-binding-protein-[Candidatus-Schekmanbacteria-bacterium] |
| WP_259170624.1 | DUF499 _2.RM.like.systems | <-HKD-DNase+SF1-SF2-Helicase <-METHYLASE <-METHYLASE <-PIN<-HTH+MPTase<-DUF3780 <-DUF499 *<-REase +35exo<-?<-?\|\|DEDD_Tnp_IS110+Transposase_20->?->DUF4096+DDE_Tnp_1+DDE_Tnp-> | 1021 | Bacteria>FCB_group>Bacteroidota/Chlorobiota_group>Rhodothermota>Rhodothermia | GCF_024807095.1 | Salinibacter_ruber | GGP42_RS07650 | 508801..511866 | 146919 | DUF499 -domain-containing-protein-[Salinibacter-ruber] |
| MBY0489937.1 | DUF499 _2.RM.like.systems | <-REase <-HKD-DNase+SF1-SF2-Helicase +i REase +CTD<-METHYLASE <-DUF499 *<-?\|\|?->?-><-MazF | 974 | Bacteria>FCB_group>Gemmatimonadota>Gemmatimonadetes>Gemmatimonadales | GCA_019752115.1 | Gemmatimonadaceae_bacterium | K2R93_08880 | 95795..98719 | 2026741 | DUF499 -domain-containing-protein-[Gemmatimonadaceae-bacterium] |
| RLB16925.1 | DUF499 _2.RM.like.systems | <-DUF499 *<-ORC-CDC6+REase <-?<-ABhydrolase | 348 | Bacteria>Myxococcota>Myxococcia>unclassified_Myxococcia | GCA_003646815.1 | Deltaproteobacteria_bacterium | DRG82_07820 | 1..1043 | 2026735 | glycosyl-transferase,-partial-[Deltaproteobacteria-bacterium] |
| MCY4488568.1 | DUF499 _2.RM.like.systems | LD-peptidase-><-?\|\|DUF499 *-> DUF3780 -> METHYLASE ->N6_MTase-> REase -> HKD-DNase+SF1-SF2-Helicase -> | 1042 | Bacteria>Myxococcota>Myxococcia>unclassified_Myxococcia | GCA_026712905.1 | Deltaproteobacteria_bacterium | OXF11_15860 | 11359..14487 | 2026735 | MAG:-DUF499 -domain-containing-protein-[Deltaproteobacteria-bacterium] |
| MCC6665145.1 | DUF499 _2.RM.like.systems | <-REase +SAVED<-HKD-DNase+SF1-SF2-Helicase <-METHYLASE <-DUF3780 <-DUF499 *<-XIS-HTH<-ParB | 1027 | Bacteria>Myxococcota>Polyangia>Polyangiales>Polyangiaceae | GCA_020849115.1 | Polyangiaceae_bacterium | IT375_15450 | 209262..212345 | 2268199 | DUF499 -domain-containing-protein-[Polyangiaceae-bacterium] |
| MDE0206328.1 | DUF499 _2.RM.like.systems | <-PAS+LuxR-HTH<-PAS+LuxR-HTH\|\|?-> DUF499 *-> DUF3780 ->OLD-ABC->?-> REase +NACHT->Ntox18->Peptidase_M78-> | 1012 | Bacteria>Nitrospinae/Tectomicrobia_group>Candidatus_Tectomicrobia>unclassified_Candidatus_Tectomicrobia | GCA_028819935.1 | Candidatus_Tectomicrobia_bacterium | OXP66_09920 | 9081..12119 | 2528274 | MAG:-DUF499 -domain-containing-protein-[Candidatus-Tectomicrobia-bacterium] |
| MCS6286386.1 | DUF499 _2.RM.like.systems | <-METHYLASE -p<-DUF3780 <-DUF499 *<-REase +35exo+Beta-Barrel+SF1-SF2-Helicase | 1044 | Bacteria>Nitrospirota>Nitrospiria>Nitrospirales>Nitrospiraceae | GCA_024998665.1 | Nitrospira_sp. | H8K08_13255 | 985..4119 | 70125 | ATP-binding-protein-[Nitrospira-sp.] |
| WP_215851339.1 | DUF499 _2.RM.like.systems | Phage_int_SAM_4+Phage_integrase->RHH-> DUF499 *-> DUF3780 ->?->?->PIN-> METHYLASE -> REase +ParB-> | 1041 | Bacteria>Pseudomonadota>Acidithiobacillia>Acidithiobacillales>Acidithiobacillaceae | GCF_031200535.1 | Acidithiobacillus_caldus | HFV02_RS10450 | 3946..7071 | 33059 | DUF499 -domain-containing-protein-[Acidithiobacillus-caldus] |
| MCZ8170923.1 | DUF499 _2.RM.like.systems | <-SMODS<-OBFOLD-NUCLEASE<-HEPN+REase <-HKD-DNase+SF1-SF2-Helicase <-METHYLASE <-DUF3780 <-DUF499 *<-CopG/RHH\|\|?-><-?<-HTH_3 | 1034 | Bacteria>Pseudomonadota>Alphaproteobacteria>Caulobacterales>Caulobacteraceae | GCF_027531485.1 | Brevundimonas_sp. | O9272_04190 | 53052..56156 | 1871086 | MAG:-DUF499 -domain-containing-protein-[Brevundimonas-sp.] |
| MCB1884749.1 | DUF499 _2.RM.like.systems | XIS-HTH->?->?->?->?-> DUF499 *-> DUF3780 -> REase -> METHYLASE -> HKD-DNase+SF1-SF2-Helicase +i REase +CTD-><-Arm-DNA-bind_2+Phage_int_SAM_3+Phage_integrase | 1009 | Bacteria>Pseudomonadota>Alphaproteobacteria>Geminicoccales>Geminicoccaceae | GCA_020445645.1 | Geminicoccaceae_bacterium | KDG89_12270 | 5302..8331 | 2448052 | ATP-binding-protein-[Geminicoccaceae-bacterium] |
| WP_178372757.1 | DUF499 _2.RM.like.systems | <-NUDIX<-?<-?<-?\|\|DUF499 *-> DUF3780 -> METHYLASE -> HKD-DNase+SF1-SF2-Helicase +i REase +CTD-> REase +PNTD1+PNTD2->PIWI->HEPN-> | 1014 | Bacteria>Pseudomonadota>Alphaproteobacteria>Hyphomicrobiales>Nitrobacteraceae | GCF_013390305.1 | Bradyrhizobium_diazoefficiens | HUW42_RS33505 | 7080156..7083200 | 1355477 | DUF499 -domain-containing-protein-[Bradyrhizobium-diazoefficiens] |
| WP_131704163.1 | DUF499 _2.RM.like.systems | <-REase \|\|RHH-> DUF499 *-> DUF3780 -> METHYLASE -> HKD-DNase+SF1-SF2-Helicase ->ParB-> REase -> | 1040 | Bacteria>Pseudomonadota>Alphaproteobacteria>Hyphomicrobiales>Rhizobiaceae | GCF_004330715.1 | Rhizobium_leguminosarum | EOH72_RS11610 | 3872..6994 | 384 | DUF499 -domain-containing-protein-[Rhizobium-leguminosarum] |
| QYK40247.1 | DUF499 _2.RM.like.systems | <-ACET<-?<-HEPN+REase <-HKD-DNase+SF1-SF2-Helicase <-METHYLASE <-DUF3780 <-DUF499 *<-CopG/RHH\|\|?-><-tRNA\|\|?->?-><-tRNA\|\|?->Peptidase_M48-> | 1034 | Bacteria>Pseudomonadota>Alphaproteobacteria>Rhodobacterales>Paracoccaceae | GCA_019454225.1 | Paracoccaceae_bacterium | KF887_12470 | 2639699..2642803 | 1904441 | MAG:-DUF499 -domain-containing-protein-[Rhodobacteraceae-bacterium] |
| AOF98012.1 | DUF499 _2.RM.like.systems | <-HIN-HTH<-HKD-DNase+SF1-SF2-Helicase +i REase +CTD<-METHYLASE -p<-DUF3780 <-DUF499 *<-REase -p<-?<-?<-OLD-ABC<-ParB+HNH | 1026 | Bacteria>Pseudomonadota>Alphaproteobacteria>Sphingomonadales>Sphingomonadaceae | GCF_001713415.1 | Sphingobium_sp._RAC03 | BSY17_1605 | 1643040..1646120 | 1843368 | hypothetical-protein-BSY17_1605-[Sphingobium-sp.-RAC03] |
| WP_237236295.1 | DUF499 _2.RM.like.systems | <-HIN-HTH<-XIS-HTH<-?<-HKD-DNase+SF1-SF2-Helicase +i REase +CTD<-METHYLASE <-DUF3780 <-DUF499 *<-REase -p<-?<-?<-OLD-ABC<-ParB+HNH | 1006 | Bacteria>Pseudomonadota>Alphaproteobacteria>Sphingomonadales>Sphingomonadaceae | GCF_001713415.1 | Sphingobium_sp._RAC03 | BSY17_RS12535 | 1643040..1646060 | 1843368 | DUF499 -domain-containing-protein-[Sphingobium-sp.-RAC03] |
| WP_054473216.1 | DUF499 _2.RM.like.systems | RHH-> DUF499 *-> DUF3780 ->?->MACRODOMAIN-> METHYLASE -> HKD-DNase+SF1-SF2-Helicase ->MORC+REase -> | 1041 | Bacteria>Pseudomonadota>Betaproteobacteria>Burkholderiales>Alcaligenaceae | GCF_001298895.1 | Achromobacter_xylosoxidans | N7677_RS16680 | 47740..50865 | 85698 | DUF499 -domain-containing-protein-[Achromobacter-xylosoxidans] |
| WP_076468429.1 | DUF499 _2.RM.like.systems | <-HAD\|\|?-><-?\|\|RHH-> DUF499 *-> DUF3780 ->?->MACRODOMAIN-> METHYLASE -> HKD-DNase+SF1-SF2-Helicase ->MORC+REase -> | 1041 | Bacteria>Pseudomonadota>Betaproteobacteria>Burkholderiales>Alcaligenaceae | GCF_001957045.1 | Achromobacter_xylosoxidans | BI147_RS15725 | 74082..77207 | 85698 | DUF499 -domain-containing-protein-[Achromobacter-xylosoxidans] |
| WP_009518701.1 | DUF499 _2.RM.like.systems | ZnR+TOPRIM+DSATPase-><-?<-?<-HTH+MPTase<-?\|\|RHH_5-> DUF499 *-> DUF3780 ->?->MACRODOMAIN-> METHYLASE -> HKD-DNase+SF1-SF2-Helicase -> REase -> | 1041 | Bacteria>Pseudomonadota>Betaproteobacteria>Burkholderiales>Comamonadaceae | GCF_001571205.1 | Hydrogenophaga | FGJ01_RS07155 | 285549..288674 | 47420 | MULTISPECIES:-DUF499 -domain-containing-protein-[Hydrogenophaga] |
| MBN9355826.1 | DUF499 _2.RM.like.systems | <-STAS<-ABC-ATPase+SbcC\|\|?->RHH_5-> DUF499 *-> DUF3780 ->?->MACRODOMAIN-> METHYLASE -> HKD-DNase+SF1-SF2-Helicase -> REase -> | 1041 | Bacteria>Pseudomonadota>Betaproteobacteria>Burkholderiales>Oxalobacteraceae | GCF_017307195.1 | Herbaspirillum_huttiense | JOI15_05180 | 142983..146108 | 863372 | DUF499 -domain-containing-protein-[Herbaspirillum-huttiense] |
| WP_276838598.1 | DUF499 _2.RM.like.systems | <-STAS<-ABC-ATPase+SbcC\|\|?->RHH_5-> DUF499 *-> DUF3780 ->?->MACRODOMAIN-> METHYLASE -> HKD-DNase+SF1-SF2-Helicase -> REase -> | 1041 | Bacteria>Pseudomonadota>Betaproteobacteria>Burkholderiales>Oxalobacteraceae | GCF_017307195.1 | Herbaspirillum_huttiense | JOI15_RS05200 | 142983..146108 | 863372 | DUF499 -domain-containing-protein-[Herbaspirillum-huttiense] |
| MBL8482095.1 | DUF499 _2.RM.like.systems | <-ZnR<-?\|\|?->Arm-DNA-bind_3+Phage_integrase-> DUF499 *-> DUF3780 ->Schlafen+HATPase+HTH-> METHYLASE -> HKD-DNase+SF1-SF2-Helicase +i REase +CTD-> REase +PNTD1+PNTD2->PIWI-> | 1008 | Bacteria>Pseudomonadota>Betaproteobacteria>Rhodocyclales>Rhodocyclaceae | GCA_016791085.1 | Rhodocyclaceae_bacterium | JNJ60_07865 | 13203..16229 | 1898103 | ATP-binding-protein-[Rhodocyclaceae-bacterium] |
| WP_218997779.1 | DUF499 _2.RM.like.systems | METHYLASE -> REase ->rve+rve_3+Mu-transpos_C->?->TetR-HTH->XIS-HTH-> DUF499 *-> DUF3780 -> METHYLASE -> HKD-DNase+SF1-SF2-Helicase ->SbcC->\|\|?->SF1-SF2-Helicase-> | 1022 | Bacteria>Pseudomonadota>Gammaproteobacteria>Alteromonadales>Shewanellaceae | GCA_019303335.1 | Shewanella_algae | KOE68_RS03920 | 10181..13249 | 38313 | DUF499 -domain-containing-protein-[Shewanella-algae] |
| ESQ16532.1 | DUF499 _2.RM.like.systems | <-HKD-DNase+SF1-SF2-Helicase +i REase +CTD<-HEPN<-PolB<-METHYLASE <-DUF3780 <-REase <-DUF499 * | 1087 | Bacteria>Pseudomonadota>Gammaproteobacteria>Chromatiales>Chromatiaceae | GCA_000495615.1 | Thiohalocapsa_sp._PB-PSB1 | N838_05685 | 9740..13003 | 1385625 | MAG:-hypothetical-protein-N838_05685-[Thiohalocapsa-sp.-PB-PSB1] |
| QQO57135.1 | DUF499 _2.RM.like.systems | DDE_Tnp_1-> DUF499 *-> REase -> DUF3780 -> METHYLASE ->PolB->HEPN-> HKD-DNase+SF1-SF2-Helicase +i REase +CTD-> | 1085 | Bacteria>Pseudomonadota>Gammaproteobacteria>Chromatiales>Chromatiaceae | GCA_016745215.1 | Thiohalocapsa_sp._PB-PSB1 | N838_31100 | 7373531..7376788 | 1385625 | MAG:-ATP-binding-protein-[Thiohalocapsa-sp.-PB-PSB1] |
| MBV5273604.1 | DUF499 _2.RM.like.systems | DUF499 *-> REase -> DUF3780 ->?->?-> METHYLASE ->STYKinase-like-> | 804 | Bacteria>Pseudomonadota>Gammaproteobacteria>Chromatiales>Chromatiaceae | GCA_019163055.1 | Lamprocystis_purpurea | JZU52_08160 | 1..2416 | 61598 | DUF499 -domain-containing-protein-[Lamprocystis-purpurea] |
| WP_211211735.1 | DUF499 _2.RM.like.systems | DUF499 *-> REase -> DUF3780 ->?->?-> METHYLASE ->STYKinase-like-> | 1034 | Bacteria>Pseudomonadota>Gammaproteobacteria>Chromatiales>Chromatiaceae | GCF_000379525.1 | Lamprocystis_purpurea | A390_RS0125930 | 13980..17084 | 61598 | DUF499 -domain-containing-protein-[Lamprocystis-purpurea] |
| WP_026199823.1 | DUF499 _2.RM.like.systems | REase -> DUF499 -p-><-HicB_lk_antitox\|\|Phd-YefM->PIN-> DUF499 *-> DUF3780 -><-HicB_lk_antitox<-HicA_toxin\|\|METHYLASE -p-> METHYLASE -> REase -> | 1035 | Bacteria>Pseudomonadota>Gammaproteobacteria>Chromatiales>Chromatiaceae | GCF_000379525.1 | Lamprocystis_purpurea | A390_RS0128810 | 112256..115363 | 61598 | DUF499 -domain-containing-protein-[Lamprocystis-purpurea] |
| WP_082673870.1 | DUF499 _2.RM.like.systems | <-ABC-ATPase+SbcC<-?\|\|FtsK+HerA+A32-> DUF499 *->?-> REase -> DUF3780 -><-?<-HIN-HTH | 1004 | Bacteria>Pseudomonadota>Gammaproteobacteria>Chromatiales>Chromatiaceae | GCF_001469165.1 | Thiohalocapsa_sp._ML1 | ML01_RS02170 | 202750..205764 | 1431688 | DUF499 -domain-containing-protein-[Thiohalocapsa-sp.-ML1] |
| WP_245969509.1 | DUF499 _2.RM.like.systems | <-ABhydrolase<-?\|\|ABhydrolase-><-?\|\|?-> DUF499 *-> REase -> DUF3780 ->?->PIN-> METHYLASE -> METHYLASE -p-> | 1016 | Bacteria>Pseudomonadota>Gammaproteobacteria>Chromatiales>Chromatiaceae | GCF_003634315.1 | Thiocapsa_rosea | BDD21_RS09910 | 2229310..2232360 | 69360 | DUF499 -domain-containing-protein-[Thiocapsa-rosea] |
| WP_245970045.1 | DUF499 _2.RM.like.systems | <-PIN<-?<-METHYLASE <-PIN<-?<-DUF3780 <-DUF499 *<-?<-REase +NACHT<-GDResolvase+HIN-HTH | 1018 | Bacteria>Pseudomonadota>Gammaproteobacteria>Chromatiales>Chromatiaceae | GCF_003634315.1 | Thiocapsa_rosea | BDD21_RS27140 | 182504..185560 | 69360 | DUF499 -domain-containing-protein-[Thiocapsa-rosea] |
| MBL1275688.1 | DUF499 _2.RM.like.systems | <-HKD-DNase+SF1-SF2-Helicase <-Schlafen+HATPase+HTH<-METHYLASE <-DUF3780 <-DUF499 *<-RHH_5<-REase \|\|SF1-SF2-Helicase->?-><-?\|\|PADR-HTH-> | 1041 | Bacteria>Pseudomonadota>Gammaproteobacteria>Chromatiales>Ectothiorhodospiraceae | GCA_002401775.2 | Ectothiorhodospiraceae_bacterium | COB30_006350 | 9914..13039 | 2026737 | DUF499 -domain-containing-protein-[Ectothiorhodospiraceae-bacterium] |
| WP_219810711.1 | DUF499 _2.RM.like.systems | HIN-HTH->HIN-HTH-><-REase <-HKD-DNase+SF1-SF2-Helicase <-METHYLASE <-DUF3780 +RAMA<-DUF499 *<-RHH<-Arm-DNA-bind_3+Phage_integrase | 1022 | Bacteria>Pseudomonadota>Gammaproteobacteria>Enterobacterales>Enterobacteriaceae | - | Enterobacter_sp._BIDMC_29 | L465_03720 | 733767..736835 | 1329841 | DUF499 -domain-containing-protein-[Enterobacter-sp.-BIDMC-29] |
| EUM08187.1 | DUF499 _2.RM.like.systems | HIN-HTH->HIN-HTH-><-REase <-HKD-DNase+SF1-SF2-Helicase <-METHYLASE <-DUF3780 +RAMA<-DUF499 *<-RHH<-Arm-DNA-bind_3+Phage_integrase | 1022 | Bacteria>Pseudomonadota>Gammaproteobacteria>Enterobacterales>Enterobacteriaceae | GCA_000534375.1 | Enterobacter_sp._BIDMC_29 | L465_03720 | 733767..736835 | 1329841 | hypothetical-protein-L465_03720-[Enterobacter-sp.-BIDMC-29] |
| WP_059294450.1 | DUF499 _2.RM.like.systems | Arm-DNA-bind_3+Phage_integrase->RHH-> DUF499 *-> DUF3780 +RAMA-> METHYLASE -> HKD-DNase+SF1-SF2-Helicase -> REase -> | 1041 | Bacteria>Pseudomonadota>Gammaproteobacteria>Enterobacterales>Enterobacteriaceae | GCF_001518515.1 | Enterobacter_hormaechei | AWI06_RS12030 | 36830..39955 | 158836 | DUF499 -domain-containing-protein-[Enterobacter-hormaechei] |
| WP_041911426.1 | DUF499 _2.RM.like.systems | HIN-HTH-><-REase <-HKD-DNase+SF1-SF2-Helicase <-METHYLASE <-DUF3780 +RAMA<-DUF499 *<-RHH<-Arm-DNA-bind_3+Phage_integrase | 1041 | Bacteria>Pseudomonadota>Gammaproteobacteria>Enterobacterales>Enterobacteriaceae | GCF_014596715.1 | Enterobacter | L465_RS18340 | 3972931..3976056 | 547 | MULTISPECIES:-DUF499 -domain-containing-protein-[Enterobacter] |
| WP_213752381.1 | DUF499 _2.RM.like.systems | wHTH(SBD)+HNH->?-> DUF499 *-> DUF3780 -> METHYLASE -> HKD-DNase+SF1-SF2-Helicase -><-REase \|\|HIN-HTH-><-Arm-DNA-bind_3+Phage_integrase | 1022 | Bacteria>Pseudomonadota>Gammaproteobacteria>Enterobacterales>Enterobacteriaceae | GCF_018446995.1 | Citrobacter | OGY13_RS10400 | 358156..361224 | 544 | DUF499 -domain-containing-protein-[Citrobacter-portucalensis] |
| HAU1604805.1 | DUF499 _2.RM.like.systems | <-RelE-ParE<-ParD_like<-?<-Y1_Tnp\|\|RHH-> DUF499 *-> DUF3780 -> METHYLASE -> HKD-DNase+SF1-SF2-Helicase ->HTH_3->RVT-> REase -> | 1041 | Bacteria>Pseudomonadota>Gammaproteobacteria>Legionellales>Legionellaceae | GCA_015989465.1 | Legionella_pneumophila | JBJ19_08155 | 5208..8333 | 446 | DUF499 -domain-containing-protein-[Legionella-pneumophila] |
| MDD5320442.1 | DUF499 _2.RM.like.systems | DUF499 *-> DUF3780 -> REase ->?->PIN-> REase -> METHYLASE -> | 1041 | Bacteria>Pseudomonadota>Gammaproteobacteria>Methylococcales>unclassified_Methylococcales | GCA_028714175.1 | Methylococcales_bacterium | PHD43_07485 | 17817..20942 | 2304002 | MAG:-DUF499 -domain-containing-protein-[Methylococcales-bacterium] |
| WP_043884792.1 | DUF499 _2.RM.like.systems | MazG-> REase +SF1-SF2-Helicase ->Arm-DNA-bind_3+Phage_integrase-><-HKD-DNase+SF1-SF2-Helicase <-METHYLASE <-DUF3780 <-DUF499 * | 1038 | Bacteria>Pseudomonadota>Gammaproteobacteria>Pseudomonadales | GCF_003836565.1 | Pseudomonadaceae | PA2G_RS09470 | 2116753..2119869 | 135621 | MULTISPECIES:-DUF499 -domain-containing-protein-[Pseudomonadaceae] |
| EAZ58601.1 | DUF499 _2.RM.like.systems | MazG-> REase +SF1-SF2-Helicase ->Arm-DNA-bind_3+Phage_integrase-><-HKD-DNase+SF1-SF2-Helicase <-METHYLASE <-DUF499 *<-?<-?<-?<-?\|\|HTH+WYL-> | 1072 | Bacteria>Pseudomonadota>Gammaproteobacteria>Pseudomonadales>Pseudomonadaceae | GCA_000152545.1 | Pseudomonas_aeruginosa_2192 | PA2G_01849 | 2116753..2119971 | 350703 | hypothetical-protein-PA2G_01849-[Pseudomonas-aeruginosa-2192] |
| MBH9417281.1 | DUF499 _2.RM.like.systems | HIN-HTH->Peptidase_M15->?->TUDOR+SF1-SF2-Helicase+i REase +CTD->?->?-> DUF499 *-> DUF3780 -> METHYLASE -> HKD-DNase+SF1-SF2-Helicase -><-Arm-DNA-bind_3+Phage_integrase<- REase +SF1-SF2-Helicase <-MazG | 1038 | Bacteria>Pseudomonadota>Gammaproteobacteria>Pseudomonadales>Pseudomonadaceae | GCA_016092775.1 | Pseudomonas_aeruginosa | GHU28_23410 | 8846..11962 | 287 | DUF499 -domain-containing-protein-[Pseudomonas-aeruginosa] |
| EIU2715573.1 | DUF499 _2.RM.like.systems | SF1-SF2-Helicase->HIN-HTH->?->TUDOR+SF1-SF2-Helicase+i REase +CTD->?->?-> DUF499 *-> DUF3780 -> METHYLASE -> HKD-DNase+SF1-SF2-Helicase -><-Arm-DNA-bind_3+Phage_integrase<- REase +SF1-SF2-Helicase <-MazG | 1072 | Bacteria>Pseudomonadota>Gammaproteobacteria>Pseudomonadales>Pseudomonadaceae | GCA_021614535.1 | Pseudomonas_aeruginosa | L4O33_002031 | 46678..49896 | 287 | DUF499 -domain-containing-protein-[Pseudomonas-aeruginosa] |
| WP_121373084.1 | DUF499 _2.RM.like.systems | REase +SF1-SF2-Helicase ->Arm-DNA-bind_3+Phage_integrase-><-HKD-DNase+SF1-SF2-Helicase <-METHYLASE <-DUF3780 <-DUF499 * | 737 | Bacteria>Pseudomonadota>Gammaproteobacteria>Pseudomonadales>Pseudomonadaceae | GCF_003630125.1 | Pseudomonas_aeruginosa | DQI18_RS19110 | 9389..11604 | 287 | DUF499 -domain-containing-protein,-partial-[Pseudomonas-aeruginosa] |
| WP_134269529.1 | DUF499 _2.RM.like.systems | MazG-> REase +SF1-SF2-Helicase ->Arm-DNA-bind_3+Phage_integrase-><-HKD-DNase+SF1-SF2-Helicase <-METHYLASE <-DUF3780 <-DUF499 *<-?<-?<-TUDOR+SF1-SF2-Helicase+i REase +CTD<-?<-Peptidase_M15<-HIN-HTH | 1038 | Bacteria>Pseudomonadota>Gammaproteobacteria>Pseudomonadales>Pseudomonadaceae | GCF_004373135.1 | Pseudomonas_aeruginosa | IPC162_RS08525 | 440785..443901 | 287 | DUF499 -domain-containing-protein-[Pseudomonas-aeruginosa] |
| WP_153986190.1 | DUF499 _2.RM.like.systems | Peptidase_M15->?->Ntox49->TUDOR+SF1-SF2-Helicase+i REase +CTD->?->?-> DUF499 *-> DUF3780 -> METHYLASE -> HKD-DNase+SF1-SF2-Helicase -><-Arm-DNA-bind_3+Phage_integrase<- REase +SF1-SF2-Helicase <-MazG | 1038 | Bacteria>Pseudomonadota>Gammaproteobacteria>Pseudomonadales>Pseudomonadaceae | GCF_009664165.1 | Pseudomonas_aeruginosa | FOZ66_RS12385 | 2584610..2587726 | 287 | DUF499 -domain-containing-protein-[Pseudomonas-aeruginosa] |
| WP_237047019.1 | DUF499 _2.RM.like.systems | MazG-> REase +SF1-SF2-Helicase ->Arm-DNA-bind_3+Phage_integrase-><-HKD-DNase+SF1-SF2-Helicase <-METHYLASE <-DUF3780 <-DUF499 * | 908 | Bacteria>Pseudomonadota>Gammaproteobacteria>Pseudomonadales>Pseudomonadaceae | GCF_009791355.1 | Pseudomonas_aeruginosa | GQF67_RS10665 | 323805..326532 | 287 | DUF499 -domain-containing-protein,-partial-[Pseudomonas-aeruginosa] |
| WP_123186567.1 | DUF499 _2.RM.like.systems | Ntox49->TUDOR+SF1-SF2-Helicase+i REase +CTD->?->?-> DUF499 *-> DUF3780 -> METHYLASE -> HKD-DNase+SF1-SF2-Helicase -><-Arm-DNA-bind_3+Phage_integrase<- REase +SF1-SF2-Helicase <-MazG | 1038 | Bacteria>Pseudomonadota>Gammaproteobacteria>Pseudomonadales>Pseudomonadaceae | GCF_013255565.1 | Pseudomonas | NLZ23_RS29065 | 7522..10638 | 286 | DUF499 -domain-containing-protein-[Pseudomonas-aeruginosa] |
| WP_206437318.1 | DUF499 _2.RM.like.systems | Peptidase_M15->?->Ntox49->TUDOR+SF1-SF2-Helicase+i REase +CTD->?->?-> DUF499 *-> DUF3780 -> METHYLASE -> HKD-DNase+SF1-SF2-Helicase -><-Arm-DNA-bind_3+Phage_integrase<- REase +SF1-SF2-Helicase <-MazG | 1072 | Bacteria>Pseudomonadota>Gammaproteobacteria>Pseudomonadales>Pseudomonadaceae | - | Pseudomonas_aeruginosa | PQC20_002029 | 43524..46742 | 287 | DUF499 -domain-containing-protein-[Pseudomonas-aeruginosa] |
| MBV1952380.1 | DUF499 _2.RM.like.systems | RHH_1-> DUF499 *-> DUF3780 -> METHYLASE -> HKD-DNase+SF1-SF2-Helicase ->SF1-SF2-Helicase-> REase -> | 1043 | Bacteria>Pseudomonadota>Gammaproteobacteria>Thiotrichales>Piscirickettsiaceae | GCA_019090295.1 | Cycloclasticus_sp. | KUG64_09355 | 786..3917 | 2024830 | DUF499 -domain-containing-protein-[Cycloclasticus-sp.] |
| MCY4011318.1 | DUF499 _2.RM.like.systems | <-HKD-DNase+SF1-SF2-Helicase +i REase +CTD<-METHYLASE -p<-METHYLASE <-DUF3780 <-DUF499 *<-REase \|\|Arm-DNA-bind_3+Phage_integrase-> | 1000 | Bacteria>Pseudomonadota>Gammaproteobacteria>unclassified_Gammaproteobacteria | GCA_026708185.1 | Gammaproteobacteria_bacterium | OXG82_01240 | 7722..10724 | 1913989 | MAG:-DUF499 -domain-containing-protein-[Gammaproteobacteria-bacterium] |
| MDE0192036.1 | DUF499 _2.RM.like.systems | <-HKD-DNase+SF1-SF2-Helicase <-REase <-N6_MTase<-METHYLASE <-DUF3780 <-DUF499 *<-RelE-ParE<-OLD-ABC<-ParB+HNH | 1021 | Bacteria>Pseudomonadota>Gammaproteobacteria>unclassified_Gammaproteobacteria | GCA_028819695.1 | Gammaproteobacteria_bacterium | OXQ90_11825 | 72946..76011 | 1913989 | MAG:-DUF499 -domain-containing-protein-[Gammaproteobacteria-bacterium] |
| MCE9614797.1 | DUF499 _2.RM.like.systems | <-HTH_3\|\|?->?->?->?->HTH-> DUF499 *-> DUF3780 -> REase ->?->DNA_methylase-> METHYLASE -> HKD-DNase+SF1-SF2-Helicase +i REase +CTD-> | 986 | Bacteria>PVC_group>Lentisphaerota>unclassified_Lentisphaerota | GCA_021413965.1 | Lentisphaerota_bacterium | K8T26_11010 | 329228..332188 | 1932692 | DUF499 -domain-containing-protein-[Lentisphaerae-bacterium] |
| MBX3412588.1 | DUF499 _2.RM.like.systems | ArdCN+MPTase->?->?->NTD+SF1-SF2-Helicase->Phage_int_SAM_1+Phage_integrase->XIS-HTH-> DUF499 *-> DUF3780 -> REase ->?-> METHYLASE ->?-> HKD-DNase+SF1-SF2-Helicase +i REase +CTD-> | 1023 | Bacteria>PVC_group>Planctomycetota>Planctomycetia>Pirellulales | GCA_019636335.1 | Pirellulales_bacterium | KF708_07865 | 38769..41840 | 2715253 | ATP-binding-protein-[Pirellulales-bacterium] |
| MCX8496484.1 | DUF499 _2.RM.like.systems | <-REase <-?\|\|HTH->MPTase->?->?-> DUF499 *-> DUF3780 -> METHYLASE -> METHYLASE -> HKD-DNase+SF1-SF2-Helicase -> REase ->HEPN-> | 1044 | Bacteria>PVC_group>Verrucomicrobiota>Verrucomicrobiae>Verrucomicrobiales | GCA_026461385.1 | Akkermansiaceae_bacterium | ORN51_09920 | 5504..8638 | 2562705 | MAG:-DUF499 -domain-containing-protein-[Akkermansiaceae-bacterium] |
| WP_172896457.1 | DUF499 _2.RM.like.systems | DUF499 *-> DUF3780 -> METHYLASE -> HKD-DNase+SF1-SF2-Helicase +i REase +CTD-> REase -> | 989 | Bacteria>Terrabacteria_group>Actinomycetota>Actinomycetes>Micromonosporales | GCF_900091445.1 | Micromonospora_echinofusca | GA0070610_RS08520 | 1858371..1861340 | 47858 | DUF499 -domain-containing-protein-[Micromonospora-echinofusca] |
| WP_260289258.1 | DUF499 _2.RM.like.systems | <-ParB<-TrwC+SF1-SF2-Helicase<-REase <-HKD-DNase+SF1-SF2-Helicase +i REase +CTD<-METHYLASE <-DUF3780 <-DUF499 *<-XIS-HTH\|\|?->ArdCN+MPTase->Primpol+PriCT1-> | 1092 | Bacteria>Terrabacteria_group>Actinomycetota>Actinomycetes>Propionibacteriales | GCF_025187725.1 | Propionibacterium_freudenreichii | EFN01_RS07350 | 14264..17542 | 1744 | DUF499 -domain-containing-protein-[Propionibacterium-freudenreichii] |
| WP_081083807.1 | DUF499 _2.RM.like.systems | XIS-HTH-> DUF499 *-> DUF3780 -> METHYLASE -> HKD-DNase+SF1-SF2-Helicase +i REase +CTD-> REase ->TrwC+SF1-SF2-Helicase->ParB-> | 1092 | Bacteria>Terrabacteria_group>Actinomycetota>Actinomycetes>Propionibacteriales | GCF_900000055.1 | Propionibacterium_freudenreichii | FAM23868_RS05705 | 570..3848 | 1744 | DUF499 -domain-containing-protein-[Propionibacterium-freudenreichii] |
| MCL6597625.1 | DUF499 _2.RM.like.systems | <-HKD-DNase+SF1-SF2-Helicase +i REase +CTD<-METHYLASE <-DUF3780 <-DUF499 *<-REase + REase <-REase <-?<-?\|\|XIS-HTH->Arm-DNA-bind_4+Phage_int_SAM_3+Phage_integrase-> | 1071 | Bacteria>Terrabacteria_group>Bacillota>Bacilli>Bacillales | GCA_023512275.1 | Alicyclobacillus_macrosporangiidus | K6T81_02655 | 38493..41708 | 392015 | DUF499 -domain-containing-protein-[Alicyclobacillus-macrosporangiidus] |
| WP_192951079.1 | DUF499 _2.RM.like.systems | TRAM+Methyltransf_31->?-> REase ->MULE-TRANSPOSASE-> DUF499 *-> DUF3780 -> METHYLASE -> HKD-DNase+SF1-SF2-Helicase -><-HTH+MPTase | 1034 | Bacteria>Terrabacteria_group>Bacillota>Bacilli>Bacillales | GCF_014890455.1 | Anoxybacillus_flavithermus | HPK01_RS05390 | 42811..45915 | 33934 | DUF499 -domain-containing-protein-[Anoxybacillus-flavithermus] |
| PHS35622.1 | DUF499 _2.RM.like.systems | <-Pkinase<-REase <-HKD-DNase+SF1-SF2-Helicase <-METHYLASE <-DUF3780 <-DUF499 *<-?<-?<-DOC+HTH<-BirA-HTH | 1030 | Bacteria>Terrabacteria_group>Bacillota>Clostridia>Eubacteriales | GCA_002733545.1 | Alkaliphilus_sp. | COA82_04085 | 77629..80721 | 1872455 | MAG:-glycosyl-transferase-[Alkaliphilus-sp.] |
| MBM7867397.1 | DUF499 _2.RM.like.systems | REase -> DUF499 *-> DUF3780 -> METHYLASE -> HKD-DNase+SF1-SF2-Helicase ->ParB+HNH-p->?->Classical-AAA-> | 1023 | Bacteria>Terrabacteria_group>Bacillota>Clostridia>Eubacteriales | GCA_016908025.1 | Heliomicrobium_gestii | JOC69_002303 | 75173..78244 | 2699 | hypothetical-protein-[Heliomicrobium-gestii] |
| WP_003361845.1 | DUF499 _2.RM.like.systems | <-PADR-HTH\|\|wHTH(SBD)+HNH->?-> DUF499 *-> DUF3780 -> METHYLASE -> HKD-DNase+SF1-SF2-Helicase -> REase +RAMA->?->Radical-SAM-> | 1031 | Bacteria>Terrabacteria_group>Bacillota>Clostridia>Eubacteriales | GCF_000020345.1 | Clostridium_botulinum | AGE29_RS02755 | 251614..254709 | 1491 | DUF499 -domain-containing-protein-[Clostridium-botulinum] |
| WP_039231429.1 | DUF499 _2.RM.like.systems | OrfB_IS605+OrfB_Zn_ribbon->SAM-methylase->?->tRNA->tRNA->tRNA->tRNA->tRNA->tRNA->?->?-> DUF499 *-> DUF3780 -> METHYLASE -> HKD-DNase+SF1-SF2-Helicase -> REase +RAMA-> | 1031 | Bacteria>Terrabacteria_group>Bacillota>Clostridia>Eubacteriales | GCF_000724265.2 | Clostridium_novyi | Z958_RS01340 | 4777..7872 | 1542 | DUF499 -domain-containing-protein-[Clostridium-novyi] |
| WP_039774266.1 | DUF499 _2.RM.like.systems | <-PADR-HTH\|\|wHTH(SBD)+HNH->?-> DUF499 *-> DUF3780 -> METHYLASE -> HKD-DNase+SF1-SF2-Helicase -> REase +RAMA->?->Radical-SAM-> | 1031 | Bacteria>Terrabacteria_group>Bacillota>Clostridia>Eubacteriales | GCF_000817935.1 | Clostridium_botulinum | T259_RS11600 | 2303984..2307079 | 1491 | DUF499 -domain-containing-protein-[Clostridium-botulinum] |
| WP_252891859.1 | DUF499 _2.RM.like.systems | DUF499 -p-> DUF499 *-> DUF499 -p-> DUF499 -p-> METHYLASE -> REase -> HKD-DNase+SF1-SF2-Helicase -p-> HKD-DNase+SF1-SF2-Helicase -> | 195 | Bacteria>Terrabacteria_group>Bacillota>Clostridia>Eubacteriales | GCF_001310995.1 | Thermoclostridium_stercorarium | JCM9324_RS16190 | 584..1171 | 1510 | DUF499 -domain-containing-protein-[Thermoclostridium-stercorarium] |
| WP_061317398.1 | DUF499 _2.RM.like.systems | <-Radical-SAM<-REase +RAMA<-HKD-DNase+SF1-SF2-Helicase <-METHYLASE -p<-METHYLASE <-DUF3780 <-DUF499 *<-?<-wHTH(SBD)+HNH\|\|PADR-HTH-> | 1031 | Bacteria>Terrabacteria_group>Bacillota>Clostridia>Eubacteriales | GCF_001573255.1 | Clostridium_botulinum | AZI51_RS06445 | 49483..52578 | 1491 | DUF499 -domain-containing-protein-[Clostridium-botulinum] |
| WP_061320311.1 | DUF499 _2.RM.like.systems | wHTH(SBD)+HNH->?-> DUF499 *-> DUF3780 -> METHYLASE -> HKD-DNase+SF1-SF2-Helicase -> REase +RAMA->?->Radical-SAM-> | 1031 | Bacteria>Terrabacteria_group>Bacillota>Clostridia>Eubacteriales | GCF_001573935.1 | Clostridium_botulinum | AZI72_RS02380 | 8952..12047 | 1491 | DUF499 -domain-containing-protein-[Clostridium-botulinum] |
| WP_065821436.1 | DUF499 _2.RM.like.systems | MULE-TRANSPOSASE-><-HD<-HKD-DNase+SF1-SF2-Helicase <-REase <-METHYLASE <-DUF3780 <-DUF499 *\|\|MULE-TRANSPOSASE-><-?<-?\|\|MULE-TRANSPOSASE-> | 1036 | Bacteria>Terrabacteria_group>Bacillota>Clostridia>Eubacteriales | GCF_001697245.1 | Thermoclostridium_stercorarium | CSTERTH_RS08760 | 2032374..2035484 | 1510 | DUF499 -domain-containing-protein-[Thermoclostridium-stercorarium] |
| WP_034844279.1 | DUF499 _2.RM.like.systems | HIN-HTH-><-?<-HIN-HTH<-HIN-HTH<-ZnR\|\|TraJ-RHH-> DUF499 *-> DUF3780 -> METHYLASE -> REase -> HKD-DNase+SF1-SF2-Helicase ->?->URI-DNase-> | 1036 | Bacteria>Terrabacteria_group>Bacillota>Clostridia>Eubacteriales | GCF_001697285.1 | Thermoclostridium_stercorarium | CSTERLE_RS00935 | 220886..223996 | 1510 | DUF499 -domain-containing-protein-[Thermoclostridium-stercorarium] |
| WP_101515073.1 | DUF499 _2.RM.like.systems | <-Radical-SAM<-?<-REase +RAMA<-HKD-DNase+SF1-SF2-Helicase <-METHYLASE <-DUF3780 <-DUF499 *<-?<-wHTH(SBD)+HNH | 1031 | Bacteria>Terrabacteria_group>Bacillota>Clostridia>Eubacteriales | GCF_002866225.1 | Clostridium_botulinum | RSJ21_RS11400 | 2254845..2257940 | 1491 | DUF499 -domain-containing-protein-[Clostridium-botulinum] |
| WP_106574724.1 | DUF499 _2.RM.like.systems | <-Radical-SAM<-REase +RAMA<-HKD-DNase+SF1-SF2-Helicase <-METHYLASE <-DUF3780 <-DUF499 *<-?<-?\|\|?->?-><-sGTP | 1031 | Bacteria>Terrabacteria_group>Bacillota>Clostridia>Eubacteriales | GCF_003014955.1 | Clostridium_botulinum | C6C12_RS10395 | 111152..114247 | 1491 | DUF499 -domain-containing-protein-[Clostridium-botulinum] |
| WP_161821648.1 | DUF499 _2.RM.like.systems | <-wHTH+SAD+HNH<-REase <-HEPN<-HKD-DNase+SF1-SF2-Helicase +i REase +CTD<-METHYLASE <-DUF3780 <-DUF499 *\|\|?-><-?<-?<-Glycos_trans_3N+Glycos_transf_3\|\|?->HAMP+HISKIN-> | 1082 | Bacteria>Terrabacteria_group>Bacillota>Clostridia>Eubacteriales | GCF_009932395.1 | Sporotomaculum_syntrophicum | SPSYN_RS06335 | 287290..290538 | 182264 | DUF499 -domain-containing-protein-[Sporotomaculum-syntrophicum] |
| WP_235920429.1 | DUF499 _2.RM.like.systems | REase -> DUF499 *-> DUF3780 -> METHYLASE -> HKD-DNase+SF1-SF2-Helicase ->ParB+HNH-p->?->Classical-AAA-> | 942 | Bacteria>Terrabacteria_group>Bacillota>Clostridia>Eubacteriales | GCF_016908025.1 | Heliomicrobium_gestii | JOC69_RS11260 | 75416..78244 | 2699 | DUF499 -domain-containing-protein-[Heliomicrobium-gestii] |
| WP_245124324.1 | DUF499 _2.RM.like.systems | sGTP-><-?<-?\|\|?->wHTH(SBD)+HNH->?-> DUF499 *-> DUF3780 -> METHYLASE -> HKD-DNase+SF1-SF2-Helicase -> REase +RAMA->Radical-SAM-> | 1031 | Bacteria>Terrabacteria_group>Bacillota>Clostridia>Eubacteriales | GCF_022936445.1 | Clostridium_botulinum | LOO64_RS12750 | 357814..360909 | 1491 | DUF499 -domain-containing-protein-[Clostridium-botulinum] |
| WP_180703528.1 | DUF499 _2.RM.like.systems | <-Phage_int_SAM_1+Phage_integrase\|\|Methyltransf_4-><-?\|\|?-> DUF499 *-> DUF3780 -> METHYLASE -> HKD-DNase+SF1-SF2-Helicase -> REase ->Phage_integrase->LexA-HTH+LexA-protease-> | 1029 | Bacteria>Terrabacteria_group>Bacillota>Clostridia>Eubacteriales | GCF_900015215.1 | Romboutsia_ilealis | CRIB_RS05530 | 1166039..1169128 | 1115758 | DUF499 -domain-containing-protein-[Romboutsia-ilealis] |
| WP_097019088.1 | DUF499 _2.RM.like.systems | <-HEPN+CBS<-HKD-DNase+SF1-SF2-Helicase <-METHYLASE <-DUF3780 <-DUF499 *<-REase <-?<-?<-?<-Methyltransf<-METHYLASE | 1029 | Bacteria>Terrabacteria_group>Bacillota>Clostridia>Halanaerobiales | GCF_900215265.1 | Orenia_metallireducens | CRN99_RS15855 | 34717..37806 | 1413210 | DUF499 -domain-containing-protein-[Orenia-metallireducens] |
| WP_075517732.1 | DUF499 _2.RM.like.systems | <-MULE-TRANSPOSASE<-?\|\|DUF499 *-> DUF3780 -> METHYLASE -> HKD-DNase+SF1-SF2-Helicase -> REase +NACHT->Mbetalac-> | 1026 | Bacteria>Terrabacteria_group>Bacillota>Clostridia>Moorellales | - | Moorella_thermoacetica | MOTE_RS11890 | 1060..4140 | 1525 | DUF499 -domain-containing-protein-[Moorella-thermoacetica] |
| WP_154407767.1 | DUF499 _2.RM.like.systems | DUF499 *-> DUF3780 -> METHYLASE -> HKD-DNase+SF1-SF2-Helicase -><-REase <-Glyoxalase\|\|ACET-> | 1040 | Bacteria>Terrabacteria_group>Bacillota>Negativicutes>Selenomonadales | GCF_009695585.1 | Anaerovibrio_slackiae | FYJ84_RS11575 | 24516..27638 | 2652309 | DUF499 -domain-containing-protein-[Anaerovibrio-slackiae] |
| GIV64890.1 | DUF499 _2.RM.like.systems | <-Peptidase<-HKD-DNase+SF1-SF2-Helicase <-METHYLASE <-DUF3780 <-DUF499 *<-REase +DUF5655\|\|Phage_int_SAM_4+Phage_integrase-> | 1029 | Bacteria>Terrabacteria_group>Chloroflexota>Anaerolineae>Anaerolineales | GCA_026003595.1 | Bellilinea_sp. | KatS3mg046_150 | 153425..156514 | 2838785 | MAG:-hypothetical-protein-KatS3mg046_150-[Bellilinea-sp.] |
| MCY3660412.1 | DUF499 _2.RM.like.systems | <-DUF499 *<-CR-ATPase8 + REase | 943 | Bacteria>Terrabacteria_group>Chloroflexota>Caldilineae>Caldilineales | GCA_026705565.1 | Caldilineaceae_bacterium | OXG36_15520 | 517..3348 | 2614250 | MAG:-DUF499 -domain-containing-protein-[Caldilineaceae-bacterium] |
| NTV99826.1 | DUF499 _2.RM.like.systems | DUF499 *-> DUF3780 -> METHYLASE -> HKD-DNase+SF1-SF2-Helicase -><-REase \|\|DDE_Tnp_1_assoc+DDE_Tnp_1->DDE_Tnp_1-> | 816 | Bacteria>Terrabacteria_group>Chloroflexota>Chloroflexia>Chloroflexales | GCA_013336555.1 | Oscillochloris_sp. | HGA19_00810 | 1512..3962 | 2547424 | DUF499 -domain-containing-protein-[Oscillochloris-sp.] |
| HCG29367.1 | DUF499 _2.RM.like.systems | REase -> DUF499 *-> DUF3780 -> METHYLASE -> HKD-DNase+SF1-SF2-Helicase ->HicA_toxin->HicB_lk_antitox-> | 1024 | Bacteria>Terrabacteria_group>Chloroflexota>unclassified_Chloroflexota | GCA_003534145.1 | Chloroflexota_bacterium | DEU95_06405 | 3683..6757 | 2026724 | hypothetical-protein-[Chloroflexi-bacterium] |
| KKZ13991.1 | DUF499 _2.RM.like.systems | HEPN-> DUF499 *-> DUF3780 -> REase ->?->ParB+HNH->OLD-ABC-> | 1040 | Bacteria>Terrabacteria_group>Cyanobacteriota/Melainabacteria_group>Cyanobacteriota>Cyanophyceae | GCA_001007635.1 | Candidatus_Synechococcus_spongiarum_15L | TQ37_02560 | 1477..4599 | 1608419 | hypothetical-protein-TQ37_02560-[Candidatus-Synechococcus-spongiarum-15L] |
| OOV26578.1 | DUF499 _2.RM.like.systems | DUF499 *-> DUF3780 -> REase ->?->ParB+HNH-> | 1040 | Bacteria>Terrabacteria_group>Cyanobacteriota/Melainabacteria_group>Cyanobacteriota>Cyanophyceae | GCA_002018045.1 | Candidatus_Synechococcus_spongiarum_LMB_bulk15M | BV61_06570 | 12460..15582 | 1943582 | hypothetical-protein-BV61_06570-[Candidatus-Synechococcus-spongiarum-LMB-bulk15M] |
| PZV10243.1 | DUF499 _2.RM.like.systems | DUF499 *-> DUF3780 -> REase +PNTD1+PNTD2->PIWI-><-MACRODOMAIN<-?\|\|METHYLASE -> | 1076 | Bacteria>Terrabacteria_group>Cyanobacteriota/Melainabacteria_group>Cyanobacteriota>Cyanophyceae | GCA_003249105.1 | Leptolyngbya_sp. | DCF32_00020 | 1234..4464 | 47254 | AAA-family-ATPase-[Leptolyngbya-sp.] |
| MXW11415.1 | DUF499 _2.RM.like.systems | <-AbrB-like\|\|?->?-> DUF499 *-> DUF3780 -> REase ->?->OLD-ABC-> METHYLASE -> | 1020 | Bacteria>Terrabacteria_group>Cyanobacteriota/Melainabacteria_group>Cyanobacteriota>Cyanophyceae | GCA_009836025.1 | Synechococcus_sp._SB0668_bin_13 | F4Z73_00765 | 8681..11743 | 2604859 | ATP-binding-protein-[Synechococcus-sp.-SB0668_bin_13] |
| MXY61915.1 | DUF499 _2.RM.like.systems | DUF499 *-> DUF3780 -> REase ->?->?->ParB+HNH->OLD-ABC-> | 1035 | Bacteria>Terrabacteria_group>Cyanobacteriota/Melainabacteria_group>Cyanobacteriota>Cyanophyceae | GCA_009837735.1 | Synechococcus_sp._SB0665_bin_28 | F4Y87_00410 | 1971..5078 | 2604856 | ATP-binding-protein-[Synechococcus-sp.-SB0665_bin_28] |
| MYG64556.1 | DUF499 _2.RM.like.systems | <-REase <-DUF3780 <-DUF499 *<-?<-?\|\|AbrB-like-> | 1025 | Bacteria>Terrabacteria_group>Cyanobacteriota/Melainabacteria_group>Cyanobacteriota>Cyanophyceae | GCA_009843455.1 | Synechococcus_sp._SB0675_bin_7 | F4158_08385 | 1344..4421 | 2604868 | ATP-binding-protein-[Synechococcus-sp.-SB0675_bin_7] |
| MYI71486.1 | DUF499 _2.RM.like.systems | DUF499 *-> DUF3780 -> REase ->?->?->OLD-ABC-> | 1020 | Bacteria>Terrabacteria_group>Cyanobacteriota/Melainabacteria_group>Cyanobacteriota>Cyanophyceae | GCA_009844645.1 | Synechococcus_sp._SB0673_bin_10 | F4099_03010 | 1152..4214 | 2604866 | ATP-binding-protein-[Synechococcus-sp.-SB0673_bin_10] |

| Protein-ID | System | Gene-Neighbourhoods | Length | Taxonomy | Assembly-ID | Strain | Locus-Tag | Genomic-coordinates | TaxID | Fasta-defline |
|---|---|---|---|---|---|---|---|---|---|---|
| -- | | | | | | | | | | |
| -- Schlafen+HATPase+wHTH | | | | | | | | | | |
| -- | | | | | | | | | | |
| WP_050460473.1 | DUF499_2.RM.like.systems | <-MarR-HTH<-HKD-DNase+SF1-SF2-Helicase+iREase+CTD<-METHYLASE<-DUF3780<-DUF499*<-Schlafen+HATPase+wHTH<-SWACOS\|\|?->RADICAL-SAM->HTH+IclR-HTH->NTP_transf+wHTH-> | 1097 | Archaea>Euryarchaeota>Stenosarchaea_group>Halobacteria>Haloferacales | GCF_001190965.1 | Haloferax_gibbonsii | ABY42_RS18700 | 86264..89557 | 35746 | DUF499-domain-containing-protein-[Haloferax-gibbonsii] |
| WP_271963542.1 | DUF499_2.RM.like.systems | <-Schlafen<-HKD-DNase+SF1-SF2-Helicase+iREase+CTD\|\|ParB+HNH-><-METHYLASE<-DUF3780<-DUF499*\|\|?->?->?-><-Phage_integrase\|\|PadR-HTH->PadR-HTH-> | 1107 | Archaea>Euryarchaeota>Stenosarchaea_group>Halobacteria>Haloferacales | GCF_028211285.1 | Halorubrum_ezzemoulense | PM025_RS15440 | 14437..17760 | 337243 | DUF499-domain-containing-protein-[Halorubrum-ezzemoulense] |
| NLA99643.1 | DUF499_2.RM.like.systems | Schlafen+HATPase+wHTH->?-><-DUF3780<-DUF499*\|\|METHYLASE-><-?<-HKD-DNase+SF1-SF2-Helicase+iREase+CTD<-?<-ORC/CDC6+HTH | 950 | Archaea>Euryarchaeota>Stenosarchaea_group>Methanomicrobia>Methanomicrobiales | GCA_012511245.1 | Methanomicrobiales_archaeon | GX837_01550 | 2570..5422 | 2184052 | DUF499-domain-containing-protein-[Methanomicrobiales-archaeon] |
| MBN1569786.1 | DUF499_2.RM.like.systems | <-wHTH+SAD+HNH<-?<-HKD-DNase+SF1-SF2-Helicase<-Schlafen+HATPase+wHTH<-METHYLASE<-DUF3780<-DUF499*<-?<-HTH+WYL | 1021 | Bacteria>Acidobacteriota>unclassified_Acidobacteriota | GCA_016929335.1 | Acidobacteriota_bacterium | JXA73_18215 | 77162..80227 | 1978231 | DUF499-domain-containing-protein-[Acidobacteria-bacterium] |
| NLT02438.1 | DUF499_2.RM.like.systems | <-ParB+HNH<-?<-HKD-DNase+SF1-SF2-Helicase<-Schlafen+HATPase+wHTH<-METHYLASE<-DUF3780<-DUF499*<-tRNA\|\|?->?->NADAR->?->Aminotran_1_2-> | 1038 | Bacteria>FCB_group>Bacteroidota/Chlorobiota_group>Bacteroidota>Bacteroidia | GCA_012718945.1 | Bacteroidales_bacterium | GXY09_01085 | 15955..19071 | 2030927 | DUF499-domain-containing-protein-[Bacteroidales-bacterium] |
| NCA71196.1 | DUF499_2.RM.like.systems | MODE-HTH+PBPII-><-DUF29\|\|DUF499*->DUF3780->Schlafen+HATPase+wHTH->METHYLASE-> | 1037 | Bacteria>FCB_group>Bacteroidota/Chlorobiota_group>Bacteroidota>Sphingobacteriia | GCA_009928635.1 | Sphingobacteriia_bacterium | EOM91_14035 | 5029..8142 | 2035813 | ATP-binding-protein-[Sphingobacteriia-bacterium] |
| MCC6769769.1 | DUF499_2.RM.like.systems | <-DUF772+DDE_Tnp_1+DDE_Tnp_1_6<-?<-?\|\|DUF499*->DUF3780->METHYLASE->Schlafen+HATPase+wHTH->HKD-DNase+SF1-SF2-Helicase+iREase+CTD-><-CHTH | 1013 | Bacteria>FCB_group>Gemmatimonadota>Gemmatimonadetes>Gemmatimonadales | GCA_020848865.1 | Gemmatimonadaceae_bacterium | IT360_01040 | 3136..6177 | 2026741 | ATP-binding-protein-[Gemmatimonadaceae-bacterium] |
| MCZ8206518.1 | DUF499_2.RM.like.systems | <-REase+NACHT\|\|DUF499*->DUF3780->METHYLASE->Schlafen+HATPase+wHTH->HKD-DNase+SF1-SF2-Helicase+iREase+CTD-> | 1013 | Bacteria>FCB_group>Gemmatimonadota>Gemmatimonadetes>Gemmatimonadales | GCA_027531815.1 | Gemmatimonas_sp. | O9271_18135 | 3551..6592 | 1962908 | MAG:-DUF499-domain-containing-protein-[Gemmatimonas-sp.] |
| MBK6307121.1 | DUF499_2.RM.like.systems | <-GDResolvase<-?\|\|?-><-HKD-DNase+SF1-SF2-Helicase+iREase+CTD<-?<-METHYLASE<-DUF499*<-Schlafen+HATPase+wHTH | 957 | Bacteria>FCB_group>Gemmatimonadota>unclassified_Gemmatimonadota | GCA_016705425.1 | Gemmatimonadota_bacterium | IPF47_15915 | 119468..122341 | 2026742 | ATP-binding-protein-[Gemmatimonadetes-bacterium] |
| MCE2900724.1 | DUF499_2.RM.like.systems | DUF499*->DUF3780->METHYLASE-p->DUF772+DDE_Tnp_1->METHYLASE-p->Schlafen+HATPase+wHTH->HKD-DNase+SF1-SF2-Helicase+iREase+CTD-> | 1013 | Bacteria>FCB_group>Gemmatimonadota>unclassified_Gemmatimonadota | GCA_021299375.1 | Gemmatimonadota_bacterium | LXA09_05735 | 68806..71847 | 2026742 | DUF499-domain-containing-protein-[Gemmatimonadetes-bacterium] |
| MCD6274302.1 | DUF499_2.RM.like.systems | DUF499*->DUF3780->METHYLASE->Schlafen-> | 1032 | Bacteria>Myxococcota>Myxococcia>unclassified_Myxococcia | GCA_021163685.1 | Deltaproteobacteria_bacterium | J7K30_16050 | 3210..6308 | 2026735 | DUF499-domain-containing-protein-[Deltaproteobacteria-bacterium] |
| MBT3366889.1 | DUF499_2.RM.like.systems | ParB+HNH+Schlafen->DUF499*->DUF3780->METHYLASE->HicB_lk_antitox->HicA_toxin->HKD-DNase+SF1-SF2-Helicase->HNH-> | 1035 | Bacteria>Nitrospinae/Tectomicrobia_group>Nitrospinota>Nitrospinia>Nitrospinales | GCA_018648125.1 | Nitrospina_sp. | HN416_07030 | 2974..6081 | 2024844 | DUF499-domain-containing-protein-[Nitrospina-sp.] |
| RSV33259.1 | DUF499_2.RM.like.systems | Phage_integrase->ZnR+TOPRIM->?->?-><-ParB+HNH\|\|?->DUF499*->DUF3780->METHYLASE-><-?\|\|HKD-DNase+SF1-SF2-Helicase+iREase+CTD-><-Schlafen<-PIWI | 1005 | Bacteria>Pseudomonadota>Alphaproteobacteria>Sphingomonadales>Sphingomonadaceae | GCA_003951715.1 | Sphingomonas_sp._ABOLH | CA237_01110 | 31273..34290 | 1985881 | DUF499-domain-containing-protein-[Sphingomonas-sp.-ABOLH] |
| WP_209445045.1 | DUF499_2.RM.like.systems | PIWI->Schlafen-><-HKD-DNase+SF1-SF2-Helicase+iREase+CTD<-?<-METHYLASE<-DUF3780<-DUF499*<-?\|\|ParB+HNH-><-?<-?<-ZnR+TOPRIM<-Phage_integrase | 1025 | Bacteria>Pseudomonadota>Alphaproteobacteria>Sphingomonadales>Sphingomonadaceae | GCF_001477495.1 | Sphingomonas_yabuuchiae | NS355_RS08100 | 26418..29495 | 172044 | DUF499-domain-containing-protein-[Sphingomonas-yabuuchiae] |
| WP_206432931.1 | DUF499_2.RM.like.systems | XIS-HTH->Phage_integrase->ZnR+TOPRIM->?->?-><-ParB+HNH\|\|DUF499*->DUF3780->METHYLASE->?->HKD-DNase+SF1-SF2-Helicase+iREase+CTD-><-Schlafen<-PIWI | 1025 | Bacteria>Pseudomonadota>Alphaproteobacteria>Sphingomonadales>Sphingomonadaceae | GCF_003951715.1 | Sphingomonas_sp._ABOLH | CA237_RS01115 | 31213..34290 | 1985881 | DUF499-domain-containing-protein-[Sphingomonas-sp.-ABOLH] |
| MBN8455770.1 | DUF499_2.RM.like.systems | <-Schlafen+HATPase+wHTH<-DUF3780<-DUF499* | 1039 | Bacteria>Pseudomonadota>Betaproteobacteria>Betaproteobacteria_incertae_sedis>Candidatus_Accumulibacter | GCA_017302415.1 | Accumulibacter_sp. | JOM31_22260 | 5561..8680 | 2053492 | ATP-binding-protein-[Candidatus-Accumulibacter-sp.] |
| MCM8580151.1 | DUF499_2.RM.like.systems | <-METHYLASE<-Schlafen+HATPase+wHTH<-DUF3780<-DUF499*<-SF1-SF2-Helicase+DUF1998+HKD-DNase+HKD-DNase<-SF1-SF2-Helicase+DUF1998+HKD-DNase+HKD-DNase | 1010 | Bacteria>Pseudomonadota>Betaproteobacteria>Betaproteobacteria_incertae_sedis>Candidatus_Accumulibacter | GCA_023806705.1 | Accumulibacter_sp. | NFV56_11820 | 5127..8159 | 2053492 | DUF499-domain-containing-protein-[Candidatus-Accumulibacter-sp.] |
| MCP5230244.1 | DUF499_2.RM.like.systems | Phage_integrase->?-><-?\|\|GDResolvase+HIN-HTH-><-URI-DNase\|\|PhdYeFM_antitox->DUF499*->DUF3780->METHYLASE->HKD-DNase+SF1-SF2-Helicase-><-Schlafen<-?\|\|ArdCN+MPTase-> | 1037 | Bacteria>Pseudomonadota>Betaproteobacteria>Betaproteobacteria_incertae_sedis>Candidatus_Accumulibacter | GCA_024235715.1 | Accumulibacter_sp. | H6941_16790 | 59686..62799 | 2053492 | ATP-binding-protein-[Candidatus-Accumulibacter-sp.] |
| MCQ1547870.1 | DUF499_2.RM.like.systems | <-Classical-AAA\|\|MetJarc-HTH-><-HKD-DNase+SF1-SF2-Helicase+iREase+CTD<-METHYLASE<-Schlafen+HATPase+wHTH<-DUF3780<-DUF499*\|\|DUF6361->HKD-DNase->SF1-SF2-Helicase-><-?<-?<-RelE-ParE | 1010 | Bacteria>Pseudomonadota>Betaproteobacteria>Betaproteobacteria_incertae_sedis>Candidatus_Accumulibacter | GCA_024380085.1 | Candidatus_Accumulibacter_phosphatis | NOF05_03365 | 13347..16379 | 327160 | DUF499-domain-containing-protein-[Candidatus-Accumulibacter-phosphatis] |
| WP_138677516.1 | DUF499_2.RM.like.systems | RelE-ParE->?->?-><-SF1-SF2-Helicase<-HKD-DNase<-DUF6361\|\|DUF499*->DUF3780->Schlafen+HATPase+wHTH->METHYLASE->HKD-DNase+SF1-SF2-Helicase+iREase+CTD-><-MetJarc-HTH\|\|Classical-AAA-> | 1010 | Bacteria>Pseudomonadota>Betaproteobacteria>Betaproteobacteria_incertae_sedis>Candidatus_Accumulibacter | GCF_005889575.1 | Candidatus_Accumulibacter_phosphatis | ACCUM_RS04320 | 32152..35184 | 327160 | DUF499-domain-containing-protein-[Candidatus-Accumulibacter-phosphatis] |
| WP_138678982.1 | DUF499_2.RM.like.systems | <-HKD-DNase+SF1-SF2-Helicase+iREase+CTD<-METHYLASE<-Schlafen+HATPase+wHTH<-DUF3780<-DUF499* | 944 | Bacteria>Pseudomonadota>Betaproteobacteria>Betaproteobacteria_incertae_sedis>Candidatus_Accumulibacter | GCF_005889575.1 | Candidatus_Accumulibacter_phosphatis | ACCUM_RS19260 | 7387..10221 | 327160 | DUF499-domain-containing-protein,-partial-[Candidatus-Accumulibacter-phosphatis] |
| WP_248595601.1 | DUF499_2.RM.like.systems | HKD-DNase->SIR2+SIR2->DUF499-p->DUF499*->DUF3780->Schlafen+HATPase+wHTH->METHYLASE->METHYLASE-p->HKD-DNase+SF1-SF2-Helicase+iREase+CTD-><-RVT | 845 | Bacteria>Pseudomonadota>Betaproteobacteria>Betaproteobacteria_incertae_sedis>Candidatus_Accumulibacter | GCF_012939955.1 | Candidatus_Accumulibacter_phosphatis | E4Q23_RS19540 | 3573..6110 | 327160 | DUF499-domain-containing-protein-[Candidatus-Accumulibacter-phosphatis] |
| WP_269498091.1 | DUF499_2.RM.like.systems | RVT->?-><-HKD-DNase+SF1-SF2-Helicase<-METHYLASE<-Schlafen+HATPase+wHTH<-DUF3780<-DUF499*<-RHH_5<-CHTH\|\|XIS-HTH->HTH_17-><-Arm-DNA-bind_3+Phage_integrase | 1041 | Bacteria>Pseudomonadota>Betaproteobacteria>Burkholderiales>Alcaligenaceae | GCF_027257175.1 | Castellaniella_sp._S9 | OOZ48_RS10600 | 328374..331499 | 2993652 | DUF499-domain-containing-protein-[Castellaniella-sp.-S9] |
| WP_212161746.1 | DUF499_2.RM.like.systems | <-REase\|\|?-><-HKD-DNase+SF1-SF2-Helicase+iREase+CTD<-METHYLASE<-Schlafen+HATPase+wHTH<-DUF3780<-DUF499*<-Methylase\|\|?-><-Radical-SAM<-MORC+DNA_gyraseB+TOPRIM | 1012 | Bacteria>Pseudomonadota>Betaproteobacteria>Burkholderiales>Burkholderiaceae | - | Burkholderia_vietnamiensis | KDW46_26565 | 46825..49863 | 60552 | DUF499-domain-containing-protein-[Burkholderia-vietnamiensis] |
| MBR8145923.1 | DUF499_2.RM.like.systems | <-REase\|\|?-><-HKD-DNase+SF1-SF2-Helicase+iREase+CTD<-METHYLASE<-Schlafen+HATPase+wHTH<-DUF3780<-DUF499*<-Methylase\|\|?-><-Radical-SAM<-MORC+DNA_gyraseB+TOPRIM | 1012 | Bacteria>Pseudomonadota>Betaproteobacteria>Burkholderiales>Burkholderiaceae | GCA_018147135.1 | Burkholderia_vietnamiensis | KDW46_26565 | 46825..49863 | 60552 | ATP-binding-protein-[Burkholderia-vietnamiensis] |
| MCA8374011.1 | DUF499_2.RM.like.systems | HIN-HTH-><-HKD-DNase+SF1-SF2-Helicase+iREase+CTD<-METHYLASE<-Schlafen+HATPase+wHTH<-DUF3780<-DUF499*<-?\|\|?->?->Pkinase-> | 1011 | Bacteria>Pseudomonadota>Betaproteobacteria>Burkholderiales>Burkholderiaceae | GCA_020420025.1 | Burkholderia_multivorans | LGM80_11600 | 411823..414858 | 87883 | DUF499-domain-containing-protein-[Burkholderia-multivorans] |
| MCO5106862.1 | DUF499_2.RM.like.systems | <-LON-AAA<-?<-?\|\|?->DUF499*->DUF3780->Schlafen+HATPase+wHTH->?->METHYLASE->HKD-DNase+SF1-SF2-Helicase+iREase+CTD-> | 1008 | Bacteria>Pseudomonadota>Betaproteobacteria>Burkholderiales>Burkholderiaceae | GCA_023954045.1 | Burkholderiaceae_bacterium | M9907_07245 | 32418..35444 | 2030806 | DUF499-domain-containing-protein-[Burkholderiaceae-bacterium] |
| WP_027788794.1 | DUF499_2.RM.like.systems | <-REase\|\|?-><-HKD-DNase+SF1-SF2-Helicase+iREase+CTD<-METHYLASE<-Schlafen+HATPase+wHTH<-DUF3780<-DUF499* | 1012 | Bacteria>Pseudomonadota>Betaproteobacteria>Burkholderiales>Burkholderiaceae | GCF_001528845.1 | Burkholderia_cepacia | BCE01_RS37115 | 24596..27634 | 292 | DUF499-domain-containing-protein-[Burkholderia-cepacia] |
| WP_088931578.1 | DUF499_2.RM.like.systems | DUF499*->DUF3780->Schlafen+HATPase+wHTH->METHYLASE->HKD-DNase+SF1-SF2-Helicase+iREase+CTD-> | 1013 | Bacteria>Pseudomonadota>Betaproteobacteria>Burkholderiales>Burkholderiaceae | GCF_003812585.1 | Burkholderia_multivorans | UA17_RS04225 | 849369..852410 | 87883 | DUF499-domain-containing-protein-[Burkholderia-multivorans] |
| WP_124581834.1 | DUF499_2.RM.like.systems | <-HKD-DNase+SF1-SF2-Helicase+iREase+CTD<-METHYLASE<-Schlafen+HATPase+wHTH<-DUF3780<-DUF499*<-Methylase\|\|?-><-Radical-SAM<-MORC+DNA_gyraseB+TOPRIM | 1012 | Bacteria>Pseudomonadota>Betaproteobacteria>Burkholderiales>Burkholderiaceae | GCF_003853875.1 | Burkholderia_contaminans | DF051_RS23135 | 9301..12339 | 488447 | DUF499-domain-containing-protein-[Burkholderia-contaminans] |
| WP_207876957.1 | DUF499_2.RM.like.systems | <-HKD-DNase+SF1-SF2-Helicase+iREase+CTD<-METHYLASE<-Schlafen+HATPase+wHTH<-DUF3780<-DUF499*<-?\|\|?->?->DEDD_Tnp_IS110+Transposase_20->SIR2-> | 1011 | Bacteria>Pseudomonadota>Betaproteobacteria>Burkholderiales>Burkholderiaceae | GCF_017378435.1 | Burkholderia_pseudomallei | J3L14_RS17850 | 4003230..4006265 | 28450 | DUF499-domain-containing-protein-[Burkholderia-pseudomallei] |
| WP_259032552.1 | DUF499_2.RM.like.systems | <-HKD-DNase+SF1-SF2-Helicase+iREase+CTD<-METHYLASE<-Schlafen+HATPase+wHTH<-DUF3780<-DUF499*<-Methylase\|\|?-><-Radical-SAM<-HIN-HTH<-?<-Phage_AlpA | 1012 | Bacteria>Pseudomonadota>Betaproteobacteria>Burkholderiales>Burkholderiaceae | GCF_024793955.1 | Burkholderia_thailandensis | NX841_RS02865 | 24753..27791 | 57975 | DUF499-domain-containing-protein-[Burkholderia-thailandensis] |
| WP_122877385.1 | DUF499_2.RM.like.systems | MORC+DNA_gyraseB+TOPRIM->Radical-SAM-><-?\|\|Methylase->DUF499*->DUF3780->Schlafen+HATPase+wHTH->METHYLASE->HKD-DNase+SF1-SF2-Helicase+iREase+CTD-><-?\|\|REase-> | 1012 | Bacteria>Pseudomonadota>Betaproteobacteria>Burkholderiales>Burkholderiaceae | GCF_900591725.1 | Burkholderia_pseudomallei | EFU85_RS04200 | 244549..247587 | 28450 | DUF499-domain-containing-protein-[Burkholderia-pseudomallei] |
| WP_173125224.1 | DUF499_2.RM.like.systems | <-HKD-DNase+SF1-SF2-Helicase+iREase+CTD<-?<-METHYLASE<-Schlafen+HATPase+wHTH<-DUF3780<-DUF499*<-?<-?<-?<-XIS-HTH<-?\|\|HTH-> | 1009 | Bacteria>Pseudomonadota>Betaproteobacteria>Burkholderiales>Burkholderiales_genera_incertae_sedis | GCF_013266755.1 | Aquabacterium_terrae | HLB44_RS18520 | 560708..563737 | 2732868 | DUF499-domain-containing-protein-[Aquabacterium-terrae] |
| MBL8328914.1 | DUF499_2.RM.like.systems | <-HKD-DNase+SF1-SF2-Helicase<-METHYLASE<-Schlafen+HATPase+wHTH<-DUF3780<-DUF499*<-RHH_1\|\|HNH->?->CopG_antitoxin-> | 1040 | Bacteria>Pseudomonadota>Betaproteobacteria>Burkholderiales>Sphaerotilaceae | GCA_016790225.1 | Rubrivivax_sp. | JNJ71_08675 | 122219..125341 | 50259 | DUF499-domain-containing-protein-[Rubrivivax-sp.] |
| MCC7099553.1 | DUF499_2.RM.like.systems | <-Schlafen<-DUF3780<-DUF499*<-RHH_1 | 1040 | Bacteria>Pseudomonadota>Betaproteobacteria>Burkholderiales>Sphaerotilaceae | GCA_020850905.1 | Rubrivivax_sp. | IT500_08190 | 1682..4804 | 50259 | DUF499-domain-containing-protein-[Rubrivivax-sp.] |
| MCW5609142.1 | DUF499_2.RM.like.systems | <-ParB+HNH<-?<-OLD-ABC<-METHYLASE<-Schlafen+HATPase+wHTH<-DUF3780<-DUF499*<-Arm-DNA-bind_3+Phage_integrase\|\|Phage_int_SAM_2+Integrase_1-><-HTH_3\|\|?-><-HTH+MPTase | 1009 | Bacteria>Pseudomonadota>Betaproteobacteria>Burkholderiales>Sphaerotilaceae | GCA_026125605.1 | Rubrivivax_sp. | KIS83_00415 | 81577..84606 | 50259 | MAG:-ATP-binding-protein-[Rubrivivax-sp.] |
| WP_009858743.1 | DUF499_2.RM.like.systems | <-HIN-HTH<-METHYLASE<-Schlafen+HATPase+wHTH<-DUF3780<-DUF499*<-RHH_1<-HTH_3\|\|Transposase_mut->SIGMA-HTH-> | 1041 | Bacteria>Pseudomonadota>Betaproteobacteria>Burkholderiales>Sphaerotilaceae | GCF_000190375.1 | Rubrivivax_benzoatilyticus | F799_RS0117815 | 6811..9936 | 316997 | DUF499-domain-containing-protein-[Rubrivivax-benzoatilyticus] |
| WP_138938924.1 | DUF499_2.RM.like.systems | <-SIGMA-HTH<-Transposase_mut\|\|HTH_3->RHH_1->DUF499*->DUF3780->Schlafen+HATPase+wHTH->METHYLASE->HEPN->ParA-Soj-PloopNTPase->ORC-CDC6-> | 1022 | Bacteria>Pseudomonadota>Betaproteobacteria>Burkholderiales>Sphaerotilaceae | GCF_011365705.1 | Rubrivivax_benzoatilyticus | G7087_RS13130 | 24930..27998 | 316997 | DUF499-domain-containing-protein-[Rubrivivax-benzoatilyticus] |
| WP_247652921.1 | DUF499_2.RM.like.systems | CopG/RHH->RHH_1->DUF499*->DUF3780->Schlafen+HATPase+wHTH->METHYLASE->HKD-DNase+SF1-SF2-Helicase-> | 1021 | Bacteria>Pseudomonadota>Betaproteobacteria>Burkholderiales>Sphaerotilaceae | - | Ideonella | KAK07_RS23525 | 27586..30651 | 36862 | MULTISPECIES:-DUF499-domain-containing-protein-[Ideonella] |
| MBI3155186.1 | DUF499_2.RM.like.systems | <-HKD-DNase+SF1-SF2-Helicase<-METHYLASE<-Schlafen+HATPase+wHTH<-DUF3780<-DUF499*<-RHH_1<-ABC-ATPase+SbcC<-PrmC_N+Methyltransf_25<-Peptidase_M20 | 1040 | Bacteria>Pseudomonadota>Betaproteobacteria>Burkholderiales>unclassified_Burkholderiales | GCA_016191995.1 | Burkholderiales_bacterium | HYZ20_07235 | 13809..16931 | 1891238 | DUF499-domain-containing-protein-[Burkholderiales-bacterium] |
| MDE2080727.1 | DUF499_2.RM.like.systems | ABC-ATPase->HAD-><-?\|\|?-><-Schlafen+HATPase+wHTH<-DUF3780<-DUF499*<-RHH_1<-?<-MORC\|\|?->?->Glycos_trans_3N+Glycos_transf_3-> | 1040 | Bacteria>Pseudomonadota>Betaproteobacteria>Burkholderiales>unclassified_Burkholderiales | GCA_028872015.1 | Burkholderiales_bacterium | KGI90_05185 | 52053..55175 | 1891238 | MAG:-DUF499-domain-containing-protein-[Burkholderiales-bacterium] |
| MBK7768857.1 | DUF499_2.RM.like.systems | <-METHYLASE-p<-Schlafen+HATPase+wHTH<-DUF3780<-DUF499*\|\|?->?-><-NTP_transf_3<-ZnR | 1005 | Bacteria>Pseudomonadota>Betaproteobacteria>Nitrosomonadales>Sterolibacteriaceae | GCA_016706645.1 | Sulfuritalea_sp. | IPI44_24195 | 5665..8682 | 2480090 | ATP-binding-protein-[Sulfuritalea-sp.] |
| MBK7421883.1 | DUF499_2.RM.like.systems | Phage_int_SAM_2+Integrase_1-><-?\|\|DUF499-p->DUF499*->DUF499-p->DUF499-p->DUF3780->Schlafen+HATPase+wHTH->Pkinase->TPR-> | 487 | Bacteria>Pseudomonadota>Betaproteobacteria>Rhodocyclales>Rhodocyclaceae | GCA_016709335.1 | Candidatus_Propionivibrio_dominans | IPJ48_01605 | 217179..218642 | 2954373 | DUF499-domain-containing-protein-[Propionivibrio-sp.] |
| MBL8482095.1 | DUF499_2.RM.like.systems | <-ZnR<-?\|\|?->Arm-DNA-bind_3+Phage_integrase->DUF499*->DUF3780->Schlafen+HATPase+wHTH->METHYLASE->HKD-DNase+SF1-SF2-Helicase+iREase+CTD->REase+PNTD1+PNTD2->PIWI-> | 1008 | Bacteria>Pseudomonadota>Betaproteobacteria>Rhodocyclales>Rhodocyclaceae | GCA_016791085.1 | Rhodocyclaceae_bacterium | JNJ60_07865 | 13203..16229 | 1898103 | ATP-binding-protein-[Rhodocyclaceae-bacterium] |
| OHC61993.1 | DUF499_2.RM.like.systems | <-HKD-DNase+SF1-SF2-Helicase+iREase+CTD<-METHYLASE<-Schlafen+HATPase+wHTH<-DUF3780<-DUF499* | 1005 | Bacteria>Pseudomonadota>Betaproteobacteria>Rhodocyclales>unclassified_Rhodocyclales | GCA_001828915.1 | Rhodocyclales_bacterium_GWA2_65_19 | A2040_16480 | 43406..46423 | 1802016 | MAG:-hypothetical-protein-A2040_16480-[Rhodocyclales-bacterium-GWA2_65_19] |
| WP_007193607.1 | DUF499_2.RM.like.systems | <-HKD-DNase+SF1-SF2-Helicase+iREase+CTD<-ParB+HNH<-METHYLASE-p<-METHYLASE<-Schlafen+HATPase+wHTH<-DUF3780<-DUF499*\|\|?->?->?->CHTH-> | 1041 | Bacteria>Pseudomonadota>Gammaproteobacteria>Chromatiales>Chromatiaceae | GCF_000223985.1 | Thiocapsa_marina | THIMADRAFT_RS13475 | 269670..272795 | 244573 | DUF499-domain-containing-protein-[Thiocapsa-marina] |
| MBL1275688.1 | DUF499_2.RM.like.systems | <-HKD-DNase+SF1-SF2-Helicase<-Schlafen+HATPase+wHTH<-METHYLASE<-DUF3780<-DUF499*<-RHH_5<-REase\|\|SF1-SF2-Helicase->?-><-?\|\|PADR-HTH-> | 1041 | Bacteria>Pseudomonadota>Gammaproteobacteria>Chromatiales>Ectothiorhodospiraceae | GCA_002401775.2 | Ectothiorhodospiraceae_bacterium | COB30_006350 | 9914..13039 | 2026737 | DUF499-domain-containing-protein-[Ectothiorhodospiraceae-bacterium] |
| WP_218806438.1 | DUF499_2.RM.like.systems | ADH_N+ADH_zinc_N-><-SF1-SF2-Helicase\|\|?->?->CopG_antitoxin->DUF499*->DUF3780->METHYLASE->Schlafen+HATPase+wHTH->?->METHYLASE->HEPN-> | 1041 | Bacteria>Pseudomonadota>Gammaproteobacteria>Methylococcales>Methylococcaceae | GCF_019285515.1 | Methylococcus_sp._Mc7 | KW115_RS14805 | 3051261..3054386 | 2860258 | DUF499-domain-containing-protein-[Methylococcus-sp.-Mc7] |
| MBG6817109.1 | DUF499_2.RM.like.systems | RHH_1->DUF499*->DUF3780->METHYLASE->Schlafen+HATPase+wHTH->METHYLASE->HKD-DNase+SF1-SF2-Helicase->MORC+REase-> | 1041 | Bacteria>Pseudomonadota>Gammaproteobacteria>Pseudomonadales>Pseudomonadaceae | GCA_015764555.1 | Pseudomonas_aeruginosa | I5J00_12285 | 138318..141443 | 287 | DUF499-domain-containing-protein-[Pseudomonas-aeruginosa] |
| WP_095598174.1 | DUF499_2.RM.like.systems | <-MORC+REase<-HKD-DNase+SF1-SF2-Helicase<-METHYLASE<-Schlafen+HATPase+wHTH<-METHYLASE<-DUF3780<-DUF499*<-RHH | 1041 | Bacteria>Pseudomonadota>Gammaproteobacteria>Pseudomonadales>Pseudomonadaceae | GCF_002286785.1 | Pseudomonas_sp._PIC25 | BZL41_RS04445 | 60362..63487 | 1958773 | DUF499-domain-containing-protein-[Pseudomonas-sp.-PIC25] |
| WP_210799995.1 | DUF499_2.RM.like.systems | RHH_1->DUF499*->DUF3780->METHYLASE->Schlafen+HATPase+wHTH->METHYLASE->HKD-DNase+SF1-SF2-Helicase->MORC+REase-> | 1041 | Bacteria>Pseudomonadota>Gammaproteobacteria>Pseudomonadales>Pseudomonadaceae | GCF_018026755.1 | Pseudomonas_aeruginosa | J5F08_RS20235 | 53337..56462 | 287 | DUF499-domain-containing-protein-[Pseudomonas-aeruginosa] |
| MCG3169754.1 | DUF499_2.RM.like.systems | <-sGTP\|\|?->?->?->?->DUF772+DDE_Tnp_1+DDE_Tnp_1_6->DUF499*->DUF3780->Schlafen+HATPase+wHTH->METHYLASE->HKD-DNase+SF1-SF2-Helicase+iREase+CTD-> | 1010 | Bacteria>Pseudomonadota>Gammaproteobacteria>Pseudomonadales>unclassified_Pseudomonadales | GCA_022072145.1 | Pseudomonadales_bacterium | CALGDGBN_01277 | 15067..18099 | 1891229 | hypothetical-protein-[Pseudomonadales-bacterium] |
| MXW50605.1 | DUF499_2.RM.like.systems | <-HicB\|\|DUF499*->DUF3780->METHYLASE->PHP+SbcC->Schlafen+HATPase+wHTH->HKD-DNase+SF1-SF2-Helicase-> | 1019 | Bacteria>Pseudomonadota>Gammaproteobacteria>unclassified_Gammaproteobacteria | GCA_009835805.1 | Gammaproteobacteria_bacterium | F4Z84_10190 | 3046..6105 | 1913989 | DUF499-domain-containing-protein-[Gammaproteobacteria-bacterium] |
| MCY4014266.1 | DUF499_2.RM.like.systems | <-HKD-DNase+SF1-SF2-Helicase<-Schlafen+HATPase+wHTH<-PHP+SbcC<-METHYLASE<-DUF3780<-DUF499* | 1023 | Bacteria>Pseudomonadota>Gammaproteobacteria>unclassified_Gammaproteobacteria | GCA_026708185.1 | Gammaproteobacteria_bacterium | OXG82_16305 | 63884..66955 | 1913989 | MAG:-DUF499-domain-containing-protein-[Gammaproteobacteria-bacterium] |
| MDE0094449.1 | DUF499_2.RM.like.systems | Schlafen+HATPase+wHTH->DUF499*->DUF499-p->DUF3780->METHYLASE->HKD-DNase+SF1-SF2-Helicase+iREase+CTD-> | 782 | Bacteria>Pseudomonadota>Gammaproteobacteria>unclassified_Gammaproteobacteria | GCA_028819195.1 | Gammaproteobacteria_bacterium | OXO49_08130 | 171410..173758 | 1913989 | MAG:-DUF499-domain-containing-protein-[Gammaproteobacteria-bacterium] |
| HAI59970.1 | DUF499_2.RM.like.systems | SF1-SF2-Helicase-><-Pkinase\|\|?-><-Synaptojanin\|\|DUF499*->DUF3780->Schlafen+HATPase+wHTH->?->METHYLASE->HKD-DNase+SF1-SF2-Helicase+iREase+CTD-> | 1013 | Bacteria>Pseudomonadota>Gammaproteobacteria>Xanthomonadales>Xanthomonadaceae | GCA_003449515.1 | Xanthomonadaceae_bacterium | DCM32_08860 | 4597..7638 | 1926873 | hypothetical-protein-[Xanthomonadaceae-bacterium] |
| MCZ8164568.1 | DUF499_2.RM.like.systems | XIS-HTH->HTH_36->?->?-><-?<-Phage_int_SAM_2+Integrase_1\|\|DUF499*->DUF3780->Schlafen+HATPase+wHTH->?->METHYLASE->HKD-DNase+SF1-SF2-Helicase+iREase+CTD-> | 1013 | Bacteria>Pseudomonadota>Gammaproteobacteria>Xanthomonadales>Xanthomonadaceae | GCA_027531545.1 | Silanimonas_sp. | O9280_02335 | 527918..530959 | 1929290 | MAG:-DUF499-domain-containing-protein-[Silanimonas-sp.] |
| WP_146472112.1 | DUF499_2.RM.like.systems | <-ABC-ATPase+SbcC\|\|?->ABhydrolase->ABC-ATPase+SbcC->?-><-?\|\|DUF499*->DUF3780->Schlafen+HATPase+wHTH->?->?->METHYLASE->HKD-DNase+SF1-SF2-Helicase+iREase+CTD-> | 1012 | Bacteria>Pseudomonadota>Gammaproteobacteria>Xanthomonadales>Xanthomonadaceae | GCF_014656335.1 | Vulcaniibacterium_thermophilum | IE210_RS02820 | 81268..84306 | 1169913 | DUF499-domain-containing-protein-[Vulcaniibacterium-thermophilum] |
| MBS0382004.1 | DUF499_2.RM.like.systems | DUF499*->DUF3780->Schlafen+HATPase+wHTH->?->METHYLASE-p-> | 889 | Bacteria>Pseudomonadota>unclassified_Pseudomonadota | GCA_018241365.1 | Pseudomonadota_bacterium | JSR56_06185 | 1..2671 | 1977087 | DUF499-domain-containing-protein-[Proteobacteria-bacterium] |
| MBS0338394.1 | DUF499_2.RM.like.systems | <-Schlafen+HATPase+wHTH<-Schlafen+HATPase+wHTH<-DUF3780<-DUF499* | 1016 | Bacteria>Pseudomonadota>unclassified_Pseudomonadota | GCA_018241645.1 | Pseudomonadota_bacterium | JSS40_16645 | 2322..5372 | 1977087 | ATP-binding-protein-[Proteobacteria-bacterium] |
| OGV20214.1 | DUF499_2.RM.like.systems | <-HKD-DNase+SF1-SF2-Helicase<-Schlafen+HATPase+wHTH<-METHYLASE<-DUF3780<-DUF499*\|\|?-><-?<-?\|\|?->STAS->ABC-ATPase+SbcC-> | 1036 | Bacteria>PVC_group>Lentisphaerota>unclassified_Lentisphaerota | GCA_001803205.1 | Lentisphaerae_bacterium_GWF2_38_69 | A2X47_06010 | 8891..12001 | 1798570 | MAG:-glycosyl-transferase-[Lentisphaerae-bacterium-GWF2_38_69] |
| WP_011117919.1 | DUF499_2.RM.like.systems | <-Schlafen<-HKD-DNase+SF1-SF2-Helicase<-TOPRIM<-OLD-ABC<-METHYLASE-p<-DUF3780<-DUF499*<-XIS-HTH | 1031 | Bacteria>PVC_group>Planctomycetota>Planctomycetia>Pirellulales | GCF_000196115.1 | Rhodopirellula_baltica | DDX19_RS10420 | 136482..139577 | 265606 | DUF499-domain-containing-protein-[Rhodopirellula-baltica] |
| WP_146459050.1 | DUF499_2.RM.like.systems | <-HKD-DNase<-?<-Schlafen<-HKD-DNase+SF1-SF2-Helicase<-METHYLASE<-DUF3780<-DUF499*<-XIS-HTH<-?<-SF1-SF2-Helicase<-?<-NERD+SF1-SF2-Helicase<-DUF2726+ZnR+TOPRIM | 1031 | Bacteria>PVC_group>Planctomycetota>Planctomycetia>Pirellulales | GCF_007860125.1 | Rubripirellula_tenax | Poly51_RS17440 | 744140..747235 | 2528015 | DUF499-domain-containing-protein-[Rubripirellula-tenax] |
| MCX5673909.1 | DUF499_2.RM.like.systems | <-REase<-REase+NACHT<-REase+NACHT\|\|DUF499*->DUF3780->Schlafen+HATPase+wHTH->METHYLASE->HKD-DNase+SF1-SF2-Helicase+iREase+CTD->?->HKD-DNase-> | 978 | Bacteria>PVC_group>Planctomycetota>unclassified_Planctomycetota | GCA_026387015.1 | Planctomycetota_bacterium | NTX87_02775 | 8015..10951 | 2026780 | MAG:-DUF499-domain-containing-protein-[Planctomycetota-bacterium] |
| MBM3879657.1 | DUF499_2.RM.like.systems | <-ABC-ATPase+SbcC<-?<-?<-?<-ABC-ATPase+SbcC\|\|DUF499*->DUF3780->Schlafen+HATPase+wHTH->?->Schlafen+HATPase+wHTH->METHYLASE->HEPN-> | 1019 | Bacteria>PVC_group>Verrucomicrobiota>unclassified_Verrucomicrobiota | GCA_016871715.1 | Verrucomicrobiota_bacterium | FJ387_08035 | 7884..10943 | 2026799 | ATP-binding-protein-[Verrucomicrobia-bacterium] |

| Protein-ID | System | Gene-Neighbourhoods | Length | Taxonomy | Assembly-ID | Strain | Locus-Tag | Genomic-coordinates | TaxID | Fasta-defline |
|---|---|---|---|---|---|---|---|---|---|---|
| MCC5848622.1 | DUF499 _2.RM.like.systems | <- Schlafen +HATPase+wHTH <- DUF3780 <- DUF499 *<-?<-?\|\|?->?->?-><-tRNA<-ABhydrolase | 1021 | Bacteria>PVC_group>Verrucomicrobiota>unclassified_Verrucomicrobiota | GCA_020831635.1 | Verrucomicrobiota_bacterium | JJU29_11045 | 2236..5301 | 2026799 | DUF499 -domain-containing-protein-[Verrucomicrobia-bacterium] |
| MCF7668420.1 | DUF499 _2.RM.like.systems | <- HKD-DNase+SF1-SF2-Helicase +iREase+CTD <-HTH+MPTase<- Schlafen +HATPase+wHTH <-SbcC+SidE<- METHYLASE <- DUF3780 <- DUF499 * | 1083 | Bacteria>PVC_group>Verrucomicrobiota>unclassified_Verrucomicrobiota | GCA_021733785.1 | Verrucomicrobiota_bacterium | K9N48_01435 | 17156..20407 | 2026799 | DUF499 -domain-containing-protein-[Verrucomicrobia-bacterium] |
| MCF7762652.1 | DUF499 _2.RM.like.systems | <- HKD-DNase+SF1-SF2-Helicase <- Schlafen +HATPase+wHTH <- METHYLASE <-wHTH+REase<- DUF3780 <- DUF499 *<-XIS-HTH\|\|SF1-SF2-Helicase-><-Glyoxalase<-?<-REC | 1027 | Bacteria>PVC_group>Verrucomicrobiota>unclassified_Verrucomicrobiota | GCA_021733945.1 | Verrucomicrobiota_bacterium | K9N62_03175 | 25626..28709 | 2026799 | DUF499 -domain-containing-protein-[Verrucomicrobia-bacterium] |
| SLM14732.1 | DUF499 _2.RM.like.systems | <- HKD-DNase+SF1-SF2-Helicase <- Schlafen +HATPase+wHTH <- METHYLASE <-OLD-ABC+TOPRIM<- DUF3780 <- DUF499 *\|\|wHTH+SAD+HNH-><-?\|\|HEPN->PolB-> | 1036 | Bacteria>Spirochaetota>Spirochaetia>Spirochaetales>environmental_samples | GCA_900163985.1 | uncultured_spirochete | SPIROBIBN47_350053 | 62416..65526 | 156406 | conserved-hypothetical-protein-[uncultured-spirochete] |
| WP_132781615.1 | DUF499 _2.RM.like.systems | <-N6_N4_MTase<-Aminotran_1_2\|\|TIR-> DUF499 *-> DUF3780 ->OLD-ABC+TOPRIM-> METHYLASE -> Schlafen +HATPase+wHTH -> HKD-DNase+SF1-SF2-Helicase -> | 1038 | Bacteria>Spirochaetota>Spirochaetia>Spirochaetales>Treponemataceae | GCF_004343725.1 | Treponema_sp._J25 | C5022_RS10535 | 15156..18272 | 2094121 | DUF499 -domain-containing-protein-[Treponema-sp.-J25] |
| MDA3832123.1 | DUF499 _2.RM.like.systems | DUF499 *-> DUF3780 -> Schlafen +HATPase+wHTH -> METHYLASE -p-> | 1006 | Bacteria>Spirochaetota>Spirochaetia>Spirochaetales>unclassified_Spirochaetales | GCA_027858175.1 | Spirochaetales_bacterium | PF495_01880 | 1..3021 | 2026792 | MAG:- DUF499 -domain-containing-protein,-partial-[Spirochaetales-bacterium] |
| MBX3108928.1 | DUF499 _2.RM.like.systems | <-HTH_3\|\|?->?->GDResolvase+HIN-HTH-><-HerA/FtsK<-?\|\| DUF499 *-> DUF3780 -> METHYLASE -> Schlafen +HATPase+wHTH -> HKD-DNase+SF1-SF2-Helicase -><-?<-DNA_methylase | 1037 | Bacteria>Terrabacteria_group>Armatimonadota>Fimbriimonadia>Fimbriimonadales | GCA_019638755.1 | Fimbriimonadaceae_bacterium | KF743_07090 | 1426545..1429658 | 2682144 | DUF499 -domain-containing-protein-[Fimbriimonadaceae-bacterium] |
| WP_023162377.1 | DUF499 _2.RM.like.systems | SIGMA-HTH->?->?->?->?->?-> DUF499 *-> DUF3780 -> METHYLASE -> HKD-DNase+SF1-SF2-Helicase -> Schlafen +HATPase+wHTH -> | 1033 | Bacteria>Terrabacteria_group>Bacillota>Clostridia>Eubacteriales | GCF_000484505.2 | Clostridium_autoethanogenum | WX72_RS00855 | 181813..184914 | 84023 | DUF499 -domain-containing-protein-[Clostridium-autoethanogenum] |
| WP_107196386.1 | DUF499 _2.RM.like.systems | SPOUT_MTase->?->?-><-Phage_integrase<-?\|\| DUF499 *-> DUF3780 -> METHYLASE -> HKD-DNase+SF1-SF2-Helicase +iREase+CTD -> Schlafen ->BirA-HTH->DOC+HTH-> | 1081 | Bacteria>Terrabacteria_group>Bacillota>Negativicutes>Veillonellales | GCF_003367905.1 | Megasphaera_stantonii | DKB62_RS10425 | 2193335..2196580 | 2144175 | DUF499 -domain-containing-protein-[Megasphaera-stantonii] |
| HEY77166.1 | DUF499 _2.RM.like.systems | PhdYeFM_antitox-><-HTH<-XIS-HTH<-GDResolvase+Recombinase+ZnR<-GDResolvase\|\|?-> DUF499 *->ASCH+ DUF3780 -> Schlafen +HATPase+wHTH ->HicB_lk_antitox-> METHYLASE ->SF1-SF2-Helicase-> HKD-DNase+SF1-SF2-Helicase +iREase+CTD -> | 1010 | Bacteria>Terrabacteria_group>Chloroflexota>Thermoflexia>unclassified_Thermoflexia | GCA_011192125.1 | Thermoflexia_bacterium | G4000_13500 | 6629..9661 | 2420334 | ATP-binding-protein-[Thermoflexia-bacterium] |
| WP_017747474.1 | DUF499 _2.RM.like.systems | <-PIN<-?<- METHYLASE <- Schlafen +TUDOR+PIWI<- DUF3780 <- DUF499 *<-?<-ABhydrolase\|\|?->MetJarc-HTH->RelE-ParE-><-REase | 1072 | Bacteria>Terrabacteria_group>Cyanobacteriota/Melainabacteria_group>Cyanobacteriota>Cyanophyceae | GCF_000346485.2 | Scytonema_hofmannii | WA1_RS47720 | 11345103..11348321 | 34078 | DUF499 -domain-containing-protein-[Scytonema-hofmannii] |
| KPA09724.1 | DUF499 _2.RM.like.systems | DUF499 *-> DUF3780 -> Schlafen +HATPase+wHTH -> METHYLASE ->ParB+HNH->OLD-ABC-> | 1021 | Bacteria>Thermodesulfobacteriota>Desulfobacteria>Desulfobacterales>Candidatus_Magnetomoraceae | GCA_001292585.1 | Candidatus_Magnetomorum_sp._HK-1 | MHK_010064 | 367..3432 | 1509431 | ATPase-AAA-[Candidatus-Magnetomorum-sp.-HK-1] |
| MBA3018061.1 | DUF499 _2.RM.like.systems | <-FtsK<-?<-?<-?\|\| DUF499 *-> DUF3780 -> METHYLASE -> Schlafen +HATPase+wHTH -> HKD-DNase+SF1-SF2-Helicase -> | 1032 | Bacteria>Thermodesulfobacteriota>Desulfobacteria>Desulfobacterales>Desulfobacteraceae | GCA_013792055.1 | Desulfobacteraceae_bacterium | FP811_07930 | 7617..10715 | 2049433 | DUF499 -domain-containing-protein-[Desulfobacteraceae-bacterium] |
| MBC8359968.1 | DUF499 _2.RM.like.systems | Schlafen +HATPase+wHTH ->BirA-HTH->?->PolB->PolB+HEPN-> DUF499 *-> | 637 | Bacteria>Thermodesulfobacteriota>Desulfobacteria>Desulfobacterales>Desulfobacterales_incertae_sedis | GCA_014382665.1 | Candidatus_Desulfatibia_profunda | H8E23_01035 | 3975..5887 | 2841695 | ATP-binding-protein-[Desulfobacterales-bacterium] |
| WP_051375312.1 | DUF499 _2.RM.like.systems | <-DDE_Tnp_1\|\| DUF499 *-> DUF499 -p-> DUF3780 -> METHYLASE -> Schlafen +HATPase+wHTH -> HKD-DNase+SF1-SF2-Helicase ->TIR-> | 790 | Bacteria>Thermodesulfobacteriota>Desulfobacteria>Desulfobacterales>Desulfosarcinaceae | GCF_028752835.1 | Desulfosarcina_sp._BuS5 | A39W_RS22855 | 2496..4868 | 933262 | DUF499 -domain-containing-protein-[Desulfosarcina-sp.-BuS5] |
| MCK9293532.1 | DUF499 _2.RM.like.systems | JAB->?-><- HKD-DNase+SF1-SF2-Helicase <- Schlafen +HATPase+wHTH <- METHYLASE <- DUF3780 <- DUF499 *<-XIS-HTH | 1027 | Bacteria>Thermodesulfobacteriota>Desulfobulbia>Desulfobulbales>Desulfobulbaceae | GCA_023230525.1 | Desulfobulbaceae_bacterium | M0P70_00480 | 103201..106284 | 2053307 | DUF499 -domain-containing-protein-[Desulfobulbaceae-bacterium] |
| MCK9175498.1 | DUF499 _2.RM.like.systems | DUF499 *-> DUF3780 -> Schlafen +HATPase+wHTH -> METHYLASE -> HKD-DNase+SF1-SF2-Helicase ->URI-DNase+RAMA-> | 1021 | Bacteria>Thermodesulfobacteriota>Desulfobulbia>Desulfobulbales>Desulfocapsaceae | GCA_023231425.1 | Desulforhopalus_sp. | M0O96_09510 | 4806..7871 | 2099376 | DUF499 -domain-containing-protein-[Desulforhopalus-sp.] |
| WP_163302422.1 | DUF499 _2.RM.like.systems | Phage_integrase-><-DUF932<- HKD-DNase+SF1-SF2-Helicase +iREase+CTD <- METHYLASE <- Schlafen +HATPase+wHTH <- DUF3780 <- DUF499 *<-?\|\|?-><-PAIREDC-HTH\|\|?->ZnR+TOPRIM+primpol+HTH-><-DUF772+DDE_Tnp_1 | 1014 | Bacteria>Thermodesulfobacteriota>Desulfovibrionia>Desulfovibrionales>Desulfovibrionaceae | GCF_013376455.1 | Desulfolutivibrio_sulfodismutans | GD606_RS12045 | 2614561..2617605 | 63561 | DUF499 -domain-containing-protein-[Desulfovibrio-sulfodismutans] |

| Protein-ID | System | Gene-Neighbourhoods | Length | Taxonomy | Assembly-ID | Strain | Locus-Tag | Genomic-coordinates | TaxID | Fasta-defline |
|---|---|---|---|---|---|---|---|---|---|---|
| -- | | | | | | | | | | |
| --HNH-linked_OLD-ABC | | | | | | | | | | |
| -- | | | | | | | | | | |
| -- OLD-ABC->HNH | | | | | | | | | | |
| --Type1.BREX | | | | | | | | | | |
| -- | | | | | | | | | | |
| AGC72568.1 | Type6.BREX | METHYLASE -><- SIGMA+OB-Fold+AAA-ATPase+Lon-Protease-BrxL <- PglZ <- HNH<- OLD-ABC <- N6- METHYLASE <-Schlafen+HATPase+wHTH<- DUF6079 *<- BrxB-BrxF- iSTAND <- BrxA-DNA-binding <- BrxE \|\|glycosylase->?->?->?-><- HNH<- OLD-ABC | 1181 | Bacteria>environmental_samples | - | uncultured_bacterium_A1Q1_fos_1870 | . | 12880..16425 | 1256554 | hypothetical-protein-[uncultured-bacterium-A1Q1_fos_1870] |
| EFG14417.1 | Type6.BREX | PRTase->?->?->tRNA->tRNA-><-?<- SIGMA+OB-Fold+AAA-ATPase+HKD-DNase-BrxL <- PglZ <- HNH<- OLD-ABC <- N6- METHYLASE <- DUF6079 -p*<- DUF6079 -p<- BrxB-BrxF- iSTAND <- BrxA-DNA-binding <- BrxE <-HTH_3<-?<-TraJ-RHH<-ZnR+ TOPRIM <-CR-ATPase4 | 762 | Bacteria>FCB_group>Bacteroidetes/Chlorobi_group>Bacteroidota>Bacteroidia>Bacteroidales | GCA_000178215.1 | Bacteroides_xylanisolvens_SD_CC_1b | CW3_0497 | 100900..103188 | 702447 | hypothetical-protein-CW3_0497-[Bacteroides-xylanisolvens-SD-CC-1b] |
| EFF57551.1 | Type6.BREX | CR-ATPase4->ZnR+ TOPRIM ->TraJ-RHH->?->HTH_3-> BrxE -> BrxA-DNA-binding -> BrxB-BrxF- iSTAND -> DUF6079 -p-> DUF6079 -p*-> N6- METHYLASE -> OLD-ABC ->HNH -> PglZ -> SIGMA+OB-Fold+AAA-ATPase+HKD-DNase-BrxL -> | 706 | Bacteria>FCB_group>Bacteroidetes/Chlorobi_group>Bacteroidota>Bacteroidia>Bacteroidales | GCA_000178295.1 | Bacteroides_xylanisolvens_SD_CC_2a | CW1_2295 | 25339..27459 | 702444 | hypothetical-protein-CW1_2295-[Bacteroides-xylanisolvens-SD-CC-2a] |
| CDM00531.1 | Type6.BREX | PRTase->?->?->tRNA->tRNA-><-?<- SIGMA+OB-Fold+AAA-ATPase+HKD-DNase-BrxL <- PglZ <- HNH<- OLD-ABC <- N6- METHYLASE <- DUF6079 -p*<- DUF6079 -p<- BrxB-BrxF- iSTAND <- BrxA-DNA-binding <- BrxE <-HTH_3<-?<-TraJ-RHH<-ZnR+ TOPRIM | 689 | Bacteria>FCB_group>Bacteroidetes/Chlorobi_group>Bacteroidota>Bacteroidia>Bacteroidales | GCA_000577955.1 | Bacteroides_xylanisolvens_SD_CC_2a | BN891_34560 | 156968..159037 | 702444 | hypothetical-protein-BN891_34560-[Bacteroides-xylanisolvens-SD-CC-2a] |
| HAM28416.1 | Type6.BREX | ChlI+Lon_C+AAA-><-RNase<-AraC-HTH\|\|?-><-?\|\|HTH_3-> BrxE -> BrxA-DNA-binding -> BrxB-BrxF- iSTAND -> DUF6079 *-> N6- METHYLASE -> OLD-ABC ->HNH -> PglZ -> SIGMA+OB-Fold+AAA-ATPase+HKD-DNase-BrxL ->?->?->35exo-> | 1218 | Bacteria>FCB_group>Bacteroidetes/Chlorobi_group>Bacteroidota>Bacteroidia>Bacteroidales | GCA_003453655.1 | Prevotella_sp. | DCP79_00675 | 66224..69880 | 59823 | BREX-system-P-loop-protein-BrxC-[Prevotella-sp.] |
| MBQ2852763.1 | Type1.BREX | AbiEi_4->AbiEii-><-Response_reg\|\|HTH_3-> BrxA-DNA-binding -> BrxB-BrxF- iSTAND -> DUF6079 *->SbcC-> N6- METHYLASE -> OLD-ABC ->HNH -> PglZ -> SIGMA+OB-Fold+AAA-ATPase+Lon-Protease-BrxL -><-?<-?<-XIS-HTH | 1215 | Bacteria>FCB_group>Bacteroidetes/Chlorobi_group>Bacteroidota>Bacteroidia>Bacteroidales | GCA_017409275.1 | Bacteroidales_bacterium | IJE76_05160 | 110084..113731 | 2030927 | hypothetical-protein-[Bacteroidales-bacterium] |
| MCI6342424.1 | Type6.BREX | DnaB+TnsC-ATPase->?-><- SIGMA+OB-Fold+AAA-ATPase+HKD-DNase-BrxL <- PglZ <- HNH<- OLD-ABC <- N6- METHYLASE <- DUF6079 *<- BrxB-BrxF- iSTAND <- BrxA-DNA-binding <- BrxE <-HTH_3 | 1217 | Bacteria>FCB_group>Bacteroidetes/Chlorobi_group>Bacteroidota>Bacteroidia>Bacteroidales | GCA_022777695.1 | Prevotella_sp. | MR788_07720 | 61686..65339 | 59823 | BREX-system-P-loop-protein-BrxC-[Prevotella-sp.] |
| MCI6336457.1 | Type1.BREX | PIWI->?-> BrxA-DNA-binding -> BrxB-BrxF- iSTAND -> DUF6079 *->ParB-> OLD-ABC ->?-> N6- METHYLASE -> OLD-ABC ->HNH -> PglZ -> SIGMA+OB-Fold+AAA-ATPase+Lon-Protease-BrxL -> | 1215 | Bacteria>FCB_group>Bacteroidetes/Chlorobi_group>Bacteroidota>Bacteroidia>Bacteroidales | GCA_022777825.1 | Prevotella_sp. | MR802_00705 | 5343..8990 | 59823 | BREX-system-P-loop-protein-BrxC-[Prevotella-sp.] |
| WP_232506819.1 | Type6.BREX | PRTase->?->?->tRNA->tRNA-><-?<- SIGMA+OB-Fold+AAA-ATPase+HKD-DNase-BrxL <- PglZ <- HNH<- OLD-ABC <- N6- METHYLASE <- DUF6079 -p*<- DUF6079 -p<- BrxB-BrxF- iSTAND <- BrxA-DNA-binding <- BrxE <-HTH_3<-?<-TraJ-RHH<-ZnR+ TOPRIM | 794 | Bacteria>FCB_group>Bacteroidetes/Chlorobi_group>Bacteroidota>Bacteroidia>Bacteroidales | GCF_000178215.1 | Bacteroides_xylanisolvens | CW3_RS0123000 | 100900..103284 | 371601 | hypothetical-protein-[Bacteroides-xylanisolvens] |
| WP_008016782.1 | Type6.BREX | <- SIGMA+OB-Fold+AAA-ATPase+HKD-DNase-BrxL <- PglZ <- HNH<- OLD-ABC <- N6- METHYLASE <- DUF6079 -p<- DUF6079 -p*<- BrxB-BrxF- iSTAND <- BrxA-DNA-binding <- BrxE <-HTH_3<-?<-TraJ-RHH<-ZnR+ TOPRIM <-CR-ATPase4 | 425 | Bacteria>FCB_group>Bacteroidetes/Chlorobi_group>Bacteroidota>Bacteroidia>Bacteroidales | GCF_000178215.1 | Bacteroides_xylanisolvens | CW3_RS26225 | 103281..104558 | 371601 | DUF6079-family-protein-[Bacteroides-xylanisolvens] |
| WP_004314649.1 | Type6.BREX | XIS-HTH->CR-ATPase4->ZnR+ TOPRIM ->TraJ-RHH->?->HTH_3-> BrxE -> BrxA-DNA-binding -> BrxB-BrxF- iSTAND -> DUF6079 -p-> DUF6079 -p*-> N6- METHYLASE -> OLD-ABC ->HNH -> PglZ -> SIGMA+OB-Fold+AAA-ATPase+HKD-DNase-BrxL -> | 515 | Bacteria>FCB_group>Bacteroidetes/Chlorobi_group>Bacteroidota>Bacteroidia>Bacteroidales | GCF_000178295.1 | Bacteroides_xylanisolvens | CW1_RS26610 | 23799..25346 | 371601 | BREX-system-P-loop-protein-BrxC-[Bacteroides-xylanisolvens] |
| WP_232505074.1 | Type6.BREX | CR-ATPase4->ZnR+ TOPRIM ->TraJ-RHH->?->?-> BrxE -> BrxA-DNA-binding -> BrxB-BrxF- iSTAND -> DUF6079 -p-> DUF6079 -p*-> N6- METHYLASE -> OLD-ABC ->HNH -> PglZ -> SIGMA+OB-Fold+AAA-ATPase+HKD-DNase-BrxL -> | 711 | Bacteria>FCB_group>Bacteroidetes/Chlorobi_group>Bacteroidota>Bacteroidia>Bacteroidales | GCF_000178295.1 | Bacteroides_xylanisolvens | CW1_RS26615 | 25324..27459 | 371601 | hypothetical-protein-[Bacteroides-xylanisolvens] |
| WP_044064719.1 | Type6.BREX | <-REase<-?<- SIGMA+OB-Fold+AAA-ATPase+HKD-DNase-BrxL <- PglZ <- HNH<- OLD-ABC <- N6- METHYLASE <- DUF6079 *<- BrxB-BrxF- iSTAND <- BrxA-DNA-binding <- BrxE <-HTH_3<-AAA<-?<-XIS-HTH<-?<-Phage_int_SAM_5+Phage_integrase | 1217 | Bacteria>FCB_group>Bacteroidetes/Chlorobi_group>Bacteroidota>Bacteroidia>Bacteroidales | GCF_000833995.1 | Prevotella_pectinovora | ST45_RS02505 | 69518..73171 | 1602169 | BREX-system-P-loop-protein-BrxC-[Prevotella-pectinovora] |
| WP_215758559.1 | Type6.BREX | Phage_int_SAM_5+Phage_integrase->?->XIS-HTH->?->AAA+->HTH_3-> BrxE -> BrxA-DNA-binding -> BrxB-BrxF- iSTAND -> DUF6079 *-> N6- METHYLASE -> OLD-ABC ->HNH -> PglZ -> SIGMA+OB-Fold+AAA-ATPase+HKD-DNase-BrxL -><-?\|\|ParA-Soj-PloopNTPase-> | 1217 | Bacteria>FCB_group>Bacteroidetes/Chlorobi_group>Bacteroidota>Bacteroidia>Bacteroidales | GCF_018789675.1 | Hallella_faecis | KM934_00340 | 78197..81850 | 2841596 | BREX-system-P-loop-protein-BrxC-[Hallella-faecis] |
| WP_008770354.1 | Type6.BREX | DDE_Tnp-><-?<- SIGMA+OB-Fold+AAA-ATPase+HKD-DNase-BrxL <- PglZ <- HNH<- OLD-ABC <- N6- METHYLASE <- DUF6079 *<- BrxB-BrxF- iSTAND <- BrxA-DNA-binding <- BrxE <-HTH_3<-?<-TraJ-RHH<-ZnR+ TOPRIM <-CR-ATPase4 | 1219 | Bacteria>FCB_group>Bacteroidetes/Chlorobi_group>Bacteroidota>Bacteroidia>Bacteroidales | GCF_902374095.1 | Bacteroides | BSAG_RS19905 | 423901..427560 | 816 | MULTISPECIES:-BREX-system-P-loop-protein-BrxC-[Bacteroides] |
| WP_273212250.1 | Type1.BREX | <-AbiEii<-wHTH+AbiEi_2<- SIGMA+OB-Fold+AAA-ATPase+Lon-Protease-BrxL <- PglZ <-Schlafen+HATPase+wHTH<-REase<- N6- METHYLASE <- HNH<- OLD-ABC <- DUF6079 *<- BrxB-BrxF- iSTAND <- BrxA-DNA-binding <-?<-?<-D5ATPase<-XIS-HTH<-?<-Arm-DNA-bind_5+Phage_int_SAM_5+Phage_integrase | 1173 | Bacteria>FCB_group>Bacteroidetes/Chlorobi_group>Bacteroidota>Cytophagia>Cytophagales | GCF_002483185.1 | Runella_zeae | B4E63_RS12460 | 307458..310979 | 94255 | BREX-system-P-loop-protein-BrxC-[Runella-zeae] |
| WP_159471214.1 | Type1.BREX | <-XIS-HTH<-AbiEii<-AbiEi_4+AbiEi_1<-REase<- SIGMA+OB-Fold+AAA-ATPase+Lon-Protease-BrxL <- PglZ <- N6- METHYLASE <- HNH<- OLD-ABC <- DUF6079 *<- BrxB-BrxF- iSTAND <- BrxA-DNA-binding \|\|CHTH->?->?->?-><-CopG/RHH | 1179 | Bacteria>FCB_group>Bacteroidetes/Chlorobi_group>Bacteroidota>Cytophagia>Cytophagales | GCF_009801115.1 | Dyadobacter_sp._3J3 | FZF08_RS17845 | 89009..92548 | 2606600 | BREX-system-P-loop-protein-BrxC-[Dyadobacter-sp.-3J3] |
| WP_141421442.1 | Type1.BREX | <-HAD+PRTase+PRTase+REase+SF1-SF2-Helicase<-?<- SIGMA+OB-Fold+AAA-ATPase+HKD-DNase-BrxL <- PglZ <-TPR+S1<-NERD+SF1-SF2-Helicase<- N6- METHYLASE <-ParB+HNH<- DUF6079 *<- BrxB-BrxF- iSTAND <- BrxA-DNA-binding <-TPR+TPR<-?<-?<-?<-Phage_int+DNA-glycosylase<- HNH<- OLD-ABC | 1165 | Bacteria>FCB_group>Bacteroidetes/Chlorobi_group>Bacteroidota>Flavobacteriia>Flavobacteriales | GCF_006546625.1 | Haloflavibacter_putidus | FKR84_RS06225 | 21354..24851 | 2576776 | BREX-system-P-loop-protein-BrxC-[Haloflavibacter-putidus] |
| MBN8706499.1 | Type1.BREX | Phage_int_SAM_4+Phage_integrase->35exo+URI-DNase->tRNA->HTH-> BrxA-DNA-binding -> BrxB-BrxF- iSTAND -> DUF6079 *-> OLD-ABC ->HNH -> N6- METHYLASE ->?->Schlafen+HATPase+wHTH-> PglZ -> SIGMA+OB-Fold+AAA-ATPase+Lon-Protease-BrxL ->SbcC-> | 1173 | Bacteria>FCB_group>Bacteroidetes/Chlorobi_group>Bacteroidota>unclassified_Bacteroidota | GCA_017303995.1 | Bacteroidota_bacterium | JOL62_11545 | 8364..11885 | 1898104 | BREX-system-P-loop-protein-BrxC-[Bacteroidota-bacterium] |
| MBS1911241.1 | Type1.BREX | <- SIGMA+OB-Fold+AAA-ATPase+Lon-Protease-BrxL <- PglZ <- HNH<- OLD-ABC <-ParB+HNH<- N6- METHYLASE <- DUF6079 *<- BrxB-BrxF- iSTAND <- BrxA-DNA-binding <-tRNA<-?\|\|?-><-?\|\|SIGMA-HTH-> | 1145 | Bacteria>FCB_group>Bacteroidetes/Chlorobi_group>Bacteroidota>unclassified_Bacteroidota | GCA_018266795.1 | Bacteroidota_bacterium | JST22_04585 | 55920..59357 | 1898104 | BREX-system-P-loop-protein-BrxC-[Bacteroidota-bacterium] |
| WP_201422032.1 | Type6.BREX | <-glycosylase<- SIGMA+OB-Fold+AAA-ATPase+Lon-Protease-BrxL <- PglZ <- HNH<- OLD-ABC <- N6- METHYLASE <- N6- METHYLASE <-Tox-RelA<- DUF6079 *<- BrxB-BrxF- iSTAND <- BrxA-DNA-binding <- BrxE <-XIS-HTH\|\|<-ACET\|\|?->?->REC+NtrC-AAA+FIS-HTH-> | 1188 | Bacteria>Myxococcota>Myxococcia>Myxococcales>Cystobacterineae>Myxococcaceae | GCF_016698685.1 | Myxococcus_xanthus | JKA73_RS29390 | 7346078..7349644 | 34 | BREX-system-P-loop-protein-BrxC-[Myxococcus-xanthus] |
| WP_267556343.1 | Type6.BREX | <-rve_3<- SIGMA+OB-Fold+AAA-ATPase+Lon-Protease-BrxL <- PglZ <- HNH<- OLD-ABC <- N6- METHYLASE <-Tox-RelA<- DUF6079 *<- BrxB-BrxF- iSTAND <- BrxA-DNA-binding <- BrxE <-XIS-HTH\|\|?-><-?<-wHTH+HNH<-?<-rve_3 | 1189 | Bacteria>Myxococcota>Myxococcia>Myxococcales>Cystobacterineae>Myxococcaceae | GCF_026626625.1 | Corallococcus_sp._BB11-1 | OV207_RS17695 | 240885..244454 | 2996783 | BREX-system-P-loop-protein-BrxC-[Corallococcus-sp.-BB11-1] |
| WP_020733584.1 | Type6.BREX | ABC-ATPase+SbcC-><- SIGMA+OB-Fold+AAA-ATPase+Lon-Protease-BrxL <- PglZ <- HNH<- OLD-ABC <- N6- METHYLASE <- N6- METHYLASE <-Schlafen+HATPase+wHTH<- DUF6079 *<- BrxB-BrxF- iSTAND <- BrxA-DNA-binding <- BrxE <-XIS-HTH\|\|?->?->ACET-> | 1182 | Bacteria>Myxococcota>Polyangia>Polyangiales>Polyangiaceae>Sorangium | GCF_000418325.1 | Sorangium_cellulosum | SCE1572_RS08150 | 2251441..2254989 | 56 | BREX-system-P-loop-protein-BrxC-[Sorangium-cellulosum] |
| MCK6536698.1 | Type1.BREX | <-TetR-HTH\|\|MerR-HTH->?->?->?-><-?\|\|XIS-HTH-> BrxA-DNA-binding -> BrxB-BrxF- iSTAND -> DUF6079 *-> N6- METHYLASE -> OLD-ABC ->HNH -> PglZ -> SIGMA+OB-Fold+AAA-ATPase+Lon-Protease-BrxL ->REase+PNTD1+PNTD2-> | 1149 | Bacteria>Myxococcota>Polyangia>Polyangiales>Polyangiaceae>unclassified_Polyangiaceae | GCA_023150915.1 | Polyangiaceae_bacterium | L6Q84_27330 | 8687..12136 | 2268199 | BREX-system-P-loop-protein-BrxC-[Polyangiaceae-bacterium] |
| MCK6527600.1 | Type6.BREX | RADICAL-SAM->?-><- SIGMA+OB-Fold+AAA-ATPase+Lon-Protease-BrxL <-?<-?<-?<- HNH<- OLD-ABC <- N6- METHYLASE <- DUF6079 *<- BrxB-BrxF- iSTAND <- BrxA-DNA-binding <- BrxE \|\|?->GDResolvase+Recombinase+ZnR-> | 1184 | Bacteria>Myxococcota>unclassified_Myxococcota | GCA_023150875.1 | Myxococcota_bacterium | L6R50_08565 | 57228..60782 | 2818507 | BREX-system-P-loop-protein-BrxC-[Myxococcota-bacterium] |
| WP_225487034.1 | Type1.BREX | <-HTH_58<-TrwC+SF1-SF2-Helicase\|\|GNAT+DDE_Tnp_1_assoc->DDE_Tnp_1_assoc->Tox-RelE->HTH_3-> wHTH+WYL+WCX -p-> BrxA-DNA-binding -> BrxB-BrxF- iSTAND -> DUF6079 *-> N6- METHYLASE -> OLD-ABC ->HNH -> PglZ -> SIGMA+OB-Fold+AAA-ATPase+Lon-Protease-BrxL -><-tRNA<-FURR-HTH | 1146 | Bacteria>Pseudomonadota>Acidithiobacillia>Acidithiobacillales>Acidithiobacillaceae>Acidithiobacillus | - | Acidithiobacillus_ferrooxidans | LDB30_08025 | 1563873..1567313 | 920 | BREX-system-P-loop-protein-BrxC-[Acidithiobacillus-ferrooxidans] |
| RWC29997.1 | Type1.BREX | SbcC+CCDC158->?->REase+PNTD1+PNTD2->PIWI-><- SIGMA+OB-Fold+AAA-ATPase+HKD-DNase-BrxL <- PglZ <- HNH<- OLD-ABC <- N6- METHYLASE <- DUF6079 *<- BrxB-BrxF- iSTAND <- BrxA-DNA-binding \|\|REase-><-?\|\|?-><-?\|\|REase->?->rve-> | 1181 | Bacteria>Pseudomonadota>Alphaproteobacteria>Hyphomicrobiales>Phyllobacteriaceae>Mesorhizobium | GCA_004017395.1 | Mesorhizobium_sp. | EOS27_14215 | 67406..70951 | 1871066 | BREX-system-P-loop-protein-BrxC-[Mesorhizobium-sp.] |
| WP_128286158.1 | Type1.BREX | ParB+HNH-><- SIGMA+OB-Fold+AAA-ATPase+HKD-DNase-BrxL <- PglZ <- HNH<- OLD-ABC <- N6- METHYLASE <- DUF6079 *<- BrxB-BrxF- iSTAND <- BrxA-DNA-binding <-RNaseH | 1181 | Bacteria>Pseudomonadota>Alphaproteobacteria>Hyphomicrobiales>Phyllobacteriaceae>Mesorhizobium | GCF_004023115.1 | Mesorhizobium_sp._M6A.T.Cr.TU.017.01.1.1 | EOB36_RS28010 | 14606..18151 | 2496774 | BREX-system-P-loop-protein-BrxC-[Mesorhizobium-sp.-M6A.T.Cr.TU.017.01.1.1] |
| WP_136091391.1 | Type1.BREX | REase+PNTD1+PNTD2->PIWI->Y1_Tnp-><- SIGMA+OB-Fold+AAA-ATPase+HKD-DNase-BrxL <- PglZ <- HNH<- OLD-ABC <- N6- METHYLASE <- DUF6079 *<- BrxB-BrxF- iSTAND <- BrxA-DNA-binding \|\|REase->?-><-?\|\|REase->rve->TnsC-ATPase->TnsC-ATPase-> | 1181 | Bacteria>Pseudomonadota>Alphaproteobacteria>Hyphomicrobiales>Phyllobacteriaceae>Mesorhizobium | GCF_004794715.1 | Mesorhizobium_sp._M8A.F.Ca.ET.207.01.1.1 | EN850_RS12645 | 56072..59617 | 2563968 | BREX-system-P-loop-protein-BrxC-[Mesorhizobium-sp.-M8A.F.Ca.ET.207.01.1.1] |
| MBF0399382.1 | Type1.BREX | BrxA-DNA-binding -> BrxB-BrxF- iSTAND -> DUF6079 *->HicB_lk_antitox->HicA_toxin-> N6- METHYLASE -> OLD-ABC ->HNH -> PglZ -> SIGMA+OB-Fold+AAA-ATPase+Lon-Protease-BrxL ->?-><-STAS | 1144 | Bacteria>Pseudomonadota>Alphaproteobacteria>Magnetococcales>unclassified_Magnetococcales | GCA_015232635.1 | Magnetococcales_bacterium | HQL90_01290 | 18375..21809 | 2026759 | BREX-system-P-loop-protein-BrxC-[Magnetococcales-bacterium] |
| QKS29579.1 | Type6.BREX | <- SIGMA+OB-Fold+AAA-ATPase+Lon-Protease-BrxL <- PglZ <- HNH<- OLD-ABC <- N6- METHYLASE <-ORC-CDC6<-ParA-Soj-PloopNTPase<-HEPN<- N6- METHYLASE <- DUF6079 *<- BrxB-BrxF- iSTAND <- BrxA-DNA-binding <- BrxE <-XIS-HTH | 1180 | Bacteria>Pseudomonadota>Betaproteobacteria>Betaproteobacteria_incertae_sedis>Candidatus_Accumulibacter | GCA_013347225.1 | Candidatus_Accumulibacter_similis | HT579_12075 | 2666587..2670129 | 2954388 | MAG:-BREX-system-P-loop-protein-BrxC-[Candidatus-Accumulibacter-similis] |
| MBK8385355.1 | Type1.BREX | <- SIGMA+OB-Fold+AAA-ATPase+Lon-Protease-BrxL <- PglZ <- N6- METHYLASE <- HNH<- OLD-ABC <- DUF6079 *<- BrxB-BrxF- iSTAND <- BrxA-DNA-binding <- BrxA-DNA-binding <-Polbeta<-wHTH+AbiEi_2 | 1154 | Bacteria>Pseudomonadota>Betaproteobacteria>Betaproteobacteria_incertae_sedis>Candidatus_Accumulibacter>unclassified_Candidatus_Accumulibacter | GCA_016711725.1 | Candidatus_Accumulibacter_sp. | IPL11_06655 | 113668..117132 | 2053492 | BREX-system-P-loop-protein-BrxC-[Candidatus-Accumulibacter-sp.] |
| MBN8436515.1 | Type1.BREX | wHTH+AbiEi_2-><-?\|\|?->wHTH+AbiEi_2->Polbeta-> BrxA-DNA-binding -> BrxA-DNA-binding -> BrxB-BrxF- iSTAND -> DUF6079 *-> N6- METHYLASE -> OLD-ABC ->HNH -> PglZ -> SIGMA+OB-Fold+AAA-ATPase+Lon-Protease-BrxL -> | 1154 | Bacteria>Pseudomonadota>Betaproteobacteria>Betaproteobacteria_incertae_sedis>Candidatus_Accumulibacter>unclassified_Candidatus_Accumulibacter | GCA_017302555.1 | Candidatus_Accumulibacter_sp. | JOL95_00470 | 105300..108764 | 2053492 | BREX-system-P-loop-protein-BrxC-[Candidatus-Accumulibacter-sp.] |
| WP_043683814.1 | Type1.BREX | DDE_Tnp_1_4->?->VirB5-><-DUF772+DDE_Tnp_1+DDE_Tnp_1_6<-?\|\| wHTH+WYL+WCX -> BrxA-DNA-binding -> BrxB-BrxF- iSTAND -> DUF6079 *->HEPN-> N6- METHYLASE -> OLD-ABC ->HNH ->DUF772+DDE_Tnp_1+DDE_Tnp_1_6->HNH-> PglZ -> SIGMA+OB-Fold+AAA-ATPase+Lon-Protease-BrxL -><-YdaS_antitoxin | 1153 | Bacteria>Pseudomonadota>Betaproteobacteria>Burkholderiales>Alcaligenaceae>Castellaniella | GCF_000612685.1 | Castellaniella_defragrans | BN940_RS13240 | 2967883..2971344 | 75697 | BREX-system-P-loop-protein-BrxC-[Castellaniella-defragrans] |
| WP_085318435.1 | Type1.BREX | <-DiS_P_DiS+Peptidase_A24<-?<-VirB11\|\|RHH_5-> BrxA-DNA-binding -> BrxB-BrxF- iSTAND -> DUF6079 *->?-> N6- METHYLASE -> OLD-ABC ->HNH -> PglZ -> SIGMA+OB-Fold+AAA-ATPase+Lon-Protease-BrxL -> | 1153 | Bacteria>Pseudomonadota>Betaproteobacteria>Burkholderiales>Alcaligenaceae>Derxia | GCF_002105195.1 | Derxia_lacustris | B9P64_RS23165 | 16077..19538 | 764842 | BREX-system-P-loop-protein-BrxC-[Derxia-lacustris] |
| WP_115063824.1 | Type1.BREX | Phage_AlpA->?->TraJ-RHH-><-REase+NACHT<-?<-RADICAL-SAM<-DUF772+DDE_Tnp_1\|\| BrxA-DNA-binding -> BrxB-BrxF- iSTAND -> DUF6079 *-> N6- METHYLASE ->DUF3644-> OLD-ABC -> TOPRIM -> OLD-ABC ->HNH -> PglZ -> SIGMA+OB-Fold+AAA-ATPase+Lon-Protease-BrxL -><-Y1_Tnp | 1155 | Bacteria>Pseudomonadota>Betaproteobacteria>Burkholderiales>Alcaligenaceae>Oligella | GCF_900454315.1 | Oligella_ureolytica | NCTC11998_02420 | 2456151..2459618 | 90244 | BREX-system-P-loop-protein-BrxC-[Oligella-ureolytica] |
| WP_108330269.1 | Type1.BREX | <- SIGMA+OB-Fold+AAA-ATPase+Lon-Protease-BrxL <- PglZ <- HNH<- OLD-ABC <- N6- METHYLASE <- REase<- OLD-ABC <- DUF6079 *<- BrxB-BrxF- iSTAND <- BrxA-DNA-binding <-PIN<-ZnR+TPR | 1154 | Bacteria>Pseudomonadota>Betaproteobacteria>Burkholderiales>Comamonadaceae>Limnohabitans | GCF_003063435.1 | Limnohabitans_sp._2KL-1 | B9Z47_RS05555 | 224189..227653 | 1100699 | BREX-system-P-loop-protein-BrxC-[Limnohabitans-sp.-2KL-1] |
| WP_108311897.1 | Type1.BREX | <-SF1-SF2-Helicase<- SIGMA+OB-Fold+AAA-ATPase+Lon-Protease-BrxL <- PglZ <- HNH<- OLD-ABC <- N6- METHYLASE <- DUF6079 *<- BrxB-BrxF- iSTAND <- BrxA-DNA-binding <-PIN<-ZnR+TPR | 1153 | Bacteria>Pseudomonadota>Betaproteobacteria>Burkholderiales>Comamonadaceae>Limnohabitans | GCF_003063455.1 | Limnohabitans_parvus | B9Z37_RS03875 | 837390..840851 | 540061 | BREX-system-P-loop-protein-BrxC-[Limnohabitans-parvus] |
| WP_101103324.1 | Type1.BREX | HTH_Tnp_1+HTH_21+rve+rve_3->RHH_5-> BrxA-DNA-binding -> BrxB-BrxF- iSTAND -> DUF6079 *->HEPN-> N6- METHYLASE -> OLD-ABC + TOPRIM ->SF1-SF2-Helicase-> OLD-ABC ->HNH ->ParB+HNH-> PglZ -> SIGMA+OB-Fold+AAA-ATPase+Lon-Protease-BrxL -> | 1153 | Bacteria>Pseudomonadota>Betaproteobacteria>Burkholderiales>Comamonadaceae>Macromonas | GCF_002837135.1 | Macromonas_bipunctata | CON76_RS10585 | 3906..7367 | 183670 | BREX-system-P-loop-protein-BrxC-[Macromonas-bipunctata] |
| MBH2043138.1 | Type1.BREX | RHH_5-> BrxA-DNA-binding -> BrxB-BrxF- iSTAND -> DUF6079 *->MarR-HTH+Polbeta->HEPN-> OLD-ABC ->HNH -> PglZ -> SIGMA+OB-Fold+AAA-ATPase+Lon-Protease-BrxL -> | 1153 | Bacteria>Pseudomonadota>Betaproteobacteria>Burkholderiales>Comamonadaceae>unclassified_Comamonadaceae | GCA_016000285.1 | Comamonadaceae_bacterium | I8H87_06315 | 1878..5339 | 1871071 | BREX-system-P-loop-protein-BrxC-[Comamonadaceae-bacterium] |
| WP_071078524.1 | Type1.BREX | Phage_integrase-><-?<- SIGMA+OB-Fold+AAA-ATPase+Lon-Protease-BrxL <- PglZ <- HNH<- OLD-ABC <- N6- METHYLASE <- DUF6079 *<- BrxB-BrxF- iSTAND <- BrxA-DNA-binding <-RHH_5<-?<-HEPN+OST-HTH | 1153 | Bacteria>Pseudomonadota>Betaproteobacteria>Burkholderiales>Oxalobacteraceae>Janthinobacterium | GCF_001854915.1 | Janthinobacterium_lividum | AKG95_RS19180 | 51113..54574 | 29581 | BREX-system-P-loop-protein-BrxC-[Janthinobacterium-lividum] |
| WP_046167751.1 | Type1.BREX | BrxA-DNA-binding -> BrxB-BrxF- iSTAND -> DUF6079 *-> N6- METHYLASE -> OLD-ABC ->HNH -> PglZ -> SIGMA+OB-Fold+AAA-ATPase+Lon-Protease-BrxL ->?->?-><-AraC-HTH+AraC-HTH | 1153 | Bacteria>Pseudomonadota>Betaproteobacteria>Neisseriales>Chromobacteriaceae>Chromobacterium_group | GCF_000971355.1 | Chromobacterium_vaccinii | VL68_RS13365 | 2032..5493 | 1108595 | BREX-system-P-loop-protein-BrxC-[Chromobacterium-vaccinii] |
| WP_096450985.1 | Type6.BREX | <- SIGMA+OB-Fold+AAA-ATPase+Lon-Protease-BrxL <- PglZ <- HNH<- OLD-ABC <- N6- METHYLASE <-ORC-CDC6<-ParA-Soj-PloopNTPase<-HEPN<- N6- METHYLASE <- DUF6079 *<- BrxB-BrxF- iSTAND <- BrxA-DNA-binding <- BrxE <-XIS-HTH<-HTH_3\|\|?-><-ParB<-?\|\|HTH_3-> | 1180 | Bacteria>Pseudomonadota>Betaproteobacteria>Rhodocyclales>Zoogloeaceae>Thauera | GCF_002354895.1 | Thauera_sp._K11 | CCZ27_RS19540 | 4450815..4454357 | 2005884 | BREX-system-P-loop-protein-BrxC-[Thauera-sp.-K11] |
| WP_148578351.1 | Type1.BREX | <-SbcC+SbcC+ABC-ATPase+SbcC\|\|?-><-HTH_3\|\|?->\|\| wHTH+WYL+WCX ->BirA-HTH-> BrxA-DNA-binding -> BrxB-BrxF- iSTAND -> DUF6079 *->HEPN-> N6- METHYLASE ->DUF932->ParB+HNH-> OLD-ABC ->HNH -> PglZ -> SIGMA+OB-Fold+AAA-ATPase+Lon-Protease-BrxL ->DUF772+DDE_Tnp_1-> | 1146 | Bacteria>Pseudomonadota>Betaproteobacteria>Rhodocyclales>Zoogloeaceae>Zoogloea | GCF_008107625.1 | Zoogloea_oleivorans | ETQ85_RS07180 | 242245..245685 | 1552750 | BREX-system-P-loop-protein-BrxC-[Zoogloea-oleivorans] |
| EJN6954191.1 | Type1.BREX | <-LexA-protease\|\|?->< wHTH+WYL+WCX \|\|ARG->?->ZnR->RHS-> BrxA-DNA-binding -> BrxB-BrxF- iSTAND -> DUF6079 *-> N6- METHYLASE -> OLD-ABC ->HNH -> PglZ ->HORMA-><-SF1-SF2-Helicase+REase<-SF1-SF2-Helicase+REase | 1157 | Bacteria>Pseudomonadota>Gammaproteobacteria>Aeromonadales>Aeromonadaceae>Aeromonas | GCA_024506355.1 | Aeromonas_hydrophila | NRD12_000732 | 206979..210452 | 644 | BREX-system-P-loop-protein-BrxC-[Aeromonas-hydrophila] |
| WP_043124789.1 | Type1.BREX | SF1-SF2-Helicase+REase->SF1-SF2-Helicase+REase-><-HORMA<- PglZ <- HNH<- OLD-ABC <- N6- METHYLASE <- DUF6079 *<- BrxB-BrxF- iSTAND <- BrxA-DNA-binding <-URI-DNase<-?\|\|LexA-protease-> | 1157 | Bacteria>Pseudomonadota>Gammaproteobacteria>Aeromonadales>Aeromonadaceae>Aeromonas | GCF_000724965.1 | Aeromonas_hydrophila | HR52_RS13935 | 3102820..3106293 | 644 | BREX-system-P-loop-protein-BrxC-[Aeromonas-hydrophila] |
| WP_113721905.1 | Type1.BREX | SF1-SF2-Helicase+REase->SF1-SF2-Helicase+REase-><-HEPN+Sel1+SEC-C<- PglZ <- HNH<- OLD-ABC <- N6- METHYLASE <- DUF6079 *<- BrxB-BrxF- iSTAND <- BrxA-DNA-binding <-RHS<-?<-ARG\|\| wHTH+WYL+WCX -><-?\|\|LexA-protease-> | 1157 | Bacteria>Pseudomonadota>Gammaproteobacteria>Aeromonadales>Aeromonadaceae>Aeromonas | GCF_003265515.1 | Aeromonas_salmonicida | CPA05_RS19945 | 20643..24116 | 645 | BREX-system-P-loop-protein-BrxC-[Aeromonas-salmonicida] |
| WP_114523382.1 | Type1.BREX | <-LexA-protease\|\|?->URI-DNase-> BrxA-DNA-binding -> BrxB-BrxF- iSTAND -> DUF6079 *-> N6- METHYLASE -> OLD-ABC ->HNH -> PglZ -> | 1157 | Bacteria>Pseudomonadota>Gammaproteobacteria>Aeromonadales>Aeromonadaceae>Aeromonas | GCF_003344485.1 | Aeromonas_sp._ARM81 | ASJ36_RS21100 | 4447..7920 | 1747384 | BREX-system-P-loop-protein-BrxC-[Aeromonas-sp.-ARM81] |
| WP_182951579.1 | Type1.BREX | <-LexA-protease\|\|?->REase+N6_ METHYLASE ->TRD+TRD->REase+SF1-SF2-Helicase->zf-ISL3+HTH_Tnp_ISL3+DDE_Tnp_ISL3->URI-DNase-> BrxA-DNA-binding -> BrxB-BrxF- iSTAND -> DUF6079 *-> N6- METHYLASE -> OLD-ABC ->HNH -> PglZ ->HIN-HTH->HTH_Tnp_1+HTH_21+rve+rve_3->HORMA-><-DDE_Tnp_1<-SF1-SF2-Helicase+REase | 1157 | Bacteria>Pseudomonadota>Gammaproteobacteria>Aeromonadales>Aeromonadaceae>Aeromonas | GCF_014168615.1 | Aeromonas_sp._WP2-W18-CRE-05 | WP2W18C05_13360 | 1426228..1429701 | 2675707 | BREX-system-P-loop-protein-BrxC-[Aeromonas-sp.-WP2-W18-CRE-05] |
| WP_203130882.1 | Type1.BREX | <- wHTH+WYL+WCX \|\|ZnR+ZnR-><-?<-LexA-protease\|\|?->URI-DNase-> BrxA-DNA-binding -> BrxB-BrxF- iSTAND -> DUF6079 -p*-> DUF6079 -p-> N6- METHYLASE -> OLD-ABC ->HNH -> PglZ ->HORMA-><-SF1-SF2-Helicase+REase<-SF1-SF2-Helicase+REase | 935 | Bacteria>Pseudomonadota>Gammaproteobacteria>Aeromonadales>Aeromonadaceae>Aeromonas | GCF_016805325.1 | Aeromonas_jandaei | GNZ06_RS06060 | 150251..153058 | 650 | BREX-system-P-loop-protein-BrxC-[Aeromonas-jandaei] |
| WP_203130883.1 | Type1.BREX | <- wHTH+WYL+WCX \|\|ZnR+ZnR-><-?<-LexA-protease\|\|?->URI-DNase-> BrxA-DNA-binding -> BrxB-BrxF- iSTAND -> DUF6079 -p-> DUF6079 -p*-> N6- METHYLASE -> OLD-ABC ->HNH -> PglZ ->HORMA-><-SF1-SF2-Helicase+REase<-SF1-SF2-Helicase+REase | 226 | Bacteria>Pseudomonadota>Gammaproteobacteria>Aeromonadales>Aeromonadaceae>Aeromonas | GCF_016805325.1 | Aeromonas_jandaei | GNZ06_RS06065 | 153058..153738 | 650 | hypothetical-protein-[Aeromonas-jandaei] |
| WP_203133261.1 | Type1.BREX | SF1-SF2-Helicase+REase->SF1-SF2-Helicase+REase-><-HORMA<- PglZ <- HNH<- OLD-ABC <- N6- METHYLASE <- DUF6079 *<- BrxB-BrxF- iSTAND <- BrxA-DNA-binding <-URI-DNase<-?\|\|LexA-protease->?-><-ZnR+ZnR\|\| wHTH+WYL+WCX -> | 1157 | Bacteria>Pseudomonadota>Gammaproteobacteria>Aeromonadales>Aeromonadaceae>Aeromonas | GCF_016805365.1 | Aeromonas_jandaei | G4224_RS08640 | 376570..380043 | 650 | BREX-system-P-loop-protein-BrxC-[Aeromonas-jandaei] |
| WP_213387487.1 | Type1.BREX | <-LexA-protease\|\|?->< wHTH+WYL+WCX \|\|ARG->?->ZnR->RHS-> BrxA-DNA-binding -> BrxB-BrxF- iSTAND -> DUF6079 *-> N6- METHYLASE -> OLD-ABC ->HNH -> PglZ ->HEPN+Sel1+SEC-C->HIN-HTH-><-SF1-SF2-Helicase+REase<-SF1-SF2-Helicase+REase | 1157 | Bacteria>Pseudomonadota>Gammaproteobacteria>Aeromonadales>Aeromonadaceae>Aeromonas | GCF_018359765.1 | Aeromonas_veronii | J4G64_RS11995 | 290568..294041 | 654 | BREX-system-P-loop-protein-BrxC-[Aeromonas-veronii] |
| WP_215805224.1 | Type1.BREX | <-HORMA<-DDE_Tnp_1<- PglZ -p<- PglZ -p<- PglZ <-HNH<- HNH<- OLD-ABC <- N6- METHYLASE <- DUF6079 *<- BrxB-BrxF- iSTAND -p<- BrxB-BrxF- iSTAND <- BrxA-DNA-binding \|\|HIN-HTH-><-URI-DNase<-?\|\|LexA-protease-> | 1157 | Bacteria>Pseudomonadota>Gammaproteobacteria>Aeromonadales>Aeromonadaceae>Aeromonas | GCF_018802365.1 | Aeromonas_jandaei | HQ398_RS10760 | 2353585..2357058 | 650 | BREX-system-P-loop-protein-BrxC-[Aeromonas-jandaei] |
| WP_216986928.1 | Type1.BREX | HDRnase-><-?<-LexA-protease\|\|ARG->ZnR->\|\|?->RHS-> BrxA-DNA-binding -> BrxB-BrxF- iSTAND -> DUF6079 *-> N6- METHYLASE -> OLD-ABC ->HNH -> PglZ ->HORMA-><-SF1-SF2-Helicase+REase<-SF1-SF2-Helicase+REase-p<-SF1-SF2-Helicase+REase-p\|\|SAM- METHYLASE -> | 1157 | Bacteria>Pseudomonadota>Gammaproteobacteria>Aeromonadales>Aeromonadaceae>Aeromonas | GCF_019048565.1 | Aeromonas_sp._FDAARGOS_1406 | I6L36_RS00705 | 137070..140543 | 2778055 | BREX-system-P-loop-protein-BrxC-[Aeromonas-sp.-FDAARGOS-1406] |
| WP_216990395.1 | Type1.BREX | DnaJ->SF1-SF2-Helicase+REase->SF1-SF2-Helicase+REase-><-HORMA<- PglZ <- HNH<- OLD-ABC <- N6- METHYLASE <- DUF6079 *<- BrxB-BrxF- iSTAND <- BrxA-DNA-binding <-RHS<-?<-?<-RNase<-DUF3584+DUF5401<-JetB<-MukF | 1157 | Bacteria>Pseudomonadota>Gammaproteobacteria>Aeromonadales>Aeromonadaceae>Aeromonas | GCF_019048705.1 | Aeromonas_sp._FDAARGOS_1409 | I6L39_RS17660 | 3801324..3804797 | 2778058 | BREX-system-P-loop-protein-BrxC-[Aeromonas-sp.-FDAARGOS-1409] |
| WP_216992842.1 | Type1.BREX | SF1-SF2-Helicase+REase->SF1-SF2-Helicase+REase-><-HORMA<- PglZ <- HNH<- OLD-ABC <- N6- METHYLASE <- DUF6079 *<- BrxB-BrxF- iSTAND <- BrxA-DNA-binding <-?\|\|LexA-protease->HTH+rve->AAA-ATPase->?-><-ZnR+ZnR\|\| wHTH+WYL+WCX -> | 1157 | Bacteria>Pseudomonadota>Gammaproteobacteria>Aeromonadales>Aeromonadaceae>Aeromonas | GCF_019048725.1 | Aeromonas_sp._FDAARGOS_1407 | I6L37_20730 | 4426496..4429969 | 2778056 | BREX-system-P-loop-protein-BrxC-[Aeromonas-sp.-FDAARGOS-1407] |
| WP_223938541.1 | Type1.BREX | <-SAM- METHYLASE <-Tnp_DNA_bind+DDE_Tnp_1+Dimer_Tnp_Tn5\|\|SF1-SF2-Helicase+REase->SF1-SF2-Helicase+REase-><-HORMA<- PglZ <- HNH<- OLD-ABC <- N6- METHYLASE <- DUF6079 *<- BrxB-BrxF- iSTAND <- BrxA-DNA-binding <-RHS | 1157 | Bacteria>Pseudomonadota>Gammaproteobacteria>Aeromonadales>Aeromonadaceae>Aeromonas | GCF_019971155.1 | Aeromonas_caviae | LDN24_RS08690 | 33954..37427 | 648 | BREX-system-P-loop-protein-BrxC-[Aeromonas-caviae] |
| WP_235462874.1 | Type1.BREX | XIS-HTH->?->?-><- wHTH+WYL+WCX \|\|ARG->?->RHS-> BrxA-DNA-binding -> BrxB-BrxF- iSTAND -> DUF6079 *-> N6- METHYLASE -> OLD-ABC ->HNH -> PglZ ->HEPN+Sel1+SEC-C-><-SF1-SF2-Helicase+REase-p | 1157 | Bacteria>Pseudomonadota>Gammaproteobacteria>Aeromonadales>Aeromonadaceae>Aeromonas | GCF_021538905.1 | Aeromonas_australiensis | EHZ86_RS12350 | 8704..12177 | 1114880 | BREX-system-P-loop-protein-BrxC-[Aeromonas-australiensis] |
| WP_241325388.1 | Type1.BREX | SF1-SF2-Helicase+REase-p->SF1-SF2-Helicase+REase-p->SF1-SF2-Helicase+REase-p-><-HEPN+Sel1+SEC-C<- PglZ <- HNH<- OLD-ABC <- N6- METHYLASE <- DUF6079 *<- BrxB-BrxF- iSTAND <- BrxA-DNA-binding <-?<-?<-?<-ABhydrolase | 1157 | Bacteria>Pseudomonadota>Gammaproteobacteria>Aeromonadales>Aeromonadaceae>Aeromonas | GCF_022549295.1 | Aeromonas_sp._MR7 | MM192_RS09020 | 75622..79095 | 2923419 | BREX-system-P-loop-protein-BrxC-[Aeromonas-sp.-MR7] |
| WP_247824022.1 | Type1.BREX | DOC+HTH-><-?<-LexA-protease\|\|?->< wHTH+WYL+WCX \|\|ZnR+ZnR->RHS-> BrxA-DNA-binding -> BrxB-BrxF- iSTAND -> DUF6079 *-> N6- METHYLASE -> OLD-ABC ->HNH -> PglZ ->HORMA-><-SF1-SF2-Helicase+REase<-SF1-SF2-Helicase+REase | 1157 | Bacteria>Pseudomonadota>Gammaproteobacteria>Aeromonadales>Aeromonadaceae>Aeromonas | GCF_023100745.1 | Aeromonas_veronii | MYF86_09055 | 1896227..1899700 | 654 | BREX-system-P-loop-protein-BrxC-[Aeromonas-veronii] |
| WP_264787731.1 | Type1.BREX | <-LexA-protease<-?\|\|RHS->ARG->\|\|RHS-> BrxA-DNA-binding -> BrxB-BrxF- iSTAND -> DUF6079 *-> N6- METHYLASE -> OLD-ABC ->HNH -> PglZ ->DDE_Tnp_1->HORMA-><-?<-zf-IS66+DDE_Tnp_IS66<-zf-IS66+DDE_Tnp_IS66 | 1157 | Bacteria>Pseudomonadota>Gammaproteobacteria>Aeromonadales>Aeromonadaceae>Aeromonas | GCF_026000055.1 | Aeromonas_caviae | OPI30_RS10860 | 2213781..2217254 | 648 | BREX-system-P-loop-protein-BrxC-[Aeromonas-caviae] |
| WP_270670318.1 | Type1.BREX | <-HTH+rve\|\|<-HEPN+Sel1+SEC-C<- PglZ <- HNH<- OLD-ABC <- N6- METHYLASE <- DUF6079 *<- BrxB-BrxF- iSTAND <- BrxA-DNA-binding <-RHS<-?\|\|<-ZnR<-ARG | 1157 | Bacteria>Pseudomonadota>Gammaproteobacteria>Aeromonadales>Aeromonadaceae>Aeromonas | GCF_027705165.1 | Aeromonas_sp._QDB04 | OKC37_RS13865 | 9577..13050 | 2990477 | BREX-system-P-loop-protein-BrxC-[Aeromonas-sp.-QDB04] |
| WP_270809144.1 | Type1.BREX | SF1-SF2-Helicase+REase-p-><-HORMA<- PglZ <- HNH<- OLD-ABC <-ParB+HNH<- N6- METHYLASE <- DUF6079 *<- BrxB-BrxF- iSTAND <- BrxA-DNA-binding <-RHS<-?\|\|<-ZnR<-ARG | 1157 | Bacteria>Pseudomonadota>Gammaproteobacteria>Aeromonadales>Aeromonadaceae>Aeromonas | GCF_027705685.1 | Aeromonas_sp._QDB08 | OKB79_RS12620 | 11744..15217 | 2990480 | BREX-system-P-loop-protein-BrxC-[Aeromonas-sp.-QDB08] |
| WP_270659881.1 | Type1.BREX | SF1-SF2-Helicase+REase->SF1-SF2-Helicase+REase-><-HEPN+Sel1+SEC-C<- PglZ <- HNH<- OLD-ABC <- N6- METHYLASE <- DUF6079 *<- BrxB-BrxF- iSTAND <- BrxA-DNA-binding <-URI-DNase<-?\|\|LexA-protease-> | 1157 | Bacteria>Pseudomonadota>Gammaproteobacteria>Aeromonadales>Aeromonadaceae>Aeromonas | GCF_027705965.1 | Aeromonas_sp._Y301-2 | OKB83_RS15925 | 143279..146752 | 2990505 | BREX-system-P-loop-protein-BrxC-[Aeromonas-sp.-Y301-2] |
| GGW57088.1 | Type1.BREX | <- wHTH+WYL+WCX -p\|\|SIR2->?-><-?\|\|HEPN-p->JAB-> BrxA-DNA-binding -> BrxB-BrxF- iSTAND -> DUF6079 *-> N6- METHYLASE -> OLD-ABC ->HNH -> PglZ ->HEPN+Sel1+SEC-C->Immank+ANK+ANK->REase-><-?<-SF1-SF2-Helicase+REase | 1114 | Bacteria>Pseudomonadota>Gammaproteobacteria>Alteromonadales>Alteromonadaceae>Alishewanella | GCA_014652375.1 | Alishewanella_tabrizica | GCM10008111_11430 | 80462..83806 | 671278 | hypothetical-protein-GCM10008111_11430-[Alishewanella-tabrizica] |
| WP_057786312.1 | Type1.BREX | SF1-SF2-Helicase-> BrxA-DNA-binding -> BrxB-BrxF- iSTAND -> DUF6079 *-> N6- METHYLASE -> OLD-ABC ->HNH ->ParB+HNH-> PglZ ->Immank+ANK+ANK->REase-><-SF1-SF2-Helicase+REase<-SF1-SF2-Helicase+REase | 1157 | Bacteria>Pseudomonadota>Gammaproteobacteria>Alteromonadales>Alteromonadaceae>Alishewanella | GCF_001441695.1 | Alishewanella_sp._WH16-1 | AAY72_RS06860 | 5881..9354 | 1651088 | BREX-system-P-loop-protein-BrxC-[Alishewanella-sp.-WH16-1] |
| WP_229796961.1 | Type1.BREX | <- wHTH+WYL+WCX -p\|\|SIR2->?-><-?\|\|HEPN-p->JAB-> BrxA-DNA-binding -> BrxB-BrxF- iSTAND -> DUF6079 *-> N6- METHYLASE -> OLD-ABC ->HNH -> PglZ ->HEPN+Sel1+SEC-C->Immank+ANK+ANK->REase-><-?<-SF1-SF2-Helicase+REase | 1125 | Bacteria>Pseudomonadota>Gammaproteobacteria>Alteromonadales>Alteromonadaceae>Alishewanella | GCF_014652375.1 | Alishewanella_tabrizica | IE070_RS05825 | 80429..83806 | 671278 | BREX-system-P-loop-protein-BrxC-[Alishewanella-tabrizica] |
| WP_089368479.1 | Type1.BREX | <-Phage_integrase<-DUF2787<-?\|\|XIS-HTH-><-?<-?\|\| wHTH+WYL+WCX -> BrxA-DNA-binding -> BrxB-BrxF- iSTAND -> DUF6079 *-> N6- METHYLASE -> OLD-ABC ->HNH -> PglZ ->NADAR->rve->AAA-ATPase->ART-DarT->TPR+TPR-> | 1169 | Bacteria>Pseudomonadota>Gammaproteobacteria>Alteromonadales>Pseudoalteromonadaceae>Pseudoalteromonas | GCF_007991255.1 | Pseudoalteromonas_nigrifaciens | PNIG_RS10730 | 2243615..2247124 | 28109 | BREX-system-P-loop-protein-BrxC-[Pseudoalteromonas-nigrifaciens] |
| WP_182752860.1 | Type1.BREX | <-Phage_integrase<-DUF2787<-?\|\|XIS-HTH-><-?\|\| wHTH+WYL+WCX -> BrxA-DNA-binding -> BrxB-BrxF- iSTAND -> DUF6079 *-> N6- METHYLASE -> OLD-ABC ->HNH -> PglZ ->NADAR->ART-DarT->TPR+TPR->ZetaToxin-> | 1159 | Bacteria>Pseudomonadota>Gammaproteobacteria>Alteromonadales>Pseudoalteromonadaceae>Pseudoalteromonas | GCF_014163875.1 | Pseudoalteromonas_sp._SR43-3 | H5088_RS10925 | 129286..132765 | 2760943 | BREX-system-P-loop-protein-BrxC-[Pseudoalteromonas-sp.-SR43-3] |
| WP_200158841.1 | Type1.BREX | <- SIGMA+OB-Fold+AAA-ATPase+Lon-Protease-BrxL <- PglZ <- HNH<- OLD-ABC <-SF1-SF2-Helicase<- OLD-ABC + TOPRIM <- N6- METHYLASE <- PIN<- OLD-ABC <- DUF6079 *<- BrxB-BrxF- iSTAND <- BrxA-DNA-binding <-GDResolvase+HIN-HTH<-ZnR | 1154 | Bacteria>Pseudomonadota>Gammaproteobacteria>Chromatiales>Chromatiaceae>Allochromatium | GCF_016584135.1 | Allochromatium_vinosum | CK029_RS16480 | 18892..22356 | 1049 | BREX-system-P-loop-protein-BrxC-[Allochromatium-vinosum] |
| WP_264346732.1 | Type1.BREX | HEPN->?->PVC-AAA->?->?->?->GreA_GreB+SF1-SF2-Helicase-> BrxA-DNA-binding -> BrxB-BrxF- iSTAND -> DUF6079 *-> N6- METHYLASE -> OLD-ABC ->HNH ->ParB+HNH-> PglZ ->HEPN+Sel1+SEC-C-><-NERD+SF1-SF2-Helicase<-?<-SF1-SF2-Helicase+REase | 1157 | Bacteria>Pseudomonadota>Gammaproteobacteria>Chromatiales>Chromatiaceae>Rheinheimera | GCF_947090785.1 | Rheinheimera_sp. | OM978_RS09960 | 2096767..2100240 | 1869214 | BREX-system-P-loop-protein-BrxC-[Rheinheimera-sp.-MM224] |
| WP_200385867.1 | Type1.BREX | DUF6079 *->?->PIN-><-?\|\| N6- METHYLASE -> OLD-ABC ->HNH -> PglZ ->AbrB-HTH->RHH-> | 1179 | Bacteria>Pseudomonadota>Gammaproteobacteria>Chromatiales>Chromatiaceae>Thiocapsa | GCF_016583985.1 | Thiocapsa_imhoffii | CKO25_RS00005 | 26..3565 | 382777 | BREX-system-P-loop-protein-BrxC-[Thiocapsa-imhoffii] |
| MCC7280699.1 | Type6.BREX | <- SIGMA+OB-Fold+AAA-ATPase+Lon-Protease-BrxL <- PglZ <- HNH<- OLD-ABC <-RHH<- N6- METHYLASE <-Schlafen+HATPase+wHTH+HTH<- DUF6079 *<- BrxB-BrxF- iSTAND <- BrxA-DNA-binding <- BrxE | 1181 | Bacteria>Pseudomonadota>Gammaproteobacteria>Chromatiales>Chromatiaceae>unclassified_Chromatiaceae | GCA_020851955.1 | Chromatiaceae_bacterium | IT487_20620 | 13992..17537 | 2049432 | BREX-system-P-loop-protein-BrxC-[Chromatiaceae-bacterium] |
| WP_238620557.1 | Type1.BREX | sGTP-><-35exo+T7DNApol\|\|?->?-> BrxA-DNA-binding -> BrxB-BrxF- iSTAND -> DUF6079 *-> N6- METHYLASE -> OLD-ABC ->HNH -> OLD-ABC ->HNH -> PglZ -> SIGMA+OB-Fold+AAA-ATPase+Lon-Protease-BrxL -><-Schlafen+HATPase+wHTH<-Schlafen+HATPase+wHTH | 1154 | Bacteria>Pseudomonadota>Gammaproteobacteria>Chromatiales>Ectothiorhodospiraceae>Ectothiorhodospira | GCF_022227785.1 | Ectothiorhodospira_sp._B14B | LQ952_00335 | 78448..81912 | 2899127 | BREX-system-P-loop-protein-BrxC-[Ectothiorhodospira-sp.-B14B] |
| QEP43020.1 | Type1.BREX | <-DDE_Tnp_1<-HTH_Tnp_1+HTH_21+rve+rve_3<-HTH_Tnp_1<- TOPRIM <- OLD-ABC <-RNase<- N6- METHYLASE <- HNH<- OLD-ABC <- DUF6079 *<- BrxB-BrxF- iSTAND <- BrxA-DNA-binding <-Phage_holin<-?<-?\|\|GDResolvase+ZnR->?->DDE_Tnp_IS66->HD+SF1-SF2-Helicase-> | 1145 | Bacteria>Pseudomonadota>Gammaproteobacteria>Chromatiales>Ectothiorhodospiraceae>unclassified_Ectothiorhodospiraceae | GCA_008375315.1 | Ectothiorhodospiraceae_bacterium_BW-2 | D5085_07760 | 1613185..1616622 | 947515 | BREX-system-P-loop-protein-BrxC-[Ectothiorhodospiraceae-bacterium-BW-2] |
| WP_132538615.1 | Type6.BREX | <-LexA-protease\|\|?-><-?<- SIGMA+OB-Fold+AAA-ATPase+Lon-Protease-BrxL <- PglZ <- HNH<- OLD-ABC <- N6- METHYLASE <-?<- DUF6079 *<- BrxB-BrxF- iSTAND <- BrxA-DNA-binding <- BrxE <-RHH_5\|\|?->?->XIS-HTH->?->ZnR+ TOPRIM +AAA-helicase+wHTH-> | 1179 | Bacteria>Pseudomonadota>Gammaproteobacteria>Gammaproteobacteria_incertae_sedis>Candidatus_Competibacteraceae>Plasticicumulans | GCF_004341245.1 | Plasticicumulans_lactativorans | EV699_RS02475 | 34554..38093 | 1133106 | BREX-system-P-loop-protein-BrxC-[Plasticicumulans-lactativorans] |
| WP_132542053.1 | Type1.BREX | <- SIGMA+OB-Fold+AAA-ATPase+Lon-Protease-BrxL <- PglZ <- HNH<- OLD-ABC <- N6- METHYLASE <-HEPN<-PolB<- DUF6079 *<- BrxB-BrxF- iSTAND <- BrxA-DNA-binding \|\|ABhydrolase-><-Acyl_transf_3<-?<-?<-?\|\|Peptidase_M3-><-ABhydrolase | 1153 | Bacteria>Pseudomonadota>Gammaproteobacteria>Gammaproteobacteria_incertae_sedis>Candidatus_Competibacteraceae>Plasticicumulans | GCF_004341245.1 | Plasticicumulans_lactativorans | EV699_RS09620 | 184945..188406 | 1133106 | BREX-system-P-loop-protein-BrxC-[Plasticicumulans-lactativorans] |
| WP_071159674.1 | Type1.BREX | <- SIGMA+OB-Fold+AAA-ATPase+Lon-Protease-BrxL <- PglZ <- HNH<- OLD-ABC <- N6- METHYLASE <- DUF6079 *<- BrxB-BrxF- iSTAND <- BrxA-DNA-binding <-XIS-HTH\|\|?-><-?<-?<-?<-?<-JAB+sGTP | 1153 | Bacteria>Pseudomonadota>Gammaproteobacteria>Methylococcales>Methylococcaceae>Methylomonas | GCF_001856095.1 | Methylomonas_sp._LWB | BJL95_RS17210 | 210511..213972 | 1905845 | BREX-system-P-loop-protein-BrxC-[Methylomonas-sp.-LWB] |
| WP_262567770.1 | Type1.BREX | <-zf-ISL3+HTH_Tnp_ISL3+DDE_Tnp_ISL3\|\|?-><-Integrase_1<- SIGMA+OB-Fold+AAA-ATPase+Lon-Protease-BrxL <- PglZ <- HNH<- OLD-ABC <- N6- METHYLASE <- DUF6079 *<- BrxB-BrxF- iSTAND <- BrxA-DNA-binding <-?<-SF1-SF2-Helicase\|\|?-><-?<-LexA-protease | 1221 | Bacteria>Pseudomonadota>Gammaproteobacteria>Oceanospirillales>Endozoicomonadaceae>Endozoicomonas | GCF_025562715.2 | Endozoicomonas_gorgoniicola | NX722_RS09445 | 2085702..2089367 | 1234144 | BREX-system-P-loop-protein-BrxC-[Endozoicomonas-gorgoniicola] |
| WP_031385055.1 | Type1.BREX | <- PglZ <- HNH<- OLD-ABC <- N6- METHYLASE -p<-HerA/FtsK<-SIR2<-REase<- N6- METHYLASE <-Schlafen+HATPase+wHTH+HTH<- DUF6079 *<- BrxB-BrxF- iSTAND <- BrxA-DNA-binding <-XIS-HTH<-MobI\|\|XIS-HTH->AAA+HTH->XIS-HTH-><-Arm-DNA-bind_3+Phage_integrase | 1193 | Bacteria>Pseudomonadota>Gammaproteobacteria>Oceanospirillales>Halomonadaceae>Halomonas | GCF_000731955.1 | Halomonas_zincidurans | HALZIN_RS0115275 | 3254805..3258386 | 1178777 | BREX-system-P-loop-protein-BrxC-[Halomonas-zincidurans] |
| WP_107181354.1 | Type1.BREX | <- SIGMA+OB-Fold+AAA-ATPase+Lon-Protease-BrxL <- PglZ <- HNH<- OLD-ABC <-PHP+SbcC<- N6- METHYLASE <- DUF6079 *<- BrxB-BrxF- iSTAND <- BrxA-DNA-binding <-XIS-HTH | 1147 | Bacteria>Pseudomonadota>Gammaproteobacteria>Oceanospirillales>Halomonadaceae>Halomonas | GCF_003030865.2 | Halomonas_sp._SL1 | C9J49_RS09500 | 199072..202515 | 2137478 | BREX-system-P-loop-protein-BrxC-[Halomonas-sp.-SL1] |
| WP_188638083.1 | Type1.BREX | <-TarH+HAMP+MA\|\|ABC-ATPase+SbcC->?->?->?->?->tRNA-> BrxA-DNA-binding -> BrxB-BrxF- iSTAND -> DUF6079 *-> N6- METHYLASE ->ParB+HNH-> OLD-ABC ->HNH -> PglZ -> SIGMA+OB-Fold+AAA-ATPase+Lon-Protease-BrxL -> | 1154 | Bacteria>Pseudomonadota>Gammaproteobacteria>Oceanospirillales>Halomonadaceae>Halomonas | GCF_014640815.1 | Halomonas_lutescens | IEW65_RS03380 | 721900..725364 | 1602943 | BREX-system-P-loop-protein-BrxC-[Halomonas-lutescens] |
| WP_081193131.1 | Type1.BREX | Peptidase_S49+Peptidase_S49-><-ArsR-HTH\|\|?->ABhydrolase-><-?\|\|?->tRNA-> BrxA-DNA-binding -> BrxB-BrxF- iSTAND -> DUF6079 *-> N6- METHYLASE ->ParB+HNH-> OLD-ABC ->HNH -> PglZ -> SIGMA+OB-Fold+AAA-ATPase+Lon-Protease-BrxL ->\|\|Tox-RelE->HTH_3-><-RelE-ParE | 1154 | Bacteria>Pseudomonadota>Gammaproteobacteria>Oceanospirillales>Halomonadaceae>Halomonas | - | Halomonas_sp._BC1 | B7L72_RS05665 | 10416..13880 | 1670448 | BREX-system-P-loop-protein-BrxC-[Halomonas-sp.-BC1] |
| WP_087719602.1 | Type1.BREX | <-ICLR-HTH<-?\|\|?->Amidohydro_1->GNTR-HTH->XIS-HTH-> BrxA-DNA-binding -> BrxB-BrxF- iSTAND -> DUF6079 *->Schlafen+HATPase+wHTH-> N6- METHYLASE ->PHP+SbcC-> OLD-ABC ->HNH -> PglZ -> SIGMA+OB-Fold+AAA-ATPase+Lon-Protease-BrxL -> | 1145 | Bacteria>Pseudomonadota>Gammaproteobacteria>Oceanospirillales>Halomonadaceae>Salinicola | GCF_002179555.1 | Salinicola_salarius | CCD76_RS11235 | 10059..13496 | 430457 | BREX-system-P-loop-protein-BrxC-[Salinicola-salarius] |
| WP_027853089.1 | Type1.BREX | <- SIGMA+OB-Fold+AAA-ATPase+Lon-Protease-BrxL <- PglZ <- HNH<- OLD-ABC <- TOPRIM <- OLD-ABC <-ParB+HNH<- N6- METHYLASE <- DUF6079 *<- BrxB-BrxF- iSTAND <- BrxA-DNA-binding <- wHTH+WYL+WCX <-TIR<-?<-?<- wHTH+WYL+WCX | 1154 | Bacteria>Pseudomonadota>Gammaproteobacteria>Oceanospirillales>Oceanospirillaceae>Marinobacterium | GCF_000428985.1 | Marinobacterium_litorale | G403_RS0105385 | 191863..195327 | 404770 | BREX-system-P-loop-protein-BrxC-[Marinobacterium-litorale] |
| WP_181736357.1 | Type1.BREX | XIS-HTH->?->?->TIR-> wHTH+WYL+WCX -> BrxA-DNA-binding -> BrxB-BrxF- iSTAND -> DUF6079 *-> N6- METHYLASE ->ParB+HNH-> OLD-ABC ->HNH -> PglZ -> SIGMA+OB-Fold+AAA-ATPase+Lon-Protease-BrxL ->ParB+HNH-> | 1154 | Bacteria>Pseudomonadota>Gammaproteobacteria>Oceanospirillales>Oceanospirillaceae>Marinobacterium | GCF_013868415.1 | Marinobacterium_sp._3-1745 | H1S06_RS00670 | 30655..34119 | 2756129 | BREX-system-P-loop-protein-BrxC-[Marinobacterium-sp.-3-1745] |
| WP_206838922.1 | Type1.BREX | MODE-HTH+PBPII-><-ADH_N+ADH_zinc_N<-?\|\|?->?-> wHTH+WYL+WCX -> BrxA-DNA-binding -> BrxB-BrxF- iSTAND -> DUF6079 *-> N6- METHYLASE ->DUF932-> OLD-ABC -> TOPRIM -> OLD-ABC ->HNH -> PglZ -> SIGMA+OB-Fold+AAA-ATPase+Lon-Protease-BrxL ->Integrase_1-> | 1154 | Bacteria>Pseudomonadota>Gammaproteobacteria>Oceanospirillales>Oceanospirillaceae>Marinobacterium | GCF_017310015.1 | Marinobacterium_georgiense | CFI10_RS02405 | 480710..484174 | 48076 | BREX-system-P-loop-protein-BrxC-[Marinobacterium-iners] |
| WP_217336191.1 | Type1.BREX | DUF772+DDE_Tnp_1+DDE_Tnp_1_6-> wHTH+WYL+WCX -> BrxA-DNA-binding -> BrxB-BrxF- iSTAND -> DUF6079 *-> N6- METHYLASE ->DUF932-> OLD-ABC ->HNH -> OLD-ABC ->HNH -> PglZ -> SIGMA+OB-Fold+AAA-ATPase+Lon-Protease-BrxL -> | 1154 | Bacteria>Pseudomonadota>Gammaproteobacteria>Oceanospirillales>Oceanospirillaceae>Marinobacterium | GCF_019061305.1 | Marinobacterium_sp._A346 | KTN04_RS15750 | 3266..6730 | 2851016 | BREX-system-P-loop-protein-BrxC-[Marinobacterium-sp.-A346] |

| Protein-ID | System | Gene-Neighbourhoods | Length | Taxonomy | Assembly-ID | Strain | Locus-Tag | Genomic-coordinates | TaxID | Fasta-defline |
|---|---|---|---|---|---|---|---|---|---|---|
| WP_119395059.1 | Type1.BREX | SF1-SF2-Helicase->HTH+HEPN->McrB-AAA->wHTH+REase+ZnR->HKD-DNase-> OLD-ABC ->HNH -> BrxA-DNA-binding -> BrxB-BrxF- iSTAND -> DUF6079 *-> N6- METHYLASE ->sGTP->sGTP->?->HerA/FtsK->HNH->?->HNH-> PglZ -> | 1155 | Bacteria>Pseudomonadota>Gammaproteobacteria>Oceanospirillales>Saccharospirillaceae>Salinibius | GCF_003577475.1 | Salinibius_halmophilus | DUW71_RS09715 | 145555..149022 | 1853216 | BREX-system-P-loop-protein-BrxC-[Salinibius-halmophilus] |
| MBC7192650.1 | Type1.BREX | <- SIGMA+OB-Fold+AAA-ATPase+Lon-Protease-BrxL <- PglZ <- HNH<- OLD-ABC <- TOPRIM <- OLD-ABC <-HTH+REase<- N6- METHYLASE <-Schlafen+HATPase+wHTH<- DUF6079 *<- BrxB-BrxF- iSTAND <- BrxA-DNA-binding <-XIS-HTH\|\|HTH_Tnp_1+HTH_21+rve+rve_3->XIS-HTH->DnaB->?-><-Phage_int_SAM_5+Phage_integrase<-sGTP | 1145 | Bacteria>Pseudomonadota>Gammaproteobacteria>Pseudomonadales>Marinobacteraceae>Marinobacter | GCA_014359475.1 | Marinobacter_sp. | H5U31_09890 | 22634..26071 | 50741 | BREX-system-P-loop-protein-BrxC-[Marinobacter-sp.] |
| WP_007349691.1 | Type5.BREX | <-Glycos_transf_2\|\|tRNA->?->?-><- SIGMA+OB-Fold+AAA-ATPase+HKD-DNase-BrxL <- PglZ <- HNH<- OLD-ABC <- N6- METHYLASE <-Schlafen+HATPase+wHTH+HTH<- DUF6079 *<- BrxB-BrxF- iSTAND <- BrxA-DNA-binding <- TUDOR+ SF1-SF2-Helicase+iREase+CTD+REase <-TPR | 1194 | Bacteria>Pseudomonadota>Gammaproteobacteria>Pseudomonadales>Marinobacteraceae>Marinobacter | GCF_000169375.1 | Marinobacter_sp._ELB17 | MELB17_RS08075 | 35418..39002 | 270374 | BREX-system-P-loop-protein-BrxC-[Marinobacter-sp.-ELB17] |
| WP_018404435.1 | Type1.BREX | OrfB_Zn_ribbon-> BrxA-DNA-binding -> BrxB-BrxF- iSTAND -> DUF6079 *->Schlafen+HATPase+wHTH+HTH-> N6- METHYLASE -> OLD-ABC ->HNH -> PglZ -> SIGMA+OB-Fold+AAA-ATPase+HKD-DNase-BrxL -> N6- METHYLASE -> | 1194 | Bacteria>Pseudomonadota>Gammaproteobacteria>Pseudomonadales>Marinobacteraceae>Marinobacter | GCF_000372805.1 | Marinobacter_gelidimuriae | A3OI_RS0111615 | 2203..5787 | 2739064 | BREX-system-P-loop-protein-BrxC-[Marinobacter-gelidimuriae] |
| WP_033898350.1 | Type1.BREX | <- wHTH+WYL+WCX \|\|?->ARG->XIS-HTH-> BrxA-DNA-binding -> BrxB-BrxF- iSTAND -> DUF6079 *-> N6- METHYLASE -p->?-><-?\|\| N6- METHYLASE -p-> OLD-ABC ->HNH -> SIGMA+OB-Fold+AAA-ATPase+HKD-DNase-BrxL -> | 1185 | Bacteria>Pseudomonadota>Gammaproteobacteria>Pseudomonadales>Pseudomonadaceae | GCF_000510895.2 | Pseudomonas | GIW75_RS00865 | 8119..11676 | 286 | MULTISPECIES:-BREX-system-P-loop-protein-BrxC-[Pseudomonas] |
| WP_064314265.1 | Type1.BREX | <-RHH\|\|XIS-HTH->?-> wHTH+WYL+WCX -> BrxA-DNA-binding -> BrxB-BrxF- iSTAND -> DUF6079 *-> N6- METHYLASE ->ParB+HNH-> OLD-ABC ->HNH -> PglZ -> SIGMA+OB-Fold+AAA-ATPase+Lon-Protease-BrxL -><-DUF4969 | 1154 | Bacteria>Pseudomonadota>Gammaproteobacteria>Pseudomonadales>Pseudomonadaceae | GCF_021601465.1 | Pseudomonas | L6D68_RS06035 | 31411..34875 | 286 | MULTISPECIES:-BREX-system-P-loop-protein-BrxC-[Pseudomonas] |
| WP_269686934.1 | Type1.BREX | SF1-SF2-Helicase->SF1-SF2-Helicase->?-><-RADICAL-SAM<-?<-MazE_antitoxin\|\| wHTH+WYL+WCX -> BrxA-DNA-binding -> BrxB-BrxF- iSTAND -> DUF6079 *-> N6- METHYLASE ->?->?-> OLD-ABC ->HNH -> PglZ -> SIGMA+OB-Fold+AAA-ATPase+Lon-Protease-BrxL -><-HTH_3\|\|PadR-HTH-> | 1223 | Bacteria>Pseudomonadota>Gammaproteobacteria>Pseudomonadales>Pseudomonadaceae>Halopseudomonas | GCF_030295685.1 | Halopseudomonas_sp._MFKK-1 | O5O48_RS10320 | 2203460..2207131 | 3015994 | BREX-system-P-loop-protein-BrxC-[Halopseudomonas-sp.-MFKK-1] |
| TXH92079.1 | Type1.BREX | <-RelE-ParE<-RHH_1<- PglZ <- HNH<- OLD-ABC <- N6- METHYLASE -p<-NERD<- OLD-ABC <- N6- METHYLASE <- DUF6079 *<- BrxB-BrxF- iSTAND <- BrxA-DNA-binding <-XIS-HTH<-?\|\| wHTH+WYL+WCX -> | 1186 | Bacteria>Pseudomonadota>Gammaproteobacteria>Pseudomonadales>Pseudomonadaceae>Pseudomonas | GCA_008015405.1 | Pseudomonas_sp. | E6Q71_01680 | 13269..16829 | 306 | BREX-system-P-loop-protein-BrxC-[Pseudomonas-sp.] |
| MBX5763959.1 | Type1.BREX | Phage_integrase-><-?<- SIGMA+OB-Fold+AAA-ATPase+Lon-Protease-BrxL <- PglZ <- HNH<- OLD-ABC <-?<-?<- N6- METHYLASE <- DUF6079 *<- BrxB-BrxF- iSTAND <- BrxA-DNA-binding <- wHTH+WYL+WCX \|\|REase+NACHT->?-><-CHTH<-?\|\|HTH_3-><-SF1-SF2-Helicase | 1223 | Bacteria>Pseudomonadota>Gammaproteobacteria>Pseudomonadales>Pseudomonadaceae>Pseudomonas | GCA_019687095.1 | Pseudomonas_aeruginosa | ISD89_03975 | 862812..866483 | 287 | BREX-system-P-loop-protein-BrxC-[Pseudomonas-aeruginosa] |
| HBN8454461.1 | Type1.BREX | XIS-HTH-><-?<-tRNA\|\|HTH_36->Phage_int_SAM_2+Integrase_1-><- SIGMA+OB-Fold+AAA-ATPase+Lon-Protease-BrxL <- PglZ <- HNH<- OLD-ABC <- N6- METHYLASE <- DUF6079 *<- BrxB-BrxF- iSTAND <- BrxA-DNA-binding <-DOC+HTH<-Arm-DNA-bind_3+Phage_integrase<-sGTP | 1146 | Bacteria>Pseudomonadota>Gammaproteobacteria>Pseudomonadales>Pseudomonadaceae>Pseudomonas | GCA_021587585.1 | Pseudomonas_aeruginosa | L3711_005718 | 39513..42953 | 287 | BREX-system-P-loop-protein-BrxC-[Pseudomonas-aeruginosa] |
| EIU3464059.1 | Type1.BREX | <-DUF3584+DUF5401<-JetB<-MukF<- SIGMA+OB-Fold+AAA-ATPase+Lon-Protease-BrxL <- PglZ <- HNH<- OLD-ABC <-ParB+HNH<- N6- METHYLASE <- DUF6079 *<- BrxB-BrxF- iSTAND <- BrxA-DNA-binding <- wHTH+WYL+WCX <-?<-ARG\|\| wHTH+WYL+WCX ->?->CR-ATPase5+REase->LexA-protease-> | 1154 | Bacteria>Pseudomonadota>Gammaproteobacteria>Pseudomonadales>Pseudomonadaceae>Pseudomonas | GCA_021644165.1 | Pseudomonas_aeruginosa | L4P80_001287 | 38375..41839 | 287 | BREX-system-P-loop-protein-BrxC-[Pseudomonas-aeruginosa] |
| HBO3110869.1 | Type1.BREX | Tox-RelE->HTH_3-><-tRNA\|\|HTH_36->Phage_int_SAM_2+Integrase_1-><- SIGMA+OB-Fold+AAA-ATPase+Lon-Protease-BrxL <- PglZ <- HNH<- OLD-ABC <- N6- METHYLASE <- DUF6079 *<- BrxB-BrxF- iSTAND <- BrxA-DNA-binding <-DOC+HTH<-Arm-DNA-bind_3+Phage_integrase<-sGTP | 1146 | Bacteria>Pseudomonadota>Gammaproteobacteria>Pseudomonadales>Pseudomonadaceae>Pseudomonas | GCA_021651555.1 | Pseudomonas_aeruginosa | L4R01_001445 | 31462..34902 | 287 | BREX-system-P-loop-protein-BrxC-[Pseudomonas-aeruginosa] |
| HBP6330513.1 | Type1.BREX | <-RHH_1\|\|XIS-HTH-><- SIGMA+OB-Fold+AAA-ATPase+Lon-Protease-BrxL <- PglZ <- HNH<- OLD-ABC <-ParB+HNH<-DUF932<- N6- METHYLASE <- DUF6079 *<- BrxB-BrxF- iSTAND <- BrxA-DNA-binding <- wHTH+WYL+WCX \|\|REase-><-Integrase_1 | 1154 | Bacteria>Pseudomonadota>Gammaproteobacteria>Pseudomonadales>Pseudomonadaceae>Pseudomonas | GCA_021710255.1 | Pseudomonas_aeruginosa | L6A37_21805 | 72354..75818 | 287 | BREX-system-P-loop-protein-BrxC-[Pseudomonas-aeruginosa] |
| HBP6726062.1 | Type1.BREX | SF1-SF2-Helicase-><-HTH_3\|\|?->CHTH-><-?\|\| wHTH+WYL+WCX -> BrxA-DNA-binding -> BrxB-BrxF- iSTAND -> DUF6079 *-> N6- METHYLASE ->?->?-> OLD-ABC ->HNH -> PglZ -> SIGMA+OB-Fold+AAA-ATPase+Lon-Protease-BrxL ->?-><-Phage_integrase | 1223 | Bacteria>Pseudomonadota>Gammaproteobacteria>Pseudomonadales>Pseudomonadaceae>Pseudomonas | GCA_021711795.1 | Pseudomonas_aeruginosa | L6B36_03955 | 73272..76943 | 287 | BREX-system-P-loop-protein-BrxC-[Pseudomonas-aeruginosa] |
| MCK9534930.1 | Type1.BREX | HTH_Tnp_1->?->LZ_Tnp_IS66+zf-IS66+DDE_Tnp_IS66+DDE_Tnp_IS66_C-><-?\|\|?-> wHTH+WYL+WCX -> BrxA-DNA-binding -> BrxB-BrxF- iSTAND -> DUF6079 *-> N6- METHYLASE ->ParB+HNH-> OLD-ABC -> TOPRIM -> OLD-ABC ->HNH -> PglZ -> SIGMA+OB-Fold+AAA-ATPase+Lon-Protease-BrxL -> | 1154 | Bacteria>Pseudomonadota>Gammaproteobacteria>Pseudomonadales>Pseudomonadaceae>Pseudomonas | GCA_023228505.1 | Pseudomonas_sp. | MOQ98_09595 | 5981..9445 | 306 | BREX-system-P-loop-protein-BrxC-[Pseudomonas-sp.] |
| CAB5551672.1 | Type1.BREX | DUF4969-><- SIGMA+OB-Fold+AAA-ATPase+Lon-Protease-BrxL <- PglZ <- HNH<- OLD-ABC <-?<-DUF932<- N6- METHYLASE <- DUF6079 *<- BrxB-BrxF- iSTAND <- BrxA-DNA-binding <- wHTH+WYL+WCX <-?<-XIS-HTH\|\|Arm-DNA-bind_3+Phage_integrase-> | 1154 | Bacteria>Pseudomonadota>Gammaproteobacteria>Pseudomonadales>Pseudomonadaceae>Pseudomonas | GCA_903684265.1 | Pseudomonas_putida | GHA_03446 | 127342..130806 | 303 | Uncharacterised-protein-[Pseudomonas-putida] |
| WP_047306816.1 | Type1.BREX | <- SIGMA+OB-Fold+AAA-ATPase+HKD-DNase-BrxL <- PglZ <- HNH<- OLD-ABC <- N6- METHYLASE -p<-RVT<- N6- METHYLASE -p<- DUF6079 *<- BrxB-BrxF- iSTAND <- BrxA-DNA-binding <-XIS-HTH<-?\|\| wHTH+WYL+WCX ->sGTP->sGTP->sGTP-> | 1185 | Bacteria>Pseudomonadota>Gammaproteobacteria>Pseudomonadales>Pseudomonadaceae>Pseudomonas | GCF_001020715.1 | Pseudomonas_fluorescens | AA061_RS32150 | 16461..20018 | 294 | BREX-system-P-loop-protein-BrxC-[Pseudomonas-fluorescens] |
| WP_095626344.1 | Type1.BREX | <- SIGMA+OB-Fold+AAA-ATPase+Lon-Protease-BrxL <- PglZ <-ParB+HNH<- HNH<- OLD-ABC <- N6- METHYLASE <- DUF6079 *<- BrxB-BrxF- iSTAND <- BrxA-DNA-binding <- wHTH+WYL+WCX <-REase\|\|ZnR-><-HIN-HTH<-ZnR | 1154 | Bacteria>Pseudomonadota>Gammaproteobacteria>Pseudomonadales>Pseudomonadaceae>Pseudomonas | GCF_002287035.1 | Pseudomonas_sp._WN033 | CK507_RS15715 | 130328..133792 | 2032629 | BREX-system-P-loop-protein-BrxC-[Pseudomonas-sp.-WN033] |
| WP_170050473.1 | Type1.BREX | <-JetB<-MukF<- SIGMA+OB-Fold+AAA-ATPase+Lon-Protease-BrxL <- PglZ <- HNH<- OLD-ABC <-ParB+HNH<-DUF932<- N6- METHYLASE <- DUF6079 *<- BrxB-BrxF- iSTAND <- BrxA-DNA-binding <- wHTH+WYL+WCX <-?<-ARG\|\| wHTH+WYL+WCX ->RHH_5->?->LexA-protease-> | 1154 | Bacteria>Pseudomonadota>Gammaproteobacteria>Pseudomonadales>Pseudomonadaceae>Pseudomonas | GCF_012985775.1 | Pseudomonas_sp._WS_5011 | HBN63_RS18875 | 35814..39278 | 2717477 | BREX-system-P-loop-protein-BrxC-[Pseudomonas-sp.-WS-5011] |
| WP_169839752.1 | Type1.BREX | DUF4969-><- SIGMA+OB-Fold+AAA-ATPase+Lon-Protease-BrxL <- PglZ <- HNH<- OLD-ABC <- N6- METHYLASE <- DUF6079 *<- BrxB-BrxF- iSTAND <- BrxA-DNA-binding <- wHTH+WYL+WCX <-wHTH+REase+ZnR<-DnaB+McrB-AAA<-Schlafen+HATPase+wHTH<-URI-DNase+RAMA<-?<-XIS-HTH | 1154 | Bacteria>Pseudomonadota>Gammaproteobacteria>Pseudomonadales>Pseudomonadaceae>Pseudomonas | GCF_012986505.1 | Pseudomonas_lundensis | HBN72_RS14625 | 129337..132801 | 86185 | BREX-system-P-loop-protein-BrxC-[Pseudomonas-lundensis] |
| WP_196169752.1 | Type1.BREX | DUF4969-><- SIGMA+OB-Fold+AAA-ATPase+Lon-Protease-BrxL <- PglZ <- HNH<- OLD-ABC <-DUF932<- N6- METHYLASE <- DUF6079 *<- BrxB-BrxF- iSTAND <- BrxA-DNA-binding <- wHTH+WYL+WCX <-?<-XIS-HTH\|\|CopG/RHH-><-MODE-HTH+PBPII | 1154 | Bacteria>Pseudomonadota>Gammaproteobacteria>Pseudomonadales>Pseudomonadaceae>Pseudomonas | GCF_015680295.1 | Pseudomonas_guariconensis | IRZ59_RS14965 | 185250..188714 | 1288410 | BREX-system-P-loop-protein-BrxC-[Pseudomonas-guariconensis] |
| WP_198849248.1 | Type1.BREX | wHTH+WYL+WCX -><- SIGMA+OB-Fold+AAA-ATPase+HKD-DNase-BrxL <- HNH<- OLD-ABC <- N6- METHYLASE -p<-HerA/FtsK<-SIR2<- N6- METHYLASE -p<- DUF6079 *<- BrxB-BrxF- iSTAND <- BrxA-DNA-binding <-XIS-HTH<-?<-tRNA<-MerR-HTH | 1186 | Bacteria>Pseudomonadota>Gammaproteobacteria>Pseudomonadales>Pseudomonadaceae>Pseudomonas | GCF_016405505.1 | Pseudomonas_sp._MF6772 | JFT60_RS22275 | 24346..27906 | 2797533 | BREX-system-P-loop-protein-BrxC-[Pseudomonas-sp.-MF6772] |
| WP_236166438.1 | Type1.BREX | DUF4969-><- SIGMA+OB-Fold+AAA-ATPase+Lon-Protease-BrxL <- PglZ <- HNH<- OLD-ABC <-ParB+HNH<- N6- METHYLASE <- DUF6079 *<- BrxB-BrxF- iSTAND <- BrxA-DNA-binding <- wHTH+WYL+WCX <-?<-XIS-HTH | 1154 | Bacteria>Pseudomonadota>Gammaproteobacteria>Pseudomonadales>Pseudomonadaceae>Pseudomonas | GCF_021601865.1 | Pseudomonas_juntendi | L6D32_RS09730 | 148894..152358 | 2666183 | BREX-system-P-loop-protein-BrxC-[Pseudomonas-juntendi] |
| WP_270945767.1 | Type1.BREX | MerR-HTH->\|\|?->?->DUF772+DDE_Tnp_1-><-?\|\| wHTH+WYL+WCX -> BrxA-DNA-binding -> BrxB-BrxF- iSTAND -> DUF6079 *-> N6- METHYLASE ->?->?-> OLD-ABC ->HNH -> PglZ -> SIGMA+OB-Fold+AAA-ATPase+Lon-Protease-BrxL ->?->HTH_3-> | 1223 | Bacteria>Pseudomonadota>Gammaproteobacteria>Pseudomonadales>Pseudomonadaceae>Pseudomonas | GCF_027856115.1 | Pseudomonas_aeruginosa | KKQ36_RS16450 | 3486481..3490152 | 287 | BREX-system-P-loop-protein-BrxC-[Pseudomonas-aeruginosa] |
| WP_053526945.1 | Type1.BREX | <-HTH_3<-?<- SIGMA+OB-Fold+AAA-ATPase+Lon-Protease-BrxL <- PglZ <- HNH<- OLD-ABC <-?<-?<- N6- METHYLASE <- DUF6079 *<- BrxB-BrxF- iSTAND <- BrxA-DNA-binding <- wHTH+WYL+WCX \|\|?->RVT->?-><-SF1-SF2-Helicase | 1223 | Bacteria>Pseudomonadota>Gammaproteobacteria>Pseudomonadales>Pseudomonadaceae>Stutzerimonas | GCF_001276475.1 | Stutzerimonas_stutzeri | ABW53_RS00980 | 14361..18032 | 316 | BREX-system-P-loop-protein-BrxC-[Stutzerimonas-stutzeri] |
| WP_125856771.1 | Type1.BREX | <-LexA-protease<- wHTH+WYL+WCX \|\|ARG->?-> wHTH+WYL+WCX -> BrxA-DNA-binding -> BrxB-BrxF- iSTAND -> DUF6079 *-> N6- METHYLASE ->ParB+HNH-> OLD-ABC ->HNH -> PglZ -> SIGMA+OB-Fold+AAA-ATPase+Lon-Protease-BrxL ->MukF->JetB->DUF3584+DUF5401-> | 1154 | Bacteria>Pseudomonadota>Gammaproteobacteria>Pseudomonadales>Pseudomonadaceae>Stutzerimonas | GCF_003940515.1 | Stutzerimonas_stutzeri | EGJ36_RS18105 | 6035..9499 | 316 | BREX-system-P-loop-protein-BrxC-[Stutzerimonas-stutzeri] |
| WP_150138110.1 | Type1.BREX | wHTH+WYL+WCX -> BrxA-DNA-binding -> BrxB-BrxF- iSTAND -> DUF6079 *-> N6- METHYLASE -> OLD-ABC ->HNH -> PglZ -> | 1159 | Bacteria>Pseudomonadota>Gammaproteobacteria>Vibrionales>Vibrionaceae>Enterovibrio | GCF_008692425.1 | Candidatus_Enterovibrio_escacola | EDM10_RS06800 | 2982..6461 | 1927127 | BREX-system-P-loop-protein-BrxC-[Candidatus-Enterovibrio-escacola] |
| HAS6390651.1 | Type1.BREX | <-HTH\|\|RNase_H-><-DUF2787<-?<-?\|\|XIS-HTH-> wHTH+WYL+WCX -> BrxA-DNA-binding -> BrxB-BrxF- iSTAND -> DUF6079 *-> N6- METHYLASE -> OLD-ABC ->HNH -> PglZ ->ZetaToxin-><-SF1-SF2-Helicase+REase<-SF1-SF2-Helicase+REase | 1157 | Bacteria>Pseudomonadota>Gammaproteobacteria>Vibrionales>Vibrionaceae>Vibrio | GCA_015777605.1 | Vibrio_vulnificus | I7251_16610 | 90517..93990 | 672 | BREX-system-P-loop-protein-BrxC-[Vibrio-vulnificus] |
| MBH1702634.1 | Type1.BREX | <-RFC-AAA+RuvB-AAA\|\|tRNA->?->AAA+wHTH-><- SIGMA+OB-Fold+AAA-ATPase+Lon-Protease-BrxL <- PglZ <- HNH<- OLD-ABC <-ParB+HNH<- N6- METHYLASE <- DUF6079 *<- BrxB-BrxF- iSTAND <- BrxA-DNA-binding <- wHTH+WYL+WCX <-?<-?<-DUF932\|\|XIS-HTH-> | 1146 | Bacteria>Pseudomonadota>Gammaproteobacteria>Xanthomonadales>Xanthomonadaceae>Stenotrophomonas | GCA_015998095.1 | Stenotrophomonas_maltophilia | I5U93_05090 | 24102..27542 | 40324 | BREX-system-P-loop-protein-BrxC-[Stenotrophomonas-maltophilia] |
| WP_125428654.1 | Type1.BREX | <-RFC-AAA+RuvB-AAA\|\|tRNA->?->AAA+wHTH-><- SIGMA+OB-Fold+AAA-ATPase+Lon-Protease-BrxL <- PglZ <- HNH<- OLD-ABC <-ParB+HNH<- N6- METHYLASE <- DUF6079 *<- BrxB-BrxF- iSTAND <- BrxA-DNA-binding <- wHTH+WYL+WCX <-?<-?<-DUF932\|\|XIS-HTH-> | 1146 | Bacteria>Pseudomonadota>Gammaproteobacteria>Xanthomonadales>Xanthomonadaceae>Stenotrophomonas | GCF_003935615.1 | Stenotrophomonas_maltophilia | EGJ89_RS09210 | 127004..130444 | 40324 | BREX-system-P-loop-protein-BrxC-[Stenotrophomonas-maltophilia] |
| ODU42543.1 | Type1.BREX | <- SIGMA+OB-Fold+AAA-ATPase+Lon-Protease-BrxL <- PglZ <-Polbeta<-MarR-HTH<- HNH<- OLD-ABC <- N6- METHYLASE <- DUF6079 *<- BrxB-BrxF- iSTAND <- BrxA-DNA-binding \|\|?-><-?\|\|HKD-DNase->?->HNH-> N6- METHYLASE -> | 1148 | Bacteria>Pseudomonadota>Gammaproteobacteria>Xanthomonadales>Xanthomonadaceae>unclassified_Xanthomonadaceae | GCA_001725645.1 | Xanthomonadaceae_bacterium_SCN_69-123 | ABS96_27585 | 191713..195159 | 1660170 | hypothetical-protein-ABS96_27585-[Xanthomonadaceae-bacterium-SCN-69-123] |
| PIQ12823.1 | Type1.BREX | <-HARE-HTH+REase<- SIGMA+OB-Fold+AAA-ATPase+Lon-Protease-BrxL <- PglZ <- HNH<- OLD-ABC <- N6- METHYLASE <- DUF6079 *<- BrxB-BrxF- iSTAND <- BrxA-DNA-binding <-SLOG+BirA-HTH<-SF1-SF2-Helicase+PRTase<- wHTH+WYL+WCX \|\|<-ParB<-ParB<-GDResolvase+PPP1-HTH | 1146 | Bacteria>Pseudomonadota>Hydrogenophilia>Hydrogenophilales>unclassified_Hydrogenophilales | GCA_002770925.1 | Hydrogenophilales_bacterium_CG18_big_fil_WC_8_21_14_2_50_58_12 | COW70_07940 | 10882..14322 | 1974036 | hypothetical-protein-COW70_07940-[Hydrogenophilales-bacterium-CG18_big_fil_WC_8_21_14_2_50_58_12] |
| NLF73010.1 | Type1.BREX | XIS-HTH-> BrxA-DNA-binding -> BrxB-BrxF- iSTAND -> DUF6079 *-> N6- METHYLASE -> OLD-ABC ->HNH -> PglZ -> SIGMA+OB-Fold+AAA-ATPase+Lon-Protease-BrxL -> | 1145 | Bacteria>PVC_group>Planctomycetota>Planctomycetia>Pirellulales>Pirellulaceae | GCA_012515235.1 | Candidatus_Anammoximicrobium_sp. | GX575_28570 | 2105..5542 | 2699742 | BREX-system-P-loop-protein-BrxC-[Candidatus-Anammoximicrobium-sp.] |
| MCC5808698.1 | Type1.BREX | Phage_integrase-><- SIGMA+OB-Fold+AAA-ATPase+HKD-DNase-BrxL <- PglZ <- HNH<- OLD-ABC <- N6- METHYLASE <- REase<- OLD-ABC <- DUF6079 *<- BrxB-BrxF- iSTAND <- BrxA-DNA-binding <-XIS-HTH | 1183 | Bacteria>PVC_group>Verrucomicrobiota>Opitutae>Opitutales>unclassified_Opitutales | GCA_020831525.1 | Opitutales_bacterium | JJU00_20390 | 12178..15729 | 2026772 | BREX-system-P-loop-protein-BrxC-[Opitutales-bacterium] |
| OPA28160.1 | Type1.BREX | PIN->REase-> OLD-ABC ->HNH ->?-> BrxA-DNA-binding -> BrxB-BrxF- iSTAND -> DUF6079 *-> N6- METHYLASE -> PglZ -> SIGMA+OB-Fold+AAA-ATPase+Lon-Protease-BrxL ->MarR-HTH->?->?->BZIP+SF1-SF2-Helicase-> | 1190 | Bacteria>Terrabacteria_group>Bacillota>Bacilli>Bacillales>Bacillaceae | GCA_002021465.1 | Bacillus_cereus | BHL47_19745 | 10940..14512 | 1396 | hypothetical-protein-BHL47_19745-[Bacillus-cereus] |
| ATI52289.1 | Type1.BREX | GDResolvase+HIN-HTH-> TOPRIM +Topoisom_bac-> OLD-ABC ->HNH ->?-> N6- METHYLASE -p->?-> BrxA-DNA-binding -> BrxB-BrxF- iSTAND -> DUF6079 *-> N6- METHYLASE -> PglZ -> SIGMA+OB-Fold+AAA-ATPase+Lon-Protease-BrxL ->HTH_Tnp_1-> | 1190 | Bacteria>Terrabacteria_group>Bacillota>Bacilli>Bacillales>Bacillaceae | GCA_002504225.1 | Bacillus_cereus | CPZ32_18870 | 3450635..3454207 | 1396 | hypothetical-protein-CPZ32_18870-[Bacillus-cereus] |
| WP_078386197.1 | Type1.BREX | PIN->REase-> OLD-ABC ->HNH ->?-> BrxA-DNA-binding -> BrxB-BrxF- iSTAND -> DUF6079 *-> N6- METHYLASE -> PglZ -> SIGMA+OB-Fold+AAA-ATPase+Lon-Protease-BrxL ->MarR-HTH->?->?->BZIP+SF1-SF2-Helicase-> | 1190 | Bacteria>Terrabacteria_group>Bacillota>Bacilli>Bacillales>Bacillaceae | GCF_002021465.1 | Bacillus_cereus | BHL47_19745 | 10940..14512 | 1396 | BREX-system-P-loop-protein-BrxC-[Bacillus-cereus] |
| NFG40378.1 | Type1.BREX | Calcineurin->?-> BrxA-DNA-binding -> BrxB-BrxF- iSTAND -> DUF6079 *-> N6- METHYLASE ->ParB+HNH-> OLD-ABC ->HNH -> PglZ -><-REase\|\| SIGMA+OB-Fold+AAA-ATPase+Lon-Protease-BrxL ->Peptidase_M42->TIR-> | 1215 | Bacteria>Terrabacteria_group>Bacillota>Clostridia>Eubacteriales>Clostridiaceae | GCA_011013445.1 | Clostridium_botulinum | FC789_04035 | 879792..883439 | 1491 | BREX-system-P-loop-protein-BrxC-[Clostridium-botulinum] |
| NFI95403.1 | Type1.BREX | <-Peptidase_M42<- SIGMA+OB-Fold+AAA-ATPase+Lon-Protease-BrxL \|\|REase-><- PglZ <- HNH<- OLD-ABC <-ParB+HNH<- N6- METHYLASE <- DUF6079 *<- BrxB-BrxF- iSTAND <- BrxA-DNA-binding | 1215 | Bacteria>Terrabacteria_group>Bacillota>Clostridia>Eubacteriales>Clostridiaceae | GCA_011014625.1 | Clostridium_botulinum | FC961_13560 | 15228..18875 | 1491 | BREX-system-P-loop-protein-BrxC-[Clostridium-botulinum] |
| MBN1067552.1 | Type1.BREX | Calcineurin->?-> BrxA-DNA-binding -> BrxB-BrxF- iSTAND -> DUF6079 *->CR-ATPase9+REase-> N6- METHYLASE -> OLD-ABC ->HNH -> PglZ -><-REase\|\| SIGMA+OB-Fold+AAA-ATPase+Lon-Protease-BrxL ->Peptidase_M42->TIR-> | 1215 | Bacteria>Terrabacteria_group>Bacillota>Clostridia>Eubacteriales>Clostridiaceae | GCA_016925955.1 | Clostridium_botulinum | DVW02_06040 | 97050..100697 | 1491 | BREX-system-P-loop-protein-BrxC-[Clostridium-botulinum] |
| WP_043662653.1 | Type1.BREX | <-wHTH+REase<-wHTH+McrB-AAA<- SIGMA+OB-Fold+AAA-ATPase+Lon-Protease-BrxL <-?<- PglZ <- HNH<- OLD-ABC <- N6- METHYLASE <- DUF6079 *<- BrxB-BrxF- iSTAND <- BrxA-DNA-binding <- HKD-DNase+SF1-SF2-Helicase +HTH+HTH+SRA<-SAM- METHYLASE <-RNase<-HNH+NERD+ZnR<-?<-?<-Amidohydro_3 | 1216 | Bacteria>Terrabacteria_group>Bacillota>Clostridia>Eubacteriales>Clostridiaceae | GCF_000785185.1 | Clostridium_butyricum | AWN73_RS08010 | 18592..22242 | 1492 | BREX-system-P-loop-protein-BrxC-[Clostridium-butyricum] |
| WP_077851369.1 | Type1.BREX | <- SIGMA+OB-Fold+AAA-ATPase+Lon-Protease-BrxL <-?<- PglZ <- HNH<- OLD-ABC <- N6- METHYLASE <- DUF6079 *<- BrxB-BrxF- iSTAND <- BrxA-DNA-binding <-?<-?<-?<-?<-?<-?<-Aminotran_1_2 | 1216 | Bacteria>Terrabacteria_group>Bacillota>Clostridia>Eubacteriales>Clostridiaceae | GCF_002006235.2 | Clostridium_felsineum | CLAUR_RS09645 | 2191072..2194722 | 36839 | BREX-system-P-loop-protein-BrxC-[Clostridium-felsineum] |
| WP_118513467.1 | Type1.BREX | <-Tox-ART-HYD2<-TIR<- PglZ <-Schlafen+HATPase+wHTH<-AbiGi+seqA+RAMA<- N6- METHYLASE <- HNH<- OLD-ABC <- DUF6079 *<- BrxB-BrxF- iSTAND <- BrxA-DNA-binding <-HTH_26<-SIGMA-HTH+SIGMA-HTH<-HISKIN<-?<-TPR+TPR+TPR | 1183 | Bacteria>Terrabacteria_group>Bacillota>Clostridia>Eubacteriales>Clostridiaceae | GCF_003477775.1 | Clostridium_sp._AF37-5AT | DW020_RS00895 | 209396..212947 | 2292007 | BREX-system-P-loop-protein-BrxC-[Clostridium-sp.-AF37-5AT] |
| WP_164949146.1 | Type1.BREX | <-HEPN<- SIGMA+OB-Fold+AAA-ATPase+Lon-Protease-BrxL <- PglZ <- HNH<- OLD-ABC <- N6- METHYLASE <- DUF6079 *<- BrxB-BrxF- iSTAND <- BrxA-DNA-binding <-?<-CR-ATPase8+REase\|\|?->?-><-?<-?\|\| METHYLASE -> | 1215 | Bacteria>Terrabacteria_group>Bacillota>Clostridia>Eubacteriales>Clostridiaceae | GCF_011065955.2 | Clostridium_sp._FP2 | G9F71_RS17965 | 3736090..3739737 | 2724481 | BREX-system-P-loop-protein-BrxC-[Clostridium-sp.-FP2] |
| WP_171786495.1 | Type1.BREX | <-D5N+D5ATPase<- SIGMA+OB-Fold+AAA-ATPase+Lon-Protease-BrxL <- PglZ <- HNH<- OLD-ABC <-PIWI<- N6- METHYLASE <- DUF6079 *<- BrxB-BrxF- iSTAND <- BrxA-DNA-binding <-?<-HTH_3<-?<-?\|\|tRNA-><-RVT | 1216 | Bacteria>Terrabacteria_group>Bacillota>Clostridia>Eubacteriales>Clostridiaceae | GCF_013149405.1 | Clostridium_beijerinckii | B0H39_RS13645 | 3014066..3017716 | 1520 | BREX-system-P-loop-protein-BrxC-[Clostridium-beijerinckii] |
| WP_171780276.1 | Type1.BREX | rve+rve_3->AAA-ATPase-><-D5N+D5ATPase<- SIGMA+OB-Fold+AAA-ATPase+Lon-Protease-BrxL <- PglZ <- HNH<- OLD-ABC <-PIWI<- N6- METHYLASE <- DUF6079 *<- BrxB-BrxF- iSTAND <- BrxA-DNA-binding <-?<-?<-HTH_3<-?<-?\|\|tRNA-><-RVT | 1216 | Bacteria>Terrabacteria_group>Bacillota>Clostridia>Eubacteriales>Clostridiaceae | GCF_014230655.1 | Clostridium_beijerinckii | HGI39_RS06505 | 40889..44539 | 1520 | BREX-system-P-loop-protein-BrxC-[Clostridium-beijerinckii] |
| WP_216171751.1 | Type1.BREX | <- SIGMA+OB-Fold+AAA-ATPase+Lon-Protease-BrxL <- PglZ <- HNH<- OLD-ABC <- N6- METHYLASE <- DUF6079 *<- BrxB-BrxF- iSTAND <- BrxA-DNA-binding \|\|Phage_int_SAM_3-><-Phage_int_SAM_1+Phage_integrase<-Phage_integrase<-Phage_integrase\|\|Phage_integrase-> | 1215 | Bacteria>Terrabacteria_group>Bacillota>Clostridia>Eubacteriales>Clostridiaceae | GCF_026650905.1 | Clostridium_estertheticum | LL037_RS21145 | 4418214..4421861 | 238834 | BREX-system-P-loop-protein-BrxC-[Clostridium-estertheticum] |
| WP_270487117.1 | Type1.BREX | <- SIGMA+OB-Fold+AAA-ATPase+Lon-Protease-BrxL <-?<-PIN<-SbcC<- PglZ <- HNH<- OLD-ABC <-TIR<- N6- METHYLASE <- DUF6079 *<- BrxB-BrxF- iSTAND <- BrxA-DNA-binding <-HTH_26<-PIN<-?<-MACRODOMAIN | 1201 | Bacteria>Terrabacteria_group>Bacillota>Clostridia>Eubacteriales>Clostridiaceae | GCF_027681635.1 | Clostridium_fessum | PGX92_RS02760 | 620693..624298 | 2126740 | BREX-system-P-loop-protein-BrxC-[Clostridium-fessum] |
| WP_115641994.1 | Type1.BREX | NTP_transf->HEPN-><- PglZ <- HNH<- OLD-ABC <- N6- METHYLASE <-?<- DUF6079 *<- BrxB-BrxF- iSTAND <- BrxA-DNA-binding \|\|HEPN-><-?<-ABC-ATPase+SbcC<-RADICAL-SAM<-?\|\|DDE_Tnp_IS240+rve_3-> | 1215 | Bacteria>Terrabacteria_group>Bacillota>Clostridia>Eubacteriales>Clostridiaceae | GCF_900461105.1 | Clostridium_putrefaciens | DY168_RS12220 | 2676627..2680274 | 99675 | BREX-system-P-loop-protein-BrxC-[Clostridium-putrefaciens] |
| TGX97927.1 | Type1.BREX | <- OLD-ABC + TOPRIM -p<- OLD-ABC + TOPRIM -p<- PglZ <- HNH<- OLD-ABC <- N6- METHYLASE <-HEPN<-ParB+HNH<- N6- METHYLASE <- DUF6079 *<- BrxB-BrxF- iSTAND <- BrxA-DNA-binding <-HTH_26<-ZnR+ZnR+ZnR<-35exo+ADA-DBD<-?<-MACRODOMAIN | 1201 | Bacteria>Terrabacteria_group>Bacillota>Clostridia>Eubacteriales>Lachnospiraceae | GCA_004793585.1 | Lachnospiraceae_bacterium | E5357_10825 | 35909..39514 | 1898203 | BREX-system-P-loop-protein-BrxC-[Lachnospiraceae-bacterium] |
| WP_059085682.1 | Type1.BREX | MACRODOMAIN->?->ZnR+ZnR+ZnR->HTH_26-> BrxA-DNA-binding -> BrxB-BrxF- iSTAND -> DUF6079 *-> N6- METHYLASE ->ParB+HNH-> OLD-ABC ->?-> OLD-ABC ->HNH -> PglZ -> | 1196 | Bacteria>Terrabacteria_group>Bacillota>Clostridia>Eubacteriales>Lachnospiraceae | GCF_001487165.1 | Blautia_massiliensis | BN3261_RS05710 | 1028364..1031954 | 1737424 | BREX-system-P-loop-protein-BrxC-[Blautia-massiliensis] |
| WP_114002767.1 | Type1.BREX | <- PglZ <- HNH<- OLD-ABC <-?<- OLD-ABC <-ParB+HNH<- N6- METHYLASE <- DUF6079 *<- BrxB-BrxF- iSTAND <- BrxA-DNA-binding <-HTH_26<-ZnR+ZnR+ZnR<-?<-MACRODOMAIN | 1196 | Bacteria>Terrabacteria_group>Bacillota>Clostridia>Eubacteriales>Lachnospiraceae | GCF_003324155.1 | Blautia_obeum | C4886_RS16030 | 22477..26067 | 40520 | BREX-system-P-loop-protein-BrxC-[Blautia-obeum] |
| WP_154780312.1 | Type1.BREX | Peptidase->?->?->MACRODOMAIN->?->ZnR+ZnR+ZnR->HTH_26-> BrxA-DNA-binding -> BrxB-BrxF- iSTAND -> DUF6079 *-> N6- METHYLASE ->RVT->?-> OLD-ABC ->HNH ->?-> PglZ ->HEPN-> SIGMA+OB-Fold+AAA-ATPase+Lon-Protease-BrxL -> | 1201 | Bacteria>Terrabacteria_group>Bacillota>Clostridia>Eubacteriales>Lachnospiraceae | GCF_009707925.1 | Blautia_luti | GKZ57_RS08760 | 74022..77627 | 89014 | BREX-system-P-loop-protein-BrxC-[Blautia-luti] |
| WP_207739085.1 | Type1.BREX | METHYLASE ->?->?->?->HTH_26-> BrxA-DNA-binding -> BrxB-BrxF- iSTAND -> DUF6079 *-> N6- METHYLASE -> OLD-ABC ->HNH -> PglZ -> SIGMA+OB-Fold+AAA-ATPase+Lon-Protease-BrxL ->?->Arm-DNA-bind_4+Phage_int_SAM_3+Phage_integrase->?->XIS-HTH+XIS-HTH-> | 1173 | Bacteria>Terrabacteria_group>Bacillota>Clostridia>Eubacteriales>Oscillospiraceae | GCF_015558275.1 | Massiliimalia_timonensis | I2G46_RS07880 | 51772..55293 | 1987501 | BREX-system-P-loop-protein-BrxC-[Massiliimalia-timonensis] |
| WP_215659170.1 | Type1.BREX | <-TIR<- SIGMA+OB-Fold+AAA-ATPase+Lon-Protease-BrxL <- PglZ <-DUF4404<-ZnR+ZnR<- N6- METHYLASE <-?<- HNH<- OLD-ABC <- DUF6079 *<- BrxB-BrxF- iSTAND <- BrxA-DNA-binding <-HTH_26<-MazE_antitoxin+SpoVT_C<-?\|\|?-><-?<-RADICAL-SAM | 1184 | Bacteria>Terrabacteria_group>Bacillota>Clostridia>Eubacteriales>Oscillospiraceae | GCF_018783905.1 | Ruminococcus_sp._MCC718 | GPK60_RS07265 | 26956..30510 | 2592649 | BREX-system-P-loop-protein-BrxC-[Ruminococcus-sp.-MCC718] |
| CCX63523.1 | Type1.BREX | Phage_integrase-><-RelE-ParE<-MazE_antitoxin<- SIGMA+OB-Fold+AAA-ATPase+Lon-Protease-BrxL <- PglZ <- HNH<- OLD-ABC <- N6- METHYLASE <- DUF6079 *<- BrxB-BrxF- iSTAND <- BrxA-DNA-binding <-XIS-HTH\|\|VirD4-TraG+VirD4-FtsK->?->?->ArdCN+MPTase-inact+LPD19->TraJ-RHH-> | 1217 | Bacteria>Terrabacteria_group>Bacillota>environmental_samples | GCA_000431495.1 | Firmicutes_bacterium_CAG:791 | BN785_00943 | 24533..28186 | 1262993 | uncharacterized-protein-BN785_00943-[Firmicutes-bacterium-CAG:791] |
| MBR3457117.1 | Type1.BREX | <- PglZ -p<- HNH<- OLD-ABC <- N6- METHYLASE -p<-?<-PIN<- N6- METHYLASE <- DUF6079 *<- BrxB-BrxF- iSTAND -p | 1215 | Bacteria>Terrabacteria_group>Bacillota>Negativicutes>Selenomonadales>Selenomonadaceae | GCA_017509225.1 | Selenomonadaceae_bacterium | IKH16_03040 | 6412..10059 | 2652289 | BREX-system-P-loop-protein-BrxC-[Selenomonadaceae-bacterium] |
| WP_037354735.1 | Type1.BREX | DDE_Tnp_1-><-?<-?<-?\|\|?-> BrxA-DNA-binding -> BrxB-BrxF- iSTAND -> DUF6079 *-> N6- METHYLASE -> OLD-ABC ->HNH ->?-> PglZ -> SIGMA+OB-Fold+AAA-ATPase+Lon-Protease-BrxL ->HTH_3-> | 1182 | Bacteria>Terrabacteria_group>Bacillota>Negativicutes>Selenomonadales>Selenomonadaceae | GCF_000701565.1 | Selenomonas_sp._FC4001 | T320_RS08130 | 15634..19182 | 1408313 | BREX-system-P-loop-protein-BrxC-[Selenomonas-sp.-FC4001] |
| RGH23082.1 | Type1.BREX | PemK_toxin->?->?-> BrxA-DNA-binding -> BrxB-BrxF- iSTAND -> DUF6079 *-> N6- METHYLASE -> OLD-ABC ->HNH -> PglZ -><-?\|\|?->?-><-ZnR | 1180 | Bacteria>Terrabacteria_group>Bacillota>unclassified_Bacillota | GCA_003461705.1 | Firmicutes_bacterium_AF12-30 | DWV72_07055 | 48691..52233 | 2292884 | BREX-system-P-loop-protein-BrxC-[Firmicutes-bacterium-AF12-30] |
| MBE9475406.1 | Type1.BREX | BrxB-BrxF- iSTAND -p-> BrxA-DNA-binding -> BrxB-BrxF- iSTAND -> DUF6079 *-> N6- METHYLASE ->REase+AIPR-> OLD-ABC ->HNH -> PglZ -> SIGMA+OB-Fold+AAA-ATPase+Lon-Protease-BrxL -> | 1145 | Bacteria>Terrabacteria_group>Chloroflexota>unclassified_Chloroflexota | GCA_015222395.1 | Chloroflexota_bacterium | IMY85_10995 | 2370..5807 | 2026724 | BREX-system-P-loop-protein-BrxC-[Chloroflexota-bacterium] |
| MBI4317923.1 | Type1.BREX | Radical_SAM->?->?->?-> BrxA-DNA-binding -> BrxB-BrxF- iSTAND -> DUF6079 *-> N6- METHYLASE -> OLD-ABC ->HNH -> PglZ -> SIGMA+OB-Fold+AAA-ATPase+Lon-Protease-BrxL -> | 1145 | Bacteria>Terrabacteria_group>Chloroflexota>unclassified_Chloroflexota | GCA_016210495.1 | Chloroflexota_bacterium | HY675_05490 | 15525..18962 | 2026724 | BREX-system-P-loop-protein-BrxC-[Chloroflexota-bacterium] |
| MBT9546703.1 | Type1.BREX | Peptidase_M2-> BrxA-DNA-binding -> BrxB-BrxF- iSTAND -> DUF6079 *-> N6- METHYLASE ->RVT-> N6- METHYLASE -p-> OLD-ABC ->HNH -> PglZ -> SIGMA+OB-Fold+AAA-ATPase+HKD-DNase-BrxL -> | 1179 | Bacteria>Terrabacteria_group>Cyanobacteriota/Melainabacteria_group>Candidatus_Sericytochromatia>unclassified_Candidatus_Sericytochromatia | GCA_018780035.1 | Candidatus_Sericytochromatia_bacterium | IV090_15030 | 94995..98534 | 2762020 | BREX-system-P-loop-protein-BrxC-[Candidatus-Sericytochromatia-bacterium] |
| WP_069067979.1 | Type1.BREX | CHTH->?->?-> BrxA-DNA-binding -> BrxB-BrxF- iSTAND -> DUF6079 *->?-> N6- METHYLASE -> OLD-ABC ->HNH -> PglZ -> SIGMA+OB-Fold+AAA-ATPase+Lon-Protease-BrxL -><-?<-HTH_23 | 1220 | Bacteria>Terrabacteria_group>Cyanobacteriota/Melainabacteria_group>Cyanobacteriota>Cyanophyceae>Nostocales | GCF_001712795.1 | Nostoc_sp._KVJ20 | A4S05_RS00980 | 24778..28440 | 457944 | BREX-system-P-loop-protein-BrxC-[Nostoc-sp.-KVJ20] |
| WP_190429845.1 | Type1.BREX | Pkinase->HTH+HKD-DNase-><- SIGMA+OB-Fold+AAA-ATPase+Lon-Protease-BrxL <- PglZ <- HNH<- OLD-ABC <- N6- METHYLASE <-?<- DUF6079 *<- BrxB-BrxF- iSTAND <- BrxA-DNA-binding \|\|ParA-Soj-PloopNTPase->CopG/RHH->?->?->Phage_integrase+Phage_integrase-><-TRPR-HTH | 1222 | Bacteria>Terrabacteria_group>Cyanobacteriota/Melainabacteria_group>Cyanobacteriota>Cyanophyceae>Oscillatoriophycideae | GCF_014695635.1 | Coleofasciculus_sp._FACHB-712 | H6F50_RS18615 | 18748..22416 | 2692789 | BREX-system-P-loop-protein-BrxC-[Coleofasciculus-sp.-FACHB-712] |
| KPQ32454.1 | Type1.BREX | BrxA-DNA-binding -> BrxB-BrxF- iSTAND -> DUF6079 *-> OLD-ABC ->NERD-> N6- METHYLASE -> OLD-ABC ->HNH -> PglZ -> SIGMA+OB-Fold+AAA-ATPase+HKD-DNase-BrxL -><-?\|\|DDE_Tnp-> | 1200 | Bacteria>Terrabacteria_group>Cyanobacteriota/Melainabacteria_group>Cyanobacteriota>Cyanophyceae>Pseudanabaenales | GCA_001314865.1 | Phormidesmis_priestleyi_Ana | HLUCCA11_21420 | 1901..5503 | 1666911 | Brex-phage-defense-system-ATPase-BrxC-[Phormidesmis-priestleyi-Ana] |
| HBH74800.1 | Type1.BREX | PIN->?-> BrxA-DNA-binding -> BrxB-BrxF- iSTAND -> DUF6079 *-> N6- METHYLASE ->SAM- METHYLASE ->ParA-Soj-PloopNTPase->?->?-> PglZ -> OLD-ABC ->HNH -> SIGMA+OB-Fold+AAA-ATPase+Lon-Protease-BrxL -> | 1163 | Bacteria>Terrabacteria_group>Cyanobacteriota/Melainabacteria_group>Cyanobacteriota>Cyanophyceae>Synechococcales | GCA_003508015.1 | Synechococcales_bacterium_UBA10510 | DDY43_15550 | 2736..6227 | 2055795 | BREX-system-P-loop-protein-BrxC-[Synechococcales-bacterium-UBA10510] |
| WP_178365089.1 | Type1.BREX | <-DDE_Tnp\|\|?->?->PRTase->RADICAL-SAM-> BrxA-DNA-binding -> BrxB-BrxF- iSTAND -> DUF6079 *-> N6- METHYLASE -> OLD-ABC ->HNH -> PglZ -> SIGMA+OB-Fold+AAA-ATPase+HKD-DNase-BrxL ->tRNA-><-?<-PAS+NtrC-AAA+FIS-HTH\|\|?->REase-> | 1189 | Bacteria>Thermodesulfobacteriota>Desulfobacteria>Desulfobacterales>Desulfobacteraceae>Desulfobacter | GCF_013389365.1 | Desulfobacter_latus | HXW94_RS01255 | 9015..12584 | 2292 | BREX-system-P-loop-protein-BrxC-[Desulfobacter-latus] |
| WP_139446530.1 | Type1.BREX | <- SIGMA+OB-Fold+AAA-ATPase+Lon-Protease-BrxL \|\|DiS_P_DiS+Peptidase_A24-><-?\|\|?-><- SIGMA+OB-Fold+AAA-ATPase+HKD-DNase-BrxL <- PglZ <- HNH<- OLD-ABC <- N6- METHYLASE <- DUF6079 *<- BrxB-BrxF- iSTAND <- BrxA-DNA-binding <-tRNA\|\|?->?-><-?<-?<-HDRnase | 1199 | Bacteria>Thermodesulfobacteriota>Desulfobacteria>Desulfobacterales>Desulfobacteraceae>Desulfobotulus | GCF_006175995.1 | Desulfobotulus_mexicanus | FIM25_RS03945 | 130812..134411 | 2586642 | BREX-system-P-loop-protein-BrxC-[Desulfobotulus-mexicanus] |
| WP_024334643.1 | Type1.BREX | <-DDE_Tnp_ISAZ013<-DDE_Tnp_ISAZ013\|\|AIPR-> BrxA-DNA-binding -> BrxB-BrxF- iSTAND -> DUF6079 *->Schlafen+HATPase+wHTH-> N6- METHYLASE ->PHP+SbcC->ParB+HNH->DUF932-> OLD-ABC ->HNH -> PglZ -> SIGMA+OB-Fold+AAA-ATPase+Lon-Protease-BrxL -> | 1144 | Bacteria>Thermodesulfobacteriota>Desulfobacteria>Desulfobacterales>Desulfobacteraceae>Desulfotignum | GCF_000421285.1 | Desulfotignum_balticum | K365_RS0111330 | 2237205..2240639 | 115781 | BREX-system-P-loop-protein-BrxC-[Desulfotignum-balticum] |
| RZB38310.1 | Type1.BREX | <-TPR+TPR<-?\|\|?->?->?-><-?\|\|?-> BrxA-DNA-binding -> BrxB-BrxF- iSTAND -> DUF6079 *-> N6- METHYLASE -p-> N6- METHYLASE -> OLD-ABC ->HNH -> PglZ -> SIGMA+OB-Fold+AAA-ATPase+HKD-DNase-BrxL -><-?<-?<-ABC-ATPase+SbcC | 1173 | Bacteria>Thermodesulfobacteriota>Desulfobacteria>Desulfobacterales>Desulfobacteraceae>unclassified_Desulfobacteraceae | GCA_004193595.1 | Desulfobacteraceae_bacterium_Eth-SRB2 | SRB2_00058 | 60586..64107 | 2528438 | hypothetical-protein-SRB2_00058-[Desulfobacteraceae-bacterium-Eth-SRB2] |
| WP_155308948.1 | Type1.BREX | VirB10->?->DnaB+HerA/FtsK->TraH->TraG_N-><-?\|\|XIS-HTH-> BrxA-DNA-binding -> BrxB-BrxF- iSTAND -> DUF6079 *-> OLD-ABC ->HNH -> N6- METHYLASE -> N6- METHYLASE ->REase+AIPR-> PglZ -> SIGMA+OB-Fold+AAA-ATPase+Lon-Protease-BrxL -><-HEPN | 1145 | Bacteria>Thermodesulfobacteriota>Desulfobacteria>Desulfobacterales>Desulfosarcinaceae>Desulfosarcina | GCF_009689005.1 | Desulfosarcina_ovata | GN112_RS03415 | 763655..767092 | 83564 | BREX-system-P-loop-protein-BrxC-[Desulfosarcina-ovata] |
| KGO34873.1 | Type1.BREX | MlaE->ABC-ATPase+SbcC->?-><-?\|\|XIS-HTH-> BrxA-DNA-binding -> BrxB-BrxF- iSTAND -> DUF6079 *-> OLD-ABC ->REase -> N6- METHYLASE ->?-> OLD-ABC ->HNH -> PglZ -> SIGMA+OB-Fold+AAA-ATPase+Lon-Protease-BrxL -> | 1146 | Bacteria>Thermodesulfobacteriota>Desulfobulbia>Desulfobulbales>Desulfobulbaceae>Desulfobulbus | GCA_000769715.1 | Desulfobulbus_sp._Tol-SR | JT06_05895 | 26559..29999 | 1536652 | hypothetical-protein-JT06_05895-[Desulfobulbus-sp.-Tol-SR] |
| WP_066605667.1 | Type1.BREX | HDRnase->?-><-?<-tRNA<-?\|\|?->?->?-> BrxA-DNA-binding -> BrxB-BrxF- iSTAND -> DUF6079 *-> N6- METHYLASE -> OLD-ABC ->HNH -> PglZ -> SIGMA+OB-Fold+AAA-ATPase+Lon-Protease-BrxL -><-RNA_ METHYLASE | 1145 | Bacteria>Thermodesulfobacteriota>Desulfovibrionia>Desulfovibrionales>Desulfomicrobiaceae>Desulfomicrobium | GCF_001553625.1 | Desulfomicrobium_orale | AXF15_RS07915 | 1725501..1728938 | 132132 | BREX-system-P-loop-protein-BrxC-[Desulfomicrobium-orale] |
| MCB6991564.1 | Type1.BREX | <-ACET<-?<-ABC-ATPase+SbcC<-ABC-ATPase+SbcC<-MarR-HTH\|\|?->HTH_26-> BrxA-DNA-binding -> BrxB-BrxF- iSTAND -> DUF6079 *-> OLD-ABC ->HNH -> N6- METHYLASE ->KAP-NTPase-> | 1183 | Bacteria>unclassified_Bacteria | GCA_020558425.1 | bacterium_210820-DFI.6.38 | LI187_13870 | 12055..15606 | 2883293 | BREX-system-P-loop-protein-BrxC-[bacterium-210820-DFI.6.38] |
| -- | | | | | | | | | | |
| -- OLD-ABC ->HNH | | | | | | | | | | |
| --Type2.BREX | | | | | | | | | | |
| -- | | | | | | | | | | |
| WP_012240290.1 | Type2.Pgl.like.BREX | PglW -> N6- METHYLASE -><-?<-?\|\|?-><-MerR-HTH<-?\|\| DUF6079 *-> OLD-ABC ->HNH -> PglZ -> BrxD-AAA -> SF1-SF2-Helicase+wHTH+Small-B-barrel+DUF1998 -><-?<-?<-?<-ZnR | 1215 | Bacteria>Myxococcota>Polyangia>Polyangiales>Polyangiaceae>Sorangium | GCF_000067165.1 | Sorangium_cellulosum | SCE_RS49865 | 10694583..10698230 | 56 | hypothetical-protein-[Sorangium-cellulosum] |
| MBN8498947.1 | Type2.Pgl.like.BREX | PglW -> N6- METHYLASE -> OLD-ABC ->HNH ->?->AbrB-HTH-> DUF6079 *-> PglZ -> BrxD-AAA -> SF1-SF2-Helicase+wHTH+Small-B-barrel+DUF1998 -> | 1215 | Bacteria>Pseudomonadota>Betaproteobacteria>Betaproteobacteria_incertae_sedis>Candidatus_Accumulibacter>unclassified_Candidatus_Accumulibacter | GCA_017302455.1 | Candidatus_Accumulibacter_sp. | J0M32_19155 | 11909..15556 | 2053492 | phage-resistance-protein-[Candidatus-Accumulibacter-sp.] |
| MBN8437012.1 | Type2.Pgl.like.BREX | HTH_26-> PglW -> N6- METHYLASE -> OLD-ABC ->HNH -> N6- METHYLASE -p-> OLD-ABC ->NERD->AbrB-HTH-> DUF6079 *-> PglZ ->PrlF_antitoxin->PIN->CR-Kinase->TPR+TPR+TPR+TPR+TPR+TPR+TPR->?->HTH->DOC+HTH-> BrxD-AAA -> | 1242 | Bacteria>Pseudomonadota>Betaproteobacteria>Betaproteobacteria_incertae_sedis>Candidatus_Accumulibacter>unclassified_Candidatus_Accumulibacter | GCA_017302555.1 | Candidatus_Accumulibacter_sp. | J0L95_03060 | 58448..62176 | 2053492 | phage-resistance-protein-[Candidatus-Accumulibacter-sp.] |
| WP_234414047.1 | Type2.Pgl.like.BREX | <-Phage_integrase<-?\|\|HTH_26-> PglW -> N6- METHYLASE ->DOC+HTH-> DUF6079 *-> PglZ -> BrxD-AAA -> SF1-SF2-Helicase+wHTH+Small-B-barrel+DUF1998 -> OLD-ABC ->HNH -><-?<-?\|\|Phage_int_SAM_1+Phage_integrase-> | 1226 | Bacteria>Pseudomonadota>Betaproteobacteria>Burkholderiales>Sphaerotilaceae>Ideonella | GCF_002198735.1 | Ideonella_sp._A_288 | EOINJALH_RS25855 | 77971..81651 | 1962181 | DUF6079-family-protein-[Ideonella-sp.-A-288] |
| GKW48594.1 | Type2.Pgl.like.BREX | <- SF1-SF2-Helicase+wHTH+Small-B-barrel+DUF1998 <- BrxD-AAA <- PglZ <- DUF6079 *<- N6- METHYLASE <- HNH<- OLD-ABC <- PglW <-XIS-HTH | 1235 | Bacteria>Pseudomonadota>Gammaproteobacteria>Oceanospirillales>Halomonadaceae | GCA_023708045.1 | Halomonas_sp._NCCP-2165 | NCCP2165_08090 | 82862..86569 | 2942298 | hypothetical-protein-NCCP2165_08090-[Halomonas-sp.-NCCP-2165] |
| WP_175072159.1 | Type2.Pgl.like.BREX | XIS-HTH-><-?\|\|?-><-?\|\|XIS-HTH-> PglW -> N6- METHYLASE -> OLD-ABC ->HNH -> DUF6079 *-> PglZ -> BrxD-AAA -> SF1-SF2-Helicase+wHTH+Small-B-barrel+DUF1998 -><-?\|\|REase-><-?<-Arm-DNA-bind_3+Phage_integrase\|\|HTH+LexA-protease-> N6-N4- METHYLASE -> | 1237 | Bacteria>Pseudomonadota>Gammaproteobacteria>Oceanospirillales>Halomonadaceae>Halomonas | GCF_013349955.1 | Halomonas_taeanensis | HUS66_RS15805 | 34399..38112 | 284577 | DUF6079-family-protein-[Halomonas-taeanensis] |
| WP_240569311.1 | Type2.Pgl.like.BREX | XIS-HTH-> PglW -> OLD-ABC ->HNH -> N6- METHYLASE -> DUF6079 *-> PglZ -> BrxD-AAA -> SF1-SF2-Helicase+wHTH+Small-B-barrel+DUF1998 ->?-> SF1-SF2-Helicase+wHTH+Small-B-barrel+DUF1998 -><-?<-GspA-ATPase | 1239 | Bacteria>Pseudomonadota>Gammaproteobacteria>Oceanospirillales>Halomonadaceae>Halomonas | GCF_022487945.1 | Halomonas_sp._EGI_63088 | MKP05_RS16485 | 45674..49393 | 2920385 | DUF6079-family-protein-[Halomonas-sp.-EGI-63088] |
| WP_251308249.1 | Type2.Pgl.like.BREX | <-EAL<-?<-GreAB-C\|\|?-><-?<- SF1-SF2-Helicase+wHTH+Small-B-barrel+DUF1998 <- BrxD-AAA <- PglZ <- DUF6079 *<- N6- METHYLASE <- HNH<- OLD-ABC <- PglW <-XIS-HTH<-?\|\|?-> wHTH+WYL+WCX ->ParB+HNH-> | 1235 | Bacteria>Pseudomonadota>Gammaproteobacteria>Oceanospirillales>Halomonadaceae>Halomonas | GCF_023708045.1 | Halomonas_sp._NCCP-2165 | NFU04_RS04185 | 82862..86569 | 2942298 | DUF6079-family-protein-[Halomonas-sp.-NCCP-2165] |
| WP_072820441.1 | Type2.Pgl.like.BREX | XIS-HTH->?->?-><-?\|\|XIS-HTH-> PglW -> OLD-ABC ->HNH -> N6- METHYLASE -> DUF6079 *-> PglZ -> BrxD-AAA -> SF1-SF2-Helicase+wHTH+Small-B-barrel+DUF1998 ->HARE-HTH+REase-><-?\|\|?->?-><-HD | 1236 | Bacteria>Pseudomonadota>Gammaproteobacteria>Oceanospirillales>Halomonadaceae>Halomonas | GCF_900128925.1 | Halomonas_ilicicola | BUA41_RS05240 | 148075..151785 | 480814 | phage-resistance-protein-[Halomonas-ilicicola] |
| WP_094873724.1 | Type2.Pgl.like.BREX | <-MULE-TRANSPOSASE<- SIGMA+OB-Fold+AAA-ATPase+Lon-Protease-BrxL <- PglZ <- HNH<- OLD-ABC <- N6- METHYLASE <- DUF6079 *<- BrxB-BrxF- iSTAND <- BrxA-DNA-binding \|\|NERD+iPkinase+Pkinase+SF1-SF2-Helicase->?-><-MULE-TRANSPOSASE<-ACET | 1142 | Bacteria>Pseudomonadota>Gammaproteobacteria>Pasteurellales>Pasteurellaceae>Gallibacterium | GCF_002263255.2 | Gallibacterium_anatis | CF557_RS03210 | 664755..668183 | 750 | BREX-system-P-loop-protein-BrxC-[Gallibacterium-anatis] |
| WP_169919065.1 | Type2.Pgl.like.BREX | <- OLD-ABC + TOPRIM <-MAD-NTD+REase<-35exo+SF1-SF2-Helicase<-?<-NERD+iPkinase+Pkinase+SF1-SF2-Helicase<-XIS-HTH<- SF1-SF2-Helicase+wHTH+Small-B-barrel+DUF1998 <- BrxD-AAA <- PglZ <- DUF6079 *<- HNH<- OLD-ABC <- N6- METHYLASE <- PglW <-tRNA<-MerR-HTH | 1231 | Bacteria>Pseudomonadota>Gammaproteobacteria>Pseudomonadales>Pseudomonadaceae>Pseudomonas | GCF_012985845.1 | Pseudomonas_sp._WS_5013 | HBN76_RS00225 | 58998..62693 | 2717475 | phage-resistance-protein-[Pseudomonas-sp.-WS-5013] |
| WP_010033155.1 | Type2.Pgl.like.BREX | PglW -> N6- METHYLASE -> OLD-ABC ->HNH -> DUF6079 *-> PglZ -> BrxD-AAA -> SF1-SF2-Helicase+wHTH+Small-B-barrel+DUF1998 -><-?<-?<-?<-?<-HTH_Tnp | 1233 | Bacteria>PVC_group>Planctomycetota>Planctomycetia>Gemmatales>Gemmataceae | GCF_003149495.1 | Gemmata_obscuriglobus | GOBSN_RS25255 | 6052898..6056599 | 114 | hypothetical-protein-[Gemmata-obscuriglobus] |
| QDU98792.1 | Type2.Pgl.like.BREX | XIS-HTH-> PglW -> N6- METHYLASE -> OLD-ABC ->HNH -> DUF6079 *-> PglZ -> BrxD-AAA -> SF1-SF2-Helicase+wHTH+Small-B-barrel+DUF1998 -> | 1225 | Bacteria>PVC_group>Planctomycetota>Planctomycetia>Pirellulales | GCA_007751035.1 | Lignipirellula_cremea | Pla8534_66660 | 8881234..8884911 | 2528010 | hypothetical-protein-Pla8534_66660-[Lignipirellula-cremea] |
| WP_105358292.1 | Type2.Pgl.like.BREX | <-SF1-SF2-Helicase\|\|?->GDResolvase+PPP1-HTH->ParB->ParB->RAMA-><- SF1-SF2-Helicase+wHTH+Small-B-barrel+DUF1998 <- BrxD-AAA <- PglZ <- DUF6079 *<- HNH<- OLD-ABC <- N6- METHYLASE <-?<-?<- PglW <-XIS-HTH | 1220 | Bacteria>PVC_group>Planctomycetota>Planctomycetia>Pirellulales>Pirellulaceae | GCF_002967725.1 | Blastopirellula_marina | C5Y98_RS24525 | 392384..396046 | 124 | phage-resistance-protein-[Blastopirellula-marina] |
| WP_145058317.1 | Type2.Pgl.like.BREX | XIS-HTH-> PglW -> N6- METHYLASE -> OLD-ABC ->HNH -> DUF6079 *-> PglZ -> BrxD-AAA -> SF1-SF2-Helicase+wHTH+Small-B-barrel+DUF1998 -><-HTH+MPTase<-?\|\|?-> N6- METHYLASE -> | 1225 | Bacteria>PVC_group>Planctomycetota>Planctomycetia>Pirellulales>Pirellulaceae | GCF_007751035.1 | Lignipirellula_cremea | Pla8534_RS33060 | 8881234..8884911 | 2528010 | phage-resistance-protein-[Lignipirellula-cremea] |
| WP_336521091.1 | Type2.Pgl.like.BREX | <- SF1-SF2-Helicase+wHTH+Small-B-barrel+DUF1998 <- BrxD-AAA <- PglZ <- DUF6079 *<-ParB+HNH<-?<- HNH<- OLD-ABC <- N6- METHYLASE <- PglW | 1216 | Bacteria>PVC_group>Planctomycetota>Planctomycetia>Planctomycetales | GCF_030622045.1 | Planctoellipticum_variicoloris | SH412_RS26695 | 6848891..6852541 | 3064265 | phage-resistance-protein-[Planctoellipticum-variicoloris] |
| WP_278468317.1 | Type2.Pgl.like.BREX | <- SF1-SF2-Helicase+wHTH+Small-B-barrel+DUF1998 <- BrxD-AAA <- PglZ <- DUF6079 *<- HNH<- OLD-ABC <- N6- METHYLASE <- PglW \|\|REase-> | 1234 | Bacteria>PVC_group>Planctomycetota>Planctomycetia>Planctomycetales | GCF_963971965.1 | Gimesia_maris | B4K77_RS12150 | 525086..528790 | 122 | phage-resistance-protein-[Gimesia-maris] |

| Protein-ID | System | Gene-Neighbourhoods | Length | Taxonomy | Assembly-ID | Strain | Locus-Tag | Genomic-coordinates | TaxID | Fasta-defline |
|---|---|---|---|---|---|---|---|---|---|---|
| MAX35346.1 | Type2.Pgl.like.BREX | <- SF1-SF2-Helicase+wHTH+Small-B-barrel+DUF1998 <-REase<- BrxD-AAA <- PglZ <- DUF6079 *<- HNH<- OLD-ABC <- N6- METHYLASE <- HNH<- OLD-ABC <- PglW | 1230 | Bacteria>PVC_group>Planctomycetota>Planctomycetia>Planctomycetales>Planctomycetaceae | GCA_002706765.1 | Gimesia_sp. | CME33_02125 | 36301..39993 | 2024833 | phage-resistance-protein-[Gimesia-sp.] |
| HBL44677.1 | Type2.Pgl.like.BREX | <- SF1-SF2-Helicase+wHTH+Small-B-barrel+DUF1998 <- SF1-SF2-Helicase+wHTH+Small-B-barrel+DUF1998 -p<-REase<- BrxD-AAA -p<- BrxD-AAA <- PglZ <- DUF6079 -p<- DUF6079 -p*<- DUF6079 -p<- HNH<- OLD-ABC <- N6- METHYLASE -p | 783 | Bacteria>PVC_group>Planctomycetota>Planctomycetia>Planctomycetales>Planctomycetaceae | GCA_003505085.1 | Planctomycetaceae_bacterium | DDZ90_14925 | 11270..13618 | 2026779 | phage-resistance-protein-[Planctomycetaceae-bacterium] |
| HBL44678.1 | Type2.Pgl.like.BREX | <- SF1-SF2-Helicase+wHTH+Small-B-barrel+DUF1998 <- SF1-SF2-Helicase+wHTH+Small-B-barrel+DUF1998 -p<-REase<- BrxD-AAA -p<- BrxD-AAA <- PglZ <- DUF6079 -p<- DUF6079 -p<- DUF6079 -p*<- HNH<- OLD-ABC <- N6- METHYLASE -p | 347 | Bacteria>PVC_group>Planctomycetota>Planctomycetia>Planctomycetales>Planctomycetaceae | GCA_003505085.1 | Planctomycetaceae_bacterium | DDZ90_14930 | 13810..14849 | 2026779 | phage-resistance-protein-[Planctomycetaceae-bacterium] |
| WP_312777488.1 | Type2.Pgl.like.BREX | <- SF1-SF2-Helicase+wHTH+Small-B-barrel+DUF1998 \|\|Phd-YefM->PIN-><- BrxD-AAA <- PglZ <- DUF6079 *<- N6- METHYLASE \|\| OLD-ABC ->HNH -><- PglW \|\|HEPN-> | 1302 | Bacteria>Terrabacteria_group>Actinomycetota>Actinomycetes>Mycobacteriales | GCF_032056685.1 | Corynebacterium_variabile | RT519_RS10550 | 105740..109648 | 1727 | DUF6079-family-protein-[Corynebacterium-variabile] |
| -- | | | | | | | | | | |
| -- OLD-ABC ->HNH | | | | | | | | | | |
| --Type3-4.BREX | | | | | | | | | | |
| -- | | | | | | | | | | |
| WP_020614466.1 | Type3.MTaseII.containing.BREX | SecA+SF1-SF2-Helicase-><-Peptidase_M23\|\|Peptidase_S26->?->?->XIS-HTH->XIS-HTH-> DUF6079 *-> OLD-ABC ->HNH -> N6-N4- METHYLASE -> TUDOR+SF1-SF2-Helicase+iREase+CTD -> PglZ -> BrxA-DNA-binding ->HIN-HTH-> | 1241 | Bacteria>Spirochaetota>Spirochaetia>Spirochaetales>Spirochaetaceae>Sediminispirochaeta | GCF_000378205.1 | Sediminispirochaeta_bajacaliforniensis | F459_RS0120160 | 20907..24632 | 148 | DUF6079-family-protein-[Sediminispirochaeta-bajacaliforniensis] |
| OQY33159.1 | Type3.MTaseII.containing.BREX | <- BrxA-DNA-binding <- PglZ <- TUDOR+SF1-SF2-Helicase+iREase+CTD <- N6-N4- METHYLASE <- HNH<- OLD-ABC <- DUF6079 *<- BrxB-BrxF- iSTAND <-AbiEi_3_N+AbiEi_3\|\|?->?->HAD+HAD->?->?-><-RADICAL-SAM | 1241 | Bacteria>Spirochaetota>Spirochaetia>Spirochaetales>Spirochaetaceae>unclassified_Spirochaetaceae | GCA_002084805.1 | Spirochaetaceae_bacterium_4572_59 | B6241_08630 | 14589..18314 | 1971727 | hypothetical-protein-B6241_08630-[Spirochaetaceae-bacterium-4572_59] |
| WP_061808755.1 | Type3.MTaseII.containing.BREX | HTH_3->?->SF1-SF2-Helicase+REase->?->?->HKD-DNase-><-CHTH\|\|REase+SF1-SF2-Helicase-> BrxB-BrxF- iSTAND -> DUF6079 *-> N6-N4- METHYLASE -> PglZ -> BrxA-DNA-binding -> TUDOR+ SF1-SF2-Helicase+iREase+CTD+REase ->ParB+HNH->?-> OLD-ABC ->HNH -> | 1226 | Bacteria>Terrabacteria_group>Bacillota>Bacilli>Bacillales>Bacillaceae | GCF_001591825.1 | Rossellomorea_vietnamensis | BVI01S_RS00200 | 36378..40058 | 218284 | DUF6079-family-protein-[Rossellomorea-vietnamensis] |
| MBR2506166.1 | Type3.MTaseII.containing.BREX | <-HTH+HTH\|\|?->HTH_36->?->?->TraJ-RHH->TraI-VirD2-RCRE->?-> BrxB-BrxF- iSTAND -> DUF6079 *-> OLD-ABC ->HNH -> N6-N4- METHYLASE ->AIPR->Schlafen+HATPase+wHTH-> TUDOR+SF1-SF2-Helicase+iREase+CTD -> PglZ -> BrxA-DNA-binding -> | 1225 | Bacteria>Terrabacteria_group>Bacillota>Bacilli>unclassified_Bacilli | GCA_017502465.1 | Bacilli_bacterium | IKB70_04475 | 8185..11862 | 1903720 | hypothetical-protein-[Bacilli-bacterium] |
| MCI8605659.1 | Type4.PglZ.containing.BREX | GGDEF-><-?<-?<-?<-tRNA\|\| OLD-ABC ->HNH ->SeqA+E3_UFM1_ligase-> PAPS-reduct + DUF4007 -><-?\|\| DUF6079 *-> iSTAND + PglZ -> SIGMA+OB-Fold+AAA-ATPase-BrxL ->ZnR-> Cysteine-Desulfurase ->ParB+HNH->VirB11+SbcC->HNH-> | 1121 | Bacteria>Terrabacteria_group>Bacillota>Clostridia>Eubacteriales>Clostridiaceae | GCA_022793615.1 | Hungatella_sp. | HFG72_02125 | 13672..17037 | 2613924 | hypothetical-protein-[Hungatella-sp.] |
| -- | | | | | | | | | | |
| -- OLD-ABC ->HNH | | | | | | | | | | |
| --BR-1 | | | | | | | | | | |
| -- | | | | | | | | | | |
| MCL4551940.1 | DUF499_1.RM.like.systems | XIS-HTH-> OLD-ABC ->HNH -> METHYLASE -> DUF499 *-> DUF499 -p->SF1-SF2-Helicase+REase-> | 336 | Archaea>TACK_group>Candidatus_Marsarchaeota>unclassified_Candidatus_Marsarchaeota | GCA_023381795.1 | Candidatus_Marsarchaeota_archaeon | M1305_00065 | 10946..11954 | 2823144 | DUF499-domain-containing-protein-[Candidatus-Marsarchaeota-archaeon] |
| MYC75109.1 | DUF499_1.RM.like.systems | <-ABC-ATPase+SbcC<-ABC-ATPase+SbcC<-ZnR<-SIGMA-HTH<-HEAT_2<- DUF499 *<-?<- METHYLASE <-?<- HNH<- OLD-ABC <- TUDOR+ SF1-SF2-Helicase+iREase+CTD+REase | 1114 | Bacteria>Bacteria_incertae_sedis>Bacteria_candidate_phyla>Candidatus_Poribacteria>unclassified_Candidatus_Poribacteria | GCA_009841255.1 | Candidatus_Poribacteria_bacterium | F4X10_04955 | 70263..73607 | 2026781 | ATP-binding-protein-[Candidatus-Poribacteria-bacterium] |
| MCG9130919.1 | DUF499_1.RM.like.systems | TUDOR+ SF1-SF2-Helicase+iREase+CTD+REase -> OLD-ABC ->HNH -> METHYLASE ->PolB->PolB+HEPN-> DUF499 *->HEAT_2->SAM-methylase->SIGMA-HTH->ZnR->ABC-ATPase+SbcC->ABC-ATPase+SbcC-> | 1114 | Bacteria>Bacteria_incertae_sedis>Bacteria_candidate_phyla>Candidatus_Poribacteria>unclassified_Candidatus_Poribacteria | GCA_022394775.1 | Candidatus_Poribacteria_bacterium | J5I95_04465 | 27738..31082 | 2026781 | ATP-binding-protein-[Candidatus-Poribacteria-bacterium] |
| MCY3550002.1 | DUF499_1.RM.like.systems | <-Polbeta<-ABC-ATPase+SbcC<-ABC-ATPase+SbcC<-ZnR<-SIGMA-HTH<-HEAT_2<- DUF499 *<-PolB<- METHYLASE <-?<- HNH<- OLD-ABC <- TUDOR+ SF1-SF2-Helicase+iREase+CTD+REase | 1122 | Bacteria>Bacteria_incertae_sedis>Bacteria_candidate_phyla>Candidatus_Poribacteria>unclassified_Candidatus_Poribacteria | GCA_026702755.1 | Candidatus_Poribacteria_bacterium | OXH39_06030 | 100519..103887 | 2026781 | MAG:-DUF499-domain-containing-protein-[Candidatus-Poribacteria-bacterium] |
| MDE0324945.1 | DUF499_1.RM.like.systems | <-HEAT_2<- DUF499 *<-PolB+HEPN<-?<- METHYLASE <-?<- HNH<- OLD-ABC | 1118 | Bacteria>Bacteria_incertae_sedis>Bacteria_candidate_phyla>Candidatus_Poribacteria>unclassified_Candidatus_Poribacteria | GCA_028820875.1 | Candidatus_Poribacteria_bacterium | OXN27_13610 | 421..3777 | 2026781 | MAG:-DUF499-domain-containing-protein-[Candidatus-Poribacteria-bacterium] |
| MBI4801911.1 | DUF499_1.RM.like.systems | STAS+HisKin->?->PIN-> OLD-ABC ->HNH -> METHYLASE -> HEPN+ DUF499 *->Glycos_trans_3N+Glycos_transf_3->Radical-SAM->Radical-SAM->?->?->RADICAL-SAM-> | 1132 | Bacteria>Elusimicrobiota>unclassified_Elusimicrobiota | GCA_016208085.1 | Elusimicrobiota_bacterium | HY796_05240 | 17514..20912 | 2030800 | ATP-binding-protein-[Elusimicrobia-bacterium] |
| WP_070725327.1 | DUF499_1.RM.like.systems | <-SF1-SF2-Helicase+REase<-?\|\| TUDOR+ SF1-SF2-Helicase+iREase+CTD+REase -> OLD-ABC ->HNH -> METHYLASE -> HEPN+ DUF499 *-><-HTH | 1110 | Bacteria>FCB_group>Bacteroidota/Chlorobiota_group>Bacteroidota>Cytophagia | GCF_001816145.1 | Hymenobacter_lapidarius | BEN47_RS03945 | 71126..74458 | 1908237 | DUF499-domain-containing-protein-[Hymenobacter-lapidarius] |
| MCB0808336.1 | DUF499_1.RM.like.systems | <- DUF499 *<- METHYLASE <- HNH<- OLD-ABC <- TUDOR+ SF1-SF2-Helicase+iREase+CTD+REase | 759 | Bacteria>FCB_group>Bacteroidota/Chlorobiota_group>Bacteroidota>Flavobacteriia | GCA_020438005.1 | Flavobacteriales_bacterium | KDB96_03570 | 1..2279 | 2021391 | ATP-binding-protein-[Flavobacteriales-bacterium] |
| NUQ24578.1 | DUF499_1.RM.like.systems | TUDOR+ SF1-SF2-Helicase+iREase+CTD+REase -> OLD-ABC ->HNH -> METHYLASE ->SF1-SF2-Helicase+REase-> HEPN+ DUF499 *->?->PolB->PolB+HEPN-><-ORC-CDC6+REase | 1095 | Bacteria>FCB_group>Bacteroidota/Chlorobiota_group>Bacteroidota>Saprospiria | GCA_013360405.1 | Saprospiraceae_bacterium | HUU34_11550 | 44219..47506 | 2202734 | DUF499-domain-containing-protein-[Saprospiraceae-bacterium] |
| WP_171805995.1 | DUF499_1.RM.like.systems | TUDOR+ SF1-SF2-Helicase+iREase+CTD+REase -> OLD-ABC ->HNH -> METHYLASE -> DUF499 *->DNA-glycosylase->CHTH->?->LZ_Tnp_IS66+zf-IS66+DDE_Tnp_IS66+DDE_Tnp_IS66_C->DUF4096+DDE_Tnp_1+DDE_Tnp_1_2-> | 942 | Bacteria>Myxococcota>Myxococcia>Myxococcales>Cystobacterineae | GCF_013155195.1 | Corallococcus_exiguus | HPP06_RS00265 | 63781..66609 | 83462 | DUF499-domain-containing-protein-[Corallococcus-exiguus] |
| WP_174489000.1 | DUF499_1.RM.like.systems | <- METHYLASE +DNA_methylase\|\|tRNA-><-DNA-glycosylase<-?<-Pkinase<-REase<- DUF499 *<- METHYLASE <- HNH<- OLD-ABC <- TUDOR+ SF1-SF2-Helicase+iREase+CTD+REase <-XIS-HTH<-PAIREDC-HTH | 943 | Bacteria>Myxococcota>Myxococcia>Myxococcales>Cystobacterineae | GCF_013336725.1 | Myxococcus_sp._CA040A | HUA75_RS36310 | 164891..167722 | 2741738 | DUF499-domain-containing-protein-[Myxococcus-sp.-CA040A] |
| KYF87240.1 | DUF499_1.RM.like.systems | <- DUF499 *<- REase<- OLD-ABC <- METHYLASE <- HNH<- OLD-ABC <- TUDOR+ SF1-SF2-Helicase+iREase+CTD+REase | 954 | Bacteria>Myxococcota>Polyangia>Polyangiales>Polyangiaceae | GCA_001589185.1 | Sorangium_cellulosum | BE17_46125 | 4063..6927 | 56 | hypothetical-protein-BE17_46125-[Sorangium-cellulosum] |
| MBZ0154723.1 | DUF499_1.RM.like.systems | PIN-> OLD-ABC ->HNH -> METHYLASE ->?->RelE-ParE-> HEPN+ DUF499 *-> | 1114 | Bacteria>Nitrospirota>Nitrospiria>Nitrospirales>Nitrospiraceae | - | Candidatus_Nitrobium_versatile | K8I29_00740 | 55978..59322 | 2884831 | DUF499-domain-containing-protein-[Alphaproteobacteria-bacterium] |
| MBU2768209.1 | DUF499_1.RM.like.systems | <- DUF499 *<- REase<- OLD-ABC <-wHTH+REase<- METHYLASE <- HNH<- OLD-ABC | 542 | Bacteria>Pseudomonadota>Acidithiobacillia>Acidithiobacillales>Acidithiobacillaceae | GCA_018853935.1 | Acidithiobacillus_ferrivorans | HAP94_19055 | 1..1626 | 160808 | ATP-binding-protein-[Acidithiobacillus-ferrivorans] |
| WP_244479882.1 | DUF499_1.RM.like.systems | TUDOR+ SF1-SF2-Helicase+iREase+CTD+REase -> OLD-ABC ->HNH -> METHYLASE -> DUF499 *->GDResolvase+HTH->ParB+HNH-><-?\|\|HTH_3-><-?\|\|REase-> | 950 | Bacteria>Pseudomonadota>Alphaproteobacteria>Hyphomicrobiales>Methylobacteriaceae | GCF_001426045.1 | Methylobacterium_sp._Leaf94 | ASG63_RS11585 | 13925..16777 | 1736250 | ATP-binding-protein-[Methylobacterium-sp.-Leaf94] |
| WP_166097056.1 | DUF499_1.RM.like.systems | MerR-HTH->tRNA-> TUDOR+ SF1-SF2-Helicase+iREase+CTD+REase -> OLD-ABC ->HNH -> METHYLASE -> DUF499 *->DUF433->PIN-><-Phage_integrase | 945 | Bacteria>Pseudomonadota>Alphaproteobacteria>Hyphomicrobiales>Nitrobacteraceae | GCF_011516635.5 | Bradyrhizobium_barranii | HAP41_RS22275 | 4684908..4687745 | 2992140 | ATP-binding-protein-[Bradyrhizobium-barranii] |
| WP_212326836.1 | DUF499_1.RM.like.systems | Phage_integrase->?->XIS-HTH->?->DnaB->HIN-HTH-><- HEPN+ DUF499 *<- METHYLASE <- HNH<- OLD-ABC <- TUDOR+ SF1-SF2-Helicase+iREase+CTD+REase | 1108 | Bacteria>Pseudomonadota>Alphaproteobacteria>Hyphomicrobiales>Nitrobacteraceae | GCF_018131105.1 | Bradyrhizobium_manausense | JQ631_RS13545 | 2838930..2842256 | 989370 | Swt1-family-HEPN-domain-containing-protein-[Bradyrhizobium-manausense] |
| WP_172236133.1 | DUF499_1.RM.like.systems | XIS-HTH->?->?->DnaB->HIN-HTH-><- HEPN+ DUF499 *<- METHYLASE <- HNH<- OLD-ABC <- TUDOR+ SF1-SF2-Helicase+iREase+CTD+REase | 1108 | Bacteria>Pseudomonadota>Alphaproteobacteria>Hyphomicrobiales>Nitrobacteraceae | GCF_902482845.1 | Bradyrhizobium | JQ619_RS32195 | 7694..11020 | 374 | MULTISPECIES:-Swt1-family-HEPN-domain-containing-protein-[Bradyrhizobium] |
| WP_116622075.1 | DUF499_1.RM.like.systems | HTH_3-> TUDOR+ SF1-SF2-Helicase+iREase+CTD+REase -> OLD-ABC ->HNH ->ASCH-> METHYLASE -> DUF499 *-><-?<-TraD\|\|MobA_MobL+SF1-SF2-helicase->XIS-HTH->?->PhdYeFM_antitox-> | 945 | Bacteria>Pseudomonadota>Alphaproteobacteria>Hyphomicrobiales>Phyllobacteriaceae | GCF_003403035.1 | Mesorhizobium_denitrificans | DY251_RS01610 | 349492..352329 | 2294114 | DUF499-domain-containing-protein-[Mesorhizobium-denitrificans] |
| WP_140734688.1 | DUF499_1.RM.like.systems | Phage_terminase-> TUDOR+ SF1-SF2-Helicase+iREase+CTD+REase -> OLD-ABC ->HNH -> METHYLASE -> DUF499 *->?->\|\|?-><-?<-?<-?\|\|Phage_integrase-> | 942 | Bacteria>Pseudomonadota>Alphaproteobacteria>Hyphomicrobiales>Phyllobacteriaceae | GCF_006442905.1 | Mesorhizobium_sp._B2-4-9 | FJ945_RS19675 | 32509..35337 | 2589940 | ATP-binding-protein-[Mesorhizobium-sp.-B2-4-9] |
| NEJ60793.1 | DUF499_1.RM.like.systems | XIS-HTH-> TUDOR+ SF1-SF2-Helicase+iREase+CTD+REase -> OLD-ABC ->HNH -> METHYLASE -> DUF499 *-><-?<-Radical-SAM | 943 | Bacteria>Pseudomonadota>Alphaproteobacteria>Hyphomicrobiales>Rhizobiaceae | GCA_010668765.1 | Rhizobium_ruizarguesonis | GUK21_31970 | 69157..71988 | 2081791 | DUF499-domain-containing-protein-[Rhizobium-leguminosarum] |
| MCJ8508804.1 | DUF499_1.RM.like.systems | <-XIS-HTH\|\|HTH_3-> TUDOR+ SF1-SF2-Helicase+iREase+CTD+REase -> OLD-ABC ->HNH -> METHYLASE -> DUF499 *->MODE-HTH+PBPII-><-ParG_RHH | 937 | Bacteria>Pseudomonadota>Alphaproteobacteria>Hyphomicrobiales>Rhizobiaceae | GCA_022968395.1 | Rhizobium_lemnae | MUU53_12875 | 44346..47159 | 1214924 | ATP-binding-protein-[Rhizobium-lemnae] |
| WP_010968623.1 | DUF499_1.RM.like.systems | HTH_3-> TUDOR+ SF1-SF2-Helicase+iREase+CTD+REase -> OLD-ABC ->HNH -> METHYLASE -> DUF499 *-><-?<-REase<-HTH_3 | 942 | Bacteria>Pseudomonadota>Alphaproteobacteria>Hyphomicrobiales>Rhizobiaceae | GCF_000006965.1 | Sinorhizobium_meliloti | J3S94_RS16455 | 3478625..3481453 | 382 | ATP-binding-protein-[Sinorhizobium-meliloti] |
| WP_012755925.1 | DUF499_1.RM.like.systems | Phage_integrase->?->?-> OLD-ABC + TOPRIM ->?-><- DUF499 *<- METHYLASE <- HNH<- OLD-ABC <- TUDOR+ SF1-SF2-Helicase+iREase+CTD+REase <-HerA/FtsK | 945 | Bacteria>Pseudomonadota>Alphaproteobacteria>Hyphomicrobiales>Rhizobiaceae | GCF_000023185.1 | Rhizobium_leguminosarum | RLEG_RS01005 | 199729..202566 | 384 | ATP-binding-protein-[Rhizobium-leguminosarum] |
| WP_069615760.1 | DUF499_1.RM.like.systems | HTH_3-> TUDOR+ SF1-SF2-Helicase+iREase+CTD+REase -> OLD-ABC ->HNH -> METHYLASE -> DUF499 *-><-?<-N6_MTase<-HTH<-?\|\|?->McrC-NTD+McrC-> | 945 | Bacteria>Pseudomonadota>Alphaproteobacteria>Hyphomicrobiales>Rhizobiaceae | GCF_001729315.1 | Rhizobium_sp._YK2 | A9Z06_RS30975 | 24492..27329 | 1860096 | ATP-binding-protein-[Rhizobium-sp.-YK2] |
| WP_130838769.1 | DUF499_1.RM.like.systems | <-REase\|\| TUDOR+ SF1-SF2-Helicase+iREase+CTD+REase -> OLD-ABC ->HNH -> METHYLASE -> DUF499 *-> | 944 | Bacteria>Pseudomonadota>Alphaproteobacteria>Hyphomicrobiales>Rhizobiaceae | GCF_004305895.1 | Rhizobium_ruizarguesonis | HA464_RS06765 | 1375987..1378821 | 2081791 | ATP-binding-protein-[Rhizobium-leguminosarum] |
| WP_164568970.1 | DUF499_1.RM.like.systems | XIS-HTH-> TUDOR+ SF1-SF2-Helicase+iREase+CTD+REase -> OLD-ABC ->HNH -> METHYLASE -> DUF499 *-><-?<-Radical-SAM | 943 | Bacteria>Pseudomonadota>Alphaproteobacteria>Hyphomicrobiales>Rhizobiaceae | GCF_010668765.1 | Rhizobium_ruizarguesonis | GUK21_31970 | 69157..71988 | 2081791 | ATP-binding-protein-[Rhizobium-leguminosarum] |
| WP_164056349.1 | DUF499_1.RM.like.systems | Arm-DNA-bind_3+Phage_integrase-><-XIS-HTH<-MobA_MobL+SF1-SF2-helicase\|\|TraD->?-><- DUF499 *<- METHYLASE <- HNH<- OLD-ABC <- TUDOR+ SF1-SF2-Helicase+iREase+CTD+REase <-HTH_3<-HigB | 944 | Bacteria>Pseudomonadota>Alphaproteobacteria>Hyphomicrobiales>Rhizobiaceae | GCF_010669145.1 | Rhizobium_oryzihabitans | G3A56_RS10085 | 1920588..1923422 | 2267833 | ATP-binding-protein-[Rhizobium-oryzihabitans] |
| WP_242223016.1 | DUF499_1.RM.like.systems | HigB->HTH_3-> TUDOR+ SF1-SF2-Helicase+iREase+CTD+REase -> OLD-ABC ->HNH -> METHYLASE -> DUF499 *-><-?<-TraD\|\|MobA_MobL+SF1-SF2-helicase-><-?\|\|XIS-HTH-> | 946 | Bacteria>Pseudomonadota>Alphaproteobacteria>Hyphomicrobiales>Rhizobiaceae | GCF_022682305.1 | Shinella_zoogloeoides | MOE34_RS09030 | 1765134..1767974 | 352475 | ATP-binding-protein-[Shinella-zoogloeoides] |
| WP_281435123.1 | DUF499_1.RM.like.systems | <-XIS-HTH\|\|HTH_3-> TUDOR+ SF1-SF2-Helicase+iREase+CTD+REase -> OLD-ABC ->HNH -> METHYLASE -> DUF499 *->MODE-HTH+PBPII-><-ParG_RHH | 943 | Bacteria>Pseudomonadota>Alphaproteobacteria>Hyphomicrobiales>Rhizobiaceae | GCF_022968395.1 | Rhizobium_lemnae | MUU53_RS12880 | 44328..47159 | 1214924 | ATP-binding-protein-[Rhizobium-lemnae] |
| WP_248132970.1 | DUF499_1.RM.like.systems | <- DUF499 *<- METHYLASE <- HNH<- OLD-ABC <- TUDOR+ SF1-SF2-Helicase+iREase+CTD+REase <-HTH_3 | 506 | Bacteria>Pseudomonadota>Alphaproteobacteria>Hyphomicrobiales>Rhizobiaceae | GCF_023119065.1 | Sinorhizobium_meliloti | J3S92_RS00005 | 1..1518 | 382 | DUF499-domain-containing-protein,-partial-[Sinorhizobium-meliloti] |
| WP_247261244.1 | DUF499_1.RM.like.systems | <-XIS-HTH\|\|HTH_3-> TUDOR+ SF1-SF2-Helicase+iREase+CTD+REase -> OLD-ABC ->HNH -> METHYLASE -> DUF499 *->MODE-HTH+PBPII-><-ParG_RHH | 937 | Bacteria>Pseudomonadota>Alphaproteobacteria>Hyphomicrobiales>Rhizobiaceae | - | Rhizobium_lemnae | MUU53_12875 | 44346..47159 | 1214924 | ATP-binding-protein-[Rhizobium-lemnae] |
| WP_200985559.1 | DUF499_1.RM.like.systems | <-VirD4-TraG+VirD4-FtsK<-?<-Classical-AAA<-PIN+OST-HTH\|\|MobA_MobL+SF1-SF2-helicase-><- DUF499 *<- METHYLASE <- HNH<- OLD-ABC <- TUDOR+ SF1-SF2-Helicase+iREase+CTD+REase <-HTH_3 | 944 | Bacteria>Pseudomonadota>Alphaproteobacteria>Hyphomicrobiales>Rhizobiaceae | - | Rhizobium_rhizogenes | JL018_RS00080 | 14669..17503 | 359 | ATP-binding-protein-[Agrobacterium-rhizogenes] |
| SDR63348.1 | DUF499_1.RM.like.systems | Phage_int_SAM_1+Phage_integrase->REase-><-?<-REase+35exo+Beta-Barrel+SF1-SF2-Helicase<- DUF499 *<- METHYLASE <- HNH<- OLD-ABC <- TUDOR+ SF1-SF2-Helicase+iREase+CTD+REase | 942 | Bacteria>Pseudomonadota>Alphaproteobacteria>Hyphomicrobiales>unclassified_Hyphomicrobiales | GCA_900103895.1 | Rhizobiales_bacterium_GAS113 | SAMN05519103_08517 | 30774..33602 | 1884352 | hypothetical-protein-SAMN05519103_08517-[Rhizobiales-bacterium-GAS113] |
| WP_013654352.1 | DUF499_1.RM.like.systems | <-PIN<-XIS-HTH\|\|?-><-ArdCN+MPTase<-HEPN<- DUF499 *<- METHYLASE <- HNH<- OLD-ABC <- TUDOR+ SF1-SF2-Helicase+iREase+CTD+REase <-HTH_3<-HigB | 944 | Bacteria>Pseudomonadota>Alphaproteobacteria>Rhodobacterales>Paracoccaceae | GCF_000192745.1 | Polymorphum_gilvum | SL003B_RS18265 | 3920112..3922946 | 991904 | DUF499-domain-containing-protein-[Polymorphum-gilvum] |
| WP_014199726.1 | DUF499_1.RM.like.systems | MODE-HTH+PBPII-><-Arm-DNA-bind_3+Phage_int_SAM_3+Phage_integrase<-?<- HEPN+ DUF499 *<-PolB+HEPN<-PolB<- METHYLASE <- HNH<- OLD-ABC <- TUDOR+ SF1-SF2-Helicase+iREase+CTD+REase | 1115 | Bacteria>Pseudomonadota>Alphaproteobacteria>Rhodospirillales>Azospirillaceae | GCF_000237365.1 | Azospirillum | GAY33_RS05480 | 11187..14534 | 191 | MULTISPECIES:-Swt1-family-HEPN-domain-containing-protein-[Azospirillum] |
| WP_209991153.1 | DUF499_1.RM.like.systems | TUDOR+ SF1-SF2-Helicase+iREase+CTD+REase -> OLD-ABC ->HNH -> METHYLASE ->?-> HEPN+ DUF499 *->?->Lon_C-> | 1114 | Bacteria>Pseudomonadota>Alphaproteobacteria>Rhodospirillales>Azospirillaceae | GCF_017876115.1 | Azospirillum_picis | J2850_RS30835 | 10080..13424 | 488438 | Swt1-family-HEPN-domain-containing-protein-[Azospirillum-picis] |
| TXH33435.1 | DUF499_1.RM.like.systems | HTH_3-> TUDOR+ SF1-SF2-Helicase+iREase+CTD+REase -> OLD-ABC ->HNH ->ASCH-> METHYLASE -> DUF499 *-><-?<-TraD\|\|MobA_MobL+SF1-SF2-helicase-> | 945 | Bacteria>Pseudomonadota>Alphaproteobacteria>Rhodospirillales>Rhodospirillaceae | GCA_008012315.1 | Rhodospirillaceae_bacterium | E6Q98_22210 | 66565..69402 | 1898112 | DUF499-domain-containing-protein-[Rhodospirillaceae-bacterium] |
| MBL8657365.1 | DUF499_1.RM.like.systems | <- DUF499 *<- METHYLASE <-ASCH<- HNH<- OLD-ABC <- TUDOR+ SF1-SF2-Helicase+iREase+CTD+REase <-HTH_3 | 944 | Bacteria>Pseudomonadota>Alphaproteobacteria>Sphingomonadales>Erythrobacteraceae | GCA_016793865.1 | Altererythrobacter_sp. | JNJ92_08435 | 1147..3981 | 1872480 | DUF499-domain-containing-protein-[Altererythrobacter-sp.] |
| RXZ34416.1 | DUF499_1.RM.like.systems | TUDOR+ SF1-SF2-Helicase+iREase+CTD+REase -> OLD-ABC ->HNH -> METHYLASE -> DUF499 *-> | 940 | Bacteria>Pseudomonadota>Alphaproteobacteria>Sphingomonadales>Sphingomonadaceae | GCA_004135605.1 | Sphingomonas_desiccabilis | EO081_01620 | 357116..359938 | 429134 | DUF499-domain-containing-protein-[Sphingomonas-desiccabilis] |
| WP_099185493.1 | DUF499_1.RM.like.systems | HTH+WYL->CopG/RHH-> TUDOR+ SF1-SF2-Helicase+iREase+CTD+REase -> OLD-ABC ->HNH -> METHYLASE -> DUF499 *-><-?\|\|LexA-protease-><-wHTH+SAD+HNH<-TrwC+SF1-SF2-helicase<-VirD4-TraG+VirD4-FtsK<-CopG/RHH | 940 | Bacteria>Pseudomonadota>Alphaproteobacteria>Sphingomonadales>Sphingomonadaceae | GCF_002723855.1 | Sphingobium | SFOMI_RS05000 | 866506..869328 | 165695 | ATP-binding-protein-[Sphingobium-fuliginis] |
| WP_130751563.1 | DUF499_1.RM.like.systems | CopG/RHH->VirD4-TraG+VirD4-FtsK->TrwC+SF1-SF2-helicase->wHTH+SAD+HNH-><-LexA-protease\|\|?-><- DUF499 *<- METHYLASE <- HNH<- OLD-ABC <- TUDOR+ SF1-SF2-Helicase+iREase+CTD+REase <-CopG/RHH<-HTH+WYL | 940 | Bacteria>Pseudomonadota>Alphaproteobacteria>Sphingomonadales>Sphingomonadaceae | GCF_004305355.1 | Sphingobium_xenophagum | MBESOW_RS00075 | 12724..15546 | 121428 | ATP-binding-protein-[Sphingobium-xenophagum] |
| WP_159367773.1 | DUF499_1.RM.like.systems | <-DDE_Tnp_IS240<-?<-LZ_Tnp_IS66+zf-IS66+DDE_Tnp_IS66+DDE_Tnp_IS66_C<-?<-HIN-HTH<-PAIREDC-HTH+HIN-HTH<- DUF499 *<- METHYLASE <- HNH<- OLD-ABC <- TUDOR+ SF1-SF2-Helicase+iREase+CTD+REase <-CopG/RHH<-HTH+WYL | 940 | Bacteria>Pseudomonadota>Alphaproteobacteria>Sphingomonadales>Sphingomonadaceae | GCF_009856545.1 | Sphingobium_yanoikuyae | GS397_RS23435 | 4919345..4922167 | 13690 | ATP-binding-protein-[Sphingobium-yanoikuyae] |
| WP_168853122.1 | DUF499_1.RM.like.systems | CopG/RHH->VirD4-TraG+VirD4-FtsK->TrwC+SF1-SF2-helicase->wHTH+SAD+HNH-><-LexA-protease\|\|?-><- DUF499 *<- METHYLASE <- HNH<- OLD-ABC <- TUDOR+ SF1-SF2-Helicase+iREase+CTD+REase <-CopG/RHH<-HTH+WYL | 940 | Bacteria>Pseudomonadota>Alphaproteobacteria>Sphingomonadales>Sphingomonadaceae | GCF_012641335.1 | Novosphingobium_sp._ERN07 | HGI47_RS12585 | 95856..98678 | 2726187 | ATP-binding-protein-[Novosphingobium-sp.-ERN07] |
| WP_206364336.1 | DUF499_1.RM.like.systems | TUDOR+ SF1-SF2-Helicase+iREase+CTD+REase -> OLD-ABC ->HNH -> METHYLASE -> DUF499 *-> | 934 | Bacteria>Pseudomonadota>Alphaproteobacteria>Sphingomonadales>Sphingomonadaceae | GCF_014196135.1 | Sphingomonas_desiccabilis | EO081_RS01620 | 357134..359938 | 429134 | ATP-binding-protein-[Sphingomonas-desiccabilis] |
| WP_260153764.1 | DUF499_1.RM.like.systems | TUDOR+ SF1-SF2-Helicase+iREase+CTD+REase -> OLD-ABC ->HNH -> METHYLASE -> DUF499 *-> | 822 | Bacteria>Pseudomonadota>Alphaproteobacteria>Sphingomonadales>Sphingomonadaceae | - | Sphingomonas_desiccabilis | GGQ81_RS01635 | 357438..359906 | 429134 | DUF499-domain-containing-protein-[Sphingomonas-desiccabilis] |
| WP_248595613.1 | DUF499_1.RM.like.systems | RelE-ParE->Phage_int_SAM_4+Phage_integrase->?-> OLD-ABC ->HNH -> METHYLASE -> DUF499 *-> DUF499 -p->Phage_int_SAM_4+Phage_integrase-> | 644 | Bacteria>Pseudomonadota>Betaproteobacteria>Betaproteobacteria_incertae_sedis>Candidatus_Accumulibacter | GCF_012939955.1 | Candidatus_Accumulibacter_phosphatis | E4Q23_RS20285 | 20006..21940 | 327160 | DUF499-domain-containing-protein-[Candidatus-Accumulibacter-phosphatis] |
| WP_169069333.1 | DUF499_1.RM.like.systems | <- HEPN+ DUF499 *<-PolB+HEPN<-PolB<- METHYLASE <- HNH<- OLD-ABC <-PemK_toxin | 1112 | Bacteria>Pseudomonadota>Betaproteobacteria>Betaproteobacteria_incertae_sedis>Candidatus_Accumulibacter | GCF_012940005.1 | Candidatus_Accumulibacter_contiguus | E4Q08_RS03235 | 1844..5182 | 2954381 | Swt1-family-HEPN-domain-containing-protein-[Candidatus-Accumulibacter-contiguus] |
| WP_217096864.1 | DUF499_1.RM.like.systems | <-ACET\|\|?->?->MarR-HTH->Phage_int_SAM_2+Integrase_1-><- DUF499 *<-Schlafen+HATPase+HTH<- METHYLASE <- METHYLASE <- HNH<- OLD-ABC <- TUDOR+ SF1-SF2-Helicase+iREase+CTD+REase | 947 | Bacteria>Pseudomonadota>Betaproteobacteria>Burkholderiales>Burkholderiaceae | GCF_019041095.1 | Burkholderia_multivorans | KTE23_RS28415 | 22880..25723 | 87883 | DUF499-domain-containing-protein-[Burkholderia-multivorans] |
| MCW5668677.1 | DUF499_1.RM.like.systems | TUDOR+ SF1-SF2-Helicase+iREase+CTD+REase -> OLD-ABC ->HNH -> METHYLASE ->?->?-> DUF499 *-><-HTH_3<-?<-HD\|\|NUDIX-> | 937 | Bacteria>Pseudomonadota>Betaproteobacteria>Burkholderiales>Comamonadaceae | GCA_026125825.1 | Hydrogenophaga_sp. | KIT86_03380 | 51119..53932 | 1904254 | MAG:-ATP-binding-protein-[Hydrogenophaga-sp.] |
| MDD2610199.1 | DUF499_1.RM.like.systems | TUDOR+ SF1-SF2-Helicase+iREase+CTD+REase -> OLD-ABC ->HNH -> METHYLASE -p-> METHYLASE ->?-> DUF499 *->?-><-?\|\|<-JAB<-UbE1+JAB<-SMODS | 951 | Bacteria>Pseudomonadota>Betaproteobacteria>Burkholderiales>Comamonadaceae | GCA_028686475.1 | Giesbergeria_sp. | PHX60_11025 | 19718..22573 | 2818473 | MAG:-DUF499-domain-containing-protein-[Giesbergeria-sp.] |
| WP_087746901.1 | DUF499_1.RM.like.systems | <-SPOUT\|\|?-><-?<-?<-?<- DUF499 *<- REase<- OLD-ABC <- METHYLASE <- HNH<- OLD-ABC <- TUDOR+ SF1-SF2-Helicase+iREase+CTD+REase | 945 | Bacteria>Pseudomonadota>Betaproteobacteria>Burkholderiales>Comamonadaceae | GCF_002176815.1 | unclassified_Acidovorax | CCX87_RS13165 | 2797139..2799976 | 2684926 | MULTISPECIES:-DUF499-domain-containing-protein-[unclassified-Acidovorax] |
| WP_055397337.1 | DUF499_1.RM.like.systems | <-ArdCN+MPTase<-?<-?<-MobA_MobL+SF1-SF2-helicase\|\|?-><-?<- DUF499 *<- REase<- OLD-ABC <- METHYLASE <- HNH<- OLD-ABC <- TUDOR+ SF1-SF2-Helicase+iREase+CTD+REase | 945 | Bacteria>Pseudomonadota>Betaproteobacteria>Burkholderiales>Comamonadaceae | GCF_017355385.1 | Acidovorax | AE621_RS12465 | 7782..10619 | 12916 | MULTISPECIES:-DUF499-domain-containing-protein-[Acidovorax] |
| WP_210568820.1 | DUF499_1.RM.like.systems | TUDOR+ SF1-SF2-Helicase+iREase+CTD+REase -> OLD-ABC ->HNH -> METHYLASE -> OLD-ABC ->REase -> DUF499 *->?->?->?-><-?\|\|SPOUT-> | 945 | Bacteria>Pseudomonadota>Betaproteobacteria>Burkholderiales>Comamonadaceae | GCF_017939685.1 | Acidovorax_sp._JG5 | J8G26_RS10065 | 96032..98869 | 2822718 | DUF499-domain-containing-protein-[Acidovorax-sp.-JG5] |
| WP_210930504.1 | DUF499_1.RM.like.systems | HTH->?->Phage_integrase->STYKinase-like-><-?\|\|DDE_Tnp_1_5+DDE_Tnp_1-><- DUF499 *<- REase<- OLD-ABC <-HEPN<- METHYLASE <- HNH<- OLD-ABC | 945 | Bacteria>Pseudomonadota>Betaproteobacteria>Burkholderiales>Comamonadaceae | GCF_018064205.1 | Comamonas_aquatica | KAQ61_RS03430 | 737208..740045 | 225991 | DUF499-domain-containing-protein-[Comamonas-aquatica] |
| MCM1512954.1 | DUF499_1.RM.like.systems | HTH_3->?->?->?-> TUDOR+ SF1-SF2-Helicase+iREase+CTD+REase -> METHYLASE -> DUF499 *-> OLD-ABC ->HNH -><-?\|\|MazE_antitoxin-> | 937 | Bacteria>Pseudomonadota>Betaproteobacteria>Burkholderiales>Oxalobacteraceae | GCA_023667225.1 | Oxalobacter_formigenes | NC112_07605 | 9124..11937 | 847 | DUF499-domain-containing-protein-[Oxalobacter-formigenes] |
| WP_182994400.1 | DUF499_1.RM.like.systems | HTH+WYL-> TUDOR+ SF1-SF2-Helicase+iREase+CTD+REase -> OLD-ABC ->HNH -> METHYLASE -> DUF499 *-><-?\|\|?-><-?<-Arm-DNA-bind_3+Phage_integrase | 941 | Bacteria>Pseudomonadota>Betaproteobacteria>Burkholderiales>Oxalobacteraceae | GCF_014171515.1 | Herbaspirillum_frisingense | G5S34_RS08630 | 1936607..1939432 | 92645 | DUF499-domain-containing-protein-[Herbaspirillum-frisingense] |
| NRT58384.1 | DUF499_1.RM.like.systems | HAD->?-><-rve_3<-ParB+HTH<-?\|\|?-><- DUF499 *<- TOPRIM <- OLD-ABC <- METHYLASE <- HNH<- OLD-ABC <- TUDOR+ SF1-SF2-Helicase+iREase+CTD+REase | 953 | Bacteria>Pseudomonadota>Betaproteobacteria>Burkholderiales>Sphaerotilaceae | GCA_013294065.1 | Leptothrix_sp._C29 | HNQ01_004152 | 18472..21333 | 2735897 | hypothetical-protein-[Leptothrix-sp.-C29] |
| WP_140628337.1 | DUF499_1.RM.like.systems | PIN+OST-HTH-><-?\|\|HTH_3-><-?<-?<-?<- DUF499 *<-?<-?<- METHYLASE <- HNH<- OLD-ABC <- TUDOR+ SF1-SF2-Helicase+iREase+CTD+REase | 936 | Bacteria>Pseudomonadota>Betaproteobacteria>Burkholderiales>Sphaerotilaceae | GCF_006386545.1 | Methylibium_rhizosphaerae | E8D38_RS09385 | 117386..120196 | 2570323 | DUF499-domain-containing-protein-[Pseudorivibacter-rhizosphaerae] |
| WP_173807403.1 | DUF499_1.RM.like.systems | HAD->?-><-ParB+HTH<-?\|\|?-><- DUF499 *<- TOPRIM <- OLD-ABC <- METHYLASE <- HNH<- OLD-ABC <- TUDOR+ SF1-SF2-Helicase+iREase+CTD+REase | 953 | Bacteria>Pseudomonadota>Betaproteobacteria>Burkholderiales>Sphaerotilaceae | GCF_013294065.1 | Leptothrix_sp._C29 | HNQ01_RS20655 | 18472..21333 | 2735897 | DUF499-domain-containing-protein-[Leptothrix-sp.-C29] |
| WP_110402335.1 | DUF499_1.RM.like.systems | <-ABhydrolase\|\|?->?-><-?<- DUF499 *<- REase<- OLD-ABC <- METHYLASE <- HNH<- OLD-ABC <- TUDOR+ SF1-SF2-Helicase+iREase+CTD+REase | 958 | Bacteria>Pseudomonadota>Betaproteobacteria>Burkholderiales>Sphaerotilaceae | - | Sphaerotilus_hippei | C7444_RS18460 | 22920..25796 | 744406 | DUF499-domain-containing-protein-[Sphaerotilus-hippei] |
| QKE41939.1 | DUF499_1.RM.like.systems | DDE_Tnp_1_assoc+DDE_Tnp_1->HTH+WYL-> TUDOR+ SF1-SF2-Helicase+iREase+CTD+REase -> OLD-ABC ->HNH ->wHTH+REase-> DUF499 *->DDE_Tnp_ISAZ013-><-?<-Hydrolase_3 | 942 | Bacteria>Pseudomonadota>Betaproteobacteria>Ferrovales>Ferrovaceae | GCA_013181415.1 | Ferrovum_myxofaciens | HO274_11940 | 2423876..2426704 | 416213 | ATP-binding-protein-[Ferrovum-myxofaciens] |
| WP_031597390.1 | DUF499_1.RM.like.systems | HTH+WYL-> TUDOR+ SF1-SF2-Helicase+iREase+CTD+REase -> OLD-ABC ->HNH -> METHYLASE -> DUF499 *->?->?->?-><-?<-?<-DUF2163+Phage_BR0599 | 944 | Bacteria>Pseudomonadota>Betaproteobacteria>Ferrovales>Ferrovaceae | GCF_000735045.1 | Ferrovum_myxofaciens | IV91_RS0107820 | 14389..17223 | 416213 | DUF499-domain-containing-protein-[Ferrovum-myxofaciens] |
| WP_273119931.1 | DUF499_1.RM.like.systems | TUDOR+ SF1-SF2-Helicase+iREase+CTD+REase -> OLD-ABC ->HNH -> METHYLASE -p-> METHYLASE -p->wHTH+REase-> DUF499 *->DDE_Tnp_ISAZ013-><-?<-Hydrolase_3 | 942 | Bacteria>Pseudomonadota>Betaproteobacteria>Ferrovales>Ferrovaceae | GCF_013181415.1 | Ferrovum_myxofaciens | HO274_RS11980 | 2423876..2426704 | 416213 | DUF499-domain-containing-protein-[Ferrovum-myxofaciens] |
| MBX3631056.1 | DUF499_1.RM.like.systems | <- DUF499 *<- METHYLASE <- HNH<- OLD-ABC <- TUDOR+ SF1-SF2-Helicase+iREase+CTD+REase <-REase+RNase | 949 | Bacteria>Pseudomonadota>Betaproteobacteria>Nitrosomonadales>Nitrosomonadaceae | GCA_019635215.1 | Nitrosomonas_sp. | KF908_14345 | 13134..15983 | 42353 | ATP-binding-protein-[Nitrosomonas-sp.] |
| MBX3668836.1 | DUF499_1.RM.like.systems | HTH_Tnp-> TUDOR+ SF1-SF2-Helicase+iREase+CTD+REase -> OLD-ABC ->HNH -> METHYLASE -> DUF499 *->?-><-Phage_lysozyme<-PIN<-?\|\|?->HTH-> | 945 | Bacteria>Pseudomonadota>Betaproteobacteria>Rhodocyclales>Rhodocyclaceae | GCA_019635235.1 | Rhodocyclaceae_bacterium | KF778_10575 | 185865..188702 | 1898103 | ATP-binding-protein-[Rhodocyclaceae-bacterium] |
| WP_018989132.1 | DUF499_1.RM.like.systems | <-PIN+PhoH<-Radical-SAM+TRAM\|\|tRNA->?-><- HEPN+ DUF499 *<- METHYLASE <- HNH<- OLD-ABC <-PIN<-?<-STAS+HisKin | 1121 | Bacteria>Pseudomonadota>Betaproteobacteria>Rhodocyclales>Rhodocyclaceae | GCF_000378245.1 | Aromatoleum_toluclasticum | F464_RS0107275 | 366039..369404 | 92003 | Swt1-family-HEPN-domain-containing-protein-[Aromatoleum-toluclasticum] |
| WP_169202998.1 | DUF499_1.RM.like.systems | <- DUF499 *<- REase<- OLD-ABC <- METHYLASE <- HNH<- OLD-ABC <- TUDOR+ SF1-SF2-Helicase+iREase+CTD+REase | 951 | Bacteria>Pseudomonadota>Betaproteobacteria>Rhodocyclales>Rhodocyclaceae | GCF_017894365.1 | Aromatoleum_bremense | GPA24_RS12870 | 12346..15201 | 76115 | DUF499-domain-containing-protein-[Aromatoleum-bremense] |
| TMW75521.1 | DUF499_1.RM.like.systems | <- DUF499 *<- METHYLASE <- HNH<- OLD-ABC <-DOC+HTH<- TUDOR+ SF1-SF2-Helicase+iREase+CTD+REase \|\|HTH_31-> | 622 | Bacteria>Pseudomonadota>Betaproteobacteria>Rhodocyclales>Zoogloeaceae | GCA_006007845.1 | Thauera_sp._UWRP | FG147_05165 | 1..1865 | 2488794 | ATP-binding-protein,-partial-[Thauera-sp.-UWRP] |
| WP_159691715.1 | DUF499_1.RM.like.systems | MODE-HTH+PBPII-><-TPR<- DUF499 *<- REase<- OLD-ABC <- METHYLASE <- HNH<- OLD-ABC <- TUDOR+ SF1-SF2-Helicase+iREase+CTD+REase | 943 | Bacteria>Pseudomonadota>Betaproteobacteria>Rhodocyclales>Zoogloeaceae | GCF_009800965.1 | Cognatazoarcus_halotolerans | GQB03_RS16405 | 71425..74256 | 2686016 | DUF499-domain-containing-protein-[Cognatazoarcus-halotolerans] |
| WP_212351537.1 | DUF499_1.RM.like.systems | <-Phage_integrase\|\|HTH+WYL-> TUDOR+ SF1-SF2-Helicase+iREase+CTD+REase -> OLD-ABC ->HNH -> METHYLASE -> DUF499 *->?-><-?<-RelA | 954 | Bacteria>Pseudomonadota>Betaproteobacteria>Rhodocyclales>Zoogloeaceae | GCF_018122735.1 | Azoarcus_sp._L1K30 | J5J83_RS19190 | 4181146..4184010 | 2820277 | DUF499-domain-containing-protein-[Azoarcus-sp.-L1K30] |
| MDA8417189.1 | DUF499_1.RM.like.systems | TUDOR+ SF1-SF2-Helicase+iREase+CTD+REase ->RVT-> OLD-ABC ->HNH -> METHYLASE ->wHTH+REase-> DUF499 *-><-?\|\|?->?->?-><-Response_reg+LuxR-HTH | 946 | Bacteria>Pseudomonadota>Betaproteobacteria>unclassified_Betaproteobacteria | GCA_028069095.1 | Betaproteobacteria_bacterium | MOZ78_09075 | 11618..14458 | 1891241 | MAG:-DUF499-domain-containing-protein-[Betaproteobacteria-bacterium] |
| QWL60397.1 | DUF499_1.RM.like.systems | URI-DNase-><-PAIREDC-HTH\|\|?-> OLD-ABC ->HNH -> METHYLASE -> DUF499 *-><-?\|\|HTH-><-RlpA<-JAB | 954 | Bacteria>Pseudomonadota>Gammaproteobacteria>Aeromonadales>Aeromonadaceae | GCA_018802265.1 | Aeromonas_jandaei | HQ400_01350 | 291564..294428 | 650 | ATP-binding-protein-[Aeromonas-jandaei] |
| WP_103858986.1 | DUF499_1.RM.like.systems | JAB->CbeA_antitoxin->RlpA-><- DUF499 *<- TOPRIM <- OLD-ABC <- METHYLASE <- HNH<- OLD-ABC <- TUDOR+ SF1-SF2-Helicase+iREase+CTD+REase | 948 | Bacteria>Pseudomonadota>Gammaproteobacteria>Aeromonadales>Aeromonadaceae | GCF_002919855.1 | Aeromonas_sp._ASNIH8 | C3418_RS17745 | 3201..6047 | 1920113 | DUF499-domain-containing-protein-[Aeromonas-sp.-ASNIH8] |
| WP_201871420.1 | DUF499_1.RM.like.systems | JAB->CbeA_antitoxin->RlpA-><- DUF499 *<- TOPRIM <- OLD-ABC <- METHYLASE <- HNH<- OLD-ABC <- TUDOR+ SF1-SF2-Helicase+iREase+CTD+REase | 948 | Bacteria>Pseudomonadota>Gammaproteobacteria>Aeromonadales>Aeromonadaceae | GCF_016728905.1 | Aeromonas_caviae | JD551_RS01070 | 83459..86305 | 648 | DUF499-domain-containing-protein-[Aeromonas-caviae] |
| WP_216991887.1 | DUF499_1.RM.like.systems | TUDOR+ SF1-SF2-Helicase+iREase+CTD+REase -> OLD-ABC ->HNH -> METHYLASE -> OLD-ABC -> TOPRIM -> DUF499 *-><-RlpA<-CbeA_antitoxin<-JAB | 948 | Bacteria>Pseudomonadota>Gammaproteobacteria>Aeromonadales>Aeromonadaceae | GCF_019048725.1 | Aeromonas_sp._FDAARGOS_1407 | I6L37_RS11490 | 2395827..2398673 | 2778056 | DUF499-domain-containing-protein-[Aeromonas-sp.-FDAARGOS-1407] |
| WP_219296379.1 | DUF499_1.RM.like.systems | JAB->RlpA-><-HTH\|\|?-><- DUF499 *<- TOPRIM <- OLD-ABC <- METHYLASE <- HNH<- OLD-ABC <- TUDOR+ SF1-SF2-Helicase+iREase+CTD+REase | 952 | Bacteria>Pseudomonadota>Gammaproteobacteria>Aeromonadales>Aeromonadaceae | GCF_019348625.1 | Aeromonas_veronii | GL270_RS12655 | 107539..110397 | 654 | DUF499-domain-containing-protein-[Aeromonas-veronii] |
| WP_223918005.1 | DUF499_1.RM.like.systems | TUDOR+ SF1-SF2-Helicase+iREase+CTD+REase -> OLD-ABC ->HNH -> METHYLASE -> OLD-ABC -> TOPRIM -> DUF499 *-><-RlpA<-CbeA_antitoxin<-JAB | 948 | Bacteria>Pseudomonadota>Gammaproteobacteria>Aeromonadales>Aeromonadaceae | GCF_019971655.1 | Aeromonas_caviae | LDN21_RS02455 | 71531..74377 | 648 | DUF499-domain-containing-protein-[Aeromonas-caviae] |
| WP_223954701.1 | DUF499_1.RM.like.systems | JAB->CbeA_antitoxin->RlpA-><- DUF499 *<- REase<- OLD-ABC <- METHYLASE <- HNH<- OLD-ABC <- TUDOR+ SF1-SF2-Helicase+iREase+CTD+REase | 946 | Bacteria>Pseudomonadota>Gammaproteobacteria>Aeromonadales>Aeromonadaceae | GCF_019972775.1 | Aeromonas_caviae | LDN68_RS17105 | 3642842..3645682 | 648 | DUF499-domain-containing-protein-[Aeromonas-caviae] |

| Protein-ID | System | Gene-Neighbourhoods | Length | Taxonomy | Assembly-ID | Strain | Locus-Tag | Genomic-coordinates | TaxID | Fasta-defline |
|---|---|---|---|---|---|---|---|---|---|---|
| WP_235462717.1 | DUF499_1.RM.like.systems | TUDOR+ SF1-SF2-Helicase+iREase+CTD+REase -> OLD-ABC->HNH -> METHYLASE -> OLD-ABC -> TOPRIM -> DUF499 *-> | 897 | Bacteria>Pseudomonadota>Gammaproteobacteria>Aeromonadales>Aeromonadaceae | GCF_021538905.1 | Aeromonas_australiensis | EHZ86_RS11610 | 15978..18671 | 1114880 | DUF499-domain-containing-protein-[Aeromonas-australiensis] |
| WP_257709589.1 | DUF499_1.RM.like.systems | TUDOR+ SF1-SF2-Helicase+iREase+CTD+REase -> OLD-ABC->HNH -> METHYLASE -> OLD-ABC -> TOPRIM -> DUF499 *-><-RlpA<-CbeA_antitoxin<-JAB<-?\|\|<-Arm-DNA-bind_3+Phage_integrase | 948 | Bacteria>Pseudomonadota>Gammaproteobacteria>Aeromonadales>Aeromonadaceae | GCF_024649925.1 | Aeromonas_caviae | NUK49_RS00060 | 12274..15120 | 648 | DUF499-domain-containing-protein-[Aeromonas-caviae] |
| WP_270668501.1 | DUF499_1.RM.like.systems | <- DUF499 *<- METHYLASE <- HNH<- OLD-ABC <-?<- TUDOR+ SF1-SF2-Helicase+iREase+CTD+REase | 954 | Bacteria>Pseudomonadota>Gammaproteobacteria>Aeromonadales>Aeromonadaceae | GCF_027705205.1 | unclassified_Aeromonas | OKB91_RS16400 | 106..2970 | 257493 | MULTISPECIES:-DUF499-domain-containing-protein-[unclassified-Aeromonas] |
| WP_101201433.1 | DUF499_1.RM.like.systems | HTH+WYL-> TUDOR+ SF1-SF2-Helicase+iREase+CTD+REase -> OLD-ABC->HNH -> METHYLASE -> DUF499 *-><-RlpA<-?\|\|?->Phage_integrase-> | 944 | Bacteria>Pseudomonadota>Gammaproteobacteria>Alteromonadales>Alteromonadaceae | GCF_002835935.1 | Paraglaciecola_sp._MB-3u-78 | CXF95_RS26560 | 32531..35365 | 2058332 | DUF499-domain-containing-protein-[Paraglaciecola-sp.-MB-3u-78] |
| WP_250655185.1 | DUF499_1.RM.like.systems | HTH+WYL-> TUDOR+ SF1-SF2-Helicase+iREase+CTD+REase -> OLD-ABC->HNH -> TUDOR+ SF1-SF2-Helicase+iREase+CTD+REase -p-> METHYLASE -> DUF499 *->SMODS->?-><-RlpA<-Phage_lysozyme | 945 | Bacteria>Pseudomonadota>Gammaproteobacteria>Alteromonadales>Alteromonadaceae | GCF_023650515.1 | Alkalimarinus_coralli | MY525_RS07060 | 1557790..1560627 | 2935863 | MULTISPECIES:-DUF499-domain-containing-protein-[unclassified-Alkalimarinus] |
| BCV35016.1 | DUF499_1.RM.like.systems | <-PRTase<-RNase\|\|?->Arm-DNA-bind_3+Phage_integrase->RlpA-><- DUF499 *<- METHYLASE <- HNH<- OLD-ABC <- TUDOR+ SF1-SF2-Helicase+iREase+CTD+REase <-HTH_3 | 943 | Bacteria>Pseudomonadota>Gammaproteobacteria>Alteromonadales>Shewanellaceae | GCA_019670565.1 | Shewanella_chilikensis | TUM17377_03440 | 374214..377045 | 558541 | hypothetical-protein-TUM17377_03440-[Shewanella-chilikensis] |
| WP_186265255.1 | DUF499_1.RM.like.systems | HTH_3-> TUDOR+ SF1-SF2-Helicase+iREase+CTD+REase -> OLD-ABC->HNH -> METHYLASE -> DUF499 *-><-RlpA<-Arm-DNA-bind_3+Phage_integrase<-?\|\|RNase->PRTase-> | 943 | Bacteria>Pseudomonadota>Gammaproteobacteria>Alteromonadales>Shewanellaceae | GCF_014263185.1 | Shewanella_algae | HU689_RS02010 | 444450..447281 | 38313 | DUF499-domain-containing-protein-[Shewanella-algae] |
| WP_115016378.1 | DUF499_1.RM.like.systems | <-JAB\|\|XIS-HTH->REase-p->REase->RlpA-><- DUF499 *<- METHYLASE <- HNH<- OLD-ABC <- TUDOR+ SF1-SF2-Helicase+iREase+CTD+REase \|\|?->DDE_Tnp_1_3-> | 943 | Bacteria>Pseudomonadota>Gammaproteobacteria>Alteromonadales>Shewanellaceae | GCF_019266125.1 | Shewanella | DX079_RS11245 | 2521097..2523928 | 22 | MULTISPECIES:-DUF499-domain-containing-protein-[Shewanella] |
| WP_219013226.1 | DUF499_1.RM.like.systems | HTH_3-> TUDOR+ SF1-SF2-Helicase+iREase+CTD+REase -> OLD-ABC->HNH -> METHYLASE -> DUF499 *-><-RlpA<-Arm-DNA-bind_3+Phage_integrase<-?\|\|RNase->PRTase-> | 943 | Bacteria>Pseudomonadota>Gammaproteobacteria>Alteromonadales>Shewanellaceae | GCF_019303275.1 | Shewanella_algae | KOE54_RS14875 | 19508..22339 | 38313 | DUF499-domain-containing-protein-[Shewanella-algae] |
| WP_079541822.1 | DUF499_1.RM.like.systems | TUDOR+ SF1-SF2-Helicase+iREase+CTD+REase -> OLD-ABC->HNH -> METHYLASE -> OLD-ABC -> TOPRIM -> DUF499 *-> | 937 | Bacteria>Pseudomonadota>Gammaproteobacteria>Cardiobacteriales>Cardiobacteriaceae | GCF_900079085.1 | Cardiobacterium_hominis | CHUV0807_RS10775 | 37520..40333 | 2718 | DUF499-domain-containing-protein-[Cardiobacterium-hominis] |
| MTW22277.1 | DUF499_1.RM.like.systems | TUDOR+ SF1-SF2-Helicase+iREase+CTD+REase -> OLD-ABC->HNH -> OLD-ABC -> TOPRIM -> DUF499 *-> | 885 | Bacteria>Pseudomonadota>Gammaproteobacteria>Chromatiales>Chromatiaceae | GCA_009720725.1 | Allochromatium_palmeri | GJ668_14455 | 38860..41517 | 231048 | DUF499-domain-containing-protein-[Allochromatium-palmeri] |
| WP_155450846.1 | DUF499_1.RM.like.systems | OLD-ABC->HNH -> METHYLASE -p-> METHYLASE -p-> OLD-ABC -> TOPRIM -> DUF499 *-> | 883 | Bacteria>Pseudomonadota>Gammaproteobacteria>Chromatiales>Chromatiaceae | GCF_009720725.1 | Allochromatium_palmeri | GJ668_RS14460 | 38866..41517 | 231048 | DUF499-domain-containing-protein-[Allochromatium-palmeri] |
| EFO1856148.1 | DUF499_1.RM.like.systems | TUDOR+ SF1-SF2-Helicase+iREase+CTD+REase -> OLD-ABC->HNH -> METHYLASE -> DUF499 *-><-HTH_Tnp_1+HTH_21+rve+rve_2<-HTH_Tnp_1+HTH_21+rve+rve_2\|\|<-?<-?\|\|XIS-HTH->DUF3987-> | 947 | Bacteria>Pseudomonadota>Gammaproteobacteria>Enterobacterales>Enterobacteriaceae | GCA_013065125.1 | Escherichia_coli | DUZ37_17320 | 27712..30555 | 562 | DUF499-domain-containing-protein-[Escherichia-coli] |
| EGQ7342643.1 | DUF499_1.RM.like.systems | RlpA-><- DUF499 *<- METHYLASE <- HNH<- OLD-ABC <- TUDOR+ SF1-SF2-Helicase+iREase+CTD+REase <-?\|\|ParB-> | 947 | Bacteria>Pseudomonadota>Gammaproteobacteria>Enterobacterales>Enterobacteriaceae | GCA_015730265.1 | Enterobacter_cloacae | I6C89_001957 | 175814..178657 | 550 | ATP-binding-protein-[Enterobacter-cloacae] |
| HAU4285776.1 | DUF499_1.RM.like.systems | TUDOR+ SF1-SF2-Helicase+iREase+CTD+REase -> OLD-ABC->HNH -> METHYLASE -> DUF499 *-><-RlpA | 947 | Bacteria>Pseudomonadota>Gammaproteobacteria>Enterobacterales>Enterobacteriaceae | GCA_016065535.1 | Citrobacter_freundii | F6W46_24070 | 10411..13254 | 546 | ATP-binding-protein-[Citrobacter-freundii] |
| HAU5791339.1 | DUF499_1.RM.like.systems | Phage_ASH->?->?-> TOPRIM +DNAG+D5N+D5ATPase+P4PRIMASE-HTH->?->RlpA-><- DUF499 *<- METHYLASE <- HNH<- OLD-ABC <- TUDOR+ SF1-SF2-Helicase+iREase+CTD+REase | 947 | Bacteria>Pseudomonadota>Gammaproteobacteria>Enterobacterales>Enterobacteriaceae | GCA_016076255.1 | Citrobacter_freundii | JD367_15405 | 174239..177082 | 546 | ATP-binding-protein-[Citrobacter-freundii] |
| HAW1058251.1 | DUF499_1.RM.like.systems | <-nSTAND1+sGTP\|\|?-> TUDOR+ SF1-SF2-Helicase+iREase+CTD+REase -> OLD-ABC->HNH -> METHYLASE -> DUF499 *-><-DEDD_Tnp_IS110+Transposase_20 | 947 | Bacteria>Pseudomonadota>Gammaproteobacteria>Enterobacterales>Enterobacteriaceae | GCA_016672425.1 | Escherichia_coli | JLK83_004317 | 23199..26042 | 562 | ATP-binding-protein-[Escherichia-coli] |
| HBC9812127.1 | DUF499_1.RM.like.systems | TUDOR+ SF1-SF2-Helicase+iREase+CTD+REase -p-> OLD-ABC->HNH -> METHYLASE -> DUF499 *-><-RlpA | 951 | Bacteria>Pseudomonadota>Gammaproteobacteria>Enterobacterales>Enterobacteriaceae | GCA_018195115.1 | Escherichia_coli | KFP54_004194 | 5317..8172 | 562 | ATP-binding-protein-[Escherichia-coli] |
| HBW9592181.1 | DUF499_1.RM.like.systems | TUDOR+ SF1-SF2-Helicase+iREase+CTD+REase -> OLD-ABC->HNH -> METHYLASE -> DUF499 *-><-RlpA | 953 | Bacteria>Pseudomonadota>Gammaproteobacteria>Enterobacterales>Enterobacteriaceae | GCA_022207645.1 | Klebsiella_pneumoniae | MFS54_26490 | 9564..12425 | 573 | DUF499-domain-containing-protein-[Klebsiella-pneumoniae] |
| HCI7559692.1 | DUF499_1.RM.like.systems | <-HIN-HTH\|\|?-> TUDOR+ SF1-SF2-Helicase+iREase+CTD+REase -> OLD-ABC->HNH -> METHYLASE -> DUF499 *-><-RlpA<-?<-Arm-DNA-bind_3+Phage_integrase<-tRNA\|\|?->?->ADH_N+ADH_zinc_N-> | 947 | Bacteria>Pseudomonadota>Gammaproteobacteria>Enterobacterales>Enterobacteriaceae | GCA_024446695.1 | Klebsiella_pneumoniae | NPW42_004570 | 42168..45011 | 573 | ATP-binding-protein-[Klebsiella-pneumoniae] |
| MCQ4415112.1 | DUF499_1.RM.like.systems | Arm-DNA-bind_3+Phage_integrase->RlpA-><- DUF499 *<- METHYLASE <- HNH<- OLD-ABC <- TUDOR+ SF1-SF2-Helicase+iREase+CTD+REase | 947 | Bacteria>Pseudomonadota>Gammaproteobacteria>Enterobacterales>Enterobacteriaceae | GCA_024451305.1 | Enterobacter_kobei | NOX26_15660 | 3292884..3295727 | 208224 | DUF499-domain-containing-protein-[Enterobacter-kobei] |
| WP_045284468.1 | DUF499_1.RM.like.systems | RlpA-><- DUF499 *<- METHYLASE <- HNH<- OLD-ABC <- TUDOR+ SF1-SF2-Helicase+iREase+CTD+REase | 947 | Bacteria>Pseudomonadota>Gammaproteobacteria>Enterobacterales>Enterobacteriaceae | GCF_000958085.1 | Enterobacter_sichuanensis | SS37_RS01295 | 119849..122692 | 2071710 | DUF499-domain-containing-protein-[Enterobacter-sichuanensis] |
| WP_086660584.1 | DUF499_1.RM.like.systems | <-nSTAND1+sGTP\|\|?-> TUDOR+ SF1-SF2-Helicase+iREase+CTD+REase -> OLD-ABC->HNH -> METHYLASE -> DUF499 *->\|\|?->CbeA_antitoxin->?-><-tRNA<-Aldedh | 947 | Bacteria>Pseudomonadota>Gammaproteobacteria>Enterobacterales>Enterobacteriaceae | GCF_014216435.1 | Escherichia_coli | GVI19_RS16185 | 90398..93241 | 562 | DUF499-domain-containing-protein-[Escherichia-coli] |
| WP_196359062.1 | DUF499_1.RM.like.systems | RlpA-><- DUF499 *<- METHYLASE <- HNH<- OLD-ABC <- TUDOR+ SF1-SF2-Helicase+iREase+CTD+REase <-?\|\|ParB-> | 947 | Bacteria>Pseudomonadota>Gammaproteobacteria>Enterobacterales>Enterobacteriaceae | GCF_015701305.1 | Enterobacter_hormaechei | I4P56_RS16415 | 165030..167873 | 158836 | DUF499-domain-containing-protein-[Enterobacter-hormaechei] |
| WP_214577408.1 | DUF499_1.RM.like.systems | RlpA-><- DUF499 *<- METHYLASE <- HNH<- OLD-ABC <- TUDOR+ SF1-SF2-Helicase+iREase+CTD+REase | 947 | Bacteria>Pseudomonadota>Gammaproteobacteria>Enterobacterales>Enterobacteriaceae | GCF_018597265.1 | Enterobacter_dykesii | KK116_RS19460 | 158324..161167 | 2797506 | DUF499-domain-containing-protein-[Enterobacter-dykesii] |
| WP_222563661.1 | DUF499_1.RM.like.systems | Arm-DNA-bind_3+Phage_integrase->RlpA-><- DUF499 *<- METHYLASE <- HNH<- OLD-ABC <- TUDOR+ SF1-SF2-Helicase+iREase+CTD+REase | 947 | Bacteria>Pseudomonadota>Gammaproteobacteria>Enterobacterales>Enterobacteriaceae | GCF_019803045.1 | Citrobacter_farmeri | Cf24236_RS16360 | 3446361..3449204 | 67824 | MULTISPECIES:-DUF499-domain-containing-protein-[Enterobacteriaceae] |
| WP_048997346.1 | DUF499_1.RM.like.systems | RlpA-><- DUF499 *<- METHYLASE <- HNH<- OLD-ABC <- TUDOR+ SF1-SF2-Helicase+iREase+CTD+REase | 947 | Bacteria>Pseudomonadota>Gammaproteobacteria>Enterobacterales>Enterobacteriaceae | GCF_021869685.1 | Enterobacter | L3068_RS02450 | 520532..523375 | 547 | MULTISPECIES:-DUF499-domain-containing-protein-[Enterobacter] |
| WP_256043565.1 | DUF499_1.RM.like.systems | Arm-DNA-bind_3+Phage_integrase->RlpA-><- DUF499 *<- METHYLASE <- HNH<- OLD-ABC <- TUDOR+ SF1-SF2-Helicase+iREase+CTD+REase | 947 | Bacteria>Pseudomonadota>Gammaproteobacteria>Enterobacterales>Enterobacteriaceae | GCF_024451305.1 | Enterobacter_kobei | NOX26_RS15660 | 3292884..3295727 | 208224 | DUF499-domain-containing-protein-[Enterobacter-kobei] |
| WP_193132801.1 | DUF499_1.RM.like.systems | HTH->?->HEPN->RlpA-><- DUF499 *<- METHYLASE <- HNH<- OLD-ABC <- TUDOR+ SF1-SF2-Helicase+iREase+CTD+REase <-?\|\|ParB-> | 942 | Bacteria>Pseudomonadota>Gammaproteobacteria>Enterobacterales>Enterobacteriaceae | GCF_032369155.1 | Enterobacter_hormaechei | CXM24_RS20100 | 4287724..4290552 | 158836 | DUF499-domain-containing-protein-[Enterobacter-hormaechei] |
| WP_086872901.1 | DUF499_1.RM.like.systems | TUDOR+ SF1-SF2-Helicase+iREase+CTD+REase -> OLD-ABC->HNH -> METHYLASE -> DUF499 *-><-RlpA<-Arm-DNA-bind_3+Phage_integrase<-tRNA\|\|REC+LuxR-HTH-> | 953 | Bacteria>Pseudomonadota>Gammaproteobacteria>Enterobacterales>Enterobacteriaceae | GCF_900184035.1 | Kosakonia_pseudosacchari | BN2591_RS15655 | 26880..29741 | 1646340 | DUF499-domain-containing-protein-[Kosakonia-pseudosacchari] |
| WP_008325059.1 | DUF499_1.RM.like.systems | TUDOR+ SF1-SF2-Helicase+iREase+CTD+REase -> OLD-ABC->HNH -> METHYLASE -> DUF499 *-><-RlpA | 947 | Bacteria>Pseudomonadota>Gammaproteobacteria>Enterobacterales | GCF_000277565.1 | Enterobacteriaceae | B397_RS01195 | 10694..13537 | 543 | MULTISPECIES:-DUF499-domain-containing-protein-[Enterobacteriaceae] |
| WP_220756270.1 | DUF499_1.RM.like.systems | TUDOR+ SF1-SF2-Helicase+iREase+CTD+REase -> OLD-ABC->HNH -> METHYLASE -> DUF499 *-><-RlpA<-RVT+HNH<-?<-ParB+HNH<-ParB+HNH | 947 | Bacteria>Pseudomonadota>Gammaproteobacteria>Enterobacterales | GCF_019655755.1 | Enterobacteriaceae | K5R72_RS10955 | 43358..46201 | 543 | DUF499-domain-containing-protein-[Escherichia-coli] |
| WP_077999168.1 | DUF499_1.RM.like.systems | JAB->RlpA-><- DUF499 *<- REase<- OLD-ABC <- METHYLASE <- HNH<- OLD-ABC <- TUDOR+ SF1-SF2-Helicase+iREase+CTD+REase | 949 | Bacteria>Pseudomonadota>Gammaproteobacteria>Enterobacterales>Hafniaceae | GCF_002006945.1 | Edwardsiella_tarda | BTN58_RS02535 | 130022..132871 | 636 | DUF499-domain-containing-protein-[Edwardsiella-tarda] |
| WP_205545050.1 | DUF499_1.RM.like.systems | <-PAIREDC-HTH+HIN-HTH\|\|?-> TUDOR+ SF1-SF2-Helicase+iREase+CTD+REase -> OLD-ABC->HNH -> METHYLASE -> DUF499 *-><-?<-HNH<-?<-DHTH+WYL\|\|nSTAND1+sGTP->HTH+WYL-> | 951 | Bacteria>Pseudomonadota>Gammaproteobacteria>Enterobacterales>Pectobacteriaceae | GCF_016949795.1 | Pectobacterium_brasiliense | H4F47_RS10595 | 47611..50466 | 180957 | DUF499-domain-containing-protein-[Pectobacterium-brasiliense] |
| SUI50248.1 | DUF499_1.RM.like.systems | GDResolvase+SIGMA-HTH->?->SF1-SF2-Helicase->RlpA-><- DUF499 *<- METHYLASE <- HNH<- OLD-ABC <- TUDOR+ SF1-SF2-Helicase+iREase+CTD+REase | 975 | Bacteria>Pseudomonadota>Gammaproteobacteria>Enterobacterales>Yersiniaceae | GCA_900457075.1 | Serratia_quinivorans | NCTC11544_01113 | 1025353..1028280 | 137545 | Protein-of-uncharacterised-function-(DUF499)-[Serratia-quinivorans] |
| CAI1753142.1 | DUF499_1.RM.like.systems | TUDOR+ SF1-SF2-Helicase+iREase+CTD+REase -> OLD-ABC->HNH -> METHYLASE -> DUF499 *-><-RlpA<-SF1-SF2-Helicase<-?<-GDResolvase+SIGMA-HTH<-?<-XIS-HTH | 975 | Bacteria>Pseudomonadota>Gammaproteobacteria>Enterobacterales>Yersiniaceae | GCA_946228205.1 | Serratia_quinivorans | SAMEA4857275_02169 | 318450..321377 | 137545 | Protein-of-uncharacterised-function-(DUF499)-[Serratia-quinivorans] |
| WP_115183101.1 | DUF499_1.RM.like.systems | <-nSTAND1+sGTP\|\|?-> TUDOR+ SF1-SF2-Helicase+iREase+CTD+REase -> OLD-ABC->HNH -> METHYLASE -> DUF499 *-><-RlpA<-SF1-SF2-Helicase<-?<-GDResolvase+SIGMA-HTH<-?<-XIS-HTH | 951 | Bacteria>Pseudomonadota>Gammaproteobacteria>Enterobacterales>Yersiniaceae | GCF_946228205.1 | Serratia | BSR03_RS13425 | 143107..145962 | 613 | MULTISPECIES:-DUF499-domain-containing-protein-[Serratia] |
| PZN71299.1 | DUF499_1.RM.like.systems | <- DUF499 *<-PIN<- METHYLASE <- HNH<- OLD-ABC <-PemK_toxin<-PrlF_antitoxin | 955 | Bacteria>Pseudomonadota>Gammaproteobacteria>Methylococcales>Methylococcales_incertae_sedis | GCA_003242955.1 | Candidatus_Methylumidiphilus_alinenensis | DM484_26720 | 13367..16234 | 2202197 | hypothetical-protein-DM484_26720-[Candidatus-Methylumidiphilus-alinenensis] |
| WP_142947823.1 | DUF499_1.RM.like.systems | Arm-DNA-bind_3+Phage_integrase-><-Phage_AlpA<-ZnR+ TOPRIM +primpol+HTH<-XIS-HTH\|\|MobI-><- DUF499 *<-Schlafen+HATPase+HTH<- METHYLASE <- METHYLASE <- HNH<- OLD-ABC <- TUDOR+ SF1-SF2-Helicase+iREase+CTD+REase | 953 | Bacteria>Pseudomonadota>Gammaproteobacteria>Oceanospirillales>Alcanivoracaceae | - | Alcanivorax_sp._DSM_26295 | FN816_RS03075 | 107194..110055 | 1938738 | DUF499-domain-containing-protein-[Alcanivorax-sp.-DSM-26295] |
| MCP5162750.1 | DUF499_1.RM.like.systems | <-ABC-ATPase+SbcC<-?<- DUF499 *<- METHYLASE <- HNH<- OLD-ABC <- TUDOR+ SF1-SF2-Helicase+iREase+CTD+REase <-HTH+WYL | 944 | Bacteria>Pseudomonadota>Gammaproteobacteria>Oceanospirillales>Hahellaceae | GCA_024234505.1 | Hahellaceae_bacterium | H7A00_13970 | 74140..76974 | 2048971 | ATP-binding-protein-[Hahellaceae-bacterium] |
| WP_239285448.1 | DUF499_1.RM.like.systems | TUDOR+ SF1-SF2-Helicase+iREase+CTD+REase -> OLD-ABC->HNH -> METHYLASE ->ParB+HNH-> DUF499 *->HTH_Tnp_1+HTH_21+rve+rve_3->?->?->AbiEi-><-CHTH<-RelE | 962 | Bacteria>Pseudomonadota>Gammaproteobacteria>Oceanospirillales>Halomonadaceae | GCF_022348135.1 | unclassified_Halomonas | MHM85_RS05495 | 207037..209925 | 2609666 | MULTISPECIES:-DUF499-domain-containing-protein-[unclassified-Halomonas] |
| WP_137094585.1 | DUF499_1.RM.like.systems | TUDOR+ SF1-SF2-Helicase+iREase+CTD+REase -> OLD-ABC->HNH -> METHYLASE -> OLD-ABC ->REase -> DUF499 *-><-MobI\|\|XIS-HTH->ZnR+ TOPRIM +primpol+HTH->Phage_AlpA-><-Arm-DNA-bind_3+Phage_integrase | 953 | Bacteria>Pseudomonadota>Gammaproteobacteria>Oceanospirillales>Halomonadaceae | GCF_028580285.1 | Halomonas | E8Q34_RS13770 | 13695..16556 | 2745 | DUF499-domain-containing-protein-[Halomonas-sp.-15WGF] |
| WP_284453091.1 | DUF499_1.RM.like.systems | Arm-DNA-bind_3+Phage_integrase-><-Phage_AlpA<-ZnR+ TOPRIM +primpol+HTH<-XIS-HTH\|\|MobI-><- DUF499 *<- REase<- OLD-ABC <- METHYLASE <- HNH<- OLD-ABC <- TUDOR+ SF1-SF2-Helicase+iREase+CTD+REase | 953 | Bacteria>Pseudomonadota>Gammaproteobacteria>Oceanospirillales>Halomonadaceae | GCF_030145725.1 | Halomonas_meridiana | KYX60_RS01235 | 106598..109459 | 29570 | DUF499-domain-containing-protein-[Halomonas-meridiana] |
| WP_072841218.1 | DUF499_1.RM.like.systems | TUDOR+ SF1-SF2-Helicase+iREase+CTD+REase -> OLD-ABC->HNH -> METHYLASE -> OLD-ABC ->REase -> DUF499 *->SMODS->?-><-RlpA\|\|?-><-HTH_26 | 942 | Bacteria>Pseudomonadota>Gammaproteobacteria>Oceanospirillales>Oceanospirillaceae | GCF_900129155.1 | Marinomonas_polaris | BUA59_RS18830 | 76459..79287 | 293552 | DUF499-domain-containing-protein-[Marinomonas-polaris] |
| WP_116685595.1 | DUF499_1.RM.like.systems | HTH+WYL-> TUDOR+ SF1-SF2-Helicase+iREase+CTD+REase -> OLD-ABC->HNH -> METHYLASE -> DUF499 *-><-?<-?\|\|?->TIR-> | 935 | Bacteria>Pseudomonadota>Gammaproteobacteria>Oceanospirillales>Oceanospirillales_incertae_sedis | GCF_003097315.1 | Pelagibaculum_spongiae | DC094_RS03025 | 690947..693754 | 2080658 | DUF499-domain-containing-protein-[Pelagibaculum-spongiae] |
| AKV97706.1 | DUF499_1.RM.like.systems | ParB->?->REase->?->DDE_Tnp_1_4->VirB4-FtsK+HerA-><- DUF499 *<- METHYLASE -p\|\|?-><- HNH<- OLD-ABC <- TUDOR+ SF1-SF2-Helicase+iREase+CTD+REase <-XIS-HTH | 951 | Bacteria>Pseudomonadota>Gammaproteobacteria>Pseudomonadales>Marinobacteraceae | GCA_001266795.1 | Marinobacter_sp._CP1 | ACP86_16945 | 3679225..3682080 | 1671721 | hypothetical-protein-ACP86_16945-[Marinobacter-sp.-CP1] |
| WP_200905073.1 | DUF499_1.RM.like.systems | REase->?->DDE_Tnp_1_4->?->VirB4-FtsK+HerA-><- DUF499 *<- METHYLASE -p\|\|?-><- METHYLASE <- HNH<- OLD-ABC <- TUDOR+ SF1-SF2-Helicase+iREase+CTD+REase | 935 | Bacteria>Pseudomonadota>Gammaproteobacteria>Pseudomonadales>Marinobacteraceae | GCF_001266795.1 | Marinobacter_sp._CP1 | ACP86_RS16950 | 3679225..3682032 | 1671721 | DUF499-domain-containing-protein-[Marinobacter-sp.-CP1] |
| WP_098424145.1 | DUF499_1.RM.like.systems | XIS-HTH-> TUDOR+ SF1-SF2-Helicase+iREase+CTD+REase -> OLD-ABC->HNH -> METHYLASE ->ParB+HNH-> DUF499 *->GDResolvase+HIN-HTH-><-?<-REase<-HNH | 946 | Bacteria>Pseudomonadota>Gammaproteobacteria>Pseudomonadales>Marinobacteraceae | GCF_002563815.1 | Marinobacter_sp._LV10R520-4 | ATG98_RS19590 | 4036541..4039381 | 1761796 | DUF499-domain-containing-protein-[Marinobacter-sp.-LV10R520-4] |
| WP_098418633.1 | DUF499_1.RM.like.systems | XIS-HTH-> TUDOR+ SF1-SF2-Helicase+iREase+CTD+REase -> OLD-ABC->HNH -> METHYLASE ->ParB+HNH-> DUF499 *->GDResolvase+HIN-HTH-><-?<-REase<-HNH<-?\|\|HIN-HTH-> | 946 | Bacteria>Pseudomonadota>Gammaproteobacteria>Pseudomonadales>Marinobacteraceae | GCF_002563885.1 | Marinobacter_sp._LV10MA510-1 | ATI45_RS05630 | 1155568..1158408 | 1415567 | DUF499-domain-containing-protein-[Marinobacter-sp.-LV10MA510-1] |
| WP_091985110.1 | DUF499_1.RM.like.systems | TUDOR+ SF1-SF2-Helicase+iREase+CTD+REase -> OLD-ABC->HNH -> METHYLASE ->MAD-NTD+EVE+McrB-AAA->wHTH+REase-> DUF499 *->SAM-methylase->HTH+HNH-><-?<-CHTH | 947 | Bacteria>Pseudomonadota>Gammaproteobacteria>Pseudomonadales>Marinobacteraceae | GCF_900115175.1 | Marinobacter_gudaonensis | BM344_RS01255 | 264147..266990 | 375760 | DUF499-domain-containing-protein-[Marinobacter-gudaonensis] |
| MBI8725677.1 | DUF499_1.RM.like.systems | TUDOR+ SF1-SF2-Helicase+iREase+CTD+REase -> OLD-ABC->HNH -> METHYLASE ->SF1-SF2-Helicase+wHTH+PRTase->?-> DUF499 *-><-RlpA<-?<-?\|\|HTH_3-><-ZnR+ZnR | 952 | Bacteria>Pseudomonadota>Gammaproteobacteria>Pseudomonadales>Pseudomonadaceae | GCA_016342885.1 | Pseudomonas_aeruginosa | JEZ69_03815 | 204270..207128 | 287 | ATP-binding-protein-[Pseudomonas-aeruginosa] |
| HCL3757805.1 | DUF499_1.RM.like.systems | TUDOR+ SF1-SF2-Helicase+iREase+CTD+REase -> OLD-ABC->HNH -> METHYLASE -> OLD-ABC -> TOPRIM -> DUF499 *-> | 667 | Bacteria>Pseudomonadota>Gammaproteobacteria>Pseudomonadales>Pseudomonadaceae | GCA_025003865.1 | Pseudomonas_aeruginosa | N2B17_005733 | 10859..12860 | 287 | ATP-binding-protein-[Pseudomonas-aeruginosa] |
| WCY08688.1 | DUF499_1.RM.like.systems | TUDOR+ SF1-SF2-Helicase+iREase+CTD+REase -> OLD-ABC->HNH ->SF1-SF2-Helicase+wHTH+PRTase->?-> DUF499 *-><-RlpA<-?<-JAB<-ParB<-?\|\|MazE_antitoxin-> | 943 | Bacteria>Pseudomonadota>Gammaproteobacteria>Pseudomonadales>Pseudomonadaceae | GCA_028609905.1 | Pseudomonas_aeruginosa | KK229_11705 | 2446793..2449624 | 287 | DUF499-domain-containing-protein-[Pseudomonas-aeruginosa] |
| WP_015271883.1 | DUF499_1.RM.like.systems | DUF932->?->ParB->JAB-><-?\|\|RlpA-><- DUF499 *<-?<-SF1-SF2-Helicase+wHTH+PRTase<- METHYLASE <- HNH<- OLD-ABC <- TUDOR+ SF1-SF2-Helicase+iREase+CTD+REase | 943 | Bacteria>Pseudomonadota>Gammaproteobacteria>Pseudomonadales>Pseudomonadaceae | GCF_000325725.1 | Pseudomonas | LU639_RS23750 | 5103653..5106484 | 286 | MULTISPECIES:-DUF499-domain-containing-protein-[Pseudomonas] |
| WP_033888172.1 | DUF499_1.RM.like.systems | TUDOR+ SF1-SF2-Helicase+iREase+CTD+REase -> OLD-ABC->HNH -> METHYLASE ->SF1-SF2-Helicase+wHTH+PRTase->?-> DUF499 *-><-RlpA<-SymE_toxin<-?<-ZNR+Pkinase+SF1-SF2-Helicase+REase | 942 | Bacteria>Pseudomonadota>Gammaproteobacteria>Pseudomonadales>Pseudomonadaceae | GCF_000506025.1 | Pseudomonas_aeruginosa | HB728_RS10975 | 2312198..2315026 | 287 | DUF499-domain-containing-protein-[Pseudomonas-aeruginosa] |
| WP_031691594.1 | DUF499_1.RM.like.systems | ZnR+ZnR-><-HTH_3\|\|?->?->RlpA-><- DUF499 *<-?<-SF1-SF2-Helicase+wHTH+PRTase<- METHYLASE <- HNH<- OLD-ABC <- TUDOR+ SF1-SF2-Helicase+iREase+CTD+REase | 954 | Bacteria>Pseudomonadota>Gammaproteobacteria>Pseudomonadales>Pseudomonadaceae | GCF_000629125.1 | Pseudomonas_aeruginosa | AJ70_RS18015 | 30417..33281 | 287 | DUF499-domain-containing-protein-[Pseudomonas-aeruginosa] |
| WP_033983017.1 | DUF499_1.RM.like.systems | TUDOR+ SF1-SF2-Helicase+iREase+CTD+REase -> OLD-ABC->HNH -> METHYLASE ->SF1-SF2-Helicase+wHTH+PRTase->?-> DUF499 *-><-RlpA<-?<-JAB<-ParB<-?\|\|MazE_antitoxin-> | 943 | Bacteria>Pseudomonadota>Gammaproteobacteria>Pseudomonadales>Pseudomonadaceae | GCF_000789805.1 | Pseudomonas_aeruginosa | NR66_RS02500 | 146772..149603 | 287 | DUF499-domain-containing-protein-[Pseudomonas-aeruginosa] |
| WP_064616121.1 | DUF499_1.RM.like.systems | HTH+WYL-> TUDOR+ SF1-SF2-Helicase+iREase+CTD+REase -> OLD-ABC->HNH -> METHYLASE -> DUF499 *-><-RlpA<-Phage_int_SAM<-Integrase_1<-?<- TOPRIM | 931 | Bacteria>Pseudomonadota>Gammaproteobacteria>Pseudomonadales>Pseudomonadaceae | GCF_001655615.1 | Pseudomonas_sp._GR_6-02 | PGR6_RS03540 | 827904..830699 | 1659194 | DUF499-domain-containing-protein-[Pseudomonas-sp.-GR-6-02] |
| WP_104882974.1 | DUF499_1.RM.like.systems | DUF932->?->ParB->JAB->RlpA-><- DUF499 *<-?<-SF1-SF2-Helicase+wHTH+PRTase<- METHYLASE <- HNH<- OLD-ABC <- TUDOR+ SF1-SF2-Helicase+iREase+CTD+REase | 943 | Bacteria>Pseudomonadota>Gammaproteobacteria>Pseudomonadales>Pseudomonadaceae | GCF_002947915.1 | Pseudomonas_sp._SWI44 | C4Q26_RS08410 | 1754987..1757818 | 2083053 | DUF499-domain-containing-protein-[Pseudomonas-sp.-SWI44] |
| WP_126634673.1 | DUF499_1.RM.like.systems | <-DDE_Tnp_1\|\|?-><-LZ_Tnp_IS66+zf-IS66+DDE_Tnp_IS66+DDE_Tnp_IS66_C<-?<-HIN-HTH\|\|RlpA-><- DUF499 *<-?<-SF1-SF2-Helicase+wHTH+PRTase<- METHYLASE <- HNH<- OLD-ABC <- TUDOR+ SF1-SF2-Helicase+iREase+CTD+REase | 943 | Bacteria>Pseudomonadota>Gammaproteobacteria>Pseudomonadales>Pseudomonadaceae | GCF_003970625.1 | Pseudomonas_aeruginosa | E5D53_RS31320 | 11116..13947 | 287 | DUF499-domain-containing-protein-[Pseudomonas-aeruginosa] |
| WP_131349395.1 | DUF499_1.RM.like.systems | TUDOR+ SF1-SF2-Helicase+iREase+CTD+REase -> OLD-ABC->HNH -> METHYLASE -> OLD-ABC ->REase -> DUF499 *-><-RlpA<-TPR<-?<-?\|\|?-><-Mbetalac | 929 | Bacteria>Pseudomonadota>Gammaproteobacteria>Pseudomonadales>Pseudomonadaceae | GCF_004327885.1 | Azotobacter_chroococcum | E0E52_RS08250 | 59995..62784 | 353 | DUF499-domain-containing-protein-[Azotobacter-chroococcum] |
| WP_135265733.1 | DUF499_1.RM.like.systems | TUDOR+ SF1-SF2-Helicase+iREase+CTD+REase -> OLD-ABC->HNH -> METHYLASE ->SF1-SF2-Helicase+wHTH+PRTase->?-> DUF499 *-><-RlpA | 943 | Bacteria>Pseudomonadota>Gammaproteobacteria>Pseudomonadales>Pseudomonadaceae | GCF_004683755.1 | Stutzerimonas_stutzeri | AK6_RS01615 | 334422..337253 | 316 | DUF499-domain-containing-protein-[Stutzerimonas-stutzeri] |
| WP_146538006.1 | DUF499_1.RM.like.systems | TUDOR+ SF1-SF2-Helicase+iREase+CTD+REase -> OLD-ABC->HNH -> METHYLASE ->SF1-SF2-Helicase+wHTH+PRTase->?-> DUF499 *-><-RlpA | 942 | Bacteria>Pseudomonadota>Gammaproteobacteria>Pseudomonadales>Pseudomonadaceae | GCF_007896855.1 | Pseudomonas_aeruginosa | FQA91_RS32330 | 37996..40824 | 287 | DUF499-domain-containing-protein-[Pseudomonas-aeruginosa] |
| WP_151883645.1 | DUF499_1.RM.like.systems | TUDOR+ SF1-SF2-Helicase+iREase+CTD+REase -> OLD-ABC->HNH -> METHYLASE -> DUF499 *-><-RlpA\|\|?-><-JAB<-ParB<-?<-DUF932 | 943 | Bacteria>Pseudomonadota>Gammaproteobacteria>Pseudomonadales>Pseudomonadaceae | GCF_009012945.1 | Pseudomonas | GB470_RS16305 | 7805..10636 | 286 | MULTISPECIES:-DUF499-domain-containing-protein-[Pseudomonas] |
| WP_165892256.1 | DUF499_1.RM.like.systems | TPR->RlpA-><- DUF499 *<-ParB+HNH<- METHYLASE <- HNH<- OLD-ABC <- TUDOR+ SF1-SF2-Helicase+iREase+CTD+REase | 933 | Bacteria>Pseudomonadota>Gammaproteobacteria>Pseudomonadales>Pseudomonadaceae | GCF_011470035.1 | Azotobacter_chroococcum | HA520_RS07840 | 19913..22714 | 353 | DUF499-domain-containing-protein-[Azotobacter-chroococcum] |
| WP_103518020.1 | DUF499_1.RM.like.systems | <-HTH+WYL\|\|?-> TUDOR+ SF1-SF2-Helicase+iREase+CTD+REase -> OLD-ABC->HNH -> METHYLASE -> DUF499 *-><-RlpA\|\|?-><-JAB<-ParB<-?<-DUF932 | 942 | Bacteria>Pseudomonadota>Gammaproteobacteria>Pseudomonadales>Pseudomonadaceae | GCF_013305665.1 | Pseudomonas | CW309_RS13950 | 90223..93051 | 286 | MULTISPECIES:-DUF499-domain-containing-protein-[Pseudomonas] |
| WP_093986953.1 | DUF499_1.RM.like.systems | TUDOR+ SF1-SF2-Helicase+iREase+CTD+REase -> OLD-ABC->HNH -> METHYLASE ->SF1-SF2-Helicase+wHTH+PRTase->?-> DUF499 *-><-RlpA<-JAB<-ParB | 943 | Bacteria>Pseudomonadota>Gammaproteobacteria>Pseudomonadales>Pseudomonadaceae | GCF_014635625.1 | Pseudomonas_fluvialis | CFY91_RS13770 | 27997..30828 | 1793966 | DUF499-domain-containing-protein-[Pseudomonas-fluvialis] |
| WP_192907661.1 | DUF499_1.RM.like.systems | TUDOR+ SF1-SF2-Helicase+iREase+CTD+REase -> OLD-ABC->HNH -> METHYLASE ->HEPN->?-> DUF499 *-><-RlpA\|\|SF1-SF2-helicase+RdgC+Beta-barrel+REase-><-REase<-Phage_int_SAM<-Integrase_1<-Arm-DNA-bind_3+Phage_integrase | 941 | Bacteria>Pseudomonadota>Gammaproteobacteria>Pseudomonadales>Pseudomonadaceae | GCF_014883875.1 | Pseudomonas_taiwanensis | ICN73_RS10045 | 2229206..2232031 | 470150 | DUF499-domain-containing-protein-[Pseudomonas-taiwanensis] |
| WP_232392569.1 | DUF499_1.RM.like.systems | DUF932->?->ParB->JAB->RlpA-><- DUF499 *<- METHYLASE <- HNH<- OLD-ABC <- TUDOR+ SF1-SF2-Helicase+iREase+CTD+REase -p<- TUDOR+ SF1-SF2-Helicase+iREase+CTD+REase <-HTH+WYL | 921 | Bacteria>Pseudomonadota>Gammaproteobacteria>Pseudomonadales>Pseudomonadaceae | GCF_021228675.1 | Pseudomonas_sp._NIBR-H-19 | LS633_RS06540 | 1376164..1378929 | 2901380 | DUF499-domain-containing-protein-[Pseudomonas-sp.-NIBR-H-19] |
| WP_232885556.1 | DUF499_1.RM.like.systems | <-HTH+WYL\|\|?-> TUDOR+ SF1-SF2-Helicase+iREase+CTD+REase -> OLD-ABC->HNH -> METHYLASE -> DUF499 *-><-RlpA\|\|?-><-JAB<-ParB<-?<-DUF932 | 943 | Bacteria>Pseudomonadota>Gammaproteobacteria>Pseudomonadales>Pseudomonadaceae | GCF_021282765.1 | Pseudomonas | QK328_RS05450 | 308623..311454 | 286 | DUF499-domain-containing-protein-[Pseudomonas-sp.-NMI795_08] |
| WP_232896897.1 | DUF499_1.RM.like.systems | DUF932->?->ParB->JAB-><-?\|\|RlpA-><- DUF499 *<-?<-SF1-SF2-Helicase+wHTH+PRTase<- METHYLASE <- HNH<- OLD-ABC <- TUDOR+ SF1-SF2-Helicase+iREase+CTD+REase | 928 | Bacteria>Pseudomonadota>Gammaproteobacteria>Pseudomonadales>Pseudomonadaceae | GCF_021283465.2 | Pseudomonas | LU687_RS25005 | 5299214..5302000 | 286 | MULTISPECIES:-DUF499-domain-containing-protein-[Pseudomonas] |
| WP_213870614.1 | DUF499_1.RM.like.systems | JAB->?-><-HTH_3\|\|?->?->RlpA-><- DUF499 *<-HEPN<- METHYLASE <- HNH<- OLD-ABC <- TUDOR+ SF1-SF2-Helicase+iREase+CTD+REase | 952 | Bacteria>Pseudomonadota>Gammaproteobacteria>Pseudomonadales>Pseudomonadaceae | GCF_029201245.1 | Pseudomonas_aeruginosa | KKT79_RS19565 | 32215..35073 | 287 | DUF499-domain-containing-protein-[Pseudomonas-aeruginosa] |
| WP_276226492.1 | DUF499_1.RM.like.systems | TUDOR+ SF1-SF2-Helicase+iREase+CTD+REase -> OLD-ABC->HNH -> METHYLASE ->SF1-SF2-Helicase+wHTH+PRTase->?-> DUF499 *-><-RlpA\|\|<- METHYLASE \|\|HTH->SbcC-> | 939 | Bacteria>Pseudomonadota>Gammaproteobacteria>Pseudomonadales>Pseudomonadaceae | GCF_029335315.1 | Pseudomonas_citronellolis | P3W75_RS06225 | 49283..52102 | 53408 | DUF499-domain-containing-protein-[Pseudomonas-citronellolis] |
| WP_128722906.1 | DUF499_1.RM.like.systems | TUDOR+ SF1-SF2-Helicase+iREase+CTD+REase -> OLD-ABC->HNH -> METHYLASE ->SF1-SF2-Helicase+wHTH+PRTase->?-> DUF499 *-><-RlpA<-SymE_toxin<-?<-ZNR+Pkinase+SF1-SF2-Helicase+REase | 942 | Bacteria>Pseudomonadota>Gammaproteobacteria>Pseudomonadales>Pseudomonadaceae | GCF_029954295.1 | Pseudomonas_aeruginosa | OLM70_RS05770 | 1186505..1189333 | 287 | DUF499-domain-containing-protein-[Pseudomonas-aeruginosa] |
| WP_285534180.1 | DUF499_1.RM.like.systems | ZnR+ZnR-><-HTH_3\|\|?->?->RlpA-><- DUF499 *<-?<-SF1-SF2-Helicase+wHTH+PRTase<- METHYLASE <- HNH<- OLD-ABC <- TUDOR+ SF1-SF2-Helicase+iREase+CTD+REase | 942 | Bacteria>Pseudomonadota>Gammaproteobacteria>Pseudomonadales>Pseudomonadaceae | GCF_030269585.1 | Stutzerimonas_stutzeri | QSJ82_RS21895 | 7754..10582 | 316 | DUF499-domain-containing-protein-[Stutzerimonas-stutzeri] |
| WP_124078188.1 | DUF499_1.RM.like.systems | ZnR+ZnR-><-HTH_3\|\|?->?->RlpA-><- DUF499 *<- TOPRIM <- OLD-ABC <- METHYLASE <- HNH<- OLD-ABC <- TUDOR+ SF1-SF2-Helicase+iREase+CTD+REase | 943 | Bacteria>Pseudomonadota>Gammaproteobacteria>Pseudomonadales>Pseudomonadaceae | GCF_033140865.1 | Pseudomonas_aeruginosa | IPC1239_RS13835 | 644387..647218 | 287 | DUF499-domain-containing-protein-[Pseudomonas-aeruginosa] |
| WP_090941615.1 | DUF499_1.RM.like.systems | TUDOR+ SF1-SF2-Helicase+iREase+CTD+REase -> OLD-ABC->HNH -> METHYLASE -> OLD-ABC ->REase -> DUF499 *-><-RlpA<-?<-?<-ZnR+ TOPRIM +D5ATPase | 929 | Bacteria>Pseudomonadota>Gammaproteobacteria>Pseudomonadales>Pseudomonadaceae | GCF_900114395.1 | Azotobacter_beijerinckii | BM240_RS15650 | 17867..20656 | 170623 | DUF499-domain-containing-protein-[Azotobacter-beijerinckii] |
| WP_280091279.1 | DUF499_1.RM.like.systems | Arm-DNA-bind_3+Phage_integrase->Phage_int_SAM-><-HNH+SAVED<-SF1-SF2-helicase+RdgC+Beta-barrel+REase\|\|RlpA-><- DUF499 *<-HEPN<- METHYLASE <- HNH<- OLD-ABC <- TUDOR+ SF1-SF2-Helicase+iREase+CTD+REase | 941 | Bacteria>Pseudomonadota>Gammaproteobacteria>Pseudomonadales>Pseudomonadaceae | GCF_947311875.1 | Pseudomonas | QME24_RS13335 | 66050..68875 | 286 | DUF499-domain-containing-protein-[Pseudomonas-juntendi] |
| WP_236481327.1 | DUF499_1.RM.like.systems | <-NERD+SF1-SF2-Helicase\|\|RlpA-><- DUF499 *<-ParB+HNH<- METHYLASE <- HNH<- OLD-ABC <- TUDOR+ SF1-SF2-Helicase+iREase+CTD+REase <-MORC+DNA_gyraseB+ TOPRIM | 946 | Bacteria>Pseudomonadota>Gammaproteobacteria>Pseudomonadales>Pseudomonadaceae | - | Pseudomonas_syringae | GIW69_RS06575 | 2105..4945 | 317 | DUF499-domain-containing-protein-[Pseudomonas-syringae] |
| WP_122622773.1 | DUF499_1.RM.like.systems | <-ParB+HNH\|\|HTH+WYL-> TUDOR+ SF1-SF2-Helicase+iREase+CTD+REase -> OLD-ABC->HNH -> METHYLASE -> DUF499 *-><-RlpA<-Phage_int_SAM<-?<- TOPRIM <-Mbetalac<-Arm-DNA-bind_3+Phage_integrase | 942 | Bacteria>Pseudomonadota>Gammaproteobacteria>Pseudomonadales>Pseudomonadaceae | - | Pseudomonas_viridiflava | EEX91_RS04620 | 101897..104725 | 33069 | DUF499-domain-containing-protein-[Pseudomonas-viridiflava] |
| WP_029406945.1 | DUF499_1.RM.like.systems | Phage_integrase-><-GDResolvase+HIN-HTH<- DUF499 *<- METHYLASE <- HNH<- OLD-ABC <- TUDOR+ SF1-SF2-Helicase+iREase+CTD+REase <-?<-HTH | 941 | Bacteria>Pseudomonadota>Gammaproteobacteria>Thiotrichales>Piscirickettsiaceae | GCF_000702325.1 | Thiomicrorhabdus_sp._Milos-T2 | BS34_RS0102315 | 537290..540115 | 90814 | DUF499-domain-containing-protein-[Thiomicrorhabdus-sp.-Milos-T2] |
| WP_020558458.1 | DUF499_1.RM.like.systems | 35exo->?->?->?-><- DUF499 *<- METHYLASE <- HNH<- OLD-ABC <- TUDOR+ SF1-SF2-Helicase+iREase+CTD+REase <-SF1-SF2-Helicase | 942 | Bacteria>Pseudomonadota>Gammaproteobacteria>Thiotrichales>Thiofilaceae | GCF_000380185.1 | Thiofilum_flexile | A3IK_RS0102855 | 95361..98189 | 125627 | DUF499-domain-containing-protein-[Thiofilum-flexile] |
| SEH06532.1 | DUF499_1.RM.like.systems | TUDOR+ SF1-SF2-Helicase+iREase+CTD+REase -> OLD-ABC->HNH -> METHYLASE -> DUF499 *->?->?->?->ParA-Soj-PloopNTPase-> | 934 | Bacteria>Pseudomonadota>Gammaproteobacteria>Thiotrichales>Thiotrichaceae | GCA_900099115.2 | Candidatus_Venteria_ishoeyi | MBHS_02395 | 35251..38055 | 1899563 | Uncharacterised-protein-[Thiotrichales-bacterium-HS_08] |
| WP_109837975.1 | DUF499_1.RM.like.systems | <-ORC-CDC6+REase\|\| TUDOR+ SF1-SF2-Helicase+iREase+CTD+REase -> OLD-ABC->HNH -> METHYLASE -> DUF499 *-> | 944 | Bacteria>Pseudomonadota>Gammaproteobacteria>Thiotrichales>Thiotrichaceae | GCF_003172875.1 | Leucothrix_pacifica | DKW60_RS12435 | 56874..59708 | 1247513 | DUF499-domain-containing-protein-[Leucothrix-pacifica] |
| WP_103920297.1 | DUF499_1.RM.like.systems | TUDOR+ SF1-SF2-Helicase+iREase+CTD+REase -> OLD-ABC->HNH -> METHYLASE -> DUF499 *->?->?->ParA-Soj-PloopNTPase-> | 934 | Bacteria>Pseudomonadota>Gammaproteobacteria>Thiotrichales>Thiotrichaceae | GCF_900099115.1 | Candidatus_Venteria_ishoeyi | MBHS_RS13590 | 35251..38055 | 1899563 | DUF499-domain-containing-protein-[Candidatus-Venteria-ishoeyi] |
| WP_078924154.1 | DUF499_1.RM.like.systems | TUDOR+ SF1-SF2-Helicase+iREase+CTD+REase -> OLD-ABC->HNH -> METHYLASE -> DUF499 *-> | 937 | Bacteria>Pseudomonadota>Gammaproteobacteria>Thiotrichales>Thiotrichaceae | GCF_900167255.1 | Thiothrix_eikelboomii | B5D50_RS18550 | 28151..30964 | 92487 | DUF499-domain-containing-protein-[Thiothrix-eikelboomii] |
| TBT35889.1 | DUF499_1.RM.like.systems | Arm-DNA-bind_3+Phage_integrase->RlpA-><- DUF499 *<- TUDOR+ SF1-SF2-Helicase+iREase+CTD+REase -p<- HNH<- OLD-ABC <-HTH+WYL<-XIS-HTH\|\|JAB-> | 940 | Bacteria>Pseudomonadota>Gammaproteobacteria>Vibrionales>Vibrionaceae | GCA_004324405.1 | Vibrio_parahaemolyticus | D5E85_08495 | 430725..433547 | 670 | ATP-binding-protein-[Vibrio-parahaemolyticus] |
| HCH6462476.1 | DUF499_1.RM.like.systems | XIS-HTH->HTH+WYL-> TUDOR+ SF1-SF2-Helicase+iREase+CTD+REase -> OLD-ABC->HNH -> METHYLASE -> DUF499 *-><-RlpA<-Arm-DNA-bind_3+Phage_integrase | 940 | Bacteria>Pseudomonadota>Gammaproteobacteria>Vibrionales>Vibrionaceae | GCA_024102875.1 | Vibrio_parahaemolyticus | NK401_003629 | 41421..44243 | 670 | ATP-binding-protein-[Vibrio-parahaemolyticus] |
| EJL6982897.1 | DUF499_1.RM.like.systems | XIS-HTH->HTH+WYL-> TUDOR+ SF1-SF2-Helicase+iREase+CTD+REase -> OLD-ABC->HNH -> METHYLASE -> DUF499 *-><-RlpA<-Arm-DNA-bind_3+Phage_integrase | 915 | Bacteria>Pseudomonadota>Gammaproteobacteria>Vibrionales>Vibrionaceae | GCA_024219855.1 | Vibrio_cholerae | NMT40_002449 | 120290..123037 | 666 | ATP-binding-protein-[Vibrio-cholerae] |
| EJR0961152.1 | DUF499_1.RM.like.systems | Arm-DNA-bind_3+Phage_integrase->RlpA-><- DUF499 *<- METHYLASE <- HNH<- OLD-ABC <- TUDOR+ SF1-SF2-Helicase+iREase+CTD+REase <-HTH+WYL<-XIS-HTH | 940 | Bacteria>Pseudomonadota>Gammaproteobacteria>Vibrionales>Vibrionaceae | GCA_024870505.1 | Vibrio_parahaemolyticus | NYK10_002560 | 278932..281754 | 670 | ATP-binding-protein-[Vibrio-parahaemolyticus] |
| WP_213964958.1 | DUF499_1.RM.like.systems | XIS-HTH->HTH+WYL-> TUDOR+ SF1-SF2-Helicase+iREase+CTD+REase -> OLD-ABC->HNH -> METHYLASE -> DUF499 *-><-RlpA<-Arm-DNA-bind_3+Phage_integrase | 940 | Bacteria>Pseudomonadota>Gammaproteobacteria>Vibrionales>Vibrionaceae | GCF_018457925.1 | Vibrio_alginolyticus | J4H38_RS12950 | 94691..97513 | 663 | DUF499-domain-containing-protein-[Vibrio-alginolyticus] |
| WP_062660547.1 | DUF499_1.RM.like.systems | HTH+WYL-> TUDOR+ SF1-SF2-Helicase+iREase+CTD+REase -> OLD-ABC->HNH -> TUDOR+ SF1-SF2-Helicase+iREase+CTD+REase -p-> METHYLASE -> DUF499 *-><-RlpA<-?\|\|?->?-><-HNH<-OBFOLD-NUCLEASE | 947 | Bacteria>Pseudomonadota>Gammaproteobacteria>Vibrionales>Vibrionaceae | GCF_900055185.1 | Grimontia_celer | GCE9029_RS00270 | 67941..70784 | 1796497 | DUF499-domain-containing-protein-[Grimontia-celer] |
| WP_254620468.1 | DUF499_1.RM.like.systems | Arm-DNA-bind_3+Phage_integrase->RlpA-><- DUF499 *<- METHYLASE <- HNH<- OLD-ABC <- TUDOR+ SF1-SF2-Helicase+iREase+CTD+REase <-HTH+WYL<-XIS-HTH | 856 | Bacteria>Pseudomonadota>Gammaproteobacteria>Vibrionales>Vibrionaceae | - | Vibrio_cholerae | KVS75_RS02420 | 551801..554371 | 666 | DUF499-domain-containing-protein-[Vibrio-cholerae] |
| MCC7256721.1 | DUF499_1.RM.like.systems | <-KAP-NTPase<-?<- DUF499 *<-KAP-NTPase<- METHYLASE <- HNH<- OLD-ABC <- TUDOR+ SF1-SF2-Helicase+iREase+CTD+REase <-PIWI | 930 | Bacteria>Pseudomonadota>Gammaproteobacteria>Xanthomonadales>Rhodanobacteraceae | GCA_020852005.1 | Dokdonella_sp. | IT467_12405 | 2246..5038 | 2291710 | ATP-binding-protein-[Dokdonella-sp.] |

| Protein-ID | System | Gene-Neighbourhoods | Length | Taxonomy | Assembly-ID | Strain | Locus-Tag | Genomic-coordinates | TaxID | Fasta-defline |
|---|---|---|---|---|---|---|---|---|---|---|
| PJK14600.1 | DUF499_1.RM.like.systems | <-DUF499*<-ParB+HNH<-METHYLASE<-HNH<-OLD-ABC<-TUDOR+SF1-SF2-Helicase+iREase+CTD+REase | 945 | Bacteria>Pseudomonadota>Gammaproteobacteria>Xanthomonadales>Xanthomonadaceae | GCA_002798295.1 | Xanthomonadaceae_bacterium_NML07-0707 | CO613_05135 | 84524..87361 | 2032574 | hypothetical-protein-CO613_05135-[Xanthomonadaceae-bacterium-NML07-0707] |
| WP_049462969.1 | DUF499_1.RM.like.systems | HTH_26->TUDOR+SF1-SF2-Helicase+iREase+CTD+REase->OLD-ABC->HNH->METHYLASE->ParB+HNH->DUF499*->?->?->?->?->TPR-> | 930 | Bacteria>Pseudomonadota>Gammaproteobacteria>Xanthomonadales>Xanthomonadaceae | GCF_001075485.1 | Stenotrophomonas_maltophilia | ADM82_RS13495 | 40138..42930 | 40324 | DUF499-domain-containing-protein-[Stenotrophomonas-maltophilia] |
| MBQ6473657.1 | DUF499_1.RM.like.systems | Schlafen+HATPase+HTH->OLD-ABC->HNH->METHYLASE->OLD-ABC->NERD->DUF499*-> | 944 | Bacteria>PVC_group>Lentisphaerota>Lentisphaeria>Victivallales | GCA_017431185.1 | Victivallales_bacterium | IJJ33_16845 | 18968..21802 | 2212475 | ATP-binding-protein-[Victivallales-bacterium] |
| WP_168963703.1 | DUF499_1.RM.like.systems | XIS-HTH-><-?<-Phage_integrase\|\|ORC-CDC6+REase->?-><-?<-DUF499*<-NERD<-OLD-ABC<-METHYLASE<-HNH<-OLD-ABC<-TUDOR+SF1-SF2-Helicase+iREase+CTD+REase | 935 | Bacteria>PVC_group>Lentisphaerota>Lentisphaeria>Victivallales | GCF_012843305.1 | Victivallis_vadensis | HF882_RS18870 | 10950..13757 | 172901 | DUF499-domain-containing-protein-[Victivallis-vadensis] |
| MBE6357833.1 | DUF499_1.RM.like.systems | ParB-HTH->?-><-?<-?<-DUF499*<-REase<-OLD-ABC<-METHYLASE<-HNH<-OLD-ABC<-TUDOR+SF1-SF2-Helicase+iREase+CTD+REase | 938 | Bacteria>PVC_group>Lentisphaerota>unclassified_Lentisphaerota | GCA_015061525.1 | Lentisphaerota_bacterium | E7057_01125 | 3177..5993 | 1932692 | ATP-binding-protein-[Lentisphaerae-bacterium] |
| MBX3408154.1 | DUF499_1.RM.like.systems | RAMA->GDResolvase+Recombinase+ZnR->ParB-><-?<-Radical-SAM<-DUF499*<-METHYLASE<-HNH<-OLD-ABC<-TUDOR+SF1-SF2-Helicase+iREase+CTD+REase | 858 | Bacteria>PVC_group>Planctomycetota>Phycisphaerae>Phycisphaerales | GCA_019636415.1 | Phycisphaeraceae_bacterium | KF869_15485 | 6654..9230 | 2026777 | ATP-binding-protein-[Phycisphaeraceae-bacterium] |
| MBX3409892.1 | DUF499_1.RM.like.systems | RAMA->GDResolvase+Recombinase+ZnR->ParB-><-?<-Radical-SAM<-DUF499*<-METHYLASE<-HNH<-OLD-ABC<-TUDOR+SF1-SF2-Helicase+iREase+CTD+REase\|\|HTH->MPTase-> | 947 | Bacteria>PVC_group>Planctomycetota>Phycisphaerae>Phycisphaerales | GCA_019636575.1 | Phycisphaeraceae_bacterium | KF859_08400 | 137437..140280 | 2026777 | ATP-binding-protein-[Phycisphaeraceae-bacterium] |
| OYP37234.1 | DUF499_1.RM.like.systems | <-wHTH+SAD+HNH<-URI-DNase+RAMA<-REase<-EVE<-REase<-DUF499*<-HNH<-OLD-ABC<-METHYLASE<-Peptidase_S8<-Classical-AAA<-TUDOR+SF1-SF2-Helicase+iREase+CTD+REase | 948 | Bacteria>PVC_group>Planctomycetota>Planctomycetia>Pirellulales | GCA_002251265.1 | Rhodopirellula_sp._MGV | CGZ80_05855 | 9695..12541 | 2023130 | hypothetical-protein-CGZ80_05855-[Rhodopirellula-sp.-MGV] |
| NLX95047.1 | DUF499_1.RM.like.systems | TPR+TPR->?->?-><-?<-PHP+SbcC<-HEPN+DUF499*<-METHYLASE<-HNH<-OLD-ABC<-TUDOR+SF1-SF2-Helicase+iREase+CTD+REase-p | 1104 | Bacteria>PVC_group>Planctomycetota>Planctomycetia>Pirellulales | GCA_012728855.1 | Rhodopirellula_sp. | GXY83_02605 | 12933..16247 | 2024855 | ATP-binding-protein-[Rhodopirellula-sp.] |
| WP_198953633.1 | DUF499_1.RM.like.systems | <-wHTH+SAD+HNH<-URI-DNase+RAMA<-REase<-EVE<-REase<-DUF499*<-HNH<-OLD-ABC<-METHYLASE<-Peptidase_S8<-Classical-AAA<-TUDOR+SF1-SF2-Helicase+iREase+CTD+REase | 951 | Bacteria>PVC_group>Planctomycetota>Planctomycetia>Pirellulales | GCF_002251265.1 | Rhodopirellula_sp._MGV | CGZ80_RS05865 | 9695..12550 | 2023130 | DUF499-domain-containing-protein-[Rhodopirellula-sp.-MGV] |
| MBX3448931.1 | DUF499_1.RM.like.systems | TUDOR+SF1-SF2-Helicase+iREase+CTD+REase->OLD-ABC->HNH->METHYLASE->OLD-ABC->REase->DUF499*->?->MORC->?->?-><-wHTH+SAD+HNH<-DNA_methylase | 948 | Bacteria>PVC_group>Planctomycetota>Planctomycetia>Planctomycetales | GCA_019636285.1 | Planctomycetaceae_bacterium | KF777_05185 | 10482..13328 | 2026779 | ATP-binding-protein-[Planctomycetaceae-bacterium] |
| MDB5334583.1 | DUF499_1.RM.like.systems | TUDOR+SF1-SF2-Helicase+iREase+CTD+REase->OLD-ABC->HNH->METHYLASE->DUF499*-> | 942 | Bacteria>PVC_group>Planctomycetota>Planctomycetia>Planctomycetales | GCA_028291605.1 | Planctomycetaceae_bacterium | JWN70_202 | 14767..17595 | 2026779 | MAG:-hypothetical-protein-JWN70_202-[Planctomycetaceae-bacterium] |
| TAH38052.1 | DUF499_1.RM.like.systems | <-wHTH+REase\|\|AbiEi_4->AbiEii-><-PIN<-ParB+HNH\|\|HKD-DNase-><-HEPN+DUF499*<-METHYLASE<-HNH<-OLD-ABC<-TUDOR+SF1-SF2-Helicase+iREase+CTD+REase<-HTH | 1098 | Bacteria>PVC_group>Planctomycetota>unclassified_Planctomycetota | GCA_004295085.1 | Planctomycetota_bacterium | EYC70_05360 | 66593..69889 | 2026780 | MAG:-ATP-binding-protein-[Planctomycetota-bacterium] |
| MCC5805073.1 | DUF499_1.RM.like.systems | OLD-ABC->HNH->METHYLASE->CHTH->RHH->PIN->HEPN+DUF499*->?->PhdYeFM_antitox->PIN-> | 1107 | Bacteria>PVC_group>Verrucomicrobiota>Opitutae>Opitutales | GCA_020831525.1 | Opitutales_bacterium | JJU00_01980 | 48883..52206 | 2026772 | DUF499-domain-containing-protein-[Opitutales-bacterium] |
| TAE90777.1 | DUF499_1.RM.like.systems | <-DUF499*<-REase<-METHYLASE<-HNH<-OLD-ABC<-TOPRIM<-OLD-ABC | 939 | Bacteria>PVC_group>Verrucomicrobiota>unclassified_Verrucomicrobiota | GCA_004293285.1 | Verrucomicrobiota_bacterium | EAZ81_08635 | 280..3099 | 2026799 | ATP-binding-protein-[Verrucomicrobia-bacterium] |
| MBI5767835.1 | DUF499_1.RM.like.systems | HTH->TUDOR+SF1-SF2-Helicase+iREase+CTD+REase->OLD-ABC->HNH->METHYLASE->DUF499*->Radical-SAM-> | 944 | Bacteria>PVC_group>Verrucomicrobiota>unclassified_Verrucomicrobiota | GCA_016235555.1 | Verrucomicrobiota_bacterium | HZA93_08560 | 101523..104357 | 2026799 | ATP-binding-protein-[Verrucomicrobia-bacterium] |
| MCX6879289.1 | DUF499_1.RM.like.systems | <-HISKIN<-DUF499*<-REase<-OLD-ABC<-METHYLASE<-HNH<-OLD-ABC<-TUDOR+SF1-SF2-Helicase+iREase+CTD+REase | 941 | Bacteria>PVC_group>Verrucomicrobiota>unclassified_Verrucomicrobiota | GCA_026397725.1 | Verrucomicrobiota_bacterium | NTW21_36620 | 9837..12662 | 2026799 | MAG:-DUF499-domain-containing-protein-[Verrucomicrobia-bacterium] |
| MCX6953048.1 | DUF499_1.RM.like.systems | TUDOR+SF1-SF2-Helicase+iREase+CTD+REase->OLD-ABC->HNH->METHYLASE->REase->DUF499*->?->?->?->ABhydrolase-> | 941 | Bacteria>PVC_group>Verrucomicrobiota>unclassified_Verrucomicrobiota | GCA_026398065.1 | Verrucomicrobiota_bacterium | NTV51_12915 | 8536..11361 | 2026799 | MAG:-DUF499-domain-containing-protein-[Verrucomicrobia-bacterium] |
| MBV5322918.1 | DUF499_1.RM.like.systems | HEPN->OLD-ABC->HNH->METHYLASE->METHYLASE->HEPN->DUF499*-> | 872 | Bacteria>Terrabacteria_group>Actinomycetota>Acidimicrobiia>Acidimicrobiales | GCA_019162935.1 | Ilumatobacteraceae_bacterium | JZU60_03690 | 12690..15310 | 2497619 | ATP-binding-protein-[Ilumatobacteraceae-bacterium] |
| WP_194396698.1 | DUF499_1.RM.like.systems | METHYLASE-><-HNH<-OLD-ABC+TOPRIM<-HEPN+DUF499*<-METHYLASE<-TUDOR+SF1-SF2-Helicase+iREase+CTD+REase<-XIS-HTH | 1130 | Bacteria>Terrabacteria_group>Actinomycetota>Actinomycetes>Micrococcales | GCF_015277815.1 | Microbacterium_atlanticum | IR212_RS15230 | 3329381..3332773 | 2782168 | DUF499-domain-containing-protein-[Microbacterium-atlanticum] |
| MBK9093501.1 | DUF499_1.RM.like.systems | TUDOR+SF1-SF2-Helicase+iREase+CTD+REase->OLD-ABC->HNH->METHYLASE->OLD-ABC->REase->DUF499*->?->ParB+HNH->?->REase-p-> | 943 | Bacteria>Terrabacteria_group>Chloroflexota>Anaerolineae>unclassified_Anaerolineae | GCA_016719785.1 | Anaerolineae_bacterium | IPM84_12145 | 210722..213553 | 2052143 | ATP-binding-protein-[Anaerolineae-bacterium] |
| MCP9859835.1 | DUF499_1.RM.like.systems | <-SIGMA-HTH<-?<-?<-DUF499*<-wHTH+SAD+HNH<-METHYLASE-p<-HNH<-OLD-ABC<-TUDOR+SF1-SF2-Helicase+iREase+CTD+REase<-LRP-HTH | 819 | Bacteria>Terrabacteria_group>Cyanobacteriota/Melainabacteria_group>Cyanobacteriota>Cyanophyceae | GCA_024345955.1 | Cyanobium_sp._Cruz-8H5 | KBZ34_12405 | 60686..63145 | 2823712 | ATP-binding-protein-[Cyanobium-sp.-Cruz-8H5] |
| WP_254953294.1 | DUF499_1.RM.like.systems | <-SIGMA-HTH<-?<-?<-DUF499*<-wHTH+SAD+HNH<-METHYLASE-p<-HNH<-OLD-ABC<-TUDOR+SF1-SF2-Helicase+iREase+CTD+REase<-LRP-HTH | 819 | Bacteria>Terrabacteria_group>Cyanobacteriota/Melainabacteria_group>Cyanobacteriota>Cyanophyceae | GCF_024345955.1 | unclassified_Cyanobium | KBZ34_12405 | 60686..63145 | 2627006 | MULTISPECIES:-DUF499-domain-containing-protein-[unclassified-Cyanobium] |
| WP_261362403.1 | DUF499_1.RM.like.systems | <-SIGMA-HTH<-?<-?<-DUF499*<-wHTH+SAD+HNH<-METHYLASE-p<-HNH<-OLD-ABC<-TUDOR+SF1-SF2-Helicase+iREase+CTD+REase<-LRP-HTH | 820 | Bacteria>Terrabacteria_group>Cyanobacteriota/Melainabacteria_group>Cyanobacteriota>Cyanophyceae | - | unclassified_Cyanobium | KBZ34_RS12460 | 60686..63148 | 2627006 | MULTISPECIES:-DUF499-domain-containing-protein-[unclassified-Cyanobium] |
| MBC2711122.1 | DUF499_1.RM.like.systems | TUDOR+SF1-SF2-Helicase+iREase+CTD+REase->?->OLD-ABC->HNH->METHYLASE->HEPN+DUF499*->?->GNTR-HTH->?->?->ABC-ATPase+SbcC->ABC-ATPase+SbcC-> | 1122 | Bacteria>Thermodesulfobacteriota>Desulfobacteria>Desulfobacterales>Desulfosarcinaceae | GCA_014237405.1 | Desulfosarcina_sp. | HGJ94_09035 | 43660..47028 | 2027861 | ATP-binding-protein-[Desulfosarcina-sp.] |
| WP_168935605.1 | DUF499_1.RM.like.systems | <-HTH_3\|\|?-><-?\|\|?->?-><-DUF499*<-REase<-OLD-ABC<-METHYLASE<-HNH<-OLD-ABC<-TUDOR+SF1-SF2-Helicase+iREase+CTD+REase | 933 | Bacteria>Thermodesulfobacteriota>Desulfovibrionia>Desulfovibrionales>Desulfovibrionaceae | GCF_012843875.1 | Desulfovibrio_piger | HF854_RS06725 | 17522..20323 | 901 | DUF499-domain-containing-protein-[Desulfovibrio-piger] |
| -- | | | | | | | | | | |
| -- OLD-ABC->HNH | | | | | | | | | | |
| --BR-2 | | | | | | | | | | |
| -- | | | | | | | | | | |
| MCY3933638.1 | DUF499_2.RM.like.systems | <-HNH<-OLD-ABC<-?<-OLD-ABC<-ParB+HNH<-DUF3780<-DUF499*<-?<-?<-SMODS\|\|HTH+MPTase-><-?<-Phd-YefM | 1021 | Bacteria>Acidobacteriota>unclassified_Acidobacteriota | GCA_026707325.1 | Acidobacteriota_bacterium | OXH70_18150 | 6925..9990 | 1978231 | MAG:-DUF499-domain-containing-protein-[Acidobacteria-bacterium] |
| MDE2976517.1 | DUF499_2.RM.like.systems | MetJarc-HTH->?->?->DUF499*->DUF3780->ParB+HNH->OLD-ABC->?->OLD-ABC->HNH-> | 1021 | Bacteria>Acidobacteriota>unclassified_Acidobacteriota | GCA_028873735.1 | Acidobacteriota_bacterium | OXU63_03230 | 1513..4578 | 1978231 | MAG:-DUF499-domain-containing-protein-[Acidobacteriota-bacterium] |
| MDE3261994.1 | DUF499_2.RM.like.systems | REase->DUF499*->DUF3780->OLD-ABC->HNH->METHYLASE->HKD-DNase+SF1-SF2-Helicase+iREase+CTD-> | 1006 | Bacteria>Acidobacteriota>unclassified_Acidobacteriota | GCA_028877265.1 | Acidobacteriota_bacterium | OYL41_08435 | 3377..6397 | 1978231 | MAG:-DUF499-domain-containing-protein-[Acidobacteriota-bacterium] |
| MCY4606377.1 | DUF499_2.RM.like.systems | OLD-ABC->HNH->DUF499*->DUF3780->?->?->?->METHYLASE->HKD-DNase+SF1-SF2-Helicase+iREase+CTD-> | 995 | Bacteria>FCB_group>Gemmatimonadota>unclassified_Gemmatimonadota | GCA_026713655.1 | Gemmatimonadota_bacterium | OXE49_19245 | 3266..6253 | 2026742 | MAG:-DUF499-domain-containing-protein-[Gemmatimonadetes-bacterium] |
| WP_198375108.1 | DUF499_2.RM.like.systems | <-HKD-DNase+SF1-SF2-Helicase+iREase+CTD<-DUF932<-METHYLASE<-HNH<-OLD-ABC<-DUF3780<-DUF499*<-DnaB<-HTH_17<-?<-Arm-DNA-bind_3+Phage_integrase\|\|Methyltransf_21->Methyltransf-> | 1033 | Bacteria>Pseudomonadota>Alphaproteobacteria>Rhodospirillales>Acetobacteraceae | GCF_016106015.1 | Roseomonas_rubea | HW577_RS05730 | 12708..15809 | 2748666 | DUF499-domain-containing-protein-[Roseomonas-rubea] |
| WP_100917768.1 | DUF499_2.RM.like.systems | Phage_int_SAM_1+Phage_integrase-><-HKD-DNase+SF1-SF2-Helicase+iREase+CTD<-HNH<-OLD-ABC<-METHYLASE<-DUF3780<-DUF499*\|\|?->?-><-?<-?<-Aminotran_4 | 1083 | Bacteria>Pseudomonadota>Gammaproteobacteria>Chromatiales>Chromatiaceae | GCF_002813775.1 | Candidatus_Thiodictyon_syntrophicum | THSYN_RS02615 | 607511..610762 | 1166950 | DUF499-domain-containing-protein-[Candidatus-Thiodictyon-syntrophicum] |
| PZN82194.1 | DUF499_2.RM.like.systems | <-METHYLASE-p<-?<-HNH<-OLD-ABC<-DUF3780<-DUF499* | 1015 | Bacteria>Pseudomonadota>Gammaproteobacteria>Methylococcales>Methylococcales_incertae_sedis | GCA_003242955.1 | Candidatus_Methylumidiphilus_alinenensis | DM484_06870 | 3894..6941 | 2202197 | hypothetical-protein-DM484_06870-[Candidatus-Methylumidiphilus-alinenensis] |
| MXY53195.1 | DUF499_2.RM.like.systems | <-TPR<-?\|\|?->?->?->DUF499*->DUF3780->OLD-ABC->HNH->METHYLASE->HKD-DNase+SF1-SF2-Helicase+iREase+CTD->OLD-ABC-> | 1009 | Bacteria>Pseudomonadota>Gammaproteobacteria>unclassified_Gammaproteobacteria | GCA_009837725.1 | Gammaproteobacteria_bacterium | F4Y86_11820 | 5670..8699 | 1913989 | ATP-binding-protein-[Gammaproteobacteria-bacterium] |
| MYB38346.1 | DUF499_2.RM.like.systems | <-OLD-ABC<-HKD-DNase+SF1-SF2-Helicase+iREase+CTD<-METHYLASE<-HNH<-OLD-ABC<-DUF3780<-DUF499* | 1009 | Bacteria>Pseudomonadota>Gammaproteobacteria>unclassified_Gammaproteobacteria | GCA_009838905.1 | Gammaproteobacteria_bacterium | F4Y26_13400 | 18076..21105 | 1913989 | DUF499-domain-containing-protein-[Gammaproteobacteria-bacterium] |
| MBK9497791.1 | DUF499_2.RM.like.systems | <-HNH<-OLD-ABC<-?<-OLD-ABC<-ParB+HNH<-DUF3780<-DUF499*<-DUF499\|\|?->?->?->Peptidase-> | 736 | Bacteria>Spirochaetota>Spirochaetia>Leptospirales>Leptospiraceae | GCA_016718635.1 | Leptospiraceae_bacterium | IPO06_00155 | 42788..44998 | 2026757 | DUF499-domain-containing-protein-[Leptospiraceae-bacterium] |
| MCY4559676.1 | DUF499_2.RM.like.systems | <-REase<-REase<-?\|\|SF1-SF2-Helicase->DUF932->XIS-HTH->DUF499*->DUF3780->OLD-ABC->HNH->METHYLASE->HKD-DNase+SF1-SF2-Helicase+iREase+CTD-> | 1013 | Bacteria>Terrabacteria_group>Chloroflexota>unclassified_Chloroflexota | GCA_026713265.1 | Chloroflexota_bacterium | OXF79_25580 | 10138..13179 | 2026724 | MAG:-DUF499-domain-containing-protein-[Chloroflexi-bacterium] |
| -- | | | | | | | | | | |
| --REase-linked_OLD-ABC | | | | | | | | | | |
| -- | | | | | | | | | | |
| -- OLD-ABC->REase | | | | | | | | | | |
| --Type1.BREX | | | | | | | | | | |
| -- | | | | | | | | | | |
| PKL50472.1 | Type1.BREX | <-MPTase<-HTH_19<-?\|\|BrxA-DNA-binding->BrxB-BrxF-iSTAND->DUF6079*->Schlafen+HATPase+wHTH+HTH->OLD-ABC->REase->N6-METHYLASE->PglZ->SIGMA+OB-Fold+AAA-ATPase+HKD-DNase-BrxL-> | 1185 | Bacteria>Bacteria_incertae_sedis>Bacteria_candidate_phyla>Candidatus_Riflebacteria>unclassified_Candidatus_Riflebacteria | GCA_002839435.1 | Candidatus_Riflebacteria_bacterium_HGW-Riflebacteria-2 | CVV42_02960 | 62937..66494 | 2013830 | MAG:-BREX-system-P-loop-protein-BrxC-[Candidatus-Riflebacteria-bacterium-HGW-Riflebacteria-2] |
| WP_187475233.1 | Type1.BREX | METHYLASE->Peptidase_M3->?->?-><-DDE_Tnp_1_4\|\|?->XIS-HTH->BrxA-DNA-binding->BrxB-BrxF-iSTAND->DUF6079*->OLD-ABC->REase->N6-METHYLASE->DrHyd->Schlafen+HATPase+wHTH->Trypsin->?->PglZ->SIGMA+OB-Fold+AAA-ATPase+HKD-DNase-BrxL-> | 1166 | Bacteria>Campylobacterota>Epsilonproteobacteria>Campylobacterales>Arcobacteraceae>Aliarcobacter | GCF_014352915.1 | Aliarcobacter_cryaerophilus | HOO34_RS05640 | 1060903..1064403 | 28198 | BREX-system-P-loop-protein-BrxC-[Aliarcobacter-cryaerophilus] |
| MBC8466717.1 | Type1.BREX | <-RAMA<-SIGMA+OB-Fold+AAA-ATPase+HKD-DNase-BrxL<-PglZ<-N6-METHYLASE<-REase<-OLD-ABC<-DUF6079*<-BrxB-BrxF-iSTAND<-BrxA-DNA-binding | 1172 | Bacteria>delta/epsilon_subdivisions>Deltaproteobacteria>unclassified_Deltaproteobacteria | GCA_014384065.1 | Deltaproteobacteria_bacterium | H8D55_02650 | 10326..13844 | 2026735 | BREX-system-P-loop-protein-BrxC-[Deltaproteobacteria-bacterium] |
| MBW1967272.1 | Type1.BREX | <-N6-METHYLASE-p<-REase<-OLD-ABC<-DUF6079*<-BrxB-BrxF-iSTAND<-BrxA-DNA-binding<-XIS-HTH | 1154 | Bacteria>delta/epsilon_subdivisions>Deltaproteobacteria>unclassified_Deltaproteobacteria | GCA_019308155.1 | Deltaproteobacteria_bacterium | JRI48_07780 | 2430..5894 | 2026735 | BREX-system-P-loop-protein-BrxC-[Deltaproteobacteria-bacterium] |
| MCC6503799.1 | Type1.BREX | <-SbcC<-Calcineurin+Calcineurin-SbcD<-SbcC+APATPase+SbcC<-SIGMA+OB-Fold+AAA-ATPase+HKD-DNase-BrxL<-PglZ<-N6-METHYLASE<-REase<-OLD-ABC<-DUF6079*<-BrxB-BrxF-iSTAND<-BrxA-DNA-binding<-XIS-HTH | 1181 | Bacteria>delta/epsilon_subdivisions>Deltaproteobacteria>unclassified_Deltaproteobacteria | GCA_020846435.1 | Deltaproteobacteria_bacterium | IT362_11790 | 16699..20244 | 2026735 | BREX-system-P-loop-protein-BrxC-[Deltaproteobacteria-bacterium] |
| MCD6532985.1 | Type1.BREX | <-PglZ<-Schlafen+HATPase+wHTH<-PHP+SbcC<-PIN<-?<-Schlafen+HATPase+wHTH<-N6-METHYLASE<-REase<-OLD-ABC<-DUF6079-p* | 691 | Bacteria>delta/epsilon_subdivisions>Deltaproteobacteria>unclassified_Deltaproteobacteria | GCA_021163445.1 | Deltaproteobacteria_bacterium | J7L25_02755 | 20972..23047 | 2026735 | BREX-system-P-loop-protein-BrxC-[Deltaproteobacteria-bacterium] |
| OQC05381.1 | Type1.BREX | BrxA-DNA-binding->BrxB-BrxF-iSTAND->DUF6079*->OLD-ABC->REase->N6-METHYLASE->PglZ->SIGMA+OB-Fold+AAA-ATPase+HKD-DNase-BrxL->HEPN-> | 1162 | Bacteria>FCB_group>Candidatus_Cloacimonetes>unclassified_Candidatus_Cloacimonetes | GCA_002070875.1 | Candidatus_Cloacimonetes_bacterium_ADurb.Bin089 | BWX76_00750 | 1812..5300 | 1852836 | MAG:-hypothetical-protein-BWX76_00750-[Candidatus-Cloacimonetes-bacterium-ADurb.Bin089] |
| MCB5271576.1 | Type1.BREX | KAP-NTPase->KAP-glycosylase->?->TatD_DNase-><-SIGMA+OB-Fold+AAA-ATPase+HKD-DNase-BrxL<-PglZ<-N6-METHYLASE<-REase<-OLD-ABC<-DUF6079*<-BrxB-BrxF-iSTAND<-BrxA-DNA-binding<-NERD+ZnR | 1171 | Bacteria>FCB_group>Candidatus_Cloacimonetes>unclassified_Candidatus_Cloacimonetes | GCA_020532545.1 | Candidatus_Cloacimonetes_bacterium | LHW56_07005 | 18816..22331 | 2030808 | BREX-system-P-loop-protein-BrxC-[Candidatus-Cloacimonadota-bacterium] |
| CDA09060.1 | Type1.BREX | <-SIGMA+OB-Fold+AAA-ATPase+Lon-Protease-BrxL<-PglZ<-CR-ATPase8+REase<-OLD-ABC+TOPRIM<-?<-?<-N6-METHYLASE<-TOPRIM<-OLD-ABC<-DUF6079*<-?<-SWIM+DUF3670+SF1-SF2-Helicase<-?<-?<-?<-SF1-SF2-Helicase | 1250 | Bacteria>Fusobacteriota>Fusobacteriia>Fusobacteriales>Fusobacteriaceae>Fusobacterium | GCA_000433695.1 | Fusobacterium_sp._CAG:649 | BN748_00884 | 23068..26820 | 1262900 | putative-uncharacterized-protein-[Fusobacterium-sp.-CAG:649] |
| WP_099987862.1 | Type1.BREX | <-BrxA-DNA-binding<-SIGMA+OB-Fold+AAA-ATPase+Lon-Protease-BrxL<-PglZ<-CR-ATPase8+REase<-OLD-ABC+TOPRIM<-?<-N6-METHYLASE<-TOPRIM<-OLD-ABC<-DUF6079*\|\|?->OrfB_IS605+OrfB_Zn_ribbon-><-URI-DNase+SF1-SF2-Helicase<-URI-DNase+SF1-SF2-Helicase<-?<-?<-Peptidase_M3_N+Peptidase_M3\|\|RNaseH-> | 1250 | Bacteria>Fusobacteriota>Fusobacteriia>Fusobacteriales>Fusobacteriaceae>Fusobacterium | GCF_002763695.1 | Fusobacterium_pseudoperiodonticum | CTM74_RS09005 | 1721055..1724807 | 2663009 | BREX-system-P-loop-protein-BrxC-[Fusobacterium-pseudoperiodonticum] |
| WP_022069638.1 | Type1.BREX | <-BrxA-DNA-binding<-SIGMA+OB-Fold+AAA-ATPase+Lon-Protease-BrxL<-PglZ<-CR-ATPase8+REase<-OLD-ABC+TOPRIM<-?<-N6-METHYLASE<-TOPRIM<-OLD-ABC<-DUF6079* | 1250 | Bacteria>Fusobacteriota>Fusobacteriia>Fusobacteriales>Fusobacteriaceae>Fusobacterium | GCF_023383545.1 | Fusobacterium_nucleatum | YWH7053_RS01985 | 23867..27619 | 851 | BREX-system-P-loop-protein-BrxC-[Fusobacterium-nucleatum] |
| WP_064218004.1 | Type1.BREX | Arm-DNA-bind_3+Phage_integrase->?->HTH_3->?->?->BrxA-DNA-binding->BrxB-BrxF-iSTAND->DUF6079*->OLD-ABC->REase->N6-METHYLASE->PglZ->SIGMA+OB-Fold+AAA-ATPase+HKD-DNase-BrxL->SbcC+SbcC-> | 1187 | Bacteria>Pseudomonadota>Acidithiobacillia>Acidithiobacillales>Acidithiobacillaceae>Acidithiobacillus | GCF_001652185.1 | Acidithiobacillus_ferrooxidans | A4H96_RS01790 | 31399..34962 | 920 | BREX-system-P-loop-protein-BrxC-[Acidithiobacillus-ferrooxidans] |
| WP_163056479.1 | Type1.BREX | XIS-HTH->?->TOPRIM->?->XIS-HTH->BrxA-DNA-binding->BrxB-BrxF-iSTAND->DUF6079*->OLD-ABC->REase->N6-METHYLASE->PglZ->SIGMA+OB-Fold+AAA-ATPase+HKD-DNase-BrxL->SbcC+SbcC->Calcineurin+Calcineurin-SbcD->SbcC-> | 1203 | Bacteria>Pseudomonadota>Acidithiobacillia>Acidithiobacillales>Acidithiobacillaceae>Acidithiobacillus | GCF_010577825.1 | Acidithiobacillus_ferrooxidans | X857_RS08730 | 38026..41637 | 920 | BREX-system-P-loop-protein-BrxC-[Acidithiobacillus-ferrooxidans] |
| WP_036280608.1 | Type1.BREX | HKD-DNase->SF1-SF2-Helicase-><-Small-B-barrel+SF1-SF2-Helicase+RdgC+REase<-SF1-SF2-Helicase+REase<-35Exo+REase\|\|wHTH+WYL+WCX->BrxA-DNA-binding->BrxB-BrxF-iSTAND->DUF6079*->OLD-ABC->REase->N6-METHYLASE->PglZ->SIGMA+OB-Fold+AAA-ATPase+HKD-DNase-BrxL->HIN-HTH->?->LZ_Tnp_IS66+zf-IS66+DDE_Tnp_IS66+DDE_Tnp_IS66_C-> | 1178 | Bacteria>Pseudomonadota>Alphaproteobacteria>Hyphomicrobiales>Methylocystaceae>Methylocystis | GCF_000188155.2 | Methylocystis_sp._ATCC_49242 | MET49242_RS03150 | 256459..259995 | 622637 | BREX-system-P-loop-protein-BrxC-[Methylocystis-sp.-ATCC-49242] |
| CAD6617282.1 | Type1.BREX | <-XIS-HTH<-Arm-DNA-bind_3+Phage_integrase-><-SIGMA+OB-Fold+AAA-ATPase+HKD-DNase-BrxL<-PglZ<-N6-METHYLASE<-REase<-OLD-ABC<-DUF6079*<-BrxB-BrxF-iSTAND<-BrxA-DNA-binding<-wHTH+WYL+WCX\|\|GDResolvase+HIN-HTH-> | 1172 | Bacteria>Pseudomonadota>Alphaproteobacteria>Hyphomicrobiales>Rhizobiaceae>Rhizobium/Agrobacterium_group | GCA_902502895.2 | Rhizobium_sp._Khangiran2 | RKHAN_03345 | 3387102..3390620 | 1435607 | BREX-system-P-loop-protein-BrxC-[Rhizobium-sp.-Khangiran2] |
| WP_004434288.1 | Type1.BREX | <-PIN<-PhdYeFM_antitox<-SIGMA+OB-Fold+AAA-ATPase+HKD-DNase-BrxL<-PglZ<-N6-METHYLASE<-REase<-OLD-ABC<-DUF6079*<-BrxB-BrxF-iSTAND<-BrxA-DNA-binding<-wHTH+WYL+WCX\|\|35Exo+REase->SF1-SF2-Helicase+REase->Small-B-barrel+SF1-SF2-Helicase+RdgC+REase-><-SF1-SF2-Helicase<-HKD-DNase | 1177 | Bacteria>Pseudomonadota>Alphaproteobacteria>Hyphomicrobiales>Rhizobiaceae>Sinorhizobium/Ensifer_group | GCF_000236945.1 | Sinorhizobium_meliloti | SM0020_RS29775 | 166866..170399 | 382 | BREX-system-P-loop-protein-BrxC-[Sinorhizobium-meliloti] |
| WP_199258165.1 | Type1.BREX | <-N6-METHYLASE-p<-REase<-OLD-ABC<-DUF6079*<-BrxB-BrxF-iSTAND<-BrxA-DNA-binding<-wHTH+WYL+WCX\|\|?->Thermonuclease->TrbM-><-?<-HTH_3 | 1173 | Bacteria>Pseudomonadota>Alphaproteobacteria>Rhodobacterales>Paracoccaceae>Paracoccus | GCF_016446475.1 | Paracoccus_binzhouensis | I9P22_RS04365 | 3344..6865 | 2796149 | BREX-system-P-loop-protein-BrxC-[Paracoccus-binzhouensis] |
| WP_106262543.1 | Type1.BREX | <-Phage_int_SAM_1+Phage_integrase-><-?<-?\|\|SKI->?-><-HEPN\|\|wHTH+WYL+WCX->BrxA-DNA-binding->BrxB-BrxF-iSTAND->DUF6079*->OLD-ABC->REase->N6-METHYLASE->PglZ->SIGMA+OB-Fold+AAA-ATPase+HKD-DNase-BrxL-><-?\|\|HTH_Tnp_1->HTH_Tnp_1+HTH_21+rve+rve_3-><-HIN-HTH | 1172 | Bacteria>Pseudomonadota>Alphaproteobacteria>Rhodobacterales>Roseobacteraceae>Donghicola | GCF_003003355.1 | Donghicola_tyrosinivorans | CLV74_RS02110 | 436162..439680 | 1652492 | BREX-system-P-loop-protein-BrxC-[Donghicola-tyrosinivorans] |
| WP_274591702.1 | Type1.BREX | XIS-HTH->wHTH+WYL+WCX->BrxA-DNA-binding->BrxB-BrxF-iSTAND->DUF6079*->OLD-ABC->REase->N6-METHYLASE->PglZ->SIGMA+OB-Fold+AAA-ATPase+HKD-DNase-BrxL-> | 1173 | Bacteria>Pseudomonadota>Alphaproteobacteria>Rhodobacterales>Roseobacteraceae>Marinovum | GCF_028807375.1 | Marinovum_sp._PR37 | PVW49_RS22970 | 3287..6808 | 3028382 | BREX-system-P-loop-protein-BrxC-[Marinovum-sp.-PR37] |
| WP_123792783.1 | Type1.BREX | <-ParB+HNH+Schlafen<-SIGMA+OB-Fold+AAA-ATPase+HKD-DNase-BrxL<-PglZ<-N6-METHYLASE<-REase<-OLD-ABC<-N6-METHYLASE<-DUF6079*<-BrxB-BrxF-iSTAND<-BrxA-DNA-binding<-HIN-HTH<-HTH_36<-?<-?<-Arm-DNA-bind_3+Phage_integrase | 1172 | Bacteria>Pseudomonadota>Alphaproteobacteria>Rhodobacterales>Roseobacteraceae>Pacificibacter | GCF_003814915.1 | Pacificibacter_maritimus | EDD53_RS08420 | 394668..398186 | 762213 | BREX-system-P-loop-protein-BrxC-[Pacificibacter-maritimus] |
| WP_150282504.1 | Type1.BREX | MPTase-><-HTH+MPTase<-?<-SIGMA+OB-Fold+AAA-ATPase+HKD-DNase-BrxL<-PglZ<-N6-METHYLASE<-REase<-OLD-ABC<-DUF6079*<-BrxB-BrxF-iSTAND<-BrxA-DNA-binding<-wHTH+WYL+WCX\|\|?->Thermonuclease->TrbM-> | 1173 | Bacteria>Pseudomonadota>Alphaproteobacteria>Rhodobacterales>Roseobacteraceae>Salipiger | GCF_008692285.1 | Salipiger_aestuarii | AL036_RS18780 | 13068..16589 | 568098 | BREX-system-P-loop-protein-BrxC-[Salipiger-aestuarii] |
| WP_085595207.1 | Type1.BREX | XIS-HTH->BrxA-DNA-binding->BrxB-BrxF-iSTAND->DUF6079*->OLD-ABC->REase->N6-METHYLASE->HTH+REase->PglZ->SIGMA+OB-Fold+AAA-ATPase+HKD-DNase-BrxL-><-SF1-SF2-Helicase | 1182 | Bacteria>Pseudomonadota>Alphaproteobacteria>Rhodospirillales>Thalassospiraceae>Thalassospira | GCF_002115825.1 | Thalassospira_sp._MCCC_1A01428 | THS27_RS23975 | 1729..5277 | 1470575 | BREX-system-P-loop-protein-BrxC-[Thalassospira-sp.-MCCC-1A01428] |
| WP_022673601.1 | Type1.BREX | HTH_3->Couple_hipA+HipA_C->HNH-><-REase+HTH+35exo+Small-B-barrel+SF1-SF2-Helicase<-SIGMA+OB-Fold+AAA-ATPase+HKD-DNase-BrxL<-PglZ<-N6-METHYLASE<-REase<-OLD-ABC<-DUF6079*<-BrxB-BrxF-iSTAND<-BrxA-DNA-binding<-wHTH+WYL+WCX<-HTH\|\|LexA-protease-><-EVE+HNH<-MobF<-VirD4-TraG+VirD4-FtsK<-TraJ-RHH | 1189 | Bacteria>Pseudomonadota>Alphaproteobacteria>Sphingomonadales>Sphingomonadaceae>Novosphingopyxis | GCF_000420305.1 | Novosphingopyxis_baekryungensis | G410_RS0110420 | 254465..258034 | 279369 | BREX-system-P-loop-protein-BrxC-[Novosphingopyxis-baekryungensis] |
| WP_021240096.1 | Type1.BREX | HKD-DNase->SF1-SF2-Helicase-><-Small-B-barrel+SF1-SF2-Helicase+RdgC+REase<-SF1-SF2-Helicase+REase<-35Exo+REase<-?\|\|wHTH+WYL+WCX->BrxA-DNA-binding->BrxB-BrxF-iSTAND->DUF6079*->OLD-ABC->REase->N6-METHYLASE->PglZ->SIGMA+OB-Fold+AAA-ATPase+HKD-DNase-BrxL-><-?<-Arm-DNA-bind_3+Phage_integrase | 1177 | Bacteria>Pseudomonadota>Alphaproteobacteria>Sphingomonadales>Sphingomonadaceae>Sphingobium | GCF_000445065.1 | Sphingobium_quisquiliarum | L288_RS41575 | 32905..36438 | 538379 | BREX-system-P-loop-protein-BrxC-[Sphingobium-quisquiliarum] |
| WP_076518288.1 | Type1.BREX | Arm-DNA-bind_3+Phage_integrase-><-SIGMA+OB-Fold+AAA-ATPase+HKD-DNase-BrxL<-PglZ<-N6-METHYLASE-p<-?<-REase<-OLD-ABC<-N6-METHYLASE<-?<-DUF6079*<-BrxB-BrxF-iSTAND<-BrxA-DNA-binding<-XIS-HTH<-Phage_int_SAM_2+Integrase_1<-Phage_rep<-DOC+HTH\|\|ACET-><-?\|\|REC+OmpR-HTH-> | 1181 | Bacteria>Pseudomonadota>Betaproteobacteria>Burkholderiales>Alcaligenaceae>Achromobacter | GCF_900156745.1 | Achromobacter_sp._MFA1_R4 | BXA00_RS09440 | 2091384..2094929 | 1881016 | BREX-system-P-loop-protein-BrxC-[Achromobacter-sp.-MFA1-R4] |
| WP_068373657.1 | Type1.BREX | <-TetR-HTH\|\|?->Phage_rep->HTH_3->Phage_int_SAM_2+Integrase_1->XIS-HTH->BrxA-DNA-binding->BrxB-BrxF-iSTAND->DUF6079*->?->N6-METHYLASE->OLD-ABC->REase->?->N6-METHYLASE-p->PglZ->SIGMA+OB-Fold+AAA-ATPase+HKD-DNase-BrxL-><-Arm-DNA-bind_3+Phage_integrase | 1176 | Bacteria>Pseudomonadota>Betaproteobacteria>Burkholderiales>Alcaligenaceae>Kerstersia | GCF_000988095.1 | Kerstersia_gyiorum | AAV32_RS14310 | 103983..107513 | 206506 | BREX-system-P-loop-protein-BrxC-[Kerstersia-gyiorum] |
| WP_132980217.1 | Type1.BREX | RelE-ParE->?->SF1-SF2-Helicase->Phage_rep->HTH_3->Phage_int_SAM_2+Integrase_1->XIS-HTH->BrxA-DNA-binding->BrxB-BrxF-iSTAND->DUF6079*->?->N6-METHYLASE->OLD-ABC->REase->?->N6-METHYLASE-p->PglZ->SIGMA+OB-Fold+AAA-ATPase+HKD-DNase-BrxL-><-Arm-DNA-bind_3+Phage_integrase | 1181 | Bacteria>Pseudomonadota>Betaproteobacteria>Burkholderiales>Alcaligenaceae>Pigmentiphaga | GCF_004352985.1 | Pigmentiphaga_sp._D-2 | EZY13_RS19720 | 185799..189344 | 1002116 | BREX-system-P-loop-protein-BrxC-[Pigmentiphaga-sp.-D-2] |
| WP_167306478.1 | Type1.BREX | <-PAS+STAS<-?<-?<-SIGMA+OB-Fold+AAA-ATPase+HKD-DNase-BrxL<-PglZ<-N6-METHYLASE<-REase<-OLD-ABC<-N6-METHYLASE<-DUF6079*<-BrxB-BrxF-iSTAND<-BrxA-DNA-binding<-XIS-HTH<-SF1-SF2-Helicase<-HKD-DNase+HKD-Rease | 1182 | Bacteria>Pseudomonadota>Betaproteobacteria>Burkholderiales>Burkholderiaceae>Paraburkholderia | GCF_900100565.1 | Paraburkholderia_steynii | BLT65_RS12240 | 289685..293233 | 1245441 | BREX-system-P-loop-protein-BrxC-[Paraburkholderia-steynii] |
| OJX16153.1 | Type1.BREX | <-SIGMA+OB-Fold+AAA-ATPase+HKD-DNase-BrxL<-PglZ<-Schlafen+HATPase+wHTH<-N6-METHYLASE-p<-?<-REase<-OLD-ABC<-N6-METHYLASE<-?<-DUF6079*<-BrxB-BrxF-iSTAND<-BrxA-DNA-binding<-Phage_int_SAM_2+Integrase_1<-Phage_rep\|\|CNMP-><-PRTase\|\|CBS+CBS->HAD+HAD-> | 1181 | Bacteria>Pseudomonadota>Betaproteobacteria>Burkholderiales>Comamonadaceae>Delftia | GCA_001898745.1 | Delftia_sp._67-8 | BGO79_00315 | 75959..79504 | 1895749 | hypothetical-protein-BGO79_00315-[Delftia-sp.-67-8] |
| WP_108330269.1 | Type1.BREX | <-SIGMA+OB-Fold+AAA-ATPase+Lon-Protease-BrxL<-PglZ<-HNH<-OLD-ABC<-N6-METHYLASE<-REase<-OLD-ABC<-DUF6079*<-BrxB-BrxF-iSTAND<-BrxA-DNA-binding<-PIN<-ZnR+TPR | 1154 | Bacteria>Pseudomonadota>Betaproteobacteria>Burkholderiales>Comamonadaceae>Limnohabitans | GCF_003063435.1 | Limnohabitans_sp._2KL-1 | B9Z47_RS05555 | 224189..227653 | 1100699 | BREX-system-P-loop-protein-BrxC-[Limnohabitans-sp.-2KL-1] |
| WP_186915942.1 | Type1.BREX | wHTH+WYL+WCX->?->?->?->wHTH+WYL+WCX-p->ANK->XIS-HTH->BrxA-DNA-binding->BrxB-BrxF-iSTAND->DUF6079*->OLD-ABC->REase->N6-METHYLASE->ParB+HNH->PglZ->SIGMA+OB-Fold+AAA-ATPase+HKD-DNase-BrxL-> | 1183 | Bacteria>Pseudomonadota>Betaproteobacteria>Burkholderiales>Oxalobacteraceae | GCF_014284115.1 | Undibacterium | H8K35_RS08325 | 157325..160876 | 401469 | MULTISPECIES:-BREX-system-P-loop-protein-BrxC-[Undibacterium] |
| WP_258819858.1 | Type1.BREX | Phage_integrase-><-?<-SbcC<-SIGMA+OB-Fold+AAA-ATPase+HKD-DNase-BrxL<-PglZ<-N6-METHYLASE-p<-REase<-OLD-ABC<-N6-METHYLASE<-DUF6079*<-BrxB-BrxF-iSTAND<-BrxA-DNA-binding<-wHTH+WYL+WCX | 1186 | Bacteria>Pseudomonadota>Betaproteobacteria>Burkholderiales>Oxalobacteraceae>Massilia_group | GCF_024753285.1 | Massilia_pinisoli | NX784_RS27510 | 42185..45745 | 1772194 | BREX-system-P-loop-protein-BrxC-[Massilia-pinisoli] |
| WP_239907373.1 | Type1.BREX | <-SIGMA+OB-Fold+AAA-ATPase+HKD-DNase-BrxL<-PglZ<-HTH+REase<-N6-METHYLASE-p<-HerA/FtsK<-SIR2<-REase<-OLD-ABC<-N6-METHYLASE<-DUF6079*<-BrxB-BrxF-iSTAND<-BrxA-DNA-binding<-wHTH+WYL+WCX<-Phage_integrase+Phage_integrase<-HTH_36<-XIS-HTH<-?<-Arm-DNA-bind_3+Phage_integrase<-sGTP | 1185 | Bacteria>Pseudomonadota>Betaproteobacteria>Neisseriales>Chromobacteriaceae>Laribacter | GCF_022394255.1 | Laribacter_hongkongensis | LH425_05855 | 38844..42401 | 168471 | BREX-system-P-loop-protein-BrxC-[Laribacter-hongkongensis] |
| WP_089417825.1 | Type1.BREX | <-SIGMA+OB-Fold+AAA-ATPase+Lon-Protease-BrxL<-PglZ<-REase+AIPR<-N6-METHYLASE<-REase<-OLD-ABC<-DUF6079*<-BrxB-BrxF-iSTAND<-BrxA-DNA-binding<-wHTH+WYL+WCX<-sGTP\|\|?->RHS+RHS+RHS+RHS->?->?-><-HTH_Tnp_1 | 1153 | Bacteria>Pseudomonadota>Betaproteobacteria>Neisseriales>Neisseriaceae>Vitreoscilla | GCF_002222655.1 | Vitreoscilla_filiformis | VITFI_RS15935 | 3397543..3401004 | 63 | BREX-system-P-loop-protein-BrxC-[Vitreoscilla-filiformis] |
| MBK7364915.1 | Type1.BREX | <-HIN-HTH+rve+rve_3\|\|tRNA-><-SIGMA+OB-Fold+AAA-ATPase+HKD-DNase-BrxL<-PglZ<-N6-METHYLASE<-REase<-OLD-ABC<-DUF6079*<-BrxB-BrxF-iSTAND<-BrxA-DNA-binding<-XIS-HTH\|\|ArdA->TRPR-HTH->?-><-DOC+HTH<-HNH<-TraI-VirD2-RCRE | 1187 | Bacteria>Pseudomonadota>Betaproteobacteria>Nitrosomonadales>Nitrosomonadaceae>Nitrosomonas | GCA_016705625.1 | Nitrosomonas_sp. | IPI97_07925 | 1671776..1675339 | 42353 | BREX-system-P-loop-protein-BrxC-[Nitrosomonas-sp.] |
| MBV6446594.1 | Type1.BREX | TraI-VirD2-RCRE->HNH->?-><-Couple_hipA+HipA_C<-TRPR-HTH<-ArdA\|\|XIS-HTH->BrxA-DNA-binding->BrxB-BrxF-iSTAND->DUF6079*->OLD-ABC->REase->N6-METHYLASE->PglZ->SIGMA+OB-Fold+AAA-ATPase+HKD-DNase-BrxL->SbcC+SbcC-><-Peptidase_S8<-BrxB-BrxF-iSTAND\|\|DOC+HTH-> | 1182 | Bacteria>Pseudomonadota>Betaproteobacteria>Nitrosomonadales>Nitrosomonadaceae>Nitrosomonas | GCA_019187415.1 | Nitrosomonas_sp. | FCKEOINB_00187 | 17095..20643 | 42353 | hypothetical-protein-[Nitrosomonas-sp.] |
| WP_205230204.1 | Type1.BREX | SbcC->?-><-SIGMA+OB-Fold+AAA-ATPase+Lon-Protease-BrxL<-PglZ<-SF1-SF2-Helicase<-OLD-ABC+TOPRIM<-N6-METHYLASE<-REase<-OLD-ABC<-DUF6079*<-BrxB-BrxF-iSTAND<-BrxA-DNA-binding<-RHH_5<-HTH_37\|\|Tn7_Tnp_TnsA_N+Tn7_Tnp_TnsA_C+rve+Mu-transpos_C->TnsC-ATPase-> | 1153 | Bacteria>Pseudomonadota>Betaproteobacteria>Rhodocyclales>Rhodocyclaceae>Azospira | GCF_905120365.1 | Azospira_oryzae | JYJ48_RS05820 | 17001..20462 | 146939 | BREX-system-P-loop-protein-BrxC-[Azospira-oryzae] |
| TXH82461.1 | Type1.BREX | PhdYeFM_antitox->PIN->BrxA-DNA-binding->BrxB-BrxF-iSTAND->DUF6079*->OLD-ABC->REase->N6-METHYLASE-p-> | 1153 | Bacteria>Pseudomonadota>Betaproteobacteria>Rhodocyclales>Zoogloeaceae>Thauera | GCA_008015255.1 | Thauera_aminoaromatica | E6Q80_15435 | 2474..5935 | 164330 | BREX-system-P-loop-protein-BrxC-[Thauera-aminoaromatica] |
| WP_107221110.1 | Type1.BREX | METHYLASE->ParA-Soj-PloopNTPase-><-SIGMA+OB-Fold+AAA-ATPase+Lon-Protease-BrxL\|\|DUF772+DDE_Tnp_1+DDE_Tnp_1_6-><-N6-METHYLASE-p<-REase<-OLD-ABC<-PIN<-MazE_antitoxin<-DUF6079*<-BrxB-BrxF-iSTAND<-BrxA-DNA-binding<-SPOUT<-RnaseHII | 1153 | Bacteria>Pseudomonadota>Betaproteobacteria>Rhodocyclales>Zoogloeaceae>Thauera | GCF_003030465.1 | Thauera_aromatica | Tharo_RS10350 | 2186270..2189731 | 59405 | BREX-system-P-loop-protein-BrxC-[Thauera-aromatica] |
| WP_247737217.1 | Type1.BREX | BrxA-DNA-binding->BrxB-BrxF-iSTAND->DUF6079*->MazE_antitoxin->PIN->OLD-ABC->REase->N6-METHYLASE-p-> | 1153 | Bacteria>Pseudomonadota>Betaproteobacteria>Rhodocyclales>Zoogloeaceae>Thauera | - | Thauera_aromatica | MX653_RS17175 | 1332..4793 | 59405 | BREX-system-P-loop-protein-BrxC-[Thauera-aromatica] |
| MBK6655412.1 | Type1.BREX | HDRnase->?-><-HTH_19\|\|?->?->PhdYeFM_antitox->PIN->BrxA-DNA-binding->BrxB-BrxF-iSTAND->DUF6079*->OLD-ABC->REase->N6-METHYLASE->ZnR->viral-capsid->?->?->TPR+REase-> | 1153 | Bacteria>Pseudomonadota>Betaproteobacteria>Rhodocyclales>Zoogloeaceae>Zoogloea | GCA_016703645.1 | Zoogloea_sp. | IPG40_16580 | 748210..751671 | 49181 | BREX-system-P-loop-protein-BrxC-[Zoogloea-sp.] |
| WP_148578091.1 | Type1.BREX | Arm-DNA-bind_2+Phage_int_SAM_3+Phage_integrase->HDRnase->?-><-HTH_19\|\|?->?->wHTH+WYL+WCX->BrxA-DNA-binding->BrxB-BrxF-iSTAND->DUF6079*->N6-METHYLASE->OLD-ABC->REase->SIR2->HerA/FtsK->N6-METHYLASE-p->HTH+REase->PglZ->SIGMA+OB-Fold+AAA-ATPase+HKD-DNase-BrxL-> | 1186 | Bacteria>Pseudomonadota>Betaproteobacteria>Rhodocyclales>Zoogloeaceae>Zoogloea | GCF_008107625.1 | Zoogloea_oleivorans | ETQ85_RS05700 | 193093..196653 | 1552750 | BREX-system-P-loop-protein-BrxC-[Zoogloea-oleivorans] |
| AUZ77883.1 | Type1.BREX | <-HIN-HTH<-SIGMA+OB-Fold+AAA-ATPase+HKD-DNase-BrxL<-PglZ<-ParB+HNH<-N6-METHYLASE<-REase<-OLD-ABC<-DUF6079*<-BrxB-BrxF-iSTAND<-BrxA-DNA-binding<-XIS-HTH\|\|<-DOC+HTH<-?<-XIS-HTH<-Arm-DNA-bind_3+Phage_integrase\|\|?-><-Sulfate_transp+STAS | 1183 | Bacteria>Pseudomonadota>Gammaproteobacteria>Aeromonadales>Aeromonadaceae>Aeromonas | GCA_002934425.1 | Aeromonas_sp._ASNIH4 | C2U40_16585 | 3601638..3605189 | 1636609 | BREX-system-P-loop-protein-BrxC-[Aeromonas-sp.-ASNIH4] |
| WP_043557612.1 | Type1.BREX | REase-><-?\|\|?-><-SIGMA+OB-Fold+AAA-ATPase+HKD-DNase-BrxL<-PglZ<-ParB+HNH<-N6-METHYLASE<-REase<-OLD-ABC<-DUF6079*<-BrxB-BrxF-iSTAND<-BrxA-DNA-binding<-XIS-HTH<-?<-AAA+HTH<-?<-XIS-HTH | 1182 | Bacteria>Pseudomonadota>Gammaproteobacteria>Aeromonadales>Aeromonadaceae>Aeromonas | GCF_000819745.1 | Aeromonas_bestiarum | BN1118_RS19650 | 69736..73284 | 105751 | BREX-system-P-loop-protein-BrxC-[Aeromonas-bestiarum] |
| WP_257706959.1 | Type1.BREX | Zn_Tnp_IS1+DDE_Tnp_IS1595-><-SIGMA+OB-Fold+AAA-ATPase+HKD-DNase-BrxL<-PglZ<-ParB+HNH<-N6-METHYLASE<-REase<-OLD-ABC<-DUF6079*<-BrxB-BrxF-iSTAND<-BrxA-DNA-binding<-XIS-HTH | 1182 | Bacteria>Pseudomonadota>Gammaproteobacteria>Aeromonadales>Aeromonadaceae>Aeromonas | GCF_024649325.1 | Aeromonas_caviae | NUK33_RS01495 | 150155..153703 | 648 | BREX-system-P-loop-protein-BrxC-[Aeromonas-caviae] |
| WP_270661186.1 | Type1.BREX | XIS-HTH->?->DOC+HTH-><-?\|\|XIS-HTH->BrxA-DNA-binding->BrxB-BrxF-iSTAND->DUF6079*->OLD-ABC->REase->N6-METHYLASE->ParB-> | 1182 | Bacteria>Pseudomonadota>Gammaproteobacteria>Aeromonadales>Aeromonadaceae>Aeromonas | GCF_027705205.1 | unclassified_Aeromonas | OKC30_RS15245 | 6003..9551 | 257493 | MULTISPECIES:-BREX-system-P-loop-protein-BrxC-[unclassified-Aeromonas] |
| WP_181014924.1 | Type1.BREX | <-HIN-HTH<-SIGMA+OB-Fold+AAA-ATPase+HKD-DNase-BrxL<-PglZ<-ParB+HNH<-N6-METHYLASE<-REase<-OLD-ABC<-DUF6079*<-BrxB-BrxF-iSTAND<-BrxA-DNA-binding<-XIS-HTH\|\|?-><-DOC+HTH<-?<-XIS-HTH\|\|?-><-Arm-DNA-bind_3+Phage_integrase | 1182 | Bacteria>Pseudomonadota>Gammaproteobacteria>Aeromonadales>Aeromonadaceae | GCF_002934425.1 | Aeromonas | C2U40_RS16580 | 3601638..3605186 | 642 | MULTISPECIES:-BREX-system-P-loop-protein-BrxC-[Aeromonas] |
| WP_006006589.1 | Type1.BREX | <-PIN<-MazE_antitoxin<-MODE-HTH+PBPII\|\|?->?->?->XIS-HTH->BrxA-DNA-binding->BrxB-BrxF-iSTAND->DUF6079*->OLD-ABC->REase->N6-METHYLASE->PglZ->SIGMA+OB-Fold+AAA-ATPase+HKD-DNase-BrxL-> | 1189 | Bacteria>Pseudomonadota>Gammaproteobacteria>Alteromonadales>Alteromonadaceae>Glaciecola | GCF_000252165.1 | Glaciecola_punicea | GPUN_RS10800 | 142812..146381 | 56804 | BREX-system-P-loop-protein-BrxC-[Glaciecola-punicea] |
| WP_136550441.1 | Type1.BREX | <-SbcC+APATPase+SbcC<-SIGMA+OB-Fold+AAA-ATPase+HKD-DNase-BrxL<-PglZ<-N6-METHYLASE<-REase<-OLD-ABC<-Schlafen+HATPase+wHTH+HTH<-DUF6079*<-BrxB-BrxF-iSTAND<-BrxA-DNA-binding<-XIS-HTH<-MobI | 1193 | Bacteria>Pseudomonadota>Gammaproteobacteria>Alteromonadales>Alteromonadaceae>Hydrocarboniclastica | GCF_004851605.1 | Hydrocarboniclastica_marina | soil367_RS18210 | 4131404..4134985 | 2259620 | BREX-system-P-loop-protein-BrxC-[Hydrocarboniclastica-marina] |

| Protein-ID | System | Gene-Neighbourhoods | Length | Taxonomy | Assembly-ID | Strain | Locus-Tag | Genomic-coordinates | TaxID | Fasta-defline |
|---|---|---|---|---|---|---|---|---|---|---|
| WP_112714387.1 | Type1.BREX | <- SIGMA+OB-Fold+AAA+ATPase+HKD-DNase-BrxL <- PglZ <-ParB+HNH<- N6- METHYLASE <- REase<- OLD-ABC <- DUF6079 *<- BrxB-BrxF- iSTAND <- BrxA-DNA-binding <-XIS-HTH\|\|?-><-OrfB_Zn_ribbon<-HTH_OrfB_IS605+OrfB_IS605+OrfB_Zn_ribbon\|\|DEDD_Tnp_IS110+Transposase_20-><-Peptidase_S51 | 1199 | Bacteria>Pseudomonadota>Gammaproteobacteria>Alteromonadales>Moritellaceae>Moritella | GCF_900465055.1 | Moritella_yayanosii | MORIYA_RS08605 | 1893116..1896715 | 69539 | BREX-system-P-loop-protein-BrxC-[Moritella-yayanosii] |
| WP_024611453.1 | Type1.BREX | <-ArdCN+MPTase<- SIGMA+OB-Fold+AAA+ATPase+HKD-DNase-BrxL <- PglZ <- N6- METHYLASE <- REase<- OLD-ABC <- DUF6079 *<- BrxB-BrxF- iSTAND <- BrxA-DNA-binding <-XIS-HTH | 1193 | Bacteria>Pseudomonadota>Gammaproteobacteria>Alteromonadales>Pseudoalteromonadaceae>Pseudoalteromonas | GCF_000498095.1 | Pseudoalteromonas_sp._TB64 | L647_RS0112855 | 131987..135568 | 1938600 | BREX-system-P-loop-protein-BrxC-[Pseudoalteromonas-sp.-TB64] |
| WP_167658112.1 | Type1.BREX | <-rve<-REase<-?<- SIGMA+OB-Fold+AAA+ATPase+HKD-DNase-BrxL <- PglZ <-ParB+HNH<- N6- METHYLASE <- REase<- OLD-ABC <- DUF6079 *<- BrxB-BrxF- iSTAND <- BrxA-DNA-binding <-XIS-HTH<-NtrC-AAA+FIS-HTH<-?<-?<-?<-?\|\|MODE-HTH+PBPII-> | 1188 | Bacteria>Pseudomonadota>Gammaproteobacteria>Alteromonadales>Pseudoalteromonadaceae>Pseudoalteromonas | GCF_011806445.1 | Pseudoalteromonas_sp._HF66 | HCZ11_RS07660 | 212811..216377 | 2721559 | BREX-system-P-loop-protein-BrxC-[Pseudoalteromonas-sp.-HF66] |
| PKG74712.1 | Type1.BREX | <- SIGMA+OB-Fold+AAA+ATPase+HKD-DNase-BrxL <- PglZ <-ParB+HNH<- N6- METHYLASE <- REase<- OLD-ABC <- DUF6079 *<- BrxB-BrxF- iSTAND <- BrxA-DNA-binding <-XIS-HTH<-?<-TnsC-ATPase<-rve+Mu-transpos_C<-REase+wHTH | 1191 | Bacteria>Pseudomonadota>Gammaproteobacteria>Alteromonadales>Shewanellaceae>Shewanella | GCA_002836135.1 | Shewanella_sp._GutCb | CXF86_11690 | 122253..125828 | 2058315 | BREX-system-P-loop-protein-BrxC-[Shewanella-sp.-GutCb] |
| WP_257985331.1 | Type1.BREX | <- SIGMA+OB-Fold+AAA+ATPase+HKD-DNase-BrxL <- PglZ <-ParB+HNH<- N6- METHYLASE <- REase<- OLD-ABC <- DUF6079 *<- BrxB-BrxF- iSTAND <- BrxA-DNA-binding <-XIS-HTH<-?<-TnsC-ATPase<-rve+Mu-transpos_C<-REase+wHTH | 1193 | Bacteria>Pseudomonadota>Gammaproteobacteria>Alteromonadales>Shewanellaceae>Shewanella | GCF_002836135.1 | Shewanella_sp._GutCb | CXF86_RS11730 | 122253..125834 | 2058315 | BREX-system-P-loop-protein-BrxC-[Shewanella-sp.-GutCb] |
| WP_112352819.1 | Type1.BREX | MerR-HTH->?-><- SIGMA+OB-Fold+AAA+ATPase+HKD-DNase-BrxL <- PglZ <- N6- METHYLASE <- REase<- OLD-ABC <- DUF6079 *<- BrxB-BrxF- iSTAND <- BrxA-DNA-binding <-XIS-HTH<-HNH+NERD+ZnR<-SAM- METHYLASE +wHTH+HNH<-?\|\|URI-DNase-><-MarR-HTH | 1197 | Bacteria>Pseudomonadota>Gammaproteobacteria>Alteromonadales>Shewanellaceae>Shewanella | GCF_900476435.1 | Shewanella_benthica | SHEWBE_RS12055 | 2634220..2637813 | 43661 | BREX-system-P-loop-protein-BrxC-[Shewanella-benthica] |
| WP_102039083.1 | Type1.BREX | <- SIGMA+OB-Fold+AAA+ATPase+HKD-DNase-BrxL <- PglZ <-ParB+HNH<- N6- METHYLASE <- REase<- OLD-ABC <- DUF6079 *<- BrxB-BrxF- iSTAND <- BrxA-DNA-binding <-XIS-HTH<-?<-TnsC-ATPase<-rve+Mu-transpos_C<-REase+wHTH | 1191 | Bacteria>Pseudomonadota>Gammaproteobacteria>Alteromonadales>Shewanellaceae>Shewanella | - | Shewanella_sp._GutCb | CXF86_11690 | 122253..125828 | 2058315 | BREX-system-P-loop-protein-BrxC-[Shewanella-sp.-GutCb] |
| WP_265366287.1 | Type1.BREX | Arm-DNA-bind_3+Phage_integrase-><-XIS-HTH<-AAA+HTH<-XIS-HTH\|\|MobI->XIS-HTH-> BrxA-DNA-binding -> BrxB-BrxF- iSTAND -> DUF6079 *->Schlafen+HATPase+wHTH+HTH-> OLD-ABC ->REase -> N6- METHYLASE ->ParB+HNH-> PglZ -> SIGMA+OB-Fold+AAA+ATPase+HKD-DNase-BrxL ->SbcC+APATPase+SbcC-> | 1193 | Bacteria>Pseudomonadota>Gammaproteobacteria>Cellvibrionales>Cellvibrionaceae>Marinimicrobium | GCF_026153455.1 | Marinimicrobium_koreense | OOT55_RS13025 | 3024461..3028042 | 306545 | BREX-system-P-loop-protein-BrxC-[Marinimicrobium-sp.-C6131] |
| WP_084199902.1 | Type1.BREX | CHTH->Calcineurin-><-?<-TnsC-ATPase<-?<-?\|\|XIS-HTH-> BrxA-DNA-binding -> BrxB-BrxF- iSTAND -> DUF6079 *-> OLD-ABC ->REase -> N6- METHYLASE -> PglZ -> SIGMA+OB-Fold+AAA+ATPase+HKD-DNase-BrxL -> | 1184 | Bacteria>Pseudomonadota>Gammaproteobacteria>Cellvibrionales>Halieaceae>Halioglobus | GCF_014652255.1 | Halioglobus_japonicus | C0029_RS05530 | 1195751..1199305 | 930805 | BREX-system-P-loop-protein-BrxC-[Halioglobus-japonicus] |
| MTW21824.1 | Type1.BREX | <-SAM- METHYLASE <- SIGMA+OB-Fold+AAA+ATPase+HKD-DNase-BrxL <- PglZ <-ParB+HNH<-Schlafen+HATPase+wHTH<-PhdYeFM_antitox<- N6- METHYLASE <- REase<- OLD-ABC <- DUF6079 *<- BrxB-BrxF- iSTAND <- BrxA-DNA-binding <-DnaB+HINT+DnaB+HINT | 1173 | Bacteria>Pseudomonadota>Gammaproteobacteria>Chromatiales>Chromatiaceae>Allochromatium | GCA_009720725.1 | Allochromatium_palmeri | GJ668_12065 | 37225..40746 | 231048 | BREX-system-P-loop-protein-BrxC-[Allochromatium-palmeri] |
| WP_186343071.1 | Type1.BREX | <-SAM- METHYLASE <- SIGMA+OB-Fold+AAA+ATPase+HKD-DNase-BrxL <- PglZ <-ParB+HNH<-Schlafen+HATPase+wHTH<-PhdYeFM_antitox<- N6- METHYLASE <- REase<- OLD-ABC <- DUF6079 *<- BrxB-BrxF- iSTAND <- BrxA-DNA-binding <-DnaB+HINT+DnaB+HINT | 1172 | Bacteria>Pseudomonadota>Gammaproteobacteria>Chromatiales>Chromatiaceae>Allochromatium | GCF_009720725.1 | Allochromatium_palmeri | GJ668_RS12065 | 37225..40743 | 231048 | BREX-system-P-loop-protein-BrxC-[Allochromatium-palmeri] |
| NEX15944.1 | Type1.BREX | <-NTP_transf<-?<- SIGMA+OB-Fold+AAA+ATPase+HKD-DNase-BrxL <- PglZ -p<- PglZ <-ParB+HNH<- N6- METHYLASE <- REase<- OLD-ABC <- DUF6079 *<- BrxB-BrxF- iSTAND <- BrxA-DNA-binding <-?<-?<-ABC-ATPase+SbcC<-?<-?\|\|PALP-> | 1173 | Bacteria>Pseudomonadota>Gammaproteobacteria>Chromatiales>Chromatiaceae>Halochromatium | GCA_010912705.1 | Halochromatium_sp. | C1943_04720 | 67001..70522 | 2049430 | BREX-system-P-loop-protein-BrxC-[Halochromatium-sp.] |
| WP_201222419.1 | Type1.BREX | <-PCMT+ METHYLASE <-?<-?<-?\|\|?->tRNA->?-> BrxA-DNA-binding -> BrxB-BrxF- iSTAND -> DUF6079 *-> OLD-ABC ->REase -> N6- METHYLASE ->Schlafen+HATPase+wHTH->ParB+HNH-> PglZ ->REase->MIT_C-> | 1174 | Bacteria>Pseudomonadota>Gammaproteobacteria>Chromatiales>Chromatiaceae>Halochromatium | GCF_016653315.1 | Halochromatium_roseum | CCR96_RS19410 | 137258..140782 | 391920 | BREX-system-P-loop-protein-BrxC-[Halochromatium-roseum] |
| WP_200250838.1 | Type1.BREX | <-ParB+HNH<- HNH<- OLD-ABC <- N6- METHYLASE <- REase<- OLD-ABC <- DUF6079 *<- BrxB-BrxF- iSTAND <- BrxA-DNA-binding | 1173 | Bacteria>Pseudomonadota>Gammaproteobacteria>Chromatiales>Chromatiaceae>Lamprobacter | GCF_016583905.1 | Lamprobacter_modestohalophilus | CK042_RS24315 | 8724..12245 | 1064514 | BREX-system-P-loop-protein-BrxC-[Lamprobacter-modestohalophilus] |
| QIG07578.1 | Type1.BREX | HTH_8->?->XIS-HTH->XIS-HTH->?->?-> BrxA-DNA-binding -> BrxB-BrxF- iSTAND -> DUF6079 *-> OLD-ABC ->REase -> N6- METHYLASE -> PglZ -> SIGMA+OB-Fold+AAA+ATPase+HKD-DNase-BrxL -><-Arm-DNA-bind_3+Phage_integrase | 1183 | Bacteria>Pseudomonadota>Gammaproteobacteria>Enterobacterales>Morganellaceae>Proteus | GCA_011046025.1 | Proteus_sp._ZN5 | GTK47_19170 | 4181966..4185517 | 2697019 | BREX-system-P-loop-protein-BrxC-[Proteus-sp.-ZN5] |
| WP_206535871.1 | Type1.BREX | HTH_8->?->XIS-HTH->XIS-HTH->?->?-> BrxA-DNA-binding -> BrxB-BrxF- iSTAND -> DUF6079 *-> OLD-ABC ->REase -> N6- METHYLASE -> PglZ -> SIGMA+OB-Fold+AAA+ATPase+HKD-DNase-BrxL -><-Arm-DNA-bind_3+Phage_integrase | 1185 | Bacteria>Pseudomonadota>Gammaproteobacteria>Enterobacterales>Morganellaceae>Proteus | GCF_011046025.1 | Proteus_sp._ZN5 | GTK47_RS19230 | 4181960..4185517 | 2697019 | BREX-system-P-loop-protein-BrxC-[Proteus-sp.-ZN5] |
| WP_272670012.1 | Type1.BREX | <-GTPase-AIG<-CopG/RHH\|\|?->XIS-HTH->?-> BrxA-DNA-binding -> BrxB-BrxF- iSTAND -> DUF6079 *-> OLD-ABC ->REase -> N6- METHYLASE -> PglZ -> SIGMA+OB-Fold+AAA+ATPase+HKD-DNase-BrxL -><-HTH_Tnp_1+HTH_21+rve+rve_3 | 1185 | Bacteria>Pseudomonadota>Gammaproteobacteria>Enterobacterales>Morganellaceae>Providencia | GCF_028480605.1 | unclassified_Providencia | PSE59_RS21125 | 8138..11695 | 2633465 | MULTISPECIES:-BREX-system-P-loop-protein-BrxC-[unclassified-Providencia] |
| MBS3952424.1 | Type1.BREX | XIS-HTH-> BrxA-DNA-binding -> BrxB-BrxF- iSTAND -> DUF6079 *-> OLD-ABC ->REase -> N6- METHYLASE ->PhdYeFM_antitox->PIN-> N6- METHYLASE -p->HTH+REase->Schlafen+HATPase+wHTH->SIR2-> | 1178 | Bacteria>Pseudomonadota>Gammaproteobacteria>Methylococcales>Methylococcaceae>Methylomicrobium | GCA_018333785.1 | Methylomicrobium_sp. | KGZ88_05695 | 2093..5629 | 2303549 | BREX-system-P-loop-protein-BrxC-[Methylomicrobium-sp.] |
| WP_096876853.1 | Type1.BREX | GDResolvase+HIN-HTH->?->XIS-HTH-> BrxA-DNA-binding -> BrxB-BrxF- iSTAND -> DUF6079 *-> OLD-ABC ->REase -> N6- METHYLASE ->HTH+REase-> PglZ -> SIGMA+OB-Fold+AAA+ATPase+HKD-DNase-BrxL ->Calcineurin-SbcD->SbcC+CCDC158->SbcC-> | 1180 | Bacteria>Pseudomonadota>Gammaproteobacteria>Methylococcales>Methylococcaceae>Methylomonas | GCF_002441955.1 | Methylomonas_koyamae | MKLM6_RS22285 | 58076..61618 | 702114 | BREX-system-P-loop-protein-BrxC-[Methylomonas-koyamae] |
| WP_004984466.1 | Type1.BREX | ZnR+ZnR->DUF932->?->?->XIS-HTH-> BrxA-DNA-binding -> BrxB-BrxF- iSTAND -> DUF6079 *-> OLD-ABC ->REase -> N6- METHYLASE -> PglZ -> SIGMA+OB-Fold+AAA+ATPase+HKD-DNase-BrxL ->?->?->ABC-ATPase+SbcC-><-ArsR-HTH | 1190 | Bacteria>Pseudomonadota>Gammaproteobacteria>Moraxellales>Moraxellaceae>Acinetobacter | GCF_000368805.1 | Acinetobacter_johnsonii | F946_RS15710 | 148915..152487 | 40214 | BREX-system-P-loop-protein-BrxC-[Acinetobacter-johnsonii] |
| WP_131214141.1 | Type1.BREX | ABC-ATPase+SbcC+ABC-ATPase+SbcC->?-><- SIGMA+OB-Fold+AAA+ATPase+HKD-DNase-BrxL <- PglZ <- N6- METHYLASE <- REase<- OLD-ABC <- DUF6079 *<- BrxB-BrxF- iSTAND <- BrxA-DNA-binding <-XIS-HTH<-XIS-HTH<-?<-?<-DUF932<-ZnR+ZnR | 1190 | Bacteria>Pseudomonadota>Gammaproteobacteria>Moraxellales>Moraxellaceae>Acinetobacter | GCF_004331285.1 | Acinetobacter_sp._ANC_3781 | E0H89_RS01980 | 72363..75935 | 2529835 | BREX-system-P-loop-protein-BrxC-[Acinetobacter-sp.-ANC-3781] |
| WP_151834043.1 | Type1.BREX | XIS-HTH-> BrxA-DNA-binding -> BrxB-BrxF- iSTAND -> DUF6079 *-> OLD-ABC ->REase -> N6- METHYLASE -> PglZ -> SIGMA+OB-Fold+AAA+ATPase+HKD-DNase-BrxL ->?->HerA/FtsK->ParB+HNH->ParB+HNH-> | 1190 | Bacteria>Pseudomonadota>Gammaproteobacteria>Moraxellales>Moraxellaceae>Acinetobacter | GCF_009012055.1 | Acinetobacter_ursingii | GBX89_RS12550 | 5375..8947 | 108980 | BREX-system-P-loop-protein-BrxC-[Acinetobacter-ursingii] |
| WP_180108075.1 | Type1.BREX | <- SIGMA+OB-Fold+AAA+ATPase+HKD-DNase-BrxL <- PglZ <- N6- METHYLASE <- REase<- OLD-ABC <- DUF6079 *<- BrxB-BrxF- iSTAND <-XIS-HTH<-?<-wHTH+HNH<-SF1-SF2-Helicase<-?<-SAM- METHYLASE \|\|?-><-ParB+HNH | 1190 | Bacteria>Pseudomonadota>Gammaproteobacteria>Moraxellales>Moraxellaceae>Acinetobacter | GCF_013419415.1 | Acinetobacter_sp._YH12085 | FTU28_RS13915 | 49013..52585 | 2601077 | BREX-system-P-loop-protein-BrxC-[Acinetobacter-sp.-YH12085] |
| WP_216936951.1 | Type1.BREX | <-HTH_3\|\|?-><-Phage_integrase<-Phage_integrase+Phage_integrase<-HTH\|\|XIS-HTH-> BrxA-DNA-binding -> BrxB-BrxF- iSTAND -> DUF6079 *-> OLD-ABC ->REase -> N6- METHYLASE -> PglZ -> SIGMA+OB-Fold+AAA+ATPase+HKD-DNase-BrxL -><-?<-REase\|\|ZnR-> | 1190 | Bacteria>Pseudomonadota>Gammaproteobacteria>Moraxellales>Moraxellaceae>Acinetobacter | GCF_019038255.1 | unclassified_Acinetobacter | KUG40_RS10660 | 10626..14198 | 196816 | MULTISPECIES:-BREX-system-P-loop-protein-BrxC-[unclassified-Acinetobacter] |
| MBL0229728.1 | Type1.BREX | <-HIN-HTH\|\|?->XIS-HTH->RelB-><-?<-?\|\| BrxA-DNA-binding -> BrxB-BrxF- iSTAND -> DUF6079 *-> OLD-ABC ->REase -> N6- METHYLASE ->PHP+SbcC->Schlafen+HATPase+wHTH-><-HTH_Tnp_1+HTH_21+rve+rve_3<-HIN-HTH<-DUF932 | 1147 | Bacteria>Pseudomonadota>Gammaproteobacteria>Moraxellales>Moraxellaceae>unclassified_Moraxellaceae | GCA_016721165.1 | Moraxellaceae_bacterium | IPP76_02595 | 6399..9842 | 1889775 | BREX-system-P-loop-protein-BrxC-[Moraxellaceae-bacterium] |
| MCH2557814.1 | Type1.BREX | Arm-DNA-bind_3+Phage_integrase-><-XIS-HTH<-AAA+HTH<-XIS-HTH\|\|MobI->XIS-HTH-> BrxA-DNA-binding -> BrxB-BrxF- iSTAND -> DUF6079 *->Schlafen+HATPase+wHTH+HTH-> OLD-ABC ->REase -> N6- METHYLASE -> PglZ -> SIGMA+OB-Fold+AAA+ATPase+HKD-DNase-BrxL ->SbcC+SbcC->Calcineurin+Calcineurin-SbcD->SbcC-> | 1193 | Bacteria>Pseudomonadota>Gammaproteobacteria>Oceanospirillales>Alcanivoracaceae>Alcanivorax | GCA_022449455.1 | Alcanivorax_sp. | MK005_11000 | 47715..51296 | 1872427 | BREX-system-P-loop-protein-BrxC-[Alcanivorax-sp.] |
| WP_103916700.1 | Type1.BREX | <- SIGMA+OB-Fold+AAA+ATPase+HKD-DNase-BrxL <-?<- PglZ <-ParB+HNH<- N6- METHYLASE <- REase<- OLD-ABC <-Schlafen+HATPase+wHTH+HTH<- DUF6079 *<- BrxB-BrxF- iSTAND <- BrxA-DNA-binding <-XIS-HTH<-?\|\| wHTH+WYL+WCX -><-XIS-HTH\|\|HIN-HTH-> | 1193 | Bacteria>Pseudomonadota>Gammaproteobacteria>Oceanospirillales>Alcanivoracaceae>Alcanivorax | GCF_900107995.1 | Alcanivorax_sp._DSM_26293 | C4S63_RS03510 | 780383..783964 | 1798238 | BREX-system-P-loop-protein-BrxC-[Alcanivorax-sp.-DSM-26293] |
| WP_228232948.1 | Type1.BREX | <-SbcC<-Calcineurin+Calcineurin-SbcD<-SbcC+SbcC<- SIGMA+OB-Fold+AAA+ATPase+HKD-DNase-BrxL <- PglZ <- N6- METHYLASE <- REase<- OLD-ABC <-Schlafen+HATPase+wHTH+HTH<- DUF6079 *<- BrxB-BrxF- iSTAND <- BrxA-DNA-binding <-XIS-HTH<-MobI\|\|XIS-HTH->AAA+HTH->XIS-HTH-><-Phage_integrase | 1193 | Bacteria>Pseudomonadota>Gammaproteobacteria>Oceanospirillales>Alcanivoracaceae>Alloalcanivorax | GCF_020782135.1 | Alloalcanivorax_marinus | LL252_RS03465 | 287759..291340 | 1177169 | BREX-system-P-loop-protein-BrxC-[Alloalcanivorax-marinus] |
| WP_008734404.1 | Type1.BREX | <- SIGMA+OB-Fold+AAA+ATPase+HKD-DNase-BrxL <- PglZ <- N6- METHYLASE <- REase<- OLD-ABC <-Schlafen+HATPase+wHTH+HTH<- DUF6079 *<- BrxB-BrxF- iSTAND <- BrxA-DNA-binding <-XIS-HTH<-sGTP<-?<-RNaseP | 1192 | Bacteria>Pseudomonadota>Gammaproteobacteria>Oceanospirillales>Alcanivoracaceae>Isoalcanivorax | GCF_000299335.2 | Isoalcanivorax_pacificus | S7S_RS18615 | 4158589..4162167 | 1306787 | BREX-system-P-loop-protein-BrxC-[Isoalcanivorax-pacificus] |
| WP_257265108.1 | Type1.BREX | XIS-HTH-> BrxA-DNA-binding -> BrxB-BrxF- iSTAND -> DUF6079 *-> OLD-ABC ->REase -> N6- METHYLASE -p-> | 1183 | Bacteria>Pseudomonadota>Gammaproteobacteria>Oceanospirillales>Endozoicomonadaceae>Endozoicomonas | GCF_024606365.1 | Endozoicomonas_sp._ONNA2 | KDK14_RS10760 | 1444..4995 | 2828741 | BREX-system-P-loop-protein-BrxC-[Endozoicomonas-sp.-ONNA2] |
| WP_019020105.1 | Type1.BREX | <-AlcRN+AraC-HTH<-?<-?<-?\|\|tRNA-><-Phage_integrase\|\|?->XIS-HTH-> BrxA-DNA-binding -> BrxB-BrxF- iSTAND -> DUF6079 *-> OLD-ABC ->REase -> N6- METHYLASE ->ParB+HNH-> PglZ ->Immank+ANK+ANK->REase-> SIGMA+OB-Fold+AAA+ATPase+HKD-DNase-BrxL -> | 1193 | Bacteria>Pseudomonadota>Gammaproteobacteria>Oceanospirillales>Halomonadaceae>Halomonas | GCF_014651575.1 | Halomonas_lutea | IE067_RS16235 | 61170..64751 | 453962 | BREX-system-P-loop-protein-BrxC-[Halomonas-lutea] |
| WP_027854332.1 | Type1.BREX | GDResolvase+HIN-HTH->?->?-><- SIGMA+OB-Fold+AAA+ATPase+HKD-DNase-BrxL <- PglZ <- N6- METHYLASE <- REase<- OLD-ABC <-Schlafen+HATPase+wHTH+HTH<- DUF6079 *<- BrxB-BrxF- iSTAND <- BrxA-DNA-binding <-XIS-HTH<-MobI\|\|XIS-HTH->AAA+HTH->XIS-HTH-><-Arm-DNA-bind_3+Phage_integrase | 1193 | Bacteria>Pseudomonadota>Gammaproteobacteria>Oceanospirillales>Oceanospirillaceae>Marinobacterium | GCF_000428985.1 | Marinobacterium_litorale | G403_RS0112740 | 74962..78543 | 404770 | BREX-system-P-loop-protein-BrxC-[Marinobacterium-litorale] |
| WP_239495510.1 | Type1.BREX | Phage_AlpA->Arm-DNA-bind_2+Phage_int_SAM_3+Phage_integrase->?-><- SIGMA+OB-Fold+AAA+ATPase+HKD-DNase-BrxL <- PglZ <-ParB+HNH<- N6- METHYLASE <- REase<- OLD-ABC <- DUF6079 *<- BrxB-BrxF- iSTAND <- BrxA-DNA-binding <-HTH_26\|\|<-HTH_Tnp_1+HTH_21+rve+rve_3<-HTH_19\|\|?->SF1-SF2-Helicase->TPR+HEPN-><- SF1-SF2-Helicase+wHTH+Small-B-barrel+DUF1998 | 1199 | Bacteria>Pseudomonadota>Gammaproteobacteria>Oceanospirillales>Oceanospirillaceae>Marinomonas | GCF_003285205.1 | Marinomonas_primoryensis | A8139_RS14980 | 3171018..3174617 | 178399 | BREX-system-P-loop-protein-BrxC-[Marinomonas-primoryensis] |
| MBE0507005.1 | Type1.BREX | BrxA-DNA-binding -> BrxB-BrxF- iSTAND -> DUF6079 *-> OLD-ABC ->REase -> N6- METHYLASE ->HTH+REase->REase-> PglZ -> SIGMA+OB-Fold+AAA+ATPase+Lon-Protease-BrxL -> | 1183 | Bacteria>Pseudomonadota>Gammaproteobacteria>Oceanospirillales>Oceanospirillaceae>Marinospirillum | GCA_014859565.1 | Marinospirillum_sp. | IBX50_09850 | 1716..5267 | 2183934 | BREX-system-P-loop-protein-BrxC-[Marinospirillum-sp.] |
| WP_022988850.1 | Type1.BREX | <- PglZ <- TOPRIM <- OLD-ABC <- HNH<- OLD-ABC <- N6- METHYLASE <- REase<- OLD-ABC <-Schlafen+HATPase+wHTH+HTH<- DUF6079 *<- BrxB-BrxF- iSTAND <- BrxA-DNA-binding <-XIS-HTH<-MobI | 1193 | Bacteria>Pseudomonadota>Gammaproteobacteria>Pseudomonadales>Marinobacteraceae>Marinobacter | GCF_000475255.1 | Marinobacter_sp._ES-1 | Q668_RS06800 | 88554..92135 | 1396858 | BREX-system-P-loop-protein-BrxC-[Marinobacter-sp.-ES-1] |
| WP_141068072.1 | Type1.BREX | Arm-DNA-bind_3+Phage_integrase-><-XIS-HTH<-AAA+HTH<-XIS-HTH\|\|MobI->XIS-HTH-> BrxA-DNA-binding -> BrxB-BrxF- iSTAND -> DUF6079 *->Schlafen+HATPase+wHTH+HTH-> OLD-ABC ->REase -> N6- METHYLASE ->ParB+HNH-> PglZ -> SIGMA+OB-Fold+AAA+ATPase+HKD-DNase-BrxL ->SbcC+APATPase+SbcC-> | 1193 | Bacteria>Pseudomonadota>Gammaproteobacteria>Pseudomonadales>Marinobacteraceae>Marinobacter | GCF_006516615.1 | Marinobacter_nauticus | FH712_RS17655 | 33566..37147 | 2743 | BREX-system-P-loop-protein-BrxC-[Marinobacter-nauticus] |
| WP_092033189.1 | Type1.BREX | Arm-DNA-bind_3+Phage_integrase-><-XIS-HTH<-AAA+HTH<-XIS-HTH\|\|MobI->XIS-HTH-> BrxA-DNA-binding -> BrxB-BrxF- iSTAND -> DUF6079 *->Schlafen+HATPase+wHTH+HTH-> OLD-ABC ->REase -> N6- METHYLASE -> PglZ -> SIGMA+OB-Fold+AAA+ATPase+HKD-DNase-BrxL ->SbcC->Calcineurin+Calcineurin-SbcD->SbcC-> | 1193 | Bacteria>Pseudomonadota>Gammaproteobacteria>Pseudomonadales>Marinobacteraceae>Marinobacter | GCF_900112835.1 | Marinobacter_sp._DSM_26671 | BM035_RS16335 | 75345..78926 | 1761793 | BREX-system-P-loop-protein-BrxC-[Marinobacter-sp.-DSM-26671] |
| WP_116918937.1 | Type1.BREX | <-XIS-HTH<-?<-?<-Arm-DNA-bind_2+Phage_int_SAM_3+Phage_integrase<-Phage_AlpA\|\|HTH_26-> BrxA-DNA-binding -> BrxB-BrxF- iSTAND -> DUF6079 *->Schlafen+HATPase+wHTH+HTH-> OLD-ABC ->REase -> N6- METHYLASE -> PglZ ->Immank+ANK+ANK->REase-> SIGMA+OB-Fold+AAA+ATPase+HKD-DNase-BrxL -> | 1193 | Bacteria>Pseudomonadota>Gammaproteobacteria>Pseudomonadales>Marinobacteraceae>Tamilnaduibacter | GCF_003096655.1 | Tamilnaduibacter_salinus | C8D92_104115 | 123558..127139 | 1484056 | BREX-system-P-loop-protein-BrxC-[Tamilnaduibacter-salinus] |
| WP_105523318.1 | Type1.BREX | <- SIGMA+OB-Fold+AAA+ATPase+HKD-DNase-BrxL <- PglZ <- N6- METHYLASE -p<-?<- REase<- OLD-ABC <- N6- METHYLASE <- DUF6079 *<- BrxB-BrxF- iSTAND <- BrxA-DNA-binding <-XIS-HTH<-ARG<-?\|\| wHTH+WYL+WCX -><-sGTP<-sGTP | 1185 | Bacteria>Pseudomonadota>Gammaproteobacteria>Pseudomonadales>Pseudomonadaceae | GCF_002975295.1 | Pseudomonas | CQ048_RS15780 | 97047..100604 | 286 | MULTISPECIES:-BREX-system-P-loop-protein-BrxC-[Pseudomonas] |
| WP_060765715.1 | Type1.BREX | <-HKD-DNase+HKD-DNase<-?\|\|\|\|<- SIGMA+OB-Fold+AAA+ATPase+HKD-DNase-BrxL <- PglZ <-ParB+HNH<- N6- METHYLASE -p<- REase<- OLD-ABC <- N6- METHYLASE -p<- DUF6079 *<- BrxB-BrxF- iSTAND <- BrxA-DNA-binding <-XIS-HTH<-?<-?<-XIS-HTH\|\|RHH-><-tRNA<-MerR-HTH | 1186 | Bacteria>Pseudomonadota>Gammaproteobacteria>Pseudomonadales>Pseudomonadaceae | GCF_029959765.1 | Pseudomonas | PFL603g_RS03160 | 383988..387548 | 286 | MULTISPECIES:-BREX-system-P-loop-protein-BrxC-[Pseudomonas] |
| HCF5867680.1 | Type1.BREX | <-ParA-Soj-PloopNTPase<- METHYLASE <-?<-?\|\|Phage_rep->Phage_int_SAM_2+Integrase_1->XIS-HTH-> BrxA-DNA-binding -> BrxB-BrxF- iSTAND -> DUF6079 *->?-> N6- METHYLASE -> OLD-ABC ->REase -> N6- METHYLASE -p->Schlafen+HATPase+wHTH-> PglZ -> SIGMA+OB-Fold+AAA+ATPase+HKD-DNase-BrxL -><-Arm-DNA-bind_3+Phage_integrase | 1181 | Bacteria>Pseudomonadota>Gammaproteobacteria>Pseudomonadales>Pseudomonadaceae>Pseudomonas | GCA_023959235.1 | Pseudomonas_aeruginosa | NIH49_002287 | 171515..175060 | 287 | BREX-system-P-loop-protein-BrxC-[Pseudomonas-aeruginosa] |
| WP_010463595.1 | Type1.BREX | <- SIGMA+OB-Fold+AAA+ATPase+HKD-DNase-BrxL <- PglZ <-ParB+HNH<- N6- METHYLASE -p<- REase<- OLD-ABC <- N6- METHYLASE -p<- DUF6079 *<- BrxB-BrxF- iSTAND <- BrxA-DNA-binding <-XIS-HTH<-ARG<-?\|\| wHTH+WYL+WCX -><-sGTP<-sGTP | 1185 | Bacteria>Pseudomonadota>Gammaproteobacteria>Pseudomonadales>Pseudomonadaceae>Pseudomonas | GCF_000257545.3 | Pseudomonas_mandelii | OU5_RS13450 | 3003724..3007281 | 75612 | BREX-system-P-loop-protein-BrxC-[Pseudomonas-mandelii] |
| WP_048385719.1 | Type1.BREX | <- SIGMA+OB-Fold+AAA+ATPase+HKD-DNase-BrxL <- PglZ <-ParB+HNH<- N6- METHYLASE -p<- REase<- OLD-ABC <- N6- METHYLASE -p<- DUF6079 *<- BrxB-BrxF- iSTAND <- BrxA-DNA-binding <-XIS-HTH<-ARG<-?\|\| wHTH+WYL+WCX -><-sGTP<-sGTP | 1185 | Bacteria>Pseudomonadota>Gammaproteobacteria>Pseudomonadales>Pseudomonadaceae>Pseudomonas | GCF_001043065.1 | Pseudomonas_helleri | TU85_RS02975 | 88982..92539 | 1608996 | BREX-system-P-loop-protein-BrxC-[Pseudomonas-helleri] |
| WP_150073350.1 | Type1.BREX | XIS-HTH->?->XIS-HTH-> BrxA-DNA-binding -> BrxB-BrxF- iSTAND -> DUF6079 *-> N6- METHYLASE -p-> OLD-ABC ->REase -> N6- METHYLASE -p->ParB+HNH-> PglZ -> SIGMA+OB-Fold+AAA+ATPase+HKD-DNase-BrxL -> | 1186 | Bacteria>Pseudomonadota>Gammaproteobacteria>Pseudomonadales>Pseudomonadaceae>Pseudomonas | GCF_008632325.1 | Pseudomonas_lactis | F3K52_RS19340 | 4156..7716 | 1615674 | BREX-system-P-loop-protein-BrxC-[Pseudomonas-lactis] |
| WP_152745526.1 | Type1.BREX | <-CopG/RHH\|\|XIS-HTH->?->XIS-HTH-> BrxA-DNA-binding -> BrxB-BrxF- iSTAND -> DUF6079 *-> OLD-ABC ->REase -> N6- METHYLASE -> PglZ -> SIGMA+OB-Fold+AAA+ATPase+Lon-Protease-BrxL ->MIT_C-> | 1183 | Bacteria>Pseudomonadota>Gammaproteobacteria>Pseudomonadales>Pseudomonadaceae>Pseudomonas | GCF_009296165.1 | Pseudomonas_kitaguniensis | F0169_RS03300 | 21652..25203 | 2607908 | BREX-system-P-loop-protein-BrxC-[Pseudomonas-kitaguniensis] |
| WP_153384138.1 | Type1.BREX | MerR-HTH->tRNA-><-RHH\|\|XIS-HTH->?->XIS-HTH-> BrxA-DNA-binding -> BrxB-BrxF- iSTAND -> DUF6079 *-> N6- METHYLASE -p-> OLD-ABC ->REase -> N6- METHYLASE -p->ParB+HNH-> PglZ -> SIGMA+OB-Fold+AAA+ATPase+HKD-DNase-BrxL ->\|\|<-?<- wHTH+WYL+WCX | 1186 | Bacteria>Pseudomonadota>Gammaproteobacteria>Pseudomonadales>Pseudomonadaceae>Pseudomonas | GCF_009600565.1 | unclassified_Pseudomonas | GHO24_RS06625 | 35218..38778 | 196821 | MULTISPECIES:-BREX-system-P-loop-protein-BrxC-[unclassified-Pseudomonas] |
| WP_177095197.1 | Type1.BREX | <- N6- METHYLASE -p<- REase<- OLD-ABC <- N6- METHYLASE <- DUF6079 *<- BrxB-BrxF- iSTAND <- BrxA-DNA-binding <-XIS-HTH<-ARG<-?\|\| wHTH+WYL+WCX -><-sGTP<-sGTP | 1185 | Bacteria>Pseudomonadota>Gammaproteobacteria>Pseudomonadales>Pseudomonadaceae>Pseudomonas | GCF_013387065.1 | Pseudomonas_reactans | HX796_RS11550 | 4461..8018 | 117680 | BREX-system-P-loop-protein-BrxC-[Pseudomonas-reactans] |
| WP_198790924.1 | Type1.BREX | <-CopG/RHH\|\|XIS-HTH->?->?->XIS-HTH-> BrxA-DNA-binding -> BrxB-BrxF- iSTAND -> DUF6079 *-> N6- METHYLASE -p-> OLD-ABC ->REase ->?-> N6- METHYLASE -p->ParB+HNH-> PglZ -> SIGMA+OB-Fold+AAA+ATPase+HKD-DNase-BrxL -> | 1185 | Bacteria>Pseudomonadota>Gammaproteobacteria>Pseudomonadales>Pseudomonadaceae>Pseudomonas | GCF_016405175.1 | Pseudomonas_sp._MF7451 | JFT65_RS10900 | 194530..198087 | 2797538 | BREX-system-P-loop-protein-BrxC-[Pseudomonas-sp.-MF7451] |
| WP_223023201.1 | Type1.BREX | <-REase\|\|DNA_ METHYLASE -><-CopG/RHH\|\|XIS-HTH->?->XIS-HTH-> BrxA-DNA-binding -> BrxB-BrxF- iSTAND -> DUF6079 *-> OLD-ABC ->REase -> N6- METHYLASE -> PglZ -> SIGMA+OB-Fold+AAA+ATPase+HKD-DNase-BrxL -><-?<-?<-?\|\|HTH_3-> | 1183 | Bacteria>Pseudomonadota>Gammaproteobacteria>Pseudomonadales>Pseudomonadaceae>Pseudomonas | GCF_019891425.1 | Pseudomonas_carnis | J1G36_06600 | 89435..92986 | 2487355 | BREX-system-P-loop-protein-BrxC-[Pseudomonas-carnis] |
| WP_236238652.1 | Type1.BREX | <-HIN-HTH<- SIGMA+OB-Fold+AAA+ATPase+HKD-DNase-BrxL <- PglZ <-ParB+HNH<- N6- METHYLASE -p<- REase<- OLD-ABC <- N6- METHYLASE -p<- DUF6079 *<- BrxB-BrxF- iSTAND <- BrxA-DNA-binding <-XIS-HTH | 1185 | Bacteria>Pseudomonadota>Gammaproteobacteria>Pseudomonadales>Pseudomonadaceae>Pseudomonas | GCF_021603345.1 | Pseudomonas_rhodesiae | L6E14_RS13730 | 266364..269921 | 76760 | BREX-system-P-loop-protein-BrxC-[Pseudomonas-rhodesiae] |
| WP_265072431.1 | Type1.BREX | <- SIGMA+OB-Fold+AAA+ATPase+HKD-DNase-BrxL <- PglZ <-ParB+HNH<- N6- METHYLASE -p<- REase<- OLD-ABC <- N6- METHYLASE -p<- DUF6079 *<- BrxB-BrxF- iSTAND <- BrxA-DNA-binding <-XIS-HTH\|\|HTH+REase-><-?<-?<-?<-Glyco_hydro_68 | 1186 | Bacteria>Pseudomonadota>Gammaproteobacteria>Pseudomonadales>Pseudomonadaceae>Pseudomonas | GCF_026016265.1 | Pseudomonas_viciae | LOY66_RS25465 | 5798638..5802198 | 2505979 | BREX-system-P-loop-protein-BrxC-[Pseudomonas-viciae] |
| MAQ23221.1 | Type1.BREX | <-Calcineurin<-Calcineurin-SbcD<- PglZ <-ParB+HNH<- N6- METHYLASE <- REase<- OLD-ABC <-Schlafen+HATPase+wHTH+HTH<- DUF6079 *<- BrxB-BrxF- iSTAND <- BrxA-DNA-binding <-XIS-HTH<-MobI\|\|XIS-HTH->AAA+HTH->XIS-HTH-><-Arm-DNA-bind_3+Phage_integrase | 1193 | Bacteria>Pseudomonadota>Gammaproteobacteria>Pseudomonadales>unclassified_Pseudomonadales | GCA_002701315.1 | Pseudomonadales_bacterium | CMK83_03295 | 18015..21596 | 1891229 | hypothetical-protein-[Pseudomonadales-bacterium] |
| MCP5344826.1 | Type1.BREX | <-PALP<- SIGMA+OB-Fold+AAA+ATPase+Lon-Protease-BrxL <- PglZ <-PHP+SbcC<- N6- METHYLASE <- REase<- OLD-ABC <- DUF6079 -p* | 834 | Bacteria>Pseudomonadota>Gammaproteobacteria>Pseudomonadales>unclassified_Pseudomonadales | GCA_024234345.1 | Pseudomonadales_bacterium | H7A05_09410 | 50993..53498 | 1891229 | BREX-system-P-loop-protein-BrxC-[Pseudomonadales-bacterium] |
| PCJ88241.1 | Type1.BREX | <-RelE-ParE<-MetJarc-HTH\|\|?-><-HKD-DNase+DUF3150<- SIGMA+OB-Fold+AAA+ATPase+HKD-DNase-BrxL <- PglZ <- N6- METHYLASE <- REase<- OLD-ABC <- DUF6079 *<- BrxB-BrxF- iSTAND <- BrxA-DNA-binding <-XIS-HTH<-?<-?\|\|RHH_5->?->?-><-HISKIN | 1177 | Bacteria>Pseudomonadota>Gammaproteobacteria>Thiotrichales>Thiotrichaceae>unclassified_Thiotrichaceae | GCA_002746365.1 | Thiotrichaceae_bacterium | COA54_02955 | 15853..19386 | 2030882 | hypothetical-protein-COA54_02955-[Thiotrichaceae-bacterium] |
| HIJ32929.1 | Type1.BREX | BrxA-DNA-binding -> BrxB-BrxF- iSTAND -> DUF6079 *-> OLD-ABC ->REase -> N6- METHYLASE -p-> | 1200 | Bacteria>Pseudomonadota>Gammaproteobacteria>unclassified_Gammaproteobacteria | GCA_013349805.1 | Gammaproteobacteria_bacterium | HPP65_00765 | 772..4374 | 1913989 | BREX-system-P-loop-protein-BrxC-[Gammaproteobacteria-bacterium] |
| WP_245395138.1 | Type1.BREX | LON-AAA->Calcineurin-SbcD->Calcineurin-><-?\|\|?->\|\| BrxA-DNA-binding -> DUF6079 -p-><- REase<- OLD-ABC <- DUF6079 *<-HDRnase\|\|<-Arm-DNA-bind_2+Phage_int_SAM_3+Phage_integrase<-Phage_AlpA\|\|HTH_26-><-?<-tRNA\|\|RADICAL-SAM->?->DDE_Tnp_1-> | 1115 | Bacteria>Pseudomonadota>Gammaproteobacteria>unclassified_Gammaproteobacteria | GCA_016097405.1 | methane-oxidizing_endosymbiont_of_Gigantopelta_aegis | JEU20_RS06805 | 1356880..1360227 | 2794938 | BREX-system-P-loop-protein-BrxC-[methane-oxidizing-endosymbiont-of-Gigantopelta-aegis] |
| WP_006647488.1 | Type1.BREX | HTH_Tnp_1+HTH_21+rve+rve_3-><- SIGMA+OB-Fold+AAA+ATPase+HKD-DNase-BrxL <- PglZ <-ParB+HNH<- N6- METHYLASE <- REase<- OLD-ABC <-HTH_Tnp_1+HTH_21+rve+rve_3<- OLD-ABC <- DUF6079 *<- BrxB-BrxF- iSTAND <- BrxA-DNA-binding <-XIS-HTH<-?<-TnsC-ATPase\|\|HTH_Tnp_1+HTH_21+rve+rve_3-><-rve+Mu-transpos_C<-REase+wHTH | 1193 | Bacteria>Pseudomonadota>Gammaproteobacteria>Vibrionales>Vibrionaceae>Photobacterium | GCF_000153325.1 | Photobacterium_sp._SKA34 | SKA34_RS21260 | 87534..91115 | 121723 | BREX-system-P-loop-protein-BrxC-[Photobacterium-sp.-SKA34] |
| ASG04983.1 | Type1.BREX | <-DeoR-HTH+ISOCOT<-DeoR-HTH+ISOCOT<-DUF2787<-?<-?\|\|XIS-HTH->XIS-HTH-> BrxA-DNA-binding -> BrxB-BrxF- iSTAND -> DUF6079 *-> OLD-ABC ->REase -> N6- METHYLASE -> PglZ -> SIGMA+OB-Fold+AAA+ATPase+HKD-DNase-BrxL -><-Arm-DNA-bind_3+Phage_integrase | 1185 | Bacteria>Pseudomonadota>Gammaproteobacteria>Vibrionales>Vibrionaceae>Vibrio | GCA_002211985.1 | Vibrio_anguillarum | CEJ46_04105 | 862780..866337 | 55601 | hypothetical-protein-CEJ46_04105-[Vibrio-anguillarum] |
| NAZ69809.1 | Type1.BREX | <- SIGMA+OB-Fold+AAA+ATPase+HKD-DNase-BrxL <- PglZ <- N6- METHYLASE <- REase<- OLD-ABC <- DUF6079 *<- BrxB-BrxF- iSTAND <- BrxA-DNA-binding <-XIS-HTH<-HNH<-?<-XIS-HTH\|\|?->DDE_Tnp_1_5+DDE_Tnp_1-> | 1185 | Bacteria>Pseudomonadota>Gammaproteobacteria>Vibrionales>Vibrionaceae>Vibrio | GCA_009906175.1 | Vibrio_toranzoniae | GL180_08045 | 10464..14021 | 1194427 | BREX-system-P-loop-protein-BrxC-[Vibrio-toranzoniae] |
| EGQ8448095.1 | Type1.BREX | <- SIGMA+OB-Fold+AAA+ATPase+HKD-DNase-BrxL <- PglZ <- N6- METHYLASE <- REase<- OLD-ABC <- DUF6079 *<- BrxB-BrxF- iSTAND <- BrxA-DNA-binding <-XIS-HTH<-HNH<-XIS-HTH\|\|?->?->DUF2787-> wHTH+WYL+WCX -> | 1185 | Bacteria>Pseudomonadota>Gammaproteobacteria>Vibrionales>Vibrionaceae>Vibrio | GCA_015775815.1 | Vibrio_alginolyticus | GTP59_14930 | 64166..67723 | 663 | BREX-system-P-loop-protein-BrxC-[Vibrio-alginolyticus] |
| MBH9740666.1 | Type1.BREX | SF1-SF2-Helicase+HTH->MukF->JetB->XIS-HTH-> BrxA-DNA-binding -> BrxB-BrxF- iSTAND -> DUF6079 *-> OLD-ABC ->REase -> N6- METHYLASE -> PglZ -> SIGMA+OB-Fold+AAA+ATPase+HKD-DNase-BrxL -> | 1185 | Bacteria>Pseudomonadota>Gammaproteobacteria>Vibrionales>Vibrionaceae>Vibrio | GCA_016094345.1 | Vibrio_navarrensis | CJ419_13180 | 152917..156474 | 29495 | hypothetical-protein-[Vibrio-navarrensis] |
| MBT0011870.1 | Type1.BREX | <- SIGMA+OB-Fold+AAA+ATPase+HKD-DNase-BrxL <- PglZ <- N6- METHYLASE <- REase<- OLD-ABC <- DUF6079 *<- BrxB-BrxF- iSTAND <- BrxA-DNA-binding <-XIS-HTH<-HNH<-XIS-HTH\|\|?->?->DUF2787-> wHTH+WYL+WCX -> | 1187 | Bacteria>Pseudomonadota>Gammaproteobacteria>Vibrionales>Vibrionaceae>Vibrio | GCA_018457755.1 | Vibrio_alginolyticus | N7I24_002738 | 64007..67570 | 663 | BREX-system-P-loop-protein-BrxC-[Vibrio-alginolyticus] |
| EIZ1010574.1 | Type1.BREX | HTH_3->Couple_hipA+HipA_C->SAM- METHYLASE -> HKD-DNase+SF1-SF2-Helicase +HTH+HTH+SRA-><- SIGMA+OB-Fold+AAA+ATPase+HKD-DNase-BrxL <- PglZ <- N6- METHYLASE <- REase<- OLD-ABC <- DUF6079 *<- BrxB-BrxF- iSTAND <- BrxA-DNA-binding <-XIS-HTH<-RNase<-DUF3584+DUF5401<-JetB<-MukF\|\| wHTH+WYL+WCX -><-TPR | 1187 | Bacteria>Pseudomonadota>Gammaproteobacteria>Vibrionales>Vibrionaceae>Vibrio | GCA_022609845.1 | Vibrio_vulnificus | MOS09_002154 | 288384..291947 | 672 | BREX-system-P-loop-protein-BrxC-[Vibrio-vulnificus] |
| HCG7393723.1 | Type1.BREX | McrB-AAA->wHTH+REase->?->HNH->?-><-HEPN\|\|XIS-HTH-> BrxA-DNA-binding -> BrxB-BrxF- iSTAND -> DUF6079 *-> OLD-ABC ->REase -> N6- METHYLASE -> PglZ -> SIGMA+OB-Fold+AAA+ATPase+HKD-DNase-BrxL ->Phage_integrase-><-ParB<-GDResolvase+HIN-HTH\|\| OLD-ABC -> | 1187 | Bacteria>Pseudomonadota>Gammaproteobacteria>Vibrionales>Vibrionaceae>Vibrio | GCA_024044075.1 | Vibrio_parahaemolyticus | NJ383_004129 | 13694..17257 | 670 | BREX-system-P-loop-protein-BrxC-[Vibrio-parahaemolyticus] |
| EJV5950193.1 | Type1.BREX | <- SIGMA+OB-Fold+AAA+ATPase+HKD-DNase-BrxL <- PglZ <- N6- METHYLASE <- REase<- OLD-ABC <- DUF6079 *<- BrxB-BrxF- iSTAND <- BrxA-DNA-binding <-XIS-HTH<-HNH<-XIS-HTH\|\|?->?->DUF2787-> wHTH+WYL+WCX -> | 1187 | Bacteria>Pseudomonadota>Gammaproteobacteria>Vibrionales>Vibrionaceae>Vibrio | GCA_025320405.1 | Vibrio_alginolyticus | N7I24_002738 | 64007..67570 | 663 | BREX-system-P-loop-protein-BrxC-[Vibrio-alginolyticus] |
| GLT15886.1 | Type1.BREX | <- wHTH+WYL+WCX \|\|MukF->JetB->DUF3584+DUF5401->RNase->XIS-HTH-> BrxA-DNA-binding -> BrxB-BrxF- iSTAND -> DUF6079 *-> OLD-ABC ->REase -> N6- METHYLASE -> PglZ -> SIGMA+OB-Fold+AAA+ATPase+HKD-DNase-BrxL ->?->HTH_3->Couple_hipA+HipA_C-> | 1186 | Bacteria>Pseudomonadota>Gammaproteobacteria>Vibrionales>Vibrionaceae>Vibrio | GCA_030161215.1 | Vibrio_algivorus | GCM10007931_28610 | 496776..500336 | 1667024 | hypothetical-protein-GCM10007931_28610-[Vibrio-algivorus] |
| WP_198300681.1 | Type1.BREX | <-DeoR-HTH+ISOCOT<-DeoR-HTH+ISOCOT<-?<-?\|\|XIS-HTH->XIS-HTH-> BrxA-DNA-binding -> BrxB-BrxF- iSTAND -> DUF6079 *-> OLD-ABC ->REase -> N6- METHYLASE -> PglZ -> SIGMA+OB-Fold+AAA+ATPase+HKD-DNase-BrxL -><-Arm-DNA-bind_3+Phage_integrase | 1187 | Bacteria>Pseudomonadota>Gammaproteobacteria>Vibrionales>Vibrionaceae>Vibrio | GCF_002211985.1 | Vibrio_anguillarum | CEJ46_RS04105 | 862774..866337 | 55601 | BREX-system-P-loop-protein-BrxC-[Vibrio-anguillarum] |
| WP_089123599.1 | Type1.BREX | <-Couple_hipA+HipA_C<-HTH_3<-?<- SIGMA+OB-Fold+AAA+ATPase+HKD-DNase-BrxL <- PglZ <- N6- METHYLASE <- REase<- OLD-ABC <- DUF6079 *<- BrxB-BrxF- iSTAND <- BrxA-DNA-binding <-XIS-HTH<-RNase<-DUF3584+DUF5401<-JetB<-MukF\|\| wHTH+WYL+WCX -> | 1184 | Bacteria>Pseudomonadota>Gammaproteobacteria>Vibrionales>Vibrionaceae>Vibrio | GCF_002218005.2 | Vibrio_algivorus | AVR_RS07075 | 1502603..1506157 | 1667024 | BREX-system-P-loop-protein-BrxC-[Vibrio-algivorus] |
| WP_202601719.1 | Type1.BREX | <- SIGMA+OB-Fold+AAA+ATPase+HKD-DNase-BrxL <- PglZ <- N6- METHYLASE <- REase<- OLD-ABC <- DUF6079 *<- BrxB-BrxF- iSTAND <- BrxA-DNA-binding <-XIS-HTH<-HNH<-SF1-SF2-Helicase+HTH<-SF1-SF2-Helicase<-?<-XIS-HTH | 1187 | Bacteria>Pseudomonadota>Gammaproteobacteria>Vibrionales>Vibrionaceae>Vibrio | GCF_009906175.1 | Vibrio_toranzoniae | GL180_RS08040 | 10464..14027 | 1194427 | BREX-system-P-loop-protein-BrxC-[Vibrio-toranzoniae] |
| WP_198106729.1 | Type1.BREX | SF1-SF2-Helicase+HTH->MukF->JetB->DUF3584+DUF5401->RNase->XIS-HTH-> BrxA-DNA-binding -> BrxB-BrxF- iSTAND -> DUF6079 *-> OLD-ABC ->REase -> N6- METHYLASE -> PglZ -> SIGMA+OB-Fold+AAA+ATPase+HKD-DNase-BrxL ->Phage_integrase-> | 1187 | Bacteria>Pseudomonadota>Gammaproteobacteria>Vibrionales>Vibrionaceae>Vibrio | GCF_016094345.1 | Vibrio_navarrensis | CJ419_RS12980 | 152911..156474 | 29495 | BREX-system-P-loop-protein-BrxC-[Vibrio-navarrensis] |
| WP_250156418.1 | Type1.BREX | <- wHTH+WYL+WCX <-DUF2787<-?<-?\|\|XIS-HTH->HNH->XIS-HTH-> BrxA-DNA-binding -> BrxB-BrxF- iSTAND -> DUF6079 *-> OLD-ABC ->REase -> N6- METHYLASE -> PglZ -> SIGMA+OB-Fold+AAA+ATPase+HKD-DNase-BrxL -> | 1185 | Bacteria>Pseudomonadota>Gammaproteobacteria>Vibrionales>Vibrionaceae>Vibrio | GCF_018457845.1 | Vibrio_alginolyticus | J4H60_RS12980 | 321648..325205 | 663 | BREX-system-P-loop-protein-BrxC-[Vibrio-alginolyticus] |
| WP_267363047.1 | Type1.BREX | <-XIS-HTH<-Arm-DNA-bind_3+Phage_int_SAM_3+Phage_integrase\|\|?->?->\|\|?-><-HEPN\|\|XIS-HTH-> BrxA-DNA-binding -> BrxB-BrxF- iSTAND -> DUF6079 *-> OLD-ABC ->REase -> N6- METHYLASE -> PglZ -> SIGMA+OB-Fold+AAA+ATPase+HKD-DNase-BrxL -> | 1187 | Bacteria>Pseudomonadota>Gammaproteobacteria>Vibrionales>Vibrionaceae>Vibrio | GCF_026537625.1 | Vibrio_parahaemolyticus | NOL37_RS24175 | 9284..12847 | 670 | BREX-system-P-loop-protein-BrxC-[Vibrio-parahaemolyticus] |
| WP_268670710.1 | Type1.BREX | MODE-HTH+PBPII-><-?\|\|XIS-HTH->?->SF1-SF2-Helicase+HTH->HNH->XIS-HTH-> BrxA-DNA-binding -> BrxB-BrxF- iSTAND -> DUF6079 *-> OLD-ABC ->REase -> N6- METHYLASE -> PglZ -> SIGMA+OB-Fold+AAA+ATPase+HKD-DNase-BrxL -><-PALP<-?<-MODE-HTH+PBPII | 1185 | Bacteria>Pseudomonadota>Gammaproteobacteria>Vibrionales>Vibrionaceae>Vibrio | GCF_026798035.1 | Vibrio_coralliirubri | OTK49_RS06670 | 229636..233193 | 1516159 | BREX-system-P-loop-protein-BrxC-[Vibrio-coralliirubri] |
| WP_274674022.1 | Type1.BREX | <-HAMP+GGDEF<-?<- SIGMA+OB-Fold+AAA+ATPase+HKD-DNase-BrxL <- PglZ <- N6- METHYLASE <- REase<- OLD-ABC <- DUF6079 *<- BrxB-BrxF- iSTAND <- BrxA-DNA-binding <-XIS-HTH | 1187 | Bacteria>Pseudomonadota>Gammaproteobacteria>Vibrionales>Vibrionaceae>Vibrio | GCF_028857845.1 | Vibrio_aestuarianus | L9W73_RS08980 | 42250..45813 | 28171 | BREX-system-P-loop-protein-BrxC-[Vibrio-aestuarianus] |
| WP_213868350.1 | Type1.BREX | <- SIGMA+OB-Fold+AAA+ATPase+HKD-DNase-BrxL <- PglZ <- N6- METHYLASE <- REase<- OLD-ABC <- DUF6079 *<- BrxB-BrxF- iSTAND <- BrxA-DNA-binding <-XIS-HTH<-HNH<-XIS-HTH\|\|?->?->DUF2787-> wHTH+WYL+WCX -> | 1187 | Bacteria>Pseudomonadota>Gammaproteobacteria>Vibrionales>Vibrionaceae>Vibrio | - | Vibrio_alginolyticus | N7I24_002738 | 64007..67570 | 663 | BREX-system-P-loop-protein-BrxC-[Vibrio-alginolyticus] |
| WP_225763940.1 | Type1.BREX | <-TRPR-HTH\|\|?->PRTase->HTH_Tnp_1+HTH_21+rve+rve_3->XIS-HTH-> BrxA-DNA-binding -> BrxB-BrxF- iSTAND -> DUF6079 *->?-> N6- METHYLASE -> OLD-ABC ->REase ->?-> N6- METHYLASE -p-> PglZ -> SIGMA+OB-Fold+AAA+ATPase+HKD-DNase-BrxL -> | 1181 | Bacteria>Pseudomonadota>Gammaproteobacteria>Xanthomonadales>Xanthomonadaceae>Stenotrophomonas | GCF_916618915.1 | Stenotrophomonas_sp._Marseille-Q4652 | LG380_RS05370 | 1171221..1174766 | 2866595 | BREX-system-P-loop-protein-BrxC-[Stenotrophomonas-sp.-Marseille-Q4652] |
| WP_218196661.1 | Type1.BREX | Arm-DNA-bind_3+Phage_integrase-><- SIGMA+OB-Fold+AAA+ATPase+HKD-DNase-BrxL <- PglZ <- N6- METHYLASE -p<-?<- REase<- OLD-ABC <- N6- METHYLASE <-?<- DUF6079 *<- BrxB-BrxF- iSTAND <- BrxA-DNA-binding <-XIS-HTH<-Phage_int_SAM_2+Integrase_1<-Phage_rep | 1182 | Bacteria>Pseudomonadota>Gammaproteobacteria>Xanthomonadales>Xanthomonadaceae>Xanthomonas | GCF_019193005.1 | Xanthomonas_euvesicatoria | KV691_RS15985 | 182541..186089 | 456327 | BREX-system-P-loop-protein-BrxC-[Xanthomonas-euvesicatoria] |
| MBU4235493.1 | Type1.BREX | <- N6- METHYLASE <- REase<- OLD-ABC <-Schlafen+HATPase+wHTH<- DUF6079 *<- BrxB-BrxF- iSTAND <- BrxA-DNA-binding <-XIS-HTH | 1147 | Bacteria>Pseudomonadota>unclassified_Pseudomonadota | GCA_018897885.1 | Pseudomonadota_bacterium | KKI01_04080 | 7059..10502 | 1977087 | BREX-system-P-loop-protein-BrxC-[Pseudomonadota-bacterium] |
| WP_145350724.1 | Type1.BREX | <-SIR2<-?<- SIGMA+OB-Fold+AAA+ATPase+HKD-DNase-BrxL <- PglZ <-REase+TPR+GreAB-C+PIN<- N6- METHYLASE <-Schlafen+HATPase+wHTH<- REase<- OLD-ABC <- DUF6079 *<- BrxB-BrxF- iSTAND <- BrxA-DNA-binding <-?\|\|TrwC+SF1-SF2-Helicase->?-><-?<-?<-ArdCN+MPTase | 1169 | Bacteria>PVC_group>Planctomycetota>Planctomycetia>Pirellulales>Pirellulaceae | GCF_007741495.1 | Roseimaritima_multifibrata | FF011L_RS05780 | 1542279..1545788 | 1930274 | BREX-system-P-loop-protein-BrxC-[Roseimaritima-multifibrata] |
| MCC5808698.1 | Type1.BREX | Phage_integrase-><- SIGMA+OB-Fold+AAA+ATPase+HKD-DNase-BrxL <- PglZ <- HNH<- OLD-ABC <- N6- METHYLASE <- REase<- OLD-ABC <- DUF6079 *<- BrxB-BrxF- iSTAND <- BrxA-DNA-binding <-XIS-HTH | 1183 | Bacteria>PVC_group>Verrucomicrobiota>Opitutae>Opitutales>unclassified_Opitutales | GCA_020831525.1 | Opitutales_bacterium | JJU00_20390 | 12178..15729 | 2026772 | BREX-system-P-loop-protein-BrxC-[Opitutales-bacterium] |
| HCF96075.1 | Type1.BREX | <- N6- METHYLASE -p<- N6- METHYLASE <- REase<- OLD-ABC <- DUF6079 -p<- DUF6079 -p*<- DUF6079 -p<- DUF6079 -p<- DUF6079 -p<- BrxB-BrxF- iSTAND <- BrxA-DNA-binding | 268 | Bacteria>PVC_group>Verrucomicrobiota>unclassified_Verrucomicrobiota | GCA_003534995.1 | Verrucomicrobiota_bacterium | DEW46_13535 | 6770..7576 | 2026799 | BREX-system-P-loop-protein-BrxC-[Verrucomicrobiota-bacterium] |
| HCF96078.1 | Type1.BREX | <- N6- METHYLASE -p<- N6- METHYLASE <- REase<- OLD-ABC <- DUF6079 -p<- DUF6079 -p<- DUF6079 -p<- DUF6079 -p<- DUF6079 -p*<- BrxB-BrxF- iSTAND <- BrxA-DNA-binding | 531 | Bacteria>PVC_group>Verrucomicrobiota>unclassified_Verrucomicrobiota | GCA_003534995.1 | Verrucomicrobiota_bacterium | DEW46_13550 | 8732..10325 | 2026799 | BREX-system-P-loop-protein-BrxC-[Verrucomicrobiota-bacterium] |
| WP_114628929.1 | Type1.BREX | <-Thymidylate_kin\|\|ParB+HNH->ParB+HNH-><-D5N+D5ATPase\|\|XIS-HTH-> BrxA-DNA-binding -> BrxB-BrxF- iSTAND -> DUF6079 *-> OLD-ABC ->REase -> N6- METHYLASE -> SIGMA+OB-Fold+AAA+ATPase+HKD-DNase-BrxL -><-AbiEii<-AbiEi_3_N+AbiEi_3 | 1173 | Bacteria>Spirochaetota>Spirochaetia>Spirochaetales>Spirochaetaceae>Oceanispirochaeta | GCF_013166645.2 | unclassified_Oceanispirochaeta | HNV09_RS06050 | 178318..181839 | 2635722 | MULTISPECIES:-BREX-system-P-loop-protein-BrxC-[unclassified-Oceanispirochaeta] |
| WP_026673780.1 | Type1.BREX | ParD_antitoxin-><- SIGMA+OB-Fold+AAA+ATPase+Lon-Protease-BrxL <- PglZ <- REase<- OLD-ABC <- N6- METHYLASE <- DUF6079 *<- BrxB-BrxF- iSTAND <- BrxA-DNA-binding <-?<-?<-HTH_31 | 1188 | Bacteria>Terrabacteria_group>Bacillota>Bacilli>Bacillales>Bacillaceae | GCF_000621445.1 | Alkalihalobacillus_bogoriensis | T323_RS0113555 | 41147..44713 | 246272 | BREX-system-P-loop-protein-BrxC-[Alkalihalobacillus-bogoriensis] |
| WP_099517056.1 | Type1.BREX | <-SIGMA-HTH<- PglZ <- REase<- OLD-ABC <- N6- METHYLASE <- DUF6079 *<- BrxB-BrxF- iSTAND <- BrxA-DNA-binding <-HTH_26<-?<-Tox-RES<-TRAM+ METHYLASE <-?<-?<-TRPR-HTH | 1196 | Bacteria>Terrabacteria_group>Bacillota>Bacilli>Bacillales>Paenibacillaceae | GCF_002741035.1 | Paenibacillus_sp._BIHB4019 | BBD42_RS03740 | 863001..866591 | 1870819 | BREX-system-P-loop-protein-BrxC-[Paenibacillus-sp.-BIHB-4019] |
| WP_217303321.1 | Type1.BREX | OLD-ABC -><-DOC<-Phd-YefM<- SIGMA+OB-Fold+AAA+ATPase+Lon-Protease-BrxL <- PglZ <- REase<- OLD-ABC <-?<- N6- METHYLASE <- DUF6079 *<- BrxB-BrxF- iSTAND <- BrxA-DNA-binding <-tRNA<-?<-?<-?<-HTH_3<-HTH_OrfB_IS605+OrfB_IS605+OrfB_Zn_ribbon | 1188 | Bacteria>Terrabacteria_group>Bacillota>Clostridia>Eubacteriales>Clostridiaceae | GCF_019056555.1 | Clostridium_sp._001 | B5S50_RS05665 | 1268220..1271786 | 1970093 | BREX-system-P-loop-protein-BrxC-[Clostridium-sp.-001] |
| TGY98248.1 | Type1.BREX | <-MarR-HTH<-?<-RhoN+RhoN+RNase+S1<-?<-?<- PglZ <- REase<- OLD-ABC <- N6- METHYLASE <- DUF6079 *<- BrxB-BrxF- iSTAND <- BrxA-DNA-binding <-?<-PRTase | 1218 | Bacteria>Terrabacteria_group>Bacillota>Clostridia>Eubacteriales>Lachnospiraceae | GCA_004793545.1 | Petralouisia_muris | E5329_00220 | 47888..51544 | 3032872 | BREX-system-P-loop-protein-BrxC-[Petralouisia-muris] |
| MCD7743820.1 | Type1.BREX | HTH_42->ABhydrolase+ABhydrolase->?->?->HTH_26-> BrxA-DNA-binding -> BrxB-BrxF- iSTAND -> DUF6079 *->?-> N6- METHYLASE -> OLD-ABC ->REase -> PglZ ->?-> SIGMA+OB-Fold+AAA+ATPase+Lon-Protease-BrxL ->MACRODOMAIN-> | 1187 | Bacteria>Terrabacteria_group>Bacillota>Clostridia>Eubacteriales>Lachnospiraceae | GCA_021199475.1 | Lachnospiraceae_bacterium | LUI13_00760 | 15415..18978 | 1898203 | BREX-system-P-loop-protein-BrxC-[Lachnospiraceae-bacterium] |
| WP_135987576.1 | Type1.BREX | <-MarR-HTH<-?<-RhoN+RhoN+RNase+S1<-?<-?<- PglZ <- REase<- OLD-ABC <- N6- METHYLASE <- DUF6079 *<- BrxB-BrxF- iSTAND <- BrxA-DNA-binding <-?<-PRTase | 1218 | Bacteria>Terrabacteria_group>Bacillota>Clostridia>Eubacteriales>Lachnospiraceae | GCF_004793545.1 | Petralouisia_muris | E5329_RS00220 | 47888..51544 | 3032872 | BREX-system-P-loop-protein-BrxC-[Petralouisia-muris] |
| MBD5149892.1 | Type1.BREX | ArsR-HTH->HTH_3-><-?<-?<-NTP_transf_3\|\|?->HTH_26-> BrxA-DNA-binding -> BrxB-BrxF- iSTAND -> DUF6079 *-> N6- METHYLASE -> OLD-ABC ->REase -> PglZ ->DDE_Tnp_1_assoc->?->?->ABC-ATPase+SbcC+ABC-ATPase+SbcC->HTH-> | 1183 | Bacteria>Terrabacteria_group>Bacillota>Clostridia>Eubacteriales>Oscillospiraceae | GCA_014799845.1 | Oscillibacter_sp. | HDT18_05880 | 26304..29855 | 1945593 | BREX-system-P-loop-protein-BrxC-[Oscillibacter-sp.] |
| UYJ11714.1 | Type1.BREX | <-CR-ATPase8+REase<-?<-CR-ATPase5+REase<-?<-?\|\|DOC+HTH-> BrxA-DNA-binding -> BrxB-BrxF- iSTAND -> DUF6079 *->?->?-> N6- METHYLASE -> OLD-ABC ->REase -> PglZ -> SIGMA+OB-Fold+AAA+ATPase+Lon-Protease-BrxL ->ParB+HNH->REase+SF1-SF2-Helicase-> | 1182 | Bacteria>Terrabacteria_group>Bacillota>Clostridia>Eubacteriales>Oscillospiraceae | GCA_025757845.1 | Oscillospiraceae_bacterium | OGM78_02700 | 567882..571430 | 2485925 | MAG:-BREX-system-P-loop-protein-BrxC-[Oscillospiraceae-bacterium] |
| WP_010247739.1 | Type1.BREX | <-Herpes_LMP1+REase<- SIGMA+OB-Fold+AAA+ATPase+Lon-Protease-BrxL <- PglZ <- REase<- OLD-ABC <- N6- METHYLASE <- DUF6079 *<- BrxB-BrxF- iSTAND <- BrxA-DNA-binding <-?<-NUDIX+DUF429<-?<-DDE_Tnp_IS66+DDE_Tnp_IS66_C<-?<-HIN-HTH<-ParB+HNH | 1183 | Bacteria>Terrabacteria_group>Bacillota>Clostridia>Eubacteriales>Oscillospiraceae | GCF_000179595.2 | Acetivibrio_cellulolyticus | ACECE_RS0212925 | 179923..183474 | 35830 | BREX-system-P-loop-protein-BrxC-[Acetivibrio-cellulolyticus] |
| WP_120485839.1 | Type1.BREX | <- SIGMA+OB-Fold+AAA+ATPase+Lon-Protease-BrxL <- PglZ <- REase<- OLD-ABC <- N6- METHYLASE <- DUF6079 *<- BrxB-BrxF- iSTAND <- BrxA-DNA-binding <-HTH_26<-JetC<-?<-HNH\|\|HTH_3->PemK_toxin->ParB-> | 1178 | Bacteria>Terrabacteria_group>Bacillota>Clostridia>Eubacteriales>Oscillospiraceae | GCF_009917605.1 | unclassified_Anaerotruncus | D5269_RS12205 | 11128..14664 | 2641626 | MULTISPECIES:-BREX-system-P-loop-protein-BrxC-[unclassified-Anaerotruncus] |
| WP_204901129.1 | Type1.BREX | SF1-SF2-Helicase-><- SIGMA+OB-Fold+AAA+ATPase+Lon-Protease-BrxL <- PglZ <- REase<- OLD-ABC <-?<-Tox-RES<- N6- METHYLASE <-SIR2<- DUF6079 *<- BrxB-BrxF- iSTAND <- BrxA-DNA-binding <-SIGMA-HTH | 1179 | Bacteria>Terrabacteria_group>Bacillota>Clostridia>Eubacteriales>Oscillospiraceae | GCF_016902575.1 | Pseudoflavonifractor_capillosus | H9X86_RS08600 | 19493..23032 | 106588 | BREX-system-P-loop-protein-BrxC-[Pseudoflavonifractor-capillosus] |

<table>
<tr><th>Protein-ID</th><th>System</th><th>Gene-Neighbourhoods</th><th>Length</th><th>Taxonomy</th><th>Assembly-ID</th><th>Strain</th><th>Locus-Tag</th><th>Genomic-coordinates</th><th>TaxID</th><th>Fasta-define</th></tr>
<tr><td>WP_099191952.1</td><td>Type1.BREX</td><td>DNA_METHYLASE->?->HHH+DNA_pol_B+RNase->TRAM+METHYLASE->NERD+HRDC->?->REase->BrxA-DNA-binding->BrxB-BrxF-iSTAND->DUF6079*->N6-METHYLASE->OLD-ABC->TOPRIM->SF1-SF2-Helicase+REase+HTH->OLD-ABC->REase->PglZ->SIGMA+OB-Fold+AAA-ATPase+Lon-Protease-BrxL-></td><td>1190</td><td>Bacteria>Terrabacteria_group>Bacillota>Clostridia>Eubacteriales>Peptostreptococcaceae</td><td>GCF_002724055.1</td><td>Tepidibacter_mesophilus</td><td>Tmeso_RS13165</td><td>34071..37643</td><td>655607</td><td>BREX-system-P-loop-protein-BrxC-[Tepidibacter-mesophilus]</td></tr>
<tr><td>MBS5426410.1</td><td>Type1.BREX</td><td><-Schlafen+HATPase+wHTH<-?||?-><-SIGMA+OB-Fold+AAA-ATPase+Lon-Protease-BrxL<-PglZ<-REase<-OLD-ABC<-N6-METHYLASE<-Schlafen+HATPase+wHTH<-DUF6079*<-BrxB-BrxF-iSTAND<-BrxA-DNA-binding<-zf-ISL3+HTH_Tnp_ISL3+DDE_Tnp_ISL3</td><td>1207</td><td>Bacteria>Terrabacteria_group>Bacillota>Negativicutes>Acidaminococcales>Acidaminococcaceae</td><td>GCA_018368575.1</td><td>Phascolarctobacterium_succinatutens</td><td>KHX39_04785</td><td>71877..75500</td><td>626940</td><td>BREX-system-P-loop-protein-BrxC-[Phascolarctobacterium-succinatutens]</td></tr>
<tr><td>MBE6098022.1</td><td>Type1.BREX</td><td><-SIGMA+OB-Fold+AAA-ATPase+Lon-Protease-BrxL<-PglZ<-REase<-OLD-ABC<-N6-METHYLASE<-DUF6079*<-BrxB-BrxF-iSTAND<-BrxA-DNA-binding<-PIN<-?<-Peptidase_M15</td><td>1220</td><td>Bacteria>Terrabacteria_group>Bacillota>Negativicutes>Selenomonadales>Selenomonadaceae</td><td>GCA_015058685.1</td><td>Schwartzia_succinivorans</td><td>E7198_09540</td><td>51689..55351</td><td>55507</td><td>BREX-system-P-loop-protein-BrxC-[Schwartzia-succinivorans]</td></tr>
<tr><td>WP_273038291.1</td><td>Type1.BREX</td><td><-SIGMA+OB-Fold+AAA-ATPase+Lon-Protease-BrxL<-PglZ<-REase<-OLD-ABC<-N6-METHYLASE<-DUF6079*<-BrxB-BrxF-iSTAND<-BrxA-DNA-binding<-PIN<-?<-Peptidase_M15</td><td>1220</td><td>Bacteria>Terrabacteria_group>Bacillota>Negativicutes>Selenomonadales>Selenomonadaceae</td><td>GCF_015058685.1</td><td>Schwartzia_succinivorans</td><td>E7198_RS09640</td><td>51689..55351</td><td>55507</td><td>BREX-system-P-loop-protein-BrxC-[Schwartzia-succinivorans]</td></tr>
<tr><td>MBE0430956.1</td><td>Type1.BREX</td><td>XIS-HTH->BrxA-DNA-binding->BrxB-BrxF-iSTAND->DUF6079*->Schlafen+HATPase+wHTH+HTH->OLD-ABC->REase->N6-METHYLASE->PglZ->p-></td><td>1193</td><td>Bacteria>Terrabacteria_group>Chloroflexota>Dehalococcoidia>unclassified_Dehalococcoidia</td><td>GCA_014859205.1</td><td>Dehalococcoidia_bacterium</td><td>IBX67_03920</td><td>8033..11614</td><td>2026734</td><td>BREX-system-P-loop-protein-BrxC-[Dehalococcoidia-bacterium]</td></tr>
<tr><td>MBX9724057.1</td><td>Type1.BREX</td><td><-SIGMA+OB-Fold+AAA-ATPase+HKD-DNase-BrxL||BrxA-DNA-binding->BrxB-BrxF-iSTAND->DUF6079*->OLD-ABC->REase->N6-METHYLASE->PglZ->p-></td><td>1175</td><td>Bacteria>Terrabacteria_group>Cyanobacteriota/Melainabacteria_group>Candidatus_Melainabacteria>Candidatus_Obscuribacterales>unclassified_Candidatus_Obscuribacterales</td><td>GCA_019747345.1</td><td>Candidatus_Obscuribacterales_bacterium</td><td>K2X81_21805</td><td>3793..7320</td><td>2364082</td><td>BREX-system-P-loop-protein-BrxC-[Candidatus-Obscuribacterales-bacterium]</td></tr>
<tr><td>BAZ90345.1</td><td>Type1.BREX</td><td><-SIGMA+OB-Fold+AAA-ATPase+HKD-DNase-BrxL<-PglZ<-HTH+REase<-N6-METHYLASE<-REase<-OLD-ABC<-PIN<-DUF433<-DUF6079*<-BrxB-BrxF-iSTAND||?->?-><-?<-?||?-><-?<-ABC-ATPase+ABC-ATPase<-NTP_transferase+MannoseP_isomer</td><td>1166</td><td>Bacteria>Terrabacteria_group>Cyanobacteriota/Melainabacteria_group>Cyanobacteriota>Cyanophyceae>Nostocales</td><td>GCA_002368135.1</td><td>Raphidiopsis_curvata_NIES-932</td><td>NIES932_18440</td><td>1957862..1961362</td><td>1973482</td><td>hypothetical-protein-NIES932_18440-[Raphidiopsis-curvata-NIES-932]</td></tr>
<tr><td>WP_057176814.1</td><td>Type1.BREX</td><td><-ACT||?->ParA-Soj-PloopNTPase->?->?->?-><-?<-?||BrxB-BrxF-iSTAND->DUF6079*->DUF433->PIN->OLD-ABC->REase->REase->N6-METHYLASE->HTH+REase->PglZ->SIGMA+OB-Fold+AAA-ATPase+HKD-DNase-BrxL-></td><td>1166</td><td>Bacteria>Terrabacteria_group>Cyanobacteriota/Melainabacteria_group>Cyanobacteriota>Cyanophyceae>Nostocales</td><td>GCF_001432185.1</td><td>Cylindrospermopsis_sp._CR12</td><td>ASL19_RS00210</td><td>43411..46911</td><td>1747196</td><td>BREX-system-P-loop-protein-BrxC-[Cylindrospermopsis-sp.-CR12]</td></tr>
<tr><td>WP_071242349.1</td><td>Type1.BREX</td><td><-ACT||?->ParA-Soj-PloopNTPase->?->?->?-><-?<-?||BrxB-BrxF-iSTAND->DUF6079*->DUF433->PIN->OLD-ABC->REase->N6-METHYLASE->HTH+REase->PglZ->SIGMA+OB-Fold+AAA-ATPase+HKD-DNase-BrxL->REase-></td><td>1166</td><td>Bacteria>Terrabacteria_group>Cyanobacteriota/Melainabacteria_group>Cyanobacteriota>Cyanophyceae>Nostocales</td><td>GCF_001858115.1</td><td>Cylindrospermopsis_raciborskii</td><td>BCV63_RS11865</td><td>45142..48642</td><td>77022</td><td>BREX-system-P-loop-protein-BrxC-[Cylindrospermopsis-raciborskii]</td></tr>
<tr><td>WP_071248143.1</td><td>Type1.BREX</td><td><-REase<-REase<-REase<-REase||?->BrxA-DNA-binding->BrxB-BrxF-iSTAND->DUF6079*->OLD-ABC->REase->N6-METHYLASE->HTH+REase->PglZ->SIGMA+OB-Fold+AAA-ATPase+HKD-DNase-BrxL-><-DsbD<-Thioredoxin</td><td>1172</td><td>Bacteria>Terrabacteria_group>Cyanobacteriota/Melainabacteria_group>Cyanobacteriota>Cyanophyceae>Nostocales</td><td>GCF_001858125.1</td><td>Cylindrospermopsis_raciborskii</td><td>CENA302_RS09780</td><td>17155..20673</td><td>77022</td><td>BREX-system-P-loop-protein-BrxC-[Cylindrospermopsis-raciborskii]</td></tr>
<tr><td>WP_102947438.1</td><td>Type1.BREX</td><td><-SIGMA+OB-Fold+AAA-ATPase+HKD-DNase-BrxL<-PglZ<-HTH+REase<-N6-METHYLASE<-REase<-OLD-ABC<-PIN<-DUF433<-DUF6079*<-BrxB-BrxF-iSTAND||?->?-><-?<-?<-?<-ParA-Soj-PloopNTPase<-?||ACT-></td><td>1166</td><td>Bacteria>Terrabacteria_group>Cyanobacteriota/Melainabacteria_group>Cyanobacteriota>Cyanophyceae>Nostocales</td><td>GCF_002893285.1</td><td>Cylindrospermopsis_raciborskii</td><td>CEP07_RS06090</td><td>15682..19182</td><td>77022</td><td>BREX-system-P-loop-protein-BrxC-[Cylindrospermopsis-raciborskii]</td></tr>
<tr><td>WP_115538370.1</td><td>Type1.BREX</td><td><-HTH+REase<-N6-METHYLASE-p<-DDE_3<-HTH_28+HTH_32+DDE_3<-N6-METHYLASE<-REase<-OLD-ABC<-PIN<-DUF433<-DUF6079*<-BrxB-BrxF-iSTAND||?->?-><-?<-?<-?<-ParA-Soj-PloopNTPase<-?||ACT-></td><td>1166</td><td>Bacteria>Terrabacteria_group>Cyanobacteriota/Melainabacteria_group>Cyanobacteriota>Cyanophyceae>Nostocales</td><td>GCF_003367075.2</td><td>Cylindrospermopsis_raciborskii</td><td>C6N34_008515</td><td>1862557..1866057</td><td>77022</td><td>BREX-system-P-loop-protein-BrxC-[Cylindrospermopsis-raciborskii]</td></tr>
<tr><td>WP_141302954.1</td><td>Type1.BREX</td><td><-PglZ||DDE_Tnp_IS1-><-N6-METHYLASE-p<-N6-METHYLASE-p<-N6-METHYLASE-p<-REase<-OLD-ABC<-PIN<-DUF433<-DUF6079*<-BrxB-BrxF-iSTAND||?->?-><-?<-?<-?<-ParA-Soj-PloopNTPase<-?||ACT-></td><td>1166</td><td>Bacteria>Terrabacteria_group>Cyanobacteriota/Melainabacteria_group>Cyanobacteriota>Cyanophyceae>Nostocales</td><td>GCF_006523545.1</td><td>Cylindrospermopsis_raciborskii</td><td>FIV49_RS01515</td><td>324659..328159</td><td>77022</td><td>BREX-system-P-loop-protein-BrxC-[Cylindrospermopsis-raciborskii]</td></tr>
<tr><td>WP_187705577.1</td><td>Type1.BREX</td><td>NTP_transferase+MannoseP_isomer->ABC-ATPase+ABC-ATPase->?-><-?||?->?-><-?<-?||BrxB-BrxF-iSTAND->DUF6079*->DUF433->PIN->OLD-ABC->REase->N6-METHYLASE->HTH+REase->PglZ->SIGMA+OB-Fold+AAA-ATPase+HKD-DNase-BrxL-></td><td>1166</td><td>Bacteria>Terrabacteria_group>Cyanobacteriota/Melainabacteria_group>Cyanobacteriota>Cyanophyceae>Nostocales</td><td>GCF_014489415.1</td><td>Cylindrospermopsis_curvispora</td><td>IAR63_RS13115</td><td>2945663..2949163</td><td>747548</td><td>BREX-system-P-loop-protein-BrxC-[Cylindrospermopsis-curvispora]</td></tr>
<tr><td>WP_236308453.1</td><td>Type1.BREX</td><td><-SIGMA+OB-Fold+AAA-ATPase+HKD-DNase-BrxL<-PglZ<-HTH+REase<-N6-METHYLASE<-REase<-OLD-ABC<-PIN<-DUF433<-DUF6079*<-BrxB-BrxF-iSTAND||?->?-><-?<-?<-?<-ParA-Soj-PloopNTPase<-?||ACT-></td><td>1166</td><td>Bacteria>Terrabacteria_group>Cyanobacteriota/Melainabacteria_group>Cyanobacteriota>Cyanophyceae>Nostocales</td><td>GCF_021650815.1</td><td>Cylindrospermopsis_raciborskii</td><td>L3I90_RS00125</td><td>24709..28209</td><td>77022</td><td>BREX-system-P-loop-protein-BrxC-[Cylindrospermopsis-raciborskii]</td></tr>
<tr><td>CCI01315.1</td><td>Type1.BREX</td><td>Pkinase+TPR+TPR+TPR->BrxB-BrxF-iSTAND->DUF6079*->OLD-ABC->REase->N6-METHYLASE->REase->?->PglZ-><-SPOUT||Peptidase_S51-></td><td>1188</td><td>Bacteria>Terrabacteria_group>Cyanobacteriota/Melainabacteria_group>Cyanobacteriota>Cyanophyceae>Oscillatoriophycideae</td><td>GCA_000312185.1</td><td>Microcystis_aeruginosa_PCC_9443</td><td>MICAC_1830011</td><td>77203..80769</td><td>1160281</td><td>conserved-hypothetical-protein-[Microcystis-aeruginosa-PCC-9443]</td></tr>
<tr><td>REJ45948.1</td><td>Type1.BREX</td><td>Pkinase+TPR+TPR+TPR->BrxB-BrxF-iSTAND->DUF6079*->OLD-ABC->REase->N6-METHYLASE->DOC+HTH->PglZ-><-SPOUT||Peptidase_S51->?-><-DDE_5+DDE_Tnp_1</td><td>1168</td><td>Bacteria>Terrabacteria_group>Cyanobacteriota/Melainabacteria_group>Cyanobacteriota>Cyanophyceae>Oscillatoriophycideae</td><td>GCA_003389435.1</td><td>Microcystis_flos-aquae_DF17</td><td>DWQ53_12245</td><td>7047..10553</td><td>2060470</td><td>BREX-system-P-loop-protein-BrxC-[Microcystis-flos-aquae-DF17]</td></tr>
<tr><td>TRU20104.1</td><td>Type1.BREX</td><td>OrfB_IS605+OrfB_Zn_ribbon-><-Pkinase<-Peptidase_M16+Peptidase_M16_C||ncRNA->?->?-><-?||?->tRNA->Pkinase+TPR+TPR+TPR->BrxB-BrxF-iSTAND->DUF6079*->OLD-ABC->REase->N6-METHYLASE->DOC+HTH->PglZ-><-SPOUT||Peptidase_S51->?->DUF488-1-></td><td>1168</td><td>Bacteria>Terrabacteria_group>Cyanobacteriota/Melainabacteria_group>Cyanobacteriota>Cyanophyceae>Oscillatoriophycideae</td><td>GCA_007095795.1</td><td>Microcystis_aeruginosa_Ma_MB_S_20031200_S102D</td><td>EWV79_18805</td><td>31642..35148</td><td>2486255</td><td>BREX-system-P-loop-protein-BrxC-[Microcystis-aeruginosa-Ma_MB_S_20031200_S102D]</td></tr>
<tr><td>NCR78979.1</td><td>Type1.BREX</td><td><-Peptidase_S51||SPOUT-><-PglZ<-?<-REase<-N6-METHYLASE<-REase<-OLD-ABC<-DUF6079*<-BrxB-BrxF-iSTAND<-Pkinase+TPR+TPR+TPR<-tRNA<-HTH_Tnp_1</td><td>1168</td><td>Bacteria>Terrabacteria_group>Cyanobacteriota/Melainabacteria_group>Cyanobacteriota>Cyanophyceae>Oscillatoriophycideae</td><td>GCA_009996105.1</td><td>Microcystis_aeruginosa_K13-10</td><td>GPI91_02970</td><td>22657..26163</td><td>2685970</td><td>BREX-system-P-loop-protein-BrxC-[Microcystis-aeruginosa-K13-10]</td></tr>
<tr><td>NCS23184.1</td><td>Type1.BREX</td><td><-Peptidase_S51||SPOUT-><-PglZ<-DOC+HTH<-?<-N6-METHYLASE<-REase<-OLD-ABC<-DUF6079*<-BrxB-BrxF-iSTAND<-Pkinase+TPR+TPR+TPR<-tRNA<-?||?-><-?<-?<-ncRNA||Peptidase_M16+Peptidase_M16_C->Pkinase-><-OrfB_IS605+OrfB_Zn_ribbon</td><td>1168</td><td>Bacteria>Terrabacteria_group>Cyanobacteriota/Melainabacteria_group>Cyanobacteriota>Cyanophyceae>Oscillatoriophycideae</td><td>GCA_009996485.1</td><td>Microcystis_aeruginosa_BS13-02</td><td>GPI96_01080</td><td>43062..46568</td><td>2685952</td><td>BREX-system-P-loop-protein-BrxC-[Microcystis-aeruginosa-BS13-02]</td></tr>
<tr><td>MBE5230557.1</td><td>Type1.BREX</td><td>Pkinase+TPR+TPR+TPR->BrxB-BrxF-iSTAND->DUF6079*->OLD-ABC->REase->N6-METHYLASE-></td><td>1168</td><td>Bacteria>Terrabacteria_group>Cyanobacteriota/Melainabacteria_group>Cyanobacteriota>Cyanophyceae>Oscillatoriophycideae</td><td>GCA_014946735.1</td><td>Microcystis_aeruginosa_PMC_728.11</td><td>INR81_16200</td><td>47803..51309</td><td>2316423</td><td>BREX-system-P-loop-protein-BrxC-[Microcystis-aeruginosa-PMC-728.11]</td></tr>
<tr><td>MCA2504292.1</td><td>Type1.BREX</td><td><-DUF488-1<-?<-Peptidase_S51||SPOUT-><-PglZ<-DOC+HTH<-N6-METHYLASE<-REase<-OLD-ABC<-DUF6079*<-BrxB-BrxF-iSTAND<-Pkinase+TPR+TPR+TPR</td><td>1168</td><td>Bacteria>Terrabacteria_group>Cyanobacteriota/Melainabacteria_group>Cyanobacteriota>Cyanophyceae>Oscillatoriophycideae</td><td>GCA_020244565.1</td><td>Microcystis_sp._M62BS1</td><td>IM561_02895</td><td>20537..24043</td><td>2771204</td><td>BREX-system-P-loop-protein-BrxC-[Microcystis-sp.-M62BS1]</td></tr>
<tr><td>WP_004266780.1</td><td>Type1.BREX</td><td><-Peptidase_S51||SPOUT-><-PglZ<-?<-REase<-N6-METHYLASE-p<-N6-METHYLASE<-REase<-OLD-ABC<-DUF6079*<-BrxB-BrxF-iSTAND<-Pkinase+TPR+TPR+TPR<-tRNA<-?<-?||?-><-?<-?<-ncRNA<-?<-DADA-Ligase</td><td>1168</td><td>Bacteria>Terrabacteria_group>Cyanobacteriota/Melainabacteria_group>Cyanobacteriota>Cyanophyceae>Oscillatoriophycideae</td><td>GCF_000312165.1</td><td>Microcystis_aeruginosa</td><td>MICAB_RS21670</td><td>36173..39679</td><td>1126</td><td>BREX-system-P-loop-protein-BrxC-[Microcystis-aeruginosa]</td></tr>
<tr><td>WP_043996480.1</td><td>Type1.BREX</td><td>Pkinase+TPR+TPR+TPR->BrxB-BrxF-iSTAND->DUF6079*->OLD-ABC->REase->N6-METHYLASE->REase->?->PglZ-><-SPOUT||Peptidase_S51-></td><td>1168</td><td>Bacteria>Terrabacteria_group>Cyanobacteriota/Melainabacteria_group>Cyanobacteriota>Cyanophyceae>Oscillatoriophycideae</td><td>GCF_000312185.1</td><td>Microcystis_aeruginosa</td><td>MICAC_RS06610</td><td>77263..80769</td><td>1126</td><td>BREX-system-P-loop-protein-BrxC-[Microcystis-aeruginosa]</td></tr>
<tr><td>WP_002790046.1</td><td>Type1.BREX</td><td>PIN-><-HicB_lk_antitox<-?||?-><-N6-METHYLASE-p<-REase<-OLD-ABC<-DUF6079-p<-DUF6079-p*<-DUF6079-p<-BrxB-BrxF-iSTAND<-?<-Pkinase+TPR+TPR+TPR</td><td>532</td><td>Bacteria>Terrabacteria_group>Cyanobacteriota/Melainabacteria_group>Cyanobacteriota>Cyanophyceae>Oscillatoriophycideae</td><td>GCF_000312225.1</td><td>Microcystis_aeruginosa</td><td>MICAF_RS30295</td><td>136943..138541</td><td>1126</td><td>BREX-system-P-loop-protein-BrxC-[Microcystis-aeruginosa]</td></tr>
<tr><td>WP_193959346.1</td><td>Type1.BREX</td><td><-PglZ<-?<-REase<-N6-METHYLASE<-REase<-OLD-ABC<-DUF6079*<-BrxB-BrxF-iSTAND</td><td>1168</td><td>Bacteria>Terrabacteria_group>Cyanobacteriota/Melainabacteria_group>Cyanobacteriota>Cyanophyceae>Oscillatoriophycideae</td><td>GCF_015207185.1</td><td>Microcystis_sp._LEGE_08355</td><td>IQ242_RS05955</td><td>8581..12087</td><td>1828687</td><td>BREX-system-P-loop-protein-BrxC-[Microcystis-sp.-LEGE-08355]</td></tr>
<tr><td>WP_271989206.1</td><td>Type1.BREX</td><td><-Peptidase_S51||SPOUT-><-PglZ<-?<-REase<-N6-METHYLASE<-REase<-OLD-ABC<-DUF6079*<-BrxB-BrxF-iSTAND<-Pkinase+TPR+TPR+TPR</td><td>1168</td><td>Bacteria>Terrabacteria_group>Cyanobacteriota/Melainabacteria_group>Cyanobacteriota>Cyanophyceae>Oscillatoriophycideae</td><td>GCF_028330905.1</td><td>Microcystis_aeruginosa</td><td>PN502_RS07790</td><td>14255..17761</td><td>1126</td><td>BREX-system-P-loop-protein-BrxC-[Microcystis-aeruginosa]</td></tr>
<tr><td>WP_272076547.1</td><td>Type1.BREX</td><td><-REase<-Pkinase<-Peptidase_M16+Peptidase_M16_C||ncRNA->?->?-><-?||?->tRNA->Pkinase+TPR+TPR+TPR->BrxB-BrxF-iSTAND->DUF6079*->OLD-ABC->REase->N6-METHYLASE->DOC+HTH->PglZ-><-SPOUT||Peptidase_S51->?->DUF488-1-></td><td>1168</td><td>Bacteria>Terrabacteria_group>Cyanobacteriota/Melainabacteria_group>Cyanobacteriota>Cyanophyceae>Oscillatoriophycideae</td><td>GCF_028331105.1</td><td>Microcystis_aeruginosa</td><td>PN487_RS04630</td><td>38433..41939</td><td>1126</td><td>BREX-system-P-loop-protein-BrxC-[Microcystis-aeruginosa]</td></tr>
<tr><td>WP_193855895.1</td><td>Type1.BREX</td><td><-Acyltransferase<-ACET<-SF1-SF2-Helicase<-?<-TRD||BrxA-DNA-binding->BrxB-BrxF-iSTAND->DUF6079*->OLD-ABC->REase->N6-METHYLASE-></td><td>1185</td><td>Bacteria>Terrabacteria_group>Cyanobacteriota/Melainabacteria_group>Cyanobacteriota>Cyanophyceae>Pseudanabaenales</td><td>GCF_015207105.1</td><td>cf._Phormidesmis_sp._LEGE_11477</td><td>IQ256_RS12585</td><td>42701..46258</td><td>1828680</td><td>BREX-system-P-loop-protein-BrxC-[cf.-Phormidesmis-sp.-LEGE-11477]</td></tr>
<tr><td>MBU6345615.1</td><td>Type1.BREX</td><td><-N6-METHYLASE-p<-REase<-OLD-ABC<-PIN<-DUF433<-DUF6079*<-BrxB-BrxF-iSTAND||?->?-><-?<-?<-?<-ParA-Soj-PloopNTPase<-?||ACT-></td><td>1166</td><td>Bacteria>Terrabacteria_group>Cyanobacteriota/Melainabacteria_group>Cyanobacteriota>unclassified_Cyanobacteriota</td><td>GCA_018969285.1</td><td>Cyanobacteria_bacterium_REEB494</td><td>KGR70_09985</td><td>4368..7868</td><td>2800107</td><td>BREX-system-P-loop-protein-BrxC-[Cyanobacteria-bacterium-REEB494]</td></tr>
<tr><td>WP_146889206.1</td><td>Type5.BREX</td><td><-HNH<-?<-ARSR-HTH<-TUDOR+SF1-SF2-Helicase+iREase+CTD+REase<-?||Peptidase_M48+TerB->SF1-SF2-Helicase+REase->BrxA-DNA-binding->BrxB-BrxF-iSTAND->DUF6079*->N6-METHYLASE->HicB->OLD-ABC->REase->PglZ+Phosphodiest->SIGMA+OB-Fold+AAA-ATPase+Lon-Protease-BrxL->SF1-SF2-Helicase->?->SF1-SF2-Helicase-></td><td>1186</td><td>Bacteria>Terrabacteria_group>Deinococcota>Deinococci>Deinococcales>Deinococcaceae</td><td>GCF_007990775.1</td><td>Deinococcus_cellulosilyticus</td><td>DC3_RS23585</td><td>23663..27223</td><td>401558</td><td>BREX-system-P-loop-protein-BrxC-[Deinococcus-cellulosilyticus]</td></tr>
<tr><td>WP_028324416.1</td><td>Type1.BREX</td><td>CHTH->N6-METHYLASE->Schlafen+HATPase+wHTH->REase+SF1-SF2-Helicase->DOC+HTH->BrxA-DNA-binding->BrxB-BrxF-iSTAND->DUF6079*->OLD-ABC->REase->N6-METHYLASE->OLD-ABC->TOPRIM->PglZ->SIGMA+OB-Fold+AAA-ATPase+Lon-Protease-BrxL->SbcC->PIN-></td><td>1145</td><td>Bacteria>Thermodesulfobacteriota>Desulfobacteria>Desulfobacterales>Desulfatirhabdiaceae>Desulfatirhabdium</td><td>GCF_000429925.1</td><td>Desulfatirhabdium_butyrativorans</td><td>G492_RS0109380</td><td>84085..87522</td><td>340467</td><td>BREX-system-P-loop-protein-BrxC-[Desulfatirhabdium-butyrativorans]</td></tr>
<tr><td>MBF0232702.1</td><td>Type1.BREX</td><td><-N6-METHYLASE<-REase<-OLD-ABC<-DUF6079*<-BrxB-BrxF-iSTAND<-BrxA-DNA-binding</td><td>1220</td><td>Bacteria>Thermodesulfobacteriota>Desulfobacteria>Desulfobacterales>Desulfobacteraceae>Desulfamplus</td><td>GCA_015231525.1</td><td>Desulfamplus_sp.</td><td>HQK65_06645</td><td>4455..8117</td><td>2006184</td><td>BREX-system-P-loop-protein-BrxC-[Desulfamplus-sp.]</td></tr>
<tr><td>WP_163354568.1</td><td>Type1.BREX</td><td><-35exo||GDResolvase+HIN-HTH-><-?<-HHH_5+SF1-SF2-Helicase||XIS-HTH->BrxA-DNA-binding->BrxB-BrxF-iSTAND->DUF6079*->OLD-ABC->REase->N6-METHYLASE->PglZ->SIGMA+OB-Fold+AAA-ATPase+HKD-DNase-BrxL-><-Zn_Tnp_IS91+Y2_Tnp</td><td>1187</td><td>Bacteria>Thermodesulfobacteriota>Desulfobacteria>Desulfobacterales>Desulfobacteraceae>Desulfobacter</td><td>-</td><td>Desulfobacter_hydrogenophilus</td><td>G3N28_17830</td><td>17749..21312</td><td>2291</td><td>BREX-system-P-loop-protein-BrxC-[Desulfobacter-hydrogenophilus]</td></tr>
<tr><td>NDY73873.1</td><td>Type1.BREX</td><td><-35exo||GDResolvase+HIN-HTH-><-?<-HHH_5+SF1-SF2-Helicase||XIS-HTH->BrxA-DNA-binding->BrxB-BrxF-iSTAND->DUF6079*->OLD-ABC->REase->N6-METHYLASE->PglZ->SIGMA+OB-Fold+AAA-ATPase+HKD-DNase-BrxL-><-Zn_Tnp_IS91+Y2_Tnp</td><td>1187</td><td>Bacteria>Thermodesulfobacteriota>Desulfobacteria>Desulfobacterales>Desulfobacteraceae>Desulfobacter</td><td>GCA_010499545.1</td><td>Desulfobacter_hydrogenophilus</td><td>G3N28_17830</td><td>17749..21312</td><td>2291</td><td>BREX-system-P-loop-protein-BrxC-[Desulfobacter-hydrogenophilus]</td></tr>
<tr><td>WP_035236115.1</td><td>Type1.BREX</td><td><-SIGMA+OB-Fold+AAA-ATPase+HKD-DNase-BrxL<-PglZ<-ParB+HNH<-N6-METHYLASE<-REase<-OLD-ABC<-DUF6079*<-BrxB-BrxF-iSTAND<-BrxA-DNA-binding<-XIS-HTH||?->?->EAL->S1+GTPase-AIG-></td><td>1189</td><td>Bacteria>Thermodesulfobacteriota>Desulfobacteria>Desulfobacterales>Desulfobacteraceae>Desulfobacter</td><td>GCF_000745975.1</td><td>Desulfobacter_vibrioformis</td><td>Q366_RS02915</td><td>253933..257502</td><td>34031</td><td>BREX-system-P-loop-protein-BrxC-[Desulfobacter-vibrioformis]</td></tr>
<tr><td>WP_111959628.1</td><td>Type1.BREX</td><td>Zn_Tnp_IS91+Y2_Tnp-><-SIGMA+OB-Fold+AAA-ATPase+HKD-DNase-BrxL<-PglZ<-N6-METHYLASE<-REase<-OLD-ABC<-DUF6079*<-BrxB-BrxF-iSTAND<-BrxA-DNA-binding<-XIS-HTH||HHH_5+SF1-SF2-Helicase->?-><-GDResolvase+HIN-HTH||35exo-></td><td>1188</td><td>Bacteria>Thermodesulfobacteriota>Desulfobacteria>Desulfobacterales>Desulfobacteraceae>Desulfobacter</td><td>GCF_004319545.1</td><td>Desulfobacter_hydrogenophilus</td><td>DO021_RS19095</td><td>11943..15509</td><td>2291</td><td>BREX-system-P-loop-protein-BrxC-[Desulfobacter-hydrogenophilus]</td></tr>
<tr><td>MCD4719518.1</td><td>Type6.BREX</td><td>BrxE->BrxA-DNA-binding->BrxB-BrxF-iSTAND->DUF6079*->OLD-ABC->REase->N6-METHYLASE->PglZ-></td><td>1208</td><td>Bacteria>Thermodesulfobacteriota>Desulfobacteria>Desulfobacterales>Desulfobacteraceae>Desulfobacula</td><td>GCA_021107415.1</td><td>Desulfobacula_sp.</td><td>K8S13_06615</td><td>2783..6409</td><td>2593537</td><td>BREX-system-P-loop-protein-BrxC-[Desulfobacula-sp.]</td></tr>
<tr><td>MBL7173047.1</td><td>Type1.BREX</td><td><-wHTH+WYL+WCX<-AbiEi_4||?->BrxA-DNA-binding->BrxB-BrxF-iSTAND->DUF6079*->OLD-ABC->REase->N6-METHYLASE->p-></td><td>1172</td><td>Bacteria>Thermodesulfobacteriota>Desulfobacteria>Desulfobacterales>Desulfobacteraceae>unclassified_Desulfobacteraceae</td><td>GCA_016782905.1</td><td>Desulfobacteraceae_bacterium</td><td>ISS68_09440</td><td>5839..9357</td><td>2049433</td><td>BREX-system-P-loop-protein-BrxC-[Desulfobacteraceae-bacterium]</td></tr>
<tr><td>CBX26795.1</td><td>Type6.BREX</td><td><-rve<-SIGMA+OB-Fold+AAA-ATPase+HKD-DNase-BrxL<-PglZ<-Schlafen+HATPase+wHTH<-N6-METHYLASE<-REase<-REase<-OLD-ABC<-DUF6079*<-BrxB-BrxF-iSTAND<-BrxA-DNA-binding<-BrxE||URI-DNase-><-tRNA<-?||?-><-PRTase<-DnaJ</td><td>1208</td><td>Bacteria>Thermodesulfobacteriota>Desulfobacteria>Desulfobacterales>Desulfobacteriaceae>Desulfobacterium</td><td>-</td><td>uncultured_Desulfobacterium_sp.</td><td>N47_A08240</td><td>98686..102312</td><td>201089</td><td>hypothetical-protein-N47_A08240-[uncultured-Desulfobacterium-sp.]</td></tr>
<tr><td>MBC8421197.1</td><td>Type1.BREX</td><td><-AbiEii<-AbiEi_4||?->BrxA-DNA-binding->BrxB-BrxF-iSTAND->DUF6079*->OLD-ABC->REase->N6-METHYLASE-></td><td>1172</td><td>Bacteria>Thermodesulfobacteriota>Desulfobacteria>Desulfobacterales>unclassified_Desulfobacterales</td><td>GCA_014382905.1</td><td>Desulfobacterales_bacterium</td><td>H8E10_21680</td><td>4515..8033</td><td>2044940</td><td>BREX-system-P-loop-protein-BrxC-[Desulfobacterales-bacterium]</td></tr>
<tr><td>KGO34873.1</td><td>Type1.BREX</td><td>MlaE->ABC-ATPase+SbcC->?-><-?||XIS-HTH->BrxA-DNA-binding->BrxB-BrxF-iSTAND->DUF6079*->OLD-ABC->REase->N6-METHYLASE->?->OLD-ABC->HNH->PglZ->SIGMA+OB-Fold+AAA-ATPase+Lon-Protease-BrxL-></td><td>1146</td><td>Bacteria>Thermodesulfobacteriota>Desulfobulbia>Desulfobulbales>Desulfobulbaceae>Desulfobulbus</td><td>GCA_000769715.1</td><td>Desulfobulbus_sp._Tol-SR</td><td>JT06_05895</td><td>26559..29999</td><td>1536652</td><td>hypothetical-protein-JT06_05895-[Desulfobulbus-sp.-Tol-SR]</td></tr>
<tr><td>WP_028584971.1</td><td>Type1.BREX</td><td><-ZnR<-SIGMA+OB-Fold+AAA-ATPase+HKD-DNase-BrxL<-PglZ<-PadR-HTH<-Schlafen+HATPase+wHTH<-N6-METHYLASE<-REase<-OLD-ABC<-DUF6079*<-BrxB-BrxF-iSTAND<-BrxA-DNA-binding<-?<-VirB11<-?<-PCMT+METHYLASE</td><td>1201</td><td>Bacteria>Thermodesulfobacteriota>Desulfobulbia>Desulfobulbales>Desulfobulbaceae>Desulfogranum</td><td>GCF_000429965.1</td><td>Desulfogranum_mediterraneum</td><td>G494_RS0113505</td><td>11738..15343</td><td>160661</td><td>BREX-system-P-loop-protein-BrxC-[Desulfogranum-mediterraneum]</td></tr>
<tr><td>MBA3007526.1</td><td>Type1.BREX</td><td>Phage_integrase->XIS-HTH->BrxA-DNA-binding->BrxB-BrxF-iSTAND->DUF6079*->Schlafen+HATPase+wHTH->OLD-ABC->REase->N6-METHYLASE->Schlafen+HATPase+wHTH->PglZ->SIGMA+OB-Fold+AAA-ATPase+Lon-Protease-BrxL-><-?||35exo+SF1-SF2-Helicase-></td><td>1147</td><td>Bacteria>Thermodesulfobacteriota>Desulfobulbia>Desulfobulbales>Desulfocapsaceae>Desulfocapsa</td><td>GCA_013792135.1</td><td>Desulfocapsa_sp.</td><td>FP810_13320</td><td>53855..57298</td><td>2066021</td><td>BREX-system-P-loop-protein-BrxC-[Desulfocapsa-sp.]</td></tr>
<tr><td>TKB07808.1</td><td>Type1.BREX</td><td><-HEPN<-SbcC<-Calcineurin+Calcineurin-SbcD<-SbcC+SbcC<-SIGMA+OB-Fold+AAA-ATPase+HKD-DNase-BrxL<-PglZ<-N6-METHYLASE<-REase<-OLD-ABC<-DUF6079*<-BrxB-BrxF-iSTAND<-BrxA-DNA-binding<-XIS-HTH<-?<-?<-?<-wHTH</td><td>1183</td><td>Bacteria>Thermodesulfobacteriota>Desulfobulbia>Desulfobulbales>Desulfocapsaceae>Desulforhopalus</td><td>GCA_005116575.1</td><td>Desulforhopalus_sp._IMCC35007</td><td>FCL48_15775</td><td>80281..83832</td><td>2569543</td><td>BREX-system-P-loop-protein-BrxC-[Desulforhopalus-sp.-IMCC35007]</td></tr>
<tr><td>WP_169309248.1</td><td>Type1.BREX</td><td><-HEPN<-SbcC<-Calcineurin+Calcineurin-SbcD<-SbcC+SbcC<-SIGMA+OB-Fold+AAA-ATPase+HKD-DNase-BrxL<-PglZ<-N6-METHYLASE<-REase<-OLD-ABC<-DUF6079*<-BrxB-BrxF-iSTAND<-BrxA-DNA-binding<-XIS-HTH<-?<-?<-?<-wHTH</td><td>1182</td><td>Bacteria>Thermodesulfobacteriota>Desulfobulbia>Desulfobulbales>Desulfocapsaceae>Desulforhopalus</td><td>GCF_005116575.1</td><td>Desulforhopalus_sp._IMCC35007</td><td>FCL48_RS15780</td><td>80281..83829</td><td>2569543</td><td>BREX-system-P-loop-protein-BrxC-[Desulforhopalus-sp.-IMCC35007]</td></tr>
<tr><td>WP_035067990.1</td><td>Type1.BREX</td><td><-35exo<-?<-SbcC+SbcC<-SIGMA+OB-Fold+AAA-ATPase+HKD-DNase-BrxL<-PglZ<-N6-METHYLASE<-REase<-OLD-ABC<-DUF6079*<-BrxB-BrxF-iSTAND<-BrxA-DNA-binding<-wHTH+WYL+WCX||RHH_1->RelE-ParE-><-?||Phd-YefM->PIN-></td><td>1182</td><td>Bacteria>Thermodesulfobacteriota>Desulfovibrionia>Desulfovibrionales>Desulfovibrionaceae>Cupidesulfovibrio</td><td>GCF_000504305.1</td><td>Cupidesulfovibrio_termitidis</td><td>DESTE_RS12450</td><td>3095808..3099356</td><td>42252</td><td>BREX-system-P-loop-protein-BrxC-[Cupidesulfovibrio-termitidis]</td></tr>
<tr><td>AAS96500.1</td><td>Type1.BREX</td><td><-GGDEF<-ZnR+HTH_Tnp_ISL3+DDE_Tnp_ISL3<-SIGMA+OB-Fold+AAA-ATPase+HKD-DNase-BrxL<-PglZ<-REase<-N6-METHYLASE<-REase<-OLD-ABC<-DUF6079*<-BrxB-BrxF-iSTAND||?-><-wHTH+WYL+WCX-p||HIN-HTH->HIN-HTH-><-?||?->35exo-></td><td>1183</td><td>Bacteria>Thermodesulfobacteriota>Desulfovibrionia>Desulfovibrionales>Desulfovibrionaceae>Desulfovibrio</td><td>GCA_000195755.1</td><td>Desulfovibrio_vulgaris_str._Hildenborough</td><td>DVU_2025</td><td>2106300..2109851</td><td>882</td><td>conserved-hypothetical-protein-[Desulfovibrio-vulgaris-str.-Hildenborough]</td></tr>
<tr><td>WP_014524353.1</td><td>Type1.BREX</td><td><-GGDEF<-ZnR+HTH_Tnp_ISL3+DDE_Tnp_ISL3<-SbcC<-SIGMA+OB-Fold+AAA-ATPase+HKD-DNase-BrxL<-PglZ<-REase<-N6-METHYLASE<-REase<-OLD-ABC<-DUF6079*<-BrxB-BrxF-iSTAND<-BrxA-DNA-binding<-wHTH+WYL+WCX-p||HIN-HTH-><-wHTH+WYL+WCX||?->35exo->?-><-RelE-ParE</td><td>1182</td><td>Bacteria>Thermodesulfobacteriota>Desulfovibrionia>Desulfovibrionales>Desulfovibrionaceae>Desulfovibrio</td><td>GCF_000195755.1</td><td>Desulfovibrio_vulgaris</td><td>DVU_RS09565</td><td>2106300..2109848</td><td>881</td><td>BREX-system-P-loop-protein-BrxC-[Desulfovibrio-vulgaris]</td></tr>
<tr><td>WP_043638458.1</td><td>Type1.BREX</td><td>NTP_transferase-><-?||GAF+GAF+HISKIN-><-?<-?||ParB+N6_N4_METHYLASE->?->BrxA-DNA-binding->BrxB-BrxF-iSTAND->DUF6079*->OLD-ABC->REase->N6-METHYLASE->PglZ->SIGMA+OB-Fold+AAA-ATPase+HKD-DNase-BrxL-></td><td>1182</td><td>Bacteria>Thermodesulfobacteriota>Desulfovibrionia>Desulfovibrionales>Desulfovibrionaceae>Desulfovibrio</td><td>GCF_000801335.2</td><td>Desulfovibrio_sp._TomC</td><td>NY78_RS16925</td><td>95476..99024</td><td>1562888</td><td>BREX-system-P-loop-protein-BrxC-[Desulfovibrio-sp.-TomC]</td></tr>
<tr><td>MCH5276204.1</td><td>Type1.BREX</td><td><-SIGMA+OB-Fold+AAA-ATPase+Lon-Protease-BrxL<-PglZ<-?<-N6-METHYLASE-p<-HTH+REase<-N6-METHYLASE<-REase<-OLD-ABC<-DUF6079*<-BrxB-BrxF-iSTAND<-BrxA-DNA-binding<-Arm-DNA-bind_3+Phage_integrase<-tRNA<-?<-?<-ABC-ATPase+SbcC</td><td>1145</td><td>Bacteria>Thermodesulfobacteriota>Desulfovibrionia>Desulfovibrionales>Desulfovibrionaceae>unclassified_Desulfovibrionaceae</td><td>GCA_022510785.1</td><td>Desulfovibrionaceae_bacterium</td><td>J1E80_00045</td><td>14777..18214</td><td>2049043</td><td>BREX-system-P-loop-protein-BrxC-[Desulfovibrionaceae-bacterium]</td></tr>
<tr><td>MBE0574417.1</td><td>Type1.BREX</td><td><-VirD4-TraG+VirD4-FtsK<-?<-?||XIS-HTH->BrxA-DNA-binding->BrxB-BrxF-iSTAND->DUF6079*->OLD-ABC->REase->N6-METHYLASE->REase->PglZ->SIGMA+OB-Fold+AAA-ATPase+HKD-DNase-BrxL->SbcC+APATPase+SbcC->Calcineurin+Calcineurin-SbcD->SbcC-></td><td>1183</td><td>Bacteria>Thermodesulfobacteriota>Desulfuromonadia>Desulfuromonadales>unclassified_Desulfuromonadales</td><td>GCA_014860225.1</td><td>Desulfuromonadales_bacterium</td><td>IH613_00815</td><td>71489..75040</td><td>2099678</td><td>BREX-system-P-loop-protein-BrxC-[Desulfuromonadales-bacterium]</td></tr>
<tr><td>MBW6509585.1</td><td>Type1.BREX</td><td>Amidohydro_1->?->?-><-Four-helical-protein||XIS-HTH->BrxA-DNA-binding->BrxB-BrxF-iSTAND->DUF6079*->Schlafen+HATPase+wHTH+HTH->OLD-ABC->REase->N6-METHYLASE->REase-></td><td>1178</td><td>Bacteria>Thermodesulfobacteriota>Desulfuromonadia>Desulfuromonadales>unclassified_Desulfuromonadales</td><td>GCA_019429255.1</td><td>Desulfuromonadales_bacterium</td><td>KOA94_08605</td><td>43514..47050</td><td>2099678</td><td>BREX-system-P-loop-protein-BrxC-[Desulfuromonadales-bacterium]</td></tr>
<tr><td>WP_207162791.1</td><td>Type1.BREX</td><td><-Peptidase_C14<-TIR<-SIGMA+OB-Fold+AAA-ATPase+HKD-DNase-BrxL<-PglZ<-TOPRIM<-OLD-ABC<-N6-METHYLASE<-REase<-OLD-ABC<-DUF6079*<-BrxB-BrxF-iSTAND<-BrxA-DNA-binding<-AbiEi_4+AbiEi_1<-ParB<-ParB<-GDResolvase+PPP1-HTH||?-><-Arm-DNA-bind_3+Phage_integrase</td><td>1175</td><td>Bacteria>Thermodesulfobacteriota>Desulfuromonadia>Geobacterales>Geobacteraceae>Geobacter</td><td>GCF_017338855.1</td><td>Geobacter_benzoatilyticus</td><td>JZM60_RS12610</td><td>2741421..2744948</td><td>2815309</td><td>BREX-system-P-loop-protein-BrxC-[Geobacter-benzoatilyticus]</td></tr>
<tr><td>MBJ6802709.1</td><td>Type1.BREX</td><td>BrxA-DNA-binding->BrxB-BrxF-iSTAND->DUF6079*->OLD-ABC->REase->N6-METHYLASE->PglZ->SIGMA+OB-Fold+AAA-ATPase+HKD-DNase-BrxL->?-><-?<-NERD+ZnR<-wHTH+SAD+HNH</td><td>1183</td><td>Bacteria>Thermodesulfobacteriota>Desulfuromonadia>Geobacterales>Geobacteraceae>Geomonas</td><td>GCA_016458235.1</td><td>Geomonas_propionica</td><td>JFN90_21475</td><td>2703..6254</td><td>2798582</td><td>BREX-system-P-loop-protein-BrxC-[Geomonas-propionica]</td></tr>
<tr><td>WP_236019113.1</td><td>Type1.BREX</td><td>BrxA-DNA-binding->BrxB-BrxF-iSTAND->DUF6079*->OLD-ABC->REase->N6-METHYLASE->PglZ->SIGMA+OB-Fold+AAA-ATPase+HKD-DNase-BrxL->?-><-?<-NERD+ZnR<-wHTH+SAD+HNH</td><td>1191</td><td>Bacteria>Thermodesulfobacteriota>Desulfuromonadia>Geobacterales>Geobacteraceae>Geomonas</td><td>GCF_016458235.1</td><td>Geomonas_propionica</td><td>JFN90_RS21475</td><td>2679..6254</td><td>2798582</td><td>BREX-system-P-loop-protein-BrxC-[Geomonas-propionica]</td></tr>
<tr><td>--<br>-- OLD-ABC->REase<br>--Type3.BREX<br>--</td><td></td><td></td><td></td><td></td><td></td><td></td><td></td><td></td><td></td><td></td></tr>
<tr><td>MBK8230678.1</td><td>Type3.MTaseII.containing.BREX</td><td><-BrxA-DNA-binding<-PglZ<-REase<-OLD-ABC<-N6-N4-METHYLASE<-HEPN<-DUF6079*<-?<-TetR-HTH||?->?->?-><-HTH_3</td><td>1232</td><td>Bacteria>Bacteria_incertae_sedis>Bacteria_candidate_phyla>Candidatus_Eisenbacteria</td><td>GCA_016712505.1</td><td>Candidatus_Eisenbacteria_bacterium</td><td>IPK72_08880</td><td>13493..17191</td><td>2212470</td><td>ATP-binding-protein-[Candidatus-Eisenbacteria-bacterium]</td></tr>
<tr><td>MBW2068238.1</td><td>Type3.MTaseII.containing.BREX</td><td><-ParA-Soj-PloopNTPase<-RADICAL-SAM||?->Peptidase_M22->?->XIS-HTH->BrxB-BrxF-iSTAND->DUF6079*->N6-N4-METHYLASE->OLD-ABC->REase->?->TUDOR+SF1-SF2-Helicase+iREase+CTD->SF1-SF2-Helicase+REase+HTH->PglZ->BrxA-DNA-binding-></td><td>1245</td><td>Bacteria>delta/epsilon_subdivisions>Deltaproteobacteria>unclassified_Deltaproteobacteria</td><td>GCA_019309115.1</td><td>Deltaproteobacteria_bacterium</td><td>JRI31_05095</td><td>91446..95183</td><td>2026735</td><td>ATP-binding-protein-[Deltaproteobacteria-bacterium]</td></tr>
<tr><td>WP_109907991.1</td><td>Type3.MTaseII.containing.BREX</td><td>TPR->?->?->?-><-HIN-HTH<-DUF772+DDE_Tnp_1+DDE_Tnp_1_6||XIS-HTH->BrxB-BrxF-iSTAND->DUF6079*->Schlafen+HATPase+wHTH->MORC->N6-N4-METHYLASE->OLD-ABC->REase->TUDOR+SF1-SF2-Helicase+iREase+CTD->PglZ->BrxA-DNA-binding-></td><td>1246</td><td>Bacteria>Pseudomonadota>Alphaproteobacteria>Rhodospirillales>Zavarziniaceae>Zavarzinia</td><td>GCF_003173035.1</td><td>Zavarzinia_aquatilis</td><td>DKG74_RS20205</td><td>21531..25271</td><td>2211142</td><td>DUF6079-family-protein-[Zavarzinia-aquatilis]</td></tr>
<tr><td>MCM8625486.1</td><td>Type3.MTaseII.containing.BREX</td><td><-BrxA-DNA-binding<-PglZ<-TUDOR+SF1-SF2-Helicase+iREase+CTD<-REase<-OLD-ABC<-N6-N4-METHYLASE<-?<-DUF6079*<-BrxB-BrxF-iSTAND<-?||NtrC-AAA+FIS-HTH->RNase->?->?->STYKIN+CR-Kinase->SF1-SF2-Helicase->HTH+HEPN-></td><td>1241</td><td>Bacteria>Pseudomonadota>Betaproteobacteria>Betaproteobacteria_incertae_sedis>Candidatus_Accumulibacter>unclassified_Candidatus_Accumulibacter</td><td>GCA_023806625.1</td><td>Candidatus_Accumulibacter_sp.</td><td>NFW05_05900</td><td>113658..117383</td><td>2053492</td><td>DUF6079-family-protein-[Candidatus-Accumulibacter-sp.]</td></tr>
<tr><td>QLQ00182.1</td><td>Type3.MTaseII.containing.BREX</td><td>Phage_lysozyme-><-BrxA-DNA-binding<-TUDOR+SF1-SF2-Helicase+iREase+CTD<-REase<-OLD-ABC<-N6-N4-METHYLASE<-OLD-ABC<-DUF6079*<-BrxB-BrxF-iSTAND||?->HerA/FtsK->N6-N4-METHYLASE-><-ZnR+TOPRIM+AAA-helicase+wHTH<-ParB<-GDResolvase+PPP1-HTH<-?<-AbiEi_4</td><td>1241</td><td>Bacteria>Pseudomonadota>Betaproteobacteria>Burkholderiales>Burkholderiaceae>unclassified_Burkholderiaceae</td><td>GCA_013693755.1</td><td>Burkholderiaceae_bacterium</td><td>HZY78_08275</td><td>1798535..1802260</td><td>2030806</td><td>ATP-binding-protein-[Burkholderiaceae-bacterium]</td></tr>
<tr><td>WP_132764525.1</td><td>Type3.MTaseII.containing.BREX</td><td>Phage_lysozyme-><-BrxA-DNA-binding<-PglZ<-TUDOR+SF1-SF2-Helicase+iREase+CTD<-REase<-OLD-ABC<-N6-N4-METHYLASE<-?<-?<-DUF6079*<-BrxB-BrxF-iSTAND<-XIS-HTH||<-?<-HIN-HTH<-HAMP+HISKIN<-REC+OmpR-HTH</td><td>1241</td><td>Bacteria>Pseudomonadota>Betaproteobacteria</td><td>GCF_026057715.1</td><td>Burkholderiales</td><td>ONS87_RS02500</td><td>493494..497219</td><td>80840</td><td>MULTISPECIES:-DUF6079-family-protein-[Burkholderiales]</td></tr>
<tr><td>WP_022652891.1</td><td>Type3.MTaseII.containing.BREX</td><td><-BrxA-DNA-binding<-PglZ<-TUDOR+SF1-SF2-Helicase+iREase+CTD<-TOPRIM<-OLD-ABC<-REase<-OLD-ABC<-N6-N4-METHYLASE<-DUF6079*<-BrxB-BrxF-iSTAND<-?<-tRNA||?->?->?->?->?->Peptidase_M50-></td><td>1246</td><td>Bacteria>Pseudomonadota>Betaproteobacteria>Neisseriales>Chromobacteriaceae>Aquaspirillum</td><td>GCF_000420525.1</td><td>Aquaspirillum_serpens</td><td>G465_RS0102410</td><td>25502..29242</td><td>190</td><td>DUF6079-family-protein-[Aquaspirillum-serpens]</td></tr>
<tr><td>AVZ79926.1</td><td>Type3.MTaseII.containing.BREX</td><td><-BrxA-DNA-binding<-PglZ<-TUDOR+SF1-SF2-Helicase+iREase+CTD<-REase<-OLD-ABC<-N6-N4-METHYLASE<-TOPRIM<-OLD-ABC<-DUF6079*<-BrxB-BrxF-iSTAND<-wHTH+WYL+WCX||?->DUF3262->TraM->TraE->TraK-></td><td>1241</td><td>Bacteria>Pseudomonadota>Betaproteobacteria>Rhodocyclales>Zoogloeaceae>unclassified_Zoogloeaceae</td><td>GCA_003054515.1</td><td>Zoogloeaceae_bacterium_Par-f-2</td><td>C3497_11130</td><td>2343564..2347289</td><td>2080469</td><td>ATPase-[Zoogloeaceae-bacterium-Par-f-2]</td></tr>
<tr><td>WP_198322148.1</td><td>Type3.MTaseII.containing.BREX</td><td><-BrxA-DNA-binding<-PglZ<-TUDOR+SF1-SF2-Helicase+iREase+CTD<-REase<-OLD-ABC<-N6-N4-METHYLASE-p<-PIN<-PhdYefM_antitox<-DUF6079*<-BrxB-BrxF-iSTAND<-RNA_METHYLASE<-Aminotran_3</td><td>1242</td><td>Bacteria>Pseudomonadota>Gammaproteobacteria>Methylococcales>Methylococcaceae>Methylococcus</td><td>GCF_016106025.1</td><td>Methylococcus_sp._BF19-07</td><td>HUK90_RS04065</td><td>110959..114687</td><td>2743472</td><td>DUF6079-family-protein-[Methylococcus-sp.-BF19-07]</td></tr>
<tr><td>WP_218808314.1</td><td>Type3.MTaseII.containing.BREX</td><td><-BrxA-DNA-binding<-PglZ<-TUDOR+SF1-SF2-Helicase+iREase+CTD<-REase<-OLD-ABC<-N6-N4-METHYLASE<-?<-PIN<-PhdYefM_antitox<-DUF6079*<-BrxB-BrxF-iSTAND<-RNA_METHYLASE<-Aminotran_3<-?<-?<-NTP_transf_3+Hexapep+Hexapep</td><td>1265</td><td>Bacteria>Pseudomonadota>Gammaproteobacteria>Methylococcales>Methylococcaceae>Methylococcus</td><td>GCF_019285515.1</td><td>Methylococcus_sp._Mc7</td><td>KW115_RS06340</td><td>1279422..1283219</td><td>2860258</td><td>DUF6079-family-protein-[Methylococcus-sp.-Mc7]</td></tr>
<tr><td>MBK9496395.1</td><td>Type3.MTaseII.containing.BREX</td><td>CRISPR_Cas6-><-AAA-ATPase<-IstB_IS21<-HTH+rve||MerR-HTH->?->BrxB-BrxF-iSTAND->DUF6079*->N6-N4-METHYLASE->OLD-ABC->REase->TUDOR+SF1-SF2-Helicase+iREase+CTD->PglZ->BrxA-DNA-binding->McrB-AAA->wHTH+REase-><-TPR+REase</td><td>1242</td><td>Bacteria>Pseudomonadota>Gammaproteobacteria>Xanthomonadales>unclassified_Xanthomonadales</td><td>GCA_016718605.1</td><td>Xanthomonadales_bacterium</td><td>IPO08_18210</td><td>258206..261934</td><td>2006849</td><td>ATP-binding-protein-[Xanthomonadales-bacterium]</td></tr>
<tr><td>MBN5005304.1</td><td>Type3.MTaseII.containing.BREX</td><td>SIGMA-HTH+SIGMA-HTH+SIGMA-HTH->tRNA->Arm-DNA-bind_3+Phage_integrase->wHTH+WYL+WCX->BrxB-BrxF-iSTAND->DUF6079*->N6-N4-METHYLASE->OLD-ABC->REase->TUDOR+SF1-SF2-Helicase+iREase+CTD->PglZ->BrxA-DNA-binding->ParB+HNH->?->DUF932-></td><td>1258</td><td>Bacteria>Pseudomonadota>Gammaproteobacteria>Xanthomonadales>Xanthomonadaceae>Stenotrophomonas</td><td>GCA_017149675.1</td><td>Stenotrophomonas_maltophilia</td><td>JY447_14725</td><td>10675..14451</td><td>40324</td><td>hypothetical-protein-[Stenotrophomonas-maltophilia]</td></tr>
<tr><td>WP_135965894.1</td><td>Type3.MTaseII.containing.BREX</td><td>METHYLASE->RNaseH->35exo-><-?||tRNA->wHTH+WYL+WCX->BrxB-BrxF-iSTAND->DUF6079*->?->N6-N4-METHYLASE->OLD-ABC->REase->TUDOR+SF1-SF2-Helicase+iREase+CTD->PglZ->BrxA-DNA-binding->ParB+HNH->DUF932-></td><td>1258</td><td>Bacteria>Pseudomonadota>Gammaproteobacteria>Xanthomonadales>Xanthomonadaceae>Stenotrophomonas</td><td>GCF_029839415.1</td><td>Stenotrophomonas_maltophilia</td><td>N5C11_RS09555</td><td>28442..32218</td><td>40324</td><td>MULTISPECIES:-DUF6079-family-protein-[Stenotrophomonas]</td></tr>
<tr><td>WP_100449517.1</td><td>Type3.MTaseII.containing.BREX</td><td>METHYLASE->RNaseH->35exo-><-?||tRNA->wHTH+WYL+WCX->BrxB-BrxF-iSTAND->DUF6079*->?->N6-N4-METHYLASE->OLD-ABC->REase->TUDOR+SF1-SF2-Helicase+iREase+CTD->PglZ->BrxA-DNA-binding->ParB+HNH->DUF932-></td><td>1258</td><td>Bacteria>Pseudomonadota>Gammaproteobacteria>Xanthomonadales>Xanthomonadaceae>Stenotrophomonas</td><td>-</td><td>Stenotrophomonas_maltophilia</td><td>B9Y88_RS03200</td><td>89921..93697</td><td>40324</td><td>DUF6079-family-protein-[Stenotrophomonas-maltophilia]</td></tr>
<tr><td>--<br>-- OLD-ABC->REase<br>--BR-1<br>--</td><td></td><td></td><td></td><td></td><td></td><td></td><td></td><td></td><td></td><td></td></tr>
<tr><td>ABD41477.1</td><td>DUF499_1.RM.like.systems</td><td>OLD-ABC->OLD-ABC->REase->METHYLASE->OLD-ABC->TOPRIM->HEPN+DUF499*-></td><td>1094</td><td>Archaea>Euryarchaeota>Stenosarchaea_group>Methanomicrobia>Methanomicrobiales</td><td>GCA_000013445.1</td><td>Methanospirillum_hungatei_JF-1</td><td>Mhun_1756</td><td>1984858..1988142</td><td>323259</td><td>conserved-hypothetical-protein-[Methanospirillum-hungatei-JF-1]</td></tr>
<tr><td>WP_204222940.1</td><td>DUF499_1.RM.like.systems</td><td>OLD-ABC->OLD-ABC->REase->METHYLASE->OLD-ABC->TOPRIM->HEPN+DUF499*-></td><td>1082</td><td>Archaea>Euryarchaeota>Stenosarchaea_group>Methanomicrobia>Methanomicrobiales</td><td>GCF_000013445.1</td><td>Methanospirillum_hungatei</td><td>MHUN_RS09170</td><td>1984894..1988142</td><td>2203</td><td>DUF499-domain-containing-protein-[Methanospirillum-hungatei]</td></tr>
<tr><td>MCB5246762.1</td><td>DUF499_1.RM.like.systems</td><td>METHYLASE->OLD-ABC->REase->DUF499*-></td><td>957</td><td>Bacteria>FCB_group>Candidatus_Cloacimonadota>unclassified_Candidatus_Cloacimonadota</td><td>GCA_020532215.1</td><td>Candidatus_Cloacimonadota_bacterium</td><td>LHW57_01885</td><td>3263..6136</td><td>2030808</td><td>DUF499-domain-containing-protein-[Candidatus-Cloacimonetes-bacterium]</td></tr>
<tr><td>MDE2733147.1</td><td>DUF499_1.RM.like.systems</td><td><-TUDOR+SF1-SF2-Helicase+iREase+CTD+REase<-HEPN+DUF499*||REase->?-><-?<-METHYLASE<-REase<-OLD-ABC</td><td>1112</td><td>Bacteria>FCB_group>Gemmatimonadota>unclassified_Gemmatimonadota</td><td>GCA_028875335.1</td><td>Gemmatimonadota_bacterium</td><td>OXI72_02050</td><td>10955..14293</td><td>2026742</td><td>MAG:-Swt1-family-HEPN-domain-containing-protein-[Gemmatimonadota-bacterium]</td></tr>
<tr><td>KYF87240.1</td><td>DUF499_1.RM.like.systems</td><td><-DUF499*<-REase<-OLD-ABC<-METHYLASE<-HNH<-OLD-ABC<-TUDOR+SF1-SF2-Helicase+iREase+CTD+REase</td><td>954</td><td>Bacteria>Myxococcota>Polyangia>Polyangiales>Polyangiaceae</td><td>GCA_001589185.1</td><td>Sorangium_cellulosum</td><td>BE17_46125</td><td>4063..6927</td><td>56</td><td>hypothetical-protein-BE17_46125-[Sorangium-cellulosum]</td></tr>
<tr><td>MBU0551512.1</td><td>DUF499_1.RM.like.systems</td><td>TUDOR+SF1-SF2-Helicase+iREase+CTD+REase-p->HTH+MPTase->PIN->METHYLASE->OLD-ABC->REase->DUF499*-><-?<-?<-RecR+TOPRIM</td><td>942</td><td>Bacteria>Myxococcota>unclassified_Myxococcota</td><td>GCA_018813065.1</td><td>Myxococcota_bacterium</td><td>KJ940_08450</td><td>6900..9728</td><td>2818507</td><td>DUF499-domain-containing-protein-[Myxococcota-bacterium]</td></tr>
<tr><td>MBU2768209.1</td><td>DUF499_1.RM.like.systems</td><td><-DUF499*<-REase<-OLD-ABC<-wHTH+REase<-METHYLASE<-HNH<-OLD-ABC</td><td>542</td><td>Bacteria>Pseudomonadota>Acidithiobacillia>Acidithiobacillales>Acidithiobacillaceae</td><td>GCA_018853935.1</td><td>Acidithiobacillus_ferrivorans</td><td>HAP94_19055</td><td>1..1626</td><td>160808</td><td>ATP-binding-protein-[Acidithiobacillus-ferrivorans]</td></tr>
<tr><td>MBF0177023.1</td><td>DUF499_1.RM.like.systems</td><td>SAM-methylase-><-?<-?<-SF1-SF2-Helicase+HRDC<-DUF499*<-REase<-OLD-ABC<-METHYLASE</td><td>950</td><td>Bacteria>Pseudomonadota>Alphaproteobacteria>Magnetococcales>unclassified_Magnetococcales</td><td>GCA_015231175.1</td><td>Magnetococcales_bacterium</td><td>HQL63_09275</td><td>22537..25389</td><td>2026759</td><td>ATP-binding-protein-[Magnetococcales-bacterium]</td></tr>
<tr><td>SBT04363.1</td><td>DUF499_1.RM.like.systems</td><td>TP_methylase-><-DUF499-p<-DUF499*<-REase<-OLD-ABC<-METHYLASE-p</td><td>671</td><td>Bacteria>Pseudomonadota>Betaproteobacteria>Betaproteobacteria_incertae_sedis>Candidatus_Accumulibacter</td><td>GCA_900089955.1</td><td>Candidatus_Accumulibacter_aalborgensis</td><td>ACCAA_130242</td><td>226085..228100</td><td>1860102</td><td>conserved-hypothetical-protein-[Candidatus-Accumulibacter-aalborgensis]</td></tr>
<tr><td>MBP6559243.1</td><td>DUF499_1.RM.like.systems</td><td><-DUF499*<-REase<-OLD-ABC<-METHYLASE-p</td><td>945</td><td>Bacteria>Pseudomonadota>Betaproteobacteria>Burkholderiales>Burkholderiaceae</td><td>GCA_017989415.1</td><td>Burkholderiaceae_bacterium</td><td>KA212_02995</td><td>512..3349</td><td>2030806</td><td>ATP-binding-protein-[Burkholderiaceae-bacterium]</td></tr>
<tr><td>WP_132962623.1</td><td>DUF499_1.RM.like.systems</td><td>GST_N+GST_C->tRNA->tRNA-><-DUF499*<-REase<-OLD-ABC<-METHYLASE<-PIN<-TUDOR+SF1-SF2-Helicase+iREase+CTD+REase||Peptidase_M24-></td><td>949</td><td>Bacteria>Pseudomonadota>Betaproteobacteria>Burkholderiales>Burkholderiales_genera_incertae_sedis</td><td>GCF_004342625.1</td><td>Tepidimonas_ignava</td><td>EDC36_RS07390</td><td>30173..33022</td><td>114249</td><td>DUF499-domain-containing-protein-[Tepidimonas-ignava]</td></tr>
<tr><td>WP_252101816.1</td><td>DUF499_1.RM.like.systems</td><td>TUDOR+SF1-SF2-Helicase+iREase+CTD+REase->Schlafen+HATPase+HTH->?->METHYLASE->OLD-ABC->REase->DUF499*-><-?<-Phage_integrase+Phage_integrase<-?<-HIN-HTH<-Phage_integrase+Phage_integrase</td><td>945</td><td>Bacteria>Pseudomonadota>Betaproteobacteria>Burkholderiales>Comamonadaceae</td><td>-</td><td>[Acidovorax]_ebreus</td><td>MRB47_RS02565</td><td>530947..533784</td><td>721785</td><td>DUF499-domain-containing-protein-[[Acidovorax]</td></tr>
<tr><td>TFI45392.1</td><td>DUF499_1.RM.like.systems</td><td><-MobA_MobL+SF1-SF2-helicase||?-><-?<-DUF499*<-REase<-OLD-ABC<-METHYLASE<-PIN<-TUDOR+SF1-SF2-Helicase+iREase+CTD+REase</td><td>945</td><td>Bacteria>Pseudomonadota>Betaproteobacteria>Burkholderiales>Comamonadaceae</td><td>-</td><td>Diaphorobacter_sp._DS2</td><td>E4093_19980</td><td>3252..6089</td><td>1302548</td><td>ATP-binding-protein-[Diaphorobacter-sp.-DS2]</td></tr>
<tr><td>MBP7545005.1</td><td>DUF499_1.RM.like.systems</td><td><-DUF499*<-REase<-OLD-ABC<-METHYLASE-p</td><td>949</td><td>Bacteria>Pseudomonadota>Betaproteobacteria>Burkholderiales>Comamonadaceae</td><td>GCA_017998835.1</td><td>Acidovorax_sp.</td><td>KA749_05805</td><td>238..3087</td><td>1872122</td><td>ATP-binding-protein-[Acidovorax-sp.]</td></tr>
<tr><td>WP_087746901.1</td><td>DUF499_1.RM.like.systems</td><td><-SPOUT||?-><-?<-?<-?<-DUF499*<-REase<-OLD-ABC<-METHYLASE<-HNH<-OLD-ABC<-TUDOR+SF1-SF2-Helicase+iREase+CTD+REase</td><td>945</td><td>Bacteria>Pseudomonadota>Betaproteobacteria>Burkholderiales>Comamonadaceae</td><td>GCF_002176815.1</td><td>unclassified_Acidovorax</td><td>CCX87_RS13165</td><td>2797139..2799976</td><td>2684926</td><td>MULTISPECIES:-DUF499-domain-containing-protein-[unclassified-Acidovorax]</td></tr>
<tr><td>WP_055397337.1</td><td>DUF499_1.RM.like.systems</td><td><-ArdCN+MPTase<-?<-?<-MobA_MobL+SF1-SF2-helicase||?-><-?<-DUF499*<-REase<-OLD-ABC<-METHYLASE<-HNH<-OLD-ABC<-TUDOR+SF1-SF2-Helicase+iREase+CTD+REase</td><td>945</td><td>Bacteria>Pseudomonadota>Betaproteobacteria>Burkholderiales>Comamonadaceae</td><td>GCF_017355385.1</td><td>Acidovorax</td><td>AE621_RS12465</td><td>7782..10619</td><td>12916</td><td>MULTISPECIES:-DUF499-domain-containing-protein-[Acidovorax]</td></tr>
<tr><td>WP_210568820.1</td><td>DUF499_1.RM.like.systems</td><td>TUDOR+SF1-SF2-Helicase+iREase+CTD+REase->OLD-ABC->HNH->METHYLASE->OLD-ABC->REase->DUF499*->?->?->?-><-?||SPOUT-></td><td>945</td><td>Bacteria>Pseudomonadota>Betaproteobacteria>Burkholderiales>Comamonadaceae</td><td>GCF_017939685.1</td><td>Acidovorax_sp._JG5</td><td>J8G26_RS10065</td><td>96032..98869</td><td>2822718</td><td>DUF499-domain-containing-protein-[Acidovorax-sp.-JG5]</td></tr>
<tr><td>WP_210930504.1</td><td>DUF499_1.RM.like.systems</td><td>HTH->?->Phage_integrase->STYKinase-like-><-?||DDE_Tnp_1_5+DDE_Tnp_1-><-DUF499*<-REase<-OLD-ABC<-HEPN<-METHYLASE<-HNH<-OLD-ABC</td><td>945</td><td>Bacteria>Pseudomonadota>Betaproteobacteria>Burkholderiales>Comamonadaceae</td><td>GCF_018064205.1</td><td>Comamonas_aquatica</td><td>KAQ61_RS03430</td><td>737208..740045</td><td>225991</td><td>DUF499-domain-containing-protein-[Comamonas-aquatica]</td></tr>
</table>

| Protein-ID | System | Gene-Neighbourhoods | Length | Taxonomy | Assembly-ID | Strain | Locus-Tag | Genomic-coordinates | TaxID | Fasta-defline |
|---|---|---|---|---|---|---|---|---|---|---|
| WP_209423136.1 | DUF499_1.RM.like.systems | SF1-SF2-Helicase+DUF1998->?-><-?<-?<-?\|\|?-><- DUF499 *<- REase<- OLD-ABC <- TOPRIM <- OLD-ABC <- METHYLASE -p<- METHYLASE | 943 | Bacteria>Pseudomonadota>Betaproteobacteria>Burkholderiales>Comamonadaceae | GCF_021249385.1 | Melaminivora_jejuensis | LVC68_RS01445 | 284242..287073 | 1267217 | DUF499-domain-containing-protein-[Melaminivora-jejuensis] |
| WP_066335883.1 | DUF499_1.RM.like.systems | <-RNase+SF1-SF2-Helicase<-?<-?<-MACRODOMAIN<- DUF499 *<- REase<- OLD-ABC <- METHYLASE <-?<- TUDOR+ SF1-SF2-Helicase+iREase+CTD+REase | 939 | Bacteria>Pseudomonadota>Betaproteobacteria>Burkholderiales>Sphaerotilaceae | GCF_001571085.1 | Azohydromonas_lata | ALA01S_RS10810 | 44580..47399 | 45677 | DUF499-domain-containing-protein-[Azohydromonas-lata] |
| WP_110402335.1 | DUF499_1.RM.like.systems | <-ABhydrolase\|\|?->?-><-?<- DUF499 *<- REase<- OLD-ABC <- METHYLASE <- HNH<- OLD-ABC <- TUDOR+ SF1-SF2-Helicase+iREase+CTD+REase | 958 | Bacteria>Pseudomonadota>Betaproteobacteria>Burkholderiales>Sphaerotilaceae | - | Sphaerotilus_hippei | C7444_RS18460 | 22920..25796 | 744406 | DUF499-domain-containing-protein-[Sphaerotilus-hippei] |
| MBK6716517.1 | DUF499_1.RM.like.systems | METHYLASE -p-> OLD-ABC ->REase -> DUF499 *-><-?<-?\|\| METHYLASE -p-> | 948 | Bacteria>Pseudomonadota>Betaproteobacteria>Burkholderiales>unclassified_Burkholderiales | GCA_016703325.1 | Burkholderiales_bacterium | IPG57_15845 | 2519..5365 | 1891238 | ATP-binding-protein-[Burkholderiales-bacterium] |
| MCZ2407521.1 | DUF499_1.RM.like.systems | Phage_integrase+Phage_integrase->?-><- DUF499 *<- REase<- OLD-ABC <- METHYLASE | 945 | Bacteria>Pseudomonadota>Betaproteobacteria>Burkholderiales>unclassified_Burkholderiales | GCA_026931285.1 | Burkholderiales_bacterium | LC097_12425 | 115108..117945 | 1891238 | MAG:-DUF499-domain-containing-protein-[Burkholderiales-bacterium] |
| WP_169202998.1 | DUF499_1.RM.like.systems | <- DUF499 *<- REase<- OLD-ABC <- METHYLASE <- HNH<- OLD-ABC <- TUDOR+ SF1-SF2-Helicase+iREase+CTD+REase | 951 | Bacteria>Pseudomonadota>Betaproteobacteria>Rhodocyclales>Rhodocyclaceae | GCF_017894365.1 | Aromatoleum_bremense | GPA24_RS12870 | 12346..15201 | 76115 | DUF499-domain-containing-protein-[Aromatoleum-bremense] |
| MDD3354816.1 | DUF499_1.RM.like.systems | PIN-><- DUF499 *<- REase<- OLD-ABC <- METHYLASE | 948 | Bacteria>Pseudomonadota>Betaproteobacteria>Rhodocyclales>Zoogloeaceae | GCA_028693495.1 | Zoogloea_sp. | PHD22_14435 | 4878..7724 | 49181 | MAG:-DUF499-domain-containing-protein-[Zoogloea-sp.] |
| WP_153168444.1 | DUF499_1.RM.like.systems | <-Radical-SAM+TRAM<-?\|\|tRNA->?->?->?-><-?<- DUF499 *<-wHTH+REase<- REase<- OLD-ABC <- METHYLASE <-HEPN<- TUDOR+ SF1-SF2-Helicase+iREase+CTD+REase | 941 | Bacteria>Pseudomonadota>Betaproteobacteria>Rhodocyclales>Zoogloeaceae | GCF_009469595.2 | Thauera_sp._2A1 | GH664_RS12700 | 85060..87885 | 2570191 | DUF499-domain-containing-protein-[Thauera-sp.-2A1] |
| WP_159691715.1 | DUF499_1.RM.like.systems | MODE-HTH+PBPII-><-TPR<- DUF499 *<- REase<- OLD-ABC <- METHYLASE <- HNH<- OLD-ABC <- TUDOR+ SF1-SF2-Helicase+iREase+CTD+REase | 943 | Bacteria>Pseudomonadota>Betaproteobacteria>Rhodocyclales>Zoogloeaceae | GCF_009800965.1 | Cognatazoarcus_halotolerans | GQB03_RS16405 | 71425..74256 | 2686016 | DUF499-domain-containing-protein-[Cognatazoarcus-halotolerans] |
| WP_223954701.1 | DUF499_1.RM.like.systems | JAB->CbeA_antitoxin->RlpA-><- DUF499 *<- REase<- OLD-ABC <- METHYLASE <- HNH<- OLD-ABC <- TUDOR+ SF1-SF2-Helicase+iREase+CTD+REase | 946 | Bacteria>Pseudomonadota>Gammaproteobacteria>Aeromonadales>Aeromonadaceae | GCF_019972775.1 | Aeromonas_caviae | LDN68_RS17105 | 3642842..3645682 | 648 | DUF499-domain-containing-protein-[Aeromonas-caviae] |
| WP_087504467.1 | DUF499_1.RM.like.systems | <-Phage_integrase\|\|HTH+WYL-> TUDOR+ SF1-SF2-Helicase+iREase+CTD+REase -> METHYLASE -> OLD-ABC ->REase -> DUF499 *-><-?<-?<-?<-?<-?\|\|REase-> | 940 | Bacteria>Pseudomonadota>Gammaproteobacteria>Alteromonadales>Echinimonadaceae | GCF_014636195.1 | Neiella_marina | CBE68_RS02320 | 109774..112596 | 508461 | DUF499-domain-containing-protein-[Neiella-marina] |
| KTG26834.1 | DUF499_1.RM.like.systems | Arm-DNA-bind_3+Phage_integrase-><-Phage_AlpA<-ZnR+ TOPRIM +primpol+HTH<-XIS-HTH\|\|MobI-><- DUF499 *<- REase<- OLD-ABC <- METHYLASE <- OLD-ABC <- TUDOR+ SF1-SF2-Helicase+iREase+CTD+REase <-HTH+WYL | 953 | Bacteria>Pseudomonadota>Gammaproteobacteria>Alteromonadales>Idiomarinaceae | GCA_001469885.1 | Idiomarina_sp._H105 | AUR68_16335 | 105470..108331 | 1766622 | hypothetical-protein-AUR68_16335-[Idiomarina-sp.-H105] |
| MCK9506376.1 | DUF499_1.RM.like.systems | OLD-ABC ->REase -> DUF499 *->Classical-AAA-><-?<-DUF932 | 951 | Bacteria>Pseudomonadota>Gammaproteobacteria>Cellvibrionales>Porticoccaceae | GCA_023228865.1 | Porticoccaceae_bacterium | MOQ95_19625 | 878..3733 | 2026782 | DUF499-domain-containing-protein-[Porticoccaceae-bacterium] |
| WP_213378539.1 | DUF499_1.RM.like.systems | MerR-HTH+GDResolvase->HTH_OrfB_IS605+OrfB_IS605+OrfB_Zn_ribbon->?-><- DUF499 *<- REase<- OLD-ABC <- METHYLASE <- TUDOR+ SF1-SF2-Helicase+iREase+CTD+REase <-HTH+WYL\|\|HTH_OrfB_IS605+OrfB_IS605+OrfB_Zn_ribbon-> | 943 | Bacteria>Pseudomonadota>Gammaproteobacteria>Chromatiales>Chromatiaceae | GCF_018409545.1 | Allochromatium_tepidum | Atep_RS10840 | 2243146..2245977 | 553982 | DUF499-domain-containing-protein-[Allochromatium-tepidum] |
| TQE99812.1 | DUF499_1.RM.like.systems | HEPN->ParA-Soj-PloopNTPase->ORC-CDC6-> METHYLASE -> OLD-ABC ->REase -> DUF499 *->?->UbE1+JAB->?->Schlafen-> | 955 | Bacteria>Pseudomonadota>Gammaproteobacteria>Chromatiales>Ectothiorhodospiraceae | GCA_006569195.1 | Spiribacter_salinus | FKY71_06670 | 12316..15183 | 1335746 | ATP-binding-protein-[Spiribacter-salinus] |
| WP_025282662.1 | DUF499_1.RM.like.systems | TraH->?->TraG_N-><-?\|\|AbiEi_4-><- DUF499 *<- REase<- OLD-ABC <- METHYLASE <-?<- TUDOR+ SF1-SF2-Helicase+iREase+CTD+REase \|\|TPR+TPR-> | 956 | Bacteria>Pseudomonadota>Gammaproteobacteria>Chromatiales>Ectothiorhodospiraceae | GCF_000583815.1 | Ectothiorhodospira_haloalkaliphila | ECTHA_RS0102365 | 117439..120309 | 421628 | DUF499-domain-containing-protein-[Ectothiorhodospira-haloalkaliphila] |
| WP_186934038.1 | DUF499_1.RM.like.systems | <- DUF499 *<- REase<- OLD-ABC <- METHYLASE <-SbcC<- TUDOR+ SF1-SF2-Helicase+iREase+CTD+REase | 946 | Bacteria>Pseudomonadota>Gammaproteobacteria>Enterobacterales>Enterobacteriaceae | GCF_014285945.1 | Klebsiella | H8M57_RS27310 | 138..2978 | 570 | MULTISPECIES:-DUF499-domain-containing-protein-[Klebsiella] |
| WP_219173436.1 | DUF499_1.RM.like.systems | HIN-HTH-> TUDOR+ SF1-SF2-Helicase+iREase+CTD+REase ->SbcC-> METHYLASE -> OLD-ABC ->REase -> DUF499 *-><-RlpA<-Arm-DNA-bind_3+Phage_integrase\|\|HIN-HTH->\|\|?->?->ParB+HNH-> | 946 | Bacteria>Pseudomonadota>Gammaproteobacteria>Enterobacterales>Enterobacteriaceae | GCF_019334445.1 | Klebsiella/Raoultella_group | LVJ43_RS03725 | 746872..749712 | 2890311 | DUF499-domain-containing-protein-[Raoultella-ornithinolytica] |
| WP_224381981.1 | DUF499_1.RM.like.systems | TUDOR+ SF1-SF2-Helicase+iREase+CTD+REase ->SbcC-> METHYLASE -> OLD-ABC ->REase -> DUF499 *-> | 350 | Bacteria>Pseudomonadota>Gammaproteobacteria>Enterobacterales>Enterobacteriaceae | GCF_020118495.1 | Klebsiella_michiganensis | FMK66_RS24360 | 60161..61211 | 1134687 | DUF499-domain-containing-protein,-partial-[Klebsiella-michiganensis] |
| WP_134906061.1 | DUF499_1.RM.like.systems | Arm-DNA-bind_3+Phage_integrase->RlpA-><- DUF499 *<- REase<- OLD-ABC <- METHYLASE <-SbcC<- TUDOR+ SF1-SF2-Helicase+iREase+CTD+REase <-HIN-HTH | 946 | Bacteria>Pseudomonadota>Gammaproteobacteria>Enterobacterales | GCF_004801595.1 | Enterobacteriaceae | I8Q62_RS26385 | 2639..5479 | 543 | MULTISPECIES:-DUF499-domain-containing-protein-[Enterobacteriaceae] |
| WP_077999168.1 | DUF499_1.RM.like.systems | JAB->RlpA-><- DUF499 *<- REase<- OLD-ABC <- METHYLASE <- HNH<- OLD-ABC <- TUDOR+ SF1-SF2-Helicase+iREase+CTD+REase | 949 | Bacteria>Pseudomonadota>Gammaproteobacteria>Enterobacterales>Hafniaceae | GCF_002006945.1 | Edwardsiella_tarda | BTN58_RS02535 | 130022..132871 | 636 | DUF499-domain-containing-protein-[Edwardsiella-tarda] |
| WP_218800673.1 | DUF499_1.RM.like.systems | <-TIR<- DUF499 *<- REase<- OLD-ABC <- METHYLASE <-PIN<- TUDOR+ SF1-SF2-Helicase+iREase+CTD+REase <-HTH_Tnp | 942 | Bacteria>Pseudomonadota>Gammaproteobacteria>Methylococcales>Methylococcaceae | GCF_019285535.1 | Methylococcus_capsulatus | M3M30_RS10970 | 2380003..2382831 | 414 | DUF499-domain-containing-protein-[Methylococcus-capsulatus] |
| WP_022949219.1 | DUF499_1.RM.like.systems | LON+AAA-> TUDOR+ SF1-SF2-Helicase+iREase+CTD+REase -> METHYLASE -> OLD-ABC ->REase -> DUF499 *->?->RADICAL-SAM->Glyco_transf_8-> | 952 | Bacteria>Pseudomonadota>Gammaproteobacteria>Methylococcales>Methylothermaceae | GCF_000421465.1 | Methylohalobius_crimeensis | H035_RS0111990 | 2436080..2438938 | 244365 | DUF499-domain-containing-protein-[Methylohalobius-crimeensis] |
| MAE39903.1 | DUF499_1.RM.like.systems | HTH_26-> TUDOR+ SF1-SF2-Helicase+iREase+CTD+REase -> METHYLASE -> OLD-ABC ->REase -> DUF499 *->?->?-><-Phage_integrase<-Phage_integrase | 941 | Bacteria>Pseudomonadota>Gammaproteobacteria>Moraxellales>Moraxellaceae | GCA_002686735.1 | Psychrobacter_sp. | CML07_03075 | 11656..14481 | 56811 | hypothetical-protein-[Psychrobacter-sp.] |
| MCC6374735.1 | DUF499_1.RM.like.systems | <- DUF499 *<- REase<- OLD-ABC <- METHYLASE -p | 878 | Bacteria>Pseudomonadota>Gammaproteobacteria>Moraxellales>Moraxellaceae | GCA_020846355.1 | Moraxellaceae_bacterium | IT465_07990 | 1..2635 | 1889775 | ATP-binding-protein-[Moraxellaceae-bacterium] |
| WP_107866351.1 | DUF499_1.RM.like.systems | METHYLASE -> OLD-ABC ->REase -> DUF499 *-><-ParB+HNH<-?<-ParB | 941 | Bacteria>Pseudomonadota>Gammaproteobacteria>Moraxellales>Moraxellaceae | GCF_003051055.1 | Agitococcus_lubricus | C8N29_RS12225 | 4668..7493 | 1077255 | DUF499-domain-containing-protein-[Agitococcus-lubricus] |
| BBI71108.2 | DUF499_1.RM.like.systems | Phage_integrase-><-Phage_AlpA<- TOPRIM +primpol<-?<-XIS-HTH<- DUF499 -p<- DUF499 *<- DUF499 -p<- REase<- OLD-ABC <-?<- METHYLASE -p<- METHYLASE -p | 386 | Bacteria>Pseudomonadota>Gammaproteobacteria>Oceanospirillales>Halomonadaceae | GCA_007163885.2 | Halomonas_axialensis | HAALTHF_10150n | 665831..666991 | 115555 | hypothetical-protein-HAALTHF_10150n-[Halomonas-axialensis] |
| MCD6438846.1 | DUF499_1.RM.like.systems | HNH-> METHYLASE -> OLD-ABC ->REase ->EVE+McrB-AAA->wHTH+REase-> DUF499 *-> | 282 | Bacteria>Pseudomonadota>Gammaproteobacteria>Oceanospirillales>Halomonadaceae | GCA_021161325.1 | Halomonas_sp. | J7L56_11360 | 10325..11171 | 1486246 | ATP-binding-protein-[Halomonas-sp.] |
| WP_197030572.1 | DUF499_1.RM.like.systems | OLD-ABC ->REase ->MAD-NTD+EVE+McrB-AAA->?->wHTH+REase->?-> DUF499 *-> DUF499 ->?-><-?\|\|?->REase-> | 264 | Bacteria>Pseudomonadota>Gammaproteobacteria>Oceanospirillales>Halomonadaceae | GCF_000568385.1 | Halomonas_sp._BC04 | Q427_RS35225 | 32151..32945 | 1403540 | DUF499-domain-containing-protein-[Halomonas-sp.-BC04] |
| WP_097789710.1 | DUF499_1.RM.like.systems | Peptidase_M20->Arm-DNA-bind_3+Phage_integrase->TPR+TPR-><-?<- DUF499 *<-wHTH+REase<-MAD-NTD+EVE+McrB-AAA<- REase<- OLD-ABC <-HEPN+CBS<- METHYLASE | 957 | Bacteria>Pseudomonadota>Gammaproteobacteria>Oceanospirillales>Halomonadaceae | GCF_002549795.1 | Halomonas_beimenensis | BEI_RS11820 | 2512898..2515771 | 475662 | DUF499-domain-containing-protein-[Halomonas-beimenensis] |
| WP_044628625.1 | DUF499_1.RM.like.systems | METHYLASE -> OLD-ABC ->REase -> DUF499 *-><-MobI\|\|XIS-HTH->ZnR+ TOPRIM +primpol+HTH->Phage_AlpA-><-Arm-DNA-bind_3+Phage_integrase | 953 | Bacteria>Pseudomonadota>Gammaproteobacteria>Oceanospirillales>Halomonadaceae | GCF_011398715.1 | Halomonas | QWE69_RS13020 | 3280..6141 | 2745 | MULTISPECIES:-DUF499-domain-containing-protein-[Halomonas] |
| WP_231663150.1 | DUF499_1.RM.like.systems | HTH+WYL-> TUDOR+ SF1-SF2-Helicase+iREase+CTD+REase -> METHYLASE -> OLD-ABC ->REase ->ParB+HNH-> DUF499 *->Primpol+D5N+D5ATPase-><-Arm-DNA-bind_3+Phage_integrase\|\|?->FAD-NAD-dep-oxidoreductase-> | 957 | Bacteria>Pseudomonadota>Gammaproteobacteria>Oceanospirillales>Halomonadaceae | GCF_021151495.1 | Halomonas_sp._IOP_31 | K7I12_RS04335 | 15378..18251 | 2876584 | DUF499-domain-containing-protein-[Halomonas-sp.-IOP_31] |
| WP_234246274.1 | DUF499_1.RM.like.systems | METHYLASE -> METHYLASE ->Schlafen+HATPase+HTH->HEPN+CBS-> OLD-ABC ->REase -> DUF499 *-> | 957 | Bacteria>Pseudomonadota>Gammaproteobacteria>Oceanospirillales>Halomonadaceae | GCF_021404345.1 | Halomonas_desiderata | HOP57_RS15495 | 85349..88222 | 52021 | DUF499-domain-containing-protein-[Halomonas-desiderata] |
| WP_239295914.1 | DUF499_1.RM.like.systems | <-HTH_Tnp_1+HTH_21+rve+rve_2<-?<- DUF499 *<-?<- REase<- OLD-ABC <- METHYLASE <-?<- TUDOR+ SF1-SF2-Helicase+iREase+CTD+REase | 958 | Bacteria>Pseudomonadota>Gammaproteobacteria>Oceanospirillales>Halomonadaceae | GCF_022348215.1 | unclassified_Halomonas | MHM38_RS03720 | 8315..11191 | 2609666 | MULTISPECIES:-DUF499-domain-containing-protein-[unclassified-Halomonas] |
| WP_253017818.1 | DUF499_1.RM.like.systems | HNH-> METHYLASE -> OLD-ABC ->REase ->EVE+McrB-AAA->wHTH+REase-> DUF499 *->?->AbiEi-><-CHTH<-RelE | 961 | Bacteria>Pseudomonadota>Gammaproteobacteria>Oceanospirillales>Halomonadaceae | GCF_024124605.1 | Halomonas_sp._CnH100-B | NH398_RS08385 | 365976..368861 | 2954490 | DUF499-domain-containing-protein-[Halomonas-sp.-CnH100-B] |
| WP_253027667.1 | DUF499_1.RM.like.systems | Phage_integrase-><-Phage_AlpA<-ZnR+ TOPRIM +primpol+HTH<-XIS-HTH\|\|MobI-><- DUF499 *<- REase<- OLD-ABC <- METHYLASE <- OLD-ABC <- TUDOR+ SF1-SF2-Helicase+iREase+CTD+REase <-HTH+WYL | 953 | Bacteria>Pseudomonadota>Gammaproteobacteria>Oceanospirillales>Halomonadaceae | GCF_024124725.1 | Halomonas_sp._Ps84H-12 | NH344_RS06635 | 49336..52197 | 2954501 | DUF499-domain-containing-protein-[Halomonas-sp.-Ps84H-12] |
| WP_137094585.1 | DUF499_1.RM.like.systems | TUDOR+ SF1-SF2-Helicase+iREase+CTD+REase -> OLD-ABC ->HNH -> METHYLASE -> OLD-ABC ->REase -> DUF499 *-><-MobI\|\|XIS-HTH->ZnR+ TOPRIM +primpol+HTH->Phage_AlpA-><-Arm-DNA-bind_3+Phage_integrase | 953 | Bacteria>Pseudomonadota>Gammaproteobacteria>Oceanospirillales>Halomonadaceae | GCF_028580285.1 | Halomonas | E8Q34_RS13770 | 13695..16556 | 2745 | DUF499-domain-containing-protein-[Halomonas-sp.-15WGF] |
| WP_284453091.1 | DUF499_1.RM.like.systems | Arm-DNA-bind_3+Phage_integrase-><-Phage_AlpA<-ZnR+ TOPRIM +primpol+HTH<-XIS-HTH\|\|MobI-><- DUF499 *<- REase<- OLD-ABC <- METHYLASE <- HNH<- OLD-ABC <- TUDOR+ SF1-SF2-Helicase+iREase+CTD+REase | 953 | Bacteria>Pseudomonadota>Gammaproteobacteria>Oceanospirillales>Halomonadaceae | GCF_030145725.1 | Halomonas_meridiana | KYX60_RS01235 | 106598..109459 | 29570 | DUF499-domain-containing-protein-[Halomonas-meridiana] |
| WP_072841218.1 | DUF499_1.RM.like.systems | TUDOR+ SF1-SF2-Helicase+iREase+CTD+REase -> OLD-ABC ->HNH -> METHYLASE -> OLD-ABC ->REase -> DUF499 *->SMODS->?-><-RlpA\|\|?->HTH_26 | 942 | Bacteria>Pseudomonadota>Gammaproteobacteria>Oceanospirillales>Oceanospirillaceae | GCF_900129155.1 | Marinomonas_polaris | BUA59_RS18830 | 76459..79287 | 293552 | DUF499-domain-containing-protein-[Marinomonas-polaris] |
| WP_230706347.1 | DUF499_1.RM.like.systems | Phage_integrase-><-Phage_AlpA<-ZnR+ TOPRIM +primpol+HTH<-XIS-HTH\|\|MobI-><- DUF499 *<- REase<- OLD-ABC \|\|\|\|?-><-?<-?<-Resolvase | 953 | Bacteria>Pseudomonadota>Gammaproteobacteria>Pseudomonadales>Marinobacteraceae | GCF_021021495.1 | Marinobacter_adhaerens | K7I10_RS06135 | 1373639..1376500 | 1033846 | DUF499-domain-containing-protein-[Marinobacter-adhaerens] |
| WP_230713447.1 | DUF499_1.RM.like.systems | Phage_integrase-><-Phage_AlpA<-ZnR+ TOPRIM +primpol+HTH<-XIS-HTH\|\|MobI-><- DUF499 *<- REase<- OLD-ABC <- METHYLASE <- OLD-ABC <- TUDOR+ SF1-SF2-Helicase+iREase+CTD+REase <-HTH+WYL | 953 | Bacteria>Pseudomonadota>Gammaproteobacteria>Pseudomonadales>Marinobacteraceae | GCF_021021515.1 | Marinobacter_shengliensis | K7I15_RS06080 | 103063..105924 | 1389223 | DUF499-domain-containing-protein-[Marinobacter-shengliensis] |
| MBG7398186.1 | DUF499_1.RM.like.systems | SF1-SF2-Helicase-><-SF1-SF2-helicase+RdgC+Beta-barrel+REase<-35exo+SF1-SF2-Helicase\|\|ABhydrolase->?->RlpA-><- DUF499 *<- REase<- OLD-ABC <- METHYLASE <-HEPN<- TUDOR+ SF1-SF2-Helicase+iREase+CTD+REase | 942 | Bacteria>Pseudomonadota>Gammaproteobacteria>Pseudomonadales>Pseudomonadaceae | GCA_015766545.1 | Pseudomonas_aeruginosa | GHU66_27420 | 38429..41257 | 287 | ATP-binding-protein-[Pseudomonas-aeruginosa] |
| HBO5114778.1 | DUF499_1.RM.like.systems | TUDOR+ SF1-SF2-Helicase+iREase+CTD+REase ->HEPN-> METHYLASE -> OLD-ABC ->REase -> DUF499 *-><-RlpA<-?<-ABhydrolase | 946 | Bacteria>Pseudomonadota>Gammaproteobacteria>Pseudomonadales>Pseudomonadaceae | GCA_021662155.1 | Pseudomonas_aeruginosa | L4W37_005387 | 12730..15570 | 287 | ATP-binding-protein-[Pseudomonas-aeruginosa] |
| MCO2346212.1 | DUF499_1.RM.like.systems | TUDOR+ SF1-SF2-Helicase+iREase+CTD+REase -><-?\|\| METHYLASE -> OLD-ABC ->REase -> DUF499 *-><-RlpA<-?\|\|35exo+SF1-SF2-Helicase-><-?<-HIN-HTH | 942 | Bacteria>Pseudomonadota>Gammaproteobacteria>Pseudomonadales>Pseudomonadaceae | GCA_023907105.1 | Pseudomonas_aeruginosa | FA362_14075 | 17744..20572 | 287 | ATP-binding-protein-[Pseudomonas-aeruginosa] |
| HCI3160560.1 | DUF499_1.RM.like.systems | ABhydrolase->?->RlpA-><- DUF499 *<- REase<- OLD-ABC <- METHYLASE <-HEPN<- TUDOR+ SF1-SF2-Helicase+iREase+CTD+REase | 942 | Bacteria>Pseudomonadota>Gammaproteobacteria>Pseudomonadales>Pseudomonadaceae | GCA_024356045.1 | Pseudomonas_aeruginosa | NOV63_004506 | 69244..72072 | 287 | ATP-binding-protein-[Pseudomonas-aeruginosa] |
| MCR7692359.1 | DUF499_1.RM.like.systems | TUDOR+ SF1-SF2-Helicase+iREase+CTD+REase ->HEPN-> METHYLASE -> OLD-ABC ->REase -> DUF499 *-> | 417 | Bacteria>Pseudomonadota>Gammaproteobacteria>Pseudomonadales>Pseudomonadaceae | GCA_024717035.1 | Pseudomonas_aeruginosa | NTA33_29240 | 17686..18936 | 287 | DUF499-domain-containing-protein-[Pseudomonas-aeruginosa] |
| WP_023835622.1 | DUF499_1.RM.like.systems | TUDOR+ SF1-SF2-Helicase+iREase+CTD+REase ->HEPN-> METHYLASE -> OLD-ABC ->REase -> DUF499 *-><-RlpA<-?<-JAB<-ParB<-?\|\|HIN-HTH-> | 944 | Bacteria>Pseudomonadota>Gammaproteobacteria>Pseudomonadales>Pseudomonadaceae | GCF_000503175.1 | Pseudomonas_aeruginosa | CF344_RS23300 | 12270..15104 | 287 | DUF499-domain-containing-protein-[Pseudomonas-aeruginosa] |
| WP_031643596.1 | DUF499_1.RM.like.systems | TUDOR+ SF1-SF2-Helicase+iREase+CTD+REase ->HEPN-> METHYLASE -> OLD-ABC ->REase -> DUF499 *-><-RlpA<-?<-ABhydrolase<-?<-?\|\|HTH_3-> | 950 | Bacteria>Pseudomonadota>Gammaproteobacteria>Pseudomonadales>Pseudomonadaceae | GCF_000629305.1 | Pseudomonas_aeruginosa | AJ61_RS31490 | 12829..15681 | 287 | DUF499-domain-containing-protein-[Pseudomonas-aeruginosa] |
| WP_049308056.1 | DUF499_1.RM.like.systems | TUDOR+ SF1-SF2-Helicase+iREase+CTD+REase ->HEPN-> METHYLASE -> OLD-ABC ->REase -> DUF499 *-><-RlpA<-?<-?\|\|SF1-SF2-helicase+RdgC+Beta-barrel+REase-><-SF1-SF2-Helicase<-HKD-DNase | 942 | Bacteria>Pseudomonadota>Gammaproteobacteria>Pseudomonadales>Pseudomonadaceae | GCF_001061475.1 | Pseudomonas_aeruginosa | ADM76_RS18680 | 17855..20683 | 287 | DUF499-domain-containing-protein-[Pseudomonas-aeruginosa] |
| WP_049271643.1 | DUF499_1.RM.like.systems | ParB->JAB->?->RlpA-><- DUF499 *<- REase<- OLD-ABC <- METHYLASE | 943 | Bacteria>Pseudomonadota>Gammaproteobacteria>Pseudomonadales>Pseudomonadaceae | GCF_001077135.1 | Pseudomonas_aeruginosa | ADL79_RS26490 | 2020..4851 | 287 | DUF499-domain-containing-protein-[Pseudomonas-aeruginosa] |
| WP_054902927.1 | DUF499_1.RM.like.systems | DUF932->?->ParB->JAB->?->RlpA-><- DUF499 *<- REase<- OLD-ABC <- METHYLASE <-HEPN<- TUDOR+ SF1-SF2-Helicase+iREase+CTD+REase \|\|HTH+WYL-> | 944 | Bacteria>Pseudomonadota>Gammaproteobacteria>Pseudomonadales>Pseudomonadaceae | GCF_001320525.1 | Pseudomonas_sp._NBRC_111131 | APH34_RS22145 | 20096..22930 | 1661046 | DUF499-domain-containing-protein-[Pseudomonas-sp.-NBRC-111131] |
| WP_074197337.1 | DUF499_1.RM.like.systems | DUF932->?->ParB->JAB->?->RlpA-><- DUF499 *<- REase<- OLD-ABC <- METHYLASE <-HEPN<- TUDOR+ SF1-SF2-Helicase+iREase+CTD+REase | 946 | Bacteria>Pseudomonadota>Gammaproteobacteria>Pseudomonadales>Pseudomonadaceae | GCF_001909445.1 | Pseudomonas_aeruginosa | BH596_RS15220 | 11473..14313 | 287 | DUF499-domain-containing-protein-[Pseudomonas-aeruginosa] |
| WP_085592307.1 | DUF499_1.RM.like.systems | MerR-HTH->tRNA-> TUDOR+ SF1-SF2-Helicase+iREase+CTD+REase -> METHYLASE -> OLD-ABC ->REase -> DUF499 *-><-RlpA<-REase+SF1-SF2-Helicase<-N6_N4_MTase\|\|MAD-NTD+HNH-> | 941 | Bacteria>Pseudomonadota>Gammaproteobacteria>Pseudomonadales>Pseudomonadaceae | GCF_002113255.1 | Pseudomonas_sp._B14(2017) | CAG34_RS21645 | 108391..111216 | 1981745 | DUF499-domain-containing-protein-[Pseudomonas-sp.-B14(2017)] |
| WP_131349395.1 | DUF499_1.RM.like.systems | TUDOR+ SF1-SF2-Helicase+iREase+CTD+REase -> OLD-ABC ->HNH -> METHYLASE -> OLD-ABC ->REase -> DUF499 *-><-RlpA<-TPR<-?<-?\|\|?-><-Mbetalac | 929 | Bacteria>Pseudomonadota>Gammaproteobacteria>Pseudomonadales>Pseudomonadaceae | GCF_004327885.1 | Azotobacter_chroococcum | EOE52_RS08250 | 59995..62784 | 353 | DUF499-domain-containing-protein-[Azotobacter-chroococcum] |
| WP_154355052.1 | DUF499_1.RM.like.systems | TUDOR+ SF1-SF2-Helicase+iREase+CTD+REase ->HEPN-> METHYLASE -> OLD-ABC ->REase -> DUF499 *-><-RlpA<-?<-JAB<-ParB<-?<-DUF932 | 942 | Bacteria>Pseudomonadota>Gammaproteobacteria>Pseudomonadales>Pseudomonadaceae | GCF_009676785.1 | Pseudomonas_aeruginosa | GJV02_RS03270 | 700187..703015 | 287 | DUF499-domain-containing-protein-[Pseudomonas-aeruginosa] |
| WP_192069117.1 | DUF499_1.RM.like.systems | Integrase_1->Phage_int_SAM->N6_N4_MTase->REase+SF1-SF2-Helicase->RlpA-><- DUF499 *<- REase<- OLD-ABC <- METHYLASE <- TUDOR+ SF1-SF2-Helicase+iREase+CTD+REase \|\|?-><-HTH_3 | 941 | Bacteria>Pseudomonadota>Gammaproteobacteria>Pseudomonadales>Pseudomonadaceae | GCF_014842375.1 | Pseudomonas_coleopterorum | IFT40_RS20700 | 77168..79993 | 1605838 | DUF499-domain-containing-protein-[Pseudomonas-coleopterorum] |
| WP_198580799.1 | DUF499_1.RM.like.systems | TUDOR+ SF1-SF2-Helicase+iREase+CTD+REase ->HEPN-> METHYLASE -> OLD-ABC ->REase -> DUF499 *-><-RlpA<-?<-JAB<-ParB<-?<-DUF932 | 944 | Bacteria>Pseudomonadota>Gammaproteobacteria>Pseudomonadales>Pseudomonadaceae | GCF_016134915.1 | Pseudomonas_aeruginosa | QMY34_RS12045 | 381175..384009 | 287 | DUF499-domain-containing-protein-[Pseudomonas-aeruginosa] |
| WP_200582036.1 | DUF499_1.RM.like.systems | TUDOR+ SF1-SF2-Helicase+iREase+CTD+REase ->HEPN-> METHYLASE -> OLD-ABC ->REase -> DUF499 *-><-RlpA<-?<-SymE_toxin<-?<-ZNR+Pkinase+SF1-SF2-Helicase+REase | 943 | Bacteria>Pseudomonadota>Gammaproteobacteria>Pseudomonadales>Pseudomonadaceae | GCF_016622185.1 | Pseudomonas_aeruginosa | GFM12_RS08425 | 1738931..1741762 | 287 | DUF499-domain-containing-protein-[Pseudomonas-aeruginosa] |
| WP_219806875.1 | DUF499_1.RM.like.systems | <-MazE_antitoxin\|\|?->ParB->JAB->?->RlpA-><- DUF499 *<- REase<- OLD-ABC <- METHYLASE <-HEPN<- TUDOR+ SF1-SF2-Helicase+iREase+CTD+REase <-Peptidase | 943 | Bacteria>Pseudomonadota>Gammaproteobacteria>Pseudomonadales>Pseudomonadaceae | GCF_019429735.1 | Pseudomonas_aeruginosa | KZ843_RS17265 | 5657..8488 | 287 | DUF499-domain-containing-protein-[Pseudomonas-aeruginosa] |
| WP_033975677.1 | DUF499_1.RM.like.systems | TUDOR+ SF1-SF2-Helicase+iREase+CTD+REase ->HEPN-> METHYLASE -> OLD-ABC ->REase -> DUF499 *-><-RlpA<-?<-SymE_toxin<-?<-ZNR+Pkinase+SF1-SF2-Helicase+REase | 943 | Bacteria>Pseudomonadota>Gammaproteobacteria>Pseudomonadales>Pseudomonadaceae | GCF_022569915.1 | Pseudomonas_aeruginosa | CGU39_RS16575 | 614189..617020 | 287 | DUF499-domain-containing-protein-[Pseudomonas-aeruginosa] |
| WP_266022655.1 | DUF499_1.RM.like.systems | TUDOR+ SF1-SF2-Helicase+iREase+CTD+REase ->HEPN-> METHYLASE -> OLD-ABC ->REase -> DUF499 *-><-RlpA<-?<-JAB<-ParB<-?<-DUF932 | 948 | Bacteria>Pseudomonadota>Gammaproteobacteria>Pseudomonadales>Pseudomonadaceae | GCF_026241065.1 | unclassified_Pseudomonas | OO409_RS22205 | 32872..35718 | 196821 | MULTISPECIES:-DUF499-domain-containing-protein-[unclassified-Pseudomonas] |
| WP_272438616.1 | DUF499_1.RM.like.systems | HKD-DNase->SF1-SF2-Helicase-><-SF1-SF2-helicase+RdgC+Beta-barrel+REase\|\|?->?->RlpA-><- DUF499 *<- REase<- OLD-ABC <- METHYLASE <-HEPN<- TUDOR+ SF1-SF2-Helicase+iREase+CTD+REase | 942 | Bacteria>Pseudomonadota>Gammaproteobacteria>Pseudomonadales>Pseudomonadaceae | GCF_028435625.1 | Pseudomonas_aeruginosa | KGZ09_RS30315 | 38429..41257 | 287 | DUF499-domain-containing-protein-[Pseudomonas-aeruginosa] |
| WP_090734260.1 | DUF499_1.RM.like.systems | TUDOR+ SF1-SF2-Helicase+iREase+CTD+REase ->?-> METHYLASE -> OLD-ABC ->REase -> DUF499 *-><-RlpA<-?\|\|PAIREDC-HTH-> | 929 | Bacteria>Pseudomonadota>Gammaproteobacteria>Pseudomonadales>Pseudomonadaceae | GCF_900108885.1 | Azotobacter_beijerinckii | BMW56_RS18070 | 16681..19470 | 170623 | DUF499-domain-containing-protein-[Azotobacter-beijerinckii] |
| WP_090941615.1 | DUF499_1.RM.like.systems | TUDOR+ SF1-SF2-Helicase+iREase+CTD+REase -> OLD-ABC ->HNH -> METHYLASE -> OLD-ABC ->REase -> DUF499 *-><-RlpA<-?<-?<-ZnR+ TOPRIM +D5ATPase | 929 | Bacteria>Pseudomonadota>Gammaproteobacteria>Pseudomonadales>Pseudomonadaceae | GCF_900114395.1 | Azotobacter_beijerinckii | BM240_RS15650 | 17867..20656 | 170623 | DUF499-domain-containing-protein-[Azotobacter-beijerinckii] |
| WP_033982644.1 | DUF499_1.RM.like.systems | ABhydrolase->?->RlpA-><- DUF499 *<- REase<- OLD-ABC <- METHYLASE <-HEPN<- TUDOR+ SF1-SF2-Helicase+iREase+CTD+REase | 942 | Bacteria>Pseudomonadota>Gammaproteobacteria>Pseudomonadales>Pseudomonadaceae | GCF_900707645.1 | Pseudomonas_aeruginosa | BUH55_RS18155 | 7471..10299 | 287 | DUF499-domain-containing-protein-[Pseudomonas-aeruginosa] |
| WP_132674023.1 | DUF499_1.RM.like.systems | DUF932->?->ParB->JAB->?->RlpA-><- DUF499 *<- REase<- OLD-ABC <- METHYLASE <-HEPN<- TUDOR+ SF1-SF2-Helicase+iREase+CTD+REase | 944 | Bacteria>Pseudomonadota>Gammaproteobacteria>Pseudomonadales>Pseudomonadaceae | - | Pseudomonas_aeruginosa | EQ623_RS00180 | 30601..33435 | 287 | DUF499-domain-containing-protein-[Pseudomonas-aeruginosa] |
| RTZ75007.1 | DUF499_1.RM.like.systems | <-NERD+SF1-SF2-Helicase<- DUF499 *<- REase<- OLD-ABC <- METHYLASE -p | 943 | Bacteria>Pseudomonadota>Gammaproteobacteria>unclassified_Gammaproteobacteria | GCA_003973075.1 | Gammaproteobacteria_bacterium | DSZ00_02995 | 2492..5323 | 1913989 | hypothetical-protein-DSZ00_02995-[Gammaproteobacteria-bacterium] |
| HFD86238.1 | DUF499_1.RM.like.systems | <- DUF499 *<- REase<- OLD-ABC <- METHYLASE -p | 692 | Bacteria>Pseudomonadota>Gammaproteobacteria>unclassified_Gammaproteobacteria | GCA_011322685.1 | Gammaproteobacteria_bacterium | ENJ35_01010 | 1..2075 | 1913989 | ATP-binding-protein-[Gammaproteobacteria-bacterium] |
| MCK9487878.1 | DUF499_1.RM.like.systems | <-SIGMA-HTH<- OLD-ABC <-?<-?<-?<- DUF499 *<- REase<- OLD-ABC <- METHYLASE <-HEPN<-PIN<-PemK_toxin | 944 | Bacteria>Pseudomonadota>Gammaproteobacteria>Xanthomonadales>unclassified_Xanthomonadales | GCA_023228845.1 | Xanthomonadales_bacterium | MOQ42_00470 | 105333..108167 | 2006849 | DUF499-domain-containing-protein-[Xanthomonadales-bacterium] |
| MCU0753612.1 | DUF499_1.RM.like.systems | <-ArsR-HTH\|\|?->?->?->?->?-><- DUF499 *<-Schlafen+HATPase+HTH<-ParB+HNH<- REase<- OLD-ABC <- METHYLASE <-HNH | 955 | Bacteria>Pseudomonadota>Gammaproteobacteria>Xanthomonadales>unclassified_Xanthomonadales | GCA_025458445.1 | Xanthomonadales_bacterium | MUE46_00615 | 13163..16030 | 2006849 | DUF499-domain-containing-protein-[Xanthomonadales-bacterium] |
| GIX36093.1 | DUF499_1.RM.like.systems | METHYLASE -p-> METHYLASE -p-> OLD-ABC ->REase -> DUF499 *-><-?<-?<-?<-?<-?<-Radical-SAM | 950 | Bacteria>Pseudomonadota>Gammaproteobacteria>Xanthomonadales>Xanthomonadaceae | GCA_026005075.1 | Xanthomonadaceae_bacterium | KatS3mg126_1872 | 4497..7349 | 1926873 | MAG:-hypothetical-protein-KatS3mg126_1872-[Xanthomonadaceae-bacterium] |
| MBU0673024.1 | DUF499_1.RM.like.systems | TUDOR+ SF1-SF2-Helicase+iREase+CTD+REase ->PIN->?-> METHYLASE -> OLD-ABC ->REase -> DUF499 *-> | 943 | Bacteria>Pseudomonadota>unclassified_Pseudomonadota | GCA_018814305.1 | Pseudomonadota_bacterium | KJ950_00090 | 19515..22346 | 1977087 | DUF499-domain-containing-protein-[Proteobacteria-bacterium] |
| MBE6357833.1 | DUF499_1.RM.like.systems | ParB-HTH->?-><-?<-?<- DUF499 *<- REase<- OLD-ABC <- METHYLASE <- HNH<- OLD-ABC <- TUDOR+ SF1-SF2-Helicase+iREase+CTD+REase | 938 | Bacteria>PVC_group>Lentisphaerota>unclassified_Lentisphaerota | GCA_015061525.1 | Lentisphaerota_bacterium | E7057_01125 | 3177..5993 | 1932692 | ATP-binding-protein-[Lentisphaerae-bacterium] |
| WP_076345234.1 | DUF499_1.RM.like.systems | Glyoxalase->?->?->?->?->tRNA-><-XIS-HTH<- DUF499 *<- REase<- OLD-ABC <- METHYLASE <- TUDOR+ SF1-SF2-Helicase+iREase+CTD+REase <-ParB | 941 | Bacteria>PVC_group>Planctomycetota>Planctomycetia>Isosphaerales | GCF_001956985.1 | Paludisphaera_borealis | BSF38_RS09985 | 2556227..2559052 | 1387353 | DUF499-domain-containing-protein-[Paludisphaera-borealis] |
| WP_250846938.1 | DUF499_1.RM.like.systems | <- DUF499 *<- REase<- OLD-ABC <- METHYLASE <- TUDOR+ SF1-SF2-Helicase+iREase+CTD+REase | 939 | Bacteria>PVC_group>Planctomycetota>Planctomycetia>Isosphaerales | GCF_011064615.1 | Aquisphaera_insulae | G6054_RS16920 | 61801..64620 | 2712864 | DUF499-domain-containing-protein-[Aquisphaera-sp.-JC669] |
| WP_230262563.1 | DUF499_1.RM.like.systems | <-RAMA<-URI-DNase+RAMA<-REase<-EVE<-REase<- DUF499 *<- REase<- OLD-ABC <- METHYLASE <-Peptidase_S8<-Classical-AAA<- TUDOR+ SF1-SF2-Helicase+iREase+CTD+REase | 950 | Bacteria>PVC_group>Planctomycetota>Planctomycetia>Pirellulales | GCF_020966635.1 | Stieleria_sp._JC731 | LOC67_RS10680 | 1300044..1302896 | 2894195 | DUF499-domain-containing-protein-[Stieleria-sp.-JC731] |
| HBI42788.1 | DUF499_1.RM.like.systems | SIGMA-HTH->?-><- DUF499 *<- REase<- OLD-ABC <- METHYLASE | 939 | Bacteria>PVC_group>Planctomycetota>Planctomycetia>Planctomycetales | GCA_003508275.1 | Planctomycetales_bacterium | DDY78_08010 | 2578..5397 | 2053591 | hypothetical-protein-[Planctomycetales-bacterium] |
| MBX3448931.1 | DUF499_1.RM.like.systems | TUDOR+ SF1-SF2-Helicase+iREase+CTD+REase -> OLD-ABC ->HNH -> METHYLASE -> OLD-ABC ->REase -> DUF499 *->?->MORC->?->?-><-wHTH+SAD+HNH<-DNA_methylase | 948 | Bacteria>PVC_group>Planctomycetota>Planctomycetia>Planctomycetales | GCA_019636285.1 | Planctomycetaceae_bacterium | KF777_05185 | 10482..13328 | 2026779 | ATP-binding-protein-[Planctomycetaceae-bacterium] |
| MCA8917196.1 | DUF499_1.RM.like.systems | XIS-HTH->?-><- DUF499 *<- REase<- OLD-ABC <- METHYLASE | 951 | Bacteria>PVC_group>Planctomycetota>unclassified_Planctomycetota | GCA_020425605.1 | Planctomycetota_bacterium | KDB90_17530 | 3869..6724 | 2026780 | ATP-binding-protein-[Planctomycetota-bacterium] |
| MCY2978566.1 | DUF499_1.RM.like.systems | <- DUF499 *<- REase<- OLD-ABC <- METHYLASE <- TUDOR+ SF1-SF2-Helicase+iREase+CTD+REase <-XIS-HTH | 968 | Bacteria>PVC_group>Planctomycetota>unclassified_Planctomycetota | GCA_026692945.1 | Planctomycetota_bacterium | NTU79_07885 | 10024..12930 | 2026780 | MAG:-ATP-binding-protein-[Planctomycetota-bacterium] |
| MCX6879289.1 | DUF499_1.RM.like.systems | <-HISKIN<- DUF499 *<- REase<- OLD-ABC <- METHYLASE <- HNH<- OLD-ABC <- TUDOR+ SF1-SF2-Helicase+iREase+CTD+REase | 941 | Bacteria>PVC_group>Verrucomicrobiota>unclassified_Verrucomicrobiota | GCA_026397725.1 | Verrucomicrobiota_bacterium | NTW21_36620 | 9837..12662 | 2026799 | MAG:-DUF499-domain-containing-protein-[Verrucomicrobia-bacterium] |
| MBK6433234.1 | DUF499_1.RM.like.systems | <- DUF499 *<- REase<- OLD-ABC <-HNH | 943 | Bacteria>Terrabacteria_group>Chloroflexota>Anaerolineae>unclassified_Anaerolineae | GCA_016704685.1 | Anaerolineae_bacterium | IPF85_21385 | 32175..35006 | 2052143 | ATP-binding-protein-[Anaerolineae-bacterium] |
| MBK9093501.1 | DUF499_1.RM.like.systems | TUDOR+ SF1-SF2-Helicase+iREase+CTD+REase -> OLD-ABC ->HNH -> METHYLASE -> OLD-ABC ->REase -> DUF499 *->?->ParB+HNH->?->REase-p-> | 943 | Bacteria>Terrabacteria_group>Chloroflexota>Anaerolineae>unclassified_Anaerolineae | GCA_016719785.1 | Anaerolineae_bacterium | IPM84_12145 | 210722..213553 | 2052143 | ATP-binding-protein-[Anaerolineae-bacterium] |
| MCL0064885.1 | DUF499_1.RM.like.systems | <-TPR+TPR+TPR+TPR+TPR+TPR+TPR<-?<- DUF499 *<-REase<- REase<- OLD-ABC <- METHYLASE | 925 | Bacteria>Terrabacteria_group>Chloroflexota>Dehalococcoidia>unclassified_Dehalococcoidia | GCA_023271695.1 | Dehalococcoidia_bacterium | M1N84_03205 | 1953..4730 | 2026734 | DUF499-domain-containing-protein-[Dehalococcoidia-bacterium] |
| QWK10277.1 | DUF499_1.RM.like.systems | <-OrfB_IS605+OrfB_Zn_ribbon<-?\|\|ABC-ATPase+SbcC-><-?<-?<-?<- HEPN+ DUF499 *<-HEPN<-PolB<- METHYLASE <- REase<- OLD-ABC + TOPRIM <- TUDOR+ SF1-SF2-Helicase+iREase+CTD+REase | 1119 | Bacteria>Terrabacteria_group>Chloroflexota>Thermoflexia>Thermoflexales | GCA_018771565.1 | Thermoflexus_hugenholtzii | KNN16_13035 | 2858619..2861978 | 1495650 | MAG:-DUF499-domain-containing-protein-[Thermoflexus-hugenholtzii] |
| MBI4322294.1 | DUF499_1.RM.like.systems | TUDOR+ SF1-SF2-Helicase+iREase+CTD+REase ->Schlafen+HATPase+HTH->?-> METHYLASE -> OLD-ABC ->REase -> DUF499 *-> | 938 | Bacteria>Terrabacteria_group>Chloroflexota>unclassified_Chloroflexota | GCA_016210495.1 | Chloroflexota_bacterium | HY675_27690 | 51598..54414 | 2026724 | ATP-binding-protein-[Chloroflexi-bacterium] |
| MCL5961065.1 | DUF499_1.RM.like.systems | TUDOR+ SF1-SF2-Helicase+iREase+CTD+REase ->Schlafen+HATPase+HTH-> METHYLASE -> OLD-ABC ->REase -> DUF499 *->MerR-HTH->?->REase->IclR-HTH-> | 939 | Bacteria>Terrabacteria_group>Chloroflexota>unclassified_Chloroflexota | GCA_023489135.1 | Chloroflexota_bacterium | M1358_17465 | 13442..16261 | 2026724 | ATP-binding-protein-[Chloroflexi-bacterium] |
| WP_229782279.1 | DUF499_1.RM.like.systems | OrfB_Zn_ribbon->?->ParB-><- DUF499 *<- REase<- OLD-ABC <- METHYLASE <- TUDOR+ SF1-SF2-Helicase+iREase+CTD+REase | 959 | Bacteria>Terrabacteria_group>Deinococcota>Deinococci>Deinococcales | GCF_014647175.1 | Deinococcus_daejeonensis | IEY59_RS18160 | 23471..26350 | 1007098 | DUF499-domain-containing-protein-[Deinococcus-daejeonensis] |
| WP_178366469.1 | DUF499_1.RM.like.systems | <- DUF499 *<- REase<- OLD-ABC <- METHYLASE <- OLD-ABC <- TUDOR+ SF1-SF2-Helicase+iREase+CTD+REase <-Radical-SAM+TRAM | 952 | Bacteria>Thermodesulfobacteriota>Desulfobacteria>Desulfobacterales>Desulfobacteraceae | GCF_013389365.1 | Desulfobacter_latus | HXW94_RS08505 | 32689..35547 | 2292 | DUF499-domain-containing-protein-[Desulfobacter-latus] |
| WP_168935605.1 | DUF499_1.RM.like.systems | <-HTH_3\|\|?-><-?\|\|?->?-><- DUF499 *<- REase<- OLD-ABC <- METHYLASE <- HNH<- OLD-ABC <- TUDOR+ SF1-SF2-Helicase+iREase+CTD+REase | 933 | Bacteria>Thermodesulfobacteriota>Desulfovibrionia>Desulfovibrionales>Desulfovibrionaceae | GCF_012843875.1 | Desulfovibrio_piger | HF854_RS06725 | 17522..20323 | 901 | DUF499-domain-containing-protein-[Desulfovibrio-piger] |

--
-- OLD-ABC ->REase
--BR-3
--

| | | | | | | | | | | |
|---|---|---|---|---|---|---|---|---|---|---|
| MCX5862619.1 | DUF499_3.PglZ.associated.systems | <- TUDOR+SF1-SF2-Helicase+iREase+CTD <- iSTAND <- METHYLASE <- REase<- OLD-ABC <- PglZ <- DUF499 * | 922 | Bacteria>Myxococcota>Myxococcia>unclassified_Myxococcia | GCA_026388625.1 | Deltaproteobacteria_bacterium | NTW27_10950 | 92147..94915 | 2026735 | MAG:-DUF499-domain-containing-protein-[Deltaproteobacteria-bacterium] |

<table>
<tr><th>Protein-ID</th><th>System</th><th>Gene-Neighbourhoods</th><th>Length</th><th>Taxonomy</th><th>Assembly-ID</th><th>Strain</th><th>Locus-Tag</th><th>Genomic-coordinates</th><th>TaxID</th><th>Fasta-defline</th></tr>
<tr><td>WP_169728956.1</td><td>DUF499_3.PglZ.associated.systems</td><td><- iSTAND <- METHYLASE <- REase<- OLD-ABC <- PglZ <- DUF499 *<-XIS-HTH<-?<-GDResolvase+Recombinase+ZnR<-URI-DNase</td><td>756</td><td>Bacteria>Thermodesulfobacteriota>Desulfobacteria>Desulfobacterales>Desulfatirhabdiaceae</td><td>GCF_000429925.1</td><td>Desulfatirhabdium_butyrativorans</td><td>G492_RS0112005</td><td>53087..55357</td><td>340467</td><td>DUF499-domain-containing-protein-[Desulfatirhabdium-butyrativorans]</td></tr>
<tr><td>MBF0302150.1</td><td>DUF499_3.PglZ.associated.systems</td><td><- iSTAND <- METHYLASE -p<- METHYLASE <- REase<- OLD-ABC <- PglZ <- DUF499 *||SF1-SF2-Helicase-></td><td>934</td><td>Bacteria>Thermodesulfobacteriota>Desulfobacteria>Desulfobacterales>Desulfobacteraceae</td><td>GCA_015232035.1</td><td>Desulfamplus_sp.</td><td>HQK73_03850</td><td>16841..19645</td><td>2006184</td><td>DUF499-domain-containing-protein-[Desulfamplus-sp.]</td></tr>
<tr><td>MDD4464448.1</td><td>DUF499_3.PglZ.associated.systems</td><td><- METHYLASE -p<- REase<- OLD-ABC <- PglZ <- DUF499 *</td><td>183</td><td>Bacteria>Thermodesulfobacteriota>Desulfobacteria>Desulfobacterales>unclassified_Desulfobacterales</td><td>-</td><td>Desulfobacterales_bacterium</td><td>PHR86_09450</td><td>3623..4176</td><td>2044940</td><td>MAG:-hypothetical-protein-PHR86_09450,-partial-[Desulfobacterales-bacterium]</td></tr>
<tr><td>MDD2316829.1</td><td>DUF499_3.PglZ.associated.systems</td><td>XIS-HTH-> DUF499 *-> PglZ -> OLD-ABC ->REase -> METHYLASE -> OLD-ABC -> TOPRIM -></td><td>951</td><td>Bacteria>Thermodesulfobacteriota>Desulfobacteria>Desulfobacterales>unclassified_Desulfobacterales</td><td>GCA_028683495.1</td><td>Desulfobacterales_bacterium</td><td>PHD57_05725</td><td>1896..4751</td><td>2044940</td><td>MAG:-hypothetical-protein-PHD57_05725-[Desulfobacterales-bacterium]</td></tr>
<tr><td>--</td><td></td><td></td><td></td><td></td><td></td><td></td><td></td><td></td><td></td><td></td></tr>
<tr><td>--PIN-linked_OLD-ABC</td><td></td><td></td><td></td><td></td><td></td><td></td><td></td><td></td><td></td><td></td></tr>
<tr><td>--</td><td></td><td></td><td></td><td></td><td></td><td></td><td></td><td></td><td></td><td></td></tr>
<tr><td>-- OLD-ABC ->PIN</td><td></td><td></td><td></td><td></td><td></td><td></td><td></td><td></td><td></td><td></td></tr>
<tr><td>--Type1.BREX</td><td></td><td></td><td></td><td></td><td></td><td></td><td></td><td></td><td></td><td></td></tr>
<tr><td>--</td><td></td><td></td><td></td><td></td><td></td><td></td><td></td><td></td><td></td><td></td></tr>
<tr><td>MBR4647494.1</td><td>Type1.BREX</td><td>MNS+CR-ATPase5-><-?||?->?->Calcineurin->SbcC+CCDC158-> BrxA-DNA-binding -> BrxB-BrxF- iSTAND -> DUF6079 *-> OLD-ABC ->PIN -> N6- METHYLASE -> PglZ -> SIGMA+OB-Fold+AAA-ATPase+Lon-Protease-BrxL ->?->?->?-> Cysteine-Desulfurase -></td><td>1174</td><td>Bacteria>FCB_group>Bacteroidetes/Chlorobi_group>Bacteroidota>Bacteroidia>Bacteroidales</td><td>GCA_017525245.1</td><td>Bacteroidales_bacterium</td><td>IKO75_10320</td><td>20247..23771</td><td>2030927</td><td>BREX-system-P-loop-protein-BrxC-[Bacteroidales-bacterium]</td></tr>
<tr><td>WP_234146092.1</td><td>Type1.BREX</td><td>Arm-DNA-bind_5+Phage_int_SAM_5+Phage_integrase-><-Schlafen+HATPase+wHTH<- SIGMA+OB-Fold+AAA-ATPase+Lon-Protease-BrxL <- PglZ <- N6- METHYLASE <- PIN<- OLD-ABC <- DUF6079 *<- BrxB-BrxF- iSTAND <- BrxA-DNA-binding <-HTH_3||?->?-> N6- METHYLASE -><-AAA+<-HTH_36<-XIS-HTH</td><td>1180</td><td>Bacteria>FCB_group>Bacteroidetes/Chlorobi_group>Bacteroidota>Bacteroidia>Bacteroidales</td><td>GCF_024760085.1</td><td>Bacteroides_uniformis</td><td>J8872_RS07820</td><td>80453..83995</td><td>820</td><td>BREX-system-P-loop-protein-BrxC-[Bacteroides-uniformis]</td></tr>
<tr><td>MCB5286841.1</td><td>Type1.BREX</td><td><-IG+IG||?->?-><-tRNA||?->?->XIS-HTH-> BrxA-DNA-binding -> BrxB-BrxF- iSTAND -> DUF6079 *-> OLD-ABC ->PIN -> N6- METHYLASE ->BirA-HTH-> PglZ -> SIGMA+OB-Fold+AAA-ATPase+Lon-Protease-BrxL -></td><td>1145</td><td>Bacteria>FCB_group>Candidatus_Cloacimonetes>unclassified_Candidatus_Cloacimonetes</td><td>GCA_020532145.1</td><td>Candidatus_Cloacimonetes_bacterium</td><td>LHW64_03450</td><td>9843..13280</td><td>2030808</td><td>BREX-system-P-loop-protein-BrxC-[Candidatus-Cloacimonadota-bacterium]</td></tr>
<tr><td>MBQ2886288.1</td><td>Type1.BREX</td><td><-wHTH+WYL<-REase<- SIGMA+OB-Fold+AAA-ATPase+Lon-Protease-BrxL <- PglZ <-Schlafen+HATPase+wHTH<- N6- METHYLASE <- PIN<- OLD-ABC <- DUF6079 *<- BrxB-BrxF- iSTAND <- BrxA-DNA-binding <-HTH_3</td><td>1180</td><td>Bacteria>Pseudomonadota>Alphaproteobacteria>unclassified_Alphaproteobacteria</td><td>GCA_017409945.1</td><td>Alphaproteobacteria_bacterium</td><td>IJE43_21410</td><td>13139..16681</td><td>1913988</td><td>BREX-system-P-loop-protein-BrxC-[Alphaproteobacteria-bacterium]</td></tr>
<tr><td>WP_076471941.1</td><td>Type6.BREX</td><td><-DUF3262<-?<-?<-?<-JetA|| wHTH+WYL+WCX -> BrxE -> BrxA-DNA-binding -> BrxB-BrxF- iSTAND -> DUF6079 *-> OLD-ABC ->PIN -> N6- METHYLASE -> N6- METHYLASE ->ParB+HNH-> PglZ -> SIGMA+OB-Fold+AAA-ATPase+HKD-DNase-BrxL -><-JAB</td><td>1207</td><td>Bacteria>Pseudomonadota>Betaproteobacteria>Burkholderiales>Alcaligenaceae>Achromobacter</td><td>GCF_001957055.1</td><td>Achromobacter_xylosoxidans</td><td>BIZ53_RS00410</td><td>81330..84953</td><td>85698</td><td>BREX-system-P-loop-protein-BrxC-[Achromobacter-xylosoxidans]</td></tr>
<tr><td>WP_141017182.1</td><td>Type6.BREX</td><td>PhdYeFM_antitox->RelE-ParE-><-DOC+HTH<-Phage_integrase<-?|| wHTH+WYL+WCX -> BrxE -> BrxA-DNA-binding -> BrxB-BrxF- iSTAND -> DUF6079 *-> OLD-ABC ->PIN -> N6- METHYLASE -> N6- METHYLASE ->?->ParB+HNH-> PglZ -> SIGMA+OB-Fold+AAA-ATPase+HKD-DNase-BrxL -><-XIS-HTH</td><td>1207</td><td>Bacteria>Pseudomonadota>Betaproteobacteria>Rhodocyclales>Zoogloeaceae>Azoarcus</td><td>GCF_006496635.1</td><td>Azoarcus_sp._DD4</td><td>CJ010_RS05935</td><td>1245819..1249442</td><td>2027405</td><td>BREX-system-P-loop-protein-BrxC-[Azoarcus-sp.-DD4]</td></tr>
<tr><td>WP_144225094.1</td><td>Type1.BREX</td><td><-Phage_integrase<-DUF2787<-?<-?||XIS-HTH->XIS-HTH-> BrxA-DNA-binding -> BrxB-BrxF- iSTAND -p-> DUF6079 *-> OLD-ABC ->PIN -> N6- METHYLASE -> PglZ -> SIGMA+OB-Fold+AAA-ATPase+Lon-Protease-BrxL ->?-><-Arm-DNA-bind_3+Phage_integrase<-?||RNasePH-></td><td>1145</td><td>Bacteria>Pseudomonadota>Gammaproteobacteria>Alteromonadales>Shewanellaceae>Shewanella</td><td>GCF_017580345.1</td><td>Shewanella_algae</td><td>I6M43_RS01845</td><td>49386..52823</td><td>38313</td><td>BREX-system-P-loop-protein-BrxC-[Shewanella-algae]</td></tr>
<tr><td>WP_200158841.1</td><td>Type1.BREX</td><td><- SIGMA+OB-Fold+AAA-ATPase+Lon-Protease-BrxL <- PglZ <- HNH<- OLD-ABC <-SF1-SF2-Helicase<- OLD-ABC + TOPRIM <- N6- METHYLASE <- PIN<- OLD-ABC <- DUF6079 *<- BrxB-BrxF- iSTAND <- BrxA-DNA-binding <-GDResolvase+HIN-HTH<-ZnR</td><td>1154</td><td>Bacteria>Pseudomonadota>Gammaproteobacteria>Chromatiales>Chromatiaceae>Allochromatium</td><td>GCF_016584135.1</td><td>Allochromatium_vinosum</td><td>CK029_RS16480</td><td>18892..22356</td><td>1049</td><td>BREX-system-P-loop-protein-BrxC-[Allochromatium-vinosum]</td></tr>
<tr><td>WP_062271501.1</td><td>Type1.BREX</td><td><- OLD-ABC <-RHH<-SF1-SF2-Helicase<- OLD-ABC + TOPRIM <- N6- METHYLASE <- PIN<- OLD-ABC <-PIN<-?<- DUF6079 *<- BrxB-BrxF- iSTAND <- BrxA-DNA-binding <-tRNA<-Aminotran_1_2||SF1-SF2-Helicase-></td><td>1153</td><td>Bacteria>Pseudomonadota>Gammaproteobacteria>Chromatiales>Chromatiaceae>Marichromatium</td><td>GCF_001583505.1</td><td>Marichromatium_gracile</td><td>AY586_RS00820</td><td>23071..26532</td><td>1048</td><td>BREX-system-P-loop-protein-BrxC-[Marichromatium-gracile]</td></tr>
<tr><td>MCK7579547.1</td><td>Type1.BREX</td><td><- OLD-ABC + TOPRIM <-?<-?<- N6- METHYLASE <- PIN<- OLD-ABC ||?-><- DUF6079 -p<- DUF6079 -p*<-?<- DUF6079 -p<- BrxB-BrxF- iSTAND <- BrxA-DNA-binding -p<-ZnR</td><td>299</td><td>Bacteria>Pseudomonadota>Gammaproteobacteria>Chromatiales>unclassified_Chromatiales</td><td>GCA_023156155.1</td><td>Chromatiales_bacterium</td><td>MZV65_30115</td><td>14491..15390</td><td>2026725</td><td>hypothetical-protein-[Chromatiales-bacterium]</td></tr>
<tr><td>MCK7579549.1</td><td>Type1.BREX</td><td><- N6- METHYLASE <- PIN<- OLD-ABC ||?-><- DUF6079 -p<- DUF6079 -p<-?<- DUF6079 -p*<- BrxB-BrxF- iSTAND <- BrxA-DNA-binding -p<-ZnR</td><td>648</td><td>Bacteria>Pseudomonadota>Gammaproteobacteria>Chromatiales>unclassified_Chromatiales</td><td>GCA_023156155.1</td><td>Chromatiales_bacterium</td><td>MZV65_30125</td><td>15595..17541</td><td>2026725</td><td>BREX-system-P-loop-protein-BrxC-[Chromatiales-bacterium]</td></tr>
<tr><td>WP_040239271.1</td><td>Type1.BREX</td><td><-sGTP<-?<-?||?->tRNA-> BrxA-DNA-binding -> BrxB-BrxF- iSTAND -> DUF6079 *-> OLD-ABC ->PIN -> N6- METHYLASE ->ParB+HNH->?->?->?-> PglZ -> SIGMA+OB-Fold+AAA-ATPase+Lon-Protease-BrxL -></td><td>1144</td><td>Bacteria>Pseudomonadota>Gammaproteobacteria>Oceanospirillales>Halomonadaceae>Chromohalobacter</td><td>GCF_000821045.2</td><td>Chromohalobacter_japonicus</td><td>BN995_RS00415</td><td>38131..41565</td><td>223900</td><td>BREX-system-P-loop-protein-BrxC-[Chromohalobacter-japonicus]</td></tr>
<tr><td>WP_189514712.1</td><td>Type1.BREX</td><td><-DUF4096+DDE_Tnp_1+DDE_Tnp_1_2<-?||?->DUF932->?->?->?-> BrxA-DNA-binding -> BrxB-BrxF- iSTAND -> DUF6079 *-> OLD-ABC ->PIN -> N6- METHYLASE ->ParB+HNH-> PglZ -> SIGMA+OB-Fold+AAA-ATPase+Lon-Protease-BrxL ->McrB-AAA->wHTH+REase-></td><td>1144</td><td>Bacteria>Pseudomonadota>Gammaproteobacteria>Oceanospirillales>Halomonadaceae>Kushneria</td><td>GCF_014652795.1</td><td>Kushneria_pakistanensis</td><td>IE123_RS02380</td><td>531699..535133</td><td>1508770</td><td>BREX-system-P-loop-protein-BrxC-[Kushneria-pakistanensis]</td></tr>
<tr><td>WP_163650581.1</td><td>Type1.BREX</td><td><- N6- METHYLASE -p<- PIN<- OLD-ABC <- DUF6079 *<- BrxB-BrxF- iSTAND <- BrxA-DNA-binding <-tRNA<-?||?->?->sGTP-></td><td>1144</td><td>Bacteria>Pseudomonadota>Gammaproteobacteria>Oceanospirillales>Halomonadaceae>Modicisalibacter</td><td>GCF_010993675.1</td><td>Modicisalibacter_sp._'Wilcox'</td><td>G7A98_RS15465</td><td>3721..7155</td><td>2679914</td><td>BREX-system-P-loop-protein-BrxC-[Modicisalibacter-sp.-'Wilcox']</td></tr>
<tr><td>WP_224416377.1</td><td>Type1.BREX</td><td><- PglZ <-?<-?<-?<- N6- METHYLASE -p<-HTH_Tnp_1+HTH_21+rve+rve_3<- N6- METHYLASE <- PIN<- OLD-ABC <- DUF6079 *<- BrxB-BrxF- iSTAND <- BrxA-DNA-binding <-tRNA<-?||?->?->sGTP-></td><td>1144</td><td>Bacteria>Pseudomonadota>Gammaproteobacteria>Oceanospirillales>Halomonadaceae>Modicisalibacter</td><td>GCF_020147965.1</td><td>Modicisalibacter_tunisiensis</td><td>KGQ90_RS13860</td><td>2977413..2980847</td><td>390637</td><td>BREX-system-P-loop-protein-BrxC-[Modicisalibacter-tunisiensis]</td></tr>
<tr><td>WP_202806908.1</td><td>Type1.BREX</td><td><- PglZ <- TOPRIM <- OLD-ABC <-SbcC<- N6- METHYLASE -p<- N6- METHYLASE <- N6- METHYLASE -p<- PIN<- OLD-ABC <- DUF6079 -p*<- DUF6079 -p<- BrxB-BrxF- iSTAND <- BrxA-DNA-binding <-XIS-HTH<-?<-HDRnase<-Phage_int_SAM<-Integrase_1||XIS-HTH-></td><td>460</td><td>Bacteria>Pseudomonadota>Gammaproteobacteria>Oceanospirillales>Oceanospirillaceae>Nitrincola</td><td>GCF_000585235.1</td><td>Nitrincola_nitratireducens</td><td>D791_RS20495</td><td>12347..13729</td><td>1229521</td><td>hypothetical-protein-[Nitrincola-nitratireducens]</td></tr>
<tr><td>WP_202806909.1</td><td>Type1.BREX</td><td><- TOPRIM <- OLD-ABC <-SbcC<- N6- METHYLASE -p<- N6- METHYLASE <- N6- METHYLASE -p<- PIN<- OLD-ABC <- DUF6079 -p<- DUF6079 -p*<- BrxB-BrxF- iSTAND <- BrxA-DNA-binding <-XIS-HTH<-?<-HDRnase<-Phage_int_SAM<-Integrase_1||XIS-HTH->Phage_integrase-></td><td>697</td><td>Bacteria>Pseudomonadota>Gammaproteobacteria>Oceanospirillales>Oceanospirillaceae>Nitrincola</td><td>GCF_000585235.1</td><td>Nitrincola_nitratireducens</td><td>D791_RS20500</td><td>13689..15782</td><td>1229521</td><td>BREX-system-P-loop-protein-BrxC-[Nitrincola-nitratireducens]</td></tr>
<tr><td>HBO4843314.1</td><td>Type6.BREX</td><td><-ACET<-YjhX_toxin||?->?-><-MODE-HTH+PBPII|| wHTH+WYL+WCX -> BrxE -> BrxA-DNA-binding -> BrxB-BrxF- iSTAND -> DUF6079 *-> OLD-ABC ->PIN -> N6- METHYLASE ->?-> PglZ -> SIGMA+OB-Fold+AAA-ATPase+HKD-DNase-BrxL -><-HDRnase||PrlF_antitoxin-></td><td>1207</td><td>Bacteria>Pseudomonadota>Gammaproteobacteria>Pseudomonadales>Pseudomonadaceae>Pseudomonas</td><td>GCA_021660595.1</td><td>Pseudomonas_aeruginosa</td><td>L4V37_002405</td><td>61513..65136</td><td>287</td><td>BREX-system-P-loop-protein-BrxC-[Pseudomonas-aeruginosa]</td></tr>
<tr><td>MBN5083090.1</td><td>Type6.BREX</td><td><-DUF932<- SIGMA+OB-Fold+AAA-ATPase+HKD-DNase-BrxL <- PglZ <-ParB+HNH<- N6- METHYLASE <- N6- METHYLASE <- PIN<- OLD-ABC <- DUF6079 *<- BrxB-BrxF- iSTAND <- BrxA-DNA-binding <- BrxE <- wHTH+WYL+WCX <-Arm-DNA-bind_3+Phage_integrase<-tRNA<-SIGMA-HTH+SIGMA-HTH+SIGMA-HTH</td><td>1207</td><td>Bacteria>Pseudomonadota>Gammaproteobacteria>Xanthomonadales>Xanthomonadaceae>Stenotrophomonas</td><td>GCA_017150495.1</td><td>Stenotrophomonas_maltophilia</td><td>JY467_05250</td><td>61681..65304</td><td>40324</td><td>BREX-system-P-loop-protein-BrxC-[Stenotrophomonas-maltophilia]</td></tr>
<tr><td>WP_004146422.1</td><td>Type6.BREX</td><td><-XIS-HTH||?->Phage_integrase->tRNA->Arm-DNA-bind_3+Phage_integrase-> wHTH+WYL+WCX -> BrxE -> BrxA-DNA-binding -> BrxB-BrxF- iSTAND -> DUF6079 *-> OLD-ABC ->PIN -> N6- METHYLASE -> N6- METHYLASE ->ParB+HNH-> PglZ -> SIGMA+OB-Fold+AAA-ATPase+HKD-DNase-BrxL ->DUF932-></td><td>1207</td><td>Bacteria>Pseudomonadota>Gammaproteobacteria>Xanthomonadales>Xanthomonadaceae>Stenotrophomonas</td><td>GCF_000355725.1</td><td>Stenotrophomonas_maltophilia</td><td>G5A34_RS07915</td><td>81973..85596</td><td>40324</td><td>BREX-system-P-loop-protein-BrxC-[Stenotrophomonas-maltophilia]</td></tr>
<tr><td>VUJ46804.1</td><td>Type6.BREX</td><td><-LexA-protease||?-><-Phage_integrase||Arm-DNA-bind_3+Phage_integrase-> wHTH+WYL+WCX -> BrxE -> BrxA-DNA-binding -> BrxB-BrxF- iSTAND -> DUF6079 *-> OLD-ABC ->PIN -> N6- METHYLASE -> N6- METHYLASE ->ParB+HNH-> PglZ -> SIGMA+OB-Fold+AAA-ATPase+HKD-DNase-BrxL ->DUF932-></td><td>1207</td><td>Bacteria>Pseudomonadota>Gammaproteobacteria>Xanthomonadales>Xanthomonadaceae>Stenotrophomonas</td><td>-</td><td>Stenotrophomonas_maltophilia</td><td>N5D45_RS14200</td><td>24687..28310</td><td>40324</td><td>hypothetical-protein-NGKLMNPD_00401-[Stenotrophomonas-maltophilia]</td></tr>
<tr><td>NBB80453.1</td><td>Type1.BREX</td><td>HTH_3-> BrxA-DNA-binding -> BrxB-BrxF- iSTAND -> DUF6079 *-> OLD-ABC ->PIN -> N6- METHYLASE -> OLD-ABC ->HNH ->ParB+HNH->Schlafen+HATPase+wHTH-> PglZ -> SIGMA+OB-Fold+AAA-ATPase+Lon-Protease-BrxL -></td><td>1147</td><td>Bacteria>PVC_group>Verrucomicrobiota>unclassified_Verrucomicrobiota</td><td>GCA_009908325.1</td><td>Verrucomicrobiota_bacterium</td><td>GVY36_13595</td><td>2227..5670</td><td>2026799</td><td>BREX-system-P-loop-protein-BrxC-[Verrucomicrobiota-bacterium]</td></tr>
<tr><td>CDD75188.1</td><td>Type1.BREX</td><td>MACRODOMAIN->?->PIN->ZnR+ZnR+ZnR->HTH_26-> BrxA-DNA-binding -> BrxB-BrxF- iSTAND -> DUF6079 *-> OLD-ABC ->PIN -> N6- METHYLASE -> OLD-ABC ->NERD->HEPN->Schlafen+HATPase+wHTH-> PglZ ->DUF4365+REase-></td><td>1200</td><td>Bacteria>Terrabacteria_group>Bacillota>Clostridia>Eubacteriales>Clostridiaceae</td><td>GCA_000436175.1</td><td>Clostridium_sp._CAG:62</td><td>BN737_01375</td><td>69547..73149</td><td>1262828</td><td>uncharacterized-protein-BN737_01375-[Clostridium-sp.-CAG:62]</td></tr>
<tr><td>WP_118744885.1</td><td>Type1.BREX</td><td>ACET->REase->?->?->?->HTH_26-> BrxA-DNA-binding -> BrxB-BrxF- iSTAND -> DUF6079 *-> OLD-ABC ->PIN -> N6- METHYLASE ->HEPN->?-> OLD-ABC ->HNH ->?->TIR-></td><td>1203</td><td>Bacteria>Terrabacteria_group>Bacillota>Clostridia>Eubacteriales>Clostridiaceae</td><td>GCF_003480745.1</td><td>Clostridium_sp._AM27-31LB</td><td>DW721_RS03885</td><td>61324..64935</td><td>2293026</td><td>BREX-system-P-loop-protein-BrxC-[Clostridium-sp.-AM27-31LB]</td></tr>
<tr><td>RGI65884.1</td><td>Type1.BREX</td><td>HTH_26-> BrxA-DNA-binding -> BrxB-BrxF- iSTAND -> DUF6079 *-> OLD-ABC ->PIN -> N6- METHYLASE ->TIR->HEPN->Schlafen+HATPase+wHTH-> PglZ ->Tox-ART-HYD3-> SIGMA+OB-Fold+AAA-ATPase+Lon-Protease-BrxL -></td><td>1196</td><td>Bacteria>Terrabacteria_group>Bacillota>Clostridia>Eubacteriales>Lachnospiraceae</td><td>GCA_003436035.1</td><td>[Eubacterium]_rectale</td><td>DXD95_13285</td><td>1619..5209</td><td>39491</td><td>BREX-system-P-loop-protein-BrxC-[[Eubacterium]</td></tr>
<tr><td>MCI7121458.1</td><td>Type1.BREX</td><td>HATPase+wHTH->?->MACRODOMAIN->?->35exo+ADA-DBD->ZnR+ZnR+ZnR->HTH_26-> BrxA-DNA-binding -> BrxB-BrxF- iSTAND -> DUF6079 *-> OLD-ABC ->PIN -> N6- METHYLASE ->?-> PglZ ->Schlafen+HATPase+wHTH-> SIGMA+OB-Fold+AAA-ATPase+Lon-Protease-BrxL ->LRR_5+LRR_8->Cas9+REC+HNH+Cas9-></td><td>1201</td><td>Bacteria>Terrabacteria_group>Bacillota>Clostridia>Eubacteriales>Lachnospiraceae</td><td>GCA_022775325.1</td><td>[Ruminococcus]_gnavus</td><td>MR941_06720</td><td>10105..13710</td><td>33038</td><td>BREX-system-P-loop-protein-BrxC-[[Ruminococcus]</td></tr>
<tr><td>WP_173865463.1</td><td>Type1.BREX</td><td><-STYKIN<-?<-LRR_5+LRR_8<- SIGMA+OB-Fold+AAA-ATPase+Lon-Protease-BrxL <-Schlafen+HATPase+wHTH<- PglZ <- N6- METHYLASE <- PIN<- OLD-ABC <- DUF6079 *<- BrxB-BrxF- iSTAND <- BrxA-DNA-binding <-HTH_26<-ZnR+ZnR+ZnR<-35exo+ADA-DBD<-?<-MACRODOMAIN<-?<-DOC+zhth</td><td>1201</td><td>Bacteria>Terrabacteria_group>Bacillota>Clostridia>Eubacteriales>Lachnospiraceae</td><td>GCF_013300965.1</td><td>Faecalicatena_fissicatena</td><td>HFN12_RS00390</td><td>75728..79333</td><td>290055</td><td>BREX-system-P-loop-protein-BrxC-[Faecalicatena-fissicatena]</td></tr>
<tr><td>WP_270645287.1</td><td>Type1.BREX</td><td>HTH_19-><- SIGMA+OB-Fold+AAA-ATPase+Lon-Protease-BrxL <-?<- PglZ <- PIN<- OLD-ABC <- OLD-ABC + TOPRIM <- N6- METHYLASE <- DUF6079 *<- BrxB-BrxF- iSTAND <- BrxA-DNA-binding <-HTH_26<-?<-?<-?<-ArdCN+MPTase-inact+LPD19<-?<- TOPRIM +Topoisom_bac</td><td>1205</td><td>Bacteria>Terrabacteria_group>Bacillota>Clostridia>Eubacteriales>Lachnospiraceae</td><td>GCF_027692785.1</td><td>Merdimonas_faecis</td><td>PG729_RS15130</td><td>28205..31822</td><td>1653435</td><td>BREX-system-P-loop-protein-BrxC-[Merdimonas-faecis]</td></tr>
<tr><td>MCI8835830.1</td><td>Type1.BREX</td><td>ABC-ATPase->ABC-ATPase+SbcC->?-> BrxA-DNA-binding -> BrxB-BrxF- iSTAND -> DUF6079 *->Schlafen+HATPase+wHTH-> N6- METHYLASE -> OLD-ABC ->PIN -> PglZ -></td><td>1205</td><td>Bacteria>Terrabacteria_group>Bacillota>Clostridia>Eubacteriales>Oscillospiraceae</td><td>GCA_022795995.1</td><td>Ruminococcus_sp.</td><td>HFH24_10725</td><td>31305..34922</td><td>41978</td><td>BREX-system-P-loop-protein-BrxC-[Ruminococcus-sp.]</td></tr>
<tr><td>MBQ3415902.1</td><td>Type1.BREX</td><td><- SIGMA+OB-Fold+AAA-ATPase+Lon-Protease-BrxL <- PglZ <-?<- PIN<- OLD-ABC <- N6- METHYLASE <-?<- DUF6079 *<- BrxB-BrxF- iSTAND <- BrxA-DNA-binding <-?<-?<-RHH_8</td><td>1216</td><td>Bacteria>Terrabacteria_group>Bacillota>Clostridia>unclassified_Clostridia</td><td>GCA_017404025.1</td><td>Clostridia_bacterium</td><td>IJH39_11340</td><td>11283..14933</td><td>2044939</td><td>BREX-system-P-loop-protein-BrxC-[Clostridia-bacterium]</td></tr>
<tr><td>WP_273698075.1</td><td>Type1.BREX</td><td><-REase<- PglZ <- N6- METHYLASE <- PIN<- OLD-ABC <- HNH<- OLD-ABC <- DUF6079 *<- BrxB-BrxF- iSTAND <- BrxA-DNA-binding <-?<-?<-?<-?<-SKI<-?||O6MGT-HTH-></td><td>1181</td><td>Bacteria>Terrabacteria_group>Bacillota>Negativicutes>Selenomonadales>Selenomonadaceae</td><td>GCF_905371815.1</td><td>Selenomonas_noxia</td><td>NW174_RS07850</td><td>36953..40498</td><td>135083</td><td>BREX-system-P-loop-protein-BrxC-[Selenomonas-noxia]</td></tr>
<tr><td>WP_273079788.1</td><td>Type1.BREX</td><td>BrxA-DNA-binding -> BrxB-BrxF- iSTAND -> DUF6079 *-> OLD-ABC ->HNH -> OLD-ABC ->PIN -> N6- METHYLASE -> PglZ ->SIR2->HerA/FtsK-></td><td>1187</td><td>Bacteria>Terrabacteria_group>Bacillota>Negativicutes>Selenomonadales>Selenomonadaceae</td><td>GCF_905371945.1</td><td>Selenomonas_felix</td><td>NW177_RS10645</td><td>2029..5592</td><td>1944634</td><td>BREX-system-P-loop-protein-BrxC-[Selenomonas-felix]</td></tr>
<tr><td>HJA77858.1</td><td>Type1.BREX</td><td>BrxB-BrxF- iSTAND -p-><-?||ACET-> BrxA-DNA-binding -> BrxB-BrxF- iSTAND -> DUF6079 *-> OLD-ABC ->PIN -> N6- METHYLASE ->PHP+SbcC-> PglZ -> SIGMA+OB-Fold+AAA-ATPase+Lon-Protease-BrxL ->SIR2-></td><td>1145</td><td>Bacteria>Thermodesulfobacteriota>Desulfovibrionia>Desulfovibrionales>Desulfovibrionaceae>Desulfovibrio</td><td>GCA_019118745.1</td><td>Candidatus_Desulfovibrio_gallistercoris</td><td>H9774_13535</td><td>7700..11137</td><td>2838533</td><td>BREX-system-P-loop-protein-BrxC-[Candidatus-Desulfovibrio-gallistercoris]</td></tr>
<tr><td>--</td><td></td><td></td><td></td><td></td><td></td><td></td><td></td><td></td><td></td><td></td></tr>
<tr><td>-- OLD-ABC ->PIN</td><td></td><td></td><td></td><td></td><td></td><td></td><td></td><td></td><td></td><td></td></tr>
<tr><td>--Type2.BREX</td><td></td><td></td><td></td><td></td><td></td><td></td><td></td><td></td><td></td><td></td></tr>
<tr><td>--</td><td></td><td></td><td></td><td></td><td></td><td></td><td></td><td></td><td></td><td></td></tr>
<tr><td>WP_266728674.1</td><td>Type2.Pgl.like.BREX</td><td>HISKIN->REC+OmpR-HTH-><-?||NTP_transf_3->?-><- SF1-SF2-Helicase+wHTH+Small-B-barrel+DUF1998 <- BrxD-AAA <- PglZ <- DUF6079 *<-?<- PIN<- OLD-ABC + TOPRIM <-ST-Hiskin<-SF1-SF2-Helicase||?->HTH_31+DUF5753->DUF397-><- N6- METHYLASE</td><td>1285</td><td>Bacteria>Terrabacteria_group>Actinomycetota>Actinomycetes>Kitasatosporales>Streptomycetaceae</td><td>GCF_026342115.1</td><td>Streptomyces_sp._NBC_00378</td><td>OOK13_RS07585</td><td>1669600..1673457</td><td>2975732</td><td>PglY-protein-[Streptomyces-sp.-NBC_00378]</td></tr>
<tr><td>WP_244805207.1</td><td>Type2.Pgl.like.BREX</td><td><- SF1-SF2-Helicase+wHTH+Small-B-barrel+DUF1998 ||?->HTH_3-><- BrxD-AAA <- PglZ <- DUF6079 *<- N6- METHYLASE <- PglW <- PIN<- OLD-ABC <-URI-DNase+SF1-SF2-Helicase<-REase+SF1-SF2-Helicase||35exo+TerB+TerB+BRCT+SIGMA-HTH-> OLD-ABC ->HNH -></td><td>1292</td><td>Bacteria>Terrabacteria_group>Actinomycetota>Actinomycetes>Mycobacteriales>Corynebacteriaceae</td><td>GCF_022869005.1</td><td>Corynebacterium_kalidii</td><td>MUN33_RS12285</td><td>327139..331017</td><td>2931982</td><td>DUF6079-family-protein-[Corynebacterium-kalidii]</td></tr>
<tr><td>WP_073883613.1</td><td>Type2.Pgl.like.BREX</td><td><- SF1-SF2-Helicase+wHTH+Small-B-barrel+DUF1998 <-SbcC<- BrxD-AAA <- PglZ <- DUF6079 *<- N6- METHYLASE <-?<- PglW <- PIN<- OLD-ABC <-REase+SF1-SF2-Helicase<-REase+SF1-SF2-Helicase|| OLD-ABC ->HNH -></td><td>1316</td><td>Bacteria>Terrabacteria_group>Actinomycetota>Actinomycetes>Mycobacteriales>Corynebacteriaceae</td><td>GCF_900015285.1</td><td>Corynebacterium_variabile</td><td>CVAR292_RS03495</td><td>9616..13566</td><td>1727</td><td>DUF6079-family-protein-[Corynebacterium-variabile]</td></tr>
<tr><td>--</td><td></td><td></td><td></td><td></td><td></td><td></td><td></td><td></td><td></td><td></td></tr>
<tr><td>-- OLD-ABC ->PIN</td><td></td><td></td><td></td><td></td><td></td><td></td><td></td><td></td><td></td><td></td></tr>
<tr><td>--Type3.BREX</td><td></td><td></td><td></td><td></td><td></td><td></td><td></td><td></td><td></td><td></td></tr>
<tr><td>--</td><td></td><td></td><td></td><td></td><td></td><td></td><td></td><td></td><td></td><td></td></tr>
<tr><td>MBF0213222.1</td><td>Type3.MTaseII.containing.BREX</td><td><- N6-N4- METHYLASE -p<-PIN<-SbcC<- N6-N4- METHYLASE <- PIN<- OLD-ABC <- DUF6079 *<- BrxB-BrxF- iSTAND <- wHTH+WYL+WCX ||?->?-><-?<-?<-MoxR-AAA</td><td>1246</td><td>Bacteria>Pseudomonadota>Alphaproteobacteria>Magnetococcales>unclassified_Magnetococcales</td><td>GCA_015231295.1</td><td>Magnetococcales_bacterium</td><td>HQM00_06620</td><td>7843..11583</td><td>2026759</td><td>ATP-binding-protein-[Magnetococcales-bacterium]</td></tr>
<tr><td>WP_106453122.1</td><td>Type3.MTaseII.containing.BREX</td><td>DOC+HTH->HTH_36->TraJ-RHH->VirB5->?->TrbL->TrbL-> wHTH+WYL+WCX -> BrxB-BrxF- iSTAND -> DUF6079 *-> OLD-ABC ->PIN ->?->MORC->Schlafen+HATPase+wHTH-> N6-N4- METHYLASE -> TUDOR+SF1-SF2-Helicase+iREase+CTD -> PglZ -> BrxA-DNA-binding -></td><td>1241</td><td>Bacteria>Pseudomonadota>Gammaproteobacteria>Aeromonadales>Aeromonadaceae>Zobellella</td><td>GCF_003012795.1</td><td>Zobellella_taiwanensis</td><td>C7I36_RS07620</td><td>82671..86396</td><td>347535</td><td>DUF6079-family-protein-[Zobellella-taiwanensis]</td></tr>
<tr><td>QVL51256.1</td><td>Type3.MTaseII.containing.BREX</td><td><- TUDOR+SF1-SF2-Helicase+iREase+CTD <- TOPRIM <- OLD-ABC <- N6-N4- METHYLASE <-HEPN<- PIN<- OLD-ABC <-PIN<-PhdYeFM_antitox<- DUF6079 *<- BrxB-BrxF- iSTAND ||?-><-?<- TOPRIM <-?<-?<-?<-?<-MoxR-AAA</td><td>1241</td><td>Bacteria>Pseudomonadota>Gammaproteobacteria>Chromatiales>Chromatiaceae>Thiocapsa</td><td>GCF_018399035.1</td><td>Thiocapsa_sp.</td><td>KFB96_03140</td><td>681179..684904</td><td>2024551</td><td>MAG:-ATP-binding-protein-[Thiocapsa-sp.]</td></tr>
<tr><td>HCF2077872.1</td><td>Type3.MTaseII.containing.BREX</td><td>DUF4969->ZnR+ZnR-><-HTH_3||?->?->HNH-> wHTH+WYL+WCX -> BrxB-BrxF- iSTAND -> DUF6079 *-> OLD-ABC ->PIN ->MORC->Schlafen+HATPase+wHTH-> N6-N4- METHYLASE -> TUDOR+SF1-SF2-Helicase+iREase+CTD -> PglZ -> BrxA-DNA-binding -></td><td>1241</td><td>Bacteria>Pseudomonadota>Gammaproteobacteria>Pseudomonadales>Pseudomonadaceae>Pseudomonas</td><td>GCA_023942995.1</td><td>Pseudomonas_aeruginosa</td><td>NH997_004587</td><td>20722..24447</td><td>287</td><td>ATP-binding-protein-[Pseudomonas-aeruginosa]</td></tr>
<tr><td>WP_249920774.1</td><td>Type3.MTaseII.containing.BREX</td><td><- BrxA-DNA-binding <- PglZ <- TUDOR+SF1-SF2-Helicase+iREase+CTD <- N6-N4- METHYLASE <-Schlafen+HATPase+wHTH<-MORC<- PIN<- OLD-ABC <- DUF6079 *<- BrxB-BrxF- iSTAND <- wHTH+WYL+WCX <-HNH<-?<-?||HTH_3-><-ZnR+ZnR<-DUF4969</td><td>1243</td><td>Bacteria>Pseudomonadota>Gammaproteobacteria>Pseudomonadales>Pseudomonadaceae>Pseudomonas</td><td>GCF_001765455.1</td><td>Pseudomonas_aeruginosa</td><td>BAW40_RS11025</td><td>50364..54095</td><td>287</td><td>DUF6079-family-protein-[Pseudomonas-aeruginosa]</td></tr>
<tr><td>WP_070341174.1</td><td>Type3.MTaseII.containing.BREX</td><td><- BrxA-DNA-binding <- PglZ <- TUDOR+SF1-SF2-Helicase+iREase+CTD <- N6-N4- METHYLASE <-Schlafen+HATPase+wHTH<-MORC<- PIN<- OLD-ABC <- DUF6079 *<- BrxB-BrxF- iSTAND <- wHTH+WYL+WCX <-HNH<-?<-?||HTH_3-><-ZnR+ZnR<-DUF4969</td><td>1241</td><td>Bacteria>Pseudomonadota>Gammaproteobacteria>Pseudomonadales>Pseudomonadaceae>Pseudomonas</td><td>GCF_001765485.1</td><td>Pseudomonas_aeruginosa</td><td>BAW37_RS11780</td><td>29962..33687</td><td>287</td><td>DUF6079-family-protein-[Pseudomonas-aeruginosa]</td></tr>
<tr><td>WP_070331909.1</td><td>Type3.MTaseII.containing.BREX</td><td><- BrxA-DNA-binding <- PglZ <- TUDOR+SF1-SF2-Helicase+iREase+CTD <- N6-N4- METHYLASE <-Schlafen+HATPase+wHTH<-MORC<- PIN<- OLD-ABC <- DUF6079 *<- BrxB-BrxF- iSTAND <- wHTH+WYL+WCX <-HNH<-?<-?||HTH_3-><-ZnR+ZnR<-DUF4969</td><td>1241</td><td>Bacteria>Pseudomonadota>Gammaproteobacteria>Pseudomonadales>Pseudomonadaceae>Pseudomonas</td><td>GCF_001766035.1</td><td>Pseudomonas_aeruginosa</td><td>BAW72_RS11835</td><td>79269..82994</td><td>287</td><td>DUF6079-family-protein-[Pseudomonas-aeruginosa]</td></tr>
<tr><td>WP_228292836.1</td><td>Type3.MTaseII.containing.BREX</td><td><- BrxA-DNA-binding <- PglZ <- TUDOR+SF1-SF2-Helicase+iREase+CTD <- N6-N4- METHYLASE <-Schlafen+HATPase+wHTH<-MORC<- PIN<- OLD-ABC <- DUF6079 *<- BrxB-BrxF- iSTAND <- wHTH+WYL+WCX <-HNH<-?<-?||HTH_3-><-ZnR+ZnR<-DUF4969</td><td>1243</td><td>Bacteria>Pseudomonadota>Gammaproteobacteria>Pseudomonadales>Pseudomonadaceae>Pseudomonas</td><td>GCF_017741985.1</td><td>Pseudomonas_aeruginosa</td><td>BAW68_RS10935</td><td>136915..140646</td><td>287</td><td>DUF6079-family-protein-[Pseudomonas-aeruginosa]</td></tr>
<tr><td>WP_031631068.1</td><td>Type3.MTaseII.containing.BREX</td><td><- BrxA-DNA-binding <- PglZ <- TUDOR+SF1-SF2-Helicase+iREase+CTD <- N6-N4- METHYLASE <-Schlafen+HATPase+wHTH<-MORC<- PIN<- OLD-ABC <- DUF6079 *<- BrxB-BrxF- iSTAND <- wHTH+WYL+WCX <-HNH<-?<-?||HTH_3-><-ZnR+ZnR<-DUF4969</td><td>1241</td><td>Bacteria>Pseudomonadota>Gammaproteobacteria>Pseudomonadales>Pseudomonadaceae>Pseudomonas</td><td>GCF_900707525.1</td><td>Pseudomonas_aeruginosa</td><td>E4Z30_RS22025</td><td>15967..19692</td><td>287</td><td>DUF6079-family-protein-[Pseudomonas-aeruginosa]</td></tr>
<tr><td>HCF94153.1</td><td>Type3.MTaseII.containing.BREX</td><td>XIS-HTH-> BrxB-BrxF- iSTAND -> DUF6079 *->Schlafen+HATPase+wHTH+HTH-> N6-N4- METHYLASE ->?-> OLD-ABC ->PIN ->?-> TUDOR+SF1-SF2-Helicase+iREase+CTD ->REase-> PglZ -></td><td>1288</td><td>Bacteria>PVC_group>Verrucomicrobiota>unclassified_Verrucomicrobiota</td><td>GCA_003534995.1</td><td>Verrucomicrobiota_bacterium</td><td>DEW46_03760</td><td>1029..4895</td><td>2026799</td><td>ATPase-[Verrucomicrobiota-bacterium]</td></tr>
<tr><td>--</td><td></td><td></td><td></td><td></td><td></td><td></td><td></td><td></td><td></td><td></td></tr>
<tr><td>-- OLD-ABC ->PIN</td><td></td><td></td><td></td><td></td><td></td><td></td><td></td><td></td><td></td><td></td></tr>
<tr><td>--BR-1</td><td></td><td></td><td></td><td></td><td></td><td></td><td></td><td></td><td></td><td></td></tr>
<tr><td>--</td><td></td><td></td><td></td><td></td><td></td><td></td><td></td><td></td><td></td><td></td></tr>
<tr><td>MBP7799276.1</td><td>DUF499_1.RM.like.systems</td><td><- HEPN+ DUF499 *<- PIN<- OLD-ABC <- METHYLASE <- TOPRIM <- OLD-ABC</td><td>1114</td><td>Bacteria>Acidobacteriota>Thermoanaerobaculia>Thermoanaerobaculales>Thermoanaerobaculaceae</td><td>GCA_018000955.1</td><td>Thermoanaerobaculaceae_bacterium</td><td>KA072_11695</td><td>22806..26150</td><td>2699758</td><td>DUF499-domain-containing-protein-[Thermoanaerobaculaceae-bacterium]</td></tr>
<tr><td>MYN64243.1</td><td>DUF499_1.RM.like.systems</td><td>OLD-ABC ->PIN -> OLD-ABC -> TOPRIM ->MetJarc-HTH->PIN-> DUF499 *-><-?||?->?->PIN-></td><td>935</td><td>Bacteria>Acidobacteriota>unclassified_Acidobacteriota</td><td>GCA_009861545.1</td><td>Acidobacteriota_bacterium</td><td>F4X11_04340</td><td>45133..47940</td><td>1978231</td><td>ATP-binding-protein-[Acidobacteria-bacterium]</td></tr>
<tr><td>MCY4659240.1</td><td>DUF499_1.RM.like.systems</td><td><-ORC-CDC6+REase<- DUF499 *<- PIN<- OLD-ABC <- METHYLASE <-ParB+HNH+Schlafen<- TOPRIM <- OLD-ABC</td><td>941</td><td>Bacteria>Acidobacteriota>unclassified_Acidobacteriota</td><td>GCA_026713985.1</td><td>Acidobacteriota_bacterium</td><td>OXF93_05425</td><td>1121..3946</td><td>1978231</td><td>MAG:-DUF499-domain-containing-protein-[Acidobacteria-bacterium]</td></tr>
<tr><td>MCL4821456.1</td><td>DUF499_1.RM.like.systems</td><td>TUDOR+ SF1-SF2-Helicase+iREase+CTD+REase -p->XIS-HTH+RVT-> METHYLASE -> OLD-ABC ->PIN -> HEPN+ DUF499 *->SF1-SF2-Helicase-><-?<-?||Pkinase-></td><td>1115</td><td>Bacteria>Acidobacteriota>Vicinamibacteria>unclassified_Vicinamibacteria</td><td>GCA_023384615.1</td><td>Vicinamibacteria_bacterium</td><td>KJ067_20160</td><td>8492..11839</td><td>2793154</td><td>DUF499-domain-containing-protein-[Vicinamibacteria-bacterium]</td></tr>
<tr><td>WP_122031531.1</td><td>DUF499_1.RM.like.systems</td><td>TUDOR+ SF1-SF2-Helicase+iREase+CTD+REase ->ORC-CDC6+REase-> METHYLASE -> OLD-ABC ->PIN -> HEPN+ DUF499 *->Radical-SAM+TRAM->PfkB->?->35exo+SF1-SF2-Helicase->?-><-NTP_transf_2</td><td>1110</td><td>Bacteria>Bacteria_incertae_sedis>Bacteria_candidate_phyla>Candidatus_Bipolaricaulota>Bipolaricaulis</td><td>GCF_900465355.1</td><td>Candidatus_Bipolaricaulis_anaerobius</td><td>BARAN1_RS05370</td><td>1116813..1120145</td><td>2026885</td><td>Swt1-family-HEPN-domain-containing-protein-[Candidatus-Bipolaricaulis-anaerobius]</td></tr>
<tr><td>MCY3594931.1</td><td>DUF499_1.RM.like.systems</td><td>Peptidase_S41-><-?<- DUF499 *<-REase<- PIN<- OLD-ABC <- METHYLASE <- TUDOR+ SF1-SF2-Helicase+iREase+CTD+REase</td><td>913</td><td>Bacteria>FCB_group>Bacteroidota/Chlorobiota_group>Bacteroidota>unclassified_Bacteroidota</td><td>GCA_026705205.1</td><td>Bacteroidota_bacterium</td><td>OXH01_06810</td><td>22578..25319</td><td>1898104</td><td>MAG:-DUF499-domain-containing-protein-[Bacteroidetes-bacterium]</td></tr>
<tr><td>MCY3614378.1</td><td>DUF499_1.RM.like.systems</td><td>SWACOS-><- DUF499 *<- PIN<- OLD-ABC <- METHYLASE -p</td><td>924</td><td>Bacteria>FCB_group>Bacteroidota/Chlorobiota_group>Bacteroidota>unclassified_Bacteroidota</td><td>GCA_026705365.1</td><td>Bacteroidota_bacterium</td><td>OXH03_04690</td><td>2086..4860</td><td>1898104</td><td>MAG:-DUF499-domain-containing-protein-[Bacteroidetes-bacterium]</td></tr>
<tr><td>MCY3630037.1</td><td>DUF499_1.RM.like.systems</td><td><- DUF499 *<- PIN<- OLD-ABC <- METHYLASE <-wHTH+REase<- TUDOR+ SF1-SF2-Helicase+iREase+CTD+REase</td><td>884</td><td>Bacteria>FCB_group>Bacteroidota/Chlorobiota_group>Bacteroidota>unclassified_Bacteroidota</td><td>GCA_026705375.1</td><td>Bacteroidota_bacterium</td><td>OXG94_08505</td><td>1..2653</td><td>1898104</td><td>MAG:-DUF499-domain-containing-protein,-partial-[Bacteroidetes-bacterium]</td></tr>
<tr><td>MCY4001546.1</td><td>DUF499_1.RM.like.systems</td><td>wHTH+REase-> METHYLASE -> OLD-ABC ->PIN -> DUF499 *-></td><td>903</td><td>Bacteria>FCB_group>Bacteroidota/Chlorobiota_group>Bacteroidota>unclassified_Bacteroidota</td><td>GCA_026708685.1</td><td>Bacteroidota_bacterium</td><td>OXF84_12180</td><td>5800..8511</td><td>1898104</td><td>MAG:-DUF499-domain-containing-protein-[Bacteroidetes-bacterium]</td></tr>
<tr><td>MDE2646784.1</td><td>DUF499_1.RM.like.systems</td><td>ABhydrolase-><- DUF499 *<- PIN<- OLD-ABC <- METHYLASE <- METHYLASE -p<-wHTH+REase<- TUDOR+ SF1-SF2-Helicase+iREase+CTD+REase</td><td>924</td><td>Bacteria>FCB_group>Bacteroidota/Chlorobiota_group>Bacteroidota>unclassified_Bacteroidota</td><td>GCA_028875465.1</td><td>Bacteroidota_bacterium</td><td>OXI05_13240</td><td>2968..5742</td><td>1898104</td><td>MAG:-DUF499-domain-containing-protein-[Bacteroidota-bacterium]</td></tr>
<tr><td>MXW15222.1</td><td>DUF499_1.RM.like.systems</td><td>TUDOR+ SF1-SF2-Helicase+iREase+CTD+REase ->wHTH+REase-> OLD-ABC ->PIN -> DUF499 *-><-ABhydrolase</td><td>924</td><td>Bacteria>FCB_group>Bacteroidota/Chlorobiota_group>Rhodothermota>Rhodothermia</td><td>GCA_009835915.1</td><td>Rhodothermaceae_bacterium</td><td>F4Z76_09955</td><td>11969..14743</td><td>2026787</td><td>ATP-binding-protein-[Rhodothermaceae-bacterium]</td></tr>
<tr><td>MXZ58767.1</td><td>DUF499_1.RM.like.systems</td><td>ABhydrolase-><- DUF499 *<- PIN<- OLD-ABC <- METHYLASE <-wHTH+REase<- TUDOR+ SF1-SF2-Helicase+iREase+CTD+REase</td><td>924</td><td>Bacteria>FCB_group>Bacteroidota/Chlorobiota_group>Rhodothermota>Rhodothermia</td><td>GCA_009837125.1</td><td>Rhodothermaceae_bacterium</td><td>F4Z16_09865</td><td>2884..5658</td><td>2026787</td><td>ATP-binding-protein-[Rhodothermaceae-bacterium]</td></tr>
<tr><td>MXX97465.1</td><td>DUF499_1.RM.like.systems</td><td>TUDOR+ SF1-SF2-Helicase+iREase+CTD+REase ->wHTH+REase-> METHYLASE -> OLD-ABC ->PIN -> DUF499 *-><-ABhydrolase</td><td>907</td><td>Bacteria>FCB_group>Bacteroidota/Chlorobiota_group>Rhodothermota>Rhodothermia</td><td>GCA_009838195.1</td><td>Rhodothermaceae_bacterium</td><td>F4Y64_07620</td><td>10395..13118</td><td>2026787</td><td>ATP-binding-protein-[Rhodothermaceae-bacterium]</td></tr>
<tr><td>MYC03829.1</td><td>DUF499_1.RM.like.systems</td><td>ABhydrolase-><- DUF499 *<- PIN<- OLD-ABC <-wHTH+REase<- TUDOR+ SF1-SF2-Helicase+iREase+CTD+REase</td><td>907</td><td>Bacteria>FCB_group>Bacteroidota/Chlorobiota_group>Rhodothermota>Rhodothermia</td><td>GCA_009840265.1</td><td>Rhodothermaceae_bacterium</td><td>F4X61_04285</td><td>3864..6587</td><td>2026787</td><td>ATP-binding-protein-[Rhodothermaceae-bacterium]</td></tr>
<tr><td>MYH11281.1</td><td>DUF499_1.RM.like.systems</td><td>TUDOR+ SF1-SF2-Helicase+iREase+CTD+REase ->wHTH+REase-> METHYLASE -> OLD-ABC ->PIN -> DUF499 *-><-ABhydrolase</td><td>924</td><td>Bacteria>FCB_group>Bacteroidota/Chlorobiota_group>Rhodothermota>Rhodothermia</td><td>GCA_009842805.1</td><td>Rhodothermaceae_bacterium</td><td>F4191_00350</td><td>10395..13169</td><td>2026787</td><td>ATP-binding-protein-[Rhodothermaceae-bacterium]</td></tr>
<tr><td>MDD9852708.1</td><td>DUF499_1.RM.like.systems</td><td>TUDOR+ SF1-SF2-Helicase+iREase+CTD+REase ->?->?-> METHYLASE -> OLD-ABC ->PIN -> HEPN+ DUF499 *-></td><td>632</td><td>Bacteria>Myxococcota>Myxococcia>unclassified_Myxococcia</td><td>GCA_028817165.1</td><td>Deltaproteobacteria_bacterium</td><td>OXU78_02000</td><td>21524..23422</td><td>2026735</td><td>MAG:-Swt1-family-HEPN-domain-containing-protein-[Deltaproteobacteria-bacterium]</td></tr>
<tr><td>MDD9860639.1</td><td>DUF499_1.RM.like.systems</td><td>SPOUT-><-LuxR-HTH||?->?->?->?-><- DUF499 *<- PIN<- OLD-ABC <- METHYLASE -p</td><td>939</td><td>Bacteria>Nitrospirota>Nitrospiria>Nitrospirales>Nitrospiraceae</td><td>GCA_028817215.1</td><td>Nitrospira_sp.</td><td>OXU40_09370</td><td>15116..17935</td><td>70125</td><td>MAG:-DUF499-domain-containing-protein-[Nitrospira-sp.]</td></tr>
<tr><td>MDD9819374.1</td><td>DUF499_1.RM.like.systems</td><td><- DUF499 *<- PIN<- OLD-ABC <- METHYLASE -p</td><td>939</td><td>Bacteria>Nitrospirota>Nitrospiria>Nitrospirales>Nitrospiraceae</td><td>GCA_028817545.1</td><td>Nitrospira_sp.</td><td>OXR07_00640</td><td>2590..5409</td><td>70125</td><td>MAG:-DUF499-domain-containing-protein-[Nitrospira-sp.]</td></tr>
<tr><td>MBI5313198.1</td><td>DUF499_1.RM.like.systems</td><td>TUDOR+ SF1-SF2-Helicase+iREase+CTD+REase ->Ntox18->Peptidase_M78-> METHYLASE -> OLD-ABC ->PIN -> HEPN+ DUF499 *->?-><-HTH</td><td>1107</td><td>Bacteria>Pseudomonadota>Alphaproteobacteria>Hyphomicrobiales>Methylocystaceae</td><td>GCA_016219665.1</td><td>Methylocystis_sp.</td><td>HZB28_08775</td><td>48507..51830</td><td>1911079</td><td>DUF499-domain-containing-protein-[Methylocystis-sp.]</td></tr>
<tr><td>WP_023715932.1</td><td>DUF499_1.RM.like.systems</td><td><-ParB<-?<-REase<- HEPN+ DUF499 *<- PIN<- OLD-ABC <- METHYLASE <- TUDOR+ SF1-SF2-Helicase+iREase+CTD+REase</td><td>1115</td><td>Bacteria>Pseudomonadota>Alphaproteobacteria>Hyphomicrobiales>Phyllobacteriaceae</td><td>GCF_000503075.1</td><td>Mesorhizobium_sp._LSHC422A00</td><td>X761_RS140575</td><td>209394..212741</td><td>1287294</td><td>DUF499-domain-containing-protein-[Mesorhizobium-sp.-LSHC422A00]</td></tr>
<tr><td>WP_135929916.1</td><td>DUF499_1.RM.like.systems</td><td>TUDOR+ SF1-SF2-Helicase+iREase+CTD+REase -> METHYLASE -> OLD-ABC ->PIN -> HEPN+ DUF499 *->?-><-?||Methylase-><-Radical-SAM<-Arm-DNA-bind_3+Phage_integrase</td><td>1110</td><td>Bacteria>Pseudomonadota>Alphaproteobacteria>Hyphomicrobiales>Phyllobacteriaceae</td><td>GCF_004791875.1</td><td>unclassified_Mesorhizobium</td><td>EN810_RS20200</td><td>141593..144925</td><td>325217</td><td>MULTISPECIES:-DUF499-domain-containing-protein-[unclassified-Mesorhizobium]</td></tr>
<tr><td>MCA3363599.1</td><td>DUF499_1.RM.like.systems</td><td>METHYLASE -> OLD-ABC ->PIN -> HEPN+ DUF499 *-></td><td>483</td><td>Bacteria>Pseudomonadota>Alphaproteobacteria>Rhodospirillales>Acetobacteraceae</td><td>GCA_020249535.1</td><td>Roseomonas_sp.</td><td>INF74_16815</td><td>3977..5424</td><td>1869190</td><td>DUF499-domain-containing-protein-[Roseomonas-sp.]</td></tr>
<tr><td>WP_180283596.1</td><td>DUF499_1.RM.like.systems</td><td><-PfkB||DeoR-HTH+ISOCOT->Transposase_31+Yae1_N-><- HEPN+ DUF499 *<- PIN<- OLD-ABC <- METHYLASE <- TUDOR+ SF1-SF2-Helicase+iREase+CTD+REase ||HTH->GDResolvase+PPP1-HTH-></td><td>1135</td><td>Bacteria>Pseudomonadota>Alphaproteobacteria>Rhodospirillales>Azospirillaceae</td><td>GCF_013423495.1</td><td>Azospirillum_oleiclasticum</td><td>HND93_RS19070</td><td>2984..6391</td><td>2735135</td><td>DUF499-domain-containing-protein-[Azospirillum-oleiclasticum]</td></tr>
<tr><td>WP_238518868.1</td><td>DUF499_1.RM.like.systems</td><td><-SF1-SF2-Helicase+DUF1998+HKD-DNase+HKD-DNase<-?<- HEPN+ DUF499 *<- PIN<- OLD-ABC <- METHYLASE <-?<-Schlafen+HATPase+HTH<- TUDOR+ SF1-SF2-Helicase+iREase+CTD+REase</td><td>1154</td><td>Bacteria>Pseudomonadota>Alphaproteobacteria>Rhodospirillales>Azospirillaceae</td><td>GCF_022214805.1</td><td>Azospirillum_doebereinerae</td><td>MCW82_RS00020</td><td>10412..13876</td><td>92933</td><td>DUF499-domain-containing-protein-[Azospirillum-doebereinerae]</td></tr>
<tr><td>SEP49898.1</td><td>DUF499_1.RM.like.systems</td><td><- HEPN+ DUF499 *<- PIN<- OLD-ABC <- METHYLASE <- TUDOR+ SF1-SF2-Helicase+iREase+CTD+REase <-HNH</td><td>1099</td><td>Bacteria>Pseudomonadota>Alphaproteobacteria>Rhodospirillales>unclassified_Rhodospirillales</td><td>GCA_900110395.1</td><td>Rhodospirillales_bacterium_URHD0017</td><td>SAMN02990966_07447</td><td>5640..8939</td><td>1380357</td><td>hypothetical-protein-SAMN02990966_07447-[Rhodospirillales-bacterium-URHD0017]</td></tr>
<tr><td>MCL6741777.1</td><td>DUF499_1.RM.like.systems</td><td>wHTH+REase-> TUDOR+ SF1-SF2-Helicase+iREase+CTD+REase -> METHYLASE -> OLD-ABC ->PIN -> HEPN+ DUF499 *-><-?<-BetaPropeller+ABhydrolase||?-><-ABhydrolase<-MODE-HTH+PBPII</td><td>1094</td><td>Bacteria>Pseudomonadota>Alphaproteobacteria>Sphingomonadales>Sphingomonadaceae</td><td>GCA_023516505.1</td><td>Sphingomonas_brevis</td><td>LZ518_11630</td><td>2249137..2252421</td><td>2908206</td><td>DUF499-domain-containing-protein-[Sphingomonas-sp.-RB56-2]</td></tr>
<tr><td>WP_283938333.1</td><td>DUF499_1.RM.like.systems</td><td>wHTH+REase-> TUDOR+ SF1-SF2-Helicase+iREase+CTD+REase -> METHYLASE -> OLD-ABC ->PIN -> HEPN+ DUF499 *-><-?<-BetaPropeller+ABhydrolase||?-><-ABhydrolase<-MODE-HTH+PBPII</td><td>1103</td><td>Bacteria>Pseudomonadota>Alphaproteobacteria>Sphingomonadales>Sphingomonadaceae</td><td>GCF_023516505.1</td><td>Sphingomonas_brevis</td><td>LZ518_RS11630</td><td>2249110..2252421</td><td>2908206</td><td>DUF499-domain-containing-protein-[Sphingomonas-brevis]</td></tr>
<tr><td>WP_249916143.1</td><td>DUF499_1.RM.like.systems</td><td>wHTH+REase-> TUDOR+ SF1-SF2-Helicase+iREase+CTD+REase -> METHYLASE -> OLD-ABC ->PIN -> HEPN+ DUF499 *-><-?<-BetaPropeller+ABhydrolase||?-><-ABhydrolase<-MODE-HTH+PBPII</td><td>1094</td><td>Bacteria>Pseudomonadota>Alphaproteobacteria>Sphingomonadales>Sphingomonadaceae</td><td>-</td><td>Sphingomonas_brevis</td><td>LZ518_11630</td><td>2249137..2252421</td><td>2908206</td><td>DUF499-domain-containing-protein-[Sphingomonas-sp.-RB56-2]</td></tr>
<tr><td>MCK6404267.1</td><td>DUF499_1.RM.like.systems</td><td><-zf-IS66+DDE_Tnp_IS66<-DDE_Tnp_1_assoc+DDE_Tnp_1<-ParB+HTH<-?||?-><- DUF499 *<- TOPRIM <- OLD-ABC <- METHYLASE <-?<- PIN<- OLD-ABC</td><td>955</td><td>Bacteria>Pseudomonadota>Betaproteobacteria>Burkholderiales>Sphaerotilaceae</td><td>GCA_023150255.1</td><td>Sphaerotilus_sulfidivorans</td><td>L6Q74_20550</td><td>5829..8696</td><td>639200</td><td>DUF499-domain-containing-protein-[Sphaerotilus-sulfidivorans]</td></tr>
<tr><td>WP_273473498.1</td><td>DUF499_1.RM.like.systems</td><td><-zf-IS66+DDE_Tnp_IS66<-DDE_Tnp_1_assoc+DDE_Tnp_1<-ParB+HTH<-?||?-><- DUF499 *<- TOPRIM <- OLD-ABC <- METHYLASE <-?<- PIN<- OLD-ABC</td><td>955</td><td>Bacteria>Pseudomonadota>Betaproteobacteria>Burkholderiales>Sphaerotilaceae</td><td>GCF_023150255.1</td><td>Sphaerotilus_sulfidivorans</td><td>L6Q74_RS20510</td><td>5829..8696</td><td>639200</td><td>DUF499-domain-containing-protein-[Sphaerotilus-sulfidivorans]</td></tr>
</table>

| Protein-ID | Domain | Assembly-ID | Strain | Taxonomy | Length | Locus-Tag | Genomic-coordinates | TaxID | Fasta-defline |
|---|---|---|---|---|---|---|---|---|---|
| --HKD-DNase+OGG-like | | | | | | | | | |
| MCC2230795.1 | HKD-DNase+OGG-like | GCA_020687485.1 | Hominifimenecus_microfluidus | Bacteria>Bacillati>Bacillota>Clostridia>Lachnospirales>Lachnospiraceae>Hominifimenecus | 477 | LKD81_07260 | 12048..13481 | 2885348 | phospholipase-D-family-protein-[Lachnospiraceae-bacterium-CLA-AA-H215] |
| MCC2230796.1 | HKD-DNase+OGG-like | GCA_020687485.1 | Hominifimenecus_microfluidus | Bacteria>Bacillati>Bacillota>Clostridia>Lachnospirales>Lachnospiraceae>Hominifimenecus | 519 | LKD81_07265 | 13692..15251 | 2885348 | cardiolipin-synthase-[Lachnospiraceae-bacterium-CLA-AA-H215] |
| WP_032410719.1 | HKD-DNase+OGG-like | GCF_004312665.1 | Enterobacterales | Bacteria>Pseudomonadati>Pseudomonadota>Gammaproteobacteria | 449 | KFB82_RS19605 | 71800..73149 | 91347 | MULTISPECIES:-phospholipase-D-like-domain-containing-protein-[Enterobacterales] |
| WP_117029420.1 | HKD-DNase+OGG-like | GCF_903971055.1 | Enterobacterales | Bacteria>Pseudomonadati>Pseudomonadota>Gammaproteobacteria | 449 | MOA98_RS25005 | 240875..242224 | 91347 | MULTISPECIES:-phospholipase-D-like-domain-containing-protein-[Klebsiella] |
| WP_000766444.1 | HKD-DNase+OGG-like | GCF_000166575.1 | Enterobacteriaceae | Bacteria>Pseudomonadati>Pseudomonadota>Gammaproteobacteria>Enterobacterales | 449 | NO030_RS20200 | 18421..19770 | 543 | MULTISPECIES:-phospholipase-D-like-domain-containing-protein-[Enterobacteriaceae] |
| WP_057055972.1 | HKD-DNase+OGG-like | GCF_014169295.1 | Enterobacteriaceae | Bacteria>Pseudomonadati>Pseudomonadota>Gammaproteobacteria>Enterobacterales | 449 | H7R61_RS03340 | 673017..674366 | 543 | MULTISPECIES:-phospholipase-D-like-domain-containing-protein-[Enterobacter-cloacae-complex] |
| WP_102104492.1 | HKD-DNase+OGG-like | GCF_029842965.1 | Enterobacteriaceae | Bacteria>Pseudomonadati>Pseudomonadota>Gammaproteobacteria>Enterobacterales | 449 | F159LOC_RS08750 | 156650..157999 | 543 | MULTISPECIES:-phospholipase-D-like-domain-containing-protein-[Enterobacteriaceae] |
| WP_192468118.1 | HKD-DNase+OGG-like | GCF_023151575.1 | Enterobacteriaceae | Bacteria>Pseudomonadati>Pseudomonadota>Gammaproteobacteria>Enterobacterales | 449 | L8T09_RS18440 | 39885..41234 | 543 | phospholipase-D-like-domain-containing-protein-[Enterobacter-asburiae] |
| WP_196010427.1 | HKD-DNase+OGG-like | GCF_029717585.1 | Enterobacteriaceae | Bacteria>Pseudomonadati>Pseudomonadota>Gammaproteobacteria>Enterobacterales | 449 | NFJ54_RS22985 | 4706502..4707851 | 543 | MULTISPECIES:-phospholipase-D-like-domain-containing-protein-[Enterobacteriaceae] |
| WP_223177675.1 | HKD-DNase+OGG-like | GCF_040365915.1 | Enterobacteriaceae | Bacteria>Pseudomonadati>Pseudomonadota>Gammaproteobacteria>Enterobacterales | 172 | LF666_RS16230 | 25352..25870 | 543 | hypothetical-protein-[Klebsiella-pneumoniae] |
| WP_023224631.1 | HKD-DNase+OGG-like | GCF_001569545.1 | Salmonella | Bacteria>Pseudomonadati>Pseudomonadota>Gammaproteobacteria>Enterobacterales>Enterobacteriaceae | 449 | NOQ95_RS21590 | 4462278..4463627 | 590 | MULTISPECIES:-phospholipase-D-like-domain-containing-protein-[Salmonella] |
| WP_040216476.1 | HKD-DNase+OGG-like | GCF_014169255.1 | Klebsiella/Raoultella_group | Bacteria>Pseudomonadati>Pseudomonadota>Gammaproteobacteria>Enterobacterales>Enterobacteriaceae | 635 | B6J71_RS18345 | 38118..40025 | 2890311 | MULTISPECIES:-hypothetical-protein-[Klebsiella] |
| WP_049088893.1 | HKD-DNase+OGG-like | GCF_003851825.1 | Klebsiella/Raoultella_group | Bacteria>Pseudomonadati>Pseudomonadota>Gammaproteobacteria>Enterobacterales>Enterobacteriaceae | 635 | KPPSC5_RS01090 | 221996..223903 | 2890311 | MULTISPECIES:-hypothetical-protein-[Klebsiella] |
| WP_063308432.1 | HKD-DNase+OGG-like | GCF_001623605.1 | Enterobacter | Bacteria>Pseudomonadati>Pseudomonadota>Gammaproteobacteria>Enterobacterales>Enterobacteriaceae | 449 | A4308_RS00225 | 47485..48834 | 547 | MULTISPECIES:-phospholipase-D-like-domain-containing-protein-[Enterobacter] |
| WP_128348266.1 | HKD-DNase+OGG-like | GCF_024607805.1 | Enterobacter | Bacteria>Pseudomonadati>Pseudomonadota>Gammaproteobacteria>Enterobacterales>Enterobacteriaceae | 449 | PVM09_RS00220 | 46465..47814 | 547 | MULTISPECIES:-phospholipase-D-like-domain-containing-protein-[Enterobacter] |
| WP_200025361.1 | HKD-DNase+OGG-like | GCF_032347795.1 | Citrobacter | Bacteria>Pseudomonadati>Pseudomonadota>Gammaproteobacteria>Enterobacterales>Enterobacteriaceae | 449 | I5388_RS15330 | 371548..372897 | 544 | phospholipase-D-like-domain-containing-protein-[Citrobacter-freundii] |
| WP_046487346.1 | HKD-DNase+OGG-like | GCF_000981805.1 | Citrobacter_amalonaticus | Bacteria>Pseudomonadati>Pseudomonadota>Gammaproteobacteria>Enterobacterales>Enterobacteriaceae>Citrobacter | 449 | F384_RS16695 | 3647649..3648998 | 35703 | phospholipase-D-like-domain-containing-protein-[Citrobacter-amalonaticus] |
| WP_192466361.1 | HKD-DNase+OGG-like | GCF_014855915.1 | Citrobacter_koseri | Bacteria>Pseudomonadati>Pseudomonadota>Gammaproteobacteria>Enterobacterales>Enterobacteriaceae>Citrobacter | 449 | FOT44_RS18765 | 327985..329334 | 545 | phospholipase-D-like-domain-containing-protein-[Citrobacter-koseri] |
| HCL6754278.1 | HKD-DNase+OGG-like | GCA_025039155.1 | Citrobacter_freundii | Bacteria>Pseudomonadati>Pseudomonadota>Gammaproteobacteria>Enterobacterales>Enterobacteriaceae>Citrobacter>Citrobacter_freundii_complex | 449 | N2R09_003048 | 102931..104280 | 546 | phosphatidylserine/phosphatidylglycerophosphate/cardiolipin-synthase-family-protein-[Citrobacter-freundii] |
| WP_135912650.1 | HKD-DNase+OGG-like | GCF_004792475.1 | Citrobacter_freundii | Bacteria>Pseudomonadati>Pseudomonadota>Gammaproteobacteria>Enterobacterales>Enterobacteriaceae>Citrobacter>Citrobacter_freundii_complex | 449 | E5284_RS20160 | 4218976..4220325 | 546 | phospholipase-D-like-domain-containing-protein-[Citrobacter-freundii] |
| WP_241511099.1 | HKD-DNase+OGG-like | GCF_022585175.1 | Citrobacter_freundii | Bacteria>Pseudomonadati>Pseudomonadota>Gammaproteobacteria>Enterobacterales>Enterobacteriaceae>Citrobacter>Citrobacter_freundii_complex | 449 | MMQ01_RS10375 | 132342..133691 | 546 | phospholipase-D-like-domain-containing-protein-[Citrobacter-freundii] |
| WP_032967173.1 | HKD-DNase+OGG-like | GCF_000409225.1 | Cronobacter_dublinensis | Bacteria>Pseudomonadati>Pseudomonadota>Gammaproteobacteria>Enterobacterales>Enterobacteriaceae>Cronobacter | 449 | AFK67_RS02765 | 582139..583488 | 413497 | phospholipase-D-like-domain-containing-protein-[Cronobacter-dublinensis] |
| WP_258133206.1 | HKD-DNase+OGG-like | GCF_002974775.1 | Cronobacter_sakazakii | Bacteria>Pseudomonadati>Pseudomonadota>Gammaproteobacteria>Enterobacterales>Enterobacteriaceae>Cronobacter | 113 | C5959_RS02050 | 318753..319094 | 28141 | hypothetical-protein-[Cronobacter-sakazakii] |
| PWG69501.1 | HKD-DNase+OGG-like | GCA_003129945.1 | Enterobacter_mori | Bacteria>Pseudomonadati>Pseudomonadota>Gammaproteobacteria>Enterobacterales>Enterobacteriaceae>Enterobacter | 449 | DEM28_19790 | 158681..160030 | 539813 | phospholipase-[Enterobacter-mori] |
| WP_058688923.1 | HKD-DNase+OGG-like | GCF_965285355.1 | Enterobacter_cloacae_complex | Bacteria>Pseudomonadati>Pseudomonadota>Gammaproteobacteria>Enterobacterales>Enterobacteriaceae>Enterobacter | 449 | ASU86_RS07695 | 173678..175027 | 354276 | phospholipase-D-like-domain-containing-protein-[Enterobacter-hormaechei] |
| WP_021240661.1 | HKD-DNase+OGG-like | GCF_000467655.2 | Enterobacter_roggenkampii | Bacteria>Pseudomonadati>Pseudomonadota>Gammaproteobacteria>Enterobacterales>Enterobacteriaceae>Enterobacter>Enterobacter_cloacae_complex | 449 | L799_RS145610 | 340693..342042 | 1812935 | phospholipase-D-like-domain-containing-protein-[Enterobacter-roggenkampii] |
| WP_095453568.1 | HKD-DNase+OGG-like | GCF_002278355.1 | Enterobacter_cloacae | Bacteria>Pseudomonadati>Pseudomonadota>Gammaproteobacteria>Enterobacterales>Enterobacteriaceae>Enterobacter>Enterobacter_cloacae_complex | 449 | CIW57_RS16440 | 149973..151322 | 550 | phospholipase-D-like-domain-containing-protein-[Enterobacter-cloacae] |
| WP_163363280.1 | HKD-DNase+OGG-like | GCF_010594505.1 | Enterobacter_cloacae | Bacteria>Pseudomonadati>Pseudomonadota>Gammaproteobacteria>Enterobacterales>Enterobacteriaceae>Enterobacter>Enterobacter_cloacae_complex | 449 | G4556_RS12880 | 5151..6500 | 550 | phospholipase-D-like-domain-containing-protein-[Enterobacter-cloacae] |
| WP_203465573.1 | HKD-DNase+OGG-like | - | Enterobacter_cloacae | Bacteria>Pseudomonadati>Pseudomonadota>Gammaproteobacteria>Enterobacterales>Enterobacteriaceae>Enterobacter>Enterobacter_cloacae_complex | 449 | GEA59_RS02595 | 541833..543182 | 550 | phospholipase-D-like-domain-containing-protein-[Enterobacter-cloacae] |
| WP_252909242.1 | HKD-DNase+OGG-like | GCF_024104775.1 | Enterobacter_roggenkampii | Bacteria>Pseudomonadati>Pseudomonadota>Gammaproteobacteria>Enterobacterales>Enterobacteriaceae>Enterobacter>Enterobacter_cloacae_complex | 395 | KGY84_RS21885 | 339476..340663 | 1812935 | phospholipase-D-like-domain-containing-protein-[Enterobacter-roggenkampii] |
| CTT27140.1 | HKD-DNase+OGG-like | GCA_001283645.1 | Escherichia_coli | Bacteria>Pseudomonadati>Pseudomonadota>Gammaproteobacteria>Enterobacterales>Enterobacteriaceae>Escherichia | 449 | ERS085343_03082 | 12186..13535 | 562 | phospholipase-D-[Escherichia-coli] |
| CTW38815.1 | HKD-DNase+OGG-like | GCA_001283765.1 | Escherichia_coli | Bacteria>Pseudomonadati>Pseudomonadota>Gammaproteobacteria>Enterobacterales>Enterobacteriaceae>Escherichia | 449 | ERS085413_04501 | 11317..12666 | 562 | phospholipase-D-[Escherichia-coli] |
| CUA46985.1 | HKD-DNase+OGG-like | GCA_001286505.1 | Escherichia_coli | Bacteria>Pseudomonadati>Pseudomonadota>Gammaproteobacteria>Enterobacterales>Enterobacteriaceae>Escherichia | 449 | ERS139261_03505 | 12108..13457 | 562 | phospholipase-D-[Escherichia-coli] |
| EEC7405254.1 | HKD-DNase+OGG-like | - | Escherichia_coli | Bacteria>Pseudomonadati>Pseudomonadota>Gammaproteobacteria>Enterobacterales>Enterobacteriaceae>Escherichia | 449 | D4L89_RS08025 | 83939..85288 | 562 | phosphatidylserine/phosphatidylglycerophosphate/cardiolipin-synthase-family-protein-[Escherichia-coli] |
| EEC9685947.1 | HKD-DNase+OGG-like | - | Escherichia_coli | Bacteria>Pseudomonadati>Pseudomonadota>Gammaproteobacteria>Enterobacterales>Enterobacteriaceae>Escherichia | 449 | D8K54_RS17555 | 11679..13028 | 562 | phosphatidylserine/phosphatidylglycerophosphate/cardiolipin-synthase-family-protein-[Escherichia-coli] |
| EED0771056.1 | HKD-DNase+OGG-like | - | Escherichia_coli | Bacteria>Pseudomonadati>Pseudomonadota>Gammaproteobacteria>Enterobacterales>Enterobacteriaceae>Escherichia | 449 | ED219_RS15250 | 13284..14633 | 562 | phosphatidylserine/phosphatidylglycerophosphate/cardiolipin-synthase-family-protein-[Escherichia-coli] |
| EED1762071.1 | HKD-DNase+OGG-like | - | Escherichia_coli | Bacteria>Pseudomonadati>Pseudomonadota>Gammaproteobacteria>Enterobacterales>Enterobacteriaceae>Escherichia | 449 | EEP34_RS13390 | 11930..13279 | 562 | phosphatidylserine/phosphatidylglycerophosphate/cardiolipin-synthase-family-protein-[Escherichia-coli] |
| EEG9579275.1 | HKD-DNase+OGG-like | - | Escherichia_coli | Bacteria>Pseudomonadati>Pseudomonadota>Gammaproteobacteria>Enterobacterales>Enterobacteriaceae>Escherichia | 449 | EB576_RS15560 | 12044..13393 | 562 | phosphatidylserine/phosphatidylglycerophosphate/cardiolipin-synthase-family-protein-[Escherichia-coli] |
| EEQ1797643.1 | HKD-DNase+OGG-like | - | Escherichia_coli | Bacteria>Pseudomonadati>Pseudomonadota>Gammaproteobacteria>Enterobacterales>Enterobacteriaceae>Escherichia | 449 | GUY43_002905 | 11295..12644 | 562 | phosphatidylserine/phosphatidylglycerophosphate/cardiolipin-synthase-family-protein-[Escherichia-coli] |
| EEQ2509740.1 | HKD-DNase+OGG-like | - | Escherichia_coli | Bacteria>Pseudomonadati>Pseudomonadota>Gammaproteobacteria>Enterobacterales>Enterobacteriaceae>Escherichia | 449 | GSE75_002863 | 13434..14783 | 562 | phosphatidylserine/phosphatidylglycerophosphate/cardiolipin-synthase-family-protein-[Escherichia-coli] |
| EEQ7780475.1 | HKD-DNase+OGG-like | - | Escherichia_coli | Bacteria>Pseudomonadati>Pseudomonadota>Gammaproteobacteria>Enterobacterales>Enterobacteriaceae>Escherichia | 449 | GLS89_002418 | 20063..21412 | 562 | phosphatidylserine/phosphatidylglycerophosphate/cardiolipin-synthase-family-protein-[Escherichia-coli] |
| EER7097200.1 | HKD-DNase+OGG-like | - | Escherichia_coli | Bacteria>Pseudomonadati>Pseudomonadota>Gammaproteobacteria>Enterobacterales>Enterobacteriaceae>Escherichia | 449 | F2Z67_003274 | 11905..13254 | 562 | phosphatidylserine/phosphatidylglycerophosphate/cardiolipin-synthase-family-protein-[Escherichia-coli] |
| EER7518153.1 | HKD-DNase+OGG-like | - | Escherichia_coli | Bacteria>Pseudomonadati>Pseudomonadota>Gammaproteobacteria>Enterobacterales>Enterobacteriaceae>Escherichia | 449 | F1Z98_003255 | 11682..13031 | 562 | phosphatidylserine/phosphatidylglycerophosphate/cardiolipin-synthase-family-protein-[Escherichia-coli] |
| EER8632683.1 | HKD-DNase+OGG-like | - | Escherichia_coli | Bacteria>Pseudomonadati>Pseudomonadota>Gammaproteobacteria>Enterobacterales>Enterobacteriaceae>Escherichia | 449 | F0E18_004252 | 11953..13302 | 562 | phosphatidylserine/phosphatidylglycerophosphate/cardiolipin-synthase-family-protein-[Escherichia-coli] |
| EES8430776.1 | HKD-DNase+OGG-like | - | Escherichia_coli | Bacteria>Pseudomonadati>Pseudomonadota>Gammaproteobacteria>Enterobacterales>Enterobacteriaceae>Escherichia | 449 | FU162_003517 | 11676..13025 | 562 | phosphatidylserine/phosphatidylglycerophosphate/cardiolipin-synthase-family-protein-[Escherichia-coli] |
| EET5967873.1 | HKD-DNase+OGG-like | - | Escherichia_coli | Bacteria>Pseudomonadati>Pseudomonadota>Gammaproteobacteria>Enterobacterales>Enterobacteriaceae>Escherichia | 449 | FE570_001652 | 11906..13255 | 562 | phosphatidylserine/phosphatidylglycerophosphate/cardiolipin-synthase-family-protein-[Escherichia-coli] |
| EEU4094851.1 | HKD-DNase+OGG-like | - | Escherichia_coli | Bacteria>Pseudomonadati>Pseudomonadota>Gammaproteobacteria>Enterobacterales>Enterobacteriaceae>Escherichia | 449 | E8H53_003145 | 11048..12397 | 562 | phosphatidylserine/phosphatidylglycerophosphate/cardiolipin-synthase-family-protein-[Escherichia-coli] |
| EEV0352860.1 | HKD-DNase+OGG-like | - | Escherichia_coli | Bacteria>Pseudomonadati>Pseudomonadota>Gammaproteobacteria>Enterobacterales>Enterobacteriaceae>Escherichia | 449 | EYE11_003103 | 12295..13644 | 562 | phosphatidylserine/phosphatidylglycerophosphate/cardiolipin-synthase-family-protein-[Escherichia-coli] |
| EEV8657619.1 | HKD-DNase+OGG-like | - | Escherichia_coli | Bacteria>Pseudomonadati>Pseudomonadota>Gammaproteobacteria>Enterobacterales>Enterobacteriaceae>Escherichia | 449 | EHJ50_001187 | 12251..13600 | 562 | phosphatidylserine/phosphatidylglycerophosphate/cardiolipin-synthase-family-protein-[Escherichia-coli] |
| EEW0980714.1 | HKD-DNase+OGG-like | - | Escherichia_coli | Bacteria>Pseudomonadati>Pseudomonadota>Gammaproteobacteria>Enterobacterales>Enterobacteriaceae>Escherichia | 449 | ED289_06200 | 123553..124902 | 562 | phosphatidylserine/phosphatidylglycerophosphate/cardiolipin-synthase-family-protein-[Escherichia-coli] |
| EEW3089666.1 | HKD-DNase+OGG-like | - | Escherichia_coli | Bacteria>Pseudomonadati>Pseudomonadota>Gammaproteobacteria>Enterobacterales>Enterobacteriaceae>Escherichia | 449 | EBA98_002643 | 27046..28395 | 562 | phosphatidylserine/phosphatidylglycerophosphate/cardiolipin-synthase-family-protein-[Escherichia-coli] |
| EEW3885156.1 | HKD-DNase+OGG-like | - | Escherichia_coli | Bacteria>Pseudomonadati>Pseudomonadota>Gammaproteobacteria>Enterobacterales>Enterobacteriaceae>Escherichia | 449 | D9Y07_001396 | 19515..20864 | 562 | phosphatidylserine/phosphatidylglycerophosphate/cardiolipin-synthase-family-protein-[Escherichia-coli] |
| EEW4281074.1 | HKD-DNase+OGG-like | - | Escherichia_coli | Bacteria>Pseudomonadati>Pseudomonadota>Gammaproteobacteria>Enterobacterales>Enterobacteriaceae>Escherichia | 449 | D9T01_23365 | 11967..13316 | 562 | phosphatidylserine/phosphatidylglycerophosphate/cardiolipin-synthase-family-protein-[Escherichia-coli] |
| EEW6030680.1 | HKD-DNase+OGG-like | - | Escherichia_coli | Bacteria>Pseudomonadati>Pseudomonadota>Gammaproteobacteria>Enterobacterales>Enterobacteriaceae>Escherichia | 449 | D7V62_16735 | 12115..13464 | 562 | phosphatidylserine/phosphatidylglycerophosphate/cardiolipin-synthase-family-protein-[Escherichia-coli] |
| EEX1719686.1 | HKD-DNase+OGG-like | - | Escherichia_coli | Bacteria>Pseudomonadati>Pseudomonadota>Gammaproteobacteria>Enterobacterales>Enterobacteriaceae>Escherichia | 449 | D3S79_003411 | 19034..20383 | 562 | phosphatidylserine/phosphatidylglycerophosphate/cardiolipin-synthase-family-protein-[Escherichia-coli] |
| EEX4954057.1 | HKD-DNase+OGG-like | - | Escherichia_coli | Bacteria>Pseudomonadati>Pseudomonadota>Gammaproteobacteria>Enterobacterales>Enterobacteriaceae>Escherichia | 449 | D0791_003837 | 12955..14304 | 562 | phosphatidylserine/phosphatidylglycerophosphate/cardiolipin-synthase-family-protein-[Escherichia-coli] |
| EEX6773449.1 | HKD-DNase+OGG-like | - | Escherichia_coli | Bacteria>Pseudomonadati>Pseudomonadota>Gammaproteobacteria>Enterobacterales>Enterobacteriaceae>Escherichia | 449 | DYL55_002407 | 11673..13022 | 562 | phosphatidylserine/phosphatidylglycerophosphate/cardiolipin-synthase-family-protein-[Escherichia-coli] |
| EEX7673120.1 | HKD-DNase+OGG-like | - | Escherichia_coli | Bacteria>Pseudomonadati>Pseudomonadota>Gammaproteobacteria>Enterobacterales>Enterobacteriaceae>Escherichia | 449 | DX034_003684 | 12179..13528 | 562 | phosphatidylserine/phosphatidylglycerophosphate/cardiolipin-synthase-family-protein-[Escherichia-coli] |
| EEY1600794.1 | HKD-DNase+OGG-like | - | Escherichia_coli | Bacteria>Pseudomonadati>Pseudomonadota>Gammaproteobacteria>Enterobacterales>Enterobacteriaceae>Escherichia | 449 | DP662_003731 | 22068..23417 | 562 | phosphatidylserine/phosphatidylglycerophosphate/cardiolipin-synthase-family-protein-[Escherichia-coli] |
| EEY2596889.1 | HKD-DNase+OGG-like | - | Escherichia_coli | Bacteria>Pseudomonadati>Pseudomonadota>Gammaproteobacteria>Enterobacterales>Enterobacteriaceae>Escherichia | 449 | E9W20_002912 | 11809..13158 | 562 | phosphatidylserine/phosphatidylglycerophosphate/cardiolipin-synthase-family-protein-[Escherichia-coli] |
| EEY5880450.1 | HKD-DNase+OGG-like | - | Escherichia_coli | Bacteria>Pseudomonadati>Pseudomonadota>Gammaproteobacteria>Enterobacterales>Enterobacteriaceae>Escherichia | 449 | EU637_10840 | 83906..85255 | 562 | phosphatidylserine/phosphatidylglycerophosphate/cardiolipin-synthase-family-protein-[Escherichia-coli] |
| EEZ3165540.1 | HKD-DNase+OGG-like | - | Escherichia_coli | Bacteria>Pseudomonadati>Pseudomonadota>Gammaproteobacteria>Enterobacterales>Enterobacteriaceae>Escherichia | 449 | DIN68_003958 | 12065..13414 | 562 | phosphatidylserine/phosphatidylglycerophosphate/cardiolipin-synthase-family-protein-[Escherichia-coli] |
| EEZ5289155.1 | HKD-DNase+OGG-like | - | Escherichia_coli | Bacteria>Pseudomonadati>Pseudomonadota>Gammaproteobacteria>Enterobacterales>Enterobacteriaceae>Escherichia | 449 | DDL85_000898 | 123498..124847 | 562 | phosphatidylserine/phosphatidylglycerophosphate/cardiolipin-synthase-family-protein-[Escherichia-coli] |
| EEZ7212410.1 | HKD-DNase+OGG-like | - | Escherichia_coli | Bacteria>Pseudomonadati>Pseudomonadota>Gammaproteobacteria>Enterobacterales>Enterobacteriaceae>Escherichia | 449 | DAB40_000424 | 14547..15896 | 562 | phosphatidylserine/phosphatidylglycerophosphate/cardiolipin-synthase-family-protein-[Escherichia-coli] |
| EEZ9156823.1 | HKD-DNase+OGG-like | - | Escherichia_coli | Bacteria>Pseudomonadati>Pseudomonadota>Gammaproteobacteria>Enterobacterales>Enterobacteriaceae>Escherichia | 449 | DDB94_002974 | 11743..13092 | 562 | phosphatidylserine/phosphatidylglycerophosphate/cardiolipin-synthase-family-protein-[Escherichia-coli] |
| EFA5158826.1 | HKD-DNase+OGG-like | - | Escherichia_coli | Bacteria>Pseudomonadati>Pseudomonadota>Gammaproteobacteria>Enterobacterales>Enterobacteriaceae>Escherichia | 449 | E2930_09995 | 133400..134749 | 562 | phosphatidylserine/phosphatidylglycerophosphate/cardiolipin-synthase-family-protein-[Escherichia-coli] |
| EFB2626045.1 | HKD-DNase+OGG-like | - | Escherichia_coli | Bacteria>Pseudomonadati>Pseudomonadota>Gammaproteobacteria>Enterobacterales>Enterobacteriaceae>Escherichia | 449 | FHL63_20505 | 137..1486 | 562 | phosphatidylserine/phosphatidylglycerophosphate/cardiolipin-synthase-family-protein-[Escherichia-coli] |
| EFD1659143.1 | HKD-DNase+OGG-like | - | Escherichia_coli | Bacteria>Pseudomonadati>Pseudomonadota>Gammaproteobacteria>Enterobacterales>Enterobacteriaceae>Escherichia | 449 | FGU32_03055 | 81935..83284 | 562 | phosphatidylserine/phosphatidylglycerophosphate/cardiolipin-synthase-family-protein-[Escherichia-coli] |
| EFD4961603.1 | HKD-DNase+OGG-like | - | Escherichia_coli | Bacteria>Pseudomonadati>Pseudomonadota>Gammaproteobacteria>Enterobacterales>Enterobacteriaceae>Escherichia | 449 | F2I36_20000 | 16252..17601 | 562 | phosphatidylserine/phosphatidylglycerophosphate/cardiolipin-synthase-family-protein-[Escherichia-coli] |
| EFE1965723.1 | HKD-DNase+OGG-like | - | Escherichia_coli | Bacteria>Pseudomonadati>Pseudomonadota>Gammaproteobacteria>Enterobacterales>Enterobacteriaceae>Escherichia | 449 | CJV81_002671 | 27045..28394 | 562 | phosphatidylserine/phosphatidylglycerophosphate/cardiolipin-synthase-family-protein-[Escherichia-coli] |
| EFF3839664.1 | HKD-DNase+OGG-like | - | Escherichia_coli | Bacteria>Pseudomonadati>Pseudomonadota>Gammaproteobacteria>Enterobacterales>Enterobacteriaceae>Escherichia | 449 | B6310_003619 | 10797..12146 | 562 | phosphatidylserine/phosphatidylglycerophosphate/cardiolipin-synthase-family-protein-[Escherichia-coli] |
| EFG3005932.1 | HKD-DNase+OGG-like | - | Escherichia_coli | Bacteria>Pseudomonadati>Pseudomonadota>Gammaproteobacteria>Enterobacterales>Enterobacteriaceae>Escherichia | 449 | BOZ89_003091 | 11402..12751 | 562 | phosphatidylserine/phosphatidylglycerophosphate/cardiolipin-synthase-family-protein-[Escherichia-coli] |
| EFG4404342.1 | HKD-DNase+OGG-like | - | Escherichia_coli | Bacteria>Pseudomonadati>Pseudomonadota>Gammaproteobacteria>Enterobacterales>Enterobacteriaceae>Escherichia | 449 | BM772_002476 | 27046..28395 | 562 | phosphatidylserine/phosphatidylglycerophosphate/cardiolipin-synthase-family-protein-[Escherichia-coli] |
| EFH0548060.1 | HKD-DNase+OGG-like | - | Escherichia_coli | Bacteria>Pseudomonadati>Pseudomonadota>Gammaproteobacteria>Enterobacterales>Enterobacteriaceae>Escherichia | 449 | BJE23_002824 | 11078..12427 | 562 | phosphatidylserine/phosphatidylglycerophosphate/cardiolipin-synthase-family-protein-[Escherichia-coli] |
| EFH4936895.1 | HKD-DNase+OGG-like | - | Escherichia_coli | Bacteria>Pseudomonadati>Pseudomonadota>Gammaproteobacteria>Enterobacterales>Enterobacteriaceae>Escherichia | 449 | F9408_05605 | 122485..123834 | 562 | phosphatidylserine/phosphatidylglycerophosphate/cardiolipin-synthase-family-protein-[Escherichia-coli] |
| EFI1580326.1 | HKD-DNase+OGG-like | - | Escherichia_coli | Bacteria>Pseudomonadati>Pseudomonadota>Gammaproteobacteria>Enterobacterales>Enterobacteriaceae>Escherichia | 449 | BFN79_003003 | 11924..13273 | 562 | phosphatidylserine/phosphatidylglycerophosphate/cardiolipin-synthase-family-protein-[Escherichia-coli] |
| EFI3320062.1 | HKD-DNase+OGG-like | - | Escherichia_coli | Bacteria>Pseudomonadati>Pseudomonadota>Gammaproteobacteria>Enterobacterales>Enterobacteriaceae>Escherichia | 449 | BCC72_001612 | 289..1638 | 562 | phosphatidylserine/phosphatidylglycerophosphate/cardiolipin-synthase-family-protein-[Escherichia-coli] |
| EFJ5167528.1 | HKD-DNase+OGG-like | - | Escherichia_coli | Bacteria>Pseudomonadati>Pseudomonadota>Gammaproteobacteria>Enterobacterales>Enterobacteriaceae>Escherichia | 449 | A5T96_003405 | 12254..13603 | 562 | phosphatidylserine/phosphatidylglycerophosphate/cardiolipin-synthase-family-protein-[Escherichia-coli] |
| EFL0544866.1 | HKD-DNase+OGG-like | - | Escherichia_coli | Bacteria>Pseudomonadati>Pseudomonadota>Gammaproteobacteria>Enterobacterales>Enterobacteriaceae>Escherichia | 449 | HIZ25_002085 | 11120..12469 | 562 | phosphatidylserine/phosphatidylglycerophosphate/cardiolipin-synthase-family-protein-[Escherichia-coli] |
| EFL4011093.1 | HKD-DNase+OGG-like | - | Escherichia_coli | Bacteria>Pseudomonadati>Pseudomonadota>Gammaproteobacteria>Enterobacterales>Enterobacteriaceae>Escherichia | 449 | AC312_001865 | 103668..105017 | 562 | phosphatidylserine/phosphatidylglycerophosphate/cardiolipin-synthase-family-protein-[Escherichia-coli] |
| EFM7055938.1 | HKD-DNase+OGG-like | - | Escherichia_coli | Bacteria>Pseudomonadati>Pseudomonadota>Gammaproteobacteria>Enterobacterales>Enterobacteriaceae>Escherichia | 449 | BAU66_002364 | 11863..13212 | 562 | phosphatidylserine/phosphatidylglycerophosphate/cardiolipin-synthase-family-protein-[Escherichia-coli] |
| EFN5734796.1 | HKD-DNase+OGG-like | - | Escherichia_coli | Bacteria>Pseudomonadati>Pseudomonadota>Gammaproteobacteria>Enterobacterales>Enterobacteriaceae>Escherichia | 449 | FQJ28_11205 | 123603..124952 | 562 | phosphatidylserine/phosphatidylglycerophosphate/cardiolipin-synthase-family-protein-[Escherichia-coli] |
| EFR2049113.1 | HKD-DNase+OGG-like | - | Escherichia_coli | Bacteria>Pseudomonadati>Pseudomonadota>Gammaproteobacteria>Enterobacterales>Enterobacteriaceae>Escherichia | 449 | HZO98_003741 | 11306..12655 | 562 | phosphatidylserine/phosphatidylglycerophosphate/cardiolipin-synthase-family-protein-[Escherichia-coli] |
| EFR9656375.1 | HKD-DNase+OGG-like | - | Escherichia_coli | Bacteria>Pseudomonadati>Pseudomonadota>Gammaproteobacteria>Enterobacterales>Enterobacteriaceae>Escherichia | 449 | H2N11_003250 | 11684..13033 | 562 | phosphatidylserine/phosphatidylglycerophosphate/cardiolipin-synthase-family-protein-[Escherichia-coli] |
| EFS8463117.1 | HKD-DNase+OGG-like | - | Escherichia_coli | Bacteria>Pseudomonadati>Pseudomonadota>Gammaproteobacteria>Enterobacterales>Enterobacteriaceae>Escherichia | 449 | HWT39_002323 | 109512..110861 | 562 | phosphatidylserine/phosphatidylglycerophosphate/cardiolipin-synthase-family-protein-[Escherichia-coli] |
| EFU0098531.1 | HKD-DNase+OGG-like | - | Escherichia_coli | Bacteria>Pseudomonadati>Pseudomonadota>Gammaproteobacteria>Enterobacterales>Enterobacteriaceae>Escherichia | 449 | HVV04_003693 | 11738..13087 | 562 | phosphatidylserine/phosphatidylglycerophosphate/cardiolipin-synthase-family-protein-[Escherichia-coli] |
| EGD5155867.1 | HKD-DNase+OGG-like | - | Escherichia_coli | Bacteria>Pseudomonadati>Pseudomonadota>Gammaproteobacteria>Enterobacterales>Enterobacteriaceae>Escherichia | 449 | DTY31_07120 | 84062..85411 | 562 | phosphatidylserine/phosphatidylglycerophosphate/cardiolipin-synthase-family-protein-[Escherichia-coli] |
| EGK3603431.1 | HKD-DNase+OGG-like | - | Escherichia_coli | Bacteria>Pseudomonadati>Pseudomonadota>Gammaproteobacteria>Enterobacterales>Enterobacteriaceae>Escherichia | 449 | IOT28_003453 | 12116..13465 | 562 | phosphatidylserine/phosphatidylglycerophosphate/cardiolipin-synthase-family-protein-[Escherichia-coli] |
| EGY5620867.1 | HKD-DNase+OGG-like | - | Escherichia_coli | Bacteria>Pseudomonadati>Pseudomonadota>Gammaproteobacteria>Enterobacterales>Enterobacteriaceae>Escherichia | 449 | JI433_002894 | 27045..28394 | 562 | phosphatidylserine/phosphatidylglycerophosphate/cardiolipin-synthase-family-protein-[Escherichia-coli] |
| EGY5636090.1 | HKD-DNase+OGG-like | - | Escherichia_coli | Bacteria>Pseudomonadati>Pseudomonadota>Gammaproteobacteria>Enterobacterales>Enterobacteriaceae>Escherichia | 449 | JI450_003026 | 12225..13574 | 562 | phosphatidylserine/phosphatidylglycerophosphate/cardiolipin-synthase-family-protein-[Escherichia-coli] |
| EGY8947060.1 | HKD-DNase+OGG-like | - | Escherichia_coli | Bacteria>Pseudomonadati>Pseudomonadota>Gammaproteobacteria>Enterobacterales>Enterobacteriaceae>Escherichia | 449 | JI613_002969 | 11000..12349 | 562 | phosphatidylserine/phosphatidylglycerophosphate/cardiolipin-synthase-family-protein-[Escherichia-coli] |
| EGY9082472.1 | HKD-DNase+OGG-like | - | Escherichia_coli | Bacteria>Pseudomonadati>Pseudomonadota>Gammaproteobacteria>Enterobacterales>Enterobacteriaceae>Escherichia | 449 | JI646_002307 | 12236..13585 | 562 | phosphatidylserine/phosphatidylglycerophosphate/cardiolipin-synthase-family-protein-[Escherichia-coli] |
| EGY9336118.1 | HKD-DNase+OGG-like | - | Escherichia_coli | Bacteria>Pseudomonadati>Pseudomonadota>Gammaproteobacteria>Enterobacterales>Enterobacteriaceae>Escherichia | 449 | HZO78_003646 | 12243..13592 | 562 | phosphatidylserine/phosphatidylglycerophosphate/cardiolipin-synthase-family-protein-[Escherichia-coli] |
| EGZ0051360.1 | HKD-DNase+OGG-like | - | Escherichia_coli | Bacteria>Pseudomonadati>Pseudomonadota>Gammaproteobacteria>Enterobacterales>Enterobacteriaceae>Escherichia | 449 | JJA24_002222 | 11213..12562 | 562 | phosphatidylserine/phosphatidylglycerophosphate/cardiolipin-synthase-family-protein-[Escherichia-coli] |
| EGZ1391644.1 | HKD-DNase+OGG-like | - | Escherichia_coli | Bacteria>Pseudomonadati>Pseudomonadota>Gammaproteobacteria>Enterobacterales>Enterobacteriaceae>Escherichia | 449 | FG160_002858 | 11078..12427 | 562 | phosphatidylserine/phosphatidylglycerophosphate/cardiolipin-synthase-family-protein-[Escherichia-coli] |
| EGZ1415764.1 | HKD-DNase+OGG-like | - | Escherichia_coli | Bacteria>Pseudomonadati>Pseudomonadota>Gammaproteobacteria>Enterobacterales>Enterobacteriaceae>Escherichia | 449 | FFU33_001574 | 144118..145467 | 562 | phosphatidylserine/phosphatidylglycerophosphate/cardiolipin-synthase-family-protein-[Escherichia-coli] |
| EGZ1559471.1 | HKD-DNase+OGG-like | - | Escherichia_coli | Bacteria>Pseudomonadati>Pseudomonadota>Gammaproteobacteria>Enterobacterales>Enterobacteriaceae>Escherichia | 449 | FAV16_003768 | 11015..12364 | 562 | phosphatidylserine/phosphatidylglycerophosphate/cardiolipin-synthase-family-protein-[Escherichia-coli] |
| EGZ1869335.1 | HKD-DNase+OGG-like | - | Escherichia_coli | Bacteria>Pseudomonadati>Pseudomonadota>Gammaproteobacteria>Enterobacterales>Enterobacteriaceae>Escherichia | 449 | EYE54_001996 | 11099..12448 | 562 | phosphatidylserine/phosphatidylglycerophosphate/cardiolipin-synthase-family-protein-[Escherichia-coli] |
| EGZ2354233.1 | HKD-DNase+OGG-like | - | Escherichia_coli | Bacteria>Pseudomonadati>Pseudomonadota>Gammaproteobacteria>Enterobacterales>Enterobacteriaceae>Escherichia | 449 | D7J47_002048 | 27046..28395 | 562 | phosphatidylserine/phosphatidylglycerophosphate/cardiolipin-synthase-family-protein-[Escherichia-coli] |
| EGZ2497687.1 | HKD-DNase+OGG-like | - | Escherichia_coli | Bacteria>Pseudomonadati>Pseudomonadota>Gammaproteobacteria>Enterobacterales>Enterobacteriaceae>Escherichia | 449 | D6W78_003635 | 11620..12969 | 562 | phosphatidylserine/phosphatidylglycerophosphate/cardiolipin-synthase-family-protein-[Escherichia-coli] |
| EGZ2674124.1 | HKD-DNase+OGG-like | - | Escherichia_coli | Bacteria>Pseudomonadati>Pseudomonadota>Gammaproteobacteria>Enterobacterales>Enterobacteriaceae>Escherichia | 449 | D1H08_002344 | 11999..13348 | 562 | phosphatidylserine/phosphatidylglycerophosphate/cardiolipin-synthase-family-protein-[Escherichia-coli] |
| EGZ2777084.1 | HKD-DNase+OGG-like | - | Escherichia_coli | Bacteria>Pseudomonadati>Pseudomonadota>Gammaproteobacteria>Enterobacterales>Enterobacteriaceae>Escherichia | 449 | DVO41_002393 | 27046..28395 | 562 | phosphatidylserine/phosphatidylglycerophosphate/cardiolipin-synthase-family-protein-[Escherichia-coli] |
| EGZ3242786.1 | HKD-DNase+OGG-like | - | Escherichia_coli | Bacteria>Pseudomonadati>Pseudomonadota>Gammaproteobacteria>Enterobacterales>Enterobacteriaceae>Escherichia | 449 | C5N70_002408 | 52340..53689 | 562 | phosphatidylserine/phosphatidylglycerophosphate/cardiolipin-synthase-family-protein-[Escherichia-coli] |
| EGZ4145356.1 | HKD-DNase+OGG-like | - | Escherichia_coli | Bacteria>Pseudomonadati>Pseudomonadota>Gammaproteobacteria>Enterobacterales>Enterobacteriaceae>Escherichia | 449 | CKY95_002789 | 11998..13347 | 562 | phosphatidylserine/phosphatidylglycerophosphate/cardiolipin-synthase-family-protein-[Escherichia-coli] |
| EGZ7370263.1 | HKD-DNase+OGG-like | - | Escherichia_coli | Bacteria>Pseudomonadati>Pseudomonadota>Gammaproteobacteria>Enterobacterales>Enterobacteriaceae>Escherichia | 449 | BAU61_002961 | 11351..12700 | 562 | phosphatidylserine/phosphatidylglycerophosphate/cardiolipin-synthase-family-protein-[Escherichia-coli] |
| EGZ7391811.1 | HKD-DNase+OGG-like | - | Escherichia_coli | Bacteria>Pseudomonadati>Pseudomonadota>Gammaproteobacteria>Enterobacterales>Enterobacteriaceae>Escherichia | 449 | BAU88_004035 | 11621..12970 | 562 | phosphatidylserine/phosphatidylglycerophosphate/cardiolipin-synthase-family-protein-[Escherichia-coli] |
| EGZ7468565.1 | HKD-DNase+OGG-like | - | Escherichia_coli | Bacteria>Pseudomonadati>Pseudomonadota>Gammaproteobacteria>Enterobacterales>Enterobacteriaceae>Escherichia | 449 | A6F02_004046 | 11506..12855 | 562 | phosphatidylserine/phosphatidylglycerophosphate/cardiolipin-synthase-family-protein-[Escherichia-coli] |
| EGZ7821801.1 | HKD-DNase+OGG-like | - | Escherichia_coli | Bacteria>Pseudomonadati>Pseudomonadota>Gammaproteobacteria>Enterobacterales>Enterobacteriaceae>Escherichia | 449 | AWC87_001494 | 10898..12247 | 562 | phosphatidylserine/phosphatidylglycerophosphate/cardiolipin-synthase-family-protein-[Escherichia-coli] |
| EGZ8838312.1 | HKD-DNase+OGG-like | - | Escherichia_coli | Bacteria>Pseudomonadati>Pseudomonadota>Gammaproteobacteria>Enterobacterales>Enterobacteriaceae>Escherichia | 449 | AT823_000497 | 21068..22417 | 562 | phosphatidylserine/phosphatidylglycerophosphate/cardiolipin-synthase-family-protein-[Escherichia-coli] |
| EGZ9024444.1 | HKD-DNase+OGG-like | - | Escherichia_coli | Bacteria>Pseudomonadati>Pseudomonadota>Gammaproteobacteria>Enterobacterales>Enterobacteriaceae>Escherichia | 449 | AFE50_004180 | 11600..12949 | 562 | phosphatidylserine/phosphatidylglycerophosphate/cardiolipin-synthase-family-protein-[Escherichia-coli] |
| EHA0949392.1 | HKD-DNase+OGG-like | - | Escherichia_coli | Bacteria>Pseudomonadati>Pseudomonadota>Gammaproteobacteria>Enterobacterales>Enterobacteriaceae>Escherichia | 449 | JI650_002934 | 10999..12348 | 562 | phosphatidylserine/phosphatidylglycerophosphate/cardiolipin-synthase-family-protein-[Escherichia-coli] |
| EHA9087560.1 | HKD-DNase+OGG-like | - | Escherichia_coli | Bacteria>Pseudomonadati>Pseudomonadota>Gammaproteobacteria>Enterobacterales>Enterobacteriaceae>Escherichia | 449 | D4094_002234 | 27048..28397 | 562 | phosphatidylserine/phosphatidylglycerophosphate/cardiolipin-synthase-family-protein-[Escherichia-coli] |

| Protein-ID | Domain | Assembly-ID | Strain | Taxonomy | Length | Locus-Tag | Genomic-coordinates | TaxID | Fasta-defline |
|---|---|---|---|---|---|---|---|---|---|
| EHC2076261.1 | HKD-DNase+OGG-like | - | Escherichia_coli | Bacteria>Pseudomonadati>Pseudomonadota>Gammaproteobacteria>Enterobacterales>Enterobacteriaceae>Escherichia | 449 | JT651_000263 | 21122..22471 | 562 | phosphatidylserine/phosphatidylglycerophosphate/cardiolipin-synthase-family-protein-[Escherichia-coli] |
| EHJ8640901.1 | HKD-DNase+OGG-like | GCA_017945505.1 | Escherichia_coli | Bacteria>Pseudomonadati>Pseudomonadota>Gammaproteobacteria>Enterobacterales>Enterobacteriaceae>Escherichia | 449 | KB366_000311 | 343310..344659 | 562 | phosphatidylserine/phosphatidylglycerophosphate/cardiolipin-synthase-family-protein-[Escherichia-coli] |
| EHK0633850.1 | HKD-DNase+OGG-like | GCA_017971565.1 | Escherichia_coli | Bacteria>Pseudomonadati>Pseudomonadota>Gammaproteobacteria>Enterobacterales>Enterobacteriaceae>Escherichia | 449 | KCG30_002482 | 123583..124932 | 562 | phosphatidylserine/phosphatidylglycerophosphate/cardiolipin-synthase-family-protein-[Escherichia-coli] |
| EHK3416140.1 | HKD-DNase+OGG-like | GCA_018020775.1 | Escherichia_coli | Bacteria>Pseudomonadati>Pseudomonadota>Gammaproteobacteria>Enterobacterales>Enterobacteriaceae>Escherichia | 449 | JYE89_003266 | 12091..13440 | 562 | phosphatidylserine/phosphatidylglycerophosphate/cardiolipin-synthase-family-protein-[Escherichia-coli] |
| EHL0363100.1 | HKD-DNase+OGG-like | GCA_018090865.1 | Escherichia_coli | Bacteria>Pseudomonadati>Pseudomonadota>Gammaproteobacteria>Enterobacterales>Enterobacteriaceae>Escherichia | 449 | D1H14_003273 | 11474..12823 | 562 | phosphatidylserine/phosphatidylglycerophosphate/cardiolipin-synthase-family-protein-[Escherichia-coli] |
| EHL0920541.1 | HKD-DNase+OGG-like | GCA_018093285.1 | Escherichia_coli | Bacteria>Pseudomonadati>Pseudomonadota>Gammaproteobacteria>Enterobacterales>Enterobacteriaceae>Escherichia | 449 | CFF49_003162 | 11014..12363 | 562 | phosphatidylserine/phosphatidylglycerophosphate/cardiolipin-synthase-family-protein-[Escherichia-coli] |
| EHL1132632.1 | HKD-DNase+OGG-like | GCA_018094045.1 | Escherichia_coli | Bacteria>Pseudomonadati>Pseudomonadota>Gammaproteobacteria>Enterobacterales>Enterobacteriaceae>Escherichia | 449 | BIB01_002908 | 10967..12316 | 562 | phosphatidylserine/phosphatidylglycerophosphate/cardiolipin-synthase-family-protein-[Escherichia-coli] |
| EHL1162652.1 | HKD-DNase+OGG-like | GCA_018094265.1 | Escherichia_coli | Bacteria>Pseudomonadati>Pseudomonadota>Gammaproteobacteria>Enterobacterales>Enterobacteriaceae>Escherichia | 449 | BE892_002475 | 27047..28396 | 562 | phosphatidylserine/phosphatidylglycerophosphate/cardiolipin-synthase-family-protein-[Escherichia-coli] |
| EHL1168109.1 | HKD-DNase+OGG-like | GCA_018094165.1 | Escherichia_coli | Bacteria>Pseudomonadati>Pseudomonadota>Gammaproteobacteria>Enterobacterales>Enterobacteriaceae>Escherichia | 449 | BFU70_002924 | 12280..13629 | 562 | phosphatidylserine/phosphatidylglycerophosphate/cardiolipin-synthase-family-protein-[Escherichia-coli] |
| EHM5922459.1 | HKD-DNase+OGG-like | GCA_018270935.1 | Escherichia_coli | Bacteria>Pseudomonadati>Pseudomonadota>Gammaproteobacteria>Enterobacterales>Enterobacteriaceae>Escherichia | 449 | KGR67_002712 | 19073..20422 | 562 | phosphatidylserine/phosphatidylglycerophosphate/cardiolipin-synthase-family-protein-[Escherichia-coli] |
| EHN3342074.1 | HKD-DNase+OGG-like | GCA_018356405.1 | Escherichia_coli | Bacteria>Pseudomonadati>Pseudomonadota>Gammaproteobacteria>Enterobacterales>Enterobacteriaceae>Escherichia | 449 | KIR71_003051 | 27046..28395 | 562 | phosphatidylserine/phosphatidylglycerophosphate/cardiolipin-synthase-family-protein-[Escherichia-coli] |
| EHQ7823794.1 | HKD-DNase+OGG-like | GCA_018975885.1 | Escherichia_coli | Bacteria>Pseudomonadati>Pseudomonadota>Gammaproteobacteria>Enterobacterales>Enterobacteriaceae>Escherichia | 449 | KSS40_002994 | 11494..12843 | 562 | phosphatidylserine/phosphatidylglycerophosphate/cardiolipin-synthase-family-protein-[Escherichia-coli] |
| EHW5636700.1 | HKD-DNase+OGG-like | GCA_019482305.1 | Escherichia_coli | Bacteria>Pseudomonadati>Pseudomonadota>Gammaproteobacteria>Enterobacterales>Enterobacteriaceae>Escherichia | 449 | K2955_003131 | 48057..49406 | 562 | phosphatidylserine/phosphatidylglycerophosphate/cardiolipin-synthase-family-protein-[Escherichia-coli] |
| EHX2153198.1 | HKD-DNase+OGG-like | GCA_019516635.1 | Escherichia_coli | Bacteria>Pseudomonadati>Pseudomonadota>Gammaproteobacteria>Enterobacterales>Enterobacteriaceae>Escherichia | 449 | K3P68_001534 | 13745..15094 | 562 | phosphatidylserine/phosphatidylglycerophosphate/cardiolipin-synthase-family-protein-[Escherichia-coli] |
| EHX8392336.1 | HKD-DNase+OGG-like | GCA_019550135.1 | Escherichia_coli | Bacteria>Pseudomonadati>Pseudomonadota>Gammaproteobacteria>Enterobacterales>Enterobacteriaceae>Escherichia | 449 | K3368_004094 | 5089..6438 | 562 | phosphatidylserine/phosphatidylglycerophosphate/cardiolipin-synthase-family-protein-[Escherichia-coli] |
| EHX8476221.1 | HKD-DNase+OGG-like | GCA_019550495.1 | Escherichia_coli | Bacteria>Pseudomonadati>Pseudomonadota>Gammaproteobacteria>Enterobacterales>Enterobacteriaceae>Escherichia | 449 | K3324_002229 | 35542..36891 | 562 | phosphatidylserine/phosphatidylglycerophosphate/cardiolipin-synthase-family-protein-[Escherichia-coli] |
| EHY5298662.1 | HKD-DNase+OGG-like | GCA_019588505.1 | Escherichia_coli | Bacteria>Pseudomonadati>Pseudomonadota>Gammaproteobacteria>Enterobacterales>Enterobacteriaceae>Escherichia | 449 | K4I57_002538 | 11296..12645 | 562 | phosphatidylserine/phosphatidylglycerophosphate/cardiolipin-synthase-family-protein-[Escherichia-coli] |
| EHY5690085.1 | HKD-DNase+OGG-like | GCA_019590565.1 | Escherichia_coli | Bacteria>Pseudomonadati>Pseudomonadota>Gammaproteobacteria>Enterobacterales>Enterobacteriaceae>Escherichia | 449 | K4J54_002563 | 27045..28394 | 562 | phosphatidylserine/phosphatidylglycerophosphate/cardiolipin-synthase-family-protein-[Escherichia-coli] |
| EHZ2564705.1 | HKD-DNase+OGG-like | GCA_019681855.1 | Escherichia_coli | Bacteria>Pseudomonadati>Pseudomonadota>Gammaproteobacteria>Enterobacterales>Enterobacteriaceae>Escherichia | 449 | K5N16_003079 | 12102..13451 | 562 | phosphatidylserine/phosphatidylglycerophosphate/cardiolipin-synthase-family-protein-[Escherichia-coli] |
| EID3638526.1 | HKD-DNase+OGG-like | GCA_020041725.1 | Escherichia_coli | Bacteria>Pseudomonadati>Pseudomonadota>Gammaproteobacteria>Enterobacterales>Enterobacteriaceae>Escherichia | 449 | LB206_001716 | 93038..94387 | 562 | phosphatidylserine/phosphatidylglycerophosphate/cardiolipin-synthase-family-protein-[Escherichia-coli] |
| EID8685853.1 | HKD-DNase+OGG-like | GCA_020094635.1 | Escherichia_coli | Bacteria>Pseudomonadati>Pseudomonadota>Gammaproteobacteria>Enterobacterales>Enterobacteriaceae>Escherichia | 449 | LCI88_002088 | 141778..143127 | 562 | phosphatidylserine/phosphatidylglycerophosphate/cardiolipin-synthase-family-protein-[Escherichia-coli] |
| EIF9349178.1 | HKD-DNase+OGG-like | GCA_020282835.1 | Escherichia_coli | Bacteria>Pseudomonadati>Pseudomonadota>Gammaproteobacteria>Enterobacterales>Enterobacteriaceae>Escherichia | 449 | LF100_002185 | 11158..12507 | 562 | phosphatidylserine/phosphatidylglycerophosphate/cardiolipin-synthase-family-protein-[Escherichia-coli] |
| EIG2625395.1 | HKD-DNase+OGG-like | GCA_020303335.1 | Escherichia_coli | Bacteria>Pseudomonadati>Pseudomonadota>Gammaproteobacteria>Enterobacterales>Enterobacteriaceae>Escherichia | 449 | GLS16_002894 | 11026..12375 | 562 | phosphatidylserine/phosphatidylglycerophosphate/cardiolipin-synthase-family-protein-[Escherichia-coli] |
| EIG7860378.1 | HKD-DNase+OGG-like | GCA_020324895.1 | Escherichia_coli | Bacteria>Pseudomonadati>Pseudomonadota>Gammaproteobacteria>Enterobacterales>Enterobacteriaceae>Escherichia | 449 | D0584_004205 | 11942..13291 | 562 | phosphatidylserine/phosphatidylglycerophosphate/cardiolipin-synthase-family-protein-[Escherichia-coli] |
| EIH7117206.1 | HKD-DNase+OGG-like | GCA_020376505.1 | Escherichia_coli | Bacteria>Pseudomonadati>Pseudomonadota>Gammaproteobacteria>Enterobacterales>Enterobacteriaceae>Escherichia | 449 | BB077_003963 | 12235..13584 | 562 | phosphatidylserine/phosphatidylglycerophosphate/cardiolipin-synthase-family-protein-[Escherichia-coli] |
| EIO0019322.1 | HKD-DNase+OGG-like | GCA_020946425.1 | Escherichia_coli | Bacteria>Pseudomonadati>Pseudomonadota>Gammaproteobacteria>Enterobacterales>Enterobacteriaceae>Escherichia | 449 | LO099_004447 | 11137..12486 | 562 | phosphatidylserine/phosphatidylglycerophosphate/cardiolipin-synthase-family-protein-[Escherichia-coli] |
| EIX9838480.1 | HKD-DNase+OGG-like | GCA_022433775.1 | Escherichia_coli | Bacteria>Pseudomonadati>Pseudomonadota>Gammaproteobacteria>Enterobacterales>Enterobacteriaceae>Escherichia | 449 | ML439_002963 | 12336..13685 | 562 | phosphatidylserine/phosphatidylglycerophosphate/cardiolipin-synthase-family-protein-[Escherichia-coli] |
| EJA4563487.1 | HKD-DNase+OGG-like | GCA_022888875.1 | Escherichia_coli | Bacteria>Pseudomonadati>Pseudomonadota>Gammaproteobacteria>Enterobacterales>Enterobacteriaceae>Escherichia | 449 | MVR17_003834 | 22390..23739 | 562 | phosphatidylserine/phosphatidylglycerophosphate/cardiolipin-synthase-family-protein-[Escherichia-coli] |
| EJH0789767.1 | HKD-DNase+OGG-like | GCA_023637565.1 | Escherichia_coli | Bacteria>Pseudomonadati>Pseudomonadota>Gammaproteobacteria>Enterobacterales>Enterobacteriaceae>Escherichia | 449 | NCI57_002422 | 12032..13381 | 562 | phosphatidylserine/phosphatidylglycerophosphate/cardiolipin-synthase-family-protein-[Escherichia-coli] |
| EJM6441913.1 | HKD-DNase+OGG-like | GCA_024340725.1 | Escherichia_coli | Bacteria>Pseudomonadati>Pseudomonadota>Gammaproteobacteria>Enterobacterales>Enterobacteriaceae>Escherichia | 449 | NOR68_003997 | 11731..13080 | 562 | phosphatidylserine/phosphatidylglycerophosphate/cardiolipin-synthase-family-protein-[Escherichia-coli] |
| EJN8021042.1 | HKD-DNase+OGG-like | GCA_024524615.1 | Escherichia_coli | Bacteria>Pseudomonadati>Pseudomonadota>Gammaproteobacteria>Enterobacterales>Enterobacteriaceae>Escherichia | 449 | NRW59_000783 | 14556..15905 | 562 | phosphatidylserine/phosphatidylglycerophosphate/cardiolipin-synthase-family-protein-[Escherichia-coli] |
| EJO9498077.1 | HKD-DNase+OGG-like | GCA_024612475.1 | Escherichia_coli | Bacteria>Pseudomonadati>Pseudomonadota>Gammaproteobacteria>Enterobacterales>Enterobacteriaceae>Escherichia | 449 | NUZ09_002879 | 20219..21568 | 562 | phosphatidylserine/phosphatidylglycerophosphate/cardiolipin-synthase-family-protein-[Escherichia-coli] |
| HAH2520955.1 | HKD-DNase+OGG-like | GCA_012530295.1 | Escherichia_coli | Bacteria>Pseudomonadati>Pseudomonadota>Gammaproteobacteria>Enterobacterales>Enterobacteriaceae>Escherichia | 449 | GF659_15760 | 83633..84982 | 562 | TPA:-phosphatidylserine/phosphatidylglycerophosphate/cardiolipin-synthase-family-protein-[Escherichia-coli] |
| HAH8430184.1 | HKD-DNase+OGG-like | GCA_012716735.1 | Escherichia_coli | Bacteria>Pseudomonadati>Pseudomonadota>Gammaproteobacteria>Enterobacterales>Enterobacteriaceae>Escherichia | 449 | HIR43_003172 | 11250..12599 | 562 | TPA:-phosphatidylserine/phosphatidylglycerophosphate/cardiolipin-synthase-family-protein-[Escherichia-coli] |
| HAH9940670.1 | HKD-DNase+OGG-like | GCA_012753595.1 | Escherichia_coli | Bacteria>Pseudomonadati>Pseudomonadota>Gammaproteobacteria>Enterobacterales>Enterobacteriaceae>Escherichia | 449 | HIF13_004048 | 22069..23418 | 562 | TPA:-phosphatidylserine/phosphatidylglycerophosphate/cardiolipin-synthase-family-protein-[Escherichia-coli] |
| HAI9090454.1 | HKD-DNase+OGG-like | GCA_012925565.1 | Escherichia_coli | Bacteria>Pseudomonadati>Pseudomonadota>Gammaproteobacteria>Enterobacterales>Enterobacteriaceae>Escherichia | 449 | HKC72_003110 | 12252..13601 | 562 | TPA:-phosphatidylserine/phosphatidylglycerophosphate/cardiolipin-synthase-family-protein-[Escherichia-coli] |
| HAJ1412082.1 | HKD-DNase+OGG-like | GCA_013000285.1 | Escherichia_coli | Bacteria>Pseudomonadati>Pseudomonadota>Gammaproteobacteria>Enterobacterales>Enterobacteriaceae>Escherichia | 449 | HMG43_22085 | 11850..13199 | 562 | TPA:-phosphatidylserine/phosphatidylglycerophosphate/cardiolipin-synthase-family-protein-[Escherichia-coli] |
| HAL9997358.1 | HKD-DNase+OGG-like | GCA_014189205.1 | Escherichia_coli | Bacteria>Pseudomonadati>Pseudomonadota>Gammaproteobacteria>Enterobacterales>Enterobacteriaceae>Escherichia | 449 | H7W48_003270 | 11954..13303 | 562 | TPA:-phosphatidylserine/phosphatidylglycerophosphate/cardiolipin-synthase-family-protein-[Escherichia-coli] |
| HAO2821708.1 | HKD-DNase+OGG-like | GCA_014823275.1 | Escherichia_coli | Bacteria>Pseudomonadati>Pseudomonadota>Gammaproteobacteria>Enterobacterales>Enterobacteriaceae>Escherichia | 449 | IHL91_001059 | 12045..13394 | 562 | phosphatidylserine/phosphatidylglycerophosphate/cardiolipin-synthase-family-protein-[Escherichia-coli] |
| HAW1448907.1 | HKD-DNase+OGG-like | GCA_016674355.1 | Escherichia_coli | Bacteria>Pseudomonadati>Pseudomonadota>Gammaproteobacteria>Enterobacterales>Enterobacteriaceae>Escherichia | 449 | JLM49_000130 | 148059..149408 | 562 | TPA:-phosphatidylserine/phosphatidylglycerophosphate/cardiolipin-synthase-family-protein-[Escherichia-coli] |
| HAX4031535.1 | HKD-DNase+OGG-like | GCA_017003755.1 | Escherichia_coli | Bacteria>Pseudomonadati>Pseudomonadota>Gammaproteobacteria>Enterobacterales>Enterobacteriaceae>Escherichia | 449 | JV397_001894 | 172911..174260 | 562 | TPA:-phosphatidylserine/phosphatidylglycerophosphate/cardiolipin-synthase-family-protein-[Escherichia-coli] |
| HAX9732777.1 | HKD-DNase+OGG-like | GCA_017073235.1 | Escherichia_coli | Bacteria>Pseudomonadati>Pseudomonadota>Gammaproteobacteria>Enterobacterales>Enterobacteriaceae>Escherichia | 449 | JS163_001167 | 116534..117883 | 562 | TPA:-phosphatidylserine/phosphatidylglycerophosphate/cardiolipin-synthase-family-protein-[Escherichia-coli] |
| HBA3375043.1 | HKD-DNase+OGG-like | GCA_017728515.1 | Escherichia_coli | Bacteria>Pseudomonadati>Pseudomonadota>Gammaproteobacteria>Enterobacterales>Enterobacteriaceae>Escherichia | 449 | J8Z50_001955 | 123835..125184 | 562 | TPA:-phosphatidylserine/phosphatidylglycerophosphate/cardiolipin-synthase-family-protein-[Escherichia-coli] |
| HBA6912642.1 | HKD-DNase+OGG-like | GCA_017762865.1 | Escherichia_coli | Bacteria>Pseudomonadati>Pseudomonadota>Gammaproteobacteria>Enterobacterales>Enterobacteriaceae>Escherichia | 449 | J1V61_002567 | 21086..22435 | 562 | TPA:-phosphatidylserine/phosphatidylglycerophosphate/cardiolipin-synthase-family-protein-[Escherichia-coli] |
| HBB4280099.1 | HKD-DNase+OGG-like | GCA_017899135.1 | Escherichia_coli | Bacteria>Pseudomonadati>Pseudomonadota>Gammaproteobacteria>Enterobacterales>Enterobacteriaceae>Escherichia | 380 | KAW34_004651 | 11884..13025 | 562 | phosphatidylserine/phosphatidylglycerophosphate/cardiolipin-synthase-family-protein-[Escherichia-coli] |
| HBD5250195.1 | HKD-DNase+OGG-like | GCA_018460085.1 | Escherichia_coli | Bacteria>Pseudomonadati>Pseudomonadota>Gammaproteobacteria>Enterobacterales>Enterobacteriaceae>Escherichia | 449 | KKL26_003780 | 18671..20020 | 562 | TPA:-phosphatidylserine/phosphatidylglycerophosphate/cardiolipin-synthase-family-protein-[Escherichia-coli] |
| HBJ0235558.1 | HKD-DNase+OGG-like | GCA_019980105.1 | Escherichia_coli | Bacteria>Pseudomonadati>Pseudomonadota>Gammaproteobacteria>Enterobacterales>Enterobacteriaceae>Escherichia | 449 | LAM28_003682 | 538..1887 | 562 | TPA:-phosphatidylserine/phosphatidylglycerophosphate/cardiolipin-synthase-family-protein-[Escherichia-coli] |
| HBM8747265.1 | HKD-DNase+OGG-like | GCA_021417505.1 | Escherichia_coli | Bacteria>Pseudomonadati>Pseudomonadota>Gammaproteobacteria>Enterobacterales>Enterobacteriaceae>Escherichia | 449 | L0V31_003264 | 52199..53548 | 562 | TPA:-phosphatidylserine/phosphatidylglycerophosphate/cardiolipin-synthase-family-protein-[Escherichia-coli] |
| HBN0573059.1 | HKD-DNase+OGG-like | GCA_021454505.1 | Escherichia_coli | Bacteria>Pseudomonadati>Pseudomonadota>Gammaproteobacteria>Enterobacterales>Enterobacteriaceae>Escherichia | 449 | L1N12_002089 | 25014..26363 | 562 | phosphatidylserine/phosphatidylglycerophosphate/cardiolipin-synthase-family-protein-[Escherichia-coli] |
| HCN4662828.1 | HKD-DNase+OGG-like | GCA_025276155.1 | Escherichia_coli | Bacteria>Pseudomonadati>Pseudomonadota>Gammaproteobacteria>Enterobacterales>Enterobacteriaceae>Escherichia | 449 | N6W02_002923 | 14522..15871 | 562 | TPA:-phosphatidylserine/phosphatidylglycerophosphate/cardiolipin-synthase-family-protein-[Escherichia-coli] |
| HCO6583164.1 | HKD-DNase+OGG-like | GCA_025412985.1 | Escherichia_coli | Bacteria>Pseudomonadati>Pseudomonadota>Gammaproteobacteria>Enterobacterales>Enterobacteriaceae>Escherichia | 449 | OBR12_001372 | 123599..124948 | 562 | TPA:-phosphatidylserine/phosphatidylglycerophosphate/cardiolipin-synthase-family-protein-[Escherichia-coli] |
| HCP1392416.1 | HKD-DNase+OGG-like | GCA_025491965.1 | Escherichia_coli | Bacteria>Pseudomonadati>Pseudomonadota>Gammaproteobacteria>Enterobacterales>Enterobacteriaceae>Escherichia | 449 | OCX38_000206 | 208288..209637 | 562 | phosphatidylserine/phosphatidylglycerophosphate/cardiolipin-synthase-family-protein-[Escherichia-coli] |
| KAA9812771.1 | HKD-DNase+OGG-like | GCA_008755545.1 | Escherichia_coli | Bacteria>Pseudomonadati>Pseudomonadota>Gammaproteobacteria>Enterobacterales>Enterobacteriaceae>Escherichia | 449 | F7F73_23805 | 11384..12733 | 562 | phosphatidylserine/phosphatidylglycerophosphate/cardiolipin-synthase-family-protein-[Escherichia-coli] |
| MBC0786091.1 | HKD-DNase+OGG-like | GCA_012382335.2 | Escherichia_coli | Bacteria>Pseudomonadati>Pseudomonadota>Gammaproteobacteria>Enterobacterales>Enterobacteriaceae>Escherichia | 449 | FSG57_002385 | 123373..124722 | 562 | phosphatidylserine/phosphatidylglycerophosphate/cardiolipin-synthase-family-protein-[Escherichia-coli] |
| WP_000766442.1 | HKD-DNase+OGG-like | GCF_000010765.1 | Escherichia_coli | Bacteria>Pseudomonadati>Pseudomonadota>Gammaproteobacteria>Enterobacterales>Enterobacteriaceae>Escherichia | 449 | MLIZ_RS15495 | 11374..12723 | 562 | phospholipase-D-like-domain-containing-protein-[Escherichia-coli] |
| WP_001695296.1 | HKD-DNase+OGG-like | GCF_000354335.2 | Escherichia_coli | Bacteria>Pseudomonadati>Pseudomonadota>Gammaproteobacteria>Enterobacterales>Enterobacteriaceae>Escherichia | 449 | EXA04_RS21895 | 4491774..4493123 | 562 | phospholipase-D-like-domain-containing-protein-[Escherichia-coli] |
| WP_001715915.1 | HKD-DNase+OGG-like | GCF_000355055.1 | Escherichia_coli | Bacteria>Pseudomonadati>Pseudomonadota>Gammaproteobacteria>Enterobacterales>Enterobacteriaceae>Escherichia | 449 | EC2850400_RS01445 | 304318..305667 | 562 | phospholipase-D-like-domain-containing-protein-[Escherichia-coli] |
| WP_024243516.1 | HKD-DNase+OGG-like | GCF_000446965.2 | Escherichia_coli | Bacteria>Pseudomonadati>Pseudomonadota>Gammaproteobacteria>Enterobacterales>Enterobacteriaceae>Escherichia | 449 | PG805_RS04715 | 97358..98707 | 562 | phospholipase-D-like-domain-containing-protein-[Escherichia-coli] |
| WP_032257289.1 | HKD-DNase+OGG-like | GCF_001286385.1 | Escherichia_coli | Bacteria>Pseudomonadati>Pseudomonadota>Gammaproteobacteria>Enterobacterales>Enterobacteriaceae>Escherichia | 449 | AOJ18_RS01765 | 49556..50905 | 562 | phospholipase-D-like-domain-containing-protein-[Escherichia-coli] |
| WP_053892088.1 | HKD-DNase+OGG-like | GCF_001285285.1 | Escherichia_coli | Bacteria>Pseudomonadati>Pseudomonadota>Gammaproteobacteria>Enterobacterales>Enterobacteriaceae>Escherichia | 449 | AOH78_RS17155 | 49511..50860 | 562 | phospholipase-D-like-domain-containing-protein-[Escherichia-coli] |
| WP_058101451.1 | HKD-DNase+OGG-like | GCF_024608215.1 | Escherichia_coli | Bacteria>Pseudomonadati>Pseudomonadota>Gammaproteobacteria>Enterobacterales>Enterobacteriaceae>Escherichia | 395 | NO078_RS04985 | 54364..55551 | 562 | phospholipase-D-like-domain-containing-protein-[Escherichia-coli] |
| WP_106883966.1 | HKD-DNase+OGG-like | GCF_003018495.1 | Escherichia_coli | Bacteria>Pseudomonadati>Pseudomonadota>Gammaproteobacteria>Enterobacterales>Enterobacteriaceae>Escherichia | 449 | C6953_RS06905 | 1314089..1315438 | 562 | phospholipase-D-like-domain-containing-protein-[Escherichia-coli] |
| WP_130090530.1 | HKD-DNase+OGG-like | GCF_021698335.1 | Escherichia_coli | Bacteria>Pseudomonadati>Pseudomonadota>Gammaproteobacteria>Enterobacterales>Enterobacteriaceae>Escherichia | 449 | IHV39_RS14505 | 25288..26637 | 562 | phospholipase-D-like-domain-containing-protein-[Escherichia-coli] |
| WP_164476771.1 | HKD-DNase+OGG-like | GCF_011045235.1 | Escherichia_coli | Bacteria>Pseudomonadati>Pseudomonadota>Gammaproteobacteria>Enterobacterales>Enterobacteriaceae>Escherichia | 449 | G6Z93_RS19900 | 4200261..4201610 | 562 | phospholipase-D-like-domain-containing-protein-[Escherichia-coli] |
| WP_169710967.1 | HKD-DNase+OGG-like | GCF_030296855.1 | Escherichia_fergusonii | Bacteria>Pseudomonadati>Pseudomonadota>Gammaproteobacteria>Enterobacterales>Enterobacteriaceae>Escherichia | 449 | H1X95_RS06235 | 193916..195265 | 564 | phospholipase-D-like-domain-containing-protein-[Escherichia-fergusonii] |
| CEK08492.1 | HKD-DNase+OGG-like | GCA_900000205.1 | Escherichia_coli_O26:H11 | Bacteria>Pseudomonadati>Pseudomonadota>Gammaproteobacteria>Enterobacterales>Enterobacteriaceae>Escherichia>Escherichia_coli | 449 | ECO26H__800028 | 25654..27003 | 244319 | Phospholipase-[Escherichia-coli-O26:H11] |
| EEZ5688040.1 | HKD-DNase+OGG-like | - | Escherichia_coli_O25 | Bacteria>Pseudomonadati>Pseudomonadota>Gammaproteobacteria>Enterobacterales>Enterobacteriaceae>Escherichia>Escherichia_coli | 449 | DBU11_001478 | 125706..127055 | 1095709 | phosphatidylserine/phosphatidylglycerophosphate/cardiolipin-synthase-family-protein-[Escherichia-coli-O25] |
| EEZ8952066.1 | HKD-DNase+OGG-like | - | Escherichia_coli_O35 | Bacteria>Pseudomonadati>Pseudomonadota>Gammaproteobacteria>Enterobacterales>Enterobacteriaceae>Escherichia>Escherichia_coli | 449 | C2D51_003187 | 21972..23321 | 2072460 | phosphatidylserine/phosphatidylglycerophosphate/cardiolipin-synthase-family-protein-[Escherichia-coli-O35] |
| EFO2091248.1 | HKD-DNase+OGG-like | - | Escherichia_coli_O54 | Bacteria>Pseudomonadati>Pseudomonadota>Gammaproteobacteria>Enterobacterales>Enterobacteriaceae>Escherichia>Escherichia_coli | 449 | DP047_19900 | 22116..23465 | 2184075 | phosphatidylserine/phosphatidylglycerophosphate/cardiolipin-synthase-family-protein-[Escherichia-coli-O54] |
| EHW21304.1 | HKD-DNase+OGG-like | GCA_000249875.2 | Escherichia_coli_DEC8E | Bacteria>Pseudomonadati>Pseudomonadota>Gammaproteobacteria>Enterobacterales>Enterobacteriaceae>Escherichia>Escherichia_coli | 395 | ECDEC8E_5313 | 136349..137536 | 868174 | hypothetical-protein-ECDEC8E_5313-[Escherichia-coli-DEC8E] |
| EHX52151.1 | HKD-DNase+OGG-like | GCA_000250315.2 | Escherichia_coli_DEC13A | Bacteria>Pseudomonadati>Pseudomonadota>Gammaproteobacteria>Enterobacterales>Enterobacteriaceae>Escherichia>Escherichia_coli | 449 | ECDEC13A_0067 | 66994..68343 | 868196 | hypothetical-protein-ECDEC13A_0067-[Escherichia-coli-DEC13A] |
| EHX55084.1 | HKD-DNase+OGG-like | GCA_000250335.2 | Escherichia_coli_DEC13B | Bacteria>Pseudomonadati>Pseudomonadota>Gammaproteobacteria>Enterobacterales>Enterobacteriaceae>Escherichia>Escherichia_coli | 449 | ECDEC13B_4626 | 90759..92108 | 868197 | hypothetical-protein-ECDEC13B_4626-[Escherichia-coli-DEC13B] |
| WP_086519896.1 | HKD-DNase+OGG-like | GCF_900775425.1 | Klebsiella | Bacteria>Pseudomonadati>Pseudomonadota>Gammaproteobacteria>Enterobacterales>Enterobacteriaceae>Klebsiella/Raoultella_group | 449 | G3W03_RS23060 | 49835..51184 | 570 | phospholipase-D-like-domain-containing-protein-[Klebsiella-pneumoniae] |
| WP_100751490.1 | HKD-DNase+OGG-like | GCF_036953065.1 | Klebsiella | Bacteria>Pseudomonadati>Pseudomonadota>Gammaproteobacteria>Enterobacterales>Enterobacteriaceae>Klebsiella/Raoultella_group | 449 | CWM56_RS26105 | 24682..26031 | 570 | phospholipase-D-like-domain-containing-protein-[Klebsiella-sp.-E-Nf3] |
| WP_101858979.1 | HKD-DNase+OGG-like | GCF_020479405.1 | Klebsiella | Bacteria>Pseudomonadati>Pseudomonadota>Gammaproteobacteria>Enterobacterales>Enterobacteriaceae>Klebsiella/Raoultella_group | 449 | PZY30_RS10250 | 12318..13667 | 570 | phospholipase-D-like-domain-containing-protein-[Klebsiella-variicola] |
| WP_168234026.1 | HKD-DNase+OGG-like | GCF_964207715.1 | Klebsiella | Bacteria>Pseudomonadati>Pseudomonadota>Gammaproteobacteria>Enterobacterales>Enterobacteriaceae>Klebsiella/Raoultella_group | 449 | HGK55_RS15755 | 327053..328402 | 570 | phospholipase-D-like-domain-containing-protein-[Klebsiella-variicola] |
| HBV8971415.1 | HKD-DNase+OGG-like | GCA_022162485.1 | Klebsiella_oxytoca | Bacteria>Pseudomonadati>Pseudomonadota>Gammaproteobacteria>Enterobacterales>Enterobacteriaceae>Klebsiella/Raoultella_group>Klebsiella | 449 | MD887_005077 | 34869..36218 | 571 | phosphatidylserine/phosphatidylglycerophosphate/cardiolipin-synthase-family-protein-[Klebsiella-oxytoca] |
| STR42112.1 | HKD-DNase+OGG-like | GCA_900451375.1 | Klebsiella_michiganensis | Bacteria>Pseudomonadati>Pseudomonadota>Gammaproteobacteria>Enterobacterales>Enterobacteriaceae>Klebsiella/Raoultella_group>Klebsiella | 294 | NCTC11694_03321 | 2635540..2636424 | 1134687 | Uncharacterised-protein-[Klebsiella-michiganensis] |
| VUT19922.1 | HKD-DNase+OGG-like | GCA_902158685.1 | Klebsiella_pasteurii | Bacteria>Pseudomonadati>Pseudomonadota>Gammaproteobacteria>Enterobacterales>Enterobacteriaceae>Klebsiella/Raoultella_group>Klebsiella | 449 | SB6424_04694 | 309704..311053 | 2587529 | hypothetical-protein-SB6424_04694-[Klebsiella-pasteurii] |
| WP_032425506.1 | HKD-DNase+OGG-like | GCF_902499505.1 | Klebsiella_pneumoniae_complex | Bacteria>Pseudomonadati>Pseudomonadota>Gammaproteobacteria>Enterobacterales>Enterobacteriaceae>Klebsiella/Raoultella_group>Klebsiella | 449 | L1C80_RS19985 | 156517..157866 | 3390273 | MULTISPECIES:-phospholipase-D-like-domain-containing-protein-[Klebsiella] |
| WP_064406994.1 | HKD-DNase+OGG-like | GCF_001598695.1 | Klebsiella_oxytoca | Bacteria>Pseudomonadati>Pseudomonadota>Gammaproteobacteria>Enterobacterales>Enterobacteriaceae>Klebsiella/Raoultella_group>Klebsiella | 449 | KO1_RS15030 | 52393..53742 | 571 | phospholipase-D-like-domain-containing-protein-[Klebsiella-oxytoca] |
| WP_109220875.1 | HKD-DNase+OGG-like | GCF_900407115.1 | Klebsiella_oxytoca | Bacteria>Pseudomonadati>Pseudomonadota>Gammaproteobacteria>Enterobacterales>Enterobacteriaceae>Klebsiella/Raoultella_group>Klebsiella | 449 | DK336_RS07870 | 57462..58811 | 571 | phospholipase-D-like-domain-containing-protein-[Klebsiella-oxytoca] |
| WP_265815518.1 | HKD-DNase+OGG-like | GCF_026224285.1 | Klebsiella_oxytoca | Bacteria>Pseudomonadati>Pseudomonadota>Gammaproteobacteria>Enterobacterales>Enterobacteriaceae>Klebsiella/Raoultella_group>Klebsiella | 449 | ORM16_RS17350 | 97096..98445 | 571 | phospholipase-D-like-domain-containing-protein-[Klebsiella-oxytoca] |
| EIV7918664.1 | HKD-DNase+OGG-like | GCA_022009395.1 | Klebsiella_pneumoniae | Bacteria>Pseudomonadati>Pseudomonadota>Gammaproteobacteria>Enterobacterales>Enterobacteriaceae>Klebsiella/Raoultella_group>Klebsiella>Klebsiella_pneumoniae_complex | 449 | MA315_005127 | 27059..28408 | 573 | phosphatidylserine/phosphatidylglycerophosphate/cardiolipin-synthase-family-protein-[Klebsiella-pneumoniae] |
| EIX9551664.1 | HKD-DNase+OGG-like | GCA_022431205.1 | Klebsiella_pneumoniae | Bacteria>Pseudomonadati>Pseudomonadota>Gammaproteobacteria>Enterobacterales>Enterobacteriaceae>Klebsiella/Raoultella_group>Klebsiella>Klebsiella_pneumoniae_complex | 449 | ML061_004134 | 29610..30959 | 573 | phosphatidylserine/phosphatidylglycerophosphate/cardiolipin-synthase-family-protein-[Klebsiella-pneumoniae] |
| GKN17803.1 | HKD-DNase+OGG-like | GCA_026006635.1 | Klebsiella_pneumoniae | Bacteria>Pseudomonadati>Pseudomonadota>Gammaproteobacteria>Enterobacterales>Enterobacteriaceae>Klebsiella/Raoultella_group>Klebsiella>Klebsiella_pneumoniae_complex | 449 | NUBL17184_44970 | 2133..3482 | 573 | hypothetical-protein-NUBL17184_44970-[Klebsiella-pneumoniae] |
| HBQ6953844.1 | HKD-DNase+OGG-like | GCA_021875055.1 | Klebsiella_pneumoniae | Bacteria>Pseudomonadati>Pseudomonadota>Gammaproteobacteria>Enterobacterales>Enterobacteriaceae>Klebsiella/Raoultella_group>Klebsiella>Klebsiella_pneumoniae_complex | 635 | L8O79_001069 | 34902..36809 | 573 | TPA:-hypothetical-protein-L8O79_001069-[Klebsiella-pneumoniae] |
| HBR1743018.1 | HKD-DNase+OGG-like | GCA_021911065.1 | Klebsiella_pneumoniae | Bacteria>Pseudomonadati>Pseudomonadota>Gammaproteobacteria>Enterobacterales>Enterobacteriaceae>Klebsiella/Raoultella_group>Klebsiella>Klebsiella_pneumoniae_complex | 449 | L9M23_004131 | 49931..51280 | 573 | phosphatidylserine/phosphatidylglycerophosphate/cardiolipin-synthase-family-protein-[Klebsiella-pneumoniae] |
| HBR4416048.1 | HKD-DNase+OGG-like | GCA_021925925.1 | Klebsiella_pneumoniae | Bacteria>Pseudomonadati>Pseudomonadota>Gammaproteobacteria>Enterobacterales>Enterobacteriaceae>Klebsiella/Raoultella_group>Klebsiella>Klebsiella_pneumoniae_complex | 449 | L9V39_001893 | 28439..29788 | 573 | phosphatidylserine/phosphatidylglycerophosphate/cardiolipin-synthase-family-protein-[Klebsiella-pneumoniae] |
| HBR6062606.1 | HKD-DNase+OGG-like | GCA_021932975.1 | Klebsiella_pneumoniae | Bacteria>Pseudomonadati>Pseudomonadota>Gammaproteobacteria>Enterobacterales>Enterobacteriaceae>Klebsiella/Raoultella_group>Klebsiella>Klebsiella_pneumoniae_complex | 449 | L9O24_005229 | 16362..17711 | 573 | phosphatidylserine/phosphatidylglycerophosphate/cardiolipin-synthase-family-protein-[Klebsiella-pneumoniae] |
| HBU3475139.1 | HKD-DNase+OGG-like | GCA_022085545.1 | Klebsiella_pneumoniae | Bacteria>Pseudomonadati>Pseudomonadota>Gammaproteobacteria>Enterobacterales>Enterobacteriaceae>Klebsiella/Raoultella_group>Klebsiella>Klebsiella_pneumoniae_complex | 449 | MCW21_004538 | 49639..50988 | 573 | TPA:-phosphatidylserine/phosphatidylglycerophosphate/cardiolipin-synthase-family-protein-[Klebsiella-pneumoniae] |
| HBV1956332.1 | HKD-DNase+OGG-like | GCA_022126105.1 | Klebsiella_pneumoniae | Bacteria>Pseudomonadati>Pseudomonadota>Gammaproteobacteria>Enterobacterales>Enterobacteriaceae>Klebsiella/Raoultella_group>Klebsiella>Klebsiella_pneumoniae_complex | 449 | MDQ85_004851 | 14654..16003 | 573 | TPA:-phosphatidylserine/phosphatidylglycerophosphate/cardiolipin-synthase-family-protein-[Klebsiella-pneumoniae] |
| HBW3476171.1 | HKD-DNase+OGG-like | GCA_022184305.1 | Klebsiella_pneumoniae | Bacteria>Pseudomonadati>Pseudomonadota>Gammaproteobacteria>Enterobacterales>Enterobacteriaceae>Klebsiella/Raoultella_group>Klebsiella>Klebsiella_pneumoniae_complex | 449 | MEZ45_004506 | 70745..72094 | 573 | TPA:-phosphatidylserine/phosphatidylglycerophosphate/cardiolipin-synthase-family-protein-[Klebsiella-pneumoniae] |
| HBW3608490.1 | HKD-DNase+OGG-like | GCA_022184865.1 | Klebsiella_pneumoniae | Bacteria>Pseudomonadati>Pseudomonadota>Gammaproteobacteria>Enterobacterales>Enterobacteriaceae>Klebsiella/Raoultella_group>Klebsiella>Klebsiella_pneumoniae_complex | 449 | MEZ81_004960 | 28404..29753 | 573 | phosphatidylserine/phosphatidylglycerophosphate/cardiolipin-synthase-family-protein-[Klebsiella-pneumoniae] |
| HBW7740803.1 | HKD-DNase+OGG-like | GCA_022200625.1 | Klebsiella_pneumoniae | Bacteria>Pseudomonadati>Pseudomonadota>Gammaproteobacteria>Enterobacterales>Enterobacteriaceae>Klebsiella/Raoultella_group>Klebsiella>Klebsiella_pneumoniae_complex | 449 | MFE66_003736 | 28420..29769 | 573 | phosphatidylserine/phosphatidylglycerophosphate/cardiolipin-synthase-family-protein-[Klebsiella-pneumoniae] |
| HBX0790081.1 | HKD-DNase+OGG-like | GCA_022212165.1 | Klebsiella_pneumoniae | Bacteria>Pseudomonadati>Pseudomonadota>Gammaproteobacteria>Enterobacterales>Enterobacteriaceae>Klebsiella/Raoultella_group>Klebsiella>Klebsiella_pneumoniae_complex | 449 | MF115_18900 | 62336..63685 | 573 | TPA:-phosphatidylserine/phosphatidylglycerophosphate/cardiolipin-synthase-family-protein-[Klebsiella-pneumoniae] |
| HBX5767908.1 | HKD-DNase+OGG-like | GCA_022248915.1 | Klebsiella_pneumoniae | Bacteria>Pseudomonadati>Pseudomonadota>Gammaproteobacteria>Enterobacterales>Enterobacteriaceae>Klebsiella/Raoultella_group>Klebsiella>Klebsiella_pneumoniae_complex | 635 | MH553_25850 | 7072..8979 | 573 | hypothetical-protein-[Klebsiella-pneumoniae] |
| HBY1687641.1 | HKD-DNase+OGG-like | GCA_022272455.1 | Klebsiella_pneumoniae | Bacteria>Pseudomonadati>Pseudomonadota>Gammaproteobacteria>Enterobacterales>Enterobacteriaceae>Klebsiella/Raoultella_group>Klebsiella>Klebsiella_pneumoniae_complex | 449 | MIQ44_23645 | 66501..67850 | 573 | TPA:-phosphatidylserine/phosphatidylglycerophosphate/cardiolipin-synthase-family-protein-[Klebsiella-pneumoniae] |
| HBZ8012779.1 | HKD-DNase+OGG-like | GCA_022500545.1 | Klebsiella_pneumoniae | Bacteria>Pseudomonadati>Pseudomonadota>Gammaproteobacteria>Enterobacterales>Enterobacteriaceae>Klebsiella/Raoultella_group>Klebsiella>Klebsiella_pneumoniae_complex | 449 | MM158_002945 | 28759..30108 | 573 | phosphatidylserine/phosphatidylglycerophosphate/cardiolipin-synthase-family-protein-[Klebsiella-pneumoniae] |
| HCA0942646.1 | HKD-DNase+OGG-like | GCA_022528725.1 | Klebsiella_pneumoniae | Bacteria>Pseudomonadati>Pseudomonadota>Gammaproteobacteria>Enterobacterales>Enterobacteriaceae>Klebsiella/Raoultella_group>Klebsiella>Klebsiella_pneumoniae_complex | 449 | MNJ11_004339 | 29609..30958 | 573 | TPA:-phosphatidylserine/phosphatidylglycerophosphate/cardiolipin-synthase-family-protein-[Klebsiella-pneumoniae] |
| HCI5816637.1 | HKD-DNase+OGG-like | GCA_024431185.1 | Klebsiella_pneumoniae | Bacteria>Pseudomonadati>Pseudomonadota>Gammaproteobacteria>Enterobacterales>Enterobacteriaceae>Klebsiella/Raoultella_group>Klebsiella>Klebsiella_pneumoniae_complex | 635 | NPN32_003715 | 22633..24540 | 573 | TPA:-hypothetical-protein-NPN32_003715-[Klebsiella-pneumoniae] |
| MBL3233192.1 | HKD-DNase+OGG-like | GCA_016755095.1 | Klebsiella_pneumoniae | Bacteria>Pseudomonadati>Pseudomonadota>Gammaproteobacteria>Enterobacterales>Enterobacteriaceae>Klebsiella/Raoultella_group>Klebsiella>Klebsiella_pneumoniae_complex | 449 | ELP31_24675 | 195..1544 | 573 | phosphatidylserine/phosphatidylglycerophosphate/cardiolipin-synthase-family-protein-[Klebsiella-pneumoniae] |
| MCP6627034.1 | HKD-DNase+OGG-like | GCA_024240675.1 | Klebsiella_pneumoniae | Bacteria>Pseudomonadati>Pseudomonadota>Gammaproteobacteria>Enterobacterales>Enterobacteriaceae>Klebsiella/Raoultella_group>Klebsiella>Klebsiella_pneumoniae_complex | 449 | NL507_17470 | 178263..179612 | 573 | phospholipase-D-like-domain-containing-protein-[Klebsiella-pneumoniae] |
| QJK24917.1 | HKD-DNase+OGG-like | GCA_012967125.1 | Klebsiella_pneumoniae | Bacteria>Pseudomonadati>Pseudomonadota>Gammaproteobacteria>Enterobacterales>Enterobacteriaceae>Klebsiella/Raoultella_group>Klebsiella>Klebsiella_pneumoniae_complex | 449 | HJX28_24020 | 4760422..4761771 | 573 | phosphatidylserine/phosphatidylglycerophosphate/cardiolipin-synthase-family-protein-[Klebsiella-pneumoniae] |
| SYD76206.1 | HKD-DNase+OGG-like | GCA_900514375.1 | Klebsiella_pneumoniae | Bacteria>Pseudomonadati>Pseudomonadota>Gammaproteobacteria>Enterobacterales>Enterobacteriaceae>Klebsiella/Raoultella_group>Klebsiella>Klebsiella_pneumoniae_complex | 395 | SAMEA3515241_04812 | 19900..21087 | 573 | Predicted-HKD-family-nuclease-[Klebsiella-pneumoniae] |
| VTO17825.1 | HKD-DNase+OGG-like | GCA_901421915.1 | Klebsiella_variicola | Bacteria>Pseudomonadati>Pseudomonadota>Gammaproteobacteria>Enterobacterales>Enterobacteriaceae>Klebsiella/Raoultella_group>Klebsiella>Klebsiella_pneumoniae_complex | 189 | NCTC9178_02475 | 2076684..2077253 | 244366 | Uncharacterised-protein-[Klebsiella-variicola] |
| VTO17828.1 | HKD-DNase+OGG-like | GCA_901421915.1 | Klebsiella_variicola | Bacteria>Pseudomonadati>Pseudomonadota>Gammaproteobacteria>Enterobacterales>Enterobacteriaceae>Klebsiella/Raoultella_group>Klebsiella>Klebsiella_pneumoniae_complex | 188 | NCTC9178_02476 | 2077250..2077816 | 244366 | Uncharacterised-protein-[Klebsiella-variicola] |
| WP_032409631.1 | HKD-DNase+OGG-like | GCF_040365185.1 | Klebsiella_pneumoniae | Bacteria>Pseudomonadati>Pseudomonadota>Gammaproteobacteria>Enterobacterales>Enterobacteriaceae>Klebsiella/Raoultella_group>Klebsiella>Klebsiella_pneumoniae_complex | 449 | PXW84_RS24995 | 24647..25996 | 573 | phospholipase-D-like-domain-containing-protein-[Klebsiella-pneumoniae] |
| WP_038432200.1 | HKD-DNase+OGG-like | GCF_902161325.1 | Klebsiella_pneumoniae | Bacteria>Pseudomonadati>Pseudomonadota>Gammaproteobacteria>Enterobacterales>Enterobacteriaceae>Klebsiella/Raoultella_group>Klebsiella>Klebsiella_pneumoniae_complex | 449 | GVM87_RS22950 | 19335..20684 | 573 | phospholipase-D-like-domain-containing-protein-[Klebsiella-pneumoniae] |
| WP_048334006.1 | HKD-DNase+OGG-like | GCF_004313665.1 | Klebsiella_pneumoniae | Bacteria>Pseudomonadati>Pseudomonadota>Gammaproteobacteria>Enterobacterales>Enterobacteriaceae>Klebsiella/Raoultella_group>Klebsiella>Klebsiella_pneumoniae_complex | 449 | KFB18_RS20820 | 24424..25773 | 573 | phospholipase-D-like-domain-containing-protein-[Klebsiella-pneumoniae] |
| WP_064171381.1 | HKD-DNase+OGG-like | GCF_900084795.1 | Klebsiella_variicola | Bacteria>Pseudomonadati>Pseudomonadota>Gammaproteobacteria>Enterobacterales>Enterobacteriaceae>Klebsiella/Raoultella_group>Klebsiella>Klebsiella_pneumoniae_complex | 449 | GJ687_RS21850 | 29927..31276 | 244366 | phospholipase-D-like-domain-containing-protein-[Klebsiella-variicola] |
| WP_085803386.1 | HKD-DNase+OGG-like | GCF_900774735.1 | Klebsiella_pneumoniae | Bacteria>Pseudomonadati>Pseudomonadota>Gammaproteobacteria>Enterobacterales>Enterobacteriaceae>Klebsiella/Raoultella_group>Klebsiella>Klebsiella_pneumoniae_complex | 449 | EF470_RS14865 | 6908..8257 | 573 | phospholipase-D-like-domain-containing-protein-[Klebsiella-pneumoniae] |
| WP_087652993.1 | HKD-DNase+OGG-like | GCF_900978435.1 | Klebsiella_variicola | Bacteria>Pseudomonadati>Pseudomonadota>Gammaproteobacteria>Enterobacterales>Enterobacteriaceae>Klebsiella/Raoultella_group>Klebsiella>Klebsiella_pneumoniae_complex | 449 | E9559_RS11295 | 74057..75406 | 244366 | phospholipase-D-like-domain-containing-protein-[Klebsiella-variicola] |

| Protein-ID | Domain | Assembly-ID | Strain | Taxonomy | Length | Locus-Tag | Genomic-coordinates | TaxID | Fasta-defline |
|---|---|---|---|---|---|---|---|---|---|
| WP_094487776.1 | HKD-DNase+OGG-like | GCF_900775555.1 | Klebsiella_pneumoniae | Bacteria>Pseudomonadati>Pseudomonadota>Gammaproteobacteria>Enterobacterales>Enterobacteriaceae>Klebsiella/Raoultella_group>Klebsiella>Klebsiella_pneumoniae_complex | 449 | CI667_RS22355 | 4596668..4598017 | 573 | phospholipase-D-like-domain-containing-protein-[Klebsiella-pneumoniae] |
| WP_110245494.1 | HKD-DNase+OGG-like | GCF_901482415.1 | Klebsiella_pneumoniae | Bacteria>Pseudomonadati>Pseudomonadota>Gammaproteobacteria>Enterobacterales>Enterobacteriaceae>Klebsiella/Raoultella_group>Klebsiella>Klebsiella_pneumoniae_complex | 449 | P3L21_RS14230 | 49707..51056 | 573 | phospholipase-D-like-domain-containing-protein-[Klebsiella-pneumoniae] |
| WP_115658223.1 | HKD-DNase+OGG-like | GCF_900498955.1 | Klebsiella_pneumoniae | Bacteria>Pseudomonadati>Pseudomonadota>Gammaproteobacteria>Enterobacterales>Enterobacteriaceae>Klebsiella/Raoultella_group>Klebsiella>Klebsiella_pneumoniae_complex | 449 | DY137_RS22240 | 313126..314475 | 573 | phospholipase-D-like-domain-containing-protein-[Klebsiella-pneumoniae] |
| WP_117041988.1 | HKD-DNase+OGG-like | GCF_900775385.1 | Klebsiella_pneumoniae | Bacteria>Pseudomonadati>Pseudomonadota>Gammaproteobacteria>Enterobacterales>Enterobacteriaceae>Klebsiella/Raoultella_group>Klebsiella>Klebsiella_pneumoniae_complex | 449 | MC274_RS12685 | 199345..200694 | 573 | phospholipase-D-like-domain-containing-protein-[Klebsiella-pneumoniae] |
| WP_117104233.1 | HKD-DNase+OGG-like | GCF_039604935.1 | Klebsiella_pneumoniae | Bacteria>Pseudomonadati>Pseudomonadota>Gammaproteobacteria>Enterobacterales>Enterobacteriaceae>Klebsiella/Raoultella_group>Klebsiella>Klebsiella_pneumoniae_complex | 449 | D2Z44_RS21360 | 226496..227845 | 573 | phospholipase-D-like-domain-containing-protein-[Klebsiella-pneumoniae] |
| WP_117133596.1 | HKD-DNase+OGG-like | GCF_964207705.1 | Klebsiella_variicola | Bacteria>Pseudomonadati>Pseudomonadota>Gammaproteobacteria>Enterobacterales>Enterobacteriaceae>Klebsiella/Raoultella_group>Klebsiella>Klebsiella_pneumoniae_complex | 449 | D2R55_RS21925 | 221402..222751 | 244366 | phospholipase-D-like-domain-containing-protein-[Klebsiella-variicola] |
| WP_131033945.1 | HKD-DNase+OGG-like | GCF_004312565.1 | Klebsiella_variicola | Bacteria>Pseudomonadati>Pseudomonadota>Gammaproteobacteria>Enterobacterales>Enterobacteriaceae>Klebsiella/Raoultella_group>Klebsiella>Klebsiella_pneumoniae_complex | 449 | EON46_RS23985 | 96811..98160 | 244366 | phospholipase-D-like-domain-containing-protein-[Klebsiella-variicola] |
| WP_169546891.1 | HKD-DNase+OGG-like | GCF_015290145.1 | Klebsiella_pneumoniae | Bacteria>Pseudomonadati>Pseudomonadota>Gammaproteobacteria>Enterobacterales>Enterobacteriaceae>Klebsiella/Raoultella_group>Klebsiella>Klebsiella_pneumoniae_complex | 449 | KQ276_RS10505 | 169567..170916 | 573 | phospholipase-D-like-domain-containing-protein-[Klebsiella-pneumoniae] |
| WP_200794275.1 | HKD-DNase+OGG-like | GCF_016635835.1 | Klebsiella_pneumoniae | Bacteria>Pseudomonadati>Pseudomonadota>Gammaproteobacteria>Enterobacterales>Enterobacteriaceae>Klebsiella/Raoultella_group>Klebsiella>Klebsiella_pneumoniae_complex | 449 | HZ910_RS23430 | 28407..29756 | 573 | phospholipase-D-like-domain-containing-protein-[Klebsiella-pneumoniae] |
| WP_223364326.1 | HKD-DNase+OGG-like | GCF_040365915.1 | Klebsiella_pneumoniae | Bacteria>Pseudomonadati>Pseudomonadota>Gammaproteobacteria>Enterobacterales>Enterobacteriaceae>Klebsiella/Raoultella_group>Klebsiella>Klebsiella_pneumoniae_complex | 276 | LF666_RS16225 | 24522..25352 | 573 | phospholipase-D-like-domain-containing-protein-[Klebsiella-pneumoniae] |
| WP_251347546.1 | HKD-DNase+OGG-like | GCF_023703955.1 | Klebsiella_pneumoniae | Bacteria>Pseudomonadati>Pseudomonadota>Gammaproteobacteria>Enterobacterales>Enterobacteriaceae>Klebsiella/Raoultella_group>Klebsiella>Klebsiella_pneumoniae_complex | 449 | KU670_RS11570 | 2412218..2413567 | 573 | phospholipase-D-like-domain-containing-protein-[Klebsiella-pneumoniae] |
| WP_278254849.1 | HKD-DNase+OGG-like | - | Klebsiella_variicola | Bacteria>Pseudomonadati>Pseudomonadota>Gammaproteobacteria>Enterobacterales>Enterobacteriaceae>Klebsiella/Raoultella_group>Klebsiella>Klebsiella_pneumoniae_complex | 622 | GSE35_RS20730 | 53238..55106 | 244366 | hypothetical-protein-[Klebsiella-variicola] |
| HBR1265007.1 | HKD-DNase+OGG-like | GCA_021909255.1 | Klebsiella_quasipneumoniae_subsp._quasipneumoniae | Bacteria>Pseudomonadati>Pseudomonadota>Gammaproteobacteria>Enterobacterales>Enterobacteriaceae>Klebsiella/Raoultella_group>Klebsiella>Klebsiella_pneumoniae_complex>Klebsiella_quasipneumoniae | 395 | L9L24_003170 | 24735..25922 | 1667327 | phosphatidylserine/phosphatidylglycerophosphate/cardiolipin-synthase-family-protein-[Klebsiella-quasipneumoniae-subsp.-quasipneumoniae] |
| HCB0434652.1 | HKD-DNase+OGG-like | GCA_023030065.1 | Klebsiella_variicola_subsp._variicola | Bacteria>Pseudomonadati>Pseudomonadota>Gammaproteobacteria>Enterobacterales>Enterobacteriaceae>Klebsiella/Raoultella_group>Klebsiella>Klebsiella_pneumoniae_complex>Klebsiella_variicola | 449 | MYM82_004340 | 30973..32322 | 2590157 | phosphatidylserine/phosphatidylglycerophosphate/cardiolipin-synthase-family-protein-[Klebsiella-variicola-subsp.-variicola] |
| WP_144838639.1 | HKD-DNase+OGG-like | GCF_007680555.1 | Leclercia_adecarboxylata | Bacteria>Pseudomonadati>Pseudomonadota>Gammaproteobacteria>Enterobacterales>Enterobacteriaceae>Leclercia | 449 | FS784_RS12395 | 131585..132934 | 83655 | phospholipase-D-like-domain-containing-protein-[Leclercia-adecarboxylata] |
| EAA8875711.1 | HKD-DNase+OGG-like | GCA_004223005.1 | Salmonella_enterica | Bacteria>Pseudomonadati>Pseudomonadota>Gammaproteobacteria>Enterobacterales>Enterobacteriaceae>Salmonella | 449 | CIU45_09655 | 395895..397244 | 28901 | phosphatidylserine/phosphatidylglycerophosphate/cardiolipin-synthase-family-protein-[Salmonella-enterica] |
| EAB2831584.1 | HKD-DNase+OGG-like | GCA_004240605.1 | Salmonella_enterica | Bacteria>Pseudomonadati>Pseudomonadota>Gammaproteobacteria>Enterobacterales>Enterobacteriaceae>Salmonella | 449 | EAV48_05300 | 171974..173323 | 28901 | phosphatidylserine/phosphatidylglycerophosphate/cardiolipin-synthase-family-protein-[Salmonella-enterica] |
| EAM3013488.1 | HKD-DNase+OGG-like | GCA_005412095.1 | Salmonella_enterica | Bacteria>Pseudomonadati>Pseudomonadota>Gammaproteobacteria>Enterobacterales>Enterobacteriaceae>Salmonella | 449 | CBJ93_01895 | 408686..410035 | 28901 | phosphatidylserine/phosphatidylglycerophosphate/cardiolipin-synthase-family-protein-[Salmonella-enterica] |
| EAM3141095.1 | HKD-DNase+OGG-like | GCA_005412435.1 | Salmonella_enterica | Bacteria>Pseudomonadati>Pseudomonadota>Gammaproteobacteria>Enterobacterales>Enterobacteriaceae>Salmonella | 449 | DMT78_01440 | 296264..297613 | 28901 | phosphatidylserine/phosphatidylglycerophosphate/cardiolipin-synthase-family-protein-[Salmonella-enterica] |
| EAT1639069.1 | HKD-DNase+OGG-like | GCA_005720135.1 | Salmonella_enterica | Bacteria>Pseudomonadati>Pseudomonadota>Gammaproteobacteria>Enterobacterales>Enterobacteriaceae>Salmonella | 449 | EKF56_11770 | 39090..40439 | 28901 | phosphatidylserine/phosphatidylglycerophosphate/cardiolipin-synthase-family-protein-[Salmonella-enterica] |
| EAT2430713.1 | HKD-DNase+OGG-like | GCA_005726115.1 | Salmonella_enterica | Bacteria>Pseudomonadati>Pseudomonadota>Gammaproteobacteria>Enterobacterales>Enterobacteriaceae>Salmonella | 449 | EI207_04355 | 191214..192563 | 28901 | phosphatidylserine/phosphatidylglycerophosphate/cardiolipin-synthase-family-protein-[Salmonella-enterica] |
| EAW3206013.1 | HKD-DNase+OGG-like | GCA_005905825.1 | Salmonella_enterica | Bacteria>Pseudomonadati>Pseudomonadota>Gammaproteobacteria>Enterobacterales>Enterobacteriaceae>Salmonella | 449 | FEL99_04720 | 221290..222639 | 28901 | phosphatidylserine/phosphatidylglycerophosphate/cardiolipin-synthase-family-protein-[Salmonella-enterica] |
| EAW9396393.1 | HKD-DNase+OGG-like | GCA_005940865.1 | Salmonella_enterica | Bacteria>Pseudomonadati>Pseudomonadota>Gammaproteobacteria>Enterobacterales>Enterobacteriaceae>Salmonella | 449 | QE38_19765 | 62208..63557 | 28901 | phosphatidylserine/phosphatidylglycerophosphate/cardiolipin-synthase-family-protein-[Salmonella-enterica] |
| EAY5508624.1 | HKD-DNase+OGG-like | GCA_006012725.1 | Salmonella_enterica | Bacteria>Pseudomonadati>Pseudomonadota>Gammaproteobacteria>Enterobacterales>Enterobacteriaceae>Salmonella | 449 | NE47_11045 | 63209..64558 | 28901 | phosphatidylserine/phosphatidylglycerophosphate/cardiolipin-synthase-family-protein-[Salmonella-enterica] |
| EAZ9311967.1 | HKD-DNase+OGG-like | GCA_006073635.1 | Salmonella_enterica | Bacteria>Pseudomonadati>Pseudomonadota>Gammaproteobacteria>Enterobacterales>Enterobacteriaceae>Salmonella | 449 | CBJ28_03055 | 172759..174108 | 28901 | phosphatidylserine/phosphatidylglycerophosphate/cardiolipin-synthase-family-protein-[Salmonella-enterica] |
| EBF4455785.1 | HKD-DNase+OGG-like | GCA_006343165.1 | Salmonella_enterica | Bacteria>Pseudomonadati>Pseudomonadota>Gammaproteobacteria>Enterobacterales>Enterobacteriaceae>Salmonella | 449 | FH443_09935 | 425918..427267 | 28901 | phosphatidylserine/phosphatidylglycerophosphate/cardiolipin-synthase-family-protein-[Salmonella-enterica] |
| EBM6187526.1 | HKD-DNase+OGG-like | GCA_006738745.1 | Salmonella_enterica | Bacteria>Pseudomonadati>Pseudomonadota>Gammaproteobacteria>Enterobacterales>Enterobacteriaceae>Salmonella | 449 | D1943_04260 | 114648..115997 | 28901 | phosphatidylserine/phosphatidylglycerophosphate/cardiolipin-synthase-family-protein-[Salmonella-enterica] |
| EBR5149899.1 | HKD-DNase+OGG-like | GCA_006958505.1 | Salmonella_enterica | Bacteria>Pseudomonadati>Pseudomonadota>Gammaproteobacteria>Enterobacterales>Enterobacteriaceae>Salmonella | 449 | B5W64_09875 | 364612..365961 | 28901 | phosphatidylserine/phosphatidylglycerophosphate/cardiolipin-synthase-family-protein-[Salmonella-enterica] |
| ECG9327781.1 | HKD-DNase+OGG-like | GCA_007700105.1 | Salmonella_enterica | Bacteria>Pseudomonadati>Pseudomonadota>Gammaproteobacteria>Enterobacterales>Enterobacteriaceae>Salmonella | 449 | FO199_03385 | 120069..121418 | 28901 | phosphatidylserine/phosphatidylglycerophosphate/cardiolipin-synthase-family-protein-[Salmonella-enterica] |
| ECJ6092452.1 | HKD-DNase+OGG-like | GCA_007834095.1 | Salmonella_enterica | Bacteria>Pseudomonadati>Pseudomonadota>Gammaproteobacteria>Enterobacterales>Enterobacteriaceae>Salmonella | 449 | GR78_06790 | 386416..387765 | 28901 | phosphatidylserine/phosphatidylglycerophosphate/cardiolipin-synthase-family-protein-[Salmonella-enterica] |
| EDB0845248.1 | HKD-DNase+OGG-like | - | Salmonella_enterica | Bacteria>Pseudomonadati>Pseudomonadota>Gammaproteobacteria>Enterobacterales>Enterobacteriaceae>Salmonella | 449 | F9C59_12705 | 156773..158122 | 28901 | phosphatidylserine/phosphatidylglycerophosphate/cardiolipin-synthase-family-protein-[Salmonella-enterica] |
| EDX0369809.1 | HKD-DNase+OGG-like | - | Salmonella_enterica | Bacteria>Pseudomonadati>Pseudomonadota>Gammaproteobacteria>Enterobacterales>Enterobacteriaceae>Salmonella | 449 | GTL21_000367 | 400612..401961 | 28901 | phosphatidylserine/phosphatidylglycerophosphate/cardiolipin-synthase-family-protein-[Salmonella-enterica] |
| EDZ0760568.1 | HKD-DNase+OGG-like | - | Salmonella_enterica | Bacteria>Pseudomonadati>Pseudomonadota>Gammaproteobacteria>Enterobacterales>Enterobacteriaceae>Salmonella | 449 | GPN34_002366 | 424464..425813 | 28901 | phosphatidylserine/phosphatidylglycerophosphate/cardiolipin-synthase-family-protein-[Salmonella-enterica] |
| EHO5648346.1 | HKD-DNase+OGG-like | GCA_018535125.1 | Salmonella_enterica | Bacteria>Pseudomonadati>Pseudomonadota>Gammaproteobacteria>Enterobacterales>Enterobacteriaceae>Salmonella | 449 | KLR08_000694 | 156773..158122 | 28901 | phosphatidylserine/phosphatidylglycerophosphate/cardiolipin-synthase-family-protein-[Salmonella-enterica] |
| EHS4662095.1 | HKD-DNase+OGG-like | GCA_019144395.1 | Salmonella_enterica | Bacteria>Pseudomonadati>Pseudomonadota>Gammaproteobacteria>Enterobacterales>Enterobacteriaceae>Salmonella | 449 | KV604_003668 | 61761..63110 | 28901 | phosphatidylserine/phosphatidylglycerophosphate/cardiolipin-synthase-family-protein-[Salmonella-enterica] |
| EHU7518558.1 | HKD-DNase+OGG-like | GCA_019318345.1 | Salmonella_enterica | Bacteria>Pseudomonadati>Pseudomonadota>Gammaproteobacteria>Enterobacterales>Enterobacteriaceae>Salmonella | 449 | KZL07_001989 | 287050..288399 | 28901 | phosphatidylserine/phosphatidylglycerophosphate/cardiolipin-synthase-family-protein-[Salmonella-enterica] |
| EHX4782369.1 | HKD-DNase+OGG-like | GCA_019533715.1 | Salmonella_enterica | Bacteria>Pseudomonadati>Pseudomonadota>Gammaproteobacteria>Enterobacterales>Enterobacteriaceae>Salmonella | 449 | K3Y02_002644 | 156596..157945 | 28901 | phosphatidylserine/phosphatidylglycerophosphate/cardiolipin-synthase-family-protein-[Salmonella-enterica] |
| EJC3638312.1 | HKD-DNase+OGG-like | GCA_023047235.1 | Salmonella_enterica | Bacteria>Pseudomonadati>Pseudomonadota>Gammaproteobacteria>Enterobacterales>Enterobacteriaceae>Salmonella | 449 | MYX88_003399 | 165171..166520 | 28901 | phosphatidylserine/phosphatidylglycerophosphate/cardiolipin-synthase-family-protein-[Salmonella-enterica] |
| MBH5277078.1 | HKD-DNase+OGG-like | GCA_016019715.1 | Salmonella_enterica | Bacteria>Pseudomonadati>Pseudomonadota>Gammaproteobacteria>Enterobacterales>Enterobacteriaceae>Salmonella | 449 | I3A81_01620 | 346852..348201 | 28901 | phosphatidylserine/phosphatidylglycerophosphate/cardiolipin-synthase-family-protein-[Salmonella-enterica] |
| WP_001523588.1 | HKD-DNase+OGG-like | GCF_000170235.1 | Salmonella_enterica | Bacteria>Pseudomonadati>Pseudomonadota>Gammaproteobacteria>Enterobacterales>Enterobacteriaceae>Salmonella | 449 | SN48_RS13455 | 32202..33551 | 28901 | phospholipase-D-like-domain-containing-protein-[Salmonella-enterica] |
| WP_071055532.1 | HKD-DNase+OGG-like | GCF_040956775.1 | Salmonella_enterica | Bacteria>Pseudomonadati>Pseudomonadota>Gammaproteobacteria>Enterobacterales>Enterobacteriaceae>Salmonella | 449 | SES26_RS19345 | 4023535..4024884 | 28901 | phospholipase-D-like-domain-containing-protein-[Salmonella-enterica] |
| WP_079808683.1 | HKD-DNase+OGG-like | GCF_002035875.1 | Salmonella_enterica | Bacteria>Pseudomonadati>Pseudomonadota>Gammaproteobacteria>Enterobacterales>Enterobacteriaceae>Salmonella | 449 | Y985_RS08700 | 108269..109618 | 28901 | phospholipase-D-like-domain-containing-protein-[Salmonella-enterica] |
| WP_079897966.1 | HKD-DNase+OGG-like | GCF_051548365.1 | Salmonella_enterica | Bacteria>Pseudomonadati>Pseudomonadota>Gammaproteobacteria>Enterobacterales>Enterobacteriaceae>Salmonella | 449 | VY62_RS13035 | 170176..171525 | 28901 | phospholipase-D-like-domain-containing-protein-[Salmonella-enterica] |
| WP_079965460.1 | HKD-DNase+OGG-like | GCF_002046745.1 | Salmonella_enterica | Bacteria>Pseudomonadati>Pseudomonadota>Gammaproteobacteria>Enterobacterales>Enterobacteriaceae>Salmonella | 449 | VX54_RS10680 | 66258..67607 | 28901 | phospholipase-D-like-domain-containing-protein-[Salmonella-enterica] |
| WP_080250289.1 | HKD-DNase+OGG-like | GCF_037083205.1 | Salmonella_enterica | Bacteria>Pseudomonadati>Pseudomonadota>Gammaproteobacteria>Enterobacterales>Enterobacteriaceae>Salmonella | 449 | RZ13_RS02325 | 178861..180210 | 28901 | phospholipase-D-like-domain-containing-protein-[Salmonella-enterica] |
| WP_086805520.1 | HKD-DNase+OGG-like | GCF_002156225.1 | Salmonella_enterica | Bacteria>Pseudomonadati>Pseudomonadota>Gammaproteobacteria>Enterobacterales>Enterobacteriaceae>Salmonella | 449 | CCL33_RS10700 | 40611..41960 | 28901 | phospholipase-D-like-domain-containing-protein-[Salmonella-enterica] |
| WP_218201155.1 | HKD-DNase+OGG-like | GCF_019173605.1 | Salmonella_enterica | Bacteria>Pseudomonadati>Pseudomonadota>Gammaproteobacteria>Enterobacterales>Enterobacteriaceae>Salmonella | 395 | KXD81_RS02490 | 10209..11396 | 28901 | phospholipase-D-like-domain-containing-protein-[Salmonella-enterica] |
| WP_235864754.1 | HKD-DNase+OGG-like | GCF_002034005.1 | Salmonella_enterica | Bacteria>Pseudomonadati>Pseudomonadota>Gammaproteobacteria>Enterobacterales>Enterobacteriaceae>Salmonella | 389 | AJH59_RS23260 | 71..1240 | 28901 | phospholipase-D-like-domain-containing-protein-[Salmonella-enterica] |
| EAB6843531.1 | HKD-DNase+OGG-like | GCA_004258725.1 | Salmonella_enterica_subsp._salamae | Bacteria>Pseudomonadati>Pseudomonadota>Gammaproteobacteria>Enterobacterales>Enterobacteriaceae>Salmonella>Salmonella_enterica | 449 | XT66_14215 | 65004..66353 | 59202 | phosphatidylserine/phosphatidylglycerophosphate/cardiolipin-synthase-family-protein-[Salmonella-enterica-subsp.-salamae] |
| EAB9078802.1 | HKD-DNase+OGG-like | GCA_004265565.1 | Salmonella_enterica_subsp._enterica | Bacteria>Pseudomonadati>Pseudomonadota>Gammaproteobacteria>Enterobacterales>Enterobacteriaceae>Salmonella>Salmonella_enterica | 449 | AAL03_01540 | 66438..67787 | 59201 | phosphatidylserine/phosphatidylglycerophosphate/cardiolipin-synthase-family-protein-[Salmonella-enterica-subsp.-enterica] |
| ECC1667305.1 | HKD-DNase+OGG-like | GCA_007463295.1 | Salmonella_enterica_subsp._salamae | Bacteria>Pseudomonadati>Pseudomonadota>Gammaproteobacteria>Enterobacterales>Enterobacteriaceae>Salmonella>Salmonella_enterica | 449 | FNN95_01285 | 281914..283263 | 59202 | phosphatidylserine/phosphatidylglycerophosphate/cardiolipin-synthase-family-protein-[Salmonella-enterica-subsp.-salamae] |
| ECD9608668.1 | HKD-DNase+OGG-like | GCA_007542335.1 | Salmonella_enterica_subsp._salamae | Bacteria>Pseudomonadati>Pseudomonadota>Gammaproteobacteria>Enterobacterales>Enterobacteriaceae>Salmonella>Salmonella_enterica | 449 | FNZ18_04590 | 270588..271937 | 59202 | phosphatidylserine/phosphatidylglycerophosphate/cardiolipin-synthase-family-protein-[Salmonella-enterica-subsp.-salamae] |
| ECG0705021.1 | HKD-DNase+OGG-like | GCA_007641515.1 | Salmonella_enterica_subsp._salamae | Bacteria>Pseudomonadati>Pseudomonadota>Gammaproteobacteria>Enterobacterales>Enterobacteriaceae>Salmonella>Salmonella_enterica | 395 | DL122_04730 | 49775..50962 | 59202 | phosphatidylserine/phosphatidylglycerophosphate/cardiolipin-synthase-family-protein-[Salmonella-enterica-subsp.-salamae] |
| EDT7223679.1 | HKD-DNase+OGG-like | - | Salmonella_enterica_subsp._enterica | Bacteria>Pseudomonadati>Pseudomonadota>Gammaproteobacteria>Enterobacterales>Enterobacteriaceae>Salmonella>Salmonella_enterica | 449 | AH268_003340 | 183089..184438 | 59201 | phosphatidylserine/phosphatidylglycerophosphate/cardiolipin-synthase-family-protein-[Salmonella-enterica-subsp.-enterica] |
| EDV5116996.1 | HKD-DNase+OGG-like | - | Salmonella_enterica_subsp._enterica | Bacteria>Pseudomonadati>Pseudomonadota>Gammaproteobacteria>Enterobacterales>Enterobacteriaceae>Salmonella>Salmonella_enterica | 449 | ZJ36_000831 | 42987..44336 | 59201 | phosphatidylserine/phosphatidylglycerophosphate/cardiolipin-synthase-family-protein-[Salmonella-enterica-subsp.-enterica] |
| EDV9109404.1 | HKD-DNase+OGG-like | - | Salmonella_enterica_subsp._enterica | Bacteria>Pseudomonadati>Pseudomonadota>Gammaproteobacteria>Enterobacterales>Enterobacteriaceae>Salmonella>Salmonella_enterica | 449 | ZK68_001606 | 66064..67413 | 59201 | phosphatidylserine/phosphatidylglycerophosphate/cardiolipin-synthase-family-protein-[Salmonella-enterica-subsp.-enterica] |
| EDW4488621.1 | HKD-DNase+OGG-like | - | Salmonella_enterica_subsp._enterica | Bacteria>Pseudomonadati>Pseudomonadota>Gammaproteobacteria>Enterobacterales>Enterobacteriaceae>Salmonella>Salmonella_enterica | 449 | YV39_000493 | 180138..181487 | 59201 | phosphatidylserine/phosphatidylglycerophosphate/cardiolipin-synthase-family-protein-[Salmonella-enterica-subsp.-enterica] |
| EGB5697875.1 | HKD-DNase+OGG-like | - | Salmonella_enterica_subsp._enterica | Bacteria>Pseudomonadati>Pseudomonadota>Gammaproteobacteria>Enterobacterales>Enterobacteriaceae>Salmonella>Salmonella_enterica | 449 | H7531_003065 | 5775..7124 | 59201 | phosphatidylserine/phosphatidylglycerophosphate/cardiolipin-synthase-family-protein-[Salmonella-enterica-subsp.-enterica] |
| AZT73897.1 | HKD-DNase+OGG-like | GCA_003994735.1 | Salmonella_enterica_subsp._enterica_serovar_Waycross | Bacteria>Pseudomonadati>Pseudomonadota>Gammaproteobacteria>Enterobacterales>Enterobacteriaceae>Salmonella>Salmonella_enterica>Salmonella_enterica_subsp._enterica | 449 | ELZ91_19550 | 3965201..3966550 | 58101 | phosphatidylserine/phosphatidylglycerophosphate/cardiolipin-synthase-family-protein-[Salmonella-enterica-subsp.-enterica-serovar-Waycross] |
| EAA1506274.1 | HKD-DNase+OGG-like | GCA_004160295.1 | Salmonella_enterica_subsp._enterica_serovar_Agama | Bacteria>Pseudomonadati>Pseudomonadota>Gammaproteobacteria>Enterobacterales>Enterobacteriaceae>Salmonella>Salmonella_enterica>Salmonella_enterica_subsp._enterica | 449 | DL104_05760 | 43293..44642 | 399581 | phosphatidylserine/phosphatidylglycerophosphate/cardiolipin-synthase-family-protein-[Salmonella-enterica-subsp.-enterica] |
| EAA5580616.1 | HKD-DNase+OGG-like | GCA_004185805.1 | Salmonella_enterica_subsp._enterica_serovar_Glostrup | Bacteria>Pseudomonadati>Pseudomonadota>Gammaproteobacteria>Enterobacterales>Enterobacteriaceae>Salmonella>Salmonella_enterica>Salmonella_enterica_subsp._enterica | 449 | DQD90_06870 | 73003..74352 | 1151180 | phosphatidylserine/phosphatidylglycerophosphate/cardiolipin-synthase-family-protein-[Salmonella-enterica-subsp.-enterica] |
| EBU7641094.1 | HKD-DNase+OGG-like | GCA_007142065.1 | Salmonella_enterica_subsp._enterica_serovar_Brevik | Bacteria>Pseudomonadati>Pseudomonadota>Gammaproteobacteria>Enterobacterales>Enterobacteriaceae>Salmonella>Salmonella_enterica>Salmonella_enterica_subsp._enterica | 449 | DK709_13745 | 73464..74813 | 2564275 | phosphatidylserine/phosphatidylglycerophosphate/cardiolipin-synthase-family-protein-[Salmonella-enterica-subsp.-enterica-serovar-Brevik] |
| EBU9347427.1 | HKD-DNase+OGG-like | GCA_007149485.1 | Salmonella_enterica_subsp._enterica_serovar_Angoda | Bacteria>Pseudomonadati>Pseudomonadota>Gammaproteobacteria>Enterobacterales>Enterobacteriaceae>Salmonella>Salmonella_enterica>Salmonella_enterica_subsp._enterica | 449 | DMU95_04660 | 151705..153054 | 2564159 | phosphatidylserine/phosphatidylglycerophosphate/cardiolipin-synthase-family-protein-[Salmonella-enterica-subsp.-enterica-serovar-Angoda] |
| EBU9775084.1 | HKD-DNase+OGG-like | GCA_007151525.1 | Salmonella_enterica_subsp._enterica_serovar_Kumasi | Bacteria>Pseudomonadati>Pseudomonadota>Gammaproteobacteria>Enterobacterales>Enterobacteriaceae>Salmonella>Salmonella_enterica>Salmonella_enterica_subsp._enterica | 138 | DM304_21675 | 17140..17553 | 2493647 | phosphatidylserine/phosphatidylglycerophosphate/cardiolipin-synthase-family-protein-[Salmonella-enterica-subsp.-enterica-serovar-Kumasi] |
| EBV2735698.1 | HKD-DNase+OGG-like | GCA_007164145.1 | Salmonella_enterica_subsp._enterica_serovar_Stourbridge | Bacteria>Pseudomonadati>Pseudomonadota>Gammaproteobacteria>Enterobacterales>Enterobacteriaceae>Salmonella>Salmonella_enterica>Salmonella_enterica_subsp._enterica | 449 | DOA65_09475 | 73010..74359 | 1160766 | phosphatidylserine/phosphatidylglycerophosphate/cardiolipin-synthase-family-protein-[Salmonella-enterica-subsp.-enterica-serovar-Stourbridge] |
| EBV4609762.1 | HKD-DNase+OGG-like | GCA_007170635.1 | Salmonella_enterica_subsp._enterica_serovar_Solt | Bacteria>Pseudomonadati>Pseudomonadota>Gammaproteobacteria>Enterobacterales>Enterobacteriaceae>Salmonella>Salmonella_enterica>Salmonella_enterica_subsp._enterica | 449 | DOX24_02205 | 17136..18485 | 2565040 | phosphatidylserine/phosphatidylglycerophosphate/cardiolipin-synthase-family-protein-[Salmonella-enterica-subsp.-enterica-serovar-Solt] |
| EBX0519403.1 | HKD-DNase+OGG-like | GCA_007236875.1 | Salmonella_enterica_subsp._enterica_serovar_Saintpaul | Bacteria>Pseudomonadati>Pseudomonadota>Gammaproteobacteria>Enterobacterales>Enterobacteriaceae>Salmonella>Salmonella_enterica>Salmonella_enterica_subsp._enterica | 449 | AHL42_03340 | 156369..157718 | 90105 | phosphatidylserine/phosphatidylglycerophosphate/cardiolipin-synthase-family-protein-[Salmonella-enterica-subsp.-enterica-serovar-Saintpaul] |
| EBY1767821.1 | HKD-DNase+OGG-like | GCA_007283275.1 | Salmonella_enterica_subsp._enterica_serovar_Georgia | Bacteria>Pseudomonadati>Pseudomonadota>Gammaproteobacteria>Enterobacterales>Enterobacteriaceae>Salmonella>Salmonella_enterica>Salmonella_enterica_subsp._enterica | 449 | DU102_16135 | 64690..66039 | 2564467 | phosphatidylserine/phosphatidylglycerophosphate/cardiolipin-synthase-family-protein-[Salmonella-enterica-subsp.-enterica-serovar-Georgia] |
| EBZ6491798.1 | HKD-DNase+OGG-like | GCA_007345015.1 | Salmonella_enterica_subsp._enterica_serovar_Dugbe | Bacteria>Pseudomonadati>Pseudomonadota>Gammaproteobacteria>Enterobacterales>Enterobacteriaceae>Salmonella>Salmonella_enterica>Salmonella_enterica_subsp._enterica | 449 | ED480_20090 | 30913..32262 | 2564376 | phosphatidylserine/phosphatidylglycerophosphate/cardiolipin-synthase-family-protein-[Salmonella-enterica-subsp.-enterica-serovar-Dugbe] |
| ECB3805270.1 | HKD-DNase+OGG-like | GCA_007422765.1 | Salmonella_enterica_subsp._enterica_serovar_Fufu | Bacteria>Pseudomonadati>Pseudomonadota>Gammaproteobacteria>Enterobacterales>Enterobacteriaceae>Salmonella>Salmonella_enterica>Salmonella_enterica_subsp._enterica | 449 | EW367_11390 | 66148..67497 | 2564449 | phosphatidylserine/phosphatidylglycerophosphate/cardiolipin-synthase-family-protein-[Salmonella-enterica-subsp.-enterica-serovar-Fufu] |
| ECD2821741.1 | HKD-DNase+OGG-like | GCA_007513645.1 | Salmonella_enterica_subsp._enterica_serovar_Fanti | Bacteria>Pseudomonadati>Pseudomonadota>Gammaproteobacteria>Enterobacterales>Enterobacteriaceae>Salmonella>Salmonella_enterica>Salmonella_enterica_subsp._enterica | 449 | EXA45_11180 | 66384..67733 | 2564418 | phosphatidylserine/phosphatidylglycerophosphate/cardiolipin-synthase-family-protein-[Salmonella-enterica-subsp.-enterica-serovar-Fanti] |
| ECF0142195.1 | HKD-DNase+OGG-like | GCA_007595955.1 | Salmonella_enterica_subsp._enterica_serovar_Monschaui | Bacteria>Pseudomonadati>Pseudomonadota>Gammaproteobacteria>Enterobacterales>Enterobacteriaceae>Salmonella>Salmonella_enterica>Salmonella_enterica_subsp._enterica | 449 | EX931_14185 | 83474..84823 | 29480 | phosphatidylserine/phosphatidylglycerophosphate/cardiolipin-synthase-family-protein-[Salmonella-enterica-subsp.-enterica-serovar-Monschaui] |
| ECG8301820.1 | HKD-DNase+OGG-like | GCA_007694625.1 | Salmonella_enterica_subsp._enterica_serovar_Glostrup | Bacteria>Pseudomonadati>Pseudomonadota>Gammaproteobacteria>Enterobacterales>Enterobacteriaceae>Salmonella>Salmonella_enterica>Salmonella_enterica_subsp._enterica | 449 | E1853_02165 | 73001..74350 | 1151180 | phosphatidylserine/phosphatidylglycerophosphate/cardiolipin-synthase-family-protein-[Salmonella-enterica-subsp.-enterica-serovar-Glostrup] |
| ECY3255621.1 | HKD-DNase+OGG-like | GCA_008731355.1 | Salmonella_enterica_subsp._enterica_serovar_Alachua | Bacteria>Pseudomonadati>Pseudomonadota>Gammaproteobacteria>Enterobacterales>Enterobacteriaceae>Salmonella>Salmonella_enterica>Salmonella_enterica_subsp._enterica | 449 | AU577_03170 | 30863..32212 | 913240 | phosphatidylserine/phosphatidylglycerophosphate/cardiolipin-synthase-family-protein-[Salmonella-enterica-subsp.-enterica-serovar-Alachua] |
| EDN6133539.1 | HKD-DNase+OGG-like | - | Salmonella_enterica_subsp._enterica_serovar_Kentucky | Bacteria>Pseudomonadati>Pseudomonadota>Gammaproteobacteria>Enterobacterales>Enterobacteriaceae>Salmonella>Salmonella_enterica>Salmonella_enterica_subsp._enterica | 449 | GJQ78_12640 | 76181..77530 | 192955 | phosphatidylserine/phosphatidylglycerophosphate/cardiolipin-synthase-family-protein-[Salmonella-enterica-subsp.-enterica-serovar-Kentucky] |
| EDQ2405067.1 | HKD-DNase+OGG-like | - | Salmonella_enterica_subsp._enterica_serovar_Saintpaul | Bacteria>Pseudomonadati>Pseudomonadota>Gammaproteobacteria>Enterobacterales>Enterobacteriaceae>Salmonella>Salmonella_enterica>Salmonella_enterica_subsp._enterica | 449 | DQ96_001229 | 156773..158122 | 90105 | phosphatidylserine/phosphatidylglycerophosphate/cardiolipin-synthase-family-protein-[Salmonella-enterica-subsp.-enterica-serovar-Saintpaul] |
| EDS3575405.1 | HKD-DNase+OGG-like | - | Salmonella_enterica_subsp._enterica_serovar_Sangera | Bacteria>Pseudomonadati>Pseudomonadota>Gammaproteobacteria>Enterobacterales>Enterobacteriaceae>Salmonella>Salmonella_enterica>Salmonella_enterica_subsp._enterica | 449 | GR179_002524 | 17202..18551 | 1173574 | phosphatidylserine/phosphatidylglycerophosphate/cardiolipin-synthase-family-protein-[Salmonella-enterica-subsp.-enterica] |
| EEE1152708.1 | HKD-DNase+OGG-like | - | Salmonella_enterica_subsp._enterica_serovar_Teko | Bacteria>Pseudomonadati>Pseudomonadota>Gammaproteobacteria>Enterobacterales>Enterobacteriaceae>Salmonella>Salmonella_enterica>Salmonella_enterica_subsp._enterica | 449 | CAE49_001969 | 319039..320388 | 2079198 | phosphatidylserine/phosphatidylglycerophosphate/cardiolipin-synthase-family-protein-[Salmonella-enterica-subsp.-enterica-serovar-Teko] |
| EEE6742968.1 | HKD-DNase+OGG-like | - | Salmonella_enterica_subsp._enterica_serovar_Westhampton | Bacteria>Pseudomonadati>Pseudomonadota>Gammaproteobacteria>Enterobacterales>Enterobacteriaceae>Salmonella>Salmonella_enterica>Salmonella_enterica_subsp._enterica | 449 | CBJ05_002026 | 422868..424217 | 356621 | phosphatidylserine/phosphatidylglycerophosphate/cardiolipin-synthase-family-protein-[Salmonella-enterica-subsp.-enterica-serovar-Westhampton] |
| EEH2707252.1 | HKD-DNase+OGG-like | - | Salmonella_enterica_subsp._enterica_serovar_Lille | Bacteria>Pseudomonadati>Pseudomonadota>Gammaproteobacteria>Enterobacterales>Enterobacteriaceae>Salmonella>Salmonella_enterica>Salmonella_enterica_subsp._enterica | 449 | GYE71_000443 | 68075..69424 | 486997 | phosphatidylserine/phosphatidylglycerophosphate/cardiolipin-synthase-family-protein-[Salmonella-enterica-subsp.-enterica] |
| EEI5362891.1 | HKD-DNase+OGG-like | - | Salmonella_enterica_subsp._enterica_serovar_Kentucky | Bacteria>Pseudomonadati>Pseudomonadota>Gammaproteobacteria>Enterobacterales>Enterobacteriaceae>Salmonella>Salmonella_enterica>Salmonella_enterica_subsp._enterica | 449 | GXM95_001569 | 187176..188525 | 192955 | phosphatidylserine/phosphatidylglycerophosphate/cardiolipin-synthase-family-protein-[Salmonella-enterica-subsp.-enterica-serovar-Kentucky] |
| EEM2455581.1 | HKD-DNase+OGG-like | - | Salmonella_enterica_subsp._enterica_serovar_Kentucky | Bacteria>Pseudomonadati>Pseudomonadota>Gammaproteobacteria>Enterobacterales>Enterobacteriaceae>Salmonella>Salmonella_enterica>Salmonella_enterica_subsp._enterica | 449 | CWG70_02805 | 187509..188858 | 192955 | phosphatidylserine/phosphatidylglycerophosphate/cardiolipin-synthase-family-protein-[Salmonella-enterica-subsp.-enterica-serovar-Kentucky] |
| EEO3564966.1 | HKD-DNase+OGG-like | - | Salmonella_enterica_subsp._enterica_serovar_Poona | Bacteria>Pseudomonadati>Pseudomonadota>Gammaproteobacteria>Enterobacterales>Enterobacteriaceae>Salmonella>Salmonella_enterica>Salmonella_enterica_subsp._enterica | 449 | DEI68_09025 | 66452..67801 | 436295 | phosphatidylserine/phosphatidylglycerophosphate/cardiolipin-synthase-family-protein-[Salmonella-enterica-subsp.-enterica-serovar-Poona] |
| EHJ7117739.1 | HKD-DNase+OGG-like | GCA_017926475.1 | Salmonella_enterica_subsp._enterica_serovar_Kentucky | Bacteria>Pseudomonadati>Pseudomonadota>Gammaproteobacteria>Enterobacterales>Enterobacteriaceae>Salmonella>Salmonella_enterica>Salmonella_enterica_subsp._enterica | 449 | KAP12_002971 | 426132..427481 | 192955 | phosphatidylserine/phosphatidylglycerophosphate/cardiolipin-synthase-family-protein-[Salmonella-enterica-subsp.-enterica-serovar-Kentucky] |
| EIJ6143998.1 | HKD-DNase+OGG-like | GCA_020548845.1 | Salmonella_enterica_subsp._enterica_serovar_4:a:- | Bacteria>Pseudomonadati>Pseudomonadota>Gammaproteobacteria>Enterobacterales>Enterobacteriaceae>Salmonella>Salmonella_enterica>Salmonella_enterica_subsp._enterica | 449 | LI678_001020 | 408794..410143 | 2885150 | phosphatidylserine/phosphatidylglycerophosphate/cardiolipin-synthase-family-protein-[Salmonella-enterica-subsp.-enterica-serovar-4:a:-] |
| UTZ95788.1 | HKD-DNase+OGG-like | GCA_024442275.1 | Salmonella_enterica_subsp._enterica_serovar_Kentucky | Bacteria>Pseudomonadati>Pseudomonadota>Gammaproteobacteria>Enterobacterales>Enterobacteriaceae>Salmonella>Salmonella_enterica>Salmonella_enterica_subsp._enterica | 449 | NOQ95_21590 | 4462278..4463627 | 192955 | phospholipase-D-like-domain-containing-protein-[Salmonella-enterica-subsp.-enterica-serovar-Kentucky] |
| ESG75021.1 | HKD-DNase+OGG-like | GCA_000487055.1 | Salmonella_enterica_subsp._enterica_serovar_Kentucky_str._ATCC_9263 | Bacteria>Pseudomonadati>Pseudomonadota>Gammaproteobacteria>Enterobacterales>Enterobacteriaceae>Salmonella>Salmonella_enterica>Salmonella_enterica_subsp._enterica>Salmonella_enterica_subsp._enterica_serovar_Kentucky | 449 | SEEK9263_16892 | 33021..34370 | 984236 | hypothetical-protein-SEEK9263_16892-[Salmonella-enterica-subsp.-enterica-serovar-Kentucky-str.-ATCC-9263] |
| ASE02934.1 | HKD-DNase+OGG-like | GCA_002208315.1 | Salmonella_enterica_subsp._enterica_serovar_Quebec_str._S-1267 | Bacteria>Pseudomonadati>Pseudomonadota>Gammaproteobacteria>Enterobacterales>Enterobacteriaceae>Salmonella>Salmonella_enterica>Salmonella_enterica_subsp._enterica>Salmonella_enterica_subsp._enterica_serovar_Quebec | 449 | LFZ40_19375 | 3798430..3799779 | 1243591 | phospholipase-[Salmonella-enterica-subsp.-enterica-serovar-Quebec-str.-S-1267] |
| EJU7772069.1 | HKD-DNase+OGG-like | GCA_025208445.1 | Salmonella_enterica_subsp._salamae_serovar_4,12:e,n,x:1,6 | Bacteria>Pseudomonadati>Pseudomonadota>Gammaproteobacteria>Enterobacterales>Enterobacteriaceae>Salmonella>Salmonella_enterica>Salmonella_enterica_subsp._salamae | 444 | KAB52_003014 | 293490..294822 | 2978379 | phosphatidylserine/phosphatidylglycerophosphate/cardiolipin-synthase-family-protein-[Salmonella-enterica-subsp.-salamae-serovar-4,12:e,n,x:1,6] |
| WP_222233104.1 | HKD-DNase+OGG-like | GCF_019795975.1 | Pantoea_sp._DY-17 | Bacteria>Pseudomonadati>Pseudomonadota>Gammaproteobacteria>Enterobacterales>Erwiniaceae>Pantoea>unclassified_Pantoea | 449 | K5Y33_RS01195 | 240373..241722 | 2871490 | phospholipase-D-like-domain-containing-protein-[Pantoea-sp.-DY-17] |
| WP_060426774.1 | HKD-DNase+OGG-like | GCF_946413725.1 | Serratia | Bacteria>Pseudomonadati>Pseudomonadota>Gammaproteobacteria>Enterobacterales>Yersiniaceae | 449 | AWX06_RS15865 | 195075..196424 | 613 | MULTISPECIES:-phospholipase-D-like-domain-containing-protein-[Serratia] |
| WP_195869498.1 | HKD-DNase+OGG-like | GCF_015645305.1 | Serratia | Bacteria>Pseudomonadati>Pseudomonadota>Gammaproteobacteria>Enterobacterales>Yersiniaceae | 449 | I1A46_RS14695 | 202218..203567 | 613 | phospholipase-D-like-domain-containing-protein-[Serratia-liquefaciens] |
| WP_112197433.1 | HKD-DNase+OGG-like | GCF_003263525.1 | Rahnella_sp._NRRL_B-41462 | Bacteria>Pseudomonadati>Pseudomonadota>Gammaproteobacteria>Enterobacterales>Yersiniaceae>Rahnella>unclassified_Rahnella | 449 | UC14_RS06700 | 45550..46899 | 1610579 | phospholipase-D-like-domain-containing-protein-[Rahnella-sp.-NRRL-B-41462] |
| WP_197785678.1 | HKD-DNase+OGG-like | GCF_016007555.1 | Serratia_ureilytica | Bacteria>Pseudomonadati>Pseudomonadota>Gammaproteobacteria>Enterobacterales>Yersiniaceae>Serratia | 449 | I5P92_RS12880 | 36736..38085 | 300181 | phospholipase-D-like-domain-containing-protein-[Serratia-ureilytica] |
| WP_212602143.1 | HKD-DNase+OGG-like | GCF_018257315.1 | Serratia_rubidaea | Bacteria>Pseudomonadati>Pseudomonadota>Gammaproteobacteria>Enterobacterales>Yersiniaceae>Serratia | 449 | JK228_RS02480 | 46464..47813 | 61652 | phospholipase-D-like-domain-containing-protein-[Serratia-rubidaea] |
| WP_244757917.1 | HKD-DNase+OGG-like | GCF_022870585.1 | Serratia_marcescens | Bacteria>Pseudomonadati>Pseudomonadota>Gammaproteobacteria>Enterobacterales>Yersiniaceae>Serratia | 449 | HW877_RS02440 | 525807..527156 | 615 | phospholipase-D-like-domain-containing-protein-[Serratia-marcescens] |
| WP_261135882.1 | HKD-DNase+OGG-like | GCF_946226775.1 | Serratia_liquefaciens | Bacteria>Pseudomonadati>Pseudomonadota>Gammaproteobacteria>Enterobacterales>Yersiniaceae>Serratia | 449 | N8M82_RS16070 | 334741..336090 | 614 | phospholipase-D-like-domain-containing-protein-[Serratia-liquefaciens] |
| WP_261427670.1 | HKD-DNase+OGG-like | GCF_946227925.1 | Serratia_proteamaculans | Bacteria>Pseudomonadati>Pseudomonadota>Gammaproteobacteria>Enterobacterales>Yersiniaceae>Serratia | 449 | OCH17_RS14315 | 369942..371291 | 28151 | phospholipase-D-like-domain-containing-protein-[Serratia-proteamaculans] |
| WP_053011793.1 | HKD-DNase+OGG-like | GCF_001244675.1 | Yersinia_intermedia | Bacteria>Pseudomonadati>Pseudomonadota>Gammaproteobacteria>Enterobacterales>Yersiniaceae>Yersinia | 449 | ALC37_RS11605 | 473965..475314 | 631 | phospholipase-D-like-domain-containing-protein-[Yersinia-intermedia] |

| Protein-ID | Domain | Assembly-ID | Strain | Taxonomy | Length | Locus-Tag | Genomic-coordinates | TaxID | Fasta-defline |
|---|---|---|---|---|---|---|---|---|---|
| --HNH(standalone) | | | | | | | | | |
| WP_160163127.1 | HNH | GCF_000337915.1 | Halorubrum_saccharovorum | Archaea>Methanobacteriati>Methanobacteriota>Stenosarchaea_group>Halobacteria>Halobacteriales>Haloferacaceae>Halorubrum | 137 | C471_RS16245 | 8082..8495 | 2248 | HNH-endonuclease-signature-motif-containing-protein-[Halorubrum-saccharovorum] |
| PHQ44029.1 | HNH | GCA_002727095.1 | Halorubrum_sp._C191 | Archaea>Methanobacteriati>Methanobacteriota>Stenosarchaea_group>Halobacteria>Halobacteriales>Haloferacaceae>Halorubrum>unclassified_Halorubrum | 146 | Z052_00925 | 29480..29920 | 1383842 | hypothetical-protein-Z052_00925-[Halorubrum-sp.-C191] |
| WP_241897867.1 | HNH | GCF_002727095.1 | Halorubrum_sp._C191 | Archaea>Methanobacteriati>Methanobacteriota>Stenosarchaea_group>Halobacteria>Halobacteriales>Haloferacaceae>Halorubrum>unclassified_Halorubrum | 52 | Z052_RS20810 | 29762..29920 | 1383842 | hypothetical-protein-[Halorubrum-sp.-C191] |
| MCW3138468.1 | HNH | GCA_025966495.1 | Candidatus_Methanophagales_archaeon | Archaea>Methanobacteriati>Methanobacteriota>Stenosarchaea_group>Methanomicrobia>Candidatus_Methanophagales>unclassified_Candidatus_Methanophagales | 197 | N2V76_08670 | 11119..11712 | 2056316 | MAG:-DUF2807-domain-containing-protein-[Methanophagales-archaeon] |
| MCL4551938.1 | HNH | GCA_023381795.1 | Candidatus_Marsarchaeota_archaeon | Archaea>Thermoproteati>Candidatus_Marsarchaeota>unclassified_Candidatus_Marsarchaeota | 215 | M1305_00055 | 7356..8003 | 2823144 | TIGR02646-family-protein-[Candidatus-Marsarchaeota-archaeon] |
| MYH43291.1 | HNH | GCA_009843995.1 | Acidimicrobiaceae_bacterium | Bacteria>Bacillati>Actinomycetota>Acidimicrobiia>Acidimicrobiales>Acidimicrobiaceae>unclassified_Acidimicrobiaceae | 157 | F4131_06255 | 178..651 | 2024894 | HNH-endonuclease-[Acidimicrobiaceae-bacterium] |
| MBV5322914.1 | HNH | GCA_019162935.1 | Ilumatobacteraceae_bacterium | Bacteria>Bacillati>Actinomycetota>Acidimicrobiia>Acidimicrobiales>Ilumatobacteraceae>unclassified_Ilumatobacteraceae | 209 | JZU60_03670 | 7566..8195 | 2497619 | TIGR02646-family-protein-[Ilumatobacteraceae-bacterium] |
| WP_100495670.1 | HNH | GCF_002802865.1 | Bifidobacterium_scaligerum | Bacteria>Bacillati>Actinomycetota>Actinomycetes>Bifidobacteriales>Bifidobacteriaceae>Bifidobacterium | 326 | CUU80_RS01810 | 440065..441045 | 2052656 | hypothetical-protein-[Bifidobacterium-scaligerum] |
| WP_162453102.1 | HNH | - | Phytoactinopolyspora_mesophila | Bacteria>Bacillati>Actinomycetota>Actinomycetes>Jiangellales>Jiangellaceae>Phytoactinopolyspora | 42 | F7O44_RS25365 | 159653..159781 | 2650750 | HNH-endonuclease-signature-motif-containing-protein-[Phytoactinopolyspora-mesophila] |
| WP_242328936.1 | HNH | GCF_022647665.1 | Streptomyces_formicae | Bacteria>Bacillati>Actinomycetota>Actinomycetes>Kitasatosporales>Streptomycetaceae>Streptomyces | 196 | J4032_RS01975 | 421866..422456 | 1616117 | hypothetical-protein-[Streptomyces-formicae] |
| WP_228479363.1 | HNH | GCF_015277815.1 | Microbacterium_atlanticum | Bacteria>Bacillati>Actinomycetota>Actinomycetes>Micrococcales>Microbacteriaceae>Microbacterium | 142 | IR212_RS15220 | 3326843..3327271 | 2782168 | hypothetical-protein-[Microbacterium-atlanticum] |
| WP_231637072.1 | HNH | GCF_001314225.1 | Microbacterium_sp._No._7 | Bacteria>Bacillati>Actinomycetota>Actinomycetes>Micrococcales>Microbacteriaceae>Microbacterium>unclassified_Microbacterium | 97 | AOA12_RS22850 | 103512..103805 | 1714373 | hypothetical-protein-[Microbacterium-sp.-No.-7] |
| MBS3179173.1 | HNH | GCA_018304855.1 | Pseudoclavibacter_sp._Marseille-Q4354 | Bacteria>Bacillati>Actinomycetota>Actinomycetes>Micrococcales>Microbacteriaceae>Pseudoclavibacter>unclassified_Pseudoclavibacter | 254 | JSQ82_10730 | 814229..814993 | 2810515 | HNH-endonuclease-[Pseudoclavibacter-sp.-Marseille-Q4354] |
| WP_249312438.1 | HNH | - | Pseudoclavibacter_sp._Marseille-Q4354 | Bacteria>Bacillati>Actinomycetota>Actinomycetes>Micrococcales>Microbacteriaceae>Pseudoclavibacter>unclassified_Pseudoclavibacter | 252 | JSQ82_RS10730 | 814229..814987 | 2810515 | HNH-endonuclease-family-protein-[Pseudoclavibacter-sp.-Marseille-Q4354] |
| WP_256336593.1 | HNH | GCF_900102895.1 | Arthrobacter_sp._ov407 | Bacteria>Bacillati>Actinomycetota>Actinomycetes>Micrococcales>Micrococcaceae>Arthrobacter>unclassified_Arthrobacter | 80 | BLS42_RS20710 | 10723..10965 | 1761748 | HNH-endonuclease-[Arthrobacter-sp.-ov407] |
| WP_022924601.1 | HNH | GCF_000421245.1 | Serinicoccus_marinus | Bacteria>Bacillati>Actinomycetota>Actinomycetes>Micrococcales>Ornithinimicrobiaceae>Serinicoccus | 292 | FU792_RS06145 | 1314861..1315739 | 247333 | HNH-endonuclease-[Serinicoccus-marinus] |
| WP_091191944.1 | HNH | GCF_900089605.1 | Micromonospora_narathiwatensis | Bacteria>Bacillati>Actinomycetota>Actinomycetes>Micromonosporales>Micromonosporaceae>Micromonospora | 210 | GA0070621_RS04935 | 1088306..1088938 | 299146 | HNH-endonuclease-family-protein-[Micromonospora-narathiwatensis] |
| WP_312777483.1 | HNH | GCF_032056685.1 | Corynebacterium_variabile | Bacteria>Bacillati>Actinomycetota>Actinomycetes>Mycobacteriales>Corynebacteriaceae>Corynebacterium | 215 | RT519_RS10565 | 114740..115387 | 1727 | retron-system-putative-HNH-endonuclease-[Corynebacterium-variabile] |
| WP_220472178.1 | HNH | GCF_014121175.1 | Tomitella_gaofuii | Bacteria>Bacillati>Actinomycetota>Actinomycetes>Mycobacteriales>Corynebacteriales_incertae_sedis>Tomitella | 300 | H7F09_RS10465 | 69179..70081 | 2760083 | HNH-endonuclease-signature-motif-containing-protein-[Tomitella-gaofuii] |
| WP_243580599.1 | HNH | GCF_022810365.1 | unclassified_Rhodococcus_(in:_high_G+C_Gram-positive_bacteria) | Bacteria>Bacillati>Actinomycetota>Actinomycetes>Mycobacteriales>Nocardiaceae>Rhodococcus | 220 | MTX38_RS18985 | 45555..46217 | 192944 | MULTISPECIES:-HNH-endonuclease-[unclassified-Rhodococcus-(in:-high-G+C-Gram-positive-bacteria)] |
| WP_317550153.1 | HNH | GCF_033042295.1 | Rhodococcus_aetherivorans | Bacteria>Bacillati>Actinomycetota>Actinomycetes>Mycobacteriales>Nocardiaceae>Rhodococcus | 159 | R3Q02_RS04850 | 125649..126128 | 191292 | HNH-endonuclease-domain-containing-protein-[Rhodococcus-aetherivorans] |
| WP_116057357.1 | HNH | - | Rhodococcus_erythropolis | Bacteria>Bacillati>Actinomycetota>Actinomycetes>Mycobacteriales>Nocardiaceae>Rhodococcus>Rhodococcus_erythropolis_group | 193 | DVG80_RS18300 | 3921625..3922206 | 1833 | HNH-endonuclease-[Rhodococcus-erythropolis] |
| MBY6706226.1 | HNH | GCA_019836145.1 | Rhodococcus_sp._BP-241 | Bacteria>Bacillati>Actinomycetota>Actinomycetes>Mycobacteriales>Nocardiaceae>Rhodococcus>unclassified_Rhodococcus_(in:_high_G+C_Gram-positive_bacteria) | 290 | HQ308_05370 | 439682..440554 | 2739441 | HNH-endonuclease-[Rhodococcus-sp.-BP-241] |
| QCQ90483.1 | HNH | GCA_005484805.1 | Rhodococcus_sp._SGAir0479 | Bacteria>Bacillati>Actinomycetota>Actinomycetes>Mycobacteriales>Nocardiaceae>Rhodococcus>unclassified_Rhodococcus_(in:_high_G+C_Gram-positive_bacteria) | 282 | E7742_04080 | 866894..867742 | 2567884 | HNH-endonuclease-[Rhodococcus-sp.-SGAir0479] |
| WP_166139840.1 | HNH | GCF_011420305.2 | Nocardioides_ochotonae | Bacteria>Bacillati>Actinomycetota>Actinomycetes>Propionibacteriales>Nocardioidaceae>Nocardioides | 299 | HBO46_RS08115 | 1677724..1678623 | 2685869 | HNH-endonuclease-[Nocardioides-ochotonae] |
| WP_202421703.1 | HNH | - | Nocardiopsis_alba | Bacteria>Bacillati>Actinomycetota>Actinomycetes>Streptosporangiales>Nocardiopsidaceae>Nocardiopsis | 235 | GTW20_RS21045 | 4835081..4835788 | 53437 | DUF5710-domain-containing-protein-[Nocardiopsis-alba] |
| MDA0169819.1 | HNH | GCA_027587225.1 | Solirubrobacter_taibaiensis | Bacteria>Bacillati>Actinomycetota>Thermoleophilia>Solirubrobacterales>Solirubrobacteraceae>Solirubrobacter | 303 | OJ998_12050 | 151996..152907 | 1395955 | HNH-endonuclease-[Solirubrobacter-taibaiensis] |
| WP_016481844.1 | HNH | GCF_000427095.1 | Chthonomonas_calidirosea | Bacteria>Bacillati>Armatimonadota>Chthonomonadia>Chthonomonadales>Chthonomonadaceae>Chthonomonas | 164 | CCALI_RS02235 | 508277..508771 | 454171 | hypothetical-protein-[Chthonomonas-calidirosea] |
| WP_075162427.1 | HNH | GCF_001457975.1 | Chthonomonas_calidirosea | Bacteria>Bacillati>Armatimonadota>Chthonomonadia>Chthonomonadales>Chthonomonadaceae>Chthonomonas | 164 | CP488_RS03520 | 845004..845498 | 454171 | hypothetical-protein-[Chthonomonas-calidirosea] |
| WP_061531640.1 | HNH | GCF_029504955.1 | Bacillus | Bacteria>Bacillati>Bacillota>Bacilli>Bacillales>Bacillaceae | 436 | M2246_RS18840 | 187580..188890 | 1386 | MULTISPECIES:-HNH-endonuclease-[Bacillus] |
| WP_151627819.1 | HNH | GCF_008923665.1 | Bacillus_luti | Bacteria>Bacillati>Bacillota>Bacilli>Bacillales>Bacillaceae>Bacillus>Bacillus_cereus_group | 204 | F8163_RS24255 | 203105..203719 | 2026191 | HNH-endonuclease-[Bacillus-luti] |
| WP_061808763.1 | HNH | GCF_001591825.1 | Rossellomorea_vietnamensis | Bacteria>Bacillati>Bacillota>Bacilli>Bacillales>Bacillaceae>Rossellomorea | 291 | BVI01S_RS00240 | 53081..53956 | 218284 | HNH-endonuclease-[Rossellomorea-vietnamensis] |
| WP_108587673.1 | HNH | GCF_003058165.1 | Paenisporosarcina_sp._OV554 | Bacteria>Bacillati>Bacillota>Bacilli>Bacillales>Caryophanaceae>Paenisporosarcina>unclassified_Paenisporosarcina | 198 | C8K15_RS17280 | 32335..32931 | 2135694 | phosphonate-C-P-lyase-system-protein-PhnH-[Paenisporosarcina-sp.-OV554] |
| WP_079914142.1 | HNH | GCF_002042965.1 | Paenibacillus_sp._32352 | Bacteria>Bacillati>Bacillota>Bacilli>Bacillales>Paenibacillaceae>Paenibacillus>unclassified_Paenibacillus | 336 | B1728_RS37075 | 171298..172308 | 1969111 | HNH-endonuclease-[Paenibacillus-sp.-32352] |
| WP_211334340.1 | HNH | GCF_003946325.1 | Companilactobacillus_furfuricola | Bacteria>Bacillati>Bacillota>Bacilli>Lactobacillales>Lactobacillaceae>Companilactobacillus | 165 | EKM78_RS12960 | 15073..15570 | 1462575 | HNH-endonuclease-family-protein-[Companilactobacillus-furfuricola] |
| GHN53706.1 | HNH | GCA_021600265.1 | Lactobacillus_delbrueckii | Bacteria>Bacillati>Bacillota>Bacilli>Lactobacillales>Lactobacillaceae>Lactobacillus | 157 | ME802_14790 | 18778..19251 | 1584 | hypothetical-protein-ME802_14790-[Lactobacillus-delbrueckii] |
| MBR2506168.1 | HNH | GCA_017502465.1 | Bacilli_bacterium | Bacteria>Bacillati>Bacillota>Bacilli>unclassified_Bacilli | 208 | IKB70_04485 | 13126..13752 | 1903720 | TIGR02646-family-protein-[Bacilli-bacterium] |
| HIT17412.1 | HNH | GCA_018716205.1 | Candidatus_Caccosoma_faecigallinarum | Bacteria>Bacillati>Bacillota>Bacillota_incertae_sedis>Candidatus_Caccosoma | 237 | IAD04_03395 | 13264..13977 | 2840720 | hypothetical-protein-[Candidatus-Caccosoma-faecigallinarum] |
| WP_077851366.1 | HNH | GCF_002006235.2 | Clostridium_felsineum | Bacteria>Bacillati>Bacillota>Clostridia>Eubacteriales>Clostridiaceae>Clostridium | 235 | CLAUR_RS09630 | 2185364..2186071 | 36839 | TIGR02646-family-protein-[Clostridium-felsineum] |
| WP_171780280.1 | HNH | GCF_014230655.1 | Clostridium_beijerinckii | Bacteria>Bacillati>Bacillota>Clostridia>Eubacteriales>Clostridiaceae>Clostridium | 235 | B0H39_RS13625 | 3006075..3006782 | 1520 | TIGR02646-family-protein-[Clostridium-beijerinckii] |
| MBN1067556.1 | HNH | GCA_016925955.1 | Clostridium_botulinum | Bacteria>Bacillati>Bacillota>Clostridia>Eubacteriales>Clostridiaceae>Clostridium | 233 | DVW02_06060 | 107164..107865 | 1491 | TIGR02646-family-protein-[Clostridium-botulinum] |
| NFG40382.1 | HNH | GCA_011013445.1 | Clostridium_botulinum | Bacteria>Bacillati>Bacillota>Clostridia>Eubacteriales>Clostridiaceae>Clostridium | 233 | FC789_04055 | 890907..891608 | 1491 | TIGR02646-family-protein-[Clostridium-botulinum] |
| NFI95399.1 | HNH | GCA_011014625.1 | Clostridium_botulinum | Bacteria>Bacillati>Bacillota>Clostridia>Eubacteriales>Clostridiaceae>Clostridium | 233 | FC961_13535 | 7131..7832 | 1491 | TIGR02646-family-protein-[Clostridium-botulinum] |
| WP_270487113.1 | HNH | GCF_027681635.1 | Clostridium_fessum | Bacteria>Bacillati>Bacillota>Clostridia>Eubacteriales>Clostridiaceae>Clostridium | 225 | PGX92_RS02740 | 613212..613889 | 2126740 | hypothetical-protein-[Clostridium-fessum] |
| WP_172556351.1 | HNH | GCF_900461105.1 | Clostridium_putrefaciens | Bacteria>Bacillati>Bacillota>Clostridia>Eubacteriales>Clostridiaceae>Clostridium | 219 | DY168_RS12195 | 2670044..2670703 | 99675 | TIGR02646-family-protein-[Clostridium-putrefaciens] |
| WP_216171754.1 | HNH | GCF_026651005.1 | Clostridium_estertheticum | Bacteria>Bacillati>Bacillota>Clostridia>Eubacteriales>Clostridiaceae>Clostridium | 213 | LL037_RS21130 | 4412312..4412953 | 238834 | TIGR02646-family-protein-[Clostridium-estertheticum] |
| WP_052188304.1 | HNH | GCF_000785185.1 | Clostridium_butyricum | Bacteria>Bacillati>Bacillota>Clostridia>Eubacteriales>Clostridiaceae>Clostridium | 223 | AWN73_RS07990 | 11406..12077 | 1492 | HNH-endonuclease-[Clostridium-butyricum] |
| SCH09454.1 | HNH | GCA_900066525.1 | uncultured_Clostridium_sp. | Bacteria>Bacillati>Bacillota>Clostridia>Eubacteriales>Clostridiaceae>Clostridium>environmental_samples | 211 | SAMEA3545335_00164 | 182450..183085 | 59620 | Predicted-restriction-endonuclease-[uncultured-Clostridium-sp.] |
| WP_118513463.1 | HNH | - | Clostridium_sp._AF37-5AT | Bacteria>Bacillati>Bacillota>Clostridia>Eubacteriales>Clostridiaceae>Clostridium>unclassified_Clostridium | 259 | DW020_RS00885 | 207314..208093 | 2292007 | hypothetical-protein-[Clostridium-sp.-AF37-5AT] |
| WP_164949143.1 | HNH | GCF_011065955.2 | Clostridium_sp._FP2 | Bacteria>Bacillati>Bacillota>Clostridia>Eubacteriales>Clostridiaceae>Clostridium>unclassified_Clostridium | 224 | G9F71_RS17950 | 3730172..3730846 | 2724481 | TIGR02646-family-protein-[Clostridium-sp.-FP2] |
| MBS6228513.1 | HNH | GCA_018374725.1 | Clostridium_sp. | Bacteria>Bacillati>Bacillota>Clostridia>Eubacteriales>Clostridiaceae>Clostridium>unclassified_Clostridium | 223 | KH509_01910 | 62343..63014 | 1506 | hypothetical-protein-[Clostridium-sp.] |
| WP_089984084.1 | HNH | GCF_001517425.1 | Clostridium_sp._C105KSO13 | Bacteria>Bacillati>Bacillota>Clostridia>Eubacteriales>Clostridiaceae>Clostridium>unclassified_Clostridium | 330 | BN3456_RS10440 | 20434..21426 | 1776045 | hypothetical-protein-[Clostridium-sp.-C105KSO13] |
| NLE25891.1 | HNH | GCA_012514215.1 | Clostridiaceae_bacterium | Bacteria>Bacillati>Bacillota>Clostridia>Eubacteriales>Clostridiaceae>unclassified_Clostridiaceae | 208 | GX625_11235 | 3116..3742 | 1898204 | hypothetical-protein-[Clostridiaceae-bacterium] |
| MCC2258800.1 | HNH | GCA_020687605.1 | Intestinimonas_aquisgranensis | Bacteria>Bacillati>Bacillota>Clostridia>Eubacteriales>Eubacteriales_incertae_sedis>Intestinimonas | 221 | LKD74_12660 | 7197..7862 | 2881257 | hypothetical-protein-[Intestinimonas-sp.-CLA-AA-H199] |
| WP_235255114.1 | HNH | - | Allofournierella_massiliensis | Bacteria>Bacillati>Bacillota>Clostridia>Eubacteriales>Oscillospiraceae>Allofournierella | 219 | JQM64_RS00365 | 76895..77554 | 1650663 | HNH-endonuclease-[Fournierella-massiliensis] |
| WP_195464811.1 | HNH | GCF_015554605.1 | Faecalispora_jeddahensis | Bacteria>Bacillati>Bacillota>Clostridia>Eubacteriales>Oscillospiraceae>Faecalispora | 239 | I2E80_RS10455 | 138786..139505 | 1414721 | hypothetical-protein-[Clostridium-jeddahense] |
| WP_195543173.1 | HNH | GCF_015558275.1 | Massiliimalia_timonensis | Bacteria>Bacillati>Bacillota>Clostridia>Eubacteriales>Oscillospiraceae>Massiliimalia | 132 | I2G46_RS07895 | 59688..60086 | 1987501 | hypothetical-protein-[Massiliimalia-timonensis] |
| MCI9347609.1 | HNH | GCA_022799735.1 | Oscillibacter_sp. | Bacteria>Bacillati>Bacillota>Clostridia>Eubacteriales>Oscillospiraceae>Oscillibacter>unclassified_Oscillibacter | 222 | HFF83_00070 | 16176..16844 | 1945593 | hypothetical-protein-[Oscillibacter-sp.] |
| MBS7204847.1 | HNH | GCA_018383015.1 | Ruminococcus_bicirculans_(ex_Wegman_et_al._2014) | Bacteria>Bacillati>Bacillota>Clostridia>Eubacteriales>Oscillospiraceae>Ruminococcus | 221 | KH048_01065 | 240475..241140 | 1160721 | hypothetical-protein-[Ruminococcus-bicirculans] |
| MBT9624602.1 | HNH | GCA_018784005.1 | Ruminococcus_bicirculans_(ex_Wegman_et_al._2014) | Bacteria>Bacillati>Bacillota>Clostridia>Eubacteriales>Oscillospiraceae>Ruminococcus | 221 | GPK54_04615 | 3058..3723 | 1160721 | hypothetical-protein-[Ruminococcus-bicirculans] |
| WP_215659168.1 | HNH | GCF_018783905.1 | Ruminococcus_sp._MCC718 | Bacteria>Bacillati>Bacillota>Clostridia>Eubacteriales>Oscillospiraceae>Ruminococcus>unclassified_Ruminococcus | 258 | GPK60_RS07255 | 24877..25653 | 2592649 | hypothetical-protein-[Ruminococcus-sp.-MCC718] |
| MBQ3756406.1 | HNH | GCA_017395085.1 | Oscillospiraceae_bacterium | Bacteria>Bacillati>Bacillota>Clostridia>Eubacteriales>Oscillospiraceae>unclassified_Oscillospiraceae | 222 | II873_04760 | 26483..27151 | 2485925 | hypothetical-protein-[Oscillospiraceae-bacterium] |
| MCD7847860.1 | HNH | GCA_021200775.1 | Oscillospiraceae_bacterium | Bacteria>Bacillati>Bacillota>Clostridia>Eubacteriales>Oscillospiraceae>unclassified_Oscillospiraceae | 221 | LUG49_07560 | 31137..31802 | 2485925 | hypothetical-protein-[Oscillospiraceae-bacterium] |
| MBE5813907.1 | HNH | GCA_015056315.1 | Clostridiales_bacterium | Bacteria>Bacillati>Bacillota>Clostridia>Eubacteriales>unclassified_Eubacteriales | 236 | E7320_01690 | 125982..126692 | 1898207 | hypothetical-protein-[Clostridiales-bacterium] |
| MCD8238625.1 | HNH | GCA_021204175.1 | Clostridiales_bacterium | Bacteria>Bacillati>Bacillota>Clostridia>Eubacteriales>unclassified_Eubacteriales | 222 | LUC92_04715 | 22006..22674 | 1898207 | hypothetical-protein-[Clostridiales-bacterium] |
| MCD7774060.1 | HNH | GCA_021199775.1 | Clostridiales_bacterium | Bacteria>Bacillati>Bacillota>Clostridia>Eubacteriales>unclassified_Eubacteriales | 221 | LUH40_00555 | 12140..12805 | 1898207 | hypothetical-protein-[Clostridiales-bacterium] |
| WP_059085688.1 | HNH | GCF_001487165.1 | Blautia | Bacteria>Bacillati>Bacillota>Clostridia>Lachnospirales>Lachnospiraceae | 225 | C4886_RS16000 | 13482..14159 | 572511 | MULTISPECIES:-hypothetical-protein-[Blautia] |
| WP_246240540.1 | HNH | GCF_009931695.1 | Anaerocolumna_sedimenticola | Bacteria>Bacillati>Bacillota>Clostridia>Lachnospirales>Lachnospiraceae>Anaerocolumna | 249 | Ana3638_RS23000 | 5164337..5165086 | 2696063 | HNH-endonuclease-[Anaerocolumna-sedimenticola] |
| WP_154780317.1 | HNH | GCF_033096465.1 | Blautia_luti | Bacteria>Bacillati>Bacillota>Clostridia>Lachnospirales>Lachnospiraceae>Blautia | 225 | GKZ57_RS08785 | 88095..88772 | 89014 | hypothetical-protein-[Blautia-luti] |
| WP_026654581.1 | HNH | GCF_000686065.1 | Butyrivibrio_sp._AE3003 | Bacteria>Bacillati>Bacillota>Clostridia>Lachnospirales>Lachnospiraceae>Butyrivibrio>unclassified_Butyrivibrio | 266 | CD00_RS0103085 | 639508..640308 | 1496721 | hypothetical-protein-[Butyrivibrio-sp.-AE3003] |
| TGX97922.1 | HNH | GCA_004793585.1 | Hominisplanchenecus_murintestinalis | Bacteria>Bacillati>Bacillota>Clostridia>Lachnospirales>Lachnospiraceae>Hominisplanchenecus | 225 | E5357_10795 | 27048..27725 | 2941517 | hypothetical-protein-E5357_10795-[Lachnospiraceae-bacterium] |
| WP_187018527.1 | HNH | GCF_014288005.1 | Hungatella_hominis | Bacteria>Bacillati>Bacillota>Clostridia>Lachnospirales>Lachnospiraceae>Hungatella | 235 | H8S75_RS00665 | 146849..147556 | 2763050 | hypothetical-protein-[Hungatella-sp.-L12] |
| MCI8605655.1 | HNH | GCA_022793615.1 | Hungatella_sp. | Bacteria>Bacillati>Bacillota>Clostridia>Lachnospirales>Lachnospiraceae>Hungatella>unclassified_Hungatella | 244 | HFG72_02105 | 7784..8518 | 2613924 | hypothetical-protein-[Hungatella-sp.] |
| MCI8605666.1 | HNH | GCA_022793615.1 | Hungatella_sp. | Bacteria>Bacillati>Bacillota>Clostridia>Lachnospirales>Lachnospiraceae>Hungatella>unclassified_Hungatella | 263 | HFG72_02160 | 24409..25200 | 2613924 | hypothetical-protein-[Hungatella-sp.] |
| MCI8664745.1 | HNH | GCA_022794315.1 | Hungatella_sp. | Bacteria>Bacillati>Bacillota>Clostridia>Lachnospirales>Lachnospiraceae>Hungatella>unclassified_Hungatella | 254 | HFG69_16140 | 109339..110103 | 2613924 | hypothetical-protein-[Hungatella-sp.] |
| WP_024292373.1 | HNH | GCF_000526995.1 | Lacrimispora_indolis | Bacteria>Bacillati>Bacillota>Clostridia>Lachnospirales>Lachnospiraceae>Lacrimispora | 265 | K401_RS0107485 | 1571627..1572424 | 69825 | hypothetical-protein-[Lacrimispora-indolis] |
| RGE15630.1 | HNH | GCA_003435545.1 | Lachnospiraceae_bacterium_OF11-28 | Bacteria>Bacillati>Bacillota>Clostridia>Lachnospirales>Lachnospiraceae>unclassified_Lachnospiraceae | 223 | DXB03_06320 | 26091..26762 | 2305244 | hypothetical-protein-DXB03_06320-[Lachnospiraceae-bacterium-OF11-28] |
| BDF48896.1 | HNH | GCA_022845955.1 | Lachnospiraceae_bacterium | Bacteria>Bacillati>Bacillota>Clostridia>Lachnospirales>Lachnospiraceae>unclassified_Lachnospiraceae | 131 | CE91St56_60190 | 6878777..6879172 | 1898203 | hypothetical-protein-CE91St56_60190-[Lachnospiraceae-bacterium] |
| BDF48897.1 | HNH | GCA_022845955.1 | Lachnospiraceae_bacterium | Bacteria>Bacillati>Bacillota>Clostridia>Lachnospirales>Lachnospiraceae>unclassified_Lachnospiraceae | 100 | CE91St56_60200 | 6879182..6879484 | 1898203 | hypothetical-protein-CE91St56_60200-[Lachnospiraceae-bacterium] |
| MBD5550162.1 | HNH | GCA_014804795.1 | Lachnospiraceae_bacterium | Bacteria>Bacillati>Bacillota>Clostridia>Lachnospirales>Lachnospiraceae>unclassified_Lachnospiraceae | 273 | HDQ96_03150 | 119935..120756 | 1898203 | hypothetical-protein-[Lachnospiraceae-bacterium] |
| MBR4981298.1 | HNH | GCA_017555455.1 | Lachnospiraceae_bacterium | Bacteria>Bacillati>Bacillota>Clostridia>Lachnospirales>Lachnospiraceae>unclassified_Lachnospiraceae | 273 | IKY94_01910 | 659..1480 | 1898203 | hypothetical-protein-[Lachnospiraceae-bacterium] |
| MCI9323600.1 | HNH | GCA_022799595.1 | Lachnospiraceae_bacterium | Bacteria>Bacillati>Bacillota>Clostridia>Lachnospirales>Lachnospiraceae>unclassified_Lachnospiraceae | 265 | HFH81_10145 | 52944..53741 | 1898203 | hypothetical-protein-[Lachnospiraceae-bacterium] |
| MCI9535142.1 | HNH | GCA_022801195.1 | Lachnospiraceae_bacterium | Bacteria>Bacillati>Bacillota>Clostridia>Lachnospirales>Lachnospiraceae>unclassified_Lachnospiraceae | 263 | HFG53_10145 | 44032..44823 | 1898203 | hypothetical-protein-[Lachnospiraceae-bacterium] |
| MST58784.1 | HNH | GCA_009695995.1 | Waltera_intestinalis | Bacteria>Bacillati>Bacillota>Clostridia>Lachnospirales>Lachnospiraceae>Waltera | 249 | FYJ59_11155 | 24862..25611 | 2606635 | hypothetical-protein-[Waltera-intestinalis] |
| WP_268878552.1 | HNH | - | Waltera_intestinalis | Bacteria>Bacillati>Bacillota>Clostridia>Lachnospirales>Lachnospiraceae>Waltera | 188 | FYJ59_RS11175 | 24862..25428 | 2606635 | HNH-endonuclease-[Waltera-intestinalis] |
| RJX18828.1 | HNH | GCA_003604985.1 | Ammonifex_sp. | Bacteria>Bacillati>Bacillota>Clostridia>Thermoanaerobacterales>Thermoanaerobacteraceae>Ammonifex>unclassified_Ammonifex | 238 | C4570_06135 | 4683..5399 | 1872493 | MAG:-HNH-endonuclease-[Ammonifex-sp.] |
| MCD8003210.1 | HNH | GCA_021202035.1 | Clostridia_bacterium | Bacteria>Bacillati>Bacillota>Clostridia>unclassified_Clostridia | 221 | LUG88_04785 | 17952..18617 | 2044939 | hypothetical-protein-[Clostridia-bacterium] |
| MBR1677454.1 | HNH | GCA_017476895.1 | Clostridia_bacterium | Bacteria>Bacillati>Bacillota>Clostridia>unclassified_Clostridia | 130 | IJ706_09120 | 32311..32703 | 2044939 | hypothetical-protein-[Clostridia-bacterium] |
| MBR1677455.1 | HNH | GCA_017476895.1 | Clostridia_bacterium | Bacteria>Bacillati>Bacillota>Clostridia>unclassified_Clostridia | 91 | IJ706_09125 | 32718..32993 | 2044939 | hypothetical-protein-[Clostridia-bacterium] |
| WP_037354741.1 | HNH | GCF_000701565.1 | Selenomonas_sp._FC4001 | Bacteria>Bacillati>Bacillota>Negativicutes>Selenomonadales>Selenomonadaceae>Selenomonas>unclassified_Selenomonas | 212 | T320_RS08145 | 24123..24761 | 1408313 | HNH-endonuclease-[Selenomonas-sp.-FC4001] |
| MBR3457111.1 | HNH | GCA_017509225.1 | Selenomonadaceae_bacterium | Bacteria>Bacillati>Bacillota>Negativicutes>Selenomonadales>Selenomonadaceae>unclassified_Selenomonadaceae | 223 | IKH16_03010 | 334..1005 | 2652289 | hypothetical-protein-[Selenomonadaceae-bacterium] |
| RGH23085.1 | HNH | GCA_003461705.1 | Firmicutes_bacterium_AF12-30 | Bacteria>Bacillati>Bacillota>unclassified_Bacillota | 244 | DWV72_07070 | 57069..57803 | 2292884 | TIGR02646-family-protein-[Firmicutes-bacterium-AF12-30] |
| CCX63519.1 | HNH | GCA_000431495.1 | Firmicutes_bacterium_CAG:791 | Bacteria>Bacillati>Bacillota>unclassified_Bacillota | 240 | BN785_00940 | 18654..19376 | 1262993 | putative-uncharacterized-protein-[Firmicutes-bacterium-CAG:791] |
| NSW92346.1 | HNH | GCA_013314855.1 | Bacillota_bacterium | Bacteria>Bacillati>Bacillota>unclassified_Bacillota | 113 | HPY74_17050 | 11362..11703 | 1879010 | hypothetical-protein-[Bacillota-bacterium] |
| MBN1136464.1 | HNH | GCA_016926595.1 | Anaerolineae_bacterium | Bacteria>Bacillati>Chloroflexota>Anaerolineae>unclassified_Anaerolineae | 44 | JXM73_07755 | 799..933 | 2052143 | HNH-endonuclease-[Anaerolineae-bacterium] |
| MBK9093497.1 | HNH | GCA_016719785.1 | Candidatus_Amarolinea_dominans | Bacteria>Bacillati>Chloroflexota>Caldilineae>Caldilineales>Candidatus_Amarolineaceae>Candidatus_Amarolinea | 287 | IPM84_12125 | 205206..206069 | 3140696 | hypothetical-protein-[Anaerolineae-bacterium] |
| MBK6433237.1 | HNH | GCA_016704685.1 | Candidatus_Amarolinea_dominans | Bacteria>Bacillati>Chloroflexota>Caldilineae>Caldilineales>Candidatus_Amarolineaceae>Candidatus_Amarolinea | 115 | IPF85_21405 | 39848..40195 | 3140696 | TIGR02646-family-protein-[Anaerolineae-bacterium] |
| MCY4559679.1 | HNH | GCA_026713265.1 | Chloroflexota_bacterium | Bacteria>Bacillati>Chloroflexota>unclassified_Chloroflexota | 250 | OXF79_25595 | 14975..15727 | 2026724 | MAG:-TIGR02646-family-protein-[Chloroflexi-bacterium] |
| MBI4317926.1 | HNH | GCA_016210495.1 | Chloroflexota_bacterium | Bacteria>Bacillati>Chloroflexota>unclassified_Chloroflexota | 199 | HY675_05505 | 24032..24631 | 2026724 | hypothetical-protein-[Chloroflexota-bacterium] |
| MBE9475410.1 | HNH | GCA_015222395.1 | Chloroflexota_bacterium | Bacteria>Bacillati>Chloroflexota>unclassified_Chloroflexota | 198 | IMY85_11015 | 12196..12792 | 2026724 | hypothetical-protein-[Chloroflexota-bacterium] |
| HEC34271.1 | HNH | GCA_011052755.1 | Chloroflexota_bacterium | Bacteria>Bacillati>Chloroflexota>unclassified_Chloroflexota | 137 | ENI37_06090 | 11763..12176 | 2026724 | HNH-endonuclease-[Chloroflexota-bacterium] |
| MBT9546708.1 | HNH | GCA_018780035.1 | Candidatus_Sericytochromatia_bacterium | Bacteria>Bacillati>Cyanobacteriota/Melainabacteria_group>Candidatus_Sericytochromatia>unclassified_Candidatus_Sericytochromatia | 210 | IV090_15055 | 106771..107403 | 2762020 | TIGR02646-family-protein-[Candidatus-Sericytochromatia-bacterium] |
| WP_190449865.1 | HNH | GCF_014695795.1 | Cyanophyceae | Bacteria>Bacillati>Cyanobacteriota/Melainabacteria_group>Cyanobacteriota | 137 | H6F86_RS16700 | 64333..64746 | 3028117 | HNH-endonuclease-signature-motif-containing-protein-[Phormidium-sp.-FACHB-592] |
| WP_190429841.1 | HNH | GCF_014695635.1 | Coleofasciculus_sp._FACHB-712 | Bacteria>Bacillati>Cyanobacteriota/Melainabacteria_group>Cyanobacteriota>Cyanophyceae>Coleofasciculales>Coleofasciculaceae>Coleofasciculus>unclassified_Coleofasciculus | 196 | H6F50_RS18595 | 11690..12280 | 2692789 | hypothetical-protein-[Coleofasciculus-sp.-FACHB-712] |
| KPQ32459.1 | HNH | GCA_001314865.1 | Phormidesmis_priestleyi_Ana | Bacteria>Bacillati>Cyanobacteriota/Melainabacteria_group>Cyanobacteriota>Cyanophyceae>Leptolyngbyales>Leptolyngbyaceae>Phormidesmis>Phormidesmis_priestleyi | 205 | HLUCCA11_21445 | 12352..12969 | 1666911 | TIGR02646-family-protein-[Phormidesmis-priestleyi-Ana] |
| WP_096691986.1 | HNH | - | unclassified_Calothrix | Bacteria>Bacillati>Cyanobacteriota/Melainabacteria_group>Cyanobacteriota>Cyanophyceae>Nostocales>Calotrichaceae>Calothrix | 175 | CDC41_RS40670 | 9224312..9224839 | 2619626 | MULTISPECIES:-group-II-intron-maturase-specific-domain-containing-protein-[unclassified-Calothrix] |
| WP_094345565.1 | HNH | GCF_003113895.1 | Nostoc | Bacteria>Bacillati>Cyanobacteriota/Melainabacteria_group>Cyanobacteriota>Cyanophyceae>Nostocales>Nostocaceae | 97 | CDC33_RS34480 | 465962..466255 | 1177 | MULTISPECIES:-HNH-endonuclease-[Nostocaceae] |
| WP_069067983.1 | HNH | GCF_001712795.1 | Nostoc_sp._KVJ20 | Bacteria>Bacillati>Cyanobacteriota/Melainabacteria_group>Cyanobacteriota>Cyanophyceae>Nostocales>Nostocaceae>Nostoc>unclassified_Nostoc | 195 | A4S05_RS01000 | 35348..35935 | 457944 | hypothetical-protein-[Nostoc-sp.-KVJ20] |
| MCU0565451.1 | HNH | GCA_025454315.1 | Oculatellaceae_cyanobacterium_Prado106 | Bacteria>Bacillati>Cyanobacteriota/Melainabacteria_group>Cyanobacteriota>Cyanophyceae>Oculatellales>Oculatellaceae>unclassified_Oculatellaceae | 133 | MUF49_02515 | 17705..18106 | 2929566 | HNH-endonuclease-[Oculatellaceae-cyanobacterium-Prado106] |
| WP_254953300.1 | HNH | GCF_024345955.1 | unclassified_Cyanobium | Bacteria>Bacillati>Cyanobacteriota/Melainabacteria_group>Cyanobacteriota>Cyanophyceae>Synechococcales>Prochlorococcaceae>Cyanobium | 226 | KBZ34_RS12475 | 64884..65564 | 2627006 | MULTISPECIES:-TIGR02646-family-protein-[unclassified-Cyanobium] |
| WP_254950525.1 | HNH | GCF_024345955.1 | unclassified_Cyanobium | Bacteria>Bacillati>Cyanobacteriota/Melainabacteria_group>Cyanobacteriota>Cyanophyceae>Synechococcales>Prochlorococcaceae>Cyanobium | 218 | KBZ34_RS03565 | 174722..175378 | 2627006 | MULTISPECIES:-HNH-endonuclease-signature-motif-containing-protein-[unclassified-Cyanobium] |
| MCP9859838.1 | HNH | GCA_024345955.1 | Cyanobium_sp._Cruz-8H5 | Bacteria>Bacillati>Cyanobacteriota/Melainabacteria_group>Cyanobacteriota>Cyanophyceae>Synechococcales>Prochlorococcaceae>Cyanobium>unclassified_Cyanobium | 226 | KBZ34_12420 | 64884..65564 | 2823712 | TIGR02646-family-protein-[Cyanobium-sp.-Cruz-8H5] |
| WP_254937140.1 | HNH | GCF_024345395.1 | Cyanobium_sp._Tous-M-B4 | Bacteria>Bacillati>Cyanobacteriota/Melainabacteria_group>Cyanobacteriota>Cyanophyceae>Synechococcales>Prochlorococcaceae>Cyanobium>unclassified_Cyanobium | 223 | KBY73_RS11010 | 148533..149204 | 2823724 | HNH-endonuclease-signature-motif-containing-protein-[Cyanobium-sp.-Tous-M-B4] |
| WP_255117820.1 | HNH | GCF_024345835.1 | Synechococcus_sp._HJ21-Hayes | Bacteria>Bacillati>Cyanobacteriota/Melainabacteria_group>Cyanobacteriota>Cyanophyceae>Synechococcales>Synechococcaceae>Synechococcus>unclassified_Synechococcus | 202 | KBZ14_RS08115 | 77630..78238 | 2823736 | HNH-endonuclease-signature-motif-containing-protein-[Synechococcus-sp.-HJ21-Hayes] |
| HBH74808.1 | HNH | GCA_003508015.1 | Synechococcales_bacterium_UBA10510 | Bacteria>Bacillati>Cyanobacteriota/Melainabacteria_group>Cyanobacteriota>Cyanophyceae>Synechococcales>unclassified_Synechococcales | 200 | DDY43_15590 | 16963..17565 | 2055795 | hypothetical-protein-[Synechococcales-bacterium-UBA10510] |
| MCG9130915.1 | HNH | GCA_022394775.1 | Candidatus_Poribacteria_bacterium | Bacteria>Bacteria_incertae_sedis>Bacteria_candidate_phyla>Candidatus_Poribacteria>unclassified_Candidatus_Poribacteria | 292 | J5I95_04445 | 22582..23460 | 2026781 | hypothetical-protein-[Candidatus-Poribacteria-bacterium] |
| MCY3550006.1 | HNH | GCA_026702755.1 | Candidatus_Poribacteria_bacterium | Bacteria>Bacteria_incertae_sedis>Bacteria_candidate_phyla>Candidatus_Poribacteria>unclassified_Candidatus_Poribacteria | 292 | OXH39_06050 | 108337..109215 | 2026781 | MAG:-hypothetical-protein-OXH39_06050-[Candidatus-Poribacteria-bacterium] |
| MDE0324950.1 | HNH | GCA_028820875.1 | Candidatus_Poribacteria_bacterium | Bacteria>Bacteria_incertae_sedis>Bacteria_candidate_phyla>Candidatus_Poribacteria>unclassified_Candidatus_Poribacteria | 292 | OXN27_13635 | 8939..9817 | 2026781 | MAG:-hypothetical-protein-OXN27_13635-[Candidatus-Poribacteria-bacterium] |
| MYC75113.1 | HNH | GCA_009841255.1 | Candidatus_Poribacteria_bacterium | Bacteria>Bacteria_incertae_sedis>Bacteria_candidate_phyla>Candidatus_Poribacteria>unclassified_Candidatus_Poribacteria | 292 | F4X10_04975 | 77950..78828 | 2026781 | hypothetical-protein-[Candidatus-Poribacteria-bacterium] |
| MYK19442.1 | HNH | GCA_009845505.1 | Candidatus_Poribacteria_bacterium | Bacteria>Bacteria_incertae_sedis>Bacteria_candidate_phyla>Candidatus_Poribacteria>unclassified_Candidatus_Poribacteria | 215 | F4055_14935 | 3290..3937 | 2026781 | HNH-endonuclease-[Candidatus-Poribacteria-bacterium] |
| MDE0555375.1 | HNH | GCA_028824855.1 | Candidatus_Poribacteria_bacterium | Bacteria>Bacteria_incertae_sedis>Bacteria_candidate_phyla>Candidatus_Poribacteria>unclassified_Candidatus_Poribacteria | 162 | OXI24_14225 | 1871..2356 | 2026781 | MAG:-HNH-endonuclease-signature-motif-containing-protein,-partial-[Candidatus-Poribacteria-bacterium] |
| MBI2566526.1 | HNH | GCA_016188575.1 | Candidatus_Schekmanbacteria_bacterium | Bacteria>Bacteria_incertae_sedis>Bacteria_candidate_phyla>Candidatus_Schekmaniibacteriota>unclassified_Candidatus_Schekmanbacteria | 204 | HYV63_05800 | 116619..117233 | 2419984 | hypothetical-protein-[Candidatus-Schekmanbacteria-bacterium] |
| AGC72576.1 | HNH | - | uncultured_bacterium_A1Q1_fos_1870 | Bacteria>environmental_samples | 220 | . | 22120..22782 | 1256554 | hypothetical-protein-[uncultured-bacterium-A1Q1_fos_1870] |
| AGC72564.1 | HNH | - | uncultured_bacterium_A1Q1_fos_1870 | Bacteria>environmental_samples | 189 | . | 5698..6267 | 1256554 | hypothetical-protein-[uncultured-bacterium-A1Q1_fos_1870] |

| Protein-ID | Domain | Assembly-ID | Strain | Taxonomy | Length | Locus-Tag | Genomic-coordinates | TaxID | Fasta-defline |
|---|---|---|---|---|---|---|---|---|---|
| WP_098973860.1 | HNH | GCF_037897565.1 | Fusobacterium_polymorphum | Bacteria>Fusobacteriati>Fusobacteriota>Fusobacteriia>Fusobacteriales>Fusobacteriaceae>Fusobacterium | 212 | CBG58_RS00375 | 91441..92079 | 76857 | HNH-endonuclease-[Fusobacterium-polymorphum] |
| WP_118127445.1 | HNH | GCF_003470285.1 | Fusobacterium_mortiferum | Bacteria>Fusobacteriati>Fusobacteriota>Fusobacteriia>Fusobacteriales>Fusobacteriaceae>Fusobacterium | 99 | DW663_RS02775 | 242176..242475 | 850 | putative-HNHc-nuclease-[Fusobacterium-mortiferum] |
| WP_194449892.1 | HNH | GCF_015277775.1 | Paludibaculum_fermentans | Bacteria>Pseudomonadati>Acidobacteriota>Terriglobia>Bryobacterales>Bryobacteraceae>Paludibaculum | 136 | IRI77_RS36755 | 9267345..9267755 | 1473598 | hypothetical-protein-[Paludibaculum-fermentans] |
| MBP7797360.1 | HNH | GCA_018000955.1 | Thermoanaerobaculaceae_bacterium | Bacteria>Pseudomonadati>Acidobacteriota>Thermoanaerobaculia>Thermoanaerobaculales>Thermoanaerobaculaceae>unclassified_Thermoanaerobaculaceae | 319 | KA072_01900 | 30614..31573 | 2699758 | hypothetical-protein-[Thermoanaerobaculaceae-bacterium] |
| KAB2964431.1 | HNH | GCA_009026165.1 | Thermoanaerobaculia_bacterium | Bacteria>Pseudomonadati>Acidobacteriota>Thermoanaerobaculia>unclassified_Thermoanaerobaculia | 187 | F9K16_04355 | 15795..16358 | 2651171 | MAG:-HNH-endonuclease-[Thermoanaerobaculia-bacterium] |
| MDE2976523.1 | HNH | GCA_028873735.1 | Acidobacteriota_bacterium | Bacteria>Pseudomonadati>Acidobacteriota>unclassified_Acidobacteriota | 254 | OXU63_03260 | 9807..10571 | 1978231 | MAG:-TIGR02646-family-protein-[Acidobacteriota-bacterium] |
| MCY3933632.1 | HNH | GCA_026707325.1 | Acidobacteriota_bacterium | Bacteria>Pseudomonadati>Acidobacteriota>unclassified_Acidobacteriota | 248 | OXH70_18120 | 788..1534 | 1978231 | MAG:-TIGR02646-family-protein-[Acidobacteria-bacterium] |
| MDE3261997.1 | HNH | GCA_028877265.1 | Acidobacteriota_bacterium | Bacteria>Pseudomonadati>Acidobacteriota>unclassified_Acidobacteriota | 230 | OYL41_08450 | 8255..8947 | 1978231 | MAG:-TIGR02646-family-protein-[Acidobacteriota-bacterium] |
| PKP62160.1 | HNH | GCA_002841055.1 | Candidatus_Atribacteria_bacterium_HGW-Atribacteria-1 | Bacteria>Pseudomonadati>Atribacterota>unclassified_Atribacterota | 208 | CVT91_00380 | 15036..15662 | 2013676 | hypothetical-protein-CVT91_00380-[Candidatus-Atribacteria-bacterium-HGW-Atribacteria-1] |
| MBI4801909.1 | HNH | GCA_016208085.1 | Elusimicrobiota_bacterium | Bacteria>Pseudomonadati>Elusimicrobiota>unclassified_Elusimicrobiota | 186 | HY796_05230 | 14032..14592 | 2030800 | TIGR02646-family-protein-[Elusimicrobiota-bacterium] |
| WP_004314653.1 | HNH | GCF_000178215.1 | Bacteroides | Bacteria>Pseudomonadati>FCB_group>Bacteroidota/Chlorobiota_group>Bacteroidota>Bacteroidia>Bacteroidales>Bacteroidaceae | 249 | CW3_RS16430 | 94885..95634 | 816 | MULTISPECIES:-hypothetical-protein-[Bacteroides] |
| CDM00528.1 | HNH | GCA_000577955.1 | Bacteroides_xylanisolvens_SD_CC_2a | Bacteria>Pseudomonadati>FCB_group>Bacteroidota/Chlorobiota_group>Bacteroidota>Bacteroidia>Bacteroidales>Bacteroidaceae>Bacteroides>Bacteroides_xylanisolvens | 249 | BN891_34530 | 150953..151702 | 702444 | hypothetical-protein-BN891_34530-[Bacteroides-xylanisolvens-SD-CC-2a] |
| EFF57554.1 | HNH | - | Bacteroides_xylanisolvens_SD_CC_2a | Bacteria>Pseudomonadati>FCB_group>Bacteroidota/Chlorobiota_group>Bacteroidota>Bacteroidia>Bacteroidales>Bacteroidaceae>Bacteroides>Bacteroides_xylanisolvens | 249 | CW1_2298 | 32725..33474 | 702444 | conserved-hypothetical-protein-TIGR02646-[Bacteroides-xylanisolvens-SD-CC-2a] |
| EFG14414.1 | HNH | - | Bacteroides_xylanisolvens_SD_CC_1b | Bacteria>Pseudomonadati>FCB_group>Bacteroidota/Chlorobiota_group>Bacteroidota>Bacteroidia>Bacteroidales>Bacteroidaceae>Bacteroides>Bacteroides_xylanisolvens | 249 | CW3_0494 | 94885..95634 | 702447 | conserved-hypothetical-protein-TIGR02646-[Bacteroides-xylanisolvens-SD-CC-1b] |
| MVZ89500.1 | HNH | GCA_009768845.1 | Microbacter_sp._ANSKLAB05 | Bacteria>Pseudomonadati>FCB_group>Bacteroidota/Chlorobiota_group>Bacteroidota>Bacteroidia>Bacteroidales>Porphyromonadaceae>Microbacter>unclassified_Microbacter | 209 | GQF49_03225 | 705168..705797 | 2684407 | HNH-endonuclease-[Microbacter-sp.-ANSKLAB05] |
| MBU0288753.1 | HNH | GCA_018789675.1 | Hallella_faecis | Bacteria>Pseudomonadati>FCB_group>Bacteroidota/Chlorobiota_group>Bacteroidota>Bacteroidia>Bacteroidales>Prevotellaceae>Hallella | 252 | KM934_00355 | 87124..87882 | 2841596 | HNH-endonuclease-[Hallella-faecis] |
| WP_044064716.1 | HNH | GCF_000833995.1 | Prevotella_pectinovora | Bacteria>Pseudomonadati>FCB_group>Bacteroidota/Chlorobiota_group>Bacteroidota>Bacteroidia>Bacteroidales>Prevotellaceae>Prevotella | 244 | ST45_RS02490 | 63495..64229 | 1602169 | HNH-endonuclease-[Prevotella-pectinovora] |
| HAM28419.1 | HNH | GCA_003453655.1 | Prevotella_sp. | Bacteria>Pseudomonadati>FCB_group>Bacteroidota/Chlorobiota_group>Bacteroidota>Bacteroidia>Bacteroidales>Prevotellaceae>Prevotella>unclassified_Prevotella | 244 | DCP79_00690 | 75154..75888 | 59823 | hypothetical-protein-[Prevotella-sp.] |
| MCI6336463.1 | HNH | GCA_022777825.1 | Prevotella_sp. | Bacteria>Pseudomonadati>FCB_group>Bacteroidota/Chlorobiota_group>Bacteroidota>Bacteroidia>Bacteroidales>Prevotellaceae>Prevotella>unclassified_Prevotella | 178 | MR802_00735 | 17186..17722 | 59823 | hypothetical-protein-[Prevotella-sp.] |
| MCI6342421.1 | HNH | GCA_022777695.1 | Prevotella_sp. | Bacteria>Pseudomonadati>FCB_group>Bacteroidota/Chlorobiota_group>Bacteroidota>Bacteroidia>Bacteroidales>Prevotellaceae>Prevotella>unclassified_Prevotella | 175 | MR788_07705 | 55864..56391 | 59823 | HNH-endonuclease-[Prevotella-sp.] |
| MBQ2852767.1 | HNH | GCA_017409275.1 | Bacteroidales_bacterium | Bacteria>Pseudomonadati>FCB_group>Bacteroidota/Chlorobiota_group>Bacteroidota>Bacteroidia>Bacteroidales>unclassified_Bacteroidales | 236 | IJE76_05180 | 121178..121888 | 2030927 | hypothetical-protein-[Bacteroidales-bacterium] |
| MCM1267120.1 | HNH | GCA_023665065.1 | Bacteroidales_bacterium | Bacteria>Pseudomonadati>FCB_group>Bacteroidota/Chlorobiota_group>Bacteroidota>Bacteroidia>Bacteroidales>unclassified_Bacteroidales | 338 | NC302_04370 | 38461..39477 | 2030927 | hypothetical-protein-[Bacteroidales-bacterium] |
| NCB25243.1 | HNH | GCA_009929165.1 | Bacteroidia_bacterium | Bacteria>Pseudomonadati>FCB_group>Bacteroidota/Chlorobiota_group>Bacteroidota>Bacteroidia>unclassified_Bacteroidia | 229 | EOM62_07180 | 2525..3214 | 2044936 | hypothetical-protein-[Bacteroidia-bacterium] |
| WP_159471210.1 | HNH | GCF_009801115.1 | Dyadobacter_sp._3J3 | Bacteria>Pseudomonadati>FCB_group>Bacteroidota/Chlorobiota_group>Bacteroidota>Cytophagia>Cytophagales>Spirosomataceae>Dyadobacter>unclassified_Dyadobacter | 210 | FZF08_RS17835 | 87107..87739 | 2606600 | hypothetical-protein-[Dyadobacter-sp.-3J3] |
| WP_273212246.1 | HNH | GCF_002483185.1 | Runella_zeae | Bacteria>Pseudomonadati>FCB_group>Bacteroidota/Chlorobiota_group>Bacteroidota>Cytophagia>Cytophagales>Spirosomataceae>Runella | 207 | B4E63_RS12450 | 305563..306186 | 94255 | hypothetical-protein-[Runella-zeae] |
| WP_141421450.1 | HNH | GCF_006546625.1 | Haloflavibacter_putidus | Bacteria>Pseudomonadati>FCB_group>Bacteroidota/Chlorobiota_group>Bacteroidota>Flavobacteriia>Flavobacteriales>Flavobacteriaceae>Haloflavibacter | 222 | FKR84_RS06270 | 31326..31994 | 2576776 | HNH-endonuclease-[Haloflavibacter-putidus] |
| WP_076662680.1 | HNH | GCF_001975575.1 | Mangrovimonas_sp._DI_80 | Bacteria>Pseudomonadati>FCB_group>Bacteroidota/Chlorobiota_group>Bacteroidota>Flavobacteriia>Flavobacteriales>Flavobacteriaceae>Mangrovimonas>unclassified_Mangrovimonas | 155 | BKM32_RS04310 | 833485..833952 | 1779330 | hypothetical-protein-[Mangrovimonas-sp.-DI-80] |
| MCB0808338.1 | HNH | GCA_020438005.1 | Flavobacteriales_bacterium | Bacteria>Pseudomonadati>FCB_group>Bacteroidota/Chlorobiota_group>Bacteroidota>Flavobacteriia>Flavobacteriales>unclassified_Flavobacteriales | 196 | KDB96_03580 | 5173..5763 | 2021391 | TIGR02646-family-protein-[Flavobacteriales-bacterium] |
| NUQ24575.1 | HNH | GCA_013360405.1 | Saprospiraceae_bacterium | Bacteria>Pseudomonadati>FCB_group>Bacteroidota/Chlorobiota_group>Bacteroidota>Saprospiria>Saprospirales>Saprospiraceae>unclassified_Saprospiraceae | 297 | HUU34_11535 | 38604..39497 | 2202734 | HNH-endonuclease-[Saprospiraceae-bacterium] |
| MCB0517808.1 | HNH | GCA_020436325.1 | Saprospiraceae_bacterium | Bacteria>Pseudomonadati>FCB_group>Bacteroidota/Chlorobiota_group>Bacteroidota>Saprospiria>Saprospirales>Saprospiraceae>unclassified_Saprospiraceae | 184 | KDD27_02655 | 16662..17216 | 2202734 | HNH-endonuclease-[Saprospiraceae-bacterium] |
| MBN8706501.1 | HNH | GCA_017303995.1 | Bacteroidota_bacterium | Bacteria>Pseudomonadati>FCB_group>Bacteroidota/Chlorobiota_group>Bacteroidota>unclassified_Bacteroidota | 203 | JOL62_11555 | 13154..13765 | 1898104 | hypothetical-protein-[Bacteroidota-bacterium] |
| MBS1911237.1 | HNH | GCA_018266795.1 | Bacteroidota_bacterium | Bacteria>Pseudomonadati>FCB_group>Bacteroidota/Chlorobiota_group>Bacteroidota>unclassified_Bacteroidota | 199 | JST22_04565 | 48108..48707 | 1898104 | hypothetical-protein-[Bacteroidota-bacterium] |
| NLW30560.1 | HNH | GCA_012728995.1 | Fibrobacter_sp. | Bacteria>Pseudomonadati>FCB_group>Fibrobacterota>Fibrobacteria>Fibrobacterales>Fibrobacteraceae>Fibrobacter>unclassified_Fibrobacter | 243 | GXY77_03785 | 993..1724 | 35828 | HNH-endonuclease-[Fibrobacter-sp.] |
| MCY4606376.1 | HNH | GCA_026713655.1 | Gemmatimonadota_bacterium | Bacteria>Pseudomonadati>FCB_group>Gemmatimonadota>unclassified_Gemmatimonadota | 258 | OXE49_19240 | 2327..3103 | 2026742 | MAG:-TIGR02646-family-protein-[Gemmatimonadetes-bacterium] |
| WP_171805993.1 | HNH | GCF_013155195.1 | Corallococcus_exiguus | Bacteria>Pseudomonadati>Myxococcota>Myxococcia>Myxococcales>Cystobacterineae>Myxococcaceae>Corallococcus | 214 | HPP06_RS00255 | 60228..60872 | 83462 | TIGR02646-family-protein-[Corallococcus-exiguus] |
| WP_267556339.1 | HNH | GCF_026626625.1 | Corallococcus_sp._BB11-1 | Bacteria>Pseudomonadati>Myxococcota>Myxococcia>Myxococcales>Cystobacterineae>Myxococcaceae>Corallococcus>unclassified_Corallococcus | 221 | OV207_RS17675 | 233935..234600 | 2996783 | hypothetical-protein-[Corallococcus-sp.-BB11-1] |
| WP_201422027.1 | HNH | GCF_016698685.1 | Myxococcus_xanthus | Bacteria>Pseudomonadati>Myxococcota>Myxococcia>Myxococcales>Cystobacterineae>Myxococcaceae>Myxococcus | 221 | JKA73_RS29365 | 7339111..7339776 | 34 | hypothetical-protein-[Myxococcus-xanthus] |
| WP_174489002.1 | HNH | GCF_013336725.1 | Myxococcus_sp._CA040A | Bacteria>Pseudomonadati>Myxococcota>Myxococcia>Myxococcales>Cystobacterineae>Myxococcaceae>Myxococcus>unclassified_Myxococcus | 211 | HUA75_RS36320 | 170634..171269 | 2741738 | TIGR02646-family-protein-[Myxococcus-sp.-CA040A] |
| HCF56395.1 | HNH | GCA_003534605.1 | Myxococcales_bacterium | Bacteria>Pseudomonadati>Myxococcota>Myxococcia>Myxococcales>unclassified_Myxococcales | 286 | DFS52_00160 | 10625..11485 | 2026763 | hypothetical-protein-[Myxococcales-bacterium] |
| MBI5504527.1 | HNH | GCA_016218925.1 | Deltaproteobacteria_bacterium | Bacteria>Pseudomonadati>Myxococcota>Myxococcia>unclassified_Myxococcia | 202 | HY899_06975 | 25058..25666 | 2026735 | HNH-endonuclease-[Deltaproteobacteria-bacterium] |
| WP_157907221.1 | HNH | GCF_000067165.1 | Sorangium_cellulosum | Bacteria>Pseudomonadati>Myxococcota>Polyangia>Polyangiales>Polyangiaceae>Sorangium | 221 | SCE_RS49870 | 10699658..10700323 | 56 | hypothetical-protein-[Sorangium-cellulosum] |
| KYF87244.1 | HNH | GCA_001589185.1 | Sorangium_cellulosum | Bacteria>Pseudomonadati>Myxococcota>Polyangia>Polyangiales>Polyangiaceae>Sorangium | 215 | BE17_46145 | 11395..12042 | 56 | hypothetical-protein-BE17_46145-[Sorangium-cellulosum] |
| WP_020733580.1 | HNH | GCF_000418325.1 | Sorangium_cellulosum | Bacteria>Pseudomonadati>Myxococcota>Polyangia>Polyangiales>Polyangiaceae>Sorangium | 189 | SCE1572_RS08125 | 2244320..2244889 | 56 | hypothetical-protein-[Sorangium-cellulosum] |
| MCK6536701.1 | HNH | GCA_023150915.1 | Polyangiaceae_bacterium | Bacteria>Pseudomonadati>Myxococcota>Polyangia>Polyangiales>Polyangiaceae>unclassified_Polyangiaceae | 217 | L6Q84_27345 | 16766..17419 | 2268199 | hypothetical-protein-[Polyangiaceae-bacterium] |
| MCK6527597.1 | HNH | GCA_023150875.1 | Myxococcota_bacterium | Bacteria>Pseudomonadati>Myxococcota>unclassified_Myxococcota | 185 | L6R50_08550 | 51103..51660 | 2818507 | hypothetical-protein-[Myxococcota-bacterium] |
| MBT3366895.1 | HNH | GCA_018648125.1 | Nitrospina_sp. | Bacteria>Pseudomonadati>Nitrospinota/Tectimicrobiota_group>Nitrospinota>Nitrospinia>Nitrospinales>Nitrospinaceae>Nitrospina>unclassified_Nitrospina | 59 | HN416_07060 | 12963..13142 | 2024844 | hypothetical-protein-[Nitrospina-sp.] |
| MBZ0154719.1 | HNH | GCA_019912415.1 | Candidatus_Nitrobium_versatile | Bacteria>Pseudomonadati>Nitrospirota>Nitrospiria>Nitrospirales>Nitrospiraceae>Candidatus_Nitrobium | 212 | K8I29_00720 | 52015..52653 | 2884831 | TIGR02646-family-protein-[Alphaproteobacteria-bacterium] |
| WP_150138113.1 | HNH | GCF_014163875.1 | Gammaproteobacteria | Bacteria>Pseudomonadati>Pseudomonadota | 216 | H5088_RS10940 | 137451..138101 | 1236 | MULTISPECIES:-TIGR02646-family-protein-[Gammaproteobacteria] |
| MBU2768214.1 | HNH | GCA_018853935.1 | Acidithiobacillus_ferrivorans | Bacteria>Pseudomonadati>Pseudomonadota>Acidithiobacillia>Acidithiobacillales>Acidithiobacillaceae>Acidithiobacillus | 238 | HAP94_19080 | 7342..8058 | 160808 | TIGR02646-family-protein-[Acidithiobacillus-ferrivorans] |
| UBU61067.1 | HNH | GCA_020221655.1 | Acidithiobacillus_ferrooxidans | Bacteria>Pseudomonadati>Pseudomonadota>Acidithiobacillia>Acidithiobacillales>Acidithiobacillaceae>Acidithiobacillus | 199 | LDB30_08040 | 1571999..1572598 | 920 | hypothetical-protein-LDB30_08040-[Acidithiobacillus-ferrooxidans] |
| WP_232474707.1 | HNH | GCF_016106015.1 | Neoroseomonas_rubea | Bacteria>Pseudomonadati>Pseudomonadota>Alphaproteobacteria>Acetobacterales>Acetobacteraceae>Neoroseomonas | 250 | HW577_RS05715 | 10173..10925 | 2748666 | TIGR02646-family-protein-[Roseomonas-rubea] |
| WP_082559273.1 | HNH | GCF_001426045.1 | Methylobacterium_sp._Leaf94 | Bacteria>Pseudomonadati>Pseudomonadota>Alphaproteobacteria>Hyphomicrobiales>Methylobacteriaceae>Methylobacterium>unclassified_Methylobacterium | 213 | ASG63_RS24160 | 10375..11016 | 1736250 | TIGR02646-family-protein-[Methylobacterium-sp.-Leaf94] |
| KFG67136.1 | HNH | GCA_000739015.1 | Microvirga_sp._BSC39 | Bacteria>Pseudomonadati>Pseudomonadota>Alphaproteobacteria>Hyphomicrobiales>Methylobacteriaceae>Microvirga>unclassified_Microvirga | 209 | JH26_23880 | 117803..118432 | 1549810 | hypothetical-protein-JH26_23880-[Microvirga-sp.-BSC39] |
| WP_036357320.1 | HNH | GCF_000739015.1 | Microvirga_sp._BSC39 | Bacteria>Pseudomonadati>Pseudomonadota>Alphaproteobacteria>Hyphomicrobiales>Methylobacteriaceae>Microvirga>unclassified_Microvirga | 209 | MA07_RS22805 | 117803..118432 | 1549810 | hypothetical-protein-[Microvirga-sp.-BSC39] |
| KFG67135.1 | HNH | GCA_000739015.1 | Microvirga_sp._BSC39 | Bacteria>Pseudomonadati>Pseudomonadota>Alphaproteobacteria>Hyphomicrobiales>Methylobacteriaceae>Microvirga>unclassified_Microvirga | 99 | JH26_23875 | 117504..117803 | 1549810 | hypothetical-protein-JH26_23875-[Microvirga-sp.-BSC39] |
| WP_036357319.1 | HNH | GCF_000739015.1 | Microvirga_sp._BSC39 | Bacteria>Pseudomonadati>Pseudomonadota>Alphaproteobacteria>Hyphomicrobiales>Methylobacteriaceae>Microvirga>unclassified_Microvirga | 99 | MA07_RS22800 | 117504..117803 | 1549810 | hypothetical-protein-[Microvirga-sp.-BSC39] |
| WP_172236131.1 | HNH | GCF_902482845.1 | Bradyrhizobium | Bacteria>Pseudomonadati>Pseudomonadota>Alphaproteobacteria>Hyphomicrobiales>Nitrobacteraceae | 205 | JQ619_RS32205 | 13915..14532 | 374 | MULTISPECIES:-TIGR02646-family-protein-[Bradyrhizobium] |
| WP_166097058.1 | HNH | GCF_011516635.5 | Bradyrhizobium_barranii | Bacteria>Pseudomonadati>Pseudomonadota>Alphaproteobacteria>Hyphomicrobiales>Nitrobacteraceae>Bradyrhizobium | 213 | HAP41_RS22265 | 4681348..4681989 | 2992140 | TIGR02646-family-protein-[Bradyrhizobium-barranii] |
| WP_212326839.1 | HNH | GCF_018131105.1 | Bradyrhizobium_manausense | Bacteria>Pseudomonadati>Pseudomonadota>Alphaproteobacteria>Hyphomicrobiales>Nitrobacteraceae>Bradyrhizobium | 205 | JQ631_RS13555 | 2845193..2845810 | 989370 | TIGR02646-family-protein-[Bradyrhizobium-manausense] |
| WP_116622074.1 | HNH | GCF_050247095.1 | Mesorhizobium_denitrificans | Bacteria>Pseudomonadati>Pseudomonadota>Alphaproteobacteria>Hyphomicrobiales>Phyllobacteriaceae>Mesorhizobium | 215 | DY251_RS01600 | 345438..346085 | 2294114 | TIGR02646-family-protein-[Mesorhizobium-denitrificans] |
| WP_244506972.1 | HNH | GCF_044005695.1 | Mesorhizobium_ciceri | Bacteria>Pseudomonadati>Pseudomonadota>Alphaproteobacteria>Hyphomicrobiales>Phyllobacteriaceae>Mesorhizobium | 228 | A4R29_RS11150 | 2458388..2459074 | 39645 | HNH-endonuclease-signature-motif-containing-protein-[Mesorhizobium-ciceri] |
| WP_140734683.1 | HNH | GCF_006442905.1 | Mesorhizobium_sp._B2-4-9 | Bacteria>Pseudomonadati>Pseudomonadota>Alphaproteobacteria>Hyphomicrobiales>Phyllobacteriaceae>Mesorhizobium>unclassified_Mesorhizobium | 213 | FJ945_RS19665 | 28978..29619 | 2589940 | TIGR02646-family-protein-[Mesorhizobium-sp.-B2-4-9] |
| WP_128286155.1 | HNH | GCF_004023115.1 | Mesorhizobium_sp._M6A.T.Cr.TU.017.01.1.1 | Bacteria>Pseudomonadati>Pseudomonadota>Alphaproteobacteria>Hyphomicrobiales>Phyllobacteriaceae>Mesorhizobium>unclassified_Mesorhizobium | 210 | EOB36_RS27995 | 9158..9790 | 2496774 | TIGR02646-family-protein-[Mesorhizobium-sp.-M6A.T.Cr.TU.017.01.1.1] |
| RWC29994.1 | HNH | GCA_004017395.1 | Mesorhizobium_sp. | Bacteria>Pseudomonadati>Pseudomonadota>Alphaproteobacteria>Hyphomicrobiales>Phyllobacteriaceae>Mesorhizobium>unclassified_Mesorhizobium | 209 | EOS27_14200 | 61973..62602 | 1871066 | TIGR02646-family-protein-[Mesorhizobium-sp.] |
| WP_136091388.1 | HNH | GCF_004794715.1 | Mesorhizobium_sp._M8A.F.Ca.ET.207.01.1.1 | Bacteria>Pseudomonadati>Pseudomonadota>Alphaproteobacteria>Hyphomicrobiales>Phyllobacteriaceae>Mesorhizobium>unclassified_Mesorhizobium | 209 | EN850_RS12630 | 50639..51268 | 2563968 | TIGR02646-family-protein-[Mesorhizobium-sp.-M8A.F.Ca.ET.207.01.1.1] |
| MCJ8508802.1 | HNH | GCA_022968395.1 | Rhizobium_lemnae | Bacteria>Pseudomonadati>Pseudomonadota>Alphaproteobacteria>Hyphomicrobiales>Rhizobiaceae>Rhizobium/Agrobacterium_group>Rhizobium | 215 | MUU53_12865 | 40793..41440 | 1214924 | TIGR02646-family-protein-[Rhizobium-lemnae] |
| WP_247261242.1 | HNH | GCF_022968395.1 | Rhizobium_lemnae | Bacteria>Pseudomonadati>Pseudomonadota>Alphaproteobacteria>Hyphomicrobiales>Rhizobiaceae>Rhizobium/Agrobacterium_group>Rhizobium | 215 | MUU53_RS12870 | 40793..41440 | 1214924 | TIGR02646-family-protein-[Rhizobium-lemnae] |
| WP_200985536.1 | HNH | - | Rhizobium_rhizogenes | Bacteria>Pseudomonadati>Pseudomonadota>Alphaproteobacteria>Hyphomicrobiales>Rhizobiaceae>Rhizobium/Agrobacterium_group>Rhizobium | 214 | JL018_RS00090 | 20430..21074 | 359 | TIGR02646-family-protein-[Agrobacterium-rhizogenes] |
| NEJ60791.1 | HNH | GCA_010668765.1 | Rhizobium_ruizarguesonis | Bacteria>Pseudomonadati>Pseudomonadota>Alphaproteobacteria>Hyphomicrobiales>Rhizobiaceae>Rhizobium/Agrobacterium_group>Rhizobium | 213 | GUK21_31960 | 65585..66226 | 2081791 | TIGR02646-family-protein-[Rhizobium-leguminosarum] |
| WP_012755927.1 | HNH | - | Rhizobium_leguminosarum | Bacteria>Pseudomonadati>Pseudomonadota>Alphaproteobacteria>Hyphomicrobiales>Rhizobiaceae>Rhizobium/Agrobacterium_group>Rhizobium | 213 | RLEG_RS01015 | 205492..206133 | 384 | TIGR02646-family-protein-[Rhizobium-leguminosarum] |
| WP_130838771.1 | HNH | GCF_004305895.1 | Rhizobium_ruizarguesonis | Bacteria>Pseudomonadati>Pseudomonadota>Alphaproteobacteria>Hyphomicrobiales>Rhizobiaceae>Rhizobium/Agrobacterium_group>Rhizobium | 212 | HA464_RS06755 | 1372422..1373060 | 2081791 | TIGR02646-family-protein-[Rhizobium-leguminosarum] |
| WP_246230969.1 | HNH | GCF_010669145.1 | Rhizobium_oryzihabitans | Bacteria>Pseudomonadati>Pseudomonadota>Alphaproteobacteria>Hyphomicrobiales>Rhizobiaceae>Rhizobium/Agrobacterium_group>Rhizobium | 208 | G3A56_RS10095 | 1926366..1926992 | 2267833 | TIGR02646-family-protein-[Rhizobium-oryzihabitans] |
| WP_069615758.1 | HNH | GCF_001729315.1 | Rhizobium_sp._YK2 | Bacteria>Pseudomonadati>Pseudomonadota>Alphaproteobacteria>Hyphomicrobiales>Rhizobiaceae>Rhizobium/Agrobacterium_group>Rhizobium>unclassified_Rhizobium | 215 | A9Z06_RS30965 | 20934..21581 | 1860096 | TIGR02646-family-protein-[Rhizobium-sp.-YK2] |
| WP_242223012.1 | HNH | GCF_022682305.1 | Shinella_zoogloeoides | Bacteria>Pseudomonadati>Pseudomonadota>Alphaproteobacteria>Hyphomicrobiales>Rhizobiaceae>Shinella | 213 | MOE34_RS09020 | 1761575..1762216 | 352475 | TIGR02646-family-protein-[Shinella-zoogloeoides] |
| WP_010968625.1 | HNH | GCF_000006965.1 | Sinorhizobium_meliloti | Bacteria>Pseudomonadati>Pseudomonadota>Alphaproteobacteria>Hyphomicrobiales>Rhizobiaceae>Sinorhizobium/Ensifer_group>Sinorhizobium | 215 | J3S92_RS00015 | 4430..5077 | 382 | TIGR02646-family-protein-[Sinorhizobium-meliloti] |
| SDR63350.1 | HNH | GCA_900103895.1 | Rhizobiales_bacterium_GAS113 | Bacteria>Pseudomonadati>Pseudomonadota>Alphaproteobacteria>Hyphomicrobiales>unclassified_Hyphomicrobiales | 218 | SAMN05519103_08519 | 36531..37187 | 1884352 | TIGR02646-family-protein-[Rhizobiales-bacterium-GAS113] |
| MBF0399387.1 | HNH | GCA_015232635.1 | Magnetococcales_bacterium | Bacteria>Pseudomonadati>Pseudomonadota>Alphaproteobacteria>Magnetococcales>unclassified_Magnetococcales | 200 | HQL90_01315 | 27036..27638 | 2026759 | hypothetical-protein-[Magnetococcales-bacterium] |
| WP_013654354.1 | HNH | GCF_000192745.1 | Polymorphum_gilvum | Bacteria>Pseudomonadati>Pseudomonadota>Alphaproteobacteria>Rhodobacterales>Paracoccaceae>Polymorphum | 213 | SL003B_RS18275 | 3925864..3926505 | 991904 | TIGR02646-family-protein-[Polymorphum-gilvum] |
| WP_211114072.1 | HNH | - | Azospirillum_picis | Bacteria>Pseudomonadati>Pseudomonadota>Alphaproteobacteria>Rhodospirillales>Azospirillaceae>Azospirillum | 263 | J2850_RS30820 | 6228..7019 | 488438 | TIGR02646-family-protein-[Azospirillum-picis] |
| WP_247891469.1 | HNH | - | Azospirillum_argentinense | Bacteria>Pseudomonadati>Pseudomonadota>Alphaproteobacteria>Rhodospirillales>Azospirillaceae>Azospirillum | 243 | GAY33_RS05500 | 18067..18798 | 2970906 | TIGR02646-family-protein-[Azospirillum-argentinense] |
| TXH33433.1 | HNH | GCA_008012315.1 | Rhodospirillaceae_bacterium | Bacteria>Pseudomonadati>Pseudomonadota>Alphaproteobacteria>Rhodospirillales>Rhodospirillaceae>unclassified_Rhodospirillaceae | 215 | E6Q98_22195 | 62511..63158 | 1898112 | TIGR02646-family-protein-[Rhodospirillaceae-bacterium] |
| MBL8657368.1 | HNH | GCA_016793865.1 | Altererythrobacter_sp. | Bacteria>Pseudomonadati>Pseudomonadota>Alphaproteobacteria>Sphingomonadales>Erythrobacteraceae>Altererythrobacter>unclassified_Altererythrobacter | 220 | JNJ92_08450 | 7386..8048 | 1872480 | TIGR02646-family-protein-[Altererythrobacter-sp.] |
| MCV0384727.1 | HNH | GCA_025698865.1 | Erythrobacter_sp. | Bacteria>Pseudomonadati>Pseudomonadota>Alphaproteobacteria>Sphingomonadales>Erythrobacteraceae>Erythrobacter/Porphyrobacter_group>Erythrobacter>unclassified_Erythrobacter | 158 | K5799_14995 | 4159..4635 | 1042 | MAG:-HNH-endonuclease-[Erythrobacter-sp.] |
| WP_099185491.1 | HNH | GCF_002723855.1 | Sphingobium | Bacteria>Pseudomonadati>Pseudomonadota>Alphaproteobacteria>Sphingomonadales>Sphingomonadaceae | 213 | SFOMI_RS04990 | 862951..863592 | 165695 | MULTISPECIES:-TIGR02646-family-protein-[Sphingobium] |
| WP_168853124.1 | HNH | GCF_012641335.1 | Novosphingobium_sp._ERN07 | Bacteria>Pseudomonadati>Pseudomonadota>Alphaproteobacteria>Sphingomonadales>Sphingomonadaceae>Novosphingobium>unclassified_Novosphingobium | 213 | HGI47_RS12595 | 101586..102227 | 2726187 | TIGR02646-family-protein-[Novosphingobium-sp.-ERN07] |
| WP_130751565.1 | HNH | GCF_004305355.1 | Sphingobium_xenophagum | Bacteria>Pseudomonadati>Pseudomonadota>Alphaproteobacteria>Sphingomonadales>Sphingomonadaceae>Sphingobium | 213 | MBESOW_RS00085 | 18469..19110 | 121428 | TIGR02646-family-protein-[Sphingobium-xenophagum] |
| WP_159367775.1 | HNH | GCF_009856545.1 | Sphingobium_yanoikuyae | Bacteria>Pseudomonadati>Pseudomonadota>Alphaproteobacteria>Sphingomonadales>Sphingomonadaceae>Sphingobium | 213 | GS397_RS23445 | 4925090..4925731 | 13690 | TIGR02646-family-protein-[Sphingobium-yanoikuyae] |
| RXZ34414.1 | HNH | GCA_004135605.1 | Sphingomonas_desiccabilis | Bacteria>Pseudomonadati>Pseudomonadota>Alphaproteobacteria>Sphingomonadales>Sphingomonadaceae>Sphingomonas | 214 | EO081_01610 | 353545..354189 | 429134 | TIGR02646-family-protein-[Sphingomonas-desiccabilis] |
| WP_129340203.1 | HNH | GCF_014196135.1 | Sphingomonas_desiccabilis | Bacteria>Pseudomonadati>Pseudomonadota>Alphaproteobacteria>Sphingomonadales>Sphingomonadaceae>Sphingomonas | 214 | GGQ81_RS01625 | 353513..354157 | 429134 | TIGR02646-family-protein-[Sphingomonas-desiccabilis] |
| WP_169068311.1 | HNH | GCF_012939955.1 | Candidatus_Accumulibacter_phosphatis | Bacteria>Pseudomonadati>Pseudomonadota>Betaproteobacteria>Betaproteobacteria_incertae_sedis>Candidatus_Accumulibacter | 238 | E4Q23_RS20275 | 16371..17087 | 327160 | TIGR02646-family-protein-[Candidatus-Accumulibacter-phosphatis] |
| MBK8578268.1 | HNH | GCA_016714935.1 | Candidatus_Accumulibacter_propinquus | Bacteria>Pseudomonadati>Pseudomonadota>Betaproteobacteria>Betaproteobacteria_incertae_sedis>Candidatus_Accumulibacter | 200 | IPN78_08385 | 111362..111964 | 2954380 | hypothetical-protein-[Candidatus-Accumulibacter-sp.] |
| QKS29572.1 | HNH | GCA_013347225.1 | Candidatus_Accumulibacter_similis | Bacteria>Pseudomonadati>Pseudomonadota>Betaproteobacteria>Betaproteobacteria_incertae_sedis>Candidatus_Accumulibacter | 191 | HT579_12040 | 2657060..2657635 | 2954388 | MAG:-hypothetical-protein-HT579_12040-[Candidatus-Accumulibacter-similis] |
| WP_248594674.1 | HNH | - | Candidatus_Accumulibacter_contiguus | Bacteria>Pseudomonadati>Pseudomonadota>Betaproteobacteria>Betaproteobacteria_incertae_sedis>Candidatus_Accumulibacter | 183 | E4Q08_RS03255 | 8732..9283 | 2954381 | TIGR02646-family-protein-[Candidatus-Accumulibacter-contiguus] |
| MBN8437007.1 | HNH | GCA_017302555.1 | Accumulibacter_sp. | Bacteria>Pseudomonadati>Pseudomonadota>Betaproteobacteria>Betaproteobacteria_incertae_sedis>Candidatus_Accumulibacter>unclassified_Candidatus_Accumulibacter | 257 | J0L95_03035 | 54389..55162 | 2053492 | TIGR02646-family-protein-[Candidatus-Accumulibacter-sp.] |
| MBN8498944.1 | HNH | GCA_017302455.1 | Accumulibacter_sp. | Bacteria>Pseudomonadati>Pseudomonadota>Betaproteobacteria>Betaproteobacteria_incertae_sedis>Candidatus_Accumulibacter>unclassified_Candidatus_Accumulibacter | 257 | J0M32_19140 | 10569..11342 | 2053492 | TIGR02646-family-protein-[Candidatus-Accumulibacter-sp.] |
| MBK8385353.1 | HNH | GCA_016711725.1 | Accumulibacter_sp. | Bacteria>Pseudomonadati>Pseudomonadota>Betaproteobacteria>Betaproteobacteria_incertae_sedis>Candidatus_Accumulibacter>unclassified_Candidatus_Accumulibacter | 200 | IPL11_06640 | 108355..108957 | 2053492 | hypothetical-protein-[Candidatus-Accumulibacter-sp.] |
| MBN8436518.1 | HNH | GCA_017302555.1 | Accumulibacter_sp. | Bacteria>Pseudomonadati>Pseudomonadota>Betaproteobacteria>Betaproteobacteria_incertae_sedis>Candidatus_Accumulibacter>unclassified_Candidatus_Accumulibacter | 200 | J0L95_00485 | 113442..114044 | 2053492 | hypothetical-protein-[Candidatus-Accumulibacter-sp.] |
| WP_043683818.1 | HNH | GCF_000612685.1 | Castellaniella_defragrans | Bacteria>Pseudomonadati>Pseudomonadota>Betaproteobacteria>Burkholderiales>Alcaligenaceae>Castellaniella | 131 | BN940_RS13260 | 2979142..2979537 | 75697 | hypothetical-protein-[Castellaniella-defragrans] |
| WP_148304966.1 | HNH | GCF_000612685.1 | Castellaniella_defragrans | Bacteria>Pseudomonadati>Pseudomonadota>Betaproteobacteria>Burkholderiales>Alcaligenaceae>Castellaniella | 123 | BN940_RS19065 | 2977146..2977517 | 75697 | hypothetical-protein-[Castellaniella-defragrans] |
| WP_085318437.1 | HNH | GCF_002105195.1 | Derxia_lacustris | Bacteria>Pseudomonadati>Pseudomonadota>Betaproteobacteria>Burkholderiales>Alcaligenaceae>Derxia | 214 | B9P64_RS23180 | 24549..25193 | 764842 | TIGR02646-family-protein-[Derxia-lacustris] |
| SUA58553.1 | HNH | GCA_900454315.1 | Oligella_ureolytica | Bacteria>Pseudomonadati>Pseudomonadota>Betaproteobacteria>Burkholderiales>Alcaligenaceae>Oligella | 199 | NCTC11998_02426 | 2467203..2467802 | 90244 | Uncharacterised-protein-[Oligella-ureolytica] |
| WP_013697446.1 | HNH | GCF_000194745.1 | Burkholderia_gladioli | Bacteria>Pseudomonadati>Pseudomonadota>Betaproteobacteria>Burkholderiales>Burkholderiaceae>Burkholderia | 178 | BGLA_RS07090 | 1588164..1588700 | 28095 | hypothetical-protein-[Burkholderia-gladioli] |
| WP_208457321.1 | HNH | GCF_902832475.1 | Burkholderia_gladioli | Bacteria>Pseudomonadati>Pseudomonadota>Betaproteobacteria>Burkholderiales>Burkholderiaceae>Burkholderia | 146 | H9G27_RS36835 | 159441..159881 | 28095 | hypothetical-protein-[Burkholderia-gladioli] |
| WP_217096868.1 | HNH | GCF_019041095.1 | Burkholderia_multivorans | Bacteria>Pseudomonadati>Pseudomonadota>Betaproteobacteria>Burkholderiales>Burkholderiaceae>Burkholderia>Burkholderia_cepacia_complex | 245 | KTE23_RS28435 | 32692..33429 | 87883 | TIGR02646-family-protein-[Burkholderia-multivorans] |
| WP_260448061.1 | HNH | GCF_003857875.1 | Burkholderia_stagnalis | Bacteria>Pseudomonadati>Pseudomonadota>Betaproteobacteria>Burkholderiales>Burkholderiaceae>Burkholderia>Burkholderia_cepacia_complex | 81 | DF113_RS32920 | 3712..3957 | 1503054 | hypothetical-protein-[Burkholderia-stagnalis] |
| WP_080401597.1 | HNH | GCF_001718355.1 | Burkholderia_ubonensis | Bacteria>Pseudomonadati>Pseudomonadota>Betaproteobacteria>Burkholderiales>Burkholderiaceae>Burkholderia>Burkholderia_cepacia_complex | 216 | WI31_RS35055 | 19653..20303 | 101571 | HNH-endonuclease-signature-motif-containing-protein-[Burkholderia-ubonensis] |
| ABO55014.1 | HNH | GCA_000016205.1 | Burkholderia_vietnamiensis_G4 | Bacteria>Pseudomonadati>Pseudomonadota>Betaproteobacteria>Burkholderiales>Burkholderiaceae>Burkholderia>Burkholderia_cepacia_complex>Burkholderia_vietnamiensis | 189 | Bcep1808_2011 | 2200494..2201063 | 269482 | conserved-hypothetical-protein-[Burkholderia-vietnamiensis-G4] |
| WP_122833863.1 | HNH | GCF_900597175.2 | Burkholderia_pseudomallei | Bacteria>Pseudomonadati>Pseudomonadota>Betaproteobacteria>Burkholderiales>Burkholderiaceae>Burkholderia>pseudomallei_group | 178 | EFT75_RS19505 | 207487..208023 | 28450 | hypothetical-protein-[Burkholderia-pseudomallei] |
| WP_201659273.1 | HNH | GCF_904848645.1 | Paraburkholderia_sabiae | Bacteria>Pseudomonadati>Pseudomonadota>Betaproteobacteria>Burkholderiales>Burkholderiaceae>Paraburkholderia | 178 | JMY25_RS35975 | 21197..21733 | 273251 | hypothetical-protein-[Paraburkholderia-sabiae] |
| WP_241974608.1 | HNH | GCF_003634175.1 | Paraburkholderia_sp._BL17N1 | Bacteria>Pseudomonadati>Pseudomonadota>Betaproteobacteria>Burkholderiales>Burkholderiaceae>Paraburkholderia>unclassified_Paraburkholderia | 187 | B0G82_RS17795 | 3928344..3928907 | 1938798 | hypothetical-protein-[Paraburkholderia-sp.-BL17N1] |
| WP_055397345.1 | HNH | GCF_017355385.1 | Acidovorax | Bacteria>Pseudomonadati>Pseudomonadota>Betaproteobacteria>Burkholderiales>Comamonadaceae | 196 | AE621_RS12485 | 15292..15882 | 12916 | MULTISPECIES:-TIGR02646-family-protein-[Acidovorax] |
| WP_087746912.1 | HNH | GCF_002176815.1 | unclassified_Acidovorax | Bacteria>Pseudomonadati>Pseudomonadota>Betaproteobacteria>Burkholderiales>Comamonadaceae>Acidovorax | 246 | CCX87_RS13185 | 2804642..2805382 | 2684926 | MULTISPECIES:-TIGR02646-family-protein-[unclassified-Acidovorax] |
| WP_086912101.1 | HNH | GCF_002157165.1 | Acidovorax_carolinensis | Bacteria>Pseudomonadati>Pseudomonadota>Betaproteobacteria>Burkholderiales>Comamonadaceae>Acidovorax | 106 | CBP33_RS08110 | 1704544..1704864 | 553814 | hypothetical-protein-[Acidovorax-carolinensis] |
| WP_094097617.1 | HNH | GCF_002157145.1 | Acidovorax_carolinensis | Bacteria>Pseudomonadati>Pseudomonadota>Betaproteobacteria>Burkholderiales>Comamonadaceae>Acidovorax | 106 | CBP34_RS06485 | 1383109..1383429 | 553814 | hypothetical-protein-[Acidovorax-carolinensis] |
| WP_210568818.1 | HNH | GCF_017939685.1 | Acidovorax_sp._JG5 | Bacteria>Pseudomonadati>Pseudomonadota>Betaproteobacteria>Burkholderiales>Comamonadaceae>Acidovorax>unclassified_Acidovorax | 246 | J8G26_RS10045 | 90626..91366 | 2822718 | TIGR02646-family-protein-[Acidovorax-sp.-JG5] |
| GAO25247.1 | HNH | GCA_000974645.1 | Alicycliphilus_sp._B1 | Bacteria>Pseudomonadati>Pseudomonadota>Betaproteobacteria>Burkholderiales>Comamonadaceae>Alicycliphilus>unclassified_Alicycliphilus | 106 | ALISP_5067 | 1537933..1538253 | 1603291 | hypothetical-protein-ALISP_5067-[Alicycliphilus-sp.-B1] |
| WP_210930509.1 | HNH | GCF_018064205.1 | Comamonas_aquatica | Bacteria>Pseudomonadati>Pseudomonadota>Betaproteobacteria>Burkholderiales>Comamonadaceae>Comamonas | 196 | KAQ61_RS03455 | 745367..745957 | 225991 | TIGR02646-family-protein-[Comamonas-aquatica] |
| WP_172207176.1 | HNH | GCF_013154915.1 | Diaphorobacter_sp._JS3050 | Bacteria>Pseudomonadati>Pseudomonadota>Betaproteobacteria>Burkholderiales>Comamonadaceae>Diaphorobacter>unclassified_Diaphorobacter | 106 | HND92_RS11870 | 2477960..2478280 | 2735554 | hypothetical-protein-[Diaphorobacter-sp.-JS3050] |
| MDD2610195.1 | HNH | GCA_028686475.1 | Giesbergeria_sp. | Bacteria>Pseudomonadati>Pseudomonadota>Betaproteobacteria>Burkholderiales>Comamonadaceae>Giesbergeria>unclassified_Giesbergeria | 197 | PHX60_11005 | 15493..16086 | 2818473 | MAG:-TIGR02646-family-protein-[Giesbergeria-sp.] |
| MCW5668673.1 | HNH | GCA_026125825.1 | Hydrogenophaga_sp. | Bacteria>Pseudomonadati>Pseudomonadota>Betaproteobacteria>Burkholderiales>Comamonadaceae>Hydrogenophaga>unclassified_Hydrogenophaga | 241 | KIT86_03360 | 44183..44908 | 1904254 | MAG:-TIGR02646-family-protein-[Hydrogenophaga-sp.] |
| WP_108311677.1 | HNH | GCF_003063455.1 | Limnohabitans_parvus | Bacteria>Pseudomonadati>Pseudomonadota>Betaproteobacteria>Burkholderiales>Comamonadaceae>Limnohabitans | 212 | B9Z37_RS03860 | 832064..832702 | 540061 | TIGR02646-family-protein-[Limnohabitans-parvus] |
| WP_108330264.1 | HNH | GCF_003063435.1 | Limnohabitans_sp._2KL-1 | Bacteria>Pseudomonadati>Pseudomonadota>Betaproteobacteria>Burkholderiales>Comamonadaceae>Limnohabitans>unclassified_Limnohabitans | 238 | B9Z47_RS05530 | 217005..217721 | 1100699 | hypothetical-protein-[Limnohabitans-sp.-2KL-1] |
| WP_101103330.1 | HNH | GCF_002837135.1 | Macromonas_bipunctata | Bacteria>Pseudomonadati>Pseudomonadota>Betaproteobacteria>Burkholderiales>Comamonadaceae>Macromonas | 228 | CON76_RS10615 | 16886..17572 | 183670 | hypothetical-protein-[Macromonas-bipunctata] |
| MBH2043142.1 | HNH | GCA_016000285.1 | Comamonadaceae_bacterium | Bacteria>Pseudomonadati>Pseudomonadota>Betaproteobacteria>Burkholderiales>Comamonadaceae>unclassified_Comamonadaceae | 218 | I8H87_06340 | 11278..11934 | 1871071 | TIGR02646-family-protein-[Comamonadaceae-bacterium] |
| WP_182994398.1 | HNH | GCF_014171515.1 | Herbaspirillum_frisingense | Bacteria>Pseudomonadati>Pseudomonadota>Betaproteobacteria>Burkholderiales>Oxalobacteraceae>Herbaspirillum | 241 | G5S34_RS08620 | 1932979..1933704 | 92645 | TIGR02646-family-protein-[Herbaspirillum-frisingense] |

| Protein-ID | Domain | Assembly-ID | Strain | Taxonomy | Length | Locus-Tag | Genomic-coordinates | TaxID | Fasta-defline |
|---|---|---|---|---|---|---|---|---|---|
| WP_081344814.1 | HNH | GCF_001854915.1 | Janthinobacterium_lividum | Bacteria>Pseudomonadati>Pseudomonadota>Betaproteobacteria>Burkholderiales>Oxalobacteraceae>Janthinobacterium | 237 | AKG95_RS19165 | 45670..46383 | 29581 | TIGR02646-family-protein-[Janthinobacterium-lividum] |
| MCM1512956.1 | HNH | GCA_023667225.1 | Oxalobacter_formigenes | Bacteria>Pseudomonadati>Pseudomonadota>Betaproteobacteria>Burkholderiales>Oxalobacteraceae>Oxalobacter | 209 | NC112_07615 | 13233..13862 | 847 | TIGR02646-family-protein-[Oxalobacter-formigenes] |
| WP_144290172.1 | HNH | GCF_002198735.1 | Ideonella_sp._A_288 | Bacteria>Pseudomonadati>Pseudomonadota>Betaproteobacteria>Burkholderiales>Sphaerotilaceae>Ideonella>unclassified_Ideonella | 209 | EOINJALH_RS31415 | 89320..89949 | 1962181 | HNH-endonuclease-[Ideonella-sp.-A-288] |
| WP_140628353.1 | HNH | GCF_006386545.1 | Methylibium_rhizosphaerae | Bacteria>Pseudomonadati>Pseudomonadota>Betaproteobacteria>Burkholderiales>Sphaerotilaceae>Methylibium | 241 | E8D38_RS09405 | 126373..127098 | 2570323 | TIGR02646-family-protein-[Pseudorivibacter-rhizosphaerae] |
| NRT58388.1 | HNH | GCA_013294065.1 | Sphaerotilus_uruguayifluvii | Bacteria>Pseudomonadati>Pseudomonadota>Betaproteobacteria>Burkholderiales>Sphaerotilaceae>Sphaerotilus | 214 | HNQ01_004156 | 26080..26724 | 2735897 | uncharacterized-protein-(TIGR02646-family)-[Leptothrix-sp.-C29] |
| WP_173807407.1 | HNH | GCF_013294065.1 | Sphaerotilus_uruguayifluvii | Bacteria>Pseudomonadati>Pseudomonadota>Betaproteobacteria>Burkholderiales>Sphaerotilaceae>Sphaerotilus | 214 | HNQ01_RS20675 | 26080..26724 | 2735897 | retron-Ec78-anti-phage-system-effector-HNH-endonuclease-PtuB-[Leptothrix-sp.-C29] |
| WP_110402339.1 | HNH | GCF_003201595.1 | Sphaerotilus_hippei | Bacteria>Pseudomonadati>Pseudomonadota>Betaproteobacteria>Burkholderiales>Sphaerotilaceae>Sphaerotilus | 196 | C7444_RS18480 | 30406..30996 | 744406 | TIGR02646-family-protein-[Sphaerotilus-hippei] |
| WP_031597388.1 | HNH | GCF_000735045.1 | Ferrovum_myxofaciens | Bacteria>Pseudomonadati>Pseudomonadota>Betaproteobacteria>Ferrovales>Ferrovaceae>Ferrovum | 241 | IV91_RS0107810 | 10764..11489 | 416213 | TIGR02646-family-protein-[Ferrovum-myxofaciens] |
| QKE41937.1 | HNH | GCA_013181415.1 | Ferrovum_myxofaciens | Bacteria>Pseudomonadati>Pseudomonadota>Betaproteobacteria>Ferrovales>Ferrovaceae>Ferrovum | 196 | HO274_11925 | 2420266..2420856 | 416213 | TIGR02646-family-protein-[Ferrovum-myxofaciens] |
| WP_273119924.1 | HNH | GCF_013181415.1 | Ferrovum_myxofaciens | Bacteria>Pseudomonadati>Pseudomonadota>Betaproteobacteria>Ferrovales>Ferrovaceae>Ferrovum | 196 | HO274_RS11960 | 2420266..2420856 | 416213 | TIGR02646-family-protein-[Ferrovum-myxofaciens] |
| MCG9054665.1 | HNH | GCA_022394275.1 | Laribacter_hongkongensis | Bacteria>Pseudomonadati>Pseudomonadota>Betaproteobacteria>Neisseriales>Aquaspirillaceae>Laribacter | 248 | LH442_01445 | 292074..292820 | 168471 | DCL-family-protein-[Laribacter-hongkongensis] |
| WP_239901857.1 | HNH | GCF_022394235.1 | Laribacter_hongkongensis | Bacteria>Pseudomonadati>Pseudomonadota>Betaproteobacteria>Neisseriales>Aquaspirillaceae>Laribacter | 248 | LH447_RS07080 | 205..951 | 168471 | DCL-family-protein-[Laribacter-hongkongensis] |
| WP_046167754.1 | HNH | GCF_000971355.1 | Chromobacterium_vaccinii | Bacteria>Pseudomonadati>Pseudomonadota>Betaproteobacteria>Neisseriales>Chromobacteriaceae>Chromobacterium | 198 | VL68_RS13380 | 10247..10843 | 1108595 | hypothetical-protein-[Chromobacterium-vaccinii] |
| WP_240067804.1 | HNH | GCF_009650905.1 | Eikenella_corrodens | Bacteria>Pseudomonadati>Pseudomonadota>Betaproteobacteria>Neisseriales>Neisseriaceae>Eikenella | 100 | GJ661_RS08710 | 25521..25823 | 539 | DCL-family-protein-[Eikenella-corrodens] |
| MBX3631058.1 | HNH | GCA_019635215.1 | Nitrosomonas_sp. | Bacteria>Pseudomonadati>Pseudomonadota>Betaproteobacteria>Nitrosomonadales>Nitrosomonadaceae>Nitrosomonas>unclassified_Nitrosomonas | 226 | KF908_14355 | 18971..19651 | 42353 | TIGR02646-family-protein-[Nitrosomonas-sp.] |
| MCW5608746.1 | HNH | GCA_026125505.1 | Nitrosomonas_sp. | Bacteria>Pseudomonadati>Pseudomonadota>Betaproteobacteria>Nitrosomonadales>Nitrosomonadaceae>Nitrosomonas>unclassified_Nitrosomonas | 226 | KIT26_11945 | 322..1002 | 42353 | MAG:-TIGR02646-family-protein-[Nitrosomonas-sp.] |
| WP_084600429.1 | HNH | GCF_000378245.1 | Aromatoleum_toluclasticum | Bacteria>Pseudomonadati>Pseudomonadota>Betaproteobacteria>Rhodocyclales>Rhodocyclaceae>Aromatoleum | 208 | F464_RS30235 | 372299..372925 | 92003 | TIGR02646-family-protein-[Aromatoleum-toluclasticum] |
| WP_169203002.1 | HNH | GCF_017894365.1 | Aromatoleum_bremense | Bacteria>Pseudomonadati>Pseudomonadota>Betaproteobacteria>Rhodocyclales>Rhodocyclaceae>Aromatoleum | 196 | GPA24_RS12890 | 19846..20436 | 76115 | TIGR02646-family-protein-[Aromatoleum-bremense] |
| MBX3668834.1 | HNH | GCA_019635235.1 | Rhodocyclaceae_bacterium | Bacteria>Pseudomonadati>Pseudomonadota>Betaproteobacteria>Rhodocyclales>Rhodocyclaceae>unclassified_Rhodocyclaceae | 270 | KF778_10565 | 182144..182956 | 1898103 | TIGR02646-family-protein-[Rhodocyclaceae-bacterium] |
| WP_096450971.1 | HNH | GCF_002354895.1 | Thauera | Bacteria>Pseudomonadati>Pseudomonadota>Betaproteobacteria>Rhodocyclales>Zoogloeaceae | 191 | CCZ27_RS19505 | 4441267..4441842 | 33057 | hypothetical-protein-[Thauera-sp.-K11] |
| WP_249116082.1 | HNH | - | Azoarcus_sp._L1K30 | Bacteria>Pseudomonadati>Pseudomonadota>Betaproteobacteria>Rhodocyclales>Zoogloeaceae>Azoarcus>unclassified_Azoarcus | 240 | J5J83_RS19180 | 4177511..4178233 | 2820277 | TIGR02646-family-protein-[Azoarcus-sp.-L1K30] |
| WP_159691723.1 | HNH | GCF_009800965.1 | Cognatazoarcus_halotolerans | Bacteria>Pseudomonadati>Pseudomonadota>Betaproteobacteria>Rhodocyclales>Zoogloeaceae>Cognatazoarcus | 196 | GQB03_RS16425 | 78888..79478 | 2686016 | TIGR02646-family-protein-[Cognatazoarcus-halotolerans] |
| TMW75523.1 | HNH | GCA_006007845.1 | Thauera_sp._UPWRP | Bacteria>Pseudomonadati>Pseudomonadota>Betaproteobacteria>Rhodocyclales>Zoogloeaceae>Thauera>unclassified_Thauera | 243 | FG147_05180 | 5146..5877 | 2488794 | TIGR02646-family-protein-[Thauera-sp.-UPWRP] |
| WP_148578356.1 | HNH | GCF_008107625.1 | Zoogloea_oleivorans | Bacteria>Pseudomonadati>Pseudomonadota>Betaproteobacteria>Rhodocyclales>Zoogloeaceae>Zoogloea | 202 | ETQ85_RS07205 | 253125..253733 | 1552750 | hypothetical-protein-[Zoogloea-oleivorans] |
| MDA8417186.1 | HNH | GCA_028069095.1 | Betaproteobacteria_bacterium | Bacteria>Pseudomonadati>Pseudomonadota>Betaproteobacteria>unclassified_Betaproteobacteria | 196 | MOZ78_09060 | 7052..7642 | 1891241 | MAG:-TIGR02646-family-protein-[Betaproteobacteria-bacterium] |
| WP_019341810.1 | HNH | GCF_000235745.1 | Pseudomonadales | Bacteria>Pseudomonadati>Pseudomonadota>Gammaproteobacteria | 100 | NAU09_RS00505 | 107782..108084 | 72274 | MULTISPECIES:-hypothetical-protein-[Pseudomonadaceae] |
| WP_033769533.1 | HNH | GCF_035013565.1 | Enterobacterales | Bacteria>Pseudomonadati>Pseudomonadota>Gammaproteobacteria | 100 | T296_RS18230 | 294271..294573 | 91347 | hypothetical-protein-[Pantoea-agglomerans] |
| WP_064355136.1 | HNH | GCF_018255815.1 | Enterobacterales | Bacteria>Pseudomonadati>Pseudomonadota>Gammaproteobacteria | 100 | KFO54_RS02635 | 549734..550036 | 91347 | MULTISPECIES:-hypothetical-protein-[Enterobacteriaceae] |
| WP_086076507.1 | HNH | GCF_902704295.1 | Enterobacterales | Bacteria>Pseudomonadati>Pseudomonadota>Gammaproteobacteria | 100 | FL632_RS17855 | 44843..45145 | 91347 | MULTISPECIES:-hypothetical-protein-[Klebsiella] |
| WP_129942180.1 | HNH | GCF_023544415.1 | Enterobacterales | Bacteria>Pseudomonadati>Pseudomonadota>Gammaproteobacteria | 100 | BSR02_RS08545 | 201652..201954 | 91347 | MULTISPECIES:-hypothetical-protein-[Enterobacterales] |
| WP_103858990.1 | HNH | GCF_019048725.1 | Aeromonas | Bacteria>Pseudomonadati>Pseudomonadota>Gammaproteobacteria>Aeromonadales>Aeromonadaceae | 239 | NUK49_RS00040 | 6879..7598 | 642 | MULTISPECIES:-TIGR02646-family-protein-[Aeromonas] |
| WP_219296385.1 | HNH | GCF_963896005.1 | Aeromonas | Bacteria>Pseudomonadati>Pseudomonadota>Gammaproteobacteria>Aeromonadales>Aeromonadaceae | 209 | EHZ86_RS11590 | 10459..11088 | 642 | MULTISPECIES:-TIGR02646-family-protein-[Aeromonas] |
| WP_182951582.1 | HNH | GCF_014168615.1 | Aeromonas | Bacteria>Pseudomonadati>Pseudomonadota>Gammaproteobacteria>Aeromonadales>Aeromonadaceae | 202 | OKB79_RS12600 | 4374..4982 | 642 | MULTISPECIES:-hypothetical-protein-[unclassified-Aeromonas] |
| WP_223954705.1 | HNH | GCF_019972775.1 | Aeromonas | Bacteria>Pseudomonadati>Pseudomonadota>Gammaproteobacteria>Aeromonadales>Aeromonadaceae | 197 | LDN68_RS17125 | 3650135..3650728 | 642 | TIGR02646-family-protein-[Aeromonas-caviae] |
| WP_228520151.1 | HNH | GCF_046226845.1 | Aeromonas | Bacteria>Pseudomonadati>Pseudomonadota>Gammaproteobacteria>Aeromonadales>Aeromonadaceae | 101 | F881_RS0123065 | 9595..9900 | 642 | MULTISPECIES:-hypothetical-protein-[Aeromonas] |
| WP_232354301.1 | HNH | GCF_963886775.1 | Aeromonas | Bacteria>Pseudomonadati>Pseudomonadota>Gammaproteobacteria>Aeromonadales>Aeromonadaceae | 101 | PYU98_RS15705 | 3393443..3393748 | 642 | MULTISPECIES:-hypothetical-protein-[Aeromonas] |
| WP_024946324.1 | HNH | GCF_026797875.1 | Aeromonas | Bacteria>Pseudomonadati>Pseudomonadota>Gammaproteobacteria>Aeromonadales>Aeromonadaceae | 100 | OTH18_RS02125 | 457537..457839 | 642 | MULTISPECIES:-hypothetical-protein-[Aeromonas] |
| WP_180911157.1 | HNH | GCF_013488005.1 | Aeromonas | Bacteria>Pseudomonadati>Pseudomonadota>Gammaproteobacteria>Aeromonadales>Aeromonadaceae | 100 | NQU91_RS15355 | 3374651..3374953 | 642 | MULTISPECIES:-hypothetical-protein-[Aeromonas] |
| MCF3097924.1 | HNH | GCA_021538905.1 | Aeromonas_australiensis | Bacteria>Pseudomonadati>Pseudomonadota>Gammaproteobacteria>Aeromonadales>Aeromonadaceae>Aeromonas | 209 | EHZ86_11620 | 10459..11088 | 1114880 | TIGR02646-family-protein-[Aeromonas-australiensis] |
| WP_201871423.1 | HNH | GCF_016728905.1 | Aeromonas_caviae | Bacteria>Pseudomonadati>Pseudomonadota>Gammaproteobacteria>Aeromonadales>Aeromonadaceae>Aeromonas | 209 | JD551_RS01090 | 91026..91655 | 648 | TIGR02646-family-protein-[Aeromonas-caviae] |
| WP_223918003.1 | HNH | GCF_019971655.1 | Aeromonas_caviae | Bacteria>Pseudomonadati>Pseudomonadota>Gammaproteobacteria>Aeromonadales>Aeromonadaceae>Aeromonas | 209 | LDN21_RS02435 | 66180..66809 | 648 | TIGR02646-family-protein-[Aeromonas-caviae] |
| EJN6954194.1 | HNH | GCA_024506355.1 | Aeromonas_hydrophila | Bacteria>Pseudomonadati>Pseudomonadota>Gammaproteobacteria>Aeromonadales>Aeromonadaceae>Aeromonas | 202 | NRD12_000735 | 215371..215979 | 644 | hypothetical-protein-[Aeromonas-hydrophila] |
| UPK56455.1 | HNH | GCA_023100745.1 | Aeromonas_veronii | Bacteria>Pseudomonadati>Pseudomonadota>Gammaproteobacteria>Aeromonadales>Aeromonadaceae>Aeromonas | 202 | MYF86_09070 | 1904534..1905142 | 654 | hypothetical-protein-MYF86_09070-[Aeromonas-veronii] |
| WP_213387490.1 | HNH | GCF_018359765.1 | Aeromonas_veronii | Bacteria>Pseudomonadati>Pseudomonadota>Gammaproteobacteria>Aeromonadales>Aeromonadaceae>Aeromonas | 202 | J4G64_RS12010 | 298878..299486 | 654 | hypothetical-protein-[Aeromonas-veronii] |
| WP_216986931.1 | HNH | GCF_019048565.1 | unclassified_Aeromonas | Bacteria>Pseudomonadati>Pseudomonadota>Gammaproteobacteria>Aeromonadales>Aeromonadaceae>Aeromonas | 202 | I6L36_RS00720 | 145359..145967 | 257493 | hypothetical-protein-[Aeromonas-sp.-FDAARGOS-1406] |
| WP_223938535.1 | HNH | GCF_019971155.1 | Aeromonas_caviae | Bacteria>Pseudomonadati>Pseudomonadota>Gammaproteobacteria>Aeromonadales>Aeromonadaceae>Aeromonas | 202 | LDN24_RS08675 | 28506..29114 | 648 | hypothetical-protein-[Aeromonas-caviae] |
| WP_235462877.1 | HNH | GCF_021538905.1 | Aeromonas_australiensis | Bacteria>Pseudomonadati>Pseudomonadota>Gammaproteobacteria>Aeromonadales>Aeromonadaceae>Aeromonas | 202 | EHZ86_RS12365 | 17062..17670 | 1114880 | hypothetical-protein-[Aeromonas-australiensis] |
| WP_264787734.1 | HNH | GCF_026000055.1 | Aeromonas_caviae | Bacteria>Pseudomonadati>Pseudomonadota>Gammaproteobacteria>Aeromonadales>Aeromonadaceae>Aeromonas | 202 | OPI30_RS10875 | 2222082..2222690 | 648 | hypothetical-protein-[Aeromonas-caviae] |
| WP_113721902.1 | HNH | GCF_963900125.1 | Aeromonas_salmonicida | Bacteria>Pseudomonadati>Pseudomonadota>Gammaproteobacteria>Aeromonadales>Aeromonadaceae>Aeromonas | 200 | CPA05_RS19930 | 15180..15782 | 645 | hypothetical-protein-[Aeromonas-salmonicida] |
| WP_080722333.1 | HNH | GCF_047780535.1 | Aeromonas_hydrophila | Bacteria>Pseudomonadati>Pseudomonadota>Gammaproteobacteria>Aeromonadales>Aeromonadaceae>Aeromonas | 199 | HR52_RS24350 | 3097366..3097965 | 644 | hypothetical-protein-[Aeromonas-hydrophila] |
| WP_203130886.1 | HNH | GCF_016805365.1 | Aeromonas_jandaei | Bacteria>Pseudomonadati>Pseudomonadota>Gammaproteobacteria>Aeromonadales>Aeromonadaceae>Aeromonas | 199 | GNZ06_RS06080 | 158573..159172 | 650 | hypothetical-protein-[Aeromonas-jandaei] |
| QWL57199.1 | HNH | GCA_018802265.1 | Aeromonas_jandaei | Bacteria>Pseudomonadati>Pseudomonadota>Gammaproteobacteria>Aeromonadales>Aeromonadaceae>Aeromonas | 197 | HQ400_01340 | 287926..288519 | 650 | TIGR02646-family-protein-[Aeromonas-jandaei] |
| WP_064338621.1 | HNH | GCF_001634325.1 | Aeromonas_veronii | Bacteria>Pseudomonadati>Pseudomonadota>Gammaproteobacteria>Aeromonadales>Aeromonadaceae>Aeromonas | 197 | WM43_RS07855 | 1782736..1783329 | 654 | TIGR02646-family-protein-[Aeromonas-veronii] |
| WP_064341059.1 | HNH | GCF_014169835.1 | Aeromonas_veronii | Bacteria>Pseudomonadati>Pseudomonadota>Gammaproteobacteria>Aeromonadales>Aeromonadaceae>Aeromonas | 197 | QBS04_RS15335 | 3278652..3279245 | 654 | TIGR02646-family-protein-[Aeromonas-veronii] |
| WP_270668499.1 | HNH | GCF_027705205.1 | unclassified_Aeromonas | Bacteria>Pseudomonadati>Pseudomonadota>Gammaproteobacteria>Aeromonadales>Aeromonadaceae>Aeromonas | 197 | OKB91_RS16410 | 6018..6611 | 257493 | MULTISPECIES:-TIGR02646-family-protein-[unclassified-Aeromonas] |
| WP_215805220.1 | HNH | GCF_018802365.1 | Aeromonas_jandaei | Bacteria>Pseudomonadati>Pseudomonadota>Gammaproteobacteria>Aeromonadales>Aeromonadaceae>Aeromonas | 122 | HQ398_RS10740 | 2348103..2348471 | 650 | hypothetical-protein-[Aeromonas-jandaei] |
| WP_215805221.1 | HNH | GCF_018802365.1 | Aeromonas_jandaei | Bacteria>Pseudomonadati>Pseudomonadota>Gammaproteobacteria>Aeromonadales>Aeromonadaceae>Aeromonas | 75 | HQ398_RS10745 | 2348484..2348711 | 650 | hypothetical-protein-[Aeromonas-jandaei] |
| WP_240332640.1 | HNH | GCF_003491245.1 | Aeromonas_hydrophila | Bacteria>Pseudomonadati>Pseudomonadota>Gammaproteobacteria>Aeromonadales>Aeromonadaceae>Aeromonas | 104 | BFW41_RS15380 | 3359202..3359516 | 644 | hypothetical-protein-[Aeromonas-hydrophila] |
| WEE24936.1 | HNH | GCA_029025785.1 | Aeromonas_hydrophila | Bacteria>Pseudomonadati>Pseudomonadota>Gammaproteobacteria>Aeromonadales>Aeromonadaceae>Aeromonas | 101 | PY771_14785 | 3200106..3200411 | 644 | hypothetical-protein-PY771_14785-[Aeromonas-hydrophila] |
| WP_235695655.1 | HNH | GCF_019348735.1 | Aeromonas_veronii | Bacteria>Pseudomonadati>Pseudomonadota>Gammaproteobacteria>Aeromonadales>Aeromonadaceae>Aeromonas | 101 | GL272_RS08120 | 388953..389258 | 654 | hypothetical-protein-[Aeromonas-veronii] |
| WP_244365009.1 | HNH | GCF_963900055.1 | Aeromonas_dhakensis | Bacteria>Pseudomonadati>Pseudomonadota>Gammaproteobacteria>Aeromonadales>Aeromonadaceae>Aeromonas | 101 | MRR68_RS06355 | 18865..19170 | 196024 | hypothetical-protein-[Aeromonas-dhakensis] |
| AXV36632.1 | HNH | GCA_003491245.1 | Aeromonas_hydrophila | Bacteria>Pseudomonadati>Pseudomonadota>Gammaproteobacteria>Aeromonadales>Aeromonadaceae>Aeromonas | 100 | BFW41_14990 | 3359202..3359504 | 644 | hypothetical-protein-BFW41_14990-[Aeromonas-hydrophila] |
| MBL0521038.1 | HNH | GCA_016730045.1 | Aeromonas_enteropelogenes | Bacteria>Pseudomonadati>Pseudomonadota>Gammaproteobacteria>Aeromonadales>Aeromonadaceae>Aeromonas | 100 | JD523_08980 | 524748..525050 | 29489 | hypothetical-protein-[Aeromonas-enteropelogenes] |
| MVG13647.1 | HNH | GCA_009749285.1 | Aeromonas_jandaei | Bacteria>Pseudomonadati>Pseudomonadota>Gammaproteobacteria>Aeromonadales>Aeromonadaceae>Aeromonas | 100 | EF849_03845 | 843671..843973 | 650 | hypothetical-protein-[Aeromonas-jandaei] |
| WP_275056627.1 | HNH | GCF_029024985.1 | Aeromonas_allosaccharophila | Bacteria>Pseudomonadati>Pseudomonadota>Gammaproteobacteria>Aeromonadales>Aeromonadaceae>Aeromonas | 127 | PYU98_RS15760 | 3409497..3409880 | 656 | HNH-endonuclease-[Aeromonas-allosaccharophila] |
| BBQ25123.1 | HNH | GCA_014168615.1 | Aeromonas_sp._WP2-W18-CRE-05 | Bacteria>Pseudomonadati>Pseudomonadota>Gammaproteobacteria>Aeromonadales>Aeromonadaceae>Aeromonas>unclassified_Aeromonas | 202 | WP2W18C05_13390 | 1434563..1435171 | 2675707 | hypothetical-protein-WP2W18C05_13390-[Aeromonas-sp.-WP2-W18-CRE-05] |
| QXC33931.1 | HNH | GCA_019048725.1 | Aeromonas_sp._FDAARGOS_1407 | Bacteria>Pseudomonadati>Pseudomonadota>Gammaproteobacteria>Aeromonadales>Aeromonadaceae>Aeromonas>unclassified_Aeromonas | 202 | I6L37_20715 | 4421054..4421662 | 2778056 | hypothetical-protein-I6L37_20715-[Aeromonas-sp.-FDAARGOS-1407] |
| WP_114523385.1 | HNH | GCF_003344485.1 | Aeromonas_sp._ARM81 | Bacteria>Pseudomonadati>Pseudomonadota>Gammaproteobacteria>Aeromonadales>Aeromonadaceae>Aeromonas>unclassified_Aeromonas | 202 | ASJ36_RS21115 | 12782..13390 | 1747384 | hypothetical-protein-[Aeromonas-sp.-ARM81] |
| WP_216990392.1 | HNH | GCF_019048705.1 | Aeromonas_sp._FDAARGOS_1409 | Bacteria>Pseudomonadati>Pseudomonadota>Gammaproteobacteria>Aeromonadales>Aeromonadaceae>Aeromonas>unclassified_Aeromonas | 202 | I6L39_RS17645 | 3795876..3796484 | 2778058 | hypothetical-protein-[Aeromonas-sp.-FDAARGOS-1409] |
| WP_241325384.1 | HNH | GCF_963899385.1 | Aeromonas_sp._MR7 | Bacteria>Pseudomonadati>Pseudomonadota>Gammaproteobacteria>Aeromonadales>Aeromonadaceae>Aeromonas>unclassified_Aeromonas | 202 | MM192_RS09005 | 70147..70755 | 2923419 | hypothetical-protein-[Aeromonas-sp.-MR7] |
| WP_270659876.1 | HNH | GCF_027705965.1 | Aeromonas_sp._Y301-2 | Bacteria>Pseudomonadati>Pseudomonadota>Gammaproteobacteria>Aeromonadales>Aeromonadaceae>Aeromonas>unclassified_Aeromonas | 202 | OKB83_RS15910 | 137804..138412 | 2990505 | hypothetical-protein-[Aeromonas-sp.-Y301-2] |
| WP_270670313.1 | HNH | GCF_027705165.1 | Aeromonas_sp._QDB04 | Bacteria>Pseudomonadati>Pseudomonadota>Gammaproteobacteria>Aeromonadales>Aeromonadaceae>Aeromonas>unclassified_Aeromonas | 202 | OKC37_RS13850 | 4129..4737 | 2990477 | hypothetical-protein-[Aeromonas-sp.-QDB04] |
| WP_108651328.1 | HNH | GCF_003072645.1 | Dongshaea_marina | Bacteria>Pseudomonadati>Pseudomonadota>Gammaproteobacteria>Aeromonadales>Aeromonadaceae>Dongshaea | 100 | DB847_RS14410 | 3042108..3042410 | 2047966 | hypothetical-protein-[Dongshaea-marina] |
| GGW57108.1 | HNH | GCA_014652375.1 | Alishewanella_tabrizica | Bacteria>Pseudomonadati>Pseudomonadota>Gammaproteobacteria>Alteromonadales>Alteromonadaceae>Alishewanella | 221 | GCM10008111_11460 | 88610..89275 | 671278 | hypothetical-protein-GCM10008111_11460-[Alishewanella-tabrizica] |
| WP_189481410.1 | HNH | GCF_014652375.1 | Alishewanella_tabrizica | Bacteria>Pseudomonadati>Pseudomonadota>Gammaproteobacteria>Alteromonadales>Alteromonadaceae>Alishewanella | 221 | IE070_RS05840 | 88610..89275 | 671278 | TIGR02646-family-protein-[Alishewanella-tabrizica] |
| WP_057786318.1 | HNH | GCF_001441695.1 | Alishewanella_sp._WH16-1 | Bacteria>Pseudomonadati>Pseudomonadota>Gammaproteobacteria>Alteromonadales>Alteromonadaceae>Alishewanella>unclassified_Alishewanella | 224 | AAY72_RS06875 | 14132..14806 | 1651088 | TIGR02646-family-protein-[Alishewanella-sp.-WH16-1] |
| WP_250655182.1 | HNH | GCF_023650515.1 | Alkalimarinus_coralli | Bacteria>Pseudomonadati>Pseudomonadota>Gammaproteobacteria>Alteromonadales>Alteromonadaceae>Alkalimarinus | 224 | MY525_RS07045 | 1553972..1554646 | 2935863 | TIGR02646-family-protein-[Alkalimarinus-coralli] |
| WP_073324022.1 | HNH | GCF_900129565.1 | Marisediminitalea_aggregata | Bacteria>Pseudomonadati>Pseudomonadota>Gammaproteobacteria>Alteromonadales>Alteromonadaceae>Marisediminitalea | 100 | BUB81_RS14485 | 47895..48197 | 634436 | hypothetical-protein-[Marisediminitalea-aggregata] |
| WP_101201431.1 | HNH | GCF_002835935.1 | Paraglaciecola_sp._MB-3u-78 | Bacteria>Pseudomonadati>Pseudomonadota>Gammaproteobacteria>Alteromonadales>Alteromonadaceae>Paraglaciecola>unclassified_Paraglaciecola | 214 | CXF95_RS26550 | 28931..29575 | 2058332 | TIGR02646-family-protein-[Paraglaciecola-sp.-MB-3u-78] |
| WP_089368482.1 | HNH | GCF_007991255.1 | Pseudoalteromonas_nigrifaciens | Bacteria>Pseudomonadati>Pseudomonadota>Gammaproteobacteria>Alteromonadales>Pseudoalteromonadaceae>Pseudoalteromonas | 216 | PNIG_RS10745 | 2251832..2252482 | 28109 | TIGR02646-family-protein-[Pseudoalteromonas-nigrifaciens] |
| WP_010557253.1 | HNH | GCF_000238335.3 | Pseudoalteromonas_marina | Bacteria>Pseudomonadati>Pseudomonadota>Gammaproteobacteria>Alteromonadales>Pseudoalteromonadaceae>Pseudoalteromonas | 100 | PMAN_RS08610 | 1880479..1880781 | 267375 | hypothetical-protein-[Pseudoalteromonas-marina] |
| WP_115016382.1 | HNH | GCF_019266125.1 | Shewanella | Bacteria>Pseudomonadati>Pseudomonadota>Gammaproteobacteria>Alteromonadales>Shewanellaceae | 208 | DX079_RS11255 | 2526923..2527549 | 22 | MULTISPECIES:-TIGR02646-family-protein-[Shewanella] |
| WP_186265253.1 | HNH | GCF_014263185.1 | Shewanella | Bacteria>Pseudomonadati>Pseudomonadota>Gammaproteobacteria>Alteromonadales>Shewanellaceae | 208 | KOE54_RS14865 | 15890..16516 | 22 | TIGR02646-family-protein-[Shewanella-algae] |
| BCV35018.1 | HNH | GCA_019670565.1 | Shewanella_chilikensis | Bacteria>Pseudomonadati>Pseudomonadota>Gammaproteobacteria>Alteromonadales>Shewanellaceae>Shewanella | 208 | TUM17377_03460 | 380034..380660 | 558541 | hypothetical-protein-TUM17377_03460-[Shewanella-chilikensis] |
| WP_257574449.1 | HNH | GCF_019303235.1 | Shewanella_algae | Bacteria>Pseudomonadati>Pseudomonadota>Gammaproteobacteria>Alteromonadales>Shewanellaceae>Shewanella | 139 | KOE59_RS06890 | 125023..125442 | 38313 | hypothetical-protein-[Shewanella-algae] |
| WP_069455071.1 | HNH | GCF_002074855.1 | Shewanella_xiamenensis | Bacteria>Pseudomonadati>Pseudomonadota>Gammaproteobacteria>Alteromonadales>Shewanellaceae>Shewanella | 100 | VOO0_RS07280 | 21174..21476 | 332186 | hypothetical-protein-[Shewanella-xiamenensis] |
| WP_259569837.1 | HNH | GCF_024999565.1 | Shewanella_baltica | Bacteria>Pseudomonadati>Pseudomonadota>Gammaproteobacteria>Alteromonadales>Shewanellaceae>Shewanella | 100 | G3445_RS08170 | 184958..185260 | 62322 | hypothetical-protein-[Shewanella-baltica] |
| WP_263191767.1 | HNH | GCF_025653705.1 | Shewanella_sp._SM87 | Bacteria>Pseudomonadati>Pseudomonadota>Gammaproteobacteria>Alteromonadales>Shewanellaceae>Shewanella>unclassified_Shewanella | 100 | L5M11_RS07345 | 28514..28816 | 2912808 | hypothetical-protein-[Shewanella-sp.-SM87] |
| WP_079541814.1 | HNH | GCF_043179105.1 | Cardiobacterium_hominis | Bacteria>Pseudomonadati>Pseudomonadota>Gammaproteobacteria>Cardiobacteriales>Cardiobacteriaceae>Cardiobacterium | 196 | CHUV0807_RS10755 | 31712..32302 | 2718 | TIGR02646-family-protein-[Cardiobacterium-hominis] |
| WP_200158834.1 | HNH | GCF_016584135.1 | Allochromatium_vinosum | Bacteria>Pseudomonadati>Pseudomonadota>Gammaproteobacteria>Chromatiales>Chromatiaceae>Allochromatium | 240 | CKO29_RS16445 | 8054..8776 | 1049 | hypothetical-protein-[Allochromatium-vinosum] |
| MTW22274.1 | HNH | GCA_009720725.1 | Allochromatium_palmeri | Bacteria>Pseudomonadati>Pseudomonadota>Gammaproteobacteria>Chromatiales>Chromatiaceae>Allochromatium | 215 | GJ668_14430 | 34395..35042 | 231048 | TIGR02646-family-protein-[Allochromatium-palmeri] |
| WP_155450842.1 | HNH | GCF_009720725.1 | Allochromatium_palmeri | Bacteria>Pseudomonadati>Pseudomonadota>Gammaproteobacteria>Chromatiales>Chromatiaceae>Allochromatium | 215 | GJ668_RS14435 | 34395..35042 | 231048 | TIGR02646-family-protein-[Allochromatium-palmeri] |
| WP_264346735.1 | HNH | GCF_947090785.1 | Rheinheimera_sp._MM224 | Bacteria>Pseudomonadati>Pseudomonadota>Gammaproteobacteria>Chromatiales>Chromatiaceae>Rheinheimera>unclassified_Rheinheimera | 224 | OM978_RS09975 | 2105073..2105747 | 3019969 | TIGR02646-family-protein-[Rheinheimera-sp.-MM224] |
| WP_132582329.1 | HNH | GCF_004345025.1 | Rheinheimera_sp._D18 | Bacteria>Pseudomonadati>Pseudomonadota>Gammaproteobacteria>Chromatiales>Chromatiaceae>Rheinheimera>unclassified_Rheinheimera | 100 | EOZ06_RS04625 | 966684..966986 | 2545632 | hypothetical-protein-[Rheinheimera-sp.-D18] |
| WP_200385873.1 | HNH | GCF_016583985.1 | Thiocapsa_imhoffii | Bacteria>Pseudomonadati>Pseudomonadota>Gammaproteobacteria>Chromatiales>Chromatiaceae>Thiocapsa | 233 | CKO25_RS00035 | 10917..11618 | 382777 | hypothetical-protein-[Thiocapsa-imhoffii] |
| WP_100917764.1 | HNH | GCF_002813775.1 | Candidatus_Thiodictyon_syntrophicum | Bacteria>Pseudomonadati>Pseudomonadota>Gammaproteobacteria>Chromatiales>Chromatiaceae>Thiodictyon | 269 | THSYN_RS02595 | 600466..601275 | 1166950 | hypothetical-protein-[Candidatus-Thiodictyon-syntrophicum] |
| MCC7280694.1 | HNH | GCA_020851955.1 | Chromatiaceae_bacterium | Bacteria>Pseudomonadati>Pseudomonadota>Gammaproteobacteria>Chromatiales>Chromatiaceae>unclassified_Chromatiaceae | 235 | IT487_20595 | 5437..6144 | 2049432 | hypothetical-protein-[Chromatiaceae-bacterium] |
| MCG5499268.1 | HNH | GCA_022227785.1 | Ectothiorhodospira_lacustris | Bacteria>Pseudomonadati>Pseudomonadota>Gammaproteobacteria>Chromatiales>Ectothiorhodospiraceae>Ectothiorhodospira | 212 | LQ952_00350 | 86792..87430 | 2899127 | TIGR02646-family-protein-[Ectothiorhodospira-sp.-B14B] |
| MCG5499270.1 | HNH | GCA_022227785.1 | Ectothiorhodospira_lacustris | Bacteria>Pseudomonadati>Pseudomonadota>Gammaproteobacteria>Chromatiales>Ectothiorhodospiraceae>Ectothiorhodospira | 202 | LQ952_00360 | 89040..89648 | 2899127 | hypothetical-protein-[Ectothiorhodospira-sp.-B14B] |
| QEP43018.1 | HNH | GCA_008375315.1 | Ectothiorhodospiraceae_bacterium_BW-2 | Bacteria>Pseudomonadati>Pseudomonadota>Gammaproteobacteria>Chromatiales>Ectothiorhodospiraceae>unclassified_Ectothiorhodospiraceae | 236 | D5085_07750 | 1611117..1611827 | 947515 | HNH-endonuclease-[Ectothiorhodospiraceae-bacterium-BW-2] |
| WP_008325055.1 | HNH | GCF_000277565.1 | Enterobacteriaceae | Bacteria>Pseudomonadati>Pseudomonadota>Gammaproteobacteria>Enterobacterales | 197 | K5R72_RS10945 | 39728..40321 | 543 | MULTISPECIES:-TIGR02646-family-protein-[Enterobacteriaceae] |
| WP_222563663.1 | HNH | GCF_019803045.1 | Enterobacteriaceae | Bacteria>Pseudomonadati>Pseudomonadota>Gammaproteobacteria>Enterobacterales | 197 | NOX26_RS15670 | 3298762..3299355 | 543 | MULTISPECIES:-TIGR02646-family-protein-[Enterobacteriaceae] |
| WP_028016410.1 | HNH | GCF_003665375.1 | Enterobacteriaceae | Bacteria>Pseudomonadati>Pseudomonadota>Gammaproteobacteria>Enterobacterales | 100 | KKT92_RS23400 | 33547..33849 | 543 | MULTISPECIES:-hypothetical-protein-[Enterobacteriaceae] |
| WP_038636812.1 | HNH | GCF_015136055.1 | Enterobacteriaceae | Bacteria>Pseudomonadati>Pseudomonadota>Gammaproteobacteria>Enterobacterales | 100 | JLF39_RS03130 | 635569..635871 | 543 | MULTISPECIES:-hypothetical-protein-[Enterobacteriaceae] |
| WP_047715696.1 | HNH | GCF_964304045.1 | Enterobacteriaceae | Bacteria>Pseudomonadati>Pseudomonadota>Gammaproteobacteria>Enterobacterales | 100 | HS092_RS16835 | 3484447..3484749 | 543 | MULTISPECIES:-hypothetical-protein-[Enterobacteriaceae] |
| WP_088569514.1 | HNH | GCF_014103065.1 | Enterobacteriaceae | Bacteria>Pseudomonadati>Pseudomonadota>Gammaproteobacteria>Enterobacterales | 100 | CEQ52_RS25765 | 14546..14848 | 543 | MULTISPECIES:-hypothetical-protein-[Enterobacteriaceae] |
| WP_048997339.1 | HNH | GCF_965179935.1 | Enterobacter | Bacteria>Pseudomonadati>Pseudomonadota>Gammaproteobacteria>Enterobacterales>Enterobacteriaceae | 197 | L3068_RS02460 | 526418..527011 | 547 | MULTISPECIES:-TIGR02646-family-protein-[Enterobacter] |
| WP_109017745.1 | HNH | GCF_029718275.1 | Citrobacter | Bacteria>Pseudomonadati>Pseudomonadota>Gammaproteobacteria>Enterobacterales>Enterobacteriaceae | 100 | OIU54_RS00770 | 156437..156739 | 544 | MULTISPECIES:-hypothetical-protein-[Citrobacter-freundii-complex] |
| WP_275380010.1 | HNH | GCF_037477975.1 | Citrobacter | Bacteria>Pseudomonadati>Pseudomonadota>Gammaproteobacteria>Enterobacterales>Enterobacteriaceae | 100 | L2101_RS11615 | 145172..145474 | 544 | hypothetical-protein-[Citrobacter-portucalensis] |
| HAU4285774.1 | HNH | GCA_016065535.1 | Citrobacter_freundii | Bacteria>Pseudomonadati>Pseudomonadota>Gammaproteobacteria>Enterobacterales>Enterobacteriaceae>Citrobacter>Citrobacter_freundii_complex | 197 | F6W46_24060 | 6765..7358 | 546 | TIGR02646-family-protein-[Citrobacter-freundii] |
| HAU5791341.1 | HNH | GCA_016076255.1 | Citrobacter_freundii | Bacteria>Pseudomonadati>Pseudomonadota>Gammaproteobacteria>Enterobacterales>Enterobacteriaceae>Citrobacter>Citrobacter_freundii_complex | 197 | JD367_15415 | 180126..180719 | 546 | TIGR02646-family-protein-[Citrobacter-freundii] |
| EIX7373167.1 | HNH | GCA_022400985.1 | Citrobacter_freundii | Bacteria>Pseudomonadati>Pseudomonadota>Gammaproteobacteria>Enterobacterales>Enterobacteriaceae>Citrobacter>Citrobacter_freundii_complex | 100 | MK620_002541 | 168306..168608 | 546 | hypothetical-protein-[Citrobacter-freundii] |
| HAT3908779.1 | HNH | GCA_015929465.1 | Citrobacter_freundii | Bacteria>Pseudomonadati>Pseudomonadota>Gammaproteobacteria>Enterobacterales>Enterobacteriaceae>Citrobacter>Citrobacter_freundii_complex | 100 | I9Y28_003031 | 165579..165881 | 546 | TPA:-hypothetical-protein-I9Y28_003031-[Citrobacter-freundii] |
| WP_204562700.1 | HNH | GCF_016893745.1 | Citrobacter_cronae | Bacteria>Pseudomonadati>Pseudomonadota>Gammaproteobacteria>Enterobacterales>Enterobacteriaceae>Citrobacter>Citrobacter_freundii_complex | 88 | JSN23_RS08095 | 1889884..1890148 | 1748967 | hypothetical-protein,-partial-[Citrobacter-cronae] |
| WP_199954356.1 | HNH | GCF_016505335.1 | Citrobacter_sp._FDAARGOS_156 | Bacteria>Pseudomonadati>Pseudomonadota>Gammaproteobacteria>Enterobacterales>Enterobacteriaceae>Citrobacter>unclassified_Citrobacter | 100 | I5510_RS10680 | 347515..347817 | 1702170 | hypothetical-protein-[Citrobacter-sp.-FDAARGOS_156] |
| WP_214577410.1 | HNH | GCF_018597265.1 | Enterobacter_dykesii | Bacteria>Pseudomonadati>Pseudomonadota>Gammaproteobacteria>Enterobacterales>Enterobacteriaceae>Enterobacter | 197 | KK116_RS19470 | 164202..164795 | 2797506 | TIGR02646-family-protein-[Enterobacter-dykesii] |
| WP_262778594.1 | HNH | GCF_025536155.1 | Enterobacter_quasiroggenkampii | Bacteria>Pseudomonadati>Pseudomonadota>Gammaproteobacteria>Enterobacterales>Enterobacteriaceae>Enterobacter | 100 | KSU18_RS16635 | 116675..116977 | 2497436 | hypothetical-protein-[Enterobacter-quasiroggenkampii] |
| MCQ4415114.1 | HNH | GCA_024451305.1 | Enterobacter_kobei | Bacteria>Pseudomonadati>Pseudomonadota>Gammaproteobacteria>Enterobacterales>Enterobacteriaceae>Enterobacter>Enterobacter_cloacae_complex | 197 | NOX26_15670 | 3298762..3299355 | 208224 | TIGR02646-family-protein-[Enterobacter-kobei] |
| WP_045284470.1 | HNH | GCF_000958085.1 | Enterobacter_sichuanensis | Bacteria>Pseudomonadati>Pseudomonadota>Gammaproteobacteria>Enterobacterales>Enterobacteriaceae>Enterobacter>Enterobacter_cloacae_complex | 197 | SS37_RS01305 | 125737..126330 | 2071710 | hypothetical-protein-[Enterobacter-sichuanensis] |
| WP_193132803.1 | HNH | GCF_032369155.1 | Enterobacter_hormaechei | Bacteria>Pseudomonadati>Pseudomonadota>Gammaproteobacteria>Enterobacterales>Enterobacteriaceae>Enterobacter>Enterobacter_cloacae_complex | 197 | CXM24_RS20110 | 4293595..4294188 | 158836 | TIGR02646-family-protein-[Enterobacter-hormaechei] |
| WP_196359064.1 | HNH | GCF_015701305.1 | Enterobacter_hormaechei | Bacteria>Pseudomonadati>Pseudomonadota>Gammaproteobacteria>Enterobacterales>Enterobacteriaceae>Enterobacter>Enterobacter_cloacae_complex | 197 | I4P56_RS16425 | 170918..171511 | 158836 | TIGR02646-family-protein-[Enterobacter-hormaechei] |
| HCA7855108.1 | HNH | GCA_023000845.1 | Enterobacter_hormaechei | Bacteria>Pseudomonadati>Pseudomonadota>Gammaproteobacteria>Enterobacterales>Enterobacteriaceae>Enterobacter>Enterobacter_cloacae_complex | 100 | MX526_001459 | 148768..149070 | 158836 | hypothetical-protein-[Enterobacter-hormaechei] |
| WP_126497378.1 | HNH | GCF_003965685.1 | Enterobacter_hormaechei | Bacteria>Pseudomonadati>Pseudomonadota>Gammaproteobacteria>Enterobacterales>Enterobacteriaceae>Enterobacter>Enterobacter_cloacae_complex | 100 | EKN63_RS10105 | 351991..352293 | 158836 | hypothetical-protein-[Enterobacter-hormaechei] |
| HCM9196133.1 | HNH | GCA_025209675.1 | Enterobacter_cloacae_subsp._dissolvens | Bacteria>Pseudomonadati>Pseudomonadota>Gammaproteobacteria>Enterobacterales>Enterobacteriaceae>Enterobacter>Enterobacter_cloacae_complex>Enterobacter_cloacae | 100 | N5K92_004836 | 4921..5223 | 69219 | TPA:-hypothetical-protein-N5K92_004836-[Enterobacter-cloacae-subsp.-dissolvens] |
| EFO1856146.1 | HNH | - | Escherichia_coli | Bacteria>Pseudomonadati>Pseudomonadota>Gammaproteobacteria>Enterobacterales>Enterobacteriaceae>Escherichia | 197 | DUZ37_17310 | 24075..24668 | 562 | TIGR02646-family-protein-[Escherichia-coli] |

| Protein-ID | Domain | Assembly-ID | Strain | Taxonomy | Length | Locus-Tag | Genomic-coordinates | TaxID | Fasta-defline |
|---|---|---|---|---|---|---|---|---|---|
| HAW1058249.1 | HNH | GCA_016672425.1 | Escherichia_coli | Bacteria>Pseudomonadati>Pseudomonadota>Gammaproteobacteria>Enterobacterales>Enterobacteriaceae>Escherichia | 197 | JLK83_004315 | 19562..20155 | 562 | TPA:-TIGR02646-family-protein-[Escherichia-coli] |
| HBC9812125.1 | HNH | GCA_018195115.1 | Escherichia_coli | Bacteria>Pseudomonadati>Pseudomonadota>Gammaproteobacteria>Enterobacterales>Enterobacteriaceae>Escherichia | 197 | KFP54_004192 | 1681..2274 | 562 | TIGR02646-family-protein-[Escherichia-coli] |
| WP_096210927.1 | HNH | GCF_014216435.1 | Escherichia_coli | Bacteria>Pseudomonadati>Pseudomonadota>Gammaproteobacteria>Enterobacterales>Enterobacteriaceae>Escherichia | 197 | GVI19_RS16175 | 86764..87357 | 562 | TIGR02646-family-protein-[Escherichia-coli] |
| HAX9835791.1 | HNH | GCA_017073735.1 | Escherichia_coli | Bacteria>Pseudomonadati>Pseudomonadota>Gammaproteobacteria>Enterobacterales>Enterobacteriaceae>Escherichia | 100 | JS210_001196 | 59973..60275 | 562 | hypothetical-protein-[Escherichia-coli] |
| MBB9297693.1 | HNH | GCA_012376355.2 | Escherichia_coli | Bacteria>Pseudomonadati>Pseudomonadota>Gammaproteobacteria>Enterobacterales>Enterobacteriaceae>Escherichia | 100 | FSI29_018545 | 13134..13436 | 562 | hypothetical-protein-[Escherichia-coli] |
| MCN9861069.1 | HNH | GCA_023889565.1 | Escherichia_coli | Bacteria>Pseudomonadati>Pseudomonadota>Gammaproteobacteria>Enterobacterales>Enterobacteriaceae>Escherichia | 100 | MLQ95_06025 | 24859..25161 | 562 | hypothetical-protein-[Escherichia-coli] |
| WP_097755478.1 | HNH | GCF_005388805.1 | Escherichia_coli | Bacteria>Pseudomonadati>Pseudomonadota>Gammaproteobacteria>Enterobacterales>Enterobacteriaceae>Escherichia | 100 | FFF01_RS10400 | 164836..165138 | 562 | hypothetical-protein-[Escherichia-coli] |
| EEU4096210.1 | HNH | - | Escherichia_coli | Bacteria>Pseudomonadati>Pseudomonadota>Gammaproteobacteria>Enterobacterales>Enterobacteriaceae>Escherichia | 40 | E8H53_004594 | 4870..4992 | 562 | HNH-endonuclease-[Escherichia-coli] |
| EEV1146574.1 | HNH | - | Escherichia_coli | Bacteria>Pseudomonadati>Pseudomonadota>Gammaproteobacteria>Enterobacterales>Enterobacteriaceae>Escherichia | 40 | EUB49_004349 | 5020..5142 | 562 | HNH-endonuclease-[Escherichia-coli] |
| EEW3089658.1 | HNH | - | Escherichia_coli | Bacteria>Pseudomonadati>Pseudomonadota>Gammaproteobacteria>Enterobacterales>Enterobacteriaceae>Escherichia | 40 | EBA98_002634 | 11004..11126 | 562 | HNH-endonuclease-[Escherichia-coli] |
| EEW3885148.1 | HNH | - | Escherichia_coli | Bacteria>Pseudomonadati>Pseudomonadota>Gammaproteobacteria>Enterobacterales>Enterobacteriaceae>Escherichia | 40 | D9Y07_001387 | 3473..3595 | 562 | HNH-endonuclease-[Escherichia-coli] |
| EEX1719678.1 | HNH | - | Escherichia_coli | Bacteria>Pseudomonadati>Pseudomonadota>Gammaproteobacteria>Enterobacterales>Enterobacteriaceae>Escherichia | 40 | D3S79_003402 | 2992..3114 | 562 | HNH-endonuclease-[Escherichia-coli] |
| EEZ2582143.1 | HNH | - | Escherichia_coli | Bacteria>Pseudomonadati>Pseudomonadota>Gammaproteobacteria>Enterobacterales>Enterobacteriaceae>Escherichia | 40 | DMH54_003963 | 5973..6095 | 562 | HNH-endonuclease-[Escherichia-coli] |
| EEZ3215395.1 | HNH | - | Escherichia_coli | Bacteria>Pseudomonadati>Pseudomonadota>Gammaproteobacteria>Enterobacterales>Enterobacteriaceae>Escherichia | 40 | DKV48_004293 | 3002..3124 | 562 | HNH-endonuclease-[Escherichia-coli] |
| EFC3446205.1 | HNH | - | Escherichia_coli | Bacteria>Pseudomonadati>Pseudomonadota>Gammaproteobacteria>Enterobacterales>Enterobacteriaceae>Escherichia | 40 | CO751_004603 | 6105..6227 | 562 | HNH-endonuclease-[Escherichia-coli] |
| EFD3459870.1 | HNH | - | Escherichia_coli | Bacteria>Pseudomonadati>Pseudomonadota>Gammaproteobacteria>Enterobacterales>Enterobacteriaceae>Escherichia | 40 | CL211_004297 | 3066..3188 | 562 | HNH-endonuclease-[Escherichia-coli] |
| EFE1965715.1 | HNH | - | Escherichia_coli | Bacteria>Pseudomonadati>Pseudomonadota>Gammaproteobacteria>Enterobacterales>Enterobacteriaceae>Escherichia | 40 | CJV81_002662 | 11003..11125 | 562 | HNH-endonuclease-[Escherichia-coli] |
| EFG4404334.1 | HNH | - | Escherichia_coli | Bacteria>Pseudomonadati>Pseudomonadota>Gammaproteobacteria>Enterobacterales>Enterobacteriaceae>Escherichia | 40 | BM772_002467 | 11004..11126 | 562 | HNH-endonuclease-[Escherichia-coli] |
| EFH0549020.1 | HNH | - | Escherichia_coli | Bacteria>Pseudomonadati>Pseudomonadota>Gammaproteobacteria>Enterobacterales>Enterobacteriaceae>Escherichia | 40 | BJE23_003832 | 36299..36421 | 562 | HNH-endonuclease-[Escherichia-coli] |
| EGY5620859.1 | HNH | - | Escherichia_coli | Bacteria>Pseudomonadati>Pseudomonadota>Gammaproteobacteria>Enterobacterales>Enterobacteriaceae>Escherichia | 40 | JI433_002885 | 11003..11125 | 562 | HNH-endonuclease-[Escherichia-coli] |
| EGY7451908.1 | HNH | - | Escherichia_coli | Bacteria>Pseudomonadati>Pseudomonadota>Gammaproteobacteria>Enterobacterales>Enterobacteriaceae>Escherichia | 40 | JI519_004382 | 6475..6597 | 562 | HNH-endonuclease-[Escherichia-coli] |
| EGY8948623.1 | HNH | - | Escherichia_coli | Bacteria>Pseudomonadati>Pseudomonadota>Gammaproteobacteria>Enterobacterales>Enterobacteriaceae>Escherichia | 40 | JI613_004640 | 4920..5042 | 562 | HNH-endonuclease-[Escherichia-coli] |
| EGY9306078.1 | HNH | - | Escherichia_coli | Bacteria>Pseudomonadati>Pseudomonadota>Gammaproteobacteria>Enterobacterales>Enterobacteriaceae>Escherichia | 40 | HZO96_004635 | 396..518 | 562 | HNH-endonuclease-[Escherichia-coli] |
| EGZ1560304.1 | HNH | - | Escherichia_coli | Bacteria>Pseudomonadati>Pseudomonadota>Gammaproteobacteria>Enterobacterales>Enterobacteriaceae>Escherichia | 40 | FAV16_004666 | 4903..5025 | 562 | HNH-endonuclease-[Escherichia-coli] |
| EGZ1871555.1 | HNH | - | Escherichia_coli | Bacteria>Pseudomonadati>Pseudomonadota>Gammaproteobacteria>Enterobacterales>Enterobacteriaceae>Escherichia | 40 | EYE54_004346 | 4819..4941 | 562 | HNH-endonuclease-[Escherichia-coli] |
| EGZ2189933.1 | HNH | - | Escherichia_coli | Bacteria>Pseudomonadati>Pseudomonadota>Gammaproteobacteria>Enterobacterales>Enterobacteriaceae>Escherichia | 40 | ED609_004470 | 5455..5577 | 562 | HNH-endonuclease-[Escherichia-coli] |
| EGZ2354225.1 | HNH | - | Escherichia_coli | Bacteria>Pseudomonadati>Pseudomonadota>Gammaproteobacteria>Enterobacterales>Enterobacteriaceae>Escherichia | 40 | D7J47_002039 | 11004..11126 | 562 | HNH-endonuclease-[Escherichia-coli] |
| EGZ2777076.1 | HNH | - | Escherichia_coli | Bacteria>Pseudomonadati>Pseudomonadota>Gammaproteobacteria>Enterobacterales>Enterobacteriaceae>Escherichia | 40 | DVO41_002384 | 11004..11126 | 562 | HNH-endonuclease-[Escherichia-coli] |
| EGZ3242778.1 | HNH | - | Escherichia_coli | Bacteria>Pseudomonadati>Pseudomonadota>Gammaproteobacteria>Enterobacterales>Enterobacteriaceae>Escherichia | 40 | C5N70_002399 | 36298..36420 | 562 | HNH-endonuclease-[Escherichia-coli] |
| EGZ5166762.1 | HNH | - | Escherichia_coli | Bacteria>Pseudomonadati>Pseudomonadota>Gammaproteobacteria>Enterobacterales>Enterobacteriaceae>Escherichia | 40 | CDS35_004371 | 5445..5567 | 562 | HNH-endonuclease-[Escherichia-coli] |
| EGZ5591351.1 | HNH | - | Escherichia_coli | Bacteria>Pseudomonadati>Pseudomonadota>Gammaproteobacteria>Enterobacterales>Enterobacteriaceae>Escherichia | 40 | BZZ56_004478 | 5692..5814 | 562 | HNH-endonuclease-[Escherichia-coli] |
| EGZ5731921.1 | HNH | - | Escherichia_coli | Bacteria>Pseudomonadati>Pseudomonadota>Gammaproteobacteria>Enterobacterales>Enterobacteriaceae>Escherichia | 40 | BL334_004675 | 6226..6348 | 562 | HNH-endonuclease-[Escherichia-coli] |
| EGZ5978146.1 | HNH | - | Escherichia_coli | Bacteria>Pseudomonadati>Pseudomonadota>Gammaproteobacteria>Enterobacterales>Enterobacteriaceae>Escherichia | 40 | BIB10_004099 | 36299..36421 | 562 | HNH-endonuclease-[Escherichia-coli] |
| EGZ8838304.1 | HNH | - | Escherichia_coli | Bacteria>Pseudomonadati>Pseudomonadota>Gammaproteobacteria>Enterobacterales>Enterobacteriaceae>Escherichia | 40 | AT823_000488 | 5026..5148 | 562 | HNH-endonuclease-[Escherichia-coli] |
| EHA9087552.1 | HNH | - | Escherichia_coli | Bacteria>Pseudomonadati>Pseudomonadota>Gammaproteobacteria>Enterobacterales>Enterobacteriaceae>Escherichia | 40 | D4094_002225 | 11006..11128 | 562 | HNH-endonuclease-[Escherichia-coli] |
| EHJ8640893.1 | HNH | GCA_017945505.1 | Escherichia_coli | Bacteria>Pseudomonadati>Pseudomonadota>Gammaproteobacteria>Enterobacterales>Enterobacteriaceae>Escherichia | 40 | KB366_000302 | 327268..327390 | 562 | HNH-endonuclease-[Escherichia-coli] |
| EHL0175575.1 | HNH | GCA_018090545.1 | Escherichia_coli | Bacteria>Pseudomonadati>Pseudomonadota>Gammaproteobacteria>Enterobacterales>Enterobacteriaceae>Escherichia | 40 | D7K57_004373 | 6621..6743 | 562 | HNH-endonuclease-[Escherichia-coli] |
| EHL0676195.1 | HNH | GCA_018092345.1 | Escherichia_coli | Bacteria>Pseudomonadati>Pseudomonadota>Gammaproteobacteria>Enterobacterales>Enterobacteriaceae>Escherichia | 40 | CP667_004248 | 6136..6258 | 562 | HNH-endonuclease-[Escherichia-coli] |
| EHL1134574.1 | HNH | GCA_018094045.1 | Escherichia_coli | Bacteria>Pseudomonadati>Pseudomonadota>Gammaproteobacteria>Enterobacterales>Enterobacteriaceae>Escherichia | 40 | BIB01_004966 | 4951..5073 | 562 | HNH-endonuclease-[Escherichia-coli] |
| EHL1162644.1 | HNH | GCA_018094265.1 | Escherichia_coli | Bacteria>Pseudomonadati>Pseudomonadota>Gammaproteobacteria>Enterobacterales>Enterobacteriaceae>Escherichia | 40 | BE892_002466 | 11005..11127 | 562 | HNH-endonuclease-[Escherichia-coli] |
| EHN3342066.1 | HNH | GCA_018356405.1 | Escherichia_coli | Bacteria>Pseudomonadati>Pseudomonadota>Gammaproteobacteria>Enterobacterales>Enterobacteriaceae>Escherichia | 40 | KIR71_003042 | 11004..11126 | 562 | HNH-endonuclease-[Escherichia-coli] |
| EHY5690077.1 | HNH | GCA_019590565.1 | Escherichia_coli | Bacteria>Pseudomonadati>Pseudomonadota>Gammaproteobacteria>Enterobacterales>Enterobacteriaceae>Escherichia | 40 | K4J54_002554 | 11003..11125 | 562 | HNH-endonuclease-[Escherichia-coli] |
| EIB0293610.1 | HNH | GCA_019848435.1 | Escherichia_coli | Bacteria>Pseudomonadati>Pseudomonadota>Gammaproteobacteria>Enterobacterales>Enterobacteriaceae>Escherichia | 40 | K7644_004615 | 5160..5282 | 562 | HNH-endonuclease-[Escherichia-coli] |
| EIG6470763.1 | HNH | GCA_020319175.1 | Escherichia_coli | Bacteria>Pseudomonadati>Pseudomonadota>Gammaproteobacteria>Enterobacterales>Enterobacteriaceae>Escherichia | 40 | EG665_005076 | 5474..5596 | 562 | HNH-endonuclease-[Escherichia-coli] |
| EIS9260704.1 | HNH | GCA_021435985.1 | Escherichia_coli | Bacteria>Pseudomonadati>Pseudomonadota>Gammaproteobacteria>Enterobacterales>Enterobacteriaceae>Escherichia | 40 | L0765_004698 | 7095..7217 | 562 | HNH-endonuclease-[Escherichia-coli] |
| EJW3212701.1 | HNH | GCA_025418235.1 | Escherichia_coli | Bacteria>Pseudomonadati>Pseudomonadota>Gammaproteobacteria>Enterobacterales>Enterobacteriaceae>Escherichia | 40 | OBZ04_004377 | 4825..4947 | 562 | HNH-endonuclease-[Escherichia-coli] |
| EEZ5786490.1 | HNH | - | Escherichia_coli_O107 | Bacteria>Pseudomonadati>Pseudomonadota>Gammaproteobacteria>Enterobacterales>Enterobacteriaceae>Escherichia>Escherichia_coli | 100 | DBU15_004379 | 650..952 | 2162915 | hypothetical-protein-[Escherichia-coli-O107] |
| POU04209.1 | HNH | GCA_002918705.1 | Escherichia_sp._ESNIH1 | Bacteria>Pseudomonadati>Pseudomonadota>Gammaproteobacteria>Enterobacterales>Enterobacteriaceae>Escherichia>unclassified_Escherichia | 224 | C3369_02320 | 483654..484328 | 1985876 | HNH-endonuclease-[Escherichia-sp.-ESNIH1] |
| WP_032738381.1 | HNH | GCF_026010935.1 | Klebsiella | Bacteria>Pseudomonadati>Pseudomonadota>Gammaproteobacteria>Enterobacterales>Enterobacteriaceae>Klebsiella/Raoultella_group | 100 | I8N79_RS22835 | 4672351..4672653 | 570 | MULTISPECIES:-hypothetical-protein-[Klebsiella] |
| WP_064279007.1 | HNH | GCF_902165215.1 | Klebsiella | Bacteria>Pseudomonadati>Pseudomonadota>Gammaproteobacteria>Enterobacterales>Enterobacteriaceae>Klebsiella/Raoultella_group | 100 | FN701_RS21065 | 23599..23901 | 570 | MULTISPECIES:-hypothetical-protein-[Klebsiella] |
| WP_064279016.1 | HNH | GCF_902165215.1 | Klebsiella | Bacteria>Pseudomonadati>Pseudomonadota>Gammaproteobacteria>Enterobacterales>Enterobacteriaceae>Klebsiella/Raoultella_group | 239 | FN701_RS21015 | 9567..10286 | 570 | MULTISPECIES:-HNH-endonuclease-signature-motif-containing-protein-[Klebsiella] |
| WP_279936014.1 | HNH | GCF_029837515.1 | Klebsiella_oxytoca | Bacteria>Pseudomonadati>Pseudomonadota>Gammaproteobacteria>Enterobacterales>Enterobacteriaceae>Klebsiella/Raoultella_group>Klebsiella | 146 | N7642_RS28030 | 6033..6475 | 571 | TIGR02646-family-protein,-partial-[Klebsiella-oxytoca] |
| HBW9592179.1 | HNH | GCA_022207645.1 | Klebsiella_pneumoniae | Bacteria>Pseudomonadati>Pseudomonadota>Gammaproteobacteria>Enterobacterales>Enterobacteriaceae>Klebsiella/Raoultella_group>Klebsiella>Klebsiella_pneumoniae_complex | 197 | MFS54_26480 | 5927..6520 | 573 | TIGR02646-family-protein-[Klebsiella-pneumoniae] |
| HCI7559690.1 | HNH | GCA_024446695.1 | Klebsiella_pneumoniae | Bacteria>Pseudomonadati>Pseudomonadota>Gammaproteobacteria>Enterobacterales>Enterobacteriaceae>Klebsiella/Raoultella_group>Klebsiella>Klebsiella_pneumoniae_complex | 197 | NPW42_004568 | 38537..39130 | 573 | TPA:-TIGR02646-family-protein-[Klebsiella-pneumoniae] |
| WP_229532554.1 | HNH | GCF_020577415.1 | Klebsiella_variicola | Bacteria>Pseudomonadati>Pseudomonadota>Gammaproteobacteria>Enterobacterales>Enterobacteriaceae>Klebsiella/Raoultella_group>Klebsiella>Klebsiella_pneumoniae_complex | 197 | KAG03_RS12060 | 337856..338449 | 244366 | TIGR02646-family-protein-[Klebsiella-variicola] |
| GKJ38609.1 | HNH | GCA_026007675.1 | Klebsiella_variicola | Bacteria>Pseudomonadati>Pseudomonadota>Gammaproteobacteria>Enterobacterales>Enterobacteriaceae>Klebsiella/Raoultella_group>Klebsiella>Klebsiella_pneumoniae_complex | 100 | NUKP24_47610 | 1893..2195 | 244366 | hypothetical-protein-NUKP24_47610-[Klebsiella-variicola] |
| HBS7446099.1 | HNH | GCA_021978815.1 | Klebsiella_pneumoniae | Bacteria>Pseudomonadati>Pseudomonadota>Gammaproteobacteria>Enterobacterales>Enterobacteriaceae>Klebsiella/Raoultella_group>Klebsiella>Klebsiella_pneumoniae_complex | 100 | MAQ81_001496 | 311323..311625 | 573 | TPA:-hypothetical-protein-MAQ81_001496-[Klebsiella-pneumoniae] |
| HBT3663668.1 | HNH | GCA_022030495.1 | Klebsiella_pneumoniae | Bacteria>Pseudomonadati>Pseudomonadota>Gammaproteobacteria>Enterobacterales>Enterobacteriaceae>Klebsiella/Raoultella_group>Klebsiella>Klebsiella_pneumoniae_complex | 100 | MBH96_005324 | 1167..1469 | 573 | hypothetical-protein-[Klebsiella-pneumoniae] |
| HBT9662841.1 | HNH | GCA_022069925.1 | Klebsiella_pneumoniae | Bacteria>Pseudomonadati>Pseudomonadota>Gammaproteobacteria>Enterobacterales>Enterobacteriaceae>Klebsiella/Raoultella_group>Klebsiella>Klebsiella_pneumoniae_complex | 100 | MCN56_004859 | 45933..46235 | 573 | hypothetical-protein-[Klebsiella-pneumoniae] |
| HBX6133590.1 | HNH | GCA_022250115.1 | Klebsiella_pneumoniae | Bacteria>Pseudomonadati>Pseudomonadota>Gammaproteobacteria>Enterobacterales>Enterobacteriaceae>Klebsiella/Raoultella_group>Klebsiella>Klebsiella_pneumoniae_complex | 100 | MH627_25380 | 19268..19570 | 573 | TPA:-hypothetical-protein-MH627_25380-[Klebsiella-pneumoniae] |
| HCM4054965.1 | HNH | GCA_025081835.1 | Klebsiella_quasipneumoniae | Bacteria>Pseudomonadati>Pseudomonadota>Gammaproteobacteria>Enterobacterales>Enterobacteriaceae>Klebsiella/Raoultella_group>Klebsiella>Klebsiella_pneumoniae_complex | 100 | N3I99_003991 | 11332..11634 | 1463165 | TPA:-hypothetical-protein-N3I99_003991-[Klebsiella-quasipneumoniae] |
| HCM5386364.1 | HNH | GCA_025094345.1 | Klebsiella_quasipneumoniae | Bacteria>Pseudomonadati>Pseudomonadota>Gammaproteobacteria>Enterobacterales>Enterobacteriaceae>Klebsiella/Raoultella_group>Klebsiella>Klebsiella_pneumoniae_complex | 100 | N3N82_003986 | 11326..11628 | 1463165 | TPA:-hypothetical-protein-N3N82_003986-[Klebsiella-quasipneumoniae] |
| MCF0347737.1 | HNH | GCA_021455545.1 | Klebsiella_pneumoniae | Bacteria>Pseudomonadati>Pseudomonadota>Gammaproteobacteria>Enterobacterales>Enterobacteriaceae>Klebsiella/Raoultella_group>Klebsiella>Klebsiella_pneumoniae_complex | 100 | L0F49_12155 | 76474..76776 | 573 | hypothetical-protein-[Klebsiella-pneumoniae] |
| MCP6235004.1 | HNH | GCA_024239175.1 | Klebsiella_pneumoniae | Bacteria>Pseudomonadati>Pseudomonadota>Gammaproteobacteria>Enterobacterales>Enterobacteriaceae>Klebsiella/Raoultella_group>Klebsiella>Klebsiella_pneumoniae_complex | 100 | NL436_15850 | 160252..160554 | 573 | hypothetical-protein-[Klebsiella-pneumoniae] |
| SBM35828.1 | HNH | GCA_900084005.1 | Klebsiella_pneumoniae | Bacteria>Pseudomonadati>Pseudomonadota>Gammaproteobacteria>Enterobacterales>Enterobacteriaceae>Klebsiella/Raoultella_group>Klebsiella>Klebsiella_pneumoniae_complex | 100 | SAMEA2273678_04061 | 177931..178233 | 573 | Uncharacterised-protein-[Klebsiella-pneumoniae] |
| WP_087867054.1 | HNH | GCF_016762175.1 | Klebsiella_pneumoniae | Bacteria>Pseudomonadati>Pseudomonadota>Gammaproteobacteria>Enterobacterales>Enterobacteriaceae>Klebsiella/Raoultella_group>Klebsiella>Klebsiella_pneumoniae_complex | 100 | H9R52_RS24870 | 54294..54596 | 573 | hypothetical-protein-[Klebsiella-pneumoniae] |
| WP_101825926.1 | HNH | GCF_002853035.1 | Klebsiella_variicola | Bacteria>Pseudomonadati>Pseudomonadota>Gammaproteobacteria>Enterobacterales>Enterobacteriaceae>Klebsiella/Raoultella_group>Klebsiella>Klebsiella_pneumoniae_complex | 100 | CLQ66_RS24370 | 14079..14381 | 244366 | hypothetical-protein-[Klebsiella-variicola] |
| WP_131026266.1 | HNH | GCF_004312745.1 | Klebsiella_variicola | Bacteria>Pseudomonadati>Pseudomonadota>Gammaproteobacteria>Enterobacterales>Enterobacteriaceae>Klebsiella/Raoultella_group>Klebsiella>Klebsiella_pneumoniae_complex | 100 | EON55_RS11895 | 66154..66456 | 244366 | hypothetical-protein-[Klebsiella-variicola] |
| WP_255089528.1 | HNH | GCF_024355505.1 | Klebsiella_pneumoniae | Bacteria>Pseudomonadati>Pseudomonadota>Gammaproteobacteria>Enterobacterales>Enterobacteriaceae>Klebsiella/Raoultella_group>Klebsiella>Klebsiella_pneumoniae_complex | 100 | L1C10_RS16390 | 20334..20636 | 573 | hypothetical-protein-[Klebsiella-pneumoniae] |
| WP_265011142.1 | HNH | GCF_026007675.1 | Klebsiella_variicola | Bacteria>Pseudomonadati>Pseudomonadota>Gammaproteobacteria>Enterobacterales>Enterobacteriaceae>Klebsiella/Raoultella_group>Klebsiella>Klebsiella_pneumoniae_complex | 100 | OPB36_RS24660 | 1893..2195 | 244366 | hypothetical-protein-[Klebsiella-variicola] |
| SBM35894.1 | HNH | GCA_900084005.1 | Klebsiella_pneumoniae | Bacteria>Pseudomonadati>Pseudomonadota>Gammaproteobacteria>Enterobacterales>Enterobacteriaceae>Klebsiella/Raoultella_group>Klebsiella>Klebsiella_pneumoniae_complex | 239 | SAMEA2273678_04070 | 191546..192265 | 573 | Uncharacterised-protein-[Klebsiella-pneumoniae] |
| WP_228299656.1 | HNH | - | Klebsiella_pneumoniae | Bacteria>Pseudomonadati>Pseudomonadota>Gammaproteobacteria>Enterobacterales>Enterobacteriaceae>Klebsiella/Raoultella_group>Klebsiella>Klebsiella_pneumoniae_complex | 126 | KFB18_RS26740 | 40067..40447 | 573 | HNH-endonuclease-signature-motif-containing-protein-[Klebsiella-pneumoniae] |
| WP_181890668.1 | HNH | GCF_902807195.1 | Raoultella_planticola | Bacteria>Pseudomonadati>Pseudomonadota>Gammaproteobacteria>Enterobacterales>Enterobacteriaceae>Klebsiella/Raoultella_group>Raoultella | 100 | HUW11_RS11180 | 74342..74644 | 575 | hypothetical-protein-[Raoultella-planticola] |
| WP_086872899.1 | HNH | GCF_900184035.1 | Kosakonia_pseudosacchari | Bacteria>Pseudomonadati>Pseudomonadota>Gammaproteobacteria>Enterobacterales>Enterobacteriaceae>Kosakonia | 197 | BN2591_RS15645 | 23247..23840 | 1646340 | TIGR02646-family-protein-[Kosakonia-pseudosacchari] |
| WP_233218411.1 | HNH | GCF_003028445.1 | Kosakonia_sp._H7A | Bacteria>Pseudomonadati>Pseudomonadota>Gammaproteobacteria>Enterobacterales>Enterobacteriaceae>Kosakonia>unclassified_Kosakonia | 101 | CWM66_RS20280 | 30474..30779 | 2054598 | hypothetical-protein-[Kosakonia-sp.-H7A] |
| WP_084884527.1 | HNH | GCF_032559815.1 | Pantoea | Bacteria>Pseudomonadati>Pseudomonadota>Gammaproteobacteria>Enterobacterales>Erwiniaceae | 100 | HA46_RS12260 | 80675..80977 | 53335 | hypothetical-protein-[Pantoea-septica] |
| WP_065650240.1 | HNH | GCF_001691555.1 | Pantoea_eucrina | Bacteria>Pseudomonadati>Pseudomonadota>Gammaproteobacteria>Enterobacterales>Erwiniaceae>Pantoea | 100 | BBA90_RS17410 | 2400360..2400662 | 472693 | hypothetical-protein-[Pantoea-eucrina] |
| WP_187500226.1 | HNH | GCF_014354155.1 | Pantoea_agglomerans | Bacteria>Pseudomonadati>Pseudomonadota>Gammaproteobacteria>Enterobacterales>Erwiniaceae>Pantoea>Pantoea_agglomerans_group | 100 | IBU41_RS09220 | 816386..816688 | 549 | hypothetical-protein-[Pantoea-agglomerans] |
| WP_077999172.1 | HNH | GCF_002006945.1 | Edwardsiella_tarda | Bacteria>Pseudomonadati>Pseudomonadota>Gammaproteobacteria>Enterobacterales>Hafniaceae>Edwardsiella | 209 | BTN58_RS02555 | 137352..137981 | 636 | TIGR02646-family-protein-[Edwardsiella-tarda] |
| WP_234625891.1 | HNH | GCF_021440185.1 | Hafnia_paralvei | Bacteria>Pseudomonadati>Pseudomonadota>Gammaproteobacteria>Enterobacterales>Hafniaceae>Hafnia | 100 | LZ651_RS15625 | 26846..27148 | 546367 | hypothetical-protein-[Hafnia-paralvei] |
| WP_111759507.1 | HNH | GCF_902387845.1 | Morganella_morganii | Bacteria>Pseudomonadati>Pseudomonadota>Gammaproteobacteria>Enterobacterales>Morganellaceae>Morganella | 100 | FX120_RS01915 | 392194..392496 | 582 | hypothetical-protein-[Morganella-morganii] |
| WP_205545048.1 | HNH | GCF_016949795.1 | Pectobacterium | Bacteria>Pseudomonadati>Pseudomonadota>Gammaproteobacteria>Enterobacterales>Pectobacteriaceae | 197 | H4F47_RS10585 | 43965..44558 | 122277 | TIGR02646-family-protein-[Pectobacterium-brasiliense] |
| WP_115183103.1 | HNH | GCF_946228205.1 | Serratia | Bacteria>Pseudomonadati>Pseudomonadota>Gammaproteobacteria>Enterobacterales>Yersiniaceae | 197 | BSR03_RS13415 | 139474..140067 | 613 | MULTISPECIES:-TIGR02646-family-protein-[Serratia] |
| WP_120162117.1 | HNH | GCF_003610915.1 | Rahnella_variigena | Bacteria>Pseudomonadati>Pseudomonadota>Gammaproteobacteria>Enterobacterales>Yersiniaceae>Rahnella | 100 | CKQ54_RS06095 | 1306751..1307053 | 574964 | hypothetical-protein-[Rahnella-variigena] |
| CAI1752883.1 | HNH | GCA_946228205.1 | Serratia_quinivorans | Bacteria>Pseudomonadati>Pseudomonadota>Gammaproteobacteria>Enterobacterales>Yersiniaceae>Serratia | 197 | SAMEA4857275_02167 | 314889..315482 | 137545 | Uncharacterised-protein-[Serratia-quinivorans] |
| SUI50262.1 | HNH | GCA_900457075.1 | Serratia_quinivorans | Bacteria>Pseudomonadati>Pseudomonadota>Gammaproteobacteria>Enterobacterales>Yersiniaceae>Serratia | 197 | NCTC11544_01115 | 1031248..1031841 | 137545 | Uncharacterised-protein-[Serratia-quinivorans] |
| WP_132542042.1 | HNH | GCF_004341245.1 | Plasticicumulans_lactativorans | Bacteria>Pseudomonadati>Pseudomonadota>Gammaproteobacteria>Gammaproteobacteria_incertae_sedis>Candidatus_Competibacteraceae>Plasticicumulans | 226 | EV699_RS09595 | 178866..179546 | 1133106 | TIGR02646-family-protein-[Plasticicumulans-lactativorans] |
| WP_132538607.1 | HNH | GCF_004341245.1 | Plasticicumulans_lactativorans | Bacteria>Pseudomonadati>Pseudomonadota>Gammaproteobacteria>Gammaproteobacteria_incertae_sedis>Candidatus_Competibacteraceae>Plasticicumulans | 220 | EV699_RS02455 | 28266..28928 | 1133106 | hypothetical-protein-[Plasticicumulans-lactativorans] |
| MBK8753024.1 | HNH | GCA_016710535.1 | Candidatus_Competibacteraceae_bacterium | Bacteria>Pseudomonadati>Pseudomonadota>Gammaproteobacteria>Gammaproteobacteria_incertae_sedis>Candidatus_Competibacteraceae>unclassified_Candidatus_Competibacteraceae | 146 | IPL99_15985 | 4382..4822 | 2053538 | hypothetical-protein-[Candidatus-Competibacteraceae-bacterium] |
| WP_080356573.1 | HNH | GCF_051982775.1 | Stenotrophomonas_maltophilia | Bacteria>Pseudomonadati>Pseudomonadota>Gammaproteobacteria>Lysobacterales>Lysobacteraceae>Stenotrophomonas>Stenotrophomonas_maltophilia_group | 238 | ADM82_RS13480 | 34768..35484 | 40324 | TIGR02646-family-protein-[Stenotrophomonas-maltophilia] |
| MBH1702630.1 | HNH | GCA_015998095.1 | Stenotrophomonas_maltophilia | Bacteria>Pseudomonadati>Pseudomonadota>Gammaproteobacteria>Lysobacterales>Lysobacteraceae>Stenotrophomonas>Stenotrophomonas_maltophilia_group | 199 | I5U93_05070 | 17033..17632 | 40324 | hypothetical-protein-[Stenotrophomonas-maltophilia] |
| WP_185752881.1 | HNH | GCF_003935615.1 | Stenotrophomonas_maltophilia | Bacteria>Pseudomonadati>Pseudomonadota>Gammaproteobacteria>Lysobacterales>Lysobacteraceae>Stenotrophomonas>Stenotrophomonas_maltophilia_group | 199 | EGJ89_RS09190 | 119935..120534 | 40324 | hypothetical-protein-[Stenotrophomonas-maltophilia] |
| PJK14603.1 | HNH | GCA_002798295.1 | Lysobacteraceae_bacterium_NML07-0707 | Bacteria>Pseudomonadati>Pseudomonadota>Gammaproteobacteria>Lysobacterales>Lysobacteraceae>unclassified_Lysobacteraceae | 243 | CO613_05150 | 91973..92704 | 2032574 | TIGR02646-family-protein-[Xanthomonadaceae-bacterium-NML07-0707] |
| ODU42591.1 | HNH | GCA_001725645.1 | Lysobacteraceae_bacterium_SCN_69-123 | Bacteria>Pseudomonadati>Pseudomonadota>Gammaproteobacteria>Lysobacterales>Lysobacteraceae>unclassified_Lysobacteraceae | 202 | ABS96_27570 | 186300..186908 | 1660170 | hypothetical-protein-ABS96_27570-[Xanthomonadaceae-bacterium-SCN-69-123] |
| MCC7256724.1 | HNH | GCA_020852005.1 | Dokdonella_sp. | Bacteria>Pseudomonadati>Pseudomonadota>Gammaproteobacteria>Lysobacterales>Rhodanobacteraceae>Dokdonella>unclassified_Dokdonella | 196 | IT467_12420 | 9856..10446 | 2291710 | TIGR02646-family-protein-[Dokdonella-sp.] |
| MCU0753618.1 | HNH | GCA_025458445.1 | Lysobacterales_bacterium | Bacteria>Pseudomonadati>Pseudomonadota>Gammaproteobacteria>Lysobacterales>unclassified_Lysobacterales | 249 | MUE46_00645 | 23650..24399 | 2006849 | TIGR02646-family-protein-[Xanthomonadales-bacterium] |
| WP_266024643.1 | HNH | GCF_026247885.1 | Methylococcus_mesophilus | Bacteria>Pseudomonadati>Pseudomonadota>Gammaproteobacteria>Methylococcales>Methylococcaceae>Methylococcus | 248 | OOT43_RS09365 | 1985087..1985833 | 2993564 | DCL-family-protein-[Methylococcus-sp.-16-5] |
| WP_071159670.1 | HNH | GCF_001856095.1 | Methylomonas_sp._LWB | Bacteria>Pseudomonadati>Pseudomonadota>Gammaproteobacteria>Methylococcales>Methylococcaceae>Methylomonas>unclassified_Methylomonas | 198 | BJL95_RS17195 | 205198..205794 | 1905845 | hypothetical-protein-[Methylomonas-sp.-LWB] |
| PZN82191.1 | HNH | GCA_003242955.1 | Candidatus_Methylumidiphilus_alinenensis | Bacteria>Pseudomonadati>Pseudomonadota>Gammaproteobacteria>Methylococcales>Methylococcales_incertae_sedis>Candidatus_Methylumidiphilus | 260 | DM484_06855 | 1041..1823 | 2202197 | TIGR02646-family-protein-[Candidatus-Methylumidiphilus-alinenensis] |
| PZN71301.1 | HNH | GCA_003242955.1 | Candidatus_Methylumidiphilus_alinenensis | Bacteria>Pseudomonadati>Pseudomonadota>Gammaproteobacteria>Methylococcales>Methylococcales_incertae_sedis>Candidatus_Methylumidiphilus | 209 | DM484_26745 | 20791..21420 | 2202197 | TIGR02646-family-protein-[Candidatus-Methylumidiphilus-alinenensis] |
| WP_225211635.1 | HNH | - | Psychrobacter_communis | Bacteria>Pseudomonadati>Pseudomonadota>Gammaproteobacteria>Moraxellales>Moraxellaceae>Psychrobacter | 138 | H9653_RS12480 | 29524..29940 | 2762238 | HNH-endonuclease-[Psychrobacter-communis] |
| WP_170968118.1 | HNH | GCF_011398715.1 | Halomonadaceae | Bacteria>Pseudomonadati>Pseudomonadota>Gammaproteobacteria>Oceanospirillales | 267 | KYX60_RS01255 | 113937..114740 | 28256 | MULTISPECIES:-TIGR02646-family-protein-[Halomonas] |
| WP_217488117.1 | HNH | GCF_013349955.1 | Halomonadaceae | Bacteria>Pseudomonadati>Pseudomonadota>Gammaproteobacteria>Oceanospirillales | 157 | HUS66_RS15800 | 33898..34371 | 28256 | hypothetical-protein-[Halomonas-taeanensis] |
| WP_142947827.1 | HNH | - | Alcanivorax_sp._DSM_26295 | Bacteria>Pseudomonadati>Pseudomonadota>Gammaproteobacteria>Oceanospirillales>Alcanivoracaceae>Alcanivorax>unclassified_Alcanivorax | 196 | FN816_RS03095 | 117050..117640 | 1938738 | TIGR02646-family-protein-[Alcanivorax-sp.-DSM-26295] |
| WP_262567767.1 | HNH | GCF_025562715.2 | Endozoicomonas_gorgoniicola | Bacteria>Pseudomonadati>Pseudomonadota>Gammaproteobacteria>Oceanospirillales>Endozoicomonadaceae>Endozoicomonas | 213 | NX722_RS09430 | 2080144..2080785 | 1234144 | TIGR02646-family-protein-[Endozoicomonas-gorgoniicola] |
| MCP5162752.1 | HNH | GCA_024234505.1 | Hahellaceae_bacterium | Bacteria>Pseudomonadati>Pseudomonadota>Gammaproteobacteria>Oceanospirillales>Hahellaceae>unclassified_Hahellaceae | 270 | H7A00_13980 | 79903..80715 | 2048971 | TIGR02646-family-protein-[Hahellaceae-bacterium] |
| MCP5206316.1 | HNH | GCA_024234475.1 | Hahellaceae_bacterium | Bacteria>Pseudomonadati>Pseudomonadota>Gammaproteobacteria>Oceanospirillales>Hahellaceae>unclassified_Hahellaceae | 216 | H7A01_03845 | 871253..871903 | 2048971 | TIGR02646-family-protein-[Hahellaceae-bacterium] |
| WP_239285454.1 | HNH | GCF_964204995.1 | unclassified_Halomonas | Bacteria>Pseudomonadati>Pseudomonadota>Gammaproteobacteria>Oceanospirillales>Halomonadaceae>Halomonas | 214 | MHM85_RS05480 | 201749..202393 | 2609666 | MULTISPECIES:-TIGR02646-family-protein-[unclassified-Halomonas] |
| WP_240569309.1 | HNH | GCF_022487945.1 | Halomonas_flagellata | Bacteria>Pseudomonadati>Pseudomonadota>Gammaproteobacteria>Oceanospirillales>Halomonadaceae>Halomonas | 209 | MKP05_RS16475 | 41279..41908 | 2920385 | TIGR02646-family-protein-[Halomonas-sp.-EGI-63088] |
| WP_253017806.1 | HNH | GCF_024124605.1 | Halomonas_sp._CnH100-B | Bacteria>Pseudomonadati>Pseudomonadota>Gammaproteobacteria>Oceanospirillales>Halomonadaceae>Halomonas>unclassified_Halomonas | 246 | NH398_RS08355 | 355737..356477 | 2954490 | TIGR02646-family-protein-[Halomonas-sp.-CnH100-B] |
| MCD6438840.1 | HNH | GCA_021161325.1 | Halomonas_sp. | Bacteria>Pseudomonadati>Pseudomonadota>Gammaproteobacteria>Oceanospirillales>Halomonadaceae>Halomonas>unclassified_Halomonas | 214 | J7L56_11330 | 181..825 | 1486246 | TIGR02646-family-protein-[Halomonas-sp.] |
| GKW48596.1 | HNH | GCA_023708045.1 | Halomonas_sp._NCCP-2165 | Bacteria>Pseudomonadati>Pseudomonadota>Gammaproteobacteria>Oceanospirillales>Halomonadaceae>Halomonas>unclassified_Halomonas | 210 | NCCP2165_08110 | 90308..90940 | 2942298 | hypothetical-protein-NCCP2165_08110-[Halomonas-sp.-NCCP-2165] |
| WP_251308251.1 | HNH | - | Halomonas_sp._NCCP-2165 | Bacteria>Pseudomonadati>Pseudomonadota>Gammaproteobacteria>Oceanospirillales>Halomonadaceae>Halomonas>unclassified_Halomonas | 210 | NFU04_RS04195 | 90308..90940 | 2942298 | TIGR02646-family-protein-[Halomonas-sp.-NCCP-2165] |
| WP_107181350.1 | HNH | GCF_003030865.2 | Halomonas_sp._SL1 | Bacteria>Pseudomonadati>Pseudomonadota>Gammaproteobacteria>Oceanospirillales>Halomonadaceae>Halomonas>unclassified_Halomonas | 202 | C9J49_RS09480 | 190914..191522 | 2137478 | hypothetical-protein-[Halomonas-sp.-SL1] |
| WP_176212840.1 | HNH | - | Halomonas_sp._BC1 | Bacteria>Pseudomonadati>Pseudomonadota>Gammaproteobacteria>Oceanospirillales>Halomonadaceae>Halomonas>unclassified_Halomonas | 199 | B7L72_RS18870 | 20494..21093 | 1670448 | hypothetical-protein-[Halomonas-sp.-BC1] |
| WP_031385047.1 | HNH | GCF_039543745.1 | Modicisalibacter_zincidurans | Bacteria>Pseudomonadati>Pseudomonadota>Gammaproteobacteria>Oceanospirillales>Halomonadaceae>Modicisalibacter | 211 | HALZIN_RS0115230 | 3242714..3243349 | 1178777 | TIGR02646-family-protein-[Halomonas-zincidurans] |
| WP_072820437.1 | HNH | GCF_900128925.1 | Modicisalibacter_ilicicola | Bacteria>Pseudomonadati>Pseudomonadota>Gammaproteobacteria>Oceanospirillales>Halomonadaceae>Modicisalibacter | 210 | BUA41_RS05230 | 143681..144313 | 480814 | TIGR02646-family-protein-[Halomonas-ilicicola] |
| WP_087719607.1 | HNH | GCF_002179555.1 | Salinicola_salarius | Bacteria>Pseudomonadati>Pseudomonadota>Gammaproteobacteria>Oceanospirillales>Halomonadaceae>Salinicola | 199 | CCD76_RS11260 | 23136..23735 | 430457 | hypothetical-protein-[Salinicola-salarius] |
| WP_188638087.1 | HNH | GCF_014640815.1 | Vreelandella_lutescens | Bacteria>Pseudomonadati>Pseudomonadota>Gammaproteobacteria>Oceanospirillales>Halomonadaceae>Vreelandella | 199 | IEW65_RS03400 | 732028..732627 | 1602943 | hypothetical-protein-[Halomonas-lutescens] |
| WP_217336195.1 | HNH | GCF_019061305.1 | Marinobacterium_weihaiense | Bacteria>Pseudomonadati>Pseudomonadota>Gammaproteobacteria>Oceanospirillales>Oceanospirillaceae>Marinobacterium | 213 | KTN04_RS15770 | 12567..13208 | 2851016 | TIGR02646-family-protein-[Marinobacterium-sp.-A346] |
| WP_206838941.1 | HNH | GCF_017310015.1 | Marinobacterium_iners | Bacteria>Pseudomonadati>Pseudomonadota>Gammaproteobacteria>Oceanospirillales>Oceanospirillaceae>Marinobacterium | 199 | CFI10_RS02435 | 491961..492560 | 48076 | hypothetical-protein-[Marinobacterium-iners] |
| WP_027853083.1 | HNH | GCF_000428985.1 | Marinobacterium_litorale | Bacteria>Pseudomonadati>Pseudomonadota>Gammaproteobacteria>Oceanospirillales>Oceanospirillaceae>Marinobacterium | 198 | G403_RS0105355 | 182836..183432 | 404770 | hypothetical-protein-[Marinobacterium-litorale] |
| WP_181736360.1 | HNH | GCF_013868415.1 | Marinobacterium_marinum | Bacteria>Pseudomonadati>Pseudomonadota>Gammaproteobacteria>Oceanospirillales>Oceanospirillaceae>Marinobacterium | 198 | H1S06_RS00690 | 40709..41305 | 2756129 | hypothetical-protein-[Marinobacterium-sp.-3-1745] |
| WP_217336197.1 | HNH | GCF_019061305.1 | Marinobacterium_weihaiense | Bacteria>Pseudomonadati>Pseudomonadota>Gammaproteobacteria>Oceanospirillales>Oceanospirillaceae>Marinobacterium | 198 | KTN04_RS15780 | 14505..15101 | 2851016 | hypothetical-protein-[Marinobacterium-sp.-A346] |
| WP_072841214.1 | HNH | GCF_900129155.1 | Marinomonas_polaris | Bacteria>Pseudomonadati>Pseudomonadota>Gammaproteobacteria>Oceanospirillales>Oceanospirillaceae>Marinomonas | 212 | BUA59_RS18810 | 71268..71906 | 293552 | TIGR02646-family-protein-[Marinomonas-polaris] |

| Protein-ID | Domain | Assembly-ID | Strain | Taxonomy | Length | Locus-Tag | Genomic-coordinates | TaxID | Fasta-defline |
|---|---|---|---|---|---|---|---|---|---|
| WP_116685594.1 | HNH | GCF_963878705.1 | Pelagibaculum_spongiae | Bacteria>Pseudomonadati>Pseudomonadota>Gammaproteobacteria>Oceanospirillales>Oceanospirillales_incertae_sedis>Pelagibaculum | 212 | DC094_RS03015 | 687339..687977 | 2080658 | TIGR02646-family-protein-[Pelagibaculum-spongiae] |
| WP_119395056.1 | HNH | GCF_003577475.1 | Salinibius_halmophilus | Bacteria>Pseudomonadati>Pseudomonadota>Gammaproteobacteria>Oceanospirillales>Saccharospirillaceae>Salinibius | 245 | DUW71_RS09700 | 143307..144044 | 1853216 | TIGR02646-family-protein-[Salinibius-halmophilus] |
| WP_233546089.1 | HNH | GCF_003577475.1 | Salinibius_halmophilus | Bacteria>Pseudomonadati>Pseudomonadota>Gammaproteobacteria>Oceanospirillales>Saccharospirillaceae>Salinibius | 239 | DUW71_RS09755 | 161742..162461 | 1853216 | HNH-endonuclease-[Salinibius-halmophilus] |
| WP_199671196.1 | HNH | GCF_003577475.1 | Salinibius_halmophilus | Bacteria>Pseudomonadati>Pseudomonadota>Gammaproteobacteria>Oceanospirillales>Saccharospirillaceae>Salinibius | 201 | DUW71_RS09745 | 160736..161341 | 1853216 | HNH/ENDO-VII-family-nuclease-[Salinibius-halmophilus] |
| NRB78695.1 | HNH | GCA_013214075.1 | Saccharospirillaceae_bacterium | Bacteria>Pseudomonadati>Pseudomonadota>Gammaproteobacteria>Oceanospirillales>Saccharospirillaceae>unclassified_Saccharospirillaceae | 242 | HRU38_08510 | 24341..25069 | 2497634 | hypothetical-protein-[Saccharospirillaceae-bacterium] |
| WP_094873718.1 | HNH | GCF_002263255.2 | Gallibacterium_anatis | Bacteria>Pseudomonadati>Pseudomonadota>Gammaproteobacteria>Pasteurellales>Pasteurellaceae>Gallibacterium | 198 | CF557_RS03195 | 659356..659952 | 750 | hypothetical-protein-[Gallibacterium-anatis] |
| WP_053526940.1 | HNH | GCF_027856115.1 | Pseudomonadaceae | Bacteria>Pseudomonadati>Pseudomonadota>Gammaproteobacteria>Pseudomonadales | 215 | KKQ36_RS16475 | 3496961..3497608 | 135621 | MULTISPECIES:-hypothetical-protein-[Pseudomonadaceae] |
| WP_126634677.1 | HNH | GCF_045795645.1 | Pseudomonadaceae | Bacteria>Pseudomonadati>Pseudomonadota>Gammaproteobacteria>Pseudomonadales | 196 | AK6_RS01595 | 327404..327994 | 135621 | MULTISPECIES:-TIGR02646-family-protein-[Pseudomonadaceae] |
| WP_091985099.1 | HNH | GCF_900115175.1 | Marinobacter_gudaonensis | Bacteria>Pseudomonadati>Pseudomonadota>Gammaproteobacteria>Pseudomonadales>Marinobacteraceae>Marinobacter | 246 | BM344_RS01235 | 255576..256316 | 375760 | TIGR02646-family-protein-[Marinobacter-gudaonensis] |
| WP_018404439.1 | HNH | GCF_000372805.1 | Marinobacter_gelidimuriae | Bacteria>Pseudomonadati>Pseudomonadota>Gammaproteobacteria>Pseudomonadales>Marinobacteraceae>Marinobacter | 211 | A3OI_RS0111635 | 12054..12689 | 2739064 | TIGR02646-family-protein-[Marinobacter-gelidimuriae] |
| WP_098418630.1 | HNH | GCF_002563885.1 | unclassified_Marinobacter | Bacteria>Pseudomonadati>Pseudomonadota>Gammaproteobacteria>Pseudomonadales>Marinobacteraceae>Marinobacter | 211 | ATG98_RS19575 | 4030631..4031266 | 83889 | MULTISPECIES:-TIGR02646-family-protein-[unclassified-Marinobacter] |
| WP_228706216.1 | HNH | GCF_002563885.1 | unclassified_Marinobacter | Bacteria>Pseudomonadati>Pseudomonadota>Gammaproteobacteria>Pseudomonadales>Marinobacteraceae>Marinobacter | 284 | ATG98_RS22615 | 4041233..4042087 | 83889 | MULTISPECIES:-HNH-endonuclease-signature-motif-containing-protein-[unclassified-Marinobacter] |
| AKV97709.1 | HNH | GCA_001266795.1 | Marinobacter_sp._CP1 | Bacteria>Pseudomonadati>Pseudomonadota>Gammaproteobacteria>Pseudomonadales>Marinobacteraceae>Marinobacter>unclassified_Marinobacter | 211 | ACP86_16965 | 3686611..3687246 | 1671721 | hypothetical-protein-ACP86_16965-[Marinobacter-sp.-CP1] |
| WP_007349687.1 | HNH | GCF_000169375.1 | Marinobacter_sp._ELB17 | Bacteria>Pseudomonadati>Pseudomonadota>Gammaproteobacteria>Pseudomonadales>Marinobacteraceae>Marinobacter>unclassified_Marinobacter | 211 | MELB17_RS08055 | 28481..29116 | 270374 | TIGR02646-family-protein-[Marinobacter-sp.-ELB17] |
| WP_053114869.1 | HNH | GCF_001266795.1 | Marinobacter_sp._CP1 | Bacteria>Pseudomonadati>Pseudomonadota>Gammaproteobacteria>Pseudomonadales>Marinobacteraceae>Marinobacter>unclassified_Marinobacter | 211 | ACP86_RS16970 | 3686611..3687246 | 1671721 | TIGR02646-family-protein-[Marinobacter-sp.-CP1] |
| MBC7192643.1 | HNH | GCA_014359475.1 | Marinobacter_sp. | Bacteria>Pseudomonadati>Pseudomonadota>Gammaproteobacteria>Pseudomonadales>Marinobacteraceae>Marinobacter>unclassified_Marinobacter | 199 | H5U31_09855 | 10380..10979 | 50741 | hypothetical-protein-[Marinobacter-sp.] |
| WP_033898359.1 | HNH | GCF_000510895.2 | Pseudomonas | Bacteria>Pseudomonadati>Pseudomonadota>Gammaproteobacteria>Pseudomonadales>Pseudomonadaceae | 215 | GIW75_RS00895 | 18648..19295 | 286 | MULTISPECIES:-TIGR02646-family-protein-[Pseudomonas] |
| WP_269686939.1 | HNH | GCF_030295685.1 | Halopseudomonas | Bacteria>Pseudomonadati>Pseudomonadota>Gammaproteobacteria>Pseudomonadales>Pseudomonadaceae | 215 | O5O48_RS10345 | 2213966..2214613 | 2901189 | hypothetical-protein-[Halopseudomonas-sp.-MFKK-1] |
| WP_015271889.1 | HNH | GCF_000325725.1 | Pseudomonas | Bacteria>Pseudomonadati>Pseudomonadota>Gammaproteobacteria>Pseudomonadales>Pseudomonadaceae | 213 | LU687_RS25025 | 5308466..5309107 | 286 | MULTISPECIES:-TIGR02646-family-protein-[Pseudomonas] |
| WP_061303570.1 | HNH | GCF_009012945.1 | Pseudomonas | Bacteria>Pseudomonadati>Pseudomonadota>Gammaproteobacteria>Pseudomonadales>Pseudomonadaceae | 213 | GB470_RS16295 | 4205..4846 | 286 | MULTISPECIES:-TIGR02646-family-protein-[Pseudomonas] |
| WP_103518018.1 | HNH | GCF_013305665.1 | Pseudomonas | Bacteria>Pseudomonadati>Pseudomonadota>Gammaproteobacteria>Pseudomonadales>Pseudomonadaceae | 213 | CW309_RS13940 | 86411..87052 | 286 | TIGR02646-family-protein-[Pseudomonas-hunanensis] |
| WP_192907657.1 | HNH | GCF_014883875.1 | Pseudomonas | Bacteria>Pseudomonadati>Pseudomonadota>Gammaproteobacteria>Pseudomonadales>Pseudomonadaceae | 196 | ICN73_RS10025 | 2224794..2225384 | 286 | TIGR02646-family-protein-[Pseudomonas-taiwanensis] |
| WP_280091281.1 | HNH | GCF_947311875.1 | Pseudomonas | Bacteria>Pseudomonadati>Pseudomonadota>Gammaproteobacteria>Pseudomonadales>Pseudomonadaceae | 196 | QME24_RS13350 | 72597..73187 | 286 | TIGR02646-family-protein-[Pseudomonas-juntendi] |
| WP_026034349.1 | HNH | GCF_000292285.1 | Pseudomonas | Bacteria>Pseudomonadati>Pseudomonadota>Gammaproteobacteria>Pseudomonadales>Pseudomonadaceae | 100 | N805_RS25720 | 5823690..5823992 | 286 | MULTISPECIES:-hypothetical-protein-[Pseudomonas] |
| WP_041931316.1 | HNH | GCF_000194805.1 | Pseudomonas | Bacteria>Pseudomonadati>Pseudomonadota>Gammaproteobacteria>Pseudomonadales>Pseudomonadaceae | 100 | PSEBR_RS03985 | 955571..955873 | 286 | hypothetical-protein-[Pseudomonas-brassicacearum] |
| WP_058410020.1 | HNH | GCF_029959515.1 | Pseudomonas | Bacteria>Pseudomonadati>Pseudomonadota>Gammaproteobacteria>Pseudomonadales>Pseudomonadaceae | 100 | AO239_RS04365 | 20812..21114 | 286 | hypothetical-protein-[Pseudomonas-sp.-ICMP-19500] |
| WP_095027078.1 | HNH | GCF_016405455.1 | Pseudomonas | Bacteria>Pseudomonadati>Pseudomonadota>Gammaproteobacteria>Pseudomonadales>Pseudomonadaceae | 100 | JFT63_RS16105 | 192851..193153 | 286 | MULTISPECIES:-hypothetical-protein-[Pseudomonas] |
| WP_186598288.1 | HNH | GCF_014269085.1 | Pseudomonas | Bacteria>Pseudomonadati>Pseudomonadota>Gammaproteobacteria>Pseudomonadales>Pseudomonadaceae | 100 | HU747_RS04825 | 445528..445830 | 286 | hypothetical-protein-[Pseudomonas-taiwanensis] |
| WP_229605959.1 | HNH | GCF_020831405.1 | Pseudomonas | Bacteria>Pseudomonadati>Pseudomonadota>Gammaproteobacteria>Pseudomonadales>Pseudomonadaceae | 100 | LMJ29_RS13080 | 80795..81097 | 286 | hypothetical-protein-[Pseudomonas-oryzagri] |
| WP_236165762.1 | HNH | GCF_021603865.1 | Pseudomonas | Bacteria>Pseudomonadati>Pseudomonadota>Gammaproteobacteria>Pseudomonadales>Pseudomonadaceae | 100 | L6D10_RS06735 | 30579..30881 | 286 | hypothetical-protein-[Pseudomonas-fulva] |
| WP_021701532.1 | HNH | GCF_000467105.1 | Aquipseudomonas_alcaligenes | Bacteria>Pseudomonadati>Pseudomonadota>Gammaproteobacteria>Pseudomonadales>Pseudomonadaceae>Aquipseudomonas | 100 | DYC96_RS14105 | 2978807..2979109 | 43263 | hypothetical-protein-[Pseudomonas-alcaligenes] |
| WP_090941607.1 | HNH | GCF_900114395.1 | Azotobacter_beijerinckii | Bacteria>Pseudomonadati>Pseudomonadota>Gammaproteobacteria>Pseudomonadales>Pseudomonadaceae>Azotobacter_group>Azotobacter | 246 | BM240_RS15630 | 12502..13242 | 170623 | TIGR02646-family-protein-[Azotobacter-beijerinckii] |
| WP_240931548.1 | HNH | GCF_011470035.1 | Azotobacter_chroococcum | Bacteria>Pseudomonadati>Pseudomonadota>Gammaproteobacteria>Pseudomonadales>Pseudomonadaceae>Azotobacter_group>Azotobacter | 233 | HA520_RS07855 | 27348..28049 | 353 | TIGR02646-family-protein-[Azotobacter-chroococcum] |
| WP_131349391.1 | HNH | GCF_004327885.1 | Azotobacter_chroococcum | Bacteria>Pseudomonadati>Pseudomonadota>Gammaproteobacteria>Pseudomonadales>Pseudomonadaceae>Azotobacter_group>Azotobacter | 196 | EOE52_RS08230 | 54780..55370 | 353 | TIGR02646-family-protein-[Azotobacter-chroococcum] |
| WP_282398727.1 | HNH | GCF_029960275.1 | Pseudomonas_paracarnis | Bacteria>Pseudomonadati>Pseudomonadota>Gammaproteobacteria>Pseudomonadales>Pseudomonadaceae>Pseudomonas | 213 | QK328_RS05440 | 305017..305658 | 2750625 | retron-Ec78-anti-phage-system-effector-HNH-endonuclease-PtuB-[Pseudomonas-paracarnis] |
| WP_064314269.1 | HNH | GCF_021601465.1 | Pseudomonas_juntendi | Bacteria>Pseudomonadati>Pseudomonadota>Gammaproteobacteria>Pseudomonadales>Pseudomonadaceae>Pseudomonas | 199 | L6D32_RS09710 | 141696..142295 | 2666183 | MULTISPECIES:-hypothetical-protein-[Pseudomonas] |
| WP_170050468.1 | HNH | GCF_012985775.1 | unclassified_Pseudomonas | Bacteria>Pseudomonadati>Pseudomonadota>Gammaproteobacteria>Pseudomonadales>Pseudomonadaceae>Pseudomonas | 199 | HBN63_RS18850 | 27616..28215 | 196821 | hypothetical-protein-[Pseudomonas-sp.-WS-5011] |
| WP_196169748.1 | HNH | GCF_015680295.1 | Pseudomonas_guariconensis | Bacteria>Pseudomonadati>Pseudomonadota>Gammaproteobacteria>Pseudomonadales>Pseudomonadaceae>Pseudomonas | 199 | IRZ59_RS14945 | 178702..179301 | 1288410 | hypothetical-protein-[Pseudomonas-guariconensis] |
| WP_165765330.1 | HNH | GCF_014635625.1 | Pseudomonas_fluvialis | Bacteria>Pseudomonadati>Pseudomonadota>Gammaproteobacteria>Pseudomonadales>Pseudomonadaceae>Pseudomonas | 196 | CFY91_RS13750 | 20988..21578 | 1793966 | TIGR02646-family-protein-[Pseudomonas-fluvialis] |
| KHO64387.1 | HNH | GCA_000806415.1 | Pseudomonas_flexibilis | Bacteria>Pseudomonadati>Pseudomonadota>Gammaproteobacteria>Pseudomonadales>Pseudomonadaceae>Pseudomonas | 100 | PT85_13865 | 365860..366162 | 706570 | hypothetical-protein-PT85_13865-[Pseudomonas-flexibilis] |
| WP_039607052.1 | HNH | GCF_900101515.1 | Pseudomonas_flexibilis | Bacteria>Pseudomonadati>Pseudomonadota>Gammaproteobacteria>Pseudomonadales>Pseudomonadaceae>Pseudomonas | 100 | BLP69_RS07670 | 85172..85474 | 706570 | hypothetical-protein-[Pseudomonas-flexibilis] |
| WP_090241715.1 | HNH | GCF_900113745.1 | Pseudomonas_guineae | Bacteria>Pseudomonadati>Pseudomonadota>Gammaproteobacteria>Pseudomonadales>Pseudomonadaceae>Pseudomonas | 100 | BMZ32_RS09375 | 95089..95391 | 425504 | hypothetical-protein-[Pseudomonas-guineae] |
| WP_188867723.1 | HNH | GCF_014646755.1 | Pseudomonas_asuensis | Bacteria>Pseudomonadati>Pseudomonadota>Gammaproteobacteria>Pseudomonadales>Pseudomonadaceae>Pseudomonas | 100 | IEX86_RS19095 | 198109..198411 | 1825787 | hypothetical-protein-[Pseudomonas-asuensis] |
| WP_244972682.1 | HNH | - | Pseudomonas_salomonii | Bacteria>Pseudomonadati>Pseudomonadota>Gammaproteobacteria>Pseudomonadales>Pseudomonadaceae>Pseudomonas | 176 | HX810_RS30890 | 502983..503513 | 191391 | DUF1524-domain-containing-protein-[Pseudomonas-salomonii] |
| WP_240357229.1 | HNH | GCF_016925675.1 | Pseudomonas_hygromyciniae | Bacteria>Pseudomonadati>Pseudomonadota>Gammaproteobacteria>Pseudomonadales>Pseudomonadaceae>Pseudomonas | 88 | JTY93_RS19450 | 4359488..4359754 | 2812000 | hypothetical-protein-[Pseudomonas-sp.-SDM007] |
| WP_079386607.1 | HNH | GCF_009905195.1 | Pseudomonas_aeruginosa | Bacteria>Pseudomonadati>Pseudomonadota>Gammaproteobacteria>Pseudomonadales>Pseudomonadaceae>Pseudomonas>Pseudomonas_aeruginosa_group | 246 | OLM70_RS05750 | 1179340..1180080 | 287 | TIGR02646-family-protein-[Pseudomonas-aeruginosa] |
| WP_146538002.1 | HNH | GCF_007896855.1 | Pseudomonas_aeruginosa | Bacteria>Pseudomonadati>Pseudomonadota>Gammaproteobacteria>Pseudomonadales>Pseudomonadaceae>Pseudomonas>Pseudomonas_aeruginosa_group | 246 | FQA91_RS32310 | 30909..31649 | 287 | TIGR02646-family-protein-[Pseudomonas-aeruginosa] |
| WP_250133304.1 | HNH | - | Pseudomonas_aeruginosa | Bacteria>Pseudomonadati>Pseudomonadota>Gammaproteobacteria>Pseudomonadales>Pseudomonadaceae>Pseudomonas>Pseudomonas_aeruginosa_group | 240 | KKT79_RS19580 | 38881..39603 | 287 | TIGR02646-family-protein-[Pseudomonas-aeruginosa] |
| HBP6726067.1 | HNH | GCA_021711795.1 | Pseudomonas_aeruginosa | Bacteria>Pseudomonadati>Pseudomonadota>Gammaproteobacteria>Pseudomonadales>Pseudomonadaceae>Pseudomonas>Pseudomonas_aeruginosa_group | 215 | L6B36_03980 | 83770..84417 | 287 | hypothetical-protein-[Pseudomonas-aeruginosa] |
| MBX5763954.1 | HNH | GCA_019687095.1 | Pseudomonas_aeruginosa | Bacteria>Pseudomonadati>Pseudomonadota>Gammaproteobacteria>Pseudomonadales>Pseudomonadaceae>Pseudomonas>Pseudomonas_aeruginosa_group | 215 | ISD89_03950 | 855351..855998 | 287 | hypothetical-protein-[Pseudomonas-aeruginosa] |
| EIU3464055.1 | HNH | GCA_021644165.1 | Pseudomonas_aeruginosa | Bacteria>Pseudomonadati>Pseudomonadota>Gammaproteobacteria>Pseudomonadales>Pseudomonadaceae>Pseudomonas>Pseudomonas_aeruginosa_group | 199 | L4P80_001283 | 31147..31746 | 287 | hypothetical-protein-[Pseudomonas-aeruginosa] |
| HBO3110866.1 | HNH | GCA_021651555.1 | Pseudomonas_aeruginosa | Bacteria>Pseudomonadati>Pseudomonadota>Gammaproteobacteria>Pseudomonadales>Pseudomonadaceae>Pseudomonas>Pseudomonas_aeruginosa_group | 199 | L4R01_001442 | 26171..26770 | 287 | hypothetical-protein-[Pseudomonas-aeruginosa] |
| HBN8454458.1 | HNH | GCA_021587585.1 | Pseudomonas_aeruginosa | Bacteria>Pseudomonadati>Pseudomonadota>Gammaproteobacteria>Pseudomonadales>Pseudomonadaceae>Pseudomonas>Pseudomonas_aeruginosa_group | 198 | L3711_005715 | 34274..34870 | 287 | hypothetical-protein-[Pseudomonas-aeruginosa] |
| HBP6330508.1 | HNH | GCA_021710255.1 | Pseudomonas_aeruginosa | Bacteria>Pseudomonadati>Pseudomonadota>Gammaproteobacteria>Pseudomonadales>Pseudomonadaceae>Pseudomonas>Pseudomonas_aeruginosa_group | 198 | L6A37_21780 | 64201..64797 | 287 | hypothetical-protein-[Pseudomonas-aeruginosa] |
| HCL3757801.1 | HNH | GCA_025003865.1 | Pseudomonas_aeruginosa | Bacteria>Pseudomonadati>Pseudomonadota>Gammaproteobacteria>Pseudomonadales>Pseudomonadaceae>Pseudomonas>Pseudomonas_aeruginosa_group | 196 | N2B17_005729 | 5599..6189 | 287 | TIGR02646-family-protein-[Pseudomonas-aeruginosa] |
| MBI8725673.1 | HNH | GCA_016342885.1 | Pseudomonas_aeruginosa | Bacteria>Pseudomonadati>Pseudomonadota>Gammaproteobacteria>Pseudomonadales>Pseudomonadaceae>Pseudomonas>Pseudomonas_aeruginosa_group | 196 | JEZ69_03795 | 197255..197845 | 287 | TIGR02646-family-protein-[Pseudomonas-aeruginosa] |
| WCY08685.1 | HNH | GCA_028609905.1 | Pseudomonas_aeruginosa | Bacteria>Pseudomonadati>Pseudomonadota>Gammaproteobacteria>Pseudomonadales>Pseudomonadaceae>Pseudomonas>Pseudomonas_aeruginosa_group | 196 | KK229_11685 | 2439782..2440372 | 287 | TIGR02646-family-protein-[Pseudomonas-aeruginosa] |
| WP_033983009.1 | HNH | GCF_049912885.1 | Pseudomonas_aeruginosa | Bacteria>Pseudomonadati>Pseudomonadota>Gammaproteobacteria>Pseudomonadales>Pseudomonadaceae>Pseudomonas>Pseudomonas_aeruginosa_group | 196 | NR66_RS02480 | 139760..140350 | 287 | TIGR02646-family-protein-[Pseudomonas-aeruginosa] |
| WP_078451164.1 | HNH | GCF_000629125.1 | Pseudomonas_aeruginosa | Bacteria>Pseudomonadati>Pseudomonadota>Gammaproteobacteria>Pseudomonadales>Pseudomonadaceae>Pseudomonas>Pseudomonas_aeruginosa_group | 196 | AJ70_RS36405 | 39706..40296 | 287 | TIGR02646-family-protein-[Pseudomonas-aeruginosa] |
| WP_161968265.1 | HNH | GCF_041698535.1 | Pseudomonas_aeruginosa | Bacteria>Pseudomonadati>Pseudomonadota>Gammaproteobacteria>Pseudomonadales>Pseudomonadaceae>Pseudomonas>Pseudomonas_aeruginosa_group | 196 | IPC1239_RS13855 | 651909..652499 | 287 | TIGR02646-family-protein-[Pseudomonas-aeruginosa] |
| WP_276226488.1 | HNH | GCF_029335315.1 | Pseudomonas_citronellolis | Bacteria>Pseudomonadati>Pseudomonadota>Gammaproteobacteria>Pseudomonadales>Pseudomonadaceae>Pseudomonas>Pseudomonas_aeruginosa_group | 196 | P3W75_RS06205 | 42252..42842 | 53408 | TIGR02646-family-protein-[Pseudomonas-citronellolis] |
| EKW7733845.1 | HNH | - | Pseudomonas_aeruginosa | Bacteria>Pseudomonadati>Pseudomonadota>Gammaproteobacteria>Pseudomonadales>Pseudomonadaceae>Pseudomonas>Pseudomonas_aeruginosa_group | 100 | PU294_000690 | 24205..24507 | 287 | hypothetical-protein-PU294_000690-[Pseudomonas-aeruginosa] |
| HCF2077869.1 | HNH | GCA_023942995.1 | Pseudomonas_aeruginosa | Bacteria>Pseudomonadati>Pseudomonadota>Gammaproteobacteria>Pseudomonadales>Pseudomonadaceae>Pseudomonas>Pseudomonas_aeruginosa_group | 100 | NH997_004584 | 18668..18970 | 287 | TPA:-hypothetical-protein-NH997_004584-[Pseudomonas-aeruginosa] |
| MBV6142866.1 | HNH | GCA_019167125.1 | Pseudomonas_aeruginosa | Bacteria>Pseudomonadati>Pseudomonadota>Gammaproteobacteria>Pseudomonadales>Pseudomonadaceae>Pseudomonas>Pseudomonas_aeruginosa_group | 100 | KUT38_04235 | 379276..379578 | 287 | hypothetical-protein-[Pseudomonas-aeruginosa] |
| MBX6244470.1 | HNH | GCA_019686455.1 | Pseudomonas_aeruginosa | Bacteria>Pseudomonadati>Pseudomonadota>Gammaproteobacteria>Pseudomonadales>Pseudomonadaceae>Pseudomonas>Pseudomonas_aeruginosa_group | 100 | ISD49_17815 | 206614..206916 | 287 | hypothetical-protein-[Pseudomonas-aeruginosa] |
| WP_031631075.1 | HNH | GCF_900707525.1 | Pseudomonas_aeruginosa | Bacteria>Pseudomonadati>Pseudomonadota>Gammaproteobacteria>Pseudomonadales>Pseudomonadaceae>Pseudomonas>Pseudomonas_aeruginosa_group | 100 | BAW40_RS11040 | 55841..56143 | 287 | hypothetical-protein-[Pseudomonas-aeruginosa] |
| WP_124182731.1 | HNH | GCF_003839865.1 | Pseudomonas_aeruginosa | Bacteria>Pseudomonadati>Pseudomonadota>Gammaproteobacteria>Pseudomonadales>Pseudomonadaceae>Pseudomonas>Pseudomonas_aeruginosa_group | 100 | IPC226_RS26595 | 205787..206089 | 287 | hypothetical-protein-[Pseudomonas-aeruginosa] |
| WP_124984782.1 | HNH | GCF_003626935.1 | Pseudomonas_aeruginosa | Bacteria>Pseudomonadati>Pseudomonadota>Gammaproteobacteria>Pseudomonadales>Pseudomonadaceae>Pseudomonas>Pseudomonas_aeruginosa_group | 100 | D7H65_RS11300 | 2311368..2311670 | 287 | hypothetical-protein-[Pseudomonas-aeruginosa] |
| HBN9708929.1 | HNH | GCA_021591785.1 | Pseudomonas_aeruginosa | Bacteria>Pseudomonadati>Pseudomonadota>Gammaproteobacteria>Pseudomonadales>Pseudomonadaceae>Pseudomonas>Pseudomonas_aeruginosa_group | 333 | L3920_001795 | 290596..291597 | 287 | hypothetical-protein-[Pseudomonas-aeruginosa] |
| WP_033991208.1 | HNH | GCF_929614685.1 | Pseudomonas_aeruginosa | Bacteria>Pseudomonadati>Pseudomonadota>Gammaproteobacteria>Pseudomonadales>Pseudomonadaceae>Pseudomonas>Pseudomonas_aeruginosa_group | 333 | MJP43_RS27235 | 14444..15445 | 287 | hypothetical-protein-[Pseudomonas-aeruginosa] |
| WP_169839749.1 | HNH | GCF_012986505.1 | Pseudomonas_lundensis | Bacteria>Pseudomonadati>Pseudomonadota>Gammaproteobacteria>Pseudomonadales>Pseudomonadaceae>Pseudomonas>Pseudomonas_chlororaphis_group | 199 | HBN72_RS14610 | 123843..124442 | 86185 | hypothetical-protein-[Pseudomonas-lundensis] |
| NNB48791.1 | HNH | GCA_012986825.1 | Pseudomonas_fragi | Bacteria>Pseudomonadati>Pseudomonadota>Gammaproteobacteria>Pseudomonadales>Pseudomonadaceae>Pseudomonas>Pseudomonas_chlororaphis_group | 100 | HBN89_05800 | 92656..92958 | 296 | hypothetical-protein-[Pseudomonas-fragi] |
| PAA10538.1 | HNH | GCA_002269625.1 | Pseudomonas_fragi | Bacteria>Pseudomonadati>Pseudomonadota>Gammaproteobacteria>Pseudomonadales>Pseudomonadaceae>Pseudomonas>Pseudomonas_chlororaphis_group | 100 | CJU78_04030 | 87742..88044 | 296 | hypothetical-protein-CJU78_04030-[Pseudomonas-fragi] |
| WP_095039827.1 | HNH | GCF_002269625.1 | Pseudomonas_fragi | Bacteria>Pseudomonadati>Pseudomonadota>Gammaproteobacteria>Pseudomonadales>Pseudomonadaceae>Pseudomonas>Pseudomonas_chlororaphis_group | 100 | CJU78_RS04030 | 87742..88044 | 296 | hypothetical-protein-[Pseudomonas-fragi] |
| WP_169883487.1 | HNH | GCF_012986805.1 | Pseudomonas_fragi | Bacteria>Pseudomonadati>Pseudomonadota>Gammaproteobacteria>Pseudomonadales>Pseudomonadaceae>Pseudomonas>Pseudomonas_chlororaphis_group | 100 | HBN91_RS17725 | 2129..2431 | 296 | hypothetical-protein-[Pseudomonas-fragi] |
| WP_169907484.1 | HNH | GCF_012986825.1 | Pseudomonas_fragi | Bacteria>Pseudomonadati>Pseudomonadota>Gammaproteobacteria>Pseudomonadales>Pseudomonadaceae>Pseudomonas>Pseudomonas_chlororaphis_group | 100 | HBN89_RS05775 | 92656..92958 | 296 | hypothetical-protein-[Pseudomonas-fragi] |
| WP_047306811.1 | HNH | GCF_001020715.1 | Pseudomonas_fluorescens | Bacteria>Pseudomonadati>Pseudomonadota>Gammaproteobacteria>Pseudomonadales>Pseudomonadaceae>Pseudomonas>Pseudomonas_fluorescens_group | 215 | AA061_RS32120 | 7871..8518 | 294 | TIGR02646-family-protein-[Pseudomonas-fluorescens] |
| WP_017337803.1 | HNH | GCF_000293885.2 | Pseudomonas_fluorescens | Bacteria>Pseudomonadati>Pseudomonadota>Gammaproteobacteria>Pseudomonadales>Pseudomonadaceae>Pseudomonas>Pseudomonas_fluorescens_group | 100 | B723_RS16850 | 3645765..3646067 | 294 | hypothetical-protein-[Pseudomonas-fluorescens] |
| CAB5551583.1 | HNH | GCA_903684265.1 | Pseudomonas_putida | Bacteria>Pseudomonadati>Pseudomonadota>Gammaproteobacteria>Pseudomonadales>Pseudomonadaceae>Pseudomonas>Pseudomonas_putida_group | 199 | GHA_03441 | 120804..121403 | 303 | Uncharacterised-protein-[Pseudomonas-putida] |
| GLO43975.1 | HNH | GCA_030159635.1 | Pseudomonas_putida | Bacteria>Pseudomonadati>Pseudomonadota>Gammaproteobacteria>Pseudomonadales>Pseudomonadaceae>Pseudomonas>Pseudomonas_putida_group | 100 | PPUN109347_05370 | 606661..606963 | 303 | hypothetical-protein-PPUN109347_05370-[Pseudomonas-putida] |
| WP_122622771.1 | HNH | - | Pseudomonas_viridiflava | Bacteria>Pseudomonadati>Pseudomonadota>Gammaproteobacteria>Pseudomonadales>Pseudomonadaceae>Pseudomonas>Pseudomonas_syringae_group | 241 | EEX91_RS04610 | 98136..98861 | 33069 | TIGR02646-family-protein-[Pseudomonas-viridiflava] |
| WP_268945569.1 | HNH | - | Pseudomonas_syringae | Bacteria>Pseudomonadati>Pseudomonadota>Gammaproteobacteria>Pseudomonadales>Pseudomonadaceae>Pseudomonas>Pseudomonas_syringae_group>Pseudomonas_syringae_group_genomosp._1 | 209 | GIW69_RS06590 | 9605..10234 | 317 | TIGR02646-family-protein-[Pseudomonas-syringae] |
| WP_104882978.1 | HNH | GCF_002947915.1 | Pseudomonas_sp._SWI44 | Bacteria>Pseudomonadati>Pseudomonadota>Gammaproteobacteria>Pseudomonadales>Pseudomonadaceae>Pseudomonas>unclassified_Pseudomonas | 246 | C4Q26_RS08430 | 1764273..1765013 | 2083053 | TIGR02646-family-protein-[Pseudomonas-sp.-SWI44] |
| WP_082920817.1 | HNH | GCF_001655615.1 | Pseudomonas_sp._GR_6-02 | Bacteria>Pseudomonadati>Pseudomonadota>Gammaproteobacteria>Pseudomonadales>Pseudomonadaceae>Pseudomonas>unclassified_Pseudomonas | 241 | PGR6_RS03530 | 824155..824880 | 1659194 | TIGR02646-family-protein-[Pseudomonas-sp.-GR-6-02] |
| WP_232392571.1 | HNH | GCF_021228675.1 | Pseudomonas_sp._NIBR-H-19 | Bacteria>Pseudomonadati>Pseudomonadota>Gammaproteobacteria>Pseudomonadales>Pseudomonadaceae>Pseudomonas>unclassified_Pseudomonas | 241 | LS633_RS06550 | 1382008..1382733 | 2901380 | TIGR02646-family-protein-[Pseudomonas-sp.-NIBR-H-19] |
| TXH92073.1 | HNH | GCA_008015405.1 | Pseudomonas_sp. | Bacteria>Pseudomonadati>Pseudomonadota>Gammaproteobacteria>Pseudomonadales>Pseudomonadaceae>Pseudomonas>unclassified_Pseudomonas | 215 | E6Q71_01645 | 5689..6336 | 306 | TIGR02646-family-protein-[Pseudomonas-sp.] |
| WP_198849242.1 | HNH | GCF_016405505.1 | Pseudomonas_sp._MF6772 | Bacteria>Pseudomonadati>Pseudomonadota>Gammaproteobacteria>Pseudomonadales>Pseudomonadaceae>Pseudomonas>unclassified_Pseudomonas | 215 | JFT60_RS22245 | 15322..15969 | 2797533 | TIGR02646-family-protein-[Pseudomonas-sp.-MF6772] |
| WP_095626341.1 | HNH | - | Pseudomonas_sp._WN033 | Bacteria>Pseudomonadati>Pseudomonadota>Gammaproteobacteria>Pseudomonadales>Pseudomonadaceae>Pseudomonas>unclassified_Pseudomonas | 212 | CK507_RS15700 | 124808..125446 | 2032629 | TIGR02646-family-protein-[Pseudomonas-sp.-WN033] |
| MCK9534936.1 | HNH | GCA_023228505.1 | Pseudomonas_sp. | Bacteria>Pseudomonadati>Pseudomonadota>Gammaproteobacteria>Pseudomonadales>Pseudomonadaceae>Pseudomonas>unclassified_Pseudomonas | 199 | MOQ98_09625 | 17797..18396 | 306 | hypothetical-protein-[Pseudomonas-sp.] |
| WP_169919066.1 | HNH | GCF_012985845.1 | Pseudomonas_sp._WS_5013 | Bacteria>Pseudomonadati>Pseudomonadota>Gammaproteobacteria>Pseudomonadales>Pseudomonadaceae>Pseudomonas>unclassified_Pseudomonas | 198 | HBN76_RS00230 | 62719..63315 | 2717475 | MULTISPECIES:-hypothetical-protein-[Pseudomonas] |
| MBL7230770.1 | HNH | GCA_016785205.1 | Pseudomonas_sp. | Bacteria>Pseudomonadati>Pseudomonadota>Gammaproteobacteria>Pseudomonadales>Pseudomonadaceae>Pseudomonas>unclassified_Pseudomonas | 100 | JJK56_22680 | 24507..24809 | 306 | hypothetical-protein-[Pseudomonas-sp.] |
| WP_008016374.1 | HNH | GCF_000282395.1 | Pseudomonas_sp._GM55 | Bacteria>Pseudomonadati>Pseudomonadota>Gammaproteobacteria>Pseudomonadales>Pseudomonadaceae>Pseudomonas>unclassified_Pseudomonas | 100 | PMI31_RS03880 | 71424..71726 | 1144333 | hypothetical-protein-[Pseudomonas-sp.-GM55] |
| WP_101152273.1 | HNH | GCF_002836195.1 | Pseudomonas_sp._Choline-3u-10 | Bacteria>Pseudomonadati>Pseudomonadota>Gammaproteobacteria>Pseudomonadales>Pseudomonadaceae>Pseudomonas>unclassified_Pseudomonas | 100 | CXF92_RS00330 | 62057..62359 | 2058311 | hypothetical-protein-[Pseudomonas-sp.-Choline-3u-10] |
| WP_218424327.1 | HNH | GCF_019218965.1 | Pseudomonas_sp._PDM31 | Bacteria>Pseudomonadati>Pseudomonadota>Gammaproteobacteria>Pseudomonadales>Pseudomonadaceae>Pseudomonas>unclassified_Pseudomonas | 100 | KW839_RS11040 | 17453..17755 | 2854778 | hypothetical-protein-[Pseudomonas-sp.-PDM31] |
| WP_125856775.1 | HNH | GCF_003940515.1 | Stutzerimonas_stutzeri | Bacteria>Pseudomonadati>Pseudomonadota>Gammaproteobacteria>Pseudomonadales>Pseudomonadaceae>Stutzerimonas>Stutzerimonas_stutzeri_group>Stutzerimonas_stutzeri_subgroup | 198 | EGJ36_RS18125 | 16149..16745 | 316 | hypothetical-protein-[Stutzerimonas-stutzeri] |
| WP_285534184.1 | HNH | GCF_030269585.1 | Stutzerimonas_stutzeri | Bacteria>Pseudomonadati>Pseudomonadota>Gammaproteobacteria>Pseudomonadales>Pseudomonadaceae>Stutzerimonas>Stutzerimonas_stutzeri_group>Stutzerimonas_stutzeri_subgroup | 196 | QSJ82_RS21915 | 16965..17555 | 316 | retron-system-putative-HNH-endonuclease-[Stutzerimonas-stutzeri] |
| MCQ2032407.1 | HNH | GCA_024397475.1 | Stutzerimonas_kunmingensis | Bacteria>Pseudomonadati>Pseudomonadota>Gammaproteobacteria>Pseudomonadales>Pseudomonadaceae>Stutzerimonas>Stutzerimonas_stutzeri_group>Stutzerimonas_stutzeri_subgroup | 100 | NAU09_00505 | 107782..108084 | 1211807 | hypothetical-protein-[Stutzerimonas-kunmingensis] |
| WP_051640643.1 | HNH | GCF_000702325.1 | Thiomicrorhabdus_sp._Milos-T2 | Bacteria>Pseudomonadati>Pseudomonadota>Gammaproteobacteria>Thiotrichales>Piscirickettsiaceae>Thiomicrorhabdus>unclassified_Thiomicrorhabdus | 222 | BS34_RS0102325 | 542991..543659 | 90814 | TIGR02646-family-protein-[Thiomicrorhabdus-sp.-Milos-T2] |
| WP_020558460.1 | HNH | GCF_000380185.1 | Thiofilum_flexile | Bacteria>Pseudomonadati>Pseudomonadota>Gammaproteobacteria>Thiotrichales>Thiofilaceae>Thiofilum | 219 | A3IK_RS0102865 | 101172..101831 | 125627 | TIGR02646-family-protein-[Thiofilum-flexile] |
| WP_109837973.1 | HNH | GCF_003172875.1 | Leucothrix_pacifica | Bacteria>Pseudomonadati>Pseudomonadota>Gammaproteobacteria>Thiotrichales>Thiotrichaceae>Leucothrix | 212 | DKW60_RS12425 | 53284..53922 | 1247513 | TIGR02646-family-protein-[Leucothrix-pacifica] |
| WP_078924152.1 | HNH | GCF_900167255.1 | Thiothrix_eikelboomii | Bacteria>Pseudomonadati>Pseudomonadota>Gammaproteobacteria>Thiotrichales>Thiotrichaceae>Thiothrix | 236 | B5D50_RS18540 | 24440..25150 | 92487 | TIGR02646-family-protein-[Thiothrix-eikelboomii] |
| SEH06530.1 | HNH | GCA_900099115.2 | Candidatus_Venteria_ishoeyi | Bacteria>Pseudomonadati>Pseudomonadota>Gammaproteobacteria>Thiotrichales>Thiotrichaceae>Venteria | 210 | MBHS_02393 | 30939..31571 | 1899563 | Uncharacterised-protein-[Thiotrichales-bacterium-HS_08] |
| WP_103920295.1 | HNH | GCF_900099115.1 | Candidatus_Venteria_ishoeyi | Bacteria>Pseudomonadati>Pseudomonadota>Gammaproteobacteria>Thiotrichales>Thiotrichaceae>Venteria | 210 | MBHS_RS13580 | 30939..31571 | 1899563 | retron-Ec78-anti-phage-system-effector-HNH-endonuclease-PtuB-[Candidatus-Venteria-ishoeyi] |
| MXY53198.1 | HNH | GCA_009837725.1 | Gammaproteobacteria_bacterium | Bacteria>Pseudomonadati>Pseudomonadota>Gammaproteobacteria>unclassified_Gammaproteobacteria | 248 | F4Y86_11835 | 10380..11126 | 1913989 | TIGR02646-family-protein-[Gammaproteobacteria-bacterium] |
| MYB38343.1 | HNH | GCA_009838905.1 | Gammaproteobacteria_bacterium | Bacteria>Pseudomonadati>Pseudomonadota>Gammaproteobacteria>unclassified_Gammaproteobacteria | 248 | F4Y26_13385 | 15649..16395 | 1913989 | TIGR02646-family-protein-[Gammaproteobacteria-bacterium] |
| MCL4555403.1 | HNH | GCA_023382035.1 | Gammaproteobacteria_bacterium | Bacteria>Pseudomonadati>Pseudomonadota>Gammaproteobacteria>unclassified_Gammaproteobacteria | 241 | M1572_00740 | 5059..5784 | 1913989 | TIGR02646-family-protein-[Gammaproteobacteria-bacterium] |
| WP_069513737.1 | HNH | GCF_963893015.1 | Vibrio | Bacteria>Pseudomonadati>Pseudomonadota>Gammaproteobacteria>Vibrionales>Vibrionaceae | 100 | J0671_RS16305 | 101661..101963 | 662 | MULTISPECIES:-hypothetical-protein-[Vibrio] |
| WP_095469008.1 | HNH | GCF_043351775.1 | Vibrio | Bacteria>Pseudomonadati>Pseudomonadota>Gammaproteobacteria>Vibrionales>Vibrionaceae | 100 | MRS66_RS05665 | 534557..534859 | 662 | MULTISPECIES:-hypothetical-protein-[Vibrio] |
| WP_123322059.1 | HNH | GCF_049311125.1 | Vibrio | Bacteria>Pseudomonadati>Pseudomonadota>Gammaproteobacteria>Vibrionales>Vibrionaceae | 100 | EGL68_RS05360 | 1113688..1113990 | 662 | MULTISPECIES:-hypothetical-protein-[Vibrio] |
| WP_124009102.1 | HNH | GCF_018458505.1 | Vibrio | Bacteria>Pseudomonadati>Pseudomonadota>Gammaproteobacteria>Vibrionales>Vibrionaceae | 100 | J4H11_RS21295 | 6596..6898 | 662 | MULTISPECIES:-hypothetical-protein-[Vibrio-harveyi-group] |
| WP_062660544.1 | HNH | GCF_900055185.1 | Grimontia_celer | Bacteria>Pseudomonadati>Pseudomonadota>Gammaproteobacteria>Vibrionales>Vibrionaceae>Grimontia | 228 | GCE9029_RS00255 | 64152..64838 | 1796497 | TIGR02646-family-protein-[Grimontia-celer] |
| WP_062660552.1 | HNH | GCF_900055185.1 | Grimontia_celer | Bacteria>Pseudomonadati>Pseudomonadota>Gammaproteobacteria>Vibrionales>Vibrionaceae>Grimontia | 353 | GCE9029_RS26695 | 74347..75408 | 1796497 | HNH-endonuclease-signature-motif-containing-protein-[Grimontia-celer] |
| EJL6982895.1 | HNH | GCA_024219855.1 | Vibrio_cholerae | Bacteria>Pseudomonadati>Pseudomonadota>Gammaproteobacteria>Vibrionales>Vibrionaceae>Vibrio | 228 | NMT40_002447 | 116358..117044 | 666 | TIGR02646-family-protein-[Vibrio-cholerae] |
| WP_217529705.1 | HNH | GCF_019093145.1 | Vibrio_cholerae | Bacteria>Pseudomonadati>Pseudomonadota>Gammaproteobacteria>Vibrionales>Vibrionaceae>Vibrio | 228 | KVS75_RS02430 | 557767..558453 | 666 | TIGR02646-family-protein-[Vibrio-cholerae] |
| HAS6423763.1 | HNH | GCA_015777845.1 | Vibrio_vulnificus | Bacteria>Pseudomonadati>Pseudomonadota>Gammaproteobacteria>Vibrionales>Vibrionaceae>Vibrio | 199 | I7251_16625 | 99196..99795 | 672 | hypothetical-protein-[Vibrio-vulnificus] |
| EGR2106519.1 | HNH | - | Vibrio_cholerae | Bacteria>Pseudomonadati>Pseudomonadota>Gammaproteobacteria>Vibrionales>Vibrionaceae>Vibrio | 100 | DYF88_13665 | 23912..24214 | 666 | hypothetical-protein-[Vibrio-cholerae] |
| EHD1694789.1 | HNH | - | Vibrio_vulnificus | Bacteria>Pseudomonadati>Pseudomonadota>Gammaproteobacteria>Vibrionales>Vibrionaceae>Vibrio | 100 | JR728_000343 | 355336..355638 | 672 | hypothetical-protein-[Vibrio-vulnificus] |
| EJS1625585.1 | HNH | GCA_024944445.1 | Vibrio_cholerae | Bacteria>Pseudomonadati>Pseudomonadota>Gammaproteobacteria>Vibrionales>Vibrionaceae>Vibrio | 100 | NXA67_002441 | 135798..136100 | 666 | hypothetical-protein-[Vibrio-cholerae] |
| HAS6049568.1 | HNH | GCA_015776375.1 | Vibrio_vulnificus | Bacteria>Pseudomonadati>Pseudomonadota>Gammaproteobacteria>Vibrionales>Vibrionaceae>Vibrio | 100 | I7138_16165 | 9390..9692 | 672 | hypothetical-protein-[Vibrio-vulnificus] |
| WP_065206905.1 | HNH | GCF_001676015.1 | Vibrio_splendidus | Bacteria>Pseudomonadati>Pseudomonadota>Gammaproteobacteria>Vibrionales>Vibrionaceae>Vibrio | 100 | A9262_RS18555 | 295571..295873 | 29497 | hypothetical-protein-[Vibrio-splendidus] |
| WP_102397779.1 | HNH | GCF_002873235.1 | Vibrio_cyclitrophicus | Bacteria>Pseudomonadati>Pseudomonadota>Gammaproteobacteria>Vibrionales>Vibrionaceae>Vibrio | 100 | BCU59_RS21815 | 100672..100974 | 47951 | hypothetical-protein-[Vibrio-cyclitrophicus] |
| WP_102530602.1 | HNH | GCF_002878515.1 | Vibrio_lentus | Bacteria>Pseudomonadati>Pseudomonadota>Gammaproteobacteria>Vibrionales>Vibrionaceae>Vibrio | 100 | BCU87_RS25080 | 3021..3323 | 136468 | hypothetical-protein-[Vibrio-lentus] |
| WP_132761836.1 | HNH | GCF_017918175.1 | Vibrio_crassostreae | Bacteria>Pseudomonadati>Pseudomonadota>Gammaproteobacteria>Vibrionales>Vibrionaceae>Vibrio | 100 | EDB51_RS02200 | 67338..67640 | 246167 | hypothetical-protein-[Vibrio-crassostreae] |
| WP_132941349.1 | HNH | GCF_004345555.1 | Vibrio_crassostreae | Bacteria>Pseudomonadati>Pseudomonadota>Gammaproteobacteria>Vibrionales>Vibrionaceae>Vibrio | 100 | EDB60_RS00570 | 111763..112065 | 246167 | hypothetical-protein-[Vibrio-crassostreae] |
| WP_194592802.1 | HNH | GCF_015351545.2 | Vibrio_vulnificus | Bacteria>Pseudomonadati>Pseudomonadota>Gammaproteobacteria>Vibrionales>Vibrionaceae>Vibrio | 100 | IRL01_RS02860 | 622801..623103 | 672 | hypothetical-protein-[Vibrio-vulnificus] |

| Protein-ID | Domain | Assembly-ID | Strain | Taxonomy | Length | Locus-Tag | Genomic-coordinates | TaxID | Fasta-defline |
|---|---|---|---|---|---|---|---|---|---|
| WP_206445941.1 | HNH | GCF_963890835.1 | Vibrio_paracholerae | Bacteria>Pseudomonadati>Pseudomonadota>Gammaproteobacteria>Vibrionales>Vibrionaceae>Vibrio | 100 | GU254_RS11795 | 76956..77258 | 650003 | hypothetical-protein-[Vibrio-paracholerae] |
| WP_230666612.1 | HNH | GCF_021014645.1 | Vibrio_cholerae | Bacteria>Pseudomonadati>Pseudomonadota>Gammaproteobacteria>Vibrionales>Vibrionaceae>Vibrio | 100 | B4944_RS12145 | 30330..30632 | 666 | hypothetical-protein-[Vibrio-cholerae] |
| WP_233934102.1 | HNH | GCF_021394995.1 | Vibrio_fluvialis | Bacteria>Pseudomonadati>Pseudomonadota>Gammaproteobacteria>Vibrionales>Vibrionaceae>Vibrio | 100 | LZT09_RS15325 | 15858..16160 | 676 | hypothetical-protein-[Vibrio-fluvialis] |
| WP_274676092.1 | HNH | GCF_028858035.1 | Vibrio_aestuarianus | Bacteria>Pseudomonadati>Pseudomonadota>Gammaproteobacteria>Vibrionales>Vibrionaceae>Vibrio | 100 | L9X51_RS11505 | 23036..23338 | 28171 | hypothetical-protein-[Vibrio-aestuarianus] |
| WP_123322072.1 | HNH | GCF_003751765.1 | Vibrio_crassostreae | Bacteria>Pseudomonadati>Pseudomonadota>Gammaproteobacteria>Vibrionales>Vibrionaceae>Vibrio | 284 | EDB73_RS01735 | 387238..388092 | 246167 | HNH-endonuclease-[Vibrio-crassostreae] |
| WP_132941361.1 | HNH | GCF_004345555.1 | Vibrio_crassostreae | Bacteria>Pseudomonadati>Pseudomonadota>Gammaproteobacteria>Vibrionales>Vibrionaceae>Vibrio | 284 | EDB60_RS00640 | 132709..133563 | 246167 | HNH-endonuclease-[Vibrio-crassostreae] |
| WP_132761838.1 | HNH | GCF_017916825.1 | Vibrio_crassostreae | Bacteria>Pseudomonadati>Pseudomonadota>Gammaproteobacteria>Vibrionales>Vibrionaceae>Vibrio | 283 | EDB51_RS02215 | 73838..74689 | 246167 | HNH-endonuclease-signature-motif-containing-protein-[Vibrio-crassostreae] |
| WP_016790627.1 | HNH | GCF_000256605.2 | Vibrio_cyclitrophicus | Bacteria>Pseudomonadati>Pseudomonadota>Gammaproteobacteria>Vibrionales>Vibrionaceae>Vibrio | 281 | OAI_RS19755 | 256434..257279 | 47951 | HNH-endonuclease-[Vibrio-cyclitrophicus] |
| EJR0961154.1 | HNH | GCA_024870505.1 | Vibrio_parahaemolyticus | Bacteria>Pseudomonadati>Pseudomonadota>Gammaproteobacteria>Vibrionales>Vibrionaceae>Vibrio>Vibrio_harveyi_group | 228 | NYK10_002562 | 284875..285561 | 670 | TIGR02646-family-protein-[Vibrio-parahaemolyticus] |
| HCH6462474.1 | HNH | GCA_024102875.1 | Vibrio_parahaemolyticus | Bacteria>Pseudomonadati>Pseudomonadota>Gammaproteobacteria>Vibrionales>Vibrionaceae>Vibrio>Vibrio_harveyi_group | 228 | NK401_003627 | 37638..38324 | 670 | TIGR02646-family-protein-[Vibrio-parahaemolyticus] |
| TBT35891.1 | HNH | GCA_004324405.1 | Vibrio_parahaemolyticus | Bacteria>Pseudomonadati>Pseudomonadota>Gammaproteobacteria>Vibrionales>Vibrionaceae>Vibrio>Vibrio_harveyi_group | 228 | D5E85_08510 | 436669..437355 | 670 | TIGR02646-family-protein-[Vibrio-parahaemolyticus] |
| WP_213964956.1 | HNH | GCF_018457925.1 | Vibrio_alginolyticus | Bacteria>Pseudomonadati>Pseudomonadota>Gammaproteobacteria>Vibrionales>Vibrionaceae>Vibrio>Vibrio_harveyi_group | 228 | J4H38_RS12940 | 90893..91579 | 663 | TIGR02646-family-protein-[Vibrio-alginolyticus] |
| EGU0166210.1 | HNH | - | Vibrio_parahaemolyticus | Bacteria>Pseudomonadati>Pseudomonadota>Gammaproteobacteria>Vibrionales>Vibrionaceae>Vibrio>Vibrio_harveyi_group | 100 | JFQ74_002024 | 202903..203205 | 670 | hypothetical-protein-[Vibrio-parahaemolyticus] |
| EHC7288056.1 | HNH | - | Vibrio_parahaemolyticus | Bacteria>Pseudomonadati>Pseudomonadota>Gammaproteobacteria>Vibrionales>Vibrionaceae>Vibrio>Vibrio_harveyi_group | 100 | JS475_001519 | 791798..792100 | 670 | hypothetical-protein-[Vibrio-parahaemolyticus] |
| EHK2855488.1 | HNH | GCA_018017035.1 | Vibrio_parahaemolyticus | Bacteria>Pseudomonadati>Pseudomonadota>Gammaproteobacteria>Vibrionales>Vibrionaceae>Vibrio>Vibrio_harveyi_group | 100 | J7X11_003962 | 22148..22450 | 670 | hypothetical-protein-[Vibrio-parahaemolyticus] |
| EHR6926064.1 | HNH | GCA_019067225.1 | Vibrio_parahaemolyticus | Bacteria>Pseudomonadati>Pseudomonadota>Gammaproteobacteria>Vibrionales>Vibrionaceae>Vibrio>Vibrio_harveyi_group | 100 | KUK83_003907 | 26038..26340 | 670 | hypothetical-protein-[Vibrio-parahaemolyticus] |
| EIA1331997.1 | HNH | GCA_019764355.1 | Vibrio_parahaemolyticus | Bacteria>Pseudomonadati>Pseudomonadota>Gammaproteobacteria>Vibrionales>Vibrionaceae>Vibrio>Vibrio_harveyi_group | 100 | K6675_002584 | 22148..22450 | 670 | hypothetical-protein-[Vibrio-parahaemolyticus] |
| EII3096493.1 | HNH | GCA_020410925.1 | Vibrio_parahaemolyticus | Bacteria>Pseudomonadati>Pseudomonadota>Gammaproteobacteria>Vibrionales>Vibrionaceae>Vibrio>Vibrio_harveyi_group | 100 | CWL41_002102 | 330808..331110 | 670 | hypothetical-protein-CWL41_002102-[Vibrio-parahaemolyticus] |
| EIN6339955.1 | HNH | GCA_020908185.1 | Vibrio_parahaemolyticus | Bacteria>Pseudomonadati>Pseudomonadota>Gammaproteobacteria>Vibrionales>Vibrionaceae>Vibrio>Vibrio_harveyi_group | 100 | LPQ31_001308 | 340557..340859 | 670 | hypothetical-protein-[Vibrio-parahaemolyticus] |
| EJB8504852.1 | HNH | GCA_022967155.1 | Vibrio_parahaemolyticus | Bacteria>Pseudomonadati>Pseudomonadota>Gammaproteobacteria>Vibrionales>Vibrionaceae>Vibrio>Vibrio_harveyi_group | 100 | MW364_003030 | 21157..21459 | 670 | hypothetical-protein-[Vibrio-parahaemolyticus] |
| EJG0684227.1 | HNH | GCA_015814245.2 | Vibrio_parahaemolyticus | Bacteria>Pseudomonadati>Pseudomonadota>Gammaproteobacteria>Vibrionales>Vibrionaceae>Vibrio>Vibrio_harveyi_group | 100 | C4G75_RS05970 | 61112..61414 | 670 | hypothetical-protein-C4G75_RS05970-[Vibrio-parahaemolyticus] |
| HBC3410544.1 | HNH | GCA_018118845.1 | Vibrio_parahaemolyticus | Bacteria>Pseudomonadati>Pseudomonadota>Gammaproteobacteria>Vibrionales>Vibrionaceae>Vibrio>Vibrio_harveyi_group | 100 | KDY13_002796 | 103990..104292 | 670 | hypothetical-protein-[Vibrio-parahaemolyticus] |
| HCE2000760.1 | HNH | GCA_023810945.1 | Vibrio_parahaemolyticus | Bacteria>Pseudomonadati>Pseudomonadota>Gammaproteobacteria>Vibrionales>Vibrionaceae>Vibrio>Vibrio_harveyi_group | 100 | NGT00_003023 | 151827..152129 | 670 | hypothetical-protein-[Vibrio-parahaemolyticus] |
| HCE5247595.1 | HNH | GCA_023836695.1 | Vibrio_parahaemolyticus | Bacteria>Pseudomonadati>Pseudomonadota>Gammaproteobacteria>Vibrionales>Vibrionaceae>Vibrio>Vibrio_harveyi_group | 100 | NG505_004030 | 107581..107883 | 670 | TPA:-hypothetical-protein-NG505_004030-[Vibrio-parahaemolyticus] |
| HCG6379161.1 | HNH | GCA_024039965.1 | Vibrio_parahaemolyticus | Bacteria>Pseudomonadati>Pseudomonadota>Gammaproteobacteria>Vibrionales>Vibrionaceae>Vibrio>Vibrio_harveyi_group | 100 | NJ203_004430 | 17336..17638 | 670 | TPA:-hypothetical-protein-NJ203_004430-[Vibrio-parahaemolyticus] |
| WP_229626011.1 | HNH | GCF_905331745.1 | Vibrio_parahaemolyticus | Bacteria>Pseudomonadati>Pseudomonadota>Gammaproteobacteria>Vibrionales>Vibrionaceae>Vibrio>Vibrio_harveyi_group | 100 | LPS00_RS14535 | 112103..112405 | 670 | hypothetical-protein-[Vibrio-parahaemolyticus] |
| WP_243191726.1 | HNH | GCF_022802175.1 | Vibrio_parahaemolyticus | Bacteria>Pseudomonadati>Pseudomonadota>Gammaproteobacteria>Vibrionales>Vibrionaceae>Vibrio>Vibrio_harveyi_group | 100 | JL838_RS10880 | 28071..28373 | 670 | hypothetical-protein-[Vibrio-parahaemolyticus] |
| WP_258672249.1 | HNH | GCF_024747115.1 | Vibrio_parahaemolyticus | Bacteria>Pseudomonadati>Pseudomonadota>Gammaproteobacteria>Vibrionales>Vibrionaceae>Vibrio>Vibrio_harveyi_group | 100 | NDJ77_RS09305 | 99424..99726 | 670 | hypothetical-protein-[Vibrio-parahaemolyticus] |
| EGU0166221.1 | HNH | - | Vibrio_parahaemolyticus | Bacteria>Pseudomonadati>Pseudomonadota>Gammaproteobacteria>Vibrionales>Vibrionaceae>Vibrio>Vibrio_harveyi_group | 282 | JFQ74_002035 | 218894..219742 | 670 | HNH-endonuclease-[Vibrio-parahaemolyticus] |
| EIN6339965.1 | HNH | GCA_020908185.1 | Vibrio_parahaemolyticus | Bacteria>Pseudomonadati>Pseudomonadota>Gammaproteobacteria>Vibrionales>Vibrionaceae>Vibrio>Vibrio_harveyi_group | 282 | LPQ31_001318 | 355953..356801 | 670 | HNH-endonuclease-[Vibrio-parahaemolyticus] |
| PIQ12820.1 | HNH | GCA_002770925.1 | Hydrogenophilales_bacterium_CG18_big_fil_WC_8_21_14_2_50_58_12 | Bacteria>Pseudomonadati>Pseudomonadota>Hydrogenophilia>Hydrogenophilales>unclassified_Hydrogenophilales | 198 | COW70_07925 | 5621..6217 | 1974036 | hypothetical-protein-COW70_07925-[Hydrogenophilales-bacterium-CG18_big_fil_WC_8_21_14_2_50_58_12] |
| MCX7197673.1 | HNH | GCA_026399745.1 | Pseudomonadota_bacterium | Bacteria>Pseudomonadati>Pseudomonadota>unclassified_Pseudomonadota | 153 | NTW37_06705 | 9299..9760 | 1977087 | MAG:-HNH-endonuclease-[Proteobacteria-bacterium] |
| MBQ6473653.1 | HNH | GCA_017431185.1 | Victivallales_bacterium | Bacteria>Pseudomonadati>PVC_group>Lentisphaerota>Lentisphaeria>Victivallales>unclassified_Victivallales | 231 | IJJ33_16825 | 13725..14420 | 2212475 | TIGR02646-family-protein-[Victivallales-bacterium] |
| WP_168963707.1 | HNH | GCF_012843305.1 | Victivallis_vadensis | Bacteria>Pseudomonadati>PVC_group>Lentisphaerota>Lentisphaeria>Victivallales>Victivallaceae>Victivallis | 260 | HF882_RS18890 | 18380..19162 | 172901 | TIGR02646-family-protein-[Victivallis-vadensis] |
| MBE6357837.1 | HNH | GCA_015061525.1 | Lentisphaerota_bacterium | Bacteria>Pseudomonadati>PVC_group>Lentisphaerota>unclassified_Lentisphaerota | 204 | E7057_01145 | 10873..11487 | 1932692 | TIGR02646-family-protein-[Lentisphaerota-bacterium] |
| MBC8553110.1 | HNH | GCA_014384865.1 | Candidatus_Brocadiales_bacterium | Bacteria>Pseudomonadati>PVC_group>Planctomycetota>Candidatus_Brocadiia>Candidatus_Brocadiales>unclassified_Candidatus_Brocadiales | 172 | H8D23_26115 | 11036..11552 | 2760054 | endonuclease-[Candidatus-Brocadiales-bacterium] |
| MBX3408156.1 | HNH | GCA_019636415.1 | Phycisphaeraceae_bacterium | Bacteria>Pseudomonadati>PVC_group>Planctomycetota>Phycisphaerae>Phycisphaerales>Phycisphaeraceae>unclassified_Phycisphaeraceae | 219 | KF869_15495 | 12412..13071 | 2026777 | TIGR02646-family-protein-[Phycisphaeraceae-bacterium] |
| MBX3409894.1 | HNH | GCA_019636575.1 | Phycisphaeraceae_bacterium | Bacteria>Pseudomonadati>PVC_group>Planctomycetota>Phycisphaerae>Phycisphaerales>Phycisphaeraceae>unclassified_Phycisphaeraceae | 219 | KF859_08410 | 143195..143854 | 2026777 | TIGR02646-family-protein-[Phycisphaeraceae-bacterium] |
| WP_010033157.1 | HNH | GCF_003149495.1 | Gemmata_obscuriglobus | Bacteria>Pseudomonadati>PVC_group>Planctomycetota>Planctomycetia>Gemmatales>Gemmataceae>Gemmata | 295 | GOBSN_RS25250 | 6051988..6052875 | 114 | hypothetical-protein-[Gemmata-obscuriglobus] |
| WP_105358294.1 | HNH | GCF_002967725.1 | Blastopirellula_marina | Bacteria>Pseudomonadati>PVC_group>Planctomycetota>Planctomycetia>Pirellulales>Pirellulaceae>Blastopirellula | 291 | C5Y98_RS24530 | 396094..396969 | 124 | hypothetical-protein-[Blastopirellula-marina] |
| NLF73013.1 | HNH | GCA_012515235.1 | Candidatus_Anammoximicrobium_sp. | Bacteria>Pseudomonadati>PVC_group>Planctomycetota>Planctomycetia>Pirellulales>Pirellulaceae>Candidatus_Anammoximicrobium>unclassified_Candidatus_Anammoximicrobium | 199 | GX575_28585 | 10565..11164 | 2699742 | hypothetical-protein-[Candidatus-Anammoximicrobium-sp.] |
| QDU98791.1 | HNH | GCA_007751035.1 | Lignipirellula_cremea | Bacteria>Pseudomonadati>PVC_group>Planctomycetota>Planctomycetia>Pirellulales>Pirellulaceae>Lignipirellula | 292 | Pla8534_66650 | 8880333..8881211 | 2528010 | hypothetical-protein-Pla8534_66650-[Lignipirellula-cremea] |
| WP_145058315.1 | HNH | GCF_007751035.1 | Lignipirellula_cremea | Bacteria>Pseudomonadati>PVC_group>Planctomycetota>Planctomycetia>Pirellulales>Pirellulaceae>Lignipirellula | 292 | Pla8534_RS33055 | 8880333..8881211 | 2528010 | hypothetical-protein-[Lignipirellula-cremea] |
| OYP37213.1 | HNH | GCA_002251265.1 | Rhodopirellula_sp._MGV | Bacteria>Pseudomonadati>PVC_group>Planctomycetota>Planctomycetia>Pirellulales>Pirellulaceae>Rhodopirellula>unclassified_Rhodopirellula | 239 | CGZ80_05860 | 12559..13278 | 2023130 | TIGR02646-family-protein-[Rhodopirellula-sp.-MGV] |
| WP_158222573.1 | HNH | GCF_002251265.1 | Rhodopirellula_sp._MGV | Bacteria>Pseudomonadati>PVC_group>Planctomycetota>Planctomycetia>Pirellulales>Pirellulaceae>Rhodopirellula>unclassified_Rhodopirellula | 220 | CGZ80_RS05870 | 12559..13221 | 2023130 | TIGR02646-family-protein-[Rhodopirellula-sp.-MGV] |
| NLX95049.1 | HNH | GCA_012728855.1 | Rhodopirellula_sp. | Bacteria>Pseudomonadati>PVC_group>Planctomycetota>Planctomycetia>Pirellulales>Pirellulaceae>Rhodopirellula>unclassified_Rhodopirellula | 213 | GXY83_02615 | 19281..19922 | 2024855 | TIGR02646-family-protein-[Rhodopirellula-sp.] |
| WP_278468319.1 | HNH | GCF_963971965.1 | Gimesia_maris | Bacteria>Pseudomonadati>PVC_group>Planctomycetota>Planctomycetia>Planctomycetales>Planctomycetaceae>Gimesia | 286 | B4K77_RS12155 | 528823..529683 | 122 | hypothetical-protein-[Gimesia-maris] |
| MAX35347.1 | HNH | GCA_002706765.1 | Gimesia_sp. | Bacteria>Pseudomonadati>PVC_group>Planctomycetota>Planctomycetia>Planctomycetales>Planctomycetaceae>Gimesia>unclassified_Gimesia | 293 | CME33_02130 | 40037..40918 | 2024833 | hypothetical-protein-[Gimesia-sp.] |
| MAX35350.1 | HNH | GCA_002706765.1 | Gimesia_sp. | Bacteria>Pseudomonadati>PVC_group>Planctomycetota>Planctomycetia>Planctomycetales>Planctomycetaceae>Gimesia>unclassified_Gimesia | 231 | CME33_02145 | 46251..46946 | 2024833 | HNH-endonuclease-[Gimesia-sp.] |
| WP_336521094.1 | HNH | GCF_030622045.1 | Planctellipticum_variicoloris | Bacteria>Pseudomonadati>PVC_group>Planctomycetota>Planctomycetia>Planctomycetales>Planctomycetaceae>Planctellipticum | 293 | SH412_RS26710 | 6854718..6855599 | 3064265 | hypothetical-protein-[Planctoellipticum-variicoloris] |
| HBL44679.1 | HNH | GCA_003505085.1 | Planctomycetaceae_bacterium | Bacteria>Pseudomonadati>PVC_group>Planctomycetota>Planctomycetia>Planctomycetales>Planctomycetaceae>unclassified_Planctomycetaceae | 293 | DDZ90_14935 | 14893..15774 | 2026779 | hypothetical-protein-[Planctomycetaceae-bacterium] |
| MDB5334581.1 | HNH | GCA_028291605.1 | Planctomycetaceae_bacterium | Bacteria>Pseudomonadati>PVC_group>Planctomycetota>Planctomycetia>Planctomycetales>Planctomycetaceae>unclassified_Planctomycetaceae | 235 | JWN70_200 | 11130..11837 | 2026779 | MAG:-Uncharacterized-protein-JWN70_200-[Planctomycetaceae-bacterium] |
| MBX3448927.1 | HNH | GCA_019636285.1 | Planctomycetaceae_bacterium | Bacteria>Pseudomonadati>PVC_group>Planctomycetota>Planctomycetia>Planctomycetales>Planctomycetaceae>unclassified_Planctomycetaceae | 216 | KF777_05165 | 5329..5979 | 2026779 | TIGR02646-family-protein-[Planctomycetaceae-bacterium] |
| TAH38054.1 | HNH | GCA_004295085.1 | Planctomycetota_bacterium | Bacteria>Pseudomonadati>PVC_group>Planctomycetota>unclassified_Planctomycetota | 210 | EYC70_05370 | 72784..73416 | 2026780 | MAG:-TIGR02646-family-protein-[Planctomycetota-bacterium] |
| MCC5808693.1 | HNH | GCA_020831525.1 | Opitutales_bacterium | Bacteria>Pseudomonadati>PVC_group>Verrucomicrobiota>Opitutia>Opitutales>unclassified_Opitutales | 211 | JJU00_20365 | 5194..5829 | 2026772 | TIGR02646-family-protein-[Opitutales-bacterium] |
| MCC5805068.1 | HNH | GCA_020831525.1 | Opitutales_bacterium | Bacteria>Pseudomonadati>PVC_group>Verrucomicrobiota>Opitutia>Opitutales>unclassified_Opitutales | 194 | JJU00_01955 | 44409..44993 | 2026772 | TIGR02646-family-protein-[Opitutales-bacterium] |
| MCX6879293.1 | HNH | GCA_026397725.1 | Verrucomicrobiota_bacterium | Bacteria>Pseudomonadati>PVC_group>Verrucomicrobiota>unclassified_Verrucomicrobiota | 275 | NTW21_36640 | 17283..18110 | 2026799 | MAG:-TIGR02646-family-protein-[Verrucomicrobia-bacterium] |
| MBI5767833.1 | HNH | GCA_016235555.1 | Verrucomicrobiota_bacterium | Bacteria>Pseudomonadati>PVC_group>Verrucomicrobiota>unclassified_Verrucomicrobiota | 238 | HZA93_08550 | 97903..98619 | 2026799 | TIGR02646-family-protein-[Verrucomicrobiota-bacterium] |
| TAE90780.1 | HNH | GCA_004293285.1 | Verrucomicrobiota_bacterium | Bacteria>Pseudomonadati>PVC_group>Verrucomicrobiota>unclassified_Verrucomicrobiota | 238 | EAZ81_08650 | 7224..7940 | 2026799 | TIGR02646-family-protein-[Verrucomicrobiota-bacterium] |
| MCX6953045.1 | HNH | GCA_026398065.1 | Verrucomicrobiota_bacterium | Bacteria>Pseudomonadati>PVC_group>Verrucomicrobiota>unclassified_Verrucomicrobiota | 199 | NTV51_12900 | 3842..4441 | 2026799 | MAG:-TIGR02646-family-protein-[Verrucomicrobia-bacterium] |
| HAO79057.1 | HNH | GCA_003455565.1 | Limisphaerales_bacterium | Bacteria>Pseudomonadati>PVC_group>Verrucomicrobiota>Verrucomicrobiia>Limisphaerales>unclassified_Limisphaerales | 207 | DCQ92_08795 | 6539..7162 | 2052186 | HNH-endonuclease-[Verrucomicrobia-subdivision-3-bacterium] |
| WP_215427526.1 | HNH | GCF_018709685.1 | Akkermansia_muciniphila | Bacteria>Pseudomonadati>PVC_group>Verrucomicrobiota>Verrucomicrobiia>Verrucomicrobiales>Akkermansiaceae>Akkermansia | 250 | HHJ06_RS03165 | 99751..100503 | 239935 | HNH-endonuclease-[Akkermansia-muciniphila] |
| WP_215427500.1 | HNH | GCF_018709685.1 | Akkermansia_muciniphila | Bacteria>Pseudomonadati>PVC_group>Verrucomicrobiota>Verrucomicrobiia>Verrucomicrobiales>Akkermansiaceae>Akkermansia | 184 | HHJ06_RS03105 | 84317..84871 | 239935 | hypothetical-protein-[Akkermansia-muciniphila] |
| MBL9159967.1 | HNH | GCA_016795385.1 | Verrucomicrobiales_bacterium | Bacteria>Pseudomonadati>PVC_group>Verrucomicrobiota>Verrucomicrobiia>Verrucomicrobiales>unclassified_Verrucomicrobiales | 204 | JNJ70_20970 | 159211..159825 | 2026801 | HNH-endonuclease-[Verrucomicrobiales-bacterium] |
| MBK9497785.1 | HNH | GCA_016718635.1 | Leptospiraceae_bacterium | Bacteria>Pseudomonadati>Spirochaetota>Spirochaetia>Leptospirales>Leptospiraceae>unclassified_Leptospiraceae | 301 | IPO06_00120 | 32165..33070 | 2026757 | hypothetical-protein-[Leptospiraceae-bacterium] |
| WP_020614468.1 | HNH | GCF_000378205.1 | Sediminispirochaeta_bajacaliforniensis | Bacteria>Pseudomonadati>Spirochaetota>Spirochaetia>Spirochaetales>Spirochaetaceae>Sediminispirochaeta | 221 | F459_RS0120170 | 25914..26579 | 148 | TIGR02646-family-protein-[Sediminispirochaeta-bajacaliforniensis] |
| OQY33157.1 | HNH | GCA_002084805.1 | Spirochaetaceae_bacterium_4572_59 | Bacteria>Pseudomonadati>Spirochaetota>Spirochaetia>Spirochaetales>Spirochaetaceae>unclassified_Spirochaetaceae | 223 | B6241_08620 | 12634..13305 | 1971727 | hypothetical-protein-B6241_08620-[Spirochaetaceae-bacterium-4572_59] |
| VBB42855.1 | HNH | GCA_900498135.1 | uncultured_Desulfatiglans_sp. | Bacteria>Pseudomonadati>Thermodesulfobacteriota>Desulfobacteria>Desulfatiglandales>Desulfatiglandaceae>Desulfatiglans>environmental_samples | 250 | TRIP_B220103 | 103706..104458 | 1748965 | hypothetical-protein-TRIP_B220103-[uncultured-Desulfatiglans-sp.] |
| WP_178365092.1 | HNH | GCF_013389365.1 | Desulfobacter_latus | Bacteria>Pseudomonadati>Thermodesulfobacteriota>Desulfobacteria>Desulfobacterales>Desulfobacteraceae>Desulfobacter | 211 | HXW94_RS01270 | 17516..18151 | 2292 | TIGR02646-family-protein-[Desulfobacter-latus] |
| WP_139446524.1 | HNH | GCF_006175995.1 | Desulfobotulus_mexicanus | Bacteria>Pseudomonadati>Thermodesulfobacteriota>Desulfobacteria>Desulfobacterales>Desulfobacteraceae>Desulfobotulus | 212 | FIM25_RS03930 | 125176..125814 | 2586642 | TIGR02646-family-protein-[Desulfobotulus-mexicanus] |
| WP_024334650.1 | HNH | GCF_000421285.1 | Desulfotignum_balticum | Bacteria>Pseudomonadati>Thermodesulfobacteriota>Desulfobacteria>Desulfobacterales>Desulfobacteraceae>Desulfotignum | 199 | K365_RS0111365 | 2253158..2253757 | 115781 | hypothetical-protein-[Desulfotignum-balticum] |
| RZB38314.1 | HNH | GCA_004193595.1 | Desulfobacteraceae_bacterium_Eth-SRB2 | Bacteria>Pseudomonadati>Thermodesulfobacteriota>Desulfobacteria>Desulfobacterales>Desulfobacteraceae>unclassified_Desulfobacteraceae | 91 | SRB2_00062 | 69028..69303 | 2528438 | hypothetical-protein-SRB2_00062-[Desulfobacteraceae-bacterium-Eth-SRB2] |
| WP_155308950.1 | HNH | GCF_009689005.1 | Desulfosarcina_ovata | Bacteria>Pseudomonadati>Thermodesulfobacteriota>Desulfobacteria>Desulfobacterales>Desulfosarcinaceae>Desulfosarcina | 235 | GN112_RS03425 | 768437..769144 | 83564 | HNH-endonuclease-[Desulfosarcina-ovata] |
| MBC2711120.1 | HNH | GCA_014237405.1 | Desulfosarcina_sp. | Bacteria>Pseudomonadati>Thermodesulfobacteriota>Desulfobacteria>Desulfobacterales>Desulfosarcinaceae>Desulfosarcina>unclassified_Desulfosarcina | 215 | HGJ94_09025 | 40117..40764 | 2027861 | TIGR02646-family-protein-[Desulfosarcina-sp.] |
| MBF0121416.1 | HNH | GCA_015228685.1 | Desulfobacterales_bacterium | Bacteria>Pseudomonadati>Thermodesulfobacteriota>Desulfobacteria>Desulfobacterales>unclassified_Desulfobacterales | 171 | HQK79_21510 | 1036..1551 | 2044940 | HNH-endonuclease-[Desulfobacterales-bacterium] |
| KGO34780.1 | HNH | GCA_000769715.1 | Desulfobulbus_sp._Tol-SR | Bacteria>Pseudomonadati>Thermodesulfobacteriota>Desulfobulbia>Desulfobulbales>Desulfobulbaceae>Desulfobulbus>unclassified_Desulfobulbus | 199 | JT06_05930 | 37799..38398 | 1536652 | hypothetical-protein-JT06_05930-[Desulfobulbus-sp.-Tol-SR] |
| WP_083517941.1 | HNH | GCF_001553625.1 | Desulfomicrobium_orale | Bacteria>Pseudomonadati>Thermodesulfobacteriota>Desulfovibrionia>Desulfovibrionales>Desulfomicrobiaceae>Desulfomicrobium | 198 | AXF15_RS13510 | 1733679..1734275 | 132132 | hypothetical-protein-[Desulfomicrobium-orale] |
| WP_168935609.1 | HNH | GCF_012843875.1 | Desulfovibrio_piger | Bacteria>Pseudomonadati>Thermodesulfobacteriota>Desulfovibrionia>Desulfovibrionales>Desulfovibrionaceae>Desulfovibrio | 261 | HF854_RS06745 | 24784..25569 | 901 | TIGR02646-family-protein-[Desulfovibrio-piger] |
| WP_143068398.1 | HNH | GCF_900109185.1 | Deinococcus_reticulitermitis | Bacteria>Thermotogati>Deinococcota>Deinococci>Deinococcales>Deinococcaceae>Deinococcus | 213 | BMY43_RS15280 | 7502..8143 | 856736 | HNH-endonuclease-domain-containing-protein-[Deinococcus-reticulitermitis] |
| WP_135230002.1 | HNH | GCF_004634215.1 | Deinococcus_fonticola | Bacteria>Thermotogati>Deinococcota>Deinococci>Deinococcales>Deinococcaceae>Deinococcus | 210 | E5Z01_RS14420 | 38158..38790 | 2528713 | HNH-endonuclease-domain-containing-protein-[Deinococcus-fonticola] |
| GGM57124.1 | HNH | GCA_014647995.1 | Deinococcus_arenae | Bacteria>Thermotogati>Deinococcota>Deinococci>Deinococcales>Deinococcaceae>Deinococcus | 196 | GCM10008956_36000 | 8919..9509 | 1452751 | hypothetical-protein-GCM10008956_36000-[Deinococcus-arenae] |
| WP_189062790.1 | HNH | GCF_014647995.1 | Deinococcus_arenae | Bacteria>Thermotogati>Deinococcota>Deinococci>Deinococcales>Deinococcaceae>Deinococcus | 196 | IEY66_RS18040 | 8919..9509 | 1452751 | hypothetical-protein-[Deinococcus-arenae] |
| WP_143721447.1 | HNH | GCF_007280555.1 | Deinococcus_detaillensis | Bacteria>Thermotogati>Deinococcota>Deinococci>Deinococcales>Deinococcaceae>Deinococcus | 194 | FNU79_RS14100 | 39148..39732 | 2592048 | HNH-endonuclease-[Deinococcus-detaillensis] |
| MBX8463531.1 | HNH | GCA_019718715.1 | Deinococcus_sp._RIT780 | Bacteria>Thermotogati>Deinococcota>Deinococci>Deinococcales>Deinococcaceae>Deinococcus>unclassified_Deinococcus | 218 | K5D85_00045 | 13885..14541 | 2870472 | hypothetical-protein-[Deinococcus-sp.-RIT780] |
| WP_221587157.1 | HNH | GCF_019718715.1 | Deinococcus_sp._RIT780 | Bacteria>Thermotogati>Deinococcota>Deinococci>Deinococcales>Deinococcaceae>Deinococcus>unclassified_Deinococcus | 218 | K5D85_RS00045 | 13885..14541 | 2870472 | hypothetical-protein-[Deinococcus-sp.-RIT780] |
| WP_244348267.1 | HNH | GCF_022846295.1 | Thermus_thermophilus | Bacteria>Thermotogati>Deinococcota>Deinococci>Thermales>Thermaceae>Thermus | 121 | MWM13_RS12125 | 30169..30534 | 274 | HNH-endonuclease-signature-motif-containing-protein-[Thermus-thermophilus] |
| WP_208321082.1 | HNH | GCF_004366815.1 | Aminivibrio_pyruvatiphilus | Bacteria>Thermotogati>Synergistota>Synergistia>Synergistales>Aminobacteriaceae>Aminivibrio | 208 | C8D99_RS04790 | 134807..135433 | 1005740 | hypothetical-protein-[Aminivibrio-pyruvatiphilus] |
| MCB6991566.1 | HNH | GCA_020558425.1 | bacterium_210820-DFI.6.38 | Bacteria>unclassified_Bacteria | 259 | LI187_13880 | 16910..17689 | 2883293 | hypothetical-protein-[bacterium-210820-DFI.6.38] |
| MCI0602496.1 | HNH | GCA_022616075.1 | bacterium | Bacteria>unclassified_Bacteria | 288 | L0156_05735 | 4120..4986 | 1869227 | HNH-endonuclease-[bacterium] |
| MBU4348831.1 | HNH | GCA_018896635.1 | bacterium | Bacteria>unclassified_Bacteria | 208 | KJ599_00735 | 11965..12591 | 1869227 | hypothetical-protein-[bacterium] |
| --ParB+HNH | | | | | | | | | |
| WP_211087438.1 | ParB+HNH | GCF_024298885.1 | Halobacteriales | Archaea>Methanobacteriati>Methanobacteriota>Stenosarchaea_group>Halobacteria | 643 | NL114_RS21515 | 121294..123225 | 2235 | MULTISPECIES:-DUF262-domain-containing-protein-[Halobacteriales] |
| WP_252490028.1 | ParB+HNH | GCF_023913515.1 | Natrialbaceae | Archaea>Methanobacteriati>Methanobacteriota>Stenosarchaea_group>Halobacteria>Halobacteriales | 709 | NED97_RS07170 | 1402106..1404235 | 1644061 | DUF262-domain-containing-protein-[Natronococcus-sp.-CG52] |
| WP_076432492.1 | ParB+HNH | GCF_900156425.1 | Haladaptatus_litoreus | Archaea>Methanobacteriati>Methanobacteriota>Stenosarchaea_group>Halobacteria>Halobacteriales>Haladaptataceae>Haladaptatus | 572 | BW974_RS20495 | 116176..117894 | 553468 | DUF262-domain-containing-protein-[Haladaptatus-litoreus] |
| WP_229123015.1 | ParB+HNH | GCF_017094465.1 | Halapricum_desulfuricans | Archaea>Methanobacteriati>Methanobacteriota>Stenosarchaea_group>Halobacteria>Halobacteriales>Haloarculaceae>Halapricum | 610 | HSEST_RS14115 | 96745..98577 | 2841257 | DUF262-domain-containing-protein-[Halapricum-desulfuricans] |
| WP_146418282.1 | ParB+HNH | GCF_902209435.1 | Haloarcula_hispanica | Archaea>Methanobacteriati>Methanobacteriota>Stenosarchaea_group>Halobacteria>Halobacteriales>Haloarculaceae>Haloarcula | 571 | FUT35_RS18480 | 3793..5508 | 51589 | DUF262-domain-containing-protein-[Haloarcula-hispanica] |
| WP_254824800.1 | ParB+HNH | GCF_024298885.1 | Haloglomus_halophilum | Archaea>Methanobacteriati>Methanobacteriota>Stenosarchaea_group>Halobacteria>Halobacteriales>Haloarculaceae>Haloglomus | 709 | NL114_RS20935 | 101597..103726 | 2962672 | DUF262-domain-containing-protein-[Haloglomus-sp.-ZY41] |
| WP_226011846.1 | ParB+HNH | GCF_020405185.1 | Halomicrobium_salinisoli | Archaea>Methanobacteriati>Methanobacteriota>Stenosarchaea_group>Halobacteria>Halobacteriales>Haloarculaceae>Halomicrobium | 694 | LE162_RS01600 | 304882..306966 | 2878391 | DUF262-domain-containing-protein-[Halomicrobium-salinisoli] |
| WP_159663621.1 | ParB+HNH | GCF_009831625.1 | Halobaculum_saliterrae | Archaea>Methanobacteriati>Methanobacteriota>Stenosarchaea_group>Halobacteria>Halobacteriales>Haloferacaceae>Halobaculum | 612 | GRX01_RS03880 | 201888..203726 | 2073113 | DUF262-domain-containing-protein-[Halobaculum-saliterrae] |
| WP_170216822.1 | ParB+HNH | GCF_004799685.1 | Halobellus_limi | Archaea>Methanobacteriati>Methanobacteriota>Stenosarchaea_group>Halobacteria>Halobacteriales>Haloferacaceae>Halobellus | 712 | DV707_RS08420 | 1698026..1700164 | 699433 | DUF262-domain-containing-protein-[Halobellus-limi] |
| WP_049916461.1 | ParB+HNH | GCF_000336795.1 | Haloferax_volcanii | Archaea>Methanobacteriati>Methanobacteriota>Stenosarchaea_group>Halobacteria>Halobacteriales>Haloferacaceae>Haloferax | 709 | C456_RS03010 | 78413..80542 | 2246 | DUF262-domain-containing-protein-[Haloferax-lucentense] |
| WP_193493616.1 | ParB+HNH | - | Haloferax_gibbonsii | Archaea>Methanobacteriati>Methanobacteriota>Stenosarchaea_group>Halobacteria>Halobacteriales>Haloferacaceae>Haloferax | 443 | HfgLR_RS17315 | 399728..401059 | 35746 | DUF1524-domain-containing-protein-[Haloferax-gibbonsii] |
| MBC9987876.1 | ParB+HNH | GCA_014530665.1 | Haloferax_sp._AS1 | Archaea>Methanobacteriati>Methanobacteriota>Stenosarchaea_group>Halobacteria>Halobacteriales>Haloferacaceae>Haloferax>unclassified_Haloferax | 643 | E4P24_16080 | 313189..315120 | 2562277 | DUF262-domain-containing-protein-[Haloferax-sp.-AS1] |
| WP_188148786.1 | ParB+HNH | GCF_014530665.1 | Haloferax_sp._AS1 | Archaea>Methanobacteriati>Methanobacteriota>Stenosarchaea_group>Halobacteria>Halobacteriales>Haloferacaceae>Haloferax>unclassified_Haloferax | 643 | E4P24_RS16215 | 313189..315120 | 2562277 | DUF262-domain-containing-protein-[Haloferax-sp.-AS1] |
| WP_271963539.1 | ParB+HNH | GCF_028211285.1 | Halorubrum_ezzemoulense | Archaea>Methanobacteriati>Methanobacteriota>Stenosarchaea_group>Halobacteria>Halobacteriales>Haloferacaceae>Halorubrum | 643 | PM025_RS15425 | 9047..10978 | 337243 | DUF262-domain-containing-HNH-endonuclease-family-protein-[Halorubrum-ezzemoulense] |
| WP_241154922.1 | ParB+HNH | GCF_003781945.1 | Halorubrum_sp._CSM-61 | Archaea>Methanobacteriati>Methanobacteriota>Stenosarchaea_group>Halobacteria>Halobacteriales>Haloferacaceae>Halorubrum>unclassified_Halorubrum | 709 | EDM04_RS13305 | 112182..114311 | 2485838 | DUF262-domain-containing-protein-[Halorubrum-sp.-CSM-61] |
| WP_207290713.1 | ParB+HNH | GCF_017352155.2 | Haloterrigena_alkaliphila | Archaea>Methanobacteriati>Methanobacteriota>Stenosarchaea_group>Halobacteria>Halobacteriales>Natrialbaceae>Haloterrigena | 616 | J0X25_RS27475 | 1780184..1782034 | 2816475 | DUF262-domain-containing-protein-[Haloterrigena-alkaliphila] |
| WP_008160992.1 | ParB+HNH | GCF_000337735.1 | Natronorubrum_sulfidifaciens | Archaea>Methanobacteriati>Methanobacteriota>Stenosarchaea_group>Halobacteria>Halobacteriales>Natrialbaceae>Natronorubrum | 563 | C495_RS06080 | 90296..91987 | 388259 | DUF262-domain-containing-protein-[Natronorubrum-sulfidifaciens] |
| WP_083687734.1 | ParB+HNH | GCF_001971705.1 | Natronorubrum_daqingense | Archaea>Methanobacteriati>Methanobacteriota>Stenosarchaea_group>Halobacteria>Halobacteriales>Natrialbaceae>Natronorubrum | 619 | BW960_RS07760 | 349709..351568 | 588898 | DUF262-domain-containing-protein-[Natronorubrum-daqingense] |
| ELY46665.1 | ParB+HNH | GCA_000337735.1 | Natronorubrum_sulfidifaciens_JCM_14089 | Archaea>Methanobacteriati>Methanobacteriota>Stenosarchaea_group>Halobacteria>Halobacteriales>Natrialbaceae>Natronorubrum>Natronorubrum_sulfidifaciens | 563 | C495_06143 | 90296..91987 | 1230460 | hypothetical-protein-C495_06143-[Natronorubrum-sulfidifaciens-JCM-14089] |
| WP_253430482.1 | ParB+HNH | GCF_024138145.1 | Natronosalvus_halobius | Archaea>Methanobacteriati>Methanobacteriota>Stenosarchaea_group>Halobacteria>Halobacteriales>Natrialbaceae>Natronosalvus | 604 | NGM15_RS10400 | 2113801..2115615 | 2953746 | DUF262-domain-containing-protein-[Halovivax-sp.-AGai3-5] |
| WP_202614607.1 | ParB+HNH | GCF_004785955.1 | Halostella_litorea | Archaea>Methanobacteriati>Methanobacteriota>Stenosarchaea_group>Halobacteria>Halobacteriales>Natronoarchaeaceae>Halostella | 709 | EYW40_RS18710 | 37177..39306 | 2528831 | DUF262-domain-containing-protein-[Halostella-litorea] |
| WP_256392870.1 | ParB+HNH | GCF_024494695.1 | Natronoarchaeum_rubrum | Archaea>Methanobacteriati>Methanobacteriota>Stenosarchaea_group>Halobacteria>Halobacteriales>Natronoarchaeaceae>Natronoarchaeum | 680 | NO931_RS13585 | 566887..568929 | 755311 | DUF262-domain-containing-protein-[Natronoarchaeum-rubrum] |
| WP_020446092.1 | ParB+HNH | GCF_000403645.1 | Salinarchaeum_sp._Harcht-Bsk1 | Archaea>Methanobacteriati>Methanobacteriota>Stenosarchaea_group>Halobacteria>Halobacteriales>Natronoarchaeaceae>Salinarchaeum>unclassified_Salinarchaeum | 705 | L593_RS06225 | 1346262..1348379 | 1333523 | DUF262-domain-containing-protein-[Salinarchaeum-sp.-Harcht-Bsk1] |
| WP_253456787.1 | ParB+HNH | GCF_024170265.1 | Methanocalculus_sp._AMF5 | Archaea>Methanobacteriati>Methanobacteriota>Stenosarchaea_group>Methanomicrobia>Methanomicrobiales>Methanocalculaceae>Methanocalculus>unclassified_Methanocalculus | 563 | J2T61_RS02240 | 77062..78753 | 1198257 | DUF262-domain-containing-protein-[Methanocalculus-sp.-AMF5] |
| RXE56038.1 | ParB+HNH | GCA_004102725.1 | Methanoculleus_taiwanensis | Archaea>Methanobacteriati>Methanobacteriota>Stenosarchaea_group>Methanomicrobia>Methanomicrobiales>Methanomicrobiaceae>Methanoculleus | 560 | ABH15_07535 | 555360..557042 | 1550565 | hypothetical-protein-ABH15_07535-[Methanoculleus-taiwanensis] |
| WP_128693763.1 | ParB+HNH | GCF_004102725.1 | Methanoculleus_taiwanensis | Archaea>Methanobacteriati>Methanobacteriota>Stenosarchaea_group>Methanomicrobia>Methanomicrobiales>Methanomicrobiaceae>Methanoculleus | 560 | ABH15_RS07620 | 555360..557042 | 1550565 | DUF262-domain-containing-protein-[Methanoculleus-taiwanensis] |
| WP_067072533.1 | ParB+HNH | GCF_001602375.1 | Methanoculleus_horonobensis | Archaea>Methanobacteriati>Methanobacteriota>Stenosarchaea_group>Methanomicrobia>Methanomicrobiales>Methanomicrobiaceae>Methanoculleus | 557 | MCUHO_RS01270 | 26683..28356 | 528314 | DUF262-domain-containing-protein-[Methanoculleus-horonobensis] |
| WP_197540525.1 | ParB+HNH | GCF_000304355.2 | Methanoculleus_bourgensis | Archaea>Methanobacteriati>Methanobacteriota>Stenosarchaea_group>Methanomicrobia>Methanomicrobiales>Methanomicrobiaceae>Methanoculleus | 547 | BN140_RS09860 | 2037935..2039578 | 83986 | DUF262-domain-containing-protein-[Methanoculleus-bourgensis] |
| CCJ36895.1 | ParB+HNH | GCA_000304355.2 | Methanoculleus_bourgensis_MS2 | Archaea>Methanobacteriati>Methanobacteriota>Stenosarchaea_group>Methanomicrobia>Methanomicrobiales>Methanomicrobiaceae>Methanoculleus>Methanoculleus_bourgensis | 557 | BN140_1972 | 2037905..2039578 | 1201294 | hypothetical-protein-BN140_1972-[Methanoculleus-bourgensis-MS2] |
| MCE5337192.1 | ParB+HNH | GCA_021371835.1 | Methanomicrobiaceae_archaeon | Archaea>Methanobacteriati>Methanobacteriota>Stenosarchaea_group>Methanomicrobia>Methanomicrobiales>Methanomicrobiaceae>unclassified_Methanomicrobiaceae | 557 | LLF90_00680 | 36892..38565 | 2716262 | DUF262-domain-containing-protein-[Methanomicrobiaceae-archaeon] |
| NLM31069.1 | ParB+HNH | GCA_012521035.1 | Methanomicrobiales_archaeon | Archaea>Methanobacteriati>Methanobacteriota>Stenosarchaea_group>Methanomicrobia>Methanomicrobiales>unclassified_Methanomicrobiales | 559 | GX216_11750 | 3128..4807 | 2184052 | DUF262-domain-containing-protein-[Methanomicrobiales-archaeon] |
| WP_200895275.1 | ParB+HNH | GCF_000979415.1 | unclassified_Methanosarcina | Archaea>Methanobacteriati>Methanobacteriota>Stenosarchaea_group>Methanomicrobia>Methanosarcinales>Methanosarcinaceae>Methanosarcina | 297 | EO97_RS19885 | 22719..23612 | 2644672 | MULTISPECIES:-DUF262-domain-containing-protein-[unclassified-Methanosarcina] |
| WP_229395642.1 | ParB+HNH | GCF_020804205.1 | Methanosarcina_sp._DH1 | Archaea>Methanobacteriati>Methanobacteriota>Stenosarchaea_group>Methanomicrobia>Methanosarcinales>Methanosarcinaceae>Methanosarcina>unclassified_Methanosarcina | 601 | FXW07_RS12035 | 181631..183436 | 2605695 | DUF262-domain-containing-protein-[Methanosarcina-sp.-DH1] |
| CAG0952182.1 | ParB+HNH | GCA_905339305.1 | Methanosarcinales_archaeon | Archaea>Methanobacteriati>Methanobacteriota>Stenosarchaea_group>Methanomicrobia>Methanosarcinales>unclassified_Methanosarcinales | 448 | METP3_00278 | 24524..25870 | 2250255 | hypothetical-protein-METP3_00278-[Methanosarcinales-archaeon] |

| Protein-ID | Domain | Assembly-ID | Strain | Taxonomy | Length | Locus-Tag | Genomic-coordinates | TaxID | Fasta-defline |
|---|---|---|---|---|---|---|---|---|---|
| MBP7066720.1 | ParB+HNH | GCA_017994095.1 | Methanothrix_sp. | Archaea>Methanobacteriati>Methanobacteriota>Stenosarchaea_group>Methanomicrobia>Methanotrichales>Methanotrichaceae>Methanothrix>unclassified_Methanothrix | 625 | KBB04_00345 | 74721..76598 | 90426 | DUF262-domain-containing-protein-[Methanothrix-sp.] |
| NHV98107.1 | ParB+HNH | GCA_011605725.1 | Nitrososphaerota_archaeon | Archaea>Thermoproteati>Nitrososphaerota>unclassified_Nitrososphaerota | 569 | HA496_00510 | 56695..58404 | 2026795 | DUF262-domain-containing-protein-[Nitrososphaerota-archaeon] |
| MXZ86863.1 | ParB+HNH | GCA_009837425.1 | Acidimicrobiia_bacterium | Bacteria>Bacillati>Actinomycetota>Acidimicrobiia>unclassified_Acidimicrobiia | 381 | F4Z02_14570 | 42500..43645 | 2080302 | DUF262-domain-containing-protein-[Acidimicrobiia-bacterium] |
| WP_109093609.1 | ParB+HNH | GCF_003123745.1 | Ancrocorticia_populi | Bacteria>Bacillati>Actinomycetota>Actinomycetes>Actinomycetales>Actinomycetaceae>Ancrocorticia | 546 | DD236_RS06705 | 264544..266184 | 2175228 | DUF262-domain-containing-protein-[Ancrocorticia-populi] |
| WP_284940341.1 | ParB+HNH | GCF_030250405.1 | Bifidobacterium_catenulatum | Bacteria>Bacillati>Actinomycetota>Actinomycetes>Bifidobacteriales>Bifidobacteriaceae>Bifidobacterium | 351 | K1513_RS02310 | 518527..519582 | 1686 | DUF262-domain-containing-protein-[Bifidobacterium-catenulatum] |
| WP_236827010.1 | ParB+HNH | GCF_021698995.1 | unclassified_Blastococcus | Bacteria>Bacillati>Actinomycetota>Actinomycetes>Geodermatophilales>Geodermatophilaceae>Blastococcus | 661 | E9565_RS02690 | 61241..63226 | 2619396 | MULTISPECIES:-DUF262-domain-containing-protein-[unclassified-Blastococcus] |
| WP_113812540.1 | ParB+HNH | GCF_003319095.1 | Blastococcus_sp._TBT05-19 | Bacteria>Bacillati>Actinomycetota>Actinomycetes>Geodermatophilales>Geodermatophilaceae>Blastococcus>unclassified_Blastococcus | 642 | DQ244_RS16240 | 127500..129428 | 2250581 | DUF262-domain-containing-protein-[Blastococcus-sp.-TBT05-19] |
| WP_269185213.1 | ParB+HNH | GCF_026967925.1 | Modestobacter_sp._VKM_Ac-2985 | Bacteria>Bacillati>Actinomycetota>Actinomycetes>Geodermatophilales>Geodermatophilaceae>Modestobacter>unclassified_Modestobacter | 555 | O2V61_RS21020 | 1075268..1076935 | 3004139 | DUF262-domain-containing-protein-[Modestobacter-sp.-VKM-Ac-2985] |
| WP_147142493.1 | ParB+HNH | GCF_039542685.1 | Stackebrandtia_albiflava | Bacteria>Bacillati>Actinomycetota>Actinomycetes>Glycomycetales>Glycomycetaceae>Stackebrandtia | 784 | LX16_RS21805 | 171589..173943 | 406432 | DUF262-domain-containing-protein-[Stackebrandtia-albiflava] |
| WP_067071157.1 | ParB+HNH | GCF_001543895.1 | Carbonactinospora_thermoautotrophica | Bacteria>Bacillati>Actinomycetota>Actinomycetes>Kitasatosporales>Carbonactinosporaceae>Carbonactinospora | 660 | TR74_RS19800 | 6627..8609 | 1469144 | DUF262-domain-containing-protein-[Carbonactinospora-thermoautotrophica] |
| WP_267593978.1 | ParB+HNH | GCF_026614415.1 | Carbonactinospora_thermoautotrophica | Bacteria>Bacillati>Actinomycetota>Actinomycetes>Kitasatosporales>Carbonactinosporaceae>Carbonactinospora | 607 | C3Y87_RS08205 | 80609..82432 | 1469144 | DUF262-domain-containing-protein-[Carbonactinospora-thermoautotrophica] |
| WP_044379852.1 | ParB+HNH | GCF_014656215.1 | Streptomyces | Bacteria>Bacillati>Actinomycetota>Actinomycetes>Kitasatosporales>Streptomycetaceae | 607 | IE311_RS17195 | 200350..202173 | 1883 | MULTISPECIES:-DUF262-domain-containing-protein-[Streptomyces] |
| WP_234526772.1 | ParB+HNH | GCF_021442415.2 | Streptomyces | Bacteria>Bacillati>Actinomycetota>Actinomycetes>Kitasatosporales>Streptomycetaceae | 605 | B0E36_RS09875 | 22176..23993 | 1883 | MULTISPECIES:-DUF262-domain-containing-protein-[unclassified-Streptomyces] |
| WP_081522442.1 | ParB+HNH | GCF_002080455.1 | Streptomyces | Bacteria>Bacillati>Actinomycetota>Actinomycetes>Kitasatosporales>Streptomycetaceae | 603 | B6R96_RS11905 | 2767602..2769413 | 1883 | DUF262-domain-containing-protein-[Streptomyces-sp.-Sge12] |
| WP_114246644.1 | ParB+HNH | GCF_003330865.1 | Streptomyces | Bacteria>Bacillati>Actinomycetota>Actinomycetes>Kitasatosporales>Streptomycetaceae | 602 | C5746_RS28315 | 6147416..6149224 | 1883 | DUF262-domain-containing-protein-[Streptomyces-atratus] |
| ROR38149.1 | ParB+HNH | GCA_003751605.1 | Kitasatospora_cineracea | Bacteria>Bacillati>Actinomycetota>Actinomycetes>Kitasatosporales>Streptomycetaceae>Kitasatospora | 606 | EDD39_6321 | 1163301..1165121 | 88074 | hypothetical-protein-EDD39_6321-[Kitasatospora-cineracea] |
| WP_123562573.1 | ParB+HNH | GCF_003751605.1 | Kitasatospora_cineracea | Bacteria>Bacillati>Actinomycetota>Actinomycetes>Kitasatosporales>Streptomycetaceae>Kitasatospora | 606 | EDD39_RS31495 | 1163301..1165121 | 88074 | DUF262-domain-containing-protein-[Kitasatospora-cineracea] |
| MCQ4043240.1 | ParB+HNH | GCA_024436055.1 | Streptantibioticus_rubrisoli | Bacteria>Bacillati>Actinomycetota>Actinomycetes>Kitasatosporales>Streptomycetaceae>Streptantibioticus | 593 | NON19_14670 | 104746..106527 | 1387313 | DUF262-domain-containing-protein-[Streptomyces-rubrisoli] |
| WP_255928150.1 | ParB+HNH | GCF_024436055.1 | Streptantibioticus_rubrisoli | Bacteria>Bacillati>Actinomycetota>Actinomycetes>Kitasatosporales>Streptomycetaceae>Streptantibioticus | 593 | NON19_RS14715 | 104746..106527 | 1387313 | DUF262-domain-containing-protein-[Streptomyces-rubrisoli] |
| WP_267842814.1 | ParB+HNH | GCF_026627185.1 | unclassified_Streptomyces | Bacteria>Bacillati>Actinomycetota>Actinomycetes>Kitasatosporales>Streptomycetaceae>Streptomyces | 607 | OTB23_RS33975 | 60657..62480 | 2593676 | MULTISPECIES:-DUF262-domain-containing-protein-[unclassified-Streptomyces] |
| WP_252540691.1 | ParB+HNH | GCF_023930305.1 | unclassified_Streptomyces | Bacteria>Bacillati>Actinomycetota>Actinomycetes>Kitasatosporales>Streptomycetaceae>Streptomyces | 605 | LRR80_RS21250 | 28296..30113 | 2593676 | DUF262-domain-containing-protein-[Streptomyces-sp.-RO-S4] |
| WP_073727346.1 | ParB+HNH | GCF_042736535.1 | unclassified_Streptomyces | Bacteria>Bacillati>Actinomycetota>Actinomycetes>Kitasatosporales>Streptomycetaceae>Streptomyces | 601 | A6A29_RS36850 | 513896..515701 | 2593676 | DUF262-domain-containing-protein-[Streptomyces-sp.-TSRI0281] |
| WTB22522.1 | ParB+HNH | GCA_045161805.1 | Streptomyces_virginiae | Bacteria>Bacillati>Actinomycetota>Actinomycetes>Kitasatosporales>Streptomycetaceae>Streptomyces | 600 | OG253_14025 | 3179133..3180935 | 1961 | DUF262-domain-containing-protein-[Streptomyces-virginiae] |
| WP_242752936.1 | ParB+HNH | GCF_022699385.1 | Streptomyces_tubbatahanensis | Bacteria>Bacillati>Actinomycetota>Actinomycetes>Kitasatosporales>Streptomycetaceae>Streptomyces | 604 | MMF93_RS18275 | 4251003..4252817 | 2923272 | DUF262-domain-containing-protein-[Streptomyces-tubbatahanensis] |
| WP_189744986.1 | ParB+HNH | GCF_014650435.1 | Streptomyces_tendae | Bacteria>Bacillati>Actinomycetota>Actinomycetes>Kitasatosporales>Streptomycetaceae>Streptomyces | 605 | IE258_RS18995 | 423042..424859 | 1932 | DUF262-domain-containing-protein-[Streptomyces-tendae] |
| WP_245226725.1 | ParB+HNH | - | Streptomyces_smyrnaeus | Bacteria>Bacillati>Actinomycetota>Actinomycetes>Kitasatosporales>Streptomycetaceae>Streptomyces | 584 | JW613_RS33025 | 38245..39999 | 1387713 | DUF262-domain-containing-protein-[Streptomyces-smyrnaeus] |
| WP_121887753.1 | ParB+HNH | GCF_003688995.1 | Streptomyces_shenzhenensis | Bacteria>Bacillati>Actinomycetota>Actinomycetes>Kitasatosporales>Streptomycetaceae>Streptomyces | 605 | CTZ28_RS03725 | 166699..168516 | 943815 | DUF262-domain-containing-protein-[Streptomyces-shenzhenensis] |
| WP_189531554.1 | ParB+HNH | GCF_014649335.1 | Streptomyces_roseolilacinus | Bacteria>Bacillati>Actinomycetota>Actinomycetes>Kitasatosporales>Streptomycetaceae>Streptomyces | 605 | IE106_RS08470 | 248436..250253 | 66904 | DUF262-domain-containing-protein-[Streptomyces-roseolilacinus] |
| WP_179083038.1 | ParB+HNH | GCF_013394065.1 | Streptomyces_rectiverticillatus | Bacteria>Bacillati>Actinomycetota>Actinomycetes>Kitasatosporales>Streptomycetaceae>Streptomyces | 605 | FGW37_RS24645 | 5632376..5634193 | 173860 | DUF262-domain-containing-protein-[Streptomyces-rectiverticillatus] |
| KOX56113.1 | ParB+HNH | GCA_001280155.1 | Streptomyces_purpurogeneiscleroticus | Bacteria>Bacillati>Actinomycetota>Actinomycetes>Kitasatosporales>Streptomycetaceae>Streptomyces | 693 | ADL19_10435 | 24420..26501 | 68259 | hypothetical-protein-ADL19_10435-[Streptomyces-purpurogeneiscleroticus] |
| WP_249836749.1 | ParB+HNH | GCF_023516615.1 | Streptomyces_panaciradicis | Bacteria>Bacillati>Actinomycetota>Actinomycetes>Kitasatosporales>Streptomycetaceae>Streptomyces | 602 | M8J74_RS14320 | 95153..96961 | 1470261 | DUF262-domain-containing-protein-[Streptomyces-panaciradicis] |
| GAA3523430.1 | ParB+HNH | GCA_039538545.1 | Streptomyces_osmaniensis | Bacteria>Bacillati>Actinomycetota>Actinomycetes>Kitasatosporales>Streptomycetaceae>Streptomyces | 602 | GCM10022295_01170 | 144208..146016 | 593134 | DUF262-domain-containing-protein-[Streptomyces-osmaniensis] |
| WP_109284044.1 | ParB+HNH | GCF_040679255.1 | Streptomyces_orinoci | Bacteria>Bacillati>Actinomycetota>Actinomycetes>Kitasatosporales>Streptomycetaceae>Streptomyces | 607 | CV755_RS31545 | 57344..59167 | 67339 | DUF262-domain-containing-protein-[Streptomyces-orinoci] |
| WP_189852663.1 | ParB+HNH | GCF_014650895.1 | Streptomyces_omiyaensis | Bacteria>Bacillati>Actinomycetota>Actinomycetes>Kitasatosporales>Streptomycetaceae>Streptomyces | 607 | IE287_RS31565 | 75478..77301 | 68247 | DUF262-domain-containing-protein-[Streptomyces-omiyaensis] |
| WP_275462593.1 | ParB+HNH | GCF_029167025.1 | Streptomyces_noursei | Bacteria>Bacillati>Actinomycetota>Actinomycetes>Kitasatosporales>Streptomycetaceae>Streptomyces | 607 | P0D74_RS11320 | 2592505..2594328 | 1971 | DUF262-domain-containing-protein-[Streptomyces-noursei] |
| WP_329144531.1 | ParB+HNH | GCF_036227585.1 | Streptomyces_niveus | Bacteria>Bacillati>Actinomycetota>Actinomycetes>Kitasatosporales>Streptomycetaceae>Streptomyces | 604 | OG275_RS09095 | 2077510..2079324 | 193462 | DUF262-domain-containing-protein-[Streptomyces-niveus] |
| WP_216339495.1 | ParB+HNH | GCF_018883605.1 | Streptomyces_niphimycinicus | Bacteria>Bacillati>Actinomycetota>Actinomycetes>Kitasatosporales>Streptomycetaceae>Streptomyces | 615 | KN815_RS01810 | 12510..14357 | 2842201 | DUF262-domain-containing-protein-[Streptomyces-sp.-4503] |
| GAA0539979.1 | ParB+HNH | GCA_039521245.1 | Streptomyces_mordarskii | Bacteria>Bacillati>Actinomycetota>Actinomycetes>Kitasatosporales>Streptomycetaceae>Streptomyces | 602 | GCM10010390_47460 | 371866..373674 | 1226758 | DUF262-domain-containing-protein-[Streptomyces-mordarskii] |
| WP_316738723.1 | ParB+HNH | GCF_032595465.1 | Streptomyces_mirabilis | Bacteria>Bacillati>Actinomycetota>Actinomycetes>Kitasatosporales>Streptomycetaceae>Streptomyces | 612 | PU648_RS58360 | 76808..78646 | 68239 | DUF262-domain-containing-protein-[Streptomyces-mirabilis] |
| WP_226071744.1 | ParB+HNH | GCF_020410765.1 | Streptomyces_marincola | Bacteria>Bacillati>Actinomycetota>Actinomycetes>Kitasatosporales>Streptomycetaceae>Streptomyces | 601 | LC193_RS04380 | 1035443..1037248 | 2878388 | DUF262-domain-containing-protein-[Streptomyces-marincola] |
| WP_150950775.1 | ParB+HNH | GCF_008806595.1 | Streptomyces_luteolifulvus | Bacteria>Bacillati>Actinomycetota>Actinomycetes>Kitasatosporales>Streptomycetaceae>Streptomyces | 602 | F7R91_RS21825 | 71641..73449 | 2615112 | DUF262-domain-containing-protein-[Streptomyces-luteolifulvus] |
| WP_330344468.1 | ParB+HNH | GCF_045161325.1 | Streptomyces_longwoodensis | Bacteria>Bacillati>Actinomycetota>Actinomycetes>Kitasatosporales>Streptomycetaceae>Streptomyces | 602 | OG547_RS11295 | 2759992..2761800 | 68231 | DUF262-domain-containing-protein-[Streptomyces-longwoodensis] |
| WP_326752423.1 | ParB+HNH | GCF_035918035.1 | Streptomyces_hirsutus | Bacteria>Bacillati>Actinomycetota>Actinomycetes>Kitasatosporales>Streptomycetaceae>Streptomyces | 601 | OIE73_RS11095 | 2423550..2425355 | 35620 | DUF262-domain-containing-protein-[Streptomyces-hirsutus] |
| WP_181660541.1 | ParB+HNH | GCF_013778455.1 | Streptomyces_himalayensis | Bacteria>Bacillati>Actinomycetota>Actinomycetes>Kitasatosporales>Streptomycetaceae>Streptomyces | 557 | H1D24_RS28265 | 54322..55995 | 2820085 | DUF262-domain-containing-protein-[Streptomyces-himalayensis] |
| WP_328775749.1 | ParB+HNH | GCF_036175285.1 | Streptomyces_goshikiensis | Bacteria>Bacillati>Actinomycetota>Actinomycetes>Kitasatosporales>Streptomycetaceae>Streptomyces | 607 | OHU17_RS09820 | 2199546..2201369 | 1942 | DUF262-domain-containing-protein-[Streptomyces-goshikiensis] |
| GAA1555955.1 | ParB+HNH | GCA_039529415.1 | Streptomyces_globosus | Bacteria>Bacillati>Actinomycetota>Actinomycetes>Kitasatosporales>Streptomycetaceae>Streptomyces | 614 | GCM10009731_07900 | 76099..77943 | 68209 | DUF262-domain-containing-protein-[Streptomyces-globosus] |
| WP_281277968.1 | ParB+HNH | - | Streptomyces_globosus | Bacteria>Bacillati>Actinomycetota>Actinomycetes>Kitasatosporales>Streptomycetaceae>Streptomyces | 601 | C0216_RS08085 | 1739027..1740832 | 68209 | DUF262-domain-containing-protein-[Streptomyces-globosus] |
| WP_105973290.1 | ParB+HNH | GCF_002954775.1 | Streptomyces_geranii | Bacteria>Bacillati>Actinomycetota>Actinomycetes>Kitasatosporales>Streptomycetaceae>Streptomyces | 604 | CW359_RS29825 | 28913..30727 | 2058923 | DUF262-domain-containing-protein-[Streptomyces-geranii] |
| WP_150231681.1 | ParB+HNH | GCF_008634015.1 | Streptomyces_filamentosus | Bacteria>Bacillati>Actinomycetota>Actinomycetes>Kitasatosporales>Streptomycetaceae>Streptomyces | 604 | CP979_RS24285 | 5255600..5257414 | 67294 | DUF262-domain-containing-protein-[Streptomyces-filamentosus] |
| WP_230991896.1 | ParB+HNH | - | Streptomyces_endocoffeicus | Bacteria>Bacillati>Actinomycetota>Actinomycetes>Kitasatosporales>Streptomycetaceae>Streptomyces | 620 | JK364_RS14440 | 436394..438256 | 2898945 | DUF262-domain-containing-protein-[Streptomyces-endocoffeicus] |
| GAA2342631.1 | ParB+HNH | GCA_039532705.1 | Streptomyces_cuspidosporus | Bacteria>Bacillati>Actinomycetota>Actinomycetes>Kitasatosporales>Streptomycetaceae>Streptomyces | 602 | GCM10010246_30000 | 60567..62375 | 66882 | DUF262-domain-containing-protein-[Streptomyces-cuspidosporus] |
| WP_184852648.1 | ParB+HNH | GCF_039540655.1 | Streptomyces_collinus | Bacteria>Bacillati>Actinomycetota>Actinomycetes>Kitasatosporales>Streptomycetaceae>Streptomyces | 606 | HNR72_RS32130 | 7199812..7201632 | 42684 | DUF262-domain-containing-protein-[Streptomyces-collinus] |
| WP_189191870.1 | ParB+HNH | GCF_014645835.1 | Streptomyces_albiflavescens | Bacteria>Bacillati>Actinomycetota>Actinomycetes>Kitasatosporales>Streptomycetaceae>Streptomyces | 602 | IEY95_RS43785 | 52011..53819 | 1623582 | DUF262-domain-containing-protein-[Streptomyces-albiflavescens] |
| GAA1254449.1 | ParB+HNH | GCA_039525965.1 | Streptomyces_yogyakartensis | Bacteria>Bacillati>Actinomycetota>Actinomycetes>Kitasatosporales>Streptomycetaceae>Streptomyces>Streptomyces_violaceusniger_group | 602 | GCM10009579_10520 | 281927..283735 | 114692 | DUF262-domain-containing-protein-[Streptomyces-yogyakartensis] |
| WP_060944828.1 | ParB+HNH | GCF_001553455.1 | Streptomyces_hygroscopicus | Bacteria>Bacillati>Actinomycetota>Actinomycetes>Kitasatosporales>Streptomycetaceae>Streptomyces>Streptomyces_violaceusniger_group | 602 | AXW62_RS14125 | 37034..38842 | 1912 | DUF262-domain-containing-protein-[Streptomyces-hygroscopicus] |
| WP_062010204.1 | ParB+HNH | - | Streptomyces_hygroscopicus | Bacteria>Bacillati>Actinomycetota>Actinomycetes>Kitasatosporales>Streptomycetaceae>Streptomyces>Streptomyces_violaceusniger_group | 602 | AXW65_RS13455 | 89520..91328 | 1912 | DUF262-domain-containing-protein-[Streptomyces-hygroscopicus] |
| WP_330811469.1 | ParB+HNH | GCF_036352935.1 | Streptomyces_asiaticus | Bacteria>Bacillati>Actinomycetota>Actinomycetes>Kitasatosporales>Streptomycetaceae>Streptomyces>Streptomyces_violaceusniger_group | 602 | V2J94_RS25875 | 107070..108878 | 114695 | DUF262-domain-containing-protein-[Streptomyces-sp.-DSM-41524] |
| WP_254711940.1 | ParB+HNH | - | Streptomyces_sp._TRM64462 | Bacteria>Bacillati>Actinomycetota>Actinomycetes>Kitasatosporales>Streptomycetaceae>Streptomyces>unclassified_Streptomyces | 594 | HTZ82_RS18010 | 67005..68789 | 2741726 | DUF262-domain-containing-protein-[Streptomyces-sp.-TRM64462] |
| WP_142151607.1 | ParB+HNH | GCF_006715395.1 | Streptomyces_sp._SLBN-31 | Bacteria>Bacillati>Actinomycetota>Actinomycetes>Kitasatosporales>Streptomycetaceae>Streptomyces>unclassified_Streptomyces | 605 | FBY22_RS32825 | 2795646..2797463 | 2768444 | DUF262-domain-containing-protein-[Streptomyces-sp.-SLBN-31] |
| WP_203660772.1 | ParB+HNH | - | Streptomyces_sp._SID14446 | Bacteria>Bacillati>Actinomycetota>Actinomycetes>Kitasatosporales>Streptomycetaceae>Streptomyces>unclassified_Streptomyces | 594 | G3I62_RS24955 | 17983..19767 | 2706072 | DUF262-domain-containing-protein-[Streptomyces-sp.-SID14446] |
| WP_268974405.1 | ParB+HNH | GCF_026898115.1 | Streptomyces_sp._S465 | Bacteria>Bacillati>Actinomycetota>Actinomycetes>Kitasatosporales>Streptomycetaceae>Streptomyces>unclassified_Streptomyces | 602 | N6H00_RS26940 | 6342672..6344480 | 2979468 | DUF262-domain-containing-protein-[Streptomyces-sp.-S465] |
| WP_037951100.1 | ParB+HNH | GCF_000590515.1 | Streptomyces_sp._PRh5 | Bacteria>Bacillati>Actinomycetota>Actinomycetes>Kitasatosporales>Streptomycetaceae>Streptomyces>unclassified_Streptomyces | 602 | Z951_RS11155 | 104144..105952 | 1158056 | DUF262-domain-containing-protein-[Streptomyces-sp.-PRh5] |
| WP_279329416.1 | ParB+HNH | GCF_029759795.1 | Streptomyces_sp._OS603R | Bacteria>Bacillati>Actinomycetota>Actinomycetes>Kitasatosporales>Streptomycetaceae>Streptomyces>unclassified_Streptomyces | 602 | P3T58_RS14570 | 160352..162160 | 3035287 | DUF262-domain-containing-protein-[Streptomyces-sp.-OS603R] |
| KPC88171.1 | ParB+HNH | GCA_001294395.1 | Streptomyces_sp._NRRL_WC-3753 | Bacteria>Bacillati>Actinomycetota>Actinomycetes>Kitasatosporales>Streptomycetaceae>Streptomyces>unclassified_Streptomyces | 605 | ADL35_06460 | 150074..151891 | 1609104 | hypothetical-protein-ADL35_06460-[Streptomyces-sp.-NRRL-WC-3753] |
| WP_266662814.1 | ParB+HNH | GCF_026340075.1 | Streptomyces_sp._NBC_01443 | Bacteria>Bacillati>Actinomycetota>Actinomycetes>Kitasatosporales>Streptomycetaceae>Streptomyces>unclassified_Streptomyces | 607 | OG755_RS09960 | 2247663..2249486 | 2903868 | DUF262-domain-containing-protein-[Streptomyces-sp.-NBC_01443] |
| WP_329194503.1 | ParB+HNH | GCF_036228565.1 | Streptomyces_sp._NBC_01435 | Bacteria>Bacillati>Actinomycetota>Actinomycetes>Kitasatosporales>Streptomycetaceae>Streptomyces>unclassified_Streptomyces | 602 | OG769_RS10330 | 2482521..2484329 | 2903865 | DUF262-domain-containing-protein-[Streptomyces-sp.-NBC_01435] |
| WP_266503380.1 | ParB+HNH | GCF_026341515.1 | Streptomyces_sp._NBC_00878 | Bacteria>Bacillati>Actinomycetota>Actinomycetes>Kitasatosporales>Streptomycetaceae>Streptomyces>unclassified_Streptomyces | 602 | OHA11_RS35710 | 8232760..8234568 | 2975854 | DUF262-domain-containing-protein-[Streptomyces-sp.-NBC_00878] |
| WP_329299359.1 | ParB+HNH | GCF_036226925.1 | Streptomyces_sp._NBC_00659 | Bacteria>Bacillati>Actinomycetota>Actinomycetes>Kitasatosporales>Streptomycetaceae>Streptomyces>unclassified_Streptomyces | 604 | OG410_RS13495 | 3119821..3121635 | 2903669 | DUF262-domain-containing-protein-[Streptomyces-sp.-NBC_00659] |
| WP_266582208.1 | ParB+HNH | GCF_026342135.1 | Streptomyces_sp._NBC_00365 | Bacteria>Bacillati>Actinomycetota>Actinomycetes>Kitasatosporales>Streptomycetaceae>Streptomyces>unclassified_Streptomyces | 605 | OOK36_RS13490 | 2967902..2969719 | 2975726 | DUF262-domain-containing-protein-[Streptomyces-sp.-NBC_00365] |
| WP_266686672.1 | ParB+HNH | GCF_026342335.1 | Streptomyces_sp._NBC_00268 | Bacteria>Bacillati>Actinomycetota>Actinomycetes>Kitasatosporales>Streptomycetaceae>Streptomyces>unclassified_Streptomyces | 604 | OOK47_RS16825 | 3632822..3634636 | 2975695 | DUF262-domain-containing-protein-[Streptomyces-sp.-NBC_00268] |
| WP_328846443.1 | ParB+HNH | GCF_036182465.1 | Streptomyces_sp._NBC_00258 | Bacteria>Bacillati>Actinomycetota>Actinomycetes>Kitasatosporales>Streptomycetaceae>Streptomyces>unclassified_Streptomyces | 602 | OG718_RS39400 | 8837652..8839460 | 2903642 | DUF262-domain-containing-protein-[Streptomyces-sp.-NBC_00258] |
| WP_326723504.1 | ParB+HNH | GCF_035917175.1 | Streptomyces_sp._NBC_00243 | Bacteria>Bacillati>Actinomycetota>Actinomycetes>Kitasatosporales>Streptomycetaceae>Streptomyces>unclassified_Streptomyces | 602 | OHT59_RS33030 | 7421468..7423276 | 2975688 | DUF262-domain-containing-protein-[Streptomyces-sp.-NBC_00243] |
| QKW04625.1 | ParB+HNH | GCA_013364175.1 | Streptomyces_sp._NA02536 | Bacteria>Bacillati>Actinomycetota>Actinomycetes>Kitasatosporales>Streptomycetaceae>Streptomyces>unclassified_Streptomyces | 665 | HUT14_10205 | 2286723..2288720 | 2742133 | DUF262-domain-containing-protein-[Streptomyces-sp.-NA02536] |
| WP_254895759.1 | ParB+HNH | GCF_013364175.1 | Streptomyces_sp._NA02536 | Bacteria>Bacillati>Actinomycetota>Actinomycetes>Kitasatosporales>Streptomycetaceae>Streptomyces>unclassified_Streptomyces | 602 | HUT14_RS10200 | 2286723..2288531 | 2742133 | DUF262-domain-containing-protein-[Streptomyces-sp.-NA02536] |
| WP_215187709.1 | ParB+HNH | GCF_018619155.1 | Streptomyces_sp._McG8 | Bacteria>Bacillati>Actinomycetota>Actinomycetes>Kitasatosporales>Streptomycetaceae>Streptomyces>unclassified_Streptomyces | 605 | HET68_RS00325 | 17350..19167 | 2725487 | DUF262-domain-containing-protein-[Streptomyces-sp.-McG8] |
| WP_166002745.1 | ParB+HNH | GCF_011193355.1 | Streptomyces_sp._JB150 | Bacteria>Bacillati>Actinomycetota>Actinomycetes>Kitasatosporales>Streptomycetaceae>Streptomyces>unclassified_Streptomyces | 605 | G7Z13_RS26590 | 5780330..5782147 | 2714844 | DUF262-domain-containing-protein-[Streptomyces-sp.-JB150] |
| WP_202426314.1 | ParB+HNH | - | Streptomyces_sp._HUCO-GS316 | Bacteria>Bacillati>Actinomycetota>Actinomycetes>Kitasatosporales>Streptomycetaceae>Streptomyces>unclassified_Streptomyces | 597 | GR925_RS33045 | 8979..10772 | 2692198 | DUF262-domain-containing-protein-[Streptomyces-sp.-HUCO-GS316] |
| UOB08694.1 | ParB+HNH | GCA_022807935.1 | Streptomyces_sp._HP-A2021 | Bacteria>Bacillati>Actinomycetota>Actinomycetes>Kitasatosporales>Streptomycetaceae>Streptomyces>unclassified_Streptomyces | 605 | MQE23_06315 | 1362799..1364616 | 2927875 | DUF262-domain-containing-protein-[Streptomyces-sp.-HP-A2021] |
| WP_235479556.1 | ParB+HNH | GCF_021545335.1 | Streptomyces_sp._GQFP | Bacteria>Bacillati>Actinomycetota>Actinomycetes>Kitasatosporales>Streptomycetaceae>Streptomyces>unclassified_Streptomyces | 604 | LUX31_RS36370 | 8059928..8061742 | 2907545 | DUF262-domain-containing-protein-[Streptomyces-sp.-GQFP] |
| WP_336050186.1 | ParB+HNH | GCF_037034225.1 | Streptomyces_sp._CA2R101 | Bacteria>Bacillati>Actinomycetota>Actinomycetes>Kitasatosporales>Streptomycetaceae>Streptomyces>unclassified_Streptomyces | 613 | WA779_RS15040 | 54033..55874 | 3120152 | DUF262-domain-containing-protein-[Streptomyces-sp.-CA2R101] |
| WP_250297838.1 | ParB+HNH | GCF_940670765.1 | Streptomyces_sp._A_4/2 | Bacteria>Bacillati>Actinomycetota>Actinomycetes>Kitasatosporales>Streptomycetaceae>Streptomyces>unclassified_Streptomyces | 602 | NBY69_RS02005 | 299214..301022 | 2934314 | DUF262-domain-containing-protein-[Streptomyces-sp.-A-4/2] |
| WP_093779308.1 | ParB+HNH | GCF_900101975.1 | Streptomyces_sp._136MFCol5.1 | Bacteria>Bacillati>Actinomycetota>Actinomycetes>Kitasatosporales>Streptomycetaceae>Streptomyces>unclassified_Streptomyces | 605 | BLQ64_RS33270 | 49162..50979 | 1172182 | DUF262-domain-containing-protein-[Streptomyces-sp.-136MFCol5.1] |
| WP_108718590.1 | ParB+HNH | GCF_003074035.1 | Miniimonas_sp._S16 | Bacteria>Bacillati>Actinomycetota>Actinomycetes>Micrococcales>Beutenbergiaceae>Miniimonas>unclassified_Miniimonas | 662 | DCS86_RS08150 | 1889293..1891281 | 2171623 | DUF262-domain-containing-protein-[Miniimonas-sp.-S16] |
| WP_152196010.1 | ParB+HNH | GCF_009193155.1 | Georgenia_subflava | Bacteria>Bacillati>Actinomycetota>Actinomycetes>Micrococcales>Bogoriellaceae>Georgenia | 354 | GD748_RS14920 | 338989..340053 | 1622177 | DUF262-domain-containing-protein-[Georgenia-subflava] |
| WP_272902429.1 | ParB+HNH | - | Brevibacterium_daeguense | Bacteria>Bacillati>Actinomycetota>Actinomycetes>Micrococcales>Brevibacteriaceae>Brevibacterium | 188 | LXN01_RS16180 | 43754..44320 | 909936 | HNH-endonuclease-family-protein-[Brevibacterium-daeguense] |
| WP_236865288.1 | ParB+HNH | GCF_039537685.1 | Brevibacterium_daeguense | Bacteria>Bacillati>Actinomycetota>Actinomycetes>Micrococcales>Brevibacteriaceae>Brevibacterium | 103 | LXN01_RS11495 | 42556..42867 | 909936 | DUF262-domain-containing-protein-[Brevibacterium-daeguense] |
| QZE24958.1 | ParB+HNH | GCA_019720815.1 | Brevibacterium_casei | Bacteria>Bacillati>Actinomycetota>Actinomycetes>Micrococcales>Brevibacteriaceae>Brevibacterium | 592 | K4X33_12610 | 2730094..2731872 | 33889 | DUF262-domain-containing-protein-[Brevibacterium-casei] |
| WP_144588168.1 | ParB+HNH | GCF_002339175.1 | Brevibacterium_casei | Bacteria>Bacillati>Actinomycetota>Actinomycetes>Micrococcales>Brevibacteriaceae>Brevibacterium | 592 | B3D11_RS04575 | 71196..72974 | 33889 | DUF262-domain-containing-protein-[Brevibacterium-casei] |
| WP_052367169.1 | ParB+HNH | GCF_000701465.1 | Paraoerskovia_marina | Bacteria>Bacillati>Actinomycetota>Actinomycetes>Micrococcales>Cellulomonadaceae>Paraoerskovia | 372 | Q343_RS14170 | 105399..106517 | 545619 | DUF262-domain-containing-protein-[Paraoerskovia-marina] |
| WP_052589427.1 | ParB+HNH | GCF_001190945.1 | Luteipulveratus_mongoliensis | Bacteria>Bacillati>Actinomycetota>Actinomycetes>Micrococcales>Dermacoccaceae>Luteipulveratus | 655 | VV02_RS01640 | 341152..343119 | 571913 | DUF262-domain-containing-protein-[Luteipulveratus-mongoliensis] |
| WP_141928903.1 | ParB+HNH | GCF_039538725.1 | Yimella_lutea | Bacteria>Bacillati>Actinomycetota>Actinomycetes>Micrococcales>Dermacoccaceae>Yimella | 653 | FB459_RS13785 | 2859962..2861923 | 587872 | DUF262-domain-containing-protein-[Yimella-lutea] |
| MBK6441772.1 | ParB+HNH | GCA_016704565.1 | Candidatus_Phosphoribacter_baldrii | Bacteria>Bacillati>Actinomycetota>Actinomycetes>Micrococcales>Dermatophilaceae>Candidatus_Phosphoribacter | 632 | IPF90_06365 | 1385305..1387203 | 2953744 | DUF262-domain-containing-protein-[Actinomycetales-bacterium] |
| WP_029212621.1 | ParB+HNH | GCF_000426385.1 | Arsenicicoccus_bolidensis | Bacteria>Bacillati>Actinomycetota>Actinomycetes>Micrococcales>Intrasporangiaceae>Arsenicicoccus | 613 | K313_RS0116905 | 17686..19527 | 229480 | DUF262-domain-containing-protein-[Arsenicicoccus-bolidensis] |
| OFE15917.1 | ParB+HNH | GCA_001758225.1 | Humibacillus_sp._DSM_29435 | Bacteria>Bacillati>Actinomycetota>Actinomycetes>Micrococcales>Intrasporangiaceae>Humibacillus>unclassified_Humibacillus | 615 | BA895_05300 | 168071..169918 | 1869167 | hypothetical-protein-BA895_05300-[Humibacillus-sp.-DSM-29435] |
| WP_070192622.1 | ParB+HNH | GCF_001758225.1 | Humibacillus_sp._DSM_29435 | Bacteria>Bacillati>Actinomycetota>Actinomycetes>Micrococcales>Intrasporangiaceae>Humibacillus>unclassified_Humibacillus | 615 | BA895_RS14650 | 168071..169918 | 1869167 | DUF262-domain-containing-protein-[Humibacillus-sp.-DSM-29435] |
| WP_034802769.1 | ParB+HNH | GCF_000576595.1 | Intrasporangium_oryzae | Bacteria>Bacillati>Actinomycetota>Actinomycetes>Micrococcales>Intrasporangiaceae>Intrasporangium | 561 | N865_RS05955 | 108543..110228 | 412687 | DUF262-domain-containing-protein-[Intrasporangium-oryzae] |
| WP_191562881.1 | ParB+HNH | GCF_904423795.1 | Janibacter_melonis | Bacteria>Bacillati>Actinomycetota>Actinomycetes>Micrococcales>Intrasporangiaceae>Janibacter | 611 | IHQ27_RS00710 | 69725..71560 | 262209 | DUF262-domain-containing-protein-[Janibacter-melonis] |
| WP_106298857.1 | ParB+HNH | GCF_003002895.1 | Knoellia_remsis | Bacteria>Bacillati>Actinomycetota>Actinomycetes>Micrococcales>Intrasporangiaceae>Knoellia | 613 | BCF74_RS15505 | 19936..21777 | 407159 | DUF262-domain-containing-protein-[Knoellia-remsis] |
| MCI1262625.1 | ParB+HNH | GCA_022641475.1 | Nostocoides_jenkinsii | Bacteria>Bacillati>Actinomycetota>Actinomycetes>Micrococcales>Intrasporangiaceae>Nostocoides | 613 | LKG20_10185 | 16696..18537 | 330834 | DUF262-domain-containing-protein-[Tetrasphaera-jenkinsii] |
| WP_159902717.1 | ParB+HNH | GCF_039541275.1 | Ornithinibacter_aureus | Bacteria>Bacillati>Actinomycetota>Actinomycetes>Micrococcales>Intrasporangiaceae>Ornithinibacter | 661 | C8E84_RS12860 | 2705167..2707152 | 622664 | DUF262-domain-containing-protein-[Ornithinibacter-aureus] |
| WP_141784691.1 | ParB+HNH | GCF_039526025.1 | Ornithinicoccus_hortensis | Bacteria>Bacillati>Actinomycetota>Actinomycetes>Micrococcales>Intrasporangiaceae>Ornithinicoccus | 615 | FB467_RS08330 | 1803570..1805417 | 82346 | DUF262-domain-containing-protein-[Ornithinicoccus-hortensis] |
| WP_164896083.1 | ParB+HNH | GCF_004025965.2 | Phycicoccus_flavus | Bacteria>Bacillati>Actinomycetota>Actinomycetes>Micrococcales>Intrasporangiaceae>Phycicoccus | 676 | EPD83_RS02110 | 94725..96755 | 2502783 | DUF262-domain-containing-protein-[Phycicoccus-flavus] |
| WP_144794199.1 | ParB+HNH | GCF_023108875.1 | Microbacterium | Bacteria>Bacillati>Actinomycetota>Actinomycetes>Micrococcales>Microbacteriaceae | 537 | MZK47_RS13290 | 4628..6241 | 33882 | MULTISPECIES:-DUF262-domain-containing-protein-[Microbacterium] |
| WP_309976838.1 | ParB+HNH | GCF_031456545.1 | Agromyces_sp._3263 | Bacteria>Bacillati>Actinomycetota>Actinomycetes>Micrococcales>Microbacteriaceae>Agromyces>unclassified_Agromyces | 766 | J2X63_RS10560 | 2298567..2300867 | 2817750 | DUF262-domain-containing-protein-[Agromyces-sp.-3263] |
| WP_309976836.1 | ParB+HNH | GCF_031456545.1 | Agromyces_sp._3263 | Bacteria>Bacillati>Actinomycetota>Actinomycetes>Micrococcales>Microbacteriaceae>Agromyces>unclassified_Agromyces | 581 | J2X63_RS10555 | 2296825..2298570 | 2817750 | DUF262-domain-containing-protein-[Agromyces-sp.-3263] |
| WBM79846.1 | ParB+HNH | GCA_027594785.1 | Cryobacterium_breve | Bacteria>Bacillati>Actinomycetota>Actinomycetes>Micrococcales>Microbacteriaceae>Cryobacterium | 625 | KIV56_17010 | 3680800..3682677 | 1259258 | DUF262-domain-containing-protein-[Cryobacterium-breve] |
| WP_281534459.1 | ParB+HNH | GCF_027594785.1 | Cryobacterium_breve | Bacteria>Bacillati>Actinomycetota>Actinomycetes>Micrococcales>Microbacteriaceae>Cryobacterium | 625 | KIV56_RS17880 | 3680800..3682677 | 1259258 | DUF262-domain-containing-protein-[Cryobacterium-breve] |
| WP_210415909.1 | ParB+HNH | GCF_007859675.1 | Humibacter_ginsenosidimutans | Bacteria>Bacillati>Actinomycetota>Actinomycetes>Micrococcales>Microbacteriaceae>Humibacter | 447 | FPZ11_RS19550 | 3801027..3802370 | 2599293 | DUF262-domain-containing-protein-[Humibacter-ginsenosidimutans] |
| WP_053352384.1 | ParB+HNH | GCF_001273845.1 | Leucobacter_musarum | Bacteria>Bacillati>Actinomycetota>Actinomycetes>Micrococcales>Microbacteriaceae>Leucobacter | 617 | CH42_RS06390 | 77567..79420 | 1930747 | DUF262-domain-containing-protein-[Leucobacter-musarum] |
| QAY61396.1 | ParB+HNH | GCA_004135285.1 | Microbacterium_protaetiae | Bacteria>Bacillati>Actinomycetota>Actinomycetes>Micrococcales>Microbacteriaceae>Microbacterium | 592 | ET475_16420 | 3520243..3522021 | 2509458 | DUF262-domain-containing-protein-[Microbacterium-protaetiae] |
| WP_129392752.1 | ParB+HNH | GCF_004135285.1 | Microbacterium_protaetiae | Bacteria>Bacillati>Actinomycetota>Actinomycetes>Micrococcales>Microbacteriaceae>Microbacterium | 592 | ET475_RS16420 | 3520243..3522021 | 2509458 | DUF262-domain-containing-protein-[Microbacterium-protaetiae] |
| MBY6061977.1 | ParB+HNH | GCA_019801485.1 | Microbacterium_esteraromaticum | Bacteria>Bacillati>Actinomycetota>Actinomycetes>Micrococcales>Microbacteriaceae>Microbacterium | 613 | KUV70_11035 | 280510..282351 | 57043 | DUF262-domain-containing-HNH-endonuclease-family-protein-[Microbacterium-esteraromaticum] |
| WP_261393917.1 | ParB+HNH | - | Microbacterium_esteraromaticum | Bacteria>Bacillati>Actinomycetota>Actinomycetes>Micrococcales>Microbacteriaceae>Microbacterium | 606 | KUV70_RS11035 | 280510..282330 | 57043 | DUF262-domain-containing-protein-[Microbacterium-esteraromaticum] |
| PPG31571.1 | ParB+HNH | GCA_002931595.1 | Rathayibacter_sp._AY2B9 | Bacteria>Bacillati>Actinomycetota>Actinomycetes>Micrococcales>Microbacteriaceae>Rathayibacter>unclassified_Rathayibacter | 853 | C5C25_07420 | 32786..35347 | 2080572 | hypothetical-protein-C5C25_07420-[Rathayibacter-sp.-AY2B9] |
| WP_144820007.1 | ParB+HNH | GCF_024655705.1 | Micrococcus | Bacteria>Bacillati>Actinomycetota>Actinomycetes>Micrococcales>Micrococcaceae | 706 | NUV33_RS00285 | 7170..9290 | 1269 | DUF262-domain-containing-protein-[Micrococcus-luteus] |
| WP_144820008.1 | ParB+HNH | GCF_024655705.1 | Micrococcus | Bacteria>Bacillati>Actinomycetota>Actinomycetes>Micrococcales>Micrococcaceae | 500 | NUV33_RS00290 | 9287..10789 | 1269 | DUF262-domain-containing-protein-[Micrococcus-luteus] |
| WP_104138816.1 | ParB+HNH | GCF_002929275.1 | Arthrobacter_sp._ZGTC131 | Bacteria>Bacillati>Actinomycetota>Actinomycetes>Micrococcales>Micrococcaceae>Arthrobacter>unclassified_Arthrobacter | 617 | CXQ90_RS09935 | 148156..150009 | 2058898 | DUF262-domain-containing-protein-[Arthrobacter-sp.-ZGTC131] |
| WP_090584039.1 | ParB+HNH | GCF_900102895.1 | Arthrobacter_sp._ov407 | Bacteria>Bacillati>Actinomycetota>Actinomycetes>Micrococcales>Micrococcaceae>Arthrobacter>unclassified_Arthrobacter | 667 | BLS42_RS19725 | 27304..29307 | 1761748 | DUF262-domain-containing-protein-[Arthrobacter-sp.-ov407] |
| WP_224048799.1 | ParB+HNH | GCF_019977335.1 | Arthrobacter_sp._NicSoilB4 | Bacteria>Bacillati>Actinomycetota>Actinomycetes>Micrococcales>Micrococcaceae>Arthrobacter>unclassified_Arthrobacter | 613 | LD013_RS04130 | 897435..899276 | 2830997 | DUF262-domain-containing-protein-[Arthrobacter-sp.-NicSoilB4] |
| BBE21860.1 | ParB+HNH | GCA_004001285.1 | Arthrobacter_sp._MN05-02 | Bacteria>Bacillati>Actinomycetota>Actinomycetes>Micrococcales>Micrococcaceae>Arthrobacter>unclassified_Arthrobacter | 713 | MN0502_07430 | 792641..794782 | 1571833 | hypothetical-protein-MN0502_07430-[Arthrobacter-sp.-MN05-02] |
| WP_262103913.1 | ParB+HNH | GCF_946892675.1 | Arthrobacter_sp._Marseille-P9274 | Bacteria>Bacillati>Actinomycetota>Actinomycetes>Micrococcales>Micrococcaceae>Arthrobacter>unclassified_Arthrobacter | 570 | OC550_RS03475 | 762664..764376 | 2866572 | DUF262-domain-containing-protein-[Arthrobacter-sp.-Marseille-P9274] |
| ALV43765.1 | ParB+HNH | GCA_001484605.1 | Pseudarthrobacter_sulfonivorans | Bacteria>Bacillati>Actinomycetota>Actinomycetes>Micrococcales>Micrococcaceae>Pseudarthrobacter | 617 | AU252_05140 | 1169113..1170966 | 121292 | hypothetical-protein-AU252_05140-[Pseudarthrobacter-sulfonivorans] |
| WP_058932759.1 | ParB+HNH | GCF_001484605.1 | Pseudarthrobacter_sulfonivorans | Bacteria>Bacillati>Actinomycetota>Actinomycetes>Micrococcales>Micrococcaceae>Pseudarthrobacter | 617 | AU252_RS05140 | 1169113..1170966 | 121292 | DUF262-domain-containing-protein-[Pseudarthrobacter-sulfonivorans] |
| QOD04366.1 | ParB+HNH | GCA_014764445.1 | Pseudarthrobacter_sp._BIM_B-2242 | Bacteria>Bacillati>Actinomycetota>Actinomycetes>Micrococcales>Micrococcaceae>Pseudarthrobacter>unclassified_Pseudarthrobacter | 699 | IDT60_04710 | 990531..992630 | 2772401 | DUF262-domain-containing-protein-[Pseudarthrobacter-sp.-BIM-B-2242] |
| WP_191081068.1 | ParB+HNH | GCF_014764445.1 | Pseudarthrobacter_sp._BIM_B-2242 | Bacteria>Bacillati>Actinomycetota>Actinomycetes>Micrococcales>Micrococcaceae>Pseudarthrobacter>unclassified_Pseudarthrobacter | 699 | IDT60_RS04710 | 990531..992630 | 2772401 | DUF262-and-DUF1524-domain-containing-protein-[Pseudarthrobacter-sp.-BIM-B-2242] |

| Protein-ID | Domain | Assembly-ID | Strain | Taxonomy | Length | Locus-Tag | Genomic-coordinates | TaxID | Fasta-defline |
|---|---|---|---|---|---|---|---|---|---|
| WP_188427985.1 | ParB+HNH | GCF_039543725.1 | Ornithinimicrobium_tianjinense | Bacteria>Bacillati>Actinomycetota>Actinomycetes>Micrococcales>Ornithinimicrobiaceae>Ornithinimicrobium | 558 | IEV65_RS02300 | 473497..475173 | 1195761 | DUF262-domain-containing-protein-[Ornithinimicrobium-tianjinense] |
| SOC53727.1 | ParB+HNH | GCA_900221015.1 | Ornithinimicrobium_cerasi | Bacteria>Bacillati>Actinomycetota>Actinomycetes>Micrococcales>Ornithinimicrobiaceae>Ornithinimicrobium | 785 | SAMN05421879_10288 | 105005..107362 | 2248773 | Protein-of-unknown-function-DUF262-[Ornithinimicrobium-cerasi] |
| WP_075958592.1 | ParB+HNH | GCF_001942405.1 | Ornithinimicrobium_sp._CNJ-824 | Bacteria>Bacillati>Actinomycetota>Actinomycetes>Micrococcales>Ornithinimicrobiaceae>Ornithinimicrobium>unclassified_Ornithinimicrobium | 592 | BJF81_RS12355 | 110569..112347 | 1904966 | DUF262-domain-containing-protein-[Ornithinimicrobium-sp.-CNJ-824] |
| WP_022924593.1 | ParB+HNH | GCF_000421245.1 | Serinicoccus_marinus | Bacteria>Bacillati>Actinomycetota>Actinomycetes>Micrococcales>Ornithinimicrobiaceae>Serinicoccus | 635 | FU792_RS06185 | 1329873..1331780 | 247333 | DUF262-domain-containing-protein-[Serinicoccus-marinus] |
| GAA4873515.1 | ParB+HNH | GCA_039544515.1 | Serinicoccus_chungangensis | Bacteria>Bacillati>Actinomycetota>Actinomycetes>Micrococcales>Ornithinimicrobiaceae>Serinicoccus | 684 | GCM10025788_09870 | 609313..611367 | 767452 | DUF262-domain-containing-protein-[Serinicoccus-chungangensis] |
| WP_139720894.1 | ParB+HNH | GCF_039544515.1 | Serinicoccus_chungangensis | Bacteria>Bacillati>Actinomycetota>Actinomycetes>Micrococcales>Ornithinimicrobiaceae>Serinicoccus | 684 | FHD63_RS05790 | 1309132..1311186 | 767452 | DUF262-domain-containing-protein-[Serinicoccus-chungangensis] |
| GAA4873519.1 | ParB+HNH | GCA_039544515.1 | Serinicoccus_chungangensis | Bacteria>Bacillati>Actinomycetota>Actinomycetes>Micrococcales>Ornithinimicrobiaceae>Serinicoccus | 443 | GCM10025788_09880 | 611369..612700 | 767452 | hypothetical-protein-GCM10025788_09880-[Serinicoccus-chungangensis] |
| WP_139720896.1 | ParB+HNH | GCF_039544515.1 | Serinicoccus_chungangensis | Bacteria>Bacillati>Actinomycetota>Actinomycetes>Micrococcales>Ornithinimicrobiaceae>Serinicoccus | 443 | FHD63_RS05795 | 1311188..1312519 | 767452 | DUF262-domain-containing-protein-[Serinicoccus-chungangensis] |
| WP_139720902.1 | ParB+HNH | GCF_039544515.1 | Serinicoccus_chungangensis | Bacteria>Bacillati>Actinomycetota>Actinomycetes>Micrococcales>Ornithinimicrobiaceae>Serinicoccus | 441 | FHD63_RS05810 | 1320410..1321735 | 767452 | DUF1524-domain-containing-protein-[Serinicoccus-chungangensis] |
| GAA4873477.1 | ParB+HNH | GCA_039544515.1 | Serinicoccus_chungangensis | Bacteria>Bacillati>Actinomycetota>Actinomycetes>Micrococcales>Ornithinimicrobiaceae>Serinicoccus | 418 | GCM10025788_09810 | 596220..597476 | 767452 | hypothetical-protein-GCM10025788_09810-[Serinicoccus-chungangensis] |
| WP_139720872.1 | ParB+HNH | GCF_039544515.1 | Serinicoccus_chungangensis | Bacteria>Bacillati>Actinomycetota>Actinomycetes>Micrococcales>Ornithinimicrobiaceae>Serinicoccus | 418 | FHD63_RS05755 | 1296039..1297295 | 767452 | DUF262-domain-containing-protein-[Serinicoccus-chungangensis] |
| SDF11604.1 | ParB+HNH | GCA_900102445.1 | Cellulosimicrobium_cellulans | Bacteria>Bacillati>Actinomycetota>Actinomycetes>Micrococcales>Promicromonosporaceae>Cellulosimicrobium | 652 | SAMN04487781_0339 | 371534..373492 | 1710 | hypothetical-protein-SAMN04487781_0339-[Cellulosimicrobium-cellulans] |
| WP_076404376.1 | ParB+HNH | GCF_900155925.1 | Cellulosimicrobium_aquatile | Bacteria>Bacillati>Actinomycetota>Actinomycetes>Micrococcales>Promicromonosporaceae>Cellulosimicrobium | 658 | BXA05_RS06495 | 512011..513987 | 1612203 | DUF262-domain-containing-protein-[Cellulosimicrobium-aquatile] |
| QAY71607.1 | ParB+HNH | GCA_004135385.1 | Xylanimonas_protaetiae | Bacteria>Bacillati>Actinomycetota>Actinomycetes>Micrococcales>Promicromonosporaceae>Xylanimonas | 613 | ET471_17510 | 3779661..3781502 | 2509457 | DUF262-domain-containing-protein-[Xylanimonas-protaetiae] |
| WP_129190459.1 | ParB+HNH | GCF_004135385.1 | Xylanimonas_protaetiae | Bacteria>Bacillati>Actinomycetota>Actinomycetes>Micrococcales>Promicromonosporaceae>Xylanimonas | 613 | ET471_RS17510 | 3779661..3781502 | 2509457 | DUF262-domain-containing-protein-[Xylanimonas-protaetiae] |
| WP_147915797.1 | ParB+HNH | GCF_008000995.1 | Ruania_zhangjianzhongii | Bacteria>Bacillati>Actinomycetota>Actinomycetes>Micrococcales>Ruaniaceae>Ruania | 651 | FU260_RS03475 | 748474..750429 | 2603206 | DUF262-domain-containing-protein-[Ruania-zhangjianzhongii] |
| WP_023560517.1 | ParB+HNH | GCF_000494755.1 | Actinoplanes_friuliensis | Bacteria>Bacillati>Actinomycetota>Actinomycetes>Micromonosporales>Micromonosporaceae>Actinoplanes | 662 | AFR_RS29715 | 6444570..6446558 | 196914 | DUF262-domain-containing-protein-[Actinoplanes-friuliensis] |
| WP_190252035.1 | ParB+HNH | GCF_014647515.1 | Dactylosporangium_sucinum | Bacteria>Bacillati>Actinomycetota>Actinomycetes>Micromonosporales>Micromonosporaceae>Dactylosporangium | 660 | IEZ31_RS23195 | 138787..140769 | 1424081 | DUF262-domain-containing-protein-[Dactylosporangium-sucinum] |
| WP_208577497.1 | ParB+HNH | GCF_017599015.1 | Micromonospora_tulbaghiae | Bacteria>Bacillati>Actinomycetota>Actinomycetes>Micromonosporales>Micromonosporaceae>Micromonospora | 562 | J5U46_RS17350 | 26522..28210 | 479978 | DUF262-domain-containing-protein-[Micromonospora-tulbaghiae] |
| WP_208576857.1 | ParB+HNH | GCF_017599015.1 | Micromonospora_tulbaghiae | Bacteria>Bacillati>Actinomycetota>Actinomycetes>Micromonosporales>Micromonosporaceae>Micromonospora | 560 | J5U46_RS08625 | 85032..86714 | 479978 | DUF262-domain-containing-protein-[Micromonospora-tulbaghiae] |
| WP_030502748.1 | ParB+HNH | - | Micromonospora_purpureochromogenes | Bacteria>Bacillati>Actinomycetota>Actinomycetes>Micromonosporales>Micromonosporaceae>Micromonospora | 560 | IH31_RS0121320 | 109320..111002 | 47872 | DUF262-domain-containing-protein-[Micromonospora-purpureochromogenes] |
| WP_088962575.1 | ParB+HNH | GCF_900091515.1 | Micromonospora_purpureochromogenes | Bacteria>Bacillati>Actinomycetota>Actinomycetes>Micromonosporales>Micromonosporaceae>Micromonospora | 552 | GA0074696_RS20365 | 4408457..4410115 | 47872 | DUF262-domain-containing-protein-[Micromonospora-purpureochromogenes] |
| WP_091191940.1 | ParB+HNH | GCF_900089605.1 | Micromonospora_narathiwatensis | Bacteria>Bacillati>Actinomycetota>Actinomycetes>Micromonosporales>Micromonosporaceae>Micromonospora | 560 | GA0070621_RS04905 | 1075488..1077170 | 299146 | DUF262-domain-containing-protein-[Micromonospora-narathiwatensis] |
| SCF44507.1 | ParB+HNH | GCA_900091555.1 | Micromonospora_mirobrigensis | Bacteria>Bacillati>Actinomycetota>Actinomycetes>Micromonosporales>Micromonosporaceae>Micromonospora | 576 | GA0070564_110101 | 117009..118739 | 262898 | Protein-of-unknown-function-DUF262-[Micromonospora-mirobrigensis] |
| WP_030271469.1 | ParB+HNH | GCF_000717585.1 | Micromonospora_globosa | Bacteria>Bacillati>Actinomycetota>Actinomycetes>Micromonosporales>Micromonosporaceae>Micromonospora | 560 | IF06_RS0114125 | 18627..20309 | 47863 | DUF262-domain-containing-protein-[Micromonospora-globosa] |
| WP_120331051.1 | ParB+HNH | GCF_003610785.1 | Micromonospora_globbae | Bacteria>Bacillati>Actinomycetota>Actinomycetes>Micromonosporales>Micromonosporaceae>Micromonospora | 560 | D7I43_RS25245 | 104862..106544 | 1894969 | DUF262-domain-containing-protein-[Micromonospora-globbae] |
| WP_088980559.1 | ParB+HNH | GCF_900091495.1 | Micromonospora_echinospora | Bacteria>Bacillati>Actinomycetota>Actinomycetes>Micromonosporales>Micromonosporaceae>Micromonospora | 580 | GA0070618_RS04845 | 1082733..1084475 | 1877 | DUF262-domain-containing-protein-[Micromonospora-echinospora] |
| WP_187693212.1 | ParB+HNH | GCF_014472405.1 | Micromonospora_chalcea | Bacteria>Bacillati>Actinomycetota>Actinomycetes>Micromonosporales>Micromonosporaceae>Micromonospora | 560 | H9X95_RS12025 | 17477..19159 | 1874 | DUF262-domain-containing-protein-[Micromonospora-chalcea] |
| WP_076473575.1 | ParB+HNH | GCF_900156065.1 | Micromonospora_avicenniae | Bacteria>Bacillati>Actinomycetota>Actinomycetes>Micromonosporales>Micromonosporaceae>Micromonospora | 568 | BXA10_RS26500 | 29275..30981 | 1198245 | DUF262-domain-containing-protein-[Micromonospora-avicenniae] |
| WP_204020359.1 | ParB+HNH | GCF_016863595.1 | Micromonospora_andamanensis | Bacteria>Bacillati>Actinomycetota>Actinomycetes>Micromonosporales>Micromonosporaceae>Micromonospora | 662 | Vwe01_RS03460 | 13595..15583 | 1287068 | DUF262-domain-containing-protein-[Micromonospora-andamanensis] |
| WP_064446383.1 | ParB+HNH | GCF_001652995.1 | Micromonospora_sp._NBRC_110037 | Bacteria>Bacillati>Actinomycetota>Actinomycetes>Micromonosporales>Micromonosporaceae>Micromonospora>unclassified_Micromonospora | 560 | A4R25_RS09720 | 2382753..2384435 | 1621261 | DUF262-domain-containing-protein-[Micromonospora-sp.-NBRC-110037] |
| WP_221380245.1 | ParB+HNH | GCF_019710575.1 | Paractinoplanes_polyasparticus | Bacteria>Bacillati>Actinomycetota>Actinomycetes>Micromonosporales>Micromonosporaceae>Paractinoplanes | 661 | KXC43_RS33635 | 34567..36552 | 2856853 | DUF262-domain-containing-protein-[Actinoplanes-polyasparticus] |
| WP_107269539.1 | ParB+HNH | GCF_003030345.1 | Plantactinospora_sp._BC1 | Bacteria>Bacillati>Actinomycetota>Actinomycetes>Micromonosporales>Micromonosporaceae>Plantactinospora>unclassified_Plantactinospora | 657 | C6361_RS28020 | 6380654..6382627 | 2108470 | DUF262-domain-containing-protein-[Plantactinospora-sp.-BC1] |
| WP_107261306.1 | ParB+HNH | GCF_003030385.1 | Plantactinospora_sp._BB1 | Bacteria>Bacillati>Actinomycetota>Actinomycetes>Micromonosporales>Micromonosporaceae>Plantactinospora>unclassified_Plantactinospora | 657 | C6W10_RS25960 | 5943871..5945844 | 2071627 | DUF262-domain-containing-protein-[Plantactinospora-sp.-BB1] |
| WP_079635264.1 | ParB+HNH | GCF_900141585.1 | Mycobacteriaceae | Bacteria>Bacillati>Actinomycetota>Actinomycetes>Mycobacteriales | 613 | B5Z43_RS22920 | 50608..52449 | 1762 | DUF262-domain-containing-protein-[Mycobacteroides-abscessus] |
| WP_259898407.1 | ParB+HNH | GCF_025152505.1 | Corynebacterium_sanguinis | Bacteria>Bacillati>Actinomycetota>Actinomycetes>Mycobacteriales>Corynebacteriaceae>Corynebacterium | 546 | M3G46_RS10580 | 275..1915 | 2594913 | DUF262-domain-containing-HNH-endonuclease-family-protein-[Corynebacterium-sanguinis] |
| WP_158726428.1 | ParB+HNH | GCF_009765205.1 | Tomitella_fengzijianii | Bacteria>Bacillati>Actinomycetota>Actinomycetes>Mycobacteriales>Corynebacteriales_incertae_sedis>Tomitella | 623 | GSQ71_RS04960 | 50262..52133 | 2597660 | DUF262-domain-containing-protein-[Tomitella-fengzijianii] |
| WP_158726436.1 | ParB+HNH | GCF_009765205.1 | Tomitella_fengzijianii | Bacteria>Bacillati>Actinomycetota>Actinomycetes>Mycobacteriales>Corynebacteriales_incertae_sedis>Tomitella | 374 | GSQ71_RS05000 | 64074..65198 | 2597660 | DUF262-domain-containing-protein-[Tomitella-fengzijianii] |
| WP_106386061.1 | ParB+HNH | GCF_002999155.1 | Dietzia | Bacteria>Bacillati>Actinomycetota>Actinomycetes>Mycobacteriales>Dietziaceae | 606 | C3V38_RS13970 | 2993285..2995105 | 37914 | MULTISPECIES:-DUF262-domain-containing-protein-[Dietzia] |
| WP_213184400.1 | ParB+HNH | GCF_018390595.1 | Dietzia_massiliensis | Bacteria>Bacillati>Actinomycetota>Actinomycetes>Mycobacteriales>Dietziaceae>Dietzia | 625 | KHA85_RS00310 | 76688..78565 | 2697499 | DUF262-domain-containing-protein-[Dietzia-massiliensis] |
| WP_213184732.1 | ParB+HNH | GCF_018390595.1 | Dietzia_massiliensis | Bacteria>Bacillati>Actinomycetota>Actinomycetes>Mycobacteriales>Dietziaceae>Dietzia | 595 | KHA85_RS02750 | 10034..11821 | 2697499 | DUF262-domain-containing-protein-[Dietzia-massiliensis] |
| WP_154827467.1 | ParB+HNH | GCF_039531565.1 | Dietzia_kunjamensis | Bacteria>Bacillati>Actinomycetota>Actinomycetes>Mycobacteriales>Dietziaceae>Dietzia | 647 | FN843_RS00165 | 46893..48836 | 322509 | DUF262-domain-containing-protein-[Dietzia-kunjamensis] |
| MCT2273889.1 | ParB+HNH | GCA_025151825.1 | Dietzia_cinnamea | Bacteria>Bacillati>Actinomycetota>Actinomycetes>Mycobacteriales>Dietziaceae>Dietzia | 647 | M3F76_05295 | 2048..3991 | 321318 | DUF262-domain-containing-HNH-endonuclease-family-protein-[Dietzia-cinnamea] |
| WP_259881458.1 | ParB+HNH | GCF_025145775.1 | Dietzia_cinnamea | Bacteria>Bacillati>Actinomycetota>Actinomycetes>Mycobacteriales>Dietziaceae>Dietzia | 647 | M3B38_RS03610 | 47057..49000 | 321318 | DUF262-domain-containing-protein-[Dietzia-cinnamea] |
| OAH45660.1 | ParB+HNH | GCA_001643175.1 | Dietzia_cinnamea | Bacteria>Bacillati>Actinomycetota>Actinomycetes>Mycobacteriales>Dietziaceae>Dietzia | 595 | AYJ66_16785 | 30876..32663 | 321318 | hypothetical-protein-AYJ66_16785-[Dietzia-cinnamea] |
| HJC59999.1 | ParB+HNH | GCA_019119875.1 | Candidatus_Dietzia_intestinigallinarum | Bacteria>Bacillati>Actinomycetota>Actinomycetes>Mycobacteriales>Dietziaceae>Dietzia | 593 | H9755_06665 | 15540..17321 | 2838537 | DUF262-domain-containing-protein-[Candidatus-Dietzia-intestinigallinarum] |
| MBB1018373.1 | ParB+HNH | GCA_014144995.1 | Dietzia_sp._DQ11-71 | Bacteria>Bacillati>Actinomycetota>Actinomycetes>Mycobacteriales>Dietziaceae>Dietzia>unclassified_Dietzia | 590 | G6023_08705 | 19070..20842 | 1630638 | DUF262-domain-containing-protein-[Dietzia-sp.-DQ11-71] |
| WP_067717289.1 | ParB+HNH | GCF_001658045.1 | Dietzia_sp._111N12-1 | Bacteria>Bacillati>Actinomycetota>Actinomycetes>Mycobacteriales>Dietziaceae>Dietzia>unclassified_Dietzia | 449 | AYO52_RS11390 | 791..2140 | 1785156 | DUF262-domain-containing-protein-[Dietzia-sp.-111N12-1] |
| WP_172812153.1 | ParB+HNH | GCF_001658045.1 | Dietzia_sp._111N12-1 | Bacteria>Bacillati>Actinomycetota>Actinomycetes>Mycobacteriales>Dietziaceae>Dietzia>unclassified_Dietzia | 267 | AYO52_RS11385 | 1..801 | 1785156 | DUF262-domain-containing-protein,-partial-[Dietzia-sp.-111N12-1] |
| WP_277242421.1 | ParB+HNH | GCF_029427115.1 | Gordonia_hongkongensis | Bacteria>Bacillati>Actinomycetota>Actinomycetes>Mycobacteriales>Gordoniaceae>Gordonia | 562 | L2299_RS00565 | 123683..125371 | 1701090 | DUF262-domain-containing-HNH-endonuclease-family-protein-[Gordonia-hongkongensis] |
| WP_279957424.1 | ParB+HNH | GCF_029853985.1 | Gordonia_alkanivorans | Bacteria>Bacillati>Actinomycetota>Actinomycetes>Mycobacteriales>Gordoniaceae>Gordonia | 600 | QEN35_RS21140 | 22604..24406 | 84096 | DUF262-domain-containing-HNH-endonuclease-family-protein-[Gordonia-alkanivorans] |
| WP_198156549.1 | ParB+HNH | GCF_001653095.1 | Hoyosella_altamirensis | Bacteria>Bacillati>Actinomycetota>Actinomycetes>Mycobacteriales>Hoyosellaceae>Hoyosella | 473 | HAL02_RS18860 | 9066..10487 | 616997 | DUF262-domain-containing-protein-[Hoyosella-altamirensis] |
| WP_198156550.1 | ParB+HNH | GCF_001653095.1 | Hoyosella_altamirensis | Bacteria>Bacillati>Actinomycetota>Actinomycetes>Mycobacteriales>Hoyosellaceae>Hoyosella | 238 | HAL02_RS24320 | 10450..11166 | 616997 | DUF1524-domain-containing-protein-[Hoyosella-altamirensis] |
| WP_073906997.1 | ParB+HNH | GCF_020884715.1 | Mycolicibacterium | Bacteria>Bacillati>Actinomycetota>Actinomycetes>Mycobacteriales>Mycobacteriaceae | 557 | MTY414_RS35930 | 9722..11395 | 1866885 | DUF262-domain-containing-protein-[Mycolicibacterium-mageritense] |
| OHV00233.1 | ParB+HNH | GCA_001854525.1 | Mycobacterium_talmoniae | Bacteria>Bacillati>Actinomycetota>Actinomycetes>Mycobacteriales>Mycobacteriaceae>Mycobacterium | 615 | BKN37_18405 | 6824..8671 | 1858794 | hypothetical-protein-BKN37_18405-[Mycobacterium-talmoniae] |
| WP_071028415.1 | ParB+HNH | GCF_001854525.1 | Mycobacterium_talmoniae | Bacteria>Bacillati>Actinomycetota>Actinomycetes>Mycobacteriales>Mycobacteriaceae>Mycobacterium | 615 | BKN37_RS18410 | 6824..8671 | 1858794 | DUF262-domain-containing-protein-[Mycobacterium-talmoniae] |
| MCQ4364654.1 | ParB+HNH | GCA_024451565.1 | Mycobacterium_gordonae | Bacteria>Bacillati>Actinomycetota>Actinomycetes>Mycobacteriales>Mycobacteriaceae>Mycobacterium | 613 | KQR54_26605 | 16328..18169 | 1778 | DUF262-domain-containing-protein-[Mycobacterium-gordonae] |
| WP_256101682.1 | ParB+HNH | GCF_024451565.1 | Mycobacterium_gordonae | Bacteria>Bacillati>Actinomycetota>Actinomycetes>Mycobacteriales>Mycobacteriaceae>Mycobacterium | 613 | KQR54_RS26620 | 16328..18169 | 1778 | DUF262-domain-containing-protein-[Mycobacterium-gordonae] |
| MCQ4364656.1 | ParB+HNH | GCA_024451565.1 | Mycobacterium_gordonae | Bacteria>Bacillati>Actinomycetota>Actinomycetes>Mycobacteriales>Mycobacteriaceae>Mycobacterium | 365 | KQR54_26615 | 21560..22657 | 1778 | DUF262-domain-containing-protein-[Mycobacterium-gordonae] |
| WP_263861824.1 | ParB+HNH | - | Mycobacterium_gordonae | Bacteria>Bacillati>Actinomycetota>Actinomycetes>Mycobacteriales>Mycobacteriaceae>Mycobacterium | 345 | KQR54_RS26630 | 21620..22657 | 1778 | DUF262-domain-containing-protein-[Mycobacterium-gordonae] |
| WP_163778794.1 | ParB+HNH | GCF_010727945.1 | Mycobacterium_cookii | Bacteria>Bacillati>Actinomycetota>Actinomycetes>Mycobacteriales>Mycobacteriaceae>Mycobacterium | 936 | G6N27_RS18570 | 3949597..3952407 | 1775 | DUF262-domain-containing-protein-[Mycobacterium-cookii] |
| WP_136246347.1 | ParB+HNH | GCF_900960235.1 | Mycobacterium_intracellulare | Bacteria>Bacillati>Actinomycetota>Actinomycetes>Mycobacteriales>Mycobacteriaceae>Mycobacterium>Mycobacterium_avium_complex_(MAC) | 615 | FAX65_RS26665 | 3044..4891 | 1767 | DUF262-domain-containing-protein-[Mycobacterium-intracellulare] |
| OMC31099.1 | ParB+HNH | GCA_001954115.1 | Mycobacterium_colombiense | Bacteria>Bacillati>Actinomycetota>Actinomycetes>Mycobacteriales>Mycobacteriaceae>Mycobacterium>Mycobacterium_avium_complex_(MAC) | 585 | A5739_14000 | 92509..94266 | 339268 | hypothetical-protein-A5739_14000-[Mycobacterium-colombiense] |
| WP_076052768.1 | ParB+HNH | GCF_001954115.1 | Mycobacterium_colombiense | Bacteria>Bacillati>Actinomycetota>Actinomycetes>Mycobacteriales>Mycobacteriaceae>Mycobacterium>Mycobacterium_avium_complex_(MAC) | 585 | A5739_RS13705 | 92509..94266 | 339268 | DUF262-domain-containing-protein-[Mycobacterium-colombiense] |
| OMC31101.1 | ParB+HNH | GCA_001954115.1 | Mycobacterium_colombiense | Bacteria>Bacillati>Actinomycetota>Actinomycetes>Mycobacteriales>Mycobacteriaceae>Mycobacterium>Mycobacterium_avium_complex_(MAC) | 558 | A5739_14020 | 103434..105110 | 339268 | hypothetical-protein-A5739_14020-[Mycobacterium-colombiense] |
| WP_076052770.1 | ParB+HNH | GCF_001954115.1 | Mycobacterium_colombiense | Bacteria>Bacillati>Actinomycetota>Actinomycetes>Mycobacteriales>Mycobacteriaceae>Mycobacterium>Mycobacterium_avium_complex_(MAC) | 558 | A5739_RS13725 | 103434..105110 | 339268 | DUF262-domain-containing-protein-[Mycobacterium-colombiense] |
| MBZ4613711.1 | ParB+HNH | GCA_020055445.1 | Mycobacterium_avium_subsp._hominissuis | Bacteria>Bacillati>Actinomycetota>Actinomycetes>Mycobacteriales>Mycobacteriaceae>Mycobacterium>Mycobacterium_avium_complex_(MAC)>Mycobacterium_avium | 559 | GBP66_23850 | 3429..5108 | 439334 | DUF262-domain-containing-protein-[Mycobacterium-avium-subsp.-hominissuis] |
| OKH62616.1 | ParB+HNH | GCA_001905305.1 | Mycobacterium_sp._SWH-M5 | Bacteria>Bacillati>Actinomycetota>Actinomycetes>Mycobacteriales>Mycobacteriaceae>Mycobacterium>unclassified_Mycobacterium | 613 | EB74_16025 | 6366..8207 | 1490483 | hypothetical-protein-EB74_16025-[Mycobacterium-sp.-SWH-M5] |
| WP_241205336.1 | ParB+HNH | GCF_022348085.1 | Mycobacterium_sp._PSTR-4-N | Bacteria>Bacillati>Actinomycetota>Actinomycetes>Mycobacteriales>Mycobacteriaceae>Mycobacterium>unclassified_Mycobacterium | 621 | MHN84_RS15065 | 173343..175208 | 2917745 | DUF262-domain-containing-protein-[Mycobacterium-sp.-PSTR-4-N] |
| WP_238886620.1 | ParB+HNH | GCF_022317005.1 | Mycobacterium_sp._IDR2000157661 | Bacteria>Bacillati>Actinomycetota>Actinomycetes>Mycobacteriales>Mycobacteriaceae>Mycobacterium>unclassified_Mycobacterium | 637 | K3G64_RS19490 | 3785522..3787435 | 2867005 | DUF262-domain-containing-protein-[Mycobacterium-sp.-IDR2000157661] |
| WP_047328959.1 | ParB+HNH | GCF_001021385.1 | Mycobacterium_sp._EPa45 | Bacteria>Bacillati>Actinomycetota>Actinomycetes>Mycobacteriales>Mycobacteriaceae>Mycobacterium>unclassified_Mycobacterium | 595 | AB431_RS04705 | 1015194..1016981 | 1545728 | DUF262-domain-containing-protein-[Mycobacterium-sp.-EPa45] |
| QEN12599.1 | ParB+HNH | GCA_008329905.1 | Mycobacterium_sp._ELW1 | Bacteria>Bacillati>Actinomycetota>Actinomycetes>Mycobacteriales>Mycobacteriaceae>Mycobacterium>unclassified_Mycobacterium | 595 | D3H54_04380 | 954977..956764 | 1547487 | DUF262-domain-containing-protein-[Mycobacterium-sp.-ELW1] |
| WP_068042709.1 | ParB+HNH | GCF_001667585.1 | Mycobacterium_sp._E2733 | Bacteria>Bacillati>Actinomycetota>Actinomycetes>Mycobacteriales>Mycobacteriaceae>Mycobacterium>unclassified_Mycobacterium | 937 | A5678_RS12480 | 52823..55636 | 1834138 | DUF262-domain-containing-protein-[Mycobacterium-sp.-E2733] |
| WP_286160126.1 | ParB+HNH | - | Mycobacterium_sp._D16Q20 | Bacteria>Bacillati>Actinomycetota>Actinomycetes>Mycobacteriales>Mycobacteriaceae>Mycobacterium>unclassified_Mycobacterium | 619 | B4386_RS22325 | 10959..12818 | 1855660 | DUF262-domain-containing-protein-[Mycobacterium-sp.-D16Q20] |
| OBB72531.1 | ParB+HNH | GCA_001665615.1 | Mycobacterium_sp._852014-52144_SCH5372336 | Bacteria>Bacillati>Actinomycetota>Actinomycetes>Mycobacteriales>Mycobacteriaceae>Mycobacterium>unclassified_Mycobacterium | 365 | A5759_18060 | 96138..97235 | 1834115 | hypothetical-protein-A5759_18060-[Mycobacterium-sp.-852014-52144_SCH5372336] |
| WP_068199840.1 | ParB+HNH | GCF_001665615.1 | Mycobacterium_sp._852014-52144_SCH5372336 | Bacteria>Bacillati>Actinomycetota>Actinomycetes>Mycobacteriales>Mycobacteriaceae>Mycobacterium>unclassified_Mycobacterium | 365 | A5759_RS17455 | 96138..97235 | 1834115 | DUF262-domain-containing-protein-[Mycobacterium-sp.-852014-52144_SCH5372336] |
| MCB0932437.1 | ParB+HNH | GCA_020438845.1 | Mycobacterium_sp. | Bacteria>Bacillati>Actinomycetota>Actinomycetes>Mycobacteriales>Mycobacteriaceae>Mycobacterium>unclassified_Mycobacterium | 613 | KDB71_11125 | 11262..13103 | 1785 | DUF262-domain-containing-protein-[Mycobacterium-sp.] |
| HRD10255.1 | ParB+HNH | GCA_035441545.1 | Mycobacterium_sp. | Bacteria>Bacillati>Actinomycetota>Actinomycetes>Mycobacteriales>Mycobacteriaceae>Mycobacterium>unclassified_Mycobacterium | 595 | PLI79_00180 | 11773..13560 | 1785 | DUF262-domain-containing-HNH-endonuclease-family-protein-[Mycobacterium-sp.] |
| OHT48138.1 | ParB+HNH | GCA_001853545.1 | Mycobacteroides_chelonae | Bacteria>Bacillati>Actinomycetota>Actinomycetes>Mycobacteriales>Mycobacteriaceae>Mycobacteroides | 613 | BKG62_21075 | 208284..210125 | 1774 | hypothetical-protein-BKG62_21075-[Mycobacteroides-chelonae] |
| WP_070930151.1 | ParB+HNH | GCF_041644775.1 | Mycobacteroides_chelonae | Bacteria>Bacillati>Actinomycetota>Actinomycetes>Mycobacteriales>Mycobacteriaceae>Mycobacteroides | 613 | BKG62_RS21075 | 208284..210125 | 1774 | DUF262-domain-containing-protein-[Mycobacteroides-chelonae] |
| WP_237140072.1 | ParB+HNH | - | Mycobacteroides_abscessus | Bacteria>Bacillati>Actinomycetota>Actinomycetes>Mycobacteriales>Mycobacteriaceae>Mycobacteroides | 607 | IUT00_RS08250 | 1655559..1657382 | 36809 | DUF262-domain-containing-protein-[Mycobacteroides-abscessus] |
| WP_079668038.1 | ParB+HNH | GCF_900141575.1 | Mycobacteroides_abscessus | Bacteria>Bacillati>Actinomycetota>Actinomycetes>Mycobacteriales>Mycobacteriaceae>Mycobacteroides | 473 | B5Y25_RS21320 | 31843..33265 | 36809 | hypothetical-protein,-partial-[Mycobacteroides-abscessus] |
| SHV22391.1 | ParB+HNH | GCA_900132885.1 | Mycobacteroides_abscessus_subsp._abscessus | Bacteria>Bacillati>Actinomycetota>Actinomycetes>Mycobacteriales>Mycobacteriaceae>Mycobacteroides>Mycobacteroides_abscessus | 613 | SAMEA2259605_05818 | 11553..13394 | 1185650 | Uncharacterized-conserved-protein-[Mycobacteroides-abscessus-subsp.-abscessus] |
| ORA83337.1 | ParB+HNH | GCA_002086285.1 | Mycolicibacter_kumamotonensis | Bacteria>Bacillati>Actinomycetota>Actinomycetes>Mycobacteriales>Mycobacteriaceae>Mycolicibacter | 365 | BST28_00060 | 19913..21010 | 354243 | hypothetical-protein-BST28_00060-[Mycolicibacter-kumamotonensis] |
| WP_133054271.1 | ParB+HNH | GCF_002086285.1 | Mycolicibacter_kumamotonensis | Bacteria>Bacillati>Actinomycetota>Actinomycetes>Mycobacteriales>Mycobacteriaceae>Mycolicibacter | 365 | BST28_RS00060 | 19913..21010 | 354243 | DUF262-domain-containing-protein-[Mycolicibacter-kumamotonensis] |
| TXI55353.1 | ParB+HNH | GCA_008015745.1 | Mycolicibacter_arupensis | Bacteria>Bacillati>Actinomycetota>Actinomycetes>Mycobacteriales>Mycobacteriaceae>Mycolicibacter | 616 | E6Q54_12725 | 10687..12537 | 342002 | DUF262-domain-containing-protein-[Mycolicibacter-arupensis] |
| WP_276761080.1 | ParB+HNH | GCF_008015745.1 | Mycolicibacter_arupensis | Bacteria>Bacillati>Actinomycetota>Actinomycetes>Mycobacteriales>Mycobacteriaceae>Mycolicibacter | 616 | E6Q54_RS12715 | 10687..12537 | 342002 | DUF262-domain-containing-protein-[Mycolicibacter-arupensis] |
| TXI48371.1 | ParB+HNH | GCA_008015745.1 | Mycolicibacter_arupensis | Bacteria>Bacillati>Actinomycetota>Actinomycetes>Mycobacteriales>Mycobacteriaceae>Mycolicibacter | 613 | E6Q54_23705 | 10295..12136 | 342002 | DUF262-domain-containing-protein-[Mycolicibacter-arupensis] |
| WP_276763636.1 | ParB+HNH | GCF_008015745.1 | Mycolicibacter_arupensis | Bacteria>Bacillati>Actinomycetota>Actinomycetes>Mycobacteriales>Mycobacteriaceae>Mycolicibacter | 613 | E6Q54_RS23750 | 10295..12136 | 342002 | DUF262-domain-containing-protein-[Mycolicibacter-arupensis] |
| WP_235881579.1 | ParB+HNH | GCF_021559935.1 | Mycolicibacterium_vanbaalenii | Bacteria>Bacillati>Actinomycetota>Actinomycetes>Mycobacteriales>Mycobacteriaceae>Mycolicibacterium | 595 | HZU38_RS11465 | 2338565..2340352 | 110539 | DUF262-domain-containing-protein-[Mycolicibacterium-vanbaalenii] |
| NTY62307.1 | ParB+HNH | GCA_013337765.1 | Mycolicibacterium_sphagni | Bacteria>Bacillati>Actinomycetota>Actinomycetes>Mycobacteriales>Mycobacteriaceae>Mycolicibacterium | 595 | FEG63_22470 | 117182..118969 | 1786 | DUF262-domain-containing-protein-[Mycolicibacterium-sphagni] |
| WP_174400024.1 | ParB+HNH | GCF_013337765.1 | Mycolicibacterium_sphagni | Bacteria>Bacillati>Actinomycetota>Actinomycetes>Mycobacteriales>Mycobacteriaceae>Mycolicibacterium | 595 | FEG63_RS22450 | 117182..118969 | 1786 | DUF262-domain-containing-protein-[Mycolicibacterium-sphagni] |
| OLP01885.1 | ParB+HNH | GCA_001942045.1 | Mycolicibacterium_porcinum | Bacteria>Bacillati>Actinomycetota>Actinomycetes>Mycobacteriales>Mycobacteriaceae>Mycolicibacterium | 595 | BVU76_13160 | 75469..77256 | 39693 | hypothetical-protein-BVU76_13160-[Mycolicibacterium-porcinum] |
| ORB28020.1 | ParB+HNH | GCA_002086815.1 | Mycolicibacterium_parafortuitum | Bacteria>Bacillati>Actinomycetota>Actinomycetes>Mycobacteriales>Mycobacteriaceae>Mycolicibacterium | 613 | BST38_21955 | 35861..37702 | 39692 | hypothetical-protein-BST38_21955-[Mycolicibacterium-parafortuitum] |
| WP_083145594.1 | ParB+HNH | GCF_002086815.1 | Mycolicibacterium_parafortuitum | Bacteria>Bacillati>Actinomycetota>Actinomycetes>Mycobacteriales>Mycobacteriaceae>Mycolicibacterium | 613 | MPP7335_RS09260 | 1904688..1906529 | 39692 | DUF262-domain-containing-protein-[Mycolicibacterium-parafortuitum] |
| WP_096312284.1 | ParB+HNH | GCF_039534765.1 | Mycolicibacterium_pallens | Bacteria>Bacillati>Actinomycetota>Actinomycetes>Mycobacteriales>Mycobacteriaceae>Mycolicibacterium | 558 | K0O64_RS24910 | 5119940..5121616 | 370524 | MULTISPECIES:-DUF262-domain-containing-protein-[Mycobacteriaceae] |
| WP_193489328.1 | ParB+HNH | GCF_039528895.1 | Mycolicibacterium_murale | Bacteria>Bacillati>Actinomycetota>Actinomycetes>Mycobacteriales>Mycobacteriaceae>Mycolicibacterium | 595 | ABD253_RS31610 | 196137..197924 | 182220 | DUF262-domain-containing-protein-[Mycolicibacterium-murale] |
| WP_267319444.1 | ParB+HNH | GCF_026501225.1 | Mycolicibacterium_mucogenicum | Bacteria>Bacillati>Actinomycetota>Actinomycetes>Mycobacteriales>Mycobacteriaceae>Mycolicibacterium | 699 | OS122_RS24295 | 492619..494718 | 56689 | DUF262-and-DUF1524-domain-containing-protein-[Mycolicibacterium-mucogenicum] |
| WP_163642249.1 | ParB+HNH | GCF_010727475.1 | Mycolicibacterium_mageritense | Bacteria>Bacillati>Actinomycetota>Actinomycetes>Mycobacteriales>Mycobacteriaceae>Mycolicibacterium | 174 | BN978_RS39980 | 1136607..1137131 | 53462 | HNH-endonuclease-family-protein-[Mycolicibacterium-mageritense] |
| WP_051578472.1 | ParB+HNH | GCF_048569395.1 | Mycolicibacterium_mageritense | Bacteria>Bacillati>Actinomycetota>Actinomycetes>Mycobacteriales>Mycobacteriaceae>Mycolicibacterium | 162 | BN978_RS38370 | 1134443..1134931 | 53462 | DUF262-domain-containing-protein-[Mycolicibacterium-mageritense] |
| WP_260941408.1 | ParB+HNH | GCF_025331195.1 | Mycolicibacterium_llatzerense | Bacteria>Bacillati>Actinomycetota>Actinomycetes>Mycobacteriales>Mycobacteriaceae>Mycolicibacterium | 387 | A7G45_RS26860 | 442296..443459 | 280871 | hypothetical-protein-[Mycolicibacterium-llatzerense] |
| MCT7366344.1 | ParB+HNH | GCA_025331195.1 | Mycolicibacterium_llatzerense | Bacteria>Bacillati>Actinomycetota>Actinomycetes>Mycobacteriales>Mycobacteriaceae>Mycolicibacterium | 218 | A7G45_26140 | 444765..445421 | 280871 | hypothetical-protein-[Mycolicibacterium-llatzerense] |
| WP_260941409.1 | ParB+HNH | GCF_039531995.1 | Mycolicibacterium_llatzerense | Bacteria>Bacillati>Actinomycetota>Actinomycetes>Mycobacteriales>Mycobacteriaceae>Mycolicibacterium | 218 | A7G45_RS26870 | 444765..445421 | 280871 | DUF262-domain-containing-protein-[Mycolicibacterium-llatzerense] |
| WP_085173432.1 | ParB+HNH | GCF_002101705.1 | Mycolicibacterium_iranicum | Bacteria>Bacillati>Actinomycetota>Actinomycetes>Mycobacteriales>Mycobacteriaceae>Mycolicibacterium | 615 | AWC12_RS07885 | 84658..86505 | 912594 | DUF262-domain-containing-protein-[Mycolicibacterium-iranicum] |
| OFJ52797.1 | ParB+HNH | GCA_001766635.1 | Mycolicibacterium_grossiae | Bacteria>Bacillati>Actinomycetota>Actinomycetes>Mycobacteriales>Mycobacteriaceae>Mycolicibacterium | 613 | BEL07_15680 | 40668..42509 | 1552759 | hypothetical-protein-BEL07_15680-[Mycobacterium-grossiae] |
| MBU8833315.1 | ParB+HNH | GCA_018524295.1 | Mycolicibacterium_goodii | Bacteria>Bacillati>Actinomycetota>Actinomycetes>Mycobacteriales>Mycobacteriaceae>Mycolicibacterium | 613 | KL858_28170 | 20953..22794 | 134601 | DUF262-domain-containing-protein-[Mycolicibacterium-goodii] |
| WP_214389767.1 | ParB+HNH | GCF_018524295.1 | Mycolicibacterium_goodii | Bacteria>Bacillati>Actinomycetota>Actinomycetes>Mycobacteriales>Mycobacteriaceae>Mycolicibacterium | 613 | KL858_RS28170 | 20953..22794 | 134601 | DUF262-domain-containing-protein-[Mycolicibacterium-goodii] |
| WP_013470724.1 | ParB+HNH | GCF_000184435.1 | Mycolicibacterium_gilvum | Bacteria>Bacillati>Actinomycetota>Actinomycetes>Mycobacteriales>Mycobacteriaceae>Mycolicibacterium | 613 | MSPYR1_RS04235 | 881724..883565 | 1804 | DUF262-domain-containing-protein-[Mycolicibacterium-gilvum] |
| WP_264058323.1 | ParB+HNH | GCF_965137455.1 | Mycolicibacterium_frederiksbergense | Bacteria>Bacillati>Actinomycetota>Actinomycetes>Mycobacteriales>Mycobacteriaceae>Mycolicibacterium | 595 | H7J08_RS31205 | 380129..381916 | 117567 | DUF262-domain-containing-HNH-endonuclease-family-protein-[Mycolicibacterium-frederiksbergense] |
| WP_128109811.1 | ParB+HNH | GCF_004014805.1 | Mycolicibacterium_elephantis | Bacteria>Bacillati>Actinomycetota>Actinomycetes>Mycobacteriales>Mycobacteriaceae>Mycolicibacterium | 614 | H7I95_RS08855 | 4649..6493 | 81858 | DUF262-domain-containing-protein-[Mycolicibacterium-elephantis] |
| OBK03535.1 | ParB+HNH | GCA_001673265.1 | Mycolicibacterium_conceptionense | Bacteria>Bacillati>Actinomycetota>Actinomycetes>Mycobacteriales>Mycobacteriaceae>Mycolicibacterium | 620 | A5639_22735 | 11306..13168 | 451644 | hypothetical-protein-A5639_22735-[Mycolicibacterium-conceptionense] |
| OMB68864.1 | ParB+HNH | GCA_001954125.1 | Mycolicibacterium_conceptionense | Bacteria>Bacillati>Actinomycetota>Actinomycetes>Mycobacteriales>Mycobacteriaceae>Mycolicibacterium | 620 | A5741_09380 | 5174..7036 | 451644 | hypothetical-protein-A5741_09380-[Mycolicibacterium-conceptionense] |
| WP_065062930.1 | ParB+HNH | GCF_001673265.1 | Mycolicibacterium_conceptionense | Bacteria>Bacillati>Actinomycetota>Actinomycetes>Mycobacteriales>Mycobacteriaceae>Mycolicibacterium | 620 | A5741_RS31965 | 5174..7036 | 451644 | DUF262-domain-containing-protein-[Mycolicibacterium-conceptionense] |
| WP_237159980.1 | ParB+HNH | GCF_002968275.1 | Mycolicibacterium_austroafricanum | Bacteria>Bacillati>Actinomycetota>Actinomycetes>Mycobacteriales>Mycobacteriaceae>Mycolicibacterium | 616 | C6A88_RS31875 | 17537..19387 | 39687 | DUF262-domain-containing-protein-[Mycolicibacterium-austroafricanum] |
| PQP39973.1 | ParB+HNH | GCA_002968275.1 | Mycolicibacterium_austroafricanum | Bacteria>Bacillati>Actinomycetota>Actinomycetes>Mycobacteriales>Mycobacteriaceae>Mycolicibacterium | 613 | C6A88_31865 | 17537..19378 | 39687 | hypothetical-protein-C6A88_31865-[Mycolicibacterium-austroafricanum] |
| MBX7452223.1 | ParB+HNH | GCA_019710595.1 | Mycolicibacterium_aurantiacum | Bacteria>Bacillati>Actinomycetota>Actinomycetes>Mycobacteriales>Mycobacteriaceae>Mycolicibacterium | 613 | GR927_29905 | 282810..284651 | 2692455 | DUF262-domain-containing-protein-[Mycolicibacterium-aurantiacum] |
| WP_163919112.1 | ParB+HNH | GCF_010731815.2 | Mycolicibacterium_arabiense | Bacteria>Bacillati>Actinomycetota>Actinomycetes>Mycobacteriales>Mycobacteriaceae>Mycolicibacterium | 613 | H5P33_RS16140 | 689244..691085 | 1286181 | DUF262-domain-containing-protein-[Mycolicibacterium-arabiense] |
| WP_239342821.1 | ParB+HNH | GCF_022348005.1 | Mycolicibacterium_sp._OfavD-34-C | Bacteria>Bacillati>Actinomycetota>Actinomycetes>Mycobacteriales>Mycobacteriaceae>Mycolicibacterium>unclassified_Mycolicibacterium | 557 | MHN82_RS23240 | 17177..18850 | 2917746 | DUF262-domain-containing-protein-[Mycolicibacterium-sp.-OfavD-34-C] |
| WP_210686517.1 | ParB+HNH | GCF_017948465.1 | Mycolicibacterium_sp._GESEQ-9 | Bacteria>Bacillati>Actinomycetota>Actinomycetes>Mycobacteriales>Mycobacteriaceae>Mycolicibacterium>unclassified_Mycolicibacterium | 595 | KCH01_RS05005 | 109633..111420 | 2812656 | DUF262-domain-containing-protein-[Mycolicibacterium-sp.-GESEQ-9] |
| WP_030536689.1 | ParB+HNH | GCF_024349805.1 | Rhodococcus | Bacteria>Bacillati>Actinomycetota>Actinomycetes>Mycobacteriales>Nocardiaceae | 558 | RE2895_RS00245 | 61484..63160 | 1827 | DUF262-domain-containing-protein-[Rhodococcus-erythropolis] |
| WP_068276442.1 | ParB+HNH | GCF_001646865.1 | Aldersonia_kunmingensis | Bacteria>Bacillati>Actinomycetota>Actinomycetes>Mycobacteriales>Nocardiaceae>Aldersonia | 558 | A8G08_RS16785 | 2386..4062 | 408066 | DUF262-domain-containing-protein-[Aldersonia-kunmingensis] |
| WP_218024102.1 | ParB+HNH | GCF_001613365.1 | Nocardia_xishanensis | Bacteria>Bacillati>Actinomycetota>Actinomycetes>Mycobacteriales>Nocardiaceae>Nocardia | 598 | NX1_RS10710 | 38563..40359 | 238964 | DUF262-domain-containing-protein-[Nocardia-xishanensis] |
| WP_245922366.1 | ParB+HNH | GCF_002933495.1 | Nocardia_nova | Bacteria>Bacillati>Actinomycetota>Actinomycetes>Mycobacteriales>Nocardiaceae>Nocardia | 180 | C5F51_RS20335 | 201566..202108 | 37330 | HNH-endonuclease-family-protein-[Nocardia-nova] |
| WP_040863303.1 | ParB+HNH | GCF_000308655.1 | Nocardia_niigatensis | Bacteria>Bacillati>Actinomycetota>Actinomycetes>Mycobacteriales>Nocardiaceae>Nocardia | 603 | ON33_RS34210 | 20886..22697 | 209249 | DUF262-domain-containing-protein-[Nocardia-niigatensis] |
| WP_228787257.1 | ParB+HNH | GCF_015478425.1 | Nocardia_farcinica | Bacteria>Bacillati>Actinomycetota>Actinomycetes>Mycobacteriales>Nocardiaceae>Nocardia | 598 | AB8M53_RS07380 | 591920..593716 | 37329 | DUF262-domain-containing-protein-[Nocardia-farcinica] |

| Protein-ID | Domain | Assembly-ID | Strain | Taxonomy | Length | Locus-Tag | Genomic-coordinates | TaxID | Fasta-defline |
|---|---|---|---|---|---|---|---|---|---|
| WP_228810787.1 | ParB+HNH | GCF_015477405.1 | Nocardia_farcinica | Bacteria>Bacillati>Actinomycetota>Actinomycetes>Mycobacteriales>Nocardiaceae>Nocardia | 598 | IU473_RS11245 | 579869..581665 | 37329 | DUF262-domain-containing-protein-[Nocardia-farcinica] |
| WP_011210978.1 | ParB+HNH | GCF_000009805.1 | Nocardia_farcinica | Bacteria>Bacillati>Actinomycetota>Actinomycetes>Mycobacteriales>Nocardiaceae>Nocardia | 556 | NFA_RS22035 | 4608933..4610603 | 37329 | DUF262-domain-containing-protein-[Nocardia-farcinica] |
| WP_056816737.1 | ParB+HNH | GCF_001427185.1 | Nocardia_sp._Root136 | Bacteria>Bacillati>Actinomycetota>Actinomycetes>Mycobacteriales>Nocardiaceae>Nocardia>unclassified_Nocardia | 602 | ASD42_RS14815 | 215950..217758 | 1736458 | DUF262-domain-containing-protein-[Nocardia-sp.-Root136] |
| WP_194806633.1 | ParB+HNH | GCF_015354405.1 | Nocardia_sp._BSTN01 | Bacteria>Bacillati>Actinomycetota>Actinomycetes>Mycobacteriales>Nocardiaceae>Nocardia>unclassified_Nocardia | 558 | IRT45_RS15955 | 109405..111081 | 2783665 | DUF262-domain-containing-protein-[Nocardia-sp.-BSTN01] |
| WP_019927710.1 | ParB+HNH | GCF_000381925.1 | Nocardia_sp._BMG111209 | Bacteria>Bacillati>Actinomycetota>Actinomycetes>Mycobacteriales>Nocardiaceae>Nocardia>unclassified_Nocardia | 623 | G361_RS0113985 | 3033721..3035592 | 1160137 | DUF262-domain-containing-protein-[Nocardia-sp.-BMG111209] |
| WP_019931230.1 | ParB+HNH | GCF_000381925.1 | Nocardia_sp._BMG111209 | Bacteria>Bacillati>Actinomycetota>Actinomycetes>Mycobacteriales>Nocardiaceae>Nocardia>unclassified_Nocardia | 600 | G361_RS0132045 | 190405..192207 | 1160137 | DUF262-domain-containing-protein-[Nocardia-sp.-BMG111209] |
| NKS26552.1 | ParB+HNH | GCA_012393535.1 | Prescottella_equi | Bacteria>Bacillati>Actinomycetota>Actinomycetes>Mycobacteriales>Nocardiaceae>Prescottella | 623 | GS505_12160 | 241068..242939 | 43767 | DUF262-domain-containing-protein-[Prescottella-equi] |
| WP_106851263.1 | ParB+HNH | GCF_003013675.1 | Prescottella_equi | Bacteria>Bacillati>Actinomycetota>Actinomycetes>Mycobacteriales>Nocardiaceae>Prescottella | 558 | C7H75_RS09235 | 1956423..1958099 | 43767 | DUF262-domain-containing-protein-[Prescottella-equi] |
| NIL92564.1 | ParB+HNH | GCA_011770045.1 | Rhodococcoides_fascians | Bacteria>Bacillati>Actinomycetota>Actinomycetes>Mycobacteriales>Nocardiaceae>Rhodococcoides | 598 | RhoFasSB10_04925 | 296..2092 | 1828 | hypothetical-protein-[Rhodococcus-fascians] |
| WP_127946805.1 | ParB+HNH | GCF_004011835.1 | Rhodococcus_spongiicola | Bacteria>Bacillati>Actinomycetota>Actinomycetes>Mycobacteriales>Nocardiaceae>Rhodococcus | 623 | EF834_RS08535 | 71248..73119 | 2487352 | DUF262-domain-containing-protein-[Rhodococcus-spongiicola] |
| WP_119699600.1 | ParB+HNH | GCF_003586525.1 | Rhodococcus_ruber | Bacteria>Bacillati>Actinomycetota>Actinomycetes>Mycobacteriales>Nocardiaceae>Rhodococcus | 563 | TN91_RS02595 | 65385..67076 | 1830 | DUF262-domain-containing-protein-[Rhodococcus-ruber] |
| WP_085470757.1 | ParB+HNH | GCF_900177695.1 | Rhodococcus_rhodochrous | Bacteria>Bacillati>Actinomycetota>Actinomycetes>Mycobacteriales>Nocardiaceae>Rhodococcus | 555 | M502_RS25915 | 17475..19142 | 1829 | DUF262-domain-containing-protein-[Rhodococcus-rhodochrous] |
| GAA4479481.1 | ParB+HNH | GCA_039540555.1 | Rhodococcus_olei | Bacteria>Bacillati>Actinomycetota>Actinomycetes>Mycobacteriales>Nocardiaceae>Rhodococcus | 403 | GCM10023094_24670 | 41995..43206 | 2161675 | hypothetical-protein-GCM10023094_24670-[Rhodococcus-olei] |
| WP_006942564.1 | ParB+HNH | GCF_000382865.1 | Rhodococcus_aetherivorans | Bacteria>Bacillati>Actinomycetota>Actinomycetes>Mycobacteriales>Nocardiaceae>Rhodococcus | 555 | EBESD8_RS20695 | 3795..5462 | 191292 | DUF262-domain-containing-protein-[Rhodococcus-aetherivorans] |
| WP_317550162.1 | ParB+HNH | GCF_033042295.1 | Rhodococcus_aetherivorans | Bacteria>Bacillati>Actinomycetota>Actinomycetes>Mycobacteriales>Nocardiaceae>Rhodococcus | 553 | R3Q02_RS04895 | 146442..148103 | 191292 | DUF262-domain-containing-protein-[Rhodococcus-aetherivorans] |
| WP_102999580.1 | ParB+HNH | GCF_002893965.1 | Rhodococcus_qingshengii | Bacteria>Bacillati>Actinomycetota>Actinomycetes>Mycobacteriales>Nocardiaceae>Rhodococcus>Rhodococcus_erythropolis_group | 562 | C1M55_RS31230 | 8437..10125 | 334542 | DUF262-domain-containing-protein-[Rhodococcus-qingshengii] |
| WP_208722092.1 | ParB+HNH | GCF_006384225.1 | Rhodococcus_qingshengii | Bacteria>Bacillati>Actinomycetota>Actinomycetes>Mycobacteriales>Nocardiaceae>Rhodococcus>Rhodococcus_erythropolis_group | 562 | IGTS8_RS29985 | 43256..44944 | 334542 | DUF262-domain-containing-protein-[Rhodococcus-qingshengii] |
| WP_269614893.1 | ParB+HNH | GCF_027270755.1 | Rhodococcus_qingshengii | Bacteria>Bacillati>Actinomycetota>Actinomycetes>Mycobacteriales>Nocardiaceae>Rhodococcus>Rhodococcus_erythropolis_group | 562 | O4271_RS30880 | 24656..26344 | 334542 | DUF262-domain-containing-HNH-endonuclease-family-protein-[Rhodococcus-qingshengii] |
| WP_231782760.1 | ParB+HNH | GCF_021165975.1 | Rhodococcus_qingshengii | Bacteria>Bacillati>Actinomycetota>Actinomycetes>Mycobacteriales>Nocardiaceae>Rhodococcus>Rhodococcus_erythropolis_group | 558 | LRQ08_RS00250 | 60792..62468 | 334542 | DUF262-domain-containing-protein-[Rhodococcus-qingshengii] |
| MCQ4152488.1 | ParB+HNH | GCA_024438075.1 | Rhodococcus_qingshengii | Bacteria>Bacillati>Actinomycetota>Actinomycetes>Mycobacteriales>Nocardiaceae>Rhodococcus>Rhodococcus_erythropolis_group | 557 | NOS52_32465 | 21162..22835 | 334542 | DUF262-domain-containing-HNH-endonuclease-family-protein-[Rhodococcus-qingshengii] |
| WP_256083184.1 | ParB+HNH | GCF_024438075.1 | Rhodococcus_qingshengii | Bacteria>Bacillati>Actinomycetota>Actinomycetes>Mycobacteriales>Nocardiaceae>Rhodococcus>Rhodococcus_erythropolis_group | 557 | NOS52_RS32460 | 21162..22835 | 334542 | DUF262-domain-containing-protein-[Rhodococcus-qingshengii] |
| WP_283569685.1 | ParB+HNH | GCF_030063715.1 | Rhodococcus_qingshengii | Bacteria>Bacillati>Actinomycetota>Actinomycetes>Mycobacteriales>Nocardiaceae>Rhodococcus>Rhodococcus_erythropolis_group | 557 | QNA24_RS33660 | 10198..11871 | 334542 | DUF262-domain-containing-HNH-endonuclease-family-protein-[Rhodococcus-qingshengii] |
| WP_102999586.1 | ParB+HNH | GCF_002893965.1 | Rhodococcus_qingshengii | Bacteria>Bacillati>Actinomycetota>Actinomycetes>Mycobacteriales>Nocardiaceae>Rhodococcus>Rhodococcus_erythropolis_group | 360 | C1M55_RS31265 | 19168..20250 | 334542 | DUF262-domain-containing-protein-[Rhodococcus-qingshengii] |
| WP_116058403.1 | ParB+HNH | GCF_016019085.1 | Rhodococcus_erythropolis | Bacteria>Bacillati>Actinomycetota>Actinomycetes>Mycobacteriales>Nocardiaceae>Rhodococcus>Rhodococcus_erythropolis_group | 562 | I3517_RS00935 | 12199..13887 | 1833 | DUF262-domain-containing-protein-[Rhodococcus-erythropolis] |
| WP_153212241.1 | ParB+HNH | GCF_009495805.1 | Rhodococcus_erythropolis | Bacteria>Bacillati>Actinomycetota>Actinomycetes>Mycobacteriales>Nocardiaceae>Rhodococcus>Rhodococcus_erythropolis_group | 562 | GEO20_RS15995 | 37197..38885 | 1833 | DUF262-domain-containing-protein-[Rhodococcus-erythropolis] |
| WP_029256837.1 | ParB+HNH | GCF_000696675.2 | Rhodococcus_erythropolis | Bacteria>Bacillati>Actinomycetota>Actinomycetes>Mycobacteriales>Nocardiaceae>Rhodococcus>Rhodococcus_erythropolis_group | 558 | H351_RS02825 | 622676..624352 | 1833 | DUF262-domain-containing-protein-[Rhodococcus-erythropolis] |
| QCQ90485.1 | ParB+HNH | GCA_005484805.1 | Rhodococcus_sp._SGAir0479 | Bacteria>Bacillati>Actinomycetota>Actinomycetes>Mycobacteriales>Nocardiaceae>Rhodococcus>unclassified_Rhodococcus_(in:_high_G+C_Gram-positive_bacteria) | 554 | E7742_04090 | 869896..871560 | 2567884 | DUF262-domain-containing-protein-[Rhodococcus-sp.-SGAir0479] |
| MBY6706233.1 | ParB+HNH | GCA_019836145.1 | Rhodococcus_sp._BP-241 | Bacteria>Bacillati>Actinomycetota>Actinomycetes>Mycobacteriales>Nocardiaceae>Rhodococcus>unclassified_Rhodococcus_(in:_high_G+C_Gram-positive_bacteria) | 591 | HQ308_05405 | 454361..456136 | 2739441 | DUF262-domain-containing-protein-[Rhodococcus-sp.-BP-241] |
| WP_228232865.1 | ParB+HNH | GCF_020782825.1 | Rhodococcus_sp._3-2 | Bacteria>Bacillati>Actinomycetota>Actinomycetes>Mycobacteriales>Nocardiaceae>Rhodococcus>unclassified_Rhodococcus_(in:_high_G+C_Gram-positive_bacteria) | 562 | LL008_RS29700 | 198395..200083 | 2890836 | DUF262-domain-containing-protein-[Rhodococcus-sp.-3-2] |
| WP_167735518.1 | ParB+HNH | GCF_004563845.1 | Rhodococcus_sp._1R11 | Bacteria>Bacillati>Actinomycetota>Actinomycetes>Mycobacteriales>Nocardiaceae>Rhodococcus>unclassified_Rhodococcus_(in:_high_G+C_Gram-positive_bacteria) | 598 | E4P29_RS16885 | 117809..119605 | 2559614 | DUF262-domain-containing-protein-[Rhodococcus-sp.-1R11] |
| OFE08175.1 | ParB+HNH | GCA_001757665.1 | Rhodococcus_sp._1139 | Bacteria>Bacillati>Actinomycetota>Actinomycetes>Mycobacteriales>Nocardiaceae>Rhodococcus>unclassified_Rhodococcus_(in:_high_G+C_Gram-positive_bacteria) | 564 | A5N83_14200 | 72618..74312 | 1833762 | hypothetical-protein-A5N83_14200-[Rhodococcus-sp.-1139] |
| WP_256261722.1 | ParB+HNH | GCF_001757665.1 | Rhodococcus_sp._1139 | Bacteria>Bacillati>Actinomycetota>Actinomycetes>Mycobacteriales>Nocardiaceae>Rhodococcus>unclassified_Rhodococcus_(in:_high_G+C_Gram-positive_bacteria) | 560 | A5N83_RS14200 | 72618..74300 | 1833762 | DUF262-domain-containing-protein-[Rhodococcus-sp.-1139] |
| WP_076482723.1 | ParB+HNH | GCF_900156495.1 | Williamsia_sterculiae | Bacteria>Bacillati>Actinomycetota>Actinomycetes>Mycobacteriales>Nocardiaceae>Williamsia | 623 | BW971_RS19025 | 28917..30788 | 1344003 | DUF262-domain-containing-protein-[Williamsia-sterculiae] |
| WP_099384689.1 | ParB+HNH | GCF_002735905.1 | Williamsia_muralis | Bacteria>Bacillati>Actinomycetota>Actinomycetes>Mycobacteriales>Nocardiaceae>Williamsia | 562 | CSW57_RS21170 | 1637083..1638771 | 85044 | DUF262-domain-containing-protein-[Williamsia-marianensis] |
| WP_020385146.1 | ParB+HNH | GCF_000372465.1 | Kribbella_catacumbae | Bacteria>Bacillati>Actinomycetota>Actinomycetes>Propionibacteriales>Kribbellaceae>Kribbella | 558 | A3ES_RS0102265 | 26360..28036 | 460086 | DUF262-domain-containing-protein-[Kribbella-catacumbae] |
| WP_153653107.1 | ParB+HNH | GCF_009649075.1 | Aeromicrobium_yanjiei | Bacteria>Bacillati>Actinomycetota>Actinomycetes>Propionibacteriales>Nocardioidaceae>Aeromicrobium | 591 | GEV26_RS10960 | 2218903..2220678 | 2662028 | DUF262-domain-containing-protein-[Aeromicrobium-yanjiei] |
| WP_124237128.1 | ParB+HNH | GCF_042681225.1 | Aeromicrobium_camelliae | Bacteria>Bacillati>Actinomycetota>Actinomycetes>Propionibacteriales>Nocardioidaceae>Aeromicrobium | 613 | EHW97_RS10555 | 81463..83304 | 1538144 | DUF262-domain-containing-protein-[Aeromicrobium-camelliae] |
| WP_166139837.1 | ParB+HNH | GCF_011420305.2 | Nocardioides_ochotonae | Bacteria>Bacillati>Actinomycetota>Actinomycetes>Propionibacteriales>Nocardioidaceae>Nocardioides | 558 | HBO46_RS08100 | 1672255..1673931 | 2685869 | DUF262-domain-containing-protein-[Nocardioides-ochotonae] |
| WP_240236716.1 | ParB+HNH | GCF_022436705.1 | Nocardioides_sp._CFH_31398 | Bacteria>Bacillati>Actinomycetota>Actinomycetes>Propionibacteriales>Nocardioidaceae>Nocardioides>unclassified_Nocardioides | 558 | MJ876_RS12950 | 79921..81597 | 2919579 | DUF262-domain-containing-protein-[Nocardioides-sp.-CFH-31398] |
| MBM0123459.1 | ParB+HNH | GCA_016803235.1 | Pimelobacter_simplex | Bacteria>Bacillati>Actinomycetota>Actinomycetes>Propionibacteriales>Nocardioidaceae>Pimelobacter | 613 | IMY96_01780 | 54582..56423 | 2045 | DUF262-domain-containing-protein-[Pimelobacter-simplex] |
| WP_203043112.1 | ParB+HNH | GCA_016803235.1 | Pimelobacter_simplex | Bacteria>Bacillati>Actinomycetota>Actinomycetes>Propionibacteriales>Nocardioidaceae>Pimelobacter | 613 | IMY96_RS01780 | 54582..56423 | 2045 | DUF262-domain-containing-protein-[Pimelobacter-simplex] |
| MAS55989.1 | ParB+HNH | GCA_002698575.1 | Pimelobacter_sp. | Bacteria>Bacillati>Actinomycetota>Actinomycetes>Propionibacteriales>Nocardioidaceae>Pimelobacter>unclassified_Pimelobacter | 619 | CMJ44_15465 | 21051..22910 | 51662 | hypothetical-protein-[Pimelobacter-sp.] |
| WP_013863117.1 | ParB+HNH | GCF_000270245.1 | Microlunatus_phosphovorus | Bacteria>Bacillati>Actinomycetota>Actinomycetes>Propionibacteriales>Propionibacteriaceae>Microlunatus | 613 | MLP_RS10780 | 2383229..2385070 | 29405 | DUF262-domain-containing-protein-[Microlunatus-phosphovorus] |
| WP_179750835.1 | ParB+HNH | GCF_013407955.1 | Microlunatus_parietis | Bacteria>Bacillati>Actinomycetota>Actinomycetes>Propionibacteriales>Propionibacteriaceae>Microlunatus | 558 | BKA15_RS11405 | 2399091..2400767 | 682979 | DUF262-domain-containing-protein-[Microlunatus-parietis] |
| MBK8462832.1 | ParB+HNH | GCA_016711325.1 | Nigerium_sp. | Bacteria>Bacillati>Actinomycetota>Actinomycetes>Propionibacteriales>Propionibacteriaceae>Nigerium>unclassified_Nigerium | 616 | IPL36_07330 | 624717..626567 | 2042655 | DUF262-domain-containing-protein-[Nigerium-sp.] |
| HQA31348.1 | ParB+HNH | GCA_035412375.1 | Propioniciclava_tarda | Bacteria>Bacillati>Actinomycetota>Actinomycetes>Propionibacteriales>Propionibacteriaceae>Propioniciclava | 660 | PKX10_08005 | 4517..6499 | 433330 | DUF262-domain-containing-HNH-endonuclease-family-protein-[Propioniciclava-tarda] |
| WP_182782742.1 | ParB+HNH | GCF_014164725.1 | Tessaracoccus_sp._MC1627 | Bacteria>Bacillati>Actinomycetota>Actinomycetes>Propionibacteriales>Propionibacteriaceae>Tessaracoccus>unclassified_Tessaracoccus | 371 | H5399_RS09130 | 138940..140055 | 2760312 | DUF262-domain-containing-protein-[Tessaracoccus-sp.-MC1627] |
| MBN9103778.1 | ParB+HNH | GCA_017306015.1 | Propionibacteriaceae_bacterium | Bacteria>Bacillati>Actinomycetota>Actinomycetes>Propionibacteriales>Propionibacteriaceae>unclassified_Propionibacteriaceae | 610 | J0I14_02125 | 374756..376588 | 2021380 | DUF262-domain-containing-protein-[Propionibacteriaceae-bacterium] |
| WP_085916423.1 | ParB+HNH | GCF_003945385.1 | Pseudonocardia | Bacteria>Bacillati>Actinomycetota>Actinomycetes>Pseudonocardiales>Pseudonocardiaceae | 633 | PSA01_RS26715 | 6642..8543 | 1847 | MULTISPECIES:-DUF262-domain-containing-protein-[Pseudonocardia] |
| WP_251785092.1 | ParB+HNH | GCF_023717215.1 | Pseudonocardia | Bacteria>Bacillati>Actinomycetota>Actinomycetes>Pseudonocardiales>Pseudonocardiaceae | 548 | ND486_RS16910 | 253170..254816 | 1847 | DUF262-domain-containing-protein-[Pseudonocardia-sp.-DR1-2] |
| GLZ41165.1 | ParB+HNH | GCA_030269645.1 | Actinokineospora_sp._NBRC_105648 | Bacteria>Bacillati>Actinomycetota>Actinomycetes>Pseudonocardiales>Pseudonocardiaceae>Actinokineospora>unclassified_Actinokineospora | 565 | Acsp05_47890 | 25012..26709 | 3032206 | hypothetical-protein-Acsp05_47890-[Actinokineospora-sp.-NBRC-105648] |
| WP_211358066.1 | ParB+HNH | GCF_006715045.1 | Amycolatopsis_cihanbeyliensis | Bacteria>Bacillati>Actinomycetota>Actinomycetes>Pseudonocardiales>Pseudonocardiaceae>Amycolatopsis | 561 | FB471_RS19220 | 4239919..4241604 | 1128664 | DUF262-domain-containing-protein-[Amycolatopsis-cihanbeyliensis] |
| WP_027930285.1 | ParB+HNH | GCF_000473265.1 | Amycolatopsis_thermoflava | Bacteria>Bacillati>Actinomycetota>Actinomycetes>Pseudonocardiales>Pseudonocardiaceae>Amycolatopsis>Amycolatopsis_methanolica_group | 609 | AMYTH_RS0110500 | 2120528..2122357 | 84480 | DUF262-domain-containing-protein-[Amycolatopsis-thermoflava] |
| WP_123687427.1 | ParB+HNH | GCF_003752125.1 | Amycolatopsis_thermoflava | Bacteria>Bacillati>Actinomycetota>Actinomycetes>Pseudonocardiales>Pseudonocardiaceae>Amycolatopsis>Amycolatopsis_methanolica_group | 608 | EDD35_RS38710 | 65665..67491 | 84480 | DUF262-domain-containing-protein-[Amycolatopsis-thermoflava] |
| MBB5890599.1 | ParB+HNH | GCA_014203355.1 | Kutzneria_kofuensis | Bacteria>Bacillati>Actinomycetota>Actinomycetes>Pseudonocardiales>Pseudonocardiaceae>Kutzneria | 602 | BJ998_001795 | 1953098..1954906 | 103725 | hypothetical-protein-[Kutzneria-kofuensis] |
| WP_281393166.1 | ParB+HNH | - | Kutzneria_kofuensis | Bacteria>Bacillati>Actinomycetota>Actinomycetes>Pseudonocardiales>Pseudonocardiaceae>Kutzneria | 595 | BJ998_RS08845 | 1953098..1954885 | 103725 | DUF262-domain-containing-protein-[Kutzneria-kofuensis] |
| SDG04274.1 | ParB+HNH | GCA_900100955.1 | Lentzea_fradiae | Bacteria>Bacillati>Actinomycetota>Actinomycetes>Pseudonocardiales>Pseudonocardiaceae>Lentzea | 762 | SAMN05216553_10538 | 53764..56052 | 200378 | Putative-DNA-binding-domain-containing-protein-[Lentzea-fradiae] |
| WP_090048718.1 | ParB+HNH | GCF_900100955.1 | Lentzea_fradiae | Bacteria>Bacillati>Actinomycetota>Actinomycetes>Pseudonocardiales>Pseudonocardiaceae>Lentzea | 762 | BLT23_RS12405 | 53764..56052 | 200378 | DUF262-domain-containing-protein-[Lentzea-fradiae] |
| GAA0926813.1 | ParB+HNH | GCA_039525585.1 | Pseudonocardia_zijingensis | Bacteria>Bacillati>Actinomycetota>Actinomycetes>Pseudonocardiales>Pseudonocardiaceae>Pseudonocardia | 773 | GCM10009559_12180 | 42766..45087 | 153376 | hypothetical-protein-GCM10009559_12180-[Pseudonocardia-zijingensis] |
| GAA4991752.1 | ParB+HNH | GCA_039544595.1 | Pseudonocardia_tropica | Bacteria>Bacillati>Actinomycetota>Actinomycetes>Pseudonocardiales>Pseudonocardiaceae>Pseudonocardia | 749 | GCM10025792_45710 | 201033..203282 | 681289 | hypothetical-protein-GCM10025792_45710-[Pseudonocardia-tropica] |
| WP_218590225.1 | ParB+HNH | GCF_019263625.1 | Pseudonocardia_oceani | Bacteria>Bacillati>Actinomycetota>Actinomycetes>Pseudonocardiales>Pseudonocardiaceae>Pseudonocardia | 633 | I4I79_RS18825 | 6840..8741 | 2792013 | DUF262-domain-containing-protein-[Pseudonocardia-oceani] |
| WP_214403720.1 | ParB+HNH | GCF_018531445.1 | Pseudonocardia_lacus | Bacteria>Bacillati>Actinomycetota>Actinomycetes>Pseudonocardiales>Pseudonocardiaceae>Pseudonocardia | 559 | KIK08_RS14825 | 9775..11454 | 2835865 | DUF262-domain-containing-protein-[Pseudonocardia-lacus] |
| WFG45379.1 | ParB+HNH | GCA_045159825.1 | Pseudonocardia_alni | Bacteria>Bacillati>Actinomycetota>Actinomycetes>Pseudonocardiales>Pseudonocardiaceae>Pseudonocardia | 752 | PaSha_20195 | 4101086..4103344 | 33907 | DUF262-domain-containing-protein-[Pseudonocardia-alni] |
| WFG45386.1 | ParB+HNH | GCA_045159825.1 | Pseudonocardia_alni | Bacteria>Bacillati>Actinomycetota>Actinomycetes>Pseudonocardiales>Pseudonocardiaceae>Pseudonocardia | 613 | PaSha_20235 | 4122642..4124483 | 33907 | DUF262-domain-containing-protein-[Pseudonocardia-alni] |
| GAA1837124.1 | ParB+HNH | GCA_039530145.1 | Pseudonocardia_alni_subsp._carboxydivorans | Bacteria>Bacillati>Actinomycetota>Actinomycetes>Pseudonocardiales>Pseudonocardiaceae>Pseudonocardia>Pseudonocardia_alni | 751 | GCM10009772_13280 | 47239..49494 | 415010 | hypothetical-protein-GCM10009772_13280-[Pseudonocardia-carboxydivorans] |
| GAA1837080.1 | ParB+HNH | GCA_039530145.1 | Pseudonocardia_alni_subsp._carboxydivorans | Bacteria>Bacillati>Actinomycetota>Actinomycetes>Pseudonocardiales>Pseudonocardiaceae>Pseudonocardia>Pseudonocardia_alni | 613 | GCM10009772_13210 | 27316..29157 | 415010 | DUF262-domain-containing-protein-[Pseudonocardia-carboxydivorans] |
| WP_251785093.1 | ParB+HNH | GCF_023717215.1 | Pseudonocardia_sp._DR1-2 | Bacteria>Bacillati>Actinomycetota>Actinomycetes>Pseudonocardiales>Pseudonocardiaceae>Pseudonocardia>unclassified_Pseudonocardia | 751 | ND486_RS16915 | 254892..257147 | 2951168 | DUF262-domain-containing-protein-[Pseudonocardia-sp.-DR1-2] |
| WP_251785085.1 | ParB+HNH | GCF_023717215.1 | Pseudonocardia_sp._DR1-2 | Bacteria>Bacillati>Actinomycetota>Actinomycetes>Pseudonocardiales>Pseudonocardiaceae>Pseudonocardia>unclassified_Pseudonocardia | 613 | ND486_RS16875 | 233448..235289 | 2951168 | DUF262-domain-containing-protein-[Pseudonocardia-sp.-DR1-2] |
| WP_190817000.1 | ParB+HNH | GCF_014697215.1 | Saccharopolyspora_pogona | Bacteria>Bacillati>Actinomycetota>Actinomycetes>Pseudonocardiales>Pseudonocardiaceae>Saccharopolyspora | 600 | DL519_RS19940 | 4375665..4377467 | 333966 | DUF262-domain-containing-protein-[Saccharopolyspora-pogona] |
| WP_077690678.1 | ParB+HNH | GCF_014927145.1 | Nocardiopsis | Bacteria>Bacillati>Actinomycetota>Actinomycetes>Streptosporangiales>Nocardiopsidaceae | 593 | NOSIN_RS11195 | 2600688..2602469 | 2013 | MULTISPECIES:-DUF262-domain-containing-protein-[Nocardiopsis] |
| WP_161111649.1 | ParB+HNH | GCF_009862005.1 | Nocardiopsis_alba | Bacteria>Bacillati>Actinomycetota>Actinomycetes>Streptosporangiales>Nocardiopsidaceae>Nocardiopsis | 593 | GTW20_RS21100 | 4854446..4856227 | 53437 | DUF262-domain-containing-protein-[Nocardiopsis-alba] |
| WP_155999757.1 | ParB+HNH | GCF_009728995.1 | Streptomonospora_sp._PA3 | Bacteria>Bacillati>Actinomycetota>Actinomycetes>Streptosporangiales>Nocardiopsidaceae>Streptomonospora>unclassified_Streptomonospora | 662 | FZ103_RS07075 | 274735..276723 | 2607326 | DUF262-domain-containing-protein-[Streptomonospora-sp.-PA3] |
| WP_223868641.1 | ParB+HNH | - | Microbispora_bryophytorum | Bacteria>Bacillati>Actinomycetota>Actinomycetes>Streptosporangiales>Streptosporangiaceae>Microbispora | 654 | IEQ31_RS35680 | 20260..22224 | 1460882 | DUF262-domain-containing-protein-[Microbispora-camponoti] |
| GHJ54459.1 | ParB+HNH | GCA_018327725.1 | Nonomuraea_sp._TT08I-71 | Bacteria>Bacillati>Actinomycetota>Actinomycetes>Streptosporangiales>Streptosporangiaceae>Nonomuraea>unclassified_Nonomuraea | 560 | Nm8I071_37660 | 3192342..3194024 | 2733864 | hypothetical-protein-Nm8I071_37660-[Nonomuraea-sp.-TT08I-71] |
| MCF6470726.1 | ParB+HNH | GCA_021699195.1 | Nonomuraea_sp._MG754425 | Bacteria>Bacillati>Actinomycetota>Actinomycetes>Streptosporangiales>Streptosporangiaceae>Nonomuraea>unclassified_Nonomuraea | 665 | FAF44_20340 | 21152..23149 | 2570319 | DUF262-domain-containing-protein-[Nonomuraea-sp.-MG754425] |
| WP_272894787.1 | ParB+HNH | - | Nonomuraea_sp._MG754425 | Bacteria>Bacillati>Actinomycetota>Actinomycetes>Streptosporangiales>Streptosporangiaceae>Nonomuraea>unclassified_Nonomuraea | 654 | FAF44_RS20635 | 21185..23149 | 2570319 | DUF262-domain-containing-protein-[Nonomuraea-sp.-MG754425] |
| WP_203946298.1 | ParB+HNH | GCF_016863335.1 | Planotetraspora_thailandica | Bacteria>Bacillati>Actinomycetota>Actinomycetes>Streptosporangiales>Streptosporangiaceae>Planotetraspora | 665 | Pth03_RS22305 | 49781..51778 | 487172 | DUF262-domain-containing-protein-[Planotetraspora-thailandica] |
| WP_203980268.1 | ParB+HNH | GCF_039527995.1 | Planotetraspora_silvatica | Bacteria>Bacillati>Actinomycetota>Actinomycetes>Streptosporangiales>Streptosporangiaceae>Planotetraspora | 665 | Psi02_RS35585 | 20184..22181 | 234614 | DUF262-domain-containing-protein-[Planotetraspora-silvatica] |
| GII94897.1 | ParB+HNH | GCA_016863435.1 | Sinosporangium_siamense | Bacteria>Bacillati>Actinomycetota>Actinomycetes>Streptosporangiales>Streptosporangiaceae>Sinosporangium | 665 | Ssi02_51280 | 165858..167855 | 1367973 | hypothetical-protein-Ssi02_51280-[Sinosporangium-siamense] |
| WP_204029816.1 | ParB+HNH | GCF_016863435.1 | Sinosporangium_siamense | Bacteria>Bacillati>Actinomycetota>Actinomycetes>Streptosporangiales>Streptosporangiaceae>Sinosporangium | 665 | Ssi02_RS24935 | 165858..167855 | 1367973 | DUF262-domain-containing-protein-[Sinosporangium-siamense] |
| WP_184983642.1 | ParB+HNH | GCF_039528315.1 | Sphaerisporangium_rubeum | Bacteria>Bacillati>Actinomycetota>Actinomycetes>Streptosporangiales>Streptosporangiaceae>Sphaerisporangium | 666 | BJ992_RS21145 | 4924075..4926075 | 321317 | DUF262-domain-containing-protein-[Sphaerisporangium-rubeum] |
| WP_246424479.1 | ParB+HNH | - | Streptosporangium_sandarakinum | Bacteria>Bacillati>Actinomycetota>Actinomycetes>Streptosporangiales>Streptosporangiaceae>Streptosporangium | 666 | HDA43_RS21560 | 649061..651061 | 1260955 | DUF262-domain-containing-protein-[Streptosporangium-sandarakinum] |
| WP_182841314.1 | ParB+HNH | GCF_039528335.1 | Actinomadura_namibiensis | Bacteria>Bacillati>Actinomycetota>Actinomycetes>Streptosporangiales>Thermomonosporaceae>Actinomadura | 576 | HNR61_RS01545 | 334066..335796 | 182080 | DUF262-domain-containing-protein-[Actinomadura-namibiensis] |
| WP_083983321.1 | ParB+HNH | GCF_001552635.1 | Actinomadura_hibisca | Bacteria>Bacillati>Actinomycetota>Actinomycetes>Streptosporangiales>Thermomonosporaceae>Actinomadura | 577 | AH6_RS22065 | 115602..117335 | 68565 | DUF262-domain-containing-protein-[Actinomadura-hibisca] |
| WP_067795476.1 | ParB+HNH | GCF_001552155.1 | Actinomadura_formosensis | Bacteria>Bacillati>Actinomycetota>Actinomycetes>Streptosporangiales>Thermomonosporaceae>Actinomadura | 577 | AF1_RS09865 | 41926..43659 | 60706 | DUF262-domain-containing-protein-[Actinomadura-formosensis] |
| MCH9729511.1 | ParB+HNH | GCA_022600605.1 | Actinomycetes_bacterium | Bacteria>Bacillati>Actinomycetota>Actinomycetes>unclassified_Actinomycetes | 616 | KOU84_07520 | 6638..8488 | 1883427 | DUF262-domain-containing-protein-[Actinomycetia-bacterium] |
| MCL4078101.1 | ParB+HNH | GCA_023371205.1 | Parvivirga_hydrogeniphila | Bacteria>Bacillati>Actinomycetota>Coriobacteriia>Anaerosomatales>Anaerosomataceae>Parvivirga | 613 | MX659_00525 | 106107..107948 | 2939460 | DUF262-domain-containing-protein-[Parvivirga-hydrogeniphila] |
| WP_267191538.1 | ParB+HNH | GCF_023371205.1 | Parvivirga_hydrogeniphila | Bacteria>Bacillati>Actinomycetota>Coriobacteriia>Anaerosomatales>Anaerosomataceae>Parvivirga | 613 | MX659_RS00525 | 106107..107948 | 2939460 | DUF262-domain-containing-protein-[Parvivirga-hydrogeniphila] |
| GAV31943.1 | ParB+HNH | GCA_001950395.1 | Coriobacteriaceae_bacterium_EMTCatB1 | Bacteria>Bacillati>Actinomycetota>Coriobacteriia>Coriobacteriales>Coriobacteriaceae>unclassified_Coriobacteriaceae | 613 | emb_1c0565 | 1377800..1379641 | 1927122 | uncharacterized-conserved-protein-[Coriobacteriaceae-bacterium-EMTCatB1] |
| WP_087190348.1 | ParB+HNH | GCF_002159895.1 | Gordonibacter | Bacteria>Bacillati>Actinomycetota>Coriobacteriia>Eggerthellales>Eggerthellaceae | 565 | B5F44_RS04020 | 73281..74978 | 644652 | MULTISPECIES:-DUF262-domain-containing-protein-[Gordonibacter] |
| MBC7265386.1 | ParB+HNH | GCA_014360605.1 | Coriobacteriia_bacterium | Bacteria>Bacillati>Actinomycetota>Coriobacteriia>unclassified_Coriobacteriia | 613 | H5T75_00170 | 38598..40439 | 2052159 | DUF262-domain-containing-protein-[Coriobacteriia-bacterium] |
| MTV27844.1 | ParB+HNH | GCA_009720505.1 | Profundirhabdus_halotolerans | Bacteria>Bacillati>Actinomycetota>Nitriliruptoria>Nitriliruptorales>Nitriliruptoraceae>Profundirhabdus | 614 | FTX61_20860 | 20318..22162 | 2604685 | DUF262-domain-containing-protein-[Nitriliruptoraceae-bacterium-ZYF776] |
| WP_217922209.1 | ParB+HNH | GCF_018141545.1 | Miltoncostaea_oceani | Bacteria>Bacillati>Actinomycetota>Thermoleophilia>Miltoncostaeales>Miltoncostaeaceae>Miltoncostaea | 779 | IU369_RS17200 | 3362197..3364536 | 2843216 | DUF262-domain-containing-protein-[Miltoncostaea-oceani] |
| MDA0169808.1 | ParB+HNH | GCA_027587225.1 | Solirubrobacter_taibaiensis | Bacteria>Bacillati>Actinomycetota>Thermoleophilia>Solirubrobacterales>Solirubrobacteraceae>Solirubrobacter | 713 | OJ998_11995 | 120158..122299 | 1395955 | DUF262-domain-containing-HNH-endonuclease-family-protein-[Solirubrobacter-taibaiensis] |
| MBJ7470369.1 | ParB+HNH | GCA_016463515.1 | Solirubrobacteraceae_bacterium | Bacteria>Bacillati>Actinomycetota>Thermoleophilia>Solirubrobacterales>Solirubrobacteraceae>unclassified_Solirubrobacteraceae | 598 | JHD16_03670 | 15164..16960 | 2586742 | DUF262-domain-containing-protein-[Solirubrobacteraceae-bacterium] |
| MBI2692602.1 | ParB+HNH | GCA_016185345.1 | Solirubrobacterales_bacterium | Bacteria>Bacillati>Actinomycetota>Thermoleophilia>Solirubrobacterales>unclassified_Solirubrobacterales | 617 | HYX29_11740 | 162349..164202 | 1909295 | DUF262-domain-containing-protein-[Solirubrobacterales-bacterium] |
| MCB0857324.1 | ParB+HNH | GCA_020438355.1 | Solirubrobacterales_bacterium | Bacteria>Bacillati>Actinomycetota>Thermoleophilia>Solirubrobacterales>unclassified_Solirubrobacterales | 592 | KDB57_04285 | 60800..62578 | 1909295 | DUF262-domain-containing-protein-[Solirubrobacterales-bacterium] |
| MDA8281343.1 | ParB+HNH | GCA_028067595.1 | Actinomycetota_bacterium | Bacteria>Bacillati>Actinomycetota>unclassified_Actinomycetota | 622 | MOZ63_13170 | 19185..21053 | 2900548 | MAG:-DUF262-domain-containing-protein-[Actinomycetota-bacterium] |
| MCA0294160.1 | ParB+HNH | GCA_020161755.1 | Actinomycetota_bacterium | Bacteria>Bacillati>Actinomycetota>unclassified_Actinomycetota | 610 | LCH96_02390 | 203663..205495 | 2900548 | DUF262-domain-containing-protein-[Actinomycetia-bacterium] |
| MDA8117684.1 | ParB+HNH | GCA_028064745.1 | Actinomycetota_bacterium | Bacteria>Bacillati>Actinomycetota>unclassified_Actinomycetota | 570 | M0000_09985 | 11130..12842 | 2900548 | MAG:-DUF262-domain-containing-protein-[Actinomycetota-bacterium] |
| MSW90816.1 | ParB+HNH | GCA_009699605.1 | Actinomycetota_bacterium | Bacteria>Bacillati>Actinomycetota>unclassified_Actinomycetota | 443 | F2812_04360 | 25587..26918 | 2900548 | DUF262-domain-containing-protein-[Actinomycetia-bacterium] |
| TAK69423.1 | ParB+HNH | GCA_004297305.1 | Actinomycetota_bacterium | Bacteria>Bacillati>Actinomycetota>unclassified_Actinomycetota | 368 | EPO13_05915 | 51536..52642 | 2900548 | DUF262-domain-containing-protein-[Actinomycetota-bacterium] |
| MBX7131091.1 | ParB+HNH | GCA_019694835.1 | Fimbriimonadaceae_bacterium | Bacteria>Bacillati>Armatimonadota>Fimbriimonadia>Fimbriimonadales>Fimbriimonadaceae>unclassified_Fimbriimonadaceae | 559 | K1X67_00280 | 63827..65506 | 2682144 | DUF262-domain-containing-protein-[Fimbriimonadaceae-bacterium] |
| WP_118637671.1 | ParB+HNH | GCF_027924705.1 | Clostridia | Bacteria>Bacillati>Bacillota | 378 | H6A32_RS13605 | 20969..22105 | 186801 | MULTISPECIES:-DUF262-domain-containing-protein-[Oscillospiraceae] |
| WP_002362822.1 | ParB+HNH | GCF_007034685.1 | Lactobacillales | Bacteria>Bacillati>Bacillota>Bacilli | 680 | AW733_RS05625 | 287629..289671 | 186826 | MULTISPECIES:-DUF4357-domain-containing-protein-[Bacteria] |
| WP_144538746.1 | ParB+HNH | - | Bacillus_sp._FJAT-22090 | Bacteria>Bacillati>Bacillota>Bacilli>Bacillales>Bacillaceae>Bacillus>unclassified_Bacillus_(in:_firmicutes) | 589 | FS503_RS11560 | 122606..124375 | 1581038 | DUF262-domain-containing-protein-[Bacillus-sp.-FJAT-22090] |
| PFG07780.1 | ParB+HNH | GCA_002563655.1 | Bacillus_sp._es.034 | Bacteria>Bacillati>Bacillota>Bacilli>Bacillales>Bacillaceae>Bacillus>unclassified_Bacillus_(in:_firmicutes) | 699 | ATG71_4689 | 4478978..4481077 | 1761763 | uncharacterized-protein-DUF1524-[Bacillus-sp.-es.034] |
| WP_251162452.1 | ParB+HNH | GCF_023713445.1 | Cytobacillus_kochii | Bacteria>Bacillati>Bacillota>Bacilli>Bacillales>Bacillaceae>Cytobacillus | 554 | M3670_RS08705 | 70967..72631 | 859143 | DUF262-domain-containing-protein-[Cytobacillus-kochii] |
| WP_066398761.1 | ParB+HNH | GCF_017874625.1 | Cytobacillus_eiseniae | Bacteria>Bacillati>Bacillota>Bacilli>Bacillales>Bacillaceae>Cytobacillus | 554 | J2Z40_RS15695 | 13743..15407 | 762947 | DUF262-domain-containing-protein-[Cytobacillus-eiseniae] |
| WP_108713782.1 | ParB+HNH | GCF_003073375.1 | unclassified_Lysinibacillus | Bacteria>Bacillati>Bacillota>Bacilli>Bacillales>Bacillaceae>Lysinibacillus | 589 | DCE79_RS14890 | 3038221..3039990 | 2636778 | MULTISPECIES:-DUF262-domain-containing-protein-[unclassified-Lysinibacillus] |
| WP_259420293.1 | ParB+HNH | GCF_024919535.1 | Lysinibacillus_sp._A4 | Bacteria>Bacillati>Bacillota>Bacilli>Bacillales>Bacillaceae>Lysinibacillus>unclassified_Lysinibacillus | 589 | NY607_RS20540 | 121506..123275 | 2976269 | DUF262-domain-containing-HNH-endonuclease-family-protein-[Lysinibacillus-sp.-A4] |
| WP_191566568.1 | ParB+HNH | GCF_904423835.1 | Metabacillus_idriensis | Bacteria>Bacillati>Bacillota>Bacilli>Bacillales>Bacillaceae>Metabacillus | 554 | IHQ32_RS17400 | 3429556..3431220 | 324768 | DUF262-domain-containing-protein-[Metabacillus-idriensis] |
| WP_166025637.1 | ParB+HNH | GCF_011393025.1 | Neobacillus_terrae | Bacteria>Bacillati>Bacillota>Bacilli>Bacillales>Bacillaceae>Neobacillus | 589 | G8761_RS08620 | 50470..52239 | 3034837 | DUF262-domain-containing-protein-[Neobacillus-terrae] |
| WP_176473356.1 | ParB+HNH | GCF_002271995.1 | Niallia_circulans | Bacteria>Bacillati>Bacillota>Bacilli>Bacillales>Bacillaceae>Niallia | 381 | CHH55_RS21980 | 43319..44464 | 1397 | DUF262-domain-containing-protein-[Niallia-circulans] |
| WP_042224730.1 | ParB+HNH | GCF_000724085.1 | Oceanobacillus_manasiensis | Bacteria>Bacillati>Bacillota>Bacilli>Bacillales>Bacillaceae>Oceanobacillus | 614 | BN985_RS16190 | 540865..542709 | 586413 | DUF262-domain-containing-protein-[Oceanobacillus-manasiensis] |
| WP_182480456.1 | ParB+HNH | GCF_003570725.1 | Peribacillus_asahii | Bacteria>Bacillati>Bacillota>Bacilli>Bacillales>Bacillaceae>Peribacillus | 362 | D1953_RS00195 | 21333..22421 | 228899 | DUF262-domain-containing-protein-[Peribacillus-asahii] |
| WP_225002272.1 | ParB+HNH | GCF_020176725.1 | Priestia_flexa | Bacteria>Bacillati>Bacillota>Bacilli>Bacillales>Bacillaceae>Priestia | 554 | LC052_RS15455 | 54606..56270 | 86664 | DUF262-domain-containing-protein-[Priestia-flexa] |
| WP_061808760.1 | ParB+HNH | GCF_001591825.1 | Rossellomorea_vietnamensis | Bacteria>Bacillati>Bacillota>Bacilli>Bacillales>Bacillaceae>Rossellomorea | 554 | BVI01S_RS00225 | 48874..50538 | 218284 | DUF262-domain-containing-protein-[Rossellomorea-vietnamensis] |
| WP_102026203.1 | ParB+HNH | GCF_900240405.1 | Salirhabdus_sp._Marseille-P4669 | Bacteria>Bacillati>Bacillota>Bacilli>Bacillales>Bacillaceae>Salirhabdus>unclassified_Salirhabdus | 554 | C1A05_RS00420 | 90106..91770 | 2042310 | DUF262-domain-containing-protein-[Salirhabdus-sp.-Marseille-P4669] |
| WP_253286656.1 | ParB+HNH | GCF_018918075.1 | Virgibacillus_sp._MSJ-26 | Bacteria>Bacillati>Bacillota>Bacilli>Bacillales>Bacillaceae>Virgibacillus>unclassified_Virgibacillus | 684 | KQI49_RS03125 | 245635..247689 | 2841522 | DUF262-and-DUF1524-domain-containing-protein-[Virgibacillus-sp.-MSJ-26] |
| WP_108587663.1 | ParB+HNH | GCF_003058165.1 | Paenisporosarcina_sp._OV554 | Bacteria>Bacillati>Bacillota>Bacilli>Bacillales>Caryophanaceae>Paenisporosarcina>unclassified_Paenisporosarcina | 691 | C8K15_RS17230 | 12439..14514 | 2135694 | DUF262-and-DUF1524-domain-containing-protein-[Paenisporosarcina-sp.-OV554] |
| WP_272441149.1 | ParB+HNH | GCF_028421645.1 | Aneurinibacillus_sp._Ricciae_BoGa-3 | Bacteria>Bacillati>Bacillota>Bacilli>Bacillales>Paenibacillaceae>Aneurinibacillus_group>Aneurinibacillus>unclassified_Aneurinibacillus | 578 | PP175_RS01555 | 306217..307953 | 3022697 | DUF262-domain-containing-HNH-endonuclease-family-protein-[Aneurinibacillus-sp.-Ricciae_BoGa-3] |
| WP_272441154.1 | ParB+HNH | GCF_028421645.1 | Aneurinibacillus_sp._Ricciae_BoGa-3 | Bacteria>Bacillati>Bacillota>Bacilli>Bacillales>Paenibacillaceae>Aneurinibacillus_group>Aneurinibacillus>unclassified_Aneurinibacillus | 348 | PP175_RS01580 | 318722..319768 | 3022697 | DUF262-domain-containing-protein-[Aneurinibacillus-sp.-Ricciae_BoGa-3] |
| WP_010494814.1 | ParB+HNH | GCF_000213315.1 | Paenibacillus_elgii | Bacteria>Bacillati>Bacillota>Bacilli>Bacillales>Paenibacillaceae>Paenibacillus | 695 | MGF69_RS0111745 | 17733..19820 | 189691 | DUF262-and-DUF1524-domain-containing-protein-[Paenibacillus-elgii] |
| WP_038596872.1 | ParB+HNH | GCF_000758525.1 | Paenibacillus_sp._FSL_H7-0357 | Bacteria>Bacillati>Bacillota>Bacilli>Bacillales>Paenibacillaceae>Paenibacillus>unclassified_Paenibacillus | 695 | H70357_RS30050 | 6825709..6827796 | 1536774 | DUF262-and-DUF1524-domain-containing-protein-[Paenibacillus-sp.-FSL-H7-0357] |

| Protein-ID | Domain | Assembly-ID | Strain | Taxonomy | Length | Locus-Tag | Genomic-coordinates | TaxID | Fasta-defline |
|---|---|---|---|---|---|---|---|---|---|
| RYL92594.1 | ParB+HNH | GCA_004153195.1 | Sporolactobacillus_sp._THM19-2 | Bacteria>Bacillati>Bacillota>Bacilli>Bacillales>Sporolactobacillaceae>Sporolactobacillus>unclassified_Sporolactobacillus | 608 | EWH91_06950 | 37418..39244 | 2511171 | DUF262-domain-containing-protein-[Sporolactobacillus-sp.-THM19-2] |
| OZT77516.1 | ParB+HNH | GCA_002265785.1 | Salinicoccus_roseus | Bacteria>Bacillati>Bacillota>Bacilli>Bacillales>Staphylococcaceae>Salinicoccus | 444 | CFN03_06145 | 224575..225909 | 45670 | hypothetical-protein-CFN03_06145-[Salinicoccus-roseus] |
| WP_179287252.1 | ParB+HNH | GCF_002265785.1 | Salinicoccus_roseus | Bacteria>Bacillati>Bacillota>Bacilli>Bacillales>Staphylococcaceae>Salinicoccus | 441 | CFN03_RS06145 | 224575..225900 | 45670 | DUF262-domain-containing-protein-[Salinicoccus-roseus] |
| WP_015638646.1 | ParB+HNH | GCF_000397165.1 | Lactobacillaceae | Bacteria>Bacillati>Bacillota>Bacilli>Lactobacillales | 595 | OSJ94_RS09285 | 18681..20468 | 33958 | MULTISPECIES:-DUF262-domain-containing-protein-[Lactobacillaceae] |
| WP_191997593.1 | ParB+HNH | - | Lactobacillaceae | Bacteria>Bacillati>Bacillota>Bacilli>Lactobacillales | 579 | FOS38_RS12520 | 1260..2999 | 33958 | MULTISPECIES:-DUF262-domain-containing-protein-[Lactobacillaceae] |
| WP_091633355.1 | ParB+HNH | GCF_900109085.1 | Alkalibacterium_gilvum | Bacteria>Bacillati>Bacillota>Bacilli>Lactobacillales>Carnobacteriaceae>Alkalibacterium | 707 | BMX83_RS03320 | 18658..20781 | 1130080 | DUF262-and-DUF1524-domain-containing-protein-[Alkalibacterium-gilvum] |
| HAJ70158.1 | ParB+HNH | GCA_003454485.1 | Alkalibacterium_sp. | Bacteria>Bacillati>Bacillota>Bacilli>Lactobacillales>Carnobacteriaceae>Alkalibacterium>unclassified_Alkalibacterium | 707 | DCO62_02900 | 48224..50347 | 1872447 | hypothetical-protein-[Alkalibacterium-sp.] |
| WP_251851840.1 | ParB+HNH | GCF_023744155.1 | Enterococcus_italicus | Bacteria>Bacillati>Bacillota>Bacilli>Lactobacillales>Enterococcaceae>Enterococcus | 597 | NE261_RS07310 | 534298..536091 | 246144 | DUF262-domain-containing-protein-[Enterococcus-italicus] |
| WP_144771936.1 | ParB+HNH | GCF_007682015.1 | Enterococcus_casseliflavus | Bacteria>Bacillati>Bacillota>Bacilli>Lactobacillales>Enterococcaceae>Enterococcus | 597 | FS854_RS14485 | 59095..60888 | 37734 | DUF262-domain-containing-protein-[Enterococcus-casseliflavus] |
| WP_156237147.1 | ParB+HNH | GCF_009734725.1 | Enterococcus_casseliflavus | Bacteria>Bacillati>Bacillota>Bacilli>Lactobacillales>Enterococcaceae>Enterococcus | 597 | EZ072_RS09900 | 39244..41037 | 37734 | DUF262-domain-containing-protein-[Enterococcus-casseliflavus] |
| WP_194194884.1 | ParB+HNH | - | Enterococcus_casseliflavus | Bacteria>Bacillati>Bacillota>Bacilli>Lactobacillales>Enterococcaceae>Enterococcus | 597 | HAX40_RS05275 | 91102..92895 | 37734 | DUF262-domain-containing-protein-[Enterococcus-casseliflavus] |
| WP_225426568.1 | ParB+HNH | GCF_003946555.1 | Companilactobacillus_hulinensis | Bacteria>Bacillati>Bacillota>Bacilli>Lactobacillales>Lactobacillaceae>Companilactobacillus | 590 | EKM66_RS06310 | 30822..32594 | 2486007 | DUF262-domain-containing-protein-[Companilactobacillus-hulinensis] |
| WP_211334341.1 | ParB+HNH | GCF_003946325.1 | Companilactobacillus_furfuricola | Bacteria>Bacillati>Bacillota>Bacilli>Lactobacillales>Lactobacillaceae>Companilactobacillus | 434 | EKM78_RS06195 | 15590..16894 | 1462575 | DUF262-domain-containing-protein-[Companilactobacillus-furfuricola] |
| WP_193820354.1 | ParB+HNH | GCF_016653515.1 | Lacticaseibacillus_rhamnosus | Bacteria>Bacillati>Bacillota>Bacilli>Lactobacillales>Lactobacillaceae>Lacticaseibacillus | 710 | JIO02_RS10950 | 2289871..2292003 | 47715 | DUF262-domain-containing-protein-[Lacticaseibacillus-rhamnosus] |
| WP_005715301.1 | ParB+HNH | GCF_000311945.1 | Lacticaseibacillus_rhamnosus | Bacteria>Bacillati>Bacillota>Bacilli>Lactobacillales>Lactobacillaceae>Lacticaseibacillus | 694 | LRHMDP2_RS10645 | 140422..142506 | 47715 | DUF262-domain-containing-protein-[Lacticaseibacillus-rhamnosus] |
| WP_014569893.1 | ParB+HNH | GCF_000026505.1 | Lacticaseibacillus_rhamnosus | Bacteria>Bacillati>Bacillota>Bacilli>Lactobacillales>Lactobacillaceae>Lacticaseibacillus | 694 | LGG_RS10070 | 2151745..2153829 | 47715 | DUF262-domain-containing-protein-[Lacticaseibacillus-rhamnosus] |
| WP_032952312.1 | ParB+HNH | GCF_925281885.1 | Lacticaseibacillus_rhamnosus | Bacteria>Bacillati>Bacillota>Bacilli>Lactobacillales>Lactobacillaceae>Lacticaseibacillus | 694 | N507_RS04235 | 882194..884278 | 47715 | DUF262-domain-containing-protein-[Lacticaseibacillus-rhamnosus] |
| WP_032957085.1 | ParB+HNH | GCF_051908385.1 | Lacticaseibacillus_rhamnosus | Bacteria>Bacillati>Bacillota>Bacilli>Lactobacillales>Lactobacillaceae>Lacticaseibacillus | 694 | FG342_RS12980 | 2553873..2555957 | 47715 | DUF262-domain-containing-protein-[Lacticaseibacillus-rhamnosus] |
| WP_048482581.1 | ParB+HNH | GCF_051123815.1 | Lacticaseibacillus_rhamnosus | Bacteria>Bacillati>Bacillota>Bacilli>Lactobacillales>Lactobacillaceae>Lacticaseibacillus | 694 | DA801_RS04650 | 54837..56921 | 47715 | DUF262-domain-containing-protein-[Lacticaseibacillus-rhamnosus] |
| WP_204129041.1 | ParB+HNH | GCF_016887865.1 | Lacticaseibacillus_rhamnosus | Bacteria>Bacillati>Bacillota>Bacilli>Lactobacillales>Lactobacillaceae>Lacticaseibacillus | 694 | JNW95_RS08710 | 94712..96796 | 47715 | DUF262-domain-containing-protein-[Lacticaseibacillus-rhamnosus] |
| WP_233026118.1 | ParB+HNH | GCF_021300595.1 | Lacticaseibacillus_rhamnosus | Bacteria>Bacillati>Bacillota>Bacilli>Lactobacillales>Lactobacillaceae>Lacticaseibacillus | 694 | LW144_RS03900 | 86049..88133 | 47715 | DUF262-domain-containing-protein-[Lacticaseibacillus-rhamnosus] |
| WP_048488334.1 | ParB+HNH | GCF_901971785.1 | Lacticaseibacillus_rhamnosus | Bacteria>Bacillati>Bacillota>Bacilli>Lactobacillales>Lactobacillaceae>Lacticaseibacillus | 595 | EFO73_RS10395 | 26549..28336 | 47715 | DUF262-domain-containing-protein-[Lacticaseibacillus-rhamnosus] |
| WP_003602970.1 | ParB+HNH | GCF_000376145.1 | Lacticaseibacillus_paracasei | Bacteria>Bacillati>Bacillota>Bacilli>Lactobacillales>Lactobacillaceae>Lacticaseibacillus | 595 | LCAUW4_RS03755 | 2281..4068 | 1597 | MULTISPECIES:-DUF262-domain-containing-protein-[Lacticaseibacillus] |
| WP_260565637.1 | ParB+HNH | GCF_025215665.1 | Lacticaseibacillus_paracasei | Bacteria>Bacillati>Bacillota>Bacilli>Lactobacillales>Lactobacillaceae>Lacticaseibacillus | 595 | EOK76_RS04800 | 1038418..1040205 | 1597 | DUF262-domain-containing-protein-[Lacticaseibacillus-paracasei] |
| WP_269189927.1 | ParB+HNH | - | Lacticaseibacillus_paracasei | Bacteria>Bacillati>Bacillota>Bacilli>Lactobacillales>Lactobacillaceae>Lacticaseibacillus | 579 | K9853_RS00475 | 15577..17316 | 1597 | DUF262-domain-containing-protein-[Lacticaseibacillus-paracasei] |
| WP_274784935.1 | ParB+HNH | GCF_028878205.1 | Lacticaseibacillus_casei | Bacteria>Bacillati>Bacillota>Bacilli>Lactobacillales>Lactobacillaceae>Lacticaseibacillus | 595 | LTZ17_RS09580 | 2717..4504 | 1582 | DUF262-domain-containing-protein-[Lacticaseibacillus-casei] |
| WP_106904929.1 | ParB+HNH | GCF_003020005.1 | Lactiplantibacillus_plantarum | Bacteria>Bacillati>Bacillota>Bacilli>Lactobacillales>Lactobacillaceae>Lactiplantibacillus | 595 | B5G53_RS16165 | 34546..36333 | 1590 | DUF262-domain-containing-protein-[Lactiplantibacillus-plantarum] |
| WP_114788346.1 | ParB+HNH | - | Lactiplantibacillus_plantarum | Bacteria>Bacillati>Bacillota>Bacilli>Lactobacillales>Lactobacillaceae>Lactiplantibacillus | 595 | DB726_RS12025 | 41382..43169 | 1590 | DUF262-domain-containing-protein-[Lactiplantibacillus-plantarum] |
| WP_128485897.1 | ParB+HNH | GCF_004101645.1 | Lactiplantibacillus_plantarum | Bacteria>Bacillati>Bacillota>Bacilli>Lactobacillales>Lactobacillaceae>Lactiplantibacillus | 595 | EQG58_RS16145 | 85802..87589 | 1590 | DUF262-domain-containing-protein-[Lactiplantibacillus-plantarum] |
| WP_128529844.1 | ParB+HNH | GCF_004078645.1 | Lactiplantibacillus_plantarum | Bacteria>Bacillati>Bacillota>Bacilli>Lactobacillales>Lactobacillaceae>Lactiplantibacillus | 595 | EQJ06_RS15705 | 51449..53236 | 1590 | DUF262-domain-containing-protein-[Lactiplantibacillus-plantarum] |
| WP_249391184.1 | ParB+HNH | GCF_964187035.1 | Lactiplantibacillus_plantarum | Bacteria>Bacillati>Bacillota>Bacilli>Lactobacillales>Lactobacillaceae>Lactiplantibacillus | 595 | M4S29_RS14370 | 17499..19286 | 1590 | DUF262-domain-containing-protein-[Lactiplantibacillus-plantarum] |
| WP_229105114.1 | ParB+HNH | - | Lactiplantibacillus_plantarum | Bacteria>Bacillati>Bacillota>Bacilli>Lactobacillales>Lactobacillaceae>Lactiplantibacillus | 579 | I2H95_RS14930 | 17853..19592 | 1590 | DUF262-domain-containing-protein-[Lactiplantibacillus-plantarum] |
| WP_188891955.1 | ParB+HNH | GCF_014656585.1 | Lactobacillus_kefiranofaciens | Bacteria>Bacillati>Bacillota>Bacilli>Lactobacillales>Lactobacillaceae>Lactobacillus | 696 | ICI50_RS03460 | 663521..665611 | 267818 | DUF262-domain-containing-protein-[Lactobacillus-kefiranofaciens] |
| WP_250825480.1 | ParB+HNH | GCF_023674385.1 | Lactobacillus_kefiranofaciens | Bacteria>Bacillati>Bacillota>Bacilli>Lactobacillales>Lactobacillaceae>Lactobacillus | 696 | MU859_RS01760 | 304782..306872 | 267818 | DUF262-domain-containing-protein-[Lactobacillus-kefiranofaciens] |
| WP_244040364.1 | ParB+HNH | GCF_014656585.1 | Lactobacillus_kefiranofaciens | Bacteria>Bacillati>Bacillota>Bacilli>Lactobacillales>Lactobacillaceae>Lactobacillus | 247 | ICI50_RS03475 | 667998..668741 | 267818 | HNH-endonuclease-family-protein-[Lactobacillus-kefiranofaciens] |
| WP_250825483.1 | ParB+HNH | - | Lactobacillus_kefiranofaciens | Bacteria>Bacillati>Bacillota>Bacilli>Lactobacillales>Lactobacillaceae>Lactobacillus | 213 | MU859_RS01775 | 309363..310004 | 267818 | HNH-endonuclease-family-protein-[Lactobacillus-kefiranofaciens] |
| MCI7713734.1 | ParB+HNH | GCA_022786405.1 | Lactobacillus_johnsonii | Bacteria>Bacillati>Bacillota>Bacilli>Lactobacillales>Lactobacillaceae>Lactobacillus | 655 | MSL49_00225 | 11483..13450 | 33959 | DUF262-domain-containing-HNH-endonuclease-family-protein-[Lactobacillus-johnsonii] |
| AUI76211.1 | ParB+HNH | GCA_002849955.1 | Lactobacillus_helveticus | Bacteria>Bacillati>Bacillota>Bacilli>Lactobacillales>Lactobacillaceae>Lactobacillus | 655 | Lh22155_05470 | 1079265..1081232 | 1587 | hypothetical-protein-Lh22155_05470-[Lactobacillus-helveticus] |
| WP_014918811.1 | ParB+HNH | GCF_000165775.1 | Lactobacillus_helveticus | Bacteria>Bacillati>Bacillota>Bacilli>Lactobacillales>Lactobacillaceae>Lactobacillus | 655 | LH5_RS02015 | 419841..421808 | 1587 | DUF262-domain-containing-protein-[Lactobacillus-helveticus] |
| WP_072748936.1 | ParB+HNH | GCF_001895375.1 | Lactobacillus_helveticus | Bacteria>Bacillati>Bacillota>Bacilli>Lactobacillales>Lactobacillaceae>Lactobacillus | 655 | BEL10_RS03680 | 27441..29408 | 1587 | DUF262-domain-containing-protein-[Lactobacillus-helveticus] |
| WP_095662163.1 | ParB+HNH | GCF_011392295.1 | Lactobacillus_helveticus | Bacteria>Bacillati>Bacillota>Bacilli>Lactobacillales>Lactobacillaceae>Lactobacillus | 655 | ERJ65_RS00210 | 12449..14416 | 1587 | DUF262-domain-containing-protein-[Lactobacillus-helveticus] |
| WP_101511689.1 | ParB+HNH | GCF_045348125.1 | Lactobacillus_helveticus | Bacteria>Bacillati>Bacillota>Bacilli>Lactobacillales>Lactobacillaceae>Lactobacillus | 655 | IMAU10149_RS03685 | 17258..19225 | 1587 | DUF262-domain-containing-protein-[Lactobacillus-helveticus] |
| WP_101812675.1 | ParB+HNH | GCF_002849955.1 | Lactobacillus_helveticus | Bacteria>Bacillati>Bacillota>Bacilli>Lactobacillales>Lactobacillaceae>Lactobacillus | 655 | Lh22155_RS05650 | 1079265..1081232 | 1587 | DUF262-domain-containing-protein-[Lactobacillus-helveticus] |
| WP_103658550.1 | ParB+HNH | GCF_015698165.1 | Lactobacillus_helveticus | Bacteria>Bacillati>Bacillota>Bacilli>Lactobacillales>Lactobacillaceae>Lactobacillus | 655 | MVU88_RS02195 | 4097..6064 | 1587 | DUF262-domain-containing-protein-[Lactobacillus-helveticus] |
| WP_172993981.1 | ParB+HNH | GCF_013280765.1 | Lactobacillus_helveticus | Bacteria>Bacillati>Bacillota>Bacilli>Lactobacillales>Lactobacillaceae>Lactobacillus | 655 | IMAU60206_RS01590 | 30415..32382 | 1587 | DUF262-domain-containing-protein-[Lactobacillus-helveticus] |
| WP_173006742.1 | ParB+HNH | GCF_013281475.1 | Lactobacillus_helveticus | Bacteria>Bacillati>Bacillota>Bacilli>Lactobacillales>Lactobacillaceae>Lactobacillus | 655 | IMAU30014_RS01870 | 30109..32076 | 1587 | DUF262-domain-containing-protein-[Lactobacillus-helveticus] |
| WP_220023634.1 | ParB+HNH | GCF_019455975.1 | Lactobacillus_helveticus | Bacteria>Bacillati>Bacillota>Bacilli>Lactobacillales>Lactobacillaceae>Lactobacillus | 655 | E0715_RS08305 | 24491..26458 | 1587 | DUF262-domain-containing-protein-[Lactobacillus-helveticus] |
| WP_220028699.1 | ParB+HNH | GCF_019455965.1 | Lactobacillus_helveticus | Bacteria>Bacillati>Bacillota>Bacilli>Lactobacillales>Lactobacillaceae>Lactobacillus | 655 | E0712_RS03730 | 170672..172639 | 1587 | DUF262-domain-containing-protein-[Lactobacillus-helveticus] |
| WP_259685555.1 | ParB+HNH | GCF_025129335.1 | Lactobacillus_helveticus | Bacteria>Bacillati>Bacillota>Bacilli>Lactobacillales>Lactobacillaceae>Lactobacillus | 655 | EFM06_RS11135 | 17850..19817 | 1587 | DUF262-domain-containing-HNH-endonuclease-family-protein-[Lactobacillus-helveticus] |
| WP_259693425.1 | ParB+HNH | GCF_025129275.1 | Lactobacillus_helveticus | Bacteria>Bacillati>Bacillota>Bacilli>Lactobacillales>Lactobacillaceae>Lactobacillus | 655 | EFM10_RS03610 | 17843..19810 | 1587 | DUF262-domain-containing-HNH-endonuclease-family-protein-[Lactobacillus-helveticus] |
| WP_014563582.1 | ParB+HNH | GCF_000189515.1 | Lactobacillus_helveticus | Bacteria>Bacillati>Bacillota>Bacilli>Lactobacillales>Lactobacillaceae>Lactobacillus | 590 | LBHH_RS05375 | 1051422..1053194 | 1587 | DUF262-domain-containing-protein-[Lactobacillus-helveticus] |
| WP_023192366.1 | ParB+HNH | GCF_000422165.1 | Lactobacillus_helveticus | Bacteria>Bacillati>Bacillota>Bacilli>Lactobacillales>Lactobacillaceae>Lactobacillus | 589 | EFP58_RS07225 | 355..2124 | 1587 | DUF262-domain-containing-protein-[Lactobacillus-helveticus] |
| WP_254261288.1 | ParB+HNH | GCF_013280955.1 | Lactobacillus_helveticus | Bacteria>Bacillati>Bacillota>Bacilli>Lactobacillales>Lactobacillaceae>Lactobacillus | 551 | IMAU50022_RS01340 | 18629..20284 | 1587 | DUF262-domain-containing-protein-[Lactobacillus-helveticus] |
| WP_238525402.1 | ParB+HNH | - | Lactobacillus_helveticus | Bacteria>Bacillati>Bacillota>Bacilli>Lactobacillales>Lactobacillaceae>Lactobacillus | 483 | LBHH_RS05450 | 1077788..1079239 | 1587 | DUF262-domain-containing-protein-[Lactobacillus-helveticus] |
| WP_233725018.1 | ParB+HNH | GCF_002849955.1 | Lactobacillus_helveticus | Bacteria>Bacillati>Bacillota>Bacilli>Lactobacillales>Lactobacillaceae>Lactobacillus | 463 | Lh22155_RS05620 | 1071155..1072546 | 1587 | DUF262-domain-containing-protein-[Lactobacillus-helveticus] |
| WP_254260692.1 | ParB+HNH | GCF_013280775.1 | Lactobacillus_helveticus | Bacteria>Bacillati>Bacillota>Bacilli>Lactobacillales>Lactobacillaceae>Lactobacillus | 343 | IMAU60211_RS09890 | 4911..5942 | 1587 | HNH-endonuclease-family-protein-[Lactobacillus-helveticus] |
| WP_172993588.1 | ParB+HNH | GCF_013280875.1 | Lactobacillus_helveticus | Bacteria>Bacillati>Bacillota>Bacilli>Lactobacillales>Lactobacillaceae>Lactobacillus | 332 | IMAU60201_RS07170 | 9882..10882 | 1587 | HNH-endonuclease-family-protein,-partial-[Lactobacillus-helveticus] |
| WP_023192223.1 | ParB+HNH | GCF_000493555.1 | Lactobacillus_helveticus | Bacteria>Bacillati>Bacillota>Bacilli>Lactobacillales>Lactobacillaceae>Lactobacillus | 274 | LHCIRMBIA104_RS13080 | 7870..8694 | 1587 | DUF262-domain-containing-protein-[Lactobacillus-helveticus] |
| WP_236656789.1 | ParB+HNH | GCF_015698145.1 | Lactobacillus_helveticus | Bacteria>Bacillati>Bacillota>Bacilli>Lactobacillales>Lactobacillaceae>Lactobacillus | 274 | EFP83_RS06600 | 15624..16448 | 1587 | DUF262-domain-containing-protein-[Lactobacillus-helveticus] |
| WP_230479753.1 | ParB+HNH | GCF_015698145.1 | Lactobacillus_helveticus | Bacteria>Bacillati>Bacillota>Bacilli>Lactobacillales>Lactobacillaceae>Lactobacillus | 270 | EFP83_RS06590 | 14483..15295 | 1587 | HNH-endonuclease-family-protein-[Lactobacillus-helveticus] |
| WP_254260691.1 | ParB+HNH | GCF_013280775.1 | Lactobacillus_helveticus | Bacteria>Bacillati>Bacillota>Bacilli>Lactobacillales>Lactobacillaceae>Lactobacillus | 203 | IMAU60211_RS09885 | 4009..4618 | 1587 | DUF262-domain-containing-protein,-partial-[Lactobacillus-helveticus] |
| WP_238525401.1 | ParB+HNH | GCF_000189515.1 | Lactobacillus_helveticus | Bacteria>Bacillati>Bacillota>Bacilli>Lactobacillales>Lactobacillaceae>Lactobacillus | 175 | LBHH_RS12575 | 1077271..1077798 | 1587 | DUF1524-domain-containing-protein-[Lactobacillus-helveticus] |
| WP_003626421.1 | ParB+HNH | GCF_000422165.1 | Lactobacillus_helveticus | Bacteria>Bacillati>Bacillota>Bacilli>Lactobacillales>Lactobacillaceae>Lactobacillus | 66 | EFP58_RS07220 | 158..358 | 1587 | DUF262-domain-containing-protein-[Lactobacillus-helveticus] |
| OUQ00068.1 | ParB+HNH | GCA_002160635.1 | Lactobacillus_gallinarum | Bacteria>Bacillati>Bacillota>Bacilli>Lactobacillales>Lactobacillaceae>Lactobacillus | 655 | B5E95_07150 | 6427..8394 | 52242 | hypothetical-protein-B5E95_07150-[Lactobacillus-gallinarum] |
| WP_087236050.1 | ParB+HNH | GCF_002160635.1 | Lactobacillus_gallinarum | Bacteria>Bacillati>Bacillota>Bacilli>Lactobacillales>Lactobacillaceae>Lactobacillus | 655 | B5E63_RS08440 | 18858..20825 | 52242 | DUF262-domain-containing-protein-[Lactobacillus-gallinarum] |
| WP_231533743.1 | ParB+HNH | GCF_001469775.1 | Lactobacillus_delbrueckii | Bacteria>Bacillati>Bacillota>Bacilli>Lactobacillales>Lactobacillaceae>Lactobacillus | 699 | AT236_RS05360 | 1073420..1075519 | 1584 | DUF262-domain-containing-protein-[Lactobacillus-delbrueckii] |
| WP_064081738.1 | ParB+HNH | GCF_925300675.1 | Lactobacillus_delbrueckii | Bacteria>Bacillati>Bacillota>Bacilli>Lactobacillales>Lactobacillaceae>Lactobacillus | 697 | NGL01_RS07835 | 543..2636 | 1584 | DUF262-domain-containing-protein-[Lactobacillus-delbrueckii] |
| WP_242073731.1 | ParB+HNH | GCF_022642945.1 | Lactobacillus_delbrueckii | Bacteria>Bacillati>Bacillota>Bacilli>Lactobacillales>Lactobacillaceae>Lactobacillus | 697 | G8B25_RS07950 | 1519789..1521882 | 1584 | DUF262-domain-containing-protein-[Lactobacillus-delbrueckii] |
| WP_231524173.1 | ParB+HNH | GCF_021134795.1 | Lactobacillus_delbrueckii | Bacteria>Bacillati>Bacillota>Bacilli>Lactobacillales>Lactobacillaceae>Lactobacillus | 695 | LOB43_RS04540 | 12465..14552 | 1584 | DUF262-domain-containing-protein-[Lactobacillus-delbrueckii] |
| WP_157660270.1 | ParB+HNH | GCF_002142575.1 | Lactobacillus_delbrueckii | Bacteria>Bacillati>Bacillota>Bacilli>Lactobacillales>Lactobacillaceae>Lactobacillus | 692 | B9N98_RS00620 | 120837..122915 | 1584 | DUF262-domain-containing-protein-[Lactobacillus-delbrueckii] |
| WP_231535726.1 | ParB+HNH | GCF_021135595.1 | Lactobacillus_delbrueckii | Bacteria>Bacillati>Bacillota>Bacilli>Lactobacillales>Lactobacillaceae>Lactobacillus | 621 | LOC07_RS06185 | 1343..3208 | 1584 | DUF262-domain-containing-protein-[Lactobacillus-delbrueckii] |
| WP_231523635.1 | ParB+HNH | GCF_021134645.1 | Lactobacillus_delbrueckii | Bacteria>Bacillati>Bacillota>Bacilli>Lactobacillales>Lactobacillaceae>Lactobacillus | 527 | O0549_RS05830 | 20640..22223 | 1584 | DUF262-domain-containing-protein-[Lactobacillus-delbrueckii] |
| WP_231115305.1 | ParB+HNH | GCF_021601025.1 | Lactobacillus_delbrueckii | Bacteria>Bacillati>Bacillota>Bacilli>Lactobacillales>Lactobacillaceae>Lactobacillus | 493 | L5981_RS02855 | 28501..29982 | 1584 | DUF1524-domain-containing-protein-[Lactobacillus-delbrueckii] |
| WP_241687507.1 | ParB+HNH | GCF_004123065.1 | Lactobacillus_delbrueckii | Bacteria>Bacillati>Bacillota>Bacilli>Lactobacillales>Lactobacillaceae>Lactobacillus | 488 | EST31_RS04475 | 25449..26915 | 1584 | DUF262-domain-containing-protein-[Lactobacillus-delbrueckii] |
| WP_241687506.1 | ParB+HNH | GCF_004123065.1 | Lactobacillus_delbrueckii | Bacteria>Bacillati>Bacillota>Bacilli>Lactobacillales>Lactobacillaceae>Lactobacillus | 220 | EST31_RS10690 | 24830..25492 | 1584 | HNH-endonuclease-family-protein-[Lactobacillus-delbrueckii] |
| WP_231523636.1 | ParB+HNH | GCF_021134645.1 | Lactobacillus_delbrueckii | Bacteria>Bacillati>Bacillota>Bacilli>Lactobacillales>Lactobacillaceae>Lactobacillus | 162 | O0549_RS05825 | 20133..20621 | 1584 | hypothetical-protein-[Lactobacillus-delbrueckii] |
| WP_232450225.1 | ParB+HNH | GCF_002142575.1 | Lactobacillus_delbrueckii | Bacteria>Bacillati>Bacillota>Bacilli>Lactobacillales>Lactobacillaceae>Lactobacillus | 162 | B9N98_RS10500 | 117380..117868 | 1584 | hypothetical-protein-[Lactobacillus-delbrueckii] |
| WP_231523634.1 | ParB+HNH | GCF_021134645.1 | Lactobacillus_delbrueckii | Bacteria>Bacillati>Bacillota>Bacilli>Lactobacillales>Lactobacillaceae>Lactobacillus | 151 | O0549_RS05840 | 23595..24050 | 1584 | hypothetical-protein-[Lactobacillus-delbrueckii] |
| GHN53707.1 | ParB+HNH | GCA_021600265.1 | Lactobacillus_delbrueckii | Bacteria>Bacillati>Bacillota>Bacilli>Lactobacillales>Lactobacillaceae>Lactobacillus | 81 | ME802_14800 | 19270..19515 | 1584 | hypothetical-protein-ME802_14800-[Lactobacillus-delbrueckii] |
| GHN53708.1 | ParB+HNH | GCA_021600265.1 | Lactobacillus_delbrueckii | Bacteria>Bacillati>Bacillota>Bacilli>Lactobacillales>Lactobacillaceae>Lactobacillus | 81 | ME802_14810 | 19499..19744 | 1584 | hypothetical-protein-ME802_14810-[Lactobacillus-delbrueckii] |
| WP_231535727.1 | ParB+HNH | GCF_021135595.1 | Lactobacillus_delbrueckii | Bacteria>Bacillati>Bacillota>Bacilli>Lactobacillales>Lactobacillaceae>Lactobacillus | 70 | LOC07_RS06190 | 3225..3437 | 1584 | hypothetical-protein-[Lactobacillus-delbrueckii] |
| MCH3996395.1 | ParB+HNH | GCA_022483225.1 | Lactobacillus_amylovorus | Bacteria>Bacillati>Bacillota>Bacilli>Lactobacillales>Lactobacillaceae>Lactobacillus | 655 | LKE67_02065 | 418977..420944 | 1604 | DUF262-domain-containing-HNH-endonuclease-family-protein-[Lactobacillus-amylovorus] |
| WP_249905372.1 | ParB+HNH | GCF_023523445.1 | Lactobacillus_amylovorus | Bacteria>Bacillati>Bacillota>Bacilli>Lactobacillales>Lactobacillaceae>Lactobacillus | 655 | LOB74_RS05640 | 1120262..1122229 | 1604 | DUF262-domain-containing-protein-[Lactobacillus-amylovorus] |
| WP_260212028.1 | ParB+HNH | GCF_903789705.1 | Lactobacillus_amylovorus | Bacteria>Bacillati>Bacillota>Bacilli>Lactobacillales>Lactobacillaceae>Lactobacillus | 655 | EFR95_RS04415 | 26031..27998 | 1604 | DUF262-domain-containing-HNH-endonuclease-family-protein-[Lactobacillus-amylovorus] |
| WP_271862778.1 | ParB+HNH | GCF_028309095.1 | Lactobacillus_amylovorus | Bacteria>Bacillati>Bacillota>Bacilli>Lactobacillales>Lactobacillaceae>Lactobacillus | 655 | ODU71_RS08755 | 27900..29867 | 1604 | DUF262-domain-containing-HNH-endonuclease-family-protein-[Lactobacillus-amylovorus] |
| WP_271873860.1 | ParB+HNH | GCF_028308865.1 | Lactobacillus_amylovorus | Bacteria>Bacillati>Bacillota>Bacilli>Lactobacillales>Lactobacillaceae>Lactobacillus | 655 | ODV08_RS03470 | 59457..61424 | 1604 | DUF262-domain-containing-HNH-endonuclease-family-protein-[Lactobacillus-amylovorus] |
| WP_271875347.1 | ParB+HNH | GCF_028308735.1 | Lactobacillus_amylovorus | Bacteria>Bacillati>Bacillota>Bacilli>Lactobacillales>Lactobacillaceae>Lactobacillus | 655 | ODV82_RS03935 | 31128..33095 | 1604 | DUF262-domain-containing-HNH-endonuclease-family-protein-[Lactobacillus-amylovorus] |
| WP_271876488.1 | ParB+HNH | GCF_028308365.1 | Lactobacillus_amylovorus | Bacteria>Bacillati>Bacillota>Bacilli>Lactobacillales>Lactobacillaceae>Lactobacillus | 655 | ODV29_RS04125 | 33803..35770 | 1604 | DUF262-domain-containing-HNH-endonuclease-family-protein-[Lactobacillus-amylovorus] |
| WP_271882931.1 | ParB+HNH | GCF_028308535.1 | Lactobacillus_amylovorus | Bacteria>Bacillati>Bacillota>Bacilli>Lactobacillales>Lactobacillaceae>Lactobacillus | 655 | ODV81_RS08100 | 5202..7169 | 1604 | DUF262-domain-containing-HNH-endonuclease-family-protein-[Lactobacillus-amylovorus] |
| WP_271897404.1 | ParB+HNH | GCF_028308945.1 | Lactobacillus_amylovorus | Bacteria>Bacillati>Bacillota>Bacilli>Lactobacillales>Lactobacillaceae>Lactobacillus | 655 | ODV12_RS07260 | 21131..23098 | 1604 | DUF262-domain-containing-HNH-endonuclease-family-protein-[Lactobacillus-amylovorus] |
| MCD5537318.1 | ParB+HNH | GCA_021135055.1 | Lactobacillus_delbrueckii_subsp._lactis | Bacteria>Bacillati>Bacillota>Bacilli>Lactobacillales>Lactobacillaceae>Lactobacillus>Lactobacillus_delbrueckii | 695 | LOB49_05215 | 25745..27832 | 29397 | DUF262-domain-containing-protein-[Lactobacillus-delbrueckii-subsp.-lactis] |
| MBT8900177.1 | ParB+HNH | GCA_018720625.1 | Lactobacillus_delbrueckii_subsp._bulgaricus | Bacteria>Bacillati>Bacillota>Bacilli>Lactobacillales>Lactobacillaceae>Lactobacillus>Lactobacillus_delbrueckii | 703 | BTI57_06445 | 13249..15360 | 1585 | hypothetical-protein-[Lactobacillus-delbrueckii-subsp.-bulgaricus] |
| MBT8901774.1 | ParB+HNH | GCA_018720695.1 | Lactobacillus_delbrueckii_subsp._bulgaricus | Bacteria>Bacillati>Bacillota>Bacilli>Lactobacillales>Lactobacillaceae>Lactobacillus>Lactobacillus_delbrueckii | 703 | BTI52_06855 | 26358..28469 | 1585 | hypothetical-protein-[Lactobacillus-delbrueckii-subsp.-bulgaricus] |
| MBT8914210.1 | ParB+HNH | GCA_018720815.1 | Lactobacillus_delbrueckii_subsp._bulgaricus | Bacteria>Bacillati>Bacillota>Bacilli>Lactobacillales>Lactobacillaceae>Lactobacillus>Lactobacillus_delbrueckii | 703 | BTI61_07590 | 625..2736 | 1585 | hypothetical-protein-[Lactobacillus-delbrueckii-subsp.-bulgaricus] |
| MBT9079753.1 | ParB+HNH | GCA_018722355.1 | Lactobacillus_delbrueckii_subsp._bulgaricus | Bacteria>Bacillati>Bacillota>Bacilli>Lactobacillales>Lactobacillaceae>Lactobacillus>Lactobacillus_delbrueckii | 697 | BTJ20_07985 | 419..2512 | 1585 | hypothetical-protein-BTJ20_07985-[Lactobacillus-delbrueckii-subsp.-bulgaricus] |
| AHI11938.1 | ParB+HNH | GCA_000525715.1 | Lactobacillus_helveticus_H9 | Bacteria>Bacillati>Bacillota>Bacilli>Lactobacillales>Lactobacillaceae>Lactobacillus>Lactobacillus_helveticus | 655 | LBH_0936 | 968554..970521 | 767456 | Protein-of-hypothetical-function-DUF262-[Lactobacillus-helveticus-H9] |
| WP_070489747.1 | ParB+HNH | GCF_001808415.1 | Lactobacillus_sp._HMSC08B12 | Bacteria>Bacillati>Bacillota>Bacilli>Lactobacillales>Lactobacillaceae>Lactobacillus>unclassified_Lactobacillus | 697 | HMPREF3168_RS09260 | 6257..8350 | 1581136 | DUF262-domain-containing-protein-[Lactobacillus-sp.-HMSC08B12] |
| WP_125647590.1 | ParB+HNH | GCF_042680365.1 | Lapidilactobacillus_gannanensis | Bacteria>Bacillati>Bacillota>Bacilli>Lactobacillales>Lactobacillaceae>Lapidilactobacillus | 595 | EKM81_RS01860 | 1151..2938 | 2486002 | DUF262-domain-containing-protein-[Lapidilactobacillus-gannanensis] |
| WP_153110822.1 | ParB+HNH | - | Lentilactobacillus_buchneri | Bacteria>Bacillati>Bacillota>Bacilli>Lactobacillales>Lactobacillaceae>Lentilactobacillus | 579 | FIU75_RS13240 | 6762..8501 | 1581 | DUF262-domain-containing-protein-[Lentilactobacillus-buchneri] |
| WP_261809682.1 | ParB+HNH | GCF_024345185.1 | Levilactobacillus_humaensis | Bacteria>Bacillati>Bacillota>Bacilli>Lactobacillales>Lactobacillaceae>Levilactobacillus | 591 | OCU14_RS02665 | 56574..58349 | 2950375 | DUF262-domain-containing-protein-[Levilactobacillus-humaensis] |
| WP_060416894.1 | ParB+HNH | GCF_001540905.1 | Levilactobacillus_brevis | Bacteria>Bacillati>Bacillota>Bacilli>Lactobacillales>Lactobacillaceae>Levilactobacillus | 562 | ABB39_RS11470 | 256..1944 | 1580 | DUF262-domain-containing-protein-[Levilactobacillus-brevis] |
| WP_021350323.1 | ParB+HNH | GCF_925284415.1 | Limosilactobacillus_fermentum | Bacteria>Bacillati>Bacillota>Bacilli>Lactobacillales>Lactobacillaceae>Limosilactobacillus | 595 | C0965_RS02340 | 485903..487690 | 1613 | DUF262-domain-containing-protein-[Limosilactobacillus-fermentum] |
| WP_269816398.1 | ParB+HNH | - | Limosilactobacillus_fermentum | Bacteria>Bacillati>Bacillota>Bacilli>Lactobacillales>Lactobacillaceae>Limosilactobacillus | 263 | LR965_RS05290 | 50..841 | 1613 | DUF262-domain-containing-protein-[Limosilactobacillus-fermentum] |
| WP_070065447.1 | ParB+HNH | GCF_001752545.1 | Oenococcus_kitaharae | Bacteria>Bacillati>Bacillota>Bacilli>Lactobacillales>Lactobacillaceae>Oenococcus | 595 | NT95_RS03525 | 1726..3513 | 336988 | DUF262-domain-containing-protein-[Oenococcus-kitaharae] |
| WP_027828093.1 | ParB+HNH | GCF_000425885.1 | Schleiferilactobacillus_harbinensis | Bacteria>Bacillati>Bacillota>Bacilli>Lactobacillales>Lactobacillaceae>Schleiferilactobacillus | 595 | FC91_RS07125 | 33363..35150 | 304207 | DUF262-domain-containing-protein-[Schleiferilactobacillus-harbinensis] |
| MCH5461953.1 | ParB+HNH | GCA_022509925.1 | Secundilactobacillus_angelensis | Bacteria>Bacillati>Bacillota>Bacilli>Lactobacillales>Lactobacillaceae>Secundilactobacillus | 691 | L3V66_04135 | 179597..181672 | 2722706 | DUF262-domain-containing-HNH-endonuclease-family-protein-[Secundilactobacillus-angelensis] |
| MCH4057842.1 | ParB+HNH | GCA_022483825.1 | Lactobacillaceae_bacterium | Bacteria>Bacillati>Bacillota>Bacilli>Lactobacillales>Lactobacillaceae>unclassified_Lactobacillaceae | 596 | LKE89_08040 | 3094..4884 | 2291323 | DUF262-domain-containing-protein-[Lactobacillaceae-bacterium] |
| WP_013403053.1 | ParB+HNH | GCF_000166355.1 | Caldicellulosiruptor_hydrothermalis | Bacteria>Bacillati>Bacillota>Bacillota_incertae_sedis>Caldicellulosiruptorales>Caldicellulosiruptoraceae>Caldicellulosiruptor | 582 | CALHY_RS05820 | 1175941..1177689 | 413888 | DUF262-domain-containing-protein-[Caldicellulosiruptor-hydrothermalis] |
| WP_015908036.1 | ParB+HNH | GCF_000022325.1 | Caldicellulosiruptor_bescii | Bacteria>Bacillati>Bacillota>Bacillota_incertae_sedis>Caldicellulosiruptorales>Caldicellulosiruptoraceae>Caldicellulosiruptor | 582 | ATHE_RS08005 | 1690566..1692314 | 31899 | DUF262-domain-containing-protein-[Caldicellulosiruptor-bescii] |
| WP_014042418.1 | ParB+HNH | GCF_000193435.2 | Caldicellulosiruptor_acetigenus | Bacteria>Bacillati>Bacillota>Bacillota_incertae_sedis>Caldicellulosiruptorales>Caldicellulosiruptoraceae>Caldicellulosiruptor | 582 | CALLA_RS05100 | 1080075..1081823 | 301953 | DUF262-domain-containing-protein-[Caldicellulosiruptor-acetigenus] |
| WP_271628808.1 | ParB+HNH | GCF_028198085.1 | Caldicellulosiruptor_sp._DIB_104C | Bacteria>Bacillati>Bacillota>Bacillota_incertae_sedis>Caldicellulosiruptorales>Caldicellulosiruptoraceae>Caldicellulosiruptor>unclassified_Caldicellulosiruptor | 582 | PGC14_RS09935 | 1999778..2001526 | 3019889 | DUF262-domain-containing-protein-[Caldicellulosiruptor-sp.-DIB-104C] |
| WP_076917457.1 | ParB+HNH | GCF_900148495.1 | Negativibacillus_massiliensis | Bacteria>Bacillati>Bacillota>Bacillota_sensu_stricto_incertae_sedis>Negativibacillus | 606 | BQ5375_RS04275 | 329162..330982 | 1871035 | DUF262-domain-containing-protein-[Negativibacillus-massiliensis] |
| WP_066893472.1 | ParB+HNH | GCF_900086595.1 | Clostridium_nigeriense | Bacteria>Bacillati>Bacillota>Clostridia>Eubacteriales>Clostridiaceae>Clostridium | 686 | BN6774_RS10765 | 2263943..2266003 | 1805470 | DUF262-domain-containing-protein-[Clostridium-nigeriense] |
| WP_161424809.1 | ParB+HNH | - | Clostridium_butyricum | Bacteria>Bacillati>Bacillota>Clostridia>Eubacteriales>Clostridiaceae>Clostridium | 393 | GND98_RS05130 | 48292..49473 | 1492 | DUF262-domain-containing-protein-[Clostridium-butyricum] |
| NFG40380.1 | ParB+HNH | GCA_011013445.1 | Clostridium_botulinum | Bacteria>Bacillati>Bacillota>Clostridia>Eubacteriales>Clostridiaceae>Clostridium | 686 | FC789_04045 | 887461..889521 | 1491 | DUF262-domain-containing-protein-[Clostridium-botulinum] |
| NFI95401.1 | ParB+HNH | GCA_011014625.1 | Clostridium_botulinum | Bacteria>Bacillati>Bacillota>Clostridia>Eubacteriales>Clostridiaceae>Clostridium | 686 | FC961_13545 | 9218..11278 | 1491 | DUF262-domain-containing-protein-[Clostridium-botulinum] |
| WP_207130135.1 | ParB+HNH | GCF_017330945.1 | Clostridium_botulinum | Bacteria>Bacillati>Bacillota>Clostridia>Eubacteriales>Clostridiaceae>Clostridium | 400 | EXQ34_RS04225 | 9472..10674 | 1491 | DUF262-domain-containing-protein-[Clostridium-botulinum] |
| NFO13002.1 | ParB+HNH | GCA_011016315.1 | Clostridium_botulinum | Bacteria>Bacillati>Bacillota>Clostridia>Eubacteriales>Clostridiaceae>Clostridium | 399 | FDB34_02040 | 205562..206761 | 1491 | DUF262-domain-containing-protein-[Clostridium-botulinum] |
| WP_226104723.1 | ParB+HNH | GCF_020443655.1 | Clostridium_algidicarnis | Bacteria>Bacillati>Bacillota>Clostridia>Eubacteriales>Clostridiaceae>Clostridium | 417 | LGK99_RS11180 | 48332..49585 | 37659 | DUF262-domain-containing-protein-[Clostridium-algidicarnis] |
| WP_185162827.1 | ParB+HNH | GCF_014212415.1 | Clostridium_algidicarnis | Bacteria>Bacillati>Bacillota>Clostridia>Eubacteriales>Clostridiaceae>Clostridium | 377 | H7D93_RS12630 | 27276..28409 | 37659 | DUF262-domain-containing-protein-[Clostridium-algidicarnis] |
| WP_252235875.1 | ParB+HNH | GCF_023846115.1 | Clostridium_sp._ZS1 | Bacteria>Bacillati>Bacillota>Clostridia>Eubacteriales>Clostridiaceae>Clostridium>unclassified_Clostridium | 399 | NG923_RS15460 | 24642..25841 | 2949989 | DUF262-domain-containing-protein-[Clostridium-sp.-ZS1] |
| WP_089984093.1 | ParB+HNH | GCF_001517425.1 | Clostridium_sp._C105KSO13 | Bacteria>Bacillati>Bacillota>Clostridia>Eubacteriales>Clostridiaceae>Clostridium>unclassified_Clostridium | 599 | BN3456_RS10490 | 36950..38749 | 1776045 | DUF262-domain-containing-protein-[Clostridium-sp.-C105KSO13] |
| HHW47354.1 | ParB+HNH | GCA_012840535.1 | Clostridiaceae_bacterium | Bacteria>Bacillati>Bacillota>Clostridia>Eubacteriales>Clostridiaceae>unclassified_Clostridiaceae | 606 | GXX14_01905 | 15527..17347 | 1898204 | DUF262-domain-containing-protein-[Clostridiaceae-bacterium] |
| WP_018214266.1 | ParB+HNH | GCF_000378805.1 | Desulfitobacterium_hafniense | Bacteria>Bacillati>Bacillota>Clostridia>Eubacteriales>Desulfitobacteriaceae>Desulfitobacterium | 374 | DESHAF_RS0122580 | 1955663..1956787 | 49338 | DUF262-domain-containing-protein-[Desulfitobacterium-hafniense] |
| WP_235255106.1 | ParB+HNH | - | Allofournierella_massiliensis | Bacteria>Bacillati>Bacillota>Clostridia>Eubacteriales>Oscillospiraceae>Allofournierella | 597 | JQM64_RS00345 | 69346..71139 | 1650663 | DUF262-domain-containing-protein-[Fournierella-massiliensis] |

| Protein-ID | Domain | Assembly-ID | Strain | Taxonomy | Length | Locus-Tag | Genomic-coordinates | TaxID | Fasta-defline |
|---|---|---|---|---|---|---|---|---|---|
| UYJ00851.1 | ParB+HNH | GCA_025757765.1 | Faecalibacterium_prausnitzii | Bacteria>Bacillati>Bacillota>Clostridia>Eubacteriales>Oscillospiraceae>Faecalibacterium | 592 | OGM65_10435 | 2164015..2165793 | 853 | MAG:-DUF262-domain-containing-protein-[Faecalibacterium-prausnitzii] |
| WP_097782658.1 | ParB+HNH | GCF_002549935.1 | Faecalibacterium_prausnitzii | Bacteria>Bacillati>Bacillota>Clostridia>Eubacteriales>Oscillospiraceae>Faecalibacterium | 592 | CGS55_RS03395 | 49739..51517 | 853 | DUF262-domain-containing-protein-[Faecalibacterium-prausnitzii] |
| WP_113992222.1 | ParB+HNH | GCF_032584275.1 | Faecalibacterium_prausnitzii | Bacteria>Bacillati>Bacillota>Clostridia>Eubacteriales>Oscillospiraceae>Faecalibacterium | 592 | DW855_RS10400 | 44122..45900 | 853 | DUF262-domain-containing-protein-[Faecalibacterium-prausnitzii] |
| WP_215675207.1 | ParB+HNH | GCF_018784645.1 | Faecalibacterium_prausnitzii | Bacteria>Bacillati>Bacillota>Clostridia>Eubacteriales>Oscillospiraceae>Faecalibacterium | 592 | GPK62_RS07885 | 50921..52699 | 853 | DUF262-domain-containing-protein-[Faecalibacterium-prausnitzii] |
| WP_158398721.1 | ParB+HNH | GCF_902167845.1 | Faecalibacterium_prausnitzii | Bacteria>Bacillati>Bacillota>Clostridia>Eubacteriales>Oscillospiraceae>Faecalibacterium | 362 | FQ260_RS07245 | 42..1130 | 853 | DUF262-domain-containing-protein-[Faecalibacterium-prausnitzii] |
| WP_275252241.1 | ParB+HNH | - | Faecalibacterium_sp._DFI.5.82 | Bacteria>Bacillati>Bacillota>Clostridia>Eubacteriales>Oscillospiraceae>Faecalibacterium>unclassified_Faecalibacterium | 217 | P0G09_RS06370 | 147510..148163 | 3031725 | DUF262-domain-containing-protein-[Faecalibacterium-sp.-DFI.5.82] |
| WP_024861709.1 | ParB+HNH | GCF_000526795.1 | Ruminococcus_flavefaciens | Bacteria>Bacillati>Bacillota>Clostridia>Eubacteriales>Oscillospiraceae>Ruminococcus | 403 | N774_RS18405 | 2999852..3001063 | 1265 | DUF262-domain-containing-protein-[Ruminococcus-flavefaciens] |
| MCR4638956.1 | ParB+HNH | GCA_024684275.1 | Ruminococcus_sp. | Bacteria>Bacillati>Bacillota>Clostridia>Eubacteriales>Oscillospiraceae>Ruminococcus>unclassified_Ruminococcus | 593 | K5677_06995 | 10152..11933 | 41978 | DUF262-domain-containing-HNH-endonuclease-family-protein-[Ruminococcus-sp.] |
| UYJ11722.1 | ParB+HNH | GCA_025757845.1 | Oscillospiraceae_bacterium | Bacteria>Bacillati>Bacillota>Clostridia>Eubacteriales>Oscillospiraceae>unclassified_Oscillospiraceae | 592 | OGM78_02740 | 582639..584417 | 2485925 | MAG:-DUF262-domain-containing-protein-[Oscillospiraceae-bacterium] |
| MBO8129692.1 | ParB+HNH | GCA_017656185.1 | Peptococcaceae_bacterium | Bacteria>Bacillati>Bacillota>Clostridia>Eubacteriales>Peptococcaceae>unclassified_Peptococcaceae | 566 | H0Z39_10955 | 3534..5234 | 2052179 | DUF262-domain-containing-protein-[Peptococcaceae-bacterium] |
| NLJ76875.1 | ParB+HNH | GCA_012518875.1 | Peptococcaceae_bacterium | Bacteria>Bacillati>Bacillota>Clostridia>Eubacteriales>Peptococcaceae>unclassified_Peptococcaceae | 551 | GX325_06420 | 256..1911 | 2052179 | DUF262-domain-containing-protein-[Peptococcaceae-bacterium] |
| MBC7075382.1 | ParB+HNH | GCA_014360155.1 | Syntrophomonadaceae_bacterium | Bacteria>Bacillati>Bacillota>Clostridia>Eubacteriales>Syntrophomonadaceae>unclassified_Syntrophomonadaceae | 603 | H5T98_04810 | 14870..16681 | 2093811 | DUF262-domain-containing-protein-[Syntrophomonadaceae-bacterium] |
| HHY76909.1 | ParB+HNH | GCA_012842725.1 | Clostridiales_bacterium | Bacteria>Bacillati>Bacillota>Clostridia>Eubacteriales>unclassified_Eubacteriales | 633 | GX498_00180 | 338..2239 | 1898207 | DUF262-domain-containing-protein-[Clostridiales-bacterium] |
| MBD9283723.1 | ParB+HNH | GCA_014848695.1 | Clostridiales_bacterium | Bacteria>Bacillati>Bacillota>Clostridia>Eubacteriales>unclassified_Eubacteriales | 597 | EGR96_02210 | 180..1973 | 1898207 | DUF262-domain-containing-protein-[Clostridiales-bacterium] |
| MCF8011658.1 | ParB+HNH | GCA_021736925.1 | Clostridiales_bacterium | Bacteria>Bacillati>Bacillota>Clostridia>Eubacteriales>unclassified_Eubacteriales | 556 | K9L56_10270 | 9791..11461 | 1898207 | DUF262-domain-containing-protein-[Clostridiales-bacterium] |
| WP_073159270.1 | ParB+HNH | GCF_900102605.1 | Halanaerobium | Bacteria>Bacillati>Bacillota>Clostridia>Halanaerobiales>Halanaerobiaceae | 571 | DER71_RS07020 | 27430..29145 | 2330 | MULTISPECIES:-DUF262-domain-containing-protein-[Halanaerobium] |
| WP_089655504.1 | ParB+HNH | GCF_900100575.1 | Halanaerobium_congolense | Bacteria>Bacillati>Bacillota>Clostridia>Halanaerobiales>Halanaerobiaceae>Halanaerobium | 581 | BLS10_RS04570 | 108612..110357 | 54121 | DUF262-domain-containing-protein-[Halanaerobium-congolense] |
| WP_134059929.1 | ParB+HNH | GCF_004366375.1 | Halanaerobium_congolense | Bacteria>Bacillati>Bacillota>Clostridia>Halanaerobiales>Halanaerobiaceae>Halanaerobium | 571 | C7954_RS10320 | 4449..6164 | 54121 | DUF262-domain-containing-protein-[Halanaerobium-congolense] |
| WP_059085684.1 | ParB+HNH | GCF_001487165.1 | Blautia | Bacteria>Bacillati>Bacillota>Clostridia>Lachnospirales>Lachnospiraceae | 398 | C4886_RS16020 | 17445..18641 | 572511 | MULTISPECIES:-DUF262-domain-containing-protein-[Blautia] |
| MCI7107134.1 | ParB+HNH | GCA_022775045.1 | Agathobacter_sp. | Bacteria>Bacillati>Bacillota>Clostridia>Lachnospirales>Lachnospiraceae>Agathobacter>unclassified_Agathobacter | 582 | MR945_04555 | 35211..36959 | 2021311 | DUF262-domain-containing-HNH-endonuclease-family-protein-[Agathobacter-sp.] |
| WP_027204788.1 | ParB+HNH | GCF_000424425.1 | Butyrivibrio_fibrisolvens | Bacteria>Bacillati>Bacillota>Clostridia>Lachnospirales>Lachnospiraceae>Butyrivibrio | 614 | G624_RS0113180 | 57715..59559 | 831 | DUF262-domain-containing-protein-[Butyrivibrio-fibrisolvens] |
| WP_027207200.1 | ParB+HNH | GCF_000423985.1 | Butyrivibrio_fibrisolvens | Bacteria>Bacillati>Bacillota>Clostridia>Lachnospirales>Lachnospiraceae>Butyrivibrio | 564 | G591_RS0114685 | 18730..20424 | 831 | DUF262-domain-containing-protein-[Butyrivibrio-fibrisolvens] |
| WP_081647853.1 | ParB+HNH | GCF_000420945.1 | Butyrivibrio_sp._VCB2006 | Bacteria>Bacillati>Bacillota>Clostridia>Lachnospirales>Lachnospiraceae>Butyrivibrio>unclassified_Butyrivibrio | 874 | G613_RS0110120 | 35992..38616 | 1280679 | DUF262-domain-containing-protein-[Butyrivibrio-sp.-VCB2006] |
| WP_029232555.1 | ParB+HNH | GCF_000420945.1 | Butyrivibrio_sp._VCB2006 | Bacteria>Bacillati>Bacillota>Clostridia>Lachnospirales>Lachnospiraceae>Butyrivibrio>unclassified_Butyrivibrio | 577 | G613_RS0110125 | 38618..40351 | 1280679 | DUF262-domain-containing-protein-[Butyrivibrio-sp.-VCB2006] |
| WP_091825294.1 | ParB+HNH | GCF_900109475.1 | Butyrivibrio_sp._ob235 | Bacteria>Bacillati>Bacillota>Clostridia>Lachnospirales>Lachnospiraceae>Butyrivibrio>unclassified_Butyrivibrio | 606 | BMW61_RS19105 | 30193..32013 | 1761780 | DUF262-domain-containing-protein-[Butyrivibrio-sp.-ob235] |
| WP_026654578.1 | ParB+HNH | GCF_000686065.1 | Butyrivibrio_sp._AE3003 | Bacteria>Bacillati>Bacillota>Clostridia>Lachnospirales>Lachnospiraceae>Butyrivibrio>unclassified_Butyrivibrio | 422 | CD00_RS0103075 | 637222..638490 | 1496721 | DUF262-domain-containing-protein-[Butyrivibrio-sp.-AE3003] |
| WP_195251431.1 | ParB+HNH | GCF_015550485.1 | Coprococcus_comes | Bacteria>Bacillati>Bacillota>Clostridia>Lachnospirales>Lachnospiraceae>Coprococcus | 378 | I2A60_RS13440 | 12101..13237 | 410072 | DUF262-domain-containing-protein-[Coprococcus-comes] |
| WP_271824187.1 | ParB+HNH | GCF_028210655.1 | Enterocloster_clostridioformis | Bacteria>Bacillati>Bacillota>Clostridia>Lachnospirales>Lachnospiraceae>Enterocloster | 606 | PMX39_RS12905 | 8226..10046 | 1531 | DUF262-domain-containing-HNH-endonuclease-family-protein-[Enterocloster-clostridioformis] |
| TGX97941.1 | ParB+HNH | GCA_004793585.1 | Hominisplanchenecus_murintestinalis | Bacteria>Bacillati>Bacillota>Clostridia>Lachnospirales>Lachnospiraceae>Hominisplanchenecus | 378 | E5357_10815 | 31113..32249 | 2941517 | DUF262-domain-containing-protein-[Lachnospiraceae-bacterium] |
| MCI8664743.1 | ParB+HNH | GCA_022794315.1 | Hungatella_sp. | Bacteria>Bacillati>Bacillota>Clostridia>Lachnospirales>Lachnospiraceae>Hungatella>unclassified_Hungatella | 461 | HFG69_16130 | 107005..108390 | 2613924 | DUF262-domain-containing-protein-[Hungatella-sp.] |
| MCI8605664.1 | ParB+HNH | GCA_022793615.1 | Hungatella_sp. | Bacteria>Bacillati>Bacillota>Clostridia>Lachnospirales>Lachnospiraceae>Hungatella>unclassified_Hungatella | 435 | HFG72_02150 | 22124..23431 | 2613924 | DUF262-domain-containing-protein-[Hungatella-sp.] |
| WP_029067209.1 | ParB+HNH | GCF_000687515.1 | Lachnobacterium_bovis | Bacteria>Bacillati>Bacillota>Clostridia>Lachnospirales>Lachnospiraceae>Lachnobacterium | 579 | CC91_RS0108450 | 388006..389745 | 140626 | DUF262-domain-containing-protein-[Lachnobacterium-bovis] |
| WP_024292375.1 | ParB+HNH | GCF_000526995.1 | Lacrimispora_indolis | Bacteria>Bacillati>Bacillota>Clostridia>Lachnospirales>Lachnospiraceae>Lacrimispora | 431 | K401_RS0107495 | 1573372..1574667 | 69825 | DUF262-domain-containing-protein-[Lacrimispora-indolis] |
| WP_118278837.1 | ParB+HNH | GCF_027662525.1 | [Ruminococcus]_lactaris | Bacteria>Bacillati>Bacillota>Clostridia>Lachnospirales>Lachnospiraceae>Mediterraneibacter | 578 | DW116_RS04320 | 5911..7647 | 46228 | DUF262-domain-containing-protein-[[Ruminococcus] |
| WP_090815506.1 | ParB+HNH | GCF_900107835.1 | Oribacterium_sp._KHPX15 | Bacteria>Bacillati>Bacillota>Clostridia>Lachnospirales>Lachnospiraceae>Oribacterium>unclassified_Oribacterium | 615 | BLV86_RS05630 | 34599..36446 | 1855342 | DUF262-domain-containing-protein-[Oribacterium-sp.-KHPX15] |
| SFG07289.1 | ParB+HNH | GCA_900113385.1 | Lachnospiraceae_bacterium_C7 | Bacteria>Bacillati>Bacillota>Clostridia>Lachnospirales>Lachnospiraceae>unclassified_Lachnospiraceae | 673 | SAMN04487761_10382 | 91756..93777 | 1792306 | protein-of-unknown-function-[Lachnospiraceae-bacterium-C7] |
| MBR3171571.1 | ParB+HNH | GCA_017514845.1 | Lachnospiraceae_bacterium | Bacteria>Bacillati>Bacillota>Clostridia>Lachnospirales>Lachnospiraceae>unclassified_Lachnospiraceae | 599 | IKF22_10055 | 4899..6698 | 1898203 | DUF262-domain-containing-protein-[Lachnospiraceae-bacterium] |
| MCI9535144.1 | ParB+HNH | GCA_022801195.1 | Lachnospiraceae_bacterium | Bacteria>Bacillati>Bacillota>Clostridia>Lachnospirales>Lachnospiraceae>unclassified_Lachnospiraceae | 458 | HFG53_10155 | 45792..47168 | 1898203 | DUF262-domain-containing-protein-[Lachnospiraceae-bacterium] |
| MCI9323602.1 | ParB+HNH | GCA_022799595.1 | Lachnospiraceae_bacterium | Bacteria>Bacillati>Bacillota>Clostridia>Lachnospirales>Lachnospiraceae>unclassified_Lachnospiraceae | 448 | HFH81_10155 | 54719..56065 | 1898203 | DUF262-domain-containing-protein-[Lachnospiraceae-bacterium] |
| MBD5550160.1 | ParB+HNH | GCA_014804795.1 | Lachnospiraceae_bacterium | Bacteria>Bacillati>Bacillota>Clostridia>Lachnospirales>Lachnospiraceae>unclassified_Lachnospiraceae | 429 | HDQ96_03140 | 117647..118936 | 1898203 | DUF262-domain-containing-protein-[Lachnospiraceae-bacterium] |
| MBR4981300.1 | ParB+HNH | GCA_017555455.1 | Lachnospiraceae_bacterium | Bacteria>Bacillati>Bacillota>Clostridia>Lachnospirales>Lachnospiraceae>unclassified_Lachnospiraceae | 429 | IKY94_01920 | 2479..3768 | 1898203 | DUF262-domain-containing-protein-[Lachnospiraceae-bacterium] |
| MCM1125307.1 | ParB+HNH | GCA_023663945.1 | Lachnospiraceae_bacterium | Bacteria>Bacillati>Bacillota>Clostridia>Lachnospirales>Lachnospiraceae>unclassified_Lachnospiraceae | 379 | NC429_02430 | 32105..33244 | 1898203 | DUF262-domain-containing-protein-[Lachnospiraceae-bacterium] |
| MBP3458812.1 | ParB+HNH | GCA_017936785.1 | Lachnospiraceae_bacterium | Bacteria>Bacillati>Bacillota>Clostridia>Lachnospirales>Lachnospiraceae>unclassified_Lachnospiraceae | 371 | J6K58_06355 | 24661..25776 | 1898203 | DUF262-domain-containing-protein-[Lachnospiraceae-bacterium] |
| WP_088552560.1 | ParB+HNH | GCF_002207765.1 | Calderihabitans_maritimus | Bacteria>Bacillati>Bacillota>Clostridia>Moorellales>Calderihabitantaceae>Calderihabitans | 561 | KKC1_RS00520 | 39283..40968 | 1246530 | DUF262-domain-containing-protein-[Calderihabitans-maritimus] |
| WP_054937990.1 | ParB+HNH | GCF_002995805.1 | Moorella_stamsii | Bacteria>Bacillati>Bacillota>Clostridia>Moorellales>Moorellaceae>Moorella | 573 | MOST_RS05230 | 70157..71878 | 1266720 | MULTISPECIES:-DUF262-domain-containing-protein-[Moorella] |
| WP_244262368.1 | ParB+HNH | - | Thermoanaerobacter_wiegelii | Bacteria>Bacillati>Bacillota>Clostridia>Thermoanaerobacterales>Thermoanaerobacteraceae>Thermoanaerobacter | 618 | THEWI_RS10755 | 2206032..2207888 | 46354 | DUF262-domain-containing-protein-[Thermoanaerobacter-wiegelii] |
| RKL62723.1 | ParB+HNH | GCA_003615265.1 | Thermoanaerobacteraceae_bacterium_SP2 | Bacteria>Bacillati>Bacillota>Clostridia>Thermoanaerobacterales>Thermoanaerobacteraceae>unclassified_Thermoanaerobacteraceae | 351 | DXT63_09720 | 5012..6067 | 2292448 | DUF262-domain-containing-protein-[Thermoanaerobacteraceae-bacterium-SP2] |
| MCF6096365.1 | ParB+HNH | GCA_021608145.1 | Thermovorax_subterraneus | Bacteria>Bacillati>Bacillota>Clostridia>Thermosediminibacterales>Thermosediminibacteraceae>Thermovorax | 563 | L1766_05000 | 42183..43874 | 496831 | DUF262-domain-containing-protein-[Thermovorax-subterraneus] |
| MBR6253003.1 | ParB+HNH | GCA_017539905.1 | Clostridia_bacterium | Bacteria>Bacillati>Bacillota>Clostridia>unclassified_Clostridia | 615 | IKR04_04100 | 185518..187365 | 2044939 | DUF262-domain-containing-protein-[Clostridia-bacterium] |
| MCR5783577.1 | ParB+HNH | GCA_024699005.1 | Clostridia_bacterium | Bacteria>Bacillati>Bacillota>Clostridia>unclassified_Clostridia | 599 | K6G90_12690 | 34544..36343 | 2044939 | DUF262-domain-containing-protein-[Clostridia-bacterium] |
| MBQ8616933.1 | ParB+HNH | GCA_017472845.1 | Clostridia_bacterium | Bacteria>Bacillati>Bacillota>Clostridia>unclassified_Clostridia | 592 | IJ418_05395 | 3678..5456 | 2044939 | DUF262-domain-containing-protein-[Clostridia-bacterium] |
| WP_270396148.1 | ParB+HNH | GCF_027659655.1 | Catenibacterium_mitsuokai | Bacteria>Bacillati>Bacillota>Erysipelotrichia>Erysipelotrichales>Coprobacillaceae>Catenibacterium | 365 | PG085_RS08460 | 8371..9468 | 100886 | DUF262-domain-containing-protein-[Catenibacterium-mitsuokai] |
| WP_132026807.1 | ParB+HNH | GCF_016767175.1 | Keratinibaculum_paraultunense | Bacteria>Bacillati>Bacillota>Tissierellia>Tissierellales>Tepidimicrobiaceae>Keratinibaculum | 684 | JL105_RS06880 | 1380734..1382788 | 1278232 | DUF262-domain-containing-protein-[Keratinibaculum-paraultunense] |
| NPV81614.1 | ParB+HNH | GCA_013178415.1 | Bacillota_bacterium | Bacteria>Bacillati>Bacillota>unclassified_Bacillota | 617 | HPY52_15355 | 5168..7021 | 1879010 | DUF262-domain-containing-protein-[Bacillota-bacterium] |
| NLG80274.1 | ParB+HNH | GCA_012516125.1 | Bacillota_bacterium | Bacteria>Bacillati>Bacillota>unclassified_Bacillota | 562 | GX492_10725 | 28147..29835 | 1879010 | DUF262-domain-containing-protein-[Bacillota-bacterium] |
| MCB9452415.1 | ParB+HNH | GCA_020634115.1 | Anaerolineaceae_bacterium | Bacteria>Bacillati>Chloroflexota>Anaerolineae>Anaerolineales>Anaerolineaceae>unclassified_Anaerolineaceae | 685 | H6672_13340 | 115647..117704 | 2024896 | DUF262-domain-containing-protein-[Anaerolineaceae-bacterium] |
| MBQ6518173.1 | ParB+HNH | GCA_017433515.1 | Anaerolineaceae_bacterium | Bacteria>Bacillati>Chloroflexota>Anaerolineae>Anaerolineales>Anaerolineaceae>unclassified_Anaerolineaceae | 597 | IJI14_05590 | 8886..10679 | 2024896 | DUF262-domain-containing-protein-[Anaerolineaceae-bacterium] |
| MBP9040765.1 | ParB+HNH | GCA_018057145.1 | Anaerolineaceae_bacterium | Bacteria>Bacillati>Chloroflexota>Anaerolineae>Anaerolineales>Anaerolineaceae>unclassified_Anaerolineaceae | 348 | KBF64_03220 | 3172..4218 | 2024896 | DUF262-domain-containing-protein-[Anaerolineaceae-bacterium] |
| MBE7435436.1 | ParB+HNH | GCA_015075725.1 | Anaerolineales_bacterium | Bacteria>Bacillati>Chloroflexota>Anaerolineae>Anaerolineales>unclassified_Anaerolineales | 837 | HS100_16090 | 3327087..3329600 | 2073117 | DUF262-domain-containing-protein-[Anaerolineales-bacterium] |
| MCB0100634.1 | ParB+HNH | GCA_020433725.1 | Anaerolineales_bacterium | Bacteria>Bacillati>Chloroflexota>Anaerolineae>Anaerolineales>unclassified_Anaerolineales | 573 | KDD74_01215 | 28279..30000 | 2073117 | DUF262-domain-containing-protein-[Anaerolineales-bacterium] |
| MCZ7550959.1 | ParB+HNH | GCA_027494025.1 | Anaerolineales_bacterium | Bacteria>Bacillati>Chloroflexota>Anaerolineae>Anaerolineales>unclassified_Anaerolineales | 542 | M5U11_17675 | 3654416..3656044 | 2073117 | MAG:-DUF262-domain-containing-protein-[Anaerolineales-bacterium] |
| NLF02864.1 | ParB+HNH | GCA_012514695.1 | Anaerolineales_bacterium | Bacteria>Bacillati>Chloroflexota>Anaerolineae>Anaerolineales>unclassified_Anaerolineales | 462 | GX601_18025 | 18056..19444 | 2073117 | DUF262-domain-containing-protein-[Anaerolineales-bacterium] |
| MBN1139290.1 | ParB+HNH | GCA_016926595.1 | Anaerolineae_bacterium | Bacteria>Bacillati>Chloroflexota>Anaerolineae>unclassified_Anaerolineae | 844 | JXM73_22125 | 57855..60389 | 2052143 | DUF262-domain-containing-protein-[Anaerolineae-bacterium] |
| MCB8944349.1 | ParB+HNH | GCA_020634415.1 | Ardenticatenaceae_bacterium | Bacteria>Bacillati>Chloroflexota>Ardenticatenia>Ardenticatenales>Ardenticatenaceae>unclassified_Ardenticatenaceae | 587 | H6658_11420 | 13018..14781 | 2614247 | DUF262-domain-containing-protein-[Ardenticatenaceae-bacterium] |
| MCC6456913.1 | ParB+HNH | GCA_020846635.1 | Caldilineaceae_bacterium | Bacteria>Bacillati>Chloroflexota>Caldilineae>Caldilineales>Caldilineaceae>unclassified_Caldilineaceae | 659 | IT328_18295 | 6286..8265 | 2614250 | DUF262-domain-containing-protein-[Caldilineaceae-bacterium] |
| MDE0078892.1 | ParB+HNH | GCA_028819025.1 | Caldilineaceae_bacterium | Bacteria>Bacillati>Chloroflexota>Caldilineae>Caldilineales>Caldilineaceae>unclassified_Caldilineaceae | 605 | OXO50_15335 | 12142..13959 | 2614250 | MAG:-DUF262-domain-containing-protein-[Caldilineaceae-bacterium] |
| MBK9096484.1 | ParB+HNH | GCA_016719785.1 | Candidatus_Amarolinea_dominans | Bacteria>Bacillati>Chloroflexota>Caldilineae>Caldilineales>Candidatus_Amarolineaceae>Candidatus_Amarolinea | 541 | IPM84_27750 | 235160..236785 | 3140696 | DUF262-domain-containing-protein-[Anaerolineae-bacterium] |
| HEX76882.1 | ParB+HNH | GCA_011170395.1 | Dehalococcoidia_bacterium | Bacteria>Bacillati>Chloroflexota>Dehalococcoidia>unclassified_Dehalococcoidia | 568 | G4O03_00465 | 90934..92640 | 2026734 | DUF262-domain-containing-protein-[Dehalococcoidia-bacterium] |
| MCL4240071.1 | ParB+HNH | GCA_023382755.1 | Dehalococcoidia_bacterium | Bacteria>Bacillati>Chloroflexota>Dehalococcoidia>unclassified_Dehalococcoidia | 565 | KJ048_01860 | 26049..27746 | 2026734 | DUF262-domain-containing-protein-[Dehalococcoidia-bacterium] |
| WP_201387655.1 | ParB+HNH | GCF_016587395.1 | Ktedonobacter_sp._SOSP1-52 | Bacteria>Bacillati>Chloroflexota>Ktedonobacteria>Ktedonobacterales>Ktedonobacteraceae>Ktedonobacter>unclassified_Ktedonobacter | 588 | JMJ16_RS30305 | 1129873..1131639 | 2778366 | DUF262-domain-containing-protein-[Ktedonobacter-sp.-SOSP1-52] |
| MCX5991387.1 | ParB+HNH | GCA_026389585.1 | Chloroflexota_bacterium | Bacteria>Bacillati>Chloroflexota>unclassified_Chloroflexota | 638 | NTZ04_03530 | 78307..80223 | 2026724 | MAG:-DUF262-domain-containing-protein-[Chloroflexi-bacterium] |
| MBU1746751.1 | ParB+HNH | GCA_018823415.1 | Chloroflexota_bacterium | Bacteria>Bacillati>Chloroflexota>unclassified_Chloroflexota | 588 | KKA73_03610 | 5287..7053 | 2026724 | DUF262-domain-containing-protein-[Chloroflexota-bacterium] |
| MBP8000884.1 | ParB+HNH | GCA_018004095.1 | Chloroflexota_bacterium | Bacteria>Bacillati>Chloroflexota>unclassified_Chloroflexota | 587 | KA338_16650 | 22901..24664 | 2026724 | DUF262-domain-containing-protein-[Chloroflexota-bacterium] |
| HDQ70820.1 | ParB+HNH | GCA_011046405.1 | Chloroflexota_bacterium | Bacteria>Bacillati>Chloroflexota>unclassified_Chloroflexota | 584 | ENN19_01850 | 1273..3027 | 2026724 | DUF262-domain-containing-protein-[Chloroflexota-bacterium] |
| MCX6033327.1 | ParB+HNH | GCA_026389845.1 | Chloroflexota_bacterium | Bacteria>Bacillati>Chloroflexota>unclassified_Chloroflexota | 578 | NT169_29100 | 5823..7559 | 2026724 | MAG:-DUF262-domain-containing-protein-[Chloroflexi-bacterium] |
| MCY3570530.1 | ParB+HNH | GCA_026705025.1 | Chloroflexota_bacterium | Bacteria>Bacillati>Chloroflexota>unclassified_Chloroflexota | 455 | OXH19_04220 | 12267..13634 | 2026724 | MAG:-DUF262-domain-containing-protein-[Chloroflexi-bacterium] |
| MCE7858271.1 | ParB+HNH | GCA_021404705.1 | Chloroflexi_bacterium_CFX2 | Bacteria>Bacillati>Chloroflexota>unclassified_Chloroflexota | 542 | DYG86_00610 | 109123..110751 | 2293708 | DUF262-domain-containing-protein-[Chloroflexi-bacterium-CFX2] |
| OJX40543.1 | ParB+HNH | GCA_001899005.1 | Chloroflexi_bacterium_44-23 | Bacteria>Bacillati>Chloroflexota>unclassified_Chloroflexota | 581 | BG078_06025 | 149773..151518 | 1895927 | hypothetical-protein-BG078_06025-[Chloroflexi-bacterium-44-23] |
| MBE9018010.1 | ParB+HNH | GCA_015206905.1 | Chroococcidiopsidales_cyanobacterium_LEGE_13417 | Bacteria>Bacillati>Cyanobacteriota/Melainabacteria_group>Cyanobacteriota>Cyanophyceae>Chroococcidiopsidales>unclassified_Chroococcidiopsidales | 557 | IQ272_18045 | 1112..2785 | 1828829 | DUF262-domain-containing-protein-[Chroococcidiopsidales-cyanobacterium-LEGE-13417] |
| WP_006100269.1 | ParB+HNH | GCF_000155555.1 | Coleofasciculus_chthonoplastes | Bacteria>Bacillati>Cyanobacteriota/Melainabacteria_group>Cyanobacteriota>Cyanophyceae>Coleofasciculales>Coleofasciculaceae>Coleofasciculus | 790 | MC7420_RS09355 | 59431..61803 | 64178 | DUF262-domain-containing-HNH-endonuclease-family-protein-[Coleofasciculus-chthonoplastes] |
| WP_242040644.1 | ParB+HNH | - | Coleofasciculus_sp._FACHB-1120 | Bacteria>Bacillati>Cyanobacteriota/Melainabacteria_group>Cyanobacteriota>Cyanophyceae>Coleofasciculales>Coleofasciculaceae>Coleofasciculus>unclassified_Coleofasciculus | 391 | H6H02_RS09845 | 181349..182524 | 2692783 | DUF262-domain-containing-protein-[Coleofasciculus-sp.-FACHB-1120] |
| WP_268243225.1 | ParB+HNH | GCF_026735145.1 | Leptolyngbya_sp._GGD | Bacteria>Bacillati>Cyanobacteriota/Melainabacteria_group>Cyanobacteriota>Cyanophyceae>Leptolyngbyales>Leptolyngbyaceae>Leptolyngbya_group>Leptolyngbya>unclassified_Leptolyngbya | 605 | OU814_RS30715 | 3753..5570 | 2997907 | DUF262-domain-containing-protein-[Leptolyngbya-sp.-GGD] |
| WP_190797463.1 | ParB+HNH | GCF_014695875.1 | Leptolyngbya_sp._FACHB-541 | Bacteria>Bacillati>Cyanobacteriota/Melainabacteria_group>Cyanobacteriota>Cyanophyceae>Leptolyngbyales>Leptolyngbyaceae>Leptolyngbya_group>Leptolyngbya>unclassified_Leptolyngbya | 597 | H6G00_RS15665 | 12008..13801 | 2692810 | DUF262-domain-containing-protein-[Leptolyngbya-sp.-FACHB-541] |
| WP_228040297.1 | ParB+HNH | GCF_015207395.1 | Nodosilinea_sp._LEGE_07088 | Bacteria>Bacillati>Cyanobacteriota/Melainabacteria_group>Cyanobacteriota>Cyanophyceae>Nodosilineales>Nodosilineaceae>Nodosilinea>unclassified_Nodosilinea | 616 | IQ254_RS06895 | 9438..11288 | 2777968 | DUF262-domain-containing-protein-[Nodosilinea-sp.-LEGE-07088] |
| BDA72469.1 | ParB+HNH | GCA_019977735.1 | Calothrix_sp._PCC_7716 | Bacteria>Bacillati>Cyanobacteriota/Melainabacteria_group>Cyanobacteriota>Cyanophyceae>Nostocales>Calotrichaceae>Calothrix>unclassified_Calothrix | 559 | CAL7716_066350 | 7615502..7617181 | 2853227 | DUF262-[Calothrix-sp.-PCC-7716] |
| MBD2467867.1 | ParB+HNH | GCA_014697455.1 | Nostoc_sp._FACHB-145 | Bacteria>Bacillati>Cyanobacteriota/Melainabacteria_group>Cyanobacteriota>Cyanophyceae>Nostocales>Nostocaceae>Nostoc>unclassified_Nostoc | 558 | H6G90_09470 | 29748..31424 | 2692836 | DUF262-domain-containing-protein-[Nostoc-sp.-FACHB-145] |
| WP_190754801.1 | ParB+HNH | GCF_014697455.1 | Nostoc_sp._FACHB-145 | Bacteria>Bacillati>Cyanobacteriota/Melainabacteria_group>Cyanobacteriota>Cyanophyceae>Nostocales>Nostocaceae>Nostoc>unclassified_Nostoc | 558 | H6G90_RS09470 | 29748..31424 | 2692836 | DUF262-domain-containing-protein-[Nostoc-sp.-FACHB-145] |
| MBD2438836.1 | ParB+HNH | GCA_014697425.1 | Nostoc_sp._FACHB-110 | Bacteria>Bacillati>Cyanobacteriota/Melainabacteria_group>Cyanobacteriota>Cyanophyceae>Nostocales>Nostocaceae>Nostoc>unclassified_Nostoc | 557 | H6G69_18785 | 67933..69606 | 2692834 | DUF262-domain-containing-protein-[Nostoc-sp.-FACHB-110] |
| WP_190690585.1 | ParB+HNH | GCF_014697425.1 | Nostoc_sp._FACHB-110 | Bacteria>Bacillati>Cyanobacteriota/Melainabacteria_group>Cyanobacteriota>Cyanophyceae>Nostocales>Nostocaceae>Nostoc>unclassified_Nostoc | 557 | H6G69_RS18790 | 67933..69606 | 2692834 | DUF262-domain-containing-protein-[Nostoc-sp.-FACHB-110] |
| WP_086759839.1 | ParB+HNH | GCF_963873765.1 | Nostoc_sp._106C | Bacteria>Bacillati>Cyanobacteriota/Melainabacteria_group>Cyanobacteriota>Cyanophyceae>Nostocales>Nostocaceae>Nostoc>unclassified_Nostoc | 745 | BV375_RS09710 | 50458..52695 | 1932667 | DUF262-domain-containing-HNH-endonuclease-family-protein-[Nostoc-sp.-106C] |
| WP_275570988.1 | ParB+HNH | GCF_029167235.1 | Kamptonema_sp._UHCC_0994 | Bacteria>Bacillati>Cyanobacteriota/Melainabacteria_group>Cyanobacteriota>Cyanophyceae>Oscillatoriophycideae>Oscillatoriales>Microcoleaceae>Kamptonema>unclassified_Kamptonema | 767 | PY364_RS28050 | 20750..23053 | 3031329 | DUF262-domain-containing-HNH-endonuclease-family-protein-[Kamptonema-sp.-UHCC-0994] |
| MCC3434394.1 | ParB+HNH | GCA_020738785.1 | Microcoleus_sp._PH2017_05_CCC_O_A | Bacteria>Bacillati>Cyanobacteriota/Melainabacteria_group>Cyanobacteriota>Cyanophyceae>Oscillatoriophycideae>Oscillatoriales>Microcoleaceae>Microcoleus>unclassified_Microcoleus | 760 | JGK20_02540 | 20715..22997 | 2798816 | DUF262-domain-containing-protein-[Microcoleus-sp.-PH2017_05_CCC_O_A] |
| OAB59994.1 | ParB+HNH | GCA_001637315.1 | Phormidium_willei_BDU_130791 | Bacteria>Bacillati>Cyanobacteriota/Melainabacteria_group>Cyanobacteriota>Cyanophyceae>Oscillatoriophycideae>Oscillatoriales>Oscillatoriaceae>Phormidium>Phormidium_willei | 759 | AY600_15265 | 8948..11227 | 399101 | hypothetical-protein-AY600_15265-[Phormidium-willei-BDU-130791] |
| KPQ40626.1 | ParB+HNH | GCA_001314905.1 | Phormidium_sp._OSCR | Bacteria>Bacillati>Cyanobacteriota/Melainabacteria_group>Cyanobacteriota>Cyanophyceae>Oscillatoriophycideae>Oscillatoriales>Oscillatoriaceae>Phormidium>unclassified_Phormidium | 758 | HLUCCO16_02260 | 111214..113490 | 1666905 | hypothetical-protein-HLUCCO16_02260-[Phormidium-sp.-OSCR] |
| WP_242033560.1 | ParB+HNH | - | Phormidium_sp._FACHB-592 | Bacteria>Bacillati>Cyanobacteriota/Melainabacteria_group>Cyanobacteriota>Cyanophyceae>Oscillatoriophycideae>Oscillatoriales>Oscillatoriaceae>Phormidium>unclassified_Phormidium | 412 | H6F86_RS16685 | 62055..63293 | 2692850 | DUF262-domain-containing-protein-[Phormidium-sp.-FACHB-592] |
| WP_253274407.1 | ParB+HNH | - | Myxosarcina_sp._GI1 | Bacteria>Bacillati>Cyanobacteriota/Melainabacteria_group>Cyanobacteriota>Cyanophyceae>Pleurocapsales>Hyellaceae>Myxosarcina>unclassified_Myxosarcina | 533 | KV40_RS27980 | 26162..27763 | 1541065 | DUF262-domain-containing-protein-[Myxosarcina-sp.-GI1] |
| WP_233424289.1 | ParB+HNH | - | Pseudanabaena_biceps | Bacteria>Bacillati>Cyanobacteriota/Melainabacteria_group>Cyanobacteriota>Cyanophyceae>Pseudanabaenales>Pseudanabaenaceae>Pseudanabaena | 570 | PSE7429_RS03415 | 26544..28256 | 927669 | DUF262-domain-containing-protein-[Pseudanabaena-biceps] |
| WP_094532955.1 | ParB+HNH | GCF_002251945.1 | Pseudanabaena_sp._SR411 | Bacteria>Bacillati>Cyanobacteriota/Melainabacteria_group>Cyanobacteriota>Cyanophyceae>Pseudanabaenales>Pseudanabaenaceae>Pseudanabaena>unclassified_Pseudanabaena | 572 | B9G53_RS13015 | 21987..23705 | 1980935 | DUF262-domain-containing-protein-[Pseudanabaena-sp.-SR411] |
| WP_201323616.1 | ParB+HNH | GCF_016584665.1 | Pseudanabaena_sp._lw0831 | Bacteria>Bacillati>Cyanobacteriota/Melainabacteria_group>Cyanobacteriota>Cyanophyceae>Pseudanabaenales>Pseudanabaenaceae>Pseudanabaena>unclassified_Pseudanabaena | 555 | APA_RS17895 | 59288..60955 | 1357935 | DUF262-domain-containing-protein-[Pseudanabaena-sp.-lw0831] |
| WP_255001870.1 | ParB+HNH | GCF_024346435.1 | Cyanobium_sp._Alchichica_3B3-8F6 | Bacteria>Bacillati>Cyanobacteriota/Melainabacteria_group>Cyanobacteriota>Cyanophyceae>Synechococcales>Prochlorococcaceae>Cyanobium>unclassified_Cyanobium | 602 | KBY65_RS13030 | 11322..13130 | 2823696 | DUF262-domain-containing-protein-[Cyanobium-sp.-Alchichica-3B3-8F6] |
| WP_255094988.1 | ParB+HNH | GCF_024345795.1 | Synechococcus_sp._J7-Johnson | Bacteria>Bacillati>Cyanobacteriota/Melainabacteria_group>Cyanobacteriota>Cyanophyceae>Synechococcales>Synechococcaceae>Synechococcus>unclassified_Synechococcus | 352 | KBY93_RS13705 | 24530..25588 | 2823737 | DUF262-domain-containing-protein-[Synechococcus-sp.-J7-Johnson] |
| MBF2075135.1 | ParB+HNH | GCA_015272325.1 | Synechococcales_cyanobacterium_C42_A2020_086 | Bacteria>Bacillati>Cyanobacteriota/Melainabacteria_group>Cyanobacteriota>Cyanophyceae>Synechococcales>unclassified_Synechococcales | 784 | IGS50_15440 | 27437..29791 | 2774344 | DUF262-domain-containing-protein-[Synechococcales-cyanobacterium-C42_A2020_086] |
| MBD1853922.1 | ParB+HNH | GCA_014695325.1 | Cyanobacteria_bacterium_FACHB-502 | Bacteria>Bacillati>Cyanobacteriota/Melainabacteria_group>Cyanobacteriota>unclassified_Cyanobacteriota | 597 | H6F87_28830 | 35208..37001 | 2692765 | DUF262-domain-containing-protein-[Cyanobacteria-bacterium-FACHB-502] |
| MBE0461500.1 | ParB+HNH | GCA_014859315.1 | Candidatus_Aminicenantes_bacterium | Bacteria>Bacteria_incertae_sedis>Bacteria_candidate_phyla>Candidatus_Aminicenantota>unclassified_Candidatus_Aminicenantes | 582 | IBX60_07690 | 1528..3276 | 2052149 | DUF262-domain-containing-protein-[Candidatus-Aminicenantes-bacterium] |
| MCF7889852.1 | ParB+HNH | GCA_021735245.1 | Candidatus_Bipolaricaulota_bacterium | Bacteria>Bacteria_incertae_sedis>Bacteria_candidate_phyla>Candidatus_Bipolaricaulota>unclassified_Candidatus_Bipolaricaulota | 613 | K9M78_01385 | 71940..73781 | 2306012 | DUF262-domain-containing-HNH-endonuclease-family-protein-[Candidatus-Bipolaricaulota-bacterium] |
| MBO7131880.1 | ParB+HNH | GCA_017645955.1 | Candidatus_Saccharibacteria_bacterium | Bacteria>Bacteria_incertae_sedis>Bacteria_candidate_phyla>Candidatus_Saccharibacteria>unclassified_Candidatus_Saccharibacteria | 610 | J6V85_01255 | 46216..48048 | 2026720 | DUF262-domain-containing-protein-[Candidatus-Saccharibacteria-bacterium] |
| WP_146981100.1 | ParB+HNH | GCF_007997015.1 | Lujinxingia_vulgaris | Bacteria>delta/epsilon_subdivisions>Deltaproteobacteria>Bradymonadales>Lujinxingiaceae>Lujinxingia | 473 | FRC98_RS09575 | 48184..49605 | 2600176 | DUF262-domain-containing-protein-[Lujinxingia-vulgaris] |
| WP_246325324.1 | ParB+HNH | GCF_010499435.1 | Dissulfurirhabdus_thermomarina | Bacteria>delta/epsilon_subdivisions>Deltaproteobacteria>Deltaproteobacteria_incertae_sedis>Dissulfurirhabdaceae>Dissulfurirhabdus | 600 | G3N55_RS03840 | 320..2122 | 1765737 | DUF262-domain-containing-protein-[Dissulfurirhabdus-thermomarina] |
| WP_231940477.1 | ParB+HNH | - | Fusobacterium_ulcerans | Bacteria>Fusobacteriati>Fusobacteriota>Fusobacteriia>Fusobacteriales>Fusobacteriaceae>Fusobacterium | 564 | FXX38_RS11825 | 328077..329771 | 861 | DUF262-domain-containing-protein-[Fusobacterium-ulcerans] |
| WP_198847743.1 | ParB+HNH | GCF_016406015.1 | Pseudomonadota | Bacteria>Pseudomonadati | 611 | K4W73_RS30445 | 388496..390331 | 1224 | MULTISPECIES:-DUF262-domain-containing-protein-[Pseudomonadota] |
| WP_180216147.1 | ParB+HNH | GCF_023101305.1 | Pseudomonadota | Bacteria>Pseudomonadati | 602 | MX621_RS22765 | 4835822..4837630 | 1224 | MULTISPECIES:-DUF262-domain-containing-protein-[Pseudomonadota] |
| WP_240685341.1 | ParB+HNH | GCF_025461165.1 | Pseudomonadota | Bacteria>Pseudomonadati | 602 | JAK52_RS02490 | 279471..281279 | 1224 | MULTISPECIES:-DUF262-domain-containing-protein-[Pseudomonadota] |
| MBD0371451.1 | ParB+HNH | GCA_014534425.1 | Pyrinomonadaceae_bacterium | Bacteria>Pseudomonadati>Acidobacteriota>Blastocatellia>Blastocatellales>Pyrinomonadaceae>unclassified_Pyrinomonadaceae | 436 | ICV60_11485 | 22399..23709 | 2283092 | DUF262-domain-containing-protein-[Pyrinomonadaceae-bacterium] |
| MBK9963537.1 | ParB+HNH | GCA_016722225.1 | Holophagales_bacterium | Bacteria>Pseudomonadati>Acidobacteriota>Holophagae>Holophagales>unclassified_Holophagales | 571 | IPP07_00995 | 221636..223351 | 2478486 | DUF262-domain-containing-protein-[Holophagales-bacterium] |
| MCY4585109.1 | ParB+HNH | GCA_026713545.1 | Bryobacterales_bacterium | Bacteria>Pseudomonadati>Acidobacteriota>Terriglobia>Bryobacterales>unclassified_Bryobacterales | 369 | OXB98_03635 | 6361..7470 | 2026791 | MAG:-DUF262-domain-containing-protein-[Bryobacterales-bacterium] |
| MBZ5523273.1 | ParB+HNH | GCA_020072595.1 | Terriglobia_bacterium | Bacteria>Pseudomonadati>Acidobacteriota>Terriglobia>unclassified_Terriglobia | 563 | LAP21_13630 | 6763..8454 | 2081523 | DUF262-domain-containing-HNH-endonuclease-family-protein-[Acidobacteriia-bacterium] |
| MCG3192291.1 | ParB+HNH | GCA_022071945.1 | Thermoanaerobaculia_bacterium | Bacteria>Pseudomonadati>Acidobacteriota>Thermoanaerobaculia>unclassified_Thermoanaerobaculia | 598 | DIJKHBIC_01529 | 8766..10562 | 2651171 | hypothetical-protein-[Thermoanaerobaculia-bacterium] |
| KAB2964426.1 | ParB+HNH | GCA_009026165.1 | Thermoanaerobaculia_bacterium | Bacteria>Pseudomonadati>Acidobacteriota>Thermoanaerobaculia>unclassified_Thermoanaerobaculia | 569 | F9K16_04330 | 5640..7349 | 2651171 | MAG:-DUF262-domain-containing-protein-[Thermoanaerobaculia-bacterium] |
| MBP7777878.1 | ParB+HNH | GCA_018000455.1 | Acidobacteriota_bacterium | Bacteria>Pseudomonadati>Acidobacteriota>unclassified_Acidobacteriota | 561 | KA371_12165 | 125775..127460 | 1978231 | DUF262-domain-containing-protein-[Acidobacteriota-bacterium] |
| MBI4902183.1 | ParB+HNH | GCA_016213775.1 | Acidobacteriota_bacterium | Bacteria>Pseudomonadati>Acidobacteriota>unclassified_Acidobacteriota | 558 | HY820_01020 | 10430..12106 | 1978231 | DUF262-domain-containing-protein-[Acidobacteriota-bacterium] |
| MBA4124437.1 | ParB+HNH | GCA_013820745.1 | Acidobacteriota_bacterium | Bacteria>Pseudomonadati>Acidobacteriota>unclassified_Acidobacteriota | 556 | H0X72_18495 | 6768..8438 | 1978231 | DUF262-domain-containing-protein-[Acidobacteriota-bacterium] |
| MCY3933636.1 | ParB+HNH | GCA_026707325.1 | Acidobacteriota_bacterium | Bacteria>Pseudomonadati>Acidobacteriota>unclassified_Acidobacteriota | 415 | OXH70_18140 | 5012..6259 | 1978231 | MAG:-DUF262-domain-containing-protein-[Acidobacteria-bacterium] |
| MDE2976519.1 | ParB+HNH | GCA_028873735.1 | Acidobacteriota_bacterium | Bacteria>Pseudomonadati>Acidobacteriota>unclassified_Acidobacteriota | 414 | OXU63_03240 | 5098..6342 | 1978231 | MAG:-DUF262-domain-containing-protein-[Acidobacteriota-bacterium] |
| MCY3932056.1 | ParB+HNH | GCA_026707325.1 | Acidobacteriota_bacterium | Bacteria>Pseudomonadati>Acidobacteriota>unclassified_Acidobacteriota | 198 | OXH70_10070 | 61692..62288 | 1978231 | MAG:-DUF262-domain-containing-protein-[Acidobacteria-bacterium] |
| MCB9367523.1 | ParB+HNH | GCA_020637445.1 | Calditrichota_bacterium | Bacteria>Pseudomonadati>Calditrichota>unclassified_Calditrichota | 455 | H6507_00205 | 53738..55105 | 2212469 | DUF262-domain-containing-protein-[Calditrichota-bacterium] |

| Protein-ID | Domain | Assembly-ID | Strain | Taxonomy | Length | Locus-Tag | Genomic-coordinates | TaxID | Fasta-defline |
|---|---|---|---|---|---|---|---|---|---|
| NPV88407.1 | ParB+HNH | GCA_013178255.1 | Coprothermobacteraceae_bacterium | Bacteria>Pseudomonadati>Coprothermobacterota>Coprothermobacteria>Coprothermobacterales>Coprothermobacteraceae>unclassified_Coprothermobacteraceae | 725 | HPY42_02625 | 515922..518099 | 2736298 | DUF262-domain-containing-protein-[Coprothermobacteraceae-bacterium] |
| SHF69827.1 | ParB+HNH | GCA_900128905.1 | Bacteroides_luti | Bacteria>Pseudomonadati>FCB_group>Bacteroidota/Chlorobiota_group>Bacteroidota>Bacteroidia>Bacteroidales>Bacteroidaceae>Bacteroides | 627 | SAMN05444405_11260 | 74922..76805 | 1297750 | Protein-of-unknown-function-[Bacteroides-luti] |
| WP_073402633.1 | ParB+HNH | GCF_900128905.1 | Bacteroides_luti | Bacteria>Pseudomonadati>FCB_group>Bacteroidota/Chlorobiota_group>Bacteroidota>Bacteroidia>Bacteroidales>Bacteroidaceae>Bacteroides | 627 | BUC21_RS11775 | 74922..76805 | 1297750 | DUF262-domain-containing-protein-[Bacteroides-luti] |
| MBS6238333.1 | ParB+HNH | GCA_018374845.1 | Bacteroides_sp. | Bacteria>Pseudomonadati>FCB_group>Bacteroidota/Chlorobiota_group>Bacteroidota>Bacteroidia>Bacteroidales>Bacteroidaceae>Bacteroides>unclassified_Bacteroides | 830 | KH897_08185 | 196442..198934 | 29523 | DUF262-domain-containing-protein-[Bacteroides-sp.] |
| MBS6238334.1 | ParB+HNH | GCA_018374845.1 | Bacteroides_sp. | Bacteria>Pseudomonadati>FCB_group>Bacteroidota/Chlorobiota_group>Bacteroidota>Bacteroidia>Bacteroidales>Bacteroidaceae>Bacteroides>unclassified_Bacteroides | 659 | KH897_08190 | 198937..200916 | 29523 | DUF262-domain-containing-protein-[Bacteroides-sp.] |
| MVZ89489.1 | ParB+HNH | GCA_009768845.1 | Microbacter_sp._ANSKLAB05 | Bacteria>Pseudomonadati>FCB_group>Bacteroidota/Chlorobiota_group>Bacteroidota>Bacteroidia>Bacteroidales>Porphyromonadaceae>Microbacter>unclassified_Microbacter | 647 | GQF49_03170 | 682634..684577 | 2684407 | DUF262-domain-containing-protein-[Microbacter-sp.-ANSKLAB05] |
| SHK68241.1 | ParB+HNH | GCA_900142525.1 | Xylanibacter_ruminicola | Bacteria>Pseudomonadati>FCB_group>Bacteroidota/Chlorobiota_group>Bacteroidota>Bacteroidia>Bacteroidales>Prevotellaceae>Xylanibacter | 632 | SAMN05216463_10951 | 64936..66834 | 839 | Protein-of-unknown-function-[Prevotella-ruminicola] |
| SHK68218.1 | ParB+HNH | GCA_900142525.1 | Xylanibacter_ruminicola | Bacteria>Pseudomonadati>FCB_group>Bacteroidota/Chlorobiota_group>Bacteroidota>Bacteroidia>Bacteroidales>Prevotellaceae>Xylanibacter | 420 | SAMN05216463_10950 | 63663..64925 | 839 | Protein-of-unknown-function-DUF262-[Prevotella-ruminicola] |
| WP_073207529.1 | ParB+HNH | GCF_900142525.1 | Xylanibacter_ruminicola | Bacteria>Pseudomonadati>FCB_group>Bacteroidota/Chlorobiota_group>Bacteroidota>Bacteroidia>Bacteroidales>Prevotellaceae>Xylanibacter | 420 | BUB86_RS07095 | 63663..64925 | 839 | DUF262-domain-containing-protein-[Prevotella-ruminicola] |
| WP_079684010.1 | ParB+HNH | GCF_900168155.1 | Parabacteroides_chartae | Bacteria>Pseudomonadati>FCB_group>Bacteroidota/Chlorobiota_group>Bacteroidota>Bacteroidia>Bacteroidales>Tannerellaceae>Parabacteroides | 750 | B5X57_RS12600 | 61517..63769 | 1037355 | DUF262-domain-containing-protein-[Parabacteroides-chartae] |
| WP_079684009.1 | ParB+HNH | GCF_900168155.1 | Parabacteroides_chartae | Bacteria>Pseudomonadati>FCB_group>Bacteroidota/Chlorobiota_group>Bacteroidota>Bacteroidia>Bacteroidales>Tannerellaceae>Parabacteroides | 583 | B5X57_RS12595 | 59794..61545 | 1037355 | DUF262-domain-containing-protein-[Parabacteroides-chartae] |
| NLT02432.1 | ParB+HNH | GCA_012718945.1 | Bacteroidales_bacterium | Bacteria>Pseudomonadati>FCB_group>Bacteroidota/Chlorobiota_group>Bacteroidota>Bacteroidia>Bacteroidales>unclassified_Bacteroidales | 832 | GXY09_01055 | 3555..6053 | 2030927 | DUF262-domain-containing-protein-[Bacteroidales-bacterium] |
| MBP6872707.1 | ParB+HNH | GCA_017991835.1 | Bacteroidales_bacterium | Bacteria>Pseudomonadati>FCB_group>Bacteroidota/Chlorobiota_group>Bacteroidota>Bacteroidia>Bacteroidales>unclassified_Bacteroidales | 553 | KBC43_11930 | 53807..55468 | 2030927 | DUF262-domain-containing-protein-[Bacteroidales-bacterium] |
| WP_075603950.1 | ParB+HNH | GCF_001660705.2 | Saccharicrinis_aurantiacus | Bacteria>Pseudomonadati>FCB_group>Bacteroidota/Chlorobiota_group>Bacteroidota>Bacteroidia>Marinilabiliales>Marinilabiliaceae>Saccharicrinis | 696 | A8B35_RS21025 | 74357..76447 | 1849719 | DUF262-domain-containing-protein-[Saccharicrinis-aurantiacus] |
| MBP8114171.1 | ParB+HNH | GCA_018005575.1 | Chitinophagaceae_bacterium | Bacteria>Pseudomonadati>FCB_group>Bacteroidota/Chlorobiota_group>Bacteroidota>Chitinophagia>Chitinophagales>Chitinophagaceae>unclassified_Chitinophagaceae | 370 | KAY50_02380 | 11021..12133 | 1869212 | DUF262-domain-containing-protein-[Chitinophagaceae-bacterium] |
| NCU04697.1 | ParB+HNH | GCA_009996555.1 | Chitinophagaceae_bacterium | Bacteria>Pseudomonadati>FCB_group>Bacteroidota/Chlorobiota_group>Bacteroidota>Chitinophagia>Chitinophagales>Chitinophagaceae>unclassified_Chitinophagaceae | 358 | GXC73_11990 | 13798..14874 | 1869212 | DUF262-domain-containing-protein-[Chitinophagaceae-bacterium] |
| MBN8676637.1 | ParB+HNH | GCA_017303475.1 | Chitinophagales_bacterium | Bacteria>Pseudomonadati>FCB_group>Bacteroidota/Chlorobiota_group>Bacteroidota>Chitinophagia>Chitinophagales>unclassified_Chitinophagales | 559 | J0M29_00340 | 97516..99195 | 2448779 | DUF262-domain-containing-protein-[Chitinophagales-bacterium] |
| MBX7108624.1 | ParB+HNH | GCA_019694975.1 | Chitinophagales_bacterium | Bacteria>Pseudomonadati>FCB_group>Bacteroidota/Chlorobiota_group>Bacteroidota>Chitinophagia>Chitinophagales>unclassified_Chitinophagales | 370 | K1X61_08275 | 477086..478198 | 2448779 | DUF262-domain-containing-protein-[Chitinophagales-bacterium] |
| WP_209329598.1 | ParB+HNH | GCF_017811155.1 | Lunatimonas_salinarum | Bacteria>Pseudomonadati>FCB_group>Bacteroidota/Chlorobiota_group>Bacteroidota>Cytophagia>Cytophagales>Cyclobacteriaceae>Lunatimonas | 565 | IAQ59_RS04520 | 168862..170559 | 1774590 | DUF262-domain-containing-protein-[Lunatimonas-salinarum] |
| WP_086597019.1 | ParB+HNH | GCF_002154225.1 | Hymenobacter_crusticola | Bacteria>Pseudomonadati>FCB_group>Bacteroidota/Chlorobiota_group>Bacteroidota>Cytophagia>Cytophagales>Hymenobacteraceae>Hymenobacter | 634 | BXP70_RS25825 | 25166..27070 | 1770526 | DUF262-domain-containing-protein-[Hymenobacter-crusticola] |
| WP_053405042.1 | ParB+HNH | GCF_001274635.1 | Persicobacter_sp._CCB-QB2 | Bacteria>Pseudomonadati>FCB_group>Bacteroidota/Chlorobiota_group>Bacteroidota>Cytophagia>Cytophagales>Persicobacteraceae>Persicobacter>unclassified_Persicobacter | 609 | NG22_RS06815 | 1496601..1498430 | 1561025 | DUF262-domain-containing-protein-[Persicobacter-sp.-CCB-QB2] |
| WP_114065480.1 | ParB+HNH | GCF_003325355.1 | Runella_rosea | Bacteria>Pseudomonadati>FCB_group>Bacteroidota/Chlorobiota_group>Bacteroidota>Cytophagia>Cytophagales>Spirosomataceae>Runella | 355 | DR864_RS02580 | 584535..585602 | 2259595 | DUF262-domain-containing-protein-[Runella-rosea] |
| WP_179314665.1 | ParB+HNH | GCF_019278425.1 | Winogradskyella | Bacteria>Pseudomonadati>FCB_group>Bacteroidota/Chlorobiota_group>Bacteroidota>Flavobacteriia>Flavobacteriales>Flavobacteriaceae | 843 | H0I32_RS04035 | 922199..924730 | 286104 | MULTISPECIES:-DUF262-domain-containing-protein-[Winogradskyella] |
| WP_179314666.1 | ParB+HNH | GCF_019278425.1 | Winogradskyella | Bacteria>Pseudomonadati>FCB_group>Bacteroidota/Chlorobiota_group>Bacteroidota>Flavobacteriia>Flavobacteriales>Flavobacteriaceae | 763 | H0I32_RS04040 | 924738..927029 | 286104 | MULTISPECIES:-DUF262-domain-containing-protein-[Winogradskyella] |
| WP_244340074.1 | ParB+HNH | GCF_022836875.1 | Flavobacterium | Bacteria>Pseudomonadati>FCB_group>Bacteroidota/Chlorobiota_group>Bacteroidota>Flavobacteriia>Flavobacteriales>Flavobacteriaceae | 564 | LZF87_RS14565 | 3204873..3206567 | 237 | MULTISPECIES:-DUF262-domain-containing-protein-[Flavobacterium] |
| WP_271393757.1 | ParB+HNH | GCF_947492035.1 | Aequorivita_sinensis | Bacteria>Pseudomonadati>FCB_group>Bacteroidota/Chlorobiota_group>Bacteroidota>Flavobacteriia>Flavobacteriales>Flavobacteriaceae>Aequorivita | 616 | POR58_RS11105 | 21894..23744 | 1382458 | DUF262-domain-containing-HNH-endonuclease-family-protein-[Aequorivita-sinensis] |
| WP_206736027.1 | ParB+HNH | GCF_011065475.1 | Aequorivita_sp._KMM_9714 | Bacteria>Pseudomonadati>FCB_group>Bacteroidota/Chlorobiota_group>Bacteroidota>Flavobacteriia>Flavobacteriales>Flavobacteriaceae>Aequorivita>unclassified_Aequorivita | 697 | G3A46_RS13580 | 28505..30598 | 2707173 | DUF262-domain-containing-protein-[Aequorivita-sp.-KMM-9714] |
| WP_103999285.1 | ParB+HNH | GCF_900108015.1 | Flavobacterium_urumqiense | Bacteria>Pseudomonadati>FCB_group>Bacteroidota/Chlorobiota_group>Bacteroidota>Flavobacteriia>Flavobacteriales>Flavobacteriaceae>Flavobacterium | 564 | C4S35_RS05870 | 202494..204188 | 935224 | DUF262-domain-containing-protein-[Flavobacterium-urumqiense] |
| WP_059014615.1 | ParB+HNH | GCF_001485515.1 | Flavobacterium_psychrophilum | Bacteria>Pseudomonadati>FCB_group>Bacteroidota/Chlorobiota_group>Bacteroidota>Flavobacteriia>Flavobacteriales>Flavobacteriaceae>Flavobacterium | 564 | FPK15_RS07985 | 1431..3125 | 96345 | DUF262-domain-containing-protein-[Flavobacterium-psychrophilum] |
| WP_131908311.1 | ParB+HNH | GCF_004349085.1 | Flavobacterium_caseinilyticum | Bacteria>Pseudomonadati>FCB_group>Bacteroidota/Chlorobiota_group>Bacteroidota>Flavobacteriia>Flavobacteriales>Flavobacteriaceae>Flavobacterium | 693 | E0F89_RS02680 | 592802..594883 | 2541732 | DUF262-domain-containing-protein-[Flavobacterium-caseinilyticum] |
| WP_089081504.1 | ParB+HNH | GCF_007990815.1 | Flavobacterium_branchiophilum | Bacteria>Pseudomonadati>FCB_group>Bacteroidota/Chlorobiota_group>Bacteroidota>Flavobacteriia>Flavobacteriales>Flavobacteriaceae>Flavobacterium | 756 | FB1_RS11085 | 6308..8578 | 55197 | DUF262-domain-containing-protein-[Flavobacterium-branchiophilum] |
| WP_089081503.1 | ParB+HNH | GCF_007990815.1 | Flavobacterium_branchiophilum | Bacteria>Pseudomonadati>FCB_group>Bacteroidota/Chlorobiota_group>Bacteroidota>Flavobacteriia>Flavobacteriales>Flavobacteriaceae>Flavobacterium | 623 | FB1_RS11090 | 8556..10427 | 55197 | DUF262-domain-containing-protein-[Flavobacterium-branchiophilum] |
| GEM55952.1 | ParB+HNH | GCA_007990815.1 | Flavobacterium_branchiophilum_NBRC_15030_=_ATCC_35035 | Bacteria>Pseudomonadati>FCB_group>Bacteroidota/Chlorobiota_group>Bacteroidota>Flavobacteriia>Flavobacteriales>Flavobacteriaceae>Flavobacterium>Flavobacterium_branchiophilum | 753 | FB1_21730 | 6308..8569 | 1218110 | hypothetical-protein-FB1_21730-[Flavobacterium-branchiophilum-NBRC-15030-=-ATCC-35035] |
| GEM55953.1 | ParB+HNH | GCA_007990815.1 | Flavobacterium_branchiophilum_NBRC_15030_=_ATCC_35035 | Bacteria>Pseudomonadati>FCB_group>Bacteroidota/Chlorobiota_group>Bacteroidota>Flavobacteriia>Flavobacteriales>Flavobacteriaceae>Flavobacterium>Flavobacterium_branchiophilum | 623 | FB1_21740 | 8556..10427 | 1218110 | hypothetical-protein-FB1_21740-[Flavobacterium-branchiophilum-NBRC-15030-=-ATCC-35035] |
| WP_027137223.1 | ParB+HNH | - | Gaetbulibacter_saemankumensis | Bacteria>Pseudomonadati>FCB_group>Bacteroidota/Chlorobiota_group>Bacteroidota>Flavobacteriia>Flavobacteriales>Flavobacteriaceae>Gaetbulibacter | 604 | H611_RS0104465 | 470812..472626 | 311208 | DUF262-domain-containing-protein-[Gaetbulibacter-saemankumensis] |
| WP_141421441.1 | ParB+HNH | GCF_006546625.1 | Haloflavibacter_putidus | Bacteria>Pseudomonadati>FCB_group>Bacteroidota/Chlorobiota_group>Bacteroidota>Flavobacteriia>Flavobacteriales>Flavobacteriaceae>Haloflavibacter | 618 | FKR84_RS06220 | 19488..21344 | 2576776 | DUF262-domain-containing-protein-[Haloflavibacter-putidus] |
| OIQ21192.1 | ParB+HNH | GCA_001874145.1 | Lacinutrix_sp._MedPE-SW | Bacteria>Pseudomonadati>FCB_group>Bacteroidota/Chlorobiota_group>Bacteroidota>Flavobacteriia>Flavobacteriales>Flavobacteriaceae>Lacinutrix>unclassified_Lacinutrix | 602 | BM549_09460 | 34259..36067 | 1860087 | MAG:-hypothetical-protein-BM549_09460-[Lacinutrix-sp.-MedPE-SW] |
| WP_076662674.1 | ParB+HNH | GCF_001975575.1 | Mangrovimonas_sp._DI_80 | Bacteria>Pseudomonadati>FCB_group>Bacteroidota/Chlorobiota_group>Bacteroidota>Flavobacteriia>Flavobacteriales>Flavobacteriaceae>Mangrovimonas>unclassified_Mangrovimonas | 691 | BKM32_RS04280 | 821782..823857 | 1779330 | DUF262-domain-containing-protein-[Mangrovimonas-sp.-DI-80] |
| WP_079511501.1 | ParB+HNH | GCF_900167935.1 | Maribacter_arcticus | Bacteria>Pseudomonadati>FCB_group>Bacteroidota/Chlorobiota_group>Bacteroidota>Flavobacteriia>Flavobacteriales>Flavobacteriaceae>Maribacter | 630 | B5X55_RS05005 | 480208..482100 | 561365 | DUF262-domain-containing-protein-[Maribacter-arcticus] |
| WP_132215842.1 | ParB+HNH | GCF_944607305.1 | Mariniflexile_fucanivorans | Bacteria>Pseudomonadati>FCB_group>Bacteroidota/Chlorobiota_group>Bacteroidota>Flavobacteriia>Flavobacteriales>Flavobacteriaceae>Mariniflexile | 618 | EV196_RS05480 | 437273..439129 | 264023 | DUF262-domain-containing-protein-[Mariniflexile-fucanivorans] |
| WP_153266435.1 | ParB+HNH | GCF_009498295.1 | Oceanihabitans_sp._IOP_32 | Bacteria>Pseudomonadati>FCB_group>Bacteroidota/Chlorobiota_group>Bacteroidota>Flavobacteriia>Flavobacteriales>Flavobacteriaceae>Oceanihabitans>unclassified_Oceanihabitans | 615 | FEZ18_RS00170 | 34870..36717 | 2529032 | DUF262-domain-containing-protein-[Oceanihabitans-sp.-IOP_32] |
| WP_090838879.1 | ParB+HNH | GCF_947496445.1 | Olleya_namhaensis | Bacteria>Pseudomonadati>FCB_group>Bacteroidota/Chlorobiota_group>Bacteroidota>Flavobacteriia>Flavobacteriales>Flavobacteriaceae>Olleya | 844 | BMZ09_RS07020 | 315006..317540 | 1144750 | DUF262-domain-containing-protein-[Olleya-namhaensis] |
| WP_090838880.1 | ParB+HNH | GCF_900114005.1 | Olleya_namhaensis | Bacteria>Pseudomonadati>FCB_group>Bacteroidota/Chlorobiota_group>Bacteroidota>Flavobacteriia>Flavobacteriales>Flavobacteriaceae>Olleya | 764 | BMZ09_RS07025 | 317545..319839 | 1144750 | DUF262-domain-containing-protein-[Olleya-namhaensis] |
| WP_272023599.1 | ParB+HNH | GCF_947496445.1 | Olleya_namhaensis | Bacteria>Pseudomonadati>FCB_group>Bacteroidota/Chlorobiota_group>Bacteroidota>Flavobacteriia>Flavobacteriales>Flavobacteriaceae>Olleya | 764 | PQG05_RS12450 | 12330..14624 | 1144750 | DUF262-domain-containing-protein-[Olleya-namhaensis] |
| WP_198639222.1 | ParB+HNH | GCF_016236975.1 | Salegentibacter_maritimus | Bacteria>Pseudomonadati>FCB_group>Bacteroidota/Chlorobiota_group>Bacteroidota>Flavobacteriia>Flavobacteriales>Flavobacteriaceae>Salegentibacter | 697 | I6U50_RS13460 | 29316..31409 | 2794347 | DUF262-domain-containing-protein-[Salegentibacter-maritimus] |
| MAM30276.1 | ParB+HNH | GCA_002693005.1 | Flavobacteriaceae_bacterium | Bacteria>Pseudomonadati>FCB_group>Bacteroidota/Chlorobiota_group>Bacteroidota>Flavobacteriia>Flavobacteriales>Flavobacteriaceae>unclassified_Flavobacteriaceae | 616 | CMC13_14760 | 18001..19851 | 1871037 | hypothetical-protein-[Flavobacteriaceae-bacterium] |
| WP_089756258.1 | ParB+HNH | GCF_900100115.1 | Chryseobacterium_soldanellicola | Bacteria>Pseudomonadati>FCB_group>Bacteroidota/Chlorobiota_group>Bacteroidota>Flavobacteriia>Flavobacteriales>Weeksellaceae>Chryseobacterium_group>Chryseobacterium | 603 | BLT04_RS13245 | 826534..828345 | 311333 | DUF262-domain-containing-protein-[Chryseobacterium-soldanellicola] |
| WP_089756267.1 | ParB+HNH | GCF_900100115.1 | Chryseobacterium_soldanellicola | Bacteria>Pseudomonadati>FCB_group>Bacteroidota/Chlorobiota_group>Bacteroidota>Flavobacteriia>Flavobacteriales>Weeksellaceae>Chryseobacterium_group>Chryseobacterium | 334 | BLT04_RS13295 | 841063..842067 | 311333 | DUF262-domain-containing-protein-[Chryseobacterium-soldanellicola] |
| QOW09783.1 | ParB+HNH | GCA_015191005.1 | Kaistella_flava_(ex_Peng_et_al._2021) | Bacteria>Pseudomonadati>FCB_group>Bacteroidota/Chlorobiota_group>Bacteroidota>Flavobacteriia>Flavobacteriales>Weeksellaceae>Chryseobacterium_group>Kaistella | 603 | Q73A0000_05080 | 1129628..1131439 | 2038776 | DUF262-domain-containing-protein-[Kaistella-flava-(ex-Peng-et-al.-2021)] |
| NUQ24409.1 | ParB+HNH | GCA_013360405.1 | Saprospiraceae_bacterium | Bacteria>Pseudomonadati>FCB_group>Bacteroidota/Chlorobiota_group>Bacteroidota>Saprospiria>Saprospirales>Saprospiraceae>unclassified_Saprospiraceae | 385 | HUU34_10675 | 17146..18303 | 2202734 | DUF262-domain-containing-protein-[Saprospiraceae-bacterium] |
| TAK42620.1 | ParB+HNH | GCA_004297645.1 | Saprospiraceae_bacterium | Bacteria>Pseudomonadati>FCB_group>Bacteroidota/Chlorobiota_group>Bacteroidota>Saprospiria>Saprospirales>Saprospiraceae>unclassified_Saprospiraceae | 179 | EPO28_07075 | 21200..21739 | 2202734 | DUF262-domain-containing-protein-[Saprospiraceae-bacterium] |
| WP_207425051.1 | ParB+HNH | GCF_017355855.1 | Pedobacter_sp._SYSU_D00535 | Bacteria>Pseudomonadati>FCB_group>Bacteroidota/Chlorobiota_group>Bacteroidota>Sphingobacteriia>Sphingobacteriales>Sphingobacteriaceae>Pedobacter>unclassified_Pedobacter | 560 | J4N23_RS00525 | 151173..152855 | 2810308 | DUF262-domain-containing-protein-[Pedobacter-sp.-SYSU-D00535] |
| OYD42849.1 | ParB+HNH | GCA_002245855.1 | Sphingobacterium_cellulitidis | Bacteria>Pseudomonadati>FCB_group>Bacteroidota/Chlorobiota_group>Bacteroidota>Sphingobacteriia>Sphingobacteriales>Sphingobacteriaceae>Sphingobacterium | 599 | CHT99_08540 | 564356..566155 | 1768011 | hypothetical-protein-CHT99_08540-[Sphingobacterium-cellulitidis] |
| WP_094282241.1 | ParB+HNH | GCF_002245765.1 | Sphingobacterium_cellulitidis | Bacteria>Pseudomonadati>FCB_group>Bacteroidota/Chlorobiota_group>Bacteroidota>Sphingobacteriia>Sphingobacteriales>Sphingobacteriaceae>Sphingobacterium | 599 | CHU00_RS15760 | 12970..14769 | 1768011 | DUF262-domain-containing-protein-[Sphingobacterium-cellulitidis] |
| MBS1776513.1 | ParB+HNH | GCA_018267925.1 | Bacteroidota_bacterium | Bacteria>Pseudomonadati>FCB_group>Bacteroidota/Chlorobiota_group>Bacteroidota>unclassified_Bacteroidota | 565 | JSS64_09570 | 73146..74843 | 1898104 | DUF262-domain-containing-protein-[Bacteroidota-bacterium] |
| MBS1911239.1 | ParB+HNH | GCA_018266795.1 | Bacteroidota_bacterium | Bacteria>Pseudomonadati>FCB_group>Bacteroidota/Chlorobiota_group>Bacteroidota>unclassified_Bacteroidota | 565 | JST22_04575 | 50695..52392 | 1898104 | DUF262-domain-containing-protein-[Bacteroidota-bacterium] |
| MBN1449252.1 | ParB+HNH | GCA_016928515.1 | Bacteroidota_bacterium | Bacteria>Pseudomonadati>FCB_group>Bacteroidota/Chlorobiota_group>Bacteroidota>unclassified_Bacteroidota | 564 | JXA28_15100 | 3198..4892 | 1898104 | DUF262-domain-containing-protein-[Bacteroidota-bacterium] |
| MBK7887719.1 | ParB+HNH | GCA_016713925.1 | Bacteroidota_bacterium | Bacteria>Pseudomonadati>FCB_group>Bacteroidota/Chlorobiota_group>Bacteroidota>unclassified_Bacteroidota | 402 | IPJ86_10630 | 607273..608481 | 1898104 | HNH-endonuclease-[Bacteroidota-bacterium] |
| WP_109646584.1 | ParB+HNH | GCF_003150675.1 | Rhodohalobacter_mucosus | Bacteria>Pseudomonadati>FCB_group>Bacteroidota/Chlorobiota_group>Balneolota>Balneolia>Balneolales>Balneolaceae>Rhodohalobacter | 649 | DDZ15_RS08070 | 62788..64737 | 2079485 | DUF262-domain-containing-protein-[Rhodohalobacter-mucosus] |
| GIV55104.1 | ParB+HNH | GCA_026003475.1 | Candidatus_Kapaibacterium_sp. | Bacteria>Pseudomonadati>FCB_group>Bacteroidota/Chlorobiota_group>Candidatus_Kapaibacteriota>Candidatus_Kapaibacteriia>Candidatus_Kapaibacteriales>Candidatus_Kapaibacteriaceae>Candidatus_Kapaibacterium>unclassified_Candidatus_Kapaibacterium | 93 | KatS3mg039_1622 | 1984219..1984500 | 2447898 | MAG:-hypothetical-protein-KatS3mg039_1622-[Candidatus-Kapabacteria-bacterium] |
| MBV5326248.1 | ParB+HNH | GCA_019162945.1 | Chlorobium_sp. | Bacteria>Pseudomonadati>FCB_group>Bacteroidota/Chlorobiota_group>Chlorobiota>Chlorobiia>Chlorobiales>Chlorobiaceae>Chlorobium/Pelodictyon_group>Chlorobium>unclassified_Chlorobium | 366 | JZU65_01225 | 67..1167 | 1095 | DUF262-domain-containing-protein-[Chlorobium-sp.] |
| WP_139626759.1 | ParB+HNH | GCF_006265245.1 | Prosthecochloris_vibrioformis | Bacteria>Pseudomonadati>FCB_group>Bacteroidota/Chlorobiota_group>Chlorobiota>Chlorobiia>Chlorobiales>Chlorobiaceae>Prosthecochloris | 573 | FGF68_RS08490 | 2656..4377 | 1098 | DUF262-domain-containing-protein-[Prosthecochloris-vibrioformis] |
| WP_068867047.1 | ParB+HNH | GCF_001687065.1 | Prosthecochloris_sp._CIB_2401 | Bacteria>Pseudomonadati>FCB_group>Bacteroidota/Chlorobiota_group>Chlorobiota>Chlorobiia>Chlorobiales>Chlorobiaceae>Prosthecochloris>unclassified_Prosthecochloris | 573 | Ptc2401_RS05060 | 1067783..1069504 | 1868325 | DUF262-domain-containing-protein-[Prosthecochloris-sp.-CIB-2401] |
| MBM2814378.1 | ParB+HNH | GCA_016844745.1 | Ignavibacteria_bacterium | Bacteria>Pseudomonadati>FCB_group>Bacteroidota/Chlorobiota_group>Ignavibacteriota>Ignavibacteria>unclassified_Ignavibacteria | 600 | HW421_1140 | 5668..7470 | 2053306 | hypothetical-protein-[Ignavibacteria-bacterium] |
| WP_251951635.1 | ParB+HNH | GCF_943908255.1 | Salinibacter_ruber | Bacteria>Pseudomonadati>FCB_group>Bacteroidota/Chlorobiota_group>Rhodothermota>Rhodothermia>Rhodothermales>Salinibacteraceae>Salinibacter | 694 | NGL47_RS11285 | 16004..18088 | 146919 | DUF262-domain-containing-protein-[Salinibacter-ruber] |
| WP_073283046.1 | ParB+HNH | GCF_900142465.1 | unclassified_Fibrobacter | Bacteria>Pseudomonadati>FCB_group>Fibrobacterota>Fibrobacteria>Fibrobacterales>Fibrobacteraceae>Fibrobacter | 599 | B7993_RS04330 | 157306..159105 | 2634177 | MULTISPECIES:-DUF262-domain-containing-protein-[unclassified-Fibrobacter] |
| SHK42541.1 | ParB+HNH | GCA_900142505.1 | Fibrobacter_sp._UWH5 | Bacteria>Pseudomonadati>FCB_group>Fibrobacterota>Fibrobacteria>Fibrobacterales>Fibrobacteraceae>Fibrobacter>unclassified_Fibrobacter | 599 | SAMN05720764_101396 | 447916..449715 | 1896211 | Protein-of-unknown-function-[Fibrobacter-sp.-UWH5] |
| WP_073156272.1 | ParB+HNH | GCF_900142505.1 | Fibrobacter_sp._UWH5 | Bacteria>Pseudomonadati>FCB_group>Fibrobacterota>Fibrobacteria>Fibrobacterales>Fibrobacteraceae>Fibrobacter>unclassified_Fibrobacter | 599 | BUB38_RS01955 | 447916..449715 | 1896211 | DUF262-domain-containing-protein-[Fibrobacter-sp.-UWH5] |
| NLW30566.1 | ParB+HNH | GCA_012728995.1 | Fibrobacter_sp. | Bacteria>Pseudomonadati>FCB_group>Fibrobacterota>Fibrobacteria>Fibrobacterales>Fibrobacteraceae>Fibrobacter>unclassified_Fibrobacter | 638 | GXY77_03815 | 12114..14030 | 35828 | DUF262-domain-containing-protein-[Fibrobacter-sp.] |
| MXZ76450.1 | ParB+HNH | GCA_009837395.1 | Gemmatimonadota_bacterium | Bacteria>Pseudomonadati>FCB_group>Gemmatimonadota>unclassified_Gemmatimonadota | 661 | F4Z03_15250 | 68354..70339 | 2026742 | DUF262-domain-containing-protein-[Gemmatimonadota-bacterium] |
| MBK8059518.1 | ParB+HNH | GCA_016713015.1 | Gemmatimonadota_bacterium | Bacteria>Pseudomonadati>FCB_group>Gemmatimonadota>unclassified_Gemmatimonadota | 623 | IPK33_17170 | 749911..751782 | 2026742 | DUF262-domain-containing-protein-[Gemmatimonadota-bacterium] |
| MYH20373.1 | ParB+HNH | GCA_009844015.1 | Gemmatimonadota_bacterium | Bacteria>Pseudomonadati>FCB_group>Gemmatimonadota>unclassified_Gemmatimonadota | 570 | F4132_14855 | 86962..88674 | 2026742 | DUF262-domain-containing-protein-[Gemmatimonadota-bacterium] |
| MDO9018001.1 | ParB+HNH | GCA_030652255.1 | Myxococcales_bacterium | Bacteria>Pseudomonadati>Myxococcota>Myxococcia>Myxococcales>unclassified_Myxococcales | 588 | Q7V43_13885 | 22438..24204 | 2026763 | DUF262-domain-containing-HNH-endonuclease-family-protein-[Myxococcales-bacterium] |
| MCE9575763.1 | ParB+HNH | GCA_021413675.1 | Deltaproteobacteria_bacterium | Bacteria>Pseudomonadati>Myxococcota>Myxococcia>unclassified_Myxococcia | 848 | K8W52_21615 | 64169..66715 | 2026735 | DUF262-domain-containing-protein-[Deltaproteobacteria-bacterium] |
| MBN1335621.1 | ParB+HNH | GCA_016927815.1 | Deltaproteobacteria_bacterium | Bacteria>Pseudomonadati>Myxococcota>Myxococcia>unclassified_Myxococcia | 637 | JXB39_06645 | 8920..10833 | 2026735 | DUF262-domain-containing-protein-[Deltaproteobacteria-bacterium] |
| MBI5504533.1 | ParB+HNH | GCA_016218925.1 | Deltaproteobacteria_bacterium | Bacteria>Pseudomonadati>Myxococcota>Myxococcia>unclassified_Myxococcia | 575 | HY899_07005 | 35041..36768 | 2026735 | DUF262-domain-containing-protein-[Deltaproteobacteria-bacterium] |
| MDE0030311.1 | ParB+HNH | GCA_028818775.1 | Deltaproteobacteria_bacterium | Bacteria>Pseudomonadati>Myxococcota>Myxococcia>unclassified_Myxococcia | 377 | OXU42_13030 | 17844..18977 | 2026735 | MAG:-DUF262-domain-containing-protein-[Deltaproteobacteria-bacterium] |
| MBA3548107.1 | ParB+HNH | GCA_013812955.1 | Nannocystis_sp. | Bacteria>Pseudomonadati>Myxococcota>Polyangia>Nannocystales>Nannocystaceae>Nannocystis>unclassified_Nannocystis | 620 | H0T76_16620 | 35431..37293 | 1962667 | DUF262-domain-containing-protein-[Nannocystis-sp.] |
| MBK8262776.1 | ParB+HNH | GCA_016712315.1 | Nannocystis_sp. | Bacteria>Pseudomonadati>Myxococcota>Polyangia>Nannocystales>Nannocystaceae>Nannocystis>unclassified_Nannocystis | 558 | IPK80_15755 | 190951..192627 | 1962667 | DUF262-domain-containing-protein-[Nannocystis-sp.] |
| MCK6512395.1 | ParB+HNH | GCA_023150835.1 | Myxococcota_bacterium | Bacteria>Pseudomonadati>Myxococcota>unclassified_Myxococcota | 598 | L6R29_20895 | 10643..12439 | 2818507 | DUF262-domain-containing-HNH-endonuclease-family-protein-[Myxococcota-bacterium] |
| NGZ08747.1 | ParB+HNH | GCA_011090425.1 | Nitrospira_sp._LK70 | Bacteria>Pseudomonadati>Nitrospirota>Nitrospiria>Nitrospirales>Nitrospiraceae>Nitrospira>unclassified_Nitrospira | 569 | CV088_05075 | 53240..54949 | 2039809 | DUF262-domain-containing-protein-[Nitrospira-sp.-LK70] |
| MBP6261840.1 | ParB+HNH | GCA_017986555.1 | Nitrospira_sp. | Bacteria>Pseudomonadati>Nitrospirota>Nitrospiria>Nitrospirales>Nitrospiraceae>Nitrospira>unclassified_Nitrospira | 570 | KA426_01220 | 6802..8514 | 70125 | DUF262-domain-containing-protein-[Nitrospira-sp.] |
| MCS6288010.1 | ParB+HNH | GCA_024998625.1 | Nitrospira_sp. | Bacteria>Pseudomonadati>Nitrospirota>Nitrospiria>Nitrospirales>Nitrospiraceae>Nitrospira>unclassified_Nitrospira | 570 | H8K10_03420 | 256106..257818 | 70125 | DUF262-domain-containing-protein-[Nitrospira-sp.] |
| MBS0184606.1 | ParB+HNH | GCA_018242685.1 | Nitrospira_sp. | Bacteria>Pseudomonadati>Nitrospirota>Nitrospiria>Nitrospirales>Nitrospiraceae>Nitrospira>unclassified_Nitrospira | 537 | JSS39_19630 | 25155..26768 | 70125 | DUF262-domain-containing-protein-[Nitrospira-sp.] |
| WP_103916695.1 | ParB+HNH | GCF_900107995.1 | Gammaproteobacteria | Bacteria>Pseudomonadati>Pseudomonadota | 626 | C4S63_RS03485 | 771549..773429 | 1236 | DUF262-domain-containing-protein-[Alcanivorax-sp.-DSM-26293] |
| WP_125941665.1 | ParB+HNH | GCF_019046925.1 | Gammaproteobacteria | Bacteria>Pseudomonadati>Pseudomonadota | 597 | I6L32_RS07275 | 1518450..1520243 | 1236 | MULTISPECIES:-DUF262-domain-containing-protein-[Gammaproteobacteria] |
| WP_000283752.1 | ParB+HNH | GCF_000022585.1 | Gammaproteobacteria | Bacteria>Pseudomonadati>Pseudomonadota | 587 | MMZ64_RS09100 | 1893927..1895690 | 1236 | MULTISPECIES:-DUF262-domain-containing-protein-[Gammaproteobacteria] |
| WP_010129446.1 | ParB+HNH | - | Gammaproteobacteria | Bacteria>Pseudomonadati>Pseudomonadota | 587 | O4N70_RS12815 | 224057..225820 | 1236 | MULTISPECIES:-DUF262-domain-containing-protein-[Gammaproteobacteria] |
| WP_102887271.1 | ParB+HNH | GCF_021431945.1 | Gammaproteobacteria | Bacteria>Pseudomonadati>Pseudomonadota | 587 | LY388_RS16335 | 2423199..2424962 | 1236 | MULTISPECIES:-DUF262-domain-containing-protein-[Gammaproteobacteria] |
| WP_111805007.1 | ParB+HNH | GCF_011045875.1 | Gammaproteobacteria | Bacteria>Pseudomonadati>Pseudomonadota | 587 | IXW67_RS19935 | 16646..18409 | 1236 | MULTISPECIES:-DUF262-domain-containing-protein-[Gammaproteobacteria] |
| WP_113603074.1 | ParB+HNH | GCF_030731885.1 | Gammaproteobacteria | Bacteria>Pseudomonadati>Pseudomonadota | 587 | DLR65_RS02200 | 58323..60086 | 1236 | DUF262-domain-containing-protein-[Vibrio-tarriae] |
| WP_119567030.1 | ParB+HNH | GCF_013602615.1 | Gammaproteobacteria | Bacteria>Pseudomonadati>Pseudomonadota | 587 | G9G97_RS25350 | 24862..26625 | 1236 | MULTISPECIES:-DUF262-domain-containing-protein-[Gammaproteobacteria] |
| WP_123161579.1 | ParB+HNH | GCF_009763665.1 | Gammaproteobacteria | Bacteria>Pseudomonadati>Pseudomonadota | 587 | GPY01_RS14955 | 2269018..2270781 | 1236 | DUF262-domain-containing-protein-[Vibrio-cholerae] |
| WP_140256558.1 | ParB+HNH | GCF_029167175.1 | Gammaproteobacteria | Bacteria>Pseudomonadati>Pseudomonadota | 587 | PY479_RS00085 | 20032..21795 | 1236 | MULTISPECIES:-DUF262-domain-containing-protein-[Gammaproteobacteria] |
| WP_206611793.1 | ParB+HNH | GCF_003721225.2 | Acidithiobacillus_sulfuriphilus | Bacteria>Pseudomonadati>Pseudomonadota>Acidithiobacillia>Acidithiobacillales>Acidithiobacillaceae>Acidithiobacillus | 616 | EC580_RS10645 | 4835..6685 | 1867749 | DUF262-domain-containing-protein-[Acidithiobacillus-sulfuriphilus] |
| WP_215855741.1 | ParB+HNH | GCF_018854635.1 | Acidithiobacillus_ferridurans | Bacteria>Pseudomonadati>Pseudomonadota>Acidithiobacillia>Acidithiobacillales>Acidithiobacillaceae>Acidithiobacillus | 602 | HF641_RS00670 | 34930..36738 | 1232575 | DUF262-domain-containing-protein-[Acidithiobacillus-ferridurans] |
| MBE7157120.1 | ParB+HNH | GCA_015069705.1 | Acetobacter_sp. | Bacteria>Pseudomonadati>Pseudomonadota>Alphaproteobacteria>Acetobacterales>Acetobacteraceae>Acetobacter>unclassified_Acetobacter | 797 | INR62_01570 | 3734..6127 | 440 | DUF262-domain-containing-protein-[Rhodospirillales-bacterium] |
| MBE7157119.1 | ParB+HNH | GCA_015069705.1 | Acetobacter_sp. | Bacteria>Pseudomonadati>Pseudomonadota>Alphaproteobacteria>Acetobacterales>Acetobacteraceae>Acetobacter>unclassified_Acetobacter | 575 | INR62_01565 | 2010..3737 | 440 | DUF262-domain-containing-protein-[Rhodospirillales-bacterium] |
| WP_201727552.1 | ParB+HNH | GCF_907164655.1 | Acidocella_sp._C78 | Bacteria>Pseudomonadati>Pseudomonadota>Alphaproteobacteria>Acetobacterales>Acidocellaceae>Acidocella>unclassified_Acidocella | 342 | ACIDOCELLA78_RS01050 | 240881..241909 | 1671486 | HNH-endonuclease-family-protein-[Acidocella-sp.-C78] |
| WP_201727551.1 | ParB+HNH | GCF_907164655.1 | Acidocella_sp._C78 | Bacteria>Pseudomonadati>Pseudomonadota>Alphaproteobacteria>Acetobacterales>Acidocellaceae>Acidocella>unclassified_Acidocella | 237 | ACIDOCELLA78_RS01040 | 238996..239709 | 1671486 | DUF262-domain-containing-protein-[Acidocella-sp.-C78] |
| WP_075822682.1 | ParB+HNH | GCF_001950415.1 | Roseomonas | Bacteria>Pseudomonadati>Pseudomonadota>Alphaproteobacteria>Acetobacterales>Roseomonadaceae | 698 | MHL39_RS17990 | 12820..14916 | 125216 | MULTISPECIES:-DUF262-and-DUF1524-domain-containing-protein-[Roseomonas] |
| QDD97340.1 | ParB+HNH | GCA_006384895.1 | Roseomonas_mucosa | Bacteria>Pseudomonadati>Pseudomonadota>Alphaproteobacteria>Acetobacterales>Roseomonadaceae>Roseomonas | 588 | ADP8_05157 | 108432..110198 | 207340 | putative-cytosolic-protein-[Roseomonas-mucosa] |
| TPG55474.1 | ParB+HNH | GCA_006438825.1 | Muricoccus_nepalensis | Bacteria>Pseudomonadati>Pseudomonadota>Alphaproteobacteria>Acetobacterales>Roseomonadaceae>Roseomonas | 662 | EAH89_14580 | 40605..42593 | 1854500 | DUF262-domain-containing-protein-[Roseomonas-nepalensis] |
| WP_140884410.1 | ParB+HNH | GCF_006438825.1 | Muricoccus_nepalensis | Bacteria>Pseudomonadati>Pseudomonadota>Alphaproteobacteria>Acetobacterales>Roseomonadaceae>Roseomonas | 662 | EAH89_RS14640 | 40605..42593 | 1854500 | DUF262-domain-containing-protein-[Roseomonas-nepalensis] |
| MCA3288091.1 | ParB+HNH | GCA_020249245.1 | Roseomonas_sp. | Bacteria>Pseudomonadati>Pseudomonadota>Alphaproteobacteria>Acetobacterales>Roseomonadaceae>Roseomonas>unclassified_Roseomonas | 647 | ING09_16235 | 131462..133405 | 1869190 | DUF262-domain-containing-protein-[Roseomonas-sp.] |
| MCA4920340.1 | ParB+HNH | GCA_020285315.1 | Roseomonas_sp. | Bacteria>Pseudomonadati>Pseudomonadota>Alphaproteobacteria>Acetobacterales>Roseomonadaceae>Roseomonas>unclassified_Roseomonas | 647 | ING82_13010 | 11625..13568 | 1869190 | DUF262-domain-containing-protein-[Roseomonas-sp.] |
| WP_254454363.1 | ParB+HNH | - | Siccirubricoccus_sp._G192 | Bacteria>Pseudomonadati>Pseudomonadota>Alphaproteobacteria>Acetobacterales>Roseomonadaceae>Siccirubricoccus>unclassified_Siccirubricoccus | 1074 | KSF81_RS28010 | 7398..10622 | 2849651 | DUF262-domain-containing-protein-[Siccirubricoccus-sp.-G192] |
| WP_217382204.1 | ParB+HNH | GCF_019084075.1 | Siccirubricoccus_sp._G192 | Bacteria>Pseudomonadati>Pseudomonadota>Alphaproteobacteria>Acetobacterales>Roseomonadaceae>Siccirubricoccus>unclassified_Siccirubricoccus | 698 | KSF81_RS28680 | 172763..174859 | 2849651 | DUF262-and-DUF1524-domain-containing-protein-[Siccirubricoccus-sp.-G192] |
| WP_031235655.1 | ParB+HNH | GCF_000495835.1 | Asticcacaulis_sp._AC402 | Bacteria>Pseudomonadati>Pseudomonadota>Alphaproteobacteria>Caulobacterales>Caulobacteraceae>Asticcacaulis>unclassified_Asticcacaulis | 596 | ABAC402_RS39740 | 113661..115451 | 1282361 | DUF262-domain-containing-protein-[Asticcacaulis-sp.-AC402] |
| KAF0185698.1 | ParB+HNH | GCA_009773335.1 | Caulobacteraceae_bacterium | Bacteria>Pseudomonadati>Pseudomonadota>Alphaproteobacteria>Caulobacterales>Caulobacteraceae>unclassified_Caulobacteraceae | 545 | FD160_507 | 45328..46965 | 1978230 | hypothetical-protein-FD160_507-[Caulobacteraceae-bacterium] |
| WP_078709665.1 | ParB+HNH | GCF_900167365.1 | Consotaella_salsifontis | Bacteria>Pseudomonadati>Pseudomonadota>Alphaproteobacteria>Hyphomicrobiales>Aurantimonadaceae>Consotaella | 562 | B5D38_RS17145 | 47980..49668 | 1365950 | DUF262-domain-containing-protein-[Consotaella-salsifontis] |
| WP_259030289.1 | ParB+HNH | GCF_024799785.1 | Brucella | Bacteria>Pseudomonadati>Pseudomonadota>Alphaproteobacteria>Hyphomicrobiales>Brucellaceae>Brucella/Ochrobactrum_group | 560 | NW321_RS01570 | 339444..341126 | 234 | DUF262-domain-containing-protein-[Brucella-anthropi] |
| UYN95210.1 | ParB+HNH | GCA_025808875.1 | Enhydrobacter_sp. | Bacteria>Pseudomonadati>Pseudomonadota>Alphaproteobacteria>Hyphomicrobiales>Hyphomicrobiales_incertae_sedis>Enhydrobacter>unclassified_Enhydrobacter | 566 | KIT25_24905 | 5024120..5025820 | 1894999 | MAG:-DUF262-domain-containing-protein-[Enhydrobacter-sp.] |
| WP_082559274.1 | ParB+HNH | GCF_001426045.1 | Methylobacterium_sp._Leaf94 | Bacteria>Pseudomonadati>Pseudomonadota>Alphaproteobacteria>Hyphomicrobiales>Methylobacteriaceae>Methylobacterium>unclassified_Methylobacterium | 620 | ASG63_RS24170 | 17928..19790 | 1736250 | DUF262-domain-containing-protein-[Methylobacterium-sp.-Leaf94] |
| WP_079537875.1 | ParB+HNH | GCF_900141755.1 | Bradyrhizobium_lablabi | Bacteria>Pseudomonadati>Pseudomonadota>Alphaproteobacteria>Hyphomicrobiales>Nitrobacteraceae>Bradyrhizobium | 635 | B5526_RS08935 | 1908947..1910854 | 722472 | DUF262-domain-containing-protein-[Bradyrhizobium-lablabi] |
| WP_213286694.1 | ParB+HNH | GCF_018398875.1 | Bradyrhizobium_sp._sGM-13 | Bacteria>Pseudomonadati>Pseudomonadota>Alphaproteobacteria>Hyphomicrobiales>Nitrobacteraceae>Bradyrhizobium>unclassified_Bradyrhizobium | 616 | KJQ96_RS11315 | 152434..154284 | 2831781 | DUF262-domain-containing-protein-[Bradyrhizobium-sp.-sGM-13] |
| MBR1240290.1 | ParB+HNH | GCA_018130325.1 | Bradyrhizobium_sp._AUGA_SZCCT0274 | Bacteria>Pseudomonadati>Pseudomonadota>Alphaproteobacteria>Hyphomicrobiales>Nitrobacteraceae>Bradyrhizobium>unclassified_Bradyrhizobium | 616 | JQ620_09150 | 68595..70445 | 2807670 | DUF262-domain-containing-protein-[Bradyrhizobium-sp.-AUGA-SZCCT0274] |
| WP_212207482.1 | ParB+HNH | GCF_018130325.1 | Bradyrhizobium_sp._AUGA_SZCCT0274 | Bacteria>Pseudomonadati>Pseudomonadota>Alphaproteobacteria>Hyphomicrobiales>Nitrobacteraceae>Bradyrhizobium>unclassified_Bradyrhizobium | 616 | JQ620_RS09155 | 68595..70445 | 2807670 | DUF262-domain-containing-protein-[Bradyrhizobium-sp.-AUGA-SZCCT0274] |
| WP_247344334.1 | ParB+HNH | GCF_023073075.1 | Bradyrhizobium_sp._61 | Bacteria>Pseudomonadati>Pseudomonadota>Alphaproteobacteria>Hyphomicrobiales>Nitrobacteraceae>Bradyrhizobium>unclassified_Bradyrhizobium | 559 | IVB46_RS33330 | 513837..515516 | 2782679 | DUF262-domain-containing-protein-[Bradyrhizobium-sp.-61] |
| CAI2932246.1 | ParB+HNH | GCA_946995915.1 | Aminobacter_niigataensis | Bacteria>Pseudomonadati>Pseudomonadota>Alphaproteobacteria>Hyphomicrobiales>Phyllobacteriaceae>Aminobacter | 560 | AMD1_0964 | 949389..951071 | 83265 | conserved-protein-of-unknown-function-[Aminobacter-niigataensis] |
| WP_066996952.1 | ParB+HNH | GCF_001672445.1 | Mesorhizobium_sp._WSM3873 | Bacteria>Pseudomonadati>Pseudomonadota>Alphaproteobacteria>Hyphomicrobiales>Phyllobacteriaceae>Mesorhizobium>unclassified_Mesorhizobium | 756 | A9K71_RS15165 | 44684..46954 | 1854056 | DUF262-domain-containing-protein-[Mesorhizobium-sp.-WSM3873] |
| WP_082980534.1 | ParB+HNH | GCF_001672445.1 | Mesorhizobium_sp._WSM3873 | Bacteria>Pseudomonadati>Pseudomonadota>Alphaproteobacteria>Hyphomicrobiales>Phyllobacteriaceae>Mesorhizobium>unclassified_Mesorhizobium | 595 | A9K71_RS15170 | 46951..48738 | 1854056 | DUF262-domain-containing-protein-[Mesorhizobium-sp.-WSM3873] |
| WP_128286153.1 | ParB+HNH | GCF_004023115.1 | Mesorhizobium_sp._M6A.T.Cr.TU.017.01.1.1 | Bacteria>Pseudomonadati>Pseudomonadota>Alphaproteobacteria>Hyphomicrobiales>Phyllobacteriaceae>Mesorhizobium>unclassified_Mesorhizobium | 652 | EOB36_RS27980 | 2477..4435 | 2496774 | DUF262-domain-containing-protein-[Mesorhizobium-sp.-M6A.T.Cr.TU.017.01.1.1] |
| WP_023809225.1 | ParB+HNH | GCF_000502875.1 | Mesorhizobium_sp._L2C084A000 | Bacteria>Pseudomonadati>Pseudomonadota>Alphaproteobacteria>Hyphomicrobiales>Phyllobacteriaceae>Mesorhizobium>unclassified_Mesorhizobium | 598 | X735_RS161790 | 855721..857517 | 1287116 | DUF262-domain-containing-protein-[Mesorhizobium-sp.-L2C084A000] |
| TIL20468.1 | ParB+HNH | GCA_004961195.1 | Mesorhizobium_sp. | Bacteria>Pseudomonadati>Pseudomonadota>Alphaproteobacteria>Hyphomicrobiales>Phyllobacteriaceae>Mesorhizobium>unclassified_Mesorhizobium | 600 | E5Y88_32430 | 12431..14233 | 1871066 | DUF262-domain-containing-protein-[Mesorhizobium-sp.] |

| Protein-ID | Domain | Assembly-ID | Strain | Taxonomy | Length | Locus-Tag | Genomic-coordinates | TaxID | Fasta-defline |
|---|---|---|---|---|---|---|---|---|---|
| RWD70284.1 | ParB+HNH | GCA_004018245.1 | Mesorhizobium_sp. | Bacteria>Pseudomonadati>Pseudomonadota>Alphaproteobacteria>Hyphomicrobiales>Phyllobacteriaceae>Mesorhizobium>unclassified_Mesorhizobium | 567 | EOS37_15305 | 32135..33838 | 1871066 | DUF262-domain-containing-protein-[Mesorhizobium-sp.] |
| WP_250810300.1 | ParB+HNH | - | Neorhizobium_tomejilense | Bacteria>Pseudomonadati>Pseudomonadota>Alphaproteobacteria>Hyphomicrobiales>Rhizobiaceae>Rhizobium/Agrobacterium_group>Neorhizobium | 637 | L1082_RS18515 | 66306..68219 | 2093828 | DUF262-domain-containing-protein-[Neorhizobium-tomejilense] |
| MBX5048872.1 | ParB+HNH | GCA_019684595.1 | Rhizobium_lentis | Bacteria>Pseudomonadati>Pseudomonadota>Alphaproteobacteria>Hyphomicrobiales>Rhizobiaceae>Rhizobium/Agrobacterium_group>Rhizobium | 599 | HJB64_19895 | 186491..188290 | 1138194 | DUF262-domain-containing-protein-[Rhizobium-lentis] |
| WP_168349958.1 | ParB+HNH | GCF_012276085.1 | Rhizobium_leguminosarum | Bacteria>Pseudomonadati>Pseudomonadota>Alphaproteobacteria>Hyphomicrobiales>Rhizobiaceae>Rhizobium/Agrobacterium_group>Rhizobium | 599 | GFM28_RS29295 | 3046..4845 | 384 | DUF262-domain-containing-protein-[Rhizobium-leguminosarum] |
| WP_127665090.1 | ParB+HNH | GCF_004001475.1 | Sinorhizobium_meliloti | Bacteria>Pseudomonadati>Pseudomonadota>Alphaproteobacteria>Hyphomicrobiales>Rhizobiaceae>Sinorhizobium/Ensifer_group>Sinorhizobium | 555 | CN231_RS17680 | 54342..56009 | 382 | DUF262-domain-containing-protein-[Sinorhizobium-meliloti] |
| WP_055658302.1 | ParB+HNH | GCF_001403915.1 | Roseibium_aggregatum | Bacteria>Pseudomonadati>Pseudomonadota>Alphaproteobacteria>Hyphomicrobiales>Stappiaceae>Roseibium | 561 | LAL4801_RS18640 | 1346278..1347963 | 187304 | DUF262-domain-containing-protein-[Roseibium-aggregatum] |
| MCW5685474.1 | ParB+HNH | GCA_026125885.1 | Pseudolabrys_sp. | Bacteria>Pseudomonadati>Pseudomonadota>Alphaproteobacteria>Hyphomicrobiales>Xanthobacteraceae>Pseudolabrys>unclassified_Pseudolabrys | 560 | KIT85_13800 | 141979..143661 | 1960880 | MAG:-DUF262-domain-containing-protein-[Pseudolabrys-sp.] |
| WP_084695238.1 | ParB+HNH | GCF_000740775.1 | Haematobacter_missouriensis | Bacteria>Pseudomonadati>Pseudomonadota>Alphaproteobacteria>Rhodobacterales>Paracoccaceae>Haematobacter | 560 | CG51_RS16235 | 14904..16586 | 366616 | DUF262-domain-containing-protein-[Haematobacter-missouriensis] |
| WP_245853284.1 | ParB+HNH | GCF_900184945.1 | Maliponia_aquimaris | Bacteria>Pseudomonadati>Pseudomonadota>Alphaproteobacteria>Rhodobacterales>Paracoccaceae>Maliponia | 642 | MAA8898_RS05740 | 28766..30694 | 1673631 | DUF262-domain-containing-protein-[Maliponia-aquimaris] |
| MCQ0971013.1 | ParB+HNH | GCA_024359525.1 | Paracoccus_albicereus | Bacteria>Pseudomonadati>Pseudomonadota>Alphaproteobacteria>Rhodobacterales>Paracoccaceae>Paracoccus | 560 | MLD63_11315 | 57027..58709 | 2922394 | DUF262-domain-containing-protein-[Paracoccus-sp.-TK19116] |
| WP_255330036.1 | ParB+HNH | GCF_024359525.1 | Paracoccus_albicereus | Bacteria>Pseudomonadati>Pseudomonadota>Alphaproteobacteria>Rhodobacterales>Paracoccaceae>Paracoccus | 560 | MLD63_RS11365 | 57027..58709 | 2922394 | DUF262-domain-containing-protein-[Paracoccus-sp.-TK19116] |
| WP_089279821.1 | ParB+HNH | GCF_900188425.1 | Antarctobacter_heliothermus | Bacteria>Pseudomonadati>Pseudomonadota>Alphaproteobacteria>Rhodobacterales>Roseobacteraceae>Antarctobacter | 650 | CHB90_RS21225 | 18802..20754 | 74033 | DUF262-domain-containing-protein-[Antarctobacter-heliothermus] |
| MAE92767.1 | ParB+HNH | GCA_002687285.1 | Pelagibaca_sp. | Bacteria>Pseudomonadati>Pseudomonadota>Alphaproteobacteria>Rhodobacterales>Roseobacteraceae>Pelagibaca>unclassified_Pelagibaca | 642 | CMI67_24860 | 12107..14035 | 1979400 | hypothetical-protein-[Pelagibaca-sp.] |
| WP_212538480.1 | ParB+HNH | GCF_018224955.1 | Thetidibacter_halocola | Bacteria>Pseudomonadati>Pseudomonadota>Alphaproteobacteria>Rhodobacterales>Roseobacteraceae>Thetidibacter | 602 | KB874_RS20745 | 24017..25825 | 2827239 | DUF262-domain-containing-protein-[Aestuariicoccus-sp.-KMU-90] |
| WP_245758748.1 | ParB+HNH | GCF_900112335.1 | Tropicimonas_isoalkanivorans | Bacteria>Pseudomonadati>Pseudomonadota>Alphaproteobacteria>Rhodobacterales>Roseobacteraceae>Tropicimonas | 642 | BM075_RS05200 | 196337..198265 | 441112 | DUF262-domain-containing-protein-[Tropicimonas-isoalkanivorans] |
| MBF9033360.1 | ParB+HNH | GCA_015689685.1 | Rhodobacterales_bacterium_HKCCE2091 | Bacteria>Pseudomonadati>Pseudomonadota>Alphaproteobacteria>Rhodobacterales>unclassified_Rhodobacterales | 641 | HKCCE2091_03840 | 303373..305298 | 2792619 | DUF262-domain-containing-protein-[Rhodobacterales-bacterium-HKCCE2091] |
| WP_145610982.1 | ParB+HNH | GCF_007828035.1 | Nitrospirillum_amazonense | Bacteria>Pseudomonadati>Pseudomonadota>Alphaproteobacteria>Rhodospirillales>Azospirillaceae>Nitrospirillum | 570 | FBZ87_RS14550 | 650768..652480 | 28077 | DUF262-domain-containing-protein-[Nitrospirillum-amazonense] |
| SUS04808.1 | ParB+HNH | GCA_900498315.1 | uncultured_Defluviicoccus_sp. | Bacteria>Pseudomonadati>Pseudomonadota>Alphaproteobacteria>Rhodospirillales>Rhodospirillaceae>Defluviicoccus>environmental_samples | 437 | DF3PB_150001 | 186..1499 | 543062 | conserved-hypothetical-protein-[uncultured-Defluviicoccus-sp.] |
| WP_027288604.1 | ParB+HNH | GCF_000515255.1 | Rhodovibrio_salinarum | Bacteria>Pseudomonadati>Pseudomonadota>Alphaproteobacteria>Rhodospirillales>Rhodovibrionaceae>Rhodovibrio | 600 | RHOSA_RS0110340 | 2234074..2235876 | 1087 | DUF262-domain-containing-protein-[Rhodovibrio-salinarum] |
| HCW69912.1 | ParB+HNH | GCA_003539675.1 | Thalassospira_lucentensis | Bacteria>Pseudomonadati>Pseudomonadota>Alphaproteobacteria>Rhodospirillales>Thalassospiraceae>Thalassospira | 602 | DHR80_22430 | 40223..42031 | 168935 | DUF262-domain-containing-protein-[Thalassospira-lucentensis] |
| TVR84539.1 | ParB+HNH | GCA_007695045.1 | Rhodospirillales_bacterium | Bacteria>Pseudomonadati>Pseudomonadota>Alphaproteobacteria>Rhodospirillales>unclassified_Rhodospirillales | 602 | EA405_01425 | 7041..8849 | 2026786 | DUF262-domain-containing-protein-[Rhodospirillales-bacterium] |
| WP_193715188.1 | ParB+HNH | GCF_015143595.1 | Sneathiella_sp._P13V-1 | Bacteria>Pseudomonadati>Pseudomonadota>Alphaproteobacteria>Sneathiellales>Sneathiellaceae>Sneathiella>unclassified_Sneathiella | 628 | GUA87_RS03850 | 783319..785205 | 2697366 | DUF262-domain-containing-protein-[Sneathiella-sp.-P13V-1] |
| WP_017183454.1 | ParB+HNH | GCF_000277525.1 | Sphingobium_xenophagum | Bacteria>Pseudomonadati>Pseudomonadota>Alphaproteobacteria>Sphingomonadales>Sphingomonadaceae>Sphingobium | 560 | QYYP_RS0113490 | 8639..10321 | 121428 | DUF262-domain-containing-protein-[Sphingobium-xenophagum] |
| MBN8436947.1 | ParB+HNH | GCA_017302555.1 | Accumulibacter_sp. | Bacteria>Pseudomonadati>Pseudomonadota>Betaproteobacteria>Betaproteobacteria_incertae_sedis>Candidatus_Accumulibacter>unclassified_Candidatus_Accumulibacter | 610 | JOL95_02710 | 181098..182930 | 2053492 | DUF262-domain-containing-protein-[Candidatus-Accumulibacter-sp.] |
| TLD46492.1 | ParB+HNH | GCA_005524045.1 | Accumulibacter_sp. | Bacteria>Pseudomonadati>Pseudomonadota>Betaproteobacteria>Betaproteobacteria_incertae_sedis>Candidatus_Accumulibacter>unclassified_Candidatus_Accumulibacter | 571 | FAZ92_01229 | 5052..6767 | 2053492 | MAG:-hypothetical-protein-FAZ92_01229-[Candidatus-Accumulibacter-sp.] |
| WP_076471949.1 | ParB+HNH | GCF_001957055.1 | Achromobacter_xylosoxidans | Bacteria>Pseudomonadati>Pseudomonadota>Betaproteobacteria>Burkholderiales>Alcaligenaceae>Achromobacter | 602 | BIZ53_RS00435 | 90601..92409 | 85698 | DUF262-domain-containing-protein-[Achromobacter-xylosoxidans] |
| WP_107316050.1 | ParB+HNH | GCF_022976495.1 | Achromobacter_xylosoxidans | Bacteria>Pseudomonadati>Pseudomonadota>Betaproteobacteria>Burkholderiales>Alcaligenaceae>Achromobacter | 602 | DBA06_RS11660 | 111976..113784 | 85698 | DUF262-domain-containing-protein-[Achromobacter-xylosoxidans] |
| WP_050447542.1 | ParB+HNH | GCF_001189595.1 | Achromobacter_spanius | Bacteria>Pseudomonadati>Pseudomonadota>Betaproteobacteria>Burkholderiales>Alcaligenaceae>Achromobacter | 562 | AFM18_RS14315 | 78326..80014 | 217203 | DUF262-domain-containing-protein-[Achromobacter-spanius] |
| WP_063584994.1 | ParB+HNH | GCF_001637105.1 | Achromobacter_ruhlandii | Bacteria>Pseudomonadati>Pseudomonadota>Betaproteobacteria>Burkholderiales>Alcaligenaceae>Achromobacter | 610 | AOT39_RS22730 | 29952..31784 | 72557 | DUF262-domain-containing-protein-[Achromobacter-ruhlandii] |
| WP_180097557.1 | ParB+HNH | GCF_902859995.1 | Achromobacter_mucicolens | Bacteria>Pseudomonadati>Pseudomonadota>Betaproteobacteria>Burkholderiales>Alcaligenaceae>Achromobacter | 602 | H0520_RS02895 | 608503..610311 | 1389922 | DUF262-domain-containing-protein-[Achromobacter-mucicolens] |
| WP_212312050.1 | ParB+HNH | GCF_018195695.1 | Achromobacter_sp._Marseille-Q0513 | Bacteria>Pseudomonadati>Pseudomonadota>Betaproteobacteria>Burkholderiales>Alcaligenaceae>Achromobacter>unclassified_Achromobacter | 620 | KDH83_RS00285 | 70103..71965 | 2829161 | DUF262-domain-containing-protein-[Achromobacter-sp.-Marseille-Q0513] |
| WP_259250632.1 | ParB+HNH | GCF_024806975.1 | Achromobacter_sp._JUb104 | Bacteria>Pseudomonadati>Pseudomonadota>Betaproteobacteria>Burkholderiales>Alcaligenaceae>Achromobacter>unclassified_Achromobacter | 602 | M2361_RS16400 | 999748..1001556 | 2940590 | DUF262-domain-containing-protein-[Achromobacter-sp.-JUb104] |
| WP_130302969.1 | ParB+HNH | GCF_004217095.1 | Advenella_incenata | Bacteria>Pseudomonadati>Pseudomonadota>Betaproteobacteria>Burkholderiales>Alcaligenaceae>Advenella | 682 | EV681_RS00045 | 12089..14137 | 267800 | DUF262-domain-containing-protein-[Advenella-incenata] |
| WP_199265312.1 | ParB+HNH | GCF_016446305.1 | Alcaligenes_faecalis | Bacteria>Pseudomonadati>Pseudomonadota>Betaproteobacteria>Burkholderiales>Alcaligenaceae>Alcaligenes | 620 | GTU60_RS02705 | 598114..599976 | 511 | DUF262-domain-containing-protein-[Alcaligenes-faecalis] |
| WP_109088942.1 | ParB+HNH | GCF_003122065.1 | Alcaligenes_faecalis | Bacteria>Pseudomonadati>Pseudomonadota>Betaproteobacteria>Burkholderiales>Alcaligenaceae>Alcaligenes | 610 | DF183_RS09475 | 791872..793704 | 511 | DUF262-domain-containing-protein-[Alcaligenes-faecalis] |
| WP_026638250.1 | ParB+HNH | GCF_000518845.1 | Bordetella_petrii | Bacteria>Pseudomonadati>Pseudomonadota>Betaproteobacteria>Burkholderiales>Alcaligenaceae>Bordetella | 610 | L636_RS0102135 | 148968..150800 | 94624 | DUF262-domain-containing-protein-[Bordetella-petrii] |
| WP_032972593.1 | ParB+HNH | GCF_006770345.1 | Bordetella_hinzii | Bacteria>Pseudomonadati>Pseudomonadota>Betaproteobacteria>Burkholderiales>Alcaligenaceae>Bordetella | 602 | CBR67_RS06395 | 1271018..1272826 | 103855 | DUF262-domain-containing-protein-[Bordetella-hinzii] |
| RIY39879.1 | ParB+HNH | GCA_003585905.1 | Neopusillimonas_maritima | Bacteria>Pseudomonadati>Pseudomonadota>Betaproteobacteria>Burkholderiales>Alcaligenaceae>Neopusillimonas | 598 | CJP73_12380 | 12521..14317 | 2026239 | type-I-restriction-endonuclease-[Pusillimonas-maritima] |
| WP_119516654.1 | ParB+HNH | GCF_003585905.1 | Neopusillimonas_maritima | Bacteria>Pseudomonadati>Pseudomonadota>Betaproteobacteria>Burkholderiales>Alcaligenaceae>Neopusillimonas | 598 | CJP73_RS12410 | 12521..14317 | 2026239 | DUF262-domain-containing-protein-[Pusillimonas-maritima] |
| WP_148812765.1 | ParB+HNH | GCF_008119665.1 | Pigmentiphaga_aceris | Bacteria>Pseudomonadati>Pseudomonadota>Betaproteobacteria>Burkholderiales>Alcaligenaceae>Pigmentiphaga | 562 | FXN63_RS03155 | 745904..747592 | 1940612 | DUF262-domain-containing-protein-[Pigmentiphaga-aceris] |
| NYT63363.1 | ParB+HNH | GCA_013416445.1 | Alcaligenaceae_bacterium | Bacteria>Pseudomonadati>Pseudomonadota>Betaproteobacteria>Burkholderiales>Alcaligenaceae>unclassified_Alcaligenaceae | 626 | HOA66_13670 | 52467..54347 | 2021368 | DUF262-domain-containing-protein-[Alcaligenaceae-bacterium] |
| WP_186043212.1 | ParB+HNH | GCF_902830235.1 | Burkholderia_gladioli | Bacteria>Pseudomonadati>Pseudomonadota>Betaproteobacteria>Burkholderiales>Burkholderiaceae>Burkholderia | 602 | H9C96_RS14810 | 6770..8578 | 28095 | DUF262-domain-containing-protein-[Burkholderia-gladioli] |
| WP_208457320.1 | ParB+HNH | GCF_902832475.1 | Burkholderia_gladioli | Bacteria>Pseudomonadati>Pseudomonadota>Betaproteobacteria>Burkholderiales>Burkholderiaceae>Burkholderia | 367 | H9G27_RS36830 | 158308..159411 | 28095 | DUF262-domain-containing-protein-[Burkholderia-gladioli] |
| WP_208457319.1 | ParB+HNH | GCF_902832475.1 | Burkholderia_gladioli | Bacteria>Pseudomonadati>Pseudomonadota>Betaproteobacteria>Burkholderiales>Burkholderiaceae>Burkholderia | 64 | H9G27_RS37215 | 158096..158290 | 28095 | DUF262-domain-containing-protein-[Burkholderia-gladioli] |
| WP_059944776.1 | ParB+HNH | GCF_001527205.1 | Burkholderia_territorii | Bacteria>Pseudomonadati>Pseudomonadota>Betaproteobacteria>Burkholderiales>Burkholderiaceae>Burkholderia>Burkholderia_cepacia_complex | 602 | WT00_RS19575 | 114095..115903 | 1503055 | DUF262-domain-containing-protein-[Burkholderia-territorii] |
| MCA8485079.1 | ParB+HNH | GCA_020421385.1 | Burkholderia_multivorans | Bacteria>Pseudomonadati>Pseudomonadota>Betaproteobacteria>Burkholderiales>Burkholderiaceae>Burkholderia>Burkholderia_cepacia_complex | 610 | LGM85_14165 | 194901..196733 | 87883 | DUF262-domain-containing-HNH-endonuclease-family-protein-[Burkholderia-multivorans] |
| WP_059936875.1 | ParB+HNH | GCF_001528605.1 | Burkholderia_multivorans | Bacteria>Pseudomonadati>Pseudomonadota>Betaproteobacteria>Burkholderiales>Burkholderiaceae>Burkholderia>Burkholderia_cepacia_complex | 602 | WK33_RS03200 | 2356..4164 | 87883 | DUF262-domain-containing-protein-[Burkholderia-multivorans] |
| WP_060225081.1 | ParB+HNH | GCF_001531525.1 | Burkholderia_cepacia | Bacteria>Pseudomonadati>Pseudomonadota>Betaproteobacteria>Burkholderiales>Burkholderiaceae>Burkholderia>Burkholderia_cepacia_complex | 610 | WL34_RS17285 | 245422..247254 | 292 | DUF262-domain-containing-protein-[Burkholderia-cepacia] |
| WP_226255242.1 | ParB+HNH | GCF_020419005.1 | Burkholderia_cepacia | Bacteria>Pseudomonadati>Pseudomonadota>Betaproteobacteria>Burkholderiales>Burkholderiaceae>Burkholderia>Burkholderia_cepacia_complex | 73 | LGM39_RS24945 | 374145..374366 | 292 | HNH-endonuclease-family-protein-[Burkholderia-cepacia] |
| WP_091908366.1 | ParB+HNH | GCF_900095735.1 | Chitinasiproducens_palmae | Bacteria>Pseudomonadati>Pseudomonadota>Betaproteobacteria>Burkholderiales>Burkholderiaceae>Chitinasiproducens | 602 | BLU54_RS11880 | 223695..225503 | 1770053 | DUF262-domain-containing-protein-[Chitinasiproducens-palmae] |
| WP_125094461.1 | ParB+HNH | GCF_003892345.1 | Lautropia_dentalis | Bacteria>Pseudomonadati>Pseudomonadota>Betaproteobacteria>Burkholderiales>Burkholderiaceae>Lautropia | 610 | EHV23_RS01860 | 487415..489247 | 2490857 | DUF262-domain-containing-protein-[Lautropia-dentalis] |
| WP_275272078.1 | ParB+HNH | GCF_029087225.1 | Limnobacter_olei | Bacteria>Pseudomonadati>Pseudomonadota>Betaproteobacteria>Burkholderiales>Burkholderiaceae>Limnobacter | 581 | PY990_RS13685 | 16226..17971 | 3031298 | DUF262-domain-containing-protein-[Limnobacter-sp.-P1] |
| MBX3589938.1 | ParB+HNH | GCA_019635275.1 | Burkholderiaceae_bacterium | Bacteria>Pseudomonadati>Pseudomonadota>Betaproteobacteria>Burkholderiales>Burkholderiaceae>unclassified_Burkholderiaceae | 621 | KF755_03425 | 46604..48469 | 2030806 | DUF262-domain-containing-protein-[Burkholderiaceae-bacterium] |
| WP_166837247.1 | ParB+HNH | GCF_011742065.1 | Aquabacterium_sp._A08 | Bacteria>Pseudomonadati>Pseudomonadota>Betaproteobacteria>Burkholderiales>Burkholderiales_genera_incertae_sedis>Aquabacterium>unclassified_Aquabacterium | 613 | HA630_RS20325 | 23194..25035 | 2718532 | DUF262-domain-containing-protein-[Aquabacterium-sp.-A08] |
| MCK6421965.1 | ParB+HNH | GCA_023150345.1 | Aquabacterium_sp. | Bacteria>Pseudomonadati>Pseudomonadota>Betaproteobacteria>Burkholderiales>Burkholderiales_genera_incertae_sedis>Aquabacterium>unclassified_Aquabacterium | 562 | L6Q73_13815 | 11728..13416 | 1872578 | DUF262-domain-containing-HNH-endonuclease-family-protein-[Aquabacterium-sp.] |
| WP_086927315.1 | ParB+HNH | GCF_002157185.1 | Acidovorax_carolinensis | Bacteria>Pseudomonadati>Pseudomonadota>Betaproteobacteria>Burkholderiales>Comamonadaceae>Acidovorax | 562 | CBP35_RS08900 | 1926449..1928137 | 553814 | DUF262-domain-containing-protein-[Acidovorax-carolinensis] |
| WP_057202650.1 | ParB+HNH | GCF_001428645.1 | Acidovorax_sp._Root217 | Bacteria>Pseudomonadati>Pseudomonadota>Betaproteobacteria>Burkholderiales>Comamonadaceae>Acidovorax>unclassified_Acidovorax | 620 | ASE31_RS15195 | 101950..103812 | 1736492 | DUF262-domain-containing-protein-[Acidovorax-sp.-Root217] |
| WP_034379105.1 | ParB+HNH | GCF_000761335.1 | Comamonas_thiooxydans | Bacteria>Pseudomonadati>Pseudomonadota>Betaproteobacteria>Burkholderiales>Comamonadaceae>Comamonas | 602 | P245_RS10555 | 80003..81811 | 363952 | DUF262-domain-containing-protein-[Comamonas-thiooxydans] |
| KEH09932.1 | ParB+HNH | GCA_000710165.1 | Delftia_tsuruhatensis | Bacteria>Pseudomonadati>Pseudomonadota>Betaproteobacteria>Burkholderiales>Comamonadaceae>Delftia | 624 | GY14_08985 | 30233..32107 | 180282 | hypothetical-protein-GY14_08985-[Delftia-tsuruhatensis] |
| WP_239478239.1 | ParB+HNH | GCF_016903055.1 | Hydrogenophaga_sp._YM1 | Bacteria>Pseudomonadati>Pseudomonadota>Betaproteobacteria>Burkholderiales>Comamonadaceae>Hydrogenophaga>unclassified_Hydrogenophaga | 624 | JNX00_RS02370 | 467896..469770 | 2806262 | DUF262-domain-containing-protein-[Hydrogenophaga-sp.-YM1] |
| QRR36378.1 | ParB+HNH | GCA_016903055.1 | Hydrogenophaga_sp._YM1 | Bacteria>Pseudomonadati>Pseudomonadota>Betaproteobacteria>Burkholderiales>Comamonadaceae>Hydrogenophaga>unclassified_Hydrogenophaga | 611 | JNX00_02370 | 467896..469731 | 2806262 | DUF262-domain-containing-protein-[Hydrogenophaga-sp.-YM1] |
| MBS3911797.1 | ParB+HNH | GCA_018333605.1 | Hydrogenophaga_sp. | Bacteria>Pseudomonadati>Pseudomonadota>Betaproteobacteria>Burkholderiales>Comamonadaceae>Hydrogenophaga>unclassified_Hydrogenophaga | 611 | KGZ70_08205 | 43861..45696 | 1904254 | DUF262-domain-containing-protein-[Hydrogenophaga-sp.] |
| WP_101103331.1 | ParB+HNH | GCF_002837135.1 | Macromonas_bipunctata | Bacteria>Pseudomonadati>Pseudomonadota>Betaproteobacteria>Burkholderiales>Comamonadaceae>Macromonas | 562 | CON76_RS10620 | 17628..19316 | 183670 | DUF262-domain-containing-protein-[Macromonas-bipunctata] |
| WP_054257716.1 | ParB+HNH | GCF_001298675.1 | Paenacidovorax_caeni | Bacteria>Pseudomonadati>Pseudomonadota>Betaproteobacteria>Burkholderiales>Comamonadaceae>Paenacidovorax | 614 | BN2503_RS16430 | 7861..9705 | 343013 | DUF262-domain-containing-protein-[Acidovorax-caeni] |
| WP_053842379.1 | ParB+HNH | GCF_001282395.1 | Paracidovorax_avenae | Bacteria>Pseudomonadati>Pseudomonadota>Betaproteobacteria>Burkholderiales>Comamonadaceae>Paracidovorax | 611 | AOG77_RS00365 | 71065..72900 | 80867 | DUF262-domain-containing-protein-[Acidovorax-avenae] |
| CDS49858.1 | ParB+HNH | GCA_000751355.1 | Polaromonas_sp._CG9_12 | Bacteria>Pseudomonadati>Pseudomonadota>Betaproteobacteria>Burkholderiales>Comamonadaceae>Polaromonas>unclassified_Polaromonas | 621 | . | 476013..477878 | 1504672 | hypothetical-protein-[Polaromonas-sp.-CG9_12] |
| WP_169420218.1 | ParB+HNH | GCF_012927085.1 | Ramlibacter_agri | Bacteria>Pseudomonadati>Pseudomonadota>Betaproteobacteria>Burkholderiales>Comamonadaceae>Ramlibacter | 562 | HHL11_RS19355 | 181880..183568 | 2728837 | DUF262-domain-containing-protein-[Ramlibacter-agri] |
| MCP5281007.1 | ParB+HNH | GCA_024235935.1 | Rhodoferax_sp. | Bacteria>Pseudomonadati>Pseudomonadota>Betaproteobacteria>Burkholderiales>Comamonadaceae>Rhodoferax>unclassified_Rhodoferax | 570 | H6930_05355 | 229728..231440 | 50421 | DUF262-domain-containing-protein-[Rhodoferax-sp.] |
| WP_045536715.1 | ParB+HNH | GCF_000828915.1 | Serpentinimonas_maccroryi | Bacteria>Pseudomonadati>Pseudomonadota>Betaproteobacteria>Burkholderiales>Comamonadaceae>Serpentinimonas | 572 | SMCB_RS09945 | 2160511..2162229 | 1458426 | DUF262-domain-containing-protein-[Serpentinimonas-maccroryi] |
| WP_095541485.1 | ParB+HNH | GCF_002285255.1 | Vandammella_animalimorsus | Bacteria>Pseudomonadati>Pseudomonadota>Betaproteobacteria>Burkholderiales>Comamonadaceae>Vandammella | 566 | CLI93_RS07375 | 86301..88001 | 2029117 | DUF262-domain-containing-protein-[Vandammella-animalimorsus] |
| WP_269747355.1 | ParB+HNH | - | Variovorax_sp._WDL1 | Bacteria>Pseudomonadati>Pseudomonadota>Betaproteobacteria>Burkholderiales>Comamonadaceae>Variovorax>unclassified_Variovorax | 617 | APY03_RS00985 | 196696..198549 | 207745 | DUF262-domain-containing-protein-[Variovorax-sp.-WDL1] |
| WP_222611183.1 | ParB+HNH | GCF_014284115.1 | Undibacterium | Bacteria>Pseudomonadati>Pseudomonadota>Betaproteobacteria>Burkholderiales>Oxalobacteraceae | 625 | H8K35_RS08345 | 166014..167891 | 401469 | MULTISPECIES:-DUF262-domain-containing-protein-[Undibacterium] |
| WP_141835415.1 | ParB+HNH | GCF_006715485.1 | Herbaspirillum_sp._SJZ107 | Bacteria>Pseudomonadati>Pseudomonadota>Betaproteobacteria>Burkholderiales>Oxalobacteraceae>Herbaspirillum>unclassified_Herbaspirillum | 620 | FBX97_RS19475 | 303665..305527 | 2572881 | DUF262-domain-containing-protein-[Herbaspirillum-sp.-SJZ107] |
| WP_152879052.1 | ParB+HNH | GCF_009295655.1 | Duganella_sp._FT27W | Bacteria>Pseudomonadati>Pseudomonadota>Betaproteobacteria>Burkholderiales>Oxalobacteraceae>Telluria_group>Duganella>unclassified_Duganella | 602 | F8197_RS00395 | 82286..84094 | 2654636 | DUF262-domain-containing-protein-[Duganella-sp.-FT27W] |
| WP_106758221.1 | ParB+HNH | GCF_003011895.2 | Massilia_glaciei | Bacteria>Pseudomonadati>Pseudomonadota>Betaproteobacteria>Burkholderiales>Oxalobacteraceae>Telluria_group>Massilia | 563 | C7C56_RS15215 | 37276..38967 | 1524097 | DUF262-domain-containing-protein-[Massilia-glaciei] |
| WP_157271681.1 | ParB+HNH | GCF_009760915.1 | Azohydromonas_aeria | Bacteria>Pseudomonadati>Pseudomonadota>Betaproteobacteria>Burkholderiales>Sphaerotilaceae>Azohydromonas | 571 | FJ694_RS32515 | 4099..5814 | 2590212 | DUF262-domain-containing-protein-[Azohydromonas-aeria] |
| MBK7616705.1 | ParB+HNH | GCA_016710065.1 | Candidatus_Ideonella_esbjergensis | Bacteria>Pseudomonadati>Pseudomonadota>Betaproteobacteria>Burkholderiales>Sphaerotilaceae>Ideonella | 621 | IPJ08_20455 | 308656..310521 | 3400289 | DUF262-domain-containing-protein-[Burkholderiales-bacterium] |
| ODU09020.1 | ParB+HNH | GCA_001724995.1 | Rubrivivax_sp._SCN_71-131 | Bacteria>Pseudomonadati>Pseudomonadota>Betaproteobacteria>Burkholderiales>Sphaerotilaceae>Rubrivivax>unclassified_Rubrivivax | 611 | ABS84_09935 | 25223..27058 | 1660138 | MAG:-hypothetical-protein-ABS84_09935-[Rubrivivax-sp.-SCN-71-131] |
| WP_130480159.1 | ParB+HNH | GCF_004216565.1 | Sphaerotilus_mobilis | Bacteria>Pseudomonadati>Pseudomonadota>Betaproteobacteria>Burkholderiales>Sphaerotilaceae>Sphaerotilus | 556 | EV685_RS01110 | 191500..193170 | 47994 | DUF262-domain-containing-protein-[Leptothrix-mobilis] |
| OGB35304.1 | ParB+HNH | GCA_001770935.1 | Burkholderiales_bacterium_RIFCSPLOWO2_12_FULL_61_40 | Bacteria>Pseudomonadati>Pseudomonadota>Betaproteobacteria>Burkholderiales>unclassified_Burkholderiales | 602 | A3F78_11680 | 41284..43092 | 1797566 | hypothetical-protein-A3F78_11680-[Burkholderiales-bacterium-RIFCSPLOWO2_12_FULL_61_40] |
| MCC6474797.1 | ParB+HNH | GCA_020846575.1 | Burkholderiales_bacterium | Bacteria>Pseudomonadati>Pseudomonadota>Betaproteobacteria>Burkholderiales>unclassified_Burkholderiales | 570 | IT514_13745 | 9149..10861 | 1891238 | DUF262-domain-containing-protein-[Burkholderiales-bacterium] |
| MCP5275504.1 | ParB+HNH | GCA_024235795.1 | Burkholderiales_bacterium | Bacteria>Pseudomonadati>Pseudomonadota>Betaproteobacteria>Burkholderiales>unclassified_Burkholderiales | 359 | H6936_11795 | 662520..663599 | 1891238 | DUF262-domain-containing-protein-[Burkholderiales-bacterium] |
| MCG9054654.1 | ParB+HNH | GCA_022394275.1 | Laribacter_hongkongensis | Bacteria>Pseudomonadati>Pseudomonadota>Betaproteobacteria>Neisseriales>Aquaspirillaceae>Laribacter | 611 | LH442_01390 | 274059..275894 | 168471 | DUF262-domain-containing-HNH-endonuclease-family-protein-[Laribacter-hongkongensis] |
| WP_239901867.1 | ParB+HNH | GCF_022394235.1 | Laribacter_hongkongensis | Bacteria>Pseudomonadati>Pseudomonadota>Betaproteobacteria>Neisseriales>Aquaspirillaceae>Laribacter | 611 | LH447_RS07135 | 17131..18966 | 168471 | DUF262-domain-containing-protein-[Laribacter-hongkongensis] |
| WP_194117487.1 | ParB+HNH | GCF_015223195.1 | Chitinilyticum_piscinae | Bacteria>Pseudomonadati>Pseudomonadota>Betaproteobacteria>Neisseriales>Chitinibacteraceae>Chitinilyticum | 602 | INR99_RS16760 | 31313..33121 | 2866724 | DUF262-domain-containing-protein-[Chitinilyticum-piscinae] |
| WP_153715430.1 | ParB+HNH | GCF_009650905.1 | Eikenella_corrodens | Bacteria>Pseudomonadati>Pseudomonadota>Betaproteobacteria>Neisseriales>Neisseriaceae>Eikenella | 602 | GJ661_RS08655 | 8176..9984 | 539 | DUF262-domain-containing-protein-[Eikenella-corrodens] |
| WP_040564281.1 | ParB+HNH | GCF_016127355.1 | Kingella_denitrificans | Bacteria>Pseudomonadati>Pseudomonadota>Betaproteobacteria>Neisseriales>Neisseriaceae>Kingella | 611 | I6I17_RS09795 | 1931337..1933172 | 502 | DUF262-domain-containing-protein-[Kingella-denitrificans] |
| ROH85755.1 | ParB+HNH | GCA_003730115.1 | Pseudomethylobacillus_aquaticus | Bacteria>Pseudomonadati>Pseudomonadota>Betaproteobacteria>Nitrosomonadales>Methylophilaceae>Pseudomethylobacillus | 610 | ED236_08395 | 63240..65072 | 2676064 | DUF262-domain-containing-protein-[Pseudomethylobacillus-aquaticus] |
| WP_123237529.1 | ParB+HNH | GCF_003730115.1 | Pseudomethylobacillus_aquaticus | Bacteria>Pseudomonadati>Pseudomonadota>Betaproteobacteria>Nitrosomonadales>Methylophilaceae>Pseudomethylobacillus | 610 | ED236_RS08395 | 63240..65072 | 2676064 | DUF262-domain-containing-protein-[Pseudomethylobacillus-aquaticus] |
| CAG1003397.1 | ParB+HNH | GCA_905339275.1 | Methylophilaceae_bacterium | Bacteria>Pseudomonadati>Pseudomonadota>Betaproteobacteria>Nitrosomonadales>Methylophilaceae>unclassified_Methylophilaceae | 570 | MTYP_03089 | 11070..12782 | 2030816 | hypothetical-protein-MTYP_03089-[Methylophilaceae-bacterium] |
| WP_181258529.1 | ParB+HNH | GCF_003051045.1 | Nitrosomonas_ureae | Bacteria>Pseudomonadati>Pseudomonadota>Betaproteobacteria>Nitrosomonadales>Nitrosomonadaceae>Nitrosomonas | 559 | C8R28_RS01180 | 3722..5401 | 44577 | DUF262-domain-containing-protein-[Nitrosomonas-ureae] |
| SDW36924.1 | ParB+HNH | GCA_900106545.1 | Nitrosomonas_communis | Bacteria>Pseudomonadati>Pseudomonadota>Betaproteobacteria>Nitrosomonadales>Nitrosomonadaceae>Nitrosomonas | 598 | SAMN05421882_100916 | 43166..44962 | 44574 | Protein-of-unknown-function-[Nitrosomonas-communis] |
| WP_074666144.1 | ParB+HNH | GCF_900106545.1 | Nitrosomonas_communis | Bacteria>Pseudomonadati>Pseudomonadota>Betaproteobacteria>Nitrosomonadales>Nitrosomonadaceae>Nitrosomonas | 598 | BLV21_RS05415 | 43166..44962 | 44574 | DUF262-domain-containing-protein-[Nitrosomonas-communis] |
| PTN11854.1 | ParB+HNH | GCA_003046585.1 | Nitrosomonas_aestuarii | Bacteria>Pseudomonadati>Pseudomonadota>Betaproteobacteria>Nitrosomonadales>Nitrosomonadaceae>Nitrosomonas | 366 | C8R11_107139 | 178532..179632 | 52441 | uncharacterized-protein-DUF262-[Nitrosomonas-aestuarii] |
| WP_074776133.1 | ParB+HNH | GCF_900107715.1 | Nitrosospira_multiformis | Bacteria>Pseudomonadati>Pseudomonadota>Betaproteobacteria>Nitrosomonadales>Nitrosomonadaceae>Nitrosospira | 570 | BLV78_RS14495 | 12908..14620 | 1231 | DUF262-domain-containing-protein-[Nitrosospira-multiformis] |
| SFW23248.1 | ParB+HNH | GCA_900119085.1 | Nitrosovibrio_sp._Nv17 | Bacteria>Pseudomonadati>Pseudomonadota>Betaproteobacteria>Nitrosomonadales>Nitrosomonadaceae>Nitrosovibrio>unclassified_Nitrosovibrio | 602 | SAMN05216414_10759 | 61745..63553 | 1855339 | Protein-of-unknown-function-[Nitrosovibrio-sp.-Nv17] |
| WP_072293618.1 | ParB+HNH | GCF_900119085.1 | Nitrosovibrio_sp._Nv17 | Bacteria>Pseudomonadati>Pseudomonadota>Betaproteobacteria>Nitrosomonadales>Nitrosomonadaceae>Nitrosovibrio>unclassified_Nitrosovibrio | 602 | BT108_RS05895 | 61745..63553 | 1855339 | DUF262-domain-containing-protein-[Nitrosovibrio-sp.-Nv17] |
| WP_008060481.1 | ParB+HNH | GCF_000214035.2 | Methyloversatilis_universalis | Bacteria>Pseudomonadati>Pseudomonadota>Betaproteobacteria>Nitrosomonadales>Sterolibacteriaceae>Methyloversatilis | 611 | METUNV1_RS07600 | 11549..13384 | 378211 | DUF262-domain-containing-protein-[Methyloversatilis-universalis] |
| WP_202636303.1 | ParB+HNH | GCF_000934545.1 | Rugosibacter_aromaticivorans | Bacteria>Pseudomonadati>Pseudomonadota>Betaproteobacteria>Nitrosomonadales>Sterolibacteriaceae>Rugosibacter | 611 | PG1C_RS05015 | 988128..989963 | 1565605 | DUF262-domain-containing-protein-[Rugosibacter-aromaticivorans] |
| MBN8780448.1 | ParB+HNH | GCA_017304205.1 | Thiobacillus_sp. | Bacteria>Pseudomonadati>Pseudomonadota>Betaproteobacteria>Nitrosomonadales>Thiobacillaceae>Thiobacillus>unclassified_Thiobacillus | 611 | J0J07_12950 | 552255..554090 | 924 | DUF262-domain-containing-protein-[Thiobacillus-sp.] |
| MBW8470841.1 | ParB+HNH | GCA_019459125.1 | Thiobacillus_sp. | Bacteria>Pseudomonadati>Pseudomonadota>Betaproteobacteria>Nitrosomonadales>Thiobacillaceae>Thiobacillus>unclassified_Thiobacillus | 596 | K0M67_21430 | 10602..12392 | 924 | DUF262-domain-containing-HNH-endonuclease-family-protein-[Thiobacillus-sp.] |
| MBK9161128.1 | ParB+HNH | GCA_016716325.1 | Nitrosomonadales_bacterium | Bacteria>Pseudomonadati>Pseudomonadota>Betaproteobacteria>Nitrosomonadales>unclassified_Nitrosomonadales | 610 | IPM27_06145 | 1287060..1288892 | 2026761 | DUF262-domain-containing-protein-[Nitrosomonadales-bacterium] |
| MBN8461446.1 | ParB+HNH | GCA_017302355.1 | Dechloromonas_sp. | Bacteria>Pseudomonadati>Pseudomonadota>Betaproteobacteria>Rhodocyclales>Azonexaceae>Dechloromonas>unclassified_Dechloromonas | 611 | J0M01_01320 | 25431..27266 | 1917218 | DUF262-domain-containing-protein-[Dechloromonas-sp.] |
| MBP6421396.1 | ParB+HNH | GCA_017988215.1 | Propionivibrio_sp. | Bacteria>Pseudomonadati>Pseudomonadota>Betaproteobacteria>Rhodocyclales>Rhodocyclaceae>Propionivibrio>unclassified_Propionivibrio | 621 | KA271_00735 | 5173..7038 | 2212460 | DUF262-domain-containing-protein-[Propionivibrio-sp.] |
| MBQ7625732.1 | ParB+HNH | GCA_017444505.1 | Rhodocyclaceae_bacterium | Bacteria>Pseudomonadati>Pseudomonadota>Betaproteobacteria>Rhodocyclales>Rhodocyclaceae>unclassified_Rhodocyclaceae | 565 | IJS87_01545 | 929..2626 | 1898103 | DUF262-domain-containing-protein-[Rhodocyclaceae-bacterium] |
| MCA3121406.1 | ParB+HNH | GCA_020248305.1 | Rhodocyclaceae_bacterium | Bacteria>Pseudomonadati>Pseudomonadota>Betaproteobacteria>Rhodocyclales>Rhodocyclaceae>unclassified_Rhodocyclaceae | 351 | ING90_02075 | 357554..358609 | 1898103 | DUF262-domain-containing-protein-[Rhodocyclaceae-bacterium] |
| WP_168986548.1 | ParB+HNH | GCF_963887635.1 | Azoarcus_taiwanensis | Bacteria>Pseudomonadati>Pseudomonadota>Betaproteobacteria>Rhodocyclales>Zoogloeaceae>Azoarcus | 571 | GPA21_RS02040 | 6948..8663 | 666964 | DUF262-domain-containing-protein-[Azoarcus-taiwanensis] |
| WP_141017188.1 | ParB+HNH | GCF_006496635.1 | Azoarcus_sp._DD4 | Bacteria>Pseudomonadati>Pseudomonadota>Betaproteobacteria>Rhodocyclales>Zoogloeaceae>Azoarcus>unclassified_Azoarcus | 620 | CJ010_RS05965 | 1255704..1257566 | 2027405 | DUF262-domain-containing-protein-[Azoarcus-sp.-DD4] |
| MBD5801784.1 | ParB+HNH | GCA_014822185.1 | Azoarcus_sp._Aa7 | Bacteria>Pseudomonadati>Pseudomonadota>Betaproteobacteria>Rhodocyclales>Zoogloeaceae>Azoarcus>unclassified_Azoarcus | 604 | AZOA_12080 | 9129..10943 | 2303328 | hypothetical-protein-[Azoarcus-sp.-Aa7] |
| WP_002936079.1 | ParB+HNH | GCF_000310125.2 | Thauera_sp._27 | Bacteria>Pseudomonadati>Pseudomonadota>Betaproteobacteria>Rhodocyclales>Zoogloeaceae>Thauera>unclassified_Thauera | 757 | B447_RS02420 | 111916..114189 | 305700 | DUF262-domain-containing-protein-[Thauera-sp.-27] |
| WP_148578354.1 | ParB+HNH | GCF_008107625.1 | Zoogloea_oleivorans | Bacteria>Pseudomonadati>Pseudomonadota>Betaproteobacteria>Rhodocyclales>Zoogloeaceae>Zoogloea | 609 | ETQ85_RS07195 | 250019..251848 | 1552750 | DUF262-domain-containing-protein-[Zoogloea-oleivorans] |
| WP_001548946.1 | ParB+HNH | GCF_000277565.1 | Enterobacterales | Bacteria>Pseudomonadati>Pseudomonadota>Gammaproteobacteria | 673 | K5R72_RS10990 | 51471..53492 | 91347 | MULTISPECIES:-DUF262-domain-containing-protein-[Enterobacteriaceae] |
| WP_117259787.1 | ParB+HNH | GCF_040061975.1 | Enterobacterales | Bacteria>Pseudomonadati>Pseudomonadota>Gammaproteobacteria | 599 | SZ66_RS15310 | 349541..351340 | 91347 | MULTISPECIES:-DUF262-domain-containing-protein-[Enterobacterales] |
| WP_021546938.1 | ParB+HNH | GCF_019973795.1 | Enterobacterales | Bacteria>Pseudomonadati>Pseudomonadota>Gammaproteobacteria | 587 | L2578_RS22545 | 3227..4990 | 91347 | MULTISPECIES:-DUF262-domain-containing-protein-[Enterobacteriaceae] |
| WP_047929055.1 | ParB+HNH | GCF_904799765.1 | Enterobacterales | Bacteria>Pseudomonadati>Pseudomonadota>Gammaproteobacteria | 587 | CDN88_RS21125 | 8959..10722 | 91347 | MULTISPECIES:-DUF262-domain-containing-protein-[Enterobacteriaceae] |
| WP_061350061.1 | ParB+HNH | GCF_004311945.1 | Enterobacterales | Bacteria>Pseudomonadati>Pseudomonadota>Gammaproteobacteria | 587 | LTT78_RS21785 | 1170..2933 | 91347 | MULTISPECIES:-DUF262-domain-containing-protein-[Enterobacteriaceae] |
| WP_079805578.1 | ParB+HNH | GCF_014168815.1 | Enterobacterales | Bacteria>Pseudomonadati>Pseudomonadota>Gammaproteobacteria | 584 | L2W90_RS09960 | 164361..166115 | 91347 | MULTISPECIES:-DUF262-domain-containing-protein-[Enterobacterales] |
| WP_094898296.1 | ParB+HNH | GCF_965118735.1 | Enterobacterales | Bacteria>Pseudomonadati>Pseudomonadota>Gammaproteobacteria | 584 | CIG23_RS11865 | 62906..64660 | 91347 | DUF262-domain-containing-protein-[Raoultella-planticola] |
| WP_120066190.1 | ParB+HNH | GCF_021440285.1 | Enterobacterales | Bacteria>Pseudomonadati>Pseudomonadota>Gammaproteobacteria | 584 | D6029_RS18470 | 9200..10954 | 91347 | MULTISPECIES:-DUF262-domain-containing-protein-[Enterobacterales] |
| WP_121868992.1 | ParB+HNH | GCF_965118295.1 | Enterobacterales | Bacteria>Pseudomonadati>Pseudomonadota>Gammaproteobacteria | 584 | EAI39_RS15065 | 82259..84013 | 91347 | MULTISPECIES:-DUF262-domain-containing-protein-[Enterobacteriaceae] |
| WP_102989249.1 | ParB+HNH | GCF_002934425.1 | Aeromonas | Bacteria>Pseudomonadati>Pseudomonadota>Gammaproteobacteria>Aeromonadales>Aeromonadaceae | 627 | C2U40_RS16560 | 3594599..3596482 | 642 | MULTISPECIES:-DUF262-domain-containing-protein-[Aeromonas] |
| WP_270809142.1 | ParB+HNH | GCF_027705685.1 | Aeromonas | Bacteria>Pseudomonadati>Pseudomonadota>Gammaproteobacteria>Aeromonadales>Aeromonadaceae | 597 | OKB79_RS12610 | 6329..8122 | 642 | DUF262-domain-containing-HNH-endonuclease-family-protein-[Aeromonas-sp.-QDB08] |
| WP_159144200.1 | ParB+HNH | GCF_009834065.1 | Aeromonas | Bacteria>Pseudomonadati>Pseudomonadota>Gammaproteobacteria>Aeromonadales>Aeromonadaceae | 584 | GRF56_RS18100 | 3931955..3933709 | 642 | DUF262-domain-containing-protein-[Aeromonas-veronii] |
| WP_139442436.1 | ParB+HNH | GCF_046082965.1 | Aeromonas | Bacteria>Pseudomonadati>Pseudomonadota>Gammaproteobacteria>Aeromonadales>Aeromonadaceae | 583 | CF108_RS03270 | 128056..129807 | 642 | DUF262-domain-containing-protein-[Aeromonas-veronii] |
| WP_201934228.1 | ParB+HNH | GCF_024347775.1 | Aeromonas | Bacteria>Pseudomonadati>Pseudomonadota>Gammaproteobacteria>Aeromonadales>Aeromonadaceae | 583 | OCX50_RS09765 | 2032316..2034067 | 642 | MULTISPECIES:-DUF262-domain-containing-protein-[Aeromonas] |

| Protein-ID | Domain | Assembly-ID | Strain | Taxonomy | Length | Locus-Tag | Genomic-coordinates | TaxID | Fasta-defline |
|---|---|---|---|---|---|---|---|---|---|
| WP_183149507.1 | ParB+HNH | GCF_014158455.1 | Aeromonas | Bacteria>Pseudomonadati>Pseudomonadota>Gammaproteobacteria>Aeromonadales>Aeromonadaceae | 581 | J4G65_RS17590 | 83647..85392 | 642 | MULTISPECIES:-DUF262-domain-containing-protein-[Aeromonas] |
| WP_069785633.1 | ParB+HNH | GCF_963870325.1 | Aeromonas | Bacteria>Pseudomonadati>Pseudomonadota>Gammaproteobacteria>Aeromonadales>Aeromonadaceae | 558 | QBS29_RS17440 | 3045221..3046897 | 642 | MULTISPECIES:-DUF262-domain-containing-protein-[Aeromonas] |
| WP_171280308.1 | ParB+HNH | GCF_963888675.1 | Aeromonas | Bacteria>Pseudomonadati>Pseudomonadota>Gammaproteobacteria>Aeromonadales>Aeromonadaceae | 558 | PVL97_RS11255 | 2522375..2524051 | 642 | MULTISPECIES:-DUF262-domain-containing-protein-[Aeromonas] |
| WP_270796653.1 | ParB+HNH | GCF_029024985.1 | Aeromonas | Bacteria>Pseudomonadati>Pseudomonadota>Gammaproteobacteria>Aeromonadales>Aeromonadaceae | 558 | OKC40_RS19410 | 2250..3926 | 642 | MULTISPECIES:-DUF262-domain-containing-HNH-endonuclease-family-protein-[Aeromonas] |
| WP_120415688.1 | ParB+HNH | GCF_003611985.1 | Aeromonas_veronii | Bacteria>Pseudomonadati>Pseudomonadota>Gammaproteobacteria>Aeromonadales>Aeromonadaceae>Aeromonas | 583 | D6R50_RS16915 | 831419..833170 | 654 | DUF262-domain-containing-protein-[Aeromonas-veronii] |
| WP_193000842.1 | ParB+HNH | GCF_014892695.1 | Aeromonas_simiae | Bacteria>Pseudomonadati>Pseudomonadota>Gammaproteobacteria>Aeromonadales>Aeromonadaceae>Aeromonas | 583 | FE240_RS10730 | 2269101..2270852 | 218936 | DUF262-domain-containing-protein-[Aeromonas-simiae] |
| WP_087756127.1 | ParB+HNH | GCF_963873515.1 | Aeromonas_salmonicida | Bacteria>Pseudomonadati>Pseudomonadota>Gammaproteobacteria>Aeromonadales>Aeromonadaceae>Aeromonas | 583 | O23A_RS10575 | 2333319..2335070 | 645 | DUF262-domain-containing-protein-[Aeromonas-salmonicida] |
| EGX6956812.1 | ParB+HNH | - | Aeromonas_hydrophila | Bacteria>Pseudomonadati>Pseudomonadota>Gammaproteobacteria>Aeromonadales>Aeromonadaceae>Aeromonas | 583 | JFQ85_000576 | 648875..650626 | 644 | DUF262-domain-containing-protein-[Aeromonas-hydrophila] |
| WP_241871098.1 | ParB+HNH | GCF_022631175.1 | Aeromonas_hydrophila | Bacteria>Pseudomonadati>Pseudomonadota>Gammaproteobacteria>Aeromonadales>Aeromonadaceae>Aeromonas | 558 | MOO40_RS17040 | 3747075..3748751 | 644 | DUF262-domain-containing-protein-[Aeromonas-hydrophila] |
| WP_010634559.1 | ParB+HNH | GCF_033096545.1 | Aeromonas_dhakensis | Bacteria>Pseudomonadati>Pseudomonadota>Gammaproteobacteria>Aeromonadales>Aeromonadaceae>Aeromonas | 599 | AQU_RS10060 | 131437..133236 | 196024 | DUF262-domain-containing-protein-[Aeromonas-dhakensis] |
| WP_095593100.1 | ParB+HNH | GCF_002285935.1 | Aeromonas_dhakensis | Bacteria>Pseudomonadati>Pseudomonadota>Gammaproteobacteria>Aeromonadales>Aeromonadaceae>Aeromonas | 583 | CK627_RS18860 | 4070173..4071924 | 196024 | DUF262-domain-containing-protein-[Aeromonas-dhakensis] |
| WP_257706955.1 | ParB+HNH | GCF_024649325.1 | Aeromonas_caviae | Bacteria>Pseudomonadati>Pseudomonadota>Gammaproteobacteria>Aeromonadales>Aeromonadaceae>Aeromonas | 627 | NUK33_RS01475 | 143161..145044 | 648 | DUF262-domain-containing-protein-[Aeromonas-caviae] |
| BDO08312.1 | ParB+HNH | GCA_024347775.1 | Aeromonas_caviae | Bacteria>Pseudomonadati>Pseudomonadota>Gammaproteobacteria>Aeromonadales>Aeromonadaceae>Aeromonas | 583 | KAM643c_18850 | 2032316..2034067 | 648 | hypothetical-protein-KAM643c_18850-[Aeromonas-caviae] |
| WP_223931721.1 | ParB+HNH | GCF_019972495.1 | Aeromonas_caviae | Bacteria>Pseudomonadati>Pseudomonadota>Gammaproteobacteria>Aeromonadales>Aeromonadaceae>Aeromonas | 583 | LDN23_RS02845 | 16378..18129 | 648 | DUF262-domain-containing-protein-[Aeromonas-caviae] |
| BBR09396.1 | ParB+HNH | GCA_014158455.1 | Aeromonas_caviae | Bacteria>Pseudomonadati>Pseudomonadota>Gammaproteobacteria>Aeromonadales>Aeromonadaceae>Aeromonas | 581 | WP3S18E02_10570 | 1124145..1125890 | 648 | hypothetical-protein-WP3S18E02_10570-[Aeromonas-caviae] |
| WP_043557609.1 | ParB+HNH | GCF_963866355.1 | Aeromonas_bestiarum | Bacteria>Pseudomonadati>Pseudomonadota>Gammaproteobacteria>Aeromonadales>Aeromonadaceae>Aeromonas | 627 | BN1118_RS19630 | 62703..64586 | 105751 | DUF262-domain-containing-protein-[Aeromonas-bestiarum] |
| MBS4697264.1 | ParB+HNH | GCA_018359965.1 | Aeromonas_allosaccharophila | Bacteria>Pseudomonadati>Pseudomonadota>Gammaproteobacteria>Aeromonadales>Aeromonadaceae>Aeromonas | 581 | J4G65_17595 | 83647..85392 | 656 | DUF262-domain-containing-protein-[Aeromonas-allosaccharophila] |
| WP_103244182.1 | ParB+HNH | GCF_963877715.1 | Aeromonas_sp._ASNIH5 | Bacteria>Pseudomonadati>Pseudomonadota>Gammaproteobacteria>Aeromonadales>Aeromonadaceae>Aeromonas>unclassified_Aeromonas | 602 | C2U30_RS17340 | 3682733..3684541 | 1758179 | DUF262-domain-containing-protein-[Aeromonas-sp.-ASNIH5] |
| AUZ76294.1 | ParB+HNH | GCA_002934425.1 | Aeromonas_sp._ASNIH4 | Bacteria>Pseudomonadati>Pseudomonadota>Gammaproteobacteria>Aeromonadales>Aeromonadaceae>Aeromonas>unclassified_Aeromonas | 627 | C2U40_16565 | 3594599..3596482 | 1636609 | hypothetical-protein-C2U40_16565-[Aeromonas-sp.-ASNIH4] |
| MCR4553829.1 | ParB+HNH | GCA_024683965.1 | Succinivibrionaceae_bacterium | Bacteria>Pseudomonadati>Pseudomonadota>Gammaproteobacteria>Aeromonadales>Succinivibrionaceae>unclassified_Succinivibrionaceae | 601 | K5752_02795 | 202..2007 | 2053620 | DUF262-domain-containing-protein-[Succinivibrionaceae-bacterium] |
| WP_057786319.1 | ParB+HNH | GCF_001441695.1 | Alishewanella_sp._WH16-1 | Bacteria>Pseudomonadati>Pseudomonadota>Gammaproteobacteria>Alteromonadales>Alteromonadaceae>Alishewanella>unclassified_Alishewanella | 557 | AAY72_RS06880 | 14816..16489 | 1651088 | DUF262-domain-containing-protein-[Alishewanella-sp.-WH16-1] |
| WP_207611652.1 | ParB+HNH | GCF_017368475.1 | Alteromonas_sp._5E99-2 | Bacteria>Pseudomonadati>Pseudomonadota>Gammaproteobacteria>Alteromonadales>Alteromonadaceae>Alteromonas/Salinimonas_group>Alteromonas>unclassified_Alteromonas | 562 | J3L16_RS04220 | 86333..88021 | 2817683 | DUF262-domain-containing-protein-[Alteromonas-sp.-5E99-2] |
| WP_166424590.1 | ParB+HNH | GCF_011378905.1 | Paraglaciecola_sp._20A4 | Bacteria>Pseudomonadati>Pseudomonadota>Gammaproteobacteria>Alteromonadales>Alteromonadaceae>Paraglaciecola>unclassified_Paraglaciecola | 408 | GQS54_RS13945 | 98453..99679 | 2687288 | DUF262-domain-containing-protein-[Paraglaciecola-sp.-20A4] |
| WP_085284215.1 | ParB+HNH | GCF_900109795.1 | Colwellia_chukchiensis | Bacteria>Pseudomonadati>Pseudomonadota>Gammaproteobacteria>Alteromonadales>Colwelliaceae>Colwellia | 641 | B6K83_RS06990 | 115928..117853 | 641665 | DUF262-domain-containing-protein-[Colwellia-chukchiensis] |
| AWB57835.1 | ParB+HNH | GCA_003061515.1 | Colwellia_sp._Arc7-D | Bacteria>Pseudomonadati>Pseudomonadota>Gammaproteobacteria>Alteromonadales>Colwelliaceae>Colwellia>unclassified_Colwellia | 643 | DBO93_09805 | 2228550..2230481 | 2161872 | DUF262-domain-containing-protein-[Colwellia-sp.-Arc7-D] |
| WP_108456182.1 | ParB+HNH | GCF_003061515.1 | Colwellia_sp._Arc7-D | Bacteria>Pseudomonadati>Pseudomonadota>Gammaproteobacteria>Alteromonadales>Colwelliaceae>Colwellia>unclassified_Colwellia | 643 | DBO93_RS09805 | 2228550..2230481 | 2161872 | DUF262-domain-containing-protein-[Colwellia-sp.-Arc7-D] |
| KXS36210.1 | ParB+HNH | GCA_001577655.1 | Idiomarina_sp._T82-3 | Bacteria>Pseudomonadati>Pseudomonadota>Gammaproteobacteria>Alteromonadales>Idiomarinaceae>Idiomarina>unclassified_Idiomarina | 598 | AWU56_325 | 95340..97136 | 1794809 | type-I-restriction-modification-system-protein-[Idiomarina-sp.-T82-3] |
| KXS36217.1 | ParB+HNH | GCA_001577655.1 | Idiomarina_sp._T82-3 | Bacteria>Pseudomonadati>Pseudomonadota>Gammaproteobacteria>Alteromonadales>Idiomarinaceae>Idiomarina>unclassified_Idiomarina | 334 | AWU56_332 | 104037..105041 | 1794809 | hypothetical-protein-AWU56_332-[Idiomarina-sp.-T82-3] |
| WP_112714379.1 | ParB+HNH | GCF_900465055.1 | Moritella_yayanosii | Bacteria>Pseudomonadati>Pseudomonadota>Gammaproteobacteria>Alteromonadales>Moritellaceae>Moritella | 635 | MORIYA_RS08585 | 1886124..1888031 | 69539 | DUF262-domain-containing-protein-[Moritella-yayanosii] |
| WP_064663365.1 | ParB+HNH | GCF_014925285.1 | Pseudoalteromonas | Bacteria>Pseudomonadati>Pseudomonadota>Gammaproteobacteria>Alteromonadales>Pseudoalteromonadaceae | 641 | PLIP_RS19225 | 625665..627590 | 53246 | MULTISPECIES:-DUF262-domain-containing-protein-[Pseudoalteromonas] |
| WP_089368386.1 | ParB+HNH | GCF_007991255.1 | Pseudoalteromonas | Bacteria>Pseudomonadati>Pseudomonadota>Gammaproteobacteria>Alteromonadales>Pseudoalteromonadaceae | 580 | PNIG_RS09945 | 2058572..2060314 | 53246 | DUF262-domain-containing-protein-[Pseudoalteromonas-nigrifaciens] |
| WP_076922915.1 | ParB+HNH | GCF_016464045.1 | unclassified_Pseudoalteromonas | Bacteria>Pseudomonadati>Pseudomonadota>Gammaproteobacteria>Alteromonadales>Pseudoalteromonadaceae>Pseudoalteromonas | 641 | JIG41_RS20345 | 10074..11999 | 194690 | MULTISPECIES:-DUF262-domain-containing-protein-[unclassified-Pseudoalteromonas] |
| WP_096038336.1 | ParB+HNH | GCF_007991235.1 | Pseudoalteromonas_tetraodonis | Bacteria>Pseudomonadati>Pseudomonadota>Gammaproteobacteria>Alteromonadales>Pseudoalteromonadaceae>Pseudoalteromonas | 596 | PTE01_RS15655 | 20935..22725 | 43659 | DUF262-domain-containing-protein-[Pseudoalteromonas-tetraodonis] |
| WP_193038369.1 | ParB+HNH | GCF_014897925.1 | Pseudoalteromonas_nigrifaciens | Bacteria>Pseudomonadati>Pseudomonadota>Gammaproteobacteria>Alteromonadales>Pseudoalteromonadaceae>Pseudoalteromonas | 562 | INT11_RS00020 | 2822..4510 | 28109 | DUF262-domain-containing-protein-[Pseudoalteromonas-nigrifaciens] |
| WP_010607318.1 | ParB+HNH | GCF_000259115.1 | Pseudoalteromonas_maricaloris | Bacteria>Pseudomonadati>Pseudomonadota>Gammaproteobacteria>Alteromonadales>Pseudoalteromonadaceae>Pseudoalteromonas | 622 | UY7_RS0121190 | 85233..87101 | 184924 | DUF262-domain-containing-protein-[Pseudoalteromonas-flavipulchra] |
| WP_105264927.1 | ParB+HNH | GCF_002964725.1 | Pseudoalteromonas_sp._T1lg76 | Bacteria>Pseudomonadati>Pseudomonadota>Gammaproteobacteria>Alteromonadales>Pseudoalteromonadaceae>Pseudoalteromonas>unclassified_Pseudoalteromonas | 647 | C2M35_RS06865 | 39384..41327 | 2077103 | DUF262-domain-containing-protein-[Pseudoalteromonas-sp.-T1lg76] |
| TMO42410.1 | ParB+HNH | GCA_005886815.1 | Pseudoalteromonas_sp._S4389 | Bacteria>Pseudomonadati>Pseudomonadota>Gammaproteobacteria>Alteromonadales>Pseudoalteromonadaceae>Pseudoalteromonas>unclassified_Pseudoalteromonas | 645 | CWC25_15735 | 17014..18951 | 579556 | DUF262-domain-containing-protein-[Pseudoalteromonas-sp.-S4389] |
| WP_054983664.1 | ParB+HNH | GCF_001400005.1 | Pseudoalteromonas_sp._P1-30 | Bacteria>Pseudomonadati>Pseudomonadota>Gammaproteobacteria>Alteromonadales>Pseudoalteromonadaceae>Pseudoalteromonas>unclassified_Pseudoalteromonas | 596 | AN395_RS16630 | 56057..57847 | 1723760 | DUF262-domain-containing-protein-[Pseudoalteromonas-sp.-P1-30] |
| WP_130052621.1 | ParB+HNH | GCF_004195145.1 | Pseudoalteromonas_sp._MEBiC_03485 | Bacteria>Pseudomonadati>Pseudomonadota>Gammaproteobacteria>Alteromonadales>Pseudoalteromonadaceae>Pseudoalteromonas>unclassified_Pseudoalteromonas | 641 | EVU92_RS18420 | 346175..348100 | 2571103 | DUF262-domain-containing-protein-[Pseudoalteromonas-sp.-MEBiC-03485] |
| WP_167658108.1 | ParB+HNH | GCF_011806445.1 | Pseudoalteromonas_sp._HF66 | Bacteria>Pseudomonadati>Pseudomonadota>Gammaproteobacteria>Alteromonadales>Pseudoalteromonadaceae>Pseudoalteromonas>unclassified_Pseudoalteromonas | 622 | HCZ11_RS07640 | 205733..207601 | 2721559 | DUF262-domain-containing-protein-[Pseudoalteromonas-sp.-HF66] |
| WP_037429575.1 | ParB+HNH | GCF_000282755.1 | Shewanella | Bacteria>Pseudomonadati>Pseudomonadota>Gammaproteobacteria>Alteromonadales>Shewanellaceae | 600 | SHEWPOL2_RS15300 | 1840..3642 | 22 | MULTISPECIES:-DUF262-domain-containing-protein-[Shewanella] |
| WP_263177452.1 | ParB+HNH | GCF_025654415.1 | unclassified_Shewanella | Bacteria>Pseudomonadati>Pseudomonadota>Gammaproteobacteria>Alteromonadales>Shewanellaceae>Shewanella | 587 | L5M16_RS21420 | 9572..11335 | 196818 | MULTISPECIES:-DUF262-domain-containing-protein-[unclassified-Shewanella] |
| WP_248084388.1 | ParB+HNH | GCF_023159095.1 | unclassified_Shewanella | Bacteria>Pseudomonadati>Pseudomonadota>Gammaproteobacteria>Alteromonadales>Shewanellaceae>Shewanella | 583 | MOD77_RS02685 | 223107..224858 | 196818 | MULTISPECIES:-DUF262-domain-containing-protein-[unclassified-Shewanella] |
| WP_224020604.1 | ParB+HNH | GCF_019973655.1 | Shewanella_xiamenensis | Bacteria>Pseudomonadati>Pseudomonadota>Gammaproteobacteria>Alteromonadales>Shewanellaceae>Shewanella | 599 | LDO18_RS12240 | 2741070..2742869 | 332186 | DUF262-domain-containing-protein-[Shewanella-xiamenensis] |
| WP_259476697.1 | ParB+HNH | GCF_024972795.1 | Shewanella_xiamenensis | Bacteria>Pseudomonadati>Pseudomonadota>Gammaproteobacteria>Alteromonadales>Shewanellaceae>Shewanella | 587 | K5M76_RS09820 | 2216196..2217959 | 332186 | DUF262-domain-containing-protein-[Shewanella-xiamenensis] |
| WP_220062215.1 | ParB+HNH | GCF_019457515.1 | Shewanella_mesophila | Bacteria>Pseudomonadati>Pseudomonadota>Gammaproteobacteria>Alteromonadales>Shewanellaceae>Shewanella | 562 | KOI73_RS17045 | 3874951..3876639 | 2864208 | DUF262-domain-containing-protein-[Shewanella-sp.-FJAT-53870] |
| WP_244899714.1 | ParB+HNH | GCF_002075795.1 | Shewanella_japonica | Bacteria>Pseudomonadati>Pseudomonadota>Gammaproteobacteria>Alteromonadales>Shewanellaceae>Shewanella | 542 | SJ2017_RS16910 | 3939955..3941583 | 93973 | DUF262-domain-containing-protein-[Shewanella-japonica] |
| WP_028764578.1 | ParB+HNH | GCF_000518705.1 | Shewanella_colwelliana | Bacteria>Pseudomonadati>Pseudomonadota>Gammaproteobacteria>Alteromonadales>Shewanellaceae>Shewanella | 562 | L876_RS0115165 | 11923..13611 | 23 | DUF262-domain-containing-protein-[Shewanella-colwelliana] |
| WP_006086466.1 | ParB+HNH | GCF_000231345.1 | Shewanella_baltica | Bacteria>Pseudomonadati>Pseudomonadota>Gammaproteobacteria>Alteromonadales>Shewanellaceae>Shewanella | 598 | SBAL625DRAFT_RS15530 | 216660..218456 | 62322 | DUF262-domain-containing-protein-[Shewanella-baltica] |
| WP_259506666.1 | ParB+HNH | GCF_024999425.1 | Shewanella_baltica | Bacteria>Pseudomonadati>Pseudomonadota>Gammaproteobacteria>Alteromonadales>Shewanellaceae>Shewanella | 562 | G3479_RS14585 | 88315..90003 | 62322 | DUF262-domain-containing-protein-[Shewanella-baltica] |
| WP_101059465.1 | ParB+HNH | GCF_017580385.1 | Shewanella_algae | Bacteria>Pseudomonadati>Pseudomonadota>Gammaproteobacteria>Alteromonadales>Shewanellaceae>Shewanella | 599 | CX652_RS12230 | 9404..11203 | 38313 | DUF262-domain-containing-protein-[Shewanella-algae] |
| WP_144191361.1 | ParB+HNH | GCF_017580415.1 | Shewanella_algae | Bacteria>Pseudomonadati>Pseudomonadota>Gammaproteobacteria>Alteromonadales>Shewanellaceae>Shewanella | 599 | I6M45_RS07405 | 503055..504854 | 38313 | DUF262-domain-containing-protein-[Shewanella-algae] |
| WP_208124437.1 | ParB+HNH | GCF_017570045.1 | Shewanella_algae | Bacteria>Pseudomonadati>Pseudomonadota>Gammaproteobacteria>Alteromonadales>Shewanellaceae>Shewanella | 599 | JKK45_RS07525 | 1716408..1718207 | 38313 | DUF262-domain-containing-protein-[Shewanella-algae] |
| WP_208223526.1 | ParB+HNH | GCF_017580425.1 | Shewanella_algae | Bacteria>Pseudomonadati>Pseudomonadota>Gammaproteobacteria>Alteromonadales>Shewanellaceae>Shewanella | 599 | I6M46_RS07455 | 8831..10630 | 38313 | DUF262-domain-containing-protein-[Shewanella-algae] |
| WP_264921056.1 | ParB+HNH | GCF_026013945.1 | Shewanella_algae | Bacteria>Pseudomonadati>Pseudomonadota>Gammaproteobacteria>Alteromonadales>Shewanellaceae>Shewanella | 599 | OLL83_RS07705 | 1710567..1712366 | 38313 | DUF262-domain-containing-HNH-endonuclease-family-protein-[Shewanella-algae] |
| WP_221058243.1 | ParB+HNH | GCF_019670685.1 | Shewanella_algae | Bacteria>Pseudomonadati>Pseudomonadota>Gammaproteobacteria>Alteromonadales>Shewanellaceae>Shewanella | 587 | K5631_RS12955 | 2924553..2926316 | 38313 | DUF262-domain-containing-protein-[Shewanella-algae] |
| WP_047539593.1 | ParB+HNH | GCF_000798835.1 | Shewanella_sp._ZOR0012 | Bacteria>Pseudomonadati>Pseudomonadota>Gammaproteobacteria>Alteromonadales>Shewanellaceae>Shewanella>unclassified_Shewanella | 600 | HT094_RS07825 | 1791231..1793033 | 1339231 | DUF262-domain-containing-protein-[Shewanella-sp.-ZOR0012] |
| WP_263145903.1 | ParB+HNH | GCF_025653395.1 | Shewanella_sp._SW24 | Bacteria>Pseudomonadati>Pseudomonadota>Gammaproteobacteria>Alteromonadales>Shewanellaceae>Shewanella>unclassified_Shewanella | 583 | L5M17_RS05795 | 183065..184816 | 2912815 | DUF262-domain-containing-protein-[Shewanella-sp.-SW24] |
| WP_055648626.1 | ParB+HNH | GCF_001308045.1 | Shewanella_sp._Sh95 | Bacteria>Pseudomonadati>Pseudomonadota>Gammaproteobacteria>Alteromonadales>Shewanellaceae>Shewanella>unclassified_Shewanella | 562 | AEA42_RS19835 | 31403..33091 | 1689868 | DUF262-domain-containing-protein-[Shewanella-sp.-Sh95] |
| WP_011622485.1 | ParB+HNH | GCF_000014685.1 | Shewanella_sp._MR-4 | Bacteria>Pseudomonadati>Pseudomonadota>Gammaproteobacteria>Alteromonadales>Shewanellaceae>Shewanella>unclassified_Shewanella | 562 | SHEWMR4_RS09040 | 2026421..2028109 | 60480 | DUF262-domain-containing-protein-[Shewanella-sp.-MR-4] |
| WP_220768074.1 | ParB+HNH | GCF_019654915.1 | Shewanella_sp._MBTL60-007 | Bacteria>Pseudomonadati>Pseudomonadota>Gammaproteobacteria>Alteromonadales>Shewanellaceae>Shewanella>unclassified_Shewanella | 587 | K5Q92_RS05660 | 58436..60199 | 2815911 | DUF262-domain-containing-protein-[Shewanella-sp.-MBTL60-007] |
| PKG74685.1 | ParB+HNH | GCA_002836135.1 | Shewanella_sp._GutCb | Bacteria>Pseudomonadati>Pseudomonadota>Gammaproteobacteria>Alteromonadales>Shewanellaceae>Shewanella>unclassified_Shewanella | 624 | CXF86_11670 | 115167..117041 | 2058315 | hypothetical-protein-CXF86_11670-[Shewanella-sp.-GutCb] |
| WP_102039064.1 | ParB+HNH | GCF_002836135.1 | Shewanella_sp._GutCb | Bacteria>Pseudomonadati>Pseudomonadota>Gammaproteobacteria>Alteromonadales>Shewanellaceae>Shewanella>unclassified_Shewanella | 624 | CXF86_RS11710 | 115167..117041 | 2058315 | DUF262-domain-containing-protein-[Shewanella-sp.-GutCb] |
| WP_011716830.1 | ParB+HNH | GCF_000203935.1 | Shewanella_sp._ANA-3 | Bacteria>Pseudomonadati>Pseudomonadota>Gammaproteobacteria>Alteromonadales>Shewanellaceae>Shewanella>unclassified_Shewanella | 562 | SHEWANA3_RS09600 | 2151348..2153036 | 94122 | DUF262-domain-containing-protein-[Shewanella-sp.-ANA-3] |
| HAY94627.1 | ParB+HNH | GCA_003486485.1 | Shewanella_sp. | Bacteria>Pseudomonadati>Pseudomonadota>Gammaproteobacteria>Alteromonadales>Shewanellaceae>Shewanella>unclassified_Shewanella | 536 | DCY70_12145 | 12012..13622 | 50422 | hypothetical-protein-[Shewanella-sp.] |
| MBV7434870.1 | ParB+HNH | GCA_019218315.1 | Cardiobacteriaceae_bacterium_TAE3-ERU3 | Bacteria>Pseudomonadati>Pseudomonadota>Gammaproteobacteria>Cardiobacteriales>Cardiobacteriaceae>unclassified_Cardiobacteriaceae | 627 | KRX19_07500 | 259213..261096 | 2856340 | DUF262-domain-containing-protein-[Cardiobacteriaceae-bacterium-TAE3-ERU3] |
| WP_265366292.1 | ParB+HNH | GCF_026153455.1 | Marinimicrobium_sp._C6131 | Bacteria>Pseudomonadati>Pseudomonadota>Gammaproteobacteria>Cellvibrionales>Cellvibrionaceae>Marinimicrobium>unclassified_Marinimicrobium | 626 | OOT55_RS13050 | 3035053..3036933 | 3022676 | DUF262-domain-containing-protein-[Marinimicrobium-sp.-C6131] |
| WP_187410608.1 | ParB+HNH | GCF_014336955.1 | Saccharophagus_sp._K07 | Bacteria>Pseudomonadati>Pseudomonadota>Gammaproteobacteria>Cellvibrionales>Cellvibrionaceae>Saccharophagus>unclassified_Saccharophagus | 610 | DWB84_RS14915 | 81474..83306 | 2283636 | DUF262-domain-containing-protein-[Saccharophagus-sp.-K07] |
| WP_043319014.1 | ParB+HNH | GCF_000708675.1 | Microbulbifer_sp._HZ11 | Bacteria>Pseudomonadati>Pseudomonadota>Gammaproteobacteria>Cellvibrionales>Microbulbiferaceae>Microbulbifer>unclassified_Microbulbifer | 564 | AU11_RS09625 | 444846..446540 | 1453501 | DUF262-domain-containing-protein-[Microbulbifer-sp.-HZ11] |
| WP_036859483.1 | ParB+HNH | GCF_000744735.1 | Porticoccus_hydrocarbonoclasticus | Bacteria>Pseudomonadati>Pseudomonadota>Gammaproteobacteria>Cellvibrionales>Porticoccaceae>Porticoccus | 627 | U740_RS05155 | 1084626..1086509 | 1073414 | DUF262-domain-containing-protein-[Porticoccus-hydrocarbonoclasticus] |
| WP_268905391.1 | ParB+HNH | GCF_026891035.1 | Dasania_phycosphaerae | Bacteria>Pseudomonadati>Pseudomonadota>Gammaproteobacteria>Cellvibrionales>Spongiibacteraceae>Dasania | 642 | OOV09_RS18290 | 14093..16021 | 2950436 | DUF262-domain-containing-HNH-endonuclease-family-protein-[Dasania-sp.-GY-19] |
| WP_273033641.1 | ParB+HNH | GCF_950061765.1 | Spongiibacter_tropicus | Bacteria>Pseudomonadati>Pseudomonadota>Gammaproteobacteria>Cellvibrionales>Spongiibacteraceae>Spongiibacter | 626 | B4C78_RS03760 | 92396..94276 | 454602 | DUF262-domain-containing-protein-[Spongiibacter-tropicus] |
| WP_190766545.1 | ParB+HNH | GCF_014705705.1 | Spongiibacter_pelagi | Bacteria>Pseudomonadati>Pseudomonadota>Gammaproteobacteria>Cellvibrionales>Spongiibacteraceae>Spongiibacter | 581 | IB286_RS13885 | 26203..27948 | 2760804 | DUF262-domain-containing-protein-[Spongiibacter-pelagi] |
| MTW21818.1 | ParB+HNH | GCA_009720725.1 | Allochromatium_palmeri | Bacteria>Pseudomonadati>Pseudomonadota>Gammaproteobacteria>Chromatiales>Chromatiaceae>Allochromatium | 573 | GJ668_12035 | 28646..30367 | 231048 | DUF262-domain-containing-protein-[Allochromatium-palmeri] |
| WP_155450397.1 | ParB+HNH | GCF_009720725.1 | Allochromatium_palmeri | Bacteria>Pseudomonadati>Pseudomonadota>Gammaproteobacteria>Chromatiales>Chromatiaceae>Allochromatium | 573 | GJ668_RS12035 | 28646..30367 | 231048 | DUF262-domain-containing-protein-[Allochromatium-palmeri] |
| WP_201222436.1 | ParB+HNH | GCF_016653315.1 | Halochromatium_roseum | Bacteria>Pseudomonadati>Pseudomonadota>Gammaproteobacteria>Chromatiales>Chromatiaceae>Halochromatium | 570 | CCR96_RS19435 | 147854..149566 | 391920 | DUF262-domain-containing-protein-[Halochromatium-roseum] |
| NEX15940.1 | ParB+HNH | GCA_010912705.1 | Halochromatium_sp. | Bacteria>Pseudomonadati>Pseudomonadota>Gammaproteobacteria>Chromatiales>Chromatiaceae>Halochromatium>unclassified_Halochromatium | 573 | C1943_04700 | 60166..61887 | 2049430 | DUF262-domain-containing-protein-[Halochromatium-sp.] |
| WP_264346736.1 | ParB+HNH | GCF_947090785.1 | Rheinheimera_sp._MM224 | Bacteria>Pseudomonadati>Pseudomonadota>Gammaproteobacteria>Chromatiales>Chromatiaceae>Rheinheimera>unclassified_Rheinheimera | 557 | OM978_RS09980 | 2105757..2107430 | 3019969 | DUF262-domain-containing-HNH-endonuclease-family-protein-[Rheinheimera-sp.-MM224] |
| WP_007193602.1 | ParB+HNH | GCF_000223985.1 | Thiocapsa_marina | Bacteria>Pseudomonadati>Pseudomonadota>Gammaproteobacteria>Chromatiales>Chromatiaceae>Thiocapsa | 620 | THIMADRAFT_RS13455 | 262509..264371 | 244573 | DUF262-domain-containing-protein-[Thiocapsa-marina] |
| WP_009148238.1 | ParB+HNH | GCF_033954835.1 | Thiorhodovibrio_frisius | Bacteria>Pseudomonadati>Pseudomonadota>Gammaproteobacteria>Chromatiales>Chromatiaceae>Thiorhodovibrio | 580 | THI970DRAFT_RS08810 | 540281..542023 | 631362 | DUF262-domain-containing-protein-[Thiorhodovibrio-sp.-970] |
| MCP5417445.1 | ParB+HNH | GCA_024235215.1 | Chromatiaceae_bacterium | Bacteria>Pseudomonadati>Pseudomonadota>Gammaproteobacteria>Chromatiales>Chromatiaceae>unclassified_Chromatiaceae | 701 | H6965_10160 | 388167..390272 | 2049432 | DUF262-domain-containing-protein-[Chromatiaceae-bacterium] |
| WP_018716652.1 | ParB+HNH | GCF_000374645.1 | Arhodomonas_aquaeolei | Bacteria>Pseudomonadati>Pseudomonadota>Gammaproteobacteria>Chromatiales>Ectothiorhodospiraceae>Arhodomonas | 562 | C516_RS0101840 | 431004..432692 | 2369 | DUF262-domain-containing-protein-[Arhodomonas-aquaeolei] |
| WP_239679774.1 | ParB+HNH | GCF_022227625.1 | Ectothiorhodospira_variabilis | Bacteria>Pseudomonadati>Pseudomonadota>Gammaproteobacteria>Chromatiales>Ectothiorhodospiraceae>Ectothiorhodospira | 615 | LPW30_RS00075 | 19364..21211 | 505694 | DUF262-domain-containing-protein-[Ectothiorhodospira-variabilis] |
| MBK5942669.1 | ParB+HNH | GCA_016653405.1 | Halorhodospira_halophila | Bacteria>Pseudomonadati>Pseudomonadota>Gammaproteobacteria>Chromatiales>Ectothiorhodospiraceae>Halorhodospira | 473 | CCR79_02080 | 99..1520 | 1053 | hypothetical-protein-[Halorhodospira-halophila] |
| WP_201168220.1 | ParB+HNH | GCF_016653405.1 | Halorhodospira_halophila | Bacteria>Pseudomonadati>Pseudomonadota>Gammaproteobacteria>Chromatiales>Ectothiorhodospiraceae>Halorhodospira | 473 | CCR79_RS02120 | 99..1520 | 1053 | DUF262-domain-containing-protein-[Halorhodospira-halophila] |
| WP_123401870.1 | ParB+HNH | GCF_003751635.1 | Inmirania_thermothiophila | Bacteria>Pseudomonadati>Pseudomonadota>Gammaproteobacteria>Chromatiales>Ectothiorhodospiraceae>Inmirania | 600 | EDC57_RS08870 | 681045..682847 | 1750597 | DUF262-domain-containing-protein-[Inmirania-thermothiophila] |
| WP_253479341.1 | ParB+HNH | GCF_024170285.1 | Natronocella_acetinitrilica | Bacteria>Pseudomonadati>Pseudomonadota>Gammaproteobacteria>Chromatiales>Ectothiorhodospiraceae>Natronocella | 615 | J2T57_RS14020 | 321053..322900 | 414046 | DUF262-domain-containing-protein-[Natronocella-acetinitrilica] |
| WP_016353356.1 | ParB+HNH | GCF_000319575.2 | Spiribacter_salinus | Bacteria>Pseudomonadati>Pseudomonadota>Gammaproteobacteria>Chromatiales>Ectothiorhodospiraceae>Spiribacter | 596 | SPISAL_RS04855 | 968997..970787 | 1335746 | DUF262-domain-containing-protein-[Spiribacter-salinus] |
| TQE98723.1 | ParB+HNH | GCA_006569195.1 | Spiribacter_salinus | Bacteria>Pseudomonadati>Pseudomonadota>Gammaproteobacteria>Chromatiales>Ectothiorhodospiraceae>Spiribacter | 237 | FKY71_12290 | 1..716 | 1335746 | HNH-endonuclease,-partial-[Spiribacter-salinus] |
| AGM41049.1 | ParB+HNH | GCA_000319575.2 | Spiribacter_salinus_M19-40 | Bacteria>Pseudomonadati>Pseudomonadota>Gammaproteobacteria>Chromatiales>Ectothiorhodospiraceae>Spiribacter>Spiribacter_salinus | 596 | SPISAL_04775 | 968997..970787 | 1260251 | hypothetical-protein-SPISAL_04775-[Spiribacter-salinus-M19-40] |
| KAF0285944.1 | ParB+HNH | GCA_009805925.1 | Spiribacter_sp._SSL99 | Bacteria>Pseudomonadati>Pseudomonadota>Gammaproteobacteria>Chromatiales>Ectothiorhodospiraceae>Spiribacter>unclassified_Spiribacter | 596 | BA899_01300 | 113776..115566 | 1866884 | hypothetical-protein-BA899_01300-[Spiribacter-sp.-SSL99] |
| WP_159299839.1 | ParB+HNH | GCF_009805925.1 | Spiribacter_sp._SSL99 | Bacteria>Pseudomonadati>Pseudomonadota>Gammaproteobacteria>Chromatiales>Ectothiorhodospiraceae>Spiribacter>unclassified_Spiribacter | 596 | BA899_RS04185 | 113776..115566 | 1866884 | DUF262-domain-containing-protein-[Spiribacter-sp.-SSL99] |
| WP_154296277.1 | ParB+HNH | GCF_009676705.1 | Spiribacter_sp._2438 | Bacteria>Pseudomonadati>Pseudomonadota>Gammaproteobacteria>Chromatiales>Ectothiorhodospiraceae>Spiribacter>unclassified_Spiribacter | 596 | GJ672_RS05605 | 1116289..1118079 | 2666185 | DUF262-domain-containing-protein-[Spiribacter-sp.-2438] |
| WP_026331450.1 | ParB+HNH | GCF_000381205.1 | Thioalkalivibrio_sp._ALE11 | Bacteria>Pseudomonadati>Pseudomonadota>Gammaproteobacteria>Chromatiales>Ectothiorhodospiraceae>Thioalkalivibrio>unclassified_Thioalkalivibrio | 585 | F620_RS0112060 | 37367..39124 | 1265494 | DUF262-domain-containing-protein-[Thioalkalivibrio-sp.-ALE11] |
| WP_019611717.1 | ParB+HNH | GCF_000381705.1 | Thioalkalivibrio_sp._AKL7 | Bacteria>Pseudomonadati>Pseudomonadota>Gammaproteobacteria>Chromatiales>Ectothiorhodospiraceae>Thioalkalivibrio>unclassified_Thioalkalivibrio | 615 | G315_RS0100965 | 229657..231504 | 1158155 | DUF262-domain-containing-protein-[Thioalkalivibrio-sp.-AKL7] |
| MCG7876674.1 | ParB+HNH | GCA_016842655.1 | Candidatus_Thiodiazotropha_taylori | Bacteria>Pseudomonadati>Pseudomonadota>Gammaproteobacteria>Chromatiales>Sedimenticolaceae>Candidatus_Thiodiazotropha | 628 | JAY78_10090 | 12616..14502 | 2792791 | DUF262-domain-containing-protein-[Candidatus-Thiodiazotropha-taylori] |
| MCG7951161.1 | ParB+HNH | GCA_016842765.1 | Candidatus_Thiodiazotropha_taylori | Bacteria>Pseudomonadati>Pseudomonadota>Gammaproteobacteria>Chromatiales>Sedimenticolaceae>Candidatus_Thiodiazotropha | 628 | JAZ09_08060 | 30134..32020 | 2792791 | DUF262-domain-containing-protein-[Candidatus-Thiodiazotropha-taylori] |
| MCG8058212.1 | ParB+HNH | GCA_016843165.1 | Candidatus_Thiodiazotropha_taylori | Bacteria>Pseudomonadati>Pseudomonadota>Gammaproteobacteria>Chromatiales>Sedimenticolaceae>Candidatus_Thiodiazotropha | 628 | JAZ21_00060 | 21313..23199 | 2792791 | MAG:-DUF262-domain-containing-protein-[Candidatus-Thiodiazotropha-taylori] |
| MCG8102613.1 | ParB+HNH | GCA_016842845.1 | Candidatus_Thiodiazotropha_taylori | Bacteria>Pseudomonadati>Pseudomonadota>Gammaproteobacteria>Chromatiales>Sedimenticolaceae>Candidatus_Thiodiazotropha | 628 | JAY70_09500 | 30164..32050 | 2792791 | MAG:-DUF262-domain-containing-protein-[Candidatus-Thiodiazotropha-taylori] |
| WP_050072999.1 | ParB+HNH | GCF_001088505.1 | Yersiniaceae | Bacteria>Pseudomonadati>Pseudomonadota>Gammaproteobacteria>Enterobacterales | 563 | ORG37_RS00060 | 15943..17634 | 1903411 | MULTISPECIES:-DUF262-domain-containing-protein-[Yersiniaceae] |
| WP_241542953.1 | ParB+HNH | GCF_963898675.1 | Morganellaceae | Bacteria>Pseudomonadati>Pseudomonadota>Gammaproteobacteria>Enterobacterales | 587 | MNY72_RS15940 | 13136..14899 | 1903414 | DUF262-domain-containing-protein-[Moellerella-wisconsensis] |
| WP_223429133.1 | ParB+HNH | GCF_036493655.1 | Enterobacteriaceae | Bacteria>Pseudomonadati>Pseudomonadota>Gammaproteobacteria>Enterobacterales | 622 | FD764_RS25530 | 391..2259 | 543 | MULTISPECIES:-DUF262-domain-containing-protein-[Enterobacteriaceae] |
| WP_001698868.1 | ParB+HNH | GCF_000340255.1 | Enterobacteriaceae | Bacteria>Pseudomonadati>Pseudomonadota>Gammaproteobacteria>Enterobacterales | 587 | C201_RS01210 | 13043..14806 | 543 | MULTISPECIES:-DUF262-domain-containing-protein-[Enterobacteriaceae] |
| WP_006788204.1 | ParB+HNH | GCF_000388555.1 | Enterobacteriaceae | Bacteria>Pseudomonadati>Pseudomonadota>Gammaproteobacteria>Enterobacterales | 587 | HKB44_RS26580 | 6233..7996 | 543 | MULTISPECIES:-DUF262-domain-containing-protein-[Enterobacteriaceae] |
| WP_064161343.1 | ParB+HNH | - | Enterobacteriaceae | Bacteria>Pseudomonadati>Pseudomonadota>Gammaproteobacteria>Enterobacterales | 587 | GVI22_RS02070 | 24590..26353 | 543 | MULTISPECIES:-DUF262-domain-containing-protein-[Enterobacteriaceae] |
| WP_224452733.1 | ParB+HNH | GCF_020162075.2 | Enterobacteriaceae | Bacteria>Pseudomonadati>Pseudomonadota>Gammaproteobacteria>Enterobacterales | 587 | KD159_RS16655 | 45212..46975 | 543 | MULTISPECIES:-DUF262-domain-containing-protein-[Enterobacteriaceae] |
| WP_038422886.1 | ParB+HNH | GCF_000632415.1 | Enterobacteriaceae | Bacteria>Pseudomonadati>Pseudomonadota>Gammaproteobacteria>Enterobacterales | 584 | J415_RS00265 | 49584..51338 | 543 | DUF262-domain-containing-protein-[Klebsiella-michiganensis] |
| WP_039268697.1 | ParB+HNH | GCF_954736865.1 | Enterobacteriaceae | Bacteria>Pseudomonadati>Pseudomonadota>Gammaproteobacteria>Enterobacterales | 584 | LI66_RS02635 | 556113..557867 | 543 | MULTISPECIES:-DUF262-domain-containing-protein-[Enterobacter] |
| WP_049245418.1 | ParB+HNH | GCF_004312085.1 | Enterobacteriaceae | Bacteria>Pseudomonadati>Pseudomonadota>Gammaproteobacteria>Enterobacterales | 584 | HUC40_RS15470 | 6747..8501 | 543 | MULTISPECIES:-DUF262-domain-containing-protein-[Enterobacteriaceae] |
| WP_097365541.1 | ParB+HNH | GCF_902158845.1 | Enterobacteriaceae | Bacteria>Pseudomonadati>Pseudomonadota>Gammaproteobacteria>Enterobacterales | 584 | FN096_RS13270 | 26193..27947 | 543 | MULTISPECIES:-DUF262-domain-containing-protein-[Enterobacteriaceae] |
| WP_224385470.1 | ParB+HNH | GCF_023153855.1 | Enterobacteriaceae | Bacteria>Pseudomonadati>Pseudomonadota>Gammaproteobacteria>Enterobacterales | 584 | L8P05_RS09270 | 352547..354301 | 543 | MULTISPECIES:-DUF262-domain-containing-protein-[Enterobacteriaceae] |
| WP_023329157.1 | ParB+HNH | GCF_021264605.1 | Enterobacteriaceae | Bacteria>Pseudomonadati>Pseudomonadota>Gammaproteobacteria>Enterobacterales | 563 | LVW29_RS13115 | 83467..85158 | 543 | DUF262-domain-containing-protein-[Klebsiella-oxytoca] |
| WP_211449380.1 | ParB+HNH | GCF_018299605.1 | Enterobacteriaceae | Bacteria>Pseudomonadati>Pseudomonadota>Gammaproteobacteria>Enterobacterales | 563 | GUY16_RS21835 | 53844..55535 | 543 | DUF262-domain-containing-protein-[Enterobacter-mori] |
| WP_008321087.1 | ParB+HNH | GCF_000277565.1 | Enterobacteriaceae | Bacteria>Pseudomonadati>Pseudomonadota>Gammaproteobacteria>Enterobacterales | 486 | K5R72_RS10995 | 53493..54953 | 543 | MULTISPECIES:-DUF262-domain-containing-protein-[Enterobacteriaceae] |
| WP_000284940.1 | ParB+HNH | GCF_901212535.1 | Salmonella | Bacteria>Pseudomonadati>Pseudomonadota>Gammaproteobacteria>Enterobacterales>Enterobacteriaceae | 584 | PQP98_RS20375 | 4152139..4153893 | 590 | DUF262-domain-containing-protein-[Salmonella-enterica] |
| WP_000283751.1 | ParB+HNH | GCF_000026225.1 | Escherichia | Bacteria>Pseudomonadati>Pseudomonadota>Gammaproteobacteria>Enterobacterales>Enterobacteriaceae | 587 | EFER_RS00090 | 18711..20474 | 561 | MULTISPECIES:-DUF262-domain-containing-protein-[Escherichia] |
| WP_164474331.1 | ParB+HNH | GCF_011045335.1 | Enterobacter | Bacteria>Pseudomonadati>Pseudomonadota>Gammaproteobacteria>Enterobacterales>Enterobacteriaceae | 599 | G5C52_RS10090 | 2126200..2127999 | 547 | DUF262-domain-containing-protein-[Enterobacter-hormaechei] |
| WP_094760993.1 | ParB+HNH | GCF_013791465.1 | Citrobacter | Bacteria>Pseudomonadati>Pseudomonadota>Gammaproteobacteria>Enterobacterales>Enterobacteriaceae | 584 | HV034_RS17815 | 3738148..3739902 | 544 | MULTISPECIES:-DUF262-domain-containing-protein-[Citrobacter] |
| WP_124023335.1 | ParB+HNH | GCF_003818135.1 | Buttiauxella_warmboldiae | Bacteria>Pseudomonadati>Pseudomonadota>Gammaproteobacteria>Enterobacterales>Enterobacteriaceae>Buttiauxella | 563 | EHN07_RS06340 | 56096..57787 | 82993 | DUF262-domain-containing-protein-[Buttiauxella-warmboldiae] |
| RJT19509.1 | ParB+HNH | GCA_003601925.1 | Buttiauxella_izardii | Bacteria>Pseudomonadati>Pseudomonadota>Gammaproteobacteria>Enterobacterales>Enterobacteriaceae>Buttiauxella | 584 | D6029_18520 | 9200..10954 | 82991 | DUF262-domain-containing-protein-[Buttiauxella-izardii] |
| CAG0284404.1 | ParB+HNH | - | Citrobacter_koseri | Bacteria>Pseudomonadati>Pseudomonadota>Gammaproteobacteria>Enterobacterales>Enterobacteriaceae>Citrobacter | 584 | AN2353V1_3686 | 3971055..3972809 | 545 | hypothetical-protein-AN2353V1_3686-[Citrobacter-koseri] |
| WP_048236248.1 | ParB+HNH | GCF_048296775.1 | Citrobacter_freundii_complex | Bacteria>Pseudomonadati>Pseudomonadota>Gammaproteobacteria>Enterobacterales>Enterobacteriaceae>Citrobacter | 584 | SK38_RS14785 | 3147680..3149434 | 1344959 | DUF262-domain-containing-protein-[Citrobacter-sp.-MGH110] |
| EGT0661409.1 | ParB+HNH | - | Citrobacter_werkmanii | Bacteria>Pseudomonadati>Pseudomonadota>Gammaproteobacteria>Enterobacterales>Enterobacteriaceae>Citrobacter>Citrobacter_freundii_complex | 584 | JAF83_001576 | 370186..371940 | 67827 | DUF262-domain-containing-protein-[Citrobacter-werkmanii] |
| MBM9568542.1 | ParB+HNH | GCA_014931625.2 | Citrobacter_sedlakii | Bacteria>Pseudomonadati>Pseudomonadota>Gammaproteobacteria>Enterobacterales>Enterobacteriaceae>Citrobacter>Citrobacter_freundii_complex | 584 | INP82_14065 | 162244..163998 | 67826 | DUF262-domain-containing-protein-[Citrobacter-sedlakii] |
| WP_179154329.1 | ParB+HNH | GCF_013395115.1 | Citrobacter_freundii | Bacteria>Pseudomonadati>Pseudomonadota>Gammaproteobacteria>Enterobacterales>Enterobacteriaceae>Citrobacter>Citrobacter_freundii_complex | 599 | HYG96_RS13010 | 246433..248232 | 546 | DUF262-domain-containing-protein-[Citrobacter-freundii] |

| Protein-ID | Domain | Assembly-ID | Strain | Taxonomy | Length | Locus-Tag | Genomic-coordinates | TaxID | Fasta-defline |
|---|---|---|---|---|---|---|---|---|---|
| WP_205688204.1 | ParB+HNH | GCF_017086505.1 | Citrobacter_freundii | Bacteria>Pseudomonadati>Pseudomonadota>Gammaproteobacteria>Enterobacterales>Enterobacteriaceae>Citrobacter>Citrobacter_freundii_complex | 599 | GHT30_RS20255 | 4257569..4259368 | 546 | DUF262-domain-containing-protein-[Citrobacter-freundii] |
| SYX30362.1 | ParB+HNH | GCA_900520375.1 | Citrobacter_freundii | Bacteria>Pseudomonadati>Pseudomonadota>Gammaproteobacteria>Enterobacterales>Enterobacteriaceae>Citrobacter>Citrobacter_freundii_complex | 584 | CFU2785_04434 | 4463267..4465021 | 546 | Uncharacterised-protein-[Citrobacter-freundii] |
| WP_200054818.1 | ParB+HNH | GCF_016504925.1 | Citrobacter_freundii | Bacteria>Pseudomonadati>Pseudomonadota>Gammaproteobacteria>Enterobacterales>Enterobacteriaceae>Citrobacter>Citrobacter_freundii_complex | 563 | GHT45_RS09250 | 244149..245840 | 546 | DUF262-domain-containing-protein-[Citrobacter-freundii] |
| OXU12832.1 | ParB+HNH | - | Citrobacter_braakii | Bacteria>Pseudomonadati>Pseudomonadota>Gammaproteobacteria>Enterobacterales>Enterobacteriaceae>Citrobacter>Citrobacter_freundii_complex | 584 | B9P84_05300 | 65999..67753 | 57706 | hypothetical-protein-B9P84_05300-[Citrobacter-braakii] |
| QLR25816.1 | ParB+HNH | GCA_013781985.1 | Citrobacter_sp._RHBSTW-01013 | Bacteria>Pseudomonadati>Pseudomonadota>Gammaproteobacteria>Enterobacterales>Enterobacteriaceae>Citrobacter>unclassified_Citrobacter | 584 | HV350_21475 | 4474175..4475929 | 2742677 | DUF262-domain-containing-protein-[Citrobacter-sp.-RHBSTW-01013] |
| MBA8131542.1 | ParB+HNH | GCA_014104375.1 | Citrobacter_sp._RHBSTW-00013 | Bacteria>Pseudomonadati>Pseudomonadota>Gammaproteobacteria>Enterobacterales>Enterobacteriaceae>Citrobacter>unclassified_Citrobacter | 584 | HV034_17815 | 3738148..3739902 | 2742628 | DUF262-domain-containing-protein-[Citrobacter-sp.-RHBSTW-00013] |
| KAB0815084.1 | ParB+HNH | GCA_008805175.1 | Cronobacter_sakazakii | Bacteria>Pseudomonadati>Pseudomonadota>Gammaproteobacteria>Enterobacterales>Enterobacteriaceae>Cronobacter | 584 | FZI24_17900 | 36110..37864 | 28141 | DUF262-domain-containing-protein-[Cronobacter-sakazakii] |
| PUV47026.1 | ParB+HNH | GCA_002532375.1 | Cronobacter_sakazakii | Bacteria>Pseudomonadati>Pseudomonadota>Gammaproteobacteria>Enterobacterales>Enterobacteriaceae>Cronobacter | 584 | CDU02_01550 | 340576..342330 | 28141 | DUF262-domain-containing-protein-[Cronobacter-sakazakii] |
| WP_097563836.1 | ParB+HNH | GCF_017845055.1 | Cronobacter_sakazakii | Bacteria>Pseudomonadati>Pseudomonadota>Gammaproteobacteria>Enterobacterales>Enterobacteriaceae>Cronobacter | 584 | KAR72_RS09240 | 215870..217624 | 28141 | DUF262-domain-containing-protein-[Cronobacter-sakazakii] |
| WP_104677903.1 | ParB+HNH | GCF_002942265.1 | Cronobacter_sakazakii | Bacteria>Pseudomonadati>Pseudomonadota>Gammaproteobacteria>Enterobacterales>Enterobacteriaceae>Cronobacter | 584 | C3D67_RS10960 | 275262..277016 | 28141 | DUF262-domain-containing-protein-[Cronobacter-sakazakii] |
| WP_126349339.1 | ParB+HNH | GCF_003965695.1 | Enterobacter_roggenkampii | Bacteria>Pseudomonadati>Pseudomonadota>Gammaproteobacteria>Enterobacterales>Enterobacteriaceae>Enterobacter>Enterobacter_cloacae_complex | 599 | EKN52_RS20265 | 30454..32253 | 1812935 | DUF262-domain-containing-protein-[Enterobacter-roggenkampii] |
| WP_129694024.1 | ParB+HNH | GCF_004138605.1 | Enterobacter_roggenkampii | Bacteria>Pseudomonadati>Pseudomonadota>Gammaproteobacteria>Enterobacterales>Enterobacteriaceae>Enterobacter>Enterobacter_cloacae_complex | 599 | D8675_RS06380 | 1080474..1082273 | 1812935 | DUF262-domain-containing-protein-[Enterobacter-roggenkampii] |
| AFP68424.1 | ParB+HNH | GCA_000286275.1 | Enterobacter_kobei | Bacteria>Pseudomonadati>Pseudomonadota>Gammaproteobacteria>Enterobacterales>Enterobacteriaceae>Enterobacter>Enterobacter_cloacae_complex | 584 | ECENHK_02655 | 579397..581151 | 208224 | hypothetical-protein-ECENHK_02655-[Enterobacter-kobei] |
| WP_014882330.1 | ParB+HNH | GCF_000286275.1 | Enterobacter_kobei | Bacteria>Pseudomonadati>Pseudomonadota>Gammaproteobacteria>Enterobacterales>Enterobacteriaceae>Enterobacter>Enterobacter_cloacae_complex | 584 | ECENHK_RS02755 | 579397..581151 | 208224 | DUF262-domain-containing-protein-[Enterobacter-kobei] |
| WP_155023645.1 | ParB+HNH | GCF_902160145.1 | Enterobacter_hormaechei | Bacteria>Pseudomonadati>Pseudomonadota>Gammaproteobacteria>Enterobacterales>Enterobacteriaceae>Enterobacter>Enterobacter_cloacae_complex | 599 | FN179_RS14120 | 338127..339926 | 158836 | DUF262-domain-containing-protein-[Enterobacter-hormaechei] |
| HBL8846882.1 | ParB+HNH | GCA_021095225.1 | Enterobacter_hormaechei | Bacteria>Pseudomonadati>Pseudomonadota>Gammaproteobacteria>Enterobacterales>Enterobacteriaceae>Enterobacter>Enterobacter_cloacae_complex | 587 | LTW48_004336 | 3288..5051 | 158836 | DUF262-domain-containing-protein-[Enterobacter-hormaechei] |
| WP_125357893.1 | ParB+HNH | GCF_003937295.2 | Enterobacter_hormaechei | Bacteria>Pseudomonadati>Pseudomonadota>Gammaproteobacteria>Enterobacterales>Enterobacteriaceae>Enterobacter>Enterobacter_cloacae_complex | 587 | EGK38_RS19820 | 4106878..4108641 | 158836 | DUF262-domain-containing-protein-[Enterobacter-hormaechei] |
| AWF33115.1 | ParB+HNH | GCA_003073995.1 | Enterobacter_hormaechei | Bacteria>Pseudomonadati>Pseudomonadota>Gammaproteobacteria>Enterobacterales>Enterobacteriaceae>Enterobacter>Enterobacter_cloacae_complex | 584 | CSC19_3710 | 3847585..3849339 | 158836 | hypothetical-protein-CSC19_3710-[Enterobacter-hormaechei] |
| MBK3145474.1 | ParB+HNH | GCA_016616795.1 | Enterobacter_hormaechei | Bacteria>Pseudomonadati>Pseudomonadota>Gammaproteobacteria>Enterobacterales>Enterobacteriaceae>Enterobacter>Enterobacter_cloacae_complex | 584 | FKK11_12160 | 277191..278945 | 158836 | DUF262-domain-containing-protein-[Enterobacter-hormaechei] |
| WP_062933983.1 | ParB+HNH | GCF_003073995.1 | Enterobacter_hormaechei | Bacteria>Pseudomonadati>Pseudomonadota>Gammaproteobacteria>Enterobacterales>Enterobacteriaceae>Enterobacter>Enterobacter_cloacae_complex | 584 | CSC19_RS19000 | 3847585..3849339 | 158836 | DUF262-domain-containing-protein-[Enterobacter-hormaechei] |
| WP_063436697.1 | ParB+HNH | GCF_001631925.1 | Enterobacter_hormaechei | Bacteria>Pseudomonadati>Pseudomonadota>Gammaproteobacteria>Enterobacterales>Enterobacteriaceae>Enterobacter>Enterobacter_cloacae_complex | 584 | A3N67_RS02930 | 132856..134610 | 158836 | DUF262-domain-containing-protein-[Enterobacter-hormaechei] |
| WP_200535482.1 | ParB+HNH | GCF_021491635.1 | Enterobacter_hormaechei | Bacteria>Pseudomonadati>Pseudomonadota>Gammaproteobacteria>Enterobacterales>Enterobacteriaceae>Enterobacter>Enterobacter_cloacae_complex | 584 | FKK11_RS12015 | 277191..278945 | 158836 | DUF262-domain-containing-protein-[Enterobacter-hormaechei] |
| WP_241512122.1 | ParB+HNH | GCF_022585395.1 | Enterobacter_hormaechei | Bacteria>Pseudomonadati>Pseudomonadota>Gammaproteobacteria>Enterobacterales>Enterobacteriaceae>Enterobacter>Enterobacter_cloacae_complex | 584 | MMO50_RS10495 | 123311..125065 | 158836 | DUF262-domain-containing-protein-[Enterobacter-hormaechei] |
| WP_110875384.1 | ParB+HNH | GCF_003207215.1 | Enterobacter_hormaechei | Bacteria>Pseudomonadati>Pseudomonadota>Gammaproteobacteria>Enterobacterales>Enterobacteriaceae>Enterobacter>Enterobacter_cloacae_complex | 563 | DOM70_RS06240 | 336023..337714 | 158836 | DUF262-domain-containing-protein-[Enterobacter-hormaechei] |
| CAF2998355.1 | ParB+HNH | - | Enterobacter_cloacae | Bacteria>Pseudomonadati>Pseudomonadota>Gammaproteobacteria>Enterobacterales>Enterobacteriaceae>Enterobacter>Enterobacter_cloacae_complex | 584 | AI2972V1_0507 | 555839..557593 | 550 | hypothetical-protein-AI2972V1_0507-[Enterobacter-cloacae] |
| MBG0523799.1 | ParB+HNH | GCA_015704905.1 | Enterobacter_cloacae | Bacteria>Pseudomonadati>Pseudomonadota>Gammaproteobacteria>Enterobacterales>Enterobacteriaceae>Enterobacter>Enterobacter_cloacae_complex | 584 | IOR24_17910 | 3772598..3774352 | 550 | DUF262-domain-containing-protein-[Enterobacter-cloacae] |
| WP_196373087.1 | ParB+HNH | GCF_015704905.1 | Enterobacter_cloacae | Bacteria>Pseudomonadati>Pseudomonadota>Gammaproteobacteria>Enterobacterales>Enterobacteriaceae>Enterobacter>Enterobacter_cloacae_complex | 584 | IOR24_RS17910 | 3772598..3774352 | 550 | DUF262-domain-containing-protein-[Enterobacter-cloacae] |
| WP_252033144.1 | ParB+HNH | GCF_023752605.1 | Enterobacter_asburiae | Bacteria>Pseudomonadati>Pseudomonadota>Gammaproteobacteria>Enterobacterales>Enterobacteriaceae>Enterobacter>Enterobacter_cloacae_complex | 563 | M8S83_RS11460 | 337440..339131 | 61645 | DUF262-domain-containing-protein-[Enterobacter-asburiae] |
| HCM9559900.1 | ParB+HNH | GCA_025214055.1 | Enterobacter_cloacae_subsp._cloacae | Bacteria>Pseudomonadati>Pseudomonadota>Gammaproteobacteria>Enterobacterales>Enterobacteriaceae>Enterobacter>Enterobacter_cloacae_complex>Enterobacter_cloacae | 599 | N5O13_002968 | 183470..185269 | 336306 | DUF262-domain-containing-protein-[Enterobacter-cloacae-subsp.-cloacae] |
| AJB80306.1 | ParB+HNH | GCA_000814225.1 | Enterobacter_hormaechei_subsp._xiangfangensis | Bacteria>Pseudomonadati>Pseudomonadota>Gammaproteobacteria>Enterobacterales>Enterobacteriaceae>Enterobacter>Enterobacter_cloacae_complex>Enterobacter_hormaechei | 584 | LI66_02635 | 556113..557867 | 1296536 | hypothetical-protein-LI66_02635-[Enterobacter-hormaechei-subsp.-xiangfangensis] |
| KZR28107.1 | ParB+HNH | GCA_001631925.1 | Enterobacter_hormaechei_subsp._steigerwaltii | Bacteria>Pseudomonadati>Pseudomonadota>Gammaproteobacteria>Enterobacterales>Enterobacteriaceae>Enterobacter>Enterobacter_cloacae_complex>Enterobacter_hormaechei | 584 | A3N67_04035 | 132856..134610 | 299766 | hypothetical-protein-A3N67_04035-[Enterobacter-hormaechei-subsp.-steigerwaltii] |
| RYA52301.1 | ParB+HNH | GCA_004136805.1 | Enterobacter_cloacae_complex_sp._GF14B | Bacteria>Pseudomonadati>Pseudomonadota>Gammaproteobacteria>Enterobacterales>Enterobacteriaceae>Enterobacter>Enterobacter_cloacae_complex>unclassified_Enterobacter_cloacae_complex | 563 | DD606_17365 | 19310..21001 | 2511982 | hypothetical-protein-DD606_17365-[Enterobacter-cloacae-complex-sp.-GF14B] |
| WP_126800554.1 | ParB+HNH | GCF_003986875.1 | Enterobacter_sp._WCHEn090032 | Bacteria>Pseudomonadati>Pseudomonadota>Gammaproteobacteria>Enterobacterales>Enterobacteriaceae>Enterobacter>unclassified_Enterobacter | 603 | EKN83_RS06690 | 47664..49475 | 2497435 | DUF262-domain-containing-protein-[Enterobacter-sp.-WCHEn090032] |
| WP_230238965.1 | ParB+HNH | GCF_020971765.1 | Enterobacter_sp._AN-K1 | Bacteria>Pseudomonadati>Pseudomonadota>Gammaproteobacteria>Enterobacterales>Enterobacteriaceae>Enterobacter>unclassified_Enterobacter | 599 | LOC22_RS13490 | 2823200..2824999 | 2894203 | DUF262-domain-containing-protein-[Enterobacter-sp.-AN-K1] |
| STN20145.1 | ParB+HNH | GCA_900450565.1 | Escherichia_fergusonii | Bacteria>Pseudomonadati>Pseudomonadota>Gammaproteobacteria>Enterobacterales>Enterobacteriaceae>Escherichia | 513 | NCTC12128_00060 | 41722..43263 | 564 | Uncharacterized-conserved-protein-[Escherichia-fergusonii] |
| STN20144.1 | ParB+HNH | GCA_900450565.1 | Escherichia_fergusonii | Bacteria>Pseudomonadati>Pseudomonadota>Gammaproteobacteria>Enterobacterales>Enterobacteriaceae>Escherichia | 91 | NCTC12128_00059 | 41499..41774 | 564 | Uncharacterized-conserved-protein-[Escherichia-fergusonii] |
| WP_249519094.1 | ParB+HNH | GCF_011077045.1 | Escherichia_coli | Bacteria>Pseudomonadati>Pseudomonadota>Gammaproteobacteria>Enterobacterales>Enterobacteriaceae>Escherichia | 622 | G6Q90_RS23630 | 18511..20379 | 562 | DUF262-domain-containing-protein-[Escherichia-coli] |
| EEW8735902.1 | ParB+HNH | - | Escherichia_coli | Bacteria>Pseudomonadati>Pseudomonadota>Gammaproteobacteria>Enterobacterales>Enterobacteriaceae>Escherichia | 587 | D5Q61_02345 | 119922..121685 | 562 | DUF262-domain-containing-protein-[Escherichia-coli] |
| EFB3218131.1 | ParB+HNH | - | Escherichia_coli | Bacteria>Pseudomonadati>Pseudomonadota>Gammaproteobacteria>Enterobacterales>Enterobacteriaceae>Escherichia | 587 | FLO28_20240 | 6268..8031 | 562 | DUF262-domain-containing-protein-[Escherichia-coli] |
| EFB6629152.1 | ParB+HNH | - | Escherichia_coli | Bacteria>Pseudomonadati>Pseudomonadota>Gammaproteobacteria>Enterobacterales>Enterobacteriaceae>Escherichia | 587 | FQF26_18835 | 75765..77528 | 562 | DUF262-domain-containing-protein-[Escherichia-coli] |
| EFH7369083.1 | ParB+HNH | - | Escherichia_coli | Bacteria>Pseudomonadati>Pseudomonadota>Gammaproteobacteria>Enterobacterales>Enterobacteriaceae>Escherichia | 587 | GKQ19_21215 | 9359..11122 | 562 | DUF262-domain-containing-protein-[Escherichia-coli] |
| EFT2808155.1 | ParB+HNH | - | Escherichia_coli | Bacteria>Pseudomonadati>Pseudomonadota>Gammaproteobacteria>Enterobacterales>Enterobacteriaceae>Escherichia | 587 | HVL53_003850 | 7139..8902 | 562 | DUF262-domain-containing-protein-[Escherichia-coli] |
| EHK3830300.1 | ParB+HNH | GCA_018024135.1 | Escherichia_coli | Bacteria>Pseudomonadati>Pseudomonadota>Gammaproteobacteria>Enterobacterales>Enterobacteriaceae>Escherichia | 587 | JR760_003916 | 6569..8332 | 562 | DUF262-domain-containing-protein-[Escherichia-coli] |
| EID8792052.1 | ParB+HNH | GCA_020095335.1 | Escherichia_coli | Bacteria>Pseudomonadati>Pseudomonadota>Gammaproteobacteria>Enterobacterales>Enterobacteriaceae>Escherichia | 587 | LCJ17_004432 | 7364..9127 | 562 | DUF262-domain-containing-protein-[Escherichia-coli] |
| EJE3980762.1 | ParB+HNH | GCA_023310495.1 | Escherichia_coli | Bacteria>Pseudomonadati>Pseudomonadota>Gammaproteobacteria>Enterobacterales>Enterobacteriaceae>Escherichia | 587 | M4B16_004239 | 9361..11124 | 562 | DUF262-domain-containing-protein-[Escherichia-coli] |
| EJN1844349.1 | ParB+HNH | GCA_024422675.1 | Escherichia_coli | Bacteria>Pseudomonadati>Pseudomonadota>Gammaproteobacteria>Enterobacterales>Enterobacteriaceae>Escherichia | 587 | NPJ45_004377 | 6950..8713 | 562 | DUF262-domain-containing-protein-[Escherichia-coli] |
| EJU9647233.1 | ParB+HNH | GCA_025224895.1 | Escherichia_coli | Bacteria>Pseudomonadati>Pseudomonadota>Gammaproteobacteria>Enterobacterales>Enterobacteriaceae>Escherichia | 587 | N5N25_003987 | 9156..10919 | 562 | DUF262-domain-containing-protein-[Escherichia-coli] |
| HAI3857479.1 | ParB+HNH | - | Escherichia_coli | Bacteria>Pseudomonadati>Pseudomonadota>Gammaproteobacteria>Enterobacterales>Enterobacteriaceae>Escherichia | 587 | HJN25_004327 | 8955..10718 | 562 | TPA:-DUF262-domain-containing-protein-[Escherichia-coli] |
| HAJ7472470.1 | ParB+HNH | GCA_013077245.1 | Escherichia_coli | Bacteria>Pseudomonadati>Pseudomonadota>Gammaproteobacteria>Enterobacterales>Enterobacteriaceae>Escherichia | 587 | HNV92_21030 | 17259..19022 | 562 | DUF262-domain-containing-protein-[Escherichia-coli] |
| HAX1741552.1 | ParB+HNH | GCA_016984655.1 | Escherichia_coli | Bacteria>Pseudomonadati>Pseudomonadota>Gammaproteobacteria>Enterobacterales>Enterobacteriaceae>Escherichia | 587 | JXS76_004486 | 2400..4163 | 562 | TPA:-DUF262-domain-containing-protein-[Escherichia-coli] |
| HBC6897267.1 | ParB+HNH | GCA_018163035.1 | Escherichia_coli | Bacteria>Pseudomonadati>Pseudomonadota>Gammaproteobacteria>Enterobacterales>Enterobacteriaceae>Escherichia | 587 | KEZ94_003916 | 8955..10718 | 562 | DUF262-domain-containing-protein-[Escherichia-coli] |
| HBE5682465.1 | ParB+HNH | GCA_018567345.1 | Escherichia_coli | Bacteria>Pseudomonadati>Pseudomonadota>Gammaproteobacteria>Enterobacterales>Enterobacteriaceae>Escherichia | 587 | KMB60_001823 | 1590..3353 | 562 | TPA:-DUF262-domain-containing-protein-[Escherichia-coli] |
| HBN0250040.1 | ParB+HNH | GCA_021454405.1 | Escherichia_coli | Bacteria>Pseudomonadati>Pseudomonadota>Gammaproteobacteria>Enterobacterales>Enterobacteriaceae>Escherichia | 587 | L1L95_003625 | 60818..62581 | 562 | DUF262-domain-containing-protein-[Escherichia-coli] |
| MCK3244658.1 | ParB+HNH | GCA_023098955.1 | Escherichia_coli | Bacteria>Pseudomonadati>Pseudomonadota>Gammaproteobacteria>Enterobacterales>Enterobacteriaceae>Escherichia | 587 | MZH07_24525 | 9382..11145 | 562 | DUF262-domain-containing-protein-[Escherichia-coli] |
| OWF25644.1 | ParB+HNH | GCA_002190095.1 | Escherichia_coli | Bacteria>Pseudomonadati>Pseudomonadota>Gammaproteobacteria>Enterobacterales>Enterobacteriaceae>Escherichia | 587 | A8M76_10405 | 2726..4489 | 562 | hypothetical-protein-A8M76_10405-[Escherichia-coli] |
| TZE05561.1 | ParB+HNH | GCA_008273905.1 | Escherichia_coli | Bacteria>Pseudomonadati>Pseudomonadota>Gammaproteobacteria>Enterobacterales>Enterobacteriaceae>Escherichia | 587 | EOI14_22610 | 13647..15410 | 562 | DUF262-domain-containing-protein-[Escherichia-coli] |
| TZF09960.1 | ParB+HNH | GCA_008273745.1 | Escherichia_coli | Bacteria>Pseudomonadati>Pseudomonadota>Gammaproteobacteria>Enterobacterales>Enterobacteriaceae>Escherichia | 587 | EOI42_22660 | 14369..16132 | 562 | DUF262-domain-containing-protein-[Escherichia-coli] |
| WP_023063794.1 | ParB+HNH | GCF_036495545.1 | Escherichia_coli | Bacteria>Pseudomonadati>Pseudomonadota>Gammaproteobacteria>Enterobacterales>Enterobacteriaceae>Escherichia | 587 | M5X65_RS22545 | 6284..8047 | 562 | DUF262-domain-containing-protein-[Escherichia-coli] |
| WP_072752449.1 | ParB+HNH | GCF_001894415.1 | Escherichia_coli | Bacteria>Pseudomonadati>Pseudomonadota>Gammaproteobacteria>Enterobacterales>Enterobacteriaceae>Escherichia | 587 | BK380_RS19180 | 21383..23146 | 562 | DUF262-domain-containing-protein-[Escherichia-coli] |
| WP_115785250.1 | ParB+HNH | GCF_029870405.1 | Escherichia_coli | Bacteria>Pseudomonadati>Pseudomonadota>Gammaproteobacteria>Enterobacterales>Enterobacteriaceae>Escherichia | 587 | OZI01_RS14445 | 43066..44829 | 562 | DUF262-domain-containing-protein-[Escherichia-coli] |
| EFD0312066.1 | ParB+HNH | - | Escherichia_coli | Bacteria>Pseudomonadati>Pseudomonadota>Gammaproteobacteria>Enterobacterales>Enterobacteriaceae>Escherichia | 584 | E6D44_20530 | 17612..19366 | 562 | DUF262-domain-containing-protein-[Escherichia-coli] |
| EFN5198233.1 | ParB+HNH | - | Escherichia_coli | Bacteria>Pseudomonadati>Pseudomonadota>Gammaproteobacteria>Enterobacterales>Enterobacteriaceae>Escherichia | 584 | FQG10_00770 | 6571..8325 | 562 | DUF262-domain-containing-protein-[Escherichia-coli] |
| EHY5030620.1 | ParB+HNH | GCA_019582855.1 | Escherichia_coli | Bacteria>Pseudomonadati>Pseudomonadota>Gammaproteobacteria>Enterobacterales>Enterobacteriaceae>Escherichia | 584 | K4I20_002755 | 18805..20559 | 562 | DUF262-domain-containing-protein-[Escherichia-coli] |
| NUC65043.1 | ParB+HNH | GCA_013342565.1 | Escherichia_coli | Bacteria>Pseudomonadati>Pseudomonadota>Gammaproteobacteria>Enterobacterales>Enterobacteriaceae>Escherichia | 584 | HUC40_15470 | 6747..8501 | 562 | DUF262-domain-containing-protein-[Escherichia-coli] |
| RLY30109.1 | ParB+HNH | GCA_003687125.1 | Escherichia_coli | Bacteria>Pseudomonadati>Pseudomonadota>Gammaproteobacteria>Enterobacterales>Enterobacteriaceae>Escherichia | 584 | EAI39_15060 | 82259..84013 | 562 | DUF262-domain-containing-protein-[Escherichia-coli] |
| WP_097448433.1 | ParB+HNH | GCF_002516105.1 | Escherichia_coli | Bacteria>Pseudomonadati>Pseudomonadota>Gammaproteobacteria>Enterobacterales>Enterobacteriaceae>Escherichia | 584 | A7Y91_RS22680 | 18883..20637 | 562 | DUF262-domain-containing-protein-[Escherichia-coli] |
| EGK3607994.1 | ParB+HNH | - | Escherichia_coli | Bacteria>Pseudomonadati>Pseudomonadota>Gammaproteobacteria>Enterobacterales>Enterobacteriaceae>Escherichia | 583 | IOT31_003183 | 95946..97697 | 562 | DUF262-domain-containing-protein-[Escherichia-coli] |
| HCO5071634.1 | ParB+HNH | GCA_025390635.1 | Escherichia_coli | Bacteria>Pseudomonadati>Pseudomonadota>Gammaproteobacteria>Enterobacterales>Enterobacteriaceae>Escherichia | 67 | N8039_004384 | 9359..9559 | 562 | DUF262-domain-containing-protein-[Escherichia-coli] |
| EEZ5714530.1 | ParB+HNH | - | Escherichia_coli_O25 | Bacteria>Pseudomonadati>Pseudomonadota>Gammaproteobacteria>Enterobacterales>Enterobacteriaceae>Escherichia>Escherichia_coli | 584 | DBU27_000027 | 23867..25621 | 1095709 | DUF262-domain-containing-protein-[Escherichia-coli-O25] |
| KEM88220.1 | ParB+HNH | GCA_000713255.1 | Escherichia_coli_6-537-08_S4_C1 | Bacteria>Pseudomonadati>Pseudomonadota>Gammaproteobacteria>Enterobacterales>Enterobacteriaceae>Escherichia>Escherichia_coli | 587 | AC92_4029 | 10031..11794 | 1444227 | hypothetical-protein-AC92_4029-[Escherichia-coli-6-537-08_S4_C1] |
| POU04215.1 | ParB+HNH | GCA_002918705.1 | Escherichia_sp._ESNIH1 | Bacteria>Pseudomonadati>Pseudomonadota>Gammaproteobacteria>Enterobacterales>Enterobacteriaceae>Escherichia>unclassified_Escherichia | 599 | C3369_02370 | 495157..496956 | 1985876 | DUF262-domain-containing-protein-[Escherichia-sp.-ESNIH1] |
| WP_117247357.1 | ParB+HNH | GCF_004313365.1 | Klebsiella | Bacteria>Pseudomonadati>Pseudomonadota>Gammaproteobacteria>Enterobacterales>Enterobacteriaceae>Klebsiella/Raoultella_group | 584 | D9K63_RS23210 | 4670731..4672485 | 570 | MULTISPECIES:-DUF262-domain-containing-protein-[Klebsiella] |
| WP_142984066.1 | ParB+HNH | GCF_902158715.1 | Klebsiella_pasteurii | Bacteria>Pseudomonadati>Pseudomonadota>Gammaproteobacteria>Enterobacterales>Enterobacteriaceae>Klebsiella/Raoultella_group>Klebsiella | 584 | FN110_RS17005 | 9198..10952 | 2587529 | DUF262-domain-containing-protein-[Klebsiella-pasteurii] |
| EGT0044781.1 | ParB+HNH | - | Klebsiella_oxytoca | Bacteria>Pseudomonadati>Pseudomonadota>Gammaproteobacteria>Enterobacterales>Enterobacteriaceae>Klebsiella/Raoultella_group>Klebsiella | 584 | I9Y61_001543 | 41234..42988 | 571 | DUF262-domain-containing-protein-[Klebsiella-oxytoca] |
| MBZ7196033.1 | ParB+HNH | GCA_020118595.1 | Klebsiella_oxytoca | Bacteria>Pseudomonadati>Pseudomonadota>Gammaproteobacteria>Enterobacterales>Enterobacteriaceae>Klebsiella/Raoultella_group>Klebsiella | 584 | FMK61_04490 | 43798..45552 | 571 | DUF262-domain-containing-protein-[Klebsiella-oxytoca] |
| WP_182012818.1 | ParB+HNH | GCF_013926025.1 | Klebsiella_michiganensis | Bacteria>Pseudomonadati>Pseudomonadota>Gammaproteobacteria>Enterobacterales>Enterobacteriaceae>Klebsiella/Raoultella_group>Klebsiella | 587 | HV264_RS11960 | 2486651..2488414 | 1134687 | DUF262-domain-containing-protein-[Klebsiella-michiganensis] |
| PLL44192.1 | ParB+HNH | GCA_002856085.1 | Klebsiella_michiganensis | Bacteria>Pseudomonadati>Pseudomonadota>Gammaproteobacteria>Enterobacterales>Enterobacteriaceae>Klebsiella/Raoultella_group>Klebsiella | 584 | CWN50_01595 | 2675..4429 | 1134687 | DUF262-domain-containing-protein-[Klebsiella-michiganensis] |
| WP_163467461.1 | ParB+HNH | GCF_963887275.1 | Klebsiella_michiganensis | Bacteria>Pseudomonadati>Pseudomonadota>Gammaproteobacteria>Enterobacterales>Enterobacteriaceae>Klebsiella/Raoultella_group>Klebsiella | 584 | G4403_RS10435 | 96386..98140 | 1134687 | DUF262-domain-containing-protein-[Klebsiella-michiganensis] |
| WP_142384532.1 | ParB+HNH | - | Klebsiella_michiganensis | Bacteria>Pseudomonadati>Pseudomonadota>Gammaproteobacteria>Enterobacterales>Enterobacteriaceae>Klebsiella/Raoultella_group>Klebsiella | 338 | CWN04_RS12965 | 19839..20852 | 1134687 | DUF262-domain-containing-protein,-partial-[Klebsiella-michiganensis] |
| WEJ89010.1 | ParB+HNH | GCA_029202365.1 | Klebsiella_huaxiensis | Bacteria>Pseudomonadati>Pseudomonadota>Gammaproteobacteria>Enterobacterales>Enterobacteriaceae>Klebsiella/Raoultella_group>Klebsiella | 584 | P0Y63_27790 | 5831494..5833248 | 2153354 | MAG:-DUF262-domain-containing-protein-[Klebsiella-huaxiensis] |
| WP_217074493.1 | ParB+HNH | GCF_019048885.1 | Klebsiella_aerogenes | Bacteria>Pseudomonadati>Pseudomonadota>Gammaproteobacteria>Enterobacterales>Enterobacteriaceae>Klebsiella/Raoultella_group>Klebsiella | 588 | KSL85_RS22225 | 4662301..4664067 | 548 | DUF262-domain-containing-protein-[Klebsiella-aerogenes] |
| QHW99330.1 | ParB+HNH | GCA_010365565.1 | Klebsiella_variicola | Bacteria>Pseudomonadati>Pseudomonadota>Gammaproteobacteria>Enterobacterales>Enterobacteriaceae>Klebsiella/Raoultella_group>Klebsiella>Klebsiella_pneumoniae_complex | 584 | GZS05_24300 | 4953510..4955264 | 244366 | DUF262-domain-containing-protein-[Klebsiella-variicola] |
| WP_163590844.1 | ParB+HNH | GCF_010602405.1 | Klebsiella_variicola | Bacteria>Pseudomonadati>Pseudomonadota>Gammaproteobacteria>Enterobacterales>Enterobacteriaceae>Klebsiella/Raoultella_group>Klebsiella>Klebsiella_pneumoniae_complex | 584 | G4552_RS17220 | 27460..29214 | 244366 | DUF262-domain-containing-protein-[Klebsiella-variicola] |
| HBQ4939448.1 | ParB+HNH | GCA_021861305.1 | Klebsiella_pneumoniae | Bacteria>Pseudomonadati>Pseudomonadota>Gammaproteobacteria>Enterobacterales>Enterobacteriaceae>Klebsiella/Raoultella_group>Klebsiella>Klebsiella_pneumoniae_complex | 603 | L7366_003902 | 47022..48833 | 573 | DUF262-domain-containing-protein-[Klebsiella-pneumoniae] |
| WP_254541267.1 | ParB+HNH | GCF_024224875.1 | Klebsiella_pneumoniae | Bacteria>Pseudomonadati>Pseudomonadota>Gammaproteobacteria>Enterobacterales>Enterobacteriaceae>Klebsiella/Raoultella_group>Klebsiella>Klebsiella_pneumoniae_complex | 603 | NLZ12_RS02815 | 589609..591420 | 573 | DUF262-domain-containing-protein-[Klebsiella-pneumoniae] |
| HBS5818884.1 | ParB+HNH | GCA_021971955.1 | Klebsiella_pneumoniae | Bacteria>Pseudomonadati>Pseudomonadota>Gammaproteobacteria>Enterobacterales>Enterobacteriaceae>Klebsiella/Raoultella_group>Klebsiella>Klebsiella_pneumoniae_complex | 599 | MAK71_003678 | 10141..11940 | 573 | TPA:-DUF262-domain-containing-protein-[Klebsiella-pneumoniae] |
| MCM6196116.1 | ParB+HNH | GCA_023741115.1 | Klebsiella_pneumoniae | Bacteria>Pseudomonadati>Pseudomonadota>Gammaproteobacteria>Enterobacterales>Enterobacteriaceae>Klebsiella/Raoultella_group>Klebsiella>Klebsiella_pneumoniae_complex | 599 | M4C47_13310 | 316332..318131 | 573 | DUF262-domain-containing-HNH-endonuclease-family-protein-[Klebsiella-pneumoniae] |
| WP_096812543.1 | ParB+HNH | GCF_009907625.1 | Klebsiella_pneumoniae | Bacteria>Pseudomonadati>Pseudomonadota>Gammaproteobacteria>Enterobacterales>Enterobacteriaceae>Klebsiella/Raoultella_group>Klebsiella>Klebsiella_pneumoniae_complex | 599 | GV742_RS00810 | 10203..12002 | 573 | DUF262-domain-containing-protein-[Klebsiella-pneumoniae] |
| HBR1749212.1 | ParB+HNH | GCA_021911085.1 | Klebsiella_pneumoniae | Bacteria>Pseudomonadati>Pseudomonadota>Gammaproteobacteria>Enterobacterales>Enterobacteriaceae>Klebsiella/Raoultella_group>Klebsiella>Klebsiella_pneumoniae_complex | 587 | L9M14_005023 | 10441..12204 | 573 | TPA:-DUF262-domain-containing-protein-[Klebsiella-pneumoniae] |
| HBY8309080.1 | ParB+HNH | - | Klebsiella_pneumoniae | Bacteria>Pseudomonadati>Pseudomonadota>Gammaproteobacteria>Enterobacterales>Enterobacteriaceae>Klebsiella/Raoultella_group>Klebsiella>Klebsiella_pneumoniae_complex | 587 | MI837_25470 | 3523..5286 | 573 | TPA:-DUF262-domain-containing-protein-[Klebsiella-pneumoniae] |
| WP_243069916.1 | ParB+HNH | GCF_022760705.1 | Klebsiella_pneumoniae | Bacteria>Pseudomonadati>Pseudomonadota>Gammaproteobacteria>Enterobacterales>Enterobacteriaceae>Klebsiella/Raoultella_group>Klebsiella>Klebsiella_pneumoniae_complex | 587 | MUG00_RS24955 | 9019..10782 | 573 | DUF262-domain-containing-protein-[Klebsiella-pneumoniae] |
| AUN71069.1 | ParB+HNH | GCA_002870985.1 | Klebsiella_pneumoniae | Bacteria>Pseudomonadati>Pseudomonadota>Gammaproteobacteria>Enterobacterales>Enterobacteriaceae>Klebsiella/Raoultella_group>Klebsiella>Klebsiella_pneumoniae_complex | 584 | C0077_01575 | 325058..326812 | 573 | DUF262-domain-containing-protein-[Klebsiella-pneumoniae] |
| EIV3098058.1 | ParB+HNH | GCA_021854435.1 | Klebsiella_pneumoniae | Bacteria>Pseudomonadati>Pseudomonadota>Gammaproteobacteria>Enterobacterales>Enterobacteriaceae>Klebsiella/Raoultella_group>Klebsiella>Klebsiella_pneumoniae_complex | 584 | L7W77_004839 | 49376..51130 | 573 | DUF262-domain-containing-protein-[Klebsiella-pneumoniae] |
| EIX9370753.1 | ParB+HNH | GCA_022430905.1 | Klebsiella_pneumoniae | Bacteria>Pseudomonadati>Pseudomonadota>Gammaproteobacteria>Enterobacterales>Enterobacteriaceae>Klebsiella/Raoultella_group>Klebsiella>Klebsiella_pneumoniae_complex | 584 | ML041_004567 | 35610..37364 | 573 | DUF262-domain-containing-protein-[Klebsiella-pneumoniae] |
| HBQ2793927.1 | ParB+HNH | GCA_021848645.1 | Klebsiella_pneumoniae | Bacteria>Pseudomonadati>Pseudomonadota>Gammaproteobacteria>Enterobacterales>Enterobacteriaceae>Klebsiella/Raoultella_group>Klebsiella>Klebsiella_pneumoniae_complex | 584 | L7V03_004585 | 35989..37743 | 573 | TPA:-DUF262-domain-containing-protein-[Klebsiella-pneumoniae] |
| MBN7733856.1 | ParB+HNH | GCA_017255495.1 | Klebsiella_pneumoniae | Bacteria>Pseudomonadati>Pseudomonadota>Gammaproteobacteria>Enterobacterales>Enterobacteriaceae>Klebsiella/Raoultella_group>Klebsiella>Klebsiella_pneumoniae_complex | 584 | JX616_18225 | 35717..37471 | 573 | DUF262-domain-containing-protein-[Klebsiella-pneumoniae] |
| QJM44417.1 | ParB+HNH | GCA_012970665.1 | Klebsiella_pneumoniae | Bacteria>Pseudomonadati>Pseudomonadota>Gammaproteobacteria>Enterobacterales>Enterobacteriaceae>Klebsiella/Raoultella_group>Klebsiella>Klebsiella_pneumoniae_complex | 584 | HJW71_23040 | 4624267..4626021 | 573 | DUF262-domain-containing-protein-[Klebsiella-pneumoniae] |
| WP_073511309.1 | ParB+HNH | GCF_965181005.1 | Klebsiella_pneumoniae | Bacteria>Pseudomonadati>Pseudomonadota>Gammaproteobacteria>Enterobacterales>Enterobacteriaceae>Klebsiella/Raoultella_group>Klebsiella>Klebsiella_pneumoniae_complex | 584 | LZ602_RS12605 | 35739..37493 | 573 | DUF262-domain-containing-protein-[Klebsiella-pneumoniae] |
| WP_117076896.1 | ParB+HNH | GCF_022229165.1 | Klebsiella_pneumoniae | Bacteria>Pseudomonadati>Pseudomonadota>Gammaproteobacteria>Enterobacterales>Enterobacteriaceae>Klebsiella/Raoultella_group>Klebsiella>Klebsiella_pneumoniae_complex | 584 | LLG23_RS24000 | 28290..30044 | 573 | DUF262-domain-containing-protein-[Klebsiella-pneumoniae] |
| MBZ6545108.1 | ParB+HNH | GCA_020115175.1 | Klebsiella_pneumoniae | Bacteria>Pseudomonadati>Pseudomonadota>Gammaproteobacteria>Enterobacterales>Enterobacteriaceae>Klebsiella/Raoultella_group>Klebsiella>Klebsiella_pneumoniae_complex | 469 | FMJ66_23205 | 23644..25052 | 573 | DUF262-domain-containing-protein-[Klebsiella-pneumoniae] |
| HBQ6039881.1 | ParB+HNH | GCA_021866875.1 | Klebsiella_pneumoniae_subsp._pneumoniae | Bacteria>Pseudomonadati>Pseudomonadota>Gammaproteobacteria>Enterobacterales>Enterobacteriaceae>Klebsiella/Raoultella_group>Klebsiella>Klebsiella_pneumoniae_complex>Klebsiella_pneumoniae | 584 | L7649_004475 | 57094..58848 | 72407 | DUF262-domain-containing-protein-[Klebsiella-pneumoniae-subsp.-pneumoniae] |
| HCD1392380.1 | ParB+HNH | GCA_023530865.1 | Klebsiella_pneumoniae_subsp._pneumoniae | Bacteria>Pseudomonadati>Pseudomonadota>Gammaproteobacteria>Enterobacterales>Enterobacteriaceae>Klebsiella/Raoultella_group>Klebsiella>Klebsiella_pneumoniae_complex>Klebsiella_pneumoniae | 584 | M9022_004644 | 21528..23282 | 72407 | DUF262-domain-containing-protein-[Klebsiella-pneumoniae-subsp.-pneumoniae] |
| HBR1784210.1 | ParB+HNH | GCA_021911385.1 | Klebsiella_quasipneumoniae_subsp._quasipneumoniae | Bacteria>Pseudomonadati>Pseudomonadota>Gammaproteobacteria>Enterobacterales>Enterobacteriaceae>Klebsiella/Raoultella_group>Klebsiella>Klebsiella_pneumoniae_complex>Klebsiella_quasipneumoniae | 584 | L9M28_003409 | 24109..25863 | 1667327 | DUF262-domain-containing-protein-[Klebsiella-quasipneumoniae-subsp.-quasipneumoniae] |
| HBS2746979.1 | ParB+HNH | GCA_021960165.1 | Klebsiella_quasipneumoniae_subsp._quasipneumoniae | Bacteria>Pseudomonadati>Pseudomonadota>Gammaproteobacteria>Enterobacterales>Enterobacteriaceae>Klebsiella/Raoultella_group>Klebsiella>Klebsiella_pneumoniae_complex>Klebsiella_quasipneumoniae | 584 | MAE70_005432 | 19342..21096 | 1667327 | DUF262-domain-containing-protein-[Klebsiella-quasipneumoniae-subsp.-quasipneumoniae] |
| MCS5967897.1 | ParB+HNH | GCA_024990005.1 | Klebsiella_variicola_subsp._variicola | Bacteria>Pseudomonadati>Pseudomonadota>Gammaproteobacteria>Enterobacterales>Enterobacteriaceae>Klebsiella/Raoultella_group>Klebsiella>Klebsiella_pneumoniae_complex>Klebsiella_variicola | 430 | LNP26_07890 | 1521219..1522511 | 2590157 | DUF262-domain-containing-protein-[Klebsiella-variicola-subsp.-variicola] |
| MCS5967896.1 | ParB+HNH | GCA_024990005.1 | Klebsiella_variicola_subsp._variicola | Bacteria>Pseudomonadati>Pseudomonadota>Gammaproteobacteria>Enterobacterales>Enterobacteriaceae>Klebsiella/Raoultella_group>Klebsiella>Klebsiella_pneumoniae_complex>Klebsiella_variicola | 120 | LNP26_07885 | 1520756..1521118 | 2590157 | DUF262-domain-containing-protein-[Klebsiella-variicola-subsp.-variicola] |
| WP_211777930.1 | ParB+HNH | GCF_018223925.1 | Klebsiella_sp._10 | Bacteria>Pseudomonadati>Pseudomonadota>Gammaproteobacteria>Enterobacterales>Enterobacteriaceae>Klebsiella/Raoultella_group>Klebsiella>unclassified_Klebsiella | 599 | KHQ26_RS17855 | 10299..12098 | 2822434 | DUF262-domain-containing-protein-[Klebsiella-sp.-10] |
| URQ56459.1 | ParB+HNH | - | Raoultella_ornithinolytica | Bacteria>Pseudomonadati>Pseudomonadota>Gammaproteobacteria>Enterobacterales>Enterobacteriaceae>Klebsiella/Raoultella_group>Raoultella | 587 | . | 83530..85293 | 54291 | Hypothetical-protein-[Raoultella-ornithinolytica] |
| EJG2380287.1 | ParB+HNH | GCA_016088515.2 | Raoultella_ornithinolytica | Bacteria>Pseudomonadati>Pseudomonadota>Gammaproteobacteria>Enterobacterales>Enterobacteriaceae>Klebsiella/Raoultella_group>Raoultella | 584 | AM381_RS08840 | 23277..25031 | 54291 | DUF262-domain-containing-protein-[Raoultella-ornithinolytica] |
| HAV2047148.1 | ParB+HNH | GCA_016445365.1 | Raoultella_ornithinolytica | Bacteria>Pseudomonadati>Pseudomonadota>Gammaproteobacteria>Enterobacterales>Enterobacteriaceae>Klebsiella/Raoultella_group>Raoultella | 584 | JHJ87_003573 | 40055..41809 | 54291 | DUF262-domain-containing-protein-[Raoultella-ornithinolytica] |
| WP_104896652.1 | ParB+HNH | GCF_021699015.1 | Raoultella_ornithinolytica | Bacteria>Pseudomonadati>Pseudomonadota>Gammaproteobacteria>Enterobacterales>Enterobacteriaceae>Klebsiella/Raoultella_group>Raoultella | 584 | INS59_RS15570 | 204139..205893 | 54291 | DUF262-domain-containing-protein-[Raoultella-ornithinolytica] |
| HAT1572919.1 | ParB+HNH | GCA_015892965.1 | Kluyvera_cryocrescens | Bacteria>Pseudomonadati>Pseudomonadota>Gammaproteobacteria>Enterobacterales>Enterobacteriaceae>Kluyvera | 587 | I8Y10_004332 | 45437..47200 | 580 | DUF262-domain-containing-protein-[Kluyvera-cryocrescens] |
| QXB20654.1 | ParB+HNH | GCA_019048185.1 | Lelliottia_amnigena | Bacteria>Pseudomonadati>Pseudomonadota>Gammaproteobacteria>Enterobacterales>Enterobacteriaceae>Lelliottia | 584 | I6L76_15760 | 3246116..3247870 | 61646 | DUF262-domain-containing-protein-[Lelliottia-amnigena] |
| WP_216980737.1 | ParB+HNH | GCF_019048185.1 | Lelliottia_amnigena | Bacteria>Pseudomonadati>Pseudomonadota>Gammaproteobacteria>Enterobacterales>Enterobacteriaceae>Lelliottia | 584 | I6L76_RS15760 | 3246116..3247870 | 61646 | DUF262-domain-containing-protein-[Lelliottia-amnigena] |
| WP_265846294.1 | ParB+HNH | GCF_026240155.1 | Plesiomonas_shigelloides | Bacteria>Pseudomonadati>Pseudomonadota>Gammaproteobacteria>Enterobacterales>Enterobacteriaceae>Plesiomonas | 583 | OQ486_RS15175 | 43545..45296 | 703 | DUF262-domain-containing-protein-[Plesiomonas-shigelloides] |
| EBT6907984.1 | ParB+HNH | GCA_007067565.1 | Salmonella_enterica | Bacteria>Pseudomonadati>Pseudomonadota>Gammaproteobacteria>Enterobacterales>Enterobacteriaceae>Salmonella | 785 | CLA26_04475 | 179034..181391 | 28901 | DUF262-domain-containing-protein-[Salmonella-enterica] |
| EDX0369818.1 | ParB+HNH | - | Salmonella_enterica | Bacteria>Pseudomonadati>Pseudomonadota>Gammaproteobacteria>Enterobacterales>Enterobacteriaceae>Salmonella | 785 | GTL21_000376 | 417534..419891 | 28901 | DUF262-domain-containing-protein-[Salmonella-enterica] |
| WP_045341600.1 | ParB+HNH | GCF_014518275.1 | Salmonella_enterica | Bacteria>Pseudomonadati>Pseudomonadota>Gammaproteobacteria>Enterobacterales>Enterobacteriaceae>Salmonella | 785 | IBX74_RS21050 | 332951..335308 | 28901 | DUF262-domain-containing-HNH-endonuclease-family-protein-[Salmonella-enterica] |
| EJV9378860.1 | ParB+HNH | GCA_025367915.1 | Salmonella_enterica | Bacteria>Pseudomonadati>Pseudomonadota>Gammaproteobacteria>Enterobacterales>Enterobacteriaceae>Salmonella | 587 | N7887_004538 | 6617..8380 | 28901 | DUF262-domain-containing-protein-[Salmonella-enterica] |
| AXD95437.1 | ParB+HNH | GCA_003325015.1 | Salmonella_enterica | Bacteria>Pseudomonadati>Pseudomonadota>Gammaproteobacteria>Enterobacterales>Enterobacteriaceae>Salmonella | 584 | CHC75_12740 | 2555942..2557696 | 28901 | DUF262-domain-containing-protein-[Salmonella-enterica] |
| EAU0181516.1 | ParB+HNH | GCA_005756755.1 | Salmonella_enterica | Bacteria>Pseudomonadati>Pseudomonadota>Gammaproteobacteria>Enterobacterales>Enterobacteriaceae>Salmonella | 584 | A4I02_04470 | 215300..217054 | 28901 | DUF262-domain-containing-protein-[Salmonella-enterica] |
| EAY5330994.1 | ParB+HNH | GCA_006011505.1 | Salmonella_enterica | Bacteria>Pseudomonadati>Pseudomonadota>Gammaproteobacteria>Enterobacterales>Enterobacteriaceae>Salmonella | 584 | LS33_08705 | 7876..9630 | 28901 | DUF262-domain-containing-protein-[Salmonella-enterica] |
| EAY8201056.1 | ParB+HNH | GCA_006024175.1 | Salmonella_enterica | Bacteria>Pseudomonadati>Pseudomonadota>Gammaproteobacteria>Enterobacterales>Enterobacteriaceae>Salmonella | 584 | ATQ84_14880 | 14778..16532 | 28901 | DUF262-domain-containing-protein-[Salmonella-enterica] |
| EBF3214240.1 | ParB+HNH | GCA_006336125.1 | Salmonella_enterica | Bacteria>Pseudomonadati>Pseudomonadota>Gammaproteobacteria>Enterobacterales>Enterobacteriaceae>Salmonella | 584 | FHK34_03270 | 186840..188594 | 28901 | DUF262-domain-containing-protein-[Salmonella-enterica] |
| EBS7807127.1 | ParB+HNH | GCA_007023485.1 | Salmonella_enterica | Bacteria>Pseudomonadati>Pseudomonadota>Gammaproteobacteria>Enterobacterales>Enterobacteriaceae>Salmonella | 584 | CD664_12545 | 47640..49394 | 28901 | DUF262-domain-containing-protein-[Salmonella-enterica] |

| Protein-ID | Domain | Assembly-ID | Strain | Taxonomy | Length | Locus-Tag | Genomic-coordinates | TaxID | Fasta-defline |
|---|---|---|---|---|---|---|---|---|---|
| ECK3339582.1 | ParB+HNH | GCA_007873945.1 | Salmonella_enterica | Bacteria>Pseudomonadati>Pseudomonadota>Gammaproteobacteria>Enterobacterales>Enterobacteriaceae>Salmonella | 584 | FRH68_09855 | 224078..225832 | 28901 | DUF262-domain-containing-protein-[Salmonella-enterica] |
| EHE5127203.1 | ParB+HNH | - | Salmonella_enterica | Bacteria>Pseudomonadati>Pseudomonadota>Gammaproteobacteria>Enterobacterales>Enterobacteriaceae>Salmonella | 584 | JMY69_002121 | 185452..187206 | 28901 | DUF262-domain-containing-protein-[Salmonella-enterica] |
| EIB7574747.1 | ParB+HNH | GCA_019905755.1 | Salmonella_enterica | Bacteria>Pseudomonadati>Pseudomonadota>Gammaproteobacteria>Enterobacterales>Enterobacteriaceae>Salmonella | 584 | K9I12_000374 | 410174..411928 | 28901 | DUF262-domain-containing-protein-[Salmonella-enterica] |
| WP_274894926.1 | ParB+HNH | GCF_028892745.1 | Salmonella_enterica | Bacteria>Pseudomonadati>Pseudomonadota>Gammaproteobacteria>Enterobacterales>Enterobacteriaceae>Salmonella | 584 | PQQ04_RS20385 | 4152352..4154106 | 28901 | DUF262-domain-containing-protein-[Salmonella-enterica] |
| ECJ2539586.1 | ParB+HNH | GCA_007808255.1 | Salmonella_enterica_subsp._salamae | Bacteria>Pseudomonadati>Pseudomonadota>Gammaproteobacteria>Enterobacterales>Enterobacteriaceae>Salmonella>Salmonella_enterica | 599 | FNI64_03980 | 379637..381436 | 59202 | DUF262-domain-containing-protein-[Salmonella-enterica-subsp.-salamae] |
| EAA4435939.1 | ParB+HNH | GCA_004179155.1 | Salmonella_enterica_subsp._salamae | Bacteria>Pseudomonadati>Pseudomonadota>Gammaproteobacteria>Enterobacterales>Enterobacteriaceae>Salmonella>Salmonella_enterica | 584 | DPE44_05510 | 75174..76928 | 59202 | DUF262-domain-containing-protein-[Salmonella-enterica-subsp.-salamae] |
| EEC5291037.1 | ParB+HNH | - | Salmonella_enterica_subsp._enterica | Bacteria>Pseudomonadati>Pseudomonadota>Gammaproteobacteria>Enterobacterales>Enterobacteriaceae>Salmonella>Salmonella_enterica | 785 | ZD54_001381 | 6469..8826 | 59201 | DUF262-domain-containing-protein-[Salmonella-enterica-subsp.-enterica] |
| EBZ8208147.1 | ParB+HNH | GCA_007351255.1 | Salmonella_enterica_subsp._enterica_serovar_Zanzibar | Bacteria>Pseudomonadati>Pseudomonadota>Gammaproteobacteria>Enterobacterales>Enterobacteriaceae>Salmonella>Salmonella_enterica>Salmonella_enterica_subsp._enterica | 584 | EGK94_05905 | 414965..416719 | 399588 | DUF262-domain-containing-protein-[Salmonella-enterica-subsp.-enterica-serovar-Zanzibar] |
| EBW8181805.1 | ParB+HNH | GCA_007227235.1 | Salmonella_enterica_subsp._enterica_serovar_Uganda | Bacteria>Pseudomonadati>Pseudomonadota>Gammaproteobacteria>Enterobacterales>Enterobacteriaceae>Salmonella>Salmonella_enterica>Salmonella_enterica_subsp._enterica | 584 | AR874_12570 | 409994..411748 | 487004 | DUF262-domain-containing-protein-[Salmonella-enterica-subsp.-enterica-serovar-Uganda] |
| EDB2044588.1 | ParB+HNH | - | Salmonella_enterica_subsp._enterica_serovar_Uganda | Bacteria>Pseudomonadati>Pseudomonadota>Gammaproteobacteria>Enterobacterales>Enterobacteriaceae>Salmonella>Salmonella_enterica>Salmonella_enterica_subsp._enterica | 584 | F9P84_01855 | 409787..411541 | 487004 | DUF262-domain-containing-protein-[Salmonella-enterica-subsp.-enterica] |
| ECD4420809.1 | ParB+HNH | GCA_007522325.1 | Salmonella_enterica_subsp._enterica_serovar_Berkeley | Bacteria>Pseudomonadati>Pseudomonadota>Gammaproteobacteria>Enterobacterales>Enterobacteriaceae>Salmonella>Salmonella_enterica>Salmonella_enterica_subsp._enterica | 584 | E0910_10435 | 11918..13672 | 1965103 | DUF262-domain-containing-protein-[Salmonella-enterica-subsp.-enterica-serovar-Berkeley] |
| APV85543.1 | ParB+HNH | GCA_000487155.2 | Salmonella_enterica_subsp._enterica_serovar_Inverness_str._ATCC_10720 | Bacteria>Pseudomonadati>Pseudomonadota>Gammaproteobacteria>Enterobacterales>Enterobacteriaceae>Salmonella>Salmonella_enterica>Salmonella_enterica_subsp._enterica>Salmonella_enterica_subsp._enterica_serovar_Inverness | 584 | SEEI0720_021520 | 4186700..4188454 | 941187 | hypothetical-protein-SEEI0720_021520-[Salmonella-enterica-subsp.-enterica-serovar-Inverness-str.-ATCC-10720] |
| EHC84521.1 | ParB+HNH | - | Salmonella_enterica_subsp._enterica_serovar_Uganda_str._R8-3404 | Bacteria>Pseudomonadati>Pseudomonadota>Gammaproteobacteria>Enterobacterales>Enterobacteriaceae>Salmonella>Salmonella_enterica>Salmonella_enterica_subsp._enterica>Salmonella_enterica_subsp._enterica_serovar_Uganda | 584 | LTSEUGA_5993 | 9012..10766 | 913083 | hypothetical-protein-LTSEUGA_5993-[Salmonella-enterica-subsp.-enterica-serovar-Uganda-str.-R8-3404] |
| MRS92358.1 | ParB+HNH | GCA_009668015.1 | Enterobacteriaceae_bacterium_RIT714 | Bacteria>Pseudomonadati>Pseudomonadota>Gammaproteobacteria>Enterobacterales>Enterobacteriaceae>unclassified_Enterobacteriaceae | 599 | GJV04_20315 | 37606..39405 | 2666193 | DUF262-domain-containing-protein-[Enterobacteriaceae-bacterium-RIT714] |
| UCB31816.1 | ParB+HNH | GCA_020342335.1 | Duffyella_gerundensis | Bacteria>Pseudomonadati>Pseudomonadota>Gammaproteobacteria>Enterobacterales>Erwiniaceae>Duffyella | 584 | J9874_02367 | 2573125..2574879 | 1619313 | hypothetical-protein-J9874_02367-[Duffyella-gerundensis] |
| WP_225700911.1 | ParB+HNH | GCF_020342335.1 | Duffyella_gerundensis | Bacteria>Pseudomonadati>Pseudomonadota>Gammaproteobacteria>Enterobacterales>Erwiniaceae>Duffyella | 584 | J9874_RS11825 | 2573125..2574879 | 1619313 | DUF262-domain-containing-protein-[Duffyella-gerundensis] |
| HCP27300.1 | ParB+HNH | GCA_003533915.1 | Pantoea_ananatis | Bacteria>Pseudomonadati>Pseudomonadota>Gammaproteobacteria>Enterobacterales>Erwiniaceae>Pantoea | 584 | DIT85_12775 | 26597..28351 | 553 | hypothetical-protein-[Pantoea-ananatis] |
| NEG80385.1 | ParB+HNH | GCA_010667945.1 | Pantoea_agglomerans | Bacteria>Pseudomonadati>Pseudomonadota>Gammaproteobacteria>Enterobacterales>Erwiniaceae>Pantoea>Pantoea_agglomerans_group | 584 | GQQ17_05155 | 298543..300297 | 549 | DUF262-domain-containing-protein-[Pantoea-agglomerans] |
| WP_163850526.1 | ParB+HNH | GCF_010667945.1 | Pantoea_agglomerans | Bacteria>Pseudomonadati>Pseudomonadota>Gammaproteobacteria>Enterobacterales>Erwiniaceae>Pantoea>Pantoea_agglomerans_group | 584 | GQQ17_RS05160 | 298543..300297 | 549 | DUF262-domain-containing-protein-[Pantoea-agglomerans] |
| WP_187492465.1 | ParB+HNH | GCF_014353895.1 | Pantoea_agglomerans | Bacteria>Pseudomonadati>Pseudomonadota>Gammaproteobacteria>Enterobacterales>Erwiniaceae>Pantoea>Pantoea_agglomerans_group | 563 | IBU46_RS01730 | 17907..19598 | 549 | DUF262-domain-containing-protein-[Pantoea-agglomerans] |
| NIE78125.1 | ParB+HNH | GCA_011752565.1 | Pantoea_sp._Ap-967 | Bacteria>Pseudomonadati>Pseudomonadota>Gammaproteobacteria>Enterobacterales>Erwiniaceae>Pantoea>unclassified_Pantoea | 624 | F3J45_27235 | 45631..47505 | 2608362 | DUF262-domain-containing-protein-[Pantoea-sp.-Ap-967] |
| WP_108479633.1 | ParB+HNH | GCF_003057415.1 | Providencia | Bacteria>Pseudomonadati>Pseudomonadota>Gammaproteobacteria>Enterobacterales>Morganellaceae | 587 | C6581_RS17860 | 43414..45177 | 586 | MULTISPECIES:-DUF262-domain-containing-protein-[Providencia] |
| MBS5193170.1 | ParB+HNH | GCA_018366325.1 | Morganella_morganii | Bacteria>Pseudomonadati>Pseudomonadota>Gammaproteobacteria>Enterobacterales>Morganellaceae>Morganella | 599 | KHY92_05170 | 13089..14888 | 582 | DUF262-domain-containing-protein-[Morganella-morganii] |
| WP_274765497.1 | ParB+HNH | GCF_028867915.1 | Morganella_morganii | Bacteria>Pseudomonadati>Pseudomonadota>Gammaproteobacteria>Enterobacterales>Morganellaceae>Morganella | 599 | PS169_RS13675 | 166001..167800 | 582 | DUF262-domain-containing-protein-[Morganella-morganii] |
| WP_081119710.1 | ParB+HNH | - | Morganella_morganii | Bacteria>Pseudomonadati>Pseudomonadota>Gammaproteobacteria>Enterobacterales>Morganellaceae>Morganella | 587 | B2H99_RS12755 | 12504..14267 | 582 | DUF262-domain-containing-protein-[Morganella-morganii] |
| WP_214179408.1 | ParB+HNH | GCF_018474625.1 | Morganella_morganii | Bacteria>Pseudomonadati>Pseudomonadota>Gammaproteobacteria>Enterobacterales>Morganellaceae>Morganella | 586 | IS084_RS18055 | 9432..11192 | 582 | DUF262-domain-containing-protein-[Morganella-morganii] |
| AUR32854.1 | ParB+HNH | GCA_002891475.1 | Morganella_morganii | Bacteria>Pseudomonadati>Pseudomonadota>Gammaproteobacteria>Enterobacterales>Morganellaceae>Morganella | 584 | C1O70_15735 | 3346076..3347830 | 582 | DUF262-domain-containing-protein-[Morganella-morganii] |
| WP_032098174.1 | ParB+HNH | GCF_002891475.1 | Morganella_morganii | Bacteria>Pseudomonadati>Pseudomonadota>Gammaproteobacteria>Enterobacterales>Morganellaceae>Morganella | 584 | C1O70_RS15735 | 3346076..3347830 | 582 | DUF262-domain-containing-protein-[Morganella-morganii] |
| HAT2164886.1 | ParB+HNH | GCA_015896365.1 | Proteus_mirabilis | Bacteria>Pseudomonadati>Pseudomonadota>Gammaproteobacteria>Enterobacterales>Morganellaceae>Proteus | 587 | I8052_002911 | 13773..15536 | 584 | DUF262-domain-containing-protein-[Proteus-mirabilis] |
| MBI6417159.1 | ParB+HNH | GCA_016238675.1 | Proteus_mirabilis | Bacteria>Pseudomonadati>Pseudomonadota>Gammaproteobacteria>Enterobacterales>Morganellaceae>Proteus | 587 | JEP37_15650 | 67275..69038 | 584 | DUF262-domain-containing-protein-[Proteus-mirabilis] |
| WP_152118570.1 | ParB+HNH | GCF_009183735.1 | Proteus_mirabilis | Bacteria>Pseudomonadati>Pseudomonadota>Gammaproteobacteria>Enterobacterales>Morganellaceae>Proteus | 587 | GBN13_RS03940 | 72761..74524 | 584 | DUF262-domain-containing-protein-[Proteus-mirabilis] |
| WP_210849184.1 | ParB+HNH | GCF_022349905.1 | Proteus_mirabilis | Bacteria>Pseudomonadati>Pseudomonadota>Gammaproteobacteria>Enterobacterales>Morganellaceae>Proteus | 587 | MJ636_RS02910 | 45610..47373 | 584 | DUF262-domain-containing-protein-[Proteus-mirabilis] |
| WP_261598776.1 | ParB+HNH | GCF_025399735.1 | Proteus_mirabilis | Bacteria>Pseudomonadati>Pseudomonadota>Gammaproteobacteria>Enterobacterales>Morganellaceae>Proteus | 587 | NMC40_RS18455 | 13817..15580 | 584 | DUF262-domain-containing-protein-[Proteus-mirabilis] |
| UVX22919.1 | ParB+HNH | - | Proteus_terrae_subsp._cibarius | Bacteria>Pseudomonadati>Pseudomonadota>Gammaproteobacteria>Enterobacterales>Morganellaceae>Proteus>Proteus_terrae | 536 | . | 70811..72421 | 626774 | hypothetical-protein-[Proteus-terrae-subsp.-cibarius] |
| EJD6671301.1 | ParB+HNH | GCA_020680725.2 | Providencia_rettgeri | Bacteria>Pseudomonadati>Pseudomonadota>Gammaproteobacteria>Enterobacterales>Morganellaceae>Providencia | 587 | MOK80_RS10545 | 477493..479256 | 587 | DUF262-domain-containing-protein-[Providencia-rettgeri] |
| MCK9997622.1 | ParB+HNH | GCA_022387885.2 | Providencia_rettgeri | Bacteria>Pseudomonadati>Pseudomonadota>Gammaproteobacteria>Enterobacterales>Morganellaceae>Providencia | 587 | MKT39_000565 | 22976..24739 | 587 | DUF262-domain-containing-protein-[Providencia-rettgeri] |
| MCL0006617.1 | ParB+HNH | GCA_022388475.2 | Providencia_rettgeri | Bacteria>Pseudomonadati>Pseudomonadota>Gammaproteobacteria>Enterobacterales>Morganellaceae>Providencia | 587 | MKT60_007085 | 62630..64393 | 587 | DUF262-domain-containing-protein-[Providencia-rettgeri] |
| MCL0017527.1 | ParB+HNH | GCA_022387615.2 | Providencia_rettgeri | Bacteria>Pseudomonadati>Pseudomonadota>Gammaproteobacteria>Enterobacterales>Morganellaceae>Providencia | 587 | MKT11_000355 | 76304..78067 | 587 | DUF262-domain-containing-protein-[Providencia-rettgeri] |
| OBY34181.1 | ParB+HNH | GCA_001680125.1 | Providencia_rettgeri | Bacteria>Pseudomonadati>Pseudomonadota>Gammaproteobacteria>Enterobacterales>Morganellaceae>Providencia | 587 | PR729_03905 | 48630..50393 | 587 | hypothetical-protein-PR729_03905-[Providencia-rettgeri] |
| WP_272668878.1 | ParB+HNH | GCF_028481005.1 | Providencia_sp._PROV151 | Bacteria>Pseudomonadati>Pseudomonadota>Gammaproteobacteria>Enterobacterales>Morganellaceae>Providencia>unclassified_Providencia | 586 | PSE52_RS00995 | 224397..226157 | 2949861 | DUF262-domain-containing-protein-[Providencia-sp.-PROV151] |
| WP_272687950.1 | ParB+HNH | GCF_028484835.1 | Providencia_sp._PROV035 | Bacteria>Pseudomonadati>Pseudomonadota>Gammaproteobacteria>Enterobacterales>Morganellaceae>Providencia>unclassified_Providencia | 587 | PSF71_RS08205 | 8721..10484 | 2949766 | DUF262-domain-containing-protein-[Providencia-sp.-PROV035] |
| WP_269723553.1 | ParB+HNH | GCF_027362595.1 | Providencia_sp._21OH12SH02B-Prov | Bacteria>Pseudomonadati>Pseudomonadota>Gammaproteobacteria>Enterobacterales>Morganellaceae>Providencia>unclassified_Providencia | 586 | O7C57_RS02365 | 498953..500713 | 3015951 | DUF262-domain-containing-protein-[Providencia-sp.-21OH12SH02B-Prov] |
| WP_211024714.1 | ParB+HNH | GCF_018094745.1 | Pectobacterium_versatile | Bacteria>Pseudomonadati>Pseudomonadota>Gammaproteobacteria>Enterobacterales>Pectobacteriaceae>Pectobacterium | 599 | F9U42_RS09285 | 84663..86462 | 2488639 | DUF262-domain-containing-protein-[Pectobacterium-versatile] |
| MCL6350166.1 | ParB+HNH | GCA_023507955.1 | Pectobacterium_polaris | Bacteria>Pseudomonadati>Pseudomonadota>Gammaproteobacteria>Enterobacterales>Pectobacteriaceae>Pectobacterium | 599 | EXT50_03105 | 77555..79354 | 2042057 | DUF262-domain-containing-protein-[Pectobacterium-polaris] |
| GKV76956.1 | ParB+HNH | GCA_024342785.1 | Pectobacterium_carotovorum_subsp._carotovorum | Bacteria>Pseudomonadati>Pseudomonadota>Gammaproteobacteria>Enterobacterales>Pectobacteriaceae>Pectobacterium>Pectobacterium_carotovorum | 599 | PEC106568_21300 | 191170..192969 | 555 | hypothetical-protein-PEC106568_21300-[Pectobacterium-carotovorum-subsp.-carotovorum] |
| WP_050135884.1 | ParB+HNH | GCF_025758435.1 | Yersinia | Bacteria>Pseudomonadati>Pseudomonadota>Gammaproteobacteria>Enterobacterales>Yersiniaceae | 587 | N4W04_RS19105 | 4128583..4130346 | 629 | MULTISPECIES:-DUF262-domain-containing-protein-[Yersinia] |
| WP_232770160.1 | ParB+HNH | GCF_021276285.1 | Rahnella_victoriana | Bacteria>Pseudomonadati>Pseudomonadota>Gammaproteobacteria>Enterobacterales>Yersiniaceae>Rahnella | 563 | J9880_RS04865 | 1046658..1048349 | 1510570 | DUF262-domain-containing-protein-[Rahnella-victoriana] |
| WP_217147152.1 | ParB+HNH | GCF_019049515.1 | Rahnella_aceris | Bacteria>Pseudomonadati>Pseudomonadota>Gammaproteobacteria>Enterobacterales>Yersiniaceae>Rahnella | 599 | J1789_RS04110 | 6533..8332 | 2703885 | DUF262-domain-containing-protein-[Rahnella-aceris] |
| WP_223536800.1 | ParB+HNH | GCF_019968705.1 | unclassified_Serratia_(in:_enterobacteria) | Bacteria>Pseudomonadati>Pseudomonadota>Gammaproteobacteria>Enterobacterales>Yersiniaceae>Serratia | 613 | KGP26_RS00395 | 102437..104278 | 2647522 | MULTISPECIES:-DUF262-domain-containing-protein-[unclassified-Serratia-(in:-enterobacteria)] |
| WP_206196475.1 | ParB+HNH | GCF_050037615.1 | Serratia_ureilytica | Bacteria>Pseudomonadati>Pseudomonadota>Gammaproteobacteria>Enterobacterales>Yersiniaceae>Serratia | 599 | JY544_RS23385 | 44457..46256 | 300181 | DUF262-domain-containing-protein-[Serratia-ureilytica] |
| WP_273866016.1 | ParB+HNH | GCF_028616515.1 | Serratia_ureilytica | Bacteria>Pseudomonadati>Pseudomonadota>Gammaproteobacteria>Enterobacterales>Yersiniaceae>Serratia | 599 | PVU68_RS17155 | 223135..224934 | 300181 | DUF262-domain-containing-protein-[Serratia-ureilytica] |
| WP_262254722.1 | ParB+HNH | GCF_946228755.1 | Serratia_proteamaculans | Bacteria>Pseudomonadati>Pseudomonadota>Gammaproteobacteria>Enterobacterales>Yersiniaceae>Serratia | 584 | OC575_RS10150 | 2137451..2139205 | 28151 | DUF262-domain-containing-protein-[Serratia-proteamaculans] |
| EGS9996489.1 | ParB+HNH | - | Serratia_marcescens | Bacteria>Pseudomonadati>Pseudomonadota>Gammaproteobacteria>Enterobacterales>Yersiniaceae>Serratia | 584 | I9Y15_003568 | 182726..184480 | 615 | DUF262-domain-containing-protein-[Serratia-marcescens] |
| HBK4794039.1 | ParB+HNH | GCA_020682225.1 | Serratia_marcescens | Bacteria>Pseudomonadati>Pseudomonadota>Gammaproteobacteria>Enterobacterales>Yersiniaceae>Serratia | 584 | LLT15_004324 | 39380..41134 | 615 | DUF262-domain-containing-protein-[Serratia-marcescens] |
| WP_260965785.1 | ParB+HNH | GCF_025336405.1 | Serratia_marcescens | Bacteria>Pseudomonadati>Pseudomonadota>Gammaproteobacteria>Enterobacterales>Yersiniaceae>Serratia | 584 | N7751_RS05100 | 1085784..1087538 | 615 | DUF262-domain-containing-protein-[Serratia-marcescens] |
| NYA44682.1 | ParB+HNH | GCA_013401715.1 | Serratia_fonticola | Bacteria>Pseudomonadati>Pseudomonadota>Gammaproteobacteria>Enterobacterales>Yersiniaceae>Serratia | 613 | HZI31_15390 | 43611..45452 | 47917 | DUF262-domain-containing-protein-[Serratia-fonticola] |
| WP_179248584.1 | ParB+HNH | GCF_013401715.1 | Serratia_fonticola | Bacteria>Pseudomonadati>Pseudomonadota>Gammaproteobacteria>Enterobacterales>Yersiniaceae>Serratia | 613 | HZI31_RS15390 | 43611..45452 | 47917 | DUF262-domain-containing-protein-[Serratia-fonticola] |
| WP_046051666.1 | ParB+HNH | GCF_000968115.1 | Yersinia_enterocolitica | Bacteria>Pseudomonadati>Pseudomonadota>Gammaproteobacteria>Enterobacterales>Yersiniaceae>Yersinia | 599 | YE5303_RS19620 | 4220568..4222367 | 630 | DUF262-domain-containing-protein-[Yersinia-enterocolitica] |
| RUQ39815.1 | ParB+HNH | GCA_003989075.1 | Candidatus_Competibacteraceae_bacterium | Bacteria>Pseudomonadati>Pseudomonadota>Gammaproteobacteria>Gammaproteobacteria_incertae_sedis>Candidatus_Competibacteraceae>unclassified_Candidatus_Competibacteraceae | 571 | EKK69_07725 | 1009..2724 | 2053538 | DUF262-domain-containing-protein-[Candidatus-Competibacteraceae-bacterium] |
| WP_075675232.1 | ParB+HNH | GCF_001939825.1 | Stenotrophomonas | Bacteria>Pseudomonadati>Pseudomonadota>Gammaproteobacteria>Lysobacterales>Lysobacteraceae | 611 | BKE09_RS06570 | 1388677..1390512 | 40323 | MULTISPECIES:-DUF262-domain-containing-protein-[Stenotrophomonas] |
| WP_135965899.1 | ParB+HNH | GCF_051986355.1 | Stenotrophomonas | Bacteria>Pseudomonadati>Pseudomonadota>Gammaproteobacteria>Lysobacterales>Lysobacteraceae | 611 | N5C11_RS09595 | 43523..45358 | 40323 | MULTISPECIES:-DUF262-domain-containing-protein-[Stenotrophomonas] |
| WP_253077539.1 | ParB+HNH | GCF_024126795.1 | Stenotrophomonas | Bacteria>Pseudomonadati>Pseudomonadota>Gammaproteobacteria>Lysobacterales>Lysobacteraceae | 611 | NJG15_RS04345 | 98312..100147 | 40323 | DUF262-domain-containing-protein-[Stenotrophomonas-maltophilia] |
| WP_049450101.1 | ParB+HNH | GCF_900766255.1 | Stenotrophomonas | Bacteria>Pseudomonadati>Pseudomonadota>Gammaproteobacteria>Lysobacterales>Lysobacteraceae | 602 | BRB91_RS07195 | 64999..66807 | 40323 | MULTISPECIES:-DUF262-domain-containing-protein-[Stenotrophomonas] |
| WP_062607675.1 | ParB+HNH | GCF_033100275.1 | Stenotrophomonas | Bacteria>Pseudomonadati>Pseudomonadota>Gammaproteobacteria>Lysobacterales>Lysobacteraceae | 602 | N5D45_RS14225 | 33901..35709 | 40323 | MULTISPECIES:-DUF262-domain-containing-protein-[Stenotrophomonas] |
| WP_070426726.1 | ParB+HNH | GCF_001806305.1 | Stenotrophomonas | Bacteria>Pseudomonadati>Pseudomonadota>Gammaproteobacteria>Lysobacterales>Lysobacteraceae | 602 | DQN96_RS04560 | 973471..975279 | 40323 | MULTISPECIES:-DUF262-domain-containing-protein-[Stenotrophomonas] |
| WP_144815142.1 | ParB+HNH | GCF_007830115.1 | Aerolutibacter_ruishenii | Bacteria>Pseudomonadati>Pseudomonadota>Gammaproteobacteria>Lysobacterales>Lysobacteraceae>Aerolutibacter | 576 | IP93_RS09865 | 3409..5139 | 686800 | DUF262-domain-containing-protein-[Lysobacter-ruishenii] |
| WP_191823355.1 | ParB+HNH | GCF_014779555.2 | Lysobacter_capsici | Bacteria>Pseudomonadati>Pseudomonadota>Gammaproteobacteria>Lysobacterales>Lysobacteraceae>Lysobacter | 611 | IEQ11_RS15885 | 3914964..3916799 | 435897 | DUF262-domain-containing-protein-[Lysobacter-capsici] |
| WP_271250639.1 | ParB+HNH | GCF_027941875.1 | Pseudoxanthomonas_mexicana | Bacteria>Pseudomonadati>Pseudomonadota>Gammaproteobacteria>Lysobacterales>Lysobacteraceae>Pseudoxanthomonas | 611 | PE064_RS03365 | 684564..686399 | 128785 | DUF262-domain-containing-protein-[Pseudoxanthomonas-mexicana] |
| WP_162376313.1 | ParB+HNH | GCF_042675825.1 | Pseudoxanthomonas_kaohsiungensis | Bacteria>Pseudomonadati>Pseudomonadota>Gammaproteobacteria>Lysobacterales>Lysobacteraceae>Pseudoxanthomonas | 607 | CSC66_RS05110 | 186199..188022 | 283923 | DUF262-domain-containing-protein-[Pseudoxanthomonas-kaohsiungensis] |
| MBH1645060.1 | ParB+HNH | GCA_015997905.1 | Stenotrophomonas_maltophilia | Bacteria>Pseudomonadati>Pseudomonadota>Gammaproteobacteria>Lysobacterales>Lysobacteraceae>Stenotrophomonas>Stenotrophomonas_maltophilia_group | 611 | I5U75_16560 | 12584..14419 | 40324 | DUF262-domain-containing-protein-[Stenotrophomonas-maltophilia] |
| MBH1712519.1 | ParB+HNH | GCA_015998125.1 | Stenotrophomonas_maltophilia | Bacteria>Pseudomonadati>Pseudomonadota>Gammaproteobacteria>Lysobacterales>Lysobacteraceae>Stenotrophomonas>Stenotrophomonas_maltophilia_group | 611 | I5U96_12285 | 13974..15809 | 40324 | DUF262-domain-containing-protein-[Stenotrophomonas-maltophilia] |
| MBN5005311.1 | ParB+HNH | GCA_017149675.1 | Stenotrophomonas_maltophilia | Bacteria>Pseudomonadati>Pseudomonadota>Gammaproteobacteria>Lysobacterales>Lysobacteraceae>Stenotrophomonas>Stenotrophomonas_maltophilia_group | 611 | JY447_14760 | 24459..26294 | 40324 | DUF262-domain-containing-protein-[Stenotrophomonas-maltophilia] |
| PZS99529.1 | ParB+HNH | GCA_003244415.1 | Stenotrophomonas_maltophilia | Bacteria>Pseudomonadati>Pseudomonadota>Gammaproteobacteria>Lysobacterales>Lysobacteraceae>Stenotrophomonas>Stenotrophomonas_maltophilia_group | 611 | A7X66_04350 | 31990..33825 | 40324 | hypothetical-protein-A7X66_04350-[Stenotrophomonas-maltophilia] |
| QNG75239.1 | ParB+HNH | GCA_014236105.1 | Stenotrophomonas_maltophilia | Bacteria>Pseudomonadati>Pseudomonadota>Gammaproteobacteria>Lysobacterales>Lysobacteraceae>Stenotrophomonas>Stenotrophomonas_maltophilia_group | 611 | EIELFIGP_04096 | 4380146..4381981 | 40324 | hypothetical-protein-EIELFIGP_04096-[Stenotrophomonas-maltophilia] |
| WP_049407014.1 | ParB+HNH | GCF_009676585.1 | Stenotrophomonas_maltophilia | Bacteria>Pseudomonadati>Pseudomonadota>Gammaproteobacteria>Lysobacterales>Lysobacteraceae>Stenotrophomonas>Stenotrophomonas_maltophilia_group | 611 | FEO88_RS04760 | 1047838..1049673 | 40324 | DUF262-domain-containing-protein-[Stenotrophomonas-maltophilia] |
| WP_099490587.1 | ParB+HNH | GCF_002739155.1 | Stenotrophomonas_maltophilia | Bacteria>Pseudomonadati>Pseudomonadota>Gammaproteobacteria>Lysobacterales>Lysobacteraceae>Stenotrophomonas>Stenotrophomonas_maltophilia_group | 611 | BRB81_RS09955 | 117740..119575 | 40324 | DUF262-domain-containing-protein-[Stenotrophomonas-maltophilia] |
| MBH1702632.1 | ParB+HNH | GCA_015998095.1 | Stenotrophomonas_maltophilia | Bacteria>Pseudomonadati>Pseudomonadota>Gammaproteobacteria>Lysobacterales>Lysobacteraceae>Stenotrophomonas>Stenotrophomonas_maltophilia_group | 609 | I5U93_05080 | 18918..20747 | 40324 | DUF262-domain-containing-protein-[Stenotrophomonas-maltophilia] |
| MBH1753228.1 | ParB+HNH | GCA_015998385.1 | Stenotrophomonas_maltophilia | Bacteria>Pseudomonadati>Pseudomonadota>Gammaproteobacteria>Lysobacterales>Lysobacteraceae>Stenotrophomonas>Stenotrophomonas_maltophilia_group | 602 | I5V37_01910 | 16595..18403 | 40324 | DUF262-domain-containing-protein-[Stenotrophomonas-maltophilia] |
| MBN5083085.1 | ParB+HNH | GCA_017150495.1 | Stenotrophomonas_maltophilia | Bacteria>Pseudomonadati>Pseudomonadota>Gammaproteobacteria>Lysobacterales>Lysobacteraceae>Stenotrophomonas>Stenotrophomonas_maltophilia_group | 602 | JY467_05225 | 54282..56090 | 40324 | DUF262-domain-containing-protein-[Stenotrophomonas-maltophilia] |
| WP_164145914.1 | ParB+HNH | GCF_010605185.1 | Stenotrophomonas_maltophilia | Bacteria>Pseudomonadati>Pseudomonadota>Gammaproteobacteria>Lysobacterales>Lysobacteraceae>Stenotrophomonas>Stenotrophomonas_maltophilia_group | 602 | G5A34_RS07940 | 91187..92995 | 40324 | DUF262-domain-containing-protein-[Stenotrophomonas-maltophilia] |
| WP_180875345.1 | ParB+HNH | GCF_013463485.1 | Stenotrophomonas_maltophilia | Bacteria>Pseudomonadati>Pseudomonadota>Gammaproteobacteria>Lysobacterales>Lysobacteraceae>Stenotrophomonas>Stenotrophomonas_maltophilia_group | 602 | D7U74_RS02185 | 21325..23133 | 40324 | DUF262-domain-containing-protein-[Stenotrophomonas-maltophilia] |
| WP_262214638.1 | ParB+HNH | - | Stenotrophomonas_maltophilia | Bacteria>Pseudomonadati>Pseudomonadota>Gammaproteobacteria>Lysobacterales>Lysobacteraceae>Stenotrophomonas>Stenotrophomonas_maltophilia_group | 602 | JAK45_RS19260 | 85683..87491 | 40324 | DUF262-domain-containing-protein-[Stenotrophomonas-maltophilia] |
| WP_229766857.1 | ParB+HNH | - | Stenotrophomonas_maltophilia | Bacteria>Pseudomonadati>Pseudomonadota>Gammaproteobacteria>Lysobacterales>Lysobacteraceae>Stenotrophomonas>Stenotrophomonas_maltophilia_group | 575 | ABW45_RS08205 | 24808..26535 | 40324 | DUF262-domain-containing-protein-[Stenotrophomonas-maltophilia] |
| WP_049462968.1 | ParB+HNH | GCF_001075485.1 | Stenotrophomonas_maltophilia | Bacteria>Pseudomonadati>Pseudomonadota>Gammaproteobacteria>Lysobacterales>Lysobacteraceae>Stenotrophomonas>Stenotrophomonas_maltophilia_group | 551 | ADM82_RS13490 | 38480..40135 | 40324 | DUF262-domain-containing-protein-[Stenotrophomonas-maltophilia] |
| WP_164085132.1 | ParB+HNH | GCF_010605045.1 | Stenotrophomonas_maltophilia | Bacteria>Pseudomonadati>Pseudomonadota>Gammaproteobacteria>Lysobacterales>Lysobacteraceae>Stenotrophomonas>Stenotrophomonas_maltophilia_group | 449 | G5A32_RS08670 | 79364..80712 | 40324 | DUF262-domain-containing-protein,-partial-[Stenotrophomonas-maltophilia] |
| WP_225591914.1 | ParB+HNH | - | Stenotrophomonas_sp._STM01 | Bacteria>Pseudomonadati>Pseudomonadota>Gammaproteobacteria>Lysobacterales>Lysobacteraceae>Stenotrophomonas>unclassified_Stenotrophomonas | 599 | IB227_RS14900 | 157154..158953 | 2769278 | DUF262-domain-containing-protein-[Stenotrophomonas-sp.-STM01] |
| WP_125428650.1 | ParB+HNH | GCF_052148605.1 | Stenotrophomonas_sp._RAC2 | Bacteria>Pseudomonadati>Pseudomonadota>Gammaproteobacteria>Lysobacterales>Lysobacteraceae>Stenotrophomonas>unclassified_Stenotrophomonas | 609 | EGJ89_RS09200 | 121820..123649 | 3064902 | DUF262-domain-containing-protein-[Stenotrophomonas-maltophilia] |
| WP_055771170.1 | ParB+HNH | GCF_001422025.1 | Stenotrophomonas_sp._Leaf70 | Bacteria>Pseudomonadati>Pseudomonadota>Gammaproteobacteria>Lysobacterales>Lysobacteraceae>Stenotrophomonas>unclassified_Stenotrophomonas | 576 | ASF01_RS13965 | 57762..59492 | 1736233 | DUF262-domain-containing-protein-[Stenotrophomonas-sp.-Leaf70] |
| WP_271679104.1 | ParB+HNH | GCF_028201335.1 | Thermomonas_mangrovi | Bacteria>Pseudomonadati>Pseudomonadota>Gammaproteobacteria>Lysobacterales>Lysobacteraceae>Thermomonas | 573 | ON759_RS04125 | 94788..96509 | 2993316 | DUF262-domain-containing-protein-[Thermomonas-sp.-2C3345] |
| PJK14601.1 | ParB+HNH | GCA_002798295.1 | Lysobacteraceae_bacterium_NML07-0707 | Bacteria>Pseudomonadati>Pseudomonadota>Gammaproteobacteria>Lysobacterales>Lysobacteraceae>unclassified_Lysobacteraceae | 566 | CO613_05140 | 87373..89073 | 2032574 | hypothetical-protein-CO613_05140-[Xanthomonadaceae-bacterium-NML07-0707] |
| MBU6247857.1 | ParB+HNH | GCA_018970895.1 | Lysobacteraceae_bacterium | Bacteria>Pseudomonadati>Pseudomonadota>Gammaproteobacteria>Lysobacterales>Lysobacteraceae>unclassified_Lysobacteraceae | 558 | KGN77_08875 | 12182..13858 | 1926873 | DUF262-domain-containing-protein-[Xanthomonadaceae-bacterium] |
| WP_005996543.1 | ParB+HNH | GCF_000192025.1 | Xanthomonas_vesicatoria | Bacteria>Pseudomonadati>Pseudomonadota>Gammaproteobacteria>Lysobacterales>Lysobacteraceae>Xanthomonas | 610 | XVE_RS19990 | 8073..9905 | 56460 | DUF262-domain-containing-protein-[Xanthomonas-vesicatoria] |
| WP_260816967.1 | ParB+HNH | GCF_025263745.1 | Xanthomonas_prunicola | Bacteria>Pseudomonadati>Pseudomonadota>Gammaproteobacteria>Lysobacterales>Lysobacteraceae>Xanthomonas | 610 | MOD45_RS16875 | 3947365..3949197 | 2053930 | DUF262-domain-containing-protein-[Xanthomonas-prunicola] |
| EGD07601.1 | ParB+HNH | - | Xanthomonas_vesicatoria_ATCC_35937 | Bacteria>Pseudomonadati>Pseudomonadota>Gammaproteobacteria>Lysobacterales>Lysobacteraceae>Xanthomonas>Xanthomonas_vesicatoria | 610 | XVE_4191 | 8073..9905 | 925775 | hypothetical-protein-XVE_4191-[Xanthomonas-vesicatoria-ATCC-35937] |
| WP_091239203.1 | ParB+HNH | GCF_900101825.1 | Aquimonas_voraii | Bacteria>Pseudomonadati>Pseudomonadota>Gammaproteobacteria>Lysobacterales>Rhodanobacteraceae>Aquimonas | 572 | BLP83_RS04260 | 983954..985672 | 265719 | DUF262-domain-containing-protein-[Aquimonas-voraii] |
| WP_017462292.1 | ParB+HNH | GCF_000334915.1 | Dyella_ginsengisoli | Bacteria>Pseudomonadati>Pseudomonadota>Gammaproteobacteria>Lysobacterales>Rhodanobacteraceae>Dyella | 610 | C693_RS0110805 | 70115..71947 | 363848 | DUF262-domain-containing-protein-[Dyella-ginsengisoli] |
| WP_017462918.1 | ParB+HNH | GCF_000334915.1 | Dyella_ginsengisoli | Bacteria>Pseudomonadati>Pseudomonadota>Gammaproteobacteria>Lysobacterales>Rhodanobacteraceae>Dyella | 602 | C693_RS0114070 | 47795..49603 | 363848 | DUF262-domain-containing-protein-[Dyella-ginsengisoli] |
| MBK8067121.1 | ParB+HNH | GCA_016713135.1 | Rhodanobacteraceae_bacterium | Bacteria>Pseudomonadati>Pseudomonadota>Gammaproteobacteria>Lysobacterales>Rhodanobacteraceae>unclassified_Rhodanobacteraceae | 614 | IPK27_05705 | 330836..332680 | 2052182 | DUF262-domain-containing-protein-[Rhodanobacteraceae-bacterium] |
| MBK7210547.1 | ParB+HNH | GCA_016707485.1 | Lysobacterales_bacterium | Bacteria>Pseudomonadati>Pseudomonadota>Gammaproteobacteria>Lysobacterales>unclassified_Lysobacterales | 706 | IPH99_09370 | 1017418..1019538 | 2006849 | DUF262-domain-containing-protein-[Xanthomonadales-bacterium] |
| MCU0753614.1 | ParB+HNH | GCA_025458445.1 | Lysobacterales_bacterium | Bacteria>Pseudomonadati>Pseudomonadota>Gammaproteobacteria>Lysobacterales>unclassified_Lysobacterales | 567 | MUE46_00625 | 17352..19055 | 2006849 | DUF262-domain-containing-HNH-endonuclease-family-protein-[Xanthomonadales-bacterium] |
| WP_266024633.1 | ParB+HNH | GCF_026247885.1 | Methylococcus_mesophilus | Bacteria>Pseudomonadati>Pseudomonadota>Gammaproteobacteria>Methylococcales>Methylococcaceae>Methylococcus | 611 | OOT43_RS09310 | 1967545..1969380 | 2993564 | DUF262-domain-containing-HNH-endonuclease-family-protein-[Methylococcus-sp.-16-5] |
| WP_005369528.1 | ParB+HNH | GCF_000214275.2 | Methylomicrobium_album | Bacteria>Pseudomonadati>Pseudomonadota>Gammaproteobacteria>Methylococcales>Methylococcaceae>Methylomicrobium | 639 | METAL_RS03100 | 739917..741836 | 39775 | DUF262-domain-containing-protein-[Methylomicrobium-album] |
| MCK4843009.1 | ParB+HNH | GCA_023134955.1 | Methylococcales_bacterium | Bacteria>Pseudomonadati>Pseudomonadota>Gammaproteobacteria>Methylococcales>unclassified_Methylococcales | 717 | KAT04_14200 | 20340..22493 | 2304002 | DUF262-domain-containing-protein-[Methylococcales-bacterium] |
| WP_011513932.1 | ParB+HNH | GCF_000013905.1 | Psychrobacter | Bacteria>Pseudomonadati>Pseudomonadota>Gammaproteobacteria>Moraxellales>Moraxellaceae | 699 | CEP87_RS02895 | 668652..670751 | 497 | DUF262-and-DUF1524-domain-containing-protein-[Psychrobacter-cryohalolentis] |
| WP_004811064.1 | ParB+HNH | GCF_017132735.1 | Acinetobacter | Bacteria>Pseudomonadati>Pseudomonadota>Gammaproteobacteria>Moraxellales>Moraxellaceae | 647 | JYB86_RS06385 | 1337383..1339326 | 469 | MULTISPECIES:-DUF262-domain-containing-protein-[Acinetobacter] |
| WP_010116437.1 | ParB+HNH | GCF_000214135.1 | Acinetobacter | Bacteria>Pseudomonadati>Pseudomonadota>Gammaproteobacteria>Moraxellales>Moraxellaceae | 647 | NVV26_RS12520 | 14682..16625 | 469 | MULTISPECIES:-DUF262-domain-containing-protein-[Acinetobacter] |
| WP_086393429.1 | ParB+HNH | GCF_015676315.1 | Acinetobacter | Bacteria>Pseudomonadati>Pseudomonadota>Gammaproteobacteria>Moraxellales>Moraxellaceae | 647 | IMO39_RS19280 | 14638..16581 | 469 | MULTISPECIES:-DUF262-domain-containing-protein-[Acinetobacter] |
| WP_005401823.1 | ParB+HNH | GCF_000162055.1 | Acinetobacter | Bacteria>Pseudomonadati>Pseudomonadota>Gammaproteobacteria>Moraxellales>Moraxellaceae | 646 | CF596_RS10755 | 9787..11727 | 469 | MULTISPECIES:-DUF262-domain-containing-protein-[Acinetobacter] |
| WP_058952664.1 | ParB+HNH | GCF_001484935.1 | Acinetobacter | Bacteria>Pseudomonadati>Pseudomonadota>Gammaproteobacteria>Moraxellales>Moraxellaceae | 646 | HHC64_RS10955 | 2266277..2268217 | 469 | MULTISPECIES:-DUF262-domain-containing-protein-[Acinetobacter] |
| WP_169025141.1 | ParB+HNH | GCF_021602685.1 | Acinetobacter | Bacteria>Pseudomonadati>Pseudomonadota>Gammaproteobacteria>Moraxellales>Moraxellaceae | 646 | L6B80_RS11010 | 21852..23792 | 469 | MULTISPECIES:-DUF262-domain-containing-protein-[Acinetobacter] |
| WP_178319336.1 | ParB+HNH | GCF_013390985.1 | Acinetobacter | Bacteria>Pseudomonadati>Pseudomonadota>Gammaproteobacteria>Moraxellales>Moraxellaceae | 646 | HX743_RS11990 | 42596..44536 | 469 | MULTISPECIES:-DUF262-domain-containing-protein-[Acinetobacter] |
| WP_002010861.1 | ParB+HNH | GCF_000516575.2 | Acinetobacter | Bacteria>Pseudomonadati>Pseudomonadota>Gammaproteobacteria>Moraxellales>Moraxellaceae | 645 | UN97_RS12390 | 253688..255625 | 469 | MULTISPECIES:-DUF262-domain-containing-protein-[Acinetobacter] |
| WP_005236277.1 | ParB+HNH | GCF_001077555.2 | Acinetobacter | Bacteria>Pseudomonadati>Pseudomonadota>Gammaproteobacteria>Moraxellales>Moraxellaceae | 645 | MSH04_RS15105 | 27071..29008 | 469 | MULTISPECIES:-DUF262-domain-containing-protein-[Acinetobacter] |
| WP_043972761.1 | ParB+HNH | GCF_000876115.1 | Acinetobacter | Bacteria>Pseudomonadati>Pseudomonadota>Gammaproteobacteria>Moraxellales>Moraxellaceae | 645 | B9Y01_RS02175 | 5180..7117 | 469 | MULTISPECIES:-DUF262-domain-containing-protein-[Acinetobacter] |
| WP_161409487.1 | ParB+HNH | GCF_030329445.1 | Acinetobacter | Bacteria>Pseudomonadati>Pseudomonadota>Gammaproteobacteria>Moraxellales>Moraxellaceae | 645 | GPS61_RS07945 | 60171..62108 | 469 | DUF262-domain-containing-protein-[Acinetobacter-haemolyticus] |
| WP_151794802.1 | ParB+HNH | GCF_008997035.1 | Acinetobacter | Bacteria>Pseudomonadati>Pseudomonadota>Gammaproteobacteria>Moraxellales>Moraxellaceae | 620 | OLN40_RS03920 | 65873..67735 | 469 | MULTISPECIES:-DUF262-domain-containing-protein-[Acinetobacter] |
| WP_108675356.1 | ParB+HNH | GCF_003072585.2 | Acinetobacter | Bacteria>Pseudomonadati>Pseudomonadota>Gammaproteobacteria>Moraxellales>Moraxellaceae | 598 | CO119_RS05580 | 1122602..1124398 | 469 | MULTISPECIES:-DUF262-domain-containing-protein-[Acinetobacter] |
| WP_005021598.1 | ParB+HNH | GCF_000214135.1 | Acinetobacter | Bacteria>Pseudomonadati>Pseudomonadota>Gammaproteobacteria>Moraxellales>Moraxellaceae | 551 | J559_RS13425 | 11628..13283 | 469 | MULTISPECIES:-HNH-endonuclease-signature-motif-containing-protein-[Acinetobacter] |
| WP_031974807.1 | ParB+HNH | GCF_963579595.1 | Acinetobacter | Bacteria>Pseudomonadati>Pseudomonadota>Gammaproteobacteria>Moraxellales>Moraxellaceae | 551 | L3V46_RS22560 | 19577..21232 | 469 | MULTISPECIES:-HNH-endonuclease-signature-motif-containing-protein-[Acinetobacter] |
| WP_104500998.1 | ParB+HNH | GCF_017132735.1 | Acinetobacter | Bacteria>Pseudomonadati>Pseudomonadota>Gammaproteobacteria>Moraxellales>Moraxellaceae | 551 | NI393_RS13275 | 17720..19375 | 469 | MULTISPECIES:-HNH-endonuclease-signature-motif-containing-protein-[Acinetobacter] |
| WP_138001002.1 | ParB+HNH | GCF_008982065.1 | Acinetobacter | Bacteria>Pseudomonadati>Pseudomonadota>Gammaproteobacteria>Moraxellales>Moraxellaceae | 551 | GBR91_RS15670 | 16909..18564 | 469 | MULTISPECIES:-HNH-endonuclease-signature-motif-containing-protein-[Acinetobacter] |
| WP_180045981.1 | ParB+HNH | GCF_030325865.1 | Acinetobacter | Bacteria>Pseudomonadati>Pseudomonadota>Gammaproteobacteria>Moraxellales>Moraxellaceae | 544 | FTU55_RS01860 | 151749..153383 | 469 | HNH-endonuclease-[Acinetobacter-sp.-YH12117] |
| WP_221260383.1 | ParB+HNH | GCF_019702905.1 | Acinetobacter | Bacteria>Pseudomonadati>Pseudomonadota>Gammaproteobacteria>Moraxellales>Moraxellaceae | 544 | MMP64_RS09465 | 137988..139622 | 469 | MULTISPECIES:-HNH-endonuclease-[unclassified-Acinetobacter] |
| WP_265466009.1 | ParB+HNH | GCF_026168575.1 | Acinetobacter | Bacteria>Pseudomonadati>Pseudomonadota>Gammaproteobacteria>Moraxellales>Moraxellaceae | 544 | OKC24_RS16320 | 5733..7367 | 469 | HNH-endonuclease-[Acinetobacter-sp.-BIT-DXN8] |
| WP_236163090.1 | ParB+HNH | GCF_021602685.1 | unclassified_Acinetobacter | Bacteria>Pseudomonadati>Pseudomonadota>Gammaproteobacteria>Moraxellales>Moraxellaceae>Acinetobacter | 544 | L6B80_RS10960 | 6768..8402 | 196816 | MULTISPECIES:-HNH-endonuclease-[unclassified-Acinetobacter] |

| Protein-ID | Domain | Assembly-ID | Strain | Taxonomy | Length | Locus-Tag | Genomic-coordinates | TaxID | Fasta-defline |
|---|---|---|---|---|---|---|---|---|---|
| WP_242052641.1 | ParB+HNH | - | unclassified_Acinetobacter | Bacteria>Pseudomonadati>Pseudomonadota>Gammaproteobacteria>Moraxellales>Moraxellaceae>Acinetobacter | 479 | IMO39_RS19800 | 3651..5090 | 196816 | MULTISPECIES:-HNH-endonuclease-signature-motif-containing-protein-[unclassified-Acinetobacter] |
| WP_153389473.1 | ParB+HNH | GCF_009601085.1 | Acinetobacter_wanghuae | Bacteria>Pseudomonadati>Pseudomonadota>Gammaproteobacteria>Moraxellales>Moraxellaceae>Acinetobacter | 645 | GHJ48_RS13150 | 12423..14360 | 2662362 | DUF262-domain-containing-protein-[Acinetobacter-wanghuae] |
| WP_005236296.1 | ParB+HNH | GCF_020099375.1 | Acinetobacter_variabilis | Bacteria>Pseudomonadati>Pseudomonadota>Gammaproteobacteria>Moraxellales>Moraxellaceae>Acinetobacter | 544 | LA331_RS04695 | 979374..981008 | 70346 | HNH-endonuclease-[Acinetobacter-variabilis] |
| WP_026438054.1 | ParB+HNH | GCF_039716135.1 | Acinetobacter_towneri | Bacteria>Pseudomonadati>Pseudomonadota>Gammaproteobacteria>Moraxellales>Moraxellaceae>Acinetobacter | 551 | H048_RS0104315 | 21764..23419 | 202956 | HNH-endonuclease-signature-motif-containing-protein-[Acinetobacter-sp.-MDS7A] |
| WP_025095056.1 | ParB+HNH | GCF_000633005.1 | Acinetobacter_soli | Bacteria>Pseudomonadati>Pseudomonadota>Gammaproteobacteria>Moraxellales>Moraxellaceae>Acinetobacter | 647 | EG58_RS0106975 | 19155..21098 | 487316 | DUF262-domain-containing-protein-[Acinetobacter-soli] |
| WP_119025215.1 | ParB+HNH | GCF_003569585.1 | Acinetobacter_soli | Bacteria>Pseudomonadati>Pseudomonadota>Gammaproteobacteria>Moraxellales>Moraxellaceae>Acinetobacter | 645 | D5I70_RS01325 | 15687..17624 | 487316 | DUF262-domain-containing-protein-[Acinetobacter-soli] |
| WP_075040291.1 | ParB+HNH | GCF_051486965.1 | Acinetobacter_radioresistens | Bacteria>Pseudomonadati>Pseudomonadota>Gammaproteobacteria>Moraxellales>Moraxellaceae>Acinetobacter | 646 | SA188iCDA_RS03915 | 61602..63542 | 40216 | DUF262-domain-containing-protein-[Acinetobacter-radioresistens] |
| WP_111281974.1 | ParB+HNH | GCF_003258335.1 | Acinetobacter_radioresistens | Bacteria>Pseudomonadati>Pseudomonadota>Gammaproteobacteria>Moraxellales>Moraxellaceae>Acinetobacter | 645 | NVV25_RS10915 | 24365..26302 | 40216 | DUF262-domain-containing-protein-[Acinetobacter-radioresistens] |
| WP_138000995.1 | ParB+HNH | GCF_005519305.1 | Acinetobacter_radioresistens | Bacteria>Pseudomonadati>Pseudomonadota>Gammaproteobacteria>Moraxellales>Moraxellaceae>Acinetobacter | 645 | E3H47_RS14115 | 1266..3203 | 40216 | DUF262-domain-containing-protein-[Acinetobacter-radioresistens] |
| WP_265761592.1 | ParB+HNH | GCF_026229485.1 | Acinetobacter_radioresistens | Bacteria>Pseudomonadati>Pseudomonadota>Gammaproteobacteria>Moraxellales>Moraxellaceae>Acinetobacter | 551 | NVV26_RS12560 | 28606..30261 | 40216 | HNH-endonuclease-signature-motif-containing-protein-[Acinetobacter-radioresistens] |
| WP_262579270.1 | ParB+HNH | GCF_050032335.1 | Acinetobacter_junii | Bacteria>Pseudomonadati>Pseudomonadota>Gammaproteobacteria>Moraxellales>Moraxellaceae>Acinetobacter | 645 | KTH64_RS13000 | 5047..6984 | 40215 | DUF262-domain-containing-HNH-endonuclease-family-protein-[Acinetobacter-junii] |
| USI88237.1 | ParB+HNH | GCA_023822225.1 | Acinetobacter_johnsonii | Bacteria>Pseudomonadati>Pseudomonadota>Gammaproteobacteria>Moraxellales>Moraxellaceae>Acinetobacter | 647 | LZ086_18675 | 17486..19429 | 40214 | DUF262-domain-containing-HNH-endonuclease-family-protein-[Acinetobacter-johnsonii] |
| WP_160255368.1 | ParB+HNH | GCF_009914395.1 | Acinetobacter_indicus | Bacteria>Pseudomonadati>Pseudomonadota>Gammaproteobacteria>Moraxellales>Moraxellaceae>Acinetobacter | 646 | GA646_RS00520 | 94832..96772 | 756892 | DUF262-domain-containing-protein-[Acinetobacter-indicus] |
| WP_241338468.1 | ParB+HNH | GCF_022549435.1 | Acinetobacter_higginsii | Bacteria>Pseudomonadati>Pseudomonadota>Gammaproteobacteria>Moraxellales>Moraxellaceae>Acinetobacter | 645 | MMP64_RS09405 | 113093..115030 | 70347 | DUF262-domain-containing-protein-[Acinetobacter-sp.-ANC-5659] |
| WP_218698251.1 | ParB+HNH | GCF_019279015.1 | Acinetobacter_harbinensis | Bacteria>Pseudomonadati>Pseudomonadota>Gammaproteobacteria>Moraxellales>Moraxellaceae>Acinetobacter | 643 | KZA13_RS08650 | 28745..30676 | 1353941 | DUF262-domain-containing-protein-[Acinetobacter-harbinensis] |
| WP_265466017.1 | ParB+HNH | GCF_026168575.1 | Acinetobacter_entericus | Bacteria>Pseudomonadati>Pseudomonadota>Gammaproteobacteria>Moraxellales>Moraxellaceae>Acinetobacter | 647 | OKC24_RS16375 | 26663..28606 | 2989714 | DUF262-domain-containing-HNH-endonuclease-family-protein-[Acinetobacter-sp.-BIT-DXN8] |
| WP_171530294.1 | ParB+HNH | GCF_013072655.1 | Acinetobacter_defluvii | Bacteria>Pseudomonadati>Pseudomonadota>Gammaproteobacteria>Moraxellales>Moraxellaceae>Acinetobacter | 646 | A7P53_RS01870 | 8893..10833 | 1871111 | DUF262-domain-containing-protein-[Acinetobacter-defluvii] |
| WP_171530301.1 | ParB+HNH | GCF_013072655.1 | Acinetobacter_defluvii | Bacteria>Pseudomonadati>Pseudomonadota>Gammaproteobacteria>Moraxellales>Moraxellaceae>Acinetobacter | 551 | A7P53_RS01910 | 22794..24449 | 1871111 | HNH-endonuclease-signature-motif-containing-protein-[Acinetobacter-defluvii] |
| WP_128202467.1 | ParB+HNH | GCF_016804045.1 | Acinetobacter_calcoaceticus/baumannii_complex | Bacteria>Pseudomonadati>Pseudomonadota>Gammaproteobacteria>Moraxellales>Moraxellaceae>Acinetobacter | 647 | NT399_RS17660 | 5240..7183 | 909768 | MULTISPECIES:-DUF262-domain-containing-protein-[Acinetobacter-calcoaceticus/baumannii-complex] |
| WP_069122564.1 | ParB+HNH | GCF_001716945.1 | Acinetobacter_pittii | Bacteria>Pseudomonadati>Pseudomonadota>Gammaproteobacteria>Moraxellales>Moraxellaceae>Acinetobacter>Acinetobacter_calcoaceticus/baumannii_complex | 647 | AXH23_RS05635 | 254795..256738 | 48296 | DUF262-domain-containing-protein-[Acinetobacter-pittii] |
| WP_099475575.1 | ParB+HNH | - | Acinetobacter_pittii | Bacteria>Pseudomonadati>Pseudomonadota>Gammaproteobacteria>Moraxellales>Moraxellaceae>Acinetobacter>Acinetobacter_calcoaceticus/baumannii_complex | 646 | CT136_RS19275 | 3175..5115 | 48296 | DUF262-domain-containing-protein-[Acinetobacter-pittii] |
| WP_274579358.1 | ParB+HNH | GCF_028774645.1 | Acinetobacter_lactucae | Bacteria>Pseudomonadati>Pseudomonadota>Gammaproteobacteria>Moraxellales>Moraxellaceae>Acinetobacter>Acinetobacter_calcoaceticus/baumannii_complex | 647 | MOO54_RS17985 | 31554..33497 | 1785128 | DUF262-domain-containing-HNH-endonuclease-family-protein-[Acinetobacter-lactucae] |
| WP_174727832.1 | ParB+HNH | GCF_013344765.1 | Acinetobacter_lactucae | Bacteria>Pseudomonadati>Pseudomonadota>Gammaproteobacteria>Moraxellales>Moraxellaceae>Acinetobacter>Acinetobacter_calcoaceticus/baumannii_complex | 645 | HUN22_RS07905 | 9011..10948 | 1785128 | DUF262-domain-containing-protein-[Acinetobacter-lactucae] |
| WP_198080384.1 | ParB+HNH | GCF_016082245.1 | Acinetobacter_calcoaceticus | Bacteria>Pseudomonadati>Pseudomonadota>Gammaproteobacteria>Moraxellales>Moraxellaceae>Acinetobacter>Acinetobacter_calcoaceticus/baumannii_complex | 647 | BL242_RS20740 | 10284..12227 | 471 | DUF262-domain-containing-protein-[Acinetobacter-calcoaceticus] |
| HAV3580161.1 | ParB+HNH | GCA_016485765.1 | Acinetobacter_baumannii | Bacteria>Pseudomonadati>Pseudomonadota>Gammaproteobacteria>Moraxellales>Moraxellaceae>Acinetobacter>Acinetobacter_calcoaceticus/baumannii_complex | 647 | JIE63_001441 | 175325..177268 | 470 | DUF262-domain-containing-protein-[Acinetobacter-baumannii] |
| WP_109426633.1 | ParB+HNH | GCF_013416255.1 | Acinetobacter_baumannii | Bacteria>Pseudomonadati>Pseudomonadota>Gammaproteobacteria>Moraxellales>Moraxellaceae>Acinetobacter>Acinetobacter_calcoaceticus/baumannii_complex | 647 | KPE97_RS15720 | 90622..92565 | 470 | DUF262-domain-containing-protein-[Acinetobacter-baumannii] |
| WP_119047912.1 | ParB+HNH | GCF_003569645.1 | Acinetobacter_baumannii | Bacteria>Pseudomonadati>Pseudomonadota>Gammaproteobacteria>Moraxellales>Moraxellaceae>Acinetobacter>Acinetobacter_calcoaceticus/baumannii_complex | 647 | D5I76_RS05705 | 63792..65735 | 470 | DUF262-domain-containing-protein-[Acinetobacter-baumannii] |
| WP_223129288.1 | ParB+HNH | GCF_019921105.1 | Acinetobacter_baumannii | Bacteria>Pseudomonadati>Pseudomonadota>Gammaproteobacteria>Moraxellales>Moraxellaceae>Acinetobacter>Acinetobacter_calcoaceticus/baumannii_complex | 647 | HLH91_RS14155 | 12528..14471 | 470 | DUF262-domain-containing-protein-[Acinetobacter-baumannii] |
| WP_228414409.1 | ParB+HNH | GCF_020810655.1 | Acinetobacter_baumannii | Bacteria>Pseudomonadati>Pseudomonadota>Gammaproteobacteria>Moraxellales>Moraxellaceae>Acinetobacter>Acinetobacter_calcoaceticus/baumannii_complex | 647 | NLHDIDDJ_RS06555 | 1359343..1361286 | 470 | DUF262-domain-containing-protein-[Acinetobacter-baumannii] |
| WP_257259084.1 | ParB+HNH | GCF_024589115.1 | Acinetobacter_baumannii | Bacteria>Pseudomonadati>Pseudomonadota>Gammaproteobacteria>Moraxellales>Moraxellaceae>Acinetobacter>Acinetobacter_calcoaceticus/baumannii_complex | 647 | L2615_RS07855 | 175325..177268 | 470 | DUF262-domain-containing-HNH-endonuclease-family-protein-[Acinetobacter-baumannii] |
| WP_005120350.1 | ParB+HNH | - | Acinetobacter_baumannii | Bacteria>Pseudomonadati>Pseudomonadota>Gammaproteobacteria>Moraxellales>Moraxellaceae>Acinetobacter>Acinetobacter_calcoaceticus/baumannii_complex | 646 | ACIN5098_RS09910 | 38092..40032 | 470 | DUF262-domain-containing-protein-[Acinetobacter-baumannii] |
| WP_185218778.1 | ParB+HNH | GCF_014218985.1 | Acinetobacter_baumannii | Bacteria>Pseudomonadati>Pseudomonadota>Gammaproteobacteria>Moraxellales>Moraxellaceae>Acinetobacter>Acinetobacter_calcoaceticus/baumannii_complex | 646 | H6W49_RS17745 | 13768..15708 | 470 | DUF262-domain-containing-protein-[Acinetobacter-baumannii] |
| BCZ16229.1 | ParB+HNH | GCA_019703285.1 | Acinetobacter_baumannii | Bacteria>Pseudomonadati>Pseudomonadota>Gammaproteobacteria>Moraxellales>Moraxellaceae>Acinetobacter>Acinetobacter_calcoaceticus/baumannii_complex | 645 | OCUAc18_37690 | 51832..53769 | 470 | hypothetical-protein-OCUAc18_37690-(plasmid)-[Acinetobacter-baumannii] |
| HAV4979220.1 | ParB+HNH | GCA_016507565.1 | Acinetobacter_baumannii | Bacteria>Pseudomonadati>Pseudomonadota>Gammaproteobacteria>Moraxellales>Moraxellaceae>Acinetobacter>Acinetobacter_calcoaceticus/baumannii_complex | 645 | JIZ05_14405 | 18184..20121 | 470 | DUF262-domain-containing-protein-[Acinetobacter-baumannii] |
| HAV6134266.1 | ParB+HNH | GCA_016514395.1 | Acinetobacter_baumannii | Bacteria>Pseudomonadati>Pseudomonadota>Gammaproteobacteria>Moraxellales>Moraxellaceae>Acinetobacter>Acinetobacter_calcoaceticus/baumannii_complex | 645 | JI295_12645 | 20097..22034 | 470 | TPA:-DUF262-domain-containing-protein-[Acinetobacter-baumannii] |
| HCA4923660.1 | ParB+HNH | GCA_022931145.1 | Acinetobacter_baumannii | Bacteria>Pseudomonadati>Pseudomonadota>Gammaproteobacteria>Moraxellales>Moraxellaceae>Acinetobacter>Acinetobacter_calcoaceticus/baumannii_complex | 645 | MW112_001900 | 185360..187297 | 470 | TPA:-DUF262-domain-containing-protein-[Acinetobacter-baumannii] |
| WP_005115114.1 | ParB+HNH | GCF_000828935.1 | Acinetobacter_baumannii | Bacteria>Pseudomonadati>Pseudomonadota>Gammaproteobacteria>Moraxellales>Moraxellaceae>Acinetobacter>Acinetobacter_calcoaceticus/baumannii_complex | 645 | G3N14_RS16460 | 5285..7222 | 470 | DUF262-domain-containing-protein-[Acinetobacter-baumannii] |
| WP_065184481.1 | ParB+HNH | GCF_040983945.1 | Acinetobacter_baumannii | Bacteria>Pseudomonadati>Pseudomonadota>Gammaproteobacteria>Moraxellales>Moraxellaceae>Acinetobacter>Acinetobacter_calcoaceticus/baumannii_complex | 645 | GKP36_RS06460 | 10582..12519 | 470 | DUF262-domain-containing-protein-[Acinetobacter-baumannii] |
| WP_032068277.1 | ParB+HNH | GCF_964300555.1 | Acinetobacter_baumannii | Bacteria>Pseudomonadati>Pseudomonadota>Gammaproteobacteria>Moraxellales>Moraxellaceae>Acinetobacter>Acinetobacter_calcoaceticus/baumannii_complex | 603 | FJU84_RS09075 | 19628..21439 | 470 | DUF262-domain-containing-protein-[Acinetobacter-baumannii] |
| WP_034686571.1 | ParB+HNH | GCF_000581035.1 | Acinetobacter_sp._869535 | Bacteria>Pseudomonadati>Pseudomonadota>Gammaproteobacteria>Moraxellales>Moraxellaceae>Acinetobacter>Acinetobacter_calcoaceticus/baumannii_complex>unclassified_Acinetobacter_calcoaceticus/baumannii_complex | 266 | J520_RS12330 | 11648..12446 | 1310621 | DUF262-domain-containing-protein,-partial-[Acinetobacter-sp.-869535] |
| WP_180045996.1 | ParB+HNH | GCF_013418865.1 | Acinetobacter_sp._YH12117 | Bacteria>Pseudomonadati>Pseudomonadota>Gammaproteobacteria>Moraxellales>Moraxellaceae>Acinetobacter>unclassified_Acinetobacter | 645 | FTU55_RS01910 | 166664..168601 | 2601104 | DUF262-domain-containing-protein-[Acinetobacter-sp.-YH12117] |
| WP_221260376.1 | ParB+HNH | GCF_019702905.1 | Acinetobacter_sp._Tol_5 | Bacteria>Pseudomonadati>Pseudomonadota>Gammaproteobacteria>Moraxellales>Moraxellaceae>Acinetobacter>unclassified_Acinetobacter | 642 | TOL5_RS12605 | 2691311..2693239 | 710648 | DUF262-domain-containing-protein-[Acinetobacter-sp.-Tol-5] |
| WP_233945505.1 | ParB+HNH | GCF_021391555.1 | Acinetobacter_sp._SH20PTE14 | Bacteria>Pseudomonadati>Pseudomonadota>Gammaproteobacteria>Moraxellales>Moraxellaceae>Acinetobacter>unclassified_Acinetobacter | 647 | LXF01_RS17025 | 111094..113037 | 2905879 | DUF262-domain-containing-protein-[Acinetobacter-sp.-SH20PTE14] |
| WP_233945512.1 | ParB+HNH | GCF_021391555.1 | Acinetobacter_sp._SH20PTE14 | Bacteria>Pseudomonadati>Pseudomonadota>Gammaproteobacteria>Moraxellales>Moraxellaceae>Acinetobacter>unclassified_Acinetobacter | 544 | LXF01_RS17075 | 127813..129447 | 2905879 | HNH-endonuclease-[Acinetobacter-sp.-SH20PTE14] |
| WP_019836723.1 | ParB+HNH | GCF_021560715.1 | Acinetobacter_sp._SCLZS86 | Bacteria>Pseudomonadati>Pseudomonadota>Gammaproteobacteria>Moraxellales>Moraxellaceae>Acinetobacter>unclassified_Acinetobacter | 646 | LZP46_RS14895 | 830..2770 | 2908637 | MULTISPECIES:-DUF262-domain-containing-protein-[unclassified-Acinetobacter] |
| WP_236095245.1 | ParB+HNH | GCF_021560715.1 | Acinetobacter_sp._SCLZS86 | Bacteria>Pseudomonadati>Pseudomonadota>Gammaproteobacteria>Moraxellales>Moraxellaceae>Acinetobacter>unclassified_Acinetobacter | 544 | LZP46_RS14945 | 16234..17868 | 2908637 | HNH-endonuclease-[Acinetobacter-sp.-SCLZS86] |
| WP_005303504.1 | ParB+HNH | GCF_000369825.1 | Acinetobacter_sp._NIPH_542 | Bacteria>Pseudomonadati>Pseudomonadota>Gammaproteobacteria>Moraxellales>Moraxellaceae>Acinetobacter>unclassified_Acinetobacter | 645 | F886_RS07840 | 472015..473952 | 1217688 | DUF262-domain-containing-protein-[Acinetobacter-sp.-NIPH-542] |
| WP_005162490.1 | ParB+HNH | GCF_000369425.1 | Acinetobacter_sp._NIPH_284 | Bacteria>Pseudomonadati>Pseudomonadota>Gammaproteobacteria>Moraxellales>Moraxellaceae>Acinetobacter>unclassified_Acinetobacter | 620 | F908_RS12555 | 1393201..1395063 | 1217704 | DUF262-domain-containing-protein-[Acinetobacter-sp.-NIPH-284] |
| WP_252322554.1 | ParB+HNH | - | Acinetobacter_sp._NBRC_110496 | Bacteria>Pseudomonadati>Pseudomonadota>Gammaproteobacteria>Moraxellales>Moraxellaceae>Acinetobacter>unclassified_Acinetobacter | 413 | Aba26S_RS21435 | 1487..2728 | 1550715 | HNH-endonuclease-signature-motif-containing-protein-[Acinetobacter-sp.-NBRC-110496] |
| ENX11169.1 | ParB+HNH | GCA_000369645.1 | Acinetobacter_sp._CIP_64.2 | Bacteria>Pseudomonadati>Pseudomonadota>Gammaproteobacteria>Moraxellales>Moraxellaceae>Acinetobacter>unclassified_Acinetobacter | 645 | F895_03862 | 20210..22147 | 1217694 | hypothetical-protein-F895_03862-[Acinetobacter-sp.-CIP-64.2] |
| WP_131273638.1 | ParB+HNH | GCF_004331195.1 | Acinetobacter_sp._ANC_4216 | Bacteria>Pseudomonadati>Pseudomonadota>Gammaproteobacteria>Moraxellales>Moraxellaceae>Acinetobacter>unclassified_Acinetobacter | 599 | E0H88_RS07235 | 74851..76650 | 2529840 | DUF262-domain-containing-protein-[Acinetobacter-sp.-ANC-4216] |
| WP_104442409.1 | ParB+HNH | GCF_002934965.1 | Acinetobacter_sp._ACNIH2 | Bacteria>Pseudomonadati>Pseudomonadota>Gammaproteobacteria>Moraxellales>Moraxellaceae>Acinetobacter>unclassified_Acinetobacter | 645 | C3F34_RS18680 | 66997..68934 | 1758189 | DUF262-domain-containing-protein-[Acinetobacter-sp.-ACNIH2] |
| WP_146164464.1 | ParB+HNH | GCF_003051055.1 | Agitococcus_lubricus | Bacteria>Pseudomonadati>Pseudomonadota>Gammaproteobacteria>Moraxellales>Moraxellaceae>Agitococcus | 818 | C8N29_RS12230 | 7583..10039 | 1077255 | DUF262-domain-containing-protein-[Agitococcus-lubricus] |
| WP_201579700.1 | ParB+HNH | GCF_904846705.1 | unclassified_Psychrobacter | Bacteria>Pseudomonadati>Pseudomonadota>Gammaproteobacteria>Moraxellales>Moraxellaceae>Psychrobacter | 593 | JMY00_RS07540 | 22723..24504 | 196806 | DUF262-domain-containing-protein-[Psychrobacter-sp.-Pi2-52] |
| WP_201542135.1 | ParB+HNH | GCF_904846515.1 | Psychrobacter_sanguinis | Bacteria>Pseudomonadati>Pseudomonadota>Gammaproteobacteria>Moraxellales>Moraxellaceae>Psychrobacter | 581 | LK453_RS13665 | 3246561..3248306 | 861445 | DUF262-domain-containing-protein-[Psychrobacter-sanguinis] |
| WP_198334110.1 | ParB+HNH | GCF_904846355.1 | Psychrobacter_namhaensis | Bacteria>Pseudomonadati>Pseudomonadota>Gammaproteobacteria>Moraxellales>Moraxellaceae>Psychrobacter | 582 | JMX36_RS06950 | 25197..26945 | 292734 | DUF262-domain-containing-protein-[Psychrobacter-namhaensis] |
| WP_201558715.1 | ParB+HNH | - | Psychrobacter_immobilis | Bacteria>Pseudomonadati>Pseudomonadota>Gammaproteobacteria>Moraxellales>Moraxellaceae>Psychrobacter | 698 | JMW74_RS07710 | 15571..17667 | 498 | DUF262-domain-containing-protein-[Psychrobacter-immobilis] |
| WP_201625910.1 | ParB+HNH | GCF_904846275.1 | Psychrobacter_immobilis | Bacteria>Pseudomonadati>Pseudomonadota>Gammaproteobacteria>Moraxellales>Moraxellaceae>Psychrobacter | 598 | JMX31_RS11860 | 6141..7937 | 498 | DUF262-domain-containing-protein-[Psychrobacter-immobilis] |
| WP_191691207.1 | ParB+HNH | GCF_014836505.1 | Psychrobacter_communis | Bacteria>Pseudomonadati>Pseudomonadota>Gammaproteobacteria>Moraxellales>Moraxellaceae>Psychrobacter | 843 | H9653_RS06025 | 34329..36860 | 2762238 | DUF262-domain-containing-protein-[Psychrobacter-communis] |
| WP_191691205.1 | ParB+HNH | GCF_014836505.1 | Psychrobacter_communis | Bacteria>Pseudomonadati>Pseudomonadota>Gammaproteobacteria>Moraxellales>Moraxellaceae>Psychrobacter | 614 | H9653_RS06020 | 32485..34329 | 2762238 | DUF262-domain-containing-protein-[Psychrobacter-communis] |
| WP_207991850.1 | ParB+HNH | GCF_017498085.1 | Psychrobacter_coccoides | Bacteria>Pseudomonadati>Pseudomonadota>Gammaproteobacteria>Moraxellales>Moraxellaceae>Psychrobacter | 694 | J3492_RS09690 | 56003..58087 | 2818440 | DUF262-domain-containing-protein-[Psychrobacter-coccoides] |
| ABE75383.1 | ParB+HNH | GCA_000013905.1 | Psychrobacter_cryohalolentis_K5 | Bacteria>Pseudomonadati>Pseudomonadota>Gammaproteobacteria>Moraxellales>Moraxellaceae>Psychrobacter>Psychrobacter_cryohalolentis | 699 | Pcryo_1606 | 1971228..1973327 | 335284 | protein-of-unknown-function-DUF1524-RloF-[Psychrobacter-cryohalolentis-K5] |
| MCG3880116.1 | ParB+HNH | GCA_022129235.1 | Psychrobacter_sp._Ps6 | Bacteria>Pseudomonadati>Pseudomonadota>Gammaproteobacteria>Moraxellales>Moraxellaceae>Psychrobacter>unclassified_Psychrobacter | 599 | I3255_11475 | 59659..61458 | 2790960 | DUF262-domain-containing-protein-[Psychrobacter-sp.-Ps6] |
| WP_238038937.1 | ParB+HNH | GCF_022129235.1 | Psychrobacter_sp._Ps6 | Bacteria>Pseudomonadati>Pseudomonadota>Gammaproteobacteria>Moraxellales>Moraxellaceae>Psychrobacter>unclassified_Psychrobacter | 599 | I3255_RS11475 | 59659..61458 | 2790960 | DUF262-domain-containing-protein-[Psychrobacter-sp.-Ps6] |
| WP_194364107.1 | ParB+HNH | GCF_015277735.1 | Psychrobacter_sp._NG254 | Bacteria>Pseudomonadati>Pseudomonadota>Gammaproteobacteria>Moraxellales>Moraxellaceae>Psychrobacter>unclassified_Psychrobacter | 680 | IP510_RS07335 | 472275..474317 | 2782003 | DUF262-domain-containing-protein-[Psychrobacter-sp.-NG254] |
| WP_194364106.1 | ParB+HNH | GCF_015277735.1 | Psychrobacter_sp._NG254 | Bacteria>Pseudomonadati>Pseudomonadota>Gammaproteobacteria>Moraxellales>Moraxellaceae>Psychrobacter>unclassified_Psychrobacter | 498 | IP510_RS07330 | 470761..472257 | 2782003 | DUF262-domain-containing-protein-[Psychrobacter-sp.-NG254] |
| WP_105245347.1 | ParB+HNH | GCF_900291405.1 | Psychrobacter_sp._Marseille-P5312 | Bacteria>Pseudomonadati>Pseudomonadota>Gammaproteobacteria>Moraxellales>Moraxellaceae>Psychrobacter>unclassified_Psychrobacter | 599 | C6W44_RS10145 | 2352659..2354458 | 2086574 | DUF262-domain-containing-protein-[Psychrobacter-sp.-Marseille-P5312] |
| WP_261869383.1 | ParB+HNH | GCF_000586435.1 | Psychrobacter_sp._JCM_18901 | Bacteria>Pseudomonadati>Pseudomonadota>Gammaproteobacteria>Moraxellales>Moraxellaceae>Psychrobacter>unclassified_Psychrobacter | 386 | TT11_RS05315 | 14905..16065 | 1298609 | DUF262-domain-containing-protein-[Psychrobacter-sp.-JCM-18901] |
| WP_089554754.1 | ParB+HNH | GCF_002224105.1 | Psychrobacter_sp._DAB_AL32B | Bacteria>Pseudomonadati>Pseudomonadota>Gammaproteobacteria>Moraxellales>Moraxellaceae>Psychrobacter>unclassified_Psychrobacter | 593 | CAN34_RS05285 | 11741..13522 | 1028414 | DUF262-domain-containing-protein-[Psychrobacter-sp.-DAB_AL32B] |
| WP_228349568.1 | ParB+HNH | GCF_016918765.2 | Flagellatimonas_centrodinii | Bacteria>Pseudomonadati>Pseudomonadota>Gammaproteobacteria>Nevskiales>Nevskiaceae>Flagellatimonas | 577 | JN531_RS14480 | 2996521..2998254 | 2806210 | DUF262-domain-containing-protein-[Flagellatimonas-centrodinii] |
| WP_035558495.1 | ParB+HNH | GCF_963977585.1 | Halomonadaceae | Bacteria>Pseudomonadati>Pseudomonadota>Gammaproteobacteria>Oceanospirillales | 581 | KO116_RS11685 | 2510000..2511745 | 28256 | DUF262-domain-containing-protein-[Halomonas-sp.-KO116] |
| WP_249692552.1 | ParB+HNH | GCF_023509475.1 | Aestuariirhabdus_haliotis | Bacteria>Pseudomonadati>Pseudomonadota>Gammaproteobacteria>Oceanospirillales>Aestuariirhabdaceae>Aestuariirhabdus | 641 | MIB40_RS07405 | 63743..65668 | 2918751 | DUF262-domain-containing-protein-[Aestuariirhabdus-haliotis] |
| WP_153499074.1 | ParB+HNH | GCF_009601165.1 | Alcanivorax_sediminis | Bacteria>Pseudomonadati>Pseudomonadota>Gammaproteobacteria>Oceanospirillales>Alcanivoracaceae>Alcanivorax | 629 | GFN93_RS03815 | 870579..872468 | 2663008 | DUF262-domain-containing-protein-[Alcanivorax-sediminis] |
| WP_161384456.1 | ParB+HNH | - | Alcanivorax_sp._ZXX171 | Bacteria>Pseudomonadati>Pseudomonadota>Gammaproteobacteria>Oceanospirillales>Alcanivoracaceae>Alcanivorax>unclassified_Alcanivorax | 582 | GN155_RS10560 | 121822..123570 | 2695858 | DUF262-domain-containing-protein-[Alcanivorax-sp.-ZXX171] |
| MBA3979211.1 | ParB+HNH | GCA_013823245.1 | Alcanivorax_sp. | Bacteria>Pseudomonadati>Pseudomonadota>Gammaproteobacteria>Oceanospirillales>Alcanivoracaceae>Alcanivorax>unclassified_Alcanivorax | 564 | C0462_01280 | 19613..21307 | 1872427 | hypothetical-protein-[Alcanivorax-sp.] |
| WP_206679220.1 | ParB+HNH | GCF_052071395.1 | Endozoicomonas_acroporae | Bacteria>Pseudomonadati>Pseudomonadota>Gammaproteobacteria>Oceanospirillales>Endozoicomonadaceae>Endozoicomonas | 383 | EO230_RS03440 | 15282..16433 | 1701104 | DUF262-domain-containing-protein-[Endozoicomonas-acroporae] |
| MRI34246.1 | ParB+HNH | GCA_009653635.1 | Endozoicomonas_sp._OPT23 | Bacteria>Pseudomonadati>Pseudomonadota>Gammaproteobacteria>Oceanospirillales>Endozoicomonadaceae>Endozoicomonas>unclassified_Endozoicomonas | 649 | EOPP23_14725 | 154603..156552 | 2072845 | hypothetical-protein-[Endozoicomonas-sp.-OPT23] |
| MCP5206315.1 | ParB+HNH | GCA_024234475.1 | Hahellaceae_bacterium | Bacteria>Pseudomonadati>Pseudomonadota>Gammaproteobacteria>Oceanospirillales>Hahellaceae>unclassified_Hahellaceae | 596 | H7A01_03840 | 869412..871202 | 2048971 | DUF262-domain-containing-protein-[Hahellaceae-bacterium] |
| WP_247619863.1 | ParB+HNH | GCF_025393375.1 | Chromohalobacter | Bacteria>Pseudomonadati>Pseudomonadota>Gammaproteobacteria>Oceanospirillales>Halomonadaceae | 570 | KZO79_RS04225 | 943387..945099 | 42054 | MULTISPECIES:-DUF262-domain-containing-protein-[Chromohalobacter] |
| WP_040239278.1 | ParB+HNH | GCF_000821045.2 | Chromohalobacter_japonicus | Bacteria>Pseudomonadati>Pseudomonadota>Gammaproteobacteria>Oceanospirillales>Halomonadaceae>Chromohalobacter | 563 | BN995_RS00435 | 46919..48610 | 223900 | DUF262-domain-containing-protein-[Chromohalobacter-japonicus] |
| WP_239285450.1 | ParB+HNH | GCF_964204995.1 | unclassified_Halomonas | Bacteria>Pseudomonadati>Pseudomonadota>Gammaproteobacteria>Oceanospirillales>Halomonadaceae>Halomonas | 561 | MHM85_RS05490 | 205349..207034 | 2609666 | MULTISPECIES:-DUF262-domain-containing-protein-[unclassified-Halomonas] |
| WP_161432255.1 | ParB+HNH | GCF_009902005.1 | Halomonas_alimentaria | Bacteria>Pseudomonadati>Pseudomonadota>Gammaproteobacteria>Oceanospirillales>Halomonadaceae>Halomonas | 617 | GRB96_RS11395 | 2391927..2393780 | 147248 | DUF262-domain-containing-protein-[Halomonas-alimentaria] |
| WP_251308258.1 | ParB+HNH | - | Halomonas_sp._NCCP-2165 | Bacteria>Pseudomonadati>Pseudomonadota>Gammaproteobacteria>Oceanospirillales>Halomonadaceae>Halomonas>unclassified_Halomonas | 566 | NFU04_RS04235 | 100099..101799 | 2942298 | DUF262-domain-containing-protein-[Halomonas-sp.-NCCP-2165] |
| WP_263591017.1 | ParB+HNH | GCF_025725685.1 | Halomonas_sp._M4R1S46 | Bacteria>Pseudomonadati>Pseudomonadota>Gammaproteobacteria>Oceanospirillales>Halomonadaceae>Halomonas>unclassified_Halomonas | 561 | OCT48_RS01530 | 315661..317346 | 2982692 | DUF262-domain-containing-protein-[Halomonas-sp.-M4R1S46] |
| WP_231663149.1 | ParB+HNH | GCF_021151495.1 | Halomonas_sp._IOP_31 | Bacteria>Pseudomonadati>Pseudomonadota>Gammaproteobacteria>Oceanospirillales>Halomonadaceae>Halomonas>unclassified_Halomonas | 562 | K7I12_RS04330 | 13690..15378 | 2876584 | DUF262-domain-containing-protein-[Halomonas-sp.-IOP_31] |
| WP_081193135.1 | ParB+HNH | - | Halomonas_sp._BC1 | Bacteria>Pseudomonadati>Pseudomonadota>Gammaproteobacteria>Oceanospirillales>Halomonadaceae>Halomonas>unclassified_Halomonas | 571 | B7L72_RS05675 | 17462..19177 | 1670448 | DUF262-domain-containing-protein-[Halomonas-sp.-BC1] |
| NQY75748.1 | ParB+HNH | GCA_013213865.1 | Halomonas_sp. | Bacteria>Pseudomonadati>Pseudomonadota>Gammaproteobacteria>Oceanospirillales>Halomonadaceae>Halomonas>unclassified_Halomonas | 581 | HRT78_01190 | 154359..156104 | 1486246 | DUF262-domain-containing-protein-[Halomonas-sp.] |
| PHR03202.1 | ParB+HNH | GCA_002733025.1 | Halomonas_sp. | Bacteria>Pseudomonadati>Pseudomonadota>Gammaproteobacteria>Oceanospirillales>Halomonadaceae>Halomonas>unclassified_Halomonas | 561 | COB32_05635 | 11430..13115 | 1486246 | hypothetical-protein-COB32_05635-[Halomonas-sp.] |
| WP_189514720.1 | ParB+HNH | GCF_014652795.1 | Kushneria_pakistanensis | Bacteria>Pseudomonadati>Pseudomonadota>Gammaproteobacteria>Oceanospirillales>Halomonadaceae>Kushneria | 561 | IE123_RS02400 | 540487..542172 | 1508770 | DUF262-domain-containing-protein-[Kushneria-pakistanensis] |
| WP_086901158.1 | ParB+HNH | GCF_002157205.1 | Kushneria_marisflavi | Bacteria>Pseudomonadati>Pseudomonadota>Gammaproteobacteria>Oceanospirillales>Halomonadaceae>Kushneria | 585 | B9H00_RS13920 | 3066053..3067810 | 157779 | DUF262-domain-containing-protein-[Kushneria-marisflavi] |
| WP_092844124.1 | ParB+HNH | GCF_900113605.1 | Modicisalibacter_xianhensis | Bacteria>Pseudomonadati>Pseudomonadota>Gammaproteobacteria>Oceanospirillales>Halomonadaceae>Modicisalibacter | 601 | BM194_RS04480 | 222713..224518 | 442341 | DUF262-domain-containing-protein-[Halomonas-xianhensis] |
| WP_019020109.1 | ParB+HNH | GCF_014651575.1 | Modicisalibacter_luteus | Bacteria>Pseudomonadati>Pseudomonadota>Gammaproteobacteria>Oceanospirillales>Halomonadaceae>Modicisalibacter | 626 | IE067_RS16255 | 70111..71991 | 453962 | DUF262-domain-containing-protein-[Halomonas-lutea] |
| WP_110669872.1 | ParB+HNH | GCF_003206155.1 | Salinicola_halophilus | Bacteria>Pseudomonadati>Pseudomonadota>Gammaproteobacteria>Oceanospirillales>Halomonadaceae>Salinicola | 615 | DOM33_RS13695 | 408739..410586 | 184065 | DUF262-domain-containing-protein-[Salinicola-halophilus] |
| WP_153843411.1 | ParB+HNH | GCF_012427705.1 | Vreelandella_piezotolerans | Bacteria>Pseudomonadati>Pseudomonadota>Gammaproteobacteria>Oceanospirillales>Halomonadaceae>Vreelandella | 570 | GYM47_RS02310 | 491579..493291 | 2609667 | DUF262-domain-containing-protein-[Halomonas-piezotolerans] |
| WP_188638085.1 | ParB+HNH | GCF_014640815.1 | Vreelandella_lutescens | Bacteria>Pseudomonadati>Pseudomonadota>Gammaproteobacteria>Oceanospirillales>Halomonadaceae>Vreelandella | 600 | IEW65_RS03390 | 728912..730714 | 1602943 | DUF262-domain-containing-protein-[Halomonas-lutescens] |
| WP_017428904.1 | ParB+HNH | GCF_000334215.1 | Vreelandella_jeotgali | Bacteria>Pseudomonadati>Pseudomonadota>Gammaproteobacteria>Oceanospirillales>Halomonadaceae>Vreelandella | 596 | C793_RS0100105 | 20311..22101 | 553386 | DUF262-domain-containing-protein-[Halomonas-jeotgali] |
| WP_179915945.1 | ParB+HNH | GCF_013415125.1 | Vreelandella_glaciei | Bacteria>Pseudomonadati>Pseudomonadota>Gammaproteobacteria>Oceanospirillales>Halomonadaceae>Vreelandella | 581 | HZS80_RS09685 | 32522..34267 | 186761 | DUF262-domain-containing-protein-[Halomonas-glaciei] |
| WP_089705406.1 | ParB+HNH | GCF_900103865.1 | Vreelandella_arcis | Bacteria>Pseudomonadati>Pseudomonadota>Gammaproteobacteria>Oceanospirillales>Halomonadaceae>Vreelandella | 626 | BLR97_RS09405 | 259382..261262 | 416873 | DUF262-domain-containing-protein-[Halomonas-arcis] |
| WP_159343436.1 | ParB+HNH | GCF_009846525.1 | Vreelandella_aquamarina | Bacteria>Pseudomonadati>Pseudomonadota>Gammaproteobacteria>Oceanospirillales>Halomonadaceae>Vreelandella | 626 | CTT34_RS16790 | 3552423..3554303 | 77097 | DUF262-domain-containing-protein-[Halomonas-meridiana] |
| WP_172515126.1 | ParB+HNH | GCF_011764625.1 | Vreelandella_aquamarina | Bacteria>Pseudomonadati>Pseudomonadota>Gammaproteobacteria>Oceanospirillales>Halomonadaceae>Vreelandella | 626 | HMEPL2_RS11735 | 2485081..2486961 | 77097 | DUF262-domain-containing-protein-[Halomonas-meridiana] |
| WP_261843644.1 | ParB+HNH | GCF_024347215.1 | Aliamphritea_ceti | Bacteria>Pseudomonadati>Pseudomonadota>Gammaproteobacteria>Oceanospirillales>Oceanospirillaceae>Aliamphritea | 587 | OCU49_RS03515 | 767878..769641 | 1524258 | DUF262-domain-containing-protein-[Aliamphritea-ceti] |
| WP_181736363.1 | ParB+HNH | GCF_013868415.1 | Marinobacterium_marinum | Bacteria>Pseudomonadati>Pseudomonadota>Gammaproteobacteria>Oceanospirillales>Oceanospirillaceae>Marinobacterium | 596 | H1S06_RS00705 | 45799..47589 | 2756129 | DUF262-domain-containing-protein-[Marinobacterium-sp.-3-1745] |
| WP_220183707.1 | ParB+HNH | GCF_013868415.1 | Marinobacterium_marinum | Bacteria>Pseudomonadati>Pseudomonadota>Gammaproteobacteria>Oceanospirillales>Oceanospirillaceae>Marinobacterium | 569 | H1S06_RS00680 | 37668..39377 | 2756129 | DUF262-domain-containing-protein-[Marinobacterium-sp.-3-1745] |
| WP_027853087.1 | ParB+HNH | GCF_000428985.1 | Marinobacterium_litorale | Bacteria>Pseudomonadati>Pseudomonadota>Gammaproteobacteria>Oceanospirillales>Oceanospirillaceae>Marinobacterium | 610 | G403_RS0105375 | 186508..188340 | 404770 | DUF262-domain-containing-protein-[Marinobacterium-litorale] |
| AWY01110.1 | ParB+HNH | GCA_003285205.1 | Marinomonas_primoryensis | Bacteria>Pseudomonadati>Pseudomonadota>Gammaproteobacteria>Oceanospirillales>Oceanospirillaceae>Marinomonas | 625 | A8139_14835 | 3164001..3165878 | 178399 | hypothetical-protein-A8139_14835-[Marinomonas-primoryensis] |
| WP_112139350.1 | ParB+HNH | GCF_003285205.1 | Marinomonas_primoryensis | Bacteria>Pseudomonadati>Pseudomonadota>Gammaproteobacteria>Oceanospirillales>Oceanospirillaceae>Marinomonas | 625 | A8139_RS14960 | 3164001..3165878 | 178399 | DUF262-domain-containing-protein-[Marinomonas-primoryensis] |
| WP_100635911.1 | ParB+HNH | GCF_002806845.1 | Marinomonas_sp._ef1 | Bacteria>Pseudomonadati>Pseudomonadota>Gammaproteobacteria>Oceanospirillales>Oceanospirillaceae>Marinomonas>unclassified_Marinomonas | 607 | CCS02_RS04325 | 405132..406955 | 2005043 | DUF262-domain-containing-protein-[Marinomonas-sp.-ef1] |
| MBE0508134.1 | ParB+HNH | GCA_014859565.1 | Marinospirillum_sp. | Bacteria>Pseudomonadati>Pseudomonadota>Gammaproteobacteria>Oceanospirillales>Oceanospirillaceae>Marinospirillum>unclassified_Marinospirillum | 562 | IBX50_15705 | 3221..4909 | 2183934 | DUF262-domain-containing-protein-[Marinospirillum-sp.] |
| QEW05879.1 | ParB+HNH | GCA_008727795.1 | Nitrincola_iocasae | Bacteria>Pseudomonadati>Pseudomonadota>Gammaproteobacteria>Oceanospirillales>Oceanospirillaceae>Nitrincola | 612 | F5I99_04905 | 1042019..1043857 | 2614693 | DUF262-domain-containing-protein-[Nitrincola-iocasae] |
| WP_151053918.1 | ParB+HNH | GCF_008727795.1 | Nitrincola_iocasae | Bacteria>Pseudomonadati>Pseudomonadota>Gammaproteobacteria>Oceanospirillales>Oceanospirillaceae>Nitrincola | 612 | F5I99_RS04905 | 1042019..1043857 | 2614693 | DUF262-domain-containing-protein-[Nitrincola-iocasae] |
| WP_260997649.1 | ParB+HNH | GCF_025345565.1 | Thalassolituus_hydrocarboniclasticus | Bacteria>Pseudomonadati>Pseudomonadota>Gammaproteobacteria>Oceanospirillales>Oceanospirillaceae>Thalassolituus | 568 | HUF19_RS16705 | 3717930..3719636 | 2742796 | DUF262-domain-containing-HNH-endonuclease-family-protein-[Thalassolituus-hydrocarbonoclasticus] |
| MAX99655.1 | ParB+HNH | GCA_002706025.1 | Oceanospirillaceae_bacterium | Bacteria>Pseudomonadati>Pseudomonadota>Gammaproteobacteria>Oceanospirillales>Oceanospirillaceae>unclassified_Oceanospirillaceae | 564 | CMI08_10730 | 5223..6917 | 1899355 | hypothetical-protein-[Oceanospirillaceae-bacterium] |
| WP_116688141.1 | ParB+HNH | GCF_963878705.1 | Pelagibaculum_spongiae | Bacteria>Pseudomonadati>Pseudomonadota>Gammaproteobacteria>Oceanospirillales>Oceanospirillales_incertae_sedis>Pelagibaculum | 580 | DC094_RS16045 | 162166..163908 | 2080658 | DUF262-domain-containing-protein-[Pelagibaculum-spongiae] |
| WP_100257383.1 | ParB+HNH | GCF_002795845.1 | Reinekea_forsetii | Bacteria>Pseudomonadati>Pseudomonadota>Gammaproteobacteria>Oceanospirillales>Saccharospirillaceae>Reinekea | 579 | REIFOR_RS09810 | 2108517..2110256 | 1336806 | DUF262-domain-containing-protein-[Reinekea-forsetii] |
| MBR9867111.1 | ParB+HNH | GCA_018223265.1 | Oceanospirillales_bacterium | Bacteria>Pseudomonadati>Pseudomonadota>Gammaproteobacteria>Oceanospirillales>unclassified_Oceanospirillales | 587 | GYB18_08410 | 19389..21152 | 1914300 | DUF262-domain-containing-protein-[Oceanospirillales-bacterium] |
| WP_023094206.1 | ParB+HNH | GCF_901482535.1 | Pseudomonadaceae | Bacteria>Pseudomonadati>Pseudomonadota>Gammaproteobacteria>Pseudomonadales | 602 | D3P44_RS15340 | 3243141..3244949 | 135621 | MULTISPECIES:-DUF262-domain-containing-protein-[Pseudomonadaceae] |
| WP_098418632.1 | ParB+HNH | GCF_002563885.1 | unclassified_Marinobacter | Bacteria>Pseudomonadati>Pseudomonadota>Gammaproteobacteria>Pseudomonadales>Marinobacteraceae>Marinobacter | 739 | ATG98_RS19585 | 4034201..4036420 | 83889 | MULTISPECIES:-DUF262-domain-containing-HNH-endonuclease-family-protein-[unclassified-Marinobacter] |
| WP_266096196.1 | ParB+HNH | - | unclassified_Marinobacter | Bacteria>Pseudomonadati>Pseudomonadota>Gammaproteobacteria>Pseudomonadales>Marinobacteraceae>Marinobacter | 452 | JIF54_RS17255 | 3698..5056 | 83889 | MULTISPECIES:-HNH-endonuclease-family-protein-[unclassified-Marinobacter] |
| WP_092021231.1 | ParB+HNH | GCF_900114775.1 | Marinobacter_zhejiangensis | Bacteria>Pseudomonadati>Pseudomonadota>Gammaproteobacteria>Pseudomonadales>Marinobacteraceae>Marinobacter | 662 | BMZ67_RS06965 | 341625..343613 | 488535 | DUF262-domain-containing-protein-[Marinobacter-zhejiangensis] |
| WP_216008128.1 | ParB+HNH | GCF_018860765.1 | Marinobacter_salexigens | Bacteria>Pseudomonadati>Pseudomonadota>Gammaproteobacteria>Pseudomonadales>Marinobacteraceae>Marinobacter | 583 | KO508_RS09740 | 164926..166677 | 1925763 | DUF262-domain-containing-protein-[Marinobacter-salexigens] |
| WP_213480156.1 | ParB+HNH | GCF_018424725.1 | Marinobacter_salarius | Bacteria>Pseudomonadati>Pseudomonadota>Gammaproteobacteria>Pseudomonadales>Marinobacteraceae>Marinobacter | 564 | DYI41_RS19375 | 7145..8839 | 1420917 | DUF262-domain-containing-protein-[Marinobacter-salarius] |
| WP_048384703.1 | ParB+HNH | GCF_001043175.1 | Marinobacter_psychrophilus | Bacteria>Pseudomonadati>Pseudomonadota>Gammaproteobacteria>Pseudomonadales>Marinobacteraceae>Marinobacter | 564 | ABA45_RS05815 | 1307760..1309454 | 330734 | DUF262-domain-containing-protein-[Marinobacter-psychrophilus] |
| WP_141068076.1 | ParB+HNH | GCF_006516615.1 | Marinobacter_nauticus | Bacteria>Pseudomonadati>Pseudomonadota>Gammaproteobacteria>Pseudomonadales>Marinobacteraceae>Marinobacter | 626 | FH712_RS17680 | 44128..46008 | 2743 | DUF262-domain-containing-protein-[Marinobacter-nauticus] |
| WP_248236649.1 | ParB+HNH | GCF_023156245.1 | Marinobacter_bryozoorum | Bacteria>Pseudomonadati>Pseudomonadota>Gammaproteobacteria>Pseudomonadales>Marinobacteraceae>Marinobacter | 564 | MLC59_RS17285 | 10298..11992 | 256324 | DUF262-domain-containing-protein-[Marinobacter-bryozoorum] |
| WP_247116149.1 | ParB+HNH | GCF_023016045.1 | Marinobacter_sp._S6332 | Bacteria>Pseudomonadati>Pseudomonadota>Gammaproteobacteria>Pseudomonadales>Marinobacteraceae>Marinobacter>unclassified_Marinobacter | 373 | MWU62_RS17905 | 13511..14632 | 2926403 | DUF262-domain-containing-protein-[Marinobacter-sp.-S6332] |

| Protein-ID | Domain | Assembly-ID | Strain | Taxonomy | Length | Locus-Tag | Genomic-coordinates | TaxID | Fasta-defline |
|---|---|---|---|---|---|---|---|---|---|
| WP_249013855.1 | ParB+HNH | GCF_023311895.1 | Marinobacter_sp._M3C | Bacteria>Pseudomonadati>Pseudomonadota>Gammaproteobacteria>Pseudomonadales>Marinobacteraceae>Marinobacter>unclassified_Marinobacter | 580 | MIH18_RS02745 | 623405..625147 | 2917715 | DUF262-domain-containing-protein-[Marinobacter-sp.-M3C] |
| WP_200371319.1 | ParB+HNH | GCF_016597815.1 | Marinobacter_sp._DY40_1A1 | Bacteria>Pseudomonadati>Pseudomonadota>Gammaproteobacteria>Pseudomonadales>Marinobacteraceae>Marinobacter>unclassified_Marinobacter | 570 | FFI61_RS14520 | 352912..354624 | 2583229 | DUF262-domain-containing-protein-[Marinobacter-sp.-DY40_1A1] |
| WP_217632574.1 | ParB+HNH | - | Marinobacter_sp._AC-23 | Bacteria>Pseudomonadati>Pseudomonadota>Gammaproteobacteria>Pseudomonadales>Marinobacteraceae>Marinobacter>unclassified_Marinobacter | 567 | BCA33_RS00495 | 114278..115981 | 1879031 | HNH-endonuclease-family-protein-[Marinobacter-sp.-AC-23] |
| MBE95562.1 | ParB+HNH | GCA_002714765.1 | Marinobacter_sp. | Bacteria>Pseudomonadati>Pseudomonadota>Gammaproteobacteria>Pseudomonadales>Marinobacteraceae>Marinobacter>unclassified_Marinobacter | 626 | CMG89_11205 | 13114..14994 | 50741 | hypothetical-protein-[Marinobacter-sp.] |
| WP_229694595.1 | ParB+HNH | GCF_014642795.1 | Pseudohongiella_nitratireducens | Bacteria>Pseudomonadati>Pseudomonadota>Gammaproteobacteria>Pseudomonadales>Pseudohongiellaceae>Pseudohongiella | 619 | IEW51_RS03955 | 201029..202888 | 1768907 | DUF262-domain-containing-protein-[Pseudohongiella-nitratireducens] |
| WP_017245160.1 | ParB+HNH | GCF_000280555.1 | Stutzerimonas | Bacteria>Pseudomonadati>Pseudomonadota>Gammaproteobacteria>Pseudomonadales>Pseudomonadaceae | 578 | A598_RS0110890 | 39262..40998 | 2901164 | DUF262-domain-containing-protein-[Stutzerimonas-stutzeri] |
| WP_017245157.1 | ParB+HNH | GCF_000280555.1 | Stutzerimonas | Bacteria>Pseudomonadati>Pseudomonadota>Gammaproteobacteria>Pseudomonadales>Pseudomonadaceae | 562 | A598_RS0110875 | 33230..34918 | 2901164 | DUF262-domain-containing-protein-[Stutzerimonas-stutzeri] |
| WP_138763792.1 | ParB+HNH | GCF_016613085.1 | Pseudomonas | Bacteria>Pseudomonadati>Pseudomonadota>Gammaproteobacteria>Pseudomonadales>Pseudomonadaceae | 757 | FHL14_RS16370 | 29806..32079 | 286 | MULTISPECIES:-DUF262-domain-containing-protein-[Pseudomonas] |
| WP_041506100.1 | ParB+HNH | GCF_000831585.1 | Pseudomonas | Bacteria>Pseudomonadati>Pseudomonadota>Gammaproteobacteria>Pseudomonadales>Pseudomonadaceae | 625 | NJC29_RS17725 | 141275..143152 | 286 | MULTISPECIES:-DUF262-domain-containing-protein-[Pseudomonas] |
| WP_150073352.1 | ParB+HNH | GCF_029959765.1 | Pseudomonas | Bacteria>Pseudomonadati>Pseudomonadota>Gammaproteobacteria>Pseudomonadales>Pseudomonadaceae | 624 | GHO24_RS06650 | 44205..46079 | 286 | MULTISPECIES:-DUF262-domain-containing-protein-[Pseudomonas] |
| WP_209953643.1 | ParB+HNH | GCF_017875115.1 | Pseudomonas | Bacteria>Pseudomonadati>Pseudomonadota>Gammaproteobacteria>Pseudomonadales>Pseudomonadaceae | 624 | J3A71_RS03880 | 849930..851804 | 286 | MULTISPECIES:-DUF262-domain-containing-protein-[Pseudomonas] |
| WP_025298036.1 | ParB+HNH | GCF_900707575.1 | Pseudomonas | Bacteria>Pseudomonadati>Pseudomonadota>Gammaproteobacteria>Pseudomonadales>Pseudomonadaceae | 623 | PMJ90_RS13005 | 2710040..2711911 | 286 | MULTISPECIES:-DUF262-domain-containing-protein-[Pseudomonas] |
| WP_019470417.1 | ParB+HNH | GCF_000292285.1 | Pseudomonas | Bacteria>Pseudomonadati>Pseudomonadota>Gammaproteobacteria>Pseudomonadales>Pseudomonadaceae | 616 | N805_RS25705 | 5817187..5819037 | 286 | MULTISPECIES:-DUF262-domain-containing-protein-[Pseudomonas] |
| WP_033986134.1 | ParB+HNH | GCF_047198885.1 | Pseudomonas | Bacteria>Pseudomonadati>Pseudomonadota>Gammaproteobacteria>Pseudomonadales>Pseudomonadaceae | 616 | AO894_RS16030 | 6235..8085 | 286 | MULTISPECIES:-DUF262-domain-containing-protein-[Pseudomonas] |
| WP_229605962.1 | ParB+HNH | GCF_020831405.1 | Pseudomonas | Bacteria>Pseudomonadati>Pseudomonadota>Gammaproteobacteria>Pseudomonadales>Pseudomonadaceae | 616 | LMJ29_RS13095 | 85750..87600 | 286 | DUF262-domain-containing-protein-[Pseudomonas-oryzagri] |
| WP_236165765.1 | ParB+HNH | GCF_021603865.1 | Pseudomonas | Bacteria>Pseudomonadati>Pseudomonadota>Gammaproteobacteria>Pseudomonadales>Pseudomonadaceae | 616 | L6D10_RS06750 | 35493..37343 | 286 | DUF262-domain-containing-protein-[Pseudomonas-fulva] |
| WP_138763793.1 | ParB+HNH | GCF_016613085.1 | Pseudomonas | Bacteria>Pseudomonadati>Pseudomonadota>Gammaproteobacteria>Pseudomonadales>Pseudomonadaceae | 600 | FHL14_RS16375 | 32076..33878 | 286 | MULTISPECIES:-DUF262-domain-containing-protein-[Pseudomonas] |
| WP_057429195.1 | ParB+HNH | GCF_963970795.1 | Pseudomonas | Bacteria>Pseudomonadati>Pseudomonadota>Gammaproteobacteria>Pseudomonadales>Pseudomonadaceae | 597 | ALO87_RS04330 | 134988..136781 | 286 | DUF262-domain-containing-protein-[Pseudomonas-syringae-group-genomosp.-3] |
| WP_095018754.1 | ParB+HNH | GCF_902329585.1 | Pseudomonas | Bacteria>Pseudomonadati>Pseudomonadota>Gammaproteobacteria>Pseudomonadales>Pseudomonadaceae | 597 | PPTH39_RS22295 | 121060..122853 | 286 | MULTISPECIES:-DUF262-domain-containing-protein-[Pseudomonas] |
| WP_021219095.1 | ParB+HNH | GCF_900707355.1 | Pseudomonas | Bacteria>Pseudomonadati>Pseudomonadota>Gammaproteobacteria>Pseudomonadales>Pseudomonadaceae | 578 | E4Z10_RS26775 | 8536..10272 | 286 | MULTISPECIES:-DUF262-domain-containing-protein-[Pseudomonas] |
| WP_023115273.1 | ParB+HNH | GCF_900707185.1 | Pseudomonas | Bacteria>Pseudomonadati>Pseudomonadota>Gammaproteobacteria>Pseudomonadales>Pseudomonadaceae | 557 | IPC175_RS29780 | 6882..8555 | 286 | MULTISPECIES:-DUF262-domain-containing-protein-[Pseudomonas] |
| WP_076583432.1 | ParB+HNH | GCF_900156545.1 | Aquipseudomonas_alcaligenes | Bacteria>Pseudomonadati>Pseudomonadota>Gammaproteobacteria>Pseudomonadales>Pseudomonadaceae>Aquipseudomonas | 624 | BW984_RS17020 | 262795..264669 | 43263 | DUF262-domain-containing-protein-[Pseudomonas-alcaligenes] |
| WP_165892257.1 | ParB+HNH | GCF_011470035.1 | Azotobacter_chroococcum | Bacteria>Pseudomonadati>Pseudomonadota>Gammaproteobacteria>Pseudomonadales>Pseudomonadaceae>Azotobacter_group>Azotobacter | 572 | HA520_RS07845 | 22711..24429 | 353 | DUF262-domain-containing-protein-[Azotobacter-chroococcum] |
| HIQ43648.1 | ParB+HNH | GCA_015661735.1 | Ectopseudomonas_oleovorans | Bacteria>Pseudomonadati>Pseudomonadota>Gammaproteobacteria>Pseudomonadales>Pseudomonadaceae>Ectopseudomonas | 572 | EYH47_12965 | 34825..36543 | 301 | DUF262-domain-containing-protein-[Pseudomonas-oleovorans] |
| WP_143505432.1 | ParB+HNH | - | Ectopseudomonas_mendocina | Bacteria>Pseudomonadati>Pseudomonadota>Gammaproteobacteria>Pseudomonadales>Pseudomonadaceae>Ectopseudomonas | 653 | EQ826_RS09470 | 205061..207022 | 300 | DUF262-domain-containing-protein-[Pseudomonas-mendocina] |
| TRO09200.1 | ParB+HNH | GCA_007049815.1 | Ectopseudomonas_mendocina | Bacteria>Pseudomonadati>Pseudomonadota>Gammaproteobacteria>Pseudomonadales>Pseudomonadaceae>Ectopseudomonas | 611 | EQ829_23240 | 32907..34742 | 300 | DUF262-domain-containing-protein-[Pseudomonas-mendocina] |
| WP_143503143.1 | ParB+HNH | GCF_007049785.1 | Ectopseudomonas_mendocina | Bacteria>Pseudomonadati>Pseudomonadota>Gammaproteobacteria>Pseudomonadales>Pseudomonadaceae>Ectopseudomonas | 611 | EQ836_RS24245 | 55905..57740 | 300 | DUF262-domain-containing-protein-[Pseudomonas-mendocina] |
| WP_003246713.1 | ParB+HNH | - | Ectopseudomonas_mendocina | Bacteria>Pseudomonadati>Pseudomonadota>Gammaproteobacteria>Pseudomonadales>Pseudomonadaceae>Ectopseudomonas | 572 | A471_RS19810 | 17606..19324 | 300 | DUF262-domain-containing-protein-[Pseudomonas-mendocina] |
| WP_143508092.1 | ParB+HNH | - | Ectopseudomonas_mendocina | Bacteria>Pseudomonadati>Pseudomonadota>Gammaproteobacteria>Pseudomonadales>Pseudomonadaceae>Ectopseudomonas | 572 | EQ828_RS12685 | 218866..220584 | 300 | DUF262-domain-containing-protein-[Pseudomonas-mendocina] |
| WP_129484087.1 | ParB+HNH | GCF_021495345.1 | Ectopseudomonas_hydrolytica | Bacteria>Pseudomonadati>Pseudomonadota>Gammaproteobacteria>Pseudomonadales>Pseudomonadaceae>Ectopseudomonas | 572 | L1F06_RS00040 | 13199..14917 | 2493633 | DUF262-domain-containing-protein-[Pseudomonas-hydrolytica] |
| WP_090214595.1 | ParB+HNH | GCF_900105885.1 | Geopseudomonas_guangdongensis | Bacteria>Pseudomonadati>Pseudomonadota>Gammaproteobacteria>Pseudomonadales>Pseudomonadaceae>Geopseudomonas | 578 | BLU22_RS11510 | 2426459..2428195 | 1245526 | DUF262-domain-containing-protein-[Pseudomonas-guangdongensis] |
| WP_104739189.1 | ParB+HNH | GCF_014641295.1 | Halopseudomonas_oceani | Bacteria>Pseudomonadati>Pseudomonadota>Gammaproteobacteria>Pseudomonadales>Pseudomonadaceae>Halopseudomonas | 625 | IEX06_RS08420 | 378212..380089 | 1708783 | DUF262-domain-containing-protein-[Halopseudomonas-oceani] |
| WP_102684490.1 | ParB+HNH | GCF_017350895.1 | unclassified_Pseudomonas | Bacteria>Pseudomonadati>Pseudomonadota>Gammaproteobacteria>Pseudomonadales>Pseudomonadaceae>Pseudomonas | 578 | J4E12_RS18515 | 6740..8476 | 196821 | MULTISPECIES:-DUF262-domain-containing-protein-[unclassified-Pseudomonas] |
| WP_092146140.1 | ParB+HNH | GCF_900102655.1 | unclassified_Pseudomonas | Bacteria>Pseudomonadati>Pseudomonadota>Gammaproteobacteria>Pseudomonadales>Pseudomonadaceae>Pseudomonas | 557 | BLP71_RS14880 | 96025..97698 | 196821 | MULTISPECIES:-DUF262-domain-containing-protein-[unclassified-Pseudomonas] |
| WP_161928860.1 | ParB+HNH | GCF_009932815.1 | Pseudomonas_yangonensis | Bacteria>Pseudomonadati>Pseudomonadota>Gammaproteobacteria>Pseudomonadales>Pseudomonadaceae>Pseudomonas | 616 | GYH25_RS19720 | 94659..96509 | 2579922 | DUF262-domain-containing-protein-[Pseudomonas-yangonensis] |
| MVW77191.1 | ParB+HNH | GCA_009763245.1 | Pseudomonas_xionganensis | Bacteria>Pseudomonadati>Pseudomonadota>Gammaproteobacteria>Pseudomonadales>Pseudomonadaceae>Pseudomonas | 578 | GJV18_17875 | 23048..24784 | 2654845 | DUF262-domain-containing-protein-[Pseudomonas-xionganensis] |
| WP_160347848.1 | ParB+HNH | GCF_009763245.1 | Pseudomonas_xionganensis | Bacteria>Pseudomonadati>Pseudomonadota>Gammaproteobacteria>Pseudomonadales>Pseudomonadaceae>Pseudomonas | 578 | GJV18_RS17870 | 23048..24784 | 2654845 | DUF262-domain-containing-protein-[Pseudomonas-xionganensis] |
| MVW77188.1 | ParB+HNH | GCA_009763245.1 | Pseudomonas_xionganensis | Bacteria>Pseudomonadati>Pseudomonadota>Gammaproteobacteria>Pseudomonadales>Pseudomonadaceae>Pseudomonas | 562 | GJV18_17860 | 17029..18717 | 2654845 | DUF262-domain-containing-protein-[Pseudomonas-xionganensis] |
| WP_160347845.1 | ParB+HNH | GCF_009763245.1 | Pseudomonas_xionganensis | Bacteria>Pseudomonadati>Pseudomonadota>Gammaproteobacteria>Pseudomonadales>Pseudomonadaceae>Pseudomonas | 562 | GJV18_RS17855 | 17029..18717 | 2654845 | DUF262-domain-containing-protein-[Pseudomonas-xionganensis] |
| WP_265072426.1 | ParB+HNH | GCF_026016265.1 | Pseudomonas_viciae | Bacteria>Pseudomonadati>Pseudomonadota>Gammaproteobacteria>Pseudomonadales>Pseudomonadaceae>Pseudomonas | 624 | LOY66_RS25440 | 5791374..5793248 | 2505979 | DUF262-domain-containing-protein-[Pseudomonas-viciae] |
| WP_186598285.1 | ParB+HNH | GCF_014269085.1 | Pseudomonas_taiwanensis | Bacteria>Pseudomonadati>Pseudomonadota>Gammaproteobacteria>Pseudomonadales>Pseudomonadaceae>Pseudomonas | 616 | HU747_RS04810 | 439025..440875 | 470150 | DUF262-domain-containing-protein-[Pseudomonas-taiwanensis] |
| NWF08287.1 | ParB+HNH | GCA_013386885.1 | Pseudomonas_salomonii | Bacteria>Pseudomonadati>Pseudomonadota>Gammaproteobacteria>Pseudomonadales>Pseudomonadaceae>Pseudomonas | 562 | HX810_11505 | 502983..504671 | 191391 | DUF262-domain-containing-protein-[Pseudomonas-salomonii] |
| WP_090347587.1 | ParB+HNH | GCF_900104805.1 | Pseudomonas_oryzae | Bacteria>Pseudomonadati>Pseudomonadota>Gammaproteobacteria>Pseudomonadales>Pseudomonadaceae>Pseudomonas | 557 | BLT78_RS03195 | 675442..677115 | 1392877 | DUF262-domain-containing-protein-[Pseudomonas-oryzae] |
| WP_150549234.1 | ParB+HNH | GCF_900682675.2 | Pseudomonas_marincola | Bacteria>Pseudomonadati>Pseudomonadota>Gammaproteobacteria>Pseudomonadales>Pseudomonadaceae>Pseudomonas | 571 | PMYSY11_RS11205 | 2410986..2412701 | 437900 | DUF262-domain-containing-protein-[Pseudomonas-marincola] |
| WP_064314267.1 | ParB+HNH | GCF_021601465.1 | Pseudomonas_juntendi | Bacteria>Pseudomonadati>Pseudomonadota>Gammaproteobacteria>Pseudomonadales>Pseudomonadaceae>Pseudomonas | 595 | L6D32_RS09720 | 143621..145408 | 2666183 | MULTISPECIES:-DUF262-domain-containing-protein-[Pseudomonas] |
| WP_205518898.1 | ParB+HNH | GCF_016925675.1 | Pseudomonas_hygromyciniae | Bacteria>Pseudomonadati>Pseudomonadota>Gammaproteobacteria>Pseudomonadales>Pseudomonadaceae>Pseudomonas | 488 | JTY93_RS19455 | 4359712..4361178 | 2812000 | DUF262-domain-containing-protein-[Pseudomonas-sp.-SDM007] |
| WP_048385711.1 | ParB+HNH | GCF_001043065.1 | Pseudomonas_helleri | Bacteria>Pseudomonadati>Pseudomonadota>Gammaproteobacteria>Pseudomonadales>Pseudomonadaceae>Pseudomonas | 624 | TU85_RS02950 | 81663..83537 | 1608996 | DUF262-domain-containing-protein-[Pseudomonas-helleri] |
| WP_153329474.1 | ParB+HNH | GCF_009600225.1 | Pseudomonas_helleri | Bacteria>Pseudomonadati>Pseudomonadota>Gammaproteobacteria>Pseudomonadales>Pseudomonadaceae>Pseudomonas | 623 | GHO39_RS16730 | 23811..25682 | 1608996 | DUF262-domain-containing-protein-[Pseudomonas-helleri] |
| WP_090241712.1 | ParB+HNH | GCF_900113745.1 | Pseudomonas_guineae | Bacteria>Pseudomonadati>Pseudomonadota>Gammaproteobacteria>Pseudomonadales>Pseudomonadaceae>Pseudomonas | 620 | BMZ32_RS09360 | 88599..90461 | 425504 | DUF262-domain-containing-protein-[Pseudomonas-guineae] |
| WP_102082560.1 | ParB+HNH | GCF_002864915.1 | Pseudomonas_guariconensis | Bacteria>Pseudomonadati>Pseudomonadota>Gammaproteobacteria>Pseudomonadales>Pseudomonadaceae>Pseudomonas | 624 | CXG53_RS23710 | 50143..52017 | 1288410 | DUF262-domain-containing-protein-[Pseudomonas-guariconensis] |
| WP_150597439.1 | ParB+HNH | GCF_039522465.1 | Pseudomonas_fluorescens_group | Bacteria>Pseudomonadati>Pseudomonadota>Gammaproteobacteria>Pseudomonadales>Pseudomonadaceae>Pseudomonas | 597 | F8N16_RS17970 | 126860..128653 | 136843 | DUF262-domain-containing-protein-[Pseudomonas-fluorescens] |
| WP_196476670.1 | ParB+HNH | GCF_015712105.2 | Pseudomonas_chaetocerotis | Bacteria>Pseudomonadati>Pseudomonadota>Gammaproteobacteria>Pseudomonadales>Pseudomonadaceae>Pseudomonas | 557 | H3221_RS24560 | 40262..41935 | 2758695 | DUF262-domain-containing-protein-[Pseudomonas-chaetocerotis] |
| WP_229999480.1 | ParB+HNH | GCF_918697895.1 | Pseudomonas_carnis | Bacteria>Pseudomonadati>Pseudomonadota>Gammaproteobacteria>Pseudomonadales>Pseudomonadaceae>Pseudomonas | 597 | LQ224_RS08495 | 117341..119134 | 2487355 | DUF262-domain-containing-protein-[Pseudomonas-carnis] |
| WP_127881956.1 | ParB+HNH | - | Pseudomonas_carnis | Bacteria>Pseudomonadati>Pseudomonadota>Gammaproteobacteria>Pseudomonadales>Pseudomonadaceae>Pseudomonas | 563 | PCSC_RS23720 | 89940..91631 | 2487355 | DUF262-domain-containing-protein-[Pseudomonas-carnis] |
| WP_013692227.1 | ParB+HNH | GCF_000194805.1 | Pseudomonas_brassicacearum | Bacteria>Pseudomonadati>Pseudomonadota>Gammaproteobacteria>Pseudomonadales>Pseudomonadaceae>Pseudomonas | 602 | PSEBR_RS03970 | 949150..950958 | 930166 | DUF262-domain-containing-protein-[Pseudomonas-brassicacearum] |
| WP_182367090.1 | ParB+HNH | GCF_014109765.1 | Pseudomonas_berkeleyensis | Bacteria>Pseudomonadati>Pseudomonadota>Gammaproteobacteria>Pseudomonadales>Pseudomonadaceae>Pseudomonas | 557 | HS968_RS15920 | 3431928..3433601 | 2726956 | DUF262-domain-containing-protein-[Pseudomonas-sp.-MSPm1] |
| WP_243243003.1 | ParB+HNH | GCF_022802935.1 | Pseudomonas_atacamensis | Bacteria>Pseudomonadati>Pseudomonadota>Gammaproteobacteria>Pseudomonadales>Pseudomonadaceae>Pseudomonas | 557 | PSD_RS21545 | 55248..56921 | 2565368 | DUF262-domain-containing-protein-[Pseudomonas-atacamensis] |
| WP_196184085.1 | ParB+HNH | GCF_015680805.1 | Pseudomonas_asiatica | Bacteria>Pseudomonadati>Pseudomonadota>Gammaproteobacteria>Pseudomonadales>Pseudomonadaceae>Pseudomonas | 595 | IR013_RS15050 | 46134..47921 | 2219225 | DUF262-domain-containing-protein-[Pseudomonas-asiatica] |
| WP_203542489.1 | ParB+HNH | GCF_016865555.1 | Pseudomonas_arcuscaelestis | Bacteria>Pseudomonadati>Pseudomonadota>Gammaproteobacteria>Pseudomonadales>Pseudomonadaceae>Pseudomonas | 610 | IIE18_RS12205 | 353387..355219 | 2710591 | DUF262-domain-containing-protein-[Pseudomonas-arcuscaelestis] |
| WP_043106444.1 | ParB+HNH | GCF_900706985.1 | Pseudomonas_aeruginosa_group | Bacteria>Pseudomonadati>Pseudomonadota>Gammaproteobacteria>Pseudomonadales>Pseudomonadaceae>Pseudomonas | 557 | NQ84_RS28345 | 8114..9787 | 136841 | MULTISPECIES:-DUF262-domain-containing-protein-[Pseudomonas-aeruginosa-group] |
| MBV5724747.1 | ParB+HNH | GCA_019166825.1 | Pseudomonas_aeruginosa | Bacteria>Pseudomonadati>Pseudomonadota>Gammaproteobacteria>Pseudomonadales>Pseudomonadaceae>Pseudomonas>Pseudomonas_aeruginosa_group | 623 | KUT37_26450 | 33969..35840 | 287 | DUF262-domain-containing-protein-[Pseudomonas-aeruginosa] |
| WP_033944478.1 | ParB+HNH | GCF_947387505.1 | Pseudomonas_aeruginosa | Bacteria>Pseudomonadati>Pseudomonadota>Gammaproteobacteria>Pseudomonadales>Pseudomonadaceae>Pseudomonas>Pseudomonas_aeruginosa_group | 623 | GPV34_RS03085 | 6814..8685 | 287 | DUF262-domain-containing-protein-[Pseudomonas-aeruginosa] |
| HBO3309318.1 | ParB+HNH | GCA_021652015.1 | Pseudomonas_aeruginosa | Bacteria>Pseudomonadati>Pseudomonadota>Gammaproteobacteria>Pseudomonadales>Pseudomonadaceae>Pseudomonas>Pseudomonas_aeruginosa_group | 616 | L4R58_004355 | 39298..41148 | 287 | TPA:-DUF262-domain-containing-protein-[Pseudomonas-aeruginosa] |
| HBP5527126.1 | ParB+HNH | GCA_021712095.1 | Pseudomonas_aeruginosa | Bacteria>Pseudomonadati>Pseudomonadota>Gammaproteobacteria>Pseudomonadales>Pseudomonadaceae>Pseudomonas>Pseudomonas_aeruginosa_group | 616 | L6B52_09525 | 12571..14421 | 287 | DUF262-domain-containing-protein-[Pseudomonas-aeruginosa] |
| KAA5679900.1 | ParB+HNH | GCA_008632665.1 | Pseudomonas_aeruginosa | Bacteria>Pseudomonadati>Pseudomonadota>Gammaproteobacteria>Pseudomonadales>Pseudomonadaceae>Pseudomonas>Pseudomonas_aeruginosa_group | 616 | F3G60_11270 | 517527..519377 | 287 | DUF262-domain-containing-protein-[Pseudomonas-aeruginosa] |
| MBG3983984.1 | ParB+HNH | GCA_015733145.1 | Pseudomonas_aeruginosa | Bacteria>Pseudomonadati>Pseudomonadota>Gammaproteobacteria>Pseudomonadales>Pseudomonadaceae>Pseudomonas>Pseudomonas_aeruginosa_group | 616 | I4919_20905 | 95739..97589 | 287 | DUF262-domain-containing-protein-[Pseudomonas-aeruginosa] |
| MCD2801265.1 | ParB+HNH | GCA_021060565.1 | Pseudomonas_aeruginosa | Bacteria>Pseudomonadati>Pseudomonadota>Gammaproteobacteria>Pseudomonadales>Pseudomonadaceae>Pseudomonas>Pseudomonas_aeruginosa_group | 611 | K4W73_30390 | 388496..390331 | 287 | DUF262-domain-containing-HNH-endonuclease-family-protein-[Pseudomonas-aeruginosa] |
| WP_058181799.1 | ParB+HNH | GCF_001453595.1 | Pseudomonas_aeruginosa | Bacteria>Pseudomonadati>Pseudomonadota>Gammaproteobacteria>Pseudomonadales>Pseudomonadaceae>Pseudomonas>Pseudomonas_aeruginosa_group | 611 | APA84_RS22725 | 29128..30963 | 287 | DUF262-domain-containing-protein-[Pseudomonas-aeruginosa] |
| WP_058184870.1 | ParB+HNH | GCF_001453775.1 | Pseudomonas_aeruginosa | Bacteria>Pseudomonadati>Pseudomonadota>Gammaproteobacteria>Pseudomonadales>Pseudomonadaceae>Pseudomonas>Pseudomonas_aeruginosa_group | 611 | APB11_RS24365 | 75383..77218 | 287 | DUF262-domain-containing-protein-[Pseudomonas-aeruginosa] |
| WP_069376992.1 | ParB+HNH | GCF_001721765.1 | Pseudomonas_aeruginosa | Bacteria>Pseudomonadati>Pseudomonadota>Gammaproteobacteria>Pseudomonadales>Pseudomonadaceae>Pseudomonas>Pseudomonas_aeruginosa_group | 611 | A6681_RS28290 | 6148692..6150527 | 287 | DUF262-domain-containing-protein-[Pseudomonas-aeruginosa] |
| WP_126152454.1 | ParB+HNH | GCF_003955475.1 | Pseudomonas_aeruginosa | Bacteria>Pseudomonadati>Pseudomonadota>Gammaproteobacteria>Pseudomonadales>Pseudomonadaceae>Pseudomonas>Pseudomonas_aeruginosa_group | 611 | EKL37_RS17265 | 57941..59776 | 287 | DUF262-domain-containing-protein-[Pseudomonas-aeruginosa] |
| MCO2467175.1 | ParB+HNH | GCA_023907445.1 | Pseudomonas_aeruginosa | Bacteria>Pseudomonadati>Pseudomonadota>Gammaproteobacteria>Pseudomonadales>Pseudomonadaceae>Pseudomonas>Pseudomonas_aeruginosa_group | 610 | FA343_15160 | 62242..64074 | 287 | DUF262-domain-containing-protein-[Pseudomonas-aeruginosa] |
| WP_023113966.1 | ParB+HNH | GCF_000481245.1 | Pseudomonas_aeruginosa | Bacteria>Pseudomonadati>Pseudomonadota>Gammaproteobacteria>Pseudomonadales>Pseudomonadaceae>Pseudomonas>Pseudomonas_aeruginosa_group | 610 | Q036_RS16820 | 2146321..2148153 | 287 | DUF262-domain-containing-protein-[Pseudomonas-aeruginosa] |
| HBN8119631.1 | ParB+HNH | GCA_021586615.1 | Pseudomonas_aeruginosa | Bacteria>Pseudomonadati>Pseudomonadota>Gammaproteobacteria>Pseudomonadales>Pseudomonadaceae>Pseudomonas>Pseudomonas_aeruginosa_group | 602 | L3655_000124 | 102873..104681 | 287 | DUF262-domain-containing-protein-[Pseudomonas-aeruginosa] |
| HCE6004666.1 | ParB+HNH | GCA_023904815.1 | Pseudomonas_aeruginosa | Bacteria>Pseudomonadati>Pseudomonadota>Gammaproteobacteria>Pseudomonadales>Pseudomonadaceae>Pseudomonas>Pseudomonas_aeruginosa_group | 602 | NHK94_000103 | 112722..114530 | 287 | TPA:-DUF262-domain-containing-protein-[Pseudomonas-aeruginosa] |
| HCF1683400.1 | ParB+HNH | GCA_023941775.1 | Pseudomonas_aeruginosa | Bacteria>Pseudomonadati>Pseudomonadota>Gammaproteobacteria>Pseudomonadales>Pseudomonadaceae>Pseudomonas>Pseudomonas_aeruginosa_group | 602 | NH867_002020 | 41066..42874 | 287 | DUF262-domain-containing-protein-[Pseudomonas-aeruginosa] |
| MBG6382133.1 | ParB+HNH | GCA_015762815.1 | Pseudomonas_aeruginosa | Bacteria>Pseudomonadati>Pseudomonadota>Gammaproteobacteria>Pseudomonadales>Pseudomonadaceae>Pseudomonas>Pseudomonas_aeruginosa_group | 602 | I5I12_28775 | 9156..10964 | 287 | DUF262-domain-containing-protein-[Pseudomonas-aeruginosa] |
| MBH3980675.1 | ParB+HNH | GCA_016011075.1 | Pseudomonas_aeruginosa | Bacteria>Pseudomonadati>Pseudomonadota>Gammaproteobacteria>Pseudomonadales>Pseudomonadaceae>Pseudomonas>Pseudomonas_aeruginosa_group | 602 | I5Q70_02590 | 513541..515349 | 287 | DUF262-domain-containing-protein-[Pseudomonas-aeruginosa] |
| MBN0299740.1 | ParB+HNH | GCA_016922145.1 | Pseudomonas_aeruginosa | Bacteria>Pseudomonadati>Pseudomonadota>Gammaproteobacteria>Pseudomonadales>Pseudomonadaceae>Pseudomonas>Pseudomonas_aeruginosa_group | 602 | JTM78_25100 | 117337..119145 | 287 | DUF262-domain-containing-protein-[Pseudomonas-aeruginosa] |
| MBN0536369.1 | ParB+HNH | GCA_016922535.1 | Pseudomonas_aeruginosa | Bacteria>Pseudomonadati>Pseudomonadota>Gammaproteobacteria>Pseudomonadales>Pseudomonadaceae>Pseudomonas>Pseudomonas_aeruginosa_group | 602 | JTM59_25820 | 117341..119149 | 287 | DUF262-domain-containing-protein-[Pseudomonas-aeruginosa] |
| WP_010792142.1 | ParB+HNH | GCF_026126725.1 | Pseudomonas_aeruginosa | Bacteria>Pseudomonadati>Pseudomonadota>Gammaproteobacteria>Pseudomonadales>Pseudomonadaceae>Pseudomonas>Pseudomonas_aeruginosa_group | 602 | K3Z93_RS02340 | 14765..16573 | 287 | DUF262-domain-containing-protein-[Pseudomonas-aeruginosa] |
| WP_033990394.1 | ParB+HNH | GCF_020907765.1 | Pseudomonas_aeruginosa | Bacteria>Pseudomonadati>Pseudomonadota>Gammaproteobacteria>Pseudomonadales>Pseudomonadaceae>Pseudomonas>Pseudomonas_aeruginosa_group | 602 | LOH47_RS10845 | 112617..114425 | 287 | DUF262-domain-containing-protein-[Pseudomonas-aeruginosa] |
| WP_034038621.1 | ParB+HNH | GCF_900147165.1 | Pseudomonas_aeruginosa | Bacteria>Pseudomonadati>Pseudomonadota>Gammaproteobacteria>Pseudomonadales>Pseudomonadaceae>Pseudomonas>Pseudomonas_aeruginosa_group | 602 | LC626_RS08360 | 492732..494540 | 287 | DUF262-domain-containing-protein-[Pseudomonas-aeruginosa] |
| WP_057380118.1 | ParB+HNH | GCF_026727755.1 | Pseudomonas_aeruginosa | Bacteria>Pseudomonadati>Pseudomonadota>Gammaproteobacteria>Pseudomonadales>Pseudomonadaceae>Pseudomonas>Pseudomonas_aeruginosa_group | 602 | PAC42_RS26740 | 5550074..5551882 | 287 | DUF262-domain-containing-protein-[Pseudomonas-aeruginosa] |
| WP_134249506.1 | ParB+HNH | GCF_004374205.1 | Pseudomonas_aeruginosa | Bacteria>Pseudomonadati>Pseudomonadota>Gammaproteobacteria>Pseudomonadales>Pseudomonadaceae>Pseudomonas>Pseudomonas_aeruginosa_group | 602 | IPC112_RS05860 | 117909..119717 | 287 | DUF262-domain-containing-protein-[Pseudomonas-aeruginosa] |
| WP_209939196.1 | ParB+HNH | GCF_017838075.1 | Pseudomonas_aeruginosa | Bacteria>Pseudomonadati>Pseudomonadota>Gammaproteobacteria>Pseudomonadales>Pseudomonadaceae>Pseudomonas>Pseudomonas_aeruginosa_group | 602 | JOA89_RS01695 | 351537..353345 | 287 | DUF262-domain-containing-protein-[Pseudomonas-aeruginosa] |
| WP_227364594.1 | ParB+HNH | GCF_020639995.1 | Pseudomonas_aeruginosa | Bacteria>Pseudomonadati>Pseudomonadota>Gammaproteobacteria>Pseudomonadales>Pseudomonadaceae>Pseudomonas>Pseudomonas_aeruginosa_group | 602 | LJG07_RS13355 | 98915..100723 | 287 | DUF262-domain-containing-protein-[Pseudomonas-aeruginosa] |
| WP_126525536.1 | ParB+HNH | GCF_003969495.1 | Pseudomonas_aeruginosa | Bacteria>Pseudomonadati>Pseudomonadota>Gammaproteobacteria>Pseudomonadales>Pseudomonadaceae>Pseudomonas>Pseudomonas_aeruginosa_group | 598 | DZA06_RS08850 | 44237..46033 | 287 | DUF262-domain-containing-protein-[Pseudomonas-aeruginosa] |
| HBO1619177.1 | ParB+HNH | GCA_021598085.1 | Pseudomonas_aeruginosa | Bacteria>Pseudomonadati>Pseudomonadota>Gammaproteobacteria>Pseudomonadales>Pseudomonadaceae>Pseudomonas>Pseudomonas_aeruginosa_group | 597 | L4F62_005710 | 11722..13515 | 287 | DUF262-domain-containing-protein-[Pseudomonas-aeruginosa] |
| EIU3464057.1 | ParB+HNH | GCA_021644165.1 | Pseudomonas_aeruginosa | Bacteria>Pseudomonadati>Pseudomonadota>Gammaproteobacteria>Pseudomonadales>Pseudomonadaceae>Pseudomonas>Pseudomonas_aeruginosa_group | 595 | L4P80_001285 | 33072..34859 | 287 | DUF262-domain-containing-protein-[Pseudomonas-aeruginosa] |
| HBP6330510.1 | ParB+HNH | GCA_021710255.1 | Pseudomonas_aeruginosa | Bacteria>Pseudomonadati>Pseudomonadota>Gammaproteobacteria>Pseudomonadales>Pseudomonadaceae>Pseudomonas>Pseudomonas_aeruginosa_group | 595 | L6A37_21790 | 66123..67910 | 287 | DUF262-domain-containing-protein-[Pseudomonas-aeruginosa] |
| WP_119519034.1 | ParB+HNH | GCF_040186885.1 | Pseudomonas_aeruginosa | Bacteria>Pseudomonadati>Pseudomonadota>Gammaproteobacteria>Pseudomonadales>Pseudomonadaceae>Pseudomonas>Pseudomonas_aeruginosa_group | 578 | AXX04_RS18770 | 16237..17973 | 287 | DUF262-domain-containing-protein-[Pseudomonas-aeruginosa] |
| MBG3917697.1 | ParB+HNH | GCA_015732935.1 | Pseudomonas_aeruginosa | Bacteria>Pseudomonadati>Pseudomonadota>Gammaproteobacteria>Pseudomonadales>Pseudomonadaceae>Pseudomonas>Pseudomonas_aeruginosa_group | 572 | I4931_21705 | 90652..92370 | 287 | DUF262-domain-containing-protein-[Pseudomonas-aeruginosa] |
| HBN9708926.1 | ParB+HNH | GCA_021591785.1 | Pseudomonas_aeruginosa | Bacteria>Pseudomonadati>Pseudomonadota>Gammaproteobacteria>Pseudomonadales>Pseudomonadaceae>Pseudomonas>Pseudomonas_aeruginosa_group | 557 | L3920_001792 | 284201..285874 | 287 | DUF262-domain-containing-protein-[Pseudomonas-aeruginosa] |
| HBO1206627.1 | ParB+HNH | GCA_021596545.1 | Pseudomonas_aeruginosa | Bacteria>Pseudomonadati>Pseudomonadota>Gammaproteobacteria>Pseudomonadales>Pseudomonadaceae>Pseudomonas>Pseudomonas_aeruginosa_group | 557 | L4E78_005913 | 18412..20085 | 287 | TPA:-DUF262-domain-containing-protein-[Pseudomonas-aeruginosa] |
| HBO6336110.1 | ParB+HNH | GCA_021670735.1 | Pseudomonas_aeruginosa | Bacteria>Pseudomonadati>Pseudomonadota>Gammaproteobacteria>Pseudomonadales>Pseudomonadaceae>Pseudomonas>Pseudomonas_aeruginosa_group | 557 | L4747_005553 | 17141..18814 | 287 | TPA:-DUF262-domain-containing-protein-[Pseudomonas-aeruginosa] |
| HCE7261579.1 | ParB+HNH | GCA_023919205.1 | Pseudomonas_aeruginosa | Bacteria>Pseudomonadati>Pseudomonadota>Gammaproteobacteria>Pseudomonadales>Pseudomonadaceae>Pseudomonas>Pseudomonas_aeruginosa_group | 557 | NHQ13_005990 | 17035..18708 | 287 | TPA:-DUF262-domain-containing-protein-[Pseudomonas-aeruginosa] |
| HCH0453165.1 | ParB+HNH | GCA_024072695.1 | Pseudomonas_aeruginosa | Bacteria>Pseudomonadati>Pseudomonadota>Gammaproteobacteria>Pseudomonadales>Pseudomonadaceae>Pseudomonas>Pseudomonas_aeruginosa_group | 557 | NKO07_001956 | 238259..239932 | 287 | TPA:-DUF262-domain-containing-protein-[Pseudomonas-aeruginosa] |
| MBG4373389.1 | ParB+HNH | GCA_015734445.1 | Pseudomonas_aeruginosa | Bacteria>Pseudomonadati>Pseudomonadota>Gammaproteobacteria>Pseudomonadales>Pseudomonadaceae>Pseudomonas>Pseudomonas_aeruginosa_group | 557 | I5F50_23110 | 18338..20011 | 287 | DUF262-domain-containing-protein-[Pseudomonas-aeruginosa] |
| MBG4395806.1 | ParB+HNH | GCA_015734545.1 | Pseudomonas_aeruginosa | Bacteria>Pseudomonadati>Pseudomonadota>Gammaproteobacteria>Pseudomonadales>Pseudomonadaceae>Pseudomonas>Pseudomonas_aeruginosa_group | 557 | GHT73_12665 | 17369..19042 | 287 | DUF262-domain-containing-protein-[Pseudomonas-aeruginosa] |
| MBG4463098.1 | ParB+HNH | GCA_015734755.1 | Pseudomonas_aeruginosa | Bacteria>Pseudomonadati>Pseudomonadota>Gammaproteobacteria>Pseudomonadales>Pseudomonadaceae>Pseudomonas>Pseudomonas_aeruginosa_group | 557 | GHU50_17070 | 20598..22271 | 287 | DUF262-domain-containing-protein-[Pseudomonas-aeruginosa] |
| MBG5216133.1 | ParB+HNH | GCA_015737185.1 | Pseudomonas_aeruginosa | Bacteria>Pseudomonadati>Pseudomonadota>Gammaproteobacteria>Pseudomonadales>Pseudomonadaceae>Pseudomonas>Pseudomonas_aeruginosa_group | 557 | GHT75_27150 | 17273..18946 | 287 | DUF262-domain-containing-protein-[Pseudomonas-aeruginosa] |
| MBI8737748.1 | ParB+HNH | GCA_016342945.1 | Pseudomonas_aeruginosa | Bacteria>Pseudomonadati>Pseudomonadota>Gammaproteobacteria>Pseudomonadales>Pseudomonadaceae>Pseudomonas>Pseudomonas_aeruginosa_group | 557 | JEZ70_02845 | 65416..67089 | 287 | DUF262-domain-containing-protein-[Pseudomonas-aeruginosa] |
| MBX5915347.1 | ParB+HNH | GCA_019688355.1 | Pseudomonas_aeruginosa | Bacteria>Pseudomonadati>Pseudomonadota>Gammaproteobacteria>Pseudomonadales>Pseudomonadaceae>Pseudomonas>Pseudomonas_aeruginosa_group | 557 | ISE50_29140 | 17147..18820 | 287 | DUF262-domain-containing-protein-[Pseudomonas-aeruginosa] |
| MBX6172273.1 | ParB+HNH | GCA_019686195.1 | Pseudomonas_aeruginosa | Bacteria>Pseudomonadati>Pseudomonadota>Gammaproteobacteria>Pseudomonadales>Pseudomonadaceae>Pseudomonas>Pseudomonas_aeruginosa_group | 557 | ISD36_12525 | 239788..241461 | 287 | DUF262-domain-containing-protein-[Pseudomonas-aeruginosa] |
| WP_043087432.1 | ParB+HNH | GCF_024300845.1 | Pseudomonas_aeruginosa | Bacteria>Pseudomonadati>Pseudomonadota>Gammaproteobacteria>Pseudomonadales>Pseudomonadaceae>Pseudomonas>Pseudomonas_aeruginosa_group | 557 | HMPREF2746_RS21460 | 4739..6412 | 287 | DUF262-domain-containing-protein-[Pseudomonas-aeruginosa] |
| WP_108110610.1 | ParB+HNH | GCF_016820125.1 | Pseudomonas_aeruginosa | Bacteria>Pseudomonadati>Pseudomonadota>Gammaproteobacteria>Pseudomonadales>Pseudomonadaceae>Pseudomonas>Pseudomonas_aeruginosa_group | 557 | JQS52_RS11335 | 2397376..2399049 | 287 | DUF262-domain-containing-protein-[Pseudomonas-aeruginosa] |
| WP_124187487.1 | ParB+HNH | GCF_003840445.1 | Pseudomonas_aeruginosa | Bacteria>Pseudomonadati>Pseudomonadota>Gammaproteobacteria>Pseudomonadales>Pseudomonadaceae>Pseudomonas>Pseudomonas_aeruginosa_group | 557 | IPC460_RS14110 | 63965..65638 | 287 | DUF262-domain-containing-protein-[Pseudomonas-aeruginosa] |
| WP_132538222.1 | ParB+HNH | - | Pseudomonas_aeruginosa | Bacteria>Pseudomonadati>Pseudomonadota>Gammaproteobacteria>Pseudomonadales>Pseudomonadaceae>Pseudomonas>Pseudomonas_aeruginosa_group | 557 | EQ548_RS01385 | 17140..18813 | 287 | DUF262-domain-containing-protein-[Pseudomonas-aeruginosa] |
| WP_134242484.1 | ParB+HNH | GCF_004370105.1 | Pseudomonas_aeruginosa | Bacteria>Pseudomonadati>Pseudomonadota>Gammaproteobacteria>Pseudomonadales>Pseudomonadaceae>Pseudomonas>Pseudomonas_aeruginosa_group | 557 | IPC1592_RS30110 | 6739..8412 | 287 | DUF262-domain-containing-protein-[Pseudomonas-aeruginosa] |
| WP_172792836.1 | ParB+HNH | GCF_013260445.1 | Pseudomonas_aeruginosa | Bacteria>Pseudomonadati>Pseudomonadota>Gammaproteobacteria>Pseudomonadales>Pseudomonadaceae>Pseudomonas>Pseudomonas_aeruginosa_group | 557 | HQN50_RS11940 | 2502881..2504554 | 287 | DUF262-domain-containing-protein-[Pseudomonas-aeruginosa] |
| WP_260092497.1 | ParB+HNH | GCF_025152895.1 | Pseudomonas_aeruginosa | Bacteria>Pseudomonadati>Pseudomonadota>Gammaproteobacteria>Pseudomonadales>Pseudomonadaceae>Pseudomonas>Pseudomonas_aeruginosa_group | 557 | MNU23_RS16415 | 101908..103581 | 287 | DUF262-domain-containing-HNH-endonuclease-family-protein-[Pseudomonas-aeruginosa] |
| EQL43540.1 | ParB+HNH | GCA_000467675.1 | Pseudomonas_aeruginosa_VRFPA03 | Bacteria>Pseudomonadati>Pseudomonadota>Gammaproteobacteria>Pseudomonadales>Pseudomonadaceae>Pseudomonas>Pseudomonas_aeruginosa_group>Pseudomonas_aeruginosa | 602 | M770_31800 | 10414..12222 | 1350465 | hypothetical-protein-M770_31800-[Pseudomonas-aeruginosa-VRFPA03] |
| NLU01328.1 | ParB+HNH | GCA_012720005.1 | Pseudomonas_lundensis | Bacteria>Pseudomonadati>Pseudomonadota>Gammaproteobacteria>Pseudomonadales>Pseudomonadaceae>Pseudomonas>Pseudomonas_chlororaphis_group | 644 | GXW88_11985 | 15338..17272 | 86185 | DUF262-domain-containing-protein-[Pseudomonas-lundensis] |
| WP_047283133.1 | ParB+HNH | GCF_018449005.1 | Pseudomonas_lundensis | Bacteria>Pseudomonadati>Pseudomonadota>Gammaproteobacteria>Pseudomonadales>Pseudomonadaceae>Pseudomonas>Pseudomonas_chlororaphis_group | 597 | KIY13_RS15520 | 3395212..3397005 | 86185 | DUF262-domain-containing-protein-[Pseudomonas-lundensis] |
| WP_094987828.1 | ParB+HNH | GCF_012986325.1 | Pseudomonas_lundensis | Bacteria>Pseudomonadati>Pseudomonadota>Gammaproteobacteria>Pseudomonadales>Pseudomonadaceae>Pseudomonas>Pseudomonas_chlororaphis_group | 597 | HBN80_RS14465 | 96550..98343 | 86185 | DUF262-domain-containing-protein-[Pseudomonas-lundensis] |
| ARQ76273.1 | ParB+HNH | GCA_002128325.1 | Pseudomonas_fragi | Bacteria>Pseudomonadati>Pseudomonadota>Gammaproteobacteria>Pseudomonadales>Pseudomonadaceae>Pseudomonas>Pseudomonas_chlororaphis_group | 597 | B6D87_19580 | 4221881..4223674 | 296 | type-I-restriction-endonuclease-[Pseudomonas-fragi] |
| MBM1201836.1 | ParB+HNH | GCA_016812215.1 | Pseudomonas_fragi | Bacteria>Pseudomonadati>Pseudomonadota>Gammaproteobacteria>Pseudomonadales>Pseudomonadaceae>Pseudomonas>Pseudomonas_chlororaphis_group | 597 | GYM96_18830 | 98258..100051 | 296 | DUF262-domain-containing-protein-[Pseudomonas-fragi] |
| PAA41062.1 | ParB+HNH | GCA_002269445.1 | Pseudomonas_fragi | Bacteria>Pseudomonadati>Pseudomonadota>Gammaproteobacteria>Pseudomonadales>Pseudomonadaceae>Pseudomonas>Pseudomonas_chlororaphis_group | 597 | CJU80_12355 | 114498..116291 | 296 | type-I-restriction-endonuclease-[Pseudomonas-fragi] |
| WP_086798824.1 | ParB+HNH | GCF_002128325.1 | Pseudomonas_fragi | Bacteria>Pseudomonadati>Pseudomonadota>Gammaproteobacteria>Pseudomonadales>Pseudomonadaceae>Pseudomonas>Pseudomonas_chlororaphis_group | 597 | B6D87_RS19600 | 4221881..4223674 | 296 | DUF262-domain-containing-protein-[Pseudomonas-fragi] |
| WP_095025596.1 | ParB+HNH | GCF_002269515.1 | Pseudomonas_fragi | Bacteria>Pseudomonadati>Pseudomonadota>Gammaproteobacteria>Pseudomonadales>Pseudomonadaceae>Pseudomonas>Pseudomonas_chlororaphis_group | 597 | CJU79_RS17665 | 24263..26056 | 296 | DUF262-domain-containing-protein-[Pseudomonas-fragi] |
| WP_203327705.1 | ParB+HNH | GCF_016812215.1 | Pseudomonas_fragi | Bacteria>Pseudomonadati>Pseudomonadota>Gammaproteobacteria>Pseudomonadales>Pseudomonadaceae>Pseudomonas>Pseudomonas_chlororaphis_group | 597 | GYM96_RS18905 | 98258..100051 | 296 | DUF262-domain-containing-protein-[Pseudomonas-fragi] |
| WP_169872739.1 | ParB+HNH | GCF_012986835.1 | Pseudomonas_fragi | Bacteria>Pseudomonadati>Pseudomonadota>Gammaproteobacteria>Pseudomonadales>Pseudomonadaceae>Pseudomonas>Pseudomonas_chlororaphis_group | 572 | HBN88_RS02015 | 42725..44443 | 296 | DUF262-domain-containing-protein-[Pseudomonas-fragi] |
| WP_155678127.1 | ParB+HNH | GCF_004919535.4 | Pseudomonas_veronii | Bacteria>Pseudomonadati>Pseudomonadota>Gammaproteobacteria>Pseudomonadales>Pseudomonadaceae>Pseudomonas>Pseudomonas_fluorescens_group | 640 | E4167_RS17845 | 3890339..3892261 | 76761 | DUF262-domain-containing-protein-[Pseudomonas-veronii] |
| QCG65908.1 | ParB+HNH | GCA_004919535.4 | Pseudomonas_veronii | Bacteria>Pseudomonadati>Pseudomonadota>Gammaproteobacteria>Pseudomonadales>Pseudomonadaceae>Pseudomonas>Pseudomonas_fluorescens_group | 597 | E4167_12605 | 2658505..2660298 | 76761 | DUF262-domain-containing-protein-[Pseudomonas-veronii] |

| Protein-ID | Domain | Assembly-ID | Strain | Taxonomy | Length | Locus-Tag | Genomic-coordinates | TaxID | Fasta-defline |
|---|---|---|---|---|---|---|---|---|---|
| WP_197678339.1 | ParB+HNH | GCF_002091795.1 | Pseudomonas_synxantha | Bacteria>Pseudomonadati>Pseudomonadota>Gammaproteobacteria>Pseudomonadales>Pseudomonadaceae>Pseudomonas>Pseudomonas_fluorescens_group | 578 | TU77_RS22290 | 186470..188206 | 47883 | DUF262-domain-containing-protein-[Pseudomonas-synxantha] |
| WP_125970725.1 | ParB+HNH | GCF_018336155.1 | Pseudomonas_rustica | Bacteria>Pseudomonadati>Pseudomonadota>Gammaproteobacteria>Pseudomonadales>Pseudomonadaceae>Pseudomonas>Pseudomonas_fluorescens_group | 597 | EJJ21_RS24480 | 5249311..5251104 | 2827099 | MULTISPECIES:-DUF262-domain-containing-protein-[Pseudomonas-fluorescens-group] |
| WP_236238648.1 | ParB+HNH | GCF_021603345.1 | Pseudomonas_rhodesiae | Bacteria>Pseudomonadati>Pseudomonadota>Gammaproteobacteria>Pseudomonadales>Pseudomonadaceae>Pseudomonas>Pseudomonas_fluorescens_group | 624 | L6E14_RS13705 | 259008..260882 | 76760 | DUF262-domain-containing-protein-[Pseudomonas-rhodesiae] |
| WP_258640414.1 | ParB+HNH | GCF_024748715.1 | Pseudomonas_moraviensis | Bacteria>Pseudomonadati>Pseudomonadota>Gammaproteobacteria>Pseudomonadales>Pseudomonadaceae>Pseudomonas>Pseudomonas_fluorescens_group | 597 | LOY57_RS03690 | 877098..878891 | 321662 | DUF262-domain-containing-protein-[Pseudomonas-moraviensis] |
| WP_010463582.1 | ParB+HNH | GCF_000257545.3 | Pseudomonas_mandelii | Bacteria>Pseudomonadati>Pseudomonadota>Gammaproteobacteria>Pseudomonadales>Pseudomonadaceae>Pseudomonas>Pseudomonas_fluorescens_group | 624 | OU5_RS13425 | 2996463..2998337 | 75612 | DUF262-domain-containing-protein-[Pseudomonas-mandelii] |
| WP_110599713.1 | ParB+HNH | - | Pseudomonas_koreensis | Bacteria>Pseudomonadati>Pseudomonadota>Gammaproteobacteria>Pseudomonadales>Pseudomonadaceae>Pseudomonas>Pseudomonas_fluorescens_group | 572 | DMX04_RS04465 | 137494..139212 | 198620 | DUF262-domain-containing-protein-[Pseudomonas-koreensis] |
| WP_060765710.1 | ParB+HNH | GCF_001542725.1 | Pseudomonas_fluorescens | Bacteria>Pseudomonadati>Pseudomonadota>Gammaproteobacteria>Pseudomonadales>Pseudomonadaceae>Pseudomonas>Pseudomonas_fluorescens_group | 624 | PFL603g_RS03135 | 376687..378561 | 294 | DUF262-domain-containing-protein-[Pseudomonas-fluorescens] |
| WP_017337800.1 | ParB+HNH | GCF_000293885.2 | Pseudomonas_fluorescens | Bacteria>Pseudomonadati>Pseudomonadota>Gammaproteobacteria>Pseudomonadales>Pseudomonadaceae>Pseudomonas>Pseudomonas_fluorescens_group | 602 | B723_RS16835 | 3639332..3641140 | 294 | DUF262-domain-containing-protein-[Pseudomonas-fluorescens] |
| WP_115284292.1 | ParB+HNH | GCF_900455465.1 | Pseudomonas_fluorescens | Bacteria>Pseudomonadati>Pseudomonadota>Gammaproteobacteria>Pseudomonadales>Pseudomonadaceae>Pseudomonas>Pseudomonas_fluorescens_group | 597 | DYC60_RS18915 | 4151510..4153303 | 294 | DUF262-domain-containing-protein-[Pseudomonas-fluorescens] |
| WP_154949569.1 | ParB+HNH | - | Pseudomonas_fluorescens | Bacteria>Pseudomonadati>Pseudomonadota>Gammaproteobacteria>Pseudomonadales>Pseudomonadaceae>Pseudomonas>Pseudomonas_fluorescens_group | 597 | F8N22_RS18765 | 166985..168778 | 294 | DUF262-domain-containing-protein-[Pseudomonas-fluorescens] |
| VVQ23885.1 | ParB+HNH | GCA_902498245.1 | Pseudomonas_fluorescens | Bacteria>Pseudomonadati>Pseudomonadota>Gammaproteobacteria>Pseudomonadales>Pseudomonadaceae>Pseudomonas>Pseudomonas_fluorescens_group | 578 | PS928_05642 | 89554..91290 | 294 | hypothetical-protein-PS928_05642-[Pseudomonas-fluorescens] |
| WP_150787877.1 | ParB+HNH | GCF_902498245.1 | Pseudomonas_fluorescens | Bacteria>Pseudomonadati>Pseudomonadota>Gammaproteobacteria>Pseudomonadales>Pseudomonadaceae>Pseudomonas>Pseudomonas_fluorescens_group | 578 | F8N86_RS29040 | 89554..91290 | 294 | DUF262-domain-containing-protein-[Pseudomonas-fluorescens] |
| TKK14315.1 | ParB+HNH | GCA_005233695.1 | Pseudomonas_fluorescens | Bacteria>Pseudomonadati>Pseudomonadota>Gammaproteobacteria>Pseudomonadales>Pseudomonadaceae>Pseudomonas>Pseudomonas_fluorescens_group | 562 | PflCFBP13510_04725 | 503970..505658 | 294 | hypothetical-protein-PflCFBP13510_04725-[Pseudomonas-fluorescens] |
| WP_137203957.1 | ParB+HNH | - | Pseudomonas_fluorescens | Bacteria>Pseudomonadati>Pseudomonadota>Gammaproteobacteria>Pseudomonadales>Pseudomonadaceae>Pseudomonas>Pseudomonas_fluorescens_group | 562 | PflCFBP13510_RS04730 | 503970..505658 | 294 | DUF262-domain-containing-protein-[Pseudomonas-fluorescens] |
| WP_032902478.1 | ParB+HNH | - | Pseudomonas_fluorescens | Bacteria>Pseudomonadati>Pseudomonadota>Gammaproteobacteria>Pseudomonadales>Pseudomonadaceae>Pseudomonas>Pseudomonas_fluorescens_group | 557 | H098_RS08865 | 7802..9475 | 294 | DUF262-domain-containing-protein-[Pseudomonas-fluorescens] |
| WP_198726224.1 | ParB+HNH | GCF_016307475.1 | Pseudomonas_corrugata | Bacteria>Pseudomonadati>Pseudomonadota>Gammaproteobacteria>Pseudomonadales>Pseudomonadaceae>Pseudomonas>Pseudomonas_fluorescens_group | 557 | YA0788_RS19990 | 40425..42098 | 47879 | DUF262-domain-containing-protein-[Pseudomonas-corrugata] |
| SBW85149.1 | ParB+HNH | GCA_900092355.1 | Pseudomonas_veronii_1YdBTEX2 | Bacteria>Pseudomonadati>Pseudomonadota>Gammaproteobacteria>Pseudomonadales>Pseudomonadaceae>Pseudomonas>Pseudomonas_fluorescens_group>Pseudomonas_veronii | 597 | PVE_P0106 | 96277..98070 | 1295141 | hypothetical-protein-PVE_P0106-[Pseudomonas-veronii-1YdBTEX2] |
| WP_180689222.1 | ParB+HNH | GCF_013425825.1 | Pseudomonas_putida | Bacteria>Pseudomonadati>Pseudomonadota>Gammaproteobacteria>Pseudomonadales>Pseudomonadaceae>Pseudomonas>Pseudomonas_putida_group | 624 | HOH12_RS03385 | 766435..768309 | 303 | DUF262-domain-containing-protein-[Pseudomonas-putida] |
| WP_112249963.1 | ParB+HNH | - | Pseudomonas_putida | Bacteria>Pseudomonadati>Pseudomonadota>Gammaproteobacteria>Pseudomonadales>Pseudomonadaceae>Pseudomonas>Pseudomonas_putida_group | 595 | GT37_RS03110 | 10130..11917 | 303 | DUF262-domain-containing-protein-[Pseudomonas-putida] |
| WP_181088910.1 | ParB+HNH | GCF_013523035.1 | Pseudomonas_plecoglossicida | Bacteria>Pseudomonadati>Pseudomonadota>Gammaproteobacteria>Pseudomonadales>Pseudomonadaceae>Pseudomonas>Pseudomonas_putida_group | 608 | G7011_RS14195 | 102628..104454 | 70775 | DUF262-domain-containing-protein-[Pseudomonas-plecoglossicida] |
| WP_181088900.1 | ParB+HNH | GCF_013523035.1 | Pseudomonas_plecoglossicida | Bacteria>Pseudomonadati>Pseudomonadota>Gammaproteobacteria>Pseudomonadales>Pseudomonadaceae>Pseudomonas>Pseudomonas_putida_group | 602 | G7011_RS14145 | 87391..89199 | 70775 | DUF262-domain-containing-protein-[Pseudomonas-plecoglossicida] |
| WP_007159174.1 | ParB+HNH | GCF_000236825.1 | Pseudomonas_oryzihabitans | Bacteria>Pseudomonadati>Pseudomonadota>Gammaproteobacteria>Pseudomonadales>Pseudomonadaceae>Pseudomonas>Pseudomonas_putida_group | 653 | PPL19_RS03895 | 146418..148379 | 47885 | DUF262-domain-containing-protein-[Pseudomonas-psychrotolerans] |
| WP_042135796.1 | ParB+HNH | GCF_000730625.1 | Pseudomonas_oryzihabitans | Bacteria>Pseudomonadati>Pseudomonadota>Gammaproteobacteria>Pseudomonadales>Pseudomonadaceae>Pseudomonas>Pseudomonas_putida_group | 624 | POR01S_RS10895 | 111539..113413 | 47885 | DUF262-domain-containing-protein-[Pseudomonas-oryzihabitans] |
| WP_236167513.1 | ParB+HNH | GCF_021602545.1 | Pseudomonas_fulva | Bacteria>Pseudomonadati>Pseudomonadota>Gammaproteobacteria>Pseudomonadales>Pseudomonadaceae>Pseudomonas>Pseudomonas_putida_group | 757 | L6D33_RS20175 | 884..3157 | 47880 | DUF262-domain-containing-protein-[Pseudomonas-fulva] |
| WP_236167514.1 | ParB+HNH | GCF_021602545.1 | Pseudomonas_fulva | Bacteria>Pseudomonadati>Pseudomonadota>Gammaproteobacteria>Pseudomonadales>Pseudomonadaceae>Pseudomonas>Pseudomonas_putida_group | 600 | L6D33_RS20180 | 3154..4956 | 47880 | DUF262-domain-containing-protein-[Pseudomonas-fulva] |
| WP_122521124.1 | ParB+HNH | GCF_900576635.1 | Pseudomonas_viridiflava | Bacteria>Pseudomonadati>Pseudomonadota>Gammaproteobacteria>Pseudomonadales>Pseudomonadaceae>Pseudomonas>Pseudomonas_syringae_group | 597 | EET02_RS10960 | 382597..384390 | 33069 | DUF262-domain-containing-protein-[Pseudomonas-viridiflava] |
| WP_122696459.1 | ParB+HNH | - | Pseudomonas_viridiflava | Bacteria>Pseudomonadati>Pseudomonadota>Gammaproteobacteria>Pseudomonadales>Pseudomonadaceae>Pseudomonas>Pseudomonas_syringae_group | 562 | EE173_RS11545 | 13106..14794 | 33069 | DUF262-domain-containing-protein-[Pseudomonas-viridiflava] |
| WP_122811393.1 | ParB+HNH | - | Pseudomonas_viridiflava | Bacteria>Pseudomonadati>Pseudomonadota>Gammaproteobacteria>Pseudomonadales>Pseudomonadaceae>Pseudomonas>Pseudomonas_syringae_group | 562 | EE220_RS21975 | 73034..74722 | 33069 | DUF262-domain-containing-protein-[Pseudomonas-viridiflava] |
| WP_163022006.1 | ParB+HNH | - | Pseudomonas_viridiflava | Bacteria>Pseudomonadati>Pseudomonadota>Gammaproteobacteria>Pseudomonadales>Pseudomonadaceae>Pseudomonas>Pseudomonas_syringae_group | 507 | EEX91_RS04590 | 90163..91686 | 33069 | DUF262-domain-containing-protein-[Pseudomonas-viridiflava] |
| WP_230691812.1 | ParB+HNH | - | Pseudomonas_syringae_group_genomosp._3 | Bacteria>Pseudomonadati>Pseudomonadota>Gammaproteobacteria>Pseudomonadales>Pseudomonadaceae>Pseudomonas>Pseudomonas_syringae_group | 578 | I4I80_RS00300 | 49986..51722 | 251701 | DUF262-domain-containing-protein-[Pseudomonas-syringae-group-genomosp.-3] |
| WP_046835094.1 | ParB+HNH | GCF_030252675.1 | Pseudomonas_syringae_group_genomosp._2 | Bacteria>Pseudomonadati>Pseudomonadota>Gammaproteobacteria>Pseudomonadales>Pseudomonadaceae>Pseudomonas>Pseudomonas_syringae_group | 572 | QO021_RS27765 | 6028529..6030247 | 251698 | DUF262-domain-containing-protein-[Pseudomonas-amygdali] |
| WP_236471268.1 | ParB+HNH | GCF_021606115.1 | Pseudomonas_syringae | Bacteria>Pseudomonadati>Pseudomonadota>Gammaproteobacteria>Pseudomonadales>Pseudomonadaceae>Pseudomonas>Pseudomonas_syringae_group>Pseudomonas_syringae_group_genomosp._1 | 597 | GIW32_RS04315 | 14710..16503 | 317 | DUF262-domain-containing-protein-[Pseudomonas-syringae] |
| WP_213582010.1 | ParB+HNH | GCF_018383465.1 | Pseudomonas_syringae | Bacteria>Pseudomonadati>Pseudomonadota>Gammaproteobacteria>Pseudomonadales>Pseudomonadaceae>Pseudomonas>Pseudomonas_syringae_group>Pseudomonas_syringae_group_genomosp._1 | 572 | KID98_RS09085 | 2030526..2032244 | 317 | DUF262-domain-containing-protein-[Pseudomonas-syringae] |
| WP_236481328.1 | ParB+HNH | - | Pseudomonas_syringae | Bacteria>Pseudomonadati>Pseudomonadota>Gammaproteobacteria>Pseudomonadales>Pseudomonadaceae>Pseudomonas>Pseudomonas_syringae_group>Pseudomonas_syringae_group_genomosp._1 | 572 | GIW69_RS06580 | 4942..6660 | 317 | DUF262-domain-containing-protein-[Pseudomonas-syringae] |
| MBF9242734.1 | ParB+HNH | GCA_015694365.1 | Pseudomonas_syringae_pv._tomato | Bacteria>Pseudomonadati>Pseudomonadota>Gammaproteobacteria>Pseudomonadales>Pseudomonadaceae>Pseudomonas>Pseudomonas_syringae_group>Pseudomonas_syringae_group_genomosp._3 | 578 | I4I80_00300 | 49986..51722 | 323 | DUF262-domain-containing-protein-[Pseudomonas-syringae-pv.-tomato] |
| NMY25072.1 | ParB+HNH | GCA_012985535.1 | Pseudomonas_sp._WS_5021 | Bacteria>Pseudomonadati>Pseudomonadota>Gammaproteobacteria>Pseudomonadales>Pseudomonadaceae>Pseudomonas>unclassified_Pseudomonas | 562 | HBO19_03700 | 802163..803851 | 2717490 | DUF262-domain-containing-protein-[Pseudomonas-sp.-WS-5021] |
| WP_170056916.1 | ParB+HNH | GCF_012985535.1 | Pseudomonas_sp._WS_5021 | Bacteria>Pseudomonadati>Pseudomonadota>Gammaproteobacteria>Pseudomonadales>Pseudomonadaceae>Pseudomonas>unclassified_Pseudomonas | 562 | HBO19_RS03690 | 802163..803851 | 2717490 | DUF262-domain-containing-protein-[Pseudomonas-sp.-WS-5021] |
| WP_170050470.1 | ParB+HNH | GCF_012985775.1 | Pseudomonas_sp._WS_5011 | Bacteria>Pseudomonadati>Pseudomonadota>Gammaproteobacteria>Pseudomonadales>Pseudomonadaceae>Pseudomonas>unclassified_Pseudomonas | 595 | HBN63_RS18860 | 29541..31328 | 2717477 | DUF262-domain-containing-protein-[Pseudomonas-sp.-WS-5011] |
| WP_095626340.1 | ParB+HNH | - | Pseudomonas_sp._WN033 | Bacteria>Pseudomonadati>Pseudomonadota>Gammaproteobacteria>Pseudomonadales>Pseudomonadaceae>Pseudomonas>unclassified_Pseudomonas | 598 | CK507_RS15695 | 122972..124768 | 2032629 | DUF262-domain-containing-protein-[Pseudomonas-sp.-WN033] |
| WP_130899397.1 | ParB+HNH | GCF_004307505.1 | Pseudomonas_sp._Sample_11 | Bacteria>Pseudomonadati>Pseudomonadota>Gammaproteobacteria>Pseudomonadales>Pseudomonadaceae>Pseudomonas>unclassified_Pseudomonas | 597 | D8U60_RS02320 | 33290..35083 | 2448261 | DUF262-domain-containing-protein-[Pseudomonas-sp.-Sample_11] |
| WP_109933368.1 | ParB+HNH | GCF_003176655.1 | Pseudomonas_sp._RW407 | Bacteria>Pseudomonadati>Pseudomonadota>Gammaproteobacteria>Pseudomonadales>Pseudomonadaceae>Pseudomonas>unclassified_Pseudomonas | 557 | DK254_RS08140 | 342758..344431 | 2202894 | DUF262-domain-containing-protein-[Pseudomonas-sp.-RW407] |
| MBY8973662.1 | ParB+HNH | GCA_019891455.1 | Pseudomonas_sp._P867 | Bacteria>Pseudomonadati>Pseudomonadota>Gammaproteobacteria>Pseudomonadales>Pseudomonadaceae>Pseudomonas>unclassified_Pseudomonas | 562 | J1G33_25095 | 7654..9342 | 2816050 | DUF262-domain-containing-protein-[Pseudomonas-sp.-P867] |
| WP_223018980.1 | ParB+HNH | GCF_019891455.1 | Pseudomonas_sp._P867 | Bacteria>Pseudomonadati>Pseudomonadota>Gammaproteobacteria>Pseudomonadales>Pseudomonadaceae>Pseudomonas>unclassified_Pseudomonas | 562 | J1G33_RS25090 | 7654..9342 | 2816050 | DUF262-domain-containing-protein-[Pseudomonas-sp.-P867] |
| WP_173406511.1 | ParB+HNH | GCF_013283895.1 | Pseudomonas_sp._MPDS | Bacteria>Pseudomonadati>Pseudomonadota>Gammaproteobacteria>Pseudomonadales>Pseudomonadaceae>Pseudomonas>unclassified_Pseudomonas | 597 | HQ912_RS04615 | 1055092..1056885 | 2762896 | DUF262-domain-containing-protein-[Pseudomonas-sp.-MPDS] |
| WP_198790928.1 | ParB+HNH | GCF_016405175.1 | Pseudomonas_sp._MF7451 | Bacteria>Pseudomonadati>Pseudomonadota>Gammaproteobacteria>Pseudomonadales>Pseudomonadaceae>Pseudomonas>unclassified_Pseudomonas | 624 | JFT65_RS10930 | 203489..205363 | 2797538 | DUF262-domain-containing-protein-[Pseudomonas-sp.-MF7451] |
| BDM24920.1 | ParB+HNH | GCA_024349685.1 | Pseudomonas_sp._LRP2-20 | Bacteria>Pseudomonadati>Pseudomonadota>Gammaproteobacteria>Pseudomonadales>Pseudomonadaceae>Pseudomonas>unclassified_Pseudomonas | 595 | KMS_R46770 | 5323154..5324941 | 2944234 | DUF262-domain-containing-HNH-endonuclease-family-protein-[Pseudomonas-sp.-LRP2-20] |
| WP_261941652.1 | ParB+HNH | GCF_024349685.1 | Pseudomonas_sp._LRP2-20 | Bacteria>Pseudomonadati>Pseudomonadota>Gammaproteobacteria>Pseudomonadales>Pseudomonadaceae>Pseudomonas>unclassified_Pseudomonas | 595 | OCX61_RS23885 | 5323154..5324941 | 2944234 | DUF262-domain-containing-protein-[Pseudomonas-sp.-LRP2-20] |
| WP_079783559.1 | ParB+HNH | - | Pseudomonas_sp._LPH1 | Bacteria>Pseudomonadati>Pseudomonadota>Gammaproteobacteria>Pseudomonadales>Pseudomonadaceae>Pseudomonas>unclassified_Pseudomonas | 557 | BHQ29_RS09375 | 2040160..2041833 | 1898684 | DUF262-domain-containing-protein-[Pseudomonas-sp.-LPH1] |
| WP_259155462.1 | ParB+HNH | GCF_024809425.1 | Pseudomonas_sp._JAI111 | Bacteria>Pseudomonadati>Pseudomonadota>Gammaproteobacteria>Pseudomonadales>Pseudomonadaceae>Pseudomonas>unclassified_Pseudomonas | 597 | HNR03_RS20410 | 280064..281857 | 2735913 | DUF262-domain-containing-protein-[Pseudomonas-sp.-JAI111] |
| WP_095181953.1 | ParB+HNH | GCF_900187595.1 | Pseudomonas_sp._Irchel_3F6 | Bacteria>Pseudomonadati>Pseudomonadota>Gammaproteobacteria>Pseudomonadales>Pseudomonadaceae>Pseudomonas>unclassified_Pseudomonas | 578 | CKW30_RS22370 | 285622..287358 | 2009003 | DUF262-domain-containing-protein-[Pseudomonas-sp.-Irchel-3F6] |
| WP_226045700.1 | ParB+HNH | GCF_020401845.1 | Pseudomonas_sp._HS-18 | Bacteria>Pseudomonadati>Pseudomonadota>Gammaproteobacteria>Pseudomonadales>Pseudomonadaceae>Pseudomonas>unclassified_Pseudomonas | 572 | LDJ84_RS00040 | 13243..14961 | 2879114 | DUF262-domain-containing-protein-[Pseudomonas-sp.-HS-18] |
| ATN11142.1 | ParB+HNH | GCA_002591235.1 | Pseudomonas_sp._FDAARGOS_380 | Bacteria>Pseudomonadati>Pseudomonadota>Gammaproteobacteria>Pseudomonadales>Pseudomonadaceae>Pseudomonas>unclassified_Pseudomonas | 562 | CRN80_16475 | 3515143..3516831 | 2018067 | hypothetical-protein-CRN80_16475-[Pseudomonas-sp.-FDAARGOS_380] |
| WP_098950599.1 | ParB+HNH | GCF_002591235.1 | Pseudomonas_sp._FDAARGOS_380 | Bacteria>Pseudomonadati>Pseudomonadota>Gammaproteobacteria>Pseudomonadales>Pseudomonadaceae>Pseudomonas>unclassified_Pseudomonas | 562 | CRN80_RS16475 | 3515143..3516831 | 2018067 | DUF262-domain-containing-protein-[Pseudomonas-sp.-FDAARGOS_380] |
| WP_192985629.1 | ParB+HNH | GCF_014892995.1 | Pseudomonas_sp._EggHat1 | Bacteria>Pseudomonadati>Pseudomonadota>Gammaproteobacteria>Pseudomonadales>Pseudomonadaceae>Pseudomonas>unclassified_Pseudomonas | 578 | INS93_RS22760 | 64074..65810 | 2761624 | DUF262-domain-containing-protein-[Pseudomonas-sp.-EggHat1] |
| WP_129388881.1 | ParB+HNH | GCF_004124275.1 | Pseudomonas_sp._DTU12.3 | Bacteria>Pseudomonadati>Pseudomonadota>Gammaproteobacteria>Pseudomonadales>Pseudomonadaceae>Pseudomonas>unclassified_Pseudomonas | 597 | C2E19_RS05080 | 1150498..1152291 | 2073078 | DUF262-domain-containing-protein-[Pseudomonas-sp.-DTU12.3] |
| WP_265980018.1 | ParB+HNH | GCF_026241005.1 | Pseudomonas_sp._DCB_BG | Bacteria>Pseudomonadati>Pseudomonadota>Gammaproteobacteria>Pseudomonadales>Pseudomonadaceae>Pseudomonas>unclassified_Pseudomonas | 623 | OO254_RS23960 | 56726..58597 | 2993595 | DUF262-domain-containing-protein-[Pseudomonas-sp.-DCB_BG] |
| WP_265969624.1 | ParB+HNH | GCF_026240895.1 | Pseudomonas_sp._DCB_AW | Bacteria>Pseudomonadati>Pseudomonadota>Gammaproteobacteria>Pseudomonadales>Pseudomonadaceae>Pseudomonas>unclassified_Pseudomonas | 563 | OO306_RS19215 | 18635..20326 | 2993596 | DUF262-domain-containing-HNH-endonuclease-family-protein-[Pseudomonas-sp.-DCB_AW] |
| WP_137823531.1 | ParB+HNH | GCF_005502555.1 | Pseudomonas_sp._D(2018) | Bacteria>Pseudomonadati>Pseudomonadota>Gammaproteobacteria>Pseudomonadales>Pseudomonadaceae>Pseudomonas>unclassified_Pseudomonas | 562 | FFS60_RS25270 | 17388..19076 | 2502238 | DUF262-domain-containing-protein-[Pseudomonas-sp.-D(2018)] |
| WP_101152279.1 | ParB+HNH | GCF_002836195.1 | Pseudomonas_sp._Choline-3u-10 | Bacteria>Pseudomonadati>Pseudomonadota>Gammaproteobacteria>Pseudomonadales>Pseudomonadaceae>Pseudomonas>unclassified_Pseudomonas | 653 | CXF92_RS00345 | 67008..68969 | 2058311 | DUF262-domain-containing-protein-[Pseudomonas-sp.-Choline-3u-10] |
| WP_231661143.1 | ParB+HNH | GCF_021147865.1 | Pseudomonas_sp._CDFA_610 | Bacteria>Pseudomonadati>Pseudomonadota>Gammaproteobacteria>Pseudomonadales>Pseudomonadaceae>Pseudomonas>unclassified_Pseudomonas | 597 | KDX39_RS16875 | 247746..249539 | 2829825 | DUF262-domain-containing-protein-[Pseudomonas-sp.-CDFA-610] |
| WP_258626728.1 | ParB+HNH | GCF_024749045.1 | Pseudomonas_sp._B21-028 | Bacteria>Pseudomonadati>Pseudomonadota>Gammaproteobacteria>Pseudomonadales>Pseudomonadaceae>Pseudomonas>unclassified_Pseudomonas | 562 | LOY35_RS21310 | 4951287..4952975 | 2895480 | DUF262-domain-containing-HNH-endonuclease-family-protein-[Pseudomonas-sp.-B21-028] |
| WP_136489734.1 | ParB+HNH | GCF_004801855.1 | Pseudomonas_sp._A-1 | Bacteria>Pseudomonadati>Pseudomonadota>Gammaproteobacteria>Pseudomonadales>Pseudomonadaceae>Pseudomonas>unclassified_Pseudomonas | 557 | E5198_RS09385 | 7282..8955 | 1821274 | DUF262-domain-containing-protein-[Pseudomonas-sp.-A-1] |
| PIB47067.1 | ParB+HNH | GCA_002742495.1 | Pseudomonas_sp._2588-5 | Bacteria>Pseudomonadati>Pseudomonadota>Gammaproteobacteria>Pseudomonadales>Pseudomonadaceae>Pseudomonas>unclassified_Pseudomonas | 562 | AOA57_22430 | 335648..337336 | 1712676 | hypothetical-protein-AOA57_22430-[Pseudomonas-sp.-2588-5] |
| WP_065945474.1 | ParB+HNH | GCF_900004795.1 | Pseudomonas_sp._24_R_17 | Bacteria>Pseudomonadati>Pseudomonadota>Gammaproteobacteria>Pseudomonadales>Pseudomonadaceae>Pseudomonas>unclassified_Pseudomonas | 557 | BFP98_RS06340 | 92444..94117 | 1844096 | DUF262-domain-containing-protein-[Pseudomonas-sp.-24-R-17] |
| WP_257780135.1 | ParB+HNH | GCF_024655625.1 | Pseudomonas_sp._18.1.10 | Bacteria>Pseudomonadati>Pseudomonadota>Gammaproteobacteria>Pseudomonadales>Pseudomonadaceae>Pseudomonas>unclassified_Pseudomonas | 578 | NUV89_RS02210 | 16908..18644 | 2969302 | DUF262-domain-containing-protein-[Pseudomonas-sp.-18.1.10] |
| WP_088513984.1 | ParB+HNH | GCF_002157495.1 | Pseudomonas_sp._1239 | Bacteria>Pseudomonadati>Pseudomonadota>Gammaproteobacteria>Pseudomonadales>Pseudomonadaceae>Pseudomonas>unclassified_Pseudomonas | 574 | B8W70_RS03335 | 11145..12869 | 1985343 | DUF262-domain-containing-protein-[Pseudomonas-sp.-1239] |
| WP_046164312.1 | ParB+HNH | GCF_000970615.1 | Pseudomonas_sp._10B238 | Bacteria>Pseudomonadati>Pseudomonadota>Gammaproteobacteria>Pseudomonadales>Pseudomonadaceae>Pseudomonas>unclassified_Pseudomonas | 624 | RT21_RS21320 | 4963..6837 | 1586417 | DUF262-domain-containing-protein-[Pseudomonas-sp.-10B238] |
| MCK9534932.1 | ParB+HNH | GCA_023228505.1 | Pseudomonas_sp. | Bacteria>Pseudomonadati>Pseudomonadota>Gammaproteobacteria>Pseudomonadales>Pseudomonadaceae>Pseudomonas>unclassified_Pseudomonas | 560 | MOQ98_09605 | 12968..14650 | 306 | DUF262-domain-containing-HNH-endonuclease-family-protein-[Pseudomonas-sp.] |
| MCQ2029881.1 | ParB+HNH | GCA_024397535.1 | Stutzerimonas_zhaodongensis | Bacteria>Pseudomonadati>Pseudomonadota>Gammaproteobacteria>Pseudomonadales>Pseudomonadaceae>Stutzerimonas | 624 | NA613_09810 | 29133..31007 | 1176257 | DUF262-domain-containing-protein-[[Pseudomonas] |
| WP_255830670.1 | ParB+HNH | GCF_024397535.1 | Stutzerimonas_zhaodongensis | Bacteria>Pseudomonadati>Pseudomonadota>Gammaproteobacteria>Pseudomonadales>Pseudomonadaceae>Stutzerimonas | 624 | NA613_RS09835 | 29133..31007 | 1176257 | DUF262-domain-containing-protein-[[Pseudomonas] |
| WP_120997714.1 | ParB+HNH | GCF_003640395.1 | Stutzerimonas_urumqiensis | Bacteria>Pseudomonadati>Pseudomonadota>Gammaproteobacteria>Pseudomonadales>Pseudomonadaceae>Stutzerimonas | 653 | D7325_RS17355 | 30815..32776 | 638269 | DUF262-domain-containing-protein-[[Pseudomonas] |
| OWG36867.1 | ParB+HNH | GCA_002193135.1 | Stutzerimonas_stutzeri | Bacteria>Pseudomonadati>Pseudomonadota>Gammaproteobacteria>Pseudomonadales>Pseudomonadaceae>Stutzerimonas>Stutzerimonas_stutzeri_group>Stutzerimonas_stutzeri_subgroup | 624 | CAQ69_18780 | 27379..29253 | 316 | hypothetical-protein-CAQ69_18780-[Stutzerimonas-stutzeri] |
| WP_125856773.1 | ParB+HNH | GCF_003940515.1 | Stutzerimonas_stutzeri | Bacteria>Pseudomonadati>Pseudomonadota>Gammaproteobacteria>Pseudomonadales>Pseudomonadaceae>Stutzerimonas>Stutzerimonas_stutzeri_group>Stutzerimonas_stutzeri_subgroup | 595 | EGJ36_RS18115 | 13036..14823 | 316 | DUF262-domain-containing-protein-[Stutzerimonas-stutzeri] |
| MCF6781035.1 | ParB+HNH | GCA_021713665.1 | Stutzerimonas_stutzeri | Bacteria>Pseudomonadati>Pseudomonadota>Gammaproteobacteria>Pseudomonadales>Pseudomonadaceae>Stutzerimonas>Stutzerimonas_stutzeri_group>Stutzerimonas_stutzeri_subgroup | 572 | L3X17_04515 | 20959..22677 | 316 | DUF262-domain-containing-HNH-endonuclease-family-protein-[Stutzerimonas-stutzeri] |
| MCF6804931.1 | ParB+HNH | GCA_021713635.1 | Stutzerimonas_stutzeri | Bacteria>Pseudomonadati>Pseudomonadota>Gammaproteobacteria>Pseudomonadales>Pseudomonadaceae>Stutzerimonas>Stutzerimonas_stutzeri_group>Stutzerimonas_stutzeri_subgroup | 572 | L3X13_09155 | 20963..22681 | 316 | DUF262-domain-containing-HNH-endonuclease-family-protein-[Stutzerimonas-stutzeri] |
| MTI93109.1 | ParB+HNH | GCA_009711815.1 | Stutzerimonas_stutzeri | Bacteria>Pseudomonadati>Pseudomonadota>Gammaproteobacteria>Pseudomonadales>Pseudomonadaceae>Stutzerimonas>Stutzerimonas_stutzeri_group>Stutzerimonas_stutzeri_subgroup | 572 | FH750_17250 | 16960..18678 | 316 | DUF262-domain-containing-protein-[Stutzerimonas-stutzeri] |
| WP_125852813.1 | ParB+HNH | GCF_003936145.1 | Stutzerimonas_stutzeri | Bacteria>Pseudomonadati>Pseudomonadota>Gammaproteobacteria>Pseudomonadales>Pseudomonadaceae>Stutzerimonas>Stutzerimonas_stutzeri_group>Stutzerimonas_stutzeri_subgroup | 572 | EGJ42_RS05715 | 50666..52384 | 316 | DUF262-domain-containing-protein-[Stutzerimonas-stutzeri] |
| WP_125867553.1 | ParB+HNH | - | Stutzerimonas_stutzeri | Bacteria>Pseudomonadati>Pseudomonadota>Gammaproteobacteria>Pseudomonadales>Pseudomonadaceae>Stutzerimonas>Stutzerimonas_stutzeri_group>Stutzerimonas_stutzeri_subgroup | 572 | EGI99_RS04495 | 992693..994411 | 316 | DUF262-domain-containing-protein-[Stutzerimonas-stutzeri] |
| WP_161612278.1 | ParB+HNH | GCF_029839225.1 | Stutzerimonas_stutzeri | Bacteria>Pseudomonadati>Pseudomonadota>Gammaproteobacteria>Pseudomonadales>Pseudomonadaceae>Stutzerimonas>Stutzerimonas_stutzeri_group>Stutzerimonas_stutzeri_subgroup | 572 | N5C32_RS13920 | 182468..184186 | 316 | DUF262-domain-containing-protein-[Stutzerimonas-stutzeri] |
| WP_283258800.1 | ParB+HNH | GCF_030056015.1 | Stutzerimonas_stutzeri | Bacteria>Pseudomonadati>Pseudomonadota>Gammaproteobacteria>Pseudomonadales>Pseudomonadaceae>Stutzerimonas>Stutzerimonas_stutzeri_group>Stutzerimonas_stutzeri_subgroup | 572 | QM290_RS03920 | 6537..8255 | 316 | DUF262-domain-containing-protein-[Stutzerimonas-stutzeri] |
| MCQ4256504.1 | ParB+HNH | GCA_024448895.1 | Stutzerimonas_stutzeri | Bacteria>Pseudomonadati>Pseudomonadota>Gammaproteobacteria>Pseudomonadales>Pseudomonadaceae>Stutzerimonas>Stutzerimonas_stutzeri_group>Stutzerimonas_stutzeri_subgroup | 557 | NA619_00620 | 140016..141689 | 316 | DUF262-domain-containing-HNH-endonuclease-family-protein-[Stutzerimonas-stutzeri] |
| WP_218422423.1 | ParB+HNH | GCF_019203165.1 | Stutzerimonas_stutzeri | Bacteria>Pseudomonadati>Pseudomonadota>Gammaproteobacteria>Pseudomonadales>Pseudomonadaceae>Stutzerimonas>Stutzerimonas_stutzeri_group>Stutzerimonas_stutzeri_subgroup | 557 | KUD99_RS07260 | 1506455..1508128 | 316 | DUF262-domain-containing-protein-[Stutzerimonas-stutzeri] |
| WP_236450579.1 | ParB+HNH | GCF_021610005.1 | Stutzerimonas_stutzeri | Bacteria>Pseudomonadati>Pseudomonadota>Gammaproteobacteria>Pseudomonadales>Pseudomonadaceae>Stutzerimonas>Stutzerimonas_stutzeri_group>Stutzerimonas_stutzeri_subgroup | 557 | L1776_RS13350 | 2921199..2922872 | 316 | DUF262-domain-containing-protein-[Stutzerimonas-stutzeri] |
| WP_255958971.1 | ParB+HNH | GCF_024448895.1 | Stutzerimonas_stutzeri | Bacteria>Pseudomonadati>Pseudomonadota>Gammaproteobacteria>Pseudomonadales>Pseudomonadaceae>Stutzerimonas>Stutzerimonas_stutzeri_group>Stutzerimonas_stutzeri_subgroup | 557 | NA619_RS00620 | 140016..141689 | 316 | DUF262-domain-containing-protein-[Stutzerimonas-stutzeri] |
| WP_023443897.1 | ParB+HNH | GCF_000495915.1 | Stutzerimonas_chloritidismutans | Bacteria>Pseudomonadati>Pseudomonadota>Gammaproteobacteria>Pseudomonadales>Pseudomonadaceae>Stutzerimonas>Stutzerimonas_stutzeri_group>Stutzerimonas_stutzeri_subgroup | 602 | F753_RS26970 | 69275..71083 | 203192 | DUF262-domain-containing-protein-[Stutzerimonas-chloritidismutans] |
| WP_053105416.1 | ParB+HNH | GCF_001267275.1 | Thiopseudomonas_alkaliphila | Bacteria>Pseudomonadati>Pseudomonadota>Gammaproteobacteria>Pseudomonadales>Pseudomonadaceae>Thiopseudomonas | 560 | AKN90_RS00625 | 130273..131955 | 1697053 | DUF262-domain-containing-protein-[Thiopseudomonas-alkaliphila] |
| MCP5171287.1 | ParB+HNH | GCA_024234435.1 | Pseudomonadales_bacterium | Bacteria>Pseudomonadati>Pseudomonadota>Gammaproteobacteria>Pseudomonadales>unclassified_Pseudomonadales | 642 | H7A02_03335 | 710533..712461 | 1891229 | DUF262-domain-containing-protein-[Pseudomonadales-bacterium] |
| MAQ23216.1 | ParB+HNH | GCA_002701315.1 | Pseudomonadales_bacterium | Bacteria>Pseudomonadati>Pseudomonadota>Gammaproteobacteria>Pseudomonadales>unclassified_Pseudomonadales | 626 | CMK83_03270 | 9124..11004 | 1891229 | hypothetical-protein-[Pseudomonadales-bacterium] |
| MAB42109.1 | ParB+HNH | GCA_002684155.1 | Pseudomonadales_bacterium | Bacteria>Pseudomonadati>Pseudomonadota>Gammaproteobacteria>Pseudomonadales>unclassified_Pseudomonadales | 567 | CMK77_07480 | 8440..10143 | 1891229 | hypothetical-protein-[Pseudomonadales-bacterium] |
| MCP5179007.1 | ParB+HNH | GCA_024234635.1 | Pseudomonadales_bacterium | Bacteria>Pseudomonadati>Pseudomonadota>Gammaproteobacteria>Pseudomonadales>unclassified_Pseudomonadales | 562 | H6994_03750 | 327114..328802 | 1891229 | DUF262-domain-containing-protein-[Pseudomonadales-bacterium] |
| MCB1656849.1 | ParB+HNH | GCA_020443205.1 | Pseudomonadales_bacterium | Bacteria>Pseudomonadati>Pseudomonadota>Gammaproteobacteria>Pseudomonadales>unclassified_Pseudomonadales | 556 | KDI39_01320 | 7644..9314 | 1891229 | DUF262-domain-containing-protein-[Pseudomonadales-bacterium] |
| WP_007146325.1 | ParB+HNH | GCF_000214595.1 | Methylophaga_aminisulfidivorans | Bacteria>Pseudomonadati>Pseudomonadota>Gammaproteobacteria>Thiotrichales>Piscirickettsiaceae>Methylophaga | 627 | MAMP_RS16280 | 81689..83572 | 230105 | DUF262-domain-containing-protein-[Methylophaga-aminisulfidivorans] |
| HEC58781.1 | ParB+HNH | GCA_011053005.1 | Methylophaga_sp. | Bacteria>Pseudomonadati>Pseudomonadota>Gammaproteobacteria>Thiotrichales>Piscirickettsiaceae>Methylophaga>unclassified_Methylophaga | 604 | ENI24_04345 | 32282..34096 | 2024840 | DUF262-domain-containing-protein-[Methylophaga-sp.] |
| CAA6806504.1 | ParB+HNH | - | uncultured_Thiotrichaceae_bacterium | Bacteria>Pseudomonadati>Pseudomonadota>Gammaproteobacteria>Thiotrichales>Thiotrichaceae>environmental_samples | 146 | HELGO_WM31108 | 1..438 | 298394 | RloF,-partial-[uncultured-Thiotrichaceae-bacterium] |
| MCY4286574.1 | ParB+HNH | GCA_026709935.1 | Thiotrichales_bacterium | Bacteria>Pseudomonadati>Pseudomonadota>Gammaproteobacteria>Thiotrichales>unclassified_Thiotrichales | 360 | OXC65_14700 | 761..1843 | 2026796 | MAG:-DUF262-domain-containing-protein-[Thiotrichales-bacterium] |
| MCB1859933.1 | ParB+HNH | GCA_020445415.1 | Gammaproteobacteria_bacterium | Bacteria>Pseudomonadati>Pseudomonadota>Gammaproteobacteria>unclassified_Gammaproteobacteria | 701 | KDI63_16795 | 9257..11362 | 1913989 | DUF262-domain-containing-protein-[Gammaproteobacteria-bacterium] |
| MBU1286068.1 | ParB+HNH | GCA_018819505.1 | Gammaproteobacteria_bacterium | Bacteria>Pseudomonadati>Pseudomonadota>Gammaproteobacteria>unclassified_Gammaproteobacteria | 669 | KJ794_21935 | 11577..13586 | 1913989 | DUF262-domain-containing-HNH-endonuclease-family-protein-[Gammaproteobacteria-bacterium] |
| MCP5421178.1 | ParB+HNH | GCA_024235155.1 | Gammaproteobacteria_bacterium | Bacteria>Pseudomonadati>Pseudomonadota>Gammaproteobacteria>unclassified_Gammaproteobacteria | 573 | H6969_11920 | 7285..9006 | 1913989 | DUF262-domain-containing-protein-[Gammaproteobacteria-bacterium] |
| NLY14763.1 | ParB+HNH | GCA_012728415.1 | Gammaproteobacteria_bacterium | Bacteria>Pseudomonadati>Pseudomonadota>Gammaproteobacteria>unclassified_Gammaproteobacteria | 566 | GXZ05_00065 | 7586..9286 | 1913989 | DUF262-domain-containing-protein-[Gammaproteobacteria-bacterium] |
| MBU1286067.1 | ParB+HNH | GCA_018819505.1 | Gammaproteobacteria_bacterium | Bacteria>Pseudomonadati>Pseudomonadota>Gammaproteobacteria>unclassified_Gammaproteobacteria | 494 | KJ794_21930 | 10096..11580 | 1913989 | DUF262-domain-containing-protein-[Gammaproteobacteria-bacterium] |
| MDD9983189.1 | ParB+HNH | GCA_028818395.1 | Gammaproteobacteria_bacterium | Bacteria>Pseudomonadati>Pseudomonadota>Gammaproteobacteria>unclassified_Gammaproteobacteria | 363 | OXU81_17830 | 10055..11146 | 1913989 | MAG:-DUF262-domain-containing-protein-[Gammaproteobacteria-bacterium] |
| WP_025525719.1 | ParB+HNH | GCF_000489535.1 | Vibrionaceae | Bacteria>Pseudomonadati>Pseudomonadota>Gammaproteobacteria>Vibrionales | 642 | FH974_RS17025 | 236142..238070 | 641 | MULTISPECIES:-DUF262-domain-containing-protein-[Vibrionaceae] |
| WP_025985247.1 | ParB+HNH | GCF_000347495.1 | Vibrio | Bacteria>Pseudomonadati>Pseudomonadota>Gammaproteobacteria>Vibrionales>Vibrionaceae | 698 | VPPCV08_RS0113755 | 237245..239341 | 662 | MULTISPECIES:-DUF262-and-DUF1524-domain-containing-protein-[Vibrio] |
| WP_074189991.1 | ParB+HNH | GCF_001909305.1 | Vibrio | Bacteria>Pseudomonadati>Pseudomonadota>Gammaproteobacteria>Vibrionales>Vibrionaceae | 642 | H058_RS04220 | 537897..539825 | 662 | MULTISPECIES:-DUF262-domain-containing-protein-[Vibrio-harveyi-group] |
| WP_102582641.1 | ParB+HNH | GCF_041470425.1 | Vibrio | Bacteria>Pseudomonadati>Pseudomonadota>Gammaproteobacteria>Vibrionales>Vibrionaceae | 641 | BCT76_RS12985 | 34975..36900 | 662 | DUF262-domain-containing-protein-[Vibrio-tasmaniensis] |
| WP_054579584.1 | ParB+HNH | GCF_014769135.1 | Vibrio | Bacteria>Pseudomonadati>Pseudomonadota>Gammaproteobacteria>Vibrionales>Vibrionaceae | 601 | NB477_RS23060 | 8402..10207 | 662 | DUF262-domain-containing-protein-[Vibrio-alginolyticus] |
| WP_065090482.1 | ParB+HNH | GCF_021725735.1 | Vibrio | Bacteria>Pseudomonadati>Pseudomonadota>Gammaproteobacteria>Vibrionales>Vibrionaceae | 601 | L3V27_RS03740 | 861234..863039 | 662 | MULTISPECIES:-DUF262-domain-containing-protein-[Vibrio] |
| WP_188047086.1 | ParB+HNH | GCF_014525155.1 | Vibrio | Bacteria>Pseudomonadati>Pseudomonadota>Gammaproteobacteria>Vibrionales>Vibrionaceae | 601 | GO186_RS04085 | 582987..584792 | 662 | DUF262-domain-containing-protein-[Vibrio-vulnificus] |
| WP_198107478.1 | ParB+HNH | GCF_963891765.1 | Vibrio | Bacteria>Pseudomonadati>Pseudomonadota>Gammaproteobacteria>Vibrionales>Vibrionaceae | 601 | O8Z82_RS06525 | 623292..625097 | 662 | MULTISPECIES:-DUF262-domain-containing-protein-[Vibrio] |
| WP_213899841.1 | ParB+HNH | GCF_018458525.1 | Vibrio | Bacteria>Pseudomonadati>Pseudomonadota>Gammaproteobacteria>Vibrionales>Vibrionaceae | 601 | J4H25_RS19075 | 17475..19280 | 662 | DUF262-domain-containing-protein-[Vibrio-alginolyticus] |
| WP_210942221.1 | ParB+HNH | GCF_963893545.1 | Vibrio | Bacteria>Pseudomonadati>Pseudomonadota>Gammaproteobacteria>Vibrionales>Vibrionaceae | 597 | GSG51_RS14600 | 16488..18281 | 662 | DUF262-domain-containing-protein-[Vibrio-paracholerae] |
| WP_025612389.1 | ParB+HNH | GCF_000489615.1 | Vibrio | Bacteria>Pseudomonadati>Pseudomonadota>Gammaproteobacteria>Vibrionales>Vibrionaceae | 562 | DX35_RS0111120 | 10322..12010 | 662 | DUF262-domain-containing-protein-[Vibrio-parahaemolyticus] |
| WP_159406355.1 | ParB+HNH | GCF_009883895.1 | Vibrio | Bacteria>Pseudomonadati>Pseudomonadota>Gammaproteobacteria>Vibrionales>Vibrionaceae | 562 | O8X55_RS08185 | 519123..520811 | 662 | DUF262-domain-containing-protein-[Vibrio-parahaemolyticus] |
| WP_130049300.1 | ParB+HNH | GCF_028765545.1 | Aliivibrio | Bacteria>Pseudomonadati>Pseudomonadota>Gammaproteobacteria>Vibrionales>Vibrionaceae | 698 | ERW57_RS17810 | 5070..7166 | 511678 | MULTISPECIES:-DUF262-and-DUF1524-domain-containing-protein-[Aliivibrio] |
| WP_236733965.1 | ParB+HNH | GCF_001690915.1 | Aliivibrio_fischeri | Bacteria>Pseudomonadati>Pseudomonadota>Gammaproteobacteria>Vibrionales>Vibrionaceae>Aliivibrio | 704 | A6E13_RS16585 | 264633..266747 | 668 | DUF262-and-DUF1524-domain-containing-protein-[Aliivibrio-fischeri] |
| WP_133416084.1 | ParB+HNH | GCF_004359415.1 | Aliivibrio_fischeri | Bacteria>Pseudomonadati>Pseudomonadota>Gammaproteobacteria>Vibrionales>Vibrionaceae>Aliivibrio | 641 | VFFQA001_RS16060 | 462614..464539 | 668 | DUF262-domain-containing-protein-[Aliivibrio-fischeri] |
| WP_155683403.1 | ParB+HNH | GCF_009727645.1 | Aliivibrio_fischeri | Bacteria>Pseudomonadati>Pseudomonadota>Gammaproteobacteria>Vibrionales>Vibrionaceae>Aliivibrio | 594 | GNP73_RS13860 | 806101..807885 | 668 | DUF262-domain-containing-protein-[Aliivibrio-fischeri] |
| WP_005419544.1 | ParB+HNH | GCF_000241785.1 | Aliivibrio_fischeri | Bacteria>Pseudomonadati>Pseudomonadota>Gammaproteobacteria>Vibrionales>Vibrionaceae>Aliivibrio | 562 | VFSR5_RS07510 | 1599325..1601013 | 668 | DUF262-domain-containing-protein-[Aliivibrio-fischeri] |
| WP_130071730.1 | ParB+HNH | GCF_004167985.1 | Aliivibrio_finisterrensis | Bacteria>Pseudomonadati>Pseudomonadota>Gammaproteobacteria>Vibrionales>Vibrionaceae>Aliivibrio | 698 | ERW52_RS16060 | 18277..20373 | 511998 | DUF262-and-DUF1524-domain-containing-protein-[Aliivibrio-finisterrensis] |
| PMH64397.1 | ParB+HNH | - | Enterovibrio_norvegicus | Bacteria>Pseudomonadati>Pseudomonadota>Gammaproteobacteria>Vibrionales>Vibrionaceae>Enterovibrio | 562 | BCU62_16185 | 19929..21617 | 188144 | hypothetical-protein-BCU62_16185-[Enterovibrio-norvegicus] |
| WP_232580806.1 | ParB+HNH | GCF_021504545.1 | Photobacterium_phosphoreum | Bacteria>Pseudomonadati>Pseudomonadota>Gammaproteobacteria>Vibrionales>Vibrionaceae>Photobacterium | 146 | GLP29_RS03820 | 144858..145298 | 659 | DUF262-domain-containing-protein-[Photobacterium-phosphoreum] |
| WP_239972149.1 | ParB+HNH | GCF_022398535.1 | Photobacterium_damselae | Bacteria>Pseudomonadati>Pseudomonadota>Gammaproteobacteria>Vibrionales>Vibrionaceae>Photobacterium | 698 | L1D45_RS07340 | 271708..273804 | 38293 | DUF262-and-DUF1524-domain-containing-protein-[Photobacterium-damselae] |
| WP_115426812.1 | ParB+HNH | GCF_003355625.1 | Photobacterium_damselae | Bacteria>Pseudomonadati>Pseudomonadota>Gammaproteobacteria>Vibrionales>Vibrionaceae>Photobacterium | 644 | DA092_RS00800 | 178693..180627 | 38293 | DUF262-domain-containing-protein-[Photobacterium-damselae] |
| WP_138222249.1 | ParB+HNH | GCF_005819835.1 | Photobacterium_damselae | Bacteria>Pseudomonadati>Pseudomonadota>Gammaproteobacteria>Vibrionales>Vibrionaceae>Photobacterium | 636 | FD717_RS16640 | 63144..65054 | 38293 | DUF262-domain-containing-protein-[Photobacterium-damselae] |
| WP_101767573.1 | ParB+HNH | GCF_027923345.1 | Photobacterium_carnosum | Bacteria>Pseudomonadati>Pseudomonadota>Gammaproteobacteria>Vibrionales>Vibrionaceae>Photobacterium | 636 | CIK00_RS03625 | 188020..189930 | 2023717 | DUF262-domain-containing-protein-[Photobacterium-carnosum] |

| Protein-ID | Domain | Assembly-ID | Strain | Taxonomy | Length | Locus-Tag | Genomic-coordinates | TaxID | Fasta-defline |
|---|---|---|---|---|---|---|---|---|---|
| WP_087820335.1 | ParB+HNH | GCF_900185565.1 | Photobacterium_aquimaris | Bacteria>Pseudomonadati>Pseudomonadota>Gammaproteobacteria>Vibrionales>Vibrionaceae>Photobacterium | 644 | PAQU9191_RS07300 | 454938..456872 | 512643 | DUF262-domain-containing-protein-[Photobacterium-aquimaris] |
| WP_006647483.1 | ParB+HNH | GCF_000153325.1 | Photobacterium_sp._SKA34 | Bacteria>Pseudomonadati>Pseudomonadota>Gammaproteobacteria>Vibrionales>Vibrionaceae>Photobacterium>unclassified_Photobacterium | 624 | SKA34_RS21225 | 79199..81073 | 121723 | DUF262-domain-containing-protein-[Photobacterium-sp.-SKA34] |
| EHK9043272.1 | ParB+HNH | GCA_018080905.1 | Vibrio_vulnificus | Bacteria>Pseudomonadati>Pseudomonadota>Gammaproteobacteria>Vibrionales>Vibrionaceae>Vibrio | 698 | KCU37_001414 | 202880..204976 | 672 | DUF262-domain-containing-protein-[Vibrio-vulnificus] |
| HAS6334001.1 | ParB+HNH | GCA_015778755.1 | Vibrio_vulnificus | Bacteria>Pseudomonadati>Pseudomonadota>Gammaproteobacteria>Vibrionales>Vibrionaceae>Vibrio | 698 | I7245_08205 | 208007..210103 | 672 | DUF262-domain-containing-protein-[Vibrio-vulnificus] |
| HAS6345175.1 | ParB+HNH | GCA_015778705.1 | Vibrio_vulnificus | Bacteria>Pseudomonadati>Pseudomonadota>Gammaproteobacteria>Vibrionales>Vibrionaceae>Vibrio | 698 | I7243_17870 | 94097..96193 | 672 | DUF262-domain-containing-protein-[Vibrio-vulnificus] |
| WP_080540647.1 | ParB+HNH | GCF_002073135.1 | Vibrio_vulnificus | Bacteria>Pseudomonadati>Pseudomonadota>Gammaproteobacteria>Vibrionales>Vibrionaceae>Vibrio | 698 | XM77_RS07285 | 66730..68826 | 672 | DUF262-and-DUF1524-domain-containing-protein-[Vibrio-vulnificus] |
| WP_206630454.1 | ParB+HNH | GCF_017291155.1 | Vibrio_vulnificus | Bacteria>Pseudomonadati>Pseudomonadota>Gammaproteobacteria>Vibrionales>Vibrionaceae>Vibrio | 698 | JOJ25_RS14680 | 84446..86542 | 672 | DUF262-and-DUF1524-domain-containing-protein-[Vibrio-vulnificus] |
| EGR0072243.1 | ParB+HNH | - | Vibrio_vulnificus | Bacteria>Pseudomonadati>Pseudomonadota>Gammaproteobacteria>Vibrionales>Vibrionaceae>Vibrio | 644 | FR269_18520 | 10261..12195 | 672 | DUF262-domain-containing-protein-[Vibrio-vulnificus] |
| EJE8545411.1 | ParB+HNH | GCA_023352275.1 | Vibrio_vulnificus | Bacteria>Pseudomonadati>Pseudomonadota>Gammaproteobacteria>Vibrionales>Vibrionaceae>Vibrio | 644 | M5238_000044 | 59550..61484 | 672 | DUF262-domain-containing-protein-[Vibrio-vulnificus] |
| HAS6362910.1 | ParB+HNH | GCA_015777565.1 | Vibrio_vulnificus | Bacteria>Pseudomonadati>Pseudomonadota>Gammaproteobacteria>Vibrionales>Vibrionaceae>Vibrio | 644 | I7230_15355 | 47044..48978 | 672 | DUF262-domain-containing-protein-[Vibrio-vulnificus] |
| WP_225482455.1 | ParB+HNH | GCF_020251885.1 | Vibrio_vulnificus | Bacteria>Pseudomonadati>Pseudomonadota>Gammaproteobacteria>Vibrionales>Vibrionaceae>Vibrio | 644 | JKP23_RS02670 | 243683..245617 | 672 | DUF262-domain-containing-protein-[Vibrio-vulnificus] |
| EHD2253059.1 | ParB+HNH | - | Vibrio_vulnificus | Bacteria>Pseudomonadati>Pseudomonadota>Gammaproteobacteria>Vibrionales>Vibrionaceae>Vibrio | 642 | JRT24_004018 | 10344..12272 | 672 | DUF262-domain-containing-protein-[Vibrio-vulnificus] |
| EHW0635154.1 | ParB+HNH | GCA_019442565.1 | Vibrio_vulnificus | Bacteria>Pseudomonadati>Pseudomonadota>Gammaproteobacteria>Vibrionales>Vibrionaceae>Vibrio | 642 | K1B29_001384 | 66187..68115 | 672 | DUF262-domain-containing-protein-[Vibrio-vulnificus] |
| HAS8502531.1 | ParB+HNH | GCA_015820675.1 | Vibrio_vulnificus | Bacteria>Pseudomonadati>Pseudomonadota>Gammaproteobacteria>Vibrionales>Vibrionaceae>Vibrio | 642 | I7764_01780 | 438105..440033 | 672 | DUF262-domain-containing-protein-[Vibrio-vulnificus] |
| WP_225454269.1 | ParB+HNH | GCF_020252125.1 | Vibrio_vulnificus | Bacteria>Pseudomonadati>Pseudomonadota>Gammaproteobacteria>Vibrionales>Vibrionaceae>Vibrio | 641 | JKP08_RS08935 | 229765..231690 | 672 | DUF262-domain-containing-protein-[Vibrio-vulnificus] |
| WP_231894801.1 | ParB+HNH | GCF_001675245.1 | Vibrio_vulnificus | Bacteria>Pseudomonadati>Pseudomonadota>Gammaproteobacteria>Vibrionales>Vibrionaceae>Vibrio | 589 | FORC17_RS08370 | 1784745..1786514 | 672 | DUF262-domain-containing-protein-[Vibrio-vulnificus] |
| HAS6187653.1 | ParB+HNH | GCA_015777055.1 | Vibrio_vulnificus | Bacteria>Pseudomonadati>Pseudomonadota>Gammaproteobacteria>Vibrionales>Vibrionaceae>Vibrio | 587 | I7170_07510 | 256191..257954 | 672 | DUF262-domain-containing-protein-[Vibrio-vulnificus] |
| HAS6259594.1 | ParB+HNH | GCA_015777345.1 | Vibrio_vulnificus | Bacteria>Pseudomonadati>Pseudomonadota>Gammaproteobacteria>Vibrionales>Vibrionaceae>Vibrio | 587 | I7226_03365 | 262133..263896 | 672 | DUF262-domain-containing-protein-[Vibrio-vulnificus] |
| HAS8442987.1 | ParB+HNH | GCA_015820485.1 | Vibrio_vulnificus | Bacteria>Pseudomonadati>Pseudomonadota>Gammaproteobacteria>Vibrionales>Vibrionaceae>Vibrio | 587 | I7740_20135 | 20733..22496 | 672 | TPA:-DUF262-domain-containing-protein-[Vibrio-vulnificus] |
| WP_224640812.1 | ParB+HNH | GCF_020169625.1 | Vibrio_vulnificus | Bacteria>Pseudomonadati>Pseudomonadota>Gammaproteobacteria>Vibrionales>Vibrionaceae>Vibrio | 587 | KUO11_RS09485 | 209508..211271 | 672 | DUF262-domain-containing-protein-[Vibrio-vulnificus] |
| HAS6310368.1 | ParB+HNH | GCA_015778905.1 | Vibrio_vulnificus | Bacteria>Pseudomonadati>Pseudomonadota>Gammaproteobacteria>Vibrionales>Vibrionaceae>Vibrio | 562 | I7207_04125 | 37892..39580 | 672 | DUF262-domain-containing-protein-[Vibrio-vulnificus] |
| WP_130200310.1 | ParB+HNH | GCF_004214625.1 | Vibrio_vulnificus | Bacteria>Pseudomonadati>Pseudomonadota>Gammaproteobacteria>Vibrionales>Vibrionaceae>Vibrio | 562 | D8T51_RS18935 | 8398..10086 | 672 | DUF262-domain-containing-protein-[Vibrio-vulnificus] |
| WP_182291543.1 | ParB+HNH | GCF_014107515.1 | Vibrio_vulnificus | Bacteria>Pseudomonadati>Pseudomonadota>Gammaproteobacteria>Vibrionales>Vibrionaceae>Vibrio | 562 | F6X00_RS18215 | 380073..381761 | 672 | DUF262-domain-containing-protein-[Vibrio-vulnificus] |
| WP_237468685.1 | ParB+HNH | - | Vibrio_stylophorae | Bacteria>Pseudomonadati>Pseudomonadota>Gammaproteobacteria>Vibrionales>Vibrionaceae>Vibrio | 487 | L9P36_RS15965 | 500453..501916 | 659351 | DUF262-domain-containing-protein-[Vibrio-stylophorae] |
| WP_237468617.1 | ParB+HNH | GCF_921293875.1 | Vibrio_stylophorae | Bacteria>Pseudomonadati>Pseudomonadota>Gammaproteobacteria>Vibrionales>Vibrionaceae>Vibrio | 182 | L9P36_RS15970 | 501880..502428 | 659351 | DUF5655-domain-containing-protein-[Vibrio-stylophorae] |
| WP_102456720.1 | ParB+HNH | GCF_002877315.1 | Vibrio_splendidus | Bacteria>Pseudomonadati>Pseudomonadota>Gammaproteobacteria>Vibrionales>Vibrionaceae>Vibrio | 735 | BCU08_RS19890 | 74911..77118 | 29497 | DUF262-domain-containing-protein-[Vibrio-splendidus] |
| WP_102456767.1 | ParB+HNH | GCF_002877315.1 | Vibrio_splendidus | Bacteria>Pseudomonadati>Pseudomonadota>Gammaproteobacteria>Vibrionales>Vibrionaceae>Vibrio | 641 | BCU08_RS20215 | 13641..15566 | 29497 | DUF262-domain-containing-protein-[Vibrio-splendidus] |
| WP_102573394.1 | ParB+HNH | GCF_963877475.1 | Vibrio_splendidus | Bacteria>Pseudomonadati>Pseudomonadota>Gammaproteobacteria>Vibrionales>Vibrionaceae>Vibrio | 641 | BCT62_RS21765 | 16955..18880 | 29497 | DUF262-domain-containing-protein-[Vibrio-splendidus] |
| WP_243579242.1 | ParB+HNH | GCF_963898895.1 | Vibrio_splendidus | Bacteria>Pseudomonadati>Pseudomonadota>Gammaproteobacteria>Vibrionales>Vibrionaceae>Vibrio | 641 | LTQ02_RS02795 | 615706..617631 | 29497 | DUF262-domain-containing-protein-[Vibrio-splendidus] |
| WP_108187617.1 | ParB+HNH | GCF_003049965.1 | Vibrio_splendidus | Bacteria>Pseudomonadati>Pseudomonadota>Gammaproteobacteria>Vibrionales>Vibrionaceae>Vibrio | 594 | CWO07_RS09235 | 48709..50493 | 29497 | DUF262-domain-containing-protein-[Vibrio-splendidus] |
| WP_100754279.1 | ParB+HNH | GCF_963876685.1 | Vibrio_salilacus | Bacteria>Pseudomonadati>Pseudomonadota>Gammaproteobacteria>Vibrionales>Vibrionaceae>Vibrio | 644 | CDZ87_RS15120 | 97..2031 | 1323749 | DUF262-domain-containing-protein-[Vibrio-salilacus] |
| WP_261872673.1 | ParB+HNH | GCF_024347075.1 | Vibrio_rarus | Bacteria>Pseudomonadati>Pseudomonadota>Gammaproteobacteria>Vibrionales>Vibrionaceae>Vibrio | 601 | OCU56_RS07800 | 1696964..1698769 | 413403 | DUF262-domain-containing-HNH-endonuclease-family-protein-[Vibrio-rarus] |
| WP_254619132.1 | ParB+HNH | - | Vibrio_metschnikovii | Bacteria>Pseudomonadati>Pseudomonadota>Gammaproteobacteria>Vibrionales>Vibrionaceae>Vibrio | 575 | KVT01_RS15800 | 21528..23255 | 28172 | DUF262-domain-containing-protein-[Vibrio-metschnikovii] |
| WP_095470309.1 | ParB+HNH | GCF_002283955.1 | Vibrio_metoecus | Bacteria>Pseudomonadati>Pseudomonadota>Gammaproteobacteria>Vibrionales>Vibrionaceae>Vibrio | 597 | CGT93_RS13880 | 66343..68136 | 1481663 | DUF262-domain-containing-protein-[Vibrio-metoecus] |
| WP_095475144.1 | ParB+HNH | GCF_002283835.1 | Vibrio_metoecus | Bacteria>Pseudomonadati>Pseudomonadota>Gammaproteobacteria>Vibrionales>Vibrionaceae>Vibrio | 597 | CGT99_RS14690 | 66584..68377 | 1481663 | DUF262-domain-containing-protein-[Vibrio-metoecus] |
| WP_102514127.1 | ParB+HNH | GCF_002877415.2 | Vibrio_lentus | Bacteria>Pseudomonadati>Pseudomonadota>Gammaproteobacteria>Vibrionales>Vibrionaceae>Vibrio | 735 | BCU13_RS17430 | 185266..187473 | 136468 | DUF262-domain-containing-protein-[Vibrio-lentus] |
| TKE90003.1 | ParB+HNH | GCA_005145785.1 | Vibrio_kanaloae | Bacteria>Pseudomonadati>Pseudomonadota>Gammaproteobacteria>Vibrionales>Vibrionaceae>Vibrio | 642 | FCV44_20765 | 13528..15456 | 170673 | DUF262-domain-containing-protein-[Vibrio-kanaloae] |
| WP_104040816.1 | ParB+HNH | GCF_002929885.1 | Vibrio_hyugaensis | Bacteria>Pseudomonadati>Pseudomonadota>Gammaproteobacteria>Vibrionales>Vibrionaceae>Vibrio | 698 | COQ55_RS11510 | 19590..21686 | 1534743 | DUF262-and-DUF1524-domain-containing-protein-[Vibrio-hyugaensis] |
| WP_029810019.1 | ParB+HNH | GCF_000490615.1 | Vibrio_harveyi_group | Bacteria>Pseudomonadati>Pseudomonadota>Gammaproteobacteria>Vibrionales>Vibrionaceae>Vibrio | 642 | DX93_RS0104670 | 14380..16308 | 717610 | DUF262-domain-containing-protein-[Vibrio-parahaemolyticus] |
| MBY8207857.1 | ParB+HNH | GCA_019854355.1 | Vibrio_fluvialis | Bacteria>Pseudomonadati>Pseudomonadota>Gammaproteobacteria>Vibrionales>Vibrionaceae>Vibrio | 699 | KW531_20825 | 10509..12608 | 676 | DUF262-and-DUF1524-domain-containing-protein-[Vibrio-fluvialis] |
| WP_100267430.1 | ParB+HNH | GCF_002796765.2 | Vibrio_fluvialis | Bacteria>Pseudomonadati>Pseudomonadota>Gammaproteobacteria>Vibrionales>Vibrionaceae>Vibrio | 587 | CV024_RS02750 | 228547..230310 | 676 | DUF262-domain-containing-protein-[Vibrio-fluvialis] |
| MBY8044955.1 | ParB+HNH | GCA_019853775.1 | Vibrio_fluvialis | Bacteria>Pseudomonadati>Pseudomonadota>Gammaproteobacteria>Vibrionales>Vibrionaceae>Vibrio | 562 | KW486_21165 | 9072..10760 | 676 | DUF262-domain-containing-HNH-endonuclease-family-protein-[Vibrio-fluvialis] |
| MBY8148859.1 | ParB+HNH | GCA_019854195.1 | Vibrio_fluvialis | Bacteria>Pseudomonadati>Pseudomonadota>Gammaproteobacteria>Vibrionales>Vibrionaceae>Vibrio | 562 | KW504_05145 | 34720..36408 | 676 | DUF262-domain-containing-HNH-endonuclease-family-protein-[Vibrio-fluvialis] |
| WP_172566990.1 | ParB+HNH | GCF_012274945.1 | Vibrio_fluvialis | Bacteria>Pseudomonadati>Pseudomonadota>Gammaproteobacteria>Vibrionales>Vibrionaceae>Vibrio | 562 | HF303_RS09160 | 441218..442906 | 676 | DUF262-domain-containing-protein-[Vibrio-fluvialis] |
| WP_102402205.1 | ParB+HNH | GCF_002873825.2 | Vibrio_cyclitrophicus | Bacteria>Pseudomonadati>Pseudomonadota>Gammaproteobacteria>Vibrionales>Vibrionaceae>Vibrio | 641 | BCV09_RS12235 | 13668..15593 | 47951 | DUF262-domain-containing-protein-[Vibrio-cyclitrophicus] |
| WP_102465506.1 | ParB+HNH | GCF_002873515.2 | Vibrio_cyclitrophicus | Bacteria>Pseudomonadati>Pseudomonadota>Gammaproteobacteria>Vibrionales>Vibrionaceae>Vibrio | 641 | BCU82_RS15545 | 45945..47870 | 47951 | DUF262-domain-containing-protein-[Vibrio-cyclitrophicus] |
| WP_016790633.1 | ParB+HNH | GCF_000256605.2 | Vibrio_cyclitrophicus | Bacteria>Pseudomonadati>Pseudomonadota>Gammaproteobacteria>Vibrionales>Vibrionaceae>Vibrio | 601 | OAI_RS19785 | 266646..268451 | 47951 | DUF262-domain-containing-protein-[Vibrio-cyclitrophicus] |
| WP_123322058.1 | ParB+HNH | GCF_003751765.1 | Vibrio_crassostreae | Bacteria>Pseudomonadati>Pseudomonadota>Gammaproteobacteria>Vibrionales>Vibrionaceae>Vibrio | 585 | EDB73_RS01665 | 366932..368689 | 246167 | DUF262-domain-containing-protein-[Vibrio-crassostreae] |
| WP_132941348.1 | ParB+HNH | GCF_004345555.1 | Vibrio_crassostreae | Bacteria>Pseudomonadati>Pseudomonadota>Gammaproteobacteria>Vibrionales>Vibrionaceae>Vibrio | 585 | EDB60_RS00565 | 109697..111454 | 246167 | DUF262-domain-containing-protein-[Vibrio-crassostreae] |
| WP_132957336.1 | ParB+HNH | GCF_963880805.1 | Vibrio_crassostreae | Bacteria>Pseudomonadati>Pseudomonadota>Gammaproteobacteria>Vibrionales>Vibrionaceae>Vibrio | 562 | EDB41_RS15390 | 74760..76448 | 246167 | DUF262-domain-containing-protein-[Vibrio-crassostreae] |
| WP_050646848.1 | ParB+HNH | GCF_001243675.1 | Vibrio_coralliirubri | Bacteria>Pseudomonadati>Pseudomonadota>Gammaproteobacteria>Vibrionales>Vibrionaceae>Vibrio | 562 | VCR4J2_RS00150 | 30603..32291 | 1516159 | DUF262-domain-containing-protein-[Vibrio-sp.-J2-4] |
| GHW31169.1 | ParB+HNH | GCA_023163105.1 | Vibrio_cholerae | Bacteria>Pseudomonadati>Pseudomonadota>Gammaproteobacteria>Vibrionales>Vibrionaceae>Vibrio | 642 | VCSRO150_3320 | 9536..11464 | 666 | hypothetical-protein-VCSRO150_3320-[Vibrio-cholerae] |
| HBC3376472.1 | ParB+HNH | GCA_018118685.1 | Vibrio_cholerae | Bacteria>Pseudomonadati>Pseudomonadota>Gammaproteobacteria>Vibrionales>Vibrionaceae>Vibrio | 642 | KDX67_000918 | 97875..99803 | 666 | DUF262-domain-containing-protein-[Vibrio-cholerae] |
| WP_148532384.1 | ParB+HNH | GCF_008084645.1 | Vibrio_cholerae | Bacteria>Pseudomonadati>Pseudomonadota>Gammaproteobacteria>Vibrionales>Vibrionaceae>Vibrio | 601 | FXE54_RS16725 | 33981..35786 | 666 | DUF262-domain-containing-protein-[Vibrio-cholerae] |
| EGR1309073.1 | ParB+HNH | - | Vibrio_cholerae | Bacteria>Pseudomonadati>Pseudomonadota>Gammaproteobacteria>Vibrionales>Vibrionaceae>Vibrio | 587 | D9A28_03780 | 153155..154918 | 666 | DUF262-domain-containing-protein-[Vibrio-cholerae] |
| EII3725634.1 | ParB+HNH | GCA_020414365.1 | Vibrio_cholerae | Bacteria>Pseudomonadati>Pseudomonadota>Gammaproteobacteria>Vibrionales>Vibrionaceae>Vibrio | 587 | LG468_000347 | 127418..129181 | 666 | DUF262-domain-containing-protein-[Vibrio-cholerae] |
| EJL6432477.1 | ParB+HNH | GCA_024216615.1 | Vibrio_cholerae | Bacteria>Pseudomonadati>Pseudomonadota>Gammaproteobacteria>Vibrionales>Vibrionaceae>Vibrio | 587 | NMR72_000937 | 125121..126884 | 666 | DUF262-domain-containing-protein-[Vibrio-cholerae] |
| HCJ6872076.1 | ParB+HNH | GCA_024621265.1 | Vibrio_cholerae | Bacteria>Pseudomonadati>Pseudomonadota>Gammaproteobacteria>Vibrionales>Vibrionaceae>Vibrio | 587 | NU929_001220 | 140922..142685 | 666 | DUF262-domain-containing-protein-[Vibrio-cholerae] |
| HCJ7299596.1 | ParB+HNH | GCA_024632775.1 | Vibrio_cholerae | Bacteria>Pseudomonadati>Pseudomonadota>Gammaproteobacteria>Vibrionales>Vibrionaceae>Vibrio | 587 | NVH34_000347 | 34034..35797 | 666 | DUF262-domain-containing-protein-[Vibrio-cholerae] |
| MVB73517.1 | ParB+HNH | GCA_009747185.1 | Vibrio_cholerae | Bacteria>Pseudomonadati>Pseudomonadota>Gammaproteobacteria>Vibrionales>Vibrionaceae>Vibrio | 587 | D6U90_04205 | 147435..149198 | 666 | DUF262-domain-containing-protein-[Vibrio-cholerae] |
| MVC74395.1 | ParB+HNH | GCA_009747245.1 | Vibrio_cholerae | Bacteria>Pseudomonadati>Pseudomonadota>Gammaproteobacteria>Vibrionales>Vibrionaceae>Vibrio | 587 | D6U22_06270 | 187837..189600 | 666 | DUF262-domain-containing-protein-[Vibrio-cholerae] |
| WP_001963846.1 | ParB+HNH | GCF_000304915.2 | Vibrio_cholerae | Bacteria>Pseudomonadati>Pseudomonadota>Gammaproteobacteria>Vibrionales>Vibrionaceae>Vibrio | 587 | VCCP1035_RS13190 | 136871..138634 | 666 | DUF262-domain-containing-protein-[Vibrio-cholerae] |
| WP_032482145.1 | ParB+HNH | GCF_000736875.1 | Vibrio_cholerae | Bacteria>Pseudomonadati>Pseudomonadota>Gammaproteobacteria>Vibrionales>Vibrionaceae>Vibrio | 587 | DN38_RS05540 | 12478..14241 | 666 | DUF262-domain-containing-protein-[Vibrio-cholerae] |
| WP_033931156.1 | ParB+HNH | GCF_000788515.1 | Vibrio_cholerae | Bacteria>Pseudomonadati>Pseudomonadota>Gammaproteobacteria>Vibrionales>Vibrionaceae>Vibrio | 587 | NH59_RS10855 | 61007..62770 | 666 | DUF262-domain-containing-protein-[Vibrio-cholerae] |
| WP_142738825.1 | ParB+HNH | GCF_006803025.1 | Vibrio_cholerae | Bacteria>Pseudomonadati>Pseudomonadota>Gammaproteobacteria>Vibrionales>Vibrionaceae>Vibrio | 587 | FLL79_RS01415 | 39715..41478 | 666 | DUF262-domain-containing-protein-[Vibrio-cholerae] |
| WP_148532769.1 | ParB+HNH | GCF_008084025.1 | Vibrio_cholerae | Bacteria>Pseudomonadati>Pseudomonadota>Gammaproteobacteria>Vibrionales>Vibrionaceae>Vibrio | 587 | FXE74_RS08215 | 131107..132870 | 666 | DUF262-domain-containing-protein-[Vibrio-cholerae] |
| WP_148536809.1 | ParB+HNH | GCF_008085615.1 | Vibrio_cholerae | Bacteria>Pseudomonadati>Pseudomonadota>Gammaproteobacteria>Vibrionales>Vibrionaceae>Vibrio | 587 | FXE25_RS13675 | 33999..35762 | 666 | DUF262-domain-containing-protein-[Vibrio-cholerae] |
| EGR0489750.1 | ParB+HNH | - | Vibrio_cholerae | Bacteria>Pseudomonadati>Pseudomonadota>Gammaproteobacteria>Vibrionales>Vibrionaceae>Vibrio | 562 | FG089_09190 | 73917..75605 | 666 | DUF262-domain-containing-protein-[Vibrio-cholerae] |
| EGR4188111.1 | ParB+HNH | - | Vibrio_cholerae | Bacteria>Pseudomonadati>Pseudomonadota>Gammaproteobacteria>Vibrionales>Vibrionaceae>Vibrio | 562 | DDM99_15765 | 14562..16250 | 666 | DUF262-domain-containing-protein-[Vibrio-cholerae] |
| WP_000041893.1 | ParB+HNH | GCF_000220785.1 | Vibrio_cholerae | Bacteria>Pseudomonadati>Pseudomonadota>Gammaproteobacteria>Vibrionales>Vibrionaceae>Vibrio | 562 | VCHE48_RS12975 | 870768..872456 | 666 | DUF262-domain-containing-protein-[Vibrio-cholerae] |
| WP_096070870.1 | ParB+HNH | GCF_002313025.1 | Vibrio_cholerae | Bacteria>Pseudomonadati>Pseudomonadota>Gammaproteobacteria>Vibrionales>Vibrionaceae>Vibrio | 562 | FXF10_RS00330 | 24180..25868 | 666 | DUF262-domain-containing-protein-[Vibrio-cholerae] |
| WP_102376358.1 | ParB+HNH | GCF_002876115.1 | Vibrio_breoganii | Bacteria>Pseudomonadati>Pseudomonadota>Gammaproteobacteria>Vibrionales>Vibrionaceae>Vibrio | 594 | BCT61_RS03010 | 194241..196025 | 553239 | DUF262-domain-containing-protein-[Vibrio-breoganii] |
| WP_040122840.1 | ParB+HNH | GCF_003031205.1 | Vibrio_anguillarum | Bacteria>Pseudomonadati>Pseudomonadota>Gammaproteobacteria>Vibrionales>Vibrionaceae>Vibrio | 601 | CEA93_RS18115 | 697442..699247 | 55601 | DUF262-domain-containing-protein-[Vibrio-anguillarum] |
| WP_088731320.1 | ParB+HNH | GCF_002212025.1 | Vibrio_anguillarum | Bacteria>Pseudomonadati>Pseudomonadota>Gammaproteobacteria>Vibrionales>Vibrionaceae>Vibrio | 587 | CEQ50_RS00055 | 13835..15598 | 55601 | DUF262-domain-containing-protein-[Vibrio-anguillarum] |
| WP_171375392.1 | ParB+HNH | GCF_013114495.1 | Vibrio_aestuarianus | Bacteria>Pseudomonadati>Pseudomonadota>Gammaproteobacteria>Vibrionales>Vibrionaceae>Vibrio | 642 | F0230_RS00335 | 86492..88420 | 28171 | DUF262-domain-containing-protein-[Vibrio-aestuarianus] |
| WP_168780464.1 | ParB+HNH | GCF_944738875.1 | Vibrio_aestuarianus | Bacteria>Pseudomonadati>Pseudomonadota>Gammaproteobacteria>Vibrionales>Vibrionaceae>Vibrio | 641 | OCX59_RS00140 | 19653..21578 | 28171 | DUF262-domain-containing-protein-[Vibrio-aestuarianus] |
| WP_253658041.1 | ParB+HNH | GCF_024181795.1 | unclassified_Vibrio | Bacteria>Pseudomonadati>Pseudomonadota>Gammaproteobacteria>Vibrionales>Vibrionaceae>Vibrio | 698 | NL644_RS04165 | 941819..943915 | 2614977 | MULTISPECIES:-DUF262-and-DUF1524-domain-containing-protein-[unclassified-Vibrio] |
| WP_285401333.1 | ParB+HNH | GCF_030248535.1 | unclassified_Vibrio | Bacteria>Pseudomonadati>Pseudomonadota>Gammaproteobacteria>Vibrionales>Vibrionaceae>Vibrio | 641 | KJB41_RS10795 | 107452..109377 | 2614977 | MULTISPECIES:-DUF262-domain-containing-HNH-endonuclease-family-protein-[unclassified-Vibrio] |
| WP_253644854.1 | ParB+HNH | GCF_024181785.1 | Vibrio_sp._Y42_MX_L11 | Bacteria>Pseudomonadati>Pseudomonadota>Gammaproteobacteria>Vibrionales>Vibrionaceae>Vibrio>unclassified_Vibrio | 644 | NL643_RS16670 | 55475..57409 | 2957765 | DUF262-domain-containing-protein-[Vibrio-sp.-Y42_MX_L11] |
| WP_206946497.1 | ParB+HNH | GCF_017313505.1 | Vibrio_sp._Vb2424 | Bacteria>Pseudomonadati>Pseudomonadota>Gammaproteobacteria>Vibrionales>Vibrionaceae>Vibrio>unclassified_Vibrio | 601 | J0679_RS01035 | 234923..236728 | 2816074 | DUF262-domain-containing-protein-[Vibrio-sp.-Vb2424] |
| WP_264912145.1 | ParB+HNH | GCF_026000495.1 | Vibrio_sp._STUT-A16 | Bacteria>Pseudomonadati>Pseudomonadota>Gammaproteobacteria>Vibrionales>Vibrionaceae>Vibrio>unclassified_Vibrio | 698 | OO778_RS24835 | 1832050..1834146 | 2976237 | DUF262-and-DUF1524-domain-containing-protein-[Vibrio-sp.-STUT-A16] |
| WP_172438074.1 | ParB+HNH | GCF_002608565.1 | Vibrio_sp._PID17_43 | Bacteria>Pseudomonadati>Pseudomonadota>Gammaproteobacteria>Vibrionales>Vibrionaceae>Vibrio>unclassified_Vibrio | 736 | AK965_RS05455 | 171999..174209 | 1583451 | DUF262-domain-containing-protein-[Vibrio-sp.-PID17_43] |
| WP_236110296.1 | ParB+HNH | GCF_021568915.1 | Vibrio_sp._McD22-P3 | Bacteria>Pseudomonadati>Pseudomonadota>Gammaproteobacteria>Vibrionales>Vibrionaceae>Vibrio>unclassified_Vibrio | 601 | L2W36_RS13795 | 201937..203742 | 2724880 | DUF262-domain-containing-protein-[Vibrio-sp.-McD22-P3] |
| GAJ78756.1 | ParB+HNH | GCA_000695055.1 | Vibrio_sp._JCM_18905 | Bacteria>Pseudomonadati>Pseudomonadota>Gammaproteobacteria>Vibrionales>Vibrionaceae>Vibrio>unclassified_Vibrio | 379 | JCM18905_4746 | 11148..12287 | 1298600 | hypothetical-protein-JCM18905_4746-[Vibrio-sp.-JCM-18905] |
| GAJ78757.1 | ParB+HNH | GCA_000695055.1 | Vibrio_sp._JCM_18905 | Bacteria>Pseudomonadati>Pseudomonadota>Gammaproteobacteria>Vibrionales>Vibrionaceae>Vibrio>unclassified_Vibrio | 227 | JCM18905_4747 | 12395..13078 | 1298600 | hypothetical-protein-JCM18905_4747-[Vibrio-sp.-JCM-18905] |
| WP_226420706.1 | ParB+HNH | GCF_020515785.1 | Vibrio_sp._E14 | Bacteria>Pseudomonadati>Pseudomonadota>Gammaproteobacteria>Vibrionales>Vibrionaceae>Vibrio>unclassified_Vibrio | 698 | LIX84_RS04100 | 949282..951378 | 2849869 | DUF262-and-DUF1524-domain-containing-protein-[Vibrio-sp.-E14] |
| PMO03879.1 | ParB+HNH | GCA_002875525.1 | Vibrio_sp._10N.222.55.C12 | Bacteria>Pseudomonadati>Pseudomonadota>Gammaproteobacteria>Vibrionales>Vibrionaceae>Vibrio>unclassified_Vibrio | 698 | BCT20_08075 | 17391..19487 | 1884470 | hypothetical-protein-BCT20_08075-[Vibrio-sp.-10N.222.55.C12] |
| EGR2744344.1 | ParB+HNH | - | Vibrio_parahaemolyticus | Bacteria>Pseudomonadati>Pseudomonadota>Gammaproteobacteria>Vibrionales>Vibrionaceae>Vibrio>Vibrio_harveyi_group | 699 | DMW33_24580 | 10381..12480 | 670 | DUF262-domain-containing-protein-[Vibrio-parahaemolyticus] |
| HBC3399175.1 | ParB+HNH | GCA_018118805.1 | Vibrio_parahaemolyticus | Bacteria>Pseudomonadati>Pseudomonadota>Gammaproteobacteria>Vibrionales>Vibrionaceae>Vibrio>Vibrio_harveyi_group | 699 | KDX79_000985 | 167941..170040 | 670 | DUF262-domain-containing-protein-[Vibrio-parahaemolyticus] |
| HCG8269542.1 | ParB+HNH | GCA_024047655.1 | Vibrio_parahaemolyticus | Bacteria>Pseudomonadati>Pseudomonadota>Gammaproteobacteria>Vibrionales>Vibrionaceae>Vibrio>Vibrio_harveyi_group | 699 | NJ590_004435 | 22930..25029 | 670 | DUF262-domain-containing-protein-[Vibrio-parahaemolyticus] |
| MCR9764760.1 | ParB+HNH | GCA_024745395.1 | Vibrio_parahaemolyticus | Bacteria>Pseudomonadati>Pseudomonadota>Gammaproteobacteria>Vibrionales>Vibrionaceae>Vibrio>Vibrio_harveyi_group | 699 | NB556_01255 | 10949..13048 | 670 | DUF262-and-DUF1524-domain-containing-protein-[Vibrio-parahaemolyticus] |
| WP_029830479.1 | ParB+HNH | GCF_000491395.1 | Vibrio_parahaemolyticus | Bacteria>Pseudomonadati>Pseudomonadota>Gammaproteobacteria>Vibrionales>Vibrionaceae>Vibrio>Vibrio_harveyi_group | 699 | DY37_RS0104870 | 9850..11949 | 670 | DUF262-and-DUF1524-domain-containing-protein-[Vibrio-parahaemolyticus] |
| WP_258517487.1 | ParB+HNH | GCF_024745395.1 | Vibrio_parahaemolyticus | Bacteria>Pseudomonadati>Pseudomonadota>Gammaproteobacteria>Vibrionales>Vibrionaceae>Vibrio>Vibrio_harveyi_group | 699 | NB556_RS01255 | 10949..13048 | 670 | DUF262-and-DUF1524-domain-containing-protein-[Vibrio-parahaemolyticus] |
| HBC3540117.1 | ParB+HNH | GCA_018119555.1 | Vibrio_parahaemolyticus | Bacteria>Pseudomonadati>Pseudomonadota>Gammaproteobacteria>Vibrionales>Vibrionaceae>Vibrio>Vibrio_harveyi_group | 698 | KDZ97_004676 | 1099..3195 | 670 | DUF262-domain-containing-protein-[Vibrio-parahaemolyticus] |
| HBK3330410.1 | ParB+HNH | GCA_020473965.1 | Vibrio_parahaemolyticus | Bacteria>Pseudomonadati>Pseudomonadota>Gammaproteobacteria>Vibrionales>Vibrionaceae>Vibrio>Vibrio_harveyi_group | 698 | KD833_003431 | 25359..27455 | 670 | DUF262-domain-containing-protein-[Vibrio-parahaemolyticus] |
| HCE2679241.1 | ParB+HNH | GCA_023813525.1 | Vibrio_parahaemolyticus | Bacteria>Pseudomonadati>Pseudomonadota>Gammaproteobacteria>Vibrionales>Vibrionaceae>Vibrio>Vibrio_harveyi_group | 698 | NGU92_000681 | 142431..144527 | 670 | DUF262-domain-containing-protein-[Vibrio-parahaemolyticus] |
| WP_228075535.1 | ParB+HNH | GCF_020740925.1 | Vibrio_parahaemolyticus | Bacteria>Pseudomonadati>Pseudomonadota>Gammaproteobacteria>Vibrionales>Vibrionaceae>Vibrio>Vibrio_harveyi_group | 698 | IB290_RS12365 | 169028..171124 | 670 | DUF262-and-DUF1524-domain-containing-protein-[Vibrio-parahaemolyticus] |
| WP_274020829.1 | ParB+HNH | GCF_028617275.1 | Vibrio_parahaemolyticus | Bacteria>Pseudomonadati>Pseudomonadota>Gammaproteobacteria>Vibrionales>Vibrionaceae>Vibrio>Vibrio_harveyi_group | 698 | O8254_RS13590 | 100736..102832 | 670 | DUF262-and-DUF1524-domain-containing-protein-[Vibrio-parahaemolyticus] |
| EHH1264394.1 | ParB+HNH | GCA_017604425.1 | Vibrio_parahaemolyticus | Bacteria>Pseudomonadati>Pseudomonadota>Gammaproteobacteria>Vibrionales>Vibrionaceae>Vibrio>Vibrio_harveyi_group | 644 | J7I02_003662 | 21682..23616 | 670 | DUF262-domain-containing-protein-[Vibrio-parahaemolyticus] |
| HCG7769726.1 | ParB+HNH | GCA_024045435.1 | Vibrio_parahaemolyticus | Bacteria>Pseudomonadati>Pseudomonadota>Gammaproteobacteria>Vibrionales>Vibrionaceae>Vibrio>Vibrio_harveyi_group | 644 | NJ450_003035 | 7028..8962 | 670 | DUF262-domain-containing-protein-[Vibrio-parahaemolyticus] |
| WP_122015996.1 | ParB+HNH | GCF_003691525.1 | Vibrio_parahaemolyticus | Bacteria>Pseudomonadati>Pseudomonadota>Gammaproteobacteria>Vibrionales>Vibrionaceae>Vibrio>Vibrio_harveyi_group | 644 | D0871_RS08255 | 1548853..1550787 | 670 | DUF262-domain-containing-protein-[Vibrio-parahaemolyticus] |
| WP_140272581.1 | ParB+HNH | GCF_006372715.1 | Vibrio_parahaemolyticus | Bacteria>Pseudomonadati>Pseudomonadota>Gammaproteobacteria>Vibrionales>Vibrionaceae>Vibrio>Vibrio_harveyi_group | 644 | CGI87_RS22515 | 10261..12195 | 670 | DUF262-domain-containing-protein-[Vibrio-parahaemolyticus] |
| WP_191116550.1 | ParB+HNH | GCF_014769095.1 | Vibrio_parahaemolyticus | Bacteria>Pseudomonadati>Pseudomonadota>Gammaproteobacteria>Vibrionales>Vibrionaceae>Vibrio>Vibrio_harveyi_group | 644 | EUV06_RS14925 | 112136..114070 | 670 | DUF262-domain-containing-protein-[Vibrio-parahaemolyticus] |
| WP_256897645.1 | ParB+HNH | GCF_024530955.1 | Vibrio_parahaemolyticus | Bacteria>Pseudomonadati>Pseudomonadota>Gammaproteobacteria>Vibrionales>Vibrionaceae>Vibrio>Vibrio_harveyi_group | 644 | H2C82_RS04840 | 216977..218911 | 670 | DUF262-domain-containing-protein-[Vibrio-parahaemolyticus] |
| EHK0048696.1 | ParB+HNH | GCA_017968185.1 | Vibrio_parahaemolyticus | Bacteria>Pseudomonadati>Pseudomonadota>Gammaproteobacteria>Vibrionales>Vibrionaceae>Vibrio>Vibrio_harveyi_group | 642 | KCA08_003290 | 80100..82028 | 670 | DUF262-domain-containing-protein-[Vibrio-parahaemolyticus] |
| EIY9800892.1 | ParB+HNH | GCA_022594155.1 | Vibrio_parahaemolyticus | Bacteria>Pseudomonadati>Pseudomonadota>Gammaproteobacteria>Vibrionales>Vibrionaceae>Vibrio>Vibrio_harveyi_group | 642 | MNO17_002469 | 357269..359197 | 670 | DUF262-domain-containing-protein-[Vibrio-parahaemolyticus] |
| EJF4458716.1 | ParB+HNH | GCA_023466975.1 | Vibrio_parahaemolyticus | Bacteria>Pseudomonadati>Pseudomonadota>Gammaproteobacteria>Vibrionales>Vibrionaceae>Vibrio>Vibrio_harveyi_group | 642 | MT303_002824 | 132655..134583 | 670 | DUF262-domain-containing-protein-[Vibrio-parahaemolyticus] |
| EJG0638706.1 | ParB+HNH | GCA_015813785.2 | Vibrio_parahaemolyticus | Bacteria>Pseudomonadati>Pseudomonadota>Gammaproteobacteria>Vibrionales>Vibrionaceae>Vibrio>Vibrio_harveyi_group | 642 | C4G37_RS13080 | 155276..157204 | 670 | DUF262-domain-containing-protein-[Vibrio-parahaemolyticus] |
| EJG0966119.1 | ParB+HNH | GCA_015813265.2 | Vibrio_parahaemolyticus | Bacteria>Pseudomonadati>Pseudomonadota>Gammaproteobacteria>Vibrionales>Vibrionaceae>Vibrio>Vibrio_harveyi_group | 642 | C4G65_RS20925 | 31743..33671 | 670 | DUF262-domain-containing-protein-[Vibrio-parahaemolyticus] |
| HBC3986620.1 | ParB+HNH | GCA_018126825.1 | Vibrio_parahaemolyticus | Bacteria>Pseudomonadati>Pseudomonadota>Gammaproteobacteria>Vibrionales>Vibrionaceae>Vibrio>Vibrio_harveyi_group | 642 | KD867_003186 | 73236..75164 | 670 | DUF262-domain-containing-protein-[Vibrio-parahaemolyticus] |
| HCG8025123.1 | ParB+HNH | GCA_024046515.1 | Vibrio_parahaemolyticus | Bacteria>Pseudomonadati>Pseudomonadota>Gammaproteobacteria>Vibrionales>Vibrionaceae>Vibrio>Vibrio_harveyi_group | 642 | NJ909_002127 | 188720..190648 | 670 | TPA:-DUF262-domain-containing-protein-[Vibrio-parahaemolyticus] |
| HCH5919842.1 | ParB+HNH | GCA_024097015.1 | Vibrio_parahaemolyticus | Bacteria>Pseudomonadati>Pseudomonadota>Gammaproteobacteria>Vibrionales>Vibrionaceae>Vibrio>Vibrio_harveyi_group | 642 | NK048_004354 | 26615..28543 | 670 | TPA:-DUF262-domain-containing-protein-[Vibrio-parahaemolyticus] |
| WP_025605142.1 | ParB+HNH | GCF_000489075.1 | Vibrio_parahaemolyticus | Bacteria>Pseudomonadati>Pseudomonadota>Gammaproteobacteria>Vibrionales>Vibrionaceae>Vibrio>Vibrio_harveyi_group | 642 | DX10_RS0106520 | 19005..20933 | 670 | DUF262-domain-containing-protein-[Vibrio-parahaemolyticus] |
| WP_069530138.1 | ParB+HNH | GCF_030994225.1 | Vibrio_parahaemolyticus | Bacteria>Pseudomonadati>Pseudomonadota>Gammaproteobacteria>Vibrionales>Vibrionaceae>Vibrio>Vibrio_harveyi_group | 642 | BBN06_RS08180 | 21246..23174 | 670 | DUF262-domain-containing-protein-[Vibrio-parahaemolyticus] |
| WP_140323903.1 | ParB+HNH | GCF_006371905.1 | Vibrio_parahaemolyticus | Bacteria>Pseudomonadati>Pseudomonadota>Gammaproteobacteria>Vibrionales>Vibrionaceae>Vibrio>Vibrio_harveyi_group | 642 | CGJ25_RS23770 | 11035..12963 | 670 | DUF262-domain-containing-protein-[Vibrio-parahaemolyticus] |
| WP_178326689.1 | ParB+HNH | - | Vibrio_parahaemolyticus | Bacteria>Pseudomonadati>Pseudomonadota>Gammaproteobacteria>Vibrionales>Vibrionaceae>Vibrio>Vibrio_harveyi_group | 642 | HX770_RS05785 | 32078..34006 | 670 | DUF262-domain-containing-protein-[Vibrio-parahaemolyticus] |
| WP_267365289.1 | ParB+HNH | GCF_026537625.1 | Vibrio_parahaemolyticus | Bacteria>Pseudomonadati>Pseudomonadota>Gammaproteobacteria>Vibrionales>Vibrionaceae>Vibrio>Vibrio_harveyi_group | 641 | NOL37_RS23790 | 26775..28700 | 670 | DUF262-domain-containing-HNH-endonuclease-family-protein-[Vibrio-parahaemolyticus] |
| EGR3003174.1 | ParB+HNH | - | Vibrio_parahaemolyticus | Bacteria>Pseudomonadati>Pseudomonadota>Gammaproteobacteria>Vibrionales>Vibrionaceae>Vibrio>Vibrio_harveyi_group | 601 | DMW06_23020 | 7547..9352 | 670 | DUF262-domain-containing-protein-[Vibrio-parahaemolyticus] |
| EGR4671253.1 | ParB+HNH | - | Vibrio_parahaemolyticus | Bacteria>Pseudomonadati>Pseudomonadota>Gammaproteobacteria>Vibrionales>Vibrionaceae>Vibrio>Vibrio_harveyi_group | 601 | C4G45_02520 | 549869..551674 | 670 | DUF262-domain-containing-protein-[Vibrio-parahaemolyticus] |
| EIU7849847.1 | ParB+HNH | GCA_021716735.1 | Vibrio_parahaemolyticus | Bacteria>Pseudomonadati>Pseudomonadota>Gammaproteobacteria>Vibrionales>Vibrionaceae>Vibrio>Vibrio_harveyi_group | 601 | L3I74_001634 | 170246..172051 | 670 | DUF262-domain-containing-protein-[Vibrio-parahaemolyticus] |
| EJE8564155.1 | ParB+HNH | GCA_023352375.1 | Vibrio_parahaemolyticus | Bacteria>Pseudomonadati>Pseudomonadota>Gammaproteobacteria>Vibrionales>Vibrionaceae>Vibrio>Vibrio_harveyi_group | 601 | M5224_000721 | 52422..54227 | 670 | DUF262-domain-containing-protein-[Vibrio-parahaemolyticus] |
| EJG0997617.1 | ParB+HNH | GCA_015813305.2 | Vibrio_parahaemolyticus | Bacteria>Pseudomonadati>Pseudomonadota>Gammaproteobacteria>Vibrionales>Vibrionaceae>Vibrio>Vibrio_harveyi_group | 601 | C4G52_RS11750 | 15991..17796 | 670 | DUF262-domain-containing-protein-[Vibrio-parahaemolyticus] |
| EJG1873895.1 | ParB+HNH | GCA_015817955.2 | Vibrio_parahaemolyticus | Bacteria>Pseudomonadati>Pseudomonadota>Gammaproteobacteria>Vibrionales>Vibrionaceae>Vibrio>Vibrio_harveyi_group | 601 | BS053_RS09080 | 165425..167230 | 670 | DUF262-domain-containing-protein-[Vibrio-parahaemolyticus] |
| HBC3481461.1 | ParB+HNH | GCA_018119205.1 | Vibrio_parahaemolyticus | Bacteria>Pseudomonadati>Pseudomonadota>Gammaproteobacteria>Vibrionales>Vibrionaceae>Vibrio>Vibrio_harveyi_group | 601 | KDY88_002918 | 72223..74028 | 670 | TPA:-DUF262-domain-containing-protein-[Vibrio-parahaemolyticus] |
| HBC3921011.1 | ParB+HNH | GCA_018126325.1 | Vibrio_parahaemolyticus | Bacteria>Pseudomonadati>Pseudomonadota>Gammaproteobacteria>Vibrionales>Vibrionaceae>Vibrio>Vibrio_harveyi_group | 601 | KD834_003322 | 45224..47029 | 670 | TPA:-DUF262-domain-containing-protein-[Vibrio-parahaemolyticus] |

| Protein-ID | Domain | Assembly-ID | Strain | Taxonomy | Length | Locus-Tag | Genomic-coordinates | TaxID | Fasta-defline |
|---|---|---|---|---|---|---|---|---|---|
| HCG5108674.1 | ParB+HNH | GCA_024034975.1 | Vibrio_parahaemolyticus | Bacteria>Pseudomonadati>Pseudomonadota>Gammaproteobacteria>Vibrionales>Vibrionaceae>Vibrio>Vibrio_harveyi_group | 601 | NJY97_002342 | 341311..343116 | 670 | DUF262-domain-containing-protein-[Vibrio-parahaemolyticus] |
| WP_228096074.1 | ParB+HNH | GCF_020741035.1 | Vibrio_parahaemolyticus | Bacteria>Pseudomonadati>Pseudomonadota>Gammaproteobacteria>Vibrionales>Vibrionaceae>Vibrio>Vibrio_harveyi_group | 601 | IB294_RS01065 | 240730..242535 | 670 | DUF262-domain-containing-protein-[Vibrio-parahaemolyticus] |
| WP_229611427.1 | ParB+HNH | GCF_905331955.1 | Vibrio_parahaemolyticus | Bacteria>Pseudomonadati>Pseudomonadota>Gammaproteobacteria>Vibrionales>Vibrionaceae>Vibrio>Vibrio_harveyi_group | 601 | LPR99_RS16750 | 25515..27320 | 670 | DUF262-domain-containing-protein-[Vibrio-parahaemolyticus] |
| EGU0166209.1 | ParB+HNH | - | Vibrio_parahaemolyticus | Bacteria>Pseudomonadati>Pseudomonadota>Gammaproteobacteria>Vibrionales>Vibrionaceae>Vibrio>Vibrio_harveyi_group | 599 | JFQ74_002023 | 200787..202586 | 670 | DUF262-domain-containing-protein-[Vibrio-parahaemolyticus] |
| EGR0983686.1 | ParB+HNH | - | Vibrio_parahaemolyticus | Bacteria>Pseudomonadati>Pseudomonadota>Gammaproteobacteria>Vibrionales>Vibrionaceae>Vibrio>Vibrio_harveyi_group | 587 | EH301_01890 | 137904..139667 | 670 | DUF262-domain-containing-protein-[Vibrio-parahaemolyticus] |
| HBC3861548.1 | ParB+HNH | GCA_018126725.1 | Vibrio_parahaemolyticus | Bacteria>Pseudomonadati>Pseudomonadota>Gammaproteobacteria>Vibrionales>Vibrionaceae>Vibrio>Vibrio_harveyi_group | 587 | KD871_001929 | 254397..256160 | 670 | DUF262-domain-containing-protein-[Vibrio-parahaemolyticus] |
| HCG7048823.1 | ParB+HNH | GCA_024042595.1 | Vibrio_parahaemolyticus | Bacteria>Pseudomonadati>Pseudomonadota>Gammaproteobacteria>Vibrionales>Vibrionaceae>Vibrio>Vibrio_harveyi_group | 587 | NJ298_000554 | 544219..545982 | 670 | DUF262-domain-containing-protein-[Vibrio-parahaemolyticus] |
| MBE3824872.1 | ParB+HNH | GCA_014918955.1 | Vibrio_parahaemolyticus | Bacteria>Pseudomonadati>Pseudomonadota>Gammaproteobacteria>Vibrionales>Vibrionaceae>Vibrio>Vibrio_harveyi_group | 587 | HJ179_16465 | 254183..255946 | 670 | DUF262-domain-containing-protein-[Vibrio-parahaemolyticus] |
| WP_114867812.1 | ParB+HNH | - | Vibrio_parahaemolyticus | Bacteria>Pseudomonadati>Pseudomonadota>Gammaproteobacteria>Vibrionales>Vibrionaceae>Vibrio>Vibrio_harveyi_group | 587 | CFH89_RS14225 | 2817702..2819465 | 670 | DUF262-domain-containing-protein-[Vibrio-parahaemolyticus] |
| WP_269665660.1 | ParB+HNH | GCF_027277905.1 | Vibrio_parahaemolyticus | Bacteria>Pseudomonadati>Pseudomonadota>Gammaproteobacteria>Vibrionales>Vibrionaceae>Vibrio>Vibrio_harveyi_group | 587 | O4M94_RS09590 | 137873..139636 | 670 | DUF262-domain-containing-protein-[Vibrio-parahaemolyticus] |
| EII3096492.1 | ParB+HNH | GCA_020410925.1 | Vibrio_parahaemolyticus | Bacteria>Pseudomonadati>Pseudomonadota>Gammaproteobacteria>Vibrionales>Vibrionaceae>Vibrio>Vibrio_harveyi_group | 585 | CWL41_002101 | 328734..330491 | 670 | DUF262-domain-containing-protein-[Vibrio-parahaemolyticus] |
| EIN6339954.1 | ParB+HNH | GCA_020908185.1 | Vibrio_parahaemolyticus | Bacteria>Pseudomonadati>Pseudomonadota>Gammaproteobacteria>Vibrionales>Vibrionaceae>Vibrio>Vibrio_harveyi_group | 585 | LPQ31_001307 | 338483..340240 | 670 | DUF262-domain-containing-protein-[Vibrio-parahaemolyticus] |
| WP_023624068.1 | ParB+HNH | GCF_003691525.1 | Vibrio_parahaemolyticus | Bacteria>Pseudomonadati>Pseudomonadota>Gammaproteobacteria>Vibrionales>Vibrionaceae>Vibrio>Vibrio_harveyi_group | 585 | D0871_RS11645 | 2300470..2302227 | 670 | DUF262-domain-containing-protein-[Vibrio-parahaemolyticus] |
| WP_274616968.1 | ParB+HNH | - | Vibrio_parahaemolyticus | Bacteria>Pseudomonadati>Pseudomonadota>Gammaproteobacteria>Vibrionales>Vibrionaceae>Vibrio>Vibrio_harveyi_group | 575 | K6U15_RS27505 | 39380..41107 | 670 | DUF262-domain-containing-HNH-endonuclease-family-protein-[Vibrio-parahaemolyticus] |
| EGQ9599055.1 | ParB+HNH | - | Vibrio_parahaemolyticus | Bacteria>Pseudomonadati>Pseudomonadota>Gammaproteobacteria>Vibrionales>Vibrionaceae>Vibrio>Vibrio_harveyi_group | 562 | F3R80_21575 | 15316..17004 | 670 | DUF262-domain-containing-protein-[Vibrio-parahaemolyticus] |
| EHW0654652.1 | ParB+HNH | GCA_019442625.1 | Vibrio_parahaemolyticus | Bacteria>Pseudomonadati>Pseudomonadota>Gammaproteobacteria>Vibrionales>Vibrionaceae>Vibrio>Vibrio_harveyi_group | 562 | K1B36_003416 | 68395..70083 | 670 | DUF262-domain-containing-protein-[Vibrio-parahaemolyticus] |
| EJO3860681.1 | ParB+HNH | GCA_024563785.1 | Vibrio_parahaemolyticus | Bacteria>Pseudomonadati>Pseudomonadota>Gammaproteobacteria>Vibrionales>Vibrionaceae>Vibrio>Vibrio_harveyi_group | 562 | NRI58_000917 | 1067404..1069092 | 670 | DUF262-domain-containing-protein-[Vibrio-parahaemolyticus] |
| HBC3940912.1 | ParB+HNH | GCA_018127045.1 | Vibrio_parahaemolyticus | Bacteria>Pseudomonadati>Pseudomonadota>Gammaproteobacteria>Vibrionales>Vibrionaceae>Vibrio>Vibrio_harveyi_group | 562 | KD880_004473 | 18670..20358 | 670 | DUF262-domain-containing-protein-[Vibrio-parahaemolyticus] |
| HCG6132050.1 | ParB+HNH | GCA_024038995.1 | Vibrio_parahaemolyticus | Bacteria>Pseudomonadati>Pseudomonadota>Gammaproteobacteria>Vibrionales>Vibrionaceae>Vibrio>Vibrio_harveyi_group | 562 | NJ068_002905 | 62571..64259 | 670 | DUF262-domain-containing-protein-[Vibrio-parahaemolyticus] |
| MBE5192783.1 | ParB+HNH | GCA_014963905.1 | Vibrio_parahaemolyticus | Bacteria>Pseudomonadati>Pseudomonadota>Gammaproteobacteria>Vibrionales>Vibrionaceae>Vibrio>Vibrio_harveyi_group | 562 | HJ051_04860 | 1086907..1088595 | 670 | DUF262-domain-containing-protein-[Vibrio-parahaemolyticus] |
| MBM4838761.1 | ParB+HNH | GCA_016876235.1 | Vibrio_parahaemolyticus | Bacteria>Pseudomonadati>Pseudomonadota>Gammaproteobacteria>Vibrionales>Vibrionaceae>Vibrio>Vibrio_harveyi_group | 562 | HY055_06520 | 133416..135104 | 670 | DUF262-domain-containing-protein-[Vibrio-parahaemolyticus] |
| MBM4875014.1 | ParB+HNH | GCA_016876415.1 | Vibrio_parahaemolyticus | Bacteria>Pseudomonadati>Pseudomonadota>Gammaproteobacteria>Vibrionales>Vibrionaceae>Vibrio>Vibrio_harveyi_group | 562 | HY045_05450 | 524639..526327 | 670 | DUF262-domain-containing-protein-[Vibrio-parahaemolyticus] |
| WP_024701191.1 | ParB+HNH | GCF_000500485.1 | Vibrio_parahaemolyticus | Bacteria>Pseudomonadati>Pseudomonadota>Gammaproteobacteria>Vibrionales>Vibrionaceae>Vibrio>Vibrio_harveyi_group | 562 | T247_RS0121010 | 31082..32770 | 670 | DUF262-domain-containing-protein-[Vibrio-parahaemolyticus] |
| WP_025796288.1 | ParB+HNH | GCF_000591555.1 | Vibrio_parahaemolyticus | Bacteria>Pseudomonadati>Pseudomonadota>Gammaproteobacteria>Vibrionales>Vibrionaceae>Vibrio>Vibrio_harveyi_group | 562 | T6V_RS20305 | 8740..10428 | 670 | DUF262-domain-containing-protein-[Vibrio-parahaemolyticus] |
| WP_116920544.1 | ParB+HNH | GCF_905331795.1 | Vibrio_parahaemolyticus | Bacteria>Pseudomonadati>Pseudomonadota>Gammaproteobacteria>Vibrionales>Vibrionaceae>Vibrio>Vibrio_harveyi_group | 562 | LPR62_RS04720 | 416291..417979 | 670 | DUF262-domain-containing-protein-[Vibrio-parahaemolyticus] |
| WP_140247447.1 | ParB+HNH | GCF_048884405.1 | Vibrio_parahaemolyticus | Bacteria>Pseudomonadati>Pseudomonadota>Gammaproteobacteria>Vibrionales>Vibrionaceae>Vibrio>Vibrio_harveyi_group | 562 | CGJ41_RS23670 | 8454..10142 | 670 | DUF262-domain-containing-protein-[Vibrio-parahaemolyticus] |
| WP_140344324.1 | ParB+HNH | GCF_006372045.1 | Vibrio_parahaemolyticus | Bacteria>Pseudomonadati>Pseudomonadota>Gammaproteobacteria>Vibrionales>Vibrionaceae>Vibrio>Vibrio_harveyi_group | 562 | CGJ19_RS21465 | 10842..12530 | 670 | DUF262-domain-containing-protein-[Vibrio-parahaemolyticus] |
| WP_005444583.1 | ParB+HNH | GCF_000259935.1 | Vibrio_harveyi | Bacteria>Pseudomonadati>Pseudomonadota>Gammaproteobacteria>Vibrionales>Vibrionaceae>Vibrio>Vibrio_harveyi_group | 642 | MUQ_RS25195 | 10938..12866 | 669 | DUF262-domain-containing-protein-[Vibrio-harveyi] |
| WP_061065629.1 | ParB+HNH | GCF_001558435.2 | Vibrio_harveyi | Bacteria>Pseudomonadati>Pseudomonadota>Gammaproteobacteria>Vibrionales>Vibrionaceae>Vibrio>Vibrio_harveyi_group | 597 | AL538_RS13480 | 2682457..2684250 | 669 | DUF262-domain-containing-protein-[Vibrio-harveyi] |
| WP_045378171.1 | ParB+HNH | GCF_000818335.1 | Vibrio_campbellii | Bacteria>Pseudomonadati>Pseudomonadota>Gammaproteobacteria>Vibrionales>Vibrionaceae>Vibrio>Vibrio_harveyi_group | 698 | TY56_RS07060 | 524452..526548 | 680 | DUF262-and-DUF1524-domain-containing-protein-[Vibrio-campbellii] |
| EGQ7646158.1 | ParB+HNH | - | Vibrio_alginolyticus | Bacteria>Pseudomonadati>Pseudomonadota>Gammaproteobacteria>Vibrionales>Vibrionaceae>Vibrio>Vibrio_harveyi_group | 699 | I2703_000230 | 292022..294121 | 663 | DUF262-domain-containing-protein-[Vibrio-alginolyticus] |
| WP_213968026.1 | ParB+HNH | GCF_018458215.1 | Vibrio_alginolyticus | Bacteria>Pseudomonadati>Pseudomonadota>Gammaproteobacteria>Vibrionales>Vibrionaceae>Vibrio>Vibrio_harveyi_group | 698 | J4H41_RS21350 | 20345..22441 | 663 | DUF262-and-DUF1524-domain-containing-protein-[Vibrio-alginolyticus] |
| WP_222133203.1 | ParB+HNH | GCF_019793245.1 | Vibrio_alginolyticus | Bacteria>Pseudomonadati>Pseudomonadota>Gammaproteobacteria>Vibrionales>Vibrionaceae>Vibrio>Vibrio_harveyi_group | 644 | K6L11_RS14315 | 34728..36662 | 663 | DUF262-domain-containing-protein-[Vibrio-alginolyticus] |
| EGR0719597.1 | ParB+HNH | - | Vibrio_alginolyticus | Bacteria>Pseudomonadati>Pseudomonadota>Gammaproteobacteria>Vibrionales>Vibrionaceae>Vibrio>Vibrio_harveyi_group | 601 | EX464_05805 | 15614..17419 | 663 | DUF262-domain-containing-protein-[Vibrio-alginolyticus] |
| EJU9535830.1 | ParB+HNH | GCA_025224335.1 | Vibrio_alginolyticus | Bacteria>Pseudomonadati>Pseudomonadota>Gammaproteobacteria>Vibrionales>Vibrionaceae>Vibrio>Vibrio_harveyi_group | 601 | N5M46_001146 | 9277..11082 | 663 | DUF262-domain-containing-protein-[Vibrio-alginolyticus] |
| WP_005376174.1 | ParB+HNH | GCF_000153505.1 | Vibrio_alginolyticus | Bacteria>Pseudomonadati>Pseudomonadota>Gammaproteobacteria>Vibrionales>Vibrionaceae>Vibrio>Vibrio_harveyi_group | 601 | V12G01_RS31825 | 1605763..1607568 | 663 | DUF262-domain-containing-protein-[Vibrio-alginolyticus] |
| WP_213874131.1 | ParB+HNH | GCF_018458545.1 | Vibrio_alginolyticus | Bacteria>Pseudomonadati>Pseudomonadota>Gammaproteobacteria>Vibrionales>Vibrionaceae>Vibrio>Vibrio_harveyi_group | 601 | J4H28_RS04445 | 1025266..1027071 | 663 | DUF262-domain-containing-protein-[Vibrio-alginolyticus] |
| WP_238967173.1 | ParB+HNH | GCF_022343145.1 | Vibrio_alginolyticus | Bacteria>Pseudomonadati>Pseudomonadota>Gammaproteobacteria>Vibrionales>Vibrionaceae>Vibrio>Vibrio_harveyi_group | 601 | K6750_RS21890 | 1372058..1373863 | 663 | DUF262-domain-containing-protein-[Vibrio-alginolyticus] |
| WP_250156567.1 | ParB+HNH | - | Vibrio_alginolyticus | Bacteria>Pseudomonadati>Pseudomonadota>Gammaproteobacteria>Vibrionales>Vibrionaceae>Vibrio>Vibrio_harveyi_group | 575 | J4H58_RS14525 | 103183..104910 | 663 | DUF262-domain-containing-protein-[Vibrio-alginolyticus] |
| WP_272166067.1 | ParB+HNH | GCF_947503745.1 | Vibrio_diabolicus | Bacteria>Pseudomonadati>Pseudomonadota>Gammaproteobacteria>Vibrionales>Vibrionaceae>Vibrio>Vibrio_harveyi_group>Vibrio_diabolicus_subgroup | 644 | PQF93_RS21005 | 22391..24325 | 50719 | DUF262-domain-containing-HNH-endonuclease-family-protein-[Vibrio-diabolicus] |
| KLI66365.1 | ParB+HNH | GCA_001013325.1 | Vibrio_vulnificus_CladeA-yb158 | Bacteria>Pseudomonadati>Pseudomonadota>Gammaproteobacteria>Vibrionales>Vibrionaceae>Vibrio>Vibrio_vulnificus | 698 | VVYB158_12210 | 20360..22456 | 1112911 | hypothetical-protein-VVYB158_12210-[Vibrio-vulnificus-CladeA-yb158] |
| PIX55558.1 | ParB+HNH | GCA_002782825.1 | Zetaproteobacteria_bacterium_CG_4_10_14_3_um_filter_54_28 | Bacteria>Pseudomonadati>Pseudomonadota>Zetaproteobacteria>unclassified_Zetaproteobacteria | 602 | COZ50_02105 | 9749..11557 | 1974114 | DUF262-domain-containing-protein-[Zetaproteobacteria-bacterium-CG_4_10_14_3_um_filter_54_28] |
| MDD4116919.1 | ParB+HNH | GCA_028701395.1 | Kiritimatiellia_bacterium | Bacteria>Pseudomonadati>PVC_group>Kiritimatiellota>Kiritimatiellia>unclassified_Kiritimatiellia | 561 | PHI39_01745 | 21696..23381 | 2212473 | MAG:-DUF262-domain-containing-protein-[Kiritimatiellae-bacterium] |
| MBX3380226.1 | ParB+HNH | GCA_019636565.1 | Phycisphaeraceae_bacterium | Bacteria>Pseudomonadati>PVC_group>Planctomycetota>Phycisphaerae>Phycisphaerales>Phycisphaeraceae>unclassified_Phycisphaeraceae | 575 | KF805_09015 | 64574..66301 | 2026777 | DUF262-domain-containing-protein-[Phycisphaeraceae-bacterium] |
| MBX3380227.1 | ParB+HNH | GCA_019636565.1 | Phycisphaeraceae_bacterium | Bacteria>Pseudomonadati>PVC_group>Planctomycetota>Phycisphaerae>Phycisphaerales>Phycisphaeraceae>unclassified_Phycisphaeraceae | 555 | KF805_09020 | 66298..67965 | 2026777 | DUF262-domain-containing-protein-[Phycisphaeraceae-bacterium] |
| MCG3126048.1 | ParB+HNH | GCA_022072445.1 | Phycisphaerae_bacterium | Bacteria>Pseudomonadati>PVC_group>Planctomycetota>Phycisphaerae>unclassified_Phycisphaerae | 591 | CHACPFDD_00876 | 42829..44604 | 2026778 | hypothetical-protein-[Phycisphaerae-bacterium] |
| TWU09955.1 | ParB+HNH | GCA_007860105.1 | Allorhodopirellula_heiligendammensis | Bacteria>Pseudomonadati>PVC_group>Planctomycetota>Planctomycetia>Pirellulales>Pirellulaceae>Allorhodopirellula | 587 | Poly21_52840 | 34939..36702 | 2714739 | hypothetical-protein-Poly21_52840-[Planctomycetes-bacterium-Poly21] |
| MCM2371084.1 | ParB+HNH | GCA_023701485.1 | Aporhodopirellula_aestuarii | Bacteria>Pseudomonadati>PVC_group>Planctomycetota>Planctomycetia>Pirellulales>Pirellulaceae>Aporhodopirellula | 568 | NB063_10730 | 36345..38051 | 2950107 | DUF262-domain-containing-protein-[Rhodopirellula-aestuarii] |
| WP_250928727.1 | ParB+HNH | GCF_023701485.1 | Aporhodopirellula_aestuarii | Bacteria>Pseudomonadati>PVC_group>Planctomycetota>Planctomycetia>Pirellulales>Pirellulaceae>Aporhodopirellula | 568 | NB063_RS10710 | 36345..38051 | 2950107 | DUF262-domain-containing-protein-[Aporhodopirellula-aestuarii] |
| WP_207398111.1 | ParB+HNH | GCF_013618625.1 | Bremerella_alba | Bacteria>Pseudomonadati>PVC_group>Planctomycetota>Planctomycetia>Pirellulales>Pirellulaceae>Bremerella | 586 | HOV93_RS18980 | 113832..115592 | 980252 | DUF262-domain-containing-protein-[Bremerella-alba] |
| MCE2752490.1 | ParB+HNH | GCA_021299695.1 | Pirellula_sp. | Bacteria>Pseudomonadati>PVC_group>Planctomycetota>Planctomycetia>Pirellulales>Pirellulaceae>Pirellula>unclassified_Pirellula | 665 | LW720_11450 | 16042..18039 | 117 | DUF262-domain-containing-protein-[Pirellula-sp.] |
| MCE2752487.1 | ParB+HNH | GCA_021299695.1 | Pirellula_sp. | Bacteria>Pseudomonadati>PVC_group>Planctomycetota>Planctomycetia>Pirellulales>Pirellulaceae>Pirellula>unclassified_Pirellula | 565 | LW720_11435 | 10442..12139 | 117 | DUF262-domain-containing-HNH-endonuclease-family-protein-[Pirellula-sp.] |
| NLX98928.1 | ParB+HNH | GCA_012728855.1 | Rhodopirellula_sp. | Bacteria>Pseudomonadati>PVC_group>Planctomycetota>Planctomycetia>Pirellulales>Pirellulaceae>Rhodopirellula>unclassified_Rhodopirellula | 651 | GXY83_22510 | 1921..3876 | 2024855 | DUF262-domain-containing-protein-[Rhodopirellula-sp.] |
| WP_145095866.1 | ParB+HNH | GCF_007753095.1 | Rosistilla_carotiformis | Bacteria>Pseudomonadati>PVC_group>Planctomycetota>Planctomycetia>Pirellulales>Pirellulaceae>Rosistilla | 586 | Poly24_RS13220 | 3651414..3653174 | 2528017 | DUF262-domain-containing-protein-[Rosistilla-carotiformis] |
| WP_336521092.1 | ParB+HNH | GCF_030622045.1 | Planctellipticum_variicoloris | Bacteria>Pseudomonadati>PVC_group>Planctomycetota>Planctomycetia>Planctomycetales>Planctomycetaceae>Planctellipticum | 437 | SH412_RS26700 | 6852575..6853891 | 3064265 | DUF262-domain-containing-protein-[Planctoellipticum-variicoloris] |
| WP_082846328.1 | ParB+HNH | GCF_001610835.1 | Planctomyces_sp._SH-PL14 | Bacteria>Pseudomonadati>PVC_group>Planctomycetota>Planctomycetia>Planctomycetales>Planctomycetaceae>Planctomyces>unclassified_Planctomyces | 439 | VT03_RS18115 | 4716741..4718060 | 1632864 | DUF262-domain-containing-protein-[Planctomyces-sp.-SH-PL14] |
| QDT36022.1 | ParB+HNH | GCA_007744515.1 | Stratiformator_vulcanicus | Bacteria>Pseudomonadati>PVC_group>Planctomycetota>Planctomycetia>Planctomycetales>Planctomycetaceae>Stratiformator | 569 | Pan189_03770 | 490987..492696 | 2527980 | hypothetical-protein-Pan189_03770-[Planctomycetes-bacterium-Pan189] |
| MCA9232426.1 | ParB+HNH | GCA_020427155.1 | Planctomycetales_bacterium | Bacteria>Pseudomonadati>PVC_group>Planctomycetota>Planctomycetia>Planctomycetales>unclassified_Planctomycetales | 590 | KDA57_17385 | 204..1976 | 2053591 | DUF262-domain-containing-protein-[Planctomycetales-bacterium] |
| MBI3271451.1 | ParB+HNH | GCA_016197205.1 | Planctomycetota_bacterium | Bacteria>Pseudomonadati>PVC_group>Planctomycetota>unclassified_Planctomycetota | 879 | HYZ53_20845 | 58408..61047 | 2026780 | DUF262-domain-containing-protein-[Planctomycetota-bacterium] |
| MCE9567307.1 | ParB+HNH | GCA_021413735.1 | Planctomycetota_bacterium | Bacteria>Pseudomonadati>PVC_group>Planctomycetota>unclassified_Planctomycetota | 658 | K8U57_35345 | 42841..44817 | 2026780 | DUF262-domain-containing-HNH-endonuclease-family-protein-[Planctomycetota-bacterium] |
| MCD6405189.1 | ParB+HNH | GCA_021160625.1 | Planctomycetota_bacterium | Bacteria>Pseudomonadati>PVC_group>Planctomycetota>unclassified_Planctomycetota | 609 | J7M19_05145 | 7306..9135 | 2026780 | DUF262-domain-containing-protein-[Planctomycetota-bacterium] |
| TAH38050.1 | ParB+HNH | GCA_004295085.1 | Planctomycetota_bacterium | Bacteria>Pseudomonadati>PVC_group>Planctomycetota>unclassified_Planctomycetota | 590 | EYC70_05350 | 62821..64593 | 2026780 | MAG:-DUF262-domain-containing-protein-[Planctomycetota-bacterium] |
| MCE9554577.1 | ParB+HNH | GCA_021413705.1 | Planctomycetota_bacterium | Bacteria>Pseudomonadati>PVC_group>Planctomycetota>unclassified_Planctomycetota | 578 | K8T91_14560 | 3274..5010 | 2026780 | DUF262-domain-containing-HNH-endonuclease-family-protein-[Planctomycetota-bacterium] |
| MCZ7645667.1 | ParB+HNH | GCA_027493525.1 | Planctomycetota_bacterium | Bacteria>Pseudomonadati>PVC_group>Planctomycetota>unclassified_Planctomycetota | 568 | M5U26_10340 | 2477243..2478949 | 2026780 | MAG:-DUF262-domain-containing-protein-[Planctomycetota-bacterium] |
| MCE9553882.1 | ParB+HNH | GCA_021413705.1 | Planctomycetota_bacterium | Bacteria>Pseudomonadati>PVC_group>Planctomycetota>unclassified_Planctomycetota | 562 | K8T91_10980 | 59270..60958 | 2026780 | DUF262-domain-containing-protein-[Planctomycetota-bacterium] |
| MBI3462534.1 | ParB+HNH | GCA_016196255.1 | Planctomycetota_bacterium | Bacteria>Pseudomonadati>PVC_group>Planctomycetota>unclassified_Planctomycetota | 545 | HY000_05660 | 1097..2734 | 2026780 | DUF262-domain-containing-protein-[Planctomycetota-bacterium] |
| MCE9567306.1 | ParB+HNH | GCA_021413735.1 | Planctomycetota_bacterium | Bacteria>Pseudomonadati>PVC_group>Planctomycetota>unclassified_Planctomycetota | 469 | K8U57_35340 | 41439..42848 | 2026780 | DUF262-domain-containing-protein-[Planctomycetota-bacterium] |
| TET34866.1 | ParB+HNH | GCA_004376375.1 | Planctomycetota_bacterium | Bacteria>Pseudomonadati>PVC_group>Planctomycetota>unclassified_Planctomycetota | 359 | E3J72_12075 | 39773..40852 | 2026780 | MAG:-DUF262-domain-containing-protein-[Planctomycetota-bacterium] |
| WP_189517052.1 | ParB+HNH | GCF_014651635.1 | Cerasicoccus_arenae | Bacteria>Pseudomonadati>PVC_group>Verrucomicrobiota>Opitutia>Puniceicoccales>Cerasicoccaceae>Cerasicoccus | 582 | JIN80_RS05845 | 23856..25604 | 424488 | DUF262-domain-containing-protein-[Cerasicoccus-arenae] |
| MBM3880419.1 | ParB+HNH | GCA_016871715.1 | Verrucomicrobiota_bacterium | Bacteria>Pseudomonadati>PVC_group>Verrucomicrobiota>unclassified_Verrucomicrobiota | 569 | FJ387_11995 | 3726..5435 | 2026799 | DUF262-domain-containing-protein-[Verrucomicrobiota-bacterium] |
| MBS0647001.1 | ParB+HNH | GCA_018239885.1 | Verrucomicrobiota_bacterium | Bacteria>Pseudomonadati>PVC_group>Verrucomicrobiota>unclassified_Verrucomicrobiota | 555 | JSR97_10500 | 7235..8902 | 2026799 | DUF262-domain-containing-protein-[Verrucomicrobiota-bacterium] |
| OQC32222.1 | ParB+HNH | GCA_002071025.1 | Verrucomicrobia_bacterium_ADurb.Bin070 | Bacteria>Pseudomonadati>PVC_group>Verrucomicrobiota>unclassified_Verrucomicrobiota | 617 | BWX70_00670 | 13544..15397 | 1852927 | hypothetical-protein-BWX70_00670-[Verrucomicrobia-bacterium-ADurb.Bin070] |
| WP_215427528.1 | ParB+HNH | GCF_018709685.1 | Akkermansia_muciniphila | Bacteria>Pseudomonadati>PVC_group>Verrucomicrobiota>Verrucomicrobiia>Verrucomicrobiales>Akkermansiaceae>Akkermansia | 689 | HHJ06_RS03170 | 100513..102582 | 239935 | DUF262-and-DUF1524-domain-containing-protein-[Akkermansia-muciniphila] |
| MBE7502031.1 | ParB+HNH | GCA_015075445.1 | Verrucomicrobiales_bacterium | Bacteria>Pseudomonadati>PVC_group>Verrucomicrobiota>Verrucomicrobiia>Verrucomicrobiales>unclassified_Verrucomicrobiales | 553 | HS113_17405 | 117722..119383 | 2026801 | DUF262-domain-containing-protein-[Verrucomicrobiales-bacterium] |
| MBE7497137.1 | ParB+HNH | GCA_015075365.1 | Verrucomicrobiaceae_bacterium | Bacteria>Pseudomonadati>PVC_group>Verrucomicrobiota>Verrucomicrobiia>Verrucomicrobiales>Verrucomicrobiaceae>unclassified_Verrucomicrobiaceae | 559 | HS117_19520 | 522193..523872 | 2026800 | DUF262-domain-containing-protein-[Verrucomicrobiaceae-bacterium] |
| WP_210413549.1 | ParB+HNH | - | Leptospira_selangorensis | Bacteria>Pseudomonadati>Spirochaetota>Spirochaetia>Leptospirales>Leptospiraceae>Leptospira | 639 | EHQ82_RS10895 | 7313..9232 | 2484982 | DUF262-domain-containing-protein-[Leptospira-selangorensis] |
| WP_238754861.1 | ParB+HNH | GCF_022267385.1 | Leptospira_bandrabouensis | Bacteria>Pseudomonadati>Spirochaetota>Spirochaetia>Leptospirales>Leptospiraceae>Leptospira | 594 | LFX13_RS16590 | 41822..43606 | 2484903 | DUF262-domain-containing-protein-[Leptospira-bandrabouensis] |
| EOQ90761.1 | ParB+HNH | GCA_000332475.2 | Leptospira_yanagawae_serovar_Saopaulo_str._Sao_Paulo_=_ATCC_700523 | Bacteria>Pseudomonadati>Spirochaetota>Spirochaetia>Leptospirales>Leptospiraceae>Leptospira>Leptospira_yanagawae>Leptospira_yanagawae_serovar_Saopaulo | 594 | LEP1GSC202_0421 | 22041..23825 | 1249483 | PF03235-family-protein-[Leptospira-yanagawae-serovar-Saopaulo-str.-Sao-Paulo-=-ATCC-700523] |
| MCB1144529.1 | ParB+HNH | GCA_020440865.1 | Leptospiraceae_bacterium | Bacteria>Pseudomonadati>Spirochaetota>Spirochaetia>Leptospirales>Leptospiraceae>unclassified_Leptospiraceae | 582 | KDK54_19900 | 6001..7749 | 2026757 | DUF262-domain-containing-protein-[Leptospiraceae-bacterium] |
| MBK9498685.1 | ParB+HNH | GCA_016718635.1 | Leptospiraceae_bacterium | Bacteria>Pseudomonadati>Spirochaetota>Spirochaetia>Leptospirales>Leptospiraceae>unclassified_Leptospiraceae | 576 | IP006_04735 | 66218..67948 | 2026757 | DUF262-domain-containing-protein-[Leptospiraceae-bacterium] |
| MBP6738377.1 | ParB+HNH | GCA_017990775.1 | Leptospiraceae_bacterium | Bacteria>Pseudomonadati>Spirochaetota>Spirochaetia>Leptospirales>Leptospiraceae>unclassified_Leptospiraceae | 574 | KA146_00230 | 6332..8056 | 2026757 | DUF262-domain-containing-protein-[Leptospiraceae-bacterium] |
| MBP6739050.1 | ParB+HNH | GCA_017990775.1 | Leptospiraceae_bacterium | Bacteria>Pseudomonadati>Spirochaetota>Spirochaetia>Leptospirales>Leptospiraceae>unclassified_Leptospiraceae | 458 | KA146_03615 | 3762..5138 | 2026757 | DUF262-domain-containing-protein-[Leptospiraceae-bacterium] |
| MBK9497789.1 | ParB+HNH | GCA_016718635.1 | Leptospiraceae_bacterium | Bacteria>Pseudomonadati>Spirochaetota>Spirochaetia>Leptospirales>Leptospiraceae>unclassified_Leptospiraceae | 456 | IP006_00145 | 40753..42123 | 2026757 | DUF262-domain-containing-protein-[Leptospiraceae-bacterium] |
| AEJ18698.1 | ParB+HNH | GCA_000219725.1 | Gracilinema_caldarium_DSM_7334 | Bacteria>Pseudomonadati>Spirochaetota>Spirochaetia>Spirochaetales>Breznakiellaceae>Gracilinema>Gracilinema_caldarium | 562 | Spica_0537 | 577999..579687 | 744872 | protein-of-unknown-function-DUF262-[Gracilinema-caldarium-DSM-7334] |
| WP_083048610.1 | ParB+HNH | GCF_002087085.1 | Marispirochaeta_aestuarii | Bacteria>Pseudomonadati>Spirochaetota>Spirochaetia>Spirochaetales>Spirochaetaceae>Marispirochaeta | 598 | B4O97_RS04135 | 256413..258209 | 1963862 | DUF262-domain-containing-protein-[Marispirochaeta-aestuarii] |
| ORC33986.1 | ParB+HNH | GCA_002087085.1 | Marispirochaeta_aestuarii | Bacteria>Pseudomonadati>Spirochaetota>Spirochaetia>Spirochaetales>Spirochaetaceae>Marispirochaeta | 568 | B4O97_13945 | 6462..8168 | 1963862 | hypothetical-protein-B4O97_13945-[Marispirochaeta-aestuarii] |
| WP_158084305.1 | ParB+HNH | GCF_002087085.1 | Marispirochaeta_aestuarii | Bacteria>Pseudomonadati>Spirochaetota>Spirochaetia>Spirochaetales>Spirochaetaceae>Marispirochaeta | 558 | B4O97_RS13945 | 6462..8138 | 1963862 | DUF262-domain-containing-protein-[Marispirochaeta-aestuarii] |
| OQY41121.1 | ParB+HNH | GCA_002084785.1 | Spirochaetaceae_bacterium_4572_7 | Bacteria>Pseudomonadati>Spirochaetota>Spirochaetia>Spirochaetales>Spirochaetaceae>unclassified_Spirochaetaceae | 750 | B6229_00045 | 12985..15237 | 1972456 | hypothetical-protein-B6229_00045-[Spirochaetaceae-bacterium-4572_7] |
| OQY41120.1 | ParB+HNH | GCA_002084785.1 | Spirochaetaceae_bacterium_4572_7 | Bacteria>Pseudomonadati>Spirochaetota>Spirochaetia>Spirochaetales>Spirochaetaceae>unclassified_Spirochaetaceae | 622 | B6229_00040 | 11127..12995 | 1972456 | hypothetical-protein-B6229_00040-[Spirochaetaceae-bacterium-4572_7] |
| MCL1994398.1 | ParB+HNH | GCA_009778155.1 | Spirochaetota_bacterium | Bacteria>Pseudomonadati>Spirochaetota>unclassified_Spirochaetota | 404 | FWG66_15750 | 25678..26892 | 2202144 | DUF262-domain-containing-protein-[Spirochaetota-bacterium] |
| OQB96943.1 | ParB+HNH | GCA_002070755.1 | Spirochaetes_bacterium_ADurb.Bin110 | Bacteria>Pseudomonadati>Spirochaetota>unclassified_Spirochaetota | 603 | BWX81_01212 | 8904..10715 | 1852908 | hypothetical-protein-BWX81_01212-[Spirochaetes-bacterium-ADurb.Bin110] |
| VBB42848.1 | ParB+HNH | GCA_900498135.1 | uncultured_Desulfatiglans_sp. | Bacteria>Pseudomonadati>Thermodesulfobacteriota>Desulfobacteria>Desulfatiglandales>Desulfatiglandaceae>Desulfatiglans>environmental_samples | 568 | TRIP_B220096 | 91510..93216 | 1748965 | conserved-hypothetical-protein-[uncultured-Desulfatiglans-sp.] |
| KPA09728.1 | ParB+HNH | GCA_001292585.1 | Candidatus_Magnetomorum_sp._HK-1 | Bacteria>Pseudomonadati>Thermodesulfobacteriota>Desulfobacteria>Desulfobacterales>Candidatus_Magnetomoraceae>Candidatus_Magnetomorum>unclassified_Candidatus_Magnetomorum | 393 | MHK_010068 | 8241..9422 | 1509431 | protein-containing-DUF262-[Candidatus-Magnetomorum-sp.-HK-1] |
| WP_035236108.1 | ParB+HNH | GCF_000745975.1 | Desulfobacter_vibrioformis | Bacteria>Pseudomonadati>Thermodesulfobacteriota>Desulfobacteria>Desulfobacterales>Desulfobacteraceae>Desulfobacter | 573 | Q366_RS02895 | 247042..248763 | 34031 | DUF262-domain-containing-protein-[Desulfobacter-vibrioformis] |
| WP_084067310.1 | ParB+HNH | GCF_900176365.1 | Desulfocicer_vacuolatum | Bacteria>Pseudomonadati>Thermodesulfobacteriota>Desulfobacteria>Desulfobacterales>Desulfobacteraceae>Desulfocicer | 415 | B9A32_RS05110 | 99213..100460 | 2298 | DUF262-domain-containing-protein-[Desulfocicer-vacuolatum] |
| WP_024334647.1 | ParB+HNH | GCF_000421285.1 | Desulfotignum_balticum | Bacteria>Pseudomonadati>Thermodesulfobacteriota>Desulfobacteria>Desulfobacterales>Desulfobacteraceae>Desulfotignum | 598 | K365_RS0111350 | 2248919..2250715 | 115781 | DUF262-domain-containing-protein-[Desulfotignum-balticum] |
| OQY49643.1 | ParB+HNH | GCA_002085465.1 | Desulfobacteraceae_bacterium_4572_89 | Bacteria>Pseudomonadati>Thermodesulfobacteriota>Desulfobacteria>Desulfobacterales>Desulfobacteraceae>unclassified_Desulfobacteraceae | 382 | B6230_07755 | 1365..2513 | 1972445 | hypothetical-protein-B6230_07755-[Desulfobacteraceae-bacterium-4572_89] |
| MBA3017258.1 | ParB+HNH | GCA_013792055.1 | Desulfobacteraceae_bacterium | Bacteria>Pseudomonadati>Thermodesulfobacteriota>Desulfobacteria>Desulfobacterales>Desulfobacteraceae>unclassified_Desulfobacteraceae | 372 | FP811_03745 | 33224..34342 | 2049433 | DUF262-domain-containing-protein-[Desulfobacteraceae-bacterium] |
| MCF8025631.1 | ParB+HNH | GCA_021737095.1 | Desulfobacteraceae_bacterium | Bacteria>Pseudomonadati>Thermodesulfobacteriota>Desulfobacteria>Desulfobacterales>Desulfobacteraceae>unclassified_Desulfobacteraceae | 173 | K9K82_09100 | 9737..10254 | 2049433 | DUF262-domain-containing-protein-[Desulfobacteraceae-bacterium] |
| PIP37813.1 | ParB+HNH | GCA_002772195.1 | Desulfobacterales_bacterium_CG23_combo_of_CG06-09_8_20_14_all_51_8 | Bacteria>Pseudomonadati>Thermodesulfobacteriota>Desulfobacteria>Desulfobacterales>unclassified_Desulfobacterales | 751 | COX19_15645 | 4765..7020 | 1973983 | hypothetical-protein-COX19_15645-[Desulfobacterales-bacterium-CG23_combo_of_CG06-09_8_20_14_all_51_8] |
| MDD5204469.1 | ParB+HNH | GCA_028713085.1 | Desulfobacterales_bacterium | Bacteria>Pseudomonadati>Thermodesulfobacteriota>Desulfobacteria>Desulfobacterales>unclassified_Desulfobacterales | 459 | PHS17_03560 | 6187..7568 | 2044940 | MAG:-HNH-endonuclease-family-protein,-partial-[Desulfobacterales-bacterium] |
| KGO34921.1 | ParB+HNH | GCA_000769715.1 | Desulfobulbus_sp._Tol-SR | Bacteria>Pseudomonadati>Thermodesulfobacteriota>Desulfobulbia>Desulfobulbales>Desulfobulbaceae>Desulfobulbus>unclassified_Desulfobulbus | 573 | JT06_05455 | 3358..5079 | 1536652 | hypothetical-protein-JT06_05455-[Desulfobulbus-sp.-Tol-SR] |
| WP_092376848.1 | ParB+HNH | GCF_900114115.1 | Desulfomicrobium_apsheronum | Bacteria>Pseudomonadati>Thermodesulfobacteriota>Desulfovibrionia>Desulfovibrionales>Desulfomicrobiaceae>Desulfomicrobium | 570 | BMZ40_RS13605 | 78007..79719 | 52560 | DUF262-domain-containing-protein-[Desulfomicrobium-apsheronum] |
| MBI4803782.1 | ParB+HNH | GCA_016208105.1 | Desulfovibrio_sp. | Bacteria>Pseudomonadati>Thermodesulfobacteriota>Desulfovibrionia>Desulfovibrionales>Desulfovibrionaceae>Desulfovibrio>unclassified_Desulfovibrio | 598 | HY795_00940 | 34580..36376 | 885 | DUF262-domain-containing-protein-[Desulfovibrio-sp.] |
| WP_243366431.1 | ParB+HNH | GCF_022808695.1 | Fundidesulfovibrio_soli | Bacteria>Pseudomonadati>Thermodesulfobacteriota>Desulfovibrionia>Desulfovibrionales>Desulfovibrionaceae>Fundidesulfovibrio | 583 | MLE18_RS00815 | 170863..172614 | 2922716 | DUF262-domain-containing-protein-[Fundidesulfovibrio-soli] |
| MBL3581667.1 | ParB+HNH | GCA_016756815.1 | Oleidesulfovibrio_alaskensis | Bacteria>Pseudomonadati>Thermodesulfobacteriota>Desulfovibrionia>Desulfovibrionales>Desulfovibrionaceae>Oleidesulfovibrio | 598 | JMJ96_04550 | 100224..102020 | 58180 | DUF262-domain-containing-protein-[Oleidesulfovibrio-alaskensis] |
| WP_273192717.1 | ParB+HNH | GCF_016756815.1 | Oleidesulfovibrio_alaskensis | Bacteria>Pseudomonadati>Thermodesulfobacteriota>Desulfovibrionia>Desulfovibrionales>Desulfovibrionaceae>Oleidesulfovibrio | 598 | JMJ96_RS04560 | 100224..102020 | 58180 | DUF262-domain-containing-HNH-endonuclease-family-protein-[Oleidesulfovibrio-alaskensis] |
| AJY71185.1 | ParB+HNH | GCA_000961035.1 | Geobacter_sulfurreducens | Bacteria>Pseudomonadati>Thermodesulfobacteriota>Desulfuromonadia>Geobacterales>Geobacteraceae>Geobacter | 573 | RW64_17295 | 3782597..3784318 | 35554 | hypothetical-protein-RW64_17295-[Geobacter-sulfurreducens] |
| RII28690.1 | ParB+HNH | GCA_003574885.1 | Geobacter_sp. | Bacteria>Pseudomonadati>Thermodesulfobacteriota>Desulfuromonadia>Geobacterales>Geobacteraceae>Geobacter>unclassified_Geobacter | 786 | CXR31_02070 | 451669..454029 | 46610 | MAG:-DUF262-domain-containing-protein-[Geobacter-sp.] |
| OPY81319.1 | ParB+HNH | GCA_002067745.1 | Syntrophus_sp._PtaU1.Bin005 | Bacteria>Pseudomonadati>Thermodesulfobacteriota>Syntrophia>Syntrophales>Syntrophaceae>Syntrophus>unclassified_Syntrophus_(in:_bacteria) | 608 | A4E70_01337 | 16347..18173 | 1811712 | MAG:-hypothetical-protein-A4E70_01337-[Syntrophus-sp.-PtaU1.Bin005] |
| WP_137424091.1 | ParB+HNH | GCF_005377625.1 | Desulfoglaeba_alkanexedens | Bacteria>Pseudomonadati>Thermodesulfobacteriota>Syntrophobacteria>Syntrophobacterales>Syntrophobacteraceae>Desulfoglaeba | 598 | FDQ92_RS08060 | 1782990..1784786 | 361111 | DUF262-domain-containing-protein-[Desulfoglaeba-alkanexedens] |
| OPX97690.1 | ParB+HNH | GCA_002067585.1 | Syntrophorhabdus_sp._PtaB.Bin027 | Bacteria>Pseudomonadati>Thermodesulfobacteriota>Syntrophorhabdia>Syntrophorhabdales>Syntrophorhabdaceae>Syntrophorhabdus>unclassified_Syntrophorhabdus | 618 | A4E59_00671 | 25320..27176 | 1811699 | MAG:-hypothetical-protein-A4E59_00671-[Syntrophorhabdus-sp.-PtaB.Bin027] |
| MDD3846876.1 | ParB+HNH | GCA_028698615.1 | Syntrophorhabdaceae_bacterium | Bacteria>Pseudomonadati>Thermodesulfobacteriota>Syntrophorhabdia>Syntrophorhabdales>Syntrophorhabdaceae>unclassified_Syntrophorhabdaceae | 368 | PHC90_11020 | 2804..3910 | 2660714 | MAG:-DUF262-domain-containing-protein-[Syntrophorhabdaceae-bacterium] |
| WP_133161747.1 | ParB+HNH | GCF_002901445.1 | Deinococcus_koreensis | Bacteria>Thermotogati>Deinococcota>Deinococci>Deinococcales>Deinococcaceae>Deinococcus | 777 | CVO96_RS20715 | 1094534..1096867 | 2054903 | DUF262-domain-containing-HNH-endonuclease-family-protein-[Deinococcus-koreensis] |
| PNY82878.1 | ParB+HNH | GCA_002901445.1 | Deinococcus_koreensis | Bacteria>Thermotogati>Deinococcota>Deinococci>Deinococcales>Deinococcaceae>Deinococcus | 449 | CVO96_05145 | 1094534..1095883 | 2054903 | hypothetical-protein-CVO96_05145-[Deinococcus-koreensis] |
| SIN70231.1 | ParB+HNH | GCA_900129645.1 | Acetomicrobium_flavidum | Bacteria>Thermotogati>Synergistota>Synergistia>Synergistales>Acetomicrobiaceae>Acetomicrobium | 587 | SAMN05444368_1316 | 1296642..1298405 | 49896 | Protein-of-unknown-function-DUF262-[Acetomicrobium-flavidum] |
| AFM22456.1 | ParB+HNH | GCA_000266925.1 | Acetomicrobium_mobile_DSM_13181 | Bacteria>Thermotogati>Synergistota>Synergistia>Synergistales>Acetomicrobiaceae>Acetomicrobium>Acetomicrobium_mobile | 592 | Anamo_1870 | 1892942..1894720 | 891968 | hypothetical-protein-Anamo_1870-[Acetomicrobium-mobile-DSM-13181] |
| WP_012870224.1 | ParB+HNH | GCF_000024905.1 | Thermanaerovibrio_acidaminovorans | Bacteria>Thermotogati>Synergistota>Synergistia>Synergistales>Synergistaceae>Thermanaerovibrio | 600 | TACI_RS07675 | 1549814..1551616 | 81462 | DUF262-domain-containing-protein-[Thermanaerovibrio-acidaminovorans] |
| WP_012870222.1 | ParB+HNH | GCF_000024905.1 | Thermanaerovibrio_acidaminovorans | Bacteria>Thermotogati>Synergistota>Synergistia>Synergistales>Synergistaceae>Thermanaerovibrio | 576 | TACI_RS07665 | 1547709..1549439 | 81462 | DUF262-domain-containing-protein-[Thermanaerovibrio-acidaminovorans] |
| WP_099772689.1 | ParB+HNH | GCF_002752615.1 | unclassified_Mesotoga | Bacteria>Thermotogati>Thermotogota>Thermotogae>Kosmotogales>Kosmotogaceae>Mesotoga | 597 | Y696_RS00845 | 34860..36653 | 1184398 | MULTISPECIES:-DUF262-domain-containing-protein-[unclassified-Mesotoga] |
| WP_181419070.1 | ParB+HNH | GCF_003228205.1 | Mesotoga_sp._TolDC | Bacteria>Thermotogati>Thermotogota>Thermotogae>Kosmotogales>Kosmotogaceae>Mesotoga>unclassified_Mesotoga | 589 | LH53_RS05210 | 23573..25342 | 1389250 | DUF262-domain-containing-protein-[Mesotoga-sp.-TolDC] |
| WP_103134079.1 | ParB+HNH | GCF_002896535.1 | Mesotoga_sp._B105.6.4 | Bacteria>Thermotogati>Thermotogota>Thermotogae>Kosmotogales>Kosmotogaceae>Mesotoga>unclassified_Mesotoga | 596 | RJ60_RS04170 | 249260..251050 | 1582224 | DUF262-domain-containing-protein-[Mesotoga-sp.-B105.6.4] |
| KUK23105.1 | ParB+HNH | GCA_001508125.1 | Thermotoga_petrophila | Bacteria>Thermotogati>Thermotogota>Thermotogae>Thermotogales>Thermotogaceae>Thermotoga | 106 | XD57_0800 | 8997..9317 | 93929 | MAG:-Uncharacterized-protein-XD57_0800,-partial-[Thermotoga-petrophila] |
| WP_004080566.1 | ParB+HNH | GCF_000008545.1 | Thermotoga_maritima | Bacteria>Thermotogati>Thermotogota>Thermotogae>Thermotogales>Thermotogaceae>Thermotoga | 584 | TM_RS05050 | 1009825..1011579 | 2336 | DUF262-domain-containing-protein-[Thermotoga-maritima] |
| ACM23762.1 | ParB+HNH | GCA_000018945.1 | Thermotoga_neapolitana_DSM_4359 | Bacteria>Thermotogati>Thermotogota>Thermotogae>Thermotogales>Thermotogaceae>Thermotoga>Thermotoga_neapolitana | 584 | CTN_1586 | 1551742..1553496 | 309803 | Putative-uncharacterized-protein-[Thermotoga-neapolitana-DSM-4359] |

| Protein-ID | Domain | Assembly-ID | Strain | Taxonomy | Length | Locus-Tag | Genomic-coordinates | TaxID | Fasta-defline |
|---|---|---|---|---|---|---|---|---|---|
| WP_236681706.1 | ParB+HNH | - | Thermotoga_sp._RQ7 | Bacteria>Thermotogati>Thermotogota>Thermotogae>Thermotogales>Thermotogaceae>Thermotoga>unclassified_Thermotoga | 332 | TRQ7_RS09825 | 1795785..1796783 | 126738 | DUF262-domain-containing-protein-[Thermotoga-sp.-RQ7] |
| WP_236681707.1 | ParB+HNH | - | Thermotoga_sp._RQ7 | Bacteria>Thermotogati>Thermotogota>Thermotogae>Thermotogales>Thermotogaceae>Thermotoga>unclassified_Thermotoga | 189 | TRQ7_RS09830 | 1796969..1797538 | 126738 | hypothetical-protein-[Thermotoga-sp.-RQ7] |
| WP_012311352.1 | ParB+HNH | GCF_000019625.1 | Thermotoga_sp._RQ2 | Bacteria>Thermotogati>Thermotogota>Thermotogae>Thermotogales>Thermotogaceae>Thermotoga>unclassified_Thermotoga | 584 | TRQ2_RS09305 | 1821347..1823101 | 126740 | DUF262-domain-containing-protein-[Thermotoga-sp.-RQ2] |
| WP_235280336.1 | ParB+HNH | GCF_000784835.1 | Thermotoga_sp._Mc24 | Bacteria>Thermotogati>Thermotogota>Thermotogae>Thermotogales>Thermotogaceae>Thermotoga>unclassified_Thermotoga | 329 | AS158_RS09330 | 75578..76567 | 1231241 | MULTISPECIES:-DUF262-domain-containing-protein-[unclassified-Thermotoga] |
| WP_241241284.1 | ParB+HNH | - | Thermotoga_sp._38H-to | Bacteria>Thermotogati>Thermotogota>Thermotogae>Thermotogales>Thermotogaceae>Thermotoga>unclassified_Thermotoga | 205 | AS158_RS09325 | 74811..75428 | 1755812 | hypothetical-protein-[Thermotoga-sp.-38H-to] |
| MBZ4661019.1 | ParB+HNH | GCA_020054995.1 | Thermotoga_sp. | Bacteria>Thermotogati>Thermotogota>Thermotogae>Thermotogales>Thermotogaceae>Thermotoga>unclassified_Thermotoga | 584 | JG779_173 | 6275..8029 | 28240 | hypothetical-protein-[Thermotoga-sp.] |
| RLJ22038.1 | ParB+HNH | GCA_003660225.1 | bacterium_endosymbiont_of_Escarpia_laminata | Bacteria>unclassified_Bacteria | 427 | DJ030_02875 | 47694..48977 | 1221601 | hypothetical-protein-DJ030_02875-[bacterium-endosymbiont-of-Escarpia-laminata] |
| MBI1292836.1 | ParB+HNH | GCA_016125005.1 | bacterium | Bacteria>unclassified_Bacteria | 558 | GC173_16600 | 18507..20183 | 1869227 | DUF262-domain-containing-protein-[bacterium] |
| MBC7327981.1 | ParB+HNH | GCA_014360465.1 | bacterium | Bacteria>unclassified_Bacteria | 462 | H5T87_07690 | 2217..3605 | 1869227 | DUF262-domain-containing-protein-[bacterium] |
| --wHTH(SBD)+HNH | | | | | | | | | |
| MCQ2357406.1 | wHTH(SBD)+HNH | GCA_024405795.1 | Methanocorpusculum_sp. | Archaea>Methanobacteriati>Methanobacteriota>Stenosarchaea_group>Methanomicrobia>Methanomicrobiales>Methanocorpusculaceae>Methanocorpusculum>unclassified_Methanocorpusculum | 385 | MJ005_01355 | 36771..37928 | 2058474 | hypothetical-protein-[Methanocorpusculum-sp.] |
| WP_113812543.1 | wHTH(SBD)+HNH | GCF_003319095.1 | Blastococcus_sp._TBT05-19 | Bacteria>Bacillati>Actinomycetota>Actinomycetes>Geodermatophilales>Geodermatophilaceae>Blastococcus>unclassified_Blastococcus | 301 | DQ244_RS16260 | 137435..138340 | 2250581 | HNH-endonuclease-[Blastococcus-sp.-TBT05-19] |
| WP_161236369.1 | wHTH(SBD)+HNH | - | unclassified_Streptomyces | Bacteria>Bacillati>Actinomycetota>Actinomycetes>Kitasatosporales>Streptomycetaceae>Streptomyces | 301 | G3I23_RS06855 | 3761..4666 | 2593676 | MULTISPECIES:-HNH-endonuclease-[unclassified-Streptomyces] |
| WP_187645672.1 | wHTH(SBD)+HNH | GCF_008271805.1 | Streptomyces_sp._TRM49041 | Bacteria>Bacillati>Actinomycetota>Actinomycetes>Kitasatosporales>Streptomycetaceae>Streptomyces>unclassified_Streptomyces | 307 | FVS66_RS24230 | 252973..253896 | 2603216 | HNH-endonuclease-[Streptomyces-sp.-TRM49041] |
| OFE15919.1 | wHTH(SBD)+HNH | GCA_001758225.1 | Humibacillus_sp._DSM_29435 | Bacteria>Bacillati>Actinomycetota>Actinomycetes>Micrococcales>Intrasporangiaceae>Humibacillus>unclassified_Humibacillus | 261 | BA895_05320 | 177643..178428 | 1869167 | hypothetical-protein-BA895_05320-[Humibacillus-sp.-DSM-29435] |
| WP_083283377.1 | wHTH(SBD)+HNH | GCF_001758225.1 | Humibacillus_sp._DSM_29435 | Bacteria>Bacillati>Actinomycetota>Actinomycetes>Micrococcales>Intrasporangiaceae>Humibacillus>unclassified_Humibacillus | 287 | BA895_RS14670 | 177643..178506 | 1869167 | HNH-endonuclease-[Humibacillus-sp.-DSM-29435] |
| WP_169735705.1 | wHTH(SBD)+HNH | GCF_000576595.1 | Intrasporangium_oryzae | Bacteria>Bacillati>Actinomycetota>Actinomycetes>Micrococcales>Intrasporangiaceae>Intrasporangium | 291 | N865_RS21625 | 98607..99482 | 412687 | HNH-endonuclease-[Intrasporangium-oryzae] |
| MCI1262618.1 | wHTH(SBD)+HNH | GCA_022641475.1 | Nostocoides_jenkinsii | Bacteria>Bacillati>Actinomycetota>Actinomycetes>Micrococcales>Intrasporangiaceae>Nostocoides | 302 | LKG20_10150 | 9582..10490 | 330834 | HNH-endonuclease-[Tetrasphaera-jenkinsii] |
| WP_224048787.1 | wHTH(SBD)+HNH | GCF_019977335.1 | Arthrobacter_sp._NicSoilB4 | Bacteria>Bacillati>Actinomycetota>Actinomycetes>Micrococcales>Micrococcaceae>Arthrobacter>unclassified_Arthrobacter | 375 | LDO13_RS04070 | 873242..874369 | 2830997 | HNH-endonuclease-[Arthrobacter-sp.-NicSoilB4] |
| WP_188427980.1 | wHTH(SBD)+HNH | GCF_039543725.1 | Ornithinimicrobium_tianjinense | Bacteria>Bacillati>Actinomycetota>Actinomycetes>Micrococcales>Ornithinimicrobiaceae>Ornithinimicrobium | 308 | IEV65_RS02270 | 463844..464770 | 1195761 | HNH-endonuclease-signature-motif-containing-protein-[Ornithinimicrobium-tianjinense] |
| SDF11501.1 | wHTH(SBD)+HNH | GCA_900102445.1 | Cellulosimicrobium_cellulans | Bacteria>Bacillati>Actinomycetota>Actinomycetes>Micrococcales>Promicromonosporaceae>Cellulosimicrobium | 310 | SAMN04487781_0334 | 365317..366249 | 1710 | putative-restriction-endonuclease-[Cellulosimicrobium-cellulans] |
| WP_076404380.1 | wHTH(SBD)+HNH | GCF_900155925.1 | Cellulosimicrobium_aquatile | Bacteria>Bacillati>Actinomycetota>Actinomycetes>Micrococcales>Promicromonosporaceae>Cellulosimicrobium | 311 | BXA05_RS06520 | 519316..520251 | 1612203 | HNH-endonuclease-[Cellulosimicrobium-aquatile] |
| WP_147915795.1 | wHTH(SBD)+HNH | GCF_008000995.1 | Ruania_zhangjianzhongii | Bacteria>Bacillati>Actinomycetota>Actinomycetes>Micrococcales>Ruaniaceae>Ruania | 297 | FU260_RS03465 | 744443..745336 | 2603206 | HNH-endonuclease-[Ruania-zhangjianzhongii] |
| WP_232070211.1 | wHTH(SBD)+HNH | GCF_010731635.1 | Mycobacterium_noviomagense | Bacteria>Bacillati>Actinomycetota>Actinomycetes>Mycobacteriales>Mycobacteriaceae>Mycobacterium | 358 | BST37_RS15660 | 45571..46647 | 459858 | HNH-endonuclease-domain-containing-protein-[Mycobacterium-noviomagense] |
| WP_267319440.1 | wHTH(SBD)+HNH | GCF_026501225.1 | Mycolicibacterium_mucogenicum | Bacteria>Bacillati>Actinomycetota>Actinomycetes>Mycobacteriales>Mycobacteriaceae>Mycolicibacterium | 368 | OS122_RS24275 | 487014..488120 | 56689 | HNH-endonuclease-domain-containing-protein-[Mycolicibacterium-mucogenicum] |
| MBM4528838.1 | wHTH(SBD)+HNH | GCA_016879355.1 | Prescottella_equi | Bacteria>Bacillati>Actinomycetota>Actinomycetes>Mycobacteriales>Nocardiaceae>Prescottella | 315 | GS563_07705 | 1638450..1639397 | 43767 | HNH-endonuclease-[Prescottella-equi] |
| WP_233281888.1 | wHTH(SBD)+HNH | GCF_003013675.1 | Prescottella_equi | Bacteria>Bacillati>Actinomycetota>Actinomycetes>Mycobacteriales>Nocardiaceae>Prescottella | 291 | C7H75_RS09215 | 1946924..1947799 | 43767 | HNH-endonuclease-signature-motif-containing-protein-[Prescottella-equi] |
| WP_260845259.1 | wHTH(SBD)+HNH | GCF_007672705.1 | Prescottella_equi | Bacteria>Bacillati>Actinomycetota>Actinomycetes>Mycobacteriales>Nocardiaceae>Prescottella | 306 | FS672_RS22140 | 115980..116900 | 43767 | HNH-endonuclease-[Prescottella-equi] |
| WP_097214069.1 | wHTH(SBD)+HNH | GCF_900215125.1 | unclassified_Rhodococcus_(in:_high_G+C_Gram-positive_bacteria) | Bacteria>Bacillati>Actinomycetota>Actinomycetes>Mycobacteriales>Nocardiaceae>Rhodococcus | 298 | C8K38_RS25085 | 95933..96829 | 192944 | MULTISPECIES:-HNH-endonuclease-[unclassified-Rhodococcus-(in:-high-G+C-Gram-positive-bacteria)] |
| WP_240236712.1 | wHTH(SBD)+HNH | GCF_022436705.1 | Nocardioides_sp._CFH_31398 | Bacteria>Bacillati>Actinomycetota>Actinomycetes>Propionibacteriales>Nocardioidaceae>Nocardioides>unclassified_Nocardioides | 256 | MJ876_RS12930 | 69103..69873 | 2919579 | HNH-endonuclease-[Nocardioides-sp.-CFH-31398] |
| WP_013863120.1 | wHTH(SBD)+HNH | GCF_000270245.1 | Microlunatus_phosphovorus | Bacteria>Bacillati>Actinomycetota>Actinomycetes>Propionibacteriales>Propionibacteriaceae>Microlunatus | 309 | MLP_RS10795 | 2389864..2390793 | 29405 | HNH-endonuclease-[Microlunatus-phosphovorus] |
| WP_184988639.1 | wHTH(SBD)+HNH | GCF_039528315.1 | Sphaerisporangium_rubeum | Bacteria>Bacillati>Actinomycetota>Actinomycetes>Streptosporangiales>Streptosporangiaceae>Sphaerisporangium | 294 | BJ992_RS21125 | 4914399..4915283 | 321317 | HNH-endonuclease-[Sphaerisporangium-rubeum] |
| PFG07770.1 | wHTH(SBD)+HNH | GCA_002563655.1 | Bacillus_sp._es.034 | Bacteria>Bacillati>Bacillota>Bacilli>Bacillales>Bacillaceae>Bacillus>unclassified_Bacillus_(in:_firmicutes) | 350 | ATG71_4679 | 4462009..4463061 | 1761763 | HNH-endonuclease-[Bacillus-sp.-es.034] |
| WP_066398786.1 | wHTH(SBD)+HNH | GCF_017874625.1 | Cytobacillus_eiseniae | Bacteria>Bacillati>Bacillota>Bacilli>Bacillales>Bacillaceae>Cytobacillus | 252 | J2Z40_RS15740 | 29456..30214 | 762947 | hypothetical-protein-[Cytobacillus-eiseniae] |
| WP_191566560.1 | wHTH(SBD)+HNH | GCF_904423835.1 | Metabacillus_idriensis | Bacteria>Bacillati>Bacillota>Bacilli>Bacillales>Bacillaceae>Metabacillus | 256 | IHQ32_RS17360 | 3414675..3415445 | 324768 | hypothetical-protein-[Metabacillus-idriensis] |
| WP_102026194.1 | wHTH(SBD)+HNH | GCF_900240405.1 | Salirhabdus_sp._Marseille-P4669 | Bacteria>Bacillati>Bacillota>Bacilli>Bacillales>Bacillaceae>Salirhabdus>unclassified_Salirhabdus | 254 | C1A05_RS00375 | 75270..76034 | 2042310 | hypothetical-protein-[Salirhabdus-sp.-Marseille-P4669] |
| EGT3599212.1 | wHTH(SBD)+HNH | - | Clostridium_perfringens | Bacteria>Bacillati>Bacillota>Clostridia>Eubacteriales>Clostridiaceae>Clostridium | 307 | FT888_03070 | 90396..91319 | 1502 | HNH-endonuclease-[Clostridium-perfringens] |
| EHK2362609.1 | wHTH(SBD)+HNH | GCA_018011955.1 | Clostridium_perfringens | Bacteria>Bacillati>Bacillota>Clostridia>Eubacteriales>Clostridiaceae>Clostridium | 304 | KCL45_000422 | 454784..455698 | 1502 | HNH-endonuclease-[Clostridium-perfringens] |
| EJT5935290.1 | wHTH(SBD)+HNH | GCA_025090725.1 | Clostridium_perfringens | Bacteria>Bacillati>Bacillota>Clostridia>Eubacteriales>Clostridiaceae>Clostridium | 225 | N2W52_000015 | 24863..25540 | 1502 | HNH-endonuclease-[Clostridium-perfringens] |
| EJT6164544.1 | wHTH(SBD)+HNH | GCA_025091785.1 | Clostridium_perfringens | Bacteria>Bacillati>Bacillota>Clostridia>Eubacteriales>Clostridiaceae>Clostridium | 307 | N2W22_000996 | 1096910..1097833 | 1502 | HNH-endonuclease-[Clostridium-perfringens] |
| EJT6169497.1 | wHTH(SBD)+HNH | GCA_025091845.1 | Clostridium_perfringens | Bacteria>Bacillati>Bacillota>Clostridia>Eubacteriales>Clostridiaceae>Clostridium | 304 | N2W53_000039 | 57754..58668 | 1502 | HNH-endonuclease-[Clostridium-perfringens] |
| EJT6540220.1 | wHTH(SBD)+HNH | GCA_025095895.1 | Clostridium_perfringens | Bacteria>Bacillati>Bacillota>Clostridia>Eubacteriales>Clostridiaceae>Clostridium | 304 | N2W31_000039 | 57677..58591 | 1502 | HNH-endonuclease-[Clostridium-perfringens] |
| HAT4275163.1 | wHTH(SBD)+HNH | GCA_015933065.1 | Clostridium_perfringens | Bacteria>Bacillati>Bacillota>Clostridia>Eubacteriales>Clostridiaceae>Clostridium | 307 | I9084_000697 | 7576..8499 | 1502 | HNH-endonuclease-[Clostridium-perfringens] |
| HCG3022576.1 | wHTH(SBD)+HNH | GCA_024023955.1 | Clostridium_perfringens | Bacteria>Bacillati>Bacillota>Clostridia>Eubacteriales>Clostridiaceae>Clostridium | 304 | NJU98_002510 | 57677..58591 | 1502 | HNH-endonuclease-[Clostridium-perfringens] |
| MBS5995889.1 | wHTH(SBD)+HNH | GCA_018372935.1 | Clostridium_perfringens | Bacteria>Bacillati>Bacillota>Clostridia>Eubacteriales>Clostridiaceae>Clostridium | 304 | KIB05_13850 | 57307..58221 | 1502 | HNH-endonuclease-[Clostridium-perfringens] |
| WP_003452094.1 | wHTH(SBD)+HNH | GCF_000171135.1 | Clostridium_perfringens | Bacteria>Bacillati>Bacillota>Clostridia>Eubacteriales>Clostridiaceae>Clostridium | 304 | KLF33_RS07620 | 1738116..1739030 | 1502 | HNH-endonuclease-signature-motif-containing-protein-[Clostridium-perfringens] |
| WP_003473427.1 | wHTH(SBD)+HNH | GCF_000172455.1 | Clostridium_perfringens | Bacteria>Bacillati>Bacillota>Clostridia>Eubacteriales>Clostridiaceae>Clostridium | 306 | CJD_RS01710 | 80543..81463 | 1502 | HNH-endonuclease-signature-motif-containing-protein-[Clostridium-perfringens] |
| WP_012720740.1 | wHTH(SBD)+HNH | GCF_000020345.1 | Clostridium_botulinum | Bacteria>Bacillati>Bacillota>Clostridia>Eubacteriales>Clostridiaceae>Clostridium | 389 | T259_RS11590 | 2302251..2303420 | 1491 | HNH-endonuclease-domain-containing-protein-[Clostridium-botulinum] |
| WP_060670437.1 | wHTH(SBD)+HNH | GCF_001304735.1 | Clostridium_perfringens | Bacteria>Bacillati>Bacillota>Clostridia>Eubacteriales>Clostridiaceae>Clostridium | 307 | FORC3_RS07380 | 1733338..1734261 | 1502 | HNH-endonuclease-signature-motif-containing-protein-[Clostridium-perfringens] |
| WP_061317399.1 | wHTH(SBD)+HNH | GCF_001573255.1 | Clostridium_botulinum | Bacteria>Bacillati>Bacillota>Clostridia>Eubacteriales>Clostridiaceae>Clostridium | 389 | AZI51_RS06455 | 53142..54311 | 1491 | HNH-endonuclease-domain-containing-protein-[Clostridium-botulinum] |
| WP_061320310.1 | wHTH(SBD)+HNH | GCF_001573935.1 | Clostridium_botulinum | Bacteria>Bacillati>Bacillota>Clostridia>Eubacteriales>Clostridiaceae>Clostridium | 389 | AZI72_RS02370 | 7228..8397 | 1491 | HNH-endonuclease-domain-containing-protein-[Clostridium-botulinum] |
| WP_111744220.1 | wHTH(SBD)+HNH | GCF_965138575.1 | Clostridium_perfringens | Bacteria>Bacillati>Bacillota>Clostridia>Eubacteriales>Clostridiaceae>Clostridium | 225 | I6G61_RS10160 | 2175932..2176609 | 1502 | HNH-endonuclease-[Clostridium-perfringens] |
| WP_125003673.1 | wHTH(SBD)+HNH | GCF_003865095.1 | Clostridium_tagluense | Bacteria>Bacillati>Bacillota>Clostridia>Eubacteriales>Clostridiaceae>Clostridium | 339 | Ctaglu_RS16525 | 89176..90195 | 360422 | HNH-endonuclease-signature-motif-containing-protein-[Clostridium-tagluense] |
| WP_142710735.1 | wHTH(SBD)+HNH | GCF_902166005.1 | Clostridium_perfringens | Bacteria>Bacillati>Bacillota>Clostridia>Eubacteriales>Clostridiaceae>Clostridium | 304 | FN719_RS05055 | 233682..234596 | 1502 | HNH-endonuclease-signature-motif-containing-protein-[Clostridium-perfringens] |
| WP_142975593.1 | wHTH(SBD)+HNH | GCF_902166025.1 | Clostridium_perfringens | Bacteria>Bacillati>Bacillota>Clostridia>Eubacteriales>Clostridiaceae>Clostridium | 307 | FN715_RS03310 | 757236..758159 | 1502 | HNH-endonuclease-signature-motif-containing-protein-[Clostridium-perfringens] |
| WP_164804799.1 | wHTH(SBD)+HNH | GCF_018135545.1 | Clostridium_perfringens | Bacteria>Bacillati>Bacillota>Clostridia>Eubacteriales>Clostridiaceae>Clostridium | 306 | G6Z22_RS06030 | 19414..20334 | 1502 | HNH-endonuclease-signature-motif-containing-protein-[Clostridium-perfringens] |
| WP_169990732.1 | wHTH(SBD)+HNH | GCF_002866225.1 | Clostridium_botulinum | Bacteria>Bacillati>Bacillota>Clostridia>Eubacteriales>Clostridiaceae>Clostridium | 389 | RSJ21_RS11410 | 2258495..2259664 | 1491 | HNH-endonuclease-domain-containing-protein-[Clostridium-botulinum] |
| WP_195381617.1 | wHTH(SBD)+HNH | GCF_051057975.1 | Clostridium_perfringens | Bacteria>Bacillati>Bacillota>Clostridia>Eubacteriales>Clostridiaceae>Clostridium | 306 | H8J86_RS06945 | 64188..65108 | 1502 | HNH-endonuclease-signature-motif-containing-protein-[Clostridium-perfringens] |
| WP_198605366.1 | wHTH(SBD)+HNH | GCF_016236125.1 | Clostridium_perfringens | Bacteria>Bacillati>Bacillota>Clostridia>Eubacteriales>Clostridiaceae>Clostridium | 306 | H8J62_RS02860 | 32312..33232 | 1502 | HNH-endonuclease-[Clostridium-perfringens] |
| WP_198616414.1 | wHTH(SBD)+HNH | GCF_016236635.1 | Clostridium_perfringens | Bacteria>Bacillati>Bacillota>Clostridia>Eubacteriales>Clostridiaceae>Clostridium | 312 | H8J95_RS08930 | 17531..18469 | 1502 | HNH-endonuclease-signature-motif-containing-protein-[Clostridium-perfringens] |
| WP_198621145.1 | wHTH(SBD)+HNH | GCF_016236545.1 | Clostridium_perfringens | Bacteria>Bacillati>Bacillota>Clostridia>Eubacteriales>Clostridiaceae>Clostridium | 307 | H8J92_RS02885 | 84739..85662 | 1502 | HNH-endonuclease-signature-motif-containing-protein-[Clostridium-perfringens] |
| WP_228572335.1 | wHTH(SBD)+HNH | GCF_020819925.1 | Clostridium_perfringens | Bacteria>Bacillati>Bacillota>Clostridia>Eubacteriales>Clostridiaceae>Clostridium | 312 | LMS36_RS03140 | 128325..129263 | 1502 | HNH-endonuclease-signature-motif-containing-protein-[Clostridium-perfringens] |
| WP_245124314.1 | wHTH(SBD)+HNH | GCF_022936445.1 | Clostridium_botulinum | Bacteria>Bacillati>Bacillota>Clostridia>Eubacteriales>Clostridiaceae>Clostridium | 389 | LOO64_RS12740 | 356090..357259 | 1491 | HNH-endonuclease-domain-containing-protein-[Clostridium-botulinum] |
| WP_207649731.1 | wHTH(SBD)+HNH | GCF_900100785.1 | Desulfosporosinus_hippei | Bacteria>Bacillati>Bacillota>Clostridia>Eubacteriales>Desulfitobacteriaceae>Desulfosporosinus | 258 | BLS16_RS05435 | 165452..166228 | 569859 | HNH-endonuclease-signature-motif-containing-protein-[Desulfosporosinus-hippei] |
| MBQ8951399.1 | wHTH(SBD)+HNH | GCA_017460785.1 | Eubacterium_sp. | Bacteria>Bacillati>Bacillota>Clostridia>Eubacteriales>Eubacteriaceae>Eubacterium>unclassified_Eubacterium | 341 | IJ065_09670 | 19487..20512 | 142586 | HNH-endonuclease-[Eubacterium-sp.] |
| MCI9127454.1 | wHTH(SBD)+HNH | GCA_022797855.1 | Eubacterium_sp. | Bacteria>Bacillati>Bacillota>Clostridia>Eubacteriales>Eubacteriaceae>Eubacterium>unclassified_Eubacterium | 336 | HFG28_09685 | 14116..15126 | 142586 | HNH-endonuclease-[Eubacterium-sp.] |
| WP_014256032.1 | wHTH(SBD)+HNH | GCF_000237085.1 | Acetivibrio_clariflavus | Bacteria>Bacillati>Bacillota>Clostridia>Eubacteriales>Oscillospiraceae>Acetivibrio | 288 | CLOCLA_RS0107460 | 1200831..1201697 | 288965 | HNH-endonuclease-signature-motif-containing-protein-[Acetivibrio-clariflavus] |
| WP_161840115.1 | wHTH(SBD)+HNH | GCF_009931695.1 | Anaerocolumna_sedimenticola | Bacteria>Bacillati>Bacillota>Clostridia>Lachnospirales>Lachnospiraceae>Anaerocolumna | 329 | Ana3638_RS23005 | 5165165..5166154 | 2696063 | HNH-endonuclease-signature-motif-containing-protein-[Anaerocolumna-sedimenticola] |
| WP_246240543.1 | wHTH(SBD)+HNH | GCF_009931695.1 | Anaerocolumna_sedimenticola | Bacteria>Bacillati>Bacillota>Clostridia>Lachnospirales>Lachnospiraceae>Anaerocolumna | 275 | Ana3638_RS23010 | 5166289..5167116 | 2696063 | HNH-endonuclease-[Anaerocolumna-sedimenticola] |
| WP_242967692.1 | wHTH(SBD)+HNH | GCF_002904165.1 | Herbinix_hemicellulosilytica | Bacteria>Bacillati>Bacillota>Clostridia>Lachnospirales>Lachnospiraceae>Herbinix | 272 | HHT355_RS13275 | 80742..81560 | 1564487 | HNH-endonuclease-signature-motif-containing-protein-[Herbinix-hemicellulosilytica] |
| WP_075865022.1 | wHTH(SBD)+HNH | GCF_001950325.1 | Carboxydothermus_islandicus | Bacteria>Bacillati>Bacillota>Clostridia>Thermoanaerobacterales>Thermoanaerobacteraceae>Carboxydothermus | 158 | ciss_RS04010 | 11595..12071 | 661089 | hypothetical-protein-[Carboxydothermus-islandicus] |
| GIV81899.1 | wHTH(SBD)+HNH | GCA_026003715.1 | Anaerolineae_bacterium | Bacteria>Bacillati>Chloroflexota>Anaerolineae>unclassified_Anaerolineae | 374 | KatS3mg051_1253 | 7938..9062 | 2052143 | MAG:-hypothetical-protein-KatS3mg051_1253-[Anaerolineae-bacterium] |
| MBP7042333.1 | wHTH(SBD)+HNH | GCA_017993775.1 | Chloroflexota_bacterium | Bacteria>Bacillati>Chloroflexota>unclassified_Chloroflexota | 374 | KBE23_06290 | 10786..11910 | 2026724 | HNH-endonuclease-[Chloroflexota-bacterium] |
| MBP8002223.1 | wHTH(SBD)+HNH | GCA_018004095.1 | Chloroflexota_bacterium | Bacteria>Bacillati>Chloroflexota>unclassified_Chloroflexota | 375 | KA338_23445 | 6725..7852 | 2026724 | HNH-endonuclease-[Chloroflexota-bacterium] |
| WP_242037122.1 | wHTH(SBD)+HNH | - | Coleofasciculus_sp._FACHB-125 | Bacteria>Bacillati>Cyanobacteriota/Melainabacteria_group>Cyanobacteriota>Cyanophyceae>Coleofasciculales>Coleofasciculaceae>Coleofasciculus>unclassified_Coleofasciculus | 168 | H6F83_RS29660 | 47714..48220 | 2692784 | HNH-endonuclease-domain-containing-protein-[Coleofasciculus-sp.-FACHB-125] |
| MBD2364928.1 | wHTH(SBD)+HNH | GCA_014697125.1 | Anabaena_minutissima_FACHB-250 | Bacteria>Bacillati>Cyanobacteriota/Melainabacteria_group>Cyanobacteriota>Cyanophyceae>Nostocales>Nostocaceae>Anabaena>Anabaena_minutissima | 460 | H6G36_27840 | 1104..2486 | 2692882 | hypothetical-protein-[Anabaena-minutissima-FACHB-250] |
| WP_221704718.1 | wHTH(SBD)+HNH | - | Desmonostoc_muscorum | Bacteria>Bacillati>Cyanobacteriota/Melainabacteria_group>Cyanobacteriota>Cyanophyceae>Nostocales>Nostocaceae>Desmonostoc | 457 | H1Q63_RS02890 | 29291..30664 | 1179 | HNH-endonuclease-domain-containing-protein-[Desmonostoc-muscorum] |
| AFZ48543.1 | wHTH(SBD)+HNH | GCA_000317655.1 | Cyanobacterium_stanieri_PCC_7202 | Bacteria>Bacillati>Cyanobacteriota/Melainabacteria_group>Cyanobacteriota>Cyanophyceae>Oscillatoriophycideae>Chroococcales>Geminocystaceae>Cyanobacterium>Cyanobacterium_stanieri | 374 | Cyast_2600 | 2858668..2859792 | 292563 | hypothetical-protein-Cyast_2600-[Cyanobacterium-stanieri-PCC-7202] |
| WP_228059573.1 | wHTH(SBD)+HNH | - | Plectonema_radiosum | Bacteria>Bacillati>Cyanobacteriota/Melainabacteria_group>Cyanobacteriota>Cyanophyceae>Oscillatoriophycideae>Oscillatoriales>Microcoleaceae>Plectonema | 172 | IQ247_RS32395 | 11530..12048 | 945768 | HNH-endonuclease-domain-containing-protein-[Plectonema-radiosum] |
| WP_204104299.1 | wHTH(SBD)+HNH | GCF_016888125.1 | Spirulina_sp._CCY15215 | Bacteria>Bacillati>Cyanobacteriota/Melainabacteria_group>Cyanobacteriota>Cyanophyceae>Spirulinales>Spirulinaceae>Spirulina>unclassified_Spirulina | 379 | JUI58_RS13390 | 6132..7271 | 2767591 | HNH-endonuclease-domain-containing-protein-[Spirulina-major] |
| WP_254953296.1 | wHTH(SBD)+HNH | GCF_024345955.1 | unclassified_Cyanobium | Bacteria>Bacillati>Cyanobacteriota/Melainabacteria_group>Cyanobacteriota>Cyanophyceae>Synechococcales>Prochlorococcaceae>Cyanobium | 307 | KBZ34_RS12465 | 63290..64213 | 2627006 | MULTISPECIES:-HNH-endonuclease-[unclassified-Cyanobium] |
| MCP9859836.1 | wHTH(SBD)+HNH | GCA_024345955.1 | Cyanobium_sp._Cruz-8H5 | Bacteria>Bacillati>Cyanobacteriota/Melainabacteria_group>Cyanobacteriota>Cyanophyceae>Synechococcales>Prochlorococcaceae>Cyanobium>unclassified_Cyanobium | 307 | KBZ34_12410 | 63290..64213 | 2823712 | HNH-endonuclease-[Cyanobium-sp.-Cruz-8H5] |
| HBH74015.1 | wHTH(SBD)+HNH | GCA_003508015.1 | Synechococcales_bacterium_UBA10510 | Bacteria>Bacillati>Cyanobacteriota/Melainabacteria_group>Cyanobacteriota>Cyanophyceae>Synechococcales>unclassified_Synechococcales | 307 | DDY43_11400 | 27310..28233 | 2055795 | restriction-endonuclease-[Synechococcales-bacterium-UBA10510] |
| WP_105482024.1 | wHTH(SBD)+HNH | GCF_002973605.1 | Abditibacterium_utsteinense | Bacteria>Pseudomonadati>Abditibacteriota>Abditibacteriia>Abditibacteriales>Abditibacteriaceae>Abditibacterium | 388 | B1R32_RS00025 | 7936..9102 | 1960156 | HNH-endonuclease-domain-containing-protein-[Abditibacterium-utsteinense] |
| MCQ2302013.1 | wHTH(SBD)+HNH | GCA_024400425.1 | Bacteroidales_bacterium | Bacteria>Pseudomonadati>FCB_group>Bacteroidota/Chlorobiota_group>Bacteroidota>Bacteroidia>Bacteroidales>unclassified_Bacteroidales | 207 | MJZ94_05240 | 11473..12096 | 2030927 | hypothetical-protein-[Bacteroidales-bacterium] |
| WP_092740287.1 | wHTH(SBD)+HNH | GCF_900107135.1 | Hymenobacter_psychrophilus | Bacteria>Pseudomonadati>FCB_group>Bacteroidota/Chlorobiota_group>Bacteroidota>Cytophagia>Cytophagales>Hymenobacteraceae>Hymenobacter | 363 | BLW21_RS11935 | 110890..111981 | 651662 | HNH-endonuclease-domain-containing-protein-[Hymenobacter-psychrophilus] |
| WP_267556350.1 | wHTH(SBD)+HNH | GCF_026626625.1 | Corallococcus_sp._BB11-1 | Bacteria>Pseudomonadati>Myxococcota>Myxococcia>Myxococcales>Cystobacterineae>Myxococcaceae>Corallococcus>unclassified_Corallococcus | 225 | OV207_RS17730 | 249123..249800 | 2996783 | hypothetical-protein-[Corallococcus-sp.-BB11-1] |
| MBP0112789.1 | wHTH(SBD)+HNH | GCA_017788805.1 | Bradyrhizobium_vignae | Bacteria>Pseudomonadati>Pseudomonadota>Alphaproteobacteria>Hyphomicrobiales>Nitrobacteraceae>Bradyrhizobium | 272 | JWS04_17190 | 16183..17001 | 1549949 | HNH-endonuclease-[Bradyrhizobium-vignae] |
| WP_240950174.1 | wHTH(SBD)+HNH | - | Novosphingobium_sp._ERN07 | Bacteria>Pseudomonadati>Pseudomonadota>Alphaproteobacteria>Sphingomonadales>Sphingomonadaceae>Novosphingobium>unclassified_Novosphingobium | 130 | HGI47_RS22675 | 94406..94798 | 2726187 | HNH-endonuclease-[Novosphingobium-sp.-ERN07] |
| WP_099185613.1 | wHTH(SBD)+HNH | GCF_002723855.1 | Sphingobium_fuliginis | Bacteria>Pseudomonadati>Pseudomonadota>Alphaproteobacteria>Sphingomonadales>Sphingomonadaceae>Sphingobium | 302 | SFOMI_RS05015 | 870388..871296 | 336203 | HNH-endonuclease-[Sphingobium-fuliginis] |
| WP_262503491.1 | wHTH(SBD)+HNH | - | Sphingobium_xenophagum | Bacteria>Pseudomonadati>Pseudomonadota>Alphaproteobacteria>Sphingomonadales>Sphingomonadaceae>Sphingobium | 306 | MBESOW_RS00060 | 10744..11664 | 121428 | HNH-endonuclease-[Sphingobium-xenophagum] |
| WP_013697448.1 | wHTH(SBD)+HNH | GCF_000194745.1 | Burkholderia_gladioli | Bacteria>Pseudomonadati>Pseudomonadota>Betaproteobacteria>Burkholderiales>Burkholderiaceae>Burkholderia | 328 | BGLA_RS38690 | 1590240..1591226 | 28095 | HNH-endonuclease-[Burkholderia-gladioli] |
| WP_194759725.1 | wHTH(SBD)+HNH | GCF_052059325.1 | Aeromonas_hydrophila | Bacteria>Pseudomonadati>Pseudomonadota>Gammaproteobacteria>Aeromonadales>Aeromonadaceae>Aeromonas | 353 | F3H83_RS12445 | 35356..36417 | 644 | HNH-endonuclease-[Aeromonas-hydrophila] |
| WP_173294706.1 | wHTH(SBD)+HNH | GCF_013248905.1 | Shewanella_sp._VB17 | Bacteria>Pseudomonadati>Pseudomonadota>Gammaproteobacteria>Alteromonadales>Shewanellaceae>Shewanella>unclassified_Shewanella | 374 | HQQ94_RS12335 | 2873602..2874726 | 2739432 | HNH-endonuclease-signature-motif-containing-protein-[Shewanella-sp.-VB17] |
| WP_000168579.1 | wHTH(SBD)+HNH | GCF_000215185.1 | Enterobacteriaceae | Bacteria>Pseudomonadati>Pseudomonadota>Gammaproteobacteria>Enterobacterales | 290 | FQ032_RS06150 | 107861..108733 | 543 | MULTISPECIES:-HNH-endonuclease-[Enterobacteriaceae] |
| WP_022646907.1 | wHTH(SBD)+HNH | GCF_010319625.1 | Enterobacter | Bacteria>Pseudomonadati>Pseudomonadota>Gammaproteobacteria>Enterobacterales>Enterobacteriaceae | 289 | NVR78_RS04755 | 314678..315547 | 547 | MULTISPECIES:-HNH-endonuclease-[Enterobacter] |
| WP_047736971.1 | wHTH(SBD)+HNH | GCF_020971765.1 | Enterobacter | Bacteria>Pseudomonadati>Pseudomonadota>Gammaproteobacteria>Enterobacterales>Enterobacteriaceae | 289 | LOC22_RS13525 | 2835125..2835994 | 547 | MULTISPECIES:-HNH-endonuclease-[Enterobacter] |
| WP_121585416.1 | wHTH(SBD)+HNH | GCF_003665575.1 | Citrobacter | Bacteria>Pseudomonadati>Pseudomonadota>Gammaproteobacteria>Enterobacterales>Enterobacteriaceae | 259 | CUC49_RS22170 | 4540093..4540872 | 544 | HNH-endonuclease-[Citrobacter-pasteurii] |
| WP_121585420.1 | wHTH(SBD)+HNH | GCF_003665575.1 | Citrobacter | Bacteria>Pseudomonadati>Pseudomonadota>Gammaproteobacteria>Enterobacterales>Enterobacteriaceae | 290 | CUC49_RS22210 | 4553848..4554720 | 544 | MULTISPECIES:-HNH-endonuclease-[Citrobacter] |
| WP_213752380.1 | wHTH(SBD)+HNH | GCF_018446995.1 | Citrobacter | Bacteria>Pseudomonadati>Pseudomonadota>Gammaproteobacteria>Enterobacterales>Enterobacteriaceae | 290 | OGY13_RS10385 | 356120..356992 | 544 | HNH-endonuclease-[Citrobacter-portucalensis] |
| HAV2024003.1 | wHTH(SBD)+HNH | GCA_016444345.1 | Citrobacter_koseri | Bacteria>Pseudomonadati>Pseudomonadota>Gammaproteobacteria>Enterobacterales>Enterobacteriaceae>Citrobacter | 259 | JHJ49_004191 | 9296..10075 | 545 | HNH-endonuclease-[Citrobacter-koseri] |
| WP_082139604.1 | wHTH(SBD)+HNH | GCF_013741395.1 | Citrobacter_freundii_complex | Bacteria>Pseudomonadati>Pseudomonadota>Gammaproteobacteria>Enterobacterales>Enterobacteriaceae>Citrobacter | 290 | SK34_RS19930 | 339859..340731 | 1344959 | MULTISPECIES:-HNH-endonuclease-[Citrobacter-freundii-complex] |
| EIX7373179.1 | wHTH(SBD)+HNH | GCA_022400985.1 | Citrobacter_freundii | Bacteria>Pseudomonadati>Pseudomonadota>Gammaproteobacteria>Enterobacterales>Enterobacteriaceae>Citrobacter>Citrobacter_freundii_complex | 290 | MK620_002554 | 186631..187503 | 546 | HNH-endonuclease-[Citrobacter-freundii] |
| HAT3908791.1 | wHTH(SBD)+HNH | GCA_015929465.1 | Citrobacter_freundii | Bacteria>Pseudomonadati>Pseudomonadota>Gammaproteobacteria>Enterobacterales>Enterobacteriaceae>Citrobacter>Citrobacter_freundii_complex | 290 | I9Y28_003044 | 183904..184776 | 546 | TPA:-HNH-endonuclease-[Citrobacter-freundii] |
| WP_048233993.1 | wHTH(SBD)+HNH | GCF_001034485.1 | Citrobacter_sp._MGH104 | Bacteria>Pseudomonadati>Pseudomonadota>Gammaproteobacteria>Enterobacterales>Enterobacteriaceae>Citrobacter>Citrobacter_freundii_complex>unclassified_Citrobacter_freundii_complex | 259 | SK34_RS19970 | 353714..354493 | 1686379 | HNH-endonuclease-[Citrobacter-sp.-MGH104] |
| KAB0815078.1 | wHTH(SBD)+HNH | GCA_008805175.1 | Cronobacter_sakazakii | Bacteria>Pseudomonadati>Pseudomonadota>Gammaproteobacteria>Enterobacterales>Enterobacteriaceae>Cronobacter | 338 | FZI24_17850 | 17041..18057 | 28141 | HNH-endonuclease-[Cronobacter-sakazakii] |
| PUV47032.1 | wHTH(SBD)+HNH | GCA_002532375.1 | Cronobacter_sakazakii | Bacteria>Pseudomonadati>Pseudomonadota>Gammaproteobacteria>Enterobacterales>Enterobacteriaceae>Cronobacter | 338 | CDU02_01585 | 353137..354153 | 28141 | HNH-endonuclease-[Cronobacter-sakazakii] |
| WP_097563840.1 | wHTH(SBD)+HNH | GCF_017845055.1 | Cronobacter_sakazakii | Bacteria>Pseudomonadati>Pseudomonadota>Gammaproteobacteria>Enterobacterales>Enterobacteriaceae>Cronobacter | 338 | KAR72_RS09210 | 204047..205063 | 28141 | HNH-endonuclease-domain-containing-protein-[Cronobacter-sakazakii] |
| WP_105536609.1 | wHTH(SBD)+HNH | GCF_002974985.1 | Cronobacter_sakazakii | Bacteria>Pseudomonadati>Pseudomonadota>Gammaproteobacteria>Enterobacterales>Enterobacteriaceae>Cronobacter | 338 | FC600_RS12540 | 208126..209142 | 28141 | HNH-endonuclease-domain-containing-protein-[Cronobacter-sakazakii] |
| WP_249324835.1 | wHTH(SBD)+HNH | GCF_018299605.1 | Enterobacter_mori | Bacteria>Pseudomonadati>Pseudomonadota>Gammaproteobacteria>Enterobacterales>Enterobacteriaceae>Enterobacter | 316 | GUY16_RS21805 | 41452..42402 | 539813 | HNH-endonuclease-[Enterobacter-mori] |
| WP_262778586.1 | wHTH(SBD)+HNH | GCF_025536155.1 | Enterobacter_quasiroggenkampii | Bacteria>Pseudomonadati>Pseudomonadota>Gammaproteobacteria>Enterobacterales>Enterobacteriaceae>Enterobacter | 289 | KSU18_RS16675 | 130946..131815 | 2497436 | HNH-endonuclease-[Enterobacter-quasiroggenkampii] |
| HCT3226641.1 | wHTH(SBD)+HNH | GCA_026310715.1 | Enterobacter_hormaechei | Bacteria>Pseudomonadati>Pseudomonadota>Gammaproteobacteria>Enterobacterales>Enterobacteriaceae>Enterobacter>Enterobacter_cloacae_complex | 289 | OTS66_001783 | 77415..78284 | 158836 | TPA:-HNH-endonuclease-[Enterobacter-hormaechei] |
| HCT3226647.1 | wHTH(SBD)+HNH | GCA_026310715.1 | Enterobacter_hormaechei | Bacteria>Pseudomonadati>Pseudomonadota>Gammaproteobacteria>Enterobacterales>Enterobacteriaceae>Enterobacter>Enterobacter_cloacae_complex | 259 | OTS66_001789 | 89566..90345 | 158836 | TPA:-HNH-endonuclease-[Enterobacter-hormaechei] |
| MBG0523805.1 | wHTH(SBD)+HNH | GCA_015704905.1 | Enterobacter_cloacae | Bacteria>Pseudomonadati>Pseudomonadota>Gammaproteobacteria>Enterobacterales>Enterobacteriaceae>Enterobacter>Enterobacter_cloacae_complex | 289 | IOR24_17945 | 3783730..3784599 | 550 | HNH-endonuclease-[Enterobacter-cloacae] |
| WP_013095399.1 | wHTH(SBD)+HNH | GCF_000025565.1 | Enterobacter_cloacae | Bacteria>Pseudomonadati>Pseudomonadota>Gammaproteobacteria>Enterobacterales>Enterobacteriaceae>Enterobacter>Enterobacter_cloacae_complex | 289 | NGL52_RS07565 | 50273..51142 | 550 | HNH-endonuclease-[Enterobacter-cloacae] |
| WP_063139202.1 | wHTH(SBD)+HNH | GCF_902160625.1 | Enterobacter_hormaechei | Bacteria>Pseudomonadati>Pseudomonadota>Gammaproteobacteria>Enterobacterales>Enterobacteriaceae>Enterobacter>Enterobacter_cloacae_complex | 289 | H0848_RS18585 | 42037..42906 | 158836 | HNH-endonuclease-[Enterobacter-hormaechei] |
| WP_196357234.1 | wHTH(SBD)+HNH | GCF_015704165.1 | Enterobacter_hormaechei | Bacteria>Pseudomonadati>Pseudomonadota>Gammaproteobacteria>Enterobacterales>Enterobacteriaceae>Enterobacter>Enterobacter_cloacae_complex | 259 | I4P33_RS13225 | 217093..217872 | 158836 | HNH-endonuclease-[Enterobacter-hormaechei] |
| WP_196357284.1 | wHTH(SBD)+HNH | GCF_015704165.1 | Enterobacter_hormaechei | Bacteria>Pseudomonadati>Pseudomonadota>Gammaproteobacteria>Enterobacterales>Enterobacteriaceae>Enterobacter>Enterobacter_cloacae_complex | 289 | I4P33_RS13265 | 230847..231716 | 158836 | HNH-endonuclease-[Enterobacter-hormaechei] |
| WP_196373092.1 | wHTH(SBD)+HNH | GCF_015704905.1 | Enterobacter_cloacae | Bacteria>Pseudomonadati>Pseudomonadota>Gammaproteobacteria>Enterobacterales>Enterobacteriaceae>Enterobacter>Enterobacter_cloacae_complex | 289 | IOR24_RS17945 | 3783730..3784599 | 550 | HNH-endonuclease-[Enterobacter-cloacae] |
| WP_251853645.1 | wHTH(SBD)+HNH | GCF_903935915.1 | Enterobacter_cloacae | Bacteria>Pseudomonadati>Pseudomonadota>Gammaproteobacteria>Enterobacterales>Enterobacteriaceae>Enterobacter>Enterobacter_cloacae_complex | 259 | NGL52_RS07605 | 64119..64898 | 550 | HNH-endonuclease-[Enterobacter-cloacae] |
| WP_252033133.1 | wHTH(SBD)+HNH | GCF_023752605.1 | Enterobacter_asburiae | Bacteria>Pseudomonadati>Pseudomonadota>Gammaproteobacteria>Enterobacterales>Enterobacteriaceae>Enterobacter>Enterobacter_cloacae_complex | 316 | M8S83_RS11435 | 326418..327368 | 61645 | HNH-endonuclease-[Enterobacter-asburiae] |
| HCM9196141.1 | wHTH(SBD)+HNH | GCA_025209675.1 | Enterobacter_cloacae_subsp._dissolvens | Bacteria>Pseudomonadati>Pseudomonadota>Gammaproteobacteria>Enterobacterales>Enterobacteriaceae>Enterobacter>Enterobacter_cloacae_complex>Enterobacter_cloacae | 289 | N5K92_004844 | 19354..20223 | 69219 | HNH-endonuclease-[Enterobacter-cloacae-subsp.-dissolvens] |
| EEU4096209.1 | wHTH(SBD)+HNH | - | Escherichia_coli | Bacteria>Pseudomonadati>Pseudomonadota>Gammaproteobacteria>Enterobacterales>Enterobacteriaceae>Escherichia | 290 | E8H53_004593 | 3916..4788 | 562 | HNH-endonuclease-[Escherichia-coli] |
| EEV1146575.1 | wHTH(SBD)+HNH | - | Escherichia_coli | Bacteria>Pseudomonadati>Pseudomonadota>Gammaproteobacteria>Enterobacterales>Enterobacteriaceae>Escherichia | 290 | EUB49_004350 | 5224..6096 | 562 | HNH-endonuclease-[Escherichia-coli] |
| EEW3089659.1 | wHTH(SBD)+HNH | - | Escherichia_coli | Bacteria>Pseudomonadati>Pseudomonadota>Gammaproteobacteria>Enterobacterales>Enterobacteriaceae>Escherichia | 290 | EBA98_002635 | 11208..12080 | 562 | HNH-endonuclease-[Escherichia-coli] |

| Protein-ID | Domain | Assembly-ID | Strain | Taxonomy | Length | Locus-Tag | Genomic-coordinates | TaxID | Fasta-defline |
|---|---|---|---|---|---|---|---|---|---|
| EEW3885149.1 | wHTH(SBD)+HNH | - | Escherichia_coli | Bacteria>Pseudomonadati>Pseudomonadota>Gammaproteobacteria>Enterobacterales>Enterobacteriaceae>Escherichia | 290 | D9Y07_001388 | 3677..4549 | 562 | HNH-endonuclease-[Escherichia-coli] |
| EEX1719679.1 | wHTH(SBD)+HNH | - | Escherichia_coli | Bacteria>Pseudomonadati>Pseudomonadota>Gammaproteobacteria>Enterobacterales>Enterobacteriaceae>Escherichia | 290 | D3S79_003403 | 3196..4068 | 562 | HNH-endonuclease-[Escherichia-coli] |
| EEZ2582142.1 | wHTH(SBD)+HNH | - | Escherichia_coli | Bacteria>Pseudomonadati>Pseudomonadota>Gammaproteobacteria>Enterobacterales>Enterobacteriaceae>Escherichia | 290 | DMH54_003962 | 5019..5891 | 562 | HNH-endonuclease-[Escherichia-coli] |
| EEZ3215396.1 | wHTH(SBD)+HNH | - | Escherichia_coli | Bacteria>Pseudomonadati>Pseudomonadota>Gammaproteobacteria>Enterobacterales>Enterobacteriaceae>Escherichia | 290 | DKV48_004294 | 3206..4078 | 562 | HNH-endonuclease-[Escherichia-coli] |
| EFC3446204.1 | wHTH(SBD)+HNH | - | Escherichia_coli | Bacteria>Pseudomonadati>Pseudomonadota>Gammaproteobacteria>Enterobacterales>Enterobacteriaceae>Escherichia | 290 | CO751_004602 | 5151..6023 | 562 | HNH-endonuclease-[Escherichia-coli] |
| EFD3459871.1 | wHTH(SBD)+HNH | - | Escherichia_coli | Bacteria>Pseudomonadati>Pseudomonadota>Gammaproteobacteria>Enterobacterales>Enterobacteriaceae>Escherichia | 290 | CL211_004298 | 3270..4142 | 562 | HNH-endonuclease-[Escherichia-coli] |
| EFE1965716.1 | wHTH(SBD)+HNH | - | Escherichia_coli | Bacteria>Pseudomonadati>Pseudomonadota>Gammaproteobacteria>Enterobacterales>Enterobacteriaceae>Escherichia | 290 | CJV81_002663 | 11207..12079 | 562 | HNH-endonuclease-[Escherichia-coli] |
| EFG4404335.1 | wHTH(SBD)+HNH | - | Escherichia_coli | Bacteria>Pseudomonadati>Pseudomonadota>Gammaproteobacteria>Enterobacterales>Enterobacteriaceae>Escherichia | 290 | BM772_002468 | 11208..12080 | 562 | HNH-endonuclease-[Escherichia-coli] |
| EFH0549021.1 | wHTH(SBD)+HNH | - | Escherichia_coli | Bacteria>Pseudomonadati>Pseudomonadota>Gammaproteobacteria>Enterobacterales>Enterobacteriaceae>Escherichia | 290 | BJE23_003833 | 36503..37375 | 562 | HNH-endonuclease-[Escherichia-coli] |
| EGY5620860.1 | wHTH(SBD)+HNH | - | Escherichia_coli | Bacteria>Pseudomonadati>Pseudomonadota>Gammaproteobacteria>Enterobacterales>Enterobacteriaceae>Escherichia | 290 | JI433_002886 | 11207..12079 | 562 | HNH-endonuclease-[Escherichia-coli] |
| EGY7451907.1 | wHTH(SBD)+HNH | - | Escherichia_coli | Bacteria>Pseudomonadati>Pseudomonadota>Gammaproteobacteria>Enterobacterales>Enterobacteriaceae>Escherichia | 290 | JI519_004381 | 5521..6393 | 562 | HNH-endonuclease-[Escherichia-coli] |
| EGY8948622.1 | wHTH(SBD)+HNH | - | Escherichia_coli | Bacteria>Pseudomonadati>Pseudomonadota>Gammaproteobacteria>Enterobacterales>Enterobacteriaceae>Escherichia | 290 | JI613_004639 | 3966..4838 | 562 | HNH-endonuclease-[Escherichia-coli] |
| EGY9306079.1 | wHTH(SBD)+HNH | - | Escherichia_coli | Bacteria>Pseudomonadati>Pseudomonadota>Gammaproteobacteria>Enterobacterales>Enterobacteriaceae>Escherichia | 290 | HZO96_004636 | 600..1472 | 562 | HNH-endonuclease-[Escherichia-coli] |
| EGZ1560303.1 | wHTH(SBD)+HNH | - | Escherichia_coli | Bacteria>Pseudomonadati>Pseudomonadota>Gammaproteobacteria>Enterobacterales>Enterobacteriaceae>Escherichia | 290 | FAV16_004665 | 3949..4821 | 562 | HNH-endonuclease-[Escherichia-coli] |
| EGZ1871554.1 | wHTH(SBD)+HNH | - | Escherichia_coli | Bacteria>Pseudomonadati>Pseudomonadota>Gammaproteobacteria>Enterobacterales>Enterobacteriaceae>Escherichia | 290 | EYE54_004345 | 3865..4737 | 562 | HNH-endonuclease-[Escherichia-coli] |
| EGZ2189932.1 | wHTH(SBD)+HNH | - | Escherichia_coli | Bacteria>Pseudomonadati>Pseudomonadota>Gammaproteobacteria>Enterobacterales>Enterobacteriaceae>Escherichia | 290 | ED609_004469 | 4501..5373 | 562 | HNH-endonuclease-[Escherichia-coli] |
| EGZ2354226.1 | wHTH(SBD)+HNH | - | Escherichia_coli | Bacteria>Pseudomonadati>Pseudomonadota>Gammaproteobacteria>Enterobacterales>Enterobacteriaceae>Escherichia | 290 | D7J47_002040 | 11208..12080 | 562 | HNH-endonuclease-[Escherichia-coli] |
| EGZ2777077.1 | wHTH(SBD)+HNH | - | Escherichia_coli | Bacteria>Pseudomonadati>Pseudomonadota>Gammaproteobacteria>Enterobacterales>Enterobacteriaceae>Escherichia | 290 | DVO41_002385 | 11208..12080 | 562 | HNH-endonuclease-[Escherichia-coli] |
| EGZ3242779.1 | wHTH(SBD)+HNH | - | Escherichia_coli | Bacteria>Pseudomonadati>Pseudomonadota>Gammaproteobacteria>Enterobacterales>Enterobacteriaceae>Escherichia | 290 | C5N70_002400 | 36502..37374 | 562 | HNH-endonuclease-[Escherichia-coli] |
| EGZ5166761.1 | wHTH(SBD)+HNH | - | Escherichia_coli | Bacteria>Pseudomonadati>Pseudomonadota>Gammaproteobacteria>Enterobacterales>Enterobacteriaceae>Escherichia | 290 | CDS35_004370 | 4491..5363 | 562 | HNH-endonuclease-[Escherichia-coli] |
| EGZ5591350.1 | wHTH(SBD)+HNH | - | Escherichia_coli | Bacteria>Pseudomonadati>Pseudomonadota>Gammaproteobacteria>Enterobacterales>Enterobacteriaceae>Escherichia | 290 | BZZ56_004477 | 4738..5610 | 562 | HNH-endonuclease-[Escherichia-coli] |
| EGZ5731920.1 | wHTH(SBD)+HNH | - | Escherichia_coli | Bacteria>Pseudomonadati>Pseudomonadota>Gammaproteobacteria>Enterobacterales>Enterobacteriaceae>Escherichia | 290 | BL334_004674 | 5272..6144 | 562 | HNH-endonuclease-[Escherichia-coli] |
| EGZ5978147.1 | wHTH(SBD)+HNH | - | Escherichia_coli | Bacteria>Pseudomonadati>Pseudomonadota>Gammaproteobacteria>Enterobacterales>Enterobacteriaceae>Escherichia | 290 | BIB10_004100 | 36503..37375 | 562 | HNH-endonuclease-[Escherichia-coli] |
| EGZ8838305.1 | wHTH(SBD)+HNH | - | Escherichia_coli | Bacteria>Pseudomonadati>Pseudomonadota>Gammaproteobacteria>Enterobacterales>Enterobacteriaceae>Escherichia | 290 | AT823_000489 | 5230..6102 | 562 | HNH-endonuclease-[Escherichia-coli] |
| EHA9087553.1 | wHTH(SBD)+HNH | - | Escherichia_coli | Bacteria>Pseudomonadati>Pseudomonadota>Gammaproteobacteria>Enterobacterales>Enterobacteriaceae>Escherichia | 290 | D4094_002226 | 11210..12082 | 562 | HNH-endonuclease-[Escherichia-coli] |
| EHJ8640894.1 | wHTH(SBD)+HNH | GCA_017945505.1 | Escherichia_coli | Bacteria>Pseudomonadati>Pseudomonadota>Gammaproteobacteria>Enterobacterales>Enterobacteriaceae>Escherichia | 290 | KB366_000303 | 327472..328344 | 562 | HNH-endonuclease-[Escherichia-coli] |
| EHL0175574.1 | wHTH(SBD)+HNH | GCA_018090545.1 | Escherichia_coli | Bacteria>Pseudomonadati>Pseudomonadota>Gammaproteobacteria>Enterobacterales>Enterobacteriaceae>Escherichia | 290 | D7K57_004372 | 5667..6539 | 562 | HNH-endonuclease-[Escherichia-coli] |
| EHL0676196.1 | wHTH(SBD)+HNH | GCA_018092345.1 | Escherichia_coli | Bacteria>Pseudomonadati>Pseudomonadota>Gammaproteobacteria>Enterobacterales>Enterobacteriaceae>Escherichia | 290 | CP667_004249 | 6340..7212 | 562 | HNH-endonuclease-[Escherichia-coli] |
| EHL1134573.1 | wHTH(SBD)+HNH | GCA_018094045.1 | Escherichia_coli | Bacteria>Pseudomonadati>Pseudomonadota>Gammaproteobacteria>Enterobacterales>Enterobacteriaceae>Escherichia | 290 | BIB01_004965 | 3997..4869 | 562 | HNH-endonuclease-[Escherichia-coli] |
| EHL1162645.1 | wHTH(SBD)+HNH | GCA_018094265.1 | Escherichia_coli | Bacteria>Pseudomonadati>Pseudomonadota>Gammaproteobacteria>Enterobacterales>Enterobacteriaceae>Escherichia | 290 | BE892_002467 | 11209..12081 | 562 | HNH-endonuclease-[Escherichia-coli] |
| EHN3342067.1 | wHTH(SBD)+HNH | GCA_018356405.1 | Escherichia_coli | Bacteria>Pseudomonadati>Pseudomonadota>Gammaproteobacteria>Enterobacterales>Enterobacteriaceae>Escherichia | 290 | KIR71_003043 | 11208..12080 | 562 | HNH-endonuclease-[Escherichia-coli] |
| EHY5690078.1 | wHTH(SBD)+HNH | GCA_019590565.1 | Escherichia_coli | Bacteria>Pseudomonadati>Pseudomonadota>Gammaproteobacteria>Enterobacterales>Enterobacteriaceae>Escherichia | 290 | K4J54_002555 | 11207..12079 | 562 | HNH-endonuclease-[Escherichia-coli] |
| EIB0293609.1 | wHTH(SBD)+HNH | GCA_019848435.1 | Escherichia_coli | Bacteria>Pseudomonadati>Pseudomonadota>Gammaproteobacteria>Enterobacterales>Enterobacteriaceae>Escherichia | 290 | K7644_004614 | 4206..5078 | 562 | HNH-endonuclease-[Escherichia-coli] |
| EIG6470762.1 | wHTH(SBD)+HNH | GCA_020319175.1 | Escherichia_coli | Bacteria>Pseudomonadati>Pseudomonadota>Gammaproteobacteria>Enterobacterales>Enterobacteriaceae>Escherichia | 290 | EG665_005075 | 4520..5392 | 562 | HNH-endonuclease-[Escherichia-coli] |
| EIS9260703.1 | wHTH(SBD)+HNH | GCA_021435985.1 | Escherichia_coli | Bacteria>Pseudomonadati>Pseudomonadota>Gammaproteobacteria>Enterobacterales>Enterobacteriaceae>Escherichia | 290 | L0765_004697 | 6141..7013 | 562 | HNH-endonuclease-[Escherichia-coli] |
| EJW3212700.1 | wHTH(SBD)+HNH | GCA_025418235.1 | Escherichia_coli | Bacteria>Pseudomonadati>Pseudomonadota>Gammaproteobacteria>Enterobacterales>Enterobacteriaceae>Escherichia | 290 | OBZ04_004376 | 3871..4743 | 562 | HNH-endonuclease-[Escherichia-coli] |
| GCK20486.1 | wHTH(SBD)+HNH | GCA_005401165.1 | Escherichia_coli | Bacteria>Pseudomonadati>Pseudomonadota>Gammaproteobacteria>Enterobacterales>Enterobacteriaceae>Escherichia | 290 | BvCmsA58A_02117 | 107761..108633 | 562 | restriction-endonuclease-[Escherichia-coli] |
| HAW1502908.1 | wHTH(SBD)+HNH | GCA_016674655.1 | Escherichia_coli | Bacteria>Pseudomonadati>Pseudomonadota>Gammaproteobacteria>Enterobacterales>Enterobacteriaceae>Escherichia | 291 | JLM52_004942 | 13933..14808 | 562 | HNH-endonuclease-[Escherichia-coli] |
| POU04221.1 | wHTH(SBD)+HNH | GCA_002918705.1 | Escherichia_sp._ESNIH1 | Bacteria>Pseudomonadati>Pseudomonadota>Gammaproteobacteria>Enterobacterales>Enterobacteriaceae>Escherichia>unclassified_Escherichia | 341 | C3369_02405 | 507201..508226 | 1985876 | HNH-endonuclease-[Escherichia-sp.-ESNIH1] |
| WP_023320239.1 | wHTH(SBD)+HNH | GCF_964207445.1 | Klebsiella | Bacteria>Pseudomonadati>Pseudomonadota>Gammaproteobacteria>Enterobacterales>Enterobacteriaceae>Klebsiella/Raoultella_group | 337 | DK336_RS07885 | 60968..61981 | 570 | MULTISPECIES:-HNH-endonuclease-domain-containing-protein-[Klebsiella] |
| WP_045394768.1 | wHTH(SBD)+HNH | GCF_018388785.1 | Klebsiella | Bacteria>Pseudomonadati>Pseudomonadota>Gammaproteobacteria>Enterobacterales>Enterobacteriaceae>Klebsiella/Raoultella_group | 289 | KSL85_RS22145 | 4640279..4641148 | 570 | MULTISPECIES:-HNH-endonuclease-[Klebsiella] |
| WEJ89003.1 | wHTH(SBD)+HNH | GCA_029202365.1 | Klebsiella_huaxiensis | Bacteria>Pseudomonadati>Pseudomonadota>Gammaproteobacteria>Enterobacterales>Enterobacteriaceae>Klebsiella/Raoultella_group>Klebsiella | 337 | P0Y63_27745 | 5815994..5817007 | 2153354 | MAG:-HNH-endonuclease-domain-containing-protein-[Klebsiella-huaxiensis] |
| WP_042944160.1 | wHTH(SBD)+HNH | GCF_902164935.1 | Klebsiella_michiganensis | Bacteria>Pseudomonadati>Pseudomonadota>Gammaproteobacteria>Enterobacterales>Enterobacteriaceae>Klebsiella/Raoultella_group>Klebsiella | 337 | C2U44_RS27580 | 5591464..5592477 | 1134687 | MULTISPECIES:-HNH-endonuclease-domain-containing-protein-[Klebsiella] |
| WP_064342114.1 | wHTH(SBD)+HNH | GCF_902159715.1 | Klebsiella_grimontii | Bacteria>Pseudomonadati>Pseudomonadota>Gammaproteobacteria>Enterobacterales>Enterobacteriaceae>Klebsiella/Raoultella_group>Klebsiella | 337 | CWN04_RS12930 | 7695..8708 | 2058152 | MULTISPECIES:-HNH-endonuclease-domain-containing-protein-[Klebsiella] |
| EIY5117088.1 | wHTH(SBD)+HNH | GCA_022503905.1 | Klebsiella_quasipneumoniae | Bacteria>Pseudomonadati>Pseudomonadota>Gammaproteobacteria>Enterobacterales>Enterobacteriaceae>Klebsiella/Raoultella_group>Klebsiella>Klebsiella_pneumoniae_complex | 289 | ML076_005002 | 34166..35035 | 1463165 | HNH-endonuclease-[Klebsiella-quasipneumoniae] |
| SBM35922.1 | wHTH(SBD)+HNH | GCA_900084005.1 | Klebsiella_pneumoniae | Bacteria>Pseudomonadati>Pseudomonadota>Gammaproteobacteria>Enterobacterales>Enterobacteriaceae>Klebsiella/Raoultella_group>Klebsiella>Klebsiella_pneumoniae_complex | 289 | SAMEA2273678_04074 | 195204..196073 | 573 | restriction-endonuclease-[Klebsiella-pneumoniae] |
| WP_155011875.1 | wHTH(SBD)+HNH | GCF_902165215.1 | Klebsiella_variicola | Bacteria>Pseudomonadati>Pseudomonadota>Gammaproteobacteria>Enterobacterales>Enterobacteriaceae>Klebsiella/Raoultella_group>Klebsiella>Klebsiella_pneumoniae_complex | 289 | FN701_RS20990 | 5759..6628 | 244366 | HNH-endonuclease-[Klebsiella-variicola] |
| WP_046886451.1 | wHTH(SBD)+HNH | GCF_048545315.1 | Leclercia_pneumoniae | Bacteria>Pseudomonadati>Pseudomonadota>Gammaproteobacteria>Enterobacterales>Enterobacteriaceae>Leclercia | 337 | OA46_RS16215 | 123824..124837 | 2815358 | HNH-endonuclease-domain-containing-protein-[Enterobacter-cloacae] |
| EAY5508614.1 | wHTH(SBD)+HNH | GCA_006012725.1 | Salmonella_enterica | Bacteria>Pseudomonadati>Pseudomonadota>Gammaproteobacteria>Enterobacterales>Enterobacteriaceae>Salmonella | 205 | NE47_10995 | 45356..45973 | 28901 | HNH-endonuclease-[Salmonella-enterica] |
| EIC8525251.1 | wHTH(SBD)+HNH | GCA_019993965.1 | Salmonella_enterica | Bacteria>Pseudomonadati>Pseudomonadota>Gammaproteobacteria>Enterobacterales>Enterobacteriaceae>Salmonella | 205 | LAR10_002211 | 26407..27024 | 28901 | HNH-endonuclease-[Salmonella-enterica] |
| WP_061423976.1 | wHTH(SBD)+HNH | GCF_004194635.3 | Salmonella_enterica | Bacteria>Pseudomonadati>Pseudomonadota>Gammaproteobacteria>Enterobacterales>Enterobacteriaceae>Salmonella | 259 | DBZ85_RS25375 | 345622..346401 | 28901 | HNH-endonuclease-[Salmonella-enterica] |
| ECJ2539583.1 | wHTH(SBD)+HNH | GCA_007808255.1 | Salmonella_enterica_subsp._salamae | Bacteria>Pseudomonadati>Pseudomonadota>Gammaproteobacteria>Enterobacterales>Enterobacteriaceae>Salmonella>Salmonella_enterica | 291 | FNI64_03960 | 372069..372944 | 59202 | HNH-endonuclease-[Salmonella-enterica-subsp.-salamae] |
| NEG80377.1 | wHTH(SBD)+HNH | GCA_010667945.1 | Pantoea_agglomerans | Bacteria>Pseudomonadati>Pseudomonadota>Gammaproteobacteria>Enterobacterales>Erwiniaceae>Pantoea>Pantoea_agglomerans_group | 337 | GQQ17_05105 | 284131..285144 | 549 | HNH-endonuclease-[Pantoea-agglomerans] |
| WP_031592601.1 | wHTH(SBD)+HNH | GCF_022966105.1 | Pantoea_agglomerans | Bacteria>Pseudomonadati>Pseudomonadota>Gammaproteobacteria>Enterobacterales>Erwiniaceae>Pantoea>Pantoea_agglomerans_group | 337 | IBU46_RS01790 | 33085..34098 | 549 | HNH-endonuclease-domain-containing-protein-[Pantoea-agglomerans] |
| WP_163850615.1 | wHTH(SBD)+HNH | GCF_010667945.1 | Pantoea_agglomerans | Bacteria>Pseudomonadati>Pseudomonadota>Gammaproteobacteria>Enterobacterales>Erwiniaceae>Pantoea>Pantoea_agglomerans_group | 337 | GQQ17_RS05110 | 284131..285144 | 549 | HNH-endonuclease-domain-containing-protein-[Pantoea-agglomerans] |
| AUR32849.1 | wHTH(SBD)+HNH | GCA_002891475.1 | Morganella_morganii | Bacteria>Pseudomonadati>Pseudomonadota>Gammaproteobacteria>Enterobacterales>Morganellaceae>Morganella | 290 | C1O70_15700 | 3338063..3338935 | 582 | restriction-endonuclease-[Morganella-morganii] |
| WP_032098169.1 | wHTH(SBD)+HNH | GCF_002891475.1 | Morganella_morganii | Bacteria>Pseudomonadati>Pseudomonadota>Gammaproteobacteria>Enterobacterales>Morganellaceae>Morganella | 290 | C1O70_RS15700 | 3338063..3338935 | 582 | HNH-endonuclease-[Morganella-morganii] |
| WP_040031756.1 | wHTH(SBD)+HNH | GCF_026723725.1 | Pectobacterium | Bacteria>Pseudomonadati>Pseudomonadota>Gammaproteobacteria>Enterobacterales>Pectobacteriaceae | 291 | OWC53_RS08245 | 1787330..1788205 | 122277 | MULTISPECIES:-HNH-endonuclease-[Pectobacterium] |
| MCL6350159.1 | wHTH(SBD)+HNH | GCA_023507955.1 | Pectobacterium_polaris | Bacteria>Pseudomonadati>Pseudomonadota>Gammaproteobacteria>Enterobacterales>Pectobacteriaceae>Pectobacterium | 281 | EXT50_03065 | 66065..66910 | 2042057 | restriction-endonuclease-[Pectobacterium-polaris] |
| WP_103199127.1 | wHTH(SBD)+HNH | GCF_002904085.1 | Pectobacterium_odoriferum | Bacteria>Pseudomonadati>Pseudomonadota>Gammaproteobacteria>Enterobacterales>Pectobacteriaceae>Pectobacterium | 291 | BV921_RS15405 | 119000..119875 | 78398 | HNH-endonuclease-[Pectobacterium-odoriferum] |
| WP_116156021.1 | wHTH(SBD)+HNH | GCF_003382585.2 | Pectobacterium_aquaticum | Bacteria>Pseudomonadati>Pseudomonadota>Gammaproteobacteria>Enterobacterales>Pectobacteriaceae>Pectobacterium | 291 | DMB81_RS10720 | 2622..3497 | 2204145 | HNH-endonuclease-[Pectobacterium-aquaticum] |
| WP_181847899.1 | wHTH(SBD)+HNH | GCF_013913645.1 | Pectobacterium_aroidearum | Bacteria>Pseudomonadati>Pseudomonadota>Gammaproteobacteria>Enterobacterales>Pectobacteriaceae>Pectobacterium | 291 | H2Y54_RS11300 | 241633..242508 | 1201031 | HNH-endonuclease-[Pectobacterium-aroidearum] |
| GKX39598.1 | wHTH(SBD)+HNH | GCA_024343445.1 | Pectobacterium_carotovorum_subsp._carotovorum | Bacteria>Pseudomonadati>Pseudomonadota>Gammaproteobacteria>Enterobacterales>Pectobacteriaceae>Pectobacterium>Pectobacterium_carotovorum | 278 | SOASR014_33370 | 129854..130690 | 555 | restriction-endonuclease-[Pectobacterium-carotovorum-subsp.-carotovorum] |
| WP_004811077.1 | wHTH(SBD)+HNH | GCF_000368465.1 | Acinetobacter_schindleri | Bacteria>Pseudomonadati>Pseudomonadota>Gammaproteobacteria>Moraxellales>Moraxellaceae>Acinetobacter | 339 | F965_RS04745 | 399897..400916 | 108981 | HNH-endonuclease-signature-motif-containing-protein-[Acinetobacter-schindleri] |
| WP_032068286.1 | wHTH(SBD)+HNH | GCF_964300555.1 | Acinetobacter_calcoaceticus/baumannii_complex | Bacteria>Pseudomonadati>Pseudomonadota>Gammaproteobacteria>Moraxellales>Moraxellaceae>Acinetobacter | 341 | OLN40_RS03890 | 55114..56139 | 909768 | MULTISPECIES:-HNH-endonuclease-domain-containing-protein-[Acinetobacter-calcoaceticus/baumannii-complex] |
| WP_075040285.1 | wHTH(SBD)+HNH | GCF_051486965.1 | Acinetobacter_radioresistens | Bacteria>Pseudomonadati>Pseudomonadota>Gammaproteobacteria>Moraxellales>Moraxellaceae>Acinetobacter | 339 | SA188iCDA_RS03885 | 51109..52128 | 40216 | HNH-endonuclease-signature-motif-containing-protein-[Acinetobacter-radioresistens] |
| WP_111281991.1 | wHTH(SBD)+HNH | GCF_003258335.1 | Acinetobacter_radioresistens | Bacteria>Pseudomonadati>Pseudomonadota>Gammaproteobacteria>Moraxellales>Moraxellaceae>Acinetobacter | 339 | NVV25_RS10875 | 10137..11156 | 40216 | HNH-endonuclease-signature-motif-containing-protein-[Acinetobacter-radioresistens] |
| WP_151794805.1 | wHTH(SBD)+HNH | GCF_008997035.1 | Acinetobacter_soli | Bacteria>Pseudomonadati>Pseudomonadota>Gammaproteobacteria>Moraxellales>Moraxellaceae>Acinetobacter | 341 | GBP95_RS04075 | 343400..344425 | 487316 | HNH-endonuclease-domain-containing-protein-[Acinetobacter-soli] |
| WP_168370059.1 | wHTH(SBD)+HNH | GCF_012278715.1 | Acinetobacter_indicus | Bacteria>Pseudomonadati>Pseudomonadota>Gammaproteobacteria>Moraxellales>Moraxellaceae>Acinetobacter | 335 | HHC64_RS10925 | 2255774..2256781 | 756892 | HNH-endonuclease-signature-motif-containing-protein-[Acinetobacter-indicus] |
| WP_171065473.1 | wHTH(SBD)+HNH | - | Acinetobacter_variabilis | Bacteria>Pseudomonadati>Pseudomonadota>Gammaproteobacteria>Moraxellales>Moraxellaceae>Acinetobacter | 339 | LA331_RS04675 | 969569..970588 | 70346 | HNH-endonuclease-domain-containing-protein-[Acinetobacter-variabilis] |
| WP_180127360.1 | wHTH(SBD)+HNH | GCF_013419465.1 | unclassified_Acinetobacter | Bacteria>Pseudomonadati>Pseudomonadota>Gammaproteobacteria>Moraxellales>Moraxellaceae>Acinetobacter | 350 | FTU25_RS07955 | 186781..187833 | 196816 | MULTISPECIES:-HNH-endonuclease-signature-motif-containing-protein-[unclassified-Acinetobacter] |
| WP_221260724.1 | wHTH(SBD)+HNH | - | Acinetobacter_higginsii | Bacteria>Pseudomonadati>Pseudomonadota>Gammaproteobacteria>Moraxellales>Moraxellaceae>Acinetobacter | 335 | TOL5_RS12635 | 2702750..2703757 | 70347 | HNH-endonuclease-signature-motif-containing-protein-[Acinetobacter-sp.-Tol-5] |
| WP_265466023.1 | wHTH(SBD)+HNH | - | Acinetobacter_entericus | Bacteria>Pseudomonadati>Pseudomonadota>Gammaproteobacteria>Moraxellales>Moraxellaceae>Acinetobacter | 339 | OKC24_RS16345 | 16135..17154 | 2989714 | HNH-endonuclease-signature-motif-containing-protein-[Acinetobacter-sp.-BIT-DXN8] |
| WP_277607251.1 | wHTH(SBD)+HNH | - | Acinetobacter_higginsii | Bacteria>Pseudomonadati>Pseudomonadota>Gammaproteobacteria>Moraxellales>Moraxellaceae>Acinetobacter | 316 | MMP64_RS09445 | 128193..129143 | 70347 | HNH-endonuclease-signature-motif-containing-protein-[Acinetobacter-sp.-ANC-5659] |
| WP_111034562.1 | wHTH(SBD)+HNH | GCF_029847455.2 | Acinetobacter_baumannii | Bacteria>Pseudomonadati>Pseudomonadota>Gammaproteobacteria>Moraxellales>Moraxellaceae>Acinetobacter>Acinetobacter_calcoaceticus/baumannii_complex | 339 | KMP96_RS12145 | 13014..14033 | 470 | HNH-endonuclease-signature-motif-containing-protein-[Acinetobacter-baumannii] |
| WP_171265253.1 | wHTH(SBD)+HNH | - | Acinetobacter_baumannii | Bacteria>Pseudomonadati>Pseudomonadota>Gammaproteobacteria>Moraxellales>Moraxellaceae>Acinetobacter>Acinetobacter_calcoaceticus/baumannii_complex | 335 | B9Y01_RS02215 | 20305..21312 | 470 | HNH-endonuclease-signature-motif-containing-protein-[Acinetobacter-baumannii] |
| WP_104852220.1 | wHTH(SBD)+HNH | GCF_002946595.1 | Acinetobacter_sp._SWBY1 | Bacteria>Pseudomonadati>Pseudomonadota>Gammaproteobacteria>Moraxellales>Moraxellaceae>Acinetobacter>unclassified_Acinetobacter | 335 | C3Y93_RS04650 | 956755..957762 | 2079596 | HNH-endonuclease-signature-motif-containing-protein-[Acinetobacter-sp.-SWBY1] |
| WP_131273643.1 | wHTH(SBD)+HNH | GCF_004331195.1 | Acinetobacter_sp._ANC_4216 | Bacteria>Pseudomonadati>Pseudomonadota>Gammaproteobacteria>Moraxellales>Moraxellaceae>Acinetobacter>unclassified_Acinetobacter | 341 | E0H88_RS07265 | 86450..87475 | 2529840 | HNH-endonuclease-domain-containing-protein-[Acinetobacter-sp.-ANC-4216] |
| WP_180049761.1 | wHTH(SBD)+HNH | GCF_013419095.1 | Acinetobacter_sp._YH12105 | Bacteria>Pseudomonadati>Pseudomonadota>Gammaproteobacteria>Moraxellales>Moraxellaceae>Acinetobacter>unclassified_Acinetobacter | 335 | FTU44_RS05470 | 209547..210554 | 2601093 | HNH-endonuclease-signature-motif-containing-protein-[Acinetobacter-sp.-YH12105] |
| WP_218949078.1 | wHTH(SBD)+HNH | GCF_013418865.1 | Acinetobacter_sp._YH12117 | Bacteria>Pseudomonadati>Pseudomonadota>Gammaproteobacteria>Moraxellales>Moraxellaceae>Acinetobacter>unclassified_Acinetobacter | 333 | FTU55_RS01880 | 156176..157177 | 2601104 | HNH-endonuclease-signature-motif-containing-protein-[Acinetobacter-sp.-YH12117] |
| WP_233945510.1 | wHTH(SBD)+HNH | GCF_021391555.1 | Acinetobacter_sp._SH20PTE14 | Bacteria>Pseudomonadati>Pseudomonadota>Gammaproteobacteria>Moraxellales>Moraxellaceae>Acinetobacter>unclassified_Acinetobacter | 339 | LXF01_RS17065 | 126138..127157 | 2905879 | HNH-endonuclease-domain-containing-protein-[Acinetobacter-sp.-SH20PTE14] |
| WP_198334114.1 | wHTH(SBD)+HNH | GCF_904846355.1 | Psychrobacter_namhaensis | Bacteria>Pseudomonadati>Pseudomonadota>Gammaproteobacteria>Moraxellales>Moraxellaceae>Psychrobacter | 353 | JMX36_RS06930 | 18926..19987 | 292734 | HNH-endonuclease-[Psychrobacter-namhaensis] |
| WP_201567151.1 | wHTH(SBD)+HNH | GCF_904846115.1 | Psychrobacter_immobilis | Bacteria>Pseudomonadati>Pseudomonadota>Gammaproteobacteria>Moraxellales>Moraxellaceae>Psychrobacter | 353 | JMW93_RS11060 | 36553..37614 | 498 | HNH-endonuclease-[Psychrobacter-immobilis] |
| SEH06871.1 | wHTH(SBD)+HNH | GCA_900099115.2 | Candidatus_Venteria_ishoeyi | Bacteria>Pseudomonadati>Pseudomonadota>Gammaproteobacteria>Thiotrichales>Thiotrichaceae>Venteria | 294 | MBHS_02737 | 10215..11099 | 1899563 | Uncharacterised-protein-[Thiotrichales-bacterium-HS_08] |
| WP_132761826.1 | wHTH(SBD)+HNH | GCF_017918175.1 | Vibrio_crassostreae | Bacteria>Pseudomonadati>Pseudomonadota>Gammaproteobacteria>Vibrionales>Vibrionaceae>Vibrio | 375 | EDB51_RS02140 | 48933..50060 | 246167 | HNH-endonuclease-[Vibrio-crassostreae] |
| OYP37208.1 | wHTH(SBD)+HNH | GCA_002251265.1 | Rhodopirellula_sp._MGV | Bacteria>Pseudomonadati>PVC_group>Planctomycetota>Planctomycetia>Pirellulales>Pirellulaceae>Rhodopirellula>unclassified_Rhodopirellula | 295 | CGZ80_05830 | 5486..6373 | 2023130 | hypothetical-protein-CGZ80_05830-[Rhodopirellula-sp.-MGV] |
| WP_094412328.1 | wHTH(SBD)+HNH | GCF_002251265.1 | Rhodopirellula_sp._MGV | Bacteria>Pseudomonadati>PVC_group>Planctomycetota>Planctomycetia>Pirellulales>Pirellulaceae>Rhodopirellula>unclassified_Rhodopirellula | 295 | CGZ80_RS05840 | 5486..6373 | 2023130 | HNH-endonuclease-[Rhodopirellula-sp.-MGV] |
| MBR4835358.1 | wHTH(SBD)+HNH | GCA_017559825.1 | Thermoguttaceae_bacterium | Bacteria>Pseudomonadati>PVC_group>Planctomycetota>Planctomycetia>Pirellulales>Thermoguttaceae>unclassified_Thermoguttaceae | 272 | IKU86_13665 | 17244..18062 | 2691551 | hypothetical-protein-[Thermoguttaceae-bacterium] |
| HBJ34345.1 | wHTH(SBD)+HNH | GCA_003506095.1 | Planctomycetaceae_bacterium | Bacteria>Pseudomonadati>PVC_group>Planctomycetota>Planctomycetia>Planctomycetales>Planctomycetaceae>unclassified_Planctomycetaceae | 299 | DDZ51_06195 | 19806..20705 | 2026779 | hypothetical-protein-[Planctomycetaceae-bacterium] |
| MBX3448936.1 | wHTH(SBD)+HNH | GCA_019636285.1 | Planctomycetaceae_bacterium | Bacteria>Pseudomonadati>PVC_group>Planctomycetota>Planctomycetia>Planctomycetales>Planctomycetaceae>unclassified_Planctomycetaceae | 253 | KF777_05210 | 18739..19500 | 2026779 | HNH-endonuclease-[Planctomycetaceae-bacterium] |
| WP_207678201.1 | wHTH(SBD)+HNH | GCF_017377395.1 | Desulfonema_magnum | Bacteria>Pseudomonadati>Thermodesulfobacteriota>Desulfobacteria>Desulfobacterales>Desulfococcaceae>Desulfonema | 169 | dnm_RS17480 | 4572213..4572722 | 45655 | hypothetical-protein-[Desulfonema-magnum] |
| KEF34341.1 | wHTH(SBD)+HNH | GCA_000708185.1 | Deinococcus_sp._RL | Bacteria>Thermotogati>Deinococcota>Deinococci>Deinococcales>Deinococcaceae>Deinococcus>unclassified_Deinococcus | 393 | RDMS_08050 | 111688..112869 | 1489678 | hypothetical-protein-RDMS_08050-[Deinococcus-sp.-RL] |
| WP_034404737.1 | wHTH(SBD)+HNH | GCF_000708185.1 | Deinococcus_sp._RL | Bacteria>Thermotogati>Deinococcota>Deinococci>Deinococcales>Deinococcaceae>Deinococcus>unclassified_Deinococcus | 393 | RDMS_RS07915 | 111688..112869 | 1489678 | HNH-endonuclease-domain-containing-protein-[Deinococcus-sp.-RL] |
| HBP9919079.1 | wHTH(SBD)+HNH | - | - | - | 290 | L7G70_002286 | 11207..12079 | - | TPA:-HNH-endonuclease-[Escherichia-coli] |
| --SAD(SRA)+wHTH(SBD)+HNH | | | | | | | | | |
| WP_251142865.1 | SAD(SRA)+wHTH+HNH | GCF_018619155.1 | Streptomyces_sp._McG8 | Bacteria>Bacillati>Actinomycetota>Actinomycetes>Kitasatosporales>Streptomycetaceae>Streptomyces>unclassified_Streptomyces | 600 | HET68_RS34365 | 43461..45263 | 2725487 | HNH-endonuclease-[Streptomyces-sp.-McG8] |
| WP_259471465.1 | SAD(SRA)+wHTH+HNH | GCF_003688995.1 | Streptomyces_shenzhenensis | Bacteria>Bacillati>Actinomycetota>Actinomycetes>Kitasatosporales>Streptomycetaceae>Streptomyces | 600 | CTZ28_RS48000 | 142156..143958 | 943815 | HNH-endonuclease-[Streptomyces-shenzhenensis] |
| WP_229892201.1 | SAD(SRA)+wHTH+HNH | GCF_014650435.1 | Streptomyces_tendae | Bacteria>Bacillati>Actinomycetota>Actinomycetes>Kitasatosporales>Streptomycetaceae>Streptomyces | 599 | IE258_RS41245 | 401388..403187 | 1932 | HNH-endonuclease-[Streptomyces-tendae] |
| WP_280669736.1 | SAD(SRA)+wHTH+HNH | GCF_029892095.1 | unclassified_Kitasatospora | Bacteria>Bacillati>Actinomycetota>Actinomycetes>Kitasatosporales>Streptomycetaceae>Kitasatospora | 595 | ABIA34_RS11525 | 276189..277976 | 2633591 | HNH-endonuclease-[Kitasatospora-sp.-MAP12-44] |
| WP_326723514.1 | SAD(SRA)+wHTH+HNH | GCF_035917175.1 | Streptomyces_sp._NBC_00243 | Bacteria>Bacillati>Actinomycetota>Actinomycetes>Kitasatosporales>Streptomycetaceae>Streptomyces>unclassified_Streptomyces | 529 | OHT59_RS33080 | 7443661..7445250 | 2975688 | HNH-endonuclease-[Streptomyces-sp.-NBC_00243] |
| WP_237519866.1 | SAD(SRA)+wHTH+HNH | GCF_009827025.1 | Streptomyces_sp._HUCO-GS316 | Bacteria>Bacillati>Actinomycetota>Actinomycetes>Kitasatosporales>Streptomycetaceae>Streptomyces>unclassified_Streptomyces | 528 | GR925_RS40065 | 31169..32755 | 2692198 | HNH-endonuclease-[Streptomyces-sp.-HUCO-GS316] |
| WP_234899917.1 | SAD(SRA)+wHTH+HNH | GCF_008634015.1 | Streptomyces_filamentosus | Bacteria>Bacillati>Actinomycetota>Actinomycetes>Kitasatosporales>Streptomycetaceae>Streptomyces | 509 | CP979_RS37310 | 5276764..5278293 | 67294 | HNH-endonuclease-[Streptomyces-filamentosus] |
| WTB22514.1 | SAD(SRA)+wHTH+HNH | GCA_045161805.1 | Streptomyces_virginiae | Bacteria>Bacillati>Actinomycetota>Actinomycetes>Kitasatosporales>Streptomycetaceae>Streptomyces | 497 | OG253_13980 | 3159029..3160522 | 1961 | HNH-endonuclease-[Streptomyces-virginiae] |
| ADL44856.1 | SAD(SRA)+wHTH+HNH | GCA_000145235.1 | Micromonospora_aurantiaca_ATCC_27029 | Bacteria>Bacillati>Actinomycetota>Actinomycetes>Micromonosporales>Micromonosporaceae>Micromonospora>Micromonospora_aurantiaca | 496 | Micau_1294 | 1338741..1340231 | 644283 | HNH-nuclease-[Micromonospora-aurantiaca-ATCC-27029] |
| WP_013284495.1 | SAD(SRA)+wHTH+HNH | GCF_000145235.1 | Micromonospora_aurantiaca | Bacteria>Bacillati>Actinomycetota>Actinomycetes>Micromonosporales>Micromonosporaceae>Micromonospora | 496 | MICAU_RS32930 | 1338741..1340231 | 47850 | HNH-endonuclease-[Micromonospora-aurantiaca] |
| WP_237291399.1 | SAD(SRA)+wHTH+HNH | GCF_002080455.1 | Streptomyces | Bacteria>Bacillati>Actinomycetota>Actinomycetes>Kitasatosporales>Streptomycetaceae | 496 | B6R96_RS38065 | 2746648..2748138 | 1883 | HNH-endonuclease-[Streptomyces-sp.-Sge12] |
| WP_266686675.1 | SAD(SRA)+wHTH+HNH | GCF_026342335.1 | Streptomyces_sp._NBC_00268 | Bacteria>Bacillati>Actinomycetota>Actinomycetes>Kitasatosporales>Streptomycetaceae>Streptomyces>unclassified_Streptomyces | 496 | OOK47_RS16840 | 3638395..3639885 | 2975695 | HNH-endonuclease-[Streptomyces-sp.-NBC_00268] |
| WP_239155547.1 | SAD(SRA)+wHTH+HNH | GCF_010548515.1 | Streptomyces_sp._SID14446 | Bacteria>Bacillati>Actinomycetota>Actinomycetes>Kitasatosporales>Streptomycetaceae>Streptomyces>unclassified_Streptomyces | 491 | G3I62_RS41730 | 12693..14168 | 2706072 | HNH-endonuclease-[Streptomyces-sp.-SID14446] |
| WP_329299362.1 | SAD(SRA)+wHTH+HNH | GCF_036226925.1 | Streptomyces_sp._NBC_00659 | Bacteria>Bacillati>Actinomycetota>Actinomycetes>Kitasatosporales>Streptomycetaceae>Streptomyces>unclassified_Streptomyces | 491 | OG410_RS13510 | 3125420..3126895 | 2903669 | HNH-endonuclease-[Streptomyces-sp.-NBC_00659] |
| WP_198033299.1 | SAD(SRA)+wHTH+HNH | GCF_023072705.1 | unclassified_Micromonospora | Bacteria>Bacillati>Actinomycetota>Actinomycetes>Micromonosporales>Micromonosporaceae>Micromonospora | 490 | FJ969_RS31395 | 50977..52449 | 2617518 | MULTISPECIES:-HNH-endonuclease-[unclassified-Micromonospora] |
| WP_198039649.1 | SAD(SRA)+wHTH+HNH | GCF_040634605.1 | Micromonospora | Bacteria>Bacillati>Actinomycetota>Actinomycetes>Micromonosporales>Micromonosporaceae | 477 | IF06_RS33885 | 41595..43028 | 1873 | HNH-endonuclease-[Micromonospora-globosa] |
| WP_250297825.1 | SAD(SRA)+wHTH+HNH | GCF_940670765.1 | Streptomyces_sp._A_4/2 | Bacteria>Bacillati>Actinomycetota>Actinomycetes>Kitasatosporales>Streptomycetaceae>Streptomyces>unclassified_Streptomyces | 464 | NBY69_RS01945 | 274529..275923 | 2934314 | HNH-endonuclease-[Streptomyces-sp.-A-4/2] |
| WP_329194509.1 | SAD(SRA)+wHTH+HNH | GCF_036228565.1 | Streptomyces_sp._NBC_01435 | Bacteria>Bacillati>Actinomycetota>Actinomycetes>Kitasatosporales>Streptomycetaceae>Streptomyces>unclassified_Streptomyces | 394 | OG769_RS10345 | 2488213..2489397 | 2903865 | HNH-endonuclease-[Streptomyces-sp.-NBC_01435] |
| WP_189531532.1 | SAD(SRA)+wHTH+HNH | GCF_014649335.1 | Streptomyces_roseolilacinus | Bacteria>Bacillati>Actinomycetota>Actinomycetes>Kitasatosporales>Streptomycetaceae>Streptomyces | 289 | IE106_RS08410 | 223028..223897 | 66904 | hypothetical-protein-[Streptomyces-roseolilacinus] |

| Protein-ID | Domain | Assembly-ID | Strain | Taxonomy | Length | Locus-Tag | Genomic-coordinates | TaxID | Fasta-defline |
|---|---|---|---|---|---|---|---|---|---|
| --PIN-EndoRNase | | | | | | | | | |
| WP_202318888.1 | PIN-EndoRNase | GCF_016757965.1 | Archaeoglobus_neptunius | Archaea>Methanobacteriati>Methanobacteriota>Archaeoglobi>Archaeoglobales>Archaeoglobaceae>Archaeoglobus | 150 | JFQ59_RS02765 | 85728..86180 | 2798580 | type-II-toxin-antitoxin-system-VapC-family-toxin-[Archaeoglobus-neptunius] |
| MCL2156396.1 | PIN-EndoRNase | GCA_009776495.1 | Methanobrevibacter_sp. | Archaea>Methanobacteriati>Methanobacteriota>Methanomada_group>Methanobacteria>Methanobacteriales>Methanobacteriaceae>Methanobrevibacter>unclassified_Methanobrevibacter | 129 | FWH54_00040 | 12552..12941 | 66852 | PIN-domain-containing-protein-[Methanobrevibacter-sp.] |
| WP_135304846.1 | PIN-EndoRNase | GCF_004681185.1 | Haloarcula_amylovorans | Archaea>Methanobacteriati>Methanobacteriota>Stenosarchaea_group>Halobacteria>Halobacteriales>Haloarculaceae>Haloarcula | 168 | E4S61_RS15385 | 198314..198820 | 2562280 | hypothetical-protein-[Halomicroarcula-amylolytica] |
| WP_229775396.1 | PIN-EndoRNase | GCF_014647415.1 | Haloarcula_sebkhae | Archaea>Methanobacteriati>Methanobacteriota>Stenosarchaea_group>Halobacteria>Halobacteriales>Haloarculaceae>Haloarcula | 189 | IEY38_RS19895 | 65277..65846 | 932660 | hypothetical-protein-[Haloarcula-sebkhae] |
| ELZ19694.1 | PIN-EndoRNase | GCA_000337455.1 | Halosimplex_carlsbadense_2-9-1 | Archaea>Methanobacteriati>Methanobacteriota>Stenosarchaea_group>Halobacteria>Halobacteriales>Haloarculaceae>Halosimplex>Halosimplex_carlsbadense | 168 | C475_22154 | 54904..55410 | 797114 | hypothetical-protein-C475_22154-[Halosimplex-carlsbadense-2-9-1] |
| WP_188786132.1 | PIN-EndoRNase | GCF_014646875.1 | Halobellus_salinus | Archaea>Methanobacteriati>Methanobacteriota>Stenosarchaea_group>Halobacteria>Halobacteriales>Haloferacaceae>Halobellus | 168 | IEX72_RS04210 | 172536..173042 | 931585 | hypothetical-protein-[Halobellus-salinus] |
| WP_240942963.1 | PIN-EndoRNase | GCF_012726105.1 | Haloferax_volcanii | Archaea>Methanobacteriati>Methanobacteriota>Stenosarchaea_group>Halobacteria>Halobacteriales>Haloferacaceae>Haloferax | 289 | GOC85_RS10650 | 2079329..2080198 | 2246 | hypothetical-protein-[Haloferax-alexandrinus] |
| WP_015911585.1 | PIN-EndoRNase | GCF_000022205.1 | Halorubrum_lacusprofundi | Archaea>Methanobacteriati>Methanobacteriota>Stenosarchaea_group>Halobacteria>Halobacteriales>Haloferacaceae>Halorubrum | 144 | HLAC_RS15640 | 432758..433192 | 2247 | PIN-domain-containing-protein-[Halorubrum-lacusprofundi] |
| WP_193308742.1 | PIN-EndoRNase | GCF_008806895.1 | Halorubrum_halophilum | Archaea>Methanobacteriati>Methanobacteriota>Stenosarchaea_group>Halobacteria>Halobacteriales>Haloferacaceae>Halorubrum | 190 | HrrH1147_RS02375 | 466547..467119 | 413816 | hypothetical-protein-[Halorubrum-halophilum] |
| WP_179261308.1 | PIN-EndoRNase | GCF_013402815.2 | Natrinema_halophilum | Archaea>Methanobacteriati>Methanobacteriota>Stenosarchaea_group>Halobacteria>Halobacteriales>Natrialbaceae>Natrinema | 144 | HYG82_RS33130 | 2544428..2544862 | 1699371 | PIN-domain-containing-protein-[Natrinema-halophilum] |
| AEN07518.1 | PIN-EndoRNase | GCA_000224475.1 | halophilic_archaeon_DL31 | Archaea>Methanobacteriati>Methanobacteriota>Stenosarchaea_group>Halobacteria>Halobacteriales>unclassified_Halobacteriales | 144 | Halar_0258 | 254914..255348 | 756883 | PilT-protein-domain-protein-[halophilic-archaeon-DL31] |
| WP_015286392.1 | PIN-EndoRNase | GCF_000327485.1 | Methanoregula_formicica | Archaea>Methanobacteriati>Methanobacteriota>Stenosarchaea_group>Methanomicrobia>Methanomicrobiales>Methanoregulaceae>Methanoregula | 153 | METFOR_RS11870 | 2387738..2388199 | 882104 | type-II-toxin-antitoxin-system-VapC-family-toxin-[Methanoregula-formicica] |
| AGB03430.1 | PIN-EndoRNase | GCA_000327485.1 | Methanoregula_formicica_SMSP | Archaea>Methanobacteriati>Methanobacteriota>Stenosarchaea_group>Methanomicrobia>Methanomicrobiales>Methanoregulaceae>Methanoregula>Methanoregula_formicica | 153 | Metfor_2430 | 2387738..2388199 | 593750 | putative-nucleic-acid-binding-protein,-contains-PIN-domain-[Methanoregula-formicica-SMSP] |
| WP_139681368.1 | PIN-EndoRNase | GCF_006274605.1 | Thermococcus_indicus | Archaea>Methanobacteriati>Methanobacteriota>Thermococci>Thermococcales>Thermococcaceae>Thermococcus | 232 | FH039_RS11125 | 2046675..2047373 | 2586643 | hypothetical-protein-[Thermococcus-indicus] |
| WP_010477370.1 | PIN-EndoRNase | GCF_000258515.1 | Thermococcus_zilligii | Archaea>Methanobacteriati>Methanobacteriota>Thermococci>Thermococcales>Thermococcaceae>Thermococcus | 238 | TZI_RS0101410 | 263245..263961 | 54076 | hypothetical-protein-[Thermococcus-zilligii] |
| WP_167893363.1 | PIN-EndoRNase | GCF_012027305.1 | Thermococcus_sp._MAR1 | Archaea>Methanobacteriati>Methanobacteriota>Thermococci>Thermococcales>Thermococcaceae>Thermococcus>unclassified_Thermococcus | 232 | E3E25_RS11230 | 190875..191573 | 1638263 | hypothetical-protein-[Thermococcus-sp.-MAR1] |
| WP_167910222.1 | PIN-EndoRNase | GCF_012027595.1 | Thermococcus_sp._Bubb.Bath | Archaea>Methanobacteriati>Methanobacteriota>Thermococci>Thermococcales>Thermococcaceae>Thermococcus>unclassified_Thermococcus | 232 | E3E29_RS06915 | 299965..300663 | 1638242 | hypothetical-protein-[Thermococcus-sp.-Bubb.Bath] |
| OYT64647.1 | PIN-EndoRNase | GCA_002255045.1 | Candidatus_Bathyarchaeota_archaeon_ex4484_205 | Archaea>Thermoproteati>Candidatus_Bathyarchaeota>unclassified_Candidatus_Bathyarchaeota | 145 | B6U74_05080 | 3652..4089 | 2012510 | hypothetical-protein-B6U74_05080-[Candidatus-Bathyarchaeota-archaeon-ex4484_205] |
| MXY61027.1 | PIN-EndoRNase | GCA_009837825.1 | Cenarchaeum_sp._SB0665_bin_23 | Archaea>Thermoproteati>Nitrososphaerota>Candidatus_Cenarchaeales>Candidatus_Cenarchaeaceae>Candidatus_Cenarchaeum>unclassified_Candidatus_Cenarchaeum | 67 | F4Y82_02790 | 7753..7956 | 2604876 | hypothetical-protein-[Cenarchaeum-sp.-SB0665_bin_23] |
| MYG32965.1 | PIN-EndoRNase | GCA_009842575.1 | Cenarchaeum_sp._SB0677_bin_16 | Archaea>Thermoproteati>Nitrososphaerota>Candidatus_Cenarchaeales>Candidatus_Cenarchaeaceae>Candidatus_Cenarchaeum>unclassified_Candidatus_Cenarchaeum | 67 | F4202_03065 | 8097..8300 | 2604883 | hypothetical-protein-[Cenarchaeum-sp.-SB0677_bin_16] |
| MYG32490.1 | PIN-EndoRNase | GCA_009842575.1 | Cenarchaeum_sp._SB0677_bin_16 | Archaea>Thermoproteati>Nitrososphaerota>Candidatus_Cenarchaeales>Candidatus_Cenarchaeaceae>Candidatus_Cenarchaeum>unclassified_Candidatus_Cenarchaeum | 99 | F4202_00560 | 1462..1761 | 2604883 | hypothetical-protein-[Cenarchaeum-sp.-SB0677_bin_16] |
| GIU72779.1 | PIN-EndoRNase | GCA_026002835.1 | Candidatus_Nitrosocaldaceae_archaeon | Archaea>Thermoproteati>Nitrososphaerota>Nitrososphaeria>Candidatus_Nitrosocaldales>Candidatus_Nitrosocaldaceae>unclassified_Candidatus_Nitrosocaldaceae | 142 | KatS3mg003_2258 | 17168..17596 | 2838776 | MAG:-hypothetical-protein-KatS3mg003_2258-[Candidatus-Nitrosocaldaceae-archaeon] |
| WP_009070039.1 | PIN-EndoRNase | GCF_000243315.1 | Metallosphaera_yellowstonensis | Archaea>Thermoproteati>Thermoproteota>Thermoprotei>Sulfolobales>Sulfolobaceae>Metallosphaera | 141 | METMK1DRAFT_RS01480 | 130772..131197 | 1111107 | type-II-toxin-antitoxin-system-VapC-family-toxin-[Metallosphaera-yellowstonensis] |
| WP_312722310.1 | PIN-EndoRNase | GCF_036842645.1 | Actinomycetes | Bacteria>Bacillati>Actinomycetota | 187 | RT527_RS09235 | 18637..19200 | 1760 | PIN-domain-containing-protein-[Mobilicoccus-sp.] |
| MDE0160528.1 | PIN-EndoRNase | GCA_028819505.1 | Acidimicrobiaceae_bacterium | Bacteria>Bacillati>Actinomycetota>Acidimicrobiia>Acidimicrobiales>Acidimicrobiaceae>unclassified_Acidimicrobiaceae | 124 | OXL98_02210 | 8739..9113 | 2024894 | MAG:-DUF5615-family-PIN-like-protein-[Acidimicrobiaceae-bacterium] |
| MDE0067325.1 | PIN-EndoRNase | GCA_028819005.1 | Acidimicrobiaceae_bacterium | Bacteria>Bacillati>Actinomycetota>Acidimicrobiia>Acidimicrobiales>Acidimicrobiaceae>unclassified_Acidimicrobiaceae | 129 | OXN44_10715 | 1423..1812 | 2024894 | MAG:-PIN-domain-containing-protein-[Acidimicrobiaceae-bacterium] |
| MDE0320764.1 | PIN-EndoRNase | GCA_028820815.1 | Acidimicrobiaceae_bacterium | Bacteria>Bacillati>Actinomycetota>Acidimicrobiia>Acidimicrobiales>Acidimicrobiaceae>unclassified_Acidimicrobiaceae | 132 | OXI97_12890 | 64070..64468 | 2024894 | MAG:-type-II-toxin-antitoxin-system-VapC-family-toxin-[Acidimicrobiaceae-bacterium] |
| MDE0160525.1 | PIN-EndoRNase | GCA_028819505.1 | Acidimicrobiaceae_bacterium | Bacteria>Bacillati>Actinomycetota>Acidimicrobiia>Acidimicrobiales>Acidimicrobiaceae>unclassified_Acidimicrobiaceae | 134 | OXL98_02195 | 5063..5467 | 2024894 | MAG:-DUF5615-family-PIN-like-protein-[Acidimicrobiaceae-bacterium] |
| MDE0495498.1 | PIN-EndoRNase | GCA_028824895.1 | Acidimicrobiaceae_bacterium | Bacteria>Bacillati>Actinomycetota>Acidimicrobiia>Acidimicrobiales>Acidimicrobiaceae>unclassified_Acidimicrobiaceae | 139 | OXH54_16265 | 40586..41005 | 2024894 | MAG:-hypothetical-protein-OXH54_16265-[Acidimicrobiaceae-bacterium] |
| MXY09520.1 | PIN-EndoRNase | GCA_009838345.1 | Acidimicrobiaceae_bacterium | Bacteria>Bacillati>Actinomycetota>Acidimicrobiia>Acidimicrobiales>Acidimicrobiaceae>unclassified_Acidimicrobiaceae | 139 | F4Y56_03040 | 75788..76207 | 2024894 | type-II-toxin-antitoxin-system-VapC-family-toxin-[Acidimicrobiaceae-bacterium] |
| MYJ29886.1 | PIN-EndoRNase | GCA_009845185.1 | Acidimicrobiaceae_bacterium | Bacteria>Bacillati>Actinomycetota>Acidimicrobiia>Acidimicrobiales>Acidimicrobiaceae>unclassified_Acidimicrobiaceae | 139 | F4072_07950 | 9991..10410 | 2024894 | type-II-toxin-antitoxin-system-VapC-family-toxin-[Acidimicrobiaceae-bacterium] |
| WP_015443643.1 | PIN-EndoRNase | GCF_000348785.1 | Ilumatobacter_coccineus | Bacteria>Bacillati>Actinomycetota>Acidimicrobiia>Acidimicrobiales>Ilumatobacteraceae>Ilumatobacter | 208 | YM304_RS20485 | 4616682..4617308 | 467094 | NYN-domain-containing-protein-[Ilumatobacter-coccineus] |
| BAN04396.1 | PIN-EndoRNase | GCA_000348785.1 | Ilumatobacter_coccineus_YM16-304 | Bacteria>Bacillati>Actinomycetota>Acidimicrobiia>Acidimicrobiales>Ilumatobacteraceae>Ilumatobacter>Ilumatobacter_coccineus | 208 | YM304_40820 | 4616682..4617308 | 1313172 | hypothetical-protein-YM304_40820-[Ilumatobacter-coccineus-YM16-304] |
| MXX42190.1 | PIN-EndoRNase | GCA_009836375.1 | Acidimicrobiales_bacterium | Bacteria>Bacillati>Actinomycetota>Acidimicrobiia>Acidimicrobiales>unclassified_Acidimicrobiales | 145 | F4Z53_03975 | 39870..40307 | 2201156 | type-II-toxin-antitoxin-system-VapC-family-toxin-[Acidimicrobiales-bacterium] |
| MCB0987058.1 | PIN-EndoRNase | GCA_020439505.1 | Acidimicrobiales_bacterium | Bacteria>Bacillati>Actinomycetota>Acidimicrobiia>Acidimicrobiales>unclassified_Acidimicrobiales | 159 | KDB09_04715 | 9470..9949 | 2201156 | DUF4411-family-protein-[Acidimicrobiales-bacterium] |
| MXW42981.1 | PIN-EndoRNase | GCA_009835645.1 | Acidimicrobiia_bacterium | Bacteria>Bacillati>Actinomycetota>Acidimicrobiia>unclassified_Acidimicrobiia | 121 | F4Z91_07835 | 12122..12487 | 2080302 | hypothetical-protein-[Acidimicrobiia-bacterium] |
| MXW42975.1 | PIN-EndoRNase | GCA_009835645.1 | Acidimicrobiia_bacterium | Bacteria>Bacillati>Actinomycetota>Acidimicrobiia>unclassified_Acidimicrobiia | 130 | F4Z91_07805 | 3154..3546 | 2080302 | type-II-toxin-antitoxin-system-VapC-family-toxin-[Acidimicrobiia-bacterium] |
| MYG93977.1 | PIN-EndoRNase | GCA_009843865.1 | Acidimicrobiia_bacterium | Bacteria>Bacillati>Actinomycetota>Acidimicrobiia>unclassified_Acidimicrobiia | 130 | F4138_03145 | 12663..13055 | 2080302 | type-II-toxin-antitoxin-system-VapC-family-toxin-[Acidimicrobiia-bacterium] |
| MXZ78954.1 | PIN-EndoRNase | GCA_009837345.1 | Acidimicrobiia_bacterium | Bacteria>Bacillati>Actinomycetota>Acidimicrobiia>unclassified_Acidimicrobiia | 138 | F4Z06_11885 | 15988..16404 | 2080302 | type-II-toxin-antitoxin-system-VapC-family-toxin-[Acidimicrobiia-bacterium] |
| MYB44034.1 | PIN-EndoRNase | GCA_009839965.1 | Acidimicrobiia_bacterium | Bacteria>Bacillati>Actinomycetota>Acidimicrobiia>unclassified_Acidimicrobiia | 152 | F4X74_03735 | 14378..14836 | 2080302 | type-II-toxin-antitoxin-system-VapC-family-toxin-[Acidimicrobiia-bacterium] |
| MYB44041.1 | PIN-EndoRNase | GCA_009839965.1 | Acidimicrobiia_bacterium | Bacteria>Bacillati>Actinomycetota>Acidimicrobiia>unclassified_Acidimicrobiia | 290 | F4X74_03770 | 26945..27817 | 2080302 | NYN-domain-containing-protein-[Acidimicrobiia-bacterium] |
| WP_249935204.1 | PIN-EndoRNase | GCF_012976645.1 | Mobiluncus_mulieris | Bacteria>Bacillati>Actinomycetota>Actinomycetes>Actinomycetales>Actinomycetaceae>Mobiluncus | 93 | HHJ77_RS04680 | 60442..60723 | 2052 | PIN-family-toxin-antitoxin-system-[Mobiluncus-mulieris] |
| WP_211583459.1 | PIN-EndoRNase | - | Mobiluncus_mulieris | Bacteria>Bacillati>Actinomycetota>Actinomycetes>Actinomycetales>Actinomycetaceae>Mobiluncus | 117 | HHJ79_RS06615 | 30791..31144 | 2052 | type-II-toxin-antitoxin-system-VapC-family-toxin-[Mobiluncus-mulieris] |
| WP_165316370.1 | PIN-EndoRNase | GCF_011038885.2 | Schaalia_sp._ZJ405 | Bacteria>Bacillati>Actinomycetota>Actinomycetes>Actinomycetales>Actinomycetaceae>Schaalia>unclassified_Schaalia | 264 | G7Y41_RS09390 | 2200915..2201709 | 2709403 | NYN-domain-containing-protein-[Schaalia-sp.-ZJ405] |
| WP_240945038.1 | PIN-EndoRNase | GCF_012932365.1 | Bifidobacterium_moraviense | Bacteria>Bacillati>Actinomycetota>Actinomycetes>Bifidobacteriales>Bifidobacteriaceae>Bifidobacterium | 198 | G1C96_RS01255 | 282893..283489 | 2675323 | PIN-domain-containing-protein-[Bifidobacterium-sp.-DSM-109958] |
| WP_169274833.1 | PIN-EndoRNase | GCF_012932365.1 | Bifidobacterium_moraviense | Bacteria>Bacillati>Actinomycetota>Actinomycetes>Bifidobacteriales>Bifidobacteriaceae>Bifidobacterium | 218 | G1C96_RS01280 | 286304..286960 | 2675323 | type-II-toxin-antitoxin-system-VapC-family-toxin-[Bifidobacterium-sp.-DSM-109958] |
| WP_066069024.1 | PIN-EndoRNase | GCF_001636565.1 | Frankia_sp._EI5c | Bacteria>Bacillati>Actinomycetota>Actinomycetes>Frankiales>Frankiaceae>Frankia>unclassified_Frankia | 154 | UG55_RS15550 | 103088..103552 | 683316 | DUF4411-family-protein-[Frankia-sp.-EI5c] |
| ROR38154.1 | PIN-EndoRNase | GCA_003751605.1 | Kitasatospora_cineracea | Bacteria>Bacillati>Actinomycetota>Actinomycetes>Kitasatosporales>Streptomycetaceae>Kitasatospora | 156 | EDD39_6326 | 1174915..1175385 | 88074 | hypothetical-protein-EDD39_6326-[Kitasatospora-cineracea] |
| WP_123562583.1 | PIN-EndoRNase | GCF_003751605.1 | Kitasatospora_cineracea | Bacteria>Bacillati>Actinomycetota>Actinomycetes>Kitasatosporales>Streptomycetaceae>Kitasatospora | 156 | EDD39_RS31520 | 1174915..1175385 | 88074 | type-II-toxin-antitoxin-system-VapC-family-toxin-[Kitasatospora-cineracea] |
| WP_073727328.1 | PIN-EndoRNase | GCF_042736535.1 | unclassified_Streptomyces | Bacteria>Bacillati>Actinomycetota>Actinomycetes>Kitasatosporales>Streptomycetaceae>Streptomyces | 124 | A6A29_RS36800 | 502309..502683 | 2593676 | type-II-toxin-antitoxin-system-VapC-family-toxin-[Streptomyces-sp.-TSRI0281] |
| WP_209310652.1 | PIN-EndoRNase | - | Streptomyces_spiramenti | Bacteria>Bacillati>Actinomycetota>Actinomycetes>Kitasatosporales>Streptomycetaceae>Streptomyces | 124 | HCJ92_RS02705 | 10983..11357 | 2720606 | PIN-domain-containing-protein-[Streptomyces-spiramenti] |
| MCT9080282.1 | PIN-EndoRNase | GCA_025399775.1 | Streptomyces_fulvoviolaceus | Bacteria>Bacillati>Actinomycetota>Actinomycetes>Kitasatosporales>Streptomycetaceae>Streptomyces | 135 | N4G65_27560 | 166..573 | 285535 | PIN-domain-nuclease-[Streptomyces-fulvoviolaceus] |
| WP_261725446.1 | PIN-EndoRNase | GCF_025399775.1 | Streptomyces_fulvoviolaceus | Bacteria>Bacillati>Actinomycetota>Actinomycetes>Kitasatosporales>Streptomycetaceae>Streptomyces | 135 | N4G65_RS27520 | 166..573 | 285535 | PIN-domain-nuclease-[Streptomyces-fulvoviolaceus] |
| WP_266738639.1 | PIN-EndoRNase | GCF_026340175.1 | Streptomyces_platensis | Bacteria>Bacillati>Actinomycetota>Actinomycetes>Kitasatosporales>Streptomycetaceae>Streptomyces | 145 | OG775_RS34860 | 28998..29435 | 58346 | hypothetical-protein-[Streptomyces-platensis] |
| WP_226071731.1 | PIN-EndoRNase | GCF_020410765.1 | Streptomyces_marincola | Bacteria>Bacillati>Actinomycetota>Actinomycetes>Kitasatosporales>Streptomycetaceae>Streptomyces | 154 | LC193_RS04360 | 1027105..1027569 | 2878388 | type-II-toxin-antitoxin-system-VapC-family-toxin-[Streptomyces-marincola] |
| WP_209310645.1 | PIN-EndoRNase | - | Streptomyces_spiramenti | Bacteria>Bacillati>Actinomycetota>Actinomycetes>Kitasatosporales>Streptomycetaceae>Streptomyces | 157 | HCJ92_RS02725 | 13076..13549 | 2720606 | PIN-domain-containing-protein-[Streptomyces-spiramenti] |
| WP_241838778.1 | PIN-EndoRNase | GCF_042736535.1 | unclassified_Streptomyces | Bacteria>Bacillati>Actinomycetota>Actinomycetes>Kitasatosporales>Streptomycetaceae>Streptomyces | 165 | A6A29_RS36830 | 505249..505746 | 2593676 | type-II-toxin-antitoxin-system-VapC-family-toxin-[Streptomyces-sp.-TSRI0281] |
| WP_250297830.1 | PIN-EndoRNase | GCF_940670765.1 | Streptomyces_sp._A_4/2 | Bacteria>Bacillati>Actinomycetota>Actinomycetes>Kitasatosporales>Streptomycetaceae>Streptomyces>unclassified_Streptomyces | 124 | NBY69_RS01965 | 288206..288580 | 2934314 | type-II-toxin-antitoxin-system-VapC-family-toxin-[Streptomyces-sp.-A-4/2] |
| WP_250297843.1 | PIN-EndoRNase | GCF_940670765.1 | Streptomyces_sp._A_4/2 | Bacteria>Bacillati>Actinomycetota>Actinomycetes>Kitasatosporales>Streptomycetaceae>Streptomyces>unclassified_Streptomyces | 124 | NBY69_RS02035 | 311930..312304 | 2934314 | type-II-toxin-antitoxin-system-VapC-family-toxin-[Streptomyces-sp.-A-4/2] |
| NEB00543.1 | PIN-EndoRNase | GCA_010548375.1 | Streptomyces_sp._SID13726 | Bacteria>Bacillati>Actinomycetota>Actinomycetes>Kitasatosporales>Streptomycetaceae>Streptomyces>unclassified_Streptomyces | 135 | G3I78_15950 | 363372..363779 | 2706058 | PIN-domain-nuclease-[Streptomyces-sp.-SID13726] |
| WP_306332897.1 | PIN-EndoRNase | GCF_030766985.1 | Streptomyces_sp._KL118A | Bacteria>Bacillati>Actinomycetota>Actinomycetes>Kitasatosporales>Streptomycetaceae>Streptomyces>unclassified_Streptomyces | 135 | RBJ68_RS04470 | 66401..66808 | 3045153 | PIN-domain-nuclease-[Streptomyces-sp.-KL118A] |
| WTB16639.1 | PIN-EndoRNase | GCA_045161795.1 | Streptomyces_sp._NBC_00829 | Bacteria>Bacillati>Actinomycetota>Actinomycetes>Kitasatosporales>Streptomycetaceae>Streptomyces>unclassified_Streptomyces | 135 | OG293_25275 | 5691557..5691964 | 2903679 | PIN-domain-nuclease-[Streptomyces-sp.-NBC_00829] |
| WP_329194495.1 | PIN-EndoRNase | GCF_036228565.1 | Streptomyces_sp._NBC_01435 | Bacteria>Bacillati>Actinomycetota>Actinomycetes>Kitasatosporales>Streptomycetaceae>Streptomyces>unclassified_Streptomyces | 160 | OG769_RS10310 | 2473921..2474403 | 2903865 | type-II-toxin-antitoxin-system-VapC-family-toxin-[Streptomyces-sp.-NBC_01435] |
| QKW00215.1 | PIN-EndoRNase | GCA_013364175.1 | Streptomyces_sp._NA02536 | Bacteria>Bacillati>Actinomycetota>Actinomycetes>Kitasatosporales>Streptomycetaceae>Streptomyces>unclassified_Streptomyces | 185 | HUT14_10185 | 2277913..2278470 | 2742133 | hypothetical-protein-HUT14_10185-[Streptomyces-sp.-NA02536] |
| WP_264176809.1 | PIN-EndoRNase | - | Agilicoccus_flavus | Bacteria>Bacillati>Actinomycetota>Actinomycetes>Micrococcales>Dermatophilaceae>Agilicoccus | 181 | ICW24_RS18610 | 14018..14563 | 2775968 | PIN-domain-containing-protein-[Agilicoccus-flavus] |
| WP_048693908.1 | PIN-EndoRNase | GCF_001050535.1 | Nostocoides_australiense | Bacteria>Bacillati>Actinomycetota>Actinomycetes>Micrococcales>Intrasporangiaceae>Nostocoides | 264 | BN11_RS02225 | 468552..469346 | 99480 | hypothetical-protein-[Tetrasphaera-australiensis] |
| WP_028130945.1 | PIN-EndoRNase | GCF_000421245.1 | Serinicoccus_marinus | Bacteria>Bacillati>Actinomycetota>Actinomycetes>Micrococcales>Ornithinimicrobiaceae>Serinicoccus | 161 | FU792_RS06155 | 1317933..1318418 | 247333 | DUF4411-family-protein-[Serinicoccus-marinus] |
| WP_147915793.1 | PIN-EndoRNase | GCF_008000995.1 | Ruania_zhangjianzhongii | Bacteria>Bacillati>Actinomycetota>Actinomycetes>Micrococcales>Ruaniaceae>Ruania | 109 | FU260_RS03455 | 743242..743571 | 2603206 | hypothetical-protein-[Ruania-zhangjianzhongii] |
| GII23649.1 | PIN-EndoRNase | GCA_016863255.1 | Planosporangium_mesophilum | Bacteria>Bacillati>Actinomycetota>Actinomycetes>Micromonosporales>Micromonosporaceae>Planosporangium | 209 | Pme01_32460 | 158013..158642 | 689768 | hypothetical-protein-Pme01_32460-[Planosporangium-mesophilum] |
| NJC84886.1 | PIN-EndoRNase | GCA_011947235.1 | Planosporangium_mesophilum | Bacteria>Bacillati>Actinomycetota>Actinomycetes>Micromonosporales>Micromonosporaceae>Planosporangium | 209 | HC030_20335 | 8130..8759 | 689768 | NYN-domain-containing-protein-[Planosporangium-mesophilum] |
| WP_168116347.1 | PIN-EndoRNase | GCF_016863255.1 | Planosporangium_mesophilum | Bacteria>Bacillati>Actinomycetota>Actinomycetes>Micromonosporales>Micromonosporaceae>Planosporangium | 209 | Pme01_RS16105 | 158013..158642 | 689768 | NYN-domain-containing-protein-[Planosporangium-mesophilum] |
| WP_312800201.1 | PIN-EndoRNase | GCF_032053705.1 | Corynebacterium_variabile | Bacteria>Bacillati>Actinomycetota>Actinomycetes>Mycobacteriales>Corynebacteriaceae>Corynebacterium | 138 | RTY08_RS06045 | 403387..403803 | 1727 | type-II-toxin-antitoxin-system-VapC-family-toxin-[Corynebacterium-variabile] |
| WP_312898985.1 | PIN-EndoRNase | GCF_032057675.1 | Corynebacterium_variabile | Bacteria>Bacillati>Actinomycetota>Actinomycetes>Mycobacteriales>Corynebacteriaceae>Corynebacterium | 138 | RTZ97_RS05785 | 83224..83640 | 1727 | type-II-toxin-antitoxin-system-VapC-family-toxin-[Corynebacterium-variabile] |
| WP_073883689.1 | PIN-EndoRNase | GCF_900015285.1 | Corynebacterium_variabile | Bacteria>Bacillati>Actinomycetota>Actinomycetes>Mycobacteriales>Corynebacteriaceae>Corynebacterium | 154 | CVAR292_RS03515 | 22356..22820 | 1727 | hypothetical-protein-[Corynebacterium-variabile] |
| WP_244805210.1 | PIN-EndoRNase | GCF_022869005.1 | Corynebacterium_kalidii | Bacteria>Bacillati>Actinomycetota>Actinomycetes>Mycobacteriales>Corynebacteriaceae>Corynebacterium | 154 | MUN33_RS12300 | 338943..339407 | 2931982 | hypothetical-protein-[Corynebacterium-kalidii] |
| WP_158726426.1 | PIN-EndoRNase | GCF_009765205.1 | Tomitella_fengzijianii | Bacteria>Bacillati>Actinomycetota>Actinomycetes>Mycobacteriales>Corynebacteriales_incertae_sedis>Tomitella | 167 | GSQ71_RS04950 | 46085..46588 | 2597660 | DUF4411-family-protein-[Tomitella-fengzijianii] |
| WP_182347796.1 | PIN-EndoRNase | GCF_014121175.1 | Tomitella_gaofuii | Bacteria>Bacillati>Actinomycetota>Actinomycetes>Mycobacteriales>Corynebacteriales_incertae_sedis>Tomitella | 167 | H7F09_RS10450 | 63107..63610 | 2760083 | DUF4411-family-protein-[Tomitella-gaofuii] |
| WP_217999516.1 | PIN-EndoRNase | - | Gordonia_aichiensis | Bacteria>Bacillati>Actinomycetota>Actinomycetes>Mycobacteriales>Gordoniaceae>Gordonia | 96 | GOACH_RS12120 | 81088..81378 | 36820 | hypothetical-protein-[Gordonia-aichiensis] |
| WP_267616416.1 | PIN-EndoRNase | GCF_026616955.1 | Gordonia_bronchialis | Bacteria>Bacillati>Actinomycetota>Actinomycetes>Mycobacteriales>Gordoniaceae>Gordonia | 126 | NOB45_RS12890 | 109304..109684 | 2054 | type-II-toxin-antitoxin-system-VapC-family-toxin-[Gordonia-bronchialis] |
| WP_226512488.1 | PIN-EndoRNase | GCF_020520305.1 | Gordonia_sp._WA4-43 | Bacteria>Bacillati>Actinomycetota>Actinomycetes>Mycobacteriales>Gordoniaceae>Gordonia>unclassified_Gordonia_(in:_high_G+C_Gram-positive_bacteria) | 340 | LEL84_RS18925 | 4166769..4167791 | 2878678 | PIN-domain-containing-protein-[Gordonia-sp.-WA4-43] |
| WP_205874820.1 | PIN-EndoRNase | GCF_017086405.1 | Mycobacterium_gordonae | Bacteria>Bacillati>Actinomycetota>Actinomycetes>Mycobacteriales>Mycobacteriaceae>Mycobacterium | 202 | JX552_RS26905 | 6272451..6273059 | 1778 | PIN-domain-containing-protein-[Mycobacterium-gordonae] |
| AFC52223.1 | PIN-EndoRNase | GCA_000276825.1 | Mycobacterium_paraintracellulare | Bacteria>Bacillati>Actinomycetota>Actinomycetes>Mycobacteriales>Mycobacteriaceae>Mycobacterium>Mycobacterium_avium_complex_(MAC) | 199 | OCQ_07100 | 705632..706231 | 1138383 | hypothetical-protein-OCQ_07100-[Mycobacterium-paraintracellulare] |
| OSC29538.1 | PIN-EndoRNase | GCA_002104735.1 | Mycobacterium_paraintracellulare | Bacteria>Bacillati>Actinomycetota>Actinomycetes>Mycobacteriales>Mycobacteriaceae>Mycobacterium>Mycobacterium_avium_complex_(MAC) | 199 | B8W68_04680 | 139241..139840 | 1138383 | PIN-domain-containing-protein-[Mycobacterium-paraintracellulare] |
| WP_014383821.1 | PIN-EndoRNase | GCF_000276825.1 | Mycobacterium_paraintracellulare | Bacteria>Bacillati>Actinomycetota>Actinomycetes>Mycobacteriales>Mycobacteriaceae>Mycobacterium>Mycobacterium_avium_complex_(MAC) | 199 | B8W68_RS04680 | 139241..139840 | 1138383 | PIN-domain-containing-protein-[Mycobacterium-paraintracellulare] |
| WP_197538264.1 | PIN-EndoRNase | GCF_000328785.1 | Mycobacterium_canetti | Bacteria>Bacillati>Actinomycetota>Actinomycetes>Mycobacteriales>Mycobacteriaceae>Mycobacterium>Mycobacterium_tuberculosis_complex | 135 | BN42_RS05715 | 1223473..1223880 | 78331 | PIN-domain-containing-protein-[Mycobacterium-canettii] |
| WP_062879809.1 | PIN-EndoRNase | GCF_001606235.1 | Mycobacteroides_abscessus | Bacteria>Bacillati>Actinomycetota>Actinomycetes>Mycobacteriales>Mycobacteriaceae>Mycobacteroides | 199 | B5431_RS13010 | 727601..728200 | 36809 | PIN-domain-containing-protein-[Mycobacteroides-abscessus] |
| WP_225506774.1 | PIN-EndoRNase | GCF_020227835.1 | Mycolicibacterium_fortuitum | Bacteria>Bacillati>Actinomycetota>Actinomycetes>Mycobacteriales>Mycobacteriaceae>Mycolicibacterium | 199 | H8Z59_RS27025 | 5483650..5484249 | 1766 | PIN-domain-containing-protein-[Mycolicibacterium-fortuitum] |
| WP_096312281.1 | PIN-EndoRNase | GCF_039534765.1 | Mycolicibacterium_pallens | Bacteria>Bacillati>Actinomycetota>Actinomycetes>Mycobacteriales>Mycobacteriaceae>Mycolicibacterium | 202 | K0O64_RS24890 | 5111894..5112502 | 370524 | MULTISPECIES:-PIN-domain-containing-protein-[Mycobacteriaceae] |
| UHJ57680.1 | PIN-EndoRNase | GCA_021249325.1 | Mycolicibacterium_fortuitum | Bacteria>Bacillati>Actinomycetota>Actinomycetes>Mycobacteriales>Mycobacteriaceae>Mycolicibacterium | 213 | LT337_13405 | 2627614..2628255 | 1766 | hypothetical-protein-LT337_13405-[Mycolicibacterium-fortuitum] |
| EHB57975.1 | PIN-EndoRNase | - | Mycolicibacterium_rhodesiae_JS60 | Bacteria>Bacillati>Actinomycetota>Actinomycetes>Mycobacteriales>Mycobacteriaceae>Mycolicibacterium>Mycolicibacterium_rhodesiae | 202 | MycrhDRAFT_0411 | 401241..401849 | 931627 | hypothetical-protein-MycrhDRAFT_0411-[Mycolicibacterium-rhodesiae-JS60] |
| WP_072804382.1 | PIN-EndoRNase | GCF_001895005.1 | Rhodococcoides_yunnanense | Bacteria>Bacillati>Actinomycetota>Actinomycetes>Mycobacteriales>Nocardiaceae>Rhodococcoides | 128 | RY1_RS14850 | 82319..82705 | 278209 | PIN-domain-containing-protein-[Rhodococcus-yunnanensis] |
| WP_176444451.1 | PIN-EndoRNase | GCF_900188125.1 | Rhodococcoides_kyotonense | Bacteria>Bacillati>Actinomycetota>Actinomycetes>Mycobacteriales>Nocardiaceae>Rhodococcoides | 128 | CHB96_RS22965 | 69045..69431 | 398843 | PIN-domain-containing-protein-[Rhodococcus-kyotonensis] |
| WP_124259268.1 | PIN-EndoRNase | GCF_003844175.1 | Rhodococcus_ruber | Bacteria>Bacillati>Actinomycetota>Actinomycetes>Mycobacteriales>Nocardiaceae>Rhodococcus | 183 | TN91_RS21985 | 2784..3335 | 1830 | PIN-domain-containing-protein-[Rhodococcus-ruber] |
| WP_227735035.1 | PIN-EndoRNase | - | Tessaracoccus_sp._OS52 | Bacteria>Bacillati>Actinomycetota>Actinomycetes>Propionibacteriales>Propionibacteriaceae>Tessaracoccus>unclassified_Tessaracoccus | 97 | LKO27_RS00960 | 222083..222376 | 2886691 | hypothetical-protein-[Tessaracoccus-sp.-OS52] |
| WP_017595715.1 | PIN-EndoRNase | GCF_000341105.1 | Nocardiopsis_potens | Bacteria>Bacillati>Actinomycetota>Actinomycetes>Streptosporangiales>Nocardiopsidaceae>Nocardiopsis | 183 | D459_RS0124850 | 21204..21755 | 1246458 | PIN-domain-containing-protein-[Nocardiopsis-potens] |
| MCH9729508.1 | PIN-EndoRNase | GCA_022600605.1 | Actinomycetes_bacterium | Bacteria>Bacillati>Actinomycetota>Actinomycetes>unclassified_Actinomycetes | 101 | KOU84_07505 | 1..307 | 1883427 | PIN-domain-containing-protein-[Actinomycetia-bacterium] |
| MCL2317099.1 | PIN-EndoRNase | GCA_009785885.1 | Actinomycetes_bacterium | Bacteria>Bacillati>Actinomycetota>Actinomycetes>unclassified_Actinomycetes | 138 | FWC46_08470 | 1888..2304 | 1883427 | PIN-domain-containing-protein-[Actinomycetia-bacterium] |
| MBR3317264.1 | PIN-EndoRNase | GCA_017510785.1 | Atopobiaceae_bacterium | Bacteria>Bacillati>Actinomycetota>Coriobacteriia>Coriobacteriales>Atopobiaceae>unclassified_Atopobiaceae | 172 | IKG21_05540 | 99178..99696 | 2485926 | DUF4411-family-protein-[Atopobiaceae-bacterium] |
| MTV27842.1 | PIN-EndoRNase | GCA_009720505.1 | Profundirhabdus_halotolerans | Bacteria>Bacillati>Actinomycetota>Nitriliruptoria>Nitriliruptorales>Nitriliruptoraceae>Profundirhabdus | 164 | FTX61_20850 | 18469..18963 | 2604685 | DUF4411-family-protein-[Nitriliruptoraceae-bacterium-ZYF776] |
| WP_219975493.1 | PIN-EndoRNase | GCF_019448335.1 | Rubrobacter_xylanophilus | Bacteria>Bacillati>Actinomycetota>Rubrobacteria>Rubrobacterales>Rubrobacteraceae>Rubrobacter | 144 | Rxycam_RS04980 | 972747..973181 | 49319 | type-II-toxin-antitoxin-system-VapC-family-toxin-[Rubrobacter-xylanophilus] |
| QIN77524.1 | PIN-EndoRNase | GCA_011492965.1 | Rubrobacter_marinus | Bacteria>Bacillati>Actinomycetota>Rubrobacteria>Rubrobacterales>Rubrobacteraceae>Rubrobacter | 150 | GBA65_02280 | 460191..460643 | 2653852 | PIN-domain-containing-protein-[Rubrobacter-marinus] |
| WP_166395206.1 | PIN-EndoRNase | GCF_011492965.1 | Rubrobacter_marinus | Bacteria>Bacillati>Actinomycetota>Rubrobacteria>Rubrobacterales>Rubrobacteraceae>Rubrobacter | 150 | GBA65_RS02345 | 460191..460643 | 2653852 | PIN-domain-containing-protein-[Rubrobacter-marinus] |
| WP_219975496.1 | PIN-EndoRNase | GCF_019448335.1 | Rubrobacter_xylanophilus | Bacteria>Bacillati>Actinomycetota>Rubrobacteria>Rubrobacterales>Rubrobacteraceae>Rubrobacter | 208 | Rxycam_RS04995 | 976328..976954 | 49319 | NYN-domain-containing-protein-[Rubrobacter-xylanophilus] |
| MBA3426228.1 | PIN-EndoRNase | GCA_013812245.1 | Rubrobacter_sp. | Bacteria>Bacillati>Actinomycetota>Rubrobacteria>Rubrobacterales>Rubrobacteraceae>Rubrobacter>unclassified_Rubrobacter | 91 | HOU04_17210 | 4604..4879 | 2036010 | hypothetical-protein-[Rubrobacter-sp.] |
| MBX6764705.1 | PIN-EndoRNase | GCA_019690035.1 | Rubrobacteraceae_bacterium | Bacteria>Bacillati>Actinomycetota>Rubrobacteria>Rubrobacterales>Rubrobacteraceae>unclassified_Rubrobacteraceae | 144 | IRY88_13655 | 1252..1686 | 2740537 | type-II-toxin-antitoxin-system-VapC-family-toxin-[Rubrobacteraceae-bacterium] |
| MCP9484918.1 | PIN-EndoRNase | GCA_024330215.1 | Candidatus_Gaiellasilicea_maunaloa | Bacteria>Bacillati>Actinomycetota>Thermoleophilia>Gaiellales>Gaiellaceae>Candidatus_Gaiellasilicea | 153 | MSC30_03580 | 45412..45873 | 2928748 | hypothetical-protein-[Candidatus-Gaiellasilicea-maunaloa] |
| UGS34695.1 | PIN-EndoRNase | GCA_021172045.1 | Capillimicrobium_parvum | Bacteria>Bacillati>Actinomycetota>Thermoleophilia>Solirubrobacterales>Capillimicrobiaceae>Capillimicrobium | 138 | DSM104329_01076 | 1080188..1080604 | 2884022 | Ribonuclease-VapC38-[Capillimicrobium-parvum] |
| WP_259314359.1 | PIN-EndoRNase | GCF_021172045.1 | Capillimicrobium_parvum | Bacteria>Bacillati>Actinomycetota>Thermoleophilia>Solirubrobacterales>Capillimicrobiaceae>Capillimicrobium | 138 | DSM104329_RS05340 | 1080188..1080604 | 2884022 | PIN-domain-containing-protein-[Capillimicrobium-parvum] |
| TXH45587.1 | PIN-EndoRNase | GCA_008012465.1 | Actinomycetota_bacterium | Bacteria>Bacillati>Actinomycetota>unclassified_Actinomycetota | 127 | E6Q90_00230 | 24057..24440 | 2900548 | type-II-toxin-antitoxin-system-VapC-family-toxin-[Actinomycetota-bacterium] |
| TXH45534.1 | PIN-EndoRNase | GCA_008012465.1 | Actinomycetota_bacterium | Bacteria>Bacillati>Actinomycetota>unclassified_Actinomycetota | 137 | E6Q90_00250 | 29089..29502 | 2900548 | PIN-domain-nuclease-[Actinomycetota-bacterium] |
| NLO26756.1 | PIN-EndoRNase | GCA_012523095.1 | Actinomycetota_bacterium | Bacteria>Bacillati>Actinomycetota>unclassified_Actinomycetota | 146 | GX113_01030 | 8665..9105 | 2900548 | type-II-toxin-antitoxin-system-VapC-family-toxin-[Actinomycetia-bacterium] |
| MBW3642894.1 | PIN-EndoRNase | GCA_019348235.1 | Actinomycetota_bacterium | Bacteria>Bacillati>Actinomycetota>unclassified_Actinomycetota | 159 | KY447_08285 | 5651..6130 | 2900548 | hypothetical-protein-[Actinomycetia-bacterium] |

| Protein-ID | Domain | Assembly-ID | Strain | Taxonomy | Length | Locus-Tag | Genomic-coordinates | TaxID | Fasta-defline |
|---|---|---|---|---|---|---|---|---|---|
| MBC7252725.1 | PIN-EndoRNase | GCA_014360685.1 | Actinomycetota_bacterium | Bacteria>Bacillati>Actinomycetota>unclassified_Actinomycetota | 230 | H5T72_01975 | 82256..82948 | 2900548 | NYN-domain-containing-protein-[Actinomycetia-bacterium] |
| NLO26758.1 | PIN-EndoRNase | GCA_012523095.1 | Actinomycetota_bacterium | Bacteria>Bacillati>Actinomycetota>unclassified_Actinomycetota | 243 | GX113_01040 | 12012..12743 | 2900548 | NYN-domain-containing-protein-[Actinomycetia-bacterium] |
| NCO42946.1 | PIN-EndoRNase | GCA_009992845.1 | Armatimonadota_bacterium | Bacteria>Bacillati>Armatimonadota>unclassified_Armatimonadota | 126 | GW892_30115 | 1179..1559 | 2033014 | hypothetical-protein-[Armatimonadota-bacterium] |
| PIY44269.1 | PIN-EndoRNase | GCA_002784925.1 | Armatimonadetes_bacterium_CG_4_10_14_3_um_filter_59_10 | Bacteria>Bacillati>Armatimonadota>unclassified_Armatimonadota | 126 | COZ05_08655 | 2964..3344 | 1973909 | hypothetical-protein-COZ05_08655-[Armatimonadetes-bacterium-CG_4_10_14_3_um_filter_59_10] |
| MBI3912158.1 | PIN-EndoRNase | GCA_016199525.1 | Armatimonadota_bacterium | Bacteria>Bacillati>Armatimonadota>unclassified_Armatimonadota | 133 | HY320_14660 | 1047..1448 | 2033014 | DUF5615-family-PIN-like-protein-[Armatimonadota-bacterium] |
| WP_259548273.1 | PIN-EndoRNase | - | Heyndrickxia_oleronia | Bacteria>Bacillati>Bacillota>Bacilli>Bacillales>Bacillaceae>Heyndrickxia | 346 | KXF32_RS03860 | 731560..732600 | 38875 | PIN-domain-containing-protein-[Heyndrickxia-oleronia] |
| WP_243459536.1 | PIN-EndoRNase | GCF_017497975.1 | Metabacillus_bambusae | Bacteria>Bacillati>Bacillota>Bacilli>Bacillales>Bacillaceae>Metabacillus | 288 | I7822_RS17990 | 34046..34912 | 2795218 | hypothetical-protein-[Metabacillus-bambusae] |
| MBO1513533.1 | PIN-EndoRNase | GCA_017497975.1 | Metabacillus_bambusae | Bacteria>Bacillati>Bacillota>Bacilli>Bacillales>Bacillaceae>Metabacillus | 306 | I7822_17980 | 33992..34912 | 2795218 | DUF4935-domain-containing-protein-[Metabacillus-bambusae] |
| HHW19262.1 | PIN-EndoRNase | GCA_012840505.1 | Candidatus_Fermentithermobacillaceae_bacterium | Bacteria>Bacillati>Bacillota>Candidatus_Fermentithermobacillia>Candidatus_Fermentithermobacillales>Candidatus_Fermentithermobacillaceae>unclassified_Candidatus_Fermentithermobacillaceae | 123 | GXX30_10290 | 61567..61938 | 3085339 | DUF5615-family-PIN-like-protein-[Bacillota-bacterium] |
| MBE3519593.1 | PIN-EndoRNase | GCA_014896535.1 | Candidatus_Fermentithermobacillaceae_bacterium | Bacteria>Bacillati>Bacillota>Candidatus_Fermentithermobacillia>Candidatus_Fermentithermobacillales>Candidatus_Fermentithermobacillaceae>unclassified_Candidatus_Fermentithermobacillaceae | 131 | IMW97_04745 | 105731..106126 | 3085339 | type-II-toxin-antitoxin-system-VapC-family-toxin-[Bacillota-bacterium] |
| HHY35074.1 | PIN-EndoRNase | GCA_012842495.1 | Candidatus_Fermentithermobacillaceae_bacterium | Bacteria>Bacillati>Bacillota>Candidatus_Fermentithermobacillia>Candidatus_Fermentithermobacillales>Candidatus_Fermentithermobacillaceae>unclassified_Candidatus_Fermentithermobacillaceae | 164 | GX510_05505 | 14248..14742 | 3085339 | DUF4411-family-protein-[Bacillota-bacterium] |
| WP_173680571.1 | PIN-EndoRNase | GCF_014230485.1 | Clostridium_tetanomorphum | Bacteria>Bacillati>Bacillota>Clostridia>Eubacteriales>Clostridiaceae>Clostridium | 162 | FHX96_RS20150 | 4171978..4172466 | 1553 | hypothetical-protein-[Clostridium-tetanomorphum] |
| WP_270807569.1 | PIN-EndoRNase | GCF_027723425.1 | Clostridium_fessum | Bacteria>Bacillati>Bacillota>Clostridia>Eubacteriales>Clostridiaceae>Clostridium | 176 | PG965_RS01485 | 325017..325547 | 2126740 | hypothetical-protein-[Clostridium-fessum] |
| WP_270487110.1 | PIN-EndoRNase | GCF_027681635.1 | Clostridium_fessum | Bacteria>Bacillati>Bacillota>Clostridia>Eubacteriales>Clostridiaceae>Clostridium | 177 | PGX92_RS02725 | 608678..609211 | 2126740 | hypothetical-protein-[Clostridium-fessum] |
| WP_231016618.1 | PIN-EndoRNase | GCF_021043395.1 | Clostridium_guangxiense | Bacteria>Bacillati>Bacillota>Clostridia>Eubacteriales>Clostridiaceae>Clostridium | 191 | LQK79_RS11215 | 16220..16795 | 1662055 | hypothetical-protein-[Clostridium-guangxiense] |
| WP_270487122.1 | PIN-EndoRNase | GCF_027681635.1 | Clostridium_fessum | Bacteria>Bacillati>Bacillota>Clostridia>Eubacteriales>Clostridiaceae>Clostridium | 218 | PG965_RS01530 | 339354..340010 | 2126740 | PIN-domain-containing-protein-[Clostridium-fessum] |
| CDD75190.1 | PIN-EndoRNase | GCA_000436175.1 | Clostridium_sp._CAG:62 | Bacteria>Bacillati>Bacillota>Clostridia>Eubacteriales>Clostridiaceae>Clostridium>unclassified_Clostridium | 164 | BN737_01377 | 74442..74936 | 1262828 | putative-uncharacterized-protein-[Clostridium-sp.-CAG:62] |
| WP_118744887.1 | PIN-EndoRNase | GCF_003480745.1 | Clostridium_sp._AM27-31LB | Bacteria>Bacillati>Bacillota>Clostridia>Eubacteriales>Clostridiaceae>Clostridium>unclassified_Clostridium | 164 | DW721_RS03895 | 66226..66720 | 2293026 | hypothetical-protein-[Clostridium-sp.-AM27-31LB] |
| CDD75183.1 | PIN-EndoRNase | GCA_000436175.1 | Clostridium_sp._CAG:62 | Bacteria>Bacillati>Bacillota>Clostridia>Eubacteriales>Clostridiaceae>Clostridium>unclassified_Clostridium | 218 | BN737_01370 | 66428..67084 | 1262828 | putative-uncharacterized-protein-[Clostridium-sp.-CAG:62] |
| WP_092487586.1 | PIN-EndoRNase | GCF_900115975.1 | Desulfoscipio_geothermicus | Bacteria>Bacillati>Bacillota>Clostridia>Eubacteriales>Desulfallaceae>Desulfoscipio | 161 | BM299_RS17675 | 5312..5797 | 39060 | hypothetical-protein-[Desulfoscipio-geothermicus] |
| WP_066668120.1 | PIN-EndoRNase | GCF_001655685.1 | Desulfotomaculum_copahuensis | Bacteria>Bacillati>Bacillota>Clostridia>Eubacteriales>Desulfotomaculaceae>Desulfotomaculum | 129 | A6M21_RS09380 | 58631..59020 | 1838280 | type-II-toxin-antitoxin-system-VapC-family-toxin-[Desulfotomaculum-copahuensis] |
| WP_227612031.1 | PIN-EndoRNase | GCF_020686905.1 | Faecalibacterium | Bacteria>Bacillati>Bacillota>Clostridia>Eubacteriales>Oscillospiraceae | 145 | LKD27_RS12635 | 20350..20787 | 216851 | DUF188-domain-containing-protein-[Faecalibacterium-sp.-CLA-AA-H283] |
| MCI8835834.1 | PIN-EndoRNase | GCA_022795995.1 | Ruminococcus_sp. | Bacteria>Bacillati>Bacillota>Clostridia>Eubacteriales>Oscillospiraceae>Ruminococcus>unclassified_Ruminococcus | 179 | HFH24_10745 | 40612..41151 | 41978 | hypothetical-protein-[Ruminococcus-sp.] |
| WP_072867393.1 | PIN-EndoRNase | GCF_900142025.1 | Desulfofundulus_thermosubterraneus | Bacteria>Bacillati>Bacillota>Clostridia>Eubacteriales>Peptococcaceae>Desulfofundulus | 129 | BUA65_RS03530 | 6882..7271 | 348840 | type-II-toxin-antitoxin-system-VapC-family-toxin-[Desulfofundulus-thermosubterraneus] |
| WP_073162466.1 | PIN-EndoRNase | GCF_900129285.1 | Desulfofundulus_australicus | Bacteria>Bacillati>Bacillota>Clostridia>Eubacteriales>Peptococcaceae>Desulfofundulus | 129 | BUB44_RS00900 | 190480..190869 | 1566 | type-II-toxin-antitoxin-system-VapC-family-toxin-[Desulfofundulus-australicus] |
| WP_027356908.1 | PIN-EndoRNase | GCF_000686645.1 | Desulfofundulus_thermocisternus | Bacteria>Bacillati>Bacillota>Clostridia>Eubacteriales>Peptococcaceae>Desulfofundulus | 137 | BR83_RS0109625 | 54525..54938 | 42471 | putative-toxin-antitoxin-system-toxin-component,-PIN-family-[Desulfofundulus-thermocisternus] |
| AEG13740.1 | PIN-EndoRNase | GCA_000214705.1 | Desulfofundulus_kuznetsovii_DSM_6115 | Bacteria>Bacillati>Bacillota>Clostridia>Eubacteriales>Peptococcaceae>Desulfofundulus>Desulfofundulus_kuznetsovii | 129 | Desku_0092 | 89328..89717 | 760568 | PilT-protein-domain-protein-[Desulfofundulus-kuznetsovii-DSM-6115] |
| NHM26611.1 | PIN-EndoRNase | GCA_011393055.1 | Desulfofundulus_sp._TPOSR | Bacteria>Bacillati>Bacillota>Clostridia>Eubacteriales>Peptococcaceae>Desulfofundulus>unclassified_Desulfofundulus | 129 | G7K71_06330 | 1303651..1304040 | 2714340 | type-II-toxin-antitoxin-system-VapC-family-toxin-[Desulfofundulus-sp.-TPOSR] |
| WP_166344336.1 | PIN-EndoRNase | GCF_011393055.1 | Desulfofundulus_sp._TPOSR | Bacteria>Bacillati>Bacillota>Clostridia>Eubacteriales>Peptococcaceae>Desulfofundulus>unclassified_Desulfofundulus | 129 | G7K71_RS06325 | 1303651..1304040 | 2714340 | type-II-toxin-antitoxin-system-VapC-family-toxin-[Desulfofundulus-sp.-TPOSR] |
| NLW38081.1 | PIN-EndoRNase | GCA_012728715.1 | Peptococcaceae_bacterium | Bacteria>Bacillati>Bacillota>Clostridia>Eubacteriales>Peptococcaceae>unclassified_Peptococcaceae | 140 | GXY90_05850 | 2521..2943 | 2052179 | PIN-domain-containing-protein-[Peptococcaceae-bacterium] |
| WP_173865461.1 | PIN-EndoRNase | GCF_013300965.1 | Lachnospiraceae | Bacteria>Bacillati>Bacillota>Clostridia>Lachnospirales | 164 | HFN12_RS00380 | 73945..74439 | 186803 | hypothetical-protein-[Faecalicatena-fissicatena] |
| RGI65886.1 | PIN-EndoRNase | GCA_003436035.1 | Agathobacter_rectalis | Bacteria>Bacillati>Bacillota>Clostridia>Lachnospirales>Lachnospiraceae>Agathobacter | 164 | DXD95_13295 | 6500..6994 | 39491 | hypothetical-protein-DXD95_13295-[[Eubacterium] |
| WP_186847748.1 | PIN-EndoRNase | GCF_014287735.1 | Coprococcus_hominis_(ex_Liu_et_al._2022) | Bacteria>Bacillati>Bacillota>Clostridia>Lachnospirales>Lachnospiraceae>Coprococcus | 316 | H8S09_RS09165 | 19867..20817 | 2763039 | PIN-domain-containing-protein-[Coprococcus-sp.-NSJ-10] |
| MCI7121460.1 | PIN-EndoRNase | GCA_022775325.1 | Mediterraneibacter_gnavus | Bacteria>Bacillati>Bacillota>Clostridia>Lachnospirales>Lachnospiraceae>Mediterraneibacter | 164 | MR941_06730 | 14999..15493 | 33038 | hypothetical-protein-[[Ruminococcus] |
| WP_270645283.1 | PIN-EndoRNase | GCF_027692785.1 | Merdimonas_faecis | Bacteria>Bacillati>Bacillota>Clostridia>Lachnospirales>Lachnospiraceae>Merdimonas | 173 | PG729_RS15110 | 21049..21570 | 1653435 | hypothetical-protein-[Merdimonas-faecis] |
| HCI18464.1 | PIN-EndoRNase | GCA_003521585.1 | Lachnospiraceae_bacterium | Bacteria>Bacillati>Bacillota>Clostridia>Lachnospirales>Lachnospiraceae>unclassified_Lachnospiraceae | 61 | DFH32_07600 | 1..185 | 1898203 | hypothetical-protein-[Lachnospiraceae-bacterium] |
| MBR4345191.1 | PIN-EndoRNase | GCA_017542245.1 | Lachnospiraceae_bacterium | Bacteria>Bacillati>Bacillota>Clostridia>Lachnospirales>Lachnospiraceae>unclassified_Lachnospiraceae | 137 | IKP88_21215 | 54900..55313 | 1898203 | putative-toxin-antitoxin-system-toxin-component,-PIN-family-[Lachnospiraceae-bacterium] |
| HCX41714.1 | PIN-EndoRNase | GCA_003539645.1 | Lachnospiraceae_bacterium | Bacteria>Bacillati>Bacillota>Clostridia>Lachnospirales>Lachnospiraceae>unclassified_Lachnospiraceae | 157 | DHH50_05150 | 197405..197878 | 1898203 | YaiI/YqxD-family-protein-[Lachnospiraceae-bacterium] |
| MBD5489948.1 | PIN-EndoRNase | GCA_014804485.1 | Lachnospiraceae_bacterium | Bacteria>Bacillati>Bacillota>Clostridia>Lachnospirales>Lachnospiraceae>unclassified_Lachnospiraceae | 168 | HDR13_14350 | 30381..30887 | 1898203 | PIN-domain-containing-protein-[Lachnospiraceae-bacterium] |
| MBQ3415898.1 | PIN-EndoRNase | GCA_017404025.1 | Clostridia_bacterium | Bacteria>Bacillati>Bacillota>Clostridia>unclassified_Clostridia | 162 | IJH39_11320 | 5232..5720 | 2044939 | hypothetical-protein-[Clostridia-bacterium] |
| MBE6098025.1 | PIN-EndoRNase | GCA_015058685.1 | Schwartzia_succinivorans | Bacteria>Bacillati>Bacillota>Negativicutes>Selenomonadales>Selenomonadaceae>Schwartzia | 129 | E7198_09555 | 56762..57151 | 55507 | type-II-toxin-antitoxin-system-VapC-family-toxin-[Schwartzia-succinivorans] |
| WP_273038295.1 | PIN-EndoRNase | GCF_015058685.1 | Schwartzia_succinivorans | Bacteria>Bacillati>Bacillota>Negativicutes>Selenomonadales>Selenomonadaceae>Schwartzia | 129 | E7198_RS09655 | 56762..57151 | 55507 | PIN-domain-containing-protein-[Schwartzia-succinivorans] |
| WP_273079792.1 | PIN-EndoRNase | GCF_905371945.1 | Selenomonas_felix | Bacteria>Bacillati>Bacillota>Negativicutes>Selenomonadales>Selenomonadaceae>Selenomonas | 163 | NW177_RS10665 | 8838..9329 | 1944634 | hypothetical-protein-[Selenomonas-felix] |
| WP_273698071.1 | PIN-EndoRNase | GCF_905371815.1 | Selenomonas_noxia | Bacteria>Bacillati>Bacillota>Negativicutes>Selenomonadales>Selenomonadaceae>Selenomonas | 163 | NW174_RS07830 | 33216..33707 | 135083 | hypothetical-protein-[Selenomonas-noxia] |
| SEH31375.1 | PIN-EndoRNase | GCA_900108415.1 | Selenomonas_sp._KH1T6 | Bacteria>Bacillati>Bacillota>Negativicutes>Selenomonadales>Selenomonadaceae>Selenomonas>unclassified_Selenomonas | 162 | SAMN05216583_1178 | 6188..6676 | 3158784 | hypothetical-protein-SAMN05216583_1178-[Selenomonas-ruminantium] |
| MBR3457115.1 | PIN-EndoRNase | GCA_017509225.1 | Selenomonadaceae_bacterium | Bacteria>Bacillati>Bacillota>Negativicutes>Selenomonadales>Selenomonadaceae>unclassified_Selenomonadaceae | 152 | IKH16_03030 | 3330..3788 | 2652289 | hypothetical-protein-[Selenomonadaceae-bacterium] |
| MCL6474637.1 | PIN-EndoRNase | GCA_023511455.1 | Bacillota_bacterium | Bacteria>Bacillati>Bacillota>unclassified_Bacillota | 87 | K6U75_06245 | 15211..15474 | 1879010 | type-II-toxin-antitoxin-system-VapC-family-toxin-[Bacillota-bacterium] |
| MBK9748866.1 | PIN-EndoRNase | GCA_016717205.1 | Candidatus_Flexifilum_breve | Bacteria>Bacillati>Chloroflexota>Anaerolineae>Aggregatilineales>Candidatus_Flexifilaceae>Candidatus_Flexifilum | 111 | IPO91_19090 | 82073..82408 | 3140694 | DUF5615-family-PIN-like-protein-[Chloroflexota-bacterium] |
| MBE2269144.1 | PIN-EndoRNase | GCA_014879865.1 | Anaerolinea_sp. | Bacteria>Bacillati>Chloroflexota>Anaerolineae>Anaerolineales>Anaerolineaceae>Anaerolinea>unclassified_Anaerolinea | 111 | IAE80_13010 | 6281..6616 | 1872519 | DUF5615-family-PIN-like-protein-[Anaerolinea-sp.] |
| WP_201777307.1 | PIN-EndoRNase | - | Flexilinea_flocculi | Bacteria>Bacillati>Chloroflexota>Anaerolineae>Anaerolineales>Anaerolineaceae>Flexilinea | 92 | ATC1_RS15000 | 815529..815807 | 1678840 | NYN-domain-containing-protein-[Flexilinea-flocculi] |
| WP_201777265.1 | PIN-EndoRNase | GCF_001192795.1 | Flexilinea_flocculi | Bacteria>Bacillati>Chloroflexota>Anaerolineae>Anaerolineales>Anaerolineaceae>Flexilinea | 115 | ATC1_RS14995 | 815163..815510 | 1678840 | hypothetical-protein-[Flexilinea-flocculi] |
| MCB9078009.1 | PIN-EndoRNase | GCA_020634955.1 | Anaerolineaceae_bacterium | Bacteria>Bacillati>Chloroflexota>Anaerolineae>Anaerolineales>Anaerolineaceae>unclassified_Anaerolineaceae | 105 | H6631_10490 | 6251..6568 | 2024896 | DUF5615-family-PIN-like-protein-[Anaerolineaceae-bacterium] |
| MBQ6343451.1 | PIN-EndoRNase | GCA_017432985.1 | Anaerolineaceae_bacterium | Bacteria>Bacillati>Chloroflexota>Anaerolineae>Anaerolineales>Anaerolineaceae>unclassified_Anaerolineaceae | 135 | IJI41_10030 | 8478..8885 | 2024896 | PIN-domain-containing-protein-[Anaerolineaceae-bacterium] |
| MCB9452418.1 | PIN-EndoRNase | GCA_020634115.1 | Anaerolineaceae_bacterium | Bacteria>Bacillati>Chloroflexota>Anaerolineae>Anaerolineales>Anaerolineaceae>unclassified_Anaerolineaceae | 142 | H6672_13355 | 121715..122143 | 2024896 | type-II-toxin-antitoxin-system-VapC-family-toxin-[Anaerolineaceae-bacterium] |
| TEU13002.1 | PIN-EndoRNase | GCA_004377365.1 | Anaerolineales_bacterium | Bacteria>Bacillati>Chloroflexota>Anaerolineae>Anaerolineales>unclassified_Anaerolineales | 123 | E3J21_19775 | 9566..9937 | 2073117 | type-II-toxin-antitoxin-system-VapC-family-toxin-[Anaerolineales-bacterium] |
| MBE7511261.1 | PIN-EndoRNase | GCA_015075625.1 | Anaerolineales_bacterium | Bacteria>Bacillati>Chloroflexota>Anaerolineae>Anaerolineales>unclassified_Anaerolineales | 140 | HS103_00395 | 90104..90526 | 2073117 | type-II-toxin-antitoxin-system-VapC-family-toxin-[Anaerolineales-bacterium] |
| MCC6958029.1 | PIN-EndoRNase | GCA_020853835.1 | Anaerolineales_bacterium | Bacteria>Bacillati>Chloroflexota>Anaerolineae>Anaerolineales>unclassified_Anaerolineales | 208 | IT316_14610 | 10397..11023 | 2073117 | NYN-domain-containing-protein-[Anaerolineales-bacterium] |
| MCK6580688.1 | PIN-EndoRNase | GCA_023151025.1 | Anaerolineae_bacterium | Bacteria>Bacillati>Chloroflexota>Anaerolineae>unclassified_Anaerolineae | 111 | L6Q98_21560 | 24971..25306 | 2052143 | DUF5615-family-PIN-like-protein-[Anaerolineae-bacterium] |
| MBL7200855.1 | PIN-EndoRNase | GCA_016783145.1 | Anaerolineae_bacterium | Bacteria>Bacillati>Chloroflexota>Anaerolineae>unclassified_Anaerolineae | 134 | ISS56_11945 | 1540..1944 | 2052143 | DUF3368-domain-containing-protein-[Anaerolineae-bacterium] |
| NPV68432.1 | PIN-EndoRNase | GCA_013178165.1 | Anaerolineae_bacterium | Bacteria>Bacillati>Chloroflexota>Anaerolineae>unclassified_Anaerolineae | 139 | HPY64_14915 | 28053..28472 | 2052143 | PIN-domain-containing-protein-[Anaerolineae-bacterium] |
| MCL4868350.1 | PIN-EndoRNase | GCA_023384415.1 | Anaerolineae_bacterium | Bacteria>Bacillati>Chloroflexota>Anaerolineae>unclassified_Anaerolineae | 143 | KJ063_05230 | 198149..198580 | 2052143 | type-II-toxin-antitoxin-system-VapC-family-toxin-[Anaerolineae-bacterium] |
| MCS7282893.1 | PIN-EndoRNase | GCA_025062415.1 | Anaerolineae_bacterium | Bacteria>Bacillati>Chloroflexota>Anaerolineae>unclassified_Anaerolineae | 146 | NZ769_06040 | 4541..4981 | 2052143 | putative-toxin-antitoxin-system-toxin-component,-PIN-family-[Anaerolineae-bacterium] |
| MCL4237390.1 | PIN-EndoRNase | GCA_023382715.1 | Anaerolineae_bacterium | Bacteria>Bacillati>Chloroflexota>Anaerolineae>unclassified_Anaerolineae | 150 | KJ047_03995 | 73215..73667 | 2052143 | PIN-domain-containing-protein-[Anaerolineae-bacterium] |
| MBN1139683.1 | PIN-EndoRNase | GCA_016926595.1 | Anaerolineae_bacterium | Bacteria>Bacillati>Chloroflexota>Anaerolineae>unclassified_Anaerolineae | 150 | JXM73_24130 | 14870..15322 | 2052143 | type-II-toxin-antitoxin-system-VapC-family-toxin-[Anaerolineae-bacterium] |
| MCR4407535.1 | PIN-EndoRNase | GCA_024653325.1 | Anaerolineae_bacterium | Bacteria>Bacillati>Chloroflexota>Anaerolineae>unclassified_Anaerolineae | 155 | NUW24_11550 | 11389..11856 | 2052143 | PIN-domain-containing-protein-[Anaerolineae-bacterium] |
| MCZ2114538.1 | PIN-EndoRNase | GCA_026954395.1 | Anaerolineae_bacterium | Bacteria>Bacillati>Chloroflexota>Anaerolineae>unclassified_Anaerolineae | 164 | LC131_12005 | 7667..8161 | 2052143 | MAG:-PIN-domain-containing-protein-[Anaerolineae-bacterium] |
| NPV86422.1 | PIN-EndoRNase | GCA_013178295.1 | Anaerolineae_bacterium | Bacteria>Bacillati>Chloroflexota>Anaerolineae>unclassified_Anaerolineae | 208 | HPY45_10495 | 180995..181621 | 2052143 | NYN-domain-containing-protein-[Anaerolineae-bacterium] |
| MCL4264801.1 | PIN-EndoRNase | GCA_023382655.1 | Anaerolineae_bacterium | Bacteria>Bacillati>Chloroflexota>Anaerolineae>unclassified_Anaerolineae | 355 | KJ069_16380 | 139386..140453 | 2052143 | NYN-domain-containing-protein-[Anaerolineae-bacterium] |
| MCZ7570816.1 | PIN-EndoRNase | GCA_027493995.1 | Ardenticatenaceae_bacterium | Bacteria>Bacillati>Chloroflexota>Ardenticatenia>Ardenticatenales>Ardenticatenaceae>unclassified_Ardenticatenaceae | 142 | M5U01_19830 | 4783054..4783482 | 2614247 | MAG:-putative-toxin-antitoxin-system-toxin-component,-PIN-family-[Ardenticatenaceae-bacterium] |
| MBK8988273.1 | PIN-EndoRNase | GCA_016716065.1 | Candidatus_Leptovillus_gracilis | Bacteria>Bacillati>Chloroflexota>Ardenticatenia>Candidatus_Promineifilales>Candidatus_Promineifilaceae>Candidatus_Leptovillus | 137 | IPM39_19765 | 143482..143895 | 3140690 | putative-toxin-antitoxin-system-toxin-component,-PIN-family-[Chloroflexota-bacterium] |
| MBK7180416.1 | PIN-EndoRNase | GCA_016707865.1 | Candidatus_Leptovillus_affinis | Bacteria>Bacillati>Chloroflexota>Ardenticatenia>Candidatus_Promineifilales>Candidatus_Promineifilaceae>Candidatus_Leptovillus | 163 | IPH82_25050 | 81687..82178 | 3140691 | DUF3368-domain-containing-protein-[Chloroflexota-bacterium] |
| MBK8988275.1 | PIN-EndoRNase | GCA_016716065.1 | Candidatus_Leptovillus_gracilis | Bacteria>Bacillati>Chloroflexota>Ardenticatenia>Candidatus_Promineifilales>Candidatus_Promineifilaceae>Candidatus_Leptovillus | 208 | IPM39_19775 | 144239..144865 | 3140690 | type-II-toxin-antitoxin-system-VapC-family-toxin-[Chloroflexota-bacterium] |
| MCC6169406.1 | PIN-EndoRNase | GCA_020845455.1 | Caldilineaceae_bacterium | Bacteria>Bacillati>Chloroflexota>Caldilineae>Caldilineales>Caldilineaceae>unclassified_Caldilineaceae | 164 | IT329_19450 | 79..573 | 2614250 | type-II-toxin-antitoxin-system-VapC-family-toxin-[Caldilineaceae-bacterium] |
| MBP7961437.1 | PIN-EndoRNase | GCA_018003515.1 | Caldilineaceae_bacterium | Bacteria>Bacillati>Chloroflexota>Caldilineae>Caldilineales>Caldilineaceae>unclassified_Caldilineaceae | 170 | KA003_05665 | 14839..15351 | 2614250 | hypothetical-protein-[Caldilineaceae-bacterium] |
| MDE0079140.1 | PIN-EndoRNase | GCA_028819025.1 | Caldilineaceae_bacterium | Bacteria>Bacillati>Chloroflexota>Caldilineae>Caldilineales>Caldilineaceae>unclassified_Caldilineaceae | 206 | OXO50_16585 | 7089..7709 | 2614250 | MAG:-NYN-domain-containing-protein-[Caldilineaceae-bacterium] |
| MCY3902334.1 | PIN-EndoRNase | GCA_026707125.1 | Caldilineaceae_bacterium | Bacteria>Bacillati>Chloroflexota>Caldilineae>Caldilineales>Caldilineaceae>unclassified_Caldilineaceae | 243 | OXF76_04140 | 56571..57302 | 2614250 | MAG:-NYN-domain-containing-protein-[Caldilineaceae-bacterium] |
| NJN66670.1 | PIN-EndoRNase | GCA_012032475.1 | Chloroflexaceae_bacterium | Bacteria>Bacillati>Chloroflexota>Chloroflexia>Chloroflexales>Chloroflexineae>Chloroflexaceae>unclassified_Chloroflexaceae | 154 | HC884_08085 | 892..1356 | 2562117 | type-II-toxin-antitoxin-system-VapC-family-toxin-[Chloroflexaceae-bacterium] |
| WP_097651204.1 | PIN-EndoRNase | GCF_002532075.1 | Candidatus_Chloroploca_asiatica | Bacteria>Bacillati>Chloroflexota>Chloroflexia>Chloroflexales>Chloroflexineae>Oscillochloridaceae>Candidatus_Chloroploca | 150 | A9Q02_RS04710 | 66661..67113 | 1506545 | type-II-toxin-antitoxin-system-VapC-family-toxin-[Candidatus-Chloroploca-asiatica] |
| WP_141508709.1 | PIN-EndoRNase | GCF_002532075.1 | Candidatus_Chloroploca_asiatica | Bacteria>Bacillati>Chloroflexota>Chloroflexia>Chloroflexales>Chloroflexineae>Oscillochloridaceae>Candidatus_Chloroploca | 155 | A9Q02_RS04725 | 69824..70291 | 1506545 | type-II-toxin-antitoxin-system-VapC-family-toxin-[Candidatus-Chloroploca-asiatica] |
| MYD51868.1 | PIN-EndoRNase | GCA_009841605.1 | Dehalococcoidia_bacterium | Bacteria>Bacillati>Chloroflexota>Dehalococcoidia>unclassified_Dehalococcoidia | 139 | F4W93_10355 | 49629..50048 | 2026734 | hypothetical-protein-[Dehalococcoidia-bacterium] |
| MBM3143890.1 | PIN-EndoRNase | GCA_016865865.1 | Chloroflexota_bacterium | Bacteria>Bacillati>Chloroflexota>unclassified_Chloroflexota | 93 | FJ010_02750 | 21137..21418 | 2026724 | NYN-domain-containing-protein-[Chloroflexota-bacterium] |
| NWF79205.1 | PIN-EndoRNase | GCA_013388805.1 | Chloroflexota_bacterium | Bacteria>Bacillati>Chloroflexota>unclassified_Chloroflexota | 104 | HXY37_04045 | 13194..13508 | 2026724 | DUF5615-family-PIN-like-protein-[Chloroflexota-bacterium] |
| MCA0453854.1 | PIN-EndoRNase | GCA_020161935.1 | Chloroflexota_bacterium | Bacteria>Bacillati>Chloroflexota>unclassified_Chloroflexota | 111 | LCI00_07770 | 56926..57261 | 2026724 | DUF5615-family-PIN-like-protein-[Chloroflexota-bacterium] |
| MBC8448147.1 | PIN-EndoRNase | GCA_014383605.1 | Chloroflexota_bacterium | Bacteria>Bacillati>Chloroflexota>unclassified_Chloroflexota | 117 | H8D78_10380 | 14088..14441 | 2026724 | DUF5615-family-PIN-like-protein-[Chloroflexota-bacterium] |
| MCY3958690.1 | PIN-EndoRNase | GCA_026707585.1 | Chloroflexota_bacterium | Bacteria>Bacillati>Chloroflexota>unclassified_Chloroflexota | 118 | OXG65_10385 | 13695..14051 | 2026724 | MAG:-DUF5615-family-PIN-like-protein-[Chloroflexi-bacterium] |
| MDE2669828.1 | PIN-EndoRNase | GCA_028875515.1 | Chloroflexota_bacterium | Bacteria>Bacillati>Chloroflexota>unclassified_Chloroflexota | 121 | OXI51_09275 | 51904..52269 | 2026724 | MAG:-DUF5615-family-PIN-like-protein-[Chloroflexota-bacterium] |
| MBI2958635.1 | PIN-EndoRNase | GCA_016189825.1 | Chloroflexota_bacterium | Bacteria>Bacillati>Chloroflexota>unclassified_Chloroflexota | 127 | HYY32_07330 | 495..878 | 2026724 | DUF5615-family-PIN-like-protein-[Chloroflexota-bacterium] |
| MBI4295303.1 | PIN-EndoRNase | GCA_016210605.1 | Chloroflexota_bacterium | Bacteria>Bacillati>Chloroflexota>unclassified_Chloroflexota | 127 | HY669_03960 | 8804..9187 | 2026724 | DUF5615-family-PIN-like-protein-[Chloroflexota-bacterium] |
| MCY4474613.1 | PIN-EndoRNase | GCA_026712855.1 | Chloroflexota_bacterium | Bacteria>Bacillati>Chloroflexota>unclassified_Chloroflexota | 130 | OXC83_04170 | 176789..177181 | 2026724 | MAG:-type-II-toxin-antitoxin-system-VapC-family-toxin-[Chloroflexi-bacterium] |
| MCX6066947.1 | PIN-EndoRNase | GCA_026389995.1 | Chloroflexota_bacterium | Bacteria>Bacillati>Chloroflexota>unclassified_Chloroflexota | 137 | NT121_14525 | 7689..8102 | 2026724 | MAG:-type-II-toxin-antitoxin-system-VapC-family-toxin-[Chloroflexi-bacterium] |
| MCC6174342.1 | PIN-EndoRNase | GCA_020845495.1 | Chloroflexota_bacterium | Bacteria>Bacillati>Chloroflexota>unclassified_Chloroflexota | 138 | IT305_03470 | 198803..199219 | 2026724 | PIN-domain-containing-protein-[Chloroflexota-bacterium] |
| MCA0453857.1 | PIN-EndoRNase | GCA_020161935.1 | Chloroflexota_bacterium | Bacteria>Bacillati>Chloroflexota>unclassified_Chloroflexota | 141 | LCI00_07785 | 60422..60847 | 2026724 | type-II-toxin-antitoxin-system-VapC-family-toxin-[Chloroflexota-bacterium] |
| MCL4487043.1 | PIN-EndoRNase | GCA_023381515.1 | Chloroflexota_bacterium | Bacteria>Bacillati>Chloroflexota>unclassified_Chloroflexota | 146 | M1570_02785 | 15169..15609 | 2026724 | putative-toxin-antitoxin-system-toxin-component,-PIN-family-[Chloroflexota-bacterium] |
| MBI2864202.1 | PIN-EndoRNase | GCA_016190625.1 | Chloroflexota_bacterium | Bacteria>Bacillati>Chloroflexota>unclassified_Chloroflexota | 149 | HYX94_06525 | 19007..19456 | 2026724 | type-II-toxin-antitoxin-system-VapC-family-toxin-[Chloroflexota-bacterium] |
| MCC6629585.1 | PIN-EndoRNase | GCA_020849725.1 | Chloroflexota_bacterium | Bacteria>Bacillati>Chloroflexota>unclassified_Chloroflexota | 155 | IT340_19575 | 35145..35612 | 2026724 | PIN-domain-containing-protein-[Chloroflexota-bacterium] |
| MBI3961041.1 | PIN-EndoRNase | GCA_016199425.1 | Chloroflexota_bacterium | Bacteria>Bacillati>Chloroflexota>unclassified_Chloroflexota | 160 | HY328_19690 | 2869..3351 | 2026724 | hypothetical-protein-[Chloroflexota-bacterium] |
| GIW20260.1 | PIN-EndoRNase | GCA_026004015.1 | Chloroflexota_bacterium | Bacteria>Bacillati>Chloroflexota>unclassified_Chloroflexota | 211 | KatS3mg065_0556 | 202064..202699 | 2026724 | MAG:-hypothetical-protein-KatS3mg065_0556-[Chloroflexota-bacterium] |
| WP_190449860.1 | PIN-EndoRNase | GCF_014695795.1 | Cyanophyceae | Bacteria>Bacillati>Cyanobacteriota/Melainabacteria_group>Cyanobacteriota | 155 | H6F86_RS16675 | 57950..58417 | 3028117 | hypothetical-protein-[Phormidium-sp.-FACHB-592] |
| WP_190676127.1 | PIN-EndoRNase | GCF_014695435.1 | Cyanophyceae | Bacteria>Bacillati>Cyanobacteriota/Melainabacteria_group>Cyanobacteriota | 161 | H6F60_RS14225 | 74063..74548 | 3028117 | hypothetical-protein-[Coleofasciculus-sp.-FACHB-129] |
| WP_257720872.1 | PIN-EndoRNase | GCF_001277295.1 | Nostocales | Bacteria>Bacillati>Cyanobacteriota/Melainabacteria_group>Cyanobacteriota>Cyanophyceae | 103 | HEQ36_RS21570 | 42804..43115 | 1161 | MULTISPECIES:-type-II-toxin-antitoxin-system-VapC-family-toxin-[Nostocales] |
| WP_094560568.1 | PIN-EndoRNase | GCF_024345455.1 | Synechococcales | Bacteria>Bacillati>Cyanobacteriota/Melainabacteria_group>Cyanobacteriota>Cyanophyceae | 132 | CJZ78_RS08340 | 53991..54389 | 1890424 | MULTISPECIES:-type-II-toxin-antitoxin-system-VapC-family-toxin-[Synechococcales] |
| WP_212661902.1 | PIN-EndoRNase | GCF_018336915.1 | Acaryochloris_marina | Bacteria>Bacillati>Cyanobacteriota/Melainabacteria_group>Cyanobacteriota>Cyanophyceae>Acaryochloridales>Acaryochloridaceae>Acaryochloris | 109 | I1H34_RS15295 | 3286571..3286900 | 155978 | DUF5615-family-PIN-like-protein-[Acaryochloris-marina] |
| WP_024125477.1 | PIN-EndoRNase | GCF_000505665.1 | Thermosynechococcus_sp._NK55a | Bacteria>Bacillati>Cyanobacteriota/Melainabacteria_group>Cyanobacteriota>Cyanophyceae>Acaryochloridales>Thermosynechococcaceae>Thermosynechococcus>unclassified_Thermosynechococcus | 128 | NK55_RS09310 | 1925678..1926064 | 1394889 | DUF5615-family-PIN-like-protein-[Thermosynechococcus-sp.-NK55a] |
| WP_024125474.1 | PIN-EndoRNase | GCF_000505665.1 | Thermosynechococcus_sp._NK55a | Bacteria>Bacillati>Cyanobacteriota/Melainabacteria_group>Cyanobacteriota>Cyanophyceae>Acaryochloridales>Thermosynechococcaceae>Thermosynechococcus>unclassified_Thermosynechococcus | 140 | NK55_RS09295 | 1922012..1922434 | 1394889 | PIN-domain-containing-protein-[Thermosynechococcus-sp.-NK55a] |
| WP_106170665.1 | PIN-EndoRNase | GCF_023546805.1 | Chroococcidiopsis | Bacteria>Bacillati>Cyanobacteriota/Melainabacteria_group>Cyanobacteriota>Cyanophyceae>Chroococcidiopsidales>Chroococcidiopsidaceae | 129 | M5J74_RS19065 | 4100525..4100914 | 54298 | MULTISPECIES:-type-II-toxin-antitoxin-system-VapC-family-toxin-[Chroococcidiopsis] |
| WP_015155781.1 | PIN-EndoRNase | GCF_000317125.1 | Chroococcidiopsis_thermalis | Bacteria>Bacillati>Cyanobacteriota/Melainabacteria_group>Cyanobacteriota>Cyanophyceae>Chroococcidiopsidales>Chroococcidiopsidaceae>Chroococcidiopsis | 129 | CHRO_RS18620 | 4310288..4310677 | 54299 | type-II-toxin-antitoxin-system-VapC-family-toxin-[Chroococcidiopsis-thermalis] |
| WP_015186348.1 | PIN-EndoRNase | GCF_000317515.1 | Allocoleopsis_franciscana | Bacteria>Bacillati>Cyanobacteriota/Melainabacteria_group>Cyanobacteriota>Cyanophyceae>Coleofasciculales>Coleofasciculaceae>Allocoleopsis | 166 | MIC7113_RS32620 | 24847..25347 | 2886352 | PIN-domain-containing-protein-[Allocoleopsis-franciscana] |
| WP_190566703.1 | PIN-EndoRNase | GCF_014696935.1 | unclassified_Desertifilum | Bacteria>Bacillati>Cyanobacteriota/Melainabacteria_group>Cyanobacteriota>Cyanophyceae>Desertifilales>Desertifilaceae>Desertifilum | 129 | H6G23_RS20100 | 69216..69605 | 2621682 | MULTISPECIES:-type-II-toxin-antitoxin-system-VapC-family-toxin-[unclassified-Desertifilum] |
| MCD8490339.1 | PIN-EndoRNase | GCA_021208125.1 | Desertifilum_sp. | Bacteria>Bacillati>Cyanobacteriota/Melainabacteria_group>Cyanobacteriota>Cyanophyceae>Desertifilales>Desertifilaceae>Desertifilum>unclassified_Desertifilum | 129 | LRZ84_27335 | 16584..16973 | 2497424 | type-II-toxin-antitoxin-system-VapC-family-toxin-[Desertifilum-sp.] |
| MCD8487849.1 | PIN-EndoRNase | GCA_021208125.1 | Desertifilum_sp. | Bacteria>Bacillati>Cyanobacteriota/Melainabacteria_group>Cyanobacteriota>Cyanophyceae>Desertifilales>Desertifilaceae>Desertifilum>unclassified_Desertifilum | 131 | LRZ84_14225 | 5090..5485 | 2497424 | type-II-toxin-antitoxin-system-VapC-family-toxin-[Desertifilum-sp.] |
| NMG59375.1 | PIN-EndoRNase | GCA_012911965.1 | Geitlerinema_sp._P-1104 | Bacteria>Bacillati>Cyanobacteriota/Melainabacteria_group>Cyanobacteriota>Cyanophyceae>Geitlerinematales>Geitlerinemataceae>Geitlerinema>unclassified_Geitlerinema | 144 | E1H12_12825 | 9281..9715 | 2546230 | PIN-domain-containing-protein-[Geitlerinema-sp.-P-1104] |
| WP_170190014.1 | PIN-EndoRNase | GCF_012911965.1 | Geitlerinema_sp._P-1104 | Bacteria>Bacillati>Cyanobacteriota/Melainabacteria_group>Cyanobacteriota>Cyanophyceae>Geitlerinematales>Geitlerinemataceae>Geitlerinema>unclassified_Geitlerinema | 144 | E1H12_RS13020 | 9281..9715 | 2546230 | type-II-toxin-antitoxin-system-VapC-family-toxin-[Geitlerinema-sp.-P-1104] |
| WAS06091.1 | PIN-EndoRNase | GCA_022206215.1 | Gloeomargaritales_cyanobacterium_VI4D9 | Bacteria>Bacillati>Cyanobacteriota/Melainabacteria_group>Cyanobacteriota>Cyanophyceae>Gloeomargaritales>unclassified_Gloeomargaritales | 105 | LQF76_04150 | 898264..898581 | 2898145 | type-II-toxin-antitoxin-system-VapC-family-toxin-[Gloeomargaritales-cyanobacterium-VI4D9] |

| Protein-ID | Domain | Assembly-ID | Strain | Taxonomy | Length | Locus-Tag | Genomic-coordinates | TaxID | Fasta-defline |
|---|---|---|---|---|---|---|---|---|---|
| WAS06088.1 | PIN-EndoRNase | GCA_022206215.1 | Gloeomargaritales_cyanobacterium_VI4D9 | Bacteria>Bacillati>Cyanobacteriota/Melainabacteria_group>Cyanobacteriota>Cyanophyceae>Gloeomargaritales>unclassified_Gloeomargaritales | 128 | LQF76_04135 | 897434..897820 | 2898145 | type-II-toxin-antitoxin-system-VapC-family-toxin-[Gloeomargaritales-cyanobacterium-VI4D9] |
| WAS06094.1 | PIN-EndoRNase | GCA_022206215.1 | Gloeomargaritales_cyanobacterium_VI4D9 | Bacteria>Bacillati>Cyanobacteriota/Melainabacteria_group>Cyanobacteriota>Cyanophyceae>Gloeomargaritales>unclassified_Gloeomargaritales | 128 | LQF76_04165 | 902589..902975 | 2898145 | type-II-toxin-antitoxin-system-VapC-family-toxin-[Gloeomargaritales-cyanobacterium-VI4D9] |
| MBC6477471.1 | PIN-EndoRNase | GCA_014323675.1 | Hormoscilla_sp._GM7CHS1pb | Bacteria>Bacillati>Cyanobacteriota/Melainabacteria_group>Cyanobacteriota>Cyanophyceae>Gomontiellales>Gomontiellaceae>Hormoscilla>unclassified_Hormoscilla | 147 | GDA56_06440 | 1108157..1108600 | 2653677 | PIN-domain-containing-protein-[Hormoscilla-sp.-GM7CHS1pb] |
| WP_121969235.1 | PIN-EndoRNase | GCF_002286735.1 | Leptolyngbya_sp._BC1307 | Bacteria>Bacillati>Cyanobacteriota/Melainabacteria_group>Cyanobacteriota>Cyanophyceae>Leptolyngbyales>Leptolyngbyaceae>Leptolyngbya_group>Leptolyngbya>unclassified_Leptolyngbya | 134 | CKL79_RS08935 | 118564..118968 | 2029589 | PIN-domain-containing-protein-[Leptolyngbya-sp.-BC1307] |
| WP_075596944.1 | PIN-EndoRNase | GCF_001939115.1 | Leptolyngbya_sp._'hensonii' | Bacteria>Bacillati>Cyanobacteriota/Melainabacteria_group>Cyanobacteriota>Cyanophyceae>Leptolyngbyales>Leptolyngbyaceae>Leptolyngbya_group>Leptolyngbya>unclassified_Leptolyngbya | 137 | BST81_RS02360 | 88011..88424 | 1922337 | putative-toxin-antitoxin-system-toxin-component,-PIN-family-[Leptolyngbya-sp.-'hensonii'] |
| PZO60754.1 | PIN-EndoRNase | GCA_003242115.1 | Phormidesmis_priestleyi | Bacteria>Bacillati>Cyanobacteriota/Melainabacteria_group>Cyanobacteriota>Cyanophyceae>Leptolyngbyales>Leptolyngbyaceae>Phormidesmis | 140 | DCF15_01595 | 26691..27113 | 268141 | twitching-motility-protein-PilT-[Phormidesmis-priestleyi] |
| PZO60795.1 | PIN-EndoRNase | GCA_003242115.1 | Phormidesmis_priestleyi | Bacteria>Bacillati>Cyanobacteriota/Melainabacteria_group>Cyanobacteriota>Cyanophyceae>Leptolyngbyales>Leptolyngbyaceae>Phormidesmis | 148 | DCF15_01400 | 23368..23814 | 268141 | VapC-toxin-family-PIN-domain-ribonuclease-[Phormidesmis-priestleyi] |
| PZO60756.1 | PIN-EndoRNase | GCA_003242115.1 | Phormidesmis_priestleyi | Bacteria>Bacillati>Cyanobacteriota/Melainabacteria_group>Cyanobacteriota>Cyanophyceae>Leptolyngbyales>Leptolyngbyaceae>Phormidesmis | 224 | DCF15_01605 | 30211..30885 | 268141 | hypothetical-protein-DCF15_01605-[Phormidesmis-priestleyi] |
| KPQ33162.1 | PIN-EndoRNase | GCA_001314865.1 | Phormidesmis_priestleyi_Ana | Bacteria>Bacillati>Cyanobacteriota/Melainabacteria_group>Cyanobacteriota>Cyanophyceae>Leptolyngbyales>Leptolyngbyaceae>Phormidesmis>Phormidesmis_priestleyi | 137 | HLUCCA11_19350 | 6692..7105 | 1666911 | toxin-antitoxin-system-Doc-family-toxin-component-[Phormidesmis-priestleyi-Ana] |
| KPQ33027.1 | PIN-EndoRNase | GCA_001314865.1 | Phormidesmis_priestleyi_Ana | Bacteria>Bacillati>Cyanobacteriota/Melainabacteria_group>Cyanobacteriota>Cyanophyceae>Leptolyngbyales>Leptolyngbyaceae>Phormidesmis>Phormidesmis_priestleyi | 139 | HLUCCA11_19905 | 21070..21489 | 1666911 | putative-nucleic-acid-binding-protein,-contains-PIN-domain-[Phormidesmis-priestleyi-Ana] |
| NJO72424.1 | PIN-EndoRNase | GCA_012033355.1 | Leptolyngbyaceae_cyanobacterium_RM1_406_9 | Bacteria>Bacillati>Cyanobacteriota/Melainabacteria_group>Cyanobacteriota>Cyanophyceae>Leptolyngbyales>Leptolyngbyaceae>unclassified_Leptolyngbyaceae | 139 | HC833_00755 | 32840..33259 | 2720434 | type-II-toxin-antitoxin-system-VapC-family-toxin-[Leptolyngbyaceae-cyanobacterium-RM1_406_9] |
| NJO66187.1 | PIN-EndoRNase | GCA_012033405.1 | Leptolyngbyaceae_cyanobacterium_RM1_405_57 | Bacteria>Bacillati>Cyanobacteriota/Melainabacteria_group>Cyanobacteriota>Cyanophyceae>Leptolyngbyales>Leptolyngbyaceae>unclassified_Leptolyngbyaceae | 165 | HC832_00945 | 5402..5899 | 2720433 | hypothetical-protein-[Leptolyngbyaceae-cyanobacterium-RM1_405_57] |
| WP_265900466.1 | PIN-EndoRNase | GCF_026240635.1 | Trichothermofontia_sichuanensis | Bacteria>Bacillati>Cyanobacteriota/Melainabacteria_group>Cyanobacteriota>Cyanophyceae>Leptolyngbyales>Trichocoleusaceae>Trichothermofontia | 138 | OOK60_RS10510 | 2480651..2481067 | 3045816 | PIN-domain-containing-protein-[Leptolyngbya-sp.-PKUAC-SCTB231] |
| WP_204138801.1 | PIN-EndoRNase | GCF_016888165.1 | Halomicronema_sp._CCY15110 | Bacteria>Bacillati>Cyanobacteriota/Melainabacteria_group>Cyanobacteriota>Cyanophyceae>Nodosilineales>Nodosilineaceae>Halomicronema>unclassified_Halomicronema | 106 | JUK32_RS10500 | 49640..49960 | 2767773 | hypothetical-protein-[Halomicronema-sp.-CCY15110] |
| WP_204138808.1 | PIN-EndoRNase | GCF_016888165.1 | Halomicronema_sp._CCY15110 | Bacteria>Bacillati>Cyanobacteriota/Melainabacteria_group>Cyanobacteriota>Cyanophyceae>Nodosilineales>Nodosilineaceae>Halomicronema>unclassified_Halomicronema | 109 | JUK32_RS10490 | 48955..49284 | 2767773 | DUF5615-family-PIN-like-protein-[Halomicronema-sp.-CCY15110] |
| WP_017302295.1 | PIN-EndoRNase | GCF_000309385.1 | Nodosilinea_nodulosa | Bacteria>Bacillati>Cyanobacteriota/Melainabacteria_group>Cyanobacteriota>Cyanophyceae>Nodosilineales>Nodosilineaceae>Nodosilinea | 71 | LEPTO7104_RS36220 | 433647..433862 | 416001 | hypothetical-protein-[Nodosilinea-nodulosa] |
| WP_017302297.1 | PIN-EndoRNase | GCF_000309385.1 | Nodosilinea_nodulosa | Bacteria>Bacillati>Cyanobacteriota/Melainabacteria_group>Cyanobacteriota>Cyanophyceae>Nodosilineales>Nodosilineaceae>Nodosilinea | 101 | LEPTO7104_RS36225 | 435026..435331 | 416001 | hypothetical-protein-[Nodosilinea-nodulosa] |
| WP_193965435.1 | PIN-EndoRNase | GCF_015207395.1 | Nodosilinea_sp._LEGE_07088 | Bacteria>Bacillati>Cyanobacteriota/Melainabacteria_group>Cyanobacteriota>Cyanophyceae>Nodosilineales>Nodosilineaceae>Nodosilinea>unclassified_Nodosilinea | 138 | IQ254_RS06905 | 12769..13185 | 2777968 | PIN-domain-containing-protein-[Nodosilinea-sp.-LEGE-07088] |
| WP_168465294.1 | PIN-EndoRNase | GCF_014696815.1 | Aphanizomenon | Bacteria>Bacillati>Cyanobacteriota/Melainabacteria_group>Cyanobacteriota>Cyanophyceae>Nostocales>Aphanizomenonaceae | 136 | FJR37_RS03345 | 1740..2150 | 1175 | MULTISPECIES:-type-II-toxin-antitoxin-system-VapC-family-toxin-[Aphanizomenon] |
| WP_096669804.1 | PIN-EndoRNase | GCF_002368115.1 | Dolichospermum_compactum | Bacteria>Bacillati>Cyanobacteriota/Melainabacteria_group>Cyanobacteriota>Cyanophyceae>Nostocales>Aphanizomenonaceae>Dolichospermum | 126 | CA730_RS19470 | 4200117..4200497 | 136073 | type-II-toxin-antitoxin-system-VapC-family-toxin-[Dolichospermum-compactum] |
| WP_236097130.1 | PIN-EndoRNase | GCF_005402965.1 | Dolichospermum_planctonicum | Bacteria>Bacillati>Cyanobacteriota/Melainabacteria_group>Cyanobacteriota>Cyanophyceae>Nostocales>Aphanizomenonaceae>Dolichospermum | 168 | FFO06_RS11410 | 65147..65653 | 136072 | DUF3368-domain-containing-protein-[Dolichospermum-planctonicum] |
| MBO1068321.1 | PIN-EndoRNase | GCA_017355655.1 | Dolichospermum_sp._DEX189 | Bacteria>Bacillati>Cyanobacteriota/Melainabacteria_group>Cyanobacteriota>Cyanophyceae>Nostocales>Aphanizomenonaceae>Dolichospermum>unclassified_Dolichospermum | 93 | HEQ13_02490 | 532804..533085 | 2717870 | type-II-toxin-antitoxin-system-VapC-family-toxin-[Dolichospermum-sp.-DEX189] |
| MBS3030301.1 | PIN-EndoRNase | GCA_018295985.1 | Dolichospermum_sp._DET66 | Bacteria>Bacillati>Cyanobacteriota/Melainabacteria_group>Cyanobacteriota>Cyanophyceae>Nostocales>Aphanizomenonaceae>Dolichospermum>unclassified_Dolichospermum | 131 | HCA26_25910 | 5405263..5405658 | 2704415 | type-II-toxin-antitoxin-system-VapC-family-toxin-[Dolichospermum-sp.-DET66] |
| WP_199301842.1 | PIN-EndoRNase | - | unclassified_Sphaerospermopsis | Bacteria>Bacillati>Cyanobacteriota/Melainabacteria_group>Cyanobacteriota>Cyanophyceae>Nostocales>Aphanizomenonaceae>Sphaerospermopsis | 164 | H5968_RS22670 | 2590..3084 | 2646443 | MULTISPECIES:-DUF3368-domain-containing-protein-[unclassified-Sphaerospermopsis] |
| WP_015130208.1 | PIN-EndoRNase | GCF_000316575.1 | Calothrix_sp._PCC_7507 | Bacteria>Bacillati>Cyanobacteriota/Melainabacteria_group>Cyanobacteriota>Cyanophyceae>Nostocales>Calotrichaceae>Calothrix>unclassified_Calothrix | 143 | CAL7507_RS19490 | 4556139..4556570 | 99598 | type-II-toxin-antitoxin-system-VapC-family-toxin-[Calothrix-sp.-PCC-7507] |
| WP_127084962.1 | PIN-EndoRNase | GCF_003991905.1 | Dulcicalothrix_desertica | Bacteria>Bacillati>Cyanobacteriota/Melainabacteria_group>Cyanobacteriota>Cyanophyceae>Nostocales>Calotrichaceae>Dulcicalothrix | 143 | DSM106972_RS34680 | 73033..73464 | 32056 | putative-toxin-antitoxin-system-toxin-component,-PIN-family-[Dulcicalothrix-desertica] |
| MBW4670335.1 | PIN-EndoRNase | GCA_019359595.1 | Cyanomargarita_calcarea_GSE-NOS-MK-12-04C | Bacteria>Bacillati>Cyanobacteriota/Melainabacteria_group>Cyanobacteriota>Cyanophyceae>Nostocales>Cyanomargaritaceae>Cyanomargarita>Cyanomargarita_calcarea | 147 | KME60_23695 | 102980..103423 | 2839659 | PIN-domain-containing-protein-[Cyanomargarita-calcarea-GSE-NOS-MK-12-04C] |
| WP_026733569.1 | PIN-EndoRNase | GCF_000517105.1 | Fischerella_sp._PCC_9605 | Bacteria>Bacillati>Cyanobacteriota/Melainabacteria_group>Cyanobacteriota>Cyanophyceae>Nostocales>Hapalosiphonaceae>Fischerella>unclassified_Fischerella | 129 | FIS9605_RS0116405 | 332928..333317 | 1173024 | type-II-toxin-antitoxin-system-VapC-family-toxin-[Fischerella-sp.-PCC-9605] |
| WP_200988281.1 | PIN-EndoRNase | GCF_015245355.1 | Anabaenopsis_elenkinii | Bacteria>Bacillati>Cyanobacteriota/Melainabacteria_group>Cyanobacteriota>Cyanophyceae>Nostocales>Nodulariaceae>Anabaenopsis | 136 | IM676_RS18815 | 4312865..4313275 | 156213 | PIN-domain-containing-protein-[Anabaenopsis-elenkinii] |
| WP_272056147.1 | PIN-EndoRNase | GCF_028330355.1 | Nodularia_spumigena | Bacteria>Bacillati>Cyanobacteriota/Melainabacteria_group>Cyanobacteriota>Cyanophyceae>Nostocales>Nodulariaceae>Nodularia | 142 | PN490_RS03015 | 8108..8536 | 70799 | PIN-domain-containing-protein-[Nodularia-spumigena] |
| WP_271941650.1 | PIN-EndoRNase | GCF_028329345.1 | Nodularia_spumigena | Bacteria>Bacillati>Cyanobacteriota/Melainabacteria_group>Cyanobacteriota>Cyanophyceae>Nostocales>Nodulariaceae>Nodularia | 165 | PN454_RS18915 | 11248..11745 | 70799 | PIN-domain-containing-protein-[Nodularia-spumigena] |
| WP_265338635.1 | PIN-EndoRNase | GCF_020702635.1 | Nodularia_sp._LEGE_04288 | Bacteria>Bacillati>Cyanobacteriota/Melainabacteria_group>Cyanobacteriota>Cyanophyceae>Nostocales>Nodulariaceae>Nodularia>unclassified_Nodularia_(in:_cyanobacteria) | 40 | IQ240_RS26575 | 4027..4149 | 1828639 | hypothetical-protein-[Nodularia-sp.-LEGE-04288] |
| OBQ20270.1 | PIN-EndoRNase | GCA_001672085.1 | Anabaena_sp._AL93 | Bacteria>Bacillati>Cyanobacteriota/Melainabacteria_group>Cyanobacteriota>Cyanophyceae>Nostocales>Nostocaceae>Anabaena>unclassified_Anabaena | 126 | AN486_07305 | 51531..51911 | 1678133 | MAG:-twitching-motility-protein-PilT-[Anabaena-sp.-AL93] |
| MCE2718754.1 | PIN-EndoRNase | GCA_021300315.1 | Anabaena_sp._49628_E55 | Bacteria>Bacillati>Cyanobacteriota/Melainabacteria_group>Cyanobacteriota>Cyanophyceae>Nostocales>Nostocaceae>Anabaena>unclassified_Anabaena | 139 | LW708_08005 | 1489..1908 | 2904988 | type-II-toxin-antitoxin-system-VapC-family-toxin-[Anabaena-sp.-49628_E55] |
| OBQ34508.1 | PIN-EndoRNase | GCA_001672075.1 | Anabaena_sp._CRKS33 | Bacteria>Bacillati>Cyanobacteriota/Melainabacteria_group>Cyanobacteriota>Cyanophyceae>Nostocales>Nostocaceae>Anabaena>unclassified_Anabaena | 139 | AN487_17935 | 3042..3461 | 1710888 | MAG:-recombinase-[Anabaena-sp.-CRKS33] |
| WP_190368029.1 | PIN-EndoRNase | GCF_014696225.1 | Anabaena_sp._FACHB-1237 | Bacteria>Bacillati>Cyanobacteriota/Melainabacteria_group>Cyanobacteriota>Cyanophyceae>Nostocales>Nostocaceae>Anabaena>unclassified_Anabaena | 139 | H6F32_RS03900 | 66981..67400 | 2692769 | type-II-toxin-antitoxin-system-VapC-family-toxin-[Anabaena-sp.-FACHB-1237] |
| WP_168507190.1 | PIN-EndoRNase | GCF_009712035.1 | Anabaena_sp._UHCC_0187 | Bacteria>Bacillati>Cyanobacteriota/Melainabacteria_group>Cyanobacteriota>Cyanophyceae>Nostocales>Nostocaceae>Anabaena>unclassified_Anabaena | 214 | FJR11_RS20810 | 8728..9372 | 2590018 | NYN-domain-containing-protein-[Anabaena-sp.-UHCC-0187] |
| WP_199349203.1 | PIN-EndoRNase | - | Nostoc_flagelliforme | Bacteria>Bacillati>Cyanobacteriota/Melainabacteria_group>Cyanobacteriota>Cyanophyceae>Nostocales>Nostocaceae>Nostoc | 125 | H6G97_RS00990 | 250962..251339 | 1306274 | type-II-toxin-antitoxin-system-VapC-family-toxin-[Nostoc-flagelliforme] |
| WP_251956296.1 | PIN-EndoRNase | GCF_023734775.1 | Nostoc_commune | Bacteria>Bacillati>Cyanobacteriota/Melainabacteria_group>Cyanobacteriota>Cyanophyceae>Nostocales>Nostocaceae>Nostoc | 128 | ANSO36C_RS22390 | 4945359..4945745 | 1178 | type-II-toxin-antitoxin-system-VapC-family-toxin-[Nostoc-commune] |
| WP_109013092.1 | PIN-EndoRNase | GCF_003113895.1 | Nostoc_commune | Bacteria>Bacillati>Cyanobacteriota/Melainabacteria_group>Cyanobacteriota>Cyanophyceae>Nostocales>Nostocaceae>Nostoc | 165 | CDC33_RS34460 | 461150..461647 | 1178 | PIN-domain-containing-protein-[Nostoc-commune] |
| WP_162398346.1 | PIN-EndoRNase | - | Nostoc_sp._B(2019) | Bacteria>Bacillati>Cyanobacteriota/Melainabacteria_group>Cyanobacteriota>Cyanophyceae>Nostocales>Nostocaceae>Nostoc>unclassified_Nostoc | 90 | GS682_RS23155 | 20864..21136 | 2690446 | hypothetical-protein-[Nostoc-sp.-B(2019)] |
| WP_224095533.1 | PIN-EndoRNase | GCF_019976755.1 | Nostoc_sp._MS1 | Bacteria>Bacillati>Cyanobacteriota/Melainabacteria_group>Cyanobacteriota>Cyanophyceae>Nostocales>Nostocaceae>Nostoc>unclassified_Nostoc | 90 | NSMS1_RS32140 | 250823..251095 | 2764711 | hypothetical-protein-[Nostoc-sp.-MS1] |
| WP_175586580.1 | PIN-EndoRNase | - | Nostoc_sp._UIC_10630 | Bacteria>Bacillati>Cyanobacteriota/Melainabacteria_group>Cyanobacteriota>Cyanophyceae>Nostocales>Nostocaceae>Nostoc>unclassified_Nostoc | 125 | G1D98_RS07800 | 7873..8250 | 2100146 | type-II-toxin-antitoxin-system-VapC-family-toxin-[Nostoc-sp.-UIC-10630] |
| MBN3957896.1 | PIN-EndoRNase | GCA_017115045.1 | Nostoc_sp._NMS8 | Bacteria>Bacillati>Cyanobacteriota/Melainabacteria_group>Cyanobacteriota>Cyanophyceae>Nostocales>Nostocaceae>Nostoc>unclassified_Nostoc | 131 | HWQ39_03285 | 14797..15192 | 2815392 | type-II-toxin-antitoxin-system-VapC-family-toxin-[Nostoc-sp.-NMS8] |
| WP_179052051.1 | PIN-EndoRNase | GCF_013393945.1 | Nostoc_sp._TCL26-01 | Bacteria>Bacillati>Cyanobacteriota/Melainabacteria_group>Cyanobacteriota>Cyanophyceae>Nostocales>Nostocaceae>Nostoc>unclassified_Nostoc | 131 | FD725_RS31170 | 81457..81852 | 2576904 | type-II-toxin-antitoxin-system-VapC-family-toxin-[Nostoc-sp.-TCL26-01] |
| WP_179052053.1 | PIN-EndoRNase | GCF_013393945.1 | Nostoc_sp._TCL26-01 | Bacteria>Bacillati>Cyanobacteriota/Melainabacteria_group>Cyanobacteriota>Cyanophyceae>Nostocales>Nostocaceae>Nostoc>unclassified_Nostoc | 136 | FD725_RS31180 | 85317..85727 | 2576904 | type-II-toxin-antitoxin-system-VapC-family-toxin-[Nostoc-sp.-TCL26-01] |
| WP_185607254.1 | PIN-EndoRNase | GCF_014222285.1 | Nostoc_sp._UCD121 | Bacteria>Bacillati>Cyanobacteriota/Melainabacteria_group>Cyanobacteriota>Cyanophyceae>Nostocales>Nostocaceae>Nostoc>unclassified_Nostoc | 165 | GNF10_RS25040 | 10265..10762 | 2681305 | PIN-domain-containing-protein-[Nostoc-sp.-UCD121] |
| WP_094343934.1 | PIN-EndoRNase | GCF_002245985.1 | Nostoc_sp._'Peltigera_membranacea_cyanobiont'_232 | Bacteria>Bacillati>Cyanobacteriota/Melainabacteria_group>Cyanobacteriota>Cyanophyceae>Nostocales>Nostocaceae>Nostoc>unclassified_Nostoc>Nostoc_cyanobionts | 128 | CDG79_RS21450 | 7436..7822 | 2014531 | type-II-toxin-antitoxin-system-VapC-family-toxin-[Nostoc-sp.-'Peltigera-membranacea-cyanobiont'-232] |
| WP_104908182.1 | PIN-EndoRNase | GCF_002949795.1 | Nostoc_sp._'Lobaria_pulmonaria_(5183)_cyanobiont' | Bacteria>Bacillati>Cyanobacteriota/Melainabacteria_group>Cyanobacteriota>Cyanophyceae>Nostocales>Nostocaceae>Nostoc>unclassified_Nostoc>Nostoc_cyanobionts | 128 | NLP_RS21705 | 4928708..4929094 | 1618022 | type-II-toxin-antitoxin-system-VapC-family-toxin-[Nostoc-sp.-'Lobaria-pulmonaria-(5183)-cyanobiont'] |
| WP_094345567.1 | PIN-EndoRNase | GCF_002245985.1 | Nostoc_sp._'Peltigera_membranacea_cyanobiont'_232 | Bacteria>Bacillati>Cyanobacteriota/Melainabacteria_group>Cyanobacteriota>Cyanophyceae>Nostocales>Nostocaceae>Nostoc>unclassified_Nostoc>Nostoc_cyanobionts | 165 | CDG79_RS31105 | 9987..10484 | 2014531 | PIN-domain-containing-protein-[Nostoc-sp.-'Peltigera-membranacea-cyanobiont'-232] |
| WP_172355885.1 | PIN-EndoRNase | GCF_013177315.1 | Thermoleptolyngbya | Bacteria>Bacillati>Cyanobacteriota/Melainabacteria_group>Cyanobacteriota>Cyanophyceae>Oculatellales>Oculatellaceae | 131 | JTM80_RS13300 | 3245865..3246260 | 2303528 | MULTISPECIES:-type-II-toxin-antitoxin-system-VapC-family-toxin-[Thermoleptolyngbya] |
| MBW4466340.1 | PIN-EndoRNase | GCA_019359155.1 | Pegethrix_bostrychoides_GSE-TBD4-15B | Bacteria>Bacillati>Cyanobacteriota/Melainabacteria_group>Cyanobacteriota>Cyanophyceae>Oculatellales>Oculatellaceae>Pegethrix>Pegethrix_bostrychoides | 165 | KME07_13020 | 95240..95737 | 2839662 | hypothetical-protein-[Pegethrix-bostrychoides-GSE-TBD4-15B] |
| WP_268611169.1 | PIN-EndoRNase | GCF_026802175.1 | Thermocoleostomius_sinensis | Bacteria>Bacillati>Cyanobacteriota/Melainabacteria_group>Cyanobacteriota>Cyanophyceae>Oculatellales>Oculatellaceae>Thermocoleostomius | 153 | OXH18_RS04215 | 968792..969253 | 3065396 | hypothetical-protein-[Leptolyngbya-sp.-PKUAC-SCTA174] |
| MBW4580242.1 | PIN-EndoRNase | GCA_019358895.1 | Tildeniella_nuda_ZEHNDER_1965/U140 | Bacteria>Bacillati>Cyanobacteriota/Melainabacteria_group>Cyanobacteriota>Cyanophyceae>Oculatellales>Oculatellaceae>Tildeniella>Tildeniella_nuda | 148 | KME42_11815 | 256958..257404 | 2136405 | PIN-domain-containing-protein-[Tildeniella-nuda-ZEHNDER-1965/U140] |
| WP_096720870.1 | PIN-EndoRNase | - | Chondrocystis_sp._NIES-4102 | Bacteria>Bacillati>Cyanobacteriota/Melainabacteria_group>Cyanobacteriota>Cyanophyceae>Oscillatoriophycideae>Chroococcales>Chroococcaceae>Chondrocystis>unclassified_Chondrocystis | 141 | CDC35_RS11855 | 2752006..2752431 | 2005460 | type-II-toxin-antitoxin-system-VapC-family-toxin-[Chondrocystis-sp.-NIES-4102] |
| PSB44270.1 | PIN-EndoRNase | GCA_003004015.1 | Cyanosarcina_cf._burmensis_CCALA_770 | Bacteria>Bacillati>Cyanobacteriota/Melainabacteria_group>Cyanobacteriota>Cyanophyceae>Oscillatoriophycideae>Chroococcales>Chroococcaceae>Cyanosarcina>Cyanosarcina_burmensis | 129 | C7B80_21185 | 8167..8556 | 2107697 | PIN-domain-nuclease-[Cyanosarcina-cf.-burmensis-CCALA-770] |
| WP_106370050.1 | PIN-EndoRNase | GCF_963877105.1 | Chlorogloea_sp._CCALA_695 | Bacteria>Bacillati>Cyanobacteriota/Melainabacteria_group>Cyanobacteriota>Cyanophyceae>Oscillatoriophycideae>Chroococcales>Entophysalidaceae>Chlorogloea>unclassified_Chlorogloea | 136 | C7B70_RS16220 | 45765..46175 | 2107693 | PIN-domain-containing-protein-[Chlorogloea-sp.-CCALA-695] |
| WP_106371308.1 | PIN-EndoRNase | GCF_963877105.1 | Chlorogloea_sp._CCALA_695 | Bacteria>Bacillati>Cyanobacteriota/Melainabacteria_group>Cyanobacteriota>Cyanophyceae>Oscillatoriophycideae>Chroococcales>Entophysalidaceae>Chlorogloea>unclassified_Chlorogloea | 136 | C7B70_RS23205 | 14179..14589 | 2107693 | PIN-domain-containing-protein-[Chlorogloea-sp.-CCALA-695] |
| AFZ48536.1 | PIN-EndoRNase | GCA_000317655.1 | Cyanobacterium_stanieri_PCC_7202 | Bacteria>Bacillati>Cyanobacteriota/Melainabacteria_group>Cyanobacteriota>Cyanophyceae>Oscillatoriophycideae>Chroococcales>Geminocystaceae>Cyanobacterium>Cyanobacterium_stanieri | 131 | Cyast_2593 | 2847340..2847735 | 292563 | PilT-protein-domain-protein-[Cyanobacterium-stanieri-PCC-7202] |
| WP_242411966.1 | PIN-EndoRNase | - | Cyanobacterium_sp._IPPAS_B-1200 | Bacteria>Bacillati>Cyanobacteriota/Melainabacteria_group>Cyanobacteriota>Cyanophyceae>Oscillatoriophycideae>Chroococcales>Geminocystaceae>Cyanobacterium>unclassified_Cyanobacterium | 135 | A5482_RS09615 | 18464..18871 | 1562720 | type-II-toxin-antitoxin-system-VapC-family-toxin-[Cyanobacterium-sp.-IPPAS-B-1200] |
| BAQ63188.1 | PIN-EndoRNase | GCA_001548095.1 | Geminocystis_sp._NIES-3708 | Bacteria>Bacillati>Cyanobacteriota/Melainabacteria_group>Cyanobacteriota>Cyanophyceae>Oscillatoriophycideae>Chroococcales>Geminocystaceae>Geminocystis>unclassified_Geminocystis | 144 | GM3708_3594 | 37456..37890 | 1615909 | VapC-toxin-protein-[Geminocystis-sp.-NIES-3708] |
| WP_066349644.1 | PIN-EndoRNase | GCF_001548095.1 | Geminocystis_sp._NIES-3708 | Bacteria>Bacillati>Cyanobacteriota/Melainabacteria_group>Cyanobacteriota>Cyanophyceae>Oscillatoriophycideae>Chroococcales>Geminocystaceae>Geminocystis>unclassified_Geminocystis | 144 | GM3708_RS17430 | 37456..37890 | 1615909 | type-II-toxin-antitoxin-system-VapC-family-toxin-[Geminocystis-sp.-NIES-3708] |
| WP_083764500.1 | PIN-EndoRNase | GCF_000019485.1 | Picosynechococcus_sp._PCC_7002 | Bacteria>Bacillati>Cyanobacteriota/Melainabacteria_group>Cyanobacteriota>Cyanophyceae>Oscillatoriophycideae>Chroococcales>Geminocystaceae>Picosynechococcus>unclassified_Picosynechococcus | 120 | CCK85_RS15775 | 114253..114615 | 32049 | MULTISPECIES:-PIN-domain-containing-protein-[unclassified-Synechococcus] |
| WP_075888514.1 | PIN-EndoRNase | GCF_001904615.1 | [Limnothrix_rosea]_IAM_M-220 | Bacteria>Bacillati>Cyanobacteriota/Melainabacteria_group>Cyanobacteriota>Cyanophyceae>Oscillatoriophycideae>Chroococcales>Geminocystaceae>Picosynechococcus>unclassified_Picosynechococcus | 134 | NIES208_RS00255 | 61911..62315 | 454133 | type-II-toxin-antitoxin-system-VapC-family-toxin-[Limnothrix-rosea] |
| WP_052349734.1 | PIN-EndoRNase | GCF_000715475.1 | Picosynechococcus_sp._NKBG042902 | Bacteria>Bacillati>Cyanobacteriota/Melainabacteria_group>Cyanobacteriota>Cyanophyceae>Oscillatoriophycideae>Chroococcales>Geminocystaceae>Picosynechococcus>unclassified_Picosynechococcus | 144 | BG2_RS15795 | 12018..12452 | 490193 | hypothetical-protein-[Synechococcus-sp.-NKBG042902] |
| WP_223210845.1 | PIN-EndoRNase | GCF_000599945.1 | Microcystis | Bacteria>Bacillati>Cyanobacteriota/Melainabacteria_group>Cyanobacteriota>Cyanophyceae>Oscillatoriophycideae>Chroococcales>Microcystaceae | 55 | PN479_RS02410 | 5404..5571 | 1125 | hypothetical-protein-[Microcystis-aeruginosa] |
| WP_002754175.1 | PIN-EndoRNase | GCF_000307995.1 | Microcystis | Bacteria>Bacillati>Cyanobacteriota/Melainabacteria_group>Cyanobacteriota>Cyanophyceae>Oscillatoriophycideae>Chroococcales>Microcystaceae | 129 | FXO09_RS20235 | 29634..30023 | 1125 | type-II-toxin-antitoxin-system-VapC-family-toxin-[Microcystis-aeruginosa] |
| WP_002748936.1 | PIN-EndoRNase | GCF_000332585.1 | Microcystis | Bacteria>Bacillati>Cyanobacteriota/Melainabacteria_group>Cyanobacteriota>Cyanophyceae>Oscillatoriophycideae>Chroococcales>Microcystaceae | 137 | BH695_RS02865 | 540647..541060 | 1125 | MULTISPECIES:-PIN-domain-containing-protein-[Microcystis] |
| WP_002777130.1 | PIN-EndoRNase | GCF_000312285.1 | Microcystis | Bacteria>Bacillati>Cyanobacteriota/Melainabacteria_group>Cyanobacteriota>Cyanophyceae>Oscillatoriophycideae>Chroococcales>Microcystaceae | 137 | PN482_RS11745 | 1108..1521 | 1125 | MULTISPECIES:-PIN-domain-containing-protein-[Microcystis] |
| WP_103113078.1 | PIN-EndoRNase | GCF_002897275.1 | Microcystis | Bacteria>Bacillati>Cyanobacteriota/Melainabacteria_group>Cyanobacteriota>Cyanophyceae>Oscillatoriophycideae>Chroococcales>Microcystaceae | 137 | H6F92_RS24360 | 19816..20229 | 1125 | MULTISPECIES:-PIN-domain-containing-protein-[Microcystis] |
| WP_002801404.1 | PIN-EndoRNase | GCF_000412595.1 | Microcystis | Bacteria>Bacillati>Cyanobacteriota/Melainabacteria_group>Cyanobacteriota>Cyanophyceae>Oscillatoriophycideae>Chroococcales>Microcystaceae | 172 | GVO69_RS15750 | 15633..16151 | 1125 | hypothetical-protein-[Microcystis-aeruginosa] |
| WP_272105043.1 | PIN-EndoRNase | GCF_032701645.1 | Microcystis | Bacteria>Bacillati>Cyanobacteriota/Melainabacteria_group>Cyanobacteriota>Cyanophyceae>Oscillatoriophycideae>Chroococcales>Microcystaceae | 172 | PN470_RS21185 | 24503..25021 | 1125 | hypothetical-protein-[Microcystis-sp.-CS-574] |
| WP_272078195.1 | PIN-EndoRNase | GCF_028331105.1 | Microcystis_aeruginosa | Bacteria>Bacillati>Cyanobacteriota/Melainabacteria_group>Cyanobacteriota>Cyanophyceae>Oscillatoriophycideae>Chroococcales>Microcystaceae>Microcystis | 70 | PN487_RS19710 | 11503..11715 | 1126 | hypothetical-protein-[Microcystis-aeruginosa] |
| WP_016515481.1 | PIN-EndoRNase | GCF_000412595.1 | Microcystis_aeruginosa | Bacteria>Bacillati>Cyanobacteriota/Melainabacteria_group>Cyanobacteriota>Cyanophyceae>Oscillatoriophycideae>Chroococcales>Microcystaceae>Microcystis | 95 | MAESPC_RS19930 | 56212..56499 | 1126 | putative-nucleic-acid-binding-protein,-contains-PIN-domain-protein-[Microcystis-aeruginosa] |
| WP_016515483.1 | PIN-EndoRNase | GCF_000412595.1 | Microcystis_aeruginosa | Bacteria>Bacillati>Cyanobacteriota/Melainabacteria_group>Cyanobacteriota>Cyanophyceae>Oscillatoriophycideae>Chroococcales>Microcystaceae>Microcystis | 129 | EZJ55_RS08100 | 1409442..1409831 | 1126 | type-II-toxin-antitoxin-system-VapC-family-toxin-[Microcystis-aeruginosa] |
| WP_151696809.1 | PIN-EndoRNase | GCF_008974145.1 | Microcystis_aeruginosa | Bacteria>Bacillati>Cyanobacteriota/Melainabacteria_group>Cyanobacteriota>Cyanophyceae>Oscillatoriophycideae>Chroococcales>Microcystaceae>Microcystis | 140 | MiAbW_RS15330 | 42676..43098 | 1126 | PIN-domain-containing-protein-[Microcystis-aeruginosa] |
| WP_271953562.1 | PIN-EndoRNase | GCF_028331205.1 | Microcystis_aeruginosa | Bacteria>Bacillati>Cyanobacteriota/Melainabacteria_group>Cyanobacteriota>Cyanophyceae>Oscillatoriophycideae>Chroococcales>Microcystaceae>Microcystis | 140 | PN437_RS04345 | 387..809 | 1126 | PIN-domain-containing-protein-[Microcystis-aeruginosa] |
| WP_002777133.1 | PIN-EndoRNase | GCF_000312205.1 | Microcystis_aeruginosa | Bacteria>Bacillati>Cyanobacteriota/Melainabacteria_group>Cyanobacteriota>Cyanophyceae>Oscillatoriophycideae>Chroococcales>Microcystaceae>Microcystis | 172 | MICAD_RS14450 | 203974..204492 | 1126 | hypothetical-protein-[Microcystis-aeruginosa] |
| WP_024971075.1 | PIN-EndoRNase | GCF_000599945.1 | Microcystis_aeruginosa | Bacteria>Bacillati>Cyanobacteriota/Melainabacteria_group>Cyanobacteriota>Cyanophyceae>Oscillatoriophycideae>Chroococcales>Microcystaceae>Microcystis | 172 | PN479_RS02400 | 3321..3839 | 1126 | hypothetical-protein-[Microcystis-aeruginosa] |
| GBE76043.1 | PIN-EndoRNase | GCA_002933835.1 | Microcystis_aeruginosa_NIES-87 | Bacteria>Bacillati>Cyanobacteriota/Melainabacteria_group>Cyanobacteriota>Cyanophyceae>Oscillatoriophycideae>Chroococcales>Microcystaceae>Microcystis>Microcystis_aeruginosa | 129 | myaer87_32700 | 114578..114967 | 449440 | hypothetical-protein-myaer87_32700-[Microcystis-aeruginosa-NIES-87] |
| TRT82539.1 | PIN-EndoRNase | GCA_007095595.1 | Microcystis_aeruginosa_Ma_AC_P_19900807_S299 | Bacteria>Bacillati>Cyanobacteriota/Melainabacteria_group>Cyanobacteriota>Cyanophyceae>Oscillatoriophycideae>Chroococcales>Microcystaceae>Microcystis>Microcystis_aeruginosa | 129 | EWV82_11190 | 9391..9780 | 2486264 | type-II-toxin-antitoxin-system-VapC-family-toxin-[Microcystis-aeruginosa-Ma_AC_P_19900807_S299] |
| REJ55683.1 | PIN-EndoRNase | GCA_003388675.1 | Microcystis_aeruginosa_DA14 | Bacteria>Bacillati>Cyanobacteriota/Melainabacteria_group>Cyanobacteriota>Cyanophyceae>Oscillatoriophycideae>Chroococcales>Microcystaceae>Microcystis>Microcystis_aeruginosa | 137 | DWQ56_15535 | 239523..239936 | 1987506 | type-II-toxin-antitoxin-system-VapC-family-toxin-[Microcystis-aeruginosa-DA14] |
| GBE76046.1 | PIN-EndoRNase | GCA_002933835.1 | Microcystis_aeruginosa_NIES-87 | Bacteria>Bacillati>Cyanobacteriota/Melainabacteria_group>Cyanobacteriota>Cyanophyceae>Oscillatoriophycideae>Chroococcales>Microcystaceae>Microcystis>Microcystis_aeruginosa | 137 | myaer87_32730 | 118572..118985 | 449440 | PIN-domain-protein-[Microcystis-aeruginosa-NIES-87] |
| REJ55685.1 | PIN-EndoRNase | GCA_003388675.1 | Microcystis_aeruginosa_DA14 | Bacteria>Bacillati>Cyanobacteriota/Melainabacteria_group>Cyanobacteriota>Cyanophyceae>Oscillatoriophycideae>Chroococcales>Microcystaceae>Microcystis>Microcystis_aeruginosa | 137 | DWQ56_15550 | 243536..243949 | 1987506 | type-II-toxin-antitoxin-system-VapC-family-toxin-[Microcystis-aeruginosa-DA14] |
| NCR40138.1 | PIN-EndoRNase | GCA_009995705.1 | Microcystis_aeruginosa_W13-11 | Bacteria>Bacillati>Cyanobacteriota/Melainabacteria_group>Cyanobacteriota>Cyanophyceae>Oscillatoriophycideae>Chroococcales>Microcystaceae>Microcystis>Microcystis_aeruginosa | 140 | GPJ21_09835 | 22240..22662 | 2685991 | PIN-domain-containing-protein-[Microcystis-aeruginosa-W13-11] |
| TRU24938.1 | PIN-EndoRNase | GCA_007095795.1 | Microcystis_aeruginosa_Ma_MB_S_20031200_S102D | Bacteria>Bacillati>Cyanobacteriota/Melainabacteria_group>Cyanobacteriota>Cyanophyceae>Oscillatoriophycideae>Chroococcales>Microcystaceae>Microcystis>Microcystis_aeruginosa | 140 | EWV79_09035 | 23872..24294 | 2486255 | PIN-domain-containing-protein-[Microcystis-aeruginosa-Ma_MB_S_20031200_S102D] |
| ARI79892.1 | PIN-EndoRNase | GCA_002095975.1 | Microcystis_aeruginosa_PCC_7806SL | Bacteria>Bacillati>Cyanobacteriota/Melainabacteria_group>Cyanobacteriota>Cyanophyceae>Oscillatoriophycideae>Chroococcales>Microcystaceae>Microcystis>Microcystis_aeruginosa>Microcystis_aeruginosa_PCC_7806 | 137 | BH695_0611 | 540647..541060 | 1903187 | hypothetical-protein-BH695_0611-[Microcystis-aeruginosa-PCC-7806SL] |
| MCA2653780.1 | PIN-EndoRNase | GCA_020245605.1 | Microcystis_sp._M061S2 | Bacteria>Bacillati>Cyanobacteriota/Melainabacteria_group>Cyanobacteriota>Cyanophyceae>Oscillatoriophycideae>Chroococcales>Microcystaceae>Microcystis>unclassified_Microcystis | 129 | IM475_05900 | 9453..9842 | 2771171 | type-II-toxin-antitoxin-system-VapC-family-toxin-[Microcystis-sp.-M061S2] |
| MCA2716393.1 | PIN-EndoRNase | GCA_020245865.1 | Microcystis_sp._M169S2 | Bacteria>Bacillati>Cyanobacteriota/Melainabacteria_group>Cyanobacteriota>Cyanophyceae>Oscillatoriophycideae>Chroococcales>Microcystaceae>Microcystis>unclassified_Microcystis | 129 | IM485_01020 | 913..1302 | 2771157 | type-II-toxin-antitoxin-system-VapC-family-toxin-[Microcystis-sp.-M169S2] |
| MCA2627576.1 | PIN-EndoRNase | GCA_020245485.1 | Microcystis_sp._M091S2 | Bacteria>Bacillati>Cyanobacteriota/Melainabacteria_group>Cyanobacteriota>Cyanophyceae>Oscillatoriophycideae>Chroococcales>Microcystaceae>Microcystis>unclassified_Microcystis | 137 | IM505_05365 | 42822..43235 | 2771177 | PIN-domain-containing-protein-[Microcystis-sp.-M091S2] |
| MCA2653776.1 | PIN-EndoRNase | GCA_020245605.1 | Microcystis_sp._M061S2 | Bacteria>Bacillati>Cyanobacteriota/Melainabacteria_group>Cyanobacteriota>Cyanophyceae>Oscillatoriophycideae>Chroococcales>Microcystaceae>Microcystis>unclassified_Microcystis | 137 | IM475_05880 | 5187..5600 | 2771171 | PIN-domain-containing-protein-[Microcystis-sp.-M061S2] |
| MCE2669066.1 | PIN-EndoRNase | GCA_021300285.1 | Microcystis_sp._49638_E5 | Bacteria>Bacillati>Cyanobacteriota/Melainabacteria_group>Cyanobacteriota>Cyanophyceae>Oscillatoriophycideae>Chroococcales>Microcystaceae>Microcystis>unclassified_Microcystis | 137 | LW728_07575 | 16217..16630 | 2904986 | PIN-domain-containing-protein-[Microcystis-sp.-49638_E5] |
| MCA2627578.1 | PIN-EndoRNase | GCA_020245485.1 | Microcystis_sp._M091S2 | Bacteria>Bacillati>Cyanobacteriota/Melainabacteria_group>Cyanobacteriota>Cyanophyceae>Oscillatoriophycideae>Chroococcales>Microcystaceae>Microcystis>unclassified_Microcystis | 172 | IM505_05375 | 44832..45350 | 2771177 | hypothetical-protein-[Microcystis-sp.-M091S2] |
| MCE2669064.1 | PIN-EndoRNase | GCA_021300285.1 | Microcystis_sp._49638_E5 | Bacteria>Bacillati>Cyanobacteriota/Melainabacteria_group>Cyanobacteriota>Cyanophyceae>Oscillatoriophycideae>Chroococcales>Microcystaceae>Microcystis>unclassified_Microcystis | 172 | LW728_07565 | 14136..14654 | 2904986 | hypothetical-protein-[Microcystis-sp.-49638_E5] |
| WP_144390570.1 | PIN-EndoRNase | GCF_000710505.1 | Planktothrix_agardhii | Bacteria>Bacillati>Cyanobacteriota/Melainabacteria_group>Cyanobacteriota>Cyanophyceae>Oscillatoriophycideae>Oscillatoriales>Microcoleaceae>Planktothrix | 58 | L2E72_RS21835 | 6490..6666 | 1160 | type-II-toxin-antitoxin-system-VapC-family-toxin-[Planktothrix-agardhii] |
| WP_254035903.1 | PIN-EndoRNase | GCF_904830935.1 | Planktothrix_agardhii | Bacteria>Bacillati>Cyanobacteriota/Melainabacteria_group>Cyanobacteriota>Cyanophyceae>Oscillatoriophycideae>Oscillatoriales>Microcoleaceae>Planktothrix | 60 | NMG88_RS22300 | 46273..46455 | 1160 | PIN-domain-containing-protein-[Planktothrix-agardhii] |
| WP_254035902.1 | PIN-EndoRNase | GCF_904830935.1 | Planktothrix_agardhii | Bacteria>Bacillati>Cyanobacteriota/Melainabacteria_group>Cyanobacteriota>Cyanophyceae>Oscillatoriophycideae>Oscillatoriales>Microcoleaceae>Planktothrix | 78 | NMG88_RS22295 | 45776..46012 | 1160 | PIN-domain-containing-protein-[Planktothrix-agardhii] |
| WP_235758593.1 | PIN-EndoRNase | - | Planktothrix_agardhii | Bacteria>Bacillati>Cyanobacteriota/Melainabacteria_group>Cyanobacteriota>Cyanophyceae>Oscillatoriophycideae>Oscillatoriales>Microcoleaceae>Planktothrix | 121 | L2E81_RS19935 | 4060..4425 | 1160 | PIN-domain-containing-protein-[Planktothrix-agardhii] |
| WP_235751991.1 | PIN-EndoRNase | GCF_015708255.1 | Planktothrix_agardhii | Bacteria>Bacillati>Cyanobacteriota/Melainabacteria_group>Cyanobacteriota>Cyanophyceae>Oscillatoriophycideae>Oscillatoriales>Microcoleaceae>Planktothrix | 132 | I4E98_RS06860 | 97..495 | 1160 | type-II-toxin-antitoxin-system-VapC-family-toxin-[Planktothrix-agardhii] |
| WP_227351550.1 | PIN-EndoRNase | GCF_904830885.1 | Planktothrix_agardhii | Bacteria>Bacillati>Cyanobacteriota/Melainabacteria_group>Cyanobacteriota>Cyanophyceae>Oscillatoriophycideae>Oscillatoriales>Microcoleaceae>Planktothrix | 139 | NMK31_RS00270 | 64130..64549 | 1160 | type-II-toxin-antitoxin-system-VapC-family-toxin-[Planktothrix-agardhii] |
| WP_227382628.1 | PIN-EndoRNase | GCF_020594575.1 | Planktothrix_agardhii | Bacteria>Bacillati>Cyanobacteriota/Melainabacteria_group>Cyanobacteriota>Cyanophyceae>Oscillatoriophycideae>Oscillatoriales>Microcoleaceae>Planktothrix | 141 | LJE16_RS21905 | 4763..5188 | 1160 | type-II-toxin-antitoxin-system-VapC-family-toxin-[Planktothrix-agardhii] |
| WP_027255662.1 | PIN-EndoRNase | GCF_963863855.1 | Planktothrix_agardhii | Bacteria>Bacillati>Cyanobacteriota/Melainabacteria_group>Cyanobacteriota>Cyanophyceae>Oscillatoriophycideae>Oscillatoriales>Microcoleaceae>Planktothrix | 147 | N316_RS0124720 | 12254..12697 | 1160 | PIN-domain-containing-protein-[Planktothrix-agardhii] |
| WP_235758974.1 | PIN-EndoRNase | - | Planktothrix_agardhii | Bacteria>Bacillati>Cyanobacteriota/Melainabacteria_group>Cyanobacteriota>Cyanophyceae>Oscillatoriophycideae>Oscillatoriales>Microcoleaceae>Planktothrix | 147 | L2E81_RS22500 | 6110..6553 | 1160 | type-II-toxin-antitoxin-system-VapC-family-toxin-[Planktothrix-agardhii] |
| WP_254034080.1 | PIN-EndoRNase | GCF_904830845.1 | Planktothrix_agardhii | Bacteria>Bacillati>Cyanobacteriota/Melainabacteria_group>Cyanobacteriota>Cyanophyceae>Oscillatoriophycideae>Oscillatoriales>Microcoleaceae>Planktothrix | 147 | NMG87_RS21730 | 23312..23755 | 1160 | type-II-toxin-antitoxin-system-VapC-family-toxin-[Planktothrix-agardhii] |
| BBD57121.1 | PIN-EndoRNase | GCA_003609755.1 | Planktothrix_agardhii_NIES-204 | Bacteria>Bacillati>Cyanobacteriota/Melainabacteria_group>Cyanobacteriota>Cyanophyceae>Oscillatoriophycideae>Oscillatoriales>Microcoleaceae>Planktothrix>Planktothrix_agardhii | 58 | NIES204_44570 | 47414..47590 | 282423 | PilT-protein-domain-protein-[Planktothrix-agardhii-NIES-204] |
| BBD57225.1 | PIN-EndoRNase | GCA_003609755.1 | Planktothrix_agardhii_NIES-204 | Bacteria>Bacillati>Cyanobacteriota/Melainabacteria_group>Cyanobacteriota>Cyanophyceae>Oscillatoriophycideae>Oscillatoriales>Microcoleaceae>Planktothrix>Planktothrix_agardhii | 147 | NIES204_45610 | 19391..19834 | 282423 | hypothetical-protein-NIES204_45610-[Planktothrix-agardhii-NIES-204] |
| MBS9772313.1 | PIN-EndoRNase | GCA_018457135.1 | Trichodesmium_erythraeum_GBRTRLIN201 | Bacteria>Bacillati>Cyanobacteriota/Melainabacteria_group>Cyanobacteriota>Cyanophyceae>Oscillatoriophycideae>Oscillatoriales>Microcoleaceae>Trichodesmium>Trichodesmium_erythraeum | 161 | J7F05_18150 | 4051..4536 | 1045867 | DUF3368-domain-containing-protein-[Trichodesmium-erythraeum-GBRTRLIN201] |
| WP_199337025.1 | PIN-EndoRNase | GCF_963890435.1 | Oscillatoria_sp._FACHB-1407 | Bacteria>Bacillati>Cyanobacteriota/Melainabacteria_group>Cyanobacteriota>Cyanophyceae>Oscillatoriophycideae>Oscillatoriales>Oscillatoriaceae>Oscillatoria>unclassified_Oscillatoria | 115 | H6G89_RS30690 | 49375..49722 | 2692847 | hypothetical-protein-[Oscillatoria-sp.-FACHB-1407] |
| OAB59990.1 | PIN-EndoRNase | GCA_001637315.1 | Phormidium_willei_BDU_130791 | Bacteria>Bacillati>Cyanobacteriota/Melainabacteria_group>Cyanobacteriota>Cyanophyceae>Oscillatoriophycideae>Oscillatoriales>Oscillatoriaceae>Phormidium>Phormidium_willei | 126 | AY600_15245 | 2934..3314 | 399101 | twitching-motility-protein-PilT-[Phormidium-willei-BDU-130791] |
| OAB60005.1 | PIN-EndoRNase | GCA_001637315.1 | Phormidium_willei_BDU_130791 | Bacteria>Bacillati>Cyanobacteriota/Melainabacteria_group>Cyanobacteriota>Cyanophyceae>Oscillatoriophycideae>Oscillatoriales>Oscillatoriaceae>Phormidium>Phormidium_willei | 133 | AY600_15325 | 25791..26192 | 399101 | PIN-domain-protein-[Phormidium-willei-BDU-130791] |
| WP_242052255.1 | PIN-EndoRNase | GCF_014697375.1 | Phormidium_sp._FACHB-1136 | Bacteria>Bacillati>Cyanobacteriota/Melainabacteria_group>Cyanobacteriota>Cyanophyceae>Oscillatoriophycideae>Oscillatoriales>Oscillatoriaceae>Phormidium>unclassified_Phormidium | 116 | H6G62_RS24130 | 4836..5186 | 2692848 | hypothetical-protein-[Phormidium-sp.-FACHB-1136] |
| KPQ40622.1 | PIN-EndoRNase | GCA_001314905.1 | Phormidium_sp._OSCR | Bacteria>Bacillati>Cyanobacteriota/Melainabacteria_group>Cyanobacteriota>Cyanophyceae>Oscillatoriophycideae>Oscillatoriales>Oscillatoriaceae>Phormidium>unclassified_Phormidium | 126 | HLUCCO16_02240 | 105336..105716 | 1666905 | hypothetical-protein-HLUCCO16_02240-[Phormidium-sp.-OSCR] |
| WP_019503189.1 | PIN-EndoRNase | GCF_000332195.1 | Pleurocapsa_sp._PCC_7319 | Bacteria>Bacillati>Cyanobacteriota/Melainabacteria_group>Cyanobacteriota>Cyanophyceae>Pleurocapsales>Hyellaceae>Pleurocapsa>unclassified_Pleurocapsa | 141 | PLEUR7319_RS0100200 | 41539..41964 | 118161 | type-II-toxin-antitoxin-system-VapC-family-toxin-[Pleurocapsa-sp.-PCC-7319] |

| Protein-ID | Domain | Assembly-ID | Strain | Taxonomy | Length | Locus-Tag | Genomic-coordinates | TaxID | Fasta-defline |
|---|---|---|---|---|---|---|---|---|---|
| WP_019504692.1 | PIN-EndoRNase | GCF_000332195.1 | Pleurocapsa_sp._PCC_7319 | Bacteria>Bacillati>Cyanobacteriota/Melainabacteria_group>Cyanobacteriota>Cyanophyceae>Pleurocapsales>Hyellaceae>Pleurocapsa>unclassified_Pleurocapsa | 155 | PLEUR7319_RS0107980 | 845766..846233 | 118161 | hypothetical-protein-[Pleurocapsa-sp.-PCC-7319] |
| WP_202950843.1 | PIN-EndoRNase | GCF_000332315.1 | Prochlorothrix_hollandica | Bacteria>Bacillati>Cyanobacteriota/Melainabacteria_group>Cyanobacteriota>Cyanophyceae>Prochlorotrichales>Prochlorotrichaceae>Prochlorothrix | 56 | PRO9006_RS36580 | 50135..50305 | 1223 | hypothetical-protein-[Prochlorothrix-hollandica] |
| WP_281006613.1 | PIN-EndoRNase | GCF_029910235.1 | Pseudanabaena | Bacteria>Bacillati>Cyanobacteriota/Melainabacteria_group>Cyanobacteriota>Cyanophyceae>Pseudanabaenales>Pseudanabaenaceae | 138 | OA858_RS18385 | 4057390..4057806 | 1152 | PIN-domain-containing-protein-[Pseudanabaena-galeata] |
| NUN66952.1 | PIN-EndoRNase | GCA_013361095.1 | Pseudanabaena_biceps | Bacteria>Bacillati>Cyanobacteriota/Melainabacteria_group>Cyanobacteriota>Cyanophyceae>Pseudanabaenales>Pseudanabaenaceae>Pseudanabaena | 194 | HCU40_19875 | 99748..100332 | 927669 | nuclease-[Pseudanabaena-biceps] |
| WP_265264587.1 | PIN-EndoRNase | GCF_026130165.1 | Spirulina_subsalsa | Bacteria>Bacillati>Cyanobacteriota/Melainabacteria_group>Cyanobacteriota>Cyanophyceae>Spirulinales>Spirulinaceae>Spirulina | 129 | K4A83_RS11010 | 16831..17220 | 54311 | type-II-toxin-antitoxin-system-VapC-family-toxin-[Spirulina-subsalsa] |
| WP_255005453.1 | PIN-EndoRNase | GCF_024346315.1 | unclassified_Cyanobium | Bacteria>Bacillati>Cyanobacteriota/Melainabacteria_group>Cyanobacteriota>Cyanophyceae>Synechococcales>Prochlorococcaceae>Cyanobium | 77 | KBZ13_RS02060 | 365258..365491 | 2627006 | type-II-toxin-antitoxin-system-VapB-family-antitoxin-[Cyanobium-sp.-ATX-6F1] |
| WP_255005454.1 | PIN-EndoRNase | GCF_024346315.1 | unclassified_Cyanobium | Bacteria>Bacillati>Cyanobacteriota/Melainabacteria_group>Cyanobacteriota>Cyanophyceae>Synechococcales>Prochlorococcaceae>Cyanobium | 133 | KBZ13_RS02065 | 365488..365889 | 2627006 | type-II-toxin-antitoxin-system-VapC-family-toxin-[Cyanobium-sp.-ATX-6F1] |
| WP_254970969.1 | PIN-EndoRNase | GCF_024346155.1 | Cyanobium_sp._Candia_9D4 | Bacteria>Bacillati>Cyanobacteriota/Melainabacteria_group>Cyanobacteriota>Cyanophyceae>Synechococcales>Prochlorococcaceae>Cyanobium>unclassified_Cyanobium | 79 | KBZ08_RS02045 | 406874..407113 | 2823707 | AbrB/MazE/SpoVT-family-DNA-binding-domain-containing-protein-[Cyanobium-sp.-Candia-9D4] |
| WP_261358787.1 | PIN-EndoRNase | - | Cyanobium_sp._ATX_6F1 | Bacteria>Bacillati>Cyanobacteriota/Melainabacteria_group>Cyanobacteriota>Cyanophyceae>Synechococcales>Prochlorococcaceae>Cyanobium>unclassified_Cyanobium | 125 | KBZ13_RS02125 | 381826..382203 | 2823702 | type-II-toxin-antitoxin-system-VapC-family-toxin-[Cyanobium-sp.-ATX-6F1] |
| WP_254967822.1 | PIN-EndoRNase | GCF_024346135.1 | Cyanobium_sp._CH-040 | Bacteria>Bacillati>Cyanobacteriota/Melainabacteria_group>Cyanobacteriota>Cyanophyceae>Synechococcales>Prochlorococcaceae>Cyanobium>unclassified_Cyanobium | 132 | KBY90_RS07060 | 4302..4700 | 2823708 | type-II-toxin-antitoxin-system-VapC-family-toxin-[Cyanobium-sp.-CH-040] |
| WP_254940708.1 | PIN-EndoRNase | GCF_024345575.1 | Cyanobium_sp._Morenito_9A2 | Bacteria>Bacillati>Cyanobacteriota/Melainabacteria_group>Cyanobacteriota>Cyanophyceae>Synechococcales>Prochlorococcaceae>Cyanobium>unclassified_Cyanobium | 132 | KBY88_RS14785 | 38262..38660 | 2823718 | PIN-domain-containing-protein-[Cyanobium-sp.-Morenito-9A2] |
| WP_254967837.1 | PIN-EndoRNase | GCF_024346135.1 | Cyanobium_sp._CH-040 | Bacteria>Bacillati>Cyanobacteriota/Melainabacteria_group>Cyanobacteriota>Cyanophyceae>Synechococcales>Prochlorococcaceae>Cyanobium>unclassified_Cyanobium | 133 | KBY90_RS07135 | 22053..22454 | 2823708 | type-II-toxin-antitoxin-system-VapC-family-toxin-[Cyanobium-sp.-CH-040] |
| WP_087068558.1 | PIN-EndoRNase | GCF_900088535.1 | Cyanobium_sp._NIES-981 | Bacteria>Bacillati>Cyanobacteriota/Melainabacteria_group>Cyanobacteriota>Cyanophyceae>Synechococcales>Prochlorococcaceae>Cyanobium>unclassified_Cyanobium | 134 | CBM981_RS11740 | 2330295..2330699 | 1851505 | PIN-domain-nuclease-[Cyanobium-sp.-NIES-981] |
| WP_254970970.1 | PIN-EndoRNase | GCF_024346155.1 | Cyanobium_sp._Candia_9D4 | Bacteria>Bacillati>Cyanobacteriota/Melainabacteria_group>Cyanobacteriota>Cyanophyceae>Synechococcales>Prochlorococcaceae>Cyanobium>unclassified_Cyanobium | 135 | KBZ08_RS02050 | 407113..407520 | 2823707 | type-II-toxin-antitoxin-system-VapC-family-toxin-[Cyanobium-sp.-Candia-9D4] |
| WP_254967818.1 | PIN-EndoRNase | GCF_024346135.1 | Cyanobium_sp._CH-040 | Bacteria>Bacillati>Cyanobacteriota/Melainabacteria_group>Cyanobacteriota>Cyanophyceae>Synechococcales>Prochlorococcaceae>Cyanobium>unclassified_Cyanobium | 138 | KBY90_RS07040 | 413..829 | 2823708 | type-II-toxin-antitoxin-system-VapC-family-toxin-[Cyanobium-sp.-CH-040] |
| WP_187147225.1 | PIN-EndoRNase | GCF_000013225.1 | unclassified_Synechococcus | Bacteria>Bacillati>Cyanobacteriota/Melainabacteria_group>Cyanobacteriota>Cyanophyceae>Synechococcales>Synechococcaceae>Synechococcus | 56 | CYB_RS15160 | 1802894..1803064 | 2626047 | DUF5615-family-PIN-like-protein-[Synechococcus-sp.-JA-2-3B'a(2-13)] |
| MBV5261738.1 | PIN-EndoRNase | GCA_019155215.1 | Synechococcus_moorigangaii_CMS01 | Bacteria>Bacillati>Cyanobacteriota/Melainabacteria_group>Cyanobacteriota>Cyanophyceae>Synechococcales>Synechococcaceae>Synechococcus>Synechococcus_moorigangaii | 142 | FLX56_25330 | 10539..10967 | 2025807 | type-II-toxin-antitoxin-system-VapC-family-toxin-[Synechococcus-moorigangaii-CMS01] |
| WP_197170380.1 | PIN-EndoRNase | GCF_015840915.1 | Synechococcus_sp._CBW1002 | Bacteria>Bacillati>Cyanobacteriota/Melainabacteria_group>Cyanobacteriota>Cyanophyceae>Synechococcales>Synechococcaceae>Synechococcus>unclassified_Synechococcus | 74 | H8F24_RS17515 | 3554469..3554693 | 1353134 | PIN-domain-containing-protein-[Synechococcus-sp.-CBW1002] |
| WP_231597942.1 | PIN-EndoRNase | GCF_015840915.1 | Synechococcus_sp._CBW1002 | Bacteria>Bacillati>Cyanobacteriota/Melainabacteria_group>Cyanobacteriota>Cyanophyceae>Synechococcales>Synechococcaceae>Synechococcus>unclassified_Synechococcus | 113 | H8F24_RS17510 | 3554146..3554487 | 1353134 | type-II-toxin-antitoxin-system-VapC-family-toxin-[Synechococcus-sp.-CBW1002] |
| WP_197170391.1 | PIN-EndoRNase | GCF_015840915.1 | Synechococcus_sp._CBW1002 | Bacteria>Bacillati>Cyanobacteriota/Melainabacteria_group>Cyanobacteriota>Cyanophyceae>Synechococcales>Synechococcaceae>Synechococcus>unclassified_Synechococcus | 132 | H8F24_RS17580 | 3574259..3574657 | 1353134 | type-II-toxin-antitoxin-system-VapC-family-toxin-[Synechococcus-sp.-CBW1002] |
| WP_094555437.1 | PIN-EndoRNase | GCF_002252625.1 | Synechococcus_sp._1G10 | Bacteria>Bacillati>Cyanobacteriota/Melainabacteria_group>Cyanobacteriota>Cyanophyceae>Synechococcales>Synechococcaceae>Synechococcus>unclassified_Synechococcus | 140 | CJZ82_RS08955 | 62417..62839 | 2025605 | type-II-toxin-antitoxin-system-VapC-family-toxin-[Synechococcus-sp.-1G10] |
| WP_254428054.1 | PIN-EndoRNase | GCF_004209775.1 | Synechococcus_sp._WH_8101 | Bacteria>Bacillati>Cyanobacteriota/Melainabacteria_group>Cyanobacteriota>Cyanophyceae>Synechococcales>Synechococcaceae>Synechococcus>unclassified_Synechococcus | 142 | SynWH8101_RS05105 | 1016251..1016679 | 59932 | type-II-toxin-antitoxin-system-VapC-family-toxin-[Synechococcus-sp.-WH-8101] |
| NJL99824.1 | PIN-EndoRNase | GCA_012031565.1 | Synechococcaceae_cyanobacterium_SM2_3_2 | Bacteria>Bacillati>Cyanobacteriota/Melainabacteria_group>Cyanobacteriota>Cyanophyceae>Synechococcales>Synechococcaceae>unclassified_Synechococcaceae | 138 | HC924_13965 | 5737..6153 | 2720471 | putative-toxin-antitoxin-system-toxin-component,-PIN-family-[Synechococcaceae-cyanobacterium-SM2_3_2] |
| HBH74796.1 | PIN-EndoRNase | GCA_003508015.1 | Synechococcales_bacterium_UBA10510 | Bacteria>Bacillati>Cyanobacteriota/Melainabacteria_group>Cyanobacteriota>Cyanophyceae>Synechococcales>unclassified_Synechococcales | 121 | DDY43_15530 | 1..367 | 2055795 | VapC-toxin-family-PIN-domain-ribonuclease-[Synechococcales-bacterium-UBA10510] |
| MBM5817553.1 | PIN-EndoRNase | GCA_016882305.1 | Cyanobacteria_bacterium_K_Offshore_surface_m2_239 | Bacteria>Bacillati>Cyanobacteriota/Melainabacteria_group>Cyanobacteriota>unclassified_Cyanobacteriota | 134 | FJ083_13515 | 1096..1500 | 2775555 | type-II-toxin-antitoxin-system-VapC-family-toxin-[Cyanobacteria-bacterium-K_Offshore_surface_m2_239] |
| MBM5817551.1 | PIN-EndoRNase | GCA_016882305.1 | Cyanobacteria_bacterium_K_Offshore_surface_m2_239 | Bacteria>Bacillati>Cyanobacteriota/Melainabacteria_group>Cyanobacteriota>unclassified_Cyanobacteriota | 145 | FJ083_13505 | 155..592 | 2775555 | type-II-toxin-antitoxin-system-VapC-family-toxin-[Cyanobacteria-bacterium-K_Offshore_surface_m2_239] |
| MCY4173477.1 | PIN-EndoRNase | GCA_026709085.1 | Cyanobacteria_bacterium_MAG_CAR3_bin_5 | Bacteria>Bacillati>Cyanobacteriota/Melainabacteria_group>Cyanobacteriota>unclassified_Cyanobacteriota | 159 | OXF25_05345 | 8085..8564 | 2775591 | MAG:-PIN-domain-containing-protein-[Cyanobacteria-bacterium-MAG-CAR3_bin_5] |
| MDA0267770.1 | PIN-EndoRNase | GCA_027592485.1 | Cyanobacteriota_bacterium | Bacteria>Bacillati>Cyanobacteriota/Melainabacteria_group>Cyanobacteriota>unclassified_Cyanobacteriota | 166 | O3A14_12620 | 2429..2929 | 2847862 | MAG:-DUF3368-domain-containing-protein-[Cyanobacteria-bacterium] |
| HBB35647.1 | PIN-EndoRNase | GCA_003488275.1 | Cyanobacteria_bacterium_UBA9273 | Bacteria>Bacillati>Cyanobacteriota/Melainabacteria_group>Cyanobacteriota>unclassified_Cyanobacteriota | 195 | DC064_28670 | 9114..9701 | 2055791 | nuclease-[Cyanobacteria-bacterium-UBA9273] |
| NIM17955.1 | PIN-EndoRNase | GCA_011771225.1 | Candidatus_Aminicenantes_bacterium | Bacteria>Bacteria_incertae_sedis>Bacteria_candidate_phyla>Candidatus_Aminicenantota>unclassified_Candidatus_Aminicenantes | 134 | GT081_38755 | 21740..22144 | 2052149 | PIN-domain-containing-protein-[Candidatus-Aminicenantes-bacterium] |
| NIM17949.1 | PIN-EndoRNase | GCA_011771225.1 | Candidatus_Aminicenantes_bacterium | Bacteria>Bacteria_incertae_sedis>Bacteria_candidate_phyla>Candidatus_Aminicenantota>unclassified_Candidatus_Aminicenantes | 160 | GT081_38725 | 14169..14651 | 2052149 | PIN-domain-containing-protein-[Candidatus-Aminicenantes-bacterium] |
| MBF6571681.1 | PIN-EndoRNase | GCA_015478625.1 | Candidatus_Binataceae_bacterium | Bacteria>Bacteria_incertae_sedis>Bacteria_candidate_phyla>Candidatus_Binatota>Candidatus_Binatia>Candidatus_Binatales>Candidatus_Binataceae>unclassified_Candidatus_Binataceae | 124 | IVW54_22800 | 2962..3336 | 2785449 | DUF5615-family-PIN-like-protein-[Candidatus-Binataceae-bacterium] |
| MBF6571347.1 | PIN-EndoRNase | GCA_015478625.1 | Candidatus_Binataceae_bacterium | Bacteria>Bacteria_incertae_sedis>Bacteria_candidate_phyla>Candidatus_Binatota>Candidatus_Binatia>Candidatus_Binatales>Candidatus_Binataceae>unclassified_Candidatus_Binataceae | 168 | IVW54_21015 | 18813..19319 | 2785449 | type-II-toxin-antitoxin-system-VapC-family-toxin-[Candidatus-Binataceae-bacterium] |
| WP_157959501.1 | PIN-EndoRNase | GCF_900465355.1 | Candidatus_Bipolaricaulis_anaerobius | Bacteria>Bacteria_incertae_sedis>Bacteria_candidate_phyla>Candidatus_Bipolaricaulota>Candidatus_Bipolaricaulia>Candidatus_Bipolaricaulales>Candidatus_Bipolaricaulaceae>Candidatus_Bipolaricaulis | 175 | BARAN1_RS05365 | 1116305..1116832 | 2026885 | hypothetical-protein-[Candidatus-Bipolaricaulis-anaerobius] |
| HAF70685.1 | PIN-EndoRNase | GCA_003445465.1 | Candidatus_Acetothermia_bacterium | Bacteria>Bacteria_incertae_sedis>Bacteria_candidate_phyla>Candidatus_Bipolaricaulota>unclassified_Candidatus_Bipolaricaulota | 149 | DCL37_04965 | 27684..28133 | 2053493 | PIN-domain-nuclease-[Candidatus-Acetothermia-bacterium] |
| KUK26948.1 | PIN-EndoRNase | GCA_001508155.1 | Acetothermia_bacterium_64_32 | Bacteria>Bacteria_incertae_sedis>Bacteria_candidate_phyla>Candidatus_Bipolaricaulota>unclassified_Candidatus_Bipolaricaulota | 149 | XD60_0837 | 43799..44248 | 1635286 | PilT-protein-like-protein-[Acetothermia-bacterium-64_32] |
| MBI1799724.1 | PIN-EndoRNase | GCA_016178685.1 | Candidatus_Eiseniibacteriota_bacterium | Bacteria>Bacteria_incertae_sedis>Bacteria_candidate_phyla>Candidatus_Eiseniibacteriota | 216 | HYR73_08555 | 13013..13663 | 2212470 | NYN-domain-containing-protein-[Candidatus-Eisenbacteria-bacterium] |
| MBZ0160825.1 | PIN-EndoRNase | GCA_019912425.1 | Candidatus_Methylomirabilis_tolerans | Bacteria>Bacteria_incertae_sedis>Bacteria_candidate_phyla>Candidatus_Methylomirabilota>Candidatus_Methylomirabilia>Candidatus_Methylomirabilales>Candidatus_Methylomirabilaceae>Candidatus_Methylomirabilis | 120 | K8G79_11935 | 3411..3773 | 3123416 | DUF5615-family-PIN-like-protein-[bacterium] |
| MYC78294.1 | PIN-EndoRNase | GCA_009841255.1 | Candidatus_Poribacteria_bacterium | Bacteria>Bacteria_incertae_sedis>Bacteria_candidate_phyla>Candidatus_Poribacteria>unclassified_Candidatus_Poribacteria | 117 | F4X10_21235 | 50666..51019 | 2026781 | DUF4411-family-protein-[Candidatus-Poribacteria-bacterium] |
| MCY3724178.1 | PIN-EndoRNase | GCA_026705955.1 | Candidatus_Poribacteria_bacterium | Bacteria>Bacteria_incertae_sedis>Bacteria_candidate_phyla>Candidatus_Poribacteria>unclassified_Candidatus_Poribacteria | 144 | OXG97_18325 | 41..475 | 2026781 | MAG:-DUF4411-family-protein-[Candidatus-Poribacteria-bacterium] |
| MDE0014208.1 | PIN-EndoRNase | GCA_028818705.1 | Candidatus_Poribacteria_bacterium | Bacteria>Bacteria_incertae_sedis>Bacteria_candidate_phyla>Candidatus_Poribacteria>unclassified_Candidatus_Poribacteria | 166 | OXU36_23895 | 10671..11171 | 2026781 | MAG:-DUF4411-family-protein-[Candidatus-Poribacteria-bacterium] |
| MDE0681303.1 | PIN-EndoRNase | GCA_028824575.1 | Candidatus_Poribacteria_bacterium | Bacteria>Bacteria_incertae_sedis>Bacteria_candidate_phyla>Candidatus_Poribacteria>unclassified_Candidatus_Poribacteria | 166 | OXI63_00190 | 2505..3005 | 2026781 | MAG:-DUF4411-family-protein-[Candidatus-Poribacteria-bacterium] |
| MBM3210710.1 | PIN-EndoRNase | GCA_016866835.1 | Candidatus_Poribacteria_bacterium | Bacteria>Bacteria_incertae_sedis>Bacteria_candidate_phyla>Candidatus_Poribacteria>unclassified_Candidatus_Poribacteria | 166 | FJZ33_00705 | 3242..3742 | 2026781 | type-II-toxin-antitoxin-system-VapC-family-toxin-[Candidatus-Poribacteria-bacterium] |
| MBI5115314.1 | PIN-EndoRNase | GCA_016215705.1 | Candidatus_Poribacteria_bacterium | Bacteria>Bacteria_incertae_sedis>Bacteria_candidate_phyla>Candidatus_Poribacteria>unclassified_Candidatus_Poribacteria | 167 | HZA56_02470 | 3625..4128 | 2026781 | DUF4411-family-protein-[Candidatus-Poribacteria-bacterium] |
| MXV83701.1 | PIN-EndoRNase | GCA_009835485.1 | Candidatus_Poribacteria_bacterium | Bacteria>Bacteria_incertae_sedis>Bacteria_candidate_phyla>Candidatus_Poribacteria>unclassified_Candidatus_Poribacteria | 224 | F4Z98_10070 | 18397..19071 | 2026781 | DUF4411-family-protein-[Candidatus-Poribacteria-bacterium] |
| MBI4865892.1 | PIN-EndoRNase | GCA_016213845.1 | Candidatus_Wallbacteria_bacterium | Bacteria>Bacteria_incertae_sedis>Bacteria_candidate_phyla>Candidatus_Walliibacteriota>unclassified_Candidatus_Wallbacteria | 133 | HY816_02975 | 110304..110705 | 2053640 | DUF5615-family-PIN-like-protein-[Candidatus-Wallbacteria-bacterium] |
| GAK56406.1 | PIN-EndoRNase | GCA_000739535.1 | Candidatus_Vecturithrix_granuli | Bacteria>Candidatus_Moduliflexota>Candidatus_Vecturitrichia>Candidatus_Vecturitrichales>Candidatus_Vecturitrichaceae>Candidatus_Vecturithrix | 166 | U27_03368 | 802091..802591 | 1499967 | hypothetical-protein-U27_03368-[Candidatus-Vecturithrix-granuli] |
| OGQ80568.1 | PIN-EndoRNase | GCA_001798595.1 | Deltaproteobacteria_bacterium_RIFCSPLOWO2_12_FULL_60_19 | Bacteria>delta/epsilon_subdivisions>Deltaproteobacteria>unclassified_Deltaproteobacteria | 172 | A3F90_01980 | 1903..2421 | 1797894 | hypothetical-protein-A3F90_01980-[Deltaproteobacteria-bacterium-RIFCSPLOWO2_12_FULL_60_19] |
| MBA3712811.1 | PIN-EndoRNase | GCA_013816505.1 | Pyrinomonadaceae_bacterium | Bacteria>Pseudomonadati>Acidobacteriota>Blastocatellia>Blastocatellales>Pyrinomonadaceae>unclassified_Pyrinomonadaceae | 115 | HOW76_10235 | 5053..5400 | 2283092 | DUF5615-family-PIN-like-protein-[Pyrinomonadaceae-bacterium] |
| GIU78632.1 | PIN-EndoRNase | GCA_026002875.1 | Bryobacteraceae_bacterium | Bacteria>Pseudomonadati>Acidobacteriota>Terriglobia>Bryobacterales>Bryobacteraceae>unclassified_Bryobacteraceae | 165 | KatS3mg005_1870 | 2217721..2218218 | 2212468 | MAG:-hypothetical-protein-KatS3mg005_1870-[Bryobacteraceae-bacterium] |
| MCS6954440.1 | PIN-EndoRNase | GCA_025055455.1 | Bryobacteraceae_bacterium | Bacteria>Pseudomonadati>Acidobacteriota>Terriglobia>Bryobacterales>Bryobacteraceae>unclassified_Bryobacteraceae | 165 | NZM33_16465 | 15316..15813 | 2212468 | DUF4411-family-protein-[Bryobacteraceae-bacterium] |
| MCY4597194.1 | PIN-EndoRNase | GCA_026713455.1 | Bryobacterales_bacterium | Bacteria>Pseudomonadati>Acidobacteriota>Terriglobia>Bryobacterales>unclassified_Bryobacterales | 136 | OXC19_20630 | 8001..8411 | 2026791 | MAG:-PIN-domain-containing-protein-[Bryobacterales-bacterium] |
| MDE0626644.1 | PIN-EndoRNase | GCA_028824065.1 | Bryobacterales_bacterium | Bacteria>Pseudomonadati>Acidobacteriota>Terriglobia>Bryobacterales>unclassified_Bryobacterales | 168 | OXH99_09620 | 919..1425 | 2026791 | MAG:-hypothetical-protein-OXH99_09620-[Bryobacterales-bacterium] |
| WP_211425995.1 | PIN-EndoRNase | GCF_018304805.1 | Chloracidobacterium_aggregatum | Bacteria>Pseudomonadati>Acidobacteriota>Terriglobia>Terriglobales>Acidobacteriaceae>Chloracidobacterium | 205 | J8C02_RS10950 | 2613815..2614432 | 2851959 | NYN-domain-containing-protein-[Chloracidobacterium-aggregatum] |
| MBP7799277.1 | PIN-EndoRNase | GCA_018000955.1 | Thermoanaerobaculaceae_bacterium | Bacteria>Pseudomonadati>Acidobacteriota>Thermoanaerobaculia>Thermoanaerobaculales>Thermoanaerobaculaceae>unclassified_Thermoanaerobaculaceae | 174 | KA072_11700 | 26354..26878 | 2699758 | hypothetical-protein-[Thermoanaerobaculaceae-bacterium] |
| RMH20599.1 | PIN-EndoRNase | GCA_003697015.1 | Acidobacteriota_bacterium | Bacteria>Pseudomonadati>Acidobacteriota>unclassified_Acidobacteriota | 115 | D6696_07820 | 1210..1557 | 1978231 | hypothetical-protein-D6696_07820-[Acidobacteriota-bacterium] |
| MBI4455128.1 | PIN-EndoRNase | GCA_016211105.1 | Acidobacteriota_bacterium | Bacteria>Pseudomonadati>Acidobacteriota>unclassified_Acidobacteriota | 121 | HY644_04430 | 207732..208097 | 1978231 | DUF5615-family-PIN-like-protein-[Acidobacteriota-bacterium] |
| MXW39172.1 | PIN-EndoRNase | GCA_009836105.1 | Acidobacteriota_bacterium | Bacteria>Pseudomonadati>Acidobacteriota>unclassified_Acidobacteriota | 124 | F4Z65_13030 | 5780..6154 | 1978231 | hypothetical-protein-[Acidobacteriota-bacterium] |
| MCI0624091.1 | PIN-EndoRNase | GCA_022615935.1 | Acidobacteriota_bacterium | Bacteria>Pseudomonadati>Acidobacteriota>unclassified_Acidobacteriota | 128 | L0387_20975 | 9774..10160 | 1978231 | type-II-toxin-antitoxin-system-VapC-family-toxin-[Acidobacteriota-bacterium] |
| QQS46396.1 | PIN-EndoRNase | GCA_016700195.1 | Acidobacteriota_bacterium | Bacteria>Pseudomonadati>Acidobacteriota>unclassified_Acidobacteriota | 128 | IPM66_21215 | 5339828..5340214 | 1978231 | type-II-toxin-antitoxin-system-VapC-family-toxin-[Acidobacteriota-bacterium] |
| MCY3931993.1 | PIN-EndoRNase | GCA_026707325.1 | Acidobacteriota_bacterium | Bacteria>Pseudomonadati>Acidobacteriota>unclassified_Acidobacteriota | 131 | OXH70_09755 | 11194..11589 | 1978231 | MAG:-type-II-toxin-antitoxin-system-VapC-family-toxin-[Acidobacteria-bacterium] |
| QQS46116.1 | PIN-EndoRNase | GCA_016700195.1 | Acidobacteriota_bacterium | Bacteria>Pseudomonadati>Acidobacteriota>unclassified_Acidobacteriota | 132 | IPM66_19705 | 4958095..4958493 | 1978231 | type-II-toxin-antitoxin-system-VapC-family-toxin-[Acidobacteriota-bacterium] |
| MYN64247.1 | PIN-EndoRNase | GCA_009861545.1 | Acidobacteriota_bacterium | Bacteria>Pseudomonadati>Acidobacteriota>unclassified_Acidobacteriota | 149 | F4X11_04360 | 50618..51067 | 1978231 | PIN-domain-containing-protein-[Acidobacteriota-bacterium] |
| MYN64242.1 | PIN-EndoRNase | GCA_009861545.1 | Acidobacteriota_bacterium | Bacteria>Pseudomonadati>Acidobacteriota>unclassified_Acidobacteriota | 151 | F4X11_04335 | 44688..45143 | 1978231 | putative-toxin-antitoxin-system-toxin-component,-PIN-family-[Acidobacteriota-bacterium] |
| MYN64238.1 | PIN-EndoRNase | GCA_009861545.1 | Acidobacteriota_bacterium | Bacteria>Pseudomonadati>Acidobacteriota>unclassified_Acidobacteriota | 155 | F4X11_04310 | 39907..40374 | 1978231 | hypothetical-protein-[Acidobacteriota-bacterium] |
| MCY4659241.1 | PIN-EndoRNase | GCA_026713985.1 | Acidobacteriota_bacterium | Bacteria>Pseudomonadati>Acidobacteriota>unclassified_Acidobacteriota | 157 | OXF93_05430 | 4082..4555 | 1978231 | MAG:-hypothetical-protein-OXF93_05430-[Acidobacteria-bacterium] |
| MCI0393079.1 | PIN-EndoRNase | GCA_022614935.1 | Acidobacteriota_bacterium | Bacteria>Pseudomonadati>Acidobacteriota>unclassified_Acidobacteriota | 160 | MOB07_30475 | 18885..19367 | 1978231 | DUF3368-domain-containing-protein-[Acidobacteriota-bacterium] |
| MBI4455136.1 | PIN-EndoRNase | GCA_016211105.1 | Acidobacteriota_bacterium | Bacteria>Pseudomonadati>Acidobacteriota>unclassified_Acidobacteriota | 173 | HY644_04470 | 214255..214776 | 1978231 | type-II-toxin-antitoxin-system-VapC-family-toxin-[Acidobacteriota-bacterium] |
| MCI0623779.1 | PIN-EndoRNase | GCA_022615935.1 | Acidobacteriota_bacterium | Bacteria>Pseudomonadati>Acidobacteriota>unclassified_Acidobacteriota | 183 | L0387_19335 | 6864..7415 | 1978231 | PIN-domain-containing-protein-[Acidobacteriota-bacterium] |
| MCL4821455.1 | PIN-EndoRNase | GCA_023384615.1 | Vicinamibacteria_bacterium | Bacteria>Pseudomonadati>Acidobacteriota>Vicinamibacteria>unclassified_Vicinamibacteria | 159 | KJ067_20155 | 7986..8465 | 2793154 | hypothetical-protein-[Vicinamibacteria-bacterium] |
| MPZ20363.1 | PIN-EndoRNase | GCA_009377605.1 | Luteitalea_sp. | Bacteria>Pseudomonadati>Acidobacteriota>Vicinamibacteria>Vicinamibacterales>Vicinamibacteraceae>Luteitalea>unclassified_Luteitalea | 206 | GEV06_20975 | 11619..12239 | 2004800 | NYN-domain-containing-protein-[Luteitalea-sp.] |
| RUM61920.1 | PIN-EndoRNase | GCA_003978555.1 | Persephonella_sp. | Bacteria>Pseudomonadati>Aquificota>Aquificia>Aquificales>Hydrogenothermaceae>Persephonella>unclassified_Persephonella | 128 | DSY59_01050 | 1587..1973 | 2060922 | MAG:-pilus-assembly-protein-[Persephonella-sp.] |
| MBI5243144.1 | PIN-EndoRNase | GCA_016212885.1 | Elusimicrobiota_bacterium | Bacteria>Pseudomonadati>Elusimicrobiota>unclassified_Elusimicrobiota | 130 | HY922_05580 | 3647..4039 | 2030800 | type-II-toxin-antitoxin-system-VapC-family-toxin-[Elusimicrobiota-bacterium] |
| MBI5243203.1 | PIN-EndoRNase | GCA_016212885.1 | Elusimicrobiota_bacterium | Bacteria>Pseudomonadati>Elusimicrobiota>unclassified_Elusimicrobiota | 175 | HY922_05875 | 4324..4851 | 2030800 | type-II-toxin-antitoxin-system-VapC-family-toxin-[Elusimicrobiota-bacterium] |
| MCX5796461.1 | PIN-EndoRNase | GCA_026388095.1 | Elusimicrobiota_bacterium | Bacteria>Pseudomonadati>Elusimicrobiota>unclassified_Elusimicrobiota | 183 | NTY77_13290 | 56839..57390 | 2030800 | MAG:-PIN-domain-containing-protein-[Elusimicrobia-bacterium] |
| MBI5243149.1 | PIN-EndoRNase | GCA_016212885.1 | Elusimicrobiota_bacterium | Bacteria>Pseudomonadati>Elusimicrobiota>unclassified_Elusimicrobiota | 186 | HY922_05605 | 14817..15377 | 2030800 | PIN-domain-containing-protein-[Elusimicrobiota-bacterium] |
| WP_234146090.1 | PIN-EndoRNase | GCF_024760085.1 | Bacteroides_uniformis | Bacteria>Pseudomonadati>FCB_group>Bacteroidota/Chlorobiota_group>Bacteroidota>Bacteroidia>Bacteroidales>Bacteroidaceae>Bacteroides | 174 | J8872_RS07810 | 78733..79257 | 820 | hypothetical-protein-[Bacteroides-uniformis] |
| WP_264909060.1 | PIN-EndoRNase | GCF_048306495.1 | Segatella | Bacteria>Pseudomonadati>FCB_group>Bacteroidota/Chlorobiota_group>Bacteroidota>Bacteroidia>Bacteroidales>Prevotellaceae | 173 | ONT11_RS09730 | 2181638..2182159 | 2974251 | hypothetical-protein-[Prevotella-copri] |
| MBR1880070.1 | PIN-EndoRNase | GCA_017478995.1 | Prevotella_sp. | Bacteria>Pseudomonadati>FCB_group>Bacteroidota/Chlorobiota_group>Bacteroidota>Bacteroidia>Bacteroidales>Prevotellaceae>Prevotella>unclassified_Prevotella | 144 | IJ804_04905 | 6147..6581 | 59823 | PIN-domain-containing-protein-[Prevotella-sp.] |
| HAH90339.1 | PIN-EndoRNase | GCA_003450425.1 | Prevotella_sp. | Bacteria>Pseudomonadati>FCB_group>Bacteroidota/Chlorobiota_group>Bacteroidota>Bacteroidia>Bacteroidales>Prevotellaceae>Prevotella>unclassified_Prevotella | 173 | DCL96_01180 | 12484..13005 | 59823 | hypothetical-protein-[Prevotella-sp.] |
| WP_264951501.1 | PIN-EndoRNase | GCF_026015505.1 | Segatella_copri | Bacteria>Pseudomonadati>FCB_group>Bacteroidota/Chlorobiota_group>Bacteroidota>Bacteroidia>Bacteroidales>Prevotellaceae>Segatella | 173 | ONT17_RS13520 | 3307399..3307920 | 165179 | hypothetical-protein-[Prevotella-copri] |
| MCZ2341132.1 | PIN-EndoRNase | GCA_026931725.1 | Bacteroidales_bacterium | Bacteria>Pseudomonadati>FCB_group>Bacteroidota/Chlorobiota_group>Bacteroidota>Bacteroidia>Bacteroidales>unclassified_Bacteroidales | 129 | LC104_04960 | 19460..19849 | 2030927 | MAG:-type-II-toxin-antitoxin-system-VapC-family-toxin-[Bacteroidales-bacterium] |
| MBR4647496.1 | PIN-EndoRNase | GCA_017525245.1 | Bacteroidales_bacterium | Bacteria>Pseudomonadati>FCB_group>Bacteroidota/Chlorobiota_group>Bacteroidota>Bacteroidia>Bacteroidales>unclassified_Bacteroidales | 174 | IK075_10330 | 24968..25492 | 2030927 | hypothetical-protein-[Bacteroidales-bacterium] |
| WP_094415987.1 | PIN-EndoRNase | GCF_002251835.1 | Flavobacterium_cyanobacteriorum | Bacteria>Pseudomonadati>FCB_group>Bacteroidota/Chlorobiota_group>Bacteroidota>Flavobacteriia>Flavobacteriales>Flavobacteriaceae>Flavobacterium | 99 | CHU92_RS12255 | 7438..7737 | 2022802 | hypothetical-protein-[Flavobacterium-cyanobacteriorum] |
| MBK7697108.1 | PIN-EndoRNase | GCA_016706925.1 | Candidatus_Vicinibacter_affinis | Bacteria>Pseudomonadati>FCB_group>Bacteroidota/Chlorobiota_group>Bacteroidota>Saprospiria>Saprospirales>Saprospiraceae>Candidatus_Vicinibacter | 99 | IPI30_23235 | 64626..64925 | 2982026 | NYN-domain-containing-protein-[Saprospiraceae-bacterium] |
| MBK7697107.1 | PIN-EndoRNase | GCA_016706925.1 | Candidatus_Vicinibacter_affinis | Bacteria>Pseudomonadati>FCB_group>Bacteroidota/Chlorobiota_group>Bacteroidota>Saprospiria>Saprospirales>Saprospiraceae>Candidatus_Vicinibacter | 110 | IPI30_23230 | 64297..64629 | 2982026 | NYN-domain-containing-protein-[Saprospiraceae-bacterium] |
| MBK8644368.1 | PIN-EndoRNase | GCA_016715705.1 | Candidatus_Vicinibacter_affinis | Bacteria>Pseudomonadati>FCB_group>Bacteroidota/Chlorobiota_group>Bacteroidota>Saprospiria>Saprospirales>Saprospiraceae>Candidatus_Vicinibacter | 209 | IPN15_19800 | 434558..435187 | 2982026 | NYN-domain-containing-protein-[Saprospiraceae-bacterium] |
| MCY3614379.1 | PIN-EndoRNase | GCA_026705365.1 | Bacteroidota_bacterium | Bacteria>Pseudomonadati>FCB_group>Bacteroidota/Chlorobiota_group>Bacteroidota>unclassified_Bacteroidota | 157 | OXH03_04695 | 4917..5390 | 1898104 | MAG:-PIN-domain-containing-protein-[Bacteroidetes-bacterium] |
| MCY3630038.1 | PIN-EndoRNase | GCA_026705375.1 | Bacteroidota_bacterium | Bacteria>Pseudomonadati>FCB_group>Bacteroidota/Chlorobiota_group>Bacteroidota>unclassified_Bacteroidota | 157 | OXG94_08510 | 2713..3186 | 1898104 | MAG:-PIN-domain-containing-protein-[Bacteroidetes-bacterium] |
| MCY4001545.1 | PIN-EndoRNase | GCA_026708685.1 | Bacteroidota_bacterium | Bacteria>Pseudomonadati>FCB_group>Bacteroidota/Chlorobiota_group>Bacteroidota>unclassified_Bacteroidota | 157 | OXF84_12175 | 4727..5200 | 1898104 | MAG:-hypothetical-protein-OXF84_12175-[Bacteroidetes-bacterium] |
| MDE2646785.1 | PIN-EndoRNase | GCA_028875465.1 | Bacteroidota_bacterium | Bacteria>Pseudomonadati>FCB_group>Bacteroidota/Chlorobiota_group>Bacteroidota>unclassified_Bacteroidota | 157 | OXI05_13245 | 5802..6275 | 1898104 | MAG:-PIN-domain-containing-protein-[Bacteroidota-bacterium] |
| MCY3594933.1 | PIN-EndoRNase | GCA_026705205.1 | Bacteroidota_bacterium | Bacteria>Pseudomonadati>FCB_group>Bacteroidota/Chlorobiota_group>Bacteroidota>unclassified_Bacteroidota | 158 | OXH01_06820 | 26538..27014 | 1898104 | MAG:-PIN-domain-containing-protein-[Bacteroidetes-bacterium] |
| WP_210512169.1 | PIN-EndoRNase | GCF_017921895.1 | Natronogracilivirga_saccharolytica | Bacteria>Pseudomonadati>FCB_group>Bacteroidota/Chlorobiota_group>Balneolota>Balneolia>Balneolales>Cyclonatronaceae>Natronogracilivirga | 183 | NATSA_RS09905 | 95553..96104 | 2812953 | hypothetical-protein-[Natronogracilivirga-saccharolytica] |
| MBK6914967.1 | PIN-EndoRNase | GCA_016709155.1 | Ignavibacteriales_bacterium | Bacteria>Pseudomonadati>FCB_group>Bacteroidota/Chlorobiota_group>Ignavibacteriota>Ignavibacteria>Ignavibacteriales>unclassified_Ignavibacteriales | 167 | IPH11_15400 | 475491..475994 | 2049428 | type-II-toxin-antitoxin-system-VapC-family-toxin-[Ignavibacteriales-bacterium] |
| MCF8267320.1 | PIN-EndoRNase | GCA_021738695.1 | Ignavibacteriales_bacterium | Bacteria>Pseudomonadati>FCB_group>Bacteroidota/Chlorobiota_group>Ignavibacteriota>Ignavibacteria>Ignavibacteriales>unclassified_Ignavibacteriales | 167 | K9I69_04495 | 4315..4818 | 2049428 | PIN-domain-containing-protein-[Ignavibacteriales-bacterium] |
| OIO18556.1 | PIN-EndoRNase | GCA_001871325.1 | Ignavibacteria_bacterium_CG1_02_37_35 | Bacteria>Pseudomonadati>FCB_group>Bacteroidota/Chlorobiota_group>Ignavibacteriota>Ignavibacteria>unclassified_Ignavibacteria | 168 | AUJ54_07710 | 3501..4007 | 1805220 | hypothetical-protein-AUJ54_07710-[Ignavibacteria-bacterium-CG1_02_37_35] |
| MXW15221.1 | PIN-EndoRNase | GCA_009835915.1 | Rhodothermaceae_bacterium | Bacteria>Pseudomonadati>FCB_group>Bacteroidota/Chlorobiota_group>Rhodothermota>Rhodothermia>Rhodothermales>Rhodothermaceae>unclassified_Rhodothermaceae | 157 | F4Z76_09950 | 11436..11909 | 2026787 | PIN-domain-containing-protein-[Rhodothermaceae-bacterium] |
| MXX97464.1 | PIN-EndoRNase | GCA_009838195.1 | Rhodothermaceae_bacterium | Bacteria>Pseudomonadati>FCB_group>Bacteroidota/Chlorobiota_group>Rhodothermota>Rhodothermia>Rhodothermales>Rhodothermaceae>unclassified_Rhodothermaceae | 157 | F4Y64_07615 | 9862..10335 | 2026787 | PIN-domain-containing-protein-[Rhodothermaceae-bacterium] |
| MXZ58768.1 | PIN-EndoRNase | GCA_009837125.1 | Rhodothermaceae_bacterium | Bacteria>Pseudomonadati>FCB_group>Bacteroidota/Chlorobiota_group>Rhodothermota>Rhodothermia>Rhodothermales>Rhodothermaceae>unclassified_Rhodothermaceae | 157 | F4Z16_09870 | 5718..6191 | 2026787 | PIN-domain-containing-protein-[Rhodothermaceae-bacterium] |
| MYC03830.1 | PIN-EndoRNase | GCA_009840265.1 | Rhodothermaceae_bacterium | Bacteria>Pseudomonadati>FCB_group>Bacteroidota/Chlorobiota_group>Rhodothermota>Rhodothermia>Rhodothermales>Rhodothermaceae>unclassified_Rhodothermaceae | 157 | F4X61_04290 | 6647..7120 | 2026787 | PIN-domain-containing-protein-[Rhodothermaceae-bacterium] |
| MYH11280.1 | PIN-EndoRNase | GCA_009842805.1 | Rhodothermaceae_bacterium | Bacteria>Pseudomonadati>FCB_group>Bacteroidota/Chlorobiota_group>Rhodothermota>Rhodothermia>Rhodothermales>Rhodothermaceae>unclassified_Rhodothermaceae | 157 | F4191_00345 | 9862..10335 | 2026787 | MAG:-PIN-domain-containing-protein-[Rhodothermaceae-bacterium] |
| WP_272503892.1 | PIN-EndoRNase | - | Salinibacter_ruber | Bacteria>Pseudomonadati>FCB_group>Bacteroidota/Chlorobiota_group>Rhodothermota>Rhodothermia>Rhodothermales>Salinibacteraceae>Salinibacter | 53 | GGP83_RS16825 | 7571..7732 | 146919 | DUF3368-domain-containing-protein-[Salinibacter-ruber] |
| WP_272592131.1 | PIN-EndoRNase | - | Salinibacter_ruber | Bacteria>Pseudomonadati>FCB_group>Bacteroidota/Chlorobiota_group>Rhodothermota>Rhodothermia>Rhodothermales>Salinibacteraceae>Salinibacter | 53 | GGP46_RS16745 | 80075..80236 | 146919 | DUF3368-domain-containing-protein-[Salinibacter-ruber] |
| WP_251930230.1 | PIN-EndoRNase | GCF_943908145.1 | Salinibacter_ruber | Bacteria>Pseudomonadati>FCB_group>Bacteroidota/Chlorobiota_group>Rhodothermota>Rhodothermia>Rhodothermales>Salinibacteraceae>Salinibacter | 159 | NGL32_RS11830 | 80320..80799 | 146919 | DUF3368-domain-containing-protein-[Salinibacter-ruber] |
| WP_259087832.1 | PIN-EndoRNase | GCF_024809725.1 | Salinibacter_ruber | Bacteria>Pseudomonadati>FCB_group>Bacteroidota/Chlorobiota_group>Rhodothermota>Rhodothermia>Rhodothermales>Salinibacteraceae>Salinibacter | 190 | GGQ07_RS07840 | 11779..12351 | 146919 | PIN-domain-containing-protein-[Salinibacter-ruber] |
| MCB5286843.1 | PIN-EndoRNase | GCA_020532145.1 | Candidatus_Cloacimonadota_bacterium | Bacteria>Pseudomonadati>FCB_group>Candidatus_Cloacimonadota>unclassified_Candidatus_Cloacimonadota | 155 | LHW64_03460 | 14595..15062 | 2030808 | hypothetical-protein-[Candidatus-Cloacimonadota-bacterium] |
| MCD4708412.1 | PIN-EndoRNase | GCA_021107435.1 | Candidatus_Sabulitectum_sp. | Bacteria>Pseudomonadati>FCB_group>Candidatus_Fermentibacterota>Candidatus_Sabulitectum>unclassified_Candidatus_Sabulitectum | 174 | K8S62_11835 | 3530..4054 | 2822438 | DUF4411-family-protein-[Candidatus-Sabulitectum-sp.] |
| MBX3180307.1 | PIN-EndoRNase | GCA_019637335.1 | Candidatus_Hydrogenedentota_bacterium | Bacteria>Pseudomonadati>FCB_group>Candidatus_Hydrogenedentota>unclassified_Candidatus_Hydrogenedentota | 126 | KF886_23395 | 8825..9205 | 2030809 | DUF5615-family-PIN-like-protein-[Candidatus-Hydrogenedentes-bacterium] |
| KAA0227460.1 | PIN-EndoRNase | GCA_008363195.1 | Candidatus_Zhuqueibacterota_bacterium | Bacteria>Pseudomonadati>FCB_group>Candidatus_Zhuqueibacterota>unclassified_Candidatus_Zhuqueibacterota | 176 | EDS67_15780 | 88151..88681 | 2172550 | type-II-toxin-antitoxin-system-VapC-family-toxin-[candidate-division-KSB1-bacterium] |
| MBO8151134.1 | PIN-EndoRNase | GCA_017656455.1 | Fidelibacterota_bacterium | Bacteria>Pseudomonadati>FCB_group>Fidelibacterota>unclassified_Fidelibacterota | 132 | HOZ30_01390 | 329345..329743 | 2026760 | type-II-toxin-antitoxin-system-VapC-family-toxin-[Candidatus-Marinimicrobia-bacterium] |
| MCY4604208.1 | PIN-EndoRNase | GCA_026713655.1 | Gemmatimonadota_bacterium | Bacteria>Pseudomonadati>FCB_group>Gemmatimonadota>unclassified_Gemmatimonadota | 120 | OXE49_08260 | 7814..8176 | 2026742 | MAG:-DUF5615-family-PIN-like-protein-[Gemmatimonadetes-bacterium] |
| MDE2653630.1 | PIN-EndoRNase | GCA_028875795.1 | Gemmatimonadota_bacterium | Bacteria>Pseudomonadati>FCB_group>Gemmatimonadota>unclassified_Gemmatimonadota | 166 | OXI71_07415 | 11198..11698 | 2026742 | MAG:-hypothetical-protein-OXI71_07415-[Gemmatimonadota-bacterium] |
| NMB74980.1 | PIN-EndoRNase | GCA_012798435.1 | Myxococcales_bacterium | Bacteria>Pseudomonadati>Myxococcota>Myxococcia>Myxococcales>unclassified_Myxococcales | 155 | GYA21_07605 | 22748..23215 | 2026763 | type-II-toxin-antitoxin-system-VapC-family-toxin-[Myxococcales-bacterium] |

| Protein-ID | Domain | Assembly-ID | Strain | Taxonomy | Length | Locus-Tag | Genomic-coordinates | TaxID | Fasta-defline |
|---|---|---|---|---|---|---|---|---|---|
| HCF56388.1 | PIN-EndoRNase | GCA_003534605.1 | Myxococcales_bacterium | Bacteria>Pseudomonadati>Myxococcota>Myxococcia>Myxococcales>unclassified_Myxococcales | 190 | DFS52_00125 | 3318..3890 | 2026763 | hypothetical-protein-[Myxococcales-bacterium] |
| MCG5052818.1 | PIN-EndoRNase | GCA_022184915.1 | Myxococcales_bacterium | Bacteria>Pseudomonadati>Myxococcota>Myxococcia>Myxococcales>unclassified_Myxococcales | 194 | KA712_07640 | 7141..7725 | 2026763 | NYN-domain-containing-protein-[Myxococcales-bacterium] |
| MBI4818897.1 | PIN-EndoRNase | GCA_016208175.1 | Deltaproteobacteria_bacterium | Bacteria>Pseudomonadati>Myxococcota>Myxococcia>unclassified_Myxococcia | 81 | HY791_21680 | 11621..11866 | 2026735 | hypothetical-protein-[Deltaproteobacteria-bacterium] |
| MBI2179303.1 | PIN-EndoRNase | GCA_016179825.1 | Deltaproteobacteria_bacterium | Bacteria>Pseudomonadati>Myxococcota>Myxococcia>unclassified_Myxococcia | 119 | HYU31_00595 | 699..1058 | 2026735 | DUF5615-family-PIN-like-protein-[Deltaproteobacteria-bacterium] |
| MBI3799268.1 | PIN-EndoRNase | GCA_016200595.1 | Deltaproteobacteria_bacterium | Bacteria>Pseudomonadati>Myxococcota>Myxococcia>unclassified_Myxococcia | 119 | HY268_20175 | 13935..14294 | 2026735 | DUF5615-family-PIN-like-protein-[Deltaproteobacteria-bacterium] |
| MBS3919359.1 | PIN-EndoRNase | GCA_018333535.1 | Deltaproteobacteria_bacterium | Bacteria>Pseudomonadati>Myxococcota>Myxococcia>unclassified_Myxococcia | 122 | KG012_10760 | 28421..28789 | 2026735 | DUF5615-family-PIN-like-protein-[Deltaproteobacteria-bacterium] |
| MBW2149118.1 | PIN-EndoRNase | GCA_019309515.1 | Deltaproteobacteria_bacterium | Bacteria>Pseudomonadati>Myxococcota>Myxococcia>unclassified_Myxococcia | 127 | JRI22_19095 | 2459..2842 | 2026735 | DUF5615-family-PIN-like-protein-[Deltaproteobacteria-bacterium] |
| MBW2309274.1 | PIN-EndoRNase | GCA_019310535.1 | Deltaproteobacteria_bacterium | Bacteria>Pseudomonadati>Myxococcota>Myxococcia>unclassified_Myxococcia | 127 | JRG73_20320 | 2998..3381 | 2026735 | DUF5615-family-PIN-like-protein-[Deltaproteobacteria-bacterium] |
| MBI5606171.1 | PIN-EndoRNase | GCA_016219225.1 | Deltaproteobacteria_bacterium | Bacteria>Pseudomonadati>Myxococcota>Myxococcia>unclassified_Myxococcia | 130 | HY879_22795 | 17420..17812 | 2026735 | type-II-toxin-antitoxin-system-VapC-family-toxin-[Deltaproteobacteria-bacterium] |
| MBI3073656.1 | PIN-EndoRNase | GCA_016192705.1 | Deltaproteobacteria_bacterium | Bacteria>Pseudomonadati>Myxococcota>Myxococcia>unclassified_Myxococcia | 132 | HYY84_16210 | 33178..33576 | 2026735 | putative-toxin-antitoxin-system-toxin-component,-PIN-family-[Deltaproteobacteria-bacterium] |
| MBI4523258.1 | PIN-EndoRNase | GCA_016210115.1 | Deltaproteobacteria_bacterium | Bacteria>Pseudomonadati>Myxococcota>Myxococcia>unclassified_Myxococcia | 133 | HY695_05525 | 15223..15624 | 2026735 | PIN-domain-containing-protein-[Deltaproteobacteria-bacterium] |
| MBI2378667.1 | PIN-EndoRNase | GCA_016183035.1 | Deltaproteobacteria_bacterium | Bacteria>Pseudomonadati>Myxococcota>Myxococcia>unclassified_Myxococcia | 136 | HYV07_32030 | 22201..22611 | 2026735 | PIN-domain-containing-protein-[Deltaproteobacteria-bacterium] |
| MBW1911013.1 | PIN-EndoRNase | GCA_019307765.1 | Deltaproteobacteria_bacterium | Bacteria>Pseudomonadati>Myxococcota>Myxococcia>unclassified_Myxococcia | 136 | JRJ11_16000 | 3826..4236 | 2026735 | DUF3368-domain-containing-protein-[Deltaproteobacteria-bacterium] |
| PXF57199.1 | PIN-EndoRNase | GCA_003194495.1 | Deltaproteobacteria_bacterium | Bacteria>Pseudomonadati>Myxococcota>Myxococcia>unclassified_Myxococcia | 136 | C4B58_10740 | 25047..25457 | 2026735 | VapC-toxin-family-PIN-domain-ribonuclease-[Deltaproteobacteria-bacterium] |
| PXF57195.1 | PIN-EndoRNase | GCA_003194495.1 | Deltaproteobacteria_bacterium | Bacteria>Pseudomonadati>Myxococcota>Myxococcia>unclassified_Myxococcia | 139 | C4B58_10715 | 19081..19500 | 2026735 | VapC-toxin-family-PIN-domain-ribonuclease-[Deltaproteobacteria-bacterium] |
| MCD6306395.1 | PIN-EndoRNase | GCA_021159525.1 | Deltaproteobacteria_bacterium | Bacteria>Pseudomonadati>Myxococcota>Myxococcia>unclassified_Myxococcia | 141 | J7M32_08920 | 1066..1491 | 2026735 | type-II-toxin-antitoxin-system-VapC-family-toxin-[Deltaproteobacteria-bacterium] |
| MBI2378673.1 | PIN-EndoRNase | GCA_016183035.1 | Deltaproteobacteria_bacterium | Bacteria>Pseudomonadati>Myxococcota>Myxococcia>unclassified_Myxococcia | 152 | HYV07_32060 | 28463..28921 | 2026735 | PIN-domain-containing-protein-[Deltaproteobacteria-bacterium] |
| MBI4814843.1 | PIN-EndoRNase | GCA_016208175.1 | Deltaproteobacteria_bacterium | Bacteria>Pseudomonadati>Myxococcota>Myxococcia>unclassified_Myxococcia | 159 | HY791_01215 | 32834..33313 | 2026735 | DUF3368-domain-containing-protein-[Deltaproteobacteria-bacterium] |
| MBW1706456.1 | PIN-EndoRNase | GCA_019306165.1 | Deltaproteobacteria_bacterium | Bacteria>Pseudomonadati>Myxococcota>Myxococcia>unclassified_Myxococcia | 166 | JRJ86_15010 | 4423..4923 | 2026735 | type-II-toxin-antitoxin-system-VapC-family-toxin-[Deltaproteobacteria-bacterium] |
| MBW2117195.1 | PIN-EndoRNase | GCA_019309305.1 | Deltaproteobacteria_bacterium | Bacteria>Pseudomonadati>Myxococcota>Myxococcia>unclassified_Myxococcia | 166 | JRH09_04650 | 4423..4923 | 2026735 | type-II-toxin-antitoxin-system-VapC-family-toxin-[Deltaproteobacteria-bacterium] |
| MBI2179304.1 | PIN-EndoRNase | GCA_016179825.1 | Deltaproteobacteria_bacterium | Bacteria>Pseudomonadati>Myxococcota>Myxococcia>unclassified_Myxococcia | 167 | HYU31_00600 | 1240..1743 | 2026735 | PIN-domain-containing-protein-[Deltaproteobacteria-bacterium] |
| MBI4527348.1 | PIN-EndoRNase | GCA_016210115.1 | Deltaproteobacteria_bacterium | Bacteria>Pseudomonadati>Myxococcota>Myxococcia>unclassified_Myxococcia | 167 | HY695_26435 | 23751..24254 | 2026735 | PIN-domain-containing-protein-[Deltaproteobacteria-bacterium] |
| MBM4322259.1 | PIN-EndoRNase | GCA_016875155.1 | Deltaproteobacteria_bacterium | Bacteria>Pseudomonadati>Myxococcota>Myxococcia>unclassified_Myxococcia | 167 | FJ115_01755 | 4317..4820 | 2026735 | type-II-toxin-antitoxin-system-VapC-family-toxin-[Deltaproteobacteria-bacterium] |
| MBN1842853.1 | PIN-EndoRNase | GCA_016930875.1 | Deltaproteobacteria_bacterium | Bacteria>Pseudomonadati>Myxococcota>Myxococcia>unclassified_Myxococcia | 168 | JW883_11310 | 19780..20286 | 2026735 | type-II-toxin-antitoxin-system-VapC-family-toxin-[Deltaproteobacteria-bacterium] |
| MBP1716770.1 | PIN-EndoRNase | GCA_017860815.1 | Deltaproteobacteria_bacterium | Bacteria>Pseudomonadati>Myxococcota>Myxococcia>unclassified_Myxococcia | 168 | H6Q43_208 | 6033..6539 | 2026735 | PilT-like-protein-[Deltaproteobacteria-bacterium] |
| MBW1895894.1 | PIN-EndoRNase | GCA_019307715.1 | Deltaproteobacteria_bacterium | Bacteria>Pseudomonadati>Myxococcota>Myxococcia>unclassified_Myxococcia | 168 | JRI47_02440 | 2946..3452 | 2026735 | type-II-toxin-antitoxin-system-VapC-family-toxin-[Deltaproteobacteria-bacterium] |
| MBI4488885.1 | PIN-EndoRNase | GCA_016210105.1 | Deltaproteobacteria_bacterium | Bacteria>Pseudomonadati>Myxococcota>Myxococcia>unclassified_Myxococcia | 171 | HY694_07345 | 21763..22278 | 2026735 | type-II-toxin-antitoxin-system-VapC-family-toxin-[Deltaproteobacteria-bacterium] |
| MBI4523253.1 | PIN-EndoRNase | GCA_016210115.1 | Deltaproteobacteria_bacterium | Bacteria>Pseudomonadati>Myxococcota>Myxococcia>unclassified_Myxococcia | 172 | HY695_05500 | 9524..10042 | 2026735 | type-II-toxin-antitoxin-system-VapC-family-toxin-[Deltaproteobacteria-bacterium] |
| MDD9852707.1 | PIN-EndoRNase | GCA_028817165.1 | Deltaproteobacteria_bacterium | Bacteria>Pseudomonadati>Myxococcota>Myxococcia>unclassified_Myxococcia | 174 | OXU78_01995 | 20972..21496 | 2026735 | MAG:-hypothetical-protein-OXU78_01995-[Deltaproteobacteria-bacterium] |
| MBI5442124.1 | PIN-EndoRNase | GCA_016218905.1 | Deltaproteobacteria_bacterium | Bacteria>Pseudomonadati>Myxococcota>Myxococcia>unclassified_Myxococcia | 187 | HY900_13045 | 3899..4462 | 2026735 | NYN-domain-containing-protein-[Deltaproteobacteria-bacterium] |
| MBN2361665.1 | PIN-EndoRNase | GCA_016934455.1 | Deltaproteobacteria_bacterium | Bacteria>Pseudomonadati>Myxococcota>Myxococcia>unclassified_Myxococcia | 215 | JXR83_19585 | 5993..6640 | 2026735 | NYN-domain-containing-protein-[Deltaproteobacteria-bacterium] |
| MBI2378660.1 | PIN-EndoRNase | GCA_016183035.1 | Deltaproteobacteria_bacterium | Bacteria>Pseudomonadati>Myxococcota>Myxococcia>unclassified_Myxococcia | 229 | HYV07_31985 | 10655..11344 | 2026735 | NYN-domain-containing-protein-[Deltaproteobacteria-bacterium] |
| MBK8255141.1 | PIN-EndoRNase | GCA_016712345.1 | Polyangiaceae_bacterium | Bacteria>Pseudomonadati>Myxococcota>Polyangia>Polyangiales>Polyangiaceae>unclassified_Polyangiaceae | 128 | IPK82_21075 | 184265..184651 | 2268199 | type-II-toxin-antitoxin-system-VapC-family-toxin-[Polyangiaceae-bacterium] |
| MBK6517539.1 | PIN-EndoRNase | GCA_016704335.1 | Polyangiaceae_bacterium | Bacteria>Pseudomonadati>Myxococcota>Polyangia>Polyangiales>Polyangiaceae>unclassified_Polyangiaceae | 146 | IPG04_26340 | 292149..292589 | 2268199 | PIN-domain-containing-protein-[Polyangiaceae-bacterium] |
| MBK8937961.1 | PIN-EndoRNase | GCA_016719945.1 | Polyangiaceae_bacterium | Bacteria>Pseudomonadati>Myxococcota>Polyangia>Polyangiales>Polyangiaceae>unclassified_Polyangiaceae | 146 | IPM79_10035 | 419015..419455 | 2268199 | PIN-domain-containing-protein-[Polyangiaceae-bacterium] |
| MCB9617768.1 | PIN-EndoRNase | GCA_020633155.1 | Sandaracinus_sp. | Bacteria>Pseudomonadati>Myxococcota>Polyangia>Polyangiales>Sandaracinaceae>Sandaracinus>unclassified_Sandaracinus | 189 | H6722_35480 | 13472..14041 | 2024858 | NYN-domain-containing-protein-[Sandaracinus-sp.] |
| MBX3269799.1 | PIN-EndoRNase | GCA_019637095.1 | Sandaracinaceae_bacterium | Bacteria>Pseudomonadati>Myxococcota>Polyangia>Polyangiales>Sandaracinaceae>unclassified_Sandaracinaceae | 218 | KF729_06040 | 110765..111421 | 2510337 | hypothetical-protein-[Sandaracinaceae-bacterium] |
| MBU0551508.1 | PIN-EndoRNase | GCA_018813065.1 | Myxococcota_bacterium | Bacteria>Pseudomonadati>Myxococcota>unclassified_Myxococcota | 169 | KJ940_08430 | 1766..2275 | 2818507 | DUF3368-domain-containing-protein-[Myxococcota-bacterium] |
| ETX04991.1 | PIN-EndoRNase | GCA_000522445.1 | Candidatus_Entotheonella_gemina | Bacteria>Pseudomonadati>Nitrospinota/Tectimicrobiota_group>Candidatus_Tectimicrobiota>Candidatus_Entotheonellia>Candidatus_Entotheonellales>Candidatus_Entotheonellaceae>Candidatus_Entotheonella | 144 | ETSY2_25595 | 2510..2944 | 1429439 | DNA-binding-protein-[Candidatus-Entotheonella-gemina] |
| WP_139077379.1 | PIN-EndoRNase | GCF_001458775.1 | Candidatus_Nitrospira_nitrificans | Bacteria>Pseudomonadati>Nitrospirota>Nitrospiria>Nitrospirales>Nitrospiraceae>Nitrospira | 106 | COMA2_RS14260 | 23073..23393 | 1742973 | hypothetical-protein-[Candidatus-Nitrospira-nitrificans] |
| WP_090899586.1 | PIN-EndoRNase | GCF_001458775.1 | Candidatus_Nitrospira_nitrificans | Bacteria>Pseudomonadati>Nitrospirota>Nitrospiria>Nitrospirales>Nitrospiraceae>Nitrospira | 183 | COMA2_RS14280 | 28817..29368 | 1742973 | PIN-domain-containing-protein-[Candidatus-Nitrospira-nitrificans] |
| MCW5796794.1 | PIN-EndoRNase | GCA_026128925.1 | Nitrospira_sp. | Bacteria>Pseudomonadati>Nitrospirota>Nitrospiria>Nitrospirales>Nitrospiraceae>Nitrospira>unclassified_Nitrospira | 125 | KIS97_20865 | 4347..4724 | 70125 | MAG:-DUF3368-domain-containing-protein-[Nitrospira-sp.] |
| MBS0166208.1 | PIN-EndoRNase | GCA_018242765.1 | Nitrospira_sp. | Bacteria>Pseudomonadati>Nitrospirota>Nitrospiria>Nitrospirales>Nitrospiraceae>Nitrospira>unclassified_Nitrospira | 160 | JSR29_09010 | 55987..56469 | 70125 | DUF3368-domain-containing-protein-[Nitrospira-sp.] |
| MDC8448551.1 | PIN-EndoRNase | GCA_028580895.1 | Nitrospira_sp. | Bacteria>Pseudomonadati>Nitrospirota>Nitrospiria>Nitrospirales>Nitrospiraceae>Nitrospira>unclassified_Nitrospira | 162 | LV473_09345 | 4622..5110 | 70125 | MAG:-DUF3368-domain-containing-protein-[Nitrospira-sp.] |
| MDD9819375.1 | PIN-EndoRNase | GCA_028817545.1 | Nitrospira_sp. | Bacteria>Pseudomonadati>Nitrospirota>Nitrospiria>Nitrospirales>Nitrospiraceae>Nitrospira>unclassified_Nitrospira | 162 | OXR07_00645 | 5480..5968 | 70125 | MAG:-type-II-toxin-antitoxin-system-VapC-family-toxin-[Nitrospira-sp.] |
| MDD9860640.1 | PIN-EndoRNase | GCA_028817215.1 | Nitrospira_sp. | Bacteria>Pseudomonadati>Nitrospirota>Nitrospiria>Nitrospirales>Nitrospiraceae>Nitrospira>unclassified_Nitrospira | 173 | OXU40_09375 | 18006..18527 | 70125 | MAG:-PIN-domain-containing-protein-[Nitrospira-sp.] |
| GIW54144.1 | PIN-EndoRNase | GCA_026004315.1 | Nitrospiraceae_bacterium | Bacteria>Pseudomonadati>Nitrospirota>Nitrospiria>Nitrospirales>Nitrospiraceae>unclassified_Nitrospiraceae | 215 | KatS3mg082_0548 | 102463..103110 | 2026770 | MAG:-hypothetical-protein-KatS3mg082_0548-[Nitrospiraceae-bacterium] |
| WP_281755191.1 | PIN-EndoRNase | GCF_027925485.1 | Thermodesulfovibrio_islandicus | Bacteria>Pseudomonadati>Nitrospirota>Thermodesulfovibrionia>Thermodesulfovibrionales>Thermodesulfovibrionaceae>Thermodesulfovibrio | 131 | QMF67_RS07345 | 1424872..1425267 | 53423 | PIN-domain-containing-protein-[Thermodesulfovibrio-islandicus] |
| RMH36428.1 | PIN-EndoRNase | GCA_003695915.1 | Nitrospirota_bacterium | Bacteria>Pseudomonadati>Nitrospirota>unclassified_Nitrospirota | 107 | D6690_05855 | 4849..5172 | 2026887 | hypothetical-protein-D6690_05855-[Nitrospirota-bacterium] |
| MCL5421043.1 | PIN-EndoRNase | GCA_023485285.1 | Nitrospirota_bacterium | Bacteria>Pseudomonadati>Nitrospirota>unclassified_Nitrospirota | 171 | M1461_01030 | 8015..8530 | 2026887 | type-II-toxin-antitoxin-system-VapC-family-toxin-[Nitrospirota-bacterium] |
| WP_070113765.1 | PIN-EndoRNase | GCF_031200535.1 | Acidithiobacillus_caldus | Bacteria>Pseudomonadati>Pseudomonadota>Acidithiobacillia>Acidithiobacillales>Acidithiobacillaceae>Acidithiobacillus | 135 | HFV02_RS10470 | 8967..9374 | 33059 | PIN-domain-containing-protein-[Acidithiobacillus-caldus] |
| WP_215870429.1 | PIN-EndoRNase | GCF_018853695.1 | Acidithiobacillus_caldus | Bacteria>Pseudomonadati>Pseudomonadota>Acidithiobacillia>Acidithiobacillales>Acidithiobacillaceae>Acidithiobacillus | 205 | HAQ02_RS11845 | 6263..6880 | 33059 | NYN-domain-containing-protein-[Acidithiobacillus-caldus] |
| WP_048844483.1 | PIN-EndoRNase | GCF_000963945.1 | Acetobacter | Bacteria>Pseudomonadati>Pseudomonadota>Alphaproteobacteria>Acetobacterales>Acetobacteraceae | 128 | AD954_RS12060 | 5501..5887 | 434 | MULTISPECIES:-type-II-toxin-antitoxin-system-VapC-family-toxin-[Acetobacter] |
| WP_045542207.1 | PIN-EndoRNase | GCF_025995375.1 | Acetobacter | Bacteria>Pseudomonadati>Pseudomonadota>Alphaproteobacteria>Acetobacterales>Acetobacteraceae | 141 | AD954_RS12095 | 19402..19827 | 434 | MULTISPECIES:-type-II-toxin-antitoxin-system-VapC-family-toxin-[Acetobacter] |
| WP_253545421.1 | PIN-EndoRNase | GCF_024158445.1 | Acetobacter_lovaniensis | Bacteria>Pseudomonadati>Pseudomonadota>Alphaproteobacteria>Acetobacterales>Acetobacteraceae>Acetobacter | 128 | NKW44_RS07970 | 15347..15733 | 104100 | type-II-toxin-antitoxin-system-VapC-family-toxin-[Acetobacter-lovaniensis] |
| WP_018307946.1 | PIN-EndoRNase | GCF_000193495.2 | Acetobacter_aceti | Bacteria>Pseudomonadati>Pseudomonadota>Alphaproteobacteria>Acetobacterales>Acetobacteraceae>Acetobacter>Acetobacter_subgen._Acetobacter | 123 | EDC15_RS12645 | 34310..34681 | 435 | type-II-toxin-antitoxin-system-VapC-family-toxin-[Acetobacter-aceti] |
| MCA7120525.1 | PIN-EndoRNase | GCA_020418435.1 | Acidibrevibacterium_fodinaquatile | Bacteria>Pseudomonadati>Pseudomonadota>Alphaproteobacteria>Acetobacterales>Acetobacteraceae>Acidibrevibacterium | 134 | LGL72_14245 | 2850906..2851310 | 1969806 | type-II-toxin-antitoxin-system-VapC-family-toxin-[Acidibrevibacterium-sp.] |
| WP_274966104.1 | PIN-EndoRNase | GCF_020418435.1 | Acidibrevibacterium_fodinaquatile | Bacteria>Pseudomonadati>Pseudomonadota>Alphaproteobacteria>Acetobacterales>Acetobacteraceae>Acidibrevibacterium | 134 | LGL72_RS14185 | 2850906..2851310 | 1969806 | type-II-toxin-antitoxin-system-VapC-family-toxin-[Acidibrevibacterium-fodinaquatile] |
| WP_012553519.1 | PIN-EndoRNase | GCF_000021325.1 | Gluconacetobacter_diazotrophicus | Bacteria>Pseudomonadati>Pseudomonadota>Alphaproteobacteria>Acetobacterales>Acetobacteraceae>Gluconacetobacter | 128 | GDIA_RS05045 | 1152993..1153379 | 33996 | type-II-toxin-antitoxin-system-VapC-family-toxin-[Gluconacetobacter-diazotrophicus] |
| OZB26687.1 | PIN-EndoRNase | GCA_002282635.1 | Acidiphilium_sp._34-64-41 | Bacteria>Pseudomonadati>Pseudomonadota>Alphaproteobacteria>Acetobacterales>Acidocellaceae>Acidiphilium>unclassified_Acidiphilium | 135 | B7X49_12100 | 13989..14396 | 1970297 | MAG:-hypothetical-protein-B7X49_12100-[Acidiphilium-sp.-34-64-41] |
| WP_019463149.1 | PIN-EndoRNase | GCF_000292225.1 | Roseomonas | Bacteria>Pseudomonadati>Pseudomonadota>Alphaproteobacteria>Acetobacterales>Roseomonadaceae | 136 | B555_RS0120320 | 802..1212 | 125216 | MULTISPECIES:-type-II-toxin-antitoxin-system-VapC-family-toxin-[Roseomonas] |
| MCA3363598.1 | PIN-EndoRNase | GCA_020249535.1 | Roseomonas_sp. | Bacteria>Pseudomonadati>Pseudomonadota>Alphaproteobacteria>Acetobacterales>Roseomonadaceae>Roseomonas>unclassified_Roseomonas | 167 | INF74_16810 | 3477..3980 | 1869190 | hypothetical-protein-[Roseomonas-sp.] |
| KAF0185694.1 | PIN-EndoRNase | GCA_009773335.1 | Caulobacteraceae_bacterium | Bacteria>Pseudomonadati>Pseudomonadota>Alphaproteobacteria>Caulobacterales>Caulobacteraceae>unclassified_Caulobacteraceae | 134 | FD160_503 | 35130..35534 | 1978230 | putative-nucleic-acid-binding-protein-contains-PIN-domain-[Caulobacteraceae-bacterium] |
| MBN8607694.1 | PIN-EndoRNase | GCA_017303355.1 | Caulobacterales_bacterium | Bacteria>Pseudomonadati>Pseudomonadota>Alphaproteobacteria>Caulobacterales>unclassified_Caulobacterales | 134 | J0L81_12325 | 133461..133865 | 2282150 | type-II-toxin-antitoxin-system-VapC-family-toxin-[Caulobacterales-bacterium] |
| MBI5313197.1 | PIN-EndoRNase | GCA_016219665.1 | Methylocystis_sp. | Bacteria>Pseudomonadati>Pseudomonadota>Alphaproteobacteria>Hyphomicrobiales>Methylocystaceae>Methylocystis>unclassified_Methylocystis | 166 | HZB28_08770 | 48010..48510 | 1911079 | hypothetical-protein-[Methylocystis-sp.] |
| WP_166097054.1 | PIN-EndoRNase | GCF_011516635.5 | Bradyrhizobium_barranii | Bacteria>Pseudomonadati>Pseudomonadota>Alphaproteobacteria>Hyphomicrobiales>Nitrobacteraceae>Bradyrhizobium | 196 | HAP41_RS22285 | 4688604..4689194 | 2992140 | hypothetical-protein-[Bradyrhizobium-barranii] |
| WP_135929915.1 | PIN-EndoRNase | GCF_004791875.1 | unclassified_Mesorhizobium | Bacteria>Pseudomonadati>Pseudomonadota>Alphaproteobacteria>Hyphomicrobiales>Phyllobacteriaceae>Mesorhizobium | 163 | EN810_RS20195 | 141105..141596 | 325217 | MULTISPECIES:-hypothetical-protein-[unclassified-Mesorhizobium] |
| WP_156941281.1 | PIN-EndoRNase | GCF_000503075.1 | Mesorhizobium_sp._LSHC422A00 | Bacteria>Pseudomonadati>Pseudomonadota>Alphaproteobacteria>Hyphomicrobiales>Phyllobacteriaceae>Mesorhizobium>unclassified_Mesorhizobium | 134 | X761_RS0135560 | 212738..213142 | 1287294 | hypothetical-protein-[Mesorhizobium-sp.-LSHC422A00] |
| RWL15102.1 | PIN-EndoRNase | GCA_004019595.1 | Mesorhizobium_sp. | Bacteria>Pseudomonadati>Pseudomonadota>Alphaproteobacteria>Hyphomicrobiales>Phyllobacteriaceae>Mesorhizobium>unclassified_Mesorhizobium | 165 | EOR57_31090 | 9658..10155 | 1871066 | PIN-domain-containing-protein-[Mesorhizobium-sp.] |
| MBF0214026.1 | PIN-EndoRNase | GCA_015231295.1 | Magnetococcales_bacterium | Bacteria>Pseudomonadati>Pseudomonadota>Alphaproteobacteria>Magnetococcales>unclassified_Magnetococcales | 136 | HQM00_10775 | 7805..8215 | 2026759 | type-II-toxin-antitoxin-system-VapC-family-toxin-[Magnetococcales-bacterium] |
| MBF0340192.1 | PIN-EndoRNase | GCA_015232245.1 | Magnetococcales_bacterium | Bacteria>Pseudomonadati>Pseudomonadota>Alphaproteobacteria>Magnetococcales>unclassified_Magnetococcales | 136 | HQL95_04420 | 2202..2612 | 2026759 | type-II-toxin-antitoxin-system-VapC-family-toxin-[Magnetococcales-bacterium] |
| MBF0143012.1 | PIN-EndoRNase | GCA_015229005.1 | Magnetococcales_bacterium | Bacteria>Pseudomonadati>Pseudomonadota>Alphaproteobacteria>Magnetococcales>unclassified_Magnetococcales | 138 | HQL59_06085 | 10893..11309 | 2026759 | type-II-toxin-antitoxin-system-VapC-family-toxin-[Magnetococcales-bacterium] |
| MBF0185781.1 | PIN-EndoRNase | GCA_015231215.1 | Magnetococcales_bacterium | Bacteria>Pseudomonadati>Pseudomonadota>Alphaproteobacteria>Magnetococcales>unclassified_Magnetococcales | 138 | HQM06_15540 | 18813..19229 | 2026759 | type-II-toxin-antitoxin-system-VapC-family-toxin-[Magnetococcales-bacterium] |
| MBF0213220.1 | PIN-EndoRNase | GCA_015231295.1 | Magnetococcales_bacterium | Bacteria>Pseudomonadati>Pseudomonadota>Alphaproteobacteria>Magnetococcales>unclassified_Magnetococcales | 170 | HQM00_06610 | 6072..6584 | 2026759 | hypothetical-protein-[Magnetococcales-bacterium] |
| MBF0213217.1 | PIN-EndoRNase | GCA_015231295.1 | Magnetococcales_bacterium | Bacteria>Pseudomonadati>Pseudomonadota>Alphaproteobacteria>Magnetococcales>unclassified_Magnetococcales | 176 | HQM00_06595 | 1048..1578 | 2026759 | hypothetical-protein-[Magnetococcales-bacterium] |
| MBF0096956.1 | PIN-EndoRNase | GCA_015228825.1 | Magnetococcales_bacterium | Bacteria>Pseudomonadati>Pseudomonadota>Alphaproteobacteria>Magnetococcales>unclassified_Magnetococcales | 180 | HQM05_06415 | 7265..7807 | 2026759 | hypothetical-protein-[Magnetococcales-bacterium] |
| MBF0143009.1 | PIN-EndoRNase | GCA_015229005.1 | Magnetococcales_bacterium | Bacteria>Pseudomonadati>Pseudomonadota>Alphaproteobacteria>Magnetococcales>unclassified_Magnetococcales | 206 | HQL59_06070 | 7096..7716 | 2026759 | NYN-domain-containing-protein-[Magnetococcales-bacterium] |
| MBF0214023.1 | PIN-EndoRNase | GCA_015231295.1 | Magnetococcales_bacterium | Bacteria>Pseudomonadati>Pseudomonadota>Alphaproteobacteria>Magnetococcales>unclassified_Magnetococcales | 206 | HQM00_10760 | 3999..4619 | 2026759 | NYN-domain-containing-protein-[Magnetococcales-bacterium] |
| MYA89894.1 | PIN-EndoRNase | GCA_009839535.1 | Boseongicola_sp._SB0662_bin_57 | Bacteria>Pseudomonadati>Pseudomonadota>Alphaproteobacteria>Rhodobacterales>Paracoccaceae>Boseongicola>unclassified_Boseongicola | 124 | F4X97_15825 | 10429..10803 | 2604843 | toxin-antitoxin-system,-toxin-component,-PIN-family-protein-[Boseongicola-sp.-SB0662_bin_57] |
| MDE0346580.1 | PIN-EndoRNase | GCA_028821095.1 | Boseongicola_sp. | Bacteria>Pseudomonadati>Pseudomonadota>Alphaproteobacteria>Rhodobacterales>Paracoccaceae>Boseongicola>unclassified_Boseongicola | 144 | OXI66_12505 | 4000..4434 | 1886600 | MAG:-PIN-domain-containing-protein-[Boseongicola-sp.] |
| WP_152828753.1 | PIN-EndoRNase | GCF_009380135.2 | Fertoeibacter_niger | Bacteria>Pseudomonadati>Pseudomonadota>Alphaproteobacteria>Rhodobacterales>Paracoccaceae>Fertoeibacter | 140 | GEU84_RS20385 | 42039..42461 | 2656921 | type-II-toxin-antitoxin-system-VapC-family-toxin-[Fertoeibacter-niger] |
| WP_180283597.1 | PIN-EndoRNase | GCF_013423495.1 | Azospirillum_oleiclasticum | Bacteria>Pseudomonadati>Pseudomonadota>Alphaproteobacteria>Rhodospirillales>Azospirillaceae>Azospirillum | 160 | HND93_RS19075 | 6407..6889 | 2735135 | hypothetical-protein-[Azospirillum-oleiclasticum] |
| WP_238518869.1 | PIN-EndoRNase | GCF_044865155.1 | Azospirillum_doebereinerae | Bacteria>Pseudomonadati>Pseudomonadota>Alphaproteobacteria>Rhodospirillales>Azospirillaceae>Azospirillum | 163 | MCW82_RS00025 | 13888..14379 | 92933 | hypothetical-protein-[Azospirillum-doebereinerae] |
| TVR79628.1 | PIN-EndoRNase | GCA_007695045.1 | Rhodospirillales_bacterium | Bacteria>Pseudomonadati>Pseudomonadota>Alphaproteobacteria>Rhodospirillales>unclassified_Rhodospirillales | 139 | EA405_12430 | 4695..5114 | 2026786 | PIN-domain-containing-protein-[Rhodospirillales-bacterium] |
| SEP49900.1 | PIN-EndoRNase | GCA_900110395.1 | Rhodospirillales_bacterium_URHD0017 | Bacteria>Pseudomonadati>Pseudomonadota>Alphaproteobacteria>Rhodospirillales>unclassified_Rhodospirillales | 163 | SAMN02990966_07448 | 8936..9427 | 1380357 | hypothetical-protein-SAMN02990966_07448-[Rhodospirillales-bacterium-URHD0017] |
| WP_098108968.1 | PIN-EndoRNase | GCF_022832385.1 | Novosphingobium | Bacteria>Pseudomonadati>Pseudomonadota>Alphaproteobacteria>Sphingomonadales>Sphingomonadaceae | 165 | B2G71_RS22850 | 13982..14479 | 165696 | MULTISPECIES:-PIN-domain-containing-protein-[unclassified-Novosphingobium] |
| WP_268740963.1 | PIN-EndoRNase | - | Novosphingobium_album_(ex_Hu_et_al._2023) | Bacteria>Pseudomonadati>Pseudomonadota>Alphaproteobacteria>Sphingomonadales>Sphingomonadaceae>Novosphingobium | 138 | MTR64_RS16255 | 44015..44431 | 2930093 | type-II-toxin-antitoxin-system-VapC-family-toxin-[Novosphingobium-sp.-B2580] |
| MCL6741776.1 | PIN-EndoRNase | GCA_023516505.1 | Sphingomonas_brevis | Bacteria>Pseudomonadati>Pseudomonadota>Alphaproteobacteria>Sphingomonadales>Sphingomonadaceae>Sphingomonas | 161 | LZ518_11625 | 2248655..2249140 | 2908206 | hypothetical-protein-[Sphingomonas-sp.-RB56-2] |
| WP_249916142.1 | PIN-EndoRNase | GCF_023516505.1 | Sphingomonas_brevis | Bacteria>Pseudomonadati>Pseudomonadota>Alphaproteobacteria>Sphingomonadales>Sphingomonadaceae>Sphingomonas | 161 | LZ518_RS11625 | 2248655..2249140 | 2908206 | hypothetical-protein-[Sphingomonas-sp.-RB56-2] |
| WP_152417294.1 | PIN-EndoRNase | GCF_000331225.1 | Sandarakinorhabdus_sp._AAP62 | Bacteria>Pseudomonadati>Pseudomonadota>Alphaproteobacteria>Sphingomonadales>Sphingosinicellaceae>Sandarakinorhabdus>unclassified_Sandarakinorhabdus | 323 | D619_RS17785 | 162051..163022 | 1248916 | hypothetical-protein-[Sandarakinorhabdus-sp.-AAP62] |
| MBQ2886286.1 | PIN-EndoRNase | GCA_017409945.1 | Alphaproteobacteria_bacterium | Bacteria>Pseudomonadati>Pseudomonadota>Alphaproteobacteria>unclassified_Alphaproteobacteria | 174 | IJE43_21400 | 11419..11943 | 1913988 | hypothetical-protein-[Alphaproteobacteria-bacterium] |
| MDD3182287.1 | PIN-EndoRNase | GCA_028691835.1 | Alphaproteobacteria_bacterium | Bacteria>Pseudomonadati>Pseudomonadota>Alphaproteobacteria>unclassified_Alphaproteobacteria | 206 | PHD48_05750 | 12525..13145 | 1913988 | MAG:-NYN-domain-containing-protein-[Alphaproteobacteria-bacterium] |
| MDD3183270.1 | PIN-EndoRNase | GCA_028691835.1 | Alphaproteobacteria_bacterium | Bacteria>Pseudomonadati>Pseudomonadota>Alphaproteobacteria>unclassified_Alphaproteobacteria | 206 | PHD48_10785 | 5314..5934 | 1913988 | MAG:-NYN-domain-containing-protein-[Alphaproteobacteria-bacterium] |
| WP_248595567.1 | PIN-EndoRNase | GCF_012939955.1 | Candidatus_Accumulibacter_phosphatis | Bacteria>Pseudomonadati>Pseudomonadota>Betaproteobacteria>Betaproteobacteria_incertae_sedis>Candidatus_Accumulibacter | 77 | E4Q23_RS18395 | 7598..7831 | 327160 | PIN-domain-containing-protein-[Candidatus-Accumulibacter-phosphatis] |
| WP_169067997.1 | PIN-EndoRNase | GCF_012939955.1 | Candidatus_Accumulibacter_phosphatis | Bacteria>Pseudomonadati>Pseudomonadota>Betaproteobacteria>Betaproteobacteria_incertae_sedis>Candidatus_Accumulibacter | 124 | E4Q23_RS18410 | 9347..9721 | 327160 | type-II-toxin-antitoxin-system-VapC-family-toxin-[Candidatus-Accumulibacter-phosphatis] |
| MBK8576829.1 | PIN-EndoRNase | GCA_016714935.1 | Candidatus_Accumulibacter_propinquus | Bacteria>Pseudomonadati>Pseudomonadota>Betaproteobacteria>Betaproteobacteria_incertae_sedis>Candidatus_Accumulibacter | 133 | IPN78_00500 | 109744..110145 | 2954380 | type-II-toxin-antitoxin-system-VapC-family-toxin-[Candidatus-Accumulibacter-sp.] |
| WP_186409052.1 | PIN-EndoRNase | GCF_900089955.1 | Candidatus_Accumulibacter_aalborgensis | Bacteria>Pseudomonadati>Pseudomonadota>Betaproteobacteria>Betaproteobacteria_incertae_sedis>Candidatus_Accumulibacter | 133 | ACCAA_RS20225 | 16345..16746 | 1860102 | type-II-toxin-antitoxin-system-VapC-family-toxin-[Candidatus-Accumulibacter-aalborgensis] |
| QLH49691.1 | PIN-EndoRNase | GCA_013414765.1 | Candidatus_Accumulibacter_cognatus | Bacteria>Pseudomonadati>Pseudomonadota>Betaproteobacteria>Betaproteobacteria_incertae_sedis>Candidatus_Accumulibacter | 133 | HWD57_07785 | 1707978..1708379 | 2954383 | MAG:-PIN-domain-containing-protein-[Candidatus-Accumulibacter-cognatus] |
| MBK8576827.1 | PIN-EndoRNase | GCA_016714935.1 | Candidatus_Accumulibacter_propinquus | Bacteria>Pseudomonadati>Pseudomonadota>Betaproteobacteria>Betaproteobacteria_incertae_sedis>Candidatus_Accumulibacter | 141 | IPN78_00490 | 106504..106929 | 2954380 | putative-toxin-antitoxin-system-toxin-component,-PIN-family-[Candidatus-Accumulibacter-sp.] |
| MBN8450446.1 | PIN-EndoRNase | GCA_017302435.1 | Candidatus_Accumulibacter_necessarius | Bacteria>Pseudomonadati>Pseudomonadota>Betaproteobacteria>Betaproteobacteria_incertae_sedis>Candidatus_Accumulibacter | 146 | J0M13_15860 | 39159..39599 | 2954386 | DUF3368-domain-containing-protein-[Candidatus-Accumulibacter-sp.] |
| WP_186409055.1 | PIN-EndoRNase | GCF_900089955.1 | Candidatus_Accumulibacter_aalborgensis | Bacteria>Pseudomonadati>Pseudomonadota>Betaproteobacteria>Betaproteobacteria_incertae_sedis>Candidatus_Accumulibacter | 161 | ACCAA_RS20240 | 19896..20381 | 1860102 | DUF3368-domain-containing-protein-[Candidatus-Accumulibacter-aalborgensis] |
| KFB77816.1 | PIN-EndoRNase | GCA_000584975.2 | Candidatus_Accumulibacter_cognatus | Bacteria>Pseudomonadati>Pseudomonadota>Betaproteobacteria>Betaproteobacteria_incertae_sedis>Candidatus_Accumulibacter | 205 | AW06_000999 | 18129..18746 | 2954383 | MAG:-6-hydroxy-3-succinoylpyridine-3-monooxygenase-HspA-[Candidatus-Accumulibacter-cognatus] |
| MCQ1548295.1 | PIN-EndoRNase | GCA_024380085.1 | Candidatus_Accumulibacter_phosphatis | Bacteria>Pseudomonadati>Pseudomonadota>Betaproteobacteria>Betaproteobacteria_incertae_sedis>Candidatus_Accumulibacter | 205 | NOF05_05605 | 13303..13920 | 327160 | NYN-domain-containing-protein-[Candidatus-Accumulibacter-phosphatis] |
| WP_034945906.1 | PIN-EndoRNase | GCF_000584975.1 | Candidatus_Accumulibacter_cognatus | Bacteria>Pseudomonadati>Pseudomonadota>Betaproteobacteria>Betaproteobacteria_incertae_sedis>Candidatus_Accumulibacter | 205 | AW06_RS03935 | 18129..18746 | 2954383 | NYN-domain-containing-protein-[Candidatus-Accumulibacter-cognatus] |
| EXI65854.1 | PIN-EndoRNase | GCA_000585015.1 | Candidatus_Accumulibacter_adjunctus | Bacteria>Pseudomonadati>Pseudomonadota>Betaproteobacteria>Betaproteobacteria_incertae_sedis>Candidatus_Accumulibacter | 206 | AW08_02879 | 55900..56520 | 1454001 | MAG:-6-hydroxy-3-succinoylpyridine-3-monooxygenase-HspA-[Candidatus-Accumulibacter-adjunctus] |
| EXI76475.1 | PIN-EndoRNase | GCA_000584995.1 | Candidatus_Accumulibacter_sp._SK-11 | Bacteria>Pseudomonadati>Pseudomonadota>Betaproteobacteria>Betaproteobacteria_incertae_sedis>Candidatus_Accumulibacter>unclassified_Candidatus_Accumulibacter | 133 | AW07_00423 | 123770..124171 | 1454000 | MAG:-tRNA(fMet)-specific-endonuclease-VapC-[Candidatus-Accumulibacter-sp.-SK-11] |
| MBK8117088.1 | PIN-EndoRNase | GCA_016712935.1 | Accumulibacter_sp. | Bacteria>Pseudomonadati>Pseudomonadota>Betaproteobacteria>Betaproteobacteria_incertae_sedis>Candidatus_Accumulibacter>unclassified_Candidatus_Accumulibacter | 133 | IPK44_22550 | 274979..275380 | 2053492 | type-II-toxin-antitoxin-system-VapC-family-toxin-[Candidatus-Accumulibacter-sp.] |
| MBK8387657.1 | PIN-EndoRNase | GCA_016711725.1 | Accumulibacter_sp. | Bacteria>Pseudomonadati>Pseudomonadota>Betaproteobacteria>Betaproteobacteria_incertae_sedis>Candidatus_Accumulibacter>unclassified_Candidatus_Accumulibacter | 133 | IPL11_19670 | 4784..5185 | 2053492 | type-II-toxin-antitoxin-system-VapC-family-toxin-[Candidatus-Accumulibacter-sp.] |
| MBN8436883.1 | PIN-EndoRNase | GCA_017302555.1 | Accumulibacter_sp. | Bacteria>Pseudomonadati>Pseudomonadota>Betaproteobacteria>Betaproteobacteria_incertae_sedis>Candidatus_Accumulibacter>unclassified_Candidatus_Accumulibacter | 133 | J0L95_02375 | 97025..97426 | 2053492 | type-II-toxin-antitoxin-system-VapC-family-toxin-[Candidatus-Accumulibacter-sp.] |
| MCM8613689.1 | PIN-EndoRNase | GCA_023806505.1 | Accumulibacter_sp. | Bacteria>Pseudomonadati>Pseudomonadota>Betaproteobacteria>Betaproteobacteria_incertae_sedis>Candidatus_Accumulibacter>unclassified_Candidatus_Accumulibacter | 133 | NFW17_16730 | 2358..2759 | 2053492 | type-II-toxin-antitoxin-system-VapC-family-toxin-[Candidatus-Accumulibacter-sp.] |
| EXI72919.1 | PIN-EndoRNase | GCA_000584995.1 | Candidatus_Accumulibacter_sp._SK-11 | Bacteria>Pseudomonadati>Pseudomonadota>Betaproteobacteria>Betaproteobacteria_incertae_sedis>Candidatus_Accumulibacter>unclassified_Candidatus_Accumulibacter | 136 | AW07_03055 | 64238..64648 | 1454000 | MAG:-putative-nucleic-acid-binding-protein-[Candidatus-Accumulibacter-sp.-SK-11] |
| HAY27741.1 | PIN-EndoRNase | GCA_003487685.1 | Accumulibacter_sp. | Bacteria>Pseudomonadati>Pseudomonadota>Betaproteobacteria>Betaproteobacteria_incertae_sedis>Candidatus_Accumulibacter>unclassified_Candidatus_Accumulibacter | 136 | DCY47_09625 | 15529..15939 | 2053492 | PIN-domain-containing-protein-[Candidatus-Accumulibacter-sp.] |
| EXI76472.1 | PIN-EndoRNase | GCA_000584995.1 | Candidatus_Accumulibacter_sp._SK-11 | Bacteria>Pseudomonadati>Pseudomonadota>Betaproteobacteria>Betaproteobacteria_incertae_sedis>Candidatus_Accumulibacter>unclassified_Candidatus_Accumulibacter | 161 | AW07_00420 | 120171..120656 | 1454000 | MAG:-hypothetical-protein-AW07_00420-[Candidatus-Accumulibacter-sp.-SK-11] |

| Protein-ID | Domain | Assembly-ID | Strain | Taxonomy | Length | Locus-Tag | Genomic-coordinates | TaxID | Fasta-defline |
|---|---|---|---|---|---|---|---|---|---|
| MBN8436886.1 | PIN-EndoRNase | GCA_017302555.1 | Accumulibacter_sp. | Bacteria>Pseudomonadati>Pseudomonadota>Betaproteobacteria>Betaproteobacteria_incertae_sedis>Candidatus_Accumulibacter>unclassified_Candidatus_Accumulibacter | 161 | J0L95_02390 | 100528..101013 | 2053492 | DUF3368-domain-containing-protein-[Candidatus-Accumulibacter-sp.] |
| MCM8638972.1 | PIN-EndoRNase | GCA_023806635.1 | Accumulibacter_sp. | Bacteria>Pseudomonadati>Pseudomonadota>Betaproteobacteria>Betaproteobacteria_incertae_sedis>Candidatus_Accumulibacter>unclassified_Candidatus_Accumulibacter | 162 | NFW30_04520 | 1202..1690 | 2053492 | DUF3368-domain-containing-protein-[Candidatus-Accumulibacter-sp.] |
| MBN8437937.1 | PIN-EndoRNase | GCA_017302555.1 | Accumulibacter_sp. | Bacteria>Pseudomonadati>Pseudomonadota>Betaproteobacteria>Betaproteobacteria_incertae_sedis>Candidatus_Accumulibacter>unclassified_Candidatus_Accumulibacter | 185 | J0L95_07820 | 85877..86434 | 2053492 | hypothetical-protein-[Candidatus-Accumulibacter-sp.] |
| MCX8099544.1 | PIN-EndoRNase | GCA_026414825.1 | Casimicrobiaceae_bacterium | Bacteria>Pseudomonadati>Pseudomonadota>Betaproteobacteria>Betaproteobacteria_incertae_sedis>Casimicrobiaceae>unclassified_Casimicrobiaceae | 74 | N3F11_11195 | 8279..8503 | 2906524 | MAG:-hypothetical-protein-N3F11_11195-[Casimicrobiaceae-bacterium] |
| WP_076471944.1 | PIN-EndoRNase | GCF_001957055.1 | Achromobacter_xylosoxidans | Bacteria>Pseudomonadati>Pseudomonadota>Betaproteobacteria>Burkholderiales>Alcaligenaceae>Achromobacter | 161 | BIZ53_RS00420 | 86280..86765 | 85698 | hypothetical-protein-[Achromobacter-xylosoxidans] |
| WP_217103122.1 | PIN-EndoRNase | GCF_019041175.1 | Burkholderia_multivorans | Bacteria>Pseudomonadati>Pseudomonadota>Betaproteobacteria>Burkholderiales>Burkholderiaceae>Burkholderia>Burkholderia_cepacia_complex | 206 | KTD10_RS22760 | 52429..53049 | 87883 | NYN-domain-containing-protein-[Burkholderia-multivorans] |
| WP_065497246.1 | PIN-EndoRNase | GCF_001685525.1 | Burkholderia_sp._CCA53 | Bacteria>Pseudomonadati>Pseudomonadota>Betaproteobacteria>Burkholderiales>Burkholderiaceae>Burkholderia>unclassified_Burkholderia | 205 | BCR55_RS29845 | 122836..123453 | 1776288 | NYN-domain-containing-protein-[Burkholderia-sp.-CCA53] |
| WP_166837249.1 | PIN-EndoRNase | GCF_011742065.1 | Aquabacterium_sp._A08 | Bacteria>Pseudomonadati>Pseudomonadota>Betaproteobacteria>Burkholderiales>Burkholderiales_genera_incertae_sedis>Aquabacterium>unclassified_Aquabacterium | 206 | HA630_RS20330 | 25174..25794 | 2718532 | NYN-domain-containing-protein-[Aquabacterium-sp.-A08] |
| WP_165902832.1 | PIN-EndoRNase | GCF_036772355.1 | Tepidimonas_ignava | Bacteria>Pseudomonadati>Pseudomonadota>Betaproteobacteria>Burkholderiales>Burkholderiales_genera_incertae_sedis>Tepidimonas | 57 | EDC36_RS13185 | 37509..37682 | 114249 | hypothetical-protein-[Tepidimonas-ignava] |
| WP_185974532.1 | PIN-EndoRNase | GCF_007556755.1 | Tepidimonas_fonticaldi | Bacteria>Pseudomonadati>Pseudomonadota>Betaproteobacteria>Burkholderiales>Burkholderiales_genera_incertae_sedis>Tepidimonas | 61 | Tfont_RS12340 | 8071..8256 | 1101373 | hypothetical-protein-[Tepidimonas-fonticaldi] |
| WP_043699923.1 | PIN-EndoRNase | GCF_020162115.1 | Tepidimonas_taiwanensis | Bacteria>Pseudomonadati>Pseudomonadota>Betaproteobacteria>Burkholderiales>Burkholderiales_genera_incertae_sedis>Tepidimonas | 205 | PL21_RS04240 | 79696..80313 | 307486 | NYN-domain-containing-protein-[Tepidimonas-taiwanensis] |
| MBN9628767.1 | PIN-EndoRNase | GCA_017307415.1 | Acidovorax_sp. | Bacteria>Pseudomonadati>Pseudomonadota>Betaproteobacteria>Burkholderiales>Comamonadaceae>Acidovorax>unclassified_Acidovorax | 133 | J0G38_21255 | 10892..11293 | 1872122 | type-II-toxin-antitoxin-system-VapC-family-toxin-[Acidovorax-sp.] |
| ODS67616.1 | PIN-EndoRNase | GCA_001724795.1 | Acidovorax_sp._SCN_65-28 | Bacteria>Pseudomonadati>Pseudomonadota>Betaproteobacteria>Burkholderiales>Comamonadaceae>Acidovorax>unclassified_Acidovorax | 133 | ABS39_19750 | 12537..12938 | 1660086 | MAG:-plasmid-maintenance-protein-[Acidovorax-sp.-SCN-65-28] |
| WP_022771793.1 | PIN-EndoRNase | GCF_000477435.1 | Candidatus_Symbiobacter_mobilis | Bacteria>Pseudomonadati>Pseudomonadota>Betaproteobacteria>Burkholderiales>Comamonadaceae>Candidatus_Symbiobacter | 133 | CENROD_RS03855 | 936144..936545 | 1436290 | type-II-toxin-antitoxin-system-VapC-family-toxin-[Candidatus-Symbiobacter-mobilis] |
| WP_022771790.1 | PIN-EndoRNase | GCF_000477435.1 | Candidatus_Symbiobacter_mobilis | Bacteria>Pseudomonadati>Pseudomonadota>Betaproteobacteria>Burkholderiales>Comamonadaceae>Candidatus_Symbiobacter | 166 | CENROD_RS03840 | 932346..932846 | 1436290 | DUF3368-domain-containing-protein-[Candidatus-Symbiobacter-mobilis] |
| WP_042419949.1 | PIN-EndoRNase | GCF_000739875.1 | Comamonas_aquatica | Bacteria>Pseudomonadati>Pseudomonadota>Betaproteobacteria>Burkholderiales>Comamonadaceae>Comamonas | 133 | CA2_RS16415 | 6338..6739 | 225991 | type-II-toxin-antitoxin-system-VapC-family-toxin-[Comamonas-aquatica] |
| WP_210931300.1 | PIN-EndoRNase | GCF_018064205.1 | Comamonas_aquatica | Bacteria>Pseudomonadati>Pseudomonadota>Betaproteobacteria>Burkholderiales>Comamonadaceae>Comamonas | 133 | KAQ61_RS08190 | 1737069..1737470 | 225991 | type-II-toxin-antitoxin-system-VapC-family-toxin-[Comamonas-aquatica] |
| WP_042427119.1 | PIN-EndoRNase | GCF_000739995.1 | Comamonas_granuli | Bacteria>Pseudomonadati>Pseudomonadota>Betaproteobacteria>Burkholderiales>Comamonadaceae>Comamonas | 206 | CGR01S_RS11730 | 77699..78319 | 290309 | NYN-domain-containing-protein-[Comamonas-granuli] |
| TFI45396.1 | PIN-EndoRNase | GCA_004563745.1 | Diaphorobacter_sp._DS2 | Bacteria>Pseudomonadati>Pseudomonadota>Betaproteobacteria>Burkholderiales>Comamonadaceae>Diaphorobacter>unclassified_Diaphorobacter | 206 | E4093_20000 | 10745..11365 | 1302548 | NYN-domain-containing-protein-[Diaphorobacter-sp.-DS2] |
| WP_127803826.1 | PIN-EndoRNase | GCF_004006415.1 | Hydrogenophaga_sp._NH-16 | Bacteria>Pseudomonadati>Pseudomonadota>Betaproteobacteria>Burkholderiales>Comamonadaceae>Hydrogenophaga>unclassified_Hydrogenophaga | 206 | DD542_RS03460 | 721590..722210 | 2184519 | NYN-domain-containing-protein-[Hydrogenophaga-sp.-NH-16] |
| WP_045536711.1 | PIN-EndoRNase | GCF_000828915.1 | Serpentinimonas_maccroryi | Bacteria>Pseudomonadati>Pseudomonadota>Betaproteobacteria>Burkholderiales>Comamonadaceae>Serpentinimonas | 206 | SMCB_RS09930 | 2155937..2156557 | 1458426 | NYN-domain-containing-protein-[Serpentinimonas-maccroryi] |
| PIQ50874.1 | PIN-EndoRNase | GCA_002786915.1 | Comamonadaceae_bacterium_CG12_big_fil_rev_8_21_14_0_65_59_15 | Bacteria>Pseudomonadati>Pseudomonadota>Betaproteobacteria>Burkholderiales>Comamonadaceae>unclassified_Comamonadaceae | 135 | COW02_17775 | 47397..47804 | 1973952 | VapC-toxin-family-PIN-domain-ribonuclease-[Comamonadaceae-bacterium-CG12_big_fil_rev_8_21_14_0_65_59_15] |
| WP_265283909.1 | PIN-EndoRNase | GCF_026127385.1 | Verminephrobacter_aporrectodeae | Bacteria>Pseudomonadati>Pseudomonadota>Betaproteobacteria>Burkholderiales>Comamonadaceae>Verminephrobacter | 172 | D8B27_RS01350 | 115939..116457 | 1110389 | NYN-domain-containing-protein-[Verminephrobacter-aporrectodeae] |
| WP_265663938.1 | PIN-EndoRNase | GCF_026192565.1 | Verminephrobacter_aporrectodeae | Bacteria>Pseudomonadati>Pseudomonadota>Betaproteobacteria>Burkholderiales>Comamonadaceae>Verminephrobacter | 172 | D8B23_RS01240 | 18577..19095 | 1110389 | NYN-domain-containing-protein-[Verminephrobacter-aporrectodeae] |
| WP_019561368.1 | PIN-EndoRNase | GCF_000381125.1 | Caldimonas_manganoxidans | Bacteria>Pseudomonadati>Pseudomonadota>Betaproteobacteria>Burkholderiales>Sphaerotilaceae>Caldimonas | 205 | F615_RS0111650 | 13153..13770 | 196015 | NYN-domain-containing-protein-[Caldimonas-manganoxidans] |
| MCK6404272.1 | PIN-EndoRNase | GCA_023150255.1 | Sphaerotilus_sulfidivorans | Bacteria>Pseudomonadati>Pseudomonadota>Betaproteobacteria>Burkholderiales>Sphaerotilaceae>Sphaerotilus | 183 | L6Q74_20575 | 14114..14665 | 639200 | hypothetical-protein-[Sphaerotilus-sulfidivorans] |
| WP_273473507.1 | PIN-EndoRNase | GCF_023150255.1 | Sphaerotilus_sulfidivorans | Bacteria>Pseudomonadati>Pseudomonadota>Betaproteobacteria>Burkholderiales>Sphaerotilaceae>Sphaerotilus | 183 | L6Q74_RS20535 | 14114..14665 | 639200 | hypothetical-protein-[Sphaerotilus-sulfidivorans] |
| MBK6594589.1 | PIN-EndoRNase | GCA_016703945.1 | Burkholderiales_bacterium | Bacteria>Pseudomonadati>Pseudomonadota>Betaproteobacteria>Burkholderiales>unclassified_Burkholderiales | 98 | IPG23_18850 | 20589..20885 | 1891238 | hypothetical-protein-[Burkholderiales-bacterium] |
| GIX27883.1 | PIN-EndoRNase | GCA_026005015.1 | Burkholderiales_bacterium | Bacteria>Pseudomonadati>Pseudomonadota>Betaproteobacteria>Burkholderiales>unclassified_Burkholderiales | 136 | KatS3mg123_1764 | 1711397..1711807 | 1891238 | MAG:-hypothetical-protein-KatS3mg123_1764-[Burkholderiales-bacterium] |
| MDE2076669.1 | PIN-EndoRNase | GCA_028864885.1 | Burkholderiales_bacterium | Bacteria>Pseudomonadati>Pseudomonadota>Betaproteobacteria>Burkholderiales>unclassified_Burkholderiales | 150 | KGI91_06290 | 40168..40620 | 1891238 | MAG:-PIN-domain-containing-protein-[Burkholderiales-bacterium] |
| MCC6611671.1 | PIN-EndoRNase | GCA_020848075.1 | Burkholderiales_bacterium | Bacteria>Pseudomonadati>Pseudomonadota>Betaproteobacteria>Burkholderiales>unclassified_Burkholderiales | 184 | IT515_18615 | 40490..41044 | 1891238 | type-II-toxin-antitoxin-system-VapC-family-toxin-[Burkholderiales-bacterium] |
| MCC6611675.1 | PIN-EndoRNase | GCA_020848075.1 | Burkholderiales_bacterium | Bacteria>Pseudomonadati>Pseudomonadota>Betaproteobacteria>Burkholderiales>unclassified_Burkholderiales | 189 | IT515_18635 | 46142..46711 | 1891238 | PIN-domain-containing-protein-[Burkholderiales-bacterium] |
| MCZ2134309.1 | PIN-EndoRNase | GCA_026932525.1 | Burkholderiales_bacterium | Bacteria>Pseudomonadati>Pseudomonadota>Betaproteobacteria>Burkholderiales>unclassified_Burkholderiales | 206 | LC098_02600 | 50153..50773 | 1891238 | MAG:-NYN-domain-containing-protein-[Burkholderiales-bacterium] |
| MCO6428557.1 | PIN-EndoRNase | GCA_024102715.1 | Nitrosomonas_communis | Bacteria>Pseudomonadati>Pseudomonadota>Betaproteobacteria>Nitrosomonadales>Nitrosomonadaceae>Nitrosomonas | 183 | J5I83_10935 | 7141..7692 | 44574 | type-II-toxin-antitoxin-system-VapC-family-toxin-[Nitrosomonas-communis] |
| MBN8774180.1 | PIN-EndoRNase | GCA_017309255.1 | Thiobacillus_sp. | Bacteria>Pseudomonadati>Pseudomonadota>Betaproteobacteria>Nitrosomonadales>Thiobacillaceae>Thiobacillus>unclassified_Thiobacillus | 144 | J0J02_08940 | 9398..9832 | 924 | type-II-toxin-antitoxin-system-VapC-family-toxin-[Thiobacillus-sp.] |
| ODV10847.1 | PIN-EndoRNase | GCA_001725535.1 | Thiobacillus_sp._SCN_64-317 | Bacteria>Pseudomonadati>Pseudomonadota>Betaproteobacteria>Nitrosomonadales>Thiobacillaceae>Thiobacillus>unclassified_Thiobacillus | 144 | ABT22_10730 | 10367..10801 | 1660143 | MAG:-hypothetical-protein-ABT22_10730-[Thiobacillus-sp.-SCN-64-317] |
| WP_011238769.1 | PIN-EndoRNase | GCF_000025965.1 | Aromatoleum_aromaticum | Bacteria>Pseudomonadati>Pseudomonadota>Betaproteobacteria>Rhodocyclales>Rhodocyclaceae>Aromatoleum | 138 | EBN1_RS14755 | 3096137..3096553 | 551760 | putative-toxin-antitoxin-system-toxin-component,-PIN-family-[Aromatoleum-aromaticum] |
| KAB2934119.1 | PIN-EndoRNase | GCA_008933825.1 | Rhodocyclaceae_bacterium | Bacteria>Pseudomonadati>Pseudomonadota>Betaproteobacteria>Rhodocyclales>Rhodocyclaceae>unclassified_Rhodocyclaceae | 167 | F9K21_11720 | 9347..9850 | 1898103 | hypothetical-protein-F9K21_11720-[Rhodocyclaceae-bacterium] |
| MBI5901624.1 | PIN-EndoRNase | GCA_016235165.1 | Rhodocyclales_bacterium | Bacteria>Pseudomonadati>Pseudomonadota>Betaproteobacteria>Rhodocyclales>unclassified_Rhodocyclales | 168 | HZB40_20685 | 24743..25249 | 2072421 | type-II-toxin-antitoxin-system-VapC-family-toxin-[Rhodocyclales-bacterium] |
| WP_211163379.1 | PIN-EndoRNase | GCF_963887635.1 | Azoarcus_taiwanensis | Bacteria>Pseudomonadati>Pseudomonadota>Betaproteobacteria>Rhodocyclales>Zoogloeaceae>Azoarcus | 206 | GPA21_RS02050 | 11569..12189 | 666964 | NYN-domain-containing-protein-[Azoarcus-taiwanensis] |
| WP_141017184.1 | PIN-EndoRNase | GCF_006496635.1 | Azoarcus_sp._DD4 | Bacteria>Pseudomonadati>Pseudomonadota>Betaproteobacteria>Rhodocyclales>Zoogloeaceae>Azoarcus>unclassified_Azoarcus | 161 | CJ010_RS05945 | 1250769..1251254 | 2027405 | hypothetical-protein-[Azoarcus-sp.-DD4] |
| WP_247725416.1 | PIN-EndoRNase | GCF_023093595.1 | Thauera_aromatica | Bacteria>Pseudomonadati>Pseudomonadota>Betaproteobacteria>Rhodocyclales>Zoogloeaceae>Thauera | 132 | MXC99_RS10590 | 101247..101645 | 59405 | PIN-domain-containing-protein-[Thauera-aromatica] |
| WP_247724879.1 | PIN-EndoRNase | GCF_023093595.1 | Thauera_aromatica | Bacteria>Pseudomonadati>Pseudomonadota>Betaproteobacteria>Rhodocyclales>Zoogloeaceae>Thauera | 133 | MXC99_RS07605 | 5791..6192 | 59405 | PIN-domain-containing-protein-[Thauera-aromatica] |
| WP_247725479.1 | PIN-EndoRNase | GCF_023093595.1 | Thauera_aromatica | Bacteria>Pseudomonadati>Pseudomonadota>Betaproteobacteria>Rhodocyclales>Zoogloeaceae>Thauera | 138 | MXC99_RS10905 | 52882..53298 | 59405 | putative-toxin-antitoxin-system-toxin-component,-PIN-family-[Thauera-aromatica] |
| TXH82459.1 | PIN-EndoRNase | GCA_008015255.1 | Thauera_aminoaromatica | Bacteria>Pseudomonadati>Pseudomonadota>Betaproteobacteria>Rhodocyclales>Zoogloeaceae>Thauera | 191 | E6Q80_15420 | 503..1078 | 164330 | PIN-domain-containing-protein-[Thauera-aminoaromatica] |
| WP_002930902.1 | PIN-EndoRNase | GCF_000310145.2 | Thauera_sp._28 | Bacteria>Pseudomonadati>Pseudomonadota>Betaproteobacteria>Rhodocyclales>Zoogloeaceae>Thauera>unclassified_Thauera | 138 | C662_RS14990 | 6777..7193 | 303682 | putative-toxin-antitoxin-system-toxin-component,-PIN-family-[Thauera-sp.-28] |
| WP_141354349.1 | PIN-EndoRNase | GCF_006539865.1 | Zoogloea_ramigera | Bacteria>Pseudomonadati>Pseudomonadota>Betaproteobacteria>Rhodocyclales>Zoogloeaceae>Zoogloea | 138 | ZRA01_RS16825 | 6115..6531 | 350 | putative-toxin-antitoxin-system-toxin-component,-PIN-family-[Zoogloea-ramigera] |
| MBK6655409.1 | PIN-EndoRNase | GCA_016703645.1 | Zoogloea_sp. | Bacteria>Pseudomonadati>Pseudomonadota>Betaproteobacteria>Rhodocyclales>Zoogloeaceae>Zoogloea>unclassified_Zoogloea | 143 | IPG40_16565 | 746382..746813 | 49181 | type-II-toxin-antitoxin-system-VapC-family-toxin-[Zoogloea-sp.] |
| MBK7848385.1 | PIN-EndoRNase | GCA_016714095.1 | Zoogloea_sp. | Bacteria>Pseudomonadati>Pseudomonadota>Betaproteobacteria>Rhodocyclales>Zoogloeaceae>Zoogloea>unclassified_Zoogloea | 143 | IPJ73_15235 | 32882..33313 | 49181 | type-II-toxin-antitoxin-system-VapC-family-toxin-[Zoogloea-sp.] |
| MCL1825810.1 | PIN-EndoRNase | GCA_009782565.1 | Betaproteobacteria_bacterium | Bacteria>Pseudomonadati>Pseudomonadota>Betaproteobacteria>unclassified_Betaproteobacteria | 137 | FWG26_07725 | 7993..8406 | 1891241 | PIN-domain-containing-protein-[Betaproteobacteria-bacterium] |
| MPZ42200.1 | PIN-EndoRNase | GCA_009377585.1 | Betaproteobacteria_bacterium | Bacteria>Pseudomonadati>Pseudomonadota>Betaproteobacteria>unclassified_Betaproteobacteria | 141 | GEV05_02120 | 144920..145345 | 1891241 | PIN-domain-containing-protein-[Betaproteobacteria-bacterium] |
| MSQ72375.1 | PIN-EndoRNase | GCA_009693435.1 | Betaproteobacteria_bacterium | Bacteria>Pseudomonadati>Pseudomonadota>Betaproteobacteria>unclassified_Betaproteobacteria | 144 | EXR27_13925 | 25096..25530 | 1891241 | PIN-domain-containing-protein-[Betaproteobacteria-bacterium] |
| OGA65438.1 | PIN-EndoRNase | GCA_001770585.1 | Betaproteobacteria_bacterium_RIFCSPLOWO2_12_FULL_65_14 | Bacteria>Pseudomonadati>Pseudomonadota>Betaproteobacteria>unclassified_Betaproteobacteria | 164 | A3G81_12445 | 3560..4054 | 1797501 | hypothetical-protein-A3G81_12445-[Betaproteobacteria-bacterium-RIFCSPLOWO2_12_FULL_65_14] |
| OGA54125.1 | PIN-EndoRNase | GCA_001772005.1 | Betaproteobacteria_bacterium_RIFCSPLOWO2_12_FULL_62_58 | Bacteria>Pseudomonadati>Pseudomonadota>Betaproteobacteria>unclassified_Betaproteobacteria | 206 | A3F74_24220 | 8518..9138 | 1797497 | NYN-domain-containing-protein-[Betaproteobacteria-bacterium-RIFCSPLOWO2_12_FULL_62_58] |
| GHU44760.1 | PIN-EndoRNase | GCA_025997095.1 | Betaproteobacteria_bacterium | Bacteria>Pseudomonadati>Pseudomonadota>Betaproteobacteria>unclassified_Betaproteobacteria | 210 | AGMMS50289_13850 | 12027..12659 | 1891241 | hypothetical-protein-AGMMS50289_13850-[Betaproteobacteria-bacterium] |
| WP_106453124.1 | PIN-EndoRNase | GCF_003012795.1 | Zobellella_taiwanensis | Bacteria>Pseudomonadati>Pseudomonadota>Gammaproteobacteria>Aeromonadales>Aeromonadaceae>Zobellella | 172 | C7I36_RS07630 | 87674..88192 | 347535 | hypothetical-protein-[Zobellella-taiwanensis] |
| WP_144225092.1 | PIN-EndoRNase | GCF_017580345.1 | Shewanella_algae | Bacteria>Pseudomonadati>Pseudomonadota>Gammaproteobacteria>Alteromonadales>Shewanellaceae>Shewanella | 153 | I6M43_RS01855 | 54166..54627 | 38313 | hypothetical-protein-[Shewanella-algae] |
| MAL94414.1 | PIN-EndoRNase | GCA_002692965.1 | Haliea_sp. | Bacteria>Pseudomonadati>Pseudomonadota>Gammaproteobacteria>Cellvibrionales>Halieaceae>Haliea>unclassified_Haliea | 179 | CME40_04980 | 35710..36249 | 1932666 | hypothetical-protein-[Haliea-sp.] |
| WP_273009214.1 | PIN-EndoRNase | GCF_002435105.1 | Spongiibacter_tropicus | Bacteria>Pseudomonadati>Pseudomonadota>Gammaproteobacteria>Cellvibrionales>Spongiibacteraceae>Spongiibacter | 206 | B3U69_RS04095 | 6155..6775 | 454602 | NYN-domain-containing-protein-[Spongiibacter-tropicus] |
| WP_009149688.1 | PIN-EndoRNase | GCF_033954835.1 | Thiorhodovibrio | Bacteria>Pseudomonadati>Pseudomonadota>Gammaproteobacteria>Chromatiales>Chromatiaceae | 129 | THI970DRAFT_RS14145 | 1749765..1750154 | 61593 | type-II-toxin-antitoxin-system-VapC-family-toxin-[Thiorhodovibrio-sp.-970] |
| WP_201065309.1 | PIN-EndoRNase | GCF_033954455.1 | Thiorhodovibrio | Bacteria>Pseudomonadati>Pseudomonadota>Gammaproteobacteria>Chromatiales>Chromatiaceae | 131 | CCR91_RS06135 | 55792..56187 | 61593 | DUF5615-family-PIN-like-protein-[Thiorhodovibrio-winogradskyi] |
| WP_201066874.1 | PIN-EndoRNase | GCF_033954455.1 | Thiorhodovibrio | Bacteria>Pseudomonadati>Pseudomonadota>Gammaproteobacteria>Chromatiales>Chromatiaceae | 134 | CCR91_RS10525 | 19743..20147 | 61593 | PIN-domain-containing-protein-[Thiorhodovibrio-winogradskyi] |
| WP_201066885.1 | PIN-EndoRNase | GCF_033954455.1 | Thiorhodovibrio | Bacteria>Pseudomonadati>Pseudomonadota>Gammaproteobacteria>Chromatiales>Chromatiaceae | 147 | CCR91_RS10555 | 24671..25114 | 61593 | PIN-domain-containing-protein-[Thiorhodovibrio-winogradskyi] |
| WP_155450781.1 | PIN-EndoRNase | GCF_009720725.1 | Allochromatium_palmeri | Bacteria>Pseudomonadati>Pseudomonadota>Gammaproteobacteria>Chromatiales>Chromatiaceae>Allochromatium | 139 | GJ668_RS14080 | 15691..16110 | 231048 | type-II-toxin-antitoxin-system-VapC-family-toxin-[Allochromatium-palmeri] |
| WP_200158839.1 | PIN-EndoRNase | GCF_016584135.1 | Allochromatium_vinosum | Bacteria>Pseudomonadati>Pseudomonadota>Gammaproteobacteria>Chromatiales>Chromatiaceae>Allochromatium | 148 | CK029_RS16470 | 17113..17559 | 1049 | hypothetical-protein-[Allochromatium-vinosum] |
| WP_201219755.1 | PIN-EndoRNase | GCF_016653315.1 | Halochromatium_roseum | Bacteria>Pseudomonadati>Pseudomonadota>Gammaproteobacteria>Chromatiales>Chromatiaceae>Halochromatium | 163 | CCR96_RS08805 | 131061..131552 | 391920 | DUF3368-domain-containing-protein-[Halochromatium-roseum] |
| WP_201218764.1 | PIN-EndoRNase | GCF_016653315.1 | Halochromatium_roseum | Bacteria>Pseudomonadati>Pseudomonadota>Gammaproteobacteria>Chromatiales>Chromatiaceae>Halochromatium | 243 | CCR96_RS05815 | 20853..21584 | 391920 | NYN-domain-containing-protein-[Halochromatium-roseum] |
| NEX15478.1 | PIN-EndoRNase | GCA_010912705.1 | Halochromatium_sp. | Bacteria>Pseudomonadati>Pseudomonadota>Gammaproteobacteria>Chromatiales>Chromatiaceae>Halochromatium>unclassified_Halochromatium | 145 | C1943_02285 | 46281..46718 | 2049430 | hypothetical-protein-[Halochromatium-sp.] |
| WP_242479450.1 | PIN-EndoRNase | GCF_016583905.1 | Lamprobacter_modestohalophilus | Bacteria>Pseudomonadati>Pseudomonadota>Gammaproteobacteria>Chromatiales>Chromatiaceae>Lamprobacter | 128 | CK042_RS16505 | 28244..28630 | 1064514 | DUF3368-domain-containing-protein-[Lamprobacter-modestohalophilus] |
| MBK1620018.1 | PIN-EndoRNase | GCA_016583905.1 | Lamprobacter_modestohalophilus | Bacteria>Pseudomonadati>Pseudomonadota>Gammaproteobacteria>Chromatiales>Chromatiaceae>Lamprobacter | 144 | CK042_16535 | 29630..30064 | 1064514 | nucleotide-binding-protein-[Lamprobacter-modestohalophilus] |
| WP_200246413.1 | PIN-EndoRNase | GCF_016583905.1 | Lamprobacter_modestohalophilus | Bacteria>Pseudomonadati>Pseudomonadota>Gammaproteobacteria>Chromatiales>Chromatiaceae>Lamprobacter | 144 | CK042_RS16520 | 29630..30064 | 1064514 | type-II-toxin-antitoxin-system-VapC-family-toxin-[Lamprobacter-modestohalophilus] |
| MBK1620015.1 | PIN-EndoRNase | GCA_016583905.1 | Lamprobacter_modestohalophilus | Bacteria>Pseudomonadati>Pseudomonadota>Gammaproteobacteria>Chromatiales>Chromatiaceae>Lamprobacter | 166 | CK042_16520 | 28244..28744 | 1064514 | DUF3368-domain-containing-protein-[Lamprobacter-modestohalophilus] |
| MBV5272981.1 | PIN-EndoRNase | GCA_019163055.1 | Lamprocystis_purpurea | Bacteria>Pseudomonadati>Pseudomonadota>Gammaproteobacteria>Chromatiales>Chromatiaceae>Lamprocystis | 135 | JZU52_04830 | 8814..9221 | 61598 | type-II-toxin-antitoxin-system-VapC-family-toxin-[Lamprocystis-purpurea] |
| MBV5274822.1 | PIN-EndoRNase | GCA_019163055.1 | Lamprocystis_purpurea | Bacteria>Pseudomonadati>Pseudomonadota>Gammaproteobacteria>Chromatiales>Chromatiaceae>Lamprocystis | 135 | JZU52_14675 | 8600..9007 | 61598 | PIN-domain-containing-protein-[Lamprocystis-purpurea] |
| WP_020503210.1 | PIN-EndoRNase | GCF_000379525.1 | Lamprocystis_purpurea | Bacteria>Pseudomonadati>Pseudomonadota>Gammaproteobacteria>Chromatiales>Chromatiaceae>Lamprocystis | 135 | A390_RS31700 | 114949..115356 | 61598 | PIN-domain-containing-protein-[Lamprocystis-purpurea] |
| WP_020506077.1 | PIN-EndoRNase | GCF_000379525.1 | Lamprocystis_purpurea | Bacteria>Pseudomonadati>Pseudomonadota>Gammaproteobacteria>Chromatiales>Chromatiaceae>Lamprocystis | 137 | A390_RS0118100 | 28028..28441 | 61598 | PIN-domain-containing-protein-[Lamprocystis-purpurea] |
| WP_020507131.1 | PIN-EndoRNase | GCF_000379525.1 | Lamprocystis_purpurea | Bacteria>Pseudomonadati>Pseudomonadota>Gammaproteobacteria>Chromatiales>Chromatiaceae>Lamprocystis | 139 | A390_RS34840 | 25879..26298 | 61598 | PIN-domain-containing-protein-[Lamprocystis-purpurea] |
| WP_062271497.1 | PIN-EndoRNase | GCF_001583505.1 | Marichromatium_gracile | Bacteria>Pseudomonadati>Pseudomonadota>Gammaproteobacteria>Chromatiales>Chromatiaceae>Marichromatium | 147 | AY586_RS00810 | 22255..22698 | 1048 | PIN-domain-containing-protein-[Marichromatium-gracile] |
| WP_062271493.1 | PIN-EndoRNase | GCF_001583505.1 | Marichromatium_gracile | Bacteria>Pseudomonadati>Pseudomonadota>Gammaproteobacteria>Chromatiales>Chromatiaceae>Marichromatium | 153 | AY586_RS00800 | 20493..20954 | 1048 | hypothetical-protein-[Marichromatium-gracile] |
| WP_011330218.1 | PIN-EndoRNase | GCF_000012805.1 | Nitrosococcus_oceani | Bacteria>Pseudomonadati>Pseudomonadota>Gammaproteobacteria>Chromatiales>Chromatiaceae>Nitrosococcus | 183 | IB75_RS00235 | 15861..16412 | 1229 | type-II-toxin-antitoxin-system-VapC-family-toxin-[Nitrosococcus-oceani] |
| WP_013219153.1 | PIN-EndoRNase | GCF_000143085.1 | Nitrosococcus_watsonii | Bacteria>Pseudomonadati>Pseudomonadota>Gammaproteobacteria>Chromatiales>Chromatiaceae>Nitrosococcus | 183 | NWAT_RS00260 | 52541..53092 | 473531 | type-II-toxin-antitoxin-system-VapC-family-toxin-[Nitrosococcus-watsonii] |
| ADJ27041.1 | PIN-EndoRNase | GCA_000143085.1 | Nitrosococcus_watsonii_C-113 | Bacteria>Pseudomonadati>Pseudomonadota>Gammaproteobacteria>Chromatiales>Chromatiaceae>Nitrosococcus>Nitrosococcus_watsonii | 183 | Nwat_0050 | 52541..53092 | 105559 | PilT-protein-domain-protein-[Nitrosococcus-watsonii-C-113] |
| WP_153975102.1 | PIN-EndoRNase | GCF_009664085.1 | Thermochromatium_tepidum | Bacteria>Pseudomonadati>Pseudomonadota>Gammaproteobacteria>Chromatiales>Chromatiaceae>Thermochromatium | 135 | E6P07_RS07885 | 1740215..1740622 | 1050 | PIN-domain-containing-protein-[Thermochromatium-tepidum] |
| WP_132976382.1 | PIN-EndoRNase | GCF_004342175.1 | Thiobaca_trueperi | Bacteria>Pseudomonadati>Pseudomonadota>Gammaproteobacteria>Chromatiales>Chromatiaceae>Thiobaca | 123 | EDC35_RS06195 | 64728..65099 | 127458 | type-II-toxin-antitoxin-system-VapC-family-toxin-[Thiobaca-trueperi] |
| WP_147431170.1 | PIN-EndoRNase | GCF_003634315.1 | Thiocapsa_rosea | Bacteria>Pseudomonadati>Pseudomonadota>Gammaproteobacteria>Chromatiales>Chromatiaceae>Thiocapsa | 48 | BDD21_RS27560 | 4631503..4631649 | 69360 | type-II-toxin-antitoxin-system-VapC-family-toxin-[Thiocapsa-rosea] |
| WP_120797043.1 | PIN-EndoRNase | GCF_003634315.1 | Thiocapsa_rosea | Bacteria>Pseudomonadati>Pseudomonadota>Gammaproteobacteria>Chromatiales>Chromatiaceae>Thiocapsa | 133 | BDD21_RS09930 | 2234233..2234634 | 69360 | type-II-toxin-antitoxin-system-VapC-family-toxin-[Thiocapsa-rosea] |
| WP_120800304.1 | PIN-EndoRNase | GCF_003634315.1 | Thiocapsa_rosea | Bacteria>Pseudomonadati>Pseudomonadota>Gammaproteobacteria>Chromatiales>Chromatiaceae>Thiocapsa | 133 | BDD21_RS27125 | 181094..181495 | 69360 | type-II-toxin-antitoxin-system-VapC-family-toxin-[Thiocapsa-rosea] |
| WP_120796607.1 | PIN-EndoRNase | GCF_003634315.1 | Thiocapsa_rosea | Bacteria>Pseudomonadati>Pseudomonadota>Gammaproteobacteria>Chromatiales>Chromatiaceae>Thiocapsa | 138 | BDD21_RS07390 | 1612856..1613272 | 69360 | putative-toxin-antitoxin-system-toxin-component,-PIN-family-[Thiocapsa-rosea] |
| WP_200385869.1 | PIN-EndoRNase | GCF_016583985.1 | Thiocapsa_imhoffii | Bacteria>Pseudomonadati>Pseudomonadota>Gammaproteobacteria>Chromatiales>Chromatiaceae>Thiocapsa | 139 | CK025_RS00015 | 4065..4484 | 382777 | PIN-domain-containing-protein-[Thiocapsa-imhoffii] |
| WP_120800302.1 | PIN-EndoRNase | GCF_003634315.1 | Thiocapsa_rosea | Bacteria>Pseudomonadati>Pseudomonadota>Gammaproteobacteria>Chromatiales>Chromatiaceae>Thiocapsa | 141 | BDD21_RS27110 | 177334..177759 | 69360 | type-II-toxin-antitoxin-system-VapC-family-toxin-[Thiocapsa-rosea] |
| QVL49521.1 | PIN-EndoRNase | GCA_018399035.1 | Thiocapsa_sp. | Bacteria>Pseudomonadati>Pseudomonadota>Gammaproteobacteria>Chromatiales>Chromatiaceae>Thiocapsa>unclassified_Thiocapsa | 141 | KFB96_03130 | 680447..680872 | 2024551 | MAG:-type-II-toxin-antitoxin-system-VapC-family-toxin-[Thiocapsa-sp.] |
| QVL48947.1 | PIN-EndoRNase | GCA_018399035.1 | Thiocapsa_sp. | Bacteria>Pseudomonadati>Pseudomonadota>Gammaproteobacteria>Chromatiales>Chromatiaceae>Thiocapsa>unclassified_Thiocapsa | 151 | KFB96_26055 | 5691334..5691789 | 2024551 | MAG:-type-II-toxin-antitoxin-system-VapC-family-toxin-[Thiocapsa-sp.] |
| QVL49519.1 | PIN-EndoRNase | GCA_018399035.1 | Thiocapsa_sp. | Bacteria>Pseudomonadati>Pseudomonadota>Gammaproteobacteria>Chromatiales>Chromatiaceae>Thiocapsa>unclassified_Thiocapsa | 169 | KFB96_03120 | 678587..679096 | 2024551 | MAG:-hypothetical-protein-KFB96_03120-[Thiocapsa-sp.] |
| WP_242470923.1 | PIN-EndoRNase | - | Thiocystis_violacea | Bacteria>Pseudomonadati>Pseudomonadota>Gammaproteobacteria>Chromatiales>Chromatiaceae>Thiocystis | 138 | CK027_RS14220 | 25762..26178 | 13725 | PIN-domain-containing-protein-[Thiocystis-violacea] |
| AUB82639.1 | PIN-EndoRNase | GCA_002813775.1 | Candidatus_Thiodictyon_syntrophicum | Bacteria>Pseudomonadati>Pseudomonadota>Gammaproteobacteria>Chromatiales>Chromatiaceae>Thiodictyon | 139 | THSYN_17945 | 4310685..4311104 | 1166950 | hypothetical-protein-THSYN_17945-[Candidatus-Thiodictyon-syntrophicum] |
| AUB85623.1 | PIN-EndoRNase | GCA_002813775.1 | Candidatus_Thiodictyon_syntrophicum | Bacteria>Pseudomonadati>Pseudomonadota>Gammaproteobacteria>Chromatiales>Chromatiaceae>Thiodictyon | 141 | THSYN_32510 | 352796..353221 | 1166950 | VapC-toxin-family-PIN-domain-ribonuclease-[Candidatus-Thiodictyon-syntrophicum] |
| WP_100923234.1 | PIN-EndoRNase | GCF_002813775.1 | Candidatus_Thiodictyon_syntrophicum | Bacteria>Pseudomonadati>Pseudomonadota>Gammaproteobacteria>Chromatiales>Chromatiaceae>Thiodictyon | 141 | THSYN_RS32785 | 352796..353221 | 1166950 | type-II-toxin-antitoxin-system-VapC-family-toxin-[Candidatus-Thiodictyon-syntrophicum] |
| AUB82642.1 | PIN-EndoRNase | GCA_002813775.1 | Candidatus_Thiodictyon_syntrophicum | Bacteria>Pseudomonadati>Pseudomonadota>Gammaproteobacteria>Chromatiales>Chromatiaceae>Thiodictyon | 168 | THSYN_17960 | 4313873..4314379 | 1166950 | hypothetical-protein-THSYN_17960-[Candidatus-Thiodictyon-syntrophicum] |
| WP_100920749.1 | PIN-EndoRNase | GCF_002813775.1 | Candidatus_Thiodictyon_syntrophicum | Bacteria>Pseudomonadati>Pseudomonadota>Gammaproteobacteria>Chromatiales>Chromatiaceae>Thiodictyon | 170 | THSYN_RS20425 | 4837429..4837941 | 1166950 | DUF3368-domain-containing-protein-[Candidatus-Thiodictyon-syntrophicum] |
| WP_015282547.1 | PIN-EndoRNase | GCF_000327045.1 | Thioflavicoccus_mobilis | Bacteria>Pseudomonadati>Pseudomonadota>Gammaproteobacteria>Chromatiales>Chromatiaceae>Thioflavicoccus | 165 | THIMO_RS17950 | 75230..75727 | 80679 | DUF3368-domain-containing-protein-[Thioflavicoccus-mobilis] |
| WP_207148030.1 | PIN-EndoRNase | - | Thiohalocapsa_halophila | Bacteria>Pseudomonadati>Pseudomonadota>Gammaproteobacteria>Chromatiales>Chromatiaceae>Thiohalocapsa | 126 | CK031_RS15105 | 7383..7763 | 69359 | type-II-toxin-antitoxin-system-VapC-family-toxin-[Thiohalocapsa-halophila] |
| WP_200239116.1 | PIN-EndoRNase | GCF_016583825.1 | Thiohalocapsa_halophila | Bacteria>Pseudomonadati>Pseudomonadota>Gammaproteobacteria>Chromatiales>Chromatiaceae>Thiohalocapsa | 133 | CK031_RS15135 | 12736..13137 | 69359 | type-II-toxin-antitoxin-system-VapC-family-toxin-[Thiohalocapsa-halophila] |
| MCG6940981.1 | PIN-EndoRNase | GCA_022340605.1 | Thiohalocapsa_sp. | Bacteria>Pseudomonadati>Pseudomonadota>Gammaproteobacteria>Chromatiales>Chromatiaceae>Thiohalocapsa>unclassified_Thiohalocapsa | 123 | LJE69_06995 | 1264..1635 | 2497641 | PIN-domain-containing-protein-[Thiohalocapsa-sp.] |
| WP_058557214.1 | PIN-EndoRNase | GCF_001469165.1 | Thiohalocapsa_sp._ML1 | Bacteria>Pseudomonadati>Pseudomonadota>Gammaproteobacteria>Chromatiales>Chromatiaceae>Thiohalocapsa>unclassified_Thiohalocapsa | 134 | ML01_RS20285 | 4014..4418 | 1431688 | PIN-domain-containing-protein-[Thiohalocapsa-sp.-ML1] |
| WP_058554454.1 | PIN-EndoRNase | GCF_001469165.1 | Thiohalocapsa_sp._ML1 | Bacteria>Pseudomonadati>Pseudomonadota>Gammaproteobacteria>Chromatiales>Chromatiaceae>Thiohalocapsa>unclassified_Thiohalocapsa | 138 | ML01_RS05775 | 165116..165532 | 1431688 | PIN-domain-containing-protein-[Thiohalocapsa-sp.-ML1] |
| WP_058554880.1 | PIN-EndoRNase | GCF_001469165.1 | Thiohalocapsa_sp._ML1 | Bacteria>Pseudomonadati>Pseudomonadota>Gammaproteobacteria>Chromatiales>Chromatiaceae>Thiohalocapsa>unclassified_Thiohalocapsa | 143 | ML01_RS08040 | 160478..160909 | 1431688 | type-II-toxin-antitoxin-system-VapC-family-toxin-[Thiohalocapsa-sp.-ML1] |
| WP_058554675.1 | PIN-EndoRNase | GCF_001469165.1 | Thiohalocapsa_sp._ML1 | Bacteria>Pseudomonadati>Pseudomonadota>Gammaproteobacteria>Chromatiales>Chromatiaceae>Thiohalocapsa>unclassified_Thiohalocapsa | 153 | ML01_RS06950 | 79867..80328 | 1431688 | type-II-toxin-antitoxin-system-VapC-family-toxin-[Thiohalocapsa-sp.-ML1] |
| WP_245232442.1 | PIN-EndoRNase | - | Thiorhodococcus_minor | Bacteria>Pseudomonadati>Pseudomonadota>Gammaproteobacteria>Chromatiales>Chromatiaceae>Thiorhodococcus | 131 | G3446_RS10710 | 62129..62524 | 57489 | PIN-domain-containing-protein-[Thiorhodococcus-minor] |
| EIC19848.1 | PIN-EndoRNase | GCA_000228725.3 | Thiorhodovibrio_frisius | Bacteria>Pseudomonadati>Pseudomonadota>Gammaproteobacteria>Chromatiales>Chromatiaceae>Thiorhodovibrio | 76 | Thi970DRAFT_03453 | 368847..369077 | 631362 | hypothetical-protein-Thi970DRAFT_03453-[Thiorhodovibrio-sp.-970] |
| WP_009150251.1 | PIN-EndoRNase | GCF_033954835.1 | Thiorhodovibrio_frisius | Bacteria>Pseudomonadati>Pseudomonadota>Gammaproteobacteria>Chromatiales>Chromatiaceae>Thiorhodovibrio | 76 | THI970DRAFT_RS16345 | 368847..369077 | 631362 | type-II-toxin-antitoxin-system-VapC-family-toxin-[Thiorhodovibrio-sp.-970] |
| WP_009148237.1 | PIN-EndoRNase | GCF_033954835.1 | Thiorhodovibrio_frisius | Bacteria>Pseudomonadati>Pseudomonadota>Gammaproteobacteria>Chromatiales>Chromatiaceae>Thiorhodovibrio | 137 | THI970DRAFT_RS08805 | 539825..540238 | 631362 | type-II-toxin-antitoxin-system-VapC-family-toxin-[Thiorhodovibrio-sp.-970] |
| WP_009146900.1 | PIN-EndoRNase | GCF_033954835.1 | Thiorhodovibrio_frisius | Bacteria>Pseudomonadati>Pseudomonadota>Gammaproteobacteria>Chromatiales>Chromatiaceae>Thiorhodovibrio | 176 | THI970DRAFT_RS02015 | 38854..39384 | 631362 | hypothetical-protein-[Thiorhodovibrio-sp.-970] |
| MCF7978331.1 | PIN-EndoRNase | GCA_021736685.1 | Chromatiaceae_bacterium | Bacteria>Pseudomonadati>Pseudomonadota>Gammaproteobacteria>Chromatiales>Chromatiaceae>unclassified_Chromatiaceae | 75 | K9L82_09980 | 9612..9839 | 2049432 | hypothetical-protein-[Chromatiaceae-bacterium] |

| Protein-ID | Domain | Assembly-ID | Strain | Taxonomy | Length | Locus-Tag | Genomic-coordinates | TaxID | Fasta-defline |
|---|---|---|---|---|---|---|---|---|---|
| MCF8016508.1 | PIN-EndoRNase | GCA_021736865.1 | Chromatiaceae_bacterium | Bacteria>Pseudomonadati>Pseudomonadota>Gammaproteobacteria>Chromatiales>Chromatiaceae>unclassified_Chromatiaceae | 75 | K9L65_12590 | 6163..6390 | 2049432 | hypothetical-protein-[Chromatiaceae-bacterium] |
| TVQ88668.1 | PIN-EndoRNase | GCA_007693535.1 | Chromatiaceae_bacterium | Bacteria>Pseudomonadati>Pseudomonadota>Gammaproteobacteria>Chromatiales>Chromatiaceae>unclassified_Chromatiaceae | 102 | EA400_08535 | 15881..16189 | 2049432 | type-II-toxin-antitoxin-system-VapC-family-toxin-[Chromatiaceae-bacterium] |
| MBP6807298.1 | PIN-EndoRNase | GCA_017991215.1 | Chromatiaceae_bacterium | Bacteria>Pseudomonadati>Pseudomonadota>Gammaproteobacteria>Chromatiales>Chromatiaceae>unclassified_Chromatiaceae | 126 | KA125_03285 | 4964..5344 | 2049432 | type-II-toxin-antitoxin-system-VapC-family-toxin-[Chromatiaceae-bacterium] |
| MCP5407918.1 | PIN-EndoRNase | GCA_024235335.1 | Chromatiaceae_bacterium | Bacteria>Pseudomonadati>Pseudomonadota>Gammaproteobacteria>Chromatiales>Chromatiaceae>unclassified_Chromatiaceae | 126 | H6963_01305 | 252576..252956 | 2049432 | type-II-toxin-antitoxin-system-VapC-family-toxin-[Chromatiaceae-bacterium] |
| MCF7978324.1 | PIN-EndoRNase | GCA_021736685.1 | Chromatiaceae_bacterium | Bacteria>Pseudomonadati>Pseudomonadota>Gammaproteobacteria>Chromatiales>Chromatiaceae>unclassified_Chromatiaceae | 130 | K9L82_09945 | 4015..4407 | 2049432 | type-II-toxin-antitoxin-system-VapC-family-toxin-[Chromatiaceae-bacterium] |
| MCF8016501.1 | PIN-EndoRNase | GCA_021736865.1 | Chromatiaceae_bacterium | Bacteria>Pseudomonadati>Pseudomonadota>Gammaproteobacteria>Chromatiales>Chromatiaceae>unclassified_Chromatiaceae | 130 | K9L65_12555 | 566..958 | 2049432 | type-II-toxin-antitoxin-system-VapC-family-toxin-[Chromatiaceae-bacterium] |
| MBK8638070.1 | PIN-EndoRNase | GCA_016714645.1 | Chromatiaceae_bacterium | Bacteria>Pseudomonadati>Pseudomonadota>Gammaproteobacteria>Chromatiales>Chromatiaceae>unclassified_Chromatiaceae | 131 | IPN92_07170 | 46573..46968 | 2049432 | type-II-toxin-antitoxin-system-VapC-family-toxin-[Chromatiaceae-bacterium] |
| MBP6733463.1 | PIN-EndoRNase | GCA_017990755.1 | Chromatiaceae_bacterium | Bacteria>Pseudomonadati>Pseudomonadota>Gammaproteobacteria>Chromatiales>Chromatiaceae>unclassified_Chromatiaceae | 138 | KA142_02010 | 6320..6736 | 2049432 | putative-toxin-antitoxin-system-toxin-component,-PIN-family-[Chromatiaceae-bacterium] |
| MCF8005385.1 | PIN-EndoRNase | GCA_021736845.1 | Chromatiaceae_bacterium | Bacteria>Pseudomonadati>Pseudomonadota>Gammaproteobacteria>Chromatiales>Chromatiaceae>unclassified_Chromatiaceae | 144 | K9L32_14510 | 317..751 | 2049432 | type-II-toxin-antitoxin-system-VapC-family-toxin-[Chromatiaceae-bacterium] |
| TVQ88666.1 | PIN-EndoRNase | GCA_007693535.1 | Chromatiaceae_bacterium | Bacteria>Pseudomonadati>Pseudomonadota>Gammaproteobacteria>Chromatiales>Chromatiaceae>unclassified_Chromatiaceae | 144 | EA400_08520 | 14321..14755 | 2049432 | type-II-toxin-antitoxin-system-VapC-family-toxin-[Chromatiaceae-bacterium] |
| TVQ85661.1 | PIN-EndoRNase | GCA_007693535.1 | Chromatiaceae_bacterium | Bacteria>Pseudomonadati>Pseudomonadota>Gammaproteobacteria>Chromatiales>Chromatiaceae>unclassified_Chromatiaceae | 150 | EA400_14990 | 2644..3096 | 2049432 | PIN-domain-containing-protein-[Chromatiaceae-bacterium] |
| MBK8639645.1 | PIN-EndoRNase | GCA_016714645.1 | Chromatiaceae_bacterium | Bacteria>Pseudomonadati>Pseudomonadota>Gammaproteobacteria>Chromatiales>Chromatiaceae>unclassified_Chromatiaceae | 158 | IPN92_15745 | 855549..856025 | 2049432 | DUF3368-domain-containing-protein-[Chromatiaceae-bacterium] |
| MBP6807677.1 | PIN-EndoRNase | GCA_017991215.1 | Chromatiaceae_bacterium | Bacteria>Pseudomonadati>Pseudomonadota>Gammaproteobacteria>Chromatiales>Chromatiaceae>unclassified_Chromatiaceae | 158 | KA125_05220 | 2260..2736 | 2049432 | DUF3368-domain-containing-protein-[Chromatiaceae-bacterium] |
| MBK8638072.1 | PIN-EndoRNase | GCA_016714645.1 | Chromatiaceae_bacterium | Bacteria>Pseudomonadati>Pseudomonadota>Gammaproteobacteria>Chromatiales>Chromatiaceae>unclassified_Chromatiaceae | 161 | IPN92_07180 | 50660..51145 | 2049432 | type-II-toxin-antitoxin-system-VapC-family-toxin-[Chromatiaceae-bacterium] |
| MCF8003662.1 | PIN-EndoRNase | GCA_021736845.1 | Chromatiaceae_bacterium | Bacteria>Pseudomonadati>Pseudomonadota>Gammaproteobacteria>Chromatiales>Chromatiaceae>unclassified_Chromatiaceae | 163 | K9L32_05560 | 4590..5081 | 2049432 | DUF3368-domain-containing-protein-[Chromatiaceae-bacterium] |
| WP_008932881.1 | PIN-EndoRNase | GCF_000225005.1 | Ectothiorhodospira_sp._PHS-1 | Bacteria>Pseudomonadati>Pseudomonadota>Gammaproteobacteria>Chromatiales>Ectothiorhodospiraceae>Ectothiorhodospira>unclassified_Ectothiorhodospira | 140 | ECTPHS_RS11295 | 34266..34688 | 519989 | type-II-toxin-antitoxin-system-VapC-family-toxin-[Ectothiorhodospira-sp.-PHS-1] |
| WP_005004505.1 | PIN-EndoRNase | GCF_000153205.1 | Nitrococcus_mobilis | Bacteria>Pseudomonadati>Pseudomonadota>Gammaproteobacteria>Chromatiales>Ectothiorhodospiraceae>Nitrococcus | 125 | NB231_RS15005 | 3306412..3306789 | 35797 | type-II-toxin-antitoxin-system-VapC-family-toxin-[Nitrococcus-mobilis] |
| WP_040664278.1 | PIN-EndoRNase | GCF_000153205.1 | Nitrococcus_mobilis | Bacteria>Pseudomonadati>Pseudomonadota>Gammaproteobacteria>Chromatiales>Ectothiorhodospiraceae>Nitrococcus | 133 | NB231_RS14965 | 3294920..3295321 | 35797 | type-II-toxin-antitoxin-system-VapC-family-toxin-[Nitrococcus-mobilis] |
| WP_019627824.1 | PIN-EndoRNase | GCF_000381845.1 | Thioalkalivibrio_sp._AKL10 | Bacteria>Pseudomonadati>Pseudomonadota>Gammaproteobacteria>Chromatiales>Ectothiorhodospiraceae>Thioalkalivibrio>unclassified_Thioalkalivibrio | 206 | G333_RS0104930 | 1026866..1027486 | 1158158 | NYN-domain-containing-protein-[Thioalkalivibrio-sp.-AKL10] |
| WP_177417919.1 | PIN-EndoRNase | - | endosymbiont_of_Lamellibrachia_barhami | Bacteria>Pseudomonadati>Pseudomonadota>Gammaproteobacteria>Chromatiales>unclassified_Chromatiales | 133 | G9Q73_RS00020 | 4366..4767 | 205975 | PIN-domain-containing-protein-[endosymbiont-of-Lamellibrachia-barhami] |
| MCK7579543.1 | PIN-EndoRNase | GCA_023156155.1 | Chromatiales_bacterium | Bacteria>Pseudomonadati>Pseudomonadota>Gammaproteobacteria>Chromatiales>unclassified_Chromatiales | 153 | MZV65_30095 | 12297..12758 | 2026725 | hypothetical-protein-[Chromatiales-bacterium] |
| MBL8253033.1 | PIN-EndoRNase | GCA_016789525.1 | Candidatus_Competibacter_sp. | Bacteria>Pseudomonadati>Pseudomonadota>Gammaproteobacteria>Gammaproteobacteria_incertae_sedis>Candidatus_Competibacteraceae>Candidatus_Competibacter>unclassified_Candidatus_Competibacter | 111 | JNJ76_05455 | 2809..3144 | 2800415 | DUF5615-family-PIN-like-protein-[Candidatus-Competibacter-sp.] |
| MBZ4194012.1 | PIN-EndoRNase | GCA_020035205.1 | Candidatus_Contendobacter_sp. | Bacteria>Pseudomonadati>Pseudomonadota>Gammaproteobacteria>Gammaproteobacteria_incertae_sedis>Candidatus_Competibacteraceae>Candidatus_Contendobacter>unclassified_Candidatus_Contendobacter | 169 | LAE24_06865 | 3441..3950 | 2651161 | PIN-domain-containing-protein-[Candidatus-Contendobacter-sp.] |
| TVR61883.1 | PIN-EndoRNase | GCA_007695245.1 | Candidatus_Competibacteraceae_bacterium | Bacteria>Pseudomonadati>Pseudomonadota>Gammaproteobacteria>Gammaproteobacteria_incertae_sedis>Candidatus_Competibacteraceae>unclassified_Candidatus_Competibacteraceae | 111 | EA420_10935 | 3389..3724 | 2053538 | hypothetical-protein-EA420_10935-[Candidatus-Competibacteraceae-bacterium] |
| RUQ39487.1 | PIN-EndoRNase | GCA_003989075.1 | Candidatus_Competibacteraceae_bacterium | Bacteria>Pseudomonadati>Pseudomonadota>Gammaproteobacteria>Gammaproteobacteria_incertae_sedis>Candidatus_Competibacteraceae>unclassified_Candidatus_Competibacteraceae | 126 | EKK69_09100 | 17805..18185 | 2053538 | type-II-toxin-antitoxin-system-VapC-family-toxin-[Candidatus-Competibacteraceae-bacterium] |
| MCB1769901.1 | PIN-EndoRNase | GCA_020445005.1 | Candidatus_Competibacteraceae_bacterium | Bacteria>Pseudomonadati>Pseudomonadota>Gammaproteobacteria>Gammaproteobacteria_incertae_sedis>Candidatus_Competibacteraceae>unclassified_Candidatus_Competibacteraceae | 136 | KDJ31_09440 | 9080..9490 | 2053538 | type-II-toxin-antitoxin-system-VapC-family-toxin-[Candidatus-Competibacteraceae-bacterium] |
| MBK8754772.1 | PIN-EndoRNase | GCA_016710535.1 | Candidatus_Competibacteraceae_bacterium | Bacteria>Pseudomonadati>Pseudomonadota>Gammaproteobacteria>Gammaproteobacteria_incertae_sedis>Candidatus_Competibacteraceae>unclassified_Candidatus_Competibacteraceae | 140 | IPL99_25420 | 33222..33644 | 2053538 | type-II-toxin-antitoxin-system-VapC-family-toxin-[Candidatus-Competibacteraceae-bacterium] |
| MCB1769896.1 | PIN-EndoRNase | GCA_020445005.1 | Candidatus_Competibacteraceae_bacterium | Bacteria>Pseudomonadati>Pseudomonadota>Gammaproteobacteria>Gammaproteobacteria_incertae_sedis>Candidatus_Competibacteraceae>unclassified_Candidatus_Competibacteraceae | 163 | KDJ31_09415 | 6644..7135 | 2053538 | DUF3368-domain-containing-protein-[Candidatus-Competibacteraceae-bacterium] |
| MBK8752265.1 | PIN-EndoRNase | GCA_016710535.1 | Candidatus_Competibacteraceae_bacterium | Bacteria>Pseudomonadati>Pseudomonadota>Gammaproteobacteria>Gammaproteobacteria_incertae_sedis>Candidatus_Competibacteraceae>unclassified_Candidatus_Competibacteraceae | 169 | IPL99_11770 | 13783..14292 | 2053538 | DUF3368-domain-containing-protein-[Candidatus-Competibacteraceae-bacterium] |
| MCY4450796.1 | PIN-EndoRNase | GCA_026712715.1 | Immundisolibacterales_bacterium | Bacteria>Pseudomonadati>Pseudomonadota>Gammaproteobacteria>Immundisolibacterales>unclassified_Immundisolibacterales | 152 | OXC01_02490 | 122516..122974 | 2764111 | MAG:-hypothetical-protein-OXC01_02490-[Immundisolibacterales-bacterium] |
| WP_062607680.1 | PIN-EndoRNase | GCF_033100275.1 | Stenotrophomonas | Bacteria>Pseudomonadati>Pseudomonadota>Gammaproteobacteria>Lysobacterales>Lysobacteraceae | 161 | G5A32_RS22625 | 3826..4311 | 40323 | MULTISPECIES:-hypothetical-protein-[Stenotrophomonas] |
| KAA2283470.1 | PIN-EndoRNase | GCA_008386465.1 | Arenimonas_fontis | Bacteria>Pseudomonadati>Pseudomonadota>Gammaproteobacteria>Lysobacterales>Lysobacteraceae>Arenimonas | 205 | F0415_12210 | 8515..9132 | 2608255 | NYN-domain-containing-protein-[Arenimonas-fontis] |
| WP_084590777.1 | PIN-EndoRNase | GCF_000747155.1 | Arenimonas_metalli | Bacteria>Pseudomonadati>Pseudomonadota>Gammaproteobacteria>Lysobacterales>Lysobacteraceae>Arenimonas | 205 | N787_RS14790 | 55533..56150 | 948077 | NYN-domain-containing-protein-[Arenimonas-metalli] |
| WP_106991541.1 | PIN-EndoRNase | GCF_003024235.1 | Arenimonas_caeni | Bacteria>Pseudomonadati>Pseudomonadota>Gammaproteobacteria>Lysobacterales>Lysobacteraceae>Arenimonas | 205 | C6N40_RS13435 | 8210..8827 | 2058085 | NYN-domain-containing-protein-[Arenimonas-caeni] |
| WP_149861503.1 | PIN-EndoRNase | GCF_008386465.1 | Arenimonas_fontis | Bacteria>Pseudomonadati>Pseudomonadota>Gammaproteobacteria>Lysobacterales>Lysobacteraceae>Arenimonas | 205 | F0415_RS12215 | 8515..9132 | 2608255 | NYN-domain-containing-protein-[Arenimonas-fontis] |
| PZO64467.1 | PIN-EndoRNase | GCA_003240695.1 | Pseudoxanthomonas_suwonensis | Bacteria>Pseudomonadati>Pseudomonadota>Gammaproteobacteria>Lysobacterales>Lysobacteraceae>Pseudoxanthomonas | 206 | DI635_05315 | 8280..8900 | 314722 | NYN-domain-containing-protein-[Pseudoxanthomonas-suwonensis] |
| MBN5083088.1 | PIN-EndoRNase | GCA_017150495.1 | Stenotrophomonas_maltophilia | Bacteria>Pseudomonadati>Pseudomonadota>Gammaproteobacteria>Lysobacterales>Lysobacteraceae>Stenotrophomonas>Stenotrophomonas_maltophilia_group | 161 | JY467_05240 | 59869..60354 | 40324 | hypothetical-protein-[Stenotrophomonas-maltophilia] |
| WP_055771179.1 | PIN-EndoRNase | GCF_001422025.1 | Stenotrophomonas_sp._Leaf70 | Bacteria>Pseudomonadati>Pseudomonadota>Gammaproteobacteria>Lysobacterales>Lysobacteraceae>Stenotrophomonas>unclassified_Stenotrophomonas | 205 | ASF01_RS13980 | 63836..64453 | 1736233 | NYN-domain-containing-protein-[Stenotrophomonas-sp.-Leaf70] |
| MBU6247861.1 | PIN-EndoRNase | GCA_018970895.1 | Lysobacteraceae_bacterium | Bacteria>Pseudomonadati>Pseudomonadota>Gammaproteobacteria>Lysobacterales>Lysobacteraceae>unclassified_Lysobacteraceae | 205 | KGN77_08895 | 17784..18401 | 1926873 | NYN-domain-containing-protein-[Xanthomonadaceae-bacterium] |
| WP_243041015.1 | PIN-EndoRNase | GCF_022760995.1 | Dyella_sedimenti | Bacteria>Pseudomonadati>Pseudomonadota>Gammaproteobacteria>Lysobacterales>Rhodanobacteraceae>Dyella | 205 | MJ605_RS08545 | 203121..203738 | 2919947 | NYN-domain-containing-protein-[Dyella-sp.-NS14] |
| WP_008437338.1 | PIN-EndoRNase | GCF_000264375.1 | Rhodanobacter_thiooxydans | Bacteria>Pseudomonadati>Pseudomonadota>Gammaproteobacteria>Lysobacterales>Rhodanobacteraceae>Rhodanobacter | 205 | RHOFW104T7_RS11980 | 31169..31786 | 416169 | NYN-domain-containing-protein-[Rhodanobacter-thiooxydans] |
| GIX32369.1 | PIN-EndoRNase | GCA_026005055.1 | Lysobacterales_bacterium | Bacteria>Pseudomonadati>Pseudomonadota>Gammaproteobacteria>Lysobacterales>unclassified_Lysobacterales | 137 | KatS3mg125_0325 | 361510..361923 | 2006849 | MAG:-hypothetical-protein-KatS3mg125_0325-[Xanthomonadales-bacterium] |
| MCK9487883.1 | PIN-EndoRNase | GCA_023228845.1 | Lysobacterales_bacterium | Bacteria>Pseudomonadati>Pseudomonadota>Gammaproteobacteria>Lysobacterales>unclassified_Lysobacterales | 206 | MOQ42_00495 | 113935..114555 | 2006849 | NYN-domain-containing-protein-[Xanthomonadales-bacterium] |
| WP_198322147.1 | PIN-EndoRNase | GCF_036864975.1 | Methylococcus | Bacteria>Pseudomonadati>Pseudomonadota>Gammaproteobacteria>Methylococcales>Methylococcaceae | 140 | HUK90_RS04055 | 110236..110658 | 413 | MULTISPECIES:-type-II-toxin-antitoxin-system-VapC-family-toxin-[Methylococcus] |
| WP_159654894.1 | PIN-EndoRNase | GCF_009828925.2 | Methylicorpusculum_oleiharenae | Bacteria>Pseudomonadati>Pseudomonadota>Gammaproteobacteria>Methylococcales>Methylococcaceae>Methylicorpusculum | 164 | GO003_RS12070 | 2676357..2676851 | 1338687 | DUF3368-domain-containing-protein-[Methylicorpusculum-oleiharenae] |
| WP_218800669.1 | PIN-EndoRNase | GCF_019285535.1 | Methylococcus_capsulatus | Bacteria>Pseudomonadati>Pseudomonadota>Gammaproteobacteria>Methylococcales>Methylococcaceae>Methylococcus | 205 | M3M30_RS10990 | 2387459..2388076 | 414 | NYN-domain-containing-protein-[Methylococcus-capsulatus] |
| WP_218808312.1 | PIN-EndoRNase | GCF_019285515.1 | Methylococcus_sp._Mc7 | Bacteria>Pseudomonadati>Pseudomonadota>Gammaproteobacteria>Methylococcales>Methylococcaceae>Methylococcus>unclassified_Methylococcus | 141 | KW115_RS06330 | 1278627..1279052 | 2860258 | type-II-toxin-antitoxin-system-VapC-family-toxin-[Methylococcus-sp.-Mc7] |
| WP_029646270.1 | PIN-EndoRNase | GCF_000527095.1 | Methylomicrobium_lacus | Bacteria>Pseudomonadati>Pseudomonadota>Gammaproteobacteria>Methylococcales>Methylococcaceae>Methylomicrobium | 142 | METLA_RS0100470 | 99916..100344 | 136992 | type-II-toxin-antitoxin-system-VapC-family-toxin-[Methylomicrobium-lacus] |
| MBS3952429.1 | PIN-EndoRNase | GCA_018333785.1 | Methylomicrobium_sp. | Bacteria>Pseudomonadati>Pseudomonadota>Gammaproteobacteria>Methylococcales>Methylococcaceae>Methylomicrobium>unclassified_Methylomicrobium | 128 | KGZ88_05720 | 11121..11507 | 2303549 | type-II-toxin-antitoxin-system-VapC-family-toxin-[Methylomicrobium-sp.] |
| MBS3952491.1 | PIN-EndoRNase | GCA_018333785.1 | Methylomicrobium_sp. | Bacteria>Pseudomonadati>Pseudomonadota>Gammaproteobacteria>Methylococcales>Methylococcaceae>Methylomicrobium>unclassified_Methylomicrobium | 155 | KGZ88_06045 | 9274..9741 | 2303549 | DUF3368-domain-containing-protein-[Methylomicrobium-sp.] |
| WP_255188306.1 | PIN-EndoRNase | GCF_024360925.2 | Methylomonas_rapida | Bacteria>Pseudomonadati>Pseudomonadota>Gammaproteobacteria>Methylococcales>Methylococcaceae>Methylomonas | 103 | NM686_RS13190 | 2813366..2813677 | 2963939 | hypothetical-protein-[Methylomonas-rapida] |
| WP_255188307.1 | PIN-EndoRNase | GCF_024360925.2 | Methylomonas_rapida | Bacteria>Pseudomonadati>Pseudomonadota>Gammaproteobacteria>Methylococcales>Methylococcaceae>Methylomonas | 132 | NM686_RS13195 | 2813957..2814355 | 2963939 | PIN-domain-containing-protein-[Methylomonas-rapida] |
| WP_192375579.1 | PIN-EndoRNase | GCF_903064715.1 | Methylomonas_albis | Bacteria>Pseudomonadati>Pseudomonadota>Gammaproteobacteria>Methylococcales>Methylococcaceae>Methylomonas | 140 | EBA_RS15665 | 3419608..3420030 | 1854563 | type-II-toxin-antitoxin-system-VapC-family-toxin-[Methylomonas-albis] |
| WP_255188313.1 | PIN-EndoRNase | GCF_024360925.2 | Methylomonas_rapida | Bacteria>Pseudomonadati>Pseudomonadota>Gammaproteobacteria>Methylococcales>Methylococcaceae>Methylomonas | 140 | NM686_RS13225 | 2824553..2824975 | 2963939 | type-II-toxin-antitoxin-system-VapC-family-toxin-[Methylomonas-rapida] |
| WP_192375576.1 | PIN-EndoRNase | GCF_903064715.1 | Methylomonas_albis | Bacteria>Pseudomonadati>Pseudomonadota>Gammaproteobacteria>Methylococcales>Methylococcaceae>Methylomonas | 327 | EBA_RS15645 | 3410321..3411304 | 1854563 | PIN-domain-containing-protein-[Methylomonas-albis] |
| WP_198942643.1 | PIN-EndoRNase | GCF_002072955.1 | Methyloprofundus_sedimenti | Bacteria>Pseudomonadati>Pseudomonadota>Gammaproteobacteria>Methylococcales>Methylococcaceae>Methyloprofundus | 96 | AU255_RS14255 | 287504..287794 | 1420851 | hypothetical-protein-[Methyloprofundus-sedimenti] |
| WP_080523590.1 | PIN-EndoRNase | GCF_002072955.1 | Methyloprofundus_sedimenti | Bacteria>Pseudomonadati>Pseudomonadota>Gammaproteobacteria>Methylococcales>Methylococcaceae>Methyloprofundus | 149 | AU255_RS14190 | 276023..276472 | 1420851 | PIN-domain-containing-protein-[Methyloprofundus-sedimenti] |
| WP_020565136.1 | PIN-EndoRNase | GCF_000372865.1 | Methylosarcina_fibrata | Bacteria>Pseudomonadati>Pseudomonadota>Gammaproteobacteria>Methylococcales>Methylococcaceae>Methylosarcina | 131 | A3OW_RS0119475 | 4141377..4141772 | 105972 | PIN-domain-nuclease-[Methylosarcina-fibrata] |
| WP_020565138.1 | PIN-EndoRNase | GCF_000372865.1 | Methylosarcina_fibrata | Bacteria>Pseudomonadati>Pseudomonadota>Gammaproteobacteria>Methylococcales>Methylococcaceae>Methylosarcina | 143 | A3OW_RS0119485 | 4142061..4142492 | 105972 | type-II-toxin-antitoxin-system-VapC-family-toxin-[Methylosarcina-fibrata] |
| WP_017842530.1 | PIN-EndoRNase | GCF_005931095.1 | Methylotuvimicrobium_buryatense | Bacteria>Pseudomonadati>Pseudomonadota>Gammaproteobacteria>Methylococcales>Methylococcaceae>Methylotuvimicrobium | 138 | EQU24_RS16470 | 3696857..3697273 | 95641 | type-II-toxin-antitoxin-system-VapC-family-toxin-[Methylotuvimicrobium-buryatense] |
| WP_017842528.1 | PIN-EndoRNase | GCF_005931095.1 | Methylotuvimicrobium_buryatense | Bacteria>Pseudomonadati>Pseudomonadota>Gammaproteobacteria>Methylococcales>Methylococcaceae>Methylotuvimicrobium | 165 | EQU24_RS16480 | 3697628..3698125 | 95641 | DUF3368-domain-containing-protein-[Methylotuvimicrobium-buryatense] |
| WP_088619245.1 | PIN-EndoRNase | GCF_002209385.1 | Methylovulum_psychrotolerans | Bacteria>Pseudomonadati>Pseudomonadota>Gammaproteobacteria>Methylococcales>Methylococcaceae>Methylovulum | 140 | KFZ76_RS21370 | 108925..109347 | 1704499 | type-II-toxin-antitoxin-system-VapC-family-toxin-[Methylovulum-psychrotolerans] |
| MCF8007299.1 | PIN-EndoRNase | GCA_021736885.1 | Methylovulum_sp. | Bacteria>Pseudomonadati>Pseudomonadota>Gammaproteobacteria>Methylococcales>Methylococcaceae>Methylovulum>unclassified_Methylovulum | 124 | K9K84_07805 | 16191..16565 | 1916980 | type-II-toxin-antitoxin-system-VapC-family-toxin-[Methylovulum-sp.] |
| TSA38616.1 | PIN-EndoRNase | GCA_007280895.1 | Methylococcaceae_bacterium | Bacteria>Pseudomonadati>Pseudomonadota>Gammaproteobacteria>Methylococcales>Methylococcaceae>unclassified_Methylococcaceae | 130 | D4R63_10695 | 6596..6988 | 1933926 | type-II-toxin-antitoxin-system-VapC-family-toxin-[Methylococcaceae-bacterium] |
| TSA38620.1 | PIN-EndoRNase | GCA_007280895.1 | Methylococcaceae_bacterium | Bacteria>Pseudomonadati>Pseudomonadota>Gammaproteobacteria>Methylococcales>Methylococcaceae>unclassified_Methylococcaceae | 143 | D4R63_10720 | 8405..8836 | 1933926 | PIN-domain-containing-protein-[Methylococcaceae-bacterium] |
| PZN71300.1 | PIN-EndoRNase | GCA_003242955.1 | Candidatus_Methylumidiphilus_alinenensis | Bacteria>Pseudomonadati>Pseudomonadota>Gammaproteobacteria>Methylococcales>Methylococcales_incertae_sedis>Candidatus_Methylumidiphilus | 139 | DM484_26725 | 16240..16659 | 2202197 | nucleotide-binding-protein-[Candidatus-Methylumidiphilus-alinenensis] |
| MSP27873.1 | PIN-EndoRNase | GCA_009692395.1 | Methylococcales_bacterium | Bacteria>Pseudomonadati>Pseudomonadota>Gammaproteobacteria>Methylococcales>unclassified_Methylococcales | 135 | EXR80_05415 | 5588..5995 | 2304002 | VapC-toxin-family-PIN-domain-ribonuclease-[Methylococcales-bacterium] |
| MDD5268637.1 | PIN-EndoRNase | GCA_028712795.1 | Methylococcales_bacterium | Bacteria>Pseudomonadati>Pseudomonadota>Gammaproteobacteria>Methylococcales>unclassified_Methylococcales | 136 | PHO08_16165 | 11476..11886 | 2304002 | MAG:-PIN-domain-containing-protein-[Methylococcales-bacterium] |
| MDD5321590.1 | PIN-EndoRNase | GCA_028714175.1 | Methylococcales_bacterium | Bacteria>Pseudomonadati>Pseudomonadota>Gammaproteobacteria>Methylococcales>unclassified_Methylococcales | 160 | PHD43_13440 | 3603..4085 | 2304002 | MAG:-DUF3368-domain-containing-protein-[Methylococcales-bacterium] |
| MDD5320446.1 | PIN-EndoRNase | GCA_028714175.1 | Methylococcales_bacterium | Bacteria>Pseudomonadati>Pseudomonadota>Gammaproteobacteria>Methylococcales>unclassified_Methylococcales | 164 | PHD43_07505 | 22797..23291 | 2304002 | MAG:-DUF3368-domain-containing-protein-[Methylococcales-bacterium] |
| MCK0538076.1 | PIN-EndoRNase | GCA_023051175.1 | Alcanivorax_quisquiliarum | Bacteria>Pseudomonadati>Pseudomonadota>Gammaproteobacteria>Oceanospirillales>Alcanivoracaceae>Alcanivorax | 206 | MU846_10180 | 130995..131615 | 2933565 | NYN-domain-containing-protein-[Alcanivorax-sp.-CY1518] |
| WP_257264232.1 | PIN-EndoRNase | GCF_024606365.1 | Endozoicomonas_sp._ONNA2 | Bacteria>Pseudomonadati>Pseudomonadota>Gammaproteobacteria>Oceanospirillales>Endozoicomonadaceae>Endozoicomonas>unclassified_Endozoicomonas | 247 | KDK14_RS06595 | 6092..6835 | 2828741 | NYN-domain-containing-protein-[Endozoicomonas-sp.-ONNA2] |
| WP_205596294.1 | PIN-EndoRNase | GCF_020147965.1 | Modicisalibacter | Bacteria>Pseudomonadati>Pseudomonadota>Gammaproteobacteria>Oceanospirillales>Halomonadaceae | 132 | KGQ90_RS13850 | 2975630..2976028 | 574347 | MULTISPECIES:-hypothetical-protein-[Modicisalibacter] |
| WP_040239275.1 | PIN-EndoRNase | GCF_000821045.2 | Chromohalobacter_japonicus | Bacteria>Pseudomonadati>Pseudomonadota>Gammaproteobacteria>Oceanospirillales>Halomonadaceae>Chromohalobacter | 152 | BN995_RS00425 | 42890..43348 | 223900 | hypothetical-protein-[Chromohalobacter-japonicus] |
| WP_163558797.1 | PIN-EndoRNase | GCF_010994375.1 | Halomonas_sp._NO4 | Bacteria>Pseudomonadati>Pseudomonadota>Gammaproteobacteria>Oceanospirillales>Halomonadaceae>Halomonas>unclassified_Halomonas | 179 | FEF15_RS08825 | 8516..9055 | 2484813 | hypothetical-protein-[Halomonas-sp.-NO4] |
| WP_223136855.1 | PIN-EndoRNase | GCF_019915905.1 | Halomonas_sp._ANAO-440 | Bacteria>Pseudomonadati>Pseudomonadota>Gammaproteobacteria>Oceanospirillales>Halomonadaceae>Halomonas>unclassified_Halomonas | 179 | KZO25_RS10135 | 177356..177895 | 2861360 | hypothetical-protein-[Halomonas-sp.-ANAO-440] |
| WP_110069961.1 | PIN-EndoRNase | GCF_003182195.1 | Halomonas_sp._A11-A | Bacteria>Pseudomonadati>Pseudomonadota>Gammaproteobacteria>Oceanospirillales>Halomonadaceae>Halomonas>unclassified_Halomonas | 180 | DER72_RS10580 | 22917..23459 | 2183985 | hypothetical-protein-[Halomonas-sp.-A11-A] |
| WP_234259722.1 | PIN-EndoRNase | GCF_021404355.1 | Halomonas_sp._MCCC_1A11062 | Bacteria>Pseudomonadati>Pseudomonadota>Gammaproteobacteria>Oceanospirillales>Halomonadaceae>Halomonas>unclassified_Halomonas | 180 | HOP58_RS04925 | 170269..170811 | 2733485 | hypothetical-protein-[Halomonas-sp.-MCCC-1A11062] |
| WP_253448040.1 | PIN-EndoRNase | GCF_024137965.1 | Halomonas_sp._Y3 | Bacteria>Pseudomonadati>Pseudomonadota>Gammaproteobacteria>Oceanospirillales>Halomonadaceae>Halomonas>unclassified_Halomonas | 180 | NHL60_RS01515 | 302877..303419 | 2956797 | hypothetical-protein-[Halomonas-sp.-Y3] |
| WP_189514716.1 | PIN-EndoRNase | GCF_014652795.1 | Kushneria_pakistanensis | Bacteria>Pseudomonadati>Pseudomonadota>Gammaproteobacteria>Oceanospirillales>Halomonadaceae>Kushneria | 153 | IE123_RS02390 | 536458..536919 | 1508770 | hypothetical-protein-[Kushneria-pakistanensis] |
| WP_086622425.1 | PIN-EndoRNase | GCF_002155145.1 | Kushneria_konosiri | Bacteria>Pseudomonadati>Pseudomonadota>Gammaproteobacteria>Oceanospirillales>Halomonadaceae>Kushneria | 180 | B9G99_RS12350 | 2664950..2665492 | 698828 | hypothetical-protein-[Kushneria-konosiri] |
| WP_147183727.1 | PIN-EndoRNase | GCF_007954585.1 | Pistricoccus_aurantiacus | Bacteria>Pseudomonadati>Pseudomonadota>Gammaproteobacteria>Oceanospirillales>Halomonadaceae>Pistricoccus | 139 | FGL86_RS05975 | 1227401..1227820 | 1883414 | hypothetical-protein-[Pistricoccus-aurantiacus] |
| WP_147183734.1 | PIN-EndoRNase | GCF_007954585.1 | Pistricoccus_aurantiacus | Bacteria>Pseudomonadati>Pseudomonadota>Gammaproteobacteria>Oceanospirillales>Halomonadaceae>Pistricoccus | 180 | FGL86_RS06010 | 1239506..1240048 | 1883414 | hypothetical-protein-[Pistricoccus-aurantiacus] |
| WP_196781581.1 | PIN-EndoRNase | GCF_012427705.1 | Vreelandella_piezotolerans | Bacteria>Pseudomonadati>Pseudomonadota>Gammaproteobacteria>Oceanospirillales>Halomonadaceae>Vreelandella | 175 | GYM47_RS05620 | 1212858..1213385 | 2609667 | hypothetical-protein-[Halomonas-piezotolerans] |
| KAE8440083.1 | PIN-EndoRNase | GCA_009660035.1 | Vreelandella_piezotolerans | Bacteria>Pseudomonadati>Pseudomonadota>Gammaproteobacteria>Oceanospirillales>Halomonadaceae>Vreelandella | 179 | F1978_02240 | 474590..475129 | 2609667 | hypothetical-protein-F1978_02240-[Halomonas-piezotolerans] |
| WP_051514666.1 | PIN-EndoRNase | GCF_000585235.1 | Nitrincola_nitratireducens | Bacteria>Pseudomonadati>Pseudomonadota>Gammaproteobacteria>Oceanospirillales>Oceanospirillaceae>Nitrincola | 157 | D791_RS17645 | 10532..11005 | 1229521 | hypothetical-protein-[Nitrincola-nitratireducens] |
| WP_200371324.1 | PIN-EndoRNase | GCF_016597815.1 | Marinobacter_sp._DY40_1A1 | Bacteria>Pseudomonadati>Pseudomonadota>Gammaproteobacteria>Pseudomonadales>Marinobacteraceae>Marinobacter>unclassified_Marinobacter | 177 | FFI61_RS14530 | 356038..356571 | 2583229 | hypothetical-protein-[Marinobacter-sp.-DY40_1A1] |
| WP_252576708.1 | PIN-EndoRNase | GCF_021495345.1 | Ectopseudomonas_hydrolytica | Bacteria>Pseudomonadati>Pseudomonadota>Gammaproteobacteria>Pseudomonadales>Pseudomonadaceae>Ectopseudomonas | 102 | L1F06_RS00030 | 9967..10275 | 2493633 | hypothetical-protein-[Pseudomonas-hydrolytica] |
| HIQ43650.1 | PIN-EndoRNase | GCA_015661735.1 | Ectopseudomonas_oleovorans | Bacteria>Pseudomonadati>Pseudomonadota>Gammaproteobacteria>Pseudomonadales>Pseudomonadaceae>Ectopseudomonas | 206 | EYH47_12975 | 39466..40086 | 301 | NYN-domain-containing-protein-[Pseudomonas-oleovorans] |
| WP_143508090.1 | PIN-EndoRNase | - | Ectopseudomonas_mendocina | Bacteria>Pseudomonadati>Pseudomonadota>Gammaproteobacteria>Pseudomonadales>Pseudomonadaceae>Ectopseudomonas | 206 | EQ828_RS12675 | 215319..215939 | 300 | NYN-domain-containing-protein-[Pseudomonas-mendocina] |
| HBO4843316.1 | PIN-EndoRNase | GCA_021660595.1 | Pseudomonas_aeruginosa | Bacteria>Pseudomonadati>Pseudomonadota>Gammaproteobacteria>Pseudomonadales>Pseudomonadaceae>Pseudomonas>Pseudomonas_aeruginosa_group | 161 | L4V37_002407 | 66463..66948 | 287 | hypothetical-protein-[Pseudomonas-aeruginosa] |
| HCF2077874.1 | PIN-EndoRNase | GCA_023942995.1 | Pseudomonas_aeruginosa | Bacteria>Pseudomonadati>Pseudomonadota>Gammaproteobacteria>Pseudomonadales>Pseudomonadaceae>Pseudomonas>Pseudomonas_aeruginosa_group | 175 | NH997_004589 | 25725..26252 | 287 | TPA:-hypothetical-protein-NH997_004589-[Pseudomonas-aeruginosa] |
| WP_071543152.1 | PIN-EndoRNase | GCF_900707525.1 | Pseudomonas_aeruginosa | Bacteria>Pseudomonadati>Pseudomonadota>Gammaproteobacteria>Pseudomonadales>Pseudomonadaceae>Pseudomonas>Pseudomonas_aeruginosa_group | 175 | BAW40_RS31235 | 48559..49086 | 287 | hypothetical-protein-[Pseudomonas-aeruginosa] |
| MBG3917695.1 | PIN-EndoRNase | GCA_015732935.1 | Pseudomonas_aeruginosa | Bacteria>Pseudomonadati>Pseudomonadota>Gammaproteobacteria>Pseudomonadales>Pseudomonadaceae>Pseudomonas>Pseudomonas_aeruginosa_group | 206 | I4931_21695 | 87108..87728 | 287 | NYN-domain-containing-protein-[Pseudomonas-aeruginosa] |
| WP_258379903.1 | PIN-EndoRNase | - | Pseudomonas_koreensis | Bacteria>Pseudomonadati>Pseudomonadota>Gammaproteobacteria>Pseudomonadales>Pseudomonadaceae>Pseudomonas>Pseudomonas_fluorescens_group | 191 | DMX04_RS04435 | 130817..131392 | 198620 | hypothetical-protein-[Pseudomonas-koreensis] |
| WP_110599717.1 | PIN-EndoRNase | - | Pseudomonas_koreensis | Bacteria>Pseudomonadati>Pseudomonadota>Gammaproteobacteria>Pseudomonadales>Pseudomonadaceae>Pseudomonas>Pseudomonas_fluorescens_group | 206 | DMX04_RS04475 | 142146..142766 | 198620 | NYN-domain-containing-protein-[Pseudomonas-koreensis] |
| WP_005744958.1 | PIN-EndoRNase | GCF_002068135.1 | Pseudomonas_syringae_group_genomosp._2 | Bacteria>Pseudomonadati>Pseudomonadota>Gammaproteobacteria>Pseudomonadales>Pseudomonadaceae>Pseudomonas>Pseudomonas_syringae_group | 206 | QO021_RS27745 | 6023779..6024399 | 251698 | NYN-domain-containing-protein-[Pseudomonas-amygdali] |
| WP_226045698.1 | PIN-EndoRNase | GCF_020401845.1 | Pseudomonas_sp._HS-18 | Bacteria>Pseudomonadati>Pseudomonadota>Gammaproteobacteria>Pseudomonadales>Pseudomonadaceae>Pseudomonas>unclassified_Pseudomonas | 102 | LDJ84_RS00030 | 10011..10319 | 2879114 | hypothetical-protein-[Pseudomonas-sp.-HS-18] |
| WP_003246710.1 | PIN-EndoRNase | GCF_025837155.1 | Pseudomonas_sp._Z8(2022) | Bacteria>Pseudomonadati>Pseudomonadota>Gammaproteobacteria>Pseudomonadales>Pseudomonadaceae>Pseudomonas>unclassified_Pseudomonas | 206 | A471_RS19800 | 14056..14676 | 2962597 | MULTISPECIES:-NYN-domain-containing-protein-[Pseudomonas] |
| WP_237730183.1 | PIN-EndoRNase | GCF_029839225.1 | Stutzerimonas_stutzeri | Bacteria>Pseudomonadati>Pseudomonadota>Gammaproteobacteria>Pseudomonadales>Pseudomonadaceae>Stutzerimonas>Stutzerimonas_stutzeri_group>Stutzerimonas_stutzeri_subgroup | 102 | N5C32_RS13910 | 179236..179544 | 316 | hypothetical-protein-[Stutzerimonas-stutzeri] |
| WP_260469074.1 | PIN-EndoRNase | - | Stutzerimonas_stutzeri | Bacteria>Pseudomonadati>Pseudomonadota>Gammaproteobacteria>Pseudomonadales>Pseudomonadaceae>Stutzerimonas>Stutzerimonas_stutzeri_group>Stutzerimonas_stutzeri_subgroup | 102 | EGI99_RS04485 | 989452..989760 | 316 | hypothetical-protein-[Stutzerimonas-stutzeri] |
| MCF6781037.1 | PIN-EndoRNase | GCA_021713665.1 | Stutzerimonas_stutzeri | Bacteria>Pseudomonadati>Pseudomonadota>Gammaproteobacteria>Pseudomonadales>Pseudomonadaceae>Stutzerimonas>Stutzerimonas_stutzeri_group>Stutzerimonas_stutzeri_subgroup | 206 | L3X17_04525 | 25607..26227 | 316 | NYN-domain-containing-protein-[Stutzerimonas-stutzeri] |
| MCF6804933.1 | PIN-EndoRNase | GCA_021713635.1 | Stutzerimonas_stutzeri | Bacteria>Pseudomonadati>Pseudomonadota>Gammaproteobacteria>Pseudomonadales>Pseudomonadaceae>Stutzerimonas>Stutzerimonas_stutzeri_group>Stutzerimonas_stutzeri_subgroup | 206 | L3X13_09165 | 25611..26231 | 316 | NYN-domain-containing-protein-[Stutzerimonas-stutzeri] |
| MTI93111.1 | PIN-EndoRNase | GCA_009711815.1 | Stutzerimonas_stutzeri | Bacteria>Pseudomonadati>Pseudomonadota>Gammaproteobacteria>Pseudomonadales>Pseudomonadaceae>Stutzerimonas>Stutzerimonas_stutzeri_group>Stutzerimonas_stutzeri_subgroup | 206 | FH750_17260 | 21608..22228 | 316 | NYN-domain-containing-protein-[Stutzerimonas-stutzeri] |
| WP_125852811.1 | PIN-EndoRNase | GCF_003936145.1 | Stutzerimonas_stutzeri | Bacteria>Pseudomonadati>Pseudomonadota>Gammaproteobacteria>Pseudomonadales>Pseudomonadaceae>Stutzerimonas>Stutzerimonas_stutzeri_group>Stutzerimonas_stutzeri_subgroup | 206 | EGJ42_RS05705 | 47110..47730 | 316 | NYN-domain-containing-protein-[Stutzerimonas-stutzeri] |
| WP_213918364.1 | PIN-EndoRNase | GCF_018457005.1 | Stutzerimonas_stutzeri | Bacteria>Pseudomonadati>Pseudomonadota>Gammaproteobacteria>Pseudomonadales>Pseudomonadaceae>Stutzerimonas>Stutzerimonas_stutzeri_group>Stutzerimonas_stutzeri_subgroup | 206 | JR047_RS05400 | 11175..11795 | 316 | NYN-domain-containing-protein-[Stutzerimonas-stutzeri] |
| WP_283258802.1 | PIN-EndoRNase | GCF_030056015.1 | Stutzerimonas_stutzeri | Bacteria>Pseudomonadati>Pseudomonadota>Gammaproteobacteria>Pseudomonadales>Pseudomonadaceae>Stutzerimonas>Stutzerimonas_stutzeri_group>Stutzerimonas_stutzeri_subgroup | 206 | QM290_RS03930 | 11179..11799 | 316 | NYN-domain-containing-protein-[Stutzerimonas-stutzeri] |
| EPL60618.1 | PIN-EndoRNase | GCA_000416345.1 | Stutzerimonas_stutzeri_B1SMN1 | Bacteria>Pseudomonadati>Pseudomonadota>Gammaproteobacteria>Pseudomonadales>Pseudomonadaceae>Stutzerimonas>Stutzerimonas_stutzeri_group>Stutzerimonas_stutzeri_subgroup>Stutzerimonas_stutzeri | 206 | B382_19955 | 13122..13742 | 1212549 | hypothetical-protein-B382_19955-[Stutzerimonas-stutzeri-B1SMN1] |
| MAS10378.1 | PIN-EndoRNase | GCA_002702145.1 | Salinisphaera_sp. | Bacteria>Pseudomonadati>Pseudomonadota>Gammaproteobacteria>Salinisphaerales>Salinisphaeraceae>Salinisphaera>unclassified_Salinisphaera | 206 | CMN26_10005 | 12818..13438 | 1914330 | NYN-domain-containing-protein-[Salinisphaera-sp.] |
| MCY4200250.1 | PIN-EndoRNase | GCA_026709225.1 | Gammaproteobacteria_bacterium | Bacteria>Pseudomonadati>Pseudomonadota>Gammaproteobacteria>unclassified_Gammaproteobacteria | 112 | OXF31_10665 | 27026..27364 | 1913989 | MAG:-NYN-domain-containing-protein-[Gammaproteobacteria-bacterium] |
| MCY4199356.1 | PIN-EndoRNase | GCA_026709225.1 | Gammaproteobacteria_bacterium | Bacteria>Pseudomonadati>Pseudomonadota>Gammaproteobacteria>unclassified_Gammaproteobacteria | 122 | OXF31_05980 | 36131..36499 | 1913989 | MAG:-DUF5615-family-PIN-like-protein-[Gammaproteobacteria-bacterium] |
| MDE0038235.1 | PIN-EndoRNase | GCA_028818795.1 | Gammaproteobacteria_bacterium | Bacteria>Pseudomonadati>Pseudomonadota>Gammaproteobacteria>unclassified_Gammaproteobacteria | 124 | OXU77_11890 | 14534..14908 | 1913989 | MAG:-type-II-toxin-antitoxin-system-VapC-family-toxin-[Gammaproteobacteria-bacterium] |
| MDE0444967.1 | PIN-EndoRNase | GCA_028821355.1 | Gammaproteobacteria_bacterium | Bacteria>Pseudomonadati>Pseudomonadota>Gammaproteobacteria>unclassified_Gammaproteobacteria | 124 | OXL38_23030 | 20021..20395 | 1913989 | MAG:-type-II-toxin-antitoxin-system-VapC-family-toxin-[Gammaproteobacteria-bacterium] |

| Protein-ID | Domain | Assembly-ID | Strain | Taxonomy | Length | Locus-Tag | Genomic-coordinates | TaxID | Fasta-defline |
|---|---|---|---|---|---|---|---|---|---|
| MCY4157212.1 | PIN-EndoRNase | GCA_026708965.1 | Gammaproteobacteria_bacterium | Bacteria>Pseudomonadati>Pseudomonadota>Gammaproteobacteria>unclassified_Gammaproteobacteria | 124 | OXF66_07250 | 51001..51375 | 1913989 | MAG:-DUF5615-family-PIN-like-protein-[Gammaproteobacteria-bacterium] |
| MCY4164746.1 | PIN-EndoRNase | GCA_026709045.1 | Gammaproteobacteria_bacterium | Bacteria>Pseudomonadati>Pseudomonadota>Gammaproteobacteria>unclassified_Gammaproteobacteria | 124 | OXF03_01145 | 156290..156664 | 1913989 | MAG:-DUF5615-family-PIN-like-protein-[Gammaproteobacteria-bacterium] |
| MCY4255935.1 | PIN-EndoRNase | GCA_026709685.1 | Gammaproteobacteria_bacterium | Bacteria>Pseudomonadati>Pseudomonadota>Gammaproteobacteria>unclassified_Gammaproteobacteria | 124 | OXE51_07465 | 8459..8833 | 1913989 | MAG:-DUF5615-family-PIN-like-protein-[Gammaproteobacteria-bacterium] |
| MCY4340925.1 | PIN-EndoRNase | GCA_026710305.1 | Gammaproteobacteria_bacterium | Bacteria>Pseudomonadati>Pseudomonadota>Gammaproteobacteria>unclassified_Gammaproteobacteria | 124 | OXE48_06115 | 66620..66994 | 1913989 | MAG:-DUF5615-family-PIN-like-protein-[Gammaproteobacteria-bacterium] |
| MCY4342455.1 | PIN-EndoRNase | GCA_026710215.1 | Gammaproteobacteria_bacterium | Bacteria>Pseudomonadati>Pseudomonadota>Gammaproteobacteria>unclassified_Gammaproteobacteria | 124 | OXE83_02625 | 5368..5742 | 1913989 | MAG:-DUF5615-family-PIN-like-protein-[Gammaproteobacteria-bacterium] |
| MBT5223357.1 | PIN-EndoRNase | GCA_018661125.1 | Gammaproteobacteria_bacterium | Bacteria>Pseudomonadati>Pseudomonadota>Gammaproteobacteria>unclassified_Gammaproteobacteria | 128 | HOM15_11175 | 5381..5767 | 1913989 | type-II-toxin-antitoxin-system-VapC-family-toxin-[Gammaproteobacteria-bacterium] |
| MDE0478450.1 | PIN-EndoRNase | GCA_028824825.1 | Gammaproteobacteria_bacterium | Bacteria>Pseudomonadati>Pseudomonadota>Gammaproteobacteria>unclassified_Gammaproteobacteria | 142 | OXI13_02370 | 8090..8518 | 1913989 | MAG:-hypothetical-protein-OXI13_02370-[Gammaproteobacteria-bacterium] |
| MXX05924.1 | PIN-EndoRNase | GCA_009836015.1 | Gammaproteobacteria_bacterium | Bacteria>Pseudomonadati>Pseudomonadota>Gammaproteobacteria>unclassified_Gammaproteobacteria | 142 | F4Z71_04115 | 93091..93519 | 1913989 | hypothetical-protein-[Gammaproteobacteria-bacterium] |
| MCY4343960.1 | PIN-EndoRNase | GCA_026710215.1 | Gammaproteobacteria_bacterium | Bacteria>Pseudomonadati>Pseudomonadota>Gammaproteobacteria>unclassified_Gammaproteobacteria | 146 | OXE83_10360 | 10859..11299 | 1913989 | MAG:-PIN-domain-containing-protein-[Gammaproteobacteria-bacterium] |
| MCY4278445.1 | PIN-EndoRNase | GCA_026709965.1 | Gammaproteobacteria_bacterium | Bacteria>Pseudomonadati>Pseudomonadota>Gammaproteobacteria>unclassified_Gammaproteobacteria | 150 | OXE81_11545 | 1989..2441 | 1913989 | MAG:-PIN-domain-containing-protein-[Gammaproteobacteria-bacterium] |
| MDE0258536.1 | PIN-EndoRNase | GCA_028820325.1 | Gammaproteobacteria_bacterium | Bacteria>Pseudomonadati>Pseudomonadota>Gammaproteobacteria>unclassified_Gammaproteobacteria | 151 | OXR82_09160 | 5131..5586 | 1913989 | MAG:-hypothetical-protein-OXR82_09160-[Gammaproteobacteria-bacterium] |
| MDE0036815.1 | PIN-EndoRNase | GCA_028818795.1 | Gammaproteobacteria_bacterium | Bacteria>Pseudomonadati>Pseudomonadota>Gammaproteobacteria>unclassified_Gammaproteobacteria | 152 | OXU77_04525 | 5609..6067 | 1913989 | MAG:-hypothetical-protein-OXU77_04525-[Gammaproteobacteria-bacterium] |
| MDD9824621.1 | PIN-EndoRNase | GCA_028817255.1 | Gammaproteobacteria_bacterium | Bacteria>Pseudomonadati>Pseudomonadota>Gammaproteobacteria>unclassified_Gammaproteobacteria | 154 | OXU43_05570 | 2999..3463 | 1913989 | MAG:-PIN-domain-containing-protein-[Gammaproteobacteria-bacterium] |
| MCY4200253.1 | PIN-EndoRNase | GCA_026709225.1 | Gammaproteobacteria_bacterium | Bacteria>Pseudomonadati>Pseudomonadota>Gammaproteobacteria>unclassified_Gammaproteobacteria | 162 | OXF31_10680 | 32109..32597 | 1913989 | MAG:-PIN-domain-containing-protein-[Gammaproteobacteria-bacterium] |
| MDE0038226.1 | PIN-EndoRNase | GCA_028818795.1 | Gammaproteobacteria_bacterium | Bacteria>Pseudomonadati>Pseudomonadota>Gammaproteobacteria>unclassified_Gammaproteobacteria | 164 | OXU77_11845 | 2012..2506 | 1913989 | MAG:-DUF4411-family-protein-[Gammaproteobacteria-bacterium] |
| MDE0444958.1 | PIN-EndoRNase | GCA_028821355.1 | Gammaproteobacteria_bacterium | Bacteria>Pseudomonadati>Pseudomonadota>Gammaproteobacteria>unclassified_Gammaproteobacteria | 164 | OXL38_22985 | 7469..7963 | 1913989 | MAG:-DUF4411-family-protein-[Gammaproteobacteria-bacterium] |
| MBU1235380.1 | PIN-EndoRNase | GCA_018819515.1 | Gammaproteobacteria_bacterium | Bacteria>Pseudomonadati>Pseudomonadota>Gammaproteobacteria>unclassified_Gammaproteobacteria | 183 | KJ788_01185 | 185326..185877 | 1913989 | PIN-domain-containing-protein-[Gammaproteobacteria-bacterium] |
| MCY4533096.1 | PIN-EndoRNase | GCA_026713185.1 | Gammaproteobacteria_bacterium | Bacteria>Pseudomonadati>Pseudomonadota>Gammaproteobacteria>unclassified_Gammaproteobacteria | 205 | OXC84_12355 | 12456..13073 | 1913989 | MAG:-NYN-domain-containing-protein-[Gammaproteobacteria-bacterium] |
| MCY4278448.1 | PIN-EndoRNase | GCA_026709965.1 | Gammaproteobacteria_bacterium | Bacteria>Pseudomonadati>Pseudomonadota>Gammaproteobacteria>unclassified_Gammaproteobacteria | 206 | OXE81_11560 | 7222..7842 | 1913989 | MAG:-NYN-domain-containing-protein-[Gammaproteobacteria-bacterium] |
| HFD80331.1 | PIN-EndoRNase | GCA_011321775.1 | Gammaproteobacteria_bacterium | Bacteria>Pseudomonadati>Pseudomonadota>Gammaproteobacteria>unclassified_Gammaproteobacteria | 265 | ENK05_08085 | 11469..12266 | 1913989 | NYN-domain-containing-protein-[Gammaproteobacteria-bacterium] |
| WP_169116167.1 | PIN-EndoRNase | GCF_012911495.1 | Tepidiphilus | Bacteria>Pseudomonadati>Pseudomonadota>Hydrogenophilia>Hydrogenophilales>Hydrogenophilaceae | 137 | GV368_RS08155 | 16419..16832 | 203470 | MULTISPECIES:-PIN-domain-containing-protein-[Tepidiphilus] |
| WP_219972356.1 | PIN-EndoRNase | - | Hydrogenophilus_thermoluteolus | Bacteria>Pseudomonadati>Pseudomonadota>Hydrogenophilia>Hydrogenophilales>Hydrogenophilaceae>Hydrogenophilus | 104 | K1I42_RS04685 | 186551..186865 | 297 | hypothetical-protein-[Hydrogenophilus-thermoluteolus] |
| WP_169115536.1 | PIN-EndoRNase | GCF_012911495.1 | Tepidiphilus_baoligensis | Bacteria>Pseudomonadati>Pseudomonadota>Hydrogenophilia>Hydrogenophilales>Hydrogenophilaceae>Tepidiphilus | 133 | GV368_RS03835 | 178464..178865 | 2698687 | PIN-domain-containing-protein-[Tepidiphilus-baoligensis] |
| WP_037989142.1 | PIN-EndoRNase | GCF_000425565.1 | Tepidiphilus_margaritifer | Bacteria>Pseudomonadati>Pseudomonadota>Hydrogenophilia>Hydrogenophilales>Hydrogenophilaceae>Tepidiphilus | 135 | H607_RS0100580 | 9135..9542 | 203471 | type-II-toxin-antitoxin-system-VapC-family-toxin-[Tepidiphilus-margaritifer] |
| WP_169116184.1 | PIN-EndoRNase | GCF_012911495.1 | Tepidiphilus_baoligensis | Bacteria>Pseudomonadati>Pseudomonadota>Hydrogenophilia>Hydrogenophilales>Hydrogenophilaceae>Tepidiphilus | 205 | GV368_RS08275 | 47276..47893 | 2698687 | NYN-domain-containing-protein-[Tepidiphilus-baoligensis] |
| WP_028873818.1 | PIN-EndoRNase | GCF_000425565.1 | Tepidiphilus_margaritifer | Bacteria>Pseudomonadati>Pseudomonadota>Hydrogenophilia>Hydrogenophilales>Hydrogenophilaceae>Tepidiphilus | 206 | H607_RS0100550 | 392..1012 | 203471 | NYN-domain-containing-protein-[Tepidiphilus-margaritifer] |
| MBS0593920.1 | PIN-EndoRNase | GCA_018240325.1 | Pseudomonadota_bacterium | Bacteria>Pseudomonadati>Pseudomonadota>unclassified_Pseudomonadota | 144 | JSR84_10680 | 68154..68588 | 1977087 | putative-toxin-antitoxin-system-toxin-component,-PIN-family-[Pseudomonadota-bacterium] |
| MBQ9395960.1 | PIN-EndoRNase | GCA_017455165.1 | Pseudomonadota_bacterium | Bacteria>Pseudomonadati>Pseudomonadota>unclassified_Pseudomonadota | 151 | IJU23_10685 | 2095..2550 | 1977087 | putative-toxin-antitoxin-system-toxin-component,-PIN-family-[Pseudomonadota-bacterium] |
| MBE9569153.1 | PIN-EndoRNase | GCA_015222845.1 | Pseudomonadota_bacterium | Bacteria>Pseudomonadati>Pseudomonadota>unclassified_Pseudomonadota | 157 | IMF11_00870 | 1..476 | 1977087 | hypothetical-protein-[Pseudomonadota-bacterium] |
| MDE3269953.1 | PIN-EndoRNase | GCA_028877195.1 | Pseudomonadota_bacterium | Bacteria>Pseudomonadati>Pseudomonadota>unclassified_Pseudomonadota | 163 | OYH77_06685 | 8239..8732 | 1977087 | MAG:-DUF4411-family-protein,-partial-[Pseudomonadota-bacterium] |
| MBU1206363.1 | PIN-EndoRNase | GCA_018819135.1 | Pseudomonadota_bacterium | Bacteria>Pseudomonadati>Pseudomonadota>unclassified_Pseudomonadota | 168 | KKH04_05480 | 21236..21742 | 1977087 | type-II-toxin-antitoxin-system-VapC-family-toxin-[Pseudomonadota-bacterium] |
| MCL1861023.1 | PIN-EndoRNase | GCA_009778425.1 | Pseudomonadota_bacterium | Bacteria>Pseudomonadati>Pseudomonadota>unclassified_Pseudomonadota | 170 | FWG52_05745 | 101..613 | 1977087 | NYN-domain-containing-protein-[Pseudomonadota-bacterium] |
| MBS0225818.1 | PIN-EndoRNase | GCA_018242465.1 | Pseudomonadota_bacterium | Bacteria>Pseudomonadati>Pseudomonadota>unclassified_Pseudomonadota | 206 | JSS25_05705 | 44505..45125 | 1977087 | NYN-domain-containing-protein-[Pseudomonadota-bacterium] |
| MBS0576262.1 | PIN-EndoRNase | GCA_018240435.1 | Pseudomonadota_bacterium | Bacteria>Pseudomonadati>Pseudomonadota>unclassified_Pseudomonadota | 206 | JSS45_07550 | 44667..45287 | 1977087 | NYN-domain-containing-protein-[Pseudomonadota-bacterium] |
| MBU0673019.1 | PIN-EndoRNase | GCA_018814305.1 | Pseudomonadota_bacterium | Bacteria>Pseudomonadati>Pseudomonadota>unclassified_Pseudomonadota | 206 | KJ950_00065 | 13431..14051 | 1977087 | NYN-domain-containing-protein-[Pseudomonadota-bacterium] |
| NCC04602.1 | PIN-EndoRNase | GCA_009929575.1 | Pseudomonadota_bacterium | Bacteria>Pseudomonadati>Pseudomonadota>unclassified_Pseudomonadota | 213 | EOM37_11310 | 13737..14378 | 1977087 | NYN-domain-containing-protein-[Pseudomonadota-bacterium] |
| MBN2145090.1 | PIN-EndoRNase | GCA_016932965.1 | Candidatus_Auribacterota_bacterium | Bacteria>Pseudomonadati>PVC_group>Candidatus_Auribacterota>unclassified_Candidatus_Auribacterota | 136 | JW774_10740 | 7829..8239 | 2093365 | hypothetical-protein-[Candidatus-Auribacterota-bacterium] |
| RJP35068.1 | PIN-EndoRNase | GCA_003598175.1 | Candidatus_Omnitrophota_bacterium | Bacteria>Pseudomonadati>PVC_group>Candidatus_Omnitrophota>unclassified_Candidatus_Omnitrophota | 162 | C4527_01145 | 2186..2674 | 2035772 | MAG:-DUF3368-domain-containing-protein-[Candidatus-Omnitrophota-bacterium] |
| MBP7275756.1 | PIN-EndoRNase | GCA_017996315.1 | Kiritimatiellia_bacterium | Bacteria>Pseudomonadati>PVC_group>Kiritimatiellota>Kiritimatiellia>unclassified_Kiritimatiellia | 221 | KBA51_06100 | 6628..7293 | 2212473 | NYN-domain-containing-protein-[Kiritimatiellia-bacterium] |
| TFH14329.1 | PIN-EndoRNase | GCA_004525935.1 | Lentisphaerales_bacterium | Bacteria>Pseudomonadati>PVC_group>Lentisphaerota>Lentisphaeria>Lentisphaerales>unclassified_Lentisphaerales | 172 | E4H02_10100 | 1143..1661 | 2448604 | PIN-domain-containing-protein-[Lentisphaerales-bacterium] |
| MBN2450532.1 | PIN-EndoRNase | GCA_016935045.1 | Lentisphaeria_bacterium | Bacteria>Pseudomonadati>PVC_group>Lentisphaerota>Lentisphaeria>unclassified_Lentisphaeria | 138 | JXR77_09080 | 1059..1475 | 2053569 | type-II-toxin-antitoxin-system-VapC-family-toxin-[Lentisphaeria-bacterium] |
| NLX24427.1 | PIN-EndoRNase | GCA_012729265.1 | Lentisphaerota_bacterium | Bacteria>Pseudomonadati>PVC_group>Lentisphaerota>unclassified_Lentisphaerota | 167 | GXY61_00485 | 4085..4588 | 1932692 | type-II-toxin-antitoxin-system-VapC-family-toxin-[Lentisphaerota-bacterium] |
| MBT5607697.1 | PIN-EndoRNase | GCA_018666605.1 | Lentisphaerota_bacterium | Bacteria>Pseudomonadati>PVC_group>Lentisphaerota>unclassified_Lentisphaerota | 351 | HOJ57_17285 | 262..1317 | 1932692 | hypothetical-protein-[Lentisphaerota-bacterium] |
| WP_052562724.1 | PIN-EndoRNase | GCF_000949635.1 | Candidatus_Brocadia | Bacteria>Pseudomonadati>PVC_group>Planctomycetota>Candidatus_Brocadiia>Candidatus_Brocadiales>Candidatus_Brocadiaceae | 139 | FBQ80_RS01145 | 29574..29993 | 380240 | PIN-domain-containing-protein-[Candidatus-Brocadia-sinica] |
| KXK32842.1 | PIN-EndoRNase | GCA_001567345.1 | Candidatus_Brocadia_sinica | Bacteria>Pseudomonadati>PVC_group>Planctomycetota>Candidatus_Brocadiia>Candidatus_Brocadiales>Candidatus_Brocadiaceae>Candidatus_Brocadia | 139 | UZ01_00264 | 129030..129449 | 795830 | PIN-domain-protein-[Candidatus-Brocadia-sinica] |
| NUN22621.1 | PIN-EndoRNase | GCA_013360885.1 | Candidatus_Jettenia_caeni | Bacteria>Pseudomonadati>PVC_group>Planctomycetota>Candidatus_Brocadiia>Candidatus_Brocadiales>Candidatus_Brocadiaceae>Candidatus_Jettenia | 124 | HUU09_04035 | 17515..17889 | 247490 | PIN-domain-containing-protein-[Candidatus-Jettenia-caeni] |
| KAA0248062.1 | PIN-EndoRNase | GCA_008363445.1 | Candidatus_Jettenia_sp._AMX1 | Bacteria>Pseudomonadati>PVC_group>Planctomycetota>Candidatus_Brocadiia>Candidatus_Brocadiales>Candidatus_Brocadiaceae>Candidatus_Jettenia>unclassified_Candidatus_Jettenia | 128 | EDM77_13490 | 6774..7160 | 2293637 | MAG:-PIN-domain-containing-protein-[Candidatus-Jettenia-sp.-AMX1] |
| MBX3391866.1 | PIN-EndoRNase | GCA_019636515.1 | Phycisphaeraceae_bacterium | Bacteria>Pseudomonadati>PVC_group>Planctomycetota>Phycisphaerae>Phycisphaerales>Phycisphaeraceae>unclassified_Phycisphaeraceae | 206 | KF787_04390 | 1042371..1042991 | 2026777 | NYN-domain-containing-protein-[Phycisphaeraceae-bacterium] |
| MCK4341361.1 | PIN-EndoRNase | GCA_023128695.1 | Phycisphaerae_bacterium | Bacteria>Pseudomonadati>PVC_group>Planctomycetota>Phycisphaerae>unclassified_Phycisphaerae | 161 | KAY37_06530 | 5585..6070 | 2026778 | PIN-domain-containing-protein-[Phycisphaerae-bacterium] |
| WP_194535995.1 | PIN-EndoRNase | GCF_013694095.1 | Thermogemmata_fonticola | Bacteria>Pseudomonadati>PVC_group>Planctomycetota>Planctomycetia>Gemmatales>Gemmataceae>Thermogemmata | 134 | H0921_RS00035 | 18849..19253 | 2755323 | type-II-toxin-antitoxin-system-VapC-family-toxin-[Thermogemmata-fonticola] |
| GIW84012.1 | PIN-EndoRNase | GCA_026004695.1 | Gemmataceae_bacterium | Bacteria>Pseudomonadati>PVC_group>Planctomycetota>Planctomycetia>Gemmatales>Gemmataceae>unclassified_Gemmataceae | 115 | KatS3mg106_525 | 15519..15866 | 2052164 | MAG:-hypothetical-protein-KatS3mg106_525-[Gemmataceae-bacterium] |
| MSR55924.1 | PIN-EndoRNase | GCA_009691845.1 | Gemmataceae_bacterium | Bacteria>Pseudomonadati>PVC_group>Planctomycetota>Planctomycetia>Gemmatales>Gemmataceae>unclassified_Gemmataceae | 122 | EXS09_22025 | 4347..4715 | 2052164 | hypothetical-protein-[Gemmataceae-bacterium] |
| MDB5306880.1 | PIN-EndoRNase | GCA_028288765.1 | Gemmataceae_bacterium | Bacteria>Pseudomonadati>PVC_group>Planctomycetota>Planctomycetia>Gemmatales>Gemmataceae>unclassified_Gemmataceae | 129 | JWO38_1082 | 730..1119 | 2052164 | MAG:-hypothetical-protein-JWO38_1082-[Gemmataceae-bacterium] |
| WP_013566220.1 | PIN-EndoRNase | GCF_000186345.1 | Isosphaera_pallida | Bacteria>Pseudomonadati>PVC_group>Planctomycetota>Planctomycetia>Isosphaerales>Isosphaeraceae>Isosphaera | 126 | ISOP_RS18000 | 4936895..4937275 | 128 | DUF5615-family-PIN-like-protein-[Isosphaera-pallida] |
| WP_015245483.1 | PIN-EndoRNase | GCF_000242455.2 | Singulisphaera_acidiphila | Bacteria>Pseudomonadati>PVC_group>Planctomycetota>Planctomycetia>Isosphaerales>Isosphaeraceae>Singulisphaera | 132 | OSM_RS0102250 | 267284..267682 | 466153 | type-II-toxin-antitoxin-system-VapC-family-toxin-[Singulisphaera-acidiphila] |
| MBV8611621.1 | PIN-EndoRNase | GCA_019233025.1 | Singulisphaera_sp. | Bacteria>Pseudomonadati>PVC_group>Planctomycetota>Planctomycetia>Isosphaerales>Isosphaeraceae>Singulisphaera>unclassified_Singulisphaera | 132 | JO034_29780 | 869..1267 | 1906241 | type-II-toxin-antitoxin-system-VapC-family-toxin-[Singulisphaera-sp.] |
| WP_230218087.1 | PIN-EndoRNase | GCF_020966755.1 | Blastopirellula_sediminis | Bacteria>Pseudomonadati>PVC_group>Planctomycetota>Planctomycetia>Pirellulales>Pirellulaceae>Blastopirellula | 163 | LOC68_RS09620 | 1471081..1471572 | 2894196 | hypothetical-protein-[Blastopirellula-sediminis] |
| WP_283433170.1 | PIN-EndoRNase | GCF_900182915.1 | Neorhodopirellula_lusitana | Bacteria>Pseudomonadati>PVC_group>Planctomycetota>Planctomycetia>Pirellulales>Pirellulaceae>Neorhodopirellula | 163 | QOL80_RS14715 | 94876..95367 | 445327 | hypothetical-protein-[Rhodopirellula-lusitana] |
| NLX99941.1 | PIN-EndoRNase | GCA_012728855.1 | Rhodopirellula_sp. | Bacteria>Pseudomonadati>PVC_group>Planctomycetota>Planctomycetia>Pirellulales>Pirellulaceae>Rhodopirellula>unclassified_Rhodopirellula | 161 | GXY83_27905 | 5838..6323 | 2024855 | DUF3368-domain-containing-protein-[Rhodopirellula-sp.] |
| WP_168215743.1 | PIN-EndoRNase | GCF_008065135.1 | Roseimaritima_ulvae | Bacteria>Pseudomonadati>PVC_group>Planctomycetota>Planctomycetia>Pirellulales>Pirellulaceae>Roseimaritima | 175 | UC8_RS27040 | 7578584..7579111 | 980254 | hypothetical-protein-[Roseimaritima-ulvae] |
| MCO6458541.1 | PIN-EndoRNase | GCA_024102795.1 | Pirellulaceae_bacterium | Bacteria>Pseudomonadati>PVC_group>Planctomycetota>Planctomycetia>Pirellulales>Pirellulaceae>unclassified_Pirellulaceae | 130 | J5I93_24815 | 23412..23804 | 2699754 | PIN-domain-containing-protein-[Pirellulaceae-bacterium] |
| MCC7337718.1 | PIN-EndoRNase | GCA_020852965.1 | Pirellulaceae_bacterium | Bacteria>Pseudomonadati>PVC_group>Planctomycetota>Planctomycetia>Pirellulales>Pirellulaceae>unclassified_Pirellulaceae | 139 | IT422_21730 | 4656..5075 | 2699754 | PIN-domain-containing-protein-[Pirellulaceae-bacterium] |
| MCC7337729.1 | PIN-EndoRNase | GCA_020852965.1 | Pirellulaceae_bacterium | Bacteria>Pseudomonadati>PVC_group>Planctomycetota>Planctomycetia>Pirellulales>Pirellulaceae>unclassified_Pirellulaceae | 143 | IT422_21785 | 31383..31814 | 2699754 | type-II-toxin-antitoxin-system-VapC-family-toxin-[Pirellulaceae-bacterium] |
| NMC20500.1 | PIN-EndoRNase | GCA_012798155.1 | Thermogutta_sp. | Bacteria>Pseudomonadati>PVC_group>Planctomycetota>Planctomycetia>Pirellulales>Thermoguttaceae>Thermogutta>unclassified_Thermogutta | 105 | GYA33_08780 | 3935..4252 | 1962930 | DUF5615-family-PIN-like-protein-[Thermogutta-sp.] |
| MCA9013094.1 | PIN-EndoRNase | GCA_020426105.1 | Planctomycetaceae_bacterium | Bacteria>Pseudomonadati>PVC_group>Planctomycetota>Planctomycetia>Planctomycetales>Planctomycetaceae>unclassified_Planctomycetaceae | 144 | KDB01_25260 | 4417..4851 | 2026779 | type-II-toxin-antitoxin-system-VapC-family-toxin-[Planctomycetaceae-bacterium] |
| MCA9076581.1 | PIN-EndoRNase | GCA_020426505.1 | Planctomycetaceae_bacterium | Bacteria>Pseudomonadati>PVC_group>Planctomycetota>Planctomycetia>Planctomycetales>Planctomycetaceae>unclassified_Planctomycetaceae | 164 | KDA93_16270 | 17276..17770 | 2026779 | hypothetical-protein-[Planctomycetaceae-bacterium] |
| HBJ35656.1 | PIN-EndoRNase | GCA_003506095.1 | Planctomycetaceae_bacterium | Bacteria>Pseudomonadati>PVC_group>Planctomycetota>Planctomycetia>Planctomycetales>Planctomycetaceae>unclassified_Planctomycetaceae | 182 | DDZ51_13110 | 6040..6588 | 2026779 | hypothetical-protein-[Planctomycetaceae-bacterium] |
| MDA8378779.1 | PIN-EndoRNase | GCA_028068345.1 | Planctomycetia_bacterium | Bacteria>Pseudomonadati>PVC_group>Planctomycetota>Planctomycetia>unclassified_Planctomycetia | 203 | MOZ50_17435 | 8575..9186 | 2052181 | MAG:-NYN-domain-containing-protein-[Planctomycetia-bacterium] |
| MCY2991649.1 | PIN-EndoRNase | GCA_026692885.1 | Planctomycetota_bacterium | Bacteria>Pseudomonadati>PVC_group>Planctomycetota>unclassified_Planctomycetota | 119 | NTY19_27815 | 5144..5503 | 2026780 | MAG:-DUF5615-family-PIN-like-protein-[Planctomycetota-bacterium] |
| MCH7685159.1 | PIN-EndoRNase | GCA_022562015.1 | Planctomycetota_bacterium | Bacteria>Pseudomonadati>PVC_group>Planctomycetota>unclassified_Planctomycetota | 126 | IH899_00505 | 2872..3252 | 2026780 | DUF5615-family-PIN-like-protein-[Planctomycetota-bacterium] |
| MBI3271454.1 | PIN-EndoRNase | GCA_016197205.1 | Planctomycetota_bacterium | Bacteria>Pseudomonadati>PVC_group>Planctomycetota>unclassified_Planctomycetota | 151 | HYZ53_20860 | 64576..65031 | 2026780 | PIN-domain-containing-protein-[Planctomycetota-bacterium] |
| MCK6447152.1 | PIN-EndoRNase | GCA_023150475.1 | Planctomycetota_bacterium | Bacteria>Pseudomonadati>PVC_group>Planctomycetota>unclassified_Planctomycetota | 155 | L6Q99_12245 | 33465..33932 | 2026780 | PIN-domain-containing-protein-[Planctomycetota-bacterium] |
| MBM4022805.1 | PIN-EndoRNase | GCA_016872955.1 | Planctomycetota_bacterium | Bacteria>Pseudomonadati>PVC_group>Planctomycetota>unclassified_Planctomycetota | 203 | FJ284_11305 | 1991..2602 | 2026780 | NYN-domain-containing-protein-[Planctomycetota-bacterium] |
| MBI3097942.1 | PIN-EndoRNase | GCA_016192525.1 | Planctomycetota_bacterium | Bacteria>Pseudomonadati>PVC_group>Planctomycetota>unclassified_Planctomycetota | 208 | HYY93_06790 | 71717..72343 | 2026780 | NYN-domain-containing-protein-[Planctomycetota-bacterium] |
| MCY2953680.1 | PIN-EndoRNase | GCA_026693085.1 | Planctomycetota_bacterium | Bacteria>Pseudomonadati>PVC_group>Planctomycetota>unclassified_Planctomycetota | 215 | NTU53_17125 | 24078..24725 | 2026780 | MAG:-NYN-domain-containing-protein-[Planctomycetota-bacterium] |
| MCZ7645663.1 | PIN-EndoRNase | GCA_027493525.1 | Planctomycetota_bacterium | Bacteria>Pseudomonadati>PVC_group>Planctomycetota>unclassified_Planctomycetota | 281 | M5U26_10320 | 2469050..2469895 | 2026780 | MAG:-PIN-domain-containing-protein-[Planctomycetota-bacterium] |
| MCE0498495.1 | PIN-EndoRNase | GCA_021272315.1 | Candidatus_Methylacidiphilales_bacterium | Bacteria>Pseudomonadati>PVC_group>Verrucomicrobiota>Methylacidiphilae>Methylacidiphilales>unclassified_Candidatus_Methylacidiphilales | 166 | LV481_11175 | 14015..14515 | 2720631 | type-II-toxin-antitoxin-system-VapC-family-toxin-[Methylacidiphilales-bacterium] |
| MCC5805072.1 | PIN-EndoRNase | GCA_020831525.1 | Opitutales_bacterium | Bacteria>Pseudomonadati>PVC_group>Verrucomicrobiota>Opitutia>Opitutales>unclassified_Opitutales | 130 | JJU00_01975 | 48463..48855 | 2026772 | type-II-toxin-antitoxin-system-VapC-family-toxin-[Opitutales-bacterium] |
| MCC5838836.1 | PIN-EndoRNase | GCA_020831585.1 | Opitutales_bacterium | Bacteria>Pseudomonadati>PVC_group>Verrucomicrobiota>Opitutia>Opitutales>unclassified_Opitutales | 134 | JJT96_01830 | 65071..65475 | 2026772 | type-II-toxin-antitoxin-system-VapC-family-toxin-[Opitutales-bacterium] |
| MCC5838831.1 | PIN-EndoRNase | GCA_020831585.1 | Opitutales_bacterium | Bacteria>Pseudomonadati>PVC_group>Verrucomicrobiota>Opitutia>Opitutales>unclassified_Opitutales | 143 | JJT96_01805 | 57831..58262 | 2026772 | PIN-domain-containing-protein-[Opitutales-bacterium] |
| MCC5805076.1 | PIN-EndoRNase | GCA_020831525.1 | Opitutales_bacterium | Bacteria>Pseudomonadati>PVC_group>Verrucomicrobiota>Opitutia>Opitutales>unclassified_Opitutales | 156 | JJU00_01995 | 57336..57806 | 2026772 | type-II-toxin-antitoxin-system-VapC-family-toxin-[Opitutales-bacterium] |
| MCH8539499.1 | PIN-EndoRNase | GCA_022565675.1 | Opitutales_bacterium | Bacteria>Pseudomonadati>PVC_group>Verrucomicrobiota>Opitutia>Opitutales>unclassified_Opitutales | 246 | LAT58_01875 | 14329..15069 | 2026772 | NYN-domain-containing-protein-[Opitutales-bacterium] |
| MCK9588327.1 | PIN-EndoRNase | GCA_023227995.1 | Terrimicrobiaceae_bacterium | Bacteria>Pseudomonadati>PVC_group>Verrucomicrobiota>Terrimicrobiia>Terrimicrobiales>Terrimicrobiaceae>unclassified_Terrimicrobiaceae | 165 | MOQ93_03060 | 7333..7830 | 2832348 | type-II-toxin-antitoxin-system-VapC-family-toxin-[Terrimicrobiaceae-bacterium] |
| MBI2947995.1 | PIN-EndoRNase | GCA_016190065.1 | Verrucomicrobiota_bacterium | Bacteria>Pseudomonadati>PVC_group>Verrucomicrobiota>unclassified_Verrucomicrobiota | 140 | HYY23_10125 | 2606..3028 | 2026799 | putative-toxin-antitoxin-system-toxin-component,-PIN-family-[Verrucomicrobiota-bacterium] |
| MCX6876432.1 | PIN-EndoRNase | GCA_026397725.1 | Verrucomicrobiota_bacterium | Bacteria>Pseudomonadati>PVC_group>Verrucomicrobiota>unclassified_Verrucomicrobiota | 141 | NTW21_21880 | 18380..18805 | 2026799 | MAG:-putative-toxin-antitoxin-system-toxin-component,-PIN-family-[Verrucomicrobia-bacterium] |
| NBB80455.1 | PIN-EndoRNase | GCA_009908325.1 | Verrucomicrobiota_bacterium | Bacteria>Pseudomonadati>PVC_group>Verrucomicrobiota>unclassified_Verrucomicrobiota | 153 | GVY36_13605 | 6997..7458 | 2026799 | hypothetical-protein-[Verrucomicrobiota-bacterium] |
| MCL4785393.1 | PIN-EndoRNase | GCA_023383755.1 | Verrucomicrobiota_bacterium | Bacteria>Pseudomonadati>PVC_group>Verrucomicrobiota>unclassified_Verrucomicrobiota | 153 | KJ070_01165 | 30998..31459 | 2026799 | DUF4411-family-protein-[Verrucomicrobiota-bacterium] |
| AKC83095.1 | PIN-EndoRNase | GCA_000972765.1 | Verrucomicrobia_bacterium_IMCC26134 | Bacteria>Pseudomonadati>PVC_group>Verrucomicrobiota>unclassified_Verrucomicrobiota | 154 | IMCC26134_10465 | 2781556..2782020 | 1637999 | hypothetical-protein-IMCC26134_10465-[Verrucomicrobia-bacterium-IMCC26134] |
| MCX6865166.1 | PIN-EndoRNase | GCA_026397695.1 | Verrucomicrobiota_bacterium | Bacteria>Pseudomonadati>PVC_group>Verrucomicrobiota>unclassified_Verrucomicrobiota | 171 | NTV46_02915 | 6519..7034 | 2026799 | MAG:-type-II-toxin-antitoxin-system-VapC-family-toxin-[Verrucomicrobia-bacterium] |
| HCF94158.1 | PIN-EndoRNase | GCA_003534995.1 | Verrucomicrobiota_bacterium | Bacteria>Pseudomonadati>PVC_group>Verrucomicrobiota>unclassified_Verrucomicrobiota | 176 | DEW46_03785 | 11311..11841 | 2026799 | hypothetical-protein-[Verrucomicrobiota-bacterium] |
| RFC50835.1 | PIN-EndoRNase | GCA_003402705.1 | Verrucomicrobiota_bacterium | Bacteria>Pseudomonadati>PVC_group>Verrucomicrobiota>unclassified_Verrucomicrobiota | 191 | DVB22_001796 | 7114..7689 | 2026799 | putative-nucleic-acid-binding-protein,-contains-PIN-domain-[Verrucomicrobiota-bacterium] |
| MCS1417416.1 | PIN-EndoRNase | GCA_024761205.1 | Limisphaerales_bacterium | Bacteria>Pseudomonadati>PVC_group>Verrucomicrobiota>Verrucomicrobiia>Limisphaerales>unclassified_Limisphaerales | 254 | M2R46_05170 | 22789..23553 | 2052186 | hypothetical-protein-[Verrucomicrobia-subdivision-3-bacterium] |
| MCW5552138.1 | PIN-EndoRNase | GCA_026125285.1 | Verrucomicrobiia_bacterium | Bacteria>Pseudomonadati>PVC_group>Verrucomicrobiota>Verrucomicrobiia>unclassified_Verrucomicrobiia | 192 | KIS67_08215 | 92150..92728 | 2499141 | MAG:-hypothetical-protein-KIS67_08215-[Verrucomicrobiae-bacterium] |
| WP_067776565.1 | PIN-EndoRNase | GCF_001683795.1 | Akkermansia_glycaniphila | Bacteria>Pseudomonadati>PVC_group>Verrucomicrobiota>Verrucomicrobiia>Verrucomicrobiales>Akkermansiaceae>Akkermansia | 175 | AC781_RS08525 | 125485..126012 | 1679444 | hypothetical-protein-[Akkermansia-glycaniphila] |
| WP_123052209.1 | PIN-EndoRNase | GCF_003716915.1 | Akkermansia_muciniphila | Bacteria>Pseudomonadati>PVC_group>Verrucomicrobiota>Verrucomicrobiia>Verrucomicrobiales>Akkermansiaceae>Akkermansia | 183 | CUB89_RS06275 | 1396403..1396954 | 239935 | type-II-toxin-antitoxin-system-VapC-family-toxin-[Akkermansia-muciniphila] |
| WP_215841668.1 | PIN-EndoRNase | GCF_018847055.1 | Akkermansia_muciniphila | Bacteria>Pseudomonadati>PVC_group>Verrucomicrobiota>Verrucomicrobiia>Verrucomicrobiales>Akkermansiaceae>Akkermansia | 183 | J5W61_RS00470 | 120213..120764 | 239935 | type-II-toxin-antitoxin-system-VapC-family-toxin-[Akkermansia-muciniphila] |
| HCN27196.1 | PIN-EndoRNase | GCA_003545395.1 | Verrucomicrobiales_bacterium | Bacteria>Pseudomonadati>PVC_group>Verrucomicrobiota>Verrucomicrobiia>Verrucomicrobiales>unclassified_Verrucomicrobiales | 139 | DIT64_00060 | 2426..2845 | 2026801 | pilus-assembly-protein-[Verrucomicrobiales-bacterium] |
| MBP7951463.1 | PIN-EndoRNase | GCA_018003315.1 | Verrucomicrobiales_bacterium | Bacteria>Pseudomonadati>PVC_group>Verrucomicrobiota>Verrucomicrobiia>Verrucomicrobiales>unclassified_Verrucomicrobiales | 193 | KA004_17610 | 10912..11493 | 2026801 | hypothetical-protein-[Verrucomicrobiales-bacterium] |
| MBL9184320.1 | PIN-EndoRNase | GCA_016795465.1 | Verrucomicrobiaceae_bacterium | Bacteria>Pseudomonadati>PVC_group>Verrucomicrobiota>Verrucomicrobiia>Verrucomicrobiales>Verrucomicrobiaceae>unclassified_Verrucomicrobiaceae | 206 | JNN17_19400 | 5718..6338 | 2026800 | NYN-domain-containing-protein-[Verrucomicrobiaceae-bacterium] |
| WP_135740497.1 | PIN-EndoRNase | GCF_004770445.1 | Leptospira_bouyouniensis | Bacteria>Pseudomonadati>Spirochaetota>Spirochaetia>Leptospirales>Leptospiraceae>Leptospira | 216 | EHQ99_RS04870 | 5107..5757 | 2484911 | hypothetical-protein-[Leptospira-bouyouniensis] |
| TGM42336.1 | PIN-EndoRNase | GCA_004770805.1 | Leptospira_biflexa | Bacteria>Pseudomonadati>Spirochaetota>Spirochaetia>Leptospirales>Leptospiraceae>Leptospira | 218 | EHQ92_16200 | 5049..5705 | 172 | hypothetical-protein-EHQ92_16200-[Leptospira-biflexa] |
| WP_135737927.1 | PIN-EndoRNase | GCF_004770325.1 | Leptospira_biflexa | Bacteria>Pseudomonadati>Spirochaetota>Spirochaetia>Leptospirales>Leptospiraceae>Leptospira | 218 | EHQ92_RS16200 | 5049..5705 | 172 | hypothetical-protein-[Leptospira-biflexa] |
| MCB1144573.1 | PIN-EndoRNase | GCA_020440865.1 | Leptospiraceae_bacterium | Bacteria>Pseudomonadati>Spirochaetota>Spirochaetia>Leptospirales>Leptospiraceae>unclassified_Leptospiraceae | 211 | KDK54_20125 | 6353..6988 | 2026757 | hypothetical-protein-[Leptospiraceae-bacterium] |
| MCB1144578.1 | PIN-EndoRNase | GCA_020440865.1 | Leptospiraceae_bacterium | Bacteria>Pseudomonadati>Spirochaetota>Spirochaetia>Leptospirales>Leptospiraceae>unclassified_Leptospiraceae | 388 | KDK54_20150 | 14119..15285 | 2026757 | DUF4935-domain-containing-protein-[Leptospiraceae-bacterium] |
| AEJ20211.1 | PIN-EndoRNase | GCA_000219725.1 | Gracilinema_caldarium_DSM_7334 | Bacteria>Pseudomonadati>Spirochaetota>Spirochaetia>Spirochaetales>Breznakiellaceae>Gracilinema>Gracilinema_caldarium | 167 | Spica_2086 | 2350090..2350593 | 744872 | hypothetical-protein-Spica_2086-[Gracilinema-caldarium-DSM-7334] |
| HAL93443.1 | PIN-EndoRNase | GCA_003452735.1 | Spirochaetaceae_bacterium | Bacteria>Pseudomonadati>Spirochaetota>Spirochaetia>Spirochaetales>Spirochaetaceae>unclassified_Spirochaetaceae | 125 | DCP56_01615 | 13893..14270 | 1898206 | VapC-toxin-family-PIN-domain-ribonuclease-[Spirochaetaceae-bacterium] |
| HAL93439.1 | PIN-EndoRNase | GCA_003452735.1 | Spirochaetaceae_bacterium | Bacteria>Pseudomonadati>Spirochaetota>Spirochaetia>Spirochaetales>Spirochaetaceae>unclassified_Spirochaetaceae | 132 | DCP56_01595 | 9118..9516 | 1898206 | VapC-toxin-family-PIN-domain-ribonuclease-[Spirochaetaceae-bacterium] |
| MDE0219723.1 | PIN-EndoRNase | GCA_028820215.1 | Spirochaetaceae_bacterium | Bacteria>Pseudomonadati>Spirochaetota>Spirochaetia>Spirochaetales>Spirochaetaceae>unclassified_Spirochaetaceae | 158 | OXJ90_10670 | 21874..22350 | 1898206 | MAG:-hypothetical-protein-OXJ90_10670-[Spirochaetaceae-bacterium] |
| MCL2137973.1 | PIN-EndoRNase | GCA_009776715.1 | Treponema_sp. | Bacteria>Pseudomonadati>Spirochaetota>Spirochaetia>Spirochaetales>Treponemataceae>Treponema>unclassified_Treponema | 136 | FWH41_00400 | 14472..14882 | 166 | putative-toxin-antitoxin-system-toxin-component,-PIN-family-[Treponema-sp.] |
| KPA14791.1 | PIN-EndoRNase | GCA_001292585.1 | Candidatus_Magnetomorum_sp._HK-1 | Bacteria>Pseudomonadati>Thermodesulfobacteriota>Desulfobacteria>Desulfobacterales>Candidatus_Magnetomoraceae>Candidatus_Magnetomorum>unclassified_Candidatus_Magnetomorum | 127 | MHK_004994 | 8253..8636 | 1509431 | twitching-motility-protein-PilT-[Candidatus-Magnetomorum-sp.-HK-1] |
| SLM29547.1 | PIN-EndoRNase | GCA_900170035.1 | Desulfamplus_magnetovallimortis | Bacteria>Pseudomonadati>Thermodesulfobacteriota>Desulfobacteria>Desulfobacterales>Desulfobacteraceae>Desulfamplus | 183 | MTBBW1_1860037 | 46213..46764 | 1246637 | PilT-like-protein-[Desulfamplus-magnetovallimortis] |
| WP_080806580.1 | PIN-EndoRNase | GCF_900170035.1 | Desulfamplus_magnetovallimortis | Bacteria>Pseudomonadati>Thermodesulfobacteriota>Desulfobacteria>Desulfobacterales>Desulfobacteraceae>Desulfamplus | 183 | MTBBW1_RS20345 | 72228..72779 | 1246637 | type-II-toxin-antitoxin-system-VapC-family-toxin-[Desulfamplus-magnetovallimortis] |
| MBF0572398.1 | PIN-EndoRNase | GCA_015233755.1 | Desulfamplus_sp. | Bacteria>Pseudomonadati>Thermodesulfobacteriota>Desulfobacteria>Desulfobacterales>Desulfobacteraceae>Desulfamplus>unclassified_Desulfamplus | 136 | HQK69_01365 | 16849..17259 | 2006184 | nucleotide-binding-protein-[Desulfamplus-sp.] |

| Protein-ID | Domain | Assembly-ID | Strain | Taxonomy | Length | Locus-Tag | Genomic-coordinates | TaxID | Fasta-defline |
|---|---|---|---|---|---|---|---|---|---|
| WP_144683392.1 | PIN-EndoRNase | GCF_007830435.1 | Desulfobotulus_alkaliphilus | Bacteria>Pseudomonadati>Thermodesulfobacteriota>Desulfobacteria>Desulfobacterales>Desulfobacteraceae>Desulfobotulus | 183 | LZ24_RS05870 | 56061..56612 | 622671 | type-II-toxin-antitoxin-system-VapC-family-toxin-[Desulfobotulus-alkaliphilus] |
| OQX20474.1 | PIN-EndoRNase | GCA_002083955.1 | Desulfobacteraceae_bacterium_IS3 | Bacteria>Pseudomonadati>Thermodesulfobacteriota>Desulfobacteria>Desulfobacterales>Desulfobacteraceae>unclassified_Desulfobacteraceae | 144 | BWK80_35710 | 1982..2416 | 1934248 | hypothetical-protein-BWK80_35710-[Desulfobacteraceae-bacterium-IS3] |
| MBL7176675.1 | PIN-EndoRNase | GCA_016782925.1 | Desulfobacteraceae_bacterium | Bacteria>Pseudomonadati>Thermodesulfobacteriota>Desulfobacteria>Desulfobacterales>Desulfobacteraceae>unclassified_Desulfobacteraceae | 161 | ISS66_12690 | 9136..9621 | 2049433 | hypothetical-protein-[Desulfobacteraceae-bacterium] |
| MCK4391112.1 | PIN-EndoRNase | GCA_023129535.1 | Desulfobacterales_bacterium | Bacteria>Pseudomonadati>Thermodesulfobacteriota>Desulfobacteria>Desulfobacterales>unclassified_Desulfobacterales | 122 | KAV83_12850 | 5144..5512 | 2044940 | DUF5615-family-PIN-like-protein-[Desulfobacterales-bacterium] |
| MBF0121418.1 | PIN-EndoRNase | GCA_015228685.1 | Desulfobacterales_bacterium | Bacteria>Pseudomonadati>Thermodesulfobacteriota>Desulfobacteria>Desulfobacterales>unclassified_Desulfobacterales | 149 | HQK79_21520 | 4571..5020 | 2044940 | type-II-toxin-antitoxin-system-VapC-family-toxin-[Desulfobacterales-bacterium] |
| MBF0121089.1 | PIN-EndoRNase | GCA_015228685.1 | Desulfobacterales_bacterium | Bacteria>Pseudomonadati>Thermodesulfobacteriota>Desulfobacteria>Desulfobacterales>unclassified_Desulfobacterales | 157 | HQK79_19840 | 16309..16782 | 2044940 | type-II-toxin-antitoxin-system-VapC-family-toxin-[Desulfobacterales-bacterium] |
| MCK4486187.1 | PIN-EndoRNase | GCA_023131115.1 | Desulfobacterales_bacterium | Bacteria>Pseudomonadati>Thermodesulfobacteriota>Desulfobacteria>Desulfobacterales>unclassified_Desulfobacterales | 168 | KAU38_05410 | 4145..4651 | 2044940 | type-II-toxin-antitoxin-system-VapC-family-toxin-[Desulfobacterales-bacterium] |
| NVM22711.1 | PIN-EndoRNase | GCA_013375395.1 | Desulfobacterales_bacterium | Bacteria>Pseudomonadati>Thermodesulfobacteriota>Desulfobacteria>Desulfobacterales>unclassified_Desulfobacterales | 168 | HWN68_13140 | 6321..6827 | 2044940 | type-II-toxin-antitoxin-system-VapC-family-toxin-[Desulfobacterales-bacterium] |
| HCF05835.1 | PIN-EndoRNase | GCA_003530935.1 | Desulfomicrobiaceae_bacterium | Bacteria>Pseudomonadati>Thermodesulfobacteriota>Desulfovibrionia>Desulfovibrionales>Desulfomicrobiaceae>unclassified_Desulfomicrobiaceae | 206 | DEU72_06330 | 6389..7009 | 2053547 | NYN-domain-containing-protein-[Desulfomicrobiaceae-bacterium] |
| MBZ4685276.1 | PIN-EndoRNase | GCA_020063575.1 | Desulfomicrobiaceae_bacterium | Bacteria>Pseudomonadati>Thermodesulfobacteriota>Desulfovibrionia>Desulfovibrionales>Desulfomicrobiaceae>unclassified_Desulfomicrobiaceae | 206 | JG774_1021 | 68442..69062 | 2053547 | hypothetical-protein-[Desulfomicrobiaceae-bacterium] |
| WP_045216070.1 | PIN-EndoRNase | GCF_000934755.1 | Desulfonatronovibrio_magnus | Bacteria>Pseudomonadati>Thermodesulfobacteriota>Desulfovibrionia>Desulfovibrionales>Desulfonatronovibrionaceae>Desulfonatronovibrio | 140 | LZ23_RS16880 | 145238..145660 | 698827 | DNA-binding-protein-[Desulfonatronovibrio-magnus] |
| HJA77860.1 | PIN-EndoRNase | GCA_019118745.1 | Candidatus_Desulfovibrio_gallistercoris | Bacteria>Pseudomonadati>Thermodesulfobacteriota>Desulfovibrionia>Desulfovibrionales>Desulfovibrionaceae>Desulfovibrio | 151 | H9774_13545 | 12498..12953 | 2838533 | hypothetical-protein-[Candidatus-Desulfovibrio-gallistercoris] |
| MCQ4862357.1 | PIN-EndoRNase | GCA_024460775.1 | Desulfovibrio_desulfuricans | Bacteria>Pseudomonadati>Thermodesulfobacteriota>Desulfovibrionia>Desulfovibrionales>Desulfovibrionaceae>Desulfovibrio | 176 | NE637_14430 | 8183..8713 | 876 | hypothetical-protein-NE637_14430-[Desulfovibrio-desulfuricans] |
| MCQ5219598.1 | PIN-EndoRNase | GCA_024463415.1 | Desulfovibrio_desulfuricans | Bacteria>Pseudomonadati>Thermodesulfobacteriota>Desulfovibrionia>Desulfovibrionales>Desulfovibrionaceae>Desulfovibrio | 176 | NE596_15670 | 6838..7368 | 876 | hypothetical-protein-NE596_15670-[Desulfovibrio-desulfuricans] |
| WP_215646939.1 | PIN-EndoRNase | GCF_020555025.1 | Desulfovibrio_desulfuricans | Bacteria>Pseudomonadati>Thermodesulfobacteriota>Desulfovibrionia>Desulfovibrionales>Desulfovibrionaceae>Desulfovibrio | 176 | NE596_RS15655 | 6838..7368 | 876 | hypothetical-protein-[Desulfovibrio-desulfuricans] |
| SBW09996.1 | PIN-EndoRNase | GCA_900079715.1 | uncultured_Desulfovibrio_sp. | Bacteria>Pseudomonadati>Thermodesulfobacteriota>Desulfovibrionia>Desulfovibrionales>Desulfovibrionaceae>Desulfovibrio>environmental_samples | 157 | KM92DES2_12849 | 2969180..2969653 | 167968 | conserved-hypothetical-protein-[uncultured-Desulfovibrio-sp.] |
| WP_165175271.1 | PIN-EndoRNase | GCF_011039085.1 | Desulfovibrio_sp._ZJ369 | Bacteria>Pseudomonadati>Thermodesulfobacteriota>Desulfovibrionia>Desulfovibrionales>Desulfovibrionaceae>Desulfovibrio>unclassified_Desulfovibrio | 157 | G7Y51_RS04975 | 87168..87641 | 2709793 | DUF4411-family-protein-[Desulfovibrio-sp.-ZJ369] |
| WP_092055232.1 | PIN-EndoRNase | GCF_900111775.1 | Desulfuromonas_acetexigens | Bacteria>Pseudomonadati>Thermodesulfobacteriota>Desulfuromonadia>Desulfuromonadales>Desulfuromonadaceae>Desulfuromonas | 175 | BQ4888_RS06400 | 8491..9018 | 38815 | hypothetical-protein-[Desulfuromonas-acetexigens] |
| MCG2740515.1 | PIN-EndoRNase | GCA_022013455.1 | Syntrophaceae_bacterium | Bacteria>Pseudomonadati>Thermodesulfobacteriota>Syntrophia>Syntrophales>Syntrophaceae>unclassified_Syntrophaceae | 169 | L6300_09815 | 4273..4782 | 2052184 | PIN-domain-containing-protein-[Syntrophaceae-bacterium] |
| WP_073037945.1 | PIN-EndoRNase | GCF_900129305.1 | Desulfacinum_infernum | Bacteria>Pseudomonadati>Thermodesulfobacteriota>Syntrophobacteria>Syntrophobacterales>Syntrophobacteraceae>Desulfacinum | 213 | BUB04_RS06065 | 35012..35653 | 35837 | NYN-domain-containing-protein-[Desulfacinum-infernum] |
| WP_221269583.1 | PIN-EndoRNase | - | Deinococcus_budaensis | Bacteria>Thermotogati>Deinococcota>Deinococci>Deinococcales>Deinococcaceae>Deinococcus | 96 | HNQ09_RS02935 | 26187..26477 | 1665626 | DUF5615-family-PIN-like-protein-[Deinococcus-budaensis] |
| WP_241191480.1 | PIN-EndoRNase | GCF_003860465.1 | Deinococcus_psychrotolerans | Bacteria>Thermotogati>Deinococcota>Deinococci>Deinococcales>Deinococcaceae>Deinococcus | 159 | EHF33_RS19825 | 22031..22510 | 2489213 | type-II-toxin-antitoxin-system-VapC-family-toxin-[Deinococcus-psychrotolerans] |
| WP_234009277.1 | PIN-EndoRNase | GCF_002953415.1 | Deinococcus_sp._NW-56 | Bacteria>Thermotogati>Deinococcota>Deinococci>Deinococcales>Deinococcaceae>Deinococcus>unclassified_Deinococcus | 157 | C3K08_RS17185 | 346356..346829 | 2080419 | type-II-toxin-antitoxin-system-VapC-family-toxin-[Deinococcus-sp.-NW-56] |
| WP_081623146.1 | PIN-EndoRNase | GCF_037481675.1 | Thermus | Bacteria>Thermotogati>Deinococcota>Deinococci>Thermales>Thermaceae | 205 | B043_RS12700 | 83848..84465 | 270 | NYN-domain-containing-protein-[Thermus-oshimai] |
| WP_240631253.1 | PIN-EndoRNase | - | Meiothermus_granaticius | Bacteria>Thermotogati>Deinococcota>Deinococci>Thermales>Thermaceae>Meiothermus | 130 | MGR01S_RS15590 | 657..1049 | 863370 | PIN-domain-containing-protein-[Meiothermus-granaticius] |
| WP_119356453.1 | PIN-EndoRNase | GCF_007990995.1 | Meiothermus_granaticius | Bacteria>Thermotogati>Deinococcota>Deinococci>Thermales>Thermaceae>Meiothermus | 136 | MGR01S_RS15610 | 8104..8514 | 863370 | type-II-toxin-antitoxin-system-VapC-family-toxin-[Meiothermus-granaticius] |
| WP_187288494.1 | PIN-EndoRNase | GCF_000183745.1 | Oceanithermus_profundus | Bacteria>Thermotogati>Deinococcota>Deinococci>Thermales>Thermaceae>Oceanithermus | 215 | OCEPR_RS12660 | 2271167..2271814 | 187137 | NYN-domain-containing-protein-[Oceanithermus-profundus] |
| WP_015065388.1 | PIN-EndoRNase | GCF_000309885.1 | Thermus_oshimai | Bacteria>Thermotogati>Deinococcota>Deinococci>Thermales>Thermaceae>Thermus | 205 | THEOS_RS12940 | 205933..206550 | 56957 | NYN-domain-containing-protein-[Thermus-oshimai] |
| WP_135343462.1 | PIN-EndoRNase | GCF_004685085.1 | Thermus_tengchongensis | Bacteria>Thermotogati>Deinococcota>Deinococci>Thermales>Thermaceae>Thermus | 205 | E0489_RS07470 | 6458..7075 | 1214928 | NYN-domain-containing-protein-[Thermus-tengchongensis] |
| MBF6594439.1 | PIN-EndoRNase | GCA_015478585.1 | Thermaceae_bacterium | Bacteria>Thermotogati>Deinococcota>Deinococci>Thermales>Thermaceae>unclassified_Thermaceae | 136 | IVW51_08325 | 20408..20818 | 2614257 | type-II-toxin-antitoxin-system-VapC-family-toxin-[Thermaceae-bacterium] |
| WP_049773133.1 | PIN-EndoRNase | GCF_000092425.1 | Truepera_radiovictrix | Bacteria>Thermotogati>Deinococcota>Deinococci>Trueperales>Trueperaceae>Truepera | 123 | TRAD_RS13555 | 2957208..2957579 | 332249 | PIN-domain-containing-protein-[Truepera-radiovictrix] |
| WP_221401669.1 | PIN-EndoRNase | GCF_000092425.1 | Truepera_radiovictrix | Bacteria>Thermotogati>Deinococcota>Deinococci>Trueperales>Trueperaceae>Truepera | 144 | TRAD_RS05940 | 1285450..1285884 | 332249 | PIN-domain-containing-protein-[Truepera-radiovictrix] |
| WP_013179176.1 | PIN-EndoRNase | GCF_000092425.1 | Truepera_radiovictrix | Bacteria>Thermotogati>Deinococcota>Deinococci>Trueperales>Trueperaceae>Truepera | 147 | TRAD_RS13540 | 2953249..2953692 | 332249 | type-II-toxin-antitoxin-system-VapC-family-toxin-[Truepera-radiovictrix] |
| MDD3916536.1 | PIN-EndoRNase | GCA_028699385.1 | Synergistaceae_bacterium | Bacteria>Thermotogati>Synergistota>Synergistia>Synergistales>Synergistaceae>unclassified_Synergistaceae | 137 | PHX00_05240 | 15016..15429 | 2053624 | MAG:-PIN-domain-containing-protein-[Synergistaceae-bacterium] |
| MBL3593371.1 | PIN-EndoRNase | GCA_016756725.1 | Synergistaceae_bacterium | Bacteria>Thermotogati>Synergistota>Synergistia>Synergistales>Synergistaceae>unclassified_Synergistaceae | 176 | JMJ93_07655 | 74940..75470 | 2053624 | hypothetical-protein-[Synergistaceae-bacterium] |
| MBK8128937.1 | PIN-EndoRNase | GCA_016712825.1 | bacterium | Bacteria>unclassified_Bacteria | 96 | IPK53_08315 | 34759..35049 | 1869227 | putative-toxin-antitoxin-system-toxin-component,-PIN-family-[bacterium] |
| MCD4760466.1 | PIN-EndoRNase | GCA_021107775.1 | bacterium | Bacteria>unclassified_Bacteria | 104 | K8R42_01065 | 1..315 | 1869227 | PIN-domain-containing-protein-[bacterium] |
| NJL57103.1 | PIN-EndoRNase | GCA_012031555.1 | bacterium | Bacteria>unclassified_Bacteria | 111 | HC928_19630 | 657..992 | 1869227 | DUF5615-family-PIN-like-protein-[bacterium] |
| MCD6327180.1 | PIN-EndoRNase | GCA_021160245.1 | bacterium | Bacteria>unclassified_Bacteria | 115 | J7M28_06470 | 14728..15075 | 1869227 | PIN-domain-containing-protein-[bacterium] |
| MCY3924844.1 | PIN-EndoRNase | GCA_026707305.1 | bacterium | Bacteria>unclassified_Bacteria | 118 | OXG52_04950 | 20662..21018 | 1869227 | MAG:-hypothetical-protein-OXG52_04950-[bacterium] |
| MCY3924846.1 | PIN-EndoRNase | GCA_026707305.1 | bacterium | Bacteria>unclassified_Bacteria | 118 | OXG52_04965 | 23555..23911 | 1869227 | MAG:-hypothetical-protein-OXG52_04965-[bacterium] |
| MCY3953253.1 | PIN-EndoRNase | GCA_026707535.1 | bacterium | Bacteria>unclassified_Bacteria | 118 | OXG69_12040 | 18969..19325 | 1869227 | MAG:-hypothetical-protein-OXG69_12040-[bacterium] |
| MCY4103456.1 | PIN-EndoRNase | GCA_026708055.1 | bacterium | Bacteria>unclassified_Bacteria | 118 | OXG55_09380 | 98828..99184 | 1869227 | MAG:-hypothetical-protein-OXG55_09380-[bacterium] |
| OQA20373.1 | PIN-EndoRNase | GCA_002069435.1 | bacterium_ADurb.Bin363 | Bacteria>unclassified_Bacteria | 119 | BWY64_00207 | 10804..11163 | 1866927 | hypothetical-protein-BWY64_00207-[bacterium-ADurb.Bin363] |
| MBE3604314.1 | PIN-EndoRNase | GCA_014896235.1 | bacterium | Bacteria>unclassified_Bacteria | 124 | IMX07_11835 | 4625..4999 | 1869227 | DUF5615-family-PIN-like-protein-[bacterium] |
| KAF0250482.1 | PIN-EndoRNase | GCA_009773835.1 | bacterium | Bacteria>unclassified_Bacteria | 125 | FD167_108 | 5464..5841 | 1869227 | PilT-domain-containing-protein-[bacterium] |
| MBN2381435.1 | PIN-EndoRNase | GCA_016934675.1 | bacterium | Bacteria>unclassified_Bacteria | 140 | JXQ70_01000 | 6566..6988 | 1869227 | PIN-domain-containing-protein-[bacterium] |
| OQA20377.1 | PIN-EndoRNase | GCA_002069435.1 | bacterium_ADurb.Bin363 | Bacteria>unclassified_Bacteria | 140 | BWY64_00211 | 12107..12529 | 1866927 | hypothetical-protein-BWY64_00211-[bacterium-ADurb.Bin363] |
| MCG3156465.1 | PIN-EndoRNase | GCA_022072305.1 | bacterium | Bacteria>unclassified_Bacteria | 141 | DKINENOH_03089 | 110058..110483 | 1869227 | hypothetical-protein-[bacterium] |
| OQA19978.1 | PIN-EndoRNase | GCA_002069435.1 | bacterium_ADurb.Bin363 | Bacteria>unclassified_Bacteria | 141 | BWY64_00496 | 6682..7107 | 1866927 | PIN-domain-protein-[bacterium-ADurb.Bin363] |
| MBX3024937.1 | PIN-EndoRNase | GCA_019637795.1 | bacterium | Bacteria>unclassified_Bacteria | 146 | KF840_08505 | 742680..743120 | 1869227 | PIN-domain-containing-protein-[bacterium] |
| MCY4163395.1 | PIN-EndoRNase | GCA_026709025.1 | bacterium | Bacteria>unclassified_Bacteria | 146 | OXE93_04175 | 5251..5691 | 1869227 | MAG:-hypothetical-protein-OXE93_04175-[bacterium] |
| MBK8128935.1 | PIN-EndoRNase | GCA_016712825.1 | bacterium | Bacteria>unclassified_Bacteria | 150 | IPK53_08305 | 33963..34415 | 1869227 | type-II-toxin-antitoxin-system-VapC-family-toxin-[bacterium] |
| NLF40199.1 | PIN-EndoRNase | GCA_012514975.1 | bacterium | Bacteria>unclassified_Bacteria | 167 | GX586_12195 | 3969..4472 | 1869227 | type-II-toxin-antitoxin-system-VapC-family-toxin-[bacterium] |
| MBI1298961.1 | PIN-EndoRNase | GCA_016124945.1 | bacterium | Bacteria>unclassified_Bacteria | 176 | GC175_28860 | 15313..15843 | 1869227 | hypothetical-protein-[bacterium] |
| GBE48763.1 | PIN-EndoRNase | GCA_002897795.1 | bacterium_BMS3Bbin12 | Bacteria>unclassified_Bacteria | 269 | BMS3Bbin12_01951 | 21476..22285 | 2005735 | NYN-domain-protein-[bacterium-BMS3Bbin12] |

| Protein-ID | Domain | Assembly-ID | Strain | Taxonomy | Length | Locus-Tag | Genomic-coordinates | TaxID | Fasta-defline |
|---|---|---|---|---|---|---|---|---|---|
| --CR-ATPase+REase | | | | | | | | | |
| WP_255153068.1 | CR-ATPase+REase | GCF_024298865.1 | Halorarius_halobius | Archaea>Methanobacteriati>Methanobacteriota>Stenosarchaea_group>Halobacteria>Halobacteriales>Haloarculaceae>Halorarius | 568 | NL112_RS22035 | 43888..45594 | 2962671 | AAA-family-ATPase-[Halosegnis-sp.-ZY10] |
| WP_235728495.1 | CR-ATPase+REase | - | Halosimplex_carlsbadense | Archaea>Methanobacteriati>Methanobacteriota>Stenosarchaea_group>Halobacteria>Halobacteriales>Haloarculaceae>Halosimplex | 570 | C475_RS21860 | 68287..69999 | 171164 | AAA-family-ATPase-[Halosimplex-carlsbadense] |
| ELZ19703.1 | CR-ATPase+REase | GCA_000337455.1 | Halosimplex_carlsbadense_2-9-1 | Archaea>Methanobacteriati>Methanobacteriota>Stenosarchaea_group>Halobacteria>Halobacteriales>Haloarculaceae>Halosimplex>Halosimplex_carlsbadense | 569 | C475_22199 | 68287..69996 | 797114 | hypothetical-protein-C475_22199-[Halosimplex-carlsbadense-2-9-1] |
| WP_257301022.1 | CR-ATPase+REase | GCF_024609245.1 | Haloarchaeobius_sp._FL176 | Archaea>Methanobacteriati>Methanobacteriota>Stenosarchaea_group>Halobacteria>Halobacteriales>Halorubellaceae>Haloarchaeobius>unclassified_Haloarchaeobius | 520 | NO978_RS18545 | 17116..18678 | 2967129 | AAA-family-ATPase-[Haloarchaeobius-sp.-FL176] |
| WP_004078480.1 | CR-ATPase+REase | GCF_000243255.1 | Methanoplanus_limicola | Archaea>Methanobacteriati>Methanobacteriota>Stenosarchaea_group>Methanomicrobia>Methanomicrobiales>Methanomicrobiaceae>Methanoplanus | 447 | METLIM_RS10775 | 2290960..2292303 | 2315 | ATP-binding-protein-[Methanoplanus-limicola] |
| WP_048067427.1 | CR-ATPase+REase | GCF_000013445.1 | Methanospirillum_hungatei | Archaea>Methanobacteriati>Methanobacteriota>Stenosarchaea_group>Methanomicrobia>Methanomicrobiales>Methanospirillaceae>Methanospirillum | 576 | MHUN_RS09815 | 2131092..2132822 | 2203 | ATP-binding-protein-[Methanospirillum-hungatei] |
| WP_044088296.1 | CR-ATPase+REase | GCF_000771685.1 | Bifidobacterium | Bacteria>Bacillati>Actinomycetota>Actinomycetes>Bifidobacteriales>Bifidobacteriaceae | 408 | EM092_RS08285 | 1523..2749 | 1678 | MULTISPECIES:-ATP-binding-protein-[Bifidobacterium] |
| WP_214374601.1 | CR-ATPase+REase | GCF_018555335.1 | Bifidobacterium | Bacteria>Bacillati>Actinomycetota>Actinomycetes>Bifidobacteriales>Bifidobacteriaceae | 408 | JS541_RS11265 | 56193..57419 | 1678 | MULTISPECIES:-ATP-binding-protein-[Bifidobacterium] |
| WP_226829912.1 | CR-ATPase+REase | GCF_011764605.1 | Bifidobacterium | Bacteria>Bacillati>Actinomycetota>Actinomycetes>Bifidobacteriales>Bifidobacteriaceae | 227 | PGQ90_RS07440 | 216716..217399 | 1678 | MULTISPECIES:-hypothetical-protein-[Bifidobacterium] |
| WP_193852558.1 | CR-ATPase+REase | GCF_003129905.1 | Bifidobacterium_callitrichidarum | Bacteria>Bacillati>Actinomycetota>Actinomycetes>Bifidobacteriales>Bifidobacteriaceae>Bifidobacterium | 431 | DF196_RS03610 | 117895..119190 | 2052941 | ATP-binding-protein-[Bifidobacterium-callitrichidarum] |
| WP_151916620.1 | CR-ATPase+REase | GCF_009078285.1 | Bifidobacterium_jacchi | Bacteria>Bacillati>Actinomycetota>Actinomycetes>Bifidobacteriales>Bifidobacteriaceae>Bifidobacterium | 424 | EHS19_RS04640 | 37982..39256 | 2490545 | ATP-binding-protein-[Bifidobacterium-jacchi] |
| WP_094668335.1 | CR-ATPase+REase | GCF_002259745.1 | Bifidobacterium_myosotis | Bacteria>Bacillati>Actinomycetota>Actinomycetes>Bifidobacteriales>Bifidobacteriaceae>Bifidobacterium | 408 | BMYO_RS10070 | 7645..8871 | 1630166 | ATP-binding-protein-[Bifidobacterium-myosotis] |
| WP_140463092.1 | CR-ATPase+REase | GCF_006408915.1 | unclassified_Bifidobacterium | Bacteria>Bacillati>Actinomycetota>Actinomycetes>Bifidobacteriales>Bifidobacteriaceae>Bifidobacterium | 408 | BW09_RS08250 | 80457..81683 | 2608897 | MULTISPECIES:-ATP-binding-protein-[unclassified-Bifidobacterium] |
| WP_150379197.1 | CR-ATPase+REase | GCF_008698155.1 | Bifidobacterium_myosotis | Bacteria>Bacillati>Actinomycetota>Actinomycetes>Bifidobacteriales>Bifidobacteriaceae>Bifidobacterium | 408 | EM091_RS06005 | 87521..88747 | 1630166 | ATP-binding-protein-[Bifidobacterium-myosotis] |
| WP_151916616.1 | CR-ATPase+REase | GCF_009078285.1 | Bifidobacterium_jacchi | Bacteria>Bacillati>Actinomycetota>Actinomycetes>Bifidobacteriales>Bifidobacteriaceae>Bifidobacterium | 408 | EHS19_RS04615 | 30489..31715 | 2490545 | ATP-binding-protein-[Bifidobacterium-jacchi] |
| WP_152233648.1 | CR-ATPase+REase | GCF_042657605.1 | Bifidobacterium_leontopitheci | Bacteria>Bacillati>Actinomycetota>Actinomycetes>Bifidobacteriales>Bifidobacteriaceae>Bifidobacterium | 407 | F7D09_RS01355 | 127889..129112 | 2650774 | ATP-binding-protein-[Bifidobacterium-leontopitheci] |
| WP_169274837.1 | CR-ATPase+REase | GCF_012932365.1 | Bifidobacterium_moraviense | Bacteria>Bacillati>Actinomycetota>Actinomycetes>Bifidobacteriales>Bifidobacteriaceae>Bifidobacterium | 407 | G1C96_RS01300 | 292352..293575 | 2675323 | ATP-binding-protein-[Bifidobacterium-sp.-DSM-109958] |
| WP_051923733.1 | CR-ATPase+REase | GCF_000771645.1 | Bifidobacterium_biavatii | Bacteria>Bacillati>Actinomycetota>Actinomycetes>Bifidobacteriales>Bifidobacteriaceae>Bifidobacterium | 405 | OU23_RS06265 | 16493..17710 | 762212 | ATP-binding-protein-[Bifidobacterium-biavatii] |
| WP_230845803.1 | CR-ATPase+REase | GCF_001275745.1 | Bifidobacterium_longum | Bacteria>Bacillati>Actinomycetota>Actinomycetes>Bifidobacteriales>Bifidobacteriaceae>Bifidobacterium | 227 | BLOI2_RS03195 | 190405..191088 | 216816 | hypothetical-protein-[Bifidobacterium-longum] |
| WP_254291659.1 | CR-ATPase+REase | GCF_019041665.1 | Bifidobacterium_longum | Bacteria>Bacillati>Actinomycetota>Actinomycetes>Bifidobacteriales>Bifidobacteriaceae>Bifidobacterium | 134 | KTF47_RS04055 | 22752..23156 | 216816 | ATP-binding-protein-[Bifidobacterium-longum] |
| MBS6134851.1 | CR-ATPase+REase | GCA_018374015.1 | Bifidobacterium_longum | Bacteria>Bacillati>Actinomycetota>Actinomycetes>Bifidobacteriales>Bifidobacteriaceae>Bifidobacterium | 104 | KH705_09810 | 15927..16241 | 216816 | ATP-binding-protein-[Bifidobacterium-longum] |
| MCI2117725.1 | CR-ATPase+REase | GCA_022653935.1 | Bifidobacterium_breve | Bacteria>Bacillati>Actinomycetota>Actinomycetes>Bifidobacteriales>Bifidobacteriaceae>Bifidobacterium | 90 | LKK17_00150 | 35149..35421 | 1685 | ATP-binding-protein-[Bifidobacterium-breve] |
| WP_269338051.1 | CR-ATPase+REase | GCF_027215525.1 | Bifidobacterium_breve | Bacteria>Bacillati>Actinomycetota>Actinomycetes>Bifidobacteriales>Bifidobacteriaceae>Bifidobacterium | 90 | O4C30_RS07890 | 18445..18717 | 1685 | ATP-binding-protein-[Bifidobacterium-breve] |
| WP_234883583.1 | CR-ATPase+REase | GCF_902167555.1 | Bifidobacterium_longum | Bacteria>Bacillati>Actinomycetota>Actinomycetes>Bifidobacteriales>Bifidobacteriaceae>Bifidobacterium | 53 | NBW58_RS09940 | 16824..16985 | 216816 | hypothetical-protein-[Bifidobacterium-longum] |
| WP_229087696.1 | CR-ATPase+REase | GCF_017312625.1 | Bifidobacterium_longum | Bacteria>Bacillati>Actinomycetota>Actinomycetes>Bifidobacteriales>Bifidobacteriaceae>Bifidobacterium | 47 | MCC10048_RS11340 | 3095..3238 | 216816 | hypothetical-protein-[Bifidobacterium-longum] |
| WP_224302002.1 | CR-ATPase+REase | GCF_020092605.1 | Streptomyces_olivaceus | Bacteria>Bacillati>Actinomycetota>Actinomycetes>Kitasatosporales>Streptomycetaceae>Streptomyces | 418 | KVH22_RS20555 | 334206..335462 | 47716 | ATP-binding-protein-[Streptomyces-olivaceus] |
| MBF0599295.1 | CR-ATPase+REase | GCA_015234195.1 | Atopobiaceae_bacterium_FL090493 | Bacteria>Bacillati>Actinomycetota>Coriobacteriia>Coriobacteriales>Atopobiaceae>unclassified_Atopobiaceae | 197 | IM763_04385 | 104776..105369 | 2661918 | DUF4143-domain-containing-protein-[Atopobiaceae-bacterium-FL090493] |
| MBR2835997.1 | CR-ATPase+REase | GCA_017515595.1 | Coriobacteriales_bacterium | Bacteria>Bacillati>Actinomycetota>Coriobacteriia>Coriobacteriales>unclassified_Coriobacteriales | 431 | IKE43_09875 | 817..2112 | 2478480 | ATP-binding-protein-[Coriobacteriales-bacterium] |
| MBR2836002.1 | CR-ATPase+REase | GCA_017515595.1 | Coriobacteriales_bacterium | Bacteria>Bacillati>Actinomycetota>Coriobacteriia>Coriobacteriales>unclassified_Coriobacteriales | 528 | IKE43_09900 | 10113..11699 | 2478480 | AAA-family-ATPase-[Coriobacteriales-bacterium] |
| WP_251231673.1 | CR-ATPase+REase | GCF_943193145.1 | Adlercreutzia_aquisgranensis | Bacteria>Bacillati>Actinomycetota>Coriobacteriia>Eggerthellales>Eggerthellaceae>Adlercreutzia | 420 | NFR41_RS09590 | 25598..26860 | 2941323 | DUF4143-domain-containing-protein-[Adlercreutzia-aquisgranensis] |
| WP_251851847.1 | CR-ATPase+REase | GCF_023744155.1 | Enterococcus_italicus | Bacteria>Bacillati>Bacillota>Bacilli>Lactobacillales>Enterococcaceae>Enterococcus | 402 | NE261_RS07345 | 545589..546797 | 246144 | ATP-binding-protein-[Enterococcus-italicus] |
| WP_257411547.1 | CR-ATPase+REase | GCF_024622885.1 | Clostridium_sporogenes | Bacteria>Bacillati>Bacillota>Clostridia>Eubacteriales>Clostridiaceae>Clostridium | 564 | NSA52_RS02855 | 12449..14143 | 1509 | ATP-binding-protein-[Clostridium-sporogenes] |
| WP_156211828.1 | CR-ATPase+REase | - | Clostridium_butyricum | Bacteria>Bacillati>Bacillota>Clostridia>Eubacteriales>Clostridiaceae>Clostridium | 517 | GND98_RS05140 | 53139..54692 | 1492 | AAA-family-ATPase-[Clostridium-butyricum] |
| WP_039258637.1 | CR-ATPase+REase | GCF_000768655.1 | Clostridium_botulinum | Bacteria>Bacillati>Bacillota>Clostridia>Eubacteriales>Clostridiaceae>Clostridium | 513 | ADU88_RS01570 | 71161..72702 | 1491 | AAA-family-ATPase-[Clostridium-botulinum] |
| MBN1067553.1 | CR-ATPase+REase | GCA_016925955.1 | Clostridium_botulinum | Bacteria>Bacillati>Bacillota>Clostridia>Eubacteriales>Clostridiaceae>Clostridium | 504 | DVW02_06045 | 100707..102221 | 1491 | AAA-family-ATPase-[Clostridium-botulinum] |
| NFO13000.1 | CR-ATPase+REase | GCA_011016315.1 | Clostridium_botulinum | Bacteria>Bacillati>Bacillota>Clostridia>Eubacteriales>Clostridiaceae>Clostridium | 504 | FDB34_02030 | 200371..201885 | 1491 | AAA-family-ATPase-[Clostridium-botulinum] |
| WP_164949150.1 | CR-ATPase+REase | GCF_011065955.2 | Clostridium_sp._FP2 | Bacteria>Bacillati>Bacillota>Clostridia>Eubacteriales>Clostridiaceae>Clostridium>unclassified_Clostridium | 566 | G9F71_RS17985 | 3741873..3743573 | 2724481 | ATP-binding-protein-[Clostridium-sp.-FP2] |
| UYJ11708.1 | CR-ATPase+REase | GCA_025757845.1 | Oscillospiraceae_bacterium | Bacteria>Bacillati>Bacillota>Clostridia>Eubacteriales>Oscillospiraceae>unclassified_Oscillospiraceae | 399 | OGM78_02665 | 561691..562890 | 2485925 | MAG:-ATP-binding-protein-[Oscillospiraceae-bacterium] |
| MBP1566398.1 | CR-ATPase+REase | GCA_017848115.1 | Oscillospiraceae_bacterium | Bacteria>Bacillati>Bacillota>Clostridia>Eubacteriales>Oscillospiraceae>unclassified_Oscillospiraceae | 563 | J6A58_11650 | 45413..47104 | 2485925 | AAA-family-ATPase-[Oscillospiraceae-bacterium] |
| UYJ11706.1 | CR-ATPase+REase | GCA_025757845.1 | Oscillospiraceae_bacterium | Bacteria>Bacillati>Bacillota>Clostridia>Eubacteriales>Oscillospiraceae>unclassified_Oscillospiraceae | 557 | OGM78_02655 | 559122..560795 | 2485925 | MAG:-ATP-binding-protein-[Oscillospiraceae-bacterium] |
| MBP1566411.1 | CR-ATPase+REase | GCA_017848115.1 | Oscillospiraceae_bacterium | Bacteria>Bacillati>Bacillota>Clostridia>Eubacteriales>Oscillospiraceae>unclassified_Oscillospiraceae | 523 | J6A58_11715 | 66769..68340 | 2485925 | AAA-family-ATPase-[Oscillospiraceae-bacterium] |
| WP_207635334.1 | CR-ATPase+REase | - | Syntrophomonas_wolfei | Bacteria>Bacillati>Bacillota>Clostridia>Eubacteriales>Syntrophomonadaceae>Syntrophomonas | 403 | SWOL_RS05000 | 1116333..1117544 | 863 | ATP-binding-protein-[Syntrophomonas-wolfei] |
| MCR4615477.1 | CR-ATPase+REase | GCA_024684435.1 | Clostridiales_bacterium | Bacteria>Bacillati>Bacillota>Clostridia>Eubacteriales>unclassified_Eubacteriales | 410 | K5756_04940 | 68551..69783 | 1898207 | ATP-binding-protein-[Clostridiales-bacterium] |
| MBD5489939.1 | CR-ATPase+REase | GCA_014804485.1 | Lachnospiraceae_bacterium | Bacteria>Bacillati>Bacillota>Clostridia>Lachnospirales>Lachnospiraceae>unclassified_Lachnospiraceae | 628 | HDR13_14305 | 14972..16858 | 1898203 | AAA-family-ATPase-[Lachnospiraceae-bacterium] |
| MBQ6380319.1 | CR-ATPase+REase | GCA_017430965.1 | Clostridia_bacterium | Bacteria>Bacillati>Bacillota>Clostridia>unclassified_Clostridia | 410 | IJJ41_01785 | 31675..32907 | 2044939 | ATP-binding-protein-[Clostridia-bacterium] |
| GJM58068.1 | CR-ATPase+REase | GCA_022834735.1 | Erysipelotrichaceae_bacterium_OPF54 | Bacteria>Bacillati>Bacillota>Erysipelotrichia>Erysipelotrichales>Erysipelotrichaceae>unclassified_Erysipelotrichaceae | 403 | EROP_17610 | 1797323..1798534 | 2894072 | ATPase-[Erysipelotrichaceae-bacterium-OPF54] |
| MBE6101271.1 | CR-ATPase+REase | GCA_015058655.1 | Selenomonas_ruminantium | Bacteria>Bacillati>Bacillota>Negativicutes>Selenomonadales>Selenomonadaceae>Selenomonas | 523 | E7200_04110 | 50874..52445 | 971 | AAA-family-ATPase-[Selenomonas-ruminantium] |
| WP_021167320.1 | CR-ATPase+REase | - | Sporomusa_ovata | Bacteria>Bacillati>Bacillota>Negativicutes>Selenomonadales>Sporomusaceae>Sporomusa | 192 | SOV_RS04390 | 939733..940311 | 2378 | DUF4143-domain-containing-protein-[Sporomusa-ovata] |
| MCY3660413.1 | CR-ATPase+REase | GCA_026705565.1 | Caldilineaceae_bacterium | Bacteria>Bacillati>Chloroflexota>Caldilineae>Caldilineales>Caldilineaceae>unclassified_Caldilineaceae | 620 | OXG36_15525 | 4594..6456 | 2614250 | MAG:-AAA-family-ATPase-[Caldilineaceae-bacterium] |
| MBS3030292.1 | CR-ATPase+REase | GCA_018295985.1 | Dolichospermum_sp._DET66 | Bacteria>Bacillati>Cyanobacteriota/Melainabacteria_group>Cyanobacteriota>Cyanophyceae>Nostocales>Aphanizomenonaceae>Dolichospermum>unclassified_Dolichospermum | 516 | HCA26_25865 | 5391839..5393389 | 2704415 | AAA-family-ATPase-[Dolichospermum-sp.-DET66] |
| WP_200988279.1 | CR-ATPase+REase | - | Anabaenopsis_elenkinii | Bacteria>Bacillati>Cyanobacteriota/Melainabacteria_group>Cyanobacteriota>Cyanophyceae>Nostocales>Nodulariaceae>Anabaenopsis | 518 | IM676_RS18800 | 4306752..4308308 | 156213 | ATP-binding-protein-[Anabaenopsis-elenkinii] |
| WP_022069631.1 | CR-ATPase+REase | GCF_023383545.1 | Fusobacterium | Bacteria>Fusobacteriati>Fusobacteriota>Fusobacteriia>Fusobacteriales>Fusobacteriaceae | 542 | YWH7053_RS01955 | 12814..14442 | 848 | AAA-family-ATPase-[Fusobacterium-nucleatum] |
| WP_032836025.1 | CR-ATPase+REase | GCF_037904605.1 | Fusobacterium | Bacteria>Fusobacteriati>Fusobacteriota>Fusobacteriia>Fusobacteriales>Fusobacteriaceae | 542 | HMPREF1501_RS01140 | 50614..52242 | 848 | AAA-family-ATPase-[Fusobacterium-sp.-OBRC1] |
| WP_032838652.1 | CR-ATPase+REase | GCF_037911025.1 | Fusobacterium | Bacteria>Fusobacteriati>Fusobacteriota>Fusobacteriia>Fusobacteriales>Fusobacteriaceae | 542 | HMPREF1498_RS02615 | 28137..29765 | 848 | AAA-family-ATPase-[Fusobacterium-sp.-CM1] |
| WP_238972700.1 | CR-ATPase+REase | GCF_037911825.1 | Fusobacterium | Bacteria>Fusobacteriati>Fusobacteriota>Fusobacteriia>Fusobacteriales>Fusobacteriaceae | 534 | K5I19_RS02540 | 142791..144395 | 848 | AAA-family-ATPase-[Fusobacterium-nucleatum] |
| WP_005980881.1 | CR-ATPase+REase | GCF_902373815.1 | Fusobacterium_ulcerans | Bacteria>Fusobacteriati>Fusobacteriota>Fusobacteriia>Fusobacteriales>Fusobacteriaceae>Fusobacterium | 547 | FXX38_RS11810 | 318647..320290 | 861 | AAA-family-ATPase-[Fusobacterium-ulcerans] |
| MCG6839740.1 | CR-ATPase+REase | GCA_022340045.1 | Fusobacterium_nucleatum | Bacteria>Fusobacteriati>Fusobacteriota>Fusobacteriia>Fusobacteriales>Fusobacteriaceae>Fusobacterium | 542 | K5I20_00255 | 54559..56187 | 851 | ATP-binding-protein-[Fusobacterium-nucleatum] |
| WP_008692044.1 | CR-ATPase+REase | GCF_000218655.1 | Fusobacterium_animalis | Bacteria>Fusobacteriati>Fusobacteriota>Fusobacteriia>Fusobacteriales>Fusobacteriaceae>Fusobacterium | 542 | HMPREF0401_RS10090 | 570389..572017 | 76859 | AAA-family-ATPase-[Fusobacterium-animalis] |
| WP_029493947.1 | CR-ATPase+REase | GCF_000455885.1 | Fusobacterium_hwasookii | Bacteria>Fusobacteriati>Fusobacteriota>Fusobacteriia>Fusobacteriales>Fusobacteriaceae>Fusobacterium | 542 | J143_RS0110830 | 58392..60020 | 1583098 | AAA-family-ATPase-[Fusobacterium-hwasookii] |
| WP_167381728.1 | CR-ATPase+REase | GCF_002763695.1 | Fusobacterium_pseudoperiodonticum | Bacteria>Fusobacteriati>Fusobacteriota>Fusobacteriia>Fusobacteriales>Fusobacteriaceae>Fusobacterium | 542 | CTM74_RS08975 | 1710113..1711741 | 2663009 | AAA-family-ATPase-[Fusobacterium-pseudoperiodonticum] |
| WP_249524179.1 | CR-ATPase+REase | GCF_023383605.1 | Fusobacterium_nucleatum | Bacteria>Fusobacteriati>Fusobacteriota>Fusobacteriia>Fusobacteriales>Fusobacteriaceae>Fusobacterium | 542 | YWH7056_RS07015 | 17440..19068 | 851 | AAA-family-ATPase-[Fusobacterium-nucleatum] |
| WP_249525220.1 | CR-ATPase+REase | GCF_023383525.1 | Fusobacterium_nucleatum | Bacteria>Fusobacteriati>Fusobacteriota>Fusobacteriia>Fusobacteriales>Fusobacteriaceae>Fusobacterium | 542 | YWH7054_RS02280 | 10456..12084 | 851 | AAA-family-ATPase-[Fusobacterium-nucleatum] |
| WP_271333588.1 | CR-ATPase+REase | - | Fusobacterium_nucleatum | Bacteria>Fusobacteriati>Fusobacteriota>Fusobacteriia>Fusobacteriales>Fusobacteriaceae>Fusobacterium | 542 | PGW91_RS03570 | 733447..735075 | 851 | AAA-family-ATPase-[Fusobacterium-nucleatum] |
| WP_118127446.1 | CR-ATPase+REase | GCF_021532115.1 | Fusobacterium_mortiferum | Bacteria>Fusobacteriati>Fusobacteriota>Fusobacteriia>Fusobacteriales>Fusobacteriaceae>Fusobacterium | 540 | DW663_RS02780 | 242532..244154 | 850 | AAA-family-ATPase-[Fusobacterium-mortiferum] |
| WP_008793545.1 | CR-ATPase+REase | GCF_902373915.1 | Fusobacterium_periodonticum | Bacteria>Fusobacteriati>Fusobacteriota>Fusobacteriia>Fusobacteriales>Fusobacteriaceae>Fusobacterium | 392 | FY302_RS03005 | 629855..631033 | 860 | AAA-family-ATPase-[Fusobacterium-periodonticum] |
| WP_100024176.1 | CR-ATPase+REase | GCF_002763625.1 | Fusobacterium_pseudoperiodonticum | Bacteria>Fusobacteriati>Fusobacteriota>Fusobacteriia>Fusobacteriales>Fusobacteriaceae>Fusobacterium | 392 | CTM72_RS01110 | 211000..212178 | 2663009 | AAA-family-ATPase-[Fusobacterium-pseudoperiodonticum] |
| MCF2639764.1 | CR-ATPase+REase | GCA_021531615.1 | Fusobacterium_varium | Bacteria>Fusobacteriati>Fusobacteriota>Fusobacteriia>Fusobacteriales>Fusobacteriaceae>Fusobacterium | 430 | I6E31_07245 | 17183..18475 | 856 | ATP-binding-protein-[Fusobacterium-varium] |
| CDA09053.1 | CR-ATPase+REase | GCA_000433695.1 | Fusobacterium_sp._CAG:649 | Bacteria>Fusobacteriati>Fusobacteriota>Fusobacteriia>Fusobacteriales>Fusobacteriaceae>Fusobacterium>unclassified_Fusobacterium | 542 | BN748_00877 | 12155..13783 | 1262900 | putative-uncharacterized-protein-[Fusobacterium-sp.-CAG:649] |
| KAA0252889.1 | CR-ATPase+REase | GCA_008363385.1 | Acidobacteriota_bacterium | Bacteria>Pseudomonadati>Acidobacteriota>unclassified_Acidobacteriota | 402 | EDX89_16495 | 10949..12157 | 1978231 | ATP-binding-protein-[Acidobacteriota-bacterium] |
| WP_013460553.1 | CR-ATPase+REase | GCF_000183725.1 | Sulfuricurvum_kujiense | Bacteria>Pseudomonadati>Campylobacterota>Epsilonproteobacteria>Campylobacterales>Sulfurimonadaceae>Sulfuricurvum | 427 | SULKU_RS08510 | 1703077..1704360 | 148813 | ATP-binding-protein-[Sulfuricurvum-kujiense] |
| WP_258972459.1 | CR-ATPase+REase | GCF_910578265.1 | Bacteroidaceae | Bacteria>Pseudomonadati>FCB_group>Bacteroidota/Chlorobiota_group>Bacteroidota>Bacteroidia>Bacteroidales | 463 | NXX32_RS11585 | 3203234..3204625 | 815 | MULTISPECIES:-ATP-binding-protein-[Bacteroidaceae] |
| WP_250702184.1 | CR-ATPase+REase | GCF_023668595.1 | Bacteroides_ovatus | Bacteria>Pseudomonadati>FCB_group>Bacteroidota/Chlorobiota_group>Bacteroidota>Bacteroidia>Bacteroidales>Bacteroidaceae>Bacteroides | 394 | NCZ26_RS08940 | 94819..96003 | 28116 | ATP-binding-protein-[Bacteroides-ovatus] |
| WP_167963297.1 | CR-ATPase+REase | GCF_012113595.1 | Bacteroides_faecium | Bacteria>Pseudomonadati>FCB_group>Bacteroidota/Chlorobiota_group>Bacteroidota>Bacteroidia>Bacteroidales>Bacteroidaceae>Bacteroides | 452 | BacF7301_RS12640 | 3538837..3540195 | 2715212 | AAA-family-ATPase-[Bacteroides-faecium] |
| MBO4331561.1 | CR-ATPase+REase | GCA_017613615.1 | Paludibacteraceae_bacterium | Bacteria>Pseudomonadati>FCB_group>Bacteroidota/Chlorobiota_group>Bacteroidota>Bacteroidia>Bacteroidales>Paludibacteraceae>unclassified_Paludibacteraceae | 520 | J5875_00145 | 40017..41579 | 2748031 | AAA-family-ATPase-[Paludibacteraceae-bacterium] |
| MBR6041784.1 | CR-ATPase+REase | GCA_017534895.1 | Paludibacteraceae_bacterium | Bacteria>Pseudomonadati>FCB_group>Bacteroidota/Chlorobiota_group>Bacteroidota>Bacteroidia>Bacteroidales>Paludibacteraceae>unclassified_Paludibacteraceae | 520 | IKP37_04025 | 44658..46220 | 2748031 | AAA-family-ATPase-[Paludibacteraceae-bacterium] |
| WP_235402039.1 | CR-ATPase+REase | GCF_021531655.1 | Leyella_stercorea | Bacteria>Pseudomonadati>FCB_group>Bacteroidota/Chlorobiota_group>Bacteroidota>Bacteroidia>Bacteroidales>Prevotellaceae>Leyella | 429 | I6E49_RS09805 | 31109..32398 | 363265 | DUF4143-domain-containing-protein-[Prevotella-stercorea] |
| WP_235392640.1 | CR-ATPase+REase | GCF_021531355.1 | Leyella_stercorea | Bacteria>Pseudomonadati>FCB_group>Bacteroidota/Chlorobiota_group>Bacteroidota>Bacteroidia>Bacteroidales>Prevotellaceae>Leyella | 529 | I6E38_RS03175 | 37413..39002 | 363265 | ATP-binding-protein-[Prevotella-stercorea] |
| WP_235392649.1 | CR-ATPase+REase | GCF_021531355.1 | Leyella_stercorea | Bacteria>Pseudomonadati>FCB_group>Bacteroidota/Chlorobiota_group>Bacteroidota>Bacteroidia>Bacteroidales>Prevotellaceae>Leyella | 519 | I6E38_RS03225 | 54779..56338 | 363265 | ATP-binding-protein-[Prevotella-stercorea] |
| MBQ7946200.1 | CR-ATPase+REase | GCA_017468435.1 | Bacteroidales_bacterium | Bacteria>Pseudomonadati>FCB_group>Bacteroidota/Chlorobiota_group>Bacteroidota>Bacteroidia>Bacteroidales>unclassified_Bacteroidales | 487 | IJ280_05755 | 7302..8765 | 2030927 | ATP-binding-protein-[Bacteroidales-bacterium] |
| MBP3741085.1 | CR-ATPase+REase | GCA_017940305.1 | Bacteroidales_bacterium | Bacteria>Pseudomonadati>FCB_group>Bacteroidota/Chlorobiota_group>Bacteroidota>Bacteroidia>Bacteroidales>unclassified_Bacteroidales | 424 | J6I41_08220 | 73623..74897 | 2030927 | ATP-binding-protein-[Bacteroidales-bacterium] |
| MBQ7062981.1 | CR-ATPase+REase | GCA_017442615.1 | Bacteroidales_bacterium | Bacteria>Pseudomonadati>FCB_group>Bacteroidota/Chlorobiota_group>Bacteroidota>Bacteroidia>Bacteroidales>unclassified_Bacteroidales | 408 | IJM88_06810 | 34452..35678 | 2030927 | ATP-binding-protein-[Bacteroidales-bacterium] |
| MBQ8703687.1 | CR-ATPase+REase | GCA_017480785.1 | Bacteroidales_bacterium | Bacteria>Pseudomonadati>FCB_group>Bacteroidota/Chlorobiota_group>Bacteroidota>Bacteroidia>Bacteroidales>unclassified_Bacteroidales | 408 | IJ524_04880 | 125488..126714 | 2030927 | ATP-binding-protein-[Bacteroidales-bacterium] |
| MBQ7238831.1 | CR-ATPase+REase | GCA_017445605.1 | Bacteroidales_bacterium | Bacteria>Pseudomonadati>FCB_group>Bacteroidota/Chlorobiota_group>Bacteroidota>Bacteroidia>Bacteroidales>unclassified_Bacteroidales | 394 | IJS20_08570 | 15528..16712 | 2030927 | ATP-binding-protein-[Bacteroidales-bacterium] |
| MBQ8703690.1 | CR-ATPase+REase | GCA_017480785.1 | Bacteroidales_bacterium | Bacteria>Pseudomonadati>FCB_group>Bacteroidota/Chlorobiota_group>Bacteroidota>Bacteroidia>Bacteroidales>unclassified_Bacteroidales | 379 | IJ524_04895 | 130269..131408 | 2030927 | ATP-binding-protein-[Bacteroidales-bacterium] |
| MBQ8703683.1 | CR-ATPase+REase | GCA_017480785.1 | Bacteroidales_bacterium | Bacteria>Pseudomonadati>FCB_group>Bacteroidota/Chlorobiota_group>Bacteroidota>Bacteroidia>Bacteroidales>unclassified_Bacteroidales | 251 | IJ524_04860 | 117187..117942 | 2030927 | AAA-family-ATPase-[Bacteroidales-bacterium] |
| MCR5064648.1 | CR-ATPase+REase | GCA_024688465.1 | Bacteroidales_bacterium | Bacteria>Pseudomonadati>FCB_group>Bacteroidota/Chlorobiota_group>Bacteroidota>Bacteroidia>Bacteroidales>unclassified_Bacteroidales | 438 | K6A67_02655 | 10017..11333 | 2030927 | AAA-family-ATPase-[Bacteroidales-bacterium] |
| MCB0520338.1 | CR-ATPase+REase | GCA_020436325.1 | Saprospiraceae_bacterium | Bacteria>Pseudomonadati>FCB_group>Bacteroidota/Chlorobiota_group>Bacteroidota>Saprospiria>Saprospirales>Saprospiraceae>unclassified_Saprospiraceae | 390 | KDD27_15475 | 19990..21162 | 2202734 | ATP-binding-protein-[Saprospiraceae-bacterium] |
| MCB0524511.1 | CR-ATPase+REase | GCA_020436385.1 | Saprospiraceae_bacterium | Bacteria>Pseudomonadati>FCB_group>Bacteroidota/Chlorobiota_group>Bacteroidota>Saprospiria>Saprospirales>Saprospiraceae>unclassified_Saprospiraceae | 375 | KDC86_08300 | 25853..26980 | 2202734 | ATP-binding-protein-[Saprospiraceae-bacterium] |
| MBF0105430.1 | CR-ATPase+REase | GCA_015228875.1 | Deltaproteobacteria_bacterium | Bacteria>Pseudomonadati>Myxococcota>Myxococcia>unclassified_Myxococcia | 417 | HQM16_08905 | 13245..14498 | 2026735 | ATP-binding-protein-[Deltaproteobacteria-bacterium] |
| NMA29240.1 | CR-ATPase+REase | GCA_012799975.1 | Burkholderiales_bacterium | Bacteria>Pseudomonadati>Pseudomonadota>Betaproteobacteria>Burkholderiales>unclassified_Burkholderiales | 451 | GX934_15905 | 3104..4459 | 1891238 | ATP-binding-protein-[Burkholderiales-bacterium] |
| WP_001255534.1 | CR-ATPase+REase | GCF_000308235.1 | Acinetobacter | Bacteria>Pseudomonadati>Pseudomonadota>Gammaproteobacteria>Moraxellales>Moraxellaceae | 410 | OYK78_RS17965 | 3414..4646 | 469 | MULTISPECIES:-ATP-binding-protein-[Acinetobacter] |
| WP_205779677.1 | CR-ATPase+REase | GCF_017132735.1 | Acinetobacter_indicus | Bacteria>Pseudomonadati>Pseudomonadota>Gammaproteobacteria>Moraxellales>Moraxellaceae>Acinetobacter | 410 | JYB86_RS06340 | 1322291..1323523 | 756892 | ATP-binding-protein-[Acinetobacter-indicus] |
| WP_206896794.1 | CR-ATPase+REase | GCF_017309445.1 | Acinetobacter_indicus | Bacteria>Pseudomonadati>Pseudomonadota>Gammaproteobacteria>Moraxellales>Moraxellaceae>Acinetobacter | 410 | JOW33_RS13770 | 2883424..2884656 | 756892 | ATP-binding-protein-[Acinetobacter-indicus] |
| WP_109291737.1 | CR-ATPase+REase | GCF_048150035.1 | Acinetobacter_baumannii | Bacteria>Pseudomonadati>Pseudomonadota>Gammaproteobacteria>Moraxellales>Moraxellaceae>Acinetobacter>Acinetobacter_calcoaceticus/baumannii_complex | 410 | DF172_RS10475 | 548..1780 | 470 | ATP-binding-protein-[Acinetobacter-baumannii] |
| WP_200291402.1 | CR-ATPase+REase | GCF_016599775.1 | Stutzerimonas_balearica | Bacteria>Pseudomonadati>Pseudomonadota>Gammaproteobacteria>Pseudomonadales>Pseudomonadaceae>Stutzerimonas>Stutzerimonas_stutzeri_group | 416 | I6H70_RS19235 | 4087280..4088530 | 74829 | ATP-binding-protein-[Stutzerimonas-balearica] |
| WP_002710596.1 | CR-ATPase+REase | GCF_000260135.1 | Thiothrix_nivea | Bacteria>Pseudomonadati>Pseudomonadota>Gammaproteobacteria>Thiotrichales>Thiotrichaceae>Thiothrix | 520 | THINI_RS21445 | 4311323..4312885 | 1031 | ATP-binding-protein-[Thiothrix-nivea] |
| MCY3837524.1 | CR-ATPase+REase | GCA_026706425.1 | Gammaproteobacteria_bacterium | Bacteria>Pseudomonadati>Pseudomonadota>Gammaproteobacteria>unclassified_Gammaproteobacteria | 591 | OXH09_02540 | 2695..4470 | 1913989 | MAG:-AAA-family-ATPase-[Gammaproteobacteria-bacterium] |
| MBL8842554.1 | CR-ATPase+REase | GCA_016795285.1 | Planctomycetota_bacterium | Bacteria>Pseudomonadati>PVC_group>Planctomycetota>unclassified_Planctomycetota | 406 | JNL90_13625 | 286184..287404 | 2026780 | ATP-binding-protein-[Planctomycetota-bacterium] |
| MCP5494231.1 | CR-ATPase+REase | GCA_024233865.1 | Leptospiraceae_bacterium | Bacteria>Pseudomonadati>Spirochaetota>Spirochaetia>Leptospirales>Leptospiraceae>unclassified_Leptospiraceae | 577 | H7A23_06710 | 329860..331593 | 2026757 | AAA-family-ATPase-[Leptospiraceae-bacterium] |
| WP_010252916.1 | CR-ATPase+REase | GCF_000297095.1 | Treponema_primitia | Bacteria>Pseudomonadati>Spirochaetota>Spirochaetia>Spirochaetales>Treponemataceae>Treponema | 425 | TPRIMZ1_RS0100065 | 13362..14639 | 88058 | ATP-binding-protein-[Treponema-primitia] |
| MBR5915379.1 | CR-ATPase+REase | GCA_017554185.1 | Spirochaetia_bacterium | Bacteria>Pseudomonadati>Spirochaetota>Spirochaetia>unclassified_Spirochaetia | 434 | IKZ57_02895 | 21477..22781 | 2053615 | ATP-binding-protein-[Spirochaetia-bacterium] |
| MBF0120840.1 | CR-ATPase+REase | GCA_015228685.1 | Desulfobacterales_bacterium | Bacteria>Pseudomonadati>Thermodesulfobacteriota>Desulfobacteria>Desulfobacterales>unclassified_Desulfobacterales | 581 | HQK79_18590 | 25345..27090 | 2044940 | AAA-family-ATPase-[Desulfobacterales-bacterium] |
| HBI14539.1 | CR-ATPase+REase | GCA_003508005.1 | Desulfobulbaceae_bacterium | Bacteria>Pseudomonadati>Thermodesulfobacteriota>Desulfobulbia>Desulfobulbales>Desulfobulbaceae>unclassified_Desulfobulbaceae | 377 | DDY20_03335 | 166172..167305 | 2053307 | hypothetical-protein-[Desulfobulbaceae-bacterium] |
| MBQ7457437.1 | CR-ATPase+REase | GCA_017450275.1 | Desulfovibrio_sp. | Bacteria>Pseudomonadati>Thermodesulfobacteriota>Desulfovibrionia>Desulfovibrionales>Desulfovibrionaceae>Desulfovibrio>unclassified_Desulfovibrio | 554 | IJS54_07520 | 10224..11888 | 885 | AAA-family-ATPase-[Desulfovibrio-sp.] |
| MCK9435713.1 | CR-ATPase+REase | GCA_023229345.1 | Synergistaceae_bacterium | Bacteria>Thermotogati>Synergistota>Synergistia>Synergistales>Synergistaceae>unclassified_Synergistaceae | 403 | MOQ12_00730 | 1728..2939 | 2053624 | ATP-binding-protein-[Synergistaceae-bacterium] |
| NCP08804.1 | CR-ATPase+REase | GCA_009993525.1 | bacterium | Bacteria>unclassified_Bacteria | 397 | GW860_07885 | 15486..16679 | 1869227 | ATP-binding-protein-[bacterium] |
| NQX01744.1 | CR-ATPase+REase | GCA_013205625.1 | bacterium | Bacteria>unclassified_Bacteria | 396 | HQ447_13900 | 5724..6914 | 1869227 | ATP-binding-protein-[bacterium] |
| MCY4630096.1 | CR-ATPase+REase | GCA_026713745.1 | bacterium | Bacteria>unclassified_Bacteria | 637 | OXE75_04315 | 6912..8825 | 1869227 | MAG:-ATP-binding-protein-[bacterium] |
| --(w)HTH+REase | | | | | | | | | |
| WP_007538666.1 | (w)HTH+REase | - | Haloferax_larsenii | Archaea>Methanobacteriati>Methanobacteriota>Stenosarchaea_group>Halobacteria>Halobacteriales>Haloferacaceae>Haloferax | 454 | C455_RS00085 | 21971..23335 | 302484 | hypothetical-protein-[Haloferax-larsenii] |
| ELZ84486.1 | (w)HTH+REase | GCA_000336955.1 | Haloferax_larsenii_JCM_13917 | Archaea>Methanobacteriati>Methanobacteriota>Stenosarchaea_group>Halobacteria>Halobacteriales>Haloferacaceae>Haloferax>Haloferax_larsenii | 454 | C455_00097 | 21971..23335 | 1227460 | hypothetical-protein-C455_00097-[Haloferax-larsenii-JCM-13917] |
| WP_144927837.1 | (w)HTH+REase | GCF_007671685.1 | Halorubrum_salsamenti | Archaea>Methanobacteriati>Methanobacteriota>Stenosarchaea_group>Halobacteria>Halobacteriales>Haloferacaceae>Halorubrum | 441 | FGL94_RS17125 | 25310..26635 | 2583990 | hypothetical-protein-[Halorubrum-salsamenti] |

| Protein-ID | Domain | Assembly-ID | Strain | Taxonomy | Length | Locus-Tag | Genomic-coordinates | TaxID | Fasta-defline |
|---|---|---|---|---|---|---|---|---|---|
| WP_273738555.1 | (w)HTH+REase | GCF_028618695.1 | Natrinema_soli | Archaea>Methanobacteriati>Methanobacteriota>Stenosarchaea_group>Halobacteria>Halobacteriales>Natrialbaceae>Natrinema | 444 | PGX04_RS11055 | 2898..4232 | 1930624 | hypothetical-protein-[Natrinema-soli] |
| WP_255191904.1 | (w)HTH+REase | GCF_024362485.1 | Natronobeatus_ordinarius | Archaea>Methanobacteriati>Methanobacteriota>Stenosarchaea_group>Halobacteria>Halobacteriales>Natrialbaceae>Natronobeatus | 449 | NMQ09_RS17710 | 3487187..3488536 | 2963433 | hypothetical-protein-[Halovivax-sp.-WLHSJ27] |
| WP_252697929.1 | (w)HTH+REase | GCF_023973145.1 | Natronosalvus_vescus | Archaea>Methanobacteriati>Methanobacteriota>Stenosarchaea_group>Halobacteria>Halobacteriales>Natrialbaceae>Natronosalvus | 305 | NGM68_RS09800 | 2042789..2043706 | 2953881 | PD-(D/E)XK-nuclease-family-protein-[Natribaculum-sp.-CGA3] |
| WP_014867865.1 | (w)HTH+REase | GCF_000304355.2 | Methanoculleus_bourgensis | Archaea>Methanobacteriati>Methanobacteriota>Stenosarchaea_group>Methanomicrobia>Methanomicrobiales>Methanomicrobiaceae>Methanoculleus | 373 | BN140_RS09850 | 2033758..2034879 | 83986 | PDDEXK-nuclease-domain-containing-protein-[Methanoculleus-bourgensis] |
| WP_256712478.1 | (w)HTH+REase | GCF_900095385.1 | Methanoculleus_chikugoensis | Archaea>Methanobacteriati>Methanobacteriota>Stenosarchaea_group>Methanomicrobia>Methanomicrobiales>Methanomicrobiaceae>Methanoculleus | 190 | L21_RS13535 | 34104..34676 | 118126 | DUF1016-N-terminal-domain-containing-protein-[Methanoculleus-chikugoensis] |
| CCJ36892.1 | (w)HTH+REase | GCA_000304355.2 | Methanoculleus_bourgensis_MS2 | Archaea>Methanobacteriati>Methanobacteriota>Stenosarchaea_group>Methanomicrobia>Methanomicrobiales>Methanomicrobiaceae>Methanoculleus>Methanoculleus_bourgensis | 373 | BN140_1969 | 2033758..2034879 | 1201294 | hypothetical-protein-BN140_1969-[Methanoculleus-bourgensis-MS2] |
| WP_004077397.1 | (w)HTH+REase | GCF_000243255.1 | Methanoplanus_limicola | Archaea>Methanobacteriati>Methanobacteriota>Stenosarchaea_group>Methanomicrobia>Methanomicrobiales>Methanomicrobiaceae>Methanoplanus | 447 | METLIM_RS07730 | 1601205..1602548 | 2315 | PDDEXK-nuclease-domain-containing-protein-[Methanoplanus-limicola] |
| WP_004078485.1 | (w)HTH+REase | GCF_000243255.1 | Methanoplanus_limicola | Archaea>Methanobacteriati>Methanobacteriota>Stenosarchaea_group>Methanomicrobia>Methanomicrobiales>Methanomicrobiaceae>Methanoplanus | 343 | METLIM_RS10800 | 2300270..2301301 | 2315 | PDDEXK-nuclease-domain-containing-protein-[Methanoplanus-limicola] |
| WP_257742636.1 | (w)HTH+REase | GCF_024662215.1 | Methanoplanus_endosymbiosus | Archaea>Methanobacteriati>Methanobacteriota>Stenosarchaea_group>Methanomicrobia>Methanomicrobiales>Methanomicrobiaceae>Methanoplanus | 312 | L6E24_RS14395 | 3082555..3083493 | 33865 | PDDEXK-nuclease-domain-containing-protein-[Methanoplanus-endosymbiosus] |
| MCE5337195.1 | (w)HTH+REase | GCA_021371835.1 | Methanomicrobiaceae_archaeon | Archaea>Methanobacteriati>Methanobacteriota>Stenosarchaea_group>Methanomicrobia>Methanomicrobiales>Methanomicrobiaceae>unclassified_Methanomicrobiaceae | 373 | LLF90_00695 | 41609..42730 | 2716262 | PDDEXK-nuclease-domain-containing-protein-[Methanomicrobiaceae-archaeon] |
| WP_214418668.1 | (w)HTH+REase | GCF_018502485.1 | Methanospirillum_purgamenti | Archaea>Methanobacteriati>Methanobacteriota>Stenosarchaea_group>Methanomicrobia>Methanomicrobiales>Methanospirillaceae>Methanospirillum | 94 | KHC33_RS10910 | 2265827..2266111 | 2834276 | hypothetical-protein-[Methanospirillum-sp.-J.3.6.1-F.2.7.3] |
| WP_011448879.1 | (w)HTH+REase | GCF_000013445.1 | Methanospirillum_hungatei | Archaea>Methanobacteriati>Methanobacteriota>Stenosarchaea_group>Methanomicrobia>Methanomicrobiales>Methanospirillaceae>Methanospirillum | 92 | MHUN_RS09885 | 2146113..2146391 | 2203 | hypothetical-protein-[Methanospirillum-hungatei] |
| MBN1165672.1 | (w)HTH+REase | GCA_016926505.1 | Methanospirillaceae_archaeon | Archaea>Methanobacteriati>Methanobacteriota>Stenosarchaea_group>Methanomicrobia>Methanomicrobiales>Methanospirillaceae>unclassified_Methanospirillaceae | 365 | JXA44_00920 | 59844..60941 | 2813030 | DUF1016-family-protein-[Methanospirillaceae-archaeon] |
| WP_082107639.1 | (w)HTH+REase | GCF_000979975.1 | Methanosarcina_sp._1.H.A.2.2 | Archaea>Methanobacteriati>Methanobacteriota>Stenosarchaea_group>Methanomicrobia>Methanosarcinales>Methanosarcinaceae>Methanosarcina>unclassified_Methanosarcina | 395 | EO93_RS11295 | 69246..70433 | 1483601 | PDDEXK-nuclease-domain-containing-protein-[Methanosarcina-sp.-1.H.A.2.2] |
| MBE3050153.1 | (w)HTH+REase | GCA_014894645.1 | Candidatus_Bathyarchaeota_archaeon | Archaea>Thermoproteati>Candidatus_Bathyarchaeota>unclassified_Candidatus_Bathyarchaeota | 380 | IMZ48_48185 | 5664..6806 | 2026714 | winged-helix-turn-helix-domain-containing-protein-[Candidatus-Bathyarchaeota-archaeon] |
| WP_118724673.1 | (w)HTH+REase | GCF_019041895.1 | Bacillota | Bacteria>Bacillati | 337 | KTG08_RS07070 | 32401..33414 | 1239 | PDDEXK-nuclease-domain-containing-protein-[[Clostridium] |
| WP_004808453.1 | (w)HTH+REase | GCF_028201595.1 | Winkia | Bacteria>Bacillati>Actinomycetota>Actinomycetes>Actinomycetales>Actinomycetaceae | 316 | PIG85_RS08965 | 1924606..1925556 | 2692118 | MULTISPECIES:-restriction-endonuclease-[Actinomycetaceae] |
| WP_128773203.1 | (w)HTH+REase | GCF_004104015.1 | Actinomyces_oricola | Bacteria>Bacillati>Actinomycetota>Actinomycetes>Actinomycetales>Actinomycetaceae>Actinomyces | 372 | EUC46_RS05560 | 23273..24391 | 206043 | PDDEXK-nuclease-domain-containing-protein-[Actinomyces-oricola] |
| WP_124851217.1 | (w)HTH+REase | GCF_003858495.1 | Buchananella_hordeovulneris | Bacteria>Bacillati>Actinomycetota>Actinomycetes>Actinomycetales>Actinomycetaceae>Buchananella | 323 | EII12_RS02015 | 57832..58803 | 52770 | PDDEXK-nuclease-domain-containing-protein-[Buchananella-hordeovulneris] |
| WP_009381275.1 | (w)HTH+REase | GCF_000314575.1 | Brevibacterium | Bacteria>Bacillati>Actinomycetota>Actinomycetes>Micrococcales>Brevibacteriaceae | 372 | I6I57_RS03280 | 758377..759495 | 1696 | MULTISPECIES:-PDDEXK-nuclease-domain-containing-protein-[Brevibacterium] |
| WP_259801018.1 | (w)HTH+REase | GCF_032085555.1 | Brevibacterium_casei | Bacteria>Bacillati>Actinomycetota>Actinomycetes>Micrococcales>Brevibacteriaceae>Brevibacterium | 348 | M3G43_RS08130 | 24647..25693 | 33889 | PDDEXK-nuclease-domain-containing-protein-[Brevibacterium-casei] |
| WP_005885098.1 | (w)HTH+REase | GCF_000178455.1 | Brevibacterium_mcbrellneri | Bacteria>Bacillati>Actinomycetota>Actinomycetes>Micrococcales>Brevibacteriaceae>Brevibacterium | 75 | HMPREF0183_RS08770 | 46836..47063 | 53363 | hypothetical-protein-[Brevibacterium-mcbrellneri] |
| WP_217494871.1 | (w)HTH+REase | GCF_005280295.1 | Brevibacterium_sp._CS2 | Bacteria>Bacillati>Actinomycetota>Actinomycetes>Micrococcales>Brevibacteriaceae>Brevibacterium>unclassified_Brevibacterium | 389 | FDF13_RS00010 | 2844..4013 | 2575923 | PDDEXK-nuclease-domain-containing-protein-[Brevibacterium-sp.-CS2] |
| WP_247619185.1 | (w)HTH+REase | GCF_023072605.1 | Brevibacterium_sp._R8603A2 | Bacteria>Bacillati>Actinomycetota>Actinomycetes>Micrococcales>Brevibacteriaceae>Brevibacterium>unclassified_Brevibacterium | 267 | MTQ12_RS14505 | 5254..6057 | 2929779 | DUF1016-N-terminal-domain-containing-protein-[Brevibacterium-sp.-R8603A2] |
| WP_204409509.1 | (w)HTH+REase | GCF_016907455.1 | Brachybacterium_muris | Bacteria>Bacillati>Actinomycetota>Actinomycetes>Micrococcales>Dermabacteraceae>Brachybacterium | 372 | JOD52_RS08680 | 1875075..1876193 | 219301 | PDDEXK-nuclease-domain-containing-protein-[Brachybacterium-muris] |
| WP_259878500.1 | (w)HTH+REase | GCF_025148185.1 | Brachybacterium_muris | Bacteria>Bacillati>Actinomycetota>Actinomycetes>Micrococcales>Dermabacteraceae>Brachybacterium | 370 | M3C58_RS02750 | 18474..19586 | 219301 | PDDEXK-nuclease-domain-containing-protein-[Brachybacterium-muris] |
| WP_249738128.1 | (w)HTH+REase | GCF_023509525.1 | Brachybacterium_equifaecis | Bacteria>Bacillati>Actinomycetota>Actinomycetes>Micrococcales>Dermabacteraceae>Brachybacterium | 348 | Bequi_RS11770 | 98537..99583 | 2910770 | PDDEXK-nuclease-domain-containing-protein-[Brachybacterium-sp.-JHP9] |
| MBK8435771.1 | (w)HTH+REase | GCA_016711265.1 | Austwickia_sp. | Bacteria>Bacillati>Actinomycetota>Actinomycetes>Micrococcales>Dermatophilaceae>Austwickia>unclassified_Austwickia | 372 | IPL37_04410 | 986282..987400 | 1872664 | DUF1016-family-protein-[Austwickia-sp.] |
| WP_274381641.1 | (w)HTH+REase | - | Segeticoccus_rhizosphaerae | Bacteria>Bacillati>Actinomycetota>Actinomycetes>Micrococcales>Intrasporangiaceae>Segeticoccus | 330 | FCN22_RS19305 | 11154..12146 | 1104777 | PDDEXK-nuclease-domain-containing-protein-[Ornithinicoccus-soli] |
| WP_124973555.1 | (w)HTH+REase | GCF_003865375.1 | Gulosibacter_macacae | Bacteria>Bacillati>Actinomycetota>Actinomycetes>Micrococcales>Microbacteriaceae>Gulosibacter | 348 | EG850_RS11345 | 28919..29965 | 2488791 | PDDEXK-nuclease-domain-containing-protein-[Gulosibacter-macacae] |
| WP_201614894.1 | (w)HTH+REase | GCF_904848685.1 | Gulosibacter_hominis | Bacteria>Bacillati>Actinomycetota>Actinomycetes>Micrococcales>Microbacteriaceae>Gulosibacter | 348 | JMX86_RS10070 | 28911..29957 | 2770504 | PDDEXK-nuclease-domain-containing-protein-[Gulosibacter-hominis] |
| WP_100810667.1 | (w)HTH+REase | GCF_002812725.1 | unclassified_Microbacterium | Bacteria>Bacillati>Actinomycetota>Actinomycetes>Micrococcales>Microbacteriaceae>Microbacterium | 138 | CGU35_RS05970 | 374577..374993 | 2609290 | MULTISPECIES:-hypothetical-protein-[unclassified-Microbacterium] |
| WP_207484744.1 | (w)HTH+REase | GCF_017355365.1 | Microbacterium_sp._SD291 | Bacteria>Bacillati>Actinomycetota>Actinomycetes>Micrococcales>Microbacteriaceae>Microbacterium>unclassified_Microbacterium | 348 | IPV10_RS13700 | 36151..37197 | 2782007 | PDDEXK-nuclease-domain-containing-protein-[Microbacterium-sp.-SD291] |
| HBS75975.1 | (w)HTH+REase | GCA_003510265.1 | Microbacterium_sp. | Bacteria>Bacillati>Actinomycetota>Actinomycetes>Micrococcales>Microbacteriaceae>Microbacterium>unclassified_Microbacterium | 138 | DEB55_16575 | 8783..9199 | 51671 | hypothetical-protein-[Microbacterium-sp.] |
| WP_104347516.1 | (w)HTH+REase | GCF_002932695.1 | Rathayibacter_sp._AY1D2 | Bacteria>Bacillati>Actinomycetota>Actinomycetes>Micrococcales>Microbacteriaceae>Rathayibacter>unclassified_Rathayibacter | 348 | C5D04_RS07565 | 15096..16142 | 2080543 | PDDEXK-nuclease-domain-containing-protein-[Rathayibacter-sp.-AY1D2] |
| WP_230672569.1 | (w)HTH+REase | GCF_920939495.1 | Rathayibacter_sp._Leaf248 | Bacteria>Bacillati>Actinomycetota>Actinomycetes>Micrococcales>Microbacteriaceae>Rathayibacter>unclassified_Rathayibacter | 184 | LRG51_RS10060 | 2082314..2082868 | 2876555 | DUF1016-N-terminal-domain-containing-protein-[Rathayibacter-sp.-Leaf248] |
| WP_230672571.1 | (w)HTH+REase | GCF_920939495.1 | Rathayibacter_sp._Leaf248 | Bacteria>Bacillati>Actinomycetota>Actinomycetes>Micrococcales>Microbacteriaceae>Rathayibacter>unclassified_Rathayibacter | 148 | LRG51_RS10065 | 2082805..2083251 | 2876555 | PDDEXK-nuclease-domain-containing-protein-[Rathayibacter-sp.-Leaf248] |
| WP_144831475.1 | (w)HTH+REase | GCF_007677595.1 | Kocuria_rhizophila | Bacteria>Bacillati>Actinomycetota>Actinomycetes>Micrococcales>Micrococcaceae>Kocuria | 372 | FS577_RS10185 | 100069..101187 | 72000 | PDDEXK-nuclease-domain-containing-protein-[Kocuria-rhizophila] |
| THE19346.1 | (w)HTH+REase | GCA_004799725.1 | Kocuria_rosea | Bacteria>Bacillati>Actinomycetota>Actinomycetes>Micrococcales>Micrococcaceae>Kocuria | 309 | E1J17_01455 | 19282..20211 | 1275 | restriction-endonuclease-[Kocuria-rosea] |
| WP_062735899.1 | (w)HTH+REase | GCF_007992155.1 | Kocuria_turfanensis | Bacteria>Bacillati>Actinomycetota>Actinomycetes>Micrococcales>Micrococcaceae>Kocuria | 309 | KTU01_RS15715 | 10551..11480 | 388357 | restriction-endonuclease-[Kocuria-turfanensis] |
| WP_112235017.1 | (w)HTH+REase | GCF_003290245.1 | Kocuria_sp._BT304 | Bacteria>Bacillati>Actinomycetota>Actinomycetes>Micrococcales>Micrococcaceae>Kocuria>unclassified_Kocuria | 372 | DNL27_RS06215 | 1380865..1381983 | 1702043 | PDDEXK-nuclease-domain-containing-protein-[Kocuria-sp.-BT304] |
| WP_052209231.1 | (w)HTH+REase | GCF_000800955.1 | Kocuria_sp._ZOR0020 | Bacteria>Bacillati>Actinomycetota>Actinomycetes>Micrococcales>Micrococcaceae>Kocuria>unclassified_Kocuria | 336 | L979_RS00025 | 6466..7476 | 1339234 | PDDEXK-nuclease-domain-containing-protein-[Kocuria-sp.-ZOR0020] |
| WP_285140307.1 | (w)HTH+REase | GCF_030226745.1 | Micrococcus_luteus | Bacteria>Bacillati>Actinomycetota>Actinomycetes>Micrococcales>Micrococcaceae>Micrococcus | 364 | QP726_RS10465 | 8548..9642 | 1270 | PDDEXK-nuclease-domain-containing-protein-[Micrococcus-luteus] |
| MBB4882888.1 | (w)HTH+REase | GCA_014204815.1 | Micrococcus_flavus | Bacteria>Bacillati>Actinomycetota>Actinomycetes>Micrococcales>Micrococcaceae>Micrococcus | 348 | BJ976_001239 | 1349697..1350743 | 384602 | putative-nuclease-of-restriction-endonuclease-like-(RecB)-superfamily-[Micrococcus-flavus] |
| WP_135027974.1 | (w)HTH+REase | GCF_014644875.1 | Micrococcus_flavus | Bacteria>Bacillati>Actinomycetota>Actinomycetes>Micrococcales>Micrococcaceae>Micrococcus | 348 | E4A47_RS01350 | 293348..294394 | 384602 | PDDEXK-nuclease-domain-containing-protein-[Micrococcus-flavus] |
| WP_141842661.1 | (w)HTH+REase | GCF_006704135.1 | Micrococcus_sp._R8502A1 | Bacteria>Bacillati>Actinomycetota>Actinomycetes>Micrococcales>Micrococcaceae>Micrococcus>unclassified_Micrococcus | 311 | FLM69_RS00650 | 7333..8268 | 2583239 | PDDEXK-nuclease-domain-containing-protein-[Micrococcus-sp.-R8502A1] |
| WP_168888197.1 | (w)HTH+REase | GCF_012641515.1 | Nesterenkonia_sedimenti | Bacteria>Bacillati>Actinomycetota>Actinomycetes>Micrococcales>Micrococcaceae>Nesterenkonia | 357 | HGQ17_RS12075 | 47236..48309 | 1463632 | PDDEXK-nuclease-domain-containing-protein-[Nesterenkonia-sedimenti] |
| WP_083956099.1 | (w)HTH+REase | GCF_001968835.1 | Tersicoccus_phoenicis | Bacteria>Bacillati>Actinomycetota>Actinomycetes>Micrococcales>Micrococcaceae>Tersicoccus | 372 | BKD30_RS07050 | 3650..4768 | 554083 | PDDEXK-nuclease-domain-containing-protein-[Tersicoccus-phoenicis] |
| MCE1178997.1 | (w)HTH+REase | GCA_021287075.1 | Micrococcales_bacterium | Bacteria>Bacillati>Actinomycetota>Actinomycetes>Micrococcales>unclassified_Micrococcales | 372 | LWW86_08205 | 121999..123117 | 2026762 | PDDEXK-nuclease-domain-containing-protein-[Micrococcales-bacterium] |
| PMC61709.1 | (w)HTH+REase | GCA_002871715.1 | Corynebacterium_xerosis | Bacteria>Bacillati>Actinomycetota>Actinomycetes>Mycobacteriales>Corynebacteriaceae>Corynebacterium | 312 | CJ204_09385 | 8702..9640 | 1725 | DUF1016-domain-containing-protein-[Corynebacterium-xerosis] |
| WP_213244942.1 | (w)HTH+REase | GCF_009914535.1 | Gordonia | Bacteria>Bacillati>Actinomycetota>Actinomycetes>Mycobacteriales>Gordoniaceae | 351 | GII33_RS19225 | 4432604..4433659 | 2053 | PDDEXK-nuclease-domain-containing-protein-[Gordonia-pseudamarae] |
| PZT93667.1 | (w)HTH+REase | GCA_003248405.1 | Gordonia_sp._(in:_high_G+C_Gram-positive_bacteria) | Bacteria>Bacillati>Actinomycetota>Actinomycetes>Mycobacteriales>Gordoniaceae>Gordonia>unclassified_Gordonia_(in:_high_G+C_Gram-positive_bacteria) | 372 | DI630_26680 | 8463..9581 | 84139 | MAG:-DUF1016-domain-containing-protein-[Gordonia-sp.-(in:-high-G+C-Gram-positive-bacteria)] |
| PZT90278.1 | (w)HTH+REase | GCA_003248405.1 | Gordonia_sp._(in:_high_G+C_Gram-positive_bacteria) | Bacteria>Bacillati>Actinomycetota>Actinomycetes>Mycobacteriales>Gordoniaceae>Gordonia>unclassified_Gordonia_(in:_high_G+C_Gram-positive_bacteria) | 304 | DI630_31240 | 13838..14752 | 84139 | MAG:-DUF1016-domain-containing-protein-[Gordonia-sp.-(in:-high-G+C-Gram-positive-bacteria)] |
| GFM17533.1 | (w)HTH+REase | GCA_017312405.1 | Mycobacterium_sp._PO1 | Bacteria>Bacillati>Actinomycetota>Actinomycetes>Mycobacteriales>Mycobacteriaceae>Mycobacterium>unclassified_Mycobacterium | 129 | PO1_contig-018-7 | 9560..9949 | 1882219 | uncharacterized-protein-PO1_contig-018-7-[Mycobacterium-sp.-PO1] |
| WP_141142700.1 | (w)HTH+REase | GCF_013138515.1 | Mycolicibacterium_fortuitum | Bacteria>Bacillati>Actinomycetota>Actinomycetes>Mycobacteriales>Mycobacteriaceae>Mycolicibacterium | 282 | FKW78_RS12985 | 27100..27948 | 1766 | restriction-endonuclease-[Mycolicibacterium-fortuitum] |
| WP_242421150.1 | (w)HTH+REase | GCF_022691685.1 | Prescottella_equi | Bacteria>Bacillati>Actinomycetota>Actinomycetes>Mycobacteriales>Nocardiaceae>Prescottella | 74 | MPC38_RS08815 | 1855447..1855671 | 43767 | hypothetical-protein-[Prescottella-equi] |
| WP_048320047.1 | (w)HTH+REase | GCF_001037935.1 | Rhodococcoides_fascians | Bacteria>Bacillati>Actinomycetota>Actinomycetes>Mycobacteriales>Nocardiaceae>Rhodococcoides | 286 | ACG96_RS20790 | 148908..149768 | 1828 | hypothetical-protein-[Rhodococcus-fascians] |
| WP_139203963.1 | (w)HTH+REase | GCF_900111805.1 | Rhodococcoides_kroppenstedtii | Bacteria>Bacillati>Actinomycetota>Actinomycetes>Mycobacteriales>Nocardiaceae>Rhodococcoides | 286 | BMZ18_RS12895 | 133585..134445 | 293050 | hypothetical-protein-[Rhodococcus-kroppenstedtii] |
| WP_094980726.1 | (w)HTH+REase | GCF_002269365.1 | Rhodococcus_pyridinivorans | Bacteria>Bacillati>Actinomycetota>Actinomycetes>Mycobacteriales>Nocardiaceae>Rhodococcus | 270 | CJU69_RS21690 | 4614893..4615705 | 103816 | DUF1016-N-terminal-domain-containing-protein-[Rhodococcus-pyridinivorans] |
| WP_203011587.1 | (w)HTH+REase | GCF_016804345.1 | Rhodococcus_ruber | Bacteria>Bacillati>Actinomycetota>Actinomycetes>Mycobacteriales>Nocardiaceae>Rhodococcus | 238 | F1734_RS10035 | 2140945..2141661 | 1830 | hypothetical-protein-[Rhodococcus-ruber] |
| WP_222635586.1 | (w)HTH+REase | GCF_019834785.1 | unclassified_Rhodococcus_(in:_high_G+C_Gram-positive_bacteria) | Bacteria>Bacillati>Actinomycetota>Actinomycetes>Mycobacteriales>Nocardiaceae>Rhodococcus | 288 | HQ363_RS22165 | 1714..2580 | 192944 | MULTISPECIES:-hypothetical-protein-[unclassified-Rhodococcus-(in:-high-G+C-Gram-positive-bacteria)] |
| OUS97314.1 | (w)HTH+REase | GCA_002165735.1 | Rhodococcus_sp._NCIMB_12038 | Bacteria>Bacillati>Actinomycetota>Actinomycetes>Mycobacteriales>Nocardiaceae>Rhodococcus>unclassified_Rhodococcus_(in:_high_G+C_Gram-positive_bacteria) | 348 | CA951_02915 | 133772..134818 | 933800 | hypothetical-protein-CA951_02915-[Rhodococcus-sp.-NCIMB-12038] |
| WP_087555440.1 | (w)HTH+REase | GCF_002165735.1 | Rhodococcus_sp._NCIMB_12038 | Bacteria>Bacillati>Actinomycetota>Actinomycetes>Mycobacteriales>Nocardiaceae>Rhodococcus>unclassified_Rhodococcus_(in:_high_G+C_Gram-positive_bacteria) | 348 | CA951_RS02930 | 133772..134818 | 933800 | PDDEXK-nuclease-domain-containing-protein-[Rhodococcus-sp.-NCIMB-12038] |
| WP_124569534.1 | (w)HTH+REase | GCF_003852455.1 | Rhodococcus_sp._KBW08 | Bacteria>Bacillati>Actinomycetota>Actinomycetes>Mycobacteriales>Nocardiaceae>Rhodococcus>unclassified_Rhodococcus_(in:_high_G+C_Gram-positive_bacteria) | 271 | DBV08_RS27765 | 154340..155155 | 2144188 | hypothetical-protein-[Rhodococcus-sp.-KBW08] |
| WP_068569933.1 | (w)HTH+REase | GCF_001575195.1 | Tsukamurella_pseudospumae | Bacteria>Bacillati>Actinomycetota>Actinomycetes>Mycobacteriales>Tsukamurellaceae>Tsukamurella | 347 | AXK60_RS03090 | 665903..666946 | 239498 | PDDEXK-nuclease-domain-containing-protein-[Tsukamurella-pseudospumae] |
| WP_210733313.1 | (w)HTH+REase | GCF_039527875.1 | Tsukamurella_spumae | Bacteria>Bacillati>Actinomycetota>Actinomycetes>Mycobacteriales>Tsukamurellaceae>Tsukamurella | 329 | HF999_RS00905 | 193428..194417 | 44753 | PDDEXK-nuclease-domain-containing-protein-[Tsukamurella-spumae] |
| WP_197466717.1 | (w)HTH+REase | GCF_001575205.1 | Tsukamurella_pseudospumae | Bacteria>Bacillati>Actinomycetota>Actinomycetes>Mycobacteriales>Tsukamurellaceae>Tsukamurella | 222 | AXK61_RS25285 | 245776..246444 | 239498 | PDDEXK-nuclease-domain-containing-protein-[Tsukamurella-pseudospumae] |
| WP_197466716.1 | (w)HTH+REase | GCF_001575205.1 | Tsukamurella_pseudospumae | Bacteria>Bacillati>Actinomycetota>Actinomycetes>Mycobacteriales>Tsukamurellaceae>Tsukamurella | 129 | AXK61_RS25280 | 245400..245789 | 239498 | PDDEXK-nuclease-domain-containing-protein-[Tsukamurella-pseudospumae] |
| WP_014846822.1 | (w)HTH+REase | GCF_000277715.1 | Arachnia_propionica | Bacteria>Bacillati>Actinomycetota>Actinomycetes>Propionibacteriales>Propionibacteriaceae>Arachnia | 367 | J5A53_RS06800 | 1505360..1506463 | 1750 | PDDEXK-nuclease-domain-containing-protein-[Arachnia-propionica] |
| WP_082793752.1 | (w)HTH+REase | GCF_001592325.1 | Arachnia_propionica | Bacteria>Bacillati>Actinomycetota>Actinomycetes>Propionibacteriales>Propionibacteriaceae>Arachnia | 367 | PP13_RS01000 | 242715..243818 | 1750 | PDDEXK-nuclease-domain-containing-protein-[Arachnia-propionica] |
| WP_126378243.1 | (w)HTH+REase | GCF_900638645.1 | Arachnia_propionica | Bacteria>Bacillati>Actinomycetota>Actinomycetes>Propionibacteriales>Propionibacteriaceae>Arachnia | 367 | EL396_RS08530 | 1940809..1941912 | 1750 | PDDEXK-nuclease-domain-containing-protein-[Arachnia-propionica] |
| WP_130874285.1 | (w)HTH+REase | GCF_900607225.1 | [Pseudopropionibacterium]_massiliense | Bacteria>Bacillati>Actinomycetota>Actinomycetes>Propionibacteriales>Propionibacteriaceae>Arachnia | 367 | E0E39_RS08100 | 1823584..1824687 | 2220000 | PDDEXK-nuclease-domain-containing-protein-[[Pseudopropionibacterium] |
| WP_211798073.1 | (w)HTH+REase | GCF_018128345.1 | Arachnia_propionica | Bacteria>Bacillati>Actinomycetota>Actinomycetes>Propionibacteriales>Propionibacteriaceae>Arachnia | 367 | J5A61_RS08585 | 1918207..1919310 | 1750 | PDDEXK-nuclease-domain-containing-protein-[Arachnia-propionica] |
| WP_278693136.1 | (w)HTH+REase | GCF_915063725.1 | Arachnia_propionica | Bacteria>Bacillati>Actinomycetota>Actinomycetes>Propionibacteriales>Propionibacteriaceae>Arachnia | 367 | QDI34_RS08810 | 14578..15681 | 1750 | PDDEXK-nuclease-domain-containing-protein-[Arachnia-propionica] |
| WP_094364308.1 | (w)HTH+REase | GCF_002250625.1 | Enemella_dayhoffiae | Bacteria>Bacillati>Actinomycetota>Actinomycetes>Propionibacteriales>Propionibacteriaceae>Enemella | 348 | CGZ93_RS11745 | 171190..172236 | 2016507 | PDDEXK-nuclease-domain-containing-protein-[Enemella-dayhoffiae] |
| WP_082034459.1 | (w)HTH+REase | GCF_902375435.1 | Nigerium_massiliense | Bacteria>Bacillati>Actinomycetota>Actinomycetes>Propionibacteriales>Propionibacteriaceae>Nigerium | 372 | BN1187_RS02665 | 262960..264078 | 1522317 | PDDEXK-nuclease-domain-containing-protein-[Nigerium-massiliense] |
| WP_239018423.1 | (w)HTH+REase | GCF_900289195.1 | Propionibacterium_ruminifibrarum | Bacteria>Bacillati>Actinomycetota>Actinomycetes>Propionibacteriales>Propionibacteriaceae>Propionibacterium | 371 | PROPJV5_RS07030 | 35561..36676 | 1962131 | PDDEXK-nuclease-domain-containing-protein-[Propionibacterium-ruminifibrarum] |
| WP_021798348.1 | (w)HTH+REase | GCF_000478805.1 | Propionibacterium_acidifaciens | Bacteria>Bacillati>Actinomycetota>Actinomycetes>Propionibacteriales>Propionibacteriaceae>Propionibacterium | 369 | P7A72_RS09155 | 70421..71530 | 556499 | PDDEXK-nuclease-domain-containing-protein-[Propionibacterium-acidifaciens] |
| WP_232012343.1 | (w)HTH+REase | GCF_900637915.1 | Propionibacterium_australiense | Bacteria>Bacillati>Actinomycetota>Actinomycetes>Propionibacteriales>Propionibacteriaceae>Propionibacterium | 331 | EL282_RS10975 | 2478240..2479235 | 119981 | PDDEXK-nuclease-domain-containing-protein-[Propionibacterium-australiense] |
| WP_242681192.1 | (w)HTH+REase | - | Tessaracoccus_sp._SD287 | Bacteria>Bacillati>Actinomycetota>Actinomycetes>Propionibacteriales>Propionibacteriaceae>Tessaracoccus>unclassified_Tessaracoccus | 327 | IPV09_RS03375 | 68338..69321 | 2782008 | PDDEXK-nuclease-domain-containing-protein-[Tessaracoccus-sp.-SD287] |
| MBK7820963.1 | (w)HTH+REase | GCA_016714275.1 | Tessaracoccus_sp. | Bacteria>Bacillati>Actinomycetota>Actinomycetes>Propionibacteriales>Propionibacteriaceae>Tessaracoccus>unclassified_Tessaracoccus | 314 | IPJ61_07770 | 1700664..1701608 | 1971211 | DUF1016-family-protein-[Tessaracoccus-sp.] |
| MBE6477655.1 | (w)HTH+REase | GCA_015062705.1 | Propionibacteriaceae_bacterium | Bacteria>Bacillati>Actinomycetota>Actinomycetes>Propionibacteriales>Propionibacteriaceae>unclassified_Propionibacteriaceae | 369 | E7Z97_06270 | 35533..36642 | 2021380 | DUF1016-domain-containing-protein-[Propionibacteriaceae-bacterium] |
| WP_221337127.1 | (w)HTH+REase | GCF_039543005.1 | Thermocatellispora_tengchongensis | Bacteria>Bacillati>Actinomycetota>Actinomycetes>Streptosporangiales>Streptosporangiaceae>Thermocatellispora | 136 | HNP84_RS34075 | 42391..42801 | 1073253 | hypothetical-protein-[Thermocatellispora-tengchongensis] |
| SFW99816.1 | (w)HTH+REase | GCA_900119625.1 | Olsenella_sp._kh2p3 | Bacteria>Bacillati>Actinomycetota>Coriobacteriia>Coriobacteriales>Atopobiaceae>Olsenella>unclassified_Olsenella | 379 | SAMN04487823_101105 | 115627..116766 | 1797112 | restriction-system-protein-[Olsenella-sp.-kh2p3] |
| MBA3474034.1 | (w)HTH+REase | GCA_013813465.1 | Rubrobacter_sp. | Bacteria>Bacillati>Actinomycetota>Rubrobacteria>Rubrobacterales>Rubrobacteraceae>Rubrobacter>unclassified_Rubrobacter | 338 | HOT57_12550 | 280..1296 | 2036010 | DUF1016-domain-containing-protein-[Rubrobacter-sp.] |
| MCA0180886.1 | (w)HTH+REase | GCA_020160805.1 | Actinomycetota_bacterium | Bacteria>Bacillati>Actinomycetota>unclassified_Actinomycetota | 372 | LCH77_15140 | 45183..46301 | 2900548 | PDDEXK-nuclease-domain-containing-protein-[Actinomycetia-bacterium] |
| NCD16599.1 | (w)HTH+REase | GCA_009930385.1 | Actinomycetota_bacterium | Bacteria>Bacillati>Actinomycetota>unclassified_Actinomycetota | 372 | EOL91_04670 | 6567..7685 | 2900548 | DUF1016-family-protein-[Actinomycetia-bacterium] |
| MCA0180525.1 | (w)HTH+REase | GCA_020160805.1 | Actinomycetota_bacterium | Bacteria>Bacillati>Actinomycetota>unclassified_Actinomycetota | 330 | LCH77_13245 | 57664..58656 | 2900548 | PDDEXK-nuclease-domain-containing-protein-[Actinomycetia-bacterium] |
| WP_271938617.1 | (w)HTH+REase | GCF_028321805.1 | Clostridia | Bacteria>Bacillati>Bacillota | 332 | PND94_RS10180 | 47295..48293 | 186801 | PDDEXK-nuclease-domain-containing-protein-[Flavonifractor-plautii] |
| WP_068488201.1 | (w)HTH+REase | GCF_001592825.1 | Planococcus_maritimus | Bacteria>Bacillati>Bacillota>Bacilli>Bacillales>Caryophanaceae>Planococcus | 301 | AY633_RS09855 | 262030..262935 | 192421 | restriction-endonuclease-[Planococcus-maritimus] |
| WP_215106218.1 | (w)HTH+REase | GCF_018613655.1 | Planococcus_sp._ISL-109 | Bacteria>Bacillati>Bacillota>Bacilli>Bacillales>Caryophanaceae>Planococcus>unclassified_Planococcus_(in:_firmicutes) | 421 | J7E21_RS07425 | 22669..23934 | 2819166 | hypothetical-protein-[Planococcus-sp.-ISL-109] |
| MBT2292696.1 | (w)HTH+REase | GCA_018614985.1 | Paenibacillus_albidus | Bacteria>Bacillati>Bacillota>Bacilli>Bacillales>Paenibacillaceae>Paenibacillus | 208 | J7E73_26910 | 20839..21465 | 2041023 | MarR-family-transcriptional-regulator-[Paenibacillus-albidus] |
| WP_091013314.1 | (w)HTH+REase | - | Paenibacillus_amylolyticus | Bacteria>Bacillati>Bacillota>Bacilli>Bacillales>Paenibacillaceae>Paenibacillus | 208 | BQ2370_RS05135 | 6066..6692 | 1451 | MarR-family-transcriptional-regulator-[Paenibacillus-amylolyticus] |
| WP_215184625.1 | (w)HTH+REase | GCF_018614985.1 | Paenibacillus_albidus | Bacteria>Bacillati>Bacillota>Bacilli>Bacillales>Paenibacillaceae>Paenibacillus | 208 | J7E73_RS26905 | 20839..21465 | 2041023 | helix-turn-helix-domain-containing-protein-[Paenibacillus-albidus] |
| MCG7380063.1 | (w)HTH+REase | GCA_022346845.1 | Paenibacillus_sp._ACRSA | Bacteria>Bacillati>Bacillota>Bacilli>Bacillales>Paenibacillaceae>Paenibacillus>unclassified_Paenibacillus | 208 | MH215_24005 | 109256..109882 | 2918211 | MarR-family-transcriptional-regulator-[Paenibacillus-sp.-ACRSA] |
| WP_056697888.1 | (w)HTH+REase | GCF_001426865.1 | Paenibacillus_sp._Root52 | Bacteria>Bacillati>Bacillota>Bacilli>Bacillales>Paenibacillaceae>Paenibacillus>unclassified_Paenibacillus | 208 | ASD24_RS16960 | 52894..53520 | 1736552 | helix-turn-helix-domain-containing-protein-[Paenibacillus-sp.-Root52] |
| WP_222215809.1 | (w)HTH+REase | GCF_019793795.1 | Paenibacillus_sp._DR312 | Bacteria>Bacillati>Bacillota>Bacilli>Bacillales>Paenibacillaceae>Paenibacillus>unclassified_Paenibacillus | 208 | K5K90_RS02620 | 600470..601096 | 2871175 | helix-turn-helix-domain-containing-protein-[Paenibacillus-sp.-DR312] |
| WP_239293648.1 | (w)HTH+REase | GCF_022346845.1 | Paenibacillus_sp._ACRSA | Bacteria>Bacillati>Bacillota>Bacilli>Bacillales>Paenibacillaceae>Paenibacillus>unclassified_Paenibacillus | 208 | MH215_RS24000 | 109256..109882 | 2918211 | helix-turn-helix-domain-containing-protein-[Paenibacillus-sp.-ACRSA] |
| WP_153832545.1 | (w)HTH+REase | GCF_009659375.1 | Fundicoccus_ignavus | Bacteria>Bacillati>Bacillota>Bacilli>Lactobacillales>Aerococcaceae>Fundicoccus | 290 | GF867_RS07775 | 4247..5119 | 2664442 | Mrr-restriction-system-protein-[Fundicoccus-ignavus] |
| WP_154059562.1 | (w)HTH+REase | GCF_901212505.1 | Enterococcus_avium | Bacteria>Bacillati>Bacillota>Bacilli>Lactobacillales>Enterococcaceae>Enterococcus | 318 | EAVMC1_RS06645 | 34255..35211 | 33945 | restriction-endonuclease-[Enterococcus-avium] |
| WP_126762074.1 | (w)HTH+REase | GCF_003987555.1 | Vagococcus_lutrae | Bacteria>Bacillati>Bacillota>Bacilli>Lactobacillales>Enterococcaceae>Vagococcus | 328 | CBF33_RS04645 | 15563..16549 | 81947 | restriction-endonuclease-[Vagococcus-lutrae] |
| WP_096584709.1 | (w)HTH+REase | GCF_002370375.1 | Latilactobacillus | Bacteria>Bacillati>Bacillota>Bacilli>Lactobacillales>Lactobacillaceae | 321 | CCX78_RS03515 | 679163..680128 | 2767885 | restriction-endonuclease-[Latilactobacillus-sakei] |
| WP_025013014.1 | (w)HTH+REase | GCF_000829055.1 | Lacticaseibacillus | Bacteria>Bacillati>Bacillota>Bacilli>Lactobacillales>Lactobacillaceae | 315 | LBCZ_RS09670 | 1981138..1982085 | 2759736 | restriction-endonuclease-[Lacticaseibacillus-casei] |
| WP_179394512.1 | (w)HTH+REase | GCF_013407595.1 | Lacticaseibacillus_absianus | Bacteria>Bacillati>Bacillota>Bacilli>Lactobacillales>Lactobacillaceae>Lacticaseibacillus | 316 | HII33_RS04100 | 963131..964081 | 2729623 | restriction-endonuclease-[Lacticaseibacillus-absianus] |
| WP_016373141.1 | (w)HTH+REase | GCF_000410335.1 | Lacticaseibacillus_paracasei | Bacteria>Bacillati>Bacillota>Bacilli>Lactobacillales>Lactobacillaceae>Lacticaseibacillus | 315 | J3H89_RS10625 | 2194349..2195296 | 1597 | restriction-endonuclease-[Lacticaseibacillus-paracasei] |
| WP_160528674.1 | (w)HTH+REase | GCF_049532865.1 | Lacticaseibacillus_paracasei | Bacteria>Bacillati>Bacillota>Bacilli>Lactobacillales>Lactobacillaceae>Lacticaseibacillus | 315 | GRZ59_RS05180 | 16728..17675 | 1597 | restriction-endonuclease-[Lacticaseibacillus-paracasei] |
| WP_235500826.1 | (w)HTH+REase | GCF_001433745.1 | Lacticaseibacillus_zeae | Bacteria>Bacillati>Bacillota>Bacilli>Lactobacillales>Lactobacillaceae>Lacticaseibacillus | 222 | FD51_RS09350 | 86307..86975 | 57037 | restriction-endonuclease-[Lacticaseibacillus-zeae] |
| WP_232521914.1 | (w)HTH+REase | GCF_003966835.1 | Lacticaseibacillus_paracasei | Bacteria>Bacillati>Bacillota>Bacilli>Lactobacillales>Lactobacillaceae>Lacticaseibacillus | 169 | EL355_RS11555 | 2234499..2235008 | 1597 | restriction-endonuclease-[Lacticaseibacillus-paracasei] |
| WP_232521915.1 | (w)HTH+REase | GCF_003966835.1 | Lacticaseibacillus_paracasei | Bacteria>Bacillati>Bacillota>Bacilli>Lactobacillales>Lactobacillaceae>Lacticaseibacillus | 152 | EL355_RS11565 | 2236043..2236501 | 1597 | winged-helix-turn-helix-domain-containing-protein-[Lacticaseibacillus-paracasei] |
| WP_235500825.1 | (w)HTH+REase | GCF_001433745.1 | Lacticaseibacillus_zeae | Bacteria>Bacillati>Bacillota>Bacilli>Lactobacillales>Lactobacillaceae>Lacticaseibacillus | 93 | FD51_RS16705 | 86029..86310 | 57037 | winged-helix-turn-helix-domain-containing-protein-[Lacticaseibacillus-zeae] |
| WP_188891969.1 | (w)HTH+REase | GCF_014656585.1 | Lactobacillus_kefiranofaciens | Bacteria>Bacillati>Bacillota>Bacilli>Lactobacillales>Lactobacillaceae>Lactobacillus | 313 | ICI50_RS03525 | 689336..690277 | 267818 | restriction-endonuclease-[Lactobacillus-kefiranofaciens] |
| WP_270447888.1 | (w)HTH+REase | GCF_027681365.1 | Lactobacillus_delbrueckii | Bacteria>Bacillati>Bacillota>Bacilli>Lactobacillales>Lactobacillaceae>Lactobacillus | 309 | PGX26_RS03400 | 67058..67987 | 1584 | restriction-endonuclease-[Lactobacillus-delbrueckii] |
| AOO75266.1 | (w)HTH+REase | GCA_001723545.1 | Latilactobacillus_curvatus | Bacteria>Bacillati>Bacillota>Bacilli>Lactobacillales>Lactobacillaceae>Latilactobacillus | 321 | LCW_03940 | 765976..766941 | 28038 | hypothetical-protein-LCW_03940-[Latilactobacillus-curvatus] |

| Protein-ID | Domain | Assembly-ID | Strain | Taxonomy | Length | Locus-Tag | Genomic-coordinates | TaxID | Fasta-defline |
|---|---|---|---|---|---|---|---|---|---|
| WP_056966479.1 | (w)HTH+REase | GCF_001435495.1 | Latilactobacillus_curvatus | Bacteria>Bacillati>Bacillota>Bacilli>Lactobacillales>Lactobacillaceae>Latilactobacillus | 321 | PSR33_RS06725 | 1320775..1321740 | 28038 | restriction-endonuclease-[Latilactobacillus-curvatus] |
| WP_085391468.1 | (w)HTH+REase | GCF_023734335.1 | Latilactobacillus_sakei | Bacteria>Bacillati>Bacillota>Bacilli>Lactobacillales>Lactobacillaceae>Latilactobacillus | 321 | A4W84_RS06325 | 1239231..1240196 | 1599 | restriction-endonuclease-[Latilactobacillus-sakei] |
| WP_099947380.1 | (w)HTH+REase | GCF_024273085.1 | Latilactobacillus_sakei | Bacteria>Bacillati>Bacillota>Bacilli>Lactobacillales>Lactobacillaceae>Latilactobacillus | 321 | LOX63_RS01790 | 356374..357339 | 1599 | restriction-endonuclease-[Latilactobacillus-sakei] |
| WP_126133402.1 | (w)HTH+REase | GCF_003957415.1 | Latilactobacillus_curvatus | Bacteria>Bacillati>Bacillota>Bacilli>Lactobacillales>Lactobacillaceae>Latilactobacillus | 321 | CYK59_RS06540 | 1240789..1241754 | 28038 | restriction-endonuclease-[Latilactobacillus-curvatus] |
| WP_265678609.1 | (w)HTH+REase | GCF_026193275.1 | Latilactobacillus_curvatus | Bacteria>Bacillati>Bacillota>Bacilli>Lactobacillales>Lactobacillaceae>Latilactobacillus | 211 | OP752_RS00185 | 38192..38827 | 28038 | restriction-endonuclease-[Latilactobacillus-curvatus] |
| WP_265678610.1 | (w)HTH+REase | GCF_026193275.1 | Latilactobacillus_curvatus | Bacteria>Bacillati>Bacillota>Bacilli>Lactobacillales>Lactobacillaceae>Latilactobacillus | 93 | OP752_RS00190 | 38876..39157 | 28038 | winged-helix-turn-helix-domain-containing-protein-[Latilactobacillus-curvatus] |
| WP_273725614.1 | (w)HTH+REase | - | Leuconostoc_mesenteroides | Bacteria>Bacillati>Bacillota>Bacilli>Lactobacillales>Lactobacillaceae>Leuconostoc | 146 | PNN51_RS08075 | 19050..19490 | 1245 | hypothetical-protein-[Leuconostoc-mesenteroides] |
| WP_270411951.1 | (w)HTH+REase | GCF_027672425.1 | Ligilactobacillus_ruminis | Bacteria>Bacillati>Bacillota>Bacilli>Lactobacillales>Lactobacillaceae>Ligilactobacillus | 333 | PGT38_RS00035 | 5311..6312 | 1623 | restriction-endonuclease-[Ligilactobacillus-ruminis] |
| WP_009555624.1 | (w)HTH+REase | GCF_000317165.1 | Ligilactobacillus_saerimneri | Bacteria>Bacillati>Bacillota>Bacilli>Lactobacillales>Lactobacillaceae>Ligilactobacillus | 303 | D271_RS07740 | 69431..70342 | 228229 | restriction-endonuclease-[Ligilactobacillus-saerimneri] |
| WP_051006310.1 | (w)HTH+REase | GCF_000255515.1 | Liquorilactobacillus_vini | Bacteria>Bacillati>Bacillota>Bacilli>Lactobacillales>Lactobacillaceae>Liquorilactobacillus | 314 | IC26_RS02645 | 2069..3013 | 238015 | restriction-endonuclease-[Liquorilactobacillus-vini] |
| WP_152261654.1 | (w)HTH+REase | GCF_009217765.1 | Schleiferilactobacillus_harbinensis | Bacteria>Bacillati>Bacillota>Bacilli>Lactobacillales>Lactobacillaceae>Schleiferilactobacillus | 299 | D1010_RS17250 | 3431330..3432229 | 304207 | restriction-endonuclease-[Schleiferilactobacillus-harbinensis] |
| WP_150392671.1 | (w)HTH+REase | GCF_008694105.1 | Schleiferilactobacillus_harbinensis | Bacteria>Bacillati>Bacillota>Bacilli>Lactobacillales>Lactobacillaceae>Schleiferilactobacillus | 197 | FMM01_RS11910 | 2401137..2401730 | 304207 | restriction-endonuclease-[Schleiferilactobacillus-harbinensis] |
| MCR0160127.1 | (w)HTH+REase | GCA_024590065.1 | [Clostridium]_innocuum | Bacteria>Bacillati>Bacillota>Clostridia>Eubacteriales>Clostridiaceae>Clostridium | 333 | MKC37_06125 | 78490..79491 | 1522 | restriction-endonuclease-[[Clostridium] |
| WP_236911587.1 | (w)HTH+REase | GCF_021655095.1 | Clostridium_sp._Cult1 | Bacteria>Bacillati>Bacillota>Clostridia>Eubacteriales>Clostridiaceae>Clostridium>unclassified_Clostridium | 318 | C3E89_RS06130 | 1251947..1252903 | 2079002 | HTH-domain-containing-protein-[Clostridium-sp.-Cult1] |
| OLA08991.1 | (w)HTH+REase | GCA_001916415.1 | Eubacterium_sp._45_250 | Bacteria>Bacillati>Bacillota>Clostridia>Eubacteriales>Eubacteriaceae>Eubacterium>unclassified_Eubacterium_(in:_firmicutes) | 331 | BHW21_06120 | 153097..154092 | 1897004 | MAG:-restriction-endonuclease-[Eubacterium-sp.-45_250] |
| UYJ00846.1 | (w)HTH+REase | GCA_025757765.1 | Faecalibacterium_prausnitzii | Bacteria>Bacillati>Bacillota>Clostridia>Eubacteriales>Oscillospiraceae>Faecalibacterium | 348 | OGM65_10410 | 2153649..2154695 | 853 | MAG:-PDDEXK-nuclease-domain-containing-protein-[Faecalibacterium-prausnitzii] |
| WP_087287646.1 | (w)HTH+REase | GCF_002160305.1 | Pseudoflavonifractor_sp._An184 | Bacteria>Bacillati>Bacillota>Clostridia>Eubacteriales>Oscillospiraceae>Pseudoflavonifractor>unclassified_Pseudoflavonifractor | 359 | B5F19_RS02660 | 19546..20625 | 1965576 | PDDEXK-nuclease-domain-containing-protein-[Pseudoflavonifractor-sp.-An184] |
| SCH98925.1 | (w)HTH+REase | GCA_900067025.1 | uncultured_Ruminococcus_sp. | Bacteria>Bacillati>Bacillota>Clostridia>Eubacteriales>Oscillospiraceae>Ruminococcus>environmental_samples | 333 | SAMEA3545436_02041 | 70605..71606 | 165186 | Restriction-endonuclease-[uncultured-Ruminococcus-sp.] |
| WP_216553760.1 | (w)HTH+REase | GCF_018917825.1 | Ruminococcus_sp._MSJ-25 | Bacteria>Bacillati>Bacillota>Clostridia>Eubacteriales>Oscillospiraceae>Ruminococcus>unclassified_Ruminococcus | 366 | KQH97_RS08280 | 263070..264170 | 2841536 | PDDEXK-nuclease-domain-containing-protein-[Ruminococcus-sp.-MSJ-25] |
| RGG58357.1 | (w)HTH+REase | GCA_003461215.1 | Ruminococcus_sp._AF19-15 | Bacteria>Bacillati>Bacillota>Clostridia>Eubacteriales>Oscillospiraceae>Ruminococcus>unclassified_Ruminococcus | 333 | DWX34_06600 | 7147..8148 | 2293157 | restriction-endonuclease-[Ruminococcus-sp.-AF19-15] |
| MBQ6611345.1 | (w)HTH+REase | GCA_017434435.1 | Oscillospiraceae_bacterium | Bacteria>Bacillati>Bacillota>Clostridia>Eubacteriales>Oscillospiraceae>unclassified_Oscillospiraceae | 355 | IJH70_13020 | 5191..6258 | 2485925 | DUF1016-family-protein-[Oscillospiraceae-bacterium] |
| MCL2053371.1 | (w)HTH+REase | GCA_009777725.1 | Oscillospiraceae_bacterium | Bacteria>Bacillati>Bacillota>Clostridia>Eubacteriales>Oscillospiraceae>unclassified_Oscillospiraceae | 342 | FWG90_02870 | 159076..160104 | 2485925 | PDDEXK-nuclease-domain-containing-protein-[Oscillospiraceae-bacterium] |
| MCD8370865.1 | (w)HTH+REase | GCA_021205425.1 | Clostridiales_bacterium | Bacteria>Bacillati>Bacillota>Clostridia>Eubacteriales>unclassified_Eubacteriales | 353 | LUC94_11215 | 9720..10781 | 1898207 | PDDEXK-nuclease-domain-containing-protein-[Clostridiales-bacterium] |
| MBS4886750.1 | (w)HTH+REase | GCA_018363765.1 | Clostridiales_bacterium | Bacteria>Bacillati>Bacillota>Clostridia>Eubacteriales>unclassified_Eubacteriales | 332 | KH027_08815 | 22821..23819 | 1898207 | DUF1016-family-protein-[Clostridiales-bacterium] |
| WP_195942872.1 | (w)HTH+REase | GCF_015668255.1 | Lachnospiraceae | Bacteria>Bacillati>Bacillota>Clostridia>Lachnospirales | 337 | LOP77_RS14950 | 16209..17222 | 186803 | MULTISPECIES:-PDDEXK-nuclease-domain-containing-protein-[Lachnospiraceae] |
| WP_262586689.1 | (w)HTH+REase | GCF_025567525.1 | Gallintestinimicrobium | Bacteria>Bacillati>Bacillota>Clostridia>Lachnospirales>Lachnospiraceae | 332 | OCV62_RS11075 | 12433..13431 | 2981633 | PDDEXK-nuclease-domain-containing-protein-[Gallintestinimicrobium-propionicum] |
| WP_021651775.1 | (w)HTH+REase | GCF_020735365.1 | Blautia | Bacteria>Bacillati>Bacillota>Clostridia>Lachnospirales>Lachnospiraceae | 333 | LK422_RS11505 | 2401200..2402201 | 572511 | MULTISPECIES:-restriction-endonuclease-[Blautia] |
| CRL42136.1 | (w)HTH+REase | GCA_001406835.1 | Agathobacter_rectalis | Bacteria>Bacillati>Bacillota>Clostridia>Lachnospirales>Lachnospiraceae>Agathobacter | 337 | T1815_00881 | 96508..97521 | 39491 | hypothetical-protein-T1815_00881-[[Eubacterium] |
| RGF12873.1 | (w)HTH+REase | GCA_003435775.1 | Blautia_sp._AM16-16B | Bacteria>Bacillati>Bacillota>Clostridia>Lachnospirales>Lachnospiraceae>Blautia>unclassified_Blautia | 337 | DW177_12640 | 18030..19043 | 2292969 | DUF1016-domain-containing-protein-[Blautia-sp.-AM16-16B] |
| WP_195509730.1 | (w)HTH+REase | GCF_015556105.1 | Coprococcus_comes | Bacteria>Bacillati>Bacillota>Clostridia>Lachnospirales>Lachnospiraceae>Coprococcus | 332 | I2F75_RS13165 | 31438..32436 | 410072 | PDDEXK-nuclease-domain-containing-protein-[Coprococcus-comes] |
| WP_147567417.1 | (w)HTH+REase | GCF_902363775.1 | Dorea_formicigenerans | Bacteria>Bacillati>Bacillota>Clostridia>Lachnospirales>Lachnospiraceae>Dorea | 333 | G4398_RS10095 | 46983..47984 | 39486 | restriction-endonuclease-[Dorea-formicigenerans] |
| RGO24551.1 | (w)HTH+REase | GCA_003439775.1 | Dorea_sp._OM02-2LB | Bacteria>Bacillati>Bacillota>Clostridia>Lachnospirales>Lachnospiraceae>Dorea>unclassified_Dorea | 337 | DXB23_09220 | 38450..39463 | 2292347 | DUF1016-domain-containing-protein-[Dorea-sp.-OM02-2LB] |
| WP_016216875.1 | (w)HTH+REase | - | Dorea_sp._5-2 | Bacteria>Bacillati>Bacillota>Clostridia>Lachnospirales>Lachnospiraceae>Dorea>unclassified_Dorea | 213 | C817_RS00460 | 111726..112367 | 1235798 | DUF1016-N-terminal-domain-containing-protein-[Dorea-sp.-5-2] |
| MCQ4758733.1 | (w)HTH+REase | GCA_024460255.1 | Enterocloster_bolteae | Bacteria>Bacillati>Bacillota>Clostridia>Lachnospirales>Lachnospiraceae>Enterocloster | 332 | NE538_26275 | 9292..10290 | 208479 | PDDEXK-nuclease-domain-containing-protein-[Enterocloster-bolteae] |
| SFG31040.1 | (w)HTH+REase | GCA_900113415.1 | Oribacterium_sp._WCC10 | Bacteria>Bacillati>Bacillota>Clostridia>Lachnospirales>Lachnospiraceae>Oribacterium>unclassified_Oribacterium | 400 | SAMN05216356_105113 | 140831..142033 | 1855343 | Predicted-nuclease-of-restriction-endonuclease-like-(RecB)-superfamily,-DUF1016-family-[Oribacterium-sp.-WCC10] |
| WP_243090451.1 | (w)HTH+REase | GCF_003468625.1 | Roseburia_inulinivorans | Bacteria>Bacillati>Bacillota>Clostridia>Lachnospirales>Lachnospiraceae>Roseburia | 97 | DW813_RS18160 | 64734..65027 | 360807 | DUF1016-N-terminal-domain-containing-protein-[Roseburia-inulinivorans] |
| EOS42582.1 | (w)HTH+REase | GCA_000403315.2 | Lachnospiraceae_bacterium_A2 | Bacteria>Bacillati>Bacillota>Clostridia>Lachnospirales>Lachnospiraceae>unclassified_Lachnospiraceae | 383 | C810_04452 | 129146..130297 | 397290 | hypothetical-protein-C810_04452-[Lachnospiraceae-bacterium-A2] |
| SFL32312.1 | (w)HTH+REase | GCA_900114405.1 | Lachnospiraceae_bacterium_KH1T2 | Bacteria>Bacillati>Bacillota>Clostridia>Lachnospirales>Lachnospiraceae>unclassified_Lachnospiraceae | 380 | SAMN05216390_1198 | 9772..10914 | 1855374 | Predicted-nuclease-of-restriction-endonuclease-like-(RecB)-superfamily,-DUF1016-family-[Lachnospiraceae-bacterium-KH1T2] |
| MCC8150844.1 | (w)HTH+REase | GCA_020860745.1 | Lachnospiraceae_bacterium | Bacteria>Bacillati>Bacillota>Clostridia>Lachnospirales>Lachnospiraceae>unclassified_Lachnospiraceae | 373 | LIO96_05185 | 4835..5956 | 1898203 | PDDEXK-nuclease-domain-containing-protein-[Lachnospiraceae-bacterium] |
| MCD7957102.1 | (w)HTH+REase | GCA_021201665.1 | Lachnospiraceae_bacterium | Bacteria>Bacillati>Bacillota>Clostridia>Lachnospirales>Lachnospiraceae>unclassified_Lachnospiraceae | 360 | LUG93_15405 | 10406..11488 | 1898203 | PDDEXK-nuclease-domain-containing-protein-[Lachnospiraceae-bacterium] |
| MBD9124681.1 | (w)HTH+REase | GCA_014845505.1 | Lachnospiraceae_bacterium | Bacteria>Bacillati>Bacillota>Clostridia>Lachnospirales>Lachnospiraceae>unclassified_Lachnospiraceae | 332 | EGP68_08010 | 65470..66468 | 1898203 | DUF1016-domain-containing-protein-[Lachnospiraceae-bacterium] |
| MCI6533900.1 | (w)HTH+REase | GCA_022782135.1 | Lachnospiraceae_bacterium | Bacteria>Bacillati>Bacillota>Clostridia>Lachnospirales>Lachnospiraceae>unclassified_Lachnospiraceae | 304 | MR444_03715 | 35741..36655 | 1898203 | Mrr-restriction-system-protein-[Lachnospiraceae-bacterium] |
| WP_271847091.1 | (w)HTH+REase | GCF_028326405.1 | Peptostreptococcus_anaerobius | Bacteria>Bacillati>Bacillota>Clostridia>Peptostreptococcales>Peptostreptococcaceae>Peptostreptococcus | 336 | PN283_RS04685 | 25466..26476 | 1261 | restriction-endonuclease-[Peptostreptococcus-anaerobius] |
| WP_122014533.1 | (w)HTH+REase | GCF_003691585.1 | Biomaibacter_acetigenes | Bacteria>Bacillati>Bacillota>Clostridia>Thermosediminibacterales>Tepidanaerobacteraceae>Biomaibacter | 304 | D2962_RS06530 | 1341506..1342420 | 2316383 | restriction-endonuclease-[Biomaibacter-acetigenes] |
| WP_256268396.1 | (w)HTH+REase | GCF_048474855.1 | Holdemanella | Bacteria>Bacillati>Bacillota>Erysipelotrichia>Erysipelotrichales>Erysipelotrichaceae | 306 | NE544_RS03855 | 25369..26289 | 1573535 | Mrr-restriction-system-protein-[Holdemanella-sp.-MSK.7.32] |
| MCQ4803589.1 | (w)HTH+REase | GCA_024460475.1 | Holdemanella_sp._MSK.7.32 | Bacteria>Bacillati>Bacillota>Erysipelotrichia>Erysipelotrichales>Erysipelotrichaceae>Holdemanella>unclassified_Holdemanella | 306 | NE544_03855 | 25369..26289 | 2965273 | Mrr-restriction-system-protein-[Holdemanella-sp.-MSK.7.32] |
| WP_006060896.1 | (w)HTH+REase | GCF_000157995.1 | Holdemania_filiformis | Bacteria>Bacillati>Bacillota>Erysipelotrichia>Erysipelotrichales>Erysipelotrichaceae>Holdemania | 333 | HOLDEFILI_RS15940 | 3161..4162 | 61171 | restriction-endonuclease-[Holdemania-filiformis] |
| GJM58072.1 | (w)HTH+REase | GCA_022834735.1 | Erysipelotrichaceae_bacterium_OPF54 | Bacteria>Bacillati>Bacillota>Erysipelotrichia>Erysipelotrichales>Erysipelotrichaceae>unclassified_Erysipelotrichaceae | 240 | EROP_17650 | 1804298..1805020 | 2894072 | hypothetical-protein-EROP_17650-[Erysipelotrichaceae-bacterium-OPF54] |
| GJM58073.1 | (w)HTH+REase | GCA_022834735.1 | Erysipelotrichaceae_bacterium_OPF54 | Bacteria>Bacillati>Bacillota>Erysipelotrichia>Erysipelotrichales>Erysipelotrichaceae>unclassified_Erysipelotrichaceae | 87 | EROP_17660 | 1805129..1805392 | 2894072 | hypothetical-protein-EROP_17660-[Erysipelotrichaceae-bacterium-OPF54] |
| HCR67834.1 | (w)HTH+REase | GCA_003533525.1 | Erysipelotrichaceae_bacterium | Bacteria>Bacillati>Bacillota>Erysipelotrichia>Erysipelotrichales>Erysipelotrichaceae>unclassified_Erysipelotrichaceae | 305 | DIV43_01650 | 12657..13574 | 2049044 | restriction-endonuclease-[Erysipelotrichaceae-bacterium] |
| WP_270440307.1 | (w)HTH+REase | - | Acidaminococcus_provencensis | Bacteria>Bacillati>Bacillota>Negativicutes>Acidaminococcales>Acidaminococcaceae>Acidaminococcus | 94 | PGX33_RS00705 | 140266..140550 | 2058289 | PDDEXK-nuclease-domain-containing-protein-[Acidaminococcus-provencensis] |
| MBP8598622.1 | (w)HTH+REase | GCA_018052705.1 | Selenomonas_sp. | Bacteria>Bacillati>Bacillota>Negativicutes>Selenomonadales>Selenomonadaceae>Selenomonas>unclassified_Selenomonas | 337 | KBI24_05325 | 17216..18229 | 2053611 | DUF1016-family-protein-[Selenomonas-sp.] |
| WP_273012937.1 | (w)HTH+REase | GCF_014872435.1 | Dialister_invisus | Bacteria>Bacillati>Bacillota>Negativicutes>Veillonellales>Veillonellaceae>Dialister | 332 | M3287_RS05165 | 3346..4344 | 218538 | PDDEXK-nuclease-domain-containing-protein-[Dialister-invisus] |
| OUL09473.1 | (w)HTH+REase | GCA_002155105.1 | Sedimentibacter_sp._SX930 | Bacteria>Bacillati>Bacillota>Tissierellia>Tissierellia_incertae_sedis>Sedimentibacter>unclassified_Sedimentibacter | 141 | B0533_04725 | 286378..286803 | 1955060 | hypothetical-protein-B0533_04725-[Sedimentibacter-sp.-SX930] |
| RHS79251.1 | (w)HTH+REase | GCA_003480065.1 | Firmicutes_bacterium_AM43-11BH | Bacteria>Bacillati>Bacillota>unclassified_Bacillota | 337 | DW928_10635 | 83687..84700 | 2292895 | DUF1016-domain-containing-protein-[Firmicutes-bacterium-AM43-11BH] |
| MBS6916849.1 | (w)HTH+REase | GCA_018380515.1 | Bacillota_bacterium | Bacteria>Bacillati>Bacillota>unclassified_Bacillota | 332 | KH272_06720 | 40366..41364 | 1879010 | DUF1016-family-protein-[Bacillota-bacterium] |
| MCI6854479.1 | (w)HTH+REase | GCA_022776865.1 | Bacillota_bacterium | Bacteria>Bacillati>Bacillota>unclassified_Bacillota | 332 | MR908_04040 | 5933..6931 | 1879010 | PDDEXK-nuclease-domain-containing-protein-[Bacillota-bacterium] |
| NPV30264.1 | (w)HTH+REase | GCA_013177945.1 | Bacillota_bacterium | Bacteria>Bacillati>Bacillota>unclassified_Bacillota | 301 | HPY58_11585 | 6524..7429 | 1879010 | restriction-endonuclease-[Bacillota-bacterium] |
| MCE7987120.1 | (w)HTH+REase | GCA_021405325.1 | Caldilinea_sp._CFX5 | Bacteria>Bacillati>Chloroflexota>Caldilineae>Caldilineales>Caldilineaceae>Caldilinea>unclassified_Caldilinea | 371 | DYG89_38610 | 42301..43416 | 2293554 | DUF1016-domain-containing-protein-[Caldilinea-sp.-CFX5] |
| MCF7634649.1 | (w)HTH+REase | GCA_021732865.1 | Dehalococcoides_mccartyi | Bacteria>Bacillati>Chloroflexota>Dehalococcoidia>Dehalococcoidales>Dehalococcoidaceae>Dehalococcoides | 259 | LLF82_000113 | 103867..104646 | 61435 | hypothetical-protein-[Dehalococcoides-mccartyi] |
| WP_227028542.1 | (w)HTH+REase | GCF_000341655.1 | Dehalococcoides_mccartyi | Bacteria>Bacillati>Chloroflexota>Dehalococcoidia>Dehalococcoidales>Dehalococcoidaceae>Dehalococcoides | 193 | DCMB_RS01620 | 325012..325593 | 61435 | hypothetical-protein-[Dehalococcoides-mccartyi] |
| MCC6178179.1 | (w)HTH+REase | GCA_020845495.1 | Chloroflexota_bacterium | Bacteria>Bacillati>Chloroflexota>unclassified_Chloroflexota | 395 | IT305_22985 | 12684..13871 | 2026724 | DUF1016-family-protein-[Chloroflexota-bacterium] |
| OGH97203.1 | (w)HTH+REase | GCA_001783265.1 | Candidatus_Margulisbacteria_bacterium_GWD2_39_127 | Bacteria>Bacillati>Cyanobacteriota/Melainabacteria_group>Candidatus_Margulisiibacteriota>unclassified_Candidatus_Margulisiibacteriota | 338 | A2X43_08535 | 62696..63712 | 1817871 | hypothetical-protein-A2X43_08535-[Candidatus-Margulisbacteria-bacterium-GWD2_39_127] |
| OGI09202.1 | (w)HTH+REase | GCA_001784485.1 | Candidatus_Margulisbacteria_bacterium_GWE2_39_32 | Bacteria>Bacillati>Cyanobacteriota/Melainabacteria_group>Candidatus_Margulisiibacteriota>unclassified_Candidatus_Margulisiibacteriota | 338 | A2X41_01340 | 12427..13443 | 1817872 | hypothetical-protein-A2X41_01340-[Candidatus-Margulisbacteria-bacterium-GWE2_39_32] |
| PZM83792.1 | (w)HTH+REase | GCA_003242895.1 | Candidatus_Margulisiibacteriota_bacterium | Bacteria>Bacillati>Cyanobacteriota/Melainabacteria_group>Candidatus_Margulisiibacteriota>unclassified_Candidatus_Margulisiibacteriota | 333 | DKM50_01280 | 7709..8710 | 2053573 | DUF1016-domain-containing-protein-[Candidatus-Margulisbacteria-bacterium] |
| WP_036047050.1 | (w)HTH+REase | GCF_002142475.1 | Leptolyngbya | Bacteria>Bacillati>Cyanobacteriota/Melainabacteria_group>Cyanobacteriota>Cyanophyceae>Leptolyngbyales>Leptolyngbyaceae>Leptolyngbya_group | 347 | MCP04_RS33165 | 392303..393346 | 47251 | PDDEXK-nuclease-domain-containing-protein-[Leptolyngbya-boryana] |
| WP_088889981.1 | (w)HTH+REase | GCF_002215035.1 | Leptolyngbya_ohadii | Bacteria>Bacillati>Cyanobacteriota/Melainabacteria_group>Cyanobacteriota>Cyanophyceae>Leptolyngbyales>Leptolyngbyaceae>Leptolyngbya_group>Leptolyngbya | 367 | CDV24_RS06820 | 194388..195491 | 1962290 | PDDEXK-nuclease-domain-containing-protein-[Leptolyngbya-ohadii] |
| WP_199323865.1 | (w)HTH+REase | - | unclassified_Leptolyngbya | Bacteria>Bacillati>Cyanobacteriota/Melainabacteria_group>Cyanobacteriota>Cyanophyceae>Leptolyngbyales>Leptolyngbyaceae>Leptolyngbya_group>Leptolyngbya | 341 | H6G61_RS21990 | 24881..25906 | 2650499 | MULTISPECIES:-PDDEXK-nuclease-domain-containing-protein-[unclassified-Leptolyngbya] |
| WP_006277873.1 | (w)HTH+REase | GCF_000175835.1 | Cylindrospermopsis_raciborskii | Bacteria>Bacillati>Cyanobacteriota/Melainabacteria_group>Cyanobacteriota>Cyanophyceae>Nostocales>Aphanizomenonaceae>Cylindrospermopsis | 349 | CRC_RS10870 | 7080..8129 | 77022 | PDDEXK-nuclease-domain-containing-protein-[Cylindrospermopsis-raciborskii] |
| WP_071248157.1 | (w)HTH+REase | GCF_001858125.1 | Cylindrospermopsis_raciborskii | Bacteria>Bacillati>Cyanobacteriota/Melainabacteria_group>Cyanobacteriota>Cyanophyceae>Nostocales>Aphanizomenonaceae>Cylindrospermopsis | 349 | CENA302_RS09800 | 25812..26861 | 77022 | PDDEXK-nuclease-domain-containing-protein-[Cylindrospermopsis-raciborskii] |
| WP_102947436.1 | (w)HTH+REase | GCF_002893285.1 | Cylindrospermopsis_raciborskii | Bacteria>Bacillati>Cyanobacteriota/Melainabacteria_group>Cyanobacteriota>Cyanophyceae>Nostocales>Aphanizomenonaceae>Cylindrospermopsis | 349 | CEP07_RS06060 | 8443..9492 | 77022 | PDDEXK-nuclease-domain-containing-protein-[Cylindrospermopsis-raciborskii] |
| WP_236308449.1 | (w)HTH+REase | GCF_021650815.1 | Cylindrospermopsis_raciborskii | Bacteria>Bacillati>Cyanobacteriota/Melainabacteria_group>Cyanobacteriota>Cyanophyceae>Nostocales>Aphanizomenonaceae>Cylindrospermopsis | 349 | L3I90_RS00095 | 17335..18384 | 77022 | PDDEXK-nuclease-domain-containing-protein-[Cylindrospermopsis-raciborskii] |
| WP_115538498.1 | (w)HTH+REase | GCF_003367075.2 | Cylindrospermopsis_raciborskii | Bacteria>Bacillati>Cyanobacteriota/Melainabacteria_group>Cyanobacteriota>Cyanophyceae>Nostocales>Aphanizomenonaceae>Cylindrospermopsis | 148 | J9B92_RS02350 | 518111..518557 | 77022 | DUF1016-N-terminal-domain-containing-protein-[Cylindrospermopsis-raciborskii] |
| WP_143042783.1 | (w)HTH+REase | GCF_001858115.1 | Cylindrospermopsis_raciborskii | Bacteria>Bacillati>Cyanobacteriota/Melainabacteria_group>Cyanobacteriota>Cyanophyceae>Nostocales>Aphanizomenonaceae>Cylindrospermopsis | 148 | BCV63_RS11895 | 54814..55260 | 77022 | DUF1016-N-terminal-domain-containing-protein-[Cylindrospermopsis-raciborskii] |
| WP_187705579.1 | (w)HTH+REase | GCF_014489415.1 | Cylindrospermopsis_curvispora | Bacteria>Bacillati>Cyanobacteriota/Melainabacteria_group>Cyanobacteriota>Cyanophyceae>Nostocales>Aphanizomenonaceae>Cylindrospermopsis | 148 | IAR63_RS13145 | 2955554..2956000 | 747548 | DUF1016-N-terminal-domain-containing-protein-[Cylindrospermopsis-curvispora] |
| MCH4905400.1 | (w)HTH+REase | GCA_022496815.1 | Cylindrospermopsis_raciborskii_CHAB3438 | Bacteria>Bacillati>Cyanobacteriota/Melainabacteria_group>Cyanobacteriota>Cyanophyceae>Nostocales>Aphanizomenonaceae>Cylindrospermopsis>Cylindrospermopsis_raciborskii | 349 | GSN00_13710 | 7003..8052 | 1480071 | DUF1016-family-protein-[Cylindrospermopsis-raciborskii-CHAB3438] |
| UJL35157.1 | (w)HTH+REase | GCA_003367075.2 | Cylindrospermopsis_raciborskii_Cr2010 | Bacteria>Bacillati>Cyanobacteriota/Melainabacteria_group>Cyanobacteriota>Cyanophyceae>Nostocales>Aphanizomenonaceae>Cylindrospermopsis>Cylindrospermopsis_raciborskii | 148 | C6N34_008470 | 1854720..1855166 | 2108055 | DUF1016-family-protein-[Cylindrospermopsis-raciborskii-Cr2010] |
| WP_057176818.1 | (w)HTH+REase | GCF_001432185.1 | Cylindrospermopsis_sp._CR12 | Bacteria>Bacillati>Cyanobacteriota/Melainabacteria_group>Cyanobacteriota>Cyanophyceae>Nostocales>Aphanizomenonaceae>Cylindrospermopsis>unclassified_Cylindrospermopsis | 148 | ASL19_RS00240 | 53320..53766 | 1747196 | DUF1016-N-terminal-domain-containing-protein-[Cylindrospermopsis-sp.-CR12] |
| BAZ90339.1 | (w)HTH+REase | GCA_002368135.1 | Raphidiopsis_curvata_NIES-932 | Bacteria>Bacillati>Cyanobacteriota/Melainabacteria_group>Cyanobacteriota>Cyanophyceae>Nostocales>Aphanizomenonaceae>Raphidiopsis>Raphidiopsis_curvata | 148 | NIES932_18380 | 1951062..1951508 | 1973482 | hypothetical-protein-NIES932_18380-[Raphidiopsis-curvata-NIES-932] |
| MBV6627481.1 | (w)HTH+REase | GCA_019192545.1 | Rivularia_sp._MS3 | Bacteria>Bacillati>Cyanobacteriota/Melainabacteria_group>Cyanobacteriota>Cyanophyceae>Nostocales>Rivulariaceae>Rivularia>unclassified_Rivularia_(in:_cyanobacteria) | 394 | KI793_31930 | 9329..10513 | 3012223 | DUF1016-family-protein-[Rivularia-sp.-(in:-cyanobacteria)] |
| QNI65419.1 | (w)HTH+REase | GCA_014280115.1 | Synechococcus_sp._A15-44 | Bacteria>Bacillati>Cyanobacteriota/Melainabacteria_group>Cyanobacteriota>Cyanophyceae>Synechococcales>Synechococcaceae>Synechococcus>unclassified_Synechococcus | 308 | SynA1544_02498 | 2017889..2018815 | 1050646 | type-IV-restriction-modification-system/-Mrr-cat-superfamily-[Synechococcus-sp.-A15-44] |
| WP_186471769.1 | (w)HTH+REase | GCF_014279955.1 | Synechococcus_sp._BMK-MC-1 | Bacteria>Bacillati>Cyanobacteriota/Melainabacteria_group>Cyanobacteriota>Cyanophyceae>Synechococcales>Synechococcaceae>Synechococcus>unclassified_Synechococcus | 308 | SynBMKMC1_RS11035 | 1954923..1955849 | 1442551 | restriction-endonuclease-[Synechococcus-sp.-BMK-MC-1] |
| WP_188020746.1 | (w)HTH+REase | GCF_014279615.1 | Synechococcus_sp._RS9907 | Bacteria>Bacillati>Cyanobacteriota/Melainabacteria_group>Cyanobacteriota>Cyanophyceae>Synechococcales>Synechococcaceae>Synechococcus>unclassified_Synechococcus | 298 | SynRS9907_RS09000 | 1630238..1631134 | 221350 | restriction-endonuclease-[Synechococcus-sp.-RS9907] |
| MCY4332865.1 | (w)HTH+REase | GCA_026710375.1 | Cyanobacteria_bacterium_MAG_CAR1_bin_15 | Bacteria>Bacillati>Cyanobacteriota/Melainabacteria_group>Cyanobacteriota>unclassified_Cyanobacteriota | 279 | OXC96_10310 | 32581..33420 | 2775589 | MAG:-hypothetical-protein-OXC96_10310-[Cyanobacteria-bacterium-MAG-CAR1_bin_15] |
| MCP6718635.1 | (w)HTH+REase | GCA_024239015.1 | Patescibacteria_group_bacterium | Bacteria>Bacteria_incertae_sedis>Bacteria_candidate_phyla>Patescibacteria_group>unclassified_Patescibacteria_group | 301 | KJI71_00200 | 46098..47003 | 2052139 | restriction-endonuclease-[Patescibacteria-group-bacterium] |
| OGR04296.1 | (w)HTH+REase | GCA_001799195.1 | Deltaproteobacteria_bacterium_RIFOXYA12_FULL_61_11 | Bacteria>delta/epsilon_subdivisions>Deltaproteobacteria>unclassified_Deltaproteobacteria | 370 | A2284_01910 | 9535..10647 | 1797896 | cytoplasmic-protein-[Deltaproteobacteria-bacterium-RIFOXYA12_FULL_61_11] |
| EFK09423.1 | (w)HTH+REase | - | delta_proteobacterium_NaphS2 | Bacteria>delta/epsilon_subdivisions>Deltaproteobacteria>unclassified_Deltaproteobacteria | 367 | NPH_3109 | 48946..50049 | 88274 | conserved-hypothetical-protein-[delta-proteobacterium-NaphS2] |
| OGP05896.1 | (w)HTH+REase | GCA_001795495.1 | Deltaproteobacteria_bacterium_GWA2_42_85 | Bacteria>delta/epsilon_subdivisions>Deltaproteobacteria>unclassified_Deltaproteobacteria | 341 | A2056_04325 | 7025..8050 | 1797803 | hypothetical-protein-A2056_04325-[Deltaproteobacteria-bacterium-GWA2_42_85] |
| OQB76676.1 | (w)HTH+REase | GCA_002070575.1 | Deltaproteobacteria_bacterium_ADurb.Bin135 | Bacteria>delta/epsilon_subdivisions>Deltaproteobacteria>unclassified_Deltaproteobacteria | 259 | BWX92_01601 | 34884..35663 | 1852867 | hypothetical-protein-BWX92_01601-[Deltaproteobacteria-bacterium-ADurb.Bin135] |
| WP_058195269.1 | (w)HTH+REase | GCF_030034615.1 | Pseudomonadota | Bacteria>Pseudomonadati | 350 | AO969_RS25395 | 24636..25688 | 1224 | MULTISPECIES:-PDDEXK-nuclease-domain-containing-protein-[Pseudomonadota] |
| WP_060853051.1 | (w)HTH+REase | GCF_001548335.1 | Pseudomonadota | Bacteria>Pseudomonadati | 350 | BT108_RS06140 | 118649..119701 | 1224 | MULTISPECIES:-PDDEXK-nuclease-domain-containing-protein-[Pseudomonadota] |
| WP_102774926.1 | (w)HTH+REase | GCF_024464665.1 | Pseudomonadota | Bacteria>Pseudomonadati | 85 | C1I89_RS23975 | 481618..481875 | 1224 | MULTISPECIES:-hypothetical-protein-[Pseudomonadota] |
| WP_141050578.1 | (w)HTH+REase | GCF_031631975.1 | Aliarcobacter | Bacteria>Pseudomonadati>Campylobacterota>Epsilonproteobacteria>Campylobacterales>Arcobacteraceae | 343 | AV936_RS08800 | 15229..16260 | 2321111 | PDDEXK-nuclease-domain-containing-protein-[Aliarcobacter-cryaerophilus] |
| WP_172658767.1 | (w)HTH+REase | GCF_013201725.1 | Arcobacter_defluvii | Bacteria>Pseudomonadati>Campylobacterota>Epsilonproteobacteria>Campylobacterales>Arcobacteraceae>Arcobacter | 333 | ADFLV_RS07195 | 1428803..1429804 | 873191 | PDDEXK-nuclease-domain-containing-protein-[Arcobacter-defluvii] |
| WP_258237385.1 | (w)HTH+REase | GCF_004116575.1 | Arcobacter_sp._AHV-9/2010 | Bacteria>Pseudomonadati>Campylobacterota>Epsilonproteobacteria>Campylobacterales>Arcobacteraceae>Arcobacter>unclassified_Arcobacter | 310 | CRU94_RS04015 | 55977..56909 | 2021861 | PDDEXK-nuclease-domain-containing-protein-[Arcobacter-sp.-CECT-9299] |
| RXJ96079.1 | (w)HTH+REase | GCA_004116575.1 | Arcobacter_sp._AHV-9/2010 | Bacteria>Pseudomonadati>Campylobacterota>Epsilonproteobacteria>Campylobacterales>Arcobacteraceae>Arcobacter>unclassified_Arcobacter | 306 | CRU94_04015 | 55989..56909 | 2021861 | hypothetical-protein-CRU94_04015-[Arcobacter-sp.-CECT-9299] |
| PHS59249.1 | (w)HTH+REase | GCA_002733155.1 | Sulfurimonas_sp. | Bacteria>Pseudomonadati>Campylobacterota>Epsilonproteobacteria>Campylobacterales>Sulfurimonadaceae>Sulfurimonas>unclassified_Sulfurimonas | 328 | COB17_00840 | 114015..115001 | 2022749 | MAG:-hypothetical-protein-COB17_00840-[Sulfurimonas-sp.] |
| MBL0059012.1 | (w)HTH+REase | GCA_016721625.1 | Elusimicrobiota_bacterium | Bacteria>Pseudomonadati>Elusimicrobiota>unclassified_Elusimicrobiota | 395 | IPP35_07870 | 1836777..1837964 | 2030800 | DUF1016-family-protein-[Elusimicrobiota-bacterium] |
| WP_008760279.1 | (w)HTH+REase | GCF_001373135.1 | Bacteroides | Bacteria>Pseudomonadati>FCB_group>Bacteroidota/Chlorobiota_group>Bacteroidota>Bacteroidia>Bacteroidales>Bacteroidaceae | 362 | HMPREF9007_RS20225 | 177175..178263 | 816 | MULTISPECIES:-PDDEXK-nuclease-domain-containing-protein-[Bacteroides] |
| WP_130041242.1 | (w)HTH+REase | GCF_052367925.1 | Bacteroides_thetaiotaomicron | Bacteria>Pseudomonadati>FCB_group>Bacteroidota/Chlorobiota_group>Bacteroidota>Bacteroidia>Bacteroidales>Bacteroidaceae>Bacteroides | 362 | EAJ10_RS04935 | 120110..121198 | 818 | PDDEXK-nuclease-domain-containing-protein-[Bacteroides-thetaiotaomicron] |
| WP_225659471.1 | (w)HTH+REase | GCF_020313445.1 | Bacteroides_thetaiotaomicron | Bacteria>Pseudomonadati>FCB_group>Bacteroidota/Chlorobiota_group>Bacteroidota>Bacteroidia>Bacteroidales>Bacteroidaceae>Bacteroides | 362 | KFX61_RS09565 | 29813..30901 | 818 | PDDEXK-nuclease-domain-containing-protein-[Bacteroides-thetaiotaomicron] |
| WP_258972449.1 | (w)HTH+REase | GCF_024793205.1 | Bacteroides_thetaiotaomicron | Bacteria>Pseudomonadati>FCB_group>Bacteroidota/Chlorobiota_group>Bacteroidota>Bacteroidia>Bacteroidales>Bacteroidaceae>Bacteroides | 362 | NXX32_RS11550 | 3192926..3194014 | 818 | PDDEXK-nuclease-domain-containing-protein-[Bacteroides-thetaiotaomicron] |
| WP_250702520.1 | (w)HTH+REase | GCF_023668595.1 | Bacteroides_ovatus | Bacteria>Pseudomonadati>FCB_group>Bacteroidota/Chlorobiota_group>Bacteroidota>Bacteroidia>Bacteroidales>Bacteroidaceae>Bacteroides | 358 | NCZ26_RS08915 | 87547..88623 | 28116 | PDDEXK-nuclease-domain-containing-protein-[Bacteroides-ovatus] |
| MBW6491486.1 | (w)HTH+REase | GCA_019429275.1 | Lentimicrobium_sp. | Bacteria>Pseudomonadati>FCB_group>Bacteroidota/Chlorobiota_group>Bacteroidota>Bacteroidia>Bacteroidales>Lentimicrobiaceae>Lentimicrobium>unclassified_Lentimicrobium | 375 | KOB15_09880 | 51375..52502 | 2034841 | DUF1016-family-protein-[Lentimicrobium-sp.] |
| WP_135984907.1 | (w)HTH+REase | GCF_004793975.1 | Lepagella_muris | Bacteria>Pseudomonadati>FCB_group>Bacteroidota/Chlorobiota_group>Bacteroidota>Bacteroidia>Bacteroidales>Muribaculaceae>Lepagella | 401 | E5331_RS10700 | 58108..59313 | 3032870 | PDDEXK-nuclease-domain-containing-protein-[Lepagella-muris] |
| MCI1742485.1 | (w)HTH+REase | GCA_022647785.1 | Prevotella_sp. | Bacteria>Pseudomonadati>FCB_group>Bacteroidota/Chlorobiota_group>Bacteroidota>Bacteroidia>Bacteroidales>Prevotellaceae>Prevotella>unclassified_Prevotella | 355 | LKI18_09140 | 33073..34140 | 59823 | PDDEXK-nuclease-domain-containing-protein-[Prevotella-sp.] |
| MCI2087043.1 | (w)HTH+REase | GCA_022653375.1 | Prevotella_sp. | Bacteria>Pseudomonadati>FCB_group>Bacteroidota/Chlorobiota_group>Bacteroidota>Bacteroidia>Bacteroidales>Prevotellaceae>Prevotella>unclassified_Prevotella | 355 | LKK10_02005 | 5013..6080 | 59823 | PDDEXK-nuclease-domain-containing-protein-[Prevotella-sp.] |
| MCB9017745.1 | (w)HTH+REase | GCA_020636675.1 | Prevotellaceae_bacterium | Bacteria>Pseudomonadati>FCB_group>Bacteroidota/Chlorobiota_group>Bacteroidota>Bacteroidia>Bacteroidales>Prevotellaceae>unclassified_Prevotellaceae | 342 | H6544_03975 | 57399..58427 | 2049047 | DUF1016-domain-containing-protein-[Prevotellaceae-bacterium] |
| MBQ9588228.1 | (w)HTH+REase | GCA_017446745.1 | Bacteroidales_bacterium | Bacteria>Pseudomonadati>FCB_group>Bacteroidota/Chlorobiota_group>Bacteroidota>Bacteroidia>Bacteroidales>unclassified_Bacteroidales | 407 | IJR26_10285 | 26230..27453 | 2030927 | DUF1016-family-protein-[Bacteroidales-bacterium] |
| MBR5399196.1 | (w)HTH+REase | GCA_017550655.1 | Bacteroidales_bacterium | Bacteria>Pseudomonadati>FCB_group>Bacteroidota/Chlorobiota_group>Bacteroidota>Bacteroidia>Bacteroidales>unclassified_Bacteroidales | 395 | IK103_05325 | 47547..48734 | 2030927 | DUF1016-family-protein-[Bacteroidales-bacterium] |
| MCC8119637.1 | (w)HTH+REase | GCA_020860515.1 | Bacteroidales_bacterium | Bacteria>Pseudomonadati>FCB_group>Bacteroidota/Chlorobiota_group>Bacteroidota>Bacteroidia>Bacteroidales>unclassified_Bacteroidales | 372 | LIP09_12945 | 28009..29127 | 2030927 | PDDEXK-nuclease-domain-containing-protein-[Bacteroidales-bacterium] |
| MBN1340542.1 | (w)HTH+REase | GCA_016927695.1 | Bacteroidales_bacterium | Bacteria>Pseudomonadati>FCB_group>Bacteroidota/Chlorobiota_group>Bacteroidota>Bacteroidia>Bacteroidales>unclassified_Bacteroidales | 367 | JXA03_14530 | 28023..29126 | 2030927 | DUF1016-family-protein-[Bacteroidales-bacterium] |
| NLA16131.1 | (w)HTH+REase | GCA_012510445.1 | Bacteroidales_bacterium | Bacteria>Pseudomonadati>FCB_group>Bacteroidota/Chlorobiota_group>Bacteroidota>Bacteroidia>Bacteroidales>unclassified_Bacteroidales | 309 | GX877_06315 | 452..1381 | 2030927 | Mrr-restriction-system-protein-[Bacteroidales-bacterium] |

| Protein-ID | Domain | Assembly-ID | Strain | Taxonomy | Length | Locus-Tag | Genomic-coordinates | TaxID | Fasta-defline |
|---|---|---|---|---|---|---|---|---|---|
| MBK9465917.1 | (w)HTH+REase | GCA_016719015.1 | Chitinophagaceae_bacterium | Bacteria>Pseudomonadati>FCB_group>Bacteroidota/Chlorobiota_group>Bacteroidota>Chitinophagia>Chitinophagales>Chitinophagaceae>unclassified_Chitinophagaceae | 360 | IPN56_14275 | 11794..12876 | 1869212 | DUF1016-family-protein-[Chitinophagaceae-bacterium] |
| MBK9661553.1 | (w)HTH+REase | GCA_016717645.1 | Chitinophagaceae_bacterium | Bacteria>Pseudomonadati>FCB_group>Bacteroidota/Chlorobiota_group>Bacteroidota>Chitinophagia>Chitinophagales>Chitinophagaceae>unclassified_Chitinophagaceae | 360 | IPO68_16920 | 8242..9324 | 1869212 | DUF1016-family-protein-[Chitinophagaceae-bacterium] |
| MCK9404660.1 | (w)HTH+REase | GCA_023229465.1 | Chitinophagaceae_bacterium | Bacteria>Pseudomonadati>FCB_group>Bacteroidota/Chlorobiota_group>Bacteroidota>Chitinophagia>Chitinophagales>Chitinophagaceae>unclassified_Chitinophagaceae | 319 | MOQ26_14805 | 2334..3293 | 1869212 | HTH-domain-containing-protein-[Chitinophagaceae-bacterium] |
| WP_019991367.1 | (w)HTH+REase | GCF_000383955.1 | Rudanella_lutea | Bacteria>Pseudomonadati>FCB_group>Bacteroidota/Chlorobiota_group>Bacteroidota>Cytophagia>Cytophagales>Cytophagaceae>Rudanella | 352 | RUDLU_RS0125940 | 44683..45741 | 451374 | PDDEXK-nuclease-domain-containing-protein-[Rudanella-lutea] |
| TAK42616.1 | (w)HTH+REase | GCA_004297645.1 | Saprospiraceae_bacterium | Bacteria>Pseudomonadati>FCB_group>Bacteroidota/Chlorobiota_group>Bacteroidota>Saprospiria>Saprospirales>Saprospiraceae>unclassified_Saprospiraceae | 348 | EPO28_07055 | 15356..16402 | 2202734 | DUF1016-domain-containing-protein-[Saprospiraceae-bacterium] |
| WP_070561015.1 | (w)HTH+REase | GCF_036805785.1 | Sphingobacterium | Bacteria>Pseudomonadati>FCB_group>Bacteroidota/Chlorobiota_group>Bacteroidota>Sphingobacteriia>Sphingobacteriales>Sphingobacteriaceae | 334 | HMPREF3127_RS01220 | 91178..92182 | 28453 | PDDEXK-nuclease-domain-containing-protein-[Sphingobacterium-sp.-HMSC13C05] |
| WP_165305704.1 | (w)HTH+REase | GCF_011064245.1 | Sphingobacterium_sp._DR205 | Bacteria>Pseudomonadati>FCB_group>Bacteroidota/Chlorobiota_group>Bacteroidota>Sphingobacteriia>Sphingobacteriales>Sphingobacteriaceae>Sphingobacterium>unclassified_Sphingobacterium | 334 | G6053_RS05220 | 1251034..1252038 | 2713573 | PDDEXK-nuclease-domain-containing-protein-[Sphingobacterium-sp.-DR205] |
| MCY3630041.1 | (w)HTH+REase | GCA_026705375.1 | Bacteroidota_bacterium | Bacteria>Pseudomonadati>FCB_group>Bacteroidota/Chlorobiota_group>Bacteroidota>unclassified_Bacteroidota | 406 | OXG94_08525 | 7517..8737 | 1898104 | MAG:-PDDEXK-nuclease-domain-containing-protein-[Bacteroidetes-bacterium] |
| MDE2646789.1 | (w)HTH+REase | GCA_028875465.1 | Bacteroidota_bacterium | Bacteria>Pseudomonadati>FCB_group>Bacteroidota/Chlorobiota_group>Bacteroidota>unclassified_Bacteroidota | 406 | OXI05_13265 | 10403..11623 | 1898104 | MAG:-PDDEXK-nuclease-domain-containing-protein-[Bacteroidota-bacterium] |
| NQT60363.1 | (w)HTH+REase | GCA_013202335.1 | Bacteroidota_bacterium | Bacteria>Pseudomonadati>FCB_group>Bacteroidota/Chlorobiota_group>Bacteroidota>unclassified_Bacteroidota | 378 | HQ557_15390 | 12411..13547 | 1898104 | DUF1016-domain-containing-protein-[Bacteroidota-bacterium] |
| MCH7965545.1 | (w)HTH+REase | GCA_022564295.1 | Bacteroidota_bacterium | Bacteria>Pseudomonadati>FCB_group>Bacteroidota/Chlorobiota_group>Bacteroidota>unclassified_Bacteroidota | 373 | IH852_16600 | 23678..24799 | 1898104 | DUF1016-family-protein-[Bacteroidota-bacterium] |
| MCY4001542.1 | (w)HTH+REase | GCA_026708685.1 | Bacteroidota_bacterium | Bacteria>Pseudomonadati>FCB_group>Bacteroidota/Chlorobiota_group>Bacteroidota>unclassified_Bacteroidota | 122 | OXF84_12160 | 1..369 | 1898104 | MAG:-PDDEXK-nuclease-domain-containing-protein,-partial-[Bacteroidetes-bacterium] |
| MBL6956531.1 | (w)HTH+REase | GCA_016776555.1 | Chlorobium_phaeobacteroides | Bacteria>Pseudomonadati>FCB_group>Bacteroidota/Chlorobiota_group>Chlorobiota>Chlorobiia>Chlorobiales>Chlorobiaceae>Chlorobium/Pelodictyon_group>Chlorobium | 349 | ISR54_06900 | 30344..31393 | 1096 | DUF1016-family-protein-[Chlorobium-phaeobacteroides] |
| WP_082678574.1 | (w)HTH+REase | GCF_001509575.1 | Chlorobium_limicola | Bacteria>Pseudomonadati>FCB_group>Bacteroidota/Chlorobiota_group>Chlorobiota>Chlorobiia>Chlorobiales>Chlorobiaceae>Chlorobium/Pelodictyon_group>Chlorobium | 349 | ASB62_RS03900 | 14348..15397 | 1092 | PDDEXK-nuclease-domain-containing-protein-[Chlorobium-limicola] |
| KAA6232799.1 | (w)HTH+REase | GCA_008634005.1 | Chlorobium_phaeovibrioides | Bacteria>Pseudomonadati>FCB_group>Bacteroidota/Chlorobiota_group>Chlorobiota>Chlorobiia>Chlorobiales>Chlorobiaceae>Chlorobium/Pelodictyon_group>Chlorobium | 349 | FP507_06800 | 1396693..1397742 | 1094 | restriction-endonuclease-[Chlorobium-phaeovibrioides] |
| WP_151419518.1 | (w)HTH+REase | GCF_008634005.1 | Chlorobium_phaeovibrioides | Bacteria>Pseudomonadati>FCB_group>Bacteroidota/Chlorobiota_group>Chlorobiota>Chlorobiia>Chlorobiales>Chlorobiaceae>Chlorobium/Pelodictyon_group>Chlorobium | 349 | FP507_RS07275 | 1396693..1397742 | 1094 | HTH-domain-containing-protein-[Chlorobium-phaeovibrioides] |
| WP_012508726.1 | (w)HTH+REase | GCF_000020645.1 | Pelodictyon_phaeoclathratiforme | Bacteria>Pseudomonadati>FCB_group>Bacteroidota/Chlorobiota_group>Chlorobiota>Chlorobiia>Chlorobiales>Chlorobiaceae>Chlorobium/Pelodictyon_group>Pelodictyon | 400 | PPHA_RS10115 | 2131061..2132263 | 34090 | PDDEXK-nuclease-domain-containing-protein-[Pelodictyon-phaeoclathratiforme] |
| WP_198166314.1 | (w)HTH+REase | GCF_002113825.1 | Prosthecochloris_sp._HL-130-GSB | Bacteria>Pseudomonadati>FCB_group>Bacteroidota/Chlorobiota_group>Chlorobiota>Chlorobiia>Chlorobiales>Chlorobiaceae>Prosthecochloris>unclassified_Prosthecochloris | 228 | B9H02_RS07515 | 1592561..1593247 | 1974213 | winged-helix-turn-helix-domain-containing-protein-[Prosthecochloris-sp.-HL-130-GSB] |
| NMW20447.1 | (w)HTH+REase | GCA_012972835.1 | Chlorobiaceae_bacterium | Bacteria>Pseudomonadati>FCB_group>Bacteroidota/Chlorobiota_group>Chlorobiota>Chlorobiia>Chlorobiales>Chlorobiaceae>unclassified_Chlorobiaceae | 375 | HKK67_02185 | 2950..4077 | 2067991 | DUF1016-family-protein-[Chlorobiaceae-bacterium] |
| MBM3423460.1 | (w)HTH+REase | GCA_016868665.1 | Chlorobiota_bacterium | Bacteria>Pseudomonadati>FCB_group>Bacteroidota/Chlorobiota_group>Chlorobiota>unclassified_Chlorobiota | 299 | FJY09_09505 | 3274..4173 | 2268192 | restriction-endonuclease-[Chlorobiota-bacterium] |
| MXW15219.1 | (w)HTH+REase | GCA_009835915.1 | Rhodothermaceae_bacterium | Bacteria>Pseudomonadati>FCB_group>Bacteroidota/Chlorobiota_group>Rhodothermota>Rhodothermia>Rhodothermales>Rhodothermaceae>unclassified_Rhodothermaceae | 406 | F4Z76_09935 | 6088..7308 | 2026787 | DUF1016-domain-containing-protein-[Rhodothermaceae-bacterium] |
| MXX97461.1 | (w)HTH+REase | GCA_009838195.1 | Rhodothermaceae_bacterium | Bacteria>Pseudomonadati>FCB_group>Bacteroidota/Chlorobiota_group>Rhodothermota>Rhodothermia>Rhodothermales>Rhodothermaceae>unclassified_Rhodothermaceae | 406 | F4Y64_07600 | 4311..5531 | 2026787 | DUF1016-domain-containing-protein-[Rhodothermaceae-bacterium] |
| MXZ58771.1 | (w)HTH+REase | GCA_009837125.1 | Rhodothermaceae_bacterium | Bacteria>Pseudomonadati>FCB_group>Bacteroidota/Chlorobiota_group>Rhodothermota>Rhodothermia>Rhodothermales>Rhodothermaceae>unclassified_Rhodothermaceae | 406 | F4Z16_09885 | 10522..11742 | 2026787 | DUF1016-domain-containing-protein-[Rhodothermaceae-bacterium] |
| MYC03832.1 | (w)HTH+REase | GCA_009840265.1 | Rhodothermaceae_bacterium | Bacteria>Pseudomonadati>FCB_group>Bacteroidota/Chlorobiota_group>Rhodothermota>Rhodothermia>Rhodothermales>Rhodothermaceae>unclassified_Rhodothermaceae | 406 | F4X61_04305 | 11248..12468 | 2026787 | DUF1016-domain-containing-protein-[Rhodothermaceae-bacterium] |
| MYH11277.1 | (w)HTH+REase | GCA_009842805.1 | Rhodothermaceae_bacterium | Bacteria>Pseudomonadati>FCB_group>Bacteroidota/Chlorobiota_group>Rhodothermota>Rhodothermia>Rhodothermales>Rhodothermaceae>unclassified_Rhodothermaceae | 406 | F4191_00330 | 4311..5531 | 2026787 | MAG:-DUF1016-domain-containing-protein-[Rhodothermaceae-bacterium] |
| MBP9037127.1 | (w)HTH+REase | GCA_018057045.1 | Candidatus_Cloacimonas_sp. | Bacteria>Pseudomonadati>FCB_group>Candidatus_Cloacimonadota>Candidatus_Cloacimonadia>Candidatus_Cloacimonadales>Candidatus_Cloacimonadaceae>Candidatus_Cloacimonas>unclassified_Candidatus_Cloacimonas | 381 | KBG38_04990 | 1471..2616 | 2053534 | DUF1016-family-protein-[Candidatus-Cloacimonas-sp.] |
| MBI2505142.1 | (w)HTH+REase | GCA_016187725.1 | Candidatus_Latescibacterota_bacterium | Bacteria>Pseudomonadati>FCB_group>Candidatus_Latescibacterota>unclassified_Candidatus_Latescibacterota | 428 | HYW07_18135 | 18791..20077 | 2053570 | DUF1016-domain-containing-protein-[Candidatus-Latescibacteria-bacterium] |
| NOQ23472.1 | (w)HTH+REase | GCA_013138965.1 | Candidatus_Aegiribacteria_sp. | Bacteria>Pseudomonadati>FCB_group>FCB_group_incertae_sedis>Candidatus_Aegiribacteria>unclassified_Candidatus_Aegiribacteria | 408 | GQ565_12610 | 16610..17836 | 2528271 | DUF1016-family-protein-[Candidatus-Aegiribacteria-sp.] |
| MBN1601228.1 | (w)HTH+REase | GCA_016929595.1 | Chitinispirillaceae_bacterium | Bacteria>Pseudomonadati>FCB_group>Fibrobacterota>Chitinispirillia>Chitinispirillales>Chitinispirillaceae>unclassified_Chitinispirillaceae | 373 | JW915_06445 | 6984..8105 | 2813024 | DUF1016-family-protein-[Chitinispirillaceae-bacterium] |
| MCY3788285.1 | (w)HTH+REase | GCA_026706505.1 | Gemmatimonadota_bacterium | Bacteria>Pseudomonadati>FCB_group>Gemmatimonadota>unclassified_Gemmatimonadota | 416 | OXH63_05800 | 248..1498 | 2026742 | MAG:-PDDEXK-nuclease-domain-containing-protein-[Gemmatimonadetes-bacterium] |
| MDE2973200.1 | (w)HTH+REase | GCA_028873825.1 | Gemmatimonadota_bacterium | Bacteria>Pseudomonadati>FCB_group>Gemmatimonadota>unclassified_Gemmatimonadota | 385 | OXU64_00550 | 4586..5743 | 2026742 | MAG:-PDDEXK-nuclease-domain-containing-protein-[Gemmatimonadota-bacterium] |
| HCF60728.1 | (w)HTH+REase | GCA_003534605.1 | Myxococcales_bacterium | Bacteria>Pseudomonadati>Myxococcota>Myxococcia>Myxococcales>unclassified_Myxococcales | 115 | DFS52_22350 | 8847..9194 | 2026763 | hypothetical-protein-[Myxococcales-bacterium] |
| NUN14057.1 | (w)HTH+REase | GCA_013360285.1 | Myxococcales_bacterium | Bacteria>Pseudomonadati>Myxococcota>Myxococcia>Myxococcales>unclassified_Myxococcales | 79 | HUU55_10535 | 580..819 | 2026763 | hypothetical-protein-[Myxococcales-bacterium] |
| MBF0105030.1 | (w)HTH+REase | GCA_015228875.1 | Deltaproteobacteria_bacterium | Bacteria>Pseudomonadati>Myxococcota>Myxococcia>unclassified_Myxococcia | 411 | HQM16_06850 | 5548..6783 | 2026735 | DUF1016-family-protein-[Deltaproteobacteria-bacterium] |
| MBM4330410.1 | (w)HTH+REase | GCA_016875165.1 | Deltaproteobacteria_bacterium | Bacteria>Pseudomonadati>Myxococcota>Myxococcia>unclassified_Myxococcia | 409 | FJ117_04140 | 57769..58998 | 2026735 | DUF1016-domain-containing-protein-[Deltaproteobacteria-bacterium] |
| MBW1705704.1 | (w)HTH+REase | GCA_019306165.1 | Deltaproteobacteria_bacterium | Bacteria>Pseudomonadati>Myxococcota>Myxococcia>unclassified_Myxococcia | 392 | JRJ86_11135 | 6672..7850 | 2026735 | DUF1016-family-protein-[Deltaproteobacteria-bacterium] |
| MBW2343656.1 | (w)HTH+REase | GCA_019310785.1 | Deltaproteobacteria_bacterium | Bacteria>Pseudomonadati>Myxococcota>Myxococcia>unclassified_Myxococcia | 392 | JRF53_06520 | 20141..21319 | 2026735 | DUF1016-family-protein-[Deltaproteobacteria-bacterium] |
| MBW1792991.1 | (w)HTH+REase | GCA_019307135.1 | Deltaproteobacteria_bacterium | Bacteria>Pseudomonadati>Myxococcota>Myxococcia>unclassified_Myxococcia | 391 | JRJ38_00925 | 7517..8692 | 2026735 | DUF1016-family-protein-[Deltaproteobacteria-bacterium] |
| MBF0105439.1 | (w)HTH+REase | GCA_015228875.1 | Deltaproteobacteria_bacterium | Bacteria>Pseudomonadati>Myxococcota>Myxococcia>unclassified_Myxococcia | 383 | HQM16_08950 | 28497..29648 | 2026735 | DUF1016-family-protein-[Deltaproteobacteria-bacterium] |
| MCP4577329.1 | (w)HTH+REase | GCA_024231575.1 | Deltaproteobacteria_bacterium | Bacteria>Pseudomonadati>Myxococcota>Myxococcia>unclassified_Myxococcia | 380 | GY846_13715 | 18134..19276 | 2026735 | DUF1016-domain-containing-protein-[Deltaproteobacteria-bacterium] |
| MBW1982274.1 | (w)HTH+REase | GCA_019308175.1 | Deltaproteobacteria_bacterium | Bacteria>Pseudomonadati>Myxococcota>Myxococcia>unclassified_Myxococcia | 378 | JRJ12_13745 | 17482..18618 | 2026735 | DUF1016-family-protein-[Deltaproteobacteria-bacterium] |
| MBS3918743.1 | (w)HTH+REase | GCA_018333535.1 | Deltaproteobacteria_bacterium | Bacteria>Pseudomonadati>Myxococcota>Myxococcia>unclassified_Myxococcia | 354 | KG012_07625 | 5516..6580 | 2026735 | DUF1016-family-protein-[Deltaproteobacteria-bacterium] |
| MBO6936773.1 | (w)HTH+REase | GCA_017644045.1 | Deltaproteobacteria_bacterium | Bacteria>Pseudomonadati>Myxococcota>Myxococcia>unclassified_Myxococcia | 317 | JJ863_17520 | 230820..231773 | 2026735 | hypothetical-protein-[Deltaproteobacteria-bacterium] |
| MBP9202685.1 | (w)HTH+REase | GCA_018058095.1 | Kofleriaceae_bacterium | Bacteria>Pseudomonadati>Myxococcota>Polyangia>Haliangiales>Kofleriaceae>unclassified_Kofleriaceae | 143 | KBG28_01785 | 80188..80619 | 2212474 | hypothetical-protein-[Kofleriaceae-bacterium] |
| MBJ71779.1 | (w)HTH+REase | GCA_002720945.1 | Sandaracinus_sp. | Bacteria>Pseudomonadati>Myxococcota>Polyangia>Polyangiales>Sandaracinaceae>Sandaracinus>unclassified_Sandaracinus | 317 | CMN31_10635 | 8090..9043 | 2024858 | hypothetical-protein-[Sandaracinus-sp.] |
| MAT26111.1 | (w)HTH+REase | GCA_002699025.1 | Sandaracinus_sp. | Bacteria>Pseudomonadati>Myxococcota>Polyangia>Polyangiales>Sandaracinaceae>Sandaracinus>unclassified_Sandaracinus | 194 | CMN29_14295 | 45521..46105 | 2024858 | hypothetical-protein-[Sandaracinus-sp.] |
| WP_040132777.1 | (w)HTH+REase | GCF_964209095.1 | Gammaproteobacteria | Bacteria>Pseudomonadati>Pseudomonadota | 289 | EXA22_RS10070 | 4788..5657 | 1236 | MULTISPECIES:-restriction-endonuclease-[Gammaproteobacteria] |
| WP_159262619.1 | (w)HTH+REase | GCF_010692865.1 | Gammaproteobacteria | Bacteria>Pseudomonadati>Pseudomonadota | 284 | QN307_RS16735 | 27639..28493 | 1236 | MULTISPECIES:-restriction-endonuclease-[Enterobacterales] |
| WP_245398277.1 | (w)HTH+REase | GCF_030388465.1 | Gammaproteobacteria | Bacteria>Pseudomonadati>Pseudomonadota | 277 | FG482_RS02540 | 521018..521851 | 1236 | MULTISPECIES:-restriction-endonuclease-[Gammaproteobacteria] |
| WP_101538051.1 | (w)HTH+REase | GCF_018854655.1 | Acidithiobacillus | Bacteria>Pseudomonadati>Pseudomonadota>Acidithiobacillia>Acidithiobacillales>Acidithiobacillaceae | 363 | HF670_RS09015 | 17221..18312 | 119977 | MULTISPECIES:-PDDEXK-nuclease-domain-containing-protein-[Acidithiobacillus] |
| MBU2839702.1 | (w)HTH+REase | GCA_018854655.1 | Acidithiobacillus_thiooxidans | Bacteria>Pseudomonadati>Pseudomonadota>Acidithiobacillia>Acidithiobacillales>Acidithiobacillaceae>Acidithiobacillus | 363 | HF670_09025 | 17221..18312 | 930 | DUF1016-domain-containing-protein-[Acidithiobacillus-thiooxidans] |
| MBU2768212.1 | (w)HTH+REase | GCA_018853935.1 | Acidithiobacillus_ferrivorans | Bacteria>Pseudomonadati>Pseudomonadota>Acidithiobacillia>Acidithiobacillales>Acidithiobacillaceae>Acidithiobacillus | 356 | HAP94_19070 | 3307..4377 | 160808 | DUF1016-domain-containing-protein-[Acidithiobacillus-ferrivorans] |
| WP_024895394.1 | (w)HTH+REase | GCF_000559045.1 | Acidithiobacillus_thiooxidans | Bacteria>Pseudomonadati>Pseudomonadota>Acidithiobacillia>Acidithiobacillales>Acidithiobacillaceae>Acidithiobacillus | 172 | A6M27_RS14975 | 5601..6119 | 930 | PDDEXK-nuclease-domain-containing-protein-[Acidithiobacillus-thiooxidans] |
| WP_127367957.1 | (w)HTH+REase | GCF_004022915.1 | unclassified_Mesorhizobium | Bacteria>Pseudomonadati>Pseudomonadota>Alphaproteobacteria>Hyphomicrobiales>Phyllobacteriaceae>Mesorhizobium | 273 | EN925_RS00850 | 4508..5329 | 325217 | MULTISPECIES:-hypothetical-protein-[unclassified-Mesorhizobium] |
| WP_233943242.1 | (w)HTH+REase | GCF_021391595.1 | Rhizobium_leguminosarum | Bacteria>Pseudomonadati>Pseudomonadota>Alphaproteobacteria>Hyphomicrobiales>Rhizobiaceae>Rhizobium/Agrobacterium_group>Rhizobium | 296 | LZK78_RS32115 | 714192..715082 | 384 | restriction-endonuclease-[Rhizobium-leguminosarum] |
| WP_234869547.1 | (w)HTH+REase | GCF_021491355.1 | Allitabrizicola_rongguiensis | Bacteria>Pseudomonadati>Pseudomonadota>Alphaproteobacteria>Rhodobacterales>Paracoccaceae>Alitabrizicola | 271 | LOV05_RS07770 | 386258..387073 | 2909234 | hypothetical-protein-[Tabrizicola-rongguiensis] |
| WP_009810040.1 | (w)HTH+REase | GCF_000152965.1 | Roseobacter_sp._MED193 | Bacteria>Pseudomonadati>Pseudomonadota>Alphaproteobacteria>Rhodobacterales>Roseobacteraceae>Roseobacter>unclassified_Roseobacter | 317 | MED193_RS12220 | 2038732..2039685 | 314262 | hypothetical-protein-[Roseobacter-sp.-MED193] |
| WP_085595214.1 | (w)HTH+REase | GCF_002115825.1 | Thalassospira_sp._MCCC_1A01428 | Bacteria>Pseudomonadati>Pseudomonadota>Alphaproteobacteria>Rhodospirillales>Thalassospiraceae>Thalassospira>unclassified_Thalassospira | 343 | THS27_RS23995 | 10524..11555 | 1470575 | PDDEXK-nuclease-domain-containing-protein-[Thalassospira-sp.-MCCC-1A01428] |
| MCL6741772.1 | (w)HTH+REase | GCA_023516505.1 | Sphingomonas_brevis | Bacteria>Pseudomonadati>Pseudomonadota>Alphaproteobacteria>Sphingomonadales>Sphingomonadaceae>Sphingomonas | 216 | LZ518_11605 | 2239874..2240524 | 2908206 | hypothetical-protein-[Sphingomonas-sp.-RB56-2] |
| WP_249916138.1 | (w)HTH+REase | GCF_023516505.1 | Sphingomonas_brevis | Bacteria>Pseudomonadati>Pseudomonadota>Alphaproteobacteria>Sphingomonadales>Sphingomonadaceae>Sphingomonas | 216 | LZ518_RS11605 | 2239874..2240524 | 2908206 | hypothetical-protein-[Sphingomonas-sp.-RB56-2] |
| WP_169068628.1 | (w)HTH+REase | GCF_907163155.1 | Candidatus_Accumulibacter | Bacteria>Pseudomonadati>Pseudomonadota>Betaproteobacteria>Betaproteobacteria_incertae_sedis | 360 | QZ648_RS13940 | 35595..36677 | 327159 | PDDEXK-nuclease-domain-containing-protein-[Candidatus-Accumulibacter-phosphatis] |
| EXI84707.1 | (w)HTH+REase | GCA_000585075.1 | Candidatus_Accumulibacter_regalis | Bacteria>Pseudomonadati>Pseudomonadota>Betaproteobacteria>Betaproteobacteria_incertae_sedis>Candidatus_Accumulibacter | 381 | AW11_03762 | 14008..15153 | 522306 | MAG:-hypothetical-protein-AW11_03762-[Candidatus-Accumulibacter-regalis] |
| MBK7952882.1 | (w)HTH+REase | GCA_016713625.1 | Candidatus_Accumulibacter_affinis | Bacteria>Pseudomonadati>Pseudomonadota>Betaproteobacteria>Betaproteobacteria_incertae_sedis>Candidatus_Accumulibacter | 376 | IPK02_02320 | 214126..215256 | 2954384 | DUF1016-family-protein-[Candidatus-Accumulibacter-sp.] |
| WP_169070209.1 | (w)HTH+REase | GCF_036625285.1 | Candidatus_Accumulibacter_contiguus | Bacteria>Pseudomonadati>Pseudomonadota>Betaproteobacteria>Betaproteobacteria_incertae_sedis>Candidatus_Accumulibacter | 348 | E4Q08_RS09490 | 44420..45466 | 2954381 | PDDEXK-nuclease-domain-containing-protein-[Candidatus-Accumulibacter-contiguus] |
| WP_248594807.1 | (w)HTH+REase | GCF_036625285.1 | Candidatus_Accumulibacter_contiguus | Bacteria>Pseudomonadati>Pseudomonadota>Betaproteobacteria>Betaproteobacteria_incertae_sedis>Candidatus_Accumulibacter | 235 | E4Q08_RS09480 | 43401..44108 | 2954381 | DUF1016-N-terminal-domain-containing-protein-[Candidatus-Accumulibacter-contiguus] |
| MBL8396549.1 | (w)HTH+REase | GCA_016789495.1 | Accumulibacter_sp. | Bacteria>Pseudomonadati>Pseudomonadota>Betaproteobacteria>Betaproteobacteria_incertae_sedis>Candidatus_Accumulibacter>unclassified_Candidatus_Accumulibacter | 364 | JNL84_00160 | 42605..43699 | 2053492 | DUF1016-family-protein-[Candidatus-Accumulibacter-sp.] |
| MCM8599229.1 | (w)HTH+REase | GCA_023806585.1 | Accumulibacter_sp. | Bacteria>Pseudomonadati>Pseudomonadota>Betaproteobacteria>Betaproteobacteria_incertae_sedis>Candidatus_Accumulibacter>unclassified_Candidatus_Accumulibacter | 346 | NFW04_11320 | 10788..11828 | 2053492 | PDDEXK-nuclease-domain-containing-protein-[Candidatus-Accumulibacter-sp.] |
| WP_125277461.1 | (w)HTH+REase | GCF_003938225.2 | Alcaligenes_aquatilis | Bacteria>Pseudomonadati>Pseudomonadota>Betaproteobacteria>Burkholderiales>Alcaligenaceae>Alcaligenes | 103 | EGK70_RS02735 | 598873..599184 | 323284 | DUF1016-N-terminal-domain-containing-protein-[Alcaligenes-aquatilis] |
| RIY39882.1 | (w)HTH+REase | GCA_003585905.1 | Neopusillimonas_maritima | Bacteria>Pseudomonadati>Pseudomonadota>Betaproteobacteria>Burkholderiales>Alcaligenaceae>Neopusillimonas | 351 | CJP73_12395 | 18716..19771 | 2026239 | restriction-endonuclease-[Pusillimonas-maritima] |
| WP_119516657.1 | (w)HTH+REase | GCF_003585905.1 | Neopusillimonas_maritima | Bacteria>Pseudomonadati>Pseudomonadota>Betaproteobacteria>Burkholderiales>Alcaligenaceae>Neopusillimonas | 351 | CJP73_RS12425 | 18716..19771 | 2026239 | HTH-domain-containing-protein-[Pusillimonas-maritima] |
| TFI47150.1 | (w)HTH+REase | GCA_004563745.1 | Diaphorobacter_sp._DS2 | Bacteria>Pseudomonadati>Pseudomonadota>Betaproteobacteria>Burkholderiales>Comamonadaceae>Diaphorobacter>unclassified_Diaphorobacter | 343 | E4O93_14240 | 7631..8662 | 1302548 | DUF1016-domain-containing-protein-[Diaphorobacter-sp.-DS2] |
| WP_104799762.1 | (w)HTH+REase | GCF_002943475.1 | Limnohabitans_sp._TS-CS-82 | Bacteria>Pseudomonadati>Pseudomonadota>Betaproteobacteria>Burkholderiales>Comamonadaceae>Limnohabitans>unclassified_Limnohabitans | 138 | C5F52_RS18135 | 192627..193043 | 2094193 | DUF1016-N-terminal-domain-containing-protein-[Limnohabitans-sp.-TS-CS-82] |
| MBK6748239.1 | (w)HTH+REase | GCA_016703145.1 | Ottowia_sp. | Bacteria>Pseudomonadati>Pseudomonadota>Betaproteobacteria>Burkholderiales>Comamonadaceae>Ottowia>unclassified_Ottowia | 136 | IPG65_18045 | 1101805..1102215 | 1898956 | DUF1016-family-protein-[Ottowia-sp.] |
| MBG6075361.1 | (w)HTH+REase | GCA_015752225.1 | Polaromonas_sp._CG_9.11 | Bacteria>Pseudomonadati>Pseudomonadota>Betaproteobacteria>Burkholderiales>Comamonadaceae>Polaromonas>unclassified_Polaromonas | 240 | IWX85_001186 | 23118..23840 | 2787730 | putative-nuclease-of-restriction-endonuclease-like-(RecB)-superfamily-[Polaromonas-sp.-CG_9.11] |
| WP_196869056.1 | (w)HTH+REase | GCF_015752225.1 | Polaromonas_sp._CG_9.11 | Bacteria>Pseudomonadati>Pseudomonadota>Betaproteobacteria>Burkholderiales>Comamonadaceae>Polaromonas>unclassified_Polaromonas | 240 | IWX85_RS05655 | 23118..23840 | 2787730 | DUF1016-N-terminal-domain-containing-protein-[Polaromonas-sp.-CG_9.11] |
| WP_265286856.1 | (w)HTH+REase | GCF_026127525.1 | Verminephrobacter_eiseniae | Bacteria>Pseudomonadati>Pseudomonadota>Betaproteobacteria>Burkholderiales>Comamonadaceae>Verminephrobacter | 373 | D5047_RS05810 | 1234353..1235474 | 364317 | PDDEXK-nuclease-domain-containing-protein-[Verminephrobacter-eiseniae] |
| MCA3175662.1 | (w)HTH+REase | GCA_020248585.1 | Burkholderiales_bacterium | Bacteria>Pseudomonadati>Pseudomonadota>Betaproteobacteria>Burkholderiales>unclassified_Burkholderiales | 356 | ING36_09015 | 724..1794 | 1891238 | DUF1016-family-protein-[Burkholderiales-bacterium] |
| MCZ2134306.1 | (w)HTH+REase | GCA_026932525.1 | Burkholderiales_bacterium | Bacteria>Pseudomonadati>Pseudomonadota>Betaproteobacteria>Burkholderiales>unclassified_Burkholderiales | 356 | LC098_02585 | 45120..46190 | 1891238 | MAG:-PDDEXK-nuclease-domain-containing-protein-[Burkholderiales-bacterium] |
| OJV49478.1 | (w)HTH+REase | GCA_001897615.1 | Burkholderiales_bacterium_68-10 | Bacteria>Pseudomonadati>Pseudomonadota>Betaproteobacteria>Burkholderiales>unclassified_Burkholderiales | 350 | BGO36_11425 | 19936..20988 | 1895730 | hypothetical-protein-BGO36_11425-[Burkholderiales-bacterium-68-10] |
| MBN8486525.1 | (w)HTH+REase | GCA_017302485.1 | Burkholderiales_bacterium | Bacteria>Pseudomonadati>Pseudomonadota>Betaproteobacteria>Burkholderiales>unclassified_Burkholderiales | 339 | J0M20_02155 | 16249..17268 | 1891238 | DUF1016-family-protein-[Burkholderiales-bacterium] |
| OJV49479.1 | (w)HTH+REase | GCA_001897615.1 | Burkholderiales_bacterium_68-10 | Bacteria>Pseudomonadati>Pseudomonadota>Betaproteobacteria>Burkholderiales>unclassified_Burkholderiales | 89 | BGO36_11430 | 20989..21258 | 1895730 | hypothetical-protein-BGO36_11430-[Burkholderiales-bacterium-68-10] |
| QKE41938.1 | (w)HTH+REase | GCA_013181415.1 | Ferrovum_myxofaciens | Bacteria>Pseudomonadati>Pseudomonadota>Betaproteobacteria>Ferrovales>Ferrovaceae>Ferrovum | 360 | HO274_11935 | 2422797..2423879 | 416213 | DUF1016-domain-containing-protein-[Ferrovum-myxofaciens] |
| WP_273119930.1 | (w)HTH+REase | GCF_013181415.1 | Ferrovum_myxofaciens | Bacteria>Pseudomonadati>Pseudomonadota>Betaproteobacteria>Ferrovales>Ferrovaceae>Ferrovum | 360 | HO274_RS11975 | 2422797..2423879 | 416213 | PDDEXK-nuclease-domain-containing-protein-[Ferrovum-myxofaciens] |
| MCG9064560.1 | (w)HTH+REase | GCA_022394255.1 | Laribacter_hongkongensis | Bacteria>Pseudomonadati>Pseudomonadota>Betaproteobacteria>Neisseriales>Aquaspirillaceae>Laribacter | 341 | LH425_05820 | 29076..30101 | 168471 | PDDEXK-nuclease-domain-containing-protein-[Laribacter-hongkongensis] |
| WP_013292861.1 | (w)HTH+REase | GCF_000145255.1 | Gallionella_capsiferriformans | Bacteria>Pseudomonadati>Pseudomonadota>Betaproteobacteria>Nitrosomonadales>Gallionellaceae>Gallionella | 381 | GALF_RS04435 | 937562..938707 | 370405 | PDDEXK-nuclease-domain-containing-protein-[Gallionella-capsiferriformans] |
| NNM68914.1 | (w)HTH+REase | GCA_013043965.1 | Gallionella_sp. | Bacteria>Pseudomonadati>Pseudomonadota>Betaproteobacteria>Nitrosomonadales>Gallionellaceae>Gallionella>unclassified_Gallionella | 356 | HKM00_02895 | 678..1748 | 2052161 | DUF1016-domain-containing-protein-[Gallionella-sp.] |
| ROH85752.1 | (w)HTH+REase | GCA_003730115.1 | Pseudomethylobacillus_aquaticus | Bacteria>Pseudomonadati>Pseudomonadota>Betaproteobacteria>Nitrosomonadales>Methylophilaceae>Pseudomethylobacillus | 313 | ED236_08380 | 57895..58836 | 2676064 | HrgA-protein-[Pseudomethylobacillus-aquaticus] |
| WP_123237526.1 | (w)HTH+REase | GCF_003730115.1 | Pseudomethylobacillus_aquaticus | Bacteria>Pseudomonadati>Pseudomonadota>Betaproteobacteria>Nitrosomonadales>Methylophilaceae>Pseudomethylobacillus | 313 | ED236_RS12655 | 57895..58836 | 2676064 | COG2958-family-protein-[Pseudomethylobacillus-aquaticus] |
| WP_204800508.1 | (w)HTH+REase | GCF_905220645.1 | Nitrosomonas_nitrosa | Bacteria>Pseudomonadati>Pseudomonadota>Betaproteobacteria>Nitrosomonadales>Nitrosomonadaceae>Nitrosomonas | 362 | NUZ5A_RS15515 | 130095..131183 | 52442 | PDDEXK-nuclease-domain-containing-protein-[Nitrosomonas-nitrosa] |
| MBX9637027.1 | (w)HTH+REase | GCA_019748355.1 | Nitrosomonas_sp. | Bacteria>Pseudomonadati>Pseudomonadota>Betaproteobacteria>Nitrosomonadales>Nitrosomonadaceae>Nitrosomonas>unclassified_Nitrosomonas | 385 | K2Q45_05695 | 9746..10903 | 42353 | PDDEXK-nuclease-domain-containing-protein-[Nitrosomonas-sp.] |
| MBK7491772.1 | (w)HTH+REase | GCA_016707165.1 | Nitrosomonas_sp. | Bacteria>Pseudomonadati>Pseudomonadota>Betaproteobacteria>Nitrosomonadales>Nitrosomonadaceae>Nitrosomonas>unclassified_Nitrosomonas | 351 | IPI17_07020 | 59819..60874 | 42353 | DUF1016-family-protein-[Nitrosomonas-sp.] |
| MCB1936155.1 | (w)HTH+REase | GCA_020446045.1 | Nitrosomonas_sp. | Bacteria>Pseudomonadati>Pseudomonadota>Betaproteobacteria>Nitrosomonadales>Nitrosomonadaceae>Nitrosomonas>unclassified_Nitrosomonas | 139 | KDF59_09450 | 1..418 | 42353 | DUF1016-domain-containing-protein-[Nitrosomonas-sp.] |
| MBK6974206.1 | (w)HTH+REase | GCA_016708905.1 | Candidatus_Methylophosphatis_roskildensis | Bacteria>Pseudomonadati>Pseudomonadota>Betaproteobacteria>Nitrosomonadales>Sterolibacteriaceae>Candidatus_Methylophosphatis | 361 | IPH26_15110 | 457566..458651 | 2899263 | DUF1016-family-protein-[Sterolibacteriaceae-bacterium] |
| MBP9798946.1 | (w)HTH+REase | GCA_018062105.1 | Sterolibacterium_sp. | Bacteria>Pseudomonadati>Pseudomonadota>Betaproteobacteria>Nitrosomonadales>Sterolibacteriaceae>Sterolibacterium>unclassified_Sterolibacterium | 350 | KBD39_02355 | 32181..33233 | 2819261 | DUF1016-family-protein-[Sterolibacterium-sp.] |
| OGS68624.1 | (w)HTH+REase | GCA_001801025.1 | Gallionellales_bacterium_GWA2_54_124 | Bacteria>Pseudomonadati>Pseudomonadota>Betaproteobacteria>Nitrosomonadales>unclassified_Nitrosomonadales | 381 | A2Z87_08885 | 87689..88834 | 1798243 | hypothetical-protein-A2Z87_08885-[Gallionellales-bacterium-GWA2_54_124] |
| MBK9161125.1 | (w)HTH+REase | GCA_016716325.1 | Nitrosomonadales_bacterium | Bacteria>Pseudomonadati>Pseudomonadota>Betaproteobacteria>Nitrosomonadales>unclassified_Nitrosomonadales | 313 | IPM27_06130 | 1281712..1282653 | 2026761 | HrgA-protein-[Nitrosomonadales-bacterium] |
| MBP6189860.1 | (w)HTH+REase | GCA_017985755.1 | Azonexus_sp. | Bacteria>Pseudomonadati>Pseudomonadota>Betaproteobacteria>Rhodocyclales>Azonexaceae>Azonexus>unclassified_Azonexus | 364 | KA440_12995 | 4874..5968 | 1872668 | DUF1016-family-protein-[Azonexus-sp.] |
| MBN8461441.1 | (w)HTH+REase | GCA_017302355.1 | Dechloromonas_sp. | Bacteria>Pseudomonadati>Pseudomonadota>Betaproteobacteria>Rhodocyclales>Azonexaceae>Dechloromonas>unclassified_Dechloromonas | 350 | J0M01_01295 | 17575..18627 | 1917218 | DUF1016-family-protein-[Dechloromonas-sp.] |
| WP_244857074.1 | (w)HTH+REase | GCF_017894365.1 | Aromatoleum_bremense | Bacteria>Pseudomonadati>Pseudomonadota>Betaproteobacteria>Rhodocyclales>Rhodocyclaceae>Aromatoleum | 355 | GPA24_RS09415 | 36485..37552 | 76115 | PDDEXK-nuclease-domain-containing-protein-[Aromatoleum-bremense] |
| OHC72336.1 | (w)HTH+REase | GCA_001828935.1 | Rhodocyclales_bacterium_RIFCSPLOWO2_02_FULL_63_24 | Bacteria>Pseudomonadati>Pseudomonadota>Betaproteobacteria>Rhodocyclales>unclassified_Rhodocyclales | 348 | A3H93_14605 | 49552..50598 | 1802018 | MAG:-hypothetical-protein-A3H93_14605-[Rhodocyclales-bacterium-RIFCSPLOWO2_02_FULL_63_24] |
| MBD5801780.1 | (w)HTH+REase | GCA_014822185.1 | Azoarcus_sp._Aa7 | Bacteria>Pseudomonadati>Pseudomonadota>Betaproteobacteria>Rhodocyclales>Zoogloeaceae>Azoarcus>unclassified_Azoarcus | 356 | AZOA_12040 | 4140..5210 | 2303328 | hypothetical-protein-[Azoarcus-sp.-Aa7] |
| WP_108973933.1 | (w)HTH+REase | GCF_003111665.1 | Parazoarcus_communis | Bacteria>Pseudomonadati>Pseudomonadota>Betaproteobacteria>Rhodocyclales>Zoogloeaceae>Parazoarcus | 340 | CEW87_RS14080 | 3083768..3084790 | 41977 | PDDEXK-nuclease-domain-containing-protein-[Parazoarcus-communis] |
| WP_068639547.1 | (w)HTH+REase | GCF_001591165.1 | Thauera_butanivorans | Bacteria>Pseudomonadati>Pseudomonadota>Betaproteobacteria>Rhodocyclales>Zoogloeaceae>Thauera | 350 | TBU01S_RS21535 | 1950..3002 | 86174 | PDDEXK-nuclease-domain-containing-protein-[Thauera-butanivorans] |
| WP_002936074.1 | (w)HTH+REase | GCF_000310125.2 | Thauera_sp._27 | Bacteria>Pseudomonadati>Pseudomonadota>Betaproteobacteria>Rhodocyclales>Zoogloeaceae>Thauera>unclassified_Thauera | 355 | B447_RS02405 | 107112..108179 | 305700 | PDDEXK-nuclease-domain-containing-protein-[Thauera-sp.-27] |
| WP_153168445.1 | (w)HTH+REase | GCF_009469595.2 | Thauera_sp._2A1 | Bacteria>Pseudomonadati>Pseudomonadota>Betaproteobacteria>Rhodocyclales>Zoogloeaceae>Thauera>unclassified_Thauera | 355 | GH664_RS12705 | 87882..88949 | 2570191 | PDDEXK-nuclease-domain-containing-protein-[Thauera-sp.-2A1] |
| WP_218028740.1 | (w)HTH+REase | GCF_006539865.1 | Zoogloea_ramigera | Bacteria>Pseudomonadati>Pseudomonadota>Betaproteobacteria>Rhodocyclales>Zoogloeaceae>Zoogloea | 360 | ZRA01_RS16850 | 11860..12942 | 350 | PDDEXK-nuclease-domain-containing-protein-[Zoogloea-ramigera] |
| WP_148578098.1 | (w)HTH+REase | GCF_008107625.1 | Zoogloea_oleivorans | Bacteria>Pseudomonadati>Pseudomonadota>Betaproteobacteria>Rhodocyclales>Zoogloeaceae>Zoogloea | 346 | ETQ85_RS05735 | 205419..206459 | 1552750 | PDDEXK-nuclease-domain-containing-protein-[Zoogloea-oleivorans] |
| MDA8417188.1 | (w)HTH+REase | GCA_028069095.1 | Betaproteobacteria_bacterium | Bacteria>Pseudomonadati>Pseudomonadota>Betaproteobacteria>unclassified_Betaproteobacteria | 356 | M0Z78_09070 | 10551..11621 | 1891241 | MAG:-PDDEXK-nuclease-domain-containing-protein-[Betaproteobacteria-bacterium] |
| WP_001272449.1 | (w)HTH+REase | GCF_000212715.2 | Enterobacterales | Bacteria>Pseudomonadati>Pseudomonadota>Gammaproteobacteria | 289 | IXW39_RS10890 | 12879..13748 | 91347 | MULTISPECIES:-restriction-endonuclease-[Enterobacterales] |
| BBR09764.1 | (w)HTH+REase | GCA_014158455.1 | Aeromonas_caviae | Bacteria>Pseudomonadati>Pseudomonadota>Gammaproteobacteria>Aeromonadales>Aeromonadaceae>Aeromonas | 350 | WP3S18E02_14250 | 1539209..1540261 | 648 | hypothetical-protein-WP3S18E02_14250-[Aeromonas-caviae] |
| WP_036957212.1 | (w)HTH+REase | GCF_913044265.1 | Pseudoalteromonas | Bacteria>Pseudomonadati>Pseudomonadota>Gammaproteobacteria>Alteromonadales>Pseudoalteromonadaceae | 289 | QL994_RS18040 | 7309..8178 | 53246 | MULTISPECIES:-restriction-endonuclease-[Pseudoalteromonas] |
| QTL37179.1 | (w)HTH+REase | GCA_017742995.1 | Pseudoalteromonas_viridis | Bacteria>Pseudomonadati>Pseudomonadota>Gammaproteobacteria>Alteromonadales>Pseudoalteromonadaceae>Pseudoalteromonas | 284 | J5X90_01840 | 426276..427130 | 339617 | restriction-endonuclease-[Pseudoalteromonas-viridis] |
| WP_020507361.1 | (w)HTH+REase | GCF_000379525.1 | Lamprocystis_purpurea | Bacteria>Pseudomonadati>Pseudomonadota>Gammaproteobacteria>Chromatiales>Chromatiaceae>Lamprocystis | 363 | A390_RS0124810 | 123477..124568 | 61598 | PDDEXK-nuclease-domain-containing-protein-[Lamprocystis-purpurea] |

| Protein-ID | Domain | Assembly-ID | Strain | Taxonomy | Length | Locus-Tag | Genomic-coordinates | TaxID | Fasta-defline |
|---|---|---|---|---|---|---|---|---|---|
| MBV5272244.1 | (w)HTH+REase | GCA_019163055.1 | Lamprocystis_purpurea | Bacteria>Pseudomonadati>Pseudomonadota>Gammaproteobacteria>Chromatiales>Chromatiaceae>Lamprocystis | 219 | JZU52_00940 | 352..1011 | 61598 | DUF1016-family-protein-[Lamprocystis-purpurea] |
| WP_223296655.1 | (w)HTH+REase | GCF_033954835.1 | Thiorhodovibrio_frisius | Bacteria>Pseudomonadati>Pseudomonadota>Gammaproteobacteria>Chromatiales>Chromatiaceae>Thiorhodovibrio | 363 | THI970DRAFT_RS02030 | 43575..44666 | 631362 | PDDEXK-nuclease-domain-containing-protein-[Thiorhodovibrio-sp.-970] |
| MCF8003109.1 | (w)HTH+REase | GCA_021736845.1 | Chromatiaceae_bacterium | Bacteria>Pseudomonadati>Pseudomonadota>Gammaproteobacteria>Chromatiales>Chromatiaceae>unclassified_Chromatiaceae | 370 | K9L32_02695 | 10753..11865 | 2049432 | PDDEXK-nuclease-domain-containing-protein-[Chromatiaceae-bacterium] |
| MBP6733465.1 | (w)HTH+REase | GCA_017990755.1 | Chromatiaceae_bacterium | Bacteria>Pseudomonadati>Pseudomonadota>Gammaproteobacteria>Chromatiales>Chromatiaceae>unclassified_Chromatiaceae | 197 | KA142_02020 | 8650..9243 | 2049432 | DUF1016-family-protein-[Chromatiaceae-bacterium] |
| WP_032437331.1 | (w)HTH+REase | GCF_900774785.1 | Enterobacteriaceae | Bacteria>Pseudomonadati>Pseudomonadota>Gammaproteobacteria>Enterobacterales | 289 | NXA99_RS21120 | 80938..81807 | 543 | MULTISPECIES:-restriction-endonuclease-[Enterobacteriaceae] |
| WP_117101577.1 | (w)HTH+REase | GCF_016651545.1 | Enterobacteriaceae | Bacteria>Pseudomonadati>Pseudomonadota>Gammaproteobacteria>Enterobacterales | 289 | M8R77_RS12275 | 169748..170617 | 543 | MULTISPECIES:-restriction-endonuclease-[Enterobacteriaceae] |
| WP_230608236.1 | (w)HTH+REase | GCF_021010575.1 | Limnobaculum_eriocheiris | Bacteria>Pseudomonadati>Pseudomonadota>Gammaproteobacteria>Enterobacterales>Budviciaceae>Limnobaculum | 289 | LPW36_RS04305 | 72106..72975 | 2897391 | restriction-endonuclease-[Limnobaculum-eriocheiris] |
| WP_199981187.1 | (w)HTH+REase | GCF_016501115.1 | Citrobacter | Bacteria>Pseudomonadati>Pseudomonadota>Gammaproteobacteria>Enterobacterales>Enterobacteriaceae | 289 | OGW14_RS13150 | 1100..1969 | 544 | restriction-endonuclease-[Citrobacter-cronae] |
| WP_232904623.1 | (w)HTH+REase | GCF_021282435.1 | Buttiauxella_sp._W03-F01 | Bacteria>Pseudomonadati>Pseudomonadota>Gammaproteobacteria>Enterobacterales>Enterobacteriaceae>Buttiauxella>unclassified_Buttiauxella | 289 | LVQ77_RS12905 | 30763..31632 | 2904524 | restriction-endonuclease-[Buttiauxella-sp.-W03-F01] |
| HAV2294928.1 | (w)HTH+REase | GCA_016445565.1 | Citrobacter_koseri | Bacteria>Pseudomonadati>Pseudomonadota>Gammaproteobacteria>Enterobacterales>Enterobacteriaceae>Citrobacter | 289 | JHK52_001286 | 169251..170120 | 545 | restriction-endonuclease-[Citrobacter-koseri] |
| HCW3139354.1 | (w)HTH+REase | GCA_026549635.1 | Citrobacter_freundii | Bacteria>Pseudomonadati>Pseudomonadota>Gammaproteobacteria>Enterobacterales>Enterobacteriaceae>Citrobacter>Citrobacter_freundii_complex | 289 | OXC88_001873 | 162424..163293 | 546 | TPA:-restriction-endonuclease-[Citrobacter-freundii] |
| MBJ8391829.1 | (w)HTH+REase | GCA_016501115.1 | Citrobacter_cronae | Bacteria>Pseudomonadati>Pseudomonadota>Gammaproteobacteria>Enterobacterales>Enterobacteriaceae>Citrobacter>Citrobacter_freundii_complex | 289 | I6M80_16475 | 8878..9747 | 1748967 | restriction-endonuclease-[Citrobacter-cronae] |
| WP_275399060.1 | (w)HTH+REase | GCF_029104325.1 | Citrobacter_portucalensis | Bacteria>Pseudomonadati>Pseudomonadota>Gammaproteobacteria>Enterobacterales>Enterobacteriaceae>Citrobacter>Citrobacter_freundii_complex | 289 | L2173_RS05360 | 163690..164559 | 1639133 | restriction-endonuclease-[Citrobacter-portucalensis] |
| WP_284106246.1 | (w)HTH+REase | GCF_030132145.1 | Citrobacter_portucalensis | Bacteria>Pseudomonadati>Pseudomonadota>Gammaproteobacteria>Enterobacterales>Enterobacteriaceae>Citrobacter>Citrobacter_freundii_complex | 289 | QL289_RS16580 | 133738..134607 | 1639133 | restriction-endonuclease-[Citrobacter-portucalensis] |
| EHN8715932.1 | (w)HTH+REase | GCA_018454885.1 | Enterobacter_hormaechei | Bacteria>Pseudomonadati>Pseudomonadota>Gammaproteobacteria>Enterobacterales>Enterobacteriaceae>Enterobacter>Enterobacter_cloacae_complex | 289 | I4A92_000739 | 160445..161314 | 158836 | restriction-endonuclease-[Enterobacter-hormaechei] |
| EKK5919803.1 | (w)HTH+REase | GCA_028182685.1 | Enterobacter_hormaechei | Bacteria>Pseudomonadati>Pseudomonadota>Gammaproteobacteria>Enterobacterales>Enterobacteriaceae>Enterobacter>Enterobacter_cloacae_complex | 289 | POC94_000680 | 171348..172217 | 158836 | restriction-endonuclease-[Enterobacter-hormaechei] |
| MCM7420476.1 | (w)HTH+REase | GCA_023751025.1 | Enterobacter_hormaechei | Bacteria>Pseudomonadati>Pseudomonadota>Gammaproteobacteria>Enterobacterales>Enterobacteriaceae>Enterobacter>Enterobacter_cloacae_complex | 289 | M8R61_11725 | 342978..343847 | 158836 | restriction-endonuclease-[Enterobacter-hormaechei] |
| VAE22461.1 | (w)HTH+REase | GCA_900558075.1 | Enterobacter_hormaechei | Bacteria>Pseudomonadati>Pseudomonadota>Gammaproteobacteria>Enterobacterales>Enterobacteriaceae>Enterobacter>Enterobacter_cloacae_complex | 289 | SAMEA2053862_03694 | 19111..19980 | 158836 | restriction-endonuclease-[Enterobacter-hormaechei] |
| WIJ55204.1 | (w)HTH+REase | GCA_030179615.1 | Enterobacter_hormaechei | Bacteria>Pseudomonadati>Pseudomonadota>Gammaproteobacteria>Enterobacterales>Enterobacteriaceae>Enterobacter>Enterobacter_cloacae_complex | 289 | OI979_06765 | 1411346..1412215 | 158836 | restriction-endonuclease-[Enterobacter-hormaechei] |
| WP_045343101.1 | (w)HTH+REase | GCF_029207635.1 | Enterobacter_hormaechei | Bacteria>Pseudomonadati>Pseudomonadota>Gammaproteobacteria>Enterobacterales>Enterobacteriaceae>Enterobacter>Enterobacter_cloacae_complex | 289 | H0762_RS10625 | 160445..161314 | 158836 | restriction-endonuclease-[Enterobacter-hormaechei] |
| HCM9136399.1 | (w)HTH+REase | GCA_025209595.1 | Enterobacter_cloacae_subsp._cloacae | Bacteria>Pseudomonadati>Pseudomonadota>Gammaproteobacteria>Enterobacterales>Enterobacteriaceae>Enterobacter>Enterobacter_cloacae_complex>Enterobacter_cloacae | 289 | N5L02_000887 | 367152..368021 | 336306 | restriction-endonuclease-[Enterobacter-cloacae-subsp.-cloacae] |
| EFA6413125.1 | (w)HTH+REase | - | Escherichia_coli | Bacteria>Pseudomonadati>Pseudomonadota>Gammaproteobacteria>Enterobacterales>Enterobacteriaceae>Escherichia | 289 | CIW06_000357 | 46724..47593 | 562 | restriction-endonuclease-[Escherichia-coli] |
| HBB8152963.1 | (w)HTH+REase | GCA_018015815.1 | Escherichia_coli | Bacteria>Pseudomonadati>Pseudomonadota>Gammaproteobacteria>Enterobacterales>Enterobacteriaceae>Escherichia | 289 | J8710_004817 | 11560..12429 | 562 | TPA:-restriction-endonuclease-[Escherichia-coli] |
| HBE2428358.1 | (w)HTH+REase | GCA_018545885.1 | Escherichia_coli | Bacteria>Pseudomonadati>Pseudomonadota>Gammaproteobacteria>Enterobacterales>Enterobacteriaceae>Escherichia | 289 | KL339_004661 | 11785..12654 | 562 | TPA:-restriction-endonuclease-[Escherichia-coli] |
| HBM8437230.1 | (w)HTH+REase | GCA_021425605.1 | Escherichia_coli | Bacteria>Pseudomonadati>Pseudomonadota>Gammaproteobacteria>Enterobacterales>Enterobacteriaceae>Escherichia | 289 | LOV60_003797 | 10813..11682 | 562 | TPA:-restriction-endonuclease-[Escherichia-coli] |
| HCW2948665.1 | (w)HTH+REase | GCA_026548625.1 | Escherichia_coli | Bacteria>Pseudomonadati>Pseudomonadota>Gammaproteobacteria>Enterobacterales>Enterobacteriaceae>Escherichia | 289 | OXB36_004611 | 9052..9921 | 562 | TPA:-restriction-endonuclease-[Escherichia-coli] |
| MBV4586518.1 | (w)HTH+REase | GCA_019145775.1 | Escherichia_coli | Bacteria>Pseudomonadati>Pseudomonadota>Gammaproteobacteria>Enterobacterales>Enterobacteriaceae>Escherichia | 289 | I2922_14320 | 176620..177489 | 562 | restriction-endonuclease-[Escherichia-coli] |
| UZF63331.1 | (w)HTH+REase | GCA_026016605.1 | Escherichia_coli | Bacteria>Pseudomonadati>Pseudomonadota>Gammaproteobacteria>Enterobacterales>Enterobacteriaceae>Escherichia | 289 | OM434_11335 | 2364185..2365054 | 562 | restriction-endonuclease-[Escherichia-coli] |
| WP_024008841.1 | (w)HTH+REase | GCF_000507645.1 | Escherichia_coli | Bacteria>Pseudomonadati>Pseudomonadota>Gammaproteobacteria>Enterobacterales>Enterobacteriaceae>Escherichia | 289 | G831_RS00470 | 57304..58173 | 562 | restriction-endonuclease-[Escherichia-coli] |
| WP_258301441.1 | (w)HTH+REase | - | Escherichia_coli | Bacteria>Pseudomonadati>Pseudomonadota>Gammaproteobacteria>Enterobacterales>Enterobacteriaceae>Escherichia | 273 | NX108_RS16790 | 1494..2315 | 562 | restriction-endonuclease-[Escherichia-coli] |
| HBC5231498.1 | (w)HTH+REase | GCA_018155305.1 | Klebsiella_oxytoca | Bacteria>Pseudomonadati>Pseudomonadota>Gammaproteobacteria>Enterobacterales>Enterobacteriaceae>Klebsiella/Raoultella_group>Klebsiella | 289 | KEU23_000987 | 15850..16719 | 571 | restriction-endonuclease-[Klebsiella-oxytoca] |
| HCD9026455.1 | (w)HTH+REase | GCA_023760145.1 | Klebsiella_oxytoca | Bacteria>Pseudomonadati>Pseudomonadota>Gammaproteobacteria>Enterobacterales>Enterobacteriaceae>Klebsiella/Raoultella_group>Klebsiella | 289 | NEF91_003866 | 35945..36814 | 571 | restriction-endonuclease-[Klebsiella-oxytoca] |
| STR56255.1 | (w)HTH+REase | GCA_900451255.1 | Klebsiella_oxytoca | Bacteria>Pseudomonadati>Pseudomonadota>Gammaproteobacteria>Enterobacterales>Enterobacteriaceae>Klebsiella/Raoultella_group>Klebsiella | 289 | NCTC11696_05220 | 5154076..5154945 | 571 | restriction-endonuclease-[Klebsiella-oxytoca] |
| WP_102217324.1 | (w)HTH+REase | GCF_004312385.1 | Klebsiella_pneumoniae_complex | Bacteria>Pseudomonadati>Pseudomonadota>Gammaproteobacteria>Enterobacterales>Enterobacteriaceae>Klebsiella/Raoultella_group>Klebsiella | 289 | HZ923_RS12920 | 27830..28699 | 3390273 | MULTISPECIES:-restriction-endonuclease-[Klebsiella] |
| WP_134882362.1 | (w)HTH+REase | GCF_015721555.1 | Klebsiella_oxytoca | Bacteria>Pseudomonadati>Pseudomonadota>Gammaproteobacteria>Enterobacterales>Enterobacteriaceae>Klebsiella/Raoultella_group>Klebsiella | 289 | QP770_RS14015 | 153140..154009 | 571 | restriction-endonuclease-[Klebsiella-oxytoca] |
| WP_227666910.1 | (w)HTH+REase | GCF_043636175.1 | Klebsiella_pneumoniae_complex | Bacteria>Pseudomonadati>Pseudomonadota>Gammaproteobacteria>Enterobacterales>Enterobacteriaceae>Klebsiella/Raoultella_group>Klebsiella | 284 | H8S48_RS22860 | 324376..325230 | 3390273 | MULTISPECIES:-restriction-endonuclease-[Klebsiella] |
| ARA42374.1 | (w)HTH+REase | GCA_002055765.1 | Klebsiella_pneumoniae | Bacteria>Pseudomonadati>Pseudomonadota>Gammaproteobacteria>Enterobacterales>Enterobacteriaceae>Klebsiella/Raoultella_group>Klebsiella>Klebsiella_pneumoniae_complex | 289 | AM364_19145 | 3773674..3774543 | 573 | restriction-endonuclease-[Klebsiella-pneumoniae] |
| EIY1898008.1 | (w)HTH+REase | GCA_022464035.1 | Klebsiella_pneumoniae | Bacteria>Pseudomonadati>Pseudomonadota>Gammaproteobacteria>Enterobacterales>Enterobacteriaceae>Klebsiella/Raoultella_group>Klebsiella>Klebsiella_pneumoniae_complex | 289 | MMH51_002459 | 46585..47454 | 573 | restriction-endonuclease-[Klebsiella-pneumoniae] |
| HBQ8501078.1 | (w)HTH+REase | GCA_021895715.1 | Klebsiella_pneumoniae | Bacteria>Pseudomonadati>Pseudomonadota>Gammaproteobacteria>Enterobacterales>Enterobacteriaceae>Klebsiella/Raoultella_group>Klebsiella>Klebsiella_pneumoniae_complex | 289 | L9A44_004849 | 46135..47004 | 573 | TPA:-restriction-endonuclease-[Klebsiella-pneumoniae] |
| HBX9967582.1 | (w)HTH+REase | GCA_022266445.1 | Klebsiella_variicola | Bacteria>Pseudomonadati>Pseudomonadota>Gammaproteobacteria>Enterobacterales>Enterobacteriaceae>Klebsiella/Raoultella_group>Klebsiella>Klebsiella_pneumoniae_complex | 289 | MIM73_23175 | 27845..28714 | 244366 | restriction-endonuclease-[Klebsiella-variicola] |
| HBY8019966.1 | (w)HTH+REase | GCA_022301155.1 | Klebsiella_pneumoniae | Bacteria>Pseudomonadati>Pseudomonadota>Gammaproteobacteria>Enterobacterales>Enterobacteriaceae>Klebsiella/Raoultella_group>Klebsiella>Klebsiella_pneumoniae_complex | 289 | MI822_22050 | 46300..47169 | 573 | TPA:-restriction-endonuclease-[Klebsiella-pneumoniae] |
| HBZ7479229.1 | (w)HTH+REase | GCA_022497945.1 | Klebsiella_pneumoniae | Bacteria>Pseudomonadati>Pseudomonadota>Gammaproteobacteria>Enterobacterales>Enterobacteriaceae>Klebsiella/Raoultella_group>Klebsiella>Klebsiella_pneumoniae_complex | 289 | MM066_002843 | 208175..209044 | 573 | TPA:-restriction-endonuclease-[Klebsiella-pneumoniae] |
| HCJ8634264.1 | (w)HTH+REase | GCA_024674695.1 | Klebsiella_pneumoniae | Bacteria>Pseudomonadati>Pseudomonadota>Gammaproteobacteria>Enterobacterales>Enterobacteriaceae>Klebsiella/Raoultella_group>Klebsiella>Klebsiella_pneumoniae_complex | 289 | NV748_003022 | 184356..185225 | 573 | TPA:-restriction-endonuclease-[Klebsiella-pneumoniae] |
| MBK1516294.1 | (w)HTH+REase | GCA_014235385.1 | Klebsiella_variicola | Bacteria>Pseudomonadati>Pseudomonadota>Gammaproteobacteria>Enterobacterales>Enterobacteriaceae>Klebsiella/Raoultella_group>Klebsiella>Klebsiella_pneumoniae_complex | 289 | H8S48_22850 | 324361..325230 | 244366 | restriction-endonuclease-[Klebsiella-variicola] |
| MBK2656981.1 | (w)HTH+REase | GCA_016614755.1 | Klebsiella_pneumoniae | Bacteria>Pseudomonadati>Pseudomonadota>Gammaproteobacteria>Enterobacterales>Enterobacteriaceae>Klebsiella/Raoultella_group>Klebsiella>Klebsiella_pneumoniae_complex | 289 | FKL12_18760 | 4488..5357 | 573 | restriction-endonuclease-[Klebsiella-pneumoniae] |
| MBK5711245.1 | (w)HTH+REase | GCA_016651545.1 | Klebsiella_pneumoniae | Bacteria>Pseudomonadati>Pseudomonadota>Gammaproteobacteria>Enterobacterales>Enterobacteriaceae>Klebsiella/Raoultella_group>Klebsiella>Klebsiella_pneumoniae_complex | 289 | JKA42_15100 | 58351..59220 | 573 | restriction-endonuclease-[Klebsiella-pneumoniae] |
| PMD76490.1 | (w)HTH+REase | GCA_002872205.1 | Klebsiella_pneumoniae | Bacteria>Pseudomonadati>Pseudomonadota>Gammaproteobacteria>Enterobacterales>Enterobacteriaceae>Klebsiella/Raoultella_group>Klebsiella>Klebsiella_pneumoniae_complex | 289 | A8A10_21910 | 142725..143594 | 573 | restriction-endonuclease-[Klebsiella-pneumoniae] |
| VTN25620.1 | (w)HTH+REase | GCA_901420815.1 | Klebsiella_pneumoniae | Bacteria>Pseudomonadati>Pseudomonadota>Gammaproteobacteria>Enterobacterales>Enterobacteriaceae>Klebsiella/Raoultella_group>Klebsiella>Klebsiella_pneumoniae_complex | 289 | NCTC9129_01962 | 1503075..1503944 | 573 | restriction-endonuclease-[Klebsiella-pneumoniae] |
| KDL80232.1 | (w)HTH+REase | GCA_000694675.1 | Klebsiella_pneumoniae_MGH_72 | Bacteria>Pseudomonadati>Pseudomonadota>Gammaproteobacteria>Enterobacterales>Enterobacteriaceae>Klebsiella/Raoultella_group>Klebsiella>Klebsiella_pneumoniae_complex>Klebsiella_pneumoniae | 289 | AD98_04568 | 195531..196400 | 1438795 | hypothetical-protein-AD98_04568-[Klebsiella-pneumoniae-MGH-72] |
| WP_285145132.1 | (w)HTH+REase | GCF_030234145.1 | Lelliottia_wanjuensis | Bacteria>Pseudomonadati>Pseudomonadota>Gammaproteobacteria>Enterobacterales>Enterobacteriaceae>Lelliottia | 289 | QQF24_RS22380 | 140960..141829 | 3050585 | MULTISPECIES:-restriction-endonuclease-[unclassified-Lelliottia] |
| ECA3794689.1 | (w)HTH+REase | GCA_007378915.1 | Salmonella_enterica_subsp._enterica_serovar_Aqua | Bacteria>Pseudomonadati>Pseudomonadota>Gammaproteobacteria>Enterobacterales>Enterobacteriaceae>Salmonella>Salmonella_enterica>Salmonella_enterica_subsp._enterica | 289 | EKG95_23295 | 23669..24538 | 1302615 | restriction-endonuclease-[Salmonella-enterica-subsp.-enterica-serovar-Aqua] |
| WP_116197727.1 | (w)HTH+REase | GCF_017920215.1 | Pantoea_ananatis | Bacteria>Pseudomonadati>Pseudomonadota>Gammaproteobacteria>Enterobacterales>Erwiniaceae>Pantoea | 289 | HG975_RS10000 | 351408..352277 | 553 | restriction-endonuclease-[Pantoea-ananatis] |
| WP_266093838.1 | (w)HTH+REase | GCF_026241965.1 | [Curtobacterium]_plantarum | Bacteria>Pseudomonadati>Pseudomonadota>Gammaproteobacteria>Enterobacterales>Erwiniaceae>Pantoea | 289 | ORG41_RS07880 | 296247..297116 | 221276 | restriction-endonuclease-[[Curtobacterium] |
| WP_282555321.1 | (w)HTH+REase | - | Providencia | Bacteria>Pseudomonadati>Pseudomonadota>Gammaproteobacteria>Enterobacterales>Morganellaceae | 273 | I6H92_RS05415 | 1214234..1215055 | 586 | restriction-endonuclease-[Providencia-stuartii] |
| MCL8535006.1 | (w)HTH+REase | GCA_023604365.1 | Proteus_mirabilis | Bacteria>Pseudomonadati>Pseudomonadota>Gammaproteobacteria>Enterobacterales>Morganellaceae>Proteus | 284 | NAE57_17185 | 8861..9715 | 584 | restriction-endonuclease-[Proteus-mirabilis] |
| MBK1419824.1 | (w)HTH+REase | GCA_016623625.1 | Providencia_stuartii | Bacteria>Pseudomonadati>Pseudomonadota>Gammaproteobacteria>Enterobacterales>Morganellaceae>Providencia | 289 | I6H92_05415 | 1214186..1215055 | 588 | restriction-endonuclease-[Providencia-stuartii] |
| WP_038193798.1 | (w)HTH+REase | GCF_963864975.1 | Xenorhabdus_bovienii | Bacteria>Pseudomonadati>Pseudomonadota>Gammaproteobacteria>Enterobacterales>Morganellaceae>Xenorhabdus | 289 | KKI95_RS10010 | 140784..141653 | 40576 | restriction-endonuclease-[Xenorhabdus-bovienii] |
| WP_060426788.1 | (w)HTH+REase | GCF_946413725.1 | Serratia | Bacteria>Pseudomonadati>Pseudomonadota>Gammaproteobacteria>Enterobacterales>Yersiniaceae | 305 | AWX06_RS15830 | 180324..181241 | 613 | MULTISPECIES:-restriction-endonuclease-[Serratia] |
| WP_095924511.1 | (w)HTH+REase | GCF_002303785.1 | Rahnella_victoriana | Bacteria>Pseudomonadati>Pseudomonadota>Gammaproteobacteria>Enterobacterales>Yersiniaceae>Rahnella | 289 | C0635_RS19370 | 4098339..4099208 | 1510570 | restriction-endonuclease-[Rahnella-victoriana] |
| MCH9867674.1 | (w)HTH+REase | GCA_022602995.1 | Serratia_marcescens | Bacteria>Pseudomonadati>Pseudomonadota>Gammaproteobacteria>Enterobacterales>Yersiniaceae>Serratia | 305 | L8N14_16495 | 90806..91723 | 615 | restriction-endonuclease-[Serratia-marcescens] |
| WP_089185253.1 | (w)HTH+REase | GCF_002220535.1 | Serratia_marcescens | Bacteria>Pseudomonadati>Pseudomonadota>Gammaproteobacteria>Enterobacterales>Yersiniaceae>Serratia | 305 | BVG90_RS23865 | 5007677..5008594 | 615 | restriction-endonuclease-[Serratia-marcescens] |
| WP_197785665.1 | (w)HTH+REase | GCF_016007555.1 | Serratia_ureilytica | Bacteria>Pseudomonadati>Pseudomonadota>Gammaproteobacteria>Enterobacterales>Yersiniaceae>Serratia | 305 | I5P92_RS12845 | 21975..22892 | 300181 | restriction-endonuclease-[Serratia-ureilytica] |
| WP_244757922.1 | (w)HTH+REase | GCF_022870585.1 | Serratia_marcescens | Bacteria>Pseudomonadati>Pseudomonadota>Gammaproteobacteria>Enterobacterales>Yersiniaceae>Serratia | 305 | HW877_RS02475 | 540995..541912 | 615 | restriction-endonuclease-[Serratia-marcescens] |
| WP_098087300.1 | (w)HTH+REase | GCF_003546905.1 | Yersinia_pseudotuberculosis | Bacteria>Pseudomonadati>Pseudomonadota>Gammaproteobacteria>Enterobacterales>Yersiniaceae>Yersinia>Yersinia_pseudotuberculosis_complex | 284 | EGX52_RS11290 | 2331335..2332189 | 633 | restriction-endonuclease-[Yersinia-pseudotuberculosis] |
| MAZ66827.1 | (w)HTH+REase | GCA_002707725.1 | Kangiellaceae_bacterium | Bacteria>Pseudomonadati>Pseudomonadota>Gammaproteobacteria>Kangiellales>Kangiellaceae>unclassified_Kangiellaceae | 358 | CMF25_06930 | 11312..12388 | 2026751 | hypothetical-protein-[Kangiellaceae-bacterium] |
| WP_083663064.1 | (w)HTH+REase | GCF_001939825.1 | Stenotrophomonas | Bacteria>Pseudomonadati>Pseudomonadota>Gammaproteobacteria>Lysobacterales>Lysobacteraceae | 110 | BKE09_RS23065 | 1396141..1396473 | 40323 | MULTISPECIES:-DUF1016-N-terminal-domain-containing-protein-[Stenotrophomonas] |
| WP_144815722.1 | (w)HTH+REase | GCF_007830115.1 | Aerolutibacter_ruishenii | Bacteria>Pseudomonadati>Pseudomonadota>Gammaproteobacteria>Lysobacterales>Lysobacteraceae>Aerolutibacter | 372 | IP93_RS11170 | 74185..75303 | 686800 | PDDEXK-nuclease-domain-containing-protein-[Lysobacter-ruishenii] |
| MBS3952431.1 | (w)HTH+REase | GCA_018333785.1 | Methylomicrobium_sp. | Bacteria>Pseudomonadati>Pseudomonadota>Gammaproteobacteria>Methylococcales>Methylococcaceae>Methylomicrobium>unclassified_Methylomicrobium | 202 | KGZ88_05730 | 11648..12256 | 2303549 | hypothetical-protein-[Methylomicrobium-sp.] |
| WP_096876856.1 | (w)HTH+REase | GCF_002441955.1 | Methylomonas_koyamae | Bacteria>Pseudomonadati>Pseudomonadota>Gammaproteobacteria>Methylococcales>Methylococcaceae>Methylomonas | 385 | MKLM6_RS22305 | 66722..67879 | 702114 | PDDEXK-nuclease-domain-containing-protein-[Methylomonas-koyamae] |
| WP_054759928.1 | (w)HTH+REase | GCF_019669905.1 | Methylomonas_koyamae | Bacteria>Pseudomonadati>Pseudomonadota>Gammaproteobacteria>Methylococcales>Methylococcaceae>Methylomonas | 365 | JCM16701_RS06925 | 44735..45832 | 702114 | PDDEXK-nuclease-domain-containing-protein-[Methylomonas-koyamae] |
| WP_172420974.1 | (w)HTH+REase | GCF_011399095.1 | Halomonas_hydrothermalis | Bacteria>Pseudomonadati>Pseudomonadota>Gammaproteobacteria>Oceanospirillales>Halomonadaceae>Halomonas | 365 | HHSLTHF2_RS10315 | 2154657..2155754 | 115561 | PDDEXK-nuclease-domain-containing-protein-[Halomonas-hydrothermalis] |
| WP_078087406.1 | (w)HTH+REase | GCF_002009255.1 | unclassified_Halomonas | Bacteria>Pseudomonadati>Pseudomonadota>Gammaproteobacteria>Oceanospirillales>Halomonadaceae>Halomonas | 137 | BV504_RS06355 | 1463670..1464083 | 2609666 | MULTISPECIES:-DUF1016-N-terminal-domain-containing-protein-[unclassified-Halomonas] |
| MBL1268351.1 | (w)HTH+REase | GCA_002733435.2 | Halomonas_sp. | Bacteria>Pseudomonadati>Pseudomonadota>Gammaproteobacteria>Oceanospirillales>Halomonadaceae>Halomonas>unclassified_Halomonas | 365 | COA87_011525 | 235105..236202 | 1486246 | DUF1016-family-protein-[Halomonas-sp.] |
| NQY75751.1 | (w)HTH+REase | GCA_013213865.1 | Halomonas_sp. | Bacteria>Pseudomonadati>Pseudomonadota>Gammaproteobacteria>Oceanospirillales>Halomonadaceae>Halomonas>unclassified_Halomonas | 365 | HRT78_01210 | 163553..164650 | 1486246 | DUF1016-domain-containing-protein-[Halomonas-sp.] |
| WP_009095649.1 | (w)HTH+REase | GCF_000236625.2 | Halomonas_sp._GFAJ-1 | Bacteria>Pseudomonadati>Pseudomonadota>Gammaproteobacteria>Oceanospirillales>Halomonadaceae>Halomonas>unclassified_Halomonas | 148 | BB497_RS11610 | 2570524..2570970 | 1118153 | DUF1016-N-terminal-domain-containing-protein-[Halomonas-sp.-GFAJ-1] |
| WP_072820807.1 | (w)HTH+REase | GCF_900128925.1 | Modicisalibacter_ilicicola | Bacteria>Pseudomonadati>Pseudomonadota>Gammaproteobacteria>Oceanospirillales>Halomonadaceae>Modicisalibacter | 388 | BUA41_RS05260 | 158426..159592 | 480814 | restriction-endonuclease-[Halomonas-ilicicola] |
| WP_083023780.1 | (w)HTH+REase | GCF_002087295.1 | Vreelandella_lionensis | Bacteria>Pseudomonadati>Pseudomonadota>Gammaproteobacteria>Oceanospirillales>Halomonadaceae>Vreelandella | 365 | B4967_RS04185 | 26547..27644 | 1144478 | PDDEXK-nuclease-domain-containing-protein-[Halomonas-lionensis] |
| WP_179915948.1 | (w)HTH+REase | GCF_013415125.1 | Vreelandella_glaciei | Bacteria>Pseudomonadati>Pseudomonadota>Gammaproteobacteria>Oceanospirillales>Halomonadaceae>Vreelandella | 365 | HZS80_RS09705 | 41716..42813 | 186761 | PDDEXK-nuclease-domain-containing-protein-[Halomonas-glaciei] |
| MBE0507009.1 | (w)HTH+REase | GCA_014859565.1 | Marinospirillum_sp. | Bacteria>Pseudomonadati>Pseudomonadota>Gammaproteobacteria>Oceanospirillales>Oceanospirillaceae>Marinospirillum>unclassified_Marinospirillum | 355 | IBX50_09870 | 10416..11483 | 2183934 | DUF1016-domain-containing-protein-[Marinospirillum-sp.] |
| WP_138986813.1 | (w)HTH+REase | GCF_008630635.1 | Neptunomonas_concharum | Bacteria>Pseudomonadati>Pseudomonadota>Gammaproteobacteria>Oceanospirillales>Oceanospirillaceae>Neptunomonas | 348 | FOU83_RS12930 | 2737096..2738142 | 1031538 | PDDEXK-nuclease-domain-containing-protein-[Neptunomonas-concharum] |
| MAD46038.1 | (w)HTH+REase | GCA_002685055.1 | Oceanospirillaceae_bacterium | Bacteria>Pseudomonadati>Pseudomonadota>Gammaproteobacteria>Oceanospirillales>Oceanospirillaceae>unclassified_Oceanospirillaceae | 348 | CMH98_13640 | 4963..6009 | 1899355 | hypothetical-protein-[Oceanospirillaceae-bacterium] |
| MAR01292.1 | (w)HTH+REase | GCA_002701445.1 | Oceanospirillaceae_bacterium | Bacteria>Pseudomonadati>Pseudomonadota>Gammaproteobacteria>Oceanospirillales>Oceanospirillaceae>unclassified_Oceanospirillaceae | 315 | CMI00_12245 | 19084..20031 | 1899355 | restriction-endonuclease-[Oceanospirillaceae-bacterium] |
| WP_274102067.1 | (w)HTH+REase | GCF_046523955.1 | Pseudomonas | Bacteria>Pseudomonadati>Pseudomonadota>Gammaproteobacteria>Pseudomonadales>Pseudomonadaceae | 303 | NPS33_RS09485 | 97996..98907 | 286 | MULTISPECIES:-restriction-endonuclease-[Pseudomonas] |
| WP_265072434.1 | (w)HTH+REase | GCF_026016265.1 | Pseudomonas_viciae | Bacteria>Pseudomonadati>Pseudomonadota>Gammaproteobacteria>Pseudomonadales>Pseudomonadaceae>Pseudomonas | 303 | LOY66_RS25485 | 5804505..5805416 | 2505979 | restriction-endonuclease-[Pseudomonas-viciae] |
| HBP0333022.1 | (w)HTH+REase | GCA_021684735.1 | Pseudomonas_aeruginosa | Bacteria>Pseudomonadati>Pseudomonadota>Gammaproteobacteria>Pseudomonadales>Pseudomonadaceae>Pseudomonas>Pseudomonas_aeruginosa_group | 350 | L5P76_005853 | 11921..12973 | 287 | TPA:-DUF1016-domain-containing-protein-[Pseudomonas-aeruginosa] |
| HBP0494142.1 | (w)HTH+REase | GCA_021685415.1 | Pseudomonas_aeruginosa | Bacteria>Pseudomonadati>Pseudomonadota>Gammaproteobacteria>Pseudomonadales>Pseudomonadaceae>Pseudomonas>Pseudomonas_aeruginosa_group | 350 | L5Q31_004991 | 20563..21615 | 287 | DUF1016-domain-containing-protein-[Pseudomonas-aeruginosa] |
| HCF1786788.1 | (w)HTH+REase | GCA_023942115.1 | Pseudomonas_aeruginosa | Bacteria>Pseudomonadati>Pseudomonadota>Gammaproteobacteria>Pseudomonadales>Pseudomonadaceae>Pseudomonas>Pseudomonas_aeruginosa_group | 350 | NH908_000696 | 73127..74179 | 287 | TPA:-DUF1016-family-protein-[Pseudomonas-aeruginosa] |
| HCF5128591.1 | (w)HTH+REase | GCA_023952375.1 | Pseudomonas_aeruginosa | Bacteria>Pseudomonadati>Pseudomonadota>Gammaproteobacteria>Pseudomonadales>Pseudomonadaceae>Pseudomonas>Pseudomonas_aeruginosa_group | 350 | NIG22_002420 | 15110..16162 | 287 | DUF1016-family-protein-[Pseudomonas-aeruginosa] |
| WP_023112382.1 | (w)HTH+REase | GCF_013912775.1 | Pseudomonas_aeruginosa | Bacteria>Pseudomonadati>Pseudomonadota>Gammaproteobacteria>Pseudomonadales>Pseudomonadaceae>Pseudomonas>Pseudomonas_aeruginosa_group | 350 | H2N73_RS00370 | 88360..89412 | 287 | PDDEXK-nuclease-domain-containing-protein-[Pseudomonas-aeruginosa] |
| RTS56727.1 | (w)HTH+REase | GCA_003968225.1 | Pseudomonas_aeruginosa | Bacteria>Pseudomonadati>Pseudomonadota>Gammaproteobacteria>Pseudomonadales>Pseudomonadaceae>Pseudomonas>Pseudomonas_aeruginosa_group | 103 | DY943_06700 | 166970..167281 | 287 | DUF1016-family-protein-[Pseudomonas-aeruginosa] |
| HBP0494143.1 | (w)HTH+REase | GCA_021685415.1 | Pseudomonas_aeruginosa | Bacteria>Pseudomonadati>Pseudomonadota>Gammaproteobacteria>Pseudomonadales>Pseudomonadaceae>Pseudomonas>Pseudomonas_aeruginosa_group | 85 | L5Q31_004992 | 21616..21873 | 287 | TPA:-hypothetical-protein-L5Q31_004992-[Pseudomonas-aeruginosa] |
| HCF1786787.1 | (w)HTH+REase | GCA_023942115.1 | Pseudomonas_aeruginosa | Bacteria>Pseudomonadati>Pseudomonadota>Gammaproteobacteria>Pseudomonadales>Pseudomonadaceae>Pseudomonas>Pseudomonas_aeruginosa_group | 85 | NH908_000695 | 72869..73126 | 287 | TPA:-hypothetical-protein-NH908_000695-[Pseudomonas-aeruginosa] |
| HCF5128590.1 | (w)HTH+REase | GCA_023952375.1 | Pseudomonas_aeruginosa | Bacteria>Pseudomonadati>Pseudomonadota>Gammaproteobacteria>Pseudomonadales>Pseudomonadaceae>Pseudomonas>Pseudomonas_aeruginosa_group | 85 | NIG22_002419 | 14852..15109 | 287 | TPA:-hypothetical-protein-NIG22_002419-[Pseudomonas-aeruginosa] |
| WP_260472515.1 | (w)HTH+REase | GCF_003968225.1 | Pseudomonas_aeruginosa | Bacteria>Pseudomonadati>Pseudomonadota>Gammaproteobacteria>Pseudomonadales>Pseudomonadaceae>Pseudomonas>Pseudomonas_aeruginosa_group | 57 | DY943_RS33040 | 167108..167281 | 287 | DUF1016-N-terminal-domain-containing-protein-[Pseudomonas-aeruginosa] |
| WP_140786431.1 | (w)HTH+REase | GCF_003626935.1 | Pseudomonas_aeruginosa | Bacteria>Pseudomonadati>Pseudomonadota>Gammaproteobacteria>Pseudomonadales>Pseudomonadaceae>Pseudomonas>Pseudomonas_aeruginosa_group | 317 | D7H65_RS11240 | 2289320..2290273 | 287 | HrgA-protein-[Pseudomonas-aeruginosa] |
| MCO2467172.1 | (w)HTH+REase | GCA_023907445.1 | Pseudomonas_aeruginosa | Bacteria>Pseudomonadati>Pseudomonadota>Gammaproteobacteria>Pseudomonadales>Pseudomonadaceae>Pseudomonas>Pseudomonas_aeruginosa_group | 313 | FA343_15145 | 56897..57838 | 287 | HrgA-protein-[Pseudomonas-aeruginosa] |
| WP_023113963.1 | (w)HTH+REase | GCF_000481245.1 | Pseudomonas_aeruginosa | Bacteria>Pseudomonadati>Pseudomonadota>Gammaproteobacteria>Pseudomonadales>Pseudomonadaceae>Pseudomonas>Pseudomonas_aeruginosa_group | 313 | Q036_RS16805 | 2140976..2141917 | 287 | COG2958-family-protein-[Pseudomonas-aeruginosa] |
| TVT71196.1 | (w)HTH+REase | GCA_007713455.1 | Pseudomonas_sp. | Bacteria>Pseudomonadati>Pseudomonadota>Gammaproteobacteria>Pseudomonadales>Pseudomonadaceae>Pseudomonas>unclassified_Pseudomonas | 350 | FHK79_07555 | 349227..350279 | 306 | DUF1016-domain-containing-protein-[Pseudomonas-sp.] |
| TVT71195.1 | (w)HTH+REase | GCA_007713455.1 | Pseudomonas_sp. | Bacteria>Pseudomonadati>Pseudomonadota>Gammaproteobacteria>Pseudomonadales>Pseudomonadaceae>Pseudomonas>unclassified_Pseudomonas | 89 | FHK79_07550 | 348957..349226 | 306 | hypothetical-protein-FHK79_07550-[Pseudomonas-sp.] |
| MCG8610924.1 | (w)HTH+REase | GCA_022361325.1 | Pseudomonadales_bacterium | Bacteria>Pseudomonadati>Pseudomonadota>Gammaproteobacteria>Pseudomonadales>unclassified_Pseudomonadales | 369 | MI864_10360 | 3664..4773 | 1891229 | PDDEXK-nuclease-domain-containing-protein-[Pseudomonadales-bacterium] |
| WP_009727410.1 | (w)HTH+REase | GCF_000349205.1 | Methylophaga_lonarensis | Bacteria>Pseudomonadati>Pseudomonadota>Gammaproteobacteria>Thiotrichales>Piscirickettsiaceae>Methylophaga | 289 | MPL1_RS12045 | 27461..28330 | 999151 | restriction-endonuclease-[Methylophaga-lonarensis] |
| MBD3820861.1 | (w)HTH+REase | GCA_014764155.1 | Thiotrichales_bacterium | Bacteria>Pseudomonadati>Pseudomonadota>Gammaproteobacteria>Thiotrichales>unclassified_Thiotrichales | 370 | IE914_01155 | 8767..9879 | 2026796 | DUF1016-domain-containing-protein-[Thiotrichales-bacterium] |
| MCP5421180.1 | (w)HTH+REase | GCA_024235155.1 | Gammaproteobacteria_bacterium | Bacteria>Pseudomonadati>Pseudomonadota>Gammaproteobacteria>unclassified_Gammaproteobacteria | 393 | H6969_11930 | 11843..13024 | 1913989 | DUF1016-domain-containing-protein-[Gammaproteobacteria-bacterium] |
| QFY90725.1 | (w)HTH+REase | GCA_009497875.2 | gamma_proteobacterium_SS-5 | Bacteria>Pseudomonadati>Pseudomonadota>Gammaproteobacteria>unclassified_Gammaproteobacteria | 368 | D5125_15350 | 2194450..2195556 | 947516 | DUF1016-family-protein-[gamma-proteobacterium-SS-5] |
| MBR9924471.1 | (w)HTH+REase | GCA_018223005.1 | Gammaproteobacteria_bacterium | Bacteria>Pseudomonadati>Pseudomonadota>Gammaproteobacteria>unclassified_Gammaproteobacteria | 365 | GYB28_05695 | 379560..380657 | 1913989 | DUF1016-domain-containing-protein-[Gammaproteobacteria-bacterium] |
| PUB90398.1 | (w)HTH+REase | GCA_003058535.1 | gamma_proteobacterium_symbiont_of_Ctena_orbiculata | Bacteria>Pseudomonadati>Pseudomonadota>Gammaproteobacteria>unclassified_Gammaproteobacteria | 284 | DBP01_07770 | 9570..10424 | 1968598 | restriction-endonuclease-[gamma-proteobacterium-symbiont-of-Ctena-orbiculata] |
| WP_087868377.1 | (w)HTH+REase | GCF_002189065.1 | Bathymodiolus_platifrons_methanotrophic_gill_symbiont | Bacteria>Pseudomonadati>Pseudomonadota>Gammaproteobacteria>unclassified_Gammaproteobacteria | 284 | CDZ78_RS01540 | 43469..44323 | 113268 | restriction-endonuclease-[Bathymodiolus-platifrons-methanotrophic-gill-symbiont] |
| EKO3646389.1 | (w)HTH+REase | GCA_028989625.1 | Vibrio_metschnikovii | Bacteria>Pseudomonadati>Pseudomonadota>Gammaproteobacteria>Vibrionales>Vibrionaceae>Vibrio | 284 | M3925_003002 | 55537..56391 | 28172 | restriction-endonuclease-[Vibrio-metschnikovii] |
| EKO3692971.1 | (w)HTH+REase | GCA_028990005.1 | Vibrio_metschnikovii | Bacteria>Pseudomonadati>Pseudomonadota>Gammaproteobacteria>Vibrionales>Vibrionaceae>Vibrio | 284 | POF02_001014 | 202006..202860 | 28172 | restriction-endonuclease-[Vibrio-metschnikovii] |
| MBY7816965.1 | (w)HTH+REase | GCA_019852895.1 | Vibrio_fluvialis | Bacteria>Pseudomonadati>Pseudomonadota>Gammaproteobacteria>Vibrionales>Vibrionaceae>Vibrio | 284 | KW416_13065 | 319539..320393 | 676 | restriction-endonuclease-[Vibrio-fluvialis] |
| MBE4228749.1 | (w)HTH+REase | GCA_014920775.1 | Vibrio_parahaemolyticus | Bacteria>Pseudomonadati>Pseudomonadota>Gammaproteobacteria>Vibrionales>Vibrionaceae>Vibrio>Vibrio_harveyi_group | 289 | HJ082_06175 | 245731..246600 | 670 | restriction-endonuclease-[Vibrio-parahaemolyticus] |
| PIY00907.1 | (w)HTH+REase | GCA_002784065.1 | Hydrogenophilales_bacterium_CG_4_10_14_3_um_filter_58_23 | Bacteria>Pseudomonadati>Pseudomonadota>Hydrogenophilia>Hydrogenophilales>unclassified_Hydrogenophilales | 258 | COZ23_06070 | 34185..34958 | 1974027 | DUF1016-domain-containing-protein,-partial-[Hydrogenophilales-bacterium-CG_4_10_14_3_um_filter_58_23] |
| PIQ12817.1 | (w)HTH+REase | GCA_002770925.1 | Hydrogenophilales_bacterium_CG18_big_fil_WC_8_21_14_2_50_58_12 | Bacteria>Pseudomonadati>Pseudomonadota>Hydrogenophilia>Hydrogenophilales>unclassified_Hydrogenophilales | 313 | COW70_07910 | 267..1208 | 1974036 | HrgA-protein-[Hydrogenophilales-bacterium-CG18_big_fil_WC_8_21_14_2_50_58_12] |

| Protein-ID | Domain | Assembly-ID | Strain | Taxonomy | Length | Locus-Tag | Genomic-coordinates | TaxID | Fasta-defline |
|---|---|---|---|---|---|---|---|---|---|
| MBS0213166.1 | (w)HTH+REase | GCA_018242445.1 | Pseudomonadota_bacterium | Bacteria>Pseudomonadati>Pseudomonadota>unclassified_Pseudomonadota | 373 | JSR26_08330 | 10216..11337 | 1977087 | DUF1016-family-protein-[Pseudomonadota-bacterium] |
| MBU2055994.1 | (w)HTH+REase | GCA_018825885.1 | Pseudomonadota_bacterium | Bacteria>Pseudomonadati>Pseudomonadota>unclassified_Pseudomonadota | 207 | KKC25_13675 | 4287..4910 | 1977087 | hypothetical-protein-[Pseudomonadota-bacterium] |
| MBR6059866.1 | (w)HTH+REase | GCA_017542865.1 | Victivallales_bacterium | Bacteria>Pseudomonadati>PVC_group>Lentisphaerota>Lentisphaeria>Victivallales>unclassified_Victivallales | 374 | IKP58_16995 | 6484..7608 | 2212475 | DUF1016-family-protein-[Victivallales-bacterium] |
| NLF19753.1 | (w)HTH+REase | GCA_012514865.1 | Lentisphaerota_bacterium | Bacteria>Pseudomonadati>PVC_group>Lentisphaerota>unclassified_Lentisphaerota | 398 | GX595_21180 | 2709..3905 | 1932692 | DUF1016-domain-containing-protein-[Lentisphaerota-bacterium] |
| OGV31173.1 | (w)HTH+REase | GCA_001804785.1 | Lentisphaerae_bacterium_GWF2_45_14 | Bacteria>Pseudomonadati>PVC_group>Lentisphaerota>unclassified_Lentisphaerota | 369 | A2020_01690 | 6036..7145 | 1798572 | MAG:-hypothetical-protein-A2020_01690-[Lentisphaerae-bacterium-GWF2_45_14] |
| TVL98973.1 | (w)HTH+REase | GCA_007618135.1 | Candidatus_Brocadia_sp._WS118 | Bacteria>Pseudomonadati>PVC_group>Planctomycetota>Candidatus_Brocadiia>Candidatus_Brocadiales>Candidatus_Brocadiaceae>Candidatus_Brocadia>unclassified_Candidatus_Brocadia | 375 | CV087_19780 | 8926..10053 | 2040625 | MAG:-DUF1016-domain-containing-protein-[Candidatus-Brocadia-sp.-WS118] |
| TVL98974.1 | (w)HTH+REase | GCA_007618135.1 | Candidatus_Brocadia_sp._WS118 | Bacteria>Pseudomonadati>PVC_group>Planctomycetota>Candidatus_Brocadiia>Candidatus_Brocadiales>Candidatus_Brocadiaceae>Candidatus_Brocadia>unclassified_Candidatus_Brocadia | 370 | CV087_19790 | 14011..15123 | 2040625 | MAG:-DUF1016-domain-containing-protein-[Candidatus-Brocadia-sp.-WS118] |
| MBN2560622.1 | (w)HTH+REase | GCA_016935885.1 | Phycisphaerae_bacterium | Bacteria>Pseudomonadati>PVC_group>Planctomycetota>Phycisphaerae>unclassified_Phycisphaerae | 372 | JXQ75_06805 | 8315..9433 | 2026778 | DUF1016-family-protein-[Phycisphaerae-bacterium] |
| MCK6484525.1 | (w)HTH+REase | GCA_023150595.1 | Phycisphaerae_bacterium | Bacteria>Pseudomonadati>PVC_group>Planctomycetota>Phycisphaerae>unclassified_Phycisphaerae | 372 | L6R00_10380 | 51083..52201 | 2026778 | PDDEXK-nuclease-domain-containing-protein-[Phycisphaerae-bacterium] |
| NQV23512.1 | (w)HTH+REase | GCA_013203845.1 | Rhodopirellula_sp. | Bacteria>Pseudomonadati>PVC_group>Planctomycetota>Planctomycetia>Pirellulales>Pirellulaceae>Rhodopirellula>unclassified_Rhodopirellula | 362 | HQ518_04000 | 17666..18754 | 2024855 | DUF1016-domain-containing-protein-[Rhodopirellula-sp.] |
| MCL4190100.1 | (w)HTH+REase | GCA_023382995.1 | Thermoguttaceae_bacterium | Bacteria>Pseudomonadati>PVC_group>Planctomycetota>Planctomycetia>Pirellulales>Thermoguttaceae>unclassified_Thermoguttaceae | 180 | KJZ87_00030 | 11572..12114 | 2691551 | hypothetical-protein-[Thermoguttaceae-bacterium] |
| MBN1912209.1 | (w)HTH+REase | GCA_016931075.1 | Pirellulales_bacterium | Bacteria>Pseudomonadati>PVC_group>Planctomycetota>Planctomycetia>Pirellulales>unclassified_Pirellulales | 389 | JW818_20985 | 3741..4910 | 2715253 | DUF1016-family-protein-[Pirellulales-bacterium] |
| MDA8378649.1 | (w)HTH+REase | GCA_028068345.1 | Planctomycetia_bacterium | Bacteria>Pseudomonadati>PVC_group>Planctomycetota>Planctomycetia>unclassified_Planctomycetia | 356 | MOZ50_16780 | 10766..11836 | 2052181 | MAG:-PDDEXK-nuclease-domain-containing-protein-[Planctomycetia-bacterium] |
| MCE9554598.1 | (w)HTH+REase | GCA_021413705.1 | Planctomycetota_bacterium | Bacteria>Pseudomonadati>PVC_group>Planctomycetota>unclassified_Planctomycetota | 410 | K8T91_14670 | 33771..35003 | 2026780 | PDDEXK-nuclease-domain-containing-protein-[Planctomycetota-bacterium] |
| MBI3462537.1 | (w)HTH+REase | GCA_016196255.1 | Planctomycetota_bacterium | Bacteria>Pseudomonadati>PVC_group>Planctomycetota>unclassified_Planctomycetota | 377 | HY000_05675 | 9177..10310 | 2026780 | DUF1016-domain-containing-protein-[Planctomycetota-bacterium] |
| GDY21842.1 | (w)HTH+REase | GCA_014193395.1 | Verrucomicrobiota_bacterium | Bacteria>Pseudomonadati>PVC_group>Verrucomicrobiota>unclassified_Verrucomicrobiota | 474 | LBMAG56_31890 | 11063..12487 | 2026799 | hypothetical-protein-LBMAG56_31890-[Verrucomicrobiota-bacterium] |
| MCF7762650.1 | (w)HTH+REase | GCA_021733945.1 | Verrucomicrobiota_bacterium | Bacteria>Pseudomonadati>PVC_group>Verrucomicrobiota>unclassified_Verrucomicrobiota | 386 | K9N62_03165 | 23882..25042 | 2026799 | PDDEXK-nuclease-domain-containing-protein-[Verrucomicrobiota-bacterium] |
| SPE51777.1 | (w)HTH+REase | GCA_900290355.1 | Verrucomicrobiota_bacterium | Bacteria>Pseudomonadati>PVC_group>Verrucomicrobiota>unclassified_Verrucomicrobiota | 358 | SBV1_1420057 | 66649..67725 | 2026799 | conserved-hypothetical-protein-[Verrucomicrobiota-bacterium] |
| PAW64465.1 | (w)HTH+REase | GCA_002304315.1 | Verrucomicrobiia_bacterium_Tous-C4TDCM | Bacteria>Pseudomonadati>PVC_group>Verrucomicrobiota>Verrucomicrobiia>unclassified_Verrucomicrobiia | 341 | B9S38_16355 | 7392..8417 | 1982332 | MAG:-hypothetical-protein-B9S38_16355-[Verrucomicrobiae-bacterium-Tous-C4TDCM] |
| MCI0539269.1 | (w)HTH+REase | GCA_022615925.1 | Verrucomicrobiales_bacterium | Bacteria>Pseudomonadati>PVC_group>Verrucomicrobiota>Verrucomicrobiia>Verrucomicrobiales>unclassified_Verrucomicrobiales | 373 | LOZ50_29025 | 7961..9082 | 2026801 | PDDEXK-nuclease-domain-containing-protein-[Verrucomicrobiales-bacterium] |
| MBL9158384.1 | (w)HTH+REase | GCA_016795385.1 | Verrucomicrobiales_bacterium | Bacteria>Pseudomonadati>PVC_group>Verrucomicrobiota>Verrucomicrobiia>Verrucomicrobiales>unclassified_Verrucomicrobiales | 363 | JNJ70_12980 | 42907..43998 | 2026801 | DUF1016-family-protein-[Verrucomicrobiales-bacterium] |
| MCP5560210.1 | (w)HTH+REase | GCA_024232965.1 | Verrucomicrobiaceae_bacterium | Bacteria>Pseudomonadati>PVC_group>Verrucomicrobiota>Verrucomicrobiia>Verrucomicrobiales>Verrucomicrobiaceae>unclassified_Verrucomicrobiaceae | 423 | H7A55_20870 | 45492..46763 | 2026800 | DUF1016-domain-containing-protein-[Verrucomicrobiaceae-bacterium] |
| MCP5560211.1 | (w)HTH+REase | GCA_024232965.1 | Verrucomicrobiaceae_bacterium | Bacteria>Pseudomonadati>PVC_group>Verrucomicrobiota>Verrucomicrobiia>Verrucomicrobiales>Verrucomicrobiaceae>unclassified_Verrucomicrobiaceae | 403 | H7A55_20875 | 46781..47992 | 2026800 | DUF1016-domain-containing-protein-[Verrucomicrobiaceae-bacterium] |
| MBK6607331.1 | (w)HTH+REase | GCA_016703885.1 | Leptospiraceae_bacterium | Bacteria>Pseudomonadati>Spirochaetota>Spirochaetia>Leptospirales>Leptospiraceae>unclassified_Leptospiraceae | 364 | IPG24_18095 | 69598..70692 | 2026757 | DUF1016-family-protein-[Leptospiraceae-bacterium] |
| WP_010252908.1 | (w)HTH+REase | GCF_000297095.1 | Treponema_primitia | Bacteria>Pseudomonadati>Spirochaetota>Spirochaetia>Spirochaetales>Treponemataceae>Treponema | 300 | TPRIMZ1_RS18015 | 2339..3241 | 88058 | PDDEXK-nuclease-domain-containing-protein-[Treponema-primitia] |
| TXT43481.1 | (w)HTH+REase | GCA_008080685.1 | Spirochaetota_bacterium | Bacteria>Pseudomonadati>Spirochaetota>unclassified_Spirochaetota | 351 | FD137_1819 | 10829..11884 | 2202144 | hypothetical-protein-FD137_1819-[Spirochaetota-bacterium] |
| VBB41623.1 | (w)HTH+REase | GCA_900498135.1 | uncultured_Desulfatiglans_sp. | Bacteria>Pseudomonadati>Thermodesulfobacteriota>Desulfobacteria>Desulfatiglandales>Desulfatiglandaceae>Desulfatiglans>environmental_samples | 268 | TRIP_B120058 | 44313..45119 | 1748965 | conserved-hypothetical-protein-[uncultured-Desulfatiglans-sp.] |
| KPA18628.1 | (w)HTH+REase | GCA_001292585.1 | Candidatus_Magnetomorum_sp._HK-1 | Bacteria>Pseudomonadati>Thermodesulfobacteriota>Desulfobacteria>Desulfobacterales>Candidatus_Magnetomoraceae>Candidatus_Magnetomorum>unclassified_Candidatus_Magnetomorum | 355 | MHK_001156 | 6317..7384 | 1509431 | protein-containing-DUF1016-[Candidatus-Magnetomorum-sp.-HK-1] |
| VEN75479.1 | (w)HTH+REase | GCA_900659855.1 | Candidatus_Desulfarcum_epimagneticum | Bacteria>Pseudomonadati>Thermodesulfobacteriota>Desulfobacteria>Desulfobacterales>Desulfobacteraceae>Candidatus_Desulfarcum | 340 | EPICR_90078 | 85491..86513 | 2487131 | conserved-hypothetical-protein-[uncultured-Desulfobacteraceae-bacterium] |
| MBL7172015.1 | (w)HTH+REase | GCA_016782905.1 | Desulfobacteraceae_bacterium | Bacteria>Pseudomonadati>Thermodesulfobacteriota>Desulfobacteria>Desulfobacterales>Desulfobacteraceae>unclassified_Desulfobacteraceae | 352 | ISS68_04190 | 387..1445 | 2049433 | DUF1016-family-protein-[Desulfobacteraceae-bacterium] |
| WP_207688116.1 | (w)HTH+REase | GCF_017377355.1 | Desulfonema_limicola | Bacteria>Pseudomonadati>Thermodesulfobacteriota>Desulfobacteria>Desulfobacterales>Desulfococcaceae>Desulfonema | 369 | dnl_RS20645 | 4843463..4844572 | 45656 | PDDEXK-nuclease-domain-containing-protein-[Desulfonema-limicola] |
| MBF0121238.1 | (w)HTH+REase | GCA_015228685.1 | Desulfobacterales_bacterium | Bacteria>Pseudomonadati>Thermodesulfobacteriota>Desulfobacteria>Desulfobacterales>unclassified_Desulfobacterales | 383 | HQK79_20585 | 7607..8758 | 2044940 | DUF1016-family-protein-[Desulfobacterales-bacterium] |
| WP_205241477.1 | (w)HTH+REase | GCF_016918545.1 | Desulfobulbus_alkaliphilus | Bacteria>Pseudomonadati>Thermodesulfobacteriota>Desulfobulbia>Desulfobulbales>Desulfobulbaceae>Desulfobulbus | 372 | JWG43_RS03765 | 159899..161017 | 869814 | PDDEXK-nuclease-domain-containing-protein-[Desulfobulbus-alkaliphilus] |
| MBL3581664.1 | (w)HTH+REase | GCA_016756815.1 | Oleidesulfovibrio_alaskensis | Bacteria>Pseudomonadati>Thermodesulfobacteriota>Desulfovibrionia>Desulfovibrionales>Desulfovibrionaceae>Oleidesulfovibrio | 351 | JMJ96_04535 | 94770..95825 | 58180 | restriction-endonuclease-[Oleidesulfovibrio-alaskensis] |
| WP_273192708.1 | (w)HTH+REase | GCF_016756815.1 | Oleidesulfovibrio_alaskensis | Bacteria>Pseudomonadati>Thermodesulfobacteriota>Desulfovibrionia>Desulfovibrionales>Desulfovibrionaceae>Oleidesulfovibrio | 351 | JMJ96_RS04545 | 94770..95825 | 58180 | HTH-domain-containing-protein-[Oleidesulfovibrio-alaskensis] |
| WP_157917346.1 | (w)HTH+REase | GCF_900217235.1 | Pseudodesulfovibrio_profundus | Bacteria>Pseudomonadati>Thermodesulfobacteriota>Desulfovibrionia>Desulfovibrionales>Desulfovibrionaceae>Pseudodesulfovibrio | 454 | DPRO_RS02645 | 536300..537664 | 57320 | hypothetical-protein-[Pseudodesulfovibrio-profundus] |
| MBN1547397.1 | (w)HTH+REase | GCA_016929155.1 | Syntrophaceae_bacterium | Bacteria>Pseudomonadati>Thermodesulfobacteriota>Syntrophia>Syntrophales>Syntrophaceae>unclassified_Syntrophaceae | 392 | JW902_12130 | 1437..2615 | 2052184 | DUF1016-family-protein-[Syntrophaceae-bacterium] |
| MBK7550236.1 | (w)HTH+REase | GCA_016706345.1 | Syntrophaceae_bacterium | Bacteria>Pseudomonadati>Thermodesulfobacteriota>Syntrophia>Syntrophales>Syntrophaceae>unclassified_Syntrophaceae | 386 | IPI61_00445 | 91190..92350 | 2052184 | DUF1016-family-protein-[Syntrophaceae-bacterium] |
| NSW87841.1 | (w)HTH+REase | GCA_013314815.1 | Syntrophobacteraceae_bacterium | Bacteria>Pseudomonadati>Thermodesulfobacteriota>Syntrophobacteria>Syntrophobacterales>Syntrophobacteraceae>unclassified_Syntrophobacteraceae | 410 | HPY84_16150 | 4527..5759 | 2052185 | DUF1016-domain-containing-protein-[Syntrophobacteraceae-bacterium] |
| WP_051409305.1 | (w)HTH+REase | GCF_000512235.1 | Syntrophorhabdus_aromaticivorans | Bacteria>Pseudomonadati>Thermodesulfobacteriota>Syntrophorhabdia>Syntrophorhabdales>Syntrophorhabdaceae>Syntrophorhabdus | 415 | SYNAR_RS22670 | 3027389..3028636 | 328301 | PDDEXK-nuclease-domain-containing-protein-[Syntrophorhabdus-aromaticivorans] |
| MBL3593016.1 | (w)HTH+REase | GCA_016756725.1 | Synergistaceae_bacterium | Bacteria>Thermotogati>Synergistota>Synergistia>Synergistales>Synergistaceae>unclassified_Synergistaceae | 351 | JMJ93_05835 | 387738..388793 | 2053624 | restriction-endonuclease-[Synergistaceae-bacterium] |
| WP_206075998.1 | (w)HTH+REase | GCF_004135675.1 | Marinitoga_lauensis | Bacteria>Thermotogati>Thermotogota>Thermotogae>Petrotogales>Petrotogaceae>Marinitoga | 158 | DJ007_RS15170 | 245453..245929 | 2201189 | restriction-endonuclease-[Marinitoga-lauensis] |
| KUO61375.1 | (w)HTH+REase | GCA_001516025.1 | bacterium_BRH_c32 | Bacteria>unclassified_Bacteria | 384 | APF79_11980 | 6657..7811 | 1734400 | hypothetical-protein-APF79_11980-[bacterium-BRH_c32] |
| MBU2648359.1 | (w)HTH+REase | GCA_018830885.1 | bacterium | Bacteria>unclassified_Bacteria | 376 | KKI24_26875 | 4170..5300 | 1869227 | DUF1016-family-protein-[bacterium] |
| NBJ82988.1 | (w)HTH+REase | GCA_009917545.1 | bacterium_1XD42-76 | Bacteria>unclassified_Bacteria | 371 | D5271_13865 | 83723..84838 | 2320881 | DUF1016-domain-containing-protein-[bacterium-1XD42-76] |
| NCP08811.1 | (w)HTH+REase | GCA_009993525.1 | bacterium | Bacteria>unclassified_Bacteria | 348 | GW860_07935 | 28175..29221 | 1869227 | MAG:-DUF1016-domain-containing-protein-[bacterium] |
| --McrC-NTD+REase | | | | | | | | | |
| WP_121538361.1 | McrC-NTD+REase | GCF_003665885.1 | Haloarcula_sp._Atlit-120R | Archaea>Methanobacteriati>Methanobacteriota>Stenosarchaea_group>Halobacteria>Halobacteriales>Haloarculaceae>Haloarcula>unclassified_Haloarcula | 428 | DVK01_RS19385 | 58857..60143 | 2282135 | hypothetical-protein-[Haloarcula-sp.-Atlit-120R] |
| WP_247995975.1 | McrC-NTD+REase | GCF_023115355.1 | Halocatena_salina | Archaea>Methanobacteriati>Methanobacteriota>Stenosarchaea_group>Halobacteria>Halobacteriales>Haloarculaceae>Halocatena | 421 | MW046_RS18820 | 25784..27049 | 2934340 | McrC-family-protein-[Halocatena-salina] |
| WP_226023956.1 | McrC-NTD+REase | GCF_020405245.1 | Halomicrobium_salinisoli | Archaea>Methanobacteriati>Methanobacteriota>Stenosarchaea_group>Halobacteria>Halobacteriales>Haloarculaceae>Halomicrobium | 437 | LE161_RS18725 | 252220..253533 | 2878391 | McrC-family-protein-[Halomicrobium-salinisoli] |
| WP_170155634.1 | McrC-NTD+REase | GCF_003610195.1 | Halopiger_aswanensis | Archaea>Methanobacteriati>Methanobacteriota>Stenosarchaea_group>Halobacteria>Halobacteriales>Natrialbaceae>Halopiger | 415 | ATJ93_RS22170 | 19079..20326 | 148449 | hypothetical-protein-[Halopiger-aswanensis] |
| KXS44099.1 | McrC-NTD+REase | GCA_001577565.1 | Methanolobus_sp._T82-4 | Archaea>Methanobacteriati>Methanobacteriota>Stenosarchaea_group>Methanomicrobia>Methanosarcinales>Methanosarcinaceae>Methanolobus>unclassified_Methanolobus | 346 | AWU59_732 | 20422..21462 | 1794908 | MAG:-McrC-protein-[Methanolobus-sp.-T82-4] |
| WP_077384435.1 | McrC-NTD+REase | GCF_017312625.1 | Bifidobacterium_longum | Bacteria>Bacillati>Actinomycetota>Actinomycetes>Bifidobacteriales>Bifidobacteriaceae>Bifidobacterium | 356 | MCC10048_RS10805 | 5134..6204 | 216816 | 5-methylcytosine-specific-restriction-endonuclease-system-specificity-protein-McrC-[Bifidobacterium-longum] |
| WP_129664649.1 | McrC-NTD+REase | GCF_004138495.1 | Phytoactinopolyspora_endophytica | Bacteria>Bacillati>Actinomycetota>Actinomycetes>Jiangellales>Jiangellaceae>Phytoactinopolyspora | 406 | ELR45_RS07140 | 289948..291168 | 1642495 | restriction-endonuclease-[Phytoactinopolyspora-endophytica] |
| WP_240981498.1 | McrC-NTD+REase | - | unclassified_Streptomyces | Bacteria>Bacillati>Actinomycetota>Actinomycetes>Kitasatosporales>Streptomycetaceae>Streptomyces | 315 | HHL18_RS29230 | 44226..45173 | 2593676 | MULTISPECIES:-hypothetical-protein-[unclassified-Streptomyces] |
| WP_146078436.1 | McrC-NTD+REase | GCF_002930935.1 | Rathayibacter_sp._AY1A2 | Bacteria>Bacillati>Actinomycetota>Actinomycetes>Micrococcales>Microbacteriaceae>Rathayibacter>unclassified_Rathayibacter | 392 | C5B93_RS04530 | 109199..110377 | 2080520 | restriction-endonuclease-[Rathayibacter-sp.-AY1A2] |
| WP_062735892.1 | McrC-NTD+REase | GCF_007992155.1 | Kocuria_turfanensis | Bacteria>Bacillati>Actinomycetota>Actinomycetes>Micrococcales>Micrococcaceae>Kocuria | 382 | KTU01_RS15755 | 30202..31350 | 388357 | hypothetical-protein-[Kocuria-turfanensis] |
| WP_006438850.1 | McrC-NTD+REase | GCF_000332995.1 | Gordonia | Bacteria>Bacillati>Actinomycetota>Actinomycetes>Mycobacteriales>Gordoniaceae | 443 | GAA2_RS11845 | 2620363..2621694 | 2053 | MULTISPECIES:-hypothetical-protein-[Gordonia] |
| GFM17531.1 | McrC-NTD+REase | GCA_017312405.1 | Mycobacterium_sp._PO1 | Bacteria>Bacillati>Actinomycetota>Actinomycetes>Mycobacteriales>Mycobacteriaceae>Mycobacterium>unclassified_Mycobacterium | 436 | PO1_contig-018-5 | 6135..7445 | 1882219 | Predicted-protein-[Mycobacterium-sp.-PO1] |
| WP_105365957.1 | McrC-NTD+REase | GCF_002968335.2 | Mycobacterium_sp._ITM-2016-00316 | Bacteria>Bacillati>Actinomycetota>Actinomycetes>Mycobacteriales>Mycobacteriaceae>Mycobacterium>unclassified_Mycobacterium | 398 | C6A86_RS28985 | 48243..49439 | 2099695 | restriction-endonuclease-[Mycobacterium-sp.-ITM-2016-00316] |
| WP_036463105.1 | McrC-NTD+REase | GCF_900101555.1 | Mycolicibacterium_neoaurum | Bacteria>Bacillati>Actinomycetota>Actinomycetes>Mycobacteriales>Mycobacteriaceae>Mycolicibacterium | 402 | BLS83_RS25995 | 148733..149941 | 1795 | restriction-endonuclease-[Mycolicibacterium-neoaurum] |
| WP_195040820.1 | McrC-NTD+REase | GCF_015477975.1 | Nocardia_cyriacigeorgica | Bacteria>Bacillati>Actinomycetota>Actinomycetes>Mycobacteriales>Nocardiaceae>Nocardia | 413 | IU440_RS19480 | 284001..285242 | 135487 | PE-PGRS-family-protein-[Nocardia-cyriacigeorgica] |
| WP_208115416.1 | McrC-NTD+REase | GCF_001612905.1 | Nocardia_ignorata | Bacteria>Bacillati>Actinomycetota>Actinomycetes>Mycobacteriales>Nocardiaceae>Nocardia | 411 | NI2_RS26635 | 82803..84038 | 145285 | hypothetical-protein-[Nocardia-ignorata] |
| NKR55759.1 | McrC-NTD+REase | GCA_012393885.1 | Prescottella_equi | Bacteria>Bacillati>Actinomycetota>Actinomycetes>Mycobacteriales>Nocardiaceae>Prescottella | 374 | GS481_17045 | 1597465..1598589 | 43767 | 5-methylcytosine-specific-restriction-endonuclease-system-specificity-protein-McrC-[Prescottella-equi] |
| WP_175278726.1 | McrC-NTD+REase | GCF_002095035.1 | Prescottella_equi | Bacteria>Bacillati>Actinomycetota>Actinomycetes>Mycobacteriales>Nocardiaceae>Prescottella | 374 | A5N78_RS18985 | 131923..133047 | 43767 | 5-methylcytosine-specific-restriction-endonuclease-system-specificity-protein-McrC-[Prescottella-equi] |
| WP_124569533.1 | McrC-NTD+REase | GCF_003852455.1 | Rhodococcus_sp._KBW08 | Bacteria>Bacillati>Actinomycetota>Actinomycetes>Mycobacteriales>Nocardiaceae>Rhodococcus>unclassified_Rhodococcus_(in:_high_G+C_Gram-positive_bacteria) | 479 | DBV08_RS27760 | 152099..153538 | 2144188 | hypothetical-protein-[Rhodococcus-sp.-KBW08] |
| MBK9741416.1 | McrC-NTD+REase | GCA_016717115.1 | Actinomycetota_bacterium | Bacteria>Bacillati>Actinomycetota>unclassified_Actinomycetota | 327 | IPO93_18505 | 41318..42301 | 2900548 | hypothetical-protein-[Actinomycetia-bacterium] |
| WP_169099407.1 | McrC-NTD+REase | GCF_012843265.1 | Bacillus_sp._DNRA2 | Bacteria>Bacillati>Bacillota>Bacilli>Bacillales>Bacillaceae>Bacillus>unclassified_Bacillus_(in:_firmicutes) | 496 | HHO41_RS13380 | 114365..115855 | 2723053 | DUF2357-domain-containing-protein-[Bacillus-sp.-DNRA2] |
| UQX55095.1 | McrC-NTD+REase | GCA_023516215.1 | Cytobacillus_pseudoceanisediminis | Bacteria>Bacillati>Bacillota>Bacilli>Bacillales>Bacillaceae>Cytobacillus | 344 | M5V91_04865 | 905566..906600 | 3051614 | 5-methylcytosine-specific-restriction-endonuclease-system-specificity-protein-McrC-[Cytobacillus-oceanisediminis] |
| WP_260358536.1 | McrC-NTD+REase | GCF_025209795.1 | Peribacillus_frigoritolerans | Bacteria>Bacillati>Bacillota>Bacilli>Bacillales>Bacillaceae>Peribacillus | 423 | NOU24_RS18810 | 755837..757108 | 450367 | McrC-family-protein-[Peribacillus-frigoritolerans] |
| WP_131508605.1 | McrC-NTD+REase | GCF_004337165.1 | Exiguobacterium_sp._SH0S1 | Bacteria>Bacillati>Bacillota>Bacilli>Bacillales>Bacillales_incertae_sedis>Bacillales_Family_XII._Incertae_Sedis>Exiguobacterium>unclassified_Exiguobacterium | 422 | EVJ20_RS13565 | 51916..53184 | 2510949 | McrC-family-protein-[Exiguobacterium-sp.-SH0S1] |
| WP_036689505.1 | McrC-NTD+REase | GCF_000517925.1 | Paenibacillus | Bacteria>Bacillati>Bacillota>Bacilli>Bacillales>Paenibacillaceae | 421 | C171_RS32045 | 181395..182660 | 44249 | MULTISPECIES:-McrC-family-protein-[Paenibacillus] |
| WP_058302017.1 | McrC-NTD+REase | GCF_902375625.1 | Gorillibacterium_timonense | Bacteria>Bacillati>Bacillota>Bacilli>Bacillales>Paenibacillaceae>Gorillibacterium | 500 | BN2684_RS09945 | 473761..475263 | 1689269 | DUF2357-domain-containing-protein-[Gorillibacterium-timonense] |
| WP_213510440.1 | McrC-NTD+REase | GCF_018403165.1 | Paenibacillus_macerans | Bacteria>Bacillati>Bacillota>Bacilli>Bacillales>Paenibacillaceae>Paenibacillus | 430 | KJS71_RS16955 | 175297..176589 | 44252 | McrC-family-protein-[Paenibacillus-macerans] |
| WP_189000187.1 | McrC-NTD+REase | GCF_014641075.1 | Paenibacillus_nasutitermitis | Bacteria>Bacillati>Bacillota>Bacilli>Bacillales>Paenibacillaceae>Paenibacillus | 423 | IEY46_RS33940 | 37738..39009 | 1652958 | hypothetical-protein-[Paenibacillus-nasutitermitis] |
| WDH83318.1 | McrC-NTD+REase | GCA_028747665.1 | Paenibacillus_urinalis | Bacteria>Bacillati>Bacillota>Bacilli>Bacillales>Paenibacillaceae>Paenibacillus | 497 | PUW23_03475 | 778763..780256 | 521520 | DUF2357-domain-containing-protein-[Paenibacillus-urinalis] |
| WP_089863715.1 | McrC-NTD+REase | GCF_900103025.1 | Clostridium | Bacteria>Bacillati>Bacillota>Clostridia>Eubacteriales>Clostridiaceae | 341 | C3E89_RS06150 | 1256280..1257305 | 1485 | MULTISPECIES:-hypothetical-protein-[Clostridium] |
| WP_245302583.1 | McrC-NTD+REase | - | Symbiobacterium_terraclitae | Bacteria>Bacillati>Bacillota>Clostridia>Eubacteriales>Symbiobacteriaceae>Symbiobacterium | 557 | J2Z79_RS10320 | 16820..18493 | 557451 | DUF2357-domain-containing-protein-[Symbiobacterium-terraclitae] |
| WP_158531734.1 | McrC-NTD+REase | GCF_003337245.1 | Halanaerobium_sp._MA284_MarDTE_T2 | Bacteria>Bacillati>Bacillota>Clostridia>Halanaerobiales>Halanaerobiaceae>Halanaerobium>unclassified_Halanaerobium | 393 | DFR78_RS07570 | 42913..44094 | 2183913 | hypothetical-protein-[Halanaerobium-sp.-MA284_MarDTE_T2] |
| RKW51251.1 | McrC-NTD+REase | GCA_003640285.1 | Lachnospiraceae_bacterium | Bacteria>Bacillati>Bacillota>Clostridia>Lachnospirales>Lachnospiraceae>unclassified_Lachnospiraceae | 348 | D8H95_17185 | 1600..2646 | 1898203 | 5-methylcytosine-specific-restriction-endonuclease-system-specificity-protein-McrC-[Lachnospiraceae-bacterium] |
| EGG92090.1 | McrC-NTD+REase | - | Lachnospiraceae_oral_taxon_107_str._F0167 | Bacteria>Bacillati>Bacillota>Clostridia>Lachnospirales>Lachnospiraceae>unclassified_Lachnospiraceae>Lachnospiraceae_bacterium_oral_taxon_500 | 348 | HMPREF0491_00117 | 41993..43039 | 575593 | hypothetical-protein-HMPREF0491_00117-[Lachnospiraceae-oral-taxon-107-str.-F0167] |
| MBP8598619.1 | McrC-NTD+REase | GCA_018052705.1 | Selenomonas_sp. | Bacteria>Bacillati>Bacillota>Negativicutes>Selenomonadales>Selenomonadaceae>Selenomonas>unclassified_Selenomonas | 345 | KBI24_05310 | 10665..11702 | 2053611 | 5-methylcytosine-specific-restriction-endonuclease-system-specificity-protein-McrC-[Selenomonas-sp.] |
| MBN1921824.1 | McrC-NTD+REase | GCA_016931315.1 | Anaerolineae_bacterium | Bacteria>Bacillati>Chloroflexota>Anaerolineae>unclassified_Anaerolineae | 531 | JW892_11300 | 4623..6218 | 2052143 | DUF2357-domain-containing-protein-[Anaerolineae-bacterium] |
| MBZ4644072.1 | McrC-NTD+REase | GCA_020054865.1 | Deferribacteraceae_bacterium | Bacteria>Pseudomonadati>Deferribacterota>Deferribacteres>Deferribacterales>Deferribacteraceae>unclassified_Deferribacteraceae | 438 | JG767_1681 | 4889..6205 | 2053543 | McrBC-superfamily-protein-[Deferribacteraceae-bacterium] |
| WP_242820838.1 | McrC-NTD+REase | GCF_020480685.1 | Deferrivibrio_essentukiensis | Bacteria>Pseudomonadati>Deferribacterota>Deferribacteres>Deferribacterales>Deferrivibrionaceae>Deferrivibrio | 438 | LF845_RS09800 | 26938..28254 | 2880922 | McrC-family-protein-[Deferrivibrio-essentukiensis] |
| WP_072366587.1 | McrC-NTD+REase | GCF_900119105.1 | Chitinophaga_sancti | Bacteria>Pseudomonadati>FCB_group>Bacteroidota/Chlorobiota_group>Bacteroidota>Chitinophagia>Chitinophagales>Chitinophagaceae>Chitinophaga | 425 | BT118_RS32570 | 12303..13580 | 1004 | restriction-endonuclease-[Chitinophaga-sancti] |
| MBL7764606.1 | McrC-NTD+REase | GCA_016786845.1 | Chitinophagaceae_bacterium | Bacteria>Pseudomonadati>FCB_group>Bacteroidota/Chlorobiota_group>Bacteroidota>Chitinophagia>Chitinophagales>Chitinophagaceae>unclassified_Chitinophagaceae | 436 | JNJ58_00805 | 56787..58097 | 1869212 | hypothetical-protein-[Chitinophagaceae-bacterium] |
| OSZ81162.1 | McrC-NTD+REase | GCA_002127295.1 | Chitinophagaceae_bacterium_IBVUCB2 | Bacteria>Pseudomonadati>FCB_group>Bacteroidota/Chlorobiota_group>Bacteroidota>Chitinophagia>Chitinophagales>Chitinophagaceae>unclassified_Chitinophagaceae | 436 | CAP36_07980 | 1801409..1802719 | 1985174 | restriction-endonuclease-[Chitinophagaceae-bacterium-IBVUCB2] |
| WP_084121636.1 | McrC-NTD+REase | GCF_900176595.1 | Aquiflexum_balticum | Bacteria>Pseudomonadati>FCB_group>Bacteroidota/Chlorobiota_group>Bacteroidota>Cytophagia>Cytophagales>Cyclobacteriaceae>Aquiflexum | 432 | B9A52_RS17160 | 4057598..4058896 | 280473 | restriction-endonuclease-[Aquiflexum-balticum] |
| WP_008625910.1 | McrC-NTD+REase | GCF_000330725.2 | Mariniradius_saccharolyticus | Bacteria>Pseudomonadati>FCB_group>Bacteroidota/Chlorobiota_group>Bacteroidota>Cytophagia>Cytophagales>Cyclobacteriaceae>Mariniradius | 432 | C943_RS07100 | 94280..95578 | 1245591 | hypothetical-protein-[Mariniradius-saccharolyticus] |
| WP_189584881.1 | McrC-NTD+REase | GCF_014652135.1 | Mongoliitalea_lutea | Bacteria>Pseudomonadati>FCB_group>Bacteroidota/Chlorobiota_group>Bacteroidota>Cytophagia>Cytophagales>Cyclobacteriaceae>Mongoliitalea | 432 | IE080_RS15865 | 11603..12901 | 849756 | restriction-endonuclease-[Mongoliitalea-lutea] |
| WP_244713432.1 | McrC-NTD+REase | GCF_022919135.1 | Hymenobacter_cellulosivorans | Bacteria>Pseudomonadati>FCB_group>Bacteroidota/Chlorobiota_group>Bacteroidota>Cytophagia>Cytophagales>Hymenobacteraceae>Hymenobacter | 408 | MUN80_RS13055 | 3072882..3074108 | 2932249 | McrC-family-protein-[Hymenobacter-sp.-5116S-27] |
| WP_122934663.1 | McrC-NTD+REase | GCF_003711225.1 | Runella_sp._SP2 | Bacteria>Pseudomonadati>FCB_group>Bacteroidota/Chlorobiota_group>Bacteroidota>Cytophagia>Cytophagales>Spirosomataceae>Runella>unclassified_Runella | 425 | DTQ70_RS30155 | 52489..53766 | 2268026 | restriction-endonuclease-[Runella-sp.-SP2] |
| WP_013769018.1 | McrC-NTD+REase | GCF_000212735.1 | Haliscomenobacter_hydrossis | Bacteria>Pseudomonadati>FCB_group>Bacteroidota/Chlorobiota_group>Bacteroidota>Saprospiria>Saprospirales>Haliscomenobacteraceae>Haliscomenobacter | 422 | HALHY_RS33435 | 20462..21730 | 2350 | 5-methylcytosine-restriction-system-component-like-protein-[Haliscomenobacter-hydrossis] |
| MCB0520335.1 | McrC-NTD+REase | GCA_020436325.1 | Saprospiraceae_bacterium | Bacteria>Pseudomonadati>FCB_group>Bacteroidota/Chlorobiota_group>Bacteroidota>Saprospiria>Saprospirales>Saprospiraceae>unclassified_Saprospiraceae | 429 | KDD27_15460 | 12996..14285 | 2202734 | restriction-endonuclease-[Saprospiraceae-bacterium] |
| KAF0128488.1 | McrC-NTD+REase | GCA_009773165.1 | Bacteroidota_bacterium | Bacteria>Pseudomonadati>FCB_group>Bacteroidota/Chlorobiota_group>Bacteroidota>unclassified_Bacteroidota | 425 | FD155_3230 | 29421..30698 | 1898104 | 5-methylcytosine-restriction-system-component-like-protein-[Bacteroidota-bacterium] |
| MCR9155227.1 | McrC-NTD+REase | GCA_024742495.1 | Bacteroidota_bacterium | Bacteria>Pseudomonadati>FCB_group>Bacteroidota/Chlorobiota_group>Bacteroidota>unclassified_Bacteroidota | 425 | NXI09_14050 | 92774..94051 | 1898104 | McrC-family-protein-[Bacteroidota-bacterium] |
| KAA6233385.1 | McrC-NTD+REase | GCA_008634005.1 | Chlorobium_phaeovibrioides | Bacteria>Pseudomonadati>FCB_group>Bacteroidota/Chlorobiota_group>Chlorobiota>Chlorobiia>Chlorobiales>Chlorobiaceae>Chlorobium/Pelodictyon_group>Chlorobium | 345 | FP507_06790 | 1393339..1394376 | 1094 | 5-methylcytosine-specific-restriction-endonuclease-system-specificity-protein-McrC-[Chlorobium-phaeovibrioides] |
| WP_164594475.1 | McrC-NTD+REase | GCF_010668835.1 | Rhizobium_ruizarguesonis | Bacteria>Pseudomonadati>Pseudomonadota>Alphaproteobacteria>Hyphomicrobiales>Rhizobiaceae>Rhizobium/Agrobacterium_group>Rhizobium | 445 | GR223_RS33850 | 47123..48460 | 2081791 | restriction-endonuclease-[Rhizobium-leguminosarum] |
| WP_066770451.1 | McrC-NTD+REase | GCF_001679665.1 | Candidatus_Viadribacter_manganicus | Bacteria>Pseudomonadati>Pseudomonadota>Alphaproteobacteria>Hyphomonadales>Hyphomonadaceae>Candidatus_Viadribacter | 411 | ATE48_RS09175 | 1766702..1767937 | 1759059 | hypothetical-protein-[Candidatus-Viadribacter-manganicus] |
| WP_254334240.1 | McrC-NTD+REase | GCF_024206475.1 | Paraburkholderia_sp._CNPSo_3274 | Bacteria>Pseudomonadati>Pseudomonadota>Betaproteobacteria>Burkholderiales>Burkholderiaceae>Paraburkholderia>unclassified_Paraburkholderia | 470 | M3I54_RS07300 | 49044..50456 | 2940932 | McrC-family-protein-[Paraburkholderia-sp.-CNPSo-3274] |
| WP_058675370.1 | McrC-NTD+REase | GCF_030324195.1 | Enterobacter_hormaechei | Bacteria>Pseudomonadati>Pseudomonadota>Gammaproteobacteria>Enterobacterales>Enterobacteriaceae>Enterobacter>Enterobacter_cloacae_complex | 348 | NUJ48_RS00180 | 43230..44276 | 158836 | 5-methylcytosine-specific-restriction-endonuclease-system-specificity-protein-McrC-[Enterobacter-hormaechei] |
| WP_123923012.1 | McrC-NTD+REase | GCF_003812145.1 | Enterobacter_roggenkampii | Bacteria>Pseudomonadati>Pseudomonadota>Gammaproteobacteria>Enterobacterales>Enterobacteriaceae>Enterobacter>Enterobacter_cloacae_complex | 348 | HV097_RS13170 | 2630081..2631127 | 1812935 | 5-methylcytosine-specific-restriction-endonuclease-system-specificity-protein-McrC-[Enterobacter-roggenkampii] |
| WP_248053568.1 | McrC-NTD+REase | GCF_023154935.1 | Enterobacter_chengduensis | Bacteria>Pseudomonadati>Pseudomonadota>Gammaproteobacteria>Enterobacterales>Enterobacteriaceae>Enterobacter>Enterobacter_cloacae_complex | 348 | L8P32_RS20990 | 20277..21323 | 2494701 | 5-methylcytosine-specific-restriction-endonuclease-system-specificity-protein-McrC-[Enterobacter-chengduensis] |
| MBK9496403.1 | McrC-NTD+REase | GCA_016718605.1 | Lysobacterales_bacterium | Bacteria>Pseudomonadati>Pseudomonadota>Gammaproteobacteria>Lysobacterales>unclassified_Lysobacterales | 379 | IPO08_18250 | 274510..275649 | 2006849 | 5-methylcytosine-specific-restriction-endonuclease-system-specificity-protein-McrC-[Xanthomonadales-bacterium] |
| WP_075107045.1 | McrC-NTD+REase | GCF_001921765.1 | Psychrobacter_sp._C_20.9 | Bacteria>Pseudomonadati>Pseudomonadota>Gammaproteobacteria>Moraxellales>Moraxellaceae>Psychrobacter>unclassified_Psychrobacter | 429 | BTW00_RS08520 | 116674..117963 | 1926477 | hypothetical-protein-[Psychrobacter-sp.-C-20.9] |
| EWH01937.1 | McrC-NTD+REase | GCA_000568385.1 | Halomonas_sp._BC04 | Bacteria>Pseudomonadati>Pseudomonadota>Gammaproteobacteria>Oceanospirillales>Halomonadaceae>Halomonas>unclassified_Halomonas | 556 | Q427_11495 | 30050..31720 | 1403540 | hypothetical-protein-Q427_11495-[Halomonas-sp.-BC04] |
| WP_043513269.1 | McrC-NTD+REase | GCF_000568385.1 | Halomonas_sp._BC04 | Bacteria>Pseudomonadati>Pseudomonadota>Gammaproteobacteria>Oceanospirillales>Halomonadaceae>Halomonas>unclassified_Halomonas | 556 | Q427_RS09395 | 30050..31720 | 1403540 | DUF2357-domain-containing-protein-[Halomonas-sp.-BC04] |
| WP_110649046.1 | McrC-NTD+REase | GCF_003206715.1 | Salinicola_peritrichatus | Bacteria>Pseudomonadati>Pseudomonadota>Gammaproteobacteria>Oceanospirillales>Halomonadaceae>Salinicola | 495 | DOM57_RS09915 | 68206..69693 | 1267424 | hypothetical-protein-[Salinicola-peritrichatus] |
| WP_036163551.1 | McrC-NTD+REase | GCF_000521805.1 | Marinomonas_ushuaiensis | Bacteria>Pseudomonadati>Pseudomonadota>Gammaproteobacteria>Oceanospirillales>Oceanospirillaceae>Marinomonas | 437 | MUS1_RS14010 | 37172..38485 | 263818 | hypothetical-protein-[Marinomonas-ushuaiensis] |
| HCG7393715.1 | McrC-NTD+REase | GCA_024044075.1 | Vibrio_parahaemolyticus | Bacteria>Pseudomonadati>Pseudomonadota>Gammaproteobacteria>Vibrionales>Vibrionaceae>Vibrio>Vibrio_harveyi_group | 450 | NJ383_004121 | 7732..9084 | 670 | McrC-family-protein-[Vibrio-parahaemolyticus] |
| WP_221091412.1 | McrC-NTD+REase | GCF_019693445.1 | Deinococcus_aquaedulcis | Bacteria>Thermotogati>Deinococcota>Deinococci>Deinococcales>Deinococcaceae>Deinococcus | 417 | KMW22_RS18005 | 23255..24508 | 2840455 | McrC-family-protein-[Deinococcus-aquaedulcis] |
| CCU85343.1 | McrC-NTD+REase | GCA_000367705.1 | Mesotoga_infera | Bacteria>Thermotogati>Thermotogota>Thermotogae>Kosmotogales>Kosmotogaceae>Mesotoga | 432 | PHOSAC3_150343 | 372184..373482 | 1236046 | conserved-hypothetical-protein-[Mesotoga-infera] |

| Protein-ID | Domain | Assembly-ID | Strain | Taxonomy | Length | Locus-Tag | Genomic-coordinates | TaxID | Fasta-defline |
|---|---|---|---|---|---|---|---|---|---|
| WP_169698875.1 | McrC-NTD+REase | GCF_900157305.1 | Mesotoga_infera | Bacteria>Thermotogati>Thermotogota>Thermotogae>Kosmotogales>Kosmotogaceae>Mesotoga | 417 | MESINF_RS05355 | 1173253..1174506 | 1236046 | McrC-family-protein-[Mesotoga-infera] |
| WP_121550382.1 | McrC-NTD+REase | GCF_003664995.1 | Mesotoga_sp._BH458_6_3_2_1 | Bacteria>Thermotogati>Thermotogota>Thermotogae>Kosmotogales>Kosmotogaceae>Mesotoga>unclassified_Mesotoga | 432 | Y697_RS03820 | 18346..19644 | 1437446 | McrC-family-protein-[Mesotoga-sp.-BH458_6_3_2_1] |
| --REase+SF2-Helicase(Type-2_BREX) | | | | | | | | | |
| WP_009070053.1 | REase+SF2-helicase | GCF_000243315.1 | Metallosphaera_yellowstonensis | Archaea>Thermoproteati>Thermoproteota>Thermoprotei>Sulfolobales>Sulfolobaceae>Metallosphaera | 595 | METMK1DRAFT_RS01500 | 137490..139277 | 1111107 | DUF2075-domain-containing-protein-[Metallosphaera-yellowstonensis] |
| EHP69860.1 | REase+SF2-helicase | GCA_000243315.1 | Metallosphaera_yellowstonensis_MK1 | Archaea>Thermoproteati>Thermoproteota>Thermoprotei>Sulfolobales>Sulfolobaceae>Metallosphaera>Metallosphaera_yellowstonensis | 595 | MetMK1DRAFT_00003620 | 137490..139277 | 671065 | hypothetical-protein-MetMK1DRAFT_00003620-[Metallosphaera-yellowstonensis-MK1] |
| WP_092520735.1 | REase+SF2-helicase | GCF_900100925.1 | Actinopolyspora_saharensis | Bacteria>Bacillati>Actinomycetota>Actinomycetes>Actinopolysporales>Actinopolysporaceae>Actinopolyspora | 647 | BLR67_RS02365 | 528261..530204 | 995062 | DUF2075-domain-containing-protein-[Actinopolyspora-saharensis] |
| WP_218861882.1 | REase+SF2-helicase | GCF_013408175.1 | Actinopolyspora_biskrensis | Bacteria>Bacillati>Actinomycetota>Actinomycetes>Actinopolysporales>Actinopolysporaceae>Actinopolyspora | 625 | FHR84_RS02725 | 611442..613319 | 1470178 | DUF2075-domain-containing-protein-[Actinopolyspora-biskrensis] |
| WP_246300197.1 | REase+SF2-helicase | - | Actinopolyspora_biskrensis | Bacteria>Bacillati>Actinomycetota>Actinomycetes>Actinopolysporales>Actinopolysporaceae>Actinopolyspora | 380 | FHR84_RS02745 | 616121..617263 | 1470178 | DUF2075-domain-containing-protein-[Actinopolyspora-biskrensis] |
| GAA2269120.1 | REase+SF2-helicase | GCA_039531935.1 | Glycomyces_scopariae | Bacteria>Bacillati>Actinomycetota>Actinomycetes>Glycomycetales>Glycomycetaceae>Glycomyces | 625 | GCM10009853_023240 | 484208..486085 | 480997 | DUF2075-domain-containing-protein-[Glycomyces-scopariae] |
| WP_033203948.1 | REase+SF2-helicase | GCF_000716445.1 | Streptomyces | Bacteria>Bacillati>Actinomycetota>Actinomycetes>Kitasatosporales>Streptomycetaceae | 689 | OO54_RS16765 | 65361..67430 | 1883 | MULTISPECIES:-DUF2075-domain-containing-protein-[Streptomyces] |
| WP_310769026.1 | REase+SF2-helicase | GCF_031583065.1 | Streptomyces | Bacteria>Bacillati>Actinomycetota>Actinomycetes>Kitasatosporales>Streptomycetaceae | 646 | Q2K21_RS09900 | 2368052..2369992 | 1883 | DUF2075-domain-containing-protein-[Streptomyces-sp.-CGMCC-4.7035] |
| WP_042495896.1 | REase+SF2-helicase | GCF_000010605.1 | Streptomyces | Bacteria>Bacillati>Actinomycetota>Actinomycetes>Kitasatosporales>Streptomycetaceae | 630 | SGR_RS07855 | 1887535..1889427 | 1883 | MULTISPECIES:-DUF2075-domain-containing-protein-[Streptomyces] |
| WP_282203637.1 | REase+SF2-helicase | GCF_948472415.1 | Kitasatospora_fiedleri | Bacteria>Bacillati>Actinomycetota>Actinomycetes>Kitasatosporales>Streptomycetaceae>Kitasatospora | 643 | QMQ26_RS31765 | 7037011..7038942 | 2991545 | DNA/RNA-helicase-domain-containing-protein-[Kitasatospora-fiedleri] |
| MCQ4043249.1 | REase+SF2-helicase | GCA_024436055.1 | Streptantibioticus_rubrisoli | Bacteria>Bacillati>Actinomycetota>Actinomycetes>Kitasatosporales>Streptomycetaceae>Streptantibioticus | 629 | NON19_14715 | 125157..127046 | 1387313 | DUF2075-domain-containing-protein-[Streptomyces-rubrisoli] |
| ARF59783.1 | REase+SF2-helicase | GCA_002082195.1 | Streptomyces_gilvosporeus | Bacteria>Bacillati>Actinomycetota>Actinomycetes>Kitasatosporales>Streptomycetaceae>Streptomyces | 705 | B1H19_30090 | 6738503..6740620 | 553510 | AAA-family-ATPase-[Streptomyces-gilvosporeus] |
| WP_044380256.1 | REase+SF2-helicase | GCF_000935185.3 | Streptomyces_noursei | Bacteria>Bacillati>Actinomycetota>Actinomycetes>Kitasatosporales>Streptomycetaceae>Streptomyces | 705 | SE95_RS29250 | 6688860..6690977 | 1971 | DUF2075-domain-containing-protein-[Streptomyces-noursei] |
| WP_077194918.1 | REase+SF2-helicase | GCF_001984445.1 | Streptomyces_lydicus | Bacteria>Bacillati>Actinomycetota>Actinomycetes>Kitasatosporales>Streptomycetaceae>Streptomyces | 705 | BXT88_RS28505 | 6368568..6370685 | 47763 | DUF2075-domain-containing-protein-[Streptomyces-lydicus] |
| WP_275467550.1 | REase+SF2-helicase | GCF_029167025.1 | Streptomyces_noursei | Bacteria>Bacillati>Actinomycetota>Actinomycetes>Kitasatosporales>Streptomycetaceae>Streptomyces | 705 | POD74_RS11345 | 2600491..2602608 | 1971 | DUF2075-domain-containing-protein-[Streptomyces-noursei] |
| WP_189596066.1 | REase+SF2-helicase | GCF_014649515.1 | Streptomyces_pilosus | Bacteria>Bacillati>Actinomycetota>Actinomycetes>Kitasatosporales>Streptomycetaceae>Streptomyces | 646 | IE141_RS07075 | 327595..329535 | 28893 | DUF2075-domain-containing-protein-[Streptomyces-pilosus] |
| WP_274043589.1 | REase+SF2-helicase | GCF_028657195.3 | Streptomyces_iconiensis | Bacteria>Bacillati>Actinomycetota>Actinomycetes>Kitasatosporales>Streptomycetaceae>Streptomyces | 646 | NMN56_RS24270 | 210329..212269 | 1384038 | DUF2075-domain-containing-protein-[Streptomyces-iconiensis] |
| WP_328932269.1 | REase+SF2-helicase | GCF_036196325.1 | unclassified_Streptomyces | Bacteria>Bacillati>Actinomycetota>Actinomycetes>Kitasatosporales>Streptomycetaceae>Streptomyces | 646 | OG537_RS11475 | 2679547..2681487 | 2593676 | DUF2075-domain-containing-protein-[Streptomyces-sp.-NBC_00228] |
| MCQ4207772.1 | REase+SF2-helicase | GCA_024448165.1 | Streptomyces_longispororuber | Bacteria>Bacillati>Actinomycetota>Actinomycetes>Kitasatosporales>Streptomycetaceae>Streptomyces | 632 | M4J07_005563 | 15807..17705 | 68230 | DUF2075-domain-containing-protein-[Streptomyces-longispororuber] |
| WP_255970746.1 | REase+SF2-helicase | GCF_024448165.1 | Streptomyces_longispororuber | Bacteria>Bacillati>Actinomycetota>Actinomycetes>Kitasatosporales>Streptomycetaceae>Streptomyces | 632 | M4J07_RS11745 | 15807..17705 | 68230 | DUF2075-domain-containing-protein-[Streptomyces-longispororuber] |
| WP_266427956.1 | REase+SF2-helicase | GCF_045207295.1 | unclassified_Streptomyces | Bacteria>Bacillati>Actinomycetota>Actinomycetes>Kitasatosporales>Streptomycetaceae>Streptomyces | 632 | OOK30_RS07465 | 1708518..1710416 | 2593676 | MULTISPECIES:-DUF2075-domain-containing-protein-[unclassified-Streptomyces] |
| ARF57882.1 | REase+SF2-helicase | GCA_002082195.1 | Streptomyces_gilvosporeus | Bacteria>Bacillati>Actinomycetota>Actinomycetes>Kitasatosporales>Streptomycetaceae>Streptomyces | 630 | B1H19_30095 | 6741067..6742959 | 553510 | ATP-binding-protein-[Streptomyces-gilvosporeus] |
| WP_258023622.1 | REase+SF2-helicase | GCF_019400985.1 | Streptomyces_bambusae | Bacteria>Bacillati>Actinomycetota>Actinomycetes>Kitasatosporales>Streptomycetaceae>Streptomyces | 628 | GPJ59_RS00100 | 27079..28965 | 1550616 | DUF2075-domain-containing-protein-[Streptomyces-bambusae] |
| WP_229907199.1 | REase+SF2-helicase | GCF_014656155.1 | Streptomyces_vinaceus | Bacteria>Bacillati>Actinomycetota>Actinomycetes>Kitasatosporales>Streptomycetaceae>Streptomyces | 627 | IE310_RS19405 | 96697..98580 | 1960 | DUF2075-domain-containing-protein-[Streptomyces-vinaceus] |
| WP_030293903.1 | REase+SF2-helicase | GCF_000717655.1 | Streptomyces_katrae | Bacteria>Bacillati>Actinomycetota>Actinomycetes>Kitasatosporales>Streptomycetaceae>Streptomyces | 623 | IF09_RS0109405 | 75863..77734 | 68223 | DUF2075-domain-containing-protein-[Streptomyces-katrae] |
| WP_044379858.1 | REase+SF2-helicase | GCF_000935185.3 | Streptomyces_noursei | Bacteria>Bacillati>Actinomycetota>Actinomycetes>Kitasatosporales>Streptomycetaceae>Streptomyces | 623 | SE95_RS29255 | 6691430..6693301 | 1971 | DUF2075-domain-containing-protein-[Streptomyces-noursei] |
| WP_077193667.1 | REase+SF2-helicase | GCF_001984445.1 | Streptomyces_lydicus | Bacteria>Bacillati>Actinomycetota>Actinomycetes>Kitasatosporales>Streptomycetaceae>Streptomyces | 623 | BXT88_RS28510 | 6371134..6373005 | 47763 | DUF2075-domain-containing-protein-[Streptomyces-lydicus] |
| WP_189852666.1 | REase+SF2-helicase | GCF_014650895.1 | Streptomyces_omiyaensis | Bacteria>Bacillati>Actinomycetota>Actinomycetes>Kitasatosporales>Streptomycetaceae>Streptomyces | 623 | IE287_RS31580 | 81143..83014 | 68247 | DUF2075-domain-containing-protein-[Streptomyces-omiyaensis] |
| WP_275462595.1 | REase+SF2-helicase | GCF_029167025.1 | Streptomyces_noursei | Bacteria>Bacillati>Actinomycetota>Actinomycetes>Kitasatosporales>Streptomycetaceae>Streptomyces | 623 | POD74_RS11335 | 2598167..2600038 | 1971 | DUF2075-domain-containing-protein-[Streptomyces-noursei] |
| WP_326669076.1 | REase+SF2-helicase | GCF_035918175.1 | Streptomyces_canus | Bacteria>Bacillati>Actinomycetota>Actinomycetes>Kitasatosporales>Streptomycetaceae>Streptomyces>Streptomyces_aurantiacus_group | 630 | OG925_RS34810 | 7587118..7589010 | 58343 | DUF2075-domain-containing-protein-[Streptomyces-canus] |
| WP_055559995.1 | REase+SF2-helicase | GCF_040683595.1 | Streptomyces_microflavus_subgroup | Bacteria>Bacillati>Actinomycetota>Actinomycetes>Kitasatosporales>Streptomycetaceae>Streptomyces>Streptomyces_griseus_group | 630 | AOK13_RS15165 | 21224..23116 | 1482601 | DUF2075-domain-containing-protein-[Streptomyces-luridiscabiei] |
| WSU80364.1 | REase+SF2-helicase | GCA_045162865.1 | Streptomyces_globisporus | Bacteria>Bacillati>Actinomycetota>Actinomycetes>Kitasatosporales>Streptomycetaceae>Streptomyces>Streptomyces_griseus_group>Streptomyces_albovinaceus_subgroup | 632 | OG215_06805 | 1566767..1568665 | 1908 | DUF2075-domain-containing-protein-[Streptomyces-globisporus] |
| MCC8482293.1 | REase+SF2-helicase | GCA_020881045.1 | Streptomyces_globisporus | Bacteria>Bacillati>Actinomycetota>Actinomycetes>Kitasatosporales>Streptomycetaceae>Streptomyces>Streptomyces_griseus_group>Streptomyces_albovinaceus_subgroup | 630 | LMJ41_31035 | 40049..41941 | 1908 | DUF2075-domain-containing-protein-[Streptomyces-globisporus] |
| WSU88909.1 | REase+SF2-helicase | GCA_045162845.1 | Streptomyces_anulatus | Bacteria>Bacillati>Actinomycetota>Actinomycetes>Kitasatosporales>Streptomycetaceae>Streptomyces>Streptomyces_griseus_group>Streptomyces_anulatus_subgroup | 630 | OG575_09630 | 2226006..2227898 | 1892 | DUF2075-domain-containing-protein-[Streptomyces-anulatus] |
| WTC74394.1 | REase+SF2-helicase | GCA_045209825.1 | Streptomyces_anulatus | Bacteria>Bacillati>Actinomycetota>Actinomycetes>Kitasatosporales>Streptomycetaceae>Streptomyces>Streptomyces_griseus_group>Streptomyces_anulatus_subgroup | 630 | OG882_30310 | 6750663..6752555 | 1892 | DUF2075-domain-containing-protein-[Streptomyces-anulatus] |
| WTZ16472.1 | REase+SF2-helicase | GCA_045040115.1 | Streptomyces_microflavus | Bacteria>Bacillati>Actinomycetota>Actinomycetes>Kitasatosporales>Streptomycetaceae>Streptomyces>Streptomyces_griseus_group>Streptomyces_microflavus_subgroup | 630 | OG270_07065 | 1562014..1563906 | 1919 | DUF2075-domain-containing-protein-[Streptomyces-microflavus] |
| GAA1254468.1 | REase+SF2-helicase | GCA_039525965.1 | Streptomyces_yogyakartensis | Bacteria>Bacillati>Actinomycetota>Actinomycetes>Kitasatosporales>Streptomycetaceae>Streptomyces>Streptomyces_violaceusniger_group | 639 | GCM10009579_10550 | 287340..289259 | 114692 | DUF2075-domain-containing-protein-[Streptomyces-yogyakartensis] |
| WP_062010224.1 | REase+SF2-helicase | GCF_001570645.1 | Streptomyces_hygroscopicus | Bacteria>Bacillati>Actinomycetota>Actinomycetes>Kitasatosporales>Streptomycetaceae>Streptomyces>Streptomyces_violaceusniger_group | 631 | AXW65_RS13470 | 95197..97092 | 1912 | DUF2075-domain-containing-protein-[Streptomyces-hygroscopicus] |
| WP_330811480.1 | REase+SF2-helicase | GCF_036352935.1 | Streptomyces_asiaticus | Bacteria>Bacillati>Actinomycetota>Actinomycetes>Kitasatosporales>Streptomycetaceae>Streptomyces>Streptomyces_violaceusniger_group | 630 | V2J94_RS25920 | 128096..129988 | 114695 | DUF2075-domain-containing-protein-[Streptomyces-sp.-DSM-41524] |
| GAA1254476.1 | REase+SF2-helicase | GCA_039525965.1 | Streptomyces_yogyakartensis | Bacteria>Bacillati>Actinomycetota>Actinomycetes>Kitasatosporales>Streptomycetaceae>Streptomyces>Streptomyces_violaceusniger_group | 629 | GCM10009579_10560 | 290175..292064 | 114692 | DUF2075-domain-containing-protein-[Streptomyces-yogyakartensis] |
| WP_062010207.1 | REase+SF2-helicase | GCF_001570645.1 | Streptomyces_hygroscopicus | Bacteria>Bacillati>Actinomycetota>Actinomycetes>Kitasatosporales>Streptomycetaceae>Streptomyces>Streptomyces_violaceusniger_group | 629 | AXW65_RS13480 | 97837..99726 | 1912 | DUF2075-domain-containing-protein-[Streptomyces-hygroscopicus] |
| WP_239764968.1 | REase+SF2-helicase | GCF_022392825.1 | Streptomyces_sp._CL12-4 | Bacteria>Bacillati>Actinomycetota>Actinomycetes>Kitasatosporales>Streptomycetaceae>Streptomyces>unclassified_Streptomyces | 704 | JRC61_RS06880 | 1561088..1563202 | 2810306 | DUF2075-domain-containing-protein-[Streptomyces-sp.-CL12-4] |
| WP_265868481.1 | REase+SF2-helicase | GCF_026236535.1 | Streptomyces_sp._SKN60 | Bacteria>Bacillati>Actinomycetota>Actinomycetes>Kitasatosporales>Streptomycetaceae>Streptomyces>unclassified_Streptomyces | 704 | KV205_RS32675 | 89455..91569 | 2855506 | DUF2075-domain-containing-protein-[Streptomyces-sp.-SKN60] |
| NEE18796.1 | REase+SF2-helicase | GCA_010550285.1 | Streptomyces_sp._SID7499 | Bacteria>Bacillati>Actinomycetota>Actinomycetes>Kitasatosporales>Streptomycetaceae>Streptomyces>unclassified_Streptomyces | 703 | G3M58_71535 | 5911..8022 | 2706086 | DUF2075-domain-containing-protein-[Streptomyces-sp.-SID7499] |
| WP_330335234.1 | REase+SF2-helicase | GCF_036346295.1 | Streptomyces_sp._NBC_00536 | Bacteria>Bacillati>Actinomycetota>Actinomycetes>Kitasatosporales>Streptomycetaceae>Streptomyces>unclassified_Streptomyces | 702 | OHS33_RS27810 | 6393780..6395888 | 2975769 | DUF2075-domain-containing-protein-[Streptomyces-sp.-NBC_00536] |
| WP_233868583.1 | REase+SF2-helicase | GCF_021390475.1 | Streptomyces_sp._ST2-7A | Bacteria>Bacillati>Actinomycetota>Actinomycetes>Kitasatosporales>Streptomycetaceae>Streptomyces>unclassified_Streptomyces | 647 | LZF96_RS09310 | 72880..74823 | 2907214 | DUF2075-domain-containing-protein-[Streptomyces-sp.-ST2-7A] |
| WP_306948863.1 | REase+SF2-helicase | GCF_030817335.1 | Streptomyces_sp._B4I13 | Bacteria>Bacillati>Actinomycetota>Actinomycetes>Kitasatosporales>Streptomycetaceae>Streptomyces>unclassified_Streptomyces | 646 | QFZ66_RS12490 | 2614135..2616075 | 3042271 | DUF2075-domain-containing-protein-[Streptomyces-sp.-B4I13] |
| WP_328677723.1 | REase+SF2-helicase | GCF_036118775.1 | Streptomyces_sp._NBC_00343 | Bacteria>Bacillati>Actinomycetota>Actinomycetes>Kitasatosporales>Streptomycetaceae>Streptomyces>unclassified_Streptomyces | 646 | OHA62_RS11620 | 2750190..2752130 | 2975719 | DUF2075-domain-containing-protein-[Streptomyces-sp.-NBC_00343] |
| WP_329248533.1 | REase+SF2-helicase | GCF_036227225.1 | Streptomyces_sp._NBC_01478 | Bacteria>Bacillati>Actinomycetota>Actinomycetes>Kitasatosporales>Streptomycetaceae>Streptomyces>unclassified_Streptomyces | 646 | OG223_RS16415 | 3651363..3653303 | 2903882 | DUF2075-domain-containing-protein-[Streptomyces-sp.-NBC_01478] |
| WSN43127.1 | REase+SF2-helicase | GCA_035983595.1 | Streptomyces_sp._NBC_01334 | Bacteria>Bacillati>Actinomycetota>Actinomycetes>Kitasatosporales>Streptomycetaceae>Streptomyces>unclassified_Streptomyces | 646 | OG736_32460 | 7223926..7225866 | 2903827 | DUF2075-domain-containing-protein-[Streptomyces-sp.-NBC_01334] |
| WP_317859885.1 | REase+SF2-helicase | GCF_033194775.1 | Streptomyces_sp._Wh19 | Bacteria>Bacillati>Actinomycetota>Actinomycetes>Kitasatosporales>Streptomycetaceae>Streptomyces>unclassified_Streptomyces | 632 | R6M67_RS01065 | 39934..41832 | 3076629 | DUF2075-domain-containing-protein-[Streptomyces-sp.-Wh19] |
| WP_136185182.1 | REase+SF2-helicase | GCF_004795255.1 | Streptomyces_sp._RKND-216 | Bacteria>Bacillati>Actinomycetota>Actinomycetes>Kitasatosporales>Streptomycetaceae>Streptomyces>unclassified_Streptomyces | 631 | E4198_RS04235 | 939867..941762 | 2562581 | DUF2075-domain-containing-protein-[Streptomyces-sp.-RKND-216] |
| NEE18795.1 | REase+SF2-helicase | GCA_010550285.1 | Streptomyces_sp._SID7499 | Bacteria>Bacillati>Actinomycetota>Actinomycetes>Kitasatosporales>Streptomycetaceae>Streptomyces>unclassified_Streptomyces | 630 | G3M58_71530 | 3870..5762 | 2706086 | DUF2075-domain-containing-protein-[Streptomyces-sp.-SID7499] |
| WP_137230277.1 | REase+SF2-helicase | GCF_005234185.1 | Streptomyces_sp._BPSDS2 | Bacteria>Bacillati>Actinomycetota>Actinomycetes>Kitasatosporales>Streptomycetaceae>Streptomyces>unclassified_Streptomyces | 630 | FD916_RS06875 | 201373..203265 | 2571021 | DUF2075-domain-containing-protein-[Streptomyces-sp.-BPSDS2] |
| WP_210930999.1 | REase+SF2-helicase | GCF_018070665.1 | Streptomyces_sp._404i | Bacteria>Bacillati>Actinomycetota>Actinomycetes>Kitasatosporales>Streptomycetaceae>Streptomyces>unclassified_Streptomyces | 630 | KBY46_RS00840 | 208989..210881 | 2824902 | DUF2075-domain-containing-protein-[Streptomyces-sp.-404i] |
| WTK77646.1 | REase+SF2-helicase | GCA_045040335.1 | Streptomyces_sp._NBC_01511 | Bacteria>Bacillati>Actinomycetota>Actinomycetes>Kitasatosporales>Streptomycetaceae>Streptomyces>unclassified_Streptomyces | 630 | OG292_07475 | 1625946..1627838 | 2903889 | DUF2075-domain-containing-protein-[Streptomyces-sp.-NBC_01511] |
| WTK93288.1 | REase+SF2-helicase | GCA_045040315.1 | Streptomyces_sp._NBC_01508 | Bacteria>Bacillati>Actinomycetota>Actinomycetes>Kitasatosporales>Streptomycetaceae>Streptomyces>unclassified_Streptomyces | 630 | OG700_07115 | 1581370..1583262 | 2903888 | DUF2075-domain-containing-protein-[Streptomyces-sp.-NBC_01508] |
| WP_067397491.1 | REase+SF2-helicase | GCF_001653115.1 | Streptomyces_sp._F-3 | Bacteria>Bacillati>Actinomycetota>Actinomycetes>Kitasatosporales>Streptomycetaceae>Streptomyces>unclassified_Streptomyces | 629 | STXM2123_RS19065 | 50855..52744 | 1840095 | DUF2075-domain-containing-protein-[Streptomyces-sp.-F-3] |
| WP_239764487.1 | REase+SF2-helicase | GCF_022392825.1 | Streptomyces_sp._CL12-4 | Bacteria>Bacillati>Actinomycetota>Actinomycetes>Kitasatosporales>Streptomycetaceae>Streptomyces>unclassified_Streptomyces | 629 | JRC61_RS06875 | 1558998..1560887 | 2810306 | DUF2075-domain-containing-protein-[Streptomyces-sp.-CL12-4] |
| WP_240805167.1 | REase+SF2-helicase | GCF_004794975.1 | Streptomyces_sp._A1547 | Bacteria>Bacillati>Actinomycetota>Actinomycetes>Kitasatosporales>Streptomycetaceae>Streptomyces>unclassified_Streptomyces | 628 | E6W17_RS04090 | 289080..290966 | 2563105 | DUF2075-domain-containing-protein-[Streptomyces-sp.-A1547] |
| WP_319723084.1 | REase+SF2-helicase | GCF_033794505.1 | Streptomyces_sp._MB09-01 | Bacteria>Bacillati>Actinomycetota>Actinomycetes>Kitasatosporales>Streptomycetaceae>Streptomyces>unclassified_Streptomyces | 628 | PV721_RS11730 | 22487..24373 | 3028666 | DUF2075-domain-containing-protein-[Streptomyces-sp.-MB09-01] |
| WP_164615105.1 | REase+SF2-helicase | GCF_010548505.1 | Streptomyces_sp._SID14515 | Bacteria>Bacillati>Actinomycetota>Actinomycetes>Kitasatosporales>Streptomycetaceae>Streptomyces>unclassified_Streptomyces | 627 | G3I54_RS15515 | 53258..55141 | 2706074 | DUF2075-domain-containing-protein-[Streptomyces-sp.-SID14515] |
| WP_265868463.1 | REase+SF2-helicase | GCF_026236535.1 | Streptomyces_sp._SKN60 | Bacteria>Bacillati>Actinomycetota>Actinomycetes>Kitasatosporales>Streptomycetaceae>Streptomyces>unclassified_Streptomyces | 621 | KV205_RS32640 | 74861..76726 | 2855506 | DUF2075-domain-containing-protein-[Streptomyces-sp.-SKN60] |
| WP_062529098.1 | REase+SF2-helicase | GCF_000971455.1 | Demequina_rhizosphaerae | Bacteria>Bacillati>Actinomycetota>Actinomycetes>Micrococcales>Demequinaceae>Demequina | 633 | AOM12_RS04435 | 339977..341878 | 1638985 | DUF2075-domain-containing-protein-[Demequina-rhizosphaerae] |
| EWT07981.1 | REase+SF2-helicase | GCA_000576575.1 | Intrasporangium_chromatireducens_Q5-1 | Bacteria>Bacillati>Actinomycetota>Actinomycetes>Micrococcales>Intrasporangiaceae>Intrasporangium>Intrasporangium_chromatireducens | 622 | N864_11735 | 62111..63979 | 584657 | ATP-binding-protein-[Intrasporangium-chromatireducens-Q5-1] |
| WP_104161522.1 | REase+SF2-helicase | GCF_002929255.1 | Arthrobacter_sp._ZGTC212 | Bacteria>Bacillati>Actinomycetota>Actinomycetes>Micrococcales>Micrococcaceae>Arthrobacter>unclassified_Arthrobacter | 556 | CXQ89_RS12645 | 72992..74662 | 2058899 | NERD-domain-containing-protein-[Arthrobacter-sp.-ZGTC212] |
| WP_022924600.1 | REase+SF2-helicase | GCF_000421245.1 | Serinicoccus_marinus | Bacteria>Bacillati>Actinomycetota>Actinomycetes>Micrococcales>Ornithinimicrobiaceae>Serinicoccus | 624 | FU792_RS06150 | 1315842..1317716 | 247333 | DUF2075-domain-containing-protein-[Serinicoccus-marinus] |
| WP_030270098.1 | REase+SF2-helicase | GCF_040634605.1 | Micromonospora | Bacteria>Bacillati>Actinomycetota>Actinomycetes>Micromonosporales>Micromonosporaceae | 672 | IF06_RS0108630 | 52144..54162 | 1873 | DUF2075-domain-containing-protein-[Micromonospora-globosa] |
| NMO57586.1 | REase+SF2-helicase | GCA_012933445.1 | Actinoplanes_sp._TBRC_11911 | Bacteria>Bacillati>Actinomycetota>Actinomycetes>Micromonosporales>Micromonosporaceae>Actinoplanes>unclassified_Actinoplanes | 644 | HH310_41275 | 20481..22415 | 2729386 | DUF2075-domain-containing-protein-[Actinoplanes-sp.-TBRC-11911] |
| WP_169181225.1 | REase+SF2-helicase | GCF_012933445.1 | Actinoplanes_sp._TBRC_11911 | Bacteria>Bacillati>Actinomycetota>Actinomycetes>Micromonosporales>Micromonosporaceae>Actinoplanes>unclassified_Actinoplanes | 644 | HH310_RS44570 | 20481..22415 | 2729386 | DNA/RNA-helicase-domain-containing-protein-[Actinoplanes-sp.-TBRC-11911] |
| WP_248294923.1 | REase+SF2-helicase | - | Actinoplanes_sp._TBRC_11911 | Bacteria>Bacillati>Actinomycetota>Actinomycetes>Micromonosporales>Micromonosporaceae>Actinoplanes>unclassified_Actinoplanes | 357 | HH310_RS44570 | 20481..21554 | 2729386 | DUF2075-domain-containing-protein-[Actinoplanes-sp.-TBRC-11911] |
| WP_013284491.1 | REase+SF2-helicase | GCF_000145235.1 | Micromonospora_aurantiaca | Bacteria>Bacillati>Actinomycetota>Actinomycetes>Micromonosporales>Micromonosporaceae>Micromonospora | 672 | MICAU_RS06465 | 1327691..1329709 | 47850 | DUF2075-domain-containing-protein-[Micromonospora-aurantiaca] |
| WP_043326015.1 | REase+SF2-helicase | GCF_023072705.1 | unclassified_Micromonospora | Bacteria>Bacillati>Actinomycetota>Actinomycetes>Micromonosporales>Micromonosporaceae>Micromonospora | 672 | FJ969_RS15355 | 60387..62405 | 2617518 | MULTISPECIES:-DUF2075-domain-containing-protein-[unclassified-Micromonospora] |
| ADL44852.1 | REase+SF2-helicase | GCA_000145235.1 | Micromonospora_aurantiaca_ATCC_27029 | Bacteria>Bacillati>Actinomycetota>Actinomycetes>Micromonosporales>Micromonosporaceae>Micromonospora>Micromonospora_aurantiaca | 672 | Micau_1290 | 1327691..1329709 | 644283 | AAA-ATPase-[Micromonospora-aurantiaca-ATCC-27029] |
| WP_007075117.1 | REase+SF2-helicase | GCF_000158815.1 | Micromonospora_sp._ATCC_39149 | Bacteria>Bacillati>Actinomycetota>Actinomycetes>Micromonosporales>Micromonosporaceae>Micromonospora>unclassified_Micromonospora | 659 | MCAG_RS21655 | 4953505..4955484 | 219305 | DUF2075-domain-containing-protein-[Micromonospora-sp.-ATCC-39149] |
| MBU2668300.1 | REase+SF2-helicase | GCA_018829635.1 | Paractinoplanes_bogorensis | Bacteria>Bacillati>Actinomycetota>Actinomycetes>Micromonosporales>Micromonosporaceae>Paractinoplanes | 644 | KOI35_32790 | 181575..183509 | 1610840 | DUF2075-domain-containing-protein-[Actinoplanes-bogorensis] |
| WP_215792564.1 | REase+SF2-helicase | GCF_018829635.1 | Paractinoplanes_bogorensis | Bacteria>Bacillati>Actinomycetota>Actinomycetes>Micromonosporales>Micromonosporaceae>Paractinoplanes | 644 | KOI35_RS32785 | 181575..183509 | 1610840 | DUF2075-domain-containing-protein-[Actinoplanes-bogorensis] |
| WP_182347798.1 | REase+SF2-helicase | GCF_014121175.1 | Tomitella_gaofuii | Bacteria>Bacillati>Actinomycetota>Actinomycetes>Mycobacteriales>Corynebacteriales_incertae_sedis>Tomitella | 625 | H7F09_RS10470 | 70207..72084 | 2760083 | DNA/RNA-helicase-domain-containing-protein-[Tomitella-gaofuii] |
| WP_247603195.1 | REase+SF2-helicase | - | Gordonia | Bacteria>Bacillati>Actinomycetota>Actinomycetes>Mycobacteriales>Gordoniaceae | 554 | HGA01_RS21745 | 18730..20394 | 2053 | MULTISPECIES:-DUF2075-domain-containing-protein-[Gordonia] |
| WP_257881111.1 | REase+SF2-helicase | GCF_002847865.1 | Gordonia_terrae | Bacteria>Bacillati>Actinomycetota>Actinomycetes>Mycobacteriales>Gordoniaceae>Gordonia | 423 | CYJ73_RS02735 | 609268..610539 | 2055 | DUF2075-domain-containing-protein-[Gordonia-terrae] |
| SLK50035.1 | REase+SF2-helicase | GCA_900132645.1 | Mycobacteroides_abscessus_subsp._abscessus | Bacteria>Bacillati>Actinomycetota>Actinomycetes>Mycobacteriales>Mycobacteriaceae>Mycobacteroides>Mycobacteroides_abscessus | 643 | SAMEA2071448_00691 | 745305..747236 | 1185650 | Uncharacterized-conserved-protein-[Mycobacteroides-abscessus-subsp.-abscessus] |
| WP_243750113.1 | REase+SF2-helicase | GCF_001612905.1 | Nocardia_ignorata | Bacteria>Bacillati>Actinomycetota>Actinomycetes>Mycobacteriales>Nocardiaceae>Nocardia | 647 | NI2_RS26685 | 107963..109906 | 145285 | DUF2075-domain-containing-protein-[Nocardia-ignorata] |
| WP_328712362.1 | REase+SF2-helicase | GCF_036150455.1 | Nocardia_salmonicida | Bacteria>Bacillati>Actinomycetota>Actinomycetes>Mycobacteriales>Nocardiaceae>Nocardia | 373 | OHA85_RS00110 | 30089..31210 | 53431 | DUF2075-domain-containing-protein-[Nocardia-salmonicida] |
| WP_056816885.1 | REase+SF2-helicase | GCF_001427185.1 | Nocardia_sp._Root136 | Bacteria>Bacillati>Actinomycetota>Actinomycetes>Mycobacteriales>Nocardiaceae>Nocardia>unclassified_Nocardia | 619 | ASD42_RS14830 | 220986..222845 | 1736458 | DUF2075-domain-containing-protein-[Nocardia-sp.-Root136] |
| MBM4487174.1 | REase+SF2-helicase | GCA_016874395.1 | Prescottella_equi | Bacteria>Bacillati>Actinomycetota>Actinomycetes>Mycobacteriales>Nocardiaceae>Prescottella | 626 | GS471_03195 | 666302..668182 | 43767 | DUF2075-domain-containing-protein-[Prescottella-equi] |
| NKR55752.1 | REase+SF2-helicase | GCA_012393885.1 | Prescottella_equi | Bacteria>Bacillati>Actinomycetota>Actinomycetes>Mycobacteriales>Nocardiaceae>Prescottella | 626 | GS481_17005 | 1581509..1583389 | 43767 | DUF2075-domain-containing-protein-[Prescottella-equi] |
| NKR97094.1 | REase+SF2-helicase | GCA_012394005.1 | Prescottella_equi | Bacteria>Bacillati>Actinomycetota>Actinomycetes>Mycobacteriales>Nocardiaceae>Prescottella | 626 | GS473_09480 | 89172..91052 | 43767 | DUF2075-domain-containing-protein-[Prescottella-equi] |
| WP_084846697.1 | REase+SF2-helicase | GCF_002094405.3 | Prescottella_equi | Bacteria>Bacillati>Actinomycetota>Actinomycetes>Mycobacteriales>Nocardiaceae>Prescottella | 626 | A6I88_RS04050 | 901364..903244 | 43767 | DUF2075-domain-containing-protein-[Prescottella-equi] |
| WP_084848439.1 | REase+SF2-helicase | GCF_002095035.1 | Prescottella_equi | Bacteria>Bacillati>Actinomycetota>Actinomycetes>Mycobacteriales>Nocardiaceae>Prescottella | 626 | A5N78_RS19030 | 147123..149003 | 43767 | DUF2075-domain-containing-protein-[Prescottella-equi] |
| EME22381.1 | REase+SF2-helicase | GCA_000341795.1 | Rhodococcus_triatomae_BKS_15-14 | Bacteria>Bacillati>Actinomycetota>Actinomycetes>Mycobacteriales>Nocardiaceae>Rhodococcus>Rhodococcus_triatomae | 598 | G419_05597 | 79751..81547 | 1278078 | hypothetical-protein-G419_05597-[Rhodococcus-triatomae-BKS-15-14] |
| NME80342.1 | REase+SF2-helicase | GCA_012844225.1 | Rhodococcus_sp._105337 | Bacteria>Bacillati>Actinomycetota>Actinomycetes>Mycobacteriales>Nocardiaceae>Rhodococcus>unclassified_Rhodococcus_(in:_high_G+C_Gram-positive_bacteria) | 623 | HF886_14725 | 22252..24123 | 2725310 | DUF2075-domain-containing-protein-[Rhodococcus-sp.-105337] |
| WP_172936833.1 | REase+SF2-helicase | - | Rhodococcus_sp._MS16 | Bacteria>Bacillati>Actinomycetota>Actinomycetes>Mycobacteriales>Nocardiaceae>Rhodococcus>unclassified_Rhodococcus_(in:_high_G+C_Gram-positive_bacteria) | 613 | FEZ60_RS20005 | 130704..132545 | 2579941 | DUF2075-domain-containing-protein-[Rhodococcus-sp.-MS16] |
| WP_240955159.1 | REase+SF2-helicase | - | Rhodococcus_sp._105337 | Bacteria>Bacillati>Actinomycetota>Actinomycetes>Mycobacteriales>Nocardiaceae>Rhodococcus>unclassified_Rhodococcus_(in:_high_G+C_Gram-positive_bacteria) | 596 | HF886_RS14725 | 22252..24042 | 2725310 | DUF2075-domain-containing-protein-[Rhodococcus-sp.-105337] |
| WP_234974498.1 | REase+SF2-helicase | - | Williamsia_sterculiae | Bacteria>Bacillati>Actinomycetota>Actinomycetes>Mycobacteriales>Nocardiaceae>Williamsia | 566 | BW971_RS19040 | 39472..41172 | 1344003 | DUF2075-domain-containing-protein-[Williamsia-sterculiae] |
| WP_224646694.1 | REase+SF2-helicase | GCF_020169415.1 | Tsukamurella_sp._M9C | Bacteria>Bacillati>Actinomycetota>Actinomycetes>Mycobacteriales>Tsukamurellaceae>Tsukamurella>unclassified_Tsukamurella | 623 | LB823_RS09375 | 55663..57534 | 2877520 | DUF2075-domain-containing-protein-[Tsukamurella-sp.-M9C] |
| WP_034307035.1 | REase+SF2-helicase | GCF_900105055.1 | Amycolatopsis_lurida | Bacteria>Bacillati>Actinomycetota>Actinomycetes>Pseudonocardiales>Pseudonocardiaceae>Amycolatopsis | 647 | BLW75_RS07215 | 435964..437907 | 31959 | DUF2075-domain-containing-protein-[Amycolatopsis-lurida] |
| WP_039917237.1 | REase+SF2-helicase | - | Amycolatopsis_azurea | Bacteria>Bacillati>Actinomycetota>Actinomycetes>Pseudonocardiales>Pseudonocardiaceae>Amycolatopsis | 632 | C791_RS14055 | 24688..26586 | 36819 | DUF2075-domain-containing-protein-[Amycolatopsis-azurea] |
| WP_141999813.1 | REase+SF2-helicase | GCF_006715045.1 | Amycolatopsis_cihanbeyliensis | Bacteria>Bacillati>Actinomycetota>Actinomycetes>Pseudonocardiales>Pseudonocardiaceae>Amycolatopsis | 631 | FB471_RS19185 | 4224064..4225959 | 1128664 | DUF2075-domain-containing-protein-[Amycolatopsis-cihanbeyliensis] |
| WP_005155548.1 | REase+SF2-helicase | GCF_000340415.1 | Amycolatopsis_azurea | Bacteria>Bacillati>Actinomycetota>Actinomycetes>Pseudonocardiales>Pseudonocardiaceae>Amycolatopsis | 578 | C791_RS14060 | 26862..28598 | 36819 | DUF2075-domain-containing-protein-[Amycolatopsis-azurea] |
| WP_205215284.1 | REase+SF2-helicase | GCF_003312875.1 | Amycolatopsis_albispora | Bacteria>Bacillati>Actinomycetota>Actinomycetes>Pseudonocardiales>Pseudonocardiaceae>Amycolatopsis | 498 | A4R43_RS09370 | 2027330..2028826 | 1804986 | DUF2075-domain-containing-protein-[Amycolatopsis-albispora] |
| KFU81960.1 | REase+SF2-helicase | GCA_000749465.3 | Amycolatopsis_lurida_NRRL_2430 | Bacteria>Bacillati>Actinomycetota>Actinomycetes>Pseudonocardiales>Pseudonocardiaceae>Amycolatopsis>Amycolatopsis_lurida | 647 | BB31_06355 | 75613..77556 | 1460371 | hypothetical-protein-BB31_06355-[Amycolatopsis-lurida-NRRL-2430] |
| WP_329787396.1 | REase+SF2-helicase | GCF_036321065.1 | Lentzea_sp._DG1S-22 | Bacteria>Bacillati>Actinomycetota>Actinomycetes>Pseudonocardiales>Pseudonocardiaceae>Lentzea>unclassified_Lentzea | 629 | V1227_RS21595 | 4731521..4733410 | 3108822 | DUF2075-domain-containing-protein-[Lentzea-sp.-DG1S-22] |
| WP_329787395.1 | REase+SF2-helicase | GCF_036321065.1 | Lentzea_sp._DG1S-22 | Bacteria>Bacillati>Actinomycetota>Actinomycetes>Pseudonocardiales>Pseudonocardiaceae>Lentzea>unclassified_Lentzea | 610 | V1227_RS21590 | 4729589..4731421 | 3108822 | DNA/RNA-helicase-domain-containing-protein-[Lentzea-sp.-DG1S-22] |
| WP_244990183.1 | REase+SF2-helicase | - | Saccharothrix_coeruleofusca | Bacteria>Bacillati>Actinomycetota>Actinomycetes>Pseudonocardiales>Pseudonocardiaceae>Saccharothrix | 611 | JOF41_RS03760 | 759739..761574 | 33919 | DUF2075-domain-containing-protein-[Saccharothrix-coeruleofusca] |
| WP_199444069.1 | REase+SF2-helicase | GCF_016464305.1 | Umezawaea_beigongshangensis | Bacteria>Bacillati>Actinomycetota>Actinomycetes>Pseudonocardiales>Pseudonocardiaceae>Umezawaea | 645 | IMZ35_RS27285 | 17107..19044 | 2780383 | DUF2075-domain-containing-protein-[Umezawaea-beigongshangensis] |
| WP_199444071.1 | REase+SF2-helicase | GCF_016464305.1 | Umezawaea_beigongshangensis | Bacteria>Bacillati>Actinomycetota>Actinomycetes>Pseudonocardiales>Pseudonocardiaceae>Umezawaea | 624 | IMZ35_RS27295 | 20592..22466 | 2780383 | DUF2075-domain-containing-protein-[Umezawaea-beigongshangensis] |
| WP_170321718.1 | REase+SF2-helicase | GCF_039525465.1 | Acrocarpospora_pleiomorpha | Bacteria>Bacillati>Actinomycetota>Actinomycetes>Streptosporangiales>Streptosporangiaceae>Acrocarpospora | 631 | Aple_RS29455 | 82344..84239 | 90975 | DUF2075-domain-containing-protein-[Acrocarpospora-pleiomorpha] |
| WP_170321719.1 | REase+SF2-helicase | GCF_009687885.1 | Acrocarpospora_pleiomorpha | Bacteria>Bacillati>Actinomycetota>Actinomycetes>Streptosporangiales>Streptosporangiaceae>Acrocarpospora | 416 | Aple_RS29515 | 112149..113397 | 90975 | DUF2075-domain-containing-protein,-partial-[Acrocarpospora-pleiomorpha] |
| WP_184941234.1 | REase+SF2-helicase | GCF_039535335.1 | Planomonospora_venezuelensis | Bacteria>Bacillati>Actinomycetota>Actinomycetes>Streptosporangiales>Streptosporangiaceae>Planomonospora | 465 | FHS22_RS12490 | 323748..325145 | 1999 | DUF2075-domain-containing-protein-[Planomonospora-venezuelensis] |
| WP_196469675.1 | REase+SF2-helicase | - | Planomonospora_sp._ID91781 | Bacteria>Bacillati>Actinomycetota>Actinomycetes>Streptosporangiales>Streptosporangiaceae>Planomonospora>unclassified_Planomonospora | 609 | HS048_RS31355 | 220666..222495 | 2738135 | DUF2075-domain-containing-protein-[Planomonospora-sp.-ID91781] |

| Protein-ID | Domain | Assembly-ID | Strain | Taxonomy | Length | Locus-Tag | Genomic-coordinates | TaxID | Fasta-defline |
|---|---|---|---|---|---|---|---|---|---|
| WP_230882460.1 | REase+SF2-helicase | - | Planomonospora_sp._ID91781 | Bacteria>Bacillati>Actinomycetota>Actinomycetes>Streptosporangiales>Streptosporangiaceae>Planomonospora>unclassified_Planomonospora | 561 | HS048_RS31350 | 218824..220509 | 2738135 | DUF2075-domain-containing-protein-[Planomonospora-sp.-ID91781] |
| GII83988.1 | REase+SF2-helicase | GCA_016863415.1 | Sphaerisporangium_siamense | Bacteria>Bacillati>Actinomycetota>Actinomycetes>Streptosporangiales>Streptosporangiaceae>Sphaerisporangium | 634 | Ssi03_19780 | 403671..405575 | 795645 | ATP-binding-protein-[Sphaerisporangium-siamense] |
| GII83987.1 | REase+SF2-helicase | GCA_016863415.1 | Sphaerisporangium_siamense | Bacteria>Bacillati>Actinomycetota>Actinomycetes>Streptosporangiales>Streptosporangiaceae>Sphaerisporangium | 628 | Ssi03_19770 | 401485..403371 | 795645 | ATP-binding-protein-[Sphaerisporangium-siamense] |
| WP_184886694.1 | REase+SF2-helicase | GCF_016863415.1 | Sphaerisporangium_siamense | Bacteria>Bacillati>Actinomycetota>Actinomycetes>Streptosporangiales>Streptosporangiaceae>Sphaerisporangium | 628 | Ssi03_RS09585 | 401485..403371 | 795645 | DUF2075-domain-containing-protein-[Sphaerisporangium-siamense] |
| WP_184886696.1 | REase+SF2-helicase | GCF_016863415.1 | Sphaerisporangium_siamense | Bacteria>Bacillati>Actinomycetota>Actinomycetes>Streptosporangiales>Streptosporangiaceae>Sphaerisporangium | 624 | Ssi03_RS09590 | 403701..405575 | 795645 | DUF2075-domain-containing-protein-[Sphaerisporangium-siamense] |
| ROO83037.1 | REase+SF2-helicase | GCA_003751225.1 | Actinocorallia_herbida | Bacteria>Bacillati>Actinomycetota>Actinomycetes>Streptosporangiales>Thermomonosporaceae>Actinocorallia | 627 | EDD29_0526 | 493765..495648 | 58109 | hypothetical-protein-EDD29_0526-[Actinocorallia-herbida] |
| WP_246053352.1 | REase+SF2-helicase | GCF_003751225.1 | Actinocorallia_herbida | Bacteria>Bacillati>Actinomycetota>Actinomycetes>Streptosporangiales>Thermomonosporaceae>Actinocorallia | 625 | EDD29_RS02570 | 493765..495642 | 58109 | DUF2075-domain-containing-protein-[Actinocorallia-herbida] |
| WP_136846138.1 | REase+SF2-helicase | GCF_005046025.1 | Parvibacter_caecicola | Bacteria>Bacillati>Actinomycetota>Coriobacteriia>Coriobacteriales>Coriobacteriaceae>Parvibacter | 567 | E5982_RS08735 | 38508..40211 | 747645 | ATP-binding-domain-containing-protein-[Parvibacter-caecicola] |
| WP_062679364.1 | REase+SF2-helicase | GCF_007197795.1 | Anoxybacillaceae | Bacteria>Bacillati>Bacillota>Bacilli>Bacillales | 644 | B4110_RS17530 | 16775..18709 | 3120669 | DUF2075-domain-containing-protein-[Parageobacillus-toebii] |
| WP_034748793.1 | REase+SF2-helicase | GCF_000513095.1 | Halalkalibacter_wakoensis | Bacteria>Bacillati>Bacillota>Bacilli>Bacillales>Bacillaceae>Halalkalibacter | 646 | JCM9140_RS16190 | 5569..7509 | 127891 | DUF2075-domain-containing-protein-[Halalkalibacter-wakoensis] |
| WP_213145788.1 | REase+SF2-helicase | GCF_018343545.2 | Neobacillus_citreus | Bacteria>Bacillati>Bacillota>Bacilli>Bacillales>Bacillaceae>Neobacillus | 647 | KHB02_RS07270 | 29001..30944 | 2833578 | DUF2075-domain-containing-protein-[Neobacillus-citreus] |
| MBR3117627.1 | REase+SF2-helicase | GCA_017514815.1 | Oceanobacillus_sp. | Bacteria>Bacillati>Bacillota>Bacilli>Bacillales>Bacillaceae>Oceanobacillus>unclassified_Oceanobacillus | 630 | IKF29_00155 | 7340..9232 | 1871622 | DUF2075-domain-containing-protein-[Oceanobacillus-sp.] |
| WP_190997464.1 | REase+SF2-helicase | GCF_014712675.1 | Peribacillus_faecalis | Bacteria>Bacillati>Bacillota>Bacilli>Bacillales>Bacillaceae>Peribacillus | 648 | IE070_RS06035 | 48023..49969 | 2772559 | DUF2075-domain-containing-protein-[Peribacillus-faecalis] |
| WP_061808753.1 | REase+SF2-helicase | GCF_001591825.1 | Rossellomorea_vietnamensis | Bacteria>Bacillati>Bacillota>Bacilli>Bacillales>Bacillaceae>Rossellomorea | 650 | BVI01S_RS00190 | 33706..35658 | 218284 | DUF2075-domain-containing-protein-[Rossellomorea-vietnamensis] |
| WP_214689545.1 | REase+SF2-helicase | GCF_031979545.1 | Exiguobacterium | Bacteria>Bacillati>Bacillota>Bacilli>Bacillales>Bacillales_incertae_sedis>Bacillales_Family_XII._Incertae_Sedis | 630 | KN071_RS16230 | 91740..93632 | 33986 | DUF2075-domain-containing-protein-[Exiguobacterium-sp.-s163] |
| WP_275060109.1 | REase+SF2-helicase | GCF_029026745.1 | Exiguobacterium_profundum | Bacteria>Bacillati>Bacillota>Bacilli>Bacillales>Bacillales_incertae_sedis>Bacillales_Family_XII._Incertae_Sedis>Exiguobacterium | 630 | OE059_RS13160 | 2535135..2537027 | 307643 | DUF2075-domain-containing-protein-[Exiguobacterium-profundum] |
| WP_214788910.1 | REase+SF2-helicase | GCF_018617975.1 | Exiguobacterium_sp._s21 | Bacteria>Bacillati>Bacillota>Bacilli>Bacillales>Bacillales_incertae_sedis>Bacillales_Family_XII._Incertae_Sedis>Exiguobacterium>unclassified_Exiguobacterium | 630 | KNP06_RS10860 | 33210..35102 | 2751244 | DUF2075-domain-containing-protein-[Exiguobacterium-sp.-s21] |
| WP_192599214.1 | REase+SF2-helicase | GCF_014873855.1 | Sporosarcina_limicola | Bacteria>Bacillati>Bacillota>Bacilli>Bacillales>Caryophanaceae>Sporosarcina | 620 | H4683_RS12780 | 31639..33501 | 34101 | DUF2075-domain-containing-protein-[Sporosarcina-limicola] |
| WP_220271116.1 | REase+SF2-helicase | GCF_003337375.1 | Paenibacillus_prosopidis | Bacteria>Bacillati>Bacillota>Bacilli>Bacillales>Paenibacillaceae>Paenibacillus | 626 | DFP97_RS01930 | 460751..462631 | 630520 | DUF2075-domain-containing-protein-[Paenibacillus-prosopidis] |
| WP_119090035.1 | REase+SF2-helicase | GCF_003570985.1 | Candidatus_Cryosericum_hinesii | Bacteria>Pseudomonadati>Caldisericota/Cryosericota_group>Candidatus_Cryosericota>Candidatus_Cryosericia>Candidatus_Cryosericales>Candidatus_Cryosericaceae>Candidatus_Cryosericum | 624 | SMC3_RS08265 | 27856..29730 | 2290915 | DUF2075-domain-containing-protein-[Candidatus-Cryosericum-hinesii] |
| WP_018709390.1 | REase+SF2-helicase | GCF_000374585.1 | Phocaeicola_barnesiae | Bacteria>Pseudomonadati>FCB_group>Bacteroidota/Chlorobiota_group>Bacteroidota>Bacteroidia>Bacteroidales>Bacteroidaceae>Phocaeicola | 790 | C510_RS0100575 | 133325..135697 | 376804 | DUF2075-domain-containing-protein-[Phocaeicola-barnesiae] |
| WP_076500067.1 | REase+SF2-helicase | GCF_900156405.1 | Shewanella_morhuae | Bacteria>Pseudomonadati>Pseudomonadota>Gammaproteobacteria>Alteromonadales>Shewanellaceae>Shewanella | 560 | BW947_RS12460 | 5812..7494 | 365591 | NERD-domain-containing-protein-[Shewanella-morhuae] |
| WP_197054384.1 | REase+SF2-helicase | GCF_000740735.1 | Nitrosococcus_oceani | Bacteria>Pseudomonadati>Pseudomonadota>Gammaproteobacteria>Chromatiales>Chromatiaceae>Nitrosococcus | 603 | HW44_RS00280 | 15829..17640 | 1229 | nuclease-related-domain-containing-DEAD/DEAH-box-helicase-[Nitrosococcus-oceani] |
| MCP5207860.1 | REase+SF2-helicase | GCA_024234475.1 | Hahellaceae_bacterium | Bacteria>Pseudomonadati>Pseudomonadota>Gammaproteobacteria>Oceanospirillales>Hahellaceae>unclassified_Hahellaceae | 523 | H7A01_11690 | 148246..149817 | 2048971 | NERD-domain-containing-protein-[Hahellaceae-bacterium] |
| WP_023122054.1 | REase+SF2-helicase | GCF_014792125.1 | Pseudomonas | Bacteria>Pseudomonadati>Pseudomonadota>Gammaproteobacteria>Pseudomonadales>Pseudomonadaceae | 618 | IPC162_RS08500 | 430689..432545 | 286 | DUF2075-domain-containing-protein-[Pseudomonas-aeruginosa] |
| WP_033939340.1 | REase+SF2-helicase | GCF_965233265.1 | Pseudomonas | Bacteria>Pseudomonadati>Pseudomonadota>Gammaproteobacteria>Pseudomonadales>Pseudomonadaceae | 618 | FOZ66_RS12410 | 2595944..2597800 | 286 | MULTISPECIES:-DUF2075-domain-containing-protein-[Pseudomonas] |
| EIU2715578.1 | REase+SF2-helicase | - | Pseudomonas_aeruginosa | Bacteria>Pseudomonadati>Pseudomonadota>Gammaproteobacteria>Pseudomonadales>Pseudomonadaceae>Pseudomonas>Pseudomonas_aeruginosa_group | 618 | L4033_002036 | 58136..59992 | 287 | DUF2075-domain-containing-protein-[Pseudomonas-aeruginosa] |
| EKL0657910.1 | REase+SF2-helicase | GCA_028366315.1 | Pseudomonas_aeruginosa | Bacteria>Pseudomonadati>Pseudomonadota>Gammaproteobacteria>Pseudomonadales>Pseudomonadaceae>Pseudomonas>Pseudomonas_aeruginosa_group | 618 | PQC20_002034 | 54960..56816 | 287 | DUF2075-domain-containing-protein-[Pseudomonas-aeruginosa] |
| HBO2945366.1 | REase+SF2-helicase | GCA_021651875.1 | Pseudomonas_aeruginosa | Bacteria>Pseudomonadati>Pseudomonadota>Gammaproteobacteria>Pseudomonadales>Pseudomonadaceae>Pseudomonas>Pseudomonas_aeruginosa_group | 618 | L4R51_003857 | 1190..3046 | 287 | TPA:-DUF2075-domain-containing-protein-[Pseudomonas-aeruginosa] |
| HBO7186454.1 | REase+SF2-helicase | GCA_021674675.1 | Pseudomonas_aeruginosa | Bacteria>Pseudomonadati>Pseudomonadota>Gammaproteobacteria>Pseudomonadales>Pseudomonadaceae>Pseudomonas>Pseudomonas_aeruginosa_group | 618 | L4968_002131 | 14567..16423 | 287 | TPA:-DUF2075-domain-containing-protein-[Pseudomonas-aeruginosa] |
| HCF1762892.1 | REase+SF2-helicase | GCA_023941915.1 | Pseudomonas_aeruginosa | Bacteria>Pseudomonadati>Pseudomonadota>Gammaproteobacteria>Pseudomonadales>Pseudomonadaceae>Pseudomonas>Pseudomonas_aeruginosa_group | 618 | NH889_002164 | 15137..16993 | 287 | TPA:-DUF2075-domain-containing-protein-[Pseudomonas-aeruginosa] |
| MBH9417286.1 | REase+SF2-helicase | GCA_016092775.1 | Pseudomonas_aeruginosa | Bacteria>Pseudomonadati>Pseudomonadota>Gammaproteobacteria>Pseudomonadales>Pseudomonadaceae>Pseudomonas>Pseudomonas_aeruginosa_group | 618 | GHU28_23435 | 20202..22058 | 287 | DUF2075-domain-containing-protein-[Pseudomonas-aeruginosa] |
| WP_004350158.1 | REase+SF2-helicase | GCF_022552585.1 | Pseudomonas_aeruginosa | Bacteria>Pseudomonadati>Pseudomonadota>Gammaproteobacteria>Pseudomonadales>Pseudomonadaceae>Pseudomonas>Pseudomonas_aeruginosa_group | 618 | PA2G_RS09445 | 2106666..2108522 | 287 | DUF2075-domain-containing-protein-[Pseudomonas-aeruginosa] |
| WP_023088233.1 | REase+SF2-helicase | GCF_000504485.1 | Pseudomonas_aeruginosa | Bacteria>Pseudomonadati>Pseudomonadota>Gammaproteobacteria>Pseudomonadales>Pseudomonadaceae>Pseudomonas>Pseudomonas_aeruginosa_group | 618 | CD037_RS27935 | 9861..11717 | 287 | DUF2075-domain-containing-protein-[Pseudomonas-aeruginosa] |
| WP_202120100.1 | REase+SF2-helicase | - | Pseudomonas_aeruginosa | Bacteria>Pseudomonadati>Pseudomonadota>Gammaproteobacteria>Pseudomonadales>Pseudomonadaceae>Pseudomonas>Pseudomonas_aeruginosa_group | 607 | GQF67_RS10640 | 313764..315587 | 287 | DUF2075-domain-containing-protein-[Pseudomonas-aeruginosa] |
| WP_162954737.1 | REase+SF2-helicase | GCF_003630125.1 | Pseudomonas_aeruginosa | Bacteria>Pseudomonadati>Pseudomonadota>Gammaproteobacteria>Pseudomonadales>Pseudomonadaceae>Pseudomonas>Pseudomonas_aeruginosa_group | 382 | DQI18_RS19085 | 1..1149 | 287 | DUF2075-domain-containing-protein,-partial-[Pseudomonas-aeruginosa] |
| EAZ58597.1 | REase+SF2-helicase | GCA_000152545.1 | Pseudomonas_aeruginosa_2192 | Bacteria>Pseudomonadati>Pseudomonadota>Gammaproteobacteria>Pseudomonadales>Pseudomonadaceae>Pseudomonas>Pseudomonas_aeruginosa_group>Pseudomonas_aeruginosa | 618 | PA2G_01845 | 2106666..2108522 | 350703 | hypothetical-protein-PA2G_01845-[Pseudomonas-aeruginosa-2192] |
| WP_274151122.1 | REase+SF2-helicase | GCF_028682495.1 | Pseudomonas_sp._25571 | Bacteria>Pseudomonadati>Pseudomonadota>Gammaproteobacteria>Pseudomonadales>Pseudomonadaceae>Pseudomonas>unclassified_Pseudomonas | 616 | NP527_RS02625 | 581562..583412 | 2967216 | 3'-5'-exonuclease-[Pseudomonas-sp.-25571] |
| WP_197573436.1 | REase+SF2-helicase | GCF_014930975.1 | Photobacterium_damselae | Bacteria>Pseudomonadati>Pseudomonadota>Gammaproteobacteria>Vibrionales>Vibrionaceae>Photobacterium | 622 | IL982_RS10225 | 2149378..2151246 | 38293 | nuclease-related-domain-containing-DEAD/DEAH-box-helicase-[Photobacterium-damselae] |
| QOQ68166.1 | REase+SF2-helicase | GCA_014930975.1 | Photobacterium_damselae_subsp._damselae | Bacteria>Pseudomonadati>Pseudomonadota>Gammaproteobacteria>Vibrionales>Vibrionaceae>Photobacterium>Photobacterium_damselae | 622 | IL982_10225 | 2149378..2151246 | 85581 | NERD-domain-containing-protein-[Photobacterium-damselae-subsp.-damselae] |
| --REase(PDDEXK) | | | | | | | | | |
| PKL69085.1 | REase | GCA_002839705.1 | Methanobacteriales_archaeon_HGW-Methanobacteriales-1 | Archaea>Methanobacteriati>Methanobacteriota>Methanomada_group>Methanobacteria>Methanobacteriales>unclassified_Methanobacteriales | 142 | CVV28_02935 | 558564..558992 | 2013815 | hypothetical-protein-CVV28_02935-[Methanobacteriales-archaeon-HGW-Methanobacteriales-1] |
| WP_136516500.1 | REase | GCF_030734185.1 | Haloferacaceae | Archaea>Methanobacteriati>Methanobacteriota>Stenosarchaea_group>Halobacteria>Halobacteriales | 172 | EA078_RS17705 | 39398..39916 | 1644056 | hypothetical-protein-[Halobellus-captivus] |
| WP_264783327.1 | REase | GCF_026008575.1 | Haladaptatus_sp._T7 | Archaea>Methanobacteriati>Methanobacteriota>Stenosarchaea_group>Halobacteria>Halobacteriales>Haladaptataceae>Haladaptatus>unclassified_Haladaptatus | 213 | OPC11_RS21300 | 65676..66317 | 2029368 | hypothetical-protein-[Haladaptatus-sp.-T7] |
| WP_158056538.1 | REase | GCF_008831545.1 | Halorussus_halophilus | Archaea>Methanobacteriati>Methanobacteriota>Stenosarchaea_group>Halobacteria>Halobacteriales>Haladaptataceae>Halorussus | 231 | F7R90_RS06990 | 1427183..1427878 | 2650975 | hypothetical-protein-[Halorussus-halophilus] |
| WP_188997402.1 | REase | GCF_014647235.1 | Haloarcula | Archaea>Methanobacteriati>Methanobacteriota>Stenosarchaea_group>Halobacteria>Halobacteriales>Haloarculaceae | 145 | EGH26_RS17090 | 339441..339878 | 2237 | group-I-intron-associated-PD-(D/E)XK-endonuclease-[Halomicroarcula-pellucida] |
| WP_247995982.1 | REase | GCF_023115355.1 | Halocatena_salina | Archaea>Methanobacteriati>Methanobacteriota>Stenosarchaea_group>Halobacteria>Halobacteriales>Haloarculaceae>Halocatena | 214 | MW046_RS18860 | 45881..46525 | 2934340 | hypothetical-protein-[Halocatena-salina] |
| WP_108383899.1 | REase | GCF_003058365.1 | Halococcoides_cellulosivorans | Archaea>Methanobacteriati>Methanobacteriota>Stenosarchaea_group>Halobacteria>Halobacteriales>Haloarculaceae>Halococcoides | 214 | HARCEL1_RS12445 | 2558093..2558737 | 1679096 | hypothetical-protein-[Halococcoides-cellulosivorans] |
| WP_158204984.1 | REase | GCF_009791395.1 | Halomarina_oriensis | Archaea>Methanobacteriati>Methanobacteriota>Stenosarchaea_group>Halobacteria>Halobacteriales>Haloarculaceae>Halomarina | 155 | GQS65_RS12505 | 11310..11777 | 671145 | group-I-intron-associated-PD-(D/E)XK-endonuclease-[Halomarina-oriensis] |
| WP_255153069.1 | REase | GCF_024298865.1 | Halorarius_halobius | Archaea>Methanobacteriati>Methanobacteriota>Stenosarchaea_group>Halobacteria>Halobacteriales>Haloarculaceae>Halorarius | 213 | NL112_RS22040 | 45662..46303 | 2962671 | hypothetical-protein-[Halosegnis-sp.-ZY10] |
| WP_092662317.1 | REase | GCF_900110215.1 | Halorientalis_persicus | Archaea>Methanobacteriati>Methanobacteriota>Stenosarchaea_group>Halobacteria>Halobacteriales>Haloarculaceae>Halorientalis | 200 | BMX17_RS17075 | 65521..66123 | 1367881 | hypothetical-protein-[Halorientalis-persicus] |
| WP_152120784.1 | REase | GCF_009184545.1 | Halosegnis_rubeus | Archaea>Methanobacteriati>Methanobacteriota>Stenosarchaea_group>Halobacteria>Halobacteriales>Haloarculaceae>Halosegnis | 113 | DMP03_RS11355 | 8349..8690 | 2212850 | hypothetical-protein-[Halosegnis-rubeus] |
| WP_188786138.1 | REase | GCF_014646875.1 | Halobellus_salinus | Archaea>Methanobacteriati>Methanobacteriota>Stenosarchaea_group>Halobacteria>Halobacteriales>Haloferacaceae>Halobellus | 236 | IEX72_RS04250 | 183284..183994 | 931585 | restriction-endonuclease-[Halobellus-salinus] |
| RDZ30297.1 | REase | - | Haloferax_sp._Atlit-48N | Archaea>Methanobacteriati>Methanobacteriota>Stenosarchaea_group>Halobacteria>Halobacteriales>Haloferacaceae>Haloferax>unclassified_Haloferax | 99 | DEQ67_14615 | 35877..36176 | 2077198 | hypothetical-protein-DEQ67_14615-[Haloferax-sp.-Atlit-48N] |
| WP_089699569.1 | REase | GCF_900103715.1 | Halogranum_gelatinilyticum | Archaea>Methanobacteriati>Methanobacteriota>Stenosarchaea_group>Halobacteria>Halobacteriales>Haloferacaceae>Halogranum | 170 | BLR57_RS16975 | 79613..80125 | 660521 | hypothetical-protein-[Halogranum-gelatinilyticum] |
| TQQ78819.1 | REase | GCA_006861665.1 | Halonotius_terrestris | Archaea>Methanobacteriati>Methanobacteriota>Stenosarchaea_group>Halobacteria>Halobacteriales>Haloferacaceae>Halonotius | 205 | EGH24_12725 | 56511..57128 | 2487750 | hypothetical-protein-EGH24_12725-[Halonotius-terrestris] |
| WP_142980510.1 | REase | GCF_006861665.1 | Halonotius_terrestris | Archaea>Methanobacteriati>Methanobacteriota>Stenosarchaea_group>Halobacteria>Halobacteriales>Haloferacaceae>Halonotius | 205 | EGH24_RS12730 | 56511..57128 | 2487750 | hypothetical-protein-[Halonotius-terrestris] |
| WP_157744194.1 | REase | GCF_002355635.1 | Halopenitus_persicus | Archaea>Methanobacteriati>Methanobacteriota>Stenosarchaea_group>Halobacteria>Halobacteriales>Haloferacaceae>Halopenitus | 162 | CPZ00_RS15325 | 1178555..1179043 | 1048396 | hypothetical-protein-[Halopenitus-persicus] |
| WP_256476700.1 | REase | GCF_023703795.1 | Haloplanus_sp._HW8-1 | Archaea>Methanobacteriati>Methanobacteriota>Stenosarchaea_group>Halobacteria>Halobacteriales>Haloferacaceae>Haloplanus>unclassified_Haloplanus | 157 | NBT82_RS06035 | 1111068..1111541 | 2949995 | group-I-intron-associated-PD-(D/E)XK-endonuclease-[Haloplanus-sp.-HW8-1] |
| WP_251330426.1 | REase | GCF_023703795.1 | Haloplanus_sp._HW8-1 | Archaea>Methanobacteriati>Methanobacteriota>Stenosarchaea_group>Halobacteria>Halobacteriales>Haloferacaceae>Haloplanus>unclassified_Haloplanus | 169 | NBT82_RS04745 | 883837..884346 | 2949995 | hypothetical-protein-[Haloplanus-sp.-HW8-1] |
| WP_080509902.1 | REase | GCF_000700025.1 | Halorubrum_saccharovorum | Archaea>Methanobacteriati>Methanobacteriota>Stenosarchaea_group>Halobacteria>Halobacteriales>Haloferacaceae>Halorubrum | 229 | FK85_RS16470 | 57145..57834 | 2248 | restriction-endonuclease-[Halorubrum-saccharovorum] |
| WP_144927810.1 | REase | GCF_007671685.1 | Halorubrum_salsamenti | Archaea>Methanobacteriati>Methanobacteriota>Stenosarchaea_group>Halobacteria>Halobacteriales>Haloferacaceae>Halorubrum | 182 | FGL94_RS17060 | 40..588 | 2583990 | restriction-endonuclease-[Halorubrum-salsamenti] |
| WP_152663304.1 | REase | GCF_000700025.1 | Halorubrum_saccharovorum | Archaea>Methanobacteriati>Methanobacteriota>Stenosarchaea_group>Halobacteria>Halobacteriales>Haloferacaceae>Halorubrum | 163 | FK85_RS17310 | 36352..36843 | 2248 | group-I-intron-associated-PD-(D/E)XK-endonuclease-[Halorubrum-saccharovorum] |
| WP_154017954.1 | REase | GCF_000513435.1 | Halorubrum_sp._AJ67 | Archaea>Methanobacteriati>Methanobacteriota>Stenosarchaea_group>Halobacteria>Halobacteriales>Haloferacaceae>Halorubrum>unclassified_Halorubrum | 52 | BN903_RS16205 | 156313..156471 | 1173487 | hypothetical-protein-[Halorubrum-sp.-AJ67] |
| WP_254767371.1 | REase | GCF_024296665.1 | Salinilacihabitans_rarus | Archaea>Methanobacteriati>Methanobacteriota>Stenosarchaea_group>Halobacteria>Halobacteriales>Natrialbaceae>Salinilacihabitans | 141 | NKG98_RS17275 | 3278985..3279410 | 2961596 | group-I-intron-associated-PD-(D/E)XK-endonuclease-[Natribaculum-sp.-AD-4] |
| WP_020446115.1 | REase | GCF_000403645.1 | Salinarchaeum_sp._Harcht-Bsk1 | Archaea>Methanobacteriati>Methanobacteriota>Stenosarchaea_group>Halobacteria>Halobacteriales>Natronoarchaeaceae>Salinarchaeum>unclassified_Salinarchaeum | 138 | L593_RS06360 | 1395476..1395892 | 1333523 | group-I-intron-associated-PD-(D/E)XK-endonuclease-[Salinarchaeum-sp.-Harcht-Bsk1] |
| WP_074369584.1 | REase | GCF_900095385.1 | Methanoculleus_chikugoensis | Archaea>Methanobacteriati>Methanobacteriota>Stenosarchaea_group>Methanomicrobia>Methanomicrobiales>Methanomicrobiaceae>Methanoculleus | 179 | L21_RS05975 | 50533..51072 | 118126 | hypothetical-protein-[Methanoculleus-chikugoensis] |
| WP_257742629.1 | REase | GCF_024662215.1 | Methanoplanus_endosymbiosus | Archaea>Methanobacteriati>Methanobacteriota>Stenosarchaea_group>Methanomicrobia>Methanomicrobiales>Methanomicrobiaceae>Methanoplanus | 76 | L6E24_RS14355 | 3069990..3070220 | 33865 | hypothetical-protein-[Methanoplanus-endosymbiosus] |
| WP_143709442.1 | REase | GCF_000013445.1 | Methanospirillum_hungatei | Archaea>Methanobacteriati>Methanobacteriota>Stenosarchaea_group>Methanomicrobia>Methanomicrobiales>Methanospirillaceae>Methanospirillum | 180 | MHUN_RS09150 | 1979149..1979691 | 2203 | hypothetical-protein-[Methanospirillum-hungatei] |
| WP_214418676.1 | REase | GCF_018502485.1 | Methanospirillum_purgamenti | Archaea>Methanobacteriati>Methanobacteriota>Stenosarchaea_group>Methanomicrobia>Methanomicrobiales>Methanospirillaceae>Methanospirillum | 209 | KHC33_RS10955 | 2276910..2277539 | 2834276 | type-I-restriction-endonuclease-[Methanospirillum-sp.-J.3.6.1-F.2.7.3] |
| ABD41473.1 | REase | GCA_000013445.1 | Methanospirillum_hungatei_JF-1 | Archaea>Methanobacteriati>Methanobacteriota>Stenosarchaea_group>Methanomicrobia>Methanomicrobiales>Methanospirillaceae>Methanospirillum>Methanospirillum_hungatei | 187 | Mhun_1752 | 1979128..1979691 | 323259 | hypothetical-protein-Mhun_1752-[Methanospirillum-hungatei-JF-1] |
| PXF61723.1 | REase | GCA_003194445.1 | Candidatus_Methanogaster_sp. | Archaea>Methanobacteriati>Methanobacteriota>Stenosarchaea_group>Methanomicrobia>Methanosarcinales>ANME-2_cluster>Candidatus_Methanogasteraceae>Candidatus_Methanogaster>unclassified_Candidatus_Methanogaster | 194 | C4B59_02385 | 70148..70732 | 3386292 | hypothetical-protein-C4B59_02385-[ANME-2-cluster-archaeon] |
| KXS44102.1 | REase | GCA_001577565.1 | Methanolobus_sp._T82-4 | Archaea>Methanobacteriati>Methanobacteriota>Stenosarchaea_group>Methanomicrobia>Methanosarcinales>Methanosarcinaceae>Methanolobus>unclassified_Methanolobus | 82 | AWU59_735 | 24863..25111 | 1794908 | MAG:-hypothetical-protein-AWU59_735-[Methanolobus-sp.-T82-4] |
| AKB47808.1 | REase | GCA_000969945.1 | Methanosarcina_sp._Kolksee | Archaea>Methanobacteriati>Methanobacteriota>Stenosarchaea_group>Methanomicrobia>Methanosarcinales>Methanosarcinaceae>Methanosarcina>unclassified_Methanosarcina | 199 | MSKOL_2031 | 2634079..2634678 | 1434099 | hypothetical-protein-MSKOL_2031-[Methanosarcina-sp.-Kolksee] |
| WP_048156128.1 | REase | GCF_000969945.1 | Methanosarcina_sp._Kolksee | Archaea>Methanobacteriati>Methanobacteriota>Stenosarchaea_group>Methanomicrobia>Methanosarcinales>Methanosarcinaceae>Methanosarcina>unclassified_Methanosarcina | 199 | MSKOL_RS10875 | 2634079..2634678 | 1434099 | hypothetical-protein-[Methanosarcina-sp.-Kolksee] |
| WP_052297567.1 | REase | GCF_000204415.1 | Methanothrix | Archaea>Methanobacteriati>Methanobacteriota>Stenosarchaea_group>Methanomicrobia>Methanotrichales>Methanotrichaceae | 80 | QVD88_RS00975 | 24552..24794 | 2222 | hypothetical-protein-[Methanothrix-soehngenii] |
| KUK43675.1 | REase | GCA_001509375.1 | Methanothrix_harundinacea | Archaea>Methanobacteriati>Methanobacteriota>Stenosarchaea_group>Methanomicrobia>Methanotrichales>Methanotrichaceae>Methanothrix | 207 | XD72_1942 | 9803..10426 | 301375 | Uncharacterized-protein-XD72_1942-[Methanothrix-harundinacea] |
| MCP1392943.1 | REase | GCA_024166385.1 | Methanothrix_harundinacea | Archaea>Methanobacteriati>Methanobacteriota>Stenosarchaea_group>Methanomicrobia>Methanotrichales>Methanotrichaceae>Methanothrix | 207 | J5I35_09330 | 3641..4264 | 301375 | hypothetical-protein-[Methanothrix-harundinacea] |
| WP_014586223.1 | REase | GCF_000235565.1 | Methanothrix_harundinacea | Archaea>Methanobacteriati>Methanobacteriota>Stenosarchaea_group>Methanomicrobia>Methanotrichales>Methanotrichaceae>Methanothrix | 205 | MHAR_RS03415 | 697534..698151 | 301375 | hypothetical-protein-[Methanothrix-harundinacea] |
| AET64038.1 | REase | GCA_000235565.1 | Methanothrix_harundinacea_6Ac | Archaea>Methanobacteriati>Methanobacteriota>Stenosarchaea_group>Methanomicrobia>Methanotrichales>Methanotrichaceae>Methanothrix>Methanothrix_harundinacea | 205 | Mhar_0660 | 697534..698151 | 1110509 | hypothetical-protein-Mhar_0660-[Methanothrix-harundinacea-6Ac] |
| MCX6669298.1 | REase | GCA_026394475.1 | Methanothrix_sp. | Archaea>Methanobacteriati>Methanobacteriota>Stenosarchaea_group>Methanomicrobia>Methanotrichales>Methanotrichaceae>Methanothrix>unclassified_Methanothrix | 192 | NTV25_05765 | 2830..3408 | 90426 | MAG:-hypothetical-protein-NTV25_05765-[Methanothrix-sp.] |
| MDD3564061.1 | REase | GCA_028695755.1 | Methanothrix_sp. | Archaea>Methanobacteriati>Methanobacteriota>Stenosarchaea_group>Methanomicrobia>Methanotrichales>Methanotrichaceae>Methanothrix>unclassified_Methanothrix | 192 | PHN90_00130 | 48615..49193 | 90426 | MAG:-hypothetical-protein-PHN90_00130-[Methanothrix-sp.] |
| NLX39270.1 | REase | GCA_012729025.1 | Methanothrix_sp. | Archaea>Methanobacteriati>Methanobacteriota>Stenosarchaea_group>Methanomicrobia>Methanotrichales>Methanotrichaceae>Methanothrix>unclassified_Methanothrix | 190 | GXY73_07145 | 4290..4862 | 90426 | hypothetical-protein-[Methanothrix-sp.] |
| MBP7068910.1 | REase | GCA_017994095.1 | Methanothrix_sp. | Archaea>Methanobacteriati>Methanobacteriota>Stenosarchaea_group>Methanomicrobia>Methanotrichales>Methanotrichaceae>Methanothrix>unclassified_Methanothrix | 169 | KBB04_11630 | 16035..16544 | 90426 | PD-(D/E)XK-nuclease-family-protein-[Methanothrix-sp.] |
| MBN1235614.1 | REase | GCA_016927055.1 | Methanotrichaceae_archaeon | Archaea>Methanobacteriati>Methanobacteriota>Stenosarchaea_group>Methanomicrobia>Methanotrichales>Methanotrichaceae>unclassified_Methanotrichaceae | 191 | JW999_06130 | 3760..4335 | 2595067 | hypothetical-protein-[Methanotrichaceae-archaeon] |
| RLI54040.1 | REase | GCA_003662775.1 | Candidatus_Thorarchaeota_archaeon | Archaea>Promethearchaeati>Candidatus_Thorarchaeota>unclassified_Candidatus_Thorarchaeota | 214 | DRP09_14175 | 3653..4297 | 2053491 | hypothetical-protein-DRP09_14175-[Candidatus-Thorarchaeota-archaeon] |
| MCD6241713.1 | REase | GCA_021158145.1 | Candidatus_Bathyarchaeota_archaeon | Archaea>Thermoproteati>Candidatus_Bathyarchaeota>unclassified_Candidatus_Bathyarchaeota | 218 | J7K27_09455 | 13299..13955 | 2026714 | hypothetical-protein-[Candidatus-Bathyarchaeota-archaeon] |
| MCC6018925.1 | REase | GCA_020833005.1 | Candidatus_Methanomethylicota_archaeon | Archaea>Thermoproteati>Candidatus_Methanomethylicota | 165 | LM601_07835 | 12481..12978 | 2250257 | hypothetical-protein-[Candidatus-Verstraetearchaeota-archaeon] |
| WP_055408539.1 | REase | GCF_001412615.1 | Pyrodictium_delaneyi | Archaea>Thermoproteati>Thermoproteota>Thermoprotei>Desulfurococcales>Pyrodictiaceae>Pyrodictium | 116 | Pyrde_RS04370 | 850468..850818 | 1273541 | YraN-family-protein-[Pyrodictium-delaneyi] |
| MCE4605484.1 | REase | GCA_015521665.1 | Desulfurococcales_archaeon | Archaea>Thermoproteati>Thermoproteota>Thermoprotei>Desulfurococcales>unclassified_Desulfurococcales | 130 | F7C08_03015 | 104235..104627 | 2480821 | hypothetical-protein-[Desulfurococcales-archaeon] |
| MCY3894572.1 | REase | GCA_026707105.1 | Acidimicrobiaceae_bacterium | Bacteria>Bacillati>Actinomycetota>Acidimicrobiia>Acidimicrobiales>Acidimicrobiaceae>unclassified_Acidimicrobiaceae | 248 | OXF65_14900 | 88..834 | 2024894 | MAG:-DUF3883-domain-containing-protein-[Acidimicrobiaceae-bacterium] |
| MDE0498680.1 | REase | GCA_028823965.1 | Acidimicrobiaceae_bacterium | Bacteria>Bacillati>Actinomycetota>Acidimicrobiia>Acidimicrobiales>Acidimicrobiaceae>unclassified_Acidimicrobiaceae | 247 | OXH86_15130 | 61972..62715 | 2024894 | MAG:-DUF3883-domain-containing-protein-[Acidimicrobiaceae-bacterium] |
| MDE0319986.1 | REase | GCA_028820815.1 | Acidimicrobiaceae_bacterium | Bacteria>Bacillati>Actinomycetota>Acidimicrobiia>Acidimicrobiales>Acidimicrobiaceae>unclassified_Acidimicrobiaceae | 215 | OXI97_08890 | 7009..7656 | 2024894 | MAG:-DUF3883-domain-containing-protein,-partial-[Acidimicrobiaceae-bacterium] |
| MCY4516056.1 | REase | GCA_026713105.1 | Acidimicrobiaceae_bacterium | Bacteria>Bacillati>Actinomycetota>Acidimicrobiia>Acidimicrobiales>Acidimicrobiaceae>unclassified_Acidimicrobiaceae | 173 | OXB99_02325 | 8756..9277 | 2024894 | MAG:-DUF3883-domain-containing-protein-[Acidimicrobiaceae-bacterium] |
| MXX43804.1 | REase | GCA_009836375.1 | Acidimicrobiales_bacterium | Bacteria>Bacillati>Actinomycetota>Acidimicrobiia>Acidimicrobiales>unclassified_Acidimicrobiales | 198 | F4Z53_12220 | 11978..12574 | 2201156 | DUF3883-domain-containing-protein-[Acidimicrobiales-bacterium] |
| MYB24417.1 | REase | GCA_009840735.1 | Acidimicrobiia_bacterium | Bacteria>Bacillati>Actinomycetota>Acidimicrobiia>unclassified_Acidimicrobiia | 217 | F4X37_04840 | 50759..51412 | 2080302 | DUF3883-domain-containing-protein-[Acidimicrobiia-bacterium] |
| MCZ7536066.1 | REase | GCA_027494135.1 | Acidimicrobiia_bacterium | Bacteria>Bacillati>Actinomycetota>Acidimicrobiia>unclassified_Acidimicrobiia | 157 | M5T61_09160 | 16538..17011 | 2080302 | MAG:-DUF3883-domain-containing-protein-[Acidimicrobiia-bacterium] |
| MCB9390231.1 | REase | GCA_020637365.1 | Acidimicrobiia_bacterium | Bacteria>Bacillati>Actinomycetota>Acidimicrobiia>unclassified_Acidimicrobiia | 173 | H6512_11915 | 836417..836938 | 2080302 | DUF4365-domain-containing-protein-[Acidimicrobiia-bacterium] |
| MBA2283089.1 | REase | GCA_013694495.1 | Acidimicrobiia_bacterium | Bacteria>Bacillati>Actinomycetota>Acidimicrobiia>unclassified_Acidimicrobiia | 156 | HOW25_17910 | 21721..22191 | 2080302 | DUF4365-domain-containing-protein-[Acidimicrobiia-bacterium] |
| WP_003788296.1 | REase | GCF_963519085.1 | Actinomyces | Bacteria>Bacillati>Actinomycetota>Actinomycetes>Actinomycetales>Actinomycetaceae | 130 | HMPREF0059_RS11105 | 112881..113273 | 1654 | very-short-patch-repair-endonuclease-[Actinomyces-viscosus] |
| WP_073452903.1 | REase | GCF_002356195.1 | Actinomyces | Bacteria>Bacillati>Actinomycetota>Actinomycetes>Actinomycetales>Actinomycetaceae | 124 | DY404_RS06905 | 1650829..1651203 | 1654 | MULTISPECIES:-very-short-patch-repair-endonuclease-[Actinomyces] |
| WP_253188752.1 | REase | GCF_001956555.1 | Actinomyces_naeslundii | Bacteria>Bacillati>Actinomycetota>Actinomycetes>Actinomycetales>Actinomycetaceae>Actinomyces | 165 | BKH25_RS11280 | 127755..128252 | 1655 | very-short-patch-repair-endonuclease-[Actinomyces-naeslundii] |
| WP_268916836.1 | REase | - | Actinomyces_israelii | Bacteria>Bacillati>Actinomycetota>Actinomycetes>Actinomycetales>Actinomycetaceae>Actinomyces | 151 | OHJ16_RS03980 | 43578..44033 | 1659 | very-short-patch-repair-endonuclease-[Actinomyces-israelii] |
| WP_039774343.1 | REase | GCF_023169925.1 | Actinomyces_oris | Bacteria>Bacillati>Actinomycetota>Actinomycetes>Actinomycetales>Actinomycetaceae>Actinomyces | 130 | AoKodu_RS00655 | 156987..157379 | 544580 | very-short-patch-repair-endonuclease-[Actinomyces-oris] |
| WP_043508281.1 | REase | GCF_000295095.1 | Actinomyces_timonensis | Bacteria>Bacillati>Actinomycetota>Actinomycetes>Actinomycetales>Actinomycetaceae>Actinomyces | 130 | A1QA_RS0104830 | 1081929..1082321 | 1288391 | very-short-patch-repair-endonuclease-[Actinomyces-timonensis] |
| WP_043539606.1 | REase | GCF_016127855.1 | Actinomyces_naeslundii | Bacteria>Bacillati>Actinomycetota>Actinomycetes>Actinomycetales>Actinomycetaceae>Actinomyces | 130 | BKH33_RS06325 | 167711..168103 | 1655 | very-short-patch-repair-endonuclease-[Actinomyces-naeslundii] |
| WP_075390149.1 | REase | GCF_902480955.1 | Actinomyces_oris | Bacteria>Bacillati>Actinomycetota>Actinomycetes>Actinomycetales>Actinomycetaceae>Actinomyces | 130 | BKH20_RS04800 | 25636..26028 | 544580 | very-short-patch-repair-endonuclease-[Actinomyces-oris] |
| WP_075403453.1 | REase | GCF_001937505.1 | Actinomyces_oris | Bacteria>Bacillati>Actinomycetota>Actinomycetes>Actinomycetales>Actinomycetaceae>Actinomyces | 130 | BKH21_RS10545 | 44927..45319 | 544580 | very-short-patch-repair-endonuclease-[Actinomyces-oris] |
| WP_075410182.1 | REase | GCF_905372775.1 | Actinomyces_naeslundii | Bacteria>Bacillati>Actinomycetota>Actinomycetes>Actinomycetales>Actinomycetaceae>Actinomyces | 130 | QDH38_RS03515 | 48353..48745 | 1655 | very-short-patch-repair-endonuclease-[Actinomyces-naeslundii] |
| WP_075412467.1 | REase | GCF_001937365.1 | Actinomyces_oris | Bacteria>Bacillati>Actinomycetota>Actinomycetes>Actinomycetales>Actinomycetaceae>Actinomyces | 130 | BKH31_RS11935 | 23947..24339 | 544580 | very-short-patch-repair-endonuclease-[Actinomyces-oris] |
| WP_151307477.1 | REase | - | Actinomyces_oris | Bacteria>Bacillati>Actinomycetota>Actinomycetes>Actinomycetales>Actinomycetaceae>Actinomyces | 130 | BKH30_RS08630 | 15390..15782 | 544580 | very-short-patch-repair-endonuclease-[Actinomyces-oris] |
| WP_271288671.1 | REase | GCF_027945475.1 | Actinomyces_oris | Bacteria>Bacillati>Actinomycetota>Actinomycetes>Actinomycetales>Actinomycetaceae>Actinomyces | 130 | PGE45_RS00600 | 145591..145983 | 544580 | very-short-patch-repair-endonuclease-[Actinomyces-oris] |
| MBE6482580.1 | REase | GCA_015062785.1 | Actinomyces_ruminicola | Bacteria>Bacillati>Actinomycetota>Actinomycetes>Actinomycetales>Actinomycetaceae>Actinomyces | 126 | E7Z94_09490 | 4123..4503 | 332524 | very-short-patch-repair-endonuclease-[Actinomyces-ruminicola] |

| Protein-ID | Domain | Assembly-ID | Strain | Taxonomy | Length | Locus-Tag | Genomic-coordinates | TaxID | Fasta-defline |
|---|---|---|---|---|---|---|---|---|---|
| WP_103061773.1 | REase | GCF_004102025.1 | Actinomyces_qiguomingii | Bacteria>Bacillati>Actinomycetota>Actinomycetes>Actinomycetales>Actinomycetaceae>Actinomyces | 126 | CWT10_RS16090 | 3841680..3842060 | 2057800 | very-short-patch-repair-endonuclease-[Actinomyces-qiguomingii] |
| BDF97833.1 | REase | GCA_023169925.1 | Actinomyces_oris_K20 | Bacteria>Bacillati>Actinomycetota>Actinomycetes>Actinomycetales>Actinomycetaceae>Actinomyces>Actinomyces_oris | 130 | AoKodu_01340 | 156987..157379 | 871541 | hypothetical-protein-AoKodu_01340-[Actinomyces-oris-K20] |
| WP_234415357.1 | REase | GCF_900323545.1 | Actinomyces_sp._Marseille-P3109 | Bacteria>Bacillati>Actinomycetota>Actinomycetes>Actinomycetales>Actinomycetaceae>Actinomyces>unclassified_Actinomyces | 169 | BQ8008_RS10560 | 2434033..2434542 | 2083009 | very-short-patch-repair-endonuclease-[Actinomyces-sp.-Marseille-P3109] |
| WP_053588277.1 | REase | GCF_001278845.1 | Actinomyces_sp._oral_taxon_414 | Bacteria>Bacillati>Actinomycetota>Actinomycetes>Actinomycetales>Actinomycetaceae>Actinomyces>unclassified_Actinomyces | 135 | AM609_RS14665 | 3667721..3668128 | 712122 | very-short-patch-repair-endonuclease-[Actinomyces-sp.-oral-taxon-414] |
| WP_220456831.1 | REase | GCF_014041995.1 | Changpingibacter_yushuensis | Bacteria>Bacillati>Actinomycetota>Actinomycetes>Actinomycetales>Actinomycetaceae>Changpingibacter | 101 | H2017_RS05150 | 1184295..1184600 | 2758440 | hypothetical-protein-[Changpingibacter-yushuensis] |
| WP_233050713.1 | REase | GCF_016405145.1 | Schaalia_cardiffensis | Bacteria>Bacillati>Actinomycetota>Actinomycetes>Actinomycetales>Actinomycetaceae>Schaalia | 143 | JFX23_RS08505 | 30346..30777 | 181487 | very-short-patch-repair-endonuclease-[Schaalia-cardiffensis] |
| WP_003826148.1 | REase | GCF_001025155.1 | Bifidobacterium | Bacteria>Bacillati>Actinomycetota>Actinomycetes>Bifidobacteriales>Bifidobacteriaceae | 54 | PG742_RS01105 | 250967..251131 | 1678 | MULTISPECIES:-hypothetical-protein-[Bifidobacterium] |
| WP_171842433.1 | REase | GCF_001685965.1 | Bifidobacterium | Bacteria>Bacillati>Actinomycetota>Actinomycetes>Bifidobacteriales>Bifidobacteriaceae | 54 | PMN34_RS05070 | 217110..217274 | 1678 | MULTISPECIES:-hypothetical-protein-[Bifidobacterium] |
| WP_168445151.1 | REase | GCF_012310535.1 | Bifidobacterium_dentium | Bacteria>Bacillati>Actinomycetota>Actinomycetes>Bifidobacteriales>Bifidobacteriaceae>Bifidobacterium | 54 | IHQ76_RS06195 | 233771..233935 | 1689 | hypothetical-protein-[Bifidobacterium-dentium] |
| WP_217054575.1 | REase | GCF_019041665.1 | Bifidobacterium_longum | Bacteria>Bacillati>Actinomycetota>Actinomycetes>Bifidobacteriales>Bifidobacteriaceae>Bifidobacterium | 81 | KTF47_RS04060 | 23190..23435 | 216816 | hypothetical-protein-[Bifidobacterium-longum] |
| WP_190971985.1 | REase | GCF_018555535.1 | Bifidobacterium_vespertilionis | Bacteria>Bacillati>Actinomycetota>Actinomycetes>Bifidobacteriales>Bifidobacteriaceae>Bifidobacterium | 50 | JS538_RS07565 | 43918..44070 | 2562524 | hypothetical-protein-[Bifidobacterium-vespertilionis] |
| WP_211244531.1 | REase | GCF_000504285.1 | Actinospica_robiniae | Bacteria>Bacillati>Actinomycetota>Actinomycetes>Catenulisporales>Actinospicaceae>Actinospica | 179 | ACTRO_RS49245 | 8510213..8510752 | 304901 | hypothetical-protein-[Actinospica-robiniae] |
| RBY97950.1 | REase | GCA_003319175.1 | Blastococcus_sp._TF02-8 | Bacteria>Bacillati>Actinomycetota>Actinomycetes>Geodermatophilales>Geodermatophilaceae>Blastococcus>unclassified_Blastococcus | 158 | DQ237_03395 | 724884..725360 | 2250574 | very-short-patch-repair-endonuclease-[Blastococcus-sp.-TF02-8] |
| WP_113797037.1 | REase | GCF_003319175.1 | Blastococcus_sp._TF02-8 | Bacteria>Bacillati>Actinomycetota>Actinomycetes>Geodermatophilales>Geodermatophilaceae>Blastococcus>unclassified_Blastococcus | 158 | DQ237_RS03395 | 724884..725360 | 2250574 | very-short-patch-repair-endonuclease-[Blastococcus-sp.-TF02-8] |
| WP_129664645.1 | REase | GCF_004138495.1 | Phytoactinopolyspora_endophytica | Bacteria>Bacillati>Actinomycetota>Actinomycetes>Jiangellales>Jiangellaceae>Phytoactinopolyspora | 129 | ELR45_RS07130 | 287164..287553 | 1642495 | hypothetical-protein-[Phytoactinopolyspora-endophytica] |
| UER54524.1 | REase | GCA_020881835.1 | Kineosporiaceae_bacterium_SCSIO_59966 | Bacteria>Bacillati>Actinomycetota>Actinomycetes>Kineosporiales>Kineosporiaceae>unclassified_Kineosporiaceae | 227 | HJG43_08210 | 1754598..1755281 | 2730919 | hypothetical-protein-HJG43_08210-[Kineosporiaceae-bacterium-SCSIO-59966] |
| WP_249836785.1 | REase | GCF_023516615.1 | Streptomyces | Bacteria>Bacillati>Actinomycetota>Actinomycetes>Kitasatosporales>Streptomycetaceae | 198 | M8J74_RS14290 | 88574..89170 | 1883 | Uma2-family-endonuclease-[Streptomyces-panacradicis] |
| WP_263456372.1 | REase | - | Streptomyces_olivaceus | Bacteria>Bacillati>Actinomycetota>Actinomycetes>Kitasatosporales>Streptomycetaceae>Streptomyces | 228 | KVH22_RS20530 | 323241..323927 | 47716 | restriction-endonuclease-[Streptomyces-olivaceus] |
| WP_255970760.1 | REase | GCF_024448165.1 | Streptomyces_longispororuber | Bacteria>Bacillati>Actinomycetota>Actinomycetes>Kitasatosporales>Streptomycetaceae>Streptomyces | 206 | M4J07_RS11785 | 28615..29235 | 68230 | restriction-endonuclease-[Streptomyces-longispororuber] |
| WP_328932263.1 | REase | GCF_036196325.1 | unclassified_Streptomyces | Bacteria>Bacillati>Actinomycetota>Actinomycetes>Kitasatosporales>Streptomycetaceae>Streptomyces | 155 | OG537_RS11445 | 2668473..2668940 | 2593676 | restriction-endonuclease-[Streptomyces-sp.-NBC_00228] |
| WP_169086800.1 | REase | GCF_012927225.1 | unclassified_Streptomyces | Bacteria>Bacillati>Actinomycetota>Actinomycetes>Kitasatosporales>Streptomycetaceae>Streptomyces | 197 | HHL18_RS29260 | 62364..62957 | 2593676 | MULTISPECIES:-Uma2-family-endonuclease-[unclassified-Streptomyces] |
| WP_234899916.1 | REase | GCF_008634015.1 | Streptomyces_filamentosus | Bacteria>Bacillati>Actinomycetota>Actinomycetes>Kitasatosporales>Streptomycetaceae>Streptomyces | 197 | CP979_RS24320 | 5267484..5268077 | 67294 | Uma2-family-endonuclease-[Streptomyces-filamentosus] |
| WP_326752425.1 | REase | GCF_035918035.1 | Streptomyces_hirsutus | Bacteria>Bacillati>Actinomycetota>Actinomycetes>Kitasatosporales>Streptomycetaceae>Streptomyces | 189 | OIE73_RS11115 | 2429413..2429982 | 35620 | Uma2-family-endonuclease-[Streptomyces-hirsutus] |
| GGQ80798.1 | REase | GCA_014649355.1 | Streptomyces_althioticus | Bacteria>Bacillati>Actinomycetota>Actinomycetes>Kitasatosporales>Streptomycetaceae>Streptomyces>Streptomyces_althioticus_group | 158 | GCM10010250_60560 | 46473..46949 | 83380 | hypothetical-protein-GCM10010250_60560-[Streptomyces-althioticus] |
| WP_174569421.1 | REase | GCF_001611795.1 | Streptomyces_qaidamensis | Bacteria>Bacillati>Actinomycetota>Actinomycetes>Kitasatosporales>Streptomycetaceae>Streptomyces>Streptomyces_aurantiacus_group | 192 | A4E84_RS11905 | 2731570..2732148 | 1783515 | Uma2-family-endonuclease-[Streptomyces-qaidamensis] |
| BCK67996.1 | REase | GCA_009811615.2 | Streptomyces_libani_subsp._rufus | Bacteria>Bacillati>Actinomycetota>Actinomycetes>Kitasatosporales>Streptomycetaceae>Streptomyces>Streptomyces_libani | 59 | Srufu_019490 | 2044030..2044209 | 249582 | hypothetical-protein-Srufu_019490-[Streptomyces-libani-subsp.-rufus] |
| WP_168711364.1 | REase | GCF_004795255.1 | Streptomyces_sp._RKND-216 | Bacteria>Bacillati>Actinomycetota>Actinomycetes>Kitasatosporales>Streptomycetaceae>Streptomyces>unclassified_Streptomyces | 209 | E4198_RS24790 | 928427..929056 | 2562581 | restriction-endonuclease-[Streptomyces-sp.-RKND-216] |
| WP_328677711.1 | REase | GCF_036118775.1 | Streptomyces_sp._NBC_00343 | Bacteria>Bacillati>Actinomycetota>Actinomycetes>Kitasatosporales>Streptomycetaceae>Streptomyces>unclassified_Streptomyces | 209 | OHA62_RS11590 | 2739108..2739737 | 2975719 | restriction-endonuclease-[Streptomyces-sp.-NBC_00343] |
| WP_081522436.1 | REase | GCF_002080455.1 | Streptomyces_sp._Sge12 | Bacteria>Bacillati>Actinomycetota>Actinomycetes>Kitasatosporales>Streptomycetaceae>Streptomyces>unclassified_Streptomyces | 197 | B6R96_RS11870 | 2756864..2757457 | 1972846 | Uma2-family-endonuclease-[Streptomyces-sp.-Sge12] |
| WP_175410412.1 | REase | GCF_013362995.1 | Streptomyces_sp._TRM64462 | Bacteria>Bacillati>Actinomycetota>Actinomycetes>Kitasatosporales>Streptomycetaceae>Streptomyces>unclassified_Streptomyces | 197 | HTZ82_RS17975 | 56187..56780 | 2741726 | Uma2-family-endonuclease-[Streptomyces-sp.-TRM64462] |
| WP_101639483.1 | REase | GCF_900169095.1 | Brevibacterium_aurantiacum | Bacteria>Bacillati>Actinomycetota>Actinomycetes>Micrococcales>Brevibacteriaceae>Brevibacterium | 125 | BAUR920_RS16210 | 7716..8093 | 273384 | very-short-patch-repair-endonuclease-[Brevibacterium-aurantiacum] |
| WP_236850604.1 | REase | GCF_001729525.1 | Brevibacterium_aurantiacum | Bacteria>Bacillati>Actinomycetota>Actinomycetes>Micrococcales>Brevibacteriaceae>Brevibacterium | 124 | CIK60_RS17595 | 19025..19399 | 273384 | DNA-mismatch-endonuclease-Vsr-[Brevibacterium-aurantiacum] |
| WP_225755642.1 | REase | GCF_916618755.1 | Actinotalea_sp._Marseille-Q4924 | Bacteria>Bacillati>Actinomycetota>Actinomycetes>Micrococcales>Cellulomonadaceae>Actinotalea>unclassified_Actinotalea | 128 | LG379_RS16290 | 261594..261980 | 2866571 | very-short-patch-repair-endonuclease-[Actinotalea-sp.-Marseille-Q4924] |
| WP_129336328.1 | REase | GCF_004123805.1 | Cellulomonas_endophytica | Bacteria>Bacillati>Actinomycetota>Actinomycetes>Micrococcales>Cellulomonadaceae>Cellulomonas | 145 | EJA20_RS01065 | 82385..82822 | 2494735 | very-short-patch-repair-endonuclease-[Cellulomonas-endophytica] |
| WP_046530911.1 | REase | - | Cellulomonas_sp._FA1 | Bacteria>Bacillati>Actinomycetota>Actinomycetes>Micrococcales>Cellulomonadaceae>Cellulomonas>unclassified_Cellulomonas | 130 | GY42_RS11855 | 2637264..2637656 | 1346710 | very-short-patch-repair-endonuclease-[Cellulomonas-sp.-FA1] |
| WP_128365192.1 | REase | - | Phycicoccus_flavus | Bacteria>Bacillati>Actinomycetota>Actinomycetes>Micrococcales>Intrasporangiaceae>Phycicoccus | 137 | EPD83_RS11940 | 17762..18175 | 2502783 | very-short-patch-repair-endonuclease-[Phycicoccus-flavus] |
| WP_157415183.1 | REase | GCF_039529235.1 | Agromyces_allii | Bacteria>Bacillati>Actinomycetota>Actinomycetes>Micrococcales>Microbacteriaceae>Agromyces | 143 | GQR16_RS11040 | 276412..276843 | 393607 | very-short-patch-repair-endonuclease-[Agromyces-allii] |
| WP_198169815.1 | REase | - | Agromyces_laixinhei | Bacteria>Bacillati>Actinomycetota>Actinomycetes>Micrococcales>Microbacteriaceae>Agromyces | 141 | GQQ90_RS09135 | 58201..58626 | 2585717 | very-short-patch-repair-endonuclease-[Agromyces-laixinhei] |
| WP_396133129.1 | REase | GCF_031456545.1 | Agromyces_sp._3263 | Bacteria>Bacillati>Actinomycetota>Actinomycetes>Micrococcales>Microbacteriaceae>Agromyces>unclassified_Agromyces | 153 | J2X63_RS18955 | 2276901..2277362 | 2817750 | winged-helix-turn-helix-domain-containing-protein-[Agromyces-sp.-3263] |
| WP_241732360.1 | REase | - | Galbitalea_soli | Bacteria>Bacillati>Actinomycetota>Actinomycetes>Micrococcales>Microbacteriaceae>Galbitalea | 205 | HNR17_RS14815 | 986141..986758 | 1268042 | DUF4263-domain-containing-protein-[Galbitalea-soli] |
| WP_268912063.1 | REase | GCF_910593785.1 | Leucobacter_soli | Bacteria>Bacillati>Actinomycetota>Actinomycetes>Micrococcales>Microbacteriaceae>Leucobacter | 134 | KXF26_RS09190 | 51069..51473 | 2812850 | very-short-patch-repair-endonuclease-[Leucobacter-soli] |
| WP_224781231.1 | REase | GCF_020096995.1 | Leucobacter_sp._Psy1 | Bacteria>Bacillati>Actinomycetota>Actinomycetes>Micrococcales>Microbacteriaceae>Leucobacter>unclassified_Leucobacter | 141 | K8P10_RS06905 | 1474745..1475170 | 2875729 | very-short-patch-repair-endonuclease-[Leucobacter-sp.-Psy1] |
| WP_156242692.1 | REase | GCF_039536885.1 | Microbacterium_oryzae | Bacteria>Bacillati>Actinomycetota>Actinomycetes>Micrococcales>Microbacteriaceae>Microbacterium | 135 | D7D94_RS11245 | 2397174..2397581 | 743009 | very-short-patch-repair-endonuclease-[Microbacterium-oryzae] |
| WP_156377208.1 | REase | GCF_001426925.1 | Microbacterium_sp._Root53 | Bacteria>Bacillati>Actinomycetota>Actinomycetes>Micrococcales>Microbacteriaceae>Microbacterium>unclassified_Microbacterium | 184 | ASD19_RS13285 | 105377..105931 | 1736553 | DUF4365-domain-containing-protein-[Microbacterium-sp.-Root53] |
| WP_282859604.1 | REase | GCF_030011915.1 | Pseudoclavibacter_helvolus | Bacteria>Bacillati>Actinomycetota>Actinomycetes>Micrococcales>Microbacteriaceae>Pseudoclavibacter | 162 | QM024_RS14950 | 8255..8743 | 255205 | DUF3883-domain-containing-protein,-partial-[Pseudoclavibacter-helvolus] |
| WP_231831074.1 | REase | GCF_021168475.1 | Pseudoclavibacter_sp._13-3 | Bacteria>Bacillati>Actinomycetota>Actinomycetes>Micrococcales>Microbacteriaceae>Pseudoclavibacter>unclassified_Pseudoclavibacter | 127 | LRX77_RS01490 | 328763..329146 | 2901228 | very-short-patch-repair-endonuclease-[Pseudoclavibacter-sp.-13-3] |
| WP_243591602.1 | REase | GCF_022815585.1 | Rathayibacter_caricis | Bacteria>Bacillati>Actinomycetota>Actinomycetes>Micrococcales>Microbacteriaceae>Rathayibacter | 146 | MT349_RS09880 | 116328..116768 | 110936 | very-short-patch-repair-endonuclease-[Rathayibacter-caricis] |
| WP_134416009.1 | REase | - | Micrococcus | Bacteria>Bacillati>Actinomycetota>Actinomycetes>Micrococcales>Micrococcaceae | 140 | KM867_RS03185 | 36892..37314 | 1269 | MULTISPECIES:-very-short-patch-repair-endonuclease-[Micrococcus] |
| WP_210233974.1 | REase | - | Arthrobacter_jiangjiafuii | Bacteria>Bacillati>Actinomycetota>Actinomycetes>Micrococcales>Micrococcaceae>Arthrobacter | 218 | J2M53_RS14470 | 603990..604646 | 2817475 | hypothetical-protein-[Arthrobacter-jiangjiafuii] |
| WP_210233976.1 | REase | GCF_017898105.1 | Arthrobacter_jiangjiafuii | Bacteria>Bacillati>Actinomycetota>Actinomycetes>Micrococcales>Micrococcaceae>Arthrobacter | 100 | J2M53_RS17170 | 608673..608975 | 2817475 | hypothetical-protein-[Arthrobacter-jiangjiafuii] |
| WP_256385956.1 | REase | GCF_003989235.1 | unclassified_Kocuria | Bacteria>Bacillati>Actinomycetota>Actinomycetes>Micrococcales>Micrococcaceae>Kocuria | 152 | D8M38_RS00605 | 17459..17917 | 2649579 | MULTISPECIES:-DNA-mismatch-endonuclease-Vsr-[unclassified-Kocuria] |
| WP_257744335.1 | REase | GCF_024655785.1 | Kocuria_rhizophila | Bacteria>Bacillati>Actinomycetota>Actinomycetes>Micrococcales>Micrococcaceae>Kocuria | 139 | NUV30_RS04795 | 27777..28196 | 72000 | very-short-patch-repair-endonuclease-[Kocuria-rhizophila] |
| WP_259919194.1 | REase | - | Kocuria_rhizophila | Bacteria>Bacillati>Actinomycetota>Actinomycetes>Micrococcales>Micrococcaceae>Kocuria | 139 | JNE43_RS06555 | 11977..12396 | 72000 | very-short-patch-repair-endonuclease-[Kocuria-rhizophila] |
| WP_234448718.1 | REase | GCF_016620465.1 | Kocuria_rhizophila | Bacteria>Bacillati>Actinomycetota>Actinomycetes>Micrococcales>Micrococcaceae>Kocuria | 97 | JHD97_RS04245 | 35454..35747 | 72000 | very-short-patch-repair-endonuclease-[Kocuria-rhizophila] |
| WP_231559476.1 | REase | GCF_000800955.1 | Kocuria_sp._ZOR0020 | Bacteria>Bacillati>Actinomycetota>Actinomycetes>Micrococcales>Micrococcaceae>Kocuria>unclassified_Kocuria | 155 | L979_RS08855 | 15245..15712 | 1339234 | very-short-patch-repair-endonuclease-[Kocuria-sp.-ZOR0020] |
| MCV7527921.1 | REase | GCA_025149725.2 | Micrococcus_luteus | Bacteria>Bacillati>Actinomycetota>Actinomycetes>Micrococcales>Micrococcaceae>Micrococcus | 146 | M3D91_010320 | 1440..1880 | 1270 | very-short-patch-repair-endonuclease-[Micrococcus-luteus] |
| WP_082067564.1 | REase | GCF_002276745.1 | Micrococcus_luteus | Bacteria>Bacillati>Actinomycetota>Actinomycetes>Micrococcales>Micrococcaceae>Micrococcus | 146 | B8W89_RS05720 | 40254..40694 | 1270 | MULTISPECIES:-very-short-patch-repair-endonuclease-[Micrococcus] |
| WP_119217963.1 | REase | - | Micrococcus_luteus | Bacteria>Bacillati>Actinomycetota>Actinomycetes>Micrococcales>Micrococcaceae>Micrococcus | 140 | C0205_RS03330 | 712131..712553 | 1270 | very-short-patch-repair-endonuclease-[Micrococcus-luteus] |
| MCV7624009.1 | REase | GCA_025145605.2 | Micrococcus_luteus | Bacteria>Bacillati>Actinomycetota>Actinomycetes>Micrococcales>Micrococcaceae>Micrococcus | 132 | M3B28_003220 | 28032..28430 | 1270 | very-short-patch-repair-endonuclease-[Micrococcus-luteus] |
| MCV7506297.1 | REase | GCA_025151425.2 | Micrococcus_luteus | Bacteria>Bacillati>Actinomycetota>Actinomycetes>Micrococcales>Micrococcaceae>Micrococcus | 126 | M3D82_001045 | 210022..210402 | 1270 | very-short-patch-repair-endonuclease-[Micrococcus-luteus] |
| WP_085387172.1 | REase | GCF_002105775.1 | Cellulosimicrobium_sp._TH-20 | Bacteria>Bacillati>Actinomycetota>Actinomycetes>Micrococcales>Promicromonosporaceae>Cellulosimicrobium>unclassified_Cellulosimicrobium | 155 | B8281_RS05550 | 1254171..1254638 | 1980001 | hypothetical-protein-[Cellulosimicrobium-sp.-TH-20] |
| WP_085387169.1 | REase | GCF_002105775.1 | Cellulosimicrobium_sp._TH-20 | Bacteria>Bacillati>Actinomycetota>Actinomycetes>Micrococcales>Promicromonosporaceae>Cellulosimicrobium>unclassified_Cellulosimicrobium | 207 | B8281_RS05535 | 1250097..1250720 | 1980001 | restriction-endonuclease-[Cellulosimicrobium-sp.-TH-20] |
| WP_239130011.1 | REase | GCF_039536125.1 | Actinoplanes_nipponensis | Bacteria>Bacillati>Actinomycetota>Actinomycetes>Micromonosporales>Micromonosporaceae>Actinoplanes | 164 | Ani05nite_RS15590 | 106320..106814 | 135950 | very-short-patch-repair-endonuclease-[Actinoplanes-nipponensis] |
| WP_023560520.1 | REase | GCF_000494755.1 | Actinoplanes_friuliensis | Bacteria>Bacillati>Actinomycetota>Actinomycetes>Micromonosporales>Micromonosporaceae>Actinoplanes | 218 | AFR_RS29730 | 6451487..6452143 | 196914 | hypothetical-protein-[Actinoplanes-friuliensis] |
| WP_208468629.1 | REase | GCF_017592555.1 | Actinoplanes_flavus | Bacteria>Bacillati>Actinomycetota>Actinomycetes>Micromonosporales>Micromonosporaceae>Actinoplanes | 134 | J5X75_RS18405 | 126236..126640 | 2820290 | hypothetical-protein-[Actinoplanes-flavus] |
| WP_203769003.1 | REase | GCF_039536125.1 | Actinoplanes_nipponensis | Bacteria>Bacillati>Actinomycetota>Actinomycetes>Micromonosporales>Micromonosporaceae>Actinoplanes | 173 | Ani05nite_RS15645 | 125958..126479 | 135950 | hypothetical-protein-[Actinoplanes-nipponensis] |
| WP_259859985.1 | REase | GCF_039533025.1 | Dactylosporangium_fulvum | Bacteria>Bacillati>Actinomycetota>Actinomycetes>Micromonosporales>Micromonosporaceae>Dactylosporangium | 167 | Dfulv_RS45325 | 9852128..9852631 | 53359 | DUF4365-domain-containing-protein-[Dactylosporangium-fulvum] |
| WP_259859981.1 | REase | GCF_039533025.1 | Dactylosporangium_fulvum | Bacteria>Bacillati>Actinomycetota>Actinomycetes>Micromonosporales>Micromonosporaceae>Dactylosporangium | 215 | Dfulv_RS45305 | 9846530..9847177 | 53359 | hypothetical-protein-[Dactylosporangium-fulvum] |
| WP_259859980.1 | REase | GCF_039533025.1 | Dactylosporangium_fulvum | Bacteria>Bacillati>Actinomycetota>Actinomycetes>Micromonosporales>Micromonosporaceae>Dactylosporangium | 192 | Dfulv_RS45300 | 9845880..9846458 | 53359 | hypothetical-protein-[Dactylosporangium-fulvum] |
| WP_203846756.1 | REase | GCF_039534735.1 | Dactylosporangium_siamense | Bacteria>Bacillati>Actinomycetota>Actinomycetes>Micromonosporales>Micromonosporaceae>Dactylosporangium | 101 | Dsi01nite_RS15105 | 107708..108013 | 685454 | Uma2-family-endonuclease-[Dactylosporangium-siamense] |
| GAA0739632.1 | REase | GCA_039521105.1 | Dactylosporangium_roseum | Bacteria>Bacillati>Actinomycetota>Actinomycetes>Micromonosporales>Micromonosporaceae>Dactylosporangium | 212 | GCM10010199_63800 | 59969..60607 | 47989 | hypothetical-protein-GCM10010199_63800-[Dactylosporangium-roseum] |
| WP_088999524.1 | REase | GCF_900091445.1 | Micromonospora_echinofusca | Bacteria>Bacillati>Actinomycetota>Actinomycetes>Micromonosporales>Micromonosporaceae>Micromonospora | 190 | GA0070610_RS08540 | 1868255..1868827 | 47858 | hypothetical-protein-[Micromonospora-echinofusca] |
| PWU48916.1 | REase | GCA_003176735.1 | Micromonospora_globispora | Bacteria>Bacillati>Actinomycetota>Actinomycetes>Micromonosporales>Micromonosporaceae>Micromonospora | 184 | DLJ46_10765 | 27482..28036 | 1450148 | hypothetical-protein-DLJ46_10765-[Micromonospora-globispora] |
| WP_154942601.1 | REase | GCF_007829925.1 | Micromonospora_palomenae | Bacteria>Bacillati>Actinomycetota>Actinomycetes>Micromonosporales>Micromonosporaceae>Micromonospora | 149 | FHX75_RS24180 | 693865..694314 | 1461247 | hypothetical-protein-[Micromonospora-palomenae] |
| WP_219849140.1 | REase | GCF_014041935.1 | Gordonia_jinghuaiqii | Bacteria>Bacillati>Actinomycetota>Actinomycetes>Mycobacteriales>Gordoniaceae>Gordonia | 150 | H1R19_RS12165 | 2739008..2739460 | 2758710 | very-short-patch-repair-endonuclease-[Gordonia-jinghuaiqii] |
| QHN29211.1 | REase | GCA_025268675.1 | Gordonia_amarae | Bacteria>Bacillati>Actinomycetota>Actinomycetes>Mycobacteriales>Gordoniaceae>Gordonia | 148 | GII32_01245 | 298954..299400 | 36821 | DNA-mismatch-endonuclease-Vsr-[Gordonia-amarae] |
| WP_085950227.1 | REase | GCF_000247715.1 | Gordonia_polyisoprenivorans | Bacteria>Bacillati>Actinomycetota>Actinomycetes>Mycobacteriales>Gordoniaceae>Gordonia | 140 | GPOL_RS26190 | 203557..203979 | 84595 | very-short-patch-repair-endonuclease-[Gordonia-polyisoprenivorans] |
| WP_244427418.1 | REase | GCF_000241345.1 | Gordonia_amarae | Bacteria>Bacillati>Actinomycetota>Actinomycetes>Mycobacteriales>Gordoniaceae>Gordonia | 130 | GII32_RS01250 | 298954..299346 | 36821 | DNA-mismatch-endonuclease-Vsr-[Gordonia-amarae] |
| QHN32993.1 | REase | GCA_025268675.1 | Gordonia_amarae | Bacteria>Bacillati>Actinomycetota>Actinomycetes>Mycobacteriales>Gordoniaceae>Gordonia | 245 | GII32_01210 | 285077..285814 | 36821 | PD-(D/E)XK-motif-protein-[Gordonia-amarae] |
| SBS79262.1 | REase | GCA_900078375.1 | uncultured_Mycobacterium_sp. | Bacteria>Bacillati>Actinomycetota>Actinomycetes>Mycobacteriales>Mycobacteriaceae>Mycobacterium>environmental_samples | 67 | MHPYR_700002 | 2465..2668 | 171292 | hypothetical-protein-MHPYR_700002-[uncultured-Mycobacterium-sp.] |
| WP_023870226.1 | REase | GCF_000829075.1 | Mycobacterium_avium | Bacteria>Bacillati>Actinomycetota>Actinomycetes>Mycobacteriales>Mycobacteriaceae>Mycobacterium>Mycobacterium_avium_complex_(MAC) | 159 | B9C26_RS25100 | 6587..7066 | 1764 | very-short-patch-repair-endonuclease-[Mycobacterium-avium] |
| ATO65159.1 | REase | GCA_002716905.4 | Mycobacterium_avium_subsp._hominissuis | Bacteria>Bacillati>Actinomycetota>Actinomycetes>Mycobacteriales>Mycobacteriaceae>Mycobacterium>Mycobacterium_avium_complex_(MAC)>Mycobacterium_avium | 141 | BEP52_09340 | 1954935..1955360 | 439334 | very-short-patch-repair-endonuclease-[Mycobacterium-avium-subsp.-hominissuis] |
| WP_286159384.1 | REase | GCF_001665535.1 | Mycobacterium_sp._852002-51961_SCH5331710 | Bacteria>Bacillati>Actinomycetota>Actinomycetes>Mycobacteriales>Mycobacteriaceae>Mycobacterium>unclassified_Mycobacterium | 159 | A5752_RS27055 | 23658..24137 | 1834105 | very-short-patch-repair-endonuclease-[Mycobacterium-sp.-852002-51961_SCH5331710] |
| WP_243841399.1 | REase | GCF_011694515.1 | Mycobacterium_sp._DL592 | Bacteria>Bacillati>Actinomycetota>Actinomycetes>Mycobacteriales>Mycobacteriaceae>Mycobacterium>unclassified_Mycobacterium | 156 | HBE64_RS20610 | 4285508..4285978 | 2675524 | very-short-patch-repair-endonuclease-[Mycobacterium-sp.-DL592] |
| WP_125079868.1 | REase | GCF_900604305.1 | Mycobacterium_sp._P7213 | Bacteria>Bacillati>Actinomycetota>Actinomycetes>Mycobacteriales>Mycobacteriaceae>Mycobacterium>unclassified_Mycobacterium | 177 | EJK91_RS20965 | 2810..3343 | 2478465 | hypothetical-protein-[Mycobacterium-sp.-P7213] |
| WP_233213235.1 | REase | GCF_002968335.2 | Mycobacterium_sp._ITM-2016-00316 | Bacteria>Bacillati>Actinomycetota>Actinomycetes>Mycobacteriales>Mycobacteriaceae>Mycobacterium>unclassified_Mycobacterium | 232 | C6A86_RS35105 | 50273..50971 | 2099695 | hypothetical-protein-[Mycobacterium-sp.-ITM-2016-00316] |
| WP_233213236.1 | REase | GCF_002968335.2 | Mycobacterium_sp._ITM-2016-00316 | Bacteria>Bacillati>Actinomycetota>Actinomycetes>Mycobacteriales>Mycobacteriaceae>Mycobacterium>unclassified_Mycobacterium | 228 | C6A86_RS28990 | 49493..50179 | 2099695 | DUF4268-domain-containing-protein-[Mycobacterium-sp.-ITM-2016-00316] |
| OBB45455.1 | REase | GCA_001665535.1 | Mycobacterium_sp._852002-51961_SCH5331710 | Bacteria>Bacillati>Actinomycetota>Actinomycetes>Mycobacteriales>Mycobacteriaceae>Mycobacterium>unclassified_Mycobacterium | 243 | A5752_02010 | 14751..15482 | 1834105 | hypothetical-protein-A5752_02010-[Mycobacterium-sp.-852002-51961_SCH5331710] |
| WP_162113232.1 | REase | GCF_010093495.1 | Mycolicibacter_kumamotonensis | Bacteria>Bacillati>Actinomycetota>Actinomycetes>Mycobacteriales>Mycobacteriaceae>Mycolicibacter | 184 | GWR20_RS22075 | 2676..3230 | 354243 | hypothetical-protein-[Mycolicibacter-kumamotonensis] |
| WP_049792839.1 | REase | GCF_008373105.1 | Mycolicibacter_arupensis | Bacteria>Bacillati>Actinomycetota>Actinomycetes>Mycobacteriales>Mycobacteriaceae>Mycolicibacter | 152 | F0402_RS09275 | 12297..12755 | 342002 | hypothetical-protein-[Mycolicibacter-arupensis] |
| WP_131813852.1 | REase | GCF_001679965.1 | Mycolicibacter_kumamotonensis | Bacteria>Bacillati>Actinomycetota>Actinomycetes>Mycobacteriales>Mycobacteriaceae>Mycolicibacter | 107 | ACT18_RS17290 | 32363..32686 | 354243 | hypothetical-protein-[Mycolicibacter-kumamotonensis] |
| WP_220096106.1 | REase | GCF_000214155.1 | Mycolicibacter_sinensis | Bacteria>Bacillati>Actinomycetota>Actinomycetes>Mycobacteriales>Mycobacteriaceae>Mycolicibacter | 107 | JDM601_RS21750 | 801940..802263 | 875328 | hypothetical-protein-[Mycolicibacter-sinensis] |
| WP_276127105.1 | REase | GCF_029270025.1 | Mycolicibacterium_septicum | Bacteria>Bacillati>Actinomycetota>Actinomycetes>Mycobacteriales>Mycobacteriaceae>Mycolicibacterium | 159 | P3H80_RS10745 | 390954..391433 | 98668 | very-short-patch-repair-endonuclease-[Mycolicibacterium-septicum] |
| WP_141142678.1 | REase | GCF_013138515.1 | Mycolicibacterium_fortuitum | Bacteria>Bacillati>Actinomycetota>Actinomycetes>Mycobacteriales>Mycobacteriaceae>Mycolicibacterium | 146 | FKW78_RS12950 | 13119..13559 | 1766 | very-short-patch-repair-endonuclease-[Mycolicibacterium-fortuitum] |
| UHJ58261.1 | REase | GCA_021249325.1 | Mycolicibacterium_fortuitum | Bacteria>Bacillati>Actinomycetota>Actinomycetes>Mycobacteriales>Mycobacteriaceae>Mycolicibacterium | 127 | LT337_13400 | 2627188..2627571 | 1766 | very-short-patch-repair-endonuclease-[Mycolicibacterium-fortuitum] |
| SDF04709.1 | REase | GCA_900101555.1 | Mycolicibacterium_neoaurum | Bacteria>Bacillati>Actinomycetota>Actinomycetes>Mycobacteriales>Mycobacteriaceae>Mycolicibacterium | 233 | SAMN04488581_5264 | 149934..150635 | 1795 | hypothetical-protein-SAMN04488581_5264-[Mycolicibacterium-neoaurum] |
| WP_234797949.1 | REase | GCF_900101555.1 | Mycolicibacterium_neoaurum | Bacteria>Bacillati>Actinomycetota>Actinomycetes>Mycobacteriales>Mycobacteriaceae>Mycolicibacterium | 233 | BLS83_RS27630 | 149934..150635 | 1795 | hypothetical-protein-[Mycolicibacterium-neoaurum] |
| WP_193043848.1 | REase | GCF_016313185.1 | Mycolicibacterium_baixiangningiae | Bacteria>Bacillati>Actinomycetota>Actinomycetes>Mycobacteriales>Mycobacteriaceae>Mycolicibacterium | 190 | I7X18_RS20545 | 4343825..4344397 | 2761578 | hypothetical-protein-[Mycolicibacterium-baixiangningiae] |
| MBU8833313.1 | REase | GCA_018524295.1 | Mycolicibacterium_goodii | Bacteria>Bacillati>Actinomycetota>Actinomycetes>Mycobacteriales>Mycobacteriaceae>Mycolicibacterium | 223 | KL858_28160 | 15993..16664 | 134601 | hypothetical-protein-[Mycolicibacterium-goodii] |
| WP_281063407.1 | REase | GCF_022691685.1 | Prescottella_equi | Bacteria>Bacillati>Actinomycetota>Actinomycetes>Mycobacteriales>Nocardiaceae>Prescottella | 142 | MPC38_RS08800 | 1850315..1850743 | 43767 | very-short-patch-repair-endonuclease-[Prescottella-equi] |
| MBY6363245.1 | REase | GCA_019834505.1 | Rhodococcoides_corynebacterioides | Bacteria>Bacillati>Actinomycetota>Actinomycetes>Mycobacteriales>Nocardiaceae>Rhodococcoides | 121 | HQ609_08990 | 105803..106168 | 53972 | hypothetical-protein-[Rhodococcus-corynebacterioides] |
| WP_269778340.1 | REase | GCF_000239135.1 | Rhodococcus_rhodochrous | Bacteria>Bacillati>Actinomycetota>Actinomycetes>Mycobacteriales>Nocardiaceae>Rhodococcus | 134 | RHODODRAFT_RS0223980 | 1455..1859 | 1829 | very-short-patch-repair-endonuclease-[Rhodococcus-rhodochrous] |
| WP_149357519.1 | REase | - | Rhodococcus_qingshengii | Bacteria>Bacillati>Actinomycetota>Actinomycetes>Mycobacteriales>Nocardiaceae>Rhodococcus>Rhodococcus_erythropolis_group | 200 | D6M20_RS01765 | 370993..371595 | 334542 | hypothetical-protein-[Rhodococcus-qingshengii] |
| WP_172936836.1 | REase | GCF_013277795.1 | Rhodococcus_sp._MS16 | Bacteria>Bacillati>Actinomycetota>Actinomycetes>Mycobacteriales>Nocardiaceae>Rhodococcus>unclassified_Rhodococcus_(in:_high_G+C_Gram-positive_bacteria) | 240 | FEZ60_RS20020 | 136898..137620 | 2579941 | PD-(D/E)XK-motif-protein-[Rhodococcus-sp.-MS16] |
| WP_232416934.1 | REase | GCF_021228355.1 | Aeromicrobium_duanguangcaii | Bacteria>Bacillati>Actinomycetota>Actinomycetes>Propionibacteriales>Nocardioidaceae>Aeromicrobium | 126 | NP095_RS05675 | 1117234..1117614 | 2968086 | MULTISPECIES:-very-short-patch-repair-endonuclease-[unclassified-Aeromicrobium] |
| WP_179517473.1 | REase | GCF_013409425.1 | Nocardioides_perillae | Bacteria>Bacillati>Actinomycetota>Actinomycetes>Propionibacteriales>Nocardioidaceae>Nocardioides | 150 | BJ989_RS06335 | 1348551..1349003 | 1119534 | DNA-mismatch-endonuclease-Vsr-[Nocardioides-perillae] |
| WP_240770227.1 | REase | GCF_004919085.1 | Nocardioides_sp._GY_10127 | Bacteria>Bacillati>Actinomycetota>Actinomycetes>Propionibacteriales>Nocardioidaceae>Nocardioides>unclassified_Nocardioides | 162 | E8D37_RS06685 | 699208..699696 | 2569762 | very-short-patch-repair-endonuclease-[Nocardioides-sp.-GY-10127] |
| WP_078920355.1 | REase | GCF_002022225.1 | Tessaracoccus | Bacteria>Bacillati>Actinomycetota>Actinomycetes>Propionibacteriales>Propionibacteriaceae | 138 | BKM78_RS14500 | 3150360..3150776 | 72763 | MULTISPECIES:-very-short-patch-repair-endonuclease-[Tessaracoccus] |
| WP_028701424.1 | REase | GCF_001975205.1 | Acidipropionibacterium_acidipropionici | Bacteria>Bacillati>Actinomycetota>Actinomycetes>Propionibacteriales>Propionibacteriaceae>Acidipropionibacterium | 127 | ASQ49_RS03465 | 781181..781564 | 1748 | very-short-patch-repair-endonuclease-[Acidipropionibacterium-acidipropionici] |
| WP_255455490.1 | REase | GCF_009707365.1 | Microlunatus_sp._Gsoil_973 | Bacteria>Bacillati>Actinomycetota>Actinomycetes>Propionibacteriales>Propionibacteriaceae>Microlunatus>unclassified_Microlunatus | 139 | GJV80_RS18455 | 3923676..3924095 | 2672569 | very-short-patch-repair-endonuclease-[Microlunatus-sp.-Gsoil-973] |
| NLI84507.1 | REase | GCA_012517405.1 | Propionibacterium_sp. | Bacteria>Bacillati>Actinomycetota>Actinomycetes>Propionibacteriales>Propionibacteriaceae>Propionibacterium>unclassified_Propionibacterium | 188 | GX440_03715 | 103450..104016 | 1977903 | hypothetical-protein-[Propionibacterium-sp.] |
| WP_182766933.1 | REase | GCF_014164735.1 | Tessaracoccus_sp._MC1756 | Bacteria>Bacillati>Actinomycetota>Actinomycetes>Propionibacteriales>Propionibacteriaceae>Tessaracoccus>unclassified_Tessaracoccus | 188 | H5393_RS12490 | 215324..215890 | 2760311 | very-short-patch-repair-endonuclease-[Tessaracoccus-sp.-MC1756] |
| WP_236808856.1 | REase | - | Amycolatopsis_albispora | Bacteria>Bacillati>Actinomycetota>Actinomycetes>Pseudonocardiales>Pseudonocardiaceae>Amycolatopsis | 116 | A4R43_RS09295 | 1993392..1993742 | 1804986 | very-short-patch-repair-endonuclease-[Amycolatopsis-albispora] |

| Protein-ID | Domain | Assembly-ID | Strain | Taxonomy | Length | Locus-Tag | Genomic-coordinates | TaxID | Fasta-defline |
|---|---|---|---|---|---|---|---|---|---|
| WP_158005352.1 | REase | GCF_900105055.1 | Amycolatopsis_lurida | Bacteria>Bacillati>Actinomycetota>Actinomycetes>Pseudonocardiales>Pseudonocardiaceae>Amycolatopsis | 187 | BLW75_RS07175 | 415362..415925 | 31959 | sce7726-family-protein-[Amycolatopsis-lurida] |
| WP_211361745.1 | REase | GCF_006717045.1 | Pseudonocardia_cypriaca | Bacteria>Bacillati>Actinomycetota>Actinomycetes>Pseudonocardiales>Pseudonocardiaceae>Pseudonocardia | 73 | FB388_RS39780 | 740326..740547 | 882449 | DUF4365-domain-containing-protein-[Pseudonocardia-cypriaca] |
| HUY47912.1 | REase | GCA_035531875.1 | Streptosporangiaceae_bacterium | Bacteria>Bacillati>Actinomycetota>Actinomycetes>Streptosporangiales>Streptosporangiaceae>unclassified_Streptosporangiaceae | 212 | VMV92_19645 | 37282..37920 | 2339230 | Uma2-family-endonuclease-[Streptosporangiaceae-bacterium] |
| EAR26281.1 | REase | GCA_000153145.1 | marine_actinobacterium_PHSC20C1 | Bacteria>Bacillati>Actinomycetota>Actinomycetes>unclassified_Actinomycetes | 118 | A20C1_10379 | 2123075..2123431 | 312284 | hypothetical-protein-A20C1_10379-[marine-actinobacterium-PHSC20C1] |
| EAR26280.1 | REase | GCA_000153145.1 | marine_actinobacterium_PHSC20C1 | Bacteria>Bacillati>Actinomycetota>Actinomycetes>unclassified_Actinomycetes | 248 | A20C1_10374 | 2122296..2123042 | 312284 | hypothetical-protein-A20C1_10374-[marine-actinobacterium-PHSC20C1] |
| PKQ27860.1 | REase | GCA_002843005.1 | Candidatus_Anoxymicrobium_japonicum | Bacteria>Bacillati>Actinomycetota>Candidatus_Geothermincolia>Candidatus_Geothermincolales>Candidatus_Anoxymicrobiaceae>Candidatus_Anoxymicrobium | 69 | CVT63_05745 | 95..304 | 2013648 | hypothetical-protein-CVT63_05745-[Candidatus-Anoxymicrobium-japonicum] |
| WP_090862517.1 | REase | GCF_052121955.1 | Atopobiaceae | Bacteria>Bacillati>Actinomycetota>Coriobacteriia>Coriobacteriales | 129 | FYJ68_RS09130 | 22876..23265 | 1643824 | MULTISPECIES:-hypothetical-protein-[Atopobiaceae] |
| RRF95313.1 | REase | GCA_003862165.1 | Coriobacteriaceae_bacterium | Bacteria>Bacillati>Actinomycetota>Coriobacteriia>Coriobacteriales>Coriobacteriaceae>unclassified_Coriobacteriaceae | 129 | DUD31_01690 | 16473..16862 | 2011094 | hypothetical-protein-DUD31_01690-[Coriobacteriaceae-bacterium] |
| GJM55554.1 | REase | GCA_022834715.1 | Granulimonas_faecalis | Bacteria>Bacillati>Actinomycetota>Coriobacteriia>Coriobacteriales>Kribbibacteriaceae>Granulimonas | 225 | ATOP_12090 | 1352361..1353038 | 2894155 | hypothetical-protein-ATOP_12090-[Granulimonas-faecalis] |
| WP_265590829.1 | REase | GCF_022834715.1 | Granulimonas_faecalis | Bacteria>Bacillati>Actinomycetota>Coriobacteriia>Coriobacteriales>Kribbibacteriaceae>Granulimonas | 206 | OR600_RS06175 | 1352361..1352981 | 2894155 | hypothetical-protein-[Granulimonas-faecalis] |
| WP_251231667.1 | REase | GCF_943193145.1 | Adlercreutzia_aquisgranensis | Bacteria>Bacillati>Actinomycetota>Coriobacteriia>Eggerthellales>Eggerthellaceae>Adlercreutzia | 180 | NFR41_RS09560 | 16642..17184 | 2941323 | DUF4365-domain-containing-protein-[Adlercreutzia-aquisgranensis] |
| MBA3474042.1 | REase | GCA_013813465.1 | Rubrobacter_sp. | Bacteria>Bacillati>Actinomycetota>Rubrobacteria>Rubrobacterales>Rubrobacteraceae>Rubrobacter>unclassified_Rubrobacter | 189 | HOT57_12590 | 7524..8093 | 2036010 | Uma2-family-endonuclease-[Rubrobacter-sp.] |
| NCD16054.1 | REase | GCA_009930385.1 | Actinomycetota_bacterium | Bacteria>Bacillati>Actinomycetota>unclassified_Actinomycetota | 162 | EOL91_01840 | 14155..14643 | 2900548 | DNA-mismatch-endonuclease-Vsr-[Actinomycetia-bacterium] |
| MBU2074165.1 | REase | GCA_018826065.1 | Actinomycetota_bacterium | Bacteria>Bacillati>Actinomycetota>unclassified_Actinomycetota | 127 | KJ938_07000 | 15202..15585 | 2900548 | very-short-patch-repair-endonuclease-[Actinomycetia-bacterium] |
| MCX6403859.1 | REase | GCA_026392535.1 | Actinomycetota_bacterium | Bacteria>Bacillati>Actinomycetota>unclassified_Actinomycetota | 241 | NT032_08215 | 1..726 | 2900548 | MAG:-DUF3883-domain-containing-protein,-partial-[Actinobacteria-bacterium] |
| NIM06026.1 | REase | GCA_011771275.1 | Armatimonadota_bacterium | Bacteria>Bacillati>Armatimonadota>unclassified_Armatimonadota | 193 | GT078_10550 | 3784..4365 | 2033014 | hypothetical-protein-[Armatimonadota-bacterium] |
| MCL6449135.1 | REase | GCA_023511535.1 | Armatimonadota_bacterium | Bacteria>Bacillati>Armatimonadota>unclassified_Armatimonadota | 187 | K6U04_13480 | 7442..8005 | 2033014 | Uma2-family-endonuclease-[Armatimonadota-bacterium] |
| MCX6379946.1 | REase | GCA_026392135.1 | Armatimonadota_bacterium | Bacteria>Bacillati>Armatimonadota>unclassified_Armatimonadota | 121 | NT023_10800 | 3044..3409 | 2033014 | MAG:-DUF559-domain-containing-protein-[Armatimonadetes-bacterium] |
| WP_088074343.1 | REase | GCF_019042735.1 | Bacillaceae | Bacteria>Bacillati>Bacillota>Bacilli>Bacillales | 201 | KS407_RS09165 | 16564..17169 | 186817 | MULTISPECIES:-competence-protein-CoiA-family-protein-[Bacillaceae] |
| WP_163151121.1 | REase | GCF_050613275.1 | Anoxybacillus | Bacteria>Bacillati>Bacillota>Bacilli>Bacillales>Anoxybacillaceae | 162 | HPK01_RS05380 | 40981..41469 | 150247 | MULTISPECIES:-hypothetical-protein-[Anoxybacillus] |
| WP_016125572.1 | REase | GCF_001635915.1 | Bacillus | Bacteria>Bacillati>Bacillota>Bacilli>Bacillales>Bacillaceae | 182 | MR162_RS10395 | 187007..187555 | 1386 | MULTISPECIES:-restriction-endonuclease-[Bacillus] |
| WP_051556526.1 | REase | GCF_000621445.1 | Alkalihalobacterium_bogoriense | Bacteria>Bacillati>Bacillota>Bacilli>Bacillales>Bacillaceae>Alkalihalobacterium | 171 | T323_RS26710 | 36055..36570 | 246272 | hypothetical-protein-[Alkalihalobacillus-bogoriensis] |
| WP_048566955.1 | REase | GCF_016483555.1 | Bacillus_cereus_group | Bacteria>Bacillati>Bacillota>Bacilli>Bacillales>Bacillaceae>Bacillus | 182 | JDS82_RS26160 | 5987..6535 | 86661 | MULTISPECIES:-restriction-endonuclease-[Bacillus-cereus-group] |
| WP_163243261.1 | REase | GCF_019915265.1 | Bacillus_aquiflavi | Bacteria>Bacillati>Bacillota>Bacilli>Bacillales>Bacillaceae>Bacillus | 102 | K6959_RS16315 | 3367364..3367672 | 2672567 | hypothetical-protein-[Bacillus-aquiflavi] |
| WP_062353361.1 | REase | GCF_001375515.1 | Bacillus_kwashiorkori | Bacteria>Bacillati>Bacillota>Bacilli>Bacillales>Bacillaceae>Bacillus | 94 | BN1242_RS13375 | 36234..36518 | 1522318 | hypothetical-protein-[Bacillus-kwashiorkori] |
| WP_062353364.1 | REase | GCF_001375515.1 | Bacillus_kwashiorkori | Bacteria>Bacillati>Bacillota>Bacilli>Bacillales>Bacillaceae>Bacillus | 159 | BN1242_RS13380 | 36558..37037 | 1522318 | DUF4263-domain-containing-protein-[Bacillus-kwashiorkori] |
| WP_136035701.1 | REase | GCF_003581585.1 | Bacillus_salacetis | Bacteria>Bacillati>Bacillota>Bacilli>Bacillales>Bacillaceae>Bacillus | 234 | D3H55_RS12375 | 1..705 | 2315464 | PD-(D/E)XK-nuclease-family-protein,-partial-[Bacillus-salacetis] |
| PGR68863.1 | REase | GCA_002576445.1 | Bacillus_cereus | Bacteria>Bacillati>Bacillota>Bacilli>Bacillales>Bacillaceae>Bacillus>Bacillus_cereus_group | 182 | COC49_05360 | 49833..50381 | 1396 | restriction-endonuclease-[Bacillus-cereus] |
| PRT22685.1 | REase | GCA_002998815.1 | Bacillus_thuringiensis | Bacteria>Bacillati>Bacillota>Bacilli>Bacillales>Bacillaceae>Bacillus>Bacillus_cereus_group | 182 | C6351_31275 | 5399..5947 | 1428 | restriction-endonuclease-[Bacillus-thuringiensis] |
| WP_098613374.1 | REase | GCF_002567495.1 | Bacillus_cereus | Bacteria>Bacillati>Bacillota>Bacilli>Bacillales>Bacillaceae>Bacillus>Bacillus_cereus_group | 182 | COK05_RS16090 | 72185..72733 | 1396 | restriction-endonuclease-[Bacillus-cereus] |
| WP_098878452.1 | REase | GCF_002576445.1 | Bacillus_cereus | Bacteria>Bacillati>Bacillota>Bacilli>Bacillales>Bacillaceae>Bacillus>Bacillus_cereus_group | 182 | COC49_RS05360 | 49833..50381 | 1396 | restriction-endonuclease-[Bacillus-cereus] |
| WP_236787486.1 | REase | GCF_021654875.1 | Bacillus_cereus | Bacteria>Bacillati>Bacillota>Bacilli>Bacillales>Bacillaceae>Bacillus>Bacillus_cereus_group | 193 | L6489_RS02830 | 536230..536811 | 1396 | hypothetical-protein-[Bacillus-cereus] |
| WP_242225829.1 | REase | GCF_022663655.1 | Bacillus_cereus_group_sp._BfR-BA-01358 | Bacteria>Bacillati>Bacillota>Bacilli>Bacillales>Bacillaceae>Bacillus>Bacillus_cereus_group>unclassified_Bacillus_cereus_group | 182 | MR078_RS05920 | 49100..49648 | 2920320 | restriction-endonuclease-[Bacillus-cereus-group-sp.-BfR-BA-01358] |
| WP_242261896.1 | REase | GCF_022662915.1 | Bacillus_cereus_group_sp._BfR-BA-01453 | Bacteria>Bacillati>Bacillota>Bacilli>Bacillales>Bacillaceae>Bacillus>Bacillus_cereus_group>unclassified_Bacillus_cereus_group | 182 | MR070_RS20040 | 135522..136070 | 2920355 | restriction-endonuclease-[Bacillus-cereus-group-sp.-BfR-BA-01453] |
| WP_229142418.1 | REase | GCF_020731355.1 | Bacillus_sp._REN16 | Bacteria>Bacillati>Bacillota>Bacilli>Bacillales>Bacillaceae>Bacillus>unclassified_Bacillus_(in:_firmicutes) | 186 | LKW28_RS21240 | 14413..14973 | 2887296 | hypothetical-protein-[Bacillus-sp.-REN16] |
| WP_210367876.1 | REase | GCF_017912555.1 | Bacillus_sp._REN3 | Bacteria>Bacillati>Bacillota>Bacilli>Bacillales>Bacillaceae>Bacillus>unclassified_Bacillus_(in:_firmicutes) | 130 | JJQ88_RS20370 | 17213..17605 | 2802440 | hypothetical-protein-[Bacillus-sp.-REN3] |
| WP_235715383.1 | REase | GCF_000513095.1 | Halalkalibacter_wakoensis | Bacteria>Bacillati>Bacillota>Bacilli>Bacillales>Bacillaceae>Halalkalibacter | 223 | JCM9140_RS24075 | 21171..21842 | 127891 | hypothetical-protein-[Halalkalibacter-wakoensis] |
| WP_213108267.1 | REase | GCF_018343555.1 | Lederbergia_citrea | Bacteria>Bacillati>Bacillota>Bacilli>Bacillales>Bacillaceae>Lederbergia | 201 | KHA89_RS14820 | 204701..205306 | 2833581 | hypothetical-protein-[Bacillus-sp.-FJAT-49731] |
| WP_121661849.1 | REase | GCF_003667825.1 | Metabacillus_litoralis | Bacteria>Bacillati>Bacillota>Bacilli>Bacillales>Bacillaceae>Metabacillus | 193 | D9842_RS06685 | 1408822..1409403 | 152268 | hypothetical-protein-[Metabacillus-litoralis] |
| WP_213145787.1 | REase | GCF_018343545.2 | Neobacillus_citreus | Bacteria>Bacillati>Bacillota>Bacilli>Bacillales>Bacillaceae>Neobacillus | 231 | KHB02_RS07275 | 31073..31768 | 2833578 | DNA/RNA-nuclease-SfsA-[Neobacillus-citreus] |
| WP_171510266.1 | REase | GCF_013138395.1 | Niallia_circulans | Bacteria>Bacillati>Bacillota>Bacilli>Bacillales>Bacillaceae>Niallia | 194 | HLK66_RS25100 | 5174489..5175073 | 1397 | hypothetical-protein-[Niallia-circulans] |
| WP_058950071.1 | REase | GCF_001485235.1 | Oceanobacillus_picturae | Bacteria>Bacillati>Bacillota>Bacilli>Bacillales>Bacillaceae>Oceanobacillus | 193 | OPHB3_RS08870 | 14605..15186 | 171693 | hypothetical-protein-[Oceanobacillus-picturae] |
| WP_272767823.1 | REase | GCF_028548275.1 | Priestia_megaterium | Bacteria>Bacillati>Bacillota>Bacilli>Bacillales>Bacillaceae>Priestia | 138 | PNF31_RS27495 | 24123..24539 | 1404 | hypothetical-protein-[Priestia-megaterium] |
| WP_257351438.1 | REase | GCF_024609785.1 | Pseudalkalibacillus_decolorationis | Bacteria>Bacillati>Bacillota>Bacilli>Bacillales>Bacillaceae>Pseudalkalibacillus | 94 | NON16_RS21320 | 24296..24580 | 163879 | hypothetical-protein-[Pseudalkalibacillus-decolorationis] |
| WP_257351437.1 | REase | GCF_024609785.1 | Pseudalkalibacillus_decolorationis | Bacteria>Bacillati>Bacillota>Bacilli>Bacillales>Bacillaceae>Pseudalkalibacillus | 118 | NON16_RS21315 | 23770..24126 | 163879 | DUF4263-domain-containing-protein-[Pseudalkalibacillus-decolorationis] |
| GGJ55451.1 | REase | GCA_014647615.1 | Virgibacillus_kapii | Bacteria>Bacillati>Bacillota>Bacilli>Bacillales>Bacillaceae>Virgibacillus | 193 | GCM10007111_17150 | 610422..611003 | 1638645 | hypothetical-protein-GCM10007111_17150-[Virgibacillus-kapii] |
| WP_088051633.1 | REase | GCF_900155625.1 | Virgibacillus_dakarensis | Bacteria>Bacillati>Bacillota>Bacilli>Bacillales>Bacillaceae>Virgibacillus | 193 | CZ382_RS12950 | 692546..693127 | 1917889 | hypothetical-protein-[Virgibacillus-dakarensis] |
| APC47775.1 | REase | GCA_001878675.1 | Virgibacillus_halodenitrificans | Bacteria>Bacillati>Bacillota>Bacilli>Bacillales>Bacillaceae>Virgibacillus | 150 | BME96_06135 | 1177224..1177676 | 1482 | hypothetical-protein-BME96_06135-[Virgibacillus-halodenitrificans] |
| WP_208650572.1 | REase | GCF_004331915.1 | Ureibacillus | Bacteria>Bacillati>Bacillota>Bacilli>Bacillales>Caryophanaceae | 190 | DKZ56_RS14275 | 2825680..2826252 | 160795 | sce7726-family-protein-[Ureibacillus-thermophilus] |
| WP_125903402.1 | REase | GCF_003944745.1 | Bhargavaea_beijingensis | Bacteria>Bacillati>Bacillota>Bacilli>Bacillales>Caryophanaceae>Bhargavaea | 193 | EJA12_RS02415 | 154611..155192 | 426756 | hypothetical-protein-[Bhargavaea-beijingensis] |
| WP_208859813.1 | REase | GCF_002082015.1 | Sporosarcina_ureae | Bacteria>Bacillati>Bacillota>Bacilli>Bacillales>Caryophanaceae>Sporosarcina | 140 | SporoP17a_RS05175 | 1038655..1039077 | 1571 | hypothetical-protein-[Sporosarcina-ureae] |
| MBO0589403.1 | REase | GCA_017347095.1 | Sporosarcina_sp._E16_8 | Bacteria>Bacillati>Bacillota>Bacilli>Bacillales>Caryophanaceae>Sporosarcina>unclassified_Sporosarcina | 234 | I2484_19085 | 10739..11443 | 2789295 | PD-(D/E)XK-nuclease-family-protein-[Sporosarcina-sp.-E16_8] |
| WP_042184807.1 | REase | GCF_000758545.1 | unclassified_Paenibacillus | Bacteria>Bacillati>Bacillota>Bacilli>Bacillales>Paenibacillaceae>Paenibacillus | 143 | H70737_RS03485 | 781705..782136 | 185978 | hypothetical-protein-[Paenibacillus-sp.-FSL-H7-0737] |
| WP_138789735.1 | REase | GCF_042666815.1 | Paenibacillus_thermoaerophilus | Bacteria>Bacillati>Bacillota>Bacilli>Bacillales>Paenibacillaceae>Paenibacillus | 146 | FE781_RS11325 | 114156..114596 | 1215385 | hypothetical-protein-[Paenibacillus-thermoaerophilus] |
| WP_216364917.1 | REase | GCF_002741035.1 | Paenibacillus_sp._BIHB_4019 | Bacteria>Bacillati>Bacillota>Bacilli>Bacillales>Paenibacillaceae>Paenibacillus>unclassified_Paenibacillus | 174 | BBD42_RS03725 | 857424..857948 | 1870819 | hypothetical-protein-[Paenibacillus-sp.-BIHB-4019] |
| WP_220482002.1 | REase | GCF_003591485.1 | Paenibacillus_sp._1011MAR3C5 | Bacteria>Bacillati>Bacillota>Bacilli>Bacillales>Paenibacillaceae>Paenibacillus>unclassified_Paenibacillus | 222 | D3P07_RS27270 | 194240..194908 | 1675787 | hypothetical-protein-[Paenibacillus-sp.-1011MAR3C5] |
| UUZ84565.1 | REase | GCA_024703795.1 | Paenibacillus_sp._P26 | Bacteria>Bacillati>Bacillota>Bacilli>Bacillales>Paenibacillaceae>Paenibacillus>unclassified_Paenibacillus | 103 | LJK88_13385 | 2156946..2157257 | 2885204 | PD-(D/E)XK-nuclease-family-protein-[Paenibacillus-sp.-P26] |
| WP_051263247.1 | REase | GCF_000430685.1 | Tuberibacillus_calidus | Bacteria>Bacillati>Bacillota>Bacilli>Bacillales>Sporolactobacillaceae>Tuberibacillus | 114 | H532_RS18440 | 20837..21181 | 340097 | Uma2-family-endonuclease-[Tuberibacillus-calidus] |
| WP_068559444.1 | REase | GCF_900067165.1 | Trichococcus | Bacteria>Bacillati>Bacillota>Bacilli>Lactobacillales>Carnobacteriaceae | 193 | BUA33_RS02200 | 209298..209879 | 82802 | MULTISPECIES:-hypothetical-protein-[Trichococcus] |
| MCG0730678.1 | REase | GCA_021893465.1 | Lactiplantibacillus_plantarum | Bacteria>Bacillati>Bacillota>Bacilli>Lactobacillales>Lactobacillaceae>Lactiplantibacillus | 69 | IMAU60171_02933 | 13157..13366 | 1590 | hypothetical-protein-[Lactiplantibacillus-plantarum] |
| WP_260308523.1 | REase | - | Lactobacillus_johnsonii | Bacteria>Bacillati>Bacillota>Bacilli>Lactobacillales>Lactobacillaceae>Lactobacillus | 64 | FC22_RS09490 | 110482..110676 | 33959 | hypothetical-protein-[Lactobacillus-johnsonii] |
| WP_150392672.1 | REase | GCF_008694105.1 | Schleiferilactobacillus_harbinensis | Bacteria>Bacillati>Bacillota>Bacilli>Lactobacillales>Lactobacillaceae>Schleiferilactobacillus | 107 | FMM01_RS11915 | 2401777..2402100 | 304207 | winged-helix-turn-helix-domain-containing-protein-[Schleiferilactobacillus-harbinensis] |
| HHW19259.1 | REase | GCA_012840505.1 | Candidatus_Fermentithermobacillaceae_bacterium | Bacteria>Bacillati>Bacillota>Candidatus_Fermentithermobacillia>Candidatus_Fermentithermobacillales>Candidatus_Fermentithermobacillaceae>unclassified_Candidatus_Fermentithermobacillaceae | 196 | GXX30_10275 | 56936..57526 | 3085339 | hypothetical-protein-[Bacillota-bacterium] |
| WP_129723502.1 | REase | GCF_004138105.1 | Xylanivirga_thermophila | Bacteria>Bacillati>Bacillota>Clostridia>Caldicoprobacterales>Xylanivirgaceae>Xylanivirga | 203 | EJN67_RS06295 | 15629..16240 | 2496273 | hypothetical-protein-[Xylanivirga-thermophila] |
| WP_073009265.1 | REase | GCF_900142075.1 | Clostridium_amylolyticum | Bacteria>Bacillati>Bacillota>Clostridia>Eubacteriales>Clostridiaceae>Clostridium | 200 | BUA82_RS13580 | 38564..39166 | 1121298 | hypothetical-protein-[Clostridium-amylolyticum] |
| WP_078025153.1 | REase | GCF_002008265.1 | Clostridium_tepidum | Bacteria>Bacillati>Bacillota>Clostridia>Eubacteriales>Clostridiaceae>Clostridium | 200 | BS637_RS12510 | 28705..29307 | 1962263 | hypothetical-protein-[Clostridium-tepidum] |
| NFG40384.1 | REase | GCA_011013445.1 | Clostridium_botulinum | Bacteria>Bacillati>Bacillota>Clostridia>Eubacteriales>Clostridiaceae>Clostridium | 246 | FC789_04065 | 894247..894987 | 1491 | hypothetical-protein-[Clostridium-botulinum] |
| NFI95397.1 | REase | GCA_011014625.1 | Clostridium_botulinum | Bacteria>Bacillati>Bacillota>Clostridia>Eubacteriales>Clostridiaceae>Clostridium | 246 | FC961_13525 | 3752..4492 | 1491 | hypothetical-protein-[Clostridium-botulinum] |
| WP_027639918.1 | REase | GCF_012844035.1 | Clostridium_cadaveris | Bacteria>Bacillati>Bacillota>Clostridia>Eubacteriales>Clostridiaceae>Clostridium | 246 | HF846_RS03745 | 121508..122248 | 1529 | competence-protein-CoiA-family-protein-[Clostridium-cadaveris] |
| WP_066893458.1 | REase | GCF_900086595.1 | Clostridium_nigeriense | Bacteria>Bacillati>Bacillota>Clostridia>Eubacteriales>Clostridiaceae>Clostridium | 246 | BN6774_RS10740 | 2256184..2256924 | 1805470 | hypothetical-protein-[Clostridium-nigeriense] |
| WP_178312093.1 | REase | GCF_013390975.1 | Clostridium_cadaveris | Bacteria>Bacillati>Bacillota>Clostridia>Eubacteriales>Clostridiaceae>Clostridium | 246 | HXZ52_RS14465 | 5607..6347 | 1529 | competence-protein-CoiA-family-protein-[Clostridium-cadaveris] |
| MBN1067558.1 | REase | GCA_016925955.1 | Clostridium_botulinum | Bacteria>Bacillati>Bacillota>Clostridia>Eubacteriales>Clostridiaceae>Clostridium | 241 | DVW02_06070 | 110509..111234 | 1491 | hypothetical-protein-[Clostridium-botulinum] |
| NFO13006.1 | REase | GCA_011016315.1 | Clostridium_botulinum | Bacteria>Bacillati>Bacillota>Clostridia>Eubacteriales>Clostridiaceae>Clostridium | 241 | FDB34_02060 | 211282..212007 | 1491 | hypothetical-protein-[Clostridium-botulinum] |
| MBX7356545.1 | REase | GCA_019705275.1 | Clostridium_chauvoei | Bacteria>Bacillati>Bacillota>Clostridia>Eubacteriales>Clostridiaceae>Clostridium | 78 | K4H38_08480 | 51591..51827 | 46867 | hypothetical-protein-K4H38_08480-[Clostridium-chauvoei] |
| WP_021876217.1 | REase | GCF_900037355.1 | Clostridium_chauvoei | Bacteria>Bacillati>Bacillota>Clostridia>Eubacteriales>Clostridiaceae>Clostridium | 78 | J9878_RS07540 | 10832..11068 | 46867 | hypothetical-protein-[Clostridium-chauvoei] |
| WP_209702454.1 | REase | GCF_017874415.1 | Clostridium_algifaecis | Bacteria>Bacillati>Bacillota>Clostridia>Eubacteriales>Clostridiaceae>Clostridium | 113 | J2Z42_RS09770 | 13924..14265 | 1472040 | hypothetical-protein-[Clostridium-algifaecis] |
| WP_257789696.1 | REase | - | Clostridium_isatidis | Bacteria>Bacillati>Bacillota>Clostridia>Eubacteriales>Clostridiaceae>Clostridium | 99 | BEN51_RS07760 | 1641110..1641409 | 182773 | DNA/RNA-nuclease-SfsA-[Clostridium-isatidis] |
| WP_207652759.1 | REase | GCF_002285495.1 | Clostridium_isatidis | Bacteria>Bacillati>Bacillota>Clostridia>Eubacteriales>Clostridiaceae>Clostridium | 175 | BEN51_RS13920 | 1641431..1641958 | 182773 | hypothetical-protein-[Clostridium-isatidis] |
| WP_267901295.1 | REase | - | Clostridium_tagluense | Bacteria>Bacillati>Bacillota>Clostridia>Eubacteriales>Clostridiaceae>Clostridium | 116 | Ctaglu_RS16465 | 71007..71357 | 360422 | DUF3883-domain-containing-protein-[Clostridium-tagluense] |
| WP_261670008.1 | REase | GCF_025398335.1 | Clostridium_sp._CX1 | Bacteria>Bacillati>Bacillota>Clostridia>Eubacteriales>Clostridiaceae>Clostridium>unclassified_Clostridium | 100 | N4T77_RS00695 | 152923..153225 | 2978346 | DNA/RNA-nuclease-SfsA-[Clostridium-sp.-CX1] |
| WP_217303317.1 | REase | GCF_019056555.1 | Clostridium_sp._001 | Bacteria>Bacillati>Bacillota>Clostridia>Eubacteriales>Clostridiaceae>Clostridium>unclassified_Clostridium | 190 | B5S50_RS05645 | 1260231..1260803 | 1970093 | hypothetical-protein-[Clostridium-sp.-001] |
| OCZ54241.1 | REase | GCA_001707455.1 | Dehalobacter_sp._TeCB1 | Bacteria>Bacillati>Bacillota>Clostridia>Eubacteriales>Desulfitobacteriaceae>Dehalobacter>unclassified_Dehalobacter | 248 | A7D23_05580 | 29884..30630 | 1843715 | hypothetical-protein-A7D23_05580-[Dehalobacter-sp.-TeCB1] |
| PSR33878.1 | REase | GCA_003023695.1 | Sulfobacillus_benefaciens | Bacteria>Bacillati>Bacillota>Clostridia>Eubacteriales>Eubacteriales_incertae_sedis>Clostridiales_Family_XVII._Incertae_Sedis>Sulfobacillus | 168 | C7B46_07755 | 6531..7037 | 453960 | hypothetical-protein-C7B46_07755-[Sulfobacillus-benefaciens] |
| MBM7867396.1 | REase | GCA_016908025.1 | Heliomicrobium_gestii | Bacteria>Bacillati>Bacillota>Clostridia>Eubacteriales>Heliobacteriaceae>Heliomicrobium | 126 | JOC69_002302 | 74027..74407 | 2699 | hypothetical-protein-[Heliomicrobium-gestii] |
| WP_161262232.1 | REase | GCF_016908025.1 | Heliomicrobium_gestii | Bacteria>Bacillati>Bacillota>Clostridia>Eubacteriales>Heliobacteriaceae>Heliomicrobium | 126 | JOC69_RS11255 | 74027..74407 | 2699 | restriction-endonuclease-[Heliomicrobium-gestii] |
| WP_010247731.1 | REase | GCF_000179595.2 | Acetivibrio_cellulolyticus | Bacteria>Bacillati>Bacillota>Clostridia>Eubacteriales>Oscillospiraceae>Acetivibrio | 173 | ACECE_RS0212910 | 174650..175171 | 35830 | hypothetical-protein-[Acetivibrio-cellulolyticus] |
| WP_120485836.1 | REase | GCF_009917605.1 | Anaerotruncus_sp._1XD42-93 | Bacteria>Bacillati>Bacillota>Clostridia>Eubacteriales>Oscillospiraceae>Anaerotruncus>unclassified_Anaerotruncus | 179 | D5269_RS12190 | 6036..6575 | 2320853 | MULTISPECIES:-hypothetical-protein-[unclassified-Anaerotruncus] |
| MBD5149895.1 | REase | GCA_014799845.1 | Oscillibacter_sp. | Bacteria>Bacillati>Bacillota>Clostridia>Eubacteriales>Oscillospiraceae>Oscillibacter>unclassified_Oscillibacter | 178 | HDT18_05895 | 34202..34738 | 1945593 | hypothetical-protein-[Oscillibacter-sp.] |
| WP_094549801.1 | REase | GCF_002252565.1 | Petroclostridium_xylanilyticum | Bacteria>Bacillati>Bacillota>Clostridia>Eubacteriales>Oscillospiraceae>Petroclostridium | 203 | CIB29_RS11445 | 327416..328027 | 1792311 | hypothetical-protein-[Petroclostridium-xylanilyticum] |
| WP_204901122.1 | REase | GCF_016902575.1 | Pseudoflavonifractor_capillosus | Bacteria>Bacillati>Bacillota>Clostridia>Eubacteriales>Oscillospiraceae>Pseudoflavonifractor | 169 | H9X86_RS08565 | 12237..12746 | 106588 | hypothetical-protein-[Pseudoflavonifractor-capillosus] |
| WP_004621800.1 | REase | GCF_000175795.2 | Ruminiclostridium_papyrosolvens | Bacteria>Bacillati>Bacillota>Clostridia>Eubacteriales>Oscillospiraceae>Ruminiclostridium | 226 | P0092_RS08145 | 1828214..1828894 | 29362 | DNA/RNA-nuclease-SfsA-[Ruminiclostridium-papyrosolvens] |
| WP_157882728.1 | REase | GCF_001697285.1 | Thermoclostridium_stercorarium | Bacteria>Bacillati>Bacillota>Clostridia>Eubacteriales>Oscillospiraceae>Thermoclostridium | 179 | CSTERLE_RS00950 | 227477..228016 | 1510 | winged-helix-turn-helix-domain-containing-protein-[Thermoclostridium-stercorarium] |
| UYJ11719.1 | REase | GCA_025757845.1 | Oscillospiraceae_bacterium | Bacteria>Bacillati>Bacillota>Clostridia>Eubacteriales>Oscillospiraceae>unclassified_Oscillospiraceae | 177 | OGM78_02725 | 577354..577887 | 2485925 | MAG:-hypothetical-protein-OGM78_02725-[Oscillospiraceae-bacterium] |
| MBQ9664280.1 | REase | GCA_017458065.1 | Oscillospiraceae_bacterium | Bacteria>Bacillati>Bacillota>Clostridia>Eubacteriales>Oscillospiraceae>unclassified_Oscillospiraceae | 142 | IJV40_14130 | 16199..16627 | 2485925 | hypothetical-protein-[Oscillospiraceae-bacterium] |
| NHM28429.1 | REase | GCA_011393055.1 | Desulfofundulus_sp._TPOSR | Bacteria>Bacillati>Bacillota>Clostridia>Eubacteriales>Peptococcaceae>Desulfofundulus>unclassified_Desulfofundulus | 238 | G7K71_15915 | 26979..27695 | 2714340 | DUF1016-family-protein-[Desulfofundulus-sp.-TPOSR] |
| KFD41491.1 | REase | GCA_000681355.2 | Peptococcaceae_bacterium_SCADC1_2_3 | Bacteria>Bacillati>Bacillota>Clostridia>Eubacteriales>Peptococcaceae>unclassified_Peptococcaceae | 227 | DK28_0208705 | 16197..16880 | 1487582 | hypothetical-protein-DK28_0208705-[Peptococcaceae-bacterium-SCADC1_2_3] |
| KFI36118.1 | REase | GCA_000740805.1 | Peptococcaceae_bacterium_SCADC1_2_3 | Bacteria>Bacillati>Bacillota>Clostridia>Eubacteriales>Peptococcaceae>unclassified_Peptococcaceae | 227 | HY00_07500 | 39185..39868 | 1487582 | hypothetical-protein-HY00_07500-[Peptococcaceae-bacterium-SCADC1_2_3] |
| NLV15839.1 | REase | GCA_012729505.1 | Syntrophomonadaceae_bacterium | Bacteria>Bacillati>Bacillota>Clostridia>Eubacteriales>Syntrophomonadaceae>unclassified_Syntrophomonadaceae | 247 | GXY50_01330 | 79467..80210 | 2093811 | hypothetical-protein-[Syntrophomonadaceae-bacterium] |
| MCR4615478.1 | REase | GCA_024684435.1 | Clostridiales_bacterium | Bacteria>Bacillati>Bacillota>Clostridia>Eubacteriales>unclassified_Eubacteriales | 126 | K5756_04945 | 70047..70427 | 1898207 | DUF559-domain-containing-protein-[Clostridiales-bacterium] |
| TYL56951.1 | REase | GCA_008123415.1 | Agathobacter_rectalis | Bacteria>Bacillati>Bacillota>Clostridia>Lachnospirales>Lachnospiraceae>Agathobacter | 152 | FYL31_13945 | 10085..10543 | 39491 | hypothetical-protein-FYL31_13945-[[Eubacterium] |
| ONI39095.1 | REase | GCA_001983455.1 | Epulopiscium_sp._SCG-B11WGA-EpuloA1 | Bacteria>Bacillati>Bacillota>Clostridia>Lachnospirales>Lachnospiraceae>Candidatus_Epulonipiscium>unclassified_Candidatus_Epulonipiscium | 115 | AN396_09490 | 76625..76972 | 1710701 | hypothetical-protein-AN396_09490-[Epulopiscium-sp.-SCG-B11WGA-EpuloA1] |
| HAB87402.1 | REase | GCA_003448155.1 | Coprococcus_sp. | Bacteria>Bacillati>Bacillota>Clostridia>Lachnospirales>Lachnospiraceae>Coprococcus>unclassified_Coprococcus | 152 | DCE60_00040 | 11819..12277 | 2049024 | hypothetical-protein-[Coprococcus-sp.] |
| MCI9500560.1 | REase | GCA_022800895.1 | Hungatella_sp. | Bacteria>Bacillati>Bacillota>Clostridia>Lachnospirales>Lachnospiraceae>Hungatella>unclassified_Hungatella | 154 | HFG76_00885 | 24329..24793 | 2613924 | hypothetical-protein-[Hungatella-sp.] |
| WP_135987573.1 | REase | GCF_004793545.1 | Petralouisia_muris | Bacteria>Bacillati>Bacillota>Clostridia>Lachnospirales>Lachnospiraceae>Petralouisia | 148 | E5329_RS00205 | 42421..42867 | 3032872 | hypothetical-protein-[Petralouisia-muris] |
| TGY98245.1 | REase | GCA_004793545.1 | Petralouisia_muris | Bacteria>Bacillati>Bacillota>Clostridia>Lachnospirales>Lachnospiraceae>Petralouisia | 134 | E5329_00205 | 42421..42825 | 3032872 | hypothetical-protein-E5329_00205-[Petralouisia-muris] |
| MCD7743823.1 | REase | GCA_021199475.1 | Lachnospiraceae_bacterium | Bacteria>Bacillati>Bacillota>Clostridia>Lachnospirales>Lachnospiraceae>unclassified_Lachnospiraceae | 178 | LUI13_00775 | 23680..24216 | 1898203 | hypothetical-protein-[Lachnospiraceae-bacterium] |
| EOS24742.1 | REase | GCA_000403215.2 | Lachnospiraceae_bacterium_A4 | Bacteria>Bacillati>Bacillota>Clostridia>Lachnospirales>Lachnospiraceae>unclassified_Lachnospiraceae | 169 | C804_05107 | 23142..23651 | 397291 | hypothetical-protein-C804_05107-[Lachnospiraceae-bacterium-A4] |
| HCI18470.1 | REase | GCA_003521585.1 | Lachnospiraceae_bacterium | Bacteria>Bacillati>Bacillota>Clostridia>Lachnospirales>Lachnospiraceae>unclassified_Lachnospiraceae | 152 | DFH32_07635 | 9145..9603 | 1898203 | hypothetical-protein-[Lachnospiraceae-bacterium] |
| HCX41708.1 | REase | GCA_003539645.1 | Lachnospiraceae_bacterium | Bacteria>Bacillati>Bacillota>Clostridia>Lachnospirales>Lachnospiraceae>unclassified_Lachnospiraceae | 152 | DHH50_05115 | 187987..188445 | 1898203 | hypothetical-protein-[Lachnospiraceae-bacterium] |
| EOS28351.1 | REase | GCA_000403275.2 | Lachnospiraceae_bacterium_28-4 | Bacteria>Bacillati>Bacillota>Clostridia>Lachnospirales>Lachnospiraceae>unclassified_Lachnospiraceae | 150 | C807_03889 | 205764..206216 | 397287 | hypothetical-protein-C807_03889-[Lachnospiraceae-bacterium-28-4] |
| MBR4606060.1 | REase | GCA_017536085.1 | Lachnospiraceae_bacterium | Bacteria>Bacillati>Bacillota>Clostridia>Lachnospirales>Lachnospiraceae>unclassified_Lachnospiraceae | 101 | IK041_07520 | 2716..3021 | 1898203 | hypothetical-protein-[Lachnospiraceae-bacterium] |
| MBR4606061.1 | REase | GCA_017536085.1 | Lachnospiraceae_bacterium | Bacteria>Bacillati>Bacillota>Clostridia>Lachnospirales>Lachnospiraceae>unclassified_Lachnospiraceae | 86 | IK041_07525 | 3092..3352 | 1898203 | hypothetical-protein-[Lachnospiraceae-bacterium] |
| MST58785.1 | REase | GCA_009695995.1 | Waltera_intestinalis | Bacteria>Bacillati>Bacillota>Clostridia>Lachnospirales>Lachnospiraceae>Waltera | 244 | FYJ59_11160 | 25629..26363 | 2606635 | DNA/RNA-nuclease-SfsA-[Waltera-intestinalis] |

| Protein-ID | Domain | Assembly-ID | Strain | Taxonomy | Length | Locus-Tag | Genomic-coordinates | TaxID | Fasta-defline |
|---|---|---|---|---|---|---|---|---|---|
| WP_073026829.1 | REase | GCF_900142105.1 | Lutispora_thermophila | Bacteria>Bacillati>Bacillota>Clostridia>Lutisporales>Lutisporaceae>Lutispora | 203 | BUB48_RS13900 | 49013..49624 | 288966 | hypothetical-protein-[Lutispora-thermophila] |
| WP_277998839.1 | REase | - | Moorella_sulfitireducens | Bacteria>Bacillati>Bacillota>Clostridia>Moorellales>Moorellaceae>Moorella | 130 | NWF31_RS04980 | 114389..114781 | 2972948 | DUF4365-domain-containing-protein-[Moorella-sp.-SLA38] |
| WP_258359421.1 | REase | GCF_024733845.1 | Moorella_sulfitireducens | Bacteria>Bacillati>Bacillota>Clostridia>Moorellales>Moorellaceae>Moorella | 163 | NWF31_RS05050 | 140298..140789 | 2972948 | hypothetical-protein-[Moorella-sp.-SLA38] |
| PHS35618.1 | REase | GCA_002733545.1 | Alkaliphilus_sp. | Bacteria>Bacillati>Bacillota>Clostridia>Peptostreptococcales>Natronincolaceae>Alkaliphilus>unclassified_Alkaliphilus | 240 | COA82_04065 | 70661..71383 | 1872455 | MAG:-hypothetical-protein-COA82_04065-[Alkaliphilus-sp.] |
| WP_180703532.1 | REase | GCF_900015215.1 | Romboutsia_ilealis | Bacteria>Bacillati>Bacillota>Clostridia>Peptostreptococcales>Peptostreptococcaceae>Romboutsia | 139 | CRIB_RS05550 | 1175401..1175820 | 1115758 | hypothetical-protein-[Romboutsia-ilealis] |
| WP_099191964.1 | REase | GCF_002724055.1 | Tepidibacter_mesophilus | Bacteria>Bacillati>Bacillota>Clostridia>Peptostreptococcales>Peptostreptococcaceae>Tepidibacter | 190 | Tmeso_RS13195 | 49090..49662 | 655607 | hypothetical-protein-[Tepidibacter-mesophilus] |
| WP_049752537.1 | REase | GCF_000018425.1 | Candidatus_Desulforudis_audaxviator | Bacteria>Bacillati>Bacillota>Clostridia>Thermoanaerobacterales>Candidatus_Desulforudaceae>Candidatus_Desulforudis | 74 | Daudx_RS02655 | 540158..540382 | 471827 | hypothetical-protein-[Candidatus-Desulforudis-audaxviator] |
| WP_094397493.1 | REase | GCF_002250075.1 | Thermoanaerobacterium_thermosaccharolyticum | Bacteria>Bacillati>Bacillota>Clostridia>Thermoanaerobacterales>Thermoanaerobacteraceae>Thermoanaerobacterium | 203 | Thert_RS08585 | 1715957..1716568 | 1517 | hypothetical-protein-[Thermoanaerobacterium-thermosaccharolyticum] |
| MBQ9252305.1 | REase | GCA_017470145.1 | Clostridia_bacterium | Bacteria>Bacillati>Bacillota>Clostridia>unclassified_Clostridia | 145 | IJ188_06700 | 54887..55324 | 2044939 | hypothetical-protein-[Clostridia-bacterium] |
| EHM91317.1 | REase | GCA_000239735.1 | Coprobacillus_sp._3_3_56FAA | Bacteria>Bacillati>Bacillota>Erysipelotrichia>Erysipelotrichales>Coprobacillaceae>Coprobacillus>unclassified_Coprobacillus | 203 | HMPREF1021_02123 | 45678..46289 | 665941 | hypothetical-protein-HMPREF1021_02123-[Coprobacillus-sp.-3_3_56FAA] |
| MCK9524535.1 | REase | GCA_023228555.1 | Limnochordia_bacterium | Bacteria>Bacillati>Bacillota>Limnochordia>unclassified_Limnochordia | 235 | MOR49_01215 | 1793..2500 | 2497621 | hypothetical-protein-[Limnochordia-bacterium] |
| MBS5426406.1 | REase | GCA_018368575.1 | Phascolarctobacterium_succinatutens | Bacteria>Bacillati>Bacillota>Negativicutes>Acidaminococcales>Acidaminococcaceae>Phascolarctobacterium | 171 | KHX39_04765 | 65071..65586 | 626940 | hypothetical-protein-[Phascolarctobacterium-succinatutens] |
| MBE6098019.1 | REase | GCA_015058685.1 | Schwartzia_succinivorans | Bacteria>Bacillati>Bacillota>Negativicutes>Selenomonadales>Selenomonadaceae>Schwartzia | 178 | E7198_09525 | 46520..47056 | 55507 | hypothetical-protein-[Schwartzia-succinivorans] |
| WP_273038288.1 | REase | GCF_015058685.1 | Schwartzia_succinivorans | Bacteria>Bacillati>Bacillota>Negativicutes>Selenomonadales>Selenomonadaceae>Schwartzia | 178 | E7198_RS09625 | 46520..47056 | 55507 | hypothetical-protein-[Schwartzia-succinivorans] |
| WP_154440319.1 | REase | GCF_018917845.1 | Tissierella_pigra | Bacteria>Bacillati>Bacillota>Tissierellia>Tissierellales>Tissierellaceae>Tissierella | 200 | KQI41_RS04670 | 944411..945013 | 2607614 | hypothetical-protein-[Tissierella-pigra] |
| OUL09465.1 | REase | GCA_002155105.1 | Sedimentibacter_sp._SX930 | Bacteria>Bacillati>Bacillota>Tissierellia>Tissierellia_incertae_sedis>Sedimentibacter>unclassified_Sedimentibacter | 193 | B0533_04685 | 274338..274919 | 1955060 | hypothetical-protein-B0533_04685-[Sedimentibacter-sp.-SX930] |
| NLD15973.1 | REase | GCA_012513395.1 | Tissierellia_bacterium | Bacteria>Bacillati>Bacillota>Tissierellia>unclassified_Tissierellia | 143 | GX666_00130 | 1..433 | 2049431 | PD-(D/E)XK-nuclease-family-protein-[Tissierellia-bacterium] |
| NPV28410.1 | REase | GCA_013177945.1 | Bacillota_bacterium | Bacteria>Bacillati>Bacillota>unclassified_Bacillota | 153 | HPY58_01895 | 9245..9706 | 1879010 | hypothetical-protein-[Bacillota-bacterium] |
| NLN07232.1 | REase | GCA_012522075.1 | Bacillota_bacterium | Bacteria>Bacillati>Bacillota>unclassified_Bacillota | 227 | GX167_06405 | 10379..11062 | 1879010 | hypothetical-protein-[Bacillota-bacterium] |
| MCB9078005.1 | REase | GCA_020634955.1 | Anaerolineaceae_bacterium | Bacteria>Bacillati>Chloroflexota>Anaerolineae>Anaerolineales>Anaerolineaceae>unclassified_Anaerolineaceae | 134 | H6631_10470 | 1298..1702 | 2024896 | GxxExxY-protein-[Anaerolineaceae-bacterium] |
| MAS36306.1 | REase | GCA_002702065.1 | Anaerolineaceae_bacterium | Bacteria>Bacillati>Chloroflexota>Anaerolineae>Anaerolineales>Anaerolineaceae>unclassified_Anaerolineaceae | 231 | CL610_20040 | 39175..39870 | 2024896 | hypothetical-protein-[Anaerolineaceae-bacterium] |
| MBE7469120.1 | REase | GCA_015075405.1 | Anaerolineales_bacterium | Bacteria>Bacillati>Chloroflexota>Anaerolineae>Anaerolineales>unclassified_Anaerolineales | 125 | HS114_08275 | 1912208..1912585 | 2073117 | GxxExxY-protein-[Anaerolineales-bacterium] |
| MCC6958030.1 | REase | GCA_020853835.1 | Anaerolineales_bacterium | Bacteria>Bacillati>Chloroflexota>Anaerolineae>Anaerolineales>unclassified_Anaerolineales | 207 | IT316_14615 | 11276..11899 | 2073117 | hypothetical-protein-[Anaerolineales-bacterium] |
| MBN1921822.1 | REase | GCA_016931315.1 | Anaerolineae_bacterium | Bacteria>Bacillati>Chloroflexota>Anaerolineae>unclassified_Anaerolineae | 196 | JW892_11290 | 1203..1793 | 2052143 | hypothetical-protein-[Anaerolineae-bacterium] |
| MCG3142364.1 | REase | GCA_022072245.1 | Anaerolineae_bacterium | Bacteria>Bacillati>Chloroflexota>Anaerolineae>unclassified_Anaerolineae | 194 | HDKAJFGB_03810 | 2087..2671 | 2052143 | hypothetical-protein-[Anaerolineae-bacterium] |
| MCD6553271.1 | REase | GCA_021162125.1 | Anaerolineae_bacterium | Bacteria>Bacillati>Chloroflexota>Anaerolineae>unclassified_Anaerolineae | 136 | J7M16_04580 | 9048..9458 | 2052143 | GxxExxY-protein-[Anaerolineae-bacterium] |
| MBL7062618.1 | REase | GCA_016783285.1 | Anaerolineae_bacterium | Bacteria>Bacillati>Chloroflexota>Anaerolineae>unclassified_Anaerolineae | 127 | ISS49_00235 | 51385..51768 | 2052143 | GxxExxY-protein-[Anaerolineae-bacterium] |
| MCL4302339.1 | REase | GCA_023382535.1 | Anaerolineae_bacterium | Bacteria>Bacillati>Chloroflexota>Anaerolineae>unclassified_Anaerolineae | 127 | KJ077_42010 | 34060..34443 | 2052143 | GxxExxY-protein-[Anaerolineae-bacterium] |
| MBN2007575.1 | REase | GCA_016931835.1 | Anaerolineae_bacterium | Bacteria>Bacillati>Chloroflexota>Anaerolineae>unclassified_Anaerolineae | 124 | JXA21_29790 | 2518..2892 | 2052143 | GxxExxY-protein-[Anaerolineae-bacterium] |
| MCK6629897.1 | REase | GCA_023151115.1 | Anaerolineae_bacterium | Bacteria>Bacillati>Chloroflexota>Anaerolineae>unclassified_Anaerolineae | 123 | L6R45_32570 | 14167..14538 | 2052143 | GxxExxY-protein-[Anaerolineae-bacterium] |
| MCQ3973385.1 | REase | GCA_024434315.1 | Anaerolineae_bacterium | Bacteria>Bacillati>Chloroflexota>Anaerolineae>unclassified_Anaerolineae | 123 | DPW09_08080 | 24106..24477 | 2052143 | GxxExxY-protein-[Anaerolineae-bacterium] |
| MCW5847518.1 | REase | GCA_026129205.1 | Anaerolineae_bacterium | Bacteria>Bacillati>Chloroflexota>Anaerolineae>unclassified_Anaerolineae | 122 | KIS95_09825 | 1636..2004 | 2052143 | MAG:-GxxExxY-protein-[Anaerolineae-bacterium] |
| MCL4300571.1 | REase | GCA_023382535.1 | Anaerolineae_bacterium | Bacteria>Bacillati>Chloroflexota>Anaerolineae>unclassified_Anaerolineae | 82 | KJ077_32845 | 81359..81607 | 2052143 | hypothetical-protein-[Anaerolineae-bacterium] |
| MBN2393687.1 | REase | GCA_016934645.1 | Anaerolineae_bacterium | Bacteria>Bacillati>Chloroflexota>Anaerolineae>unclassified_Anaerolineae | 210 | JXR84_23345 | 13178..13810 | 2052143 | hypothetical-protein-[Anaerolineae-bacterium] |
| MCL4299605.1 | REase | GCA_023382535.1 | Anaerolineae_bacterium | Bacteria>Bacillati>Chloroflexota>Anaerolineae>unclassified_Anaerolineae | 199 | KJ077_27965 | 96323..96922 | 2052143 | hypothetical-protein-[Anaerolineae-bacterium] |
| MCB0191468.1 | REase | GCA_020433805.1 | Anaerolineae_bacterium | Bacteria>Bacillati>Chloroflexota>Anaerolineae>unclassified_Anaerolineae | 123 | KDJ65_05945 | 15102..15473 | 2052143 | NERD-domain-containing-protein-[Anaerolineae-bacterium] |
| MBZ0275670.1 | REase | GCA_019912915.1 | Anaerolineae_bacterium | Bacteria>Bacillati>Chloroflexota>Anaerolineae>unclassified_Anaerolineae | 240 | K8I60_05980 | 10211..10931 | 2052143 | NERD-domain-containing-protein-[Anaerolineae-bacterium] |
| MCZ7570814.1 | REase | GCA_027493995.1 | Ardenticatenaceae_bacterium | Bacteria>Bacillati>Chloroflexota>Ardenticatenia>Ardenticatenales>Ardenticatenaceae>unclassified_Ardenticatenaceae | 125 | M5U01_19820 | 4779069..4779446 | 2614247 | MAG:-DUF559-domain-containing-protein-[Ardenticatenaceae-bacterium] |
| MCZ7573955.1 | REase | GCA_027493995.1 | Ardenticatenaceae_bacterium | Bacteria>Bacillati>Chloroflexota>Ardenticatenia>Ardenticatenales>Ardenticatenaceae>unclassified_Ardenticatenaceae | 241 | M5U01_35840 | 8599510..8600235 | 2614247 | MAG:-DUF3883-domain-containing-protein-[Ardenticatenaceae-bacterium] |
| MBK8904529.1 | REase | GCA_016710325.1 | Candidatus_Leptofilum_proximum | Bacteria>Bacillati>Chloroflexota>Ardenticatenia>Candidatus_Promineifilales>Candidatus_Promineifilaceae>Candidatus_Leptofilum | 153 | IPM53_25345 | 3770823..3771284 | 3140689 | hypothetical-protein-[Anaerolineaceae-bacterium] |
| GIV69802.1 | REase | GCA_026003635.1 | Caldilinea_sp. | Bacteria>Bacillati>Chloroflexota>Caldilineae>Caldilineales>Caldilineaceae>Caldilinea>unclassified_Caldilinea | 141 | KatS3mg048_2664 | 135486..135911 | 2293560 | MAG:-hypothetical-protein-KatS3mg048_2664-[Caldilinea-sp.] |
| MBK6433235.1 | REase | GCA_016704685.1 | Candidatus_Amarolinea_dominans | Bacteria>Bacillati>Chloroflexota>Caldilineae>Caldilineales>Candidatus_Amarolineaceae>Candidatus_Amarolinea | 176 | IPF85_21390 | 35089..35619 | 3140696 | hypothetical-protein-[Anaerolineae-bacterium] |
| MBK9093500.1 | REase | GCA_016719785.1 | Candidatus_Amarolinea_dominans | Bacteria>Bacillati>Chloroflexota>Caldilineae>Caldilineales>Candidatus_Amarolineaceae>Candidatus_Amarolinea | 176 | IPM84_12140 | 210109..210639 | 3140696 | hypothetical-protein-[Anaerolineae-bacterium] |
| MBK6429950.1 | REase | GCA_016704685.1 | Candidatus_Amarolinea_dominans | Bacteria>Bacillati>Chloroflexota>Caldilineae>Caldilineales>Candidatus_Amarolineaceae>Candidatus_Amarolinea | 133 | IPF85_04120 | 225299..225700 | 3140696 | GxxExxY-protein-[Anaerolineae-bacterium] |
| MBK9093505.1 | REase | GCA_016719785.1 | Candidatus_Amarolinea_dominans | Bacteria>Bacillati>Chloroflexota>Caldilineae>Caldilineales>Candidatus_Amarolineaceae>Candidatus_Amarolinea | 68 | IPM84_12170 | 219237..219443 | 3140696 | hypothetical-protein-[Anaerolineae-bacterium] |
| MCS6883597.1 | REase | GCA_025057155.1 | Chloroflexaceae_bacterium | Bacteria>Bacillati>Chloroflexota>Chloroflexia>Chloroflexales>Chloroflexineae>Chloroflexaceae>unclassified_Chloroflexaceae | 125 | NZU74_19895 | 13339..13716 | 2562117 | GxxExxY-protein-[Chloroflexaceae-bacterium] |
| MCS7288798.1 | REase | GCA_025062325.1 | Roseiflexus_sp. | Bacteria>Bacillati>Chloroflexota>Chloroflexia>Chloroflexales>Roseiflexineae>Roseiflexaceae>Roseiflexus>unclassified_Roseiflexus | 137 | NZ699_06670 | 886..1299 | 2562120 | GxxExxY-protein-[Roseiflexus-sp.] |
| WP_011955033.1 | REase | GCF_000016665.1 | Roseiflexus_sp._RS-1 | Bacteria>Bacillati>Chloroflexota>Chloroflexia>Chloroflexales>Roseiflexineae>Roseiflexaceae>Roseiflexus>unclassified_Roseiflexus | 129 | ROSERS_RS01215 | 296561..296950 | 357808 | GxxExxY-protein-[Roseiflexus-sp.-RS-1] |
| MBE0430959.1 | REase | GCA_014859205.1 | Dehalococcoidia_bacterium | Bacteria>Bacillati>Chloroflexota>Dehalococcoidia>unclassified_Dehalococcoidia | 165 | IBX67_03935 | 14614..15111 | 2026734 | hypothetical-protein-[Dehalococcoidia-bacterium] |
| MCL0064886.1 | REase | GCA_023271695.1 | Dehalococcoidia_bacterium | Bacteria>Bacillati>Chloroflexota>Dehalococcoidia>unclassified_Dehalococcoidia | 206 | M1N84_03210 | 4742..5362 | 2026734 | hypothetical-protein-[Dehalococcoidia-bacterium] |
| MCL0064887.1 | REase | GCA_023271695.1 | Dehalococcoidia_bacterium | Bacteria>Bacillati>Chloroflexota>Dehalococcoidia>unclassified_Dehalococcoidia | 245 | M1N84_03215 | 5302..6039 | 2026734 | Uma2-family-endonuclease-[Dehalococcoidia-bacterium] |
| GHO59278.1 | REase | GCA_016587375.1 | Ktedonobacter_robiniae | Bacteria>Bacillati>Chloroflexota>Ktedonobacteria>Ktedonobacterales>Ktedonobacteraceae>Ktedonobacter | 149 | KSB_77530 | 1137381..1137830 | 2778365 | hypothetical-protein-KSB_77530-[Ktedonobacter-robiniae] |
| MBK6562035.1 | REase | GCA_016704105.1 | Candidatus_Amarobacter_glycogenicus | Bacteria>Bacillati>Chloroflexota>Tepidiformia>Tepidiformales>Tepidiformaceae>Candidatus_Amarobacter | 167 | IPG14_10960 | 179..682 | 3140699 | restriction-endonuclease-[Dehalococcoidia-bacterium] |
| MBN1889999.1 | REase | GCA_016931085.1 | Thermoflexales_bacterium | Bacteria>Bacillati>Chloroflexota>Thermoflexia>Thermoflexales>unclassified_Thermoflexales | 194 | JW850_18530 | 392..976 | 2497639 | hypothetical-protein-[Thermoflexales-bacterium] |
| MBU1747545.1 | REase | GCA_018823415.1 | Chloroflexota_bacterium | Bacteria>Bacillati>Chloroflexota>unclassified_Chloroflexota | 197 | KKA73_07645 | 1325..1918 | 2026724 | hypothetical-protein-[Chloroflexota-bacterium] |
| MCL5950400.1 | REase | GCA_023489485.1 | Chloroflexota_bacterium | Bacteria>Bacillati>Chloroflexota>unclassified_Chloroflexota | 194 | M1132_01525 | 23483..24067 | 2026724 | hypothetical-protein-[Chloroflexota-bacterium] |
| MBI4322293.1 | REase | GCA_016210495.1 | Chloroflexota_bacterium | Bacteria>Bacillati>Chloroflexota>unclassified_Chloroflexota | 170 | HY675_27685 | 50858..51370 | 2026724 | hypothetical-protein-[Chloroflexota-bacterium] |
| MCL5961064.1 | REase | GCA_023489135.1 | Chloroflexota_bacterium | Bacteria>Bacillati>Chloroflexota>unclassified_Chloroflexota | 170 | M1358_17460 | 12787..13299 | 2026724 | hypothetical-protein-[Chloroflexota-bacterium] |
| MCL5961068.1 | REase | GCA_023489135.1 | Chloroflexota_bacterium | Bacteria>Bacillati>Chloroflexota>unclassified_Chloroflexota | 138 | M1358_17480 | 18068..18484 | 2026724 | GxxExxY-protein-[Chloroflexota-bacterium] |
| MBI1882243.1 | REase | GCA_016178315.1 | Chloroflexota_bacterium | Bacteria>Bacillati>Chloroflexota>unclassified_Chloroflexota | 73 | HYR94_29065 | 1329..1550 | 2026724 | GxxExxY-protein-[Chloroflexota-bacterium] |
| MDE2668028.1 | REase | GCA_028875515.1 | Chloroflexota_bacterium | Bacteria>Bacillati>Chloroflexota>unclassified_Chloroflexota | 139 | OXI51_00040 | 12379..12798 | 2026724 | MAG:-DNA-mismatch-endonuclease-Vsr-[Chloroflexota-bacterium] |
| HCG29366.1 | REase | GCA_003534145.1 | Chloroflexota_bacterium | Bacteria>Bacillati>Chloroflexota>unclassified_Chloroflexota | 161 | DEU95_06400 | 3107..3592 | 2026724 | Uma2-family-endonuclease-[Chloroflexota-bacterium] |
| MBI3979240.1 | REase | GCA_016199325.1 | Chloroflexota_bacterium | Bacteria>Bacillati>Chloroflexota>unclassified_Chloroflexota | 130 | HY331_13740 | 15461..15853 | 2026724 | endonuclease-domain-containing-protein-[Chloroflexota-bacterium] |
| MDC0253129.1 | REase | GCA_028346115.1 | Chloroflexota_bacterium | Bacteria>Bacillati>Chloroflexota>unclassified_Chloroflexota | 218 | OAK52_03090 | 1..659 | 2026724 | DUF3883-domain-containing-protein,-partial-[Chloroflexi-bacterium] |
| MBI2863769.1 | REase | GCA_016190625.1 | Chloroflexota_bacterium | Bacteria>Bacillati>Chloroflexota>unclassified_Chloroflexota | 164 | HYX94_04325 | 24466..24960 | 2026724 | restriction-endonuclease-[Chloroflexota-bacterium] |
| MCC6178176.1 | REase | GCA_020845495.1 | Chloroflexota_bacterium | Bacteria>Bacillati>Chloroflexota>unclassified_Chloroflexota | 134 | IT305_22970 | 5468..5872 | 2026724 | restriction-endonuclease-[Chloroflexota-bacterium] |
| KAA3643968.1 | REase | GCA_008501685.1 | Chloroflexota_bacterium | Bacteria>Bacillati>Chloroflexota>unclassified_Chloroflexota | 224 | DWQ07_17785 | 196075..196749 | 2026724 | DUF91-domain-containing-protein-[Chloroflexota-bacterium] |
| MBC8450529.1 | REase | GCA_014383605.1 | Chloroflexota_bacterium | Bacteria>Bacillati>Chloroflexota>unclassified_Chloroflexota | 214 | H8D78_22575 | 11137..11781 | 2026724 | hypothetical-protein-[Chloroflexota-bacterium] |
| MBC8450555.1 | REase | GCA_014383605.1 | Chloroflexota_bacterium | Bacteria>Bacillati>Chloroflexota>unclassified_Chloroflexota | 213 | H8D78_22705 | 1210..1851 | 2026724 | hypothetical-protein-[Chloroflexota-bacterium] |
| MCC6567533.1 | REase | GCA_020848335.1 | Chloroflexota_bacterium | Bacteria>Bacillati>Chloroflexota>unclassified_Chloroflexota | 234 | IT298_17385 | 19495..20199 | 2026724 | hypothetical-protein-[Chloroflexota-bacterium] |
| MCX6066941.1 | REase | GCA_026389995.1 | Chloroflexota_bacterium | Bacteria>Bacillati>Chloroflexota>unclassified_Chloroflexota | 165 | NT121_14495 | 2773..3270 | 2026724 | MAG:-hypothetical-protein-NT121_14495-[Chloroflexi-bacterium] |
| MCL4459199.1 | REase | GCA_023382375.1 | Chloroflexota_bacterium | Bacteria>Bacillati>Chloroflexota>unclassified_Chloroflexota | 158 | M1577_04750 | 53955..54431 | 2026724 | hypothetical-protein-[Chloroflexota-bacterium] |
| OGH97201.1 | REase | GCA_001783265.1 | Candidatus_Margulisbacteria_bacterium_GWD2_39_127 | Bacteria>Bacillati>Cyanobacteriota/Melainabacteria_group>Candidatus_Margulisiibacteriota>unclassified_Candidatus_Margulisiibacteriota | 123 | A2X43_08525 | 58555..58926 | 1817871 | GxxExxY-protein-[Candidatus-Margulisbacteria-bacterium-GWD2_39_127] |
| OGI09204.1 | REase | GCA_001784485.1 | Candidatus_Margulisbacteria_bacterium_GWE2_39_32 | Bacteria>Bacillati>Cyanobacteriota/Melainabacteria_group>Candidatus_Margulisiibacteriota>unclassified_Candidatus_Margulisiibacteriota | 123 | A2X41_01350 | 17213..17584 | 1817872 | GxxExxY-protein-[Candidatus-Margulisbacteria-bacterium-GWE2_39_32] |
| PZM83736.1 | REase | GCA_003242895.1 | Candidatus_Margulisiibacteriota_bacterium | Bacteria>Bacillati>Cyanobacteriota/Melainabacteria_group>Candidatus_Margulisiibacteriota>unclassified_Candidatus_Margulisiibacteriota | 123 | DKM50_01290 | 12546..12917 | 2053573 | GxxExxY-protein-[Candidatus-Margulisbacteria-bacterium] |
| MBX9724059.1 | REase | GCA_019747345.1 | Candidatus_Obscuribacterales_bacterium | Bacteria>Bacillati>Cyanobacteriota/Melainabacteria_group>Candidatus_Melainabacteria>Candidatus_Obscuribacterales>unclassified_Candidatus_Obscuribacterales | 166 | K2X81_21815 | 8444..8944 | 2364082 | hypothetical-protein-[Candidatus-Obscuribacterales-bacterium] |
| WP_190667191.1 | REase | GCF_014695285.1 | Cyanophyceae | Bacteria>Bacillati>Cyanobacteriota/Melainabacteria_group>Cyanobacteriota | 188 | H6F54_RS11875 | 19734..20300 | 3028117 | hypothetical-protein-[Coleofasciculus-sp.-FACHB-501] |
| WP_190676131.1 | REase | GCF_014695435.1 | Cyanophyceae | Bacteria>Bacillati>Cyanobacteriota/Melainabacteria_group>Cyanobacteriota | 212 | H6F60_RS14240 | 78766..79404 | 3028117 | Uma2-family-endonuclease-[Coleofasciculus-sp.-FACHB-129] |
| WP_190769517.1 | REase | GCF_014696105.1 | Cyanophyceae | Bacteria>Bacillati>Cyanobacteriota/Melainabacteria_group>Cyanobacteriota | 208 | H6F67_RS25780 | 68395..69021 | 3028117 | hypothetical-protein-[Microcoleus-sp.-FACHB-1515] |
| WP_190447190.1 | REase | GCF_014695795.1 | Cyanophyceae | Bacteria>Bacillati>Cyanobacteriota/Melainabacteria_group>Cyanobacteriota | 207 | H6F86_RS05940 | 29653..30276 | 3028117 | hypothetical-protein-[Phormidium-sp.-FACHB-592] |
| WP_041444102.1 | REase | GCF_000019485.1 | Cyanophyceae | Bacteria>Bacillati>Cyanobacteriota/Melainabacteria_group>Cyanobacteriota | 121 | CCK85_RS15770 | 113713..114078 | 3028117 | MULTISPECIES:-DUF559-domain-containing-protein-[unclassified-Synechococcus] |
| WP_227788784.1 | REase | GCF_963896525.1 | Nostocales | Bacteria>Bacillati>Cyanobacteriota/Melainabacteria_group>Cyanobacteriota>Cyanophyceae | 220 | IQ240_RS03240 | 98212..98874 | 1161 | Uma2-family-endonuclease-[Nodularia-sp.-LEGE-04288] |
| WP_096669724.1 | REase | GCF_002368115.1 | Nostocales | Bacteria>Bacillati>Cyanobacteriota/Melainabacteria_group>Cyanobacteriota>Cyanophyceae | 232 | HEQ36_RS21595 | 53400..54098 | 1161 | MULTISPECIES:-type-I-restriction-endonuclease-[Dolichospermum] |
| NJR56032.1 | REase | GCA_012034865.1 | Acaryochloris_sp._CRU_2_0 | Bacteria>Bacillati>Cyanobacteriota/Melainabacteria_group>Cyanobacteriota>Cyanophyceae>Acaryochloridales>Acaryochloridaceae>Acaryochloris>unclassified_Acaryochloris | 205 | HC768_16560 | 10778..11395 | 2720402 | hypothetical-protein-[Acaryochloris-sp.-CRU_2_0] |
| WP_250683157.1 | REase | GCF_023650955.1 | Thermosynechococcus_sp._HN-54 | Bacteria>Bacillati>Cyanobacteriota/Melainabacteria_group>Cyanobacteriota>Cyanophyceae>Acaryochloridales>Thermosynechococcaceae>Thermosynechococcus>unclassified_Thermosynechococcus | 190 | NBE99_RS03720 | 774962..775534 | 2933959 | Uma2-family-endonuclease-[Thermosynechococcus-sp.-HN-54] |
| WJI23190.1 | REase | GCA_030322965.1 | Thermosynechococcus_sp._B0 | Bacteria>Bacillati>Cyanobacteriota/Melainabacteria_group>Cyanobacteriota>Cyanophyceae>Acaryochloridales>Thermosynechococcaceae>Thermosynechococcus>unclassified_Thermosynechococcus | 83 | MZ909_08105 | 1639208..1639459 | 2937284 | Uma2-family-endonuclease-[Thermosynechococcus-sp.-B0] |
| WP_015155780.1 | REase | GCF_000317125.1 | Chroococcidiopsis_thermalis | Bacteria>Bacillati>Cyanobacteriota/Melainabacteria_group>Cyanobacteriota>Cyanophyceae>Chroococcidiopsidales>Chroococcidiopsidaceae>Chroococcidiopsis | 211 | CHRO_RS18615 | 4309621..4310256 | 54299 | Uma2-family-endonuclease-[Chroococcidiopsis-thermalis] |
| WP_250012762.1 | REase | GCF_023546805.1 | Chroococcidiopsis_sp._CCNUC1 | Bacteria>Bacillati>Cyanobacteriota/Melainabacteria_group>Cyanobacteriota>Cyanophyceae>Chroococcidiopsidales>Chroococcidiopsidaceae>Chroococcidiopsis>unclassified_Chroococcidiopsis | 211 | M5J74_RS19070 | 4100946..4101581 | 2653189 | Uma2-family-endonuclease-[Chroococcidiopsis-sp.-CCNUC1] |
| WP_190566693.1 | REase | GCF_014696935.1 | unclassified_Desertifilum | Bacteria>Bacillati>Cyanobacteriota/Melainabacteria_group>Cyanobacteriota>Cyanophyceae>Desertifilales>Desertifilaceae>Desertifilum | 199 | H6G23_RS20075 | 63892..64491 | 2621682 | MULTISPECIES:-hypothetical-protein-[unclassified-Desertifilum] |
| MCD8487853.1 | REase | GCA_021208125.1 | Desertifilum_sp. | Bacteria>Bacillati>Cyanobacteriota/Melainabacteria_group>Cyanobacteriota>Cyanophyceae>Desertifilales>Desertifilaceae>Desertifilum>unclassified_Desertifilum | 199 | LRZ84_14245 | 9269..9868 | 2497424 | hypothetical-protein-[Desertifilum-sp.] |
| WP_218082164.1 | REase | GCF_018389385.1 | Anthocerotibacter_panamensis | Bacteria>Bacillati>Cyanobacteriota/Melainabacteria_group>Cyanobacteriota>Cyanophyceae>Gloeobacterales>Anthocerotibacteraceae>Anthocerotibacter | 194 | IL331_RS05740 | 1254811..1255395 | 2857077 | Uma2-family-endonuclease-[Anthocerotibacter-panamensis] |
| WP_230840688.1 | REase | GCF_021018745.1 | Gloeobacter_morelensis | Bacteria>Bacillati>Cyanobacteriota/Melainabacteria_group>Cyanobacteriota>Cyanophyceae>Gloeobacterales>Gloeobacteraceae>Gloeobacter | 216 | ISF26_RS17940 | 3738568..3739218 | 2907343 | Uma2-family-endonuclease-[Gloeobacter-morelensis] |
| OUC12870.1 | REase | GCA_002148405.1 | Alkalinema_sp._CACIAM_70d | Bacteria>Bacillati>Cyanobacteriota/Melainabacteria_group>Cyanobacteriota>Cyanophyceae>Leptolyngbyales>Leptolyngbyaceae>Alkalinema>unclassified_Alkalinema | 214 | B0A82_20150 | 19818..20462 | 1934309 | MAG:-hypothetical-protein-B0A82_20150-[Alkalinema-sp.-CACIAM-70d] |
| WP_190576381.1 | REase | GCF_014697025.1 | Alkalinema_sp._FACHB-956 | Bacteria>Bacillati>Cyanobacteriota/Melainabacteria_group>Cyanobacteriota>Cyanophyceae>Leptolyngbyales>Leptolyngbyaceae>Alkalinema>unclassified_Alkalinema | 212 | H6G21_RS22820 | 41359..41997 | 2692768 | Uma2-family-endonuclease-[Alkalinema-sp.-FACHB-956] |
| WP_233748350.1 | REase | GCF_021379005.1 | Leptodesmis_sichuanensis | Bacteria>Bacillati>Cyanobacteriota/Melainabacteria_group>Cyanobacteriota>Cyanophyceae>Leptolyngbyales>Leptolyngbyaceae>Leptodesmis | 190 | KIK02_RS09470 | 2042497..2043069 | 2906798 | Uma2-family-endonuclease-[Leptodesmis-sichuanensis] |
| WP_233748349.1 | REase | GCF_021379005.1 | Leptodesmis_sichuanensis | Bacteria>Bacillati>Cyanobacteriota/Melainabacteria_group>Cyanobacteriota>Cyanophyceae>Leptolyngbyales>Leptolyngbyaceae>Leptodesmis | 215 | KIK02_RS09465 | 2041834..2042481 | 2906798 | Uma2-family-endonuclease-[Leptodesmis-sichuanensis] |
| WP_190375079.1 | REase | GCF_014696055.1 | unclassified_Leptolyngbya | Bacteria>Bacillati>Cyanobacteriota/Melainabacteria_group>Cyanobacteriota>Cyanophyceae>Leptolyngbyales>Leptolyngbyaceae>Leptolyngbya_group>Leptolyngbya | 216 | H6G01_RS00350 | 18471..19121 | 2650499 | type-I-restriction-enzyme-HsdR-N-terminal-domain-containing-protein-[Leptolyngbya-sp.-FACHB-17] |
| WP_088889976.1 | REase | GCF_002215035.1 | Leptolyngbya_ohadii | Bacteria>Bacillati>Cyanobacteriota/Melainabacteria_group>Cyanobacteriota>Cyanophyceae>Leptolyngbyales>Leptolyngbyaceae>Leptolyngbya_group>Leptolyngbya | 166 | CDV24_RS06795 | 189285..189785 | 1962290 | DUF4058-family-protein-[Leptolyngbya-ohadii] |
| OAB62771.1 | REase | GCA_001637395.1 | Leptolyngbya_valderiana_BDU_20041 | Bacteria>Bacillati>Cyanobacteriota/Melainabacteria_group>Cyanobacteriota>Cyanophyceae>Leptolyngbyales>Leptolyngbyaceae>Leptolyngbya_group>Leptolyngbya>Leptolyngbya_valderiana | 189 | AY599_12435 | 19105..19674 | 322866 | hypothetical-protein-AY599_12435-[Leptolyngbya-valderiana-BDU-20041] |
| WP_075596947.1 | REase | GCF_001939115.1 | Leptolyngbya_sp._'hensonii' | Bacteria>Bacillati>Cyanobacteriota/Melainabacteria_group>Cyanobacteriota>Cyanophyceae>Leptolyngbyales>Leptolyngbyaceae>Leptolyngbya_group>Leptolyngbya>unclassified_Leptolyngbya | 187 | BST81_RS02375 | 91590..92153 | 1922337 | Uma2-family-endonuclease-[Leptolyngbya-sp.-'hensonii'] |
| NEQ47452.1 | REase | GCA_010671905.1 | Leptolyngbya_sp._SIOISBB | Bacteria>Bacillati>Cyanobacteriota/Melainabacteria_group>Cyanobacteriota>Cyanophyceae>Leptolyngbyales>Leptolyngbyaceae>Leptolyngbya_group>Leptolyngbya>unclassified_Leptolyngbya | 216 | F6K00_29495 | 6893778..6894428 | 2607771 | Uma2-family-endonuclease-[Leptolyngbya-sp.-SIOISBB] |
| WP_231948149.1 | REase | GCF_001650195.1 | Phormidesmis_priestleyi | Bacteria>Bacillati>Cyanobacteriota/Melainabacteria_group>Cyanobacteriota>Cyanophyceae>Leptolyngbyales>Leptolyngbyaceae>Phormidesmis | 221 | A8707_RS19250 | 12362..13027 | 268141 | Uma2-family-endonuclease-[Phormidesmis-priestleyi] |
| WP_073075159.1 | REase | GCF_001895925.1 | Phormidesmis_priestleyi | Bacteria>Bacillati>Cyanobacteriota/Melainabacteria_group>Cyanobacteriota>Cyanophyceae>Leptolyngbyales>Leptolyngbyaceae>Phormidesmis | 207 | C7B65_RS24720 | 8054..8677 | 268141 | hypothetical-protein-[Phormidesmis-priestleyi] |
| WP_073075158.1 | REase | GCF_001895925.1 | Phormidesmis_priestleyi | Bacteria>Bacillati>Cyanobacteriota/Melainabacteria_group>Cyanobacteriota>Cyanophyceae>Leptolyngbyales>Leptolyngbyaceae>Phormidesmis | 207 | C7B65_RS24725 | 8683..9306 | 268141 | restriction-endonuclease-subunit-R-[Phormidesmis-priestleyi] |
| WP_193855897.1 | REase | GCF_015207105.1 | cf._Phormidesmis_sp._LEGE_11477 | Bacteria>Bacillati>Cyanobacteriota/Melainabacteria_group>Cyanobacteriota>Cyanophyceae>Leptolyngbyales>Leptolyngbyaceae>Phormidesmis>unclassified_Phormidesmis | 173 | IQ256_RS12595 | 47340..47861 | 1828680 | hypothetical-protein-[cf.-Phormidesmis-sp.-LEGE-11477] |
| EKQ69981.1 | REase | GCA_000309945.1 | Leptolyngbyaceae_cyanobacterium_JSC-12 | Bacteria>Bacillati>Cyanobacteriota/Melainabacteria_group>Cyanobacteriota>Cyanophyceae>Leptolyngbyales>Leptolyngbyaceae>unclassified_Leptolyngbyaceae | 124 | OsccyDRAFT_0236 | 260688..261062 | 864702 | hypothetical-protein-OsccyDRAFT_0236-[Leptolyngbyaceae-cyanobacterium-JSC-12] |
| NJO76739.1 | REase | GCA_012033355.1 | Leptolyngbyaceae_cyanobacterium_RM1_406_9 | Bacteria>Bacillati>Cyanobacteriota/Melainabacteria_group>Cyanobacteriota>Cyanophyceae>Leptolyngbyales>Leptolyngbyaceae>unclassified_Leptolyngbyaceae | 216 | HC833_25185 | 550..1200 | 2720434 | Uma2-family-endonuclease-[Leptolyngbyaceae-cyanobacterium-RM1_406_9] |
| EKQ69985.1 | REase | GCA_000309945.1 | Leptolyngbyaceae_cyanobacterium_JSC-12 | Bacteria>Bacillati>Cyanobacteriota/Melainabacteria_group>Cyanobacteriota>Cyanophyceae>Leptolyngbyales>Leptolyngbyaceae>unclassified_Leptolyngbyaceae | 215 | OsccyDRAFT_0241 | 267844..268491 | 864702 | hypothetical-protein-OsccyDRAFT_0241-[Leptolyngbyaceae-cyanobacterium-JSC-12] |
| WP_169613740.1 | REase | GCF_012911975.1 | Nodosilinea_sp._P-1105 | Bacteria>Bacillati>Cyanobacteriota/Melainabacteria_group>Cyanobacteriota>Cyanophyceae>Nodosilineales>Nodosilineaceae>Nodosilinea>unclassified_Nodosilinea | 208 | E1H13_RS12445 | 29516..30142 | 2546229 | hypothetical-protein-[Nodosilinea-sp.-P-1105] |
| WP_194058628.1 | REase | GCF_015207525.1 | Nodosilinea_sp._LEGE_06152 | Bacteria>Bacillati>Cyanobacteriota/Melainabacteria_group>Cyanobacteriota>Cyanophyceae>Nodosilineales>Nodosilineaceae>Nodosilinea>unclassified_Nodosilinea | 138 | IQ265_RS24910 | 15649..16065 | 2777966 | XisH-family-protein-[Nodosilinea-sp.-LEGE-06152] |
| WP_218140785.1 | REase | GCF_014489415.1 | Cylindrospermopsis | Bacteria>Bacillati>Cyanobacteriota/Melainabacteria_group>Cyanobacteriota>Cyanophyceae>Nostocales>Aphanizomenonaceae | 171 | IAR63_RS13135 | 2951491..2952006 | 77021 | MULTISPECIES:-hypothetical-protein-[Cylindrospermopsis] |
| WP_200959782.1 | REase | GCF_018139025.1 | Cylindrospermopsis | Bacteria>Bacillati>Cyanobacteriota/Melainabacteria_group>Cyanobacteriota>Cyanophyceae>Nostocales>Aphanizomenonaceae | 67 | J9B92_RS02360 | 522181..522384 | 77021 | MULTISPECIES:-hypothetical-protein-[Cylindrospermopsis] |
| WP_200959781.1 | REase | GCF_036409495.1 | Cylindrospermopsis | Bacteria>Bacillati>Cyanobacteriota/Melainabacteria_group>Cyanobacteriota>Cyanophyceae>Nostocales>Aphanizomenonaceae | 64 | ASL19_RS17985 | 49235..49429 | 77021 | hypothetical-protein-[Cylindrospermopsis-sp.-CR12] |

| Protein-ID | Domain | Assembly-ID | Strain | Taxonomy | Length | Locus-Tag | Genomic-coordinates | TaxID | Fasta-defline |
|---|---|---|---|---|---|---|---|---|---|
| WP_057177240.1 | REase | - | Cylindrospermopsis | Bacteria>Bacillati>Cyanobacteriota/Melainabacteria_group>Cyanobacteriota>Cyanophyceae>Nostocales>Aphanizomenonaceae | 193 | J9B92_RS12740 | 2848075..2848656 | 77021 | MULTISPECIES:-Uma2-family-endonuclease-[Cylindrospermopsis] |
| WP_039200352.1 | REase | GCF_014696815.1 | Aphanizomenon_flos-aquae | Bacteria>Bacillati>Cyanobacteriota/Melainabacteria_group>Cyanobacteriota>Cyanophyceae>Nostocales>Aphanizomenonaceae>Aphanizomenon | 200 | H6F99_RS24995 | 10839..11441 | 1176 | hypothetical-protein-[Aphanizomenon-flos-aquae] |
| WP_039200354.1 | REase | GCF_014696815.1 | Aphanizomenon_flos-aquae | Bacteria>Bacillati>Cyanobacteriota/Melainabacteria_group>Cyanobacteriota>Cyanophyceae>Nostocales>Aphanizomenonaceae>Aphanizomenon | 164 | H6F99_RS24985 | 9771..10265 | 1176 | DUF4365-domain-containing-protein-[Aphanizomenon-flos-aquae] |
| WP_039201599.1 | REase | GCF_000789435.1 | Aphanizomenon_flos-aquae | Bacteria>Bacillati>Cyanobacteriota/Melainabacteria_group>Cyanobacteriota>Cyanophyceae>Nostocales>Aphanizomenonaceae>Aphanizomenon | 232 | OA07_RS10595 | 19660..20358 | 1176 | hypothetical-protein-[Aphanizomenon-flos-aquae] |
| MBO1045990.1 | REase | GCA_001593825.2 | Aphanizomenon_flos-aquae_UKL13-PB | Bacteria>Bacillati>Cyanobacteriota/Melainabacteria_group>Cyanobacteriota>Cyanophyceae>Nostocales>Aphanizomenonaceae>Aphanizomenon>Aphanizomenon_flos-aquae | 202 | A2T80_019125 | 77087..77695 | 1803587 | Uma2-family-endonuclease-[Aphanizomenon-flos-aquae-UKL13-PB] |
| WP_168466098.1 | REase | GCF_009712065.1 | Aphanizomenon_sp._UHCC_0183 | Bacteria>Bacillati>Cyanobacteriota/Melainabacteria_group>Cyanobacteriota>Cyanophyceae>Nostocales>Aphanizomenonaceae>Aphanizomenon>unclassified_Aphanizomenon | 200 | FJR37_RS11795 | 55253..55855 | 2590028 | hypothetical-protein-[Aphanizomenon-sp.-UHCC-0183] |
| WP_141302950.1 | REase | GCF_006523545.1 | Cylindrospermopsis_raciborskii | Bacteria>Bacillati>Cyanobacteriota/Melainabacteria_group>Cyanobacteriota>Cyanophyceae>Nostocales>Aphanizomenonaceae>Cylindrospermopsis | 178 | FIV49_RS01495 | 321816..322352 | 77022 | hypothetical-protein-[Cylindrospermopsis-raciborskii] |
| WP_236308451.1 | REase | GCF_021650815.1 | Cylindrospermopsis_raciborskii | Bacteria>Bacillati>Cyanobacteriota/Melainabacteria_group>Cyanobacteriota>Cyanophyceae>Nostocales>Aphanizomenonaceae>Cylindrospermopsis | 178 | L3I90_RS00105 | 21866..22402 | 77022 | hypothetical-protein-[Cylindrospermopsis-raciborskii] |
| WP_071248148.1 | REase | GCF_001858125.1 | Cylindrospermopsis_raciborskii | Bacteria>Bacillati>Cyanobacteriota/Melainabacteria_group>Cyanobacteriota>Cyanophyceae>Nostocales>Aphanizomenonaceae>Cylindrospermopsis | 165 | CENA302_RS09790 | 21788..22285 | 77022 | hypothetical-protein-[Cylindrospermopsis-raciborskii] |
| WP_211942099.1 | REase | GCF_018139025.1 | Cylindrospermopsis_raciborskii | Bacteria>Bacillati>Cyanobacteriota/Melainabacteria_group>Cyanobacteriota>Cyanophyceae>Nostocales>Aphanizomenonaceae>Cylindrospermopsis | 72 | J9B92_RS02365 | 522394..522612 | 77022 | hypothetical-protein-[Cylindrospermopsis-raciborskii] |
| WP_040009385.1 | REase | - | Cylindrospermopsis_raciborskii | Bacteria>Bacillati>Cyanobacteriota/Melainabacteria_group>Cyanobacteriota>Cyanophyceae>Nostocales>Aphanizomenonaceae>Cylindrospermopsis | 193 | CRC_RS10850 | 1076..1657 | 77022 | Uma2-family-endonuclease-[Cylindrospermopsis-raciborskii] |
| WP_071242354.1 | REase | GCF_001858115.1 | Cylindrospermopsis_raciborskii | Bacteria>Bacillati>Cyanobacteriota/Melainabacteria_group>Cyanobacteriota>Cyanophyceae>Nostocales>Aphanizomenonaceae>Cylindrospermopsis | 193 | BCV63_RS11915 | 60687..61268 | 77022 | Uma2-family-endonuclease-[Cylindrospermopsis-raciborskii] |
| UJL35161.1 | REase | GCA_003367075.2 | Cylindrospermopsis_raciborskii_Cr2010 | Bacteria>Bacillati>Cyanobacteriota/Melainabacteria_group>Cyanobacteriota>Cyanophyceae>Nostocales>Aphanizomenonaceae>Cylindrospermopsis>Cylindrospermopsis_raciborskii | 178 | C6N34_008495 | 1859697..1860233 | 2108055 | hypothetical-protein-C6N34_008495-[Cylindrospermopsis-raciborskii-Cr2010] |
| WP_168360491.1 | REase | GCF_009712075.2 | Dolichospermum_planctonicum | Bacteria>Bacillati>Cyanobacteriota/Melainabacteria_group>Cyanobacteriota>Cyanophyceae>Nostocales>Aphanizomenonaceae>Dolichospermum | 194 | FJR41_RS21620 | 27411..27995 | 136072 | hypothetical-protein-[Dolichospermum-planctonicum] |
| WP_193849070.1 | REase | GCF_015207785.1 | Dolichospermum_flos-aquae | Bacteria>Bacillati>Cyanobacteriota/Melainabacteria_group>Cyanobacteriota>Cyanophyceae>Nostocales>Aphanizomenonaceae>Dolichospermum | 194 | IQ222_RS12490 | 18594..19178 | 1166 | hypothetical-protein-[Dolichospermum-flos-aquae] |
| WP_271835225.1 | REase | GCF_028330815.1 | Dolichospermum_lemmermannii | Bacteria>Bacillati>Cyanobacteriota/Melainabacteria_group>Cyanobacteriota>Cyanophyceae>Nostocales>Aphanizomenonaceae>Dolichospermum | 200 | PN450_RS01670 | 4663..5265 | 54295 | hypothetical-protein-[Dolichospermum-lemmermannii] |
| MBS3030298.1 | REase | GCA_018295985.1 | Dolichospermum_sp._DET66 | Bacteria>Bacillati>Cyanobacteriota/Melainabacteria_group>Cyanobacteriota>Cyanophyceae>Nostocales>Aphanizomenonaceae>Dolichospermum>unclassified_Dolichospermum | 200 | HCA26_25895 | 5399793..5400395 | 2704415 | hypothetical-protein-[Dolichospermum-sp.-DET66] |
| BAZ90341.1 | REase | GCA_002368135.1 | Raphidiopsis_curvata_NIES-932 | Bacteria>Bacillati>Cyanobacteriota/Melainabacteria_group>Cyanobacteriota>Cyanophyceae>Nostocales>Aphanizomenonaceae>Raphidiopsis>Raphidiopsis_curvata | 178 | NIES932_18400 | 1955002..1955538 | 1973482 | hypothetical-protein-NIES932_18400-[Raphidiopsis-curvata-NIES-932] |
| MBW4670337.1 | REase | GCA_019359595.1 | Cyanomargarita_calcarea_GSE-NOS-MK-12-04C | Bacteria>Bacillati>Cyanobacteriota/Melainabacteria_group>Cyanobacteriota>Cyanophyceae>Nostocales>Cyanomargaritaceae>Cyanomargarita>Cyanomargarita_calcarea | 207 | KME60_23705 | 104766..105389 | 2839659 | Uma2-family-endonuclease-[Cyanomargarita-calcarea-GSE-NOS-MK-12-04C] |
| MBW4670328.1 | REase | GCA_019359595.1 | Cyanomargarita_calcarea_GSE-NOS-MK-12-04C | Bacteria>Bacillati>Cyanobacteriota/Melainabacteria_group>Cyanobacteriota>Cyanophyceae>Nostocales>Cyanomargaritaceae>Cyanomargarita>Cyanomargarita_calcarea | 196 | KME60_23660 | 94785..95375 | 2839659 | Uma2-family-endonuclease-[Cyanomargarita-calcarea-GSE-NOS-MK-12-04C] |
| MBW4588293.1 | REase | GCA_019359795.1 | Aetokthonos_hydrillicola_CCALA_1050 | Bacteria>Bacillati>Cyanobacteriota/Melainabacteria_group>Cyanobacteriota>Cyanophyceae>Nostocales>Hapalosiphonaceae>Aetokthonos>Aetokthonos_hydrillicola | 201 | KME33_24305 | 45877..46482 | 2710484 | Uma2-family-endonuclease-[Aetokthonos-hydrillicola-CCALA-1050] |
| WP_009454386.1 | REase | GCF_000231365.1 | Fischerella_thermalis | Bacteria>Bacillati>Cyanobacteriota/Melainabacteria_group>Cyanobacteriota>Cyanophyceae>Nostocales>Hapalosiphonaceae>Fischerella | 194 | FJSC11DRAFT_RS02255 | 573512..574096 | 372787 | hypothetical-protein-[Fischerella-thermalis] |
| WP_102148298.1 | REase | GCF_002870755.1 | Fischerella_thermalis | Bacteria>Bacillati>Cyanobacteriota/Melainabacteria_group>Cyanobacteriota>Cyanophyceae>Nostocales>Hapalosiphonaceae>Fischerella | 194 | CEN42_RS03270 | 11352..11936 | 372787 | hypothetical-protein-[Fischerella-thermalis] |
| EHC19654.1 | REase | - | Fischerella_thermalis_JSC-11 | Bacteria>Bacillati>Cyanobacteriota/Melainabacteria_group>Cyanobacteriota>Cyanophyceae>Nostocales>Hapalosiphonaceae>Fischerella>Fischerella_thermalis | 194 | FJSC11DRAFT_0471 | 573512..574096 | 741277 | hypothetical-protein-FJSC11DRAFT_0471-[Fischerella-thermalis-JSC-11] |
| PMB36921.1 | REase | GCA_002870755.1 | Fischerella_thermalis_CCMEE_5208 | Bacteria>Bacillati>Cyanobacteriota/Melainabacteria_group>Cyanobacteriota>Cyanophyceae>Nostocales>Hapalosiphonaceae>Fischerella>Fischerella_thermalis | 194 | CEN42_03250 | 11352..11936 | 2019669 | hypothetical-protein-CEN42_03250-[Fischerella-thermalis-CCMEE-5208] |
| WP_197036065.1 | REase | GCF_000517105.1 | Fischerella_sp._PCC_9605 | Bacteria>Bacillati>Cyanobacteriota/Melainabacteria_group>Cyanobacteriota>Cyanophyceae>Nostocales>Hapalosiphonaceae>Fischerella>unclassified_Fischerella | 190 | FIS9605_RS37260 | 336523..337095 | 1173024 | endonuclease-domain-containing-protein-[Fischerella-sp.-PCC-9605] |
| WP_200988280.1 | REase | GCF_015245355.1 | Anabaenopsis_elenkinii | Bacteria>Bacillati>Cyanobacteriota/Melainabacteria_group>Cyanobacteriota>Cyanophyceae>Nostocales>Nodulariaceae>Anabaenopsis | 203 | IM676_RS18810 | 4312126..4312737 | 156213 | Uma2-family-endonuclease-[Anabaenopsis-elenkinii] |
| WP_271941656.1 | REase | GCF_028329345.1 | Nodularia_spumigena | Bacteria>Bacillati>Cyanobacteriota/Melainabacteria_group>Cyanobacteriota>Cyanophyceae>Nostocales>Nodulariaceae>Nodularia | 220 | PN454_RS18925 | 14854..15516 | 70799 | Uma2-family-endonuclease-[Nodularia-spumigena] |
| WP_280650399.1 | REase | GCF_963920845.1 | Umezakia_ovalisporum | Bacteria>Bacillati>Cyanobacteriota/Melainabacteria_group>Cyanobacteriota>Cyanophyceae>Nostocales>Nodulariaceae>Umezakia | 194 | NWP27_RS05685 | 17034..17618 | 75695 | hypothetical-protein-[Umezakia-ovalisporum] |
| WP_280656505.1 | REase | GCF_029891765.1 | Umezakia_ovalisporum | Bacteria>Bacillati>Cyanobacteriota/Melainabacteria_group>Cyanobacteriota>Cyanophyceae>Nostocales>Nodulariaceae>Umezakia | 194 | NWP15_RS06370 | 22065..22649 | 75695 | hypothetical-protein-[Umezakia-ovalisporum] |
| WP_280692696.1 | REase | GCF_029891755.1 | Umezakia_ovalisporum | Bacteria>Bacillati>Cyanobacteriota/Melainabacteria_group>Cyanobacteriota>Cyanophyceae>Nostocales>Nodulariaceae>Umezakia | 194 | NWP23_RS06510 | 33044..33628 | 75695 | hypothetical-protein-[Umezakia-ovalisporum] |
| WP_280699382.1 | REase | GCF_029891685.1 | Umezakia_ovalisporum | Bacteria>Bacillati>Cyanobacteriota/Melainabacteria_group>Cyanobacteriota>Cyanophyceae>Nostocales>Nodulariaceae>Umezakia | 194 | NWP25_RS10980 | 21860..22444 | 75695 | hypothetical-protein-[Umezakia-ovalisporum] |
| WP_280657421.1 | REase | GCF_029891765.1 | Umezakia_ovalisporum | Bacteria>Bacillati>Cyanobacteriota/Melainabacteria_group>Cyanobacteriota>Cyanophyceae>Nostocales>Nodulariaceae>Umezakia | 220 | NWP26_RS18495 | 4760..5422 | 75695 | Uma2-family-endonuclease-[Umezakia-ovalisporum] |
| WP_190371459.1 | REase | GCF_014696335.1 | unclassified_Anabaena | Bacteria>Bacillati>Cyanobacteriota/Melainabacteria_group>Cyanobacteriota>Cyanophyceae>Nostocales>Nostocaceae>Anabaena | 194 | H6F39_RS17010 | 3393..3977 | 2619674 | MULTISPECIES:-hypothetical-protein-[unclassified-Anabaena] |
| WP_168507186.1 | REase | GCF_009712035.1 | Anabaena_sp._UHCC_0187 | Bacteria>Bacillati>Cyanobacteriota/Melainabacteria_group>Cyanobacteriota>Cyanophyceae>Nostocales>Nostocaceae>Anabaena>unclassified_Anabaena | 194 | FJR11_RS20800 | 5068..5652 | 2590018 | hypothetical-protein-[Anabaena-sp.-UHCC-0187] |
| WP_193913745.1 | REase | GCF_015207005.2 | Desmonostoc_muscorum | Bacteria>Bacillati>Cyanobacteriota/Melainabacteria_group>Cyanobacteriota>Cyanophyceae>Nostocales>Nostocaceae>Desmonostoc | 138 | IQ276_RS32455 | 7897257..7897673 | 1179 | XisH-family-protein-[Desmonostoc-muscorum] |
| WP_012411894.1 | REase | GCF_000020025.1 | Nostoc_punctiforme | Bacteria>Bacillati>Cyanobacteriota/Melainabacteria_group>Cyanobacteriota>Cyanophyceae>Nostocales>Nostocaceae>Nostoc | 206 | NPUN_RS28580 | 6975426..6976046 | 272131 | Uma2-family-endonuclease-[Nostoc-punctiforme] |
| WP_251958559.1 | REase | GCF_023734775.1 | Nostoc_commune | Bacteria>Bacillati>Cyanobacteriota/Melainabacteria_group>Cyanobacteriota>Cyanophyceae>Nostocales>Nostocaceae>Nostoc | 206 | ANSO36C_RS04430 | 989990..990610 | 1178 | Uma2-family-endonuclease-[Nostoc-commune] |
| WP_100904447.1 | REase | GCF_963875825.1 | Nostoc_flagelliforme | Bacteria>Bacillati>Cyanobacteriota/Melainabacteria_group>Cyanobacteriota>Cyanophyceae>Nostocales>Nostocaceae>Nostoc | 200 | COO91_RS47970 | 749536..750138 | 1306274 | Uma2-family-endonuclease-[Nostoc-flagelliforme] |
| WP_181929828.1 | REase | GCF_014023275.1 | Nostoc_edaphicum | Bacteria>Bacillati>Cyanobacteriota/Melainabacteria_group>Cyanobacteriota>Cyanophyceae>Nostocales>Nostocaceae>Nostoc | 154 | HUN01_RS33855 | 7494323..7494787 | 264686 | DUF3883-domain-containing-protein-[Nostoc-edaphicum] |
| WP_012411907.1 | REase | GCF_000020025.1 | Nostoc_punctiforme | Bacteria>Bacillati>Cyanobacteriota/Melainabacteria_group>Cyanobacteriota>Cyanophyceae>Nostocales>Nostocaceae>Nostoc | 138 | NPUN_RS28645 | 6994929..6995345 | 272131 | XisH-family-protein-[Nostoc-punctiforme] |
| MBW4429555.1 | REase | GCA_019359245.1 | Nostoc_desertorum_CM1-VF14 | Bacteria>Bacillati>Cyanobacteriota/Melainabacteria_group>Cyanobacteriota>Cyanophyceae>Nostocales>Nostocaceae>Nostoc>Nostoc_desertorum | 188 | KME50_35635 | 5380..5946 | 287862 | hypothetical-protein-[Nostoc-desertorum-CM1-VF14] |
| MBW4455561.1 | REase | GCA_019359215.1 | Nostoc_indistinguendum_CM1-VF10 | Bacteria>Bacillati>Cyanobacteriota/Melainabacteria_group>Cyanobacteriota>Cyanophyceae>Nostocales>Nostocaceae>Nostoc>Nostoc_indistinguendum | 200 | KME55_24200 | 26172..26774 | 287863 | Uma2-family-endonuclease-[Nostoc-indistinguendum-CM1-VF10] |
| MBI1240692.1 | REase | GCA_016124525.1 | Nostoc_sp._RI_552 | Bacteria>Bacillati>Cyanobacteriota/Melainabacteria_group>Cyanobacteriota>Cyanophyceae>Nostocales>Nostocaceae>Nostoc>unclassified_Nostoc | 194 | GC195_03650 | 35801..36385 | 2654185 | hypothetical-protein-[Nostoc-sp.-RI_552] |
| WP_138504564.1 | REase | GCF_005869855.1 | Nostoc_sp._PA-18-2419 | Bacteria>Bacillati>Cyanobacteriota/Melainabacteria_group>Cyanobacteriota>Cyanophyceae>Nostocales>Nostocaceae>Nostoc>unclassified_Nostoc | 194 | FCR06_RS25150 | 44040..44624 | 2575443 | hypothetical-protein-[Nostoc-sp.-PA-18-2419] |
| WP_162398345.1 | REase | - | Nostoc_sp._B(2019) | Bacteria>Bacillati>Cyanobacteriota/Melainabacteria_group>Cyanobacteriota>Cyanophyceae>Nostocales>Nostocaceae>Nostoc>unclassified_Nostoc | 211 | GS682_RS23150 | 20125..20760 | 2690446 | Uma2-family-endonuclease-[Nostoc-sp.-B(2019)] |
| WP_194144966.1 | REase | GCF_015206985.1 | Nostoc_sp._LEGE_12447 | Bacteria>Bacillati>Cyanobacteriota/Melainabacteria_group>Cyanobacteriota>Cyanophyceae>Nostocales>Nostocaceae>Nostoc>unclassified_Nostoc | 138 | IQ274_RS34150 | 5942..6358 | 1828640 | XisH-family-protein-[Nostoc-sp.-LEGE-12447] |
| WP_104909972.1 | REase | GCF_002949795.1 | Nostoc_sp._'Lobaria_pulmonaria_(5183)_cyanobiont' | Bacteria>Bacillati>Cyanobacteriota/Melainabacteria_group>Cyanobacteriota>Cyanophyceae>Nostocales>Nostocaceae>Nostoc>unclassified_Nostoc>Nostoc_cyanobionts | 128 | NLP_RS21720 | 4933071..4933457 | 1618022 | endonuclease-domain-containing-protein-[Nostoc-sp.-'Lobaria-pulmonaria-(5183)-cyanobiont'] |
| MBW4559184.1 | REase | GCA_019358915.1 | Trichormus_sp._ATA11-4-KO1 | Bacteria>Bacillati>Cyanobacteriota/Melainabacteria_group>Cyanobacteriota>Cyanophyceae>Nostocales>Nostocaceae>Trichormus>unclassified_Trichormus | 208 | KME59_25355 | 1670..2296 | 2835091 | type-I-restriction-endonuclease-subunit-R-[Trichormus-sp.-ATA11-4-KO1] |
| MBV6627483.1 | REase | GCA_019192545.1 | Rivularia_sp._MS3 | Bacteria>Bacillati>Cyanobacteriota/Melainabacteria_group>Cyanobacteriota>Cyanophyceae>Nostocales>Rivulariaceae>Rivularia>unclassified_Rivularia_(in:_cyanobacteria) | 214 | KI793_31945 | 15275..15919 | 3012223 | Uma2-family-endonuclease-[Rivularia-sp.-(in:-cyanobacteria)] |
| MBV6627479.1 | REase | GCA_019192545.1 | Rivularia_sp._MS3 | Bacteria>Bacillati>Cyanobacteriota/Melainabacteria_group>Cyanobacteriota>Cyanophyceae>Nostocales>Rivulariaceae>Rivularia>unclassified_Rivularia_(in:_cyanobacteria) | 212 | KI793_31920 | 7396..8034 | 3012223 | endonuclease-domain-containing-protein-[Rivularia-sp.-(in:-cyanobacteria)] |
| KAB8333556.1 | REase | GCA_000828085.3 | Scytonema_tolypothrichoides_VB-61278 | Bacteria>Bacillati>Cyanobacteriota/Melainabacteria_group>Cyanobacteriota>Cyanophyceae>Nostocales>Scytonemataceae>Scytonema>Scytonema_tolypothrichoides | 193 | SD80_012875 | 26196..26777 | 1233231 | YraN-family-protein-[Scytonema-tolypothrichoides-VB-61278] |
| WP_198940959.1 | REase | GCF_963872035.1 | Scytonema_sp._HK-05 | Bacteria>Bacillati>Cyanobacteriota/Melainabacteria_group>Cyanobacteriota>Cyanophyceae>Nostocales>Scytonemataceae>Scytonema>unclassified_Scytonema | 235 | NIES2130_RS44050 | 79554..80261 | 1137095 | endonuclease-domain-containing-protein-[Scytonema-sp.-HK-05] |
| MBR8836032.1 | REase | GCA_016903675.1 | Stigonema_ocellatum_SAG_48.90_=_DSM_106950 | Bacteria>Bacillati>Cyanobacteriota/Melainabacteria_group>Cyanobacteriota>Cyanophyceae>Nostocales>Stigonemataceae>Stigonema>Stigonema_ocellatum | 212 | DSM106950_18915 | 24970..25608 | 221294 | Uma2-family-endonuclease-[Stigonema-ocellatum-SAG-48.90-=-DSM-106950] |
| WP_045875056.1 | REase | GCF_963860465.1 | unclassified_Tolypothrix | Bacteria>Bacillati>Cyanobacteriota/Melainabacteria_group>Cyanobacteriota>Cyanophyceae>Nostocales>Tolypothrichaceae>Tolypothrix | 200 | FDUTEX481_RS39720 | 32242..32844 | 2649714 | MULTISPECIES:-Uma2-family-endonuclease-[unclassified-Tolypothrix] |
| WP_045875058.1 | REase | GCF_963860465.1 | unclassified_Tolypothrix | Bacteria>Bacillati>Cyanobacteriota/Melainabacteria_group>Cyanobacteriota>Cyanophyceae>Nostocales>Tolypothrichaceae>Tolypothrix | 137 | FDUTEX481_RS39730 | 33258..33671 | 2649714 | MULTISPECIES:-XisH-family-protein-[unclassified-Tolypothrix] |
| KAB8314076.1 | REase | GCA_000828075.3 | Tolypothrix_campylonemoides_VB511288 | Bacteria>Bacillati>Cyanobacteriota/Melainabacteria_group>Cyanobacteriota>Cyanophyceae>Nostocales>Tolypothrichaceae>Tolypothrix>Tolypothrix_campylonemoides | 215 | SD81_039395 | 93798..94445 | 1245935 | Uma2-family-endonuclease-[Tolypothrix-campylonemoides-VB511288] |
| MBW4580239.1 | REase | GCA_019358895.1 | Tildeniella_nuda_ZEHNDER_1965/U140 | Bacteria>Bacillati>Cyanobacteriota/Melainabacteria_group>Cyanobacteriota>Cyanophyceae>Oculatellales>Oculatellaceae>Tildeniella>Tildeniella_nuda | 203 | KME42_11790 | 250494..251105 | 2136405 | hypothetical-protein-[Tildeniella-nuda-ZEHNDER-1965/U140] |
| WP_096720876.1 | REase | - | Chondrocystis_sp._NIES-4102 | Bacteria>Bacillati>Cyanobacteriota/Melainabacteria_group>Cyanobacteriota>Cyanophyceae>Oscillatoriophycideae>Chroococcales>Chroococcaceae>Chondrocystis>unclassified_Chondrocystis | 246 | CDC35_RS11870 | 2756207..2756947 | 2005460 | DUF4365-domain-containing-protein-[Chondrocystis-sp.-NIES-4102] |
| PSB44269.1 | REase | GCA_003004015.1 | Cyanosarcina_cf._burmensis_CCALA_770 | Bacteria>Bacillati>Cyanobacteriota/Melainabacteria_group>Cyanobacteriota>Cyanophyceae>Oscillatoriophycideae>Chroococcales>Chroococcaceae>Cyanosarcina>Cyanosarcina_burmensis | 211 | C7B80_21180 | 7500..8135 | 2107697 | Uma2-family-endonuclease-[Cyanosarcina-cf.-burmensis-CCALA-770] |
| WP_106371306.1 | REase | GCF_963877105.1 | Chlorogloea_sp._CCALA_695 | Bacteria>Bacillati>Cyanobacteriota/Melainabacteria_group>Cyanobacteriota>Cyanophyceae>Oscillatoriophycideae>Chroococcales>Entophysalidaceae>Chlorogloea>unclassified_Chlorogloea | 188 | C7B70_RS23195 | 10473..11039 | 2107693 | hypothetical-protein-[Chlorogloea-sp.-CCALA-695] |
| WP_065715518.1 | REase | GCF_001693255.1 | Picosynechococcus_sp._PCC_7003 | Bacteria>Bacillati>Cyanobacteriota/Melainabacteria_group>Cyanobacteriota>Cyanophyceae>Oscillatoriophycideae>Chroococcales>Geminocystaceae>Picosynechococcus>unclassified_Picosynechococcus | 121 | AWQ21_RS15090 | 99532..99897 | 374981 | endonuclease-domain-containing-protein-[Synechococcus-sp.-PCC-7003] |
| WP_004266777.1 | REase | GCF_000312165.1 | Microcystis | Bacteria>Bacillati>Cyanobacteriota/Melainabacteria_group>Cyanobacteriota>Cyanophyceae>Oscillatoriophycideae>Chroococcales>Microcystaceae | 176 | MICAB_RS21660 | 34508..35038 | 1125 | hypothetical-protein-[Microcystis-aeruginosa] |
| WP_002766562.1 | REase | GCF_000312185.1 | Microcystis | Bacteria>Bacillati>Cyanobacteriota/Melainabacteria_group>Cyanobacteriota>Cyanophyceae>Oscillatoriophycideae>Chroococcales>Microcystaceae | 161 | PN487_RS04640 | 43075..43560 | 1125 | MULTISPECIES:-hypothetical-protein-[Microcystis] |
| WP_004162513.1 | REase | GCF_026222535.1 | Microcystis | Bacteria>Bacillati>Cyanobacteriota/Melainabacteria_group>Cyanobacteriota>Cyanophyceae>Oscillatoriophycideae>Chroococcales>Microcystaceae | 166 | H6F92_RS24325 | 6779..7279 | 1125 | MULTISPECIES:-hypothetical-protein-[Microcystis] |
| WP_002754172.1 | REase | GCF_003206625.1 | Microcystis | Bacteria>Bacillati>Cyanobacteriota/Melainabacteria_group>Cyanobacteriota>Cyanophyceae>Oscillatoriophycideae>Chroococcales>Microcystaceae | 126 | PN479_RS02425 | 9141..9521 | 1125 | MULTISPECIES:-GxxExxY-protein-[Microcystis] |
| WP_190359817.1 | REase | GCF_026222535.1 | Microcystis | Bacteria>Bacillati>Cyanobacteriota/Melainabacteria_group>Cyanobacteriota>Cyanophyceae>Oscillatoriophycideae>Chroococcales>Microcystaceae | 126 | H6F92_RS24375 | 23574..23954 | 1125 | MULTISPECIES:-GxxExxY-protein-[Microcystis] |
| WP_002766565.1 | REase | GCF_042134685.1 | Microcystis | Bacteria>Bacillati>Cyanobacteriota/Melainabacteria_group>Cyanobacteriota>Cyanophyceae>Oscillatoriophycideae>Chroococcales>Microcystaceae | 139 | PN502_RS07770 | 8503..8922 | 1125 | MULTISPECIES:-XisH-family-protein-[Microcystis] |
| WP_002790041.1 | REase | GCF_000312225.1 | Microcystis_aeruginosa | Bacteria>Bacillati>Cyanobacteriota/Melainabacteria_group>Cyanobacteriota>Cyanophyceae>Oscillatoriophycideae>Chroococcales>Microcystaceae>Microcystis | 161 | MICAF_RS22370 | 133752..134237 | 1126 | hypothetical-protein-[Microcystis-aeruginosa] |
| WP_271989202.1 | REase | GCF_028330905.1 | Microcystis_aeruginosa | Bacteria>Bacillati>Cyanobacteriota/Melainabacteria_group>Cyanobacteriota>Cyanophyceae>Oscillatoriophycideae>Chroococcales>Microcystaceae>Microcystis | 161 | PN502_RS07780 | 12635..13120 | 1126 | hypothetical-protein-[Microcystis-aeruginosa] |
| WP_002748928.1 | REase | GCF_000332585.1 | Microcystis_aeruginosa | Bacteria>Bacillati>Cyanobacteriota/Melainabacteria_group>Cyanobacteriota>Cyanophyceae>Oscillatoriophycideae>Chroococcales>Microcystaceae>Microcystis | 99 | BH695_RS28120 | 535681..535980 | 1126 | GxxExxY-protein-[Microcystis-aeruginosa] |
| WP_002748929.1 | REase | GCF_000332585.1 | Microcystis_aeruginosa | Bacteria>Bacillati>Cyanobacteriota/Melainabacteria_group>Cyanobacteriota>Cyanophyceae>Oscillatoriophycideae>Chroococcales>Microcystaceae>Microcystis | 49 | BH695_RS28125 | 536043..536192 | 1126 | GxxExxY-protein-[Microcystis-aeruginosa] |
| WP_150976540.1 | REase | GCF_028331105.1 | Microcystis_aeruginosa | Bacteria>Bacillati>Cyanobacteriota/Melainabacteria_group>Cyanobacteriota>Cyanophyceae>Oscillatoriophycideae>Chroococcales>Microcystaceae>Microcystis | 196 | PN487_RS04575 | 28074..28664 | 1126 | Uma2-family-endonuclease-[Microcystis-aeruginosa] |
| TRU20106.1 | REase | GCA_007095795.1 | Microcystis_aeruginosa_Ma_MB_S_20031200_S102D | Bacteria>Bacillati>Cyanobacteriota/Melainabacteria_group>Cyanobacteriota>Cyanophyceae>Oscillatoriophycideae>Chroococcales>Microcystaceae>Microcystis>Microcystis_aeruginosa | 176 | EWV79_18815 | 36284..36814 | 2486255 | hypothetical-protein-EWV79_18815-[Microcystis-aeruginosa-Ma_MB_S_20031200_S102D] |
| CCI01317.1 | REase | GCA_000312185.1 | Microcystis_aeruginosa_PCC_9443 | Bacteria>Bacillati>Cyanobacteriota/Melainabacteria_group>Cyanobacteriota>Cyanophyceae>Oscillatoriophycideae>Chroococcales>Microcystaceae>Microcystis>Microcystis_aeruginosa | 161 | MICAC_1830013 | 81905..82390 | 1160281 | conserved-hypothetical-protein-[Microcystis-aeruginosa-PCC-9443] |
| MBE5230559.1 | REase | GCA_014946735.1 | Microcystis_aeruginosa_PMC_728.11 | Bacteria>Bacillati>Cyanobacteriota/Melainabacteria_group>Cyanobacteriota>Cyanophyceae>Oscillatoriophycideae>Chroococcales>Microcystaceae>Microcystis>Microcystis_aeruginosa | 161 | INR81_16210 | 52445..52930 | 2316423 | hypothetical-protein-[Microcystis-aeruginosa-PMC-728.11] |
| NCR78977.1 | REase | GCA_009996105.1 | Microcystis_aeruginosa_K13-10 | Bacteria>Bacillati>Cyanobacteriota/Melainabacteria_group>Cyanobacteriota>Cyanophyceae>Oscillatoriophycideae>Chroococcales>Microcystaceae>Microcystis>Microcystis_aeruginosa | 161 | GPI91_02960 | 21037..21522 | 2685970 | hypothetical-protein-[Microcystis-aeruginosa-K13-10] |
| NCS23182.1 | REase | GCA_009996485.1 | Microcystis_aeruginosa_BS13-02 | Bacteria>Bacillati>Cyanobacteriota/Melainabacteria_group>Cyanobacteriota>Cyanophyceae>Oscillatoriophycideae>Chroococcales>Microcystaceae>Microcystis>Microcystis_aeruginosa | 161 | GPI96_01070 | 41441..41926 | 2685952 | hypothetical-protein-[Microcystis-aeruginosa-BS13-02] |
| NCR40135.1 | REase | GCA_009995705.1 | Microcystis_aeruginosa_W13-11 | Bacteria>Bacillati>Cyanobacteriota/Melainabacteria_group>Cyanobacteriota>Cyanophyceae>Oscillatoriophycideae>Chroococcales>Microcystaceae>Microcystis>Microcystis_aeruginosa | 126 | GPJ21_09820 | 18418..18798 | 2685991 | GxxExxY-protein-[Microcystis-aeruginosa-W13-11] |
| TRU24935.1 | REase | GCA_007095795.1 | Microcystis_aeruginosa_Ma_MB_S_20031200_S102D | Bacteria>Bacillati>Cyanobacteriota/Melainabacteria_group>Cyanobacteriota>Cyanophyceae>Oscillatoriophycideae>Chroococcales>Microcystaceae>Microcystis>Microcystis_aeruginosa | 126 | EWV79_09020 | 20015..20395 | 2486255 | GxxExxY-protein-[Microcystis-aeruginosa-Ma_MB_S_20031200_S102D] |
| CCI01319.1 | REase | GCA_000312185.1 | Microcystis_aeruginosa_PCC_9443 | Bacteria>Bacillati>Cyanobacteriota/Melainabacteria_group>Cyanobacteriota>Cyanophyceae>Oscillatoriophycideae>Chroococcales>Microcystaceae>Microcystis>Microcystis_aeruginosa | 139 | MICAC_1830015 | 86094..86513 | 1160281 | XisH-protein-[Microcystis-aeruginosa-PCC-9443] |
| NCR78975.1 | REase | GCA_009996105.1 | Microcystis_aeruginosa_K13-10 | Bacteria>Bacillati>Cyanobacteriota/Melainabacteria_group>Cyanobacteriota>Cyanophyceae>Oscillatoriophycideae>Chroococcales>Microcystaceae>Microcystis>Microcystis_aeruginosa | 139 | GPI91_02950 | 16905..17324 | 2685970 | fatty-acid-oxidation-protein-subunit-alpha-[Microcystis-aeruginosa-K13-10] |
| ARI79888.1 | REase | GCA_002095975.1 | Microcystis_aeruginosa_PCC_7806SL | Bacteria>Bacillati>Cyanobacteriota/Melainabacteria_group>Cyanobacteriota>Cyanophyceae>Oscillatoriophycideae>Chroococcales>Microcystaceae>Microcystis>Microcystis_aeruginosa>Microcystis_aeruginosa_PCC_7806 | 99 | BH695_0607 | 535681..535980 | 1903187 | hypothetical-protein-BH695_0607-[Microcystis-aeruginosa-PCC-7806SL] |
| ARI79889.1 | REase | GCA_002095975.1 | Microcystis_aeruginosa_PCC_7806SL | Bacteria>Bacillati>Cyanobacteriota/Melainabacteria_group>Cyanobacteriota>Cyanophyceae>Oscillatoriophycideae>Chroococcales>Microcystaceae>Microcystis>Microcystis_aeruginosa>Microcystis_aeruginosa_PCC_7806 | 49 | BH695_0608 | 536043..536192 | 1903187 | hypothetical-protein-BH695_0608-[Microcystis-aeruginosa-PCC-7806SL] |
| REJ45950.1 | REase | GCA_003389435.1 | Microcystis_flos-aquae_DF17 | Bacteria>Bacillati>Cyanobacteriota/Melainabacteria_group>Cyanobacteriota>Cyanophyceae>Oscillatoriophycideae>Chroococcales>Microcystaceae>Microcystis>Microcystis_flos-aquae | 161 | DWQ53_12255 | 11688..12173 | 2060470 | hypothetical-protein-DWQ53_12255-[Microcystis-flos-aquae-DF17] |
| MCA2509362.1 | REase | GCA_020244605.1 | Microcystis_sp._M60BS1 | Bacteria>Bacillati>Cyanobacteriota/Melainabacteria_group>Cyanobacteriota>Cyanophyceae>Oscillatoriophycideae>Chroococcales>Microcystaceae>Microcystis>unclassified_Microcystis | 161 | IM561_02885 | 18916..19401 | 2771202 | hypothetical-protein-[Microcystis-sp.-M60BS1] |
| WP_232306173.1 | REase | GCF_000297435.1 | Microcystis_sp._T1-4 | Bacteria>Bacillati>Cyanobacteriota/Melainabacteria_group>Cyanobacteriota>Cyanophyceae>Oscillatoriophycideae>Chroococcales>Microcystaceae>Microcystis>unclassified_Microcystis | 136 | MICAI_RS09615 | 1..411 | 1160279 | GxxExxY-protein,-partial-[Microcystis-sp.-T1-4] |
| MCA2627573.1 | REase | GCA_020245485.1 | Microcystis_sp._M091S2 | Bacteria>Bacillati>Cyanobacteriota/Melainabacteria_group>Cyanobacteriota>Cyanophyceae>Oscillatoriophycideae>Chroococcales>Microcystaceae>Microcystis>unclassified_Microcystis | 126 | IM505_05350 | 39106..39486 | 2771177 | GxxExxY-protein-[Microcystis-sp.-M091S2] |
| MCE2669069.1 | REase | GCA_021300285.1 | Microcystis_sp._49638_E5 | Bacteria>Bacillati>Cyanobacteriota/Melainabacteria_group>Cyanobacteriota>Cyanophyceae>Oscillatoriophycideae>Chroococcales>Microcystaceae>Microcystis>unclassified_Microcystis | 126 | LW728_07590 | 19966..20346 | 2904986 | GxxExxY-protein-[Microcystis-sp.-49638_E5] |
| WP_026788983.1 | REase | GCF_963863855.1 | Planktothrix | Bacteria>Bacillati>Cyanobacteriota/Melainabacteria_group>Cyanobacteriota>Cyanophyceae>Oscillatoriophycideae>Oscillatoriales>Microcoleaceae | 201 | N316_RS0113735 | 332915..333520 | 54304 | MULTISPECIES:-Uma2-family-endonuclease-[Planktothrix] |
| WP_026794457.1 | REase | GCF_000464725.1 | Planktothrix | Bacteria>Bacillati>Cyanobacteriota/Melainabacteria_group>Cyanobacteriota>Cyanophyceae>Oscillatoriophycideae>Oscillatoriales>Microcoleaceae | 201 | NMG75_RS16750 | 3653801..3654406 | 54304 | MULTISPECIES:-Uma2-family-endonuclease-[Planktothrix] |
| WP_026797171.1 | REase | GCF_904830885.1 | Planktothrix | Bacteria>Bacillati>Cyanobacteriota/Melainabacteria_group>Cyanobacteriota>Cyanophyceae>Oscillatoriophycideae>Oscillatoriales>Microcoleaceae | 201 | NMK31_RS00285 | 68837..69442 | 54304 | MULTISPECIES:-Uma2-family-endonuclease-[Planktothrix] |
| WP_026786972.1 | REase | GCF_903969095.1 | Planktothrix | Bacteria>Bacillati>Cyanobacteriota/Melainabacteria_group>Cyanobacteriota>Cyanophyceae>Oscillatoriophycideae>Oscillatoriales>Microcoleaceae | 191 | NMG75_RS11280 | 2465262..2465837 | 54304 | MULTISPECIES:-Uma2-family-endonuclease-[Planktothrix] |
| WP_026793810.1 | REase | GCF_000464725.1 | Planktothrix | Bacteria>Bacillati>Cyanobacteriota/Melainabacteria_group>Cyanobacteriota>Cyanophyceae>Oscillatoriophycideae>Oscillatoriales>Microcoleaceae | 191 | N317_RS0104210 | 880671..881246 | 54304 | MULTISPECIES:-Uma2-family-endonuclease-[Planktothrix] |
| WP_026789286.1 | REase | GCF_005402765.1 | Planktothrix | Bacteria>Bacillati>Cyanobacteriota/Melainabacteria_group>Cyanobacteriota>Cyanophyceae>Oscillatoriophycideae>Oscillatoriales>Microcoleaceae | 188 | PLAN_RS00580 | 95299..95865 | 54304 | MULTISPECIES:-Uma2-family-endonuclease-[Planktothrix] |
| WP_026788985.1 | REase | GCF_000464725.1 | Planktothrix | Bacteria>Bacillati>Cyanobacteriota/Melainabacteria_group>Cyanobacteriota>Cyanophyceae>Oscillatoriophycideae>Oscillatoriales>Microcoleaceae | 214 | NMK31_RS00295 | 70110..70754 | 54304 | MULTISPECIES:-Uma2-family-endonuclease-[Planktothrix] |
| WP_026798704.1 | REase | GCF_005402765.1 | Planktothrix | Bacteria>Bacillati>Cyanobacteriota/Melainabacteria_group>Cyanobacteriota>Cyanophyceae>Oscillatoriophycideae>Oscillatoriales>Microcoleaceae | 220 | FLZ02_RS17795 | 10326..10988 | 54304 | MULTISPECIES:-hypothetical-protein-[Planktothrix] |
| WP_036831800.1 | REase | GCF_000464725.1 | Planktothrix | Bacteria>Bacillati>Cyanobacteriota/Melainabacteria_group>Cyanobacteriota>Cyanophyceae>Oscillatoriophycideae>Oscillatoriales>Microcoleaceae | 217 | N316_RS0113750 | 337830..338483 | 54304 | MULTISPECIES:-hypothetical-protein-[Planktothrix] |
| WP_026788984.1 | REase | GCF_000464725.1 | Planktothrix | Bacteria>Bacillati>Cyanobacteriota/Melainabacteria_group>Cyanobacteriota>Cyanophyceae>Oscillatoriophycideae>Oscillatoriales>Microcoleaceae | 200 | NMG75_RS16745 | 3653209..3653811 | 54304 | MULTISPECIES:-hypothetical-protein-[Planktothrix] |
| WP_026797172.1 | REase | GCF_005402765.1 | Planktothrix | Bacteria>Bacillati>Cyanobacteriota/Melainabacteria_group>Cyanobacteriota>Cyanophyceae>Oscillatoriophycideae>Oscillatoriales>Microcoleaceae | 200 | L2E81_RS19960 | 6325..6927 | 54304 | MULTISPECIES:-hypothetical-protein-[Planktothrix] |
| WP_036831804.1 | REase | GCF_000464725.1 | Planktothrix | Bacteria>Bacillati>Cyanobacteriota/Melainabacteria_group>Cyanobacteriota>Cyanophyceae>Oscillatoriophycideae>Oscillatoriales>Microcoleaceae | 206 | NMK38_RS22710 | 83088..83708 | 54304 | MULTISPECIES:-type-I-restriction-endonuclease-[Planktothrix] |
| WP_043937983.1 | REase | GCF_021554825.1 | Planktothrix | Bacteria>Bacillati>Cyanobacteriota/Melainabacteria_group>Cyanobacteriota>Cyanophyceae>Oscillatoriophycideae>Oscillatoriales>Microcoleaceae | 206 | L2E72_RS21845 | 10339..10959 | 54304 | MULTISPECIES:-hypothetical-protein-[Planktothrix] |
| WP_043938824.1 | REase | GCF_020594525.1 | Planktothrix | Bacteria>Bacillati>Cyanobacteriota/Melainabacteria_group>Cyanobacteriota>Cyanophyceae>Oscillatoriophycideae>Oscillatoriales>Microcoleaceae | 206 | LJE15_RS21480 | 4330..4950 | 54304 | MULTISPECIES:-type-I-restriction-enzyme-HsdR-N-terminal-domain-containing-protein-[Planktothrix] |
| WP_272033241.1 | REase | GCF_028330895.1 | Kamptonema_animale | Bacteria>Bacillati>Cyanobacteriota/Melainabacteria_group>Cyanobacteriota>Cyanophyceae>Oscillatoriophycideae>Oscillatoriales>Microcoleaceae>Kamptonema | 201 | PN499_RS16770 | 5270..5875 | 92934 | Uma2-family-endonuclease-[Kamptonema-animale] |
| WP_272033245.1 | REase | GCF_028330895.1 | Kamptonema_animale | Bacteria>Bacillati>Cyanobacteriota/Melainabacteria_group>Cyanobacteriota>Cyanophyceae>Oscillatoriophycideae>Oscillatoriales>Microcoleaceae>Kamptonema | 218 | PN499_RS16790 | 9712..10368 | 92934 | Uma2-family-endonuclease-[Kamptonema-animale] |
| WP_023065262.1 | REase | GCF_000478195.2 | Lyngbya_aestuarii | Bacteria>Bacillati>Cyanobacteriota/Melainabacteria_group>Cyanobacteriota>Cyanophyceae>Oscillatoriophycideae>Oscillatoriales>Microcoleaceae>Lyngbya | 215 | M595_RS06065 | 75804..76451 | 118322 | hypothetical-protein-[Lyngbya-aestuarii] |
| EGK85304.1 | REase | - | Microcoleus_vaginatus_FGP-2 | Bacteria>Bacillati>Cyanobacteriota/Melainabacteria_group>Cyanobacteriota>Cyanophyceae>Oscillatoriophycideae>Oscillatoriales>Microcoleaceae>Microcoleus>Microcoleus_vaginatus | 195 | MicvaDRAFT_4686 | 59549..60136 | 756067 | hypothetical-protein-MicvaDRAFT_4686-[Microcoleus-vaginatus-FGP-2] |
| WP_192216666.1 | REase | GCF_014849525.1 | Microcoleus_sp._FACHB-SPT15 | Bacteria>Bacillati>Cyanobacteriota/Melainabacteria_group>Cyanobacteriota>Cyanophyceae>Oscillatoriophycideae>Oscillatoriales>Microcoleaceae>Microcoleus>unclassified_Microcoleus | 210 | H6F98_RS00240 | 51410..52042 | 2692830 | restriction-endonuclease-subunit-R-[Microcoleus-sp.-FACHB-SPT15] |
| WP_192216665.1 | REase | GCF_014849525.1 | Microcoleus_sp._FACHB-SPT15 | Bacteria>Bacillati>Cyanobacteriota/Melainabacteria_group>Cyanobacteriota>Cyanophyceae>Oscillatoriophycideae>Oscillatoriales>Microcoleaceae>Microcoleus>unclassified_Microcoleus | 209 | H6F98_RS00235 | 50770..51399 | 2692830 | type-I-restriction-endonuclease-[Microcoleus-sp.-FACHB-SPT15] |
| WP_254175249.1 | REase | GCF_904830925.1 | Planktothrix_pseudagardhii | Bacteria>Bacillati>Cyanobacteriota/Melainabacteria_group>Cyanobacteriota>Cyanophyceae>Oscillatoriophycideae>Oscillatoriales>Microcoleaceae>Planktothrix | 194 | NMK36_RS28465 | 8803..9387 | 132604 | hypothetical-protein-[Planktothrix-pseudagardhii] |
| CAD5939277.1 | REase | GCA_904830785.1 | Planktothrix_rubescens | Bacteria>Bacillati>Cyanobacteriota/Melainabacteria_group>Cyanobacteriota>Cyanophyceae>Oscillatoriophycideae>Oscillatoriales>Microcoleaceae>Planktothrix | 201 | NO108_02190 | 2318397..2319002 | 59512 | putative-protein-sll0925-[Planktothrix-rubescens] |
| CAH2571598.1 | REase | GCA_937024135.1 | Planktothrix_rubescens | Bacteria>Bacillati>Cyanobacteriota/Melainabacteria_group>Cyanobacteriota>Cyanophyceae>Oscillatoriophycideae>Oscillatoriales>Microcoleaceae>Planktothrix | 201 | PRNO82_00996 | 1098147..1098752 | 59512 | putative-protein-sll0925-[Planktothrix-rubescens] |
| WP_141294971.1 | REase | GCF_005402765.1 | Planktothrix_agardhii | Bacteria>Bacillati>Cyanobacteriota/Melainabacteria_group>Cyanobacteriota>Cyanophyceae>Oscillatoriophycideae>Oscillatoriales>Microcoleaceae>Planktothrix | 201 | FLZ02_RS17810 | 15309..15914 | 1160 | Uma2-family-endonuclease-[Planktothrix-agardhii] |

| Protein-ID | Domain | Assembly-ID | Strain | Taxonomy | Length | Locus-Tag | Genomic-coordinates | TaxID | Fasta-defline |
|---|---|---|---|---|---|---|---|---|---|
| WP_254035906.1 | REase | GCF_904830935.1 | Planktothrix_agardhii | Bacteria>Bacillati>Cyanobacteriota/Melainabacteria_group>Cyanobacteriota>Cyanophyceae>Oscillatoriophycideae>Oscillatoriales>Microcoleaceae>Planktothrix | 201 | NMG88_RS22315 | 50114..50719 | 1160 | Uma2-family-endonuclease-[Planktothrix-agardhii] |
| CAD5929349.1 | REase | GCA_904830855.1 | Planktothrix_agardhii | Bacteria>Bacillati>Cyanobacteriota/Melainabacteria_group>Cyanobacteriota>Cyanophyceae>Oscillatoriophycideae>Oscillatoriales>Microcoleaceae>Planktothrix | 191 | PANO66_01239 | 1439029..1439604 | 1160 | hypothetical-protein-PANO66_01239-[Planktothrix-agardhii] |
| WP_042151510.1 | REase | GCF_000710505.1 | Planktothrix_agardhii | Bacteria>Bacillati>Cyanobacteriota/Melainabacteria_group>Cyanobacteriota>Cyanophyceae>Oscillatoriophycideae>Oscillatoriales>Microcoleaceae>Planktothrix | 191 | NMG87_RS10525 | 2385914..2386489 | 1160 | Uma2-family-endonuclease-[Planktothrix-agardhii] |
| WP_235750930.1 | REase | GCF_015708255.1 | Planktothrix_agardhii | Bacteria>Bacillati>Cyanobacteriota/Melainabacteria_group>Cyanobacteriota>Cyanophyceae>Oscillatoriophycideae>Oscillatoriales>Microcoleaceae>Planktothrix | 191 | NMG84_RS06220 | 1439029..1439604 | 1160 | Uma2-family-endonuclease-[Planktothrix-agardhii] |
| WP_254035077.1 | REase | GCF_904830935.1 | Planktothrix_agardhii | Bacteria>Bacillati>Cyanobacteriota/Melainabacteria_group>Cyanobacteriota>Cyanophyceae>Oscillatoriophycideae>Oscillatoriales>Microcoleaceae>Planktothrix | 191 | NMG88_RS02410 | 524582..525157 | 1160 | Uma2-family-endonuclease-[Planktothrix-agardhii] |
| CAD5939246.1 | REase | GCA_904830785.1 | Planktothrix_rubescens | Bacteria>Bacillati>Cyanobacteriota/Melainabacteria_group>Cyanobacteriota>Cyanophyceae>Oscillatoriophycideae>Oscillatoriales>Microcoleaceae>Planktothrix | 214 | NO108_02188 | 2317092..2317736 | 59512 | hypothetical-protein-NO108_02188-[Planktothrix-rubescens] |
| CAD5939329.1 | REase | GCA_904830785.1 | Planktothrix_rubescens | Bacteria>Bacillati>Cyanobacteriota/Melainabacteria_group>Cyanobacteriota>Cyanophyceae>Oscillatoriophycideae>Oscillatoriales>Microcoleaceae>Planktothrix | 220 | NO108_02193 | 2323323..2323985 | 59512 | hypothetical-protein-NO108_02193-[Planktothrix-rubescens] |
| CAH2571595.1 | REase | GCA_937024135.1 | Planktothrix_rubescens | Bacteria>Bacillati>Cyanobacteriota/Melainabacteria_group>Cyanobacteriota>Cyanophyceae>Oscillatoriophycideae>Oscillatoriales>Microcoleaceae>Planktothrix | 220 | PRNO82_00993 | 1093767..1094429 | 59512 | hypothetical-protein-PRNO82_00993-[Planktothrix-rubescens] |
| CAD5939263.1 | REase | GCA_904830785.1 | Planktothrix_rubescens | Bacteria>Bacillati>Cyanobacteriota/Melainabacteria_group>Cyanobacteriota>Cyanophyceae>Oscillatoriophycideae>Oscillatoriales>Microcoleaceae>Planktothrix | 200 | NO108_02189 | 2317805..2318407 | 59512 | hypothetical-protein-NO108_02189-[Planktothrix-rubescens] |
| CAH2571599.1 | REase | GCA_937024135.1 | Planktothrix_rubescens | Bacteria>Bacillati>Cyanobacteriota/Melainabacteria_group>Cyanobacteriota>Cyanophyceae>Oscillatoriophycideae>Oscillatoriales>Microcoleaceae>Planktothrix | 200 | PRNO82_00997 | 1098742..1099344 | 59512 | hypothetical-protein-PRNO82_00997-[Planktothrix-rubescens] |
| WP_235753327.1 | REase | GCF_904830885.1 | Planktothrix_agardhii | Bacteria>Bacillati>Cyanobacteriota/Melainabacteria_group>Cyanobacteriota>Cyanophyceae>Oscillatoriophycideae>Oscillatoriales>Microcoleaceae>Planktothrix | 200 | NMK31_RS00290 | 69432..70034 | 1160 | hypothetical-protein-[Planktothrix-agardhii] |
| WP_254035907.1 | REase | GCF_904830935.1 | Planktothrix_agardhii | Bacteria>Bacillati>Cyanobacteriota/Melainabacteria_group>Cyanobacteriota>Cyanophyceae>Oscillatoriophycideae>Oscillatoriales>Microcoleaceae>Planktothrix | 200 | NMG88_RS22320 | 50709..51311 | 1160 | hypothetical-protein-[Planktothrix-agardhii] |
| CAD5939292.1 | REase | GCA_904830785.1 | Planktothrix_rubescens | Bacteria>Bacillati>Cyanobacteriota/Melainabacteria_group>Cyanobacteriota>Cyanophyceae>Oscillatoriophycideae>Oscillatoriales>Microcoleaceae>Planktothrix | 206 | NO108_02191 | 2319010..2319630 | 59512 | Putative-Restriction-endonuclease-EcoRI-[Planktothrix-rubescens] |
| CAH2571597.1 | REase | GCA_937024135.1 | Planktothrix_rubescens | Bacteria>Bacillati>Cyanobacteriota/Melainabacteria_group>Cyanobacteriota>Cyanophyceae>Oscillatoriophycideae>Oscillatoriales>Microcoleaceae>Planktothrix | 206 | PRNO82_00995 | 1097519..1098139 | 59512 | hypothetical-protein-PRNO82_00995-[Planktothrix-rubescens] |
| WP_036842261.1 | REase | GCF_903969095.1 | Planktothrix_agardhii | Bacteria>Bacillati>Cyanobacteriota/Melainabacteria_group>Cyanobacteriota>Cyanophyceae>Oscillatoriophycideae>Oscillatoriales>Microcoleaceae>Planktothrix | 206 | NMG75_RS16755 | 3654414..3655034 | 1160 | hypothetical-protein-[Planktothrix-agardhii] |
| WP_141294969.1 | REase | GCF_005402765.1 | Planktothrix_agardhii | Bacteria>Bacillati>Cyanobacteriota/Melainabacteria_group>Cyanobacteriota>Cyanophyceae>Oscillatoriophycideae>Oscillatoriales>Microcoleaceae>Planktothrix | 206 | NMK31_RS00280 | 68209..68829 | 1160 | type-I-restriction-enzyme-HsdR-N-terminal-domain-containing-protein-[Planktothrix-agardhii] |
| WP_254035905.1 | REase | GCF_904830935.1 | Planktothrix_agardhii | Bacteria>Bacillati>Cyanobacteriota/Melainabacteria_group>Cyanobacteriota>Cyanophyceae>Oscillatoriophycideae>Oscillatoriales>Microcoleaceae>Planktothrix | 206 | NMG88_RS22310 | 49486..50106 | 1160 | type-I-restriction-enzyme-HsdR-N-terminal-domain-containing-protein-[Planktothrix-agardhii] |
| WP_235758597.1 | REase | - | Planktothrix_agardhii | Bacteria>Bacillati>Cyanobacteriota/Melainabacteria_group>Cyanobacteriota>Cyanophyceae>Oscillatoriophycideae>Oscillatoriales>Microcoleaceae>Planktothrix | 184 | L2E81_RS19950 | 5168..5722 | 1160 | type-I-restriction-enzyme-HsdR-N-terminal-domain-containing-protein-[Planktothrix-agardhii] |
| WP_264479403.1 | REase | - | Planktothrix_agardhii | Bacteria>Bacillati>Cyanobacteriota/Melainabacteria_group>Cyanobacteriota>Cyanophyceae>Oscillatoriophycideae>Oscillatoriales>Microcoleaceae>Planktothrix | 185 | LJE15_RS21525 | 22808..23365 | 1160 | Rpn-family-recombination-promoting-nuclease/putative-transposase-[Planktothrix-agardhii] |
| BBD57118.1 | REase | GCA_003609755.1 | Planktothrix_agardhii_NIES-204 | Bacteria>Bacillati>Cyanobacteriota/Melainabacteria_group>Cyanobacteriota>Cyanophyceae>Oscillatoriophycideae>Oscillatoriales>Microcoleaceae>Planktothrix>Planktothrix_agardhii | 201 | NIES204_44540 | 43081..43686 | 282423 | hypothetical-protein-NIES204_44540-[Planktothrix-agardhii-NIES-204] |
| MCP9293154.1 | REase | - | Planktothrix_agardhii_LY1 | Bacteria>Bacillati>Cyanobacteriota/Melainabacteria_group>Cyanobacteriota>Cyanophyceae>Oscillatoriophycideae>Oscillatoriales>Microcoleaceae>Planktothrix>Planktothrix_agardhii | 191 | NDM07_00480 | 16212..16787 | 2950590 | Uma2-family-endonuclease-[Planktothrix-agardhii-LY1] |
| BBD57117.1 | REase | GCA_003609755.1 | Planktothrix_agardhii_NIES-204 | Bacteria>Bacillati>Cyanobacteriota/Melainabacteria_group>Cyanobacteriota>Cyanophyceae>Oscillatoriophycideae>Oscillatoriales>Microcoleaceae>Planktothrix>Planktothrix_agardhii | 200 | NIES204_44530 | 42489..43091 | 282423 | hypothetical-protein-NIES204_44530-[Planktothrix-agardhii-NIES-204] |
| BBD57119.1 | REase | GCA_003609755.1 | Planktothrix_agardhii_NIES-204 | Bacteria>Bacillati>Cyanobacteriota/Melainabacteria_group>Cyanobacteriota>Cyanophyceae>Oscillatoriophycideae>Oscillatoriales>Microcoleaceae>Planktothrix>Planktothrix_agardhii | 206 | NIES204_44550 | 43694..44314 | 282423 | hypothetical-protein-NIES204_44550-[Planktothrix-agardhii-NIES-204] |
| BBD57223.1 | REase | GCA_003609755.1 | Planktothrix_agardhii_NIES-204 | Bacteria>Bacillati>Cyanobacteriota/Melainabacteria_group>Cyanobacteriota>Cyanophyceae>Oscillatoriophycideae>Oscillatoriales>Microcoleaceae>Planktothrix>Planktothrix_agardhii | 138 | NIES204_45590 | 18560..18976 | 282423 | FdxN-element-excision-controlling-factor-protein-[Planktothrix-agardhii-NIES-204] |
| MBS9772312.1 | REase | GCA_018457135.1 | Trichodesmium_erythraeum_GBRTRLIN201 | Bacteria>Bacillati>Cyanobacteriota/Melainabacteria_group>Cyanobacteriota>Cyanophyceae>Oscillatoriophycideae>Oscillatoriales>Microcoleaceae>Trichodesmium>Trichodesmium_erythraeum | 192 | J7F05_18145 | 3463..4041 | 1045867 | hypothetical-protein-[Trichodesmium-erythraeum-GBRTRLIN201] |
| WP_285965026.1 | REase | - | Oscillatoria_laete-virens | Bacteria>Bacillati>Cyanobacteriota/Melainabacteria_group>Cyanobacteriota>Cyanophyceae>Oscillatoriophycideae>Oscillatoriales>Oscillatoriaceae>Oscillatoria | 249 | QQ056_RS17140 | 56728..57477 | 1565316 | restriction-endonuclease-[Oscillatoria-laete-virens] |
| WP_026099031.1 | REase | GCF_000332335.1 | Oscillatoria_sp._PCC_10802 | Bacteria>Bacillati>Cyanobacteriota/Melainabacteria_group>Cyanobacteriota>Cyanophyceae>Oscillatoriophycideae>Oscillatoriales>Oscillatoriaceae>Oscillatoria>unclassified_Oscillatoria | 220 | OSC10802_RS0133325 | 645017..645679 | 1173028 | Uma2-family-endonuclease-[Oscillatoria-sp.-PCC-10802] |
| WP_190577358.1 | REase | GCF_963890155.1 | Phormidium_tenue | Bacteria>Bacillati>Cyanobacteriota/Melainabacteria_group>Cyanobacteriota>Cyanophyceae>Oscillatoriophycideae>Oscillatoriales>Oscillatoriaceae>Phormidium | 206 | H6G05_RS06515 | 61090..61710 | 126344 | hypothetical-protein-[Pseudanabaena-sp.-FACHB-1050] |
| MCC5898058.1 | REase | GCA_020832215.1 | Phormidium_sp._BM_Day4_Bin.17 | Bacteria>Bacillati>Cyanobacteriota/Melainabacteria_group>Cyanobacteriota>Cyanophyceae>Oscillatoriophycideae>Oscillatoriales>Oscillatoriaceae>Phormidium>unclassified_Phormidium | 216 | JJU32_09095 | 32387..33037 | 2813767 | Uma2-family-endonuclease-[Phormidium-sp.-BM_Day4_Bin.17] |
| WP_200906364.1 | REase | GCF_000972705.2 | Limnoraphis_robusta | Bacteria>Bacillati>Cyanobacteriota/Melainabacteria_group>Cyanobacteriota>Cyanophyceae>Oscillatoriophycideae>Oscillatoriales>Sirenicapillariaceae>Limnoraphis | 215 | WN50_RS27090 | 905..1552 | 1118279 | restriction-endonuclease-subunit-R-[Limnoraphis-robusta] |
| WP_017711369.1 | REase | GCF_000332315.1 | Prochlorothrix_hollandica | Bacteria>Bacillati>Cyanobacteriota/Melainabacteria_group>Cyanobacteriota>Cyanophyceae>Prochlorotrichales>Prochlorotrichaceae>Prochlorothrix | 216 | PRO9006_RS0103840 | 891171..891821 | 1223 | type-I-restriction-enzyme-HsdR-N-terminal-domain-containing-protein-[Prochlorothrix-hollandica] |
| WP_281006616.1 | REase | GCF_029910235.1 | Pseudanabaena | Bacteria>Bacillati>Cyanobacteriota/Melainabacteria_group>Cyanobacteriota>Cyanophyceae>Pseudanabaenales>Pseudanabaenaceae | 206 | OA858_RS18400 | 4059167..4059787 | 1152 | Uma2-family-endonuclease-[Pseudanabaena-galeata] |
| WP_281006615.1 | REase | GCF_029910235.1 | Pseudanabaena | Bacteria>Bacillati>Cyanobacteriota/Melainabacteria_group>Cyanobacteriota>Cyanophyceae>Pseudanabaenales>Pseudanabaenaceae | 212 | OA858_RS18395 | 4058466..4059104 | 1152 | Uma2-family-endonuclease-[Pseudanabaena-galeata] |
| WP_281009994.1 | REase | GCF_029910235.1 | Pseudanabaena | Bacteria>Bacillati>Cyanobacteriota/Melainabacteria_group>Cyanobacteriota>Cyanophyceae>Pseudanabaenales>Pseudanabaenaceae | 207 | OA858_RS25965 | 130338..130961 | 1152 | hypothetical-protein-[Pseudanabaena-galeata] |
| WP_281006614.1 | REase | GCF_029910235.1 | Pseudanabaena | Bacteria>Bacillati>Cyanobacteriota/Melainabacteria_group>Cyanobacteriota>Cyanophyceae>Pseudanabaenales>Pseudanabaenaceae | 201 | OA858_RS18390 | 4057854..4058459 | 1152 | hypothetical-protein-[Pseudanabaena-galeata] |
| WP_009625706.1 | REase | GCF_000332215.1 | Pseudanabaena | Bacteria>Bacillati>Cyanobacteriota/Melainabacteria_group>Cyanobacteriota>Cyanophyceae>Pseudanabaenales>Pseudanabaenaceae | 137 | PSE7429_RS03655 | 17413..17826 | 1152 | MULTISPECIES:-element-excision-factor-XisH-family-protein-[Pseudanabaena] |
| WP_190351070.1 | REase | GCF_014696345.1 | Pseudanabaena_cinerea | Bacteria>Bacillati>Cyanobacteriota/Melainabacteria_group>Cyanobacteriota>Cyanophyceae>Pseudanabaenales>Pseudanabaenaceae>Pseudanabaena | 202 | H6F44_RS11365 | 48103..48711 | 2661616 | Uma2-family-endonuclease-[Pseudanabaena-sp.-FACHB-1277] |
| WP_169366111.1 | REase | GCF_012863495.1 | Pseudanabaena_yagii | Bacteria>Bacillati>Cyanobacteriota/Melainabacteria_group>Cyanobacteriota>Cyanophyceae>Pseudanabaenales>Pseudanabaenaceae>Pseudanabaena | 201 | HC246_RS24860 | 12242..12847 | 2661615 | Uma2-family-endonuclease-[Pseudanabaena-yagii] |
| WP_169366107.1 | REase | GCF_012863495.1 | Pseudanabaena_yagii | Bacteria>Bacillati>Cyanobacteriota/Melainabacteria_group>Cyanobacteriota>Cyanophyceae>Pseudanabaenales>Pseudanabaenaceae>Pseudanabaena | 217 | HC246_RS24840 | 9679..10332 | 2661615 | Uma2-family-endonuclease-[Pseudanabaena-yagii] |
| WP_169366106.1 | REase | GCF_012863495.1 | Pseudanabaena_yagii | Bacteria>Bacillati>Cyanobacteriota/Melainabacteria_group>Cyanobacteriota>Cyanophyceae>Pseudanabaenales>Pseudanabaenaceae>Pseudanabaena | 192 | HC246_RS24835 | 9078..9656 | 2661615 | Uma2-family-endonuclease-[Pseudanabaena-yagii] |
| WP_169366108.1 | REase | GCF_012863495.1 | Pseudanabaena_yagii | Bacteria>Bacillati>Cyanobacteriota/Melainabacteria_group>Cyanobacteriota>Cyanophyceae>Pseudanabaenales>Pseudanabaenaceae>Pseudanabaena | 191 | HC246_RS24845 | 10374..10949 | 2661615 | Uma2-family-endonuclease-[Pseudanabaena-yagii] |
| WP_190351064.1 | REase | GCF_014696345.1 | Pseudanabaena_cinerea | Bacteria>Bacillati>Cyanobacteriota/Melainabacteria_group>Cyanobacteriota>Cyanophyceae>Pseudanabaenales>Pseudanabaenaceae>Pseudanabaena | 138 | H6F44_RS11330 | 40375..40791 | 2661616 | XisH-family-protein-[Pseudanabaena-sp.-FACHB-1277] |
| MCE2977895.1 | REase | GCA_021298775.1 | Pseudanabaena_sp._CoA8_M7 | Bacteria>Bacillati>Cyanobacteriota/Melainabacteria_group>Cyanobacteriota>Cyanophyceae>Pseudanabaenales>Pseudanabaenaceae>Pseudanabaena>unclassified_Pseudanabaena | 206 | LW851_16620 | 3075..3695 | 2904992 | Uma2-family-endonuclease-[Pseudanabaena-sp.-CoA8_M7] |
| MCA6503990.1 | REase | GCA_020403835.1 | Pseudanabaena_sp._M090S1SP2A07QC | Bacteria>Bacillati>Cyanobacteriota/Melainabacteria_group>Cyanobacteriota>Cyanophyceae>Pseudanabaenales>Pseudanabaenaceae>Pseudanabaena>unclassified_Pseudanabaena | 205 | IM580_15725 | 3513..4130 | 2771254 | Uma2-family-endonuclease-[Pseudanabaena-sp.-M090S1SP2A07QC] |
| MCA6581760.1 | REase | GCA_020404925.1 | Pseudanabaena_sp._M34BS1SP1A06MG | Bacteria>Bacillati>Cyanobacteriota/Melainabacteria_group>Cyanobacteriota>Cyanophyceae>Pseudanabaenales>Pseudanabaenaceae>Pseudanabaena>unclassified_Pseudanabaena | 205 | IM536_06935 | 3560..4177 | 2771240 | Uma2-family-endonuclease-[Pseudanabaena-sp.-M34BS1SP1A06MG] |
| MCA6597114.1 | REase | GCA_020405005.1 | Pseudanabaena_sp._M046S1SP1A06QC | Bacteria>Bacillati>Cyanobacteriota/Melainabacteria_group>Cyanobacteriota>Cyanophyceae>Pseudanabaenales>Pseudanabaenaceae>Pseudanabaena>unclassified_Pseudanabaena | 205 | IM539_12620 | 36410..37027 | 2771236 | Uma2-family-endonuclease-[Pseudanabaena-sp.-M046S1SP1A06QC] |
| MCA6600379.1 | REase | GCA_020404965.1 | Pseudanabaena_sp._M57BS1SP1A06MG | Bacteria>Bacillati>Cyanobacteriota/Melainabacteria_group>Cyanobacteriota>Cyanophyceae>Pseudanabaenales>Pseudanabaenaceae>Pseudanabaena>unclassified_Pseudanabaena | 205 | IM537_09265 | 7265..7882 | 2771237 | Uma2-family-endonuclease-[Pseudanabaena-sp.-M57BS1SP1A06MG] |
| MCA6605400.1 | REase | GCA_020405035.1 | Pseudanabaena_sp._M007S1SP1A06QC | Bacteria>Bacillati>Cyanobacteriota/Melainabacteria_group>Cyanobacteriota>Cyanophyceae>Pseudanabaenales>Pseudanabaenaceae>Pseudanabaena>unclassified_Pseudanabaena | 205 | IM533_13240 | 2153..2770 | 2771232 | Uma2-family-endonuclease-[Pseudanabaena-sp.-M007S1SP1A06QC] |
| MCA6597118.1 | REase | GCA_020405005.1 | Pseudanabaena_sp._M046S1SP1A06QC | Bacteria>Bacillati>Cyanobacteriota/Melainabacteria_group>Cyanobacteriota>Cyanophyceae>Pseudanabaenales>Pseudanabaenaceae>Pseudanabaena>unclassified_Pseudanabaena | 221 | IM539_12645 | 41749..42414 | 2771236 | Uma2-family-endonuclease-[Pseudanabaena-sp.-M046S1SP1A06QC] |
| MCA6597119.1 | REase | GCA_020405005.1 | Pseudanabaena_sp._M046S1SP1A06QC | Bacteria>Bacillati>Cyanobacteriota/Melainabacteria_group>Cyanobacteriota>Cyanophyceae>Pseudanabaenales>Pseudanabaenaceae>Pseudanabaena>unclassified_Pseudanabaena | 215 | IM539_12650 | 42435..43082 | 2771236 | Uma2-family-endonuclease-[Pseudanabaena-sp.-M046S1SP1A06QC] |
| MCA6503989.1 | REase | GCA_020403835.1 | Pseudanabaena_sp._M090S1SP2A07QC | Bacteria>Bacillati>Cyanobacteriota/Melainabacteria_group>Cyanobacteriota>Cyanophyceae>Pseudanabaenales>Pseudanabaenaceae>Pseudanabaena>unclassified_Pseudanabaena | 212 | IM580_15720 | 2797..3435 | 2771254 | Uma2-family-endonuclease-[Pseudanabaena-sp.-M090S1SP2A07QC] |
| MCA6581761.1 | REase | GCA_020404925.1 | Pseudanabaena_sp._M34BS1SP1A06MG | Bacteria>Bacillati>Cyanobacteriota/Melainabacteria_group>Cyanobacteriota>Cyanophyceae>Pseudanabaenales>Pseudanabaenaceae>Pseudanabaena>unclassified_Pseudanabaena | 212 | IM536_06940 | 4255..4893 | 2771240 | Uma2-family-endonuclease-[Pseudanabaena-sp.-M34BS1SP1A06MG] |
| MCA6597115.1 | REase | GCA_020405005.1 | Pseudanabaena_sp._M046S1SP1A06QC | Bacteria>Bacillati>Cyanobacteriota/Melainabacteria_group>Cyanobacteriota>Cyanophyceae>Pseudanabaenales>Pseudanabaenaceae>Pseudanabaena>unclassified_Pseudanabaena | 212 | IM539_12625 | 37105..37743 | 2771236 | Uma2-family-endonuclease-[Pseudanabaena-sp.-M046S1SP1A06QC] |
| MCA6600380.1 | REase | GCA_020404965.1 | Pseudanabaena_sp._M57BS1SP1A06MG | Bacteria>Bacillati>Cyanobacteriota/Melainabacteria_group>Cyanobacteriota>Cyanophyceae>Pseudanabaenales>Pseudanabaenaceae>Pseudanabaena>unclassified_Pseudanabaena | 212 | IM537_09270 | 7960..8598 | 2771237 | Uma2-family-endonuclease-[Pseudanabaena-sp.-M57BS1SP1A06MG] |
| MCA6605399.1 | REase | GCA_020405035.1 | Pseudanabaena_sp._M007S1SP1A06QC | Bacteria>Bacillati>Cyanobacteriota/Melainabacteria_group>Cyanobacteriota>Cyanophyceae>Pseudanabaenales>Pseudanabaenaceae>Pseudanabaena>unclassified_Pseudanabaena | 212 | IM533_13235 | 1437..2075 | 2771232 | Uma2-family-endonuclease-[Pseudanabaena-sp.-M007S1SP1A06QC] |
| MCE2977894.1 | REase | GCA_021298775.1 | Pseudanabaena_sp._CoA8_M7 | Bacteria>Bacillati>Cyanobacteriota/Melainabacteria_group>Cyanobacteriota>Cyanophyceae>Pseudanabaenales>Pseudanabaenaceae>Pseudanabaena>unclassified_Pseudanabaena | 212 | LW851_16615 | 2362..3000 | 2904992 | Uma2-family-endonuclease-[Pseudanabaena-sp.-CoA8_M7] |
| MCA6503988.1 | REase | GCA_020403835.1 | Pseudanabaena_sp._M090S1SP2A07QC | Bacteria>Bacillati>Cyanobacteriota/Melainabacteria_group>Cyanobacteriota>Cyanophyceae>Pseudanabaenales>Pseudanabaenaceae>Pseudanabaena>unclassified_Pseudanabaena | 201 | IM580_15715 | 2185..2790 | 2771254 | hypothetical-protein-[Pseudanabaena-sp.-M090S1SP2A07QC] |
| MCA6581762.1 | REase | GCA_020404925.1 | Pseudanabaena_sp._M34BS1SP1A06MG | Bacteria>Bacillati>Cyanobacteriota/Melainabacteria_group>Cyanobacteriota>Cyanophyceae>Pseudanabaenales>Pseudanabaenaceae>Pseudanabaena>unclassified_Pseudanabaena | 201 | IM536_06945 | 4900..5505 | 2771240 | hypothetical-protein-[Pseudanabaena-sp.-M34BS1SP1A06MG] |
| MCA6597116.1 | REase | GCA_020405005.1 | Pseudanabaena_sp._M046S1SP1A06QC | Bacteria>Bacillati>Cyanobacteriota/Melainabacteria_group>Cyanobacteriota>Cyanophyceae>Pseudanabaenales>Pseudanabaenaceae>Pseudanabaena>unclassified_Pseudanabaena | 201 | IM539_12630 | 37750..38355 | 2771236 | hypothetical-protein-[Pseudanabaena-sp.-M046S1SP1A06QC] |
| MCA6600381.1 | REase | GCA_020404965.1 | Pseudanabaena_sp._M57BS1SP1A06MG | Bacteria>Bacillati>Cyanobacteriota/Melainabacteria_group>Cyanobacteriota>Cyanophyceae>Pseudanabaenales>Pseudanabaenaceae>Pseudanabaena>unclassified_Pseudanabaena | 201 | IM537_09275 | 8605..9210 | 2771237 | hypothetical-protein-[Pseudanabaena-sp.-M57BS1SP1A06MG] |
| MCA6605398.1 | REase | GCA_020405035.1 | Pseudanabaena_sp._M007S1SP1A06QC | Bacteria>Bacillati>Cyanobacteriota/Melainabacteria_group>Cyanobacteriota>Cyanophyceae>Pseudanabaenales>Pseudanabaenaceae>Pseudanabaena>unclassified_Pseudanabaena | 201 | IM533_13230 | 825..1430 | 2771232 | hypothetical-protein-[Pseudanabaena-sp.-M007S1SP1A06QC] |
| MCE2977893.1 | REase | GCA_021298775.1 | Pseudanabaena_sp._CoA8_M7 | Bacteria>Bacillati>Cyanobacteriota/Melainabacteria_group>Cyanobacteriota>Cyanophyceae>Pseudanabaenales>Pseudanabaenaceae>Pseudanabaena>unclassified_Pseudanabaena | 201 | LW851_16610 | 1750..2355 | 2904992 | hypothetical-protein-[Pseudanabaena-sp.-CoA8_M7] |
| MCL1489588.1 | REase | GCA_023336205.1 | Pseudanabaena_sp._Salubria-1 | Bacteria>Bacillati>Cyanobacteriota/Melainabacteria_group>Cyanobacteriota>Cyanophyceae>Pseudanabaenales>Pseudanabaenaceae>Pseudanabaena>unclassified_Pseudanabaena | 239 | M1G31_02375 | 44348..45067 | 2938918 | Uma2-family-endonuclease-[Pseudanabaena-sp.-Salubria-1] |
| WP_201323618.1 | REase | GCF_016584665.1 | Pseudanabaena_sp._lw0831 | Bacteria>Bacillati>Cyanobacteriota/Melainabacteria_group>Cyanobacteriota>Cyanophyceae>Pseudanabaenales>Pseudanabaenaceae>Pseudanabaena>unclassified_Pseudanabaena | 202 | APA_RS17905 | 63964..64572 | 1357935 | Uma2-family-endonuclease-[Pseudanabaena-sp.-lw0831] |
| WP_254923794.1 | REase | GCF_002251945.1 | Pseudanabaena_sp._SR411 | Bacteria>Bacillati>Cyanobacteriota/Melainabacteria_group>Cyanobacteriota>Cyanophyceae>Pseudanabaenales>Pseudanabaenaceae>Pseudanabaena>unclassified_Pseudanabaena | 118 | B9G53_RS28190 | 16143..16499 | 1980935 | Uma2-family-endonuclease-[Pseudanabaena-sp.-SR411] |
| WP_217901666.1 | REase | GCF_002251945.1 | Pseudanabaena_sp._SR411 | Bacteria>Bacillati>Cyanobacteriota/Melainabacteria_group>Cyanobacteriota>Cyanophyceae>Pseudanabaenales>Pseudanabaenaceae>Pseudanabaena>unclassified_Pseudanabaena | 77 | B9G53_RS28185 | 15903..16136 | 1980935 | Uma2-family-endonuclease-[Pseudanabaena-sp.-SR411] |
| WP_204104296.1 | REase | GCF_016888125.1 | Spirulina_sp._CCY15215 | Bacteria>Bacillati>Cyanobacteriota/Melainabacteria_group>Cyanobacteriota>Cyanophyceae>Spirulinales>Spirulinaceae>Spirulina>unclassified_Spirulina | 203 | JUI58_RS13375 | 1720..2331 | 2767591 | Uma2-family-endonuclease-[Spirulina-major] |
| WP_204104301.1 | REase | GCF_016888125.1 | Spirulina_sp._CCY15215 | Bacteria>Bacillati>Cyanobacteriota/Melainabacteria_group>Cyanobacteriota>Cyanophyceae>Spirulinales>Spirulinaceae>Spirulina>unclassified_Spirulina | 219 | JUI58_RS13400 | 7972..8631 | 2767591 | restriction-endonuclease-subunit-R-[Spirulina-major] |
| MBW4553109.1 | REase | GCA_019359715.1 | Aphanocapsa_sp._GSE-SYN-MK-11-07L | Bacteria>Bacillati>Cyanobacteriota/Melainabacteria_group>Cyanobacteriota>Cyanophyceae>Synechococcales>Merismopediaceae>Aphanocapsa>unclassified_Aphanocapsa | 206 | KME35_18655 | 5230..5850 | 2835086 | Uma2-family-endonuclease-[Aphanocapsa-sp.-GSE-SYN-MK-11-07L] |
| MBW4553279.1 | REase | GCA_019359715.1 | Aphanocapsa_sp._GSE-SYN-MK-11-07L | Bacteria>Bacillati>Cyanobacteriota/Melainabacteria_group>Cyanobacteriota>Cyanophyceae>Synechococcales>Merismopediaceae>Aphanocapsa>unclassified_Aphanocapsa | 216 | KME35_19525 | 7338..7988 | 2835086 | restriction-endonuclease-subunit-R-[Aphanocapsa-sp.-GSE-SYN-MK-11-07L] |
| MBW4553280.1 | REase | GCA_019359715.1 | Aphanocapsa_sp._GSE-SYN-MK-11-07L | Bacteria>Bacillati>Cyanobacteriota/Melainabacteria_group>Cyanobacteriota>Cyanophyceae>Synechococcales>Merismopediaceae>Aphanocapsa>unclassified_Aphanocapsa | 211 | KME35_19530 | 8045..8680 | 2835086 | restriction-endonuclease-subunit-R-[Aphanocapsa-sp.-GSE-SYN-MK-11-07L] |
| WP_106289189.1 | REase | GCF_963876965.1 | Merismopedia_glauca | Bacteria>Bacillati>Cyanobacteriota/Melainabacteria_group>Cyanobacteriota>Cyanophyceae>Synechococcales>Merismopediaceae>Merismopedia | 189 | C7B64_RS13510 | 4288..4857 | 292586 | hypothetical-protein-[Merismopedia-glauca] |
| WP_254950527.1 | REase | GCF_024345955.1 | unclassified_Cyanobium | Bacteria>Bacillati>Cyanobacteriota/Melainabacteria_group>Cyanobacteriota>Cyanophyceae>Synechococcales>Prochlorococcaceae>Cyanobium | 173 | KBZ34_RS03570 | 175399..175920 | 2627006 | MULTISPECIES:-hypothetical-protein-[unclassified-Cyanobium] |
| WP_277867660.1 | REase | GCF_029582805.1 | Candidatus_Synechococcus_calcipolaris | Bacteria>Bacillati>Cyanobacteriota/Melainabacteria_group>Cyanobacteriota>Cyanophyceae>Synechococcales>Synechococcaceae>Synechococcus | 96 | L3556_RS12440 | 2891..3181 | 1522304 | DUF2887-domain-containing-protein-[Candidatus-Synechococcus-calcipolaris] |
| KKZ13996.1 | REase | GCA_001007635.1 | Candidatus_Synechococcus_spongiarum_15L | Bacteria>Bacillati>Cyanobacteriota/Melainabacteria_group>Cyanobacteriota>Cyanophyceae>Synechococcales>Synechococcaceae>Synechococcus>Candidatus_Synechococcus_spongiarum | 168 | TQ37_02570 | 5061..5567 | 1608419 | hypothetical-protein-TQ37_02570-[Candidatus-Synechococcus-spongiarum-15L] |
| OOV26580.1 | REase | GCA_002018045.1 | Candidatus_Synechococcus_spongiarum_LMB_bulk15M | Bacteria>Bacillati>Cyanobacteriota/Melainabacteria_group>Cyanobacteriota>Cyanophyceae>Synechococcales>Synechococcaceae>Synechococcus>Candidatus_Synechococcus_spongiarum | 109 | BV61_06580 | 16032..16361 | 1943582 | hypothetical-protein-BV61_06580-[Candidatus-Synechococcus-spongiarum-LMB-bulk15M] |
| WP_247214941.1 | REase | GCF_963899605.1 | Synechococcus_sp._C9 | Bacteria>Bacillati>Cyanobacteriota/Melainabacteria_group>Cyanobacteriota>Cyanophyceae>Synechococcales>Synechococcaceae>Synechococcus>unclassified_Synechococcus | 192 | MLD66_RS00205 | 44196..44774 | 102119 | Uma2-family-endonuclease-[Synechococcus-sp.-C9] |
| WP_235278693.1 | REase | GCF_021533835.1 | Synechococcus_sp._Nb3U1 | Bacteria>Bacillati>Cyanobacteriota/Melainabacteria_group>Cyanobacteriota>Cyanophyceae>Synechococcales>Synechococcaceae>Synechococcus>unclassified_Synechococcus | 190 | L1047_RS09915 | 2084626..2085198 | 1914529 | Uma2-family-endonuclease-[Synechococcus-sp.-Nb3U1] |
| WP_247214942.1 | REase | GCF_963899605.1 | Synechococcus_sp._C9 | Bacteria>Bacillati>Cyanobacteriota/Melainabacteria_group>Cyanobacteriota>Cyanophyceae>Synechococcales>Synechococcaceae>Synechococcus>unclassified_Synechococcus | 190 | MLD66_RS00210 | 44796..45368 | 102119 | Uma2-family-endonuclease-[Synechococcus-sp.-C9] |
| WP_247214940.1 | REase | GCF_963899605.1 | Synechococcus_sp._C9 | Bacteria>Bacillati>Cyanobacteriota/Melainabacteria_group>Cyanobacteriota>Cyanophyceae>Synechococcales>Synechococcaceae>Synechococcus>unclassified_Synechococcus | 217 | MLD66_RS00200 | 43531..44184 | 102119 | Uma2-family-endonuclease-[Synechococcus-sp.-C9] |
| WP_235278694.1 | REase | GCF_021533835.1 | Synechococcus_sp._Nb3U1 | Bacteria>Bacillati>Cyanobacteriota/Melainabacteria_group>Cyanobacteriota>Cyanophyceae>Synechococcales>Synechococcaceae>Synechococcus>unclassified_Synechococcus | 215 | L1047_RS09920 | 2085214..2085861 | 1914529 | Uma2-family-endonuclease-[Synechococcus-sp.-Nb3U1] |
| MXW11417.1 | REase | GCA_009836025.1 | Synechococcus_sp._SB0668_bin_13 | Bacteria>Bacillati>Cyanobacteriota/Melainabacteria_group>Cyanobacteriota>Cyanophyceae>Synechococcales>Synechococcaceae>Synechococcus>unclassified_Synechococcus | 165 | F4Z73_00775 | 12196..12693 | 2604859 | DUF4365-domain-containing-protein-[Synechococcus-sp.-SB0668_bin_13] |
| MXY61917.1 | REase | GCA_009837735.1 | Synechococcus_sp._SB0665_bin_28 | Bacteria>Bacillati>Cyanobacteriota/Melainabacteria_group>Cyanobacteriota>Cyanophyceae>Synechococcales>Synechococcaceae>Synechococcus>unclassified_Synechococcus | 165 | F4Y87_00420 | 5531..6028 | 2604856 | DUF4365-domain-containing-protein-[Synechococcus-sp.-SB0665_bin_28] |
| MYG64554.1 | REase | GCA_009843455.1 | Synechococcus_sp._SB0675_bin_7 | Bacteria>Bacillati>Cyanobacteriota/Melainabacteria_group>Cyanobacteriota>Cyanophyceae>Synechococcales>Synechococcaceae>Synechococcus>unclassified_Synechococcus | 165 | F4158_08375 | 394..891 | 2604868 | DUF4365-domain-containing-protein-[Synechococcus-sp.-SB0675_bin_7] |
| MYI71488.1 | REase | GCA_009844645.1 | Synechococcus_sp._SB0673_bin_10 | Bacteria>Bacillati>Cyanobacteriota/Melainabacteria_group>Cyanobacteriota>Cyanophyceae>Synechococcales>Synechococcaceae>Synechococcus>unclassified_Synechococcus | 165 | F4099_03020 | 4667..5164 | 2604866 | DUF4365-domain-containing-protein-[Synechococcus-sp.-SB0673_bin_10] |
| WP_255117819.1 | REase | GCF_024345835.1 | Synechococcus_sp._HJ21-Hayes | Bacteria>Bacillati>Cyanobacteriota/Melainabacteria_group>Cyanobacteriota>Cyanophyceae>Synechococcales>Synechococcaceae>Synechococcus>unclassified_Synechococcus | 173 | KBZ14_RS08110 | 77040..77561 | 2823736 | hypothetical-protein-[Synechococcus-sp.-HJ21-Hayes] |
| MBF2075129.1 | REase | GCA_015272325.1 | Synechococcales_cyanobacterium_C42_A2020_086 | Bacteria>Bacillati>Cyanobacteriota/Melainabacteria_group>Cyanobacteriota>Cyanophyceae>Synechococcales>unclassified_Synechococcales | 200 | IGS50_15410 | 13538..14140 | 2774344 | Uma2-family-endonuclease-[Synechococcales-cyanobacterium-C42_A2020_086] |
| MBF2075142.1 | REase | GCA_015272325.1 | Synechococcales_cyanobacterium_C42_A2020_086 | Bacteria>Bacillati>Cyanobacteriota/Melainabacteria_group>Cyanobacteriota>Cyanophyceae>Synechococcales>unclassified_Synechococcales | 197 | IGS50_15475 | 38132..38725 | 2774344 | Uma2-family-endonuclease-[Synechococcales-cyanobacterium-C42_A2020_086] |
| MBF2000220.1 | REase | GCA_015272295.1 | Synechococcales_cyanobacterium_M58_A2018_015 | Bacteria>Bacillati>Cyanobacteriota/Melainabacteria_group>Cyanobacteriota>Cyanophyceae>Synechococcales>unclassified_Synechococcales | 190 | IGS38_05835 | 42152..42724 | 2774348 | Uma2-family-endonuclease-[Synechococcales-cyanobacterium-M58_A2018_015] |
| MBF2075188.1 | REase | GCA_015272325.1 | Synechococcales_cyanobacterium_C42_A2020_086 | Bacteria>Bacillati>Cyanobacteriota/Melainabacteria_group>Cyanobacteriota>Cyanophyceae>Synechococcales>unclassified_Synechococcales | 190 | IGS50_15715 | 90180..90752 | 2774344 | Uma2-family-endonuclease-[Synechococcales-cyanobacterium-C42_A2020_086] |
| MBF2075190.1 | REase | GCA_015272325.1 | Synechococcales_cyanobacterium_C42_A2020_086 | Bacteria>Bacillati>Cyanobacteriota/Melainabacteria_group>Cyanobacteriota>Cyanophyceae>Synechococcales>unclassified_Synechococcales | 190 | IGS50_15725 | 91568..92140 | 2774344 | Uma2-family-endonuclease-[Synechococcales-cyanobacterium-C42_A2020_086] |
| MBF2075187.1 | REase | GCA_015272325.1 | Synechococcales_cyanobacterium_C42_A2020_086 | Bacteria>Bacillati>Cyanobacteriota/Melainabacteria_group>Cyanobacteriota>Cyanophyceae>Synechococcales>unclassified_Synechococcales | 214 | IGS50_15710 | 89448..90092 | 2774344 | Uma2-family-endonuclease-[Synechococcales-cyanobacterium-C42_A2020_086] |
| MBF2000221.1 | REase | GCA_015272295.1 | Synechococcales_cyanobacterium_M58_A2018_015 | Bacteria>Bacillati>Cyanobacteriota/Melainabacteria_group>Cyanobacteriota>Cyanophyceae>Synechococcales>unclassified_Synechococcales | 209 | IGS38_05840 | 42808..43437 | 2774348 | restriction-endonuclease-subunit-R-[Synechococcales-cyanobacterium-M58_A2018_015] |
| MBF2075189.1 | REase | GCA_015272325.1 | Synechococcales_cyanobacterium_C42_A2020_086 | Bacteria>Bacillati>Cyanobacteriota/Melainabacteria_group>Cyanobacteriota>Cyanophyceae>Synechococcales>unclassified_Synechococcales | 209 | IGS50_15720 | 90855..91484 | 2774344 | restriction-endonuclease-subunit-R-[Synechococcales-cyanobacterium-C42_A2020_086] |
| MBF2075143.1 | REase | GCA_015272325.1 | Synechococcales_cyanobacterium_C42_A2020_086 | Bacteria>Bacillati>Cyanobacteriota/Melainabacteria_group>Cyanobacteriota>Cyanophyceae>Synechococcales>unclassified_Synechococcales | 69 | IGS50_15480 | 38955..39164 | 2774344 | hypothetical-protein-[Synechococcales-cyanobacterium-C42_A2020_086] |
| MBF2075125.1 | REase | GCA_015272325.1 | Synechococcales_cyanobacterium_C42_A2020_086 | Bacteria>Bacillati>Cyanobacteriota/Melainabacteria_group>Cyanobacteriota>Cyanophyceae>Synechococcales>unclassified_Synechococcales | 119 | IGS50_15390 | 7095..7454 | 2774344 | Uma2-family-endonuclease-[Synechococcales-cyanobacterium-C42_A2020_086] |
| WP_244349765.1 | REase | GCF_022848905.1 | Thermostichus_vulcanus | Bacteria>Bacillati>Cyanobacteriota/Melainabacteria_group>Cyanobacteriota>Cyanophyceae>Thermostichales>Thermostichaceae>Thermostichus | 190 | JX360_RS06135 | 26731..27303 | 32053 | Uma2-family-endonuclease-[Thermostichus-vulcanus] |
| WP_279611291.1 | REase | - | Thermostichus_vulcanus | Bacteria>Bacillati>Cyanobacteriota/Melainabacteria_group>Cyanobacteriota>Cyanophyceae>Thermostichales>Thermostichaceae>Thermostichus | 212 | JX360_RS06140 | 27374..28012 | 32053 | Uma2-family-endonuclease-[Thermostichus-vulcanus] |
| MBU6345611.1 | REase | GCA_018969285.1 | Cyanobacteria_bacterium_REEB494 | Bacteria>Bacillati>Cyanobacteriota/Melainabacteria_group>Cyanobacteriota>unclassified_Cyanobacteriota | 178 | KGR70_09965 | 1508..2044 | 2800107 | hypothetical-protein-[Cyanobacteria-bacterium-REEB494] |
| MDA0267771.1 | REase | GCA_027592485.1 | Cyanobacteriota_bacterium | Bacteria>Bacillati>Cyanobacteriota/Melainabacteria_group>Cyanobacteriota>unclassified_Cyanobacteriota | 189 | O3A14_12625 | 2926..3495 | 2847862 | MAG:-hypothetical-protein-O3A14_12625-[Cyanobacteria-bacterium] |
| HBB33195.1 | REase | GCA_003488275.1 | Cyanobacteria_bacterium_UBA9273 | Bacteria>Bacillati>Cyanobacteriota/Melainabacteria_group>Cyanobacteriota>unclassified_Cyanobacteriota | 202 | DC064_15695 | 5715..6323 | 2055791 | hypothetical-protein-[Cyanobacteria-bacterium-UBA9273] |
| MBU6346400.1 | REase | GCA_018969285.1 | Cyanobacteria_bacterium_REEB494 | Bacteria>Bacillati>Cyanobacteriota/Melainabacteria_group>Cyanobacteriota>unclassified_Cyanobacteriota | 193 | KGR70_14040 | 15180..15761 | 2800107 | Uma2-family-endonuclease-[Cyanobacteria-bacterium-REEB494] |
| OWY65327.1 | REase | GCA_002213405.1 | cyanobacterium_TDX16 | Bacteria>Bacillati>Cyanobacteriota/Melainabacteria_group>Cyanobacteriota>unclassified_Cyanobacteriota | 211 | B7486_42515 | 13093..13728 | 1503470 | hypothetical-protein-B7486_42515-[cyanobacterium-TDX16] |
| MBJ7900528.1 | REase | GCA_016471375.1 | Cyanobacteria_bacterium_RI_101 | Bacteria>Bacillati>Cyanobacteriota/Melainabacteria_group>Cyanobacteriota>unclassified_Cyanobacteriota | 210 | GC158_11545 | 13472..14104 | 2775578 | restriction-endonuclease-subunit-R-[Cyanobacteria-bacterium-RI_101] |
| MCY4331634.1 | REase | GCA_026710375.1 | Cyanobacteria_bacterium_MAG_CAR1_bin_15 | Bacteria>Bacillati>Cyanobacteriota/Melainabacteria_group>Cyanobacteriota>unclassified_Cyanobacteriota | 165 | OXC96_03835 | 57714..58211 | 2775589 | MAG:-DUF4365-domain-containing-protein-[Cyanobacteria-bacterium-MAG-CAR1_bin_15] |
| MDE0648518.1 | REase | GCA_028824215.1 | Cyanobacteria_bacterium_MAG_IRC4_bin_6 | Bacteria>Bacillati>Cyanobacteriota/Melainabacteria_group>Cyanobacteriota>unclassified_Cyanobacteriota | 165 | OXI08_10895 | 11748..12245 | 2775598 | MAG:-DUF4365-domain-containing-protein-[Cyanobacteria-bacterium-MAG-IRC4_bin_6] |
| MCY3653789.1 | REase | GCA_026705665.1 | Cyanobacteria_bacterium_MAG_IRC1_bin_28 | Bacteria>Bacillati>Cyanobacteriota/Melainabacteria_group>Cyanobacteriota>unclassified_Cyanobacteriota | 101 | OXG70_01965 | 3281..3586 | 2775596 | MAG:-DUF4365-domain-containing-protein-[Cyanobacteria-bacterium-MAG-IRC1_bin_28] |
| MCY4360133.1 | REase | GCA_026710605.1 | Cyanobacteria_bacterium_MAG_APA_bin_95 | Bacteria>Bacillati>Cyanobacteriota/Melainabacteria_group>Cyanobacteriota>unclassified_Cyanobacteriota | 62 | OXC47_05660 | 1..186 | 2775588 | MAG:-DUF4365-domain-containing-protein,-partial-[Cyanobacteria-bacterium-MAG-APA_bin_95] |
| MCX6842953.1 | REase | GCA_026397595.1 | candidate_division_WOR-3_bacterium | Bacteria>Bacteria_incertae_sedis>Bacteria_candidate_phyla>candidate_division_WOR-3 | 161 | NTX53_11800 | 9669..10154 | 2052148 | MAG:-DUF3883-domain-containing-protein-[candidate-division-WOR-3-bacterium] |
| MCX7973588.1 | REase | GCA_026416195.1 | Candidatus_Aminicenantes_bacterium | Bacteria>Bacteria_incertae_sedis>Bacteria_candidate_phyla>Candidatus_Aminicenantota>unclassified_Candidatus_Aminicenantes | 223 | N3B16_03710 | 4181..4852 | 2052149 | MAG:-restriction-endonuclease-[Candidatus-Aminicenantes-bacterium] |
| MCD6320865.1 | REase | GCA_021160365.1 | Candidatus_Bipolaricaulota_bacterium | Bacteria>Bacteria_incertae_sedis>Bacteria_candidate_phyla>Candidatus_Bipolaricaulota>unclassified_Candidatus_Bipolaricaulota | 143 | J7L84_01270 | 9984..10415 | 2306012 | hypothetical-protein-[Candidatus-Bipolaricaulota-bacterium] |
| MBK8230674.1 | REase | GCA_016712505.1 | Candidatus_Eiseniibacteriota_bacterium | Bacteria>Bacteria_incertae_sedis>Bacteria_candidate_phyla>Candidatus_Eiseniibacteriota | 164 | IPK72_08860 | 7258..7752 | 2212470 | hypothetical-protein-[Candidatus-Eisenbacteria-bacterium] |
| MBI4536773.1 | REase | GCA_016209735.1 | Candidatus_Methylomirabilota_bacterium | Bacteria>Bacteria_incertae_sedis>Bacteria_candidate_phyla>Candidatus_Methylomirabilota>unclassified_Candidatus_Methylomirabilota | 136 | HY712_02315 | 59494..59904 | 2072417 | GxxExxY-protein-[candidate-division-NC10-bacterium] |

| Protein-ID | Domain | Assembly-ID | Strain | Taxonomy | Length | Locus-Tag | Genomic-coordinates | TaxID | Fasta-defline |
|---|---|---|---|---|---|---|---|---|---|
| MDE0423421.1 | REase | GCA_028820845.1 | Candidatus_Poribacteria_bacterium | Bacteria>Bacteria_incertae_sedis>Bacteria_candidate_phyla>Candidatus_Poribacteria>unclassified_Candidatus_Poribacteria | 87 | OXN25_00985 | 13738..14001 | 2026781 | MAG:-hypothetical-protein-OXN25_00985-[Candidatus-Poribacteria-bacterium] |
| MCY3724181.1 | REase | GCA_026705955.1 | Candidatus_Poribacteria_bacterium | Bacteria>Bacteria_incertae_sedis>Bacteria_candidate_phyla>Candidatus_Poribacteria>unclassified_Candidatus_Poribacteria | 74 | OXG97_18340 | 4961..5185 | 2026781 | MAG:-hypothetical-protein-OXG97_18340-[Candidatus-Poribacteria-bacterium] |
| MDE0014211.1 | REase | GCA_028818705.1 | Candidatus_Poribacteria_bacterium | Bacteria>Bacteria_incertae_sedis>Bacteria_candidate_phyla>Candidatus_Poribacteria>unclassified_Candidatus_Poribacteria | 74 | OXU36_23910 | 15592..15816 | 2026781 | MAG:-hypothetical-protein-OXU36_23910-[Candidatus-Poribacteria-bacterium] |
| MXV83697.1 | REase | GCA_009835485.1 | Candidatus_Poribacteria_bacterium | Bacteria>Bacteria_incertae_sedis>Bacteria_candidate_phyla>Candidatus_Poribacteria>unclassified_Candidatus_Poribacteria | 74 | F4Z98_10050 | 14363..14587 | 2026781 | hypothetical-protein-[Candidatus-Poribacteria-bacterium] |
| MXV83698.1 | REase | GCA_009835485.1 | Candidatus_Poribacteria_bacterium | Bacteria>Bacteria_incertae_sedis>Bacteria_candidate_phyla>Candidatus_Poribacteria>unclassified_Candidatus_Poribacteria | 74 | F4Z98_10055 | 14607..14831 | 2026781 | hypothetical-protein-[Candidatus-Poribacteria-bacterium] |
| MXY28221.1 | REase | GCA_009838315.1 | Candidatus_Poribacteria_bacterium | Bacteria>Bacteria_incertae_sedis>Bacteria_candidate_phyla>Candidatus_Poribacteria>unclassified_Candidatus_Poribacteria | 74 | F4Y59_08700 | 3707..3931 | 2026781 | hypothetical-protein-[Candidatus-Poribacteria-bacterium] |
| MYC78291.1 | REase | GCA_009841255.1 | Candidatus_Poribacteria_bacterium | Bacteria>Bacteria_incertae_sedis>Bacteria_candidate_phyla>Candidatus_Poribacteria>unclassified_Candidatus_Poribacteria | 74 | F4X10_21220 | 46625..46849 | 2026781 | hypothetical-protein-[Candidatus-Poribacteria-bacterium] |
| MYK17051.1 | REase | GCA_009845505.1 | Candidatus_Poribacteria_bacterium | Bacteria>Bacteria_incertae_sedis>Bacteria_candidate_phyla>Candidatus_Poribacteria>unclassified_Candidatus_Poribacteria | 74 | F4055_02635 | 6247..6471 | 2026781 | hypothetical-protein-[Candidatus-Poribacteria-bacterium] |
| MYC78290.1 | REase | GCA_009841255.1 | Candidatus_Poribacteria_bacterium | Bacteria>Bacteria_incertae_sedis>Bacteria_candidate_phyla>Candidatus_Poribacteria>unclassified_Candidatus_Poribacteria | 70 | F4X10_21215 | 46393..46605 | 2026781 | hypothetical-protein-[Candidatus-Poribacteria-bacterium] |
| MXV83699.1 | REase | GCA_009835485.1 | Candidatus_Poribacteria_bacterium | Bacteria>Bacteria_incertae_sedis>Bacteria_candidate_phyla>Candidatus_Poribacteria>unclassified_Candidatus_Poribacteria | 85 | F4Z98_10060 | 14869..15126 | 2026781 | hypothetical-protein-[Candidatus-Poribacteria-bacterium] |
| MYC78292.1 | REase | GCA_009841255.1 | Candidatus_Poribacteria_bacterium | Bacteria>Bacteria_incertae_sedis>Bacteria_candidate_phyla>Candidatus_Poribacteria>unclassified_Candidatus_Poribacteria | 85 | F4X10_21225 | 46887..47144 | 2026781 | hypothetical-protein-[Candidatus-Poribacteria-bacterium] |
| PKL50475.1 | REase | GCA_002839435.1 | Candidatus_Riflebacteria_bacterium_HGW-Riflebacteria-2 | Bacteria>Bacteria_incertae_sedis>Bacteria_candidate_phyla>Candidatus_Rifleibacteriota>unclassified_Candidatus_Rifleibacteriota | 178 | CVV42_02975 | 69525..70061 | 2013830 | MAG:-hypothetical-protein-CVV42_02975-[Candidatus-Riflebacteria-bacterium-HGW-Riflebacteria-2] |
| MBI2568436.1 | REase | GCA_016188575.1 | Candidatus_Schekmanbacteria_bacterium | Bacteria>Bacteria_incertae_sedis>Bacteria_candidate_phyla>Candidatus_Schekmaniibacteriota>unclassified_Candidatus_Schekmanbacteria | 140 | HYV63_15555 | 57635..58057 | 2419984 | GxxExxY-protein-[Candidatus-Schekmanbacteria-bacterium] |
| MBI2568431.1 | REase | GCA_016188575.1 | Candidatus_Schekmanbacteria_bacterium | Bacteria>Bacteria_incertae_sedis>Bacteria_candidate_phyla>Candidatus_Schekmaniibacteriota>unclassified_Candidatus_Schekmanbacteria | 180 | HYV63_15530 | 50257..50799 | 2419984 | Uma2-family-endonuclease-[Candidatus-Schekmanbacteria-bacterium] |
| MBI2571113.1 | REase | GCA_016188575.1 | Candidatus_Schekmanbacteria_bacterium | Bacteria>Bacteria_incertae_sedis>Bacteria_candidate_phyla>Candidatus_Schekmaniibacteriota>unclassified_Candidatus_Schekmanbacteria | 209 | HYV63_29255 | 5540..6169 | 2419984 | hypothetical-protein-[Candidatus-Schekmanbacteria-bacterium] |
| MBI2571109.1 | REase | GCA_016188575.1 | Candidatus_Schekmanbacteria_bacterium | Bacteria>Bacteria_incertae_sedis>Bacteria_candidate_phyla>Candidatus_Schekmaniibacteriota>unclassified_Candidatus_Schekmanbacteria | 195 | HYV63_29235 | 2226..2813 | 2419984 | DUF3883-domain-containing-protein-[Candidatus-Schekmanbacteria-bacterium] |
| MDE2100039.1 | REase | GCA_028872085.1 | Patescibacteria_group_bacterium | Bacteria>Bacteria_incertae_sedis>Bacteria_candidate_phyla>Patescibacteria_group>unclassified_Patescibacteria_group | 195 | KGL39_22485 | 12788..13375 | 2052139 | MAG:-hypothetical-protein-KGL39_22485-[Patescibacteria-group-bacterium] |
| MCD6532983.1 | REase | GCA_021163445.1 | Candidatus_Tharpella_aukensis | Bacteria>Candidatus_Tharpellota>Candidatus_Tharpellia>Candidatus_Tharpellales>Candidatus_Tharpellaceae>Candidatus_Tharpella | 176 | J7L25_02745 | 19229..19759 | 2954148 | hypothetical-protein-[Deltaproteobacteria-bacterium] |
| WP_163298133.1 | REase | GCF_010499435.1 | Dissulfurirhabdus_thermomarina | Bacteria>delta/epsilon_subdivisions>Deltaproteobacteria>Deltaproteobacteria_incertae_sedis>Dissulfurirhabdaceae>Dissulfurirhabdus | 127 | G3N55_RS03855 | 5164..5547 | 1765737 | GxxExxY-protein-[Dissulfurirhabdus-thermomarina] |
| MBF0239770.1 | REase | GCA_015231535.1 | SAR324_cluster_bacterium | Bacteria>delta/epsilon_subdivisions>Deltaproteobacteria>Deltaproteobacteria_incertae_sedis>SAR324_cluster | 126 | HQM12_18885 | 7606..7986 | 2024889 | GxxExxY-protein-[SAR324-cluster-bacterium] |
| MBF0239763.1 | REase | GCA_015231535.1 | SAR324_cluster_bacterium | Bacteria>delta/epsilon_subdivisions>Deltaproteobacteria>Deltaproteobacteria_incertae_sedis>SAR324_cluster | 121 | HQM12_18850 | 542..907 | 2024889 | GxxExxY-protein-[SAR324-cluster-bacterium] |
| WP_197735415.1 | REase | GCF_007990525.1 | Leptotrichia_hofstadii | Bacteria>Fusobacteriati>Fusobacteriota>Fusobacteriia>Fusobacteriales>Leptotrichiaceae>Leptotrichia | 215 | FVE77_RS00760 | 174628..175275 | 157688 | DUF4263-domain-containing-protein-[Leptotrichia-hofstadii] |
| WP_036069828.1 | REase | GCF_000469525.1 | Leptotrichia_sp._oral_taxon_225 | Bacteria>Fusobacteriati>Fusobacteriota>Fusobacteriia>Fusobacteriales>Leptotrichiaceae>Leptotrichia>unclassified_Leptotrichia | 209 | HMPREF9108_RS00955 | 37544..38173 | 671213 | DUF4263-domain-containing-protein-[Leptotrichia-sp.-oral-taxon-225] |
| MBA2524267.1 | REase | GCA_013696625.1 | Pyrinomonadaceae_bacterium | Bacteria>Pseudomonadati>Acidobacteriota>Blastocatellia>Blastocatellales>Pyrinomonadaceae>unclassified_Pyrinomonadaceae | 194 | HOV18_00590 | 4712..5296 | 2283092 | hypothetical-protein-[Pyrinomonadaceae-bacterium] |
| MBZ5490312.1 | REase | GCA_020072645.1 | Terriglobia_bacterium | Bacteria>Pseudomonadati>Acidobacteriota>Terriglobia>unclassified_Terriglobia | 195 | LAO76_05195 | 368403..368990 | 2081523 | hypothetical-protein-[Acidobacteriia-bacterium] |
| KAB2964424.1 | REase | GCA_009026165.1 | Thermoanaerobaculia_bacterium | Bacteria>Pseudomonadati>Acidobacteriota>Thermoanaerobaculia>unclassified_Thermoanaerobaculia | 117 | F9K16_04320 | 1721..2074 | 2651171 | MAG:-hypothetical-protein-F9K16_04320-[Thermoanaerobaculia-bacterium] |
| MBN1568513.1 | REase | GCA_016929335.1 | Acidobacteriota_bacterium | Bacteria>Pseudomonadati>Acidobacteriota>unclassified_Acidobacteriota | 206 | JXA73_11760 | 12423..13043 | 1978231 | hypothetical-protein-[Acidobacteriota-bacterium] |
| PYV70337.1 | REase | GCA_003225375.1 | Acidobacteriota_bacterium | Bacteria>Pseudomonadati>Acidobacteriota>unclassified_Acidobacteriota | 195 | DMG97_19570 | 8288..8875 | 1978231 | hypothetical-protein-DMG97_19570-[Acidobacteriota-bacterium] |
| MCI0624093.1 | REase | GCA_022615935.1 | Acidobacteriota_bacterium | Bacteria>Pseudomonadati>Acidobacteriota>unclassified_Acidobacteriota | 146 | L0387_20990 | 11448..11888 | 1978231 | GxxExxY-protein-[Acidobacteriota-bacterium] |
| MBI4750000.1 | REase | GCA_016208495.1 | Acidobacteriota_bacterium | Bacteria>Pseudomonadati>Acidobacteriota>unclassified_Acidobacteriota | 125 | HY774_16055 | 64984..65361 | 1978231 | GxxExxY-protein-[Acidobacteriota-bacterium] |
| RMG44916.1 | REase | GCA_003696865.1 | Acidobacteriota_bacterium | Bacteria>Pseudomonadati>Acidobacteriota>unclassified_Acidobacteriota | 123 | D6723_18890 | 1032..1403 | 1978231 | GxxExxY-protein-[Acidobacteriota-bacterium] |
| HBL27436.1 | REase | GCA_003504885.1 | Acidobacteriota_bacterium | Bacteria>Pseudomonadati>Acidobacteriota>unclassified_Acidobacteriota | 229 | DD490_11430 | 3714..4401 | 1978231 | hypothetical-protein-[Acidobacteriota-bacterium] |
| HBL27435.1 | REase | GCA_003504885.1 | Acidobacteriota_bacterium | Bacteria>Pseudomonadati>Acidobacteriota>unclassified_Acidobacteriota | 89 | DD490_11425 | 3208..3477 | 1978231 | hypothetical-protein-[Acidobacteriota-bacterium] |
| MBI2833276.1 | REase | GCA_016190995.1 | Acidobacteriota_bacterium | Bacteria>Pseudomonadati>Acidobacteriota>unclassified_Acidobacteriota | 72 | HYX76_02485 | 2923..3141 | 1978231 | hypothetical-protein-[Acidobacteriota-bacterium] |
| MCC6130032.1 | REase | GCA_020845555.1 | Acidobacteriota_bacterium | Bacteria>Pseudomonadati>Acidobacteriota>unclassified_Acidobacteriota | 202 | IT186_08885 | 30302..30910 | 1978231 | hypothetical-protein-[Acidobacteriota-bacterium] |
| OFW00072.1 | REase | GCA_001767075.1 | Acidobacteria_bacterium_RIFCSPLOWO2_12_FULL_54_10 | Bacteria>Pseudomonadati>Acidobacteriota>unclassified_Acidobacteriota | 93 | A3F68_05140 | 24428..24709 | 1797186 | hypothetical-protein-A3F68_05140-[Acidobacteria-bacterium-RIFCSPLOWO2_12_FULL_54_10] |
| MBS1828813.1 | REase | GCA_018268935.1 | Acidobacteriota_bacterium | Bacteria>Pseudomonadati>Acidobacteriota>unclassified_Acidobacteriota | 163 | JST93_26155 | 37167..37658 | 1978231 | DUF4365-domain-containing-protein-[Acidobacteriota-bacterium] |
| MCB9093208.1 | REase | GCA_020635155.1 | Halobacteriovoraceae_bacterium | Bacteria>Pseudomonadati>Bdellovibrionota>Bacteriovoracia>Bacteriovoracales>Halobacteriovoraceae>unclassified_Halobacteriovoraceae | 236 | H6620_11675 | 20587..21297 | 2026745 | hypothetical-protein-[Halobacteriovoraceae-bacterium] |
| MCR4433940.1 | REase | GCA_024655865.1 | Caldisericota_bacterium | Bacteria>Pseudomonadati>Caldisericota/Cryosericota_group>Caldisericota>unclassified_Caldisericota | 202 | NUV70_07820 | 11887..12495 | 2052147 | hypothetical-protein-[Caldisericota-bacterium] |
| KAA3613618.1 | REase | GCA_008501795.1 | Calditrichota_bacterium | Bacteria>Pseudomonadati>Calditrichota>unclassified_Calditrichota | 193 | DWQ03_12330 | 122075..122656 | 2212469 | hypothetical-protein-DWQ03_12330-[Calditrichota-bacterium] |
| WP_187475235.1 | REase | GCF_014352915.1 | Aliarcobacter_cryaerophilus | Bacteria>Pseudomonadati>Campylobacterota>Epsilonproteobacteria>Campylobacterales>Arcobacteraceae>Aliarcobacter | 166 | HOO34_RS05650 | 1065545..1066045 | 28198 | hypothetical-protein-[Aliarcobacter-cryaerophilus] |
| WP_117713563.1 | REase | GCF_015558535.1 | Bacteroidales | Bacteria>Pseudomonadati>FCB_group>Bacteroidota/Chlorobiota_group>Bacteroidota>Bacteroidia | 204 | QH473_RS20235 | 25019..25633 | 171549 | MULTISPECIES:-hypothetical-protein-[Bacteroidales] |
| KAB5317655.1 | REase | GCA_009025145.1 | Bacteroides_stercoris | Bacteria>Pseudomonadati>FCB_group>Bacteroidota/Chlorobiota_group>Bacteroidota>Bacteroidia>Bacteroidales>Bacteroidaceae>Bacteroides | 204 | F9949_11580 | 59848..60462 | 46506 | hypothetical-protein-F9949_11580-[Bacteroides-stercoris] |
| RHC75804.1 | REase | GCA_003468465.1 | Bacteroides_uniformis | Bacteria>Pseudomonadati>FCB_group>Bacteroidota/Chlorobiota_group>Bacteroidota>Bacteroidia>Bacteroidales>Bacteroidaceae>Bacteroides | 204 | DW831_02870 | 24690..25304 | 820 | hypothetical-protein-DW831_02870-[Bacteroides-uniformis] |
| MBD5285712.1 | REase | GCA_014801525.1 | Bacteroides_sp. | Bacteria>Pseudomonadati>FCB_group>Bacteroidota/Chlorobiota_group>Bacteroidota>Bacteroidia>Bacteroidales>Bacteroidaceae>Bacteroides>unclassified_Bacteroides | 240 | HDS29_05485 | 13386..14108 | 29523 | hypothetical-protein-[Bacteroides-sp.] |
| WP_255395961.1 | REase | - | Bacteroides_sp._An51A | Bacteria>Pseudomonadati>FCB_group>Bacteroidota/Chlorobiota_group>Bacteroidota>Bacteroidia>Bacteroidales>Bacteroidaceae>Bacteroides>unclassified_Bacteroides | 86 | B5G04_RS20440 | 138803..139063 | 1965640 | DUF4143-domain-containing-protein-[Bacteroides-sp.-An51A] |
| WP_022053560.1 | REase | GCF_944319775.1 | Phocaeicola_plebeius | Bacteria>Pseudomonadati>FCB_group>Bacteroidota/Chlorobiota_group>Bacteroidota>Bacteroidia>Bacteroidales>Bacteroidaceae>Phocaeicola | 221 | H6A37_RS09800 | 45246..45911 | 310297 | DUF2461-domain-containing-protein-[Phocaeicola-plebeius] |
| HBQ57057.1 | REase | GCA_003511405.1 | Porphyromonadaceae_bacterium | Bacteria>Pseudomonadati>FCB_group>Bacteroidota/Chlorobiota_group>Bacteroidota>Bacteroidia>Bacteroidales>Porphyromonadaceae>unclassified_Porphyromonadaceae | 149 | DD786_07910 | 34568..35019 | 2049046 | hypothetical-protein-[Porphyromonadaceae-bacterium] |
| WP_094434340.1 | REase | GCF_002251185.1 | unclassified_Prevotella | Bacteria>Pseudomonadati>FCB_group>Bacteroidota/Chlorobiota_group>Bacteroidota>Bacteroidia>Bacteroidales>Prevotellaceae>Prevotella | 177 | CIK90_RS10275 | 65058..65591 | 2638335 | MULTISPECIES:-very-short-patch-repair-endonuclease-[unclassified-Prevotella] |
| WP_042520320.1 | REase | GCF_000834015.1 | Prevotella_pectinovora | Bacteria>Pseudomonadati>FCB_group>Bacteroidota/Chlorobiota_group>Bacteroidota>Bacteroidia>Bacteroidales>Prevotellaceae>Prevotella | 201 | ST42_RS12145 | 13631..14236 | 1602169 | DUF559-domain-containing-protein-[Prevotella-pectinovora] |
| OYP42210.1 | REase | GCA_002251295.1 | Prevotella_sp._P5-50 | Bacteria>Pseudomonadati>FCB_group>Bacteroidota/Chlorobiota_group>Bacteroidota>Bacteroidia>Bacteroidales>Prevotellaceae>Prevotella>unclassified_Prevotella | 177 | CIK88_02530 | 803..1336 | 2024217 | very-short-patch-repair-endonuclease-[Prevotella-sp.-P5-50] |
| WP_144007315.1 | REase | GCF_002251345.1 | Prevotella_sp._P4-119 | Bacteria>Pseudomonadati>FCB_group>Bacteroidota/Chlorobiota_group>Bacteroidota>Bacteroidia>Bacteroidales>Prevotellaceae>Prevotella>unclassified_Prevotella | 238 | CIK89_RS00880 | 32258..32974 | 2024218 | restriction-endonuclease-[Prevotella-sp.-P4-119] |
| MCZ2341284.1 | REase | GCA_026931725.1 | Bacteroidales_bacterium | Bacteria>Pseudomonadati>FCB_group>Bacteroidota/Chlorobiota_group>Bacteroidota>Bacteroidia>Bacteroidales>unclassified_Bacteroidales | 125 | LC104_05740 | 15663..16040 | 2030927 | GxxExxY-protein-[Bacteroidales-bacterium] |
| PLW98638.1 | REase | GCA_002869405.1 | Marinilabiliales_bacterium | Bacteria>Pseudomonadati>FCB_group>Bacteroidota/Chlorobiota_group>Bacteroidota>Bacteroidia>Marinilabiliales>unclassified_Marinilabiliales | 89 | C0593_05030 | 16348..16617 | 2053303 | MAG:-hypothetical-protein-C0593_05030-[Marinilabiliales-bacterium] |
| NCB25655.1 | REase | GCA_009929165.1 | Bacteroidia_bacterium | Bacteria>Pseudomonadati>FCB_group>Bacteroidota/Chlorobiota_group>Bacteroidota>Bacteroidia>unclassified_Bacteroidia | 247 | EOM62_09320 | 8223..8966 | 2044936 | hypothetical-protein-[Bacteroidia-bacterium] |
| WP_100629837.1 | REase | GCF_002807035.1 | Algoriphagus_formosus | Bacteria>Pseudomonadati>FCB_group>Bacteroidota/Chlorobiota_group>Bacteroidota>Cytophagia>Cytophagales>Cyclobacteriaceae>Algoriphagus | 223 | CDH51_RS17545 | 467444..468115 | 2007308 | endonuclease-NucS-[Algoriphagus-formosus] |
| MBK7652068.1 | REase | GCA_016709845.1 | Flammeovirgaceae_bacterium | Bacteria>Pseudomonadati>FCB_group>Bacteroidota/Chlorobiota_group>Bacteroidota>Cytophagia>Cytophagales>Flammeovirgaceae>unclassified_Flammeovirgaceae | 182 | IPJ20_16975 | 20233..20781 | 2026740 | hypothetical-protein-[Flammeovirgaceae-bacterium] |
| WP_035559962.1 | REase | GCF_000715495.1 | Hymenobacter_sp._IS2118 | Bacteria>Pseudomonadati>FCB_group>Bacteroidota/Chlorobiota_group>Bacteroidota>Cytophagia>Cytophagales>Hymenobacteraceae>Hymenobacter>unclassified_Hymenobacter | 228 | GM79_RS04570 | 22469..23155 | 1505605 | hypothetical-protein-[Hymenobacter-sp.-IS2118] |
| WP_183972375.1 | REase | GCF_014195535.1 | Runella_defluvii | Bacteria>Pseudomonadati>FCB_group>Bacteroidota/Chlorobiota_group>Bacteroidota>Cytophagia>Cytophagales>Spirosomataceae>Runella | 234 | FHS57_RS08100 | 115127..115831 | 370973 | hypothetical-protein-[Runella-defluvii] |
| MBC7388091.1 | REase | GCA_014376535.1 | Cytophagales_bacterium | Bacteria>Pseudomonadati>FCB_group>Bacteroidota/Chlorobiota_group>Bacteroidota>Cytophagia>Cytophagales>unclassified_Cytophagales | 233 | H7329_02665 | 2490..3191 | 2053541 | hypothetical-protein-[Opitutaceae-bacterium] |
| WP_024482430.1 | REase | GCF_000468595.1 | Cellulophaga_baltica | Bacteria>Pseudomonadati>FCB_group>Bacteroidota/Chlorobiota_group>Bacteroidota>Flavobacteriia>Flavobacteriales>Flavobacteriaceae>Cellulophaga | 228 | M665_RS21275 | 87366..88052 | 76594 | DUF3883-domain-containing-protein-[Cellulophaga-baltica] |
| WP_133532122.1 | REase | GCF_004362665.1 | Flavobacterium_dankookense | Bacteria>Pseudomonadati>FCB_group>Bacteroidota/Chlorobiota_group>Bacteroidota>Flavobacteriia>Flavobacteriales>Flavobacteriaceae>Flavobacterium | 137 | BC748_RS03770 | 562774..563187 | 706186 | hypothetical-protein-[Flavobacterium-dankookense] |
| WP_152604996.1 | REase | GCF_000759025.1 | Phaeodactylibacter_xiamenensis | Bacteria>Pseudomonadati>FCB_group>Bacteroidota/Chlorobiota_group>Bacteroidota>Saprospiria>Saprospirales>Haliscomenobacteraceae>Phaeodactylibacter | 211 | IX84_RS10895 | 20063..20698 | 1524460 | hypothetical-protein-[Phaeodactylibacter-xiamenensis] |
| WP_147165914.1 | REase | GCF_007993045.1 | Phaeodactylibacter_luteus | Bacteria>Pseudomonadati>FCB_group>Bacteroidota/Chlorobiota_group>Bacteroidota>Saprospiria>Saprospirales>Haliscomenobacteraceae>Phaeodactylibacter | 86 | FRY97_RS02855 | 79617..79877 | 1564516 | hypothetical-protein-[Phaeodactylibacter-luteus] |
| KAA6232804.1 | REase | GCA_008634005.1 | Chlorobium_phaeovibrioides | Bacteria>Pseudomonadati>FCB_group>Bacteroidota/Chlorobiota_group>Chlorobiota>Chlorobiia>Chlorobiales>Chlorobiaceae>Chlorobium/Pelodictyon_group>Chlorobium | 181 | FP507_06830 | 1407640..1408185 | 1094 | hypothetical-protein-FP507_06830-[Chlorobium-phaeovibrioides] |
| WP_151419522.1 | REase | GCF_008634005.1 | Chlorobium_phaeovibrioides | Bacteria>Pseudomonadati>FCB_group>Bacteroidota/Chlorobiota_group>Chlorobiota>Chlorobiia>Chlorobiales>Chlorobiaceae>Chlorobium/Pelodictyon_group>Chlorobium | 181 | FP507_RS07305 | 1407640..1408185 | 1094 | hypothetical-protein-[Chlorobium-phaeovibrioides] |
| WP_198166313.1 | REase | GCF_002113825.1 | Prosthecochloris_sp._HL-130-GSB | Bacteria>Pseudomonadati>FCB_group>Bacteroidota/Chlorobiota_group>Chlorobiota>Chlorobiia>Chlorobiales>Chlorobiaceae>Prosthecochloris>unclassified_Prosthecochloris | 126 | B9H02_RS07500 | 1590598..1590978 | 1974213 | restriction-endonuclease-[Prosthecochloris-sp.-HL-130-GSB] |
| MBL7989348.1 | REase | GCA_016788565.1 | Chlorobiota_bacterium | Bacteria>Pseudomonadati>FCB_group>Bacteroidota/Chlorobiota_group>Chlorobiota>unclassified_Chlorobiota | 124 | JNJ94_14900 | 62075..62449 | 2268192 | YraN-family-protein-[Chlorobiota-bacterium] |
| MCX6133012.1 | REase | GCA_026390595.1 | Ignavibacteriales_bacterium | Bacteria>Pseudomonadati>FCB_group>Bacteroidota/Chlorobiota_group>Ignavibacteriota>Ignavibacteria>Ignavibacteriales>unclassified_Ignavibacteriales | 191 | NTU47_04270 | 81903..82478 | 2049428 | MAG:-hypothetical-protein-NTU47_04270-[Ignavibacteriales-bacterium] |
| WP_251951692.1 | REase | GCF_943908255.1 | Salinibacter_ruber | Bacteria>Pseudomonadati>FCB_group>Bacteroidota/Chlorobiota_group>Rhodothermota>Rhodothermia>Rhodothermales>Salinibacteraceae>Salinibacter | 167 | NGL47_RS12310 | 29790..30293 | 146919 | hypothetical-protein-[Salinibacter-ruber] |
| WP_259094360.1 | REase | GCF_024809025.1 | Salinibacter_ruber | Bacteria>Pseudomonadati>FCB_group>Bacteroidota/Chlorobiota_group>Rhodothermota>Rhodothermia>Rhodothermales>Salinibacteraceae>Salinibacter | 167 | GGP87_RS12330 | 38300..38803 | 146919 | hypothetical-protein-[Salinibacter-ruber] |
| OQC05383.1 | REase | GCA_002070875.1 | Candidatus_Cloacimonetes_bacterium_ADurb.Bin089 | Bacteria>Pseudomonadati>FCB_group>Candidatus_Cloacimonadota>unclassified_Candidatus_Cloacimonadota | 172 | BWX76_00752 | 6342..6860 | 1852836 | MAG:-hypothetical-protein-BWX76_00752-[Candidatus-Cloacimonetes-bacterium-ADurb.Bin089] |
| MCB5271574.1 | REase | GCA_020532545.1 | Candidatus_Cloacimonadota_bacterium | Bacteria>Pseudomonadati>FCB_group>Candidatus_Cloacimonadota>unclassified_Candidatus_Cloacimonadota | 165 | LHW56_06995 | 17204..17701 | 2030808 | hypothetical-protein-[Candidatus-Cloacimonadota-bacterium] |
| MCB5246761.1 | REase | GCA_020532215.1 | Candidatus_Cloacimonadota_bacterium | Bacteria>Pseudomonadati>FCB_group>Candidatus_Cloacimonadota>unclassified_Candidatus_Cloacimonadota | 169 | LHW57_01880 | 2739..3248 | 2030808 | hypothetical-protein-[Candidatus-Cloacimonadota-bacterium] |
| MCB5253273.1 | REase | GCA_020532345.1 | Candidatus_Cloacimonadota_bacterium | Bacteria>Pseudomonadati>FCB_group>Candidatus_Cloacimonadota>unclassified_Candidatus_Cloacimonadota | 214 | LHW51_09910 | 3067..3711 | 2030808 | DUF4263-domain-containing-protein-[Candidatus-Cloacimonadota-bacterium] |
| MCD6099132.1 | REase | GCA_021158485.1 | Fidelibacterota_bacterium | Bacteria>Pseudomonadati>FCB_group>Fidelibacterota>unclassified_Fidelibacterota | 102 | J7K33_00960 | 101..409 | 2026760 | hypothetical-protein-[Candidatus-Marinimicrobia-bacterium] |
| MYC87146.1 | REase | GCA_009841015.1 | Candidatus_Palauibacter_denitrificans | Bacteria>Pseudomonadati>FCB_group>Gemmatimonadota>Gemmatimonadia>Candidatus_Palauibacterales>Candidatus_Palauibacteraceae>Candidatus_Palauibacter | 198 | F4X22_02770 | 9673..10269 | 3056734 | hypothetical-protein-[Gemmatimonadales-bacterium] |
| MDE2877863.1 | REase | GCA_028874535.1 | Candidatus_Palauibacter_soopunensis | Bacteria>Pseudomonadati>FCB_group>Gemmatimonadota>Gemmatimonadia>Candidatus_Palauibacterales>Candidatus_Palauibacteraceae>Candidatus_Palauibacter | 141 | OXO55_03045 | 12342..12767 | 3056739 | MAG:-very-short-patch-repair-endonuclease-[Gemmatimonadota-bacterium] |
| MYE70562.1 | REase | GCA_009841855.1 | Gemmatimonadota_bacterium | Bacteria>Pseudomonadati>FCB_group>Gemmatimonadota>unclassified_Gemmatimonadota | 198 | F4237_11005 | 2095..2691 | 2026742 | hypothetical-protein-[Gemmatimonadota-bacterium] |
| MDE2981418.1 | REase | GCA_028873815.1 | Gemmatimonadota_bacterium | Bacteria>Pseudomonadati>FCB_group>Gemmatimonadota>unclassified_Gemmatimonadota | 177 | OXU74_09505 | 41729..42262 | 2026742 | MAG:-hypothetical-protein-OXU74_09505-[Gemmatimonadota-bacterium] |
| MYE95019.1 | REase | GCA_009841825.1 | Gemmatimonadota_bacterium | Bacteria>Pseudomonadati>FCB_group>Gemmatimonadota>unclassified_Gemmatimonadota | 147 | F4238_16845 | 3333..3776 | 2026742 | DNA-mismatch-endonuclease-Vsr-[Gemmatimonadota-bacterium] |
| MCY3788284.1 | REase | GCA_026706505.1 | Gemmatimonadota_bacterium | Bacteria>Pseudomonadati>FCB_group>Gemmatimonadota>unclassified_Gemmatimonadota | 76 | OXH63_05795 | 1..232 | 2026742 | MAG:-DUF3883-domain-containing-protein,-partial-[Gemmatimonadetes-bacterium] |
| MDE2733152.1 | REase | GCA_028875335.1 | Gemmatimonadota_bacterium | Bacteria>Pseudomonadati>FCB_group>Gemmatimonadota>unclassified_Gemmatimonadota | 155 | OXI72_02075 | 18906..19373 | 2026742 | MAG:-hypothetical-protein-OXI72_02075-[Gemmatimonadota-bacterium] |
| MDE2733148.1 | REase | GCA_028875335.1 | Gemmatimonadota_bacterium | Bacteria>Pseudomonadati>FCB_group>Gemmatimonadota>unclassified_Gemmatimonadota | 84 | OXI72_02055 | 14531..14785 | 2026742 | MAG:-hypothetical-protein-OXI72_02055-[Gemmatimonadota-bacterium] |
| WP_012098802.1 | REase | GCF_000017505.1 | Anaeromyxobacter_sp._Fw109-5 | Bacteria>Pseudomonadati>Myxococcota>Myxococcia>Myxococcales>Cystobacterineae>Anaeromyxobacteraceae>Anaeromyxobacter>unclassified_Anaeromyxobacter | 197 | ANAE109_RS20465 | 4651303..4651896 | 404589 | hypothetical-protein-[Anaeromyxobacter-sp.-Fw109-5] |
| WP_121761666.1 | REase | GCF_003668955.1 | Corallococcus_sp._AB038B | Bacteria>Pseudomonadati>Myxococcota>Myxococcia>Myxococcales>Cystobacterineae>Myxococcaceae>Corallococcus>unclassified_Corallococcus | 249 | D7Y04_RS31660 | 367717..368466 | 2316718 | Tox-REase-5-domain-containing-protein-[Corallococcus-sp.-AB038B] |
| MCA3011285.1 | REase | GCA_020247085.1 | Myxococcaceae_bacterium | Bacteria>Pseudomonadati>Myxococcota>Myxococcia>Myxococcales>Cystobacterineae>Myxococcaceae>unclassified_Myxococcaceae | 154 | INH41_02685 | 11669..12130 | 2484252 | nuclease-[Myxococcaceae-bacterium] |
| MBI3184664.1 | REase | GCA_016191815.1 | Myxococcales_bacterium | Bacteria>Pseudomonadati>Myxococcota>Myxococcia>Myxococcales>unclassified_Myxococcales | 197 | HYZ28_21215 | 49915..50508 | 2026763 | hypothetical-protein-[Myxococcales-bacterium] |
| MBL9006088.1 | REase | GCA_016794505.1 | Myxococcales_bacterium | Bacteria>Pseudomonadati>Myxococcota>Myxococcia>Myxococcales>unclassified_Myxococcales | 197 | JNJ46_17675 | 14207..14800 | 2026763 | hypothetical-protein-[Myxococcales-bacterium] |
| MBI3184671.1 | REase | GCA_016191815.1 | Myxococcales_bacterium | Bacteria>Pseudomonadati>Myxococcota>Myxococcia>Myxococcales>unclassified_Myxococcales | 168 | HYZ28_21250 | 57266..57772 | 2026763 | very-short-patch-repair-endonuclease-[Myxococcales-bacterium] |
| MBK7583790.1 | REase | GCA_016706225.1 | Myxococcales_bacterium | Bacteria>Pseudomonadati>Myxococcota>Myxococcia>Myxococcales>unclassified_Myxococcales | 149 | IPI67_26800 | 454364..454813 | 2026763 | hypothetical-protein-[Myxococcales-bacterium] |
| MCC6503797.1 | REase | GCA_020846435.1 | Deltaproteobacteria_bacterium | Bacteria>Pseudomonadati>Myxococcota>Myxococcia>unclassified_Myxococcia | 179 | IT362_11780 | 15102..15641 | 2026735 | hypothetical-protein-[Deltaproteobacteria-bacterium] |
| MBW1967270.1 | REase | GCA_019308155.1 | Deltaproteobacteria_bacterium | Bacteria>Pseudomonadati>Myxococcota>Myxococcia>unclassified_Myxococcia | 177 | JRI48_07770 | 849..1382 | 2026735 | hypothetical-protein-[Deltaproteobacteria-bacterium] |
| MBW2068241.1 | REase | GCA_019309115.1 | Deltaproteobacteria_bacterium | Bacteria>Pseudomonadati>Myxococcota>Myxococcia>unclassified_Myxococcia | 168 | JRI31_05110 | 99826..100332 | 2026735 | hypothetical-protein-[Deltaproteobacteria-bacterium] |
| MBC8466715.1 | REase | GCA_014384065.1 | Deltaproteobacteria_bacterium | Bacteria>Pseudomonadati>Myxococcota>Myxococcia>unclassified_Myxococcia | 166 | H8D55_02640 | 8723..9223 | 2026735 | hypothetical-protein-[Deltaproteobacteria-bacterium] |
| MBI4514359.1 | REase | GCA_016210005.1 | Deltaproteobacteria_bacterium | Bacteria>Pseudomonadati>Myxococcota>Myxococcia>unclassified_Myxococcia | 143 | HY699_00885 | 119059..119490 | 2026735 | GxxExxY-protein-[Deltaproteobacteria-bacterium] |
| HBA39937.1 | REase | GCA_003490035.1 | Deltaproteobacteria_bacterium | Bacteria>Pseudomonadati>Myxococcota>Myxococcia>unclassified_Myxococcia | 121 | DCZ05_09405 | 9259..9624 | 2026735 | GxxExxY-protein-[Deltaproteobacteria-bacterium] |
| MBI4525151.1 | REase | GCA_016210115.1 | Deltaproteobacteria_bacterium | Bacteria>Pseudomonadati>Myxococcota>Myxococcia>unclassified_Myxococcia | 121 | HY695_15210 | 17307..17672 | 2026735 | GxxExxY-protein-[Deltaproteobacteria-bacterium] |
| HBA39935.1 | REase | GCA_003490035.1 | Deltaproteobacteria_bacterium | Bacteria>Pseudomonadati>Myxococcota>Myxococcia>unclassified_Myxococcia | 65 | DCZ05_09395 | 5831..6028 | 2026735 | hypothetical-protein-[Deltaproteobacteria-bacterium] |
| MBI5446428.1 | REase | GCA_016218905.1 | Deltaproteobacteria_bacterium | Bacteria>Pseudomonadati>Myxococcota>Myxococcia>unclassified_Myxococcia | 190 | HY900_35090 | 18402..18974 | 2026735 | Uma2-family-endonuclease-[Deltaproteobacteria-bacterium] |
| MBI4519049.1 | REase | GCA_016210005.1 | Deltaproteobacteria_bacterium | Bacteria>Pseudomonadati>Myxococcota>Myxococcia>unclassified_Myxococcia | 151 | HY699_24935 | 62742..63197 | 2026735 | endonuclease-domain-containing-protein-[Deltaproteobacteria-bacterium] |
| MBI3249563.1 | REase | GCA_016197285.1 | Deltaproteobacteria_bacterium | Bacteria>Pseudomonadati>Myxococcota>Myxococcia>unclassified_Myxococcia | 124 | HYZ50_23925 | 2200..2574 | 2026735 | endonuclease-domain-containing-protein-[Deltaproteobacteria-bacterium] |
| MCX5862616.1 | REase | GCA_026388625.1 | Deltaproteobacteria_bacterium | Bacteria>Pseudomonadati>Myxococcota>Myxococcia>unclassified_Myxococcia | 183 | NTW27_10935 | 89220..89771 | 2026735 | MAG:-hypothetical-protein-NTW27_10935-[Deltaproteobacteria-bacterium] |
| MCL5406609.1 | REase | GCA_023484975.1 | Deltaproteobacteria_bacterium | Bacteria>Pseudomonadati>Myxococcota>Myxococcia>unclassified_Myxococcia | 155 | M1398_07820 | 2000..2467 | 2026735 | hypothetical-protein-[Deltaproteobacteria-bacterium] |
| MBK9368676.1 | REase | GCA_016719515.1 | Deltaproteobacteria_bacterium | Bacteria>Pseudomonadati>Myxococcota>Myxococcia>unclassified_Myxococcia | 199 | IPN01_20605 | 111880..112479 | 2026735 | hypothetical-protein-[Deltaproteobacteria-bacterium] |
| MCY4488572.1 | REase | GCA_026712905.1 | Deltaproteobacteria_bacterium | Bacteria>Pseudomonadati>Myxococcota>Myxococcia>unclassified_Myxococcia | 220 | OXF11_15880 | 21622..22284 | 2026735 | MAG:-hypothetical-protein-OXF11_15880-[Deltaproteobacteria-bacterium] |
| WP_050434137.1 | REase | GCF_001189295.1 | Chondromyces_crocatus | Bacteria>Pseudomonadati>Myxococcota>Polyangia>Polyangiales>Polyangiaceae>Chondromyces | 197 | CMC5_RS33065 | 9155927..9156520 | 52 | hypothetical-protein-[Chondromyces-crocatus] |
| KYF87241.1 | REase | GCA_001589185.1 | Sorangium_cellulosum | Bacteria>Pseudomonadati>Myxococcota>Polyangia>Polyangiales>Polyangiaceae>Sorangium | 168 | BE17_46130 | 6924..7430 | 56 | cysteinyl-tRNA-synthetase-[Sorangium-cellulosum] |
| MBX3129294.1 | REase | GCA_019637295.1 | Polyangiaceae_bacterium | Bacteria>Pseudomonadati>Myxococcota>Polyangia>Polyangiales>Polyangiaceae>unclassified_Polyangiaceae | 197 | KF718_21430 | 15804..16397 | 2268199 | hypothetical-protein-[Polyangiaceae-bacterium] |
| HRG94675.1 | REase | GCA_035439005.1 | Polyangiaceae_bacterium | Bacteria>Pseudomonadati>Myxococcota>Polyangia>Polyangiales>Polyangiaceae>unclassified_Polyangiaceae | 124 | PLR99_00410 | 21782..22156 | 2268199 | GxxExxY-protein-[Polyangiaceae-bacterium] |
| MBU0551511.1 | REase | GCA_018813065.1 | Myxococcota_bacterium | Bacteria>Pseudomonadati>Myxococcota>unclassified_Myxococcota | 175 | KJ940_08445 | 6363..6890 | 2818507 | hypothetical-protein-[Myxococcota-bacterium] |
| MCK6514471.1 | REase | GCA_023150775.1 | Myxococcota_bacterium | Bacteria>Pseudomonadati>Myxococcota>unclassified_Myxococcota | 220 | L6R46_05385 | 10155..10817 | 2818507 | hypothetical-protein-[Myxococcota-bacterium] |
| MCP9446718.1 | REase | GCA_024330945.1 | Nitrospira_sp. | Bacteria>Pseudomonadati>Nitrospirota>Nitrospiria>Nitrospirales>Nitrospiraceae>Nitrospira>unclassified_Nitrospira | 123 | NNA22_03995 | 12291..12662 | 70125 | GxxExxY-protein-[Nitrospira-sp.] |

| Protein-ID | Domain | Assembly-ID | Strain | Taxonomy | Length | Locus-Tag | Genomic-coordinates | TaxID | Fasta-defline |
|---|---|---|---|---|---|---|---|---|---|
| MBS0151033.1 | REase | GCA_018242725.1 | Nitrospira_sp. | Bacteria>Pseudomonadati>Nitrospirota>Nitrospiria>Nitrospirales>Nitrospiraceae>Nitrospira>unclassified_Nitrospira | 181 | JSR31_08835 | 29725..30270 | 70125 | hypothetical-protein-[Nitrospira-sp.] |
| MBX3319016.1 | REase | GCA_019636895.1 | Nitrospira_sp. | Bacteria>Pseudomonadati>Nitrospirota>Nitrospiria>Nitrospirales>Nitrospiraceae>Nitrospira>unclassified_Nitrospira | 149 | KF890_03985 | 55676..56125 | 70125 | PD-(D/E)XK-nuclease-family-protein-[Nitrospira-sp.] |
| OIP61732.1 | REase | GCA_001873285.1 | Nitrospirae_bacterium_CG2_30_53_67 | Bacteria>Pseudomonadati>Nitrospirota>unclassified_Nitrospirota | 138 | AUK29_09780 | 4097..4513 | 1805278 | GxxExxY-protein-[Nitrospirae-bacterium-CG2_30_53_67] |
| RMH36430.1 | REase | GCA_003695915.1 | Nitrospirota_bacterium | Bacteria>Pseudomonadati>Nitrospirota>unclassified_Nitrospirota | 121 | D6690_05865 | 5774..6139 | 2026887 | GxxExxY-protein-[Nitrospirota-bacterium] |
| PIV85848.1 | REase | GCA_002781445.1 | Nitrospirae_bacterium_CG17_big_fil_post_rev_8_21_14_2_50_50_9 | Bacteria>Pseudomonadati>Nitrospirota>unclassified_Nitrospirota | 114 | COW52_00065 | 4158..4501 | 1974070 | GxxExxY-protein,-partial-[Nitrospirae-bacterium-CG17_big_fil_post_rev_8_21_14_2_50_50_9] |
| WP_103916697.1 | REase | GCF_900107995.1 | Gammaproteobacteria | Bacteria>Pseudomonadati>Pseudomonadota | 177 | C4S63_RS03495 | 776940..777473 | 1236 | hypothetical-protein-[Alcanivorax-sp.-DSM-26293] |
| WP_228232946.1 | REase | GCF_020782135.1 | Gammaproteobacteria | Bacteria>Pseudomonadati>Pseudomonadota | 165 | LL252_RS03450 | 284262..284759 | 1236 | hypothetical-protein-[Alloalcanivorax-marinus] |
| WP_064218006.1 | REase | GCF_035795835.1 | Acidithiobacillus | Bacteria>Pseudomonadati>Pseudomonadota>Acidithiobacillia>Acidithiobacillales>Acidithiobacillaceae | 179 | A4H96_RS01800 | 36020..36559 | 119977 | hypothetical-protein-[Acidithiobacillus-ferrooxidans] |
| WP_163056483.1 | REase | GCF_010577825.1 | Acidithiobacillus | Bacteria>Pseudomonadati>Pseudomonadota>Acidithiobacillia>Acidithiobacillales>Acidithiobacillaceae | 165 | X857_RS08740 | 42752..43249 | 119977 | hypothetical-protein-[Acidithiobacillus-ferrooxidans] |
| MBU2768210.1 | REase | GCA_018853935.1 | Acidithiobacillus_ferrivorans | Bacteria>Pseudomonadati>Pseudomonadota>Acidithiobacillia>Acidithiobacillales>Acidithiobacillaceae>Acidithiobacillus | 171 | HAP94_19060 | 1631..2146 | 160808 | hypothetical-protein-[Acidithiobacillus-ferrivorans] |
| UBU62552.1 | REase | GCA_020221655.1 | Acidithiobacillus_ferrooxidans | Bacteria>Pseudomonadati>Pseudomonadota>Acidithiobacillia>Acidithiobacillales>Acidithiobacillaceae>Acidithiobacillus | 113 | LDB30_00730 | 151088..151429 | 920 | hypothetical-protein-LDB30_00730-[Acidithiobacillus-ferrooxidans] |
| WP_199258163.1 | REase | GCF_016446475.1 | Rhodobacterales | Bacteria>Pseudomonadati>Pseudomonadota>Alphaproteobacteria | 166 | I9P22_RS04355 | 1666..2166 | 204455 | hypothetical-protein-[Paracoccus-binzhouensis] |
| WP_036280613.1 | REase | GCF_000188155.2 | Methylocystis_sp._ATCC_49242 | Bacteria>Pseudomonadati>Pseudomonadota>Alphaproteobacteria>Hyphomicrobiales>Methylocystaceae>Methylocystis>unclassified_Methylocystis | 166 | MET49242_RS03160 | 261161..261661 | 622637 | hypothetical-protein-[Methylocystis-sp.-ATCC-49242] |
| CDX58770.1 | REase | GCA_000824765.1 | Mesorhizobium_plurifarium | Bacteria>Pseudomonadati>Pseudomonadota>Alphaproteobacteria>Hyphomicrobiales>Phyllobacteriaceae>Mesorhizobium | 164 | MPL3365_30300 | 302879..303373 | 69974 | conserved-hypothetical-protein-[Mesorhizobium-plurifarium] |
| TPI12232.1 | REase | GCA_006439355.1 | Mesorhizobium_sp._B4-1-3 | Bacteria>Pseudomonadati>Pseudomonadota>Alphaproteobacteria>Hyphomicrobiales>Phyllobacteriaceae>Mesorhizobium>unclassified_Mesorhizobium | 217 | FJW06_17690 | 66695..67348 | 2589889 | hypothetical-protein-FJW06_17690-[Mesorhizobium-sp.-B4-1-3] |
| WP_140753512.1 | REase | GCF_006439355.1 | Mesorhizobium_sp._B4-1-3 | Bacteria>Pseudomonadati>Pseudomonadota>Alphaproteobacteria>Hyphomicrobiales>Phyllobacteriaceae>Mesorhizobium>unclassified_Mesorhizobium | 217 | FJW06_RS17690 | 66695..67348 | 2589889 | hypothetical-protein-[Mesorhizobium-sp.-B4-1-3] |
| WP_197504235.1 | REase | GCF_001672445.1 | Mesorhizobium_sp._WSM3873 | Bacteria>Pseudomonadati>Pseudomonadota>Alphaproteobacteria>Hyphomicrobiales>Phyllobacteriaceae>Mesorhizobium>unclassified_Mesorhizobium | 164 | A9K71_RS15160 | 43730..44224 | 1854056 | hypothetical-protein-[Mesorhizobium-sp.-WSM3873] |
| RWB26426.1 | REase | GCA_004016585.1 | Mesorhizobium_sp. | Bacteria>Pseudomonadati>Pseudomonadota>Alphaproteobacteria>Hyphomicrobiales>Phyllobacteriaceae>Mesorhizobium>unclassified_Mesorhizobium | 238 | EOQ43_30890 | 28853..29569 | 1871066 | hypothetical-protein-EOQ43_30890-[Mesorhizobium-sp.] |
| RWE30569.1 | REase | GCA_004017865.1 | Mesorhizobium_sp. | Bacteria>Pseudomonadati>Pseudomonadota>Alphaproteobacteria>Hyphomicrobiales>Phyllobacteriaceae>Mesorhizobium>unclassified_Mesorhizobium | 240 | EOS77_19505 | 7087..7809 | 1871066 | hypothetical-protein-EOS77_19505-[Mesorhizobium-sp.] |
| WP_154722160.1 | REase | GCF_900492205.1 | Ciceribacter_sp._T2.26MG-112.2 | Bacteria>Pseudomonadati>Pseudomonadota>Alphaproteobacteria>Hyphomicrobiales>Rhizobiaceae>Ciceribacter>unclassified_Ciceribacter | 73 | DYZ14_RS19095 | 24599..24820 | 3137154 | restriction-endonuclease-[Ciceribacter-naphthalenivorans] |
| CAD6617274.1 | REase | - | Rhizobium_sp._Khangiran2 | Bacteria>Pseudomonadati>Pseudomonadota>Alphaproteobacteria>Hyphomicrobiales>Rhizobiaceae>Rhizobium/Agrobacterium_group>Rhizobium>unclassified_Rhizobium | 167 | RKHAN_03343 | 3385415..3385918 | 1435607 | hypothetical-protein-RKHAN_03343-[Rhizobium-sp.-Khangiran2] |
| WP_149748053.1 | REase | GCF_900156005.1 | Rhizobium_sp._RU35A | Bacteria>Pseudomonadati>Pseudomonadota>Alphaproteobacteria>Hyphomicrobiales>Rhizobiaceae>Rhizobium/Agrobacterium_group>Rhizobium>unclassified_Rhizobium | 112 | F3H95_RS25730 | 2141..2479 | 1907414 | restriction-endonuclease-[Rhizobium-sp.-RU35A] |
| WP_004434283.1 | REase | GCF_000236945.1 | Sinorhizobium_meliloti | Bacteria>Pseudomonadati>Pseudomonadota>Alphaproteobacteria>Hyphomicrobiales>Rhizobiaceae>Sinorhizobium/Ensifer_group>Sinorhizobium | 166 | SM0020_RS29785 | 165197..165697 | 382 | hypothetical-protein-[Sinorhizobium-meliloti] |
| WP_238947175.1 | REase | GCF_900094735.1 | Saliniramus_fredricksonii | Bacteria>Pseudomonadati>Pseudomonadota>Alphaproteobacteria>Hyphomicrobiales>Salinarimonadaceae>Saliniramus | 135 | GA0071312_RS10320 | 789964..790371 | 1653334 | endonuclease-domain-containing-protein-[Saliniramus-fredricksonii] |
| MCZ7659372.1 | REase | GCA_027493405.1 | Xanthobacteraceae_bacterium | Bacteria>Pseudomonadati>Pseudomonadota>Alphaproteobacteria>Hyphomicrobiales>Xanthobacteraceae>unclassified_Xanthobacteraceae | 234 | M5U07_16620 | 16411..17115 | 2021379 | MAG:-hypothetical-protein-M5U07_16620-[Xanthobacteraceae-bacterium] |
| MBF0177024.1 | REase | GCA_015231175.1 | Magnetococcales_bacterium | Bacteria>Pseudomonadati>Pseudomonadota>Alphaproteobacteria>Magnetococcales>unclassified_Magnetococcales | 175 | HQL63_09280 | 25405..25932 | 2026759 | hypothetical-protein-[Magnetococcales-bacterium] |
| WP_274591704.1 | REase | GCF_041429805.1 | Marinovum | Bacteria>Pseudomonadati>Pseudomonadota>Alphaproteobacteria>Rhodobacterales>Roseobacteraceae | 166 | PVW49_RS22980 | 7992..8492 | 367771 | hypothetical-protein-[Marinovum-sp.-PR37] |
| WP_245888437.1 | REase | GCF_003003355.1 | Donghicola_tyrosinivorans | Bacteria>Pseudomonadati>Pseudomonadota>Alphaproteobacteria>Rhodobacterales>Roseobacteraceae>Donghicola | 163 | CLV74_RS02120 | 440873..441364 | 1652492 | hypothetical-protein-[Donghicola-tyrosinivorans] |
| WP_123792781.1 | REase | GCF_003814915.1 | Pacificibacter_maritimus | Bacteria>Pseudomonadati>Pseudomonadota>Alphaproteobacteria>Rhodobacterales>Roseobacteraceae>Pacificibacter | 172 | EDD53_RS08400 | 388941..389459 | 762213 | hypothetical-protein-[Pacificibacter-maritimus] |
| MAE92772.1 | REase | GCA_002687285.1 | Pelagibaca_sp. | Bacteria>Pseudomonadati>Pseudomonadota>Alphaproteobacteria>Rhodobacterales>Roseobacteraceae>Pelagibaca>unclassified_Pelagibaca | 115 | CMI67_24885 | 21629..21974 | 1979400 | hypothetical-protein-[Pelagibaca-sp.] |
| WP_170518617.1 | REase | GCF_013032825.1 | Ruegeria_atlantica | Bacteria>Pseudomonadati>Pseudomonadota>Alphaproteobacteria>Rhodobacterales>Roseobacteraceae>Ruegeria | 208 | GOW75_RS18380 | 61885..62511 | 81569 | hypothetical-protein-[Ruegeria-atlantica] |
| WP_150282502.1 | REase | GCF_008692285.1 | Salipiger_aestuarii | Bacteria>Pseudomonadati>Pseudomonadota>Alphaproteobacteria>Rhodobacterales>Roseobacteraceae>Salipiger | 166 | AL036_RS18770 | 11390..11890 | 568098 | hypothetical-protein-[Salipiger-aestuarii] |
| PZR07945.1 | REase | GCA_003243305.1 | Azospirillum_brasilense | Bacteria>Pseudomonadati>Pseudomonadota>Alphaproteobacteria>Rhodospirillales>Azospirillaceae>Azospirillum | 238 | DI532_22865 | 11769..12485 | 192 | hypothetical-protein-DI532_22865-[Azospirillum-brasilense] |
| WP_085595211.1 | REase | GCF_002115825.1 | Thalassospira_sp._MCCC_1A01428 | Bacteria>Pseudomonadati>Pseudomonadota>Alphaproteobacteria>Rhodospirillales>Thalassospiraceae>Thalassospira>unclassified_Thalassospira | 178 | THS27_RS23985 | 6449..6985 | 1470575 | hypothetical-protein-[Thalassospira-sp.-MCCC-1A01428] |
| WP_133251653.1 | REase | GCF_003173035.1 | Zavarzinia_aquatilis | Bacteria>Pseudomonadati>Pseudomonadota>Alphaproteobacteria>Rhodospirillales>Zavarziniaceae>Zavarzinia | 169 | DKG74_RS21545 | 32845..33354 | 2211142 | hypothetical-protein-[Zavarzinia-aquatilis] |
| WP_022673599.1 | REase | GCF_000420305.1 | Novosphingopyxis_baekryungensis | Bacteria>Pseudomonadati>Pseudomonadota>Alphaproteobacteria>Sphingomonadales>Sphingomonadaceae>Novosphingopyxis | 167 | G410_RS0110410 | 252793..253296 | 279369 | hypothetical-protein-[Novosphingopyxis-baekryungensis] |
| WP_021240098.1 | REase | GCF_000445065.1 | Sphingobium_quisquiliarum | Bacteria>Pseudomonadati>Pseudomonadota>Alphaproteobacteria>Sphingomonadales>Sphingomonadaceae>Sphingobium | 166 | L288_RS41585 | 37607..38107 | 538379 | hypothetical-protein-[Sphingobium-quisquiliarum] |
| MDA8776762.1 | REase | GCA_028101585.1 | Alphaproteobacteria_bacterium | Bacteria>Pseudomonadati>Pseudomonadota>Alphaproteobacteria>unclassified_Alphaproteobacteria | 213 | N9N58_04655 | 20380..21021 | 1913988 | hypothetical-protein-N9N58_04655-[Alphaproteobacteria-bacterium] |
| SBT04364.1 | REase | GCA_900089955.1 | Candidatus_Accumulibacter_aalborgensis | Bacteria>Pseudomonadati>Pseudomonadota>Betaproteobacteria>Betaproteobacteria_incertae_sedis>Candidatus_Accumulibacter | 181 | ACCAA_130243 | 228097..228642 | 1860102 | conserved-hypothetical-protein-[Candidatus-Accumulibacter-aalborgensis] |
| SBT07047.1 | REase | GCA_900089955.1 | Candidatus_Accumulibacter_aalborgensis | Bacteria>Pseudomonadati>Pseudomonadota>Betaproteobacteria>Betaproteobacteria_incertae_sedis>Candidatus_Accumulibacter | 135 | ACCAA_40013 | 16295..16702 | 1860102 | conserved-hypothetical-protein-[Candidatus-Accumulibacter-aalborgensis] |
| MCM8625482.1 | REase | GCA_023806625.1 | Accumulibacter_sp. | Bacteria>Pseudomonadati>Pseudomonadota>Betaproteobacteria>Betaproteobacteria_incertae_sedis>Candidatus_Accumulibacter>unclassified_Candidatus_Accumulibacter | 169 | NFW05_05880 | 107989..108498 | 2053492 | hypothetical-protein-[Candidatus-Accumulibacter-sp.] |
| WP_076518284.1 | REase | GCF_900156745.1 | Achromobacter_sp._MFA1_R4 | Bacteria>Pseudomonadati>Pseudomonadota>Betaproteobacteria>Burkholderiales>Alcaligenaceae>Achromobacter>unclassified_Achromobacter | 184 | BXA00_RS09420 | 2087000..2087554 | 1881016 | hypothetical-protein-[Achromobacter-sp.-MFA1-R4] |
| WP_068373670.1 | REase | GCF_000988095.1 | Kerstersia_gyiorum | Bacteria>Pseudomonadati>Pseudomonadota>Betaproteobacteria>Burkholderiales>Alcaligenaceae>Kerstersia | 180 | AAV32_RS14330 | 111385..111927 | 206506 | hypothetical-protein-[Kerstersia-gyiorum] |
| WP_132980220.1 | REase | GCF_004352985.1 | Pigmentiphaga_sp._D-2 | Bacteria>Pseudomonadati>Pseudomonadota>Betaproteobacteria>Burkholderiales>Alcaligenaceae>Pigmentiphaga>unclassified_Pigmentiphaga | 178 | EZY13_RS19740 | 193174..193710 | 1002116 | hypothetical-protein-[Pigmentiphaga-sp.-D-2] |
| WP_186144237.1 | REase | GCF_902831945.1 | Burkholderia_gladioli | Bacteria>Pseudomonadati>Pseudomonadota>Betaproteobacteria>Burkholderiales>Burkholderiaceae>Burkholderia | 208 | H9E34_RS32230 | 54000..54626 | 28095 | DUF3644-domain-containing-protein-[Burkholderia-gladioli] |
| WP_091778013.1 | REase | GCF_900100565.1 | Paraburkholderia_steynii | Bacteria>Pseudomonadati>Pseudomonadota>Betaproteobacteria>Burkholderiales>Burkholderiaceae>Paraburkholderia | 174 | BLT65_RS12225 | 286172..286696 | 1245441 | hypothetical-protein-[Paraburkholderia-steynii] |
| MBN6206493.1 | REase | GCA_017162415.1 | Ralstonia_pickettii | Bacteria>Pseudomonadati>Pseudomonadota>Betaproteobacteria>Burkholderiales>Burkholderiaceae>Ralstonia | 193 | JYK21_08510 | 17085..17666 | 329 | hypothetical-protein-[Ralstonia-pickettii] |
| QLQ00178.1 | REase | GCA_013693755.1 | Burkholderiaceae_bacterium | Bacteria>Pseudomonadati>Pseudomonadota>Betaproteobacteria>Burkholderiales>Burkholderiaceae>unclassified_Burkholderiaceae | 169 | HZY78_08250 | 1792423..1792932 | 2030806 | hypothetical-protein-HZY78_08250-[Burkholderiaceae-bacterium] |
| MBP6559244.1 | REase | GCA_017989415.1 | Burkholderiaceae_bacterium | Bacteria>Pseudomonadati>Pseudomonadota>Betaproteobacteria>Burkholderiales>Burkholderiaceae>unclassified_Burkholderiaceae | 175 | KA212_03000 | 3362..3889 | 2030806 | hypothetical-protein-[Burkholderiaceae-bacterium] |
| WP_180684521.1 | REase | GCF_013432195.1 | Tepidicella_baoligensis | Bacteria>Pseudomonadati>Pseudomonadota>Betaproteobacteria>Burkholderiales>Burkholderiales_genera_incertae_sedis>Tepidicella | 194 | HZS63_RS13530 | 8315..8899 | 2707016 | hypothetical-protein-[Tepidicella-baoligensis] |
| WP_143969783.1 | REase | GCF_007556755.1 | Tepidimonas_fonticaldi | Bacteria>Pseudomonadati>Pseudomonadota>Betaproteobacteria>Burkholderiales>Burkholderiales_genera_incertae_sedis>Tepidimonas | 194 | Tfont_RS13260 | 4455..5039 | 1101373 | hypothetical-protein-[Tepidimonas-fonticaldi] |
| WP_132962624.1 | REase | GCF_036772355.1 | Tepidimonas_ignava | Bacteria>Pseudomonadati>Pseudomonadota>Betaproteobacteria>Burkholderiales>Burkholderiales_genera_incertae_sedis>Tepidimonas | 166 | EDC36_RS07395 | 33033..33533 | 114249 | hypothetical-protein-[Tepidimonas-ignava] |
| WP_055397339.1 | REase | GCF_017355385.1 | Acidovorax | Bacteria>Pseudomonadati>Pseudomonadota>Betaproteobacteria>Burkholderiales>Comamonadaceae | 175 | AE621_RS12470 | 10632..11159 | 12916 | MULTISPECIES:-hypothetical-protein-[Acidovorax] |
| WP_087746903.1 | REase | GCF_002176815.1 | unclassified_Acidovorax | Bacteria>Pseudomonadati>Pseudomonadota>Betaproteobacteria>Burkholderiales>Comamonadaceae>Acidovorax | 175 | J8G26_RS10060 | 95492..96019 | 2684926 | MULTISPECIES:-hypothetical-protein-[unclassified-Acidovorax] |
| MBP7545006.1 | REase | GCA_017998835.1 | Acidovorax_sp. | Bacteria>Pseudomonadati>Pseudomonadota>Betaproteobacteria>Burkholderiales>Comamonadaceae>Acidovorax>unclassified_Acidovorax | 211 | KA749_05810 | 3084..3719 | 1872122 | hypothetical-protein-[Acidovorax-sp.] |
| WP_210930505.1 | REase | GCF_018064205.1 | Comamonas_aquatica | Bacteria>Pseudomonadati>Pseudomonadota>Betaproteobacteria>Burkholderiales>Comamonadaceae>Comamonas | 211 | KAQ61_RS03435 | 740042..740677 | 225991 | hypothetical-protein-[Comamonas-aquatica] |
| OJX16149.1 | REase | GCA_001898745.1 | Delftia_sp._67-8 | Bacteria>Pseudomonadati>Pseudomonadota>Betaproteobacteria>Burkholderiales>Comamonadaceae>Delftia>unclassified_Delftia | 184 | BGO79_00295 | 71575..72129 | 1895749 | hypothetical-protein-BGO79_00295-[Delftia-sp.-67-8] |
| TFI45393.1 | REase | GCA_004563745.1 | Diaphorobacter_sp._DS2 | Bacteria>Pseudomonadati>Pseudomonadota>Betaproteobacteria>Burkholderiales>Comamonadaceae>Diaphorobacter>unclassified_Diaphorobacter | 174 | E4O93_19985 | 6102..6626 | 1302548 | hypothetical-protein-E4O93_19985-[Diaphorobacter-sp.-DS2] |
| WP_252101815.1 | REase | GCF_022807955.1 | Diaphorobacter_sp._LI3 | Bacteria>Pseudomonadati>Pseudomonadota>Betaproteobacteria>Burkholderiales>Comamonadaceae>Diaphorobacter>unclassified_Diaphorobacter | 174 | MRB47_RS02560 | 530410..530934 | 2952886 | hypothetical-protein-[[Acidovorax] |
| WP_108330267.1 | REase | GCF_003063435.1 | Limnohabitans_sp._2KL-1 | Bacteria>Pseudomonadati>Pseudomonadota>Betaproteobacteria>Burkholderiales>Comamonadaceae>Limnohabitans>unclassified_Limnohabitans | 175 | B9Z47_RS05545 | 222475..223002 | 1100699 | hypothetical-protein-[Limnohabitans-sp.-2KL-1] |
| WP_209423135.1 | REase | GCF_021249385.1 | Melaminivora_jejuensis | Bacteria>Pseudomonadati>Pseudomonadota>Betaproteobacteria>Burkholderiales>Comamonadaceae>Melaminivora | 174 | LVC68_RS01450 | 287079..287603 | 1267217 | hypothetical-protein-[Melaminivora-jejuensis] |
| WP_265286860.1 | REase | GCF_026127525.1 | Verminephrobacter_eiseniae | Bacteria>Pseudomonadati>Pseudomonadota>Betaproteobacteria>Burkholderiales>Comamonadaceae>Verminephrobacter | 166 | D5047_RS05830 | 1242339..1242839 | 364317 | hypothetical-protein-[Verminephrobacter-eiseniae] |
| WP_265286859.1 | REase | GCF_026127525.1 | Verminephrobacter_eiseniae | Bacteria>Pseudomonadati>Pseudomonadota>Betaproteobacteria>Burkholderiales>Comamonadaceae>Verminephrobacter | 119 | D5047_RS05825 | 1242011..1242370 | 364317 | hypothetical-protein-[Verminephrobacter-eiseniae] |
| WP_230404447.1 | REase | GCF_014284115.1 | Undibacterium | Bacteria>Pseudomonadati>Pseudomonadota>Betaproteobacteria>Burkholderiales>Oxalobacteraceae | 134 | H8K35_RS08335 | 162083..162487 | 401469 | MULTISPECIES:-hypothetical-protein-[Undibacterium] |
| WP_258819855.1 | REase | GCF_024753285.1 | Massilia_pinisoli | Bacteria>Pseudomonadati>Pseudomonadota>Betaproteobacteria>Burkholderiales>Oxalobacteraceae>Telluria_group>Massilia | 177 | NX784_RS27495 | 38696..39229 | 1772194 | hypothetical-protein-[Massilia-pinisoli] |
| UUZ52275.1 | REase | GCA_024703915.1 | Massilia_sp._H-1 | Bacteria>Pseudomonadati>Pseudomonadota>Betaproteobacteria>Burkholderiales>Oxalobacteraceae>Telluria_group>Massilia>unclassified_Massilia | 215 | LP419_21970 | 3093881..3094528 | 2895566 | hypothetical-protein-LP419_21970-[Massilia-sp.-H-1] |
| UUZ46677.1 | REase | GCA_024703935.1 | Massilia_sp._B-10 | Bacteria>Pseudomonadati>Pseudomonadota>Betaproteobacteria>Burkholderiales>Oxalobacteraceae>Telluria_group>Massilia>unclassified_Massilia | 87 | LP420_22505 | 3092593..3092856 | 2895567 | hypothetical-protein-LP420_22505-[Massilia-sp.-B-10] |
| WP_157118906.1 | REase | GCF_001571085.1 | Azohydromonas_lata | Bacteria>Pseudomonadati>Pseudomonadota>Betaproteobacteria>Burkholderiales>Sphaerotilaceae>Azohydromonas | 208 | ALA01S_RS32795 | 47396..48022 | 45677 | hypothetical-protein-[Azohydromonas-lata] |
| WP_132764538.1 | REase | GCF_026057715.1 | Caldimonas_thermodepolymerans | Bacteria>Pseudomonadati>Pseudomonadota>Betaproteobacteria>Burkholderiales>Sphaerotilaceae>Caldimonas | 169 | ONS87_RS02475 | 487637..488146 | 215580 | hypothetical-protein-[Caldimonas-thermodepolymerans] |
| WP_110402336.1 | REase | GCF_003201595.1 | Sphaerotilus_hippei | Bacteria>Pseudomonadati>Pseudomonadota>Betaproteobacteria>Burkholderiales>Sphaerotilaceae>Sphaerotilus | 204 | C7444_RS18465 | 25793..26407 | 744406 | hypothetical-protein-[Sphaerotilus-hippei] |
| MCZ2407522.1 | REase | GCA_026931285.1 | Burkholderiales_bacterium | Bacteria>Pseudomonadati>Pseudomonadota>Betaproteobacteria>Burkholderiales>unclassified_Burkholderiales | 173 | LC097_12430 | 117951..118472 | 1891238 | MAG:-hypothetical-protein-LC097_12430-[Burkholderiales-bacterium] |
| MBK6716516.1 | REase | GCA_016703325.1 | Burkholderiales_bacterium | Bacteria>Pseudomonadati>Pseudomonadota>Betaproteobacteria>Burkholderiales>unclassified_Burkholderiales | 161 | IPG57_15840 | 1898..2383 | 1891238 | hypothetical-protein-[Burkholderiales-bacterium] |
| MCP5275502.1 | REase | GCA_024235795.1 | Burkholderiales_bacterium | Bacteria>Pseudomonadati>Pseudomonadota>Betaproteobacteria>Burkholderiales>unclassified_Burkholderiales | 180 | H6936_11785 | 658397..658939 | 1891238 | hypothetical-protein-[Burkholderiales-bacterium] |
| MCP5275802.1 | REase | GCA_024235795.1 | Burkholderiales_bacterium | Bacteria>Pseudomonadati>Pseudomonadota>Betaproteobacteria>Burkholderiales>unclassified_Burkholderiales | 180 | H6936_13320 | 1021650..1022192 | 1891238 | hypothetical-protein-[Burkholderiales-bacterium] |
| NMA29239.1 | REase | GCA_012799975.1 | Burkholderiales_bacterium | Bacteria>Pseudomonadati>Pseudomonadota>Betaproteobacteria>Burkholderiales>unclassified_Burkholderiales | 235 | GX934_15900 | 2305..3012 | 1891238 | Uma2-family-endonuclease-[Burkholderiales-bacterium] |
| WP_022652888.1 | REase | GCF_000420525.1 | Aquaspirillum_serpens | Bacteria>Pseudomonadati>Pseudomonadota>Betaproteobacteria>Neisseriales>Aquaspirillaceae>Aquaspirillum | 173 | G465_RS13155 | 21128..21649 | 190 | hypothetical-protein-[Aquaspirillum-serpens] |
| MCG9064564.1 | REase | GCA_022394255.1 | Laribacter_hongkongensis | Bacteria>Pseudomonadati>Pseudomonadota>Betaproteobacteria>Neisseriales>Aquaspirillaceae>Laribacter | 173 | LH425_05840 | 35355..35876 | 168471 | hypothetical-protein-[Laribacter-hongkongensis] |
| WP_089417823.1 | REase | GCF_002222655.1 | Vitreoscilla_filiformis | Bacteria>Pseudomonadati>Pseudomonadota>Betaproteobacteria>Neisseriales>Neisseriaceae>Vitreoscilla | 175 | VITFI_RS15925 | 3395960..3396487 | 63 | hypothetical-protein-[Vitreoscilla-filiformis] |
| MBV6446596.1 | REase | GCA_019187415.1 | Nitrosomonas_sp. | Bacteria>Pseudomonadati>Pseudomonadota>Betaproteobacteria>Nitrosomonadales>Nitrosomonadaceae>Nitrosomonas>unclassified_Nitrosomonas | 179 | FCKEOINB_00189 | 21812..22351 | 42353 | hypothetical-protein-[Nitrosomonas-sp.] |
| MBK7364913.1 | REase | GCA_016705625.1 | Nitrosomonas_sp. | Bacteria>Pseudomonadati>Pseudomonadota>Betaproteobacteria>Nitrosomonadales>Nitrosomonadaceae>Nitrosomonas>unclassified_Nitrosomonas | 165 | IPI97_07915 | 1670119..1670616 | 42353 | hypothetical-protein-[Nitrosomonas-sp.] |
| MBX3639965.1 | REase | GCA_019635175.1 | Nitrosomonas_sp. | Bacteria>Pseudomonadati>Pseudomonadota>Betaproteobacteria>Nitrosomonadales>Nitrosomonadaceae>Nitrosomonas>unclassified_Nitrosomonas | 130 | KF888_05550 | 115576..115968 | 42353 | GxxExxY-protein-[Nitrosomonas-sp.] |
| MCB1936157.1 | REase | GCA_020446045.1 | Nitrosomonas_sp. | Bacteria>Pseudomonadati>Pseudomonadota>Betaproteobacteria>Nitrosomonadales>Nitrosomonadaceae>Nitrosomonas>unclassified_Nitrosomonas | 180 | KDF59_09460 | 3904..4446 | 42353 | hypothetical-protein-[Nitrosomonas-sp.] |
| MBP9798944.1 | REase | GCA_018062105.1 | Sterolibacterium_sp. | Bacteria>Pseudomonadati>Pseudomonadota>Betaproteobacteria>Nitrosomonadales>Sterolibacteriaceae>Sterolibacterium>unclassified_Sterolibacterium | 188 | KBD39_02345 | 28806..29372 | 2819261 | endonuclease-domain-containing-protein-[Sterolibacterium-sp.] |
| MBN8462735.1 | REase | GCA_017302355.1 | Dechloromonas_sp. | Bacteria>Pseudomonadati>Pseudomonadota>Betaproteobacteria>Rhodocyclales>Azonexaceae>Dechloromonas>unclassified_Dechloromonas | 144 | J0M01_07955 | 8264..8698 | 1917218 | endonuclease-domain-containing-protein-[Dechloromonas-sp.] |
| WP_169202999.1 | REase | GCF_017894365.1 | Aromatoleum_bremense | Bacteria>Pseudomonadati>Pseudomonadota>Betaproteobacteria>Rhodocyclales>Rhodocyclaceae>Aromatoleum | 170 | GPA24_RS12875 | 15290..15802 | 76115 | hypothetical-protein-[Aromatoleum-bremense] |
| WP_205230202.1 | REase | GCF_905120365.1 | Azospira_oryzae | Bacteria>Pseudomonadati>Pseudomonadota>Betaproteobacteria>Rhodocyclales>Rhodocyclaceae>Azospira | 175 | JYJ48_RS05810 | 15418..15945 | 146939 | hypothetical-protein-[Azospira-oryzae] |
| MBQ7625734.1 | REase | GCA_017444505.1 | Rhodocyclaceae_bacterium | Bacteria>Pseudomonadati>Pseudomonadota>Betaproteobacteria>Rhodocyclales>Rhodocyclaceae>unclassified_Rhodocyclaceae | 140 | IJS87_01555 | 5510..5932 | 1898103 | YraN-family-protein-[Rhodocyclaceae-bacterium] |
| WP_159691717.1 | REase | GCF_009800965.1 | Cognatazoarcus_halotolerans | Bacteria>Pseudomonadati>Pseudomonadota>Betaproteobacteria>Rhodocyclales>Zoogloeaceae>Cognatazoarcus | 170 | GQB03_RS16410 | 74418..74930 | 2686016 | hypothetical-protein-[Cognatazoarcus-halotolerans] |
| TXH82466.1 | REase | GCA_008015255.1 | Thauera_aminoaromatica | Bacteria>Pseudomonadati>Pseudomonadota>Betaproteobacteria>Rhodocyclales>Zoogloeaceae>Thauera | 175 | E6Q80_15445 | 6991..7518 | 164330 | hypothetical-protein-E6Q80_15445-[Thauera-aminoaromatica] |
| WP_107221106.1 | REase | GCF_003030465.1 | Thauera_aromatica | Bacteria>Pseudomonadati>Pseudomonadota>Betaproteobacteria>Rhodocyclales>Zoogloeaceae>Thauera | 175 | Tharo_RS10330 | 2183943..2184470 | 59405 | hypothetical-protein-[Thauera-aromatica] |
| WP_247737220.1 | REase | GCF_023093675.1 | Thauera_aromatica | Bacteria>Pseudomonadati>Pseudomonadota>Betaproteobacteria>Rhodocyclales>Zoogloeaceae>Thauera | 135 | MX653_RS17195 | 6720..7127 | 59405 | hypothetical-protein-[Thauera-aromatica] |
| WP_153168446.1 | REase | GCF_009469595.2 | Thauera_sp._2A1 | Bacteria>Pseudomonadati>Pseudomonadota>Betaproteobacteria>Rhodocyclales>Zoogloeaceae>Thauera>unclassified_Thauera | 170 | GH664_RS12710 | 88974..89486 | 2570191 | hypothetical-protein-[Thauera-sp.-2A1] |
| AVZ79921.1 | REase | GCA_003054515.1 | Zoogloeaceae_bacteirum_Par-f-2 | Bacteria>Pseudomonadati>Pseudomonadota>Betaproteobacteria>Rhodocyclales>Zoogloeaceae>unclassified_Zoogloeaceae | 169 | C3497_11105 | 2337467..2337976 | 2080469 | hypothetical-protein-C3497_11105-[Zoogloeaceae-bacteirum-Par-f-2] |
| WP_148578094.1 | REase | GCF_008107625.1 | Zoogloea_oleivorans | Bacteria>Pseudomonadati>Pseudomonadota>Betaproteobacteria>Rhodocyclales>Zoogloeaceae>Zoogloea | 171 | ETQ85_RS05715 | 199630..200145 | 1552750 | hypothetical-protein-[Zoogloea-oleivorans] |
| MBK6655414.1 | REase | GCA_016703645.1 | Zoogloea_sp. | Bacteria>Pseudomonadati>Pseudomonadota>Betaproteobacteria>Rhodocyclales>Zoogloeaceae>Zoogloea>unclassified_Zoogloea | 175 | IPG40_16590 | 752727..753254 | 49181 | hypothetical-protein-[Zoogloea-sp.] |
| MBK7848389.1 | REase | GCA_016714095.1 | Zoogloea_sp. | Bacteria>Pseudomonadati>Pseudomonadota>Betaproteobacteria>Rhodocyclales>Zoogloeaceae>Zoogloea>unclassified_Zoogloea | 175 | IPJ73_15260 | 39228..39755 | 49181 | hypothetical-protein-[Zoogloea-sp.] |
| MDD3354817.1 | REase | GCA_028693495.1 | Zoogloea_sp. | Bacteria>Pseudomonadati>Pseudomonadota>Betaproteobacteria>Rhodocyclales>Zoogloeaceae>Zoogloea>unclassified_Zoogloea | 168 | PHD22_14440 | 7748..8254 | 49181 | MAG:-hypothetical-protein-PHD22_14440-[Zoogloea-sp.] |
| MBI5922416.1 | REase | GCA_016234745.1 | Betaproteobacteria_bacterium | Bacteria>Pseudomonadati>Pseudomonadota>Betaproteobacteria>unclassified_Betaproteobacteria | 194 | HY847_12345 | 42292..42876 | 1891241 | hypothetical-protein-[Betaproteobacteria-bacterium] |
| OGA55363.1 | REase | GCA_001772005.1 | Betaproteobacteria_bacterium_RIFCSPLOWO2_12_FULL_62_58 | Bacteria>Pseudomonadati>Pseudomonadota>Betaproteobacteria>unclassified_Betaproteobacteria | 192 | A3F74_07160 | 3167..3745 | 1797497 | hypothetical-protein-A3F74_07160-[Betaproteobacteria-bacterium-RIFCSPLOWO2_12_FULL_62_58] |
| WP_284453092.1 | REase | GCF_048595985.1 | Oceanospirillales | Bacteria>Pseudomonadati>Pseudomonadota>Gammaproteobacteria | 169 | KYX60_RS01240 | 109464..109973 | 135619 | MULTISPECIES:-hypothetical-protein-[Oceanospirillales] |
| WP_096459282.1 | REase | GCF_002355415.1 | Sulfurifustis_variabilis | Bacteria>Pseudomonadati>Pseudomonadota>Gammaproteobacteria>Acidiferrobacterales>Acidiferrobacteraceae>Sulfurifustis | 194 | SVA_RS04280 | 892160..892744 | 1675686 | hypothetical-protein-[Sulfurifustis-variabilis] |
| WP_102989253.1 | REase | GCF_002934425.1 | Aeromonas | Bacteria>Pseudomonadati>Pseudomonadota>Gammaproteobacteria>Aeromonadales>Aeromonadaceae | 180 | OKC30_RS15255 | 10606..11148 | 642 | MULTISPECIES:-hypothetical-protein-[Aeromonas] |
| WP_257706957.1 | REase | GCF_024649325.1 | Aeromonas_caviae | Bacteria>Pseudomonadati>Pseudomonadota>Gammaproteobacteria>Aeromonadales>Aeromonadaceae>Aeromonas | 180 | NUK33_RS01485 | 148558..149100 | 648 | hypothetical-protein-[Aeromonas-caviae] |
| WP_043557610.1 | REase | GCF_963866355.1 | Aeromonas_bestiarum | Bacteria>Pseudomonadati>Pseudomonadota>Gammaproteobacteria>Aeromonadales>Aeromonadaceae>Aeromonas | 179 | BN1118_RS19640 | 68088..68627 | 105751 | hypothetical-protein-[Aeromonas-bestiarum] |
| WP_223954702.1 | REase | GCF_019972775.1 | Aeromonas_caviae | Bacteria>Pseudomonadati>Pseudomonadota>Gammaproteobacteria>Aeromonadales>Aeromonadaceae>Aeromonas | 169 | LDN68_RS17110 | 3645690..3646199 | 648 | hypothetical-protein-[Aeromonas-caviae] |
| WP_080741110.1 | REase | GCF_963864655.1 | Aeromonas_bestiarum | Bacteria>Pseudomonadati>Pseudomonadota>Gammaproteobacteria>Aeromonadales>Aeromonadaceae>Aeromonas | 193 | BN1118_RS21425 | 54874..55455 | 105751 | sce7726-family-protein-[Aeromonas-bestiarum] |
| AUZ76296.1 | REase | GCA_002934425.1 | Aeromonas_sp._ASNIH4 | Bacteria>Pseudomonadati>Pseudomonadota>Gammaproteobacteria>Aeromonadales>Aeromonadaceae>Aeromonas>unclassified_Aeromonas | 180 | C2U40_16575 | 3600041..3600583 | 1636609 | hypothetical-protein-C2U40_16575-[Aeromonas-sp.-ASNIH4] |
| WP_006006593.1 | REase | GCF_000252165.1 | Glaciecola_punicea | Bacteria>Pseudomonadati>Pseudomonadota>Gammaproteobacteria>Alteromonadales>Alteromonadaceae>Glaciecola | 167 | GPUN_RS10810 | 147490..147993 | 56804 | hypothetical-protein-[Glaciecola-punicea] |
| WP_136550438.1 | REase | GCF_004851605.1 | Hydrocarboniclastica_marina | Bacteria>Pseudomonadati>Pseudomonadota>Gammaproteobacteria>Alteromonadales>Alteromonadaceae>Hydrocarboniclastica | 165 | soil367_RS18195 | 4127907..4128404 | 2259620 | hypothetical-protein-[Hydrocarboniclastica-marina] |
| WP_229744614.1 | REase | GCF_014636195.1 | Neiella_marina | Bacteria>Pseudomonadati>Pseudomonadota>Gammaproteobacteria>Alteromonadales>Echinimonadaceae>Neiella | 174 | CBE68_RS02315 | 109246..109770 | 508461 | hypothetical-protein-[Neiella-marina] |
| WP_087504473.1 | REase | GCF_014636195.1 | Neiella_marina | Bacteria>Pseudomonadati>Pseudomonadota>Gammaproteobacteria>Alteromonadales>Echinimonadaceae>Neiella | 130 | CBE68_RS02350 | 118383..118775 | 508461 | group-I-intron-associated-PD-(D/E)XK-endonuclease-[Neiella-marina] |
| KTG26835.1 | REase | GCA_001469885.1 | Idiomarina_sp._H105 | Bacteria>Pseudomonadati>Pseudomonadota>Gammaproteobacteria>Alteromonadales>Idiomarinaceae>Idiomarina>unclassified_Idiomarina | 169 | AUR68_16340 | 108336..108845 | 1766622 | hypothetical-protein-AUR68_16340-[Idiomarina-sp.-H105] |
| WP_112714383.1 | REase | GCF_900465055.1 | Moritella_yayanosii | Bacteria>Pseudomonadati>Pseudomonadota>Gammaproteobacteria>Alteromonadales>Moritellaceae>Moritella | 175 | MORIYA_RS08595 | 1891534..1892061 | 69539 | hypothetical-protein-[Moritella-yayanosii] |

| Protein-ID | Domain | Assembly-ID | Strain | Taxonomy | Length | Locus-Tag | Genomic-coordinates | TaxID | Fasta-defline |
|---|---|---|---|---|---|---|---|---|---|
| WP_024611451.1 | REase | GCF_000498095.1 | Pseudoalteromonas_sp._TB64 | Bacteria>Pseudomonadati>Pseudomonadota>Gammaproteobacteria>Alteromonadales>Pseudoalteromonadaceae>Pseudoalteromonas>unclassified_Pseudoalteromonas | 175 | L647_RS0112845 | 130348..130875 | 1938600 | hypothetical-protein-[Pseudoalteromonas-sp.-TB64] |
| WP_167658110.1 | REase | GCF_011806445.1 | Pseudoalteromonas_sp._HF66 | Bacteria>Pseudomonadati>Pseudomonadota>Gammaproteobacteria>Alteromonadales>Pseudoalteromonadaceae>Pseudoalteromonas>unclassified_Pseudoalteromonas | 175 | HCZ11_RS07650 | 211172..211699 | 2721559 | hypothetical-protein-[Pseudoalteromonas-sp.-HF66] |
| WP_254869125.1 | REase | - | Pseudoalteromonas_sp._HF66 | Bacteria>Pseudomonadati>Pseudomonadota>Gammaproteobacteria>Alteromonadales>Pseudoalteromonadaceae>Pseudoalteromonas>unclassified_Pseudoalteromonas | 185 | HCZ11_RS07620 | 198331..198888 | 2721559 | TnsA-endonuclease-N-terminal-domain-containing-protein-[Pseudoalteromonas-sp.-HF66] |
| WP_112352817.1 | REase | GCF_900476435.1 | Shewanella_benthica | Bacteria>Pseudomonadati>Pseudomonadota>Gammaproteobacteria>Alteromonadales>Shewanellaceae>Shewanella | 170 | SHEWBE_RS12045 | 2632653..2633165 | 43661 | hypothetical-protein-[Shewanella-benthica] |
| VEE64284.1 | REase | GCA_900636665.1 | Shewanella_putrefaciens | Bacteria>Pseudomonadati>Pseudomonadota>Gammaproteobacteria>Alteromonadales>Shewanellaceae>Shewanella | 203 | NCTC12093_04127 | 4663708..4664319 | 24 | Uncharacterised-protein-[Shewanella-putrefaciens] |
| WP_126513129.1 | REase | GCF_024999165.1 | Shewanella_baltica | Bacteria>Pseudomonadati>Pseudomonadota>Gammaproteobacteria>Alteromonadales>Shewanellaceae>Shewanella | 203 | G7A68_RS06635 | 11602..12213 | 62322 | MULTISPECIES:-hypothetical-protein-[Shewanella] |
| WP_218997782.1 | REase | GCF_019303335.1 | Shewanella_algae | Bacteria>Pseudomonadati>Pseudomonadota>Gammaproteobacteria>Alteromonadales>Shewanellaceae>Shewanella | 208 | K0E68_RS03895 | 5595..6221 | 38313 | TnsA-endonuclease-N-terminal-domain-containing-protein-[Shewanella-algae] |
| PKG74687.1 | REase | GCA_002836135.1 | Shewanella_sp._GutCb | Bacteria>Pseudomonadati>Pseudomonadota>Gammaproteobacteria>Alteromonadales>Shewanellaceae>Shewanella>unclassified_Shewanella | 165 | CXF86_11680 | 120638..121135 | 2058315 | hypothetical-protein-CXF86_11680-[Shewanella-sp.-GutCb] |
| WP_102039066.1 | REase | GCF_002836135.1 | Shewanella_sp._GutCb | Bacteria>Pseudomonadati>Pseudomonadota>Gammaproteobacteria>Alteromonadales>Shewanellaceae>Shewanella>unclassified_Shewanella | 165 | CXF86_RS11720 | 120638..121135 | 2058315 | hypothetical-protein-[Shewanella-sp.-GutCb] |
| WP_265366290.1 | REase | GCF_963971615.1 | Marinimicrobium_sp._C6131 | Bacteria>Pseudomonadati>Pseudomonadota>Gammaproteobacteria>Cellvibrionales>Cellvibrionaceae>Marinimicrobium>unclassified_Marinimicrobium | 177 | OOT55_RS13040 | 3030991..3031524 | 3022676 | hypothetical-protein-[Marinimicrobium-sp.-C6131] |
| WP_084199898.1 | REase | GCF_014652255.1 | Halioglobus_japonicus | Bacteria>Pseudomonadati>Pseudomonadota>Gammaproteobacteria>Cellvibrionales>Halieaceae>Halioglobus | 166 | C0029_RS05540 | 1200414..1200914 | 930805 | hypothetical-protein-[Halioglobus-japonicus] |
| WP_148062604.1 | REase | GCF_008064635.1 | Parahaliea_aestuarii | Bacteria>Pseudomonadati>Pseudomonadota>Gammaproteobacteria>Cellvibrionales>Halieaceae>Parahaliea | 168 | FVW59_RS02295 | 509254..509760 | 1852021 | hypothetical-protein-[Parahaliea-aestuarii] |
| MCK9506375.1 | REase | GCA_023228865.1 | Porticoccaceae_bacterium | Bacteria>Pseudomonadati>Pseudomonadota>Gammaproteobacteria>Cellvibrionales>Porticoccaceae>unclassified_Porticoccaceae | 172 | MOQ95_19620 | 352..870 | 2026782 | hypothetical-protein-[Porticoccaceae-bacterium] |
| MTW21822.1 | REase | GCA_009720725.1 | Allochromatium_palmeri | Bacteria>Pseudomonadati>Pseudomonadota>Gammaproteobacteria>Chromatiales>Chromatiaceae>Allochromatium | 179 | GJ668_12055 | 35628..36167 | 231048 | hypothetical-protein-[Allochromatium-palmeri] |
| WP_155450401.1 | REase | GCF_009720725.1 | Allochromatium_palmeri | Bacteria>Pseudomonadati>Pseudomonadota>Gammaproteobacteria>Chromatiales>Chromatiaceae>Allochromatium | 179 | GJ668_RS12055 | 35628..36167 | 231048 | hypothetical-protein-[Allochromatium-palmeri] |
| WP_213378540.1 | REase | GCF_018409545.1 | Allochromatium_tepidum | Bacteria>Pseudomonadati>Pseudomonadota>Gammaproteobacteria>Chromatiales>Chromatiaceae>Allochromatium | 165 | Atep_RS10845 | 2245991..2246488 | 553982 | hypothetical-protein-[Allochromatium-tepidum] |
| WP_155450779.1 | REase | GCF_009720725.1 | Allochromatium_palmeri | Bacteria>Pseudomonadati>Pseudomonadota>Gammaproteobacteria>Chromatiales>Chromatiaceae>Allochromatium | 199 | GJ668_RS14070 | 11378..11977 | 231048 | Uma2-family-endonuclease-[Allochromatium-palmeri] |
| WP_201222424.1 | REase | GCF_016653315.1 | Halochromatium_roseum | Bacteria>Pseudomonadati>Pseudomonadota>Gammaproteobacteria>Chromatiales>Chromatiaceae>Halochromatium | 165 | CCR96_RS19420 | 141831..142328 | 391920 | hypothetical-protein-[Halochromatium-roseum] |
| WP_201222440.1 | REase | GCF_016653315.1 | Halochromatium_roseum | Bacteria>Pseudomonadati>Pseudomonadota>Gammaproteobacteria>Chromatiales>Chromatiaceae>Halochromatium | 125 | CCR96_RS19445 | 152229..152606 | 391920 | GxxExxY-protein-[Halochromatium-roseum] |
| NEX15942.1 | REase | GCA_010912705.1 | Halochromatium_sp. | Bacteria>Pseudomonadati>Pseudomonadota>Gammaproteobacteria>Chromatiales>Chromatiaceae>Halochromatium>unclassified_Halochromatium | 179 | C1943_04710 | 65404..65943 | 2049430 | hypothetical-protein-[Halochromatium-sp.] |
| WP_200250818.1 | REase | GCF_016583905.1 | Lamprobacter_modestohalophilus | Bacteria>Pseudomonadati>Pseudomonadota>Gammaproteobacteria>Chromatiales>Chromatiaceae>Lamprobacter | 180 | CK042_RS24305 | 7124..7666 | 1064514 | hypothetical-protein-[Lamprobacter-modestohalophilus] |
| MBV5273605.1 | REase | GCA_019163055.1 | Lamprocystis_purpurea | Bacteria>Pseudomonadati>Pseudomonadota>Gammaproteobacteria>Chromatiales>Chromatiaceae>Lamprocystis | 127 | JZU52_08165 | 2609..2992 | 61598 | GxxExxY-protein-[Lamprocystis-purpurea] |
| WP_020507574.1 | REase | GCF_000379525.1 | Lamprocystis_purpurea | Bacteria>Pseudomonadati>Pseudomonadota>Gammaproteobacteria>Chromatiales>Chromatiaceae>Lamprocystis | 127 | A390_RS0125935 | 17277..17660 | 61598 | GxxExxY-protein-[Lamprocystis-purpurea] |
| WP_020508116.1 | REase | GCF_000379525.1 | Lamprocystis_purpurea | Bacteria>Pseudomonadati>Pseudomonadota>Gammaproteobacteria>Chromatiales>Chromatiaceae>Lamprocystis | 127 | A390_RS0128785 | 109487..109870 | 61598 | GxxExxY-protein-[Lamprocystis-purpurea] |
| WP_020508126.1 | REase | GCF_000379525.1 | Lamprocystis_purpurea | Bacteria>Pseudomonadati>Pseudomonadota>Gammaproteobacteria>Chromatiales>Chromatiaceae>Lamprocystis | 127 | A390_RS0128835 | 121211..121594 | 61598 | GxxExxY-protein-[Lamprocystis-purpurea] |
| WP_020506083.1 | REase | GCF_000379525.1 | Lamprocystis_purpurea | Bacteria>Pseudomonadati>Pseudomonadota>Gammaproteobacteria>Chromatiales>Chromatiaceae>Lamprocystis | 234 | A390_RS0118130 | 33553..34257 | 61598 | Uma2-family-endonuclease-[Lamprocystis-purpurea] |
| MBV5272243.1 | REase | GCA_019163055.1 | Lamprocystis_purpurea | Bacteria>Pseudomonadati>Pseudomonadota>Gammaproteobacteria>Chromatiales>Chromatiaceae>Lamprocystis | 34 | JZU52_00935 | 196..300 | 61598 | hypothetical-protein-[Lamprocystis-purpurea] |
| WP_011330216.1 | REase | - | Nitrosococcus_oceani | Bacteria>Pseudomonadati>Pseudomonadota>Gammaproteobacteria>Chromatiales>Chromatiaceae>Nitrosococcus | 119 | IB75_RS00215 | 11407..11766 | 1229 | PDDEXK-nuclease-domain-containing-protein-[Nitrosococcus-oceani] |
| WP_050754200.1 | REase | GCF_000012805.1 | Nitrosococcus_oceani | Bacteria>Pseudomonadati>Pseudomonadota>Gammaproteobacteria>Chromatiales>Chromatiaceae>Nitrosococcus | 106 | NOC_RS00510 | 59672..59992 | 1229 | PDDEXK-nuclease-domain-containing-protein-[Nitrosococcus-oceani] |
| GEM21690.1 | REase | GCA_007990675.1 | Nitrosococcus_oceani | Bacteria>Pseudomonadati>Pseudomonadota>Gammaproteobacteria>Chromatiales>Chromatiaceae>Nitrosococcus | 98 | NONS58_31400 | 11576..11872 | 1229 | hypothetical-protein-NONS58_31400-[Nitrosococcus-oceani] |
| EDZ65771.1 | REase | - | Nitrosococcus_oceani_AFC27 | Bacteria>Pseudomonadati>Pseudomonadota>Gammaproteobacteria>Chromatiales>Chromatiaceae>Nitrosococcus>Nitrosococcus_oceani | 98 | NOC27_2451 | 576968..577264 | 473788 | hypothetical-protein-NOC27_2451-[Nitrosococcus-oceani-AFC27] |
| WP_120797040.1 | REase | GCF_003634315.1 | Thiocapsa_rosea | Bacteria>Pseudomonadati>Pseudomonadota>Gammaproteobacteria>Chromatiales>Chromatiaceae>Thiocapsa | 129 | BDD21_RS09915 | 2232638..2233027 | 69360 | GxxExxY-protein-[Thiocapsa-rosea] |
| WP_120798743.1 | REase | GCF_003634315.1 | Thiocapsa_rosea | Bacteria>Pseudomonadati>Pseudomonadota>Gammaproteobacteria>Chromatiales>Chromatiaceae>Thiocapsa | 200 | BDD21_RS20535 | 4626041..4626643 | 69360 | Uma2-family-endonuclease-[Thiocapsa-rosea] |
| QVL48758.1 | REase | GCA_018399035.1 | Thiocapsa_sp. | Bacteria>Pseudomonadati>Pseudomonadota>Gammaproteobacteria>Chromatiales>Chromatiaceae>Thiocapsa>unclassified_Thiocapsa | 200 | KFB96_24960 | 5463976..5464578 | 2024551 | MAG:-Uma2-family-endonuclease-[Thiocapsa-sp.] |
| WP_200333074.1 | REase | GCF_016583565.1 | Thiocystis_violacea | Bacteria>Pseudomonadati>Pseudomonadota>Gammaproteobacteria>Chromatiales>Chromatiaceae>Thiocystis | 120 | CKO23_RS17310 | 13880..14242 | 13725 | endonuclease-domain-containing-protein-[Thiocystis-violacea] |
| ESQ16531.1 | REase | GCA_000495615.1 | Thiohalocapsa_sp._PB-PSB1 | Bacteria>Pseudomonadati>Pseudomonadota>Gammaproteobacteria>Chromatiales>Chromatiaceae>Thiohalocapsa>unclassified_Thiohalocapsa | 135 | N838_05680 | 9332..9739 | 1385625 | MAG:-hypothetical-protein-N838_05680-[Thiohalocapsa-sp.-PB-PSB1] |
| QQO57136.1 | REase | GCA_016745215.1 | Thiohalocapsa_sp._PB-PSB1 | Bacteria>Pseudomonadati>Pseudomonadota>Gammaproteobacteria>Chromatiales>Chromatiaceae>Thiohalocapsa>unclassified_Thiohalocapsa | 123 | N838_31105 | 7376825..7377196 | 1385625 | MAG:-GxxExxY-protein-[Thiohalocapsa-sp.-PB-PSB1] |
| WP_058553752.1 | REase | GCF_001469165.1 | Thiohalocapsa_sp._ML1 | Bacteria>Pseudomonadati>Pseudomonadota>Gammaproteobacteria>Chromatiales>Chromatiaceae>Thiohalocapsa>unclassified_Thiohalocapsa | 123 | ML01_RS02180 | 206192..206563 | 1431688 | GxxExxY-protein-[Thiohalocapsa-sp.-ML1] |
| WP_025282663.1 | REase | GCF_000583815.1 | Ectothiorhodospira_haloalkaliphila | Bacteria>Pseudomonadati>Pseudomonadota>Gammaproteobacteria>Chromatiales>Ectothiorhodospiraceae>Ectothiorhodospira | 178 | ECTHA_RS0102370 | 120349..120885 | 421628 | hypothetical-protein-[Ectothiorhodospira-haloalkaliphila] |
| TQE99817.1 | REase | GCA_006569195.1 | Spiribacter_salinus | Bacteria>Pseudomonadati>Pseudomonadota>Gammaproteobacteria>Chromatiales>Ectothiorhodospiraceae>Spiribacter | 168 | FKY71_06665 | 11782..12288 | 1335746 | hypothetical-protein-FKY71_06665-[Spiribacter-salinus] |
| WP_134906062.1 | REase | GCF_004801595.1 | Enterobacteriaceae | Bacteria>Pseudomonadati>Pseudomonadota>Gammaproteobacteria>Enterobacterales | 169 | FMK66_RS24355 | 59646..60155 | 543 | MULTISPECIES:-hypothetical-protein-[Enterobacteriaceae] |
| WP_028016401.1 | REase | GCF_014168915.1 | Enterobacteriaceae | Bacteria>Pseudomonadati>Pseudomonadota>Gammaproteobacteria>Enterobacterales | 240 | KKT92_RS26065 | 4498..5220 | 543 | MULTISPECIES:-hypothetical-protein-[Enterobacteriaceae] |
| WP_223239723.1 | REase | GCF_014596715.2 | Enterobacter | Bacteria>Pseudomonadati>Pseudomonadota>Gammaproteobacteria>Enterobacterales>Enterobacteriaceae | 113 | L465_RS18320 | 3965515..3965856 | 547 | MULTISPECIES:-winged-helix-turn-helix-domain-containing-protein-[Enterobacter] |
| WP_284143398.1 | REase | GCF_018446995.1 | Citrobacter | Bacteria>Pseudomonadati>Pseudomonadota>Gammaproteobacteria>Enterobacterales>Enterobacteriaceae | 146 | OGY13_RS10420 | 367590..368030 | 544 | MULTISPECIES:-restriction-endonuclease-[Citrobacter] |
| WP_121528279.1 | REase | GCF_003665375.1 | Enterobacter_kobei | Bacteria>Pseudomonadati>Pseudomonadota>Gammaproteobacteria>Enterobacterales>Enterobacteriaceae>Enterobacter>Enterobacter_cloacae_complex | 240 | FDW86_RS03785 | 39391..40113 | 208224 | MULTISPECIES:-hypothetical-protein-[Enterobacteriaceae] |
| WP_234784063.1 | REase | GCF_001518515.1 | Enterobacter_hormaechei | Bacteria>Pseudomonadati>Pseudomonadota>Gammaproteobacteria>Enterobacterales>Enterobacteriaceae>Enterobacter>Enterobacter_cloacae_complex | 90 | AWI06_RS12050 | 46992..47264 | 158836 | winged-helix-turn-helix-domain-containing-protein-[Enterobacter-hormaechei] |
| WP_196373091.1 | REase | GCF_015704905.1 | Enterobacter_cloacae | Bacteria>Pseudomonadati>Pseudomonadota>Gammaproteobacteria>Enterobacterales>Enterobacteriaceae>Enterobacter>Enterobacter_cloacae_complex | 171 | IOR24_RS17940 | 3782465..3782980 | 550 | hypothetical-protein-[Enterobacter-cloacae] |
| EUM08183.1 | REase | GCA_000534375.1 | Enterobacter_sp._BIDMC_29 | Bacteria>Pseudomonadati>Pseudomonadota>Gammaproteobacteria>Enterobacterales>Enterobacteriaceae>Enterobacter>Enterobacter_cloacae_complex>unclassified_Enterobacter_cloacae_complex | 108 | L465_03716 | 726351..726677 | 1329841 | hypothetical-protein-L465_03716-[Enterobacter-sp.-BIDMC-29] |
| EKU2197732.1 | REase | - | Escherichia_coli | Bacteria>Pseudomonadati>Pseudomonadota>Gammaproteobacteria>Enterobacterales>Enterobacteriaceae>Escherichia | 244 | LRR03_002280 | 227488..228222 | 562 | restriction-endonuclease,-partial-[Escherichia-coli] |
| HAO0324355.1 | REase | GCA_014771515.1 | Escherichia_coli | Bacteria>Pseudomonadati>Pseudomonadota>Gammaproteobacteria>Enterobacterales>Enterobacteriaceae>Escherichia | 184 | IGZ64_004241 | 4267..4821 | 562 | hypothetical-protein-[Escherichia-coli] |
| HBT9662832.1 | REase | GCA_022069925.1 | Klebsiella_pneumoniae | Bacteria>Pseudomonadati>Pseudomonadota>Gammaproteobacteria>Enterobacterales>Enterobacteriaceae>Klebsiella/Raoultella_group>Klebsiella>Klebsiella_pneumoniae_complex | 240 | MCN56_004850 | 30717..31439 | 573 | hypothetical-protein-[Klebsiella-pneumoniae] |
| WP_077999169.1 | REase | GCF_002006945.1 | Edwardsiella_tarda | Bacteria>Pseudomonadati>Pseudomonadota>Gammaproteobacteria>Enterobacterales>Hafniaceae>Edwardsiella | 169 | BTN58_RS02540 | 132877..133386 | 636 | hypothetical-protein-[Edwardsiella-tarda] |
| WP_267964441.1 | REase | - | Hafnia_paralvei | Bacteria>Pseudomonadati>Pseudomonadota>Gammaproteobacteria>Enterobacterales>Hafniaceae>Hafnia | 223 | LZ651_RS15670 | 41641..42312 | 546367 | hypothetical-protein-[Hafnia-paralvei] |
| WP_272670016.1 | REase | GCF_028480605.1 | Providencia | Bacteria>Pseudomonadati>Pseudomonadota>Gammaproteobacteria>Enterobacterales>Morganellaceae | 169 | PSE59_RS21135 | 12744..13253 | 586 | MULTISPECIES:-hypothetical-protein-[unclassified-Providencia] |
| QIG07334.1 | REase | GCA_011046025.1 | Proteus_sp._ZN5 | Bacteria>Pseudomonadati>Pseudomonadota>Gammaproteobacteria>Enterobacterales>Morganellaceae>Proteus>unclassified_Proteus_(in:_enterobacteria) | 167 | GTK47_19180 | 4186625..4187128 | 2697019 | hypothetical-protein-GTK47_19180-[Proteus-sp.-ZN5] |
| WP_165126220.1 | REase | GCF_011046025.1 | Proteus_sp._ZN5 | Bacteria>Pseudomonadati>Pseudomonadota>Gammaproteobacteria>Enterobacterales>Morganellaceae>Proteus>unclassified_Proteus_(in:_enterobacteria) | 167 | GTK47_RS19240 | 4186625..4187128 | 2697019 | hypothetical-protein-[Proteus-sp.-ZN5] |
| WP_247666643.1 | REase | GCF_018094745.1 | Pectobacterium_versatile | Bacteria>Pseudomonadati>Pseudomonadota>Gammaproteobacteria>Enterobacterales>Pectobacteriaceae>Pectobacterium | 156 | F9U42_RS09250 | 74411..74881 | 2488639 | hypothetical-protein-[Pectobacterium-versatile] |
| HAU1604811.1 | REase | GCA_015989465.1 | Legionella_pneumophila | Bacteria>Pseudomonadati>Pseudomonadota>Gammaproteobacteria>Legionellales>Legionellaceae>Legionella | 161 | JBJ19_08185 | 16885..17370 | 446 | hypothetical-protein-[Legionella-pneumophila] |
| WP_135965898.1 | REase | GCF_051986355.1 | Stenotrophomonas | Bacteria>Pseudomonadati>Pseudomonadota>Gammaproteobacteria>Lysobacterales>Lysobacteraceae | 169 | N5C11_RS09575 | 37376..37885 | 40323 | MULTISPECIES:-hypothetical-protein-[Stenotrophomonas] |
| MBN5005307.1 | REase | GCA_017149675.1 | Stenotrophomonas_maltophilia | Bacteria>Pseudomonadati>Pseudomonadota>Gammaproteobacteria>Lysobacterales>Lysobacteraceae>Stenotrophomonas>Stenotrophomonas_maltophilia_group | 169 | JY447_14740 | 18387..18896 | 40324 | hypothetical-protein-[Stenotrophomonas-maltophilia] |
| WP_225763944.1 | REase | GCF_916618915.1 | Stenotrophomonas_sp._Marseille-Q4652 | Bacteria>Pseudomonadati>Pseudomonadota>Gammaproteobacteria>Lysobacterales>Lysobacteraceae>Stenotrophomonas>unclassified_Stenotrophomonas | 184 | LG380_RS05390 | 1178596..1179150 | 2866595 | hypothetical-protein-[Stenotrophomonas-sp.-Marseille-Q4652] |
| GIX36092.1 | REase | GCA_026005075.1 | Lysobacteraceae_bacterium | Bacteria>Pseudomonadati>Pseudomonadota>Gammaproteobacteria>Lysobacterales>Lysobacteraceae>unclassified_Lysobacteraceae | 167 | KatS3mg126_1871 | 3978..4481 | 1926873 | MAG:-hypothetical-protein-KatS3mg126_1871-[Xanthomonadaceae-bacterium] |
| WP_218196657.1 | REase | GCF_019193005.1 | Xanthomonas_euvesicatoria | Bacteria>Pseudomonadati>Pseudomonadota>Gammaproteobacteria>Lysobacterales>Lysobacteraceae>Xanthomonas | 180 | KV691_RS15965 | 178163..178705 | 456327 | hypothetical-protein-[Xanthomonas-euvesicatoria] |
| MBK9496398.1 | REase | GCA_016718605.1 | Lysobacterales_bacterium | Bacteria>Pseudomonadati>Pseudomonadota>Gammaproteobacteria>Lysobacterales>unclassified_Lysobacterales | 169 | IPO08_18225 | 265872..266381 | 2006849 | hypothetical-protein-[Xanthomonadales-bacterium] |
| MCK9487879.1 | REase | GCA_023228845.1 | Lysobacterales_bacterium | Bacteria>Pseudomonadati>Pseudomonadota>Gammaproteobacteria>Lysobacterales>unclassified_Lysobacterales | 170 | MOQ42_00475 | 108262..108774 | 2006849 | hypothetical-protein-[Xanthomonadales-bacterium] |
| MCU0753615.1 | REase | GCA_025458445.1 | Lysobacterales_bacterium | Bacteria>Pseudomonadati>Pseudomonadota>Gammaproteobacteria>Lysobacterales>unclassified_Lysobacterales | 164 | MUE46_00630 | 19205..19699 | 2006849 | hypothetical-protein-[Xanthomonadales-bacterium] |
| GIX32364.1 | REase | GCA_026005055.1 | Lysobacterales_bacterium | Bacteria>Pseudomonadati>Pseudomonadota>Gammaproteobacteria>Lysobacterales>unclassified_Lysobacterales | 101 | KatS3mg125_0320 | 357443..357748 | 2006849 | MAG:-hypothetical-protein-KatS3mg125_0320-[Xanthomonadales-bacterium] |
| WP_156881497.1 | REase | GCF_017832375.1 | Methylococcaceae | Bacteria>Pseudomonadati>Pseudomonadota>Gammaproteobacteria>Methylococcales | 194 | QMX71_RS08990 | 1893028..1893612 | 403 | hypothetical-protein-[Methylocaldum-sp.-RMAD-M] |
| WP_198322143.1 | REase | GCF_036864975.1 | Methylococcus | Bacteria>Pseudomonadati>Pseudomonadota>Gammaproteobacteria>Methylococcales>Methylococcaceae | 169 | HUK90_RS04035 | 105923..106432 | 413 | MULTISPECIES:-hypothetical-protein-[Methylococcus] |
| WP_218808308.1 | REase | GCF_019285515.1 | unclassified_Methylococcus | Bacteria>Pseudomonadati>Pseudomonadota>Gammaproteobacteria>Methylococcales>Methylococcaceae>Methylococcus | 169 | KW115_RS06310 | 1273772..1274281 | 2618889 | hypothetical-protein-[Methylococcus-sp.-Mc7] |
| WP_218796636.1 | REase | GCF_019285555.1 | Methylococcus_capsulatus | Bacteria>Pseudomonadati>Pseudomonadota>Gammaproteobacteria>Methylococcales>Methylococcaceae>Methylococcus | 194 | KW114_RS08335 | 1841159..1841743 | 414 | hypothetical-protein-[Methylococcus-capsulatus] |
| WP_218797030.1 | REase | GCF_019285555.1 | Methylococcus_capsulatus | Bacteria>Pseudomonadati>Pseudomonadota>Gammaproteobacteria>Methylococcales>Methylococcaceae>Methylococcus | 194 | KW114_RS10925 | 2380646..2381230 | 414 | hypothetical-protein-[Methylococcus-capsulatus] |
| WP_218800672.1 | REase | GCF_019285535.1 | Methylococcus_capsulatus | Bacteria>Pseudomonadati>Pseudomonadota>Gammaproteobacteria>Methylococcales>Methylococcaceae>Methylococcus | 164 | M3M30_RS10975 | 2382854..2383348 | 414 | hypothetical-protein-[Methylococcus-capsulatus] |
| MBS3952426.1 | REase | GCA_018333785.1 | Methylomicrobium_sp. | Bacteria>Pseudomonadati>Pseudomonadota>Gammaproteobacteria>Methylococcales>Methylococcaceae>Methylomicrobium>unclassified_Methylomicrobium | 178 | KGZ88_05705 | 6768..7304 | 2303549 | hypothetical-protein-[Methylomicrobium-sp.] |
| WP_096876855.1 | REase | GCF_002441955.1 | Methylomonas_koyamae | Bacteria>Pseudomonadati>Pseudomonadota>Gammaproteobacteria>Methylococcales>Methylococcaceae>Methylomonas | 165 | MKLM6_RS22295 | 62733..63230 | 702114 | hypothetical-protein-[Methylomonas-koyamae] |
| WP_256360604.1 | REase | GCF_001312005.1 | Methylomonas_koyamae | Bacteria>Pseudomonadati>Pseudomonadota>Gammaproteobacteria>Methylococcales>Methylococcaceae>Methylomonas | 209 | JCM16701_RS30325 | 43518..44147 | 702114 | DUF4263-domain-containing-protein-[Methylomonas-koyamae] |
| WP_256360605.1 | REase | GCF_001312005.1 | Methylomonas_koyamae | Bacteria>Pseudomonadati>Pseudomonadota>Gammaproteobacteria>Methylococcales>Methylococcaceae>Methylomonas | 190 | JCM16701_RS30330 | 44170..44742 | 702114 | hypothetical-protein-[Methylomonas-koyamae] |
| MCF7971354.1 | REase | GCA_021736705.1 | Methylococcaceae_bacterium | Bacteria>Pseudomonadati>Pseudomonadota>Gammaproteobacteria>Methylococcales>Methylococcaceae>unclassified_Methylococcaceae | 124 | K9L22_09355 | 2632..3006 | 1933926 | GxxExxY-protein-[Methylococcaceae-bacterium] |
| WP_022949218.1 | REase | GCF_000421465.1 | Methylohalobius_crimeensis | Bacteria>Pseudomonadati>Pseudomonadota>Gammaproteobacteria>Methylococcales>Methylothermaceae>Methylohalobius | 170 | H035_RS0111985 | 2435563..2436075 | 244365 | hypothetical-protein-[Methylohalobius-crimeensis] |
| MDD5320444.1 | REase | GCA_028714175.1 | Methylococcales_bacterium | Bacteria>Pseudomonadati>Pseudomonadota>Gammaproteobacteria>Methylococcales>unclassified_Methylococcales | 204 | PHD43_07495 | 21616..22230 | 2304002 | MAG:-Uma2-family-endonuclease-[Methylococcales-bacterium] |
| MDD5320447.1 | REase | GCA_028714175.1 | Methylococcales_bacterium | Bacteria>Pseudomonadati>Pseudomonadota>Gammaproteobacteria>Methylococcales>unclassified_Methylococcales | 116 | PHD43_07510 | 23451..23801 | 2304002 | MAG:-DUF559-domain-containing-protein-[Methylococcales-bacterium] |
| WP_151834045.1 | REase | GCF_009012055.1 | Acinetobacter | Bacteria>Pseudomonadati>Pseudomonadota>Gammaproteobacteria>Moraxellales>Moraxellaceae | 171 | GBX89_RS12560 | 9996..10511 | 469 | hypothetical-protein-[Acinetobacter-ursingii] |
| WP_004984468.1 | REase | GCF_034183535.1 | Acinetobacter | Bacteria>Pseudomonadati>Pseudomonadota>Gammaproteobacteria>Moraxellales>Moraxellaceae | 167 | EOH89_RS01970 | 70751..71254 | 469 | MULTISPECIES:-hypothetical-protein-[Acinetobacter] |
| WP_216936953.1 | REase | GCF_019038255.1 | unclassified_Acinetobacter | Bacteria>Pseudomonadati>Pseudomonadota>Gammaproteobacteria>Moraxellales>Moraxellaceae>Acinetobacter | 167 | KUG40_RS10670 | 15307..15810 | 196816 | MULTISPECIES:-hypothetical-protein-[unclassified-Acinetobacter] |
| WP_180108073.1 | REase | GCF_013419415.1 | Acinetobacter_sp._YH12085 | Bacteria>Pseudomonadati>Pseudomonadota>Gammaproteobacteria>Moraxellales>Moraxellaceae>Acinetobacter>unclassified_Acinetobacter | 171 | FTU28_RS13905 | 47449..47964 | 2601077 | hypothetical-protein-[Acinetobacter-sp.-YH12085] |
| WP_107866350.1 | REase | GCF_003051055.1 | Agitococcus_lubricus | Bacteria>Pseudomonadati>Pseudomonadota>Gammaproteobacteria>Moraxellales>Moraxellaceae>Agitococcus | 164 | C8N29_RS12220 | 4174..4668 | 1077255 | hypothetical-protein-[Agitococcus-lubricus] |
| MAE39902.1 | REase | GCA_002686735.1 | Psychrobacter_sp. | Bacteria>Pseudomonadati>Pseudomonadota>Gammaproteobacteria>Moraxellales>Moraxellaceae>Psychrobacter>unclassified_Psychrobacter | 172 | CML07_03070 | 11135..11653 | 56811 | hypothetical-protein-[Psychrobacter-sp.] |
| WP_075107051.1 | REase | GCF_001921765.1 | Psychrobacter_sp._C_20.9 | Bacteria>Pseudomonadati>Pseudomonadota>Gammaproteobacteria>Moraxellales>Moraxellaceae>Psychrobacter>unclassified_Psychrobacter | 128 | BTW00_RS08555 | 130069..130455 | 1926477 | hypothetical-protein-[Psychrobacter-sp.-C-20.9] |
| WP_256714040.1 | REase | GCF_027919975.1 | Psychrobacter_sp._ANT_H3 | Bacteria>Pseudomonadati>Pseudomonadota>Gammaproteobacteria>Moraxellales>Moraxellaceae>Psychrobacter>unclassified_Psychrobacter | 173 | BTW00_RS08500 | 111960..112481 | 3019444 | MULTISPECIES:-Holliday-junction-resolvase-like-protein-[unclassified-Psychrobacter] |
| MBL0229730.1 | REase | GCA_016721165.1 | Moraxellaceae_bacterium | Bacteria>Pseudomonadati>Pseudomonadota>Gammaproteobacteria>Moraxellales>Moraxellaceae>unclassified_Moraxellaceae | 174 | IPP76_02605 | 10892..11416 | 1889775 | hypothetical-protein-[Moraxellaceae-bacterium] |
| MCC6374736.1 | REase | GCA_020846355.1 | Moraxellaceae_bacterium | Bacteria>Pseudomonadati>Pseudomonadota>Gammaproteobacteria>Moraxellales>Moraxellaceae>unclassified_Moraxellaceae | 172 | IT465_07995 | 2648..3166 | 1889775 | hypothetical-protein-[Moraxellaceae-bacterium] |
| WP_044628626.1 | REase | GCF_011398715.1 | Halomonadaceae | Bacteria>Pseudomonadati>Pseudomonadota>Gammaproteobacteria>Oceanospirillales | 169 | NH344_RS06640 | 52202..52711 | 28256 | MULTISPECIES:-hypothetical-protein-[Halomonas] |
| MCH2557817.1 | REase | GCA_022449455.1 | Alcanivorax_sp. | Bacteria>Pseudomonadati>Pseudomonadota>Gammaproteobacteria>Oceanospirillales>Alcanivoracaceae>Alcanivorax>unclassified_Alcanivorax | 177 | MK005_11015 | 54206..54739 | 1872427 | hypothetical-protein-[Alcanivorax-sp.] |
| WP_008734410.1 | REase | GCF_000299335.2 | Isoalcanivorax_pacificus | Bacteria>Pseudomonadati>Pseudomonadota>Gammaproteobacteria>Oceanospirillales>Alcanivoracaceae>Isoalcanivorax | 177 | S7S_RS18600 | 4155157..4155690 | 1306787 | hypothetical-protein-[Isoalcanivorax-pacificus] |
| WP_257265110.1 | REase | GCF_024606365.1 | Endozoicomonas_sp._ONNA2 | Bacteria>Pseudomonadati>Pseudomonadota>Gammaproteobacteria>Oceanospirillales>Endozoicomonadaceae>Endozoicomonas>unclassified_Endozoicomonas | 178 | KDK14_RS10770 | 6059..6595 | 2828741 | hypothetical-protein-[Endozoicomonas-sp.-ONNA2] |
| WP_206617984.1 | REase | GCF_004014695.1 | Hahella_sp._KA22 | Bacteria>Pseudomonadati>Pseudomonadota>Gammaproteobacteria>Oceanospirillales>Hahellaceae>Hahella>unclassified_Hahella | 168 | EUZ85_RS02360 | 526258..526764 | 1628392 | hypothetical-protein-[Hahella-sp.-KA22] |
| WP_234246273.1 | REase | GCF_021404345.1 | Billgrantia_desiderata | Bacteria>Pseudomonadati>Pseudomonadota>Gammaproteobacteria>Oceanospirillales>Halomonadaceae>Billgrantia | 173 | HOP57_RS15490 | 84831..85352 | 52021 | hypothetical-protein-[Halomonas-desiderata] |
| WP_097789713.1 | REase | GCF_039522625.1 | Halomonas_beimenensis | Bacteria>Pseudomonadati>Pseudomonadota>Gammaproteobacteria>Oceanospirillales>Halomonadaceae>Halomonas | 173 | BEI_RS11835 | 2520751..2521272 | 475662 | hypothetical-protein-[Halomonas-beimenensis] |
| WP_239295916.1 | REase | GCF_022348215.1 | unclassified_Halomonas | Bacteria>Pseudomonadati>Pseudomonadota>Gammaproteobacteria>Oceanospirillales>Halomonadaceae>Halomonas | 173 | MHM38_RS03730 | 11494..12015 | 2609666 | MULTISPECIES:-hypothetical-protein-[unclassified-Halomonas] |
| WP_275437442.1 | REase | - | unclassified_Halomonas | Bacteria>Pseudomonadati>Pseudomonadota>Gammaproteobacteria>Oceanospirillales>Halomonadaceae>Halomonas | 196 | MHM35_RS17080 | 124070..124660 | 2609666 | MULTISPECIES:-DUF4263-domain-containing-protein-[unclassified-Halomonas] |
| WP_231663148.1 | REase | GCF_021151495.1 | Halomonas_sp._IOP_31 | Bacteria>Pseudomonadati>Pseudomonadota>Gammaproteobacteria>Oceanospirillales>Halomonadaceae>Halomonas>unclassified_Halomonas | 201 | K7I12_RS04325 | 13092..13697 | 2876584 | hypothetical-protein-[Halomonas-sp.-IOP_31] |
| MCD6438843.1 | REase | GCA_021161325.1 | Halomonas_sp. | Bacteria>Pseudomonadati>Pseudomonadota>Gammaproteobacteria>Oceanospirillales>Halomonadaceae>Halomonas>unclassified_Halomonas | 198 | J7L56_11345 | 4780..5376 | 1486246 | hypothetical-protein-[Halomonas-sp.] |
| WP_253017812.1 | REase | GCF_024124605.1 | Halomonas_sp._CnH100-B | Bacteria>Pseudomonadati>Pseudomonadota>Gammaproteobacteria>Oceanospirillales>Halomonadaceae>Halomonas>unclassified_Halomonas | 198 | NH398_RS08370 | 360431..361027 | 2954490 | hypothetical-protein-[Halomonas-sp.-CnH100-B] |
| EWH01934.1 | REase | GCA_000568385.1 | Halomonas_sp._BC04 | Bacteria>Pseudomonadati>Pseudomonadota>Gammaproteobacteria>Oceanospirillales>Halomonadaceae>Halomonas>unclassified_Halomonas | 173 | Q427_11480 | 26663..27184 | 1403540 | cysteinyl-tRNA-synthetase-[Halomonas-sp.-BC04] |
| WP_043513265.1 | REase | GCF_000568385.1 | Halomonas_sp._BC04 | Bacteria>Pseudomonadati>Pseudomonadota>Gammaproteobacteria>Oceanospirillales>Halomonadaceae>Halomonas>unclassified_Halomonas | 173 | Q427_RS09380 | 26663..27184 | 1403540 | hypothetical-protein-[Halomonas-sp.-BC04] |
| WP_019020107.1 | REase | GCF_014651575.1 | Modicisalibacter_luteus | Bacteria>Pseudomonadati>Pseudomonadota>Gammaproteobacteria>Oceanospirillales>Halomonadaceae>Modicisalibacter | 165 | IE067_RS16245 | 66074..66571 | 453962 | hypothetical-protein-[Halomonas-lutea] |
| WP_019020112.1 | REase | GCF_014651575.1 | Modicisalibacter_luteus | Bacteria>Pseudomonadati>Pseudomonadota>Gammaproteobacteria>Oceanospirillales>Halomonadaceae>Modicisalibacter | 168 | IE067_RS16270 | 75663..76169 | 453962 | hypothetical-protein-[Halomonas-lutea] |
| BBI71110.2 | REase | GCA_007163885.2 | Vreelandella_aquamarina | Bacteria>Pseudomonadati>Pseudomonadota>Gammaproteobacteria>Oceanospirillales>Halomonadaceae>Vreelandella | 169 | HAALTHF_10170n | 667177..667686 | 77097 | hypothetical-protein-HAALTHF_10170n-[Halomonas-axialensis] |
| WP_027854329.1 | REase | GCF_000428985.1 | Marinobacterium_litorale | Bacteria>Pseudomonadati>Pseudomonadota>Gammaproteobacteria>Oceanospirillales>Oceanospirillaceae>Marinobacterium | 165 | G403_RS0112725 | 71465..71962 | 404770 | hypothetical-protein-[Marinobacterium-litorale] |
| AWY01112.1 | REase | GCA_003285205.1 | Marinomonas_primoryensis | Bacteria>Pseudomonadati>Pseudomonadota>Gammaproteobacteria>Oceanospirillales>Oceanospirillaceae>Marinomonas | 166 | A8139_14845 | 3169406..3169906 | 178399 | hypothetical-protein-A8139_14845-[Marinomonas-primoryensis] |
| WP_112139354.1 | REase | GCF_003285205.1 | Marinomonas_primoryensis | Bacteria>Pseudomonadati>Pseudomonadota>Gammaproteobacteria>Oceanospirillales>Oceanospirillaceae>Marinomonas | 166 | A8139_RS14970 | 3169406..3169906 | 178399 | hypothetical-protein-[Marinomonas-primoryensis] |
| WP_072841217.1 | REase | GCF_900129155.1 | Marinomonas_polaris | Bacteria>Pseudomonadati>Pseudomonadota>Gammaproteobacteria>Oceanospirillales>Oceanospirillaceae>Marinomonas | 170 | BUA59_RS18825 | 75942..76454 | 293552 | hypothetical-protein-[Marinomonas-polaris] |
| WP_205113930.1 | REase | GCF_016906065.1 | Marinomonas_foliarum | Bacteria>Pseudomonadati>Pseudomonadota>Gammaproteobacteria>Oceanospirillales>Oceanospirillaceae>Marinomonas | 180 | JSY38_RS15145 | 3251763..3252305 | 491950 | hypothetical-protein-[Marinomonas-foliarum] |

| Protein-ID | Domain | Assembly-ID | Strain | Taxonomy | Length | Locus-Tag | Genomic-coordinates | TaxID | Fasta-defline |
|---|---|---|---|---|---|---|---|---|---|
| MBE0507007.1 | REase | GCA_014859565.1 | Marinospirillum_sp. | Bacteria>Pseudomonadati>Pseudomonadota>Gammaproteobacteria>Oceanospirillales>Oceanospirillaceae>Marinospirillum>unclassified_Marinospirillum | 175 | IBX50_09860 | 6316..6843 | 2183934 | hypothetical-protein-[Marinospirillum-sp.] |
| MAR01285.1 | REase | GCA_002701445.1 | Oceanospirillaceae_bacterium | Bacteria>Pseudomonadati>Pseudomonadota>Gammaproteobacteria>Oceanospirillales>Oceanospirillaceae>unclassified_Oceanospirillaceae | 168 | CMI00_12210 | 6091..6597 | 1899355 | hypothetical-protein-[Oceanospirillaceae-bacterium] |
| WP_141068074.1 | REase | GCF_006516615.1 | Marinobacter_nauticus | Bacteria>Pseudomonadati>Pseudomonadota>Gammaproteobacteria>Pseudomonadales>Marinobacteraceae>Marinobacter | 177 | FH712_RS17670 | 40096..40629 | 2743 | hypothetical-protein-[Marinobacter-nauticus] |
| WP_230706348.1 | REase | GCF_021021495.1 | Marinobacter_adhaerens | Bacteria>Pseudomonadati>Pseudomonadota>Gammaproteobacteria>Pseudomonadales>Marinobacteraceae>Marinobacter | 169 | K7I10_RS06140 | 1376505..1377014 | 1033846 | hypothetical-protein-[Marinobacter-adhaerens] |
| WP_230713449.1 | REase | GCF_021021515.1 | Marinobacter_shengliensis | Bacteria>Pseudomonadati>Pseudomonadota>Gammaproteobacteria>Pseudomonadales>Marinobacteraceae>Marinobacter | 169 | K7I15_RS06085 | 105929..106438 | 1389223 | hypothetical-protein-[Marinobacter-shengliensis] |
| WP_092033194.1 | REase | GCF_900112835.1 | Marinobacter_sp._DSM_26671 | Bacteria>Pseudomonadati>Pseudomonadota>Gammaproteobacteria>Pseudomonadales>Marinobacteraceae>Marinobacter>unclassified_Marinobacter | 177 | BM035_RS16350 | 81875..82408 | 1761793 | hypothetical-protein-[Marinobacter-sp.-DSM-26671] |
| WP_022988847.1 | REase | GCF_000475255.1 | Marinobacter_sp._ES-1 | Bacteria>Pseudomonadati>Pseudomonadota>Gammaproteobacteria>Pseudomonadales>Marinobacteraceae>Marinobacter>unclassified_Marinobacter | 165 | Q668_RS06785 | 85057..85554 | 1396858 | hypothetical-protein-[Marinobacter-sp.-ES-1] |
| WP_022990213.1 | REase | GCF_000475255.1 | Marinobacter_sp._ES-1 | Bacteria>Pseudomonadati>Pseudomonadota>Gammaproteobacteria>Pseudomonadales>Marinobacteraceae>Marinobacter>unclassified_Marinobacter | 168 | Q668_RS13825 | 19942..20448 | 1396858 | hypothetical-protein-[Marinobacter-sp.-ES-1] |
| WP_247116144.1 | REase | GCF_023016045.1 | Marinobacter_sp._S6332 | Bacteria>Pseudomonadati>Pseudomonadota>Gammaproteobacteria>Pseudomonadales>Marinobacteraceae>Marinobacter>unclassified_Marinobacter | 168 | MWU62_RS17880 | 7247..7753 | 2926403 | hypothetical-protein-[Marinobacter-sp.-S6332] |
| PVY76887.1 | REase | GCA_003096655.1 | Tamilnaduibacter_salinus | Bacteria>Pseudomonadati>Pseudomonadota>Gammaproteobacteria>Pseudomonadales>Marinobacteraceae>Tamilnaduibacter | 165 | C8D92_104118 | 130139..130636 | 1484056 | hypothetical-protein-C8D92_104118-[Tamilnaduibacter-salinus] |
| PVY76891.1 | REase | GCA_003096655.1 | Tamilnaduibacter_salinus | Bacteria>Pseudomonadati>Pseudomonadota>Gammaproteobacteria>Pseudomonadales>Marinobacteraceae>Tamilnaduibacter | 168 | C8D92_104122 | 137770..138276 | 1484056 | hypothetical-protein-C8D92_104122-[Tamilnaduibacter-salinus] |
| WP_105523315.1 | REase | GCF_002975295.1 | Pseudomonas | Bacteria>Pseudomonadati>Pseudomonadota>Gammaproteobacteria>Pseudomonadales>Pseudomonadaceae | 175 | CQ048_RS15765 | 93460..93987 | 286 | MULTISPECIES:-hypothetical-protein-[Pseudomonas] |
| WP_060765712.1 | REase | GCF_044218865.1 | Pseudomonas | Bacteria>Pseudomonadati>Pseudomonadota>Gammaproteobacteria>Pseudomonadales>Pseudomonadaceae | 174 | GHO24_RS06640 | 41803..42327 | 286 | MULTISPECIES:-hypothetical-protein-[Pseudomonas] |
| WP_152745530.1 | REase | GCF_019891425.1 | Pseudomonas | Bacteria>Pseudomonadati>Pseudomonadota>Gammaproteobacteria>Pseudomonadales>Pseudomonadaceae | 165 | F0169_RS03310 | 26345..26842 | 286 | MULTISPECIES:-hypothetical-protein-[Pseudomonas] |
| WP_074867510.1 | REase | GCF_900109735.1 | Atopomonas_hussainii | Bacteria>Pseudomonadati>Pseudomonadota>Gammaproteobacteria>Pseudomonadales>Pseudomonadaceae>Atopomonas | 244 | BMX80_RS09310 | 105767..106501 | 1429083 | hypothetical-protein-[Atopomonas-hussainii] |
| WP_090734257.1 | REase | GCF_900108885.1 | Azotobacter_beijerinckii | Bacteria>Pseudomonadati>Pseudomonadota>Gammaproteobacteria>Pseudomonadales>Pseudomonadaceae>Azotobacter_group>Azotobacter | 191 | BMW56_RS18065 | 16003..16578 | 170623 | hypothetical-protein-[Azotobacter-beijerinckii] |
| WP_090941613.1 | REase | GCF_900114395.1 | Azotobacter_beijerinckii | Bacteria>Pseudomonadati>Pseudomonadota>Gammaproteobacteria>Pseudomonadales>Pseudomonadaceae>Azotobacter_group>Azotobacter | 191 | BM240_RS15645 | 17190..17765 | 170623 | hypothetical-protein-[Azotobacter-beijerinckii] |
| WP_131349394.1 | REase | GCF_004327885.1 | Azotobacter_chroococcum | Bacteria>Pseudomonadati>Pseudomonadota>Gammaproteobacteria>Pseudomonadales>Pseudomonadaceae>Azotobacter_group>Azotobacter | 191 | E0E52_RS08245 | 59318..59893 | 353 | hypothetical-protein-[Azotobacter-chroococcum] |
| WP_265072428.1 | REase | GCF_026016265.1 | Pseudomonas_viciae | Bacteria>Pseudomonadati>Pseudomonadota>Gammaproteobacteria>Pseudomonadales>Pseudomonadaceae>Pseudomonas | 179 | LOY66_RS25450 | 5795170..5795709 | 2505979 | hypothetical-protein-[Pseudomonas-viciae] |
| WP_048385713.1 | REase | GCF_001043065.1 | Pseudomonas_helleri | Bacteria>Pseudomonadati>Pseudomonadota>Gammaproteobacteria>Pseudomonadales>Pseudomonadaceae>Pseudomonas | 174 | TU85_RS02960 | 85433..85957 | 1608996 | hypothetical-protein-[Pseudomonas-helleri] |
| WP_177095191.1 | REase | GCF_013387065.1 | Pseudomonas_reactans | Bacteria>Pseudomonadati>Pseudomonadota>Gammaproteobacteria>Pseudomonadales>Pseudomonadaceae>Pseudomonas | 174 | HX796_RS11535 | 882..1406 | 117680 | hypothetical-protein-[Pseudomonas-reactans] |
| MBY8951564.1 | REase | GCA_019891425.1 | Pseudomonas_carnis | Bacteria>Pseudomonadati>Pseudomonadota>Gammaproteobacteria>Pseudomonadales>Pseudomonadaceae>Pseudomonas | 165 | J1G36_06610 | 94128..94625 | 2487355 | hypothetical-protein-[Pseudomonas-carnis] |
| WP_023835621.1 | REase | GCF_000503175.1 | Pseudomonas_aeruginosa_group | Bacteria>Pseudomonadati>Pseudomonadota>Gammaproteobacteria>Pseudomonadales>Pseudomonadaceae>Pseudomonas | 193 | QMY34_RS12040 | 380490..381071 | 136841 | hypothetical-protein-[Pseudomonas-aeruginosa] |
| WP_192069116.1 | REase | GCF_014842375.1 | Pseudomonas_coleopterorum | Bacteria>Pseudomonadati>Pseudomonadota>Gammaproteobacteria>Pseudomonadales>Pseudomonadaceae>Pseudomonas | 193 | IFT40_RS20705 | 80095..80676 | 1605838 | hypothetical-protein-[Pseudomonas-coleopterorum] |
| WP_266022657.1 | REase | GCF_026241065.1 | unclassified_Pseudomonas | Bacteria>Pseudomonadati>Pseudomonadota>Gammaproteobacteria>Pseudomonadales>Pseudomonadaceae>Pseudomonas | 173 | OO409_RS22200 | 32343..32864 | 196821 | MULTISPECIES:-hypothetical-protein-[unclassified-Pseudomonas] |
| MBY8951554.1 | REase | GCA_019891425.1 | Pseudomonas_carnis | Bacteria>Pseudomonadati>Pseudomonadota>Gammaproteobacteria>Pseudomonadales>Pseudomonadaceae>Pseudomonas | 152 | J1G36_06555 | 81563..82021 | 2487355 | DNA-mismatch-endonuclease-Vsr-[Pseudomonas-carnis] |
| HCF5867684.1 | REase | GCA_023959235.1 | Pseudomonas_aeruginosa | Bacteria>Pseudomonadati>Pseudomonadota>Gammaproteobacteria>Pseudomonadales>Pseudomonadaceae>Pseudomonas>Pseudomonas_aeruginosa_group | 180 | NIH49_002291 | 178317..178859 | 287 | hypothetical-protein-[Pseudomonas-aeruginosa] |
| HBO5114777.1 | REase | GCA_021662155.1 | Pseudomonas_aeruginosa | Bacteria>Pseudomonadati>Pseudomonadota>Gammaproteobacteria>Pseudomonadales>Pseudomonadaceae>Pseudomonas>Pseudomonas_aeruginosa_group | 193 | L4W37_005386 | 12045..12626 | 287 | hypothetical-protein-[Pseudomonas-aeruginosa] |
| WP_031643595.1 | REase | GCF_046449655.1 | Pseudomonas_aeruginosa | Bacteria>Pseudomonadati>Pseudomonadota>Gammaproteobacteria>Pseudomonadales>Pseudomonadaceae>Pseudomonas>Pseudomonas_aeruginosa_group | 193 | AJ61_RS31485 | 12144..12725 | 287 | hypothetical-protein-[Pseudomonas-aeruginosa] |
| WP_033975674.1 | REase | GCF_022569915.1 | Pseudomonas_aeruginosa | Bacteria>Pseudomonadati>Pseudomonadota>Gammaproteobacteria>Pseudomonadales>Pseudomonadaceae>Pseudomonas>Pseudomonas_aeruginosa_group | 193 | CGU39_RS16570 | 613504..614085 | 287 | hypothetical-protein-[Pseudomonas-aeruginosa] |
| MCR7692358.1 | REase | GCA_024717035.1 | Pseudomonas_aeruginosa | Bacteria>Pseudomonadati>Pseudomonadota>Gammaproteobacteria>Pseudomonadales>Pseudomonadaceae>Pseudomonas>Pseudomonas_aeruginosa_group | 173 | NTA33_29235 | 17157..17678 | 287 | hypothetical-protein-NTA33_29235-[Pseudomonas-aeruginosa] |
| WP_223723729.1 | REase | GCF_900707645.1 | Pseudomonas_aeruginosa | Bacteria>Pseudomonadati>Pseudomonadota>Gammaproteobacteria>Pseudomonadales>Pseudomonadaceae>Pseudomonas>Pseudomonas_aeruginosa_group | 173 | KGZ09_RS30320 | 41265..41786 | 287 | hypothetical-protein-[Pseudomonas-aeruginosa] |
| WP_234464498.1 | REase | GCF_050243135.1 | Pseudomonas_aeruginosa | Bacteria>Pseudomonadati>Pseudomonadota>Gammaproteobacteria>Pseudomonadales>Pseudomonadaceae>Pseudomonas>Pseudomonas_aeruginosa_group | 173 | GFM12_RS08420 | 1738402..1738923 | 287 | hypothetical-protein-[Pseudomonas-aeruginosa] |
| WP_258055199.1 | REase | - | Pseudomonas_aeruginosa | Bacteria>Pseudomonadati>Pseudomonadota>Gammaproteobacteria>Pseudomonadales>Pseudomonadaceae>Pseudomonas>Pseudomonas_aeruginosa_group | 173 | KZ843_RS17270 | 8496..9017 | 287 | hypothetical-protein-[Pseudomonas-aeruginosa] |
| HCI3160561.1 | REase | GCA_024356045.1 | Pseudomonas_aeruginosa | Bacteria>Pseudomonadati>Pseudomonadota>Gammaproteobacteria>Pseudomonadales>Pseudomonadaceae>Pseudomonas>Pseudomonas_aeruginosa_group | 170 | NOV63_004507 | 72080..72592 | 287 | TPA:-hypothetical-protein-NOV63_004507-[Pseudomonas-aeruginosa] |
| MBG7398187.1 | REase | GCA_015766545.1 | Pseudomonas_aeruginosa | Bacteria>Pseudomonadati>Pseudomonadota>Gammaproteobacteria>Pseudomonadales>Pseudomonadaceae>Pseudomonas>Pseudomonas_aeruginosa_group | 170 | GHU66_27425 | 41265..41777 | 287 | hypothetical-protein-GHU66_27425-[Pseudomonas-aeruginosa] |
| MCO2346211.1 | REase | GCA_023907105.1 | Pseudomonas_aeruginosa | Bacteria>Pseudomonadati>Pseudomonadota>Gammaproteobacteria>Pseudomonadales>Pseudomonadaceae>Pseudomonas>Pseudomonas_aeruginosa_group | 170 | FA362_14070 | 17224..17736 | 287 | hypothetical-protein-FA362_14070-[Pseudomonas-aeruginosa] |
| HBO3135815.1 | REase | GCA_021651475.1 | Pseudomonas_aeruginosa | Bacteria>Pseudomonadati>Pseudomonadota>Gammaproteobacteria>Pseudomonadales>Pseudomonadaceae>Pseudomonas>Pseudomonas_aeruginosa_group | 182 | L4Q96_001361 | 38645..39193 | 287 | hypothetical-protein-[Pseudomonas-aeruginosa] |
| WP_230091451.1 | REase | - | Pseudomonas_aeruginosa | Bacteria>Pseudomonadati>Pseudomonadota>Gammaproteobacteria>Pseudomonadales>Pseudomonadaceae>Pseudomonas>Pseudomonas_aeruginosa_group | 202 | K3Z95_RS29445 | 6818..7426 | 287 | DUF4263-domain-containing-protein-[Pseudomonas-aeruginosa] |
| WP_010463589.1 | REase | GCF_000257545.3 | Pseudomonas_mandelii | Bacteria>Pseudomonadati>Pseudomonadota>Gammaproteobacteria>Pseudomonadales>Pseudomonadaceae>Pseudomonas>Pseudomonas_fluorescens_group | 179 | OU5_RS13435 | 3000259..3000798 | 75612 | hypothetical-protein-[Pseudomonas-mandelii] |
| WP_236238650.1 | REase | GCF_021603345.1 | Pseudomonas_rhodesiae | Bacteria>Pseudomonadati>Pseudomonadota>Gammaproteobacteria>Pseudomonadales>Pseudomonadaceae>Pseudomonas>Pseudomonas_fluorescens_group | 175 | L6E14_RS13715 | 262805..263332 | 76760 | hypothetical-protein-[Pseudomonas-rhodesiae] |
| WP_198790927.1 | REase | GCF_016405175.1 | Pseudomonas_sp._MF7451 | Bacteria>Pseudomonadati>Pseudomonadota>Gammaproteobacteria>Pseudomonadales>Pseudomonadaceae>Pseudomonas>unclassified_Pseudomonas | 179 | JFT65_RS10915 | 201013..201552 | 2797538 | hypothetical-protein-[Pseudomonas-sp.-MF7451] |
| WP_054902928.1 | REase | GCF_001320525.1 | Pseudomonas_sp._NBRC_111131 | Bacteria>Pseudomonadati>Pseudomonadota>Gammaproteobacteria>Pseudomonadales>Pseudomonadaceae>Pseudomonas>unclassified_Pseudomonas | 193 | APH34_RS22150 | 23034..23615 | 1661046 | hypothetical-protein-[Pseudomonas-sp.-NBRC-111131] |
| WP_256573571.1 | REase | GCF_002113255.1 | Pseudomonas_sp._B14(2017) | Bacteria>Pseudomonadati>Pseudomonadota>Gammaproteobacteria>Pseudomonadales>Pseudomonadaceae>Pseudomonas>unclassified_Pseudomonas | 173 | CAG34_RS21640 | 107862..108383 | 1981745 | hypothetical-protein-[Pseudomonas-sp.-B14(2017)] |
| MAQ23218.1 | REase | GCA_002701315.1 | Pseudomonadales_bacterium | Bacteria>Pseudomonadati>Pseudomonadota>Gammaproteobacteria>Pseudomonadales>unclassified_Pseudomonadales | 177 | CMK83_03280 | 14533..15066 | 1891229 | hypothetical-protein-[Pseudomonadales-bacterium] |
| MCP5344824.1 | REase | GCA_024234345.1 | Pseudomonadales_bacterium | Bacteria>Pseudomonadati>Pseudomonadota>Gammaproteobacteria>Pseudomonadales>unclassified_Pseudomonadales | 165 | H7A05_09400 | 49453..49950 | 1891229 | hypothetical-protein-[Pseudomonadales-bacterium] |
| WP_203168600.1 | REase | GCF_016801985.1 | Steroidobacter_gossypii | Bacteria>Pseudomonadati>Pseudomonadota>Gammaproteobacteria>Steroidobacterales>Steroidobacteraceae>Steroidobacter | 195 | JM946_RS17395 | 833077..833664 | 2805490 | hypothetical-protein-[Steroidobacter-gossypii] |
| MBV1952385.1 | REase | GCA_019090295.1 | Cycloclasticus_sp. | Bacteria>Pseudomonadati>Pseudomonadota>Gammaproteobacteria>Thiotrichales>Piscirickettsiaceae>Cycloclasticus>unclassified_Cycloclasticus | 94 | KUG64_09380 | 13242..13524 | 2024830 | winged-helix-turn-helix-domain-containing-protein-[Cycloclasticus-sp.] |
| CAA6806528.1 | REase | - | uncultured_Thiotrichaceae_bacterium | Bacteria>Pseudomonadati>Pseudomonadota>Gammaproteobacteria>Thiotrichales>Thiotrichaceae>environmental_samples | 181 | HELGO_WM31112 | 4763..5308 | 298394 | Unknown-protein-[uncultured-Thiotrichaceae-bacterium] |
| WP_210225667.1 | REase | GCF_017901155.1 | Candidatus_Thiothrix_anitrata | Bacteria>Pseudomonadati>Pseudomonadota>Gammaproteobacteria>Thiotrichales>Thiotrichaceae>Thiothrix | 148 | J8380_RS10855 | 2104247..2104693 | 2823902 | DUF559-domain-containing-protein-[Candidatus-Thiothrix-anitrata] |
| PCJ88239.1 | REase | GCA_002746365.1 | Thiotrichaceae_bacterium | Bacteria>Pseudomonadati>Pseudomonadota>Gammaproteobacteria>Thiotrichales>Thiotrichaceae>unclassified_Thiotrichaceae | 165 | COA54_02945 | 14232..14729 | 2030882 | hypothetical-protein-COA54_02945-[Thiotrichaceae-bacterium] |
| HIJ32931.1 | REase | GCA_013349805.1 | Gammaproteobacteria_bacterium | Bacteria>Pseudomonadati>Pseudomonadota>Gammaproteobacteria>unclassified_Gammaproteobacteria | 171 | HPP65_00775 | 5425..5940 | 1913989 | hypothetical-protein-[Gammaproteobacteria-bacterium] |
| WP_198244462.1 | REase | GCF_016097405.1 | methane-oxidizing_endosymbiont_of_Gigantopelta_aegis | Bacteria>Pseudomonadati>Pseudomonadota>Gammaproteobacteria>unclassified_Gammaproteobacteria | 165 | JEU20_RS06795 | 1355340..1355837 | 2794938 | hypothetical-protein-[methane-oxidizing-endosymbiont-of-Gigantopelta-aegis] |
| MYA16810.1 | REase | GCA_009836885.1 | Gammaproteobacteria_bacterium | Bacteria>Pseudomonadati>Pseudomonadota>Gammaproteobacteria>unclassified_Gammaproteobacteria | 198 | F4Z28_08610 | 8042..8638 | 1913989 | hypothetical-protein-[Gammaproteobacteria-bacterium] |
| HFD86239.1 | REase | GCA_011322685.1 | Gammaproteobacteria_bacterium | Bacteria>Pseudomonadati>Pseudomonadota>Gammaproteobacteria>unclassified_Gammaproteobacteria | 171 | ENJ35_01015 | 2083..2598 | 1913989 | hypothetical-protein-[Gammaproteobacteria-bacterium] |
| RTZ75008.1 | REase | GCA_003973075.1 | Gammaproteobacteria_bacterium | Bacteria>Pseudomonadati>Pseudomonadota>Gammaproteobacteria>unclassified_Gammaproteobacteria | 171 | DSZ00_03000 | 5320..5835 | 1913989 | hypothetical-protein-DSZ00_03000-[Gammaproteobacteria-bacterium] |
| QFY90723.1 | REase | GCA_009497875.2 | gamma_proteobacterium_SS-5 | Bacteria>Pseudomonadati>Pseudomonadota>Gammaproteobacteria>unclassified_Gammaproteobacteria | 133 | D5125_15340 | 2190255..2190656 | 947516 | GxxExxY-protein-[gamma-proteobacterium-SS-5] |
| QFY90727.1 | REase | GCA_009497875.2 | gamma_proteobacterium_SS-5 | Bacteria>Pseudomonadati>Pseudomonadota>Gammaproteobacteria>unclassified_Gammaproteobacteria | 127 | D5125_15360 | 2198257..2198640 | 947516 | GxxExxY-protein-[gamma-proteobacterium-SS-5] |
| MBT4196265.1 | REase | GCA_018651435.1 | Gammaproteobacteria_bacterium | Bacteria>Pseudomonadati>Pseudomonadota>Gammaproteobacteria>unclassified_Gammaproteobacteria | 169 | HOE12_18210 | 19771..20280 | 1913989 | hypothetical-protein-[Gammaproteobacteria-bacterium] |
| MDE0192032.1 | REase | GCA_028819695.1 | Gammaproteobacteria_bacterium | Bacteria>Pseudomonadati>Pseudomonadota>Gammaproteobacteria>unclassified_Gammaproteobacteria | 218 | OXQ90_11805 | 65495..66151 | 1913989 | MAG:-hypothetical-protein-OXQ90_11805-[Gammaproteobacteria-bacterium] |
| WP_006647485.1 | REase | GCF_000153325.1 | Photobacterium_sp._SKA34 | Bacteria>Pseudomonadati>Pseudomonadota>Gammaproteobacteria>Vibrionales>Vibrionaceae>Photobacterium>unclassified_Photobacterium | 174 | SKA34_RS21235 | 84637..85161 | 121723 | hypothetical-protein-[Photobacterium-sp.-SKA34] |
| EIZ1010572.1 | REase | GCA_022609845.1 | Vibrio_vulnificus | Bacteria>Pseudomonadati>Pseudomonadota>Gammaproteobacteria>Vibrionales>Vibrionaceae>Vibrio | 174 | MOS09_002152 | 286748..287272 | 672 | hypothetical-protein-[Vibrio-vulnificus] |
| NAZ69807.1 | REase | GCA_009906175.1 | Vibrio_toranzoniae | Bacteria>Pseudomonadati>Pseudomonadota>Gammaproteobacteria>Vibrionales>Vibrionaceae>Vibrio | 174 | GL180_08035 | 8828..9352 | 1194427 | hypothetical-protein-[Vibrio-toranzoniae] |
| WP_161749220.1 | REase | GCF_009906175.1 | Vibrio_toranzoniae | Bacteria>Pseudomonadati>Pseudomonadota>Gammaproteobacteria>Vibrionales>Vibrionaceae>Vibrio | 174 | GL180_RS08030 | 8828..9352 | 1194427 | hypothetical-protein-[Vibrio-toranzoniae] |
| ASG03055.1 | REase | GCA_002211985.1 | Vibrio_anguillarum | Bacteria>Pseudomonadati>Pseudomonadota>Gammaproteobacteria>Vibrionales>Vibrionaceae>Vibrio | 171 | CEJ46_04115 | 867383..867898 | 55601 | hypothetical-protein-CEJ46_04115-[Vibrio-anguillarum] |
| GLT15888.1 | REase | GCA_030161215.1 | Vibrio_algivorus | Bacteria>Pseudomonadati>Pseudomonadota>Gammaproteobacteria>Vibrionales>Vibrionaceae>Vibrio | 171 | GCM10007931_28630 | 501382..501897 | 1667024 | hypothetical-protein-GCM10007931_28630-[Vibrio-algivorus] |
| WP_088720999.1 | REase | GCF_002211985.1 | Vibrio_anguillarum | Bacteria>Pseudomonadati>Pseudomonadota>Gammaproteobacteria>Vibrionales>Vibrionaceae>Vibrio | 171 | CEJ46_RS04115 | 867383..867898 | 55601 | hypothetical-protein-[Vibrio-anguillarum] |
| WP_089123109.1 | REase | GCF_030161215.1 | Vibrio_algivorus | Bacteria>Pseudomonadati>Pseudomonadota>Gammaproteobacteria>Vibrionales>Vibrionaceae>Vibrio | 171 | AVR_RS07065 | 1501042..1501557 | 1667024 | hypothetical-protein-[Vibrio-algivorus] |
| WP_274674020.1 | REase | GCF_028857845.1 | Vibrio_aestuarianus | Bacteria>Pseudomonadati>Pseudomonadota>Gammaproteobacteria>Vibrionales>Vibrionaceae>Vibrio | 171 | L9W73_RS08970 | 40692..41207 | 28171 | hypothetical-protein-[Vibrio-aestuarianus] |
| MBH9740668.1 | REase | GCA_016094345.1 | Vibrio_navarrensis | Bacteria>Pseudomonadati>Pseudomonadota>Gammaproteobacteria>Vibrionales>Vibrionaceae>Vibrio | 169 | CJ419_13190 | 157579..158088 | 29495 | hypothetical-protein-[Vibrio-navarrensis] |
| WP_198106731.1 | REase | GCF_963891765.1 | Vibrio_navarrensis | Bacteria>Pseudomonadati>Pseudomonadota>Gammaproteobacteria>Vibrionales>Vibrionaceae>Vibrio | 169 | CJ419_RS12990 | 157579..158088 | 29495 | hypothetical-protein-[Vibrio-navarrensis] |
| WP_268670712.1 | REase | GCF_026798035.1 | Vibrio_coralliirubri | Bacteria>Pseudomonadati>Pseudomonadota>Gammaproteobacteria>Vibrionales>Vibrionaceae>Vibrio | 169 | OTK49_RS06680 | 234242..234751 | 1516159 | hypothetical-protein-[Vibrio-coralliirubri] |
| WP_132941354.1 | REase | GCF_004345555.1 | Vibrio_crassostreae | Bacteria>Pseudomonadati>Pseudomonadota>Gammaproteobacteria>Vibrionales>Vibrionaceae>Vibrio | 239 | EDB60_RS00595 | 118303..119022 | 246167 | hypothetical-protein-[Vibrio-crassostreae] |
| HCG7393725.1 | REase | GCA_024044075.1 | Vibrio_parahaemolyticus | Bacteria>Pseudomonadati>Pseudomonadota>Gammaproteobacteria>Vibrionales>Vibrionaceae>Vibrio>Vibrio_harveyi_group | 169 | NJ383_004131 | 18362..18871 | 670 | hypothetical-protein-[Vibrio-parahaemolyticus] |
| WP_267363051.1 | REase | GCF_026537625.1 | Vibrio_parahaemolyticus | Bacteria>Pseudomonadati>Pseudomonadota>Gammaproteobacteria>Vibrionales>Vibrionaceae>Vibrio>Vibrio_harveyi_group | 169 | NOL37_RS24185 | 13952..14461 | 670 | hypothetical-protein-[Vibrio-parahaemolyticus] |
| EGQ8448093.1 | REase | - | Vibrio_alginolyticus | Bacteria>Pseudomonadati>Pseudomonadota>Gammaproteobacteria>Vibrionales>Vibrionaceae>Vibrio>Vibrio_harveyi_group | 168 | GTP59_14920 | 62554..63060 | 663 | hypothetical-protein-[Vibrio-alginolyticus] |
| EJV5950191.1 | REase | GCA_025320405.1 | Vibrio_alginolyticus | Bacteria>Pseudomonadati>Pseudomonadota>Gammaproteobacteria>Vibrionales>Vibrionaceae>Vibrio>Vibrio_harveyi_group | 168 | N7I24_002736 | 62395..62901 | 663 | hypothetical-protein-[Vibrio-alginolyticus] |
| WP_213868352.1 | REase | GCF_018457845.1 | Vibrio_alginolyticus | Bacteria>Pseudomonadati>Pseudomonadota>Gammaproteobacteria>Vibrionales>Vibrionaceae>Vibrio>Vibrio_harveyi_group | 168 | J4H60_RS12990 | 326311..326817 | 663 | hypothetical-protein-[Vibrio-alginolyticus] |
| WP_182037016.1 | REase | GCF_903986725.1 | Vibrio_diabolicus | Bacteria>Pseudomonadati>Pseudomonadota>Gammaproteobacteria>Vibrionales>Vibrionaceae>Vibrio>Vibrio_harveyi_group>Vibrio_diabolicus_subgroup | 239 | H7A60_RS18195 | 124501..125220 | 50719 | hypothetical-protein-[Vibrio-diabolicus] |
| MBU4235490.1 | REase | GCA_018897885.1 | Pseudomonadota_bacterium | Bacteria>Pseudomonadati>Pseudomonadota>unclassified_Pseudomonadota | 175 | KKI01_04065 | 3553..4080 | 1977087 | MAG:-hypothetical-protein-KKI01_04065-[Proteobacteria-bacterium] |
| MCG8418452.1 | REase | GCA_022360315.1 | Pseudomonadota_bacterium | Bacteria>Pseudomonadati>Pseudomonadota>unclassified_Pseudomonadota | 193 | MJE77_10970 | 3825..4406 | 1977087 | hypothetical-protein-[Pseudomonadota-bacterium] |
| MBU0673023.1 | REase | GCA_018814305.1 | Pseudomonadota_bacterium | Bacteria>Pseudomonadati>Pseudomonadota>unclassified_Pseudomonadota | 174 | KJ950_00085 | 18973..19497 | 1977087 | hypothetical-protein-[Pseudomonadota-bacterium] |
| MDE3269949.1 | REase | GCA_028877195.1 | Pseudomonadota_bacterium | Bacteria>Pseudomonadati>Pseudomonadota>unclassified_Pseudomonadota | 172 | OYH77_06665 | 687..1205 | 1977087 | MAG:-hypothetical-protein-OYH77_06665-[Pseudomonadota-bacterium] |
| MBU1208888.1 | REase | GCA_018819135.1 | Pseudomonadota_bacterium | Bacteria>Pseudomonadati>Pseudomonadota>unclassified_Pseudomonadota | 67 | KKH04_18505 | 10090..10293 | 1977087 | hypothetical-protein-[Pseudomonadota-bacterium] |
| MBK7497324.1 | REase | GCA_016706995.1 | Candidatus_Omnitrophota_bacterium | Bacteria>Pseudomonadati>PVC_group>Candidatus_Omnitrophota>unclassified_Candidatus_Omnitrophota | 125 | IPI28_18705 | 42728..43105 | 2035772 | GxxExxY-protein-[Candidatus-Omnitrophota-bacterium] |
| OGW72083.1 | REase | GCA_001803975.1 | Omnitrophica_bacterium_GWA2_41_15 | Bacteria>Pseudomonadati>PVC_group>Candidatus_Omnitrophota>unclassified_Candidatus_Omnitrophota | 132 | A2047_02000 | 18561..18959 | 1801814 | MAG:-hypothetical-protein-A2047_02000-[Omnitrophica-bacterium-GWA2_41_15] |
| OGW72077.1 | REase | GCA_001803975.1 | Omnitrophica_bacterium_GWA2_41_15 | Bacteria>Pseudomonadati>PVC_group>Candidatus_Omnitrophota>unclassified_Candidatus_Omnitrophota | 219 | A2047_01970 | 8607..9266 | 1801814 | MAG:-hypothetical-protein-A2047_01970-[Omnitrophica-bacterium-GWA2_41_15] |
| MBN2450530.1 | REase | GCA_016935045.1 | Lentisphaeria_bacterium | Bacteria>Pseudomonadati>PVC_group>Lentisphaerota>Lentisphaeria>unclassified_Lentisphaeria | 126 | JXR77_09070 | 75..455 | 2053569 | GxxExxY-protein-[Lentisphaeria-bacterium] |
| MBE6357834.1 | REase | GCA_015061525.1 | Lentisphaerota_bacterium | Bacteria>Pseudomonadati>PVC_group>Lentisphaerota>unclassified_Lentisphaerota | 174 | E7057_01130 | 6012..6536 | 1932692 | hypothetical-protein-[Lentisphaerota-bacterium] |
| NLN03022.1 | REase | GCA_012521875.1 | Lentisphaerota_bacterium | Bacteria>Pseudomonadati>PVC_group>Lentisphaerota>unclassified_Lentisphaerota | 123 | GX174_14145 | 1356..1727 | 1932692 | GxxExxY-protein-[Lentisphaerota-bacterium] |
| MCE9614799.1 | REase | GCA_021413965.1 | Lentisphaerota_bacterium | Bacteria>Pseudomonadati>PVC_group>Lentisphaerota>unclassified_Lentisphaerota | 182 | K8T26_11020 | 332614..333162 | 1932692 | very-short-patch-repair-endonuclease-[Lentisphaerota-bacterium] |
| TVL98634.1 | REase | GCA_007618135.1 | Candidatus_Brocadia_sp._WS118 | Bacteria>Pseudomonadati>PVC_group>Planctomycetota>Candidatus_Brocadiia>Candidatus_Brocadiales>Candidatus_Brocadiaceae>Candidatus_Brocadia>unclassified_Candidatus_Brocadia | 192 | CV087_21010 | 11824..12402 | 2040625 | MAG:-hypothetical-protein-CV087_21010-[Candidatus-Brocadia-sp.-WS118] |
| WP_096895082.1 | REase | GCF_002443295.1 | Candidatus_Scalindua_japonica | Bacteria>Pseudomonadati>PVC_group>Planctomycetota>Candidatus_Brocadiia>Candidatus_Brocadiales>Candidatus_Scalinduaceae>Candidatus_Scalindua | 127 | SCALIN_RS13985 | 120951..121334 | 1284222 | GxxExxY-protein-[Candidatus-Scalindua-japonica] |
| MAO24326.1 | REase | GCA_002696165.1 | Phycisphaerae_bacterium | Bacteria>Pseudomonadati>PVC_group>Planctomycetota>Phycisphaerae>unclassified_Phycisphaerae | 212 | CMJ25_26545 | 12322..12960 | 2026778 | hypothetical-protein-[Phycisphaerae-bacterium] |
| MCK6484532.1 | REase | GCA_023150595.1 | Phycisphaerae_bacterium | Bacteria>Pseudomonadati>PVC_group>Planctomycetota>Phycisphaerae>unclassified_Phycisphaerae | 213 | L6R00_10415 | 64336..64977 | 2026778 | very-short-patch-repair-endonuclease-[Phycisphaerae-bacterium] |
| WP_210660198.1 | REase | GCF_017939745.1 | Gemmata_palustris | Bacteria>Pseudomonadati>PVC_group>Planctomycetota>Planctomycetia>Gemmatales>Gemmataceae>Gemmata | 194 | J8F10_RS30115 | 7263768..7264352 | 2822762 | hypothetical-protein-[Gemmata-palustris] |
| MSR55923.1 | REase | GCA_009691845.1 | Gemmataceae_bacterium | Bacteria>Pseudomonadati>PVC_group>Planctomycetota>Planctomycetia>Gemmatales>Gemmataceae>unclassified_Gemmataceae | 198 | EXS09_22020 | 3754..4350 | 2052164 | hypothetical-protein-[Gemmataceae-bacterium] |
| WP_165226835.1 | REase | GCF_011064615.1 | Aquisphaera_insulae | Bacteria>Pseudomonadati>PVC_group>Planctomycetota>Planctomycetia>Isosphaerales>Isosphaeraceae>Aquisphaera | 171 | G6054_RS16925 | 64632..65147 | 2712864 | hypothetical-protein-[Aquisphaera-sp.-JC669] |
| WP_076345236.1 | REase | GCF_001956985.1 | Paludisphaera_borealis | Bacteria>Pseudomonadati>PVC_group>Planctomycetota>Planctomycetia>Isosphaerales>Isosphaeraceae>Paludisphaera | 168 | BSF38_RS09990 | 2559069..2559575 | 1387353 | hypothetical-protein-[Paludisphaera-borealis] |
| WP_230262564.1 | REase | GCF_020966635.1 | Pirellulaceae | Bacteria>Pseudomonadati>PVC_group>Planctomycetota>Planctomycetia>Pirellulales | 193 | LOC67_RS10685 | 1302921..1303502 | 2691357 | hypothetical-protein-[Stieleria-sp.-JC731] |
| WP_230262560.1 | REase | GCF_020966635.1 | Pirellulaceae | Bacteria>Pseudomonadati>PVC_group>Planctomycetota>Planctomycetia>Pirellulales | 222 | LOC67_RS10665 | 1297676..1298344 | 2691357 | hypothetical-protein-[Stieleria-sp.-JC731] |
| WP_146436435.1 | REase | GCF_007859755.1 | Blastopirellula_retiformator | Bacteria>Pseudomonadati>PVC_group>Planctomycetota>Planctomycetia>Pirellulales>Pirellulaceae>Blastopirellula | 196 | Enr8_RS23485 | 391191..391781 | 2527970 | hypothetical-protein-[Blastopirellula-retiformator] |
| WP_146679084.1 | REase | GCF_001610875.1 | Pirellula_sp._SH-Sr6A | Bacteria>Pseudomonadati>PVC_group>Planctomycetota>Planctomycetia>Pirellulales>Pirellulaceae>Pirellula>unclassified_Pirellula | 197 | VN12_RS23065 | 5950953..5951546 | 1632865 | hypothetical-protein-[Pirellula-sp.-SH-Sr6A] |
| NLX99943.1 | REase | GCA_012728855.1 | Rhodopirellula_sp. | Bacteria>Pseudomonadati>PVC_group>Planctomycetota>Planctomycetia>Pirellulales>Pirellulaceae>Rhodopirellula>unclassified_Rhodopirellula | 204 | GXY83_27915 | 6702..7316 | 2024855 | hypothetical-protein-[Rhodopirellula-sp.] |
| NLX99946.1 | REase | GCA_012728855.1 | Rhodopirellula_sp. | Bacteria>Pseudomonadati>PVC_group>Planctomycetota>Planctomycetia>Pirellulales>Pirellulaceae>Rhodopirellula>unclassified_Rhodopirellula | 125 | GXY83_27930 | 11099..11476 | 2024855 | GxxExxY-protein-[Rhodopirellula-sp.] |
| NLX98933.1 | REase | GCA_012728855.1 | Rhodopirellula_sp. | Bacteria>Pseudomonadati>PVC_group>Planctomycetota>Planctomycetia>Pirellulales>Pirellulaceae>Rhodopirellula>unclassified_Rhodopirellula | 138 | GXY83_22535 | 11784..12200 | 2024855 | DNA-mismatch-endonuclease-Vsr-[Rhodopirellula-sp.] |
| WP_218933034.1 | REase | GCF_007741495.1 | Roseimaritima_multifibrata | Bacteria>Pseudomonadati>PVC_group>Planctomycetota>Planctomycetia>Pirellulales>Pirellulaceae>Roseimaritima | 176 | FF011L_RS05770 | 1540598..1541128 | 1930274 | hypothetical-protein-[Roseimaritima-multifibrata] |
| WP_261366875.1 | REase | - | Stieleria_sp._JC731 | Bacteria>Pseudomonadati>PVC_group>Planctomycetota>Planctomycetia>Pirellulales>Pirellulaceae>Stieleria>unclassified_Stieleria | 158 | LOC67_RS27335 | 1299206..1299682 | 2894195 | DUF3883-domain-containing-protein-[Stieleria-sp.-JC731] |
| GIW94554.1 | REase | GCA_026004775.1 | Pirellulaceae_bacterium | Bacteria>Pseudomonadati>PVC_group>Planctomycetota>Planctomycetia>Pirellulales>Pirellulaceae>unclassified_Pirellulaceae | 122 | KatS3mg110_2595 | 16146..16514 | 2699754 | MAG:-hypothetical-protein-KatS3mg110_2595-[Pirellulaceae-bacterium] |
| MBX3412590.1 | REase | GCA_019636335.1 | Pirellulales_bacterium | Bacteria>Pseudomonadati>PVC_group>Planctomycetota>Planctomycetia>Pirellulales>unclassified_Pirellulales | 174 | KF708_07875 | 42300..42824 | 2715253 | DUF4365-domain-containing-protein-[Pirellulales-bacterium] |
| WP_197998735.1 | REase | GCF_007748195.1 | Gimesia_aquarii | Bacteria>Pseudomonadati>PVC_group>Planctomycetota>Planctomycetia>Planctomycetales>Planctomycetaceae>Gimesia | 179 | V144x_RS03470 | 841377..841916 | 2527964 | hypothetical-protein-[Gimesia-aquarii] |
| WP_145201098.1 | REase | GCF_007744255.1 | Thalassoglobus_polymorphus | Bacteria>Pseudomonadati>PVC_group>Planctomycetota>Planctomycetia>Planctomycetales>Planctomycetaceae>Thalassoglobus | 198 | Mal48_RS14945 | 4187367..4187963 | 2527994 | hypothetical-protein-[Thalassoglobus-polymorphus] |
| MCA9067328.1 | REase | GCA_020426385.1 | Planctomycetaceae_bacterium | Bacteria>Pseudomonadati>PVC_group>Planctomycetota>Planctomycetia>Planctomycetales>Planctomycetaceae>unclassified_Planctomycetaceae | 194 | KDA84_00290 | 5005..5589 | 2026779 | hypothetical-protein-[Planctomycetaceae-bacterium] |

| Protein-ID | Domain | Assembly-ID | Strain | Taxonomy | Length | Locus-Tag | Genomic-coordinates | TaxID | Fasta-defline |
|---|---|---|---|---|---|---|---|---|---|
| MBX3448930.1 | REase | GCA_019636285.1 | Planctomycetaceae_bacterium | Bacteria>Pseudomonadati>PVC_group>Planctomycetota>Planctomycetia>Planctomycetales>Planctomycetaceae>unclassified_Planctomycetaceae | 169 | KF777_05180 | 9957..10466 | 2026779 | hypothetical-protein-[Planctomycetaceae-bacterium] |
| GIX04292.1 | REase | GCA_026004855.1 | Planctomycetaceae_bacterium | Bacteria>Pseudomonadati>PVC_group>Planctomycetota>Planctomycetia>Planctomycetales>Planctomycetaceae>unclassified_Planctomycetaceae | 145 | KatS3mg114_0161 | 186303..186740 | 2026779 | MAG:-hypothetical-protein-KatS3mg114_0161-[Planctomycetaceae-bacterium] |
| MCA9067327.1 | REase | GCA_020426385.1 | Planctomycetaceae_bacterium | Bacteria>Pseudomonadati>PVC_group>Planctomycetota>Planctomycetia>Planctomycetales>Planctomycetaceae>unclassified_Planctomycetaceae | 229 | KDA84_00285 | 4035..4724 | 2026779 | hypothetical-protein-[Planctomycetaceae-bacterium] |
| MCB9950011.1 | REase | GCA_020638955.1 | Planctomycetaceae_bacterium | Bacteria>Pseudomonadati>PVC_group>Planctomycetota>Planctomycetia>Planctomycetales>Planctomycetaceae>unclassified_Planctomycetaceae | 201 | H6824_03430 | 871684..872289 | 2026779 | hypothetical-protein-[Planctomycetaceae-bacterium] |
| HBI42789.1 | REase | GCA_003508275.1 | Planctomycetales_bacterium | Bacteria>Pseudomonadati>PVC_group>Planctomycetota>Planctomycetia>Planctomycetales>unclassified_Planctomycetales | 169 | DDY78_08015 | 5409..5918 | 2053591 | hypothetical-protein-[Planctomycetales-bacterium] |
| MCA9231623.1 | REase | GCA_020427155.1 | Planctomycetales_bacterium | Bacteria>Pseudomonadati>PVC_group>Planctomycetota>Planctomycetia>Planctomycetales>unclassified_Planctomycetales | 104 | KDA57_13310 | 1..313 | 2053591 | hypothetical-protein-[Planctomycetales-bacterium] |
| MCA9231632.1 | REase | GCA_020427155.1 | Planctomycetales_bacterium | Bacteria>Pseudomonadati>PVC_group>Planctomycetota>Planctomycetia>Planctomycetales>unclassified_Planctomycetales | 166 | KDA57_13355 | 17329..17829 | 2053591 | DUF91-domain-containing-protein-[Planctomycetales-bacterium] |
| MBI2825940.1 | REase | GCA_016191115.1 | Planctomycetia_bacterium | Bacteria>Pseudomonadati>PVC_group>Planctomycetota>Planctomycetia>unclassified_Planctomycetia | 135 | HYX69_14750 | 231260..231667 | 2052181 | DNA-mismatch-endonuclease-Vsr-[Planctomycetia-bacterium] |
| MBI5364694.1 | REase | GCA_016220035.1 | Planctomycetota_bacterium | Bacteria>Pseudomonadati>PVC_group>Planctomycetota>unclassified_Planctomycetota | 197 | HZA53_16075 | 12201..12794 | 2026780 | hypothetical-protein-[Planctomycetota-bacterium] |
| MCY2928836.1 | REase | GCA_026693065.1 | Planctomycetota_bacterium | Bacteria>Pseudomonadati>PVC_group>Planctomycetota>unclassified_Planctomycetota | 196 | NTV86_04955 | 19634..20224 | 2026780 | MAG:-hypothetical-protein-NTV86_04955-[Planctomycetota-bacterium] |
| MCE9567304.1 | REase | GCA_021413735.1 | Planctomycetota_bacterium | Bacteria>Pseudomonadati>PVC_group>Planctomycetota>unclassified_Planctomycetota | 195 | K8U57_35330 | 36561..37148 | 2026780 | hypothetical-protein-[Planctomycetota-bacterium] |
| RIK66095.1 | REase | GCA_003576905.1 | Planctomycetota_bacterium | Bacteria>Pseudomonadati>PVC_group>Planctomycetota>unclassified_Planctomycetota | 195 | DCC65_10755 | 37717..38304 | 2026780 | MAG:-hypothetical-protein-DCC65_10755-[Planctomycetota-bacterium] |
| MBE7491677.1 | REase | GCA_015075515.1 | Planctomycetota_bacterium | Bacteria>Pseudomonadati>PVC_group>Planctomycetota>unclassified_Planctomycetota | 193 | HS108_07995 | 573790..574371 | 2026780 | hypothetical-protein-[Planctomycetota-bacterium] |
| MBN8597232.1 | REase | GCA_017303555.1 | Planctomycetota_bacterium | Bacteria>Pseudomonadati>PVC_group>Planctomycetota>unclassified_Planctomycetota | 190 | J0L78_06120 | 34956..35528 | 2026780 | hypothetical-protein-[Planctomycetota-bacterium] |
| MBN8644245.1 | REase | GCA_017303565.1 | Planctomycetota_bacterium | Bacteria>Pseudomonadati>PVC_group>Planctomycetota>unclassified_Planctomycetota | 183 | J0L61_03270 | 7202..7753 | 2026780 | hypothetical-protein-[Planctomycetota-bacterium] |
| MBI1914949.1 | REase | GCA_016177925.1 | Planctomycetota_bacterium | Bacteria>Pseudomonadati>PVC_group>Planctomycetota>unclassified_Planctomycetota | 145 | HYS12_09470 | 8243..8680 | 2026780 | hypothetical-protein-[Planctomycetota-bacterium] |
| MCY2978567.1 | REase | GCA_026692945.1 | Planctomycetota_bacterium | Bacteria>Pseudomonadati>PVC_group>Planctomycetota>unclassified_Planctomycetota | 182 | NTU79_07890 | 12947..13495 | 2026780 | MAG:-hypothetical-protein-NTU79_07890-[Planctomycetota-bacterium] |
| MCA8917197.1 | REase | GCA_020425605.1 | Planctomycetota_bacterium | Bacteria>Pseudomonadati>PVC_group>Planctomycetota>unclassified_Planctomycetota | 170 | KDB90_17535 | 6813..7325 | 2026780 | hypothetical-protein-[Planctomycetota-bacterium] |
| MBP7751893.1 | REase | GCA_018000135.1 | Planctomycetota_bacterium | Bacteria>Pseudomonadati>PVC_group>Planctomycetota>unclassified_Planctomycetota | 136 | KA323_14490 | 49465..49875 | 2026780 | GxxExxY-protein-[Planctomycetota-bacterium] |
| NMD35174.1 | REase | GCA_012797405.1 | Planctomycetota_bacterium | Bacteria>Pseudomonadati>PVC_group>Planctomycetota>unclassified_Planctomycetota | 136 | GYA73_04755 | 3219..3629 | 2026780 | GxxExxY-protein-[Planctomycetota-bacterium] |
| MCY2991653.1 | REase | GCA_026692885.1 | Planctomycetota_bacterium | Bacteria>Pseudomonadati>PVC_group>Planctomycetota>unclassified_Planctomycetota | 125 | NTY19_27835 | 9513..9890 | 2026780 | MAG:-GxxExxY-protein-[Planctomycetota-bacterium] |
| MBI2191312.1 | REase | GCA_016179685.1 | Planctomycetota_bacterium | Bacteria>Pseudomonadati>PVC_group>Planctomycetota>unclassified_Planctomycetota | 123 | HYU36_04940 | 75087..75458 | 2026780 | GxxExxY-protein-[Planctomycetota-bacterium] |
| MCK6472709.1 | REase | GCA_023150555.1 | Planctomycetota_bacterium | Bacteria>Pseudomonadati>PVC_group>Planctomycetota>unclassified_Planctomycetota | 201 | L6R28_13275 | 14450..15055 | 2026780 | hypothetical-protein-[Planctomycetota-bacterium] |
| MCH8343170.1 | REase | GCA_022568135.1 | Planctomycetota_bacterium | Bacteria>Pseudomonadati>PVC_group>Planctomycetota>unclassified_Planctomycetota | 201 | IH983_04195 | 76878..77483 | 2026780 | hypothetical-protein-[Planctomycetota-bacterium] |
| WP_148205080.1 | REase | GCF_000019665.1 | Candidatus_Methylacidiphilum_infernorum | Bacteria>Pseudomonadati>PVC_group>Verrucomicrobiota>Methylacidiphilae>Methylacidiphilales>Methylacidiphilaceae>Methylacidiphilum_(ex_Ratnadevi_et_al._2023) | 119 | MINF_RS11085 | 328953..329312 | 511746 | hypothetical-protein-[Methylacidiphilum-infernorum] |
| RRJ96055.1 | REase | GCA_001653825.2 | Opitutaceae_bacterium_TAV4 | Bacteria>Pseudomonadati>PVC_group>Verrucomicrobiota>Opitutia>Opitutales>Opitutaceae>unclassified_Opitutaceae | 138 | Ga0100231_019085 | 2738561..2738977 | 278959 | DNA-mismatch-endonuclease-Vsr-[Opitutaceae-bacterium-TAV4] |
| RRK00200.1 | REase | GCA_001653815.2 | Opitutaceae_bacterium_TAV3 | Bacteria>Pseudomonadati>PVC_group>Verrucomicrobiota>Opitutia>Opitutales>Opitutaceae>unclassified_Opitutaceae | 138 | Ga0100230_019760 | 2820507..2820923 | 278958 | DNA-mismatch-endonuclease-Vsr-[Opitutaceae-bacterium-TAV3] |
| MCC5808696.1 | REase | GCA_020831525.1 | Opitutales_bacterium | Bacteria>Pseudomonadati>PVC_group>Verrucomicrobiota>Opitutia>Opitutales>unclassified_Opitutales | 168 | JJU00_20380 | 10620..11126 | 2026772 | hypothetical-protein-[Opitutales-bacterium] |
| MBE2203051.1 | REase | GCA_014879575.1 | Chthoniobacterales_bacterium | Bacteria>Pseudomonadati>PVC_group>Verrucomicrobiota>Spartobacteria>Chthoniobacterales>unclassified_Chthoniobacterales | 177 | IAE94_01750 | 16182..16715 | 2201154 | hypothetical-protein-[Chthoniobacterales-bacterium] |
| HCF96072.1 | REase | GCA_003534995.1 | Verrucomicrobiota_bacterium | Bacteria>Pseudomonadati>PVC_group>Verrucomicrobiota>unclassified_Verrucomicrobiota | 165 | DEW46_13520 | 4643..5140 | 2026799 | hypothetical-protein-[Verrucomicrobiota-bacterium] |
| MBI2925505.1 | REase | GCA_016190045.1 | Verrucomicrobiota_bacterium | Bacteria>Pseudomonadati>PVC_group>Verrucomicrobiota>unclassified_Verrucomicrobiota | 194 | HYY24_07355 | 11604..12188 | 2026799 | hypothetical-protein-[Verrucomicrobiota-bacterium] |
| MCX6880520.1 | REase | GCA_026397725.1 | Verrucomicrobiota_bacterium | Bacteria>Pseudomonadati>PVC_group>Verrucomicrobiota>unclassified_Verrucomicrobiota | 192 | NTW21_42950 | 2077..2655 | 2026799 | MAG:-hypothetical-protein-NTW21_42950-[Verrucomicrobia-bacterium] |
| MCX6879290.1 | REase | GCA_026397725.1 | Verrucomicrobiota_bacterium | Bacteria>Pseudomonadati>PVC_group>Verrucomicrobiota>unclassified_Verrucomicrobiota | 182 | NTW21_36625 | 12676..13224 | 2026799 | MAG:-hypothetical-protein-NTW21_36625-[Verrucomicrobia-bacterium] |
| MBM3880844.1 | REase | GCA_016871715.1 | Verrucomicrobiota_bacterium | Bacteria>Pseudomonadati>PVC_group>Verrucomicrobiota>unclassified_Verrucomicrobiota | 149 | FJ387_14195 | 23115..23564 | 2026799 | GxxExxY-protein-[Verrucomicrobiota-bacterium] |
| MBM3880409.1 | REase | GCA_016871715.1 | Verrucomicrobiota_bacterium | Bacteria>Pseudomonadati>PVC_group>Verrucomicrobiota>unclassified_Verrucomicrobiota | 130 | FJ387_11945 | 20674..21066 | 2026799 | GxxExxY-protein-[Verrucomicrobiota-bacterium] |
| HCF94161.1 | REase | GCA_003534995.1 | Verrucomicrobiota_bacterium | Bacteria>Pseudomonadati>PVC_group>Verrucomicrobiota>unclassified_Verrucomicrobiota | 127 | DEW46_03800 | 15411..15794 | 2026799 | GxxExxY-protein-[Verrucomicrobiota-bacterium] |
| MCU0784166.1 | REase | GCA_025457745.1 | Verrucomicrobiota_bacterium | Bacteria>Pseudomonadati>PVC_group>Verrucomicrobiota>unclassified_Verrucomicrobiota | 212 | MUF81_09015 | 1..639 | 2026799 | DUF3883-domain-containing-protein-[Verrucomicrobiota-bacterium] |
| WP_205880900.1 | REase | GCF_011044775.1 | Limisphaera_ngatamarikiensis | Bacteria>Pseudomonadati>PVC_group>Verrucomicrobiota>Verrucomicrobiia>Limisphaerales>Limisphaeraceae>Limisphaera | 121 | G4L39_RS09175 | 328454..328819 | 1324935 | GxxExxY-protein-[Limisphaera-ngatamarikiensis] |
| MBX3732316.1 | REase | GCA_019634635.1 | Verrucomicrobiia_bacterium | Bacteria>Pseudomonadati>PVC_group>Verrucomicrobiota>Verrucomicrobiia>unclassified_Verrucomicrobiia | 130 | KF791_06945 | 141979..142371 | 2499141 | GxxExxY-protein-[Verrucomicrobiae-bacterium] |
| MCX8496478.1 | REase | GCA_026461385.1 | Akkermansiaceae_bacterium | Bacteria>Pseudomonadati>PVC_group>Verrucomicrobiota>Verrucomicrobiia>Verrucomicrobiales>Akkermansiaceae>unclassified_Akkermansiaceae | 248 | ORN51_09890 | 1..744 | 2562705 | MAG:-DUF2800-domain-containing-protein,-partial-[Akkermansiaceae-bacterium] |
| MBE7497141.1 | REase | GCA_015075365.1 | Verrucomicrobiaceae_bacterium | Bacteria>Pseudomonadati>PVC_group>Verrucomicrobiota>Verrucomicrobiia>Verrucomicrobiales>Verrucomicrobiaceae>unclassified_Verrucomicrobiaceae | 193 | HS117_19540 | 528896..529477 | 2026800 | hypothetical-protein-[Verrucomicrobiaceae-bacterium] |
| MCB1193038.1 | REase | GCA_020440835.1 | Leptospiraceae_bacterium | Bacteria>Pseudomonadati>Spirochaetota>Spirochaetia>Leptospirales>Leptospiraceae>unclassified_Leptospiraceae | 148 | KDK90_21520 | 1382..1828 | 2026757 | PD-(D/E)XK-nuclease-family-transposase-[Leptospiraceae-bacterium] |
| MCB1193034.1 | REase | GCA_020440835.1 | Leptospiraceae_bacterium | Bacteria>Pseudomonadati>Spirochaetota>Spirochaetia>Leptospirales>Leptospiraceae>unclassified_Leptospiraceae | 121 | KDK90_21500 | 42..407 | 2026757 | hypothetical-protein-[Leptospiraceae-bacterium] |
| MCB1193037.1 | REase | GCA_020440835.1 | Leptospiraceae_bacterium | Bacteria>Pseudomonadati>Spirochaetota>Spirochaetia>Leptospirales>Leptospiraceae>unclassified_Leptospiraceae | 69 | KDK90_21515 | 1189..1398 | 2026757 | hypothetical-protein-[Leptospiraceae-bacterium] |
| WP_114628931.1 | REase | GCF_013166645.2 | unclassified_Oceanispirochaeta | Bacteria>Pseudomonadati>Spirochaetota>Spirochaetia>Spirochaetales>Spirochaetaceae>Oceanispirochaeta | 166 | HNV09_RS06060 | 182960..183460 | 2635722 | MULTISPECIES:-hypothetical-protein-[unclassified-Oceanispirochaeta] |
| OQY41123.1 | REase | GCA_002084785.1 | Spirochaetaceae_bacterium_4572_7 | Bacteria>Pseudomonadati>Spirochaetota>Spirochaetia>Spirochaetales>Spirochaetaceae>unclassified_Spirochaetaceae | 61 | B6229_00005 | 4..189 | 1972456 | hypothetical-protein-B6229_00005-[Spirochaetaceae-bacterium-4572_7] |
| KPA15640.1 | REase | GCA_001292585.1 | Candidatus_Magnetomorum_sp._HK-1 | Bacteria>Pseudomonadati>Thermodesulfobacteriota>Desulfobacteria>Desulfobacterales>Candidatus_Magnetomoraceae>Candidatus_Magnetomorum>unclassified_Candidatus_Magnetomorum | 128 | MHK_004159 | 12891..13277 | 1509431 | protein-containing-DUF1703-[Candidatus-Magnetomorum-sp.-HK-1] |
| WP_051328026.1 | REase | GCF_000429925.1 | Desulfatirhabdium_butyrativorans | Bacteria>Pseudomonadati>Thermodesulfobacteriota>Desulfobacteria>Desulfobacterales>Desulfatirhabdiaceae>Desulfatirhabdium | 135 | G492_RS23695 | 88565..88972 | 340467 | hypothetical-protein-[Desulfatirhabdium-butyrativorans] |
| WP_028324797.1 | REase | GCF_000429925.1 | Desulfatirhabdium_butyrativorans | Bacteria>Pseudomonadati>Thermodesulfobacteriota>Desulfobacteria>Desulfobacterales>Desulfatirhabdiaceae>Desulfatirhabdium | 183 | G492_RS0111990 | 50167..50718 | 340467 | hypothetical-protein-[Desulfatirhabdium-butyrativorans] |
| MBF0232700.1 | REase | GCA_015231525.1 | Desulfamplus_sp. | Bacteria>Pseudomonadati>Thermodesulfobacteriota>Desulfobacteria>Desulfobacterales>Desulfobacteraceae>Desulfamplus>unclassified_Desulfamplus | 167 | HQK65_06635 | 2918..3421 | 2006184 | hypothetical-protein-[Desulfamplus-sp.] |
| MBF0572397.1 | REase | GCA_015233755.1 | Desulfamplus_sp. | Bacteria>Pseudomonadati>Thermodesulfobacteriota>Desulfobacteria>Desulfobacterales>Desulfobacteraceae>Desulfamplus>unclassified_Desulfamplus | 206 | HQK69_01360 | 16189..16809 | 2006184 | Uma2-family-endonuclease-[Desulfamplus-sp.] |
| MBF0302147.1 | REase | GCA_015232035.1 | Desulfamplus_sp. | Bacteria>Pseudomonadati>Thermodesulfobacteriota>Desulfobacteria>Desulfobacterales>Desulfobacteraceae>Desulfamplus>unclassified_Desulfamplus | 184 | HQK73_03835 | 13914..14468 | 2006184 | hypothetical-protein-[Desulfamplus-sp.] |
| NDY73875.1 | REase | GCA_010499545.1 | Desulfobacter_hydrogenophilus | Bacteria>Pseudomonadati>Thermodesulfobacteriota>Desulfobacteria>Desulfobacterales>Desulfobacteraceae>Desulfobacter | 179 | G3N28_17840 | 22481..23020 | 2291 | hypothetical-protein-[Desulfobacter-hydrogenophilus] |
| WP_111959624.1 | REase | GCF_004319545.1 | Desulfobacter_hydrogenophilus | Bacteria>Pseudomonadati>Thermodesulfobacteriota>Desulfobacteria>Desulfobacterales>Desulfobacteraceae>Desulfobacter | 179 | DO021_RS19085 | 10235..10774 | 2291 | hypothetical-protein-[Desulfobacter-hydrogenophilus] |
| WP_035236112.1 | REase | GCF_000745975.1 | Desulfobacter_vibrioformis | Bacteria>Pseudomonadati>Thermodesulfobacteriota>Desulfobacteria>Desulfobacterales>Desulfobacteraceae>Desulfobacter | 165 | Q366_RS02905 | 252321..252818 | 34031 | hypothetical-protein-[Desulfobacter-vibrioformis] |
| WP_178366470.1 | REase | GCF_013389365.1 | Desulfobacter_latus | Bacteria>Pseudomonadati>Thermodesulfobacteriota>Desulfobacteria>Desulfobacterales>Desulfobacteraceae>Desulfobacter | 172 | HXW94_RS08510 | 35549..36067 | 2292 | hypothetical-protein-[Desulfobacter-latus] |
| MCD4719520.1 | REase | GCA_021107415.1 | Desulfobacula_sp. | Bacteria>Pseudomonadati>Thermodesulfobacteriota>Desulfobacteria>Desulfobacterales>Desulfobacteraceae>Desulfobacula>unclassified_Desulfobacula | 160 | K8S13_06625 | 7471..7953 | 2593537 | hypothetical-protein-[Desulfobacula-sp.] |
| MBL7173049.1 | REase | GCA_016782905.1 | Desulfobacteraceae_bacterium | Bacteria>Pseudomonadati>Thermodesulfobacteriota>Desulfobacteria>Desulfobacterales>Desulfobacteraceae>unclassified_Desulfobacteraceae | 161 | ISS68_09450 | 10460..10945 | 2049433 | hypothetical-protein-[Desulfobacteraceae-bacterium] |
| MBC8432174.1 | REase | GCA_014383215.1 | Candidatus_Desulfatibia_vada | Bacteria>Pseudomonadati>Thermodesulfobacteriota>Desulfobacteria>Desulfobacterales>Desulfobacterales_incertae_sedis>Candidatus_Desulfatibia | 193 | H8D96_09650 | 5808..6389 | 2841696 | hypothetical-protein-[Desulfobacterales-bacterium] |
| CBX26792.1 | REase | - | uncultured_Desulfobacterium_sp. | Bacteria>Pseudomonadati>Thermodesulfobacteriota>Desulfobacteria>Desulfobacterales>Desulfobacteriaceae>Desulfobacterium>environmental_samples | 98 | N47_A08210 | 97151..97447 | 201089 | hypothetical-protein-N47_A08210-[uncultured-Desulfobacterium-sp.] |
| CBX26793.1 | REase | - | uncultured_Desulfobacterium_sp. | Bacteria>Pseudomonadati>Thermodesulfobacteriota>Desulfobacteria>Desulfobacterales>Desulfobacteriaceae>Desulfobacterium>environmental_samples | 64 | N47_A08220 | 97440..97634 | 201089 | hypothetical-protein-N47_A08220-[uncultured-Desulfobacterium-sp.] |
| MBC8421199.1 | REase | GCA_014382905.1 | Desulfobacterales_bacterium | Bacteria>Pseudomonadati>Thermodesulfobacteriota>Desulfobacteria>Desulfobacterales>unclassified_Desulfobacterales | 161 | H8E10_21690 | 9136..9621 | 2044940 | hypothetical-protein-[Desulfobacterales-bacterium] |
| MDD2316832.1 | REase | GCA_028683495.1 | Desulfobacterales_bacterium | Bacteria>Pseudomonadati>Thermodesulfobacteriota>Desulfobacteria>Desulfobacterales>unclassified_Desulfobacterales | 185 | PHD57_05740 | 7106..7663 | 2044940 | MAG:-hypothetical-protein-PHD57_05740-[Desulfobacterales-bacterium] |
| MDD4464445.1 | REase | GCA_028705335.1 | Desulfobacterales_bacterium | Bacteria>Pseudomonadati>Thermodesulfobacteriota>Desulfobacteria>Desulfobacterales>unclassified_Desulfobacterales | 185 | PHR86_09435 | 711..1268 | 2044940 | MAG:-hypothetical-protein-PHR86_09435-[Desulfobacterales-bacterium] |
| WP_240193566.1 | REase | GCF_016918545.1 | Desulfobulbus_alkaliphilus | Bacteria>Pseudomonadati>Thermodesulfobacteriota>Desulfobulbia>Desulfobulbales>Desulfobulbaceae>Desulfobulbus | 127 | JWG43_RS12485 | 41365..41748 | 869814 | GxxExxY-protein-[Desulfobulbus-alkaliphilus] |
| WP_205243276.1 | REase | GCF_016918545.1 | Desulfobulbus_alkaliphilus | Bacteria>Pseudomonadati>Thermodesulfobacteriota>Desulfobulbia>Desulfobulbales>Desulfobulbaceae>Desulfobulbus | 126 | JWG43_RS12515 | 53319..53699 | 869814 | GxxExxY-protein-[Desulfobulbus-alkaliphilus] |
| WP_205228198.1 | REase | GCF_016919065.1 | Desulfobulbus_rhabdoformis | Bacteria>Pseudomonadati>Thermodesulfobacteriota>Desulfobulbia>Desulfobulbales>Desulfobulbaceae>Desulfobulbus | 183 | JWJ90_RS15880 | 28266..28817 | 34032 | hypothetical-protein-[Desulfobulbus-rhabdoformis] |
| KGO34777.1 | REase | GCA_000769715.1 | Desulfobulbus_sp._Tol-SR | Bacteria>Pseudomonadati>Thermodesulfobacteriota>Desulfobulbia>Desulfobulbales>Desulfobulbaceae>Desulfobulbus>unclassified_Desulfobulbus | 175 | JT06_05905 | 31048..31575 | 1536652 | hypothetical-protein-JT06_05905-[Desulfobulbus-sp.-Tol-SR] |
| WP_028584969.1 | REase | GCF_000429965.1 | Desulfogranum_mediterraneum | Bacteria>Pseudomonadati>Thermodesulfobacteriota>Desulfobulbia>Desulfobulbales>Desulfobulbaceae>Desulfogranum | 166 | G494_RS0113495 | 10130..10630 | 160661 | hypothetical-protein-[Desulfogranum-mediterraneum] |
| MBA3007529.1 | REase | GCA_013792135.1 | Desulfocapsa_sp. | Bacteria>Pseudomonadati>Thermodesulfobacteriota>Desulfobulbia>Desulfobulbales>Desulfocapsaceae>Desulfocapsa>unclassified_Desulfocapsa | 175 | FP810_13335 | 60277..60804 | 2066021 | hypothetical-protein-[Desulfocapsa-sp.] |
| HBH32879.1 | REase | GCA_003499505.1 | Desulfofustis_sp. | Bacteria>Pseudomonadati>Thermodesulfobacteriota>Desulfobulbia>Desulfobulbales>Desulfocapsaceae>Desulfofustis>unclassified_Desulfofustis | 112 | DDX81_13460 | 4975..5313 | 2053546 | GxxExxY-protein-[Desulfofustis-sp.] |
| MBL0380704.1 | REase | GCA_016722945.1 | Desulfofustis_sp._PB-SRB1 | Bacteria>Pseudomonadati>Thermodesulfobacteriota>Desulfobulbia>Desulfobulbales>Desulfocapsaceae>Desulfofustis>unclassified_Desulfofustis | 112 | JKP90_13805 | 379359..379697 | 1385624 | GxxExxY-protein-[Desulfofustis-sp.-PB-SRB1] |
| WP_205217415.1 | REase | GCF_016918505.1 | Desulforhopalus_vacuolatus | Bacteria>Pseudomonadati>Thermodesulfobacteriota>Desulfobulbia>Desulfobulbales>Desulfocapsaceae>Desulforhopalus | 75 | JWG39_RS00660 | 161909..162136 | 40414 | GxxExxY-protein-[Desulforhopalus-vacuolatus] |
| TKB07806.1 | REase | GCA_005116575.1 | Desulforhopalus_sp._IMCC35007 | Bacteria>Pseudomonadati>Thermodesulfobacteriota>Desulfobulbia>Desulfobulbales>Desulfocapsaceae>Desulforhopalus>unclassified_Desulforhopalus | 179 | FCL48_15765 | 78684..79223 | 2569543 | hypothetical-protein-FCL48_15765-[Desulforhopalus-sp.-IMCC35007] |
| WP_136873372.1 | REase | GCF_005116575.1 | Desulforhopalus_sp._IMCC35007 | Bacteria>Pseudomonadati>Thermodesulfobacteriota>Desulfobulbia>Desulfobulbales>Desulfocapsaceae>Desulforhopalus>unclassified_Desulforhopalus | 179 | FCL48_RS15770 | 78684..79223 | 2569543 | hypothetical-protein-[Desulforhopalus-sp.-IMCC35007] |
| WP_027370875.1 | REase | GCF_000620765.1 | Desulfovermiculus_halophilus | Bacteria>Pseudomonadati>Thermodesulfobacteriota>Desulfovibrionia>Desulfovibrionales>Desulfohalobiaceae>Desulfovermiculus | 123 | N902_RS0110330 | 25931..26302 | 339722 | GxxExxY-protein-[Desulfovermiculus-halophilus] |
| WP_168935606.1 | REase | GCF_012843875.1 | Desulfovibrio_piger | Bacteria>Pseudomonadati>Thermodesulfobacteriota>Desulfovibrionia>Desulfovibrionales>Desulfovibrionaceae>Desulfovibrio | 172 | HF854_RS06730 | 20320..20838 | 901 | hypothetical-protein-[Desulfovibrio-piger] |
| WP_053062253.1 | REase | GCF_000801335.2 | Desulfovibrio_sp._TomC | Bacteria>Pseudomonadati>Thermodesulfobacteriota>Desulfovibrionia>Desulfovibrionales>Desulfovibrionaceae>Desulfovibrio>unclassified_Desulfovibrio | 177 | NY78_RS16935 | 100081..100614 | 1562888 | hypothetical-protein-[Desulfovibrio-sp.-TomC] |
| MBQ9405900.1 | REase | GCA_017454855.1 | Desulfovibrio_sp. | Bacteria>Pseudomonadati>Thermodesulfobacteriota>Desulfovibrionia>Desulfovibrionales>Desulfovibrionaceae>Desulfovibrio>unclassified_Desulfovibrio | 106 | IJU37_04060 | 8069..8389 | 885 | hypothetical-protein-[Desulfovibrio-sp.] |
| WP_010939305.1 | REase | GCF_000195755.1 | Nitratidesulfovibrio_vulgaris | Bacteria>Pseudomonadati>Thermodesulfobacteriota>Desulfovibrionia>Desulfovibrionales>Desulfovibrionaceae>Nitratidesulfovibrio | 179 | DVU_RS09555 | 2104703..2105242 | 881 | hypothetical-protein-[Desulfovibrio-vulgaris] |
| WP_035067988.1 | REase | GCF_000504305.1 | Nitratidesulfovibrio_termitidis | Bacteria>Pseudomonadati>Thermodesulfobacteriota>Desulfovibrionia>Desulfovibrionales>Desulfovibrionaceae>Nitratidesulfovibrio | 179 | DESTE_RS12440 | 3094211..3094750 | 42252 | hypothetical-protein-[Cupidesulfovibrio-termitidis] |
| WP_010939303.1 | REase | GCF_000195755.1 | Nitratidesulfovibrio_vulgaris | Bacteria>Pseudomonadati>Thermodesulfobacteriota>Desulfovibrionia>Desulfovibrionales>Desulfovibrionaceae>Nitratidesulfovibrio | 155 | DVU_RS09545 | 2100755..2101222 | 881 | GxxExxY-protein-[Desulfovibrio-vulgaris] |
| AAS96498.1 | REase | GCA_000195755.1 | Nitratidesulfovibrio_vulgaris_str._Hildenborough | Bacteria>Pseudomonadati>Thermodesulfobacteriota>Desulfovibrionia>Desulfovibrionales>Desulfovibrionaceae>Nitratidesulfovibrio>Nitratidesulfovibrio_vulgaris | 179 | DVU_2023 | 2104703..2105242 | 882 | hypothetical-protein-DVU_2023-[Desulfovibrio-vulgaris-str.-Hildenborough] |
| AAS96496.1 | REase | GCA_000195755.1 | Nitratidesulfovibrio_vulgaris_str._Hildenborough | Bacteria>Pseudomonadati>Thermodesulfobacteriota>Desulfovibrionia>Desulfovibrionales>Desulfovibrionaceae>Nitratidesulfovibrio>Nitratidesulfovibrio_vulgaris | 155 | DVU_2021 | 2100755..2101222 | 882 | hypothetical-protein-DVU_2021-[Desulfovibrio-vulgaris-str.-Hildenborough] |
| MCH5276202.1 | REase | GCA_022510785.1 | Desulfovibrionaceae_bacterium | Bacteria>Pseudomonadati>Thermodesulfobacteriota>Desulfovibrionia>Desulfovibrionales>Desulfovibrionaceae>unclassified_Desulfovibrionaceae | 171 | J1E80_00035 | 13218..13733 | 2049043 | hypothetical-protein-[Desulfovibrionaceae-bacterium] |
| MBE0574419.1 | REase | GCA_014860225.1 | Desulfuromonadales_bacterium | Bacteria>Pseudomonadati>Thermodesulfobacteriota>Desulfuromonadia>Desulfuromonadales>unclassified_Desulfuromonadales | 179 | IH613_00825 | 76098..76637 | 2099678 | hypothetical-protein-[Desulfuromonadales-bacterium] |
| MBW6509588.1 | REase | GCA_019429255.1 | Desulfuromonadales_bacterium | Bacteria>Pseudomonadati>Thermodesulfobacteriota>Desulfuromonadia>Desulfuromonadales>unclassified_Desulfuromonadales | 179 | K0A94_08620 | 49870..50409 | 2099678 | hypothetical-protein-[Desulfuromonadales-bacterium] |
| MBW6509590.1 | REase | GCA_019429255.1 | Desulfuromonadales_bacterium | Bacteria>Pseudomonadati>Thermodesulfobacteriota>Desulfuromonadia>Desulfuromonadales>unclassified_Desulfuromonadales | 80 | K0A94_08630 | 54192..54432 | 2099678 | GxxExxY-protein-[Desulfuromonadales-bacterium] |
| WP_207162789.1 | REase | GCF_017338855.1 | Geobacter_benzoatilyticus | Bacteria>Pseudomonadati>Thermodesulfobacteriota>Desulfuromonadia>Geobacterales>Geobacteraceae>Geobacter | 176 | JZM60_RS12600 | 2739837..2740367 | 2815309 | hypothetical-protein-[Geobacter-benzoatilyticus] |
| MBJ6802711.1 | REase | GCA_016458235.1 | Geomonas_propionica | Bacteria>Pseudomonadati>Thermodesulfobacteriota>Desulfuromonadia>Geobacterales>Geobacteraceae>Geomonas | 176 | JFN90_21485 | 7311..7841 | 2798582 | hypothetical-protein-[Geomonas-propionica] |
| WP_199397181.1 | REase | GCF_016458235.1 | Geomonas_propionica | Bacteria>Pseudomonadati>Thermodesulfobacteriota>Desulfuromonadia>Geobacterales>Geobacteraceae>Geomonas | 176 | JFN90_RS21485 | 7311..7841 | 2798582 | hypothetical-protein-[Geomonas-propionica] |
| WP_258932063.1 | REase | GCF_022603395.1 | Desulfoferrobacter_suflitae | Bacteria>Pseudomonadati>Thermodesulfobacteriota>Syntrophobacteria>Syntrophobacterales>Syntrophobacteraceae>Desulfoferrobacter | 134 | K1V27_RS20435 | 53351..53755 | 2865782 | GxxExxY-protein-[Desulfoferrobacter-suflitae] |
| NSW87842.1 | REase | GCA_013314815.1 | Syntrophobacteraceae_bacterium | Bacteria>Pseudomonadati>Thermodesulfobacteriota>Syntrophobacteria>Syntrophobacterales>Syntrophobacteraceae>unclassified_Syntrophobacteraceae | 138 | HPY84_16155 | 5756..6172 | 2052185 | DUF3883-domain-containing-protein-[Syntrophobacteraceae-bacterium] |
| WP_146889217.1 | REase | GCF_007990775.1 | Deinococcus_cellulosilyticus | Bacteria>Thermotogati>Deinococcota>Deinococci>Deinococcales>Deinococcaceae>Deinococcus | 173 | DC3_RS23605 | 32435..32956 | 401558 | hypothetical-protein-[Deinococcus-cellulosilyticus] |
| WP_189059187.1 | REase | GCF_014647175.1 | Deinococcus_daejeonensis | Bacteria>Thermotogati>Deinococcota>Deinococci>Deinococcales>Deinococcaceae>Deinococcus | 175 | IEY59_RS18165 | 26347..26874 | 1007098 | hypothetical-protein-[Deinococcus-daejeonensis] |
| WP_268906348.1 | REase | - | Deinococcus_aquaedulcis | Bacteria>Thermotogati>Deinococcota>Deinococci>Deinococcales>Deinococcaceae>Deinococcus | 177 | KMW22_RS19495 | 3294..3827 | 2840455 | restriction-endonuclease-[Deinococcus-aquaedulcis] |
| WP_126188141.1 | REase | GCF_003958985.1 | Thermus_scotoductus | Bacteria>Thermotogati>Deinococcota>Deinococci>Thermales>Thermaceae>Thermus | 194 | CSW50_RS14440 | 7734..8318 | 37636 | hypothetical-protein-[Thermus-scotoductus] |
| GHV28798.1 | REase | GCA_026002695.1 | Synergistales_bacterium | Bacteria>Thermotogati>Synergistota>Synergistia>Synergistales>unclassified_Synergistales | 153 | FACS1894167_06810 | 13617..14078 | 2268201 | hypothetical-protein-FACS1894167_06810-[Synergistales-bacterium] |
| CCU85344.1 | REase | GCA_000367705.1 | Mesotoga_infera | Bacteria>Thermotogati>Thermotogota>Thermotogae>Kosmotogales>Kosmotogaceae>Mesotoga | 220 | PHOSAC3_150344 | 373487..374149 | 1236046 | exported-hypothetical-protein-[Mesotoga-infera] |
| WP_121550381.1 | REase | GCF_003664995.1 | Mesotoga_sp._BH458_6_3_2_1 | Bacteria>Thermotogati>Thermotogota>Thermotogae>Kosmotogales>Kosmotogaceae>Mesotoga>unclassified_Mesotoga | 220 | Y697_RS03815 | 17670..18332 | 1437446 | hypothetical-protein-[Mesotoga-sp.-BH458_6_3_2_1] |
| WP_206075997.1 | REase | GCF_004135675.1 | Marinitoga_lauensis | Bacteria>Thermotogati>Thermotogota>Thermotogae>Petrotogales>Petrotogaceae>Marinitoga | 98 | DJ007_RS15165 | 245124..245420 | 2201189 | winged-helix-turn-helix-domain-containing-protein-[Marinitoga-lauensis] |
| MCD6238357.1 | REase | GCA_021158015.1 | Thermotogota_bacterium | Bacteria>Thermotogati>Thermotogota>unclassified_Thermotogota | 223 | J7K51_03360 | 3196..3867 | 2053689 | hypothetical-protein-[Thermotogota-bacterium] |
| MCR9296474.1 | REase | GCA_024743355.1 | bacterium | Bacteria>unclassified_Bacteria | 109 | NXI32_27505 | 4793..5123 | 1869227 | hypothetical-protein-[bacterium] |
| KAF0250484.1 | REase | GCA_009773835.1 | bacterium | Bacteria>unclassified_Bacteria | 203 | FD167_110 | 8624..9235 | 1869227 | hypothetical-protein-FD167_110-[bacterium] |
| MCY4630091.1 | REase | GCA_026713745.1 | bacterium | Bacteria>unclassified_Bacteria | 233 | OXE75_04290 | 1..703 | 1869227 | MAG:-DUF3883-domain-containing-protein,-partial-[bacterium] |
| MCE5247404.1 | REase | GCA_021372775.1 | bacterium | Bacteria>unclassified_Bacteria | 139 | LLG88_10875 | 5076..5495 | 1869227 | YraN-family-protein-[bacterium] |
| MCY4630099.1 | REase | GCA_026713745.1 | bacterium | Bacteria>unclassified_Bacteria | 198 | OXE75_04330 | 10057..10653 | 1869227 | MAG:-DUF4143-domain-containing-protein-[bacterium] |
| MBS2037454.1 | REase | GCA_018266105.1 | bacterium | Bacteria>unclassified_Bacteria | 186 | JST97_20870 | 13873..14433 | 1869227 | Rpn-family-recombination-promoting-nuclease/putative-transposase-[bacterium] |
| MBS2037453.1 | REase | GCA_018266105.1 | bacterium | Bacteria>unclassified_Bacteria | 95 | JST97_20865 | 13589..13876 | 1869227 | hypothetical-protein-[bacterium] |
| --SF1-SF2-Helicase+REase | | | | | | | | | |
| MBR3112540.1 | SF2-Helicase+REase | GCA_017545145.1 | Methanobrevibacter_sp. | Archaea>Methanobacteriati>Methanobacteriota>Methanomada_group>Methanobacteria>Methanobacteriales>Methanobacteriaceae>Methanobrevibacter>unclassified_Methanobrevibacter | 1002 | IKH29_02360 | 41440..44448 | 66852 | ATP-dependent-helicase-[Methanobrevibacter-sp.] |
| WP_122106901.1 | SF2-Helicase+REase | GCF_042657325.1 | Haloferacaceae | Archaea>Methanobacteriati>Methanobacteriota>Stenosarchaea_group>Halobacteria>Halobacteriales | 1212 | EAO78_RS16640 | 8559..12197 | 1644056 | UvrD-helicase-domain-containing-protein-[Halobellus-captivus] |
| WP_188997381.1 | SF2-Helicase+REase | GCF_014647235.1 | Haloarcula_pellucida | Archaea>Methanobacteriati>Methanobacteriota>Stenosarchaea_group>Halobacteria>Halobacteriales>Haloarculaceae>Haloarcula | 1212 | EGH26_RS17035 | 310188..313826 | 1427151 | UvrD-helicase-domain-containing-protein-[Halomicroarcula-pellucida] |

| Protein-ID | Domain | Assembly-ID | Strain | Taxonomy | Length | Locus-Tag | Genomic-coordinates | TaxID | Fasta-defline |
|---|---|---|---|---|---|---|---|---|---|
| WP_121538357.1 | SF2-Helicase+REase | GCF_003665885.1 | Haloarcula_sp._Atlit-120R | Archaea>Methanobacteriati>Methanobacteriota>Stenosarchaea_group>Halobacteria>Halobacteriales>Haloarculaceae>Haloarcula>unclassified_Haloarcula | 1212 | DVK01_RS19365 | 50913..54551 | 2282135 | UvrD-helicase-domain-containing-protein-[Haloarcula-sp.-Atlit-120R] |
| WP_247995976.1 | SF2-Helicase+REase | GCF_023115355.1 | Halocatena_salina | Archaea>Methanobacteriati>Methanobacteriota>Stenosarchaea_group>Halobacteria>Halobacteriales>Haloarculaceae>Halocatena | 1213 | MW046_RS18830 | 29790..33431 | 2934340 | UvrD-helicase-domain-containing-protein-[Halocatena-salina] |
| WP_247994820.1 | SF2-Helicase+REase | GCF_023115355.1 | Halocatena_salina | Archaea>Methanobacteriati>Methanobacteriota>Stenosarchaea_group>Halobacteria>Halobacteriales>Haloarculaceae>Halocatena | 954 | MW046_RS14190 | 274551..277415 | 2934340 | UvrD-helicase-domain-containing-protein-[Halocatena-salina] |
| WP_158203515.1 | SF2-Helicase+REase | GCF_009791395.1 | Halomarina_oriensis | Archaea>Methanobacteriati>Methanobacteriota>Stenosarchaea_group>Halobacteria>Halobacteriales>Haloarculaceae>Halomarina | 1259 | GQS65_RS04680 | 72240..76019 | 671145 | UvrD-helicase-domain-containing-protein-[Halomarina-oriensis] |
| WP_254547352.1 | SF2-Helicase+REase | GCF_024228315.1 | Halomarina_pelagica | Archaea>Methanobacteriati>Methanobacteriota>Stenosarchaea_group>Halobacteria>Halobacteriales>Haloarculaceae>Halomarina | 953 | NKI68_RS22900 | 71540..74401 | 2961599 | ATP-dependent-DNA-helicase-[Halomarina-sp.-BND7] |
| WP_226023612.1 | SF2-Helicase+REase | GCF_020405245.1 | Halomicrobium_salinisoli | Archaea>Methanobacteriati>Methanobacteriota>Stenosarchaea_group>Halobacteria>Halobacteriales>Haloarculaceae>Halomicrobium | 1211 | LE161_RS18790 | 275149..278784 | 2878391 | UvrD-helicase-domain-containing-protein-[Halomicrobium-salinisoli] |
| WP_018258692.1 | SF2-Helicase+REase | GCF_000379085.1 | Halomicrobium_katesii | Archaea>Methanobacteriati>Methanobacteriota>Stenosarchaea_group>Halobacteria>Halobacteriales>Haloarculaceae>Halomicrobium | 959 | HALKA_RS0114590 | 2769353..2772232 | 437163 | ATP-dependent-DNA-helicase-[Halomicrobium-katesii] |
| WP_135666508.1 | SF2-Helicase+REase | GCF_900880625.1 | Halorhabdus_rudnickae | Archaea>Methanobacteriati>Methanobacteriota>Stenosarchaea_group>Halobacteria>Halobacteriales>Haloarculaceae>Halorhabdus | 960 | BN2694_RS13570 | 47033..49915 | 1775544 | ATP-dependent-DNA-helicase-[Halorhabdus-rudnickae] |
| WP_152120788.1 | SF2-Helicase+REase | GCF_009184545.1 | Halosegnis_rubeus | Archaea>Methanobacteriati>Methanobacteriota>Stenosarchaea_group>Halobacteria>Halobacteriales>Haloarculaceae>Halosegnis | 1211 | DMP03_RS11375 | 16693..20328 | 2212850 | UvrD-helicase-domain-containing-protein-[Halosegnis-rubeus] |
| WP_267638592.1 | SF2-Helicase+REase | GCF_026586665.1 | Halovenus_rubra | Archaea>Methanobacteriati>Methanobacteriota>Stenosarchaea_group>Halobacteria>Halobacteriales>Haloarculaceae>Halovenus | 1238 | N5098_RS14455 | 63426..67142 | 869890 | UvrD-helicase-domain-containing-protein-[Halovenus-rubra] |
| WP_058984829.1 | SF2-Helicase+REase | GCF_001485535.1 | Halobacterium | Archaea>Methanobacteriati>Methanobacteriota>Stenosarchaea_group>Halobacteria>Halobacteriales>Halobacteriaceae | 1211 | AVZ66_RS14260 | 133971..137606 | 2239 | MULTISPECIES:-UvrD-helicase-domain-containing-protein-[Halobacterium] |
| WP_123076492.1 | SF2-Helicase+REase | GCF_003721175.2 | Halarchaeum_sp._CBA1220 | Archaea>Methanobacteriati>Methanobacteriota>Stenosarchaea_group>Halobacteria>Halobacteriales>Halobacteriaceae>Halarchaeum>unclassified_Halarchaeum | 950 | EFA46_RS09105 | 1693693..1696545 | 1853682 | ATP-dependent-DNA-helicase-[Halarchaeum-sp.-CBA1220] |
| WP_232700620.1 | SF2-Helicase+REase | GCF_021249345.1 | Halobacterium_wangiae | Archaea>Methanobacteriati>Methanobacteriota>Stenosarchaea_group>Halobacteria>Halobacteriales>Halobacteriaceae>Halobacterium | 956 | LT965_RS09810 | 1840423..1843293 | 2902623 | ATP-dependent-DNA-helicase-[Halobacterium-sp.-Gai3-17] |
| WP_244705661.1 | SF2-Helicase+REase | GCF_039522505.1 | Halococcus_dombrowskii | Archaea>Methanobacteriati>Methanobacteriota>Stenosarchaea_group>Halobacteria>Halobacteriales>Halococcaceae>Halococcus | 967 | MUK72_RS16080 | 359799..362702 | 179637 | ATP-dependent-DNA-helicase-[Halococcus-dombrowskii] |
| WP_007538676.1 | SF2-Helicase+REase | - | Haloferax_larsenii | Archaea>Methanobacteriati>Methanobacteriota>Stenosarchaea_group>Halobacteria>Halobacteriales>Haloferacaceae>Haloferax | 1144 | C455_RS00130 | 35946..39380 | 302484 | ATP-dependent-DNA-helicase-[Haloferax-larsenii] |
| ELZ84494.1 | SF2-Helicase+REase | GCA_000336955.1 | Haloferax_larsenii_JCM_13917 | Archaea>Methanobacteriati>Methanobacteriota>Stenosarchaea_group>Halobacteria>Halobacteriales>Haloferacaceae>Haloferax>Haloferax_larsenii | 1144 | C455_00137 | 35946..39380 | 1227460 | UvrD/REP-helicase-[Haloferax-larsenii-JCM-13917] |
| TQQ78809.1 | SF2-Helicase+REase | GCA_006861665.1 | Halonotius_terrestris | Archaea>Methanobacteriati>Methanobacteriota>Stenosarchaea_group>Halobacteria>Halobacteriales>Haloferacaceae>Halonotius | 1212 | EGH24_12670 | 28499..32137 | 2487750 | helicase-[Halonotius-terrestris] |
| WP_142980500.1 | SF2-Helicase+REase | GCF_006861665.1 | Halonotius_terrestris | Archaea>Methanobacteriati>Methanobacteriota>Stenosarchaea_group>Halobacteria>Halobacteriales>Haloferacaceae>Halonotius | 1212 | EGH24_RS12675 | 28499..32137 | 2487750 | UvrD-helicase-domain-containing-protein-[Halonotius-terrestris] |
| WP_096390026.1 | SF2-Helicase+REase | GCF_002355635.1 | Halopenitus_persicus | Archaea>Methanobacteriati>Methanobacteriota>Stenosarchaea_group>Halobacteria>Halobacteriales>Haloferacaceae>Halopenitus | 1213 | CPZ00_RS05725 | 1188460..1192101 | 1048396 | UvrD-helicase-domain-containing-protein-[Halopenitus-persicus] |
| WP_099255987.1 | SF2-Helicase+REase | GCF_002727125.1 | Halorubrum_persicum | Archaea>Methanobacteriati>Methanobacteriota>Stenosarchaea_group>Halobacteria>Halobacteriales>Haloferacaceae>Halorubrum | 1179 | DJ69_RS12990 | 74136..77675 | 1383844 | UvrD-helicase-domain-containing-protein-[Halorubrum-persicum] |
| WP_271958990.1 | SF2-Helicase+REase | GCF_028211185.1 | Halorubrum_ezzemoulense | Archaea>Methanobacteriati>Methanobacteriota>Stenosarchaea_group>Halobacteria>Halobacteriales>Haloferacaceae>Halorubrum | 1165 | PM020_RS15265 | 30354..33851 | 337243 | UvrD-helicase-domain-containing-protein-[Halorubrum-ezzemoulense] |
| WP_014053205.1 | SF2-Helicase+REase | GCF_000022205.1 | Halorubrum_lacusprofundi | Archaea>Methanobacteriati>Methanobacteriota>Stenosarchaea_group>Halobacteria>Halobacteriales>Haloferacaceae>Halorubrum | 948 | HLAC_RS15665 | 442652..445498 | 2247 | ATP-dependent-helicase-[Halorubrum-lacusprofundi] |
| WP_136591375.1 | SF2-Helicase+REase | GCF_004916935.1 | Salinigranum_halophilum | Archaea>Methanobacteriati>Methanobacteriota>Stenosarchaea_group>Halobacteria>Halobacteriales>Haloferacaceae>Salinigranum | 945 | E6C73_RS15440 | 218028..220865 | 2565931 | UvrD-helicase-domain-containing-protein-[Salinigranum-halophilum] |
| WP_256532743.1 | SF2-Helicase+REase | GCF_024498195.1 | Halovivax_cerinus | Archaea>Methanobacteriati>Methanobacteriota>Stenosarchaea_group>Halobacteria>Halobacteriales>Natrialbaceae>Halovivax | 952 | NO366_RS02530 | 527467..530325 | 1487865 | ATP-dependent-DNA-helicase-[Halovivax-cerinus] |
| QCC61763.1 | SF2-Helicase+REase | GCA_004799625.1 | Natrinema_thermotolerans | Archaea>Methanobacteriati>Methanobacteriota>Stenosarchaea_group>Halobacteria>Halobacteriales>Natrialbaceae>Natrinema | 1210 | DVR14_24795 | 913649..917281 | 121872 | helicase-[Natrinema-thermotolerans] |
| WP_254767386.1 | SF2-Helicase+REase | GCF_024296665.1 | Salinilacihabitans_rarus | Archaea>Methanobacteriati>Methanobacteriota>Stenosarchaea_group>Halobacteria>Halobacteriales>Natrialbaceae>Salinilacihabitans | 959 | NKG98_RS17350 | 3305151..3308030 | 2961596 | ATP-dependent-DNA-helicase-[Natribaculum-sp.-AD-4] |
| WP_049893917.1 | SF2-Helicase+REase | GCF_000403645.1 | Salinarchaeum_sp._Harcht-Bsk1 | Archaea>Methanobacteriati>Methanobacteriota>Stenosarchaea_group>Halobacteria>Halobacteriales>Natronoarchaeaceae>Salinarchaeum>unclassified_Salinarchaeum | 982 | L593_RS06280 | 1362444..1365392 | 1333523 | UvrD-helicase-domain-containing-protein-[Salinarchaeum-sp.-Harcht-Bsk1] |
| AEN07512.1 | SF2-Helicase+REase | GCA_000224475.1 | halophilic_archaeon_DL31 | Archaea>Methanobacteriati>Methanobacteriota>Stenosarchaea_group>Halobacteria>Halobacteriales>unclassified_Halobacteriales | 948 | Halar_0252 | 242608..245454 | 756883 | UvrD/REP-helicase-[halophilic-archaeon-DL31] |
| WP_270096520.1 | SF2-Helicase+REase | GCF_027594145.1 | Methanococcoides_alaskense | Archaea>Methanobacteriati>Methanobacteriota>Stenosarchaea_group>Methanomicrobia>Methanosarcinales>Methanosarcinaceae>Methanococcoides | 1020 | PBV53_RS07445 | 717252..720314 | 325778 | UvrD-helicase-domain-containing-protein-[Methanococcoides-alaskense] |
| MDD3564057.1 | SF2-Helicase+REase | GCA_028695755.1 | Methanothrix_sp. | Archaea>Methanobacteriati>Methanobacteriota>Stenosarchaea_group>Methanomicrobia>Methanotrichales>Methanotrichaceae>Methanothrix>unclassified_Methanothrix | 996 | PHN90_00110 | 37395..40385 | 90426 | MAG:-UvrD-helicase-domain-containing-protein-[Methanothrix-sp.] |
| MBT4336381.1 | SF2-Helicase+REase | GCA_018652365.1 | Candidatus_Woesearchaeota_archaeon | Archaea>Nanobdellati>Candidatus_Woesearchaeota>unclassified_Candidatus_Woesearchaeota | 1058 | HOD66_03390 | 39076..42252 | 2026803 | ATP-dependent-helicase-[Candidatus-Woesearchaeota-archaeon] |
| MBT4469964.1 | SF2-Helicase+REase | GCA_018656325.1 | Candidatus_Woesearchaeota_archaeon | Archaea>Nanobdellati>Candidatus_Woesearchaeota>unclassified_Candidatus_Woesearchaeota | 1058 | HOC62_06270 | 47625..50801 | 2026803 | ATP-dependent-helicase-[Candidatus-Woesearchaeota-archaeon] |
| MBT6744312.1 | SF2-Helicase+REase | GCA_018696595.1 | Candidatus_Woesearchaeota_archaeon | Archaea>Nanobdellati>Candidatus_Woesearchaeota>unclassified_Candidatus_Woesearchaeota | 1058 | HOA70_03225 | 39076..42252 | 2026803 | ATP-dependent-helicase-[Candidatus-Woesearchaeota-archaeon] |
| MBT6646598.1 | SF2-Helicase+REase | GCA_018697615.1 | Nitrososphaerota_archaeon | Archaea>Thermoproteati>Nitrososphaerota>unclassified_Nitrososphaerota | 896 | HOB51_03660 | 1100..3790 | 2026795 | ATP-dependent-helicase-[Nitrososphaerota-archaeon] |
| WP_129664631.1 | SF2-Helicase+REase | GCF_004138495.1 | Phytoactinopolyspora_endophytica | Bacteria>Bacillati>Actinomycetota>Actinomycetes>Jiangellales>Jiangellaceae>Phytoactinopolyspora | 1063 | ELR45_RS07085 | 270471..273662 | 1642495 | UvrD-helicase-domain-containing-protein-[Phytoactinopolyspora-endophytica] |
| WP_055898782.1 | SF2-Helicase+REase | GCF_001425665.1 | Yonghaparkia_sp._Root332 | Bacteria>Bacillati>Actinomycetota>Actinomycetes>Micrococcales>Microbacteriaceae>Microcella>unclassified_Microcella | 1097 | ASC54_RS10005 | 492897..496190 | 1736516 | UvrD-helicase-domain-containing-protein-[Yonghaparkia-sp.-Root332] |
| OBB45423.1 | SF2-Helicase+REase | GCA_001665535.1 | Mycobacterium_sp._852002-51961_SCH5331710 | Bacteria>Bacillati>Actinomycetota>Actinomycetes>Mycobacteriales>Mycobacteriaceae>Mycobacterium>unclassified_Mycobacterium | 1104 | A5752_02045 | 28601..31915 | 1834105 | hypothetical-protein-A5752_02045-[Mycobacterium-sp.-852002-51961_SCH5331710] |
| WP_067083708.1 | SF2-Helicase+REase | GCF_001665535.1 | Mycobacterium_sp._852002-51961_SCH5331710 | Bacteria>Bacillati>Actinomycetota>Actinomycetes>Mycobacteriales>Mycobacteriaceae>Mycobacterium>unclassified_Mycobacterium | 1104 | A5752_RS07220 | 28601..31915 | 1834105 | UvrD-helicase-domain-containing-protein-[Mycobacterium-sp.-852002-51961_SCH5331710] |
| WP_235612276.1 | SF2-Helicase+REase | - | Metalysinibacillus_jejuensis | Bacteria>Bacillati>Bacillota>Bacilli>Bacillales>Caryophanaceae>Metalysinibacillus | 683 | DDE80_RS04480 | 52453..54504 | 914327 | ATP-dependent-helicase-[Metalysinibacillus-jejuensis] |
| WP_036689476.1 | SF2-Helicase+REase | GCF_000517925.1 | Paenibacillus | Bacteria>Bacillati>Bacillota>Bacilli>Bacillales>Paenibacillaceae | 1080 | C171_RS31990 | 157394..160636 | 44249 | MULTISPECIES:-UvrD-helicase-domain-containing-protein-[Paenibacillus] |
| WP_023554879.1 | SF2-Helicase+REase | GCF_000503775.1 | Brevibacillus_panacihumi | Bacteria>Bacillati>Bacillota>Bacilli>Bacillales>Paenibacillaceae>Brevibacillus | 1120 | T458_RS33505 | 835646..839008 | 497735 | UvrD-helicase-domain-containing-protein-[Brevibacillus-panacihumi] |
| WP_221842509.1 | SF2-Helicase+REase | GCF_019749055.1 | Brevibacillus_agri | Bacteria>Bacillati>Bacillota>Bacilli>Bacillales>Paenibacillaceae>Brevibacillus | 1117 | H7K32_RS04475 | 4379..7732 | 51101 | UvrD-helicase-domain-containing-protein-[Brevibacillus-agri] |
| WP_170184340.1 | SF2-Helicase+REase | GCF_006540045.1 | Brevibacillus_centrosporus | Bacteria>Bacillati>Bacillota>Bacilli>Bacillales>Paenibacillaceae>Brevibacillus | 1073 | BCE02nite_RS06975 | 47535..50756 | 54910 | UvrD-helicase-domain-containing-protein-[Brevibacillus-centrosporus] |
| WP_058302026.1 | SF2-Helicase+REase | GCF_902375625.1 | Gorillibacterium_timonense | Bacteria>Bacillati>Bacillota>Bacilli>Bacillales>Paenibacillaceae>Gorillibacterium | 1099 | BN2684_RS09990 | 491385..494684 | 1689269 | UvrD-helicase-domain-containing-protein-[Gorillibacterium-timonense] |
| WP_213510442.1 | SF2-Helicase+REase | GCF_018403165.1 | Paenibacillus_macerans | Bacteria>Bacillati>Bacillota>Bacilli>Bacillales>Paenibacillaceae>Paenibacillus | 1154 | KJS71_RS16975 | 180221..183685 | 44252 | UvrD-helicase-domain-containing-protein-[Paenibacillus-macerans] |
| WP_085168715.1 | SF2-Helicase+REase | GCF_900177285.1 | Paenibacillus_barengoltzii | Bacteria>Bacillati>Bacillota>Bacilli>Bacillales>Paenibacillaceae>Paenibacillus | 1120 | B9N82_RS00030 | 7208..10570 | 343517 | UvrD-helicase-domain-containing-protein-[Paenibacillus-barengoltzii] |
| WP_054819686.1 | SF2-Helicase+REase | GCF_024519685.1 | unclassified_Paenibacillus | Bacteria>Bacillati>Bacillota>Bacilli>Bacillales>Paenibacillaceae>Paenibacillus | 1117 | NR121_RS23300 | 11368..14721 | 185978 | MULTISPECIES:-UvrD-helicase-domain-containing-protein-[unclassified-Paenibacillus] |
| WDH83309.1 | SF2-Helicase+REase | GCA_028747665.1 | Paenibacillus_urinalis | Bacteria>Bacillati>Bacillota>Bacilli>Bacillales>Paenibacillaceae>Paenibacillus | 1099 | PUW23_03430 | 759545..762844 | 521520 | UvrD-helicase-domain-containing-protein-[Paenibacillus-urinalis] |
| WP_025333199.1 | SF2-Helicase+REase | GCF_000612505.1 | Paenibacillus_sabinae | Bacteria>Bacillati>Bacillota>Bacilli>Bacillales>Paenibacillaceae>Paenibacillus | 1099 | PSAB_RS03430 | 740320..743619 | 365617 | UvrD-helicase-domain-containing-protein-[Paenibacillus-sabinae] |
| WP_189000165.1 | SF2-Helicase+REase | GCF_014641075.1 | Paenibacillus_nasutitermitis | Bacteria>Bacillati>Bacillota>Bacilli>Bacillales>Paenibacillaceae>Paenibacillus | 1099 | IEY46_RS33885 | 7831..11130 | 1652958 | UvrD-helicase-domain-containing-protein-[Paenibacillus-nasutitermitis] |
| WP_076118195.1 | SF2-Helicase+REase | GCF_001954425.1 | Paenibacillus_odorifer | Bacteria>Bacillati>Bacillota>Bacilli>Bacillales>Paenibacillaceae>Paenibacillus | 1081 | BSK52_RS07885 | 306341..309586 | 189426 | UvrD-helicase-domain-containing-protein-[Paenibacillus-odorifer] |
| WP_042210528.1 | SF2-Helicase+REase | GCF_000758665.1 | Paenibacillus_borealis | Bacteria>Bacillati>Bacillota>Bacilli>Bacillales>Paenibacillaceae>Paenibacillus | 1080 | PBOR_RS03710 | 846211..849453 | 160799 | UvrD-helicase-domain-containing-protein-[Paenibacillus-borealis] |
| WP_209876423.1 | SF2-Helicase+REase | GCF_017874735.1 | Paenibacillus_silagei | Bacteria>Bacillati>Bacillota>Bacilli>Bacillales>Paenibacillaceae>Paenibacillus | 1079 | J2Z70_RS20885 | 36507..39746 | 1670801 | UvrD-helicase-domain-containing-protein-[Paenibacillus-silagei] |
| WP_090826231.1 | SF2-Helicase+REase | GCF_900103825.1 | Paenibacillus_sp._yr247 | Bacteria>Bacillati>Bacillota>Bacilli>Bacillales>Paenibacillaceae>Paenibacillus>unclassified_Paenibacillus | 1101 | BLT91_RS25100 | 5678..8983 | 1761880 | UvrD-helicase-domain-containing-protein-[Paenibacillus-sp.-yr247] |
| WP_060645787.1 | SF2-Helicase+REase | GCF_001427935.1 | Paenibacillus_sp._Soil724D2 | Bacteria>Bacillati>Bacillota>Bacilli>Bacillales>Paenibacillaceae>Paenibacillus>unclassified_Paenibacillus | 1099 | ASG85_RS07185 | 112013..115312 | 1736392 | UvrD-helicase-domain-containing-protein-[Paenibacillus-sp.-Soil724D2] |
| UUZ84564.1 | SF2-Helicase+REase | GCA_024703795.1 | Paenibacillus_sp._P26 | Bacteria>Bacillati>Bacillota>Bacilli>Bacillales>Paenibacillaceae>Paenibacillus>unclassified_Paenibacillus | 1029 | LJK88_13380 | 2153903..2156992 | 2885204 | UvrD-helicase-domain-containing-protein-[Paenibacillus-sp.-P26] |
| WP_181742024.1 | SF2-Helicase+REase | GCF_013868055.1 | Thermoactinomyces_mirandus | Bacteria>Bacillati>Bacillota>Bacilli>Bacillales>Thermoactinomycetaceae>Thermoactinomyces | 854 | H2C83_RS15080 | 11210..13774 | 2756294 | ATP-dependent-helicase-[Thermoactinomyces-mirandus] |
| WP_068559455.1 | SF2-Helicase+REase | GCF_900129415.1 | Trichococcus_flocculiformis | Bacteria>Bacillati>Bacillota>Bacilli>Lactobacillales>Carnobacteriaceae>Trichococcus | 1269 | BUA33_RS02165 | 192950..196759 | 82803 | helicase-exonuclease-AddAB-subunit-AddA-[Trichococcus-flocculiformis] |
| WP_018248935.1 | SF2-Helicase+REase | GCF_000379025.1 | Orenia_marismortui | Bacteria>Bacillati>Bacillota>Clostridia>Halanaerobiales>Halobacteroidaceae>Orenia | 1163 | OREMA_RS0108955 | 141992..145483 | 46469 | UvrD-helicase-domain-containing-protein-[Orenia-marismortui] |
| MCR5727522.1 | SF2-Helicase+REase | GCA_024699435.1 | Lachnospiraceae_bacterium | Bacteria>Bacillati>Bacillota>Clostridia>Lachnospirales>Lachnospiraceae>unclassified_Lachnospiraceae | 1263 | K6G24_08655 | 22956..26747 | 1898203 | UvrD-helicase-domain-containing-protein-[Lachnospiraceae-bacterium] |
| OUL09461.1 | SF2-Helicase+REase | GCA_002155105.1 | Sedimentibacter_sp._SX930 | Bacteria>Bacillati>Bacillota>Tissierellia>Tissierellia_incertae_sedis>Sedimentibacter>unclassified_Sedimentibacter | 1269 | B0533_04660 | 260288..264097 | 1955060 | helicase-exonuclease-AddAB-subunit-AddA-[Sedimentibacter-sp.-SX930] |
| MTI83940.1 | SF2-Helicase+REase | GCA_009711705.1 | Bacillota_bacterium | Bacteria>Bacillati>Bacillota>unclassified_Bacillota | 626 | FH756_08530 | 80323..82203 | 1879010 | hypothetical-protein-[Bacillota-bacterium] |
| MCX6032086.1 | SF2-Helicase+REase | GCA_026389845.1 | Chloroflexota_bacterium | Bacteria>Bacillati>Chloroflexota>unclassified_Chloroflexota | 1135 | NT169_22665 | 1185..4592 | 2026724 | MAG:-UvrD-helicase-domain-containing-protein-[Chloroflexi-bacterium] |
| MCL4488351.1 | SF2-Helicase+REase | GCA_023381515.1 | Chloroflexota_bacterium | Bacteria>Bacillati>Chloroflexota>unclassified_Chloroflexota | 990 | M1570_09515 | 97337..100309 | 2026724 | ATP-dependent-helicase-[Chloroflexota-bacterium] |
| MCL5952143.1 | SF2-Helicase+REase | GCA_023489485.1 | Chloroflexota_bacterium | Bacteria>Bacillati>Chloroflexota>unclassified_Chloroflexota | 990 | M1132_10570 | 61529..64501 | 2026724 | ATP-dependent-helicase-[Chloroflexota-bacterium] |
| WP_096597411.1 | SF2-Helicase+REase | GCF_963875285.1 | Calothrix_sp._NIES-2100 | Bacteria>Bacillati>Cyanobacteriota/Melainabacteria_group>Cyanobacteriota>Cyanophyceae>Nostocales>Calotrichaceae>Calothrix>unclassified_Calothrix | 1274 | CA734_RS02090 | 548127..551951 | 1954172 | BREX-3-system-P-loop-containing-protein-BrxF-[Calothrix-sp.-NIES-2100] |
| MTJ55917.1 | SF2-Helicase+REase | GCA_009712085.1 | Anabaena_sp._UHCC_0253 | Bacteria>Bacillati>Cyanobacteriota/Melainabacteria_group>Cyanobacteriota>Cyanophyceae>Nostocales>Nostocaceae>Anabaena>unclassified_Anabaena | 1230 | FJR38_26295 | 53858..57552 | 2590019 | BREX-3-system-P-loop-containing-protein-BrxF-[Anabaena-sp.-UHCC-0253] |
| WP_190937389.1 | SF2-Helicase+REase | GCF_014698055.1 | Nostoc_sp._FACHB-133 | Bacteria>Bacillati>Cyanobacteriota/Melainabacteria_group>Cyanobacteriota>Cyanophyceae>Nostocales>Nostocaceae>Nostoc>unclassified_Nostoc | 1284 | H6G86_RS30120 | 37621..41475 | 2692835 | BREX-3-system-P-loop-containing-protein-BrxF-[Nostoc-sp.-FACHB-133] |
| WP_096683472.1 | SF2-Helicase+REase | - | Nostoc_sp._NIES-2111 | Bacteria>Bacillati>Cyanobacteriota/Melainabacteria_group>Cyanobacteriota>Cyanophyceae>Nostocales>Nostocaceae>Nostoc>unclassified_Nostoc | 1281 | CA737_RS32350 | 63598..67443 | 1973475 | BREX-3-system-P-loop-containing-protein-BrxF-[Nostoc-sp.-NIES-2111] |
| MBR8836035.1 | SF2-Helicase+REase | GCA_016903675.1 | Stigonema_ocellatum_SAG_48.90_=_DSM_106950 | Bacteria>Bacillati>Cyanobacteriota/Melainabacteria_group>Cyanobacteriota>Cyanophyceae>Nostocales>Stigonemataceae>Stigonema>Stigonema_ocellatum | 1092 | DSM106950_18930 | 28853..32131 | 221294 | UvrD-helicase-domain-containing-protein-[Stigonema-ocellatum-SAG-48.90-=-DSM-106950] |
| MYC78299.1 | SF2-Helicase+REase | GCA_009841255.1 | Candidatus_Poribacteria_bacterium | Bacteria>Bacteria_incertae_sedis>Bacteria_candidate_phyla>Candidatus_Poribacteria>unclassified_Candidatus_Poribacteria | 1169 | F4X10_21265 | 57545..61054 | 2026781 | AAA-family-ATPase-[Candidatus-Poribacteria-bacterium] |
| MDE0423413.1 | SF2-Helicase+REase | GCA_028820845.1 | Candidatus_Poribacteria_bacterium | Bacteria>Bacteria_incertae_sedis>Bacteria_candidate_phyla>Candidatus_Poribacteria>unclassified_Candidatus_Poribacteria | 1162 | OXN25_00945 | 238..3726 | 2026781 | MAG:-UvrD-helicase-domain-containing-protein-[Candidatus-Poribacteria-bacterium] |
| MYB95137.1 | SF2-Helicase+REase | GCA_009838805.1 | Candidatus_Poribacteria_bacterium | Bacteria>Bacteria_incertae_sedis>Bacteria_candidate_phyla>Candidatus_Poribacteria>unclassified_Candidatus_Poribacteria | 1149 | F4Y43_14890 | 9310..12759 | 2026781 | AAA-family-ATPase-[Candidatus-Poribacteria-bacterium] |
| MBZ5659869.1 | SF2-Helicase+REase | GCA_020073485.1 | Terriglobia_bacterium | Bacteria>Pseudomonadati>Acidobacteriota>Terriglobia>unclassified_Terriglobia | 1114 | LAO08_05620 | 155639..158983 | 2081523 | UvrD-helicase-domain-containing-protein-[Acidobacteriia-bacterium] |
| MCR9155220.1 | SF2-Helicase+REase | GCA_024742495.1 | Bacteroidota_bacterium | Bacteria>Pseudomonadati>FCB_group>Bacteroidota/Chlorobiota_group>Bacteroidota>unclassified_Bacteroidota | 1012 | NXI09_14015 | 78225..81263 | 1898104 | UvrD-helicase-domain-containing-protein-[Bacteroidota-bacterium] |
| MDE2877865.1 | SF2-Helicase+REase | GCA_028874535.1 | Candidatus_Palauibacter_soopunensis | Bacteria>Pseudomonadati>FCB_group>Gemmatimonadota>Gemmatimonadia>Candidatus_Palauibacterales>Candidatus_Palauibacteraceae>Candidatus_Palauibacter | 1015 | OXO55_03055 | 14208..17255 | 3056739 | MAG:-ATP-dependent-DNA-helicase-[Gemmatimonadota-bacterium] |
| MCC7380509.1 | SF2-Helicase+REase | GCA_020850435.1 | Deltaproteobacteria_bacterium | Bacteria>Pseudomonadati>Myxococcota>Myxococcia>unclassified_Myxococcia | 1244 | IT384_01665 | 14441..18175 | 2026735 | UvrD-helicase-domain-containing-protein-[Deltaproteobacteria-bacterium] |
| WP_229825718.1 | SF2-Helicase+REase | GCF_003967075.1 | Alphaproteobacteria | Bacteria>Pseudomonadati>Pseudomonadota | 1065 | SAMIE_RS17595 | 3639688..3642885 | 28211 | MULTISPECIES:-UvrD-helicase-domain-containing-protein-[Alphaproteobacteria] |
| WP_086466631.1 | SF2-Helicase+REase | GCF_002148795.1 | Alphaproteobacteria | Bacteria>Pseudomonadati>Pseudomonadota | 1054 | BKM74_RS15590 | 46794..49958 | 28211 | UvrD-helicase-domain-containing-protein-[Oceanibaculum-nanhaiense] |
| MAD36289.1 | SF2-Helicase+REase | GCA_002684955.1 | Tistrella_sp. | Bacteria>Pseudomonadati>Pseudomonadota>Alphaproteobacteria>Geminicoccales>Geminicoccaceae>Tistrella>unclassified_Tistrella | 1054 | CM028_06260 | 13848..17012 | 2024861 | DNA-helicase-UvrD-[Tistrella-sp.] |
| MBA78727.1 | SF2-Helicase+REase | GCA_002709065.1 | Tistrella_sp. | Bacteria>Pseudomonadati>Pseudomonadota>Alphaproteobacteria>Geminicoccales>Geminicoccaceae>Tistrella>unclassified_Tistrella | 1054 | CM030_25965 | 13848..17012 | 2024861 | DNA-helicase-UvrD-[Tistrella-sp.] |
| WP_210251079.1 | SF2-Helicase+REase | GCF_008932705.1 | Brucella_anthropi | Bacteria>Pseudomonadati>Pseudomonadota>Alphaproteobacteria>Hyphomicrobiales>Brucellaceae>Brucella/Ochrobactrum_group>Brucella | 1065 | F9L04_RS18310 | 28867..32064 | 529 | UvrD-helicase-domain-containing-protein-[Brucella-anthropi] |
| WP_210207307.1 | SF2-Helicase+REase | GCF_041929885.1 | Pseudochrobactrum_asaccharolyticum | Bacteria>Pseudomonadati>Pseudomonadota>Alphaproteobacteria>Hyphomicrobiales>Brucellaceae>Pseudochrobactrum | 1066 | DFR47_RS11495 | 56038..59238 | 354351 | UvrD-helicase-domain-containing-protein-[Pseudochrobactrum-asaccharolyticum] |
| WP_075657539.1 | SF2-Helicase+REase | GCF_001939785.1 | Pseudochrobactrum_sp._B5 | Bacteria>Pseudomonadati>Pseudomonadota>Alphaproteobacteria>Hyphomicrobiales>Brucellaceae>Pseudochrobactrum>unclassified_Pseudochrobactrum | 1068 | BKD14_RS17995 | 70215..73421 | 1289478 | UvrD-helicase-domain-containing-protein-[Pseudochrobactrum-sp.-B5] |
| WP_266332115.1 | SF2-Helicase+REase | GCF_026343955.1 | Kaistia_defluvii | Bacteria>Pseudomonadati>Pseudomonadota>Alphaproteobacteria>Hyphomicrobiales>Kaistiaceae>Kaistia | 1066 | OSH10_RS12630 | 356034..359234 | 410841 | UvrD-helicase-domain-containing-protein-[Kaistia-defluvii] |
| WP_036280598.1 | SF2-Helicase+REase | GCF_000188155.2 | Methylocystis_sp._ATCC_49242 | Bacteria>Pseudomonadati>Pseudomonadota>Alphaproteobacteria>Hyphomicrobiales>Methylocystaceae>Methylocystis>unclassified_Methylocystis | 1130 | MET49242_RS03125 | 247961..251353 | 622637 | UvrD-helicase-domain-containing-protein-[Methylocystis-sp.-ATCC-49242] |
| WP_210205443.1 | SF2-Helicase+REase | GCF_003217325.1 | Rhodopseudomonas_faecalis | Bacteria>Pseudomonadati>Pseudomonadota>Alphaproteobacteria>Hyphomicrobiales>Nitrobacteraceae>Rhodopseudomonas | 1066 | BJ122_RS08520 | 77560..80760 | 99655 | UvrD-helicase-domain-containing-protein-[Rhodopseudomonas-faecalis] |
| BCG83578.1 | SF2-Helicase+REase | GCA_016756495.1 | Mesorhizobium_sp._113-3-3 | Bacteria>Pseudomonadati>Pseudomonadota>Alphaproteobacteria>Hyphomicrobiales>Phyllobacteriaceae>Mesorhizobium>unclassified_Mesorhizobium | 1130 | MesoLj113b_71200 | 309856..313248 | 2744516 | DNA-helicase-[Mesorhizobium-sp.-113-3-3] |
| WP_009450518.1 | SF2-Helicase+REase | GCF_000300515.1 | Nitratireductor_indicus | Bacteria>Pseudomonadati>Pseudomonadota>Alphaproteobacteria>Hyphomicrobiales>Phyllobacteriaceae>Nitratireductor | 1066 | NA8A_RS11820 | 123469..126669 | 721133 | UvrD-helicase-domain-containing-protein-[Nitratireductor-indicus] |
| WP_154722154.1 | SF2-Helicase+REase | GCF_900492205.1 | Ciceribacter_sp._T2.26MG-112.2 | Bacteria>Pseudomonadati>Pseudomonadota>Alphaproteobacteria>Hyphomicrobiales>Rhizobiaceae>Ciceribacter>unclassified_Ciceribacter | 1131 | DYZ14_RS19050 | 4203..7598 | 3137154 | UvrD-helicase-domain-containing-protein-[Ciceribacter-naphthalenivorans] |
| WP_205910046.1 | SF2-Helicase+REase | GCF_014202175.1 | Rhizobium_rosettiformans | Bacteria>Pseudomonadati>Pseudomonadota>Alphaproteobacteria>Hyphomicrobiales>Rhizobiaceae>Rhizobium/Agrobacterium_group>Rhizobium | 1066 | FAA86_RS19530 | 50984..54184 | 1368430 | UvrD-helicase-domain-containing-protein-[Rhizobium-rosettiformans] |
| WP_004434307.1 | SF2-Helicase+REase | GCF_000236945.1 | Sinorhizobium_meliloti | Bacteria>Pseudomonadati>Pseudomonadota>Alphaproteobacteria>Hyphomicrobiales>Rhizobiaceae>Sinorhizobium/Ensifer_group>Sinorhizobium | 1131 | SM0020_RS29750 | 175508..178903 | 382 | UvrD-helicase-domain-containing-protein-[Sinorhizobium-meliloti] |
| WP_131389743.1 | SF2-Helicase+REase | - | Haematobacter_massiliensis | Bacteria>Pseudomonadati>Pseudomonadota>Alphaproteobacteria>Rhodobacterales>Paracoccaceae>Haematobacter | 1067 | HmaOT1_RS15935 | 79589..82792 | 195105 | UvrD-helicase-domain-containing-protein-[Haematobacter-massiliensis] |
| WP_199257816.1 | SF2-Helicase+REase | GCF_016446475.1 | Paracoccus_binzhouensis | Bacteria>Pseudomonadati>Pseudomonadota>Alphaproteobacteria>Rhodobacterales>Paracoccaceae>Paracoccus | 1067 | I9P22_RS02425 | 9724..12927 | 2796149 | UvrD-helicase-domain-containing-protein-[Paracoccus-binzhouensis] |
| MBL4843419.1 | SF2-Helicase+REase | GCA_016765055.1 | Thalassospira_sp. | Bacteria>Pseudomonadati>Pseudomonadota>Alphaproteobacteria>Rhodospirillales>Thalassospiraceae>Thalassospira>unclassified_Thalassospira | 1066 | JKY47_21610 | 6472..9672 | 1912094 | UvrD-helicase-domain-containing-protein-[Thalassospira-sp.] |
| WP_183952295.1 | SF2-Helicase+REase | GCF_014196255.1 | Sphingomonas | Bacteria>Pseudomonadati>Pseudomonadota>Alphaproteobacteria>Sphingomonadales>Sphingomonadaceae | 1128 | GGR48_RS13060 | 18013..21399 | 13687 | UvrD-helicase-domain-containing-protein-[Sphingomonas-pseudosanguinis] |
| WP_069709533.1 | SF2-Helicase+REase | - | Novosphingobium_resinovorum | Bacteria>Pseudomonadati>Pseudomonadota>Alphaproteobacteria>Sphingomonadales>Sphingomonadaceae>Novosphingobium | 1068 | BES08_RS24100 | 1538606..1541812 | 158500 | UvrD-helicase-domain-containing-protein-[Novosphingobium-resinovorum] |
| WP_031304998.1 | SF2-Helicase+REase | GCF_000445065.1 | Sphingobium_quisquiliarum | Bacteria>Pseudomonadati>Pseudomonadota>Alphaproteobacteria>Sphingomonadales>Sphingomonadaceae>Sphingobium | 1132 | L288_RS41550 | 24400..27798 | 538379 | UvrD-helicase-domain-containing-protein-[Sphingobium-quisquiliarum] |
| WP_100206933.1 | SF2-Helicase+REase | GCF_000224695.1 | Sphingobium_yanoikuyae | Bacteria>Pseudomonadati>Pseudomonadota>Alphaproteobacteria>Sphingomonadales>Sphingomonadaceae>Sphingobium | 1066 | SYA_RS0104840 | 52698..55898 | 13690 | UvrD-helicase-domain-containing-protein-[Sphingobium-yanoikuyae] |
| MBB3880221.1 | SF2-Helicase+REase | GCA_014196255.1 | Sphingomonas_pseudosanguinis | Bacteria>Pseudomonadati>Pseudomonadota>Alphaproteobacteria>Sphingomonadales>Sphingomonadaceae>Sphingomonas | 1128 | GGR48_002664 | 18013..21399 | 413712 | exodeoxyribonuclease-5-[Sphingomonas-pseudosanguinis] |
| WP_066478873.1 | SF2-Helicase+REase | GCF_001555785.1 | unclassified_Sphingomonas | Bacteria>Pseudomonadati>Pseudomonadota>Alphaproteobacteria>Sphingomonadales>Sphingomonadaceae>Sphingomonas | 1068 | AX667_RS15635 | 56777..59983 | 196159 | MULTISPECIES:-UvrD-helicase-domain-containing-protein-[unclassified-Sphingomonas] |
| MCK9506954.1 | SF2-Helicase+REase | GCA_023228545.1 | Pigmentiphaga_sp. | Bacteria>Pseudomonadati>Pseudomonadota>Betaproteobacteria>Burkholderiales>Alcaligenaceae>Pigmentiphaga>unclassified_Pigmentiphaga | 1066 | MOQ54_00805 | 14061..17261 | 1977564 | UvrD-helicase-domain-containing-protein-[Pigmentiphaga-sp.] |
| MBP7846544.1 | SF2-Helicase+REase | GCA_018001735.1 | Burkholderiales_bacterium | Bacteria>Pseudomonadati>Pseudomonadota>Betaproteobacteria>Burkholderiales>unclassified_Burkholderiales | 1180 | KAZ94_00250 | 54115..57657 | 1891238 | UvrD-helicase-domain-containing-protein-[Burkholderiales-bacterium] |
| WP_167854939.1 | SF2-Helicase+REase | GCF_004745945.1 | Mangrovimicrobium_sediminis | Bacteria>Pseudomonadati>Pseudomonadota>Gammaproteobacteria>Cellvibrionales>Halieaceae>Mangrovimicrobium | 1045 | E4634_RS16050 | 170375..173512 | 2562682 | UvrD-helicase-domain-containing-protein-[Haliea-sp.-SAOS-164] |
| WP_019531411.1 | SF2-Helicase+REase | GCF_000373485.1 | Dasania_marina | Bacteria>Pseudomonadati>Pseudomonadota>Gammaproteobacteria>Cellvibrionales>Spongiibacteraceae>Dasania | 1057 | B067_RS0117595 | 6786..9959 | 471499 | UvrD-helicase-domain-containing-protein-[Dasania-marina] |
| MCG7876678.1 | SF2-Helicase+REase | GCA_016842655.1 | Candidatus_Thiodiazotropha_taylori | Bacteria>Pseudomonadati>Pseudomonadota>Gammaproteobacteria>Chromatiales>Sedimenticolaceae>Candidatus_Thiodiazotropha | 1057 | JAY78_10110 | 21929..25102 | 2792791 | UvrD-helicase-domain-containing-protein-[Candidatus-Thiodiazotropha-taylori] |
| MCG7951165.1 | SF2-Helicase+REase | GCA_016842765.1 | Candidatus_Thiodiazotropha_taylori | Bacteria>Pseudomonadati>Pseudomonadota>Gammaproteobacteria>Chromatiales>Sedimenticolaceae>Candidatus_Thiodiazotropha | 1057 | JAZ09_08080 | 39447..42620 | 2792791 | UvrD-helicase-domain-containing-protein-[Candidatus-Thiodiazotropha-taylori] |
| MCG8058208.1 | SF2-Helicase+REase | GCA_016843165.1 | Candidatus_Thiodiazotropha_taylori | Bacteria>Pseudomonadati>Pseudomonadota>Gammaproteobacteria>Chromatiales>Sedimenticolaceae>Candidatus_Thiodiazotropha | 1057 | JAZ21_00040 | 10713..13886 | 2792791 | UvrD-helicase-domain-containing-protein-[Candidatus-Thiodiazotropha-taylori] |
| MCG8102617.1 | SF2-Helicase+REase | GCA_016842845.1 | Candidatus_Thiodiazotropha_taylori | Bacteria>Pseudomonadati>Pseudomonadota>Gammaproteobacteria>Chromatiales>Sedimenticolaceae>Candidatus_Thiodiazotropha | 1057 | JAY70_09520 | 39477..42650 | 2792791 | UvrD-helicase-domain-containing-protein-[Candidatus-Thiodiazotropha-taylori] |
| WP_248566597.1 | SF2-Helicase+REase | GCF_023195885.1 | Methylonatrum_kenyense | Bacteria>Pseudomonadati>Pseudomonadota>Gammaproteobacteria>Gammaproteobacteria_incertae_sedis>Methylonatrum | 1239 | M0534_RS01305 | 270460..274179 | 455253 | exodeoxyribonuclease-V-subunit-beta-[Methylonatrum-kenyense] |
| WP_070087174.1 | SF2-Helicase+REase | GCF_001753915.1 | Pseudomonas_sp._NBRC_111122 | Bacteria>Pseudomonadati>Pseudomonadota>Gammaproteobacteria>Pseudomonadales>Pseudomonadaceae>Pseudomonas>unclassified_Pseudomonas | 1224 | BJQ15_RS15350 | 30585..34259 | 1661037 | exodeoxyribonuclease-V-subunit-beta-[Pseudomonas-sp.-NBRC-111122] |
| MBK8482430.1 | SF2-Helicase+REase | GCA_016711215.1 | Pseudomonadota_bacterium | Bacteria>Pseudomonadati>Pseudomonadota>unclassified_Pseudomonadota | 648 | IPL40_14905 | 137288..139234 | 1977087 | ATP-dependent-helicase-[Pseudomonadota-bacterium] |
| MCE9567299.1 | SF2-Helicase+REase | GCA_021413735.1 | Planctomycetota_bacterium | Bacteria>Pseudomonadati>PVC_group>Planctomycetota>unclassified_Planctomycetota | 1133 | K8U57_35300 | 22215..25616 | 2026780 | UvrD-helicase-domain-containing-protein-[Planctomycetota-bacterium] |

| Protein-ID | Domain | Assembly-ID | Strain | Taxonomy | Length | Locus-Tag | Genomic-coordinates | TaxID | Fasta-defline |
|---|---|---|---|---|---|---|---|---|---|
| WP_143068399.1 | SF2-Helicase+REase | GCF_900109185.1 | Deinococcus_reticulitermitis | Bacteria>Thermotogati>Deinococcota>Deinococci>Deinococcales>Deinococcaceae>Deinococcus | 1093 | BMY43_RS15305 | 16708..19989 | 856736 | UvrD-helicase-domain-containing-protein-[Deinococcus-reticulitermitis] |
| WP_221091404.1 | SF2-Helicase+REase | GCF_019693445.1 | Deinococcus_aquaedulcis | Bacteria>Thermotogati>Deinococcota>Deinococci>Deinococcales>Deinococcaceae>Deinococcus | 1075 | KMW22_RS17965 | 5163..8390 | 2840455 | UvrD-helicase-domain-containing-protein-[Deinococcus-aquaedulcis] |
| MBI1293580.1 | SF2-Helicase+REase | GCA_016124945.1 | bacterium | Bacteria>unclassified_Bacteria | 1194 | GC175_01310 | 56882..60466 | 1869227 | AAA-family-ATPase-[bacterium] |
| --SF1-SF2-Helicase+REase | | | | | | | | | |
| WP_014998455.1 | SF2-Helicase+REase | - | Gammaproteobacteria | Bacteria>Pseudomonadati>Pseudomonadota | 660 | AMBLS11_RS09790 | 2286088..2288070 | 1236 | glutathione-synthase-[Alteromonas-macleodii] |
| WP_014998456.1 | SF2-Helicase+REase | - | Gammaproteobacteria | Bacteria>Pseudomonadati>Pseudomonadota | 618 | AMBLS11_RS09795 | 2288077..2289933 | 1236 | DEAD/DEAH-box-helicase-family-protein-[Alteromonas-macleodii] |
| WP_096812540.1 | SF2-Helicase+REase | GCF_017086505.1 | Enterobacterales | Bacteria>Pseudomonadati>Pseudomonadota>Gammaproteobacteria | 665 | NLZ12_RS02830 | 595411..597408 | 91347 | MULTISPECIES:-glutathione-synthase-[Enterobacteriaceae] |
| WP_158149589.1 | SF2-Helicase+REase | GCF_014353895.1 | Enterobacterales | Bacteria>Pseudomonadati>Pseudomonadota>Gammaproteobacteria | 665 | IBU46_RS01715 | 11792..13789 | 91347 | MULTISPECIES:-DEAD/DEAH-box-helicase-family-protein-[Enterobacterales] |
| WP_158149590.1 | SF2-Helicase+REase | GCF_051124655.1 | Enterobacterales | Bacteria>Pseudomonadati>Pseudomonadota>Gammaproteobacteria | 620 | GSQ69_RS16265 | 3485911..3487773 | 91347 | MULTISPECIES:-DEAD/DEAH-box-helicase-family-protein-[Enterobacterales] |
| WP_201744910.1 | SF2-Helicase+REase | GCF_009907625.1 | Enterobacterales | Bacteria>Pseudomonadati>Pseudomonadota>Gammaproteobacteria | 604 | GV742_RS00820 | 14110..15924 | 91347 | DEAD/DEAH-box-helicase-family-protein-[Klebsiella-pneumoniae] |
| WP_042889125.1 | SF2-Helicase+REase | GCF_033096545.1 | Aeromonas_dhakensis | Bacteria>Pseudomonadati>Pseudomonadota>Gammaproteobacteria>Aeromonadales>Aeromonadaceae>Aeromonas | 663 | AQU_RS10045 | 125393..127384 | 196024 | MULTISPECIES:-glutathione-synthase-[Aeromonas] |
| WP_010634557.1 | SF2-Helicase+REase | GCF_033096545.1 | Aeromonas_dhakensis | Bacteria>Pseudomonadati>Pseudomonadota>Gammaproteobacteria>Aeromonadales>Aeromonadaceae>Aeromonas | 618 | AQU_RS10050 | 127400..129256 | 196024 | DEAD/DEAH-box-helicase-family-protein-[Aeromonas-dhakensis] |
| WP_242698654.1 | SF2-Helicase+REase | GCF_017368475.1 | Alteromonas_sp._5E99-2 | Bacteria>Pseudomonadati>Pseudomonadota>Gammaproteobacteria>Alteromonadales>Alteromonadaceae>Alteromonas/Salinimonas_group>Alteromonas>unclassified_Alteromonas | 682 | J3L16_RS04235 | 91996..94044 | 2817683 | glutathione-synthase-[Alteromonas-sp.-5E99-2] |
| WP_100972005.1 | SF2-Helicase+REase | GCF_002831605.1 | Alteromonas_sp._MB-3u-76 | Bacteria>Pseudomonadati>Pseudomonadota>Gammaproteobacteria>Alteromonadales>Alteromonadaceae>Alteromonas/Salinimonas_group>Alteromonas>unclassified_Alteromonas | 660 | CW735_RS10435 | 2324796..2326778 | 2058133 | glutathione-synthase-[Alteromonas-sp.-MB-3u-76] |
| WP_207611654.1 | SF2-Helicase+REase | GCF_017368475.1 | Alteromonas_sp._5E99-2 | Bacteria>Pseudomonadati>Pseudomonadota>Gammaproteobacteria>Alteromonadales>Alteromonadaceae>Alteromonas/Salinimonas_group>Alteromonas>unclassified_Alteromonas | 618 | J3L16_RS04230 | 90136..91992 | 2817683 | DEAD/DEAH-box-helicase-family-protein-[Alteromonas-sp.-5E99-2] |
| WP_198520969.1 | SF2-Helicase+REase | GCF_002831605.1 | Alteromonas_sp._MB-3u-76 | Bacteria>Pseudomonadati>Pseudomonadota>Gammaproteobacteria>Alteromonadales>Alteromonadaceae>Alteromonas/Salinimonas_group>Alteromonas>unclassified_Alteromonas | 345 | CW735_RS10440 | 2326785..2327822 | 2058133 | DEAD/DEAH-box-helicase-family-protein-[Alteromonas-sp.-MB-3u-76] |
| WP_198520970.1 | SF2-Helicase+REase | GCF_002831605.1 | Alteromonas_sp._MB-3u-76 | Bacteria>Pseudomonadati>Pseudomonadota>Gammaproteobacteria>Alteromonadales>Alteromonadaceae>Alteromonas/Salinimonas_group>Alteromonas>unclassified_Alteromonas | 297 | CW735_RS10455 | 2330365..2331258 | 2058133 | hypothetical-protein-[Alteromonas-sp.-MB-3u-76] |
| WP_076500053.1 | SF2-Helicase+REase | GCF_900156405.1 | Shewanella_morhuae | Bacteria>Pseudomonadati>Pseudomonadota>Gammaproteobacteria>Alteromonadales>Shewanellaceae>Shewanella | 666 | BW947_RS12450 | 1834..3834 | 365591 | DEAD/DEAH-box-helicase-family-protein-[Shewanella-morhuae] |
| WP_076500065.1 | SF2-Helicase+REase | GCF_900156405.1 | Shewanella_morhuae | Bacteria>Pseudomonadati>Pseudomonadota>Gammaproteobacteria>Alteromonadales>Shewanellaceae>Shewanella | 619 | BW947_RS12455 | 3875..5734 | 365591 | DEAD/DEAH-box-helicase-family-protein-[Shewanella-morhuae] |
| WP_173294713.1 | SF2-Helicase+REase | GCF_013248905.1 | Shewanella_sp._VB17 | Bacteria>Pseudomonadati>Pseudomonadota>Gammaproteobacteria>Alteromonadales>Shewanellaceae>Shewanella>unclassified_Shewanella | 664 | HQQ94_RS12370 | 2887581..2889575 | 2739432 | DEAD/DEAH-box-helicase-family-protein-[Shewanella-sp.-VB17] |
| WP_173294712.1 | SF2-Helicase+REase | GCF_013248905.1 | Shewanella_sp._VB17 | Bacteria>Pseudomonadati>Pseudomonadota>Gammaproteobacteria>Alteromonadales>Shewanellaceae>Shewanella>unclassified_Shewanella | 617 | HQQ94_RS12365 | 2885667..2887520 | 2739432 | DEAD/DEAH-box-helicase-family-protein-[Shewanella-sp.-VB17] |
| WP_064489887.1 | SF2-Helicase+REase | GCF_902160145.1 | Enterobacteriaceae | Bacteria>Pseudomonadati>Pseudomonadota>Gammaproteobacteria>Enterobacterales | 665 | EHJ08_RS11780 | 217960..219957 | 543 | MULTISPECIES:-hypothetical-protein-[Enterobacteriaceae] |
| WP_211449374.1 | SF2-Helicase+REase | GCF_018299605.1 | Enterobacteriaceae | Bacteria>Pseudomonadati>Pseudomonadota>Gammaproteobacteria>Enterobacterales | 665 | GUY16_RS21820 | 47729..49726 | 543 | DEAD/DEAH-box-helicase-family-protein-[Enterobacter-mori] |
| WP_200895767.1 | SF2-Helicase+REase | GCF_902160145.1 | Enterobacteriaceae | Bacteria>Pseudomonadati>Pseudomonadota>Gammaproteobacteria>Enterobacterales | 604 | FN179_RS14110 | 334205..336019 | 543 | MULTISPECIES:-DEAD/DEAH-box-helicase-family-protein-[Enterobacteriaceae] |
| WP_207621934.1 | SF2-Helicase+REase | GCF_011045335.1 | Enterobacter | Bacteria>Pseudomonadati>Pseudomonadota>Gammaproteobacteria>Enterobacterales>Enterobacteriaceae | 604 | G5C52_RS10080 | 2122278..2124092 | 547 | DEAD/DEAH-box-helicase-family-protein-[Enterobacter-hormaechei] |
| WP_124023332.1 | SF2-Helicase+REase | GCF_003818135.1 | Buttiauxella_warmboldiae | Bacteria>Pseudomonadati>Pseudomonadota>Gammaproteobacteria>Enterobacterales>Enterobacteriaceae>Buttiauxella | 665 | EHN07_RS06325 | 49982..51979 | 82993 | DEAD/DEAH-box-helicase-family-protein-[Buttiauxella-warmboldiae] |
| WP_124023333.1 | SF2-Helicase+REase | GCF_003818135.1 | Buttiauxella_warmboldiae | Bacteria>Pseudomonadati>Pseudomonadota>Gammaproteobacteria>Enterobacterales>Enterobacteriaceae>Buttiauxella | 620 | EHN07_RS06330 | 52027..53889 | 82993 | DEAD/DEAH-box-helicase-family-protein-[Buttiauxella-warmboldiae] |
| WP_200054821.1 | SF2-Helicase+REase | GCF_016504925.1 | Citrobacter_freundii | Bacteria>Pseudomonadati>Pseudomonadota>Gammaproteobacteria>Enterobacterales>Enterobacteriaceae>Citrobacter>Citrobacter_freundii_complex | 665 | GHT45_RS09265 | 249958..251955 | 546 | DEAD/DEAH-box-helicase-family-protein-[Citrobacter-freundii] |
| WP_200054820.1 | SF2-Helicase+REase | GCF_016504925.1 | Citrobacter_freundii | Bacteria>Pseudomonadati>Pseudomonadota>Gammaproteobacteria>Enterobacterales>Enterobacteriaceae>Citrobacter>Citrobacter_freundii_complex | 620 | GHT45_RS09260 | 248048..249910 | 546 | DEAD/DEAH-box-helicase-family-protein-[Citrobacter-freundii] |
| WP_200143508.1 | SF2-Helicase+REase | GCF_013395115.1 | Citrobacter_freundii | Bacteria>Pseudomonadati>Pseudomonadota>Gammaproteobacteria>Enterobacterales>Enterobacteriaceae>Citrobacter>Citrobacter_freundii_complex | 604 | HYG96_RS13020 | 250340..252154 | 546 | DEAD/DEAH-box-helicase-family-protein-[Citrobacter-freundii] |
| WP_202622921.1 | SF2-Helicase+REase | GCF_009938355.1 | Cronobacter_sakazakii | Bacteria>Pseudomonadati>Pseudomonadota>Gammaproteobacteria>Enterobacterales>Enterobacteriaceae>Cronobacter | 604 | EHJ08_RS11775 | 216064..217878 | 28141 | DEAD/DEAH-box-helicase-family-protein-[Cronobacter-sakazakii] |
| WP_211449376.1 | SF2-Helicase+REase | GCF_018299605.1 | Enterobacter_mori | Bacteria>Pseudomonadati>Pseudomonadota>Gammaproteobacteria>Enterobacterales>Enterobacteriaceae>Enterobacter | 620 | GUY16_RS21825 | 49774..51636 | 539813 | DEAD/DEAH-box-helicase-family-protein-[Enterobacter-mori] |
| WP_126349342.1 | SF2-Helicase+REase | GCF_004138605.1 | Enterobacter_roggenkampii | Bacteria>Pseudomonadati>Pseudomonadota>Gammaproteobacteria>Enterobacterales>Enterobacteriaceae>Enterobacter>Enterobacter_cloacae_complex | 665 | D8675_RS06365 | 1074473..1076470 | 1812935 | glutathione-synthase-[Enterobacter-roggenkampii] |
| WP_164474328.1 | SF2-Helicase+REase | GCF_011045335.1 | Enterobacter_hormaechei | Bacteria>Pseudomonadati>Pseudomonadota>Gammaproteobacteria>Enterobacterales>Enterobacteriaceae>Enterobacter>Enterobacter_cloacae_complex | 665 | G5C52_RS10075 | 2120199..2122196 | 158836 | glutathione-synthase-[Enterobacter-hormaechei] |
| WP_252033138.1 | SF2-Helicase+REase | GCF_023752605.1 | Enterobacter_asburiae | Bacteria>Pseudomonadati>Pseudomonadota>Gammaproteobacteria>Enterobacterales>Enterobacteriaceae>Enterobacter>Enterobacter_cloacae_complex | 665 | M8S83_RS11445 | 331326..333323 | 61645 | hypothetical-protein-[Enterobacter-asburiae] |
| WP_252033140.1 | SF2-Helicase+REase | GCF_023752605.1 | Enterobacter_asburiae | Bacteria>Pseudomonadati>Pseudomonadota>Gammaproteobacteria>Enterobacterales>Enterobacteriaceae>Enterobacter>Enterobacter_cloacae_complex | 620 | M8S83_RS11450 | 333371..335233 | 61645 | DEAD/DEAH-box-helicase-family-protein-[Enterobacter-asburiae] |
| WP_198680144.1 | SF2-Helicase+REase | GCF_004138605.1 | Enterobacter_roggenkampii | Bacteria>Pseudomonadati>Pseudomonadota>Gammaproteobacteria>Enterobacterales>Enterobacteriaceae>Enterobacter>Enterobacter_cloacae_complex | 604 | D8675_RS06370 | 1076552..1078366 | 1812935 | DEAD/DEAH-box-helicase-family-protein-[Enterobacter-roggenkampii] |
| WP_126800551.1 | SF2-Helicase+REase | GCF_003986875.1 | Enterobacter_sp._WCHEn090032 | Bacteria>Pseudomonadati>Pseudomonadota>Gammaproteobacteria>Enterobacterales>Enterobacteriaceae>Enterobacter>unclassified_Enterobacter | 665 | EKN83_RS06675 | 41676..43673 | 2497435 | glutathione-synthase-[Enterobacter-sp.-WCHEn090032] |
| WP_221885391.1 | SF2-Helicase+REase | GCF_003986875.1 | Enterobacter_sp._WCHEn090032 | Bacteria>Pseudomonadati>Pseudomonadota>Gammaproteobacteria>Enterobacterales>Enterobacteriaceae>Enterobacter>unclassified_Enterobacter | 604 | EKN83_RS06680 | 43755..45569 | 2497435 | DEAD/DEAH-box-helicase-family-protein-[Enterobacter-sp.-WCHEn090032] |
| POU04212.1 | SF2-Helicase+REase | GCA_002918705.1 | Escherichia_sp._ESNIH1 | Bacteria>Pseudomonadati>Pseudomonadota>Gammaproteobacteria>Enterobacterales>Enterobacteriaceae>Escherichia>unclassified_Escherichia | 665 | C3369_02355 | 489156..491153 | 1985876 | glutathione-synthase-[Escherichia-sp.-ESNIH1] |
| POU04213.1 | SF2-Helicase+REase | GCA_002918705.1 | Escherichia_sp._ESNIH1 | Bacteria>Pseudomonadati>Pseudomonadota>Gammaproteobacteria>Enterobacterales>Enterobacteriaceae>Escherichia>unclassified_Escherichia | 618 | C3369_02360 | 491193..493049 | 1985876 | glutathione-synthase-[Escherichia-sp.-ESNIH1] |
| WP_023329160.1 | SF2-Helicase+REase | GCF_021264605.1 | Klebsiella_oxytoca | Bacteria>Pseudomonadati>Pseudomonadota>Gammaproteobacteria>Enterobacterales>Enterobacteriaceae>Klebsiella/Raoultella_group>Klebsiella | 665 | LVW29_RS13130 | 89276..91273 | 571 | DEAD/DEAH-box-helicase-family-protein-[Klebsiella-oxytoca] |
| WP_023329159.1 | SF2-Helicase+REase | GCF_021264605.1 | Klebsiella_oxytoca | Bacteria>Pseudomonadati>Pseudomonadota>Gammaproteobacteria>Enterobacterales>Enterobacteriaceae>Klebsiella/Raoultella_group>Klebsiella | 620 | LVW29_RS13125 | 87366..89228 | 571 | DEAD/DEAH-box-helicase-family-protein-[Klebsiella-oxytoca] |
| HBQ4939445.1 | SF2-Helicase+REase | GCA_021861305.1 | Klebsiella_pneumoniae | Bacteria>Pseudomonadati>Pseudomonadota>Gammaproteobacteria>Enterobacterales>Enterobacteriaceae>Klebsiella/Raoultella_group>Klebsiella>Klebsiella_pneumoniae_complex | 665 | L7366_003899 | 41034..43031 | 573 | TPA:-hypothetical-protein-L7366_003899-[Klebsiella-pneumoniae] |
| HBS5818887.1 | SF2-Helicase+REase | GCA_021971955.1 | Klebsiella_pneumoniae | Bacteria>Pseudomonadati>Pseudomonadota>Gammaproteobacteria>Enterobacterales>Enterobacteriaceae>Klebsiella/Raoultella_group>Klebsiella>Klebsiella_pneumoniae_complex | 665 | MAK71_003681 | 15944..17941 | 573 | TPA:-hypothetical-protein-MAK71_003681-[Klebsiella-pneumoniae] |
| HBQ4939446.1 | SF2-Helicase+REase | GCA_021861305.1 | Klebsiella_pneumoniae | Bacteria>Pseudomonadati>Pseudomonadota>Gammaproteobacteria>Enterobacterales>Enterobacteriaceae>Klebsiella/Raoultella_group>Klebsiella>Klebsiella_pneumoniae_complex | 604 | L7366_003900 | 43113..44927 | 573 | DEAD/DEAH-box-helicase-family-protein-[Klebsiella-pneumoniae] |
| HBS5818886.1 | SF2-Helicase+REase | GCA_021971955.1 | Klebsiella_pneumoniae | Bacteria>Pseudomonadati>Pseudomonadota>Gammaproteobacteria>Enterobacterales>Enterobacteriaceae>Klebsiella/Raoultella_group>Klebsiella>Klebsiella_pneumoniae_complex | 604 | MAK71_003680 | 14048..15862 | 573 | DEAD/DEAH-box-helicase-family-protein-[Klebsiella-pneumoniae] |
| ECJ2539584.1 | SF2-Helicase+REase | GCA_007808255.1 | Salmonella_enterica_subsp._salamae | Bacteria>Pseudomonadati>Pseudomonadota>Gammaproteobacteria>Enterobacterales>Enterobacteriaceae>Salmonella>Salmonella_enterica | 618 | FNI64_03970 | 375682..377538 | 59202 | glutathione-synthase-[Salmonella-enterica-subsp.-salamae] |
| WP_272844020.1 | SF2-Helicase+REase | GCF_028555165.1 | Pantoea_ananatis | Bacteria>Pseudomonadati>Pseudomonadota>Gammaproteobacteria>Enterobacterales>Erwiniaceae>Pantoea | 665 | SZ66_RS15325 | 355344..357341 | 553 | hypothetical-protein-[Pantoea-ananatis] |
| WP_272844019.1 | SF2-Helicase+REase | GCF_028555165.1 | Pantoea_ananatis | Bacteria>Pseudomonadati>Pseudomonadota>Gammaproteobacteria>Enterobacterales>Erwiniaceae>Pantoea | 604 | SZ66_RS15320 | 353448..355262 | 553 | DEAD/DEAH-box-helicase-family-protein-[Pantoea-ananatis] |
| WP_187492463.1 | SF2-Helicase+REase | GCF_014353895.1 | Pantoea_agglomerans | Bacteria>Pseudomonadati>Pseudomonadota>Gammaproteobacteria>Enterobacterales>Erwiniaceae>Pantoea>Pantoea_agglomerans_group | 620 | IBU46_RS01720 | 13837..15699 | 549 | DEAD/DEAH-box-helicase-family-protein-[Pantoea-agglomerans] |
| WP_191918832.1 | SF2-Helicase+REase | GCF_014838975.1 | Pantoea_agglomerans | Bacteria>Pseudomonadati>Pseudomonadota>Gammaproteobacteria>Enterobacterales>Erwiniaceae>Pantoea>Pantoea_agglomerans_group | 620 | IFU26_RS12710 | 281262..283124 | 549 | DEAD/DEAH-box-helicase-family-protein-[Pantoea-agglomerans] |
| WP_199771974.1 | SF2-Helicase+REase | GCF_014838975.1 | Pantoea_agglomerans | Bacteria>Pseudomonadati>Pseudomonadota>Gammaproteobacteria>Enterobacterales>Erwiniaceae>Pantoea>Pantoea_agglomerans_group | 521 | IFU26_RS12705 | 279217..280782 | 549 | DEAD/DEAH-box-helicase-family-protein-[Pantoea-agglomerans] |
| MCL6350163.1 | SF2-Helicase+REase | GCA_023507955.1 | Pectobacterium_polaris | Bacteria>Pseudomonadati>Pseudomonadota>Gammaproteobacteria>Enterobacterales>Pectobacteriaceae>Pectobacterium | 665 | EXT50_03090 | 71554..73551 | 2042057 | glutathione-synthase-[Pectobacterium-polaris] |
| WP_211024707.1 | SF2-Helicase+REase | GCF_018094745.1 | Pectobacterium_versatile | Bacteria>Pseudomonadati>Pseudomonadota>Gammaproteobacteria>Enterobacterales>Pectobacteriaceae>Pectobacterium | 665 | F9U42_RS09270 | 78663..80660 | 2488639 | hypothetical-protein-[Pectobacterium-versatile] |
| MCL6350164.1 | SF2-Helicase+REase | GCA_023507955.1 | Pectobacterium_polaris | Bacteria>Pseudomonadati>Pseudomonadota>Gammaproteobacteria>Enterobacterales>Pectobacteriaceae>Pectobacterium | 618 | EXT50_03095 | 73591..75447 | 2042057 | glutathione-synthase-[Pectobacterium-polaris] |
| WP_211024710.1 | SF2-Helicase+REase | GCF_018094745.1 | Pectobacterium_versatile | Bacteria>Pseudomonadati>Pseudomonadota>Gammaproteobacteria>Enterobacterales>Pectobacteriaceae>Pectobacterium | 604 | F9U42_RS09275 | 80742..82556 | 2488639 | DEAD/DEAH-box-helicase-family-protein-[Pectobacterium-versatile] |
| GKV76953.1 | SF2-Helicase+REase | GCA_024342785.1 | Pectobacterium_carotovorum_subsp._carotovorum | Bacteria>Pseudomonadati>Pseudomonadota>Gammaproteobacteria>Enterobacterales>Pectobacteriaceae>Pectobacterium>Pectobacterium_carotovorum | 665 | PEC106568_21270 | 185170..187167 | 555 | hypothetical-protein-PEC106568_21270-[Pectobacterium-carotovorum-subsp.-carotovorum] |
| GKV76954.1 | SF2-Helicase+REase | GCA_024342785.1 | Pectobacterium_carotovorum_subsp._carotovorum | Bacteria>Pseudomonadati>Pseudomonadota>Gammaproteobacteria>Enterobacterales>Pectobacteriaceae>Pectobacterium>Pectobacterium_carotovorum | 602 | PEC106568_21280 | 187207..189015 | 555 | hypothetical-protein-PEC106568_21280-[Pectobacterium-carotovorum-subsp.-carotovorum] |
| WP_232770157.1 | SF2-Helicase+REase | GCF_021276285.1 | Rahnella_victoriana | Bacteria>Pseudomonadati>Pseudomonadota>Gammaproteobacteria>Enterobacterales>Yersiniaceae>Rahnella | 665 | J9880_RS04850 | 1040544..1042541 | 1510570 | hypothetical-protein-[Rahnella-victoriana] |
| WP_266057115.1 | SF2-Helicase+REase | GCF_026241995.1 | Rahnella_perminowiae | Bacteria>Pseudomonadati>Pseudomonadota>Gammaproteobacteria>Enterobacterales>Yersiniaceae>Rahnella | 665 | ORG37_RS00045 | 9829..11826 | 2816244 | DEAD/DEAH-box-helicase-family-protein-[Rahnella-perminowiae] |
| WP_232770158.1 | SF2-Helicase+REase | GCF_021276285.1 | Rahnella_victoriana | Bacteria>Pseudomonadati>Pseudomonadota>Gammaproteobacteria>Enterobacterales>Yersiniaceae>Rahnella | 620 | J9880_RS04855 | 1042589..1044451 | 1510570 | DEAD/DEAH-box-helicase-family-protein-[Rahnella-victoriana] |
| WP_266057117.1 | SF2-Helicase+REase | GCF_026241995.1 | Rahnella_perminowiae | Bacteria>Pseudomonadati>Pseudomonadota>Gammaproteobacteria>Enterobacterales>Yersiniaceae>Rahnella | 620 | ORG37_RS00050 | 11874..13736 | 2816244 | DEAD/DEAH-box-helicase-family-protein-[Rahnella-perminowiae] |
| NYA44688.1 | SF2-Helicase+REase | GCA_013401715.1 | Serratia_fonticola | Bacteria>Pseudomonadati>Pseudomonadota>Gammaproteobacteria>Enterobacterales>Yersiniaceae>Serratia | 665 | HZI31_15420 | 55915..57912 | 47917 | DEAD/DEAH-box-helicase-family-protein-[Serratia-fonticola] |
| WP_179248589.1 | SF2-Helicase+REase | GCF_013401715.1 | Serratia_fonticola | Bacteria>Pseudomonadati>Pseudomonadota>Gammaproteobacteria>Enterobacterales>Yersiniaceae>Serratia | 665 | HZI31_RS15420 | 55915..57912 | 47917 | DEAD/DEAH-box-helicase-family-protein-[Serratia-fonticola] |
| WP_206196478.1 | SF2-Helicase+REase | GCF_050037615.1 | Serratia_ureilytica | Bacteria>Pseudomonadati>Pseudomonadota>Gammaproteobacteria>Enterobacterales>Yersiniaceae>Serratia | 665 | PVU68_RS17170 | 228923..230920 | 300181 | DEAD/DEAH-box-helicase-family-protein-[Serratia-ureilytica] |
| WP_223536785.1 | SF2-Helicase+REase | GCF_019968705.1 | unclassified_Serratia_(in:_enterobacteria) | Bacteria>Pseudomonadati>Pseudomonadota>Gammaproteobacteria>Enterobacterales>Yersiniaceae>Serratia | 665 | KGP26_RS00365 | 89995..91992 | 2647522 | MULTISPECIES:-DEAD/DEAH-box-helicase-family-protein-[unclassified-Serratia-(in:-enterobacteria)] |
| NYA44687.1 | SF2-Helicase+REase | GCA_013401715.1 | Serratia_fonticola | Bacteria>Pseudomonadati>Pseudomonadota>Gammaproteobacteria>Enterobacterales>Yersiniaceae>Serratia | 618 | HZI31_15415 | 54041..55897 | 47917 | glutathione-synthase-[Serratia-fonticola] |
| WP_179248588.1 | SF2-Helicase+REase | GCF_013401715.1 | Serratia_fonticola | Bacteria>Pseudomonadati>Pseudomonadota>Gammaproteobacteria>Enterobacterales>Yersiniaceae>Serratia | 618 | HZI31_RS15415 | 54041..55897 | 47917 | glutathione-synthase-[Serratia-fonticola] |
| WP_223536788.1 | SF2-Helicase+REase | GCF_019968705.1 | unclassified_Serratia_(in:_enterobacteria) | Bacteria>Pseudomonadati>Pseudomonadota>Gammaproteobacteria>Enterobacterales>Yersiniaceae>Serratia | 618 | KGP26_RS00370 | 92010..93866 | 2647522 | MULTISPECIES:-hypothetical-protein-[unclassified-Serratia-(in:-enterobacteria)] |
| WP_273866018.1 | SF2-Helicase+REase | GCF_028616515.1 | Serratia_ureilytica | Bacteria>Pseudomonadati>Pseudomonadota>Gammaproteobacteria>Enterobacterales>Yersiniaceae>Serratia | 618 | PVU68_RS17165 | 227042..228898 | 300181 | DEAD/DEAH-box-helicase-family-protein-[Serratia-ureilytica] |
| WP_206196477.1 | SF2-Helicase+REase | GCF_050037615.1 | Serratia_ureilytica | Bacteria>Pseudomonadati>Pseudomonadota>Gammaproteobacteria>Enterobacterales>Yersiniaceae>Serratia | 603 | JY544_RS23395 | 48364..50175 | 300181 | DEAD/DEAH-box-helicase-family-protein-[Serratia-ureilytica] |
| WP_046051663.1 | SF2-Helicase+REase | GCF_000968115.1 | Yersinia_enterocolitica | Bacteria>Pseudomonadati>Pseudomonadota>Gammaproteobacteria>Enterobacterales>Yersiniaceae>Yersinia | 665 | YE5303_RS19605 | 4214573..4216570 | 630 | glutathione-synthase-[Yersinia-enterocolitica] |
| WP_197541081.1 | SF2-Helicase+REase | GCF_000968115.1 | Yersinia_enterocolitica | Bacteria>Pseudomonadati>Pseudomonadota>Gammaproteobacteria>Enterobacterales>Yersiniaceae>Yersinia | 604 | YE5303_RS19610 | 4216652..4218466 | 630 | DEAD/DEAH-box-helicase-family-protein-[Yersinia-enterocolitica] |
| MCP5206312.1 | SF2-Helicase+REase | GCA_024234475.1 | Hahellaceae_bacterium | Bacteria>Pseudomonadati>Pseudomonadota>Gammaproteobacteria>Oceanospirillales>Hahellaceae>unclassified_Hahellaceae | 659 | H7A01_03825 | 863410..865389 | 2048971 | glutathione-synthase-[Hahellaceae-bacterium] |
| MCP5206313.1 | SF2-Helicase+REase | GCA_024234475.1 | Hahellaceae_bacterium | Bacteria>Pseudomonadati>Pseudomonadota>Gammaproteobacteria>Oceanospirillales>Hahellaceae>unclassified_Hahellaceae | 620 | H7A01_03830 | 865396..867258 | 2048971 | glutathione-synthase-[Hahellaceae-bacterium] |
| WP_113594179.1 | SF2-Helicase+REase | GCF_043352035.1 | Vibrio | Bacteria>Pseudomonadati>Pseudomonadota>Gammaproteobacteria>Vibrionales>Vibrionaceae | 665 | DLR66_RS03560 | 59188..61185 | 662 | DEAD/DEAH-box-helicase-family-protein-[Vibrio-paracholerae] |
| WP_113594180.1 | SF2-Helicase+REase | GCF_043352035.1 | Vibrio | Bacteria>Pseudomonadati>Pseudomonadota>Gammaproteobacteria>Vibrionales>Vibrionaceae | 618 | DLR66_RS03565 | 61235..63091 | 662 | DEAD/DEAH-box-helicase-family-protein-[Vibrio-paracholerae] |
| WP_197573427.1 | SF2-Helicase+REase | - | Photobacterium_damselae | Bacteria>Pseudomonadati>Pseudomonadota>Gammaproteobacteria>Vibrionales>Vibrionaceae>Photobacterium | 665 | IL982_RS10165 | 2130787..2132784 | 38293 | DEAD/DEAH-box-helicase-family-protein-[Photobacterium-damselae] |
| WP_197573428.1 | SF2-Helicase+REase | GCF_014930975.1 | Photobacterium_damselae | Bacteria>Pseudomonadati>Pseudomonadota>Gammaproteobacteria>Vibrionales>Vibrionaceae>Photobacterium | 618 | IL982_RS10170 | 2132834..2134690 | 38293 | DEAD/DEAH-box-helicase-family-protein-[Photobacterium-damselae] |
| QQQ68155.1 | SF2-Helicase+REase | GCA_014930975.1 | Photobacterium_damselae_subsp._damselae | Bacteria>Pseudomonadati>Pseudomonadota>Gammaproteobacteria>Vibrionales>Vibrionaceae>Photobacterium>Photobacterium_damselae | 665 | IL982_10165 | 2130787..2132784 | 85581 | DEAD/DEAH-box-helicase-family-protein-[Photobacterium-damselae-subsp.-damselae] |
| QQQ68156.1 | SF2-Helicase+REase | GCA_014930975.1 | Photobacterium_damselae_subsp._damselae | Bacteria>Pseudomonadati>Pseudomonadota>Gammaproteobacteria>Vibrionales>Vibrionaceae>Photobacterium>Photobacterium_damselae | 618 | IL982_10170 | 2132834..2134690 | 85581 | DEAD/DEAH-box-helicase-family-protein-[Photobacterium-damselae-subsp.-damselae] |
| WP_077485097.1 | SF2-Helicase+REase | GCF_001995745.1 | Salinivibrio_sp._ML323 | Bacteria>Pseudomonadati>Pseudomonadota>Gammaproteobacteria>Vibrionales>Vibrionaceae>Salinivibrio>unclassified_Salinivibrio | 666 | BZG13_RS03585 | 138176..140176 | 1909474 | DEAD/DEAH-box-helicase-family-protein-[Salinivibrio-sp.-ML323] |
| WP_077485095.1 | SF2-Helicase+REase | GCF_001995745.1 | Salinivibrio_sp._ML323 | Bacteria>Pseudomonadati>Pseudomonadota>Gammaproteobacteria>Vibrionales>Vibrionaceae>Salinivibrio>unclassified_Salinivibrio | 620 | BZG13_RS03580 | 136272..138134 | 1909474 | DEAD/DEAH-box-helicase-family-protein-[Salinivibrio-sp.-ML323] |
| EGR0286972.1 | SF2-Helicase+REase | - | Vibrio_cholerae | Bacteria>Pseudomonadati>Pseudomonadota>Gammaproteobacteria>Vibrionales>Vibrionaceae>Vibrio | 665 | FK206_11670 | 73999..75996 | 666 | glutathione-synthase-[Vibrio-cholerae] |
| EGR2430333.1 | SF2-Helicase+REase | - | Vibrio_cholerae | Bacteria>Pseudomonadati>Pseudomonadota>Gammaproteobacteria>Vibrionales>Vibrionaceae>Vibrio | 665 | DYB83_10170 | 23876..25873 | 666 | glutathione-synthase-[Vibrio-cholerae] |
| EJL6578338.1 | SF2-Helicase+REase | GCA_024217525.1 | Vibrio_cholerae | Bacteria>Pseudomonadati>Pseudomonadota>Gammaproteobacteria>Vibrionales>Vibrionaceae>Vibrio | 665 | NMS50_002624 | 60079..62076 | 666 | DEAD/DEAH-box-helicase-family-protein-[Vibrio-cholerae] |
| WP_045588646.1 | SF2-Helicase+REase | - | Vibrio_vulnificus | Bacteria>Pseudomonadati>Pseudomonadota>Gammaproteobacteria>Vibrionales>Vibrionaceae>Vibrio | 665 | WG98_RS00480 | 104479..106476 | 672 | DEAD/DEAH-box-helicase-family-protein-[Vibrio-vulnificus] |
| EGR0286973.1 | SF2-Helicase+REase | - | Vibrio_cholerae | Bacteria>Pseudomonadati>Pseudomonadota>Gammaproteobacteria>Vibrionales>Vibrionaceae>Vibrio | 618 | FK206_11675 | 76046..77902 | 666 | glutathione-synthase-[Vibrio-cholerae] |
| EGR2430332.1 | SF2-Helicase+REase | - | Vibrio_cholerae | Bacteria>Pseudomonadati>Pseudomonadota>Gammaproteobacteria>Vibrionales>Vibrionaceae>Vibrio | 618 | DYB83_10165 | 21970..23826 | 666 | glutathione-synthase-[Vibrio-cholerae] |
| EJL6578339.1 | SF2-Helicase+REase | GCA_024217525.1 | Vibrio_cholerae | Bacteria>Pseudomonadati>Pseudomonadota>Gammaproteobacteria>Vibrionales>Vibrionaceae>Vibrio | 618 | NMS50_002625 | 62127..63983 | 666 | DEAD/DEAH-box-helicase-family-protein-[Vibrio-cholerae] |
| WP_045588645.1 | SF2-Helicase+REase | GCF_000959745.1 | Vibrio_vulnificus | Bacteria>Pseudomonadati>Pseudomonadota>Gammaproteobacteria>Vibrionales>Vibrionaceae>Vibrio | 618 | WG98_RS00475 | 102573..104429 | 672 | DEAD/DEAH-box-helicase-family-protein-[Vibrio-vulnificus] |
| EGR1144242.1 | SF2-Helicase+REase | - | Vibrio_parahaemolyticus | Bacteria>Pseudomonadati>Pseudomonadota>Gammaproteobacteria>Vibrionales>Vibrionaceae>Vibrio>Vibrio_harveyi_group | 665 | EC562_15475 | 29788..31785 | 670 | glutathione-synthase-[Vibrio-parahaemolyticus] |
| EGR1548460.1 | SF2-Helicase+REase | - | Vibrio_parahaemolyticus | Bacteria>Pseudomonadati>Pseudomonadota>Gammaproteobacteria>Vibrionales>Vibrionaceae>Vibrio>Vibrio_harveyi_group | 665 | D6Y11_16920 | 149989..151986 | 670 | glutathione-synthase-[Vibrio-parahaemolyticus] |
| EGR3302286.1 | SF2-Helicase+REase | - | Vibrio_parahaemolyticus | Bacteria>Pseudomonadati>Pseudomonadota>Gammaproteobacteria>Vibrionales>Vibrionaceae>Vibrio>Vibrio_harveyi_group | 665 | DMJ21_13160 | 20822..22819 | 670 | glutathione-synthase-[Vibrio-parahaemolyticus] |
| EHH2507625.1 | SF2-Helicase+REase | GCA_017637525.1 | Vibrio_parahaemolyticus | Bacteria>Pseudomonadati>Pseudomonadota>Gammaproteobacteria>Vibrionales>Vibrionaceae>Vibrio>Vibrio_harveyi_group | 665 | J7X10_003788 | 98287..100284 | 670 | DEAD/DEAH-box-helicase-family-protein-[Vibrio-parahaemolyticus] |
| EIS4855662.1 | SF2-Helicase+REase | GCA_021362915.1 | Vibrio_parahaemolyticus | Bacteria>Pseudomonadati>Pseudomonadota>Gammaproteobacteria>Vibrionales>Vibrionaceae>Vibrio>Vibrio_harveyi_group | 665 | LZX80_001788 | 106155..108152 | 670 | DEAD/DEAH-box-helicase-family-protein-[Vibrio-parahaemolyticus] |
| EJC1210123.1 | SF2-Helicase+REase | GCA_022997325.1 | Vibrio_parahaemolyticus | Bacteria>Pseudomonadati>Pseudomonadota>Gammaproteobacteria>Vibrionales>Vibrionaceae>Vibrio>Vibrio_harveyi_group | 665 | MX350_002600 | 18440..20437 | 670 | DEAD/DEAH-box-helicase-family-protein-[Vibrio-parahaemolyticus] |
| EJG0981287.1 | SF2-Helicase+REase | GCA_015813375.2 | Vibrio_parahaemolyticus | Bacteria>Pseudomonadati>Pseudomonadota>Gammaproteobacteria>Vibrionales>Vibrionaceae>Vibrio>Vibrio_harveyi_group | 665 | C4G85_RS03470 | 765418..767415 | 670 | DEAD/DEAH-box-helicase-family-protein-[Vibrio-parahaemolyticus] |
| HCM0920437.1 | SF2-Helicase+REase | GCA_025068375.1 | Vibrio_parahaemolyticus | Bacteria>Pseudomonadati>Pseudomonadota>Gammaproteobacteria>Vibrionales>Vibrionaceae>Vibrio>Vibrio_harveyi_group | 665 | N2828_003375 | 106156..108153 | 670 | TPA:-DEAD/DEAH-box-helicase-family-protein-[Vibrio-parahaemolyticus] |
| ODY54471.1 | SF2-Helicase+REase | GCA_001727265.1 | Vibrio_parahaemolyticus | Bacteria>Pseudomonadati>Pseudomonadota>Gammaproteobacteria>Vibrionales>Vibrionaceae>Vibrio>Vibrio_harveyi_group | 665 | BBM25_06270 | 16171..18168 | 670 | glutathione-synthase-[Vibrio-parahaemolyticus] |
| WP_053816404.1 | SF2-Helicase+REase | - | Vibrio_parahaemolyticus | Bacteria>Pseudomonadati>Pseudomonadota>Gammaproteobacteria>Vibrionales>Vibrionaceae>Vibrio>Vibrio_harveyi_group | 665 | BED52_RS16155 | 16177..18174 | 670 | DEAD/DEAH-box-helicase-family-protein-[Vibrio-parahaemolyticus] |
| WP_062854379.1 | SF2-Helicase+REase | - | Vibrio_parahaemolyticus | Bacteria>Pseudomonadati>Pseudomonadota>Gammaproteobacteria>Vibrionales>Vibrionaceae>Vibrio>Vibrio_harveyi_group | 665 | AWQ17_RS02405 | 106563..108560 | 670 | DEAD/DEAH-box-helicase-family-protein-[Vibrio-parahaemolyticus] |
| WP_069490030.1 | SF2-Helicase+REase | - | Vibrio_parahaemolyticus | Bacteria>Pseudomonadati>Pseudomonadota>Gammaproteobacteria>Vibrionales>Vibrionaceae>Vibrio>Vibrio_harveyi_group | 665 | BBM25_RS06270 | 16171..18168 | 670 | DEAD/DEAH-box-helicase-family-protein-[Vibrio-parahaemolyticus] |
| WP_102780437.1 | SF2-Helicase+REase | - | Vibrio_parahaemolyticus | Bacteria>Pseudomonadati>Pseudomonadota>Gammaproteobacteria>Vibrionales>Vibrionaceae>Vibrio>Vibrio_harveyi_group | 665 | CSR03_RS22380 | 134357..136354 | 670 | DEAD/DEAH-box-helicase-family-protein-[Vibrio-parahaemolyticus] |
| WP_104411095.1 | SF2-Helicase+REase | GCF_001558435.2 | Vibrio_harveyi | Bacteria>Pseudomonadati>Pseudomonadota>Gammaproteobacteria>Vibrionales>Vibrionaceae>Vibrio>Vibrio_harveyi_group | 665 | AL538_RS13495 | 2688246..2690243 | 669 | DEAD/DEAH-box-helicase-family-protein-[Vibrio-harveyi] |
| EGR1144243.1 | SF2-Helicase+REase | - | Vibrio_parahaemolyticus | Bacteria>Pseudomonadati>Pseudomonadota>Gammaproteobacteria>Vibrionales>Vibrionaceae>Vibrio>Vibrio_harveyi_group | 618 | EC562_15480 | 31835..33691 | 670 | glutathione-synthase-[Vibrio-parahaemolyticus] |
| EGR1548461.1 | SF2-Helicase+REase | - | Vibrio_parahaemolyticus | Bacteria>Pseudomonadati>Pseudomonadota>Gammaproteobacteria>Vibrionales>Vibrionaceae>Vibrio>Vibrio_harveyi_group | 618 | D6Y11_16925 | 152036..153892 | 670 | glutathione-synthase-[Vibrio-parahaemolyticus] |
| EGR3302285.1 | SF2-Helicase+REase | - | Vibrio_parahaemolyticus | Bacteria>Pseudomonadati>Pseudomonadota>Gammaproteobacteria>Vibrionales>Vibrionaceae>Vibrio>Vibrio_harveyi_group | 618 | DMJ21_13155 | 18916..20772 | 670 | glutathione-synthase-[Vibrio-parahaemolyticus] |
| EHH2507626.1 | SF2-Helicase+REase | GCA_017637525.1 | Vibrio_parahaemolyticus | Bacteria>Pseudomonadati>Pseudomonadota>Gammaproteobacteria>Vibrionales>Vibrionaceae>Vibrio>Vibrio_harveyi_group | 618 | J7X10_003789 | 100334..102190 | 670 | DEAD/DEAH-box-helicase-family-protein-[Vibrio-parahaemolyticus] |
| EIS4855661.1 | SF2-Helicase+REase | GCA_021362915.1 | Vibrio_parahaemolyticus | Bacteria>Pseudomonadati>Pseudomonadota>Gammaproteobacteria>Vibrionales>Vibrionaceae>Vibrio>Vibrio_harveyi_group | 618 | LZX80_001787 | 104249..106105 | 670 | DEAD/DEAH-box-helicase-family-protein-[Vibrio-parahaemolyticus] |
| EJC1210122.1 | SF2-Helicase+REase | GCA_022997325.1 | Vibrio_parahaemolyticus | Bacteria>Pseudomonadati>Pseudomonadota>Gammaproteobacteria>Vibrionales>Vibrionaceae>Vibrio>Vibrio_harveyi_group | 618 | MX350_002599 | 16534..18390 | 670 | DEAD/DEAH-box-helicase-family-protein-[Vibrio-parahaemolyticus] |
| EJG0981288.1 | SF2-Helicase+REase | GCA_015813375.2 | Vibrio_parahaemolyticus | Bacteria>Pseudomonadati>Pseudomonadota>Gammaproteobacteria>Vibrionales>Vibrionaceae>Vibrio>Vibrio_harveyi_group | 618 | C4G85_RS03475 | 767465..769321 | 670 | DEAD/DEAH-box-helicase-family-protein-[Vibrio-parahaemolyticus] |

| Protein-ID | Domain | Assembly-ID | Strain | Taxonomy | Length | Locus-Tag | Genomic-coordinates | TaxID | Fasta-defline |
|---|---|---|---|---|---|---|---|---|---|
| HCM0920436.1 | SF2-Helicase+REase | GCA_025068375.1 | Vibrio_parahaemolyticus | Bacteria>Pseudomonadati>Pseudomonadota>Gammaproteobacteria>Vibrionales>Vibrionaceae>Vibrio>Vibrio_harveyi_group | 618 | N2828_003374 | 104250..106106 | 670 | TPA:-DEAD/DEAH-box-helicase-family-protein-[Vibrio-parahaemolyticus] |
| ODY54470.1 | SF2-Helicase+REase | GCA_001727265.1 | Vibrio_parahaemolyticus | Bacteria>Pseudomonadati>Pseudomonadota>Gammaproteobacteria>Vibrionales>Vibrionaceae>Vibrio>Vibrio_harveyi_group | 618 | BBM25_06265 | 14265..16121 | 670 | glutathione-synthase-[Vibrio-parahaemolyticus] |
| WP_061065631.1 | SF2-Helicase+REase | GCF_001558435.2 | Vibrio_harveyi | Bacteria>Pseudomonadati>Pseudomonadota>Gammaproteobacteria>Vibrionales>Vibrionaceae>Vibrio>Vibrio_harveyi_group | 618 | AL538_RS13490 | 2686350..2688206 | 669 | DEAD/DEAH-box-helicase-family-protein-[Vibrio-harveyi] |
| WP_062854378.1 | SF2-Helicase+REase | GCF_001608855.1 | Vibrio_parahaemolyticus | Bacteria>Pseudomonadati>Pseudomonadota>Gammaproteobacteria>Vibrionales>Vibrionaceae>Vibrio>Vibrio_harveyi_group | 618 | AWQ17_RS02400 | 104657..106513 | 670 | DEAD/DEAH-box-helicase-family-protein-[Vibrio-parahaemolyticus] |
| WP_065970500.1 | SF2-Helicase+REase | GCF_001704915.1 | Vibrio_parahaemolyticus | Bacteria>Pseudomonadati>Pseudomonadota>Gammaproteobacteria>Vibrionales>Vibrionaceae>Vibrio>Vibrio_harveyi_group | 618 | BED52_RS16150 | 14271..16127 | 670 | DEAD/DEAH-box-helicase-family-protein-[Vibrio-parahaemolyticus] |
| WP_069490029.1 | SF2-Helicase+REase | GCF_001727265.1 | Vibrio_parahaemolyticus | Bacteria>Pseudomonadati>Pseudomonadota>Gammaproteobacteria>Vibrionales>Vibrionaceae>Vibrio>Vibrio_harveyi_group | 618 | BBM25_RS06265 | 14265..16121 | 670 | DEAD/DEAH-box-helicase-family-protein-[Vibrio-parahaemolyticus] |
| WP_102780436.1 | SF2-Helicase+REase | GCF_030238805.1 | Vibrio_parahaemolyticus | Bacteria>Pseudomonadati>Pseudomonadota>Gammaproteobacteria>Vibrionales>Vibrionaceae>Vibrio>Vibrio_harveyi_group | 618 | CSR03_RS22375 | 132451..134307 | 670 | DEAD/DEAH-box-helicase-family-protein-[Vibrio-parahaemolyticus] |
| --TUDOR+SF1-SF2-Helicase+iREase+CTD+REase(BrxHII-Helicase) | | | | | | | | | |
| SLK50046.1 | TUDOR+SF1-SF2-Helicase+iREase+CTD+REase | GCA_900132645.1 | Mycobacteroides_abscessus_subsp._abscessus | Bacteria>Bacillati>Actinomycetota>Actinomycetes>Mycobacteriales>Mycobacteriaceae>Mycobacteroides>Mycobacteroides_abscessus | 1061 | SAMEA2071448_00692 | 747503..750688 | 1185650 | helicase-domain-containing-protein-[Mycobacteroides-abscessus-subsp.-abscessus] |
| OUS97319.1 | TUDOR+SF1-SF2-Helicase+iREase+CTD+REase | GCA_002165735.1 | Rhodococcus_sp._NCIMB_12038 | Bacteria>Bacillati>Actinomycetota>Actinomycetes>Mycobacteriales>Nocardiaceae>Rhodococcus>unclassified_Rhodococcus_(in:_high_G+C_Gram-positive_bacteria) | 1136 | CA951_02940 | 143247..146657 | 933800 | helicase-[Rhodococcus-sp.-NCIMB-12038] |
| WP_087555445.1 | TUDOR+SF1-SF2-Helicase+iREase+CTD+REase | GCF_002165735.1 | Rhodococcus_sp._NCIMB_12038 | Bacteria>Bacillati>Actinomycetota>Actinomycetes>Mycobacteriales>Nocardiaceae>Rhodococcus>unclassified_Rhodococcus_(in:_high_G+C_Gram-positive_bacteria) | 1136 | CA951_RS02955 | 143247..146657 | 933800 | helicase-related-protein-[Rhodococcus-sp.-NCIMB-12038] |
| REB77447.1 | TUDOR+SF1-SF2-Helicase+iREase+CTD+REase | GCA_003384685.1 | Cutibacterium_acnes | Bacteria>Bacillati>Actinomycetota>Actinomycetes>Propionibacteriales>Propionibacteriaceae>Cutibacterium | 1059 | CP883_05565 | 1474..4653 | 1747 | helicase-[Cutibacterium-acnes] |
| WP_029421467.1 | TUDOR+SF1-SF2-Helicase+iREase+CTD+REase | GCF_000702485.1 | Alicyclobacillus_macrosporangiidus | Bacteria>Bacillati>Bacillota>Bacilli>Bacillales>Alicyclobacillaceae>Alicyclobacillus | 1039 | N687_RS0108590 | 1716398..1719517 | 392015 | helicase-related-protein,-partial-[Alicyclobacillus-macrosporangiidus] |
| WP_241772001.1 | TUDOR+SF1-SF2-Helicase+iREase+CTD+REase | GCF_002243665.1 | Geobacillus | Bacteria>Bacillati>Bacillota>Bacilli>Bacillales>Anoxybacillaceae | 934 | GT3921_RS12985 | 2553566..2556370 | 129337 | MULTISPECIES:-helicase-related-protein-[Geobacillus] |
| WP_192949324.1 | TUDOR+SF1-SF2-Helicase+iREase+CTD+REase | GCF_014890335.1 | Anoxybacillus_flavithermus | Bacteria>Bacillati>Bacillota>Bacilli>Bacillales>Anoxybacillaceae>Anoxybacillus | 1066 | HPK03_RS06765 | 4369..7569 | 33934 | helicase-related-protein-[Anoxybacillus-flavithermus] |
| WP_077428555.1 | TUDOR+SF1-SF2-Helicase+iREase+CTD+REase | GCF_001996285.1 | Anoxybacillus_kestanbolensis | Bacteria>Bacillati>Bacillota>Bacilli>Bacillales>Anoxybacillaceae>Anoxybacillus | 1062 | BO219_RS04420 | 404253..407441 | 227476 | helicase-related-protein-[Anoxybacillus-karvacharensis] |
| MBB3906531.1 | TUDOR+SF1-SF2-Helicase+iREase+CTD+REase | GCA_014196195.1 | Anoxybacteroides_rupiense | Bacteria>Bacillati>Bacillota>Bacilli>Bacillales>Anoxybacillaceae>Anoxybacteroides | 1066 | GGQ95_000808 | 70191..73391 | 311460 | superfamily-II-DNA-or-RNA-helicase-[Anoxybacillus-rupiensis] |
| WP_275191610.1 | TUDOR+SF1-SF2-Helicase+iREase+CTD+REase | GCF_029077325.1 | Anoxybacteroides_rupiense | Bacteria>Bacillati>Bacillota>Bacilli>Bacillales>Anoxybacillaceae>Anoxybacteroides | 1066 | PNH38_RS00025 | 5268..8468 | 311460 | helicase-related-protein-[Anoxybacillus-rupiensis] |
| KZE90473.1 | TUDOR+SF1-SF2-Helicase+iREase+CTD+REase | GCA_001620045.1 | Geobacillus_stearothermophilus | Bacteria>Bacillati>Bacillota>Bacilli>Bacillales>Anoxybacillaceae>Geobacillus | 1067 | AVP43_03290 | 64849..68052 | 1422 | RNA-polymerase-associated-protein-RapA-[Geobacillus-stearothermophilus] |
| WP_160269227.1 | TUDOR+SF1-SF2-Helicase+iREase+CTD+REase | - | Geobacillus_stearothermophilus | Bacteria>Bacillati>Bacillota>Bacilli>Bacillales>Anoxybacillaceae>Geobacillus | 1066 | EPB69_RS06780 | 1340591..1343791 | 1422 | helicase-related-protein-[Geobacillus-stearothermophilus] |
| ASS85707.1 | TUDOR+SF1-SF2-Helicase+iREase+CTD+REase | GCA_002243605.1 | Geobacillus_lituanicus | Bacteria>Bacillati>Bacillota>Bacilli>Bacillales>Anoxybacillaceae>Geobacillus>Geobacillus_thermoleovorans_group | 1062 | GLN3_00255 | 54367..57555 | 169283 | helicase-[Geobacillus-lituanicus] |
| AKM18688.1 | TUDOR+SF1-SF2-Helicase+iREase+CTD+REase | GCA_001028085.1 | Geobacillus_sp._12AMOR1 | Bacteria>Bacillati>Bacillota>Bacilli>Bacillales>Anoxybacillaceae>Geobacillus>unclassified_Geobacillus | 1067 | GARCT_01395 | 1376573..1379776 | 1629723 | RNA-polymerase-associated-protein-RapA-[Geobacillus-sp.-12AMOR1] |
| WP_062679362.1 | TUDOR+SF1-SF2-Helicase+iREase+CTD+REase | GCF_001587455.1 | Parageobacillus_toebii | Bacteria>Bacillati>Bacillota>Bacilli>Bacillales>Anoxybacillaceae>Parageobacillus | 1066 | B4110_RS17520 | 13106..16306 | 153151 | helicase-related-protein-[Parageobacillus-toebii] |
| WP_003250548.1 | TUDOR+SF1-SF2-Helicase+iREase+CTD+REase | GCF_000258725.1 | Parageobacillus_thermoglucosidasius | Bacteria>Bacillati>Bacillota>Bacilli>Bacillales>Anoxybacillaceae>Parageobacillus | 1058 | GT20_RS09305 | 1827658..1830834 | 1426 | helicase-related-protein-[Parageobacillus-thermoglucosidasius] |
| WP_132948759.1 | TUDOR+SF1-SF2-Helicase+iREase+CTD+REase | GCF_004341205.1 | Thermolongibacillus_altinsuensis | Bacteria>Bacillati>Bacillota>Bacilli>Bacillales>Anoxybacillaceae>Thermolongibacillus | 1062 | EDD69_RS09650 | 69704..72892 | 575256 | helicase-related-protein-[Thermolongibacillus-altinsuensis] |
| WP_076874147.1 | TUDOR+SF1-SF2-Helicase+iREase+CTD+REase | GCF_039778265.1 | Bacillus | Bacteria>Bacillati>Bacillota>Bacilli>Bacillales>Bacillaceae | 1071 | BACERE00188_RS21760 | 59288..62503 | 1386 | MULTISPECIES:-helicase-related-protein-[Bacillus-cereus-group] |
| WP_098349133.1 | TUDOR+SF1-SF2-Helicase+iREase+CTD+REase | GCF_050171705.1 | Bacillus | Bacteria>Bacillati>Bacillota>Bacilli>Bacillales>Bacillaceae | 1066 | CN915_RS11650 | 28985..32185 | 1386 | MULTISPECIES:-helicase-related-protein-[Bacillus] |
| WP_229143296.1 | TUDOR+SF1-SF2-Helicase+iREase+CTD+REase | GCF_020686245.1 | Bacillus | Bacteria>Bacillati>Bacillota>Bacilli>Bacillales>Bacillaceae | 1063 | LKL84_RS22395 | 83475..86666 | 1386 | helicase-related-protein-[Bacillus-paranthracis] |
| WP_212956140.1 | TUDOR+SF1-SF2-Helicase+iREase+CTD+REase | GCF_018332475.1 | Bacillus | Bacteria>Bacillati>Bacillota>Bacilli>Bacillales>Bacillaceae | 1061 | KI369_RS12450 | 29180..32365 | 1386 | MULTISPECIES:-helicase-related-protein-[Bacillus] |
| WP_231419118.1 | TUDOR+SF1-SF2-Helicase+iREase+CTD+REase | GCF_030166655.1 | Pontibacillus | Bacteria>Bacillati>Bacillota>Bacilli>Bacillales>Bacillaceae | 1059 | LQ226_RS16655 | 53061..56240 | 289201 | helicase-related-protein-[Pontibacillus-sp.-HN14] |
| WP_083979171.1 | TUDOR+SF1-SF2-Helicase+iREase+CTD+REase | GCF_000948175.1 | Aeribacillus_alveayuensis | Bacteria>Bacillati>Bacillota>Bacilli>Bacillales>Bacillaceae>Aeribacillus | 1066 | LG50_RS18050 | 13546..16746 | 279215 | helicase-related-protein-[Bacillus-alveayuensis] |
| WP_146817803.1 | TUDOR+SF1-SF2-Helicase+iREase+CTD+REase | GCF_007991275.1 | Alkalibacillus_haloalkaliphilus | Bacteria>Bacillati>Bacillota>Bacilli>Bacillales>Bacillaceae>Alkalibacillus | 1068 | AHA02nite_RS12510 | 55575..58781 | 94136 | helicase-related-protein-[Alkalibacillus-haloalkaliphilus] |
| WP_242290715.1 | TUDOR+SF1-SF2-Helicase+iREase+CTD+REase | GCF_026684175.1 | Bacillus_cereus_group | Bacteria>Bacillati>Bacillota>Bacilli>Bacillales>Bacillaceae>Bacillus | 1071 | OU819_RS04795 | 940609..943824 | 86661 | MULTISPECIES:-helicase-related-protein-[Bacillus-cereus-group] |
| WP_003349670.1 | TUDOR+SF1-SF2-Helicase+iREase+CTD+REase | GCF_000262735.1 | Bacillus_methanolicus | Bacteria>Bacillati>Bacillota>Bacilli>Bacillales>Bacillaceae>Bacillus | 1066 | MGA3_RS15515 | 49774..52974 | 1471 | helicase-related-protein-[Bacillus-methanolicus] |
| WP_248893730.1 | TUDOR+SF1-SF2-Helicase+iREase+CTD+REase | GCF_023278615.1 | Bacillus_methanolicus | Bacteria>Bacillati>Bacillota>Bacilli>Bacillales>Bacillaceae>Bacillus | 1066 | C0971_RS07460 | 1491713..1494913 | 1471 | helicase-related-protein-[Bacillus-methanolicus] |
| WP_075982593.1 | TUDOR+SF1-SF2-Helicase+iREase+CTD+REase | - | Bacillus_massilionigeriensis | Bacteria>Bacillati>Bacillota>Bacilli>Bacillales>Bacillaceae>Bacillus | 1064 | FX071_RS15920 | 3233225..3236419 | 1805475 | helicase-related-protein-[Bacillus-massilionigeriensis] |
| WP_242320711.1 | TUDOR+SF1-SF2-Helicase+iREase+CTD+REase | GCF_022663735.1 | Bacillus_cereus_group | Bacteria>Bacillati>Bacillota>Bacilli>Bacillales>Bacillaceae>Bacillus | 1063 | MR162_RS10440 | 204592..207783 | 86661 | helicase-related-protein-[Bacillus-cereus-group-sp.-BfR-BA-01354] |
| WP_100332648.1 | TUDOR+SF1-SF2-Helicase+iREase+CTD+REase | - | Bacillus_xiapuensis | Bacteria>Bacillati>Bacillota>Bacilli>Bacillales>Bacillaceae>Bacillus | 1062 | CEF20_RS14900 | 662159..665347 | 2014075 | helicase-related-protein-[Bacillus-xiapuensis] |
| WP_073518922.1 | TUDOR+SF1-SF2-Helicase+iREase+CTD+REase | GCF_900094815.1 | Bacillus_cereus_group | Bacteria>Bacillati>Bacillota>Bacilli>Bacillales>Bacillaceae>Bacillus | 1061 | CCY53_RS04740 | 220032..223217 | 86661 | MULTISPECIES:-helicase-related-protein-[Bacillus-cereus-group] |
| WP_075981297.1 | TUDOR+SF1-SF2-Helicase+iREase+CTD+REase | GCF_902386855.1 | Bacillus_massilionigeriensis | Bacteria>Bacillati>Bacillota>Bacilli>Bacillales>Bacillaceae>Bacillus | 1061 | BQ6100_RS08705 | 1746691..1749876 | 1805475 | helicase-related-protein-[Bacillus-massilionigeriensis] |
| WP_098878443.1 | TUDOR+SF1-SF2-Helicase+iREase+CTD+REase | GCF_016483555.1 | Bacillus_cereus_group | Bacteria>Bacillati>Bacillota>Bacilli>Bacillales>Bacillaceae>Bacillus | 1061 | JDS82_RS26205 | 23551..26736 | 86661 | MULTISPECIES:-helicase-related-protein-[Bacillus-cereus-group] |
| WP_236801617.1 | TUDOR+SF1-SF2-Helicase+iREase+CTD+REase | GCF_021655215.1 | Bacillus_cereus_group | Bacteria>Bacillati>Bacillota>Bacilli>Bacillales>Bacillaceae>Bacillus | 1061 | L6484_RS04925 | 999407..1002592 | 86661 | helicase-related-protein-[Bacillus-cereus] |
| WP_226516203.1 | TUDOR+SF1-SF2-Helicase+iREase+CTD+REase | GCF_020519665.1 | Bacillus_shivajii | Bacteria>Bacillati>Bacillota>Bacilli>Bacillales>Bacillaceae>Bacillus | 1059 | LGQ02_RS20925 | 4376871..4380050 | 1983719 | helicase-related-protein-[Bacillus-shivajii] |
| WP_167560816.1 | TUDOR+SF1-SF2-Helicase+iREase+CTD+REase | GCF_012030395.1 | Bacillus_subtilis_group | Bacteria>Bacillati>Bacillota>Bacilli>Bacillales>Bacillaceae>Bacillus | 1057 | MSS26_RS04975 | 994002..997175 | 653685 | MULTISPECIES:-helicase-related-protein-[Bacillus-subtilis-group] |
| WP_184528629.1 | TUDOR+SF1-SF2-Helicase+iREase+CTD+REase | - | Bacillus_benzoevorans | Bacteria>Bacillati>Bacillota>Bacilli>Bacillales>Bacillaceae>Bacillus | 1057 | HNR53_RS18160 | 57580..60753 | 1456 | helicase-related-protein-[Bacillus-benzoevorans] |
| PES78866.1 | TUDOR+SF1-SF2-Helicase+iREase+CTD+REase | GCA_002558905.1 | Bacillus_anthracis | Bacteria>Bacillati>Bacillota>Bacilli>Bacillales>Bacillaceae>Bacillus>Bacillus_cereus_group | 1071 | CN504_21540 | 18656..21871 | 1392 | helicase-[Bacillus-anthracis] |
| WP_202283224.1 | TUDOR+SF1-SF2-Helicase+iREase+CTD+REase | GCF_016757895.1 | Bacillus_cereus | Bacteria>Bacillati>Bacillota>Bacilli>Bacillales>Bacillaceae>Bacillus>Bacillus_cereus_group | 1071 | IRV09_RS02760 | 77901..81116 | 1396 | helicase-related-protein-[Bacillus-cereus] |
| PED47020.1 | TUDOR+SF1-SF2-Helicase+iREase+CTD+REase | GCA_002552505.1 | Bacillus_cereus | Bacteria>Bacillati>Bacillota>Bacilli>Bacillales>Bacillaceae>Bacillus>Bacillus_cereus_group | 1066 | CON49_25645 | 19358..22558 | 1396 | helicase-[Bacillus-cereus] |
| PFA00388.1 | TUDOR+SF1-SF2-Helicase+iREase+CTD+REase | GCA_002561255.1 | Bacillus_cereus | Bacteria>Bacillati>Bacillota>Bacilli>Bacillales>Bacillaceae>Bacillus>Bacillus_cereus_group | 1066 | CN375_05465 | 25128..28328 | 1396 | helicase-[Bacillus-cereus] |
| PFT90553.1 | TUDOR+SF1-SF2-Helicase+iREase+CTD+REase | GCA_002568375.1 | Bacillus_cereus | Bacteria>Bacillati>Bacillota>Bacilli>Bacillales>Bacillaceae>Bacillus>Bacillus_cereus_group | 1066 | COK66_16910 | 25462..28662 | 1396 | helicase-[Bacillus-cereus] |
| PGQ77676.1 | TUDOR+SF1-SF2-Helicase+iREase+CTD+REase | GCA_002576945.1 | Bacillus_anthracis | Bacteria>Bacillati>Bacillota>Bacilli>Bacillales>Bacillaceae>Bacillus>Bacillus_cereus_group | 1066 | COA15_22140 | 371839..375039 | 1392 | helicase-[Bacillus-anthracis] |
| WP_001994590.1 | TUDOR+SF1-SF2-Helicase+iREase+CTD+REase | GCF_000832865.1 | Bacillus_cereus | Bacteria>Bacillati>Bacillota>Bacilli>Bacillales>Bacillaceae>Bacillus>Bacillus_cereus_group | 1066 | AK40_RS29435 | 4981255..4984455 | 1396 | helicase-related-protein-[Bacillus-cereus] |
| WP_098606670.1 | TUDOR+SF1-SF2-Helicase+iREase+CTD+REase | GCF_002575835.1 | Bacillus_cereus | Bacteria>Bacillati>Bacillota>Bacilli>Bacillales>Bacillaceae>Bacillus>Bacillus_cereus_group | 1066 | COA03_RS10385 | 77009..80209 | 1396 | helicase-related-protein-[Bacillus-cereus] |
| WP_131246815.1 | TUDOR+SF1-SF2-Helicase+iREase+CTD+REase | GCF_004330345.1 | Bacillus_mycoides | Bacteria>Bacillati>Bacillota>Bacilli>Bacillales>Bacillaceae>Bacillus>Bacillus_cereus_group | 1066 | EOM27_RS18710 | 94607..97807 | 1405 | helicase-related-protein-[Bacillus-mycoides] |
| PRT22694.1 | TUDOR+SF1-SF2-Helicase+iREase+CTD+REase | GCA_002998815.1 | Bacillus_thuringiensis | Bacteria>Bacillati>Bacillota>Bacilli>Bacillales>Bacillaceae>Bacillus>Bacillus_cereus_group | 1064 | C6351_31320 | 23044..26238 | 1428 | helicase-[Bacillus-thuringiensis] |
| WP_144565373.1 | TUDOR+SF1-SF2-Helicase+iREase+CTD+REase | - | Bacillus_mobilis | Bacteria>Bacillati>Bacillota>Bacilli>Bacillales>Bacillaceae>Bacillus>Bacillus_cereus_group | 1064 | FS794_RS23145 | 76329..79523 | 2026190 | helicase-related-protein-[Bacillus-mobilis] |
| PEY26879.1 | TUDOR+SF1-SF2-Helicase+iREase+CTD+REase | GCA_002559605.1 | Bacillus_anthracis | Bacteria>Bacillati>Bacillota>Bacilli>Bacillales>Bacillaceae>Bacillus>Bacillus_cereus_group | 1063 | CN340_12740 | 60581..63772 | 1392 | helicase-[Bacillus-anthracis] |
| WP_098147713.1 | TUDOR+SF1-SF2-Helicase+iREase+CTD+REase | GCF_002556085.1 | Bacillus_wiedmannii | Bacteria>Bacillati>Bacillota>Bacilli>Bacillales>Bacillaceae>Bacillus>Bacillus_cereus_group | 1063 | CN685_RS18705 | 16324..19515 | 1890302 | helicase-related-protein-[Bacillus-wiedmannii] |
| WP_233723730.1 | TUDOR+SF1-SF2-Helicase+iREase+CTD+REase | GCF_021655055.1 | Bacillus_cereus | Bacteria>Bacillati>Bacillota>Bacilli>Bacillales>Bacillaceae>Bacillus>Bacillus_cereus_group | 1063 | L6497_RS04800 | 959297..962488 | 1396 | helicase-related-protein-[Bacillus-cereus] |
| WP_098613364.1 | TUDOR+SF1-SF2-Helicase+iREase+CTD+REase | GCF_002567495.1 | Bacillus_cereus | Bacteria>Bacillati>Bacillota>Bacilli>Bacillales>Bacillaceae>Bacillus>Bacillus_cereus_group | 1062 | COK05_RS16040 | 48438..51626 | 1396 | helicase-related-protein-[Bacillus-cereus] |
| PGR68854.1 | TUDOR+SF1-SF2-Helicase+iREase+CTD+REase | GCA_002576445.1 | Bacillus_cereus | Bacteria>Bacillati>Bacillota>Bacilli>Bacillales>Bacillaceae>Bacillus>Bacillus_cereus_group | 1061 | COC49_05315 | 29641..32826 | 1396 | helicase-[Bacillus-cereus] |
| WP_002116632.1 | TUDOR+SF1-SF2-Helicase+iREase+CTD+REase | - | Bacillus_cereus | Bacteria>Bacillati>Bacillota>Bacilli>Bacillales>Bacillaceae>Bacillus>Bacillus_cereus_group | 1061 | IEQ_RS03965 | 801060..804245 | 1396 | helicase-related-protein-[Bacillus-cereus] |
| WP_098016183.1 | TUDOR+SF1-SF2-Helicase+iREase+CTD+REase | GCF_002551735.1 | Bacillus_toyonensis | Bacteria>Bacillati>Bacillota>Bacilli>Bacillales>Bacillaceae>Bacillus>Bacillus_cereus_group | 1061 | JDS69_RS23075 | 93170..96355 | 155322 | MULTISPECIES:-helicase-related-protein-[Bacillus-cereus-group] |
| WP_215562749.1 | TUDOR+SF1-SF2-Helicase+iREase+CTD+REase | GCF_018741585.1 | Bacillus_wiedmannii | Bacteria>Bacillati>Bacillota>Bacilli>Bacillales>Bacillaceae>Bacillus>Bacillus_cereus_group | 1061 | EXW48_RS05035 | 1027350..1030535 | 1890302 | helicase-related-protein-[Bacillus-wiedmannii] |
| WP_098359291.1 | TUDOR+SF1-SF2-Helicase+iREase+CTD+REase | GCF_002561295.1 | Bacillus_cereus | Bacteria>Bacillati>Bacillota>Bacilli>Bacillales>Bacillaceae>Bacillus>Bacillus_cereus_group | 1060 | CN373_RS10785 | 71583..74765 | 1396 | helicase-related-protein-[Bacillus-cereus] |
| EDX56282.1 | TUDOR+SF1-SF2-Helicase+iREase+CTD+REase | - | Bacillus_cereus_W | Bacteria>Bacillati>Bacillota>Bacilli>Bacillales>Bacillaceae>Bacillus>Bacillus_cereus_group>Bacillus_cereus | 1063 | BCW_0926 | 64027..67218 | 405917 | helicase-domain-protein-[Bacillus-cereus-W] |
| WP_270691667.1 | TUDOR+SF1-SF2-Helicase+iREase+CTD+REase | GCF_027715825.1 | Bacillus_cereus_group_sp._Bc061 | Bacteria>Bacillati>Bacillota>Bacilli>Bacillales>Bacillaceae>Bacillus>Bacillus_cereus_group>unclassified_Bacillus_cereus_group | 1066 | PDQ40_RS23445 | 102285..105485 | 3018117 | helicase-related-protein-[Bacillus-cereus-group-sp.-Bc061] |
| WP_242261889.1 | TUDOR+SF1-SF2-Helicase+iREase+CTD+REase | GCF_022662915.1 | Bacillus_cereus_group_sp._BfR-BA-01453 | Bacteria>Bacillati>Bacillota>Bacilli>Bacillales>Bacillaceae>Bacillus>Bacillus_cereus_group>unclassified_Bacillus_cereus_group | 1063 | MR070_RS19995 | 115308..118499 | 2920355 | helicase-related-protein-[Bacillus-cereus-group-sp.-BfR-BA-01453] |
| WP_242225821.1 | TUDOR+SF1-SF2-Helicase+iREase+CTD+REase | GCF_022663655.1 | Bacillus_cereus_group_sp._BfR-BA-01358 | Bacteria>Bacillati>Bacillota>Bacilli>Bacillales>Bacillaceae>Bacillus>Bacillus_cereus_group>unclassified_Bacillus_cereus_group | 1061 | MR078_RS05875 | 28869..32054 | 2920320 | helicase-related-protein-[Bacillus-cereus-group-sp.-BfR-BA-01358] |
| WP_161986549.1 | TUDOR+SF1-SF2-Helicase+iREase+CTD+REase | - | Bacillus_subtilis | Bacteria>Bacillati>Bacillota>Bacilli>Bacillales>Bacillaceae>Bacillus>Bacillus_subtilis_group | 1061 | GPJ55_RS17645 | 3388929..3392114 | 1423 | helicase-related-protein-[Bacillus-subtilis] |
| WP_268328402.1 | TUDOR+SF1-SF2-Helicase+iREase+CTD+REase | - | Bacillus_inaquosorum | Bacteria>Bacillati>Bacillota>Bacilli>Bacillales>Bacillaceae>Bacillus>Bacillus_subtilis_group | 1061 | MOD47_RS12350 | 100317..103502 | 483913 | helicase-related-protein-[Bacillus-inaquosorum] |
| WP_268298779.1 | TUDOR+SF1-SF2-Helicase+iREase+CTD+REase | GCF_026789485.1 | Bacillus_inaquosorum | Bacteria>Bacillati>Bacillota>Bacilli>Bacillales>Bacillaceae>Bacillus>Bacillus_subtilis_group | 1057 | MOB75_RS13805 | 106363..109536 | 483913 | helicase-related-protein-[Bacillus-inaquosorum] |
| WP_152155469.1 | TUDOR+SF1-SF2-Helicase+iREase+CTD+REase | GCF_009183765.1 | Bacillus_sp._B3-WWTP-C-10-D-3 | Bacteria>Bacillati>Bacillota>Bacilli>Bacillales>Bacillaceae>Bacillus>unclassified_Bacillus_(in:_firmicutes) | 1071 | GBN83_RS14850 | 13070..16285 | 2653217 | helicase-related-protein-[Bacillus-sp.-B3-WWTP-C-10-D-3] |
| WP_098172536.1 | TUDOR+SF1-SF2-Helicase+iREase+CTD+REase | GCF_002555445.1 | Bacillus_sp._AFS017336 | Bacteria>Bacillati>Bacillota>Bacilli>Bacillales>Bacillaceae>Bacillus>unclassified_Bacillus_(in:_firmicutes) | 1070 | CN601_RS23850 | 73244..76456 | 2033489 | helicase-related-protein-[Bacillus-sp.-AFS017336] |
| WP_169099395.1 | TUDOR+SF1-SF2-Helicase+iREase+CTD+REase | GCF_012843265.1 | Bacillus_sp._DNRA2 | Bacteria>Bacillati>Bacillota>Bacilli>Bacillales>Bacillaceae>Bacillus>unclassified_Bacillus_(in:_firmicutes) | 1068 | HH041_RS13350 | 100339..103545 | 2723053 | helicase-related-protein-[Bacillus-sp.-DNRA2] |
| WP_196478726.1 | TUDOR+SF1-SF2-Helicase+iREase+CTD+REase | GCF_015712745.1 | Bacillus_sp._SRB3LM | Bacteria>Bacillati>Bacillota>Bacilli>Bacillales>Bacillaceae>Bacillus>unclassified_Bacillus_(in:_firmicutes) | 1066 | F3I02_RS30510 | 651096..654296 | 2608689 | helicase-related-protein-[Bacillus-sp.-SRB3LM] |
| WP_208731625.1 | TUDOR+SF1-SF2-Helicase+iREase+CTD+REase | GCF_003355115.1 | Bacillus_sp._COPE52 | Bacteria>Bacillati>Bacillota>Bacilli>Bacillales>Bacillaceae>Bacillus>unclassified_Bacillus_(in:_firmicutes) | 1066 | DPQ31_RS04915 | 993445..996645 | 2233998 | helicase-related-protein-[Bacillus-sp.-COPE52] |
| WP_077114045.1 | TUDOR+SF1-SF2-Helicase+iREase+CTD+REase | - | Bacillus_sp._VT-16-64 | Bacteria>Bacillati>Bacillota>Bacilli>Bacillales>Bacillaceae>Bacillus>unclassified_Bacillus_(in:_firmicutes) | 1065 | BLX87_RS22745 | 13642..16839 | 1917019 | helicase-related-protein-[Bacillus-sp.-VT-16-64] |
| WP_001165451.1 | TUDOR+SF1-SF2-Helicase+iREase+CTD+REase | GCF_038008665.1 | Bacillus_sp._FSL_R10-2201 | Bacteria>Bacillati>Bacillota>Bacilli>Bacillales>Bacillaceae>Bacillus>unclassified_Bacillus_(in:_firmicutes) | 1061 | ICU_RS21125 | 4131237..4134422 | 2954657 | helicase-related-protein-[Bacillus-cereus] |
| OHR66992.1 | TUDOR+SF1-SF2-Helicase+iREase+CTD+REase | GCA_001838545.1 | Bacillus_sp._HMSC76G11 | Bacteria>Bacillati>Bacillota>Bacilli>Bacillales>Bacillaceae>Bacillus>unclassified_Bacillus_(in:_firmicutes) | 1060 | HMPREF3291_01060 | 79940..83122 | 1608910 | helicase-[Bacillus-sp.-HMSC76G11] |
| WP_026590662.1 | TUDOR+SF1-SF2-Helicase+iREase+CTD+REase | GCF_000482345.1 | Bacillus_sp._UNC437CL72CviS29 | Bacteria>Bacillati>Bacillota>Bacilli>Bacillales>Bacillaceae>Bacillus>unclassified_Bacillus_(in:_firmicutes) | 1060 | M014_RS0104545 | 303205..306387 | 1340430 | helicase-related-protein-[Bacillus-sp.-UNC437CL72CviS29] |
| WP_048718299.1 | TUDOR+SF1-SF2-Helicase+iREase+CTD+REase | GCF_001076885.1 | Bacillus_sp._522_BSPC | Bacteria>Bacillati>Bacillota>Bacilli>Bacillales>Bacillaceae>Bacillus>unclassified_Bacillus_(in:_firmicutes) | 1059 | ADJ83_RS15705 | 27226..30405 | 1579338 | helicase-related-protein-[Bacillus-sp.-522_BSPC] |
| WP_214723421.1 | TUDOR+SF1-SF2-Helicase+iREase+CTD+REase | GCF_018613175.1 | Bacillus_sp._ISL-47 | Bacteria>Bacillati>Bacillota>Bacilli>Bacillales>Bacillaceae>Bacillus>unclassified_Bacillus_(in:_firmicutes) | 1058 | J7I93_RS03720 | 66383..69559 | 2819130 | helicase-related-protein-[Bacillus-sp.-ISL-47] |
| PFG07774.1 | TUDOR+SF1-SF2-Helicase+iREase+CTD+REase | GCA_002563655.1 | Bacillus_sp._es.034 | Bacteria>Bacillati>Bacillota>Bacilli>Bacillales>Bacillaceae>Bacillus>unclassified_Bacillus_(in:_firmicutes) | 1057 | ATG71_4683 | 4464377..4467550 | 1761763 | SNF2-domain-containing-protein-[Bacillus-sp.-es.034] |
| WP_061570115.1 | TUDOR+SF1-SF2-Helicase+iREase+CTD+REase | GCF_001587535.1 | Caldibacillus_debilis | Bacteria>Bacillati>Bacillota>Bacilli>Bacillales>Bacillaceae>Caldibacillus | 1064 | B4135_RS15865 | 32183..35377 | 301148 | helicase-related-protein-[Caldibacillus-debilis] |
| WP_066398762.1 | TUDOR+SF1-SF2-Helicase+iREase+CTD+REase | GCF_017874625.1 | Cytobacillus_eiseniae | Bacteria>Bacillati>Bacillota>Bacilli>Bacillales>Bacillaceae>Cytobacillus | 1072 | J2Z40_RS15700 | 15428..18646 | 762947 | helicase-related-protein-[Cytobacillus-eiseniae] |
| WP_251162453.1 | TUDOR+SF1-SF2-Helicase+iREase+CTD+REase | GCF_023713445.1 | Cytobacillus_kochii | Bacteria>Bacillati>Bacillota>Bacilli>Bacillales>Bacillaceae>Cytobacillus | 1072 | M3670_RS08700 | 67711..70929 | 859143 | helicase-related-protein-[Cytobacillus-kochii] |
| WP_019382824.1 | TUDOR+SF1-SF2-Helicase+iREase+CTD+REase | GCF_000294775.2 | Cytobacillus_oceanisediminis | Bacteria>Bacillati>Bacillota>Bacilli>Bacillales>Bacillaceae>Cytobacillus | 1061 | A361_RS26930 | 5405931..5409116 | 665099 | helicase-related-protein-[Cytobacillus-oceanisediminis] |
| WP_144543243.1 | TUDOR+SF1-SF2-Helicase+iREase+CTD+REase | GCF_965137975.1 | Cytobacillus_oceanisediminis | Bacteria>Bacillati>Bacillota>Bacilli>Bacillales>Bacillaceae>Cytobacillus | 1059 | IQ19_RS15300 | 222508..225687 | 665099 | helicase-related-protein-[Cytobacillus-oceanisediminis] |
| WP_224936973.1 | TUDOR+SF1-SF2-Helicase+iREase+CTD+REase | GCF_020171845.1 | Cytobacillus_kochii | Bacteria>Bacillati>Bacillota>Bacilli>Bacillales>Bacillaceae>Cytobacillus | 1059 | LCM23_RS10840 | 34981..38160 | 859143 | helicase-related-protein-[Cytobacillus-kochii] |
| WP_241504154.1 | TUDOR+SF1-SF2-Helicase+iREase+CTD+REase | - | Cytobacillus_horneckiae | Bacteria>Bacillati>Bacillota>Bacilli>Bacillales>Bacillaceae>Cytobacillus | 963 | CWS20_RS08130 | 37106..39997 | 549687 | helicase-related-protein-[Cytobacillus-horneckiae] |
| MBU9710541.1 | TUDOR+SF1-SF2-Helicase+iREase+CTD+REase | GCA_019042215.1 | Evansella_tamaricis | Bacteria>Bacillati>Bacillota>Bacilli>Bacillales>Bacillaceae>Evansella | 1058 | KS419_02140 | 56388..59564 | 2069301 | DEAD/DEAH-box-helicase-[Evansella-tamaricis] |
| WP_078596328.1 | TUDOR+SF1-SF2-Helicase+iREase+CTD+REase | GCF_002019695.1 | Evansella_clarkii | Bacteria>Bacillati>Bacillota>Bacilli>Bacillales>Bacillaceae>Evansella | 1058 | BK582_RS17770 | 3793494..3796670 | 79879 | helicase-related-protein-[Evansella-clarkii] |
| WP_066159873.1 | TUDOR+SF1-SF2-Helicase+iREase+CTD+REase | GCF_001591945.1 | Halalkalibacter_krulwichiae | Bacteria>Bacillati>Bacillota>Bacilli>Bacillales>Bacillaceae>Halalkalibacter | 1061 | BKR01S_RS20130 | 4275..7460 | 199441 | helicase-related-protein-[Halalkalibacter-krulwichiae] |
| ASF41328.1 | TUDOR+SF1-SF2-Helicase+iREase+CTD+REase | GCA_002209425.1 | Halobacillus_halophilus | Bacteria>Bacillati>Bacillota>Bacilli>Bacillales>Bacillaceae>Halobacillus | 1059 | CEH05_20020 | 4049973..4053152 | 1570 | helicase-[Halobacillus-halophilus] |
| WP_014645299.1 | TUDOR+SF1-SF2-Helicase+iREase+CTD+REase | GCF_000284515.1 | Halobacillus_halophilus | Bacteria>Bacillati>Bacillota>Bacilli>Bacillales>Bacillaceae>Halobacillus | 1059 | HBHAL_RS19735 | 4050129..4053308 | 1570 | helicase-related-protein-[Halobacillus-halophilus] |
| WP_163530745.1 | TUDOR+SF1-SF2-Helicase+iREase+CTD+REase | GCF_902726645.1 | Halobacillus_ihumii | Bacteria>Bacillati>Bacillota>Bacilli>Bacillales>Bacillaceae>Halobacillus | 1057 | G6R08_RS20450 | 4091164..4094337 | 2686092 | helicase-related-protein-[Halobacillus-ihumii] |
| WP_019720356.1 | TUDOR+SF1-SF2-Helicase+iREase+CTD+REase | GCF_000223155.1 | Heyndrickxia_coagulans | Bacteria>Bacillati>Bacillota>Bacilli>Bacillales>Bacillaceae>Heyndrickxia | 1062 | IOE_RS0101960 | 11003..14191 | 1398 | helicase-related-protein-[Weizmannia-coagulans] |
| WP_258922095.1 | TUDOR+SF1-SF2-Helicase+iREase+CTD+REase | GCF_024758645.1 | Heyndrickxia_coagulans | Bacteria>Bacillati>Bacillota>Bacilli>Bacillales>Bacillaceae>Heyndrickxia | 1062 | L1879_RS05755 | 1216247..1219435 | 1398 | DEAD/DEAH-box-helicase-[Weizmannia-coagulans] |
| WP_202780866.1 | TUDOR+SF1-SF2-Helicase+iREase+CTD+REase | - | Heyndrickxia_vini | Bacteria>Bacillati>Bacillota>Bacilli>Bacillales>Bacillaceae>Heyndrickxia | 1057 | I5776_RS19075 | 3827988..3831161 | 1476025 | helicase-related-protein-[Heyndrickxia-vini] |
| WP_251339429.1 | TUDOR+SF1-SF2-Helicase+iREase+CTD+REase | GCF_023714665.1 | Heyndrickxia_oleronia | Bacteria>Bacillati>Bacillota>Bacilli>Bacillales>Bacillaceae>Heyndrickxia | 1057 | M3589_RS18015 | 378340..381513 | 38875 | helicase-related-protein-[Heyndrickxia-oleronia] |
| WP_257961951.1 | TUDOR+SF1-SF2-Helicase+iREase+CTD+REase | GCF_024705655.1 | Lysinibacillus_capsici | Bacteria>Bacillati>Bacillota>Bacilli>Bacillales>Bacillaceae>Lysinibacillus | 1063 | NUA17_RS16990 | 45221..48412 | 2115968 | DEAD/DEAH-box-helicase-[Lysinibacillus-capsici] |
| WP_126294750.1 | TUDOR+SF1-SF2-Helicase+iREase+CTD+REase | GCF_003966145.1 | Lysinibacillus_telephonicus | Bacteria>Bacillati>Bacillota>Bacilli>Bacillales>Bacillaceae>Lysinibacillus | 1059 | EKG35_RS12255 | 10052..13231 | 1714840 | helicase-related-protein-[Lysinibacillus-telephonicus] |
| WP_269897761.1 | TUDOR+SF1-SF2-Helicase+iREase+CTD+REase | GCF_027557635.1 | Lysinibacillus_sp._JK80 | Bacteria>Bacillati>Bacillota>Bacilli>Bacillales>Bacillaceae>Lysinibacillus>unclassified_Lysinibacillus | 1057 | HXV90_RS18585 | 3696089..3699262 | 2749809 | helicase-related-protein-[Lysinibacillus-sp.-JK80] |
| WP_221874925.1 | TUDOR+SF1-SF2-Helicase+iREase+CTD+REase | GCF_019749255.1 | Mesobacillus_maritimus | Bacteria>Bacillati>Bacillota>Bacilli>Bacillales>Bacillaceae>Mesobacillus | 1057 | H0185_RS18260 | 56665..59838 | 1643336 | helicase-related-protein-[Mesobacillus-maritimus] |
| WP_191566567.1 | TUDOR+SF1-SF2-Helicase+iREase+CTD+REase | GCF_904423835.1 | Metabacillus_idriensis | Bacteria>Bacillati>Bacillota>Bacilli>Bacillales>Bacillaceae>Metabacillus | 1072 | IHQ32_RS17395 | 3426300..3429518 | 324768 | helicase-related-protein-[Metabacillus-idriensis] |
| WP_251625218.1 | TUDOR+SF1-SF2-Helicase+iREase+CTD+REase | GCF_023712125.1 | Metabacillus_idriensis | Bacteria>Bacillati>Bacillota>Bacilli>Bacillales>Bacillaceae>Metabacillus | 1060 | M4D55_RS07625 | 56135..59317 | 324768 | helicase-related-protein-[Metabacillus-idriensis] |
| WP_155112637.1 | TUDOR+SF1-SF2-Helicase+iREase+CTD+REase | GCF_009711125.1 | Metabacillus_mangrovi | Bacteria>Bacillati>Bacillota>Bacilli>Bacillales>Bacillaceae>Metabacillus | 1059 | GKZ89_RS11895 | 37167..40346 | 1491830 | helicase-related-protein-[Metabacillus-mangrovi] |
| WP_213145786.1 | TUDOR+SF1-SF2-Helicase+iREase+CTD+REase | GCF_018343545.2 | Neobacillus_citreus | Bacteria>Bacillati>Bacillota>Bacilli>Bacillales>Bacillaceae>Neobacillus | 1068 | KHB02_RS07280 | 31957..35163 | 2833578 | helicase-related-protein-[Neobacillus-citreus] |
| WP_191274565.1 | TUDOR+SF1-SF2-Helicase+iREase+CTD+REase | GCF_014656545.1 | Neobacillus_kokaensis | Bacteria>Bacillati>Bacillota>Bacilli>Bacillales>Bacillaceae>Neobacillus | 1060 | IE222_RS16225 | 105561..108743 | 2759023 | helicase-related-protein-[Neobacillus-kokaensis] |
| PAD23519.1 | TUDOR+SF1-SF2-Helicase+iREase+CTD+REase | GCA_002271635.1 | Niallia_circulans | Bacteria>Bacillati>Bacillota>Bacilli>Bacillales>Bacillaceae>Niallia | 1061 | CHH62_21845 | 13572..16757 | 1397 | helicase-[Niallia-circulans] |
| WP_095259400.1 | TUDOR+SF1-SF2-Helicase+iREase+CTD+REase | GCF_002271635.1 | Niallia_circulans | Bacteria>Bacillati>Bacillota>Bacilli>Bacillales>Bacillaceae>Niallia | 1061 | CHH62_RS21855 | 13572..16757 | 1397 | helicase-related-protein-[Niallia-circulans] |
| MCM3032234.1 | TUDOR+SF1-SF2-Helicase+iREase+CTD+REase | GCA_023714395.1 | Niallia_sp._MER_6 | Bacteria>Bacillati>Bacillota>Bacilli>Bacillales>Bacillaceae>Niallia>unclassified_Niallia | 1057 | M3600_17140 | 103946..107119 | 2939567 | SNF2-related-protein-[Niallia-sp.-MER-6] |

| Protein-ID | Domain | Assembly-ID | Strain | Taxonomy | Length | Locus-Tag | Genomic-coordinates | TaxID | Fasta-defline |
|---|---|---|---|---|---|---|---|---|---|
| WP_188383788.1 | TUDOR+SF1-SF2-Helicase+iREase+CTD+REase | GCF_014637405.1 | Ornithinibacillus_halotolerans | Bacteria>Bacillati>Bacillota>Bacilli>Bacillales>Bacillaceae>Ornithinibacillus | 1053 | IEV35_RS05930 | 23199..26360 | 1274357 | helicase-related-protein-[Ornithinibacillus-halotolerans] |
| WP_026908396.1 | TUDOR+SF1-SF2-Helicase+iREase+CTD+REase | GCF_000482485.1 | Paucisalibacillus_globulus | Bacteria>Bacillati>Bacillota>Bacilli>Bacillales>Bacillaceae>Paucisalibacillus | 1053 | K332_RS0116110 | 145318..148479 | 351095 | helicase-related-protein-[Paucisalibacillus-globulus] |
| WP_190997465.1 | TUDOR+SF1-SF2-Helicase+iREase+CTD+REase | GCF_014712675.1 | Peribacillus_faecalis | Bacteria>Bacillati>Bacillota>Bacilli>Bacillales>Bacillaceae>Peribacillus | 1070 | IEO70_RS06040 | 50099..53311 | 2772559 | helicase-related-protein-[Peribacillus-faecalis] |
| WP_119115136.1 | TUDOR+SF1-SF2-Helicase+iREase+CTD+REase | GCF_003570725.1 | Peribacillus_asahii | Bacteria>Bacillati>Bacillota>Bacilli>Bacillales>Bacillaceae>Peribacillus | 1069 | D1953_RS00155 | 5880..9089 | 228899 | helicase-related-protein-[Peribacillus-asahii] |
| WP_117327460.1 | TUDOR+SF1-SF2-Helicase+iREase+CTD+REase | GCF_003429095.1 | Peribacillus_saganii | Bacteria>Bacillati>Bacillota>Bacilli>Bacillales>Bacillaceae>Peribacillus | 1057 | D0469_RS14505 | 24876..28049 | 2303992 | helicase-related-protein-[Peribacillus-saganii] |
| WP_260358525.1 | TUDOR+SF1-SF2-Helicase+iREase+CTD+REase | GCF_025209795.1 | Peribacillus_frigoritolerans | Bacteria>Bacillati>Bacillota>Bacilli>Bacillales>Bacillaceae>Peribacillus | 1056 | NOU24_RS18770 | 738298..741468 | 450367 | DEAD/DEAH-box-helicase-[Peribacillus-frigoritolerans] |
| WP_225002271.1 | TUDOR+SF1-SF2-Helicase+iREase+CTD+REase | GCF_020176725.1 | Priestia_flexa | Bacteria>Bacillati>Bacillota>Bacilli>Bacillales>Bacillaceae>Priestia | 1072 | LC052_RS15450 | 51342..54560 | 86664 | helicase-related-protein-[Priestia-flexa] |
| MQR86866.1 | TUDOR+SF1-SF2-Helicase+iREase+CTD+REase | GCA_009497105.1 | Priestia_megaterium | Bacteria>Bacillati>Bacillota>Bacilli>Bacillales>Bacillaceae>Priestia | 1064 | GFV16_13180 | 21489..24683 | 1404 | DEAD/DEAH-box-helicase-[Priestia-megaterium] |
| WP_272767824.1 | TUDOR+SF1-SF2-Helicase+iREase+CTD+REase | GCF_028548275.1 | Priestia_megaterium | Bacteria>Bacillati>Bacillota>Bacilli>Bacillales>Bacillaceae>Priestia | 1062 | PNF31_RS27500 | 24827..28015 | 1404 | helicase-related-protein-[Priestia-megaterium] |
| WP_098794045.1 | TUDOR+SF1-SF2-Helicase+iREase+CTD+REase | GCF_002576235.1 | Priestia_megaterium | Bacteria>Bacillati>Bacillota>Bacilli>Bacillales>Bacillaceae>Priestia | 1060 | COA18_RS28035 | 1950790..1953972 | 1404 | helicase-related-protein-[Priestia-megaterium] |
| WP_188386546.1 | TUDOR+SF1-SF2-Helicase+iREase+CTD+REase | GCF_014638355.1 | Priestia_taiwanensis | Bacteria>Bacillati>Bacillota>Bacilli>Bacillales>Bacillaceae>Priestia | 1057 | IEV41_RS00685 | 129855..133028 | 1347902 | helicase-related-protein-[Priestia-taiwanensis] |
| MCM3358658.1 | TUDOR+SF1-SF2-Helicase+iREase+CTD+REase | GCA_023713305.1 | Psychrobacillus_sp._MER_TA_171 | Bacteria>Bacillati>Bacillota>Bacilli>Bacillales>Bacillaceae>Psychrobacillus>unclassified_Psychrobacillus | 1071 | M3627_11590 | 15799..19014 | 2939577 | SNF2-related-protein-[Psychrobacillus-sp.-MER-TA-171] |
| WP_136831111.1 | TUDOR+SF1-SF2-Helicase+iREase+CTD+REase | GCF_005116465.1 | Robertmurraya_kyonggiensis | Bacteria>Bacillati>Bacillota>Bacilli>Bacillales>Bacillaceae>Robertmurraya | 1057 | FA727_RS11525 | 61609..64782 | 1037680 | helicase-related-protein-[Robertmurraya-kyonggiensis] |
| WP_061808759.1 | TUDOR+SF1-SF2-Helicase+iREase+CTD+REase | GCF_001591825.1 | Rossellomorea_vietnamensis | Bacteria>Bacillati>Bacillota>Bacilli>Bacillales>Bacillaceae>Rossellomorea | 1072 | BVI01S_RS00220 | 45618..48836 | 218284 | helicase-related-protein-[Rossellomorea-vietnamensis] |
| WP_205168138.1 | TUDOR+SF1-SF2-Helicase+iREase+CTD+REase | GCF_016908495.1 | Rossellomorea_pakistanensis | Bacteria>Bacillati>Bacillota>Bacilli>Bacillales>Bacillaceae>Rossellomorea | 1057 | JOC86_RS02200 | 432486..435659 | 992288 | helicase-related-protein-[Bacillus-pakistanensis] |
| WP_174496254.1 | TUDOR+SF1-SF2-Helicase+iREase+CTD+REase | GCF_902809875.1 | Salirhabdus_euzebyi | Bacteria>Bacillati>Bacillota>Bacilli>Bacillales>Bacillaceae>Salirhabdus | 1070 | HNQ94_RS00290 | 59642..62854 | 394506 | helicase-related-protein-[Salirhabdus-euzebyi] |
| WP_102026202.1 | TUDOR+SF1-SF2-Helicase+iREase+CTD+REase | GCF_900240405.1 | Salirhabdus_sp._Marseille-P4669 | Bacteria>Bacillati>Bacillota>Bacilli>Bacillales>Bacillaceae>Salirhabdus>unclassified_Salirhabdus | 1072 | C1A05_RS00415 | 86868..90086 | 2042310 | helicase-related-protein-[Salirhabdus-sp.-Marseille-P4669] |
| WP_093855903.1 | TUDOR+SF1-SF2-Helicase+iREase+CTD+REase | GCF_007989445.1 | Tenuibacillus_multivorans | Bacteria>Bacillati>Bacillota>Bacilli>Bacillales>Bacillaceae>Tenuibacillus | 1059 | BLP77_RS04455 | 344715..347894 | 237069 | helicase-related-protein-[Tenuibacillus-multivorans] |
| MBO8173522.1 | TUDOR+SF1-SF2-Helicase+iREase+CTD+REase | GCA_017656255.1 | Bacillaceae_bacterium | Bacteria>Bacillati>Bacillota>Bacilli>Bacillales>Bacillaceae>unclassified_Bacillaceae | 1077 | HOZ33_16770 | 2418..5651 | 1889774 | DEAD/DEAH-box-helicase-family-protein-[Bacillaceae-bacterium] |
| WP_099159041.1 | TUDOR+SF1-SF2-Helicase+iREase+CTD+REase | GCF_900187325.1 | Virgibacillus_ndiopensis | Bacteria>Bacillati>Bacillota>Bacilli>Bacillales>Bacillaceae>Virgibacillus | 1059 | CS937_RS13690 | 185420..188599 | 2004408 | helicase-related-protein-[Virgibacillus-ndiopensis] |
| STO11904.1 | TUDOR+SF1-SF2-Helicase+iREase+CTD+REase | GCA_900450595.1 | [Flavobacterium]_thermophilum | Bacteria>Bacillati>Bacillota>Bacilli>Bacillales>Bacillales_incertae_sedis | 786 | NCTC11526_00570 | 556011..558371 | 1414643 | RNA-polymerase-associated-protein-rapA-[[Flavobacterium] |
| WP_026826558.1 | TUDOR+SF1-SF2-Helicase+iREase+CTD+REase | GCF_000620845.1 | Exiguobacterium_marinum | Bacteria>Bacillati>Bacillota>Bacilli>Bacillales>Bacillales_incertae_sedis>Bacillales_Family_XII._Incertae_Sedis>Exiguobacterium | 1057 | P400_RS0112685 | 2395166..2398339 | 273528 | helicase-related-protein-[Exiguobacterium-marinum] |
| WP_026834101.1 | TUDOR+SF1-SF2-Helicase+iREase+CTD+REase | GCF_000620825.1 | Exiguobacterium_undae | Bacteria>Bacillati>Bacillota>Bacilli>Bacillales>Bacillales_incertae_sedis>Bacillales_Family_XII._Incertae_Sedis>Exiguobacterium | 1052 | P399_RS0115205 | 2855250..2858408 | 169177 | helicase-related-protein-[Exiguobacterium-undae] |
| WP_275060108.1 | TUDOR+SF1-SF2-Helicase+iREase+CTD+REase | GCF_029026745.1 | Exiguobacterium_profundum | Bacteria>Bacillati>Bacillota>Bacilli>Bacillales>Bacillales_incertae_sedis>Bacillales_Family_XII._Incertae_Sedis>Exiguobacterium | 1047 | OE059_RS13155 | 2531985..2535128 | 307643 | helicase-related-protein-[Exiguobacterium-profundum] |
| WP_131508627.1 | TUDOR+SF1-SF2-Helicase+iREase+CTD+REase | GCF_004337165.1 | Exiguobacterium_sp._SH0S1 | Bacteria>Bacillati>Bacillota>Bacilli>Bacillales>Bacillales_incertae_sedis>Bacillales_Family_XII._Incertae_Sedis>Exiguobacterium>unclassified_Exiguobacterium | 1059 | EVJ20_RS13625 | 71778..74957 | 2510949 | helicase-related-protein-[Exiguobacterium-sp.-SH0S1] |
| WP_047391759.1 | TUDOR+SF1-SF2-Helicase+iREase+CTD+REase | GCF_000798935.2 | Exiguobacterium_sp._ZWU0009 | Bacteria>Bacillati>Bacillota>Bacilli>Bacillales>Bacillales_incertae_sedis>Bacillales_Family_XII._Incertae_Sedis>Exiguobacterium>unclassified_Exiguobacterium | 1054 | L985_RS01905 | 339807..342971 | 1224749 | helicase-related-protein-[Exiguobacterium-sp.-ZWU0009] |
| WP_078145976.1 | TUDOR+SF1-SF2-Helicase+iREase+CTD+REase | GCF_002009685.1 | Exiguobacterium_sp._HVEsp1 | Bacteria>Bacillati>Bacillota>Bacilli>Bacillales>Bacillales_incertae_sedis>Bacillales_Family_XII._Incertae_Sedis>Exiguobacterium>unclassified_Exiguobacterium | 1054 | BV910_RS00645 | 132490..135654 | 1934003 | helicase-related-protein-[Exiguobacterium-sp.-HVEsp1] |
| WP_214805984.1 | TUDOR+SF1-SF2-Helicase+iREase+CTD+REase | GCF_018618875.1 | Exiguobacterium_sp._s46 | Bacteria>Bacillati>Bacillota>Bacilli>Bacillales>Bacillales_incertae_sedis>Bacillales_Family_XII._Incertae_Sedis>Exiguobacterium>unclassified_Exiguobacterium | 1054 | KNQ08_RS12540 | 183729..186893 | 2751200 | helicase-related-protein-[Exiguobacterium-sp.-s46] |
| WP_214788913.1 | TUDOR+SF1-SF2-Helicase+iREase+CTD+REase | GCF_018617975.1 | Exiguobacterium_sp._s21 | Bacteria>Bacillati>Bacillota>Bacilli>Bacillales>Bacillales_incertae_sedis>Bacillales_Family_XII._Incertae_Sedis>Exiguobacterium>unclassified_Exiguobacterium | 1048 | KNP06_RS10865 | 35109..38255 | 2751244 | helicase-related-protein-[Exiguobacterium-sp.-s21] |
| WP_214689543.1 | TUDOR+SF1-SF2-Helicase+iREase+CTD+REase | GCF_018617135.1 | Exiguobacterium_sp._s163 | Bacteria>Bacillati>Bacillota>Bacilli>Bacillales>Bacillales_incertae_sedis>Bacillales_Family_XII._Incertae_Sedis>Exiguobacterium>unclassified_Exiguobacterium | 1047 | KNO71_RS16225 | 88590..91733 | 2751287 | helicase-related-protein-[Exiguobacterium-sp.-s163] |
| WP_271306169.1 | TUDOR+SF1-SF2-Helicase+iREase+CTD+REase | GCF_027947425.1 | Exiguobacterium_sp._MMG028 | Bacteria>Bacillati>Bacillota>Bacilli>Bacillales>Bacillales_incertae_sedis>Bacillales_Family_XII._Incertae_Sedis>Exiguobacterium>unclassified_Exiguobacterium | 1047 | PJK55_RS13680 | 72986..76129 | 3021979 | helicase-related-protein-[Exiguobacterium-sp.-MMG028] |
| WP_066463085.1 | TUDOR+SF1-SF2-Helicase+iREase+CTD+REase | GCF_001700325.1 | Caryophanon_latum | Bacteria>Bacillati>Bacillota>Bacilli>Bacillales>Caryophanaceae>Caryophanon | 1057 | A6K76_RS05960 | 13764..16937 | 33977 | helicase-related-protein-[Caryophanon-latum] |
| WP_019416003.1 | TUDOR+SF1-SF2-Helicase+iREase+CTD+REase | GCF_000286315.1 | Paenisporosarcina_sp._TG20 | Bacteria>Bacillati>Bacillota>Bacilli>Bacillales>Caryophanaceae>Paenisporosarcina>unclassified_Paenisporosarcina | 1063 | B371_RS0119470 | 37382..40573 | 1211706 | helicase-related-protein-[Paenisporosarcina-sp.-TG20] |
| WP_108587669.1 | TUDOR+SF1-SF2-Helicase+iREase+CTD+REase | GCF_003058165.1 | Paenisporosarcina_sp._OV554 | Bacteria>Bacillati>Bacillota>Bacilli>Bacillales>Caryophanaceae>Paenisporosarcina>unclassified_Paenisporosarcina | 1058 | C8K15_RS17260 | 25791..28967 | 2135694 | helicase-related-protein-[Paenisporosarcina-sp.-OV554] |
| WP_065525140.1 | TUDOR+SF1-SF2-Helicase+iREase+CTD+REase | GCF_001687665.2 | Planococcus_donghaensis | Bacteria>Bacillati>Bacillota>Bacilli>Bacillales>Caryophanaceae>Planococcus | 1061 | BCM40_RS15160 | 3092323..3095508 | 414778 | helicase-related-protein-[Planococcus-donghaensis] |
| WP_149457008.1 | TUDOR+SF1-SF2-Helicase+iREase+CTD+REase | GCF_008369335.1 | Planococcus_sp._ANT_H30 | Bacteria>Bacillati>Bacillota>Bacilli>Bacillales>Caryophanaceae>Planococcus>unclassified_Planococcus_(in:_firmicutes) | 1063 | FQ085_RS14945 | 117189..120380 | 2597347 | helicase-related-protein-[Planococcus-sp.-ANT_H30] |
| WP_215106196.1 | TUDOR+SF1-SF2-Helicase+iREase+CTD+REase | GCF_018613655.1 | Planococcus_sp._ISL-109 | Bacteria>Bacillati>Bacillota>Bacilli>Bacillales>Caryophanaceae>Planococcus>unclassified_Planococcus_(in:_firmicutes) | 1063 | J7E21_RS07360 | 1271..4462 | 2819166 | helicase-related-protein-[Planococcus-sp.-ISL-109] |
| WP_274307979.1 | TUDOR+SF1-SF2-Helicase+iREase+CTD+REase | GCF_028747805.1 | Solibacillus_daqui | Bacteria>Bacillati>Bacillota>Bacilli>Bacillales>Caryophanaceae>Solibacillus | 1057 | O7776_RS16095 | 3277153..3280326 | 2912187 | helicase-related-protein-[Solibacillus-sp.-ZS111008] |
| WP_192599215.1 | TUDOR+SF1-SF2-Helicase+iREase+CTD+REase | GCF_014873855.1 | Sporosarcina_limicola | Bacteria>Bacillati>Bacillota>Bacilli>Bacillales>Caryophanaceae>Sporosarcina | 1065 | H4683_RS12785 | 34027..37224 | 34101 | helicase-related-protein-[Sporosarcina-limicola] |
| WP_107933595.1 | TUDOR+SF1-SF2-Helicase+iREase+CTD+REase | GCF_003049615.1 | Ureibacillus_chungkukjangi | Bacteria>Bacillati>Bacillota>Bacilli>Bacillales>Caryophanaceae>Ureibacillus | 1057 | C9J33_RS05730 | 60046..63219 | 1202712 | helicase-related-protein-[Ureibacillus-chungkukjangi] |
| WP_173598484.1 | TUDOR+SF1-SF2-Helicase+iREase+CTD+REase | GCF_013284375.2 | Brevibacillus | Bacteria>Bacillati>Bacillota>Bacilli>Bacillales>Paenibacillaceae | 1072 | HP435_RS06025 | 1191250..1194468 | 55080 | MULTISPECIES:-helicase-related-protein-[Brevibacillus] |
| WP_272441151.1 | TUDOR+SF1-SF2-Helicase+iREase+CTD+REase | GCF_028421645.1 | Aneurinibacillus_sp._Ricciae_BoGa-3 | Bacteria>Bacillati>Bacillota>Bacilli>Bacillales>Paenibacillaceae>Aneurinibacillus_group>Aneurinibacillus>unclassified_Aneurinibacillus | 1064 | PP175_RS01565 | 310603..313797 | 3022697 | helicase-related-protein-[Aneurinibacillus-sp.-Ricciae_BoGa-3] |
| WP_023554873.1 | TUDOR+SF1-SF2-Helicase+iREase+CTD+REase | GCF_000503775.1 | Brevibacillus_panacihumi | Bacteria>Bacillati>Bacillota>Bacilli>Bacillales>Paenibacillaceae>Brevibacillus | 1069 | T458_RS33480 | 822575..825784 | 497735 | helicase-related-protein-[Brevibacillus-panacihumi] |
| WP_122960159.1 | TUDOR+SF1-SF2-Helicase+iREase+CTD+REase | GCF_006540045.1 | Brevibacillus_centrosporus | Bacteria>Bacillati>Bacillota>Bacilli>Bacillales>Paenibacillaceae>Brevibacillus | 1069 | BCE02nite_RS06945 | 34603..37812 | 54910 | helicase-related-protein-[Brevibacillus-centrosporus] |
| WP_198826833.1 | TUDOR+SF1-SF2-Helicase+iREase+CTD+REase | GCF_016406105.1 | Brevibacillus_composti | Bacteria>Bacillati>Bacillota>Bacilli>Bacillales>Paenibacillaceae>Brevibacillus | 1069 | KDJ56_RS14940 | 2930975..2934184 | 2796470 | helicase-related-protein-[Brevibacillus-composti] |
| WP_221842521.1 | TUDOR+SF1-SF2-Helicase+iREase+CTD+REase | GCF_019749055.1 | Brevibacillus_agri | Bacteria>Bacillati>Bacillota>Bacilli>Bacillales>Paenibacillaceae>Brevibacillus | 1069 | H7K32_RS04505 | 17499..20708 | 51101 | helicase-related-protein-[Brevibacillus-agri] |
| GIO15802.1 | TUDOR+SF1-SF2-Helicase+iREase+CTD+REase | GCA_018333175.1 | Cohnella_xylanilytica | Bacteria>Bacillati>Bacillota>Bacilli>Bacillales>Paenibacillaceae>Cohnella | 1069 | J19TS2_53570 | 47856..51065 | 557555 | helicase-[Cohnella-xylanilytica] |
| WP_212962146.1 | TUDOR+SF1-SF2-Helicase+iREase+CTD+REase | GCF_018333175.1 | Cohnella_xylanilytica | Bacteria>Bacillati>Bacillota>Bacilli>Bacillales>Paenibacillaceae>Cohnella | 1069 | KI376_RS26585 | 47856..51065 | 557555 | helicase-related-protein-[Cohnella-xylanilytica] |
| WP_058302020.1 | TUDOR+SF1-SF2-Helicase+iREase+CTD+REase | GCF_902375625.1 | Gorillibacterium_timonense | Bacteria>Bacillati>Bacillota>Bacilli>Bacillales>Paenibacillaceae>Gorillibacterium | 1073 | BN2684_RS09960 | 478383..481604 | 1689269 | helicase-related-protein-[Gorillibacterium-timonense] |
| WP_114378276.1 | TUDOR+SF1-SF2-Helicase+iREase+CTD+REase | GCF_003337375.1 | Paenibacillus_prosopidis | Bacteria>Bacillati>Bacillota>Bacilli>Bacillales>Paenibacillaceae>Paenibacillus | 1075 | DFP97_RS01975 | 476454..479681 | 630520 | helicase-related-protein-[Paenibacillus-prosopidis] |
| WP_085168726.1 | TUDOR+SF1-SF2-Helicase+iREase+CTD+REase | GCF_900177285.1 | Paenibacillus_barengoltzii | Bacteria>Bacillati>Bacillota>Bacilli>Bacillales>Paenibacillaceae>Paenibacillus | 1074 | B9N82_RS00060 | 20416..23640 | 343517 | helicase-related-protein-[Paenibacillus-barengoltzii] |
| WDH83315.1 | TUDOR+SF1-SF2-Helicase+iREase+CTD+REase | GCA_028747665.1 | Paenibacillus_urinalis | Bacteria>Bacillati>Bacillota>Bacilli>Bacillales>Paenibacillaceae>Paenibacillus | 1073 | PUW23_03460 | 772509..775730 | 521520 | helicase-related-protein-[Paenibacillus-urinalis] |
| WP_189000177.1 | TUDOR+SF1-SF2-Helicase+iREase+CTD+REase | GCF_014641075.1 | Paenibacillus_nasutitermitis | Bacteria>Bacillati>Bacillota>Bacilli>Bacillales>Paenibacillaceae>Paenibacillus | 1073 | IEY46_RS33915 | 20853..24074 | 1652958 | helicase-related-protein-[Paenibacillus-nasutitermitis] |
| WP_042184805.1 | TUDOR+SF1-SF2-Helicase+iREase+CTD+REase | GCF_000758545.1 | unclassified_Paenibacillus | Bacteria>Bacillati>Bacillota>Bacilli>Bacillales>Paenibacillaceae>Paenibacillus | 1070 | H70737_RS03480 | 778468..781680 | 185978 | helicase-related-protein-[Paenibacillus-sp.-FSL-H7-0737] |
| WP_076118183.1 | TUDOR+SF1-SF2-Helicase+iREase+CTD+REase | GCF_001954425.1 | Paenibacillus_odorifer | Bacteria>Bacillati>Bacillota>Bacilli>Bacillales>Paenibacillaceae>Paenibacillus | 1069 | BSK52_RS07855 | 293313..296522 | 189426 | helicase-related-protein-[Paenibacillus-odorifer] |
| WP_138789734.1 | TUDOR+SF1-SF2-Helicase+iREase+CTD+REase | GCF_042666815.1 | Paenibacillus_thermoaerophilus | Bacteria>Bacillati>Bacillota>Bacilli>Bacillales>Paenibacillaceae>Paenibacillus | 1069 | FE781_RS11320 | 110815..114024 | 1215385 | helicase-related-protein-[Paenibacillus-thermoaerophilus] |
| WP_209876433.1 | TUDOR+SF1-SF2-Helicase+iREase+CTD+REase | GCF_017874735.1 | Paenibacillus_silagei | Bacteria>Bacillati>Bacillota>Bacilli>Bacillales>Paenibacillaceae>Paenibacillus | 1069 | J2Z70_RS20915 | 49633..52842 | 1670801 | helicase-related-protein-[Paenibacillus-silagei] |
| WP_089526504.1 | TUDOR+SF1-SF2-Helicase+iREase+CTD+REase | - | Paenibacillus_herberti | Bacteria>Bacillati>Bacillota>Bacilli>Bacillales>Paenibacillaceae>Paenibacillus | 1060 | CGZ75_RS19645 | 28390..31572 | 1619309 | helicase-related-protein-[Paenibacillus-herberti] |
| UUZ84571.1 | TUDOR+SF1-SF2-Helicase+iREase+CTD+REase | GCA_024703795.1 | Paenibacillus_sp._P26 | Bacteria>Bacillati>Bacillota>Bacilli>Bacillales>Paenibacillaceae>Paenibacillus>unclassified_Paenibacillus | 1069 | LJK88_13420 | 2167024..2170233 | 2885204 | DEAD/DEAH-box-helicase-[Paenibacillus-sp.-P26] |
| WP_090826268.1 | TUDOR+SF1-SF2-Helicase+iREase+CTD+REase | GCF_900103825.1 | Paenibacillus_sp._yr247 | Bacteria>Bacillati>Bacillota>Bacilli>Bacillales>Paenibacillaceae>Paenibacillus>unclassified_Paenibacillus | 1069 | BLT91_RS25170 | 28223..31432 | 1761880 | helicase-related-protein-[Paenibacillus-sp.-yr247] |
| WP_256832014.1 | TUDOR+SF1-SF2-Helicase+iREase+CTD+REase | GCF_024519685.1 | Paenibacillus_sp._Pae15 | Bacteria>Bacillati>Bacillota>Bacilli>Bacillales>Paenibacillaceae>Paenibacillus>unclassified_Paenibacillus | 1069 | NR121_RS23330 | 24581..27790 | 2926018 | MULTISPECIES:-helicase-related-protein-[unclassified-Paenibacillus] |
| WP_092984056.1 | TUDOR+SF1-SF2-Helicase+iREase+CTD+REase | GCF_900101075.1 | Lacicoccus_qingdaonensis | Bacteria>Bacillati>Bacillota>Bacilli>Bacillales>Salinicoccaceae>Lacicoccus | 1055 | BLR15_RS01725 | 43758..46925 | 576118 | helicase-related-protein-[Salinicoccus-qingdaonensis] |
| WP_271398739.1 | TUDOR+SF1-SF2-Helicase+iREase+CTD+REase | GCF_947502135.1 | Salinicoccus_roseus | Bacteria>Bacillati>Bacillota>Bacilli>Bacillales>Staphylococcaceae>Salinicoccus | 1054 | POR62_RS08095 | 192780..195944 | 45670 | helicase-related-protein-[Salinicoccus-roseus] |
| WP_106450123.1 | TUDOR+SF1-SF2-Helicase+iREase+CTD+REase | GCF_003008695.1 | Trichococcus_alkaliphilus | Bacteria>Bacillati>Bacillota>Bacilli>Lactobacillales>Carnobacteriaceae>Trichococcus | 1058 | C7R73_RS08670 | 59649..62825 | 2052943 | helicase-related-protein-[Trichococcus-alkaliphilus] |
| KKE80054.1 | TUDOR+SF1-SF2-Helicase+iREase+CTD+REase | GCA_000974285.1 | Bacilli_bacterium_VT-13-104 | Bacteria>Bacillati>Bacillota>Bacilli>unclassified_Bacilli | 1062 | WH51_03815 | 57115..60303 | 1637974 | helicase-[Bacilli-bacterium-VT-13-104] |
| WP_207652760.1 | TUDOR+SF1-SF2-Helicase+iREase+CTD+REase | GCF_002285495.1 | Clostridium_isatidis | Bacteria>Bacillati>Bacillota>Clostridia>Eubacteriales>Clostridiaceae>Clostridium | 882 | BEN51_RS07765 | 1642000..1644648 | 182773 | helicase-related-protein-[Clostridium-isatidis] |
| WP_185649260.1 | TUDOR+SF1-SF2-Helicase+iREase+CTD+REase | GCF_014231265.1 | Clostridium_sp._DJ247 | Bacteria>Bacillati>Bacillota>Clostridia>Eubacteriales>Clostridiaceae>Clostridium>unclassified_Clostridium | 1073 | HGI79_RS03430 | 31207..34428 | 2726188 | helicase-related-protein-[Clostridium-sp.-DJ247] |
| WP_261669786.1 | TUDOR+SF1-SF2-Helicase+iREase+CTD+REase | GCF_025398335.1 | Clostridium_sp._CX1 | Bacteria>Bacillati>Bacillota>Clostridia>Eubacteriales>Clostridiaceae>Clostridium>unclassified_Clostridium | 1073 | N4T77_RS00690 | 149683..152904 | 2978346 | DEAD/DEAH-box-helicase-[Clostridium-sp.-CX1] |
| WP_164950929.1 | TUDOR+SF1-SF2-Helicase+iREase+CTD+REase | GCF_011065955.2 | Clostridium_sp._FP2 | Bacteria>Bacillati>Bacillota>Clostridia>Eubacteriales>Clostridiaceae>Clostridium>unclassified_Clostridium | 1069 | G9F71_RS16290 | 3386323..3389532 | 2724481 | helicase-related-protein-[Clostridium-sp.-FP2] |
| KJS50637.1 | TUDOR+SF1-SF2-Helicase+iREase+CTD+REase | GCA_000961805.1 | Peptococcaceae_bacterium_BRH_c23 | Bacteria>Bacillati>Bacillota>Clostridia>Eubacteriales>Peptococcaceae>unclassified_Peptococcaceae | 1059 | VR66_01640 | 9912..13091 | 1629714 | helicase-[Peptococcaceae-bacterium-BRH_c23] |
| WP_198136465.1 | TUDOR+SF1-SF2-Helicase+iREase+CTD+REase | - | Marinobacter_sp._ELB17 | Bacteria>Pseudomonadati>Pseudomonadota>Gammaproteobacteria>Pseudomonadales>Marinobacteraceae>Marinobacter>unclassified_Marinobacter | 875 | MELB17_RS08090 | 40463..43090 | 270374 | helicase-related-protein-[Marinobacter-sp.-ELB17] |
| WP_146889188.1 | TUDOR+SF1-SF2-Helicase+iREase+CTD+REase | GCF_007990775.1 | Deinococcus_cellulosilyticus | Bacteria>Thermotogati>Deinococcota>Deinococci>Deinococcales>Deinococcaceae>Deinococcus | 925 | DC3_RS23555 | 10504..13281 | 401558 | helicase-related-protein-[Deinococcus-cellulosilyticus] |
| WP_189062801.1 | TUDOR+SF1-SF2-Helicase+iREase+CTD+REase | GCF_014647995.1 | Deinococcus_arenae | Bacteria>Thermotogati>Deinococcota>Deinococci>Deinococcales>Deinococcaceae>Deinococcus | 892 | IEY66_RS18105 | 30777..33455 | 1452751 | DEAD/DEAH-box-helicase-[Deinococcus-arenae] |
| --TUDOR+SF1-SF2-Helicase+iREase+CTD+REase-p(BrxHII-Helicase) | | | | | | | | | |
| WP_052127348.1 | TUDOR+SF1-SF2-Helicase+iREase+CTD+REase | GCF_000775615.1 | Pontibacillus_litoralis | Bacteria>Bacillati>Bacillota>Bacilli>Bacillales>Bacillaceae>Pontibacillus | 409 | N784_RS17010 | 27254..28483 | 516703 | DEAD/DEAH-box-helicase-[Pontibacillus-litoralis] |
| WP_235715380.1 | TUDOR+SF1-SF2-Helicase+iREase+CTD+REase | GCF_000513095.1 | Halalkalibacter_wakoensis | Bacteria>Bacillati>Bacillota>Bacilli>Bacillales>Bacillaceae>Halalkalibacter | 389 | JCM9140_RS24060 | 18620..19789 | 127891 | DEAD/DEAH-box-helicase-[Halalkalibacter-wakoensis] |
| WP_052127349.1 | TUDOR+SF1-SF2-Helicase+iREase+CTD+REase | GCF_000775615.1 | Pontibacillus_litoralis | Bacteria>Bacillati>Bacillota>Bacilli>Bacillales>Bacillaceae>Pontibacillus | 378 | N784_RS17015 | 28495..29631 | 516703 | C-terminal-helicase-domain-containing-protein-[Pontibacillus-litoralis] |
| STO11903.1 | TUDOR+SF1-SF2-Helicase+iREase+CTD+REase | GCA_900450595.1 | [Flavobacterium]_thermophilum | Bacteria>Bacillati>Bacillota>Bacilli>Bacillales>Bacillales_incertae_sedis | 292 | NCTC11526_00569 | 555167..556045 | 1414643 | RNA-polymerase-associated-protein-rapA-[[Flavobacterium] |
| WP_235715382.1 | TUDOR+SF1-SF2-Helicase+iREase+CTD+REase | GCF_000513095.1 | Halalkalibacter_wakoensis | Bacteria>Bacillati>Bacillota>Bacilli>Bacillales>Bacillaceae>Halalkalibacter | 241 | JCM9140_RS24070 | 20458..21183 | 127891 | hypothetical-protein-[Halalkalibacter-wakoensis] |
| WP_235715381.1 | TUDOR+SF1-SF2-Helicase+iREase+CTD+REase | GCF_000513095.1 | Halalkalibacter_wakoensis | Bacteria>Bacillati>Bacillota>Bacilli>Bacillales>Bacillaceae>Halalkalibacter | 203 | JCM9140_RS24065 | 19850..20461 | 127891 | C-terminal-helicase-domain-containing-protein-[Halalkalibacter-wakoensis] |

| Protein-ID | Domain | Assembly-ID | Strain | Taxonomy | Length | Locus-Tag | Genomic-coordinates | TaxID | Fasta-defline |
|---|---|---|---|---|---|---|---|---|---|
| --Schlafen+HATPase+wHTH | | | | | | | | | |
| CBH38390.1 | Schlafen+HATPase+wHTH | - | uncultured_archaeon | Archaea>environmental_samples | 93 | BSM_18670 | 1756647..1756928 | 115547 | hypothetical-protein-BSM_18670-[uncultured-archaeon] |
| WP_234298125.1 | Schlafen+HATPase+wHTH | GCF_021431925.1 | Halobaculum_sp._CBA1158 | Archaea>Methanobacteriati>Methanobacteriota>Stenosarchaea_group>Halobacteria>Halobacteriales>Haloferacaceae>Halobaculum>unclassified_Halobaculum | 359 | Hbl1158_RS00465 | 104827..105906 | 2904243 | ATP-binding-protein-[Halobaculum-sp.-CBA1158] |
| WP_082220747.1 | Schlafen+HATPase+wHTH | GCF_001190965.1 | Haloferax_gibbonsii | Archaea>Methanobacteriati>Methanobacteriota>Stenosarchaea_group>Halobacteria>Halobacteriales>Haloferacaceae>Haloferax | 486 | ABY42_RS19260 | 90285..91745 | 35746 | ATP-binding-protein-[Haloferax-gibbonsii] |
| WP_271963537.1 | Schlafen+HATPase+wHTH | GCF_028211285.1 | Halorubrum_ezzemoulense | Archaea>Methanobacteriati>Methanobacteriota>Stenosarchaea_group>Halobacteria>Halobacteriales>Haloferacaceae>Halorubrum | 457 | PM025_RS15415 | 3279..4652 | 337243 | ATP-binding-protein-[Halorubrum-ezzemoulense] |
| WP_121821042.1 | Schlafen+HATPase+wHTH | GCF_003675855.1 | Halostella_salina | Archaea>Methanobacteriati>Methanobacteriota>Stenosarchaea_group>Halobacteria>Halobacteriales>Natronoarchaeaceae>Halostella | 457 | D8896_RS05290 | 400778..402151 | 1547897 | ATP-binding-protein-[Halostella-salina] |
| WP_253460966.1 | Schlafen+HATPase+wHTH | GCF_024170265.1 | Methanocalculus_sp._AMF5 | Archaea>Methanobacteriati>Methanobacteriota>Stenosarchaea_group>Methanomicrobia>Methanomicrobiales>Methanocalculaceae>Methanocalculus>unclassified_Methanocalculus | 117 | J2T61_RS09760 | 9453..9806 | 1198257 | putative-DNA-binding-domain-containing-protein-[Methanocalculus-sp.-AMF5] |
| WP_167815998.1 | Schlafen+HATPase+wHTH | - | Methanocorpusculum_sp._GPch4 | Archaea>Methanobacteriati>Methanobacteriota>Stenosarchaea_group>Methanomicrobia>Methanomicrobiales>Methanocorpusculaceae>Methanocorpusculum>unclassified_Methanocorpusculum | 539 | EYA77_RS04445 | 9275..10894 | 2527877 | ATP-binding-protein-[Methanocorpusculum-sp.-GPch4] |
| WP_167815996.1 | Schlafen+HATPase+wHTH | - | Methanocorpusculum_sp._GPch4 | Archaea>Methanobacteriati>Methanobacteriota>Stenosarchaea_group>Methanomicrobia>Methanomicrobiales>Methanocorpusculaceae>Methanocorpusculum>unclassified_Methanocorpusculum | 402 | EYA77_RS04435 | 7063..8271 | 2527877 | ATP-binding-protein-[Methanocorpusculum-sp.-GPch4] |
| WP_167815511.1 | Schlafen+HATPase+wHTH | GCF_012062385.1 | Methanocorpusculum_sp._GPch4 | Archaea>Methanobacteriati>Methanobacteriota>Stenosarchaea_group>Methanomicrobia>Methanomicrobiales>Methanocorpusculaceae>Methanocorpusculum>unclassified_Methanocorpusculum | 496 | EYA77_RS00160 | 13490..14980 | 2527877 | helix-turn-helix-domain-containing-protein-[Methanocorpusculum-sp.-GPch4] |
| WP_048152689.1 | Schlafen+HATPase+wHTH | GCF_000784355.1 | Methanolacinia_paynteri | Archaea>Methanobacteriati>Methanobacteriota>Stenosarchaea_group>Methanomicrobia>Methanomicrobiales>Methanomicrobiaceae>Methanolacinia | 481 | METPAY_RS11485 | 104392..105837 | 230356 | helix-turn-helix-domain-containing-protein-[Methanolacinia-paynteri] |
| WP_048152681.1 | Schlafen+HATPase+wHTH | GCF_000784355.1 | Methanolacinia_paynteri | Archaea>Methanobacteriati>Methanobacteriota>Stenosarchaea_group>Methanomicrobia>Methanomicrobiales>Methanomicrobiaceae>Methanolacinia | 444 | METPAY_RS11450 | 94619..95953 | 230356 | ATP-binding-protein-[Methanolacinia-paynteri] |
| WP_209691004.1 | Schlafen+HATPase+wHTH | GCF_017875315.1 | Methanomicrobium_sp._W14 | Archaea>Methanobacteriati>Methanobacteriota>Stenosarchaea_group>Methanomicrobia>Methanomicrobiales>Methanomicrobiaceae>Methanomicrobium>unclassified_Methanomicrobium | 604 | J2128_RS09595 | 63555..65369 | 2817839 | putative-DNA-binding-domain-containing-protein-[Methanomicrobium-sp.-W14] |
| WP_256440668.1 | Schlafen+HATPase+wHTH | - | Methanospirillum_purgamenti | Archaea>Methanobacteriati>Methanobacteriota>Stenosarchaea_group>Methanomicrobia>Methanomicrobiales>Methanospirillaceae>Methanospirillum | 494 | KHC33_RS10915 | 2266104..2267588 | 2834276 | helix-turn-helix-domain-containing-protein-[Methanospirillum-sp.-J.3.6.1-F.2.7.3] |
| PWR75053.1 | Schlafen+HATPase+wHTH | GCA_003173335.1 | Methanospirillum_stamsii | Archaea>Methanobacteriati>Methanobacteriota>Stenosarchaea_group>Methanomicrobia>Methanomicrobiales>Methanospirillaceae>Methanospirillum | 419 | DLD82_07500 | 182202..183461 | 1277351 | transcriptional-regulator-[Methanospirillum-stamsii] |
| MCA1916606.1 | Schlafen+HATPase+wHTH | GCA_020216685.1 | Methanospirillum_hungatei | Archaea>Methanobacteriati>Methanobacteriota>Stenosarchaea_group>Methanomicrobia>Methanomicrobiales>Methanospirillaceae>Methanospirillum | 485 | LDL35_09430 | 1277..2734 | 2203 | helix-turn-helix-domain-containing-protein-[Methanospirillum-hungatei] |
| NLA99640.1 | Schlafen+HATPase+wHTH | GCA_012511245.1 | Methanomicrobiales_archaeon | Archaea>Methanobacteriati>Methanobacteriota>Stenosarchaea_group>Methanomicrobia>Methanomicrobiales>unclassified_Methanomicrobiales | 320 | GX837_01535 | 1..965 | 2184052 | winged-helix-turn-helix-transcriptional-regulator-[Methanomicrobiales-archaeon] |
| QN042139.1 | Schlafen+HATPase+wHTH | - | Candidatus_Methanogaster_sp._ANME-2c_ERB4 | Archaea>Methanobacteriati>Methanobacteriota>Stenosarchaea_group>Methanomicrobia>Methanosarcinales>ANME-2_cluster>Candidatus_Methanogasteraceae>Candidatus_Methanogaster>unclassified_Candidatus_Methanogaster | 492 | CAOPPJJI_00005 | 5856..7334 | 2759911 | hypothetical-protein-CAOPPJJI_00005-[Methanosarcinales-archaeon-ANME-2c-ERB4] |
| MCE8425366.1 | Schlafen+HATPase+wHTH | GCA_021409295.1 | Candidatus_Methanoperedens_sp. | Archaea>Methanobacteriati>Methanobacteriota>Stenosarchaea_group>Methanomicrobia>Methanosarcinales>ANME-2_cluster>Candidatus_Methanoperedenaceae>Candidatus_Methanoperedens>unclassified_Candidatus_Methanoperedens | 496 | J5U17_06260 | 936..2426 | 2032688 | putative-DNA-binding-domain-containing-protein-[Candidatus-Methanoperedens-sp.] |
| NPE28181.1 | Schlafen+HATPase+wHTH | GCA_013166765.1 | Methanococcoides_sp._SA1 | Archaea>Methanobacteriati>Methanobacteriota>Stenosarchaea_group>Methanomicrobia>Methanosarcinales>Methanosarcinaceae>Methanococcoides>unclassified_Methanococcoides | 498 | HNV12_09460 | 9338..10834 | 2735871 | AAA-family-ATPase-[Methanococcoides-sp.-SA1] |
| MCD4820924.1 | Schlafen+HATPase+wHTH | GCA_021108275.1 | Methanococcoides_sp. | Archaea>Methanobacteriati>Methanobacteriota>Stenosarchaea_group>Methanomicrobia>Methanosarcinales>Methanosarcinaceae>Methanococcoides>unclassified_Methanococcoides | 493 | K8R11_02350 | 102754..104235 | 1966350 | helix-turn-helix-domain-containing-protein-[Methanococcoides-sp.] |
| WP_123137089.1 | Schlafen+HATPase+wHTH | GCF_003722075.1 | Methanohalophilus_sp._RSK | Archaea>Methanobacteriati>Methanobacteriota>Stenosarchaea_group>Methanomicrobia>Methanosarcinales>Methanosarcinaceae>Methanohalophilus>unclassified_Methanohalophilus | 489 | EFE42_RS09710 | 41145..42614 | 2485783 | helix-turn-helix-domain-containing-protein-[Methanohalophilus-sp.-RSK] |
| OBZ35402.1 | Schlafen+HATPase+wHTH | GCA_001683555.1 | Methanohalophilus_sp._DAL1 | Archaea>Methanobacteriati>Methanobacteriota>Stenosarchaea_group>Methanomicrobia>Methanosarcinales>Methanosarcinaceae>Methanohalophilus>unclassified_Methanohalophilus | 493 | A9957_07290 | 6756..8237 | 1864608 | MAG:-AAA-family-ATPase-[Methanohalophilus-sp.-DAL1] |
| WP_256623628.1 | Schlafen+HATPase+wHTH | GCF_024500045.1 | Methanolobus_chelungpuianus | Archaea>Methanobacteriati>Methanobacteriota>Stenosarchaea_group>Methanomicrobia>Methanosarcinales>Methanosarcinaceae>Methanolobus | 494 | PV02_RS11885 | 322732..324216 | 502115 | helix-turn-helix-domain-containing-protein-[Methanolobus-chelungpuianus] |
| WP_164997806.1 | Schlafen+HATPase+wHTH | GCF_002243045.1 | Methanolobus_psychrotolerans | Archaea>Methanobacteriati>Methanobacteriota>Stenosarchaea_group>Methanomicrobia>Methanosarcinales>Methanosarcinaceae>Methanolobus | 484 | BB781_RS06875 | 24712..26166 | 1874706 | helix-turn-helix-domain-containing-protein-[Methanolobus-psychrotolerans] |
| WP_091932404.1 | Schlafen+HATPase+wHTH | GCF_900114835.1 | Methanolobus_profundi | Archaea>Methanobacteriati>Methanobacteriota>Stenosarchaea_group>Methanomicrobia>Methanosarcinales>Methanosarcinaceae>Methanolobus | 510 | BM008_RS01875 | 364617..366149 | 487685 | helix-turn-helix-domain-containing-protein-[Methanolobus-profundi] |
| AFV22362.1 | Schlafen+HATPase+wHTH | GCA_000306725.1 | Methanolobus_psychrophilus_R15 | Archaea>Methanobacteriati>Methanobacteriota>Stenosarchaea_group>Methanomicrobia>Methanosarcinales>Methanosarcinaceae>Methanolobus>Methanolobus_psychrophilus | 490 | Mpsy_0149 | 152920..154392 | 1094980 | ATP-dependent-DNA-helicase-[Methanolobus-psychrophilus-R15] |
| KXS44095.1 | Schlafen+HATPase+wHTH | GCA_001577565.1 | Methanolobus_sp._T82-4 | Archaea>Methanobacteriati>Methanobacteriota>Stenosarchaea_group>Methanomicrobia>Methanosarcinales>Methanosarcinaceae>Methanolobus>unclassified_Methanolobus | 506 | AWU59_728 | 12661..14181 | 1794908 | MAG:-putative-transcriptional-regulator-with-HTH-domain-[Methanolobus-sp.-T82-4] |
| WP_245527567.1 | Schlafen+HATPase+wHTH | GCF_000217995.1 | Methanosalsum_zhilinae | Archaea>Methanobacteriati>Methanobacteriota>Stenosarchaea_group>Methanomicrobia>Methanosarcinales>Methanosarcinaceae>Methanosalsum | 492 | MZHIL_RS04750 | 1024766..1026244 | 39669 | helix-turn-helix-domain-containing-protein-[Methanosalsum-zhilinae] |
| WP_148705387.1 | Schlafen+HATPase+wHTH | GCF_000970125.1 | Methanosarcina_siciliae | Archaea>Methanobacteriati>Methanobacteriota>Stenosarchaea_group>Methanomicrobia>Methanosarcinales>Methanosarcinaceae>Methanosarcina | 618 | MSSIH_RS10185 | 2390255..2392111 | 38027 | putative-DNA-binding-domain-containing-protein-[Methanosarcina-siciliae] |
| WP_048041738.1 | Schlafen+HATPase+wHTH | GCF_048285915.1 | Methanosarcina_mazei | Archaea>Methanobacteriati>Methanobacteriota>Stenosarchaea_group>Methanomicrobia>Methanosarcinales>Methanosarcinaceae>Methanosarcina | 610 | DU31_RS16155 | 65558..67390 | 2209 | putative-DNA-binding-domain-containing-protein-[Methanosarcina-mazei] |
| WP_015410903.1 | Schlafen+HATPase+wHTH | GCF_000007065.1 | Methanosarcina_mazei | Archaea>Methanobacteriati>Methanobacteriota>Stenosarchaea_group>Methanomicrobia>Methanosarcinales>Methanosarcinaceae>Methanosarcina | 47 | FQU78_RS06245 | 1443445..1443588 | 2209 | ATP-dependent-DNA-helicase-[Methanosarcina-mazei] |
| WP_015410902.1 | Schlafen+HATPase+wHTH | GCF_000007065.1 | Methanosarcina_mazei | Archaea>Methanobacteriati>Methanobacteriota>Stenosarchaea_group>Methanomicrobia>Methanosarcinales>Methanosarcinaceae>Methanosarcina | 53 | FQU78_RS06250 | 1443585..1443746 | 2209 | putative-DNA-binding-domain-containing-protein-[Methanosarcina-mazei] |
| KKG05627.1 | Schlafen+HATPase+wHTH | GCA_000979375.1 | Methanosarcina_mazei | Archaea>Methanobacteriati>Methanobacteriota>Stenosarchaea_group>Methanomicrobia>Methanosarcinales>Methanosarcinaceae>Methanosarcina | 53 | DU40_09120 | 244209..244370 | 2209 | transcriptional-regulator-[Methanosarcina-mazei] |
| AKB55568.1 | Schlafen+HATPase+wHTH | GCA_000970025.1 | Methanosarcina_barkeri_MS | Archaea>Methanobacteriati>Methanobacteriota>Stenosarchaea_group>Methanomicrobia>Methanosarcinales>Methanosarcinaceae>Methanosarcina>Methanosarcina_barkeri | 44 | MSBRM_2570 | 3395415..3395549 | 1434108 | ATP-dependent-DNA-helicase-[Methanosarcina-barkeri-MS] |
| AKB74627.1 | Schlafen+HATPase+wHTH | GCA_000970265.1 | Methanosarcina_lacustris_Z-7289 | Archaea>Methanobacteriati>Methanobacteriota>Stenosarchaea_group>Methanomicrobia>Methanosarcinales>Methanosarcinaceae>Methanosarcina>Methanosarcina_lacustris | 602 | MSLAZ_1366 | 1712723..1714531 | 1434111 | ATP-dependent-DNA-helicase-[Methanosarcina-lacustris-Z-7289] |
| WP_229393449.1 | Schlafen+HATPase+wHTH | GCF_020804205.1 | Methanosarcina_sp._DH1 | Archaea>Methanobacteriati>Methanobacteriota>Stenosarchaea_group>Methanomicrobia>Methanosarcinales>Methanosarcinaceae>Methanosarcina>unclassified_Methanosarcina | 473 | FXW07_RS01535 | 49329..50750 | 2605695 | ATP-binding-protein-[Methanosarcina-sp.-DH1] |
| WP_197073947.1 | Schlafen+HATPase+wHTH | GCF_000969965.1 | Methanosarcina_sp._WWM596 | Archaea>Methanobacteriati>Methanobacteriota>Stenosarchaea_group>Methanomicrobia>Methanosarcinales>Methanosarcinaceae>Methanosarcina>unclassified_Methanosarcina | 626 | MSWHS_RS14785 | 3349304..3351184 | 1434103 | putative-DNA-binding-domain-containing-protein-[Methanosarcina-sp.-WWM596] |
| WP_048162605.1 | Schlafen+HATPase+wHTH | GCF_000979975.1 | Methanosarcina_sp._1.H.A.2.2 | Archaea>Methanobacteriati>Methanobacteriota>Stenosarchaea_group>Methanomicrobia>Methanosarcinales>Methanosarcinaceae>Methanosarcina>unclassified_Methanosarcina | 456 | EO93_RS11310 | 72718..74088 | 1483601 | helix-turn-helix-domain-containing-protein-[Methanosarcina-sp.-1.H.A.2.2] |
| HHV24728.1 | Schlafen+HATPase+wHTH | GCA_012839505.1 | Methanosarcina_sp. | Archaea>Methanobacteriati>Methanobacteriota>Stenosarchaea_group>Methanomicrobia>Methanosarcinales>Methanosarcinaceae>Methanosarcina>unclassified_Methanosarcina | 606 | GXX65_09475 | 16096..17916 | 2213 | transcriptional-regulator-[Methanosarcina-sp.] |
| MCG7853931.1 | Schlafen+HATPase+wHTH | GCA_022353215.1 | Methanosarcinaceae_archaeon | Archaea>Methanobacteriati>Methanobacteriota>Stenosarchaea_group>Methanomicrobia>Methanosarcinales>Methanosarcinaceae>unclassified_Methanosarcinaceae | 397 | MIO92_15525 | 5500..6693 | 1945595 | putative-DNA-binding-domain-containing-protein-[Methanosarcinaceae-archaeon] |
| NYT10422.1 | Schlafen+HATPase+wHTH | GCA_013415605.1 | Methanosarcinales_archaeon | Archaea>Methanobacteriati>Methanobacteriota>Stenosarchaea_group>Methanomicrobia>Methanosarcinales>unclassified_Methanosarcinales | 606 | GKC09_10920 | 6672..8492 | 2250255 | AAA-family-ATPase-[Methanosarcinales-archaeon] |
| WP_014586221.1 | Schlafen+HATPase+wHTH | GCF_000235565.1 | Methanothrix_harundinacea | Archaea>Methanobacteriati>Methanobacteriota>Stenosarchaea_group>Methanomicrobia>Methanotrichales>Methanotrichaceae>Methanothrix | 565 | MHAR_RS03405 | 692672..694369 | 301375 | ATP-binding-protein-[Methanothrix-harundinacea] |
| NLJ23754.1 | Schlafen+HATPase+wHTH | GCA_012517695.1 | Methanothrix_soehngenii | Archaea>Methanobacteriati>Methanobacteriota>Stenosarchaea_group>Methanomicrobia>Methanotrichales>Methanotrichaceae>Methanothrix | 607 | GX426_11725 | 7201..9024 | 2223 | AAA-family-ATPase-[Methanothrix-soehngenii] |
| AET64036.1 | Schlafen+HATPase+wHTH | GCA_000235565.1 | Methanothrix_harundinacea_6Ac | Archaea>Methanobacteriati>Methanobacteriota>Stenosarchaea_group>Methanomicrobia>Methanotrichales>Methanotrichaceae>Methanothrix>Methanothrix_harundinacea | 565 | Mhar_0658 | 692672..694369 | 1110509 | Divergent-AAA-domain-family-[Methanothrix-harundinacea-6Ac] |
| OPX79393.1 | Schlafen+HATPase+wHTH | GCA_002067795.1 | Methanosaeta_sp._PtaB.Bin039 | Archaea>Methanobacteriati>Methanobacteriota>Stenosarchaea_group>Methanomicrobia>Methanotrichales>Methanotrichaceae>Methanothrix>unclassified_Methanothrix | 606 | A4E45_00672 | 44332..46152 | 1811685 | MAG:-Divergent-AAA-domain-protein-[Methanosaeta-sp.-PtaB.Bin039] |
| MBN1235618.1 | Schlafen+HATPase+wHTH | GCA_016927055.1 | Methanotrichaceae_archaeon | Archaea>Methanobacteriati>Methanobacteriota>Stenosarchaea_group>Methanomicrobia>Methanotrichales>Methanotrichaceae>unclassified_Methanotrichaceae | 81 | JW999_06150 | 9282..9527 | 2595067 | DeoR-family-transcriptional-regulator-[Methanotrichaceae-archaeon] |
| MBU4220829.1 | Schlafen+HATPase+wHTH | GCA_018898145.1 | Methanobacteriota_archaeon | Archaea>Methanobacteriati>Methanobacteriota>unclassified_Methanobacteriota | 501 | KKA10_04275 | 15168..16673 | 2026739 | helix-turn-helix-domain-containing-protein-[Euryarchaeota-archaeon] |
| MBU4220827.1 | Schlafen+HATPase+wHTH | GCA_018898145.1 | Methanobacteriota_archaeon | Archaea>Methanobacteriati>Methanobacteriota>unclassified_Methanobacteriota | 50 | KKA10_04265 | 14139..14291 | 2026739 | ATP-binding-protein-[Euryarchaeota-archaeon] |
| MXX20695.1 | Schlafen+HATPase+wHTH | GCA_009836545.1 | Cenarchaeum_sp._SB0667_bin_13 | Archaea>Thermoproteati>Nitrososphaerota>Candidatus_Cenarchaeales>Candidatus_Cenarchaeaceae>Candidatus_Cenarchaeum>unclassified_Candidatus_Cenarchaeum | 228 | F4Z46_02775 | 7459..8145 | 2604878 | DUF2196-domain-containing-protein-[Cenarchaeum-sp.-SB0667_bin_13] |
| MCC7078888.1 | Schlafen+HATPase+wHTH | GCA_020850945.1 | Acidimicrobiia_bacterium | Bacteria>Bacillati>Actinomycetota>Acidimicrobiia>unclassified_Acidimicrobiia | 563 | IT198_17365 | 35692..37383 | 2080302 | putative-DNA-binding-domain-containing-protein-[Acidimicrobiia-bacterium] |
| MCC7078884.1 | Schlafen+HATPase+wHTH | GCA_020850945.1 | Acidimicrobiia_bacterium | Bacteria>Bacillati>Actinomycetota>Acidimicrobiia>unclassified_Acidimicrobiia | 577 | IT198_17345 | 27127..28860 | 2080302 | putative-DNA-binding-domain-containing-protein-[Acidimicrobiia-bacterium] |
| WP_041834944.1 | Schlafen+HATPase+wHTH | GCF_000015025.1 | Acidothermus_cellulolyticus | Bacteria>Bacillati>Actinomycetota>Actinomycetes>Acidothermales>Acidothermaceae>Acidothermus | 469 | ACEL_RS04200 | 900176..901585 | 28049 | ATP-binding-protein-[Acidothermus-cellulolyticus] |
| WP_166855912.1 | Schlafen+HATPase+wHTH | GCF_014595995.2 | Actinomyces | Bacteria>Bacillati>Actinomycetota>Actinomycetes>Actinomycetales>Actinomycetaceae | 537 | HCR20_RS03225 | 138508..140121 | 1654 | MULTISPECIES:-ATP-binding-protein-[Actinomyces] |
| WP_075371845.1 | Schlafen+HATPase+wHTH | GCF_031296635.1 | Actinomyces | Bacteria>Bacillati>Actinomycetota>Actinomycetes>Actinomycetales>Actinomycetaceae | 498 | BKH23_RS02490 | 275969..277465 | 1654 | helix-turn-helix-domain-containing-protein-[Actinomyces-oris] |
| WP_205647225.1 | Schlafen+HATPase+wHTH | GCF_003073475.1 | Actinobaculum_sp._313 | Bacteria>Bacillati>Actinomycetota>Actinomycetes>Actinomycetales>Actinomycetaceae>Actinobaculum>unclassified_Actinobaculum | 149 | DDD63_RS12530 | 1988600..1989049 | 2495645 | hypothetical-protein-[Actinobaculum-sp.-313] |
| WP_205647223.1 | Schlafen+HATPase+wHTH | GCF_003073475.1 | Actinobaculum_sp._313 | Bacteria>Bacillati>Actinomycetota>Actinomycetes>Actinomycetales>Actinomycetaceae>Actinobaculum>unclassified_Actinobaculum | 257 | DDD63_RS12525 | 1987764..1988537 | 2495645 | ATP-binding-protein-[Actinobaculum-sp.-313] |
| WP_205647219.1 | Schlafen+HATPase+wHTH | GCF_003073475.1 | Actinobaculum_sp._313 | Bacteria>Bacillati>Actinomycetota>Actinomycetes>Actinomycetales>Actinomycetaceae>Actinobaculum>unclassified_Actinobaculum | 72 | DDD63_RS12510 | 1976349..1976567 | 2495645 | hypothetical-protein-[Actinobaculum-sp.-313] |
| WP_126145885.1 | Schlafen+HATPase+wHTH | GCF_003957265.1 | Actinobaculum_sp._352 | Bacteria>Bacillati>Actinomycetota>Actinomycetes>Actinomycetales>Actinomycetaceae>Actinobaculum>unclassified_Actinobaculum | 428 | EKN07_RS06080 | 225133..226419 | 2490946 | ATP-binding-protein-[Actinobaculum-sp.-352] |
| WP_271289306.1 | Schlafen+HATPase+wHTH | GCF_027945475.1 | Actinomyces_oris | Bacteria>Bacillati>Actinomycetota>Actinomycetes>Actinomycetales>Actinomycetaceae>Actinomyces | 498 | PGE45_RS05455 | 1256274..1257770 | 544580 | helix-turn-helix-domain-containing-protein-[Actinomyces-oris] |
| WP_245572171.1 | Schlafen+HATPase+wHTH | GCF_000429265.1 | Actinomyces_gerencseriae | Bacteria>Bacillati>Actinomycetota>Actinomycetes>Actinomycetales>Actinomycetaceae>Actinomyces | 333 | G448_RS14995 | 216209..217210 | 52769 | ATP-binding-protein-[Actinomyces-gerencseriae] |
| WP_141407053.1 | Schlafen+HATPase+wHTH | GCF_016127955.1 | Actinomyces_oris | Bacteria>Bacillati>Actinomycetota>Actinomycetes>Actinomycetales>Actinomycetaceae>Actinomyces | 498 | I6I08_RS06250 | 1505513..1507009 | 544580 | helix-turn-helix-domain-containing-protein-[Actinomyces-oris] |
| WP_136193202.1 | Schlafen+HATPase+wHTH | GCF_004796555.1 | Actinomyces_procaprae | Bacteria>Bacillati>Actinomycetota>Actinomycetes>Actinomycetales>Actinomycetaceae>Actinomyces | 523 | FAQ94_RS07515 | 95665..97236 | 2560010 | ATP-binding-protein-[Actinomyces-procaprae] |
| WP_127841207.1 | Schlafen+HATPase+wHTH | GCF_004010955.1 | Actinomyces_wuliandei | Bacteria>Bacillati>Actinomycetota>Actinomycetes>Actinomycetales>Actinomycetaceae>Actinomyces | 510 | CWS50_RS00395 | 88028..89560 | 2057743 | helix-turn-helix-domain-containing-protein-[Actinomyces-wuliandei] |
| WP_126382189.1 | Schlafen+HATPase+wHTH | GCF_900637165.1 | Actinomyces_howellii | Bacteria>Bacillati>Actinomycetota>Actinomycetes>Actinomycetales>Actinomycetaceae>Actinomyces | 481 | GYB74_RS04540 | 51847..53292 | 52771 | helix-turn-helix-domain-containing-protein-[Actinomyces-howellii] |
| WP_076066521.1 | Schlafen+HATPase+wHTH | GCF_001956485.1 | Actinomyces_naeslundii | Bacteria>Bacillati>Actinomycetota>Actinomycetes>Actinomycetales>Actinomycetaceae>Actinomyces | 498 | BKH36_RS03210 | 67088..68584 | 1655 | helix-turn-helix-domain-containing-protein-[Actinomyces-naeslundii] |
| WP_075550930.1 | Schlafen+HATPase+wHTH | GCF_001937415.1 | Actinomyces_oris | Bacteria>Bacillati>Actinomycetota>Actinomycetes>Actinomycetales>Actinomycetaceae>Actinomyces | 498 | BKH26_RS09130 | 60776..62272 | 544580 | helix-turn-helix-domain-containing-protein-[Actinomyces-oris] |
| WP_075411482.1 | Schlafen+HATPase+wHTH | GCF_001937365.1 | Actinomyces_oris | Bacteria>Bacillati>Actinomycetota>Actinomycetes>Actinomycetales>Actinomycetaceae>Actinomyces | 502 | BKH31_RS05945 | 47548..49056 | 544580 | helix-turn-helix-domain-containing-protein-[Actinomyces-oris] |
| WP_075400904.1 | Schlafen+HATPase+wHTH | GCF_001937425.1 | Actinomyces_oris | Bacteria>Bacillati>Actinomycetota>Actinomycetes>Actinomycetales>Actinomycetaceae>Actinomyces | 498 | BKH24_RS01010 | 85079..86575 | 544580 | helix-turn-helix-domain-containing-protein-[Actinomyces-oris] |
| WP_075392851.1 | Schlafen+HATPase+wHTH | GCF_001937725.1 | Actinomyces_oris | Bacteria>Bacillati>Actinomycetota>Actinomycetes>Actinomycetales>Actinomycetaceae>Actinomyces | 498 | BKH22_RS09300 | 64165..65661 | 544580 | helix-turn-helix-domain-containing-protein-[Actinomyces-oris] |
| WP_075374662.1 | Schlafen+HATPase+wHTH | GCF_001937535.1 | Actinomyces_oris | Bacteria>Bacillati>Actinomycetota>Actinomycetes>Actinomycetales>Actinomycetaceae>Actinomyces | 502 | BKH18_RS06070 | 88967..90475 | 544580 | helix-turn-helix-domain-containing-protein-[Actinomyces-oris] |
| QQC40837.1 | Schlafen+HATPase+wHTH | GCA_016127955.1 | Actinomyces_oris | Bacteria>Bacillati>Actinomycetota>Actinomycetes>Actinomycetales>Actinomycetaceae>Actinomyces | 498 | I6I08_06250 | 1505513..1507009 | 544580 | putative-DNA-binding-domain-containing-protein-[Actinomyces-oris] |
| OLO77536.1 | Schlafen+HATPase+wHTH | GCA_001937535.1 | Actinomyces_oris | Bacteria>Bacillati>Actinomycetota>Actinomycetes>Actinomycetales>Actinomycetaceae>Actinomyces | 502 | BKH18_06075 | 88967..90475 | 544580 | AAA-family-ATPase-[Actinomyces-oris] |
| OLO63824.1 | Schlafen+HATPase+wHTH | GCA_001937485.1 | Actinomyces_oris | Bacteria>Bacillati>Actinomycetota>Actinomycetes>Actinomycetales>Actinomycetaceae>Actinomyces | 498 | BKH23_02490 | 275969..277465 | 544580 | AAA-family-ATPase-[Actinomyces-oris] |
| OLO62961.1 | Schlafen+HATPase+wHTH | GCA_001937725.1 | Actinomyces_oris | Bacteria>Bacillati>Actinomycetota>Actinomycetes>Actinomycetales>Actinomycetaceae>Actinomyces | 498 | BKH22_09290 | 64165..65661 | 544580 | AAA-family-ATPase-[Actinomyces-oris] |
| OLO62857.1 | Schlafen+HATPase+wHTH | GCA_001937425.1 | Actinomyces_oris | Bacteria>Bacillati>Actinomycetota>Actinomycetes>Actinomycetales>Actinomycetaceae>Actinomyces | 498 | BKH24_01010 | 85079..86575 | 544580 | AAA-family-ATPase-[Actinomyces-oris] |
| OLO54915.1 | Schlafen+HATPase+wHTH | GCA_001937415.1 | Actinomyces_oris | Bacteria>Bacillati>Actinomycetota>Actinomycetes>Actinomycetales>Actinomycetaceae>Actinomyces | 498 | BKH26_09150 | 60776..62272 | 544580 | AAA-family-ATPase-[Actinomyces-oris] |
| OLO46897.1 | Schlafen+HATPase+wHTH | GCA_001937365.1 | Actinomyces_oris | Bacteria>Bacillati>Actinomycetota>Actinomycetes>Actinomycetales>Actinomycetaceae>Actinomyces | 502 | BKH31_05945 | 47548..49056 | 544580 | AAA-family-ATPase-[Actinomyces-oris] |
| WP_119835305.1 | Schlafen+HATPase+wHTH | GCF_003591215.1 | Actinomyces_sp._2119 | Bacteria>Bacillati>Actinomycetota>Actinomycetes>Actinomycetales>Actinomycetaceae>Actinomyces>unclassified_Actinomyces | 523 | D4740_RS02100 | 58629..60200 | 2321393 | ATP-binding-protein-[Actinomyces-sp.-2119] |
| RJF43787.1 | Schlafen+HATPase+wHTH | GCA_003591215.1 | Actinomyces_sp._2119 | Bacteria>Bacillati>Actinomycetota>Actinomycetes>Actinomycetales>Actinomycetaceae>Actinomyces>unclassified_Actinomyces | 523 | D4740_02100 | 58629..60200 | 2321393 | AAA-family-ATPase-[Actinomyces-sp.-2119] |
| WP_109093612.1 | Schlafen+HATPase+wHTH | GCF_003123745.1 | Ancrocorticia_populi | Bacteria>Bacillati>Actinomycetota>Actinomycetes>Actinomycetales>Actinomycetaceae>Ancrocorticia | 493 | DD236_RS06720 | 270839..272320 | 2175228 | helix-turn-helix-domain-containing-protein-[Ancrocorticia-populi] |
| WP_262508935.1 | Schlafen+HATPase+wHTH | - | Schaalia_sp._JY-X159 | Bacteria>Bacillati>Actinomycetota>Actinomycetes>Actinomycetales>Actinomycetaceae>Schaalia>unclassified_Schaalia | 366 | IDL90_RS00785 | 180752..181852 | 2758575 | putative-DNA-binding-domain-containing-protein-[Schaalia-sp.-JY-X159] |
| WP_271768976.1 | Schlafen+HATPase+wHTH | GCF_028314485.1 | Bifidobacterium | Bacteria>Bacillati>Actinomycetota>Actinomycetes>Bifidobacteriales>Bifidobacteriaceae | 512 | PMN34_RS05100 | 229872..231410 | 1678 | ATP-binding-protein-[Bifidobacterium-pseudocatenulatum] |
| WP_270271487.1 | Schlafen+HATPase+wHTH | GCF_027679525.1 | Bifidobacterium | Bacteria>Bacillati>Actinomycetota>Actinomycetes>Bifidobacteriales>Bifidobacteriaceae | 498 | PGW19_RS08950 | 6325..7821 | 1678 | ATP-binding-protein-[Bifidobacterium-adolescentis] |
| WP_232780029.1 | Schlafen+HATPase+wHTH | GCF_001264045.1 | Bifidobacterium | Bacteria>Bacillati>Actinomycetota>Actinomycetes>Bifidobacteriales>Bifidobacteriaceae | 17 | CE167_RS12725 | 23533..23586 | 1678 | MULTISPECIES:-ATP-binding-protein-[Bifidobacterium] |
| WP_230845796.1 | Schlafen+HATPase+wHTH | GCF_001264045.1 | Bifidobacterium | Bacteria>Bacillati>Actinomycetota>Actinomycetes>Bifidobacteriales>Bifidobacteriaceae | 99 | CE167_RS03775 | 23599..23898 | 1678 | MULTISPECIES:-ATP-binding-protein-[Bifidobacterium] |
| WP_214374606.1 | Schlafen+HATPase+wHTH | GCF_018555335.1 | Bifidobacterium | Bacteria>Bacillati>Actinomycetota>Actinomycetes>Bifidobacteriales>Bifidobacteriaceae | 498 | JS541_RS11290 | 64457..65953 | 1678 | MULTISPECIES:-ATP-binding-protein-[Bifidobacterium] |
| WP_117664459.1 | Schlafen+HATPase+wHTH | GCF_020525765.1 | Bifidobacterium | Bacteria>Bacillati>Actinomycetota>Actinomycetes>Bifidobacteriales>Bifidobacteriaceae | 512 | DXC45_RS01955 | 409419..410957 | 1678 | ATP-binding-protein-[Bifidobacterium-pseudocatenulatum] |
| WP_116463717.1 | Schlafen+HATPase+wHTH | GCF_008086305.1 | Bifidobacterium | Bacteria>Bacillati>Actinomycetota>Actinomycetes>Bifidobacteriales>Bifidobacteriaceae | 512 | PWA56_RS02170 | 496349..497887 | 1678 | ATP-binding-protein-[Bifidobacterium-longum] |
| WP_085380383.1 | Schlafen+HATPase+wHTH | GCF_039063575.1 | Bifidobacterium | Bacteria>Bacillati>Actinomycetota>Actinomycetes>Bifidobacteriales>Bifidobacteriaceae | 512 | LMG10734_RS05585 | 480253..481791 | 1678 | ATP-binding-protein-[Bifidobacterium-adolescentis] |
| WP_065465683.1 | Schlafen+HATPase+wHTH | GCF_001685805.1 | Bifidobacterium | Bacteria>Bacillati>Actinomycetota>Actinomycetes>Bifidobacteriales>Bifidobacteriaceae | 83 | BAD00_RS10870 | 108092..108343 | 1678 | hypothetical-protein-[Bifidobacterium-breve] |
| WP_052828893.1 | Schlafen+HATPase+wHTH | GCF_001685865.1 | Bifidobacterium | Bacteria>Bacillati>Actinomycetota>Actinomycetes>Bifidobacteriales>Bifidobacteriaceae | 100 | NRBB57_RS05020 | 1139740..1140042 | 1678 | MULTISPECIES:-hypothetical-protein-[Bifidobacterium] |
| WP_052792438.1 | Schlafen+HATPase+wHTH | GCF_001264105.1 | Bifidobacterium | Bacteria>Bacillati>Actinomycetota>Actinomycetes>Bifidobacteriales>Bifidobacteriaceae | 512 | FPG44_RS02600 | 440514..442052 | 1678 | ATP-binding-protein-[Bifidobacterium-breve] |
| WP_029679906.1 | Schlafen+HATPase+wHTH | GCF_000421385.1 | Bifidobacterium | Bacteria>Bacillati>Actinomycetota>Actinomycetes>Bifidobacteriales>Bifidobacteriaceae | 512 | BFS26_RS01860 | 9218..10756 | 1678 | ATP-binding-protein-[Bifidobacterium-longum] |
| WP_017143369.1 | Schlafen+HATPase+wHTH | GCF_000273525.1 | Bifidobacterium | Bacteria>Bacillati>Actinomycetota>Actinomycetes>Bifidobacteriales>Bifidobacteriaceae | 508 | I2D91_RS01790 | 434591..436117 | 1678 | MULTISPECIES:-ATP-binding-protein-[Bifidobacterium] |
| WP_270836346.1 | Schlafen+HATPase+wHTH | GCF_027669765.1 | Bifidobacterium_bifidum | Bacteria>Bacillati>Actinomycetota>Actinomycetes>Bifidobacteriales>Bifidobacteriaceae>Bifidobacterium | 498 | PHA16_RS02390 | 597809..599305 | 1681 | ATP-binding-protein-[Bifidobacterium-bifidum] |
| WP_270561191.1 | Schlafen+HATPase+wHTH | GCF_027698005.1 | Bifidobacterium_dentium | Bacteria>Bacillati>Actinomycetota>Actinomycetes>Bifidobacteriales>Bifidobacteriaceae>Bifidobacterium | 515 | PG823_RS09675 | 51017..52564 | 1689 | ATP-binding-protein-[Bifidobacterium-dentium] |
| WP_270559046.1 | Schlafen+HATPase+wHTH | GCF_027698785.1 | Bifidobacterium_pseudocatenulatum | Bacteria>Bacillati>Actinomycetota>Actinomycetes>Bifidobacteriales>Bifidobacteriaceae>Bifidobacterium | 498 | PG820_RS05630 | 108303..109799 | 28026 | ATP-binding-protein-[Bifidobacterium-pseudocatenulatum] |
| WP_270558498.1 | Schlafen+HATPase+wHTH | GCF_027690155.1 | Bifidobacterium_adolescentis | Bacteria>Bacillati>Actinomycetota>Actinomycetes>Bifidobacteriales>Bifidobacteriaceae>Bifidobacterium | 512 | PG057_RS05005 | 451562..453100 | 1680 | ATP-binding-protein-[Bifidobacterium-adolescentis] |
| WP_270557624.1 | Schlafen+HATPase+wHTH | GCF_027697805.1 | Bifidobacterium_pseudocatenulatum | Bacteria>Bacillati>Actinomycetota>Actinomycetes>Bifidobacteriales>Bifidobacteriaceae>Bifidobacterium | 498 | PG776_RS01100 | 267369..268865 | 28026 | ATP-binding-protein-[Bifidobacterium-pseudocatenulatum] |
| WP_270554216.1 | Schlafen+HATPase+wHTH | GCF_027695195.1 | Bifidobacterium_adolescentis | Bacteria>Bacillati>Actinomycetota>Actinomycetes>Bifidobacteriales>Bifidobacteriaceae>Bifidobacterium | 512 | PG742_RS01135 | 261359..262897 | 1680 | ATP-binding-protein-[Bifidobacterium-adolescentis] |
| WP_270547724.1 | Schlafen+HATPase+wHTH | GCF_027724105.1 | Bifidobacterium_bifidum | Bacteria>Bacillati>Actinomycetota>Actinomycetes>Bifidobacteriales>Bifidobacteriaceae>Bifidobacterium | 506 | PG972_RS08460 | 5563..7083 | 1681 | ATP-binding-protein-[Bifidobacterium-bifidum] |
| WP_270447027.1 | Schlafen+HATPase+wHTH | GCF_027682665.1 | Bifidobacterium_pseudocatenulatum | Bacteria>Bacillati>Actinomycetota>Actinomycetes>Bifidobacteriales>Bifidobacteriaceae>Bifidobacterium | 526 | PGX68_RS01410 | 358002..359582 | 28026 | ATP-binding-protein-[Bifidobacterium-pseudocatenulatum] |
| WP_270307606.1 | Schlafen+HATPase+wHTH | GCF_027664445.1 | Bifidobacterium_adolescentis | Bacteria>Bacillati>Actinomycetota>Actinomycetes>Bifidobacteriales>Bifidobacteriaceae>Bifidobacterium | 498 | PGQ86_RS05885 | 1344174..1345670 | 1680 | ATP-binding-protein-[Bifidobacterium-adolescentis] |
| WP_270294255.1 | Schlafen+HATPase+wHTH | GCF_027683145.1 | Bifidobacterium_adolescentis | Bacteria>Bacillati>Actinomycetota>Actinomycetes>Bifidobacteriales>Bifidobacteriaceae>Bifidobacterium | 498 | PGX62_RS02085 | 460390..461886 | 1680 | ATP-binding-protein-[Bifidobacterium-adolescentis] |
| WP_270271793.1 | Schlafen+HATPase+wHTH | GCF_027667785.1 | Bifidobacterium_bifidum | Bacteria>Bacillati>Actinomycetota>Actinomycetes>Bifidobacteriales>Bifidobacteriaceae>Bifidobacterium | 498 | PGS22_RS05360 | 136218..137714 | 1681 | ATP-binding-protein-[Bifidobacterium-bifidum] |
| WP_270200769.1 | Schlafen+HATPase+wHTH | GCF_027671795.1 | Bifidobacterium_pseudocatenulatum | Bacteria>Bacillati>Actinomycetota>Actinomycetes>Bifidobacteriales>Bifidobacteriaceae>Bifidobacterium | 498 | PGT11_RS00285 | 75856..77352 | 28026 | ATP-binding-protein-[Bifidobacterium-pseudocatenulatum] |
| WP_270199958.1 | Schlafen+HATPase+wHTH | GCF_027675485.1 | Bifidobacterium_pseudocatenulatum | Bacteria>Bacillati>Actinomycetota>Actinomycetes>Bifidobacteriales>Bifidobacteriaceae>Bifidobacterium | 498 | PGV18_RS06425 | 50419..51915 | 28026 | ATP-binding-protein-[Bifidobacterium-pseudocatenulatum] |
| WP_269374249.1 | Schlafen+HATPase+wHTH | GCF_027214485.1 | Bifidobacterium_breve | Bacteria>Bacillati>Actinomycetota>Actinomycetes>Bifidobacteriales>Bifidobacteriaceae>Bifidobacterium | 506 | O4C76_RS09005 | 66475..67995 | 1685 | ATP-binding-protein-[Bifidobacterium-breve] |
| WP_269355162.1 | Schlafen+HATPase+wHTH | GCF_027215185.1 | Bifidobacterium_breve | Bacteria>Bacillati>Actinomycetota>Actinomycetes>Bifidobacteriales>Bifidobacteriaceae>Bifidobacterium | 512 | O4C27_RS08980 | 58825..60363 | 1685 | ATP-binding-protein-[Bifidobacterium-breve] |
| WP_259298131.1 | Schlafen+HATPase+wHTH | GCF_027687815.1 | Bifidobacterium_bifidum | Bacteria>Bacillati>Actinomycetota>Actinomycetes>Bifidobacteriales>Bifidobacteriaceae>Bifidobacterium | 520 | PG427_RS02275 | 564350..565912 | 1681 | ATP-binding-protein-[Bifidobacterium-bifidum] |
| WP_242673999.1 | Schlafen+HATPase+wHTH | GCF_004334635.1 | Bifidobacterium_longum | Bacteria>Bacillati>Actinomycetota>Actinomycetes>Bifidobacteriales>Bifidobacteriaceae>Bifidobacterium | 399 | MCC10126_RS01970 | 1..1196 | 216816 | ATP-binding-protein,-partial-[Bifidobacterium-longum] |
| WP_242075536.1 | Schlafen+HATPase+wHTH | GCF_022643745.1 | Bifidobacterium_longum | Bacteria>Bacillati>Actinomycetota>Actinomycetes>Bifidobacteriales>Bifidobacteriaceae>Bifidobacterium | 514 | G8B15_RS07015 | 1534363..1535907 | 216816 | ATP-binding-protein-[Bifidobacterium-longum] |

| Protein-ID | Domain | Assembly-ID | Strain | Taxonomy | Length | Locus-Tag | Genomic-coordinates | TaxID | Fasta-defline |
|---|---|---|---|---|---|---|---|---|---|
| WP_241735162.1 | Schlafen+HATPase+wHTH | GCF_015703835.1 | Bifidobacterium_dentium | Bacteria>Bacillati>Actinomycetota>Actinomycetes>Bifidobacteriales>Bifidobacteriaceae>Bifidobacterium | 495 | IHQ82_RS06070 | 188812..190299 | 1689 | ATP-binding-protein-[Bifidobacterium-dentium] |
| WP_240945039.1 | Schlafen+HATPase+wHTH | GCF_012932365.1 | Bifidobacterium_moraviense | Bacteria>Bacillati>Actinomycetota>Actinomycetes>Bifidobacteriales>Bifidobacteriaceae>Bifidobacterium | 496 | G1C96_RS01310 | 297279..298769 | 2675323 | ATP-binding-protein-[Bifidobacterium-sp.-DSM-109958] |
| WP_240833136.1 | Schlafen+HATPase+wHTH | - | Bifidobacterium_pseudocatenulatum | Bacteria>Bacillati>Actinomycetota>Actinomycetes>Bifidobacteriales>Bifidobacteriaceae>Bifidobacterium | 512 | LF914_RS07620 | 47658..49196 | 28026 | ATP-binding-protein-[Bifidobacterium-pseudocatenulatum] |
| WP_237970586.1 | Schlafen+HATPase+wHTH | GCF_022136875.1 | Bifidobacterium_longum | Bacteria>Bacillati>Actinomycetota>Actinomycetes>Bifidobacteriales>Bifidobacteriaceae>Bifidobacterium | 498 | LOO84_RS04705 | 91005..92501 | 216816 | ATP-binding-protein-[Bifidobacterium-longum] |
| WP_237919302.1 | Schlafen+HATPase+wHTH | GCF_022135125.1 | Bifidobacterium_pseudocatenulatum | Bacteria>Bacillati>Actinomycetota>Actinomycetes>Bifidobacteriales>Bifidobacteriaceae>Bifidobacterium | 498 | LOM88_RS00220 | 63601..65097 | 28026 | ATP-binding-protein-[Bifidobacterium-pseudocatenulatum] |
| WP_233432575.1 | Schlafen+HATPase+wHTH | GCF_003952825.1 | Bifidobacterium_pseudocatenulatum | Bacteria>Bacillati>Actinomycetota>Actinomycetes>Bifidobacteriales>Bifidobacteriaceae>Bifidobacterium | 499 | CWS99_RS05755 | 1333282..1334781 | 28026 | ATP-binding-protein-[Bifidobacterium-pseudocatenulatum] |
| WP_230803846.1 | Schlafen+HATPase+wHTH | GCF_001189355.1 | Bifidobacterium_breve | Bacteria>Bacillati>Actinomycetota>Actinomycetes>Bifidobacteriales>Bifidobacteriaceae>Bifidobacterium | 46 | BBRI4_RS11740 | 131838..131978 | 1685 | ATP-binding-protein-[Bifidobacterium-breve] |
| WP_230803845.1 | Schlafen+HATPase+wHTH | GCF_001189355.1 | Bifidobacterium_breve | Bacteria>Bacillati>Actinomycetota>Actinomycetes>Bifidobacteriales>Bifidobacteriaceae>Bifidobacterium | 99 | BBRI4_RS05540 | 131526..131825 | 1685 | ATP-binding-protein-[Bifidobacterium-breve] |
| WP_226650297.1 | Schlafen+HATPase+wHTH | GCF_020526585.1 | Bifidobacterium_pseudocatenulatum | Bacteria>Bacillati>Actinomycetota>Actinomycetes>Bifidobacteriales>Bifidobacteriaceae>Bifidobacterium | 512 | KZP10_RS05680 | 1058652..1060190 | 28026 | ATP-binding-protein-[Bifidobacterium-pseudocatenulatum] |
| WP_223616840.1 | Schlafen+HATPase+wHTH | GCF_014898135.1 | Bifidobacterium_longum | Bacteria>Bacillati>Actinomycetota>Actinomycetes>Bifidobacteriales>Bifidobacteriaceae>Bifidobacterium | 44 | BL5934_RS10255 | 984865..984999 | 216816 | ATP-binding-protein-[Bifidobacterium-longum] |
| WP_206690743.1 | Schlafen+HATPase+wHTH | - | Bifidobacterium_dentium | Bacteria>Bacillati>Actinomycetota>Actinomycetes>Bifidobacteriales>Bifidobacteriaceae>Bifidobacterium | 494 | IHQ76_RS06165 | 222107..223591 | 1689 | ATP-binding-protein-[Bifidobacterium-dentium] |
| WP_204464692.1 | Schlafen+HATPase+wHTH | GCF_016899675.1 | Bifidobacterium_pullorum | Bacteria>Bacillati>Actinomycetota>Actinomycetes>Bifidobacteriales>Bifidobacteriaceae>Bifidobacterium | 528 | H6A68_RS03855 | 159008..160594 | 78448 | ATP-binding-protein-[Bifidobacterium-pullorum] |
| WP_204389269.1 | Schlafen+HATPase+wHTH | GCF_016900615.1 | Bifidobacterium_pullorum | Bacteria>Bacillati>Actinomycetota>Actinomycetes>Bifidobacteriales>Bifidobacteriaceae>Bifidobacterium | 529 | H6A14_RS02690 | 180930..182519 | 78448 | ATP-binding-protein-[Bifidobacterium-pullorum] |
| WP_202574829.1 | Schlafen+HATPase+wHTH | GCF_016759645.1 | Bifidobacterium_longum | Bacteria>Bacillati>Actinomycetota>Actinomycetes>Bifidobacteriales>Bifidobacteriaceae>Bifidobacterium | 514 | JL737_RS05455 | 19693..21237 | 216816 | ATP-binding-protein-[Bifidobacterium-longum] |
| WP_200406518.1 | Schlafen+HATPase+wHTH | GCF_014898255.1 | Bifidobacterium_longum | Bacteria>Bacillati>Actinomycetota>Actinomycetes>Bifidobacteriales>Bifidobacteriaceae>Bifidobacterium | 512 | BL7055_RS03255 | 799800..801338 | 216816 | ATP-binding-protein-[Bifidobacterium-longum] |
| WP_196330422.1 | Schlafen+HATPase+wHTH | GCF_017743195.1 | Bifidobacterium_dentium | Bacteria>Bacillati>Actinomycetota>Actinomycetes>Bifidobacteriales>Bifidobacteriaceae>Bifidobacterium | 499 | KZX63_RS03550 | 173428..174927 | 1689 | ATP-binding-protein-[Bifidobacterium-dentium] |
| WP_195938987.1 | Schlafen+HATPase+wHTH | GCF_015667675.1 | Bifidobacterium_pseudocatenulatum | Bacteria>Bacillati>Actinomycetota>Actinomycetes>Bifidobacteriales>Bifidobacteriaceae>Bifidobacterium | 512 | I4U64_RS06645 | 382441..383979 | 28026 | ATP-binding-protein-[Bifidobacterium-pseudocatenulatum] |
| WP_195936828.1 | Schlafen+HATPase+wHTH | GCF_015667555.1 | Bifidobacterium_bifidum | Bacteria>Bacillati>Actinomycetota>Actinomycetes>Bifidobacteriales>Bifidobacteriaceae>Bifidobacterium | 512 | I4U71_RS07220 | 6507..8045 | 1681 | ATP-binding-protein-[Bifidobacterium-bifidum] |
| WP_195482032.1 | Schlafen+HATPase+wHTH | GCF_964239665.1 | Bifidobacterium_adolescentis | Bacteria>Bacillati>Actinomycetota>Actinomycetes>Bifidobacteriales>Bifidobacteriaceae>Bifidobacterium | 512 | I2E65_RS01550 | 353857..355395 | 1680 | ATP-binding-protein-[Bifidobacterium-adolescentis] |
| WP_195451641.1 | Schlafen+HATPase+wHTH | GCF_015555105.1 | Bifidobacterium_dentium | Bacteria>Bacillati>Actinomycetota>Actinomycetes>Bifidobacteriales>Bifidobacteriaceae>Bifidobacterium | 501 | I2E42_RS03160 | 220932..222437 | 1689 | ATP-binding-protein-[Bifidobacterium-dentium] |
| WP_193530921.1 | Schlafen+HATPase+wHTH | GCF_014983005.1 | Bifidobacterium_pullorum | Bacteria>Bacillati>Actinomycetota>Actinomycetes>Bifidobacteriales>Bifidobacteriaceae>Bifidobacterium | 528 | INF27_RS02910 | 193033..194619 | 78448 | ATP-binding-protein-[Bifidobacterium-pullorum] |
| WP_168445145.1 | Schlafen+HATPase+wHTH | GCF_012310535.1 | Bifidobacterium_dentium | Bacteria>Bacillati>Actinomycetota>Actinomycetes>Bifidobacteriales>Bifidobacteriaceae>Bifidobacterium | 483 | PG133_RS04060 | 123876..125327 | 1689 | ATP-binding-protein-[Bifidobacterium-dentium] |
| WP_163196472.1 | Schlafen+HATPase+wHTH | GCF_010667645.1 | Bifidobacterium_platyrrhinorum | Bacteria>Bacillati>Actinomycetota>Actinomycetes>Bifidobacteriales>Bifidobacteriaceae>Bifidobacterium | 511 | GFD21_RS03295 | 752164..753699 | 2661628 | ATP-binding-protein-[Bifidobacterium-platyrrhinorum] |
| WP_163113199.1 | Schlafen+HATPase+wHTH | GCF_011029205.1 | Bifidobacterium_bifidum | Bacteria>Bacillati>Actinomycetota>Actinomycetes>Bifidobacteriales>Bifidobacteriaceae>Bifidobacterium | 512 | G5T23_RS07335 | 35029..36567 | 1681 | ATP-binding-protein-[Bifidobacterium-bifidum] |
| WP_154050439.1 | Schlafen+HATPase+wHTH | GCF_901212475.1 | Bifidobacterium_longum | Bacteria>Bacillati>Actinomycetota>Actinomycetes>Bifidobacteriales>Bifidobacteriaceae>Bifidobacterium | 498 | FGK88_RS06820 | 96541..98037 | 216816 | ATP-binding-protein-[Bifidobacterium-longum] |
| WP_152233653.1 | Schlafen+HATPase+wHTH | GCF_042657605.1 | Bifidobacterium_leontopitheci | Bacteria>Bacillati>Actinomycetota>Actinomycetes>Bifidobacteriales>Bifidobacteriaceae>Bifidobacterium | 502 | F7D09_RS01385 | 135870..137378 | 2650774 | ATP-binding-protein-[Bifidobacterium-leontopitheci] |
| WP_142421437.1 | Schlafen+HATPase+wHTH | GCF_006828885.1 | Bifidobacterium_pseudocatenulatum | Bacteria>Bacillati>Actinomycetota>Actinomycetes>Bifidobacteriales>Bifidobacteriaceae>Bifidobacterium | 512 | GH753_RS08030 | 1975570..1977108 | 28026 | ATP-binding-protein-[Bifidobacterium-pseudocatenulatum] |
| WP_131220404.1 | Schlafen+HATPase+wHTH | GCF_004334345.1 | Bifidobacterium_longum | Bacteria>Bacillati>Actinomycetota>Actinomycetes>Bifidobacteriales>Bifidobacteriaceae>Bifidobacterium | 512 | MCC10040_RS02095 | 109705..111243 | 216816 | ATP-binding-protein-[Bifidobacterium-longum] |
| WP_131216918.1 | Schlafen+HATPase+wHTH | GCF_004334155.1 | Bifidobacterium_longum | Bacteria>Bacillati>Actinomycetota>Actinomycetes>Bifidobacteriales>Bifidobacteriaceae>Bifidobacterium | 498 | MCC10077_RS10230 | 11036..12532 | 216816 | ATP-binding-protein-[Bifidobacterium-longum] |
| WP_129237749.1 | Schlafen+HATPase+wHTH | GCF_004135085.1 | Bifidobacterium_pullorum | Bacteria>Bacillati>Actinomycetota>Actinomycetes>Bifidobacteriales>Bifidobacteriaceae>Bifidobacterium | 542 | ESN35_RS07325 | 1673348..1674976 | 78448 | ATP-binding-protein-[Bifidobacterium-pullorum] |
| WP_124879773.1 | Schlafen+HATPase+wHTH | GCF_003860285.1 | Bifidobacterium_breve | Bacteria>Bacillati>Actinomycetota>Actinomycetes>Bifidobacteriales>Bifidobacteriaceae>Bifidobacterium | 526 | EH245_RS01915 | 439112..440692 | 1685 | ATP-binding-protein-[Bifidobacterium-breve] |
| WP_118279741.1 | Schlafen+HATPase+wHTH | GCF_003472685.1 | Bifidobacterium_pseudocatenulatum | Bacteria>Bacillati>Actinomycetota>Actinomycetes>Bifidobacteriales>Bifidobacteriaceae>Bifidobacterium | 512 | DW102_RS06330 | 107754..109292 | 28026 | ATP-binding-protein-[Bifidobacterium-pseudocatenulatum] |
| WP_118269381.1 | Schlafen+HATPase+wHTH | GCF_003473145.1 | Bifidobacterium_bifidum | Bacteria>Bacillati>Actinomycetota>Actinomycetes>Bifidobacteriales>Bifidobacteriaceae>Bifidobacterium | 512 | DW145_RS03290 | 280647..282185 | 1681 | ATP-binding-protein-[Bifidobacterium-bifidum] |
| WP_117807084.1 | Schlafen+HATPase+wHTH | GCF_003457765.1 | Bifidobacterium_adolescentis | Bacteria>Bacillati>Actinomycetota>Actinomycetes>Bifidobacteriales>Bifidobacteriaceae>Bifidobacterium | 531 | DWZ00_RS05070 | 1168372..1169967 | 1680 | ATP-binding-protein-[Bifidobacterium-adolescentis] |
| WP_117759614.1 | Schlafen+HATPase+wHTH | GCF_009832825.1 | Bifidobacterium_adolescentis | Bacteria>Bacillati>Actinomycetota>Actinomycetes>Bifidobacteriales>Bifidobacteriaceae>Bifidobacterium | 503 | DWX79_RS04600 | 1016428..1017939 | 1680 | ATP-binding-protein-[Bifidobacterium-adolescentis] |
| WP_117658185.1 | Schlafen+HATPase+wHTH | GCF_003436575.1 | Bifidobacterium_bifidum | Bacteria>Bacillati>Actinomycetota>Actinomycetes>Bifidobacteriales>Bifidobacteriaceae>Bifidobacterium | 512 | DXD34_RS01720 | 359106..360644 | 1681 | ATP-binding-protein-[Bifidobacterium-bifidum] |
| WP_117656281.1 | Schlafen+HATPase+wHTH | GCF_003437075.1 | Bifidobacterium_pseudocatenulatum | Bacteria>Bacillati>Actinomycetota>Actinomycetes>Bifidobacteriales>Bifidobacteriaceae>Bifidobacterium | 512 | DXC75_RS03270 | 108141..109679 | 28026 | ATP-binding-protein-[Bifidobacterium-pseudocatenulatum] |
| WP_115955256.1 | Schlafen+HATPase+wHTH | GCF_003390735.1 | Bifidobacterium_bifidum | Bacteria>Bacillati>Actinomycetota>Actinomycetes>Bifidobacteriales>Bifidobacteriaceae>Bifidobacterium | 514 | CJD48_RS06200 | 1410303..1411847 | 1681 | ATP-binding-protein-[Bifidobacterium-bifidum] |
| WP_101629207.1 | Schlafen+HATPase+wHTH | GCF_002861445.1 | Bifidobacterium_longum | Bacteria>Bacillati>Actinomycetota>Actinomycetes>Bifidobacteriales>Bifidobacteriaceae>Bifidobacterium | 512 | CYJ39_RS09180 | 62057..63595 | 216816 | ATP-binding-protein-[Bifidobacterium-longum] |
| WP_101617894.1 | Schlafen+HATPase+wHTH | GCF_002860355.1 | Bifidobacterium_margollesii | Bacteria>Bacillati>Actinomycetota>Actinomycetes>Bifidobacteriales>Bifidobacteriaceae>Bifidobacterium | 494 | Uis1B_RS09875 | 31243..32727 | 2020964 | ATP-binding-protein-[Bifidobacterium-margollesii] |
| WP_101025549.1 | Schlafen+HATPase+wHTH | GCF_002832945.1 | Bifidobacterium_longum | Bacteria>Bacillati>Actinomycetota>Actinomycetes>Bifidobacteriales>Bifidobacteriaceae>Bifidobacterium | 512 | DPC6317_RS01870 | 221195..222733 | 216816 | ATP-binding-protein-[Bifidobacterium-longum] |
| WP_094668338.1 | Schlafen+HATPase+wHTH | GCF_002259745.1 | Bifidobacterium_myosotis | Bacteria>Bacillati>Actinomycetota>Actinomycetes>Bifidobacteriales>Bifidobacteriaceae>Bifidobacterium | 498 | BMYO_RS10085 | 13171..14667 | 1630166 | putative-DNA-binding-domain-containing-protein-[Bifidobacterium-myosotis] |
| WP_080822639.1 | Schlafen+HATPase+wHTH | - | Bifidobacterium_dentium | Bacteria>Bacillati>Actinomycetota>Actinomycetes>Bifidobacteriales>Bifidobacteriaceae>Bifidobacterium | 499 | J7M36_RS06035 | 1415985..1417484 | 1689 | ATP-binding-protein-[Bifidobacterium-dentium] |
| WP_077383006.1 | Schlafen+HATPase+wHTH | GCF_900157155.1 | Bifidobacterium_longum | Bacteria>Bacillati>Actinomycetota>Actinomycetes>Bifidobacteriales>Bifidobacteriaceae>Bifidobacterium | 512 | BQ8631_RS08400 | 38997..40535 | 216816 | ATP-binding-protein-[Bifidobacterium-longum] |
| WP_070099217.1 | Schlafen+HATPase+wHTH | GCF_001406455.1 | Bifidobacterium_adolescentis | Bacteria>Bacillati>Actinomycetota>Actinomycetes>Bifidobacteriales>Bifidobacteriaceae>Bifidobacterium | 247 | ARB73_RS01785 | 429602..430345 | 1680 | ATP-binding-protein-[Bifidobacterium-adolescentis] |
| WP_065453249.1 | Schlafen+HATPase+wHTH | GCF_001686185.1 | Bifidobacterium_longum | Bacteria>Bacillati>Actinomycetota>Actinomycetes>Bifidobacteriales>Bifidobacteriaceae>Bifidobacterium | 512 | BAD19_RS10585 | 403665..405203 | 216816 | ATP-binding-protein-[Bifidobacterium-longum] |
| WP_065435303.1 | Schlafen+HATPase+wHTH | GCF_001686025.1 | Bifidobacterium_pseudocatenulatum | Bacteria>Bacillati>Actinomycetota>Actinomycetes>Bifidobacteriales>Bifidobacteriaceae>Bifidobacterium | 512 | BAD11_RS06855 | 37585..39123 | 28026 | ATP-binding-protein-[Bifidobacterium-pseudocatenulatum] |
| WP_053824997.1 | Schlafen+HATPase+wHTH | GCF_001281345.1 | Bifidobacterium_bifidum | Bacteria>Bacillati>Actinomycetota>Actinomycetes>Bifidobacteriales>Bifidobacteriaceae>Bifidobacterium | 512 | RY70_RS04440 | 1123418..1124956 | 1681 | ATP-binding-protein-[Bifidobacterium-bifidum] |
| WP_047298568.1 | Schlafen+HATPase+wHTH | GCF_039063425.1 | Bifidobacterium_bifidum | Bacteria>Bacillati>Actinomycetota>Actinomycetes>Bifidobacteriales>Bifidobacteriaceae>Bifidobacterium | 512 | LMG11583_RS03355 | 7885..9423 | 1681 | ATP-binding-protein-[Bifidobacterium-bifidum] |
| WP_046999383.1 | Schlafen+HATPase+wHTH | GCF_001010915.1 | Bifidobacterium_adolescentis | Bacteria>Bacillati>Actinomycetota>Actinomycetes>Bifidobacteriales>Bifidobacteriaceae>Bifidobacterium | 554 | AAX71_RS05475 | 51998..53662 | 1680 | ATP-binding-protein-[Bifidobacterium-adolescentis] |
| WP_041080198.1 | Schlafen+HATPase+wHTH | GCF_000829295.1 | Bifidobacterium_longum | Bacteria>Bacillati>Actinomycetota>Actinomycetes>Bifidobacteriales>Bifidobacteriaceae>Bifidobacterium | 512 | BL105A_RS01830 | 427651..429189 | 216816 | ATP-binding-protein-[Bifidobacterium-longum] |
| WP_038426292.1 | Schlafen+HATPase+wHTH | GCF_000772485.1 | Bifidobacterium_longum | Bacteria>Bacillati>Actinomycetota>Actinomycetes>Bifidobacteriales>Bifidobacteriaceae>Bifidobacterium | 512 | DWZ18_RS09040 | 43374..44912 | 216816 | ATP-binding-protein-[Bifidobacterium-longum] |
| WP_033509372.1 | Schlafen+HATPase+wHTH | GCF_000770965.1 | Bifidobacterium_pullorum | Bacteria>Bacillati>Actinomycetota>Actinomycetes>Bifidobacteriales>Bifidobacteriaceae>Bifidobacterium | 528 | OU28_RS04635 | 13893..15479 | 78448 | ATP-binding-protein-[Bifidobacterium-pullorum] |
| WP_033494227.1 | Schlafen+HATPase+wHTH | GCF_000771645.1 | Bifidobacterium_biavatii | Bacteria>Bacillati>Actinomycetota>Actinomycetes>Bifidobacteriales>Bifidobacteriaceae>Bifidobacterium | 502 | OU23_RS06255 | 9230..10738 | 762212 | ATP-binding-protein-[Bifidobacterium-biavatii] |
| WP_032745986.1 | Schlafen+HATPase+wHTH | GCF_000730035.1 | Bifidobacterium_longum | Bacteria>Bacillati>Actinomycetota>Actinomycetes>Bifidobacteriales>Bifidobacteriaceae>Bifidobacterium | 512 | BL171B_RS06790 | 269674..271212 | 216816 | ATP-binding-protein-[Bifidobacterium-longum] |
| WP_032738775.1 | Schlafen+HATPase+wHTH | GCF_000730055.1 | Bifidobacterium_longum | Bacteria>Bacillati>Actinomycetota>Actinomycetes>Bifidobacteriales>Bifidobacteriaceae>Bifidobacterium | 514 | BL71B_RS01475 | 374972..376516 | 216816 | ATP-binding-protein-[Bifidobacterium-longum] |
| WP_032735731.1 | Schlafen+HATPase+wHTH | GCF_000730105.1 | Bifidobacterium_longum | Bacteria>Bacillati>Actinomycetota>Actinomycetes>Bifidobacteriales>Bifidobacteriaceae>Bifidobacterium | 512 | BL15B_RS01730 | 300246..301784 | 216816 | ATP-binding-protein-[Bifidobacterium-longum] |
| WP_032684172.1 | Schlafen+HATPase+wHTH | GCF_022701235.1 | Bifidobacterium_longum | Bacteria>Bacillati>Actinomycetota>Actinomycetes>Bifidobacteriales>Bifidobacteriaceae>Bifidobacterium | 514 | LMY38_RS03495 | 748653..750197 | 216816 | ATP-binding-protein-[Bifidobacterium-longum] |
| WP_026646059.1 | Schlafen+HATPase+wHTH | GCF_000687635.1 | Bifidobacterium_ruminantium | Bacteria>Bacillati>Actinomycetota>Actinomycetes>Bifidobacteriales>Bifidobacteriaceae>Bifidobacterium | 512 | T534_RS0102100 | 11217..12755 | 78346 | ATP-binding-protein-[Bifidobacterium-ruminantium] |
| WP_003826163.1 | Schlafen+HATPase+wHTH | GCF_001025155.1 | Bifidobacterium_angulatum | Bacteria>Bacillati>Actinomycetota>Actinomycetes>Bifidobacteriales>Bifidobacteriaceae>Bifidobacterium | 510 | BBAG_RS03490 | 874539..876071 | 1683 | ATP-binding-protein-[Bifidobacterium-angulatum] |
| WP_003819011.1 | Schlafen+HATPase+wHTH | GCF_000300215.1 | Bifidobacterium_bifidum | Bacteria>Bacillati>Actinomycetota>Actinomycetes>Bifidobacteriales>Bifidobacteriaceae>Bifidobacterium | 512 | IF1010_RS06535 | 90896..92434 | 1681 | ATP-binding-protein-[Bifidobacterium-bifidum] |
| PLS30249.1 | Schlafen+HATPase+wHTH | GCA_002860355.1 | Bifidobacterium_margollesii | Bacteria>Bacillati>Actinomycetota>Actinomycetes>Bifidobacteriales>Bifidobacteriaceae>Bifidobacterium | 494 | Uis1B_1919 | 31243..32727 | 2020964 | ATP-binding-protein-[Bifidobacterium-margollesii] |
| PKC83368.1 | Schlafen+HATPase+wHTH | GCA_002832945.1 | Bifidobacterium_longum | Bacteria>Bacillati>Actinomycetota>Actinomycetes>Bifidobacteriales>Bifidobacteriaceae>Bifidobacterium | 100 | DPC6317_0881 | 245684..245986 | 216816 | ATP-binding-protein-[Bifidobacterium-longum] |
| MCK6131819.1 | Schlafen+HATPase+wHTH | GCA_023147995.1 | Bifidobacterium_dentium | Bacteria>Bacillati>Actinomycetota>Actinomycetes>Bifidobacteriales>Bifidobacteriaceae>Bifidobacterium | 499 | KZX63_03540 | 173428..174927 | 1689 | putative-DNA-binding-domain-containing-protein-[Bifidobacterium-dentium] |
| MBS7047479.1 | Schlafen+HATPase+wHTH | GCA_018381815.1 | Bifidobacterium_adolescentis | Bacteria>Bacillati>Actinomycetota>Actinomycetes>Bifidobacteriales>Bifidobacteriaceae>Bifidobacterium | 512 | KH096_03865 | 352519..354057 | 1680 | putative-DNA-binding-domain-containing-protein-[Bifidobacterium-adolescentis] |
| MBS6717592.1 | Schlafen+HATPase+wHTH | GCA_018378875.1 | Bifidobacterium_longum | Bacteria>Bacillati>Actinomycetota>Actinomycetes>Bifidobacteriales>Bifidobacteriaceae>Bifidobacterium | 495 | KH250_10255 | 14892..16379 | 216816 | putative-DNA-binding-domain-containing-protein-[Bifidobacterium-longum] |
| KFJ01340.1 | Schlafen+HATPase+wHTH | GCA_000741785.1 | Bifidobacterium_stellenboschense | Bacteria>Bacillati>Actinomycetota>Actinomycetes>Bifidobacteriales>Bifidobacteriaceae>Bifidobacterium | 506 | BSTEL_1470 | 71515..73035 | 762211 | putative-transcriptional-regulator-containing-an-HTH-domain-Bifidobacterium-stellenboschense] |
| KAB5930821.1 | Schlafen+HATPase+wHTH | GCA_009094485.1 | Bifidobacterium_adolescentis | Bacteria>Bacillati>Actinomycetota>Actinomycetes>Bifidobacteriales>Bifidobacteriaceae>Bifidobacterium | 247 | GA595_06020 | 102821..103564 | 1680 | hypothetical-protein-GA595_06020-[Bifidobacterium-adolescentis] |
| KAB5743302.1 | Schlafen+HATPase+wHTH | GCA_009075895.1 | Bifidobacterium_adolescentis | Bacteria>Bacillati>Actinomycetota>Actinomycetes>Bifidobacteriales>Bifidobacteriaceae>Bifidobacterium | 512 | GA749_07765 | 114194..115732 | 1680 | ATP-binding-protein-[Bifidobacterium-adolescentis] |
| MBE5064981.1 | Schlafen+HATPase+wHTH | GCA_014983005.1 | Bifidobacterium_pullorum_subsp._saeculare | Bacteria>Bacillati>Actinomycetota>Actinomycetes>Bifidobacteriales>Bifidobacteriaceae>Bifidobacterium>Bifidobacterium_pullorum | 528 | INF27_02910 | 193033..194619 | 78257 | putative-DNA-binding-domain-containing-protein-[Bifidobacterium-pullorum-subsp.-saeculare] |
| WP_140556585.1 | Schlafen+HATPase+wHTH | GCF_006408715.1 | Bifidobacterium_sp._UTCIF-39 | Bacteria>Bacillati>Actinomycetota>Actinomycetes>Bifidobacteriales>Bifidobacteriaceae>Bifidobacterium>unclassified_Bifidobacterium | 497 | EP30_RS04690 | 97150..98643 | 1465359 | ATP-binding-protein-[Bifidobacterium-sp.-UTCIF-39] |
| GDZ58124.1 | Schlafen+HATPase+wHTH | GCA_018327225.1 | Bifidobacteriaceae_bacterium_MCC01967 | Bacteria>Bacillati>Actinomycetota>Actinomycetes>Bifidobacteriales>Bifidobacteriaceae>unclassified_Bifidobacteriaceae_(miscellaneous) | 512 | MCC01967_09800 | 136231..137769 | 2569520 | ATPase-AAA-[Bifidobacteriaceae-bacterium-MCC01967] |
| GDY95794.1 | Schlafen+HATPase+wHTH | GCA_018326565.1 | Bifidobacteriaceae_bacterium_MCC01943 | Bacteria>Bacillati>Actinomycetota>Actinomycetes>Bifidobacteriales>Bifidobacteriaceae>unclassified_Bifidobacteriaceae_(miscellaneous) | 503 | MCC01943_10810 | 13678..15189 | 2569488 | ATPase-AAA-[Bifidobacteriaceae-bacterium-MCC01943] |
| GDY90544.1 | Schlafen+HATPase+wHTH | GCA_018326505.1 | Bifidobacteriaceae_bacterium_MCC01971 | Bacteria>Bacillati>Actinomycetota>Actinomycetes>Bifidobacteriales>Bifidobacteriaceae>unclassified_Bifidobacteriaceae_(miscellaneous) | 512 | MCC01971_21620 | 7290..8828 | 2569485 | ATPase-AAA-[Bifidobacteriaceae-bacterium-MCC01971] |
| SBW20210.1 | Schlafen+HATPase+wHTH | GCA_900067225.1 | Candidatus_Protofrankia_californiensis | Bacteria>Bacillati>Actinomycetota>Actinomycetes>Frankiales>Frankiaceae>Protofrankia | 657 | FDG2_1624 | 6133..8106 | 1839754 | hypothetical-protein-FDG2_1624-[Candidatus-Frankia-californiensis] |
| WP_147251545.1 | Schlafen+HATPase+wHTH | GCF_003319095.1 | Blastococcus_sp._TBT05-19 | Bacteria>Bacillati>Actinomycetota>Actinomycetes>Geodermatophilales>Geodermatophilaceae>Blastococcus>unclassified_Blastococcus | 566 | DQ244_RS16230 | 123051..124751 | 2250581 | ATP-binding-protein-[Blastococcus-sp.-TBT05-19] |
| WP_263976855.1 | Schlafen+HATPase+wHTH | - | Streptomyces_sp._NK08204 | Bacteria>Bacillati>Actinomycetota>Actinomycetes>Kitasatosporales>Streptomycetaceae>Streptomyces>unclassified_Streptomyces | 405 | K7G59_RS27415 | 99281..100498 | 2873260 | ATP-binding-protein-[Streptomyces-sp.-NK08204] |
| WP_260463287.1 | Schlafen+HATPase+wHTH | - | Streptomyces_sp._TRM72054 | Bacteria>Bacillati>Actinomycetota>Actinomycetes>Kitasatosporales>Streptomycetaceae>Streptomyces>unclassified_Streptomyces | 437 | K4749_RS28635 | 571..1884 | 2870562 | ATP-binding-protein-[Streptomyces-sp.-TRM72054] |
| WP_239157865.1 | Schlafen+HATPase+wHTH | - | Streptomyces_sp._SID13726 | Bacteria>Bacillati>Actinomycetota>Actinomycetes>Kitasatosporales>Streptomycetaceae>Streptomyces>unclassified_Streptomyces | 409 | G3I78_RS15975 | 374620..375849 | 2706058 | ATP-binding-protein-[Streptomyces-sp.-SID13726] |
| WP_218514037.1 | Schlafen+HATPase+wHTH | GCF_019219825.1 | Streptomyces_sp._TRM70350 | Bacteria>Bacillati>Actinomycetota>Actinomycetes>Kitasatosporales>Streptomycetaceae>Streptomyces>unclassified_Streptomyces | 436 | KWI83_RS10495 | 178166..179476 | 2856165 | ATP-binding-protein-[Streptomyces-sp.-TRM70350] |
| WP_214957742.1 | Schlafen+HATPase+wHTH | GCF_018614195.1 | Streptomyces_sp._ISL-43 | Bacteria>Bacillati>Actinomycetota>Actinomycetes>Kitasatosporales>Streptomycetaceae>Streptomyces>unclassified_Streptomyces | 392 | J7F03_RS37995 | 17860..19038 | 2819183 | ATP-binding-protein-[Streptomyces-sp.-ISL-43] |
| WP_166682463.1 | Schlafen+HATPase+wHTH | - | Streptomyces_sp._846.5 | Bacteria>Bacillati>Actinomycetota>Actinomycetes>Kitasatosporales>Streptomycetaceae>Streptomyces>unclassified_Streptomyces | 507 | EDD99_RS20225 | 4425338..4426861 | 2485156 | ATP-binding-protein-[Streptomyces-sp.-846.5] |
| WP_078852152.1 | Schlafen+HATPase+wHTH | GCF_000719505.1 | Streptomyces_sp._NRRL_B-3229 | Bacteria>Bacillati>Actinomycetota>Actinomycetes>Kitasatosporales>Streptomycetaceae>Streptomyces>unclassified_Streptomyces | 446 | IF54_RS0129565 | 217723..219063 | 1463836 | ATP-binding-protein-[Streptomyces-sp.-NRRL-B-3229] |
| NEB00547.1 | Schlafen+HATPase+wHTH | GCA_010548375.1 | Streptomyces_sp._SID13726 | Bacteria>Bacillati>Actinomycetota>Actinomycetes>Kitasatosporales>Streptomycetaceae>Streptomyces>unclassified_Streptomyces | 446 | G3I78_15975 | 374509..375849 | 2706058 | ATP-binding-protein-[Streptomyces-sp.-SID13726] |
| MCP3817896.1 | Schlafen+HATPase+wHTH | GCA_024213665.1 | Streptomyces_sp._A3M-1-3 | Bacteria>Bacillati>Actinomycetota>Actinomycetes>Kitasatosporales>Streptomycetaceae>Streptomyces>unclassified_Streptomyces | 508 | NLX86_07035 | 35031..36557 | 2962044 | ATP-binding-protein-[Streptomyces-sp.-A3M-1-3] |
| MBX9397442.1 | Schlafen+HATPase+wHTH | GCA_019733235.1 | Streptomyces_sp._TRM72054 | Bacteria>Bacillati>Actinomycetota>Actinomycetes>Kitasatosporales>Streptomycetaceae>Streptomyces>unclassified_Streptomyces | 448 | K4749_28620 | 571..1917 | 2870562 | ATP-binding-protein-[Streptomyces-sp.-TRM72054] |
| MBV7696076.1 | Schlafen+HATPase+wHTH | GCA_019219825.1 | Streptomyces_sp._TRM70350 | Bacteria>Bacillati>Actinomycetota>Actinomycetes>Kitasatosporales>Streptomycetaceae>Streptomyces>unclassified_Streptomyces | 436 | KWI83_10495 | 178166..179476 | 2856165 | ATP-binding-protein-[Streptomyces-sp.-TRM70350] |
| WP_159096450.1 | Schlafen+HATPase+wHTH | GCF_003074035.1 | Miniimonas_sp._S16 | Bacteria>Bacillati>Actinomycetota>Actinomycetes>Micrococcales>Beutenbergiaceae>Miniimonas>unclassified_Miniimonas | 611 | DCS86_RS08170 | 1895917..1897752 | 2171623 | ATP-binding-protein-[Miniimonas-sp.-S16] |
| WP_282959228.1 | Schlafen+HATPase+wHTH | - | Cellulomonas_fimi | Bacteria>Bacillati>Actinomycetota>Actinomycetes>Micrococcales>Cellulomonadaceae>Cellulomonas | 560 | HIR71_RS02960 | 32032..33714 | 1708 | ATP-binding-protein-[Cellulomonas-fimi] |
| MBK6299926.1 | Schlafen+HATPase+wHTH | GCA_016705565.1 | Candidatus_Phosphoribacter_hodrii | Bacteria>Bacillati>Actinomycetota>Actinomycetes>Micrococcales>Dermatophilaceae>Candidatus_Phosphoribacter | 582 | IPF40_02345 | 523261..525009 | 2953743 | putative-DNA-binding-domain-containing-protein-[Actinomycetales-bacterium] |
| QAY59793.1 | Schlafen+HATPase+wHTH | GCA_004135285.1 | Microbacterium_protaetiae | Bacteria>Bacillati>Actinomycetota>Actinomycetes>Micrococcales>Microbacteriaceae>Microbacterium | 480 | ET475_07175 | 1535915..1537357 | 2509458 | AAA-family-ATPase-[Microbacterium-protaetiae] |
| WP_028245013.1 | Schlafen+HATPase+wHTH | GCF_000423505.1 | Pseudoclavibacter_soli | Bacteria>Bacillati>Actinomycetota>Actinomycetes>Micrococcales>Microbacteriaceae>Pseudoclavibacter | 493 | G557_RS0103835 | 199917..201398 | 452623 | helix-turn-helix-domain-containing-protein-[Pseudoclavibacter-soli] |
| WP_202129909.1 | Schlafen+HATPase+wHTH | GCF_004011115.1 | Rathayibacter_iranicus | Bacteria>Bacillati>Actinomycetota>Actinomycetes>Micrococcales>Microbacteriaceae>Rathayibacter | 481 | C7V51_RS01490 | 316540..317985 | 59737 | helix-turn-helix-domain-containing-protein-[Rathayibacter-iranicus] |
| WP_237187174.1 | Schlafen+HATPase+wHTH | GCF_928381705.1 | Rothia_nasimurium | Bacteria>Bacillati>Actinomycetota>Actinomycetes>Micrococcales>Micrococcaceae>Rothia | 501 | L6762_RS10425 | 35412..36917 | 85336 | ATP-binding-protein-[Rothia-nasimurium] |
| WP_142113244.1 | Schlafen+HATPase+wHTH | GCF_039533185.1 | Rarobacter_incanus | Bacteria>Bacillati>Actinomycetota>Actinomycetes>Micrococcales>Rarobacteraceae>Rarobacter | 493 | FB389_RS10075 | 2430099..2431580 | 153494 | helix-turn-helix-domain-containing-protein-[Rarobacter-incanus] |
| WP_238488472.1 | Schlafen+HATPase+wHTH | - | Actinoplanes_flavus | Bacteria>Bacillati>Actinomycetota>Actinomycetes>Micromonosporales>Micromonosporaceae>Actinoplanes | 640 | J5X75_RS18435 | 132681..134603 | 2820290 | ATP-binding-protein-[Actinoplanes-flavus] |
| WP_109591758.1 | Schlafen+HATPase+wHTH | GCF_016862475.1 | Actinoplanes_xinjiangensis | Bacteria>Bacillati>Actinomycetota>Actinomycetes>Micromonosporales>Micromonosporaceae>Actinoplanes | 400 | Axi01nite_RS08925 | 153578..154780 | 512350 | ATP-binding-protein-[Actinoplanes-xinjiangensis] |
| PWK49467.1 | Schlafen+HATPase+wHTH | GCA_003148685.1 | Actinoplanes_xinjiangensis | Bacteria>Bacillati>Actinomycetota>Actinomycetes>Micromonosporales>Micromonosporaceae>Actinoplanes | 399 | BC793_104140 | 153746..154948 | 512350 | ATP-dependent-DNA-helicase-RecG-[Actinoplanes-xinjiangensis] |
| WP_285664603.1 | Schlafen+HATPase+wHTH | GCF_030269765.1 | Actinorhabdospora_filicis | Bacteria>Bacillati>Actinomycetota>Actinomycetes>Micromonosporales>Micromonosporaceae>Actinorhabdospora | 394 | QSK10_RS20920 | 119686..120870 | 1785913 | ATP-binding-protein-[Actinorhabdospora-filicis] |
| WP_111242815.1 | Schlafen+HATPase+wHTH | GCF_018326245.1 | Micromonospora_endophytica | Bacteria>Bacillati>Actinomycetota>Actinomycetes>Micromonosporales>Micromonosporaceae>Micromonospora | 457 | D3H59_RS20845 | 11119..12492 | 515350 | ATP-binding-protein-[Micromonospora-endophytica] |
| WP_080793630.1 | Schlafen+HATPase+wHTH | GCF_900169525.1 | Corynebacterium_pacaense | Bacteria>Bacillati>Actinomycetota>Actinomycetes>Mycobacteriales>Corynebacteriaceae>Corynebacterium | 494 | B7E02_RS05015 | 1046279..1047763 | 1816684 | helix-turn-helix-domain-containing-protein-[Corynebacterium-pacaense] |
| WP_244897480.1 | Schlafen+HATPase+wHTH | GCF_025823205.1 | Mycobacterium_alsense | Bacteria>Bacillati>Actinomycetota>Actinomycetes>Mycobacteriales>Mycobacteriaceae>Mycobacterium | 400 | BST11_RS03255 | 99431..100633 | 324058 | ATP-binding-protein-[Mycobacterium-alsense] |
| WP_204801724.1 | Schlafen+HATPase+wHTH | GCF_905219495.1 | Mycobacterium_riyadhense | Bacteria>Bacillati>Actinomycetota>Actinomycetes>Mycobacteriales>Mycobacteriaceae>Mycobacterium | 416 | JX354_RS03515 | 788445..789695 | 486698 | ATP-binding-protein-[Mycobacterium-riyadhense] |
| OOK63785.1 | Schlafen+HATPase+wHTH | GCA_002003585.1 | Mycobacterium_kansasii | Bacteria>Bacillati>Actinomycetota>Actinomycetes>Mycobacteriales>Mycobacteriaceae>Mycobacterium | 417 | BZL29_8432 | 2189..3442 | 1768 | divergent-AAA-domain-protein-[Mycobacterium-kansasii] |
| WP_041784791.1 | Schlafen+HATPase+wHTH | GCF_000276825.1 | Mycobacterium_paraintracellulare | Bacteria>Bacillati>Actinomycetota>Actinomycetes>Mycobacteriales>Mycobacteriaceae>Mycobacterium>Mycobacterium_avium_complex_(MAC) | 416 | B8W68_RS04655 | 127227..128477 | 1138383 | putative-DNA-binding-domain-containing-protein-[Mycobacterium-paraintracellulare] |
| WP_009974959.1 | Schlafen+HATPase+wHTH | GCF_000174035.1 | Mycobacterium_avium | Bacteria>Bacillati>Actinomycetota>Actinomycetes>Mycobacteriales>Mycobacteriaceae>Mycobacterium>Mycobacterium_avium_complex_(MAC) | 415 | LTQ57_RS17665 | 3794008..3795255 | 1764 | ATP-binding-protein-[Mycobacterium-avium] |
| OSC29535.1 | Schlafen+HATPase+wHTH | GCA_002104735.1 | Mycobacterium_paraintracellulare | Bacteria>Bacillati>Actinomycetota>Actinomycetes>Mycobacteriales>Mycobacteriaceae>Mycobacterium>Mycobacterium_avium_complex_(MAC) | 416 | B8W68_04655 | 127227..128477 | 1138383 | transcriptional-regulator-[Mycobacterium-paraintracellulare] |
| AYJ06364.1 | Schlafen+HATPase+wHTH | GCA_003640565.1 | Mycobacterium_avium | Bacteria>Bacillati>Actinomycetota>Actinomycetes>Mycobacteriales>Mycobacteriaceae>Mycobacterium>Mycobacterium_avium_complex_(MAC) | 415 | DBO90_17290 | 3697962..3699209 | 1764 | ATP-binding-protein-[Mycobacterium-avium] |
| AFC52218.1 | Schlafen+HATPase+wHTH | GCA_000276825.1 | Mycobacterium_paraintracellulare | Bacteria>Bacillati>Actinomycetota>Actinomycetes>Mycobacteriales>Mycobacteriaceae>Mycobacterium>Mycobacterium_avium_complex_(MAC) | 357 | OCQ_07050 | 693618..694691 | 1138383 | hypothetical-protein-OCQ_07050-[Mycobacterium-paraintracellulare] |
| WP_062879805.1 | Schlafen+HATPase+wHTH | GCF_001606235.1 | Mycobacteroides_abscessus | Bacteria>Bacillati>Actinomycetota>Actinomycetes>Mycobacteriales>Mycobacteriaceae>Mycobacteroides | 416 | B5431_RS12985 | 715590..716840 | 36809 | ATP-binding-protein-[Mycobacteroides-abscessus] |
| WP_276761076.1 | Schlafen+HATPase+wHTH | GCF_008015745.1 | Mycolicibacter_arupensis | Bacteria>Bacillati>Actinomycetota>Actinomycetes>Mycobacteriales>Mycobacteriaceae>Mycolicibacter | 405 | E6Q54_RS12705 | 6089..7306 | 342002 | ATP-binding-protein-[Mycolicibacter-arupensis] |
| TXI55352.1 | Schlafen+HATPase+wHTH | GCA_008015745.1 | Mycolicibacter_arupensis | Bacteria>Bacillati>Actinomycetota>Actinomycetes>Mycobacteriales>Mycobacteriaceae>Mycolicibacter | 417 | E6Q54_12715 | 6089..7342 | 342002 | ATP-binding-protein-[Mycolicibacter-arupensis] |
| WP_225506780.1 | Schlafen+HATPase+wHTH | GCF_020227835.1 | Mycolicibacterium_fortuitum | Bacteria>Bacillati>Actinomycetota>Actinomycetes>Mycobacteriales>Mycobacteriaceae>Mycolicibacterium | 416 | H8Z59_RS27045 | 5493892..5495142 | 1766 | ATP-binding-protein-[Mycolicibacterium-fortuitum] |

| Protein-ID | Domain | Assembly-ID | Strain | Taxonomy | Length | Locus-Tag | Genomic-coordinates | TaxID | Fasta-defline |
|---|---|---|---|---|---|---|---|---|---|
| WP_216868093.1 | Schlafen+HATPase+wHTH | - | Mycolicibacterium_goodii | Bacteria>Bacillati>Actinomycetota>Actinomycetes>Mycobacteriales>Mycobacteriaceae>Mycolicibacterium | 399 | KL858_RS23370 | 10654..11853 | 134601 | ATP-binding-protein-[Mycolicibacterium-goodii] |
| EHB57970.1 | Schlafen+HATPase+wHTH | - | Mycolicibacterium_rhodesiae_JS60 | Bacteria>Bacillati>Actinomycetota>Actinomycetes>Mycobacteriales>Mycobacteriaceae>Mycolicibacterium>Mycolicibacterium_rhodesiae | 416 | MycrhDRAFT_0406 | 389359..390609 | 931627 | AAA-4-family-protein-[Mycolicibacterium-rhodesiae-JS60] |
| WP_067710519.1 | Schlafen+HATPase+wHTH | GCF_001613325.1 | Nocardia_yamanashiensis | Bacteria>Bacillati>Actinomycetota>Actinomycetes>Mycobacteriales>Nocardiaceae>Nocardia | 457 | NY1_RS09385 | 73591..74964 | 209247 | ATP-binding-protein-[Nocardia-yamanashiensis] |
| WP_036920885.1 | Schlafen+HATPase+wHTH | GCF_000423465.1 | Propionicicella_superfundia | Bacteria>Bacillati>Actinomycetota>Actinomycetes>Propionibacteriales>Nocardioidaceae>Propionicicella | 480 | G555_RS0102875 | 93041..94483 | 348582 | helix-turn-helix-domain-containing-protein-[Propionicicella-superfundia] |
| WP_097828213.1 | Schlafen+HATPase+wHTH | GCF_004011075.1 | Acidipropionibacterium_jensenii | Bacteria>Bacillati>Actinomycetota>Actinomycetes>Propionibacteriales>Propionibacteriaceae>Acidipropionibacterium | 507 | COZ11_RS09895 | 2240716..2242239 | 1749 | helix-turn-helix-domain-containing-protein-[Acidipropionibacterium-jensenii] |
| MCI1746864.1 | Schlafen+HATPase+wHTH | GCA_022647765.1 | Acidipropionibacterium_sp. | Bacteria>Bacillati>Actinomycetota>Actinomycetes>Propionibacteriales>Propionibacteriaceae>Acidipropionibacterium>unclassified_Acidipropionibacterium | 111 | LKI24_01395 | 278856..279191 | 1912218 | hypothetical-protein-[Acidipropionibacterium-sp.] |
| MCI1746861.1 | Schlafen+HATPase+wHTH | GCA_022647765.1 | Acidipropionibacterium_sp. | Bacteria>Bacillati>Actinomycetota>Actinomycetes>Propionibacteriales>Propionibacteriaceae>Acidipropionibacterium>unclassified_Acidipropionibacterium | 73 | LKI24_01380 | 273824..274045 | 1912218 | ATP-binding-protein-[Acidipropionibacterium-sp.] |
| WP_130873233.1 | Schlafen+HATPase+wHTH | GCF_900607225.1 | [Pseudopropionibacterium]_massiliense | Bacteria>Bacillati>Actinomycetota>Actinomycetes>Propionibacteriales>Propionibacteriaceae>Arachnia | 493 | E0E39_RS01885 | 445564..447045 | 2220000 | helix-turn-helix-domain-containing-protein-[[Pseudopropionibacterium] |
| WP_041792944.1 | Schlafen+HATPase+wHTH | GCF_000270245.1 | Microlunatus_phosphovorus | Bacteria>Bacillati>Actinomycetota>Actinomycetes>Propionibacteriales>Propionibacteriaceae>Microlunatus | 493 | MLP_RS18420 | 4115570..4117051 | 29405 | ATP-binding-protein-[Microlunatus-phosphovorus] |
| BAK36827.1 | Schlafen+HATPase+wHTH | GCA_000270245.1 | Microlunatus_phosphovorus_NM-1 | Bacteria>Bacillati>Actinomycetota>Actinomycetes>Propionibacteriales>Propionibacteriaceae>Microlunatus>Microlunatus_phosphovorus | 481 | MLP_38130 | 4115606..4117051 | 1032480 | hypothetical-protein-MLP_38130-[Microlunatus-phosphovorus-NM-1] |
| WP_055346075.1 | Schlafen+HATPase+wHTH | GCF_001369055.1 | Propionibacterium_freudenreichii | Bacteria>Bacillati>Actinomycetota>Actinomycetes>Propionibacteriales>Propionibacteriaceae>Propionibacterium | 494 | PFCIRM516_RS05515 | 56734..58218 | 1744 | helix-turn-helix-domain-containing-protein-[Propionibacterium-freudenreichii] |
| WP_048768822.1 | Schlafen+HATPase+wHTH | GCF_001050955.1 | Propionibacterium_freudenreichii | Bacteria>Bacillati>Actinomycetota>Actinomycetes>Propionibacteriales>Propionibacteriaceae>Propionibacterium | 493 | PFCIRM1025_RS01865 | 41950..43431 | 1744 | helix-turn-helix-domain-containing-protein-[Propionibacterium-freudenreichii] |
| WP_028706414.1 | Schlafen+HATPase+wHTH | GCF_000426605.1 | Propionibacterium_acidifaciens | Bacteria>Bacillati>Actinomycetota>Actinomycetes>Propionibacteriales>Propionibacteriaceae>Propionibacterium | 112 | K336_RS0104620 | 37569..37907 | 556499 | hypothetical-protein-[Propionibacterium-acidifaciens] |
| WP_182817602.1 | Schlafen+HATPase+wHTH | GCF_014164675.1 | Propioniciclava_sp._MC1683 | Bacteria>Bacillati>Actinomycetota>Actinomycetes>Propionibacteriales>Propionibacteriaceae>Propioniciclava>unclassified_Propioniciclava | 578 | H5397_RS11635 | 15066..16802 | 2760309 | ATP-binding-protein-[Propioniciclava-sp.-MC1683] |
| WP_073189911.1 | Schlafen+HATPase+wHTH | GCF_900141915.1 | Tessaracoccus_bendigoensis | Bacteria>Bacillati>Actinomycetota>Actinomycetes>Propionibacteriales>Propionibacteriaceae>Tessaracoccus | 493 | BUC15_RS15355 | 8875..10356 | 72764 | helix-turn-helix-domain-containing-protein-[Tessaracoccus-bendigoensis] |
| MBV9142992.1 | Schlafen+HATPase+wHTH | GCA_019247305.1 | Pseudonocardiales_bacterium | Bacteria>Bacillati>Actinomycetota>Actinomycetes>Pseudonocardiales>unclassified_Pseudonocardiales | 383 | JO115_19115 | 7513..8664 | 2201155 | putative-DNA-binding-domain-containing-protein-[Pseudonocardiales-bacterium] |
| WP_237450000.1 | Schlafen+HATPase+wHTH | GCF_009862005.1 | Nocardiopsis_alba | Bacteria>Bacillati>Actinomycetota>Actinomycetes>Streptosporangiales>Nocardiopsidaceae>Nocardiopsis | 377 | GTW20_RS21065 | 4837475..4838608 | 53437 | ATP-binding-protein-[Nocardiopsis-alba] |
| WP_176226660.1 | Schlafen+HATPase+wHTH | GCF_013281895.1 | Candidatus_Hakubella_thermalkaliphila | Bacteria>Bacillati>Actinomycetota>Actinomycetota_incertae_sedis>Candidatus_Hakubellales>Candidatus_Hakubellaceae>Candidatus_Hakubella | 408 | HW437_RS01605 | 3890..5116 | 2754717 | ATP-binding-protein-[Candidatus-Hakubanella-thermoalkaliphilus] |
| GFP42120.1 | Schlafen+HATPase+wHTH | GCA_013282075.1 | Candidatus_Hakubella_thermalkaliphila | Bacteria>Bacillati>Actinomycetota>Actinomycetota_incertae_sedis>Candidatus_Hakubellales>Candidatus_Hakubellaceae>Candidatus_Hakubella | 408 | HKBW3C_01246 | 1806..3032 | 2754717 | ATP-dependent-DNA-helicase-RecG-[Candidatus-Hakubanella-thermoalkaliphilus] |
| NCO65435.1 | Schlafen+HATPase+wHTH | GCA_009993245.1 | Candidatus_Aquicultor_secundus | Bacteria>Bacillati>Actinomycetota>Candidatus_Aquicultoria>Candidatus_Aquicultorales>Candidatus_Aquicultoraceae>Candidatus_Aquicultor | 555 | GW874_02330 | 10246..11913 | 1973895 | ATPase-[Solirubrobacter-sp.] |
| PKQ27867.1 | Schlafen+HATPase+wHTH | GCA_002843005.1 | Candidatus_Anoxymicrobium_japonicum | Bacteria>Bacillati>Actinomycetota>Candidatus_Geothermincolia>Candidatus_Geothermincolales>Candidatus_Anoxymicrobiaceae>Candidatus_Anoxymicrobium | 535 | CVT63_05755 | 4241..5848 | 2013648 | transcriptional-regulator-[Candidatus-Anoxymicrobium-japonicum] |
| WP_204559879.1 | Schlafen+HATPase+wHTH | GCF_016902615.1 | Collinsella_tanakaei | Bacteria>Bacillati>Actinomycetota>Coriobacteriia>Coriobacteriales>Coriobacteriaceae>Collinsella | 542 | H6A21_RS01770 | 48309..49937 | 626935 | putative-DNA-binding-domain-containing-protein-[Collinsella-tanakaei] |
| WP_204537515.1 | Schlafen+HATPase+wHTH | - | Collinsella_tanakaei | Bacteria>Bacillati>Actinomycetota>Coriobacteriia>Coriobacteriales>Coriobacteriaceae>Collinsella | 546 | H6A33_RS00130 | 27333..28973 | 626935 | putative-DNA-binding-domain-containing-protein-[Collinsella-tanakaei] |
| WP_195443569.1 | Schlafen+HATPase+wHTH | GCF_015555045.1 | Collinsella_aerofaciens | Bacteria>Bacillati>Actinomycetota>Coriobacteriia>Coriobacteriales>Coriobacteriaceae>Collinsella | 500 | I2E45_RS00485 | 69150..70652 | 74426 | ATP-binding-protein-[Collinsella-aerofaciens] |
| WP_273384075.1 | Schlafen+HATPase+wHTH | GCF_944320695.1 | Enorma_phocaeensis | Bacteria>Bacillati>Actinomycetota>Coriobacteriia>Coriobacteriales>Coriobacteriaceae>Enorma | 574 | PUQ31_RS06685 | 22660..24384 | 1871019 | ATP-binding-protein-[Enorma-phocaeensis] |
| WP_195347159.1 | Schlafen+HATPase+wHTH | GCF_015551405.1 | Eggerthella_lenta | Bacteria>Bacillati>Actinomycetota>Coriobacteriia>Eggerthellales>Eggerthellaceae>Eggerthella | 500 | I2D50_RS06380 | 51333..52835 | 84112 | ATP-binding-protein-[Eggerthella-lenta] |
| WP_273888495.1 | Schlafen+HATPase+wHTH | GCF_028617085.1 | Rubrobacter_naiadicus | Bacteria>Bacillati>Actinomycetota>Rubrobacteria>Rubrobacterales>Rubrobacteraceae>Rubrobacter | 467 | PJB25_RS10025 | 45865..47268 | 1392641 | ATP-binding-protein-[Rubrobacter-naiadicus] |
| NLE73127.1 | Schlafen+HATPase+wHTH | GCA_012514635.1 | Actinomycetota_bacterium | Bacteria>Bacillati>Actinomycetota>unclassified_Actinomycetota | 210 | GX604_00620 | 12000..12632 | 2900548 | ATP-binding-protein-[Actinomycetia-bacterium] |
| MCA0378132.1 | Schlafen+HATPase+wHTH | GCA_020161315.1 | Actinomycetota_bacterium | Bacteria>Bacillati>Actinomycetota>unclassified_Actinomycetota | 493 | LCH36_01650 | 73078..74559 | 2900548 | helix-turn-helix-domain-containing-protein-[Actinomycetia-bacterium] |
| MBU4174458.1 | Schlafen+HATPase+wHTH | GCA_018898375.1 | Actinomycetota_bacterium | Bacteria>Bacillati>Actinomycetota>unclassified_Actinomycetota | 521 | KJ686_06085 | 24999..26564 | 2900548 | putative-DNA-binding-domain-containing-protein-[Actinomycetia-bacterium] |
| MBS3974430.1 | Schlafen+HATPase+wHTH | GCA_018334075.1 | Actinomycetota_bacterium | Bacteria>Bacillati>Actinomycetota>unclassified_Actinomycetota | 494 | KGZ89_06135 | 1159..2643 | 2900548 | putative-DNA-binding-domain-containing-protein-[Actinomycetia-bacterium] |
| MCC7230623.1 | Schlafen+HATPase+wHTH | GCA_020851885.1 | Fimbriimonadaceae_bacterium | Bacteria>Bacillati>Armatimonadota>Fimbriimonadia>Fimbriimonadales>Fimbriimonadaceae>unclassified_Fimbriimonadaceae | 489 | IT203_09545 | 12264..13733 | 2682144 | putative-DNA-binding-domain-containing-protein-[Fimbriimonadaceae-bacterium] |
| MBX3108931.1 | Schlafen+HATPase+wHTH | GCA_019638755.1 | Fimbriimonadaceae_bacterium | Bacteria>Bacillati>Armatimonadota>Fimbriimonadia>Fimbriimonadales>Fimbriimonadaceae>unclassified_Fimbriimonadaceae | 635 | KF743_07105 | 1433602..1435509 | 2682144 | putative-DNA-binding-domain-containing-protein-[Fimbriimonadaceae-bacterium] |
| MCX7689023.1 | Schlafen+HATPase+wHTH | GCA_026418845.1 | Fimbriimonadales_bacterium | Bacteria>Bacillati>Armatimonadota>Fimbriimonadia>Fimbriimonadales>unclassified_Fimbriimonadales | 119 | N2045_13755 | 1276..1635 | 2682145 | MAG:-hypothetical-protein-N2045_13755-[Fimbriimonadales-bacterium] |
| GIV04316.1 | Schlafen+HATPase+wHTH | GCA_026003055.1 | Fimbriimonadales_bacterium | Bacteria>Bacillati>Armatimonadota>Fimbriimonadia>Fimbriimonadales>unclassified_Fimbriimonadales | 453 | KatS3mg015_3146 | 8658..10019 | 2682145 | MAG:-hypothetical-protein-KatS3mg015_3146-[Fimbriimonadales-bacterium] |
| GIV02393.1 | Schlafen+HATPase+wHTH | GCA_026003055.1 | Fimbriimonadales_bacterium | Bacteria>Bacillati>Armatimonadota>Fimbriimonadia>Fimbriimonadales>unclassified_Fimbriimonadales | 1098 | KatS3mg015_1223 | 1245957..1249253 | 2682145 | MAG:-hypothetical-protein-KatS3mg015_1223-[Fimbriimonadales-bacterium] |
| MCX6379002.1 | Schlafen+HATPase+wHTH | GCA_026392135.1 | Armatimonadota_bacterium | Bacteria>Bacillati>Armatimonadota>unclassified_Armatimonadota | 644 | NT023_05915 | 1872..3806 | 2033014 | MAG:-putative-DNA-binding-domain-containing-protein-[Armatimonadetes-bacterium] |
| GIV15261.1 | Schlafen+HATPase+wHTH | GCA_026003175.1 | Armatimonadota_bacterium | Bacteria>Bacillati>Armatimonadota>unclassified_Armatimonadota | 285 | KatS3mg022_0696 | 789652..790509 | 2033014 | MAG:-hypothetical-protein-KatS3mg022_0696-[Armatimonadota-bacterium] |
| GIV15260.1 | Schlafen+HATPase+wHTH | GCA_026003175.1 | Armatimonadota_bacterium | Bacteria>Bacillati>Armatimonadota>unclassified_Armatimonadota | 531 | KatS3mg022_0695 | 787929..789524 | 2033014 | MAG:-ATP-dependent-DNA-helicase-RecG-[Armatimonadota-bacterium] |
| WP_195379254.1 | Schlafen+HATPase+wHTH | GCF_015551645.1 | Clostridia | Bacteria>Bacillati>Bacillota | 490 | GPK86_RS03015 | 9151..10623 | 186801 | MULTISPECIES:-ATP-binding-protein-[Eubacteriales] |
| WP_181968743.1 | Schlafen+HATPase+wHTH | GCF_048410155.1 | Clostridia | Bacteria>Bacillati>Bacillota | 492 | DW835_RS14820 | 29658..31136 | 186801 | putative-DNA-binding-domain-containing-protein-[Clostridium-sp.-AM34-9AC] |
| WP_173773125.1 | Schlafen+HATPase+wHTH | GCF_934875305.1 | Clostridia | Bacteria>Bacillati>Bacillota | 490 | G5A19_RS02630 | 11172..12644 | 186801 | MULTISPECIES:-ATP-binding-protein-[Lachnospiraceae] |
| WP_119247637.1 | Schlafen+HATPase+wHTH | GCF_020687485.1 | Clostridia | Bacteria>Bacillati>Bacillota | 490 | PGV47_RS07595 | 50541..52013 | 186801 | ATP-binding-protein-[Dorea-longicatena] |
| WP_238988114.1 | Schlafen+HATPase+wHTH | GCF_965137035.1 | Filibacter_tadaridae | Bacteria>Bacillati>Bacillota>Bacilli>Bacillales>Caryophanaceae>Filibacter | 559 | FILTAD_RS01275 | 9677..11356 | 2483811 | putative-DNA-binding-domain-containing-protein-[Filibacter-tadaridae] |
| MBK0347251.1 | Schlafen+HATPase+wHTH | GCA_016522115.1 | Aerococcaceae_bacterium_zg-1578 | Bacteria>Bacillati>Bacillota>Bacilli>Lactobacillales>Aerococcaceae>unclassified_Aerococcaceae | 563 | JDW15_01200 | 59498..61189 | 2796350 | putative-DNA-binding-domain-containing-protein-[Aerococcaceae-bacterium-zg-1578] |
| WP_236150041.1 | Schlafen+HATPase+wHTH | GCF_021600205.1 | Lactobacillus_delbrueckii | Bacteria>Bacillati>Bacillota>Bacilli>Lactobacillales>Lactobacillaceae>Lactobacillus | 536 | L5979_RS00550 | 17580..19190 | 1584 | ATP-binding-protein-[Lactobacillus-delbrueckii] |
| WP_231545264.1 | Schlafen+HATPase+wHTH | GCF_021134235.1 | Lactobacillus_delbrueckii | Bacteria>Bacillati>Bacillota>Bacilli>Lactobacillales>Lactobacillaceae>Lactobacillus | 536 | LOB65_RS04185 | 36259..37869 | 1584 | ATP-binding-protein-[Lactobacillus-delbrueckii] |
| WP_137638547.1 | Schlafen+HATPase+wHTH | GCF_005405165.1 | Loigolactobacillus_binensis | Bacteria>Bacillati>Bacillota>Bacilli>Lactobacillales>Lactobacillaceae>Loigolactobacillus | 482 | FFM78_RS11405 | 32576..34024 | 2559922 | putative-DNA-binding-domain-containing-protein-[Loigolactobacillus-binensis] |
| MBR2506171.1 | Schlafen+HATPase+wHTH | GCA_017502465.1 | Bacilli_bacterium | Bacteria>Bacillati>Bacillota>Bacilli>unclassified_Bacilli | 489 | IKB70_04500 | 18181..19650 | 1903720 | putative-DNA-binding-domain-containing-protein-[Bacilli-bacterium] |
| WP_052671500.1 | Schlafen+HATPase+wHTH | GCF_026914285.1 | Caldicellulosiruptor_naganoensis | Bacteria>Bacillati>Bacillota>Bacillota_incertae_sedis>Caldicellulosiruptorales>Caldicellulosiruptoraceae>Caldicellulosiruptor | 178 | OTJ99_RS09130 | 1820715..1821251 | 29324 | hypothetical-protein-[Caldicellulosiruptor-naganoensis] |
| WP_021658831.1 | Schlafen+HATPase+wHTH | GCF_900129515.1 | Eubacteriales | Bacteria>Bacillati>Bacillota>Clostridia | 480 | BUB64_RS00450 | 92469..93911 | 186802 | putative-DNA-binding-domain-containing-protein-[Bittarella-massiliensis] |
| HIZ73451.1 | Schlafen+HATPase+wHTH | GCA_019116755.1 | Candidatus_Gallimonas_intestinavium | Bacteria>Bacillati>Bacillota>Clostridia>Clostridia_incertae_sedis>Candidatus_Gallimonas | 470 | H9964_07700 | 16890..18302 | 2838603 | helix-turn-helix-domain-containing-protein-[Candidatus-Gallimonas-intestinavium] |
| WP_138373970.1 | Schlafen+HATPase+wHTH | - | Oscillospiraceae | Bacteria>Bacillati>Bacillota>Clostridia>Eubacteriales | 484 | I2E95_RS16300 | 8052..9506 | 216572 | MULTISPECIES:-ATP-binding-protein-[Oscillospiraceae] |
| WP_263041863.1 | Schlafen+HATPase+wHTH | GCF_025630755.1 | Acutalibacter_sp._LFL-21 | Bacteria>Bacillati>Bacillota>Clostridia>Eubacteriales>Acutalibacteraceae>Acutalibacter>unclassified_Acutalibacter | 480 | ODJ80_RS09360 | 221526..222968 | 2983399 | putative-DNA-binding-domain-containing-protein-[Acutalibacter-sp.-LFL-21] |
| WP_253205986.1 | Schlafen+HATPase+wHTH | GCF_018861855.1 | Clostridium_estertheticum | Bacteria>Bacillati>Bacillota>Clostridia>Eubacteriales>Clostridiaceae>Clostridium | 262 | KPL49_RS19715 | 55746..56534 | 238834 | ATP-binding-protein-[Clostridium-estertheticum] |
| WP_216137996.1 | Schlafen+HATPase+wHTH | GCF_018861855.1 | Clostridium_estertheticum | Bacteria>Bacillati>Bacillota>Clostridia>Eubacteriales>Clostridiaceae>Clostridium | 134 | KPL49_RS19705 | 54885..55289 | 238834 | ATP-binding-protein-[Clostridium-estertheticum] |
| WP_207727448.1 | Schlafen+HATPase+wHTH | GCF_015351765.1 | Clostridium_vitabionis | Bacteria>Bacillati>Bacillota>Clostridia>Eubacteriales>Clostridiaceae>Clostridium | 502 | ISF27_RS11795 | 56532..58040 | 2784388 | ATP-binding-protein-[Clostridium-vitabionis] |
| WP_117821977.1 | Schlafen+HATPase+wHTH | GCF_003461085.1 | unclassified_Clostridium | Bacteria>Bacillati>Bacillota>Clostridia>Eubacteriales>Clostridiaceae>Clostridium | 493 | DW959_RS12380 | 52848..54329 | 2614128 | MULTISPECIES:-putative-DNA-binding-domain-containing-protein-[unclassified-Clostridium] |
| WP_023162373.1 | Schlafen+HATPase+wHTH | GCF_000484505.2 | Clostridium_autoethanogenum | Bacteria>Bacillati>Bacillota>Clostridia>Eubacteriales>Clostridiaceae>Clostridium | 480 | WX72_RS00875 | 191587..193029 | 84023 | ATP-binding-protein-[Clostridium-autoethanogenum] |
| WP_195540870.1 | Schlafen+HATPase+wHTH | - | Clostridium_sp._D43t1_170807_D5 | Bacteria>Bacillati>Bacillota>Clostridia>Eubacteriales>Clostridiaceae>Clostridium>unclassified_Clostridium | 490 | I2G39_RS02175 | 459239..460711 | 2787150 | ATP-binding-protein-[Clostridium-sp.-D43t1_170807_D5] |
| WP_118742505.1 | Schlafen+HATPase+wHTH | GCF_003480665.1 | Clostridium_sp._AM28-20LB | Bacteria>Bacillati>Bacillota>Clostridia>Eubacteriales>Clostridiaceae>Clostridium>unclassified_Clostridium | 490 | DW739_RS01400 | 299294..300766 | 2293027 | ATP-binding-protein-[Clostridium-sp.-AM28-20LB] |
| WP_118740656.1 | Schlafen+HATPase+wHTH | - | Clostridium_sp._AM32-2 | Bacteria>Bacillati>Bacillota>Clostridia>Eubacteriales>Clostridiaceae>Clostridium>unclassified_Clostridium | 481 | DW807_RS07240 | 10746..12191 | 2293029 | ATP-binding-protein-[Clostridium-sp.-AM32-2] |
| WP_118740425.1 | Schlafen+HATPase+wHTH | - | Clostridium_sp._AM32-2 | Bacteria>Bacillati>Bacillota>Clostridia>Eubacteriales>Clostridiaceae>Clostridium>unclassified_Clostridium | 490 | DW807_RS02850 | 172516..173988 | 2293029 | ATP-binding-protein-[Clostridium-sp.-AM32-2] |
| WP_118670888.1 | Schlafen+HATPase+wHTH | - | Clostridium_sp._OM08-29 | Bacteria>Bacillati>Bacillota>Clostridia>Eubacteriales>Clostridiaceae>Clostridium>unclassified_Clostridium | 492 | DXC24_RS05255 | 328227..329705 | 2293049 | putative-DNA-binding-domain-containing-protein-[Clostridium-sp.-OM08-29] |
| WP_118513559.1 | Schlafen+HATPase+wHTH | - | Clostridium_sp._AF37-5AT | Bacteria>Bacillati>Bacillota>Clostridia>Eubacteriales>Clostridiaceae>Clostridium>unclassified_Clostridium | 497 | DW020_RS00870 | 201215..202708 | 2292007 | ATP-binding-protein-[Clostridium-sp.-AF37-5AT] |
| WP_118320919.1 | Schlafen+HATPase+wHTH | - | Clostridium_sp._AM22-16AC | Bacteria>Bacillati>Bacillota>Clostridia>Eubacteriales>Clostridiaceae>Clostridium>unclassified_Clostridium | 500 | DW260_RS00820 | 187013..188515 | 2292009 | ATP-binding-protein-[Clostridium-sp.-AM22-16AC] |
| WP_103227374.1 | Schlafen+HATPase+wHTH | GCF_002899675.1 | Clostridium_sp._chh4-2 | Bacteria>Bacillati>Bacillota>Clostridia>Eubacteriales>Clostridiaceae>Clostridium>unclassified_Clostridium | 489 | C0033_RS08170 | 114748..116217 | 2067550 | putative-DNA-binding-domain-containing-protein-[Clostridium-sp.-chh4-2] |
| RHU61445.1 | Schlafen+HATPase+wHTH | GCA_003481275.1 | Clostridium_sp._TF08-15 | Bacteria>Bacillati>Bacillota>Clostridia>Eubacteriales>Clostridiaceae>Clostridium>unclassified_Clostridium | 496 | DXC82_15775 | 25119..26609 | 2293052 | hypothetical-protein-DXC82_15775-[Clostridium-sp.-TF08-15] |
| RGF28777.1 | Schlafen+HATPase+wHTH | GCA_003435925.1 | Clostridium_sp._AF46-9NS | Bacteria>Bacillati>Bacillota>Clostridia>Eubacteriales>Clostridiaceae>Clostridium>unclassified_Clostridium | 499 | DW081_14970 | 27170..28669 | 2293020 | hypothetical-protein-DW081_14970-[Clostridium-sp.-AF46-9NS] |
| MBS5468803.1 | Schlafen+HATPase+wHTH | GCA_018368355.1 | Clostridium_sp. | Bacteria>Bacillati>Bacillota>Clostridia>Eubacteriales>Clostridiaceae>Clostridium>unclassified_Clostridium | 498 | KHX69_08295 | 44236..45732 | 1506 | putative-DNA-binding-domain-containing-protein-[Clostridium-sp.] |
| CDD75195.1 | Schlafen+HATPase+wHTH | GCA_000436175.1 | Clostridium_sp._CAG:62 | Bacteria>Bacillati>Bacillota>Clostridia>Eubacteriales>Clostridiaceae>Clostridium>unclassified_Clostridium | 492 | BN737_01382 | 82165..83643 | 1262828 | hypothetical-conserved-protein-[Clostridium-sp.-CAG:62] |
| WP_216574653.1 | Schlafen+HATPase+wHTH | GCF_018918265.1 | Falcatimonas_sp._MSJ-15 | Bacteria>Bacillati>Bacillota>Clostridia>Eubacteriales>Clostridiaceae>Falcatimonas>unclassified_Falcatimonas | 485 | KQI85_RS03190 | 196350..197807 | 2841515 | ATP-binding-protein-[Falcatimonas-sp.-MSJ-15] |
| NBI69483.1 | Schlafen+HATPase+wHTH | GCA_009911305.1 | Clostridiaceae_bacterium | Bacteria>Bacillati>Bacillota>Clostridia>Eubacteriales>Clostridiaceae>unclassified_Clostridiaceae | 544 | D3Z50_00060 | 14669..16303 | 1898204 | winged-helix-turn-helix-transcriptional-regulator-[Clostridiaceae-bacterium] |
| NBH73086.1 | Schlafen+HATPase+wHTH | GCA_009911255.1 | Clostridiaceae_bacterium | Bacteria>Bacillati>Bacillota>Clostridia>Eubacteriales>Clostridiaceae>unclassified_Clostridiaceae | 444 | D3Z51_13875 | 21591..22925 | 1898204 | AAA-family-ATPase-[Clostridiaceae-bacterium] |
| NBH73081.1 | Schlafen+HATPase+wHTH | GCA_009911255.1 | Clostridiaceae_bacterium | Bacteria>Bacillati>Bacillota>Clostridia>Eubacteriales>Clostridiaceae>unclassified_Clostridiaceae | 490 | D3Z51_13840 | 11145..12617 | 1898204 | winged-helix-turn-helix-transcriptional-regulator-[Clostridiaceae-bacterium] |
| NBH35977.1 | Schlafen+HATPase+wHTH | GCA_009911065.1 | Clostridiaceae_bacterium | Bacteria>Bacillati>Bacillota>Clostridia>Eubacteriales>Clostridiaceae>unclassified_Clostridiaceae | 495 | D3Z58_21190 | 13216..14703 | 1898204 | winged-helix-turn-helix-transcriptional-regulator-[Clostridiaceae-bacterium] |
| NBH17645.1 | Schlafen+HATPase+wHTH | GCA_009911105.1 | Clostridiaceae_bacterium | Bacteria>Bacillati>Bacillota>Clostridia>Eubacteriales>Clostridiaceae>unclassified_Clostridiaceae | 495 | D3Z55_09220 | 49066..50553 | 1898204 | winged-helix-turn-helix-transcriptional-regulator-[Clostridiaceae-bacterium] |
| WP_018214268.1 | Schlafen+HATPase+wHTH | GCF_000378805.1 | Desulfitobacterium_hafniense | Bacteria>Bacillati>Bacillota>Clostridia>Eubacteriales>Desulfitobacteriaceae>Desulfitobacterium | 302 | DESHAF_RS0122590 | 1958480..1959388 | 49338 | hypothetical-protein-[Desulfitobacterium-hafniense] |
| WP_186842514.1 | Schlafen+HATPase+wHTH | GCF_014284475.1 | Acetobacterium_fimetarium | Bacteria>Bacillati>Bacillota>Clostridia>Eubacteriales>Eubacteriaceae>Acetobacterium | 110 | GH808_RS09320 | 36502..36834 | 52691 | ATP-binding-protein-[Acetobacterium-fimetarium] |
| WP_055289376.1 | Schlafen+HATPase+wHTH | GCF_045002895.1 | Eubacterium_ramulus | Bacteria>Bacillati>Bacillota>Clostridia>Eubacteriales>Eubacteriaceae>Eubacterium | 490 | ARA08_RS03305 | 198113..199585 | 39490 | ATP-binding-protein-[Eubacterium-ramulus] |
| MCI5510138.1 | Schlafen+HATPase+wHTH | GCA_022764165.1 | Eubacterium_sp. | Bacteria>Bacillati>Bacillota>Clostridia>Eubacteriales>Eubacteriaceae>Eubacterium>unclassified_Eubacterium | 512 | MR436_06810 | 18116..19654 | 142586 | putative-DNA-binding-domain-containing-protein-[Eubacterium-sp.] |
| MBS6586979.1 | Schlafen+HATPase+wHTH | GCA_018377805.1 | Eubacterium_sp. | Bacteria>Bacillati>Bacillota>Clostridia>Eubacteriales>Eubacteriaceae>Eubacterium>unclassified_Eubacterium | 475 | KH345_12605 | 11244..12671 | 142586 | putative-DNA-binding-domain-containing-protein-[Eubacterium-sp.] |
| OLA08996.1 | Schlafen+HATPase+wHTH | GCA_001916415.1 | Eubacterium_sp._45_250 | Bacteria>Bacillati>Bacillota>Clostridia>Eubacteriales>Eubacteriaceae>Eubacterium>unclassified_Eubacterium_(in:_firmicutes) | 536 | BHW21_06145 | 160759..162369 | 1897004 | MAG:-transcriptional-regulator-[Eubacterium-sp.-45_250] |
| MCH4073092.1 | Schlafen+HATPase+wHTH | GCA_022483875.1 | Pseudoramibacter_sp. | Bacteria>Bacillati>Bacillota>Clostridia>Eubacteriales>Eubacteriaceae>Pseudoramibacter>unclassified_Pseudoramibacter | 246 | LKF02_09125 | 1850160..1850900 | 2034862 | putative-DNA-binding-domain-containing-protein-[Pseudoramibacter-sp.] |
| MCH4073091.1 | Schlafen+HATPase+wHTH | GCA_022483875.1 | Pseudoramibacter_sp. | Bacteria>Bacillati>Bacillota>Clostridia>Eubacteriales>Eubacteriaceae>Pseudoramibacter>unclassified_Pseudoramibacter | 281 | LKF02_09120 | 1849286..1850131 | 2034862 | hypothetical-protein-[Pseudoramibacter-sp.] |
| WP_087184152.1 | Schlafen+HATPase+wHTH | - | Gemmiger_sp._An87 | Bacteria>Bacillati>Bacillota>Clostridia>Eubacteriales>Eubacteriales_incertae_sedis>Gemmiger>unclassified_Gemmiger | 411 | B5G38_RS07665 | 41305..42540 | 1965662 | ATP-binding-protein-[Gemmiger-sp.-An87] |
| WP_227612047.1 | Schlafen+HATPase+wHTH | GCF_020686905.1 | Faecalibacterium | Bacteria>Bacillati>Bacillota>Clostridia>Eubacteriales>Oscillospiraceae | 537 | LKD27_RS12795 | 11329..12942 | 216851 | ATP-binding-protein-[Faecalibacterium-sp.-CLA-AA-H283] |
| WP_014256027.1 | Schlafen+HATPase+wHTH | GCF_000237085.1 | Acetivibrio_clariflavus | Bacteria>Bacillati>Bacillota>Clostridia>Eubacteriales>Oscillospiraceae>Acetivibrio | 386 | CLOCLA_RS0107435 | 1191060..1192220 | 288965 | ATP-binding-protein-[Acetivibrio-clariflavus] |
| OUP64013.1 | Schlafen+HATPase+wHTH | GCA_002160415.1 | Drancourtella_sp._An177 | Bacteria>Bacillati>Bacillota>Clostridia>Eubacteriales>Oscillospiraceae>Drancourtella>unclassified_Drancourtella | 505 | B5F13_08875 | 7940..9457 | 1965573 | ATP-binding-protein-[Drancourtella-sp.-An177] |
| WP_097800075.1 | Schlafen+HATPase+wHTH | GCF_002549775.1 | Faecalibacterium_prausnitzii | Bacteria>Bacillati>Bacillota>Clostridia>Eubacteriales>Oscillospiraceae>Faecalibacterium | 536 | CGS49_RS00530 | 109901..111511 | 853 | ATP-binding-protein-[Faecalibacterium-prausnitzii] |
| WP_055192164.1 | Schlafen+HATPase+wHTH | - | Faecalibacterium_prausnitzii | Bacteria>Bacillati>Bacillota>Clostridia>Eubacteriales>Oscillospiraceae>Faecalibacterium | 537 | FXW87_RS12985 | 9391..11004 | 853 | ATP-binding-protein-[Faecalibacterium-prausnitzii] |
| WP_243413740.1 | Schlafen+HATPase+wHTH | - | Pseudoflavonifractor_phocaeensis | Bacteria>Bacillati>Bacillota>Clostridia>Eubacteriales>Oscillospiraceae>Pseudoflavonifractor | 543 | H6B27_RS11845 | 7404..9035 | 1870988 | putative-DNA-binding-domain-containing-protein-[Pseudoflavonifractor-phocaeensis] |
| WP_207752431.1 | Schlafen+HATPase+wHTH | - | Pseudoflavonifractor_capillosus | Bacteria>Bacillati>Bacillota>Clostridia>Eubacteriales>Oscillospiraceae>Pseudoflavonifractor | 548 | H9X87_RS10175 | 7906..9552 | 106588 | ATP-binding-protein-[Pseudoflavonifractor-capillosus] |
| WP_206020568.1 | Schlafen+HATPase+wHTH | - | Pseudoflavonifractor_sp._SW1122 | Bacteria>Bacillati>Bacillota>Clostridia>Eubacteriales>Oscillospiraceae>Pseudoflavonifractor>unclassified_Pseudoflavonifractor | 548 | E0L15_RS04970 | 45928..47574 | 2530044 | ATP-binding-protein-[Pseudoflavonifractor-sp.-SW1122] |
| MBT9624609.1 | Schlafen+HATPase+wHTH | GCA_018784005.1 | Ruminococcus_bicirculans_(ex_Wegman_et_al._2014) | Bacteria>Bacillati>Bacillota>Clostridia>Eubacteriales>Oscillospiraceae>Ruminococcus | 68 | GPK54_04650 | 15571..15777 | 1160721 | hypothetical-protein-[Ruminococcus-bicirculans] |
| MBS7204854.1 | Schlafen+HATPase+wHTH | GCA_018383015.1 | Ruminococcus_bicirculans_(ex_Wegman_et_al._2014) | Bacteria>Bacillati>Bacillota>Clostridia>Eubacteriales>Oscillospiraceae>Ruminococcus | 68 | KH048_01100 | 252997..253203 | 1160721 | putative-DNA-binding-domain-containing-protein-[Ruminococcus-bicirculans] |
| WP_195982639.1 | Schlafen+HATPase+wHTH | GCF_015669355.1 | Ruminococcus_sp._1001275B_160808_F8 | Bacteria>Bacillati>Bacillota>Clostridia>Eubacteriales>Oscillospiraceae>Ruminococcus>unclassified_Ruminococcus | 490 | I4V83_RS01655 | 32137..33609 | 2787131 | ATP-binding-protein-[Ruminococcus-sp.-1001275B_160808_F8] |
| RHO19185.1 | Schlafen+HATPase+wHTH | GCA_003473745.1 | Ruminococcus_sp._AM18-44 | Bacteria>Bacillati>Bacillota>Clostridia>Eubacteriales>Oscillospiraceae>Ruminococcus>unclassified_Ruminococcus | 495 | DW225_05615 | 142573..144060 | 2293186 | transcriptional-regulator-[Ruminococcus-sp.-AM18-44] |
| MCI8835831.1 | Schlafen+HATPase+wHTH | GCA_022795995.1 | Ruminococcus_sp. | Bacteria>Bacillati>Bacillota>Clostridia>Eubacteriales>Oscillospiraceae>Ruminococcus>unclassified_Ruminococcus | 278 | HFH24_10730 | 34882..35718 | 41978 | hypothetical-protein-[Ruminococcus-sp.] |
| RJV95621.1 | Schlafen+HATPase+wHTH | GCA_003603335.1 | Subdoligranulum_sp._AF14-43 | Bacteria>Bacillati>Bacillota>Clostridia>Eubacteriales>Oscillospiraceae>Subdoligranulum>unclassified_Subdoligranulum | 411 | DWW15_14720 | 32398..33633 | 2302959 | ATP-binding-protein-[Subdoligranulum-sp.-AF14-43] |
| UYJ36606.1 | Schlafen+HATPase+wHTH | GCA_025758005.1 | Oscillospiraceae_bacterium | Bacteria>Bacillati>Bacillota>Clostridia>Eubacteriales>Oscillospiraceae>unclassified_Oscillospiraceae | 490 | OGM10_10395 | 2207593..2209065 | 2485925 | MAG:-putative-DNA-binding-domain-containing-protein-[Oscillospiraceae-bacterium] |
| UYJ05870.1 | Schlafen+HATPase+wHTH | GCA_025757805.1 | Oscillospiraceae_bacterium | Bacteria>Bacillati>Bacillota>Clostridia>Eubacteriales>Oscillospiraceae>unclassified_Oscillospiraceae | 536 | OGM79_08415 | 1757029..1758639 | 2485925 | MAG:-putative-DNA-binding-domain-containing-protein-[Oscillospiraceae-bacterium] |
| MCL1790895.1 | Schlafen+HATPase+wHTH | GCA_009778595.1 | Peptococcaceae_bacterium | Bacteria>Bacillati>Bacillota>Clostridia>Eubacteriales>Peptococcaceae>unclassified_Peptococcaceae | 486 | FWG40_05985 | 8120..9580 | 2052179 | putative-DNA-binding-domain-containing-protein-[Peptococcaceae-bacterium] |
| MBE6111692.1 | Schlafen+HATPase+wHTH | GCA_015058785.1 | Peptococcaceae_bacterium | Bacteria>Bacillati>Bacillota>Clostridia>Eubacteriales>Peptococcaceae>unclassified_Peptococcaceae | 511 | E7195_01570 | 63362..64897 | 2052179 | transcriptional-regulator-[Peptococcaceae-bacterium] |
| WP_251617133.1 | Schlafen+HATPase+wHTH | GCF_943193045.1 | Pumilibacter_muris | Bacteria>Bacillati>Bacillota>Clostridia>Eubacteriales>Pumilibacteraceae>Pumilibacter | 476 | NFR62_RS07855 | 63238..64668 | 2941510 | putative-DNA-binding-domain-containing-protein-[Pumilibacter-muris] |
| OKZ97586.1 | Schlafen+HATPase+wHTH | GCA_001915795.1 | Clostridiales_bacterium_42_27 | Bacteria>Bacillati>Bacillota>Clostridia>Eubacteriales>unclassified_Eubacteriales | 536 | BHV90_19775 | 11391..13001 | 1897045 | transcriptional-regulator-[Clostridiales-bacterium-42_27] |
| OAD88352.1 | Schlafen+HATPase+wHTH | GCA_001641065.1 | Clostridiales_bacterium_KLE1615 | Bacteria>Bacillati>Bacillota>Clostridia>Eubacteriales>unclassified_Eubacteriales | 495 | HMPREF2738_01641 | 72966..74453 | 1715004 | divergent-AAA-domain-protein-[Clostridiales-bacterium-KLE1615] |
| NLO81517.1 | Schlafen+HATPase+wHTH | GCA_012523455.1 | Clostridiales_bacterium | Bacteria>Bacillati>Bacillota>Clostridia>Eubacteriales>unclassified_Eubacteriales | 330 | GX094_00410 | 100..1092 | 1898207 | histidine-kinase-[Clostridiales-bacterium] |
| MBS5313237.1 | Schlafen+HATPase+wHTH | GCA_018367215.1 | Clostridiales_bacterium | Bacteria>Bacillati>Bacillota>Clostridia>Eubacteriales>unclassified_Eubacteriales | 497 | KHY53_10220 | 18543..20036 | 1898207 | putative-DNA-binding-domain-containing-protein-[Clostridiales-bacterium] |
| MBS5052654.1 | Schlafen+HATPase+wHTH | GCA_018365235.1 | Clostridiales_bacterium | Bacteria>Bacillati>Bacillota>Clostridia>Eubacteriales>unclassified_Eubacteriales | 490 | KHZ49_10875 | 17895..19367 | 1898207 | putative-DNA-binding-domain-containing-protein-[Clostridiales-bacterium] |
| WP_206706677.1 | Schlafen+HATPase+wHTH | GCF_017301615.1 | Koleobacter_methoxysyntrophicus | Bacteria>Bacillati>Bacillota>Clostridia>Koleobacterales>Koleobacteraceae>Koleobacter | 387 | H0A61_RS08420 | 1758085..1759248 | 2751313 | putative-DNA-binding-domain-containing-protein-[Koleobacter-methoxysyntrophicus] |
| QSQ09318.1 | Schlafen+HATPase+wHTH | GCA_017301615.1 | Koleobacter_methoxysyntrophicus | Bacteria>Bacillati>Bacillota>Clostridia>Koleobacterales>Koleobacteraceae>Koleobacter | 387 | H0A61_01679 | 1758085..1759248 | 2751313 | hypothetical-protein-H0A61_01679-[Koleobacter-methoxysyntrophicus] |
| WP_173865458.1 | Schlafen+HATPase+wHTH | GCF_013300965.1 | Lachnospiraceae | Bacteria>Bacillati>Bacillota>Clostridia>Lachnospirales | 572 | HFN12_RS00365 | 65804..67522 | 186803 | ATP-binding-protein-[Faecalicatena-fissicatena] |

| Protein-ID | Domain | Assembly-ID | Strain | Taxonomy | Length | Locus-Tag | Genomic-coordinates | TaxID | Fasta-defline |
|---|---|---|---|---|---|---|---|---|---|
| WP_173770659.1 | Schlafen+HATPase+wHTH | GCF_013302375.1 | Lachnospiraceae | Bacteria>Bacillati>Bacillota>Clostridia>Lachnospirales | 490 | G5A01_RS01775 | 128018..129490 | 186803 | ATP-binding-protein-[Blautia-faecis] |
| WP_118447463.1 | Schlafen+HATPase+wHTH | GCF_020560115.1 | Lachnospiraceae | Bacteria>Bacillati>Bacillota>Clostridia>Lachnospirales | 490 | LI126_RS00430 | 90784..92256 | 186803 | MULTISPECIES:-ATP-binding-protein-[Lachnospiraceae] |
| WP_117761887.1 | Schlafen+HATPase+wHTH | GCF_902364755.1 | Lachnospiraceae | Bacteria>Bacillati>Bacillota>Clostridia>Lachnospirales | 490 | FXX70_RS12960 | 70902..72374 | 186803 | MULTISPECIES:-ATP-binding-protein-[Lachnospiraceae] |
| WP_117739790.1 | Schlafen+HATPase+wHTH | GCF_013301605.1 | Lachnospiraceae | Bacteria>Bacillati>Bacillota>Clostridia>Lachnospirales | 490 | G4434_RS12245 | 47868..49340 | 186803 | MULTISPECIES:-ATP-binding-protein-[Lachnospiraceae] |
| WP_055268219.1 | Schlafen+HATPase+wHTH | GCF_001406335.1 | Lachnospiraceae | Bacteria>Bacillati>Bacillota>Clostridia>Lachnospirales | 515 | DWX26_RS14820 | 12083..13630 | 186803 | MULTISPECIES:-ATP-binding-protein-[Lachnospiraceae] |
| WP_072849723.1 | Schlafen+HATPase+wHTH | GCF_900142265.1 | Anaerotignum_lactatifermentans | Bacteria>Bacillati>Bacillota>Clostridia>Lachnospirales>Anaerotignaceae>Anaerotignum | 483 | BUB35_RS04890 | 29719..31170 | 160404 | putative-DNA-binding-domain-containing-protein-[Anaerotignum-lactatifermentans] |
| WP_138345912.1 | Schlafen+HATPase+wHTH | GCF_015669935.1 | Roseburia | Bacteria>Bacillati>Bacillota>Clostridia>Lachnospirales>Lachnospiraceae | 512 | FGQ84_RS06840 | 98396..99934 | 841 | MULTISPECIES:-ATP-binding-protein-[Roseburia] |
| WP_102289723.1 | Schlafen+HATPase+wHTH | GCF_042853085.1 | Eisenbergiella | Bacteria>Bacillati>Bacillota>Clostridia>Lachnospirales>Lachnospiraceae | 489 | C1733_RS26390 | 2132722..2134191 | 1432051 | ATP-binding-protein-[Eisenbergiella-massiliensis] |
| WP_055656464.1 | Schlafen+HATPase+wHTH | GCF_902362405.1 | Hungatella | Bacteria>Bacillati>Bacillota>Clostridia>Lachnospirales>Lachnospiraceae | 487 | FY393_RS15405 | 230173..231636 | 1649459 | ATP-binding-protein-[Hungatella-hathewayi] |
| TYL56935.1 | Schlafen+HATPase+wHTH | GCA_008123415.1 | Agathobacter_rectalis | Bacteria>Bacillati>Bacillota>Clostridia>Lachnospirales>Lachnospiraceae>Agathobacter | 492 | FYL31_13950 | 10610..12088 | 39491 | hypothetical-protein-FYL31_13950-[[Eubacterium] |
| RHD97633.1 | Schlafen+HATPase+wHTH | GCA_003468965.1 | Agathobacter_rectalis | Bacteria>Bacillati>Bacillota>Clostridia>Lachnospirales>Lachnospiraceae>Agathobacter | 512 | DW775_02185 | 12242..13780 | 39491 | ATP-binding-protein-[[Eubacterium] |
| RHA07189.1 | Schlafen+HATPase+wHTH | GCA_003466745.1 | Agathobacter_rectalis | Bacteria>Bacillati>Bacillota>Clostridia>Lachnospirales>Lachnospiraceae>Agathobacter | 512 | DW951_01500 | 319810..321348 | 39491 | ATP-binding-protein-[[Eubacterium] |
| RGZ95796.1 | Schlafen+HATPase+wHTH | GCA_003467085.1 | Agathobacter_rectalis | Bacteria>Bacillati>Bacillota>Clostridia>Lachnospirales>Lachnospiraceae>Agathobacter | 512 | DW967_00050 | 11396..12934 | 39491 | ATP-binding-protein-[[Eubacterium] |
| RGI65890.1 | Schlafen+HATPase+wHTH | GCA_003436035.1 | Agathobacter_rectalis | Bacteria>Bacillati>Bacillota>Clostridia>Lachnospirales>Lachnospiraceae>Agathobacter | 496 | DXD95_13315 | 13291..14781 | 39491 | hypothetical-protein-DXD95_13315-[[Eubacterium] |
| MCG4813358.1 | Schlafen+HATPase+wHTH | GCA_022136565.1 | Agathobacter_rectalis | Bacteria>Bacillati>Bacillota>Clostridia>Lachnospirales>Lachnospiraceae>Agathobacter | 512 | LON19_09305 | 53183..54721 | 39491 | putative-DNA-binding-domain-containing-protein-[[Eubacterium] |
| CUM83620.1 | Schlafen+HATPase+wHTH | GCA_001406375.1 | Agathobacter_rectalis | Bacteria>Bacillati>Bacillota>Clostridia>Lachnospirales>Lachnospiraceae>Agathobacter | 498 | ERS852580_00749 | 5019..6515 | 39491 | Divergent-AAA-domain-[[Eubacterium] |
| WP_158358551.1 | Schlafen+HATPase+wHTH | GCF_025567195.1 | Alitiscatomonas_aceti | Bacteria>Bacillati>Bacillota>Clostridia>Lachnospirales>Lachnospiraceae>Alitiscatomonas | 494 | OCV69_RS08265 | 61993..63477 | 2981724 | ATP-binding-protein-[Alitiscatomonas-aceti] |
| WP_075672065.1 | Schlafen+HATPase+wHTH | GCF_001940245.1 | Anaerostipes_sp._992a | Bacteria>Bacillati>Bacillota>Clostridia>Lachnospirales>Lachnospiraceae>Anaerostipes>unclassified_Anaerostipes | 484 | BHF69_RS12300 | 119788..121242 | 1261637 | ATP-binding-protein-[Anaerostipes-sp.-992a] |
| WP_227239828.1 | Schlafen+HATPase+wHTH | GCF_020563965.1 | Blautia_glucerasea | Bacteria>Bacillati>Bacillota>Clostridia>Lachnospirales>Lachnospiraceae>Blautia | 496 | LI273_RS06150 | 25864..27354 | 536633 | putative-DNA-binding-domain-containing-protein-[Blautia-glucerasea] |
| WP_118198125.1 | Schlafen+HATPase+wHTH | GCF_040095635.1 | Blautia_obeum | Bacteria>Bacillati>Bacillota>Clostridia>Lachnospirales>Lachnospiraceae>Blautia | 490 | DW272_RS15825 | 17148..18620 | 40520 | ATP-binding-protein-[Blautia-obeum] |
| WP_118051564.1 | Schlafen+HATPase+wHTH | GCF_003464645.1 | Blautia_obeum | Bacteria>Bacillati>Bacillota>Clostridia>Lachnospirales>Lachnospiraceae>Blautia | 493 | DWW07_RS08235 | 42815..44296 | 40520 | ATP-binding-protein-[Blautia-obeum] |
| WP_118012966.1 | Schlafen+HATPase+wHTH | GCF_013299585.1 | Blautia_obeum | Bacteria>Bacillati>Bacillota>Clostridia>Lachnospirales>Lachnospiraceae>Blautia | 490 | G4467_RS13385 | 22863..24335 | 40520 | ATP-binding-protein-[Blautia-obeum] |
| WP_065540594.1 | Schlafen+HATPase+wHTH | GCF_001689125.2 | Blautia_pseudococcoides | Bacteria>Bacillati>Bacillota>Clostridia>Lachnospirales>Lachnospiraceae>Blautia | 493 | A4V09_RS00340 | 83492..84973 | 1796616 | ATP-binding-protein-[Blautia-pseudococcoides] |
| SCG99591.1 | Schlafen+HATPase+wHTH | GCA_900066205.1 | uncultured_Blautia_sp. | Bacteria>Bacillati>Bacillota>Clostridia>Lachnospirales>Lachnospiraceae>Blautia>environmental_samples | 490 | SAMEA3545256_00114 | 132444..133916 | 765821 | Divergent-AAA-domain-[uncultured-Blautia-sp.] |
| WP_265190758.1 | Schlafen+HATPase+wHTH | GCF_020553125.1 | Blautia_sp._MSK20_18 | Bacteria>Bacillati>Bacillota>Clostridia>Lachnospirales>Lachnospiraceae>Blautia>unclassified_Blautia | 490 | LI091_RS01850 | 111326..112798 | 2883186 | ATP-binding-protein-[Blautia-sp.-MSK20_18] |
| WP_087273655.1 | Schlafen+HATPase+wHTH | GCF_002161645.1 | Blautia_sp._An46 | Bacteria>Bacillati>Bacillota>Clostridia>Lachnospirales>Lachnospiraceae>Blautia>unclassified_Blautia | 504 | B5G00_RS13685 | 77126..78640 | 1965636 | ATP-binding-protein-[Blautia-sp.-An46] |
| RHS91604.1 | Schlafen+HATPase+wHTH | GCA_003480245.1 | Blautia_sp._AM42-2 | Bacteria>Bacillati>Bacillota>Clostridia>Lachnospirales>Lachnospiraceae>Blautia>unclassified_Blautia | 501 | DW915_10725 | 12699..14204 | 2292976 | ATP-binding-protein-[Blautia-sp.-AM42-2] |
| MCB7506156.1 | Schlafen+HATPase+wHTH | GCA_020553125.1 | Blautia_sp._MSK20_18 | Bacteria>Bacillati>Bacillota>Clostridia>Lachnospirales>Lachnospiraceae>Blautia>unclassified_Blautia | 490 | LI091_01860 | 111326..112798 | 2883186 | putative-DNA-binding-domain-containing-protein-[Blautia-sp.-MSK20_18] |
| WP_252197944.1 | Schlafen+HATPase+wHTH | GCF_018785175.1 | Coprococcus_catus | Bacteria>Bacillati>Bacillota>Clostridia>Lachnospirales>Lachnospiraceae>Coprococcus | 231 | GPK82_RS00925 | 46540..47235 | 116085 | ATP-binding-protein-[Coprococcus-catus] |
| WP_173794828.1 | Schlafen+HATPase+wHTH | GCF_013301435.1 | Coprococcus_eutactus | Bacteria>Bacillati>Bacillota>Clostridia>Lachnospirales>Lachnospiraceae>Coprococcus | 490 | G4458_RS07860 | 100974..102446 | 33043 | ATP-binding-protein-[Coprococcus-eutactus] |
| MBU5249606.1 | Schlafen+HATPase+wHTH | GCA_018916865.1 | Coprococcus_comes | Bacteria>Bacillati>Bacillota>Clostridia>Lachnospirales>Lachnospiraceae>Coprococcus | 417 | KQH93_07485 | 19577..20830 | 410072 | putative-DNA-binding-domain-containing-protein-[Coprococcus-comes] |
| WP_118348304.1 | Schlafen+HATPase+wHTH | GCF_003458885.1 | Coprococcus_sp._AF21-14LB | Bacteria>Bacillati>Bacillota>Clostridia>Lachnospirales>Lachnospiraceae>Coprococcus>unclassified_Coprococcus | 490 | DWX73_RS07795 | 4806..6278 | 2292231 | ATP-binding-protein-[Coprococcus-sp.-AF21-14LB] |
| HAB87403.1 | Schlafen+HATPase+wHTH | GCA_003448155.1 | Coprococcus_sp. | Bacteria>Bacillati>Bacillota>Clostridia>Lachnospirales>Lachnospiraceae>Coprococcus>unclassified_Coprococcus | 492 | DCE60_00045 | 12344..13822 | 2049024 | hypothetical-protein-[Coprococcus-sp.] |
| WP_168932963.1 | Schlafen+HATPase+wHTH | GCF_012843825.1 | Dorea_formicigenerans | Bacteria>Bacillati>Bacillota>Clostridia>Lachnospirales>Lachnospiraceae>Dorea | 488 | HF855_RS01320 | 78330..79796 | 39486 | ATP-binding-protein-[Dorea-formicigenerans] |
| MCQ4892852.1 | Schlafen+HATPase+wHTH | GCA_024461355.1 | Dorea_longicatena | Bacteria>Bacillati>Bacillota>Clostridia>Lachnospirales>Lachnospiraceae>Dorea | 490 | NE607_07120 | 29998..31470 | 88431 | putative-DNA-binding-domain-containing-protein-[Dorea-longicatena] |
| WP_016216876.1 | Schlafen+HATPase+wHTH | - | Dorea_sp._5-2 | Bacteria>Bacillati>Bacillota>Clostridia>Lachnospirales>Lachnospiraceae>Dorea>unclassified_Dorea | 495 | C817_RS00465 | 112831..114318 | 1235798 | ATP-binding-protein-[Dorea-sp.-5-2] |
| MCI9239448.1 | Schlafen+HATPase+wHTH | GCA_022798835.1 | Dorea_sp. | Bacteria>Bacillati>Bacillota>Clostridia>Lachnospirales>Lachnospiraceae>Dorea>unclassified_Dorea | 490 | HFH07_13940 | 12480..13952 | 2040332 | winged-helix-turn-helix-transcriptional-regulator-[Dorea-sp.] |
| HIV92254.1 | Schlafen+HATPase+wHTH | GCA_019113175.1 | Candidatus_Eisenbergiella_stercoravium | Bacteria>Bacillati>Bacillota>Clostridia>Lachnospirales>Lachnospiraceae>Eisenbergiella | 241 | H9744_11350 | 95079..95804 | 2838556 | hypothetical-protein-[Candidatus-Eisenbergiella-stercoravium] |
| HIV92253.1 | Schlafen+HATPase+wHTH | GCA_019113175.1 | Candidatus_Eisenbergiella_stercoravium | Bacteria>Bacillati>Bacillota>Clostridia>Lachnospirales>Lachnospiraceae>Eisenbergiella | 284 | H9744_11345 | 94198..95052 | 2838556 | putative-DNA-binding-domain-containing-protein-[Candidatus-Eisenbergiella-stercoravium] |
| WP_227178522.1 | Schlafen+HATPase+wHTH | GCF_020555685.1 | Enterocloster_bolteae | Bacteria>Bacillati>Bacillota>Clostridia>Lachnospirales>Lachnospiraceae>Enterocloster | 416 | LI018_RS10110 | 529026..530276 | 208479 | ATP-binding-protein-[Enterocloster-bolteae] |
| WP_002585539.1 | Schlafen+HATPase+wHTH | GCF_013304275.1 | Enterocloster_clostridioformis | Bacteria>Bacillati>Bacillota>Clostridia>Lachnospirales>Lachnospiraceae>Enterocloster | 505 | HMPREF1088_RS04960 | 1090430..1091947 | 1531 | putative-DNA-binding-domain-containing-protein-[Enterocloster-clostridioformis] |
| WP_002578400.1 | Schlafen+HATPase+wHTH | GCF_034129265.1 | Enterocloster_bolteae | Bacteria>Bacillati>Bacillota>Clostridia>Lachnospirales>Lachnospiraceae>Enterocloster | 490 | HMPREF1095_RS19875 | 1370866..1372338 | 208479 | ATP-binding-protein-[Enterocloster-bolteae] |
| HJC65845.1 | Schlafen+HATPase+wHTH | GCA_019119955.1 | Candidatus_Enterocloster_excrementigallinarum | Bacteria>Bacillati>Bacillota>Clostridia>Lachnospirales>Lachnospiraceae>Enterocloster | 535 | H9931_03875 | 76307..77914 | 2838558 | putative-DNA-binding-domain-containing-protein-[Candidatus-Enterocloster-excrementigallinarum] |
| WP_270422338.1 | Schlafen+HATPase+wHTH | GCF_027661365.1 | Fusicatenibacter_saccharivorans | Bacteria>Bacillati>Bacillota>Clostridia>Lachnospirales>Lachnospiraceae>Fusicatenibacter | 490 | PGQ16_RS03055 | 263009..264481 | 1150298 | ATP-binding-protein-[Fusicatenibacter-saccharivorans] |
| WP_238033439.1 | Schlafen+HATPase+wHTH | GCF_022136435.1 | Fusicatenibacter_saccharivorans | Bacteria>Bacillati>Bacillota>Clostridia>Lachnospirales>Lachnospiraceae>Fusicatenibacter | 490 | LON21_RS13030 | 72225..73697 | 1150298 | ATP-binding-protein-[Fusicatenibacter-saccharivorans] |
| WP_173830347.1 | Schlafen+HATPase+wHTH | GCF_013301105.1 | Fusicatenibacter_saccharivorans | Bacteria>Bacillati>Bacillota>Clostridia>Lachnospirales>Lachnospiraceae>Fusicatenibacter | 490 | G5B05_RS14765 | 12289..13761 | 1150298 | ATP-binding-protein-[Fusicatenibacter-saccharivorans] |
| MBN2952030.1 | Schlafen+HATPase+wHTH | GCA_016939215.1 | Fusicatenibacter_saccharivorans | Bacteria>Bacillati>Bacillota>Clostridia>Lachnospirales>Lachnospiraceae>Fusicatenibacter | 515 | JTJ23_00200 | 11160..12707 | 1150298 | putative-DNA-binding-domain-containing-protein-[Fusicatenibacter-saccharivorans] |
| MCC2230799.1 | Schlafen+HATPase+wHTH | GCA_020687485.1 | Hominifimenecus_microfluidus | Bacteria>Bacillati>Bacillota>Clostridia>Lachnospirales>Lachnospiraceae>Hominifimenecus | 506 | LKD81_07280 | 20227..21747 | 2885348 | putative-DNA-binding-domain-containing-protein-[Lachnospiraceae-bacterium-CLA-AA-H215] |
| MCC2229912.1 | Schlafen+HATPase+wHTH | GCA_020687485.1 | Hominifimenecus_microfluidus | Bacteria>Bacillati>Bacillota>Clostridia>Lachnospirales>Lachnospiraceae>Hominifimenecus | 490 | LKD81_02690 | 35436..36908 | 2885348 | putative-DNA-binding-domain-containing-protein-[Lachnospiraceae-bacterium-CLA-AA-H215] |
| MCI8726397.1 | Schlafen+HATPase+wHTH | GCA_022794715.1 | Hungatella_sp. | Bacteria>Bacillati>Bacillota>Clostridia>Lachnospirales>Lachnospiraceae>Hungatella>unclassified_Hungatella | 514 | HFG66_09825 | 46445..47989 | 2613924 | ATP-binding-protein-[Hungatella-sp.] |
| WP_279221900.1 | Schlafen+HATPase+wHTH | - | Lachnoanaerobaculum_orale | Bacteria>Bacillati>Bacillota>Clostridia>Lachnospirales>Lachnospiraceae>Lachnoanaerobaculum | 487 | EHW90_RS07140 | 1505845..1507308 | 979627 | putative-DNA-binding-domain-containing-protein-[Lachnoanaerobaculum-orale] |
| WP_278320431.1 | Schlafen+HATPase+wHTH | - | Lachnoanaerobaculum_saburreum | Bacteria>Bacillati>Bacillota>Clostridia>Lachnospirales>Lachnospiraceae>Lachnoanaerobaculum | 483 | HMPREF1866_RS12100 | 16985..18436 | 467210 | putative-DNA-binding-domain-containing-protein-[Lachnoanaerobaculum-saburreum] |
| WP_087163150.1 | Schlafen+HATPase+wHTH | GCF_002160525.1 | Lachnoclostridium_sp._An169 | Bacteria>Bacillati>Bacillota>Clostridia>Lachnospirales>Lachnospiraceae>Lachnoclostridium>unclassified_Lachnoclostridium | 484 | B5F07_RS20930 | 8980..10434 | 1965569 | ATP-binding-protein-[Lachnoclostridium-sp.-An169] |
| WP_118390635.1 | Schlafen+HATPase+wHTH | GCF_003474935.1 | Lachnospira_eligens | Bacteria>Bacillati>Bacillota>Clostridia>Lachnospirales>Lachnospiraceae>Lachnospira | 491 | DWZ79_RS02885 | 269066..270541 | 39485 | putative-DNA-binding-domain-containing-protein-[Lachnospira-eligens] |
| WP_055287911.1 | Schlafen+HATPase+wHTH | GCF_902362785.1 | Lachnospira_eligens | Bacteria>Bacillati>Bacillota>Clostridia>Lachnospirales>Lachnospiraceae>Lachnospira | 492 | FXW70_RS12170 | 160761..162239 | 39485 | putative-DNA-binding-domain-containing-protein-[Lachnospira-eligens] |
| WP_270395913.1 | Schlafen+HATPase+wHTH | GCF_027666095.1 | Mediterraneibacter_massiliensis | Bacteria>Bacillati>Bacillota>Clostridia>Lachnospirales>Lachnospiraceae>Mediterraneibacter | 508 | PGR65_RS11765 | 24216..25742 | 1720300 | ATP-binding-protein-[Mediterraneibacter-massiliensis] |
| MCI7121464.1 | Schlafen+HATPase+wHTH | GCA_022775325.1 | Mediterraneibacter_gnavus | Bacteria>Bacillati>Bacillota>Clostridia>Lachnospirales>Lachnospiraceae>Mediterraneibacter | 488 | MR941_06750 | 23481..24947 | 33038 | putative-DNA-binding-domain-containing-protein-[[Ruminococcus] |
| WP_109627593.1 | Schlafen+HATPase+wHTH | GCF_024622195.1 | Murimonas_intestini | Bacteria>Bacillati>Bacillota>Clostridia>Lachnospirales>Lachnospiraceae>Murimonas | 572 | NSB30_RS07530 | 14292..16010 | 1337051 | ATP-binding-protein-[Murimonas-intestini] |
| EHL12233.1 | Schlafen+HATPase+wHTH | GCA_000238075.1 | Oribacterium_asaccharolyticum_ACB7 | Bacteria>Bacillati>Bacillota>Clostridia>Lachnospirales>Lachnospiraceae>Oribacterium>Oribacterium_asaccharolyticum | 526 | HMPREF9624_00540 | 309852..311432 | 796944 | hypothetical-protein-HMPREF9624_00540-[Oribacterium-asaccharolyticum-ACB7] |
| MBP3802958.1 | Schlafen+HATPase+wHTH | GCA_017940955.1 | Oribacterium_sp. | Bacteria>Bacillati>Bacillota>Clostridia>Lachnospirales>Lachnospiraceae>Oribacterium>unclassified_Oribacterium | 513 | J6I76_03535 | 17461..19002 | 1969407 | putative-DNA-binding-domain-containing-protein-[Oribacterium-sp.] |
| HCS67501.1 | Schlafen+HATPase+wHTH | GCA_003531095.1 | Oribacterium_sp. | Bacteria>Bacillati>Bacillota>Clostridia>Lachnospirales>Lachnospiraceae>Oribacterium>unclassified_Oribacterium | 488 | DIW34_04715 | 2688..4154 | 1969407 | ATP-binding-protein-[Oribacterium-sp.] |
| WP_227249251.1 | Schlafen+HATPase+wHTH | GCF_020557615.1 | Roseburia_faecis | Bacteria>Bacillati>Bacillota>Clostridia>Lachnospirales>Lachnospiraceae>Roseburia | 512 | LIZ84_RS08425 | 36789..38327 | 301302 | ATP-binding-protein-[Roseburia-faecis] |
| WP_195441361.1 | Schlafen+HATPase+wHTH | GCF_015554785.1 | Roseburia_intestinalis | Bacteria>Bacillati>Bacillota>Clostridia>Lachnospirales>Lachnospiraceae>Roseburia | 490 | I2F09_RS06820 | 80525..81997 | 166486 | ATP-binding-protein-[Roseburia-intestinalis] |
| CUO05308.1 | Schlafen+HATPase+wHTH | GCA_001405875.1 | Roseburia_hominis | Bacteria>Bacillati>Bacillota>Clostridia>Lachnospirales>Lachnospiraceae>Roseburia | 515 | ERS852391_01278 | 66061..67608 | 301301 | Divergent-AAA-domain-[Roseburia-hominis] |
| WP_075720908.1 | Schlafen+HATPase+wHTH | GCF_001940225.2 | Roseburia_sp._499 | Bacteria>Bacillati>Bacillota>Clostridia>Lachnospirales>Lachnospiraceae>Roseburia>unclassified_Roseburia | 491 | BIV20_RS10265 | 360860..362335 | 1261634 | putative-DNA-binding-domain-containing-protein-[Roseburia-sp.-499] |
| HCU04408.1 | Schlafen+HATPase+wHTH | GCA_003543095.1 | Roseburia_sp. | Bacteria>Bacillati>Bacillota>Clostridia>Lachnospirales>Lachnospiraceae>Roseburia>unclassified_Roseburia | 515 | DF667_12375 | 35988..37535 | 2049040 | ATP-binding-protein-[Roseburia-sp.] |
| HBA06131.1 | Schlafen+HATPase+wHTH | GCA_003491515.1 | Roseburia_sp. | Bacteria>Bacillati>Bacillota>Clostridia>Lachnospirales>Lachnospiraceae>Roseburia>unclassified_Roseburia | 484 | DC452_03340 | 21443..22897 | 2049040 | ATP-binding-protein-[Roseburia-sp.] |
| WP_181475443.1 | Schlafen+HATPase+wHTH | GCF_013694215.1 | Sellimonas_intestinalis | Bacteria>Bacillati>Bacillota>Clostridia>Lachnospirales>Lachnospiraceae>Sellimonas | 490 | H1A12_RS00270 | 59586..61058 | 1653434 | ATP-binding-protein-[Sellimonas-intestinalis] |
| WP_249326645.1 | Schlafen+HATPase+wHTH | GCF_014337175.1 | Simiaoa_sunii | Bacteria>Bacillati>Bacillota>Clostridia>Lachnospirales>Lachnospiraceae>Simiaoa | 486 | H9Q77_RS04490 | 965385..966845 | 2763672 | ATP-binding-protein-[Simiaoa-sunii] |
| QNM03387.1 | Schlafen+HATPase+wHTH | GCA_014337175.1 | Simiaoa_sunii | Bacteria>Bacillati>Bacillota>Clostridia>Lachnospirales>Lachnospiraceae>Simiaoa | 486 | H9Q77_04495 | 965385..966845 | 2763672 | putative-DNA-binding-domain-containing-protein-[Simiaoa-sunii] |
| MBP3664795.1 | Schlafen+HATPase+wHTH | GCA_017939005.1 | Tyzzerella_sp. | Bacteria>Bacillati>Bacillota>Clostridia>Lachnospirales>Lachnospiraceae>Tyzzerella>unclassified_Tyzzerella | 547 | J6J03_06500 | 2581..4224 | 2053632 | putative-DNA-binding-domain-containing-protein-[Tyzzerella-sp.] |
| SEQ84548.1 | Schlafen+HATPase+wHTH | GCA_900110635.1 | Lachnospiraceae_bacterium_NE2001 | Bacteria>Bacillati>Bacillota>Clostridia>Lachnospirales>Lachnospiraceae>unclassified_Lachnospiraceae | 571 | SAMN02910369_02527 | 32199..33914 | 1520823 | ATP-dependent-DNA-helicase-RecG-[Lachnospiraceae-bacterium-NE2001] |
| SEP83412.1 | Schlafen+HATPase+wHTH | GCA_900110975.1 | Lachnospiraceae_bacterium_RM5 | Bacteria>Bacillati>Bacillota>Clostridia>Lachnospirales>Lachnospiraceae>unclassified_Lachnospiraceae | 582 | SAMN02910289_00628 | 8610..10358 | 1200721 | ATP-dependent-DNA-helicase-RecG-[Lachnospiraceae-bacterium-RM5] |
| RKW57358.1 | Schlafen+HATPase+wHTH | GCA_003640285.1 | Lachnospiraceae_bacterium | Bacteria>Bacillati>Bacillota>Clostridia>Lachnospirales>Lachnospiraceae>unclassified_Lachnospiraceae | 465 | D8H95_03150 | 331..1728 | 1898203 | AAA-family-ATPase-[Lachnospiraceae-bacterium] |
| MCI7595224.1 | Schlafen+HATPase+wHTH | GCA_022786115.1 | Lachnospiraceae_bacterium | Bacteria>Bacillati>Bacillota>Clostridia>Lachnospirales>Lachnospiraceae>unclassified_Lachnospiraceae | 494 | MSS92_03240 | 19536..21020 | 1898203 | putative-DNA-binding-domain-containing-protein-[Lachnospiraceae-bacterium] |
| MCI6977810.1 | Schlafen+HATPase+wHTH | GCA_022775645.1 | Lachnospiraceae_bacterium | Bacteria>Bacillati>Bacillota>Clostridia>Lachnospirales>Lachnospiraceae>unclassified_Lachnospiraceae | 614 | MR930_04165 | 192508..194352 | 1898203 | putative-DNA-binding-domain-containing-protein-[Lachnospiraceae-bacterium] |
| MCH3986387.1 | Schlafen+HATPase+wHTH | GCA_022483185.1 | Lachnospiraceae_bacterium | Bacteria>Bacillati>Bacillota>Clostridia>Lachnospirales>Lachnospiraceae>unclassified_Lachnospiraceae | 517 | LKE69_00335 | 73744..75297 | 1898203 | putative-DNA-binding-domain-containing-protein-[Lachnospiraceae-bacterium] |
| MCD8362905.1 | Schlafen+HATPase+wHTH | GCA_021205385.1 | Lachnospiraceae_bacterium | Bacteria>Bacillati>Bacillota>Clostridia>Lachnospirales>Lachnospiraceae>unclassified_Lachnospiraceae | 455 | LUC98_08100 | 28661..30028 | 1898203 | putative-DNA-binding-domain-containing-protein-[Lachnospiraceae-bacterium] |
| MCD8362903.1 | Schlafen+HATPase+wHTH | GCA_021205385.1 | Lachnospiraceae_bacterium | Bacteria>Bacillati>Bacillota>Clostridia>Lachnospirales>Lachnospiraceae>unclassified_Lachnospiraceae | 545 | LUC98_08090 | 23232..24869 | 1898203 | putative-DNA-binding-domain-containing-protein-[Lachnospiraceae-bacterium] |
| MCD7763961.1 | Schlafen+HATPase+wHTH | GCA_021199655.1 | Lachnospiraceae_bacterium | Bacteria>Bacillati>Bacillota>Clostridia>Lachnospirales>Lachnospiraceae>unclassified_Lachnospiraceae | 328 | LUI14_12365 | 53357..54343 | 1898203 | winged-helix-turn-helix-transcriptional-regulator-[Lachnospiraceae-bacterium] |
| MCD7763960.1 | Schlafen+HATPase+wHTH | GCA_021199655.1 | Lachnospiraceae_bacterium | Bacteria>Bacillati>Bacillota>Clostridia>Lachnospirales>Lachnospiraceae>unclassified_Lachnospiraceae | 61 | LUI14_12360 | 53171..53356 | 1898203 | hypothetical-protein-[Lachnospiraceae-bacterium] |
| MCD7763959.1 | Schlafen+HATPase+wHTH | GCA_021199655.1 | Lachnospiraceae_bacterium | Bacteria>Bacillati>Bacillota>Clostridia>Lachnospirales>Lachnospiraceae>unclassified_Lachnospiraceae | 51 | LUI14_12355 | 53040..53195 | 1898203 | ATP-binding-protein-[Lachnospiraceae-bacterium] |
| MCD7762790.1 | Schlafen+HATPase+wHTH | GCA_021199655.1 | Lachnospiraceae_bacterium | Bacteria>Bacillati>Bacillota>Clostridia>Lachnospirales>Lachnospiraceae>unclassified_Lachnospiraceae | 467 | LUI14_06250 | 11743..13146 | 1898203 | putative-DNA-binding-domain-containing-protein-[Lachnospiraceae-bacterium] |
| MCD7717397.1 | Schlafen+HATPase+wHTH | GCA_021199235.1 | Lachnospiraceae_bacterium | Bacteria>Bacillati>Bacillota>Clostridia>Lachnospirales>Lachnospiraceae>unclassified_Lachnospiraceae | 474 | LUI39_13315 | 24577..26001 | 1898203 | putative-DNA-binding-domain-containing-protein-[Lachnospiraceae-bacterium] |
| MBR4345200.1 | Schlafen+HATPase+wHTH | GCA_017542245.1 | Lachnospiraceae_bacterium | Bacteria>Bacillati>Bacillota>Clostridia>Lachnospirales>Lachnospiraceae>unclassified_Lachnospiraceae | 493 | IKP88_21260 | 68645..70126 | 1898203 | putative-DNA-binding-domain-containing-protein-[Lachnospiraceae-bacterium] |
| MBR3507819.1 | Schlafen+HATPase+wHTH | GCA_017527205.1 | Lachnospiraceae_bacterium | Bacteria>Bacillati>Bacillota>Clostridia>Lachnospirales>Lachnospiraceae>unclassified_Lachnospiraceae | 565 | IKO03_03375 | 48115..49812 | 1898203 | putative-DNA-binding-domain-containing-protein-[Lachnospiraceae-bacterium] |
| MBQ9134966.1 | Schlafen+HATPase+wHTH | GCA_017463605.1 | Lachnospiraceae_bacterium | Bacteria>Bacillati>Bacillota>Clostridia>Lachnospirales>Lachnospiraceae>unclassified_Lachnospiraceae | 379 | IJX66_02585 | 19819..20958 | 1898203 | ATP-binding-protein-[Lachnospiraceae-bacterium] |
| HCX41709.1 | Schlafen+HATPase+wHTH | GCA_003539645.1 | Lachnospiraceae_bacterium | Bacteria>Bacillati>Bacillota>Clostridia>Lachnospirales>Lachnospiraceae>unclassified_Lachnospiraceae | 492 | DHH50_05120 | 188512..189990 | 1898203 | hypothetical-protein-[Lachnospiraceae-bacterium] |
| HCI18469.1 | Schlafen+HATPase+wHTH | GCA_003521585.1 | Lachnospiraceae_bacterium | Bacteria>Bacillati>Bacillota>Clostridia>Lachnospirales>Lachnospiraceae>unclassified_Lachnospiraceae | 492 | DFH32_07630 | 7600..9078 | 1898203 | hypothetical-protein-[Lachnospiraceae-bacterium] |
| GFI28170.1 | Schlafen+HATPase+wHTH | GCA_011959925.1 | Lachnospiraceae_bacterium | Bacteria>Bacillati>Bacillota>Clostridia>Lachnospirales>Lachnospiraceae>unclassified_Lachnospiraceae | 502 | IMSAGC012_03301 | 17215..18723 | 1898203 | hypothetical-protein-IMSAGC012_03301-[Lachnospiraceae-bacterium] |
| EOS37356.1 | Schlafen+HATPase+wHTH | GCA_000403295.2 | Lachnospiraceae_bacterium_M18-1 | Bacteria>Bacillati>Bacillota>Clostridia>Lachnospirales>Lachnospiraceae>unclassified_Lachnospiraceae | 498 | C808_03334 | 113918..115414 | 1235792 | hypothetical-protein-C808_03334-[Lachnospiraceae-bacterium-M18-1] |
| EGN43138.1 | Schlafen+HATPase+wHTH | - | Lachnospiraceae_bacterium_1_1_57FAA | Bacteria>Bacillati>Bacillota>Clostridia>Lachnospirales>Lachnospiraceae>unclassified_Lachnospiraceae | 490 | HMPREF0990_02343 | 38706..40178 | 658081 | hypothetical-protein-HMPREF0990_02343-[Lachnospiraceae-bacterium-1_1_57FAA] |
| EFV19665.1 | Schlafen+HATPase+wHTH | - | Lachnospiraceae_bacterium_8_1_57FAA | Bacteria>Bacillati>Bacillota>Clostridia>Lachnospirales>Lachnospiraceae>unclassified_Lachnospiraceae | 491 | HMPREF1026_01175 | 40881..42356 | 665951 | ATP-dependent-DNA-helicase-[Lachnospiraceae-bacterium-8_1_57FAA] |
| EHO53371.1 | Schlafen+HATPase+wHTH | GCA_000242315.1 | Lachnospiraceae_bacterium_oral_taxon_082_str._F0431 | Bacteria>Bacillati>Bacillota>Clostridia>Lachnospirales>Lachnospiraceae>unclassified_Lachnospiraceae>Lachnospiraceae_bacterium_oral_taxon_082 | 490 | HMPREF9099_00897 | 86092..87564 | 861454 | divergent-AAA-domain-protein-[Lachnospiraceae-bacterium-oral-taxon-082-str.-F0431] |
| WP_118373614.1 | Schlafen+HATPase+wHTH | GCF_014337155.1 | Wujia_chipingensis | Bacteria>Bacillati>Bacillota>Clostridia>Lachnospirales>Lachnospiraceae>Wujia | 500 | H9Q76_RS05880 | 1244053..1245555 | 2763670 | MULTISPECIES:-ATP-binding-protein-[Eubacteriales] |
| WP_073026966.1 | Schlafen+HATPase+wHTH | GCF_900142105.1 | Lutispora_thermophila | Bacteria>Bacillati>Bacillota>Clostridia>Lutisporales>Lutisporaceae>Lutispora | 386 | BUB48_RS14285 | 19722..20882 | 288966 | ATP-binding-protein-[Lutispora-thermophila] |
| WP_156275500.1 | Schlafen+HATPase+wHTH | GCF_009735625.1 | Moorella_glycerini | Bacteria>Bacillati>Bacillota>Clostridia>Moorellales>Moorellaceae>Moorella | 172 | MGLY_RS15900 | 3178069..3178587 | 55779 | ATP-binding-protein-[Moorella-glycerini] |
| WP_013361255.1 | Schlafen+HATPase+wHTH | GCF_000196455.1 | Acetoanaerobium_sticklandii | Bacteria>Bacillati>Bacillota>Clostridia>Peptostreptococcales>Filifactoraceae>Acetoanaerobium | 435 | CLOST_RS05405 | 1188968..1190275 | 1511 | putative-DNA-binding-domain-containing-protein-[Acetoanaerobium-sticklandii] |
| NCB93721.1 | Schlafen+HATPase+wHTH | GCA_009929525.1 | Clostridia_bacterium | Bacteria>Bacillati>Bacillota>Clostridia>unclassified_Clostridia | 470 | EOM40_14360 | 13777..15189 | 2044939 | ATP-binding-protein-[Clostridia-bacterium] |
| MCR5783581.1 | Schlafen+HATPase+wHTH | GCA_024699005.1 | Clostridia_bacterium | Bacteria>Bacillati>Bacillota>Clostridia>unclassified_Clostridia | 415 | K6G90_12710 | 42661..43908 | 2044939 | ATP-binding-protein-[Clostridia-bacterium] |
| MCR5783580.1 | Schlafen+HATPase+wHTH | GCA_024699005.1 | Clostridia_bacterium | Bacteria>Bacillati>Bacillota>Clostridia>unclassified_Clostridia | 487 | K6G90_12705 | 41048..42511 | 2044939 | putative-DNA-binding-domain-containing-protein-[Clostridia-bacterium] |
| MBO5955455.1 | Schlafen+HATPase+wHTH | GCA_017631935.1 | Clostridia_bacterium | Bacteria>Bacillati>Bacillota>Clostridia>unclassified_Clostridia | 518 | J6Q10_01485 | 1908..3464 | 2044939 | putative-DNA-binding-domain-containing-protein-[Clostridia-bacterium] |
| GHU63464.1 | Schlafen+HATPase+wHTH | GCA_025997195.1 | Clostridia_bacterium | Bacteria>Bacillati>Bacillota>Clostridia>unclassified_Clostridia | 524 | AGMMS49983_06970 | 214..1788 | 2044939 | hypothetical-protein-AGMMS49983_06970-[Clostridia-bacterium] |
| WP_167849985.1 | Schlafen+HATPase+wHTH | GCF_900240265.1 | Galactobacillus_timonensis | Bacteria>Bacillati>Bacillota>Erysipelotrichia>Erysipelotrichales>Erysipelotrichaceae>Galactobacillus | 501 | C1714_RS08220 | 1729471..1730976 | 2041840 | putative-DNA-binding-domain-containing-protein-[Galactobacillus-timonensis] |
| WP_102343497.1 | Schlafen+HATPase+wHTH | GCF_900240265.1 | Galactobacillus_timonensis | Bacteria>Bacillati>Bacillota>Erysipelotrichia>Erysipelotrichales>Erysipelotrichaceae>Galactobacillus | 500 | C1714_RS12170 | 358244..359746 | 2041840 | ATP-binding-protein-[Galactobacillus-timonensis] |
| WP_229030590.1 | Schlafen+HATPase+wHTH | GCF_015547685.1 | Holdemanella_biformis | Bacteria>Bacillati>Bacillota>Erysipelotrichia>Erysipelotrichales>Erysipelotrichaceae>Holdemanella | 557 | I1941_RS05770 | 5616..7289 | 1735 | putative-DNA-binding-domain-containing-protein-[Holdemanella-biformis] |
| WP_256268394.1 | Schlafen+HATPase+wHTH | GCF_024460475.1 | Holdemanella_sp._MSK.7.32 | Bacteria>Bacillati>Bacillota>Erysipelotrichia>Erysipelotrichales>Erysipelotrichaceae>Holdemanella>unclassified_Holdemanella | 557 | NE544_RS03845 | 20797..22470 | 2965273 | putative-DNA-binding-domain-containing-protein-[Holdemanella-sp.-MSK.7.32] |
| MCQ4803587.1 | Schlafen+HATPase+wHTH | GCA_024460475.1 | Holdemanella_sp._MSK.7.32 | Bacteria>Bacillati>Bacillota>Erysipelotrichia>Erysipelotrichales>Erysipelotrichaceae>Holdemanella>unclassified_Holdemanella | 557 | NE544_03845 | 20797..22470 | 2965273 | putative-DNA-binding-domain-containing-protein-[Holdemanella-sp.-MSK.7.32] |
| MCH4019938.1 | Schlafen+HATPase+wHTH | GCA_022483385.1 | Erysipelotrichaceae_bacterium | Bacteria>Bacillati>Bacillota>Erysipelotrichia>Erysipelotrichales>Erysipelotrichaceae>unclassified_Erysipelotrichaceae | 513 | LKE61_03585 | 750230..751771 | 2049044 | putative-DNA-binding-domain-containing-protein-[Erysipelotrichaceae-bacterium] |
| MCH4019928.1 | Schlafen+HATPase+wHTH | GCA_022483385.1 | Erysipelotrichaceae_bacterium | Bacteria>Bacillati>Bacillota>Erysipelotrichia>Erysipelotrichales>Erysipelotrichaceae>unclassified_Erysipelotrichaceae | 161 | LKE61_03535 | 735932..736417 | 2049044 | hypothetical-protein-[Erysipelotrichaceae-bacterium] |
| MCH4019927.1 | Schlafen+HATPase+wHTH | GCA_022483385.1 | Erysipelotrichaceae_bacterium | Bacteria>Bacillati>Bacillota>Erysipelotrichia>Erysipelotrichales>Erysipelotrichaceae>unclassified_Erysipelotrichaceae | 214 | LKE61_03530 | 735230..735874 | 2049044 | ATP-binding-protein-[Erysipelotrichaceae-bacterium] |
| HCR67832.1 | Schlafen+HATPase+wHTH | GCA_003533525.1 | Erysipelotrichaceae_bacterium | Bacteria>Bacillati>Bacillota>Erysipelotrichia>Erysipelotrichales>Erysipelotrichaceae>unclassified_Erysipelotrichaceae | 564 | DIV43_01640 | 8063..9757 | 2049044 | transcriptional-regulator-[Erysipelotrichaceae-bacterium] |
| NCB38811.1 | Schlafen+HATPase+wHTH | GCA_009928855.1 | Erysipelotrichia_bacterium | Bacteria>Bacillati>Bacillota>Erysipelotrichia>unclassified_Erysipelotrichia | 215 | EOM80_08580 | 1..645 | 2184014 | ATP-binding-protein-[Erysipelotrichia-bacterium] |

| Protein-ID | Domain | Assembly-ID | Strain | Taxonomy | Length | Locus-Tag | Genomic-coordinates | TaxID | Fasta-defline |
|---|---|---|---|---|---|---|---|---|---|
| MBS5426409.1 | Schlafen+HATPase+wHTH | GCA_018368575.1 | Phascolarctobacterium_succinatutens | Bacteria>Bacillati>Bacillota>Negativicutes>Acidaminococcales>Acidaminococcaceae>Phascolarctobacterium | 500 | KHX39_04780 | 70286..71788 | 626940 | putative-DNA-binding-domain-containing-protein-[Phascolarctobacterium-succinatutens] |
| MBS5426401.1 | Schlafen+HATPase+wHTH | GCA_018368575.1 | Phascolarctobacterium_succinatutens | Bacteria>Bacillati>Bacillota>Negativicutes>Acidaminococcales>Acidaminococcaceae>Phascolarctobacterium | 333 | KHX39_04740 | 57580..58581 | 626940 | putative-DNA-binding-domain-containing-protein-[Phascolarctobacterium-succinatutens] |
| PWM79772.1 | Schlafen+HATPase+wHTH | GCA_003150755.1 | Phascolarctobacterium_sp. | Bacteria>Bacillati>Bacillota>Negativicutes>Acidaminococcales>Acidaminococcaceae>Phascolarctobacterium>unclassified_Phascolarctobacterium | 572 | DBY32_02650 | 555051..556769 | 2049039 | MAG:-transcriptional-regulator-[Phascolarctobacterium-sp.] |
| MBQ7883791.1 | Schlafen+HATPase+wHTH | GCA_017467905.1 | Phascolarctobacterium_sp. | Bacteria>Bacillati>Bacillota>Negativicutes>Acidaminococcales>Acidaminococcaceae>Phascolarctobacterium>unclassified_Phascolarctobacterium | 488 | IJ320_05495 | 22180..23646 | 2049039 | putative-DNA-binding-domain-containing-protein-[Phascolarctobacterium-sp.] |
| MBR6862442.1 | Schlafen+HATPase+wHTH | GCA_017531245.1 | Acidaminococcaceae_bacterium | Bacteria>Bacillati>Bacillota>Negativicutes>Acidaminococcales>Acidaminococcaceae>unclassified_Acidaminococcaceae | 466 | IKM73_14090 | 192158..193558 | 2049052 | putative-DNA-binding-domain-containing-protein-[Acidaminococcaceae-bacterium] |
| WP_013740679.1 | Schlafen+HATPase+wHTH | GCF_000208405.1 | Selenomonas_sputigena | Bacteria>Bacillati>Bacillota>Negativicutes>Selenomonadales>Selenomonadaceae>Selenomonas | 490 | SELSP_RS03630 | 829034..830506 | 69823 | ATP-binding-protein-[Selenomonas-sputigena] |
| EEX76835.1 | Schlafen+HATPase+wHTH | - | Selenomonas_sputigena_ATCC_35185 | Bacteria>Bacillati>Bacillota>Negativicutes>Selenomonadales>Selenomonadaceae>Selenomonas>Selenomonas_sputigena | 482 | SELSPUOL_01792 | 550278..551726 | 546271 | divergent-AAA-domain-protein-[Selenomonas-sputigena-ATCC-35185] |
| NCD09235.1 | Schlafen+HATPase+wHTH | GCA_009930275.1 | Negativicutes_bacterium | Bacteria>Bacillati>Bacillota>Negativicutes>unclassified_Negativicutes | 508 | EOL98_07390 | 6970..8496 | 2499140 | transcriptional-regulator-[Negativicutes-bacterium] |
| WP_248289427.1 | Schlafen+HATPase+wHTH | - | Megasphaera_elsdenii | Bacteria>Bacillati>Bacillota>Negativicutes>Veillonellales>Veillonellaceae>Megasphaera | 488 | HG933_RS10940 | 7781..9247 | 907 | putative-DNA-binding-domain-containing-protein-[Megasphaera-elsdenii] |
| WP_198643526.1 | Schlafen+HATPase+wHTH | GCF_003367905.1 | Megasphaera_stantonii | Bacteria>Bacillati>Bacillota>Negativicutes>Veillonellales>Veillonellaceae>Megasphaera | 159 | DKB62_RS12730 | 2204775..2205254 | 2144175 | ATP-binding-protein-[Megasphaera-stantonii] |
| WP_091652331.1 | Schlafen+HATPase+wHTH | GCF_900103535.1 | Megasphaera_paucivorans | Bacteria>Bacillati>Bacillota>Negativicutes>Veillonellales>Veillonellaceae>Megasphaera | 484 | BLR11_RS11765 | 7303..8757 | 349095 | ATP-binding-protein-[Megasphaera-paucivorans] |
| HJE83012.1 | Schlafen+HATPase+wHTH | GCA_020742115.1 | Megasphaera_stantonii | Bacteria>Bacillati>Bacillota>Negativicutes>Veillonellales>Veillonellaceae>Megasphaera | 503 | K8V17_06230 | 4425..5936 | 2144175 | putative-DNA-binding-domain-containing-protein-[Megasphaera-stantonii] |
| MCH4217196.1 | Schlafen+HATPase+wHTH | GCA_022486845.1 | Megasphaera_sp. | Bacteria>Bacillati>Bacillota>Negativicutes>Veillonellales>Veillonellaceae>Megasphaera>unclassified_Megasphaera | 448 | LKF74_01380 | 15807..17153 | 2023260 | putative-DNA-binding-domain-containing-protein-[Megasphaera-sp.] |
| RHT37486.1 | Schlafen+HATPase+wHTH | GCA_003480425.1 | Firmicutes_bacterium_AM31-12AC | Bacteria>Bacillati>Bacillota>unclassified_Bacillota | 484 | DW790_07835 | 19121..20575 | 2292892 | ATP-binding-protein-[Firmicutes-bacterium-AM31-12AC] |
| RHP99070.1 | Schlafen+HATPase+wHTH | GCA_003478205.1 | Firmicutes_bacterium_AM59-13 | Bacteria>Bacillati>Bacillota>unclassified_Bacillota | 486 | DXA19_01145 | 52777..54237 | 2292897 | winged-helix-turn-helix-transcriptional-regulator-[Firmicutes-bacterium-AM59-13] |
| NPV80541.1 | Schlafen+HATPase+wHTH | GCA_013178415.1 | Bacillota_bacterium | Bacteria>Bacillati>Bacillota>unclassified_Bacillota | 556 | HPY52_09750 | 53503..55173 | 1879010 | transcriptional-regulator-[Bacillota-bacterium] |
| NLG83535.1 | Schlafen+HATPase+wHTH | GCA_012516115.1 | Bacillota_bacterium | Bacteria>Bacillati>Bacillota>unclassified_Bacillota | 544 | GX493_02755 | 7307..8941 | 1879010 | transcriptional-regulator-[Bacillota-bacterium] |
| MCI8646069.1 | Schlafen+HATPase+wHTH | GCA_022794115.1 | Bacillota_bacterium | Bacteria>Bacillati>Bacillota>unclassified_Bacillota | 565 | HFE76_04525 | 2507..4204 | 1879010 | transcriptional-regulator-[Bacillota-bacterium] |
| MCI8284117.1 | Schlafen+HATPase+wHTH | GCA_022790955.1 | Bacillota_bacterium | Bacteria>Bacillati>Bacillota>unclassified_Bacillota | 481 | HFE90_02455 | 67197..68642 | 1879010 | winged-helix-turn-helix-transcriptional-regulator-[Bacillota-bacterium] |
| MBA1336323.1 | Schlafen+HATPase+wHTH | GCA_013540605.1 | Bacillota_bacterium | Bacteria>Bacillati>Bacillota>unclassified_Bacillota | 219 | HPY66_2758 | 60295..60954 | 1879010 | Transcriptional-regulator-[Bacillota-bacterium] |
| MBA1336322.1 | Schlafen+HATPase+wHTH | GCA_013540605.1 | Bacillota_bacterium | Bacteria>Bacillati>Bacillota>unclassified_Bacillota | 190 | HPY66_2757 | 59720..60292 | 1879010 | Transcriptional-regulator-[Bacillota-bacterium] |
| HCT36710.1 | Schlafen+HATPase+wHTH | GCA_003543125.1 | Bacillota_bacterium | Bacteria>Bacillati>Bacillota>unclassified_Bacillota | 535 | DF292_06740 | 32234..33841 | 1879010 | transcriptional-regulator-[Bacillota-bacterium] |
| MCB9452417.1 | Schlafen+HATPase+wHTH | GCA_020634115.1 | Anaerolineaceae_bacterium | Bacteria>Bacillati>Chloroflexota>Anaerolineae>Anaerolineales>Anaerolineaceae>unclassified_Anaerolineaceae | 344 | H6672_13350 | 120617..121651 | 2024896 | ATP-binding-protein-[Anaerolineaceae-bacterium] |
| MBQ6518177.1 | Schlafen+HATPase+wHTH | GCA_017433515.1 | Anaerolineaceae_bacterium | Bacteria>Bacillati>Chloroflexota>Anaerolineae>Anaerolineales>Anaerolineaceae>unclassified_Anaerolineaceae | 552 | IJI14_05610 | 17481..19139 | 2024896 | putative-DNA-binding-domain-containing-protein-[Anaerolineaceae-bacterium] |
| MAS36309.1 | Schlafen+HATPase+wHTH | GCA_002702065.1 | Anaerolineaceae_bacterium | Bacteria>Bacillati>Chloroflexota>Anaerolineae>Anaerolineales>Anaerolineaceae>unclassified_Anaerolineaceae | 312 | CL610_20055 | 43123..44061 | 2024896 | hypothetical-protein-[Anaerolineaceae-bacterium] |
| MCJ7529876.1 | Schlafen+HATPase+wHTH | GCA_022865645.1 | Anaerolineales_bacterium | Bacteria>Bacillati>Chloroflexota>Anaerolineae>Anaerolineales>unclassified_Anaerolineales | 202 | MUO64_02440 | 9119..9727 | 2073117 | ATP-binding-protein-[Anaerolineales-bacterium] |
| MCC6192469.1 | Schlafen+HATPase+wHTH | GCA_020845955.1 | Anaerolineales_bacterium | Bacteria>Bacillati>Chloroflexota>Anaerolineae>Anaerolineales>unclassified_Anaerolineales | 287 | IT318_25865 | 17320..18183 | 2073117 | putative-DNA-binding-domain-containing-protein-[Anaerolineales-bacterium] |
| MBN1136468.1 | Schlafen+HATPase+wHTH | GCA_016926595.1 | Anaerolineae_bacterium | Bacteria>Bacillati>Chloroflexota>Anaerolineae>unclassified_Anaerolineae | 603 | JXM73_07775 | 6740..8551 | 2052143 | putative-DNA-binding-domain-containing-protein-[Anaerolineae-bacterium] |
| MBL7065774.1 | Schlafen+HATPase+wHTH | GCA_016783285.1 | Anaerolineae_bacterium | Bacteria>Bacillati>Chloroflexota>Anaerolineae>unclassified_Anaerolineae | 473 | ISS49_16480 | 8622..10043 | 2052143 | putative-DNA-binding-domain-containing-protein-[Anaerolineae-bacterium] |
| MBL7062827.1 | Schlafen+HATPase+wHTH | GCA_016783285.1 | Anaerolineae_bacterium | Bacteria>Bacillati>Chloroflexota>Anaerolineae>unclassified_Anaerolineae | 564 | ISS49_01300 | 4913..6607 | 2052143 | putative-DNA-binding-domain-containing-protein-[Anaerolineae-bacterium] |
| CAG1014957.1 | Schlafen+HATPase+wHTH | GCA_905339325.1 | Anaerolineae_bacterium | Bacteria>Bacillati>Chloroflexota>Anaerolineae>unclassified_Anaerolineae | 412 | ANRL4_05359 | 9455..10693 | 2052143 | hypothetical-protein-ANRL4_05359-[Anaerolineae-bacterium] |
| MCE7987216.1 | Schlafen+HATPase+wHTH | GCA_021405325.1 | Caldilinea_sp._CFX5 | Bacteria>Bacillati>Chloroflexota>Caldilineae>Caldilineales>Caldilineaceae>Caldilinea>unclassified_Caldilinea | 543 | DYG89_39095 | 112517..114148 | 2293554 | transcriptional-regulator-[Caldilinea-sp.-CFX5] |
| GIV69800.1 | Schlafen+HATPase+wHTH | GCA_026003635.1 | Caldilinea_sp. | Bacteria>Bacillati>Chloroflexota>Caldilineae>Caldilineales>Caldilineaceae>Caldilinea>unclassified_Caldilinea | 172 | KatS3mg048_2662 | 131185..131703 | 2293560 | MAG:-hypothetical-protein-KatS3mg048_2662-[Caldilinea-sp.] |
| MBK9093185.1 | Schlafen+HATPase+wHTH | GCA_016719785.1 | Candidatus_Amarolinea_dominans | Bacteria>Bacillati>Chloroflexota>Caldilineae>Caldilineales>Candidatus_Amarolineaceae>Candidatus_Amarolinea | 402 | IPM84_10450 | 136562..137770 | 3140696 | putative-DNA-binding-domain-containing-protein-[Anaerolineae-bacterium] |
| MBK9093183.1 | Schlafen+HATPase+wHTH | GCA_016719785.1 | Candidatus_Amarolinea_dominans | Bacteria>Bacillati>Chloroflexota>Caldilineae>Caldilineales>Candidatus_Amarolineaceae>Candidatus_Amarolinea | 552 | IPM84_10440 | 134754..136412 | 3140696 | putative-DNA-binding-domain-containing-protein-[Anaerolineae-bacterium] |
| MBK6429949.1 | Schlafen+HATPase+wHTH | GCA_016704685.1 | Candidatus_Amarolinea_dominans | Bacteria>Bacillati>Chloroflexota>Caldilineae>Caldilineales>Candidatus_Amarolineaceae>Candidatus_Amarolinea | 552 | IPF85_04105 | 219667..221325 | 3140696 | putative-DNA-binding-domain-containing-protein-[Anaerolineae-bacterium] |
| MBK6429947.1 | Schlafen+HATPase+wHTH | GCA_016704685.1 | Candidatus_Amarolinea_dominans | Bacteria>Bacillati>Chloroflexota>Caldilineae>Caldilineales>Candidatus_Amarolineaceae>Candidatus_Amarolinea | 402 | IPF85_04095 | 218309..219517 | 3140696 | putative-DNA-binding-domain-containing-protein-[Anaerolineae-bacterium] |
| WP_129628420.1 | Schlafen+HATPase+wHTH | GCF_004138135.1 | Candidatus_Oscillochloris_fontis | Bacteria>Bacillati>Chloroflexota>Chloroflexia>Chloroflexales>Chloroflexineae>Oscillochloridaceae>Oscillochloris | 548 | EKD03_RS05545 | 113585..115231 | 2496868 | putative-DNA-binding-domain-containing-protein-[Candidatus-Oscillochloris-fontis] |
| WP_012120126.1 | Schlafen+HATPase+wHTH | GCF_000017805.1 | Roseiflexus_castenholzii | Bacteria>Bacillati>Chloroflexota>Chloroflexia>Chloroflexales>Roseiflexineae>Roseiflexaceae>Roseiflexus | 386 | RCAS_RS08220 | 2069909..2071069 | 120962 | putative-DNA-binding-domain-containing-protein-[Roseiflexus-castenholzii] |
| MBE0430957.1 | Schlafen+HATPase+wHTH | GCA_014859205.1 | Dehalococcoidia_bacterium | Bacteria>Bacillati>Chloroflexota>Dehalococcoidia>unclassified_Dehalococcoidia | 556 | IBX67_03925 | 11655..13325 | 2026734 | putative-DNA-binding-domain-containing-protein-[Dehalococcoidia-bacterium] |
| HEY77168.1 | Schlafen+HATPase+wHTH | GCA_011192125.1 | Thermoflexia_bacterium | Bacteria>Bacillati>Chloroflexota>Thermoflexia>unclassified_Thermoflexia | 568 | G4O00_13510 | 10565..12271 | 2420334 | AAA-family-ATPase-[Thermoflexia-bacterium] |
| PKO06215.1 | Schlafen+HATPase+wHTH | GCA_002842045.1 | Chloroflexi_bacterium_HGW-Chloroflexi-3 | Bacteria>Bacillati>Chloroflexota>unclassified_Chloroflexota | 623 | CVU41_08915 | 241094..242965 | 2013726 | AAA-family-ATPase-[Chloroflexi-bacterium-HGW-Chloroflexi-3] |
| MDE2823074.1 | Schlafen+HATPase+wHTH | GCA_028874805.1 | Chloroflexota_bacterium | Bacteria>Bacillati>Chloroflexota>unclassified_Chloroflexota | 412 | OXK79_06175 | 66..1304 | 2026724 | MAG:-putative-DNA-binding-domain-containing-protein-[Chloroflexota-bacterium] |
| MCL5961061.1 | Schlafen+HATPase+wHTH | GCA_023489135.1 | Chloroflexota_bacterium | Bacteria>Bacillati>Chloroflexota>unclassified_Chloroflexota | 488 | M1358_17445 | 6846..8312 | 2026724 | putative-DNA-binding-domain-containing-protein-[Chloroflexota-bacterium] |
| MCL5107379.1 | Schlafen+HATPase+wHTH | GCA_023473795.1 | Chloroflexota_bacterium | Bacteria>Bacillati>Chloroflexota>unclassified_Chloroflexota | 344 | M1401_00600 | 23352..24386 | 2026724 | ATP-binding-protein-[Chloroflexota-bacterium] |
| MCL4370921.1 | Schlafen+HATPase+wHTH | GCA_023380955.1 | Chloroflexota_bacterium | Bacteria>Bacillati>Chloroflexota>unclassified_Chloroflexota | 400 | M1380_08450 | 7321..8523 | 2026724 | putative-DNA-binding-domain-containing-protein-[Chloroflexota-bacterium] |
| MCC6176200.1 | Schlafen+HATPase+wHTH | GCA_020845495.1 | Chloroflexota_bacterium | Bacteria>Bacillati>Chloroflexota>unclassified_Chloroflexota | 487 | IT305_12915 | 7430..8893 | 2026724 | putative-DNA-binding-domain-containing-protein-[Chloroflexota-bacterium] |
| MCC6174338.1 | Schlafen+HATPase+wHTH | GCA_020845495.1 | Chloroflexota_bacterium | Bacteria>Bacillati>Chloroflexota>unclassified_Chloroflexota | 77 | IT305_03450 | 193686..193919 | 2026724 | ATP-binding-protein-[Chloroflexota-bacterium] |
| MBI4332980.1 | Schlafen+HATPase+wHTH | GCA_016210565.1 | Chloroflexota_bacterium | Bacteria>Bacillati>Chloroflexota>unclassified_Chloroflexota | 546 | HY673_17070 | 16471..18111 | 2026724 | putative-DNA-binding-domain-containing-protein-[Chloroflexota-bacterium] |
| MBI4322289.1 | Schlafen+HATPase+wHTH | GCA_016210495.1 | Chloroflexota_bacterium | Bacteria>Bacillati>Chloroflexota>unclassified_Chloroflexota | 394 | HY675_27665 | 43988..45172 | 2026724 | putative-DNA-binding-domain-containing-protein-[Chloroflexota-bacterium] |
| HAF11545.1 | Schlafen+HATPase+wHTH | GCA_003446795.1 | Chloroflexota_bacterium | Bacteria>Bacillati>Chloroflexota>unclassified_Chloroflexota | 599 | DCK98_15895 | 10763..12562 | 2026724 | transcriptional-regulator-[Chloroflexota-bacterium] |
| WP_057176915.1 | Schlafen+HATPase+wHTH | GCF_014489415.1 | Cylindrospermopsis | Bacteria>Bacillati>Cyanobacteriota/Melainabacteria_group>Cyanobacteriota>Cyanophyceae>Nostocales>Aphanizomenonaceae | 387 | J9B92_RS13870 | 3107017..3108180 | 77021 | MULTISPECIES:-putative-DNA-binding-domain-containing-protein-[Cylindrospermopsis] |
| WP_240869621.1 | Schlafen+HATPase+wHTH | GCF_022496815.1 | Cylindrospermopsis_raciborskii | Bacteria>Bacillati>Cyanobacteriota/Melainabacteria_group>Cyanobacteriota>Cyanophyceae>Nostocales>Aphanizomenonaceae>Cylindrospermopsis | 387 | GSN00_RS16365 | 89743..90906 | 77022 | putative-DNA-binding-domain-containing-protein-[Cylindrospermopsis-raciborskii] |
| WP_236308181.1 | Schlafen+HATPase+wHTH | GCF_021650815.1 | Cylindrospermopsis_raciborskii | Bacteria>Bacillati>Cyanobacteriota/Melainabacteria_group>Cyanobacteriota>Cyanophyceae>Nostocales>Aphanizomenonaceae>Cylindrospermopsis | 387 | L3I90_RS11080 | 2540711..2541874 | 77022 | putative-DNA-binding-domain-containing-protein-[Cylindrospermopsis-raciborskii] |
| WP_127084956.1 | Schlafen+HATPase+wHTH | GCF_003991905.1 | Dulcicalothrix_desertica | Bacteria>Bacillati>Cyanobacteriota/Melainabacteria_group>Cyanobacteriota>Cyanophyceae>Nostocales>Calotrichaceae>Dulcicalothrix | 471 | DSM106972_RS34650 | 67317..68732 | 32056 | ATP-binding-protein-[Dulcicalothrix-desertica] |
| MBR8836039.1 | Schlafen+HATPase+wHTH | GCA_016903675.1 | Stigonema_ocellatum_SAG_48.90_=_DSM_106950 | Bacteria>Bacillati>Cyanobacteriota/Melainabacteria_group>Cyanobacteriota>Cyanophyceae>Nostocales>Stigonemataceae>Stigonema>Stigonema_ocellatum | 163 | DSM106950_18950 | 36438..36929 | 221294 | hypothetical-protein-[Stigonema-ocellatum-SAG-48.90-=-DSM-106950] |
| MBD2549132.1 | Schlafen+HATPase+wHTH | GCA_014698445.1 | Microcystis_elabens_FACHB-917 | Bacteria>Bacillati>Cyanobacteriota/Melainabacteria_group>Cyanobacteriota>Cyanophyceae>Oscillatoriophycideae>Chroococcales>Microcystaceae>Microcystis>Microcystis_elabens | 458 | H6G65_05955 | 18587..19963 | 2692898 | ArsR-family-transcriptional-regulator-[Microcystis-elabens-FACHB-917] |
| MBD2549131.1 | Schlafen+HATPase+wHTH | GCA_014698445.1 | Microcystis_elabens_FACHB-917 | Bacteria>Bacillati>Cyanobacteriota/Melainabacteria_group>Cyanobacteriota>Cyanophyceae>Oscillatoriophycideae>Chroococcales>Microcystaceae>Microcystis>Microcystis_elabens | 118 | H6G65_05950 | 18292..18648 | 2692898 | ATP-binding-protein-[Microcystis-elabens-FACHB-917] |
| MBD2549126.1 | Schlafen+HATPase+wHTH | GCA_014698445.1 | Microcystis_elabens_FACHB-917 | Bacteria>Bacillati>Cyanobacteriota/Melainabacteria_group>Cyanobacteriota>Cyanophyceae>Oscillatoriophycideae>Chroococcales>Microcystaceae>Microcystis>Microcystis_elabens | 632 | H6G65_05925 | 10452..12350 | 2692898 | putative-DNA-binding-domain-containing-protein-[Microcystis-elabens-FACHB-917] |
| WP_261354022.1 | Schlafen+HATPase+wHTH | - | Cyanobium_sp._L1E-Cus | Bacteria>Bacillati>Cyanobacteriota/Melainabacteria_group>Cyanobacteriota>Cyanophyceae>Synechococcales>Prochlorococcaceae>Cyanobium>unclassified_Cyanobium | 558 | KBZ22_RS09735 | 46632..48308 | 2823714 | putative-DNA-binding-domain-containing-protein-[Cyanobium-sp.-L1E-Cus] |
| WP_255021235.1 | Schlafen+HATPase+wHTH | GCF_024345755.1 | Cyanobium_sp._L1E-Cus | Bacteria>Bacillati>Cyanobacteriota/Melainabacteria_group>Cyanobacteriota>Cyanophyceae>Synechococcales>Prochlorococcaceae>Cyanobium>unclassified_Cyanobium | 485 | KBZ22_RS04725 | 35691..37148 | 2823714 | helix-turn-helix-domain-containing-protein-[Cyanobium-sp.-L1E-Cus] |
| WP_254995748.1 | Schlafen+HATPase+wHTH | GCF_024346415.1 | Cyanobium_sp._Aljojuca_7D2 | Bacteria>Bacillati>Cyanobacteriota/Melainabacteria_group>Cyanobacteriota>Cyanophyceae>Synechococcales>Prochlorococcaceae>Cyanobium>unclassified_Cyanobium | 478 | KBY57_RS10255 | 27502..28938 | 2823698 | helix-turn-helix-domain-containing-protein-[Cyanobium-sp.-Aljojuca-7D2] |
| WP_255118715.1 | Schlafen+HATPase+wHTH | GCF_024345775.1 | unclassified_Synechococcus | Bacteria>Bacillati>Cyanobacteriota/Melainabacteria_group>Cyanobacteriota>Cyanophyceae>Synechococcales>Synechococcaceae>Synechococcus | 622 | KBY80_RS12345 | 10062..11930 | 2626047 | MULTISPECIES:-ATP-binding-protein-[unclassified-Synechococcus] |
| WP_255112872.1 | Schlafen+HATPase+wHTH | GCF_024345735.1 | Synechococcus_lacustris | Bacteria>Bacillati>Cyanobacteriota/Melainabacteria_group>Cyanobacteriota>Cyanophyceae>Synechococcales>Synechococcaceae>Synechococcus | 590 | KBY64_RS06865 | 17388..19160 | 2116544 | ATP-binding-protein-[Synechococcus-lacustris] |
| WP_197211781.1 | Schlafen+HATPase+wHTH | GCF_015840715.1 | Synechococcus_sp._CBW1004 | Bacteria>Bacillati>Cyanobacteriota/Melainabacteria_group>Cyanobacteriota>Cyanophyceae>Synechococcales>Synechococcaceae>Synechococcus>unclassified_Synechococcus | 558 | H8F25_RS01955 | 418559..420235 | 1353136 | ATP-binding-protein-[Synechococcus-sp.-CBW1004] |
| WP_186514741.1 | Schlafen+HATPase+wHTH | GCF_014280115.1 | Synechococcus_sp._A15-44 | Bacteria>Bacillati>Cyanobacteriota/Melainabacteria_group>Cyanobacteriota>Cyanophyceae>Synechococcales>Synechococcaceae>Synechococcus>unclassified_Synechococcus | 556 | SynA1544_RS02770 | 509713..511383 | 1050646 | putative-DNA-binding-domain-containing-protein-[Synechococcus-sp.-A15-44] |
| MCK9492971.1 | Schlafen+HATPase+wHTH | GCA_023228755.1 | Acholeplasmataceae_bacterium | Bacteria>Bacillati>Mycoplasmatota>Mollicutes>Acholeplasmatales>Acholeplasmataceae>unclassified_Acholeplasmataceae | 564 | MOQ00_01240 | 11714..13408 | 1898209 | putative-DNA-binding-domain-containing-protein-[Acholeplasmataceae-bacterium] |
| MBN2056797.1 | Schlafen+HATPase+wHTH | GCA_016932295.1 | candidate_division_CSSED10-310_bacterium | Bacteria>Bacteria_incertae_sedis>Bacteria_candidate_phyla>candidate_division_CSSED10-310 | 513 | JW905_17850 | 7543..9084 | 2855610 | putative-DNA-binding-domain-containing-protein-[bacterium] |
| MBN2055104.1 | Schlafen+HATPase+wHTH | GCA_016932295.1 | candidate_division_CSSED10-310_bacterium | Bacteria>Bacteria_incertae_sedis>Bacteria_candidate_phyla>candidate_division_CSSED10-310 | 296 | JW905_09285 | 12211..13101 | 2855610 | hypothetical-protein-[bacterium] |
| MCX6842106.1 | Schlafen+HATPase+wHTH | GCA_026397595.1 | candidate_division_WOR-3_bacterium | Bacteria>Bacteria_incertae_sedis>Bacteria_candidate_phyla>candidate_division_WOR-3 | 394 | NTX53_07500 | 6801..7985 | 2052148 | MAG:-putative-DNA-binding-domain-containing-protein-[candidate-division-WOR-3-bacterium] |
| OQB57506.1 | Schlafen+HATPase+wHTH | GCA_002070465.1 | Candidatus_Aminicenantes_bacterium_ADurb.Bin147 | Bacteria>Bacteria_incertae_sedis>Bacteria_candidate_phyla>Candidatus_Aminicenantota>unclassified_Candidatus_Aminicenantes | 546 | BWX98_01439 | 12325..13965 | 1852831 | Divergent-AAA-domain-protein-[Candidatus-Aminicenantes-bacterium-ADurb.Bin147] |
| MCX6557424.1 | Schlafen+HATPase+wHTH | GCA_026393795.1 | Candidatus_Aminicenantes_bacterium | Bacteria>Bacteria_incertae_sedis>Bacteria_candidate_phyla>Candidatus_Aminicenantota>unclassified_Candidatus_Aminicenantes | 448 | NTW95_08370 | 4688..6034 | 2052149 | MAG:-ATP-binding-protein-[Candidatus-Aminicenantes-bacterium] |
| MCD6192838.1 | Schlafen+HATPase+wHTH | GCA_021157535.1 | Candidatus_Aminicenantes_bacterium | Bacteria>Bacteria_incertae_sedis>Bacteria_candidate_phyla>Candidatus_Aminicenantota>unclassified_Candidatus_Aminicenantes | 486 | J7L26_05135 | 27036..28496 | 2052149 | putative-DNA-binding-domain-containing-protein-[Candidatus-Aminicenantes-bacterium] |
| MDD5263472.1 | Schlafen+HATPase+wHTH | GCA_028713635.1 | Candidatus_Bipolaricaulis_sp. | Bacteria>Bacteria_incertae_sedis>Bacteria_candidate_phyla>Candidatus_Bipolaricaulota>Candidatus_Bipolaricaulia>Candidatus_Bipolaricaulales>Candidatus_Bipolaricaulaceae>Candidatus_Bipolaricaulis>unclassified_Candidatus_Bipolaricaulis | 552 | PHU43_01375 | 39040..40698 | 2814139 | MAG:-ATP-binding-protein-[Candidatus-Bipolaricaulis-sp.] |
| MCK4303760.1 | Schlafen+HATPase+wHTH | GCA_023128935.1 | Candidatus_Eiseniibacteriota_bacterium | Bacteria>Bacteria_incertae_sedis>Bacteria_candidate_phyla>Candidatus_Eiseniibacteriota | 547 | KAY24_05945 | 8461..10104 | 2212470 | putative-DNA-binding-domain-containing-protein-[Candidatus-Eisenbacteria-sp.] |
| MCC6649977.1 | Schlafen+HATPase+wHTH | GCA_020849095.1 | Candidatus_Eiseniibacteriota_bacterium | Bacteria>Bacteria_incertae_sedis>Bacteria_candidate_phyla>Candidatus_Eiseniibacteriota | 417 | IT348_02375 | 4271..5524 | 2212470 | putative-DNA-binding-domain-containing-protein-[Candidatus-Eisenbacteria-bacterium] |
| WP_259099625.1 | Schlafen+HATPase+wHTH | GCF_030520105.1 | Candidatus_Fervidibacter_sacchari | Bacteria>Bacteria_incertae_sedis>Bacteria_candidate_phyla>Candidatus_Fervidibacterota>Candidatus_Fervidibacter | 565 | M2350_RS13980 | 278135..279832 | 1448929 | ATP-binding-protein-[Candidatus-Fervidibacter-sacchari] |
| PWB47244.1 | Schlafen+HATPase+wHTH | GCA_003104975.1 | Candidatus_Methylomirabilota_bacterium | Bacteria>Bacteria_incertae_sedis>Bacteria_candidate_phyla>Candidatus_Methylomirabilota>unclassified_Candidatus_Methylomirabilota | 537 | C3F12_04510 | 179420..181033 | 2072417 | transcriptional-regulator-[candidate-division-NC10-bacterium] |
| MCD6507088.1 | Schlafen+HATPase+wHTH | GCA_021162805.1 | Candidatus_Poribacteria_bacterium | Bacteria>Bacteria_incertae_sedis>Bacteria_candidate_phyla>Candidatus_Poribacteria>unclassified_Candidatus_Poribacteria | 98 | J7M22_10750 | 21956..22252 | 2026781 | ATP-binding-protein-[Candidatus-Poribacteria-bacterium] |
| MCD6506889.1 | Schlafen+HATPase+wHTH | GCA_021162805.1 | Candidatus_Poribacteria_bacterium | Bacteria>Bacteria_incertae_sedis>Bacteria_candidate_phyla>Candidatus_Poribacteria>unclassified_Candidatus_Poribacteria | 458 | J7M22_09735 | 2383..3759 | 2026781 | putative-DNA-binding-domain-containing-protein-[Candidatus-Poribacteria-bacterium] |
| PKL50473.1 | Schlafen+HATPase+wHTH | GCA_002839435.1 | Candidatus_Riflebacteria_bacterium_HGW-Riflebacteria-2 | Bacteria>Bacteria_incertae_sedis>Bacteria_candidate_phyla>Candidatus_Rifleibacteriota>unclassified_Candidatus_Rifleibacteriota | 557 | CVV42_02965 | 66504..68177 | 2013830 | MAG:-ATPase-[Candidatus-Riflebacteria-bacterium-HGW-Riflebacteria-2] |
| OGL14731.1 | Schlafen+HATPase+wHTH | GCA_001790335.1 | Candidatus_Rokubacteria_bacterium_RIFCSPLOWO2_12_FULL_71_22 | Bacteria>Bacteria_incertae_sedis>Bacteria_candidate_phyla>Candidatus_Rokuibacteriota>unclassified_Candidatus_Rokuibacteriota | 411 | A3F92_03750 | 15936..17171 | 1802110 | MAG:-hypothetical-protein-A3F92_03750-[Candidatus-Rokubacteria-bacterium-RIFCSPLOWO2_12_FULL_71_22] |
| MBI2216637.1 | Schlafen+HATPase+wHTH | GCA_016184125.1 | Candidatus_Rokuibacteriota_bacterium | Bacteria>Bacteria_incertae_sedis>Bacteria_candidate_phyla>Candidatus_Rokuibacteriota>unclassified_Candidatus_Rokuibacteriota | 417 | HYU51_05005 | 38416..39669 | 2053607 | putative-DNA-binding-domain-containing-protein-[Candidatus-Rokubacteria-bacterium] |
| MCX7047274.1 | Schlafen+HATPase+wHTH | GCA_026398575.1 | Candidatus_Sumerlaeota_bacterium | Bacteria>Bacteria_incertae_sedis>Bacteria_candidate_phyla>Candidatus_Sumerlaeota>unclassified_Candidatus_Sumerlaeota | 281 | NTX50_17530 | 14320..15165 | 2107588 | MAG:-hypothetical-protein-NTX50_17530-[Candidatus-Sumerlaeota-bacterium] |
| MCX7047271.1 | Schlafen+HATPase+wHTH | GCA_026398575.1 | Candidatus_Sumerlaeota_bacterium | Bacteria>Bacteria_incertae_sedis>Bacteria_candidate_phyla>Candidatus_Sumerlaeota>unclassified_Candidatus_Sumerlaeota | 442 | NTX50_17515 | 9168..10496 | 2107588 | MAG:-putative-DNA-binding-domain-containing-protein-[Candidatus-Sumerlaeota-bacterium] |
| MBN1883900.1 | Schlafen+HATPase+wHTH | GCA_016931295.1 | Candidatus_Krumholzibacteriota_bacterium | Bacteria>Candidatus_Krumholzibacteriota>unclassified_Candidatus_Krumholzibacteriota | 554 | JW876_00080 | 22290..23954 | 2725313 | putative-DNA-binding-domain-containing-protein-[Candidatus-Krumholzibacteriota-bacterium] |
| MCD6532981.1 | Schlafen+HATPase+wHTH | GCA_021163445.1 | Candidatus_Tharpella_aukensis | Bacteria>Candidatus_Tharpellota>Candidatus_Tharpellia>Candidatus_Tharpellales>Candidatus_Tharpellaceae>Candidatus_Tharpella | 413 | J7L25_02735 | 14394..15635 | 2954148 | putative-DNA-binding-domain-containing-protein-[Deltaproteobacteria-bacterium] |
| MCD6532977.1 | Schlafen+HATPase+wHTH | GCA_021163445.1 | Candidatus_Tharpella_aukensis | Bacteria>Candidatus_Tharpellota>Candidatus_Tharpellia>Candidatus_Tharpellales>Candidatus_Tharpellaceae>Candidatus_Tharpella | 417 | J7L25_02715 | 9829..11082 | 2954148 | putative-DNA-binding-domain-containing-protein-[Deltaproteobacteria-bacterium] |
| MBN1944773.1 | Schlafen+HATPase+wHTH | GCA_016931535.1 | Bradymonadales_bacterium | Bacteria>delta/epsilon_subdivisions>Deltaproteobacteria>Bradymonadales>unclassified_Bradymonadales | 571 | JW797_03810 | 2913..4628 | 2099667 | putative-DNA-binding-domain-containing-protein-[Bradymonadales-bacterium] |
| WP_237889220.1 | Schlafen+HATPase+wHTH | GCF_022026155.1 | Dissulfurimicrobium_hydrothermale | Bacteria>delta/epsilon_subdivisions>Deltaproteobacteria>Deltaproteobacteria_incertae_sedis>Dissulfurimicrobium | 558 | LGS26_RS03270 | 699756..701432 | 1750598 | putative-DNA-binding-domain-containing-protein-[Dissulfurimicrobium-hydrothermale] |
| PKN28383.1 | Schlafen+HATPase+wHTH | GCA_002840495.1 | Deltaproteobacteria_bacterium_HGW-Deltaproteobacteria-21 | Bacteria>delta/epsilon_subdivisions>Deltaproteobacteria>unclassified_Deltaproteobacteria | 569 | CVU64_13610 | 11985..13694 | 2013749 | AAA-family-ATPase-[Deltaproteobacteria-bacterium-HGW-Deltaproteobacteria-21] |
| PJB39409.1 | Schlafen+HATPase+wHTH | GCA_002789955.1 | Deltaproteobacteria_bacterium_CG_4_9_14_3_um_filter_63_12 | Bacteria>delta/epsilon_subdivisions>Deltaproteobacteria>unclassified_Deltaproteobacteria | 488 | CO108_17310 | 5719..7185 | 1973967 | transcriptional-regulator-[Deltaproteobacteria-bacterium-CG_4_9_14_3_um_filter_63_12] |
| PIX22659.1 | Schlafen+HATPase+wHTH | GCA_002783725.1 | Deltaproteobacteria_bacterium_CG_4_8_14_3_um_filter_45_9 | Bacteria>delta/epsilon_subdivisions>Deltaproteobacteria>unclassified_Deltaproteobacteria | 572 | COZ69_10540 | 1325..3043 | 1973962 | AAA-family-ATPase-[Deltaproteobacteria-bacterium-CG_4_8_14_3_um_filter_45_9] |
| PIV21311.1 | Schlafen+HATPase+wHTH | GCA_002780715.1 | Deltaproteobacteria_bacterium_CG03_land_8_20_14_0_80_45_14 | Bacteria>delta/epsilon_subdivisions>Deltaproteobacteria>unclassified_Deltaproteobacteria | 572 | COS40_08015 | 8064..9782 | 1973969 | AAA-family-ATPase-[Deltaproteobacteria-bacterium-CG03_land_8_20_14_0_80_45_14] |
| PIV21171.1 | Schlafen+HATPase+wHTH | GCA_002780715.1 | Deltaproteobacteria_bacterium_CG03_land_8_20_14_0_80_45_14 | Bacteria>delta/epsilon_subdivisions>Deltaproteobacteria>unclassified_Deltaproteobacteria | 549 | COS40_08735 | 5247..6896 | 1973969 | ATPase-[Deltaproteobacteria-bacterium-CG03_land_8_20_14_0_80_45_14] |
| OGR05788.1 | Schlafen+HATPase+wHTH | GCA_001799225.1 | Deltaproteobacteria_bacterium_RIFOXYD12_FULL_50_9 | Bacteria>delta/epsilon_subdivisions>Deltaproteobacteria>unclassified_Deltaproteobacteria | 377 | A2511_04575 | 5874..7004 | 1797902 | transcriptional-regulator,-partial-[Deltaproteobacteria-bacterium-RIFOXYD12_FULL_50_9] |
| OGR04293.1 | Schlafen+HATPase+wHTH | GCA_001799195.1 | Deltaproteobacteria_bacterium_RIFOXYA12_FULL_61_11 | Bacteria>delta/epsilon_subdivisions>Deltaproteobacteria>unclassified_Deltaproteobacteria | 419 | A2284_01890 | 2151..3410 | 1797896 | transcriptional-regulator-[Deltaproteobacteria-bacterium-RIFOXYA12_FULL_61_11] |
| OGQ88898.1 | Schlafen+HATPase+wHTH | GCA_001799285.1 | Deltaproteobacteria_bacterium_RIFOXYD12_FULL_56_24 | Bacteria>delta/epsilon_subdivisions>Deltaproteobacteria>unclassified_Deltaproteobacteria | 502 | A2512_08465 | 5480..6988 | 1797905 | AAA-family-ATPase-[Deltaproteobacteria-bacterium-RIFOXYD12_FULL_56_24] |
| OGP84698.1 | Schlafen+HATPase+wHTH | GCA_001797945.1 | Deltaproteobacteria_bacterium_RBG_16_58_17 | Bacteria>delta/epsilon_subdivisions>Deltaproteobacteria>unclassified_Deltaproteobacteria | 557 | A2V87_08865 | 22473..24146 | 1797845 | ATPase-[Deltaproteobacteria-bacterium-RBG_16_58_17] |
| EFK08780.1 | Schlafen+HATPase+wHTH | - | delta_proteobacterium_NaphS2 | Bacteria>delta/epsilon_subdivisions>Deltaproteobacteria>unclassified_Deltaproteobacteria | 511 | NPH_0524 | 25396..26931 | 88274 | divergent-AAA-domain-protein-[delta-proteobacterium-NaphS2] |
| CAG1064744.1 | Schlafen+HATPase+wHTH | GCA_905339405.1 | uncultured_bacterium | Bacteria>environmental_samples | 556 | BAC1_00309 | 131134..132804 | 77133 | hypothetical-protein-BAC1_00309-[uncultured-bacterium] |
| AGC72567.1 | Schlafen+HATPase+wHTH | - | uncultured_bacterium_A1Q1_fos_1870 | Bacteria>environmental_samples | 455 | . | 11516..12883 | 1256554 | ATP-dependent-DNA-helicase-[uncultured-bacterium-A1Q1_fos_1870] |
| WP_238972696.1 | Schlafen+HATPase+wHTH | GCF_037911825.1 | Fusobacterium | Bacteria>Fusobacteriati>Fusobacteriota>Fusobacteriia>Fusobacteriales>Fusobacteriaceae | 513 | K5I19_RS02520 | 133879..135420 | 848 | helix-turn-helix-domain-containing-protein-[Fusobacterium-nucleatum] |
| WP_032838657.1 | Schlafen+HATPase+wHTH | GCF_037911025.1 | Fusobacterium | Bacteria>Fusobacteriati>Fusobacteriota>Fusobacteriia>Fusobacteriales>Fusobacteriaceae | 517 | HMPREF1498_RS02600 | 36410..37963 | 848 | helix-turn-helix-domain-containing-protein-[Fusobacterium-sp.-CM1] |
| WP_032836034.1 | Schlafen+HATPase+wHTH | GCF_045159995.1 | Fusobacterium | Bacteria>Fusobacteriati>Fusobacteriota>Fusobacteriia>Fusobacteriales>Fusobacteriaceae | 516 | HMPREF1501_RS01115 | 59747..61297 | 848 | helix-turn-helix-domain-containing-protein-[Fusobacterium-sp.-OBRC1] |
| WP_274179668.1 | Schlafen+HATPase+wHTH | GCF_037904935.1 | Fusobacterium_polymorphum | Bacteria>Fusobacteriati>Fusobacteriota>Fusobacteriia>Fusobacteriales>Fusobacteriaceae>Fusobacterium | 443 | PSC67_RS07235 | 1403877..1405208 | 76857 | putative-DNA-binding-domain-containing-protein-[Fusobacterium-nucleatum] |
| WP_274179667.1 | Schlafen+HATPase+wHTH | - | Fusobacterium_nucleatum | Bacteria>Fusobacteriati>Fusobacteriota>Fusobacteriia>Fusobacteriales>Fusobacteriaceae>Fusobacterium | 294 | PSC67_RS07230 | 1402968..1403852 | 851 | putative-DNA-binding-domain-containing-protein-[Fusobacterium-nucleatum] |
| WP_274179666.1 | Schlafen+HATPase+wHTH | - | Fusobacterium_nucleatum | Bacteria>Fusobacteriati>Fusobacteriota>Fusobacteriia>Fusobacteriales>Fusobacteriaceae>Fusobacterium | 228 | PSC67_RS07225 | 1402300..1402986 | 851 | ATP-binding-protein-[Fusobacterium-nucleatum] |
| WP_271333591.1 | Schlafen+HATPase+wHTH | - | Fusobacterium_nucleatum | Bacteria>Fusobacteriati>Fusobacteriota>Fusobacteriia>Fusobacteriales>Fusobacteriaceae>Fusobacterium | 443 | PGW91_RS03585 | 740526..741857 | 851 | putative-DNA-binding-domain-containing-protein-[Fusobacterium-nucleatum] |
| WP_220302804.1 | Schlafen+HATPase+wHTH | GCF_019552045.1 | Fusobacterium_polymorphum | Bacteria>Fusobacteriati>Fusobacteriota>Fusobacteriia>Fusobacteriales>Fusobacteriaceae>Fusobacterium | 521 | JY397_RS08315 | 1737405..1738970 | 76857 | helix-turn-helix-domain-containing-protein-[Fusobacterium-polymorphum] |
| WP_220297606.1 | Schlafen+HATPase+wHTH | GCF_019552085.1 | Fusobacterium_animalis | Bacteria>Fusobacteriati>Fusobacteriota>Fusobacteriia>Fusobacteriales>Fusobacteriaceae>Fusobacterium | 516 | JY398_RS09450 | 1982061..1983611 | 76859 | helix-turn-helix-domain-containing-protein-[Fusobacterium-animalis] |
| WP_220282654.1 | Schlafen+HATPase+wHTH | GCF_019552065.1 | Fusobacterium_polymorphum | Bacteria>Fusobacteriati>Fusobacteriota>Fusobacteriia>Fusobacteriales>Fusobacteriaceae>Fusobacterium | 516 | JY402_RS06725 | 1355065..1356615 | 76857 | helix-turn-helix-domain-containing-protein-[Fusobacterium-polymorphum] |
| WP_158411767.1 | Schlafen+HATPase+wHTH | GCF_037905245.1 | Fusobacterium_animalis | Bacteria>Fusobacteriati>Fusobacteriota>Fusobacteriia>Fusobacteriales>Fusobacteriaceae>Fusobacterium | 521 | RN90_RS06270 | 1294486..1296051 | 76859 | helix-turn-helix-domain-containing-protein-[Fusobacterium-animalis] |

| Protein-ID | Domain | Assembly-ID | Strain | Taxonomy | Length | Locus-Tag | Genomic-coordinates | TaxID | Fasta-defline |
|---|---|---|---|---|---|---|---|---|---|
| WP_158411766.1 | Schlafen+HATPase+wHTH | GCF_019552085.1 | Fusobacterium_animalis | Bacteria>Fusobacteriati>Fusobacteriota>Fusobacteriia>Fusobacteriales>Fusobacteriaceae>Fusobacterium | 443 | JY398_RS09455 | 1983636..1984967 | 76859 | putative-DNA-binding-domain-containing-protein-[Fusobacterium-animalis] |
| WP_099989145.1 | Schlafen+HATPase+wHTH | GCF_002763735.1 | Fusobacterium_pseudoperiodonticum | Bacteria>Fusobacteriati>Fusobacteriota>Fusobacteriia>Fusobacteriales>Fusobacteriaceae>Fusobacterium | 516 | CTM78_RS04800 | 950893..952443 | 2663009 | helix-turn-helix-domain-containing-protein-[Fusobacterium-pseudoperiodonticum] |
| WP_098997102.1 | Schlafen+HATPase+wHTH | GCF_037905975.1 | Fusobacterium_polymorphum | Bacteria>Fusobacteriati>Fusobacteriota>Fusobacteriia>Fusobacteriales>Fusobacteriaceae>Fusobacterium | 443 | CBG56_RS03305 | 138232..139563 | 76857 | MULTISPECIES:-putative-DNA-binding-domain-containing-protein-[Fusobacterium] |
| WP_098997101.1 | Schlafen+HATPase+wHTH | GCF_037905975.1 | Fusobacterium_polymorphum | Bacteria>Fusobacteriati>Fusobacteriota>Fusobacteriia>Fusobacteriales>Fusobacteriaceae>Fusobacterium | 516 | CBG56_RS03300 | 136657..138207 | 76857 | MULTISPECIES:-helix-turn-helix-domain-containing-protein-[Fusobacterium] |
| WP_035500770.1 | Schlafen+HATPase+wHTH | GCF_902373895.1 | Fusobacterium_gonidiaformans | Bacteria>Fusobacteriati>Fusobacteriota>Fusobacteriia>Fusobacteriales>Fusobacteriaceae>Fusobacterium | 442 | FY288_RS06320 | 246401..247729 | 849 | ATP-binding-protein-[Fusobacterium-gonidiaformans] |
| WP_029493950.1 | Schlafen+HATPase+wHTH | GCF_000455885.1 | Fusobacterium_hwasookii | Bacteria>Fusobacteriati>Fusobacteriota>Fusobacteriia>Fusobacteriales>Fusobacteriaceae>Fusobacterium | 514 | J143_RS0110845 | 66677..68221 | 1583098 | helix-turn-helix-domain-containing-protein-[Fusobacterium-hwasookii] |
| WP_008793541.1 | Schlafen+HATPase+wHTH | GCF_902373915.1 | Fusobacterium_periodonticum | Bacteria>Fusobacteriati>Fusobacteriota>Fusobacteriia>Fusobacteriales>Fusobacteriaceae>Fusobacterium | 443 | FY302_RS02990 | 623301..624632 | 860 | putative-DNA-binding-domain-containing-protein-[Fusobacterium-periodonticum] |
| WP_005903406.1 | Schlafen+HATPase+wHTH | GCF_000178895.1 | Fusobacterium_nucleatum | Bacteria>Fusobacteriati>Fusobacteriota>Fusobacteriia>Fusobacteriales>Fusobacteriaceae>Fusobacterium | 443 | C4N14_RS08060 | 1661659..1662990 | 851 | putative-DNA-binding-domain-containing-protein-[Fusobacterium-nucleatum] |
| MCG6839744.1 | Schlafen+HATPase+wHTH | GCA_022340045.1 | Fusobacterium_nucleatum | Bacteria>Fusobacteriati>Fusobacteriota>Fusobacteriia>Fusobacteriales>Fusobacteriaceae>Fusobacterium | 518 | K5I20_00275 | 63703..65259 | 851 | helix-turn-helix-domain-containing-protein-[Fusobacterium-nucleatum] |
| ALF25295.1 | Schlafen+HATPase+wHTH | GCA_001296185.1 | Fusobacterium_nucleatum_subsp._nucleatum | Bacteria>Fusobacteriati>Fusobacteriota>Fusobacteriia>Fusobacteriales>Fusobacteriaceae>Fusobacterium>Fusobacterium_nucleatum | 443 | RN95_02165 | 461111..462442 | 76856 | AAA-family-ATPase-[Fusobacterium-nucleatum-subsp.-nucleatum] |
| WP_070766195.1 | Schlafen+HATPase+wHTH | GCF_001815715.1 | Fusobacterium_sp._HMSC064B11 | Bacteria>Fusobacteriati>Fusobacteriota>Fusobacteriia>Fusobacteriales>Fusobacteriaceae>Fusobacterium>unclassified_Fusobacterium | 516 | HMPREF3051_RS04315 | 257771..259321 | 1739543 | helix-turn-helix-domain-containing-protein-[Fusobacterium-sp.-HMSC064B11] |
| WP_032846246.1 | Schlafen+HATPase+wHTH | GCF_000524395.1 | Fusobacterium_sp._CM22 | Bacteria>Fusobacteriati>Fusobacteriota>Fusobacteriia>Fusobacteriales>Fusobacteriaceae>Fusobacterium>unclassified_Fusobacterium | 516 | HMPREF1500_RS09205 | 120696..122246 | 936563 | helix-turn-helix-domain-containing-protein-[Fusobacterium-sp.-CM22] |
| MBA3712807.1 | Schlafen+HATPase+wHTH | GCA_013816505.1 | Pyrinomonadaceae_bacterium | Bacteria>Pseudomonadati>Acidobacteriota>Blastocatellia>Blastocatellales>Pyrinomonadaceae>unclassified_Pyrinomonadaceae | 401 | HOW76_10215 | 1..1206 | 2283092 | transcriptional-regulator-[Pyrinomonadaceae-bacterium] |
| MCI0490106.1 | Schlafen+HATPase+wHTH | GCA_022615215.1 | Blastocatellia_bacterium | Bacteria>Pseudomonadati>Acidobacteriota>Blastocatellia>unclassified_Blastocatellia | 125 | L0229_26230 | 11487..11860 | 2052146 | ATP-binding-protein-[Blastocatellia-bacterium] |
| WP_005031449.1 | Schlafen+HATPase+wHTH | - | Holophaga_foetida | Bacteria>Pseudomonadati>Acidobacteriota>Holophagae>Holophagales>Holophagaceae>Holophaga | 220 | HOLFODRAFT_RS19140 | 857531..858193 | 35839 | ATP-binding-protein-[Holophaga-foetida] |
| MCC6263217.1 | Schlafen+HATPase+wHTH | GCA_020847315.1 | Bryobacterales_bacterium | Bacteria>Pseudomonadati>Acidobacteriota>Terriglobia>Bryobacterales>unclassified_Bryobacterales | 487 | IT169_06535 | 3450..4913 | 2026791 | putative-DNA-binding-domain-containing-protein-[Bryobacterales-bacterium] |
| MBK9166731.1 | Schlafen+HATPase+wHTH | GCA_016716385.1 | Bryobacterales_bacterium | Bacteria>Pseudomonadati>Acidobacteriota>Terriglobia>Bryobacterales>unclassified_Bryobacterales | 545 | IPM24_04625 | 389959..391596 | 2026791 | putative-DNA-binding-domain-containing-protein-[Bryobacterales-bacterium] |
| MBZ5499149.1 | Schlafen+HATPase+wHTH | GCA_020072565.1 | Terriglobia_bacterium | Bacteria>Pseudomonadati>Acidobacteriota>Terriglobia>unclassified_Terriglobia | 413 | LAP85_22345 | 20168..21409 | 2081523 | putative-DNA-binding-domain-containing-protein-[Acidobacteriia-bacterium] |
| MBZ5495690.1 | Schlafen+HATPase+wHTH | GCA_020072565.1 | Terriglobia_bacterium | Bacteria>Pseudomonadati>Acidobacteriota>Terriglobia>unclassified_Terriglobia | 578 | LAP85_04760 | 207846..209582 | 2081523 | putative-DNA-binding-domain-containing-protein-[Acidobacteriia-bacterium] |
| MBP7799659.1 | Schlafen+HATPase+wHTH | GCA_018000955.1 | Thermoanaerobaculaceae_bacterium | Bacteria>Pseudomonadati>Acidobacteriota>Thermoanaerobaculia>Thermoanaerobaculales>Thermoanaerobaculaceae>unclassified_Thermoanaerobaculaceae | 656 | KA072_13650 | 13528..15498 | 2699758 | putative-DNA-binding-domain-containing-protein-[Thermoanaerobaculaceae-bacterium] |
| MCG3192276.1 | Schlafen+HATPase+wHTH | GCA_022071945.1 | Thermoanaerobaculia_bacterium | Bacteria>Pseudomonadati>Acidobacteriota>Thermoanaerobaculia>unclassified_Thermoanaerobaculia | 406 | DIJKHBIC_01514 | 139672..140892 | 2651171 | hypothetical-protein-[Thermoanaerobaculia-bacterium] |
| RIK10618.1 | Schlafen+HATPase+wHTH | GCA_003577105.1 | Acidobacteriota_bacterium | Bacteria>Pseudomonadati>Acidobacteriota>unclassified_Acidobacteriota | 213 | DCC49_02895 | 63696..64337 | 1978231 | hypothetical-protein-DCC49_02895-[Acidobacteriota-bacterium] |
| OQC41889.1 | Schlafen+HATPase+wHTH | GCA_002071135.1 | Acidobacteria_bacterium_ADurb.Bin051 | Bacteria>Pseudomonadati>Acidobacteriota>unclassified_Acidobacteriota | 545 | BWX64_00586 | 13010..14647 | 1852786 | Divergent-AAA-domain-protein-[Acidobacteria-bacterium-ADurb.Bin051] |
| OFW39747.1 | Schlafen+HATPase+wHTH | GCA_001767515.1 | Acidobacteria_bacterium_RIFCSPLOWO2_12_FULL_67_14 | Bacteria>Pseudomonadati>Acidobacteriota>unclassified_Acidobacteriota | 548 | A3F70_01780 | 15280..16926 | 1797192 | hypothetical-protein-A3F70_01780-[Acidobacteria-bacterium-RIFCSPLOWO2_12_FULL_67_14] |
| OFW17301.1 | Schlafen+HATPase+wHTH | GCA_001767195.1 | Acidobacteria_bacterium_RIFCSPLOWO2_02_FULL_67_21 | Bacteria>Pseudomonadati>Acidobacteriota>unclassified_Acidobacteriota | 112 | A3H29_10500 | 6882..7220 | 1797183 | hypothetical-protein-A3H29_10500-[Acidobacteria-bacterium-RIFCSPLOWO2_02_FULL_67_21] |
| ODS52400.1 | Schlafen+HATPase+wHTH | GCA_001724025.1 | Acidobacteria_bacterium_SCN_69-37 | Bacteria>Pseudomonadati>Acidobacteriota>unclassified_Acidobacteriota | 537 | ABS36_17615 | 159239..160852 | 1660160 | transcriptional-regulator-[Acidobacteria-bacterium-SCN-69-37] |
| NLV30312.1 | Schlafen+HATPase+wHTH | GCA_012729555.1 | Acidobacteriota_bacterium | Bacteria>Pseudomonadati>Acidobacteriota>unclassified_Acidobacteriota | 551 | GXY47_04075 | 31112..32767 | 1978231 | ATPase-[Acidobacteriota-bacterium] |
| MYN68208.1 | Schlafen+HATPase+wHTH | GCA_009861545.1 | Acidobacteriota_bacterium | Bacteria>Pseudomonadati>Acidobacteriota>unclassified_Acidobacteriota | 430 | F4X11_24855 | 48818..50110 | 1978231 | hypothetical-protein-[Acidobacteriota-bacterium] |
| MYN66474.1 | Schlafen+HATPase+wHTH | GCA_009861545.1 | Acidobacteriota_bacterium | Bacteria>Pseudomonadati>Acidobacteriota>unclassified_Acidobacteriota | 624 | F4X11_15810 | 56757..58631 | 1978231 | AAA-family-ATPase-[Acidobacteriota-bacterium] |
| MYN66278.1 | Schlafen+HATPase+wHTH | GCA_009861545.1 | Acidobacteriota_bacterium | Bacteria>Pseudomonadati>Acidobacteriota>unclassified_Acidobacteriota | 421 | F4X11_14805 | 24059..25324 | 1978231 | transcriptional-regulator-[Acidobacteriota-bacterium] |
| MYN66272.1 | Schlafen+HATPase+wHTH | GCA_009861545.1 | Acidobacteriota_bacterium | Bacteria>Pseudomonadati>Acidobacteriota>unclassified_Acidobacteriota | 430 | F4X11_14775 | 8644..9936 | 1978231 | transcriptional-regulator-[Acidobacteriota-bacterium] |
| MCY3932057.1 | Schlafen+HATPase+wHTH | GCA_026707325.1 | Acidobacteriota_bacterium | Bacteria>Pseudomonadati>Acidobacteriota>unclassified_Acidobacteriota | 506 | OXH70_10075 | 62285..63805 | 1978231 | MAG:-putative-DNA-binding-domain-containing-protein-[Acidobacteria-bacterium] |
| MCI0393107.1 | Schlafen+HATPase+wHTH | GCA_022614935.1 | Acidobacteriota_bacterium | Bacteria>Pseudomonadati>Acidobacteriota>unclassified_Acidobacteriota | 540 | MOB07_30615 | 10418..12040 | 1978231 | putative-DNA-binding-domain-containing-protein-[Acidobacteriota-bacterium] |
| MCC6131983.1 | Schlafen+HATPase+wHTH | GCA_020845555.1 | Acidobacteriota_bacterium | Bacteria>Pseudomonadati>Acidobacteriota>unclassified_Acidobacteriota | 406 | IT186_18835 | 5928..7148 | 1978231 | putative-DNA-binding-domain-containing-protein-[Acidobacteriota-bacterium] |
| MBP7569109.1 | Schlafen+HATPase+wHTH | GCA_017998655.1 | Acidobacteriota_bacterium | Bacteria>Pseudomonadati>Acidobacteriota>unclassified_Acidobacteriota | 56 | KBA95_03505 | 25483..25653 | 1978231 | hypothetical-protein-[Acidobacteriota-bacterium] |
| MBN1569783.1 | Schlafen+HATPase+wHTH | GCA_016929335.1 | Acidobacteriota_bacterium | Bacteria>Pseudomonadati>Acidobacteriota>unclassified_Acidobacteriota | 485 | JXA73_18200 | 72084..73541 | 1978231 | putative-DNA-binding-domain-containing-protein-[Acidobacteriota-bacterium] |
| MBI4454520.1 | Schlafen+HATPase+wHTH | GCA_016211105.1 | Acidobacteriota_bacterium | Bacteria>Pseudomonadati>Acidobacteriota>unclassified_Acidobacteriota | 397 | HY644_01330 | 42457..43650 | 1978231 | putative-DNA-binding-domain-containing-protein-[Acidobacteriota-bacterium] |
| MBI3951046.1 | Schlafen+HATPase+wHTH | GCA_016199695.1 | Acidobacteriota_bacterium | Bacteria>Pseudomonadati>Acidobacteriota>unclassified_Acidobacteriota | 381 | HY314_11400 | 5562..6707 | 1978231 | putative-DNA-binding-domain-containing-protein-[Acidobacteriota-bacterium] |
| MBI2837084.1 | Schlafen+HATPase+wHTH | GCA_016191015.1 | Acidobacteriota_bacterium | Bacteria>Pseudomonadati>Acidobacteriota>unclassified_Acidobacteriota | 414 | HYX75_02125 | 11360..12604 | 1978231 | putative-DNA-binding-domain-containing-protein-[Acidobacteriota-bacterium] |
| MBI2837081.1 | Schlafen+HATPase+wHTH | GCA_016191015.1 | Acidobacteriota_bacterium | Bacteria>Pseudomonadati>Acidobacteriota>unclassified_Acidobacteriota | 414 | HYX75_02110 | 5176..6420 | 1978231 | putative-DNA-binding-domain-containing-protein-[Acidobacteriota-bacterium] |
| MBI1792205.1 | Schlafen+HATPase+wHTH | GCA_016178945.1 | Acidobacteriota_bacterium | Bacteria>Pseudomonadati>Acidobacteriota>unclassified_Acidobacteriota | 553 | HYR60_32175 | 3618..5279 | 1978231 | putative-DNA-binding-domain-containing-protein-[Acidobacteriota-bacterium] |
| MBC7217417.1 | Schlafen+HATPase+wHTH | GCA_014359405.1 | Candidatus_Caldatribacterium_sp. | Bacteria>Pseudomonadati>Atribacterota>Atribacteria>Atribacterales>Candidatus_Caldatribacteriaceae>Candidatus_Caldatribacterium>unclassified_Candidatus_Caldatribacterium | 173 | H5U36_04500 | 1983..2504 | 2282143 | putative-DNA-binding-domain-containing-protein-[Candidatus-Caldatribacterium-sp.] |
| MBC7217416.1 | Schlafen+HATPase+wHTH | GCA_014359405.1 | Candidatus_Caldatribacterium_sp. | Bacteria>Pseudomonadati>Atribacterota>Atribacteria>Atribacterales>Candidatus_Caldatribacteriaceae>Candidatus_Caldatribacterium>unclassified_Candidatus_Caldatribacterium | 572 | H5U36_04495 | 174..1892 | 2282143 | putative-DNA-binding-domain-containing-protein-[Candidatus-Caldatribacterium-sp.] |
| MBE3120496.1 | Schlafen+HATPase+wHTH | GCA_014894575.1 | Candidatus_Atribacteria_bacterium | Bacteria>Pseudomonadati>Atribacterota>Atribacteria>unclassified_Atribacteria | 351 | IMZ50_17305 | 10650..11705 | 2053509 | transcriptional-regulator-[Candidatus-Atribacteria-bacterium] |
| NLI41208.1 | Schlafen+HATPase+wHTH | GCA_012517765.1 | Caldisericales_bacterium | Bacteria>Pseudomonadati>Caldisericota/Cryosericota_group>Caldisericota>Caldisericia>Caldisericales>unclassified_Caldisericales | 386 | GX421_08530 | 10219..11379 | 2614251 | ATP-dependent-DNA-helicase-RecG-[Caldisericales-bacterium] |
| NLI41206.1 | Schlafen+HATPase+wHTH | GCA_012517765.1 | Caldisericales_bacterium | Bacteria>Pseudomonadati>Caldisericota/Cryosericota_group>Caldisericota>Caldisericia>Caldisericales>unclassified_Caldisericales | 569 | GX421_08520 | 5207..6916 | 2614251 | AAA-family-ATPase-[Caldisericales-bacterium] |
| WP_260968569.1 | Schlafen+HATPase+wHTH | GCF_025335945.1 | Aliarcobacter_cryaerophilus | Bacteria>Pseudomonadati>Campylobacterota>Epsilonproteobacteria>Campylobacterales>Arcobacteraceae>Aliarcobacter | 323 | N5S77_RS10410 | 47027..47998 | 28198 | putative-DNA-binding-domain-containing-protein-[Aliarcobacter-cryaerophilus] |
| WP_260968568.1 | Schlafen+HATPase+wHTH | GCF_025335945.1 | Aliarcobacter_cryaerophilus | Bacteria>Pseudomonadati>Campylobacterota>Epsilonproteobacteria>Campylobacterales>Arcobacteraceae>Aliarcobacter | 498 | N5S77_RS10405 | 45425..46921 | 28198 | putative-DNA-binding-domain-containing-protein-[Aliarcobacter-cryaerophilus] |
| WP_260949069.1 | Schlafen+HATPase+wHTH | GCF_025334565.1 | Aliarcobacter_cryaerophilus | Bacteria>Pseudomonadati>Campylobacterota>Epsilonproteobacteria>Campylobacterales>Arcobacteraceae>Aliarcobacter | 494 | N5T87_RS09100 | 12470..13954 | 28198 | putative-DNA-binding-domain-containing-protein-[Aliarcobacter-cryaerophilus] |
| WP_260949068.1 | Schlafen+HATPase+wHTH | GCF_025334565.1 | Aliarcobacter_cryaerophilus | Bacteria>Pseudomonadati>Campylobacterota>Epsilonproteobacteria>Campylobacterales>Arcobacteraceae>Aliarcobacter | 445 | N5T87_RS09095 | 11027..12364 | 28198 | ATP-binding-protein-[Aliarcobacter-cryaerophilus] |
| WP_260916971.1 | Schlafen+HATPase+wHTH | GCF_025335525.1 | Aliarcobacter_butzleri | Bacteria>Pseudomonadati>Campylobacterota>Epsilonproteobacteria>Campylobacterales>Arcobacteraceae>Aliarcobacter | 447 | N5U17_RS09635 | 11787..13130 | 28197 | putative-DNA-binding-domain-containing-protein-[Aliarcobacter-butzleri] |
| WP_152054586.1 | Schlafen+HATPase+wHTH | GCF_902500815.1 | Aliarcobacter_butzleri | Bacteria>Pseudomonadati>Campylobacterota>Epsilonproteobacteria>Campylobacterales>Arcobacteraceae>Aliarcobacter | 494 | GDI74_RS00950 | 18786..20270 | 28197 | putative-DNA-binding-domain-containing-protein-[Aliarcobacter-butzleri] |
| WP_046997146.1 | Schlafen+HATPase+wHTH | GCF_001010585.1 | Aliarcobacter_butzleri | Bacteria>Pseudomonadati>Campylobacterota>Epsilonproteobacteria>Campylobacterales>Arcobacteraceae>Aliarcobacter | 572 | AA20_RS09515 | 20098..21816 | 28197 | putative-DNA-binding-domain-containing-protein-[Aliarcobacter-butzleri] |
| WP_226069226.1 | Schlafen+HATPase+wHTH | GCF_020410885.1 | Sulfurimonas_sp._SWIR-19 | Bacteria>Pseudomonadati>Campylobacterota>Epsilonproteobacteria>Campylobacterales>Sulfurimonadaceae>Sulfurimonas>unclassified_Sulfurimonas | 553 | LCX93_RS05515 | 1066893..1068554 | 2878390 | ATP-binding-protein-[Sulfurimonas-sp.-SWIR-19] |
| HIP19034.1 | Schlafen+HATPase+wHTH | GCA_015663065.1 | Sulfurovum_sp. | Bacteria>Pseudomonadati>Campylobacterota>Epsilonproteobacteria>Campylobacterales>Sulfurovaceae>Sulfurovum>unclassified_Sulfurovum | 599 | EYG78_06845 | 22172..23971 | 1969726 | transcriptional-regulator-[Sulfurovum-sp.] |
| WP_204430313.1 | Schlafen+HATPase+wHTH | GCF_016899805.1 | Bacteroidales | Bacteria>Pseudomonadati>FCB_group>Bacteroidota/Chlorobiota_group>Bacteroidota>Bacteroidia | 533 | NW211_RS06355 | 159605..161206 | 171549 | MULTISPECIES:-putative-DNA-binding-domain-containing-protein-[Bacteroidales] |
| WP_172774767.1 | Schlafen+HATPase+wHTH | GCF_963928575.1 | Bacteroidales | Bacteria>Pseudomonadati>FCB_group>Bacteroidota/Chlorobiota_group>Bacteroidota>Bacteroidia | 533 | KSX11_RS12370 | 129014..130615 | 171549 | MULTISPECIES:-putative-DNA-binding-domain-containing-protein-[Bacteroidaceae] |
| WP_135993584.1 | Schlafen+HATPase+wHTH | GCF_948695785.1 | Bacteroidales | Bacteria>Pseudomonadati>FCB_group>Bacteroidota/Chlorobiota_group>Bacteroidota>Bacteroidia | 551 | E5330_RS10955 | 10725..12380 | 171549 | ATP-binding-protein-[Muribaculum-intestinale] |
| WP_057098609.1 | Schlafen+HATPase+wHTH | GCF_017309675.2 | Bacteroidales | Bacteria>Pseudomonadati>FCB_group>Bacteroidota/Chlorobiota_group>Bacteroidota>Bacteroidia | 545 | DWX60_RS07010 | 7814..9451 | 171549 | MULTISPECIES:-ATP-binding-protein-[Bacteroidales] |
| WP_005795642.1 | Schlafen+HATPase+wHTH | GCF_902652125.1 | Bacteroidales | Bacteria>Pseudomonadati>FCB_group>Bacteroidota/Chlorobiota_group>Bacteroidota>Bacteroidia | 525 | BLS71_RS01415 | 100504..102081 | 171549 | MULTISPECIES:-ATP-binding-protein-[Bacteroidales] |
| WP_119962090.1 | Schlafen+HATPase+wHTH | GCF_042848865.1 | Bacteroidaceae | Bacteria>Pseudomonadati>FCB_group>Bacteroidota/Chlorobiota_group>Bacteroidota>Bacteroidia>Bacteroidales | 482 | DWY57_RS16345 | 99725..101173 | 815 | MULTISPECIES:-ATP-binding-protein-[unclassified-Bacteroides] |
| WP_259011404.1 | Schlafen+HATPase+wHTH | GCF_024791585.1 | Bacteroides | Bacteria>Pseudomonadati>FCB_group>Bacteroidota/Chlorobiota_group>Bacteroidota>Bacteroidia>Bacteroidales>Bacteroidaceae | 550 | NXX54_RS14170 | 3712315..3713967 | 816 | putative-DNA-binding-domain-containing-protein-[Bacteroides-sp.-BFG-638] |
| WP_195661288.1 | Schlafen+HATPase+wHTH | GCF_938030555.1 | Bacteroides | Bacteria>Pseudomonadati>FCB_group>Bacteroidota/Chlorobiota_group>Bacteroidota>Bacteroidia>Bacteroidales>Bacteroidaceae | 532 | KQP55_RS20555 | 5446861..5448459 | 816 | putative-DNA-binding-domain-containing-protein-[Bacteroides-nordii] |
| WP_154043603.1 | Schlafen+HATPase+wHTH | GCF_946497445.1 | Bacteroides | Bacteria>Pseudomonadati>FCB_group>Bacteroidota/Chlorobiota_group>Bacteroidota>Bacteroidia>Bacteroidales>Bacteroidaceae | 492 | I6E24_RS07130 | 5180..6658 | 816 | MULTISPECIES:-ATP-binding-protein-[Bacteroides] |
| WP_117985930.1 | Schlafen+HATPase+wHTH | GCF_026966235.1 | Bacteroides | Bacteria>Pseudomonadati>FCB_group>Bacteroidota/Chlorobiota_group>Bacteroidota>Bacteroidia>Bacteroidales>Bacteroidaceae | 544 | O1446_RS12900 | 59682..61316 | 816 | MULTISPECIES:-putative-DNA-binding-domain-containing-protein-[Bacteroides] |
| WP_115473025.1 | Schlafen+HATPase+wHTH | GCF_019733435.1 | Bacteroides | Bacteria>Pseudomonadati>FCB_group>Bacteroidota/Chlorobiota_group>Bacteroidota>Bacteroidia>Bacteroidales>Bacteroidaceae | 533 | I2F01_RS18140 | 81350..82951 | 816 | MULTISPECIES:-putative-DNA-binding-domain-containing-protein-[Bacteroides] |
| WP_032543212.1 | Schlafen+HATPase+wHTH | GCF_000710375.2 | Bacteroides | Bacteria>Pseudomonadati>FCB_group>Bacteroidota/Chlorobiota_group>Bacteroidota>Bacteroidia>Bacteroidales>Bacteroidaceae | 537 | PDN68_RS18435 | 94907..96520 | 816 | putative-DNA-binding-domain-containing-protein-[Bacteroides-fragilis] |
| WP_012055869.1 | Schlafen+HATPase+wHTH | GCF_000012825.1 | Phocaeicola | Bacteria>Pseudomonadati>FCB_group>Bacteroidota/Chlorobiota_group>Bacteroidota>Bacteroidia>Bacteroidales>Bacteroidaceae | 525 | I6I55_RS09880 | 2555879..2557456 | 909656 | MULTISPECIES:-putative-DNA-binding-domain-containing-protein-[Phocaeicola] |
| WP_009000712.1 | Schlafen+HATPase+wHTH | GCF_025147085.1 | Bacteroides | Bacteria>Pseudomonadati>FCB_group>Bacteroidota/Chlorobiota_group>Bacteroidota>Bacteroidia>Bacteroidales>Bacteroidaceae | 532 | NQ505_RS12890 | 3658131..3659729 | 816 | MULTISPECIES:-putative-DNA-binding-domain-containing-protein-[Bacteroides] |
| WP_004319890.1 | Schlafen+HATPase+wHTH | GCF_042850695.1 | Bacteroides | Bacteria>Pseudomonadati>FCB_group>Bacteroidota/Chlorobiota_group>Bacteroidota>Bacteroidia>Bacteroidales>Bacteroidaceae | 532 | KOG75_RS22175 | 11875..13473 | 816 | MULTISPECIES:-putative-DNA-binding-domain-containing-protein-[Bacteroides] |
| WP_274224869.1 | Schlafen+HATPase+wHTH | GCF_028743275.1 | Bacteroides_fragilis | Bacteria>Pseudomonadati>FCB_group>Bacteroidota/Chlorobiota_group>Bacteroidota>Bacteroidia>Bacteroidales>Bacteroidaceae>Bacteroides | 545 | PT255_RS11115 | 2679584..2681221 | 817 | ATP-binding-protein-[Bacteroides-fragilis] |
| WP_254780821.1 | Schlafen+HATPase+wHTH | GCF_900102645.1 | Bacteroides_ovatus | Bacteria>Pseudomonadati>FCB_group>Bacteroidota/Chlorobiota_group>Bacteroidota>Bacteroidia>Bacteroidales>Bacteroidaceae>Bacteroides | 506 | BLR14_RS08945 | 63258..64778 | 28116 | ATP-binding-protein-[Bacteroides-ovatus] |
| WP_250716302.1 | Schlafen+HATPase+wHTH | GCF_023654185.1 | Bacteroides_fragilis | Bacteria>Pseudomonadati>FCB_group>Bacteroidota/Chlorobiota_group>Bacteroidota>Bacteroidia>Bacteroidales>Bacteroidaceae>Bacteroides | 574 | J8L04_RS14235 | 63784..65508 | 817 | putative-DNA-binding-domain-containing-protein-[Bacteroides-fragilis] |
| WP_234146086.1 | Schlafen+HATPase+wHTH | GCF_024760085.1 | Bacteroides_uniformis | Bacteria>Pseudomonadati>FCB_group>Bacteroidota/Chlorobiota_group>Bacteroidota>Bacteroidia>Bacteroidales>Bacteroidaceae>Bacteroides | 339 | J8872_RS07790 | 69681..70700 | 820 | ATP-binding-protein-[Bacteroides-uniformis] |
| WP_227088014.1 | Schlafen+HATPase+wHTH | GCF_020560105.2 | Bacteroides_cellulosilyticus | Bacteria>Pseudomonadati>FCB_group>Bacteroidota/Chlorobiota_group>Bacteroidota>Bacteroidia>Bacteroidales>Bacteroidaceae>Bacteroides | 627 | LI190_RS20865 | 48425..50308 | 246787 | putative-DNA-binding-domain-containing-protein-[Bacteroides-cellulosilyticus] |
| WP_217725666.1 | Schlafen+HATPase+wHTH | GCF_019128915.1 | unclassified_Bacteroides | Bacteria>Pseudomonadati>FCB_group>Bacteroidota/Chlorobiota_group>Bacteroidota>Bacteroidia>Bacteroidales>Bacteroidaceae>Bacteroides | 532 | KSX87_RS09990 | 221887..223485 | 2646097 | MULTISPECIES:-putative-DNA-binding-domain-containing-protein-[unclassified-Bacteroides] |
| WP_215683074.1 | Schlafen+HATPase+wHTH | GCF_018785945.1 | Bacteroides_salyersiae | Bacteria>Pseudomonadati>FCB_group>Bacteroidota/Chlorobiota_group>Bacteroidota>Bacteroidia>Bacteroidales>Bacteroidaceae>Bacteroides | 482 | GPL06_RS15300 | 22756..24204 | 291644 | ATP-binding-protein-[Bacteroides-salyersiae] |
| WP_204447181.1 | Schlafen+HATPase+wHTH | GCF_016902245.1 | Bacteroides_caecigallinarum | Bacteria>Pseudomonadati>FCB_group>Bacteroidota/Chlorobiota_group>Bacteroidota>Bacteroidia>Bacteroidales>Bacteroidaceae>Bacteroides | 525 | H6A66_RS14130 | 13710..15287 | 1411144 | putative-DNA-binding-domain-containing-protein-[Bacteroides-caecigallinarum] |
| WP_195571586.1 | Schlafen+HATPase+wHTH | GCF_947649005.1 | Bacteroides_nordii | Bacteria>Pseudomonadati>FCB_group>Bacteroidota/Chlorobiota_group>Bacteroidota>Bacteroidia>Bacteroidales>Bacteroidaceae>Bacteroides | 532 | I2G99_RS18050 | 22748..24346 | 291645 | putative-DNA-binding-domain-containing-protein-[Bacteroides-nordii] |
| WP_195207458.1 | Schlafen+HATPase+wHTH | GCF_015547015.1 | Bacteroides_clarus | Bacteria>Pseudomonadati>FCB_group>Bacteroidota/Chlorobiota_group>Bacteroidota>Bacteroidia>Bacteroidales>Bacteroidaceae>Bacteroides | 482 | I1P93_RS11580 | 59154..60602 | 626929 | ATP-binding-protein-[Bacteroides-clarus] |
| WP_177608042.1 | Schlafen+HATPase+wHTH | GCF_028200085.1 | Bacteroides_xylanisolvens | Bacteria>Pseudomonadati>FCB_group>Bacteroidota/Chlorobiota_group>Bacteroidota>Bacteroidia>Bacteroidales>Bacteroidaceae>Bacteroides | 515 | PL493_RS08925 | 30483..32030 | 371601 | putative-DNA-binding-domain-containing-protein-[Bacteroides-xylanisolvens] |
| WP_154042814.1 | Schlafen+HATPase+wHTH | GCF_900066445.1 | Bacteroides_cellulolyticus | Bacteria>Pseudomonadati>FCB_group>Bacteroidota/Chlorobiota_group>Bacteroidota>Bacteroidia>Bacteroidales>Bacteroidaceae>Bacteroides | 576 | OCV82_RS01760 | 407827..409557 | 2981780 | putative-DNA-binding-domain-containing-protein-[Bacteroides-cellulolyticus] |
| WP_120489890.1 | Schlafen+HATPase+wHTH | GCF_019733535.1 | Bacteroides_ovatus | Bacteria>Pseudomonadati>FCB_group>Bacteroidota/Chlorobiota_group>Bacteroidota>Bacteroidia>Bacteroidales>Bacteroidaceae>Bacteroides | 532 | G7086_RS03175 | 94720..96318 | 28116 | putative-DNA-binding-domain-containing-protein-[Bacteroides-ovatus] |
| WP_074557119.1 | Schlafen+HATPase+wHTH | GCF_900102645.1 | Bacteroides_ovatus | Bacteria>Pseudomonadati>FCB_group>Bacteroidota/Chlorobiota_group>Bacteroidota>Bacteroidia>Bacteroidales>Bacteroidaceae>Bacteroides | 532 | BLR14_RS05855 | 116004..117602 | 28116 | putative-DNA-binding-domain-containing-protein-[Bacteroides-ovatus] |
| WP_074557112.1 | Schlafen+HATPase+wHTH | GCF_900102645.1 | Bacteroides_ovatus | Bacteria>Pseudomonadati>FCB_group>Bacteroidota/Chlorobiota_group>Bacteroidota>Bacteroidia>Bacteroidales>Bacteroidaceae>Bacteroides | 513 | BLR14_RS05820 | 102914..104455 | 28116 | ATP-binding-protein-[Bacteroides-ovatus] |
| WP_073402631.1 | Schlafen+HATPase+wHTH | GCF_900128905.1 | Bacteroides_luti | Bacteria>Pseudomonadati>FCB_group>Bacteroidota/Chlorobiota_group>Bacteroidota>Bacteroidia>Bacteroidales>Bacteroidaceae>Bacteroides | 635 | BUC21_RS11770 | 72863..74770 | 1297750 | ATP-binding-protein-[Bacteroides-luti] |
| WP_071148142.1 | Schlafen+HATPase+wHTH | GCF_900108345.1 | Bacteroides_ndongoniae | Bacteria>Pseudomonadati>FCB_group>Bacteroidota/Chlorobiota_group>Bacteroidota>Bacteroidia>Bacteroidales>Bacteroidaceae>Bacteroides | 482 | BQ5591_RS01355 | 256319..257767 | 1903262 | ATP-binding-protein-[Bacteroides-ndongoniae] |
| WP_060407658.1 | Schlafen+HATPase+wHTH | GCF_902387305.1 | Bacteroides_cellulosilyticus | Bacteria>Pseudomonadati>FCB_group>Bacteroidota/Chlorobiota_group>Bacteroidota>Bacteroidia>Bacteroidales>Bacteroidaceae>Bacteroides | 529 | FYC30_RS03940 | 67647..69236 | 246787 | putative-DNA-binding-domain-containing-protein-[Bacteroides-cellulosilyticus] |
| WP_060386339.1 | Schlafen+HATPase+wHTH | GCF_020091485.1 | Bacteroides_stercoris | Bacteria>Pseudomonadati>FCB_group>Bacteroidota/Chlorobiota_group>Bacteroidota>Bacteroidia>Bacteroidales>Bacteroidaceae>Bacteroides | 545 | K6V31_RS06885 | 1547023..1548660 | 46506 | ATP-binding-protein-[Bacteroides-stercoris] |
| WP_057099391.1 | Schlafen+HATPase+wHTH | GCF_932186025.1 | Bacteroides_acidifaciens | Bacteria>Pseudomonadati>FCB_group>Bacteroidota/Chlorobiota_group>Bacteroidota>Bacteroidia>Bacteroidales>Bacteroidaceae>Bacteroides | 482 | QDP01_RS07400 | 35225..36673 | 85831 | ATP-binding-protein-[Bacteroides-acidifaciens] |
| WP_032560930.1 | Schlafen+HATPase+wHTH | GCF_029872615.1 | Bacteroides_fragilis | Bacteria>Pseudomonadati>FCB_group>Bacteroidota/Chlorobiota_group>Bacteroidota>Bacteroidia>Bacteroidales>Bacteroidaceae>Bacteroides | 533 | M068_RS14685 | 119346..120947 | 817 | putative-DNA-binding-domain-containing-protein-[Bacteroides-fragilis] |
| WP_016268712.1 | Schlafen+HATPase+wHTH | GCF_000403155.2 | Bacteroides_thetaiotaomicron | Bacteria>Pseudomonadati>FCB_group>Bacteroidota/Chlorobiota_group>Bacteroidota>Bacteroidia>Bacteroidales>Bacteroidaceae>Bacteroides | 482 | C799_RS09500 | 3543661..3545109 | 818 | ATP-binding-protein-[Bacteroides-thetaiotaomicron] |
| WP_005930498.1 | Schlafen+HATPase+wHTH | GCF_902374995.1 | Bacteroides_salyersiae | Bacteria>Pseudomonadati>FCB_group>Bacteroidota/Chlorobiota_group>Bacteroidota>Bacteroidia>Bacteroidales>Bacteroidaceae>Bacteroides | 482 | NXY20_RS18865 | 4714320..4715768 | 291644 | ATP-binding-protein-[Bacteroides-salyersiae] |
| SHF69789.1 | Schlafen+HATPase+wHTH | GCA_900128905.1 | Bacteroides_luti | Bacteria>Pseudomonadati>FCB_group>Bacteroidota/Chlorobiota_group>Bacteroidota>Bacteroidia>Bacteroidales>Bacteroidaceae>Bacteroides | 635 | SAMN05444405_11259 | 72863..74770 | 1297750 | ATP-dependent-DNA-helicase-RecG-[Bacteroides-luti] |
| MCF2581766.1 | Schlafen+HATPase+wHTH | GCA_021530995.1 | Bacteroides_caecigallinarum | Bacteria>Pseudomonadati>FCB_group>Bacteroidota/Chlorobiota_group>Bacteroidota>Bacteroidia>Bacteroidales>Bacteroidaceae>Bacteroides | 492 | I6E24_07130 | 5180..6658 | 1411144 | putative-DNA-binding-domain-containing-protein-[Bacteroides-caecigallinarum] |
| KAA4743088.1 | Schlafen+HATPase+wHTH | GCA_008569225.1 | Bacteroides_fragilis | Bacteria>Pseudomonadati>FCB_group>Bacteroidota/Chlorobiota_group>Bacteroidota>Bacteroidia>Bacteroidales>Bacteroidaceae>Bacteroides | 533 | F3B36_09975 | 285383..286984 | 817 | AAA-family-ATPase-[Bacteroides-fragilis] |
| KAA3824843.1 | Schlafen+HATPase+wHTH | GCA_008565655.1 | Bacteroides_ovatus | Bacteria>Pseudomonadati>FCB_group>Bacteroidota/Chlorobiota_group>Bacteroidota>Bacteroidia>Bacteroidales>Bacteroidaceae>Bacteroides | 532 | F3F58_10395 | 9679..11277 | 28116 | AAA-family-ATPase-[Bacteroides-ovatus] |
| SCH71649.1 | Schlafen+HATPase+wHTH | GCA_900066445.1 | uncultured_Bacteroides_sp. | Bacteria>Pseudomonadati>FCB_group>Bacteroidota/Chlorobiota_group>Bacteroidota>Bacteroidia>Bacteroidales>Bacteroidaceae>Bacteroides>environmental_samples | 492 | SAMEA3545309_01209 | 46108..47586 | 162156 | Divergent-AAA-domain-[uncultured-Bacteroides-sp.] |
| SCH13796.1 | Schlafen+HATPase+wHTH | GCA_900066445.1 | uncultured_Bacteroides_sp. | Bacteria>Pseudomonadati>FCB_group>Bacteroidota/Chlorobiota_group>Bacteroidota>Bacteroidia>Bacteroidales>Bacteroidaceae>Bacteroides>environmental_samples | 576 | SAMEA3545309_00362 | 407827..409557 | 162156 | Divergent-AAA-domain-[uncultured-Bacteroides-sp.] |
| WP_258979870.1 | Schlafen+HATPase+wHTH | GCF_024793605.1 | Bacteroides_sp._BFG-637 | Bacteria>Pseudomonadati>FCB_group>Bacteroidota/Chlorobiota_group>Bacteroidota>Bacteroidia>Bacteroidales>Bacteroidaceae>Bacteroides>unclassified_Bacteroides | 549 | NXX38_RS14325 | 3712341..3713990 | 2972764 | putative-DNA-binding-domain-containing-protein-[Bacteroides-sp.-BFG-637] |
| WP_002561969.1 | Schlafen+HATPase+wHTH | GCF_000382465.1 | Bacteroides_sp._HPS0048 | Bacteria>Pseudomonadati>FCB_group>Bacteroidota/Chlorobiota_group>Bacteroidota>Bacteroidia>Bacteroidales>Bacteroidaceae>Bacteroides>unclassified_Bacteroides | 532 | HMPREF1214_RS19605 | 474137..475735 | 1078089 | putative-DNA-binding-domain-containing-protein-[Bacteroides-sp.-HPS0048] |
| OKZ20117.1 | Schlafen+HATPase+wHTH | GCA_001915545.1 | Bacteroides_sp._43_108 | Bacteria>Pseudomonadati>FCB_group>Bacteroidota/Chlorobiota_group>Bacteroidota>Bacteroidia>Bacteroidales>Bacteroidaceae>Bacteroides>unclassified_Bacteroides | 635 | BHV77_10970 | 32815..34722 | 1896974 | MAG:-hypothetical-protein-BHV77_10970-[Bacteroides-sp.-43_108] |
| OKZ19961.1 | Schlafen+HATPase+wHTH | GCA_001915545.1 | Bacteroides_sp._43_108 | Bacteria>Pseudomonadati>FCB_group>Bacteroidota/Chlorobiota_group>Bacteroidota>Bacteroidia>Bacteroidales>Bacteroidaceae>Bacteroides>unclassified_Bacteroides | 545 | BHV77_11435 | 21084..22721 | 1896974 | MAG:-hypothetical-protein-BHV77_11435-[Bacteroides-sp.-43_108] |
| OKZ04614.1 | Schlafen+HATPase+wHTH | GCA_001915435.1 | Bacteroides_sp._41_26 | Bacteria>Pseudomonadati>FCB_group>Bacteroidota/Chlorobiota_group>Bacteroidota>Bacteroidia>Bacteroidales>Bacteroidaceae>Bacteroides>unclassified_Bacteroides | 532 | BHV71_10455 | 77711..79309 | 1896973 | MAG:-AAA-family-ATPase-[Bacteroides-sp.-41_26] |
| MCD8261447.1 | Schlafen+HATPase+wHTH | GCA_021204375.1 | Bacteroides_sp. | Bacteria>Pseudomonadati>FCB_group>Bacteroidota/Chlorobiota_group>Bacteroidota>Bacteroidia>Bacteroidales>Bacteroidaceae>Bacteroides>unclassified_Bacteroides | 268 | LUD15_08075 | 471..1277 | 29523 | putative-DNA-binding-domain-containing-protein-[Bacteroides-sp.] |
| MBS6238332.1 | Schlafen+HATPase+wHTH | GCA_018374845.1 | Bacteroides_sp. | Bacteria>Pseudomonadati>FCB_group>Bacteroidota/Chlorobiota_group>Bacteroidota>Bacteroidia>Bacteroidales>Bacteroidaceae>Bacteroides>unclassified_Bacteroides | 526 | KH897_08180 | 194812..196392 | 29523 | putative-DNA-binding-domain-containing-protein-[Bacteroides-sp.] |
| MBQ8602357.1 | Schlafen+HATPase+wHTH | GCA_017472925.1 | Bacteroides_sp. | Bacteria>Pseudomonadati>FCB_group>Bacteroidota/Chlorobiota_group>Bacteroidota>Bacteroidia>Bacteroidales>Bacteroidaceae>Bacteroides>unclassified_Bacteroides | 581 | IJ413_07395 | 185801..187546 | 29523 | putative-DNA-binding-domain-containing-protein-[Bacteroides-sp.] |
| MBD5419380.1 | Schlafen+HATPase+wHTH | GCA_014803765.1 | Bacteroides_sp. | Bacteria>Pseudomonadati>FCB_group>Bacteroidota/Chlorobiota_group>Bacteroidota>Bacteroidia>Bacteroidales>Bacteroidaceae>Bacteroides>unclassified_Bacteroides | 562 | HDR48_05030 | 8700..10388 | 29523 | AAA-family-ATPase-[Bacteroides-sp.] |
| MBD5349665.1 | Schlafen+HATPase+wHTH | GCA_014802205.1 | Bacteroides_sp. | Bacteria>Pseudomonadati>FCB_group>Bacteroidota/Chlorobiota_group>Bacteroidota>Bacteroidia>Bacteroidales>Bacteroidaceae>Bacteroides>unclassified_Bacteroides | 514 | HDR89_02110 | 29735..31279 | 29523 | AAA-family-ATPase-[Bacteroides-sp.] |
| MBD5304901.1 | Schlafen+HATPase+wHTH | GCA_014801805.1 | Bacteroides_sp. | Bacteria>Pseudomonadati>FCB_group>Bacteroidota/Chlorobiota_group>Bacteroidota>Bacteroidia>Bacteroidales>Bacteroidaceae>Bacteroides>unclassified_Bacteroides | 509 | HDS12_01215 | 22953..24482 | 29523 | AAA-family-ATPase-[Bacteroides-sp.] |
| MBD5304892.1 | Schlafen+HATPase+wHTH | GCA_014801805.1 | Bacteroides_sp. | Bacteria>Pseudomonadati>FCB_group>Bacteroidota/Chlorobiota_group>Bacteroidota>Bacteroidia>Bacteroidales>Bacteroidaceae>Bacteroides>unclassified_Bacteroides | 324 | HDS12_01170 | 6743..7717 | 29523 | hypothetical-protein-[Bacteroides-sp.] |
| MBD5263516.1 | Schlafen+HATPase+wHTH | GCA_014801365.1 | Bacteroides_sp. | Bacteria>Pseudomonadati>FCB_group>Bacteroidota/Chlorobiota_group>Bacteroidota>Bacteroidia>Bacteroidales>Bacteroidaceae>Bacteroides>unclassified_Bacteroides | 548 | HDS39_05530 | 27295..28941 | 29523 | ATP-dependent-DNA-helicase-[Bacteroides-sp.] |
| MBD5262114.1 | Schlafen+HATPase+wHTH | GCA_014801315.1 | Bacteroides_sp. | Bacteria>Pseudomonadati>FCB_group>Bacteroidota/Chlorobiota_group>Bacteroidota>Bacteroidia>Bacteroidales>Bacteroidaceae>Bacteroides>unclassified_Bacteroides | 540 | HDS38_08305 | 55600..57222 | 29523 | MarR-family-transcriptional-regulator-[Bacteroides-sp.] |

| Protein-ID | Domain | Assembly-ID | Strain | Taxonomy | Length | Locus-Tag | Genomic-coordinates | TaxID | Fasta-defline |
|---|---|---|---|---|---|---|---|---|---|
| EEZ19145.1 | Schlafen+HATPase+wHTH | - | Bacteroides_sp._3_1_33FAA | Bacteria>Pseudomonadati>FCB_group>Bacteroidota/Chlorobiota_group>Bacteroidota>Bacteroidia>Bacteroidales>Bacteroidaceae>Bacteroides>unclassified_Bacteroides | 545 | HMPREF0105_4545 | 20251..21888 | 457391 | divergent-AAA-domain-protein-[Bacteroides-sp.-3_1_33FAA] |
| CDD83750.1 | Schlafen+HATPase+wHTH | GCA_000435095.1 | Bacteroides_sp._CAG:462 | Bacteria>Pseudomonadati>FCB_group>Bacteroidota/Chlorobiota_group>Bacteroidota>Bacteroidia>Bacteroidales>Bacteroidaceae>Bacteroides>unclassified_Bacteroides | 451 | BN666_00391 | 11198..12553 | 1262740 | putative-transcriptional-regulator-[Bacteroides-sp.-CAG:462] |
| CDD81789.1 | Schlafen+HATPase+wHTH | GCA_000435095.1 | Bacteroides_sp._CAG:462 | Bacteria>Pseudomonadati>FCB_group>Bacteroidota/Chlorobiota_group>Bacteroidota>Bacteroidia>Bacteroidales>Bacteroidaceae>Bacteroides>unclassified_Bacteroides | 533 | BN666_01220 | 40969..42570 | 1262740 | uncharacterized-protein-BN666_01220-[Bacteroides-sp.-CAG:462] |
| WP_270226500.1 | Schlafen+HATPase+wHTH | GCF_052363755.1 | Phocaeicola_massiliensis | Bacteria>Pseudomonadati>FCB_group>Bacteroidota/Chlorobiota_group>Bacteroidota>Bacteroidia>Bacteroidales>Bacteroidaceae>Phocaeicola | 511 | PGQ49_RS05950 | 195698..197233 | 204516 | putative-DNA-binding-domain-containing-protein-[Phocaeicola-massiliensis] |
| WP_214462060.1 | Schlafen+HATPase+wHTH | GCF_018588025.1 | Phocaeicola_dorei | Bacteria>Pseudomonadati>FCB_group>Bacteroidota/Chlorobiota_group>Bacteroidota>Bacteroidia>Bacteroidales>Bacteroidaceae>Phocaeicola | 458 | JQN06_RS10450 | 612683..614059 | 357276 | MULTISPECIES:-ATP-binding-protein-[Bacteroidaceae] |
| WP_204506198.1 | Schlafen+HATPase+wHTH | GCF_963927365.1 | Phocaeicola_plebeius | Bacteria>Pseudomonadati>FCB_group>Bacteroidota/Chlorobiota_group>Bacteroidota>Bacteroidia>Bacteroidales>Bacteroidaceae>Phocaeicola | 628 | H6A37_RS09765 | 34318..36204 | 310297 | putative-DNA-binding-domain-containing-protein-[Phocaeicola-plebeius] |
| WP_181213823.1 | Schlafen+HATPase+wHTH | GCF_013618865.1 | Phocaeicola_faecicola | Bacteria>Pseudomonadati>FCB_group>Bacteroidota/Chlorobiota_group>Bacteroidota>Bacteroidia>Bacteroidales>Bacteroidaceae>Phocaeicola | 545 | H3001_RS00980 | 107368..109005 | 2739389 | ATP-binding-protein-[Phocaeicola-faecicola] |
| WP_024993538.1 | Schlafen+HATPase+wHTH | GCF_000613805.1 | Phocaeicola_paurosaccharolyticus | Bacteria>Pseudomonadati>FCB_group>Bacteroidota/Chlorobiota_group>Bacteroidota>Bacteroidia>Bacteroidales>Bacteroidaceae>Phocaeicola | 531 | JCM15092_RS04005 | 48048..49643 | 732242 | putative-DNA-binding-domain-containing-protein-[Phocaeicola-paurosaccharolyticus] |
| NBH65792.1 | Schlafen+HATPase+wHTH | GCA_009911015.1 | Phocaeicola_sartorii | Bacteria>Pseudomonadati>FCB_group>Bacteroidota/Chlorobiota_group>Bacteroidota>Bacteroidia>Bacteroidales>Bacteroidaceae>Phocaeicola | 482 | D3Z46_05555 | 63489..64937 | 671267 | transcriptional-regulator-[Phocaeicola-sartorii] |
| MCQ2071600.1 | Schlafen+HATPase+wHTH | GCA_024398655.1 | Bacteroidaceae_bacterium | Bacteria>Pseudomonadati>FCB_group>Bacteroidota/Chlorobiota_group>Bacteroidota>Bacteroidia>Bacteroidales>Bacteroidaceae>unclassified_Bacteroidaceae | 449 | MJY96_00545 | 115045..116394 | 2212467 | putative-DNA-binding-domain-containing-protein-[Bacteroidaceae-bacterium] |
| MBR4380324.1 | Schlafen+HATPase+wHTH | GCA_017534535.1 | Bacteroidaceae_bacterium | Bacteria>Pseudomonadati>FCB_group>Bacteroidota/Chlorobiota_group>Bacteroidota>Bacteroidia>Bacteroidales>Bacteroidaceae>unclassified_Bacteroidaceae | 550 | IKP48_03620 | 22663..24315 | 2212467 | putative-DNA-binding-domain-containing-protein-[Bacteroidaceae-bacterium] |
| MBQ8866659.1 | Schlafen+HATPase+wHTH | GCA_017464815.1 | Bacteroidaceae_bacterium | Bacteria>Pseudomonadati>FCB_group>Bacteroidota/Chlorobiota_group>Bacteroidota>Bacteroidia>Bacteroidales>Bacteroidaceae>unclassified_Bacteroidaceae | 625 | IJ013_02910 | 81852..83729 | 2212467 | putative-DNA-binding-domain-containing-protein-[Bacteroidaceae-bacterium] |
| MBQ0129307.1 | Schlafen+HATPase+wHTH | GCA_018065885.1 | Candidatus_Limimorpha_equi | Bacteria>Pseudomonadati>FCB_group>Bacteroidota/Chlorobiota_group>Bacteroidota>Bacteroidia>Bacteroidales>Bacteroidales_incertae_sedis>Candidatus_Limimorpha | 552 | KBT57_05660 | 8977..10635 | 3083112 | putative-DNA-binding-domain-containing-protein-[bacterium] |
| MBQ0129302.1 | Schlafen+HATPase+wHTH | GCA_018065885.1 | Candidatus_Limimorpha_equi | Bacteria>Pseudomonadati>FCB_group>Bacteroidota/Chlorobiota_group>Bacteroidota>Bacteroidia>Bacteroidales>Bacteroidales_incertae_sedis>Candidatus_Limimorpha | 554 | KBT57_05635 | 2665..4329 | 3083112 | putative-DNA-binding-domain-containing-protein-[bacterium] |
| MBP9995127.1 | Schlafen+HATPase+wHTH | GCA_018064605.1 | Candidatus_Limimorpha_caballi | Bacteria>Pseudomonadati>FCB_group>Bacteroidota/Chlorobiota_group>Bacteroidota>Bacteroidia>Bacteroidales>Bacteroidales_incertae_sedis>Candidatus_Limimorpha | 235 | KBT67_09370 | 69936..70643 | 3083111 | putative-DNA-binding-domain-containing-protein-[bacterium] |
| CEA16539.1 | Schlafen+HATPase+wHTH | GCA_000953535.1 | Fermentimonas_caenicola | Bacteria>Pseudomonadati>FCB_group>Bacteroidota/Chlorobiota_group>Bacteroidota>Bacteroidia>Bacteroidales>Dysgonomonadaceae>Fermentimonas | 488 | ING2E5B_1795 | 2021744..2023210 | 1562970 | divergent-AAA-domain-protein-[Fermentimonas-caenicola] |
| WP_076930953.1 | Schlafen+HATPase+wHTH | GCF_900095135.1 | Proteiniphilum_saccharofermentans | Bacteria>Pseudomonadati>FCB_group>Bacteroidota/Chlorobiota_group>Bacteroidota>Bacteroidia>Bacteroidales>Dysgonomonadaceae>Proteiniphilum | 508 | PSM36_RS11105 | 2838823..2840349 | 1642647 | ATP-binding-protein-[Proteiniphilum-saccharofermentans] |
| WP_172278704.1 | Schlafen+HATPase+wHTH | GCF_013166635.1 | Lentimicrobium_sp._S6 | Bacteria>Pseudomonadati>FCB_group>Bacteroidota/Chlorobiota_group>Bacteroidota>Bacteroidia>Bacteroidales>Lentimicrobiaceae>Lentimicrobium>unclassified_Lentimicrobium | 539 | HNS40_RS06020 | 49599..51218 | 2735872 | putative-DNA-binding-domain-containing-protein-[Lentimicrobium-sp.-S6] |
| ROT18777.1 | Schlafen+HATPase+wHTH | GCA_003833095.1 | Muribaculaceae_bacterium_Isolate-114_(HZI) | Bacteria>Pseudomonadati>FCB_group>Bacteroidota/Chlorobiota_group>Bacteroidota>Bacteroidia>Bacteroidales>Muribaculaceae>unclassified_Muribaculaceae | 538 | EEL53_11900 | 17683..19299 | 2486475 | AAA-family-ATPase-[Muribaculaceae-bacterium-Isolate-114-(HZI)] |
| ROS89079.1 | Schlafen+HATPase+wHTH | GCA_003762635.1 | Muribaculaceae_bacterium_Isolate-043_(Harlan) | Bacteria>Pseudomonadati>FCB_group>Bacteroidota/Chlorobiota_group>Bacteroidota>Bacteroidia>Bacteroidales>Muribaculaceae>unclassified_Muribaculaceae | 559 | EEL36_13375 | 11362..13041 | 2486463 | transcriptional-regulator-[Muribaculaceae-bacterium-Isolate-043-(Harlan)] |
| MCF0219892.1 | Schlafen+HATPase+wHTH | GCA_021443165.1 | Muribaculaceae_bacterium | Bacteria>Pseudomonadati>FCB_group>Bacteroidota/Chlorobiota_group>Bacteroidota>Bacteroidia>Bacteroidales>Muribaculaceae>unclassified_Muribaculaceae | 572 | HUK14_08930 | 80411..82129 | 2498093 | putative-DNA-binding-domain-containing-protein-[Muribaculaceae-bacterium] |
| WP_195241295.1 | Schlafen+HATPase+wHTH | GCF_958425225.1 | Odoribacter_laneus | Bacteria>Pseudomonadati>FCB_group>Bacteroidota/Chlorobiota_group>Bacteroidota>Bacteroidia>Bacteroidales>Odoribacteraceae>Odoribacter | 482 | I2A63_RS11765 | 53256..54704 | 626933 | ATP-binding-protein-[Odoribacter-laneus] |
| WP_235417295.1 | Schlafen+HATPase+wHTH | GCF_021531515.1 | Leyella_stercorea | Bacteria>Pseudomonadati>FCB_group>Bacteroidota/Chlorobiota_group>Bacteroidota>Bacteroidia>Bacteroidales>Prevotellaceae>Leyella | 640 | I6E41_RS10115 | 8357..10279 | 363265 | ATP-binding-protein-[Prevotella-stercorea] |
| WP_205111297.1 | Schlafen+HATPase+wHTH | GCF_016899955.1 | Marseilla_massiliensis | Bacteria>Pseudomonadati>FCB_group>Bacteroidota/Chlorobiota_group>Bacteroidota>Bacteroidia>Bacteroidales>Prevotellaceae>Marseilla | 533 | H6B30_RS13035 | 31469..33070 | 1841864 | putative-DNA-binding-domain-containing-protein-[Marseilla-massiliensis] |
| WP_205109175.1 | Schlafen+HATPase+wHTH | GCF_016899955.1 | Marseilla_massiliensis | Bacteria>Pseudomonadati>FCB_group>Bacteroidota/Chlorobiota_group>Bacteroidota>Bacteroidia>Bacteroidales>Prevotellaceae>Marseilla | 462 | H6B30_RS07335 | 44826..46214 | 1841864 | ATP-binding-protein-[Marseilla-massiliensis] |
| WP_240048173.1 | Schlafen+HATPase+wHTH | GCF_008016795.1 | Prevotella_brunnea | Bacteria>Pseudomonadati>FCB_group>Bacteroidota/Chlorobiota_group>Bacteroidota>Bacteroidia>Bacteroidales>Prevotellaceae>Prevotella | 252 | ETF27_RS11545 | 5374..6132 | 2508867 | putative-DNA-binding-domain-containing-protein-[Prevotella-brunnea] |
| WP_240048172.1 | Schlafen+HATPase+wHTH | GCF_008016795.1 | Prevotella_brunnea | Bacteria>Pseudomonadati>FCB_group>Bacteroidota/Chlorobiota_group>Bacteroidota>Bacteroidia>Bacteroidales>Prevotellaceae>Prevotella | 213 | ETF27_RS11540 | 4693..5334 | 2508867 | ATP-binding-protein-[Prevotella-brunnea] |
| WP_125105110.1 | Schlafen+HATPase+wHTH | GCF_900625065.1 | Prevotella_marseillensis | Bacteria>Pseudomonadati>FCB_group>Bacteroidota/Chlorobiota_group>Bacteroidota>Bacteroidia>Bacteroidales>Prevotellaceae>Prevotella | 533 | EJL04_RS12215 | 3051786..3053387 | 2479840 | putative-DNA-binding-domain-containing-protein-[Prevotella-marseillensis] |
| WP_077197248.1 | Schlafen+HATPase+wHTH | GCF_900155655.1 | Prevotella_ihumii | Bacteria>Pseudomonadati>FCB_group>Bacteroidota/Chlorobiota_group>Bacteroidota>Bacteroidia>Bacteroidales>Prevotellaceae>Prevotella | 630 | CZ367_RS13310 | 3068815..3070707 | 1917878 | putative-DNA-binding-domain-containing-protein-[Prevotella-ihumii] |
| WP_094391619.1 | Schlafen+HATPase+wHTH | GCF_002251345.1 | Prevotella_sp._P4-119 | Bacteria>Pseudomonadati>FCB_group>Bacteroidota/Chlorobiota_group>Bacteroidota>Bacteroidia>Bacteroidales>Prevotellaceae>Prevotella>unclassified_Prevotella | 544 | CIK89_RS12090 | 10265..11899 | 2024218 | ATP-binding-protein-[Prevotella-sp.-P4-119] |
| MCI7017341.1 | Schlafen+HATPase+wHTH | GCA_022775925.1 | Prevotella_sp. | Bacteria>Pseudomonadati>FCB_group>Bacteroidota/Chlorobiota_group>Bacteroidota>Bacteroidia>Bacteroidales>Prevotellaceae>Prevotella>unclassified_Prevotella | 472 | MR923_09630 | 15581..16999 | 59823 | putative-DNA-binding-domain-containing-protein-[Prevotella-sp.] |
| MCI6403341.1 | Schlafen+HATPase+wHTH | GCA_022777225.1 | Prevotella_sp. | Bacteria>Pseudomonadati>FCB_group>Bacteroidota/Chlorobiota_group>Bacteroidota>Bacteroidia>Bacteroidales>Prevotellaceae>Prevotella>unclassified_Prevotella | 681 | MR820_04570 | 23976..26021 | 59823 | putative-DNA-binding-domain-containing-protein-[Prevotella-sp.] |
| MCI2087044.1 | Schlafen+HATPase+wHTH | GCA_022653375.1 | Prevotella_sp. | Bacteria>Pseudomonadati>FCB_group>Bacteroidota/Chlorobiota_group>Bacteroidota>Bacteroidia>Bacteroidales>Prevotellaceae>Prevotella>unclassified_Prevotella | 511 | LKK10_02010 | 6259..7794 | 59823 | putative-DNA-binding-domain-containing-protein-[Prevotella-sp.] |
| MCI1742486.1 | Schlafen+HATPase+wHTH | GCA_022647785.1 | Prevotella_sp. | Bacteria>Pseudomonadati>FCB_group>Bacteroidota/Chlorobiota_group>Bacteroidota>Bacteroidia>Bacteroidales>Prevotellaceae>Prevotella>unclassified_Prevotella | 511 | LKI18_09145 | 34319..35854 | 59823 | putative-DNA-binding-domain-containing-protein-[Prevotella-sp.] |
| MCH4018980.1 | Schlafen+HATPase+wHTH | GCA_022483425.1 | Prevotella_sp. | Bacteria>Pseudomonadati>FCB_group>Bacteroidota/Chlorobiota_group>Bacteroidota>Bacteroidia>Bacteroidales>Prevotellaceae>Prevotella>unclassified_Prevotella | 518 | LKE74_11140 | 2800971..2802527 | 59823 | putative-DNA-binding-domain-containing-protein-[Prevotella-sp.] |
| MBR6936555.1 | Schlafen+HATPase+wHTH | GCA_017519645.1 | Prevotella_sp. | Bacteria>Pseudomonadati>FCB_group>Bacteroidota/Chlorobiota_group>Bacteroidota>Bacteroidia>Bacteroidales>Prevotellaceae>Prevotella>unclassified_Prevotella | 547 | IKH63_03125 | 119002..120645 | 59823 | putative-DNA-binding-domain-containing-protein-[Prevotella-sp.] |
| MBR2237164.1 | Schlafen+HATPase+wHTH | GCA_017475265.1 | Prevotella_sp. | Bacteria>Pseudomonadati>FCB_group>Bacteroidota/Chlorobiota_group>Bacteroidota>Bacteroidia>Bacteroidales>Prevotellaceae>Prevotella>unclassified_Prevotella | 547 | IJ887_04715 | 38672..40315 | 59823 | putative-DNA-binding-domain-containing-protein-[Prevotella-sp.] |
| MBO5155855.1 | Schlafen+HATPase+wHTH | GCA_017622935.1 | Prevotella_sp. | Bacteria>Pseudomonadati>FCB_group>Bacteroidota/Chlorobiota_group>Bacteroidota>Bacteroidia>Bacteroidales>Prevotellaceae>Prevotella>unclassified_Prevotella | 582 | J6C05_01770 | 47591..49339 | 59823 | putative-DNA-binding-domain-containing-protein-[Prevotella-sp.] |
| WP_264965399.1 | Schlafen+HATPase+wHTH | GCF_026015685.1 | Segatella_copri | Bacteria>Pseudomonadati>FCB_group>Bacteroidota/Chlorobiota_group>Bacteroidota>Bacteroidia>Bacteroidales>Prevotellaceae>Segatella | 484 | ONT16_RS03145 | 724771..726225 | 165179 | ATP-binding-protein-[Prevotella-copri] |
| WP_262301576.1 | Schlafen+HATPase+wHTH | GCF_025460925.1 | Segatella_hominis | Bacteria>Pseudomonadati>FCB_group>Bacteroidota/Chlorobiota_group>Bacteroidota>Bacteroidia>Bacteroidales>Prevotellaceae>Segatella | 555 | OCW41_RS08890 | 94878..96545 | 2518605 | ATP-binding-protein-[Prevotella-hominis] |
| WP_252760929.1 | Schlafen+HATPase+wHTH | GCF_024054555.1 | Segatella_cerevisiae | Bacteria>Pseudomonadati>FCB_group>Bacteroidota/Chlorobiota_group>Bacteroidota>Bacteroidia>Bacteroidales>Prevotellaceae>Segatella | 514 | NG821_RS06915 | 13689..15233 | 2053716 | ATP-binding-protein-[Prevotella-cerevisiae] |
| WP_233339203.1 | Schlafen+HATPase+wHTH | GCF_021344195.1 | Segatella_copri | Bacteria>Pseudomonadati>FCB_group>Bacteroidota/Chlorobiota_group>Bacteroidota>Bacteroidia>Bacteroidales>Prevotellaceae>Segatella | 564 | LYY06_RS08400 | 29274..30968 | 165179 | putative-DNA-binding-domain-containing-protein-[Prevotella-copri] |
| WP_203049166.1 | Schlafen+HATPase+wHTH | GCF_016803255.1 | Segatella_copri | Bacteria>Pseudomonadati>FCB_group>Bacteroidota/Chlorobiota_group>Bacteroidota>Bacteroidia>Bacteroidales>Prevotellaceae>Segatella | 564 | F8Q89_RS01970 | 79583..81277 | 165179 | putative-DNA-binding-domain-containing-protein-[Prevotella-copri] |
| WP_117695277.1 | Schlafen+HATPase+wHTH | GCF_003437315.1 | Segatella_copri | Bacteria>Pseudomonadati>FCB_group>Bacteroidota/Chlorobiota_group>Bacteroidota>Bacteroidia>Bacteroidales>Prevotellaceae>Segatella | 564 | DXC61_RS12010 | 21019..22713 | 165179 | putative-DNA-binding-domain-containing-protein-[Prevotella-copri] |
| MCB9017747.1 | Schlafen+HATPase+wHTH | GCA_020636675.1 | Prevotellaceae_bacterium | Bacteria>Pseudomonadati>FCB_group>Bacteroidota/Chlorobiota_group>Bacteroidota>Bacteroidia>Bacteroidales>Prevotellaceae>unclassified_Prevotellaceae | 448 | H6544_03990 | 61342..62688 | 2049047 | putative-DNA-binding-domain-containing-protein-[Prevotellaceae-bacterium] |
| WP_074760631.1 | Schlafen+HATPase+wHTH | GCF_900107775.1 | Xylanibacter_ruminicola | Bacteria>Pseudomonadati>FCB_group>Bacteroidota/Chlorobiota_group>Bacteroidota>Bacteroidia>Bacteroidales>Prevotellaceae>Xylanibacter | 477 | BLU97_RS05630 | 376119..377552 | 839 | ATP-binding-protein-[Prevotella-ruminicola] |
| NBJ07684.1 | Schlafen+HATPase+wHTH | GCA_009911575.1 | Alistipes_sp._Z76 | Bacteria>Pseudomonadati>FCB_group>Bacteroidota/Chlorobiota_group>Bacteroidota>Bacteroidia>Bacteroidales>Rikenellaceae>Alistipes>unclassified_Alistipes | 536 | D1647_16100 | 24782..26392 | 2304565 | AAA-family-ATPase-[Alistipes-sp.-Z76] |
| HCV16202.1 | Schlafen+HATPase+wHTH | GCA_003542255.1 | Rikenellaceae_bacterium | Bacteria>Pseudomonadati>FCB_group>Bacteroidota/Chlorobiota_group>Bacteroidota>Bacteroidia>Bacteroidales>Rikenellaceae>unclassified_Rikenellaceae | 560 | DF637_07705 | 10396..12078 | 2049048 | hypothetical-protein-[Rikenellaceae-bacterium] |
| WP_122342694.1 | Schlafen+HATPase+wHTH | GCF_039750875.1 | Parabacteroides | Bacteria>Pseudomonadati>FCB_group>Bacteroidota/Chlorobiota_group>Bacteroidota>Bacteroidia>Bacteroidales>Tannerellaceae | 545 | DWZ81_RS06105 | 95184..96821 | 375288 | ATP-binding-protein-[Parabacteroides-merdae] |
| WP_158543337.1 | Schlafen+HATPase+wHTH | GCF_014385205.1 | Parabacteroides_acidifaciens | Bacteria>Pseudomonadati>FCB_group>Bacteroidota/Chlorobiota_group>Bacteroidota>Bacteroidia>Bacteroidales>Tannerellaceae>Parabacteroides | 552 | H8784_RS05815 | 83070..84728 | 2290935 | putative-DNA-binding-domain-containing-protein-[Parabacteroides-acidifaciens] |
| WP_122345558.1 | Schlafen+HATPase+wHTH | GCF_938033175.1 | Parabacteroides_sp._AF17-28 | Bacteria>Pseudomonadati>FCB_group>Bacteroidota/Chlorobiota_group>Bacteroidota>Bacteroidia>Bacteroidales>Tannerellaceae>Parabacteroides>unclassified_Parabacteroides | 552 | DWW90_RS15100 | 30159..31817 | 2292241 | putative-DNA-binding-domain-containing-protein-[Parabacteroides-sp.-AF17-28] |
| RHR54762.1 | Schlafen+HATPase+wHTH | GCA_003479205.1 | Parabacteroides_sp._AF17-28 | Bacteria>Pseudomonadati>FCB_group>Bacteroidota/Chlorobiota_group>Bacteroidota>Bacteroidia>Bacteroidales>Tannerellaceae>Parabacteroides>unclassified_Parabacteroides | 552 | DWW90_15095 | 30159..31817 | 2292241 | AAA-family-ATPase-[Parabacteroides-sp.-AF17-28] |
| MCC8143015.1 | Schlafen+HATPase+wHTH | GCA_020860665.1 | Tannerellaceae_bacterium | Bacteria>Pseudomonadati>FCB_group>Bacteroidota/Chlorobiota_group>Bacteroidota>Bacteroidia>Bacteroidales>Tannerellaceae>unclassified_Tannerellaceae | 270 | LIO97_03245 | 3976..4788 | 2785930 | putative-DNA-binding-domain-containing-protein-[Tannerellaceae-bacterium] |
| NLT02435.1 | Schlafen+HATPase+wHTH | GCA_012718945.1 | Bacteroidales_bacterium | Bacteria>Pseudomonadati>FCB_group>Bacteroidota/Chlorobiota_group>Bacteroidota>Bacteroidia>Bacteroidales>unclassified_Bacteroidales | 463 | GXY09_01070 | 10920..12311 | 2030927 | ArsR-family-transcriptional-regulator-[Bacteroidales-bacterium] |
| MCZ2341282.1 | Schlafen+HATPase+wHTH | GCA_026931725.1 | Bacteroidales_bacterium | Bacteria>Pseudomonadati>FCB_group>Bacteroidota/Chlorobiota_group>Bacteroidota>Bacteroidia>Bacteroidales>unclassified_Bacteroidales | 563 | LC104_05730 | 10068..11759 | 2030927 | putative-DNA-binding-domain-containing-protein-[Bacteroidales-bacterium] |
| MCQ2306875.1 | Schlafen+HATPase+wHTH | GCA_024400305.1 | Bacteroidales_bacterium | Bacteria>Pseudomonadati>FCB_group>Bacteroidota/Chlorobiota_group>Bacteroidota>Bacteroidia>Bacteroidales>unclassified_Bacteroidales | 465 | MJ000_04830 | 41046..42443 | 2030927 | putative-DNA-binding-domain-containing-protein-[Bacteroidales-bacterium] |
| MCK9303765.1 | Schlafen+HATPase+wHTH | GCA_023230265.1 | Bacteroidales_bacterium | Bacteria>Pseudomonadati>FCB_group>Bacteroidota/Chlorobiota_group>Bacteroidota>Bacteroidia>Bacteroidales>unclassified_Bacteroidales | 486 | MOP27_00020 | 6627..8087 | 2030927 | putative-DNA-binding-domain-containing-protein-[Bacteroidales-bacterium] |
| MCK9303762.1 | Schlafen+HATPase+wHTH | GCA_023230265.1 | Bacteroidales_bacterium | Bacteria>Pseudomonadati>FCB_group>Bacteroidota/Chlorobiota_group>Bacteroidota>Bacteroidia>Bacteroidales>unclassified_Bacteroidales | 548 | MOP27_00005 | 398..2044 | 2030927 | putative-DNA-binding-domain-containing-protein-[Bacteroidales-bacterium] |
| MCK4344534.1 | Schlafen+HATPase+wHTH | GCA_023128755.1 | Bacteroidales_bacterium | Bacteria>Pseudomonadati>FCB_group>Bacteroidota/Chlorobiota_group>Bacteroidota>Bacteroidia>Bacteroidales>unclassified_Bacteroidales | 504 | KAX05_04550 | 4982..6496 | 2030927 | putative-DNA-binding-domain-containing-protein-[Bacteroidales-bacterium] |
| MCC8119634.1 | Schlafen+HATPase+wHTH | GCA_020860515.1 | Bacteroidales_bacterium | Bacteria>Pseudomonadati>FCB_group>Bacteroidota/Chlorobiota_group>Bacteroidota>Bacteroidia>Bacteroidales>unclassified_Bacteroidales | 447 | LIP09_12930 | 21374..22714 | 2030927 | putative-DNA-binding-domain-containing-protein-[Bacteroidales-bacterium] |
| MCC8119630.1 | Schlafen+HATPase+wHTH | GCA_020860515.1 | Bacteroidales_bacterium | Bacteria>Pseudomonadati>FCB_group>Bacteroidota/Chlorobiota_group>Bacteroidota>Bacteroidia>Bacteroidales>unclassified_Bacteroidales | 562 | LIP09_12910 | 17241..18929 | 2030927 | putative-DNA-binding-domain-containing-protein-[Bacteroidales-bacterium] |
| MBR0038051.1 | Schlafen+HATPase+wHTH | GCA_017449065.1 | Bacteroidales_bacterium | Bacteria>Pseudomonadati>FCB_group>Bacteroidota/Chlorobiota_group>Bacteroidota>Bacteroidia>Bacteroidales>unclassified_Bacteroidales | 495 | IJP70_10510 | 34563..36050 | 2030927 | putative-DNA-binding-domain-containing-protein-[Bacteroidales-bacterium] |
| MBQ7946202.1 | Schlafen+HATPase+wHTH | GCA_017468435.1 | Bacteroidales_bacterium | Bacteria>Pseudomonadati>FCB_group>Bacteroidota/Chlorobiota_group>Bacteroidota>Bacteroidia>Bacteroidales>unclassified_Bacteroidales | 581 | IJ280_05765 | 11440..13185 | 2030927 | putative-DNA-binding-domain-containing-protein-[Bacteroidales-bacterium] |
| MBQ6275499.1 | Schlafen+HATPase+wHTH | GCA_017428605.1 | Bacteroidales_bacterium | Bacteria>Pseudomonadati>FCB_group>Bacteroidota/Chlorobiota_group>Bacteroidota>Bacteroidia>Bacteroidales>unclassified_Bacteroidales | 466 | IJK62_02180 | 72495..73895 | 2030927 | putative-DNA-binding-domain-containing-protein-[Bacteroidales-bacterium] |
| MBQ4549128.1 | Schlafen+HATPase+wHTH | GCA_017399385.1 | Bacteroidales_bacterium | Bacteria>Pseudomonadati>FCB_group>Bacteroidota/Chlorobiota_group>Bacteroidota>Bacteroidia>Bacteroidales>unclassified_Bacteroidales | 581 | IJA42_08120 | 148450..150195 | 2030927 | putative-DNA-binding-domain-containing-protein-[Bacteroidales-bacterium] |
| WP_062122669.1 | Schlafen+HATPase+wHTH | GCF_000974365.1 | Geofilum_rubicundum | Bacteria>Pseudomonadati>FCB_group>Bacteroidota/Chlorobiota_group>Bacteroidota>Bacteroidia>Marinilabiliales>Marinilabiliaceae>Geofilum | 483 | JCM15548_RS04450 | 148418..149869 | 472113 | ATP-binding-protein-[Geofilum-rubicundum] |
| WP_163710194.1 | Schlafen+HATPase+wHTH | GCF_010882175.1 | Mangrovibacterium_lignilyticum | Bacteria>Pseudomonadati>FCB_group>Bacteroidota/Chlorobiota_group>Bacteroidota>Bacteroidia>Marinilabiliales>Prolixibacteraceae>Mangrovibacterium | 531 | GJV22_RS06150 | 1522600..1524195 | 2668052 | putative-DNA-binding-domain-containing-protein-[Mangrovibacterium-lignilyticum] |
| WP_107823953.1 | Schlafen+HATPase+wHTH | GCF_963677905.1 | Mangrovibacterium_marinum | Bacteria>Pseudomonadati>FCB_group>Bacteroidota/Chlorobiota_group>Bacteroidota>Bacteroidia>Marinilabiliales>Prolixibacteraceae>Mangrovibacterium | 541 | C8N47_RS17930 | 9920..11545 | 1639118 | putative-DNA-binding-domain-containing-protein-[Mangrovibacterium-marinum] |
| MCK9411441.1 | Schlafen+HATPase+wHTH | GCA_023229575.1 | Prolixibacteraceae_bacterium | Bacteria>Pseudomonadati>FCB_group>Bacteroidota/Chlorobiota_group>Bacteroidota>Bacteroidia>Marinilabiliales>Prolixibacteraceae>unclassified_Prolixibacteraceae | 498 | MOQ53_04010 | 32167..33663 | 2053594 | hypothetical-protein-[Prolixibacteraceae-bacterium] |
| MCF8362393.1 | Schlafen+HATPase+wHTH | GCA_021740345.1 | Prolixibacteraceae_bacterium | Bacteria>Pseudomonadati>FCB_group>Bacteroidota/Chlorobiota_group>Bacteroidota>Bacteroidia>Marinilabiliales>Prolixibacteraceae>unclassified_Prolixibacteraceae | 89 | K9G70_07190 | 16513..16782 | 2053594 | ATP-binding-protein-[Prolixibacteraceae-bacterium] |
| MCF8362386.1 | Schlafen+HATPase+wHTH | GCA_021740345.1 | Prolixibacteraceae_bacterium | Bacteria>Pseudomonadati>FCB_group>Bacteroidota/Chlorobiota_group>Bacteroidota>Bacteroidia>Marinilabiliales>Prolixibacteraceae>unclassified_Prolixibacteraceae | 541 | K9G70_07155 | 6004..7629 | 2053594 | putative-DNA-binding-domain-containing-protein-[Prolixibacteraceae-bacterium] |
| GIV23746.1 | Schlafen+HATPase+wHTH | GCA_026003215.1 | Bacteroidia_bacterium | Bacteria>Pseudomonadati>FCB_group>Bacteroidota/Chlorobiota_group>Bacteroidota>Bacteroidia>unclassified_Bacteroidia | 567 | KatS3mg025_1405 | 1611564..1613267 | 2044936 | MAG:-ATPase-AAA-[Bacteroidia-bacterium] |
| WP_255036054.1 | Schlafen+HATPase+wHTH | GCF_024343775.1 | Lacihabitans_soyangensis | Bacteria>Pseudomonadati>FCB_group>Bacteroidota/Chlorobiota_group>Bacteroidota>Cytophagia>Cytophagales>Leadbetterellaceae>Lacihabitans | 250 | EGI31_RS04910 | 54315..55067 | 869394 | ArsR-family-transcriptional-regulator-[Lacihabitans-soyangensis] |
| WP_138994215.1 | Schlafen+HATPase+wHTH | GCF_006149005.1 | Larkinella_sp._C7 | Bacteria>Pseudomonadati>FCB_group>Bacteroidota/Chlorobiota_group>Bacteroidota>Cytophagia>Cytophagales>Spirosomataceae>Larkinella>unclassified_Larkinella | 537 | FH871_RS25325 | 22482..24095 | 2576607 | putative-DNA-binding-domain-containing-protein-[Larkinella-sp.-C7] |
| WP_273212241.1 | Schlafen+HATPase+wHTH | GCF_002483185.1 | Runella_zeae | Bacteria>Pseudomonadati>FCB_group>Bacteroidota/Chlorobiota_group>Bacteroidota>Cytophagia>Cytophagales>Spirosomataceae>Runella | 460 | B4E63_RS12435 | 299561..300943 | 94255 | helix-turn-helix-domain-containing-protein-[Runella-zeae] |
| WP_013929904.1 | Schlafen+HATPase+wHTH | GCF_000218895.1 | Runella_slithyformis | Bacteria>Pseudomonadati>FCB_group>Bacteroidota/Chlorobiota_group>Bacteroidota>Cytophagia>Cytophagales>Spirosomataceae>Runella | 454 | RUNSL_RS20910 | 4926952..4928316 | 106 | helix-turn-helix-domain-containing-protein-[Runella-slithyformis] |
| MBW7869166.1 | Schlafen+HATPase+wHTH | GCA_019455205.1 | Brumimicrobium_sp. | Bacteria>Pseudomonadati>FCB_group>Bacteroidota/Chlorobiota_group>Bacteroidota>Flavobacteriia>Flavobacteriales>Crocinitomicaceae>Brumimicrobium>unclassified_Brumimicrobium | 542 | H3C31_12690 | 9754..11382 | 2029867 | putative-DNA-binding-domain-containing-protein-[Brumimicrobium-sp.] |
| WP_026451334.1 | Schlafen+HATPase+wHTH | GCF_000429125.1 | Aequorivita_capsosiphonis | Bacteria>Pseudomonadati>FCB_group>Bacteroidota/Chlorobiota_group>Bacteroidota>Flavobacteriia>Flavobacteriales>Flavobacteriaceae>Aequorivita | 534 | G440_RS0110865 | 111778..113382 | 487317 | putative-DNA-binding-domain-containing-protein-[Aequorivita-capsosiphonis] |
| MBQ0768323.1 | Schlafen+HATPase+wHTH | GCA_018067115.1 | Bizionia_sp. | Bacteria>Pseudomonadati>FCB_group>Bacteroidota/Chlorobiota_group>Bacteroidota>Flavobacteriia>Flavobacteriales>Flavobacteriaceae>Bizionia>unclassified_Bizionia | 489 | KBT58_03465 | 2447..3916 | 1954480 | putative-DNA-binding-domain-containing-protein-[Bizionia-sp.] |
| WP_094415996.1 | Schlafen+HATPase+wHTH | GCF_002251835.1 | Flavobacterium_cyanobacteriorum | Bacteria>Pseudomonadati>FCB_group>Bacteroidota/Chlorobiota_group>Bacteroidota>Flavobacteriia>Flavobacteriales>Flavobacteriaceae>Flavobacterium | 564 | CHU92_RS12280 | 16626..18320 | 2022802 | putative-DNA-binding-domain-containing-protein-[Flavobacterium-cyanobacteriorum] |
| WP_199600305.1 | Schlafen+HATPase+wHTH | GCF_016469035.1 | Gelidibacter_salicanalis | Bacteria>Pseudomonadati>FCB_group>Bacteroidota/Chlorobiota_group>Bacteroidota>Flavobacteriia>Flavobacteriales>Flavobacteriaceae>Gelidibacter | 538 | JEM65_RS13230 | 168631..170247 | 291193 | putative-DNA-binding-domain-containing-protein-[Gelidibacter-salicanalis] |
| WP_129016552.1 | Schlafen+HATPase+wHTH | GCF_004115975.1 | Gelidibacter_gilvus | Bacteria>Pseudomonadati>FCB_group>Bacteroidota/Chlorobiota_group>Bacteroidota>Flavobacteriia>Flavobacteriales>Flavobacteriaceae>Gelidibacter | 476 | ESZ48_RS06595 | 15835..17265 | 59602 | putative-DNA-binding-domain-containing-protein-[Gelidibacter-gilvus] |
| WP_034059319.1 | Schlafen+HATPase+wHTH | GCF_000797445.1 | Lacinutrix_jangbogonensis | Bacteria>Pseudomonadati>FCB_group>Bacteroidota/Chlorobiota_group>Bacteroidota>Flavobacteriia>Flavobacteriales>Flavobacteriaceae>Lacinutrix | 533 | RH82_RS07095 | 66110..67711 | 1469557 | putative-DNA-binding-domain-containing-protein-[Lacinutrix-jangbogonensis] |
| NJY61424.1 | Schlafen+HATPase+wHTH | GCA_012038135.1 | Salinimicrobium_nanhaiense | Bacteria>Pseudomonadati>FCB_group>Bacteroidota/Chlorobiota_group>Bacteroidota>Flavobacteriia>Flavobacteriales>Flavobacteriaceae>Salinimicrobium | 568 | HC174_01465 | 305887..307593 | 2722701 | hypothetical-protein-[Salinimicrobium-nanhaiense] |
| WP_093139958.1 | Schlafen+HATPase+wHTH | GCF_014651275.1 | Ulvibacter_litoralis | Bacteria>Pseudomonadati>FCB_group>Bacteroidota/Chlorobiota_group>Bacteroidota>Flavobacteriia>Flavobacteriales>Flavobacteriaceae>Ulvibacter | 476 | IE050_RS03675 | 815060..816490 | 227084 | putative-DNA-binding-domain-containing-protein-[Ulvibacter-litoralis] |
| MAL23062.1 | Schlafen+HATPase+wHTH | GCA_002694765.1 | Xanthomarina_sp. | Bacteria>Pseudomonadati>FCB_group>Bacteroidota/Chlorobiota_group>Bacteroidota>Flavobacteriia>Flavobacteriales>Flavobacteriaceae>Xanthomarina>unclassified_Xanthomarina | 537 | CMP04_07450 | 105703..107316 | 1931211 | hypothetical-protein-[Xanthomarina-sp.] |
| ROI04328.1 | Schlafen+HATPase+wHTH | GCA_003729975.1 | Kaistella_haifensis | Bacteria>Pseudomonadati>FCB_group>Bacteroidota/Chlorobiota_group>Bacteroidota>Flavobacteriia>Flavobacteriales>Weeksellaceae>Chryseobacterium_group>Kaistella | 446 | EGH90_10155 | 60304..61644 | 421525 | winged-helix-turn-helix-transcriptional-regulator-[Kaistella-haifensis] |
| WP_028298623.1 | Schlafen+HATPase+wHTH | GCF_000427965.1 | Olivibacter_sitiensis | Bacteria>Pseudomonadati>FCB_group>Bacteroidota/Chlorobiota_group>Bacteroidota>Sphingobacteriia>Sphingobacteriales>Sphingobacteriaceae>Olivibacter | 487 | A37S_RS0120645 | 14300..15763 | 376470 | helix-turn-helix-domain-containing-protein-[Olivibacter-sitiensis] |
| NLJ08469.1 | Schlafen+HATPase+wHTH | GCA_012517435.1 | Sphingobacteriales_bacterium | Bacteria>Pseudomonadati>FCB_group>Bacteroidota/Chlorobiota_group>Bacteroidota>Sphingobacteriia>Sphingobacteriales>unclassified_Sphingobacteriales | 580 | GX437_12470 | 12404..14146 | 2044944 | transcriptional-regulator-[Sphingobacteriales-bacterium] |
| NCA71198.1 | Schlafen+HATPase+wHTH | GCA_009928635.1 | Sphingobacteriia_bacterium | Bacteria>Pseudomonadati>FCB_group>Bacteroidota/Chlorobiota_group>Bacteroidota>Sphingobacteriia>unclassified_Sphingobacteriia | 575 | EOM91_14045 | 8635..10362 | 2035813 | AAA-family-ATPase-[Sphingobacteriia-bacterium] |
| OQC38487.1 | Schlafen+HATPase+wHTH | GCA_002071145.1 | Bacteroidetes_bacterium_ADurb.Bin041 | Bacteria>Pseudomonadati>FCB_group>Bacteroidota/Chlorobiota_group>Bacteroidota>unclassified_Bacteroidota | 442 | BWX63_00122 | 10224..11552 | 1852801 | Divergent-AAA-domain-protein-[Bacteroidetes-bacterium-ADurb.Bin041] |
| NUQ82572.1 | Schlafen+HATPase+wHTH | GCA_013360915.1 | Bacteroidota_bacterium | Bacteria>Pseudomonadati>FCB_group>Bacteroidota/Chlorobiota_group>Bacteroidota>unclassified_Bacteroidota | 511 | HUU10_13240 | 54941..56476 | 1898104 | putative-DNA-binding-domain-containing-protein-[Bacteroidota-bacterium] |
| MQY79008.1 | Schlafen+HATPase+wHTH | GCA_009619225.1 | Bacteroidota_bacterium | Bacteria>Pseudomonadati>FCB_group>Bacteroidota/Chlorobiota_group>Bacteroidota>unclassified_Bacteroidota | 451 | GH151_07415 | 3265..4620 | 1898104 | transcriptional-regulator-[Bacteroidota-bacterium] |
| MCB0536187.1 | Schlafen+HATPase+wHTH | GCA_020436205.1 | Bacteroidota_bacterium | Bacteria>Pseudomonadati>FCB_group>Bacteroidota/Chlorobiota_group>Bacteroidota>unclassified_Bacteroidota | 474 | KDE33_01545 | 12119..13543 | 1898104 | putative-DNA-binding-domain-containing-protein-[Bacteroidota-bacterium] |
| MBN8706504.1 | Schlafen+HATPase+wHTH | GCA_017303995.1 | Bacteroidota_bacterium | Bacteria>Pseudomonadati>FCB_group>Bacteroidota/Chlorobiota_group>Bacteroidota>unclassified_Bacteroidota | 461 | JOL62_11570 | 17736..19121 | 1898104 | putative-DNA-binding-domain-containing-protein-[Bacteroidota-bacterium] |
| MBK9503942.1 | Schlafen+HATPase+wHTH | GCA_016718695.1 | Bacteroidota_bacterium | Bacteria>Pseudomonadati>FCB_group>Bacteroidota/Chlorobiota_group>Bacteroidota>unclassified_Bacteroidota | 474 | IPO03_00790 | 164501..165925 | 1898104 | putative-DNA-binding-domain-containing-protein-[Bacteroidota-bacterium] |
| MBK9455377.1 | Schlafen+HATPase+wHTH | GCA_016718365.1 | Bacteroidota_bacterium | Bacteria>Pseudomonadati>FCB_group>Bacteroidota/Chlorobiota_group>Bacteroidota>unclassified_Bacteroidota | 474 | IPO24_07200 | 116038..117462 | 1898104 | putative-DNA-binding-domain-containing-protein-[Bacteroidota-bacterium] |
| KPP94732.1 | Schlafen+HATPase+wHTH | GCA_001314545.1 | Bacteroidetes_bacterium_HLUCCA01 | Bacteria>Pseudomonadati>FCB_group>Bacteroidota/Chlorobiota_group>Bacteroidota>unclassified_Bacteroidota | 557 | HLUCCA01_08735 | 182744..184417 | 1666909 | ATP-dependent-DNA-helicase-[Bacteroidetes-bacterium-HLUCCA01] |
| KAF0128490.1 | Schlafen+HATPase+wHTH | GCA_009773165.1 | Bacteroidota_bacterium | Bacteria>Pseudomonadati>FCB_group>Bacteroidota/Chlorobiota_group>Bacteroidota>unclassified_Bacteroidota | 545 | FD155_3232 | 33823..35460 | 1898104 | transcriptional-regulator-with-an-ATP-binding-domain-[Bacteroidota-bacterium] |
| HEX04757.1 | Schlafen+HATPase+wHTH | GCA_011043795.1 | Bacteroidota_bacterium | Bacteria>Pseudomonadati>FCB_group>Bacteroidota/Chlorobiota_group>Bacteroidota>unclassified_Bacteroidota | 492 | ENH10_06320 | 3404..4882 | 1898104 | AAA-family-ATPase-[Bacteroidota-bacterium] |
| BCX00055.1 | Schlafen+HATPase+wHTH | GCA_025998875.1 | Bacteroidota_bacterium | Bacteria>Pseudomonadati>FCB_group>Bacteroidota/Chlorobiota_group>Bacteroidota>unclassified_Bacteroidota | 568 | KatS3mg041_0101 | 112144..113850 | 1898104 | MAG:-ATPase-AAA-[Bacteroidota-bacterium] |
| WP_265763891.1 | Schlafen+HATPase+wHTH | GCF_026229185.1 | Fodinibius_salsisoli | Bacteria>Pseudomonadati>FCB_group>Bacteroidota/Chlorobiota_group>Balneolota>Balneolia>Balneolales>Balneolaceae>Fodinibius | 564 | J6I44_RS00110 | 26693..28387 | 2820877 | putative-DNA-binding-domain-containing-protein-[Aliifodinibius-sp.-1BSP15-2V2] |
| MCC5907047.1 | Schlafen+HATPase+wHTH | GCA_020832285.1 | Balneolaceae_bacterium | Bacteria>Pseudomonadati>FCB_group>Bacteroidota/Chlorobiota_group>Balneolota>Balneolia>Balneolales>Balneolaceae>unclassified_Balneolaceae | 564 | JJU13_12605 | 38234..39928 | 2053516 | putative-DNA-binding-domain-containing-protein-[Balneolaceae-bacterium] |
| GIV55099.1 | Schlafen+HATPase+wHTH | GCA_026003475.1 | Candidatus_Kapaibacterium_sp. | Bacteria>Pseudomonadati>FCB_group>Bacteroidota/Chlorobiota_group>Candidatus_Kapaibacteriota>Candidatus_Kapaibacteriia>Candidatus_Kapaibacteriales>Candidatus_Kapaibacteriaceae>Candidatus_Kapaibacterium>unclassified_Candidatus_Kapaibacterium | 139 | KatS3mg039_1617 | 1976475..1976894 | 2447898 | MAG:-hypothetical-protein-KatS3mg039_1617-[Candidatus-Kapabacteria-bacterium] |
| WP_076791023.1 | Schlafen+HATPase+wHTH | GCF_001975465.1 | Chlorobium_sp._KB01 | Bacteria>Pseudomonadati>FCB_group>Bacteroidota/Chlorobiota_group>Chlorobiota>Chlorobiia>Chlorobiales>Chlorobiaceae>Chlorobium/Pelodictyon_group>Chlorobium>unclassified_Chlorobium | 556 | BOO20_RS04860 | 10633..12303 | 1917528 | ATP-binding-protein-[Chlorobium-sp.-KB01] |
| WP_076789314.1 | Schlafen+HATPase+wHTH | - | Chlorobium_sp._KB01 | Bacteria>Pseudomonadati>FCB_group>Bacteroidota/Chlorobiota_group>Chlorobiota>Chlorobiia>Chlorobiales>Chlorobiaceae>Chlorobium/Pelodictyon_group>Chlorobium>unclassified_Chlorobium | 512 | BOO20_RS00740 | 12441..13979 | 1917528 | putative-DNA-binding-domain-containing-protein-[Chlorobium-sp.-KB01] |
| MCF8215292.1 | Schlafen+HATPase+wHTH | GCA_021738525.1 | Chlorobium_sp. | Bacteria>Pseudomonadati>FCB_group>Bacteroidota/Chlorobiota_group>Chlorobiota>Chlorobiia>Chlorobiales>Chlorobiaceae>Chlorobium/Pelodictyon_group>Chlorobium>unclassified_Chlorobium | 413 | K9I96_00270 | 70799..72040 | 1095 | putative-DNA-binding-domain-containing-protein-[Chlorobium-sp.] |
| MCF8215289.1 | Schlafen+HATPase+wHTH | GCA_021738525.1 | Chlorobium_sp. | Bacteria>Pseudomonadati>FCB_group>Bacteroidota/Chlorobiota_group>Chlorobiota>Chlorobiia>Chlorobiales>Chlorobiaceae>Chlorobium/Pelodictyon_group>Chlorobium>unclassified_Chlorobium | 570 | K9I96_00255 | 62382..64094 | 1095 | putative-DNA-binding-domain-containing-protein-[Chlorobium-sp.] |
| MBV5329908.1 | Schlafen+HATPase+wHTH | GCA_019162945.1 | Chlorobium_sp. | Bacteria>Pseudomonadati>FCB_group>Bacteroidota/Chlorobiota_group>Chlorobiota>Chlorobiia>Chlorobiales>Chlorobiaceae>Chlorobium/Pelodictyon_group>Chlorobium>unclassified_Chlorobium | 610 | JZU65_20150 | 8643..10475 | 1095 | putative-DNA-binding-domain-containing-protein-[Chlorobium-sp.] |
| MBV5329668.1 | Schlafen+HATPase+wHTH | GCA_019162945.1 | Chlorobium_sp. | Bacteria>Pseudomonadati>FCB_group>Bacteroidota/Chlorobiota_group>Chlorobiota>Chlorobiia>Chlorobiales>Chlorobiaceae>Chlorobium/Pelodictyon_group>Chlorobium>unclassified_Chlorobium | 574 | JZU65_18930 | 702..2426 | 1095 | putative-DNA-binding-domain-containing-protein-[Chlorobium-sp.] |
| MBV5326251.1 | Schlafen+HATPase+wHTH | GCA_019162945.1 | Chlorobium_sp. | Bacteria>Pseudomonadati>FCB_group>Bacteroidota/Chlorobiota_group>Chlorobiota>Chlorobiia>Chlorobiales>Chlorobiaceae>Chlorobium/Pelodictyon_group>Chlorobium>unclassified_Chlorobium | 556 | JZU65_01245 | 7784..9454 | 1095 | putative-DNA-binding-domain-containing-protein-[Chlorobium-sp.] |
| MBV5326245.1 | Schlafen+HATPase+wHTH | GCA_019162945.1 | Chlorobium_sp. | Bacteria>Pseudomonadati>FCB_group>Bacteroidota/Chlorobiota_group>Chlorobiota>Chlorobiia>Chlorobiales>Chlorobiaceae>Chlorobium/Pelodictyon_group>Chlorobium>unclassified_Chlorobium | 509 | JZU65_01210 | 5981..7510 | 1095 | putative-DNA-binding-domain-containing-protein-[Chlorobium-sp.] |
| KZK74370.1 | Schlafen+HATPase+wHTH | GCA_001622165.1 | Pelodictyon_luteolum | Bacteria>Pseudomonadati>FCB_group>Bacteroidota/Chlorobiota_group>Chlorobiota>Chlorobiia>Chlorobiales>Chlorobiaceae>Chlorobium/Pelodictyon_group>Pelodictyon | 588 | A3K90_02900 | 30631..32397 | 1100 | transcriptional-regulator-[Pelodictyon-luteolum] |

| Protein-ID | Domain | Assembly-ID | Strain | Taxonomy | Length | Locus-Tag | Genomic-coordinates | TaxID | Fasta-defline |
|---|---|---|---|---|---|---|---|---|---|
| WP_139627001.1 | Schlafen+HATPase+wHTH | GCF_006265245.1 | Prosthecochloris_vibrioformis | Bacteria>Pseudomonadati>FCB_group>Bacteroidota/Chlorobiota_group>Chlorobiota>Chlorobiia>Chlorobiales>Chlorobiaceae>Prosthecochloris | 570 | FGF68_RS10430 | 13031..14743 | 1098 | ATP-binding-protein-[Prosthecochloris-vibrioformis] |
| WP_139626988.1 | Schlafen+HATPase+wHTH | - | Prosthecochloris_vibrioformis | Bacteria>Pseudomonadati>FCB_group>Bacteroidota/Chlorobiota_group>Chlorobiota>Chlorobiia>Chlorobiales>Chlorobiaceae>Prosthecochloris | 418 | FGF68_RS10375 | 20345..21601 | 1098 | ATP-binding-protein-[Prosthecochloris-vibrioformis] |
| WP_139626760.1 | Schlafen+HATPase+wHTH | GCF_006265245.1 | Prosthecochloris_vibrioformis | Bacteria>Pseudomonadati>FCB_group>Bacteroidota/Chlorobiota_group>Chlorobiota>Chlorobiia>Chlorobiales>Chlorobiaceae>Prosthecochloris | 556 | FGF68_RS08500 | 8136..9806 | 1098 | ATP-binding-protein-[Prosthecochloris-vibrioformis] |
| TNJ34103.1 | Schlafen+HATPase+wHTH | GCA_006265245.1 | Prosthecochloris_vibrioformis | Bacteria>Pseudomonadati>FCB_group>Bacteroidota/Chlorobiota_group>Chlorobiota>Chlorobiia>Chlorobiales>Chlorobiaceae>Prosthecochloris | 418 | FGF68_10380 | 20345..21601 | 1098 | transcriptional-regulator-[Prosthecochloris-vibrioformis] |
| TNJ34083.1 | Schlafen+HATPase+wHTH | GCA_006265245.1 | Prosthecochloris_vibrioformis | Bacteria>Pseudomonadati>FCB_group>Bacteroidota/Chlorobiota_group>Chlorobiota>Chlorobiia>Chlorobiales>Chlorobiaceae>Prosthecochloris | 570 | FGF68_10435 | 13031..14743 | 1098 | AAA-family-ATPase-[Prosthecochloris-vibrioformis] |
| WP_068867400.1 | Schlafen+HATPase+wHTH | GCF_001687065.1 | Prosthecochloris_sp._CIB_2401 | Bacteria>Pseudomonadati>FCB_group>Bacteroidota/Chlorobiota_group>Chlorobiota>Chlorobiia>Chlorobiales>Chlorobiaceae>Prosthecochloris>unclassified_Prosthecochloris | 572 | Ptc2401_RS06775 | 1429170..1430888 | 1868325 | ATP-binding-protein-[Prosthecochloris-sp.-CIB-2401] |
| WP_068867397.1 | Schlafen+HATPase+wHTH | GCF_001687065.1 | Prosthecochloris_sp._CIB_2401 | Bacteria>Pseudomonadati>FCB_group>Bacteroidota/Chlorobiota_group>Chlorobiota>Chlorobiia>Chlorobiales>Chlorobiaceae>Prosthecochloris>unclassified_Prosthecochloris | 413 | Ptc2401_RS06760 | 1421645..1422886 | 1868325 | putative-DNA-binding-domain-containing-protein-[Prosthecochloris-sp.-CIB-2401] |
| WP_068867045.1 | Schlafen+HATPase+wHTH | GCF_001687065.1 | Prosthecochloris_sp._CIB_2401 | Bacteria>Pseudomonadati>FCB_group>Bacteroidota/Chlorobiota_group>Chlorobiota>Chlorobiia>Chlorobiales>Chlorobiaceae>Prosthecochloris>unclassified_Prosthecochloris | 556 | Ptc2401_RS05040 | 1060879..1062549 | 1868325 | ATP-binding-protein-[Prosthecochloris-sp.-CIB-2401] |
| ANT65174.1 | Schlafen+HATPase+wHTH | GCA_001687065.1 | Prosthecochloris_sp._CIB_2401 | Bacteria>Pseudomonadati>FCB_group>Bacteroidota/Chlorobiota_group>Chlorobiota>Chlorobiia>Chlorobiales>Chlorobiaceae>Prosthecochloris>unclassified_Prosthecochloris | 572 | Ptc2401_01410 | 1429170..1430888 | 1868325 | Divergent-AAA-domain-protein-[Prosthecochloris-sp.-CIB-2401] |
| ANT65171.1 | Schlafen+HATPase+wHTH | GCA_001687065.1 | Prosthecochloris_sp._CIB_2401 | Bacteria>Pseudomonadati>FCB_group>Bacteroidota/Chlorobiota_group>Chlorobiota>Chlorobiia>Chlorobiales>Chlorobiaceae>Prosthecochloris>unclassified_Prosthecochloris | 413 | Ptc2401_01407 | 1421645..1422886 | 1868325 | Divergent-AAA-domain-protein-[Prosthecochloris-sp.-CIB-2401] |
| NMW20011.1 | Schlafen+HATPase+wHTH | GCA_012972825.1 | Chlorobiaceae_bacterium | Bacteria>Pseudomonadati>FCB_group>Bacteroidota/Chlorobiota_group>Chlorobiota>Chlorobiia>Chlorobiales>Chlorobiaceae>unclassified_Chlorobiaceae | 57 | HKK66_13580 | 7757..7926 | 2067991 | ATP-binding-protein-[Chlorobiaceae-bacterium] |
| NMW20008.1 | Schlafen+HATPase+wHTH | GCA_012972825.1 | Chlorobiaceae_bacterium | Bacteria>Pseudomonadati>FCB_group>Bacteroidota/Chlorobiota_group>Chlorobiota>Chlorobiia>Chlorobiales>Chlorobiaceae>unclassified_Chlorobiaceae | 556 | HKK66_13565 | 3493..5163 | 2067991 | ATPase-[Chlorobiaceae-bacterium] |
| MBM3423464.1 | Schlafen+HATPase+wHTH | GCA_016868665.1 | Chlorobiota_bacterium | Bacteria>Pseudomonadati>FCB_group>Bacteroidota/Chlorobiota_group>Chlorobiota>unclassified_Chlorobiota | 555 | FJY09_09525 | 8993..10660 | 2268192 | putative-DNA-binding-domain-containing-protein-[Chlorobiota-bacterium] |
| MBV6512082.1 | Schlafen+HATPase+wHTH | GCA_019187025.1 | Ignavibacteriaceae_bacterium | Bacteria>Pseudomonadati>FCB_group>Bacteroidota/Chlorobiota_group>Ignavibacteriota>Ignavibacteria>Ignavibacteriales>Ignavibacteriaceae>unclassified_Ignavibacteriaceae | 562 | FMNOHCHN_01572 | 149177..150865 | 2651163 | hypothetical-protein-[Ignavibacteriaceae-bacterium] |
| QOJ30045.1 | Schlafen+HATPase+wHTH | GCA_015709675.1 | Ignavibacteriales_bacterium | Bacteria>Pseudomonadati>FCB_group>Bacteroidota/Chlorobiota_group>Ignavibacteriota>Ignavibacteria>Ignavibacteriales>unclassified_Ignavibacteriales | 562 | HRU80_14680 | 3722806..3724494 | 2049428 | putative-DNA-binding-domain-containing-protein-[Ignavibacteriales-bacterium] |
| NLO12464.1 | Schlafen+HATPase+wHTH | GCA_012522935.1 | Candidatus_Cloacimonadota_bacterium | Bacteria>Pseudomonadati>FCB_group>Candidatus_Cloacimonadota>unclassified_Candidatus_Cloacimonadota | 637 | GX122_08620 | 3623..5536 | 2030808 | hypothetical-protein-[Candidatus-Cloacimonadota-bacterium] |
| MCK9335091.1 | Schlafen+HATPase+wHTH | GCA_023230025.1 | Candidatus_Cloacimonadota_bacterium | Bacteria>Pseudomonadati>FCB_group>Candidatus_Cloacimonadota>unclassified_Candidatus_Cloacimonadota | 610 | MOQ99_07260 | 3894..5726 | 2030808 | putative-DNA-binding-domain-containing-protein-[Candidatus-Cloacimonadota-bacterium] |
| MCB5288063.1 | Schlafen+HATPase+wHTH | GCA_020532145.1 | Candidatus_Cloacimonadota_bacterium | Bacteria>Pseudomonadati>FCB_group>Candidatus_Cloacimonadota>unclassified_Candidatus_Cloacimonadota | 453 | LHW64_09700 | 5217..6578 | 2030808 | putative-DNA-binding-domain-containing-protein-[Candidatus-Cloacimonetes-bacterium] |
| MCB5262480.1 | Schlafen+HATPase+wHTH | GCA_020532425.1 | Candidatus_Cloacimonadota_bacterium | Bacteria>Pseudomonadati>FCB_group>Candidatus_Cloacimonadota>unclassified_Candidatus_Cloacimonadota | 453 | LHW53_07105 | 2488..3849 | 2030808 | putative-DNA-binding-domain-containing-protein-[Candidatus-Cloacimonadota-bacterium] |
| MCB5253473.1 | Schlafen+HATPase+wHTH | GCA_020532345.1 | Candidatus_Cloacimonadota_bacterium | Bacteria>Pseudomonadati>FCB_group>Candidatus_Cloacimonadota>unclassified_Candidatus_Cloacimonadota | 631 | LHW51_10920 | 2273..4168 | 2030808 | putative-DNA-binding-domain-containing-protein-[Candidatus-Cloacimonadota-bacterium] |
| MCB5228481.1 | Schlafen+HATPase+wHTH | GCA_020532325.1 | Candidatus_Cloacimonadota_bacterium | Bacteria>Pseudomonadati>FCB_group>Candidatus_Cloacimonadota>unclassified_Candidatus_Cloacimonadota | 637 | LHW44_00175 | 40689..42602 | 2030808 | putative-DNA-binding-domain-containing-protein-[Candidatus-Cloacimonadota-bacterium] |
| NLO34314.1 | Schlafen+HATPase+wHTH | GCA_012523015.1 | Candidatus_Hydrogenedentota_bacterium | Bacteria>Pseudomonadati>FCB_group>Candidatus_Hydrogenedentota>unclassified_Candidatus_Hydrogenedentota | 502 | GX117_13345 | 1284..2792 | 2030809 | ATP-dependent-DNA-helicase-RecG-[Candidatus-Hydrogenedentes-bacterium] |
| MBW7865070.1 | Schlafen+HATPase+wHTH | GCA_019455225.1 | Candidatus_Hydrogenedentota_bacterium | Bacteria>Pseudomonadati>FCB_group>Candidatus_Hydrogenedentota>unclassified_Candidatus_Hydrogenedentota | 417 | H3C30_11750 | 26768..28020 | 2030809 | putative-DNA-binding-domain-containing-protein-[Candidatus-Hydrogenedentes-bacterium] |
| RKY59332.1 | Schlafen+HATPase+wHTH | GCA_003645885.1 | Candidatus_Latescibacterota_bacterium | Bacteria>Pseudomonadati>FCB_group>Candidatus_Latescibacterota>unclassified_Candidatus_Latescibacterota | 460 | DRP94_03925 | 9652..11034 | 2053570 | hypothetical-protein-DRP94_03925-[Candidatus-Latescibacteria-bacterium] |
| KMQ51970.1 | Schlafen+HATPase+wHTH | GCA_001045525.1 | Chitinispirillum_alkaliphilum | Bacteria>Pseudomonadati>FCB_group>Fibrobacterota>Chitinispirillia>Chitinispirillales>Chitinispirillaceae>Chitinispirillum | 523 | CHISP_1225 | 135862..137433 | 1008392 | ATPase-AAA-[Chitinispirillum-alkaliphilum] |
| KMQ51968.1 | Schlafen+HATPase+wHTH | GCA_001045525.1 | Chitinispirillum_alkaliphilum | Bacteria>Pseudomonadati>FCB_group>Fibrobacterota>Chitinispirillia>Chitinispirillales>Chitinispirillaceae>Chitinispirillum | 592 | CHISP_1223 | 131232..133010 | 1008392 | ATPase-AAA-[Chitinispirillum-alkaliphilum] |
| WP_073156277.1 | Schlafen+HATPase+wHTH | GCF_900142465.1 | unclassified_Fibrobacter | Bacteria>Pseudomonadati>FCB_group>Fibrobacterota>Fibrobacteria>Fibrobacterales>Fibrobacteraceae>Fibrobacter | 561 | BUB73_RS01550 | 359953..361638 | 2634177 | MULTISPECIES:-putative-DNA-binding-domain-containing-protein-[unclassified-Fibrobacter] |
| WP_088660224.1 | Schlafen+HATPase+wHTH | GCF_002210425.1 | Fibrobacter_sp._UWB2 | Bacteria>Pseudomonadati>FCB_group>Fibrobacterota>Fibrobacteria>Fibrobacterales>Fibrobacteraceae>Fibrobacter>unclassified_Fibrobacter | 533 | B7982_RS07620 | 360277..361878 | 1964358 | putative-DNA-binding-domain-containing-protein-[Fibrobacter-sp.-UWB2] |
| WP_088658679.1 | Schlafen+HATPase+wHTH | GCF_002210355.1 | Fibrobacter_sp._UWH3 | Bacteria>Pseudomonadati>FCB_group>Fibrobacterota>Fibrobacteria>Fibrobacterales>Fibrobacteraceae>Fibrobacter>unclassified_Fibrobacter | 561 | B7993_RS04340 | 162664..164349 | 1964353 | putative-DNA-binding-domain-containing-protein-[Fibrobacter-sp.-UWH3] |
| WP_072979078.1 | Schlafen+HATPase+wHTH | GCF_900142475.1 | Fibrobacter_sp._UWH4 | Bacteria>Pseudomonadati>FCB_group>Fibrobacterota>Fibrobacteria>Fibrobacterales>Fibrobacteraceae>Fibrobacter>unclassified_Fibrobacter | 533 | BUA93_RS10240 | 94530..96131 | 1896210 | putative-DNA-binding-domain-containing-protein-[Fibrobacter-sp.-UWH4] |
| SHK42617.1 | Schlafen+HATPase+wHTH | GCA_900142505.1 | Fibrobacter_sp._UWH5 | Bacteria>Pseudomonadati>FCB_group>Fibrobacterota>Fibrobacteria>Fibrobacterales>Fibrobacteraceae>Fibrobacter>unclassified_Fibrobacter | 561 | SAMN05720764_101398 | 453269..454954 | 1896211 | ATP-dependent-DNA-helicase-RecG-[Fibrobacter-sp.-UWH5] |
| OWV23331.1 | Schlafen+HATPase+wHTH | GCA_002210425.1 | Fibrobacter_sp._UWB2 | Bacteria>Pseudomonadati>FCB_group>Fibrobacterota>Fibrobacteria>Fibrobacterales>Fibrobacteraceae>Fibrobacter>unclassified_Fibrobacter | 533 | B7982_07635 | 360277..361878 | 1964358 | transcriptional-regulator-[Fibrobacter-sp.-UWB2] |
| NLW30642.1 | Schlafen+HATPase+wHTH | GCA_012728995.1 | Fibrobacter_sp. | Bacteria>Pseudomonadati>FCB_group>Fibrobacterota>Fibrobacteria>Fibrobacterales>Fibrobacteraceae>Fibrobacter>unclassified_Fibrobacter | 556 | GXY77_04200 | 6013..7683 | 35828 | ATPase-[Fibrobacter-sp.] |
| QQS04456.1 | Schlafen+HATPase+wHTH | GCA_016699655.1 | Fibrobacterota_bacterium | Bacteria>Pseudomonadati>FCB_group>Fibrobacterota>unclassified_Fibrobacterota | 613 | IPK50_19525 | 4747479..4749320 | 2052160 | putative-DNA-binding-domain-containing-protein-[Fibrobacterota-bacterium] |
| MCF7823052.1 | Schlafen+HATPase+wHTH | GCA_021734325.1 | Fidelibacterota_bacterium | Bacteria>Pseudomonadati>FCB_group>Fidelibacterota>unclassified_Fidelibacterota | 500 | K9N35_02675 | 87566..89068 | 2026760 | putative-DNA-binding-domain-containing-protein-[Candidatus-Marinimicrobia-bacterium] |
| MCZ8206521.1 | Schlafen+HATPase+wHTH | GCA_027531815.1 | Gemmatimonas_sp. | Bacteria>Pseudomonadati>FCB_group>Gemmatimonadota>Gemmatimonadia>Gemmatimonadales>Gemmatimonadaceae>Gemmatimonas>unclassified_Gemmatimonas | 520 | O9271_18150 | 10067..11629 | 1962908 | MAG:-putative-DNA-binding-domain-containing-protein-[Gemmatimonas-sp.] |
| MCC6769772.1 | Schlafen+HATPase+wHTH | GCA_020848865.1 | Gemmatimonadaceae_bacterium | Bacteria>Pseudomonadati>FCB_group>Gemmatimonadota>Gemmatimonadia>Gemmatimonadales>Gemmatimonadaceae>unclassified_Gemmatimonadaceae | 520 | IT360_01055 | 9676..11238 | 2026741 | putative-DNA-binding-domain-containing-protein-[Gemmatimonadaceae-bacterium] |
| ODT03615.1 | Schlafen+HATPase+wHTH | GCA_001724275.1 | Gemmatimonadetes_bacterium_SCN_70-22 | Bacteria>Pseudomonadati>FCB_group>Gemmatimonadota>unclassified_Gemmatimonadota | 539 | ABS52_08390 | 5688..7307 | 1660163 | transcriptional-regulator-[Gemmatimonadetes-bacterium-SCN-70-22] |
| MYH51653.1 | Schlafen+HATPase+wHTH | GCA_009843805.1 | Gemmatimonadota_bacterium | Bacteria>Pseudomonadati>FCB_group>Gemmatimonadota>unclassified_Gemmatimonadota | 628 | F4139_01750 | 17848..19734 | 2026742 | hypothetical-protein-[Gemmatimonadota-bacterium] |
| MYG34811.1 | Schlafen+HATPase+wHTH | GCA_009842515.1 | Gemmatimonadota_bacterium | Bacteria>Pseudomonadati>FCB_group>Gemmatimonadota>unclassified_Gemmatimonadota | 471 | F4205_04905 | 11126..12541 | 2026742 | AAA-family-ATPase-[Gemmatimonadota-bacterium] |
| MCE2900729.1 | Schlafen+HATPase+wHTH | GCA_021299375.1 | Gemmatimonadota_bacterium | Bacteria>Pseudomonadati>FCB_group>Gemmatimonadota>unclassified_Gemmatimonadota | 520 | LXA09_05765 | 76513..78075 | 2026742 | putative-DNA-binding-domain-containing-protein-[Gemmatimonadota-bacterium] |
| MBK6307122.1 | Schlafen+HATPase+wHTH | GCA_016705425.1 | Gemmatimonadota_bacterium | Bacteria>Pseudomonadati>FCB_group>Gemmatimonadota>unclassified_Gemmatimonadota | 453 | IPF47_15920 | 122410..123771 | 2026742 | putative-DNA-binding-domain-containing-protein-[Gemmatimonadota-bacterium] |
| MCB9507833.1 | Schlafen+HATPase+wHTH | GCA_020633615.1 | Myxococcales_bacterium | Bacteria>Pseudomonadati>Myxococcota>Myxococcia>Myxococcales>unclassified_Myxococcales | 296 | H6697_09240 | 12897..13787 | 2026763 | ATP-binding-protein-[Myxococcales-bacterium] |
| HCF56385.1 | Schlafen+HATPase+wHTH | GCA_003534605.1 | Myxococcales_bacterium | Bacteria>Pseudomonadati>Myxococcota>Myxococcia>Myxococcales>unclassified_Myxococcales | 552 | DFS52_00110 | 1..1657 | 2026763 | AAA-family-ATPase-[Myxococcales-bacterium] |
| TDB40154.1 | Schlafen+HATPase+wHTH | GCA_004348915.1 | Deltaproteobacteria_bacterium | Bacteria>Pseudomonadati>Myxococcota>Myxococcia>unclassified_Myxococcia | 556 | D9V46_00540 | 115664..117334 | 2026735 | ATPase-[Deltaproteobacteria-bacterium] |
| RLC12741.1 | Schlafen+HATPase+wHTH | GCA_003647435.1 | Deltaproteobacteria_bacterium | Bacteria>Pseudomonadati>Myxococcota>Myxococcia>unclassified_Myxococcia | 426 | DRH43_00750 | 1946..3226 | 2026735 | transcriptional-regulator-[Deltaproteobacteria-bacterium] |
| PXF60417.1 | Schlafen+HATPase+wHTH | GCA_003194495.1 | Deltaproteobacteria_bacterium | Bacteria>Pseudomonadati>Myxococcota>Myxococcia>unclassified_Myxococcia | 468 | C4B58_00695 | 167873..169279 | 2026735 | ATP-dependent-DNA-helicase-RecG-[Deltaproteobacteria-bacterium] |
| PXF55794.1 | Schlafen+HATPase+wHTH | GCA_003194485.1 | Deltaproteobacteria_bacterium | Bacteria>Pseudomonadati>Myxococcota>Myxococcia>unclassified_Myxococcia | 394 | C4B57_01980 | 116661..117845 | 2026735 | transcriptional-regulator-[Deltaproteobacteria-bacterium] |
| PXF54129.1 | Schlafen+HATPase+wHTH | GCA_003194485.1 | Deltaproteobacteria_bacterium | Bacteria>Pseudomonadati>Myxococcota>Myxococcia>unclassified_Myxococcia | 480 | C4B57_08245 | 20902..22344 | 2026735 | hypothetical-protein-C4B57_08245-[Deltaproteobacteria-bacterium] |
| NLF25484.1 | Schlafen+HATPase+wHTH | GCA_012514925.1 | Deltaproteobacteria_bacterium | Bacteria>Pseudomonadati>Myxococcota>Myxococcia>unclassified_Myxococcia | 402 | GX589_07490 | 21920..23128 | 2026735 | transcriptional-regulator-[Deltaproteobacteria-bacterium] |
| MDD9854178.1 | Schlafen+HATPase+wHTH | GCA_028817165.1 | Deltaproteobacteria_bacterium | Bacteria>Pseudomonadati>Myxococcota>Myxococcia>unclassified_Myxococcia | 565 | OXU78_09550 | 9108..10805 | 2026735 | MAG:-putative-DNA-binding-domain-containing-protein-[Deltaproteobacteria-bacterium] |
| MCP4577331.1 | Schlafen+HATPase+wHTH | GCA_024231575.1 | Deltaproteobacteria_bacterium | Bacteria>Pseudomonadati>Myxococcota>Myxococcia>unclassified_Myxococcia | 501 | GY846_13725 | 23145..24650 | 2026735 | AAA-family-ATPase-[Deltaproteobacteria-bacterium] |
| MCK9263308.1 | Schlafen+HATPase+wHTH | GCA_023230665.1 | Deltaproteobacteria_bacterium | Bacteria>Pseudomonadati>Myxococcota>Myxococcia>unclassified_Myxococcia | 556 | MOR18_05790 | 13854..15524 | 2026735 | putative-DNA-binding-domain-containing-protein-[Deltaproteobacteria-bacterium] |
| MCD6274305.1 | Schlafen+HATPase+wHTH | GCA_021163685.1 | Deltaproteobacteria_bacterium | Bacteria>Pseudomonadati>Myxococcota>Myxococcia>unclassified_Myxococcia | 223 | J7K30_16065 | 9878..10546 | 2026735 | ATP-binding-protein-[Deltaproteobacteria-bacterium] |
| MCD6261583.1 | Schlafen+HATPase+wHTH | GCA_021163815.1 | Deltaproteobacteria_bacterium | Bacteria>Pseudomonadati>Myxococcota>Myxococcia>unclassified_Myxococcia | 630 | J7J52_00985 | 66713..68605 | 2026735 | putative-DNA-binding-domain-containing-protein-[Deltaproteobacteria-bacterium] |
| MCB9731979.1 | Schlafen+HATPase+wHTH | GCA_020632715.1 | Deltaproteobacteria_bacterium | Bacteria>Pseudomonadati>Myxococcota>Myxococcia>unclassified_Myxococcia | 587 | H6745_05020 | 1398944..1400707 | 2026735 | putative-DNA-binding-domain-containing-protein-[Deltaproteobacteria-bacterium] |
| MBW2342103.1 | Schlafen+HATPase+wHTH | GCA_019310725.1 | Deltaproteobacteria_bacterium | Bacteria>Pseudomonadati>Myxococcota>Myxococcia>unclassified_Myxococcia | 551 | JRF50_17475 | 405..2060 | 2026735 | putative-DNA-binding-domain-containing-protein-[Deltaproteobacteria-bacterium] |
| MBW2077698.1 | Schlafen+HATPase+wHTH | GCA_019309065.1 | Deltaproteobacteria_bacterium | Bacteria>Pseudomonadati>Myxococcota>Myxococcia>unclassified_Myxococcia | 544 | JRI71_09175 | 883..2517 | 2026735 | putative-DNA-binding-domain-containing-protein-[Deltaproteobacteria-bacterium] |
| MBW1949833.1 | Schlafen+HATPase+wHTH | GCA_019308065.1 | Deltaproteobacteria_bacterium | Bacteria>Pseudomonadati>Myxococcota>Myxococcia>unclassified_Myxococcia | 541 | JRJ18_09845 | 27506..29131 | 2026735 | putative-DNA-binding-domain-containing-protein-[Deltaproteobacteria-bacterium] |
| MBW1938724.1 | Schlafen+HATPase+wHTH | GCA_019307905.1 | Deltaproteobacteria_bacterium | Bacteria>Pseudomonadati>Myxococcota>Myxococcia>unclassified_Myxococcia | 184 | JRI67_08180 | 1..552 | 2026735 | ATP-binding-protein-[Deltaproteobacteria-bacterium] |
| MBW1935300.1 | Schlafen+HATPase+wHTH | GCA_019307915.1 | Deltaproteobacteria_bacterium | Bacteria>Pseudomonadati>Myxococcota>Myxococcia>unclassified_Myxococcia | 558 | JRI84_07090 | 5346..7022 | 2026735 | putative-DNA-binding-domain-containing-protein-[Deltaproteobacteria-bacterium] |
| MBW1922076.1 | Schlafen+HATPase+wHTH | GCA_019307945.1 | Deltaproteobacteria_bacterium | Bacteria>Pseudomonadati>Myxococcota>Myxococcia>unclassified_Myxococcia | 541 | JRJ35_01255 | 82050..83675 | 2026735 | putative-DNA-binding-domain-containing-protein-[Deltaproteobacteria-bacterium] |
| MBW1717202.1 | Schlafen+HATPase+wHTH | GCA_019306255.1 | Deltaproteobacteria_bacterium | Bacteria>Pseudomonadati>Myxococcota>Myxococcia>unclassified_Myxococcia | 551 | JRJ77_15500 | 1380..3035 | 2026735 | putative-DNA-binding-domain-containing-protein-[Deltaproteobacteria-bacterium] |
| MBS3918751.1 | Schlafen+HATPase+wHTH | GCA_018333535.1 | Deltaproteobacteria_bacterium | Bacteria>Pseudomonadati>Myxococcota>Myxococcia>unclassified_Myxococcia | 575 | KG012_07665 | 18841..20568 | 2026735 | putative-DNA-binding-domain-containing-protein-[Deltaproteobacteria-bacterium] |
| MBN2361668.1 | Schlafen+HATPase+wHTH | GCA_016934455.1 | Deltaproteobacteria_bacterium | Bacteria>Pseudomonadati>Myxococcota>Myxococcia>unclassified_Myxococcia | 433 | JXR83_19600 | 12323..13624 | 2026735 | ATP-binding-protein-[Deltaproteobacteria-bacterium] |
| MBM4341421.1 | Schlafen+HATPase+wHTH | GCA_016875195.1 | Deltaproteobacteria_bacterium | Bacteria>Pseudomonadati>Myxococcota>Myxococcia>unclassified_Myxococcia | 551 | FJ110_17990 | 1501..3156 | 2026735 | hypothetical-protein-[Deltaproteobacteria-bacterium] |
| MBK9368671.1 | Schlafen+HATPase+wHTH | GCA_016719515.1 | Deltaproteobacteria_bacterium | Bacteria>Pseudomonadati>Myxococcota>Myxococcia>unclassified_Myxococcia | 640 | IPN01_20580 | 105232..107154 | 2026735 | hypothetical-protein-[Deltaproteobacteria-bacterium] |
| MBI5606166.1 | Schlafen+HATPase+wHTH | GCA_016219225.1 | Deltaproteobacteria_bacterium | Bacteria>Pseudomonadati>Myxococcota>Myxococcia>unclassified_Myxococcia | 456 | HY879_22770 | 10603..11973 | 2026735 | putative-DNA-binding-domain-containing-protein-[Deltaproteobacteria-bacterium] |
| MBI4818895.1 | Schlafen+HATPase+wHTH | GCA_016208175.1 | Deltaproteobacteria_bacterium | Bacteria>Pseudomonadati>Myxococcota>Myxococcia>unclassified_Myxococcia | 459 | HY791_21670 | 5869..7248 | 2026735 | ATPase-[Deltaproteobacteria-bacterium] |
| MBI4818893.1 | Schlafen+HATPase+wHTH | GCA_016208175.1 | Deltaproteobacteria_bacterium | Bacteria>Pseudomonadati>Myxococcota>Myxococcia>unclassified_Myxococcia | 636 | HY791_21660 | 2735..4645 | 2026735 | putative-DNA-binding-domain-containing-protein-[Deltaproteobacteria-bacterium] |
| MBI4764649.1 | Schlafen+HATPase+wHTH | GCA_016208255.1 | Deltaproteobacteria_bacterium | Bacteria>Pseudomonadati>Myxococcota>Myxococcia>unclassified_Myxococcia | 570 | HY787_08595 | 12943..14655 | 2026735 | putative-DNA-binding-domain-containing-protein-[Deltaproteobacteria-bacterium] |
| MBI4527345.1 | Schlafen+HATPase+wHTH | GCA_016210115.1 | Deltaproteobacteria_bacterium | Bacteria>Pseudomonadati>Myxococcota>Myxococcia>unclassified_Myxococcia | 412 | HY695_26420 | 19077..20315 | 2026735 | putative-DNA-binding-domain-containing-protein-[Deltaproteobacteria-bacterium] |
| MBI2378663.1 | Schlafen+HATPase+wHTH | GCA_016183035.1 | Deltaproteobacteria_bacterium | Bacteria>Pseudomonadati>Myxococcota>Myxococcia>unclassified_Myxococcia | 649 | HYV07_32005 | 15558..17507 | 2026735 | putative-DNA-binding-domain-containing-protein-[Deltaproteobacteria-bacterium] |
| MBF0104317.1 | Schlafen+HATPase+wHTH | GCA_015228875.1 | Deltaproteobacteria_bacterium | Bacteria>Pseudomonadati>Myxococcota>Myxococcia>unclassified_Myxococcia | 469 | HQM16_03210 | 35316..36725 | 2026735 | putative-DNA-binding-domain-containing-protein-[Deltaproteobacteria-bacterium] |
| MBC8466730.1 | Schlafen+HATPase+wHTH | GCA_014384065.1 | Deltaproteobacteria_bacterium | Bacteria>Pseudomonadati>Myxococcota>Myxococcia>unclassified_Myxococcia | 82 | H8D55_02715 | 4854..5102 | 2026735 | hypothetical-protein-[Deltaproteobacteria-bacterium] |
| MAG32378.1 | Schlafen+HATPase+wHTH | GCA_002688255.1 | Deltaproteobacteria_bacterium | Bacteria>Pseudomonadati>Myxococcota>Myxococcia>unclassified_Myxococcia | 411 | CL908_15965 | 3246..4481 | 2026735 | ATP-dependent-DNA-helicase-RecG-[Deltaproteobacteria-bacterium] |
| GIW47704.1 | Schlafen+HATPase+wHTH | GCA_026004235.1 | Deltaproteobacteria_bacterium | Bacteria>Pseudomonadati>Myxococcota>Myxococcia>unclassified_Myxococcia | 438 | KatS3mg078_1581 | 295704..297020 | 2026735 | MAG:-ATP-dependent-DNA-helicase-[Deltaproteobacteria-bacterium] |
| WP_276076301.1 | Schlafen+HATPase+wHTH | - | Polyangium_mundeleinium | Bacteria>Pseudomonadati>Myxococcota>Polyangia>Polyangiales>Polyangiaceae>Polyangium | 190 | POL67_RS52610 | 1417738..1418310 | 2995306 | ATP-binding-protein-[Polyangium-sp.-rjm3] |
| WP_129354455.1 | Schlafen+HATPase+wHTH | GCF_004135735.1 | Sorangium_cellulosum | Bacteria>Pseudomonadati>Myxococcota>Polyangia>Polyangiales>Polyangiaceae>Sorangium | 543 | SOCEGT47_RS35860 | 9456142..9457773 | 56 | ATP-binding-protein-[Sorangium-cellulosum] |
| WP_020733583.1 | Schlafen+HATPase+wHTH | GCF_000418325.1 | Sorangium_cellulosum | Bacteria>Pseudomonadati>Myxococcota>Polyangia>Polyangiales>Polyangiaceae>Sorangium | 476 | SCE1572_RS08145 | 2250014..2251444 | 56 | ATP-binding-protein-[Sorangium-cellulosum] |
| MCL2723924.1 | Schlafen+HATPase+wHTH | GCA_009781005.1 | Polyangiaceae_bacterium | Bacteria>Pseudomonadati>Myxococcota>Polyangia>Polyangiales>Polyangiaceae>unclassified_Polyangiaceae | 413 | FWD69_05745 | 10819..12060 | 2268199 | putative-DNA-binding-domain-containing-protein-[Polyangiaceae-bacterium] |
| MCB9617766.1 | Schlafen+HATPase+wHTH | GCA_020633155.1 | Sandaracinus_sp. | Bacteria>Pseudomonadati>Myxococcota>Polyangia>Polyangiales>Sandaracinaceae>Sandaracinus>unclassified_Sandaracinus | 563 | H6722_35470 | 8184..9875 | 2024858 | putative-DNA-binding-domain-containing-protein-[Sandaracinus-sp.] |
| MCK6514474.1 | Schlafen+HATPase+wHTH | GCA_023150775.1 | Myxococcota_bacterium | Bacteria>Pseudomonadati>Myxococcota>unclassified_Myxococcota | 640 | L6R46_05400 | 15152..17074 | 2818507 | hypothetical-protein-[Myxococcota-bacterium] |
| MBP7125455.1 | Schlafen+HATPase+wHTH | GCA_017994695.1 | Myxococcota_bacterium | Bacteria>Pseudomonadati>Myxococcota>unclassified_Myxococcota | 544 | KBD49_03705 | 57522..59156 | 2818507 | putative-DNA-binding-domain-containing-protein-[Myxococcota-bacterium] |
| MDE0208008.1 | Schlafen+HATPase+wHTH | GCA_028819935.1 | Candidatus_Tectimicrobiota_bacterium | Bacteria>Pseudomonadati>Nitrospinota/Tectimicrobiota_group>Candidatus_Tectimicrobiota | 431 | OXP66_18480 | 4767..6062 | 2528274 | MAG:-putative-DNA-binding-domain-containing-protein-[Candidatus-Tectomicrobia-bacterium] |
| OGW17479.1 | Schlafen+HATPase+wHTH | GCA_001803565.1 | Nitrospinae_bacterium_RIFCSPLOWO2_12_FULL_45_22 | Bacteria>Pseudomonadati>Nitrospinota/Tectimicrobiota_group>Nitrospinota>unclassified_Nitrospinota | 550 | A3G93_06405 | 1967..3619 | 1801687 | hypothetical-protein-A3G93_06405-[Nitrospinae-bacterium-RIFCSPLOWO2_12_FULL_45_22] |
| MBZ0154910.1 | Schlafen+HATPase+wHTH | GCA_019912415.1 | Candidatus_Nitrobium_versatile | Bacteria>Pseudomonadati>Nitrospirota>Nitrospiria>Nitrospirales>Nitrospiraceae>Candidatus_Nitrobium | 560 | K8I29_01685 | 63962..65644 | 2884831 | putative-DNA-binding-domain-containing-protein-[Alphaproteobacteria-bacterium] |
| MCP9446720.1 | Schlafen+HATPase+wHTH | GCA_024330945.1 | Nitrospira_sp. | Bacteria>Pseudomonadati>Nitrospirota>Nitrospiria>Nitrospirales>Nitrospiraceae>Nitrospira>unclassified_Nitrospira | 563 | NNA22_04005 | 16292..17983 | 70125 | putative-DNA-binding-domain-containing-protein-[Nitrospira-sp.] |
| GIW56392.1 | Schlafen+HATPase+wHTH | GCA_026004315.1 | Nitrospiraceae_bacterium | Bacteria>Pseudomonadati>Nitrospirota>Nitrospiria>Nitrospirales>Nitrospiraceae>unclassified_Nitrospiraceae | 308 | KatS3mg082_2796 | 5711..6637 | 2026770 | MAG:-hypothetical-protein-KatS3mg082_2796-[Nitrospiraceae-bacterium] |
| GIW56390.1 | Schlafen+HATPase+wHTH | GCA_026004315.1 | Nitrospiraceae_bacterium | Bacteria>Pseudomonadati>Nitrospirota>Nitrospiria>Nitrospirales>Nitrospiraceae>unclassified_Nitrospiraceae | 174 | KatS3mg082_2794 | 4892..5416 | 2026770 | MAG:-hypothetical-protein-KatS3mg082_2794-[Nitrospiraceae-bacterium] |
| MCC6543786.1 | Schlafen+HATPase+wHTH | GCA_020846775.1 | Nitrospirota_bacterium | Bacteria>Pseudomonadati>Nitrospirota>unclassified_Nitrospirota | 220 | IT392_04705 | 8249..8911 | 2026887 | putative-DNA-binding-domain-containing-protein-[Nitrospirota-bacterium] |
| MCC6543782.1 | Schlafen+HATPase+wHTH | GCA_020846775.1 | Nitrospirota_bacterium | Bacteria>Pseudomonadati>Nitrospirota>unclassified_Nitrospirota | 500 | IT392_04685 | 606..2108 | 2026887 | putative-DNA-binding-domain-containing-protein-[Nitrospirota-bacterium] |
| MBI5192440.1 | Schlafen+HATPase+wHTH | GCA_016212215.1 | Nitrospirota_bacterium | Bacteria>Pseudomonadati>Nitrospirota>unclassified_Nitrospirota | 500 | HZA08_03230 | 27332..28834 | 2026887 | putative-DNA-binding-domain-containing-protein-[Nitrospirota-bacterium] |
| MBI5192437.1 | Schlafen+HATPase+wHTH | GCA_016212215.1 | Nitrospirota_bacterium | Bacteria>Pseudomonadati>Nitrospirota>unclassified_Nitrospirota | 383 | HZA08_03215 | 19294..20445 | 2026887 | putative-DNA-binding-domain-containing-protein-[Nitrospirota-bacterium] |
| WP_103916699.1 | Schlafen+HATPase+wHTH | GCF_900107995.1 | Gammaproteobacteria | Bacteria>Pseudomonadati>Pseudomonadota | 556 | C4S63_RS03505 | 778672..780342 | 1236 | ATP-binding-protein-[Alcanivorax-sp.-DSM-26293] |
| WP_065846590.1 | Schlafen+HATPase+wHTH | GCF_011762225.1 | Alphaproteobacteria | Bacteria>Pseudomonadati>Pseudomonadota | 618 | FHS82_RS15745 | 29226..31082 | 28211 | MULTISPECIES:-putative-DNA-binding-domain-containing-protein-[Alphaproteobacteria] |
| WP_027854331.1 | Schlafen+HATPase+wHTH | GCF_000428985.1 | Gammaproteobacteria | Bacteria>Pseudomonadati>Pseudomonadota | 556 | LL252_RS03460 | 286048..287718 | 1236 | MULTISPECIES:-ATP-binding-protein-[Oceanospirillales] |
| WP_101538035.1 | Schlafen+HATPase+wHTH | GCF_018854655.1 | Acidithiobacillus | Bacteria>Pseudomonadati>Pseudomonadota>Acidithiobacillia>Acidithiobacillales>Acidithiobacillaceae | 556 | HF670_RS09005 | 11852..13522 | 119977 | MULTISPECIES:-ATP-binding-protein-[Acidithiobacillus] |
| WP_271780038.1 | Schlafen+HATPase+wHTH | GCF_028215975.1 | Acidithiobacillus_ferriphilus | Bacteria>Pseudomonadati>Pseudomonadota>Acidithiobacillia>Acidithiobacillales>Acidithiobacillaceae>Acidithiobacillus | 477 | PJU76_RS00605 | 124911..126344 | 1689834 | helix-turn-helix-domain-containing-protein-[Acidithiobacillus-ferriphilus] |
| WP_226844045.1 | Schlafen+HATPase+wHTH | GCF_018854875.1 | Acidithiobacillus_ferriphilus | Bacteria>Pseudomonadati>Pseudomonadota>Acidithiobacillia>Acidithiobacillales>Acidithiobacillaceae>Acidithiobacillus | 554 | HF928_RS01640 | 8371..10035 | 1689834 | ATP-binding-protein-[Acidithiobacillus-ferriphilus] |
| WP_215854620.1 | Schlafen+HATPase+wHTH | - | Acidithiobacillus_ferrooxidans | Bacteria>Pseudomonadati>Pseudomonadota>Acidithiobacillia>Acidithiobacillales>Acidithiobacillaceae>Acidithiobacillus | 521 | HF290_RS06275 | 9935..11500 | 920 | ATP-binding-protein-[Acidithiobacillus-ferrooxidans] |
| WP_054609109.1 | Schlafen+HATPase+wHTH | GCF_020887015.1 | Acidithiobacillus_ferrooxidans | Bacteria>Pseudomonadati>Pseudomonadota>Acidithiobacillia>Acidithiobacillales>Acidithiobacillaceae>Acidithiobacillus | 486 | EC577_RS04330 | 14418..15878 | 920 | helix-turn-helix-domain-containing-protein-[Acidithiobacillus-ferrooxidans] |
| WP_035192520.1 | Schlafen+HATPase+wHTH | GCF_000750615.1 | Acidithiobacillus_ferrivorans | Bacteria>Pseudomonadati>Pseudomonadota>Acidithiobacillia>Acidithiobacillales>Acidithiobacillaceae>Acidithiobacillus | 554 | AFERRI_RS08535 | 126753..128417 | 160808 | ATP-binding-protein-[Acidithiobacillus-ferrivorans] |
| WP_014029719.1 | Schlafen+HATPase+wHTH | GCF_000214095.2 | Acidithiobacillus_ferrivorans | Bacteria>Pseudomonadati>Pseudomonadota>Acidithiobacillia>Acidithiobacillales>Acidithiobacillaceae>Acidithiobacillus | 482 | ACIFE_RS11325 | 2368053..2369501 | 160808 | helix-turn-helix-domain-containing-protein-[Acidithiobacillus-ferrivorans] |
| MBU2839700.1 | Schlafen+HATPase+wHTH | GCA_018854655.1 | Acidithiobacillus_thiooxidans | Bacteria>Pseudomonadati>Pseudomonadota>Acidithiobacillia>Acidithiobacillales>Acidithiobacillaceae>Acidithiobacillus | 556 | HF670_09015 | 11852..13522 | 930 | ATPase-[Acidithiobacillus-thiooxidans] |
| MBN6740321.1 | Schlafen+HATPase+wHTH | GCA_017165985.1 | Acidithiobacillus_sp._MC6.1 | Bacteria>Pseudomonadati>Pseudomonadota>Acidithiobacillia>Acidithiobacillales>Acidithiobacillaceae>Acidithiobacillus>unclassified_Acidithiobacillus | 477 | JKG47_07205 | 16868..18301 | 2801580 | putative-DNA-binding-domain-containing-protein-[Acidithiobacillus-sp.-MC6.1] |
| WP_183267041.1 | Schlafen+HATPase+wHTH | GCF_014201825.1 | Acidocella_aromatica | Bacteria>Pseudomonadati>Pseudomonadota>Alphaproteobacteria>Acetobacterales>Acidocellaceae>Acidocella | 625 | HNP71_RS11365 | 81552..83429 | 1303579 | putative-DNA-binding-domain-containing-protein-[Acidocella-aromatica] |
| WP_227076190.1 | Schlafen+HATPase+wHTH | GCF_020567705.1 | Acidocella_sp._KAb_2-4 | Bacteria>Pseudomonadati>Pseudomonadota>Alphaproteobacteria>Acetobacterales>Acidocellaceae>Acidocella>unclassified_Acidocella | 620 | LI953_RS11305 | 1266..3128 | 2885158 | putative-DNA-binding-domain-containing-protein-[Acidocella-sp.-KAb-2-4] |
| WP_201727594.1 | Schlafen+HATPase+wHTH | GCF_907164655.1 | Acidocella_sp._C78 | Bacteria>Pseudomonadati>Pseudomonadota>Alphaproteobacteria>Acetobacterales>Acidocellaceae>Acidocella>unclassified_Acidocella | 616 | ACIDOCELLA78_RS32370 | 300458..302308 | 1671486 | putative-DNA-binding-domain-containing-protein-[Acidocella-sp.-C78] |
| OYV51916.1 | Schlafen+HATPase+wHTH | GCA_002255785.1 | Acidocella_sp._20-58-15 | Bacteria>Pseudomonadati>Pseudomonadota>Alphaproteobacteria>Acetobacterales>Acidocellaceae>Acidocella>unclassified_Acidocella | 622 | B7Z77_03060 | 4663..6531 | 1970302 | MAG:-AAA-family-ATPase-[Acidocella-sp.-20-58-15] |
| WP_149814262.1 | Schlafen+HATPase+wHTH | GCF_008386565.1 | Teichococcus_oryzae | Bacteria>Pseudomonadati>Pseudomonadota>Alphaproteobacteria>Acetobacterales>Roseomonadaceae>Roseomonas | 625 | F0Q34_RS20620 | 4801..6678 | 1608942 | putative-DNA-binding-domain-containing-protein-[Roseomonas-oryzae] |
| QDD97345.1 | Schlafen+HATPase+wHTH | GCA_006384895.1 | Roseomonas_mucosa | Bacteria>Pseudomonadati>Pseudomonadota>Alphaproteobacteria>Acetobacterales>Roseomonadaceae>Roseomonas | 624 | ADP8_05162 | 118670..120544 | 207340 | DNA-binding-protein-[Roseomonas-mucosa] |
| KAA2211336.1 | Schlafen+HATPase+wHTH | GCA_008386565.1 | Teichococcus_oryzae | Bacteria>Pseudomonadati>Pseudomonadota>Alphaproteobacteria>Acetobacterales>Roseomonadaceae>Roseomonas | 625 | F0Q34_20615 | 4801..6678 | 1608942 | AAA-family-ATPase-[Roseomonas-oryzae] |

| Protein-ID | Domain | Assembly-ID | Strain | Taxonomy | Length | Locus-Tag | Genomic-coordinates | TaxID | Fasta-defline |
|---|---|---|---|---|---|---|---|---|---|
| MCA3394954.1 | Schlafen+HATPase+wHTH | GCA_020249805.1 | Roseomonas_sp. | Bacteria>Pseudomonadati>Pseudomonadota>Alphaproteobacteria>Acetobacterales>Roseomonadaceae>Roseomonas>unclassified_Roseomonas | 622 | INF85_04835 | 21476..23344 | 1869190 | putative-DNA-binding-domain-containing-protein-[Roseomonas-sp.] |
| MCB1832522.1 | Schlafen+HATPase+wHTH | GCA_020445435.1 | Geminicoccaceae_bacterium | Bacteria>Pseudomonadati>Pseudomonadota>Alphaproteobacteria>Geminicoccales>Geminicoccaceae>unclassified_Geminicoccaceae | 621 | KDH19_03610 | 15741..17606 | 2448052 | putative-DNA-binding-domain-containing-protein-[Geminicoccaceae-bacterium] |
| WP_055036839.1 | Schlafen+HATPase+wHTH | GCF_001548155.2 | Blastochloris_viridis | Bacteria>Pseudomonadati>Pseudomonadota>Alphaproteobacteria>Hyphomicrobiales>Blastochloridaceae>Blastochloris | 622 | BV133_RS16115 | 3698206..3700074 | 1079 | putative-DNA-binding-domain-containing-protein-[Blastochloris-viridis] |
| WP_207488590.1 | Schlafen+HATPase+wHTH | GCF_017355465.1 | Brucella_pituitosa | Bacteria>Pseudomonadati>Pseudomonadota>Alphaproteobacteria>Hyphomicrobiales>Brucellaceae>Brucella/Ochrobactrum_group>Brucella | 621 | IPV26_RS10120 | 78648..80513 | 571256 | putative-DNA-binding-domain-containing-protein-[Brucella-pituitosa] |
| GLI95562.1 | Schlafen+HATPase+wHTH | GCA_027923385.1 | Methylocystis_echinoides | Bacteria>Pseudomonadati>Pseudomonadota>Alphaproteobacteria>Hyphomicrobiales>Methylocystaceae>Methylocystis | 625 | LMG27198_45540 | 62257..64134 | 29468 | hypothetical-protein-LMG27198_45540-[Methylocystis-echinoides] |
| WP_264623140.1 | Schlafen+HATPase+wHTH | GCF_025960955.1 | Bradyrhizobium_elkanii | Bacteria>Pseudomonadati>Pseudomonadota>Alphaproteobacteria>Hyphomicrobiales>Nitrobacteraceae>Bradyrhizobium | 620 | M2210_RS06795 | 1520073..1521935 | 29448 | putative-DNA-binding-domain-containing-protein-[Bradyrhizobium-elkanii] |
| WP_012110065.1 | Schlafen+HATPase+wHTH | GCF_000017565.1 | Parvibaculum_lavamentivorans | Bacteria>Pseudomonadati>Pseudomonadota>Alphaproteobacteria>Hyphomicrobiales>Parvibaculaceae>Parvibaculum | 620 | PLAV_RS18040 | 3800884..3802746 | 256618 | putative-DNA-binding-domain-containing-protein-[Parvibaculum-lavamentivorans] |
| WP_135398843.1 | Schlafen+HATPase+wHTH | GCF_014200255.1 | Rhizobium/Agrobacterium_group | Bacteria>Pseudomonadati>Pseudomonadota>Alphaproteobacteria>Hyphomicrobiales>Rhizobiaceae | 621 | GGD52_RS24615 | 159840..161705 | 227290 | putative-DNA-binding-domain-containing-protein-[Agrobacterium-tumefaciens] |
| WP_115156596.1 | Schlafen+HATPase+wHTH | GCF_004329915.1 | Rhizobium_leguminosarum | Bacteria>Pseudomonadati>Pseudomonadota>Alphaproteobacteria>Hyphomicrobiales>Rhizobiaceae>Rhizobium/Agrobacterium_group>Rhizobium | 621 | E0H64_RS12785 | 333223..335088 | 384 | putative-DNA-binding-domain-containing-protein-[Rhizobium-leguminosarum] |
| WP_004111592.1 | Schlafen+HATPase+wHTH | GCF_000359745.1 | Rhizobium_freirei | Bacteria>Pseudomonadati>Pseudomonadota>Alphaproteobacteria>Hyphomicrobiales>Rhizobiaceae>Rhizobium/Agrobacterium_group>Rhizobium | 621 | RHSP_RS07180 | 16028..17893 | 1353277 | putative-DNA-binding-domain-containing-protein-[Rhizobium-freirei] |
| WP_216330251.1 | Schlafen+HATPase+wHTH | GCF_018873185.1 | Rhizobium_sp._X9 | Bacteria>Pseudomonadati>Pseudomonadota>Alphaproteobacteria>Hyphomicrobiales>Rhizobiaceae>Rhizobium/Agrobacterium_group>Rhizobium>unclassified_Rhizobium | 621 | JZY08_RS11875 | 2436647..2438512 | 2815360 | putative-DNA-binding-domain-containing-protein-[Rhizobium-sp.-X9] |
| TGE86677.1 | Schlafen+HATPase+wHTH | GCA_004766515.1 | Rhizobium_sp._SEMIA_4032 | Bacteria>Pseudomonadati>Pseudomonadota>Alphaproteobacteria>Hyphomicrobiales>Rhizobiaceae>Rhizobium/Agrobacterium_group>Rhizobium>unclassified_Rhizobium | 621 | C9418_21780 | 39555..41420 | 2137762 | AAA-family-ATPase-[Rhizobium-sp.-SEMIA-4032] |
| MBX3567503.1 | Schlafen+HATPase+wHTH | GCA_019635715.1 | Rhizobiaceae_bacterium | Bacteria>Pseudomonadati>Pseudomonadota>Alphaproteobacteria>Hyphomicrobiales>Rhizobiaceae>unclassified_Rhizobiaceae | 621 | KF914_05555 | 66915..68780 | 1913961 | putative-DNA-binding-domain-containing-protein-[Rhizobiaceae-bacterium] |
| MCC0042684.1 | Schlafen+HATPase+wHTH | GCA_020637975.1 | Rhodobiaceae_bacterium | Bacteria>Pseudomonadati>Pseudomonadota>Alphaproteobacteria>Hyphomicrobiales>Rhodobiaceae>unclassified_Rhodobiaceae | 611 | H6880_10965 | 1..1833 | 2026785 | putative-DNA-binding-domain-containing-protein-[Rhodobiaceae-bacterium] |
| WP_085440104.1 | Schlafen+HATPase+wHTH | GCF_002109495.1 | Magnetofaba_australis | Bacteria>Pseudomonadati>Pseudomonadota>Alphaproteobacteria>Magnetococcales>Magnetococcaceae>Magnetofaba | 621 | MAIT1_RS00705 | 11710..13575 | 1472297 | putative-DNA-binding-domain-containing-protein-[Magnetofaba-australis] |
| OSM08493.1 | Schlafen+HATPase+wHTH | GCA_002109495.1 | Magnetofaba_australis_IT-1 | Bacteria>Pseudomonadati>Pseudomonadota>Alphaproteobacteria>Magnetococcales>Magnetococcaceae>Magnetofaba>Magnetofaba_australis | 621 | MAIT1_04665 | 11710..13575 | 1434232 | putative-AAA-ATPase-[Magnetofaba-australis-IT-1] |
| MBF0308897.1 | Schlafen+HATPase+wHTH | GCA_015231925.1 | Magnetococcales_bacterium | Bacteria>Pseudomonadati>Pseudomonadota>Alphaproteobacteria>Magnetococcales>unclassified_Magnetococcales | 557 | HQL56_05150 | 10603..12276 | 2026759 | putative-DNA-binding-domain-containing-protein-[Magnetococcales-bacterium] |
| MBF0181805.1 | Schlafen+HATPase+wHTH | GCA_015231235.1 | Magnetococcales_bacterium | Bacteria>Pseudomonadati>Pseudomonadota>Alphaproteobacteria>Magnetococcales>unclassified_Magnetococcales | 557 | HQM03_17445 | 4635..6308 | 2026759 | putative-DNA-binding-domain-containing-protein-[Magnetococcales-bacterium] |
| MBF0133381.1 | Schlafen+HATPase+wHTH | GCA_015228925.1 | Magnetococcales_bacterium | Bacteria>Pseudomonadati>Pseudomonadota>Alphaproteobacteria>Magnetococcales>unclassified_Magnetococcales | 645 | HQL75_12435 | 17731..19668 | 2026759 | putative-DNA-binding-domain-containing-protein-[Magnetococcales-bacterium] |
| MBF0097997.1 | Schlafen+HATPase+wHTH | GCA_015228825.1 | Magnetococcales_bacterium | Bacteria>Pseudomonadati>Pseudomonadota>Alphaproteobacteria>Magnetococcales>unclassified_Magnetococcales | 428 | HQM05_11740 | 2688..3974 | 2026759 | putative-DNA-binding-domain-containing-protein-[Magnetococcales-bacterium] |
| MBF0096958.1 | Schlafen+HATPase+wHTH | GCA_015228825.1 | Magnetococcales_bacterium | Bacteria>Pseudomonadati>Pseudomonadota>Alphaproteobacteria>Magnetococcales>unclassified_Magnetococcales | 420 | HQM05_06425 | 9314..10576 | 2026759 | putative-DNA-binding-domain-containing-protein-[Magnetococcales-bacterium] |
| MCY4460276.1 | Schlafen+HATPase+wHTH | GCA_026712665.1 | Albidovulum_sp. | Bacteria>Pseudomonadati>Pseudomonadota>Alphaproteobacteria>Rhodobacterales>Paracoccaceae>Albidovulum>unclassified_Albidovulum | 583 | OXC26_07755 | 4058..5809 | 1872424 | MAG:-putative-DNA-binding-domain-containing-protein-[Albidovulum-sp.] |
| WP_092891912.1 | Schlafen+HATPase+wHTH | GCF_014681765.1 | Roseicitreum_antarcticum | Bacteria>Pseudomonadati>Pseudomonadota>Alphaproteobacteria>Rhodobacterales>Paracoccaceae>Roseicitreum | 622 | H9529_RS17205 | 39518..41386 | 564137 | putative-DNA-binding-domain-containing-protein-[Roseicitreum-antarcticum] |
| WP_158270883.1 | Schlafen+HATPase+wHTH | GCF_003072065.1 | Thalassorhabdomicrobium_marinisediminis | Bacteria>Pseudomonadati>Pseudomonadota>Alphaproteobacteria>Rhodobacterales>Paracoccaceae>Thalassorhabdomicrobium | 1065 | DC363_RS14215 | 62143..65340 | 2170577 | putative-DNA-binding-domain-containing-protein-[Thalassorhabdomicrobium-marinisediminis] |
| MCA3449868.1 | Schlafen+HATPase+wHTH | GCA_020250135.1 | Rhodobacter_sp. | Bacteria>Pseudomonadati>Pseudomonadota>Alphaproteobacteria>Rhodobacterales>Rhodobacter_group>Rhodobacter>unclassified_Rhodobacter | 622 | INF92_05755 | 9598..11466 | 1062 | putative-DNA-binding-domain-containing-protein-[Rhodobacter-sp.] |
| WP_274389678.1 | Schlafen+HATPase+wHTH | - | Azospirillum_doebereinerae | Bacteria>Pseudomonadati>Pseudomonadota>Alphaproteobacteria>Rhodospirillales>Azospirillaceae>Azospirillum | 387 | MCW82_RS00045 | 19842..21005 | 92933 | ATP-binding-protein-[Azospirillum-doebereinerae] |
| MYK58413.1 | Schlafen+HATPase+wHTH | GCA_009846065.1 | Rhodospirillaceae_bacterium | Bacteria>Pseudomonadati>Pseudomonadota>Alphaproteobacteria>Rhodospirillales>Rhodospirillaceae>unclassified_Rhodospirillaceae | 632 | F4027_07315 | 3507..5405 | 1898112 | AAA-family-ATPase-[Rhodospirillaceae-bacterium] |
| MXY42113.1 | Schlafen+HATPase+wHTH | GCA_009838215.1 | Rhodospirillaceae_bacterium | Bacteria>Pseudomonadati>Pseudomonadota>Alphaproteobacteria>Rhodospirillales>Rhodospirillaceae>unclassified_Rhodospirillaceae | 632 | F4Y62_20205 | 49977..51875 | 1898112 | AAA-family-ATPase-[Rhodospirillaceae-bacterium] |
| MBK8211734.1 | Schlafen+HATPase+wHTH | GCA_016712375.1 | Rhodospirillales_bacterium | Bacteria>Pseudomonadati>Pseudomonadota>Alphaproteobacteria>Rhodospirillales>unclassified_Rhodospirillales | 74 | IPK78_19065 | 30458..30682 | 2026786 | ATP-binding-protein-[Rhodospirillales-bacterium] |
| WP_109907992.1 | Schlafen+HATPase+wHTH | GCF_003173035.1 | Zavarzinia_aquatilis | Bacteria>Pseudomonadati>Pseudomonadota>Alphaproteobacteria>Rhodospirillales>Zavarziniaceae>Zavarzinia | 620 | DKG74_RS20210 | 25276..27138 | 2211142 | putative-DNA-binding-domain-containing-protein-[Zavarzinia-aquatilis] |
| WP_058745240.1 | Schlafen+HATPase+wHTH | GCF_001477495.1 | Sphingomonas_yabuuchiae | Bacteria>Pseudomonadati>Pseudomonadota>Alphaproteobacteria>Sphingomonadales>Sphingomonadaceae>Sphingomonas | 273 | NS355_RS08075 | 18468..19289 | 172044 | ATP-binding-protein-[Sphingomonas-yabuuchiae] |
| WP_126012415.1 | Schlafen+HATPase+wHTH | GCF_003951715.1 | Sphingomonas_sp._ABOLH | Bacteria>Pseudomonadati>Pseudomonadota>Alphaproteobacteria>Sphingomonadales>Sphingomonadaceae>Sphingomonas>unclassified_Sphingomonas | 273 | CA237_RS01140 | 41419..42240 | 1985881 | ATP-binding-protein-[Sphingomonas-sp.-ABOLH] |
| RSV33120.1 | Schlafen+HATPase+wHTH | GCA_003951715.1 | Sphingomonas_sp._ABOLH | Bacteria>Pseudomonadati>Pseudomonadota>Alphaproteobacteria>Sphingomonadales>Sphingomonadaceae>Sphingomonas>unclassified_Sphingomonas | 273 | CA237_01135 | 41419..42240 | 1985881 | ATP-binding-protein-[Sphingomonas-sp.-ABOLH] |
| MBQ2886284.1 | Schlafen+HATPase+wHTH | GCA_017409945.1 | Alphaproteobacteria_bacterium | Bacteria>Pseudomonadati>Pseudomonadota>Alphaproteobacteria>unclassified_Alphaproteobacteria | 498 | IJE43_21390 | 6414..7910 | 1913988 | putative-DNA-binding-domain-containing-protein-[Alphaproteobacteria-bacterium] |
| WP_169068192.1 | Schlafen+HATPase+wHTH | GCF_012939955.1 | Candidatus_Accumulibacter_phosphatis | Bacteria>Pseudomonadati>Pseudomonadota>Betaproteobacteria>Betaproteobacteria_incertae_sedis>Candidatus_Accumulibacter | 578 | E4Q23_RS19550 | 6606..8342 | 327160 | ATP-binding-protein-[Candidatus-Accumulibacter-phosphatis] |
| WP_138677518.1 | Schlafen+HATPase+wHTH | GCF_005889575.1 | Candidatus_Accumulibacter_phosphatis | Bacteria>Pseudomonadati>Pseudomonadota>Betaproteobacteria>Betaproteobacteria_incertae_sedis>Candidatus_Accumulibacter | 578 | ACCUM_RS19250 | 5145..6881 | 327160 | ATP-binding-protein-[Candidatus-Accumulibacter-phosphatis] |
| WP_034945900.1 | Schlafen+HATPase+wHTH | GCF_000584975.1 | Candidatus_Accumulibacter_cognatus | Bacteria>Pseudomonadati>Pseudomonadota>Betaproteobacteria>Betaproteobacteria_incertae_sedis>Candidatus_Accumulibacter | 422 | AW06_RS03925 | 13440..14708 | 2954383 | ATP-binding-protein-[Candidatus-Accumulibacter-cognatus] |
| MCQ1548297.1 | Schlafen+HATPase+wHTH | GCA_024380085.1 | Candidatus_Accumulibacter_phosphatis | Bacteria>Pseudomonadati>Pseudomonadota>Betaproteobacteria>Betaproteobacteria_incertae_sedis>Candidatus_Accumulibacter | 422 | NOF05_05615 | 17308..18576 | 327160 | putative-DNA-binding-domain-containing-protein-[Candidatus-Accumulibacter-phosphatis] |
| MCQ1547868.1 | Schlafen+HATPase+wHTH | GCA_024380085.1 | Candidatus_Accumulibacter_phosphatis | Bacteria>Pseudomonadati>Pseudomonadota>Betaproteobacteria>Betaproteobacteria_incertae_sedis>Candidatus_Accumulibacter | 579 | NOF05_03355 | 11102..12841 | 327160 | putative-DNA-binding-domain-containing-protein-[Candidatus-Accumulibacter-phosphatis] |
| KFB77814.1 | Schlafen+HATPase+wHTH | GCA_000584975.2 | Candidatus_Accumulibacter_cognatus | Bacteria>Pseudomonadati>Pseudomonadota>Betaproteobacteria>Betaproteobacteria_incertae_sedis>Candidatus_Accumulibacter | 422 | AW06_000997 | 13440..14708 | 2954383 | MAG:-Divergent-AAA-domain-protein-[Candidatus-Accumulibacter-cognatus] |
| KFB72160.1 | Schlafen+HATPase+wHTH | GCA_000585035.2 | Candidatus_Accumulibacter_phosphatis | Bacteria>Pseudomonadati>Pseudomonadota>Betaproteobacteria>Betaproteobacteria_incertae_sedis>Candidatus_Accumulibacter | 621 | AW09_002652 | 2826..4691 | 327160 | MAG:-Divergent-AAA-domain-protein-[Candidatus-Accumulibacter-phosphatis] |
| KFB72159.1 | Schlafen+HATPase+wHTH | GCA_000585035.2 | Candidatus_Accumulibacter_phosphatis | Bacteria>Pseudomonadati>Pseudomonadota>Betaproteobacteria>Betaproteobacteria_incertae_sedis>Candidatus_Accumulibacter | 504 | AW09_002651 | 1199..2713 | 327160 | MAG:-Divergent-AAA-domain-protein-[Candidatus-Accumulibacter-phosphatis] |
| KFB70632.1 | Schlafen+HATPase+wHTH | GCA_000585035.2 | Candidatus_Accumulibacter_phosphatis | Bacteria>Pseudomonadati>Pseudomonadota>Betaproteobacteria>Betaproteobacteria_incertae_sedis>Candidatus_Accumulibacter | 153 | AW09_004272 | 3888..4349 | 327160 | MAG:-Divergent-AAA-domain-protein-[Candidatus-Accumulibacter-phosphatis] |
| KFB70631.1 | Schlafen+HATPase+wHTH | GCA_000585035.2 | Candidatus_Accumulibacter_phosphatis | Bacteria>Pseudomonadati>Pseudomonadota>Betaproteobacteria>Betaproteobacteria_incertae_sedis>Candidatus_Accumulibacter | 224 | AW09_004271 | 3268..3942 | 327160 | MAG:-hypothetical-protein-AW09_004271-[Candidatus-Accumulibacter-phosphatis] |
| MCP5230248.1 | Schlafen+HATPase+wHTH | GCA_024235715.1 | Accumulibacter_sp. | Bacteria>Pseudomonadati>Pseudomonadota>Betaproteobacteria>Betaproteobacteria_incertae_sedis>Candidatus_Accumulibacter>unclassified_Candidatus_Accumulibacter | 343 | H6941_16810 | 69230..70261 | 2053492 | ATP-binding-protein-[Candidatus-Accumulibacter-sp.] |
| MCM8626699.1 | Schlafen+HATPase+wHTH | GCA_023806625.1 | Accumulibacter_sp. | Bacteria>Pseudomonadati>Pseudomonadota>Betaproteobacteria>Betaproteobacteria_incertae_sedis>Candidatus_Accumulibacter>unclassified_Candidatus_Accumulibacter | 615 | NFW05_12170 | 8747..10594 | 2053492 | putative-DNA-binding-domain-containing-protein-[Candidatus-Accumulibacter-sp.] |
| MCM8600114.1 | Schlafen+HATPase+wHTH | GCA_023806585.1 | Accumulibacter_sp. | Bacteria>Pseudomonadati>Pseudomonadota>Betaproteobacteria>Betaproteobacteria_incertae_sedis>Candidatus_Accumulibacter>unclassified_Candidatus_Accumulibacter | 423 | NFW04_15900 | 32618..33889 | 2053492 | putative-DNA-binding-domain-containing-protein-[Candidatus-Accumulibacter-sp.] |
| MCM8580149.1 | Schlafen+HATPase+wHTH | GCA_023806705.1 | Accumulibacter_sp. | Bacteria>Pseudomonadati>Pseudomonadota>Betaproteobacteria>Betaproteobacteria_incertae_sedis>Candidatus_Accumulibacter>unclassified_Candidatus_Accumulibacter | 578 | NFV56_11810 | 2885..4621 | 2053492 | putative-DNA-binding-domain-containing-protein-[Candidatus-Accumulibacter-sp.] |
| MBN8455768.1 | Schlafen+HATPase+wHTH | GCA_017302415.1 | Accumulibacter_sp. | Bacteria>Pseudomonadati>Pseudomonadota>Betaproteobacteria>Betaproteobacteria_incertae_sedis>Candidatus_Accumulibacter>unclassified_Candidatus_Accumulibacter | 653 | J0M31_22250 | 2758..4719 | 2053492 | putative-DNA-binding-domain-containing-protein-[Candidatus-Accumulibacter-sp.] |
| WP_156860855.1 | Schlafen+HATPase+wHTH | GCF_009746125.1 | Casimicrobium_huifangae | Bacteria>Pseudomonadati>Pseudomonadota>Betaproteobacteria>Betaproteobacteria_incertae_sedis>Casimicrobiaceae>Casimicrobium | 596 | FKL89_RS00875 | 186951..188741 | 2591109 | ATP-binding-protein-[Casimicrobium-huifangae] |
| WP_194770509.1 | Schlafen+HATPase+wHTH | GCF_032085695.1 | Comamonadaceae | Bacteria>Pseudomonadati>Pseudomonadota>Betaproteobacteria>Burkholderiales | 596 | ISG32_RS08030 | 62237..64027 | 80864 | ATP-binding-protein-[Diaphorobacter-caeni] |
| WP_238920070.1 | Schlafen+HATPase+wHTH | GCF_051295355.1 | Achromobacter_xylosoxidans | Bacteria>Pseudomonadati>Pseudomonadota>Betaproteobacteria>Burkholderiales>Alcaligenaceae>Achromobacter | 621 | L7A49_RS17315 | 4545..6410 | 85698 | putative-DNA-binding-domain-containing-protein-[Achromobacter-xylosoxidans] |
| WP_175180615.1 | Schlafen+HATPase+wHTH | GCF_902859735.1 | Achromobacter_aegrifaciens | Bacteria>Pseudomonadati>Pseudomonadota>Betaproteobacteria>Burkholderiales>Alcaligenaceae>Achromobacter | 621 | HV766_RS04065 | 906168..908033 | 1287736 | putative-DNA-binding-domain-containing-protein-[Achromobacter-aegrifaciens] |
| WP_125283001.1 | Schlafen+HATPase+wHTH | GCF_003939895.1 | Achromobacter_denitrificans | Bacteria>Pseudomonadati>Pseudomonadota>Betaproteobacteria>Burkholderiales>Alcaligenaceae>Achromobacter | 596 | EGU64_RS02480 | 232540..234330 | 32002 | ATP-binding-protein-[Achromobacter-denitrificans] |
| WP_200235201.1 | Schlafen+HATPase+wHTH | GCF_016595155.1 | Advenella_mandrilli | Bacteria>Pseudomonadati>Pseudomonadota>Betaproteobacteria>Burkholderiales>Alcaligenaceae>Advenella | 506 | JHL22_RS06615 | 193357..194877 | 2800330 | helix-turn-helix-domain-containing-protein-[Advenella-mandrilli] |
| WP_121737850.1 | Schlafen+HATPase+wHTH | GCF_003671915.1 | Alcaligenes_aquatilis | Bacteria>Pseudomonadati>Pseudomonadota>Betaproteobacteria>Burkholderiales>Alcaligenaceae>Alcaligenes | 497 | D3M96_RS02100 | 468929..470422 | 323284 | helix-turn-helix-domain-containing-protein-[Alcaligenes-aquatilis] |
| WP_226349065.1 | Schlafen+HATPase+wHTH | GCF_020496585.1 | Alcaligenes_sp._13f | Bacteria>Pseudomonadati>Pseudomonadota>Betaproteobacteria>Burkholderiales>Alcaligenaceae>Alcaligenes>unclassified_Alcaligenes | 596 | KOE80_RS10210 | 33411..35201 | 2841924 | ATP-binding-protein-[Alcaligenes-sp.-13f] |
| MCB4322565.1 | Schlafen+HATPase+wHTH | GCA_020496585.1 | Alcaligenes_sp._13f | Bacteria>Pseudomonadati>Pseudomonadota>Betaproteobacteria>Burkholderiales>Alcaligenaceae>Alcaligenes>unclassified_Alcaligenes | 590 | KOE80_10180 | 33429..35201 | 2841924 | putative-DNA-binding-domain-containing-protein-[Alcaligenes-sp.-13f] |
| WP_094830674.1 | Schlafen+HATPase+wHTH | GCF_002261435.1 | Bordetella_genomosp._1 | Bacteria>Pseudomonadati>Pseudomonadota>Betaproteobacteria>Burkholderiales>Alcaligenaceae>Bordetella | 596 | CAL27_RS02670 | 575602..577392 | 1395607 | ATP-binding-protein-[Bordetella-genomosp.-1] |
| WP_033459550.1 | Schlafen+HATPase+wHTH | GCF_000690475.1 | Bordetella_bronchiseptica | Bacteria>Pseudomonadati>Pseudomonadota>Betaproteobacteria>Burkholderiales>Alcaligenaceae>Bordetella | 424 | EL288_RS19655 | 4196754..4198028 | 518 | ATP-binding-protein-[Bordetella-bronchiseptica] |
| OZI68386.1 | Schlafen+HATPase+wHTH | GCA_002261435.1 | Bordetella_genomosp._1 | Bacteria>Pseudomonadati>Pseudomonadota>Betaproteobacteria>Burkholderiales>Alcaligenaceae>Bordetella | 596 | CAL27_02675 | 575602..577392 | 1395607 | transcriptional-regulator-[Bordetella-genomosp.-1] |
| WP_066458436.1 | Schlafen+HATPase+wHTH | GCF_001592225.1 | Castellaniella_caeni | Bacteria>Pseudomonadati>Pseudomonadota>Betaproteobacteria>Burkholderiales>Alcaligenaceae>Castellaniella | 558 | CUB94_RS15505 | 5714..7390 | 266123 | putative-DNA-binding-domain-containing-protein-[Castellaniella-caeni] |
| WP_269498088.1 | Schlafen+HATPase+wHTH | GCF_027257175.1 | Castellaniella_sp._S9 | Bacteria>Pseudomonadati>Pseudomonadota>Betaproteobacteria>Burkholderiales>Alcaligenaceae>Castellaniella>unclassified_Castellaniella | 640 | O0Z48_RS10590 | 325826..327748 | 2993652 | ATP-binding-protein-[Castellaniella-sp.-S9] |
| WP_166406660.1 | Schlafen+HATPase+wHTH | GCF_011290505.1 | Paenalcaligenes_suwonensis | Bacteria>Pseudomonadati>Pseudomonadota>Betaproteobacteria>Burkholderiales>Alcaligenaceae>Paenalcaligenes | 490 | GY634_RS00675 | 158381..159853 | 1202713 | helix-turn-helix-domain-containing-protein-[Paenalcaligenes-suwonensis] |
| HBT33985.1 | Schlafen+HATPase+wHTH | GCA_003509985.1 | Pusillimonas_sp. | Bacteria>Pseudomonadati>Pseudomonadota>Betaproteobacteria>Burkholderiales>Alcaligenaceae>Pusillimonas>unclassified_Pusillimonas | 617 | DEB15_14755 | 21785..23638 | 3040095 | transcriptional-regulator-[Pusillimonas-sp.-(ex-Stolz-et-al.-2005)] |
| NYT60763.1 | Schlafen+HATPase+wHTH | GCA_013416445.1 | Alcaligenaceae_bacterium | Bacteria>Pseudomonadati>Pseudomonadota>Betaproteobacteria>Burkholderiales>Alcaligenaceae>unclassified_Alcaligenaceae | 416 | H0A66_00260 | 60907..62157 | 2021368 | putative-DNA-binding-domain-containing-protein-[Alcaligenaceae-bacterium] |
| NYT60500.1 | Schlafen+HATPase+wHTH | GCA_013416405.1 | Alcaligenaceae_bacterium | Bacteria>Pseudomonadati>Pseudomonadota>Betaproteobacteria>Burkholderiales>Alcaligenaceae>unclassified_Alcaligenaceae | 480 | H0A65_16380 | 6668..8110 | 2021368 | putative-DNA-binding-domain-containing-protein-[Alcaligenaceae-bacterium] |
| WP_088931580.1 | Schlafen+HATPase+wHTH | GCF_003812585.1 | Burkholderia | Bacteria>Pseudomonadati>Pseudomonadota>Betaproteobacteria>Burkholderiales>Burkholderiaceae | 580 | J3L14_RS17840 | 4000976..4002718 | 32008 | MULTISPECIES:-ATP-binding-protein-[Burkholderia] |
| WP_027788792.1 | Schlafen+HATPase+wHTH | GCF_001528845.1 | Burkholderia | Bacteria>Pseudomonadati>Pseudomonadota>Betaproteobacteria>Burkholderiales>Burkholderiaceae | 577 | EFU85_RS04210 | 248100..249833 | 32008 | MULTISPECIES:-ATP-binding-protein-[Burkholderia] |
| WP_264838732.1 | Schlafen+HATPase+wHTH | GCF_025995855.1 | Burkholderia_cenocepacia | Bacteria>Pseudomonadati>Pseudomonadota>Betaproteobacteria>Burkholderiales>Burkholderiaceae>Burkholderia>Burkholderia_cepacia_complex | 596 | K6Y76_RS22775 | 70319..72109 | 95486 | ATP-binding-protein-[Burkholderia-cenocepacia] |
| WP_217096865.1 | Schlafen+HATPase+wHTH | GCF_019041095.1 | Burkholderia_multivorans | Bacteria>Pseudomonadati>Pseudomonadota>Betaproteobacteria>Burkholderiales>Burkholderiaceae>Burkholderia>Burkholderia_cepacia_complex | 424 | KTE23_RS28420 | 25720..26994 | 87883 | ATP-binding-protein-[Burkholderia-multivorans] |
| WP_124581832.1 | Schlafen+HATPase+wHTH | GCF_003853875.1 | Burkholderia_contaminans | Bacteria>Pseudomonadati>Pseudomonadota>Betaproteobacteria>Burkholderiales>Burkholderiaceae>Burkholderia>Burkholderia_cepacia_complex | 577 | DF051_RS23125 | 7055..8788 | 488447 | ATP-binding-protein-[Burkholderia-contaminans] |
| WP_105831309.1 | Schlafen+HATPase+wHTH | GCF_902829415.1 | Burkholderia_ambifaria | Bacteria>Pseudomonadati>Pseudomonadota>Betaproteobacteria>Burkholderiales>Burkholderiaceae>Burkholderia>Burkholderia_cepacia_complex | 618 | HWW14_RS24140 | 139475..141331 | 152480 | putative-DNA-binding-domain-containing-protein-[Burkholderia-ambifaria] |
| QQJ97247.1 | Schlafen+HATPase+wHTH | GCA_016458785.1 | Burkholderia_ambifaria | Bacteria>Pseudomonadati>Pseudomonadota>Betaproteobacteria>Burkholderiales>Burkholderiaceae>Burkholderia>Burkholderia_cepacia_complex | 618 | JG536_00770 | 178890..180746 | 152480 | putative-DNA-binding-domain-containing-protein-[Burkholderia-ambifaria] |
| PRG07804.1 | Schlafen+HATPase+wHTH | GCA_002981655.1 | Burkholderia_ambifaria | Bacteria>Pseudomonadati>Pseudomonadota>Betaproteobacteria>Burkholderiales>Burkholderiaceae>Burkholderia>Burkholderia_cepacia_complex | 618 | C6Q14_09300 | 44442..46298 | 152480 | AAA-family-ATPase-[Burkholderia-ambifaria] |
| MCA8374009.1 | Schlafen+HATPase+wHTH | GCA_020420025.1 | Burkholderia_multivorans | Bacteria>Pseudomonadati>Pseudomonadota>Betaproteobacteria>Burkholderiales>Burkholderiaceae>Burkholderia>Burkholderia_cepacia_complex | 580 | LGM80_11590 | 409569..411311 | 87883 | putative-DNA-binding-domain-containing-protein-[Burkholderia-multivorans] |
| MBR8145921.1 | Schlafen+HATPase+wHTH | GCA_018147135.1 | Burkholderia_vietnamiensis | Bacteria>Pseudomonadati>Pseudomonadota>Betaproteobacteria>Burkholderiales>Burkholderiaceae>Burkholderia>Burkholderia_cepacia_complex | 577 | KDW46_26555 | 44579..46312 | 60552 | putative-DNA-binding-domain-containing-protein-[Burkholderia-vietnamiensis] |
| WP_259032550.1 | Schlafen+HATPase+wHTH | GCF_024793955.1 | Burkholderia_thailandensis | Bacteria>Pseudomonadati>Pseudomonadota>Betaproteobacteria>Burkholderiales>Burkholderiaceae>Burkholderia>pseudomallei_group | 580 | NX841_RS02855 | 22499..24241 | 57975 | ATP-binding-protein-[Burkholderia-thailandensis] |
| WP_122634119.1 | Schlafen+HATPase+wHTH | GCF_900596605.1 | Burkholderia_pseudomallei | Bacteria>Pseudomonadati>Pseudomonadota>Betaproteobacteria>Burkholderiales>Burkholderiaceae>Burkholderia>pseudomallei_group | 625 | EFR82_RS35450 | 8982..10859 | 28450 | putative-DNA-binding-domain-containing-protein-[Burkholderia-pseudomallei] |
| WP_124920777.1 | Schlafen+HATPase+wHTH | GCF_003857805.1 | Burkholderia_sp._Bp9099 | Bacteria>Pseudomonadati>Pseudomonadota>Betaproteobacteria>Burkholderiales>Burkholderiaceae>Burkholderia>unclassified_Burkholderia | 618 | DIE17_RS07330 | 176899..178755 | 2184568 | putative-DNA-binding-domain-containing-protein-[Burkholderia-sp.-Bp9099] |
| WP_065497251.1 | Schlafen+HATPase+wHTH | GCF_001685525.1 | Burkholderia_sp._CCA53 | Bacteria>Pseudomonadati>Pseudomonadota>Betaproteobacteria>Burkholderiales>Burkholderiaceae>Burkholderia>unclassified_Burkholderia | 423 | BCR55_RS29350 | 25002..26273 | 1776288 | ATP-binding-protein-[Burkholderia-sp.-CCA53] |
| RQZ49992.1 | Schlafen+HATPase+wHTH | GCA_003857805.1 | Burkholderia_sp._Bp9099 | Bacteria>Pseudomonadati>Pseudomonadota>Betaproteobacteria>Burkholderiales>Burkholderiaceae>Burkholderia>unclassified_Burkholderia | 618 | DIE17_07280 | 176899..178755 | 2184568 | AAA-family-ATPase-[Burkholderia-sp.-Bp9099] |
| WP_183716834.1 | Schlafen+HATPase+wHTH | GCF_014191085.1 | Paraburkholderia_tropica | Bacteria>Pseudomonadati>Pseudomonadota>Betaproteobacteria>Burkholderiales>Burkholderiaceae>Paraburkholderia | 416 | FHX57_RS30225 | 121474..122724 | 92647 | ATP-binding-protein-[Paraburkholderia-tropica] |
| NKJ47218.1 | Schlafen+HATPase+wHTH | GCA_012275575.1 | Paraburkholderia_sp._SG-MS1 | Bacteria>Pseudomonadati>Pseudomonadota>Betaproteobacteria>Burkholderiales>Burkholderiaceae>Paraburkholderia>unclassified_Paraburkholderia | 416 | CIC12_10780 | 81576..82826 | 2023741 | transcriptional-regulator-[Burkholderia-sp.-SG-MS1] |
| MCO5109350.1 | Schlafen+HATPase+wHTH | GCA_023954095.1 | Burkholderiaceae_bacterium | Bacteria>Pseudomonadati>Pseudomonadota>Betaproteobacteria>Burkholderiales>Burkholderiaceae>unclassified_Burkholderiaceae | 621 | M9929_00720 | 29811..31676 | 2030806 | putative-DNA-binding-domain-containing-protein-[Burkholderiaceae-bacterium] |
| MCO5106864.1 | Schlafen+HATPase+wHTH | GCA_023954045.1 | Burkholderiaceae_bacterium | Bacteria>Pseudomonadati>Pseudomonadota>Betaproteobacteria>Burkholderiales>Burkholderiaceae>unclassified_Burkholderiaceae | 580 | M9907_07255 | 35951..37693 | 2030806 | putative-DNA-binding-domain-containing-protein-[Burkholderiaceae-bacterium] |
| MBX7228948.1 | Schlafen+HATPase+wHTH | GCA_019695155.1 | Burkholderiaceae_bacterium | Bacteria>Pseudomonadati>Pseudomonadota>Betaproteobacteria>Burkholderiales>Burkholderiaceae>unclassified_Burkholderiaceae | 621 | K1X48_05030 | 17464..19329 | 2030806 | putative-DNA-binding-domain-containing-protein-[Burkholderiaceae-bacterium] |
| MBX3589941.1 | Schlafen+HATPase+wHTH | GCA_019635275.1 | Burkholderiaceae_bacterium | Bacteria>Pseudomonadati>Pseudomonadota>Betaproteobacteria>Burkholderiales>Burkholderiaceae>unclassified_Burkholderiaceae | 624 | KF755_03440 | 54134..56008 | 2030806 | putative-DNA-binding-domain-containing-protein-[Burkholderiaceae-bacterium] |
| MBK8767856.1 | Schlafen+HATPase+wHTH | GCA_016710525.1 | Burkholderiaceae_bacterium | Bacteria>Pseudomonadati>Pseudomonadota>Betaproteobacteria>Burkholderiales>Burkholderiaceae>unclassified_Burkholderiaceae | 152 | IPM01_25905 | 28876..29334 | 2030806 | hypothetical-protein-[Burkholderiaceae-bacterium] |
| MBK8767855.1 | Schlafen+HATPase+wHTH | GCA_016710525.1 | Burkholderiaceae_bacterium | Bacteria>Pseudomonadati>Pseudomonadota>Betaproteobacteria>Burkholderiales>Burkholderiaceae>unclassified_Burkholderiaceae | 208 | IPM01_25900 | 28277..28903 | 2030806 | putative-DNA-binding-domain-containing-protein-[Burkholderiaceae-bacterium] |
| WP_043699938.1 | Schlafen+HATPase+wHTH | GCF_020162115.1 | Tepidimonas_taiwanensis | Bacteria>Pseudomonadati>Pseudomonadota>Betaproteobacteria>Burkholderiales>Burkholderiales_genera_incertae_sedis>Tepidimonas | 423 | PL21_RS04265 | 87791..89062 | 307486 | ATP-binding-protein-[Tepidimonas-taiwanensis] |
| OJX16146.1 | Schlafen+HATPase+wHTH | GCA_001898745.1 | Delftia_sp._67-8 | Bacteria>Pseudomonadati>Pseudomonadota>Betaproteobacteria>Burkholderiales>Comamonadaceae>Delftia>unclassified_Delftia | 491 | BGO79_00280 | 68094..69569 | 1895749 | AAA-family-ATPase-[Delftia-sp.-67-8] |
| MBF5004204.1 | Schlafen+HATPase+wHTH | GCA_015354245.1 | Diaphorobacter_caeni | Bacteria>Pseudomonadati>Pseudomonadota>Betaproteobacteria>Burkholderiales>Comamonadaceae>Diaphorobacter | 596 | ISG32_08025 | 62237..64027 | 2784387 | putative-DNA-binding-domain-containing-protein-[Diaphorobacter-caeni] |
| WP_252101811.1 | Schlafen+HATPase+wHTH | GCF_022807955.1 | Diaphorobacter_sp._LI3 | Bacteria>Pseudomonadati>Pseudomonadota>Betaproteobacteria>Burkholderiales>Comamonadaceae>Diaphorobacter>unclassified_Diaphorobacter | 394 | MRB47_RS02540 | 524634..525818 | 2952886 | ATP-binding-protein-[[Acidovorax] |
| WP_233254876.1 | Schlafen+HATPase+wHTH | GCF_003063315.1 | Limnohabitans_sp._T6-5 | Bacteria>Pseudomonadati>Pseudomonadota>Betaproteobacteria>Burkholderiales>Comamonadaceae>Limnohabitans>unclassified_Limnohabitans | 478 | B9Z51_RS07565 | 411104..412540 | 1100724 | helix-turn-helix-domain-containing-protein-[Limnohabitans-sp.-T6-5] |
| MBY4597993.1 | Schlafen+HATPase+wHTH | GCA_019793415.1 | Ottowia_caeni | Bacteria>Pseudomonadati>Pseudomonadota>Betaproteobacteria>Burkholderiales>Comamonadaceae>Ottowia | 400 | K3217_20950 | 95066..96268 | 2870339 | putative-DNA-binding-domain-containing-protein-[Ottowia-caeni] |
| PIQ50871.1 | Schlafen+HATPase+wHTH | GCA_002786915.1 | Comamonadaceae_bacterium_CG12_big_fil_rev_8_21_14_0_65_59_15 | Bacteria>Pseudomonadati>Pseudomonadota>Betaproteobacteria>Burkholderiales>Comamonadaceae>unclassified_Comamonadaceae | 413 | COW02_17760 | 39824..41065 | 1973952 | transcriptional-regulator-[Comamonadaceae-bacterium-CG12_big_fil_rev_8_21_14_0_65_59_15] |
| WP_265289360.1 | Schlafen+HATPase+wHTH | GCF_026127525.1 | Verminephrobacter_eiseniae | Bacteria>Pseudomonadati>Pseudomonadota>Betaproteobacteria>Burkholderiales>Comamonadaceae>Verminephrobacter | 148 | D5047_RS20575 | 4396424..4396870 | 364317 | ATP-binding-protein-[Verminephrobacter-eiseniae] |
| WP_265289359.1 | Schlafen+HATPase+wHTH | GCF_026127525.1 | Verminephrobacter_eiseniae | Bacteria>Pseudomonadati>Pseudomonadota>Betaproteobacteria>Burkholderiales>Comamonadaceae>Verminephrobacter | 263 | D5047_RS20570 | 4395599..4396390 | 364317 | putative-DNA-binding-domain-containing-protein-[Verminephrobacter-eiseniae] |
| WP_232287681.1 | Schlafen+HATPase+wHTH | GCF_000015565.1 | Verminephrobacter_eiseniae | Bacteria>Pseudomonadati>Pseudomonadota>Betaproteobacteria>Burkholderiales>Comamonadaceae>Verminephrobacter | 439 | D8B29_RS12170 | 4558..5877 | 364317 | ATP-binding-protein-[Verminephrobacter-eiseniae] |
| WP_050467361.1 | Schlafen+HATPase+wHTH | GCF_001571165.1 | Herbaspirillum_chlorophenolicum | Bacteria>Pseudomonadati>Pseudomonadota>Betaproteobacteria>Burkholderiales>Oxalobacteraceae>Herbaspirillum | 596 | HCH01S_RS12910 | 73661..75451 | 211589 | ATP-binding-protein-[Herbaspirillum-chlorophenolicum] |
| WP_210803129.1 | Schlafen+HATPase+wHTH | GCF_018069755.1 | Ideonella | Bacteria>Pseudomonadati>Pseudomonadota>Betaproteobacteria>Burkholderiales>Sphaerotilaceae | 651 | KAK07_RS23535 | 31333..33288 | 36862 | MULTISPECIES:-ATP-binding-protein-[Ideonella] |
| WP_281915168.1 | Schlafen+HATPase+wHTH | GCF_937957785.1 | Caldimonas_thermodepolymerans | Bacteria>Pseudomonadati>Pseudomonadota>Betaproteobacteria>Burkholderiales>Sphaerotilaceae>Caldimonas | 423 | QMG53_RS04180 | 4323..5594 | 215580 | ATP-binding-protein-[Caldimonas-thermodepolymerans] |
| WP_062194126.1 | Schlafen+HATPase+wHTH | GCF_001592165.1 | Caldimonas_taiwanensis | Bacteria>Pseudomonadati>Pseudomonadota>Betaproteobacteria>Burkholderiales>Sphaerotilaceae>Caldimonas | 590 | CTA01S_RS07710 | 5880..7652 | 307483 | putative-DNA-binding-domain-containing-protein-[Caldimonas-taiwanensis] |
| WP_246312356.1 | Schlafen+HATPase+wHTH | - | Pseudaquabacterium_terrae | Bacteria>Pseudomonadati>Pseudomonadota>Betaproteobacteria>Burkholderiales>Sphaerotilaceae>Pseudaquabacterium | 586 | HLB44_RS18510 | 558415..560175 | 2732868 | ATP-binding-protein-[Aquabacterium-terrae] |
| WP_009858745.1 | Schlafen+HATPase+wHTH | GCF_000190375.1 | Rubrivivax_benzoatilyticus | Bacteria>Pseudomonadati>Pseudomonadota>Betaproteobacteria>Burkholderiales>Sphaerotilaceae>Rubrivivax | 651 | G7087_RS13140 | 28662..30617 | 316997 | ATP-binding-protein-[Rubrivivax-benzoatilyticus] |
| ODU09014.1 | Schlafen+HATPase+wHTH | GCA_001724995.1 | Rubrivivax_sp._SCN_71-131 | Bacteria>Pseudomonadati>Pseudomonadota>Betaproteobacteria>Burkholderiales>Sphaerotilaceae>Rubrivivax>unclassified_Rubrivivax | 615 | ABS84_09900 | 13053..14900 | 1660138 | MAG:-AAA-family-ATPase-[Rubrivivax-sp.-SCN-71-131] |
| MCW5609140.1 | Schlafen+HATPase+wHTH | GCA_026125605.1 | Rubrivivax_sp. | Bacteria>Pseudomonadati>Pseudomonadota>Betaproteobacteria>Burkholderiales>Sphaerotilaceae>Rubrivivax>unclassified_Rubrivivax | 582 | KIS83_00405 | 79293..81041 | 50259 | MAG:-putative-DNA-binding-domain-containing-protein-[Rubrivivax-sp.] |
| MCC7099551.1 | Schlafen+HATPase+wHTH | GCA_020850905.1 | Rubrivivax_sp. | Bacteria>Pseudomonadati>Pseudomonadota>Betaproteobacteria>Burkholderiales>Sphaerotilaceae>Rubrivivax>unclassified_Rubrivivax | 335 | IT500_08180 | 1..1006 | 50259 | putative-DNA-binding-domain-containing-protein-[Rubrivivax-sp.] |
| MBL8328912.1 | Schlafen+HATPase+wHTH | GCA_016790225.1 | Rubrivivax_sp. | Bacteria>Pseudomonadati>Pseudomonadota>Betaproteobacteria>Burkholderiales>Sphaerotilaceae>Rubrivivax>unclassified_Rubrivivax | 651 | JNJ71_08665 | 119496..121451 | 50259 | putative-DNA-binding-domain-containing-protein-[Rubrivivax-sp.] |
| RIK89377.1 | Schlafen+HATPase+wHTH | GCA_003577305.1 | Burkholderiales_bacterium | Bacteria>Pseudomonadati>Pseudomonadota>Betaproteobacteria>Burkholderiales>unclassified_Burkholderiales | 400 | DCC70_08100 | 19942..21144 | 1891238 | transcriptional-regulator-[Burkholderiales-bacterium] |
| MDE2080725.1 | Schlafen+HATPase+wHTH | GCA_028872015.1 | Burkholderiales_bacterium | Bacteria>Pseudomonadati>Pseudomonadota>Betaproteobacteria>Burkholderiales>unclassified_Burkholderiales | 661 | KGI90_05175 | 49392..51377 | 1891238 | MAG:-putative-DNA-binding-domain-containing-protein-[Burkholderiales-bacterium] |
| MCP5292582.1 | Schlafen+HATPase+wHTH | GCA_024235775.1 | Burkholderiales_bacterium | Bacteria>Pseudomonadati>Pseudomonadota>Betaproteobacteria>Burkholderiales>unclassified_Burkholderiales | 344 | H6938_08645 | 45850..46884 | 1891238 | putative-DNA-binding-domain-containing-protein-[Burkholderiales-bacterium] |

| Protein-ID | Domain | Assembly-ID | Strain | Taxonomy | Length | Locus-Tag | Genomic-coordinates | TaxID | Fasta-defline |
|---|---|---|---|---|---|---|---|---|---|
| MCP5252672.1 | Schlafen+HATPase+wHTH | GCA_024235735.1 | Burkholderiales_bacterium | Bacteria>Pseudomonadati>Pseudomonadota>Betaproteobacteria>Burkholderiales>unclassified_Burkholderiales | 344 | H6939_13340 | 20055..21089 | 1891238 | putative-DNA-binding-domain-containing-protein-[Burkholderiales-bacterium] |
| MBL8529537.1 | Schlafen+HATPase+wHTH | GCA_016791285.1 | Burkholderiales_bacterium | Bacteria>Pseudomonadati>Pseudomonadota>Betaproteobacteria>Burkholderiales>unclassified_Burkholderiales | 401 | JNL68_17800 | 1999..3204 | 1891238 | putative-DNA-binding-domain-containing-protein-[Burkholderiales-bacterium] |
| MBI3155184.1 | Schlafen+HATPase+wHTH | GCA_016191995.1 | Burkholderiales_bacterium | Bacteria>Pseudomonadati>Pseudomonadota>Betaproteobacteria>Burkholderiales>unclassified_Burkholderiales | 651 | HYZ20_07225 | 11157..13112 | 1891238 | putative-DNA-binding-domain-containing-protein-[Burkholderiales-bacterium] |
| WP_244505731.1 | Schlafen+HATPase+wHTH | GCF_900106545.1 | Nitrosomonas_communis | Bacteria>Pseudomonadati>Pseudomonadota>Betaproteobacteria>Nitrosomonadales>Nitrosomonadaceae>Nitrosomonas | 275 | BLV21_RS18395 | 51829..52656 | 44574 | ATP-binding-protein-[Nitrosomonas-communis] |
| WP_218142481.1 | Schlafen+HATPase+wHTH | GCF_900111055.1 | Nitrosomonas_ureae | Bacteria>Pseudomonadati>Pseudomonadota>Betaproteobacteria>Nitrosomonadales>Nitrosomonadaceae>Nitrosomonas | 604 | BMW48_RS11205 | 16341..18155 | 44577 | ATP-binding-protein-[Nitrosomonas-ureae] |
| WP_211752263.1 | Schlafen+HATPase+wHTH | GCF_900106545.1 | Nitrosomonas_communis | Bacteria>Pseudomonadati>Pseudomonadota>Betaproteobacteria>Nitrosomonadales>Nitrosomonadaceae>Nitrosomonas | 297 | BLV21_RS18400 | 52604..53497 | 44574 | ATP-binding-protein-[Nitrosomonas-communis] |
| WP_204800509.1 | Schlafen+HATPase+wHTH | GCF_905220645.1 | Nitrosomonas_nitrosa | Bacteria>Pseudomonadati>Pseudomonadota>Betaproteobacteria>Nitrosomonadales>Nitrosomonadaceae>Nitrosomonas | 555 | NUZ5A_RS15525 | 135463..137130 | 52442 | ATP-binding-protein-[Nitrosomonas-nitrosa] |
| WP_107790391.1 | Schlafen+HATPase+wHTH | GCF_003051105.1 | Nitrosomonas_nitrosa | Bacteria>Pseudomonadati>Pseudomonadota>Betaproteobacteria>Nitrosomonadales>Nitrosomonadaceae>Nitrosomonas | 555 | C8R30_RS13780 | 16886..18553 | 52442 | ATP-binding-protein-[Nitrosomonas-nitrosa] |
| SDW37021.1 | Schlafen+HATPase+wHTH | GCA_900106545.1 | Nitrosomonas_communis | Bacteria>Pseudomonadati>Pseudomonadota>Betaproteobacteria>Nitrosomonadales>Nitrosomonadaceae>Nitrosomonas | 297 | SAMN05421882_100920 | 52604..53497 | 44574 | ATP-dependent-DNA-helicase-RecG-[Nitrosomonas-communis] |
| SDW36999.1 | Schlafen+HATPase+wHTH | GCA_900106545.1 | Nitrosomonas_communis | Bacteria>Pseudomonadati>Pseudomonadota>Betaproteobacteria>Nitrosomonadales>Nitrosomonadaceae>Nitrosomonas | 275 | SAMN05421882_100919 | 51829..52656 | 44574 | ATP-dependent-DNA-helicase-RecG-[Nitrosomonas-communis] |
| MCO6428552.1 | Schlafen+HATPase+wHTH | GCA_024102715.1 | Nitrosomonas_communis | Bacteria>Pseudomonadati>Pseudomonadota>Betaproteobacteria>Nitrosomonadales>Nitrosomonadaceae>Nitrosomonas | 402 | J5I83_10910 | 877..2085 | 44574 | putative-DNA-binding-domain-containing-protein-[Nitrosomonas-communis] |
| MBK7768855.1 | Schlafen+HATPase+wHTH | GCA_016706645.1 | Sulfuritalea_sp. | Bacteria>Pseudomonadati>Pseudomonadota>Betaproteobacteria>Nitrosomonadales>Sterolibacteriaceae>Sulfuritalea>unclassified_Sulfuritalea | 565 | IPI44_24185 | 3477..5174 | 2480090 | putative-DNA-binding-domain-containing-protein-[Sulfuritalea-sp.] |
| MBK7665763.1 | Schlafen+HATPase+wHTH | GCA_016709795.1 | Sterolibacteriaceae_bacterium | Bacteria>Pseudomonadati>Pseudomonadota>Betaproteobacteria>Nitrosomonadales>Sterolibacteriaceae>unclassified_Sterolibacteriaceae | 452 | IPJ21_19920 | 4298834..4300192 | 2682146 | putative-DNA-binding-domain-containing-protein-[Sterolibacteriaceae-bacterium] |
| ODU50881.1 | Schlafen+HATPase+wHTH | GCA_001725135.1 | Thiobacillus_sp._SCN_63-374 | Bacteria>Pseudomonadati>Pseudomonadota>Betaproteobacteria>Nitrosomonadales>Thiobacillaceae>Thiobacillus>unclassified_Thiobacillus | 394 | ABS92_01660 | 18626..19810 | 1660149 | MAG:-transcriptional-regulator-[Thiobacillus-sp.-SCN-63-374] |
| MCX7674149.1 | Schlafen+HATPase+wHTH | GCA_026419045.1 | Thiobacillaceae_bacterium | Bacteria>Pseudomonadati>Pseudomonadota>Betaproteobacteria>Nitrosomonadales>Thiobacillaceae>unclassified_Thiobacillaceae | 424 | N2Z63_11250 | 13639..14913 | 2528409 | MAG:-putative-DNA-binding-domain-containing-protein-[Thiobacillaceae-bacterium] |
| MBI3902212.1 | Schlafen+HATPase+wHTH | GCA_016199835.1 | Nitrosomonadales_bacterium | Bacteria>Pseudomonadati>Pseudomonadota>Betaproteobacteria>Nitrosomonadales>unclassified_Nitrosomonadales | 530 | HY306_04620 | 115107..116699 | 2026761 | putative-DNA-binding-domain-containing-protein-[Nitrosomonadales-bacterium] |
| MBI1887189.1 | Schlafen+HATPase+wHTH | GCA_016177795.1 | Nitrosomonadales_bacterium | Bacteria>Pseudomonadati>Pseudomonadota>Betaproteobacteria>Nitrosomonadales>unclassified_Nitrosomonadales | 530 | HYS19_02285 | 83944..85536 | 2026761 | putative-DNA-binding-domain-containing-protein-[Nitrosomonadales-bacterium] |
| MBN8461438.1 | Schlafen+HATPase+wHTH | GCA_017302355.1 | Dechloromonas_sp. | Bacteria>Pseudomonadati>Pseudomonadota>Betaproteobacteria>Rhodocyclales>Azonexaceae>Dechloromonas>unclassified_Dechloromonas | 622 | J0M01_01280 | 12060..13928 | 1917218 | putative-DNA-binding-domain-containing-protein-[Dechloromonas-sp.] |
| MBK7421887.1 | Schlafen+HATPase+wHTH | GCA_016709335.1 | Candidatus_Propionivibrio_dominans | Bacteria>Pseudomonadati>Pseudomonadota>Betaproteobacteria>Rhodocyclales>Rhodocyclaceae>Propionivibrio | 577 | IPJ48_01625 | 220429..222162 | 2954373 | putative-DNA-binding-domain-containing-protein-[Propionivibrio-sp.] |
| MCX8086912.1 | Schlafen+HATPase+wHTH | GCA_026415155.1 | Rhodocyclaceae_bacterium | Bacteria>Pseudomonadati>Pseudomonadota>Betaproteobacteria>Rhodocyclales>Rhodocyclaceae>unclassified_Rhodocyclaceae | 423 | N3C63_08435 | 27368..28639 | 1898103 | MAG:-putative-DNA-binding-domain-containing-protein-[Rhodocyclaceae-bacterium] |
| MCA3140846.1 | Schlafen+HATPase+wHTH | GCA_020248385.1 | Rhodocyclaceae_bacterium | Bacteria>Pseudomonadati>Pseudomonadota>Betaproteobacteria>Rhodocyclales>Rhodocyclaceae>unclassified_Rhodocyclaceae | 395 | ING80_01380 | 48973..50160 | 1898103 | putative-DNA-binding-domain-containing-protein-[Rhodocyclaceae-bacterium] |
| MBL8482097.1 | Schlafen+HATPase+wHTH | GCA_016791085.1 | Rhodocyclaceae_bacterium | Bacteria>Pseudomonadati>Pseudomonadota>Betaproteobacteria>Rhodocyclales>Rhodocyclaceae>unclassified_Rhodocyclaceae | 581 | JNJ60_07875 | 16737..18482 | 1898103 | putative-DNA-binding-domain-containing-protein-[Rhodocyclaceae-bacterium] |
| AUM01339.1 | Schlafen+HATPase+wHTH | GCA_002863805.1 | Rhodocyclaceae_bacterium | Bacteria>Pseudomonadati>Pseudomonadota>Betaproteobacteria>Rhodocyclales>Rhodocyclaceae>unclassified_Rhodocyclaceae | 590 | B4966_15120 | 3169702..3171474 | 1898103 | transcriptional-regulator-[Rhodocyclaceae-bacterium] |
| OHC61991.1 | Schlafen+HATPase+wHTH | GCA_001828915.1 | Rhodocyclales_bacterium_GWA2_65_19 | Bacteria>Pseudomonadati>Pseudomonadota>Betaproteobacteria>Rhodocyclales>unclassified_Rhodocyclales | 562 | A2040_16470 | 41234..42922 | 1802016 | MAG:-transcriptional-regulator-[Rhodocyclales-bacterium-GWA2_65_19] |
| WP_015434798.1 | Schlafen+HATPase+wHTH | GCF_000349945.1 | Azoarcus_sp._KH32C | Bacteria>Pseudomonadati>Pseudomonadota>Betaproteobacteria>Rhodocyclales>Zoogloeaceae>Azoarcus>unclassified_Azoarcus | 415 | AZKH_RS05725 | 1274770..1276017 | 748247 | ATP-binding-protein-[Azoarcus-sp.-KH32C] |
| OGA55365.1 | Schlafen+HATPase+wHTH | GCA_001772005.1 | Betaproteobacteria_bacterium_RIFCSPLOWO2_12_FULL_62_58 | Bacteria>Pseudomonadati>Pseudomonadota>Betaproteobacteria>unclassified_Betaproteobacteria | 548 | A3F74_07170 | 7046..8692 | 1797497 | transcriptional-regulator-[Betaproteobacteria-bacterium-RIFCSPLOWO2_12_FULL_62_58] |
| OGA30957.1 | Schlafen+HATPase+wHTH | GCA_001771975.1 | Betaproteobacteria_bacterium_RIFCSPLOWO2_02_FULL_68_150 | Bacteria>Pseudomonadati>Pseudomonadota>Betaproteobacteria>unclassified_Betaproteobacteria | 409 | A3I65_01675 | 18902..20131 | 1797493 | hypothetical-protein-A3I65_01675-[Betaproteobacteria-bacterium-RIFCSPLOWO2_02_FULL_68_150] |
| MBI5922419.1 | Schlafen+HATPase+wHTH | GCA_016234745.1 | Betaproteobacteria_bacterium | Bacteria>Pseudomonadati>Pseudomonadota>Betaproteobacteria>unclassified_Betaproteobacteria | 401 | HY847_12360 | 46902..48107 | 1891241 | putative-DNA-binding-domain-containing-protein-[Betaproteobacteria-bacterium] |
| MBI3917897.1 | Schlafen+HATPase+wHTH | GCA_016199545.1 | Betaproteobacteria_bacterium | Bacteria>Pseudomonadati>Pseudomonadota>Betaproteobacteria>unclassified_Betaproteobacteria | 410 | HY322_12920 | 21972..23204 | 1891241 | putative-DNA-binding-domain-containing-protein-[Betaproteobacteria-bacterium] |
| GHU44751.1 | Schlafen+HATPase+wHTH | GCA_025997095.1 | Betaproteobacteria_bacterium | Bacteria>Pseudomonadati>Pseudomonadota>Betaproteobacteria>unclassified_Betaproteobacteria | 423 | AGMMS50289_13820 | 5092..6363 | 1891241 | ATPase-AAA-[Betaproteobacteria-bacterium] |
| GHU42410.1 | Schlafen+HATPase+wHTH | GCA_025997095.1 | Betaproteobacteria_bacterium | Bacteria>Pseudomonadati>Pseudomonadota>Betaproteobacteria>unclassified_Betaproteobacteria | 562 | AGMMS50289_07070 | 1446..3134 | 1891241 | DNA-binding-protein-[Betaproteobacteria-bacterium] |
| WP_106453127.1 | Schlafen+HATPase+wHTH | GCF_003012795.1 | Zobellella_taiwanensis | Bacteria>Pseudomonadati>Pseudomonadota>Gammaproteobacteria>Aeromonadales>Aeromonadaceae>Zobellella | 624 | C7I36_RS07645 | 90337..92211 | 347535 | putative-DNA-binding-domain-containing-protein-[Zobellella-taiwanensis] |
| WP_219938191.1 | Schlafen+HATPase+wHTH | GCF_019448115.1 | Succinivibrio_faecicola | Bacteria>Pseudomonadati>Pseudomonadota>Gammaproteobacteria>Aeromonadales>Succinivibrionaceae>Succinivibrio | 473 | J5V48_RS08695 | 9731..11152 | 2820300 | putative-DNA-binding-domain-containing-protein-[Succinivibrio-faecicola] |
| WP_020742687.1 | Schlafen+HATPase+wHTH | GCF_000439515.1 | Alteromonas_mediterranea | Bacteria>Pseudomonadati>Pseudomonadota>Gammaproteobacteria>Alteromonadales>Alteromonadaceae>Alteromonas/Salinimonas_group>Alteromonas | 342 | I876_RS01275 | 287880..288908 | 314275 | ATP-binding-protein-[Alteromonas-mediterranea] |
| WP_136550440.1 | Schlafen+HATPase+wHTH | GCF_004851605.1 | Hydrocarboniclastica_marina | Bacteria>Pseudomonadati>Pseudomonadota>Gammaproteobacteria>Alteromonadales>Alteromonadaceae>Hydrocarboniclastica | 556 | soil367_RS18205 | 4129693..4131363 | 2259620 | ATP-binding-protein-[Hydrocarboniclastica-marina] |
| NQY33781.1 | Schlafen+HATPase+wHTH | GCA_013216015.1 | Alteromonadaceae_bacterium | Bacteria>Pseudomonadati>Pseudomonadota>Gammaproteobacteria>Alteromonadales>Alteromonadaceae>unclassified_Alteromonadaceae | 390 | HRT37_02200 | 99434..100606 | 1916082 | putative-DNA-binding-domain-containing-protein-[Alteromonadaceae-bacterium] |
| WP_105264931.1 | Schlafen+HATPase+wHTH | GCF_002964725.1 | Pseudoalteromonas_sp._T1lg76 | Bacteria>Pseudomonadati>Pseudomonadota>Gammaproteobacteria>Alteromonadales>Pseudoalteromonadaceae>Pseudoalteromonas>unclassified_Pseudoalteromonas | 510 | C2M35_RS06885 | 46939..48471 | 2077103 | ATP-binding-protein-[Pseudoalteromonas-sp.-T1lg76] |
| WP_105190027.1 | Schlafen+HATPase+wHTH | GCF_002964785.1 | Pseudoalteromonas_sp._T1lg48 | Bacteria>Pseudomonadati>Pseudomonadota>Gammaproteobacteria>Alteromonadales>Pseudoalteromonadaceae>Pseudoalteromonas>unclassified_Pseudoalteromonas | 222 | C2M29_RS12215 | 655088..655756 | 2077100 | ATP-binding-protein-[Pseudoalteromonas-sp.-T1lg48] |
| MBU2886740.1 | Schlafen+HATPase+wHTH | GCA_018860575.1 | Gilvimarinus_agarilyticus | Bacteria>Pseudomonadati>Pseudomonadota>Gammaproteobacteria>Cellvibrionales>Cellvibrionaceae>Gilvimarinus | 631 | KO507_13280 | 319738..321633 | 679259 | putative-DNA-binding-domain-containing-protein-[Gilvimarinus-agarilyticus] |
| WP_265366288.1 | Schlafen+HATPase+wHTH | GCF_026153455.1 | Marinimicrobium_sp._C6131 | Bacteria>Pseudomonadati>Pseudomonadota>Gammaproteobacteria>Cellvibrionales>Cellvibrionaceae>Marinimicrobium>unclassified_Marinimicrobium | 556 | OOT55_RS13030 | 3028083..3029753 | 3022676 | ATP-binding-protein-[Marinimicrobium-sp.-C6131] |
| WP_182173098.1 | Schlafen+HATPase+wHTH | GCF_014077625.1 | Sediminihaliea_albiluteola | Bacteria>Pseudomonadati>Pseudomonadota>Gammaproteobacteria>Cellvibrionales>Halieaceae>Sediminihaliea | 423 | H2508_RS10785 | 847473..848744 | 2758564 | ATP-binding-protein-[Sediminihaliea-albiluteola] |
| WP_266072740.1 | Schlafen+HATPase+wHTH | GCF_026241615.1 | Microbulbifer_thermotolerans | Bacteria>Pseudomonadati>Pseudomonadota>Gammaproteobacteria>Cellvibrionales>Microbulbiferaceae>Microbulbifer | 622 | OQJ46_RS08135 | 154604..156472 | 252514 | putative-DNA-binding-domain-containing-protein-[Microbulbifer-thermotolerans] |
| WP_043319018.1 | Schlafen+HATPase+wHTH | GCF_000708675.1 | Microbulbifer_sp._HZ11 | Bacteria>Pseudomonadati>Pseudomonadota>Gammaproteobacteria>Cellvibrionales>Microbulbiferaceae>Microbulbifer>unclassified_Microbulbifer | 573 | AU11_RS09635 | 450140..451861 | 1453501 | ATP-binding-protein-[Microbulbifer-sp.-HZ11] |
| MAT52641.1 | Schlafen+HATPase+wHTH | GCA_002702725.1 | Porticoccaceae_bacterium | Bacteria>Pseudomonadati>Pseudomonadota>Gammaproteobacteria>Cellvibrionales>Porticoccaceae>unclassified_Porticoccaceae | 423 | CMK32_15790 | 16274..17545 | 2026782 | transcriptional-regulator-[Porticoccaceae-bacterium] |
| MAT50825.1 | Schlafen+HATPase+wHTH | GCA_002702725.1 | Porticoccaceae_bacterium | Bacteria>Pseudomonadati>Pseudomonadota>Gammaproteobacteria>Cellvibrionales>Porticoccaceae>unclassified_Porticoccaceae | 556 | CMK32_06555 | 6249..7919 | 2026782 | ATPase-[Porticoccaceae-bacterium] |
| WP_268905393.1 | Schlafen+HATPase+wHTH | GCF_026891035.1 | Dasania_phycosphaerae | Bacteria>Pseudomonadati>Pseudomonadota>Gammaproteobacteria>Cellvibrionales>Spongiibacteraceae>Dasania | 482 | OOV09_RS18300 | 19644..21092 | 2950436 | helix-turn-helix-domain-containing-protein-[Dasania-sp.-GY-19] |
| WP_273033647.1 | Schlafen+HATPase+wHTH | GCF_950061765.1 | Spongiibacter_tropicus | Bacteria>Pseudomonadati>Pseudomonadota>Gammaproteobacteria>Cellvibrionales>Spongiibacteraceae>Spongiibacter | 557 | B4C78_RS03790 | 101082..102755 | 454602 | ATP-binding-protein-[Spongiibacter-tropicus] |
| WP_201222434.1 | Schlafen+HATPase+wHTH | GCF_016653315.1 | Halochromatium_roseum | Bacteria>Pseudomonadati>Pseudomonadota>Gammaproteobacteria>Chromatiales>Chromatiaceae>Halochromatium | 492 | CCR96_RS19430 | 146293..147771 | 391920 | helix-turn-helix-domain-containing-protein-[Halochromatium-roseum] |
| WP_020507128.1 | Schlafen+HATPase+wHTH | GCF_000379525.1 | Lamprocystis_purpurea | Bacteria>Pseudomonadati>Pseudomonadota>Gammaproteobacteria>Chromatiales>Chromatiaceae>Lamprocystis | 553 | A390_RS0123540 | 21065..22726 | 61598 | ATP-binding-protein-[Lamprocystis-purpurea] |
| WP_020505852.1 | Schlafen+HATPase+wHTH | GCF_000379525.1 | Lamprocystis_purpurea | Bacteria>Pseudomonadati>Pseudomonadota>Gammaproteobacteria>Chromatiales>Chromatiaceae>Lamprocystis | 605 | A390_RS0116915 | 26082..27899 | 61598 | putative-DNA-binding-domain-containing-protein-[Lamprocystis-purpurea] |
| WP_020505455.1 | Schlafen+HATPase+wHTH | GCF_000379525.1 | Lamprocystis_purpurea | Bacteria>Pseudomonadati>Pseudomonadota>Gammaproteobacteria>Chromatiales>Chromatiaceae>Lamprocystis | 497 | A390_RS0114865 | 130649..132142 | 61598 | putative-DNA-binding-domain-containing-protein-[Lamprocystis-purpurea] |
| WP_002813715.1 | Schlafen+HATPase+wHTH | GCF_000012805.1 | Nitrosococcus_oceani | Bacteria>Pseudomonadati>Pseudomonadota>Gammaproteobacteria>Chromatiales>Chromatiaceae>Nitrosococcus | 555 | IB75_RS00210 | 9673..11340 | 1229 | crosslink-repair-DNA-glycosylase-YcaQ-family-protein-[Nitrosococcus-oceani] |
| GEM21689.1 | Schlafen+HATPase+wHTH | GCA_007990675.1 | Nitrosococcus_oceani | Bacteria>Pseudomonadati>Pseudomonadota>Gammaproteobacteria>Chromatiales>Chromatiaceae>Nitrosococcus | 555 | NONS58_31390 | 9779..11446 | 1229 | ATPase-[Nitrosococcus-oceani] |
| EDZ66368.1 | Schlafen+HATPase+wHTH | - | Nitrosococcus_oceani_AFC27 | Bacteria>Pseudomonadati>Pseudomonadota>Gammaproteobacteria>Chromatiales>Chromatiaceae>Nitrosococcus>Nitrosococcus_oceani | 555 | NOC27_3048 | 575216..576883 | 473788 | Divergent-AAA-domain-family-[Nitrosococcus-oceani-AFC27] |
| WP_120796692.1 | Schlafen+HATPase+wHTH | GCF_003634315.1 | Thiocapsa_rosea | Bacteria>Pseudomonadati>Pseudomonadota>Gammaproteobacteria>Chromatiales>Chromatiaceae>Thiocapsa | 425 | BDD21_RS07915 | 1737621..1738898 | 69360 | ATP-binding-protein-[Thiocapsa-rosea] |
| WP_007193605.1 | Schlafen+HATPase+wHTH | GCF_000223985.1 | Thiocapsa_marina | Bacteria>Pseudomonadati>Pseudomonadota>Gammaproteobacteria>Chromatiales>Chromatiaceae>Thiocapsa | 574 | THIMADRAFT_RS13465 | 267431..269155 | 244573 | ATP-binding-protein-[Thiocapsa-marina] |
| WP_200249823.1 | Schlafen+HATPase+wHTH | GCF_016583935.1 | Thiococcus_pfennigii | Bacteria>Pseudomonadati>Pseudomonadota>Gammaproteobacteria>Chromatiales>Chromatiaceae>Thiococcus | 505 | CKO22_RS03180 | 30758..32275 | 1057 | putative-DNA-binding-domain-containing-protein-[Thiococcus-pfennigii] |
| WP_200373983.1 | Schlafen+HATPase+wHTH | GCF_016583575.1 | Thiocystis_violacea | Bacteria>Pseudomonadati>Pseudomonadota>Gammaproteobacteria>Chromatiales>Chromatiaceae>Thiocystis | 424 | CKO27_RS06080 | 51943..53217 | 13725 | ATP-binding-protein-[Thiocystis-violacea] |
| WP_014777945.1 | Schlafen+HATPase+wHTH | GCF_000227745.2 | Thiocystis_violascens | Bacteria>Pseudomonadati>Pseudomonadota>Gammaproteobacteria>Chromatiales>Chromatiaceae>Thiocystis | 424 | THIVI_RS07135 | 1610103..1611377 | 73141 | ATP-binding-protein-[Thiocystis-violascens] |
| AUB82635.1 | Schlafen+HATPase+wHTH | GCA_002813775.1 | Candidatus_Thiodictyon_syntrophicum | Bacteria>Pseudomonadati>Pseudomonadota>Gammaproteobacteria>Chromatiales>Chromatiaceae>Thiodictyon | 553 | THSYN_17925 | 4305856..4307517 | 1166950 | ATPase-[Candidatus-Thiodictyon-syntrophicum] |
| MBK1629588.1 | Schlafen+HATPase+wHTH | GCA_016583825.1 | Thiohalocapsa_halophila | Bacteria>Pseudomonadati>Pseudomonadota>Gammaproteobacteria>Chromatiales>Chromatiaceae>Thiohalocapsa | 425 | CKO31_02300 | 112945..114222 | 69359 | hypothetical-protein-[Thiohalocapsa-halophila] |
| WP_058554451.1 | Schlafen+HATPase+wHTH | GCF_001469165.1 | Thiohalocapsa_sp._ML1 | Bacteria>Pseudomonadati>Pseudomonadota>Gammaproteobacteria>Chromatiales>Chromatiaceae>Thiohalocapsa>unclassified_Thiohalocapsa | 428 | ML01_RS05760 | 161556..162842 | 1431688 | ATP-binding-protein-[Thiohalocapsa-sp.-ML1] |
| WP_009150249.1 | Schlafen+HATPase+wHTH | GCF_033954835.1 | Thiorhodovibrio_frisius | Bacteria>Pseudomonadati>Pseudomonadota>Gammaproteobacteria>Chromatiales>Chromatiaceae>Thiorhodovibrio | 433 | THI970DRAFT_RS16335 | 364126..365427 | 631362 | ATP-binding-protein-[Thiorhodovibrio-sp.-970] |
| EIC19846.1 | Schlafen+HATPase+wHTH | GCA_000228725.3 | Thiorhodovibrio_frisius | Bacteria>Pseudomonadati>Pseudomonadota>Gammaproteobacteria>Chromatiales>Chromatiaceae>Thiorhodovibrio | 433 | Thi970DRAFT_03451 | 364126..365427 | 631362 | putative-transcriptional-regulator-with-HTH-domain-[Thiorhodovibrio-sp.-970] |
| MCP5408974.1 | Schlafen+HATPase+wHTH | GCA_024235335.1 | Chromatiaceae_bacterium | Bacteria>Pseudomonadati>Pseudomonadota>Gammaproteobacteria>Chromatiales>Chromatiaceae>unclassified_Chromatiaceae | 586 | H6963_06740 | 89402..91162 | 2049432 | putative-DNA-binding-domain-containing-protein-[Chromatiaceae-bacterium] |
| MCC7280698.1 | Schlafen+HATPase+wHTH | GCA_020851955.1 | Chromatiaceae_bacterium | Bacteria>Pseudomonadati>Pseudomonadota>Gammaproteobacteria>Chromatiales>Chromatiaceae>unclassified_Chromatiaceae | 553 | IT487_20615 | 12322..13983 | 2049432 | putative-DNA-binding-domain-containing-protein-[Chromatiaceae-bacterium] |
| MBP8282510.1 | Schlafen+HATPase+wHTH | GCA_018007135.1 | Chromatiaceae_bacterium | Bacteria>Pseudomonadati>Pseudomonadota>Gammaproteobacteria>Chromatiales>Chromatiaceae>unclassified_Chromatiaceae | 591 | KAX46_01115 | 5508..7283 | 2049432 | putative-DNA-binding-domain-containing-protein-[Chromatiaceae-bacterium] |
| MBP6806942.1 | Schlafen+HATPase+wHTH | GCA_017991215.1 | Chromatiaceae_bacterium | Bacteria>Pseudomonadati>Pseudomonadota>Gammaproteobacteria>Chromatiales>Chromatiaceae>unclassified_Chromatiaceae | 591 | KA125_01455 | 7317..9092 | 2049432 | putative-DNA-binding-domain-containing-protein-[Chromatiaceae-bacterium] |
| MBP6733684.1 | Schlafen+HATPase+wHTH | GCA_017990755.1 | Chromatiaceae_bacterium | Bacteria>Pseudomonadati>Pseudomonadota>Gammaproteobacteria>Chromatiales>Chromatiaceae>unclassified_Chromatiaceae | 553 | KA142_03155 | 6116..7777 | 2049432 | putative-DNA-binding-domain-containing-protein-[Chromatiaceae-bacterium] |
| MBP6733367.1 | Schlafen+HATPase+wHTH | GCA_017990755.1 | Chromatiaceae_bacterium | Bacteria>Pseudomonadati>Pseudomonadota>Gammaproteobacteria>Chromatiales>Chromatiaceae>unclassified_Chromatiaceae | 591 | KA142_01505 | 3767..5542 | 2049432 | putative-DNA-binding-domain-containing-protein-[Chromatiaceae-bacterium] |
| MBP6582542.1 | Schlafen+HATPase+wHTH | GCA_017989555.1 | Chromatiaceae_bacterium | Bacteria>Pseudomonadati>Pseudomonadota>Gammaproteobacteria>Chromatiales>Chromatiaceae>unclassified_Chromatiaceae | 591 | KA204_03585 | 1481..3256 | 2049432 | putative-DNA-binding-domain-containing-protein-[Chromatiaceae-bacterium] |
| MBK8637140.1 | Schlafen+HATPase+wHTH | GCA_016714645.1 | Chromatiaceae_bacterium | Bacteria>Pseudomonadati>Pseudomonadota>Gammaproteobacteria>Chromatiales>Chromatiaceae>unclassified_Chromatiaceae | 591 | IPN92_02260 | 480553..482328 | 2049432 | putative-DNA-binding-domain-containing-protein-[Chromatiaceae-bacterium] |
| WP_018716650.1 | Schlafen+HATPase+wHTH | GCF_000374645.1 | Arhodomonas_aquaeolei | Bacteria>Pseudomonadati>Pseudomonadota>Gammaproteobacteria>Chromatiales>Ectothiorhodospiraceae>Arhodomonas | 570 | C516_RS0101830 | 425707..427419 | 2369 | ATP-binding-protein-[Arhodomonas-aquaeolei] |
| WP_238627720.1 | Schlafen+HATPase+wHTH | GCF_022227695.1 | unclassified_Ectothiorhodospira | Bacteria>Pseudomonadati>Pseudomonadota>Gammaproteobacteria>Chromatiales>Ectothiorhodospiraceae>Ectothiorhodospira | 556 | LPW28_RS03065 | 42130..43800 | 2684909 | MULTISPECIES:-ATP-binding-protein-[unclassified-Ectothiorhodospira] |
| WP_238621056.1 | Schlafen+HATPase+wHTH | GCF_022227785.1 | Ectothiorhodospira_lacustris | Bacteria>Pseudomonadati>Pseudomonadota>Gammaproteobacteria>Chromatiales>Ectothiorhodospiraceae>Ectothiorhodospira | 610 | LQ952_RS02360 | 41815..43647 | 2899127 | putative-DNA-binding-domain-containing-protein-[Ectothiorhodospira-sp.-B14B] |
| MCG5499274.1 | Schlafen+HATPase+wHTH | GCA_022227785.1 | Ectothiorhodospira_lacustris | Bacteria>Pseudomonadati>Pseudomonadota>Gammaproteobacteria>Chromatiales>Ectothiorhodospiraceae>Ectothiorhodospira | 215 | LQ952_00380 | 94664..95311 | 2899127 | ATP-binding-protein-[Ectothiorhodospira-sp.-B14B] |
| MCG5499273.1 | Schlafen+HATPase+wHTH | GCA_022227785.1 | Ectothiorhodospira_lacustris | Bacteria>Pseudomonadati>Pseudomonadota>Gammaproteobacteria>Chromatiales>Ectothiorhodospiraceae>Ectothiorhodospira | 237 | LQ952_00375 | 94044..94757 | 2899127 | hypothetical-protein-[Ectothiorhodospira-sp.-B14B] |
| WP_063464067.1 | Schlafen+HATPase+wHTH | GCF_001632845.1 | Ectothiorhodospira_sp._BSL-9 | Bacteria>Pseudomonadati>Pseudomonadota>Gammaproteobacteria>Chromatiales>Ectothiorhodospiraceae>Ectothiorhodospira>unclassified_Ectothiorhodospira | 556 | ECTOBSL9_RS04505 | 918420..920090 | 1442136 | ATP-binding-protein-[Ectothiorhodospira-sp.-BSL-9] |
| WP_008932924.1 | Schlafen+HATPase+wHTH | GCF_000225005.1 | Ectothiorhodospira_sp._PHS-1 | Bacteria>Pseudomonadati>Pseudomonadota>Gammaproteobacteria>Chromatiales>Ectothiorhodospiraceae>Ectothiorhodospira>unclassified_Ectothiorhodospira | 615 | ECTPHS_RS11495 | 4823..6670 | 519989 | putative-DNA-binding-domain-containing-protein-[Ectothiorhodospira-sp.-PHS-1] |
| WP_123401499.1 | Schlafen+HATPase+wHTH | GCF_003751635.1 | Inmirania_thermothiophila | Bacteria>Pseudomonadati>Pseudomonadota>Gammaproteobacteria>Chromatiales>Ectothiorhodospiraceae>Inmirania | 467 | EDC57_RS08845 | 669087..670490 | 1750597 | ATP-binding-protein-[Inmirania-thermothiophila] |
| TQE98799.1 | Schlafen+HATPase+wHTH | GCA_006569195.1 | Spiribacter_salinus | Bacteria>Pseudomonadati>Pseudomonadota>Gammaproteobacteria>Chromatiales>Ectothiorhodospiraceae>Spiribacter | 477 | FKY71_11935 | 5411..6844 | 1335746 | AAA-family-ATPase-[Spiribacter-salinus] |
| QEP44634.1 | Schlafen+HATPase+wHTH | GCA_008375315.1 | Ectothiorhodospiraceae_bacterium_BW-2 | Bacteria>Pseudomonadati>Pseudomonadota>Gammaproteobacteria>Chromatiales>Ectothiorhodospiraceae>unclassified_Ectothiorhodospiraceae | 418 | D5085_16735 | 3572413..3573669 | 947515 | transcriptional-regulator-[Ectothiorhodospiraceae-bacterium-BW-2] |
| MBL1275685.1 | Schlafen+HATPase+wHTH | GCA_002401775.2 | Ectothiorhodospiraceae_bacterium | Bacteria>Pseudomonadati>Pseudomonadota>Gammaproteobacteria>Chromatiales>Ectothiorhodospiraceae>unclassified_Ectothiorhodospiraceae | 629 | COB30_006335 | 4341..6230 | 2026737 | putative-DNA-binding-domain-containing-protein-[Ectothiorhodospiraceae-bacterium] |
| MCG8047519.1 | Schlafen+HATPase+wHTH | GCA_016843075.1 | Candidatus_Thiodiazotropha_taylori | Bacteria>Pseudomonadati>Pseudomonadota>Gammaproteobacteria>Chromatiales>Sedimenticolaceae>Candidatus_Thiodiazotropha | 420 | JAY66_17840 | 33202..34464 | 2792791 | putative-DNA-binding-domain-containing-protein-[Candidatus-Thiodiazotropha-taylori] |
| MCG7872688.1 | Schlafen+HATPase+wHTH | GCA_016848925.1 | Candidatus_Thiodiazotropha_lotti | Bacteria>Pseudomonadati>Pseudomonadota>Gammaproteobacteria>Chromatiales>Sedimenticolaceae>Candidatus_Thiodiazotropha | 414 | JAZ11_11360 | 110353..111597 | 2792787 | putative-DNA-binding-domain-containing-protein-[Candidatus-Thiodiazotropha-lotti] |
| MCG7872378.1 | Schlafen+HATPase+wHTH | GCA_016848925.1 | Candidatus_Thiodiazotropha_lotti | Bacteria>Pseudomonadati>Pseudomonadota>Gammaproteobacteria>Chromatiales>Sedimenticolaceae>Candidatus_Thiodiazotropha | 556 | JAZ11_09795 | 177210..178880 | 2792787 | putative-DNA-binding-domain-containing-protein-[Candidatus-Thiodiazotropha-lotti] |
| WP_029131816.1 | Schlafen+HATPase+wHTH | GCF_000428045.1 | Sedimenticola_selenatireducens | Bacteria>Pseudomonadati>Pseudomonadota>Gammaproteobacteria>Chromatiales>Sedimenticolaceae>Sedimenticola | 420 | A3GO_RS0101020 | 206179..207441 | 191960 | ATP-binding-protein-[Sedimenticola-selenatireducens] |
| PLY12605.1 | Schlafen+HATPase+wHTH | GCA_002869585.1 | Sedimenticola_sp. | Bacteria>Pseudomonadati>Pseudomonadota>Gammaproteobacteria>Chromatiales>Sedimenticolaceae>Sedimenticola>unclassified_Sedimenticola | 513 | C0631_16520 | 5263..6804 | 1940285 | MAG:-AAA-family-ATPase-[Sedimenticola-sp.] |
| WP_079805586.1 | Schlafen+HATPase+wHTH | GCF_016903075.1 | Salmonella_enterica | Bacteria>Pseudomonadati>Pseudomonadota>Gammaproteobacteria>Enterobacterales>Enterobacteriaceae>Salmonella | 199 | AJH70_RS01345 | 296291..296890 | 28901 | ATP-binding-protein-[Salmonella-enterica] |
| WP_169248418.1 | Schlafen+HATPase+wHTH | GCF_012939995.1 | Candidatus_Competibacter_phosphatis | Bacteria>Pseudomonadati>Pseudomonadota>Gammaproteobacteria>Gammaproteobacteria_incertae_sedis>Candidatus_Competibacteraceae>Candidatus_Competibacter | 91 | E4P82_RS08100 | 11272..11547 | 221280 | ATP-binding-protein-[Candidatus-Competibacter-phosphatis] |
| MBT4668299.1 | Schlafen+HATPase+wHTH | GCA_018657745.1 | Candidatus_Ruthturnera_sp. | Bacteria>Pseudomonadati>Pseudomonadota>Gammaproteobacteria>Gammaproteobacteria_incertae_sedis>Candidatus_Pseudothioglobaceae>Candidatus_Ruthturnera>unclassified_Candidatus_Ruthturnera | 622 | HOC17_02015 | 5676..7544 | 2735566 | AAA-family-ATPase-[Candidatus-Ruthturnera-sp.] |
| WP_248566605.1 | Schlafen+HATPase+wHTH | GCF_023195885.1 | Methylonatrum_kenyense | Bacteria>Pseudomonadati>Pseudomonadota>Gammaproteobacteria>Gammaproteobacteria_incertae_sedis>Methylonatrum | 573 | M0534_RS01345 | 289014..290735 | 455253 | ATP-binding-protein-[Methylonatrum-kenyense] |
| WP_075675237.1 | Schlafen+HATPase+wHTH | GCF_001939825.1 | Stenotrophomonas | Bacteria>Pseudomonadati>Pseudomonadota>Gammaproteobacteria>Lysobacterales>Lysobacteraceae | 622 | BKE09_RS06605 | 1400104..1401972 | 40323 | MULTISPECIES:-putative-DNA-binding-domain-containing-protein-[Stenotrophomonas] |
| WP_033836878.1 | Schlafen+HATPase+wHTH | GCF_000454525.1 | Xanthomonas | Bacteria>Pseudomonadati>Pseudomonadota>Gammaproteobacteria>Lysobacterales>Lysobacteraceae | 626 | XMLX20_RS19130 | 4480176..4482056 | 338 | putative-DNA-binding-domain-containing-protein-[Xanthomonas-citri] |
| MCZ8164570.1 | Schlafen+HATPase+wHTH | GCA_027531545.1 | Silanimonas_sp. | Bacteria>Pseudomonadati>Pseudomonadota>Gammaproteobacteria>Lysobacterales>Lysobacteraceae>Silanimonas>unclassified_Silanimonas | 579 | O9280_02345 | 531432..533171 | 1929290 | MAG:-putative-DNA-binding-domain-containing-protein-[Silanimonas-sp.] |
| HAI59972.1 | Schlafen+HATPase+wHTH | GCA_003449515.1 | Lysobacteraceae_bacterium | Bacteria>Pseudomonadati>Pseudomonadota>Gammaproteobacteria>Lysobacterales>Lysobacteraceae>unclassified_Lysobacteraceae | 575 | DCM32_08870 | 8123..9850 | 1926873 | AAA-family-ATPase-[Xanthomonadaceae-bacterium] |
| WP_146472110.1 | Schlafen+HATPase+wHTH | GCF_014656335.1 | Vulcaniibacterium_thermophilum | Bacteria>Pseudomonadati>Pseudomonadota>Gammaproteobacteria>Lysobacterales>Lysobacteraceae>Vulcaniibacterium | 575 | IE210_RS02830 | 84789..86516 | 1169913 | ATP-binding-protein-[Vulcaniibacterium-thermophilum] |
| WP_230434263.1 | Schlafen+HATPase+wHTH | GCF_020990895.1 | Xanthomonas_melonis | Bacteria>Pseudomonadati>Pseudomonadota>Gammaproteobacteria>Lysobacterales>Lysobacteraceae>Xanthomonas | 621 | JWH05_RS03150 | 11233..13098 | 56456 | putative-DNA-binding-domain-containing-protein-[Xanthomonas-melonis] |
| WP_106891667.1 | Schlafen+HATPase+wHTH | GCF_003015185.1 | Ahniella_affigens | Bacteria>Pseudomonadati>Pseudomonadota>Gammaproteobacteria>Lysobacterales>Rhodanobacteraceae>Ahniella | 665 | C7S18_RS11310 | 2965833..2967830 | 2021234 | ATP-binding-protein-[Ahniella-affigens] |
| WP_043696848.1 | Schlafen+HATPase+wHTH | GCF_000800155.1 | Luteibacter_sp._9133 | Bacteria>Pseudomonadati>Pseudomonadota>Gammaproteobacteria>Lysobacterales>Rhodanobacteraceae>Luteibacter>unclassified_Luteibacter | 424 | FA87_RS19040 | 4370528..4371802 | 1500891 | ATP-binding-protein-[Luteibacter-sp.-9133] |
| TAN08688.1 | Schlafen+HATPase+wHTH | GCA_004299555.1 | Rhodanobacteraceae_bacterium | Bacteria>Pseudomonadati>Pseudomonadota>Gammaproteobacteria>Lysobacterales>Rhodanobacteraceae>unclassified_Rhodanobacteraceae | 423 | EPN38_01840 | 143430..144701 | 2052182 | transcriptional-regulator-[Rhodanobacteraceae-bacterium] |
| MBL8299345.1 | Schlafen+HATPase+wHTH | GCA_016790075.1 | Rhodanobacteraceae_bacterium | Bacteria>Pseudomonadati>Pseudomonadota>Gammaproteobacteria>Lysobacterales>Rhodanobacteraceae>unclassified_Rhodanobacteraceae | 624 | JNN30_13485 | 3001..4875 | 2052182 | putative-DNA-binding-domain-containing-protein-[Rhodanobacteraceae-bacterium] |
| MCU0753613.1 | Schlafen+HATPase+wHTH | GCA_025458445.1 | Lysobacterales_bacterium | Bacteria>Pseudomonadati>Pseudomonadota>Gammaproteobacteria>Lysobacterales>unclassified_Lysobacterales | 442 | MUE46_00620 | 16027..17355 | 2006849 | putative-DNA-binding-domain-containing-protein-[Xanthomonadales-bacterium] |
| WP_143341539.1 | Schlafen+HATPase+wHTH | - | Crenothrix_polyspora | Bacteria>Pseudomonadati>Pseudomonadota>Gammaproteobacteria>Methylococcales>Crenotrichaceae>Crenothrix | 136 | CCY91_RS08550 | 1..409 | 360316 | ATP-binding-protein,-partial-[Crenothrix-polyspora] |
| WP_174627170.1 | Schlafen+HATPase+wHTH | GCF_902806695.1 | Candidatus_Methylobacter_favarea | Bacteria>Pseudomonadati>Pseudomonadota>Gammaproteobacteria>Methylococcales>Methylococcaceae>Methylobacter | 250 | HUX64_RS15755 | 13746..14498 | 2707345 | ATP-binding-protein-[Candidatus-Methylobacter-favarea] |
| WP_174625957.1 | Schlafen+HATPase+wHTH | GCF_902806695.1 | Candidatus_Methylobacter_favarea | Bacteria>Pseudomonadati>Pseudomonadota>Gammaproteobacteria>Methylococcales>Methylococcaceae>Methylobacter | 416 | HUX64_RS09310 | 23834..25084 | 2707345 | putative-DNA-binding-domain-containing-protein-[Candidatus-Methylobacter-favarea] |
| MCF7967520.1 | Schlafen+HATPase+wHTH | GCA_021736725.1 | Methylobacter_tundripaludum | Bacteria>Pseudomonadati>Pseudomonadota>Gammaproteobacteria>Methylococcales>Methylococcaceae>Methylobacter | 582 | K9L79_18565 | 6826..8574 | 173365 | putative-DNA-binding-domain-containing-protein-[Methylobacter-tundripaludum] |
| WP_249385183.1 | Schlafen+HATPase+wHTH | GCF_023370095.1 | Methylococcus_capsulatus | Bacteria>Pseudomonadati>Pseudomonadota>Gammaproteobacteria>Methylococcales>Methylococcaceae>Methylococcus | 590 | M3M30_RS10795 | 2336833..2338605 | 414 | putative-DNA-binding-domain-containing-protein-[Methylococcus-capsulatus] |
| WP_218812146.1 | Schlafen+HATPase+wHTH | GCF_019285535.1 | Methylococcus_capsulatus | Bacteria>Pseudomonadati>Pseudomonadota>Gammaproteobacteria>Methylococcales>Methylococcaceae>Methylococcus | 564 | KW112_RS09050 | 1953892..1955586 | 414 | putative-DNA-binding-domain-containing-protein-[Methylococcus-capsulatus] |
| WP_218800689.1 | Schlafen+HATPase+wHTH | GCF_019285575.1 | Methylococcus_capsulatus | Bacteria>Pseudomonadati>Pseudomonadota>Gammaproteobacteria>Methylococcales>Methylococcaceae>Methylococcus | 590 | KW113_RS00575 | 124484..126256 | 414 | putative-DNA-binding-domain-containing-protein-[Methylococcus-capsulatus] |
| WP_169603381.1 | Schlafen+HATPase+wHTH | GCF_012769535.1 | Methylococcus_geothermalis | Bacteria>Pseudomonadati>Pseudomonadota>Gammaproteobacteria>Methylococcales>Methylococcaceae>Methylococcus | 399 | GNH96_RS09075 | 1902719..1903918 | 2681310 | ATP-binding-protein-[Methylococcus-geothermalis] |
| WP_017365773.1 | Schlafen+HATPase+wHTH | GCF_000297615.1 | Methylococcus_capsulatus | Bacteria>Pseudomonadati>Pseudomonadota>Gammaproteobacteria>Methylococcales>Methylococcaceae>Methylococcus | 423 | QMX71_RS07550 | 1584664..1585935 | 414 | ATP-binding-protein-[Methylococcus-capsulatus] |
| WP_010962199.1 | Schlafen+HATPase+wHTH | GCF_000008325.1 | Methylococcus_capsulatus | Bacteria>Pseudomonadati>Pseudomonadota>Gammaproteobacteria>Methylococcales>Methylococcaceae>Methylococcus | 423 | MCA_RS14765 | 3192837..3194108 | 414 | ATP-binding-protein-[Methylococcus-capsulatus] |

| Protein-ID | Domain | Assembly-ID | Strain | Taxonomy | Length | Locus-Tag | Genomic-coordinates | TaxID | Fasta-defline |
|---|---|---|---|---|---|---|---|---|---|
| WP_218806441.1 | Schlafen+HATPase+wHTH | GCF_019285515.1 | Methylococcus_sp._Mc7 | Bacteria>Pseudomonadati>Pseudomonadota>Gammaproteobacteria>Methylococcales>Methylococcaceae>Methylococcus>unclassified_Methylococcus | 433 | KW115_RS14820 | 3057910..3059211 | 2860258 | ATP-binding-protein-[Methylococcus-sp.-Mc7] |
| WP_024296679.1 | Schlafen+HATPase+wHTH | GCF_000527095.1 | Methylomicrobium_lacus | Bacteria>Pseudomonadati>Pseudomonadota>Gammaproteobacteria>Methylococcales>Methylococcaceae>Methylomicrobium | 414 | METLA_RS0100485 | 106840..108084 | 136992 | putative-DNA-binding-domain-containing-protein-[Methylomicrobium-lacus] |
| WP_215479512.1 | Schlafen+HATPase+wHTH | GCF_018722605.1 | Methylovulum_psychrotolerans | Bacteria>Pseudomonadati>Pseudomonadota>Gammaproteobacteria>Methylococcales>Methylococcaceae>Methylovulum | 414 | KFZ76_RS21395 | 116044..117288 | 1704499 | ATP-binding-protein-[Methylovulum-psychrotolerans] |
| WP_103975693.1 | Schlafen+HATPase+wHTH | GCF_002923755.1 | Methylovulum_psychrotolerans | Bacteria>Pseudomonadati>Pseudomonadota>Gammaproteobacteria>Methylococcales>Methylococcaceae>Methylovulum | 414 | AADEFJLK_RS21310 | 17185..18429 | 1704499 | ATP-binding-protein-[Methylovulum-psychrotolerans] |
| WP_088619242.1 | Schlafen+HATPase+wHTH | GCF_002209385.1 | Methylovulum_psychrotolerans | Bacteria>Pseudomonadati>Pseudomonadota>Gammaproteobacteria>Methylococcales>Methylococcaceae>Methylovulum | 414 | CEK71_RS09940 | 2181392..2182636 | 1704499 | ATP-binding-protein-[Methylovulum-psychrotolerans] |
| PZN70751.1 | Schlafen+HATPase+wHTH | GCA_003242955.1 | Candidatus_Methylumidiphilus_alinenensis | Bacteria>Pseudomonadati>Pseudomonadota>Gammaproteobacteria>Methylococcales>Methylococcales_incertae_sedis>Candidatus_Methylumidiphilus | 410 | DM484_28020 | 22783..24015 | 2202197 | transcriptional-regulator-[Candidatus-Methylumidiphilus-alinenensis] |
| PZN70745.1 | Schlafen+HATPase+wHTH | GCA_003242955.1 | Candidatus_Methylumidiphilus_alinenensis | Bacteria>Pseudomonadati>Pseudomonadota>Gammaproteobacteria>Methylococcales>Methylococcales_incertae_sedis>Candidatus_Methylumidiphilus | 387 | DM484_28005 | 16485..17648 | 2202197 | ATP-dependent-DNA-helicase-RecG-[Candidatus-Methylumidiphilus-alinenensis] |
| WP_241878430.1 | Schlafen+HATPase+wHTH | - | Psychrobacter_raelei | Bacteria>Pseudomonadati>Pseudomonadota>Gammaproteobacteria>Moraxellales>Moraxellaceae>Psychrobacter | 54 | MN210_RS12760 | 2347496..2347660 | 2565531 | ATP-binding-protein-[Psychrobacter-sp.-PraFG1] |
| WP_241878429.1 | Schlafen+HATPase+wHTH | - | Psychrobacter_raelei | Bacteria>Pseudomonadati>Pseudomonadota>Gammaproteobacteria>Moraxellales>Moraxellaceae>Psychrobacter | 102 | MN210_RS12755 | 2347093..2347401 | 2565531 | ATP-binding-protein-[Psychrobacter-sp.-PraFG1] |
| WP_241878427.1 | Schlafen+HATPase+wHTH | - | Psychrobacter_raelei | Bacteria>Pseudomonadati>Pseudomonadota>Gammaproteobacteria>Moraxellales>Moraxellaceae>Psychrobacter | 174 | MN210_RS12745 | 2346028..2346552 | 2565531 | hypothetical-protein-[Psychrobacter-sp.-PraFG1] |
| MBC7752531.1 | Schlafen+HATPase+wHTH | GCA_014378805.1 | Moraxellaceae_bacterium | Bacteria>Pseudomonadati>Pseudomonadota>Gammaproteobacteria>Moraxellales>Moraxellaceae>unclassified_Moraxellaceae | 613 | H7Z73_12615 | 14251..16092 | 1889775 | putative-DNA-binding-domain-containing-protein-[Candidatus-Saccharibacteria-bacterium] |
| WP_179915498.1 | Schlafen+HATPase+wHTH | GCF_947331475.1 | Halomonadaceae | Bacteria>Pseudomonadati>Pseudomonadota>Gammaproteobacteria>Oceanospirillales | 570 | HZS80_RS06350 | 2297..4009 | 28256 | ATP-binding-protein-[Halomonas-glaciei] |
| WP_159177831.1 | Schlafen+HATPase+wHTH | GCF_902506335.1 | Halomonadaceae | Bacteria>Pseudomonadati>Pseudomonadota>Gammaproteobacteria>Oceanospirillales | 556 | JMX33_RS24535 | 5307878..5309548 | 28256 | MULTISPECIES:-ATP-binding-protein-[unclassified-Halomonas] |
| WP_062359356.1 | Schlafen+HATPase+wHTH | GCF_020782315.1 | Halomonadaceae | Bacteria>Pseudomonadati>Pseudomonadota>Gammaproteobacteria>Oceanospirillales | 493 | BIS08_RS08530 | 3846..5327 | 28256 | MULTISPECIES:-helix-turn-helix-domain-containing-protein-[Halomonas] |
| WP_041158391.1 | Schlafen+HATPase+wHTH | GCF_900099955.1 | Halomonadaceae | Bacteria>Pseudomonadati>Pseudomonadota>Gammaproteobacteria>Oceanospirillales | 493 | BLR10_RS07710 | 74987..76468 | 28256 | MULTISPECIES:-helix-turn-helix-domain-containing-protein-[Halomonas] |
| WP_008928707.1 | Schlafen+HATPase+wHTH | GCF_000300995.1 | Alcanivorax_hongdengensis | Bacteria>Pseudomonadati>Pseudomonadota>Gammaproteobacteria>Oceanospirillales>Alcanivoracaceae>Alcanivorax | 600 | A11A3_RS07655 | 71886..73688 | 519051 | putative-DNA-binding-domain-containing-protein-[Alcanivorax-hongdengensis] |
| WP_161382581.1 | Schlafen+HATPase+wHTH | - | Alcanivorax_sp._ZXX171 | Bacteria>Pseudomonadati>Pseudomonadota>Gammaproteobacteria>Oceanospirillales>Alcanivoracaceae>Alcanivorax>unclassified_Alcanivorax | 557 | GN155_RS00645 | 136057..137730 | 2695858 | ATP-binding-protein-[Alcanivorax-sp.-ZXX171] |
| WP_142947824.1 | Schlafen+HATPase+wHTH | - | Alcanivorax_sp._DSM_26295 | Bacteria>Pseudomonadati>Pseudomonadota>Gammaproteobacteria>Oceanospirillales>Alcanivoracaceae>Alcanivorax>unclassified_Alcanivorax | 423 | FN816_RS03080 | 110052..111323 | 1938738 | ATP-binding-protein-[Alcanivorax-sp.-DSM-26295] |
| MCH2557815.1 | Schlafen+HATPase+wHTH | GCA_022449455.1 | Alcanivorax_sp. | Bacteria>Pseudomonadati>Pseudomonadota>Gammaproteobacteria>Oceanospirillales>Alcanivoracaceae>Alcanivorax>unclassified_Alcanivorax | 556 | MK005_11005 | 51337..53007 | 1872427 | putative-DNA-binding-domain-containing-protein-[Alcanivorax-sp.] |
| MCH2557382.1 | Schlafen+HATPase+wHTH | GCA_022449455.1 | Alcanivorax_sp. | Bacteria>Pseudomonadati>Pseudomonadota>Gammaproteobacteria>Oceanospirillales>Alcanivoracaceae>Alcanivorax>unclassified_Alcanivorax | 594 | MK005_08785 | 134515..136299 | 1872427 | putative-DNA-binding-domain-containing-protein-[Alcanivorax-sp.] |
| MBA3979208.1 | Schlafen+HATPase+wHTH | GCA_013823245.1 | Alcanivorax_sp. | Bacteria>Pseudomonadati>Pseudomonadota>Gammaproteobacteria>Oceanospirillales>Alcanivoracaceae>Alcanivorax>unclassified_Alcanivorax | 573 | C0462_01265 | 11463..13184 | 1872427 | AAA-family-ATPase-[Alcanivorax-sp.] |
| MAD69916.1 | Schlafen+HATPase+wHTH | GCA_002685555.1 | Alcanivorax_sp. | Bacteria>Pseudomonadati>Pseudomonadota>Gammaproteobacteria>Oceanospirillales>Alcanivoracaceae>Alcanivorax>unclassified_Alcanivorax | 557 | CL538_05265 | 29888..31561 | 1872427 | ATPase-[Alcanivorax-sp.] |
| KYZ87034.1 | Schlafen+HATPase+wHTH | GCA_001611695.1 | Alcanivorax_sp._KX64203 | Bacteria>Pseudomonadati>Pseudomonadota>Gammaproteobacteria>Oceanospirillales>Alcanivoracaceae>Alcanivorax>unclassified_Alcanivorax | 573 | A3Q32_03165 | 121165..122886 | 1817793 | AAA-family-ATPase-[Alcanivorax-sp.-KX64203] |
| HIK75732.1 | Schlafen+HATPase+wHTH | GCA_015660495.1 | Alcanivorax_sp. | Bacteria>Pseudomonadati>Pseudomonadota>Gammaproteobacteria>Oceanospirillales>Alcanivoracaceae>Alcanivorax>unclassified_Alcanivorax | 557 | EYG00_13685 | 16642..18315 | 1872427 | ATPase-[Alcanivorax-sp.] |
| WP_262731201.1 | Schlafen+HATPase+wHTH | GCF_025532125.1 | Alloalcanivorax_marinus | Bacteria>Pseudomonadati>Pseudomonadota>Gammaproteobacteria>Oceanospirillales>Alcanivoracaceae>Alloalcanivorax | 578 | B27N_RS13555 | 4296..6032 | 1177169 | ATP-binding-protein-[Alloalcanivorax-marinus] |
| WP_204429960.1 | Schlafen+HATPase+wHTH | GCF_016906305.1 | Alloalcanivorax_marinus | Bacteria>Pseudomonadati>Pseudomonadota>Gammaproteobacteria>Oceanospirillales>Alcanivoracaceae>Alloalcanivorax | 557 | IM760_RS12905 | 14651..16324 | 1177169 | ATP-binding-protein-[Alloalcanivorax-marinus] |
| WP_008734406.1 | Schlafen+HATPase+wHTH | GCF_000299335.2 | Isoalcanivorax_pacificus | Bacteria>Pseudomonadati>Pseudomonadota>Gammaproteobacteria>Oceanospirillales>Alcanivoracaceae>Isoalcanivorax | 556 | S7S_RS18610 | 4156889..4158559 | 1306787 | ATP-binding-protein-[Isoalcanivorax-pacificus] |
| MCP5207855.1 | Schlafen+HATPase+wHTH | GCA_024234475.1 | Hahellaceae_bacterium | Bacteria>Pseudomonadati>Pseudomonadota>Gammaproteobacteria>Oceanospirillales>Hahellaceae>unclassified_Hahellaceae | 612 | H7A01_11665 | 137768..139606 | 2048971 | putative-DNA-binding-domain-containing-protein-[Hahellaceae-bacterium] |
| WP_234246267.1 | Schlafen+HATPase+wHTH | GCF_021404345.1 | Billgrantia_desiderata | Bacteria>Pseudomonadati>Pseudomonadota>Gammaproteobacteria>Oceanospirillales>Halomonadaceae>Billgrantia | 423 | HOP57_RS15475 | 81676..82947 | 52021 | ATP-binding-protein-[Halomonas-desiderata] |
| WP_103969333.1 | Schlafen+HATPase+wHTH | GCF_900108215.1 | Billgrantia_desiderata | Bacteria>Pseudomonadati>Pseudomonadota>Gammaproteobacteria>Oceanospirillales>Halomonadaceae>Billgrantia | 556 | C4S71_RS12470 | 91479..93149 | 52021 | ATP-binding-protein-[Halomonas-desiderata] |
| WP_239296661.1 | Schlafen+HATPase+wHTH | GCF_022348215.1 | unclassified_Halomonas | Bacteria>Pseudomonadati>Pseudomonadota>Gammaproteobacteria>Oceanospirillales>Halomonadaceae>Halomonas | 557 | MHM35_RS10405 | 119755..121428 | 2609666 | MULTISPECIES:-ATP-binding-protein-[unclassified-Halomonas] |
| WP_192538996.1 | Schlafen+HATPase+wHTH | GCF_052353025.1 | Halomonas_colorata | Bacteria>Pseudomonadati>Pseudomonadota>Gammaproteobacteria>Oceanospirillales>Halomonadaceae>Halomonas | 493 | EI547_RS13720 | 38803..40284 | 2742615 | helix-turn-helix-domain-containing-protein-[Halomonas-sp.-FME20] |
| WP_183333608.1 | Schlafen+HATPase+wHTH | GCF_014193375.1 | Halomonas_campaniensis | Bacteria>Pseudomonadati>Pseudomonadota>Gammaproteobacteria>Oceanospirillales>Halomonadaceae>Halomonas | 558 | BDK63_RS15590 | 5681..7357 | 213554 | ATP-binding-protein-[Halomonas-campaniensis] |
| WP_136248425.1 | Schlafen+HATPase+wHTH | GCF_004798905.1 | Halomonas_borealis | Bacteria>Pseudomonadati>Pseudomonadota>Gammaproteobacteria>Oceanospirillales>Halomonadaceae>Halomonas | 594 | ETC07_RS10260 | 54628..56412 | 2508710 | putative-DNA-binding-domain-containing-protein-[Halomonas-borealis] |
| WP_133481654.1 | Schlafen+HATPase+wHTH | GCF_004361885.1 | Halomonas_ventosae | Bacteria>Pseudomonadati>Pseudomonadota>Gammaproteobacteria>Oceanospirillales>Halomonadaceae>Halomonas | 572 | DFO68_RS02355 | 100319..102037 | 229007 | putative-DNA-binding-domain-containing-protein-[Halomonas-ventosae] |
| WP_133481546.1 | Schlafen+HATPase+wHTH | GCF_004361885.1 | Halomonas_ventosae | Bacteria>Pseudomonadati>Pseudomonadota>Gammaproteobacteria>Oceanospirillales>Halomonadaceae>Halomonas | 557 | DFO68_RS01980 | 15884..17557 | 229007 | ATP-binding-protein-[Halomonas-ventosae] |
| WP_101147208.1 | Schlafen+HATPase+wHTH | GCF_037070225.1 | unclassified_Halomonas | Bacteria>Pseudomonadati>Pseudomonadota>Gammaproteobacteria>Oceanospirillales>Halomonadaceae>Halomonas | 485 | CXF87_RS15355 | 104796..106253 | 2609666 | helix-turn-helix-domain-containing-protein-[Halomonas-sp.-MES3-P3E] |
| WP_078087407.1 | Schlafen+HATPase+wHTH | GCF_002009255.1 | unclassified_Halomonas | Bacteria>Pseudomonadati>Pseudomonadota>Gammaproteobacteria>Oceanospirillales>Halomonadaceae>Halomonas | 501 | BV504_RS06360 | 1464309..1465814 | 2609666 | MULTISPECIES:-helix-turn-helix-domain-containing-protein-[unclassified-Halomonas] |
| WP_027962107.1 | Schlafen+HATPase+wHTH | GCF_000620045.1 | Halomonas_halodenitrificans | Bacteria>Pseudomonadati>Pseudomonadota>Gammaproteobacteria>Oceanospirillales>Halomonadaceae>Halomonas | 570 | Q358_RS0111240 | 17625..19337 | 28252 | ATP-binding-protein-[Halomonas-halodenitrificans] |
| WP_263591022.1 | Schlafen+HATPase+wHTH | GCF_025725685.1 | Halomonas_sp._M4R1S46 | Bacteria>Pseudomonadati>Pseudomonadota>Gammaproteobacteria>Oceanospirillales>Halomonadaceae>Halomonas>unclassified_Halomonas | 570 | OCT48_RS01555 | 325360..327072 | 2982692 | ATP-binding-protein-[Halomonas-sp.-M4R1S46] |
| WP_232090423.1 | Schlafen+HATPase+wHTH | GCF_014931605.1 | Halomonas_sp._MCCC_1A13316 | Bacteria>Pseudomonadati>Pseudomonadota>Gammaproteobacteria>Oceanospirillales>Halomonadaceae>Halomonas>unclassified_Halomonas | 611 | HNO52_RS20465 | 4396482..4398317 | 2733487 | putative-DNA-binding-domain-containing-protein-[Halomonas-sp.-MCCC-1A13316] |
| WP_231664594.1 | Schlafen+HATPase+wHTH | GCF_021151495.1 | Halomonas_sp._IOP_31 | Bacteria>Pseudomonadati>Pseudomonadota>Gammaproteobacteria>Oceanospirillales>Halomonadaceae>Halomonas>unclassified_Halomonas | 573 | K7I12_RS15020 | 24759..26480 | 2876584 | ATP-binding-protein-[Halomonas-sp.-IOP_31] |
| WP_218314906.1 | Schlafen+HATPase+wHTH | GCF_019211725.1 | Halomonas_sp._18071143 | Bacteria>Pseudomonadati>Pseudomonadota>Gammaproteobacteria>Oceanospirillales>Halomonadaceae>Halomonas>unclassified_Halomonas | 493 | KVY02_RS07720 | 1665037..1666518 | 2855441 | helix-turn-helix-domain-containing-protein-[Halomonas-sp.-18071143] |
| WP_169956448.1 | Schlafen+HATPase+wHTH | GCF_013000985.1 | Halomonas_sp._PGE1 | Bacteria>Pseudomonadati>Pseudomonadota>Gammaproteobacteria>Oceanospirillales>Halomonadaceae>Halomonas>unclassified_Halomonas | 557 | HIR79_RS02115 | 425464..427137 | 2730360 | ATP-binding-protein-[Halomonas-sp.-PGE1] |
| WP_163558375.1 | Schlafen+HATPase+wHTH | GCF_010994375.1 | Halomonas_sp._NO4 | Bacteria>Pseudomonadati>Pseudomonadota>Gammaproteobacteria>Oceanospirillales>Halomonadaceae>Halomonas>unclassified_Halomonas | 570 | FEF15_RS06630 | 14156..15868 | 2484813 | ATP-binding-protein-[Halomonas-sp.-NO4] |
| WP_081191887.1 | Schlafen+HATPase+wHTH | - | Halomonas_sp._BC1 | Bacteria>Pseudomonadati>Pseudomonadota>Gammaproteobacteria>Oceanospirillales>Halomonadaceae>Halomonas>unclassified_Halomonas | 497 | B7L72_RS00100 | 23119..24612 | 1670448 | helix-turn-helix-domain-containing-protein-[Halomonas-sp.-BC1] |
| WP_062374365.1 | Schlafen+HATPase+wHTH | GCF_001596805.1 | Halomonas_sp._KX33721 | Bacteria>Pseudomonadati>Pseudomonadota>Gammaproteobacteria>Oceanospirillales>Halomonadaceae>Halomonas>unclassified_Halomonas | 492 | A3220_RS08355 | 68236..69714 | 1819251 | helix-turn-helix-domain-containing-protein-[Halomonas-sp.-KX33721] |
| WP_048306247.1 | Schlafen+HATPase+wHTH | GCF_001039575.1 | Halomonas_sp._PR-M31 | Bacteria>Pseudomonadati>Pseudomonadota>Gammaproteobacteria>Oceanospirillales>Halomonadaceae>Halomonas>unclassified_Halomonas | 557 | CH74_RS04340 | 1160..2833 | 1471202 | ATP-binding-protein-[Halomonas-sp.-PR-M31] |
| WP_009095652.1 | Schlafen+HATPase+wHTH | GCF_000236625.2 | Halomonas_sp._GFAJ-1 | Bacteria>Pseudomonadati>Pseudomonadota>Gammaproteobacteria>Oceanospirillales>Halomonadaceae>Halomonas>unclassified_Halomonas | 493 | BB497_RS11615 | 2571174..2572655 | 1118153 | helix-turn-helix-domain-containing-protein-[Halomonas-sp.-GFAJ-1] |
| WP_008956848.1 | Schlafen+HATPase+wHTH | GCF_000235725.1 | Halomonas_sp._HAL1 | Bacteria>Pseudomonadati>Pseudomonadota>Gammaproteobacteria>Oceanospirillales>Halomonadaceae>Halomonas>unclassified_Halomonas | 485 | HAL1_RS04475 | 16902..18359 | 550984 | helix-turn-helix-domain-containing-protein-[Halomonas-sp.-HAL1] |
| PHR03207.1 | Schlafen+HATPase+wHTH | GCA_002733025.1 | Halomonas_sp. | Bacteria>Pseudomonadati>Pseudomonadota>Gammaproteobacteria>Oceanospirillales>Halomonadaceae>Halomonas>unclassified_Halomonas | 572 | COB32_05660 | 21094..22812 | 1486246 | AAA-family-ATPase-[Halomonas-sp.] |
| PHR03142.1 | Schlafen+HATPase+wHTH | GCA_002733025.1 | Halomonas_sp. | Bacteria>Pseudomonadati>Pseudomonadota>Gammaproteobacteria>Oceanospirillales>Halomonadaceae>Halomonas>unclassified_Halomonas | 493 | COB32_06230 | 69204..70685 | 1486246 | AAA-family-ATPase-[Halomonas-sp.] |
| HBQ05709.1 | Schlafen+HATPase+wHTH | GCA_003529085.1 | Halomonas_sp. | Bacteria>Pseudomonadati>Pseudomonadota>Gammaproteobacteria>Oceanospirillales>Halomonadaceae>Halomonas>unclassified_Halomonas | 493 | DD676_03535 | 658..2139 | 1486246 | AAA-family-ATPase-[Halomonas-sp.] |
| WP_031385054.1 | Schlafen+HATPase+wHTH | GCF_039543745.1 | Modicisalibacter_zincidurans | Bacteria>Pseudomonadati>Pseudomonadota>Gammaproteobacteria>Oceanospirillales>Halomonadaceae>Modicisalibacter | 557 | HALZIN_RS0115270 | 3253091..3254764 | 1178777 | ATP-binding-protein-[Halomonas-zincidurans] |
| WP_110649044.1 | Schlafen+HATPase+wHTH | GCF_003206715.1 | Salinicola_peritrichatus | Bacteria>Pseudomonadati>Pseudomonadota>Gammaproteobacteria>Oceanospirillales>Halomonadaceae>Salinicola | 573 | DOM57_RS09905 | 62848..64569 | 1267424 | ATP-binding-protein-[Salinicola-peritrichatus] |
| WP_087719603.1 | Schlafen+HATPase+wHTH | GCF_002179555.1 | Salinicola_salarius | Bacteria>Pseudomonadati>Pseudomonadota>Gammaproteobacteria>Oceanospirillales>Halomonadaceae>Salinicola | 578 | CCD76_RS11240 | 13673..15409 | 430457 | ATP-binding-protein-[Salinicola-salarius] |
| WP_193059741.1 | Schlafen+HATPase+wHTH | GCF_052103235.1 | Vreelandella_alkaliphila | Bacteria>Pseudomonadati>Pseudomonadota>Gammaproteobacteria>Oceanospirillales>Halomonadaceae>Vreelandella | 497 | INT28_RS17785 | 7229..8722 | 272774 | MULTISPECIES:-helix-turn-helix-domain-containing-protein-[unclassified-Halomonas] |
| WP_180091725.1 | Schlafen+HATPase+wHTH | GCF_013416325.1 | Vreelandella_sedimenti | Bacteria>Pseudomonadati>Pseudomonadota>Gammaproteobacteria>Oceanospirillales>Halomonadaceae>Vreelandella | 487 | HZU72_RS10345 | 62247..63710 | 2729618 | helix-turn-helix-domain-containing-protein-[Halomonas-sedimenti] |
| WP_172515132.1 | Schlafen+HATPase+wHTH | GCF_011764625.1 | Vreelandella_aquamarina | Bacteria>Pseudomonadati>Pseudomonadota>Gammaproteobacteria>Oceanospirillales>Halomonadaceae>Vreelandella | 557 | HMEPL2_RS11775 | 2495138..2496811 | 77097 | ATP-binding-protein-[Halomonas-meridiana] |
| WP_172514442.1 | Schlafen+HATPase+wHTH | GCF_011764625.1 | Vreelandella_aquamarina | Bacteria>Pseudomonadati>Pseudomonadota>Gammaproteobacteria>Oceanospirillales>Halomonadaceae>Vreelandella | 493 | HMEPL2_RS01775 | 386385..387866 | 77097 | helix-turn-helix-domain-containing-protein-[Halomonas-meridiana] |
| WP_159343440.1 | Schlafen+HATPase+wHTH | GCF_009846525.1 | Vreelandella_aquamarina | Bacteria>Pseudomonadati>Pseudomonadota>Gammaproteobacteria>Oceanospirillales>Halomonadaceae>Vreelandella | 556 | CTT34_RS16810 | 3561382..3563052 | 77097 | ATP-binding-protein-[Halomonas-meridiana] |
| WP_154045156.1 | Schlafen+HATPase+wHTH | GCF_900142895.1 | Vreelandella_subglaciescola | Bacteria>Pseudomonadati>Pseudomonadota>Gammaproteobacteria>Oceanospirillales>Halomonadaceae>Vreelandella | 57 | B5495_RS14505 | 136446..136619 | 29571 | hypothetical-protein-[Halomonas-subglaciescola] |
| WP_153844140.1 | Schlafen+HATPase+wHTH | GCF_012427705.1 | Vreelandella_piezotolerans | Bacteria>Pseudomonadati>Pseudomonadota>Gammaproteobacteria>Oceanospirillales>Halomonadaceae>Vreelandella | 493 | GYM47_RS09715 | 2109126..2110607 | 2609667 | helix-turn-helix-domain-containing-protein-[Halomonas-piezotolerans] |
| WP_125748593.1 | Schlafen+HATPase+wHTH | GCF_003950215.1 | Vreelandella_venusta | Bacteria>Pseudomonadati>Pseudomonadota>Gammaproteobacteria>Oceanospirillales>Halomonadaceae>Vreelandella | 493 | DDR56_RS08115 | 81674..83155 | 44935 | helix-turn-helix-domain-containing-protein-[Halomonas-venusta] |
| WP_089675691.1 | Schlafen+HATPase+wHTH | GCF_900110265.1 | Vreelandella_aquamarina | Bacteria>Pseudomonadati>Pseudomonadota>Gammaproteobacteria>Oceanospirillales>Halomonadaceae>Vreelandella | 493 | BMY49_RS12180 | 11220..12701 | 77097 | helix-turn-helix-domain-containing-protein-[Halomonas-aquamarina] |
| WP_074210657.1 | Schlafen+HATPase+wHTH | GCF_006540125.1 | Vreelandella_aquamarina | Bacteria>Pseudomonadati>Pseudomonadota>Gammaproteobacteria>Oceanospirillales>Halomonadaceae>Vreelandella | 497 | HME01_RS04160 | 110866..112359 | 77097 | helix-turn-helix-domain-containing-protein-[Halomonas-meridiana] |
| MAX99658.1 | Schlafen+HATPase+wHTH | GCA_002706025.1 | Oceanospirillaceae_bacterium | Bacteria>Pseudomonadati>Pseudomonadota>Gammaproteobacteria>Oceanospirillales>Oceanospirillaceae>unclassified_Oceanospirillaceae | 573 | CMI08_10745 | 13309..15030 | 1899355 | AAA-family-ATPase-[Oceanospirillaceae-bacterium] |
| RLP52822.1 | Schlafen+HATPase+wHTH | GCA_003667185.1 | Ketobacter_sp. | Bacteria>Pseudomonadati>Pseudomonadota>Gammaproteobacteria>Pseudomonadales>Ketobacteraceae>Ketobacter>unclassified_Ketobacter | 556 | D6160_18815 | 19125..20795 | 2083498 | MAG:-ATPase-[Ketobacter-sp.] |
| WP_023009220.1 | Schlafen+HATPase+wHTH | GCF_000475375.1 | Marinobacter | Bacteria>Pseudomonadati>Pseudomonadota>Gammaproteobacteria>Pseudomonadales>Marinobacteraceae | 415 | FHK81_RS16390 | 8957..10204 | 2742 | MULTISPECIES:-putative-DNA-binding-domain-containing-protein-[Marinobacter] |
| WP_022992170.1 | Schlafen+HATPase+wHTH | GCF_000475355.1 | Marinobacter | Bacteria>Pseudomonadati>Pseudomonadota>Gammaproteobacteria>Pseudomonadales>Marinobacteraceae | 557 | DET50_RS14955 | 44264..45937 | 2742 | MULTISPECIES:-ATP-binding-protein-[Marinobacter] |
| WP_273135108.1 | Schlafen+HATPase+wHTH | GCF_007713625.1 | Marinobacter_vinifirmus | Bacteria>Pseudomonadati>Pseudomonadota>Gammaproteobacteria>Pseudomonadales>Marinobacteraceae>Marinobacter | 569 | FHK81_RS16405 | 16431..18140 | 355591 | ATP-binding-protein-[Marinobacter-vinifirmus] |
| WP_248236651.1 | Schlafen+HATPase+wHTH | GCF_023156245.1 | Marinobacter_bryozoorum | Bacteria>Pseudomonadati>Pseudomonadota>Gammaproteobacteria>Pseudomonadales>Marinobacteraceae>Marinobacter | 573 | MLC59_RS17295 | 15555..17276 | 256324 | ATP-binding-protein-[Marinobacter-bryozoorum] |
| WP_248185655.1 | Schlafen+HATPase+wHTH | GCF_023156235.1 | Marinobacter_goseongensis | Bacteria>Pseudomonadati>Pseudomonadota>Gammaproteobacteria>Pseudomonadales>Marinobacteraceae>Marinobacter | 556 | MLC70_RS17475 | 1153..2823 | 453838 | ATP-binding-protein-[Marinobacter-goseongensis] |
| WP_141068073.1 | Schlafen+HATPase+wHTH | GCF_006516615.1 | Marinobacter_nauticus | Bacteria>Pseudomonadati>Pseudomonadota>Gammaproteobacteria>Pseudomonadales>Marinobacteraceae>Marinobacter | 556 | FH712_RS17660 | 37188..38858 | 2743 | ATP-binding-protein-[Marinobacter-nauticus] |
| WP_113879481.1 | Schlafen+HATPase+wHTH | GCF_003315555.1 | Marinobacter_nauticus | Bacteria>Pseudomonadati>Pseudomonadota>Gammaproteobacteria>Pseudomonadales>Marinobacteraceae>Marinobacter | 557 | DET64_RS06860 | 2780..4453 | 2743 | ATP-binding-protein-[Marinobacter-nauticus] |
| WP_104416051.1 | Schlafen+HATPase+wHTH | GCF_002934485.1 | Marinobacter_persicus | Bacteria>Pseudomonadati>Pseudomonadota>Gammaproteobacteria>Pseudomonadales>Marinobacteraceae>Marinobacter | 557 | BY455_RS08985 | 68961..70634 | 930118 | ATP-binding-protein-[Marinobacter-persicus] |
| WP_048384701.1 | Schlafen+HATPase+wHTH | GCF_001043175.1 | Marinobacter_psychrophilus | Bacteria>Pseudomonadati>Pseudomonadota>Gammaproteobacteria>Pseudomonadales>Marinobacteraceae>Marinobacter | 569 | ABA45_RS05805 | 1302435..1304144 | 330734 | ATP-binding-protein-[Marinobacter-psychrophilus] |
| WP_018404436.1 | Schlafen+HATPase+wHTH | GCF_000372805.1 | Marinobacter_gelidimuriae | Bacteria>Pseudomonadati>Pseudomonadota>Gammaproteobacteria>Pseudomonadales>Marinobacteraceae>Marinobacter | 450 | A3OI_RS0111620 | 5827..7179 | 2739064 | ATP-binding-protein-[Marinobacter-gelidimuriae] |
| WP_018403234.1 | Schlafen+HATPase+wHTH | GCF_000372805.1 | Marinobacter_gelidimuriae | Bacteria>Pseudomonadati>Pseudomonadota>Gammaproteobacteria>Pseudomonadales>Marinobacteraceae>Marinobacter | 555 | A3OI_RS0105070 | 18144..19811 | 2739064 | ATP-binding-protein-[Marinobacter-gelidimuriae] |
| WP_011784102.1 | Schlafen+HATPase+wHTH | GCF_000015365.1 | Marinobacter_nauticus | Bacteria>Pseudomonadati>Pseudomonadota>Gammaproteobacteria>Pseudomonadales>Marinobacteraceae>Marinobacter | 556 | MAQU_RS02810 | 632570..634240 | 2743 | ATP-binding-protein-[Marinobacter-nauticus] |
| TVT30738.1 | Schlafen+HATPase+wHTH | GCA_007713625.1 | Marinobacter_vinifirmus | Bacteria>Pseudomonadati>Pseudomonadota>Gammaproteobacteria>Pseudomonadales>Marinobacteraceae>Marinobacter | 569 | FHK81_16385 | 16431..18140 | 355591 | AAA-family-ATPase-[Marinobacter-vinifirmus] |
| TVT30735.1 | Schlafen+HATPase+wHTH | GCA_007713625.1 | Marinobacter_vinifirmus | Bacteria>Pseudomonadati>Pseudomonadota>Gammaproteobacteria>Pseudomonadales>Marinobacteraceae>Marinobacter | 415 | FHK81_16370 | 8957..10204 | 355591 | transcriptional-regulator-[Marinobacter-vinifirmus] |
| BBJ05052.1 | Schlafen+HATPase+wHTH | - | Marinobacter_nauticus | Bacteria>Pseudomonadati>Pseudomonadota>Gammaproteobacteria>Pseudomonadales>Marinobacteraceae>Marinobacter | 557 | YBY_29010 | 3189229..3190902 | 2743 | DNA-binding-protein-[Marinobacter-nauticus] |
| EHJ03978.1 | Schlafen+HATPase+wHTH | GCA_000235625.2 | Marinobacter_manganoxydans_MnI7-9 | Bacteria>Pseudomonadati>Pseudomonadota>Gammaproteobacteria>Pseudomonadales>Marinobacteraceae>Marinobacter>Marinobacter_manganoxydans | 557 | KYE_13695 | 45895..47568 | 1094979 | putative-transcriptional-regulator-[Marinobacter-manganoxydans-MnI7-9] |
| WP_247116151.1 | Schlafen+HATPase+wHTH | GCF_023016045.1 | Marinobacter_sp._S6332 | Bacteria>Pseudomonadati>Pseudomonadota>Gammaproteobacteria>Pseudomonadales>Marinobacteraceae>Marinobacter>unclassified_Marinobacter | 557 | MWU62_RS17915 | 18202..19875 | 2926403 | ATP-binding-protein-[Marinobacter-sp.-S6332] |
| WP_200203487.1 | Schlafen+HATPase+wHTH | GCF_016597775.1 | Marinobacter_sp._1-3A | Bacteria>Pseudomonadati>Pseudomonadota>Gammaproteobacteria>Pseudomonadales>Marinobacteraceae>Marinobacter>unclassified_Marinobacter | 394 | FE848_RS12600 | 143442..144626 | 2582920 | ATP-binding-protein-[Marinobacter-sp.-1-3A] |
| WP_200203478.1 | Schlafen+HATPase+wHTH | GCF_016597775.1 | Marinobacter_sp._1-3A | Bacteria>Pseudomonadati>Pseudomonadota>Gammaproteobacteria>Pseudomonadales>Marinobacteraceae>Marinobacter>unclassified_Marinobacter | 423 | FE848_RS12580 | 135470..136741 | 2582920 | ATP-binding-protein-[Marinobacter-sp.-1-3A] |
| WP_157133264.1 | Schlafen+HATPase+wHTH | GCF_000169375.1 | Marinobacter_sp._ELB17 | Bacteria>Pseudomonadati>Pseudomonadota>Gammaproteobacteria>Pseudomonadales>Marinobacteraceae>Marinobacter>unclassified_Marinobacter | 450 | MELB17_RS08070 | 34011..35363 | 270374 | ATP-binding-protein-[Marinobacter-sp.-ELB17] |
| WP_092033191.1 | Schlafen+HATPase+wHTH | GCF_900112835.1 | Marinobacter_sp._DSM_26671 | Bacteria>Pseudomonadati>Pseudomonadota>Gammaproteobacteria>Pseudomonadales>Marinobacteraceae>Marinobacter>unclassified_Marinobacter | 556 | BM035_RS16340 | 78967..80637 | 1761793 | ATP-binding-protein-[Marinobacter-sp.-DSM-26671] |
| WP_023011722.1 | Schlafen+HATPase+wHTH | GCF_000475315.1 | Marinobacter_sp._EN3 | Bacteria>Pseudomonadati>Pseudomonadota>Gammaproteobacteria>Pseudomonadales>Marinobacteraceae>Marinobacter>unclassified_Marinobacter | 556 | Q675_RS32880 | 60453..62123 | 1397533 | ATP-binding-protein-[Marinobacter-sp.-EN3] |
| WP_023008351.1 | Schlafen+HATPase+wHTH | GCF_000475375.1 | Marinobacter_sp._EVN1 | Bacteria>Pseudomonadati>Pseudomonadota>Gammaproteobacteria>Pseudomonadales>Marinobacteraceae>Marinobacter>unclassified_Marinobacter | 557 | Q674_RS26675 | 51059..52732 | 1397532 | ATP-binding-protein-[Marinobacter-sp.-EVN1] |
| WP_023007676.1 | Schlafen+HATPase+wHTH | GCF_000475375.1 | Marinobacter_sp._EVN1 | Bacteria>Pseudomonadati>Pseudomonadota>Gammaproteobacteria>Pseudomonadales>Marinobacteraceae>Marinobacter>unclassified_Marinobacter | 573 | Q674_RS21830 | 59916..61637 | 1397532 | ATP-binding-protein-[Marinobacter-sp.-EVN1] |
| WP_022990204.1 | Schlafen+HATPase+wHTH | GCF_000475255.1 | Marinobacter_sp._ES-1 | Bacteria>Pseudomonadati>Pseudomonadota>Gammaproteobacteria>Pseudomonadales>Marinobacteraceae>Marinobacter>unclassified_Marinobacter | 557 | Q668_RS13785 | 6161..7834 | 1396858 | ATP-binding-protein-[Marinobacter-sp.-ES-1] |
| WP_022988849.1 | Schlafen+HATPase+wHTH | GCF_000475255.1 | Marinobacter_sp._ES-1 | Bacteria>Pseudomonadati>Pseudomonadota>Gammaproteobacteria>Pseudomonadales>Marinobacteraceae>Marinobacter>unclassified_Marinobacter | 556 | Q668_RS06795 | 86843..88513 | 1396858 | ATP-binding-protein-[Marinobacter-sp.-ES-1] |
| WP_007349109.1 | Schlafen+HATPase+wHTH | GCF_000169375.1 | Marinobacter_sp._ELB17 | Bacteria>Pseudomonadati>Pseudomonadota>Gammaproteobacteria>Pseudomonadales>Marinobacteraceae>Marinobacter>unclassified_Marinobacter | 569 | MELB17_RS05275 | 140723..142432 | 270374 | ATP-binding-protein-[Marinobacter-sp.-ELB17] |
| MBE95558.1 | Schlafen+HATPase+wHTH | GCA_002714765.1 | Marinobacter_sp. | Bacteria>Pseudomonadati>Pseudomonadota>Gammaproteobacteria>Pseudomonadales>Marinobacteraceae>Marinobacter>unclassified_Marinobacter | 556 | CMG89_11185 | 4318..5988 | 50741 | ATPase-[Marinobacter-sp.] |
| MBC7192649.1 | Schlafen+HATPase+wHTH | GCA_014359475.1 | Marinobacter_sp. | Bacteria>Pseudomonadati>Pseudomonadota>Gammaproteobacteria>Pseudomonadales>Marinobacteraceae>Marinobacter>unclassified_Marinobacter | 573 | H5U31_09885 | 20895..22616 | 50741 | putative-DNA-binding-domain-containing-protein-[Marinobacter-sp.] |
| MAH31342.1 | Schlafen+HATPase+wHTH | GCA_002690365.1 | Marinobacter_sp. | Bacteria>Pseudomonadati>Pseudomonadota>Gammaproteobacteria>Pseudomonadales>Marinobacteraceae>Marinobacter>unclassified_Marinobacter | 446 | CMG78_04660 | 1..1337 | 50741 | ATPase-[Marinobacter-sp.] |
| KXS53694.1 | Schlafen+HATPase+wHTH | GCA_001577575.1 | Marinobacter_sp._T13-3 | Bacteria>Pseudomonadati>Pseudomonadota>Gammaproteobacteria>Pseudomonadales>Marinobacteraceae>Marinobacter>unclassified_Marinobacter | 403 | AWU57_1915 | 5834..7045 | 1794906 | ATP-dependent-DNA-helicase-RecG-[Marinobacter-sp.-T13-3] |
| WP_095610361.1 | Schlafen+HATPase+wHTH | GCF_002287355.1 | Tamilnaduibacter_salinus | Bacteria>Pseudomonadati>Pseudomonadota>Gammaproteobacteria>Pseudomonadales>Marinobacteraceae>Tamilnaduibacter | 591 | CF392_RS04955 | 49210..50985 | 1484056 | putative-DNA-binding-domain-containing-protein-[Tamilnaduibacter-salinus] |
| PVY76885.1 | Schlafen+HATPase+wHTH | GCA_003096655.1 | Tamilnaduibacter_salinus | Bacteria>Pseudomonadati>Pseudomonadota>Gammaproteobacteria>Pseudomonadales>Marinobacteraceae>Tamilnaduibacter | 556 | C8D92_104116 | 127180..128850 | 1484056 | ATP-dependent-DNA-helicase-RecG-[Tamilnaduibacter-salinus] |
| WP_050481610.1 | Schlafen+HATPase+wHTH | GCF_001181125.1 | Pseudomonas | Bacteria>Pseudomonadati>Pseudomonadota>Gammaproteobacteria>Pseudomonadales>Pseudomonadaceae | 596 | J4H77_RS25745 | 180297..182087 | 286 | ATP-binding-protein-[Pseudomonas-aeruginosa] |
| WP_034126878.1 | Schlafen+HATPase+wHTH | GCF_030505435.1 | Pseudomonas | Bacteria>Pseudomonadati>Pseudomonadota>Gammaproteobacteria>Pseudomonadales>Pseudomonadaceae | 416 | FHL14_RS24035 | 12990..14240 | 286 | MULTISPECIES:-ATP-binding-protein-[Pseudomonas] |
| WP_152390157.1 | Schlafen+HATPase+wHTH | GCF_009363155.1 | Azotobacter_salinestris | Bacteria>Pseudomonadati>Pseudomonadota>Gammaproteobacteria>Pseudomonadales>Pseudomonadaceae>Azotobacter_group>Azotobacter | 487 | GCU53_RS24420 | 149021..150484 | 69964 | helix-turn-helix-domain-containing-protein-[Azotobacter-salinestris] |
| WP_188867718.1 | Schlafen+HATPase+wHTH | GCF_014646755.1 | Pseudomonas_asuensis | Bacteria>Pseudomonadati>Pseudomonadota>Gammaproteobacteria>Pseudomonadales>Pseudomonadaceae>Pseudomonas | 624 | IEX86_RS19070 | 188697..190571 | 1825787 | putative-DNA-binding-domain-containing-protein-[Pseudomonas-asuensis] |
| WP_181127553.1 | Schlafen+HATPase+wHTH | GCF_013522615.1 | Pseudomonas_carnis | Bacteria>Pseudomonadati>Pseudomonadota>Gammaproteobacteria>Pseudomonadales>Pseudomonadaceae>Pseudomonas | 596 | G7029_RS19455 | 120814..122604 | 2487355 | putative-DNA-binding-domain-containing-protein-[Pseudomonas-carnis] |
| MBA1269663.1 | Schlafen+HATPase+wHTH | GCA_013522615.1 | Pseudomonas_carnis | Bacteria>Pseudomonadati>Pseudomonadota>Gammaproteobacteria>Pseudomonadales>Pseudomonadaceae>Pseudomonas | 596 | G7029_19545 | 120814..122604 | 2487355 | transcriptional-regulator-[Pseudomonas-carnis] |
| WP_247258977.1 | Schlafen+HATPase+wHTH | GCF_032334355.1 | Pseudomonas_aeruginosa | Bacteria>Pseudomonadati>Pseudomonadota>Gammaproteobacteria>Pseudomonadales>Pseudomonadaceae>Pseudomonas>Pseudomonas_aeruginosa_group | 596 | MX104_RS19080 | 61127..62917 | 287 | ATP-binding-protein-[Pseudomonas-aeruginosa] |
| WP_210799983.1 | Schlafen+HATPase+wHTH | GCF_018026755.1 | Pseudomonas_aeruginosa | Bacteria>Pseudomonadati>Pseudomonadota>Gammaproteobacteria>Pseudomonadales>Pseudomonadaceae>Pseudomonas>Pseudomonas_aeruginosa_group | 423 | J5F08_RS20250 | 59962..61233 | 287 | ATP-binding-protein-[Pseudomonas-aeruginosa] |
| WP_172907836.1 | Schlafen+HATPase+wHTH | GCF_013276295.1 | Pseudomonas_aeruginosa | Bacteria>Pseudomonadati>Pseudomonadota>Gammaproteobacteria>Pseudomonadales>Pseudomonadaceae>Pseudomonas>Pseudomonas_aeruginosa_group | 596 | GWK58_RS24970 | 34230..36020 | 287 | ATP-binding-protein-[Pseudomonas-aeruginosa] |
| WP_150035386.1 | Schlafen+HATPase+wHTH | GCF_038431945.1 | Pseudomonas_aeruginosa | Bacteria>Pseudomonadati>Pseudomonadota>Gammaproteobacteria>Pseudomonadales>Pseudomonadaceae>Pseudomonas>Pseudomonas_aeruginosa_group | 394 | NFY03_RS10230 | 2210174..2211358 | 287 | ATP-binding-protein-[Pseudomonas-aeruginosa] |
| WP_140786436.1 | Schlafen+HATPase+wHTH | GCF_003626935.1 | Pseudomonas_aeruginosa | Bacteria>Pseudomonadati>Pseudomonadota>Gammaproteobacteria>Pseudomonadales>Pseudomonadaceae>Pseudomonas>Pseudomonas_aeruginosa_group | 624 | D7H65_RS11275 | 2301885..2303759 | 287 | putative-DNA-binding-domain-containing-protein-[Pseudomonas-aeruginosa] |
| WP_124988196.1 | Schlafen+HATPase+wHTH | GCF_003863815.1 | Pseudomonas_aeruginosa | Bacteria>Pseudomonadati>Pseudomonadota>Gammaproteobacteria>Pseudomonadales>Pseudomonadaceae>Pseudomonas>Pseudomonas_aeruginosa_group | 596 | EIM11_RS16890 | 67831..69621 | 287 | ATP-binding-protein-[Pseudomonas-aeruginosa] |

| Protein-ID | Domain | Assembly-ID | Strain | Taxonomy | Length | Locus-Tag | Genomic-coordinates | TaxID | Fasta-defline |
|---|---|---|---|---|---|---|---|---|---|
| WP_124984785.1 | Schlafen+HATPase+wHTH | GCF_003863685.1 | Pseudomonas_aeruginosa | Bacteria>Pseudomonadati>Pseudomonadota>Gammaproteobacteria>Pseudomonadales>Pseudomonadaceae>Pseudomonas>Pseudomonas_aeruginosa_group | 613 | EIM22_RS03575 | 280209..282050 | 287 | putative-DNA-binding-domain-containing-protein-[Pseudomonas-aeruginosa] |
| WP_124182727.1 | Schlafen+HATPase+wHTH | GCF_003839865.1 | Pseudomonas_aeruginosa | Bacteria>Pseudomonadati>Pseudomonadota>Gammaproteobacteria>Pseudomonadales>Pseudomonadaceae>Pseudomonas>Pseudomonas_aeruginosa_group | 613 | IPC226_RS26575 | 198459..200300 | 287 | putative-DNA-binding-domain-containing-protein-[Pseudomonas-aeruginosa] |
| WP_094226378.1 | Schlafen+HATPase+wHTH | GCF_002242855.1 | Pseudomonas_aeruginosa | Bacteria>Pseudomonadati>Pseudomonadota>Gammaproteobacteria>Pseudomonadales>Pseudomonadaceae>Pseudomonas>Pseudomonas_aeruginosa_group | 596 | ACG91_RS23755 | 36198..37988 | 287 | ATP-binding-protein-[Pseudomonas-aeruginosa] |
| WP_084308436.1 | Schlafen+HATPase+wHTH | GCF_002091575.1 | Pseudomonas_flavescens | Bacteria>Pseudomonadati>Pseudomonadota>Gammaproteobacteria>Pseudomonadales>Pseudomonadaceae>Pseudomonas>Pseudomonas_aeruginosa_group | 481 | PFL01S_RS23710 | 9708..11153 | 29435 | helix-turn-helix-domain-containing-protein-[Pseudomonas-flavescens] |
| WP_070330118.1 | Schlafen+HATPase+wHTH | GCF_001766125.1 | Pseudomonas_aeruginosa | Bacteria>Pseudomonadati>Pseudomonadota>Gammaproteobacteria>Pseudomonadales>Pseudomonadaceae>Pseudomonas>Pseudomonas_aeruginosa_group | 624 | BAW37_RS11765 | 24084..25958 | 287 | putative-DNA-binding-domain-containing-protein-[Pseudomonas-aeruginosa] |
| WP_033991606.1 | Schlafen+HATPase+wHTH | GCF_000795045.1 | Pseudomonas_aeruginosa | Bacteria>Pseudomonadati>Pseudomonadota>Gammaproteobacteria>Pseudomonadales>Pseudomonadaceae>Pseudomonas>Pseudomonas_aeruginosa_group | 596 | NR30_RS22670 | 82892..84682 | 287 | putative-DNA-binding-domain-containing-protein-[Pseudomonas-aeruginosa] |
| WP_033979002.1 | Schlafen+HATPase+wHTH | GCF_025799685.1 | Pseudomonas_aeruginosa | Bacteria>Pseudomonadati>Pseudomonadota>Gammaproteobacteria>Pseudomonadales>Pseudomonadaceae>Pseudomonas>Pseudomonas_aeruginosa_group | 394 | CJT50_RS28670 | 27187..28371 | 287 | ATP-binding-protein-[Pseudomonas-aeruginosa] |
| WP_031631061.1 | Schlafen+HATPase+wHTH | GCF_900707525.1 | Pseudomonas_aeruginosa | Bacteria>Pseudomonadati>Pseudomonadota>Gammaproteobacteria>Pseudomonadales>Pseudomonadaceae>Pseudomonas>Pseudomonas_aeruginosa_group | 624 | BAW40_RS11010 | 44486..46360 | 287 | putative-DNA-binding-domain-containing-protein-[Pseudomonas-aeruginosa] |
| WP_022580060.1 | Schlafen+HATPase+wHTH | GCF_020990445.1 | Pseudomonas_aeruginosa | Bacteria>Pseudomonadati>Pseudomonadota>Gammaproteobacteria>Pseudomonadales>Pseudomonadaceae>Pseudomonas>Pseudomonas_aeruginosa_group | 596 | BM548_RS18935 | 108510..110300 | 287 | ATP-binding-protein-[Pseudomonas-aeruginosa] |
| RRI70513.1 | Schlafen+HATPase+wHTH | GCA_003863585.1 | Pseudomonas_aeruginosa | Bacteria>Pseudomonadati>Pseudomonadota>Gammaproteobacteria>Pseudomonadales>Pseudomonadaceae>Pseudomonas>Pseudomonas_aeruginosa_group | 596 | EIM10_18035 | 73037..74827 | 287 | transcriptional-regulator-[Pseudomonas-aeruginosa] |
| MCO3958046.1 | Schlafen+HATPase+wHTH | GCA_023912735.1 | Pseudomonas_aeruginosa | Bacteria>Pseudomonadati>Pseudomonadota>Gammaproteobacteria>Pseudomonadales>Pseudomonadaceae>Pseudomonas>Pseudomonas_aeruginosa_group | 596 | FA108_20860 | 45044..46834 | 287 | transcriptional-regulator-[Pseudomonas-aeruginosa] |
| MCO2926701.1 | Schlafen+HATPase+wHTH | GCA_023909145.1 | Pseudomonas_aeruginosa | Bacteria>Pseudomonadati>Pseudomonadota>Gammaproteobacteria>Pseudomonadales>Pseudomonadaceae>Pseudomonas>Pseudomonas_aeruginosa_group | 394 | FA260_26695 | 12340..13524 | 287 | transcriptional-regulator-[Pseudomonas-aeruginosa] |
| MCO1977060.1 | Schlafen+HATPase+wHTH | GCA_023905865.1 | Pseudomonas_aeruginosa | Bacteria>Pseudomonadati>Pseudomonadota>Gammaproteobacteria>Pseudomonadales>Pseudomonadaceae>Pseudomonas>Pseudomonas_aeruginosa_group | 596 | FA422_11945 | 13757..15547 | 287 | transcriptional-regulator-[Pseudomonas-aeruginosa] |
| MBX6030766.1 | Schlafen+HATPase+wHTH | GCA_019688625.1 | Pseudomonas_aeruginosa | Bacteria>Pseudomonadati>Pseudomonadota>Gammaproteobacteria>Pseudomonadales>Pseudomonadaceae>Pseudomonas>Pseudomonas_aeruginosa_group | 602 | ISE67_08705 | 279006..280814 | 287 | putative-DNA-binding-domain-containing-protein-[Pseudomonas-aeruginosa] |
| MBO3071650.1 | Schlafen+HATPase+wHTH | GCA_017582945.1 | Pseudomonas_aeruginosa | Bacteria>Pseudomonadati>Pseudomonadota>Gammaproteobacteria>Pseudomonadales>Pseudomonadaceae>Pseudomonas>Pseudomonas_aeruginosa_group | 596 | J4H77_25750 | 180297..182087 | 287 | putative-DNA-binding-domain-containing-protein-[Pseudomonas-aeruginosa] |
| MBG6817112.1 | Schlafen+HATPase+wHTH | GCA_015764555.1 | Pseudomonas_aeruginosa | Bacteria>Pseudomonadati>Pseudomonadota>Gammaproteobacteria>Pseudomonadales>Pseudomonadaceae>Pseudomonas>Pseudomonas_aeruginosa_group | 423 | I5J00_12300 | 144943..146214 | 287 | putative-DNA-binding-domain-containing-protein-[Pseudomonas-aeruginosa] |
| HCG0728346.1 | Schlafen+HATPase+wHTH | GCA_024002515.1 | Pseudomonas_aeruginosa | Bacteria>Pseudomonadati>Pseudomonadota>Gammaproteobacteria>Pseudomonadales>Pseudomonadaceae>Pseudomonas>Pseudomonas_aeruginosa_group | 596 | NJJ48_005269 | 24797..26587 | 287 | putative-DNA-binding-domain-containing-protein-[Pseudomonas-aeruginosa] |
| HCF5867686.1 | Schlafen+HATPase+wHTH | GCA_023959235.1 | Pseudomonas_aeruginosa | Bacteria>Pseudomonadati>Pseudomonadota>Gammaproteobacteria>Pseudomonadales>Pseudomonadaceae>Pseudomonas>Pseudomonas_aeruginosa_group | 495 | NIH49_002294 | 180860..182347 | 287 | putative-DNA-binding-domain-containing-protein-[Pseudomonas-aeruginosa] |
| HCF2077876.1 | Schlafen+HATPase+wHTH | GCA_023942995.1 | Pseudomonas_aeruginosa | Bacteria>Pseudomonadati>Pseudomonadota>Gammaproteobacteria>Pseudomonadales>Pseudomonadaceae>Pseudomonas>Pseudomonas_aeruginosa_group | 624 | NH997_004591 | 28451..30325 | 287 | TPA:-putative-DNA-binding-domain-containing-protein-[Pseudomonas-aeruginosa] |
| HCF1890566.1 | Schlafen+HATPase+wHTH | GCA_023942415.1 | Pseudomonas_aeruginosa | Bacteria>Pseudomonadati>Pseudomonadota>Gammaproteobacteria>Pseudomonadales>Pseudomonadaceae>Pseudomonas>Pseudomonas_aeruginosa_group | 600 | NH920_001200 | 109588..111390 | 287 | putative-DNA-binding-domain-containing-protein-[Pseudomonas-aeruginosa] |
| WP_239610956.1 | Schlafen+HATPase+wHTH | GCF_022355455.1 | Pseudomonas_sp._KSR10 | Bacteria>Pseudomonadati>Pseudomonadota>Gammaproteobacteria>Pseudomonadales>Pseudomonadaceae>Pseudomonas>unclassified_Pseudomonas | 596 | MCB86_RS11355 | 517829..519619 | 2916654 | putative-DNA-binding-domain-containing-protein-[Pseudomonas-sp.-KSR10] |
| WP_220803979.1 | Schlafen+HATPase+wHTH | GCF_019656335.1 | Pseudomonas_sp._NCCP-436 | Bacteria>Pseudomonadati>Pseudomonadota>Gammaproteobacteria>Pseudomonadales>Pseudomonadaceae>Pseudomonas>unclassified_Pseudomonas | 624 | NCCP436_RS10640 | 95972..97846 | 2842481 | putative-DNA-binding-domain-containing-protein-[Pseudomonas-sp.-NCCP-436] |
| WP_218424331.1 | Schlafen+HATPase+wHTH | GCF_019218965.1 | Pseudomonas_sp._PDM31 | Bacteria>Pseudomonadati>Pseudomonadota>Gammaproteobacteria>Pseudomonadales>Pseudomonadaceae>Pseudomonas>unclassified_Pseudomonas | 585 | KW839_RS11060 | 23326..25083 | 2854778 | putative-DNA-binding-domain-containing-protein-[Pseudomonas-sp.-PDM31] |
| WP_095598171.1 | Schlafen+HATPase+wHTH | GCF_002286785.1 | Pseudomonas_sp._PIC25 | Bacteria>Pseudomonadati>Pseudomonadota>Gammaproteobacteria>Pseudomonadales>Pseudomonadaceae>Pseudomonas>unclassified_Pseudomonas | 423 | BZL41_RS04430 | 55591..56862 | 1958773 | ATP-binding-protein-[Pseudomonas-sp.-PIC25] |
| HBC00736.1 | Schlafen+HATPase+wHTH | GCA_003488145.1 | Pseudomonas_sp. | Bacteria>Pseudomonadati>Pseudomonadota>Gammaproteobacteria>Pseudomonadales>Pseudomonadaceae>Pseudomonas>unclassified_Pseudomonas | 416 | DC032_09175 | 14261..15511 | 306 | transcriptional-regulator-[Pseudomonas-sp.] |
| MBX9914997.1 | Schlafen+HATPase+wHTH | GCA_019745275.1 | Pseudomonadaceae_bacterium | Bacteria>Pseudomonadati>Pseudomonadota>Gammaproteobacteria>Pseudomonadales>Pseudomonadaceae>unclassified_Pseudomonadaceae | 481 | K2Y25_13975 | 3551..4996 | 1914538 | helix-turn-helix-domain-containing-protein-[Pseudomonadaceae-bacterium] |
| MCP5320936.1 | Schlafen+HATPase+wHTH | GCA_024234165.1 | Pseudomonadales_bacterium | Bacteria>Pseudomonadati>Pseudomonadota>Gammaproteobacteria>Pseudomonadales>unclassified_Pseudomonadales | 419 | H7A12_08960 | 731464..732723 | 1891229 | putative-DNA-binding-domain-containing-protein-[Pseudomonadales-bacterium] |
| MCP5320935.1 | Schlafen+HATPase+wHTH | GCA_024234165.1 | Pseudomonadales_bacterium | Bacteria>Pseudomonadati>Pseudomonadota>Gammaproteobacteria>Pseudomonadales>unclassified_Pseudomonadales | 400 | H7A12_08955 | 730248..731450 | 1891229 | putative-DNA-binding-domain-containing-protein-[Pseudomonadales-bacterium] |
| MCG3169756.1 | Schlafen+HATPase+wHTH | GCA_022072145.1 | Pseudomonadales_bacterium | Bacteria>Pseudomonadati>Pseudomonadota>Gammaproteobacteria>Pseudomonadales>unclassified_Pseudomonadales | 575 | CALGDGBN_01279 | 18583..20310 | 1891229 | hypothetical-protein-[Pseudomonadales-bacterium] |
| MAQ23220.1 | Schlafen+HATPase+wHTH | GCA_002701315.1 | Pseudomonadales_bacterium | Bacteria>Pseudomonadati>Pseudomonadota>Gammaproteobacteria>Pseudomonadales>unclassified_Pseudomonadales | 556 | CMK83_03290 | 16304..17974 | 1891229 | ATPase-[Pseudomonadales-bacterium] |
| MBL1259970.1 | Schlafen+HATPase+wHTH | GCA_002401765.2 | Thiotrichaceae_bacterium | Bacteria>Pseudomonadati>Pseudomonadota>Gammaproteobacteria>Thiotrichales>Thiotrichaceae>unclassified_Thiotrichaceae | 482 | COB33_005500 | 41862..43310 | 2030882 | putative-DNA-binding-domain-containing-protein-[Thiotrichaceae-bacterium] |
| MBT3348499.1 | Schlafen+HATPase+wHTH | GCA_018648495.1 | Thiotrichales_bacterium | Bacteria>Pseudomonadati>Pseudomonadota>Gammaproteobacteria>Thiotrichales>unclassified_Thiotrichales | 470 | HN398_08690 | 10630..12042 | 2026796 | AAA-family-ATPase-[Thiotrichales-bacterium] |
| MBL1294160.1 | Schlafen+HATPase+wHTH | GCA_002401375.2 | Thiotrichales_bacterium | Bacteria>Pseudomonadati>Pseudomonadota>Gammaproteobacteria>Thiotrichales>unclassified_Thiotrichales | 609 | COB61_009835 | 13669..15498 | 2026796 | putative-DNA-binding-domain-containing-protein-[Thiotrichales-bacterium] |
| TAK54304.1 | Schlafen+HATPase+wHTH | GCA_004297335.1 | Gammaproteobacteria_bacterium | Bacteria>Pseudomonadati>Pseudomonadota>Gammaproteobacteria>unclassified_Gammaproteobacteria | 544 | EP025_07510 | 6762..8396 | 1913989 | transcriptional-regulator-[Gammaproteobacteria-bacterium] |
| PKM04363.1 | Schlafen+HATPase+wHTH | GCA_002839965.1 | Gammaproteobacteria_bacterium_HGW-Gammaproteobacteria-6 | Bacteria>Pseudomonadati>Pseudomonadota>Gammaproteobacteria>unclassified_Gammaproteobacteria | 624 | CVV16_05335 | 27579..29453 | 2013802 | AAA-family-ATPase-[Gammaproteobacteria-bacterium-HGW-Gammaproteobacteria-6] |
| NCA89501.1 | Schlafen+HATPase+wHTH | GCA_009928645.1 | Gammaproteobacteria_bacterium | Bacteria>Pseudomonadati>Pseudomonadota>Gammaproteobacteria>unclassified_Gammaproteobacteria | 425 | EOM92_11495 | 3357..4634 | 1913989 | hypothetical-protein-[Gammaproteobacteria-bacterium] |
| MXW50609.1 | Schlafen+HATPase+wHTH | GCA_009835805.1 | Gammaproteobacteria_bacterium | Bacteria>Pseudomonadati>Pseudomonadota>Gammaproteobacteria>unclassified_Gammaproteobacteria | 235 | F4Z84_10215 | 13336..14043 | 1913989 | hypothetical-protein-[Gammaproteobacteria-bacterium] |
| MDE0453906.1 | Schlafen+HATPase+wHTH | GCA_028821145.1 | Gammaproteobacteria_bacterium | Bacteria>Pseudomonadati>Pseudomonadota>Gammaproteobacteria>unclassified_Gammaproteobacteria | 139 | OXJ63_01200 | 17970..18389 | 1913989 | MAG:-ATP-binding-protein-[Gammaproteobacteria-bacterium] |
| MDE0453905.1 | Schlafen+HATPase+wHTH | GCA_028821145.1 | Gammaproteobacteria_bacterium | Bacteria>Pseudomonadati>Pseudomonadota>Gammaproteobacteria>unclassified_Gammaproteobacteria | 119 | OXJ63_01195 | 17614..17973 | 1913989 | MAG:-hypothetical-protein-OXJ63_01195-[Gammaproteobacteria-bacterium] |
| MDE0129994.1 | Schlafen+HATPase+wHTH | GCA_028819355.1 | Gammaproteobacteria_bacterium | Bacteria>Pseudomonadati>Pseudomonadota>Gammaproteobacteria>unclassified_Gammaproteobacteria | 464 | OXQ86_10570 | 56636..58030 | 1913989 | MAG:-putative-DNA-binding-domain-containing-protein-[Gammaproteobacteria-bacterium] |
| MDE0094448.1 | Schlafen+HATPase+wHTH | GCA_028819195.1 | Gammaproteobacteria_bacterium | Bacteria>Pseudomonadati>Pseudomonadota>Gammaproteobacteria>unclassified_Gammaproteobacteria | 446 | OXO49_08125 | 169936..171276 | 1913989 | MAG:-hypothetical-protein-OXO49_08125-[Gammaproteobacteria-bacterium] |
| MDD9817279.1 | Schlafen+HATPase+wHTH | GCA_028817495.1 | Gammaproteobacteria_bacterium | Bacteria>Pseudomonadati>Pseudomonadota>Gammaproteobacteria>unclassified_Gammaproteobacteria | 134 | OXU61_03950 | 8849..9253 | 1913989 | MAG:-hypothetical-protein-OXU61_03950-[Gammaproteobacteria-bacterium] |
| MCZ0943261.1 | Schlafen+HATPase+wHTH | GCA_026914145.1 | Gammaproteobacteria_bacterium | Bacteria>Pseudomonadati>Pseudomonadota>Gammaproteobacteria>unclassified_Gammaproteobacteria | 419 | OXJ53_09410 | 5886..7145 | 1913989 | MAG:-putative-DNA-binding-domain-containing-protein-[Gammaproteobacteria-bacterium] |
| MCY4343958.1 | Schlafen+HATPase+wHTH | GCA_026710215.1 | Gammaproteobacteria_bacterium | Bacteria>Pseudomonadati>Pseudomonadota>Gammaproteobacteria>unclassified_Gammaproteobacteria | 412 | OXE83_10350 | 9268..10506 | 1913989 | MAG:-putative-DNA-binding-domain-containing-protein-[Gammaproteobacteria-bacterium] |
| MCY4014262.1 | Schlafen+HATPase+wHTH | GCA_026708185.1 | Gammaproteobacteria_bacterium | Bacteria>Pseudomonadati>Pseudomonadota>Gammaproteobacteria>unclassified_Gammaproteobacteria | 415 | OXG82_16285 | 56194..57441 | 1913989 | MAG:-putative-DNA-binding-domain-containing-protein-[Gammaproteobacteria-bacterium] |
| MCP5127471.1 | Schlafen+HATPase+wHTH | GCA_024235095.1 | Gammaproteobacteria_bacterium | Bacteria>Pseudomonadati>Pseudomonadota>Gammaproteobacteria>unclassified_Gammaproteobacteria | 562 | H6973_18110 | 273817..275505 | 1913989 | putative-DNA-binding-domain-containing-protein-[Gammaproteobacteria-bacterium] |
| MCP4410537.1 | Schlafen+HATPase+wHTH | GCA_024222475.1 | Gammaproteobacteria_bacterium | Bacteria>Pseudomonadati>Pseudomonadota>Gammaproteobacteria>unclassified_Gammaproteobacteria | 414 | GY807_22895 | 9510..10754 | 1913989 | transcriptional-regulator-[Gammaproteobacteria-bacterium] |
| MCG3203222.1 | Schlafen+HATPase+wHTH | GCA_022071965.1 | Gammaproteobacteria_bacterium | Bacteria>Pseudomonadati>Pseudomonadota>Gammaproteobacteria>unclassified_Gammaproteobacteria | 409 | NFCOHLIN_03155 | 13176..14405 | 1913989 | hypothetical-protein-[Gammaproteobacteria-bacterium] |
| MCG3145053.1 | Schlafen+HATPase+wHTH | GCA_022072255.1 | Gammaproteobacteria_bacterium | Bacteria>Pseudomonadati>Pseudomonadota>Gammaproteobacteria>unclassified_Gammaproteobacteria | 550 | HONDAALG_02594 | 46002..47654 | 1913989 | hypothetical-protein-[Gammaproteobacteria-bacterium] |
| MBU1223665.1 | Schlafen+HATPase+wHTH | GCA_018819475.1 | Gammaproteobacteria_bacterium | Bacteria>Pseudomonadati>Pseudomonadota>Gammaproteobacteria>unclassified_Gammaproteobacteria | 591 | KKI05_07415 | 72790..74565 | 1913989 | putative-DNA-binding-domain-containing-protein-[Gammaproteobacteria-bacterium] |
| MBU0782762.1 | Schlafen+HATPase+wHTH | GCA_018815355.1 | Gammaproteobacteria_bacterium | Bacteria>Pseudomonadati>Pseudomonadota>Gammaproteobacteria>unclassified_Gammaproteobacteria | 596 | KJ979_02535 | 69233..71023 | 1913989 | putative-DNA-binding-domain-containing-protein-[Gammaproteobacteria-bacterium] |
| HFD80328.1 | Schlafen+HATPase+wHTH | GCA_011321775.1 | Gammaproteobacteria_bacterium | Bacteria>Pseudomonadati>Pseudomonadota>Gammaproteobacteria>unclassified_Gammaproteobacteria | 424 | ENK05_08070 | 3784..5058 | 1913989 | transcriptional-regulator-[Gammaproteobacteria-bacterium] |
| WP_130049302.1 | Schlafen+HATPase+wHTH | GCF_028765545.1 | Aliivibrio | Bacteria>Pseudomonadati>Pseudomonadota>Gammaproteobacteria>Vibrionales>Vibrionaceae | 527 | ERW52_RS16075 | 25292..26875 | 511678 | MULTISPECIES:-putative-DNA-binding-domain-containing-protein-[Aliivibrio] |
| WP_108693862.1 | Schlafen+HATPase+wHTH | GCF_043106745.1 | Vibrio | Bacteria>Pseudomonadati>Pseudomonadota>Gammaproteobacteria>Vibrionales>Vibrionaceae | 410 | DC364_RS00740 | 66326..67558 | 662 | ATP-binding-protein-[Vibrio-vulnificus] |
| WP_006071669.1 | Schlafen+HATPase+wHTH | GCF_000181535.1 | Vibrio | Bacteria>Pseudomonadati>Pseudomonadota>Gammaproteobacteria>Vibrionales>Vibrionaceae | 410 | VSAK1_RS11970 | 46943..48175 | 662 | MULTISPECIES:-ATP-binding-protein-[Vibrio] |
| WP_133416082.1 | Schlafen+HATPase+wHTH | GCF_004359415.1 | Aliivibrio_fischeri | Bacteria>Pseudomonadati>Pseudomonadota>Gammaproteobacteria>Vibrionales>Vibrionaceae>Aliivibrio | 527 | VFFQA001_RS16050 | 457262..458845 | 668 | putative-DNA-binding-domain-containing-protein-[Aliivibrio-fischeri] |
| WP_065641603.1 | Schlafen+HATPase+wHTH | GCF_001690915.1 | Aliivibrio_fischeri | Bacteria>Pseudomonadati>Pseudomonadota>Gammaproteobacteria>Vibrionales>Vibrionaceae>Aliivibrio | 527 | A6E13_RS16570 | 257332..258915 | 668 | putative-DNA-binding-domain-containing-protein-[Aliivibrio-fischeri] |
| WP_215796805.1 | Schlafen+HATPase+wHTH | GCF_001989775.2 | Vibrio_anguillarum | Bacteria>Pseudomonadati>Pseudomonadota>Gammaproteobacteria>Vibrionales>Vibrionaceae>Vibrio | 410 | PN39_RS13095 | 13827..15059 | 55601 | ATP-binding-protein-[Vibrio-anguillarum] |
| WP_065091079.1 | Schlafen+HATPase+wHTH | GCF_001675245.1 | Vibrio_vulnificus | Bacteria>Pseudomonadati>Pseudomonadota>Gammaproteobacteria>Vibrionales>Vibrionaceae>Vibrio | 410 | FORC17_RS18325 | 769912..771144 | 672 | ATP-binding-protein-[Vibrio-vulnificus] |
| QPL56155.1 | Schlafen+HATPase+wHTH | GCA_015767675.1 | Vibrio_navarrensis | Bacteria>Pseudomonadati>Pseudomonadota>Gammaproteobacteria>Vibrionales>Vibrionaceae>Vibrio | 410 | I3X05_18840 | 389161..390393 | 29495 | putative-DNA-binding-domain-containing-protein-[Vibrio-navarrensis] |
| MBY7947237.1 | Schlafen+HATPase+wHTH | GCA_019853335.1 | Vibrio_fluvialis | Bacteria>Pseudomonadati>Pseudomonadota>Gammaproteobacteria>Vibrionales>Vibrionaceae>Vibrio | 410 | KW461_01965 | 446799..448031 | 676 | putative-DNA-binding-domain-containing-protein-[Vibrio-fluvialis] |
| HAS6096893.1 | Schlafen+HATPase+wHTH | GCA_015776315.1 | Vibrio_vulnificus | Bacteria>Pseudomonadati>Pseudomonadota>Gammaproteobacteria>Vibrionales>Vibrionaceae>Vibrio | 410 | I7145_19880 | 58286..59518 | 672 | transcriptional-regulator-[Vibrio-vulnificus] |
| EHZ2755063.1 | Schlafen+HATPase+wHTH | GCA_019682715.1 | Vibrio_vulnificus | Bacteria>Pseudomonadati>Pseudomonadota>Gammaproteobacteria>Vibrionales>Vibrionaceae>Vibrio | 410 | K5N53_003180 | 44735..45967 | 672 | putative-DNA-binding-domain-containing-protein-[Vibrio-vulnificus] |
| WP_159408513.1 | Schlafen+HATPase+wHTH | GCF_009883875.1 | Vibrio_parahaemolyticus | Bacteria>Pseudomonadati>Pseudomonadota>Gammaproteobacteria>Vibrionales>Vibrionaceae>Vibrio>Vibrio_harveyi_group | 410 | EHC69_RS27025 | 6662..7894 | 670 | ATP-binding-protein-[Vibrio-parahaemolyticus] |
| EJE8525236.1 | Schlafen+HATPase+wHTH | GCA_023352175.1 | Vibrio_parahaemolyticus | Bacteria>Pseudomonadati>Pseudomonadota>Gammaproteobacteria>Vibrionales>Vibrionaceae>Vibrio>Vibrio_harveyi_group | 410 | M5229_003410 | 72427..73659 | 670 | putative-DNA-binding-domain-containing-protein-[Vibrio-parahaemolyticus] |
| EJE8514350.1 | Schlafen+HATPase+wHTH | GCA_023352155.1 | Vibrio_parahaemolyticus | Bacteria>Pseudomonadati>Pseudomonadota>Gammaproteobacteria>Vibrionales>Vibrionaceae>Vibrio>Vibrio_harveyi_group | 410 | M5244_002605 | 64635..65867 | 670 | putative-DNA-binding-domain-containing-protein-[Vibrio-parahaemolyticus] |
| WP_272245954.1 | Schlafen+HATPase+wHTH | GCF_028446755.1 | Vibrio_europaeus | Bacteria>Pseudomonadati>Pseudomonadota>Gammaproteobacteria>Vibrionales>Vibrionaceae>Vibrio>Vibrio_oreintalis_group | 410 | OPW38_RS04405 | 13209..14441 | 300876 | ATP-binding-protein-[Vibrio-europaeus] |
| WP_217125924.1 | Schlafen+HATPase+wHTH | GCF_019049855.1 | Hydrogenophilus_thiooxidans | Bacteria>Pseudomonadati>Pseudomonadota>Hydrogenophilia>Hydrogenophilales>Hydrogenophilaceae>Hydrogenophilus | 571 | KUL93_RS04630 | 91008..92723 | 2820326 | ATP-binding-protein-[Hydrogenophilus-thiooxidans] |
| BBD77931.1 | Schlafen+HATPase+wHTH | GCA_003574215.1 | Hydrogenophilus_thermoluteolus | Bacteria>Pseudomonadati>Pseudomonadota>Hydrogenophilia>Hydrogenophilales>Hydrogenophilaceae>Hydrogenophilus | 571 | HPTL_1673 | 1773735..1775450 | 297 | AAA-family-ATPase-[Hydrogenophilus-thermoluteolus] |
| WP_142810717.1 | Schlafen+HATPase+wHTH | GCF_006980705.1 | Tepidiphilus_olei | Bacteria>Pseudomonadati>Pseudomonadota>Hydrogenophilia>Hydrogenophilales>Hydrogenophilaceae>Tepidiphilus | 602 | EPD80_RS07345 | 164029..165837 | 2502184 | putative-DNA-binding-domain-containing-protein-[Tepidiphilus-olei] |
| MCX5809779.1 | Schlafen+HATPase+wHTH | GCA_026388245.1 | Pseudomonadota_bacterium | Bacteria>Pseudomonadati>Pseudomonadota>unclassified_Pseudomonadota | 561 | NTX36_10490 | 7359..9044 | 1977087 | MAG:-putative-DNA-binding-domain-containing-protein-[Proteobacteria-bacterium] |
| MCA0325061.1 | Schlafen+HATPase+wHTH | GCA_020161215.1 | Pseudomonadota_bacterium | Bacteria>Pseudomonadati>Pseudomonadota>unclassified_Pseudomonadota | 621 | LCH89_05595 | 5312..7177 | 1977087 | putative-DNA-binding-domain-containing-protein-[Pseudomonadota-bacterium] |
| MBU4294443.1 | Schlafen+HATPase+wHTH | GCA_018897105.1 | Pseudomonadota_bacterium | Bacteria>Pseudomonadati>Pseudomonadota>unclassified_Pseudomonadota | 471 | KJ862_00975 | 74025..75440 | 1977087 | putative-DNA-binding-domain-containing-protein-[Pseudomonadota-bacterium] |
| MBU4235492.1 | Schlafen+HATPase+wHTH | GCA_018897885.1 | Pseudomonadota_bacterium | Bacteria>Pseudomonadati>Pseudomonadota>unclassified_Pseudomonadota | 570 | KKI01_04075 | 5334..7046 | 1977087 | MAG:-putative-DNA-binding-domain-containing-protein-[Proteobacteria-bacterium] |
| MBU1744599.1 | Schlafen+HATPase+wHTH | GCA_018822765.1 | Pseudomonadota_bacterium | Bacteria>Pseudomonadati>Pseudomonadota>unclassified_Pseudomonadota | 488 | KJ649_06865 | 11268..12734 | 1977087 | putative-DNA-binding-domain-containing-protein-[Pseudomonadota-bacterium] |
| MBU0967617.1 | Schlafen+HATPase+wHTH | GCA_018816555.1 | Pseudomonadota_bacterium | Bacteria>Pseudomonadati>Pseudomonadota>unclassified_Pseudomonadota | 464 | KKA54_14690 | 26282..27676 | 1977087 | putative-DNA-binding-domain-containing-protein-[Pseudomonadota-bacterium] |
| MBU0673652.1 | Schlafen+HATPase+wHTH | GCA_018814305.1 | Pseudomonadota_bacterium | Bacteria>Pseudomonadati>Pseudomonadota>unclassified_Pseudomonadota | 506 | KJ950_03305 | 52414..53934 | 1977087 | putative-DNA-binding-domain-containing-protein-[Pseudomonadota-bacterium] |
| MBS0577370.1 | Schlafen+HATPase+wHTH | GCA_018240435.1 | Pseudomonadota_bacterium | Bacteria>Pseudomonadati>Pseudomonadota>unclassified_Pseudomonadota | 615 | JSS45_13270 | 6043..7890 | 1977087 | putative-DNA-binding-domain-containing-protein-[Pseudomonadota-bacterium] |
| MBS0382006.1 | Schlafen+HATPase+wHTH | GCA_018241365.1 | Pseudomonadota_bacterium | Bacteria>Pseudomonadati>Pseudomonadota>unclassified_Pseudomonadota | 572 | JSR56_06195 | 3151..4869 | 1977087 | putative-DNA-binding-domain-containing-protein-[Pseudomonadota-bacterium] |
| MBS0338392.1 | Schlafen+HATPase+wHTH | GCA_018241645.1 | Pseudomonadota_bacterium | Bacteria>Pseudomonadati>Pseudomonadota>unclassified_Pseudomonadota | 284 | JSS40_16635 | 993..1843 | 1977087 | putative-DNA-binding-domain-containing-protein-[Pseudomonadota-bacterium] |
| MBS0338391.1 | Schlafen+HATPase+wHTH | GCA_018241645.1 | Pseudomonadota_bacterium | Bacteria>Pseudomonadati>Pseudomonadota>unclassified_Pseudomonadota | 238 | JSS40_16630 | 149..865 | 1977087 | AAA-family-ATPase-[Pseudomonadota-bacterium] |
| MBK6849140.1 | Schlafen+HATPase+wHTH | GCA_016702545.1 | Pseudomonadota_bacterium | Bacteria>Pseudomonadati>Pseudomonadota>unclassified_Pseudomonadota | 420 | IPG96_16950 | 58473..59735 | 1977087 | putative-DNA-binding-domain-containing-protein-[Pseudomonadota-bacterium] |
| KAA0218650.1 | Schlafen+HATPase+wHTH | GCA_008363105.1 | Pseudomonadota_bacterium | Bacteria>Pseudomonadati>Pseudomonadota>unclassified_Pseudomonadota | 400 | EDM78_03980 | 21898..23100 | 1977087 | transcriptional-regulator-[Pseudomonadota-bacterium] |
| WP_053322518.1 | Schlafen+HATPase+wHTH | GCF_001273615.1 | Mariprofundus_ferrooxydans | Bacteria>Pseudomonadati>Pseudomonadota>Zetaproteobacteria>Mariprofundales>Mariprofundaceae>Mariprofundus | 464 | AL013_RS02420 | 93887..95281 | 314344 | helix-turn-helix-domain-containing-protein-[Mariprofundus-ferrooxydans] |
| MCG2712033.1 | Schlafen+HATPase+wHTH | GCA_022013205.1 | Candidatus_Omnitrophota_bacterium | Bacteria>Pseudomonadati>PVC_group>Candidatus_Omnitrophota>unclassified_Candidatus_Omnitrophota | 85 | L6416_06885 | 1..256 | 2035772 | ATP-binding-protein-[Candidatus-Omnitrophota-bacterium] |
| MBU4477604.1 | Schlafen+HATPase+wHTH | GCA_018895275.1 | Candidatus_Omnitrophota_bacterium | Bacteria>Pseudomonadati>PVC_group>Candidatus_Omnitrophota>unclassified_Candidatus_Omnitrophota | 512 | KKH34_00705 | 4430..5968 | 2035772 | putative-DNA-binding-domain-containing-protein-[Candidatus-Omnitrophota-bacterium] |
| MBU1810878.1 | Schlafen+HATPase+wHTH | GCA_018823915.1 | Candidatus_Omnitrophota_bacterium | Bacteria>Pseudomonadati>PVC_group>Candidatus_Omnitrophota>unclassified_Candidatus_Omnitrophota | 487 | KJ629_07040 | 1922..3385 | 2035772 | putative-DNA-binding-domain-containing-protein-[Candidatus-Omnitrophica-bacterium] |
| MBK7497320.1 | Schlafen+HATPase+wHTH | GCA_016706995.1 | Candidatus_Omnitrophota_bacterium | Bacteria>Pseudomonadati>PVC_group>Candidatus_Omnitrophota>unclassified_Candidatus_Omnitrophota | 597 | IPI28_18685 | 34969..36762 | 2035772 | putative-DNA-binding-domain-containing-protein-[Candidatus-Omnitrophota-bacterium] |
| MBQ9739435.1 | Schlafen+HATPase+wHTH | GCA_017457135.1 | Kiritimatiellia_bacterium | Bacteria>Pseudomonadati>PVC_group>Kiritimatiellota>Kiritimatiellia>unclassified_Kiritimatiellia | 580 | IJV91_00710 | 8684..10426 | 2212473 | putative-DNA-binding-domain-containing-protein-[Kiritimatiellia-bacterium] |
| MBQ6141561.1 | Schlafen+HATPase+wHTH | GCA_017432735.1 | Kiritimatiellia_bacterium | Bacteria>Pseudomonadati>PVC_group>Kiritimatiellota>Kiritimatiellia>unclassified_Kiritimatiellia | 517 | IJI54_09770 | 21926..23479 | 2212473 | putative-DNA-binding-domain-containing-protein-[Kiritimatiellia-bacterium] |
| MBN1673170.1 | Schlafen+HATPase+wHTH | GCA_016929905.1 | Kiritimatiellia_bacterium | Bacteria>Pseudomonadati>PVC_group>Kiritimatiellota>Kiritimatiellia>unclassified_Kiritimatiellia | 564 | JXR37_19150 | 47764..49458 | 2212473 | putative-DNA-binding-domain-containing-protein-[Kiritimatiellia-bacterium] |
| TFH14333.1 | Schlafen+HATPase+wHTH | GCA_004525935.1 | Lentisphaerales_bacterium | Bacteria>Pseudomonadati>PVC_group>Lentisphaerota>Lentisphaeria>Lentisphaerales>unclassified_Lentisphaerales | 365 | E4H02_10095 | 1..1094 | 2448604 | transcriptional-regulator,-partial-[Lentisphaerales-bacterium] |
| MBQ6473651.1 | Schlafen+HATPase+wHTH | GCA_017431185.1 | Victivallales_bacterium | Bacteria>Pseudomonadati>PVC_group>Lentisphaerota>Lentisphaeria>Victivallales>unclassified_Victivallales | 525 | IJJ33_16815 | 10750..12327 | 2212475 | putative-DNA-binding-domain-containing-protein-[Victivallales-bacterium] |
| OGV20212.1 | Schlafen+HATPase+wHTH | GCA_001803205.1 | Lentisphaerae_bacterium_GWF2_38_69 | Bacteria>Pseudomonadati>PVC_group>Lentisphaerota>unclassified_Lentisphaerota | 561 | A2X47_05995 | 3551..5236 | 1798570 | MAG:-hypothetical-protein-A2X47_05995-[Lentisphaerae-bacterium-GWF2_38_69] |
| NLF62185.1 | Schlafen+HATPase+wHTH | GCA_012515215.1 | Lentisphaerota_bacterium | Bacteria>Pseudomonadati>PVC_group>Lentisphaerota>unclassified_Lentisphaerota | 556 | GX574_13660 | 38303..39973 | 1932692 | ATPase-[Lentisphaerota-bacterium] |
| NLF19758.1 | Schlafen+HATPase+wHTH | GCA_012514865.1 | Lentisphaerota_bacterium | Bacteria>Pseudomonadati>PVC_group>Lentisphaerota>unclassified_Lentisphaerota | 527 | GX595_21210 | 8884..10467 | 1932692 | AAA-family-ATPase-[Lentisphaerota-bacterium] |
| NLE53430.1 | Schlafen+HATPase+wHTH | GCA_012514365.1 | Lentisphaerota_bacterium | Bacteria>Pseudomonadati>PVC_group>Lentisphaerota>unclassified_Lentisphaerota | 535 | GX617_00710 | 1404..3011 | 1932692 | transcriptional-regulator-[Lentisphaerota-bacterium] |
| MBT5606088.1 | Schlafen+HATPase+wHTH | GCA_018666605.1 | Lentisphaerota_bacterium | Bacteria>Pseudomonadati>PVC_group>Lentisphaerota>unclassified_Lentisphaerota | 533 | HOJ57_09115 | 4936..6537 | 1932692 | transcriptional-regulator-[Lentisphaerota-bacterium] |
| MBT4818761.1 | Schlafen+HATPase+wHTH | GCA_018664235.1 | Lentisphaerota_bacterium | Bacteria>Pseudomonadati>PVC_group>Lentisphaerota>unclassified_Lentisphaerota | 617 | HON70_23835 | 6714..8567 | 1932692 | AAA-family-ATPase-[Lentisphaerota-bacterium] |
| MBT3379342.1 | Schlafen+HATPase+wHTH | GCA_018648355.1 | Lentisphaerota_bacterium | Bacteria>Pseudomonadati>PVC_group>Lentisphaerota>unclassified_Lentisphaerota | 617 | HN406_27325 | 10998..12851 | 1932692 | AAA-family-ATPase-[Lentisphaerota-bacterium] |
| MBT3379338.1 | Schlafen+HATPase+wHTH | GCA_018648355.1 | Lentisphaerota_bacterium | Bacteria>Pseudomonadati>PVC_group>Lentisphaerota>unclassified_Lentisphaerota | 533 | HN406_27305 | 4818..6419 | 1932692 | transcriptional-regulator-[Lentisphaerota-bacterium] |
| WP_096895083.1 | Schlafen+HATPase+wHTH | GCF_002443295.1 | Candidatus_Scalindua_japonica | Bacteria>Pseudomonadati>PVC_group>Planctomycetota>Candidatus_Brocadiia>Candidatus_Brocadiales>Candidatus_Scalinduaceae>Candidatus_Scalindua | 585 | SCALIN_RS13990 | 121450..123207 | 1284222 | putative-DNA-binding-domain-containing-protein-[Candidatus-Scalindua-japonica] |
| MBT6049513.1 | Schlafen+HATPase+wHTH | GCA_018697755.1 | Candidatus_Scalindua_sp. | Bacteria>Pseudomonadati>PVC_group>Planctomycetota>Candidatus_Brocadiia>Candidatus_Brocadiales>Candidatus_Scalinduaceae>Candidatus_Scalindua>unclassified_Candidatus_Scalindua | 602 | HOG49_22165 | 15702..17510 | 2268189 | transcriptional-regulator-[Candidatus-Scalindua-sp.] |
| MBT6049511.1 | Schlafen+HATPase+wHTH | GCA_018697755.1 | Candidatus_Scalindua_sp. | Bacteria>Pseudomonadati>PVC_group>Planctomycetota>Candidatus_Brocadiia>Candidatus_Brocadiales>Candidatus_Scalinduaceae>Candidatus_Scalindua>unclassified_Candidatus_Scalindua | 587 | HOG49_22155 | 10971..12734 | 2268189 | AAA-family-ATPase-[Candidatus-Scalindua-sp.] |
| MBC8551962.1 | Schlafen+HATPase+wHTH | GCA_014384865.1 | Candidatus_Brocadiales_bacterium | Bacteria>Pseudomonadati>PVC_group>Planctomycetota>Candidatus_Brocadiia>Candidatus_Brocadiales>unclassified_Candidatus_Brocadiales | 489 | H8D23_20130 | 5538..7007 | 2760054 | putative-DNA-binding-domain-containing-protein-[Candidatus-Brocadiales-bacterium] |
| NQV34260.1 | Schlafen+HATPase+wHTH | GCA_013203945.1 | Phycisphaeraceae_bacterium | Bacteria>Pseudomonadati>PVC_group>Planctomycetota>Phycisphaerae>Phycisphaerales>Phycisphaeraceae>unclassified_Phycisphaeraceae | 538 | HQ515_16315 | 4389..6005 | 2026777 | putative-DNA-binding-domain-containing-protein-[Phycisphaeraceae-bacterium] |
| MCC6908124.1 | Schlafen+HATPase+wHTH | GCA_020852515.1 | Phycisphaerales_bacterium | Bacteria>Pseudomonadati>PVC_group>Planctomycetota>Phycisphaerae>Phycisphaerales>unclassified_Phycisphaerales | 546 | IT430_09305 | 96145..97785 | 2052180 | putative-DNA-binding-domain-containing-protein-[Phycisphaerales-bacterium] |
| MBK9119767.1 | Schlafen+HATPase+wHTH | GCA_016716475.1 | Phycisphaerales_bacterium | Bacteria>Pseudomonadati>PVC_group>Planctomycetota>Phycisphaerae>Phycisphaerales>unclassified_Phycisphaerales | 546 | IPM18_09240 | 372988..374628 | 2052180 | putative-DNA-binding-domain-containing-protein-[Phycisphaerales-bacterium] |
| QDV91809.1 | Schlafen+HATPase+wHTH | GCA_007753915.1 | Phycisphaerae_bacterium_RAS2 | Bacteria>Pseudomonadati>PVC_group>Planctomycetota>Phycisphaerae>unclassified_Phycisphaerae | 430 | RAS2_29150 | 3438110..3439402 | 2528035 | Divergent-AAA-domain-protein-[Phycisphaerae-bacterium-RAS2] |
| MBW7904852.1 | Schlafen+HATPase+wHTH | GCA_019454965.1 | Phycisphaerae_bacterium | Bacteria>Pseudomonadati>PVC_group>Planctomycetota>Phycisphaerae>unclassified_Phycisphaerae | 548 | H3C42_04700 | 496357..498003 | 2026778 | putative-DNA-binding-domain-containing-protein-[Phycisphaerae-bacterium] |
| MBI9018592.1 | Schlafen+HATPase+wHTH | GCA_016342495.1 | Phycisphaerae_bacterium | Bacteria>Pseudomonadati>PVC_group>Planctomycetota>Phycisphaerae>unclassified_Phycisphaerae | 555 | JEZ07_15170 | 9088..10755 | 2026778 | putative-DNA-binding-domain-containing-protein-[Phycisphaerae-bacterium] |
| GJQ27843.1 | Schlafen+HATPase+wHTH | GCA_022180675.1 | Phycisphaerae_bacterium | Bacteria>Pseudomonadati>PVC_group>Planctomycetota>Phycisphaerae>unclassified_Phycisphaerae | 457 | HBSAPP02_28750 | 72712..74085 | 2026778 | hypothetical-protein-HBSAPP02_28750-[Phycisphaerae-bacterium] |
| WP_250928723.1 | Schlafen+HATPase+wHTH | GCF_023701485.1 | Aporhodopirellula_aestuarii | Bacteria>Pseudomonadati>PVC_group>Planctomycetota>Planctomycetia>Pirellulales>Pirellulaceae>Aporhodopirellula | 570 | NB063_RS10690 | 29793..31505 | 2950107 | RNA-binding-domain-containing-protein-[Aporhodopirellula-aestuarii] |

| Protein-ID | Domain | Assembly-ID | Strain | Taxonomy | Length | Locus-Tag | Genomic-coordinates | TaxID | Fasta-defline |
|---|---|---|---|---|---|---|---|---|---|
| MCM2371080.1 | Schlafen+HATPase+wHTH | GCA_023701485.1 | Aporhodopirellula_aestuarii | Bacteria>Pseudomonadati>PVC_group>Planctomycetota>Planctomycetia>Pirellulales>Pirellulaceae>Aporhodopirellula | 570 | NB063_10710 | 29793..31505 | 2950107 | putative-DNA-binding-domain-containing-protein-[Rhodopirellula-aestuarii] |
| NLF73680.1 | Schlafen+HATPase+wHTH | GCA_012515235.1 | Candidatus_Anammoximicrobium_sp. | Bacteria>Pseudomonadati>PVC_group>Planctomycetota>Planctomycetia>Pirellulales>Pirellulaceae>Candidatus_Anammoximicrobium>unclassified_Candidatus_Anammoximicrobium | 490 | GX575_31965 | 11899..13371 | 2699742 | hypothetical-protein-[Candidatus-Anammoximicrobium-sp.] |
| WP_146600639.1 | Schlafen+HATPase+wHTH | GCF_007860185.1 | Novipirellula_aureliae | Bacteria>Pseudomonadati>PVC_group>Planctomycetota>Planctomycetia>Pirellulales>Pirellulaceae>Novipirellula | 441 | Q31b_RS15745 | 224828..226153 | 2527966 | ATP-binding-protein-[Novipirellula-aureliae] |
| WP_011117925.1 | Schlafen+HATPase+wHTH | GCF_000196115.1 | Rhodopirellula_baltica | Bacteria>Pseudomonadati>PVC_group>Planctomycetota>Planctomycetia>Pirellulales>Pirellulaceae>Rhodopirellula | 316 | DDX19_RS10390 | 127277..128227 | 265606 | ATP-binding-protein-[Rhodopirellula-baltica] |
| WP_274403449.1 | Schlafen+HATPase+wHTH | GCF_028768465.1 | Rhodopirellula_sp._P2 | Bacteria>Pseudomonadati>PVC_group>Planctomycetota>Planctomycetia>Pirellulales>Pirellulaceae>Rhodopirellula>unclassified_Rhodopirellula | 397 | PSR62_RS13055 | 3698716..3699909 | 2127060 | ATP-binding-protein-[Rhodopirellula-sp.-P2] |
| WP_274403444.1 | Schlafen+HATPase+wHTH | GCF_028768465.1 | Rhodopirellula_sp._P2 | Bacteria>Pseudomonadati>PVC_group>Planctomycetota>Planctomycetia>Pirellulales>Pirellulaceae>Rhodopirellula>unclassified_Rhodopirellula | 424 | PSR62_RS13030 | 3690026..3691300 | 2127060 | ATP-binding-protein-[Rhodopirellula-sp.-P2] |
| WP_182870883.1 | Schlafen+HATPase+wHTH | GCF_014154935.1 | Rhodopirellula_sp._JC639 | Bacteria>Pseudomonadati>PVC_group>Planctomycetota>Planctomycetia>Pirellulales>Pirellulaceae>Rhodopirellula>unclassified_Rhodopirellula | 570 | H1B37_RS29505 | 18783..20495 | 2755585 | RNA-binding-domain-containing-protein-[Rhodopirellula-sp.-JC639] |
| WP_153558878.1 | Schlafen+HATPase+wHTH | GCF_009618275.1 | Roseimaritima_sediminicola | Bacteria>Pseudomonadati>PVC_group>Planctomycetota>Planctomycetia>Pirellulales>Pirellulaceae>Roseimaritima | 537 | GE027_RS19395 | 36895..38508 | 2662066 | ATP-binding-protein-[Roseimaritima-sediminicola] |
| WP_145350722.1 | Schlafen+HATPase+wHTH | GCF_007741495.1 | Roseimaritima_multifibrata | Bacteria>Pseudomonadati>PVC_group>Planctomycetota>Planctomycetia>Pirellulales>Pirellulaceae>Roseimaritima | 539 | FF011L_RS05765 | 1538938..1540557 | 1930274 | ATP-binding-protein-[Roseimaritima-multifibrata] |
| WP_146459046.1 | Schlafen+HATPase+wHTH | GCF_007860125.1 | Rubripirellula_tenax | Bacteria>Pseudomonadati>PVC_group>Planctomycetota>Planctomycetia>Pirellulales>Pirellulaceae>Rubripirellula | 316 | Poly51_RS17420 | 736882..737832 | 2528015 | ATP-binding-protein-[Rubripirellula-tenax] |
| GIW99317.1 | Schlafen+HATPase+wHTH | GCA_026004795.1 | Pirellulaceae_bacterium | Bacteria>Pseudomonadati>PVC_group>Planctomycetota>Planctomycetia>Pirellulales>Pirellulaceae>unclassified_Pirellulaceae | 438 | KatS3mg111_2650 | 703569..704885 | 2699754 | MAG:-ATP-dependent-DNA-helicase-[Pirellulaceae-bacterium] |
| MCL6502994.1 | Schlafen+HATPase+wHTH | GCA_023511675.1 | Pirellulales_bacterium | Bacteria>Pseudomonadati>PVC_group>Planctomycetota>Planctomycetia>Pirellulales>unclassified_Pirellulales | 375 | K6T86_09945 | 18632..19759 | 2715253 | helix-turn-helix-domain-containing-protein-[Pirellulales-bacterium] |
| WP_145451935.1 | Schlafen+HATPase+wHTH | GCF_007748155.1 | Gimesia_panareensis | Bacteria>Pseudomonadati>PVC_group>Planctomycetota>Planctomycetia>Planctomycetales>Planctomycetaceae>Gimesia | 542 | Enr10x_RS26155 | 6959632..6961260 | 2527978 | ATP-binding-protein-[Gimesia-panareensis] |
| WP_145380204.1 | Schlafen+HATPase+wHTH | GCF_007747995.1 | Symmachiella_dynata | Bacteria>Pseudomonadati>PVC_group>Planctomycetota>Planctomycetia>Planctomycetales>Planctomycetaceae>Symmachiella | 580 | Mal52_RS28000 | 7357514..7359256 | 2527995 | putative-DNA-binding-domain-containing-protein-[Symmachiella-dynata] |
| HBJ35662.1 | Schlafen+HATPase+wHTH | GCA_003506095.1 | Planctomycetaceae_bacterium | Bacteria>Pseudomonadati>PVC_group>Planctomycetota>Planctomycetia>Planctomycetales>Planctomycetaceae>unclassified_Planctomycetaceae | 539 | DDZ51_13140 | 14863..16482 | 2026779 | transcriptional-regulator-[Planctomycetaceae-bacterium] |
| MCA9126639.1 | Schlafen+HATPase+wHTH | GCA_020426625.1 | Planctomycetales_bacterium | Bacteria>Pseudomonadati>PVC_group>Planctomycetota>Planctomycetia>Planctomycetales>unclassified_Planctomycetales | 397 | KDB22_06130 | 11354..12547 | 2053591 | putative-DNA-binding-domain-containing-protein-[Planctomycetales-bacterium] |
| GHT10311.1 | Schlafen+HATPase+wHTH | GCA_026001785.1 | Planctomycetales_bacterium | Bacteria>Pseudomonadati>PVC_group>Planctomycetota>Planctomycetia>Planctomycetales>unclassified_Planctomycetales | 161 | FACS1894170_02210 | 19459..19944 | 2053591 | hypothetical-protein-FACS1894170_02210-[Planctomycetales-bacterium] |
| QDU83129.1 | Schlafen+HATPase+wHTH | GCA_007750655.1 | Rohdeia_mirabilis | Bacteria>Pseudomonadati>PVC_group>Planctomycetota>Planctomycetia>Planctomycetia_incertae_sedis>Rohdeia | 402 | Pla163_02260 | 286903..288111 | 2528008 | Divergent-AAA-domain-protein-[Planctomycetes-bacterium-Pla163] |
| MBE7458018.1 | Schlafen+HATPase+wHTH | GCA_015075705.1 | Planctomycetia_bacterium | Bacteria>Pseudomonadati>PVC_group>Planctomycetota>Planctomycetia>unclassified_Planctomycetia | 547 | HS102_15575 | 51689..53332 | 2052181 | putative-DNA-binding-domain-containing-protein-[Planctomycetia-bacterium] |
| PIE22548.1 | Schlafen+HATPase+wHTH | GCA_002748355.1 | Planctomycetota_bacterium | Bacteria>Pseudomonadati>PVC_group>Planctomycetota>unclassified_Planctomycetota | 426 | CSA62_11860 | 34716..35996 | 2026780 | MAG:-transcriptional-regulator-[Planctomycetota-bacterium] |
| OQZ06363.1 | Schlafen+HATPase+wHTH | GCA_002050205.1 | Planctomycetes_bacterium_UTPLA1 | Bacteria>Pseudomonadati>PVC_group>Planctomycetota>unclassified_Planctomycetota | 624 | B6D36_05370 | 940..2814 | 1961112 | hypothetical-protein-B6D36_05370-[Planctomycetes-bacterium-UTPLA1] |
| MCX5673911.1 | Schlafen+HATPase+wHTH | GCA_026387015.1 | Planctomycetota_bacterium | Bacteria>Pseudomonadati>PVC_group>Planctomycetota>unclassified_Planctomycetota | 561 | NTX87_02785 | 11402..13087 | 2026780 | MAG:-putative-DNA-binding-domain-containing-protein-[Planctomycetota-bacterium] |
| MCC6785234.1 | Schlafen+HATPase+wHTH | GCA_020848735.1 | Planctomycetota_bacterium | Bacteria>Pseudomonadati>PVC_group>Planctomycetota>unclassified_Planctomycetota | 429 | IT457_20460 | 14581..15870 | 2026780 | putative-DNA-binding-domain-containing-protein-[Planctomycetota-bacterium] |
| MBW8002867.1 | Schlafen+HATPase+wHTH | GCA_019456255.1 | Planctomycetota_bacterium | Bacteria>Pseudomonadati>PVC_group>Planctomycetota>unclassified_Planctomycetota | 473 | FVQ80_12740 | 6645..8066 | 2026780 | HTH-domain-containing-protein-[Planctomycetota-bacterium] |
| MBM4038870.1 | Schlafen+HATPase+wHTH | GCA_016872855.1 | Planctomycetota_bacterium | Bacteria>Pseudomonadati>PVC_group>Planctomycetota>unclassified_Planctomycetota | 453 | FJ290_10175 | 10205..11566 | 2026780 | hypothetical-protein-[Planctomycetota-bacterium] |
| MBL4771869.1 | Schlafen+HATPase+wHTH | GCA_016764635.1 | Planctomycetota_bacterium | Bacteria>Pseudomonadati>PVC_group>Planctomycetota>unclassified_Planctomycetota | 368 | JKY61_12195 | 1..1106 | 2026780 | putative-DNA-binding-domain-containing-protein-[Planctomycetota-bacterium] |
| MBK9385765.1 | Schlafen+HATPase+wHTH | GCA_016719295.1 | Planctomycetota_bacterium | Bacteria>Pseudomonadati>PVC_group>Planctomycetota>unclassified_Planctomycetota | 562 | IPN34_13225 | 3522890..3524578 | 2026780 | putative-DNA-binding-domain-containing-protein-[Planctomycetota-bacterium] |
| MBI4882200.1 | Schlafen+HATPase+wHTH | GCA_016213895.1 | Planctomycetota_bacterium | Bacteria>Pseudomonadati>PVC_group>Planctomycetota>unclassified_Planctomycetota | 614 | HY812_21440 | 863..2707 | 2026780 | putative-DNA-binding-domain-containing-protein-[Planctomycetota-bacterium] |
| MBI4586091.1 | Schlafen+HATPase+wHTH | GCA_016209605.1 | Planctomycetota_bacterium | Bacteria>Pseudomonadati>PVC_group>Planctomycetota>unclassified_Planctomycetota | 558 | HY717_18940 | 12917..14593 | 2026780 | putative-DNA-binding-domain-containing-protein-[Planctomycetota-bacterium] |
| MBI4584107.1 | Schlafen+HATPase+wHTH | GCA_016209605.1 | Planctomycetota_bacterium | Bacteria>Pseudomonadati>PVC_group>Planctomycetota>unclassified_Planctomycetota | 257 | HY717_08800 | 42660..43433 | 2026780 | hypothetical-protein-[Planctomycetota-bacterium] |
| MBI2193009.1 | Schlafen+HATPase+wHTH | GCA_016179685.1 | Planctomycetota_bacterium | Bacteria>Pseudomonadati>PVC_group>Planctomycetota>unclassified_Planctomycetota | 453 | HYU36_13605 | 47988..49349 | 2026780 | putative-DNA-binding-domain-containing-protein-[Planctomycetota-bacterium] |
| MBI2191316.1 | Schlafen+HATPase+wHTH | GCA_016179685.1 | Planctomycetota_bacterium | Bacteria>Pseudomonadati>PVC_group>Planctomycetota>unclassified_Planctomycetota | 361 | HYU36_04960 | 79705..80790 | 2026780 | hypothetical-protein-[Planctomycetota-bacterium] |
| KAA3609102.1 | Schlafen+HATPase+wHTH | GCA_008501825.1 | Planctomycetota_bacterium | Bacteria>Pseudomonadati>PVC_group>Planctomycetota>unclassified_Planctomycetota | 430 | DWQ01_09785 | 13608..14900 | 2026780 | MAG:-ATP-dependent-DNA-helicase-RecG-[Planctomycetota-bacterium] |
| KAA3609100.1 | Schlafen+HATPase+wHTH | GCA_008501825.1 | Planctomycetota_bacterium | Bacteria>Pseudomonadati>PVC_group>Planctomycetota>unclassified_Planctomycetota | 546 | DWQ01_09770 | 6243..7883 | 2026780 | MAG:-transcriptional-regulator-[Planctomycetota-bacterium] |
| MCE0498494.1 | Schlafen+HATPase+wHTH | GCA_021272315.1 | Candidatus_Methylacidiphilales_bacterium | Bacteria>Pseudomonadati>PVC_group>Verrucomicrobiota>Methylacidiphilae>Methylacidiphilales>unclassified_Candidatus_Methylacidiphilales | 329 | LV481_11170 | 13029..14018 | 2720631 | putative-DNA-binding-domain-containing-protein-[Methylacidiphilales-bacterium] |
| MBL9185850.1 | Schlafen+HATPase+wHTH | GCA_016794785.1 | Opitutaceae_bacterium | Bacteria>Pseudomonadati>PVC_group>Verrucomicrobiota>Opitutia>Opitutales>Opitutaceae>unclassified_Opitutaceae | 619 | JNK23_00085 | 27681..29540 | 2006848 | putative-DNA-binding-domain-containing-protein-[Opitutaceae-bacterium] |
| MBT62429.1 | Schlafen+HATPase+wHTH | GCA_002729725.1 | Puniceicoccaceae_bacterium | Bacteria>Pseudomonadati>PVC_group>Verrucomicrobiota>Opitutia>Puniceicoccales>Puniceicoccaceae>unclassified_Puniceicoccaceae | 492 | CML13_04365 | 13592..15070 | 2026784 | AAA-family-ATPase-[Puniceicoccaceae-bacterium] |
| NBB80460.1 | Schlafen+HATPase+wHTH | GCA_009908325.1 | Verrucomicrobiota_bacterium | Bacteria>Pseudomonadati>PVC_group>Verrucomicrobiota>unclassified_Verrucomicrobiota | 493 | GVY36_13630 | 14787..16268 | 2026799 | winged-helix-turn-helix-transcriptional-regulator-[Verrucomicrobiota-bacterium] |
| MCF7762648.1 | Schlafen+HATPase+wHTH | GCA_021733945.1 | Verrucomicrobiota_bacterium | Bacteria>Pseudomonadati>PVC_group>Verrucomicrobiota>unclassified_Verrucomicrobiota | 567 | K9N62_03155 | 19161..20864 | 2026799 | putative-DNA-binding-domain-containing-protein-[Verrucomicrobiota-bacterium] |
| MCF7707717.1 | Schlafen+HATPase+wHTH | GCA_021733835.1 | Verrucomicrobiota_bacterium | Bacteria>Pseudomonadati>PVC_group>Verrucomicrobiota>unclassified_Verrucomicrobiota | 535 | K9N52_02310 | 35957..37564 | 2026799 | putative-DNA-binding-domain-containing-protein-[Verrucomicrobiota-bacterium] |
| MCF7668416.1 | Schlafen+HATPase+wHTH | GCA_021733785.1 | Verrucomicrobiota_bacterium | Bacteria>Pseudomonadati>PVC_group>Verrucomicrobiota>unclassified_Verrucomicrobiota | 383 | K9N48_01415 | 8950..10101 | 2026799 | putative-DNA-binding-domain-containing-protein-[Verrucomicrobiota-bacterium] |
| MCC5848620.1 | Schlafen+HATPase+wHTH | GCA_020831635.1 | Verrucomicrobiota_bacterium | Bacteria>Pseudomonadati>PVC_group>Verrucomicrobiota>unclassified_Verrucomicrobiota | 542 | JJU29_11035 | 1..1626 | 2026799 | putative-DNA-binding-domain-containing-protein-[Verrucomicrobiota-bacterium] |
| MCC5847924.1 | Schlafen+HATPase+wHTH | GCA_020831635.1 | Verrucomicrobiota_bacterium | Bacteria>Pseudomonadati>PVC_group>Verrucomicrobiota>unclassified_Verrucomicrobiota | 573 | JJU29_07495 | 20277..21998 | 2026799 | putative-DNA-binding-domain-containing-protein-[Verrucomicrobiota-bacterium] |
| MBP7946077.1 | Schlafen+HATPase+wHTH | GCA_018003215.1 | Verrucomicrobiota_bacterium | Bacteria>Pseudomonadati>PVC_group>Verrucomicrobiota>unclassified_Verrucomicrobiota | 229 | KA191_08405 | 3770..4459 | 2026799 | putative-DNA-binding-domain-containing-protein-[Verrucomicrobiota-bacterium] |
| MBN9688599.1 | Schlafen+HATPase+wHTH | GCA_017309115.1 | Verrucomicrobiota_bacterium | Bacteria>Pseudomonadati>PVC_group>Verrucomicrobiota>unclassified_Verrucomicrobiota | 412 | J0M24_00040 | 8530..9768 | 2026799 | putative-DNA-binding-domain-containing-protein-[Verrucomicrobiota-bacterium] |
| MBM3879661.1 | Schlafen+HATPase+wHTH | GCA_016871715.1 | Verrucomicrobiota_bacterium | Bacteria>Pseudomonadati>PVC_group>Verrucomicrobiota>unclassified_Verrucomicrobiota | 573 | FJ387_08055 | 13759..15480 | 2026799 | hypothetical-protein-[Verrucomicrobiota-bacterium] |
| MBM3879659.1 | Schlafen+HATPase+wHTH | GCA_016871715.1 | Verrucomicrobiota_bacterium | Bacteria>Pseudomonadati>PVC_group>Verrucomicrobiota>unclassified_Verrucomicrobiota | 396 | FJ387_08045 | 11515..12705 | 2026799 | transcriptional-regulator-[Verrucomicrobiota-bacterium] |
| MBI4023513.1 | Schlafen+HATPase+wHTH | GCA_016198745.1 | Verrucomicrobiota_bacterium | Bacteria>Pseudomonadati>PVC_group>Verrucomicrobiota>unclassified_Verrucomicrobiota | 587 | HY360_00935 | 15403..17166 | 2026799 | putative-DNA-binding-domain-containing-protein-[Verrucomicrobiota-bacterium] |
| KAB2639240.1 | Schlafen+HATPase+wHTH | GCA_008932935.1 | Verrucomicrobiota_bacterium | Bacteria>Pseudomonadati>PVC_group>Verrucomicrobiota>unclassified_Verrucomicrobiota | 547 | DVB25_06020 | 6622..8265 | 2026799 | transcriptional-regulator-[Verrucomicrobiota-bacterium] |
| HCF94154.1 | Schlafen+HATPase+wHTH | GCA_003534995.1 | Verrucomicrobiota_bacterium | Bacteria>Pseudomonadati>PVC_group>Verrucomicrobiota>unclassified_Verrucomicrobiota | 554 | DEW46_03765 | 4903..6567 | 2026799 | ATPase-[Verrucomicrobiota-bacterium] |
| MCP5535437.1 | Schlafen+HATPase+wHTH | GCA_024233115.1 | Akkermansiaceae_bacterium | Bacteria>Pseudomonadati>PVC_group>Verrucomicrobiota>Verrucomicrobiia>Verrucomicrobiales>Akkermansiaceae>unclassified_Akkermansiaceae | 464 | H7A51_04285 | 27334..28728 | 2562705 | putative-DNA-binding-domain-containing-protein-[Akkermansiaceae-bacterium] |
| MBX3742595.1 | Schlafen+HATPase+wHTH | GCA_019634595.1 | Akkermansiaceae_bacterium | Bacteria>Pseudomonadati>PVC_group>Verrucomicrobiota>Verrucomicrobiia>Verrucomicrobiales>Akkermansiaceae>unclassified_Akkermansiaceae | 628 | KF712_16530 | 318441..320327 | 2562705 | putative-DNA-binding-domain-containing-protein-[Akkermansiaceae-bacterium] |
| WP_142661085.1 | Schlafen+HATPase+wHTH | GCF_902143375.2 | Methylacidimicrobium_tartarophylax | Bacteria>Pseudomonadati>PVC_group>Verrucomicrobiota>Verrucomicrobiota_incertae_sedis>Methylacidimicrobium | 415 | MAMT_RS11065 | 1216..2463 | 1041768 | ATP-binding-protein-[Methylacidimicrobium-tartarophylax] |
| SLM14722.1 | Schlafen+HATPase+wHTH | GCA_900163985.1 | uncultured_spirochete | Bacteria>Pseudomonadati>Spirochaetota>Spirochaetia>Spirochaetales>environmental_samples | 479 | SPIROBIBN47_350049 | 55782..57221 | 156406 | ATPase-AAA-[uncultured-spirochete] |
| WP_117329278.1 | Schlafen+HATPase+wHTH | GCF_003429665.1 | Sphaerochaeta_halotolerans | Bacteria>Pseudomonadati>Spirochaetota>Spirochaetia>Spirochaetales>Sphaerochaetaceae>Sphaerochaeta | 592 | DYP60_RS02425 | 187573..189351 | 2293840 | putative-DNA-binding-domain-containing-protein-[Sphaerochaeta-halotolerans] |
| RFU95879.1 | Schlafen+HATPase+wHTH | GCA_003429665.1 | Sphaerochaeta_halotolerans | Bacteria>Pseudomonadati>Spirochaetota>Spirochaetia>Spirochaetales>Sphaerochaetaceae>Sphaerochaeta | 592 | DYP60_02430 | 187573..189351 | 2293840 | AAA-family-ATPase-[Sphaerochaeta-halotolerans] |
| MCH3907671.1 | Schlafen+HATPase+wHTH | GCA_022482495.1 | Sphaerochaeta_sp. | Bacteria>Pseudomonadati>Spirochaetota>Spirochaetia>Spirochaetales>Sphaerochaetaceae>Sphaerochaeta>unclassified_Sphaerochaeta | 507 | LKE28_05355 | 260813..262336 | 1972642 | putative-DNA-binding-domain-containing-protein-[Sphaerochaeta-sp.] |
| MBO7485538.1 | Schlafen+HATPase+wHTH | GCA_017649985.1 | Spirochaetaceae_bacterium | Bacteria>Pseudomonadati>Spirochaetota>Spirochaetia>Spirochaetales>Spirochaetaceae>unclassified_Spirochaetaceae | 491 | J6T84_05705 | 6402..7877 | 1898206 | putative-DNA-binding-domain-containing-protein-[Spirochaetaceae-bacterium] |
| WP_010252933.1 | Schlafen+HATPase+wHTH | GCF_000297095.1 | Treponema_primitia | Bacteria>Pseudomonadati>Spirochaetota>Spirochaetia>Spirochaetales>Treponemataceae>Treponema | 484 | TPRIMZ1_RS18035 | 28004..29458 | 88058 | putative-DNA-binding-domain-containing-protein-[Treponema-primitia] |
| WP_132781621.1 | Schlafen+HATPase+wHTH | GCF_004343725.1 | Treponema_sp._J25 | Bacteria>Pseudomonadati>Spirochaetota>Spirochaetia>Spirochaetales>Treponemataceae>Treponema>unclassified_Treponema | 483 | C5O22_RS10555 | 23459..24910 | 2094121 | putative-DNA-binding-domain-containing-protein-[Treponema-sp.-J25] |
| MCR5123680.1 | Schlafen+HATPase+wHTH | GCA_024697755.1 | Treponema_sp. | Bacteria>Pseudomonadati>Spirochaetota>Spirochaetia>Spirochaetales>Treponemataceae>Treponema>unclassified_Treponema | 559 | K6B43_00610 | 3982..5661 | 166 | putative-DNA-binding-domain-containing-protein-[Treponema-sp.] |
| MBQ5908483.1 | Schlafen+HATPase+wHTH | GCA_017381365.1 | Treponema_sp. | Bacteria>Pseudomonadati>Spirochaetota>Spirochaetia>Spirochaetales>Treponemataceae>Treponema>unclassified_Treponema | 560 | IIW80_10585 | 8765..10447 | 166 | putative-DNA-binding-domain-containing-protein-[Treponema-sp.] |
| MDD6486201.1 | Schlafen+HATPase+wHTH | GCA_029227075.1 | Spirochaetales_bacterium | Bacteria>Pseudomonadati>Spirochaetota>Spirochaetia>Spirochaetales>unclassified_Spirochaetales | 513 | PUF61_04580 | 320383..321924 | 2026792 | MAG:-putative-DNA-binding-domain-containing-protein-[Spirochaetales-bacterium] |
| MDA3832125.1 | Schlafen+HATPase+wHTH | GCA_027858175.1 | Spirochaetales_bacterium | Bacteria>Pseudomonadati>Spirochaetota>Spirochaetia>Spirochaetales>unclassified_Spirochaetales | 489 | PF495_01890 | 3602..5071 | 2026792 | MAG:-putative-DNA-binding-domain-containing-protein-[Spirochaetales-bacterium] |
| MBR5915374.1 | Schlafen+HATPase+wHTH | GCA_017554185.1 | Spirochaetia_bacterium | Bacteria>Pseudomonadati>Spirochaetota>Spirochaetia>unclassified_Spirochaetia | 414 | IKZ57_02870 | 12015..13259 | 2053615 | ATP-binding-protein-[Spirochaetia-bacterium] |
| GHT84085.1 | Schlafen+HATPase+wHTH | GCA_026002105.1 | Spirochaetia_bacterium | Bacteria>Pseudomonadati>Spirochaetota>Spirochaetia>unclassified_Spirochaetia | 400 | FACS1894137_06700 | 7502..8704 | 2053615 | hypothetical-protein-FACS1894137_06700-[Spirochaetia-bacterium] |
| GHT84066.1 | Schlafen+HATPase+wHTH | GCA_026002105.1 | Spirochaetia_bacterium | Bacteria>Pseudomonadati>Spirochaetota>Spirochaetia>unclassified_Spirochaetia | 400 | FACS1894137_06650 | 2570..3772 | 2053615 | hypothetical-protein-FACS1894137_06650-[Spirochaetia-bacterium] |
| TXT43485.1 | Schlafen+HATPase+wHTH | GCA_008080685.1 | Spirochaetota_bacterium | Bacteria>Pseudomonadati>Spirochaetota>unclassified_Spirochaetota | 150 | FD137_1823 | 15566..16018 | 2202144 | hypothetical-protein-FD137_1823-[Spirochaetota-bacterium] |
| OQB94578.1 | Schlafen+HATPase+wHTH | GCA_002070755.1 | Spirochaetes_bacterium_ADurb.Bin110 | Bacteria>Pseudomonadati>Spirochaetota>unclassified_Spirochaetota | 560 | BWX81_02437 | 20207..21889 | 1852908 | Divergent-AAA-domain-protein-[Spirochaetes-bacterium-ADurb.Bin110] |
| MDD3581158.1 | Schlafen+HATPase+wHTH | GCA_028695935.1 | Desulfobacca_sp. | Bacteria>Pseudomonadati>Thermodesulfobacteriota>Desulfobaccia>Desulfobaccales>Desulfobaccaceae>Desulfobacca>unclassified_Desulfobacca | 564 | PHW74_09075 | 10906..12600 | 2067990 | MAG:-ATP-binding-protein-[Desulfobacca-sp.] |
| WP_028321660.1 | Schlafen+HATPase+wHTH | GCF_000422285.1 | Desulfatiglans_anilini | Bacteria>Pseudomonadati>Thermodesulfobacteriota>Desulfobacteria>Desulfatiglandales>Desulfatiglandaceae>Desulfatiglans | 518 | H567_RS0112515 | 314..1870 | 90728 | putative-DNA-binding-domain-containing-protein-[Desulfatiglans-anilini] |
| KPA14789.1 | Schlafen+HATPase+wHTH | GCA_001292585.1 | Candidatus_Magnetomorum_sp._HK-1 | Bacteria>Pseudomonadati>Thermodesulfobacteriota>Desulfobacteria>Desulfobacterales>Candidatus_Magnetomoraceae>Candidatus_Magnetomorum>unclassified_Candidatus_Magnetomorum | 522 | MHK_004992 | 2929..4497 | 1509431 | ATPase-AAA-[Candidatus-Magnetomorum-sp.-HK-1] |
| KPA09726.1 | Schlafen+HATPase+wHTH | GCA_001292585.1 | Candidatus_Magnetomorum_sp._HK-1 | Bacteria>Pseudomonadati>Thermodesulfobacteriota>Desulfobacteria>Desulfobacterales>Candidatus_Magnetomoraceae>Candidatus_Magnetomorum>unclassified_Candidatus_Magnetomorum | 551 | MHK_010066 | 4020..5675 | 1509431 | divergent-AAA-domain-protein-[Candidatus-Magnetomorum-sp.-HK-1] |
| WP_073478765.1 | Schlafen+HATPase+wHTH | GCF_900142135.1 | Desulfatibacillum_alkenivorans | Bacteria>Pseudomonadati>Thermodesulfobacteriota>Desulfobacteria>Desulfobacterales>Desulfatibacillaceae>Desulfatibacillum | 569 | BUE18_RS24410 | 21925..23634 | 259354 | ATP-binding-protein-[Desulfatibacillum-alkenivorans] |
| WP_028324411.1 | Schlafen+HATPase+wHTH | GCF_000429925.1 | Desulfatirhabdium_butyrativorans | Bacteria>Pseudomonadati>Thermodesulfobacteriota>Desulfobacteria>Desulfobacterales>Desulfatirhabdiaceae>Desulfatirhabdium | 478 | G492_RS0109350 | 76406..77842 | 340467 | putative-DNA-binding-domain-containing-protein-[Desulfatirhabdium-butyrativorans] |
| WP_087881809.1 | Schlafen+HATPase+wHTH | GCF_900170035.1 | Desulfamplus_magnetovallimortis | Bacteria>Pseudomonadati>Thermodesulfobacteriota>Desulfobacteria>Desulfobacterales>Desulfobacteraceae>Desulfamplus | 613 | MTBBW1_RS20325 | 66277..68118 | 1246637 | putative-DNA-binding-domain-containing-protein-[Desulfamplus-magnetovallimortis] |
| SLM29543.1 | Schlafen+HATPase+wHTH | GCA_900170035.1 | Desulfamplus_magnetovallimortis | Bacteria>Pseudomonadati>Thermodesulfobacteriota>Desulfobacteria>Desulfobacterales>Desulfobacteraceae>Desulfamplus | 613 | MTBBW1_1860033 | 40262..42103 | 1246637 | putative-transcriptional-regulator-[Desulfamplus-magnetovallimortis] |
| HBT88576.1 | Schlafen+HATPase+wHTH | GCA_003510365.1 | Desulfobacter_sp. | Bacteria>Pseudomonadati>Thermodesulfobacteriota>Desulfobacteria>Desulfobacterales>Desulfobacteraceae>Desulfobacter>unclassified_Desulfobacter | 470 | DEB50_08950 | 32924..34336 | 2294 | AAA-family-ATPase-[Desulfobacter-sp.] |
| HBT88456.1 | Schlafen+HATPase+wHTH | GCA_003510365.1 | Desulfobacter_sp. | Bacteria>Pseudomonadati>Thermodesulfobacteriota>Desulfobacteria>Desulfobacterales>Desulfobacteraceae>Desulfobacter>unclassified_Desulfobacter | 613 | DEB50_08325 | 31493..33334 | 2294 | transcriptional-regulator-[Desulfobacter-sp.] |
| WP_024334644.1 | Schlafen+HATPase+wHTH | GCF_000421285.1 | Desulfotignum_balticum | Bacteria>Pseudomonadati>Thermodesulfobacteriota>Desulfobacteria>Desulfobacterales>Desulfobacteraceae>Desulfotignum | 574 | K365_RS0111335 | 2240636..2242360 | 115781 | ATP-binding-protein-[Desulfotignum-balticum] |
| RZB36553.1 | Schlafen+HATPase+wHTH | GCA_004193595.1 | Desulfobacteraceae_bacterium_Eth-SRB2 | Bacteria>Pseudomonadati>Thermodesulfobacteriota>Desulfobacteria>Desulfobacterales>Desulfobacteraceae>unclassified_Desulfobacteraceae | 419 | SRB2_02049 | 65458..66717 | 2528438 | hypothetical-protein-SRB2_02049-[Desulfobacteraceae-bacterium-Eth-SRB2] |
| RJR29736.1 | Schlafen+HATPase+wHTH | GCA_003599615.1 | Desulfobacteraceae_bacterium | Bacteria>Pseudomonadati>Thermodesulfobacteriota>Desulfobacteria>Desulfobacterales>Desulfobacteraceae>unclassified_Desulfobacteraceae | 569 | C4576_34880 | 11173..12882 | 2049433 | AAA-family-ATPase-[Desulfobacteraceae-bacterium] |
| MCP3943394.1 | Schlafen+HATPase+wHTH | GCA_024226935.1 | Desulfobacteraceae_bacterium | Bacteria>Pseudomonadati>Thermodesulfobacteriota>Desulfobacteria>Desulfobacterales>Desulfobacteraceae>unclassified_Desulfobacteraceae | 382 | GY710_18180 | 5918..7066 | 2049433 | ATP-dependent-DNA-helicase-RecG-[Desulfobacteraceae-bacterium] |
| MBL7204917.1 | Schlafen+HATPase+wHTH | GCA_016783005.1 | Desulfobacteraceae_bacterium | Bacteria>Pseudomonadati>Thermodesulfobacteriota>Desulfobacteria>Desulfobacterales>Desulfobacteraceae>unclassified_Desulfobacteraceae | 127 | ISS63_11395 | 12335..12718 | 2049433 | ATP-binding-protein-[Desulfobacteraceae-bacterium] |
| MBL7204911.1 | Schlafen+HATPase+wHTH | GCA_016783005.1 | Desulfobacteraceae_bacterium | Bacteria>Pseudomonadati>Thermodesulfobacteriota>Desulfobacteria>Desulfobacterales>Desulfobacteraceae>unclassified_Desulfobacteraceae | 434 | ISS63_11365 | 2683..3987 | 2049433 | putative-DNA-binding-domain-containing-protein-[Desulfobacteraceae-bacterium] |
| MBA3018064.1 | Schlafen+HATPase+wHTH | GCA_013792055.1 | Desulfobacteraceae_bacterium | Bacteria>Pseudomonadati>Thermodesulfobacteriota>Desulfobacteria>Desulfobacterales>Desulfobacteraceae>unclassified_Desulfobacteraceae | 483 | FP811_07945 | 14452..15903 | 2049433 | AAA-family-ATPase-[Desulfobacteraceae-bacterium] |
| HDR14880.1 | Schlafen+HATPase+wHTH | GCA_011048855.1 | Desulfobacteraceae_bacterium | Bacteria>Pseudomonadati>Thermodesulfobacteriota>Desulfobacteria>Desulfobacterales>Desulfobacteraceae>unclassified_Desulfobacteraceae | 507 | ENN79_05215 | 5052..6575 | 2049433 | winged-helix-turn-helix-transcriptional-regulator-[Desulfobacteraceae-bacterium] |
| MBC8360092.1 | Schlafen+HATPase+wHTH | GCA_014382665.1 | Candidatus_Desulfatibia_profunda | Bacteria>Pseudomonadati>Thermodesulfobacteriota>Desulfobacteria>Desulfobacterales>Desulfobacterales_incertae_sedis>Candidatus_Desulfatibia | 502 | H8E23_01675 | 3237..4745 | 2841695 | putative-DNA-binding-domain-containing-protein-[Desulfobacterales-bacterium] |
| MBC8359963.1 | Schlafen+HATPase+wHTH | GCA_014382665.1 | Candidatus_Desulfatibia_profunda | Bacteria>Pseudomonadati>Thermodesulfobacteriota>Desulfobacteria>Desulfobacterales>Desulfobacterales_incertae_sedis>Candidatus_Desulfatibia | 88 | H8E23_01010 | 699..965 | 2841695 | hypothetical-protein-[Desulfobacterales-bacterium] |
| CBX26789.1 | Schlafen+HATPase+wHTH | - | uncultured_Desulfobacterium_sp. | Bacteria>Pseudomonadati>Thermodesulfobacteriota>Desulfobacteria>Desulfobacterales>Desulfobacteriaceae>Desulfobacterium>environmental_samples | 463 | N47_A08180 | 91023..92414 | 201089 | hypothetical-protein-N47_A08180-[uncultured-Desulfobacterium-sp.] |
| WP_207688114.1 | Schlafen+HATPase+wHTH | GCF_017377355.1 | Desulfonema_limicola | Bacteria>Pseudomonadati>Thermodesulfobacteriota>Desulfobacteria>Desulfobacterales>Desulfococcaceae>Desulfonema | 483 | dnl_RS20635 | 4838306..4839757 | 45656 | helix-turn-helix-domain-containing-protein-[Desulfonema-limicola] |
| WP_083453313.1 | Schlafen+HATPase+wHTH | GCF_001293685.1 | Desulfatitalea_tepidiphila | Bacteria>Pseudomonadati>Thermodesulfobacteriota>Desulfobacteria>Desulfobacterales>Desulfosarcinaceae>Desulfatitalea | 401 | DFT_RS04415 | 1016484..1017689 | 1185843 | ATP-binding-protein-[Desulfatitalea-tepidiphila] |
| WP_027355265.1 | Schlafen+HATPase+wHTH | GCF_028752835.1 | Desulfosarcina_sp._BuS5 | Bacteria>Pseudomonadati>Thermodesulfobacteriota>Desulfobacteria>Desulfobacterales>Desulfosarcinaceae>Desulfosarcina>unclassified_Desulfosarcina | 485 | A39W_RS0118880 | 9162..10619 | 933262 | ATP-binding-protein-[Desulfosarcina-sp.-BuS5] |
| MCF6289520.1 | Schlafen+HATPase+wHTH | GCA_021647905.1 | Desulfobacterales_bacterium | Bacteria>Pseudomonadati>Thermodesulfobacteriota>Desulfobacteria>Desulfobacterales>unclassified_Desulfobacterales | 559 | L3J03_00735 | 79204..80883 | 2044940 | putative-DNA-binding-domain-containing-protein-[Desulfobacterales-bacterium] |
| MBN1931787.1 | Schlafen+HATPase+wHTH | GCA_016931455.1 | Desulfobacterales_bacterium | Bacteria>Pseudomonadati>Thermodesulfobacteriota>Desulfobacteria>Desulfobacterales>unclassified_Desulfobacterales | 482 | JW786_09295 | 22907..24355 | 2044940 | putative-DNA-binding-domain-containing-protein-[Desulfobacterales-bacterium] |
| MBF0121239.1 | Schlafen+HATPase+wHTH | GCA_015228685.1 | Desulfobacterales_bacterium | Bacteria>Pseudomonadati>Thermodesulfobacteriota>Desulfobacteria>Desulfobacterales>unclassified_Desulfobacterales | 489 | HQK79_20590 | 8755..10224 | 2044940 | putative-DNA-binding-domain-containing-protein-[Desulfobacterales-bacterium] |
| MBC8417233.1 | Schlafen+HATPase+wHTH | GCA_014382905.1 | Desulfobacterales_bacterium | Bacteria>Pseudomonadati>Thermodesulfobacteriota>Desulfobacteria>Desulfobacterales>unclassified_Desulfobacterales | 563 | H8E10_01395 | 9500..11191 | 2044940 | putative-DNA-binding-domain-containing-protein-[Desulfobacterales-bacterium] |
| WP_205243272.1 | Schlafen+HATPase+wHTH | GCF_016918545.1 | Desulfobulbus_alkaliphilus | Bacteria>Pseudomonadati>Thermodesulfobacteriota>Desulfobulbia>Desulfobulbales>Desulfobulbaceae>Desulfobulbus | 499 | JWG43_RS12495 | 44101..45600 | 869814 | ATP-binding-protein-[Desulfobulbus-alkaliphilus] |
| KGO32871.1 | Schlafen+HATPase+wHTH | GCA_000769715.1 | Desulfobulbus_sp._Tol-SR | Bacteria>Pseudomonadati>Thermodesulfobacteriota>Desulfobulbia>Desulfobulbales>Desulfobulbaceae>Desulfobulbus>unclassified_Desulfobulbus | 385 | JT06_17865 | 10047..11204 | 1536652 | ATP-dependent-DNA-helicase-RecG-[Desulfobulbus-sp.-Tol-SR] |
| KGO32868.1 | Schlafen+HATPase+wHTH | GCA_000769715.1 | Desulfobulbus_sp._Tol-SR | Bacteria>Pseudomonadati>Thermodesulfobacteriota>Desulfobulbia>Desulfobulbales>Desulfobulbaceae>Desulfobulbus>unclassified_Desulfobulbus | 418 | JT06_17850 | 6580..7836 | 1536652 | transcriptional-regulator-[Desulfobulbus-sp.-Tol-SR] |
| WP_208597680.1 | Schlafen+HATPase+wHTH | - | Desulfogranum_mediterraneum | Bacteria>Pseudomonadati>Thermodesulfobacteriota>Desulfobulbia>Desulfobulbales>Desulfobulbaceae>Desulfogranum | 407 | G494_RS23495 | 5401..6624 | 160661 | helix-turn-helix-domain-containing-protein-[Desulfogranum-mediterraneum] |
| WP_013164350.1 | Schlafen+HATPase+wHTH | GCF_000092205.1 | Desulfurivibrio_alkaliphilus | Bacteria>Pseudomonadati>Thermodesulfobacteriota>Desulfobulbia>Desulfobulbales>Desulfobulbaceae>Desulfurivibrio | 574 | DAAHT2_RS10980 | 2522796..2524520 | 427923 | ATP-binding-protein-[Desulfurivibrio-alkaliphilus] |
| RJX32578.1 | Schlafen+HATPase+wHTH | GCA_003605035.1 | Desulfurivibrio_sp. | Bacteria>Pseudomonadati>Thermodesulfobacteriota>Desulfobulbia>Desulfobulbales>Desulfobulbaceae>Desulfurivibrio>unclassified_Desulfurivibrio | 557 | C4531_06355 | 10193..11866 | 2093369 | MAG:-transcriptional-regulator-[Desulfurivibrio-sp.] |
| MCK9293529.1 | Schlafen+HATPase+wHTH | GCA_023230525.1 | Desulfobulbaceae_bacterium | Bacteria>Pseudomonadati>Thermodesulfobacteriota>Desulfobulbia>Desulfobulbales>Desulfobulbaceae>unclassified_Desulfobulbaceae | 504 | M0P70_00465 | 98240..99754 | 2053307 | putative-DNA-binding-domain-containing-protein-[Desulfobulbaceae-bacterium] |
| MCJ7602134.1 | Schlafen+HATPase+wHTH | GCA_022866315.1 | Desulfobulbaceae_bacterium | Bacteria>Pseudomonadati>Thermodesulfobacteriota>Desulfobulbia>Desulfobulbales>Desulfobulbaceae>unclassified_Desulfobulbaceae | 558 | MUO63_11645 | 10326..12002 | 2053307 | putative-DNA-binding-domain-containing-protein-[Desulfobulbaceae-bacterium] |
| HHD64078.1 | Schlafen+HATPase+wHTH | GCA_011372665.1 | Desulfobulbaceae_bacterium | Bacteria>Pseudomonadati>Thermodesulfobacteriota>Desulfobulbia>Desulfobulbales>Desulfobulbaceae>unclassified_Desulfobulbaceae | 385 | ENK96_06930 | 6874..8031 | 2053307 | ATP-dependent-DNA-helicase-RecG-[Desulfobulbaceae-bacterium] |
| MBA3007531.1 | Schlafen+HATPase+wHTH | GCA_013792135.1 | Desulfocapsa_sp. | Bacteria>Pseudomonadati>Thermodesulfobacteriota>Desulfobulbia>Desulfobulbales>Desulfocapsaceae>Desulfocapsa>unclassified_Desulfocapsa | 512 | FP810_13345 | 64268..65806 | 2066021 | winged-helix-turn-helix-transcriptional-regulator-[Desulfocapsa-sp.] |
| MBA3007527.1 | Schlafen+HATPase+wHTH | GCA_013792135.1 | Desulfocapsa_sp. | Bacteria>Pseudomonadati>Thermodesulfobacteriota>Desulfobulbia>Desulfobulbales>Desulfocapsaceae>Desulfocapsa>unclassified_Desulfocapsa | 570 | FP810_13325 | 57311..59023 | 2066021 | AAA-family-ATPase-[Desulfocapsa-sp.] |
| MBL0380706.1 | Schlafen+HATPase+wHTH | GCA_016722945.1 | Desulfofustis_sp._PB-SRB1 | Bacteria>Pseudomonadati>Thermodesulfobacteriota>Desulfobulbia>Desulfobulbales>Desulfocapsaceae>Desulfofustis>unclassified_Desulfofustis | 503 | JKP90_13815 | 382681..384192 | 1385624 | putative-DNA-binding-domain-containing-protein-[Desulfofustis-sp.-PB-SRB1] |
| HBH32877.1 | Schlafen+HATPase+wHTH | GCA_003499505.1 | Desulfofustis_sp. | Bacteria>Pseudomonadati>Thermodesulfobacteriota>Desulfobulbia>Desulfobulbales>Desulfocapsaceae>Desulfofustis>unclassified_Desulfofustis | 503 | DDX81_13450 | 480..1991 | 2053546 | AAA-family-ATPase-[Desulfofustis-sp.] |
| WP_205217416.1 | Schlafen+HATPase+wHTH | GCF_016918505.1 | Desulforhopalus_vacuolatus | Bacteria>Pseudomonadati>Thermodesulfobacteriota>Desulfobulbia>Desulfobulbales>Desulfocapsaceae>Desulforhopalus | 612 | JWG39_RS00665 | 162176..164014 | 40414 | putative-DNA-binding-domain-containing-protein-[Desulforhopalus-vacuolatus] |
| MCK9175500.1 | Schlafen+HATPase+wHTH | GCA_023231425.1 | Desulforhopalus_sp. | Bacteria>Pseudomonadati>Thermodesulfobacteriota>Desulfobulbia>Desulfobulbales>Desulfocapsaceae>Desulforhopalus>unclassified_Desulforhopalus | 500 | MOO96_09520 | 8461..9963 | 2099376 | putative-DNA-binding-domain-containing-protein-[Desulforhopalus-sp.] |
| WP_027370871.1 | Schlafen+HATPase+wHTH | GCF_000620765.1 | Desulfovermiculus_halophilus | Bacteria>Pseudomonadati>Thermodesulfobacteriota>Desulfovibrionia>Desulfovibrionales>Desulfohalobiaceae>Desulfovermiculus | 487 | N902_RS0110310 | 21151..22614 | 339722 | helix-turn-helix-domain-containing-protein-[Desulfovermiculus-halophilus] |

| Protein-ID | Domain | Assembly-ID | Strain | Taxonomy | Length | Locus-Tag | Genomic-coordinates | TaxID | Fasta-defline |
|---|---|---|---|---|---|---|---|---|---|
| WP_192624166.1 | Schlafen+HATPase+wHTH | GCF_014873765.1 | Desulfomicrobium_macestii | Bacteria>Pseudomonadati>Thermodesulfobacteriota>Desulfovibrionia>Desulfovibrionales>Desulfomicrobiaceae>Desulfomicrobium | 416 | H4684_RS13775 | 78146..79396 | 90731 | putative-DNA-binding-domain-containing-protein-[Desulfomicrobium-macestii] |
| MBC7355962.1 | Schlafen+HATPase+wHTH | GCA_014359945.1 | Desulfomicrobiaceae_bacterium | Bacteria>Pseudomonadati>Thermodesulfobacteriota>Desulfovibrionia>Desulfovibrionales>Desulfomicrobiaceae>unclassified_Desulfomicrobiaceae | 423 | H5U09_07165 | 23802..25073 | 2053547 | putative-DNA-binding-domain-containing-protein-[Desulfomicrobiaceae-bacterium] |
| WP_107736625.1 | Schlafen+HATPase+wHTH | GCF_003046795.1 | Desulfonatronum_sp._SC1 | Bacteria>Pseudomonadati>Thermodesulfobacteriota>Desulfovibrionia>Desulfovibrionales>Desulfonatronaceae>Desulfonatronum>unclassified_Desulfonatronum | 385 | C6366_RS06770 | 78197..79354 | 2109626 | putative-DNA-binding-domain-containing-protein-[Desulfonatronum-sp.-SC1] |
| WP_013514584.1 | Schlafen+HATPase+wHTH | GCF_000176915.2 | Pseudodesulfovibrio | Bacteria>Pseudomonadati>Thermodesulfobacteriota>Desulfovibrionia>Desulfovibrionales>Desulfovibrionaceae | 612 | DAES_RS08290 | 1826351..1828189 | 2035811 | putative-DNA-binding-domain-containing-protein-[Pseudodesulfovibrio-aespoeensis] |
| WP_163302424.1 | Schlafen+HATPase+wHTH | GCF_013376455.1 | Desulfolutivibrio_sulfodismutans | Bacteria>Pseudomonadati>Thermodesulfobacteriota>Desulfovibrionia>Desulfovibrionales>Desulfovibrionaceae>Desulfolutivibrio | 538 | GD606_RS12035 | 2612460..2614076 | 63561 | ATP-binding-protein-[Desulfovibrio-sulfodismutans] |
| MCM0755308.1 | Schlafen+HATPase+wHTH | GCA_023660105.1 | Desulfovibrio_aminophilus | Bacteria>Pseudomonadati>Thermodesulfobacteriota>Desulfovibrionia>Desulfovibrionales>Desulfovibrionaceae>Desulfovibrio | 591 | M7784_08625 | 70674..72449 | 81425 | putative-DNA-binding-domain-containing-protein-[Desulfovibrio-aminophilus] |
| MCG2733857.1 | Schlafen+HATPase+wHTH | GCA_022013435.1 | Pseudodesulfovibrio_aespoeensis | Bacteria>Pseudomonadati>Thermodesulfobacteriota>Desulfovibrionia>Desulfovibrionales>Desulfovibrionaceae>Pseudodesulfovibrio | 612 | L6357_12310 | 9635..11473 | 182210 | putative-DNA-binding-domain-containing-protein-[Pseudodesulfovibrio-aespoeensis] |
| MBW6511063.1 | Schlafen+HATPase+wHTH | GCA_019429445.1 | Desulfuromonadaceae_bacterium | Bacteria>Pseudomonadati>Thermodesulfobacteriota>Desulfuromonadia>Desulfuromonadales>Desulfuromonadaceae>unclassified_Desulfuromonadaceae | 616 | K0A93_02950 | 86332..88182 | 2099677 | putative-DNA-binding-domain-containing-protein-[Desulfuromonadaceae-bacterium] |
| AJF05948.1 | Schlafen+HATPase+wHTH | GCA_000827125.1 | Geoalkalibacter_subterraneus | Bacteria>Pseudomonadati>Thermodesulfobacteriota>Desulfuromonadia>Desulfuromonadales>Geoalkalibacteraceae>Geoalkalibacter | 592 | GSUB_04360 | 971090..972868 | 483547 | transcriptional-regulator-[Geoalkalibacter-subterraneus] |
| MBW6509586.1 | Schlafen+HATPase+wHTH | GCA_019429255.1 | Desulfuromonadales_bacterium | Bacteria>Pseudomonadati>Thermodesulfobacteriota>Desulfuromonadia>Desulfuromonadales>unclassified_Desulfuromonadales | 557 | K0A94_08610 | 47060..48733 | 2099678 | putative-DNA-binding-domain-containing-protein-[Desulfuromonadales-bacterium] |
| WP_067619280.1 | Schlafen+HATPase+wHTH | GCF_001687335.1 | Dissulfuribacter_thermophilus | Bacteria>Pseudomonadati>Thermodesulfobacteriota>Dissulfuribacteria>Dissulfuribacterales>Dissulfuribacteraceae>Dissulfuribacter | 96 | DBT_RS08705 | 74198..74488 | 1156395 | ATP-binding-protein-[Dissulfuribacter-thermophilus] |
| WP_011416812.1 | Schlafen+HATPase+wHTH | GCF_000013405.1 | Syntrophus_aciditrophicus | Bacteria>Pseudomonadati>Thermodesulfobacteriota>Syntrophia>Syntrophales>Syntrophaceae>Syntrophus | 413 | SYN_RS04270 | 936821..938062 | 316277 | putative-DNA-binding-domain-containing-protein-[Syntrophus-aciditrophicus] |
| OPY81707.1 | Schlafen+HATPase+wHTH | GCA_002067745.1 | Syntrophus_sp._PtaU1.Bin005 | Bacteria>Pseudomonadati>Thermodesulfobacteriota>Syntrophia>Syntrophales>Syntrophaceae>Syntrophus>unclassified_Syntrophus_(in:_bacteria) | 556 | A4E70_01073 | 16337..18007 | 1811712 | MAG:-Divergent-AAA-domain-protein-[Syntrophus-sp.-PtaU1.Bin005] |
| MBS3905933.1 | Schlafen+HATPase+wHTH | GCA_018333715.1 | Syntrophaceae_bacterium | Bacteria>Pseudomonadati>Thermodesulfobacteriota>Syntrophia>Syntrophales>Syntrophaceae>unclassified_Syntrophaceae | 404 | KGZ49_02745 | 11027..12241 | 2052184 | putative-DNA-binding-domain-containing-protein-[Syntrophaceae-bacterium] |
| MCK9528167.1 | Schlafen+HATPase+wHTH | GCA_023228425.1 | Syntrophales_bacterium | Bacteria>Pseudomonadati>Thermodesulfobacteriota>Syntrophia>Syntrophales>unclassified_Syntrophales | 390 | M0Q23_05895 | 61318..62490 | 2928017 | putative-DNA-binding-domain-containing-protein-[Syntrophales-bacterium] |
| MCK9230426.1 | Schlafen+HATPase+wHTH | GCA_023230845.1 | Syntrophales_bacterium | Bacteria>Pseudomonadati>Thermodesulfobacteriota>Syntrophia>Syntrophales>unclassified_Syntrophales | 390 | M0P18_07090 | 6323..7495 | 2928017 | putative-DNA-binding-domain-containing-protein-[Syntrophales-bacterium] |
| WP_246041626.1 | Schlafen+HATPase+wHTH | GCF_005377625.1 | Desulfoglaeba_alkanexedens | Bacteria>Pseudomonadati>Thermodesulfobacteriota>Syntrophobacteria>Syntrophobacterales>Syntrophobacteraceae>Desulfoglaeba | 319 | FDQ92_RS08000 | 1769219..1770178 | 361111 | ATP-binding-protein-[Desulfoglaeba-alkanexedens] |
| WP_246041625.1 | Schlafen+HATPase+wHTH | GCF_005377625.1 | Desulfoglaeba_alkanexedens | Bacteria>Pseudomonadati>Thermodesulfobacteriota>Syntrophobacteria>Syntrophobacterales>Syntrophobacteraceae>Desulfoglaeba | 74 | FDQ92_RS15690 | 1768922..1769146 | 361111 | ATP-binding-protein-[Desulfoglaeba-alkanexedens] |
| WP_137424078.1 | Schlafen+HATPase+wHTH | GCF_005377625.1 | Desulfoglaeba_alkanexedens | Bacteria>Pseudomonadati>Thermodesulfobacteriota>Syntrophobacteria>Syntrophobacterales>Syntrophobacteraceae>Desulfoglaeba | 513 | FDQ92_RS07975 | 1760760..1762301 | 361111 | putative-DNA-binding-domain-containing-protein-[Desulfoglaeba-alkanexedens] |
| WP_244082337.1 | Schlafen+HATPase+wHTH | GCF_022836495.1 | Desulforhabdus_sp._TSK | Bacteria>Pseudomonadati>Thermodesulfobacteriota>Syntrophobacteria>Syntrophobacterales>Syntrophobacteraceae>Desulforhabdus>unclassified_Desulforhabdus | 596 | MWC30_RS01310 | 289844..291634 | 2925014 | ATP-binding-protein-[Desulforhabdus-sp.-TSK] |
| WP_123290344.1 | Schlafen+HATPase+wHTH | GCF_003751385.1 | Desulfosoma_caldarium | Bacteria>Pseudomonadati>Thermodesulfobacteriota>Syntrophobacteria>Syntrophobacterales>Syntrophobacteraceae>Desulfosoma | 557 | EDC27_RS09310 | 250601..252274 | 610254 | ATP-binding-protein-[Desulfosoma-caldarium] |
| OHE16644.1 | Schlafen+HATPase+wHTH | GCA_001830835.1 | Syntrophobacterales_bacterium_GWC2_56_13 | Bacteria>Pseudomonadati>Thermodesulfobacteriota>Syntrophobacteria>Syntrophobacterales>unclassified_Syntrophobacterales | 557 | A2X96_09770 | 14302..15975 | 1802287 | ATPase-[Syntrophobacterales-bacterium-GWC2_56_13] |
| WP_189066311.1 | Schlafen+HATPase+wHTH | GCF_014648115.1 | Deinococcus_seoulensis | Bacteria>Thermotogati>Deinococcota>Deinococci>Deinococcales>Deinococcaceae>Deinococcus | 569 | IEY70_RS17435 | 25174..26883 | 1837379 | putative-DNA-binding-domain-containing-protein-[Deinococcus-seoulensis] |
| WP_129117972.1 | Schlafen+HATPase+wHTH | GCF_030161515.1 | Deinococcus_metallilatus | Bacteria>Thermotogati>Deinococcota>Deinococci>Deinococcales>Deinococcaceae>Deinococcus | 570 | QQK08_RS18430 | 9052..10764 | 1211322 | ATP-binding-protein-[Deinococcus-metallilatus] |
| WP_107138390.1 | Schlafen+HATPase+wHTH | GCF_003028415.1 | Deinococcus_arcticus | Bacteria>Thermotogati>Deinococcota>Deinococci>Deinococcales>Deinococcaceae>Deinococcus | 567 | C8263_RS12100 | 74137..75840 | 2136176 | ATP-binding-protein-[Deinococcus-arcticus] |
| WP_253581494.1 | Schlafen+HATPase+wHTH | GCF_024171495.1 | Deinococcus_sp._HSC-46F16 | Bacteria>Thermotogati>Deinococcota>Deinococci>Deinococcales>Deinococcaceae>Deinococcus>unclassified_Deinococcus | 578 | L1280_RS07640 | 445252..446988 | 2910968 | ATP-binding-protein-[Deinococcus-sp.-HSC-46F16] |
| WP_239048769.1 | Schlafen+HATPase+wHTH | GCF_022344005.1 | Deinococcus_sp._KNUC1210 | Bacteria>Thermotogati>Deinococcota>Deinococci>Deinococcales>Deinococcaceae>Deinococcus>unclassified_Deinococcus | 584 | MF271_RS10570 | 1559961..1561715 | 2917691 | ATP-binding-protein-[Deinococcus-sp.-KNUC1210] |
| WP_278359297.1 | Schlafen+HATPase+wHTH | GCF_002378905.1 | Thermus_thermophilus | Bacteria>Thermotogati>Deinococcota>Deinococci>Thermales>Thermaceae>Thermus | 464 | B3N72_RS05310 | 19140..20534 | 274 | ATP-binding-protein,-partial-[Thermus-thermophilus] |
| WP_244347429.1 | Schlafen+HATPase+wHTH | GCF_016629545.1 | Thermus_thermophilus | Bacteria>Thermotogati>Deinococcota>Deinococci>Thermales>Thermaceae>Thermus | 548 | JMH49_RS05550 | 1041471..1043117 | 274 | ATP-binding-protein-[Thermus-thermophilus] |
| WP_224065206.1 | Schlafen+HATPase+wHTH | GCF_000091545.1 | Thermus_thermophilus | Bacteria>Thermotogati>Deinococcota>Deinococci>Thermales>Thermaceae>Thermus | 428 | QUE39_RS05190 | 967399..968685 | 274 | ATP-binding-protein-[Thermus-thermophilus] |
| WP_126188070.1 | Schlafen+HATPase+wHTH | GCF_003958985.1 | Thermus_scotoductus | Bacteria>Thermotogati>Deinococcota>Deinococci>Thermales>Thermaceae>Thermus | 589 | CSW41_RS02880 | 9776..11545 | 37636 | ATP-binding-protein-[Thermus-scotoductus] |
| WP_011228380.1 | Schlafen+HATPase+wHTH | GCF_000091545.1 | Thermus_thermophilus | Bacteria>Thermotogati>Deinococcota>Deinococci>Thermales>Thermaceae>Thermus | 121 | QUE39_RS05200 | 969991..970356 | 274 | ATP-binding-protein-[Thermus-thermophilus] |
| BDG19120.1 | Schlafen+HATPase+wHTH | GCA_022846275.1 | Thermus_thermophilus | Bacteria>Thermotogati>Deinococcota>Deinococci>Thermales>Thermaceae>Thermus | 563 | TthSNM11_13230 | 1193476..1195167 | 274 | ATPase-AAA-[Thermus-thermophilus] |
| BCP66161.1 | Schlafen+HATPase+wHTH | GCA_016629545.1 | Thermus_thermophilus | Bacteria>Thermotogati>Deinococcota>Deinococci>Thermales>Thermaceae>Thermus | 563 | TthHB5018_10950 | 1041471..1043162 | 274 | ATPase-AAA-[Thermus-thermophilus] |
| WP_286878964.1 | Schlafen+HATPase+wHTH | GCF_003450455.1 | Thermus_sp. | Bacteria>Thermotogati>Deinococcota>Deinococci>Thermales>Thermaceae>Thermus>unclassified_Thermus | 463 | DCM56_RS07485 | 1..1394 | 275 | ATP-binding-protein,-partial-[Thermus-sp.] |
| MBF6594437.1 | Schlafen+HATPase+wHTH | GCA_015478585.1 | Thermaceae_bacterium | Bacteria>Thermotogati>Deinococcota>Deinococci>Thermales>Thermaceae>unclassified_Thermaceae | 114 | IVW51_08315 | 17105..17449 | 2614257 | ATP-binding-protein-[Thermaceae-bacterium] |
| MCY4354896.1 | Schlafen+HATPase+wHTH | GCA_026710445.1 | Truepera_sp. | Bacteria>Thermotogati>Deinococcota>Deinococci>Trueperales>Trueperaceae>Truepera>unclassified_Truepera | 409 | OXC09_08990 | 24626..25855 | 2478917 | MAG:-putative-DNA-binding-domain-containing-protein-[Truepera-sp.] |
| WP_236101000.1 | Schlafen+HATPase+wHTH | GCF_021568795.1 | Dethiosulfovibrio_faecalis | Bacteria>Thermotogati>Synergistota>Synergistia>Synergistales>Dethiosulfovibrionaceae>Dethiosulfovibrio | 602 | L2W58_RS00515 | 90972..92780 | 2720018 | putative-DNA-binding-domain-containing-protein-[Dethiosulfovibrio-faecalis] |
| WP_005662438.1 | Schlafen+HATPase+wHTH | GCF_000172975.1 | Dethiosulfovibrio_peptidovorans | Bacteria>Thermotogati>Synergistota>Synergistia>Synergistales>Dethiosulfovibrionaceae>Dethiosulfovibrio | 555 | DPEP_RS12010 | 2483560..2485227 | 47055 | ATP-binding-protein-[Dethiosulfovibrio-peptidovorans] |
| WP_005662434.1 | Schlafen+HATPase+wHTH | GCF_000172975.1 | Dethiosulfovibrio_peptidovorans | Bacteria>Thermotogati>Synergistota>Synergistia>Synergistales>Dethiosulfovibrionaceae>Dethiosulfovibrio | 435 | DPEP_RS12000 | 2479209..2480516 | 47055 | putative-DNA-binding-domain-containing-protein-[Dethiosulfovibrio-peptidovorans] |
| WP_006299686.1 | Schlafen+HATPase+wHTH | GCF_000165795.1 | Aminomonas_paucivorans | Bacteria>Thermotogati>Synergistota>Synergistia>Synergistales>Synergistaceae>Aminomonas | 557 | APAU_RS00535 | 114747..116420 | 81412 | ATP-binding-protein-[Aminomonas-paucivorans] |
| WP_034444781.1 | Schlafen+HATPase+wHTH | GCF_000585335.1 | Cloacibacillus_evryensis | Bacteria>Thermotogati>Synergistota>Synergistia>Synergistales>Synergistaceae>Cloacibacillus | 517 | CLOEV_RS14980 | 3363667..3365220 | 508460 | winged-helix-turn-helix-transcriptional-regulator-[Cloacibacillus-evryensis] |
| MBL3593012.1 | Schlafen+HATPase+wHTH | GCA_016756725.1 | Synergistaceae_bacterium | Bacteria>Thermotogati>Synergistota>Synergistia>Synergistales>Synergistaceae>unclassified_Synergistaceae | 573 | JMJ93_05815 | 378042..379763 | 2053624 | putative-DNA-binding-domain-containing-protein-[Synergistaceae-bacterium] |
| RKJ48597.1 | Schlafen+HATPase+wHTH | GCA_003612585.1 | bacterium_1XD42-54 | Bacteria>unclassified_Bacteria | 490 | D7Y05_12375 | 22210..23682 | 2320110 | winged-helix-turn-helix-transcriptional-regulator-[bacterium-1XD42-54] |
| RKJ05435.1 | Schlafen+HATPase+wHTH | GCA_003611905.1 | bacterium_D16-54 | Bacteria>unclassified_Bacteria | 504 | D7X87_08495 | 75409..76923 | 2320104 | ATP-binding-protein-[bacterium-D16-54] |
| NCP08808.1 | Schlafen+HATPase+wHTH | GCA_009993525.1 | bacterium | Bacteria>unclassified_Bacteria | 556 | GW860_07905 | 21741..23411 | 1869227 | ATPase-[bacterium] |
| NBK91897.1 | Schlafen+HATPase+wHTH | GCA_009917555.1 | bacterium_1XD21-13 | Bacteria>unclassified_Bacteria | 506 | D5278_07875 | 178256..179776 | 2320888 | HTH-domain-containing-protein-[bacterium-1XD21-13] |
| MDE0416422.1 | Schlafen+HATPase+wHTH | GCA_028821455.1 | bacterium | Bacteria>unclassified_Bacteria | 464 | OXI95_05725 | 22198..23592 | 1869227 | MAG:-putative-DNA-binding-domain-containing-protein-[bacterium] |
| MDD3925573.1 | Schlafen+HATPase+wHTH | GCA_028699365.1 | bacterium | Bacteria>unclassified_Bacteria | 553 | PHT33_02825 | 8072..9733 | 1869227 | MAG:-ATP-binding-protein-[bacterium] |
| MCP4405115.1 | Schlafen+HATPase+wHTH | GCA_024222835.1 | bacterium | Bacteria>unclassified_Bacteria | 456 | GY801_48415 | 1627..2997 | 1869227 | AAA-family-ATPase-[bacterium] |
| MCG3156462.1 | Schlafen+HATPase+wHTH | GCA_022072305.1 | bacterium | Bacteria>unclassified_Bacteria | 545 | DKINENOH_03086 | 103205..104842 | 1869227 | hypothetical-protein-[bacterium] |
| MBN2373592.1 | Schlafen+HATPase+wHTH | GCA_016934595.1 | bacterium | Bacteria>unclassified_Bacteria | 556 | JXL19_07390 | 9382..11052 | 1869227 | putative-DNA-binding-domain-containing-protein-[bacterium] |
| MBI9030470.1 | Schlafen+HATPase+wHTH | GCA_016342615.1 | bacterium | Bacteria>unclassified_Bacteria | 559 | JEZ13_00500 | 15860..17539 | 1869227 | putative-DNA-binding-domain-containing-protein-[bacterium] |
| KU061372.1 | Schlafen+HATPase+wHTH | GCA_001516025.1 | bacterium_BRH_c32 | Bacteria>unclassified_Bacteria | 320 | APF79_11965 | 1062..2024 | 1734400 | hypothetical-protein-APF79_11965-[bacterium-BRH_c32] |
| GBE48766.1 | Schlafen+HATPase+wHTH | GCA_002897795.1 | bacterium_BMS3Bbin12 | Bacteria>unclassified_Bacteria | 424 | BMS3Bbin12_01954 | 28742..30016 | 2005735 | divergent-AAA-domain-protein-[bacterium-BMS3Bbin12] |
| GBD39151.1 | Schlafen+HATPase+wHTH | GCA_002898375.1 | bacterium_HR37 | Bacteria>unclassified_Bacteria | 47 | HRbin37_01418 | 75241..75384 | 2035432 | hypothetical-protein-HRbin37_01418-[bacterium-HR37] |

| Protein-ID | Domain | Assembly-ID | Strain | Taxonomy | Length | Locus-Tag | Genomic-coordinates | TaxID | Fasta-defline |
|---|---|---|---|---|---|---|---|---|---|
| --OLD-ABC | | | | | | | | | |
| RLI77771.1 | OLD-ABC | GCA_003662925.1 | Archaeoglobales_archaeon | Archaea>Methanobacteriati>Methanobacteriota>Archaeoglobi>Archaeoglobales>unclassified_Archaeoglobales | 443 | DRP04_10880 | 886..2217 | 2250258 | recombinase-RecF-[Archaeoglobales-archaeon] |
| WP_214400398.1 | OLD-ABC | GCF_902827225.1 | Methanocaldococcus_lauensis | Archaea>Methanobacteriati>Methanobacteriota>Methanomada_group>Methanococci>Methanococcales>Methanocaldococcaceae>Methanocaldococcus | 369 | KMP69_RS02585 | 458374..459483 | 2546128 | AAA-family-ATPase-[Methanocaldococcus-lauensis] |
| MCW3134163.1 | OLD-ABC | GCA_025966595.1 | Candidatus_Methanophagales_archaeon | Archaea>Methanobacteriati>Methanobacteriota>Stenosarchaea_group>Methanomicrobia>Candidatus_Methanophagales>unclassified_Candidatus_Methanophagales | 322 | N2V78_07550 | 1774..2742 | 2056316 | MAG:-AAA-family-ATPase-[Methanophagales-archaeon] |
| MCW3129354.1 | OLD-ABC | GCA_025966515.1 | Candidatus_Methanophagales_archaeon | Archaea>Methanobacteriati>Methanobacteriota>Stenosarchaea_group>Methanomicrobia>Candidatus_Methanophagales>unclassified_Candidatus_Methanophagales | 321 | N2V75_04555 | 2217..3182 | 2056316 | MAG:-AAA-family-ATPase-[Methanophagales-archaeon] |
| MBR1368479.1 | OLD-ABC | GCA_018132105.1 | Methanocalculus_chunghsingensis | Archaea>Methanobacteriati>Methanobacteriota>Stenosarchaea_group>Methanomicrobia>Methanomicrobiales>Methanocalculaceae>Methanocalculus | 365 | RJ53_02750 | 14757..15854 | 156457 | chromosome-segregation-protein-SMC-[Methanocalculus-chunghsingensis] |
| WP_211530107.1 | OLD-ABC | GCF_018132105.1 | Methanocalculus_chunghsingensis | Archaea>Methanobacteriati>Methanobacteriota>Stenosarchaea_group>Methanomicrobia>Methanomicrobiales>Methanocalculaceae>Methanocalculus | 365 | RJ53_RS02815 | 14757..15854 | 156457 | AAA-family-ATPase-[Methanocalculus-chunghsingensis] |
| WP_253488283.1 | OLD-ABC | GCF_024170505.1 | Methanocalculus_alkaliphilus | Archaea>Methanobacteriati>Methanobacteriota>Stenosarchaea_group>Methanomicrobia>Methanomicrobiales>Methanocalculaceae>Methanocalculus | 365 | J2T58_RS06400 | 5147..6244 | 768730 | AAA-family-ATPase-[Methanocalculus-alkaliphilus] |
| WP_167816000.1 | OLD-ABC | - | Methanocorpusculum_sp._GPch4 | Archaea>Methanobacteriati>Methanobacteriota>Stenosarchaea_group>Methanomicrobia>Methanomicrobiales>Methanocorpusculaceae>Methanocorpusculum>unclassified_Methanocorpusculum | 391 | EYA77_RS04455 | 14161..15336 | 2527877 | AAA-family-ATPase-[Methanocorpusculum-sp.-GPch4] |
| WP_011448871.1 | OLD-ABC | GCF_000013445.1 | Methanospirillum | Archaea>Methanobacteriati>Methanobacteriota>Stenosarchaea_group>Methanomicrobia>Methanomicrobiales>Methanospirillaceae | 389 | MHUN_RS09835 | 2134488..2135657 | 2202 | ATP-binding-protein-[Methanospirillum-hungatei] |
| WP_011448735.1 | OLD-ABC | GCF_000013445.1 | Methanospirillum_hungatei | Archaea>Methanobacteriati>Methanobacteriota>Stenosarchaea_group>Methanomicrobia>Methanomicrobiales>Methanospirillaceae>Methanospirillum | 515 | MHUN_RS09140 | 1975816..1977363 | 2203 | AAA-family-ATPase-[Methanospirillum-hungatei] |
| WP_011448736.1 | OLD-ABC | GCF_000013445.1 | Methanospirillum_hungatei | Archaea>Methanobacteriati>Methanobacteriota>Stenosarchaea_group>Methanomicrobia>Methanomicrobiales>Methanospirillaceae>Methanospirillum | 513 | MHUN_RS09145 | 1977584..1979125 | 2203 | AAA-family-ATPase-[Methanospirillum-hungatei] |
| WP_048067947.1 | OLD-ABC | - | Methanospirillum_hungatei | Archaea>Methanobacteriati>Methanobacteriota>Stenosarchaea_group>Methanomicrobia>Methanomicrobiales>Methanospirillaceae>Methanospirillum | 403 | MHUN_RS09160 | 1982921..1984132 | 2203 | AAA-family-ATPase-[Methanospirillum-hungatei] |
| WP_011448107.1 | OLD-ABC | GCF_000013445.1 | Methanospirillum_hungatei | Archaea>Methanobacteriati>Methanobacteriota>Stenosarchaea_group>Methanomicrobia>Methanomicrobiales>Methanospirillaceae>Methanospirillum | 369 | MHUN_RS05650 | 1218862..1219971 | 2203 | AAA-family-ATPase-[Methanospirillum-hungatei] |
| ABD41471.1 | OLD-ABC | GCA_000013445.1 | Methanospirillum_hungatei_JF-1 | Archaea>Methanobacteriati>Methanobacteriota>Stenosarchaea_group>Methanomicrobia>Methanomicrobiales>Methanospirillaceae>Methanospirillum>Methanospirillum_hungatei | 515 | Mhun_1750 | 1975816..1977363 | 323259 | ATPase-[Methanospirillum-hungatei-JF-1] |
| ABD41472.1 | OLD-ABC | GCA_000013445.1 | Methanospirillum_hungatei_JF-1 | Archaea>Methanobacteriati>Methanobacteriota>Stenosarchaea_group>Methanomicrobia>Methanomicrobiales>Methanospirillaceae>Methanospirillum>Methanospirillum_hungatei | 513 | Mhun_1751 | 1977584..1979125 | 323259 | conserved-hypothetical-protein-[Methanospirillum-hungatei-JF-1] |
| ABD41475.1 | OLD-ABC | GCA_000013445.1 | Methanospirillum_hungatei_JF-1 | Archaea>Methanobacteriati>Methanobacteriota>Stenosarchaea_group>Methanomicrobia>Methanomicrobiales>Methanospirillaceae>Methanospirillum>Methanospirillum_hungatei | 405 | Mhun_1754 | 1982915..1984132 | 323259 | conserved-hypothetical-protein-[Methanospirillum-hungatei-JF-1] |
| ABD40829.1 | OLD-ABC | GCA_000013445.1 | Methanospirillum_hungatei_JF-1 | Archaea>Methanobacteriati>Methanobacteriota>Stenosarchaea_group>Methanomicrobia>Methanomicrobiales>Methanospirillaceae>Methanospirillum>Methanospirillum_hungatei | 369 | Mhun_1080 | 1218862..1219971 | 323259 | conserved-hypothetical-protein-[Methanospirillum-hungatei-JF-1] |
| NLL10043.1 | OLD-ABC | GCA_012520015.1 | Methanomicrobiales_archaeon | Archaea>Methanobacteriati>Methanobacteriota>Stenosarchaea_group>Methanomicrobia>Methanomicrobiales>unclassified_Methanomicrobiales | 389 | GX268_03945 | 628..1797 | 2184052 | AAA-family-ATPase-[Methanomicrobiales-archaeon] |
| QNO47557.1 | OLD-ABC | - | Candidatus_Methanogaster_sp._ANME-2c_ERB4 | Archaea>Methanobacteriati>Methanobacteriota>Stenosarchaea_group>Methanomicrobia>Methanosarcinales>ANME-2_cluster>Candidatus_Methanogasteraceae>Candidatus_Methanogaster>unclassified_Candidatus_Methanogaster | 329 | GGGHDLIA_00047 | 31690..32679 | 2759911 | DNA-replication-and-repair-protein-RecF-[Methanosarcinales-archaeon-ANME-2c-ERB4] |
| OFV65719.1 | OLD-ABC | GCA_001766825.1 | Candidatus_Syntrophoarchaeum_butanivorans | Archaea>Methanobacteriati>Methanobacteriota>Stenosarchaea_group>Methanomicrobia>Methanosarcinales>ANME-2_cluster>Candidatus_Syntrophoarchaeum | 322 | SBU_001302 | 3839..4807 | 1839936 | hypothetical-protein-SBU_001302-[Candidatus-Syntrophoarchaeum-butanivorans] |
| WP_270096525.1 | OLD-ABC | GCF_027594145.1 | Methanococcoides_alaskense | Archaea>Methanobacteriati>Methanobacteriota>Stenosarchaea_group>Methanomicrobia>Methanosarcinales>Methanosarcinaceae>Methanococcoides | 418 | PBV53_RS07470 | 730568..731824 | 325778 | ATP-binding-protein-[Methanococcoides-alaskense] |
| NPE28178.1 | OLD-ABC | GCA_013166765.1 | Methanococcoides_sp._SA1 | Archaea>Methanobacteriati>Methanobacteriota>Stenosarchaea_group>Methanomicrobia>Methanosarcinales>Methanosarcinaceae>Methanococcoides>unclassified_Methanococcoides | 420 | HNV12_09445 | 5461..6723 | 2735871 | ATP-binding-protein-[Methanococcoides-sp.-SA1] |
| WP_091932399.1 | OLD-ABC | GCF_900114835.1 | Methanolobus_profundi | Archaea>Methanobacteriati>Methanobacteriota>Stenosarchaea_group>Methanomicrobia>Methanosarcinales>Methanosarcinaceae>Methanolobus | 420 | BM008_RS01865 | 361506..362768 | 487685 | ATP-binding-protein-[Methanolobus-profundi] |
| KXS44092.1 | OLD-ABC | GCA_001577565.1 | Methanolobus_sp._T82-4 | Archaea>Methanobacteriati>Methanobacteriota>Stenosarchaea_group>Methanomicrobia>Methanosarcinales>Methanosarcinaceae>Methanolobus>unclassified_Methanolobus | 420 | AWU59_725 | 7322..8584 | 1794908 | MAG:-putative-ATPase-[Methanolobus-sp.-T82-4] |
| WP_048159311.1 | OLD-ABC | GCF_000969965.1 | Methanosarcina_sp._WWM596 | Archaea>Methanobacteriati>Methanobacteriota>Stenosarchaea_group>Methanomicrobia>Methanosarcinales>Methanosarcinaceae>Methanosarcina>unclassified_Methanosarcina | 430 | MSWHS_RS14810 | 3358572..3359864 | 1434103 | AAA-family-ATPase-[Methanosarcina-sp.-WWM596] |
| WP_273278521.1 | OLD-ABC | GCF_913757535.1 | Methanothrix_soehngenii | Archaea>Methanobacteriati>Methanobacteriota>Stenosarchaea_group>Methanomicrobia>Methanotrichales>Methanotrichaceae>Methanothrix | 381 | NW278_RS08470 | 1665125..1666270 | 2223 | AAA-family-ATPase-[Methanothrix-soehngenii] |
| MBK7385732.1 | OLD-ABC | GCA_016706325.1 | Methanothrix_sp. | Archaea>Methanobacteriati>Methanobacteriota>Stenosarchaea_group>Methanomicrobia>Methanotrichales>Methanotrichaceae>Methanothrix>unclassified_Methanothrix | 381 | IPI63_02965 | 572346..573491 | 90426 | AAA-family-ATPase-[Methanothrix-sp.] |
| MBA3046424.1 | OLD-ABC | GCA_013791785.1 | Methanobacteriota_archaeon | Archaea>Methanobacteriati>Methanobacteriota>unclassified_Methanobacteriota | 412 | FP824_09465 | 31777..33015 | 2026739 | ATP-binding-protein-[Euryarchaeota-archaeon] |
| OQA55743.1 | OLD-ABC | GCA_002069575.1 | Euryarchaeota_archaeon_ADurb.Bin294 | Archaea>Methanobacteriati>Methanobacteriota>unclassified_Methanobacteriota | 401 | BWY45_02070 | 2515..3720 | 1852878 | DNA-replication-and-repair-protein-RecF-[Euryarchaeota-archaeon-ADurb.Bin294] |
| MBU4220746.1 | OLD-ABC | GCA_018898145.1 | Methanobacteriota_archaeon | Archaea>Methanobacteriati>Methanobacteriota>unclassified_Methanobacteriota | 335 | KKA10_03845 | 4714..5721 | 2026739 | AAA-family-ATPase-[Euryarchaeota-archaeon] |
| RLF71963.1 | OLD-ABC | GCA_003663625.1 | Thermoplasmata_archaeon | Archaea>Methanobacteriati>Thermoplasmatota>Thermoplasmata>unclassified_Thermoplasmata | 327 | DRN40_01340 | 2565..3548 | 1906666 | MAG:-hypothetical-protein-DRN40_01340-[Thermoplasmata-archaeon] |
| MXW95268.1 | OLD-ABC | GCA_009836165.1 | Acidimicrobiaceae_bacterium | Bacteria>Bacillati>Actinomycetota>Acidimicrobiia>Acidimicrobiales>Acidimicrobiaceae>unclassified_Acidimicrobiaceae | 432 | F4Z64_07150 | 276..1574 | 2024894 | AAA-family-ATPase-[Acidimicrobiaceae-bacterium] |
| MYH43042.1 | OLD-ABC | GCA_009843995.1 | Acidimicrobiaceae_bacterium | Bacteria>Bacillati>Actinomycetota>Acidimicrobiia>Acidimicrobiales>Acidimicrobiaceae>unclassified_Acidimicrobiaceae | 432 | F4131_04995 | 20992..22290 | 2024894 | AAA-family-ATPase-[Acidimicrobiaceae-bacterium] |
| MCY3892498.1 | OLD-ABC | GCA_026707105.1 | Acidimicrobiaceae_bacterium | Bacteria>Bacillati>Actinomycetota>Acidimicrobiia>Acidimicrobiales>Acidimicrobiaceae>unclassified_Acidimicrobiaceae | 423 | OXF65_04285 | 1210..2481 | 2024894 | MAG:-AAA-family-ATPase-[Acidimicrobiaceae-bacterium] |
| MXW97852.1 | OLD-ABC | GCA_009835735.1 | Acidimicrobiaceae_bacterium | Bacteria>Bacillati>Actinomycetota>Acidimicrobiia>Acidimicrobiales>Acidimicrobiaceae>unclassified_Acidimicrobiaceae | 352 | F4Z89_04465 | 10206..11264 | 2024894 | AAA-family-ATPase-[Acidimicrobiaceae-bacterium] |
| MBF0967809.1 | OLD-ABC | GCA_015259445.1 | Actinomyces_sp. | Bacteria>Bacillati>Actinomycetota>Actinomycetes>Actinomycetales>Actinomycetaceae>Actinomyces>unclassified_Actinomyces | 427 | HXK08_00800 | 8005..9288 | 29317 | AAA-family-ATPase-[Actinomyces-sp.] |
| OFQ22231.1 | OLD-ABC | GCA_001813985.1 | Actinomyces_sp._HMSC062G12 | Bacteria>Bacillati>Actinomycetota>Actinomycetes>Actinomycetales>Actinomycetaceae>Actinomyces>unclassified_Actinomyces | 423 | HMPREF2946_04755 | 255101..256372 | 1739450 | chromosome-segregation-protein-SMC-[Actinomyces-sp.-HMSC062G12] |
| MBS6968674.1 | OLD-ABC | GCA_018381115.1 | Actinomyces_sp. | Bacteria>Bacillati>Actinomycetota>Actinomycetes>Actinomycetales>Actinomycetaceae>Actinomyces>unclassified_Actinomyces | 406 | KH140_06975 | 8612..9832 | 29317 | AAA-family-ATPase-[Actinomyces-sp.] |
| AOS47541.1 | OLD-ABC | GCA_001746855.1 | Pauljensenia_hongkongensis | Bacteria>Bacillati>Actinomycetota>Actinomycetes>Actinomycetales>Actinomycetaceae>Pauljensenia | 384 | BH719_06530 | 1591596..1592750 | 178339 | hypothetical-protein-BH719_06530-[Pauljensenia-hongkongensis] |
| WP_009744024.1 | OLD-ABC | GCF_001746855.1 | Pauljensenia_hongkongensis | Bacteria>Bacillati>Actinomycetota>Actinomycetes>Actinomycetales>Actinomycetaceae>Pauljensenia | 384 | BH719_RS06530 | 1591596..1592750 | 178339 | ATP-binding-protein-[Pauljensenia-hongkongensis] |
| WP_227411047.1 | OLD-ABC | GCF_016900615.1 | Bifidobacterium_pullorum | Bacteria>Bacillati>Actinomycetota>Actinomycetes>Bifidobacteriales>Bifidobacteriaceae>Bifidobacterium | 444 | INF27_RS02870 | 180456..181790 | 78448 | ATP-binding-protein-[Bifidobacterium-pullorum] |
| WP_238556889.1 | OLD-ABC | GCF_000770965.1 | Bifidobacterium_pullorum | Bacteria>Bacillati>Actinomycetota>Actinomycetes>Bifidobacteriales>Bifidobacteriaceae>Bifidobacterium | 444 | OU28_RS04670 | 28509..29843 | 78448 | ATP-binding-protein-[Bifidobacterium-pullorum] |
| PJM72968.1 | OLD-ABC | GCA_002802875.1 | Bifidobacterium_primatium | Bacteria>Bacillati>Actinomycetota>Actinomycetes>Bifidobacteriales>Bifidobacteriaceae>Bifidobacterium | 437 | CS006_06860 | 164999..166312 | 2045438 | hypothetical-protein-CS006_06860-[Bifidobacterium-primatium] |
| WP_100511071.1 | OLD-ABC | GCF_002802875.1 | Bifidobacterium_primatium | Bacteria>Bacillati>Actinomycetota>Actinomycetes>Bifidobacteriales>Bifidobacteriaceae>Bifidobacterium | 437 | CS006_RS06840 | 164999..166312 | 2045438 | ATP-binding-protein-[Bifidobacterium-primatium] |
| WP_248824781.1 | OLD-ABC | GCF_023242945.1 | Frankia_umida | Bacteria>Bacillati>Actinomycetota>Actinomycetes>Frankiales>Frankiaceae>Frankia | 250 | MXD59_RS12255 | 78998..79750 | 573489 | ABC-transporter-ATP-binding-protein-[Frankia-sp.-Ag45/Mut15] |
| WP_131786017.1 | OLD-ABC | GCF_900634745.1 | Protofrankia_symbiont_of_Coriaria_ruscifolia | Bacteria>Bacillati>Actinomycetota>Actinomycetes>Frankiales>Frankiaceae>Protofrankia>unclassified_Protofrankia | 434 | CV1CTNOD_RS11005 | 7836..9140 | 1306542 | ATP-binding-protein-[Frankia-symbiont-of-Coriaria-ruscifolia] |
| WP_161173777.1 | OLD-ABC | GCF_039533845.1 | Streptomyces | Bacteria>Bacillati>Actinomycetota>Actinomycetes>Kitasatosporales>Streptomycetaceae | 470 | BJ962_RS25975 | 2064081..2065493 | 1883 | MULTISPECIES:-AAA-family-ATPase-[Streptomyces] |
| WP_260146672.1 | OLD-ABC | GCF_044743135.1 | Streptomyces | Bacteria>Bacillati>Actinomycetota>Actinomycetes>Kitasatosporales>Streptomycetaceae | 364 | EDD95_RS00800 | 186912..188006 | 1883 | AAA-family-ATPase-[Streptomyces-sp.-CEV-2-1] |
| WP_245226712.1 | OLD-ABC | GCF_017676345.1 | Streptomyces_smyrnaeus | Bacteria>Bacillati>Actinomycetota>Actinomycetes>Kitasatosporales>Streptomycetaceae>Streptomyces | 566 | JW613_RS32970 | 13291..14991 | 1387713 | AAA-family-ATPase-[Streptomyces-smyrnaeus] |
| WP_077964580.1 | OLD-ABC | GCF_009296025.1 | Streptomyces_tsukubensis | Bacteria>Bacillati>Actinomycetota>Actinomycetes>Kitasatosporales>Streptomycetaceae>Streptomyces | 471 | GBW32_RS05165 | 1159237..1160652 | 83656 | AAA-family-ATPase-[Streptomyces-tsukubensis] |
| WP_086823273.1 | OLD-ABC | GCF_040683955.1 | unclassified_Streptomyces | Bacteria>Bacillati>Actinomycetota>Actinomycetes>Kitasatosporales>Streptomycetaceae>Streptomyces | 471 | CCN56_RS03170 | 26382..27797 | 2593676 | AAA-family-ATPase-[Streptomyces-sp.-NRRL-B-24572] |
| WP_105973300.1 | OLD-ABC | GCF_002954775.1 | Streptomyces_geranii | Bacteria>Bacillati>Actinomycetota>Actinomycetes>Kitasatosporales>Streptomycetaceae>Streptomyces | 471 | CW359_RS29880 | 51723..53138 | 2058923 | AAA-family-ATPase-[Streptomyces-geranii] |
| WP_043500387.1 | OLD-ABC | GCF_000761215.1 | Streptomyces_glaucescens | Bacteria>Bacillati>Actinomycetota>Actinomycetes>Kitasatosporales>Streptomycetaceae>Streptomyces | 470 | SGLAU_RS10265 | 2377064..2378476 | 1907 | AAA-family-ATPase-[Streptomyces-glaucescens] |
| WP_189233273.1 | OLD-ABC | GCF_014648815.1 | Streptomyces_flaveolus | Bacteria>Bacillati>Actinomycetota>Actinomycetes>Kitasatosporales>Streptomycetaceae>Streptomyces | 374 | IEZ05_RS29290 | 130569..131693 | 67297 | AAA-family-ATPase-[Streptomyces-flaveolus] |
| WP_241787659.1 | OLD-ABC | GCF_000590515.1 | Streptomyces_sp._PRh5 | Bacteria>Bacillati>Actinomycetota>Actinomycetes>Kitasatosporales>Streptomycetaceae>Streptomyces>unclassified_Streptomyces | 529 | Z951_RS11095 | 76555..78144 | 1158056 | ATP-binding-protein-[Streptomyces-sp.-PRh5] |
| WP_215167462.1 | OLD-ABC | GCF_018614415.1 | Streptomyces_sp._ISL-12 | Bacteria>Bacillati>Actinomycetota>Actinomycetes>Kitasatosporales>Streptomycetaceae>Streptomyces>unclassified_Streptomyces | 474 | J7I94_RS14410 | 81063..82487 | 2819177 | AAA-family-ATPase-[Streptomyces-sp.-ISL-12] |
| WP_235479567.1 | OLD-ABC | GCF_021545335.1 | Streptomyces_sp._GQFP | Bacteria>Bacillati>Actinomycetota>Actinomycetes>Kitasatosporales>Streptomycetaceae>Streptomyces>unclassified_Streptomyces | 471 | LUX31_RS36425 | 8082628..8084043 | 2907545 | AAA-family-ATPase-[Streptomyces-sp.-GQFP] |
| WP_266686661.1 | OLD-ABC | GCF_026342335.1 | Streptomyces_sp._NBC_00268 | Bacteria>Bacillati>Actinomycetota>Actinomycetes>Kitasatosporales>Streptomycetaceae>Streptomyces>unclassified_Streptomyces | 469 | OOK47_RS16770 | 3610666..3612075 | 2975695 | AAA-family-ATPase-[Streptomyces-sp.-NBC_00268] |
| WP_215167469.1 | OLD-ABC | GCF_018614415.1 | Streptomyces_sp._ISL-12 | Bacteria>Bacillati>Actinomycetota>Actinomycetes>Kitasatosporales>Streptomycetaceae>Streptomyces>unclassified_Streptomyces | 450 | J7I94_RS14460 | 92830..94182 | 2819177 | ATP-binding-protein-[Streptomyces-sp.-ISL-12] |
| WP_102640597.1 | OLD-ABC | GCF_002879675.3 | Streptomyces_sp._SMS_SU21 | Bacteria>Bacillati>Actinomycetota>Actinomycetes>Kitasatosporales>Streptomycetaceae>Streptomyces>unclassified_Streptomyces | 374 | C1C94_RS24650 | 1679214..1680338 | 2069440 | AAA-family-ATPase-[Streptomyces-sp.-SMS_SU21] |
| WP_214971903.1 | OLD-ABC | GCF_018614615.1 | Streptomyces_sp._ISL-100 | Bacteria>Bacillati>Actinomycetota>Actinomycetes>Kitasatosporales>Streptomycetaceae>Streptomyces>unclassified_Streptomyces | 372 | J7E89_RS10770 | 53054..54172 | 2819173 | AAA-family-ATPase-[Streptomyces-sp.-ISL-100] |
| MBX3116432.1 | OLD-ABC | GCA_019638875.1 | Cryobacterium_sp. | Bacteria>Bacillati>Actinomycetota>Actinomycetes>Micrococcales>Microbacteriaceae>Cryobacterium>unclassified_Cryobacterium | 385 | KF808_03690 | 31942..33099 | 1926290 | AAA-family-ATPase-[Cryobacterium-sp.] |
| WP_090804560.1 | OLD-ABC | GCF_016862535.1 | Asanoa_ishikariensis | Bacteria>Bacillati>Actinomycetota>Actinomycetes>Micromonosporales>Micromonosporaceae>Asanoa | 420 | BLV39_RS40685 | 268343..269605 | 137265 | AAA-family-ATPase-[Asanoa-ishikariensis] |
| WP_172894358.1 | OLD-ABC | GCF_900091515.1 | Micromonospora_purpureochromogenes | Bacteria>Bacillati>Actinomycetota>Actinomycetes>Micromonosporales>Micromonosporaceae>Micromonospora | 430 | GA0074696_RS20355 | 4406211..4407503 | 47872 | DUF3696-domain-containing-protein-[Micromonospora-purpureochromogenes] |
| WP_076473573.1 | OLD-ABC | GCF_900156065.1 | Micromonospora_avicenniae | Bacteria>Bacillati>Actinomycetota>Actinomycetes>Micromonosporales>Micromonosporaceae>Micromonospora | 419 | BXA10_RS26490 | 27161..28420 | 1198245 | AAA-family-ATPase-[Micromonospora-avicenniae] |
| WP_208577499.1 | OLD-ABC | GCF_017599015.1 | Micromonospora_tulbaghiae | Bacteria>Bacillati>Actinomycetota>Actinomycetes>Micromonosporales>Micromonosporaceae>Micromonospora | 419 | J5U46_RS17360 | 29064..30323 | 479978 | AAA-family-ATPase-[Micromonospora-tulbaghiae] |
| SCF44517.1 | OLD-ABC | GCA_900091555.1 | Micromonospora_mirobrigensis | Bacteria>Bacillati>Actinomycetota>Actinomycetes>Micromonosporales>Micromonosporaceae>Micromonospora | 417 | GA0070564_110103 | 119581..120834 | 262898 | Predicted-ATPase-[Micromonospora-mirobrigensis] |
| WP_207786237.1 | OLD-ABC | GCF_003176745.1 | Micromonospora_globispora | Bacteria>Bacillati>Actinomycetota>Actinomycetes>Micromonosporales>Micromonosporaceae>Micromonospora | 399 | DLE60_RS16065 | 5554..6753 | 1450148 | AAA-family-ATPase-[Micromonospora-globispora] |
| WP_179783412.1 | OLD-ABC | - | Micromonospora_jinlongensis | Bacteria>Bacillati>Actinomycetota>Actinomycetes>Micromonosporales>Micromonosporaceae>Micromonospora | 394 | HNR22_RS05340 | 1261877..1263061 | 1287877 | AAA-family-ATPase-[Micromonospora-jinlongensis] |
| WP_203807953.1 | OLD-ABC | GCF_016862435.1 | Paractinoplanes_tereljensis | Bacteria>Bacillati>Actinomycetota>Actinomycetes>Micromonosporales>Micromonosporaceae>Paractinoplanes | 434 | Ate02nite_RS20665 | 73547..74851 | 571912 | AAA-family-ATPase-[Actinoplanes-tereljensis] |
| WP_270679269.1 | OLD-ABC | GCF_027706745.1 | Nocardiopsis_suaedae | Bacteria>Bacillati>Actinomycetota>Actinomycetes>Streptosporangiales>Nocardiopsidaceae>Nocardiopsis | 402 | O4U47_RS19135 | 28300..29508 | 3018444 | AAA-family-ATPase-[Nocardiopsis-sp.-LSu2-4] |
| WP_182841316.1 | OLD-ABC | GCF_039528335.1 | Actinomadura_namibiensis | Bacteria>Bacillati>Actinomycetota>Actinomycetes>Streptosporangiales>Thermomonosporaceae>Actinomadura | 433 | HNR61_RS01555 | 336773..338074 | 182080 | DUF3696-domain-containing-protein-[Actinomadura-namibiensis] |
| WP_225961000.1 | OLD-ABC | GCF_014873935.1 | Actinomadura_algeriensis | Bacteria>Bacillati>Actinomycetota>Actinomycetes>Streptosporangiales>Thermomonosporaceae>Actinomadura | 432 | H4W34_RS03570 | 877065..878363 | 1679523 | ATP-binding-protein-[Actinomadura-algeriensis] |
| WP_067482820.1 | OLD-ABC | GCF_001552635.1 | Actinomadura_hibisca | Bacteria>Bacillati>Actinomycetota>Actinomycetes>Streptosporangiales>Thermomonosporaceae>Actinomadura | 420 | AH6_RS22055 | 113363..114625 | 68565 | DUF3696-domain-containing-protein-[Actinomadura-hibisca] |
| TMR42117.1 | OLD-ABC | GCA_005889745.1 | Actinomadura_geliboluensis | Bacteria>Bacillati>Actinomycetota>Actinomycetes>Streptosporangiales>Thermomonosporaceae>Actinomadura | 419 | ETD96_02090 | 29210..30469 | 882440 | ATP-binding-protein-[Actinomadura-geliboluensis] |
| WP_242884439.1 | OLD-ABC | GCF_022751195.1 | Actinomadura_litoris | Bacteria>Bacillati>Actinomycetota>Actinomycetes>Streptosporangiales>Thermomonosporaceae>Actinomadura | 419 | K2F48_RS09345 | 532385..533644 | 2678616 | ATP-binding-protein-[Actinomadura-litoris] |
| WP_084262716.1 | OLD-ABC | GCF_001552155.1 | Actinomadura_formosensis | Bacteria>Bacillati>Actinomycetota>Actinomycetes>Streptosporangiales>Thermomonosporaceae>Actinomadura | 417 | AF1_RS33145 | 44630..45883 | 60706 | DUF3696-domain-containing-protein-[Actinomadura-formosensis] |
| WP_176233421.1 | OLD-ABC | GCF_013281955.1 | Candidatus_Hakubella_thermalkaliphila | Bacteria>Bacillati>Actinomycetota>Actinomycetota_incertae_sedis>Candidatus_Hakubellales>Candidatus_Hakubellaceae>Candidatus_Hakubella | 322 | HW440_RS05450 | 8342..9310 | 2754717 | AAA-family-ATPase-[Candidatus-Hakubanella-thermoalkaliphilus] |
| GIU94954.1 | OLD-ABC | GCA_026002995.1 | Gaiellaceae_bacterium | Bacteria>Bacillati>Actinomycetota>Thermoleophilia>Gaiellales>Gaiellaceae>unclassified_Gaiellaceae | 386 | KatS3mg012_1411 | 1399223..1400383 | 2838783 | MAG:-chromosome-segregation-protein-SMC-[Gaiellaceae-bacterium] |
| MDA8117681.1 | OLD-ABC | GCA_028064745.1 | Actinomycetota_bacterium | Bacteria>Bacillati>Actinomycetota>unclassified_Actinomycetota | 391 | M0000_09970 | 6006..7181 | 2900548 | MAG:-AAA-family-ATPase-[Actinomycetota-bacterium] |
| NLH98008.1 | OLD-ABC | GCA_012517255.1 | Chthonomonadales_bacterium | Bacteria>Bacillati>Armatimonadota>Chthonomonadia>Chthonomonadales>unclassified_Chthonomonadales | 391 | GX446_00790 | 18863..20038 | 2282151 | AAA-family-ATPase-[Chthonomonadales-bacterium] |
| NCO91846.1 | OLD-ABC | GCA_009993145.1 | Armatimonadota_bacterium | Bacteria>Bacillati>Armatimonadota>unclassified_Armatimonadota | 419 | GW880_11045 | 709..1968 | 2033014 | ATP-binding-protein-[Armatimonadota-bacterium] |
| NDK16276.1 | OLD-ABC | GCA_010119115.1 | Armatimonadota_bacterium | Bacteria>Bacillati>Armatimonadota>unclassified_Armatimonadota | 419 | GW911_29970 | 619..1878 | 2033014 | ATP-binding-protein-[Armatimonadota-bacterium] |
| PIU89758.1 | OLD-ABC | GCA_002779825.1 | Armatimonadetes_bacterium_CG06_land_8_20_14_3_00_66_21 | Bacteria>Bacillati>Armatimonadota>unclassified_Armatimonadota | 419 | COS65_27520 | 1363..2622 | 1973915 | hypothetical-protein-COS65_27520-[Armatimonadetes-bacterium-CG06_land_8_20_14_3_00_66_21] |
| PIZ50109.1 | OLD-ABC | GCA_002791775.1 | Armatimonadetes_bacterium_CG_4_10_14_0_8_um_filter_66_14 | Bacteria>Bacillati>Armatimonadota>unclassified_Armatimonadota | 419 | COY42_02305 | 1383..2642 | 1973908 | hypothetical-protein-COY42_02305-[Armatimonadetes-bacterium-CG_4_10_14_0_8_um_filter_66_14] |
| PJB73373.1 | OLD-ABC | GCA_002789735.1 | Armatimonadetes_bacterium_CG_4_9_14_3_um_filter_66_14 | Bacteria>Bacillati>Armatimonadota>unclassified_Armatimonadota | 419 | CO096_06115 | 35919..37178 | 1973914 | hypothetical-protein-CO096_06115-[Armatimonadetes-bacterium-CG_4_9_14_3_um_filter_66_14] |
| OIO95803.1 | OLD-ABC | GCA_001871675.1 | Armatimonadetes_bacterium_CG2_30_66_41 | Bacteria>Bacillati>Armatimonadota>unclassified_Armatimonadota | 407 | AUJ96_25905 | 9242..10465 | 1805028 | hypothetical-protein-AUJ96_25905-[Armatimonadetes-bacterium-CG2_30_66_41] |
| AQQ52246.1 | OLD-ABC | GCA_001999905.1 | Planococcus_lenghuensis | Bacteria>Bacillati>Bacillota>Bacilli>Bacillales>Caryophanaceae>Planococcus | 380 | B0X71_03380 | 629580..630722 | 2213202 | hypothetical-protein-B0X71_03380-[Planococcus-lenghuensis] |
| WP_251420886.1 | OLD-ABC | GCF_943192955.1 | Lactococcus | Bacteria>Bacillati>Bacillota>Bacilli>Lactobacillales>Streptococcaceae | 406 | NFR64_RS02415 | 37670..38890 | 1357 | MULTISPECIES:-ATP-binding-protein-[Lactococcus] |
| WP_058211093.1 | OLD-ABC | GCF_001456725.1 | Lactococcus_lactis | Bacteria>Bacillati>Bacillota>Bacilli>Lactobacillales>Streptococcaceae>Lactococcus | 406 | KF134_RS08795 | 319683..320903 | 1358 | ATP-binding-protein-[Lactococcus-lactis] |
| WP_191672700.1 | OLD-ABC | GCF_014830225.1 | Lactococcus_petauri | Bacteria>Bacillati>Bacillota>Bacilli>Lactobacillales>Streptococcaceae>Lactococcus | 406 | DXX96_RS03200 | 87819..89039 | 1940789 | ATP-binding-protein-[Lactococcus-petauri] |
| WP_164950926.1 | OLD-ABC | GCF_011065955.2 | Clostridium_sp._FP2 | Bacteria>Bacillati>Bacillota>Clostridia>Eubacteriales>Clostridiaceae>Clostridium>unclassified_Clostridium | 364 | G9F71_RS16305 | 3392945..3394039 | 2724481 | ATP-binding-protein-[Clostridium-sp.-FP2] |
| WP_035381185.1 | OLD-ABC | GCF_000601455.1 | Fervidicella_metallireducens | Bacteria>Bacillati>Bacillota>Clostridia>Eubacteriales>Clostridiaceae>Fervidicella | 304 | Q428_RS11720 | 10693..11607 | 655338 | ATP-binding-protein-[Fervidicella-metallireducens] |
| KGP76496.1 | OLD-ABC | GCA_000770645.1 | Desulfosporosinus_sp._Tol-M | Bacteria>Bacillati>Bacillota>Clostridia>Eubacteriales>Desulfitobacteriaceae>Desulfosporosinus>unclassified_Desulfosporosinus | 425 | JT05_04445 | 11924..13201 | 1536651 | ATPase-[Desulfosporosinus-sp.-Tol-M] |
| KGP76499.1 | OLD-ABC | GCA_000770645.1 | Desulfosporosinus_sp._Tol-M | Bacteria>Bacillati>Bacillota>Clostridia>Eubacteriales>Desulfitobacteriaceae>Desulfosporosinus>unclassified_Desulfosporosinus | 406 | JT05_04460 | 14883..16103 | 1536651 | ATPase-[Desulfosporosinus-sp.-Tol-M] |
| WP_091728839.1 | OLD-ABC | GCF_900107485.1 | Proteiniborus_ethanoligenes | Bacteria>Bacillati>Bacillota>Clostridia>Eubacteriales>Eubacteriales_incertae_sedis>Proteiniborus | 345 | BLV37_RS06305 | 20200..21237 | 415015 | AAA-family-ATPase-[Proteiniborus-ethanoligenes] |
| WP_073163777.1 | OLD-ABC | GCF_900129285.1 | Desulfofundulus_australicus | Bacteria>Bacillati>Bacillota>Clostridia>Eubacteriales>Peptococcaceae>Desulfofundulus | 362 | BUB44_RS05260 | 45292..46380 | 1566 | ATP-binding-protein-[Desulfofundulus-australicus] |
| SHE91870.1 | OLD-ABC | GCA_900129285.1 | Desulfofundulus_australicus_DSM_11792 | Bacteria>Bacillati>Bacillota>Clostridia>Eubacteriales>Peptococcaceae>Desulfofundulus>Desulfofundulus_australicus | 362 | SAMN02745218_01066 | 45292..46380 | 1121425 | ATPase/GTPase,-AAA15-family-[Desulfofundulus-australicus-DSM-11792] |
| WP_270389661.1 | OLD-ABC | GCF_027661925.1 | Blautia_massiliensis_(ex_Durand_et_al._2017) | Bacteria>Bacillati>Bacillota>Clostridia>Lachnospirales>Lachnospiraceae>Blautia | 406 | PGQ23_RS05925 | 21980..23200 | 1737424 | AAA-family-ATPase-[Blautia-massiliensis] |
| TGX97923.1 | OLD-ABC | GCA_004793585.1 | Hominisplanchenecus_murintestinalis | Bacteria>Bacillati>Bacillota>Clostridia>Lachnospirales>Lachnospiraceae>Hominisplanchenecus | 438 | E5357_10800 | 27729..29045 | 2941517 | ATP-binding-protein-[Lachnospiraceae-bacterium] |
| WP_243126266.1 | OLD-ABC | GCF_015550395.1 | [Clostridium]_hylemonae | Bacteria>Bacillati>Bacillota>Clostridia>Lachnospirales>Lachnospiraceae>Lachnoclostridium | 423 | I2C33_RS01895 | 49278..50549 | 89153 | ATP-binding-protein-[[Clostridium] |
| WP_270645284.1 | OLD-ABC | GCF_027692785.1 | Merdimonas_faecis | Bacteria>Bacillati>Bacillota>Clostridia>Lachnospirales>Lachnospiraceae>Merdimonas | 392 | PG729_RS15115 | 21578..22756 | 1653435 | AAA-family-ATPase-[Merdimonas-faecis] |
| MBR4608210.1 | OLD-ABC | GCA_017536085.1 | Lachnospiraceae_bacterium | Bacteria>Bacillati>Bacillota>Clostridia>Lachnospirales>Lachnospiraceae>unclassified_Lachnospiraceae | 399 | IK041_18535 | 24995..26194 | 1898203 | AAA-family-ATPase-[Lachnospiraceae-bacterium] |
| NBJ03625.1 | OLD-ABC | GCA_009911175.1 | Lachnospiraceae_bacterium | Bacteria>Bacillati>Bacillota>Clostridia>Lachnospirales>Lachnospiraceae>unclassified_Lachnospiraceae | 361 | D3Z62_26950 | 6066..7151 | 1898203 | hypothetical-protein-[Lachnospiraceae-bacterium] |
| NBH98985.1 | OLD-ABC | GCA_009911465.1 | Lachnospiraceae_bacterium | Bacteria>Bacillati>Bacillota>Clostridia>Lachnospirales>Lachnospiraceae>unclassified_Lachnospiraceae | 338 | D3Z44_12650 | 18259..19275 | 1898203 | ATP-binding-protein-[Lachnospiraceae-bacterium] |
| WP_255228677.1 | OLD-ABC | GCF_024363565.1 | Lutispora_saccharofermentans | Bacteria>Bacillati>Bacillota>Clostridia>Lutisporales>Lutisporaceae>Lutispora | 373 | LJD61_RS16660 | 79023..80144 | 3024236 | AAA-family-ATPase-[Lutispora-saccharofermentans] |
| WP_081358601.1 | OLD-ABC | GCF_001874065.1 | Moorella_thermoacetica | Bacteria>Bacillati>Bacillota>Clostridia>Moorellales>Moorellaceae>Moorella | 417 | MOOR_RS04470 | 138519..139772 | 1525 | ATP-binding-protein-[Moorella-thermoacetica] |
| WP_099191962.1 | OLD-ABC | GCF_002724055.1 | Tepidibacter_mesophilus | Bacteria>Bacillati>Bacillota>Clostridia>Peptostreptococcales>Peptostreptococcaceae>Tepidibacter | 431 | Tmeso_RS13190 | 47802..49097 | 655607 | AAA-family-ATPase-[Tepidibacter-mesophilus] |
| WP_278281470.1 | OLD-ABC | GCF_002724055.1 | Tepidibacter_mesophilus | Bacteria>Bacillati>Bacillota>Clostridia>Peptostreptococcales>Peptostreptococcaceae>Tepidibacter | 345 | Tmeso_RS13175 | 41344..42381 | 655607 | AAA-family-ATPase-[Tepidibacter-mesophilus] |
| MCR4419672.1 | OLD-ABC | GCA_024653205.1 | Clostridia_bacterium | Bacteria>Bacillati>Bacillota>Clostridia>unclassified_Clostridia | 415 | NUV99_05990 | 81760..83007 | 2044939 | AAA-family-ATPase-[Clostridia-bacterium] |
| MBP3656730.1 | OLD-ABC | GCA_017938795.1 | Clostridia_bacterium | Bacteria>Bacillati>Bacillota>Clostridia>unclassified_Clostridia | 366 | J6K32_08550 | 20590..21690 | 2044939 | AAA-family-ATPase-[Clostridia-bacterium] |
| NCB76907.1 | OLD-ABC | GCA_009929485.1 | Negativicutes_bacterium | Bacteria>Bacillati>Bacillota>Negativicutes>unclassified_Negativicutes | 375 | EOM42_07425 | 8743..9870 | 2499140 | chromosome-segregation-protein-SMC-[Negativicutes-bacterium] |

| Protein-ID | Domain | Assembly-ID | Strain | Taxonomy | Length | Locus-Tag | Genomic-coordinates | TaxID | Fasta-defline |
|---|---|---|---|---|---|---|---|---|---|
| WP_072743895.1 | OLD-ABC | GCF_900130025.1 | Sporanaerobacter_acetigenes | Bacteria>Bacillati>Bacillota>Tissierellia>Tissierellales>Sporanaerobacteraceae>Sporanaerobacter | 345 | BUA21_RS05950 | 80504..81541 | 165813 | AAA-family-ATPase-[Sporanaerobacter-acetigenes] |
| WP_054870026.1 | OLD-ABC | GCF_001316435.1 | Caloranaerobacter_sp._TR13 | Bacteria>Bacillati>Bacillota>Tissierellia>Tissierellales>Thermohalobacteraceae>Caloranaerobacter>unclassified_Caloranaerobacter | 344 | TR13x_RS01055 | 230238..231272 | 1302151 | ATP-binding-protein-[Caloranaerobacter-sp.-TR13] |
| MBU4541892.1 | OLD-ABC | GCA_018903255.1 | Bacillota_bacterium | Bacteria>Bacillati>Bacillota>unclassified_Bacillota | 379 | KJ774_11805 | 1364..2503 | 1879010 | AAA-family-ATPase-[Bacillota-bacterium] |
| MBT9164710.1 | OLD-ABC | GCA_018725485.1 | Candidatus_Lithacetigena_glycinireducens | Bacteria>Bacillati>Candidatus_Lithacetigenota>Candidatus_Lithacetigena | 322 | DDT23_00714 | 4336..5304 | 2986055 | DNA-replication-and-repair-protein-RecF-[candidate-division-WS2-bacterium] |
| MBQ6502301.1 | OLD-ABC | GCA_017432685.1 | Flexilinea_sp. | Bacteria>Bacillati>Chloroflexota>Anaerolineae>Anaerolineales>Anaerolineaceae>Flexilinea>unclassified_Flexilinea | 421 | IJI57_00130 | 12618..13883 | 2017390 | ATP/GTP-binding-protein-[Flexilinea-sp.] |
| WP_075070886.1 | OLD-ABC | GCF_001050255.2 | Levilinea_saccharolytica | Bacteria>Bacillati>Chloroflexota>Anaerolineae>Anaerolineales>Anaerolineaceae>Levilinea | 438 | LSAC_RS18190 | 3297508..3298824 | 229921 | ATP-binding-protein-[Levilinea-saccharolytica] |
| WP_062416898.1 | OLD-ABC | GCF_001050255.2 | Levilinea_saccharolytica | Bacteria>Bacillati>Chloroflexota>Anaerolineae>Anaerolineales>Anaerolineaceae>Levilinea | 367 | LSAC_RS05155 | 237702..238805 | 229921 | AAA-family-ATPase-[Levilinea-saccharolytica] |
| MBN1317034.1 | OLD-ABC | GCA_016927355.1 | Anaerolineales_bacterium | Bacteria>Bacillati>Chloroflexota>Anaerolineae>Anaerolineales>unclassified_Anaerolineales | 414 | JXA42_16250 | 32829..34073 | 2073117 | AAA-family-ATPase-[Anaerolineales-bacterium] |
| MCL4868348.1 | OLD-ABC | GCA_023384415.1 | Anaerolineae_bacterium | Bacteria>Bacillati>Chloroflexota>Anaerolineae>unclassified_Anaerolineae | 436 | KJ063_05220 | 195789..197099 | 2052143 | AAA-family-ATPase-[Anaerolineae-bacterium] |
| MBN2007577.1 | OLD-ABC | GCA_016931835.1 | Anaerolineae_bacterium | Bacteria>Bacillati>Chloroflexota>Anaerolineae>unclassified_Anaerolineae | 433 | JXA21_29800 | 5927..7228 | 2052143 | ATP-binding-protein-[Anaerolineae-bacterium] |
| MCX7854725.1 | OLD-ABC | GCA_026417235.1 | Anaerolineae_bacterium | Bacteria>Bacillati>Chloroflexota>Anaerolineae>unclassified_Anaerolineae | 428 | N2556_01890 | 6200..7486 | 2052143 | MAG:-AAA-family-ATPase-[Anaerolineae-bacterium] |
| MCZ7568049.1 | OLD-ABC | GCA_027493995.1 | Ardenticatenaceae_bacterium | Bacteria>Bacillati>Chloroflexota>Ardenticatenia>Ardenticatenales>Ardenticatenaceae>unclassified_Ardenticatenaceae | 405 | M5U01_05520 | 1315906..1317123 | 2614247 | MAG:-AAA-family-ATPase-[Ardenticatenaceae-bacterium] |
| MCB0025414.1 | OLD-ABC | GCA_020433165.1 | Caldilinea_sp. | Bacteria>Bacillati>Chloroflexota>Caldilineae>Caldilineales>Caldilineaceae>Caldilinea>unclassified_Caldilinea | 458 | KDD91_20420 | 1044..2420 | 2293560 | AAA-family-ATPase-[Caldilinea-sp.] |
| MCB9118708.1 | OLD-ABC | GCA_020634765.1 | Caldilineaceae_bacterium | Bacteria>Bacillati>Chloroflexota>Caldilineae>Caldilineales>Caldilineaceae>unclassified_Caldilineaceae | 458 | H6640_03145 | 16245..17621 | 2614250 | AAA-family-ATPase-[Caldilineaceae-bacterium] |
| MCB9124681.1 | OLD-ABC | GCA_020634715.1 | Caldilineaceae_bacterium | Bacteria>Bacillati>Chloroflexota>Caldilineae>Caldilineales>Caldilineaceae>unclassified_Caldilineaceae | 458 | H6643_09245 | 36450..37826 | 2614250 | AAA-family-ATPase-[Caldilineaceae-bacterium] |
| MDE0631427.1 | OLD-ABC | GCA_028824195.1 | Caldilineaceae_bacterium | Bacteria>Bacillati>Chloroflexota>Caldilineae>Caldilineales>Caldilineaceae>unclassified_Caldilineaceae | 426 | OXH73_07980 | 23227..24507 | 2614250 | MAG:-AAA-family-ATPase-[Caldilineaceae-bacterium] |
| MXX27766.1 | OLD-ABC | GCA_009835815.1 | Caldilineaceae_bacterium_SB0668_bin_21 | Bacteria>Bacillati>Chloroflexota>Caldilineae>Caldilineales>Caldilineaceae>unclassified_Caldilineaceae | 426 | F4Z82_20255 | 25298..26578 | 2605264 | AAA-family-ATPase-[Caldilineaceae-bacterium-SB0668_bin_21] |
| MDE0200336.1 | OLD-ABC | GCA_028819675.1 | Caldilineaceae_bacterium | Bacteria>Bacillati>Chloroflexota>Caldilineae>Caldilineales>Caldilineaceae>unclassified_Caldilineaceae | 399 | OXK78_19175 | 3204..4403 | 2614250 | MAG:-ATP-binding-protein-[Caldilineaceae-bacterium] |
| MCY4117635.1 | OLD-ABC | GCA_026708105.1 | Caldilineaceae_bacterium | Bacteria>Bacillati>Chloroflexota>Caldilineae>Caldilineales>Caldilineaceae>unclassified_Caldilineaceae | 391 | OXF55_12180 | 32654..33829 | 2614250 | MAG:-ATP-binding-protein-[Caldilineaceae-bacterium] |
| MDE0431261.1 | OLD-ABC | GCA_028821555.1 | Caldilineaceae_bacterium | Bacteria>Bacillati>Chloroflexota>Caldilineae>Caldilineales>Caldilineaceae>unclassified_Caldilineaceae | 391 | OXH98_15975 | 80968..82143 | 2614250 | MAG:-ATP-binding-protein-[Caldilineaceae-bacterium] |
| MDE0464746.1 | OLD-ABC | GCA_028821465.1 | Caldilineaceae_bacterium | Bacteria>Bacillati>Chloroflexota>Caldilineae>Caldilineales>Caldilineaceae>unclassified_Caldilineaceae | 391 | OXH93_20175 | 4354..5529 | 2614250 | MAG:-AAA-family-ATPase-[Caldilineaceae-bacterium] |
| MCY4079346.1 | OLD-ABC | GCA_026707715.1 | Caldilineaceae_bacterium | Bacteria>Bacillati>Chloroflexota>Caldilineae>Caldilineales>Caldilineaceae>unclassified_Caldilineaceae | 370 | OXF54_03830 | 14677..15789 | 2614250 | MAG:-AAA-family-ATPase-[Caldilineaceae-bacterium] |
| MXX28117.1 | OLD-ABC | GCA_009835815.1 | Caldilineaceae_bacterium_SB0668_bin_21 | Bacteria>Bacillati>Chloroflexota>Caldilineae>Caldilineales>Caldilineaceae>unclassified_Caldilineaceae | 370 | F4Z82_22085 | 4095..5207 | 2605264 | AAA-family-ATPase-[Caldilineaceae-bacterium-SB0668_bin_21] |
| MBK9096489.1 | OLD-ABC | GCA_016719785.1 | Candidatus_Amarolinea_dominans | Bacteria>Bacillati>Chloroflexota>Caldilineae>Caldilineales>Candidatus_Amarolineaceae>Candidatus_Amarolinea | 386 | IPM84_27775 | 243762..244922 | 3140696 | AAA-family-ATPase-[Anaerolineae-bacterium] |
| MBK9093482.1 | OLD-ABC | GCA_016719785.1 | Candidatus_Amarolinea_dominans | Bacteria>Bacillati>Chloroflexota>Caldilineae>Caldilineales>Candidatus_Amarolineaceae>Candidatus_Amarolinea | 365 | IPM84_12050 | 185120..186217 | 3140696 | AAA-family-ATPase-[Anaerolineae-bacterium] |
| MBK6429945.1 | OLD-ABC | GCA_016704685.1 | Candidatus_Amarolinea_dominans | Bacteria>Bacillati>Chloroflexota>Caldilineae>Caldilineales>Candidatus_Amarolineaceae>Candidatus_Amarolinea | 360 | IPF85_04080 | 212815..213897 | 3140696 | AAA-family-ATPase-[Anaerolineae-bacterium] |
| MBK9093188.1 | OLD-ABC | GCA_016719785.1 | Candidatus_Amarolinea_dominans | Bacteria>Bacillati>Chloroflexota>Caldilineae>Caldilineales>Candidatus_Amarolineaceae>Candidatus_Amarolinea | 360 | IPM84_10465 | 142254..143336 | 3140696 | AAA-family-ATPase-[Anaerolineae-bacterium] |
| WP_129673426.1 | OLD-ABC | GCF_004138165.1 | Candidatus_Chloroploca_sp._Khr17 | Bacteria>Bacillati>Chloroflexota>Chloroflexia>Chloroflexales>Chloroflexineae>Oscillochloridaceae>Candidatus_Chloroploca>unclassified_Candidatus_Chloroploca | 422 | EKD05_RS13135 | 7143..8411 | 2496869 | ATP-binding-protein-[Candidatus-Chloroploca-sp.-Khr17] |
| WP_200808051.1 | OLD-ABC | - | Thermoflexus_hugenholtzii | Bacteria>Bacillati>Chloroflexota>Thermoflexia>Thermoflexales>Thermoflexaceae>Thermoflexus | 373 | CFB18_RS02800 | 21767..22888 | 1495650 | AAA-family-ATPase-[Thermoflexus-hugenholtzii] |
| HHY55157.1 | OLD-ABC | GCA_012842785.1 | Chloroflexota_bacterium | Bacteria>Bacillati>Chloroflexota>unclassified_Chloroflexota | 477 | GYA08_06940 | 23032..24465 | 2026724 | AAA-family-ATPase-[Chloroflexota-bacterium] |
| MBM3127992.1 | OLD-ABC | GCA_016865935.1 | Chloroflexota_bacterium | Bacteria>Bacillati>Chloroflexota>unclassified_Chloroflexota | 437 | FJ009_05070 | 19987..21300 | 2026724 | ATP-binding-protein-[Chloroflexota-bacterium] |
| MBI2863487.1 | OLD-ABC | GCA_016190625.1 | Chloroflexota_bacterium | Bacteria>Bacillati>Chloroflexota>unclassified_Chloroflexota | 429 | HYX94_02875 | 4988..6277 | 2026724 | AAA-family-ATPase-[Chloroflexota-bacterium] |
| OGN72034.1 | OLD-ABC | GCA_001796335.1 | Chloroflexi_bacterium_GWB2_49_20 | Bacteria>Bacillati>Chloroflexota>unclassified_Chloroflexota | 427 | A2X25_12865 | 143944..145227 | 1797612 | hypothetical-protein-A2X25_12865-[Chloroflexi-bacterium-GWB2_49_20] |
| MXZ89879.1 | OLD-ABC | GCA_009837355.1 | Chloroflexota_bacterium | Bacteria>Bacillati>Chloroflexota>unclassified_Chloroflexota | 426 | F4Z05_00150 | 22280..23560 | 2026724 | AAA-family-ATPase-[Chloroflexota-bacterium] |
| MYD49643.1 | OLD-ABC | GCA_009841575.1 | Chloroflexota_bacterium | Bacteria>Bacillati>Chloroflexota>unclassified_Chloroflexota | 426 | F4W95_14355 | 9405..10685 | 2026724 | AAA-family-ATPase-[Chloroflexota-bacterium] |
| MBI4327841.1 | OLD-ABC | GCA_016210465.1 | Chloroflexota_bacterium | Bacteria>Bacillati>Chloroflexota>unclassified_Chloroflexota | 418 | HY674_21635 | 2729..3985 | 2026724 | AAA-family-ATPase-[Chloroflexota-bacterium] |
| MCE7937790.1 | OLD-ABC | GCA_021404695.1 | Chloroflexi_bacterium_CFX6 | Bacteria>Bacillati>Chloroflexota>unclassified_Chloroflexota | 401 | DYG90_04155 | 3871..5076 | 2293710 | hypothetical-protein-[Chloroflexi-bacterium-CFX6] |
| MYC29510.1 | OLD-ABC | GCA_009840135.1 | Chloroflexota_bacterium | Bacteria>Bacillati>Chloroflexota>unclassified_Chloroflexota | 249 | F4X65_05405 | 6361..7110 | 2026724 | ATP-binding-protein-[Chloroflexota-bacterium] |
| PZV08802.1 | OLD-ABC | GCA_003249105.1 | Leptolyngbya_sp. | Bacteria>Bacillati>Cyanobacteriota/Melainabacteria_group>Cyanobacteriota>Cyanophyceae>Leptolyngbyales>Leptolyngbyaceae>Leptolyngbya_group>Leptolyngbya>unclassified_Leptolyngbya | 410 | DCF32_03940 | 19980..21212 | 47254 | chromosome-segregation-protein-SMC-[Leptolyngbya-sp.] |
| WP_127084965.1 | OLD-ABC | GCF_003991905.1 | Dulcicalothrix_desertica | Bacteria>Bacillati>Cyanobacteriota/Melainabacteria_group>Cyanobacteriota>Cyanophyceae>Nostocales>Calotrichaceae>Dulcicalothrix | 414 | DSM106972_RS34695 | 74210..75454 | 32056 | ATP-binding-protein-[Dulcicalothrix-desertica] |
| WP_127084955.1 | OLD-ABC | GCF_003991905.1 | Dulcicalothrix_desertica | Bacteria>Bacillati>Cyanobacteriota/Melainabacteria_group>Cyanobacteriota>Cyanophyceae>Nostocales>Calotrichaceae>Dulcicalothrix | 362 | DSM106972_RS34645 | 65604..66692 | 32056 | AAA-family-ATPase-[Dulcicalothrix-desertica] |
| MBR8836045.1 | OLD-ABC | GCA_016903675.1 | Stigonema_ocellatum_SAG_48.90_=_DSM_106950 | Bacteria>Bacillati>Cyanobacteriota/Melainabacteria_group>Cyanobacteriota>Cyanophyceae>Nostocales>Stigonemataceae>Stigonema>Stigonema_ocellatum | 408 | DSM106950_18980 | 44596..45822 | 221294 | AAA-family-ATPase-[Stigonema-ocellatum-SAG-48.90-=-DSM-106950] |
| MBR8836040.1 | OLD-ABC | GCA_016903675.1 | Stigonema_ocellatum_SAG_48.90_=_DSM_106950 | Bacteria>Bacillati>Cyanobacteriota/Melainabacteria_group>Cyanobacteriota>Cyanophyceae>Nostocales>Stigonemataceae>Stigonema>Stigonema_ocellatum | 345 | DSM106950_18955 | 37134..38171 | 221294 | ATP-binding-protein-[Stigonema-ocellatum-SAG-48.90-=-DSM-106950] |
| MCA6584619.1 | OLD-ABC | GCA_020404925.1 | Pseudanabaena_sp._M34BS1SP1A06MG | Bacteria>Bacillati>Cyanobacteriota/Melainabacteria_group>Cyanobacteriota>Cyanophyceae>Pseudanabaenales>Pseudanabaenaceae>Pseudanabaena>unclassified_Pseudanabaena | 522 | IM536_21885 | 12713..14281 | 2771240 | AAA-family-ATPase-[Pseudanabaena-sp.-M34BS1SP1A06MG] |
| MCA6593931.1 | OLD-ABC | GCA_020404945.1 | Pseudanabaena_sp._M38BS1SP1A06MG | Bacteria>Bacillati>Cyanobacteriota/Melainabacteria_group>Cyanobacteriota>Cyanophyceae>Pseudanabaenales>Pseudanabaenaceae>Pseudanabaena>unclassified_Pseudanabaena | 522 | IM535_17915 | 12741..14309 | 2771238 | AAA-family-ATPase-[Pseudanabaena-sp.-M38BS1SP1A06MG] |
| MCA6597705.1 | OLD-ABC | GCA_020405005.1 | Pseudanabaena_sp._M046S1SP1A06QC | Bacteria>Bacillati>Cyanobacteriota/Melainabacteria_group>Cyanobacteriota>Cyanophyceae>Pseudanabaenales>Pseudanabaenaceae>Pseudanabaena>unclassified_Pseudanabaena | 522 | IM539_15745 | 12705..14273 | 2771236 | AAA-family-ATPase-[Pseudanabaena-sp.-M046S1SP1A06QC] |
| MCF8140152.1 | OLD-ABC | GCA_021737825.1 | Cyanobium_usitatum_Tobar12.5m-G36 | Bacteria>Bacillati>Cyanobacteriota/Melainabacteria_group>Cyanobacteriota>Cyanophyceae>Synechococcales>Prochlorococcaceae>Cyanobium>Cyanobium_usitatum | 390 | K9J75_03470 | 35944..37116 | 2874848 | AAA-family-ATPase-[Cyanobium-usitatum-Tobar12.5m-G36] |
| MCX6842952.1 | OLD-ABC | GCA_026397595.1 | candidate_division_WOR-3_bacterium | Bacteria>Bacteria_incertae_sedis>Bacteria_candidate_phyla>candidate_division_WOR-3 | 370 | NTX53_11795 | 8555..9667 | 2052148 | MAG:-AAA-family-ATPase-[candidate-division-WOR-3-bacterium] |
| MCG2812994.1 | OLD-ABC | GCA_022013915.1 | Candidatus_Aminicenantes_bacterium | Bacteria>Bacteria_incertae_sedis>Bacteria_candidate_phyla>Candidatus_Aminicenantota>unclassified_Candidatus_Aminicenantes | 400 | L6428_16310 | 3541..4743 | 2052149 | AAA-family-ATPase-[Candidatus-Aminicenantes-bacterium] |
| QAA75942.1 | OLD-ABC | GCA_004102645.1 | Candidatus_Bipolaricaulis_sibiricus | Bacteria>Bacteria_incertae_sedis>Bacteria_candidate_phyla>Candidatus_Bipolaricaulota>Candidatus_Bipolaricaulia>Candidatus_Bipolaricaulales>Candidatus_Bipolaricaulaceae>Candidatus_Bipolaricaulis | 396 | BIP78_0174 | 166151..167341 | 2501609 | hypothetical-protein-BIP78_0174-[Candidatus-Bipolaricaulis-sibiricus] |
| MDD5559006.1 | OLD-ABC | GCA_028716865.1 | Candidatus_Methylomirabilis_sp. | Bacteria>Bacteria_incertae_sedis>Bacteria_candidate_phyla>Candidatus_Methylomirabilota>Candidatus_Methylomirabilia>Candidatus_Methylomirabilales>Candidatus_Methylomirabilaceae>Candidatus_Methylomirabilis>unclassified_Candidatus_Methylomirabilis | 387 | PHV01_05030 | 51086..52249 | 2032687 | MAG:-AAA-family-ATPase-[Candidatus-Methylomirabilis-sp.] |
| MDE2058887.1 | OLD-ABC | GCA_028864815.1 | Candidatus_Methylomirabilota_bacterium | Bacteria>Bacteria_incertae_sedis>Bacteria_candidate_phyla>Candidatus_Methylomirabilota>unclassified_Candidatus_Methylomirabilota | 434 | KGJ27_04060 | 13099..14403 | 2072417 | MAG:-ATP-binding-protein-[candidate-division-NC10-bacterium] |
| MBI4536778.1 | OLD-ABC | GCA_016209735.1 | Candidatus_Methylomirabilota_bacterium | Bacteria>Bacteria_incertae_sedis>Bacteria_candidate_phyla>Candidatus_Methylomirabilota>unclassified_Candidatus_Methylomirabilota | 412 | HY712_02340 | 65565..66803 | 2072417 | ATP-binding-protein-[candidate-division-NC10-bacterium] |
| MDE2058201.1 | OLD-ABC | GCA_028864815.1 | Candidatus_Methylomirabilota_bacterium | Bacteria>Bacteria_incertae_sedis>Bacteria_candidate_phyla>Candidatus_Methylomirabilota>unclassified_Candidatus_Methylomirabilota | 386 | KGJ27_00515 | 27734..28894 | 2072417 | MAG:-AAA-family-ATPase-[candidate-division-NC10-bacterium] |
| MDE0183889.1 | OLD-ABC | GCA_028819845.1 | Candidatus_Poribacteria_bacterium | Bacteria>Bacteria_incertae_sedis>Bacteria_candidate_phyla>Candidatus_Poribacteria>unclassified_Candidatus_Poribacteria | 499 | OXP71_00365 | 80049..81548 | 2026781 | MAG:-AAA-family-ATPase-[Candidatus-Poribacteria-bacterium] |
| MDE0183898.1 | OLD-ABC | GCA_028819845.1 | Candidatus_Poribacteria_bacterium | Bacteria>Bacteria_incertae_sedis>Bacteria_candidate_phyla>Candidatus_Poribacteria>unclassified_Candidatus_Poribacteria | 423 | OXP71_00410 | 95947..97218 | 2026781 | MAG:-AAA-family-ATPase-[Candidatus-Poribacteria-bacterium] |
| MBF0500208.1 | OLD-ABC | GCA_015233155.1 | Candidatus_Rifleibacteriota_bacterium | Bacteria>Bacteria_incertae_sedis>Bacteria_candidate_phyla>Candidatus_Rifleibacteriota>unclassified_Candidatus_Rifleibacteriota | 369 | HQM09_08745 | 71133..72242 | 2053586 | AAA-family-ATPase-[Candidatus-Riflebacteria-bacterium] |
| MBI2567005.1 | OLD-ABC | GCA_016188575.1 | Candidatus_Schekmanbacteria_bacterium | Bacteria>Bacteria_incertae_sedis>Bacteria_candidate_phyla>Candidatus_Schekmaniibacteriota>unclassified_Candidatus_Schekmanbacteria | 394 | HYV63_08255 | 60103..61287 | 2419984 | AAA-family-ATPase-[Candidatus-Schekmanbacteria-bacterium] |
| MBN1883902.1 | OLD-ABC | GCA_016931295.1 | Candidatus_Krumholzibacteriota_bacterium | Bacteria>Candidatus_Krumholzibacteriota>unclassified_Candidatus_Krumholzibacteriota | 391 | JW876_00090 | 26952..28127 | 2725313 | AAA-family-ATPase-[Candidatus-Krumholzibacteriota-bacterium] |
| PJB45677.1 | OLD-ABC | GCA_002789955.1 | Deltaproteobacteria_bacterium_CG_4_9_14_3_um_filter_63_12 | Bacteria>delta/epsilon_subdivisions>Deltaproteobacteria>unclassified_Deltaproteobacteria | 423 | CO108_06835 | 11430..12701 | 1973967 | ATPase-[Deltaproteobacteria-bacterium-CG_4_9_14_3_um_filter_63_12] |
| OGQ97434.1 | OLD-ABC | GCA_001799275.1 | Deltaproteobacteria_bacterium_RIFOXYD12_FULL_57_12 | Bacteria>delta/epsilon_subdivisions>Deltaproteobacteria>unclassified_Deltaproteobacteria | 422 | A2521_08595 | 18315..19583 | 1797906 | RloA-protein-[Deltaproteobacteria-bacterium-RIFOXYD12_FULL_57_12] |
| EFK06062.1 | OLD-ABC | - | delta_proteobacterium_NaphS2 | Bacteria>delta/epsilon_subdivisions>Deltaproteobacteria>unclassified_Deltaproteobacteria | 370 | NPH_2625 | 41309..42421 | 88274 | RecF/RecN/SMC-N-terminal-domain-protein-[delta-proteobacterium-NaphS2] |
| WP_008700609.1 | OLD-ABC | GCF_000273625.1 | Fusobacterium | Bacteria>Fusobacteriati>Fusobacteriota>Fusobacteriia>Fusobacteriales>Fusobacteriaceae | 370 | J141_RS0104315 | 26351..27463 | 848 | MULTISPECIES:-ATP-binding-protein-[Fusobacterium] |
| WP_022069637.1 | OLD-ABC | GCF_023383545.1 | Fusobacterium | Bacteria>Fusobacteriati>Fusobacteriota>Fusobacteriia>Fusobacteriales>Fusobacteriaceae | 370 | YWH7199_RS06530 | 21537..22649 | 848 | ATP-binding-protein-[Fusobacterium-nucleatum] |
| WP_032836037.1 | OLD-ABC | GCF_045159995.1 | Fusobacterium | Bacteria>Fusobacteriati>Fusobacteriota>Fusobacteriia>Fusobacteriales>Fusobacteriaceae | 370 | HMPREF1501_RS01105 | 62221..63333 | 848 | ATP-binding-protein-[Fusobacterium-sp.-OBRC1] |
| WP_032841324.1 | OLD-ABC | GCF_024140205.1 | Fusobacterium | Bacteria>Fusobacteriati>Fusobacteriota>Fusobacteriia>Fusobacteriales>Fusobacteriaceae | 370 | BKN39_RS05490 | 9678..10790 | 848 | MULTISPECIES:-ATP-binding-protein-[Fusobacterium] |
| WP_238972694.1 | OLD-ABC | GCF_037911825.1 | Fusobacterium | Bacteria>Fusobacteriati>Fusobacteriota>Fusobacteriia>Fusobacteriales>Fusobacteriaceae | 370 | K5I19_RS02510 | 131829..132941 | 848 | ATP-binding-protein-[Fusobacterium-nucleatum] |
| WP_005903410.1 | OLD-ABC | GCF_000178895.1 | Fusobacterium_nucleatum | Bacteria>Fusobacteriati>Fusobacteriota>Fusobacteriia>Fusobacteriales>Fusobacteriaceae>Fusobacterium | 370 | C4N14_RS08070 | 1666126..1667238 | 851 | ATP-binding-protein-[Fusobacterium-nucleatum] |
| WP_008692036.1 | OLD-ABC | GCF_000218655.1 | Fusobacterium_animalis | Bacteria>Fusobacteriati>Fusobacteriota>Fusobacteriia>Fusobacteriales>Fusobacteriaceae>Fusobacterium | 370 | HMPREF0401_RS10115 | 561221..562333 | 76859 | ATP-binding-protein-[Fusobacterium-animalis] |
| WP_008797394.1 | OLD-ABC | GCF_000163915.2 | Fusobacterium_vincentii | Bacteria>Fusobacteriati>Fusobacteriota>Fusobacteriia>Fusobacteriales>Fusobacteriaceae>Fusobacterium | 370 | HMPREF0405_RS05635 | 1191673..1192785 | 155615 | ATP-binding-protein-[Fusobacterium-vincentii] |
| WP_023036665.1 | OLD-ABC | GCF_048313155.1 | Fusobacterium_polymorphum | Bacteria>Fusobacteriati>Fusobacteriota>Fusobacteriia>Fusobacteriales>Fusobacteriaceae>Fusobacterium | 370 | HMPREF1767_RS09530 | 32157..33269 | 76857 | AAA-family-ATPase-[Fusobacterium-nucleatum] |
| WP_060676306.1 | OLD-ABC | GCF_001296085.1 | Fusobacterium_animalis | Bacteria>Fusobacteriati>Fusobacteriota>Fusobacteriia>Fusobacteriales>Fusobacteriaceae>Fusobacterium | 370 | RN98_RS07265 | 1460186..1461298 | 76859 | ATP-binding-protein-[Fusobacterium-animalis] |
| WP_088387743.1 | OLD-ABC | GCF_002204435.1 | Fusobacterium_polymorphum | Bacteria>Fusobacteriati>Fusobacteriota>Fusobacteriia>Fusobacteriales>Fusobacteriaceae>Fusobacterium | 370 | CA839_RS02155 | 470997..472109 | 76857 | ATP-binding-protein-[Fusobacterium-polymorphum] |
| WP_098702329.1 | OLD-ABC | GCF_002573625.1 | Fusobacterium_polymorphum | Bacteria>Fusobacteriati>Fusobacteriota>Fusobacteriia>Fusobacteriales>Fusobacteriaceae>Fusobacterium | 370 | RN96_RS03475 | 673419..674531 | 76857 | ATP-binding-protein-[Fusobacterium-polymorphum] |
| WP_098998445.1 | OLD-ABC | GCF_037897565.1 | Fusobacterium_polymorphum | Bacteria>Fusobacteriati>Fusobacteriota>Fusobacteriia>Fusobacteriales>Fusobacteriaceae>Fusobacterium | 370 | CBG58_RS00345 | 81538..82650 | 76857 | AAA-family-ATPase-[Fusobacterium-polymorphum] |
| WP_099959236.1 | OLD-ABC | GCF_002761955.1 | Fusobacterium_pseudoperiodonticum | Bacteria>Fusobacteriati>Fusobacteriota>Fusobacteriia>Fusobacteriales>Fusobacteriaceae>Fusobacterium | 370 | CTM71_RS09950 | 1965937..1967049 | 2663009 | ATP-binding-protein-[Fusobacterium-pseudoperiodonticum] |
| WP_099972120.1 | OLD-ABC | GCF_963546905.1 | Fusobacterium_pseudoperiodonticum | Bacteria>Fusobacteriati>Fusobacteriota>Fusobacteriia>Fusobacteriales>Fusobacteriaceae>Fusobacterium | 370 | CTM69_RS07780 | 479995..481107 | 2663009 | ATP-binding-protein-[Fusobacterium-pseudoperiodonticum] |
| WP_099987861.1 | OLD-ABC | GCF_002763695.1 | Fusobacterium_pseudoperiodonticum | Bacteria>Fusobacteriati>Fusobacteriota>Fusobacteriia>Fusobacteriales>Fusobacteriaceae>Fusobacterium | 370 | CTM74_RS09000 | 1719799..1720911 | 2663009 | ATP-binding-protein-[Fusobacterium-pseudoperiodonticum] |
| WP_100024173.1 | OLD-ABC | GCF_002763625.1 | Fusobacterium_pseudoperiodonticum | Bacteria>Fusobacteriati>Fusobacteriota>Fusobacteriia>Fusobacteriales>Fusobacteriaceae>Fusobacterium | 370 | CTM72_RS01090 | 203772..204884 | 2663009 | ATP-binding-protein-[Fusobacterium-pseudoperiodonticum] |
| WP_124794842.1 | OLD-ABC | GCF_003859915.1 | Fusobacterium_canifelinum | Bacteria>Fusobacteriati>Fusobacteriota>Fusobacteriia>Fusobacteriales>Fusobacteriaceae>Fusobacterium | 370 | EII27_RS00785 | 176919..178031 | 285729 | ATP-binding-protein-[Fusobacterium-canifelinum] |
| WP_158410488.1 | OLD-ABC | GCF_050219895.1 | Fusobacterium_animalis | Bacteria>Fusobacteriati>Fusobacteriota>Fusobacteriia>Fusobacteriales>Fusobacteriaceae>Fusobacterium | 370 | CI111_RS06865 | 5293..6405 | 76859 | ATP-binding-protein-[Fusobacterium-animalis] |
| WP_220302802.1 | OLD-ABC | GCF_019552045.1 | Fusobacterium_polymorphum | Bacteria>Fusobacteriati>Fusobacteriota>Fusobacteriia>Fusobacteriales>Fusobacteriaceae>Fusobacterium | 370 | JY397_RS08305 | 1735382..1736494 | 76857 | ATP-binding-protein-[Fusobacterium-polymorphum] |
| WP_220308656.1 | OLD-ABC | GCF_019552125.1 | Fusobacterium_animalis | Bacteria>Fusobacteriati>Fusobacteriota>Fusobacteriia>Fusobacteriales>Fusobacteriaceae>Fusobacterium | 370 | JY401_RS04545 | 953553..954665 | 76859 | ATP-binding-protein-[Fusobacterium-animalis] |
| WP_265152269.1 | OLD-ABC | GCF_026089295.1 | Fusobacterium_simiae | Bacteria>Fusobacteriati>Fusobacteriota>Fusobacteriia>Fusobacteriales>Fusobacteriaceae>Fusobacterium | 370 | OCK72_RS06715 | 33963..35075 | 855 | ATP-binding-protein-[Fusobacterium-simiae] |
| ALF25293.1 | OLD-ABC | GCA_001296185.1 | Fusobacterium_nucleatum_subsp._nucleatum | Bacteria>Fusobacteriati>Fusobacteriota>Fusobacteriia>Fusobacteriales>Fusobacteriaceae>Fusobacterium>Fusobacterium_nucleatum | 370 | RN95_02155 | 459089..460201 | 76856 | chromosome-segregation-protein-SMC-[Fusobacterium-nucleatum-subsp.-nucleatum] |
| CDA09059.1 | OLD-ABC | GCA_000433695.1 | Fusobacterium_sp._CAG:649 | Bacteria>Fusobacteriati>Fusobacteriota>Fusobacteriia>Fusobacteriales>Fusobacteriaceae>Fusobacterium>unclassified_Fusobacterium | 370 | BN748_00883 | 21814..22926 | 1262900 | putative-uncharacterized-protein-[Fusobacterium-sp.-CAG:649] |
| WP_032846250.1 | OLD-ABC | GCF_000524395.1 | Fusobacterium_sp._CM22 | Bacteria>Fusobacteriati>Fusobacteriota>Fusobacteriia>Fusobacteriales>Fusobacteriaceae>Fusobacterium>unclassified_Fusobacterium | 370 | HMPREF1500_RS09195 | 123181..124293 | 936563 | ATP-binding-protein-[Fusobacterium-sp.-CM22] |
| WP_070766193.1 | OLD-ABC | GCF_001815715.1 | Fusobacterium_sp._HMSC064B11 | Bacteria>Fusobacteriati>Fusobacteriota>Fusobacteriia>Fusobacteriales>Fusobacteriaceae>Fusobacterium>unclassified_Fusobacterium | 370 | HMPREF3051_RS04305 | 255735..256847 | 1739543 | ATP-binding-protein-[Fusobacterium-sp.-HMSC064B11] |
| WP_108080575.1 | OLD-ABC | GCF_019285575.1 | Pseudomonadota | Bacteria>Pseudomonadati | 386 | KW113_RS00560 | 119898..121058 | 1224 | MULTISPECIES:-AAA-family-ATPase-[Pseudomonadota] |
| QQR76219.1 | OLD-ABC | GCA_016699405.1 | Holophagales_bacterium | Bacteria>Pseudomonadati>Acidobacteriota>Holophagae>Holophagales>unclassified_Holophagales | 391 | IPJ17_18300 | 4433549..4434724 | 2478486 | AAA-family-ATPase-[Holophagales-bacterium] |
| MCY4597198.1 | OLD-ABC | GCA_026713455.1 | Bryobacterales_bacterium | Bacteria>Pseudomonadati>Acidobacteriota>Terriglobia>Bryobacterales>unclassified_Bryobacterales | 364 | OXC19_20650 | 12389..13483 | 2026791 | MAG:-AAA-family-ATPase-[Bryobacterales-bacterium] |
| MXY68844.1 | OLD-ABC | GCA_009838515.1 | Terriglobia_bacterium | Bacteria>Pseudomonadati>Acidobacteriota>Terriglobia>unclassified_Terriglobia | 367 | F4Y47_09830 | 224752..225855 | 2081523 | AAA-family-ATPase-[Acidobacteriia-bacterium] |
| MYK11748.1 | OLD-ABC | GCA_009845805.1 | Terriglobia_bacterium | Bacteria>Pseudomonadati>Acidobacteriota>Terriglobia>unclassified_Terriglobia | 367 | F4041_19175 | 3011..4114 | 2081523 | AAA-family-ATPase-[Acidobacteriia-bacterium] |
| MBP7799278.1 | OLD-ABC | GCA_018000955.1 | Thermoanaerobaculaceae_bacterium | Bacteria>Pseudomonadati>Acidobacteriota>Thermoanaerobaculia>Thermoanaerobaculales>Thermoanaerobaculaceae>unclassified_Thermoanaerobaculaceae | 450 | KA072_11705 | 26875..28227 | 2699758 | DUF3696-domain-containing-protein-[Thermoanaerobaculaceae-bacterium] |
| MCU0292533.1 | OLD-ABC | GCA_025456225.1 | Thermoanaerobaculaceae_bacterium | Bacteria>Pseudomonadati>Acidobacteriota>Thermoanaerobaculia>Thermoanaerobaculales>Thermoanaerobaculaceae>unclassified_Thermoanaerobaculaceae | 420 | MUF10_11200 | 9187..10449 | 2699758 | ATP-binding-protein-[Thermoanaerobaculaceae-bacterium] |
| MBP7799237.1 | OLD-ABC | GCA_018000955.1 | Thermoanaerobaculaceae_bacterium | Bacteria>Pseudomonadati>Acidobacteriota>Thermoanaerobaculia>Thermoanaerobaculales>Thermoanaerobaculaceae>unclassified_Thermoanaerobaculaceae | 410 | KA072_11465 | 12494..13726 | 2699758 | AAA-family-ATPase-[Thermoanaerobaculaceae-bacterium] |
| MBP7799281.1 | OLD-ABC | GCA_018000955.1 | Thermoanaerobaculaceae_bacterium | Bacteria>Pseudomonadati>Acidobacteriota>Thermoanaerobaculia>Thermoanaerobaculales>Thermoanaerobaculaceae>unclassified_Thermoanaerobaculaceae | 386 | KA072_11720 | 31790..32950 | 2699758 | AAA-family-ATPase-[Thermoanaerobaculaceae-bacterium] |
| MCG3190934.1 | OLD-ABC | GCA_022071945.1 | Thermoanaerobaculia_bacterium | Bacteria>Pseudomonadati>Acidobacteriota>Thermoanaerobaculia>unclassified_Thermoanaerobaculia | 422 | DIJKHBIC_00157 | 192662..193930 | 2651171 | hypothetical-protein-[Thermoanaerobaculia-bacterium] |
| MCG3192296.1 | OLD-ABC | GCA_022071945.1 | Thermoanaerobaculia_bacterium | Bacteria>Pseudomonadati>Acidobacteriota>Thermoanaerobaculia>unclassified_Thermoanaerobaculia | 398 | DIJKHBIC_01534 | 16761..17957 | 2651171 | hypothetical-protein-[Thermoanaerobaculia-bacterium] |
| MCG3192272.1 | OLD-ABC | GCA_022071945.1 | Thermoanaerobaculia_bacterium | Bacteria>Pseudomonadati>Acidobacteriota>Thermoanaerobaculia>unclassified_Thermoanaerobaculia | 387 | DIJKHBIC_01510 | 132245..133408 | 2651171 | ATP/GTP-phosphatase-[Thermoanaerobaculia-bacterium] |
| MCY4659242.1 | OLD-ABC | GCA_026713985.1 | Acidobacteriota_bacterium | Bacteria>Pseudomonadati>Acidobacteriota>unclassified_Acidobacteriota | 452 | OXF93_05435 | 4557..5915 | 1978231 | MAG:-AAA-family-ATPase-[Acidobacteria-bacterium] |
| MCY3965605.1 | OLD-ABC | GCA_026707545.1 | Acidobacteriota_bacterium | Bacteria>Pseudomonadati>Acidobacteriota>unclassified_Acidobacteriota | 451 | OXG83_11250 | 2827626..2828981 | 1978231 | MAG:-ATP-binding-protein-[Acidobacteria-bacterium] |
| MYN67295.1 | OLD-ABC | GCA_009861545.1 | Acidobacteriota_bacterium | Bacteria>Pseudomonadati>Acidobacteriota>unclassified_Acidobacteriota | 440 | F4X11_20075 | 75908..77230 | 1978231 | AAA-family-ATPase-[Acidobacteriota-bacterium] |
| MYN68211.1 | OLD-ABC | GCA_009861545.1 | Acidobacteriota_bacterium | Bacteria>Pseudomonadati>Acidobacteriota>unclassified_Acidobacteriota | 434 | F4X11_24870 | 54606..55910 | 1978231 | AAA-family-ATPase-[Acidobacteriota-bacterium] |
| MYN67292.1 | OLD-ABC | GCA_009861545.1 | Acidobacteriota_bacterium | Bacteria>Pseudomonadati>Acidobacteriota>unclassified_Acidobacteriota | 433 | F4X11_20055 | 69974..71275 | 1978231 | AAA-family-ATPase-[Acidobacteriota-bacterium] |
| MCC6130027.1 | OLD-ABC | GCA_020845555.1 | Acidobacteriota_bacterium | Bacteria>Pseudomonadati>Acidobacteriota>unclassified_Acidobacteriota | 398 | IT186_08860 | 23656..24852 | 1978231 | AAA-family-ATPase-[Acidobacteriota-bacterium] |
| MBI4902185.1 | OLD-ABC | GCA_016213775.1 | Acidobacteriota_bacterium | Bacteria>Pseudomonadati>Acidobacteriota>unclassified_Acidobacteriota | 397 | HY820_01030 | 15729..16922 | 1978231 | AAA-family-ATPase-[Acidobacteriota-bacterium] |
| MCY4659246.1 | OLD-ABC | GCA_026713985.1 | Acidobacteriota_bacterium | Bacteria>Pseudomonadati>Acidobacteriota>unclassified_Acidobacteriota | 385 | OXF93_05455 | 11686..12843 | 1978231 | MAG:-ATP-binding-protein-[Acidobacteria-bacterium] |
| MCY3965600.1 | OLD-ABC | GCA_026707545.1 | Acidobacteriota_bacterium | Bacteria>Pseudomonadati>Acidobacteriota>unclassified_Acidobacteriota | 378 | OXG83_11225 | 2820800..2821936 | 1978231 | MAG:-AAA-family-ATPase-[Acidobacteria-bacterium] |
| MCC6131987.1 | OLD-ABC | GCA_020845555.1 | Acidobacteriota_bacterium | Bacteria>Pseudomonadati>Acidobacteriota>unclassified_Acidobacteriota | 376 | IT186_18855 | 13412..14542 | 1978231 | AAA-family-ATPase-[Acidobacteriota-bacterium] |
| MDE2979140.1 | OLD-ABC | GCA_028873735.1 | Acidobacteriota_bacterium | Bacteria>Pseudomonadati>Acidobacteriota>unclassified_Acidobacteriota | 375 | OXU63_16710 | 11412..12539 | 1978231 | MAG:-AAA-family-ATPase-[Acidobacteriota-bacterium] |
| MYN66270.1 | OLD-ABC | GCA_009861545.1 | Acidobacteriota_bacterium | Bacteria>Pseudomonadati>Acidobacteriota>unclassified_Acidobacteriota | 369 | F4X11_14765 | 6823..7932 | 1978231 | ATP-binding-protein-[Acidobacteriota-bacterium] |
| MBI4676022.1 | OLD-ABC | GCA_016209035.1 | Elusimicrobiota_bacterium | Bacteria>Pseudomonadati>Elusimicrobiota>unclassified_Elusimicrobiota | 411 | HY748_00380 | 77545..78780 | 2030800 | ATP-binding-protein-[Elusimicrobiota-bacterium] |

| Protein-ID | Domain | Assembly-ID | Strain | Taxonomy | Length | Locus-Tag | Genomic-coordinates | TaxID | Fasta-defline |
|---|---|---|---|---|---|---|---|---|---|
| MBI4676020.1 | OLD-ABC | GCA_016209035.1 | Elusimicrobiota_bacterium | Bacteria>Pseudomonadati>Elusimicrobiota>unclassified_Elusimicrobiota | 398 | HY748_00370 | 75692..76888 | 2030800 | AAA-family-ATPase-[Elusimicrobiota-bacterium] |
| MBI5244006.1 | OLD-ABC | GCA_016212885.1 | Elusimicrobiota_bacterium | Bacteria>Pseudomonadati>Elusimicrobiota>unclassified_Elusimicrobiota | 340 | HY922_10075 | 1..1024 | 2030800 | ATP-binding-protein-[Elusimicrobiota-bacterium] |
| WP_172774763.1 | OLD-ABC | GCF_963929675.1 | Bacteroidales | Bacteria>Pseudomonadati>FCB_group>Bacteroidota/Chlorobiota_group>Bacteroidota>Bacteroidia | 404 | HKQ53_RS16465 | 73401..74615 | 171549 | MULTISPECIES:-ATP-binding-protein-[Bacteroidaceae] |
| WP_217745554.1 | OLD-ABC | GCF_019127365.1 | Prevotellaceae | Bacteria>Pseudomonadati>FCB_group>Bacteroidota/Chlorobiota_group>Bacteroidota>Bacteroidia>Bacteroidales | 239 | KSW85_RS06180 | 44745..45464 | 171552 | AAA-family-ATPase-[Prevotella-copri] |
| WP_148330684.1 | OLD-ABC | GCF_025567125.1 | Phocaeicola | Bacteria>Pseudomonadati>FCB_group>Bacteroidota/Chlorobiota_group>Bacteroidota>Bacteroidia>Bacteroidales>Bacteroidaceae | 239 | H6B28_RS04780 | 130362..131081 | 909656 | MULTISPECIES:-AAA-family-ATPase-[Bacteroidaceae] |
| WP_072542575.1 | OLD-ABC | GCF_900128455.1 | Bacteroides_mediterraneensis | Bacteria>Pseudomonadati>FCB_group>Bacteroidota/Chlorobiota_group>Bacteroidota>Bacteroidia>Bacteroidales>Bacteroidaceae>Bacteroides | 239 | BQ7480_RS08755 | 2128307..2129026 | 1841856 | AAA-family-ATPase-[Bacteroides-mediterraneensis] |
| WP_072542582.1 | OLD-ABC | GCF_900128455.1 | Bacteroides_mediterraneensis | Bacteria>Pseudomonadati>FCB_group>Bacteroidota/Chlorobiota_group>Bacteroidota>Bacteroidia>Bacteroidales>Bacteroidaceae>Bacteroides | 235 | BQ7480_RS08805 | 2138529..2139236 | 1841856 | ABC-transporter-ATP-binding-protein-[Bacteroides-mediterraneensis] |
| CDD33289.1 | OLD-ABC | GCA_000436055.1 | Bacteroides_sp._CAG:714 | Bacteria>Pseudomonadati>FCB_group>Bacteroidota/Chlorobiota_group>Bacteroidota>Bacteroidia>Bacteroidales>Bacteroidaceae>Bacteroides>unclassified_Bacteroides | 239 | BN762_00980 | 2895..3614 | 1262749 | putative-uncharacterized-protein-[Bacteroides-sp.-CAG:714] |
| CDD50094.1 | OLD-ABC | GCA_000435415.1 | Bacteroides_sp._CAG:875 | Bacteria>Pseudomonadati>FCB_group>Bacteroidota/Chlorobiota_group>Bacteroidota>Bacteroidia>Bacteroidales>Bacteroidaceae>Bacteroides>unclassified_Bacteroides | 235 | BN800_01979 | 11275..11982 | 1262752 | putative-uncharacterized-protein-[Bacteroides-sp.-CAG:875] |
| CDD50105.1 | OLD-ABC | GCA_000435415.1 | Bacteroides_sp._CAG:875 | Bacteria>Pseudomonadati>FCB_group>Bacteroidota/Chlorobiota_group>Bacteroidota>Bacteroidia>Bacteroidales>Bacteroidaceae>Bacteroides>unclassified_Bacteroides | 148 | BN800_01990 | 21865..22311 | 1262752 | putative-uncharacterized-protein-[Bacteroides-sp.-CAG:875] |
| HJF82220.1 | OLD-ABC | GCA_020742805.1 | Phocaeicola_plebeius | Bacteria>Pseudomonadati>FCB_group>Bacteroidota/Chlorobiota_group>Bacteroidota>Bacteroidia>Bacteroidales>Bacteroidaceae>Phocaeicola | 239 | K8V40_11350 | 7719..8438 | 310297 | AAA-family-ATPase-[Phocaeicola-plebeius] |
| WP_118441919.1 | OLD-ABC | GCF_003474535.1 | Phocaeicola_plebeius | Bacteria>Pseudomonadati>FCB_group>Bacteroidota/Chlorobiota_group>Bacteroidota>Bacteroidia>Bacteroidales>Bacteroidaceae>Phocaeicola | 239 | DW041_RS13380 | 8176..8895 | 310297 | AAA-family-ATPase-[Phocaeicola-plebeius] |
| MCR4768679.1 | OLD-ABC | GCA_024682395.1 | Bacteroidaceae_bacterium | Bacteria>Pseudomonadati>FCB_group>Bacteroidota/Chlorobiota_group>Bacteroidota>Bacteroidia>Bacteroidales>Bacteroidaceae>unclassified_Bacteroidaceae | 383 | K5874_00455 | 1481..2632 | 2212467 | ATP-binding-protein-[Bacteroidaceae-bacterium] |
| HAH91107.1 | OLD-ABC | GCA_003450425.1 | Prevotella_sp. | Bacteria>Pseudomonadati>FCB_group>Bacteroidota/Chlorobiota_group>Bacteroidota>Bacteroidia>Bacteroidales>Prevotellaceae>Prevotella>unclassified_Prevotella | 239 | DCL96_05235 | 11412..12131 | 59823 | hypothetical-protein-[Prevotella-sp.] |
| MBN2915287.1 | OLD-ABC | GCA_016939035.1 | Prevotella_sp. | Bacteria>Pseudomonadati>FCB_group>Bacteroidota/Chlorobiota_group>Bacteroidota>Bacteroidia>Bacteroidales>Prevotellaceae>Prevotella>unclassified_Prevotella | 239 | JTJ26_04210 | 24276..24995 | 59823 | AAA-family-ATPase-[Prevotella-sp.] |
| WP_094881846.1 | OLD-ABC | GCF_002265625.1 | Prevotella_sp._885 | Bacteria>Pseudomonadati>FCB_group>Bacteroidota/Chlorobiota_group>Bacteroidota>Bacteroidia>Bacteroidales>Prevotellaceae>Prevotella>unclassified_Prevotella | 239 | CHL74_RS14185 | 3364..4083 | 2022527 | AAA-family-ATPase-[Prevotella-sp.-885] |
| MCI6336759.1 | OLD-ABC | GCA_022777825.1 | Prevotella_sp. | Bacteria>Pseudomonadati>FCB_group>Bacteroidota/Chlorobiota_group>Bacteroidota>Bacteroidia>Bacteroidales>Prevotellaceae>Prevotella>unclassified_Prevotella | 238 | MR802_02265 | 2646..3362 | 59823 | ATP-binding-protein-[Prevotella-sp.] |
| MCI6195663.1 | OLD-ABC | GCA_022773545.1 | Prevotella_sp. | Bacteria>Pseudomonadati>FCB_group>Bacteroidota/Chlorobiota_group>Bacteroidota>Bacteroidia>Bacteroidales>Prevotellaceae>Prevotella>unclassified_Prevotella | 236 | MR670_00300 | 48847..49557 | 59823 | ATP-binding-protein-[Prevotella-sp.] |
| WP_118139865.1 | OLD-ABC | GCF_003465445.1 | Segatella_copri | Bacteria>Pseudomonadati>FCB_group>Bacteroidota/Chlorobiota_group>Bacteroidota>Bacteroidia>Bacteroidales>Prevotellaceae>Segatella | 239 | DWV60_RS07150 | 35503..36222 | 165179 | AAA-family-ATPase-[Prevotella-copri] |
| MCL2726875.1 | OLD-ABC | GCA_009781275.1 | Bacteroidales_bacterium | Bacteria>Pseudomonadati>FCB_group>Bacteroidota/Chlorobiota_group>Bacteroidota>Bacteroidia>Bacteroidales>unclassified_Bacteroidales | 428 | FWD56_00660 | 17757..19043 | 2030927 | AAA-family-ATPase-[Bacteroidales-bacterium] |
| MCQ2228647.1 | OLD-ABC | GCA_024401335.1 | Bacteroidales_bacterium | Bacteria>Pseudomonadati>FCB_group>Bacteroidota/Chlorobiota_group>Bacteroidota>Bacteroidia>Bacteroidales>unclassified_Bacteroidales | 405 | MJZ13_02745 | 170864..172081 | 2030927 | ATP-binding-protein-[Bacteroidales-bacterium] |
| MBD5181933.1 | OLD-ABC | GCA_014800235.1 | Bacteroidales_bacterium | Bacteria>Pseudomonadati>FCB_group>Bacteroidota/Chlorobiota_group>Bacteroidota>Bacteroidia>Bacteroidales>unclassified_Bacteroidales | 356 | HDS98_05220 | 27934..29004 | 2030927 | AAA-family-ATPase-[Bacteroidales-bacterium] |
| WP_188169947.1 | OLD-ABC | GCF_014534745.1 | Flammeovirga_sp._EKP202 | Bacteria>Pseudomonadati>FCB_group>Bacteroidota/Chlorobiota_group>Bacteroidota>Cytophagia>Cytophagales>Flammeovirgaceae>Flammeovirga>unclassified_Flammeovirga | 373 | IC221_RS16080 | 60639..61760 | 2770592 | ATP-binding-protein-[Flammeovirga-sp.-EKP202] |
| MBX7052696.1 | OLD-ABC | GCA_019694805.1 | Flavobacteriales_bacterium | Bacteria>Pseudomonadati>FCB_group>Bacteroidota/Chlorobiota_group>Bacteroidota>Flavobacteriia>Flavobacteriales>unclassified_Flavobacteriales | 397 | K1X54_11720 | 17246..18439 | 2021391 | AAA-family-ATPase-[Flavobacteriales-bacterium] |
| WP_020534909.1 | OLD-ABC | GCF_000379805.1 | Lewinella_cohaerens | Bacteria>Pseudomonadati>FCB_group>Bacteroidota/Chlorobiota_group>Bacteroidota>Saprospiria>Saprospirales>Lewinellaceae>Lewinella | 711 | A3EU_RS0104375 | 124062..126197 | 70995 | AAA-family-ATPase-[Lewinella-cohaerens] |
| MCB9053306.1 | OLD-ABC | GCA_020636405.1 | Lewinellaceae_bacterium | Bacteria>Pseudomonadati>FCB_group>Bacteroidota/Chlorobiota_group>Bacteroidota>Saprospiria>Saprospirales>Lewinellaceae>unclassified_Lewinellaceae | 454 | H6556_28045 | 308951..310315 | 2760807 | ATP-binding-protein-[Lewinellaceae-bacterium] |
| MCB9053320.1 | OLD-ABC | GCA_020636405.1 | Lewinellaceae_bacterium | Bacteria>Pseudomonadati>FCB_group>Bacteroidota/Chlorobiota_group>Bacteroidota>Saprospiria>Saprospirales>Lewinellaceae>unclassified_Lewinellaceae | 419 | H6556_28115 | 339350..340609 | 2760807 | ATP-binding-protein-[Lewinellaceae-bacterium] |
| MBK8920121.1 | OLD-ABC | GCA_016719895.1 | Saprospirales_bacterium | Bacteria>Pseudomonadati>FCB_group>Bacteroidota/Chlorobiota_group>Bacteroidota>Saprospiria>Saprospirales>unclassified_Saprospirales | 393 | IPM81_01195 | 287970..289151 | 2026790 | AAA-family-ATPase-[Saprospirales-bacterium] |
| MCC5934983.1 | OLD-ABC | GCA_020832405.1 | Balneolales_bacterium | Bacteria>Pseudomonadati>FCB_group>Bacteroidota/Chlorobiota_group>Balneolota>Balneolia>Balneolales>unclassified_Balneolales | 427 | JJU35_12125 | 8329..9612 | 2497613 | AAA-family-ATPase-[Balneolales-bacterium] |
| MBV5328347.1 | OLD-ABC | GCA_019162945.1 | Chlorobium_sp. | Bacteria>Pseudomonadati>FCB_group>Bacteroidota/Chlorobiota_group>Chlorobiota>Chlorobiia>Chlorobiales>Chlorobiaceae>Chlorobium/Pelodictyon_group>Chlorobium>unclassified_Chlorobium | 377 | JZU65_12030 | 5382..6515 | 1095 | AAA-family-ATPase-[Chlorobium-sp.] |
| TNJ34082.1 | OLD-ABC | GCA_006265245.1 | Prosthecochloris_vibrioformis | Bacteria>Pseudomonadati>FCB_group>Bacteroidota/Chlorobiota_group>Chlorobiota>Chlorobiia>Chlorobiales>Chlorobiaceae>Prosthecochloris | 724 | FGF68_10425 | 7233..9407 | 1098 | hypothetical-protein-FGF68_10425-[Prosthecochloris-vibrioformis] |
| MBK7265053.1 | OLD-ABC | GCA_016707285.1 | Ignavibacteriales_bacterium | Bacteria>Pseudomonadati>FCB_group>Bacteroidota/Chlorobiota_group>Ignavibacteriota>Ignavibacteria>Ignavibacteriales>unclassified_Ignavibacteriales | 435 | IPI12_01700 | 359423..360730 | 2049428 | ATP-binding-protein-[Ignavibacteriales-bacterium] |
| MCB5285392.1 | OLD-ABC | GCA_020532085.1 | Candidatus_Cloacimonadota_bacterium | Bacteria>Pseudomonadati>FCB_group>Candidatus_Cloacimonadota>unclassified_Candidatus_Cloacimonadota | 396 | LHW45_07375 | 27124..28314 | 2030808 | AAA-family-ATPase-[Candidatus-Cloacimonadota-bacterium] |
| MCB5285396.1 | OLD-ABC | GCA_020532085.1 | Candidatus_Cloacimonadota_bacterium | Bacteria>Pseudomonadati>FCB_group>Candidatus_Cloacimonadota>unclassified_Candidatus_Cloacimonadota | 382 | LHW45_07395 | 32698..33846 | 2030808 | AAA-family-ATPase-[Candidatus-Cloacimonadota-bacterium] |
| MBP7719543.1 | OLD-ABC | GCA_017999735.1 | Candidatus_Fermentibacter_sp. | Bacteria>Pseudomonadati>FCB_group>Candidatus_Fermentibacterota>Candidatus_Fermentibacteria_(class)>Candidatus_Fermentibacterales>Candidatus_Fermentibacteraceae>Candidatus_Fermentibacter>unclassified_Candidatus_Fermentibacter | 391 | KA108_02780 | 12685..13860 | 2779163 | AAA-family-ATPase-[Candidatus-Fermentibacter-sp.] |
| NOQ23477.1 | OLD-ABC | GCA_013138965.1 | Candidatus_Aegiribacteria_sp. | Bacteria>Pseudomonadati>FCB_group>FCB_group_incertae_sedis>Candidatus_Aegiribacteria>unclassified_Candidatus_Aegiribacteria | 601 | GQ565_12635 | 23308..25113 | 2528271 | ATP-binding-cassette-domain-containing-protein-[Candidatus-Aegiribacteria-sp.] |
| NOQ23478.1 | OLD-ABC | GCA_013138965.1 | Candidatus_Aegiribacteria_sp. | Bacteria>Pseudomonadati>FCB_group>FCB_group_incertae_sedis>Candidatus_Aegiribacteria>unclassified_Candidatus_Aegiribacteria | 595 | GQ565_12640 | 25110..26897 | 2528271 | ATP-binding-cassette-domain-containing-protein-[Candidatus-Aegiribacteria-sp.] |
| NLD91454.1 | OLD-ABC | GCA_012513915.1 | Fibrobacter_sp. | Bacteria>Pseudomonadati>FCB_group>Fibrobacterota>Fibrobacteria>Fibrobacterales>Fibrobacteraceae>Fibrobacter>unclassified_Fibrobacter | 416 | GX639_02175 | 59484..60734 | 35828 | AAA-family-ATPase-[Fibrobacter-sp.] |
| MYA34701.1 | OLD-ABC | GCA_009838765.1 | Gemmatimonadales_bacterium | Bacteria>Pseudomonadati>FCB_group>Gemmatimonadota>Gemmatimonadia>Gemmatimonadales>unclassified_Gemmatimonadales | 380 | F4Y33_13785 | 66577..67719 | 2448054 | AAA-family-ATPase-[Gemmatimonadales-bacterium] |
| MXX77942.1 | OLD-ABC | GCA_009836785.1 | Gemmatimonadales_bacterium | Bacteria>Pseudomonadati>FCB_group>Gemmatimonadota>Gemmatimonadia>Gemmatimonadales>unclassified_Gemmatimonadales | 211 | F4Z33_02935 | 1..638 | 2448054 | AAA-family-ATPase-[Gemmatimonadales-bacterium] |
| MXX38676.1 | OLD-ABC | GCA_009835785.1 | Gemmatimonadota_bacterium | Bacteria>Pseudomonadati>FCB_group>Gemmatimonadota>unclassified_Gemmatimonadota | 436 | F4Z85_11550 | 4574..5884 | 2026742 | AAA-family-ATPase-[Gemmatimonadota-bacterium] |
| MDE2723060.1 | OLD-ABC | GCA_028875455.1 | Gemmatimonadota_bacterium | Bacteria>Pseudomonadati>FCB_group>Gemmatimonadota>unclassified_Gemmatimonadota | 414 | OXI59_06760 | 2548..3792 | 2026742 | MAG:-AAA-family-ATPase-[Gemmatimonadota-bacterium] |
| MYD59774.1 | OLD-ABC | GCA_009841665.1 | Gemmatimonadota_bacterium | Bacteria>Pseudomonadati>FCB_group>Gemmatimonadota>unclassified_Gemmatimonadota | 413 | F4W91_01915 | 33198..34439 | 2026742 | AAA-family-ATPase-[Gemmatimonadota-bacterium] |
| MDE2654308.1 | OLD-ABC | GCA_028875795.1 | Gemmatimonadota_bacterium | Bacteria>Pseudomonadati>FCB_group>Gemmatimonadota>unclassified_Gemmatimonadota | 391 | OXI71_10910 | 87162..88337 | 2026742 | MAG:-AAA-family-ATPase-[Gemmatimonadota-bacterium] |
| MBX6362424.1 | OLD-ABC | GCA_019689775.1 | Gemmatimonadota_bacterium | Bacteria>Pseudomonadati>FCB_group>Gemmatimonadota>unclassified_Gemmatimonadota | 383 | IRZ00_01040 | 21478..22629 | 2026742 | AAA-family-ATPase-[Gemmatimonadota-bacterium] |
| MYB67638.1 | OLD-ABC | GCA_009840005.1 | Gemmatimonadota_bacterium | Bacteria>Pseudomonadati>FCB_group>Gemmatimonadota>unclassified_Gemmatimonadota | 374 | F4X75_03965 | 10601..11725 | 2026742 | AAA-family-ATPase-[Gemmatimonadota-bacterium] |
| MCY4603565.1 | OLD-ABC | GCA_026713655.1 | Gemmatimonadota_bacterium | Bacteria>Pseudomonadati>FCB_group>Gemmatimonadota>unclassified_Gemmatimonadota | 368 | OXE49_04975 | 4907..6013 | 2026742 | MAG:-ATP-binding-protein-[Gemmatimonadetes-bacterium] |
| MXX38673.1 | OLD-ABC | GCA_009835785.1 | Gemmatimonadota_bacterium | Bacteria>Pseudomonadati>FCB_group>Gemmatimonadota>unclassified_Gemmatimonadota | 205 | F4Z85_11535 | 1..620 | 2026742 | ATP-binding-protein-[Gemmatimonadota-bacterium] |
| MCB9544271.1 | OLD-ABC | GCA_020633435.1 | Myxococcales_bacterium | Bacteria>Pseudomonadati>Myxococcota>Myxococcia>Myxococcales>unclassified_Myxococcales | 483 | H6706_00080 | 20003..21454 | 2026763 | AAA-family-ATPase-[Myxococcales-bacterium] |
| HCF56390.1 | OLD-ABC | GCA_003534605.1 | Myxococcales_bacterium | Bacteria>Pseudomonadati>Myxococcota>Myxococcia>Myxococcales>unclassified_Myxococcales | 405 | DFS52_00135 | 5014..6231 | 2026763 | ATPase-[Myxococcales-bacterium] |
| MCG5052486.1 | OLD-ABC | GCA_022184915.1 | Myxococcales_bacterium | Bacteria>Pseudomonadati>Myxococcota>Myxococcia>Myxococcales>unclassified_Myxococcales | 399 | KA712_05965 | 1444932..1446131 | 2026763 | AAA-family-ATPase-[Myxococcales-bacterium] |
| MBL9045674.1 | OLD-ABC | GCA_016794465.1 | Myxococcales_bacterium | Bacteria>Pseudomonadati>Myxococcota>Myxococcia>Myxococcales>unclassified_Myxococcales | 391 | JNM83_28955 | 7822..8997 | 2026763 | AAA-family-ATPase-[Myxococcales-bacterium] |
| MAG32377.1 | OLD-ABC | GCA_002688255.1 | Deltaproteobacteria_bacterium | Bacteria>Pseudomonadati>Myxococcota>Myxococcia>unclassified_Myxococcia | 852 | CL908_15960 | 677..3235 | 2026735 | phosphoesterase-[Deltaproteobacteria-bacterium] |
| MBO6936775.1 | OLD-ABC | GCA_017644045.1 | Deltaproteobacteria_bacterium | Bacteria>Pseudomonadati>Myxococcota>Myxococcia>unclassified_Myxococcia | 448 | JJ863_17530 | 235451..236797 | 2026735 | AAA-family-ATPase-[Deltaproteobacteria-bacterium] |
| MBN1905587.1 | OLD-ABC | GCA_016931115.1 | Deltaproteobacteria_bacterium | Bacteria>Pseudomonadati>Myxococcota>Myxococcia>unclassified_Myxococcia | 447 | JW927_10860 | 36938..38281 | 2026735 | ATP-binding-protein-[Deltaproteobacteria-bacterium] |
| MBI3795504.1 | OLD-ABC | GCA_016200595.1 | Deltaproteobacteria_bacterium | Bacteria>Pseudomonadati>Myxococcota>Myxococcia>unclassified_Myxococcia | 425 | HY268_00840 | 6048..7325 | 2026735 | AAA-family-ATPase-[Deltaproteobacteria-bacterium] |
| MDD9873631.1 | OLD-ABC | GCA_028817295.1 | Deltaproteobacteria_bacterium | Bacteria>Pseudomonadati>Myxococcota>Myxococcia>unclassified_Myxococcia | 424 | OXU65_09080 | 7582..8856 | 2026735 | MAG:-AAA-family-ATPase-[Deltaproteobacteria-bacterium] |
| MCC6811535.1 | OLD-ABC | GCA_020848545.1 | Deltaproteobacteria_bacterium | Bacteria>Pseudomonadati>Myxococcota>Myxococcia>unclassified_Myxococcia | 418 | IT381_29160 | 80328..81584 | 2026735 | ATP-binding-protein-[Deltaproteobacteria-bacterium] |
| PXF54132.1 | OLD-ABC | GCA_003194485.1 | Deltaproteobacteria_bacterium | Bacteria>Pseudomonadati>Myxococcota>Myxococcia>unclassified_Myxococcia | 413 | C4B57_08260 | 23603..24844 | 2026735 | ATP-binding-protein-[Deltaproteobacteria-bacterium] |
| MBW1922687.1 | OLD-ABC | GCA_019307945.1 | Deltaproteobacteria_bacterium | Bacteria>Pseudomonadati>Myxococcota>Myxococcia>unclassified_Myxococcia | 411 | JRJ35_04375 | 3766..5001 | 2026735 | AAA-family-ATPase-[Deltaproteobacteria-bacterium] |
| RLC08400.1 | OLD-ABC | GCA_003647435.1 | Deltaproteobacteria_bacterium | Bacteria>Pseudomonadati>Myxococcota>Myxococcia>unclassified_Myxococcia | 411 | DRH43_10125 | 1671..2906 | 2026735 | hypothetical-protein-DRH43_10125-[Deltaproteobacteria-bacterium] |
| MCC6811538.1 | OLD-ABC | GCA_020848545.1 | Deltaproteobacteria_bacterium | Bacteria>Pseudomonadati>Myxococcota>Myxococcia>unclassified_Myxococcia | 410 | IT381_29175 | 86545..87777 | 2026735 | ATP-binding-protein-[Deltaproteobacteria-bacterium] |
| MBM4396667.1 | OLD-ABC | GCA_016875475.1 | Deltaproteobacteria_bacterium | Bacteria>Pseudomonadati>Myxococcota>Myxococcia>unclassified_Myxococcia | 404 | FJ087_13385 | 3779..4993 | 2026735 | AAA-family-ATPase-[Deltaproteobacteria-bacterium] |
| MBT3255201.1 | OLD-ABC | GCA_018644785.1 | Deltaproteobacteria_bacterium | Bacteria>Pseudomonadati>Myxococcota>Myxococcia>unclassified_Myxococcia | 402 | HN366_01910 | 14018..15226 | 2026735 | AAA-family-ATPase-[Deltaproteobacteria-bacterium] |
| MBI4706187.1 | OLD-ABC | GCA_016209145.1 | Deltaproteobacteria_bacterium | Bacteria>Pseudomonadati>Myxococcota>Myxococcia>unclassified_Myxococcia | 397 | HY744_34255 | 7364..8557 | 2026735 | AAA-family-ATPase-[Deltaproteobacteria-bacterium] |
| MAG32383.1 | OLD-ABC | GCA_002688255.1 | Deltaproteobacteria_bacterium | Bacteria>Pseudomonadati>Myxococcota>Myxococcia>unclassified_Myxococcia | 396 | CL908_15990 | 9781..10971 | 2026735 | chromosome-segregation-protein-SMC-[Deltaproteobacteria-bacterium] |
| MDD3553581.1 | OLD-ABC | GCA_028695735.1 | Deltaproteobacteria_bacterium | Bacteria>Pseudomonadati>Myxococcota>Myxococcia>unclassified_Myxococcia | 391 | PHC35_03565 | 34179..35354 | 2026735 | MAG:-AAA-family-ATPase-[Deltaproteobacteria-bacterium] |
| PXF57193.1 | OLD-ABC | GCA_003194495.1 | Deltaproteobacteria_bacterium | Bacteria>Pseudomonadati>Myxococcota>Myxococcia>unclassified_Myxococcia | 390 | C4B58_10705 | 13983..15155 | 2026735 | hypothetical-protein-C4B58_10705-[Deltaproteobacteria-bacterium] |
| TRZ75308.1 | OLD-ABC | GCA_007279985.1 | Deltaproteobacteria_bacterium | Bacteria>Pseudomonadati>Myxococcota>Myxococcia>unclassified_Myxococcia | 389 | D4R93_05180 | 2303..3472 | 2026735 | chromosome-segregation-protein-SMC-[Deltaproteobacteria-bacterium] |
| NLN59463.1 | OLD-ABC | GCA_012522415.1 | Deltaproteobacteria_bacterium | Bacteria>Pseudomonadati>Myxococcota>Myxococcia>unclassified_Myxococcia | 379 | GX147_01905 | 36251..37390 | 2026735 | AAA-family-ATPase-[Deltaproteobacteria-bacterium] |
| MBI4706264.1 | OLD-ABC | GCA_016209145.1 | Deltaproteobacteria_bacterium | Bacteria>Pseudomonadati>Myxococcota>Myxococcia>unclassified_Myxococcia | 377 | HY744_34640 | 16015..17148 | 2026735 | AAA-family-ATPase-[Deltaproteobacteria-bacterium] |
| MBI4818892.1 | OLD-ABC | GCA_016208175.1 | Deltaproteobacteria_bacterium | Bacteria>Pseudomonadati>Myxococcota>Myxococcia>unclassified_Myxococcia | 375 | HY791_21655 | 1495..2622 | 2026735 | AAA-family-ATPase-[Deltaproteobacteria-bacterium] |
| MBW1911018.1 | OLD-ABC | GCA_019307765.1 | Deltaproteobacteria_bacterium | Bacteria>Pseudomonadati>Myxococcota>Myxococcia>unclassified_Myxococcia | 190 | JRJ11_16025 | 8387..8960 | 2026735 | ATP-binding-protein-[Deltaproteobacteria-bacterium] |
| MCX5910050.1 | OLD-ABC | GCA_026388955.1 | Deltaproteobacteria_bacterium | Bacteria>Pseudomonadati>Myxococcota>Myxococcia>unclassified_Myxococcia | 126 | NTY64_23455 | 1..382 | 2026735 | MAG:-AAA-family-ATPase,-partial-[Deltaproteobacteria-bacterium] |
| MBK8262771.1 | OLD-ABC | GCA_016712315.1 | Nannocystis_sp. | Bacteria>Pseudomonadati>Myxococcota>Polyangia>Nannocystales>Nannocystaceae>Nannocystis>unclassified_Nannocystis | 423 | IPK80_15730 | 180500..181771 | 1962667 | AAA-family-ATPase-[Nannocystis-sp.] |
| WP_129573701.1 | OLD-ABC | GCF_004135755.1 | Sorangium | Bacteria>Pseudomonadati>Myxococcota>Polyangia>Polyangiales>Polyangiaceae | 389 | SOCE836_RS08210 | 2235280..2236449 | 39643 | MULTISPECIES:-AAA-family-ATPase-[Sorangium] |
| WP_129345943.1 | OLD-ABC | GCF_004135735.1 | Sorangium_cellulosum | Bacteria>Pseudomonadati>Myxococcota>Polyangia>Polyangiales>Polyangiaceae>Sorangium | 389 | SOCEGT47_RS05515 | 1455987..1457156 | 56 | AAA-family-ATPase-[Sorangium-cellulosum] |
| MBK8937972.1 | OLD-ABC | GCA_016719945.1 | Polyangiaceae_bacterium | Bacteria>Pseudomonadati>Myxococcota>Polyangia>Polyangiales>Polyangiaceae>unclassified_Polyangiaceae | 550 | IPM79_10090 | 430896..432548 | 2268199 | AAA-family-ATPase-[Polyangiaceae-bacterium] |
| MBK6517527.1 | OLD-ABC | GCA_016704335.1 | Polyangiaceae_bacterium | Bacteria>Pseudomonadati>Myxococcota>Polyangia>Polyangiales>Polyangiaceae>unclassified_Polyangiaceae | 412 | IPG04_26280 | 279060..280298 | 2268199 | AAA-family-ATPase-[Polyangiaceae-bacterium] |
| MBN2191666.1 | OLD-ABC | GCA_016933275.1 | Polyangiaceae_bacterium | Bacteria>Pseudomonadati>Myxococcota>Polyangia>Polyangiales>Polyangiaceae>unclassified_Polyangiaceae | 407 | JW751_02525 | 18812..20035 | 2268199 | AAA-family-ATPase-[Polyangiaceae-bacterium] |
| MCU0671309.1 | OLD-ABC | GCA_025453555.1 | Myxococcota_bacterium | Bacteria>Pseudomonadati>Myxococcota>unclassified_Myxococcota | 424 | MUE69_00805 | 253..1527 | 2818507 | AAA-family-ATPase-[Myxococcota-bacterium] |
| MBP7125457.1 | OLD-ABC | GCA_017994695.1 | Myxococcota_bacterium | Bacteria>Pseudomonadati>Myxococcota>unclassified_Myxococcota | 405 | KBD49_03715 | 62279..63496 | 2818507 | AAA-family-ATPase-[Myxococcota-bacterium] |
| MCK6575222.1 | OLD-ABC | GCA_023150895.1 | Myxococcota_bacterium | Bacteria>Pseudomonadati>Myxococcota>unclassified_Myxococcota | 401 | L6V77_29500 | 974..2179 | 2818507 | ATP-binding-protein-[Myxococcota-bacterium] |
| MCK6575225.1 | OLD-ABC | GCA_023150895.1 | Myxococcota_bacterium | Bacteria>Pseudomonadati>Myxococcota>unclassified_Myxococcota | 397 | L6V77_29515 | 7319..8512 | 2818507 | AAA-family-ATPase-[Myxococcota-bacterium] |
| MCS6797216.1 | OLD-ABC | GCA_025054695.1 | Myxococcota_bacterium | Bacteria>Pseudomonadati>Myxococcota>unclassified_Myxococcota | 380 | NZ898_01580 | 57207..58349 | 2818507 | AAA-family-ATPase-[Myxococcota-bacterium] |
| MCC6543976.1 | OLD-ABC | GCA_020846775.1 | Nitrospirota_bacterium | Bacteria>Pseudomonadati>Nitrospirota>unclassified_Nitrospirota | 393 | IT392_05660 | 169358..170539 | 2026887 | AAA-family-ATPase-[Nitrospirota-bacterium] |
| PIX82645.1 | OLD-ABC | GCA_002783945.1 | Nitrospirae_bacterium_CG_4_10_14_3_um_filter_70_108 | Bacteria>Pseudomonadati>Nitrospirota>unclassified_Nitrospirota | 391 | COZ33_09555 | 2411..3586 | 1974054 | chromosome-segregation-protein-SMC-[Nitrospirae-bacterium-CG_4_10_14_3_um_filter_70_108] |
| WP_000595812.1 | OLD-ABC | GCF_000299535.2 | Gammaproteobacteria | Bacteria>Pseudomonadati>Pseudomonadota | 389 | B934_RS0100875 | 9222..10391 | 1236 | MULTISPECIES:-AAA-family-ATPase-[Gammaproteobacteria] |
| WP_014998459.1 | OLD-ABC | - | Gammaproteobacteria | Bacteria>Pseudomonadati>Pseudomonadota | 370 | AMBLS11_RS09810 | 2292738..2293850 | 1236 | AAA-family-ATPase-[Alteromonas-macleodii] |
| WP_022992172.1 | OLD-ABC | GCF_000475355.1 | Gammaproteobacteria | Bacteria>Pseudomonadati>Pseudomonadota | 370 | DF068_RS01960 | 9199..10311 | 1236 | MULTISPECIES:-AAA-family-ATPase-[Gammaproteobacteria] |
| WP_239296669.1 | OLD-ABC | GCF_002311495.1 | Gammaproteobacteria | Bacteria>Pseudomonadati>Pseudomonadota | 370 | MHM35_RS10435 | 129920..131032 | 1236 | MULTISPECIES:-AAA-family-ATPase-[Gammaproteobacteria] |
| WP_123104886.1 | OLD-ABC | GCF_003721225.2 | Acidithiobacillus_sulfuriphilus | Bacteria>Pseudomonadati>Pseudomonadota>Acidithiobacillia>Acidithiobacillales>Acidithiobacillaceae>Acidithiobacillus | 414 | EC580_RS10655 | 7337..8581 | 1867749 | AAA-family-ATPase-[Acidithiobacillus-sulfuriphilus] |
| WP_014029721.1 | OLD-ABC | GCF_000214095.2 | Acidithiobacillus_ferrivorans | Bacteria>Pseudomonadati>Pseudomonadota>Acidithiobacillia>Acidithiobacillales>Acidithiobacillaceae>Acidithiobacillus | 388 | ACIFE_RS11335 | 2371294..2372460 | 160808 | AAA-family-ATPase-[Acidithiobacillus-ferrivorans] |
| WP_218285164.1 | OLD-ABC | GCF_019203965.1 | Elioraea_tepida | Bacteria>Pseudomonadati>Pseudomonadota>Alphaproteobacteria>Acetobacterales>Elioraeaceae>Elioraea | 391 | K0353_RS13030 | 2712857..2714032 | 2843330 | AAA-family-ATPase-[Elioraea-tepida] |
| MCB1832524.1 | OLD-ABC | GCA_020445435.1 | Geminicoccaceae_bacterium | Bacteria>Pseudomonadati>Pseudomonadota>Alphaproteobacteria>Geminicoccales>Geminicoccaceae>unclassified_Geminicoccaceae | 392 | KDH19_03620 | 18234..19412 | 2448052 | AAA-family-ATPase-[Geminicoccaceae-bacterium] |
| VXB71457.1 | OLD-ABC | GCA_902506225.1 | Enhydrobacter_sp._8BJ | Bacteria>Pseudomonadati>Pseudomonadota>Alphaproteobacteria>Hyphomicrobiales>Hyphomicrobiales_incertae_sedis>Enhydrobacter>unclassified_Enhydrobacter | 399 | ENHYD8BJ_80435 | 465199..466398 | 2653137 | Predicted-ATPase-[Enhydrobacter-sp.-8BJ] |
| WP_012755928.1 | OLD-ABC | - | Rhizobium_leguminosarum | Bacteria>Pseudomonadati>Pseudomonadota>Alphaproteobacteria>Hyphomicrobiales>Rhizobiaceae>Rhizobium/Agrobacterium_group>Rhizobium | 442 | RLEG_RS01020 | 206135..207463 | 384 | AAA-family-ATPase-[Rhizobium-leguminosarum] |
| WP_115156543.1 | OLD-ABC | GCF_004329915.1 | Rhizobium_leguminosarum | Bacteria>Pseudomonadati>Pseudomonadota>Alphaproteobacteria>Hyphomicrobiales>Rhizobiaceae>Rhizobium/Agrobacterium_group>Rhizobium | 397 | EOH64_RS12775 | 331402..332595 | 384 | AAA-family-ATPase-[Rhizobium-leguminosarum] |
| MBF0181807.1 | OLD-ABC | GCA_015231235.1 | Magnetococcales_bacterium | Bacteria>Pseudomonadati>Pseudomonadota>Alphaproteobacteria>Magnetococcales>unclassified_Magnetococcales | 413 | HQM03_17455 | 9241..10482 | 2026759 | AAA-family-ATPase-[Magnetococcales-bacterium] |
| MBF0262093.1 | OLD-ABC | GCA_015231775.1 | Magnetococcales_bacterium | Bacteria>Pseudomonadati>Pseudomonadota>Alphaproteobacteria>Magnetococcales>unclassified_Magnetococcales | 384 | HQL97_09705 | 6713..7867 | 2026759 | AAA-family-ATPase-[Magnetococcales-bacterium] |
| OOY26629.1 | OLD-ABC | GCA_002020275.1 | Thioclava_sp._L04-15 | Bacteria>Pseudomonadati>Pseudomonadota>Alphaproteobacteria>Rhodobacterales>Paracoccaceae>Thioclava>unclassified_Thioclava | 392 | BMI90_16685 | 41074..42252 | 1915318 | chromosome-segregation-protein-SMC-[Thioclava-sp.-L04-15] |
| WP_078530621.1 | OLD-ABC | GCF_002020275.1 | Thioclava_sp._L04-15 | Bacteria>Pseudomonadati>Pseudomonadota>Alphaproteobacteria>Rhodobacterales>Paracoccaceae>Thioclava>unclassified_Thioclava | 392 | BMI90_RS16615 | 41074..42252 | 1915318 | AAA-family-ATPase-[Thioclava-sp.-L04-15] |
| QCO03444.1 | OLD-ABC | GCA_005222185.1 | Azospirillum_argentinense | Bacteria>Pseudomonadati>Pseudomonadota>Alphaproteobacteria>Rhodospirillales>Azospirillaceae>Azospirillum | 398 | D3867_15340 | 175185..176381 | 2970906 | chromosome-segregation-protein-SMC-[Azospirillum-argentinense] |
| MDE0384391.1 | OLD-ABC | GCA_028821395.1 | Defluviicoccus_sp. | Bacteria>Pseudomonadati>Pseudomonadota>Alphaproteobacteria>Rhodospirillales>Rhodospirillaceae>Defluviicoccus>unclassified_Defluviicoccus | 416 | OXI22_10945 | 41075..42325 | 2682137 | MAG:-AAA-family-ATPase-[Defluviicoccus-sp.] |
| MDA8051499.1 | OLD-ABC | GCA_028063655.1 | Rhodospirillales_bacterium | Bacteria>Pseudomonadati>Pseudomonadota>Alphaproteobacteria>Rhodospirillales>unclassified_Rhodospirillales | 387 | M0002_16110 | 3586..4749 | 2026786 | MAG:-AAA-family-ATPase-[Rhodospirillales-bacterium] |
| WP_083199111.1 | OLD-ABC | GCF_001699185.1 | Sphingobium_sp._Ndbn-10 | Bacteria>Pseudomonadati>Pseudomonadota>Alphaproteobacteria>Sphingomonadales>Sphingomonadaceae>Sphingobium>unclassified_Sphingobium | 397 | FHS82_RS15735 | 27405..28598 | 1667223 | MULTISPECIES:-AAA-family-ATPase-[Alphaproteobacteria] |
| MCY3753477.1 | OLD-ABC | GCA_026706245.1 | Alphaproteobacteria_bacterium | Bacteria>Pseudomonadati>Pseudomonadota>Alphaproteobacteria>unclassified_Alphaproteobacteria | 420 | OXG99_05230 | 11190..12452 | 1913988 | MAG:-AAA-family-ATPase-[Alphaproteobacteria-bacterium] |
| MCB9746495.1 | OLD-ABC | GCA_020632815.1 | Alphaproteobacteria_bacterium | Bacteria>Pseudomonadati>Pseudomonadota>Alphaproteobacteria>unclassified_Alphaproteobacteria | 396 | H6740_28225 | 9411..10601 | 1913988 | AAA-family-ATPase-[Alphaproteobacteria-bacterium] |
| MCB9797658.1 | OLD-ABC | GCA_020632765.1 | Alphaproteobacteria_bacterium | Bacteria>Pseudomonadati>Pseudomonadota>Alphaproteobacteria>unclassified_Alphaproteobacteria | 396 | H6741_33630 | 14549..15739 | 1913988 | AAA-family-ATPase-[Alphaproteobacteria-bacterium] |
| WP_046536493.1 | OLD-ABC | GCF_035331625.1 | Candidatus_Accumulibacter | Bacteria>Pseudomonadati>Pseudomonadota>Betaproteobacteria>Betaproteobacteria_incertae_sedis | 373 | ACCUM_RS00300 | 70770..71891 | 327159 | ATP-binding-protein-[Candidatus-Accumulibacter-phosphatis] |
| QLH49278.1 | OLD-ABC | GCA_013414765.1 | Candidatus_Accumulibacter_cognatus | Bacteria>Pseudomonadati>Pseudomonadota>Betaproteobacteria>Betaproteobacteria_incertae_sedis>Candidatus_Accumulibacter | 443 | HWD57_05380 | 1165134..1166465 | 2954383 | MAG:-ATP-binding-protein-[Candidatus-Accumulibacter-cognatus] |
| KFB72162.1 | OLD-ABC | GCA_000585035.2 | Candidatus_Accumulibacter_phosphatis | Bacteria>Pseudomonadati>Pseudomonadota>Betaproteobacteria>Betaproteobacteria_incertae_sedis>Candidatus_Accumulibacter | 409 | AW09_002654 | 7484..8713 | 327160 | MAG:-putative-ATPase-[Candidatus-Accumulibacter-phosphatis] |
| KFB72164.1 | OLD-ABC | GCA_000585035.2 | Candidatus_Accumulibacter_phosphatis | Bacteria>Pseudomonadati>Pseudomonadota>Betaproteobacteria>Betaproteobacteria_incertae_sedis>Candidatus_Accumulibacter | 407 | AW09_002656 | 9519..10742 | 327160 | MAG:-putative-ATPase-[Candidatus-Accumulibacter-phosphatis] |

| Protein-ID | Domain | Assembly-ID | Strain | Taxonomy | Length | Locus-Tag | Genomic-coordinates | TaxID | Fasta-defline |
|---|---|---|---|---|---|---|---|---|---|
| RDE49750.1 | OLD-ABC | GCA_003332265.1 | Candidatus_Accumulibacter_meliphilus | Bacteria>Pseudomonadati>Pseudomonadota>Betaproteobacteria>Betaproteobacteria_incertae_sedis>Candidatus_Accumulibacter | 390 | DVS81_14915 | 16403..17575 | 2211374 | MAG:-ATP-binding-cassette-domain-containing-protein-[Candidatus-Accumulibacter-meliphilus] |
| QLH49280.1 | OLD-ABC | GCA_013414765.1 | Candidatus_Accumulibacter_cognatus | Bacteria>Pseudomonadati>Pseudomonadota>Betaproteobacteria>Betaproteobacteria_incertae_sedis>Candidatus_Accumulibacter | 373 | HWD57_05395 | 1167393..1168514 | 2954383 | MAG:-AAA-family-ATPase-[Candidatus-Accumulibacter-cognatus] |
| WP_248594811.1 | OLD-ABC | - | Candidatus_Accumulibacter_contiguus | Bacteria>Pseudomonadati>Pseudomonadota>Betaproteobacteria>Betaproteobacteria_incertae_sedis>Candidatus_Accumulibacter | 164 | E4Q08_RS24600 | 42164..42658 | 2954381 | AAA-family-ATPase-[Candidatus-Accumulibacter-contiguus] |
| WP_248594805.1 | OLD-ABC | GCF_012940005.1 | Candidatus_Accumulibacter_contiguus | Bacteria>Pseudomonadati>Pseudomonadota>Betaproteobacteria>Betaproteobacteria_incertae_sedis>Candidatus_Accumulibacter | 98 | E4Q08_RS24590 | 41454..41750 | 2954381 | AAA-family-ATPase-[Candidatus-Accumulibacter-contiguus] |
| MBN8451982.1 | OLD-ABC | GCA_017302415.1 | Accumulibacter_sp. | Bacteria>Pseudomonadati>Pseudomonadota>Betaproteobacteria>Betaproteobacteria_incertae_sedis>Candidatus_Accumulibacter>unclassified_Candidatus_Accumulibacter | 415 | J0M31_02660 | 91295..92542 | 2053492 | AAA-family-ATPase-[Candidatus-Accumulibacter-sp.] |
| MBN8517035.1 | OLD-ABC | GCA_017302385.1 | Accumulibacter_sp. | Bacteria>Pseudomonadati>Pseudomonadota>Betaproteobacteria>Betaproteobacteria_incertae_sedis>Candidatus_Accumulibacter>unclassified_Candidatus_Accumulibacter | 373 | J0L71_04010 | 50397..51518 | 2053492 | AAA-family-ATPase-[Candidatus-Accumulibacter-sp.] |
| TLD46495.1 | OLD-ABC | GCA_005524045.1 | Accumulibacter_sp. | Bacteria>Pseudomonadati>Pseudomonadota>Betaproteobacteria>Betaproteobacteria_incertae_sedis>Candidatus_Accumulibacter>unclassified_Candidatus_Accumulibacter | 356 | FAZ92_01232 | 10502..11572 | 2053492 | MAG:-hypothetical-protein-FAZ92_01232-[Candidatus-Accumulibacter-sp.] |
| MCX8099548.1 | OLD-ABC | GCA_026414825.1 | Casimicrobiaceae_bacterium | Bacteria>Pseudomonadati>Pseudomonadota>Betaproteobacteria>Betaproteobacteria_incertae_sedis>Casimicrobiaceae>unclassified_Casimicrobiaceae | 378 | N3F11_11215 | 12612..13748 | 2906524 | MAG:-AAA-family-ATPase-[Casimicrobiaceae-bacterium] |
| WP_026120980.1 | OLD-ABC | GCF_000341045.1 | Burkholderiaceae | Bacteria>Pseudomonadati>Pseudomonadota>Betaproteobacteria>Burkholderiales | 392 | SY87_RS33405 | 10555..11733 | 119060 | MULTISPECIES:-AAA-family-ATPase-[Burkholderiaceae] |
| WP_194770511.1 | OLD-ABC | GCF_032085695.1 | Comamonadaceae | Bacteria>Pseudomonadati>Pseudomonadota>Betaproteobacteria>Burkholderiales | 392 | ISG32_RS08055 | 70839..72017 | 80864 | AAA-family-ATPase-[Diaphorobacter-caeni] |
| MCH1991584.1 | OLD-ABC | GCA_022444305.1 | Achromobacter_xylosoxidans | Bacteria>Pseudomonadati>Pseudomonadota>Betaproteobacteria>Burkholderiales>Alcaligenaceae>Achromobacter | 409 | L7Q78_02310 | 20553..21782 | 85698 | AAA-family-ATPase-[Achromobacter-xylosoxidans] |
| SUA58551.1 | OLD-ABC | GCA_900454315.1 | Oligella_ureolytica | Bacteria>Pseudomonadati>Pseudomonadota>Betaproteobacteria>Burkholderiales>Alcaligenaceae>Oligella | 422 | NCTC11998_02425 | 2465938..2467206 | 90244 | Predicted-ATP-binding-protein-involved-in-virulence-[Oligella-ureolytica] |
| SUA58546.1 | OLD-ABC | GCA_900454315.1 | Oligella_ureolytica | Bacteria>Pseudomonadati>Pseudomonadota>Betaproteobacteria>Burkholderiales>Alcaligenaceae>Oligella | 368 | NCTC11998_02423 | 2464092..2465198 | 90244 | Predicted-ATPase-[Oligella-ureolytica] |
| NYT60766.1 | OLD-ABC | GCA_013416445.1 | Alcaligenaceae_bacterium | Bacteria>Pseudomonadati>Pseudomonadota>Betaproteobacteria>Burkholderiales>Alcaligenaceae>unclassified_Alcaligenaceae | 375 | H0A66_00275 | 65509..66636 | 2021368 | AAA-family-ATPase-[Alcaligenaceae-bacterium] |
| WP_071038001.1 | OLD-ABC | GCF_001854305.1 | Cupriavidus | Bacteria>Pseudomonadati>Pseudomonadota>Betaproteobacteria>Burkholderiales>Burkholderiaceae | 392 | BKK79_RS15560 | 3508150..3509328 | 106589 | MULTISPECIES:-AAA-family-ATPase-[Cupriavidus] |
| KIX33652.1 | OLD-ABC | GCA_000932075.1 | Burkholderia_pseudomallei | Bacteria>Pseudomonadati>Pseudomonadota>Betaproteobacteria>Burkholderiales>Burkholderiaceae>Burkholderia>pseudomallei_group | 392 | SY87_33355 | 10555..11733 | 28450 | chromosome-segregation-protein-SMC-[Burkholderia-pseudomallei] |
| AOY93049.1 | OLD-ABC | GCA_001854305.1 | Cupriavidus_sp._USMAA2-4 | Bacteria>Pseudomonadati>Pseudomonadota>Betaproteobacteria>Burkholderiales>Burkholderiaceae>Cupriavidus>unclassified_Cupriavidus | 392 | BKK79_15565 | 3508150..3509328 | 876364 | chromosome-segregation-protein-SMC-[Cupriavidus-sp.-USMAA2-4] |
| WP_133608525.1 | OLD-ABC | GCF_004362855.1 | Aquabacterium_commune | Bacteria>Pseudomonadati>Pseudomonadota>Betaproteobacteria>Burkholderiales>Burkholderiales_genera_incertae_sedis>Aquabacterium | 385 | EV672_RS09260 | 191505..192662 | 70586 | AAA-family-ATPase-[Aquabacterium-commune] |
| WP_143969786.1 | OLD-ABC | GCF_007556755.1 | Tepidimonas_fonticaldi | Bacteria>Pseudomonadati>Pseudomonadota>Betaproteobacteria>Burkholderiales>Burkholderiales_genera_incertae_sedis>Tepidimonas | 436 | Tfont_RS13275 | 8599..9909 | 1101373 | AAA-family-ATPase-[Tepidimonas-fonticaldi] |
| WP_043699929.1 | OLD-ABC | GCF_020162115.1 | Tepidimonas_taiwanensis | Bacteria>Pseudomonadati>Pseudomonadota>Betaproteobacteria>Burkholderiales>Burkholderiales_genera_incertae_sedis>Tepidimonas | 390 | PL21_RS04250 | 83228..84400 | 307486 | AAA-family-ATPase-[Tepidimonas-taiwanensis] |
| WP_143969663.1 | OLD-ABC | GCF_007556755.1 | Tepidimonas_fonticaldi | Bacteria>Pseudomonadati>Pseudomonadota>Betaproteobacteria>Burkholderiales>Burkholderiales_genera_incertae_sedis>Tepidimonas | 390 | Tfont_RS12350 | 11154..12326 | 1101373 | AAA-family-ATPase-[Tepidimonas-fonticaldi] |
| WP_054075978.1 | OLD-ABC | GCF_001294075.1 | Comamonas | Bacteria>Pseudomonadati>Pseudomonadota>Betaproteobacteria>Burkholderiales>Comamonadaceae | 397 | KF712C_RS25800 | 22904..24097 | 283 | MULTISPECIES:-ATP-binding-protein-[Comamonas] |
| WP_272568618.1 | OLD-ABC | GCF_030159035.1 | unclassified_Acidovorax | Bacteria>Pseudomonadati>Pseudomonadota>Betaproteobacteria>Burkholderiales>Comamonadaceae>Acidovorax | 409 | M5C96_RS08960 | 1968943..1970172 | 2684926 | ATP-binding-protein-[Acidovorax-sp.-GBBC-1281] |
| WP_094097621.1 | OLD-ABC | GCF_002157145.1 | Acidovorax_carolinensis | Bacteria>Pseudomonadati>Pseudomonadota>Betaproteobacteria>Burkholderiales>Comamonadaceae>Acidovorax | 383 | CBP34_RS06515 | 1389250..1390401 | 553814 | ATP-binding-protein-[Acidovorax-carolinensis] |
| WP_008906816.1 | OLD-ABC | GCF_000238595.1 | Acidovorax_sp._NO-1 | Bacteria>Pseudomonadati>Pseudomonadota>Betaproteobacteria>Burkholderiales>Comamonadaceae>Acidovorax>unclassified_Acidovorax | 401 | KYG_RS20345 | 14100..15305 | 512030 | AAA-family-ATPase-[Acidovorax-sp.-NO-1] |
| GAO25244.1 | OLD-ABC | GCA_000974645.1 | Alicycliphilus_sp._B1 | Bacteria>Pseudomonadati>Pseudomonadota>Betaproteobacteria>Burkholderiales>Comamonadaceae>Alicycliphilus>unclassified_Alicycliphilus | 393 | ALISP_5064 | 1532000..1533181 | 1603291 | hypothetical-protein-ALISP_5064-[Alicycliphilus-sp.-B1] |
| WP_091817121.1 | OLD-ABC | GCF_900110225.1 | Brachymonas_denitrificans | Bacteria>Pseudomonadati>Pseudomonadota>Betaproteobacteria>Burkholderiales>Comamonadaceae>Brachymonas | 409 | BMW28_RS09510 | 73013..74242 | 28220 | AAA-family-ATPase-[Brachymonas-denitrificans] |
| SEN73476.1 | OLD-ABC | GCA_900110225.1 | Brachymonas_denitrificans_DSM_15123 | Bacteria>Pseudomonadati>Pseudomonadota>Betaproteobacteria>Burkholderiales>Comamonadaceae>Brachymonas>Brachymonas_denitrificans | 409 | SAMN02745977_01906 | 73013..74242 | 1121117 | Predicted-ATPase-[Brachymonas-denitrificans-DSM-15123] |
| MBF5004209.1 | OLD-ABC | GCA_015354245.1 | Diaphorobacter_caeni | Bacteria>Pseudomonadati>Pseudomonadota>Betaproteobacteria>Burkholderiales>Comamonadaceae>Diaphorobacter | 392 | ISG32_08050 | 70839..72017 | 2784387 | AAA-family-ATPase-[Diaphorobacter-caeni] |
| QIL80932.1 | OLD-ABC | GCA_011305995.1 | Diaphorobacter_sp._HDW4A | Bacteria>Pseudomonadati>Pseudomonadota>Betaproteobacteria>Burkholderiales>Comamonadaceae>Diaphorobacter>unclassified_Diaphorobacter | 356 | G7047_14255 | 3128568..3129638 | 2714924 | AAA-family-ATPase-[Diaphorobacter-sp.-HDW4A] |
| WP_166306533.1 | OLD-ABC | GCF_011305995.1 | Diaphorobacter_sp._HDW4A | Bacteria>Pseudomonadati>Pseudomonadota>Betaproteobacteria>Burkholderiales>Comamonadaceae>Diaphorobacter>unclassified_Diaphorobacter | 356 | G7047_RS14250 | 3128568..3129638 | 2714924 | AAA-family-ATPase-[Diaphorobacter-sp.-HDW4A] |
| WP_104799764.1 | OLD-ABC | GCF_002943475.1 | Limnohabitans_sp._TS-CS-82 | Bacteria>Pseudomonadati>Pseudomonadota>Betaproteobacteria>Burkholderiales>Comamonadaceae>Limnohabitans>unclassified_Limnohabitans | 402 | C5F52_RS18145 | 193789..194997 | 2094193 | AAA-family-ATPase-[Limnohabitans-sp.-TS-CS-82] |
| WP_108373343.1 | OLD-ABC | GCF_003063315.1 | Limnohabitans_sp._T6-5 | Bacteria>Pseudomonadati>Pseudomonadota>Betaproteobacteria>Burkholderiales>Comamonadaceae>Limnohabitans>unclassified_Limnohabitans | 402 | B9Z51_RS07545 | 404520..405728 | 1100724 | ATP-binding-protein-[Limnohabitans-sp.-T6-5] |
| WP_108373346.1 | OLD-ABC | GCF_003063315.1 | Limnohabitans_sp._T6-5 | Bacteria>Pseudomonadati>Pseudomonadota>Betaproteobacteria>Burkholderiales>Comamonadaceae>Limnohabitans>unclassified_Limnohabitans | 383 | B9Z51_RS07555 | 406390..407541 | 1100724 | ATP-binding-protein-[Limnohabitans-sp.-T6-5] |
| WP_108373336.1 | OLD-ABC | GCF_003063315.1 | Limnohabitans_sp._T6-5 | Bacteria>Pseudomonadati>Pseudomonadota>Betaproteobacteria>Burkholderiales>Comamonadaceae>Limnohabitans>unclassified_Limnohabitans | 372 | B9Z51_RS07525 | 401095..402213 | 1100724 | AAA-family-ATPase-[Limnohabitans-sp.-T6-5] |
| WP_101103329.1 | OLD-ABC | GCF_002837135.1 | Macromonas_bipunctata | Bacteria>Pseudomonadati>Pseudomonadota>Betaproteobacteria>Burkholderiales>Comamonadaceae>Macromonas | 443 | CON76_RS10610 | 15558..16889 | 183670 | AAA-family-ATPase-[Macromonas-bipunctata] |
| WP_209423131.1 | OLD-ABC | GCF_021249385.1 | Melaminivora_jejuensis | Bacteria>Pseudomonadati>Pseudomonadota>Betaproteobacteria>Burkholderiales>Comamonadaceae>Melaminivora | 376 | LVC68_RS01465 | 289326..290456 | 1267217 | ATP-binding-protein-[Melaminivora-jejuensis] |
| WP_209423134.1 | OLD-ABC | GCF_021249385.1 | Melaminivora_jejuensis | Bacteria>Pseudomonadati>Pseudomonadota>Betaproteobacteria>Burkholderiales>Comamonadaceae>Melaminivora | 352 | LVC68_RS01455 | 287590..288648 | 1267217 | AAA-family-ATPase-[Melaminivora-jejuensis] |
| WP_011797937.1 | OLD-ABC | GCF_000015505.1 | Polaromonas_naphthalenivorans | Bacteria>Pseudomonadati>Pseudomonadota>Betaproteobacteria>Burkholderiales>Comamonadaceae>Polaromonas | 355 | PNAP_RS21290 | 181245..182312 | 216465 | ATP-binding-protein-[Polaromonas-naphthalenivorans] |
| WP_265288962.1 | OLD-ABC | GCF_026127525.1 | Verminephrobacter_eiseniae | Bacteria>Pseudomonadati>Pseudomonadota>Betaproteobacteria>Burkholderiales>Comamonadaceae>Verminephrobacter | 407 | D5047_RS18235 | 3907259..3908482 | 364317 | ATP-binding-protein-[Verminephrobacter-eiseniae] |
| WP_096234090.1 | OLD-ABC | GCF_002327145.1 | Janthinobacterium_svalbardensis | Bacteria>Pseudomonadati>Pseudomonadota>Betaproteobacteria>Burkholderiales>Oxalobacteraceae>Janthinobacterium | 370 | CNX70_RS06890 | 1571908..1573020 | 368607 | ATP-binding-protein-[Janthinobacterium-svalbardensis] |
| WP_173807408.1 | OLD-ABC | GCF_013294065.1 | Sphaerotilus_uruguayifluvii | Bacteria>Pseudomonadati>Pseudomonadota>Betaproteobacteria>Burkholderiales>Sphaerotilaceae>Sphaerotilus | 477 | HNQ01_RS20680 | 26711..28144 | 2735897 | AAA-family-ATPase-[Leptothrix-sp.-C29] |
| NRT58389.1 | OLD-ABC | GCA_013294065.1 | Sphaerotilus_uruguayifluvii | Bacteria>Pseudomonadati>Pseudomonadota>Betaproteobacteria>Burkholderiales>Sphaerotilaceae>Sphaerotilus | 475 | HNQ01_004157 | 26711..28138 | 2735897 | putative-ATP-binding-protein-involved-in-virulence-[Leptothrix-sp.-C29] |
| MCK6404273.1 | OLD-ABC | GCA_023150255.1 | Sphaerotilus_sulfidivorans | Bacteria>Pseudomonadati>Pseudomonadota>Betaproteobacteria>Burkholderiales>Sphaerotilaceae>Sphaerotilus | 471 | L6Q74_20580 | 14665..16080 | 639200 | DUF3696-domain-containing-protein-[Sphaerotilus-sulfidivorans] |
| WP_273473509.1 | OLD-ABC | GCF_023150255.1 | Sphaerotilus_sulfidivorans | Bacteria>Pseudomonadati>Pseudomonadota>Betaproteobacteria>Burkholderiales>Sphaerotilaceae>Sphaerotilus | 471 | L6Q74_RS20540 | 14665..16080 | 639200 | DUF3696-domain-containing-protein-[Sphaerotilus-sulfidivorans] |
| WP_251969635.1 | OLD-ABC | GCF_023734135.1 | Sphaerotilus_microaerophilus | Bacteria>Pseudomonadati>Pseudomonadota>Betaproteobacteria>Burkholderiales>Sphaerotilaceae>Sphaerotilus | 417 | NGK70_RS16740 | 3931158..3932411 | 2914710 | ATP-binding-protein-[Sphaerotilus-sp.-FB-5] |
| MCK6404269.1 | OLD-ABC | GCA_023150255.1 | Sphaerotilus_sulfidivorans | Bacteria>Pseudomonadati>Pseudomonadota>Betaproteobacteria>Burkholderiales>Sphaerotilaceae>Sphaerotilus | 396 | L6Q74_20560 | 9327..10517 | 639200 | AAA-family-ATPase-[Sphaerotilus-sulfidivorans] |
| NRT58386.1 | OLD-ABC | GCA_013294065.1 | Sphaerotilus_uruguayifluvii | Bacteria>Pseudomonadati>Pseudomonadota>Betaproteobacteria>Burkholderiales>Sphaerotilaceae>Sphaerotilus | 396 | HNQ01_004154 | 21964..23154 | 2735897 | putative-ATPase-[Leptothrix-sp.-C29] |
| WP_173807405.1 | OLD-ABC | GCF_013294065.1 | Sphaerotilus_uruguayifluvii | Bacteria>Pseudomonadati>Pseudomonadota>Betaproteobacteria>Burkholderiales>Sphaerotilaceae>Sphaerotilus | 396 | HNQ01_RS20665 | 21964..23154 | 2735897 | AAA-family-ATPase-[Leptothrix-sp.-C29] |
| WP_273473501.1 | OLD-ABC | GCF_023150255.1 | Sphaerotilus_sulfidivorans | Bacteria>Pseudomonadati>Pseudomonadota>Betaproteobacteria>Burkholderiales>Sphaerotilaceae>Sphaerotilus | 396 | L6Q74_RS20520 | 9327..10517 | 639200 | AAA-family-ATPase-[Sphaerotilus-sulfidivorans] |
| MCZ2419567.1 | OLD-ABC | GCA_026931465.1 | Burkholderiales_bacterium | Bacteria>Pseudomonadati>Pseudomonadota>Betaproteobacteria>Burkholderiales>unclassified_Burkholderiales | 444 | LC123_06990 | 89564..90898 | 1891238 | MAG:-ATP-binding-protein-[Burkholderiales-bacterium] |
| MBE0627111.1 | OLD-ABC | GCA_014860355.1 | Burkholderiales_bacterium | Bacteria>Pseudomonadati>Pseudomonadota>Betaproteobacteria>Burkholderiales>unclassified_Burkholderiales | 432 | IH606_20115 | 3272..4570 | 1891238 | AAA-family-ATPase-[Burkholderiales-bacterium] |
| MBK6715974.1 | OLD-ABC | GCA_016703325.1 | Burkholderiales_bacterium | Bacteria>Pseudomonadati>Pseudomonadota>Betaproteobacteria>Burkholderiales>unclassified_Burkholderiales | 429 | IPG57_12980 | 78921..80210 | 1891238 | AAA-family-ATPase-[Burkholderiales-bacterium] |
| MBX9598969.1 | OLD-ABC | GCA_019748155.1 | Burkholderiales_bacterium | Bacteria>Pseudomonadati>Pseudomonadota>Betaproteobacteria>Burkholderiales>unclassified_Burkholderiales | 425 | K2X04_10425 | 10608..11885 | 1891238 | ATP-binding-protein-[Burkholderiales-bacterium] |
| MBP7846548.1 | OLD-ABC | GCA_018001735.1 | Burkholderiales_bacterium | Bacteria>Pseudomonadati>Pseudomonadota>Betaproteobacteria>Burkholderiales>unclassified_Burkholderiales | 413 | KAZ94_00270 | 63037..64278 | 1891238 | ATP-binding-protein-[Burkholderiales-bacterium] |
| NMA26874.1 | OLD-ABC | GCA_012799975.1 | Burkholderiales_bacterium | Bacteria>Pseudomonadati>Pseudomonadota>Betaproteobacteria>Burkholderiales>unclassified_Burkholderiales | 391 | GX934_03850 | 828..2003 | 1891238 | AAA-family-ATPase-[Burkholderiales-bacterium] |
| QKE41380.1 | OLD-ABC | GCA_013181415.1 | Ferrovum_myxofaciens | Bacteria>Pseudomonadati>Pseudomonadota>Betaproteobacteria>Ferrovales>Ferrovaceae>Ferrovum | 405 | HO274_08660 | 1791569..1792786 | 416213 | ATP-binding-protein-[Ferrovum-myxofaciens] |
| WP_273118819.1 | OLD-ABC | GCF_013181415.1 | Ferrovum_myxofaciens | Bacteria>Pseudomonadati>Pseudomonadota>Betaproteobacteria>Ferrovales>Ferrovaceae>Ferrovum | 405 | HO274_RS08690 | 1791569..1792786 | 416213 | ATP-binding-protein-[Ferrovum-myxofaciens] |
| NDU85857.1 | OLD-ABC | GCA_010436015.1 | Ferrovum_sp. | Bacteria>Pseudomonadati>Pseudomonadota>Betaproteobacteria>Ferrovales>Ferrovaceae>Ferrovum>unclassified_Ferrovum | 405 | G3H99_04500 | 12902..14119 | 2609467 | ATP-binding-protein-[Ferrovum-sp.] |
| WP_022652887.1 | OLD-ABC | GCF_000420525.1 | Aquaspirillum_serpens | Bacteria>Pseudomonadati>Pseudomonadota>Betaproteobacteria>Neisseriales>Aquaspirillaceae>Aquaspirillum | 409 | G465_RS0102390 | 19898..21127 | 190 | AAA-family-ATPase-[Aquaspirillum-serpens] |
| WP_022652889.1 | OLD-ABC | GCF_000420525.1 | Aquaspirillum_serpens | Bacteria>Pseudomonadati>Pseudomonadota>Betaproteobacteria>Neisseriales>Aquaspirillaceae>Aquaspirillum | 344 | G465_RS0102400 | 21646..22680 | 190 | ATP-binding-protein-[Aquaspirillum-serpens] |
| MCG9052524.1 | OLD-ABC | GCA_022394235.1 | Laribacter_hongkongensis | Bacteria>Pseudomonadati>Pseudomonadota>Betaproteobacteria>Neisseriales>Aquaspirillaceae>Laribacter | 410 | LH447_05310 | 11045..12277 | 168471 | AAA-family-ATPase-[Laribacter-hongkongensis] |
| MCG9059782.1 | OLD-ABC | GCA_022394335.1 | Laribacter_hongkongensis | Bacteria>Pseudomonadati>Pseudomonadota>Betaproteobacteria>Neisseriales>Aquaspirillaceae>Laribacter | 410 | LH452_12740 | 24089..25321 | 168471 | AAA-family-ATPase-[Laribacter-hongkongensis] |
| WP_239891834.1 | OLD-ABC | GCF_022394335.1 | Laribacter_hongkongensis | Bacteria>Pseudomonadati>Pseudomonadota>Betaproteobacteria>Neisseriales>Aquaspirillaceae>Laribacter | 410 | LH452_RS12730 | 24089..25321 | 168471 | ATP-binding-protein-[Laribacter-hongkongensis] |
| WP_239900337.1 | OLD-ABC | GCF_022394235.1 | Laribacter_hongkongensis | Bacteria>Pseudomonadati>Pseudomonadota>Betaproteobacteria>Neisseriales>Aquaspirillaceae>Laribacter | 410 | LH442_RS13350 | 12283..13515 | 168471 | AAA-family-ATPase-[Laribacter-hongkongensis] |
| MCG9000249.1 | OLD-ABC | GCA_022393975.1 | Laribacter_hongkongensis | Bacteria>Pseudomonadati>Pseudomonadota>Betaproteobacteria>Neisseriales>Aquaspirillaceae>Laribacter | 407 | LH454_02015 | 64985..66208 | 168471 | AAA-family-ATPase-[Laribacter-hongkongensis] |
| WP_047248549.1 | OLD-ABC | GCF_001020585.1 | Chromobacterium_subtsugae | Bacteria>Pseudomonadati>Pseudomonadota>Betaproteobacteria>Neisseriales>Chromobacteriaceae>Chromobacterium | 351 | AA094_RS04300 | 50478..51533 | 251747 | ATP-binding-protein-[Chromobacterium-subtsugae] |
| WP_013292864.1 | OLD-ABC | GCF_000145255.1 | Gallionella_capsiferriformans | Bacteria>Pseudomonadati>Pseudomonadota>Betaproteobacteria>Nitrosomonadales>Gallionellaceae>Gallionella | 394 | GALF_RS04450 | 940834..942018 | 370405 | AAA-family-ATPase-[Gallionella-capsiferriformans] |
| MBP9798941.1 | OLD-ABC | GCA_018062105.1 | Sterolibacterium_sp. | Bacteria>Pseudomonadati>Pseudomonadota>Betaproteobacteria>Nitrosomonadales>Sterolibacteriaceae>Sterolibacterium>unclassified_Sterolibacterium | 392 | KBD39_02330 | 25765..26943 | 2819261 | AAA-family-ATPase-[Sterolibacterium-sp.] |
| NOT17076.1 | OLD-ABC | GCA_013140765.1 | Sulfuriferula_sp. | Bacteria>Pseudomonadati>Pseudomonadota>Betaproteobacteria>Nitrosomonadales>Sulfuricellaceae>Sulfuriferula>unclassified_Sulfuriferula | 390 | HOP20_03265 | 14766..15938 | 2025307 | AAA-family-ATPase-[Sulfuriferula-sp.] |
| MCX7674153.1 | OLD-ABC | GCA_026419045.1 | Thiobacillaceae_bacterium | Bacteria>Pseudomonadati>Pseudomonadota>Betaproteobacteria>Nitrosomonadales>Thiobacillaceae>unclassified_Thiobacillaceae | 389 | N2Z63_11270 | 18734..19903 | 2528409 | MAG:-AAA-family-ATPase-[Thiobacillaceae-bacterium] |
| MBI5658238.1 | OLD-ABC | GCA_016223025.1 | Nitrosomonadales_bacterium | Bacteria>Pseudomonadati>Pseudomonadota>Betaproteobacteria>Nitrosomonadales>unclassified_Nitrosomonadales | 428 | HZC43_01520 | 58253..59539 | 2026761 | ATP-binding-protein-[Nitrosomonadales-bacterium] |
| MBI1887196.1 | OLD-ABC | GCA_016177795.1 | Nitrosomonadales_bacterium | Bacteria>Pseudomonadati>Pseudomonadota>Betaproteobacteria>Nitrosomonadales>unclassified_Nitrosomonadales | 424 | HYS19_02320 | 94090..95364 | 2026761 | ATP-binding-protein-[Nitrosomonadales-bacterium] |
| MBI3902219.1 | OLD-ABC | GCA_016199835.1 | Nitrosomonadales_bacterium | Bacteria>Pseudomonadati>Pseudomonadota>Betaproteobacteria>Nitrosomonadales>unclassified_Nitrosomonadales | 424 | HY306_04655 | 126809..128083 | 2026761 | ATP-binding-protein-[Nitrosomonadales-bacterium] |
| WP_153146308.1 | OLD-ABC | GCF_009469615.1 | Dechloromonas_sp._H13 | Bacteria>Pseudomonadati>Pseudomonadota>Betaproteobacteria>Rhodocyclales>Azonexaceae>Dechloromonas>unclassified_Dechloromonas | 391 | GH660_RS08860 | 102513..103688 | 2570193 | AAA-family-ATPase-[Dechloromonas-sp.-H13] |
| WP_011238772.1 | OLD-ABC | GCF_000025965.1 | Aromatoleum_aromaticum | Bacteria>Pseudomonadati>Pseudomonadota>Betaproteobacteria>Rhodocyclales>Rhodocyclaceae>Aromatoleum | 430 | EBN1_RS14775 | 3100992..3102284 | 551760 | ATP-binding-protein-[Aromatoleum-aromaticum] |
| MCK0506217.1 | OLD-ABC | GCA_012910705.2 | Aromatoleum_anaerobium | Bacteria>Pseudomonadati>Pseudomonadota>Betaproteobacteria>Rhodocyclales>Rhodocyclaceae>Aromatoleum | 386 | GO606_004915 | 97058..98218 | 182180 | AAA-family-ATPase-[Aromatoleum-anaerobium] |
| WP_169117245.1 | OLD-ABC | GCF_012910705.2 | Aromatoleum_anaerobium | Bacteria>Pseudomonadati>Pseudomonadota>Betaproteobacteria>Rhodocyclales>Rhodocyclaceae>Aromatoleum | 386 | GO606_RS04915 | 97058..98218 | 182180 | ATP-binding-protein-[Aromatoleum-anaerobium] |
| MCK0506219.1 | OLD-ABC | GCA_012910705.2 | Aromatoleum_anaerobium | Bacteria>Pseudomonadati>Pseudomonadota>Betaproteobacteria>Rhodocyclales>Rhodocyclaceae>Aromatoleum | 351 | GO606_004925 | 99082..100137 | 182180 | AAA-family-ATPase-[Aromatoleum-anaerobium] |
| WP_011238774.1 | OLD-ABC | GCF_000025965.1 | Aromatoleum_aromaticum | Bacteria>Pseudomonadati>Pseudomonadota>Betaproteobacteria>Rhodocyclales>Rhodocyclaceae>Aromatoleum | 351 | EBN1_RS14785 | 3102934..3103989 | 551760 | ATP-binding-protein-[Aromatoleum-aromaticum] |
| WP_169117247.1 | OLD-ABC | GCF_012910705.2 | Aromatoleum_anaerobium | Bacteria>Pseudomonadati>Pseudomonadota>Betaproteobacteria>Rhodocyclales>Rhodocyclaceae>Aromatoleum | 351 | GO606_RS04925 | 99082..100137 | 182180 | ATP-binding-protein-[Aromatoleum-anaerobium] |
| WP_011238776.1 | OLD-ABC | GCF_000025965.1 | Aromatoleum_aromaticum | Bacteria>Pseudomonadati>Pseudomonadota>Betaproteobacteria>Rhodocyclales>Rhodocyclaceae>Aromatoleum | 56 | EBN1_RS14795 | 3104853..3105023 | 551760 | AAA-family-ATPase-[Aromatoleum-aromaticum] |
| WP_205230203.1 | OLD-ABC | GCF_905120365.1 | Azospira_oryzae | Bacteria>Pseudomonadati>Pseudomonadota>Betaproteobacteria>Rhodocyclales>Rhodocyclaceae>Azospira | 347 | JYJ48_RS05815 | 15942..16985 | 146939 | ATP-binding-protein-[Azospira-oryzae] |
| WP_153111721.1 | OLD-ABC | GCF_009469695.1 | Propionivibrio_limicola | Bacteria>Pseudomonadati>Pseudomonadota>Betaproteobacteria>Rhodocyclales>Rhodocyclaceae>Propionivibrio | 409 | GH659_RS12240 | 4428..5657 | 167645 | AAA-family-ATPase-[Propionivibrio-limicola] |
| TRZ91912.1 | OLD-ABC | GCA_007280105.1 | Rhodocyclaceae_bacterium | Bacteria>Pseudomonadati>Pseudomonadota>Betaproteobacteria>Rhodocyclales>Rhodocyclaceae>unclassified_Rhodocyclaceae | 418 | D4R84_13320 | 5510..6766 | 1898103 | ATPase-[Rhodocyclaceae-bacterium] |
| AUM01647.1 | OLD-ABC | GCA_002863805.1 | Rhodocyclaceae_bacterium | Bacteria>Pseudomonadati>Pseudomonadota>Betaproteobacteria>Rhodocyclales>Rhodocyclaceae>unclassified_Rhodocyclaceae | 391 | B4966_15135 | 3174879..3176054 | 1898103 | chromosome-segregation-protein-SMC-[Rhodocyclaceae-bacterium] |
| MCX8086915.1 | OLD-ABC | GCA_026415155.1 | Rhodocyclaceae_bacterium | Bacteria>Pseudomonadati>Pseudomonadota>Betaproteobacteria>Rhodocyclales>Rhodocyclaceae>unclassified_Rhodocyclaceae | 390 | N3C63_08450 | 32026..33198 | 1898103 | MAG:-AAA-family-ATPase-[Rhodocyclaceae-bacterium] |
| TRZ91910.1 | OLD-ABC | GCA_007280105.1 | Rhodocyclaceae_bacterium | Bacteria>Pseudomonadati>Pseudomonadota>Betaproteobacteria>Rhodocyclales>Rhodocyclaceae>unclassified_Rhodocyclaceae | 388 | D4R84_13305 | 1677..2843 | 1898103 | DUF2813-domain-containing-protein-[Rhodocyclaceae-bacterium] |
| CAH1748216.1 | OLD-ABC | GCA_932126575.1 | Thauera_humireducens | Bacteria>Pseudomonadati>Pseudomonadota>Betaproteobacteria>Rhodocyclales>Zoogloeaceae>Thauera | 386 | C4PIVTH_3076 | 3149823..3150983 | 1134435 | ATP-binding-cassette-domain-containing-protein-[Thauera-sp.-PIV-1] |
| WP_247725476.1 | OLD-ABC | GCF_023093595.1 | Thauera_aromatica | Bacteria>Pseudomonadati>Pseudomonadota>Betaproteobacteria>Rhodocyclales>Zoogloeaceae>Thauera | 386 | MXC99_RS10890 | 47791..48951 | 59405 | AAA-family-ATPase-[Thauera-aromatica] |
| AVZ79925.1 | OLD-ABC | GCA_003054515.1 | Zoogloeaceae_bacteirum_Par-f-2 | Bacteria>Pseudomonadati>Pseudomonadota>Betaproteobacteria>Rhodocyclales>Zoogloeaceae>unclassified_Zoogloeaceae | 386 | C3497_11125 | 2342392..2343552 | 2080469 | chromosome-segregation-protein-SMC-[Zoogloeaceae-bacteirum-Par-f-2] |
| AVZ79922.1 | OLD-ABC | GCA_003054515.1 | Zoogloeaceae_bacteirum_Par-f-2 | Bacteria>Pseudomonadati>Pseudomonadota>Betaproteobacteria>Rhodocyclales>Zoogloeaceae>unclassified_Zoogloeaceae | 344 | C3497_11110 | 2337973..2339007 | 2080469 | AAA-family-ATPase-[Zoogloeaceae-bacteirum-Par-f-2] |
| GHT91735.1 | OLD-ABC | GCA_026000515.1 | Betaproteobacteria_bacterium | Bacteria>Pseudomonadati>Pseudomonadota>Betaproteobacteria>unclassified_Betaproteobacteria | 414 | FACS1894116_00780 | 35809..37053 | 1891241 | ATPase-[Betaproteobacteria-bacterium] |
| MBI3917899.1 | OLD-ABC | GCA_016199545.1 | Betaproteobacteria_bacterium | Bacteria>Pseudomonadati>Pseudomonadota>Betaproteobacteria>unclassified_Betaproteobacteria | 414 | HY322_12930 | 26291..27535 | 1891241 | AAA-family-ATPase-[Betaproteobacteria-bacterium] |
| MBI3372695.1 | OLD-ABC | GCA_016195915.1 | Betaproteobacteria_bacterium | Bacteria>Pseudomonadati>Pseudomonadota>Betaproteobacteria>unclassified_Betaproteobacteria | 406 | HY017_13200 | 56284..57504 | 1891241 | AAA-family-ATPase-[Betaproteobacteria-bacterium] |
| MBI3917895.1 | OLD-ABC | GCA_016199545.1 | Betaproteobacteria_bacterium | Bacteria>Pseudomonadati>Pseudomonadota>Betaproteobacteria>unclassified_Betaproteobacteria | 379 | HY322_12910 | 17681..18820 | 1891241 | AAA-family-ATPase-[Betaproteobacteria-bacterium] |
| WP_121581754.1 | OLD-ABC | GCF_963882275.1 | Enterobacterales | Bacteria>Pseudomonadati>Pseudomonadota>Gammaproteobacteria | 373 | I1A46_RS14715 | 208936..210057 | 91347 | MULTISPECIES:-AAA-family-ATPase-[Enterobacterales] |
| WP_112020504.1 | OLD-ABC | GCF_013624605.1 | Enterobacterales | Bacteria>Pseudomonadati>Pseudomonadota>Gammaproteobacteria | 347 | HV094_RS19975 | 4134976..4136019 | 91347 | MULTISPECIES:-ATP-binding-protein-[Enterobacterales] |
| WP_129328056.1 | OLD-ABC | GCF_036708225.1 | Enterobacterales | Bacteria>Pseudomonadati>Pseudomonadota>Gammaproteobacteria | 347 | DD604_RS20955 | 25033..26076 | 91347 | ATP-binding-protein-[Enterobacter-cloacae] |
| WP_219296388.1 | OLD-ABC | GCF_963896005.1 | Aeromonas | Bacteria>Pseudomonadati>Pseudomonadota>Gammaproteobacteria>Aeromonadales>Aeromonadaceae | 489 | EHZ86_RS11585 | 8997..10466 | 642 | MULTISPECIES:-AAA-family-ATPase-[Aeromonas] |
| WP_103858991.1 | OLD-ABC | GCF_019048725.1 | Aeromonas | Bacteria>Pseudomonadati>Pseudomonadota>Gammaproteobacteria>Aeromonadales>Aeromonadaceae | 473 | NUK49_RS00035 | 5461..6882 | 642 | MULTISPECIES:-AAA-family-ATPase-[Aeromonas] |
| WP_059169684.1 | OLD-ABC | GCF_963869605.1 | Aeromonas | Bacteria>Pseudomonadati>Pseudomonadota>Gammaproteobacteria>Aeromonadales>Aeromonadaceae | 425 | VO70_RS19200 | 17587..18864 | 642 | AAA-family-ATPase-[Aeromonas-salmonicida] |
| WP_182788685.1 | OLD-ABC | GCF_014161425.1 | Aeromonas | Bacteria>Pseudomonadati>Pseudomonadota>Gammaproteobacteria>Aeromonadales>Aeromonadaceae | 425 | OKB81_RS14900 | 6743..8020 | 642 | MULTISPECIES:-AAA-family-ATPase-[Aeromonas] |
| WP_139392566.1 | OLD-ABC | GCF_021654355.1 | Aeromonas | Bacteria>Pseudomonadati>Pseudomonadota>Gammaproteobacteria>Aeromonadales>Aeromonadaceae | 415 | OKC32_RS09695 | 8679..9926 | 642 | MULTISPECIES:-ATP-binding-protein-[Aeromonas] |
| WP_183149505.1 | OLD-ABC | GCF_014158455.1 | Aeromonas | Bacteria>Pseudomonadati>Pseudomonadota>Gammaproteobacteria>Aeromonadales>Aeromonadaceae | 415 | H7X73_RS05540 | 1122097..1123344 | 642 | ATP-binding-protein-[Aeromonas-caviae] |
| WP_050664997.1 | OLD-ABC | GCF_000820105.1 | Aeromonas | Bacteria>Pseudomonadati>Pseudomonadota>Gammaproteobacteria>Aeromonadales>Aeromonadaceae | 414 | BN1133_RS02620 | 42301..43545 | 642 | MULTISPECIES:-ATP-binding-protein-[Aeromonas] |
| WP_160837715.1 | OLD-ABC | GCF_013085725.1 | Aeromonas | Bacteria>Pseudomonadati>Pseudomonadota>Gammaproteobacteria>Aeromonadales>Aeromonadaceae | 414 | NUK45_RS19505 | 2352..3596 | 642 | MULTISPECIES:-ATP-binding-protein-[Aeromonas] |
| WP_234605105.1 | OLD-ABC | GCF_026173065.1 | Aeromonas | Bacteria>Pseudomonadati>Pseudomonadota>Gammaproteobacteria>Aeromonadales>Aeromonadaceae | 414 | ORM39_RS00830 | 20212..21456 | 642 | ATP-binding-protein-[Aeromonas-salmonicida] |
| WP_265455064.1 | OLD-ABC | GCF_049175675.1 | Aeromonas | Bacteria>Pseudomonadati>Pseudomonadota>Gammaproteobacteria>Aeromonadales>Aeromonadaceae | 414 | ORM31_RS04465 | 131039..132283 | 642 | ATP-binding-protein-[Aeromonas-salmonicida] |
| WP_064341061.1 | OLD-ABC | GCF_014169835.1 | Aeromonas | Bacteria>Pseudomonadati>Pseudomonadota>Gammaproteobacteria>Aeromonadales>Aeromonadaceae | 395 | EHZ86_RS11600 | 14064..15251 | 642 | MULTISPECIES:-AAA-family-ATPase-[Aeromonas] |
| WP_103858988.1 | OLD-ABC | GCF_024649925.1 | Aeromonas | Bacteria>Pseudomonadati>Pseudomonadota>Gammaproteobacteria>Aeromonadales>Aeromonadaceae | 395 | NUK49_RS00050 | 10530..11717 | 642 | MULTISPECIES:-AAA-family-ATPase-[Aeromonas] |
| WP_216991886.1 | OLD-ABC | GCF_019048725.1 | Aeromonas | Bacteria>Pseudomonadati>Pseudomonadota>Gammaproteobacteria>Aeromonadales>Aeromonadaceae | 395 | I6L37_RS11480 | 2394083..2395270 | 642 | AAA-family-ATPase-[Aeromonas-sp.-FDAARGOS-1407] |
| WP_139392560.1 | OLD-ABC | GCF_018359965.1 | Aeromonas | Bacteria>Pseudomonadati>Pseudomonadota>Gammaproteobacteria>Aeromonadales>Aeromonadaceae | 349 | OKC32_RS09725 | 18368..19417 | 642 | MULTISPECIES:-ATP-binding-protein-[Aeromonas] |

| Protein-ID | Domain | Assembly-ID | Strain | Taxonomy | Length | Locus-Tag | Genomic-coordinates | TaxID | Fasta-defline |
|---|---|---|---|---|---|---|---|---|---|
| WP_201871425.1 | OLD-ABC | GCF_016728905.1 | Aeromonas_caviae | Bacteria>Pseudomonadati>Pseudomonadota>Gammaproteobacteria>Aeromonadales>Aeromonadaceae>Aeromonas | 489 | JD551_RS01095 | 91648..93117 | 648 | AAA-family-ATPase-[Aeromonas-caviae] |
| WP_223918002.1 | OLD-ABC | GCF_019971655.1 | Aeromonas_caviae | Bacteria>Pseudomonadati>Pseudomonadota>Gammaproteobacteria>Aeromonadales>Aeromonadaceae>Aeromonas | 487 | LDN21_RS02430 | 64724..66187 | 648 | AAA-family-ATPase-[Aeromonas-caviae] |
| WP_042011557.1 | OLD-ABC | GCF_000819885.1 | Aeromonas_fluvialis | Bacteria>Pseudomonadati>Pseudomonadota>Gammaproteobacteria>Aeromonadales>Aeromonadaceae>Aeromonas | 425 | BN1125_RS07640 | 78037..79314 | 591962 | AAA-family-ATPase-[Aeromonas-fluvialis] |
| WP_128297389.1 | OLD-ABC | GCF_004024195.1 | Aeromonas_caviae | Bacteria>Pseudomonadati>Pseudomonadota>Gammaproteobacteria>Aeromonadales>Aeromonadaceae>Aeromonas | 425 | DN612_RS19425 | 6171..7448 | 648 | ATP-binding-protein-[Aeromonas-caviae] |
| WP_190283729.1 | OLD-ABC | GCF_014679585.1 | Aeromonas_hydrophila | Bacteria>Pseudomonadati>Pseudomonadota>Gammaproteobacteria>Aeromonadales>Aeromonadaceae>Aeromonas | 425 | IFA22_RS17955 | 3926752..3928029 | 644 | ATP-binding-protein-[Aeromonas-hydrophila] |
| WP_270793602.1 | OLD-ABC | GCF_027705385.1 | unclassified_Aeromonas | Bacteria>Pseudomonadati>Pseudomonadota>Gammaproteobacteria>Aeromonadales>Aeromonadaceae>Aeromonas | 418 | OKB76_RS13250 | 5656..6912 | 257493 | MULTISPECIES:-ATP-binding-protein-[unclassified-Aeromonas] |
| WP_265783522.1 | OLD-ABC | GCF_026230585.1 | Aeromonas_veronii | Bacteria>Pseudomonadati>Pseudomonadota>Gammaproteobacteria>Aeromonadales>Aeromonadaceae>Aeromonas | 417 | OD810_RS03315 | 106354..107607 | 654 | AAA-family-ATPase-[Aeromonas-veronii] |
| BBR09394.1 | OLD-ABC | GCA_014158455.1 | Aeromonas_caviae | Bacteria>Pseudomonadati>Pseudomonadota>Gammaproteobacteria>Aeromonadales>Aeromonadaceae>Aeromonas | 415 | WP3S18E02_10550 | 1122097..1123344 | 648 | hypothetical-protein-WP3S18E02_10550-[Aeromonas-caviae] |
| MBS4697266.1 | OLD-ABC | GCA_018359965.1 | Aeromonas_allosaccharophila | Bacteria>Pseudomonadati>Pseudomonadota>Gammaproteobacteria>Aeromonadales>Aeromonadaceae>Aeromonas | 415 | J4G65_17605 | 86193..87440 | 656 | AAA-family-ATPase-[Aeromonas-allosaccharophila] |
| WP_157666800.1 | OLD-ABC | GCF_963873515.1 | Aeromonas_salmonicida | Bacteria>Pseudomonadati>Pseudomonadota>Gammaproteobacteria>Aeromonadales>Aeromonadaceae>Aeromonas | 415 | O23A_RS10585 | 2335871..2337118 | 645 | ATP-binding-protein-[Aeromonas-salmonicida] |
| WP_193000846.1 | OLD-ABC | GCF_014892695.1 | Aeromonas_simiae | Bacteria>Pseudomonadati>Pseudomonadota>Gammaproteobacteria>Aeromonadales>Aeromonadaceae>Aeromonas | 415 | FE240_RS10740 | 2271650..2272897 | 218936 | ATP-binding-protein-[Aeromonas-simiae] |
| WP_201932779.1 | OLD-ABC | GCF_016729935.1 | Aeromonas_caviae | Bacteria>Pseudomonadati>Pseudomonadota>Gammaproteobacteria>Aeromonadales>Aeromonadaceae>Aeromonas | 415 | JD546_RS01990 | 63444..64691 | 648 | ATP-binding-protein-[Aeromonas-caviae] |
| WP_202047007.1 | OLD-ABC | GCF_016730065.1 | Aeromonas_media | Bacteria>Pseudomonadati>Pseudomonadota>Gammaproteobacteria>Aeromonadales>Aeromonadaceae>Aeromonas | 415 | JD497_RS12370 | 529961..531208 | 651 | ATP-binding-protein-[Aeromonas-media] |
| WP_213393397.1 | OLD-ABC | GCF_018359965.1 | Aeromonas_allosaccharophila | Bacteria>Pseudomonadati>Pseudomonadota>Gammaproteobacteria>Aeromonadales>Aeromonadaceae>Aeromonas | 415 | J4G65_RS17600 | 86193..87440 | 656 | ATP-binding-protein-[Aeromonas-allosaccharophila] |
| WP_257712003.1 | OLD-ABC | GCF_024649525.1 | Aeromonas_caviae | Bacteria>Pseudomonadati>Pseudomonadota>Gammaproteobacteria>Aeromonadales>Aeromonadaceae>Aeromonas | 415 | NUK31_RS14225 | 16246..17493 | 648 | ATP-binding-protein-[Aeromonas-caviae] |
| MBW3845326.1 | OLD-ABC | GCA_019348915.1 | Aeromonas_hydrophila | Bacteria>Pseudomonadati>Pseudomonadota>Gammaproteobacteria>Aeromonadales>Aeromonadaceae>Aeromonas | 414 | GPV00_14060 | 285143..286387 | 644 | AAA-family-ATPase-[Aeromonas-hydrophila] |
| WP_076362313.1 | OLD-ABC | GCF_900155975.1 | Aeromonas_hydrophila | Bacteria>Pseudomonadati>Pseudomonadota>Gammaproteobacteria>Aeromonadales>Aeromonadaceae>Aeromonas | 414 | BW963_RS17625 | 138910..140154 | 644 | ATP-binding-protein-[Aeromonas-hydrophila] |
| WP_101617900.1 | OLD-ABC | GCF_002850695.3 | Aeromonas_hydrophila | Bacteria>Pseudomonadati>Pseudomonadota>Gammaproteobacteria>Aeromonadales>Aeromonadaceae>Aeromonas | 414 | C1A23_RS07650 | 1218856..1220100 | 644 | ATP-binding-protein-[Aeromonas-hydrophila] |
| WP_123173886.1 | OLD-ABC | GCF_003722175.1 | Aeromonas_veronii | Bacteria>Pseudomonadati>Pseudomonadota>Gammaproteobacteria>Aeromonadales>Aeromonadaceae>Aeromonas | 414 | EFI48_RS19255 | 4191765..4193009 | 654 | ATP-binding-protein-[Aeromonas-veronii] |
| WP_201944275.1 | OLD-ABC | GCF_016729595.1 | Aeromonas_veronii | Bacteria>Pseudomonadati>Pseudomonadota>Gammaproteobacteria>Aeromonadales>Aeromonadaceae>Aeromonas | 414 | JD494_RS16540 | 21357..22601 | 654 | ATP-binding-protein-[Aeromonas-veronii] |
| WP_219254718.1 | OLD-ABC | GCF_038524185.1 | Aeromonas_hydrophila | Bacteria>Pseudomonadati>Pseudomonadota>Gammaproteobacteria>Aeromonadales>Aeromonadaceae>Aeromonas | 414 | GPV00_RS13995 | 285143..286387 | 644 | ATP-binding-protein-[Aeromonas-hydrophila] |
| WP_234578018.1 | OLD-ABC | GCF_963898355.1 | Aeromonas_salmonicida | Bacteria>Pseudomonadati>Pseudomonadota>Gammaproteobacteria>Aeromonadales>Aeromonadaceae>Aeromonas | 414 | LZ632_RS15265 | 22065..23309 | 645 | ATP-binding-protein-[Aeromonas-salmonicida] |
| WP_236274796.1 | OLD-ABC | GCF_021609985.1 | Aeromonas_caviae | Bacteria>Pseudomonadati>Pseudomonadota>Gammaproteobacteria>Aeromonadales>Aeromonadaceae>Aeromonas | 414 | L1871_RS17020 | 3636805..3638049 | 648 | ATP-binding-protein-[Aeromonas-caviae] |
| WP_265776109.1 | OLD-ABC | GCF_049168655.1 | Aeromonas_veronii | Bacteria>Pseudomonadati>Pseudomonadota>Gammaproteobacteria>Aeromonadales>Aeromonadaceae>Aeromonas | 414 | OD804_RS12720 | 2588..3832 | 654 | ATP-binding-protein-[Aeromonas-veronii] |
| WP_275056846.1 | OLD-ABC | GCF_029024985.1 | Aeromonas_allosaccharophila | Bacteria>Pseudomonadati>Pseudomonadota>Gammaproteobacteria>Aeromonadales>Aeromonadaceae>Aeromonas | 414 | PYU98_RS18275 | 3927977..3929221 | 656 | ATP-binding-protein-[Aeromonas-allosaccharophila] |
| MCF3097926.1 | OLD-ABC | GCA_021538905.1 | Aeromonas_australiensis | Bacteria>Pseudomonadati>Pseudomonadota>Gammaproteobacteria>Aeromonadales>Aeromonadaceae>Aeromonas | 395 | EHZ86_11630 | 14064..15251 | 1114880 | chromosome-segregation-protein-SMC-[Aeromonas-australiensis] |
| WP_201871421.1 | OLD-ABC | GCF_016728905.1 | Aeromonas_caviae | Bacteria>Pseudomonadati>Pseudomonadota>Gammaproteobacteria>Aeromonadales>Aeromonadaceae>Aeromonas | 395 | JD551_RS01080 | 86862..88049 | 648 | AAA-family-ATPase-[Aeromonas-caviae] |
| WP_264787582.1 | OLD-ABC | GCF_026000055.1 | Aeromonas_caviae | Bacteria>Pseudomonadati>Pseudomonadota>Gammaproteobacteria>Aeromonadales>Aeromonadaceae>Aeromonas | 395 | OPI30_RS08270 | 1645083..1646270 | 648 | AAA-family-ATPase-[Aeromonas-caviae] |
| WP_270698349.1 | OLD-ABC | GCF_027705185.1 | unclassified_Aeromonas | Bacteria>Pseudomonadati>Pseudomonadota>Gammaproteobacteria>Aeromonadales>Aeromonadaceae>Aeromonas | 395 | OKC33_RS16310 | 7300..8487 | 257493 | MULTISPECIES:-AAA-family-ATPase-[unclassified-Aeromonas] |
| WP_270793606.1 | OLD-ABC | GCF_027705385.1 | unclassified_Aeromonas | Bacteria>Pseudomonadati>Pseudomonadota>Gammaproteobacteria>Aeromonadales>Aeromonadaceae>Aeromonas | 374 | OKB76_RS13270 | 11844..12968 | 257493 | MULTISPECIES:-ATP-binding-protein-[unclassified-Aeromonas] |
| BBR09398.1 | OLD-ABC | GCA_014158455.1 | Aeromonas_caviae | Bacteria>Pseudomonadati>Pseudomonadota>Gammaproteobacteria>Aeromonadales>Aeromonadaceae>Aeromonas | 353 | WP3S18E02_10590 | 1129744..1130805 | 648 | hypothetical-protein-WP3S18E02_10590-[Aeromonas-caviae] |
| WP_042011551.1 | OLD-ABC | GCF_000819885.1 | Aeromonas_fluvialis | Bacteria>Pseudomonadati>Pseudomonadota>Gammaproteobacteria>Aeromonadales>Aeromonadaceae>Aeromonas | 353 | BN1125_RS07615 | 70371..71432 | 591962 | AAA-family-ATPase-[Aeromonas-fluvialis] |
| WP_159144195.1 | OLD-ABC | GCF_009834065.1 | Aeromonas_veronii | Bacteria>Pseudomonadati>Pseudomonadota>Gammaproteobacteria>Aeromonadales>Aeromonadaceae>Aeromonas | 353 | GRF56_RS18075 | 3924779..3925840 | 654 | AAA-family-ATPase-[Aeromonas-veronii] |
| WP_183149509.1 | OLD-ABC | GCF_014158455.1 | Aeromonas_caviae | Bacteria>Pseudomonadati>Pseudomonadota>Gammaproteobacteria>Aeromonadales>Aeromonadaceae>Aeromonas | 353 | H7X73_RS05560 | 1129744..1130805 | 648 | AAA-family-ATPase-[Aeromonas-caviae] |
| WP_265783517.1 | OLD-ABC | GCF_026230585.1 | Aeromonas_veronii | Bacteria>Pseudomonadati>Pseudomonadota>Gammaproteobacteria>Aeromonadales>Aeromonadaceae>Aeromonas | 350 | OD810_RS03280 | 99076..100128 | 654 | AAA-family-ATPase-[Aeromonas-veronii] |
| MBS4697262.1 | OLD-ABC | GCA_018359965.1 | Aeromonas_allosaccharophila | Bacteria>Pseudomonadati>Pseudomonadota>Gammaproteobacteria>Aeromonadales>Aeromonadaceae>Aeromonas | 349 | J4G65_17585 | 78744..79793 | 656 | ATP-binding-protein-[Aeromonas-allosaccharophila] |
| MBW3845320.1 | OLD-ABC | GCA_019348915.1 | Aeromonas_hydrophila | Bacteria>Pseudomonadati>Pseudomonadota>Gammaproteobacteria>Aeromonadales>Aeromonadaceae>Aeromonas | 349 | GPV00_14030 | 275950..276999 | 644 | AAA-family-ATPase-[Aeromonas-hydrophila] |
| WP_202047004.1 | OLD-ABC | GCF_016730065.1 | Aeromonas_media | Bacteria>Pseudomonadati>Pseudomonadota>Gammaproteobacteria>Aeromonadales>Aeromonadaceae>Aeromonas | 349 | JD497_RS12355 | 524267..525316 | 651 | ATP-binding-protein-[Aeromonas-media] |
| WP_219254708.1 | OLD-ABC | GCF_038524185.1 | Aeromonas_hydrophila | Bacteria>Pseudomonadati>Pseudomonadota>Gammaproteobacteria>Aeromonadales>Aeromonadaceae>Aeromonas | 349 | GPV00_RS13965 | 275950..276999 | 644 | ATP-binding-protein-[Aeromonas-hydrophila] |
| WP_081250393.1 | OLD-ABC | GCF_001634325.1 | Aeromonas_veronii | Bacteria>Pseudomonadati>Pseudomonadota>Gammaproteobacteria>Aeromonadales>Aeromonadaceae>Aeromonas | 260 | WM43_RS07835 | 1777357..1778139 | 654 | AAA-family-ATPase-[Aeromonas-veronii] |
| WP_279465182.1 | OLD-ABC | GCF_029773565.1 | Aeromonas_veronii | Bacteria>Pseudomonadati>Pseudomonadota>Gammaproteobacteria>Aeromonadales>Aeromonadaceae>Aeromonas | 231 | QBS04_RS15315 | 3274574..3275269 | 654 | AAA-family-ATPase-[Aeromonas-veronii] |
| WP_279465185.1 | OLD-ABC | GCF_029773565.1 | Aeromonas_veronii | Bacteria>Pseudomonadati>Pseudomonadota>Gammaproteobacteria>Aeromonadales>Aeromonadaceae>Aeromonas | 148 | QBS04_RS15320 | 3275229..3275675 | 654 | AAA-family-ATPase-[Aeromonas-veronii] |
| WP_064338619.1 | OLD-ABC | GCF_001634325.1 | Aeromonas_veronii | Bacteria>Pseudomonadati>Pseudomonadota>Gammaproteobacteria>Aeromonadales>Aeromonadaceae>Aeromonas | 146 | WM43_RS07845 | 1779326..1779766 | 654 | AAA-family-ATPase-[Aeromonas-veronii] |
| WP_244812701.1 | OLD-ABC | GCF_022869435.1 | Aeromonas_sp._LsrichE-8G | Bacteria>Pseudomonadati>Pseudomonadota>Gammaproteobacteria>Aeromonadales>Aeromonadaceae>Aeromonas>unclassified_Aeromonas | 395 | MUW49_RS10485 | 1342..2529 | 2932048 | AAA-family-ATPase-[Aeromonas-sp.-LsrichE-8G] |
| WP_163133168.1 | OLD-ABC | GCF_900537085.1 | Agarivorans_sp._Alg241-V36 | Bacteria>Pseudomonadati>Pseudomonadota>Gammaproteobacteria>Alteromonadales>Alteromonadaceae>Agarivorans>unclassified_Agarivorans | 367 | G6R11_RS11260 | 297414..298517 | 2305992 | AAA-family-ATPase-[Agarivorans-sp.-Alg241-V36] |
| WP_232368259.1 | OLD-ABC | GCF_002831605.1 | Alteromonas_sp._MB-3u-76 | Bacteria>Pseudomonadati>Pseudomonadota>Gammaproteobacteria>Alteromonadales>Alteromonadaceae>Alteromonas/Salinimonas_group>Alteromonas>unclassified_Alteromonas | 350 | CW735_RS10470 | 2334062..2335114 | 2058133 | AAA-family-ATPase-[Alteromonas-sp.-MB-3u-76] |
| WP_146799745.1 | OLD-ABC | GCF_007954305.1 | Colwellia_hornerae | Bacteria>Pseudomonadati>Pseudomonadota>Gammaproteobacteria>Alteromonadales>Colwelliaceae>Colwellia | 370 | ESZ26_RS11975 | 111206..112318 | 89402 | AAA-family-ATPase-[Colwellia-hornerae] |
| WP_165723735.1 | OLD-ABC | GCF_943789085.1 | Pseudoalteromonas | Bacteria>Pseudomonadati>Pseudomonadota>Gammaproteobacteria>Alteromonadales>Pseudoalteromonadaceae | 389 | QDP19_RS17635 | 3064190..3065359 | 53246 | MULTISPECIES:-AAA-family-ATPase-[Pseudoalteromonas] |
| WP_130146691.1 | OLD-ABC | GCF_004208955.1 | Pseudoalteromonas_sp._CO325X | Bacteria>Pseudomonadati>Pseudomonadota>Gammaproteobacteria>Alteromonadales>Pseudoalteromonadaceae>Pseudoalteromonas>unclassified_Pseudoalteromonas | 389 | EXT46_RS08645 | 69579..70748 | 1777262 | AAA-family-ATPase-[Pseudoalteromonas-sp.-CO325X] |
| WP_243888584.1 | OLD-ABC | GCF_017580535.1 | Shewanella_algae | Bacteria>Pseudomonadati>Pseudomonadota>Gammaproteobacteria>Alteromonadales>Shewanellaceae>Shewanella | 390 | I6M49_RS22465 | 7195..8367 | 38313 | AAA-family-ATPase-[Shewanella-algae] |
| WP_006086468.1 | OLD-ABC | GCF_000231345.1 | Shewanella_baltica | Bacteria>Pseudomonadati>Pseudomonadota>Gammaproteobacteria>Alteromonadales>Shewanellaceae>Shewanella | 389 | SBAL625DRAFT_RS15535 | 218449..219618 | 62322 | AAA-family-ATPase-[Shewanella-baltica] |
| WP_208251008.1 | OLD-ABC | GCF_017580085.1 | Shewanella_algae | Bacteria>Pseudomonadati>Pseudomonadota>Gammaproteobacteria>Alteromonadales>Shewanellaceae>Shewanella | 389 | I6M29_RS02110 | 13101..14270 | 38313 | AAA-family-ATPase-[Shewanella-algae] |
| WP_076500075.1 | OLD-ABC | GCF_900156405.1 | Shewanella_morhuae | Bacteria>Pseudomonadati>Pseudomonadota>Gammaproteobacteria>Alteromonadales>Shewanellaceae>Shewanella | 370 | BW947_RS12480 | 15725..16837 | 365591 | AAA-family-ATPase-[Shewanella-morhuae] |
| WP_261732936.1 | OLD-ABC | GCF_025398315.1 | Shewanella_phaeophyticola | Bacteria>Pseudomonadati>Pseudomonadota>Gammaproteobacteria>Alteromonadales>Shewanellaceae>Shewanella | 271 | N4T56_RS08710 | 1861739..1862554 | 2978345 | hypothetical-protein-[Shewanella-sp.-KJ10-1] |
| WP_179123630.1 | OLD-ABC | GCF_043179105.1 | Cardiobacterium_hominis | Bacteria>Pseudomonadati>Pseudomonadota>Gammaproteobacteria>Cardiobacteriales>Cardiobacteriaceae>Cardiobacterium | 477 | CHUV0807_RS10750 | 30286..31719 | 2718 | AAA-family-ATPase-[Cardiobacterium-hominis] |
| WP_079541818.1 | OLD-ABC | GCF_043179105.1 | Cardiobacterium_hominis | Bacteria>Pseudomonadati>Pseudomonadota>Gammaproteobacteria>Cardiobacteriales>Cardiobacteriaceae>Cardiobacterium | 398 | CHUV0807_RS10765 | 35775..36971 | 2718 | AAA-family-ATPase-[Cardiobacterium-hominis] |
| WP_096087094.1 | OLD-ABC | GCF_002312815.1 | Agaribacterium_haliotis | Bacteria>Pseudomonadati>Pseudomonadota>Gammaproteobacteria>Cellvibrionales>Cellvibrionaceae>Agaribacterium | 389 | CFK45_RS14060 | 10811..11980 | 2013869 | AAA-family-ATPase-[Agaribacterium-haliotis] |
| WP_040392246.1 | OLD-ABC | GCF_000263355.1 | Cellvibrio_sp._BR | Bacteria>Pseudomonadati>Pseudomonadota>Gammaproteobacteria>Cellvibrionales>Cellvibrionaceae>Cellvibrio>unclassified_Cellvibrio | 396 | O59_RS11280 | 229054..230244 | 1134474 | ATP-binding-protein-[Cellvibrio-sp.-BR] |
| WP_054114050.1 | OLD-ABC | GCF_001296225.1 | Marinagarivorans_algicola | Bacteria>Pseudomonadati>Pseudomonadota>Gammaproteobacteria>Cellvibrionales>Cellvibrionaceae>Marinagarivorans | 389 | AC526_RS14085 | 29706..30875 | 1513270 | AAA-family-ATPase-[Marinagarivorans-algicola] |
| WP_011469232.1 | OLD-ABC | GCF_000013665.1 | Saccharophagus_degradans | Bacteria>Pseudomonadati>Pseudomonadota>Gammaproteobacteria>Cellvibrionales>Cellvibrionaceae>Saccharophagus | 370 | SDE_RS14415 | 3487626..3488738 | 86304 | AAA-family-ATPase-[Saccharophagus-degradans] |
| WP_135445663.1 | OLD-ABC | GCF_004745945.1 | Mangrovimicrobium_sediminis | Bacteria>Pseudomonadati>Pseudomonadota>Gammaproteobacteria>Cellvibrionales>Halieaceae>Mangrovimicrobium | 394 | E4634_RS16025 | 157649..158833 | 2562682 | AAA-family-ATPase-[Haliea-sp.-SAOS-164] |
| MTW22273.1 | OLD-ABC | GCA_009720725.1 | Allochromatium_palmeri | Bacteria>Pseudomonadati>Pseudomonadota>Gammaproteobacteria>Chromatiales>Chromatiaceae>Allochromatium | 477 | GJ668_14425 | 32960..34393 | 231048 | AAA-family-ATPase-[Allochromatium-palmeri] |
| WP_246194909.1 | OLD-ABC | - | Allochromatium_palmeri | Bacteria>Pseudomonadati>Pseudomonadota>Gammaproteobacteria>Chromatiales>Chromatiaceae>Allochromatium | 475 | GJ668_RS14430 | 32966..34393 | 231048 | AAA-family-ATPase-[Allochromatium-palmeri] |
| WP_200158840.1 | OLD-ABC | GCF_016584135.1 | Allochromatium_vinosum | Bacteria>Pseudomonadati>Pseudomonadota>Gammaproteobacteria>Chromatiales>Chromatiaceae>Allochromatium | 436 | CKO29_RS16475 | 17571..18881 | 1049 | ATP-binding-protein-[Allochromatium-vinosum] |
| WP_200158835.1 | OLD-ABC | GCF_016584135.1 | Allochromatium_vinosum | Bacteria>Pseudomonadati>Pseudomonadota>Gammaproteobacteria>Chromatiales>Chromatiaceae>Allochromatium | 423 | CKO29_RS16450 | 8773..10044 | 1049 | AAA-family-ATPase-[Allochromatium-vinosum] |
| WP_176976534.1 | OLD-ABC | - | Allochromatium_humboldtianum | Bacteria>Pseudomonadati>Pseudomonadota>Gammaproteobacteria>Chromatiales>Chromatiaceae>Allochromatium | 415 | HW932_RS10985 | 216610..217857 | 504901 | ATP-binding-protein-[Allochromatium-humboldtianum] |
| BCU06241.1 | OLD-ABC | GCA_018409545.1 | Allochromatium_tepidum | Bacteria>Pseudomonadati>Pseudomonadota>Gammaproteobacteria>Chromatiales>Chromatiaceae>Allochromatium | 407 | Atep_09180 | 997805..999028 | 553982 | ATPase-[Allochromatium-tepidum] |
| WP_213380488.1 | OLD-ABC | GCF_018409545.1 | Allochromatium_tepidum | Bacteria>Pseudomonadati>Pseudomonadota>Gammaproteobacteria>Chromatiales>Chromatiaceae>Allochromatium | 407 | Atep_RS04620 | 997805..999028 | 553982 | ATP-binding-protein-[Allochromatium-tepidum] |
| MTW22275.1 | OLD-ABC | GCA_009720725.1 | Allochromatium_palmeri | Bacteria>Pseudomonadati>Pseudomonadota>Gammaproteobacteria>Chromatiales>Chromatiaceae>Allochromatium | 396 | GJ668_14445 | 37105..38295 | 231048 | AAA-family-ATPase-[Allochromatium-palmeri] |
| WP_155450843.1 | OLD-ABC | GCF_009720725.1 | Allochromatium_palmeri | Bacteria>Pseudomonadati>Pseudomonadota>Gammaproteobacteria>Chromatiales>Chromatiaceae>Allochromatium | 396 | GJ668_RS14450 | 37105..38295 | 231048 | AAA-family-ATPase-[Allochromatium-palmeri] |
| WP_201220074.1 | OLD-ABC | GCF_016653315.1 | Halochromatium_roseum | Bacteria>Pseudomonadati>Pseudomonadota>Gammaproteobacteria>Chromatiales>Chromatiaceae>Halochromatium | 366 | CCR96_RS10450 | 42726..43826 | 391920 | AAA-family-ATPase-[Halochromatium-roseum] |
| WP_274601171.1 | OLD-ABC | - | Halochromatium_roseum | Bacteria>Pseudomonadati>Pseudomonadota>Gammaproteobacteria>Chromatiales>Chromatiaceae>Halochromatium | 200 | CCR96_RS25200 | 11114..11716 | 391920 | ATP-binding-protein-[Halochromatium-roseum] |
| WP_211211740.1 | OLD-ABC | GCF_000379525.1 | Lamprocystis_purpurea | Bacteria>Pseudomonadati>Pseudomonadota>Gammaproteobacteria>Chromatiales>Chromatiaceae>Lamprocystis | 478 | A390_RS0126770 | 73468..74904 | 61598 | ATP-binding-protein-[Lamprocystis-purpurea] |
| WP_020505850.1 | OLD-ABC | GCF_000379525.1 | Lamprocystis_purpurea | Bacteria>Pseudomonadati>Pseudomonadota>Gammaproteobacteria>Chromatiales>Chromatiaceae>Lamprocystis | 427 | A390_RS0116905 | 22041..23324 | 61598 | ATP-binding-protein-[Lamprocystis-purpurea] |
| WP_020507364.1 | OLD-ABC | GCF_000379525.1 | Lamprocystis_purpurea | Bacteria>Pseudomonadati>Pseudomonadota>Gammaproteobacteria>Chromatiales>Chromatiaceae>Lamprocystis | 369 | A390_RS0124825 | 127648..128757 | 61598 | AAA-family-ATPase-[Lamprocystis-purpurea] |
| WP_062271495.1 | OLD-ABC | GCF_001583505.1 | Marichromatium_gracile | Bacteria>Pseudomonadati>Pseudomonadota>Gammaproteobacteria>Chromatiales>Chromatiaceae>Marichromatium | 436 | AY586_RS00805 | 20951..22261 | 1048 | ATP-binding-protein-[Marichromatium-gracile] |
| WP_062271483.1 | OLD-ABC | GCF_001583505.1 | Marichromatium_gracile | Bacteria>Pseudomonadati>Pseudomonadota>Gammaproteobacteria>Chromatiales>Chromatiaceae>Marichromatium | 422 | AY586_RS00775 | 11833..13101 | 1048 | AAA-family-ATPase-[Marichromatium-gracile] |
| WP_200387141.1 | OLD-ABC | GCF_016583985.1 | Thiocapsa_imhoffii | Bacteria>Pseudomonadati>Pseudomonadota>Gammaproteobacteria>Chromatiales>Chromatiaceae>Thiocapsa | 461 | CKO25_RS06625 | 41749..43134 | 382777 | AAA-family-ATPase-[Thiocapsa-imhoffii] |
| QVL48950.1 | OLD-ABC | GCA_018399035.1 | Thiocapsa_sp. | Bacteria>Pseudomonadati>Pseudomonadota>Gammaproteobacteria>Chromatiales>Chromatiaceae>Thiocapsa>unclassified_Thiocapsa | 460 | KFB96_26075 | 5693748..5695130 | 2024551 | MAG:-AAA-family-ATPase-[Thiocapsa-sp.] |
| QVL49520.1 | OLD-ABC | GCA_018399035.1 | Thiocapsa_sp. | Bacteria>Pseudomonadati>Pseudomonadota>Gammaproteobacteria>Chromatiales>Chromatiaceae>Thiocapsa>unclassified_Thiocapsa | 429 | KFB96_03125 | 679089..680378 | 2024551 | MAG:-AAA-family-ATPase-[Thiocapsa-sp.] |
| QVL49517.1 | OLD-ABC | GCA_018399035.1 | Thiocapsa_sp. | Bacteria>Pseudomonadati>Pseudomonadota>Gammaproteobacteria>Chromatiales>Chromatiaceae>Thiocapsa>unclassified_Thiocapsa | 408 | KFB96_03105 | 672899..674125 | 2024551 | MAG:-AAA-family-ATPase-[Thiocapsa-sp.] |
| QVL47239.1 | OLD-ABC | GCA_018399035.1 | Thiocapsa_sp. | Bacteria>Pseudomonadati>Pseudomonadota>Gammaproteobacteria>Chromatiales>Chromatiaceae>Thiocapsa>unclassified_Thiocapsa | 389 | KFB96_16120 | 3546852..3548021 | 2024551 | MAG:-AAA-family-ATPase-[Thiocapsa-sp.] |
| QVL47944.1 | OLD-ABC | GCA_018399035.1 | Thiocapsa_sp. | Bacteria>Pseudomonadati>Pseudomonadota>Gammaproteobacteria>Chromatiales>Chromatiaceae>Thiocapsa>unclassified_Thiocapsa | 384 | KFB96_20140 | 4421746..4422900 | 2024551 | MAG:-AAA-family-ATPase-[Thiocapsa-sp.] |
| QVL49211.1 | OLD-ABC | GCA_018399035.1 | Thiocapsa_sp. | Bacteria>Pseudomonadati>Pseudomonadota>Gammaproteobacteria>Chromatiales>Chromatiaceae>Thiocapsa>unclassified_Thiocapsa | 374 | KFB96_01385 | 296233..297357 | 2024551 | MAG:-AAA-family-ATPase-[Thiocapsa-sp.] |
| WP_200375564.1 | OLD-ABC | GCF_016583575.1 | Thiocystis_violacea | Bacteria>Pseudomonadati>Pseudomonadota>Gammaproteobacteria>Chromatiales>Chromatiaceae>Thiocystis | 383 | CKO27_RS14225 | 26209..27360 | 13725 | ATP-binding-protein-[Thiocystis-violacea] |
| WP_216644669.1 | OLD-ABC | GCF_002813775.1 | Candidatus_Thiodictyon_syntrophicum | Bacteria>Pseudomonadati>Pseudomonadota>Gammaproteobacteria>Chromatiales>Chromatiaceae>Thiodictyon | 466 | THSYN_RS03430 | 786524..787924 | 1166950 | ATP-binding-protein-[Candidatus-Thiodictyon-syntrophicum] |
| WP_100919680.1 | OLD-ABC | GCF_002813775.1 | Candidatus_Thiodictyon_syntrophicum | Bacteria>Pseudomonadati>Pseudomonadota>Gammaproteobacteria>Chromatiales>Chromatiaceae>Thiodictyon | 453 | THSYN_RS13915 | 3311132..3312493 | 1166950 | AAA-family-ATPase-[Candidatus-Thiodictyon-syntrophicum] |
| AUB82575.1 | OLD-ABC | GCA_002813775.1 | Candidatus_Thiodictyon_syntrophicum | Bacteria>Pseudomonadati>Pseudomonadota>Gammaproteobacteria>Chromatiales>Chromatiaceae>Thiodictyon | 438 | THSYN_17575 | 4230137..4231453 | 1166950 | recombinase-RecF-[Candidatus-Thiodictyon-syntrophicum] |
| WP_100920297.1 | OLD-ABC | GCF_002813775.1 | Candidatus_Thiodictyon_syntrophicum | Bacteria>Pseudomonadati>Pseudomonadota>Gammaproteobacteria>Chromatiales>Chromatiaceae>Thiodictyon | 438 | THSYN_RS17710 | 4230137..4231453 | 1166950 | AAA-family-ATPase-[Candidatus-Thiodictyon-syntrophicum] |
| WP_150092065.1 | OLD-ABC | GCF_008632335.1 | Thiohalocapsa_marina | Bacteria>Pseudomonadati>Pseudomonadota>Gammaproteobacteria>Chromatiales>Chromatiaceae>Thiohalocapsa | 407 | F2Q65_RS07745 | 161649..162872 | 424902 | ATP-binding-protein-[Thiohalocapsa-marina] |
| WP_058557219.1 | OLD-ABC | GCF_001469165.1 | Thiohalocapsa_sp._ML1 | Bacteria>Pseudomonadati>Pseudomonadota>Gammaproteobacteria>Chromatiales>Chromatiaceae>Thiohalocapsa>unclassified_Thiohalocapsa | 391 | ML01_RS20310 | 8834..10009 | 1431688 | ATP-binding-protein-[Thiohalocapsa-sp.-ML1] |
| WP_164452827.1 | OLD-ABC | GCF_010820565.1 | Thiorhodococcus_minor | Bacteria>Pseudomonadati>Pseudomonadota>Gammaproteobacteria>Chromatiales>Chromatiaceae>Thiorhodococcus | 414 | G3446_RS10730 | 65078..66322 | 57489 | AAA-family-ATPase-[Thiorhodococcus-minor] |
| MBK8637139.1 | OLD-ABC | GCA_016714645.1 | Chromatiaceae_bacterium | Bacteria>Pseudomonadati>Pseudomonadota>Gammaproteobacteria>Chromatiales>Chromatiaceae>unclassified_Chromatiaceae | 423 | IPN92_02250 | 476459..477730 | 2049432 | AAA-family-ATPase-[Chromatiaceae-bacterium] |
| MBP8282508.1 | OLD-ABC | GCA_018007135.1 | Chromatiaceae_bacterium | Bacteria>Pseudomonadati>Pseudomonadota>Gammaproteobacteria>Chromatiales>Chromatiaceae>unclassified_Chromatiaceae | 410 | KAX46_01105 | 1402..2634 | 2049432 | AAA-family-ATPase-[Chromatiaceae-bacterium] |
| MBP6582545.1 | OLD-ABC | GCA_017989555.1 | Chromatiaceae_bacterium | Bacteria>Pseudomonadati>Pseudomonadota>Gammaproteobacteria>Chromatiales>Chromatiaceae>unclassified_Chromatiaceae | 387 | KA204_03600 | 4049..5212 | 2049432 | AAA-family-ATPase-[Chromatiaceae-bacterium] |
| MBP6733370.1 | OLD-ABC | GCA_017990755.1 | Chromatiaceae_bacterium | Bacteria>Pseudomonadati>Pseudomonadota>Gammaproteobacteria>Chromatiales>Chromatiaceae>unclassified_Chromatiaceae | 387 | KA142_01520 | 6335..7498 | 2049432 | AAA-family-ATPase-[Chromatiaceae-bacterium] |
| MBP6806940.1 | OLD-ABC | GCA_017991215.1 | Chromatiaceae_bacterium | Bacteria>Pseudomonadati>Pseudomonadota>Gammaproteobacteria>Chromatiales>Chromatiaceae>unclassified_Chromatiaceae | 387 | KA125_01445 | 5377..6540 | 2049432 | AAA-family-ATPase-[Chromatiaceae-bacterium] |
| MBP8282513.1 | OLD-ABC | GCA_018007135.1 | Chromatiaceae_bacterium | Bacteria>Pseudomonadati>Pseudomonadota>Gammaproteobacteria>Chromatiales>Chromatiaceae>unclassified_Chromatiaceae | 387 | KAX46_01130 | 8076..9239 | 2049432 | AAA-family-ATPase-[Chromatiaceae-bacterium] |
| MBP6733365.1 | OLD-ABC | GCA_017990755.1 | Chromatiaceae_bacterium | Bacteria>Pseudomonadati>Pseudomonadota>Gammaproteobacteria>Chromatiales>Chromatiaceae>unclassified_Chromatiaceae | 298 | KA142_01495 | 1..893 | 2049432 | AAA-family-ATPase-[Chromatiaceae-bacterium] |
| WP_238621058.1 | OLD-ABC | GCF_022227785.1 | Ectothiorhodospira_lacustris | Bacteria>Pseudomonadati>Pseudomonadota>Gammaproteobacteria>Chromatiales>Ectothiorhodospiraceae>Ectothiorhodospira | 410 | LQ952_RS02370 | 44306..45538 | 2899127 | AAA-family-ATPase-[Ectothiorhodospira-sp.-B14B] |
| WP_238617953.1 | OLD-ABC | GCF_022227765.1 | Ectothiorhodospira_shaposhnikovii | Bacteria>Pseudomonadati>Pseudomonadota>Gammaproteobacteria>Chromatiales>Ectothiorhodospiraceae>Ectothiorhodospira | 397 | LQD11_RS13160 | 106472..107665 | 1054 | ATP-binding-protein-[Ectothiorhodospira-shaposhnikovii] |
| WP_090207820.1 | OLD-ABC | GCF_900110965.1 | Ectothiorhodospira_magna | Bacteria>Pseudomonadati>Pseudomonadota>Gammaproteobacteria>Chromatiales>Ectothiorhodospiraceae>Ectothiorhodospira | 371 | BMX62_RS09260 | 44635..45750 | 867345 | AAA-family-ATPase-[Ectothiorhodospira-magna] |
| WP_238627726.1 | OLD-ABC | GCF_022227695.1 | unclassified_Ectothiorhodospira | Bacteria>Pseudomonadati>Pseudomonadota>Gammaproteobacteria>Chromatiales>Ectothiorhodospiraceae>Ectothiorhodospira | 370 | LPW28_RS03035 | 34125..35237 | 2684909 | MULTISPECIES:-AAA-family-ATPase-[unclassified-Ectothiorhodospira] |
| WP_005002339.1 | OLD-ABC | GCF_000153205.1 | Nitrococcus_mobilis | Bacteria>Pseudomonadati>Pseudomonadota>Gammaproteobacteria>Chromatiales>Ectothiorhodospiraceae>Nitrococcus | 409 | NB231_RS09975 | 2188011..2189240 | 35797 | AAA-family-ATPase-[Nitrococcus-mobilis] |
| MBF2761738.1 | OLD-ABC | GCA_015278165.1 | Ectothiorhodospiraceae_bacterium_AqS1 | Bacteria>Pseudomonadati>Pseudomonadota>Gammaproteobacteria>Chromatiales>Ectothiorhodospiraceae>unclassified_Ectothiorhodospiraceae | 465 | ISN28_16005 | 817..2214 | 1725326 | AAA-family-ATPase-[Ectothiorhodospiraceae-bacterium-AqS1] |
| MBF2761744.1 | OLD-ABC | GCA_015278165.1 | Ectothiorhodospiraceae_bacterium_AqS1 | Bacteria>Pseudomonadati>Pseudomonadota>Gammaproteobacteria>Chromatiales>Ectothiorhodospiraceae>unclassified_Ectothiorhodospiraceae | 453 | ISN28_16035 | 10058..11419 | 1725326 | AAA-family-ATPase-[Ectothiorhodospiraceae-bacterium-AqS1] |
| QEP43019.1 | OLD-ABC | GCA_008375315.1 | Ectothiorhodospiraceae_bacterium_BW-2 | Bacteria>Pseudomonadati>Pseudomonadota>Gammaproteobacteria>Chromatiales>Ectothiorhodospiraceae>unclassified_Ectothiorhodospiraceae | 448 | D5085_07755 | 1611828..1613174 | 947515 | AAA-family-ATPase-[Ectothiorhodospiraceae-bacterium-BW-2] |
| QEP43015.1 | OLD-ABC | GCA_008375315.1 | Ectothiorhodospiraceae_bacterium_BW-2 | Bacteria>Pseudomonadati>Pseudomonadota>Gammaproteobacteria>Chromatiales>Ectothiorhodospiraceae>unclassified_Ectothiorhodospiraceae | 356 | D5085_07735 | 1605667..1606737 | 947515 | chromosome-segregation-protein-SMC-[Ectothiorhodospiraceae-bacterium-BW-2] |
| PLY12610.1 | OLD-ABC | GCA_002869585.1 | Sedimenticola_sp. | Bacteria>Pseudomonadati>Pseudomonadota>Gammaproteobacteria>Chromatiales>Sedimenticolaceae>Sedimenticola>unclassified_Sedimenticola | 430 | C0631_16545 | 12697..13989 | 1940285 | MAG:-ATPase-[Sedimenticola-sp.] |
| WP_177417921.1 | OLD-ABC | GCF_011947365.2 | endosymbiont_of_Lamellibrachia_barhami | Bacteria>Pseudomonadati>Pseudomonadota>Gammaproteobacteria>Chromatiales>unclassified_Chromatiales | 384 | G9Q73_RS00030 | 8403..9557 | 205975 | ATP-binding-protein-[endosymbiont-of-Lamellibrachia-barhami] |
| WP_006812021.1 | OLD-ABC | GCF_013635755.1 | Enterobacteriaceae | Bacteria>Pseudomonadati>Pseudomonadota>Gammaproteobacteria>Enterobacterales | 373 | K4X45_RS16770 | 79469..80590 | 543 | MULTISPECIES:-AAA-family-ATPase-[Enterobacteriaceae] |
| WP_117108866.1 | OLD-ABC | GCF_903684385.1 | Enterobacteriaceae | Bacteria>Pseudomonadati>Pseudomonadota>Gammaproteobacteria>Enterobacterales | 373 | GQQ45_RS24735 | 5626..6747 | 543 | MULTISPECIES:-AAA-family-ATPase-[Enterobacteriaceae] |
| WP_222691533.1 | OLD-ABC | GCF_019841255.1 | Enterobacteriaceae | Bacteria>Pseudomonadati>Pseudomonadota>Gammaproteobacteria>Enterobacterales | 373 | K7A35_RS15415 | 8801..9922 | 543 | MULTISPECIES:-AAA-family-ATPase-[Enterobacteriaceae] |

| Protein-ID | Domain | Assembly-ID | Strain | Taxonomy | Length | Locus-Tag | Genomic-coordinates | TaxID | Fasta-defline |
|---|---|---|---|---|---|---|---|---|---|
| WP_210849182.1 | OLD-ABC | GCF_022349905.1 | Morganellaceae | Bacteria>Pseudomonadati>Pseudomonadota>Gammaproteobacteria>Enterobacterales | 372 | MJ636_RS02900 | 40844..41962 | 1903414 | AAA-family-ATPase-[Proteus-mirabilis] |
| WP_023157615.1 | OLD-ABC | GCF_033235525.1 | Enterobacteriaceae | Bacteria>Pseudomonadati>Pseudomonadota>Gammaproteobacteria>Enterobacterales | 363 | M4W13_RS12110 | 114837..115928 | 543 | MULTISPECIES:-ATP-binding-protein-[Enterobacteriaceae] |
| WP_057055968.1 | OLD-ABC | GCF_014169295.1 | Enterobacteriaceae | Bacteria>Pseudomonadati>Pseudomonadota>Gammaproteobacteria>Enterobacterales | 361 | H7R61_RS03360 | 680005..681090 | 543 | MULTISPECIES:-AAA-family-ATPase-[Enterobacter-cloacae-complex] |
| WP_112781822.1 | OLD-ABC | GCF_902387755.1 | Enterobacteriaceae | Bacteria>Pseudomonadati>Pseudomonadota>Gammaproteobacteria>Enterobacterales | 361 | FYC20_RS23055 | 4738..5823 | 543 | MULTISPECIES:-ATP-binding-protein-[Enterobacteriaceae] |
| WP_163363284.1 | OLD-ABC | GCF_046630575.1 | Enterobacteriaceae | Bacteria>Pseudomonadati>Pseudomonadota>Gammaproteobacteria>Enterobacterales | 360 | G4556_RS12900 | 12137..13219 | 543 | ATP-binding-protein-[Enterobacter-cloacae] |
| WP_215723023.1 | OLD-ABC | GCF_019351595.1 | Enterobacter | Bacteria>Pseudomonadati>Pseudomonadota>Gammaproteobacteria>Enterobacterales>Enterobacteriaceae | 623 | JW319_RS20525 | 167495..169366 | 547 | MULTISPECIES:-AAA-family-ATPase-[Enterobacter] |
| WP_000371966.1 | OLD-ABC | GCF_000264195.1 | Escherichia | Bacteria>Pseudomonadati>Pseudomonadota>Gammaproteobacteria>Enterobacterales>Enterobacteriaceae | 373 | PP93_RS24140 | 21908..23029 | 561 | MULTISPECIES:-AAA-family-ATPase-[Escherichia] |
| WP_023224627.1 | OLD-ABC | GCF_001569545.1 | Salmonella | Bacteria>Pseudomonadati>Pseudomonadota>Gammaproteobacteria>Enterobacterales>Enterobacteriaceae | 373 | NOQ95_RS21570 | 4455794..4456915 | 590 | MULTISPECIES:-AAA-family-ATPase-[Salmonella] |
| WP_001526178.1 | OLD-ABC | GCF_000006945.2 | Salmonella | Bacteria>Pseudomonadati>Pseudomonadota>Gammaproteobacteria>Enterobacterales>Enterobacteriaceae | 363 | AJ382_RS05880 | 3940..5031 | 590 | MULTISPECIES:-ATP-binding-protein-[Salmonella] |
| WP_080162867.1 | OLD-ABC | GCF_046569485.1 | Salmonella | Bacteria>Pseudomonadati>Pseudomonadota>Gammaproteobacteria>Enterobacterales>Enterobacteriaceae | 363 | SO66_RS04065 | 166135..167226 | 590 | ATP-binding-protein-[Salmonella-enterica] |
| HAU4366731.1 | OLD-ABC | GCA_016081455.1 | Citrobacter_amalonaticus | Bacteria>Pseudomonadati>Pseudomonadota>Gammaproteobacteria>Enterobacterales>Enterobacteriaceae>Citrobacter | 363 | F6W26_02165 | 128108..129199 | 35703 | ATP-binding-protein-[Citrobacter-amalonaticus] |
| WP_044264601.1 | OLD-ABC | GCF_000731055.1 | Citrobacter_amalonaticus | Bacteria>Pseudomonadati>Pseudomonadota>Gammaproteobacteria>Enterobacterales>Enterobacteriaceae>Citrobacter | 363 | DQ02_RS05325 | 405918..407009 | 35703 | ATP-binding-protein-[Citrobacter-amalonaticus] |
| WP_149691148.1 | OLD-ABC | GCF_008373795.1 | Citrobacter_pasteurii | Bacteria>Pseudomonadati>Pseudomonadota>Gammaproteobacteria>Enterobacterales>Enterobacteriaceae>Citrobacter | 347 | DXF85_RS00940 | 188678..189721 | 1563222 | ATP-binding-protein-[Citrobacter-pasteurii] |
| HBC2890665.1 | OLD-ABC | GCA_018089985.1 | Citrobacter_freundii | Bacteria>Pseudomonadati>Pseudomonadota>Gammaproteobacteria>Enterobacterales>Enterobacteriaceae>Citrobacter>Citrobacter_freundii_complex | 373 | JD321_004757 | 8768..9889 | 546 | AAA-family-ATPase-[Citrobacter-freundii] |
| WP_203457739.1 | OLD-ABC | GCF_016859695.1 | Citrobacter_braakii | Bacteria>Pseudomonadati>Pseudomonadota>Gammaproteobacteria>Enterobacterales>Enterobacteriaceae>Citrobacter>Citrobacter_freundii_complex | 365 | GF710_RS09105 | 92351..93448 | 57706 | ATP-binding-protein-[Citrobacter-braakii] |
| WP_241511103.1 | OLD-ABC | GCF_022585175.1 | Citrobacter_freundii | Bacteria>Pseudomonadati>Pseudomonadota>Gammaproteobacteria>Enterobacterales>Enterobacteriaceae>Citrobacter>Citrobacter_freundii_complex | 361 | MMQ01_RS10395 | 139301..140386 | 546 | ATP-binding-protein-[Citrobacter-freundii] |
| WP_269506676.1 | OLD-ABC | GCF_027277125.1 | Cronobacter_sakazakii | Bacteria>Pseudomonadati>Pseudomonadota>Gammaproteobacteria>Enterobacterales>Enterobacteriaceae>Cronobacter | 363 | O6D31_RS12685 | 371466..372557 | 28141 | AAA-family-ATPase-[Cronobacter-sakazakii] |
| WP_032634488.1 | OLD-ABC | GCF_019974415.1 | Enterobacter_cloacae_complex | Bacteria>Pseudomonadati>Pseudomonadota>Gammaproteobacteria>Enterobacterales>Enterobacteriaceae>Enterobacter | 373 | CI630_RS14015 | 9258..10379 | 354276 | MULTISPECIES:-AAA-family-ATPase-[Enterobacter-cloacae-complex] |
| WP_058675374.1 | OLD-ABC | GCF_030324195.1 | Enterobacter_cloacae_complex | Bacteria>Pseudomonadati>Pseudomonadota>Gammaproteobacteria>Enterobacterales>Enterobacteriaceae>Enterobacter | 364 | NUJ48_RS00160 | 36500..37594 | 354276 | ATP-binding-protein-[Enterobacter-hormaechei] |
| WP_257246764.1 | OLD-ABC | GCF_024607805.1 | unclassified_Enterobacter | Bacteria>Pseudomonadati>Pseudomonadota>Gammaproteobacteria>Enterobacterales>Enterobacteriaceae>Enterobacter | 363 | NR293_RS15375 | 77110..78201 | 2608935 | MULTISPECIES:-ATP-binding-protein-[unclassified-Enterobacter] |
| WP_219280524.1 | OLD-ABC | GCF_019351595.1 | Enterobacter_cloacae_complex | Bacteria>Pseudomonadati>Pseudomonadota>Gammaproteobacteria>Enterobacterales>Enterobacteriaceae>Enterobacter | 360 | L8Q49_RS17930 | 121580..122662 | 354276 | MULTISPECIES:-ATP-binding-protein-[Enterobacter-cloacae-complex] |
| WP_253179318.1 | OLD-ABC | GCF_024128585.1 | Enterobacter_mori | Bacteria>Pseudomonadati>Pseudomonadota>Gammaproteobacteria>Enterobacterales>Enterobacteriaceae>Enterobacter | 360 | NJA11_RS17920 | 127235..128317 | 539813 | ATP-binding-protein-[Enterobacter-mori] |
| WP_100165589.1 | OLD-ABC | GCF_002785605.1 | Enterobacter_mori | Bacteria>Pseudomonadati>Pseudomonadota>Gammaproteobacteria>Enterobacterales>Enterobacteriaceae>Enterobacter | 346 | B9Q20_RS04875 | 371005..372045 | 539813 | ATP-binding-protein-[Enterobacter-mori] |
| WP_059444839.1 | OLD-ABC | GCF_051561185.1 | Enterobacter_hormaechei | Bacteria>Pseudomonadati>Pseudomonadota>Gammaproteobacteria>Enterobacterales>Enterobacteriaceae>Enterobacter>Enterobacter_cloacae_complex | 373 | DDA68_RS23720 | 8825..9946 | 158836 | MULTISPECIES:-AAA-family-ATPase-[Enterobacter-cloacae-complex] |
| WP_248053565.1 | OLD-ABC | GCF_023154935.1 | Enterobacter_chengduensis | Bacteria>Pseudomonadati>Pseudomonadota>Gammaproteobacteria>Enterobacterales>Enterobacteriaceae>Enterobacter>Enterobacter_cloacae_complex | 373 | L8P32_RS20970 | 13774..14895 | 2494701 | AAA-family-ATPase-[Enterobacter-chengduensis] |
| WP_265188115.1 | OLD-ABC | GCF_027920365.1 | Enterobacter_hormaechei | Bacteria>Pseudomonadati>Pseudomonadota>Gammaproteobacteria>Enterobacterales>Enterobacteriaceae>Enterobacter>Enterobacter_cloacae_complex | 373 | PBS85_RS02760 | 576407..577528 | 158836 | AAA-family-ATPase-[Enterobacter-hormaechei] |
| WP_274416988.1 | OLD-ABC | GCF_028765355.1 | Enterobacter_kobei | Bacteria>Pseudomonadati>Pseudomonadota>Gammaproteobacteria>Enterobacterales>Enterobacteriaceae>Enterobacter>Enterobacter_cloacae_complex | 364 | PVM09_RS00200 | 39748..40842 | 208224 | ATP-binding-protein-[Enterobacter-kobei] |
| WP_063851022.1 | OLD-ABC | GCF_902160625.1 | Enterobacter_hormaechei | Bacteria>Pseudomonadati>Pseudomonadota>Gammaproteobacteria>Enterobacterales>Enterobacteriaceae>Enterobacter>Enterobacter_cloacae_complex | 363 | H0848_RS18565 | 35077..36168 | 158836 | ATP-binding-protein-[Enterobacter-hormaechei] |
| WP_095453565.1 | OLD-ABC | GCF_050972325.1 | Enterobacter_cloacae | Bacteria>Pseudomonadati>Pseudomonadota>Gammaproteobacteria>Enterobacterales>Enterobacteriaceae>Enterobacter>Enterobacter_cloacae_complex | 363 | CIW57_RS16420 | 143271..144362 | 550 | ATP-binding-protein-[Enterobacter-cloacae] |
| MCE1563406.1 | OLD-ABC | GCA_021285765.1 | Enterobacter_hormaechei | Bacteria>Pseudomonadati>Pseudomonadota>Gammaproteobacteria>Enterobacterales>Enterobacteriaceae>Enterobacter>Enterobacter_cloacae_complex | 347 | LWT81_04620 | 333091..334134 | 158836 | ATP-binding-protein-[Enterobacter-hormaechei] |
| WP_032621385.1 | OLD-ABC | GCF_023734955.1 | Enterobacter_hormaechei | Bacteria>Pseudomonadati>Pseudomonadota>Gammaproteobacteria>Enterobacterales>Enterobacteriaceae>Enterobacter>Enterobacter_cloacae_complex | 347 | NVR78_RS04775 | 321444..322487 | 158836 | ATP-binding-protein-[Enterobacter-hormaechei] |
| WP_069597814.1 | OLD-ABC | GCF_001729805.1 | Enterobacter_roggenkampii | Bacteria>Pseudomonadati>Pseudomonadota>Gammaproteobacteria>Enterobacterales>Enterobacteriaceae>Enterobacter>Enterobacter_cloacae_complex | 347 | BFV67_RS02550 | 550719..551762 | 1812935 | ATP-binding-protein-[Enterobacter-roggenkampii] |
| WP_110108111.1 | OLD-ABC | GCF_003186415.1 | Enterobacter_hormaechei | Bacteria>Pseudomonadati>Pseudomonadota>Gammaproteobacteria>Enterobacterales>Enterobacteriaceae>Enterobacter>Enterobacter_cloacae_complex | 347 | BET69_RS02800 | 567175..568218 | 158836 | ATP-binding-protein-[Enterobacter-hormaechei] |
| WP_123923009.1 | OLD-ABC | GCF_003812145.1 | Enterobacter_roggenkampii | Bacteria>Pseudomonadati>Pseudomonadota>Gammaproteobacteria>Enterobacterales>Enterobacteriaceae>Enterobacter>Enterobacter_cloacae_complex | 347 | HV097_RS13190 | 2636779..2637822 | 1812935 | ATP-binding-protein-[Enterobacter-roggenkampii] |
| MCI9498941.1 | OLD-ABC | GCA_022800955.1 | Enterobacter_hormaechei_subsp._steigerwaltii | Bacteria>Pseudomonadati>Pseudomonadota>Gammaproteobacteria>Enterobacterales>Enterobacteriaceae>Enterobacter>Enterobacter_cloacae_complex>Enterobacter_hormaechei | 373 | HFE69_15010 | 338041..339162 | 299766 | AAA-family-ATPase-[Enterobacter-hormaechei-subsp.-steigerwaltii] |
| MBT2005128.1 | OLD-ABC | GCA_018597745.1 | Enterobacter_hormaechei_subsp._xiangfangensis | Bacteria>Pseudomonadati>Pseudomonadota>Gammaproteobacteria>Enterobacterales>Enterobacteriaceae>Enterobacter>Enterobacter_cloacae_complex>Enterobacter_hormaechei | 360 | KKZ39_07405 | 335890..336972 | 1296536 | AAA-family-ATPase-[Enterobacter-hormaechei-subsp.-xiangfangensis] |
| WP_182062525.1 | OLD-ABC | GCF_013927005.1 | Enterobacter_sp._RHBSTW-00175 | Bacteria>Pseudomonadati>Pseudomonadota>Gammaproteobacteria>Enterobacterales>Enterobacteriaceae>Enterobacter>unclassified_Enterobacter | 373 | HV107_RS06445 | 1375985..1377106 | 2742639 | AAA-family-ATPase-[Enterobacter-sp.-RHBSTW-00175] |
| EFA6389816.1 | OLD-ABC | - | Escherichia_coli | Bacteria>Pseudomonadati>Pseudomonadota>Gammaproteobacteria>Enterobacterales>Enterobacteriaceae>Escherichia | 373 | CIW09_001011 | 347124..348245 | 562 | AAA-family-ATPase-[Escherichia-coli] |
| EFC0631245.1 | OLD-ABC | - | Escherichia_coli | Bacteria>Pseudomonadati>Pseudomonadota>Gammaproteobacteria>Enterobacterales>Enterobacteriaceae>Escherichia | 373 | FA817_13720 | 293873..294994 | 562 | ABC-transporter-ATP-binding-protein-[Escherichia-coli] |
| HAG7757104.1 | OLD-ABC | GCA_012297605.1 | Escherichia_coli | Bacteria>Pseudomonadati>Pseudomonadota>Gammaproteobacteria>Enterobacterales>Enterobacteriaceae>Escherichia | 373 | HGR18_09665 | 347079..348200 | 562 | TPA:-AAA-family-ATPase-[Escherichia-coli] |
| HAX4880265.1 | OLD-ABC | GCA_017016975.1 | Escherichia_coli | Bacteria>Pseudomonadati>Pseudomonadota>Gammaproteobacteria>Enterobacterales>Enterobacteriaceae>Escherichia | 373 | JZF83_001470 | 104426..105547 | 562 | TPA:-AAA-family-ATPase-[Escherichia-coli] |
| HCJ9453758.1 | OLD-ABC | GCA_024682335.1 | Escherichia_coli | Bacteria>Pseudomonadati>Pseudomonadota>Gammaproteobacteria>Enterobacterales>Enterobacteriaceae>Escherichia | 373 | NV934_004006 | 16566..17687 | 562 | TPA:-AAA-family-ATPase-[Escherichia-coli] |
| WP_001552309.1 | OLD-ABC | GCF_902505225.1 | Escherichia_coli | Bacteria>Pseudomonadati>Pseudomonadota>Gammaproteobacteria>Enterobacterales>Enterobacteriaceae>Escherichia | 373 | GDK13_RS03585 | 261189..262310 | 562 | AAA-family-ATPase-[Escherichia-coli] |
| WP_038989006.1 | OLD-ABC | GCF_000812725.1 | Escherichia_coli | Bacteria>Pseudomonadati>Pseudomonadota>Gammaproteobacteria>Enterobacterales>Enterobacteriaceae>Escherichia | 373 | RK29_RS16975 | 10219..11340 | 562 | AAA-family-ATPase-[Escherichia-coli] |
| WP_050858671.1 | OLD-ABC | GCF_030295055.1 | Escherichia_coli | Bacteria>Pseudomonadati>Pseudomonadota>Gammaproteobacteria>Enterobacterales>Enterobacteriaceae>Escherichia | 373 | H5E40_RS01435 | 286061..287182 | 562 | AAA-family-ATPase-[Escherichia-coli] |
| WP_052920516.1 | OLD-ABC | GCF_001262905.1 | Escherichia_coli | Bacteria>Pseudomonadati>Pseudomonadota>Gammaproteobacteria>Enterobacterales>Enterobacteriaceae>Escherichia | 373 | AGA26_RS17900 | 9108..10229 | 562 | AAA-family-ATPase-[Escherichia-coli] |
| WP_088209950.1 | OLD-ABC | GCF_943733315.1 | Escherichia_coli | Bacteria>Pseudomonadati>Pseudomonadota>Gammaproteobacteria>Enterobacterales>Enterobacteriaceae>Escherichia | 373 | LCR13_RS23615 | 143962..145083 | 562 | AAA-family-ATPase-[Escherichia-coli] |
| WP_089663050.1 | OLD-ABC | GCF_022489725.1 | Escherichia_coli | Bacteria>Pseudomonadati>Pseudomonadota>Gammaproteobacteria>Enterobacterales>Enterobacteriaceae>Escherichia | 373 | ML322_RS02975 | 182312..183433 | 562 | AAA-family-ATPase-[Escherichia-coli] |
| WP_115723903.1 | OLD-ABC | GCF_029542125.1 | Escherichia_coli | Bacteria>Pseudomonadati>Pseudomonadota>Gammaproteobacteria>Enterobacterales>Enterobacteriaceae>Escherichia | 373 | OM087_RS11610 | 2414839..2415960 | 562 | AAA-family-ATPase-[Escherichia-coli] |
| MCN5345818.1 | OLD-ABC | GCA_023869305.1 | Escherichia_coli | Bacteria>Pseudomonadati>Pseudomonadota>Gammaproteobacteria>Enterobacterales>Enterobacteriaceae>Escherichia | 363 | MLU97_13820 | 77114..78205 | 562 | ATP-binding-protein-[Escherichia-coli] |
| WP_060616118.1 | OLD-ABC | GCF_034719175.1 | Escherichia_coli | Bacteria>Pseudomonadati>Pseudomonadota>Gammaproteobacteria>Enterobacterales>Enterobacteriaceae>Escherichia | 363 | DT570_RS09700 | 5950..7041 | 562 | ATP-binding-protein-[Escherichia-coli] |
| WP_196588767.1 | OLD-ABC | GCF_024917295.1 | Klebsiella | Bacteria>Pseudomonadati>Pseudomonadota>Gammaproteobacteria>Enterobacterales>Enterobacteriaceae>Klebsiella/Raoultella_group | 363 | M5R35_RS24850 | 5272617..5273708 | 570 | MULTISPECIES:-ATP-binding-protein-[Klebsiella] |
| WP_154062668.1 | OLD-ABC | GCF_902159535.1 | Klebsiella | Bacteria>Pseudomonadati>Pseudomonadota>Gammaproteobacteria>Enterobacterales>Enterobacteriaceae>Klebsiella/Raoultella_group | 347 | FN635_RS25510 | 5349597..5350640 | 570 | ATP-binding-protein-[Klebsiella-michiganensis] |
| WP_264975993.1 | OLD-ABC | GCF_026008315.1 | Klebsiella_pneumoniae_complex | Bacteria>Pseudomonadati>Pseudomonadota>Gammaproteobacteria>Enterobacterales>Enterobacteriaceae>Klebsiella/Raoultella_group>Klebsiella | 362 | OPC12_RS09045 | 180981..182069 | 3390273 | AAA-family-ATPase-[Klebsiella-quasipneumoniae] |
| WP_182409422.1 | OLD-ABC | GCF_014129405.1 | Klebsiella_michiganensis | Bacteria>Pseudomonadati>Pseudomonadota>Gammaproteobacteria>Enterobacterales>Enterobacteriaceae>Klebsiella/Raoultella_group>Klebsiella | 347 | H0753_RS11140 | 93396..94439 | 1134687 | ATP-binding-protein-[Klebsiella-michiganensis] |
| WP_217074482.1 | OLD-ABC | GCF_019048885.1 | Klebsiella_aerogenes | Bacteria>Pseudomonadati>Pseudomonadota>Gammaproteobacteria>Enterobacterales>Enterobacteriaceae>Klebsiella/Raoultella_group>Klebsiella | 347 | KSL85_RS22175 | 4648192..4649235 | 548 | ATP-binding-protein-[Klebsiella-aerogenes] |
| WP_224260389.1 | OLD-ABC | GCF_026224455.1 | Klebsiella_michiganensis | Bacteria>Pseudomonadati>Pseudomonadota>Gammaproteobacteria>Enterobacterales>Enterobacteriaceae>Klebsiella/Raoultella_group>Klebsiella | 347 | ORL69_RS11450 | 325400..326443 | 1134687 | ATP-binding-protein-[Klebsiella-michiganensis] |
| EIV3902454.1 | OLD-ABC | GCA_021871195.1 | Klebsiella_pneumoniae | Bacteria>Pseudomonadati>Pseudomonadota>Gammaproteobacteria>Enterobacterales>Enterobacteriaceae>Klebsiella/Raoultella_group>Klebsiella>Klebsiella_pneumoniae_complex | 373 | L8E75_001737 | 321193..322314 | 573 | AAA-family-ATPase-[Klebsiella-pneumoniae] |
| EIY5117084.1 | OLD-ABC | GCA_022503905.1 | Klebsiella_quasipneumoniae | Bacteria>Pseudomonadati>Pseudomonadota>Gammaproteobacteria>Enterobacterales>Enterobacteriaceae>Klebsiella/Raoultella_group>Klebsiella>Klebsiella_pneumoniae_complex | 373 | ML076_004998 | 27279..28400 | 1463165 | AAA-family-ATPase-[Klebsiella-quasipneumoniae] |
| HBR1207025.1 | OLD-ABC | GCA_021909055.1 | Klebsiella_pneumoniae | Bacteria>Pseudomonadati>Pseudomonadota>Gammaproteobacteria>Enterobacterales>Enterobacteriaceae>Klebsiella/Raoultella_group>Klebsiella>Klebsiella_pneumoniae_complex | 373 | L9K92_004099 | 87794..88915 | 573 | AAA-family-ATPase-[Klebsiella-pneumoniae] |
| HBX5876369.1 | OLD-ABC | GCA_022249175.1 | Klebsiella_pneumoniae | Bacteria>Pseudomonadati>Pseudomonadota>Gammaproteobacteria>Enterobacterales>Enterobacteriaceae>Klebsiella/Raoultella_group>Klebsiella>Klebsiella_pneumoniae_complex | 373 | MH570_26225 | 9298..10419 | 573 | TPA:-ABC-transporter-ATP-binding-protein-[Klebsiella-pneumoniae] |
| WP_048999821.1 | OLD-ABC | GCF_040365915.1 | Klebsiella_pneumoniae | Bacteria>Pseudomonadati>Pseudomonadota>Gammaproteobacteria>Enterobacterales>Enterobacteriaceae>Klebsiella/Raoultella_group>Klebsiella>Klebsiella_pneumoniae_complex | 373 | LF666_RS16205 | 18032..19153 | 573 | AAA-family-ATPase-[Klebsiella-pneumoniae] |
| WP_135732749.1 | OLD-ABC | GCF_900775595.1 | Klebsiella_quasipneumoniae | Bacteria>Pseudomonadati>Pseudomonadota>Gammaproteobacteria>Enterobacterales>Enterobacteriaceae>Klebsiella/Raoultella_group>Klebsiella>Klebsiella_pneumoniae_complex | 373 | ICC30_RS25230 | 19005..20126 | 1463165 | AAA-family-ATPase-[Klebsiella-quasipneumoniae] |
| WP_117268634.1 | OLD-ABC | GCF_900509065.1 | Klebsiella_pneumoniae | Bacteria>Pseudomonadati>Pseudomonadota>Gammaproteobacteria>Enterobacterales>Enterobacteriaceae>Klebsiella/Raoultella_group>Klebsiella>Klebsiella_pneumoniae_complex | 363 | D2728_RS15285 | 263426..264517 | 573 | ATP-binding-protein-[Klebsiella-pneumoniae] |
| GKP62589.1 | OLD-ABC | GCA_026008315.1 | Klebsiella_quasipneumoniae | Bacteria>Pseudomonadati>Pseudomonadota>Gammaproteobacteria>Enterobacterales>Enterobacteriaceae>Klebsiella/Raoultella_group>Klebsiella>Klebsiella_pneumoniae_complex | 362 | NUKP47_17650 | 180981..182069 | 1463165 | hypothetical-protein-NUKP47_17650-[Klebsiella-quasipneumoniae] |
| HBR1207680.1 | OLD-ABC | GCA_021909055.1 | Klebsiella_pneumoniae | Bacteria>Pseudomonadati>Pseudomonadota>Gammaproteobacteria>Enterobacterales>Enterobacteriaceae>Klebsiella/Raoultella_group>Klebsiella>Klebsiella_pneumoniae_complex | 360 | L9K92_004775 | 35902..36984 | 573 | AAA-family-ATPase-[Klebsiella-pneumoniae] |
| HBS6654370.1 | OLD-ABC | GCA_021975495.1 | Klebsiella_pneumoniae | Bacteria>Pseudomonadati>Pseudomonadota>Gammaproteobacteria>Enterobacterales>Enterobacteriaceae>Klebsiella/Raoultella_group>Klebsiella>Klebsiella_pneumoniae_complex | 360 | MAN10_004112 | 48795..49877 | 573 | TPA:-AAA-family-ATPase-[Klebsiella-pneumoniae] |
| HBS7652076.1 | OLD-ABC | GCA_021979555.1 | Klebsiella_pneumoniae | Bacteria>Pseudomonadati>Pseudomonadota>Gammaproteobacteria>Enterobacterales>Enterobacteriaceae>Klebsiella/Raoultella_group>Klebsiella>Klebsiella_pneumoniae_complex | 360 | MAR21_002850 | 324843..325925 | 573 | TPA:-AAA-family-ATPase-[Klebsiella-pneumoniae] |
| HBS9599070.1 | OLD-ABC | GCA_021986995.1 | Klebsiella_pneumoniae | Bacteria>Pseudomonadati>Pseudomonadota>Gammaproteobacteria>Enterobacterales>Enterobacteriaceae>Klebsiella/Raoultella_group>Klebsiella>Klebsiella_pneumoniae_complex | 360 | MAW98_001664 | 48794..49876 | 573 | TPA:-AAA-family-ATPase-[Klebsiella-pneumoniae] |
| HBX7751795.1 | OLD-ABC | GCA_022258155.1 | Klebsiella_pneumoniae | Bacteria>Pseudomonadati>Pseudomonadota>Gammaproteobacteria>Enterobacterales>Enterobacteriaceae>Klebsiella/Raoultella_group>Klebsiella>Klebsiella_pneumoniae_complex | 360 | MIG23_15240 | 10044..11126 | 573 | TPA:-ATP-binding-protein-[Klebsiella-pneumoniae] |
| HCF8566125.1 | OLD-ABC | GCA_023980665.1 | Klebsiella_pneumoniae | Bacteria>Pseudomonadati>Pseudomonadota>Gammaproteobacteria>Enterobacterales>Enterobacteriaceae>Klebsiella/Raoultella_group>Klebsiella>Klebsiella_pneumoniae_complex | 360 | NIV99_001796 | 27527..28609 | 573 | TPA:-AAA-family-ATPase-[Klebsiella-pneumoniae] |
| TWY76109.1 | OLD-ABC | GCA_007955295.1 | Klebsiella_pneumoniae | Bacteria>Pseudomonadati>Pseudomonadota>Gammaproteobacteria>Enterobacterales>Enterobacteriaceae>Klebsiella/Raoultella_group>Klebsiella>Klebsiella_pneumoniae_complex | 360 | FTO41_25010 | 27512..28594 | 573 | AAA-family-ATPase-[Klebsiella-pneumoniae] |
| WP_040153451.1 | OLD-ABC | GCF_900775295.1 | Klebsiella_pneumoniae | Bacteria>Pseudomonadati>Pseudomonadota>Gammaproteobacteria>Enterobacterales>Enterobacteriaceae>Klebsiella/Raoultella_group>Klebsiella>Klebsiella_pneumoniae_complex | 360 | JEK00_RS18430 | 15007..16089 | 573 | ATP-binding-protein-[Klebsiella-pneumoniae] |
| WP_048289719.1 | OLD-ABC | GCF_934247105.1 | Klebsiella_pneumoniae | Bacteria>Pseudomonadati>Pseudomonadota>Gammaproteobacteria>Enterobacterales>Enterobacteriaceae>Klebsiella/Raoultella_group>Klebsiella>Klebsiella_pneumoniae_complex | 360 | A8T29_RS25600 | 20785..21867 | 573 | ATP-binding-protein-[Klebsiella-pneumoniae] |
| WP_064143070.1 | OLD-ABC | GCF_900085185.1 | Klebsiella_variicola | Bacteria>Pseudomonadati>Pseudomonadota>Gammaproteobacteria>Enterobacterales>Enterobacteriaceae>Klebsiella/Raoultella_group>Klebsiella>Klebsiella_pneumoniae_complex | 347 | A8S52_RS23965 | 27132..28175 | 244366 | ATP-binding-protein-[Klebsiella-variicola] |
| WP_104443229.1 | OLD-ABC | GCF_018156605.1 | Klebsiella_pneumoniae | Bacteria>Pseudomonadati>Pseudomonadota>Gammaproteobacteria>Enterobacterales>Enterobacteriaceae>Klebsiella/Raoultella_group>Klebsiella>Klebsiella_pneumoniae_complex | 347 | J5D66_RS24895 | 9662..10705 | 573 | ATP-binding-protein-[Klebsiella-pneumoniae] |
| WP_223370573.1 | OLD-ABC | GCF_019927485.1 | Klebsiella_pneumoniae | Bacteria>Pseudomonadati>Pseudomonadota>Gammaproteobacteria>Enterobacterales>Enterobacteriaceae>Klebsiella/Raoultella_group>Klebsiella>Klebsiella_pneumoniae_complex | 347 | PXW47_RS20400 | 332043..333086 | 573 | ATP-binding-protein-[Klebsiella-pneumoniae] |
| HCB0569861.1 | OLD-ABC | GCA_023030625.1 | Klebsiella_quasipneumoniae_subsp._similipneumoniae | Bacteria>Pseudomonadati>Pseudomonadota>Gammaproteobacteria>Enterobacterales>Enterobacteriaceae>Klebsiella/Raoultella_group>Klebsiella>Klebsiella_pneumoniae_complex>Klebsiella_quasipneumoniae | 414 | MYL58_003258 | 40940..42184 | 1463164 | ATP-binding-protein-[Klebsiella-quasipneumoniae-subsp.-similipneumoniae] |
| HCI5938463.1 | OLD-ABC | GCA_024431715.1 | Klebsiella_quasipneumoniae_subsp._quasipneumoniae | Bacteria>Pseudomonadati>Pseudomonadota>Gammaproteobacteria>Enterobacterales>Enterobacteriaceae>Klebsiella/Raoultella_group>Klebsiella>Klebsiella_pneumoniae_complex>Klebsiella_quasipneumoniae | 373 | NPN45_004674 | 9491..10612 | 1667327 | AAA-family-ATPase-[Klebsiella-quasipneumoniae-subsp.-quasipneumoniae] |
| HCI4589556.1 | OLD-ABC | GCA_024379455.1 | Klebsiella_quasipneumoniae_subsp._quasipneumoniae | Bacteria>Pseudomonadati>Pseudomonadota>Gammaproteobacteria>Enterobacterales>Enterobacteriaceae>Klebsiella/Raoultella_group>Klebsiella>Klebsiella_pneumoniae_complex>Klebsiella_quasipneumoniae | 360 | NO899_004584 | 23537..24619 | 1667327 | AAA-family-ATPase-[Klebsiella-quasipneumoniae-subsp.-quasipneumoniae] |
| WP_102104496.1 | OLD-ABC | GCF_002870005.1 | unclassified_Lelliottia | Bacteria>Pseudomonadati>Pseudomonadota>Gammaproteobacteria>Enterobacterales>Enterobacteriaceae>Lelliottia | 373 | F159LOC_RS08770 | 163368..164489 | 2642424 | MULTISPECIES:-AAA-family-ATPase-[unclassified-Lelliottia] |
| WP_233672192.1 | OLD-ABC | GCF_021368255.1 | Plesiomonas_sp._PI-19 | Bacteria>Pseudomonadati>Pseudomonadota>Gammaproteobacteria>Enterobacterales>Enterobacteriaceae>Plesiomonas>unclassified_Plesiomonas | 419 | LQ901_RS09595 | 2144225..2145484 | 2898798 | AAA-family-ATPase-[Plesiomonas-sp.-PI-19] |
| WP_233672187.1 | OLD-ABC | GCF_021368255.1 | Plesiomonas_sp._PI-19 | Bacteria>Pseudomonadati>Pseudomonadota>Gammaproteobacteria>Enterobacterales>Enterobacteriaceae>Plesiomonas>unclassified_Plesiomonas | 353 | LQ901_RS09570 | 2137995..2139056 | 2898798 | ATP-binding-protein-[Plesiomonas-sp.-PI-19] |
| EAB2013164.1 | OLD-ABC | GCA_004236885.1 | Salmonella_enterica | Bacteria>Pseudomonadati>Pseudomonadota>Gammaproteobacteria>Enterobacterales>Enterobacteriaceae>Salmonella | 373 | D7J64_23100 | 12448..13569 | 28901 | ABC-transporter-ATP-binding-protein-[Salmonella-enterica] |
| EAP9950626.1 | OLD-ABC | GCA_005576875.1 | Salmonella_enterica | Bacteria>Pseudomonadati>Pseudomonadota>Gammaproteobacteria>Enterobacterales>Enterobacteriaceae>Salmonella | 373 | D3G27_06345 | 183429..184550 | 28901 | ABC-transporter-ATP-binding-protein-[Salmonella-enterica] |
| EAW3206017.1 | OLD-ABC | GCA_005905825.1 | Salmonella_enterica | Bacteria>Pseudomonadati>Pseudomonadota>Gammaproteobacteria>Enterobacterales>Enterobacteriaceae>Salmonella | 373 | FEL99_04740 | 228003..229124 | 28901 | ABC-transporter-ATP-binding-protein-[Salmonella-enterica] |
| EAW9396389.1 | OLD-ABC | GCA_005940865.1 | Salmonella_enterica | Bacteria>Pseudomonadati>Pseudomonadota>Gammaproteobacteria>Enterobacterales>Enterobacteriaceae>Salmonella | 373 | QE38_19745 | 55725..56846 | 28901 | ABC-transporter-ATP-binding-protein-[Salmonella-enterica] |
| EAY5508620.1 | OLD-ABC | GCA_006012725.1 | Salmonella_enterica | Bacteria>Pseudomonadati>Pseudomonadota>Gammaproteobacteria>Enterobacterales>Enterobacteriaceae>Salmonella | 373 | NE47_11025 | 56711..57832 | 28901 | ABC-transporter-ATP-binding-protein-[Salmonella-enterica] |
| EBF4455781.1 | OLD-ABC | GCA_006343165.1 | Salmonella_enterica | Bacteria>Pseudomonadati>Pseudomonadota>Gammaproteobacteria>Enterobacterales>Enterobacteriaceae>Salmonella | 373 | FH443_09915 | 419434..420555 | 28901 | ABC-transporter-ATP-binding-protein-[Salmonella-enterica] |
| EBH5838353.1 | OLD-ABC | GCA_006503215.1 | Salmonella_enterica | Bacteria>Pseudomonadati>Pseudomonadota>Gammaproteobacteria>Enterobacterales>Enterobacteriaceae>Salmonella | 373 | CQL58_06365 | 300375..301496 | 28901 | ABC-transporter-ATP-binding-protein-[Salmonella-enterica] |
| EBR5149895.1 | OLD-ABC | GCA_006958505.1 | Salmonella_enterica | Bacteria>Pseudomonadati>Pseudomonadota>Gammaproteobacteria>Enterobacterales>Enterobacteriaceae>Salmonella | 373 | B5W64_09855 | 358128..359249 | 28901 | AAA-family-ATPase-[Salmonella-enterica] |
| EDZ0760564.1 | OLD-ABC | - | Salmonella_enterica | Bacteria>Pseudomonadati>Pseudomonadota>Gammaproteobacteria>Enterobacterales>Enterobacteriaceae>Salmonella | 373 | GPN34_002362 | 417966..419087 | 28901 | AAA-family-ATPase-[Salmonella-enterica] |
| EIC8525257.1 | OLD-ABC | GCA_019993965.1 | Salmonella_enterica | Bacteria>Pseudomonadati>Pseudomonadota>Gammaproteobacteria>Enterobacterales>Enterobacteriaceae>Salmonella | 373 | LAR10_002217 | 37725..38846 | 28901 | AAA-family-ATPase-[Salmonella-enterica] |
| WP_064013331.1 | OLD-ABC | GCF_002208315.1 | Salmonella_enterica | Bacteria>Pseudomonadati>Pseudomonadota>Gammaproteobacteria>Enterobacterales>Enterobacteriaceae>Salmonella | 373 | CE137_RS21060 | 4103091..4104212 | 28901 | AAA-family-ATPase-[Salmonella-enterica] |
| EAA3385005.1 | OLD-ABC | GCA_004174495.1 | Salmonella_enterica | Bacteria>Pseudomonadati>Pseudomonadota>Gammaproteobacteria>Enterobacterales>Enterobacteriaceae>Salmonella | 363 | DMV54_10720 | 139271..140362 | 28901 | ATP-binding-protein-[Salmonella-enterica] |
| EAA8317756.1 | OLD-ABC | GCA_004221325.1 | Salmonella_enterica | Bacteria>Pseudomonadati>Pseudomonadota>Gammaproteobacteria>Enterobacterales>Enterobacteriaceae>Salmonella | 363 | AOS09_04975 | 169971..171062 | 28901 | ATP-binding-cassette-domain-containing-protein-[Salmonella-enterica] |
| EAA8875707.1 | OLD-ABC | GCA_004223005.1 | Salmonella_enterica | Bacteria>Pseudomonadati>Pseudomonadota>Gammaproteobacteria>Enterobacterales>Enterobacteriaceae>Salmonella | 363 | CIU45_09635 | 389188..390279 | 28901 | ATP-binding-cassette-domain-containing-protein-[Salmonella-enterica] |
| EAA9319738.1 | OLD-ABC | GCA_004226405.1 | Salmonella_enterica | Bacteria>Pseudomonadati>Pseudomonadota>Gammaproteobacteria>Enterobacterales>Enterobacteriaceae>Salmonella | 363 | CRQ39_09910 | 174734..175825 | 28901 | ATP-binding-protein-[Salmonella-enterica] |
| EAB1768965.1 | OLD-ABC | GCA_004236005.1 | Salmonella_enterica | Bacteria>Pseudomonadati>Pseudomonadota>Gammaproteobacteria>Enterobacterales>Enterobacteriaceae>Salmonella | 363 | EIS13_11230 | 25633..26724 | 28901 | ATP-binding-protein-[Salmonella-enterica] |
| EAB2910640.1 | OLD-ABC | GCA_004240965.1 | Salmonella_enterica | Bacteria>Pseudomonadati>Pseudomonadota>Gammaproteobacteria>Enterobacterales>Enterobacteriaceae>Salmonella | 363 | EHO38_04045 | 127794..128885 | 28901 | ATP-binding-protein-[Salmonella-enterica] |
| EAM5877056.1 | OLD-ABC | GCA_005424775.1 | Salmonella_enterica | Bacteria>Pseudomonadati>Pseudomonadota>Gammaproteobacteria>Enterobacterales>Enterobacteriaceae>Salmonella | 363 | EPF42_07735 | 341543..342634 | 28901 | ATP-binding-protein-[Salmonella-enterica] |
| EAM6081347.1 | OLD-ABC | GCA_005424535.1 | Salmonella_enterica | Bacteria>Pseudomonadati>Pseudomonadota>Gammaproteobacteria>Enterobacterales>Enterobacteriaceae>Salmonella | 363 | ERR94_15340 | 12620..13711 | 28901 | ATP-binding-protein-[Salmonella-enterica] |
| EAO8797693.1 | OLD-ABC | GCA_005530015.1 | Salmonella_enterica | Bacteria>Pseudomonadati>Pseudomonadota>Gammaproteobacteria>Enterobacterales>Enterobacteriaceae>Salmonella | 363 | E0391_14545 | 33236..34327 | 28901 | ATP-binding-protein-[Salmonella-enterica] |
| EAP0128503.1 | OLD-ABC | GCA_005535515.1 | Salmonella_enterica | Bacteria>Pseudomonadati>Pseudomonadota>Gammaproteobacteria>Enterobacterales>Enterobacteriaceae>Salmonella | 363 | DKP73_00825 | 168721..169812 | 28901 | hypothetical-protein-DKP73_00825-[Salmonella-enterica] |
| EAP0644641.1 | OLD-ABC | GCA_005538295.1 | Salmonella_enterica | Bacteria>Pseudomonadati>Pseudomonadota>Gammaproteobacteria>Enterobacterales>Enterobacteriaceae>Salmonella | 363 | C6277_11595 | 103249..104340 | 28901 | ATP-binding-protein-[Salmonella-enterica] |
| EAQ0497029.1 | OLD-ABC | GCA_005582475.1 | Salmonella_enterica | Bacteria>Pseudomonadati>Pseudomonadota>Gammaproteobacteria>Enterobacterales>Enterobacteriaceae>Salmonella | 363 | EW683_07765 | 44247..45338 | 28901 | ATP-binding-protein-[Salmonella-enterica] |
| EAQ7293577.1 | OLD-ABC | GCA_005612115.1 | Salmonella_enterica | Bacteria>Pseudomonadati>Pseudomonadota>Gammaproteobacteria>Enterobacterales>Enterobacteriaceae>Salmonella | 363 | CRQ46_08455 | 414076..415167 | 28901 | ATP-binding-protein-[Salmonella-enterica] |
| EAS9365412.1 | OLD-ABC | GCA_005712675.1 | Salmonella_enterica | Bacteria>Pseudomonadati>Pseudomonadota>Gammaproteobacteria>Enterobacterales>Enterobacteriaceae>Salmonella | 363 | EPA30_12280 | 178206..179297 | 28901 | ATP-binding-protein-[Salmonella-enterica] |
| EAT1838143.1 | OLD-ABC | GCA_005717435.1 | Salmonella_enterica | Bacteria>Pseudomonadati>Pseudomonadota>Gammaproteobacteria>Enterobacterales>Enterobacteriaceae>Salmonella | 363 | EJR21_08325 | 110349..111440 | 28901 | ATP-binding-protein-[Salmonella-enterica] |
| EAT2430709.1 | OLD-ABC | GCA_005726115.1 | Salmonella_enterica | Bacteria>Pseudomonadati>Pseudomonadota>Gammaproteobacteria>Enterobacterales>Enterobacteriaceae>Salmonella | 363 | EI207_04335 | 184506..185597 | 28901 | ATP-binding-protein-[Salmonella-enterica] |
| EAU0116269.1 | OLD-ABC | GCA_005756415.1 | Salmonella_enterica | Bacteria>Pseudomonadati>Pseudomonadota>Gammaproteobacteria>Enterobacterales>Enterobacteriaceae>Salmonella | 363 | ASR58_09440 | 33248..34339 | 28901 | ATP-binding-cassette-domain-containing-protein-[Salmonella-enterica] |
| EAU0152693.1 | OLD-ABC | GCA_005756695.1 | Salmonella_enterica | Bacteria>Pseudomonadati>Pseudomonadota>Gammaproteobacteria>Enterobacterales>Enterobacteriaceae>Salmonella | 363 | AZG01_01875 | 398095..399186 | 28901 | ATP-binding-cassette-domain-containing-protein-[Salmonella-enterica] |
| EAV3792366.1 | OLD-ABC | GCA_005840905.1 | Salmonella_enterica | Bacteria>Pseudomonadati>Pseudomonadota>Gammaproteobacteria>Enterobacterales>Enterobacteriaceae>Salmonella | 363 | E2X64_08245 | 32971..34062 | 28901 | ATP-binding-protein-[Salmonella-enterica] |

| Protein-ID | Domain | Assembly-ID | Strain | Taxonomy | Length | Locus-Tag | Genomic-coordinates | TaxID | Fasta-defline |
|---|---|---|---|---|---|---|---|---|---|
| EAV4850772.1 | OLD-ABC | GCA_005847515.1 | Salmonella_enterica | Bacteria>Pseudomonadati>Pseudomonadota>Gammaproteobacteria>Enterobacterales>Enterobacteriaceae>Salmonella | 363 | EGE73_10380 | 32216..33307 | 28901 | ATP-binding-protein-[Salmonella-enterica] |
| EAW5830199.1 | OLD-ABC | GCA_005917905.1 | Salmonella_enterica | Bacteria>Pseudomonadati>Pseudomonadota>Gammaproteobacteria>Enterobacterales>Enterobacteriaceae>Salmonella | 363 | FF074_06115 | 178076..179167 | 28901 | ATP-binding-protein-[Salmonella-enterica] |
| EAX4456712.1 | OLD-ABC | GCA_005963445.1 | Salmonella_enterica | Bacteria>Pseudomonadati>Pseudomonadota>Gammaproteobacteria>Enterobacterales>Enterobacteriaceae>Salmonella | 363 | AC467_14830 | 25402..26493 | 28901 | ATP-binding-protein-[Salmonella-enterica] |
| EAZ5773632.1 | OLD-ABC | GCA_006057755.1 | Salmonella_enterica | Bacteria>Pseudomonadati>Pseudomonadota>Gammaproteobacteria>Enterobacterales>Enterobacteriaceae>Salmonella | 363 | CCZ51_07195 | 103250..104341 | 28901 | ATP-binding-protein-[Salmonella-enterica] |
| EBB0684613.1 | OLD-ABC | GCA_006125875.1 | Salmonella_enterica | Bacteria>Pseudomonadati>Pseudomonadota>Gammaproteobacteria>Enterobacterales>Enterobacteriaceae>Salmonella | 363 | CG859_01600 | 340876..341967 | 28901 | ATP-binding-protein-[Salmonella-enterica] |
| EBB1941186.1 | OLD-ABC | GCA_006131775.1 | Salmonella_enterica | Bacteria>Pseudomonadati>Pseudomonadota>Gammaproteobacteria>Enterobacterales>Enterobacteriaceae>Salmonella | 363 | CJP21_07740 | 46043..47134 | 28901 | ATP-binding-protein-[Salmonella-enterica] |
| EBC0774149.1 | OLD-ABC | GCA_006172705.1 | Salmonella_enterica | Bacteria>Pseudomonadati>Pseudomonadota>Gammaproteobacteria>Enterobacterales>Enterobacteriaceae>Salmonella | 363 | CS696_13035 | 18729..19820 | 28901 | ATP-binding-cassette-domain-containing-protein-[Salmonella-enterica] |
| EBC4250053.1 | OLD-ABC | GCA_006188785.1 | Salmonella_enterica | Bacteria>Pseudomonadati>Pseudomonadota>Gammaproteobacteria>Enterobacterales>Enterobacteriaceae>Salmonella | 363 | C2B00_07700 | 221213..222304 | 28901 | ATP-binding-protein-[Salmonella-enterica] |
| EBD9210542.1 | OLD-ABC | GCA_006272795.1 | Salmonella_enterica | Bacteria>Pseudomonadati>Pseudomonadota>Gammaproteobacteria>Enterobacterales>Enterobacteriaceae>Salmonella | 363 | DA198_05745 | 182188..183279 | 28901 | ATP-binding-protein-[Salmonella-enterica] |
| EBE1252191.1 | OLD-ABC | GCA_006282055.1 | Salmonella_enterica | Bacteria>Pseudomonadati>Pseudomonadota>Gammaproteobacteria>Enterobacterales>Enterobacteriaceae>Salmonella | 363 | DDQ39_02650 | 131781..132872 | 28901 | ATP-binding-protein-[Salmonella-enterica] |
| EBE5553330.1 | OLD-ABC | GCA_006301715.1 | Salmonella_enterica | Bacteria>Pseudomonadati>Pseudomonadota>Gammaproteobacteria>Enterobacterales>Enterobacteriaceae>Salmonella | 363 | CP597_01390 | 305117..306208 | 28901 | ATP-binding-protein-[Salmonella-enterica] |
| EBE9623450.1 | OLD-ABC | GCA_006320265.1 | Salmonella_enterica | Bacteria>Pseudomonadati>Pseudomonadota>Gammaproteobacteria>Enterobacterales>Enterobacteriaceae>Salmonella | 363 | DA967_03435 | 178327..179418 | 28901 | ATP-binding-protein-[Salmonella-enterica] |
| EBG3411775.1 | OLD-ABC | GCA_006423935.1 | Salmonella_enterica | Bacteria>Pseudomonadati>Pseudomonadota>Gammaproteobacteria>Enterobacterales>Enterobacteriaceae>Salmonella | 363 | FIW39_06215 | 178392..179483 | 28901 | ATP-binding-protein-[Salmonella-enterica] |
| EBH0811991.1 | OLD-ABC | GCA_006475165.1 | Salmonella_enterica | Bacteria>Pseudomonadati>Pseudomonadota>Gammaproteobacteria>Enterobacterales>Enterobacteriaceae>Salmonella | 363 | FJ896_02680 | 178296..179387 | 28901 | ATP-binding-protein-[Salmonella-enterica] |
| EBH4970958.1 | OLD-ABC | GCA_006499395.1 | Salmonella_enterica | Bacteria>Pseudomonadati>Pseudomonadota>Gammaproteobacteria>Enterobacterales>Enterobacteriaceae>Salmonella | 363 | CKR35_08935 | 177711..178802 | 28901 | ATP-binding-protein-[Salmonella-enterica] |
| EBI8027932.1 | OLD-ABC | GCA_006571625.1 | Salmonella_enterica | Bacteria>Pseudomonadati>Pseudomonadota>Gammaproteobacteria>Enterobacterales>Enterobacteriaceae>Salmonella | 363 | CHM23_07400 | 178283..179374 | 28901 | ATP-binding-protein-[Salmonella-enterica] |
| EBK7108452.1 | OLD-ABC | GCA_006653645.1 | Salmonella_enterica | Bacteria>Pseudomonadati>Pseudomonadota>Gammaproteobacteria>Enterobacterales>Enterobacteriaceae>Salmonella | 363 | DSC93_06695 | 178293..179384 | 28901 | ATP-binding-protein-[Salmonella-enterica] |
| EBK9871841.1 | OLD-ABC | GCA_006668805.1 | Salmonella_enterica | Bacteria>Pseudomonadati>Pseudomonadota>Gammaproteobacteria>Enterobacterales>Enterobacteriaceae>Salmonella | 363 | DYD89_19160 | 33248..34339 | 28901 | ATP-binding-cassette-domain-containing-protein-[Salmonella-enterica] |
| EBL1260731.1 | OLD-ABC | GCA_006672445.1 | Salmonella_enterica | Bacteria>Pseudomonadati>Pseudomonadota>Gammaproteobacteria>Enterobacterales>Enterobacteriaceae>Salmonella | 363 | D0790_07480 | 178494..179585 | 28901 | ATP-binding-protein-[Salmonella-enterica] |
| EBL2857390.1 | OLD-ABC | GCA_006679365.1 | Salmonella_enterica | Bacteria>Pseudomonadati>Pseudomonadota>Gammaproteobacteria>Enterobacterales>Enterobacteriaceae>Salmonella | 363 | D3Q95_12490 | 127577..128668 | 28901 | ATP-binding-protein-[Salmonella-enterica] |
| EBL7150922.1 | OLD-ABC | GCA_006698245.1 | Salmonella_enterica | Bacteria>Pseudomonadati>Pseudomonadota>Gammaproteobacteria>Enterobacterales>Enterobacteriaceae>Salmonella | 363 | CFL44_11220 | 75362..76453 | 28901 | ATP-binding-protein-[Salmonella-enterica] |
| EBR4438817.1 | OLD-ABC | GCA_006955825.1 | Salmonella_enterica | Bacteria>Pseudomonadati>Pseudomonadota>Gammaproteobacteria>Enterobacterales>Enterobacteriaceae>Salmonella | 363 | BSE18_11495 | 103250..104341 | 28901 | AAA-family-ATPase-[Salmonella-enterica] |
| EBT4520740.1 | OLD-ABC | GCA_007056145.1 | Salmonella_enterica | Bacteria>Pseudomonadati>Pseudomonadota>Gammaproteobacteria>Enterobacterales>Enterobacteriaceae>Salmonella | 363 | CKQ08_17670 | 103250..104341 | 28901 | AAA-family-ATPase-[Salmonella-enterica] |
| EBU3750856.1 | OLD-ABC | GCA_007101645.1 | Salmonella_enterica | Bacteria>Pseudomonadati>Pseudomonadota>Gammaproteobacteria>Enterobacterales>Enterobacteriaceae>Salmonella | 363 | CVX24_10935 | 33245..34336 | 28901 | AAA-family-ATPase-[Salmonella-enterica] |
| ECH3013708.1 | OLD-ABC | GCA_007716945.1 | Salmonella_enterica | Bacteria>Pseudomonadati>Pseudomonadota>Gammaproteobacteria>Enterobacterales>Enterobacteriaceae>Salmonella | 363 | FOV18_07410 | 338284..339375 | 28901 | AAA-family-ATPase-[Salmonella-enterica] |
| ECJ6092448.1 | OLD-ABC | GCA_007834095.1 | Salmonella_enterica | Bacteria>Pseudomonadati>Pseudomonadota>Gammaproteobacteria>Enterobacterales>Enterobacteriaceae>Salmonella | 363 | GR78_06770 | 379708..380799 | 28901 | ATP-binding-protein-[Salmonella-enterica] |
| ECO5284316.1 | OLD-ABC | GCA_008153185.1 | Salmonella_enterica | Bacteria>Pseudomonadati>Pseudomonadota>Gammaproteobacteria>Enterobacterales>Enterobacteriaceae>Salmonella | 363 | FYE58_11855 | 405315..406406 | 28901 | AAA-family-ATPase-[Salmonella-enterica] |
| EDC2377953.1 | OLD-ABC | - | Salmonella_enterica | Bacteria>Pseudomonadati>Pseudomonadota>Gammaproteobacteria>Enterobacterales>Enterobacteriaceae>Salmonella | 363 | GAA61_09315 | 33248..34339 | 28901 | AAA-family-ATPase-[Salmonella-enterica] |
| EDH2237972.1 | OLD-ABC | - | Salmonella_enterica | Bacteria>Pseudomonadati>Pseudomonadota>Gammaproteobacteria>Enterobacterales>Enterobacteriaceae>Salmonella | 363 | GC580_06320 | 177877..178968 | 28901 | AAA-family-ATPase-[Salmonella-enterica] |
| EDI9669679.1 | OLD-ABC | - | Salmonella_enterica | Bacteria>Pseudomonadati>Pseudomonadota>Gammaproteobacteria>Enterobacterales>Enterobacteriaceae>Salmonella | 363 | GEV75_12815 | 46032..47123 | 28901 | ATP-binding-protein-[Salmonella-enterica] |
| EDL6685016.1 | OLD-ABC | - | Salmonella_enterica | Bacteria>Pseudomonadati>Pseudomonadota>Gammaproteobacteria>Enterobacterales>Enterobacteriaceae>Salmonella | 363 | CXC07_02680 | 178277..179368 | 28901 | ATP-binding-protein-[Salmonella-enterica] |
| EDX0369805.1 | OLD-ABC | - | Salmonella_enterica | Bacteria>Pseudomonadati>Pseudomonadota>Gammaproteobacteria>Enterobacterales>Enterobacteriaceae>Salmonella | 363 | GTL21_000363 | 393904..394995 | 28901 | AAA-family-ATPase-[Salmonella-enterica] |
| EEB8473038.1 | OLD-ABC | - | Salmonella_enterica | Bacteria>Pseudomonadati>Pseudomonadota>Gammaproteobacteria>Enterobacterales>Enterobacteriaceae>Salmonella | 363 | GN471_06055 | 178082..179173 | 28901 | AAA-family-ATPase-[Salmonella-enterica] |
| EEH0007755.1 | OLD-ABC | - | Salmonella_enterica | Bacteria>Pseudomonadati>Pseudomonadota>Gammaproteobacteria>Enterobacterales>Enterobacteriaceae>Salmonella | 363 | G3P26_001652 | 364157..365248 | 28901 | AAA-family-ATPase-[Salmonella-enterica] |
| EEH3158634.1 | OLD-ABC | - | Salmonella_enterica | Bacteria>Pseudomonadati>Pseudomonadota>Gammaproteobacteria>Enterobacterales>Enterobacteriaceae>Salmonella | 363 | GZB32_002494 | 103250..104341 | 28901 | AAA-family-ATPase-[Salmonella-enterica] |
| EEI0202359.1 | OLD-ABC | - | Salmonella_enterica | Bacteria>Pseudomonadati>Pseudomonadota>Gammaproteobacteria>Enterobacterales>Enterobacteriaceae>Salmonella | 363 | GNM15_17110 | 103250..104341 | 28901 | AAA-family-ATPase-[Salmonella-enterica] |
| EEL0135867.1 | OLD-ABC | - | Salmonella_enterica | Bacteria>Pseudomonadati>Pseudomonadota>Gammaproteobacteria>Enterobacterales>Enterobacteriaceae>Salmonella | 363 | G4L63_001978 | 121306..122397 | 28901 | AAA-family-ATPase-[Salmonella-enterica] |
| EEP5746979.1 | OLD-ABC | - | Salmonella_enterica | Bacteria>Pseudomonadati>Pseudomonadota>Gammaproteobacteria>Enterobacterales>Enterobacteriaceae>Salmonella | 363 | HBB29_002273 | 103250..104341 | 28901 | AAA-family-ATPase-[Salmonella-enterica] |
| EFP5236619.1 | OLD-ABC | - | Salmonella_enterica | Bacteria>Pseudomonadati>Pseudomonadota>Gammaproteobacteria>Enterobacterales>Enterobacteriaceae>Salmonella | 363 | H1F05_002029 | 177712..178803 | 28901 | AAA-family-ATPase-[Salmonella-enterica] |
| EFQ1578019.1 | OLD-ABC | - | Salmonella_enterica | Bacteria>Pseudomonadati>Pseudomonadota>Gammaproteobacteria>Enterobacterales>Enterobacteriaceae>Salmonella | 363 | H1L81_001834 | 46036..47127 | 28901 | AAA-family-ATPase-[Salmonella-enterica] |
| EFQ9845220.1 | OLD-ABC | - | Salmonella_enterica | Bacteria>Pseudomonadati>Pseudomonadota>Gammaproteobacteria>Enterobacterales>Enterobacteriaceae>Salmonella | 363 | HZL63_001577 | 145203..146294 | 28901 | AAA-family-ATPase-[Salmonella-enterica] |
| EFR0716067.1 | OLD-ABC | - | Salmonella_enterica | Bacteria>Pseudomonadati>Pseudomonadota>Gammaproteobacteria>Enterobacterales>Enterobacteriaceae>Salmonella | 363 | HYQ91_004638 | 5887..6978 | 28901 | AAA-family-ATPase-[Salmonella-enterica] |
| EFS7669870.1 | OLD-ABC | - | Salmonella_enterica | Bacteria>Pseudomonadati>Pseudomonadota>Gammaproteobacteria>Enterobacterales>Enterobacteriaceae>Salmonella | 363 | HWZ11_001255 | 6962..8053 | 28901 | AAA-family-ATPase-[Salmonella-enterica] |
| EFU4335448.1 | OLD-ABC | - | Salmonella_enterica | Bacteria>Pseudomonadati>Pseudomonadota>Gammaproteobacteria>Enterobacterales>Enterobacteriaceae>Salmonella | 363 | HUC58_001815 | 177713..178804 | 28901 | AAA-family-ATPase-[Salmonella-enterica] |
| EGC3767044.1 | OLD-ABC | - | Salmonella_enterica | Bacteria>Pseudomonadati>Pseudomonadota>Gammaproteobacteria>Enterobacterales>Enterobacteriaceae>Salmonella | 363 | H9A68_002332 | 46036..47127 | 28901 | AAA-family-ATPase-[Salmonella-enterica] |
| EGD1472165.1 | OLD-ABC | - | Salmonella_enterica | Bacteria>Pseudomonadati>Pseudomonadota>Gammaproteobacteria>Enterobacterales>Enterobacteriaceae>Salmonella | 363 | IAV68_002848 | 103250..104341 | 28901 | AAA-family-ATPase-[Salmonella-enterica] |
| EGF1249455.1 | OLD-ABC | - | Salmonella_enterica | Bacteria>Pseudomonadati>Pseudomonadota>Gammaproteobacteria>Enterobacterales>Enterobacteriaceae>Salmonella | 363 | IB037_001323 | 178283..179374 | 28901 | AAA-family-ATPase-[Salmonella-enterica] |
| EGG4120338.1 | OLD-ABC | - | Salmonella_enterica | Bacteria>Pseudomonadati>Pseudomonadota>Gammaproteobacteria>Enterobacterales>Enterobacteriaceae>Salmonella | 363 | SI35_20750 | 164689..165780 | 28901 | AAA-family-ATPase-[Salmonella-enterica] |
| EGH1765316.1 | OLD-ABC | - | Salmonella_enterica | Bacteria>Pseudomonadati>Pseudomonadota>Gammaproteobacteria>Enterobacterales>Enterobacteriaceae>Salmonella | 363 | ID459_000691 | 178051..179142 | 28901 | AAA-family-ATPase-[Salmonella-enterica] |
| EGH8508055.1 | OLD-ABC | - | Salmonella_enterica | Bacteria>Pseudomonadati>Pseudomonadota>Gammaproteobacteria>Enterobacterales>Enterobacteriaceae>Salmonella | 363 | IFE76_001738 | 33457..34548 | 28901 | AAA-family-ATPase-[Salmonella-enterica] |
| EGL7108186.1 | OLD-ABC | - | Salmonella_enterica | Bacteria>Pseudomonadati>Pseudomonadota>Gammaproteobacteria>Enterobacterales>Enterobacteriaceae>Salmonella | 363 | IMT80_001263 | 178293..179384 | 28901 | AAA-family-ATPase-[Salmonella-enterica] |
| EGP1242557.1 | OLD-ABC | - | Salmonella_enterica | Bacteria>Pseudomonadati>Pseudomonadota>Gammaproteobacteria>Enterobacterales>Enterobacteriaceae>Salmonella | 363 | IV995_001917 | 67504..68595 | 28901 | AAA-family-ATPase-[Salmonella-enterica] |
| EGT1190056.1 | OLD-ABC | - | Salmonella_enterica | Bacteria>Pseudomonadati>Pseudomonadota>Gammaproteobacteria>Enterobacterales>Enterobacteriaceae>Salmonella | 363 | I3G58_002373 | 103250..104341 | 28901 | AAA-family-ATPase-[Salmonella-enterica] |
| EGU3680635.1 | OLD-ABC | - | Salmonella_enterica | Bacteria>Pseudomonadati>Pseudomonadota>Gammaproteobacteria>Enterobacterales>Enterobacteriaceae>Salmonella | 363 | H0R40_002397 | 34760..35851 | 28901 | AAA-family-ATPase-[Salmonella-enterica] |
| EGU5502541.1 | OLD-ABC | - | Salmonella_enterica | Bacteria>Pseudomonadati>Pseudomonadota>Gammaproteobacteria>Enterobacterales>Enterobacteriaceae>Salmonella | 363 | HGK88_001889 | 338482..339573 | 28901 | AAA-family-ATPase-[Salmonella-enterica] |
| EGV8384250.1 | OLD-ABC | - | Salmonella_enterica | Bacteria>Pseudomonadati>Pseudomonadota>Gammaproteobacteria>Enterobacterales>Enterobacteriaceae>Salmonella | 363 | EHN43_00840 | 178285..179376 | 28901 | ATP-binding-protein-[Salmonella-enterica] |
| EHD7670520.1 | OLD-ABC | - | Salmonella_enterica | Bacteria>Pseudomonadati>Pseudomonadota>Gammaproteobacteria>Enterobacterales>Enterobacteriaceae>Salmonella | 363 | JQD35_001756 | 177946..179037 | 28901 | AAA-family-ATPase-[Salmonella-enterica] |
| EHH9023836.1 | OLD-ABC | GCA_017748765.1 | Salmonella_enterica | Bacteria>Pseudomonadati>Pseudomonadota>Gammaproteobacteria>Enterobacterales>Enterobacteriaceae>Salmonella | 363 | J8439_003631 | 25681..26772 | 28901 | AAA-family-ATPase-[Salmonella-enterica] |
| EHI1197050.1 | OLD-ABC | GCA_017785955.1 | Salmonella_enterica | Bacteria>Pseudomonadati>Pseudomonadota>Gammaproteobacteria>Enterobacterales>Enterobacteriaceae>Salmonella | 363 | J8551_000468 | 103250..104341 | 28901 | AAA-family-ATPase-[Salmonella-enterica] |
| EHJ0877871.1 | OLD-ABC | GCA_017846115.1 | Salmonella_enterica | Bacteria>Pseudomonadati>Pseudomonadota>Gammaproteobacteria>Enterobacterales>Enterobacteriaceae>Salmonella | 363 | KAB51_002699 | 25489..26580 | 28901 | AAA-family-ATPase-[Salmonella-enterica] |
| EHN4465872.1 | OLD-ABC | GCA_018387865.1 | Salmonella_enterica | Bacteria>Pseudomonadati>Pseudomonadota>Gammaproteobacteria>Enterobacterales>Enterobacteriaceae>Salmonella | 363 | KIZ21_001042 | 343586..344677 | 28901 | AAA-family-ATPase-[Salmonella-enterica] |
| EHO5627403.1 | OLD-ABC | GCA_018535225.1 | Salmonella_enterica | Bacteria>Pseudomonadati>Pseudomonadota>Gammaproteobacteria>Enterobacterales>Enterobacteriaceae>Salmonella | 363 | KLQ90_002309 | 38131..39222 | 28901 | AAA-family-ATPase-[Salmonella-enterica] |
| EHQ7935669.1 | OLD-ABC | GCA_018976465.1 | Salmonella_enterica | Bacteria>Pseudomonadati>Pseudomonadota>Gammaproteobacteria>Enterobacterales>Enterobacteriaceae>Salmonella | 363 | KSM73_002315 | 103250..104341 | 28901 | AAA-family-ATPase-[Salmonella-enterica] |
| EHT6941868.1 | OLD-ABC | GCA_019250575.1 | Salmonella_enterica | Bacteria>Pseudomonadati>Pseudomonadota>Gammaproteobacteria>Enterobacterales>Enterobacteriaceae>Salmonella | 363 | KYH33_001189 | 178282..179373 | 28901 | AAA-family-ATPase-[Salmonella-enterica] |
| EHU5044812.1 | OLD-ABC | GCA_019291225.1 | Salmonella_enterica | Bacteria>Pseudomonadati>Pseudomonadota>Gammaproteobacteria>Enterobacterales>Enterobacteriaceae>Salmonella | 363 | KY920_001889 | 94919..96010 | 28901 | AAA-family-ATPase-[Salmonella-enterica] |
| EHU5815176.1 | OLD-ABC | GCA_019298295.1 | Salmonella_enterica | Bacteria>Pseudomonadati>Pseudomonadota>Gammaproteobacteria>Enterobacterales>Enterobacteriaceae>Salmonella | 363 | KZE80_000351 | 391709..392800 | 28901 | AAA-family-ATPase-[Salmonella-enterica] |
| EHZ3783990.1 | OLD-ABC | GCA_019697575.1 | Salmonella_enterica | Bacteria>Pseudomonadati>Pseudomonadota>Gammaproteobacteria>Enterobacterales>Enterobacteriaceae>Salmonella | 363 | K5U30_001274 | 178283..179374 | 28901 | AAA-family-ATPase-[Salmonella-enterica] |
| EHZ9862032.1 | OLD-ABC | GCA_019744275.1 | Salmonella_enterica | Bacteria>Pseudomonadati>Pseudomonadota>Gammaproteobacteria>Enterobacterales>Enterobacteriaceae>Salmonella | 363 | K6Z50_000358 | 403255..404346 | 28901 | AAA-family-ATPase-[Salmonella-enterica] |
| EIA3194371.1 | OLD-ABC | GCA_019788365.1 | Salmonella_enterica | Bacteria>Pseudomonadati>Pseudomonadota>Gammaproteobacteria>Enterobacterales>Enterobacteriaceae>Salmonella | 363 | K7D38_001959 | 182187..183278 | 28901 | AAA-family-ATPase-[Salmonella-enterica] |
| EIA3335813.1 | OLD-ABC | GCA_019789135.1 | Salmonella_enterica | Bacteria>Pseudomonadati>Pseudomonadota>Gammaproteobacteria>Enterobacterales>Enterobacteriaceae>Salmonella | 363 | K7D78_001300 | 407493..408584 | 28901 | AAA-family-ATPase-[Salmonella-enterica] |
| EIB1064376.1 | OLD-ABC | GCA_019855255.1 | Salmonella_enterica | Bacteria>Pseudomonadati>Pseudomonadota>Gammaproteobacteria>Enterobacterales>Enterobacteriaceae>Salmonella | 363 | K7907_001963 | 103250..104341 | 28901 | AAA-family-ATPase-[Salmonella-enterica] |
| EIC4097434.1 | OLD-ABC | GCA_019951695.1 | Salmonella_enterica | Bacteria>Pseudomonadati>Pseudomonadota>Gammaproteobacteria>Enterobacterales>Enterobacteriaceae>Salmonella | 363 | K9796_003005 | 29522..30613 | 28901 | AAA-family-ATPase-[Salmonella-enterica] |
| EIF9427097.1 | OLD-ABC | GCA_020283115.1 | Salmonella_enterica | Bacteria>Pseudomonadati>Pseudomonadota>Gammaproteobacteria>Enterobacterales>Enterobacteriaceae>Salmonella | 363 | HW154_002105 | 33248..34339 | 28901 | AAA-family-ATPase-[Salmonella-enterica] |
| EIF9623262.1 | OLD-ABC | GCA_020284415.1 | Salmonella_enterica | Bacteria>Pseudomonadati>Pseudomonadota>Gammaproteobacteria>Enterobacterales>Enterobacteriaceae>Salmonella | 363 | HQ981_002277 | 33248..34339 | 28901 | AAA-family-ATPase-[Salmonella-enterica] |
| EIJ0643396.1 | OLD-ABC | GCA_020492735.1 | Salmonella_enterica | Bacteria>Pseudomonadati>Pseudomonadota>Gammaproteobacteria>Enterobacterales>Enterobacteriaceae>Salmonella | 363 | LH177_001937 | 67484..68575 | 28901 | AAA-family-ATPase-[Salmonella-enterica] |
| EIJ2988434.1 | OLD-ABC | GCA_020513955.1 | Salmonella_enterica | Bacteria>Pseudomonadati>Pseudomonadota>Gammaproteobacteria>Enterobacterales>Enterobacteriaceae>Salmonella | 363 | LIS40_000163 | 178283..179374 | 28901 | AAA-family-ATPase-[Salmonella-enterica] |
| EIM3792265.1 | OLD-ABC | GCA_020780095.1 | Salmonella_enterica | Bacteria>Pseudomonadati>Pseudomonadota>Gammaproteobacteria>Enterobacterales>Enterobacteriaceae>Salmonella | 363 | LM721_003556 | 34712..35803 | 28901 | AAA-family-ATPase-[Salmonella-enterica] |
| EIO7765745.1 | OLD-ABC | GCA_020997775.1 | Salmonella_enterica | Bacteria>Pseudomonadati>Pseudomonadota>Gammaproteobacteria>Enterobacterales>Enterobacteriaceae>Salmonella | 363 | LQR80_000701 | 178283..179374 | 28901 | AAA-family-ATPase-[Salmonella-enterica] |
| EIQ4281223.1 | OLD-ABC | GCA_021189835.1 | Salmonella_enterica | Bacteria>Pseudomonadati>Pseudomonadota>Gammaproteobacteria>Enterobacterales>Enterobacteriaceae>Salmonella | 363 | LU454_001107 | 110194..111285 | 28901 | AAA-family-ATPase-[Salmonella-enterica] |
| EIR8875061.1 | OLD-ABC | GCA_021305935.1 | Salmonella_enterica | Bacteria>Pseudomonadati>Pseudomonadota>Gammaproteobacteria>Enterobacterales>Enterobacteriaceae>Salmonella | 363 | LXE31_003494 | 10143..11234 | 28901 | AAA-family-ATPase-[Salmonella-enterica] |
| EJB4514210.1 | OLD-ABC | GCA_022945175.1 | Salmonella_enterica | Bacteria>Pseudomonadati>Pseudomonadota>Gammaproteobacteria>Enterobacterales>Enterobacteriaceae>Salmonella | 363 | MT088_002308 | 387504..388595 | 28901 | AAA-family-ATPase-[Salmonella-enterica] |
| EJF4683725.1 | OLD-ABC | GCA_023476405.1 | Salmonella_enterica | Bacteria>Pseudomonadati>Pseudomonadota>Gammaproteobacteria>Enterobacterales>Enterobacteriaceae>Salmonella | 363 | M8N25_002791 | 103250..104341 | 28901 | AAA-family-ATPase-[Salmonella-enterica] |
| EJF5860348.1 | OLD-ABC | GCA_023495345.1 | Salmonella_enterica | Bacteria>Pseudomonadati>Pseudomonadota>Gammaproteobacteria>Enterobacterales>Enterobacteriaceae>Salmonella | 363 | M8X04_001883 | 177878..178969 | 28901 | AAA-family-ATPase-[Salmonella-enterica] |
| EJN9121678.1 | OLD-ABC | GCA_024535515.1 | Salmonella_enterica | Bacteria>Pseudomonadati>Pseudomonadota>Gammaproteobacteria>Enterobacterales>Enterobacteriaceae>Salmonella | 363 | NR521_001805 | 178293..179384 | 28901 | AAA-family-ATPase-[Salmonella-enterica] |
| EJS8182322.1 | OLD-ABC | GCA_024988615.1 | Salmonella_enterica | Bacteria>Pseudomonadati>Pseudomonadota>Gammaproteobacteria>Enterobacterales>Enterobacteriaceae>Salmonella | 363 | NWJ04_001332 | 33248..34339 | 28901 | AAA-family-ATPase-[Salmonella-enterica] |
| HAC8921683.1 | OLD-ABC | GCA_010188005.1 | Salmonella_enterica | Bacteria>Pseudomonadati>Pseudomonadota>Gammaproteobacteria>Enterobacterales>Enterobacteriaceae>Salmonella | 363 | G0J09_01280 | 53817..54908 | 28901 | TPA_asm:-ATP-binding-protein-[Salmonella-enterica] |
| HAF0812801.1 | OLD-ABC | GCA_011125695.1 | Salmonella_enterica | Bacteria>Pseudomonadati>Pseudomonadota>Gammaproteobacteria>Enterobacterales>Enterobacteriaceae>Salmonella | 363 | G9G17_001781 | 103250..104341 | 28901 | TPA:-AAA-family-ATPase-[Salmonella-enterica] |
| HAF9991658.1 | OLD-ABC | GCA_011779145.1 | Salmonella_enterica | Bacteria>Pseudomonadati>Pseudomonadota>Gammaproteobacteria>Enterobacterales>Enterobacteriaceae>Salmonella | 363 | G8Q07_002650 | 178108..179199 | 28901 | TPA:-AAA-family-ATPase-[Salmonella-enterica] |
| HAG2842214.1 | OLD-ABC | GCA_011791385.1 | Salmonella_enterica | Bacteria>Pseudomonadati>Pseudomonadota>Gammaproteobacteria>Enterobacterales>Enterobacteriaceae>Salmonella | 363 | G8Z41_002590 | 110159..111250 | 28901 | TPA:-AAA-family-ATPase-[Salmonella-enterica] |
| HAG5552354.1 | OLD-ABC | GCA_011798725.1 | Salmonella_enterica | Bacteria>Pseudomonadati>Pseudomonadota>Gammaproteobacteria>Enterobacterales>Enterobacteriaceae>Salmonella | 363 | G8Q25_001510 | 181956..183047 | 28901 | TPA:-AAA-family-ATPase-[Salmonella-enterica] |
| HAK8810849.1 | OLD-ABC | GCA_013646795.1 | Salmonella_enterica | Bacteria>Pseudomonadati>Pseudomonadota>Gammaproteobacteria>Enterobacterales>Enterobacteriaceae>Salmonella | 363 | H2X42_001964 | 165729..166820 | 28901 | AAA-family-ATPase-[Salmonella-enterica] |
| MCM9949029.1 | OLD-ABC | GCA_023843835.1 | Salmonella_enterica | Bacteria>Pseudomonadati>Pseudomonadota>Gammaproteobacteria>Enterobacterales>Enterobacteriaceae>Salmonella | 363 | M5G05_07270 | 343509..344600 | 28901 | AAA-family-ATPase-[Salmonella-enterica] |
| MDG46365.1 | OLD-ABC | GCA_003733245.1 | Salmonella_enterica | Bacteria>Pseudomonadati>Pseudomonadota>Gammaproteobacteria>Enterobacterales>Enterobacteriaceae>Salmonella | 363 | EB289_07930 | 46036..47127 | 28901 | ATP-binding-protein-[Salmonella-enterica] |
| WP_000683870.1 | OLD-ABC | GCF_967854935.1 | Salmonella_enterica | Bacteria>Pseudomonadati>Pseudomonadota>Gammaproteobacteria>Enterobacterales>Enterobacteriaceae>Salmonella | 363 | J3T40_RS14915 | 387754..388845 | 28901 | ATP-binding-protein-[Salmonella-enterica] |
| WP_023217631.1 | OLD-ABC | GCF_000412855.1 | Salmonella_enterica | Bacteria>Pseudomonadati>Pseudomonadota>Gammaproteobacteria>Enterobacterales>Enterobacteriaceae>Salmonella | 363 | GX95_RS20915 | 4062446..4063537 | 28901 | ATP-binding-protein-[Salmonella-enterica] |
| WP_023258576.1 | OLD-ABC | GCF_027943645.1 | Salmonella_enterica | Bacteria>Pseudomonadati>Pseudomonadota>Gammaproteobacteria>Enterobacterales>Enterobacteriaceae>Salmonella | 363 | F1E41_RS17170 | 158038..159129 | 28901 | ATP-binding-protein-[Salmonella-enterica] |
| WP_039590206.1 | OLD-ABC | GCF_943119195.1 | Salmonella_enterica | Bacteria>Pseudomonadati>Pseudomonadota>Gammaproteobacteria>Enterobacterales>Enterobacteriaceae>Salmonella | 363 | OCU40_RS05655 | 1162687..1163778 | 28901 | ATP-binding-protein-[Salmonella-enterica] |
| WP_058114202.1 | OLD-ABC | GCF_003324795.1 | Salmonella_enterica | Bacteria>Pseudomonadati>Pseudomonadota>Gammaproteobacteria>Enterobacterales>Enterobacteriaceae>Salmonella | 363 | DOE60_RS06780 | 1371694..1372785 | 28901 | ATP-binding-protein-[Salmonella-enterica] |
| WP_061584063.1 | OLD-ABC | GCF_016519005.1 | Salmonella_enterica | Bacteria>Pseudomonadati>Pseudomonadota>Gammaproteobacteria>Enterobacterales>Enterobacteriaceae>Salmonella | 363 | C5I04_RS23485 | 341565..342656 | 28901 | ATP-binding-protein-[Salmonella-enterica] |
| WP_079808686.1 | OLD-ABC | GCF_002035875.1 | Salmonella_enterica | Bacteria>Pseudomonadati>Pseudomonadota>Gammaproteobacteria>Enterobacterales>Enterobacteriaceae>Salmonella | 363 | Y985_RS08720 | 115235..116326 | 28901 | ATP-binding-protein-[Salmonella-enterica] |
| WP_079902378.1 | OLD-ABC | GCF_003325275.1 | Salmonella_enterica | Bacteria>Pseudomonadati>Pseudomonadota>Gammaproteobacteria>Enterobacterales>Enterobacteriaceae>Salmonella | 363 | CHE40_RS23170 | 4538709..4539800 | 28901 | ATP-binding-protein-[Salmonella-enterica] |
| WP_080205564.1 | OLD-ABC | GCF_943114325.1 | Salmonella_enterica | Bacteria>Pseudomonadati>Pseudomonadota>Gammaproteobacteria>Enterobacterales>Enterobacteriaceae>Salmonella | 363 | OCU44_RS00995 | 219375..220466 | 28901 | ATP-binding-protein-[Salmonella-enterica] |
| WP_218201153.1 | OLD-ABC | GCF_019173605.1 | Salmonella_enterica | Bacteria>Pseudomonadati>Pseudomonadota>Gammaproteobacteria>Enterobacterales>Enterobacteriaceae>Salmonella | 363 | KXD81_RS02510 | 17174..18265 | 28901 | ATP-binding-protein-[Salmonella-enterica] |
| WP_267272755.1 | OLD-ABC | GCF_026460795.1 | Salmonella_enterica | Bacteria>Pseudomonadati>Pseudomonadota>Gammaproteobacteria>Enterobacterales>Enterobacteriaceae>Salmonella | 363 | OWO74_RS14185 | 6545..7636 | 28901 | ATP-binding-protein-[Salmonella-enterica] |
| EAA7604356.1 | OLD-ABC | GCA_004218255.1 | Salmonella_enterica | Bacteria>Pseudomonadati>Pseudomonadota>Gammaproteobacteria>Enterobacterales>Enterobacteriaceae>Salmonella | 362 | ADQ66_03565 | 137820..138908 | 28901 | ATP-binding-protein-[Salmonella-enterica] |
| EAB2831580.1 | OLD-ABC | GCA_004240605.1 | Salmonella_enterica | Bacteria>Pseudomonadati>Pseudomonadota>Gammaproteobacteria>Enterobacterales>Enterobacteriaceae>Salmonella | 362 | EAV48_05280 | 165268..166356 | 28901 | ATP-binding-protein-[Salmonella-enterica] |
| EAM3013484.1 | OLD-ABC | GCA_005412095.1 | Salmonella_enterica | Bacteria>Pseudomonadati>Pseudomonadota>Gammaproteobacteria>Enterobacterales>Enterobacteriaceae>Salmonella | 362 | CBJ93_01875 | 401981..403069 | 28901 | ATP-binding-protein-[Salmonella-enterica] |
| EAM5567468.1 | OLD-ABC | GCA_005423155.1 | Salmonella_enterica | Bacteria>Pseudomonadati>Pseudomonadota>Gammaproteobacteria>Enterobacterales>Enterobacteriaceae>Salmonella | 362 | EK556_21770 | 6291..7379 | 28901 | ATP-binding-protein-[Salmonella-enterica] |
| EAO2684616.1 | OLD-ABC | GCA_005496785.1 | Salmonella_enterica | Bacteria>Pseudomonadati>Pseudomonadota>Gammaproteobacteria>Enterobacterales>Enterobacteriaceae>Salmonella | 362 | E2X65_05660 | 74894..75982 | 28901 | ATP-binding-protein-[Salmonella-enterica] |
| EAQ6130545.1 | OLD-ABC | GCA_005607075.1 | Salmonella_enterica | Bacteria>Pseudomonadati>Pseudomonadota>Gammaproteobacteria>Enterobacterales>Enterobacteriaceae>Salmonella | 362 | AXA91_18740 | 69719..70807 | 28901 | ATP-binding-protein-[Salmonella-enterica] |
| EAT1639065.1 | OLD-ABC | GCA_005720135.1 | Salmonella_enterica | Bacteria>Pseudomonadati>Pseudomonadota>Gammaproteobacteria>Enterobacterales>Enterobacteriaceae>Salmonella | 362 | EKF56_11750 | 32385..33473 | 28901 | ATP-binding-protein-[Salmonella-enterica] |
| EAW1006426.1 | OLD-ABC | GCA_005898145.1 | Salmonella_enterica | Bacteria>Pseudomonadati>Pseudomonadota>Gammaproteobacteria>Enterobacterales>Enterobacteriaceae>Salmonella | 362 | EA086_11670 | 164425..165513 | 28901 | ATP-binding-protein-[Salmonella-enterica] |
| EAW8145857.1 | OLD-ABC | GCA_005934015.1 | Salmonella_enterica | Bacteria>Pseudomonadati>Pseudomonadota>Gammaproteobacteria>Enterobacterales>Enterobacteriaceae>Salmonella | 362 | HM10_03835 | 170860..171948 | 28901 | ATP-binding-protein-[Salmonella-enterica] |

| Protein-ID | Domain | Assembly-ID | Strain | Taxonomy | Length | Locus-Tag | Genomic-coordinates | TaxID | Fasta-defline |
|---|---|---|---|---|---|---|---|---|---|
| EAW9404813.1 | OLD-ABC | GCA_005940875.1 | Salmonella_enterica | Bacteria>Pseudomonadati>Pseudomonadota>Gammaproteobacteria>Enterobacterales>Enterobacteriaceae>Salmonella | 362 | QD58_16780 | 32734..33822 | 28901 | ATP-binding-protein-[Salmonella-enterica] |
| EAX8471829.1 | OLD-ABC | GCA_005980785.1 | Salmonella_enterica | Bacteria>Pseudomonadati>Pseudomonadota>Gammaproteobacteria>Enterobacterales>Enterobacteriaceae>Salmonella | 362 | AH421_04150 | 253882..254970 | 28901 | ATP-binding-protein-[Salmonella-enterica] |
| EAZ9311963.1 | OLD-ABC | GCA_006073635.1 | Salmonella_enterica | Bacteria>Pseudomonadati>Pseudomonadota>Gammaproteobacteria>Enterobacterales>Enterobacteriaceae>Salmonella | 362 | CBJ28_03035 | 166054..167142 | 28901 | ATP-binding-protein-[Salmonella-enterica] |
| EBA4497719.1 | OLD-ABC | GCA_006097275.1 | Salmonella_enterica | Bacteria>Pseudomonadati>Pseudomonadota>Gammaproteobacteria>Enterobacterales>Enterobacteriaceae>Salmonella | 362 | A7G53_12030 | 216246..217334 | 28901 | ATP-binding-protein-[Salmonella-enterica] |
| EBB3832831.1 | OLD-ABC | GCA_006140415.1 | Salmonella_enterica | Bacteria>Pseudomonadati>Pseudomonadota>Gammaproteobacteria>Enterobacterales>Enterobacteriaceae>Salmonella | 362 | CJ140_05530 | 157103..158191 | 28901 | ATP-binding-protein-[Salmonella-enterica] |
| EBF0840958.1 | OLD-ABC | GCA_006325425.1 | Salmonella_enterica | Bacteria>Pseudomonadati>Pseudomonadota>Gammaproteobacteria>Enterobacterales>Enterobacteriaceae>Salmonella | 362 | DDQ86_07960 | 157635..158723 | 28901 | ATP-binding-protein-[Salmonella-enterica] |
| EBK5803037.1 | OLD-ABC | GCA_006648525.1 | Salmonella_enterica | Bacteria>Pseudomonadati>Pseudomonadota>Gammaproteobacteria>Enterobacterales>Enterobacteriaceae>Salmonella | 362 | DPM88_04075 | 165782..166870 | 28901 | ATP-binding-protein-[Salmonella-enterica] |
| EBM6187522.1 | OLD-ABC | GCA_006738745.1 | Salmonella_enterica | Bacteria>Pseudomonadati>Pseudomonadota>Gammaproteobacteria>Enterobacterales>Enterobacteriaceae>Salmonella | 362 | D1943_04240 | 107943..109031 | 28901 | ATP-binding-protein-[Salmonella-enterica] |
| EBS7702176.1 | OLD-ABC | GCA_007022165.1 | Salmonella_enterica | Bacteria>Pseudomonadati>Pseudomonadota>Gammaproteobacteria>Enterobacterales>Enterobacteriaceae>Salmonella | 362 | CD603_08300 | 59570..60658 | 28901 | ATP-binding-protein-[Salmonella-enterica] |
| ECG9327785.1 | OLD-ABC | GCA_007700105.1 | Salmonella_enterica | Bacteria>Pseudomonadati>Pseudomonadota>Gammaproteobacteria>Enterobacterales>Enterobacteriaceae>Salmonella | 362 | FO199_03405 | 127035..128123 | 28901 | ATP-binding-protein-[Salmonella-enterica] |
| ECQ9432770.1 | OLD-ABC | GCA_008312495.1 | Salmonella_enterica | Bacteria>Pseudomonadati>Pseudomonadota>Gammaproteobacteria>Enterobacterales>Enterobacteriaceae>Salmonella | 362 | F0371_05175 | 10174..11262 | 28901 | ATP-binding-protein-[Salmonella-enterica] |
| EDL7421470.1 | OLD-ABC | - | Salmonella_enterica | Bacteria>Pseudomonadati>Pseudomonadota>Gammaproteobacteria>Enterobacterales>Enterobacteriaceae>Salmonella | 362 | CU840_19775 | 18381..19469 | 28901 | ATP-binding-protein-[Salmonella-enterica] |
| EDV2155934.1 | OLD-ABC | - | Salmonella_enterica | Bacteria>Pseudomonadati>Pseudomonadota>Gammaproteobacteria>Enterobacterales>Enterobacteriaceae>Salmonella | 362 | JR65_004157 | 32460..33548 | 28901 | ATP-binding-protein-[Salmonella-enterica] |
| EDW7722490.1 | OLD-ABC | - | Salmonella_enterica | Bacteria>Pseudomonadati>Pseudomonadota>Gammaproteobacteria>Enterobacterales>Enterobacteriaceae>Salmonella | 362 | GTR06_001873 | 405276..406364 | 28901 | ATP-binding-protein-[Salmonella-enterica] |
| EEL7097505.1 | OLD-ABC | - | Salmonella_enterica | Bacteria>Pseudomonadati>Pseudomonadota>Gammaproteobacteria>Enterobacterales>Enterobacteriaceae>Salmonella | 362 | GKH84_08585 | 166250..167338 | 28901 | AAA-family-ATPase-[Salmonella-enterica] |
| EEP2127994.1 | OLD-ABC | - | Salmonella_enterica | Bacteria>Pseudomonadati>Pseudomonadota>Gammaproteobacteria>Enterobacterales>Enterobacteriaceae>Salmonella | 362 | HA780_002412 | 118632..119720 | 28901 | ATP-binding-protein-[Salmonella-enterica] |
| EFS3421934.1 | OLD-ABC | - | Salmonella_enterica | Bacteria>Pseudomonadati>Pseudomonadota>Gammaproteobacteria>Enterobacterales>Enterobacteriaceae>Salmonella | 362 | HX201_001607 | 157504..158592 | 28901 | ATP-binding-protein-[Salmonella-enterica] |
| EFS5965940.1 | OLD-ABC | - | Salmonella_enterica | Bacteria>Pseudomonadati>Pseudomonadota>Gammaproteobacteria>Enterobacterales>Enterobacteriaceae>Salmonella | 362 | HW654_000974 | 202683..203771 | 28901 | ATP-binding-protein-[Salmonella-enterica] |
| EGB5523464.1 | OLD-ABC | - | Salmonella_enterica | Bacteria>Pseudomonadati>Pseudomonadota>Gammaproteobacteria>Enterobacterales>Enterobacteriaceae>Salmonella | 362 | H7074_002106 | 166418..167506 | 28901 | ATP-binding-protein-[Salmonella-enterica] |
| EGH2238323.1 | OLD-ABC | - | Salmonella_enterica | Bacteria>Pseudomonadati>Pseudomonadota>Gammaproteobacteria>Enterobacterales>Enterobacteriaceae>Salmonella | 362 | ID764_000356 | 398181..399269 | 28901 | ATP-binding-protein-[Salmonella-enterica] |
| EGI8941864.1 | OLD-ABC | - | Salmonella_enterica | Bacteria>Pseudomonadati>Pseudomonadota>Gammaproteobacteria>Enterobacterales>Enterobacteriaceae>Salmonella | 362 | IG687_002355 | 280365..281453 | 28901 | ATP-binding-protein-[Salmonella-enterica] |
| EGK3406184.1 | OLD-ABC | - | Salmonella_enterica | Bacteria>Pseudomonadati>Pseudomonadota>Gammaproteobacteria>Enterobacterales>Enterobacteriaceae>Salmonella | 362 | IOS83_004391 | 20389..21477 | 28901 | ATP-binding-protein-[Salmonella-enterica] |
| EHA2523075.1 | OLD-ABC | - | Salmonella_enterica | Bacteria>Pseudomonadati>Pseudomonadota>Gammaproteobacteria>Enterobacterales>Enterobacteriaceae>Salmonella | 362 | JKI82_003471 | 173255..174343 | 28901 | ATP-binding-protein-[Salmonella-enterica] |
| EHL7445943.1 | OLD-ABC | GCA_018171815.1 | Salmonella_enterica | Bacteria>Pseudomonadati>Pseudomonadota>Gammaproteobacteria>Enterobacterales>Enterobacteriaceae>Salmonella | 362 | KEH66_000996 | 393337..394425 | 28901 | ATP-binding-protein-[Salmonella-enterica] |
| EHO5101498.1 | OLD-ABC | GCA_018532585.1 | Salmonella_enterica | Bacteria>Pseudomonadati>Pseudomonadota>Gammaproteobacteria>Enterobacterales>Enterobacteriaceae>Salmonella | 362 | KLP74_002743 | 7607..8695 | 28901 | ATP-binding-protein-[Salmonella-enterica] |
| EHS4662099.1 | OLD-ABC | GCA_019144395.1 | Salmonella_enterica | Bacteria>Pseudomonadati>Pseudomonadota>Gammaproteobacteria>Enterobacterales>Enterobacteriaceae>Salmonella | 362 | KV604_003672 | 68727..69815 | 28901 | ATP-binding-protein-[Salmonella-enterica] |
| EHT1034283.1 | OLD-ABC | GCA_019186465.1 | Salmonella_enterica | Bacteria>Pseudomonadati>Pseudomonadota>Gammaproteobacteria>Enterobacterales>Enterobacteriaceae>Salmonella | 362 | KW750_001286 | 156481..157569 | 28901 | ATP-binding-protein-[Salmonella-enterica] |
| EHU7518554.1 | OLD-ABC | GCA_019318345.1 | Salmonella_enterica | Bacteria>Pseudomonadati>Pseudomonadota>Gammaproteobacteria>Enterobacterales>Enterobacteriaceae>Salmonella | 362 | KZL07_001985 | 280345..281433 | 28901 | ATP-binding-protein-[Salmonella-enterica] |
| EHV2030461.1 | OLD-ABC | GCA_019356775.1 | Salmonella_enterica | Bacteria>Pseudomonadati>Pseudomonadota>Gammaproteobacteria>Enterobacterales>Enterobacteriaceae>Salmonella | 362 | KZ138_001533 | 397060..398148 | 28901 | ATP-binding-protein-[Salmonella-enterica] |
| EHW6237843.1 | OLD-ABC | GCA_019485205.1 | Salmonella_enterica | Bacteria>Pseudomonadati>Pseudomonadota>Gammaproteobacteria>Enterobacterales>Enterobacteriaceae>Salmonella | 362 | K2495_002339 | 38118..39206 | 28901 | ATP-binding-protein-[Salmonella-enterica] |
| EIU6597716.1 | OLD-ABC | GCA_021692515.1 | Salmonella_enterica | Bacteria>Pseudomonadati>Pseudomonadota>Gammaproteobacteria>Enterobacterales>Enterobacteriaceae>Salmonella | 362 | L5W02_001570 | 384621..385709 | 28901 | ATP-binding-protein-[Salmonella-enterica] |
| EJC3638308.1 | OLD-ABC | GCA_023047235.1 | Salmonella_enterica | Bacteria>Pseudomonadati>Pseudomonadota>Gammaproteobacteria>Enterobacterales>Enterobacteriaceae>Salmonella | 362 | MYX88_003395 | 158467..159555 | 28901 | ATP-binding-protein-[Salmonella-enterica] |
| EJK0228890.1 | OLD-ABC | GCA_024068175.1 | Salmonella_enterica | Bacteria>Pseudomonadati>Pseudomonadota>Gammaproteobacteria>Enterobacterales>Enterobacteriaceae>Salmonella | 362 | NKK90_001817 | 318361..319449 | 28901 | ATP-binding-protein-[Salmonella-enterica] |
| EJM7985885.1 | OLD-ABC | GCA_024373005.1 | Salmonella_enterica | Bacteria>Pseudomonadati>Pseudomonadota>Gammaproteobacteria>Enterobacterales>Enterobacteriaceae>Salmonella | 362 | NO486_001114 | 376450..377538 | 28901 | ATP-binding-protein-[Salmonella-enterica] |
| EJO5979499.1 | OLD-ABC | GCA_024573615.1 | Salmonella_enterica | Bacteria>Pseudomonadati>Pseudomonadota>Gammaproteobacteria>Enterobacterales>Enterobacteriaceae>Salmonella | 362 | NRR85_002627 | 166418..167506 | 28901 | ATP-binding-protein-[Salmonella-enterica] |
| HAK8092290.1 | OLD-ABC | GCA_013643495.1 | Salmonella_enterica | Bacteria>Pseudomonadati>Pseudomonadota>Gammaproteobacteria>Enterobacterales>Enterobacteriaceae>Salmonella | 362 | H2V91_002194 | 160459..161547 | 28901 | ATP-binding-protein-[Salmonella-enterica] |
| MBH5277082.1 | OLD-ABC | GCA_016019715.1 | Salmonella_enterica | Bacteria>Pseudomonadati>Pseudomonadota>Gammaproteobacteria>Enterobacterales>Enterobacteriaceae>Salmonella | 362 | I3A81_01640 | 353819..354907 | 28901 | ATP-binding-protein-[Salmonella-enterica] |
| WP_061380449.1 | OLD-ABC | GCF_001570285.1 | Salmonella_enterica | Bacteria>Pseudomonadati>Pseudomonadota>Gammaproteobacteria>Enterobacterales>Enterobacteriaceae>Salmonella | 362 | AW153_RS18925 | 165720..166808 | 28901 | ATP-binding-protein-[Salmonella-enterica] |
| WP_071055529.1 | OLD-ABC | GCF_040956775.1 | Salmonella_enterica | Bacteria>Pseudomonadati>Pseudomonadota>Gammaproteobacteria>Enterobacterales>Enterobacteriaceae>Salmonella | 362 | SES26_RS19325 | 4016830..4017918 | 28901 | ATP-binding-protein-[Salmonella-enterica] |
| WP_076736166.1 | OLD-ABC | GCF_001975445.1 | Salmonella_enterica | Bacteria>Pseudomonadati>Pseudomonadota>Gammaproteobacteria>Enterobacterales>Enterobacteriaceae>Salmonella | 362 | LFZ24_RS24370 | 4838272..4839360 | 28901 | ATP-binding-protein-[Salmonella-enterica] |
| WP_076937573.1 | OLD-ABC | GCF_001975325.1 | Salmonella_enterica | Bacteria>Pseudomonadati>Pseudomonadota>Gammaproteobacteria>Enterobacterales>Enterobacteriaceae>Salmonella | 362 | LFZ3_RS17415 | 3413749..3414837 | 28901 | ATP-binding-protein-[Salmonella-enterica] |
| WP_079944400.1 | OLD-ABC | GCF_050412455.1 | Salmonella_enterica | Bacteria>Pseudomonadati>Pseudomonadota>Gammaproteobacteria>Enterobacterales>Enterobacteriaceae>Salmonella | 362 | VX95_RS09550 | 158742..159830 | 28901 | ATP-binding-protein-[Salmonella-enterica] |
| WP_079965459.1 | OLD-ABC | GCF_002046745.1 | Salmonella_enterica | Bacteria>Pseudomonadati>Pseudomonadota>Gammaproteobacteria>Enterobacterales>Enterobacteriaceae>Salmonella | 362 | VX54_RS10660 | 59553..60641 | 28901 | ATP-binding-protein-[Salmonella-enterica] |
| WP_080090699.1 | OLD-ABC | GCF_002234755.2 | Salmonella_enterica | Bacteria>Pseudomonadati>Pseudomonadota>Gammaproteobacteria>Enterobacterales>Enterobacteriaceae>Salmonella | 362 | CHD02_RS05515 | 1132756..1133844 | 28901 | ATP-binding-protein-[Salmonella-enterica] |
| WP_080166836.1 | OLD-ABC | GCF_002208255.1 | Salmonella_enterica | Bacteria>Pseudomonadati>Pseudomonadota>Gammaproteobacteria>Enterobacterales>Enterobacteriaceae>Salmonella | 362 | LFZ35_RS21100 | 4069349..4070437 | 28901 | ATP-binding-protein-[Salmonella-enterica] |
| WP_080213760.1 | OLD-ABC | GCF_025176825.1 | Salmonella_enterica | Bacteria>Pseudomonadati>Pseudomonadota>Gammaproteobacteria>Enterobacterales>Enterobacteriaceae>Salmonella | 362 | AS04_RS02260 | 51762..52850 | 28901 | ATP-binding-protein-[Salmonella-enterica] |
| WP_080250287.1 | OLD-ABC | GCF_037083205.1 | Salmonella_enterica | Bacteria>Pseudomonadati>Pseudomonadota>Gammaproteobacteria>Enterobacterales>Enterobacteriaceae>Salmonella | 362 | RZ13_RS02305 | 172156..173244 | 28901 | ATP-binding-protein-[Salmonella-enterica] |
| WP_094887771.1 | OLD-ABC | GCF_040281485.1 | Salmonella_enterica | Bacteria>Pseudomonadati>Pseudomonadota>Gammaproteobacteria>Enterobacterales>Enterobacteriaceae>Salmonella | 362 | CCO43_RS23245 | 8064..9152 | 28901 | ATP-binding-protein-[Salmonella-enterica] |
| WP_094914770.1 | OLD-ABC | GCF_903930985.1 | Salmonella_enterica | Bacteria>Pseudomonadati>Pseudomonadota>Gammaproteobacteria>Enterobacterales>Enterobacteriaceae>Salmonella | 362 | DQN83_RS20190 | 3954041..3955129 | 28901 | ATP-binding-protein-[Salmonella-enterica] |
| WP_094946978.1 | OLD-ABC | GCF_002266315.1 | Salmonella_enterica | Bacteria>Pseudomonadati>Pseudomonadota>Gammaproteobacteria>Enterobacterales>Enterobacteriaceae>Salmonella | 362 | CCO36_RS01890 | 394343..395431 | 28901 | ATP-binding-protein-[Salmonella-enterica] |
| WP_094964981.1 | OLD-ABC | GCF_002266055.1 | Salmonella_enterica | Bacteria>Pseudomonadati>Pseudomonadota>Gammaproteobacteria>Enterobacterales>Enterobacteriaceae>Salmonella | 362 | CCO49_RS14345 | 406443..407531 | 28901 | ATP-binding-protein-[Salmonella-enterica] |
| WP_111769568.1 | OLD-ABC | GCF_051417595.1 | Salmonella_enterica | Bacteria>Pseudomonadati>Pseudomonadota>Gammaproteobacteria>Enterobacterales>Enterobacteriaceae>Salmonella | 362 | R537_RS06020 | 149636..150724 | 28901 | ATP-binding-protein-[Salmonella-enterica] |
| WP_140237023.1 | OLD-ABC | GCF_006379925.2 | Salmonella_enterica | Bacteria>Pseudomonadati>Pseudomonadota>Gammaproteobacteria>Enterobacterales>Enterobacteriaceae>Salmonella | 362 | FG677_RS21920 | 10220..11308 | 28901 | ATP-binding-protein-[Salmonella-enterica] |
| WP_183072253.1 | OLD-ABC | GCF_903931105.1 | Salmonella_enterica | Bacteria>Pseudomonadati>Pseudomonadota>Gammaproteobacteria>Enterobacterales>Enterobacteriaceae>Salmonella | 362 | H7T17_RS07620 | 61535..62623 | 28901 | ATP-binding-protein-[Salmonella-enterica] |
| ECI8263213.1 | OLD-ABC | GCA_007791395.1 | Salmonella_enterica_subsp._salamae | Bacteria>Pseudomonadati>Pseudomonadota>Gammaproteobacteria>Enterobacterales>Enterobacteriaceae>Salmonella>Salmonella_enterica | 373 | S481_13085 | 129323..130444 | 59202 | ABC-transporter-ATP-binding-protein-[Salmonella-enterica-subsp.-salamae] |
| EDT7223675.1 | OLD-ABC | - | Salmonella_enterica_subsp._enterica | Bacteria>Pseudomonadati>Pseudomonadota>Gammaproteobacteria>Enterobacterales>Enterobacteriaceae>Salmonella>Salmonella_enterica | 373 | AH268_003336 | 176605..177726 | 59201 | AAA-family-ATPase-[Salmonella-enterica-subsp.-enterica] |
| EGB5697879.1 | OLD-ABC | - | Salmonella_enterica_subsp._enterica | Bacteria>Pseudomonadati>Pseudomonadota>Gammaproteobacteria>Enterobacterales>Enterobacteriaceae>Salmonella>Salmonella_enterica | 373 | H7531_003069 | 12487..13608 | 59201 | AAA-family-ATPase-[Salmonella-enterica-subsp.-enterica] |
| EAA5451233.1 | OLD-ABC | GCA_004184965.1 | Salmonella_enterica_subsp._enterica | Bacteria>Pseudomonadati>Pseudomonadota>Gammaproteobacteria>Enterobacterales>Enterobacteriaceae>Salmonella>Salmonella_enterica | 363 | DQC86_00810 | 167889..168980 | 59201 | ATP-binding-protein-[Salmonella-enterica-subsp.-enterica] |
| EAA9627984.1 | OLD-ABC | GCA_004227565.1 | Salmonella_enterica_subsp._enterica | Bacteria>Pseudomonadati>Pseudomonadota>Gammaproteobacteria>Enterobacterales>Enterobacteriaceae>Salmonella>Salmonella_enterica | 363 | DV01_07265 | 178283..179374 | 59201 | ATP-binding-protein-[Salmonella-enterica-subsp.-enterica] |
| EAB6843527.1 | OLD-ABC | GCA_004258725.1 | Salmonella_enterica_subsp._salamae | Bacteria>Pseudomonadati>Pseudomonadota>Gammaproteobacteria>Enterobacterales>Enterobacteriaceae>Salmonella>Salmonella_enterica | 363 | XT66_14195 | 58296..59387 | 59202 | ATP-binding-protein-[Salmonella-enterica-subsp.-salamae] |
| EBG5655089.1 | OLD-ABC | GCA_006430455.1 | Salmonella_enterica_subsp._enterica | Bacteria>Pseudomonadati>Pseudomonadota>Gammaproteobacteria>Enterobacterales>Enterobacteriaceae>Salmonella>Salmonella_enterica | 363 | FI236_11965 | 103250..104341 | 59201 | ATP-binding-protein-[Salmonella-enterica-subsp.-enterica] |
| ECC0743074.1 | OLD-ABC | GCA_007457215.1 | Salmonella_enterica_subsp._enterica | Bacteria>Pseudomonadati>Pseudomonadota>Gammaproteobacteria>Enterobacterales>Enterobacteriaceae>Salmonella>Salmonella_enterica | 363 | FMV06_07385 | 103250..104341 | 59201 | AAA-family-ATPase-[Salmonella-enterica-subsp.-enterica] |
| ECC1667301.1 | OLD-ABC | GCA_007463295.1 | Salmonella_enterica_subsp._salamae | Bacteria>Pseudomonadati>Pseudomonadota>Gammaproteobacteria>Enterobacterales>Enterobacteriaceae>Salmonella>Salmonella_enterica | 363 | FNN95_01265 | 275206..276297 | 59202 | ATP-binding-protein-[Salmonella-enterica-subsp.-salamae] |
| ECC3171799.1 | OLD-ABC | GCA_007472855.1 | Salmonella_enterica_subsp._enterica | Bacteria>Pseudomonadati>Pseudomonadota>Gammaproteobacteria>Enterobacterales>Enterobacteriaceae>Salmonella>Salmonella_enterica | 363 | R914_02605 | 84971..86062 | 59201 | ATP-binding-protein-[Salmonella-enterica-subsp.-enterica] |
| ECD1705585.1 | OLD-ABC | GCA_007509785.1 | Salmonella_enterica_subsp._enterica | Bacteria>Pseudomonadati>Pseudomonadota>Gammaproteobacteria>Enterobacterales>Enterobacteriaceae>Salmonella>Salmonella_enterica | 363 | EWD17_08940 | 103250..104341 | 59201 | ATP-binding-protein-[Salmonella-enterica-subsp.-enterica] |
| ECD9608664.1 | OLD-ABC | GCA_007542335.1 | Salmonella_enterica_subsp._salamae | Bacteria>Pseudomonadati>Pseudomonadota>Gammaproteobacteria>Enterobacterales>Enterobacteriaceae>Salmonella>Salmonella_enterica | 363 | FNZ18_04570 | 263880..264971 | 59202 | AAA-family-ATPase-[Salmonella-enterica-subsp.-salamae] |
| ECG0705017.1 | OLD-ABC | GCA_007641515.1 | Salmonella_enterica_subsp._salamae | Bacteria>Pseudomonadati>Pseudomonadota>Gammaproteobacteria>Enterobacterales>Enterobacteriaceae>Salmonella>Salmonella_enterica | 363 | DL122_04710 | 42906..43997 | 59202 | ATP-binding-protein-[Salmonella-enterica-subsp.-salamae] |
| ECG1893119.1 | OLD-ABC | GCA_007650535.1 | Salmonella_enterica_subsp._enterica | Bacteria>Pseudomonadati>Pseudomonadota>Gammaproteobacteria>Enterobacterales>Enterobacteriaceae>Salmonella>Salmonella_enterica | 363 | EI460_05115 | 163000..164091 | 59201 | ATP-binding-protein-[Salmonella-enterica-subsp.-enterica] |
| ECG5099191.1 | OLD-ABC | GCA_007668905.1 | Salmonella_enterica_subsp._enterica | Bacteria>Pseudomonadati>Pseudomonadota>Gammaproteobacteria>Enterobacterales>Enterobacteriaceae>Salmonella>Salmonella_enterica | 363 | E0547_20055 | 67276..68367 | 59201 | ATP-binding-protein-[Salmonella-enterica-subsp.-enterica] |
| ECI6492269.1 | OLD-ABC | GCA_007781245.1 | Salmonella_enterica_subsp._enterica | Bacteria>Pseudomonadati>Pseudomonadota>Gammaproteobacteria>Enterobacterales>Enterobacteriaceae>Salmonella>Salmonella_enterica | 363 | EIL18_20705 | 63157..64248 | 59201 | ATP-binding-protein-[Salmonella-enterica-subsp.-enterica] |
| ECI6612373.1 | OLD-ABC | GCA_007781585.1 | Salmonella_enterica_subsp._enterica | Bacteria>Pseudomonadati>Pseudomonadota>Gammaproteobacteria>Enterobacterales>Enterobacteriaceae>Salmonella>Salmonella_enterica | 363 | EU408_19965 | 70110..71201 | 59201 | ATP-binding-protein-[Salmonella-enterica-subsp.-enterica] |
| ECI7333734.1 | OLD-ABC | GCA_007783945.1 | Salmonella_enterica_subsp._enterica | Bacteria>Pseudomonadati>Pseudomonadota>Gammaproteobacteria>Enterobacterales>Enterobacteriaceae>Salmonella>Salmonella_enterica | 363 | EZ611_10905 | 103250..104341 | 59201 | ATP-binding-protein-[Salmonella-enterica-subsp.-enterica] |
| ECR2528672.1 | OLD-ABC | GCA_008341925.1 | Salmonella_enterica_subsp._enterica | Bacteria>Pseudomonadati>Pseudomonadota>Gammaproteobacteria>Enterobacterales>Enterobacteriaceae>Salmonella>Salmonella_enterica | 363 | F1J42_15855 | 103250..104341 | 59201 | AAA-family-ATPase-[Salmonella-enterica-subsp.-enterica] |
| EDU7470135.1 | OLD-ABC | - | Salmonella_enterica_subsp._enterica | Bacteria>Pseudomonadati>Pseudomonadota>Gammaproteobacteria>Enterobacterales>Enterobacteriaceae>Salmonella>Salmonella_enterica | 363 | CQO18_002338 | 103250..104341 | 59201 | AAA-family-ATPase-[Salmonella-enterica-subsp.-enterica] |
| EDU9161210.1 | OLD-ABC | - | Salmonella_enterica_subsp._enterica | Bacteria>Pseudomonadati>Pseudomonadota>Gammaproteobacteria>Enterobacterales>Enterobacteriaceae>Salmonella>Salmonella_enterica | 363 | IW54_004327 | 6766..7857 | 59201 | AAA-family-ATPase-[Salmonella-enterica-subsp.-enterica] |
| EDU9624697.1 | OLD-ABC | - | Salmonella_enterica_subsp._enterica | Bacteria>Pseudomonadati>Pseudomonadota>Gammaproteobacteria>Enterobacterales>Enterobacteriaceae>Salmonella>Salmonella_enterica | 363 | XW65_002879 | 67169..68260 | 59201 | AAA-family-ATPase-[Salmonella-enterica-subsp.-enterica] |
| EDV4604929.1 | OLD-ABC | - | Salmonella_enterica_subsp._enterica | Bacteria>Pseudomonadati>Pseudomonadota>Gammaproteobacteria>Enterobacterales>Enterobacteriaceae>Salmonella>Salmonella_enterica | 363 | ZP62_000971 | 67608..68699 | 59201 | AAA-family-ATPase-[Salmonella-enterica-subsp.-enterica] |
| EDZ7084222.1 | OLD-ABC | - | Salmonella_enterica_subsp._enterica | Bacteria>Pseudomonadati>Pseudomonadota>Gammaproteobacteria>Enterobacterales>Enterobacteriaceae>Salmonella>Salmonella_enterica | 363 | GSO86_003558 | 10179..11270 | 59201 | AAA-family-ATPase-[Salmonella-enterica] |
| EEC1070528.1 | OLD-ABC | - | Salmonella_enterica_subsp._enterica | Bacteria>Pseudomonadati>Pseudomonadota>Gammaproteobacteria>Enterobacterales>Enterobacteriaceae>Salmonella>Salmonella_enterica | 363 | AH545_000140 | 147353..148444 | 59201 | AAA-family-ATPase-[Salmonella-enterica-subsp.-enterica] |
| HAT0208694.1 | OLD-ABC | GCA_015882745.1 | Salmonella_enterica_subsp._enterica | Bacteria>Pseudomonadati>Pseudomonadota>Gammaproteobacteria>Enterobacterales>Enterobacteriaceae>Salmonella>Salmonella_enterica | 363 | I8Q75_000458 | 70251..71342 | 59201 | TPA:-AAA-family-ATPase-[Salmonella-enterica-subsp.-enterica] |
| HAV0565025.1 | OLD-ABC | GCA_016249375.1 | Salmonella_enterica_subsp._enterica | Bacteria>Pseudomonadati>Pseudomonadota>Gammaproteobacteria>Enterobacterales>Enterobacteriaceae>Salmonella>Salmonella_enterica | 363 | JF300_000746 | 84559..85650 | 59201 | TPA:-AAA-family-ATPase-[Salmonella-enterica-subsp.-enterica] |
| SUG73764.1 | OLD-ABC | GCA_900456435.1 | Salmonella_enterica_subsp._enterica | Bacteria>Pseudomonadati>Pseudomonadota>Gammaproteobacteria>Enterobacterales>Enterobacteriaceae>Salmonella>Salmonella_enterica | 363 | NCTC5798_05053 | 4081615..4082706 | 59201 | spermidine/putrescine-ABC-transporter-ATPase-[Salmonella-enterica-subsp.-enterica] |
| EAB6416575.1 | OLD-ABC | GCA_004256685.1 | Salmonella_enterica_subsp._enterica | Bacteria>Pseudomonadati>Pseudomonadota>Gammaproteobacteria>Enterobacterales>Enterobacteriaceae>Salmonella>Salmonella_enterica | 362 | D6S13_10920 | 59661..60749 | 59201 | ATP-binding-protein-[Salmonella-enterica-subsp.-enterica] |
| EAB9078798.1 | OLD-ABC | GCA_004265565.1 | Salmonella_enterica_subsp._enterica | Bacteria>Pseudomonadati>Pseudomonadota>Gammaproteobacteria>Enterobacterales>Enterobacteriaceae>Salmonella>Salmonella_enterica | 362 | AAL03_01520 | 59733..60821 | 59201 | ATP-binding-protein-[Salmonella-enterica-subsp.-enterica] |
| EAC1236047.1 | OLD-ABC | GCA_004276135.1 | Salmonella_enterica_subsp._enterica | Bacteria>Pseudomonadati>Pseudomonadota>Gammaproteobacteria>Enterobacterales>Enterobacteriaceae>Salmonella>Salmonella_enterica | 362 | DTG84_00820 | 165298..166386 | 59201 | ATP-binding-protein-[Salmonella-enterica-subsp.-enterica] |
| ECG1483124.1 | OLD-ABC | GCA_007649655.1 | Salmonella_enterica_subsp._enterica | Bacteria>Pseudomonadati>Pseudomonadota>Gammaproteobacteria>Enterobacterales>Enterobacteriaceae>Salmonella>Salmonella_enterica | 362 | DT144_20830 | 59780..60868 | 59201 | ATP-binding-protein-[Salmonella-enterica-subsp.-enterica] |
| ECI3716682.1 | OLD-ABC | GCA_007768905.1 | Salmonella_enterica_subsp._enterica | Bacteria>Pseudomonadati>Pseudomonadota>Gammaproteobacteria>Enterobacterales>Enterobacteriaceae>Salmonella>Salmonella_enterica | 362 | DK077_11330 | 51767..52855 | 59201 | ATP-binding-protein-[Salmonella-enterica-subsp.-enterica] |
| ECI6683603.1 | OLD-ABC | GCA_007781825.1 | Salmonella_enterica_subsp._enterica | Bacteria>Pseudomonadati>Pseudomonadota>Gammaproteobacteria>Enterobacterales>Enterobacteriaceae>Salmonella>Salmonella_enterica | 362 | EHE65_20735 | 59471..60559 | 59201 | ATP-binding-protein-[Salmonella-enterica-subsp.-enterica] |
| EDS5421360.1 | OLD-ABC | - | Salmonella_enterica_subsp._enterica | Bacteria>Pseudomonadati>Pseudomonadota>Gammaproteobacteria>Enterobacterales>Enterobacteriaceae>Salmonella>Salmonella_enterica | 362 | ZH17_000158 | 165611..166699 | 59201 | ATP-binding-protein-[Salmonella-enterica-subsp.-enterica] |
| EDT2838944.1 | OLD-ABC | - | Salmonella_enterica_subsp._enterica | Bacteria>Pseudomonadati>Pseudomonadota>Gammaproteobacteria>Enterobacterales>Enterobacteriaceae>Salmonella>Salmonella_enterica | 362 | S958_003411 | 10003..11091 | 59201 | ATP-binding-protein-[Salmonella-enterica-subsp.-enterica] |
| EDV5116992.1 | OLD-ABC | - | Salmonella_enterica_subsp._enterica | Bacteria>Pseudomonadati>Pseudomonadota>Gammaproteobacteria>Enterobacterales>Enterobacteriaceae>Salmonella>Salmonella_enterica | 362 | ZJ36_000827 | 36282..37370 | 59201 | ATP-binding-protein-[Salmonella-enterica-subsp.-enterica] |
| EDV9109400.1 | OLD-ABC | - | Salmonella_enterica_subsp._enterica | Bacteria>Pseudomonadati>Pseudomonadota>Gammaproteobacteria>Enterobacterales>Enterobacteriaceae>Salmonella>Salmonella_enterica | 362 | ZK68_001602 | 59359..60447 | 59201 | ATP-binding-protein-[Salmonella-enterica-subsp.-enterica] |
| EDW0696247.1 | OLD-ABC | - | Salmonella_enterica_subsp._enterica | Bacteria>Pseudomonadati>Pseudomonadota>Gammaproteobacteria>Enterobacterales>Enterobacteriaceae>Salmonella>Salmonella_enterica | 362 | KV01_000537 | 157729..158817 | 59201 | ATP-binding-protein-[Salmonella-enterica-subsp.-enterica] |
| EDW4488617.1 | OLD-ABC | - | Salmonella_enterica_subsp._enterica | Bacteria>Pseudomonadati>Pseudomonadota>Gammaproteobacteria>Enterobacterales>Enterobacteriaceae>Salmonella>Salmonella_enterica | 362 | YV39_000489 | 173433..174521 | 59201 | ATP-binding-protein-[Salmonella-enterica-subsp.-enterica] |
| EEC0292873.1 | OLD-ABC | - | Salmonella_enterica_subsp._enterica | Bacteria>Pseudomonadati>Pseudomonadota>Gammaproteobacteria>Enterobacterales>Enterobacteriaceae>Salmonella>Salmonella_enterica | 362 | YV76_000181 | 193750..194838 | 59201 | ATP-binding-protein-[Salmonella-enterica-subsp.-enterica] |
| EEJ3565317.1 | OLD-ABC | - | Salmonella_enterica_subsp._enterica | Bacteria>Pseudomonadati>Pseudomonadota>Gammaproteobacteria>Enterobacterales>Enterobacteriaceae>Salmonella>Salmonella_enterica | 362 | AA437_001846 | 133073..134161 | 59201 | ATP-binding-protein-[Salmonella-enterica-subsp.-enterica] |
| EEJ7235530.1 | OLD-ABC | - | Salmonella_enterica_subsp._salamae | Bacteria>Pseudomonadati>Pseudomonadota>Gammaproteobacteria>Enterobacterales>Enterobacteriaceae>Salmonella>Salmonella_enterica | 362 | S483_003521 | 159760..160848 | 59202 | ATP-binding-protein-[Salmonella-enterica-subsp.-salamae] |
| EHA8894995.1 | OLD-ABC | - | Salmonella_enterica_subsp._enterica | Bacteria>Pseudomonadati>Pseudomonadota>Gammaproteobacteria>Enterobacterales>Enterobacteriaceae>Salmonella>Salmonella_enterica | 362 | JHW15_001256 | 51741..52829 | 59201 | ATP-binding-protein-[Salmonella-enterica-subsp.-enterica] |
| MLQ35366.1 | OLD-ABC | GCA_003910355.1 | Salmonella_enterica_subsp._enterica | Bacteria>Pseudomonadati>Pseudomonadota>Gammaproteobacteria>Enterobacterales>Enterobacteriaceae>Salmonella>Salmonella_enterica | 362 | DRM12_04730 | 257119..258207 | 59201 | ATP-binding-protein-[Salmonella-enterica-subsp.-enterica] |
| AZT73893.1 | OLD-ABC | GCA_003994735.1 | Salmonella_enterica_subsp._enterica_serovar_Waycross | Bacteria>Pseudomonadati>Pseudomonadota>Gammaproteobacteria>Enterobacterales>Enterobacteriaceae>Salmonella>Salmonella_enterica>Salmonella_enterica_subsp._enterica | 373 | ELZ91_19530 | 3958717..3959838 | 58101 | ABC-transporter-ATP-binding-protein-[Salmonella-enterica-subsp.-enterica-serovar-Waycross] |
| EBU9347431.1 | OLD-ABC | GCA_007149485.1 | Salmonella_enterica_subsp._enterica_serovar_Angoda | Bacteria>Pseudomonadati>Pseudomonadota>Gammaproteobacteria>Enterobacterales>Enterobacteriaceae>Salmonella>Salmonella_enterica>Salmonella_enterica_subsp._enterica | 373 | DMU95_04680 | 158417..159538 | 2564159 | ABC-transporter-ATP-binding-protein-[Salmonella-enterica-subsp.-enterica-serovar-Angoda] |
| ECF0142191.1 | OLD-ABC | GCA_007595955.1 | Salmonella_enterica_subsp._enterica_serovar_Monschaui | Bacteria>Pseudomonadati>Pseudomonadota>Gammaproteobacteria>Enterobacterales>Enterobacteriaceae>Salmonella>Salmonella_enterica>Salmonella_enterica_subsp._enterica | 373 | EX931_14165 | 76995..78116 | 29480 | ABC-transporter-ATP-binding-protein-[Salmonella-enterica-subsp.-enterica-serovar-Monschaui] |
| ECY3255617.1 | OLD-ABC | GCA_008731355.1 | Salmonella_enterica_subsp._enterica_serovar_Alachua | Bacteria>Pseudomonadati>Pseudomonadota>Gammaproteobacteria>Enterobacterales>Enterobacteriaceae>Salmonella>Salmonella_enterica>Salmonella_enterica_subsp._enterica | 373 | AU577_03150 | 24378..25499 | 913240 | AAA-family-ATPase-[Salmonella-enterica-subsp.-enterica-serovar-Alachua] |
| EDN6133535.1 | OLD-ABC | - | Salmonella_enterica_subsp._enterica_serovar_Kentucky | Bacteria>Pseudomonadati>Pseudomonadota>Gammaproteobacteria>Enterobacterales>Enterobacteriaceae>Salmonella>Salmonella_enterica>Salmonella_enterica_subsp._enterica | 373 | GJQ78_12620 | 69697..70818 | 192955 | AAA-family-ATPase-[Salmonella-enterica-subsp.-enterica-serovar-Kentucky] |
| EDS3575401.1 | OLD-ABC | - | Salmonella_enterica_subsp._enterica_serovar_Sangera | Bacteria>Pseudomonadati>Pseudomonadota>Gammaproteobacteria>Enterobacterales>Enterobacteriaceae>Salmonella>Salmonella_enterica>Salmonella_enterica_subsp._enterica | 373 | GR179_002520 | 10718..11839 | 1173574 | AAA-family-ATPase-[Salmonella-enterica-subsp.-enterica] |
| EEI5362887.1 | OLD-ABC | - | Salmonella_enterica_subsp._enterica_serovar_Kentucky | Bacteria>Pseudomonadati>Pseudomonadota>Gammaproteobacteria>Enterobacterales>Enterobacteriaceae>Salmonella>Salmonella_enterica>Salmonella_enterica_subsp._enterica | 373 | GXM95_001565 | 180692..181813 | 192955 | AAA-family-ATPase-[Salmonella-enterica-subsp.-enterica-serovar-Kentucky] |
| EEM2455577.1 | OLD-ABC | - | Salmonella_enterica_subsp._enterica_serovar_Kentucky | Bacteria>Pseudomonadati>Pseudomonadota>Gammaproteobacteria>Enterobacterales>Enterobacteriaceae>Salmonella>Salmonella_enterica>Salmonella_enterica_subsp._enterica | 373 | CWG70_02785 | 181025..182146 | 192955 | AAA-family-ATPase-[Salmonella-enterica-subsp.-enterica-serovar-Kentucky] |
| EHJ7117735.1 | OLD-ABC | GCA_017926475.1 | Salmonella_enterica_subsp._enterica_serovar_Kentucky | Bacteria>Pseudomonadati>Pseudomonadota>Gammaproteobacteria>Enterobacterales>Enterobacteriaceae>Salmonella>Salmonella_enterica>Salmonella_enterica_subsp._enterica | 373 | KAP12_002967 | 419648..420769 | 192955 | AAA-family-ATPase-[Salmonella-enterica-subsp.-enterica-serovar-Kentucky] |
| UTZ95784.1 | OLD-ABC | GCA_024442275.1 | Salmonella_enterica_subsp._enterica_serovar_Kentucky | Bacteria>Pseudomonadati>Pseudomonadota>Gammaproteobacteria>Enterobacterales>Enterobacteriaceae>Salmonella>Salmonella_enterica>Salmonella_enterica_subsp._enterica | 373 | NOQ95_21570 | 4455794..4456915 | 192955 | AAA-family-ATPase-[Salmonella-enterica-subsp.-enterica-serovar-Kentucky] |
| EAA3988070.1 | OLD-ABC | GCA_004177075.1 | Salmonella_enterica_subsp._enterica_serovar_Typhimurium | Bacteria>Pseudomonadati>Pseudomonadota>Gammaproteobacteria>Enterobacterales>Enterobacteriaceae>Salmonella>Salmonella_enterica>Salmonella_enterica_subsp._enterica | 363 | DO557_10285 | 127864..128955 | 90371 | ATP-binding-protein-[Salmonella-enterica-subsp.-enterica] |
| EAA4842520.1 | OLD-ABC | GCA_004183005.1 | Salmonella_enterica_subsp._enterica_serovar_Typhimurium | Bacteria>Pseudomonadati>Pseudomonadota>Gammaproteobacteria>Enterobacterales>Enterobacteriaceae>Salmonella>Salmonella_enterica>Salmonella_enterica_subsp._enterica | 363 | DPP49_10715 | 67480..68571 | 90371 | ATP-binding-protein-[Salmonella-enterica-subsp.-enterica] |
| EAA6168128.1 | OLD-ABC | GCA_004188635.1 | Salmonella_enterica_subsp._enterica_serovar_Ohio | Bacteria>Pseudomonadati>Pseudomonadota>Gammaproteobacteria>Enterobacterales>Enterobacteriaceae>Salmonella>Salmonella_enterica>Salmonella_enterica_subsp._enterica | 363 | DQZ68_14455 | 19842..20933 | 117541 | ATP-binding-protein-[Salmonella-enterica-subsp.-enterica] |
| EAA7109617.1 | OLD-ABC | GCA_004192185.1 | Salmonella_enterica_subsp._enterica_serovar_Ouagadougou | Bacteria>Pseudomonadati>Pseudomonadota>Gammaproteobacteria>Enterobacterales>Enterobacteriaceae>Salmonella>Salmonella_enterica>Salmonella_enterica_subsp._enterica | 363 | DSE94_16640 | 59503..60594 | 2564899 | ATP-binding-protein-[Salmonella-enterica-subsp.-enterica] |

| Protein-ID | Domain | Assembly-ID | Strain | Taxonomy | Length | Locus-Tag | Genomic-coordinates | TaxID | Fasta-defline |
|---|---|---|---|---|---|---|---|---|---|
| EAA7310019.1 | OLD-ABC | GCA_004192645.1 | Salmonella_enterica_subsp._enterica_serovar_Duesseldorf | Bacteria>Pseudomonadati>Pseudomonadota>Gammaproteobacteria>Enterobacterales>Enterobacteriaceae>Salmonella>Salmonella_enterica>Salmonella_enterica_subsp._enterica | 363 | TL61_10435 | 163357..164448 | 322832 | ATP-binding-cassette-domain-containing-protein-[Salmonella-enterica-subsp.-enterica] |
| EAA8523363.1 | OLD-ABC | GCA_004222265.1 | Salmonella_enterica_subsp._enterica_serovar_Cerro | Bacteria>Pseudomonadati>Pseudomonadota>Gammaproteobacteria>Enterobacterales>Enterobacteriaceae>Salmonella>Salmonella_enterica>Salmonella_enterica_subsp._enterica | 363 | ADQ53_00825 | 168355..169446 | 340188 | ATP-binding-protein-[Salmonella-enterica-subsp.-enterica] |
| EAB9374477.1 | OLD-ABC | GCA_004270415.1 | Salmonella_enterica_subsp._enterica_serovar_Llandoff | Bacteria>Pseudomonadati>Pseudomonadota>Gammaproteobacteria>Enterobacterales>Enterobacteriaceae>Salmonella>Salmonella_enterica>Salmonella_enterica_subsp._enterica | 363 | D7N03_15210 | 68047..69138 | 2564701 | ATP-binding-protein-[Salmonella-enterica-subsp.-enterica] |
| EAC0120239.1 | OLD-ABC | GCA_004271765.1 | Salmonella_enterica_subsp._enterica_serovar_Typhimurium | Bacteria>Pseudomonadati>Pseudomonadota>Gammaproteobacteria>Enterobacterales>Enterobacteriaceae>Salmonella>Salmonella_enterica>Salmonella_enterica_subsp._enterica | 363 | EBO89_06935 | 65555..66646 | 90371 | ATP-binding-protein-[Salmonella-enterica-subsp.-enterica] |
| EAC1009597.1 | OLD-ABC | GCA_004275715.1 | Salmonella_enterica_subsp._enterica_serovar_Jangwani | Bacteria>Pseudomonadati>Pseudomonadota>Gammaproteobacteria>Enterobacterales>Enterobacteriaceae>Salmonella>Salmonella_enterica>Salmonella_enterica_subsp._enterica | 363 | YR14_13810 | 157771..158862 | 2564577 | ATP-binding-cassette-domain-containing-protein-[Salmonella-enterica-subsp.-enterica] |
| EAO4363251.1 | OLD-ABC | GCA_005505735.1 | Salmonella_enterica_subsp._enterica_serovar_Bere | Bacteria>Pseudomonadati>Pseudomonadota>Gammaproteobacteria>Enterobacterales>Enterobacteriaceae>Salmonella>Salmonella_enterica>Salmonella_enterica_subsp._enterica | 363 | E5197_09140 | 182722..183813 | 2024271 | ATP-binding-protein-[Salmonella-enterica] |
| EBB4495257.1 | OLD-ABC | GCA_006143655.1 | Salmonella_enterica_subsp._enterica_serovar_Typhimurium | Bacteria>Pseudomonadati>Pseudomonadota>Gammaproteobacteria>Enterobacterales>Enterobacteriaceae>Salmonella>Salmonella_enterica>Salmonella_enterica_subsp._enterica | 363 | CET45_10240 | 178282..179373 | 90371 | ATP-binding-protein-[Salmonella-enterica-subsp.-enterica-serovar-Typhimurium] |
| EBL4178055.1 | OLD-ABC | GCA_006685525.1 | Salmonella_enterica_subsp._enterica_serovar_Typhimurium | Bacteria>Pseudomonadati>Pseudomonadota>Gammaproteobacteria>Enterobacterales>Enterobacteriaceae>Salmonella>Salmonella_enterica>Salmonella_enterica_subsp._enterica | 363 | E5L78_02835 | 25528..26619 | 90371 | AAA-family-ATPase-[Salmonella-enterica-subsp.-enterica-serovar-Typhimurium] |
| EBL6294789.1 | OLD-ABC | GCA_006694585.1 | Salmonella_enterica_subsp._enterica_serovar_Typhimurium | Bacteria>Pseudomonadati>Pseudomonadota>Gammaproteobacteria>Enterobacterales>Enterobacteriaceae>Salmonella>Salmonella_enterica>Salmonella_enterica_subsp._enterica | 363 | AC864_13985 | 176999..178090 | 90371 | AAA-family-ATPase-[Salmonella-enterica-subsp.-enterica-serovar-Typhimurium] |
| EBR9945163.1 | OLD-ABC | GCA_006983125.1 | Salmonella_enterica_subsp._enterica_serovar_Krefeld | Bacteria>Pseudomonadati>Pseudomonadota>Gammaproteobacteria>Enterobacterales>Enterobacteriaceae>Salmonella>Salmonella_enterica>Salmonella_enterica_subsp._enterica | 363 | DTF03_11155 | 160770..161861 | 1242084 | ATP-binding-protein-[Salmonella-enterica-subsp.-enterica-serovar-Krefeld] |
| EBS4429084.1 | OLD-ABC | GCA_007008305.1 | Salmonella_enterica_subsp._enterica_serovar_Poona | Bacteria>Pseudomonadati>Pseudomonadota>Gammaproteobacteria>Enterobacterales>Enterobacteriaceae>Salmonella>Salmonella_enterica>Salmonella_enterica_subsp._enterica | 363 | DQJ97_07375 | 33258..34349 | 436295 | ATP-binding-cassette-domain-containing-protein-[Salmonella-enterica-subsp.-enterica-serovar-Poona] |
| EBU6515331.1 | OLD-ABC | GCA_007137465.1 | Salmonella_enterica_subsp._enterica_serovar_Typhimurium | Bacteria>Pseudomonadati>Pseudomonadota>Gammaproteobacteria>Enterobacterales>Enterobacteriaceae>Salmonella>Salmonella_enterica>Salmonella_enterica_subsp._enterica | 363 | DKJ21_01480 | 12178..13269 | 90371 | hypothetical-protein-DKJ21_01480-[Salmonella-enterica-subsp.-enterica-serovar-Typhimurium] |
| EBU6749121.1 | OLD-ABC | GCA_007137985.1 | Salmonella_enterica_subsp._enterica_serovar_Typhimurium | Bacteria>Pseudomonadati>Pseudomonadota>Gammaproteobacteria>Enterobacterales>Enterobacteriaceae>Salmonella>Salmonella_enterica>Salmonella_enterica_subsp._enterica | 363 | DKS55_01045 | 6350..7441 | 90371 | hypothetical-protein-DKS55_01045-[Salmonella-enterica-subsp.-enterica-serovar-Typhimurium] |
| EBU9878491.1 | OLD-ABC | GCA_007152025.1 | Salmonella_enterica_subsp._enterica_serovar_Typhimurium | Bacteria>Pseudomonadati>Pseudomonadota>Gammaproteobacteria>Enterobacterales>Enterobacteriaceae>Salmonella>Salmonella_enterica>Salmonella_enterica_subsp._enterica | 363 | DM361_04540 | 67579..68670 | 90371 | hypothetical-protein-DM361_04540-[Salmonella-enterica-subsp.-enterica-serovar-Typhimurium] |
| EBV0268832.1 | OLD-ABC | GCA_007152865.1 | Salmonella_enterica_subsp._enterica_serovar_Typhimurium | Bacteria>Pseudomonadati>Pseudomonadota>Gammaproteobacteria>Enterobacterales>Enterobacteriaceae>Salmonella>Salmonella_enterica>Salmonella_enterica_subsp._enterica | 363 | DNH74_05355 | 67365..68456 | 90371 | hypothetical-protein-DNH74_05355-[Salmonella-enterica-subsp.-enterica-serovar-Typhimurium] |
| EBV5450233.1 | OLD-ABC | GCA_007175875.1 | Salmonella_enterica_subsp._enterica_serovar_Typhimurium | Bacteria>Pseudomonadati>Pseudomonadota>Gammaproteobacteria>Enterobacterales>Enterobacteriaceae>Salmonella>Salmonella_enterica>Salmonella_enterica_subsp._enterica | 363 | DO810_06280 | 67583..68674 | 90371 | ATP-binding-protein-[Salmonella-enterica-subsp.-enterica-serovar-Typhimurium] |
| EBV7110756.1 | OLD-ABC | GCA_007180535.1 | Salmonella_enterica_subsp._enterica_serovar_Ohio | Bacteria>Pseudomonadati>Pseudomonadota>Gammaproteobacteria>Enterobacterales>Enterobacteriaceae>Salmonella>Salmonella_enterica>Salmonella_enterica_subsp._enterica | 363 | YY60_02800 | 162067..163158 | 117541 | ATP-binding-protein-[Salmonella-enterica-subsp.-enterica-serovar-Ohio] |
| EBW1953458.1 | OLD-ABC | GCA_007201375.1 | Salmonella_enterica_subsp._enterica_serovar_Cerro | Bacteria>Pseudomonadati>Pseudomonadota>Gammaproteobacteria>Enterobacterales>Enterobacteriaceae>Salmonella>Salmonella_enterica>Salmonella_enterica_subsp._enterica | 363 | C3N96_11420 | 51423..52514 | 340188 | ATP-binding-protein-[Salmonella-enterica-subsp.-enterica-serovar-Cerro] |
| EBW2731418.1 | OLD-ABC | GCA_007205375.1 | Salmonella_enterica_subsp._enterica_serovar_Typhimurium | Bacteria>Pseudomonadati>Pseudomonadota>Gammaproteobacteria>Enterobacterales>Enterobacteriaceae>Salmonella>Salmonella_enterica>Salmonella_enterica_subsp._enterica | 363 | DPA65_05345 | 178299..179390 | 90371 | ATP-binding-protein-[Salmonella-enterica-subsp.-enterica-serovar-Typhimurium] |
| EBW3295110.1 | OLD-ABC | GCA_007207035.1 | Salmonella_enterica_subsp._enterica_serovar_Cerro | Bacteria>Pseudomonadati>Pseudomonadota>Gammaproteobacteria>Enterobacterales>Enterobacteriaceae>Salmonella>Salmonella_enterica>Salmonella_enterica_subsp._enterica | 363 | DPE09_02370 | 51422..52513 | 340188 | ATP-binding-protein-[Salmonella-enterica-subsp.-enterica-serovar-Cerro] |
| EBY1767817.1 | OLD-ABC | GCA_007283275.1 | Salmonella_enterica_subsp._enterica_serovar_Georgia | Bacteria>Pseudomonadati>Pseudomonadota>Gammaproteobacteria>Enterobacterales>Enterobacteriaceae>Salmonella>Salmonella_enterica>Salmonella_enterica_subsp._enterica | 363 | DU102_16115 | 57982..59073 | 2564467 | ATP-binding-protein-[Salmonella-enterica-subsp.-enterica-serovar-Georgia] |
| EBZ6440066.1 | OLD-ABC | GCA_007344875.1 | Salmonella_enterica_subsp._enterica_serovar_Typhimurium | Bacteria>Pseudomonadati>Pseudomonadota>Gammaproteobacteria>Enterobacterales>Enterobacteriaceae>Salmonella>Salmonella_enterica>Salmonella_enterica_subsp._enterica | 363 | ED482_12675 | 6419..7510 | 90371 | ATP-binding-protein-[Salmonella-enterica-subsp.-enterica-serovar-Typhimurium] |
| ECF4176039.1 | OLD-ABC | GCA_007613175.1 | Salmonella_enterica_subsp._enterica_serovar_Typhimurium | Bacteria>Pseudomonadati>Pseudomonadota>Gammaproteobacteria>Enterobacterales>Enterobacteriaceae>Salmonella>Salmonella_enterica>Salmonella_enterica_subsp._enterica | 363 | E5U03_12945 | 103250..104341 | 90371 | ATP-binding-protein-[Salmonella-enterica-subsp.-enterica-serovar-Typhimurium] |
| ECI2073656.1 | OLD-ABC | GCA_007762095.1 | Salmonella_enterica_subsp._enterica_serovar_Kentucky | Bacteria>Pseudomonadati>Pseudomonadota>Gammaproteobacteria>Enterobacterales>Enterobacteriaceae>Salmonella>Salmonella_enterica>Salmonella_enterica_subsp._enterica | 363 | AUD11_09535 | 103250..104341 | 192955 | AAA-family-ATPase-[Salmonella-enterica-subsp.-enterica-serovar-Kentucky] |
| ECK5773202.1 | OLD-ABC | GCA_007886645.1 | Salmonella_enterica_subsp._enterica_serovar_Typhimurium | Bacteria>Pseudomonadati>Pseudomonadota>Gammaproteobacteria>Enterobacterales>Enterobacteriaceae>Salmonella>Salmonella_enterica>Salmonella_enterica_subsp._enterica | 363 | AXS32_00825 | 179941..181032 | 90371 | AAA-family-ATPase-[Salmonella-enterica-subsp.-enterica-serovar-Typhimurium] |
| ECM3029730.1 | OLD-ABC | GCA_008020915.1 | Salmonella_enterica_subsp._enterica_serovar_Typhimurium | Bacteria>Pseudomonadati>Pseudomonadota>Gammaproteobacteria>Enterobacterales>Enterobacteriaceae>Salmonella>Salmonella_enterica>Salmonella_enterica_subsp._enterica | 363 | QB31_08415 | 67591..68682 | 90371 | AAA-family-ATPase-[Salmonella-enterica-subsp.-enterica-serovar-Typhimurium] |
| ECM3363967.1 | OLD-ABC | GCA_008021815.1 | Salmonella_enterica_subsp._enterica_serovar_Typhimurium | Bacteria>Pseudomonadati>Pseudomonadota>Gammaproteobacteria>Enterobacterales>Enterobacteriaceae>Salmonella>Salmonella_enterica>Salmonella_enterica_subsp._enterica | 363 | PW85_07010 | 103250..104341 | 90371 | AAA-family-ATPase-[Salmonella-enterica-subsp.-enterica-serovar-Typhimurium] |
| ECN0902566.1 | OLD-ABC | GCA_008061715.1 | Salmonella_enterica_subsp._enterica_serovar_Typhimurium | Bacteria>Pseudomonadati>Pseudomonadota>Gammaproteobacteria>Enterobacterales>Enterobacteriaceae>Salmonella>Salmonella_enterica>Salmonella_enterica_subsp._enterica | 363 | YR82_02415 | 67366..68457 | 90371 | AAA-family-ATPase-[Salmonella-enterica-subsp.-enterica-serovar-Typhimurium] |
| ECO0332140.1 | OLD-ABC | GCA_008112305.1 | Salmonella_enterica_subsp._enterica_serovar_Typhimurium | Bacteria>Pseudomonadati>Pseudomonadota>Gammaproteobacteria>Enterobacterales>Enterobacteriaceae>Salmonella>Salmonella_enterica>Salmonella_enterica_subsp._enterica | 363 | ZS66_00465 | 103250..104341 | 90371 | AAA-family-ATPase-[Salmonella-enterica-subsp.-enterica-serovar-Typhimurium] |
| ECU0288389.1 | OLD-ABC | GCA_008492025.1 | Salmonella_enterica_subsp._enterica_serovar_Typhimurium | Bacteria>Pseudomonadati>Pseudomonadota>Gammaproteobacteria>Enterobacterales>Enterobacteriaceae>Salmonella>Salmonella_enterica>Salmonella_enterica_subsp._enterica | 363 | C7B88_00720 | 147389..148480 | 90371 | ATP-binding-protein-[Salmonella-enterica-subsp.-enterica-serovar-Typhimurium] |
| ECU9351202.1 | OLD-ABC | GCA_008530195.1 | Salmonella_enterica_subsp._enterica_serovar_Typhimurium | Bacteria>Pseudomonadati>Pseudomonadota>Gammaproteobacteria>Enterobacterales>Enterobacteriaceae>Salmonella>Salmonella_enterica>Salmonella_enterica_subsp._enterica | 363 | CIH13_12050 | 178393..179484 | 90371 | ATP-binding-protein-[Salmonella-enterica-subsp.-enterica-serovar-Typhimurium] |
| ECV7778491.1 | OLD-ABC | GCA_008576505.1 | Salmonella_enterica_subsp._enterica_serovar_Typhimurium | Bacteria>Pseudomonadati>Pseudomonadota>Gammaproteobacteria>Enterobacterales>Enterobacteriaceae>Salmonella>Salmonella_enterica>Salmonella_enterica_subsp._enterica | 363 | ZX52_02600 | 67366..68457 | 90371 | AAA-family-ATPase-[Salmonella-enterica-subsp.-enterica-serovar-Typhimurium] |
| ECY8229163.1 | OLD-ABC | GCA_008752395.1 | Salmonella_enterica_subsp._enterica_serovar_Stanley | Bacteria>Pseudomonadati>Pseudomonadota>Gammaproteobacteria>Enterobacterales>Enterobacteriaceae>Salmonella>Salmonella_enterica>Salmonella_enterica_subsp._enterica | 363 | AU695_08290 | 168031..169122 | 192953 | AAA-family-ATPase-[Salmonella-enterica-subsp.-enterica-serovar-Stanley] |
| ECZ9372115.1 | OLD-ABC | GCA_008834045.1 | Salmonella_enterica_subsp._enterica_serovar_Typhimurium | Bacteria>Pseudomonadati>Pseudomonadota>Gammaproteobacteria>Enterobacterales>Enterobacteriaceae>Salmonella>Salmonella_enterica>Salmonella_enterica_subsp._enterica | 363 | A3U95_10235 | 103250..104341 | 90371 | AAA-family-ATPase-[Salmonella-enterica-subsp.-enterica-serovar-Typhimurium] |
| EDA5898038.1 | OLD-ABC | GCA_008864045.1 | Salmonella_enterica_subsp._enterica_serovar_Poona | Bacteria>Pseudomonadati>Pseudomonadota>Gammaproteobacteria>Enterobacterales>Enterobacteriaceae>Salmonella>Salmonella_enterica>Salmonella_enterica_subsp._enterica | 363 | F9X73_05945 | 33248..34339 | 436295 | AAA-family-ATPase-[Salmonella-enterica-subsp.-enterica] |
| EDA7420102.1 | OLD-ABC | - | Salmonella_enterica_subsp._enterica_serovar_Typhimurium | Bacteria>Pseudomonadati>Pseudomonadota>Gammaproteobacteria>Enterobacterales>Enterobacteriaceae>Salmonella>Salmonella_enterica>Salmonella_enterica_subsp._enterica | 363 | A3U84_08770 | 67583..68674 | 90371 | AAA-family-ATPase-[Salmonella-enterica-subsp.-enterica-serovar-Typhimurium] |
| EDB9656970.1 | OLD-ABC | - | Salmonella_enterica_subsp._enterica_serovar_Typhimurium | Bacteria>Pseudomonadati>Pseudomonadota>Gammaproteobacteria>Enterobacterales>Enterobacteriaceae>Salmonella>Salmonella_enterica>Salmonella_enterica_subsp._enterica | 363 | BEO39_09555 | 34819..35910 | 90371 | AAA-family-ATPase-[Salmonella-enterica-subsp.-enterica-serovar-Typhimurium] |
| EDE2293302.1 | OLD-ABC | - | Salmonella_enterica_subsp._enterica_serovar_Typhimurium | Bacteria>Pseudomonadati>Pseudomonadota>Gammaproteobacteria>Enterobacterales>Enterobacteriaceae>Salmonella>Salmonella_enterica>Salmonella_enterica_subsp._enterica | 363 | GBE85_07245 | 103250..104341 | 90371 | AAA-family-ATPase-[Salmonella-enterica] |
| EDI5281732.1 | OLD-ABC | - | Salmonella_enterica_subsp._enterica_serovar_Typhimurium | Bacteria>Pseudomonadati>Pseudomonadota>Gammaproteobacteria>Enterobacterales>Enterobacteriaceae>Salmonella>Salmonella_enterica>Salmonella_enterica_subsp._enterica | 363 | CEM90_10185 | 178182..179273 | 90371 | ATP-binding-protein-[Salmonella-enterica-subsp.-enterica-serovar-Typhimurium] |
| EDL1929629.1 | OLD-ABC | - | Salmonella_enterica_subsp._enterica_serovar_Typhimurium | Bacteria>Pseudomonadati>Pseudomonadota>Gammaproteobacteria>Enterobacterales>Enterobacteriaceae>Salmonella>Salmonella_enterica>Salmonella_enterica_subsp._enterica | 363 | CQD92_11440 | 103250..104341 | 90371 | ATP-binding-protein-[Salmonella-enterica-subsp.-enterica-serovar-Typhimurium] |
| EDL2092521.1 | OLD-ABC | - | Salmonella_enterica_subsp._enterica_serovar_Typhimurium | Bacteria>Pseudomonadati>Pseudomonadota>Gammaproteobacteria>Enterobacterales>Enterobacteriaceae>Salmonella>Salmonella_enterica>Salmonella_enterica_subsp._enterica | 363 | CQF25_02845 | 67599..68690 | 90371 | ATP-binding-protein-[Salmonella-enterica-subsp.-enterica-serovar-Typhimurium] |
| EDM4126307.1 | OLD-ABC | - | Salmonella_enterica_subsp._enterica_serovar_Typhimurium | Bacteria>Pseudomonadati>Pseudomonadota>Gammaproteobacteria>Enterobacterales>Enterobacteriaceae>Salmonella>Salmonella_enterica>Salmonella_enterica_subsp._enterica | 363 | CR810_06745 | 179942..181033 | 90371 | ATP-binding-protein-[Salmonella-enterica-subsp.-enterica-serovar-Typhimurium] |
| EDM6640282.1 | OLD-ABC | - | Salmonella_enterica_subsp._enterica_serovar_Typhimurium | Bacteria>Pseudomonadati>Pseudomonadota>Gammaproteobacteria>Enterobacterales>Enterobacteriaceae>Salmonella>Salmonella_enterica>Salmonella_enterica_subsp._enterica | 363 | CUM00_00820 | 178281..179372 | 90371 | ATP-binding-protein-[Salmonella-enterica-subsp.-enterica-serovar-Typhimurium] |
| EDT5820086.1 | OLD-ABC | - | Salmonella_enterica_subsp._enterica_serovar_4,[5],12:i:- | Bacteria>Pseudomonadati>Pseudomonadota>Gammaproteobacteria>Enterobacterales>Enterobacteriaceae>Salmonella>Salmonella_enterica>Salmonella_enterica_subsp._enterica | 363 | B7921_002248 | 103250..104341 | 440524 | AAA-family-ATPase-[Salmonella-enterica-subsp.-enterica-serovar-4,[5] |
| EDU3766994.1 | OLD-ABC | - | Salmonella_enterica_subsp._enterica_serovar_Minnesota | Bacteria>Pseudomonadati>Pseudomonadota>Gammaproteobacteria>Enterobacterales>Enterobacteriaceae>Salmonella>Salmonella_enterica>Salmonella_enterica_subsp._enterica | 363 | C4O09_000359 | 398123..399214 | 70803 | AAA-family-ATPase-[Salmonella-enterica-subsp.-enterica-serovar-Minnesota] |
| EDV2418010.1 | OLD-ABC | - | Salmonella_enterica_subsp._enterica_serovar_Ohio | Bacteria>Pseudomonadati>Pseudomonadota>Gammaproteobacteria>Enterobacterales>Enterobacteriaceae>Salmonella>Salmonella_enterica>Salmonella_enterica_subsp._enterica | 363 | CQ222_000231 | 220938..222029 | 117541 | AAA-family-ATPase-[Salmonella-enterica-subsp.-enterica-serovar-Ohio] |
| EDW2626494.1 | OLD-ABC | - | Salmonella_enterica_subsp._enterica_serovar_Sangalkam | Bacteria>Pseudomonadati>Pseudomonadota>Gammaproteobacteria>Enterobacterales>Enterobacteriaceae>Salmonella>Salmonella_enterica>Salmonella_enterica_subsp._enterica | 363 | XX60_002384 | 164061..165152 | 2564992 | AAA-family-ATPase-[Salmonella-enterica-subsp.-enterica] |
| EDX7895851.1 | OLD-ABC | - | Salmonella_enterica_subsp._enterica_serovar_4,[5],12:i:- | Bacteria>Pseudomonadati>Pseudomonadota>Gammaproteobacteria>Enterobacterales>Enterobacteriaceae>Salmonella>Salmonella_enterica>Salmonella_enterica_subsp._enterica | 363 | C2Y98_001770 | 46026..47117 | 440524 | AAA-family-ATPase-[Salmonella-enterica-subsp.-enterica-serovar-4,[5] |
| EEE6742964.1 | OLD-ABC | - | Salmonella_enterica_subsp._enterica_serovar_Westhampton | Bacteria>Pseudomonadati>Pseudomonadota>Gammaproteobacteria>Enterobacterales>Enterobacteriaceae>Salmonella>Salmonella_enterica>Salmonella_enterica_subsp._enterica | 363 | CBJ05_002022 | 416159..417250 | 356621 | AAA-family-ATPase-[Salmonella-enterica-subsp.-enterica-serovar-Westhampton] |
| EEH9995096.1 | OLD-ABC | - | Salmonella_enterica_subsp._enterica_serovar_4,[5],12:i:- | Bacteria>Pseudomonadati>Pseudomonadota>Gammaproteobacteria>Enterobacterales>Enterobacteriaceae>Salmonella>Salmonella_enterica>Salmonella_enterica_subsp._enterica | 363 | G7H85_000161 | 177783..178874 | 440524 | AAA-family-ATPase-[Salmonella-enterica-subsp.-enterica-serovar-4,[5] |
| EEJ7298182.1 | OLD-ABC | - | Salmonella_enterica_subsp._enterica_serovar_Livingstone | Bacteria>Pseudomonadati>Pseudomonadota>Gammaproteobacteria>Enterobacterales>Enterobacteriaceae>Salmonella>Salmonella_enterica>Salmonella_enterica_subsp._enterica | 363 | DV24_002303 | 307305..308396 | 260680 | AAA-family-ATPase-[Salmonella-enterica-subsp.-enterica-serovar-Livingstone] |
| EEJ9771968.1 | OLD-ABC | - | Salmonella_enterica_subsp._enterica_serovar_Typhimurium | Bacteria>Pseudomonadati>Pseudomonadota>Gammaproteobacteria>Enterobacterales>Enterobacteriaceae>Salmonella>Salmonella_enterica>Salmonella_enterica_subsp._enterica | 363 | HL79_09640 | 178295..179386 | 90371 | AAA-family-ATPase-[Salmonella-enterica-subsp.-enterica-serovar-Typhimurium] |
| EEK0538629.1 | OLD-ABC | - | Salmonella_enterica_subsp._enterica_serovar_Typhimurium | Bacteria>Pseudomonadati>Pseudomonadota>Gammaproteobacteria>Enterobacterales>Enterobacteriaceae>Salmonella>Salmonella_enterica>Salmonella_enterica_subsp._enterica | 363 | IH96_05500 | 145209..146300 | 90371 | AAA-family-ATPase-[Salmonella-enterica-subsp.-enterica-serovar-Typhimurium] |
| EEP8318665.1 | OLD-ABC | - | Salmonella_enterica_subsp._enterica_serovar_Livingstone | Bacteria>Pseudomonadati>Pseudomonadota>Gammaproteobacteria>Enterobacterales>Enterobacteriaceae>Salmonella>Salmonella_enterica>Salmonella_enterica_subsp._enterica | 363 | HCM12_000956 | 59771..60862 | 260680 | AAA-family-ATPase-[Salmonella-enterica-subsp.-enterica] |
| EEP9788443.1 | OLD-ABC | - | Salmonella_enterica_subsp._enterica_serovar_Saintpaul | Bacteria>Pseudomonadati>Pseudomonadota>Gammaproteobacteria>Enterobacterales>Enterobacteriaceae>Salmonella>Salmonella_enterica>Salmonella_enterica_subsp._enterica | 363 | HCP67_002107 | 72280..73371 | 90105 | AAA-family-ATPase-[Salmonella-enterica-subsp.-enterica-serovar-Saintpaul] |
| EGA9157242.1 | OLD-ABC | - | Salmonella_enterica_subsp._enterica_serovar_Kirkee | Bacteria>Pseudomonadati>Pseudomonadota>Gammaproteobacteria>Enterobacterales>Enterobacteriaceae>Salmonella>Salmonella_enterica>Salmonella_enterica_subsp._enterica | 363 | H6M98_002633 | 59665..60756 | 2564624 | AAA-family-ATPase-[Salmonella-enterica-subsp.-enterica-serovar-Kirkee] |
| EGT0819115.1 | OLD-ABC | - | Salmonella_enterica_subsp._enterica_serovar_Ohio | Bacteria>Pseudomonadati>Pseudomonadota>Gammaproteobacteria>Enterobacterales>Enterobacteriaceae>Salmonella>Salmonella_enterica>Salmonella_enterica_subsp._enterica | 363 | I3I53_003216 | 20307..21398 | 117541 | AAA-family-ATPase-[Salmonella-enterica-subsp.-enterica-serovar-Ohio] |
| EGZ8568477.1 | OLD-ABC | - | Salmonella_enterica_subsp._enterica_serovar_Typhimurium | Bacteria>Pseudomonadati>Pseudomonadota>Gammaproteobacteria>Enterobacterales>Enterobacteriaceae>Salmonella>Salmonella_enterica>Salmonella_enterica_subsp._enterica | 363 | JJM06_003394 | 19949..21040 | 90371 | AAA-family-ATPase-[Salmonella-enterica-subsp.-enterica-serovar-Typhimurium] |
| EHC3781323.1 | OLD-ABC | - | Salmonella_enterica_subsp._enterica_serovar_Typhimurium | Bacteria>Pseudomonadati>Pseudomonadota>Gammaproteobacteria>Enterobacterales>Enterobacteriaceae>Salmonella>Salmonella_enterica>Salmonella_enterica_subsp._enterica | 363 | JS923_000752 | 180408..181499 | 90371 | AAA-family-ATPase-[Salmonella-enterica-subsp.-enterica-serovar-Typhimurium] |
| EHD9459647.1 | OLD-ABC | - | Salmonella_enterica_subsp._enterica_serovar_Typhimurium | Bacteria>Pseudomonadati>Pseudomonadota>Gammaproteobacteria>Enterobacterales>Enterobacteriaceae>Salmonella>Salmonella_enterica>Salmonella_enterica_subsp._enterica | 363 | JYL48_000533 | 178296..179387 | 90371 | AAA-family-ATPase-[Salmonella-enterica-subsp.-enterica-serovar-Typhimurium] |
| EHJ1242126.1 | OLD-ABC | GCA_017851875.1 | Salmonella_enterica_subsp._enterica_serovar_Typhimurium | Bacteria>Pseudomonadati>Pseudomonadota>Gammaproteobacteria>Enterobacterales>Enterobacteriaceae>Salmonella>Salmonella_enterica>Salmonella_enterica_subsp._enterica | 363 | KAC06_001464 | 76544..77635 | 90371 | AAA-family-ATPase-[Salmonella-enterica-subsp.-enterica-serovar-Typhimurium] |
| EHN4525915.1 | OLD-ABC | GCA_018388245.1 | Salmonella_enterica_subsp._enterica_serovar_Ohio | Bacteria>Pseudomonadati>Pseudomonadota>Gammaproteobacteria>Enterobacterales>Enterobacteriaceae>Salmonella>Salmonella_enterica>Salmonella_enterica_subsp._enterica | 363 | KI018_001083 | 99383..100474 | 117541 | AAA-family-ATPase-[Salmonella-enterica-subsp.-enterica-serovar-Ohio] |
| EHS3976102.1 | OLD-ABC | GCA_019140515.1 | Salmonella_enterica_subsp._enterica_serovar_Poona | Bacteria>Pseudomonadati>Pseudomonadota>Gammaproteobacteria>Enterobacterales>Enterobacteriaceae>Salmonella>Salmonella_enterica>Salmonella_enterica_subsp._enterica | 363 | KV430_003534 | 9340..10431 | 436295 | AAA-family-ATPase-[Salmonella-enterica-subsp.-enterica-serovar-Poona] |
| EIE3342238.1 | OLD-ABC | GCA_020155375.1 | Salmonella_enterica_subsp._enterica_serovar_Typhimurium | Bacteria>Pseudomonadati>Pseudomonadota>Gammaproteobacteria>Enterobacterales>Enterobacteriaceae>Salmonella>Salmonella_enterica>Salmonella_enterica_subsp._enterica | 363 | LDC84_002300 | 67573..68664 | 90371 | AAA-family-ATPase-[Salmonella-enterica-subsp.-enterica-serovar-Typhimurium] |
| EII7462486.1 | OLD-ABC | GCA_020462505.1 | Salmonella_enterica_subsp._enterica_serovar_Typhimurium | Bacteria>Pseudomonadati>Pseudomonadota>Gammaproteobacteria>Enterobacterales>Enterobacteriaceae>Salmonella>Salmonella_enterica>Salmonella_enterica_subsp._enterica | 363 | LHR57_000984 | 6322..7413 | 90371 | AAA-family-ATPase-[Salmonella-enterica-subsp.-enterica-serovar-Typhimurium] |
| EIJ0372101.1 | OLD-ABC | GCA_020490715.1 | Salmonella_enterica_subsp._enterica_serovar_Typhimurium | Bacteria>Pseudomonadati>Pseudomonadota>Gammaproteobacteria>Enterobacterales>Enterobacteriaceae>Salmonella>Salmonella_enterica>Salmonella_enterica_subsp._enterica | 363 | LH081_001561 | 178283..179374 | 90371 | AAA-family-ATPase-[Salmonella-enterica-subsp.-enterica-serovar-Typhimurium] |
| EJN2869330.1 | OLD-ABC | GCA_024442015.1 | Salmonella_enterica_subsp._enterica_serovar_Derby | Bacteria>Pseudomonadati>Pseudomonadota>Gammaproteobacteria>Enterobacterales>Enterobacteriaceae>Salmonella>Salmonella_enterica>Salmonella_enterica_subsp._enterica | 363 | NPT71_003580 | 172382..173473 | 28144 | AAA-family-ATPase-[Salmonella-enterica-subsp.-enterica-serovar-Derby] |
| EJW7283566.1 | OLD-ABC | GCA_025474975.1 | Salmonella_enterica_subsp._enterica_serovar_Typhimurium | Bacteria>Pseudomonadati>Pseudomonadota>Gammaproteobacteria>Enterobacterales>Enterobacteriaceae>Salmonella>Salmonella_enterica>Salmonella_enterica_subsp._enterica | 363 | OCR56_001319 | 76453..77544 | 90371 | AAA-family-ATPase-[Salmonella-enterica-subsp.-enterica-serovar-Typhimurium] |
| HAD0448192.1 | OLD-ABC | GCA_010193865.1 | Salmonella_enterica_subsp._enterica_serovar_Typhimurium | Bacteria>Pseudomonadati>Pseudomonadota>Gammaproteobacteria>Enterobacterales>Enterobacteriaceae>Salmonella>Salmonella_enterica>Salmonella_enterica_subsp._enterica | 363 | GON57_00700 | 145212..146303 | 90371 | TPA_asm:-AAA-family-ATPase-[Salmonella-enterica-subsp.-enterica-serovar-Typhimurium] |
| HAD1283636.1 | OLD-ABC | GCA_010197825.1 | Salmonella_enterica_subsp._enterica_serovar_Typhimurium | Bacteria>Pseudomonadati>Pseudomonadota>Gammaproteobacteria>Enterobacterales>Enterobacteriaceae>Salmonella>Salmonella_enterica>Salmonella_enterica_subsp._enterica | 363 | GTH76_16765 | 103250..104341 | 90371 | TPA_asm:-AAA-family-ATPase-[Salmonella-enterica-subsp.-enterica-serovar-Typhimurium] |
| HAE3790825.1 | OLD-ABC | GCA_010485735.1 | Salmonella_enterica_subsp._enterica_serovar_Typhimurium | Bacteria>Pseudomonadati>Pseudomonadota>Gammaproteobacteria>Enterobacterales>Enterobacteriaceae>Salmonella>Salmonella_enterica>Salmonella_enterica_subsp._enterica | 363 | G4A23_000145 | 154066..155157 | 90371 | TPA_asm:-AAA-family-ATPase-[Salmonella-enterica-subsp.-enterica-serovar-Typhimurium] |
| HAS9574205.1 | OLD-ABC | GCA_015871885.1 | Salmonella_enterica_subsp._enterica_serovar_Typhimurium | Bacteria>Pseudomonadati>Pseudomonadota>Gammaproteobacteria>Enterobacterales>Enterobacteriaceae>Salmonella>Salmonella_enterica>Salmonella_enterica_subsp._enterica | 363 | I8M64_002287 | 103251..104342 | 90371 | TPA:-AAA-family-ATPase-[Salmonella-enterica-subsp.-enterica-serovar-Typhimurium] |
| HBJ3502653.1 | OLD-ABC | GCA_020048005.1 | Salmonella_enterica_subsp._enterica_serovar_Typhimurium | Bacteria>Pseudomonadati>Pseudomonadota>Gammaproteobacteria>Enterobacterales>Enterobacteriaceae>Salmonella>Salmonella_enterica>Salmonella_enterica_subsp._enterica | 363 | LB282_001659 | 178067..179158 | 90371 | TPA:-AAA-family-ATPase-[Salmonella-enterica-subsp.-enterica-serovar-Typhimurium] |
| HBL9882171.1 | OLD-ABC | GCA_021103155.1 | Salmonella_enterica_subsp._enterica_serovar_Jangwani | Bacteria>Pseudomonadati>Pseudomonadota>Gammaproteobacteria>Enterobacterales>Enterobacteriaceae>Salmonella>Salmonella_enterica>Salmonella_enterica_subsp._enterica | 363 | LT029_003492 | 162575..163666 | 2564577 | TPA:-AAA-family-ATPase-[Salmonella-enterica-subsp.-enterica-serovar-Jangwani] |
| MBT0981087.1 | OLD-ABC | GCA_018531305.1 | Salmonella_enterica_subsp._enterica_serovar_Typhimurium | Bacteria>Pseudomonadati>Pseudomonadota>Gammaproteobacteria>Enterobacterales>Enterobacteriaceae>Salmonella>Salmonella_enterica>Salmonella_enterica_subsp._enterica | 363 | KJ032_08030 | 178313..179404 | 90371 | AAA-family-ATPase-[Salmonella-enterica-subsp.-enterica-serovar-Typhimurium] |
| MBZ4898655.1 | OLD-ABC | GCA_020070625.1 | Salmonella_enterica_subsp._enterica_serovar_Typhimurium | Bacteria>Pseudomonadati>Pseudomonadota>Gammaproteobacteria>Enterobacterales>Enterobacteriaceae>Salmonella>Salmonella_enterica>Salmonella_enterica_subsp._enterica | 363 | HU117_01205 | 54134..55225 | 90371 | AAA-family-ATPase-[Salmonella-enterica-subsp.-enterica-serovar-Typhimurium] |
| MCL8965754.1 | OLD-ABC | GCA_023608175.1 | Salmonella_enterica_subsp._enterica_serovar_Enteritidis | Bacteria>Pseudomonadati>Pseudomonadota>Gammaproteobacteria>Enterobacterales>Enterobacteriaceae>Salmonella>Salmonella_enterica>Salmonella_enterica_subsp._enterica | 363 | MU637_15185 | 177913..179004 | 149539 | AAA-family-ATPase-[Salmonella-enterica-subsp.-enterica-serovar-Enteritidis] |
| OFB20552.1 | OLD-ABC | GCA_001756205.1 | Salmonella_enterica_subsp._enterica_serovar_Typhimurium | Bacteria>Pseudomonadati>Pseudomonadota>Gammaproteobacteria>Enterobacterales>Enterobacteriaceae>Salmonella>Salmonella_enterica>Salmonella_enterica_subsp._enterica | 363 | BG487_15365 | 7329..8420 | 90371 | hypothetical-protein-BG487_15365-[Salmonella-enterica-subsp.-enterica-serovar-Typhimurium] |
| QSG23650.1 | OLD-ABC | GCA_017094585.1 | Salmonella_enterica_subsp._enterica_serovar_Typhimurium | Bacteria>Pseudomonadati>Pseudomonadota>Gammaproteobacteria>Enterobacterales>Enterobacteriaceae>Salmonella>Salmonella_enterica>Salmonella_enterica_subsp._enterica | 363 | IRY58_23105 | 4744884..4745975 | 90371 | ATP-binding-protein-[Salmonella-enterica-subsp.-enterica-serovar-Typhimurium] |
| SUH94772.1 | OLD-ABC | GCA_900456925.1 | Salmonella_enterica_subsp._enterica_serovar_Typhimurium | Bacteria>Pseudomonadati>Pseudomonadota>Gammaproteobacteria>Enterobacterales>Enterobacteriaceae>Salmonella>Salmonella_enterica>Salmonella_enterica_subsp._enterica | 363 | NCTC13347_04215 | 4180362..4181453 | 90371 | spermidine/putrescine-ABC-transporter-ATPase-[Salmonella-enterica-subsp.-enterica-serovar-Typhimurium] |
| EAA1506270.1 | OLD-ABC | GCA_004160295.1 | Salmonella_enterica_subsp._enterica_serovar_Agama | Bacteria>Pseudomonadati>Pseudomonadota>Gammaproteobacteria>Enterobacterales>Enterobacteriaceae>Salmonella>Salmonella_enterica>Salmonella_enterica_subsp._enterica | 362 | DL104_05740 | 36588..37676 | 399581 | ATP-binding-protein-[Salmonella-enterica-subsp.-enterica] |
| EAA5580612.1 | OLD-ABC | GCA_004185805.1 | Salmonella_enterica_subsp._enterica_serovar_Glostrup | Bacteria>Pseudomonadati>Pseudomonadota>Gammaproteobacteria>Enterobacterales>Enterobacteriaceae>Salmonella>Salmonella_enterica>Salmonella_enterica_subsp._enterica | 362 | DQD90_06850 | 66298..67386 | 1151180 | ATP-binding-protein-[Salmonella-enterica-subsp.-enterica] |
| EAA5642508.1 | OLD-ABC | GCA_004185395.1 | Salmonella_enterica_subsp._enterica_serovar_Javiana | Bacteria>Pseudomonadati>Pseudomonadota>Gammaproteobacteria>Enterobacterales>Enterobacteriaceae>Salmonella>Salmonella_enterica>Salmonella_enterica_subsp._enterica | 362 | DQJ71_10400 | 156434..157522 | 363569 | ATP-binding-protein-[Salmonella-enterica-subsp.-enterica] |
| EAA6071872.1 | OLD-ABC | GCA_004187475.1 | Salmonella_enterica_subsp._enterica_serovar_Corvallis | Bacteria>Pseudomonadati>Pseudomonadota>Gammaproteobacteria>Enterobacterales>Enterobacteriaceae>Salmonella>Salmonella_enterica>Salmonella_enterica_subsp._enterica | 362 | DQZ25_06120 | 165625..166713 | 593905 | ATP-binding-protein-[Salmonella-enterica-subsp.-enterica] |
| EAB6380445.1 | OLD-ABC | GCA_004255205.1 | Salmonella_enterica_subsp._enterica_serovar_Emek | Bacteria>Pseudomonadati>Pseudomonadota>Gammaproteobacteria>Enterobacterales>Enterobacteriaceae>Salmonella>Salmonella_enterica>Salmonella_enterica_subsp._enterica | 362 | CWG97_20710 | 25333..26421 | 286784 | ATP-binding-protein-[Salmonella-enterica-subsp.-enterica] |
| EAB7776928.1 | OLD-ABC | GCA_004262645.1 | Salmonella_enterica_subsp._enterica_serovar_Cannstatt | Bacteria>Pseudomonadati>Pseudomonadota>Gammaproteobacteria>Enterobacterales>Enterobacteriaceae>Salmonella>Salmonella_enterica>Salmonella_enterica_subsp._enterica | 362 | D6413_08725 | 59697..60785 | 2564307 | ATP-binding-protein-[Salmonella-enterica-subsp.-enterica] |
| EBF8311519.1 | OLD-ABC | GCA_006394955.1 | Salmonella_enterica_subsp._enterica_serovar_Tamberma | Bacteria>Pseudomonadati>Pseudomonadota>Gammaproteobacteria>Enterobacterales>Enterobacteriaceae>Salmonella>Salmonella_enterica>Salmonella_enterica_subsp._enterica | 362 | FGE90_18590 | 60583..61671 | 2565079 | ATP-binding-protein-[Salmonella-enterica-subsp.-enterica] |
| EBG0770114.1 | OLD-ABC | GCA_006406395.1 | Salmonella_enterica_subsp._enterica_serovar_Derby | Bacteria>Pseudomonadati>Pseudomonadota>Gammaproteobacteria>Enterobacterales>Enterobacteriaceae>Salmonella>Salmonella_enterica>Salmonella_enterica_subsp._enterica | 362 | FH890_08335 | 10425..11513 | 28144 | ATP-binding-protein-[Salmonella-enterica] |
| EBU7641090.1 | OLD-ABC | GCA_007142065.1 | Salmonella_enterica_subsp._enterica_serovar_Brevik | Bacteria>Pseudomonadati>Pseudomonadota>Gammaproteobacteria>Enterobacterales>Enterobacteriaceae>Salmonella>Salmonella_enterica>Salmonella_enterica_subsp._enterica | 362 | DK709_13725 | 66759..67847 | 2564275 | ATP-binding-protein-[Salmonella-enterica-subsp.-enterica-serovar-Brevik] |
| EBU8677658.1 | OLD-ABC | GCA_007146445.1 | Salmonella_enterica_subsp._enterica_serovar_Parkroyal | Bacteria>Pseudomonadati>Pseudomonadota>Gammaproteobacteria>Enterobacterales>Enterobacteriaceae>Salmonella>Salmonella_enterica>Salmonella_enterica_subsp._enterica | 362 | DLR30_20705 | 35212..36300 | 2564909 | hypothetical-protein-[Salmonella-enterica-subsp.-enterica-serovar-Parkroyal] |
| EBU9775080.1 | OLD-ABC | GCA_007151525.1 | Salmonella_enterica_subsp._enterica_serovar_Kumasi | Bacteria>Pseudomonadati>Pseudomonadota>Gammaproteobacteria>Enterobacterales>Enterobacteriaceae>Salmonella>Salmonella_enterica>Salmonella_enterica_subsp._enterica | 362 | DM304_21655 | 10435..11523 | 2493647 | ATP-binding-protein-[Salmonella-enterica-subsp.-enterica-serovar-Kumasi] |
| EBV2735694.1 | OLD-ABC | GCA_007164145.1 | Salmonella_enterica_subsp._enterica_serovar_Stourbridge | Bacteria>Pseudomonadati>Pseudomonadota>Gammaproteobacteria>Enterobacterales>Enterobacteriaceae>Salmonella>Salmonella_enterica>Salmonella_enterica_subsp._enterica | 362 | DOA65_09455 | 66305..67393 | 1160766 | ATP-binding-protein-[Salmonella-enterica-subsp.-enterica-serovar-Stourbridge] |
| EBV4609758.1 | OLD-ABC | GCA_007170635.1 | Salmonella_enterica_subsp._enterica_serovar_Solt | Bacteria>Pseudomonadati>Pseudomonadota>Gammaproteobacteria>Enterobacterales>Enterobacteriaceae>Salmonella>Salmonella_enterica>Salmonella_enterica_subsp._enterica | 362 | DOX24_02185 | 10431..11519 | 2565040 | ATP-binding-protein-[Salmonella-enterica-subsp.-enterica-serovar-Solt] |
| EBV4672045.1 | OLD-ABC | GCA_007170935.1 | Salmonella_enterica_subsp._enterica_serovar_Tamberma | Bacteria>Pseudomonadati>Pseudomonadota>Gammaproteobacteria>Enterobacterales>Enterobacteriaceae>Salmonella>Salmonella_enterica>Salmonella_enterica_subsp._enterica | 362 | DOW22_12530 | 156539..157627 | 2565079 | ATP-binding-protein-[Salmonella-enterica-subsp.-enterica-serovar-Tamberma] |
| EBW5570926.1 | OLD-ABC | GCA_007216555.1 | Salmonella_enterica_subsp._enterica_serovar_Emek | Bacteria>Pseudomonadati>Pseudomonadota>Gammaproteobacteria>Enterobacterales>Enterobacteriaceae>Salmonella>Salmonella_enterica>Salmonella_enterica_subsp._enterica | 362 | DPY54_19335 | 25325..26413 | 286784 | ATP-binding-protein-[Salmonella-enterica-subsp.-enterica-serovar-Emek] |
| EBW8394990.1 | OLD-ABC | GCA_007228915.1 | Salmonella_enterica_subsp._enterica_serovar_Florida | Bacteria>Pseudomonadati>Pseudomonadota>Gammaproteobacteria>Enterobacterales>Enterobacteriaceae>Salmonella>Salmonella_enterica>Salmonella_enterica_subsp._enterica | 362 | A3X38_19925 | 10432..11520 | 2564436 | ATP-binding-protein-[Salmonella-enterica-subsp.-enterica-serovar-Florida] |
| EBX4839079.1 | OLD-ABC | GCA_007256215.1 | Salmonella_enterica_subsp._enterica_serovar_Tamberma | Bacteria>Pseudomonadati>Pseudomonadota>Gammaproteobacteria>Enterobacterales>Enterobacteriaceae>Salmonella>Salmonella_enterica>Salmonella_enterica_subsp._enterica | 362 | DSB42_14910 | 170932..172020 | 2565079 | ATP-binding-protein-[Salmonella-enterica-subsp.-enterica-serovar-Tamberma] |
| EBX6018291.1 | OLD-ABC | GCA_007258805.1 | Salmonella_enterica_subsp._enterica_serovar_Dortmund | Bacteria>Pseudomonadati>Pseudomonadota>Gammaproteobacteria>Enterobacterales>Enterobacteriaceae>Salmonella>Salmonella_enterica>Salmonella_enterica_subsp._enterica | 362 | DSR74_22235 | 7229..8317 | 2564368 | ATP-binding-protein-[Salmonella-enterica-subsp.-enterica-serovar-Dortmund] |
| EBX6302553.1 | OLD-ABC | GCA_007260765.1 | Salmonella_enterica_subsp._enterica_serovar_Enugu | Bacteria>Pseudomonadati>Pseudomonadota>Gammaproteobacteria>Enterobacterales>Enterobacteriaceae>Salmonella>Salmonella_enterica>Salmonella_enterica_subsp._enterica | 362 | DS191_04280 | 33166..34254 | 2564400 | ATP-binding-protein-[Salmonella-enterica-subsp.-enterica-serovar-Enugu] |
| EBX8836174.1 | OLD-ABC | GCA_007270555.1 | Salmonella_enterica_subsp._enterica_serovar_Urbana | Bacteria>Pseudomonadati>Pseudomonadota>Gammaproteobacteria>Enterobacterales>Enterobacteriaceae>Salmonella>Salmonella_enterica>Salmonella_enterica_subsp._enterica | 362 | DTU76_05380 | 59590..60678 | 260677 | ATP-binding-protein-[Salmonella-enterica-subsp.-enterica-serovar-Urbana] |
| EBY1390422.1 | OLD-ABC | GCA_007281935.1 | Salmonella_enterica_subsp._enterica_serovar_Wagenia | Bacteria>Pseudomonadati>Pseudomonadota>Gammaproteobacteria>Enterobacterales>Enterobacteriaceae>Salmonella>Salmonella_enterica>Salmonella_enterica_subsp._enterica | 362 | DUW06_08895 | 148074..149162 | 322833 | ATP-binding-protein-[Salmonella-enterica-subsp.-enterica-serovar-Wagenia] |
| EBY2665622.1 | OLD-ABC | GCA_007287115.1 | Salmonella_enterica_subsp._enterica_serovar_Teko | Bacteria>Pseudomonadati>Pseudomonadota>Gammaproteobacteria>Enterobacterales>Enterobacteriaceae>Salmonella>Salmonella_enterica>Salmonella_enterica_subsp._enterica | 362 | DVF23_02325 | 24480..25568 | 2079198 | ATP-binding-protein-[Salmonella-enterica-subsp.-enterica-serovar-Teko] |
| EBZ6491794.1 | OLD-ABC | GCA_007345015.1 | Salmonella_enterica_subsp._enterica_serovar_Dugbe | Bacteria>Pseudomonadati>Pseudomonadota>Gammaproteobacteria>Enterobacterales>Enterobacteriaceae>Salmonella>Salmonella_enterica>Salmonella_enterica_subsp._enterica | 362 | ED480_20070 | 24208..25296 | 2564376 | ATP-binding-protein-[Salmonella-enterica-subsp.-enterica-serovar-Dugbe] |
| ECA8969864.1 | OLD-ABC | GCA_007399965.1 | Salmonella_enterica_subsp._enterica_serovar_Omuna | Bacteria>Pseudomonadati>Pseudomonadota>Gammaproteobacteria>Enterobacterales>Enterobacteriaceae>Salmonella>Salmonella_enterica>Salmonella_enterica_subsp._enterica | 362 | ETB55_05735 | 65111..66199 | 2564887 | ATP-binding-protein-[Salmonella-enterica-subsp.-enterica-serovar-Omuna] |
| ECB3805266.1 | OLD-ABC | GCA_007422765.1 | Salmonella_enterica_subsp._enterica_serovar_Fufu | Bacteria>Pseudomonadati>Pseudomonadota>Gammaproteobacteria>Enterobacterales>Enterobacteriaceae>Salmonella>Salmonella_enterica>Salmonella_enterica_subsp._enterica | 362 | EW367_11370 | 59443..60531 | 2564449 | ATP-binding-protein-[Salmonella-enterica-subsp.-enterica-serovar-Fufu] |
| ECD2821737.1 | OLD-ABC | GCA_007513645.1 | Salmonella_enterica_subsp._enterica_serovar_Fanti | Bacteria>Pseudomonadati>Pseudomonadota>Gammaproteobacteria>Enterobacterales>Enterobacteriaceae>Salmonella>Salmonella_enterica>Salmonella_enterica_subsp._enterica | 362 | EXA45_11160 | 59679..60767 | 2564418 | ATP-binding-protein-[Salmonella-enterica-subsp.-enterica-serovar-Fanti] |
| ECD3895176.1 | OLD-ABC | GCA_007520185.1 | Salmonella_enterica_subsp._enterica_serovar_Gloucester | Bacteria>Pseudomonadati>Pseudomonadota>Gammaproteobacteria>Enterobacterales>Enterobacteriaceae>Salmonella>Salmonella_enterica>Salmonella_enterica_subsp._enterica | 362 | EOU70_05960 | 172118..173206 | 2564474 | ATP-binding-protein-[Salmonella-enterica-subsp.-enterica-serovar-Gloucester] |
| ECE0383739.1 | OLD-ABC | GCA_007548695.1 | Salmonella_enterica_subsp._enterica_serovar_Aba | Bacteria>Pseudomonadati>Pseudomonadota>Gammaproteobacteria>Enterobacterales>Enterobacteriaceae>Salmonella>Salmonella_enterica>Salmonella_enterica_subsp._enterica | 362 | XZ45_12020 | 33056..34144 | 2564103 | ATP-binding-protein-[Salmonella-enterica-subsp.-enterica] |
| ECG3781075.1 | OLD-ABC | GCA_007660485.1 | Salmonella_enterica_subsp._enterica_serovar_Breukelen | Bacteria>Pseudomonadati>Pseudomonadota>Gammaproteobacteria>Enterobacterales>Enterobacteriaceae>Salmonella>Salmonella_enterica>Salmonella_enterica_subsp._enterica | 362 | EXA83_17830 | 59472..60560 | 2564274 | ATP-binding-protein-[Salmonella-enterica-subsp.-enterica-serovar-Breukelen] |
| ECG8301816.1 | OLD-ABC | GCA_007694625.1 | Salmonella_enterica_subsp._enterica_serovar_Glostrup | Bacteria>Pseudomonadati>Pseudomonadota>Gammaproteobacteria>Enterobacterales>Enterobacteriaceae>Salmonella>Salmonella_enterica>Salmonella_enterica_subsp._enterica | 362 | E1853_02145 | 66296..67384 | 1151180 | ATP-binding-protein-[Salmonella-enterica-subsp.-enterica-serovar-Glostrup] |
| ECT8081316.1 | OLD-ABC | GCA_008481645.1 | Salmonella_enterica_subsp._enterica_serovar_Carrau | Bacteria>Pseudomonadati>Pseudomonadota>Gammaproteobacteria>Enterobacterales>Enterobacteriaceae>Salmonella>Salmonella_enterica>Salmonella_enterica_subsp._enterica | 362 | BLY96_09870 | 157455..158543 | 1160739 | ATP-binding-protein-[Salmonella-enterica-subsp.-enterica-serovar-Carrau] |
| EDH5631747.1 | OLD-ABC | - | Salmonella_enterica_subsp._enterica_serovar_Claibornei | Bacteria>Pseudomonadati>Pseudomonadota>Gammaproteobacteria>Enterobacterales>Enterobacteriaceae>Salmonella>Salmonella_enterica>Salmonella_enterica_subsp._enterica | 362 | CB172_15090 | 165779..166867 | 2572709 | ATP-binding-protein-[Salmonella-enterica-subsp.-enterica-serovar-Claibornei] |
| EDR5087376.1 | OLD-ABC | - | Salmonella_enterica_subsp._enterica_serovar_Kingston | Bacteria>Pseudomonadati>Pseudomonadota>Gammaproteobacteria>Enterobacterales>Enterobacteriaceae>Salmonella>Salmonella_enterica>Salmonella_enterica_subsp._enterica | 362 | GRZ33_003683 | 33511..34599 | 2161663 | ATP-binding-protein-[Salmonella-enterica-subsp.-enterica] |

| Protein-ID | Domain | Assembly-ID | Strain | Taxonomy | Length | Locus-Tag | Genomic-coordinates | TaxID | Fasta-defline |
|---|---|---|---|---|---|---|---|---|---|
| EDS5116134.1 | OLD-ABC | - | Salmonella_enterica_subsp._enterica_serovar_Agoueve | Bacteria>Pseudomonadati>Pseudomonadota>Gammaproteobacteria>Enterobacterales>Enterobacteriaceae>Salmonella>Salmonella_enterica>Salmonella_enterica_subsp._enterica | 362 | CBJ63_000887 | 172397..173485 | 1505961 | ATP-binding-protein-[Salmonella-enterica-subsp.-enterica-serovar-Agoueve] |
| EDT1456937.1 | OLD-ABC | - | Salmonella_enterica_subsp._enterica_serovar_Ramatgan | Bacteria>Pseudomonadati>Pseudomonadota>Gammaproteobacteria>Enterobacterales>Enterobacteriaceae>Salmonella>Salmonella_enterica>Salmonella_enterica_subsp._enterica | 362 | CE894_001672 | 160524..161612 | 2564942 | ATP-binding-protein-[Salmonella-enterica-subsp.-enterica-serovar-Ramatgan] |
| EDT7239826.1 | OLD-ABC | - | Salmonella_enterica_subsp._enterica_serovar_Warragul | Bacteria>Pseudomonadati>Pseudomonadota>Gammaproteobacteria>Enterobacterales>Enterobacteriaceae>Salmonella>Salmonella_enterica>Salmonella_enterica_subsp._enterica | 362 | AHZ96_001969 | 100743..101831 | 2565180 | ATP-binding-protein-[Salmonella-enterica-subsp.-enterica] |
| EDV9644206.1 | OLD-ABC | - | Salmonella_enterica_subsp._enterica_serovar_Carno | Bacteria>Pseudomonadati>Pseudomonadota>Gammaproteobacteria>Enterobacterales>Enterobacteriaceae>Salmonella>Salmonella_enterica>Salmonella_enterica_subsp._enterica | 362 | HM25_003999 | 171877..172965 | 2024268 | ATP-binding-protein-[Salmonella-enterica-subsp.-enterica-serovar-Carno] |
| EED3636777.1 | OLD-ABC | - | Salmonella_enterica_subsp._enterica_serovar_Sundsvall | Bacteria>Pseudomonadati>Pseudomonadota>Gammaproteobacteria>Enterobacterales>Enterobacteriaceae>Salmonella>Salmonella_enterica>Salmonella_enterica_subsp._enterica | 362 | S719_003283 | 164940..166028 | 2024273 | ATP-binding-protein-[Salmonella-enterica-subsp.-enterica] |
| EEE1152712.1 | OLD-ABC | - | Salmonella_enterica_subsp._enterica_serovar_Teko | Bacteria>Pseudomonadati>Pseudomonadota>Gammaproteobacteria>Enterobacterales>Enterobacteriaceae>Salmonella>Salmonella_enterica>Salmonella_enterica_subsp._enterica | 362 | CAE49_001973 | 326005..327093 | 2079198 | ATP-binding-protein-[Salmonella-enterica-subsp.-enterica-serovar-Teko] |
| EEH2707248.1 | OLD-ABC | - | Salmonella_enterica_subsp._enterica_serovar_Lille | Bacteria>Pseudomonadati>Pseudomonadota>Gammaproteobacteria>Enterobacterales>Enterobacteriaceae>Salmonella>Salmonella_enterica>Salmonella_enterica_subsp._enterica | 362 | GYE71_000439 | 61370..62458 | 486997 | ATP-binding-protein-[Salmonella-enterica-subsp.-enterica] |
| EEO3564962.1 | OLD-ABC | - | Salmonella_enterica_subsp._enterica_serovar_Poona | Bacteria>Pseudomonadati>Pseudomonadota>Gammaproteobacteria>Enterobacterales>Enterobacteriaceae>Salmonella>Salmonella_enterica>Salmonella_enterica_subsp._enterica | 362 | DEI68_09005 | 59747..60835 | 436295 | AAA-family-ATPase-[Salmonella-enterica-subsp.-enterica-serovar-Poona] |
| EGH9456390.1 | OLD-ABC | - | Salmonella_enterica_subsp._enterica_serovar_Leeuwarden | Bacteria>Pseudomonadati>Pseudomonadota>Gammaproteobacteria>Enterobacterales>Enterobacteriaceae>Salmonella>Salmonella_enterica>Salmonella_enterica_subsp._enterica | 362 | HKU92_000349 | 396698..397786 | 2564678 | ATP-binding-protein-[Salmonella-enterica-subsp.-enterica-serovar-Leeuwarden] |
| EHO4506762.1 | OLD-ABC | GCA_018524125.1 | Salmonella_enterica_subsp._enterica_serovar_Carrau | Bacteria>Pseudomonadati>Pseudomonadota>Gammaproteobacteria>Enterobacterales>Enterobacteriaceae>Salmonella>Salmonella_enterica>Salmonella_enterica_subsp._enterica | 362 | KD666_001068 | 156641..157729 | 1160739 | ATP-binding-protein-[Salmonella-enterica-subsp.-enterica-serovar-Carrau] |
| EHP4799942.1 | OLD-ABC | GCA_018776965.1 | Salmonella_enterica_subsp._enterica_serovar_Sangera | Bacteria>Pseudomonadati>Pseudomonadota>Gammaproteobacteria>Enterobacterales>Enterobacteriaceae>Salmonella>Salmonella_enterica>Salmonella_enterica_subsp._enterica | 362 | KM018_002017 | 52035..53123 | 1173574 | ATP-binding-protein-[Salmonella-enterica-subsp.-enterica-serovar-Sangera] |
| EHS5456437.1 | OLD-ABC | GCA_019151305.1 | Salmonella_enterica_subsp._enterica_serovar_Corvallis | Bacteria>Pseudomonadati>Pseudomonadota>Gammaproteobacteria>Enterobacterales>Enterobacteriaceae>Salmonella>Salmonella_enterica>Salmonella_enterica_subsp._enterica | 362 | KV802_001388 | 172559..173647 | 593905 | ATP-binding-protein-[Salmonella-enterica-subsp.-enterica-serovar-Corvallis] |
| EIJ6143994.1 | OLD-ABC | GCA_020548845.1 | Salmonella_enterica_subsp._enterica_serovar_4:a:- | Bacteria>Pseudomonadati>Pseudomonadota>Gammaproteobacteria>Enterobacterales>Enterobacteriaceae>Salmonella>Salmonella_enterica>Salmonella_enterica_subsp._enterica | 362 | LI678_001016 | 402089..403177 | 2885150 | ATP-binding-protein-[Salmonella-enterica-subsp.-enterica-serovar-4:a:-] |
| EIR7569904.1 | OLD-ABC | GCA_021281445.1 | Salmonella_enterica_subsp._enterica_serovar_Agbeni | Bacteria>Pseudomonadati>Pseudomonadota>Gammaproteobacteria>Enterobacterales>Enterobacteriaceae>Salmonella>Salmonella_enterica>Salmonella_enterica_subsp._enterica | 362 | LXI61_003346 | 25540..26628 | 1967642 | ATP-binding-protein-[Salmonella-enterica-subsp.-enterica-serovar-Agbeni] |
| HCB4841066.1 | OLD-ABC | GCA_023082205.1 | Salmonella_enterica_subsp._enterica_serovar_Bredeney | Bacteria>Pseudomonadati>Pseudomonadota>Gammaproteobacteria>Enterobacterales>Enterobacteriaceae>Salmonella>Salmonella_enterica>Salmonella_enterica_subsp._enterica | 362 | MOF44_000353 | 397764..398852 | 134047 | ATP-binding-protein-[Salmonella-enterica-subsp.-enterica-serovar-Bredeney] |
| MIX28206.1 | OLD-ABC | GCA_003876625.1 | Salmonella_enterica_subsp._enterica_serovar_Livingstone | Bacteria>Pseudomonadati>Pseudomonadota>Gammaproteobacteria>Enterobacterales>Enterobacteriaceae>Salmonella>Salmonella_enterica>Salmonella_enterica_subsp._enterica | 362 | CRQ34_16020 | 166543..167631 | 260680 | ATP-binding-protein-[Salmonella-enterica-subsp.-enterica-serovar-Livingstone] |
| SUI30168.1 | OLD-ABC | GCA_900456825.1 | Salmonella_enterica_subsp._enterica_serovar_Derby | Bacteria>Pseudomonadati>Pseudomonadota>Gammaproteobacteria>Enterobacterales>Enterobacteriaceae>Salmonella>Salmonella_enterica>Salmonella_enterica_subsp._enterica | 362 | NCTC5722_04072 | 4069831..4070919 | 28144 | spermidine/putrescine-ABC-transporter-ATPase-[Salmonella-enterica-subsp.-enterica-serovar-Derby] |
| ESG75017.1 | OLD-ABC | GCA_000487055.1 | Salmonella_enterica_subsp._enterica_serovar_Kentucky_str._ATCC_9263 | Bacteria>Pseudomonadati>Pseudomonadota>Gammaproteobacteria>Enterobacterales>Enterobacteriaceae>Salmonella>Salmonella_enterica>Salmonella_enterica_subsp._enterica>Salmonella_enterica_subsp._enterica_serovar_Kentucky | 373 | SEEK9263_16872 | 26537..27658 | 984236 | RecF/RecN/SMC-N-domain,-putative-[Salmonella-enterica-subsp.-enterica-serovar-Kentucky-str.-ATCC-9263] |
| EDH9995455.1 | OLD-ABC | - | Salmonella_enterica_subsp._enterica_serovar_Java | Bacteria>Pseudomonadati>Pseudomonadota>Gammaproteobacteria>Enterobacterales>Enterobacteriaceae>Salmonella>Salmonella_enterica>Salmonella_enterica_subsp._enterica>Salmonella_enterica_subsp._enterica_serovar_Paratyphi_B | 363 | CC627_22215 | 6216..7307 | 224729 | ATP-binding-protein-[Salmonella-enterica-subsp.-enterica-serovar-Java] |
| ASE02938.1 | OLD-ABC | GCA_002208315.1 | Salmonella_enterica_subsp._enterica_serovar_Quebec_str._S-1267 | Bacteria>Pseudomonadati>Pseudomonadota>Gammaproteobacteria>Enterobacterales>Enterobacteriaceae>Salmonella>Salmonella_enterica>Salmonella_enterica_subsp._enterica>Salmonella_enterica_subsp._enterica_serovar_Quebec | 373 | LFZ40_19395 | 3805142..3806263 | 1243591 | ABC-transporter-ATP-binding-protein-[Salmonella-enterica-subsp.-enterica-serovar-Quebec-str.-S-1267] |
| ECS4740323.1 | OLD-ABC | GCA_008423865.1 | Salmonella_enterica_subsp._enterica_serovar_Typhimurium_var._5- | Bacteria>Pseudomonadati>Pseudomonadota>Gammaproteobacteria>Enterobacterales>Enterobacteriaceae>Salmonella>Salmonella_enterica>Salmonella_enterica_subsp._enterica>Salmonella_enterica_subsp._enterica_serovar_Typhimurium | 363 | A9R80_15950 | 81184..82275 | 1620419 | ATP-binding-protein-[Salmonella-enterica-subsp.-enterica-serovar-Typhimurium-var.-5-] |
| EDV6600964.1 | OLD-ABC | - | Salmonella_enterica_subsp._enterica_serovar_Typhimurium_var._5- | Bacteria>Pseudomonadati>Pseudomonadota>Gammaproteobacteria>Enterobacterales>Enterobacteriaceae>Salmonella>Salmonella_enterica>Salmonella_enterica_subsp._enterica>Salmonella_enterica_subsp._enterica_serovar_Typhimurium | 363 | BH535_001150 | 76529..77620 | 1620419 | AAA-family-ATPase-[Salmonella-enterica-subsp.-enterica-serovar-Typhimurium-var.-5-] |
| HCM2417896.1 | OLD-ABC | GCA_025074775.1 | Salmonella_enterica_subsp._enterica_serovar_Typhimurium_var._monophasic_4,[5],12:i:- | Bacteria>Pseudomonadati>Pseudomonadota>Gammaproteobacteria>Enterobacterales>Enterobacteriaceae>Salmonella>Salmonella_enterica>Salmonella_enterica_subsp._enterica>Salmonella_enterica_subsp._enterica_serovar_Typhimurium | 363 | N3D22_002714 | 103250..104341 | 1461749 | TPA:-AAA-family-ATPase-[Salmonella-enterica-subsp.-enterica-serovar-Typhimurium-var.-monophasic-4,[5] |
| EJU7772065.1 | OLD-ABC | GCA_025208445.1 | Salmonella_enterica_subsp._salamae_serovar_4,12:e,n,x:1,6 | Bacteria>Pseudomonadati>Pseudomonadota>Gammaproteobacteria>Enterobacterales>Enterobacteriaceae>Salmonella>Salmonella_enterica>Salmonella_enterica_subsp._salamae | 363 | KAB52_003010 | 286782..287873 | 2978379 | ATP-binding-protein-[Salmonella-enterica-subsp.-salamae-serovar-4,12:e,n,x:1,6] |
| VUC90728.1 | OLD-ABC | GCA_902109495.1 | Salmonella_sp._NCTC_11881 | Bacteria>Pseudomonadati>Pseudomonadota>Gammaproteobacteria>Enterobacterales>Enterobacteriaceae>Salmonella>unclassified_Salmonella | 192 | NCTC11881_05567 | 4066314..4066892 | 2583579 | spermidine/putrescine-ABC-transporter-ATPase-[Salmonella-sp.-NCTC-11881] |
| VUC90727.1 | OLD-ABC | GCA_902109495.1 | Salmonella_sp._NCTC_11881 | Bacteria>Pseudomonadati>Pseudomonadota>Gammaproteobacteria>Enterobacterales>Enterobacteriaceae>Salmonella>unclassified_Salmonella | 149 | NCTC11881_05566 | 4065800..4066249 | 2583579 | spermidine/putrescine-ABC-transporter-ATPase-[Salmonella-sp.-NCTC-11881] |
| WP_007890772.1 | OLD-ABC | GCF_000282675.1 | Pantoea | Bacteria>Pseudomonadati>Pseudomonadota>Gammaproteobacteria>Enterobacterales>Erwiniaceae | 369 | PMI17_RS16955 | 199455..200564 | 53335 | MULTISPECIES:-AAA-family-ATPase-[unclassified-Pantoea] |
| WP_137241125.1 | OLD-ABC | GCF_005233495.2 | Pantoea_agglomerans | Bacteria>Pseudomonadati>Pseudomonadota>Gammaproteobacteria>Enterobacterales>Erwiniaceae>Pantoea>Pantoea_agglomerans_group | 366 | PagCFBP13505_RS08750 | 48187..49287 | 549 | ATP-binding-protein-[Pantoea-agglomerans] |
| WP_154209856.1 | OLD-ABC | GCF_009668105.1 | Pantoea_agglomerans | Bacteria>Pseudomonadati>Pseudomonadota>Gammaproteobacteria>Enterobacterales>Erwiniaceae>Pantoea>Pantoea_agglomerans_group | 347 | GJV05_RS04770 | 294878..295921 | 549 | ATP-binding-protein-[Pantoea-agglomerans] |
| WP_245115612.1 | OLD-ABC | GCF_022966105.1 | Pantoea_agglomerans | Bacteria>Pseudomonadati>Pseudomonadota>Gammaproteobacteria>Enterobacterales>Erwiniaceae>Pantoea>Pantoea_agglomerans_group | 347 | LZ609_RS16065 | 2639883..2640926 | 549 | ATP-binding-protein-[Pantoea-agglomerans] |
| WP_007890765.1 | OLD-ABC | GCF_000282675.1 | Pantoea_sp._GM01 | Bacteria>Pseudomonadati>Pseudomonadota>Gammaproteobacteria>Enterobacterales>Erwiniaceae>Pantoea>unclassified_Pantoea | 365 | PMI17_RS16915 | 189695..190792 | 1144320 | ATP-binding-protein-[Pantoea-sp.-GM01] |
| WP_222233108.1 | OLD-ABC | GCF_019795975.1 | Pantoea_sp._DY-17 | Bacteria>Pseudomonadati>Pseudomonadota>Gammaproteobacteria>Enterobacterales>Erwiniaceae>Pantoea>unclassified_Pantoea | 347 | K5Y33_RS01215 | 247361..248404 | 2871490 | ATP-binding-protein-[Pantoea-sp.-DY-17] |
| WP_226228487.1 | OLD-ABC | - | Hafnia_paralvei | Bacteria>Pseudomonadati>Pseudomonadota>Gammaproteobacteria>Enterobacterales>Hafniaceae>Hafnia | 366 | LA328_RS17870 | 3840625..3841725 | 546367 | ATP-binding-protein-[Hafnia-paralvei] |
| WP_265497330.1 | OLD-ABC | GCF_026173365.1 | Providencia | Bacteria>Pseudomonadati>Pseudomonadota>Gammaproteobacteria>Enterobacterales>Morganellaceae | 389 | ORM28_RS01810 | 13050..14219 | 586 | AAA-family-ATPase-[Providencia-rustigianii] |
| WP_241501908.1 | OLD-ABC | GCF_963899015.1 | Moellerella_wisconsensis | Bacteria>Pseudomonadati>Pseudomonadota>Gammaproteobacteria>Enterobacterales>Morganellaceae>Moellerella | 389 | MNY66_RS17050 | 200720..201889 | 158849 | AAA-family-ATPase-[Moellerella-wisconsensis] |
| WP_241543890.1 | OLD-ABC | GCF_963899445.1 | Moellerella_wisconsensis | Bacteria>Pseudomonadati>Pseudomonadota>Gammaproteobacteria>Enterobacterales>Morganellaceae>Moellerella | 389 | MNY64_RS16760 | 124500..125669 | 158849 | AAA-family-ATPase-[Moellerella-wisconsensis] |
| WP_209036129.1 | OLD-ABC | GCF_017693395.1 | Proteus_mirabilis | Bacteria>Pseudomonadati>Pseudomonadota>Gammaproteobacteria>Enterobacterales>Morganellaceae>Proteus | 389 | IGB04_RS13135 | 86297..87466 | 584 | AAA-family-ATPase-[Proteus-mirabilis] |
| MBI6417157.1 | OLD-ABC | GCA_016238675.1 | Proteus_mirabilis | Bacteria>Pseudomonadati>Pseudomonadota>Gammaproteobacteria>Enterobacterales>Morganellaceae>Proteus | 372 | JEP37_15640 | 62602..63720 | 584 | AAA-family-ATPase-[Proteus-mirabilis] |
| HAT2164888.1 | OLD-ABC | GCA_015896365.1 | Proteus_mirabilis | Bacteria>Pseudomonadati>Pseudomonadota>Gammaproteobacteria>Enterobacterales>Morganellaceae>Proteus | 371 | I8052_002913 | 19202..20317 | 584 | AAA-family-ATPase-[Proteus-mirabilis] |
| WP_137022262.1 | OLD-ABC | GCF_005157325.1 | Providencia_heimbachae | Bacteria>Pseudomonadati>Pseudomonadota>Gammaproteobacteria>Enterobacterales>Morganellaceae>Providencia | 365 | C9446_RS17900 | 3861004..3862101 | 333962 | AAA-family-ATPase-[Providencia-heimbachae] |
| WP_272668880.1 | OLD-ABC | GCF_028481005.1 | Providencia_sp._PROV151 | Bacteria>Pseudomonadati>Pseudomonadota>Gammaproteobacteria>Enterobacterales>Morganellaceae>Providencia>unclassified_Providencia | 372 | PSE52_RS01005 | 229805..230923 | 2949861 | AAA-family-ATPase-[Providencia-sp.-PROV151] |
| WP_045967685.1 | OLD-ABC | GCF_963866745.1 | Xenorhabdus_doucetiae | Bacteria>Pseudomonadati>Pseudomonadota>Gammaproteobacteria>Enterobacterales>Morganellaceae>Xenorhabdus | 372 | LY16_RS12975 | 19370..20488 | 351671 | AAA-family-ATPase-[Xenorhabdus-doucetiae] |
| WP_103199122.1 | OLD-ABC | GCF_002904085.1 | Pectobacterium_odoriferum | Bacteria>Pseudomonadati>Pseudomonadota>Gammaproteobacteria>Enterobacterales>Pectobacteriaceae>Pectobacterium | 366 | BV921_RS15380 | 109168..110268 | 78398 | ATP-binding-protein-[Pectobacterium-odoriferum] |
| WP_005974428.1 | OLD-ABC | GCF_000291725.1 | Pectobacterium_wasabiae | Bacteria>Pseudomonadati>Pseudomonadota>Gammaproteobacteria>Enterobacterales>Pectobacteriaceae>Pectobacterium | 364 | Y17_RS17680 | 17513..18607 | 55208 | ATP-binding-protein-[Pectobacterium-wasabiae] |
| WP_181847883.1 | OLD-ABC | GCF_013913645.1 | Pectobacterium_aroidearum | Bacteria>Pseudomonadati>Pseudomonadota>Gammaproteobacteria>Enterobacterales>Pectobacteriaceae>Pectobacterium | 364 | H2Y54_RS11275 | 231836..232930 | 1201031 | AAA-family-ATPase-[Pectobacterium-aroidearum] |
| WP_116155989.1 | OLD-ABC | GCF_003382585.2 | Pectobacterium_aquaticum | Bacteria>Pseudomonadati>Pseudomonadota>Gammaproteobacteria>Enterobacterales>Pectobacteriaceae>Pectobacterium | 363 | DMB81_RS10745 | 12237..13328 | 2204145 | ATP-binding-protein-[Pectobacterium-aquaticum] |
| WP_268198600.1 | OLD-ABC | GCF_026723725.1 | Pectobacterium_brasiliense | Bacteria>Pseudomonadati>Pseudomonadota>Gammaproteobacteria>Enterobacterales>Pectobacteriaceae>Pectobacterium | 350 | OWC53_RS08220 | 1777594..1778646 | 180957 | ATP-binding-protein-[Pectobacterium-brasiliense] |
| WP_103160991.1 | OLD-ABC | GCF_002904045.1 | Pectobacterium_odoriferum | Bacteria>Pseudomonadati>Pseudomonadota>Gammaproteobacteria>Enterobacterales>Pectobacteriaceae>Pectobacterium | 347 | BVY05_RS19620 | 81306..82349 | 78398 | ATP-binding-protein-[Pectobacterium-odoriferum] |
| GKX39593.1 | OLD-ABC | GCA_024343445.1 | Pectobacterium_carotovorum_subsp._carotovorum | Bacteria>Pseudomonadati>Pseudomonadota>Gammaproteobacteria>Enterobacterales>Pectobacteriaceae>Pectobacterium>Pectobacterium_carotovorum | 350 | SOASR014_33320 | 120112..121164 | 555 | hypothetical-protein-SOASR014_33320-[Pectobacterium-carotovorum-subsp.-carotovorum] |
| WP_112197429.1 | OLD-ABC | GCF_003263525.1 | Rahnella_sp._NRRL_B-41462 | Bacteria>Pseudomonadati>Pseudomonadota>Gammaproteobacteria>Enterobacterales>Yersiniaceae>Rahnella>unclassified_Rahnella | 362 | UC14_RS06680 | 38820..39908 | 1610579 | ATP-binding-protein-[Rahnella-sp.-NRRL-B-41462] |
| HBE9089662.1 | OLD-ABC | GCA_018627075.1 | Serratia_fonticola | Bacteria>Pseudomonadati>Pseudomonadota>Gammaproteobacteria>Enterobacterales>Yersiniaceae>Serratia | 347 | KNG84_001686 | 41955..42998 | 47917 | AAA-family-ATPase-[Serratia-fonticola] |
| WP_038251185.1 | OLD-ABC | GCF_000173755.1 | Yersinia_ruckeri | Bacteria>Pseudomonadati>Pseudomonadota>Gammaproteobacteria>Enterobacterales>Yersiniaceae>Yersinia | 373 | YRUCK0001_RS14090 | 22639..23760 | 29486 | AAA-family-ATPase-[Yersinia-ruckeri] |
| WP_053011789.1 | OLD-ABC | GCF_001244675.1 | Yersinia_intermedia | Bacteria>Pseudomonadati>Pseudomonadota>Gammaproteobacteria>Enterobacterales>Yersiniaceae>Yersinia | 366 | ALC37_RS11585 | 467242..468342 | 631 | ATP-binding-protein-[Yersinia-intermedia] |
| WP_050124852.1 | OLD-ABC | GCF_001115185.1 | Yersinia_aleksiciae | Bacteria>Pseudomonadati>Pseudomonadota>Gammaproteobacteria>Enterobacterales>Yersiniaceae>Yersinia | 360 | AEB18_RS00875 | 223975..225057 | 263819 | ATP-binding-protein-[Yersinia-aleksiciae] |
| WP_145532657.1 | OLD-ABC | GCF_902170345.1 | Yersinia_alsatica | Bacteria>Pseudomonadati>Pseudomonadota>Gammaproteobacteria>Enterobacterales>Yersiniaceae>Yersinia | 360 | FTB07_RS01250 | 29974..31056 | 2890317 | ATP-binding-protein-[Yersinia-alsatica] |
| WP_145580443.1 | OLD-ABC | GCF_902170795.1 | Yersinia_mollaretii | Bacteria>Pseudomonadati>Pseudomonadota>Gammaproteobacteria>Enterobacterales>Yersiniaceae>Yersinia | 360 | FTB33_RS16515 | 14062..15144 | 33060 | ATP-binding-protein-[Yersinia-mollaretii] |
| WP_145931272.1 | OLD-ABC | GCF_902168565.1 | Yersinia_bercovieri | Bacteria>Pseudomonadati>Pseudomonadota>Gammaproteobacteria>Enterobacterales>Yersiniaceae>Yersinia | 347 | FS908_RS01785 | 60994..62037 | 634 | ATP-binding-protein-[Yersinia-bercovieri] |
| QDW34851.1 | OLD-ABC | GCA_005937895.2 | Yersinia_sp._KBS0713 | Bacteria>Pseudomonadati>Pseudomonadota>Gammaproteobacteria>Enterobacterales>Yersiniaceae>Yersinia>unclassified_Yersinia_(in:_enterobacteria) | 361 | FFE93_018520 | 3900944..3902029 | 1179669 | AAA-family-ATPase-[Yersinia-sp.-KBS0713] |
| BAL53119.1 | OLD-ABC | - | uncultured_Gammaproteobacteria_bacterium | Bacteria>Pseudomonadati>Pseudomonadota>Gammaproteobacteria>environmental_samples | 435 | HGMM_F04D09C16 | 13995..15302 | 86473 | hypothetical-conserved-protein-[uncultured-Gammaproteobacteria-bacterium] |
| WP_132538609.1 | OLD-ABC | GCF_004341245.1 | Plasticicumulans_lactativorans | Bacteria>Pseudomonadati>Pseudomonadota>Gammaproteobacteria>Gammaproteobacteria_incertae_sedis>Candidatus_Competibacteraceae>Plasticicumulans | 426 | EV699_RS02460 | 28925..30205 | 1133106 | ATP-binding-protein-[Plasticicumulans-lactativorans] |
| MBK8754675.1 | OLD-ABC | GCA_016710535.1 | Candidatus_Competibacteraceae_bacterium | Bacteria>Pseudomonadati>Pseudomonadota>Gammaproteobacteria>Gammaproteobacteria_incertae_sedis>Candidatus_Competibacteraceae>unclassified_Candidatus_Competibacteraceae | 412 | IPL99_24900 | 43000..44238 | 2053538 | AAA-family-ATPase-[Candidatus-Competibacteraceae-bacterium] |
| MBK8754769.1 | OLD-ABC | GCA_016710535.1 | Candidatus_Competibacteraceae_bacterium | Bacteria>Pseudomonadati>Pseudomonadota>Gammaproteobacteria>Gammaproteobacteria_incertae_sedis>Candidatus_Competibacteraceae>unclassified_Candidatus_Competibacteraceae | 391 | IPL99_25405 | 31031..32206 | 2053538 | AAA-family-ATPase-[Candidatus-Competibacteraceae-bacterium] |
| WP_248566602.1 | OLD-ABC | GCF_023195885.1 | Methylonatrum_kenyense | Bacteria>Pseudomonadati>Pseudomonadota>Gammaproteobacteria>Gammaproteobacteria_incertae_sedis>Methylonatrum | 361 | M0534_RS01330 | 281124..282209 | 455253 | AAA-family-ATPase-[Methylonatrum-kenyense] |
| MAZ66825.1 | OLD-ABC | GCA_002707725.1 | Kangiellaceae_bacterium | Bacteria>Pseudomonadati>Pseudomonadota>Gammaproteobacteria>Kangiellales>Kangiellaceae>unclassified_Kangiellaceae | 371 | CMF25_06920 | 6447..7562 | 2026751 | ABC-transporter-ATP-binding-protein-[Kangiellaceae-bacterium] |
| WP_054661689.1 | OLD-ABC | GCF_002739155.1 | Stenotrophomonas | Bacteria>Pseudomonadati>Pseudomonadota>Gammaproteobacteria>Lysobacterales>Lysobacteraceae | 409 | BRB81_RS09930 | 110201..111430 | 40323 | MULTISPECIES:-AAA-family-ATPase-[Stenotrophomonas] |
| PZO64465.1 | OLD-ABC | GCA_003240695.1 | Pseudoxanthomonas_suwonensis | Bacteria>Pseudomonadati>Pseudomonadota>Gammaproteobacteria>Lysobacterales>Lysobacteraceae>Pseudoxanthomonas | 377 | DI635_05305 | 4159..5292 | 314722 | hypothetical-protein-DI635_05305-[Pseudoxanthomonas-suwonensis] |
| ALJ26655.1 | OLD-ABC | GCA_001314305.1 | Stenotrophomonas_acidaminiphila | Bacteria>Pseudomonadati>Pseudomonadota>Gammaproteobacteria>Lysobacterales>Lysobacteraceae>Stenotrophomonas | 409 | AOT14_01940 | 237843..239072 | 128780 | ATPase-[Stenotrophomonas-acidaminiphila] |
| MTI74444.1 | OLD-ABC | GCA_009711865.1 | Stenotrophomonas_sp. | Bacteria>Pseudomonadati>Pseudomonadota>Gammaproteobacteria>Lysobacterales>Lysobacteraceae>Stenotrophomonas>unclassified_Stenotrophomonas | 421 | FH747_12435 | 112267..113532 | 69392 | ATP-binding-protein-[Stenotrophomonas-sp.] |
| WP_230434265.1 | OLD-ABC | GCF_020990895.1 | Xanthomonas_melonis | Bacteria>Pseudomonadati>Pseudomonadota>Gammaproteobacteria>Lysobacterales>Lysobacteraceae>Xanthomonas | 394 | JWH05_RS03160 | 13722..14906 | 56456 | AAA-family-ATPase-[Xanthomonas-melonis] |
| WP_259393301.1 | OLD-ABC | - | Dokdonella_fugitiva | Bacteria>Pseudomonadati>Pseudomonadota>Gammaproteobacteria>Lysobacterales>Rhodanobacteraceae>Dokdonella | 390 | FHW13_RS05505 | 302296..303468 | 328517 | AAA-family-ATPase-[Dokdonella-fugitiva] |
| WP_262965892.1 | OLD-ABC | GCF_025583945.1 | Methylobacter_psychrophilus | Bacteria>Pseudomonadati>Pseudomonadota>Gammaproteobacteria>Methylococcales>Methylococcaceae>Methylobacter | 381 | N7974_RS12250 | 22374..23519 | 96941 | ATP-binding-protein-[Methylobacter-psychrophilus] |
| WP_031438513.1 | OLD-ABC | GCF_000733835.1 | Methylobacter_tundripaludum | Bacteria>Pseudomonadati>Pseudomonadota>Gammaproteobacteria>Methylococcales>Methylococcaceae>Methylobacter | 380 | T449_RS0121495 | 4647389..4648531 | 173365 | ATP-binding-protein-[Methylobacter-tundripaludum] |
| WP_218796633.1 | OLD-ABC | GCF_019285555.1 | Methylococcus_capsulatus | Bacteria>Pseudomonadati>Pseudomonadota>Gammaproteobacteria>Methylococcales>Methylococcaceae>Methylococcus | 436 | KW114_RS08320 | 1836292..1837602 | 414 | AAA-family-ATPase-[Methylococcus-capsulatus] |
| WP_255188311.1 | OLD-ABC | GCF_024360925.2 | Methylomonas_rapida | Bacteria>Pseudomonadati>Pseudomonadota>Gammaproteobacteria>Methylococcales>Methylococcaceae>Methylomonas | 386 | NM686_RS13215 | 2819819..2820979 | 2963939 | ATP-binding-protein-[Methylomonas-rapida] |
| WP_255187195.1 | OLD-ABC | GCF_024360925.2 | Methylomonas_rapida | Bacteria>Pseudomonadati>Pseudomonadota>Gammaproteobacteria>Methylococcales>Methylococcaceae>Methylomonas | 356 | NM686_RS07165 | 1512858..1513928 | 2963939 | AAA-family-ATPase-[Methylomonas-rapida] |
| WP_255189601.1 | OLD-ABC | GCF_024360925.2 | Methylomonas_rapida | Bacteria>Pseudomonadati>Pseudomonadota>Gammaproteobacteria>Methylococcales>Methylococcaceae>Methylomonas | 351 | NM686_RS20135 | 4254635..4255690 | 2963939 | ATP-binding-protein-[Methylomonas-rapida] |
| MBS3964851.1 | OLD-ABC | GCA_018333915.1 | Methylomonas_sp. | Bacteria>Pseudomonadati>Pseudomonadota>Gammaproteobacteria>Methylococcales>Methylococcaceae>Methylomonas>unclassified_Methylomonas | 357 | KGZ80_10230 | 14549..15622 | 418 | ATP-binding-protein-[Methylomonas-sp.] |
| NOT11399.1 | OLD-ABC | GCA_013140705.1 | Methylococcaceae_bacterium | Bacteria>Pseudomonadati>Pseudomonadota>Gammaproteobacteria>Methylococcales>Methylococcaceae>unclassified_Methylococcaceae | 427 | HOP23_06145 | 40843..42126 | 1933926 | ATP-binding-protein-[Methylococcaceae-bacterium] |
| NOT11402.1 | OLD-ABC | GCA_013140705.1 | Methylococcaceae_bacterium | Bacteria>Pseudomonadati>Pseudomonadota>Gammaproteobacteria>Methylococcales>Methylococcaceae>unclassified_Methylococcaceae | 394 | HOP23_06160 | 45753..46937 | 1933926 | AAA-family-ATPase-[Methylococcaceae-bacterium] |
| WP_108675355.1 | OLD-ABC | GCF_003072585.2 | Acinetobacter | Bacteria>Pseudomonadati>Pseudomonadota>Gammaproteobacteria>Moraxellales>Moraxellaceae | 389 | C0119_RS05585 | 1124391..1125560 | 469 | MULTISPECIES:-AAA-family-ATPase-[Acinetobacter] |
| WP_087528530.1 | OLD-ABC | GCF_002165305.2 | Acinetobacter_sp._WCHA55 | Bacteria>Pseudomonadati>Pseudomonadota>Gammaproteobacteria>Moraxellales>Moraxellaceae>Acinetobacter>unclassified_Acinetobacter | 375 | CDG62_RS00805 | 30673..31800 | 2004646 | ATP-binding-protein-[Acinetobacter-sp.-WCHA55] |
| WP_107866911.1 | OLD-ABC | GCF_003051055.1 | Agitococcus_lubricus | Bacteria>Pseudomonadati>Pseudomonadota>Gammaproteobacteria>Moraxellales>Moraxellaceae>Agitococcus | 372 | C8N29_RS15155 | 15245..16363 | 1077255 | ATP-binding-protein-[Agitococcus-lubricus] |
| OLF41365.1 | OLD-ABC | GCA_001921755.1 | Psychrobacter_sp._Rd_27.2 | Bacteria>Pseudomonadati>Pseudomonadota>Gammaproteobacteria>Moraxellales>Moraxellaceae>Psychrobacter>unclassified_Psychrobacter | 369 | BTV99_04800 | 271798..272907 | 1926479 | recombinase-RecF-[Psychrobacter-sp.-Rd-27.2] |
| WP_075101226.1 | OLD-ABC | GCF_001921755.1 | Psychrobacter_sp._Rd_27.2 | Bacteria>Pseudomonadati>Pseudomonadota>Gammaproteobacteria>Moraxellales>Moraxellaceae>Psychrobacter>unclassified_Psychrobacter | 369 | BTV99_RS04795 | 271798..272907 | 1926479 | AAA-family-ATPase-[Psychrobacter-sp.-Rd-27.2] |
| MCP5177611.1 | OLD-ABC | GCA_024234655.1 | Moraxellaceae_bacterium | Bacteria>Pseudomonadati>Pseudomonadota>Gammaproteobacteria>Moraxellales>Moraxellaceae>unclassified_Moraxellaceae | 394 | H6997_08680 | 40843..42027 | 1889775 | AAA-family-ATPase-[Moraxellaceae-bacterium] |
| TQC95076.1 | OLD-ABC | GCA_006546595.1 | Moraxellaceae_bacterium_AER2_44_116 | Bacteria>Pseudomonadati>Pseudomonadota>Gammaproteobacteria>Moraxellales>Moraxellaceae>unclassified_Moraxellaceae | 386 | FK216_14940 | 7376..8536 | 2591038 | AAA-family-ATPase-[Moraxellaceae-bacterium-AER2_44_116] |
| WP_093286148.1 | OLD-ABC | GCF_900111015.1 | Solimonas_aquatica | Bacteria>Pseudomonadati>Pseudomonadota>Gammaproteobacteria>Nevskiales>Nevskiaceae>Solimonas | 356 | BMY12_RS11540 | 201684..202754 | 489703 | AAA-family-ATPase-[Solimonas-aquatica] |
| WP_041158394.1 | OLD-ABC | GCF_900099955.1 | Halomonadaceae | Bacteria>Pseudomonadati>Pseudomonadota>Gammaproteobacteria>Oceanospirillales | 372 | BLR10_RS07680 | 63903..65021 | 28256 | MULTISPECIES:-ATP-binding-protein-[Halomonas] |
| HIK75726.1 | OLD-ABC | GCA_015660495.1 | Alcanivorax_sp. | Bacteria>Pseudomonadati>Pseudomonadota>Gammaproteobacteria>Oceanospirillales>Alcanivoracaceae>Alcanivorax>unclassified_Alcanivorax | 370 | EYG00_13655 | 7038..8150 | 1872427 | hypothetical-protein-[Alcanivorax-sp.] |
| MAD69922.1 | OLD-ABC | GCA_002685555.1 | Alcanivorax_sp. | Bacteria>Pseudomonadati>Pseudomonadota>Gammaproteobacteria>Oceanospirillales>Alcanivoracaceae>Alcanivorax>unclassified_Alcanivorax | 370 | CL538_05295 | 40053..41165 | 1872427 | hypothetical-protein-[Alcanivorax-sp.] |
| MBE8233360.1 | OLD-ABC | GCA_015163765.1 | Endozoicomonadaceae_bacterium | Bacteria>Pseudomonadati>Pseudomonadota>Gammaproteobacteria>Oceanospirillales>Endozoicomonadaceae>unclassified_Endozoicomonadaceae | 393 | HAW67_06450 | 9243..10424 | 2590577 | AAA-family-ATPase-[Endozoicomonadaceae-bacterium] |
| WP_011397284.1 | OLD-ABC | GCF_000012985.1 | Hahella_chejuensis | Bacteria>Pseudomonadati>Pseudomonadota>Gammaproteobacteria>Oceanospirillales>Hahellaceae>Hahella | 386 | HCH_RS15595 | 3601492..3602652 | 158327 | AAA-family-ATPase-[Hahella-chejuensis] |
| WP_133481542.1 | OLD-ABC | GCF_004361885.1 | Halomonas_ventosae | Bacteria>Pseudomonadati>Pseudomonadota>Gammaproteobacteria>Oceanospirillales>Halomonadaceae>Halomonas | 426 | DFO68_RS01970 | 11011..12291 | 229007 | AAA-family-ATPase-[Halomonas-ventosae] |
| WP_101147212.1 | OLD-ABC | GCF_037070215.1 | unclassified_Halomonas | Bacteria>Pseudomonadati>Pseudomonadota>Gammaproteobacteria>Oceanospirillales>Halomonadaceae>Halomonas | 356 | CXF87_RS15375 | 111848..112918 | 2609666 | ATP-binding-protein-[Halomonas-sp.-MES3-P3E] |
| WP_172420969.1 | OLD-ABC | GCF_011399095.1 | Halomonas_hydrothermalis | Bacteria>Pseudomonadati>Pseudomonadota>Gammaproteobacteria>Oceanospirillales>Halomonadaceae>Halomonas | 356 | HHSLTHF2_RS10285 | 2146810..2147880 | 115561 | ATP-binding-protein-[Halomonas-hydrothermalis] |
| WP_081191893.1 | OLD-ABC | - | Halomonas_sp._BC1 | Bacteria>Pseudomonadati>Pseudomonadota>Gammaproteobacteria>Oceanospirillales>Halomonadaceae>Halomonas>unclassified_Halomonas | 394 | B7L72_RS00120 | 28903..30087 | 1670448 | ATP-binding-protein-[Halomonas-sp.-BC1] |
| WP_218314909.1 | OLD-ABC | GCF_019211725.1 | Halomonas_sp._18071143 | Bacteria>Pseudomonadati>Pseudomonadota>Gammaproteobacteria>Oceanospirillales>Halomonadaceae>Halomonas>unclassified_Halomonas | 394 | KVY02_RS07740 | 1670800..1671984 | 2855441 | ATP-binding-protein-[Halomonas-sp.-18071143] |
| WP_169956452.1 | OLD-ABC | GCF_013000985.1 | Halomonas_sp._PGE1 | Bacteria>Pseudomonadati>Pseudomonadota>Gammaproteobacteria>Oceanospirillales>Halomonadaceae>Halomonas>unclassified_Halomonas | 370 | HIR79_RS02125 | 430733..431845 | 2730360 | AAA-family-ATPase-[Halomonas-sp.-PGE1] |
| WP_138801526.1 | OLD-ABC | GCF_005937175.1 | Halomonas_sp._ATBC28 | Bacteria>Pseudomonadati>Pseudomonadota>Gammaproteobacteria>Oceanospirillales>Halomonadaceae>Halomonas>unclassified_Halomonas | 366 | E0L35_RS22995 | 8544..9644 | 2545264 | AAA-family-ATPase-[Halomonas-sp.-ATBC28] |
| WP_231664591.1 | OLD-ABC | GCF_021151495.1 | Halomonas_sp._IOP_31 | Bacteria>Pseudomonadati>Pseudomonadota>Gammaproteobacteria>Oceanospirillales>Halomonadaceae>Halomonas>unclassified_Halomonas | 361 | K7I12_RS15005 | 16780..17865 | 2876584 | AAA-family-ATPase-[Halomonas-sp.-IOP_31] |
| WP_062374374.1 | OLD-ABC | GCF_001596805.1 | Halomonas_sp._KX33721 | Bacteria>Pseudomonadati>Pseudomonadota>Gammaproteobacteria>Oceanospirillales>Halomonadaceae>Halomonas>unclassified_Halomonas | 356 | A3220_RS08375 | 75327..76397 | 1819251 | ATP-binding-protein-[Halomonas-sp.-KX33721] |
| WP_193059743.1 | OLD-ABC | GCF_052263255.1 | Vreelandella_alkaliphila | Bacteria>Pseudomonadati>Pseudomonadota>Gammaproteobacteria>Oceanospirillales>Halomonadaceae>Vreelandella | 398 | INT28_RS17805 | 13007..14203 | 272774 | MULTISPECIES:-ATP-binding-protein-[unclassified-Halomonas] |
| WP_079550436.1 | OLD-ABC | GCF_900142895.1 | Vreelandella_subglaciescola | Bacteria>Pseudomonadati>Pseudomonadota>Gammaproteobacteria>Oceanospirillales>Halomonadaceae>Vreelandella | 394 | B5495_RS00620 | 140551..141735 | 29571 | ATP-binding-protein-[Halomonas-subglaciescola] |
| WP_027853084.1 | OLD-ABC | GCF_000428985.1 | Marinobacterium_litorale | Bacteria>Pseudomonadati>Pseudomonadota>Gammaproteobacteria>Oceanospirillales>Oceanospirillaceae>Marinobacterium | 425 | G403_RS0105360 | 183432..184709 | 404770 | ATP-binding-protein-[Marinobacterium-litorale] |
| WP_277987740.1 | OLD-ABC | GCF_017310015.1 | Marinobacterium_iners | Bacteria>Pseudomonadati>Pseudomonadota>Gammaproteobacteria>Oceanospirillales>Oceanospirillaceae>Marinobacterium | 425 | CFI10_RS02430 | 490682..491959 | 48076 | ATP-binding-protein-[Marinobacterium-iners] |
| WP_254579408.1 | OLD-ABC | GCF_024267675.1 | Marinobacterium_sedimentorum | Bacteria>Pseudomonadati>Pseudomonadota>Gammaproteobacteria>Oceanospirillales>Oceanospirillaceae>Marinobacterium | 412 | MRS74_RS06190 | 158050..159288 | 2927804 | AAA-family-ATPase-[Marinobacterium-sedimentorum] |
| WP_027853086.1 | OLD-ABC | GCF_000428985.1 | Marinobacterium_litorale | Bacteria>Pseudomonadati>Pseudomonadota>Gammaproteobacteria>Oceanospirillales>Oceanospirillaceae>Marinobacterium | 367 | G403_RS0105370 | 185386..186489 | 404770 | AAA-family-ATPase-[Marinobacterium-litorale] |
| WP_206838936.1 | OLD-ABC | GCF_017310015.1 | Marinobacterium_iners | Bacteria>Pseudomonadati>Pseudomonadota>Gammaproteobacteria>Oceanospirillales>Oceanospirillaceae>Marinobacterium | 367 | CFI10_RS02420 | 488902..490005 | 48076 | AAA-family-ATPase-[Marinobacterium-iners] |
| WP_067206356.1 | OLD-ABC | GCF_900089755.1 | Marinomonas_aquimarina | Bacteria>Pseudomonadati>Pseudomonadota>Gammaproteobacteria>Oceanospirillales>Oceanospirillaceae>Marinomonas | 389 | MAQ5080_RS04930 | 82750..83919 | 295068 | AAA-family-ATPase-[Marinomonas-aquimarina] |
| WP_226752735.1 | OLD-ABC | GCF_020532605.1 | Marinomonas_algarum | Bacteria>Pseudomonadati>Pseudomonadota>Gammaproteobacteria>Oceanospirillales>Oceanospirillaceae>Marinomonas | 389 | LG368_RS00220 | 45245..46414 | 2883105 | AAA-family-ATPase-[Marinomonas-algarum] |
| WP_036514530.1 | OLD-ABC | GCF_000585235.1 | Nitrincola_nitratireducens | Bacteria>Pseudomonadati>Pseudomonadota>Gammaproteobacteria>Oceanospirillales>Oceanospirillaceae>Nitrincola | 445 | D791_RS17650 | 11002..12339 | 1229521 | ATP-binding-protein-[Nitrincola-nitratireducens] |
| QEW05877.1 | OLD-ABC | GCA_008727795.1 | Nitrincola_iocasae | Bacteria>Pseudomonadati>Pseudomonadota>Gammaproteobacteria>Oceanospirillales>Oceanospirillaceae>Nitrincola | 412 | F5I99_04895 | 1040107..1041345 | 2614693 | AAA-family-ATPase-[Nitrincola-iocasae] |
| WP_151053916.1 | OLD-ABC | GCF_008727795.1 | Nitrincola_iocasae | Bacteria>Pseudomonadati>Pseudomonadota>Gammaproteobacteria>Oceanospirillales>Oceanospirillaceae>Nitrincola | 412 | F5I99_RS04895 | 1040107..1041345 | 2614693 | AAA-family-ATPase-[Nitrincola-iocasae] |

| Protein-ID | Domain | Assembly-ID | Strain | Taxonomy | Length | Locus-Tag | Genomic-coordinates | TaxID | Fasta-defline |
|---|---|---|---|---|---|---|---|---|---|
| WP_202806903.1 | OLD-ABC | GCF_000585235.1 | Nitrincola_nitratireducens | Bacteria>Pseudomonadati>Pseudomonadota>Gammaproteobacteria>Oceanospirillales>Oceanospirillaceae>Nitrincola | 257 | D791_RS17635 | 5914..6687 | 1229521 | AAA-family-ATPase-[Nitrincola-nitratireducens] |
| WP_260997648.1 | OLD-ABC | GCF_025345565.1 | Thalassolituus_hydrocarboniclasticus | Bacteria>Pseudomonadati>Pseudomonadota>Gammaproteobacteria>Oceanospirillales>Oceanospirillaceae>Thalassolituus | 367 | HUF19_RS16700 | 3716744..3717847 | 2742796 | AAA-family-ATPase-[Thalassolituus-hydrocarbonoclasticus] |
| MBM94538.1 | OLD-ABC | GCA_002723385.1 | Oceanospirillaceae_bacterium | Bacteria>Pseudomonadati>Pseudomonadota>Gammaproteobacteria>Oceanospirillales>Oceanospirillaceae>unclassified_Oceanospirillaceae | 389 | CMI09_01630 | 46663..47832 | 1899355 | chromosome-segregation-protein-SMC-[Oceanospirillaceae-bacterium] |
| MBM94535.1 | OLD-ABC | GCA_002723385.1 | Oceanospirillaceae_bacterium | Bacteria>Pseudomonadati>Pseudomonadota>Gammaproteobacteria>Oceanospirillales>Oceanospirillaceae>unclassified_Oceanospirillaceae | 371 | CMI09_01615 | 41058..42173 | 1899355 | ABC-transporter-ATP-binding-protein-[Oceanospirillaceae-bacterium] |
| WP_241504074.1 | OLD-ABC | GCF_963878705.1 | Pelagibaculum_spongiae | Bacteria>Pseudomonadati>Pseudomonadota>Gammaproteobacteria>Oceanospirillales>Oceanospirillales_incertae_sedis>Pelagibaculum | 401 | DC094_RS16065 | 169346..170551 | 2080658 | AAA-family-ATPase-[Pelagibaculum-spongiae] |
| WP_116688143.1 | OLD-ABC | GCF_963878705.1 | Pelagibaculum_spongiae | Bacteria>Pseudomonadati>Pseudomonadota>Gammaproteobacteria>Oceanospirillales>Oceanospirillales_incertae_sedis>Pelagibaculum | 379 | DC094_RS16055 | 167611..168750 | 2080658 | AAA-family-ATPase-[Pelagibaculum-spongiae] |
| MBR9867109.1 | OLD-ABC | GCA_018223265.1 | Oceanospirillales_bacterium | Bacteria>Pseudomonadati>Pseudomonadota>Gammaproteobacteria>Oceanospirillales>unclassified_Oceanospirillales | 371 | GYB18_08400 | 14593..15708 | 1914300 | AAA-family-ATPase-[Oceanospirillales-bacterium] |
| WP_057391036.1 | OLD-ABC | GCF_905479555.1 | Pseudomonadaceae | Bacteria>Pseudomonadati>Pseudomonadota>Gammaproteobacteria>Pseudomonadales | 367 | ACP89_RS12345 | 14716..15819 | 135621 | MULTISPECIES:-AAA-family-ATPase-[Pseudomonas-aeruginosa-group] |
| WP_022988845.1 | OLD-ABC | GCF_000475255.1 | Marinobacter | Bacteria>Pseudomonadati>Pseudomonadota>Gammaproteobacteria>Pseudomonadales>Marinobacteraceae | 426 | DET50_RS14965 | 49539..50819 | 2742 | MULTISPECIES:-AAA-family-ATPase-[Marinobacter] |
| WP_023008349.1 | OLD-ABC | GCF_000475375.1 | unclassified_Marinobacter | Bacteria>Pseudomonadati>Pseudomonadota>Gammaproteobacteria>Pseudomonadales>Marinobacteraceae>Marinobacter | 426 | Q674_RS26665 | 46186..47466 | 83889 | AAA-family-ATPase-[Marinobacter-sp.-EVN1] |
| WP_092021232.1 | OLD-ABC | GCF_900114775.1 | Marinobacter_zhejiangensis | Bacteria>Pseudomonadati>Pseudomonadota>Gammaproteobacteria>Pseudomonadales>Marinobacteraceae>Marinobacter | 376 | BMZ67_RS06970 | 343610..344740 | 488535 | ATP-binding-protein-[Marinobacter-zhejiangensis] |
| BBJ05057.1 | OLD-ABC | - | Marinobacter_nauticus | Bacteria>Pseudomonadati>Pseudomonadota>Gammaproteobacteria>Pseudomonadales>Marinobacteraceae>Marinobacter | 370 | YBY_29060 | 3197860..3198972 | 2743 | chromosome-segregation-protein-SMC-[Marinobacter-nauticus] |
| WP_011784106.1 | OLD-ABC | GCF_000015365.1 | Marinobacter_nauticus | Bacteria>Pseudomonadati>Pseudomonadota>Gammaproteobacteria>Pseudomonadales>Marinobacteraceae>Marinobacter | 370 | MAQU_RS02830 | 641372..642484 | 2743 | AAA-family-ATPase-[Marinobacter-nauticus] |
| EHJ03980.1 | OLD-ABC | GCA_000235625.2 | Marinobacter_manganoxydans_MnI7-9 | Bacteria>Pseudomonadati>Pseudomonadota>Gammaproteobacteria>Pseudomonadales>Marinobacteraceae>Marinobacter>Marinobacter_manganoxydans | 370 | KYE_13705 | 51134..52246 | 1094979 | hypothetical-protein-KYE_13705-[Marinobacter-manganoxydans-MnI7-9] |
| MBC7192644.1 | OLD-ABC | GCA_014359475.1 | Marinobacter_sp. | Bacteria>Pseudomonadati>Pseudomonadota>Gammaproteobacteria>Pseudomonadales>Marinobacteraceae>Marinobacter>unclassified_Marinobacter | 427 | H5U31_09860 | 10976..12259 | 50741 | ATP-binding-protein-[Marinobacter-sp.] |
| WP_022988848.1 | OLD-ABC | GCF_000475255.1 | Marinobacter_sp._ES-1 | Bacteria>Pseudomonadati>Pseudomonadota>Gammaproteobacteria>Pseudomonadales>Marinobacteraceae>Marinobacter>unclassified_Marinobacter | 388 | Q668_RS06790 | 85551..86717 | 1396858 | ATP-binding-protein-[Marinobacter-sp.-ES-1] |
| WP_022988843.1 | OLD-ABC | GCF_000475255.1 | Marinobacter_sp._ES-1 | Bacteria>Pseudomonadati>Pseudomonadota>Gammaproteobacteria>Pseudomonadales>Marinobacteraceae>Marinobacter>unclassified_Marinobacter | 370 | Q668_RS06765 | 78420..79532 | 1396858 | AAA-family-ATPase-[Marinobacter-sp.-ES-1] |
| MBC7192646.1 | OLD-ABC | GCA_014359475.1 | Marinobacter_sp. | Bacteria>Pseudomonadati>Pseudomonadota>Gammaproteobacteria>Pseudomonadales>Marinobacteraceae>Marinobacter>unclassified_Marinobacter | 361 | H5U31_09870 | 12977..14062 | 50741 | AAA-family-ATPase-[Marinobacter-sp.] |
| KXS54421.1 | OLD-ABC | GCA_001577575.1 | Marinobacter_sp._T13-3 | Bacteria>Pseudomonadati>Pseudomonadota>Gammaproteobacteria>Pseudomonadales>Marinobacteraceae>Marinobacter>unclassified_Marinobacter | 360 | AWU57_1203 | 13015..14097 | 1794906 | SMC-domain-containing-protein-[Marinobacter-sp.-T13-3] |
| WP_152390155.1 | OLD-ABC | GCF_009363155.1 | Azotobacter_salinestris | Bacteria>Pseudomonadati>Pseudomonadota>Gammaproteobacteria>Pseudomonadales>Pseudomonadaceae>Azotobacter_group>Azotobacter | 389 | GCU53_RS24410 | 144185..145354 | 69964 | ATP-binding-protein-[Azotobacter-salinestris] |
| WP_206408614.1 | OLD-ABC | GCF_041378955.2 | Ectopseudomonas_oleovorans | Bacteria>Pseudomonadati>Pseudomonadota>Gammaproteobacteria>Pseudomonadales>Pseudomonadaceae>Ectopseudomonas | 424 | ACP89_RS12330 | 10959..12233 | 301 | AAA-family-ATPase-[Pseudomonas-oleovorans] |
| MBA1269668.1 | OLD-ABC | GCA_013522615.1 | Pseudomonas_carnis | Bacteria>Pseudomonadati>Pseudomonadota>Gammaproteobacteria>Pseudomonadales>Pseudomonadaceae>Pseudomonas | 392 | G7029_19570 | 129428..130606 | 2487355 | AAA-family-ATPase-[Pseudomonas-carnis] |
| WP_181127556.1 | OLD-ABC | GCF_013522615.1 | Pseudomonas_carnis | Bacteria>Pseudomonadati>Pseudomonadota>Gammaproteobacteria>Pseudomonadales>Pseudomonadaceae>Pseudomonas | 392 | G7029_RS19480 | 129428..130606 | 2487355 | AAA-family-ATPase-[Pseudomonas-carnis] |
| WP_161892247.1 | OLD-ABC | GCF_009932375.1 | Pseudomonas_juntendi | Bacteria>Pseudomonadati>Pseudomonadota>Gammaproteobacteria>Pseudomonadales>Pseudomonadaceae>Pseudomonas | 367 | GYA94_RS10095 | 83602..84705 | 2666183 | AAA-family-ATPase-[Pseudomonas-juntendi] |
| WP_204881709.1 | OLD-ABC | GCF_900682675.2 | Pseudomonas_marincola | Bacteria>Pseudomonadati>Pseudomonadota>Gammaproteobacteria>Pseudomonadales>Pseudomonadaceae>Pseudomonas | 367 | PMYSY11_RS11200 | 2409762..2410865 | 437900 | AAA-family-ATPase-[Pseudomonas-marincola] |
| WP_229999478.1 | OLD-ABC | GCF_918697895.1 | Pseudomonas_carnis | Bacteria>Pseudomonadati>Pseudomonadota>Gammaproteobacteria>Pseudomonadales>Pseudomonadaceae>Pseudomonas | 367 | LQ224_RS08500 | 119151..120254 | 2487355 | AAA-family-ATPase-[Pseudomonas-carnis] |
| HCL3757800.1 | OLD-ABC | GCA_025003865.1 | Pseudomonas_aeruginosa | Bacteria>Pseudomonadati>Pseudomonadota>Gammaproteobacteria>Pseudomonadales>Pseudomonadaceae>Pseudomonas>Pseudomonas_aeruginosa_group | 436 | N2B17_005728 | 4296..5606 | 287 | AAA-family-ATPase-[Pseudomonas-aeruginosa] |
| WP_124078195.1 | OLD-ABC | GCF_033140865.1 | Pseudomonas_aeruginosa | Bacteria>Pseudomonadati>Pseudomonadota>Gammaproteobacteria>Pseudomonadales>Pseudomonadaceae>Pseudomonas>Pseudomonas_aeruginosa_group | 436 | IPC1239_RS13860 | 652492..653802 | 287 | AAA-family-ATPase-[Pseudomonas-aeruginosa] |
| MBX6030768.1 | OLD-ABC | GCA_019688625.1 | Pseudomonas_aeruginosa | Bacteria>Pseudomonadati>Pseudomonadota>Gammaproteobacteria>Pseudomonadales>Pseudomonadaceae>Pseudomonas>Pseudomonas_aeruginosa_group | 410 | ISE67_08715 | 281473..282705 | 287 | AAA-family-ATPase-[Pseudomonas-aeruginosa] |
| RRI70511.1 | OLD-ABC | GCA_003863585.1 | Pseudomonas_aeruginosa | Bacteria>Pseudomonadati>Pseudomonadota>Gammaproteobacteria>Pseudomonadales>Pseudomonadaceae>Pseudomonas>Pseudomonas_aeruginosa_group | 409 | EIM10_18025 | 71149..72378 | 287 | ATPase-[Pseudomonas-aeruginosa] |
| WP_124988194.1 | OLD-ABC | GCF_003863815.1 | Pseudomonas_aeruginosa | Bacteria>Pseudomonadati>Pseudomonadota>Gammaproteobacteria>Pseudomonadales>Pseudomonadaceae>Pseudomonas>Pseudomonas_aeruginosa_group | 409 | EIM11_RS16900 | 70280..71509 | 287 | AAA-family-ATPase-[Pseudomonas-aeruginosa] |
| WP_247258976.1 | OLD-ABC | GCF_032334355.1 | Pseudomonas_aeruginosa | Bacteria>Pseudomonadati>Pseudomonadota>Gammaproteobacteria>Pseudomonadales>Pseudomonadaceae>Pseudomonas>Pseudomonas_aeruginosa_group | 409 | MX104_RS19090 | 63576..64805 | 287 | AAA-family-ATPase-[Pseudomonas-aeruginosa] |
| WP_270932597.1 | OLD-ABC | GCF_027856215.1 | Pseudomonas_aeruginosa | Bacteria>Pseudomonadati>Pseudomonadota>Gammaproteobacteria>Pseudomonadales>Pseudomonadaceae>Pseudomonas>Pseudomonas_aeruginosa_group | 409 | JR315_RS22290 | 17080..18309 | 287 | AAA-family-ATPase-[Pseudomonas-aeruginosa] |
| HCK4430922.1 | OLD-ABC | GCA_024830635.1 | Pseudomonas_aeruginosa | Bacteria>Pseudomonadati>Pseudomonadota>Gammaproteobacteria>Pseudomonadales>Pseudomonadaceae>Pseudomonas>Pseudomonas_aeruginosa_group | 396 | N0G72_005467 | 44394..45584 | 287 | AAA-family-ATPase-[Pseudomonas-aeruginosa] |
| HCL3757803.1 | OLD-ABC | GCA_025003865.1 | Pseudomonas_aeruginosa | Bacteria>Pseudomonadati>Pseudomonadota>Gammaproteobacteria>Pseudomonadales>Pseudomonadaceae>Pseudomonas>Pseudomonas_aeruginosa_group | 396 | N2B17_005731 | 9107..10297 | 287 | AAA-family-ATPase-[Pseudomonas-aeruginosa] |
| WP_124078190.1 | OLD-ABC | GCF_038431925.1 | Pseudomonas_aeruginosa | Bacteria>Pseudomonadati>Pseudomonadota>Gammaproteobacteria>Pseudomonadales>Pseudomonadaceae>Pseudomonas>Pseudomonas_aeruginosa_group | 396 | EQH74_RS32320 | 3883..5073 | 287 | AAA-family-ATPase-[Pseudomonas-aeruginosa] |
| HBO1294565.1 | OLD-ABC | GCA_021597005.1 | Pseudomonas_aeruginosa | Bacteria>Pseudomonadati>Pseudomonadota>Gammaproteobacteria>Pseudomonadales>Pseudomonadaceae>Pseudomonas>Pseudomonas_aeruginosa_group | 392 | L4E86_006701 | 4118..5296 | 287 | AAA-family-ATPase-[Pseudomonas-aeruginosa] |
| HBO7921139.1 | OLD-ABC | GCA_021678155.1 | Pseudomonas_aeruginosa | Bacteria>Pseudomonadati>Pseudomonadota>Gammaproteobacteria>Pseudomonadales>Pseudomonadaceae>Pseudomonas>Pseudomonas_aeruginosa_group | 388 | L5C37_005433 | 26046..27212 | 287 | AAA-family-ATPase-[Pseudomonas-aeruginosa] |
| MBG4863330.1 | OLD-ABC | GCA_015735985.1 | Pseudomonas_aeruginosa | Bacteria>Pseudomonadati>Pseudomonadota>Gammaproteobacteria>Pseudomonadales>Pseudomonadaceae>Pseudomonas>Pseudomonas_aeruginosa_group | 388 | GHU23_29000 | 11016..12182 | 287 | AAA-family-ATPase-[Pseudomonas-aeruginosa] |
| MBG6730510.1 | OLD-ABC | GCA_015764285.1 | Pseudomonas_aeruginosa | Bacteria>Pseudomonadati>Pseudomonadota>Gammaproteobacteria>Pseudomonadales>Pseudomonadaceae>Pseudomonas>Pseudomonas_aeruginosa_group | 388 | I5I92_11345 | 13151..14317 | 287 | AAA-family-ATPase-[Pseudomonas-aeruginosa] |
| WP_094776637.1 | OLD-ABC | GCF_002239565.1 | Pseudomonas_aeruginosa | Bacteria>Pseudomonadati>Pseudomonadota>Gammaproteobacteria>Pseudomonadales>Pseudomonadaceae>Pseudomonas>Pseudomonas_aeruginosa_group | 388 | CF337_RS09115 | 70603..71769 | 287 | AAA-family-ATPase-[Pseudomonas-aeruginosa] |
| HBO7921134.1 | OLD-ABC | GCA_021678155.1 | Pseudomonas_aeruginosa | Bacteria>Pseudomonadati>Pseudomonadota>Gammaproteobacteria>Pseudomonadales>Pseudomonadaceae>Pseudomonas>Pseudomonas_aeruginosa_group | 380 | L5C37_005428 | 19102..20244 | 287 | AAA-family-ATPase-[Pseudomonas-aeruginosa] |
| MBG4863327.1 | OLD-ABC | GCA_015735985.1 | Pseudomonas_aeruginosa | Bacteria>Pseudomonadati>Pseudomonadota>Gammaproteobacteria>Pseudomonadales>Pseudomonadaceae>Pseudomonas>Pseudomonas_aeruginosa_group | 379 | GHU23_28980 | 7165..8304 | 287 | AAA-family-ATPase-[Pseudomonas-aeruginosa] |
| WP_124127281.1 | OLD-ABC | GCF_022551795.1 | Pseudomonas_aeruginosa | Bacteria>Pseudomonadati>Pseudomonadota>Gammaproteobacteria>Pseudomonadales>Pseudomonadaceae>Pseudomonas>Pseudomonas_aeruginosa_group | 371 | IPC776_RS25335 | 80658..81773 | 287 | AAA-family-ATPase-[Pseudomonas-aeruginosa] |
| HBO1619176.1 | OLD-ABC | GCA_021598085.1 | Pseudomonas_aeruginosa | Bacteria>Pseudomonadati>Pseudomonadota>Gammaproteobacteria>Pseudomonadales>Pseudomonadaceae>Pseudomonas>Pseudomonas_aeruginosa_group | 367 | L4F62_005709 | 10602..11705 | 287 | TPA:-AAA-family-ATPase-[Pseudomonas-aeruginosa] |
| WP_023127682.1 | OLD-ABC | GCF_025696785.1 | Pseudomonas_aeruginosa | Bacteria>Pseudomonadati>Pseudomonadota>Gammaproteobacteria>Pseudomonadales>Pseudomonadaceae>Pseudomonas>Pseudomonas_aeruginosa_group | 367 | KUC66_RS22990 | 86144..87247 | 287 | AAA-family-ATPase-[Pseudomonas-aeruginosa] |
| ARQ76272.1 | OLD-ABC | GCA_002128325.1 | Pseudomonas_fragi | Bacteria>Pseudomonadati>Pseudomonadota>Gammaproteobacteria>Pseudomonadales>Pseudomonadaceae>Pseudomonas>Pseudomonas_chlororaphis_group | 367 | B6D87_19575 | 4220761..4221864 | 296 | recombinase-RecF-[Pseudomonas-fragi] |
| WP_047283132.1 | OLD-ABC | GCF_018449005.1 | Pseudomonas_lundensis | Bacteria>Pseudomonadati>Pseudomonadota>Gammaproteobacteria>Pseudomonadales>Pseudomonadaceae>Pseudomonas>Pseudomonas_chlororaphis_group | 367 | KIY13_RS15515 | 3394092..3395195 | 86185 | AAA-family-ATPase-[Pseudomonas-lundensis] |
| WP_086798822.1 | OLD-ABC | GCF_002128325.1 | Pseudomonas_fragi | Bacteria>Pseudomonadati>Pseudomonadota>Gammaproteobacteria>Pseudomonadales>Pseudomonadaceae>Pseudomonas>Pseudomonas_chlororaphis_group | 367 | B6D87_RS19595 | 4220761..4221864 | 296 | AAA-family-ATPase-[Pseudomonas-fragi] |
| WP_094987829.1 | OLD-ABC | GCF_012986325.1 | Pseudomonas_lundensis | Bacteria>Pseudomonadati>Pseudomonadota>Gammaproteobacteria>Pseudomonadales>Pseudomonadaceae>Pseudomonas>Pseudomonas_chlororaphis_group | 367 | HBN80_RS14460 | 95430..96533 | 86185 | AAA-family-ATPase-[Pseudomonas-lundensis] |
| WP_106118417.1 | OLD-ABC | GCF_003001555.1 | Pseudomonas_fluorescens | Bacteria>Pseudomonadati>Pseudomonadota>Gammaproteobacteria>Pseudomonadales>Pseudomonadaceae>Pseudomonas>Pseudomonas_fluorescens_group | 395 | C7A10_RS23715 | 134436..135623 | 294 | ATP-binding-protein-[Pseudomonas-fluorescens] |
| WP_261753791.1 | OLD-ABC | GCF_025397875.1 | Pseudomonas_veronii | Bacteria>Pseudomonadati>Pseudomonadota>Gammaproteobacteria>Pseudomonadales>Pseudomonadaceae>Pseudomonas>Pseudomonas_fluorescens_group | 371 | MN113_RS13515 | 503654..504769 | 76761 | AAA-family-ATPase-[Pseudomonas-veronii] |
| WP_236361677.1 | OLD-ABC | GCF_021605645.1 | Pseudomonas_gessardii | Bacteria>Pseudomonadati>Pseudomonadota>Gammaproteobacteria>Pseudomonadales>Pseudomonadaceae>Pseudomonas>Pseudomonas_fluorescens_group | 367 | GIW56_RS19745 | 20153..21256 | 78544 | AAA-family-ATPase-[Pseudomonas-gessardii] |
| CAB5576333.1 | OLD-ABC | GCA_903684465.1 | Pseudomonas_putida | Bacteria>Pseudomonadati>Pseudomonadota>Gammaproteobacteria>Pseudomonadales>Pseudomonadaceae>Pseudomonas>Pseudomonas_putida_group | 371 | GHA_00149 | 170658..171773 | 303 | recombination-protein-F-[Pseudomonas-putida] |
| WP_122701525.1 | OLD-ABC | - | Pseudomonas_viridiflava | Bacteria>Pseudomonadati>Pseudomonadota>Gammaproteobacteria>Pseudomonadales>Pseudomonadaceae>Pseudomonas>Pseudomonas_syringae_group | 371 | EFA20_RS09795 | 79405..80520 | 33069 | AAA-family-ATPase-[Pseudomonas-viridiflava] |
| MCK9534935.1 | OLD-ABC | GCA_023228505.1 | Pseudomonas_sp. | Bacteria>Pseudomonadati>Pseudomonadota>Gammaproteobacteria>Pseudomonadales>Pseudomonadaceae>Pseudomonas>unclassified_Pseudomonas | 421 | M0Q98_09620 | 16535..17800 | 306 | AAA-family-ATPase-[Pseudomonas-sp.] |
| TXH86521.1 | OLD-ABC | GCA_008015405.1 | Pseudomonas_sp. | Bacteria>Pseudomonadati>Pseudomonadota>Gammaproteobacteria>Pseudomonadales>Pseudomonadaceae>Pseudomonas>unclassified_Pseudomonas | 396 | E6Q71_06455 | 4160..5350 | 306 | chromosome-segregation-protein-SMC-[Pseudomonas-sp.] |
| WP_222018160.1 | OLD-ABC | GCF_019775695.1 | Pseudomonas_sp._AO-1 | Bacteria>Pseudomonadati>Pseudomonadota>Gammaproteobacteria>Pseudomonadales>Pseudomonadaceae>Pseudomonas>unclassified_Pseudomonas | 395 | KVQ82_RS04460 | 957855..959042 | 2855434 | ATP-binding-protein-[Pseudomonas-sp.-AO-1] |
| WP_273917577.1 | OLD-ABC | GCF_028656895.1 | Pseudomonas_sp._TNT2022_ID1044 | Bacteria>Pseudomonadati>Pseudomonadota>Gammaproteobacteria>Pseudomonadales>Pseudomonadaceae>Pseudomonas>unclassified_Pseudomonas | 395 | M5G20_RS01565 | 346273..347460 | 2942636 | ATP-binding-protein-[Pseudomonas-sp.-TNT2022-ID1044] |
| MCK9534933.1 | OLD-ABC | GCA_023228505.1 | Pseudomonas_sp. | Bacteria>Pseudomonadati>Pseudomonadota>Gammaproteobacteria>Pseudomonadales>Pseudomonadaceae>Pseudomonas>unclassified_Pseudomonas | 369 | M0Q98_09610 | 14665..15774 | 306 | AAA-family-ATPase-[Pseudomonas-sp.] |
| MBB59701.1 | OLD-ABC | GCA_002709345.1 | Pseudomonas_sp. | Bacteria>Pseudomonadati>Pseudomonadota>Gammaproteobacteria>Pseudomonadales>Pseudomonadaceae>Pseudomonas>unclassified_Pseudomonas | 367 | CML00_00830 | 32389..33492 | 306 | recombinase-RecF-[Pseudomonas-sp.] |
| MCQ4319997.1 | OLD-ABC | GCA_024448935.1 | Stutzerimonas_stutzeri | Bacteria>Pseudomonadati>Pseudomonadota>Gammaproteobacteria>Pseudomonadales>Pseudomonadaceae>Stutzerimonas>Stutzerimonas_stutzeri_group>Stutzerimonas_stutzeri_subgroup | 395 | NA637_07220 | 21377..22564 | 316 | AAA-family-ATPase-[Stutzerimonas-stutzeri] |
| PAO90331.1 | OLD-ABC | GCA_002282995.1 | Stutzerimonas_stutzeri | Bacteria>Pseudomonadati>Pseudomonadota>Gammaproteobacteria>Pseudomonadales>Pseudomonadaceae>Stutzerimonas>Stutzerimonas_stutzeri_group>Stutzerimonas_stutzeri_subgroup | 367 | BV581_21205 | 14138..15241 | 316 | recombinase-RecF-[Stutzerimonas-stutzeri] |
| WP_053105418.1 | OLD-ABC | GCF_001267275.1 | Thiopseudomonas_alkaliphila | Bacteria>Pseudomonadati>Pseudomonadota>Gammaproteobacteria>Pseudomonadales>Pseudomonadaceae>Thiopseudomonas | 369 | AKN90_RS00630 | 131969..133078 | 1697053 | AAA-family-ATPase-[Thiopseudomonas-alkaliphila] |
| MCP5320939.1 | OLD-ABC | GCA_024234165.1 | Pseudomonadales_bacterium | Bacteria>Pseudomonadati>Pseudomonadota>Gammaproteobacteria>Pseudomonadales>unclassified_Pseudomonadales | 405 | H7A12_08975 | 736820..738037 | 1891229 | AAA-family-ATPase-[Pseudomonadales-bacterium] |
| MCG8614352.1 | OLD-ABC | GCA_022361325.1 | Pseudomonadales_bacterium | Bacteria>Pseudomonadati>Pseudomonadota>Gammaproteobacteria>Pseudomonadales>unclassified_Pseudomonadales | 395 | MI864_27895 | 12593..13780 | 1891229 | AAA-family-ATPase-[Pseudomonadales-bacterium] |
| MAB42108.1 | OLD-ABC | GCA_002684155.1 | Pseudomonadales_bacterium | Bacteria>Pseudomonadati>Pseudomonadota>Gammaproteobacteria>Pseudomonadales>unclassified_Pseudomonadales | 367 | CMK77_07475 | 7236..8339 | 1891229 | recombinase-RecF-[Pseudomonadales-bacterium] |
| MAS10376.1 | OLD-ABC | GCA_002702145.1 | Salinisphaera_sp. | Bacteria>Pseudomonadati>Pseudomonadota>Gammaproteobacteria>Salinisphaerales>Salinisphaeraceae>Salinisphaera>unclassified_Salinisphaera | 396 | CMN26_09995 | 8648..9838 | 1914330 | chromosome-segregation-protein-SMC-[Salinisphaera-sp.] |
| WP_022954804.1 | OLD-ABC | GCF_000419525.1 | Leucothrix_mucor | Bacteria>Pseudomonadati>Pseudomonadota>Gammaproteobacteria>Thiotrichales>Thiotrichaceae>Leucothrix | 384 | LEUMU_RS0123790 | 72582..73736 | 45248 | ATP-binding-protein-[Leucothrix-mucor] |
| OQX14375.1 | OLD-ABC | GCA_002083875.1 | Thiothrix_lacustris | Bacteria>Pseudomonadati>Pseudomonadota>Gammaproteobacteria>Thiotrichales>Thiotrichaceae>Thiothrix | 556 | BWK73_09680 | 15042..16712 | 525917 | hypothetical-protein-BWK73_09680-[Thiothrix-lacustris] |
| WP_002710593.1 | OLD-ABC | GCF_000260135.1 | Thiothrix_nivea | Bacteria>Pseudomonadati>Pseudomonadota>Gammaproteobacteria>Thiotrichales>Thiotrichaceae>Thiothrix | 421 | THINI_RS21430 | 4305656..4306921 | 1031 | ATP-binding-protein-[Thiothrix-nivea] |
| WP_210226004.1 | OLD-ABC | GCF_017901155.1 | Candidatus_Thiothrix_anitrata | Bacteria>Pseudomonadati>Pseudomonadota>Gammaproteobacteria>Thiotrichales>Thiotrichaceae>Thiothrix | 414 | J8380_RS12840 | 2551203..2552447 | 2823902 | AAA-family-ATPase-[Candidatus-Thiothrix-anitrata] |
| QLQ31993.1 | OLD-ABC | GCA_013693955.1 | Candidatus_Thiothrix_singaporensis | Bacteria>Pseudomonadati>Pseudomonadota>Gammaproteobacteria>Thiotrichales>Thiotrichaceae>Thiothrix | 279 | HZT40_10785 | 2138880..2139719 | 2799669 | AAA-family-ATPase-[Candidatus-Thiothrix-singaporensis] |
| QLQ31994.1 | OLD-ABC | GCA_013693955.1 | Candidatus_Thiothrix_singaporensis | Bacteria>Pseudomonadati>Pseudomonadota>Gammaproteobacteria>Thiotrichales>Thiotrichaceae>Thiothrix | 193 | HZT40_10790 | 2139753..2140334 | 2799669 | ATP-binding-protein-[Candidatus-Thiothrix-singaporensis] |
| MCY4286577.1 | OLD-ABC | GCA_026709935.1 | Thiotrichales_bacterium | Bacteria>Pseudomonadati>Pseudomonadota>Gammaproteobacteria>Thiotrichales>unclassified_Thiotrichales | 418 | OXC65_14715 | 5874..7130 | 2026796 | MAG:-AAA-family-ATPase-[Thiotrichales-bacterium] |
| MDD9824622.1 | OLD-ABC | GCA_028817255.1 | Gammaproteobacteria_bacterium | Bacteria>Pseudomonadati>Pseudomonadota>Gammaproteobacteria>unclassified_Gammaproteobacteria | 449 | OXU43_05575 | 3464..4813 | 1913989 | MAG:-AAA-family-ATPase-[Gammaproteobacteria-bacterium] |
| MDD9817277.1 | OLD-ABC | GCA_028817495.1 | Gammaproteobacteria_bacterium | Bacteria>Pseudomonadati>Pseudomonadota>Gammaproteobacteria>unclassified_Gammaproteobacteria | 423 | OXU61_03940 | 4376..5647 | 1913989 | MAG:-ATP-binding-protein-[Gammaproteobacteria-bacterium] |
| MDD9864537.1 | OLD-ABC | GCA_028817305.1 | Gammaproteobacteria_bacterium | Bacteria>Pseudomonadati>Pseudomonadota>Gammaproteobacteria>unclassified_Gammaproteobacteria | 423 | OXU54_08195 | 2422..3693 | 1913989 | MAG:-ATP-binding-protein-[Gammaproteobacteria-bacterium] |
| MXY90449.1 | OLD-ABC | GCA_009837675.1 | Gammaproteobacteria_bacterium | Bacteria>Pseudomonadati>Pseudomonadota>Gammaproteobacteria>unclassified_Gammaproteobacteria | 416 | F4Y89_07915 | 38138..39388 | 1913989 | ATP-binding-protein-[Gammaproteobacteria-bacterium] |
| NPU94900.1 | OLD-ABC | GCA_013178375.1 | Gammaproteobacteria_bacterium | Bacteria>Pseudomonadati>Pseudomonadota>Gammaproteobacteria>unclassified_Gammaproteobacteria | 414 | HPY82_23595 | 10988..12232 | 1913989 | AAA-family-ATPase-[Gammaproteobacteria-bacterium] |
| MDE0038693.1 | OLD-ABC | GCA_028818795.1 | Gammaproteobacteria_bacterium | Bacteria>Pseudomonadati>Pseudomonadota>Gammaproteobacteria>unclassified_Gammaproteobacteria | 410 | OXU77_14240 | 13954..15186 | 1913989 | MAG:-AAA-family-ATPase-[Gammaproteobacteria-bacterium] |
| MCZ0943263.1 | OLD-ABC | GCA_026914145.1 | Gammaproteobacteria_bacterium | Bacteria>Pseudomonadati>Pseudomonadota>Gammaproteobacteria>unclassified_Gammaproteobacteria | 393 | OXJ53_09420 | 7844..9025 | 1913989 | MAG:-AAA-family-ATPase-[Gammaproteobacteria-bacterium] |
| MDE0073869.1 | OLD-ABC | GCA_028819075.1 | Gammaproteobacteria_bacterium | Bacteria>Pseudomonadati>Pseudomonadota>Gammaproteobacteria>unclassified_Gammaproteobacteria | 391 | OXU32_07795 | 1554..2729 | 1913989 | MAG:-AAA-family-ATPase-[Gammaproteobacteria-bacterium] |
| TAK54306.1 | OLD-ABC | GCA_004297335.1 | Gammaproteobacteria_bacterium | Bacteria>Pseudomonadati>Pseudomonadota>Gammaproteobacteria>unclassified_Gammaproteobacteria | 391 | EPO25_07520 | 11392..12567 | 1913989 | chromosome-segregation-protein-SMC-[Gammaproteobacteria-bacterium] |
| MCP5451259.1 | OLD-ABC | GCA_024235115.1 | Gammaproteobacteria_bacterium | Bacteria>Pseudomonadati>Pseudomonadota>Gammaproteobacteria>unclassified_Gammaproteobacteria | 369 | H6972_12090 | 49273..50382 | 1913989 | AAA-family-ATPase-[Gammaproteobacteria-bacterium] |
| NLY14762.1 | OLD-ABC | GCA_012728415.1 | Gammaproteobacteria_bacterium | Bacteria>Pseudomonadati>Pseudomonadota>Gammaproteobacteria>unclassified_Gammaproteobacteria | 367 | GXZ05_00060 | 6385..7488 | 1913989 | AAA-family-ATPase-[Gammaproteobacteria-bacterium] |
| MCP4410535.1 | OLD-ABC | GCA_024222475.1 | Gammaproteobacteria_bacterium | Bacteria>Pseudomonadati>Pseudomonadota>Gammaproteobacteria>unclassified_Gammaproteobacteria | 365 | GY807_22885 | 7754..8851 | 1913989 | AAA-family-ATPase-[Gammaproteobacteria-bacterium] |
| MDE0157708.1 | OLD-ABC | GCA_028819495.1 | Gammaproteobacteria_bacterium | Bacteria>Pseudomonadati>Pseudomonadota>Gammaproteobacteria>unclassified_Gammaproteobacteria | 359 | OXS28_19160 | 1774..2853 | 1913989 | MAG:-AAA-family-ATPase-[Gammaproteobacteria-bacterium] |
| MDE0512494.1 | OLD-ABC | GCA_028823865.1 | Gammaproteobacteria_bacterium | Bacteria>Pseudomonadati>Pseudomonadota>Gammaproteobacteria>unclassified_Gammaproteobacteria | 359 | OXI88_11980 | 9085..10164 | 1913989 | MAG:-AAA-family-ATPase-[Gammaproteobacteria-bacterium] |
| WP_249373669.1 | OLD-ABC | GCF_013048605.1 | Vibrio | Bacteria>Pseudomonadati>Pseudomonadota>Gammaproteobacteria>Vibrionales>Vibrionaceae | 390 | FKN13_RS12345 | 68830..70002 | 662 | AAA-family-ATPase-[Vibrio-sp.-2-2(9)] |
| WP_085609971.1 | OLD-ABC | GCF_032463505.1 | Vibrio | Bacteria>Pseudomonadati>Pseudomonadota>Gammaproteobacteria>Vibrionales>Vibrionaceae | 389 | B5C31_RS21490 | 49690..50859 | 662 | MULTISPECIES:-AAA-family-ATPase-[Vibrio] |
| WP_170894126.1 | OLD-ABC | GCF_033441435.1 | Vibrio | Bacteria>Pseudomonadati>Pseudomonadota>Gammaproteobacteria>Vibrionales>Vibrionaceae | 389 | G3220_RS19470 | 65242..66411 | 662 | MULTISPECIES:-AAA-family-ATPase-[Vibrio] |
| WP_113594183.1 | OLD-ABC | GCF_043352035.1 | Vibrio | Bacteria>Pseudomonadati>Pseudomonadota>Gammaproteobacteria>Vibrionales>Vibrionaceae | 361 | DLR66_RS03580 | 65885..66970 | 662 | AAA-family-ATPase-[Vibrio-paracholerae] |
| WP_054775222.1 | OLD-ABC | GCF_014647395.1 | Aliivibrio_fischeri | Bacteria>Pseudomonadati>Pseudomonadota>Gammaproteobacteria>Vibrionales>Vibrionaceae>Aliivibrio | 367 | AVFI_RS07670 | 1641346..1642449 | 668 | AAA-family-ATPase-[Aliivibrio-fischeri] |
| WP_190395655.1 | OLD-ABC | GCF_014694495.1 | Aliivibrio_sp._S10_S31 | Bacteria>Pseudomonadati>Pseudomonadota>Gammaproteobacteria>Vibrionales>Vibrionaceae>Aliivibrio>unclassified_Aliivibrio | 367 | HC728_RS13690 | 64953..66056 | 2720224 | AAA-family-ATPase-[Aliivibrio-sp.-S10_S31] |
| PLC56103.1 | OLD-ABC | GCA_002849605.1 | Photobacterium_carnosum | Bacteria>Pseudomonadati>Pseudomonadota>Gammaproteobacteria>Vibrionales>Vibrionaceae>Photobacterium | 393 | CIK00_20180 | 16095..17276 | 2023717 | chromosome-segregation-protein-SMC-[Photobacterium-carnosum] |
| WP_176285152.1 | OLD-ABC | GCF_013371045.1 | Photobacterium_damselae | Bacteria>Pseudomonadati>Pseudomonadota>Gammaproteobacteria>Vibrionales>Vibrionaceae>Photobacterium | 367 | HU831_RS17615 | 64099..65202 | 38293 | AAA-family-ATPase-[Photobacterium-damselae] |
| WP_197573431.1 | OLD-ABC | GCF_014930975.1 | Photobacterium_damselae | Bacteria>Pseudomonadati>Pseudomonadota>Gammaproteobacteria>Vibrionales>Vibrionaceae>Photobacterium | 367 | IL982_RS10185 | 2137484..2138587 | 38293 | AAA-family-ATPase-[Photobacterium-damselae] |
| CAQ34954.1 | OLD-ABC | - | Photobacterium_damselae_subsp._piscicida | Bacteria>Pseudomonadati>Pseudomonadota>Gammaproteobacteria>Vibrionales>Vibrionaceae>Photobacterium>Photobacterium_damselae | 389 | . | 24853..26022 | 38294 | hypothetical-protein-[Photobacterium-damselae-subsp.-piscicida] |
| QOQ68159.1 | OLD-ABC | GCA_014930975.1 | Photobacterium_damselae_subsp._damselae | Bacteria>Pseudomonadati>Pseudomonadota>Gammaproteobacteria>Vibrionales>Vibrionaceae>Photobacterium>Photobacterium_damselae | 367 | IL982_10185 | 2137484..2138587 | 85581 | AAA-family-ATPase-[Photobacterium-damselae-subsp.-damselae] |
| WP_077485089.1 | OLD-ABC | GCF_001995745.1 | Salinivibrio_sp._ML323 | Bacteria>Pseudomonadati>Pseudomonadota>Gammaproteobacteria>Vibrionales>Vibrionaceae>Salinivibrio>unclassified_Salinivibrio | 367 | BZG13_RS03565 | 132357..133460 | 1909474 | AAA-family-ATPase-[Salinivibrio-sp.-ML323] |
| TNF18457.1 | OLD-ABC | GCA_006226665.1 | Vibrionaceae_bacterium | Bacteria>Pseudomonadati>Pseudomonadota>Gammaproteobacteria>Vibrionales>Vibrionaceae>unclassified_Vibrionaceae | 370 | EP325_05035 | 58751..59863 | 1889773 | recombinase-RecF-[Vibrionaceae-bacterium] |
| WP_054541289.1 | OLD-ABC | GCF_963869775.1 | Vibrio_splendidus | Bacteria>Pseudomonadati>Pseudomonadota>Gammaproteobacteria>Vibrionales>Vibrionaceae>Vibrio | 390 | AN167_RS02940 | 111171..112343 | 29497 | AAA-family-ATPase-[Vibrio-splendidus] |
| EGQ8649254.1 | OLD-ABC | - | Vibrio_cholerae | Bacteria>Pseudomonadati>Pseudomonadota>Gammaproteobacteria>Vibrionales>Vibrionaceae>Vibrio | 389 | GNS53_08425 | 56009..57178 | 666 | AAA-family-ATPase-[Vibrio-cholerae] |
| EJB8347939.1 | OLD-ABC | GCA_022966375.1 | Vibrio_cholerae | Bacteria>Pseudomonadati>Pseudomonadota>Gammaproteobacteria>Vibrionales>Vibrionaceae>Vibrio | 389 | MW328_000281 | 39415..40584 | 666 | AAA-family-ATPase-[Vibrio-cholerae] |
| EJB8581803.1 | OLD-ABC | GCA_022967455.1 | Vibrio_cholerae | Bacteria>Pseudomonadati>Pseudomonadota>Gammaproteobacteria>Vibrionales>Vibrionaceae>Vibrio | 389 | MW361_003122 | 33790..34959 | 666 | AAA-family-ATPase-[Vibrio-cholerae] |
| EJK2098983.1 | OLD-ABC | GCA_024103575.1 | Vibrio_cholerae | Bacteria>Pseudomonadati>Pseudomonadota>Gammaproteobacteria>Vibrionales>Vibrionaceae>Vibrio | 389 | NK421_001792 | 212889..214058 | 666 | AAA-family-ATPase-[Vibrio-cholerae] |
| WP_274695776.1 | OLD-ABC | GCF_028858205.1 | Vibrio_aestuarianus | Bacteria>Pseudomonadati>Pseudomonadota>Gammaproteobacteria>Vibrionales>Vibrionaceae>Vibrio | 389 | MEG05_RS14950 | 3091712..3092881 | 28171 | AAA-family-ATPase-[Vibrio-aestuarianus] |
| WP_261872480.1 | OLD-ABC | GCF_024347075.1 | Vibrio_rarus | Bacteria>Pseudomonadati>Pseudomonadota>Gammaproteobacteria>Vibrionales>Vibrionaceae>Vibrio | 380 | OCU56_RS06795 | 1449381..1450523 | 413403 | ATP-binding-protein-[Vibrio-rarus] |
| WP_108187612.1 | OLD-ABC | GCF_003049965.1 | Vibrio_splendidus | Bacteria>Pseudomonadati>Pseudomonadota>Gammaproteobacteria>Vibrionales>Vibrionaceae>Vibrio | 373 | CWO07_RS09205 | 39355..40476 | 29497 | ATP-binding-protein-[Vibrio-splendidus] |
| EII3725632.1 | OLD-ABC | GCA_020414365.1 | Vibrio_cholerae | Bacteria>Pseudomonadati>Pseudomonadota>Gammaproteobacteria>Vibrionales>Vibrionaceae>Vibrio | 372 | LG468_000345 | 122625..123743 | 666 | AAA-family-ATPase-[Vibrio-cholerae] |
| WP_221886920.1 | OLD-ABC | GCF_006803025.1 | Vibrio_cholerae | Bacteria>Pseudomonadati>Pseudomonadota>Gammaproteobacteria>Vibrionales>Vibrionaceae>Vibrio | 372 | FLL79_RS01425 | 45143..46261 | 666 | AAA-family-ATPase-[Vibrio-cholerae] |
| HAS6259592.1 | OLD-ABC | GCA_015777345.1 | Vibrio_vulnificus | Bacteria>Pseudomonadati>Pseudomonadota>Gammaproteobacteria>Vibrionales>Vibrionaceae>Vibrio | 371 | I7226_03355 | 257341..258456 | 672 | AAA-family-ATPase-[Vibrio-vulnificus] |

| Protein-ID | Domain | Assembly-ID | Strain | Taxonomy | Length | Locus-Tag | Genomic-coordinates | TaxID | Fasta-defline |
|---|---|---|---|---|---|---|---|---|---|
| WP_032482147.1 | OLD-ABC | GCF_000736875.1 | Vibrio_cholerae | Bacteria>Pseudomonadati>Pseudomonadota>Gammaproteobacteria>Vibrionales>Vibrionaceae>Vibrio | 371 | DN38_RS05530 | 17931..19046 | 666 | AAA-family-ATPase-[Vibrio-cholerae] |
| WP_102376354.1 | OLD-ABC | GCF_002876115.1 | Vibrio_breoganii | Bacteria>Pseudomonadati>Pseudomonadota>Gammaproteobacteria>Vibrionales>Vibrionaceae>Vibrio | 370 | BCT61_RS02990 | 187744..188856 | 553239 | ATP-binding-protein-[Vibrio-breoganii] |
| WP_045588642.1 | OLD-ABC | GCF_000959745.1 | Vibrio_vulnificus | Bacteria>Pseudomonadati>Pseudomonadota>Gammaproteobacteria>Vibrionales>Vibrionaceae>Vibrio | 367 | WG98_RS00460 | 98676..99779 | 672 | AAA-family-ATPase-[Vibrio-vulnificus] |
| WP_210447444.1 | OLD-ABC | GCF_017917465.1 | Vibrio_crassostreae | Bacteria>Pseudomonadati>Pseudomonadota>Gammaproteobacteria>Vibrionales>Vibrionaceae>Vibrio | 367 | KB624_RS13670 | 49392..50495 | 246167 | AAA-family-ATPase-[Vibrio-crassostreae] |
| EGR0286976.1 | OLD-ABC | - | Vibrio_cholerae | Bacteria>Pseudomonadati>Pseudomonadota>Gammaproteobacteria>Vibrionales>Vibrionaceae>Vibrio | 361 | FK206_11690 | 80696..81781 | 666 | recombinase-RecF-[Vibrio-cholerae] |
| EGR2430329.1 | OLD-ABC | - | Vibrio_cholerae | Bacteria>Pseudomonadati>Pseudomonadota>Gammaproteobacteria>Vibrionales>Vibrionaceae>Vibrio | 361 | DYB83_10150 | 18091..19176 | 666 | recombinase-RecF-[Vibrio-cholerae] |
| EJL6578342.1 | OLD-ABC | GCA_024217525.1 | Vibrio_cholerae | Bacteria>Pseudomonadati>Pseudomonadota>Gammaproteobacteria>Vibrionales>Vibrionaceae>Vibrio | 361 | NMS50_002628 | 66777..67862 | 666 | AAA-family-ATPase-[Vibrio-cholerae] |
| AAL59730.1 | OLD-ABC | - | Vibrio_cholerae | Bacteria>Pseudomonadati>Pseudomonadota>Gammaproteobacteria>Vibrionales>Vibrionaceae>Vibrio | 250 | . | 32488..33240 | 666 | unknown-[Vibrio-cholerae] |
| AAL59731.1 | OLD-ABC | - | Vibrio_cholerae | Bacteria>Pseudomonadati>Pseudomonadota>Gammaproteobacteria>Vibrionales>Vibrionaceae>Vibrio | 160 | . | 32037..32519 | 666 | unknown-[Vibrio-cholerae] |
| EJE4207361.1 | OLD-ABC | GCA_023313315.1 | Vibrio_parahaemolyticus | Bacteria>Pseudomonadati>Pseudomonadota>Gammaproteobacteria>Vibrionales>Vibrionaceae>Vibrio>Vibrio_harveyi_group | 389 | M3899_001772 | 265929..267098 | 670 | AAA-family-ATPase-[Vibrio-parahaemolyticus] |
| HCE3362111.1 | OLD-ABC | GCA_023821665.1 | Vibrio_parahaemolyticus | Bacteria>Pseudomonadati>Pseudomonadota>Gammaproteobacteria>Vibrionales>Vibrionaceae>Vibrio>Vibrio_harveyi_group | 389 | NG589_000243 | 271279..272448 | 670 | AAA-family-ATPase-[Vibrio-parahaemolyticus] |
| MBE3881971.1 | OLD-ABC | GCA_014919165.1 | Vibrio_parahaemolyticus | Bacteria>Pseudomonadati>Pseudomonadota>Gammaproteobacteria>Vibrionales>Vibrionaceae>Vibrio>Vibrio_harveyi_group | 389 | HJ168_09870 | 229941..231110 | 670 | AAA-family-ATPase-[Vibrio-parahaemolyticus] |
| WP_134990669.1 | OLD-ABC | - | Vibrio_parahaemolyticus | Bacteria>Pseudomonadati>Pseudomonadota>Gammaproteobacteria>Vibrionales>Vibrionaceae>Vibrio>Vibrio_harveyi_group | 389 | DA442_RS13695 | 2689942..2691111 | 670 | AAA-family-ATPase-[Vibrio-parahaemolyticus] |
| WP_274056406.1 | OLD-ABC | GCF_028616275.1 | Vibrio_parahaemolyticus | Bacteria>Pseudomonadati>Pseudomonadota>Gammaproteobacteria>Vibrionales>Vibrionaceae>Vibrio>Vibrio_harveyi_group | 389 | O8051_RS20175 | 63263..64432 | 670 | AAA-family-ATPase-[Vibrio-parahaemolyticus] |
| MBE3824870.1 | OLD-ABC | GCA_014918955.1 | Vibrio_parahaemolyticus | Bacteria>Pseudomonadati>Pseudomonadota>Gammaproteobacteria>Vibrionales>Vibrionaceae>Vibrio>Vibrio_harveyi_group | 371 | HJ179_16455 | 249375..250490 | 670 | AAA-family-ATPase-[Vibrio-parahaemolyticus] |
| EGR1144246.1 | OLD-ABC | - | Vibrio_parahaemolyticus | Bacteria>Pseudomonadati>Pseudomonadota>Gammaproteobacteria>Vibrionales>Vibrionaceae>Vibrio>Vibrio_harveyi_group | 367 | EC562_15495 | 36485..37588 | 670 | recombinase-RecF-[Vibrio-parahaemolyticus] |
| EII2403785.1 | OLD-ABC | GCA_020407245.1 | Vibrio_parahaemolyticus | Bacteria>Pseudomonadati>Pseudomonadota>Gammaproteobacteria>Vibrionales>Vibrionaceae>Vibrio>Vibrio_harveyi_group | 367 | LG159_002758 | 112268..113371 | 670 | AAA-family-ATPase-[Vibrio-parahaemolyticus] |
| EIS4855658.1 | OLD-ABC | GCA_021362915.1 | Vibrio_parahaemolyticus | Bacteria>Pseudomonadati>Pseudomonadota>Gammaproteobacteria>Vibrionales>Vibrionaceae>Vibrio>Vibrio_harveyi_group | 367 | LZX80_001784 | 100352..101455 | 670 | AAA-family-ATPase-[Vibrio-parahaemolyticus] |
| EJG0936609.1 | OLD-ABC | GCA_015814185.2 | Vibrio_parahaemolyticus | Bacteria>Pseudomonadati>Pseudomonadota>Gammaproteobacteria>Vibrionales>Vibrionaceae>Vibrio>Vibrio_harveyi_group | 367 | C4G97_RS14665 | 113720..114823 | 670 | AAA-family-ATPase-[Vibrio-parahaemolyticus] |
| HCM0920433.1 | OLD-ABC | GCA_025068375.1 | Vibrio_parahaemolyticus | Bacteria>Pseudomonadati>Pseudomonadota>Gammaproteobacteria>Vibrionales>Vibrionaceae>Vibrio>Vibrio_harveyi_group | 367 | N2828_003371 | 100353..101456 | 670 | TPA:-AAA-family-ATPase-[Vibrio-parahaemolyticus] |
| KOY19347.1 | OLD-ABC | GCA_001280635.1 | Vibrio_parahaemolyticus | Bacteria>Pseudomonadati>Pseudomonadota>Gammaproteobacteria>Vibrionales>Vibrionaceae>Vibrio>Vibrio_harveyi_group | 367 | ACX06_21950 | 162888..163991 | 670 | recombinase-RecF-[Vibrio-parahaemolyticus] |
| WP_053816408.1 | OLD-ABC | GCF_001280635.1 | Vibrio_parahaemolyticus | Bacteria>Pseudomonadati>Pseudomonadota>Gammaproteobacteria>Vibrionales>Vibrionaceae>Vibrio>Vibrio_harveyi_group | 367 | ACX06_RS21950 | 162888..163991 | 670 | AAA-family-ATPase-[Vibrio-parahaemolyticus] |
| WP_062854375.1 | OLD-ABC | GCF_001608855.1 | Vibrio_parahaemolyticus | Bacteria>Pseudomonadati>Pseudomonadota>Gammaproteobacteria>Vibrionales>Vibrionaceae>Vibrio>Vibrio_harveyi_group | 367 | AWQ17_RS02385 | 100760..101863 | 670 | AAA-family-ATPase-[Vibrio-parahaemolyticus] |
| WP_238812497.1 | OLD-ABC | GCF_022290755.1 | Vibrio_parahaemolyticus | Bacteria>Pseudomonadati>Pseudomonadota>Gammaproteobacteria>Vibrionales>Vibrionaceae>Vibrio>Vibrio_harveyi_group | 367 | K6U44_RS14240 | 14759..15862 | 670 | AAA-family-ATPase-[Vibrio-parahaemolyticus] |
| EGR1548464.1 | OLD-ABC | - | Vibrio_parahaemolyticus | Bacteria>Pseudomonadati>Pseudomonadota>Gammaproteobacteria>Vibrionales>Vibrionaceae>Vibrio>Vibrio_harveyi_group | 361 | D6Y11_16940 | 156686..157771 | 670 | recombinase-RecF-[Vibrio-parahaemolyticus] |
| EGR3302282.1 | OLD-ABC | - | Vibrio_parahaemolyticus | Bacteria>Pseudomonadati>Pseudomonadota>Gammaproteobacteria>Vibrionales>Vibrionaceae>Vibrio>Vibrio_harveyi_group | 361 | DMJ21_13140 | 15037..16122 | 670 | recombinase-RecF-[Vibrio-parahaemolyticus] |
| EHH2507629.1 | OLD-ABC | GCA_017637525.1 | Vibrio_parahaemolyticus | Bacteria>Pseudomonadati>Pseudomonadota>Gammaproteobacteria>Vibrionales>Vibrionaceae>Vibrio>Vibrio_harveyi_group | 361 | J7X10_003792 | 104977..106062 | 670 | AAA-family-ATPase-[Vibrio-parahaemolyticus] |
| EJC1210119.1 | OLD-ABC | GCA_022997325.1 | Vibrio_parahaemolyticus | Bacteria>Pseudomonadati>Pseudomonadota>Gammaproteobacteria>Vibrionales>Vibrionaceae>Vibrio>Vibrio_harveyi_group | 361 | MX350_002596 | 12655..13740 | 670 | AAA-family-ATPase-[Vibrio-parahaemolyticus] |
| EJG0981291.1 | OLD-ABC | GCA_015813375.2 | Vibrio_parahaemolyticus | Bacteria>Pseudomonadati>Pseudomonadota>Gammaproteobacteria>Vibrionales>Vibrionaceae>Vibrio>Vibrio_harveyi_group | 361 | C4G85_RS03490 | 772108..773193 | 670 | AAA-family-ATPase-[Vibrio-parahaemolyticus] |
| ODY54467.1 | OLD-ABC | GCA_001727265.1 | Vibrio_parahaemolyticus | Bacteria>Pseudomonadati>Pseudomonadota>Gammaproteobacteria>Vibrionales>Vibrionaceae>Vibrio>Vibrio_harveyi_group | 361 | BBM25_06250 | 10393..11478 | 670 | recombinase-RecF-[Vibrio-parahaemolyticus] |
| WP_029846592.1 | OLD-ABC | GCF_000491375.1 | Vibrio_parahaemolyticus | Bacteria>Pseudomonadati>Pseudomonadota>Gammaproteobacteria>Vibrionales>Vibrionaceae>Vibrio>Vibrio_harveyi_group | 361 | PO77_RS19440 | 163290..164375 | 670 | AAA-family-ATPase-[Vibrio-parahaemolyticus] |
| WP_065970496.1 | OLD-ABC | GCF_001704915.1 | Vibrio_parahaemolyticus | Bacteria>Pseudomonadati>Pseudomonadota>Gammaproteobacteria>Vibrionales>Vibrionaceae>Vibrio>Vibrio_harveyi_group | 361 | BBM25_RS06250 | 10393..11478 | 670 | AAA-family-ATPase-[Vibrio-parahaemolyticus] |
| WP_102780433.1 | OLD-ABC | GCF_030238805.1 | Vibrio_parahaemolyticus | Bacteria>Pseudomonadati>Pseudomonadota>Gammaproteobacteria>Vibrionales>Vibrionaceae>Vibrio>Vibrio_harveyi_group | 361 | CSR03_RS22360 | 128572..129657 | 670 | AAA-family-ATPase-[Vibrio-parahaemolyticus] |
| WP_104969988.1 | OLD-ABC | GCF_002953335.1 | Vibrio_diabolicus | Bacteria>Pseudomonadati>Pseudomonadota>Gammaproteobacteria>Vibrionales>Vibrionaceae>Vibrio>Vibrio_harveyi_group>Vibrio_diabolicus_subgroup | 389 | AL552_RS16290 | 3105919..3107088 | 50719 | AAA-family-ATPase-[Vibrio-diabolicus] |
| OEE71516.1 | OLD-ABC | GCA_000287075.2 | Vibrio_ordalii_FF-167 | Bacteria>Pseudomonadati>Pseudomonadota>Gammaproteobacteria>Vibrionales>Vibrionaceae>Vibrio>Vibrio_ordalii | 389 | A1QQ_08600 | 36353..37522 | 617131 | chromosome-segregation-protein-SMC-[Vibrio-ordalii-FF-167] |
| BBD77929.1 | OLD-ABC | GCA_003574215.1 | Hydrogenophilus_thermoluteolus | Bacteria>Pseudomonadati>Pseudomonadota>Hydrogenophilia>Hydrogenophilales>Hydrogenophilaceae>Hydrogenophilus | 406 | HPTL_1671 | 1769280..1770500 | 297 | ATPase-[Hydrogenophilus-thermoluteolus] |
| NPU98922.1 | OLD-ABC | GCA_013177805.1 | Candidatus_Omnitrophota_bacterium | Bacteria>Pseudomonadati>PVC_group>Candidatus_Omnitrophota>unclassified_Candidatus_Omnitrophota | 382 | HPY51_17135 | 16402..17550 | 2035772 | AAA-family-ATPase-[Candidatus-Omnitrophota-bacterium] |
| MCF7790537.1 | OLD-ABC | GCA_021733985.1 | Victivallales_bacterium | Bacteria>Pseudomonadati>PVC_group>Lentisphaerota>Lentisphaeria>Victivallales>unclassified_Victivallales | 431 | K9M56_00940 | 7720..9015 | 2212475 | ATP-binding-protein-[Victivallales-bacterium] |
| NLF62181.1 | OLD-ABC | GCA_012515215.1 | Lentisphaerota_bacterium | Bacteria>Pseudomonadati>PVC_group>Lentisphaerota>unclassified_Lentisphaerota | 387 | GX574_13640 | 34965..36128 | 1932692 | AAA-family-ATPase-[Lentisphaerota-bacterium] |
| OQC12435.1 | OLD-ABC | GCA_002070945.1 | Lentisphaerae_bacterium_ADurb.Bin082 | Bacteria>Pseudomonadati>PVC_group>Lentisphaerota>unclassified_Lentisphaerota | 372 | BWX73_02928 | 51991..53109 | 1852899 | MAG:-hypothetical-protein-BWX73_02928-[Lentisphaerae-bacterium-ADurb.Bin082] |
| MCK6483013.1 | OLD-ABC | GCA_023150595.1 | Phycisphaerae_bacterium | Bacteria>Pseudomonadati>PVC_group>Planctomycetota>Phycisphaerae>unclassified_Phycisphaerae | 420 | L6R00_02625 | 135046..136308 | 2026778 | AAA-family-ATPase-[Phycisphaerae-bacterium] |
| MBP7937489.1 | OLD-ABC | GCA_018003015.1 | Phycisphaerae_bacterium | Bacteria>Pseudomonadati>PVC_group>Planctomycetota>Phycisphaerae>unclassified_Phycisphaerae | 403 | KA354_22845 | 8337..9548 | 2026778 | AAA-family-ATPase-[Phycisphaerae-bacterium] |
| MBY0527365.1 | OLD-ABC | GCA_019752375.1 | Gemmataceae_bacterium | Bacteria>Pseudomonadati>PVC_group>Planctomycetota>Planctomycetia>Gemmatales>Gemmataceae>unclassified_Gemmataceae | 411 | K2R98_28475 | 31603..32838 | 2052164 | AAA-family-ATPase-[Gemmataceae-bacterium] |
| GIW78773.1 | OLD-ABC | GCA_026004675.1 | Gemmatales_bacterium | Bacteria>Pseudomonadati>PVC_group>Planctomycetota>Planctomycetia>Gemmatales>unclassified_Gemmatales | 417 | KatS3mg105_0580 | 674698..675951 | 2838789 | MAG:-ATPase-[Gemmatales-bacterium] |
| WP_011117922.1 | OLD-ABC | GCF_000196115.1 | Rhodopirellula_baltica | Bacteria>Pseudomonadati>PVC_group>Planctomycetota>Planctomycetia>Pirellulales>Pirellulaceae>Rhodopirellula | 392 | DDX19_RS10405 | 131622..132800 | 265606 | ATP-binding-protein-[Rhodopirellula-baltica] |
| WP_145167964.1 | OLD-ABC | GCF_007741535.1 | Rubripirellula_lacrimiformis | Bacteria>Pseudomonadati>PVC_group>Planctomycetota>Planctomycetia>Pirellulales>Pirellulaceae>Rubripirellula | 422 | K227x_RS02720 | 778821..780089 | 1930273 | ATP-binding-protein-[Rubripirellula-lacrimiformis] |
| NLY99306.1 | OLD-ABC | GCA_012744395.1 | Pirellulaceae_bacterium | Bacteria>Pseudomonadati>PVC_group>Planctomycetota>Planctomycetia>Pirellulales>Pirellulaceae>unclassified_Pirellulaceae | 425 | GXY25_02080 | 6526..7803 | 2699754 | ATP-binding-protein-[Pirellulaceae-bacterium] |
| GIW94557.1 | OLD-ABC | GCA_026004775.1 | Pirellulaceae_bacterium | Bacteria>Pseudomonadati>PVC_group>Planctomycetota>Planctomycetia>Pirellulales>Pirellulaceae>unclassified_Pirellulaceae | 411 | KatS3mg110_2598 | 17615..18850 | 2699754 | MAG:-DNA-recombination-protein-RecF-[Pirellulaceae-bacterium] |
| MDD4270714.1 | OLD-ABC | GCA_028703175.1 | Thermoguttaceae_bacterium | Bacteria>Pseudomonadati>PVC_group>Planctomycetota>Planctomycetia>Pirellulales>Thermoguttaceae>unclassified_Thermoguttaceae | 425 | PHN77_21015 | 1872..3149 | 2691551 | MAG:-ATP-binding-protein-[Thermoguttaceae-bacterium] |
| WP_075094279.1 | OLD-ABC | GCF_001610835.1 | Planctomyces_sp._SH-PL14 | Bacteria>Pseudomonadati>PVC_group>Planctomycetota>Planctomycetia>Planctomycetales>Planctomycetaceae>Planctomyces>unclassified_Planctomyces | 395 | VT03_RS18105 | 4714663..4715850 | 1632864 | ATP-binding-protein-[Planctomyces-sp.-SH-PL14] |
| MCA9232430.1 | OLD-ABC | GCA_020427155.1 | Planctomycetales_bacterium | Bacteria>Pseudomonadati>PVC_group>Planctomycetota>Planctomycetia>Planctomycetales>unclassified_Planctomycetales | 429 | KDA57_17405 | 8535..9824 | 2053591 | AAA-family-ATPase-[Planctomycetales-bacterium] |
| OJW08009.1 | OLD-ABC | GCA_001898385.1 | Planctomycetales_bacterium_71-10 | Bacteria>Pseudomonadati>PVC_group>Planctomycetota>Planctomycetia>Planctomycetales>unclassified_Planctomycetales | 409 | BGO49_20685 | 21933..23162 | 1895807 | MAG:-hypothetical-protein-BGO49_20685-[Planctomycetales-bacterium-71-10] |
| MCA9126643.1 | OLD-ABC | GCA_020426625.1 | Planctomycetales_bacterium | Bacteria>Pseudomonadati>PVC_group>Planctomycetota>Planctomycetia>Planctomycetales>unclassified_Planctomycetales | 390 | KDB22_06150 | 17482..18654 | 2053591 | AAA-family-ATPase-[Planctomycetales-bacterium] |
| HBI47074.1 | OLD-ABC | GCA_003508275.1 | Planctomycetales_bacterium | Bacteria>Pseudomonadati>PVC_group>Planctomycetota>Planctomycetia>Planctomycetales>unclassified_Planctomycetales | 352 | DDY78_30105 | 28605..29663 | 2053591 | hypothetical-protein-[Planctomycetales-bacterium] |
| QDU83132.1 | OLD-ABC | GCA_007750655.1 | Rohdeia_mirabilis | Bacteria>Pseudomonadati>PVC_group>Planctomycetota>Planctomycetia>Planctomycetia_incertae_sedis>Rohdeia | 391 | Pla163_02290 | 292115..293290 | 2528008 | recombination-protein-F-[Planctomycetes-bacterium-Pla163] |
| GIW70821.1 | OLD-ABC | GCA_026004615.1 | Planctomycetota_bacterium | Bacteria>Pseudomonadati>PVC_group>Planctomycetota>unclassified_Planctomycetota | 738 | KatS3mg102_0363 | 455154..457370 | 2026780 | MAG:-hypothetical-protein-KatS3mg102_0363-[Planctomycetota-bacterium] |
| MCG3135899.1 | OLD-ABC | GCA_022072415.1 | Planctomycetota_bacterium | Bacteria>Pseudomonadati>PVC_group>Planctomycetota>unclassified_Planctomycetota | 433 | HMLKMBBP_03675 | 10480..11781 | 2026780 | hypothetical-protein-[Planctomycetota-bacterium] |
| MBI4586096.1 | OLD-ABC | GCA_016209605.1 | Planctomycetota_bacterium | Bacteria>Pseudomonadati>PVC_group>Planctomycetota>unclassified_Planctomycetota | 431 | HY717_18965 | 23795..25090 | 2026780 | AAA-family-ATPase-[Planctomycetota-bacterium] |
| NQW48681.1 | OLD-ABC | GCA_013205375.1 | Planctomycetota_bacterium | Bacteria>Pseudomonadati>PVC_group>Planctomycetota>unclassified_Planctomycetota | 425 | HQ464_13050 | 9589..10866 | 2026780 | ATP-binding-protein-[Planctomycetota-bacterium] |
| MBM4083199.1 | OLD-ABC | GCA_016873055.1 | Planctomycetota_bacterium | Bacteria>Pseudomonadati>PVC_group>Planctomycetota>unclassified_Planctomycetota | 418 | FJ272_00240 | 3268..4524 | 2026780 | ATP-binding-protein-[Planctomycetota-bacterium] |
| MBW3538716.1 | OLD-ABC | GCA_019347905.1 | Planctomycetota_bacterium | Bacteria>Pseudomonadati>PVC_group>Planctomycetota>unclassified_Planctomycetota | 418 | KY476_00450 | 27328..28584 | 2026780 | AAA-family-ATPase-[Planctomycetota-bacterium] |
| MBI4584117.1 | OLD-ABC | GCA_016209605.1 | Planctomycetota_bacterium | Bacteria>Pseudomonadati>PVC_group>Planctomycetota>unclassified_Planctomycetota | 411 | HY717_08850 | 11846..13081 | 2026780 | AAA-family-ATPase-[Planctomycetota-bacterium] |
| MBK6942985.1 | OLD-ABC | GCA_016709145.1 | Planctomycetota_bacterium | Bacteria>Pseudomonadati>PVC_group>Planctomycetota>unclassified_Planctomycetota | 410 | IPH13_22670 | 260927..262159 | 2026780 | AAA-family-ATPase-[Planctomycetota-bacterium] |
| RIK67120.1 | OLD-ABC | GCA_003576905.1 | Planctomycetota_bacterium | Bacteria>Pseudomonadati>PVC_group>Planctomycetota>unclassified_Planctomycetota | 402 | DCC65_07830 | 148465..149673 | 2026780 | MAG:-chromosome-segregation-protein-SMC-[Planctomycetota-bacterium] |
| MBI1826655.1 | OLD-ABC | GCA_016178855.1 | Planctomycetota_bacterium | Bacteria>Pseudomonadati>PVC_group>Planctomycetota>unclassified_Planctomycetota | 400 | HYR83_09760 | 17970..19172 | 2026780 | AAA-family-ATPase-[Planctomycetota-bacterium] |
| OQA03648.1 | OLD-ABC | GCA_002068635.1 | Planctomycetes_bacterium_ADurb.Bin401 | Bacteria>Pseudomonadati>PVC_group>Planctomycetota>unclassified_Planctomycetota | 365 | BWY69_00333 | 10849..11946 | 1852904 | hypothetical-protein-BWY69_00333-[Planctomycetes-bacterium-ADurb.Bin401] |
| AHF94668.1 | OLD-ABC | GCA_000242935.3 | Opitutaceae_bacterium_TAV5 | Bacteria>Pseudomonadati>PVC_group>Verrucomicrobiota>Opitutia>Opitutales>Opitutaceae>unclassified_Opitutaceae | 379 | OPIT5_23240 | 5475665..5476804 | 794903 | hypothetical-protein-OPIT5_23240-[Opitutaceae-bacterium-TAV5] |
| MCH8539502.1 | OLD-ABC | GCA_022565675.1 | Opitutales_bacterium | Bacteria>Pseudomonadati>PVC_group>Verrucomicrobiota>Opitutia>Opitutales>unclassified_Opitutales | 438 | LAT58_01890 | 19989..21305 | 2026772 | ATP-binding-protein-[Opitutales-bacterium] |
| MBE2203058.1 | OLD-ABC | GCA_014879575.1 | Chthoniobacterales_bacterium | Bacteria>Pseudomonadati>PVC_group>Verrucomicrobiota>Spartobacteria>Chthoniobacterales>unclassified_Chthoniobacterales | 428 | IAE94_01785 | 25833..27119 | 2201154 | AAA-family-ATPase-[Chthoniobacterales-bacterium] |
| TAE90781.1 | OLD-ABC | GCA_004293285.1 | Verrucomicrobiota_bacterium | Bacteria>Pseudomonadati>PVC_group>Verrucomicrobiota>unclassified_Verrucomicrobiota | 461 | EAZ81_08655 | 7943..9328 | 2026799 | ATP-binding-protein-[Verrucomicrobiota-bacterium] |
| MBP7947573.1 | OLD-ABC | GCA_018003215.1 | Verrucomicrobiota_bacterium | Bacteria>Pseudomonadati>PVC_group>Verrucomicrobiota>unclassified_Verrucomicrobiota | 444 | KA191_16105 | 5119..6453 | 2026799 | AAA-family-ATPase-[Verrucomicrobiota-bacterium] |
| MBP7947740.1 | OLD-ABC | GCA_018003215.1 | Verrucomicrobiota_bacterium | Bacteria>Pseudomonadati>PVC_group>Verrucomicrobiota>unclassified_Verrucomicrobiota | 436 | KA191_16995 | 7935..9245 | 2026799 | AAA-family-ATPase-[Verrucomicrobiota-bacterium] |
| NMD22051.1 | OLD-ABC | GCA_012797305.1 | Verrucomicrobiota_bacterium | Bacteria>Pseudomonadati>PVC_group>Verrucomicrobiota>unclassified_Verrucomicrobiota | 436 | GYA76_17660 | 3765..5075 | 2026799 | AAA-family-ATPase-[Verrucomicrobiota-bacterium] |
| MCF7707713.1 | OLD-ABC | GCA_021733835.1 | Verrucomicrobiota_bacterium | Bacteria>Pseudomonadati>PVC_group>Verrucomicrobiota>unclassified_Verrucomicrobiota | 428 | K9N52_02290 | 30097..31383 | 2026799 | AAA-family-ATPase-[Verrucomicrobiota-bacterium] |
| MCX6966133.1 | OLD-ABC | GCA_026398255.1 | Verrucomicrobiota_bacterium | Bacteria>Pseudomonadati>PVC_group>Verrucomicrobiota>unclassified_Verrucomicrobiota | 428 | NTW41_12410 | 32667..33953 | 2026799 | MAG:-AAA-family-ATPase-[Verrucomicrobia-bacterium] |
| TAE90783.1 | OLD-ABC | GCA_004293285.1 | Verrucomicrobiota_bacterium | Bacteria>Pseudomonadati>PVC_group>Verrucomicrobiota>unclassified_Verrucomicrobiota | 428 | EAZ81_08665 | 10074..11360 | 2026799 | hypothetical-protein-EAZ81_08665-[Verrucomicrobiota-bacterium] |
| MBP7947742.1 | OLD-ABC | GCA_018003215.1 | Verrucomicrobiota_bacterium | Bacteria>Pseudomonadati>PVC_group>Verrucomicrobiota>unclassified_Verrucomicrobiota | 423 | KA191_17005 | 9922..11193 | 2026799 | ATP-binding-protein-[Verrucomicrobiota-bacterium] |
| NLH73613.1 | OLD-ABC | GCA_012517065.1 | Verrucomicrobiota_bacterium | Bacteria>Pseudomonadati>PVC_group>Verrucomicrobiota>unclassified_Verrucomicrobiota | 423 | GX456_11215 | 19862..21133 | 2026799 | ATP-binding-protein-[Verrucomicrobiota-bacterium] |
| NMD22049.1 | OLD-ABC | GCA_012797305.1 | Verrucomicrobiota_bacterium | Bacteria>Pseudomonadati>PVC_group>Verrucomicrobiota>unclassified_Verrucomicrobiota | 423 | GYA76_17650 | 1817..3088 | 2026799 | AAA-family-ATPase-[Verrucomicrobiota-bacterium] |
| MBU6399158.1 | OLD-ABC | GCA_018969165.1 | Verrucomicrobiota_bacterium | Bacteria>Pseudomonadati>PVC_group>Verrucomicrobiota>unclassified_Verrucomicrobiota | 408 | KGS61_02470 | 872..2098 | 2026799 | AAA-family-ATPase-[Verrucomicrobiota-bacterium] |
| MBI4624649.1 | OLD-ABC | GCA_016209255.1 | Verrucomicrobiota_bacterium | Bacteria>Pseudomonadati>PVC_group>Verrucomicrobiota>unclassified_Verrucomicrobiota | 406 | HY736_15700 | 13312..14532 | 2026799 | AAA-family-ATPase-[Verrucomicrobiota-bacterium] |
| MBM3880422.1 | OLD-ABC | GCA_016871715.1 | Verrucomicrobiota_bacterium | Bacteria>Pseudomonadati>PVC_group>Verrucomicrobiota>unclassified_Verrucomicrobiota | 403 | FJ387_12010 | 9151..10362 | 2026799 | chromosome-segregation-protein-SMC-[Verrucomicrobiota-bacterium] |
| MBM3880204.1 | OLD-ABC | GCA_016871715.1 | Verrucomicrobiota_bacterium | Bacteria>Pseudomonadati>PVC_group>Verrucomicrobiota>unclassified_Verrucomicrobiota | 400 | FJ387_10875 | 26021..27223 | 2026799 | hypothetical-protein-[Verrucomicrobiota-bacterium] |
| PWU13196.1 | OLD-ABC | GCA_003176045.1 | Verrucomicrobiota_bacterium | Bacteria>Pseudomonadati>PVC_group>Verrucomicrobiota>unclassified_Verrucomicrobiota | 397 | C5B50_19750 | 17212..18405 | 2026799 | hypothetical-protein-C5B50_19750-[Verrucomicrobiota-bacterium] |
| MCF7668201.1 | OLD-ABC | GCA_021733785.1 | Verrucomicrobiota_bacterium | Bacteria>Pseudomonadati>PVC_group>Verrucomicrobiota>unclassified_Verrucomicrobiota | 391 | K9N48_00285 | 27794..28969 | 2026799 | AAA-family-ATPase-[Verrucomicrobiota-bacterium] |
| MCC5848507.1 | OLD-ABC | GCA_020831635.1 | Verrucomicrobiota_bacterium | Bacteria>Pseudomonadati>PVC_group>Verrucomicrobiota>unclassified_Verrucomicrobiota | 389 | JJU29_10455 | 23301..24470 | 2026799 | AAA-family-ATPase-[Verrucomicrobiota-bacterium] |
| RME93003.1 | OLD-ABC | GCA_003695015.1 | Verrucomicrobiota_bacterium | Bacteria>Pseudomonadati>PVC_group>Verrucomicrobiota>unclassified_Verrucomicrobiota | 387 | D6766_09040 | 7488..8651 | 2026799 | chromosome-segregation-protein-SMC-[Verrucomicrobiota-bacterium] |
| MBU4201310.1 | OLD-ABC | GCA_018898455.1 | Verrucomicrobiota_bacterium | Bacteria>Pseudomonadati>PVC_group>Verrucomicrobiota>unclassified_Verrucomicrobiota | 329 | KJ919_15490 | 17510..18499 | 2026799 | ATP-binding-protein-[Verrucomicrobiota-bacterium] |
| WP_165106902.1 | OLD-ABC | GCF_011044775.1 | Limisphaera_ngatamarikiensis | Bacteria>Pseudomonadati>PVC_group>Verrucomicrobiota>Verrucomicrobiia>Limisphaerales>Limisphaeraceae>Limisphaera | 415 | G4L39_RS06745 | 85970..87217 | 1324935 | ATP-binding-protein-[Limisphaera-ngatamarikiensis] |
| MBE7502032.1 | OLD-ABC | GCA_015075445.1 | Verrucomicrobiales_bacterium | Bacteria>Pseudomonadati>PVC_group>Verrucomicrobiota>Verrucomicrobiia>Verrucomicrobiales>unclassified_Verrucomicrobiales | 429 | HS113_17410 | 119505..120794 | 2026801 | ATP-binding-protein-[Verrucomicrobiales-bacterium] |
| MBE7503147.1 | OLD-ABC | GCA_015075445.1 | Verrucomicrobiales_bacterium | Bacteria>Pseudomonadati>PVC_group>Verrucomicrobiota>Verrucomicrobiia>Verrucomicrobiales>unclassified_Verrucomicrobiales | 423 | HS113_23255 | 53847..55118 | 2026801 | ATP-binding-protein-[Verrucomicrobiales-bacterium] |
| MCI0536693.1 | OLD-ABC | GCA_022615925.1 | Verrucomicrobiales_bacterium | Bacteria>Pseudomonadati>PVC_group>Verrucomicrobiota>Verrucomicrobiia>Verrucomicrobiales>unclassified_Verrucomicrobiales | 403 | LOZ50_15860 | 6901..8112 | 2026801 | AAA-family-ATPase-[Verrucomicrobiales-bacterium] |
| MBO7485542.1 | OLD-ABC | GCA_017649985.1 | Spirochaetaceae_bacterium | Bacteria>Pseudomonadati>Spirochaetota>Spirochaetia>Spirochaetales>Spirochaetaceae>unclassified_Spirochaetaceae | 432 | J6T84_05725 | 13218..14516 | 1898206 | AAA-family-ATPase-[Spirochaetaceae-bacterium] |
| MDE0447653.1 | OLD-ABC | GCA_028821475.1 | Spirochaetaceae_bacterium | Bacteria>Pseudomonadati>Spirochaetota>Spirochaetia>Spirochaetales>Spirochaetaceae>unclassified_Spirochaetaceae | 427 | OXH96_13350 | 28729..30012 | 1898206 | MAG:-AAA-family-ATPase-[Spirochaetaceae-bacterium] |
| MDD9989003.1 | OLD-ABC | GCA_028818385.1 | Spirochaetaceae_bacterium | Bacteria>Pseudomonadati>Spirochaetota>Spirochaetia>Spirochaetales>Spirochaetaceae>unclassified_Spirochaetaceae | 414 | OXQ31_22220 | 6989..8233 | 1898206 | MAG:-AAA-family-ATPase-[Spirochaetaceae-bacterium] |
| OQY41117.1 | OLD-ABC | GCA_002084785.1 | Spirochaetaceae_bacterium_4572_7 | Bacteria>Pseudomonadati>Spirochaetota>Spirochaetia>Spirochaetales>Spirochaetaceae>unclassified_Spirochaetaceae | 395 | B6229_00025 | 5654..6841 | 1972456 | chromosome-segregation-protein-SMC-[Spirochaetaceae-bacterium-4572_7] |
| GHU24471.1 | OLD-ABC | GCA_026002265.1 | Spirochaetia_bacterium | Bacteria>Pseudomonadati>Spirochaetota>Spirochaetia>unclassified_Spirochaetia | 386 | FACS1894164_11190 | 4618..5778 | 2053615 | chromosome-segregation-protein-SMC-[Spirochaetia-bacterium] |
| GHV83240.1 | OLD-ABC | GCA_025997755.1 | Spirochaetia_bacterium | Bacteria>Pseudomonadati>Spirochaetota>Spirochaetia>unclassified_Spirochaetia | 383 | AGMMS50212_05800 | 4436..5587 | 2053615 | chromosome-segregation-protein-SMC-[Spirochaetia-bacterium] |
| OQB96946.1 | OLD-ABC | GCA_002070755.1 | Spirochaetes_bacterium_ADurb.Bin110 | Bacteria>Pseudomonadati>Spirochaetota>unclassified_Spirochaetota | 445 | BWX81_01215 | 14279..15616 | 1852908 | hypothetical-protein-BWX81_01215-[Spirochaetes-bacterium-ADurb.Bin110] |
| TXT44738.1 | OLD-ABC | GCA_008080685.1 | Spirochaetota_bacterium | Bacteria>Pseudomonadati>Spirochaetota>unclassified_Spirochaetota | 407 | FD137_1513 | 8876..10099 | 2202144 | hypothetical-protein-FD137_1513-[Spirochaetota-bacterium] |
| HEC67881.1 | OLD-ABC | GCA_011052795.1 | Candidatus_Desulfofervidus_auxilii | Bacteria>Pseudomonadati>Thermodesulfobacteriota>Candidatus_Desulfofervidia>Candidatus_Desulfofervidales>Candidatus_Desulfofervidaceae>Candidatus_Desulfofervidus | 339 | ENI35_03595 | 7556..8575 | 1621989 | hypothetical-protein-[Candidatus-Desulfofervidus-auxilii] |
| WP_083610987.1 | OLD-ABC | GCF_900142135.1 | Desulfatibacillum_alkenivorans | Bacteria>Pseudomonadati>Thermodesulfobacteriota>Desulfobacteria>Desulfobacterales>Desulfatibacillaceae>Desulfatibacillum | 370 | BUE18_RS12855 | 64568..65680 | 259354 | AAA-family-ATPase-[Desulfatibacillum-alkenivorans] |
| WP_028324162.1 | OLD-ABC | GCF_000429925.1 | Desulfatirhabdium_butyrativorans | Bacteria>Pseudomonadati>Thermodesulfobacteriota>Desulfobacteria>Desulfobacterales>Desulfatirhabdiaceae>Desulfatirhabdium | 385 | G492_RS0107600 | 72633..73790 | 340467 | ATP-binding-protein-[Desulfatirhabdium-butyrativorans] |
| WP_028324419.1 | OLD-ABC | GCF_000429925.1 | Desulfatirhabdium_butyrativorans | Bacteria>Pseudomonadati>Thermodesulfobacteriota>Desulfobacteria>Desulfobacterales>Desulfatirhabdiaceae>Desulfatirhabdium | 359 | G492_RS0109400 | 92639..93718 | 340467 | AAA-family-ATPase-[Desulfatirhabdium-butyrativorans] |
| WP_028324417.1 | OLD-ABC | GCF_000429925.1 | Desulfatirhabdium_butyrativorans | Bacteria>Pseudomonadati>Thermodesulfobacteriota>Desulfobacteria>Desulfobacterales>Desulfatirhabdiaceae>Desulfatirhabdium | 344 | G492_RS0109385 | 87534..88568 | 340467 | ATP-binding-protein-[Desulfatirhabdium-butyrativorans] |
| WP_080806588.1 | OLD-ABC | GCF_900170035.1 | Desulfamplus_magnetovallimortis | Bacteria>Pseudomonadati>Thermodesulfobacteriota>Desulfobacteria>Desulfobacterales>Desulfobacteraceae>Desulfamplus | 430 | MTBBW1_RS20355 | 75974..77266 | 1246637 | AAA-family-ATPase-[Desulfamplus-magnetovallimortis] |
| MBF0468019.1 | OLD-ABC | GCA_015232935.1 | Desulfamplus_sp. | Bacteria>Pseudomonadati>Thermodesulfobacteriota>Desulfobacteria>Desulfobacterales>Desulfobacteraceae>Desulfamplus>unclassified_Desulfamplus | 417 | HQK61_03905 | 3584..4837 | 2006184 | AAA-family-ATPase-[Desulfamplus-sp.] |
| HBT88454.1 | OLD-ABC | GCA_003510365.1 | Desulfobacter_sp. | Bacteria>Pseudomonadati>Thermodesulfobacteriota>Desulfobacteria>Desulfobacterales>Desulfobacteraceae>Desulfobacter>unclassified_Desulfobacter | 412 | DEB50_08315 | 27339..28577 | 2294 | ATPase-[Desulfobacter-sp.] |
| MCF8089783.1 | OLD-ABC | GCA_021737465.1 | Desulfotignum_sp. | Bacteria>Pseudomonadati>Thermodesulfobacteriota>Desulfobacteria>Desulfobacterales>Desulfobacteraceae>Desulfotignum>unclassified_Desulfotignum | 376 | K9L23_17200 | 6676..7806 | 2049434 | AAA-family-ATPase-[Desulfotignum-sp.] |
| MCF8089778.1 | OLD-ABC | GCA_021737465.1 | Desulfotignum_sp. | Bacteria>Pseudomonadati>Thermodesulfobacteriota>Desulfobacteria>Desulfobacterales>Desulfobacteraceae>Desulfotignum>unclassified_Desulfotignum | 79 | K9L23_17175 | 1..242 | 2049434 | hypothetical-protein-[Desulfotignum-sp.] |
| MBC8432680.1 | OLD-ABC | GCA_014383215.1 | Candidatus_Desulfatibia_vada | Bacteria>Pseudomonadati>Thermodesulfobacteriota>Desulfobacteria>Desulfobacterales>Desulfobacterales_incertae_sedis>Candidatus_Desulfatibia | 396 | H8D96_12280 | 10629..11819 | 2841696 | AAA-family-ATPase-[Desulfobacterales-bacterium] |
| WP_207678193.1 | OLD-ABC | GCF_017377395.1 | Desulfonema_magnum | Bacteria>Pseudomonadati>Thermodesulfobacteriota>Desulfobacteria>Desulfobacterales>Desulfococcaceae>Desulfonema | 450 | dnm_RS17440 | 4555075..4556427 | 45655 | ATP-binding-protein-[Desulfonema-magnum] |
| WP_155316143.1 | OLD-ABC | GCF_009688945.1 | Desulfosarcina_alkanivorans | Bacteria>Pseudomonadati>Thermodesulfobacteriota>Desulfobacteria>Desulfobacterales>Desulfosarcinaceae>Desulfosarcina | 395 | DSCA_RS09160 | 2077141..2078328 | 571177 | AAA-family-ATPase-[Desulfosarcina-alkanivorans] |

| Protein-ID | Domain | Assembly-ID | Strain | Taxonomy | Length | Locus-Tag | Genomic-coordinates | TaxID | Fasta-defline |
|---|---|---|---|---|---|---|---|---|---|
| MBC2710214.1 | OLD-ABC | GCA_014237405.1 | Desulfosarcina_sp. | Bacteria>Pseudomonadati>Thermodesulfobacteriota>Desulfobacteria>Desulfobacterales>Desulfosarcinaceae>Desulfosarcina>unclassified_Desulfosarcina | 394 | HGJ94_04185 | 16177..17361 | 2027861 | AAA-family-ATPase-[Desulfosarcina-sp.] |
| MDD2316831.1 | OLD-ABC | GCA_028683495.1 | Desulfobacterales_bacterium | Bacteria>Pseudomonadati>Thermodesulfobacteriota>Desulfobacteria>Desulfobacterales>unclassified_Desulfobacterales | 463 | PHD57_05735 | 5715..7106 | 2044940 | MAG:-ATP-binding-protein-[Desulfobacterales-bacterium] |
| MBF0101463.1 | OLD-ABC | GCA_015228675.1 | Desulfobacterales_bacterium | Bacteria>Pseudomonadati>Thermodesulfobacteriota>Desulfobacteria>Desulfobacterales>unclassified_Desulfobacterales | 433 | HQK77_11195 | 8064..9365 | 2044940 | AAA-family-ATPase-[Desulfobacterales-bacterium] |
| MBL0712137.1 | OLD-ABC | GCA_016735415.1 | Desulfobacterales_bacterium | Bacteria>Pseudomonadati>Thermodesulfobacteriota>Desulfobacteria>Desulfobacterales>unclassified_Desulfobacterales | 430 | JJV98_00420 | 9447..10739 | 2044940 | AAA-family-ATPase-[Desulfosarcina-sp.] |
| MDD2316834.1 | OLD-ABC | GCA_028683495.1 | Desulfobacterales_bacterium | Bacteria>Pseudomonadati>Thermodesulfobacteriota>Desulfobacteria>Desulfobacterales>unclassified_Desulfobacterales | 395 | PHD57_05750 | 11232..12419 | 2044940 | MAG:-AAA-family-ATPase-[Desulfobacterales-bacterium] |
| MCG2778951.1 | OLD-ABC | GCA_022013615.1 | Desulfobacterales_bacterium | Bacteria>Pseudomonadati>Thermodesulfobacteriota>Desulfobacteria>Desulfobacterales>unclassified_Desulfobacterales | 381 | L6406_25020 | 7138..8283 | 2044940 | AAA-family-ATPase-[Desulfobacterales-bacterium] |
| MCU0558986.1 | OLD-ABC | GCA_025454255.1 | Desulfobacterales_bacterium | Bacteria>Pseudomonadati>Thermodesulfobacteriota>Desulfobacteria>Desulfobacterales>unclassified_Desulfobacterales | 378 | MUD16_02175 | 70142..71278 | 2044940 | AAA-family-ATPase-[Desulfobacterales-bacterium] |
| MBF0101461.1 | OLD-ABC | GCA_015228675.1 | Desulfobacterales_bacterium | Bacteria>Pseudomonadati>Thermodesulfobacteriota>Desulfobacteria>Desulfobacterales>unclassified_Desulfobacterales | 357 | HQK77_11185 | 6291..7364 | 2044940 | AAA-family-ATPase-[Desulfobacterales-bacterium] |
| WP_028319486.1 | OLD-ABC | GCF_000621145.1 | Desulfobulbus_elongatus | Bacteria>Pseudomonadati>Thermodesulfobacteriota>Desulfobulbia>Desulfobulbales>Desulfobulbaceae>Desulfobulbus | 384 | Q362_RS0116025 | 7316..8470 | 53332 | ATP-binding-protein-[Desulfobulbus-elongatus] |
| WP_205241968.1 | OLD-ABC | GCF_016918545.1 | Desulfobulbus_alkaliphilus | Bacteria>Pseudomonadati>Thermodesulfobacteriota>Desulfobulbia>Desulfobulbales>Desulfobulbaceae>Desulfobulbus | 369 | JWG43_RS06290 | 124152..125261 | 869814 | AAA-family-ATPase-[Desulfobulbus-alkaliphilus] |
| WP_218915011.1 | OLD-ABC | - | Desulfurivibrio_alkaliphilus | Bacteria>Pseudomonadati>Thermodesulfobacteriota>Desulfobulbia>Desulfobulbales>Desulfobulbaceae>Desulfurivibrio | 332 | DAAHT2_RS10970 | 2517995..2518993 | 427923 | AAA-family-ATPase-[Desulfurivibrio-alkaliphilus] |
| MBE0584990.1 | OLD-ABC | GCA_014860245.1 | Desulfofustis_sp. | Bacteria>Pseudomonadati>Thermodesulfobacteriota>Desulfobulbia>Desulfobulbales>Desulfocapsaceae>Desulfofustis>unclassified_Desulfofustis | 395 | IH612_14685 | 1438..2625 | 2053546 | AAA-family-ATPase-[Desulfofustis-sp.] |
| BDV01249.1 | OLD-ABC | GCA_027925145.1 | Thermodesulfomicrobium_sp._WS | Bacteria>Pseudomonadati>Thermodesulfobacteriota>Desulfovibrionia>Desulfovibrionales>Desulfomicrobiaceae>Thermodesulfomicrobium>unclassified_Thermodesulfomicrobium | 414 | TDMWS_13340 | 1434825..1436069 | 3004129 | ATPase-[Thermodesulfomicrobium-sp.-WS] |
| HCF05838.1 | OLD-ABC | GCA_003530935.1 | Desulfomicrobiaceae_bacterium | Bacteria>Pseudomonadati>Thermodesulfobacteriota>Desulfovibrionia>Desulfovibrionales>Desulfomicrobiaceae>unclassified_Desulfomicrobiaceae | 390 | DEU72_06345 | 11002..12174 | 2053547 | chromosome-segregation-protein-SMC-[Desulfomicrobiaceae-bacterium] |
| MBZ4685273.1 | OLD-ABC | GCA_020063575.1 | Desulfomicrobiaceae_bacterium | Bacteria>Pseudomonadati>Thermodesulfobacteriota>Desulfovibrionia>Desulfovibrionales>Desulfomicrobiaceae>unclassified_Desulfomicrobiaceae | 390 | JG774_1018 | 63277..64449 | 2053547 | recombination-protein-[Desulfomicrobiaceae-bacterium] |
| WP_008869233.1 | OLD-ABC | GCF_000174435.1 | Desulfonatronospira | Bacteria>Pseudomonadati>Thermodesulfobacteriota>Desulfovibrionia>Desulfovibrionales>Desulfonatronovibrionaceae | 371 | DTHIO_RS04845 | 1065096..1066211 | 488937 | AAA-family-ATPase-[Desulfonatronospira-thiodismutans] |
| WP_045217260.1 | OLD-ABC | GCF_000934755.1 | Desulfonatronovibrio_magnus | Bacteria>Pseudomonadati>Thermodesulfobacteriota>Desulfovibrionia>Desulfovibrionales>Desulfonatronovibrionaceae>Desulfonatronovibrio | 430 | LZ23_RS20535 | 19411..20703 | 698827 | AAA-family-ATPase-[Desulfonatronovibrio-magnus] |
| WP_270570096.1 | OLD-ABC | - | Bilophila_wadsworthia | Bacteria>Pseudomonadati>Thermodesulfobacteriota>Desulfovibrionia>Desulfovibrionales>Desulfovibrionaceae>Bilophila | 430 | PG042_RS02160 | 64811..66103 | 35833 | AAA-family-ATPase-[Bilophila-wadsworthia] |
| MCI7405012.1 | OLD-ABC | GCA_022783735.1 | Desulfovibrio_piger | Bacteria>Pseudomonadati>Thermodesulfobacteriota>Desulfovibrionia>Desulfovibrionales>Desulfovibrionaceae>Desulfovibrio | 362 | MSS64_01030 | 30305..31393 | 901 | AAA-family-ATPase-[Desulfovibrio-piger] |
| PLX83268.1 | OLD-ABC | GCA_002868845.1 | Desulfuromonas_sp. | Bacteria>Pseudomonadati>Thermodesulfobacteriota>Desulfuromonadia>Desulfuromonadales>Desulfuromonadaceae>Desulfuromonas>unclassified_Desulfuromonas | 451 | C0617_12105 | 152495..153850 | 892 | MAG:-abortive-phage-resistance-protein-[Desulfuromonas-sp.] |
| WP_207162787.1 | OLD-ABC | GCF_017338855.1 | Geobacter_benzoatilyticus | Bacteria>Pseudomonadati>Thermodesulfobacteriota>Desulfuromonadia>Geobacterales>Geobacteraceae>Geobacter | 447 | JZM60_RS12590 | 2734827..2736170 | 2815309 | AAA-family-ATPase-[Geobacter-benzoatilyticus] |
| WP_207162790.1 | OLD-ABC | GCF_017338855.1 | Geobacter_benzoatilyticus | Bacteria>Pseudomonadati>Thermodesulfobacteriota>Desulfuromonadia>Geobacterales>Geobacteraceae>Geobacter | 354 | JZM60_RS12605 | 2740360..2741424 | 2815309 | AAA-family-ATPase-[Geobacter-benzoatilyticus] |
| WP_011415946.1 | OLD-ABC | - | Syntrophus_aciditrophicus | Bacteria>Pseudomonadati>Thermodesulfobacteriota>Syntrophia>Syntrophales>Syntrophaceae>Syntrophus | 397 | SYN_RS00160 | 34900..36093 | 316277 | AAA-family-ATPase-[Syntrophus-aciditrophicus] |
| ABC75911.1 | OLD-ABC | GCA_000013405.1 | Syntrophus_aciditrophicus_SB | Bacteria>Pseudomonadati>Thermodesulfobacteriota>Syntrophia>Syntrophales>Syntrophaceae>Syntrophus>Syntrophus_aciditrophicus | 397 | SYN_00010 | 34900..36093 | 56780 | ATPase-[Syntrophus-aciditrophicus-SB] |
| OPY81709.1 | OLD-ABC | GCA_002067745.1 | Syntrophus_sp._PtaU1.Bin005 | Bacteria>Pseudomonadati>Thermodesulfobacteriota>Syntrophia>Syntrophales>Syntrophaceae>Syntrophus>unclassified_Syntrophus_(in:_bacteria) | 448 | A4E70_01075 | 21802..23148 | 1811712 | MAG:-hypothetical-protein-A4E70_01075-[Syntrophus-sp.-PtaU1.Bin005] |
| OIP91482.1 | OLD-ABC | GCA_001873745.1 | Syntrophaceae_bacterium_CG2_30_58_14 | Bacteria>Pseudomonadati>Thermodesulfobacteriota>Syntrophia>Syntrophales>Syntrophaceae>unclassified_Syntrophaceae | 395 | AUK26_06765 | 1351..2538 | 1805378 | hypothetical-protein-AUK26_06765-[Syntrophaceae-bacterium-CG2_30_58_14] |
| MCK9230424.1 | OLD-ABC | GCA_023230845.1 | Syntrophales_bacterium | Bacteria>Pseudomonadati>Thermodesulfobacteriota>Syntrophia>Syntrophales>unclassified_Syntrophales | 391 | MOP18_07080 | 2310..3485 | 2928017 | AAA-family-ATPase-[Syntrophales-bacterium] |
| MCK9528169.1 | OLD-ABC | GCA_023228425.1 | Syntrophales_bacterium | Bacteria>Pseudomonadati>Thermodesulfobacteriota>Syntrophia>Syntrophales>unclassified_Syntrophales | 391 | MOQ23_05905 | 65328..66503 | 2928017 | AAA-family-ATPase-[Syntrophales-bacterium] |
| WP_170180246.1 | OLD-ABC | GCF_005377625.1 | Desulfoglaeba_alkanexedens | Bacteria>Pseudomonadati>Thermodesulfobacteriota>Syntrophobacteria>Syntrophobacterales>Syntrophobacteraceae>Desulfoglaeba | 411 | FDQ92_RS07900 | 1744238..1745473 | 361111 | ATP-binding-protein-[Desulfoglaeba-alkanexedens] |
| WP_137423743.1 | OLD-ABC | GCF_005377625.1 | Desulfoglaeba_alkanexedens | Bacteria>Pseudomonadati>Thermodesulfobacteriota>Syntrophobacteria>Syntrophobacterales>Syntrophobacteraceae>Desulfoglaeba | 377 | FDQ92_RS06000 | 1283157..1284290 | 361111 | ATP-binding-protein-[Desulfoglaeba-alkanexedens] |
| WP_170161765.1 | OLD-ABC | GCF_003751385.1 | Desulfosoma_caldarium | Bacteria>Pseudomonadati>Thermodesulfobacteriota>Syntrophobacteria>Syntrophobacterales>Syntrophobacteraceae>Desulfosoma | 412 | EDC27_RS10780 | 103541..104779 | 610254 | AAA-family-ATPase-[Desulfosoma-caldarium] |
| WP_123290641.1 | OLD-ABC | GCF_003751385.1 | Desulfosoma_caldarium | Bacteria>Pseudomonadati>Thermodesulfobacteriota>Syntrophobacteria>Syntrophobacterales>Syntrophobacteraceae>Desulfosoma | 383 | EDC27_RS10765 | 98379..99530 | 610254 | ATP-binding-protein-[Desulfosoma-caldarium] |
| MBW6486651.1 | OLD-ABC | GCA_019429105.1 | Syntrophobacterales_bacterium | Bacteria>Pseudomonadati>Thermodesulfobacteriota>Syntrophobacteria>Syntrophobacterales>unclassified_Syntrophobacterales | 396 | K0B01_10940 | 9621..10811 | 2282155 | AAA-family-ATPase-[Syntrophobacterales-bacterium] |
| PIV05608.1 | OLD-ABC | GCA_002779905.1 | Syntrophobacterales_bacterium_CG03_land_8_20_14_0_80_58_14 | Bacteria>Pseudomonadati>Thermodesulfobacteriota>Syntrophobacteria>Syntrophobacterales>unclassified_Syntrophobacterales | 364 | COS57_06990 | 10224..11318 | 1974100 | chromosome-segregation-protein-SMC-[Syntrophobacterales-bacterium-CG03_land_8_20_14_0_80_58_14] |
| WP_028893188.1 | OLD-ABC | GCF_000512235.1 | Syntrophorhabdus_aromaticivorans | Bacteria>Pseudomonadati>Thermodesulfobacteriota>Syntrophorhabdia>Syntrophorhabdales>Syntrophorhabdaceae>Syntrophorhabdus | 384 | SYNAR_RS0100635 | 127027..128181 | 328301 | ATP-binding-protein-[Syntrophorhabdus-aromaticivorans] |
| OPY73254.1 | OLD-ABC | GCA_002067415.1 | Syntrophorhabdus_sp._PtaU1.Bin002 | Bacteria>Pseudomonadati>Thermodesulfobacteriota>Syntrophorhabdia>Syntrophorhabdales>Syntrophorhabdaceae>Syntrophorhabdus>unclassified_Syntrophorhabdus | 399 | A4E62_00570 | 21790..22989 | 1811702 | MAG:-recombination-protein-F-[Syntrophorhabdus-sp.-PtaU1.Bin002] |
| MBX3143488.1 | OLD-ABC | GCA_019637455.1 | Trueperaceae_bacterium | Bacteria>Thermotogati>Deinococcota>Deinococci>Trueperales>Trueperaceae>unclassified_Trueperaceae | 394 | KF813_07020 | 10141..11325 | 2026798 | AAA-family-ATPase-[Trueperaceae-bacterium] |
| MCY3637188.1 | OLD-ABC | GCA_026705625.1 | bacterium | Bacteria>unclassified_Bacteria | 422 | OXH23_16425 | 23192..24460 | 1869227 | MAG:-AAA-family-ATPase-[bacterium] |
| MCP4662277.1 | OLD-ABC | GCA_024231315.1 | bacterium | Bacteria>unclassified_Bacteria | 398 | GY856_43300 | 5871..7067 | 1869227 | AAA-family-ATPase-[bacterium] |
| KAF0250485.1 | OLD-ABC | GCA_009773835.1 | bacterium | Bacteria>unclassified_Bacteria | 392 | FD167_111 | 9238..10416 | 1869227 | SMC-domain-containing-protein-[bacterium] |
| MBD5384947.1 | OLD-ABC | GCA_014802625.1 | bacterium | Bacteria>unclassified_Bacteria | 356 | HDR69_00775 | 101087..102157 | 1869227 | AAA-family-ATPase-[bacterium] |
| MBU1261808.1 | OLD-ABC | GCA_018819855.1 | bacterium | Bacteria>unclassified_Bacteria | 331 | KJ640_02505 | 1954..2949 | 1869227 | AAA-family-ATPase-[bacterium] |
| MBU1598554.1 | OLD-ABC | GCA_018822205.1 | bacterium | Bacteria>unclassified_Bacteria | 331 | KKG61_00340 | 1882..2877 | 1869227 | AAA-family-ATPase-[bacterium] |
| MBU2462378.1 | OLD-ABC | GCA_018828965.1 | bacterium | Bacteria>unclassified_Bacteria | 331 | KKH65_05845 | 1292..2287 | 1869227 | AAA-family-ATPase-[bacterium] |
| --OLD-ABC+TOPRIM | | | | | | | | | |
| WP_247995984.1 | OLD-ABC+TOPRIM | GCF_023115355.1 | Halocatena_salina | Archaea>Methanobacteriati>Methanobacteriota>Stenosarchaea_group>Halobacteria>Halobacteriales>Haloarculaceae>Halocatena | 554 | MW046_RS18870 | 50837..52501 | 2934340 | AAA-family-ATPase-[Halocatena-salina] |
| WP_049941287.1 | OLD-ABC+TOPRIM | GCF_000023945.1 | Halorhabdus_utahensis | Archaea>Methanobacteriati>Methanobacteriota>Stenosarchaea_group>Halobacteria>Halobacteriales>Haloarculaceae>Halorhabdus | 531 | HUTA_RS09360 | 1929947..1931542 | 146826 | AAA-family-ATPase-[Halorhabdus-utahensis] |
| WP_049892747.1 | OLD-ABC+TOPRIM | GCF_000069025.1 | Halobacterium_salinarum | Archaea>Methanobacteriati>Methanobacteriota>Stenosarchaea_group>Halobacteria>Halobacteriales>Halobacteriaceae>Halobacterium | 394 | OE_RS13025 | 224681..225865 | 2242 | AAA-family-ATPase-[Halobacterium-salinarum] |
| WP_074878701.1 | OLD-ABC+TOPRIM | GCF_900115675.1 | Halolamina | Archaea>Methanobacteriati>Methanobacteriota>Stenosarchaea_group>Halobacteria>Halobacteriales>Haloferacaceae | 551 | BMZ83_RS11270 | 74546..76201 | 1075397 | MULTISPECIES:-AAA-family-ATPase-[Halolamina] |
| WP_103992126.1 | OLD-ABC+TOPRIM | GCF_004799685.1 | Halobellus_limi | Archaea>Methanobacteriati>Methanobacteriota>Stenosarchaea_group>Halobacteria>Halobacteriales>Haloferacaceae>Halobellus | 533 | DV707_RS08445 | 1709400..1711001 | 699433 | AAA-family-ATPase-[Halobellus-limi] |
| SFP79062.1 | OLD-ABC+TOPRIM | GCA_900115675.1 | Halolamina_pelagica | Archaea>Methanobacteriati>Methanobacteriota>Stenosarchaea_group>Halobacteria>Halobacteriales>Haloferacaceae>Halolamina | 551 | SAMN05216277_10863 | 74546..76201 | 699431 | putative-ATP-dependent-endonuclease-of-the-OLD-family-[Halolamina-pelagica] |
| WP_096396321.1 | OLD-ABC+TOPRIM | GCF_002355655.1 | Halorubrum_trapanicum | Archaea>Methanobacteriati>Methanobacteriota>Stenosarchaea_group>Halobacteria>Halobacteriales>Haloferacaceae>Halorubrum | 554 | CPZ01_RS13865 | 9233..10897 | 29284 | AAA-family-ATPase-[Halorubrum-trapanicum] |
| WP_008446378.1 | OLD-ABC+TOPRIM | GCF_000336875.1 | Halorubrum_californiense | Archaea>Methanobacteriati>Methanobacteriota>Stenosarchaea_group>Halobacteria>Halobacteriales>Haloferacaceae>Halorubrum | 553 | C463_RS17220 | 33206..34867 | 416585 | AAA-family-ATPase-[Halorubrum-californiense] |
| ELZ39025.1 | OLD-ABC+TOPRIM | GCA_000336875.1 | Halorubrum_californiense_DSM_19288 | Archaea>Methanobacteriati>Methanobacteriota>Stenosarchaea_group>Halobacteria>Halobacteriales>Haloferacaceae>Halorubrum>Halorubrum_californiense | 553 | C463_17715 | 33206..34867 | 1227465 | ATP-dependent-endonuclease-of-the-OLD-family--like-protein-[Halorubrum-californiense-DSM-19288] |
| QCC61759.1 | OLD-ABC+TOPRIM | GCA_004799625.1 | Natrinema_thermotolerans | Archaea>Methanobacteriati>Methanobacteriota>Stenosarchaea_group>Halobacteria>Halobacteriales>Natrialbaceae>Natrinema | 551 | DVR14_24765 | 899438..901093 | 121872 | ATP-dependent-endonuclease-[Natrinema-thermotolerans] |
| WP_049967091.1 | OLD-ABC+TOPRIM | GCF_031165565.1 | Natrinema_thermotolerans | Archaea>Methanobacteriati>Methanobacteriota>Stenosarchaea_group>Halobacteria>Halobacteriales>Natrialbaceae>Natrinema | 551 | HL44_RS19245 | 136338..137993 | 121872 | AAA-family-ATPase-[Natrinema-thermotolerans] |
| MBR1368471.1 | OLD-ABC+TOPRIM | GCA_018132105.1 | Methanocalculus_chunghsingensis | Archaea>Methanobacteriati>Methanobacteriota>Stenosarchaea_group>Methanomicrobia>Methanomicrobiales>Methanocalculaceae>Methanocalculus | 631 | RJ53_02710 | 548..2443 | 156457 | hypothetical-protein-[Methanocalculus-chunghsingensis] |
| WP_211530099.1 | OLD-ABC+TOPRIM | GCF_018132105.1 | Methanocalculus_chunghsingensis | Archaea>Methanobacteriati>Methanobacteriota>Stenosarchaea_group>Methanomicrobia>Methanomicrobiales>Methanocalculaceae>Methanocalculus | 631 | RJ53_RS02775 | 548..2443 | 156457 | AAA-family-ATPase-[Methanocalculus-chunghsingensis] |
| QN042141.1 | OLD-ABC+TOPRIM | - | Candidatus_Methanogaster_sp._ANME-2c_ERB4 | Archaea>Methanobacteriati>Methanobacteriota>Stenosarchaea_group>Methanomicrobia>Methanosarcinales>ANME-2_cluster>Candidatus_Methanogasteraceae>Candidatus_Methanogaster>unclassified_Candidatus_Methanogaster | 638 | CAOPPJJI_00007 | 9298..11214 | 2759911 | DNA-replication-and-repair-protein-RecF-[Methanosarcinales-archaeon-ANME-2c-ERB4] |
| WP_135605608.1 | OLD-ABC+TOPRIM | GCF_900774285.1 | Methanococcoides_sp._NM1 | Archaea>Methanobacteriati>Methanobacteriota>Stenosarchaea_group>Methanomicrobia>Methanosarcinales>Methanosarcinaceae>Methanococcoides>unclassified_Methanococcoides | 661 | E7X56_RS05455 | 149497..151482 | 1201013 | AAA-family-ATPase-[Methanococcoides-sp.-NM1] |
| WP_209616530.1 | OLD-ABC+TOPRIM | GCF_017873415.1 | Methanolobus_bombayensis | Archaea>Methanobacteriati>Methanobacteriota>Stenosarchaea_group>Methanomicrobia>Methanosarcinales>Methanosarcinaceae>Methanolobus | 605 | J2746_RS03795 | 327937..329754 | 38023 | DUF2813-domain-containing-protein-[Methanolobus-bombayensis] |
| WP_013898591.1 | OLD-ABC+TOPRIM | GCF_000217995.1 | Methanosalsum_zhilinae | Archaea>Methanobacteriati>Methanobacteriota>Stenosarchaea_group>Methanomicrobia>Methanosarcinales>Methanosarcinaceae>Methanosalsum | 605 | MZHIL_RS06575 | 1406392..1408209 | 39669 | DUF2813-domain-containing-protein-[Methanosalsum-zhilinae] |
| WP_048126485.1 | OLD-ABC+TOPRIM | GCF_000970265.1 | Methanosarcina_lacustris | Archaea>Methanobacteriati>Methanobacteriota>Stenosarchaea_group>Methanomicrobia>Methanosarcinales>Methanosarcinaceae>Methanosarcina | 670 | MSLAZ_RS10140 | 2461327..2463339 | 170861 | AAA-family-ATPase-[Methanosarcina-lacustris] |
| WP_148705389.1 | OLD-ABC+TOPRIM | GCF_000970125.1 | Methanosarcina_siciliae | Archaea>Methanobacteriati>Methanobacteriota>Stenosarchaea_group>Methanomicrobia>Methanosarcinales>Methanosarcinaceae>Methanosarcina | 612 | MSSIH_RS10195 | 2394975..2396813 | 38027 | ATP-binding-protein-[Methanosarcina-siciliae] |
| WP_082108016.1 | OLD-ABC+TOPRIM | GCF_000979385.1 | Methanosarcina_sp._2.H.A.1B.4 | Archaea>Methanobacteriati>Methanobacteriota>Stenosarchaea_group>Methanomicrobia>Methanosarcinales>Methanosarcinaceae>Methanosarcina>unclassified_Methanosarcina | 692 | EO92_RS09690 | 20413..22491 | 1483600 | AAA-family-ATPase-[Methanosarcina-sp.-2.H.A.1B.4] |
| HII01699.1 | OLD-ABC+TOPRIM | GCA_013329415.1 | Methanosarcinaceae_archaeon | Archaea>Methanobacteriati>Methanobacteriota>Stenosarchaea_group>Methanomicrobia>Methanosarcinales>Methanosarcinaceae>unclassified_Methanosarcinaceae | 507 | HA351_08655 | 3631..5153 | 1945595 | AAA-family-ATPase-[Methanosarcinaceae-archaeon] |
| WP_148695185.1 | OLD-ABC+TOPRIM | GCF_002906215.1 | Candidatus_Nitrosocaldus | Archaea>Thermoproteati>Nitrososphaerota>Nitrososphaeria>Candidatus_Nitrosocaldales>Candidatus_Nitrosocaldaceae | 432 | CRG89_RS04295 | 832286..833584 | 498374 | MULTISPECIES:-ATP-binding-protein-[Candidatus-Nitrosocaldus] |
| WP_256266338.1 | OLD-ABC+TOPRIM | GCF_048469655.1 | Bacillota | Bacteria>Bacillati | 546 | NE589_RS05465 | 114357..115997 | 1239 | AAA-family-ATPase-[Faecalibacterium-prausnitzii] |
| WP_115955246.1 | OLD-ABC+TOPRIM | GCF_003390735.1 | Bifidobacterium_bifidum | Bacteria>Bacillati>Actinomycetota>Actinomycetes>Bifidobacteriales>Bifidobacteriaceae>Bifidobacterium | 794 | CJD48_RS06175 | 1400854..1403238 | 1681 | AAA-family-ATPase-[Bifidobacterium-bifidum] |
| WP_072727062.1 | OLD-ABC+TOPRIM | GCF_001895165.1 | Bifidobacterium_lemurum | Bacteria>Bacillati>Actinomycetota>Actinomycetes>Bifidobacteriales>Bifidobacteriaceae>Bifidobacterium | 652 | BRTL_RS11115 | 285..2243 | 1603886 | AAA-family-ATPase-[Bifidobacterium-lemurum] |
| WP_033203933.1 | OLD-ABC+TOPRIM | GCF_000716445.1 | Streptomyces | Bacteria>Bacillati>Actinomycetota>Actinomycetes>Kitasatosporales>Streptomycetaceae | 642 | OO54_RS16725 | 48913..50841 | 1883 | MULTISPECIES:-AAA-family-ATPase-[Streptomyces] |
| WP_303656580.1 | OLD-ABC+TOPRIM | - | Streptantibioticus_ferralitis | Bacteria>Bacillati>Actinomycetota>Actinomycetes>Kitasatosporales>Streptomycetaceae>Streptantibioticus | 710 | P2L57_RS22730 | 145706..147838 | 236510 | AAA-family-ATPase-[Streptantibioticus-ferralitis] |
| WP_150501935.1 | OLD-ABC+TOPRIM | GCF_008705255.1 | Streptomyces_venezuelae | Bacteria>Bacillati>Actinomycetota>Actinomycetes>Kitasatosporales>Streptomycetaceae>Streptomyces | 642 | DEJ45_RS06875 | 1623990..1625918 | 54571 | AAA-family-ATPase-[Streptomyces-venezuelae] |
| WP_137992475.1 | OLD-ABC+TOPRIM | GCF_005517195.1 | Streptomyces_vilmorinianum | Bacteria>Bacillati>Actinomycetota>Actinomycetes>Kitasatosporales>Streptomycetaceae>Streptomyces | 641 | FDM97_RS23470 | 5022743..5024668 | 3051092 | AAA-family-ATPase-[Streptomyces-exfoliatus] |
| WP_256259967.1 | OLD-ABC+TOPRIM | GCF_045286965.1 | Streptomyces_atratus | Bacteria>Bacillati>Actinomycetota>Actinomycetes>Kitasatosporales>Streptomycetaceae>Streptomyces | 635 | BT364_RS17780 | 152182..154089 | 1893 | AAA-family-ATPase-[Streptomyces-atratus] |
| WP_266728679.1 | OLD-ABC+TOPRIM | GCF_036077095.1 | unclassified_Streptomyces | Bacteria>Bacillati>Actinomycetota>Actinomycetes>Kitasatosporales>Streptomycetaceae>Streptomyces | 635 | OOK13_RS07600 | 1676224..1678131 | 2593676 | AAA-family-ATPase-[Streptomyces-sp.-NBC_00378] |
| WP_136324452.1 | OLD-ABC+TOPRIM | GCF_004795105.1 | Streptomyces_sp._A0642 | Bacteria>Bacillati>Actinomycetota>Actinomycetes>Kitasatosporales>Streptomycetaceae>Streptomyces>unclassified_Streptomyces | 640 | E6R60_RS01240 | 299216..301138 | 2563100 | ATP-dependent-endonuclease-[Streptomyces-sp.-A0642] |
| WP_266868154.1 | OLD-ABC+TOPRIM | GCF_026340305.1 | Streptomyces_sp._NBC_01381 | Bacteria>Bacillati>Actinomycetota>Actinomycetes>Kitasatosporales>Streptomycetaceae>Streptomyces>unclassified_Streptomyces | 637 | OG453_RS15055 | 3219730..3221643 | 2903845 | AAA-family-ATPase-[Streptomyces-sp.-NBC_01381] |
| WP_282599422.1 | OLD-ABC+TOPRIM | - | Brevibacterium_sp._R8603A2 | Bacteria>Bacillati>Actinomycetota>Actinomycetes>Micrococcales>Brevibacteriaceae>Brevibacterium>unclassified_Brevibacterium | 363 | MTQ12_RS15090 | 51..1142 | 2929779 | AAA-family-ATPase-[Brevibacterium-sp.-R8603A2] |
| WP_152667291.1 | OLD-ABC+TOPRIM | GCF_000986825.1 | Cellulomonas_sp._FA1 | Bacteria>Bacillati>Actinomycetota>Actinomycetes>Micrococcales>Cellulomonadaceae>Cellulomonas>unclassified_Cellulomonas | 579 | GY42_RS11865 | 2639173..2640912 | 1346710 | AAA-family-ATPase-[Cellulomonas-sp.-FA1] |
| WP_204409505.1 | OLD-ABC+TOPRIM | GCF_016907455.1 | Brachybacterium_muris | Bacteria>Bacillati>Actinomycetota>Actinomycetes>Micrococcales>Dermabacteraceae>Brachybacterium | 580 | JOD52_RS08670 | 1869857..1871599 | 219301 | AAA-family-ATPase-[Brachybacterium-muris] |
| WP_040162129.1 | OLD-ABC+TOPRIM | GCF_902375455.1 | Mobilicoccus_massiliensis | Bacteria>Bacillati>Actinomycetota>Actinomycetes>Micrococcales>Dermatophilaceae>Mobilicoccus | 619 | FX004_RS16840 | 302043..303902 | 1522310 | AAA-family-ATPase-[Mobilicoccus-massiliensis] |
| WP_194396697.1 | OLD-ABC+TOPRIM | GCF_015277815.1 | Microbacterium_atlanticum | Bacteria>Bacillati>Actinomycetota>Actinomycetes>Micrococcales>Microbacteriaceae>Microbacterium | 672 | IR212_RS15225 | 3327295..3329313 | 2782168 | AAA-family-ATPase-[Microbacterium-atlanticum] |
| WP_248539434.1 | OLD-ABC+TOPRIM | GCF_023195655.1 | Microbacterium_aurugineum | Bacteria>Bacillati>Actinomycetota>Actinomycetes>Micrococcales>Microbacteriaceae>Microbacterium | 672 | M0721_RS02795 | 590481..592499 | 2851642 | AAA-family-ATPase-[Microbacterium-sp.-SSW1-7] |
| WP_255162090.1 | OLD-ABC+TOPRIM | GCF_024362265.1 | Microbacterium_maritypicum | Bacteria>Bacillati>Actinomycetota>Actinomycetes>Micrococcales>Microbacteriaceae>Microbacterium | 585 | NMQ05_RS13055 | 2676192..2677949 | 33918 | AAA-family-ATPase-[Microbacterium-liquefaciens] |
| MBY6061969.1 | OLD-ABC+TOPRIM | GCA_019801485.1 | Microbacterium_esteraromaticum | Bacteria>Bacillati>Actinomycetota>Actinomycetes>Micrococcales>Microbacteriaceae>Microbacterium | 499 | KUV70_10995 | 266075..267574 | 57043 | AAA-family-ATPase-[Microbacterium-esteraromaticum] |
| WP_222446680.1 | OLD-ABC+TOPRIM | GCF_019801485.1 | Microbacterium_esteraromaticum | Bacteria>Bacillati>Actinomycetota>Actinomycetes>Micrococcales>Microbacteriaceae>Microbacterium | 499 | KUV70_RS10995 | 266075..267574 | 57043 | AAA-family-ATPase-[Microbacterium-esteraromaticum] |
| PZU44170.1 | OLD-ABC+TOPRIM | GCA_003248655.1 | Microbacterium_sp. | Bacteria>Bacillati>Actinomycetota>Actinomycetes>Micrococcales>Microbacteriaceae>Microbacterium>unclassified_Microbacterium | 680 | DI566_12715 | 10370..12412 | 51671 | OLD-family-endonuclease-[Microbacterium-sp.] |
| WP_054682989.1 | OLD-ABC+TOPRIM | GCF_001314225.1 | Microbacterium_sp._No._7 | Bacteria>Bacillati>Actinomycetota>Actinomycetes>Micrococcales>Microbacteriaceae>Microbacterium>unclassified_Microbacterium | 672 | AOA12_RS21695 | 101494..103512 | 1714373 | AAA-family-ATPase-[Microbacterium-sp.-No.-7] |
| WP_274500271.1 | OLD-ABC+TOPRIM | GCF_028765365.1 | Plantibacter_flavus | Bacteria>Bacillati>Actinomycetota>Actinomycetes>Micrococcales>Microbacteriaceae>Plantibacter | 575 | PUY80_RS13435 | 279662..281389 | 150123 | ATP-binding-protein-[Plantibacter-flavus] |
| WP_144831469.1 | OLD-ABC+TOPRIM | GCF_007677595.1 | Kocuria_rhizophila | Bacteria>Bacillati>Actinomycetota>Actinomycetes>Micrococcales>Micrococcaceae>Kocuria | 617 | FS577_RS10175 | 94741..96594 | 72000 | AAA-family-ATPase-[Kocuria-rhizophila] |
| WP_112235019.1 | OLD-ABC+TOPRIM | GCF_003290245.1 | Kocuria_sp._BT304 | Bacteria>Bacillati>Actinomycetota>Actinomycetes>Micrococcales>Micrococcaceae>Kocuria>unclassified_Kocuria | 614 | DNL27_RS06225 | 1385458..1387302 | 1702043 | AAA-family-ATPase-[Kocuria-sp.-BT304] |
| WP_285140309.1 | OLD-ABC+TOPRIM | GCF_030226745.1 | Micrococcus_luteus | Bacteria>Bacillati>Actinomycetota>Actinomycetes>Micrococcales>Micrococcaceae>Micrococcus | 613 | QP726_RS10475 | 13117..14958 | 1270 | AAA-family-ATPase-[Micrococcus-luteus] |
| WP_141842657.1 | OLD-ABC+TOPRIM | GCF_006704135.1 | Micrococcus_sp._R8502A1 | Bacteria>Bacillati>Actinomycetota>Actinomycetes>Micrococcales>Micrococcaceae>Micrococcus>unclassified_Micrococcus | 613 | FLM69_RS00640 | 2017..3858 | 2583239 | AAA-family-ATPase-[Micrococcus-sp.-R8502A1] |
| WP_172894356.1 | OLD-ABC+TOPRIM | GCF_900091515.1 | Micromonospora_purpureochromogenes | Bacteria>Bacillati>Actinomycetota>Actinomycetes>Micromonosporales>Micromonosporaceae>Micromonospora | 624 | GA0074696_RS20315 | 4388136..4390010 | 47872 | AAA-family-ATPase-[Micromonospora-purpureochromogenes] |
| WP_278145174.1 | OLD-ABC+TOPRIM | GCF_029626165.1 | Micromonospora_sp._WMMD967 | Bacteria>Bacillati>Actinomycetota>Actinomycetes>Micromonosporales>Micromonosporaceae>Micromonospora>unclassified_Micromonospora | 576 | O7631_RS17590 | 3787766..3789496 | 3016101 | AAA-family-ATPase-[Micromonospora-sp.-WMMD967] |
| WP_182347792.1 | OLD-ABC+TOPRIM | GCF_014121175.1 | Tomitella_gaofuii | Bacteria>Bacillati>Actinomycetota>Actinomycetes>Mycobacteriales>Corynebacteriales_incertae_sedis>Tomitella | 539 | H7F09_RS10430 | 51352..52971 | 2760083 | AAA-family-ATPase-[Tomitella-gaofuii] |
| WP_213249147.1 | OLD-ABC+TOPRIM | GCF_009914535.1 | Gordonia | Bacteria>Bacillati>Actinomycetota>Actinomycetes>Mycobacteriales>Gordoniaceae | 647 | IDH24_RS11705 | 150297..152240 | 2053 | AAA-family-ATPase-[Gordonia-pseudamarae] |
| PZT93669.1 | OLD-ABC+TOPRIM | GCA_003248405.1 | Gordonia_sp._(in:_high_G+C_Gram-positive_bacteria) | Bacteria>Bacillati>Actinomycetota>Actinomycetes>Mycobacteriales>Gordoniaceae>Gordonia>unclassified_Gordonia_(in:_high_G+C_Gram-positive_bacteria) | 617 | DI630_26690 | 13056..14909 | 84139 | MAG:-hypothetical-protein-DI630_26690-[Gordonia-sp.-(in:-high-G+C-Gram-positive-bacteria)] |
| WP_082948830.1 | OLD-ABC+TOPRIM | GCF_001668625.1 | Mycobacterium_sp._1274756.6 | Bacteria>Bacillati>Actinomycetota>Actinomycetes>Mycobacteriales>Mycobacteriaceae>Mycobacterium>unclassified_Mycobacterium | 567 | A5643_RS17710 | 88341..90044 | 1834076 | ATP-binding-protein-[Mycobacterium-sp.-1274756.6] |
| WP_186322401.1 | OLD-ABC+TOPRIM | GCF_007672705.1 | Prescottella_equi | Bacteria>Bacillati>Actinomycetota>Actinomycetes>Mycobacteriales>Nocardiaceae>Prescottella | 635 | FS672_RS22175 | 129243..131150 | 43767 | AAA-family-ATPase-[Prescottella-equi] |
| WP_222632850.1 | OLD-ABC+TOPRIM | GCF_019836125.1 | unclassified_Rhodococcus_(in:_high_G+C_Gram-positive_bacteria) | Bacteria>Bacillati>Actinomycetota>Actinomycetes>Mycobacteriales>Nocardiaceae>Rhodococcus | 726 | HQ302_RS17045 | 234344..236524 | 192944 | MULTISPECIES:-AAA-family-ATPase-[unclassified-Rhodococcus-(in:-high-G+C-Gram-positive-bacteria)] |
| WP_068111683.1 | OLD-ABC+TOPRIM | GCF_001653335.1 | Nocardioides_dokdonensis | Bacteria>Bacillati>Actinomycetota>Actinomycetes>Propionibacteriales>Nocardioidaceae>Nocardioides | 616 | I601_RS15625 | 3305858..3307708 | 450734 | AAA-family-ATPase-[Nocardioides-dokdonensis] |
| WP_204918872.1 | OLD-ABC+TOPRIM | GCF_039531735.1 | Microlunatus_panaciterrae | Bacteria>Bacillati>Actinomycetota>Actinomycetes>Propionibacteriales>Propionibacteriaceae>Microlunatus | 617 | JOE57_RS13915 | 3038890..3040743 | 400768 | AAA-family-ATPase-[Microlunatus-panaciterrae] |
| WP_119162164.1 | OLD-ABC+TOPRIM | GCF_900637915.1 | Propionibacterium_australiense | Bacteria>Bacillati>Actinomycetota>Actinomycetes>Propionibacteriales>Propionibacteriaceae>Propionibacterium | 539 | EL282_RS10955 | 2472279..2473898 | 119981 | AAA-family-ATPase-[Propionibacterium-australiense] |
| MBK7820966.1 | OLD-ABC+TOPRIM | GCA_016714275.1 | Tessaracoccus_sp. | Bacteria>Bacillati>Actinomycetota>Actinomycetes>Propionibacteriales>Propionibacteriaceae>Tessaracoccus>unclassified_Tessaracoccus | 617 | IPJ61_07785 | 1706244..1708097 | 1971211 | AAA-family-ATPase-[Tessaracoccus-sp.] |
| WP_182784907.1 | OLD-ABC+TOPRIM | GCF_014164725.1 | Tessaracoccus_sp._MC1627 | Bacteria>Bacillati>Actinomycetota>Actinomycetes>Propionibacteriales>Propionibacteriaceae>Tessaracoccus>unclassified_Tessaracoccus | 617 | H5399_RS17215 | 2448..4301 | 2760312 | AAA-family-ATPase-[Tessaracoccus-sp.-MC1627] |
| WP_182799527.1 | OLD-ABC+TOPRIM | GCF_014164775.1 | Tessaracoccus_sp._MC1679 | Bacteria>Bacillati>Actinomycetota>Actinomycetes>Propionibacteriales>Propionibacteriaceae>Tessaracoccus>unclassified_Tessaracoccus | 617 | H5398_RS07165 | 387926..389779 | 2760313 | AAA-family-ATPase-[Tessaracoccus-sp.-MC1679] |
| GII83977.1 | OLD-ABC+TOPRIM | GCA_016863415.1 | Sphaerisporangium_siamense | Bacteria>Bacillati>Actinomycetota>Actinomycetes>Streptosporangiales>Streptosporangiaceae>Sphaerisporangium | 619 | Ssi03_19670 | 378108..379967 | 795645 | hypothetical-protein-Ssi03_19670-[Sphaerisporangium-siamense] |
| WP_184886677.1 | OLD-ABC+TOPRIM | GCF_016863415.1 | Sphaerisporangium_siamense | Bacteria>Bacillati>Actinomycetota>Actinomycetes>Streptosporangiales>Streptosporangiaceae>Sphaerisporangium | 619 | Ssi03_RS09535 | 378108..379967 | 795645 | hypothetical-protein-[Sphaerisporangium-siamense] |
| WP_067479642.1 | OLD-ABC+TOPRIM | GCF_001552635.1 | Actinomadura_hibisca | Bacteria>Bacillati>Actinomycetota>Actinomycetes>Streptosporangiales>Thermomonosporaceae>Actinomadura | 607 | AH6_RS14635 | 24951..26774 | 68565 | AAA-family-ATPase-[Actinomadura-hibisca] |
| KUK48425.1 | OLD-ABC+TOPRIM | GCA_001508275.1 | Actinobacteria_bacterium_66_15 | Bacteria>Bacillati>Actinomycetota>Actinomycetes>unclassified_Actinomycetes | 615 | XD74_0978 | 13428..15275 | 1635289 | ATP-dependent-endonuclease-of-the-OLD-family-like-protein-[Actinobacteria-bacterium-66_15] |
| MBM6989333.1 | OLD-ABC+TOPRIM | GCA_016901295.1 | Parafannyhessea_umbonata | Bacteria>Bacillati>Actinomycetota>Coriobacteriia>Coriobacteriales>Atopobiaceae>Parafannyhessea | 578 | I3I99_07015 | 42883..44619 | 604330 | ATP-binding-protein-[Parafannyhessea-umbonata] |

| Protein-ID | Domain | Assembly-ID | Strain | Taxonomy | Length | Locus-Tag | Genomic-coordinates | TaxID | Fasta-defline |
|---|---|---|---|---|---|---|---|---|---|
| WP_186297390.1 | OLD-ABC+TOPRIM | GCF_007655325.1 | Enorma_shizhengliae | Bacteria>Bacillati>Actinomycetota>Coriobacteriia>Coriobacteriales>Coriobacteriaceae>Enorma | 548 | GJE22_RS07720 | 30048..31694 | 2606615 | AAA-family-ATPase-[Enorma-shizhengliae] |
| MTV27837.1 | OLD-ABC+TOPRIM | GCA_009720505.1 | Profundirhabdus_halotolerans | Bacteria>Bacillati>Actinomycetota>Nitriliruptoria>Nitriliruptorales>Nitriliruptoraceae>Profundirhabdus | 509 | FTX61_20825 | 5719..7248 | 2604685 | AAA-family-ATPase-[Nitriliruptoraceae-bacterium-ZYF776] |
| MSW49532.1 | OLD-ABC+TOPRIM | GCA_009699505.1 | Actinomycetota_bacterium | Bacteria>Bacillati>Actinomycetota>unclassified_Actinomycetota | 781 | F2817_01405 | 11591..13936 | 2900548 | AAA-family-ATPase-[Actinomycetia-bacterium] |
| MCL4554257.1 | OLD-ABC+TOPRIM | GCA_023382015.1 | Actinomycetota_bacterium | Bacteria>Bacillati>Actinomycetota>unclassified_Actinomycetota | 569 | M1565_03865 | 1205..2914 | 2900548 | AAA-family-ATPase-[Actinomycetota-bacterium] |
| MDA8047539.1 | OLD-ABC+TOPRIM | GCA_028063615.1 | Actinomycetota_bacterium | Bacteria>Bacillati>Actinomycetota>unclassified_Actinomycetota | 566 | MOZ30_20290 | 9381..11081 | 2900548 | MAG:-AAA-family-ATPase-[Actinomycetota-bacterium] |
| MCA0180523.1 | OLD-ABC+TOPRIM | GCA_020160805.1 | Actinomycetota_bacterium | Bacteria>Bacillati>Actinomycetota>unclassified_Actinomycetota | 529 | LCH77_13230 | 52339..53928 | 2900548 | AAA-family-ATPase-[Actinomycetia-bacterium] |
| GIV04314.1 | OLD-ABC+TOPRIM | GCA_026003055.1 | Fimbriimonadales_bacterium | Bacteria>Bacillati>Armatimonadota>Fimbriimonadia>Fimbriimonadales>unclassified_Fimbriimonadales | 580 | KatS3mg015_3144 | 6555..8297 | 2682145 | MAG:-ATPase-AAA-[Fimbriimonadales-bacterium] |
| MBN1458233.1 | OLD-ABC+TOPRIM | GCA_016928475.1 | Armatimonadota_bacterium | Bacteria>Bacillati>Armatimonadota>unclassified_Armatimonadota | 560 | JXA57_01770 | 19522..21204 | 2033014 | AAA-family-ATPase-[Armatimonadota-bacterium] |
| WP_117466977.1 | OLD-ABC+TOPRIM | GCF_048410155.1 | Clostridia | Bacteria>Bacillati>Bacillota | 554 | DW835_RS14830 | 33880..35544 | 186801 | AAA-family-ATPase-[Clostridium-sp.-AM34-9AC] |
| WP_002578397.1 | OLD-ABC+TOPRIM | GCF_020687485.1 | Clostridia | Bacteria>Bacillati>Bacillota | 546 | PGV47_RS07610 | 56826..58466 | 186801 | MULTISPECIES:-AAA-family-ATPase-[Lachnospiraceae] |
| WP_023554872.1 | OLD-ABC+TOPRIM | GCF_900177285.1 | Paenibacillaceae | Bacteria>Bacillati>Bacillota>Bacilli>Bacillales | 642 | B9N82_RS00065 | 23862..25790 | 186822 | MULTISPECIES:-AAA-family-ATPase-[Paenibacillaceae] |
| MBB3906529.1 | OLD-ABC+TOPRIM | GCA_014196195.1 | Anoxybacteroides_rupiense | Bacteria>Bacillati>Bacillota>Bacilli>Bacillales>Anoxybacillaceae>Anoxybacteroides | 676 | GGQ95_000806 | 67155..69185 | 311460 | putative-ATP-dependent-endonuclease-of-OLD-family-[Anoxybacillus-rupiensis] |
| WP_275191609.1 | OLD-ABC+TOPRIM | GCF_029077325.1 | Anoxybacteroides_rupiense | Bacteria>Bacillati>Bacillota>Bacilli>Bacillales>Anoxybacillaceae>Anoxybacteroides | 676 | PNH38_RS00015 | 2227..4257 | 311460 | AAA-family-ATPase-[Anoxybacillus-rupiensis] |
| WP_015863622.1 | OLD-ABC+TOPRIM | GCF_004341205.1 | Thermolongibacillus_altinsuensis | Bacteria>Bacillati>Bacillota>Bacilli>Bacillales>Anoxybacillaceae>Thermolongibacillus | 634 | EDD69_RS09655 | 73226..75130 | 575256 | ATP-binding-protein-[Thermolongibacillus-altinsuensis] |
| WP_066159863.1 | OLD-ABC+TOPRIM | GCF_001591945.1 | Halalkalibacter_krulwichiae | Bacteria>Bacillati>Bacillota>Bacilli>Bacillales>Bacillaceae>Halalkalibacter | 636 | BKR01S_RS20115 | 1..1910 | 199441 | AAA-family-ATPase,-partial-[Halalkalibacter-krulwichiae] |
| WP_259420298.1 | OLD-ABC+TOPRIM | GCF_024919535.1 | Lysinibacillus_sp._A4 | Bacteria>Bacillati>Bacillota>Bacilli>Bacillales>Bacillaceae>Lysinibacillus>unclassified_Lysinibacillus | 599 | NY607_RS20565 | 131420..133219 | 2976269 | AAA-family-ATPase-[Lysinibacillus-sp.-A4] |
| WP_272767822.1 | OLD-ABC+TOPRIM | GCF_028548275.1 | Priestia_megaterium | Bacteria>Bacillati>Bacillota>Bacilli>Bacillales>Bacillaceae>Priestia | 683 | PNF31_RS27490 | 21362..23413 | 1404 | AAA-family-ATPase-[Priestia-megaterium] |
| WP_221842523.1 | OLD-ABC+TOPRIM | GCF_019749055.1 | Brevibacillus_agri | Bacteria>Bacillati>Bacillota>Bacilli>Bacillales>Paenibacillaceae>Brevibacillus | 606 | H7K32_RS04510 | 20730..22550 | 51101 | AAA-family-ATPase-[Brevibacillus-agri] |
| WP_198826832.1 | OLD-ABC+TOPRIM | GCF_016406105.1 | Brevibacillus_composti | Bacteria>Bacillati>Bacillota>Bacilli>Bacillales>Paenibacillaceae>Brevibacillus | 585 | KDJ56_RS14935 | 2929196..2930953 | 2796470 | AAA-family-ATPase-[Brevibacillus-composti] |
| GIO15801.1 | OLD-ABC+TOPRIM | GCA_018333175.1 | Cohnella_xylanilytica | Bacteria>Bacillati>Bacillota>Bacilli>Bacillales>Paenibacillaceae>Cohnella | 642 | J19TS2_53560 | 45691..47619 | 557555 | ATP-dependent-endonuclease-[Cohnella-xylanilytica] |
| WP_212962145.1 | OLD-ABC+TOPRIM | GCF_018333175.1 | Cohnella_xylanilytica | Bacteria>Bacillati>Bacillota>Bacilli>Bacillales>Paenibacillaceae>Cohnella | 642 | KI376_RS26580 | 45691..47619 | 557555 | AAA-family-ATPase-[Cohnella-xylanilytica] |
| WP_256832015.1 | OLD-ABC+TOPRIM | GCF_024519685.1 | unclassified_Paenibacillus | Bacteria>Bacillati>Bacillota>Bacilli>Bacillales>Paenibacillaceae>Paenibacillus | 606 | NR121_RS23335 | 27812..29632 | 185978 | MULTISPECIES:-AAA-family-ATPase-[unclassified-Paenibacillus] |
| UUZ84572.1 | OLD-ABC+TOPRIM | GCA_024703795.1 | Paenibacillus_sp._P26 | Bacteria>Bacillati>Bacillota>Bacilli>Bacillales>Paenibacillaceae>Paenibacillus>unclassified_Paenibacillus | 606 | LJK88_13425 | 2170255..2172075 | 2885204 | AAA-family-ATPase-[Paenibacillus-sp.-P26] |
| WP_198143286.1 | OLD-ABC+TOPRIM | GCF_900067165.1 | Trichococcus | Bacteria>Bacillati>Bacillota>Bacilli>Lactobacillales>Carnobacteriaceae | 621 | BUA33_RS02180 | 202473..204338 | 82802 | MULTISPECIES:-AAA-family-ATPase-[Trichococcus] |
| MBD8998783.1 | OLD-ABC+TOPRIM | GCA_014847545.1 | Ligilactobacillus_ruminis | Bacteria>Bacillati>Bacillota>Bacilli>Lactobacillales>Lactobacillaceae>Ligilactobacillus | 554 | EGQ92_00785 | 8957..10621 | 1623 | hypothetical-protein-[Ligilactobacillus-ruminis] |
| WP_013403047.1 | OLD-ABC+TOPRIM | GCF_000022325.1 | Caldicellulosiruptor | Bacteria>Bacillati>Bacillota>Bacillota_incertae_sedis>Caldicellulosiruptorales>Caldicellulosiruptoraceae | 685 | ATHE_RS08035 | 1703965..1706022 | 44000 | MULTISPECIES:-AAA-family-ATPase-[Caldicellulosiruptor] |
| WP_256861317.1 | OLD-ABC+TOPRIM | GCF_000022325.1 | Caldicellulosiruptor_bescii | Bacteria>Bacillati>Bacillota>Bacillota_incertae_sedis>Caldicellulosiruptorales>Caldicellulosiruptoraceae>Caldicellulosiruptor | 647 | ATHE_RS08000 | 1688349..1690292 | 31899 | AAA-family-ATPase-[Caldicellulosiruptor-bescii] |
| WP_013430105.1 | OLD-ABC+TOPRIM | GCF_000166775.1 | Caldicellulosiruptor_kronotskyensis | Bacteria>Bacillati>Bacillota>Bacillota_incertae_sedis>Caldicellulosiruptorales>Caldicellulosiruptoraceae>Caldicellulosiruptor | 631 | CALKRO_RS05675 | 1292393..1294288 | 413889 | AAA-family-ATPase-[Caldicellulosiruptor-kronotskyensis] |
| WP_231133369.1 | OLD-ABC+TOPRIM | GCF_013664685.1 | Capillibacterium_thermochitinicola | Bacteria>Bacillati>Bacillota>Bacillota_sensu_stricto_incertae_sedis>Capillibacterium | 488 | G5B42_RS08305 | 140567..142032 | 2699427 | AAA-family-ATPase,-partial-[Capillibacterium-thermochitinicola] |
| WP_138373958.1 | OLD-ABC+TOPRIM | GCF_015554965.1 | Oscillospiraceae | Bacteria>Bacillati>Bacillota>Clostridia>Eubacteriales | 546 | I2E95_RS16285 | 1602..3242 | 216572 | MULTISPECIES:-AAA-family-ATPase-[Oscillospiraceae] |
| WP_117821975.1 | OLD-ABC+TOPRIM | GCF_003461085.1 | unclassified_Clostridium | Bacteria>Bacillati>Bacillota>Clostridia>Eubacteriales>Clostridiaceae>Clostridium | 554 | DW959_RS12390 | 57083..58747 | 2614128 | MULTISPECIES:-AAA-family-ATPase-[unclassified-Clostridium] |
| WP_072897338.1 | OLD-ABC+TOPRIM | GCF_900129365.1 | Clostridium_fallax | Bacteria>Bacillati>Bacillota>Clostridia>Eubacteriales>Clostridiaceae>Clostridium | 394 | BUA70_RS11135 | 22443..23627 | 1533 | AAA-family-ATPase-[Clostridium-fallax] |
| RGF28775.1 | OLD-ABC+TOPRIM | GCA_003435925.1 | Clostridium_sp._AF46-9NS | Bacteria>Bacillati>Bacillota>Clostridia>Eubacteriales>Clostridiaceae>Clostridium>unclassified_Clostridium | 554 | DW081_14960 | 22762..24426 | 2293020 | hypothetical-protein-DW081_14960-[Clostridium-sp.-AF46-9NS] |
| RHU61443.1 | OLD-ABC+TOPRIM | GCA_003481275.1 | Clostridium_sp._TF08-15 | Bacteria>Bacillati>Bacillota>Clostridia>Eubacteriales>Clostridiaceae>Clostridium>unclassified_Clostridium | 554 | DXC82_15765 | 20709..22373 | 2293052 | hypothetical-protein-DXC82_15765-[Clostridium-sp.-TF08-15] |
| WP_118650261.1 | OLD-ABC+TOPRIM | - | Clostridium_sp._OF09-10 | Bacteria>Bacillati>Bacillota>Clostridia>Eubacteriales>Clostridiaceae>Clostridium>unclassified_Clostridium | 546 | DXA91_RS01345 | 58223..59863 | 2292309 | AAA-family-ATPase-[Clostridium-sp.-OF09-10] |
| NLE27046.1 | OLD-ABC+TOPRIM | GCA_012514215.1 | Clostridiaceae_bacterium | Bacteria>Bacillati>Bacillota>Clostridia>Eubacteriales>Clostridiaceae>unclassified_Clostridiaceae | 597 | GX625_17230 | 33463..35256 | 1898204 | AAA-family-ATPase-[Clostridiaceae-bacterium] |
| WP_015260692.1 | OLD-ABC+TOPRIM | GCF_000243135.2 | Desulfitobacterium_dichloroeliminans | Bacteria>Bacillati>Bacillota>Clostridia>Eubacteriales>Desulfitobacteriaceae>Desulfitobacterium | 678 | DESDI_RS00610 | 125574..127610 | 233055 | AAA-family-ATPase-[Desulfitobacterium-dichloroeliminans] |
| WP_015942907.1 | OLD-ABC+TOPRIM | GCF_000021925.1 | Desulfitobacterium_hafniense | Bacteria>Bacillati>Bacillota>Clostridia>Eubacteriales>Desulfitobacteriaceae>Desulfitobacterium | 601 | DPCES_RS04015 | 858180..859985 | 49338 | AAA-family-ATPase-[Desulfitobacterium-hafniense] |
| MCI5510141.1 | OLD-ABC+TOPRIM | GCA_022764165.1 | Eubacterium_sp. | Bacteria>Bacillati>Bacillota>Clostridia>Eubacteriales>Eubacteriaceae>Eubacterium>unclassified_Eubacterium | 546 | MR436_06825 | 24511..26151 | 142586 | AAA-family-ATPase-[Eubacterium-sp.] |
| MCQ4886537.1 | OLD-ABC+TOPRIM | GCA_024460915.1 | Faecalibacterium_prausnitzii | Bacteria>Bacillati>Bacillota>Clostridia>Eubacteriales>Oscillospiraceae>Faecalibacterium | 546 | NE589_05465 | 114357..115997 | 853 | AAA-family-ATPase-[Faecalibacterium-prausnitzii] |
| NJP39663.1 | OLD-ABC+TOPRIM | GCA_012034405.1 | Oscillospiraceae_bacterium_HV4-5-C5C | Bacteria>Bacillati>Bacillota>Clostridia>Eubacteriales>Oscillospiraceae>unclassified_Oscillospiraceae | 521 | HCH52_01140 | 258737..260302 | 2721777 | AAA-family-ATPase-[Oscillospiraceae-bacterium-HV4-5-C5C] |
| WP_259498353.1 | OLD-ABC+TOPRIM | GCF_024958695.1 | Desulfofundulus_thermocisternus | Bacteria>Bacillati>Bacillota>Clostridia>Eubacteriales>Peptococcaceae>Desulfofundulus | 576 | NZJ93_RS02600 | 108546..110276 | 42471 | AAA-family-ATPase-[Desulfofundulus-thermocisternus] |
| WP_013175443.1 | OLD-ABC+TOPRIM | GCF_000092405.1 | Syntrophothermus_lipocalidus | Bacteria>Bacillati>Bacillota>Clostridia>Eubacteriales>Syntrophomonadaceae>Syntrophothermus | 536 | SLIP_RS06290 | 1304459..1306069 | 86170 | AAA-family-ATPase-[Syntrophothermus-lipocalidus] |
| NLI53237.1 | OLD-ABC+TOPRIM | GCA_012517905.1 | Clostridiales_bacterium | Bacteria>Bacillati>Bacillota>Clostridia>Eubacteriales>unclassified_Eubacteriales | 576 | GX417_02815 | 12764..14494 | 1898207 | ATP-binding-protein-[Clostridiales-bacterium] |
| WP_108141014.1 | OLD-ABC+TOPRIM | GCF_003053565.1 | Halanaerobium | Bacteria>Bacillati>Bacillota>Clostridia>Halanaerobiales>Halanaerobiaceae | 593 | C8C79_RS10210 | 10391..12172 | 2330 | MULTISPECIES:-AAA-family-ATPase-[Halanaerobium] |
| WP_005488048.1 | OLD-ABC+TOPRIM | GCF_000350165.1 | Halanaerobium_saccharolyticum | Bacteria>Bacillati>Bacillota>Clostridia>Halanaerobiales>Halanaerobiaceae>Halanaerobium | 598 | HSACCH_RS04035 | 791923..793719 | 43595 | AAA-family-ATPase-[Halanaerobium-saccharolyticum] |
| WP_055268225.1 | OLD-ABC+TOPRIM | GCF_001406335.1 | Lachnospiraceae | Bacteria>Bacillati>Bacillota>Clostridia>Lachnospirales | 546 | LON21_RS13015 | 65772..67412 | 186803 | MULTISPECIES:-AAA-family-ATPase-[Lachnospiraceae] |
| WP_270389666.1 | OLD-ABC+TOPRIM | GCF_027661925.1 | Blautia_massiliensis_(ex_Durand_et_al._2017) | Bacteria>Bacillati>Bacillota>Clostridia>Lachnospirales>Lachnospiraceae>Blautia | 485 | PGQ23_RS05940 | 26462..27919 | 1737424 | AAA-family-ATPase-[Blautia-massiliensis] |
| RHS91607.1 | OLD-ABC+TOPRIM | GCA_003480245.1 | Blautia_sp._AM42-2 | Bacteria>Bacillati>Bacillota>Clostridia>Lachnospirales>Lachnospiraceae>Blautia>unclassified_Blautia | 546 | DW915_10740 | 19014..20654 | 2292976 | ATP-dependent-endonuclease-[Blautia-sp.-AM42-2] |
| WP_073386263.1 | OLD-ABC+TOPRIM | GCF_900129945.1 | Butyrivibrio_fibrisolvens | Bacteria>Bacillati>Bacillota>Clostridia>Lachnospirales>Lachnospiraceae>Butyrivibrio | 626 | BUC27_RS05960 | 12364..14244 | 831 | AAA-family-ATPase-[Butyrivibrio-fibrisolvens] |
| WP_026495924.1 | OLD-ABC+TOPRIM | GCF_000424265.1 | Butyrivibrio_sp._WCD3002 | Bacteria>Bacillati>Bacillota>Clostridia>Lachnospirales>Lachnospiraceae>Butyrivibrio>unclassified_Butyrivibrio | 571 | G608_RS0110200 | 175564..177279 | 1280676 | AAA-family-ATPase-[Butyrivibrio-sp.-WCD3002] |
| MCQ4892855.1 | OLD-ABC+TOPRIM | GCA_024461355.1 | Dorea_longicatena | Bacteria>Bacillati>Bacillota>Clostridia>Lachnospirales>Lachnospiraceae>Dorea | 546 | NE607_07135 | 36283..37923 | 88431 | AAA-family-ATPase-[Dorea-longicatena] |
| WP_168932960.1 | OLD-ABC+TOPRIM | GCF_012843825.1 | Dorea_formicigenerans | Bacteria>Bacillati>Bacillota>Clostridia>Lachnospirales>Lachnospiraceae>Dorea | 546 | HF855_RS01305 | 71878..73518 | 39486 | AAA-family-ATPase-[Dorea-formicigenerans] |
| WP_118740422.1 | OLD-ABC+TOPRIM | GCF_027661365.1 | Fusicatenibacter_saccharivorans | Bacteria>Bacillati>Bacillota>Clostridia>Lachnospirales>Lachnospiraceae>Fusicatenibacter | 546 | PGQ16_RS03070 | 269294..270934 | 1150298 | MULTISPECIES:-AAA-family-ATPase-[Eubacteriales] |
| MCC2229915.1 | OLD-ABC+TOPRIM | GCA_020687485.1 | Hominifimenecus_microfluidus | Bacteria>Bacillati>Bacillota>Clostridia>Lachnospirales>Lachnospiraceae>Hominifimenecus | 546 | LKD81_02705 | 41721..43361 | 2885348 | AAA-family-ATPase-[Lachnospiraceae-bacterium-CLA-AA-H215] |
| WP_087163138.1 | OLD-ABC+TOPRIM | GCF_002160525.1 | Lachnoclostridium_sp._An169 | Bacteria>Bacillati>Bacillota>Clostridia>Lachnospirales>Lachnospiraceae>Lachnoclostridium>unclassified_Lachnoclostridium | 546 | B5F07_RS20915 | 2530..4170 | 1965569 | AAA-family-ATPase-[Lachnoclostridium-sp.-An169] |
| WP_055287909.1 | OLD-ABC+TOPRIM | GCF_902362785.1 | Lachnospira_eligens | Bacteria>Bacillati>Bacillota>Clostridia>Lachnospirales>Lachnospiraceae>Lachnospira | 485 | FXW70_RS12160 | 156564..158021 | 39485 | AAA-family-ATPase-[Lachnospira-eligens] |
| WP_117549495.1 | OLD-ABC+TOPRIM | GCF_015560805.1 | Mediterraneibacter_glycyrrhizinilyticus | Bacteria>Bacillati>Bacillota>Clostridia>Lachnospirales>Lachnospiraceae>Mediterraneibacter | 610 | I2H87_RS09330 | 37402..39234 | 342942 | ATP-binding-protein-[Mediterraneibacter-glycyrrhizinilyticus] |
| WP_227165166.1 | OLD-ABC+TOPRIM | GCF_020560115.1 | Mediterraneibacter_sp._210702-DFI.5.30 | Bacteria>Bacillati>Bacillota>Clostridia>Lachnospirales>Lachnospiraceae>Mediterraneibacter>unclassified_Mediterraneibacter | 610 | LI126_RS08120 | 86505..88337 | 2883232 | ATP-binding-protein-[Mediterraneibacter-sp.-210702-DFI.5.30] |
| WP_270645285.1 | OLD-ABC+TOPRIM | GCF_027692785.1 | Merdimonas_faecis | Bacteria>Bacillati>Bacillota>Clostridia>Lachnospirales>Lachnospiraceae>Merdimonas | 554 | PG729_RS15120 | 22931..24595 | 1653435 | AAA-family-ATPase-[Merdimonas-faecis] |
| RGC74029.1 | OLD-ABC+TOPRIM | GCA_003434385.1 | Lachnospiraceae_bacterium_AM23-2LB | Bacteria>Bacillati>Bacillota>Clostridia>Lachnospirales>Lachnospiraceae>unclassified_Lachnospiraceae | 610 | DW655_01910 | 398968..400800 | 2302964 | hypothetical-protein-DW655_01910-[Lachnospiraceae-bacterium-AM23-2LB] |
| HCI18467.1 | OLD-ABC+TOPRIM | GCA_003521585.1 | Lachnospiraceae_bacterium | Bacteria>Bacillati>Bacillota>Clostridia>Lachnospirales>Lachnospiraceae>unclassified_Lachnospiraceae | 554 | DFH32_07620 | 3182..4846 | 1898203 | hypothetical-protein-[Lachnospiraceae-bacterium] |
| HCX41711.1 | OLD-ABC+TOPRIM | GCA_003539645.1 | Lachnospiraceae_bacterium | Bacteria>Bacillati>Bacillota>Clostridia>Lachnospirales>Lachnospiraceae>unclassified_Lachnospiraceae | 554 | DHH50_05130 | 192744..194408 | 1898203 | hypothetical-protein-[Lachnospiraceae-bacterium] |
| NBH98988.1 | OLD-ABC+TOPRIM | GCA_009911465.1 | Lachnospiraceae_bacterium | Bacteria>Bacillati>Bacillota>Clostridia>Lachnospirales>Lachnospiraceae>unclassified_Lachnospiraceae | 486 | D3Z44_12665 | 22369..23829 | 1898203 | ATP-dependent-endonuclease-[Lachnospiraceae-bacterium] |
| WP_012301640.1 | OLD-ABC+TOPRIM | GCF_000018425.1 | Candidatus_Desulforudis_audaxviator | Bacteria>Bacillati>Bacillota>Clostridia>Thermoanaerobacterales>Candidatus_Desulforudaceae>Candidatus_Desulforudis | 663 | Daudx_RS02670 | 544581..546572 | 471827 | ATP-dependent-endonuclease-[Candidatus-Desulforudis-audaxviator] |
| WP_115793168.1 | OLD-ABC+TOPRIM | GCF_003368535.1 | Ammonifex_thiophilus | Bacteria>Bacillati>Bacillota>Clostridia>Thermoanaerobacterales>Thermoanaerobacteraceae>Ammonifex | 578 | DXX99_RS09070 | 9186..10922 | 444093 | AAA-family-ATPase-[Ammonifex-thiophilus] |
| MBQ6961344.1 | OLD-ABC+TOPRIM | GCA_017448975.1 | Clostridia_bacterium | Bacteria>Bacillati>Bacillota>Clostridia>unclassified_Clostridia | 644 | IJP78_10250 | 11544..13478 | 2044939 | AAA-family-ATPase-[Clostridia-bacterium] |
| QRN85662.1 | OLD-ABC+TOPRIM | GCA_016887505.1 | Clostridia_bacterium | Bacteria>Bacillati>Bacillota>Clostridia>unclassified_Clostridia | 644 | JR334_12050 | 2553096..2555030 | 2044939 | AAA-family-ATPase-[Clostridia-bacterium] |
| WP_166083464.1 | OLD-ABC+TOPRIM | GCF_031215075.1 | Erysipelothrix | Bacteria>Bacillati>Bacillota>Erysipelotrichia>Erysipelotrichales>Erysipelotrichaceae | 662 | GON62_RS10695 | 1627..3615 | 1647 | AAA-family-ATPase-[Erysipelothrix-anatis] |
| MCH4019941.1 | OLD-ABC+TOPRIM | GCA_022483385.1 | Erysipelotrichaceae_bacterium | Bacteria>Bacillati>Bacillota>Erysipelotrichia>Erysipelotrichales>Erysipelotrichaceae>unclassified_Erysipelotrichaceae | 574 | LKE61_03600 | 756489..758213 | 2049044 | AAA-family-ATPase-[Erysipelotrichaceae-bacterium] |
| MBE3584145.1 | OLD-ABC+TOPRIM | GCA_014896305.1 | Limnochordaceae_bacterium | Bacteria>Bacillati>Bacillota>Limnochordia>Limnochordales>Limnochordaceae>unclassified_Limnochordaceae | 622 | IMX01_08545 | 23162..25030 | 2773236 | ATP-dependent-endonuclease-[Limnochordaceae-bacterium] |
| NLN06910.1 | OLD-ABC+TOPRIM | GCA_012522075.1 | Bacillota_bacterium | Bacteria>Bacillati>Bacillota>unclassified_Bacillota | 613 | GX167_04750 | 8784..10625 | 1879010 | AAA-family-ATPase-[Bacillota-bacterium] |
| NPV89871.1 | OLD-ABC+TOPRIM | GCA_013178275.1 | Bacillota_bacterium | Bacteria>Bacillati>Bacillota>unclassified_Bacillota | 583 | HPY50_03735 | 183328..185079 | 1879010 | AAA-family-ATPase-[Bacillota-bacterium] |
| HBL50376.1 | OLD-ABC+TOPRIM | GCA_003504715.1 | Bacillota_bacterium | Bacteria>Bacillati>Bacillota>unclassified_Bacillota | 579 | DDZ65_09070 | 2351..4090 | 1879010 | hypothetical-protein-[Bacillota-bacterium] |
| WP_062416904.1 | OLD-ABC+TOPRIM | GCF_001050255.2 | Levilinea_saccharolytica | Bacteria>Bacillati>Chloroflexota>Anaerolineae>Anaerolineales>Anaerolineaceae>Levilinea | 657 | LSAC_RS05195 | 249259..251232 | 229921 | AAA-family-ATPase-[Levilinea-saccharolytica] |
| MBN1920384.1 | OLD-ABC+TOPRIM | GCA_016931315.1 | Anaerolineae_bacterium | Bacteria>Bacillati>Chloroflexota>Anaerolineae>unclassified_Anaerolineae | 584 | JW892_04015 | 5196..6950 | 2052143 | AAA-family-ATPase-[Anaerolineae-bacterium] |
| MBN1991943.1 | OLD-ABC+TOPRIM | GCA_016931895.1 | Anaerolineae_bacterium | Bacteria>Bacillati>Chloroflexota>Anaerolineae>unclassified_Anaerolineae | 579 | JW953_04520 | 59190..60929 | 2052143 | AAA-family-ATPase-[Anaerolineae-bacterium] |
| MDE0143356.1 | OLD-ABC+TOPRIM | GCA_028819395.1 | Caldilineaceae_bacterium | Bacteria>Bacillati>Chloroflexota>Caldilineae>Caldilineales>Caldilineaceae>unclassified_Caldilineaceae | 659 | OXN19_17920 | 6755..8734 | 2614250 | MAG:-ATP-dependent-endonuclease-[Caldilineaceae-bacterium] |
| MDE0312850.1 | OLD-ABC+TOPRIM | GCA_028820765.1 | Caldilineaceae_bacterium | Bacteria>Bacillati>Chloroflexota>Caldilineae>Caldilineales>Caldilineaceae>unclassified_Caldilineaceae | 658 | OXI52_11345 | 12313..14289 | 2614250 | MAG:-ATP-dependent-endonuclease-[Caldilineaceae-bacterium] |
| MYD91966.1 | OLD-ABC+TOPRIM | GCA_009839295.1 | Caldilineaceae_bacterium_SB0662_bin_9 | Bacteria>Bacillati>Chloroflexota>Caldilineae>Caldilineales>Caldilineaceae>unclassified_Caldilineaceae | 653 | F4Y08_16825 | 85275..87236 | 2605258 | AAA-family-ATPase-[Caldilineaceae-bacterium-SB0662_bin_9] |
| MBK9093476.1 | OLD-ABC+TOPRIM | GCA_016719785.1 | Candidatus_Amarolinea_dominans | Bacteria>Bacillati>Chloroflexota>Caldilineae>Caldilineales>Candidatus_Amarolineaceae>Candidatus_Amarolinea | 621 | IPM84_12020 | 175442..177307 | 3140696 | AAA-family-ATPase-[Anaerolineae-bacterium] |
| NUQ36642.1 | OLD-ABC+TOPRIM | GCA_013360575.1 | Caldilineales_bacterium | Bacteria>Bacillati>Chloroflexota>Caldilineae>Caldilineales>unclassified_Caldilineales | 577 | HUU23_02370 | 1331..3064 | 2306055 | AAA-family-ATPase-[Caldilineales-bacterium] |
| PFG73565.1 | OLD-ABC+TOPRIM | GCA_002563855.1 | Tepidiforma_thermophila | Bacteria>Bacillati>Chloroflexota>Tepidiformia>Tepidiformales>Tepidiformaceae>Tepidiforma | 572 | A9A59_0765 | 807601..809319 | 2761530 | putative-ATP-dependent-endonuclease-of-OLD-family-[Chloroflexi-bacterium-G233] |
| WP_098503016.1 | OLD-ABC+TOPRIM | GCF_002563855.1 | Tepidiforma_thermophila | Bacteria>Bacillati>Chloroflexota>Tepidiformia>Tepidiformales>Tepidiformaceae>Tepidiforma | 572 | A9A59_RS03815 | 807601..809319 | 2761530 | AAA-family-ATPase-[Tepidiforma-thermophila] |
| QWK10282.1 | OLD-ABC+TOPRIM | GCA_018771565.1 | Thermoflexus_hugenholtzii | Bacteria>Bacillati>Chloroflexota>Thermoflexia>Thermoflexales>Thermoflexaceae>Thermoflexus | 530 | KNN16_13060 | 2867651..2869243 | 1495650 | MAG:-AAA-family-ATPase-[Thermoflexus-hugenholtzii] |
| MCL4466050.1 | OLD-ABC+TOPRIM | GCA_023382155.1 | Chloroflexota_bacterium | Bacteria>Bacillati>Chloroflexota>unclassified_Chloroflexota | 674 | M1389_08485 | 4502..6526 | 2026724 | ATP-dependent-endonuclease-[Chloroflexota-bacterium] |
| NJD66805.1 | OLD-ABC+TOPRIM | GCA_012027015.1 | Chloroflexota_bacterium | Bacteria>Bacillati>Chloroflexota>unclassified_Chloroflexota | 646 | FIB00_16455 | 3519..5459 | 2026724 | DUF2813-domain-containing-protein-[Chloroflexota-bacterium] |
| MCC6178175.1 | OLD-ABC+TOPRIM | GCA_020845495.1 | Chloroflexota_bacterium | Bacteria>Bacillati>Chloroflexota>unclassified_Chloroflexota | 623 | IT305_22965 | 3562..5433 | 2026724 | AAA-family-ATPase-[Chloroflexota-bacterium] |
| MCQ3935717.1 | OLD-ABC+TOPRIM | GCA_024434165.1 | Chloroflexota_bacterium | Bacteria>Bacillati>Chloroflexota>unclassified_Chloroflexota | 595 | DPW18_01595 | 25215..27002 | 2026724 | ATP-dependent-endonuclease-[Chloroflexota-bacterium] |
| HHY55160.1 | OLD-ABC+TOPRIM | GCA_012842785.1 | Chloroflexota_bacterium | Bacteria>Bacillati>Chloroflexota>unclassified_Chloroflexota | 574 | GYA08_06955 | 28200..29924 | 2026724 | AAA-family-ATPase-[Chloroflexota-bacterium] |
| WP_232321576.1 | OLD-ABC+TOPRIM | GCF_963870475.1 | Picosynechococcus_sp._PCC_7117 | Bacteria>Bacillati>Cyanobacteriota/Melainabacteria_group>Cyanobacteriota>Cyanophyceae>Oscillatoriophycideae>Chroococcales>Geminocystaceae>Picosynechococcus>unclassified_Picosynechococcus | 660 | AWQ22_RS15035 | 26802..28784 | 195498 | ATP-binding-protein-[Synechococcus-sp.-PCC-7117] |
| HDN19514.1 | OLD-ABC+TOPRIM | GCA_011042375.1 | Candidatus_Acetothermia_bacterium | Bacteria>Bacteria_incertae_sedis>Bacteria_candidate_phyla>Candidatus_Bipolaricaulota>unclassified_Candidatus_Bipolaricaulota | 574 | ENF46_00255 | 5029..6753 | 2053493 | AAA-family-ATPase-[Candidatus-Acetothermia-bacterium] |
| WP_022069632.1 | OLD-ABC+TOPRIM | GCF_023383545.1 | Fusobacterium | Bacteria>Fusobacteriati>Fusobacteriota>Fusobacteriia>Fusobacteriales>Fusobacteriaceae | 606 | YWH7053_RS01960 | 14485..16305 | 848 | AAA-family-ATPase-[Fusobacterium-nucleatum] |
| WP_008692042.1 | OLD-ABC+TOPRIM | GCF_000218655.1 | Fusobacterium_animalis | Bacteria>Fusobacteriati>Fusobacteriota>Fusobacteriia>Fusobacteriales>Fusobacteriaceae>Fusobacterium | 606 | HMPREF0401_RS10095 | 568527..570347 | 76859 | AAA-family-ATPase-[Fusobacterium-animalis] |
| WP_099987857.1 | OLD-ABC+TOPRIM | GCF_002763695.1 | Fusobacterium_pseudoperiodonticum | Bacteria>Fusobacteriati>Fusobacteriota>Fusobacteriia>Fusobacteriales>Fusobacteriaceae>Fusobacterium | 606 | CTM74_RS08980 | 1711783..1713603 | 2663009 | AAA-family-ATPase-[Fusobacterium-pseudoperiodonticum] |
| WP_124794850.1 | OLD-ABC+TOPRIM | GCF_003859915.1 | Fusobacterium_canifelinum | Bacteria>Fusobacteriati>Fusobacteriota>Fusobacteriia>Fusobacteriales>Fusobacteriaceae>Fusobacterium | 606 | EII27_RS00805 | 184241..186061 | 285729 | AAA-family-ATPase-[Fusobacterium-canifelinum] |
| WP_005903403.1 | OLD-ABC+TOPRIM | GCF_000178895.1 | Fusobacterium_nucleatum | Bacteria>Fusobacteriati>Fusobacteriota>Fusobacteriia>Fusobacteriales>Fusobacteriaceae>Fusobacterium | 602 | C4N14_RS08045 | 1654507..1656315 | 851 | AAA-family-ATPase-[Fusobacterium-nucleatum] |
| WP_220308652.1 | OLD-ABC+TOPRIM | GCF_019552125.1 | Fusobacterium_animalis | Bacteria>Fusobacteriati>Fusobacteriota>Fusobacteriia>Fusobacteriales>Fusobacteriaceae>Fusobacterium | 602 | JY401_RS04525 | 945442..947250 | 76859 | AAA-family-ATPase-[Fusobacterium-animalis] |
| CDA09054.1 | OLD-ABC+TOPRIM | GCA_000433695.1 | Fusobacterium_sp._CAG:649 | Bacteria>Fusobacteriati>Fusobacteriota>Fusobacteriia>Fusobacteriales>Fusobacteriaceae>Fusobacterium>unclassified_Fusobacterium | 606 | BN748_00878 | 13826..15646 | 1262900 | putative-uncharacterized-protein-[Fusobacterium-sp.-CAG:649] |
| VTX48848.1 | OLD-ABC+TOPRIM | GCA_901873485.1 | uncultured_Leptotrichia_sp. | Bacteria>Fusobacteriati>Fusobacteriota>Fusobacteriia>Fusobacteriales>Leptotrichiaceae>Leptotrichia>environmental_samples | 564 | FPIDKHFM_01497 | 18744..20438 | 159271 | DNA-replication-and-repair-protein-RecF-[uncultured-Leptotrichia-sp.] |
| MBQ7256252.1 | OLD-ABC+TOPRIM | GCA_017445025.1 | Abditibacteriota_bacterium | Bacteria>Pseudomonadati>Abditibacteriota>unclassified_Abditibacteriota | 540 | IJS60_00985 | 30858..32480 | 2783868 | AAA-family-ATPase-[Abditibacteriota-bacterium] |
| MBK9963540.1 | OLD-ABC+TOPRIM | GCA_016722225.1 | Holophagales_bacterium | Bacteria>Pseudomonadati>Acidobacteriota>Holophagae>Holophagales>unclassified_Holophagales | 605 | IPP07_01010 | 230293..232110 | 2478486 | AAA-family-ATPase-[Holophagales-bacterium] |
| MBP7799235.1 | OLD-ABC+TOPRIM | GCA_018000955.1 | Thermoanaerobaculaceae_bacterium | Bacteria>Pseudomonadati>Acidobacteriota>Thermoanaerobaculia>Thermoanaerobaculales>Thermoanaerobaculaceae>unclassified_Thermoanaerobaculaceae | 656 | KA072_11455 | 7499..9469 | 2699758 | AAA-family-ATPase-[Thermoanaerobaculaceae-bacterium] |
| MCK6683890.1 | OLD-ABC+TOPRIM | GCA_023151375.1 | Thermoanaerobaculia_bacterium | Bacteria>Pseudomonadati>Acidobacteriota>Thermoanaerobaculia>unclassified_Thermoanaerobaculia | 502 | L6R30_15930 | 16630..18138 | 2651171 | AAA-family-ATPase-[Thermoanaerobaculia-bacterium] |
| MBS1816797.1 | OLD-ABC+TOPRIM | GCA_018268955.1 | Acidobacteriota_bacterium | Bacteria>Pseudomonadati>Acidobacteriota>unclassified_Acidobacteriota | 685 | JSU08_02610 | 191430..193487 | 1978231 | AAA-family-ATPase-[Acidobacteriota-bacterium] |
| MCY4077552.1 | OLD-ABC+TOPRIM | GCA_026707615.1 | Acidobacteriota_bacterium | Bacteria>Pseudomonadati>Acidobacteriota>unclassified_Acidobacteriota | 666 | OXH04_19215 | 7410..9410 | 1978231 | MAG:-AAA-family-ATPase-[Acidobacteria-bacterium] |
| MBI1792209.1 | OLD-ABC+TOPRIM | GCA_016178945.1 | Acidobacteriota_bacterium | Bacteria>Pseudomonadati>Acidobacteriota>unclassified_Acidobacteriota | 587 | HYR60_32195 | 11911..13674 | 1978231 | AAA-family-ATPase-[Acidobacteriota-bacterium] |
| MBS1819351.1 | OLD-ABC+TOPRIM | GCA_018268955.1 | Acidobacteriota_bacterium | Bacteria>Pseudomonadati>Acidobacteriota>unclassified_Acidobacteriota | 577 | JSU08_15560 | 74437..76170 | 1978231 | AAA-family-ATPase-[Acidobacteriota-bacterium] |
| WP_226069223.1 | OLD-ABC+TOPRIM | GCF_020410885.1 | Sulfurimonas_sp._SWIR-19 | Bacteria>Pseudomonadati>Campylobacterota>Epsilonproteobacteria>Campylobacterales>Sulfurimonadaceae>Sulfurimonas>unclassified_Sulfurimonas | 590 | LCX93_RS05500 | 1063110..1064882 | 2878390 | AAA-family-ATPase-[Sulfurimonas-sp.-SWIR-19] |
| HIP19031.1 | OLD-ABC+TOPRIM | GCA_015663065.1 | Sulfurovum_sp. | Bacteria>Pseudomonadati>Campylobacterota>Epsilonproteobacteria>Campylobacterales>Sulfurovaceae>Sulfurovum>unclassified_Sulfurovum | 569 | EYG78_06830 | 17252..18961 | 1969726 | DUF2813-domain-containing-protein-[Sulfurovum-sp.] |
| MCX5796463.1 | OLD-ABC+TOPRIM | GCA_026388095.1 | Elusimicrobiota_bacterium | Bacteria>Pseudomonadati>Elusimicrobiota>unclassified_Elusimicrobiota | 609 | NTY77_13300 | 60262..62091 | 2030800 | MAG:-AAA-family-ATPase-[Elusimicrobia-bacterium] |
| WP_217718388.1 | OLD-ABC+TOPRIM | GCF_963928575.1 | Bacteroidales | Bacteria>Pseudomonadati>FCB_group>Bacteroidota/Chlorobiota_group>Bacteroidota>Bacteroidia | 673 | KSX11_RS12360 | 125296..127317 | 171549 | AAA-family-ATPase-[Bacteroides-uniformis] |
| WP_117985928.1 | OLD-ABC+TOPRIM | GCF_026966235.1 | Bacteroides | Bacteria>Pseudomonadati>FCB_group>Bacteroidota/Chlorobiota_group>Bacteroidota>Bacteroidia>Bacteroidales>Bacteroidaceae | 673 | O1446_RS12910 | 63013..65034 | 816 | MULTISPECIES:-AAA-family-ATPase-[Bacteroides] |

| Protein-ID | Domain | Assembly-ID | Strain | Taxonomy | Length | Locus-Tag | Genomic-coordinates | TaxID | Fasta-defline |
|---|---|---|---|---|---|---|---|---|---|
| WP_002561971.1 | OLD-ABC+TOPRIM | GCF_000382465.1 | Bacteroides_sp._HPS0048 | Bacteria>Pseudomonadati>FCB_group>Bacteroidota/Chlorobiota_group>Bacteroidota>Bacteroidia>Bacteroidales>Bacteroidaceae>Bacteroides>unclassified_Bacteroides | 678 | HMPREF1214_RS19615 | 477429..479465 | 1078089 | AAA-family-ATPase-[Bacteroides-sp.-HPS0048] |
| WP_172774765.1 | OLD-ABC+TOPRIM | GCF_012976275.1 | Phocaeicola_vulgatus | Bacteria>Pseudomonadati>FCB_group>Bacteroidota/Chlorobiota_group>Bacteroidota>Bacteroidia>Bacteroidales>Bacteroidaceae>Phocaeicola | 673 | HKQ53_RS16525 | 86650..88671 | 821 | AAA-family-ATPase-[Phocaeicola-vulgatus] |
| MBQ3604729.1 | OLD-ABC+TOPRIM | GCA_017399915.1 | Muribaculaceae_bacterium | Bacteria>Pseudomonadati>FCB_group>Bacteroidota/Chlorobiota_group>Bacteroidota>Bacteroidia>Bacteroidales>Muribaculaceae>unclassified_Muribaculaceae | 604 | II990_00545 | 529..2343 | 2498093 | AAA-family-ATPase-[Muribaculaceae-bacterium] |
| MCH4147880.1 | OLD-ABC+TOPRIM | GCA_022484605.1 | Prevotella_sp. | Bacteria>Pseudomonadati>FCB_group>Bacteroidota/Chlorobiota_group>Bacteroidota>Bacteroidia>Bacteroidales>Prevotellaceae>Prevotella>unclassified_Prevotella | 534 | LKF33_05580 | 27366..28970 | 59823 | AAA-family-ATPase-[Prevotella-sp.] |
| WP_079684007.1 | OLD-ABC+TOPRIM | GCF_900168155.1 | Parabacteroides_chartae | Bacteria>Pseudomonadati>FCB_group>Bacteroidota/Chlorobiota_group>Bacteroidota>Bacteroidia>Bacteroidales>Tannerellaceae>Parabacteroides | 585 | B5X57_RS12585 | 55868..57625 | 1037355 | AAA-family-ATPase-[Parabacteroides-chartae] |
| MBQ7712011.1 | OLD-ABC+TOPRIM | GCA_017449975.1 | Bacteroidales_bacterium | Bacteria>Pseudomonadati>FCB_group>Bacteroidota/Chlorobiota_group>Bacteroidota>Bacteroidia>Bacteroidales>unclassified_Bacteroidales | 682 | IJT39_09340 | 46789..48837 | 2030927 | AAA-family-ATPase-[Bacteroidales-bacterium] |
| MBR5214689.1 | OLD-ABC+TOPRIM | GCA_017549605.1 | Bacteroidales_bacterium | Bacteria>Pseudomonadati>FCB_group>Bacteroidota/Chlorobiota_group>Bacteroidota>Bacteroidia>Bacteroidales>unclassified_Bacteroidales | 577 | IKV80_00520 | 5167..6900 | 2030927 | AAA-family-ATPase-[Bacteroidales-bacterium] |
| WP_234861367.1 | OLD-ABC+TOPRIM | GCF_021491425.1 | Mariniradius_sediminis | Bacteria>Pseudomonadati>FCB_group>Bacteroidota/Chlorobiota_group>Bacteroidota>Cytophagia>Cytophagales>Cyclobacteriaceae>Mariniradius | 550 | LOU89_RS09875 | 258951..260603 | 2909237 | AAA-family-ATPase-[Mariniradius-sp.-RY-2] |
| WP_188169953.1 | OLD-ABC+TOPRIM | GCF_014534745.1 | Flammeovirga_sp._EKP202 | Bacteria>Pseudomonadati>FCB_group>Bacteroidota/Chlorobiota_group>Bacteroidota>Cytophagia>Cytophagales>Flammeovirgaceae>Flammeovirga>unclassified_Flammeovirga | 605 | IC221_RS16115 | 78227..80044 | 2770592 | AAA-family-ATPase-[Flammeovirga-sp.-EKP202] |
| WP_201982529.1 | OLD-ABC+TOPRIM | GCF_016734815.1 | Hymenobacter_rubidus | Bacteria>Pseudomonadati>FCB_group>Bacteroidota/Chlorobiota_group>Bacteroidota>Cytophagia>Cytophagales>Hymenobacteraceae>Hymenobacter | 611 | JKL48_RS11470 | 11186..13021 | 1441626 | AAA-family-ATPase-[Hymenobacter-rubidus] |
| WP_194850139.1 | OLD-ABC+TOPRIM | GCF_015356755.1 | Nonlabens_antarcticus | Bacteria>Pseudomonadati>FCB_group>Bacteroidota/Chlorobiota_group>Bacteroidota>Flavobacteriia>Flavobacteriales>Flavobacteriaceae>Nonlabens | 643 | IQ282_RS01315 | 61986..63917 | 392714 | AAA-family-ATPase-[Nonlabens-antarcticus] |
| HEX04761.1 | OLD-ABC+TOPRIM | GCA_011043795.1 | Bacteroidota_bacterium | Bacteria>Pseudomonadati>FCB_group>Bacteroidota/Chlorobiota_group>Bacteroidota>unclassified_Bacteroidota | 595 | ENH10_06340 | 10456..12240 | 1898104 | hypothetical-protein-[Bacteroidota-bacterium] |
| KZK74368.1 | OLD-ABC+TOPRIM | GCA_001622165.1 | Pelodictyon_luteolum | Bacteria>Pseudomonadati>FCB_group>Bacteroidota/Chlorobiota_group>Chlorobiota>Chlorobiia>Chlorobiales>Chlorobiaceae>Chlorobium/Pelodictyon_group>Pelodictyon | 571 | A3K90_02890 | 26133..27848 | 1100 | hypothetical-protein-A3K90_02890-[Pelodictyon-luteolum] |
| BBM69557.1 | OLD-ABC+TOPRIM | GCA_009936255.1 | Rhodothermus_marinus | Bacteria>Pseudomonadati>FCB_group>Bacteroidota/Chlorobiota_group>Rhodothermota>Rhodothermia>Rhodothermales>Rhodothermaceae>Rhodothermus | 617 | RmaAA213_14030 | 1626804..1628657 | 29549 | hypothetical-protein-RmaAA213_14030-[Rhodothermus-marinus] |
| WP_161541047.1 | OLD-ABC+TOPRIM | GCF_009936255.1 | Rhodothermus_marinus | Bacteria>Pseudomonadati>FCB_group>Bacteroidota/Chlorobiota_group>Rhodothermota>Rhodothermia>Rhodothermales>Rhodothermaceae>Rhodothermus | 617 | Rma1338_RS07055 | 1626804..1628657 | 29549 | ATP-binding-protein-[Rhodothermus-marinus] |
| WP_259084848.1 | OLD-ABC+TOPRIM | GCF_024809025.1 | Salinibacter_ruber | Bacteria>Pseudomonadati>FCB_group>Bacteroidota/Chlorobiota_group>Rhodothermota>Rhodothermia>Rhodothermales>Salinibacteraceae>Salinibacter | 722 | GGP87_RS12300 | 25845..28013 | 146919 | AAA-family-ATPase-[Salinibacter-ruber] |
| WP_103016188.1 | OLD-ABC+TOPRIM | GCF_002894575.1 | Salinibacter_ruber | Bacteria>Pseudomonadati>FCB_group>Bacteroidota/Chlorobiota_group>Rhodothermota>Rhodothermia>Rhodothermales>Salinibacteraceae>Salinibacter | 654 | B7R92_RS05360 | 17617..19581 | 146919 | AAA-family-ATPase-[Salinibacter-ruber] |
| WP_251951629.1 | OLD-ABC+TOPRIM | GCF_943908255.1 | Salinibacter_ruber | Bacteria>Pseudomonadati>FCB_group>Bacteroidota/Chlorobiota_group>Rhodothermota>Rhodothermia>Rhodothermales>Salinibacteraceae>Salinibacter | 653 | NGL47_RS11260 | 7327..9288 | 146919 | AAA-family-ATPase-[Salinibacter-ruber] |
| WP_259182575.1 | OLD-ABC+TOPRIM | GCF_024806795.1 | Salinibacter_ruber | Bacteria>Pseudomonadati>FCB_group>Bacteroidota/Chlorobiota_group>Rhodothermota>Rhodothermia>Rhodothermales>Salinibacteraceae>Salinibacter | 653 | GGQ19_RS12655 | 75305..77266 | 146919 | AAA-family-ATPase-[Salinibacter-ruber] |
| WP_259195585.1 | OLD-ABC+TOPRIM | GCF_024809635.1 | Salinibacter_ruber | Bacteria>Pseudomonadati>FCB_group>Bacteroidota/Chlorobiota_group>Rhodothermota>Rhodothermia>Rhodothermales>Salinibacteraceae>Salinibacter | 644 | GGP50_RS04145 | 18699..20633 | 146919 | ATP-dependent-endonuclease-[Salinibacter-ruber] |
| MCS3668711.1 | OLD-ABC+TOPRIM | GCA_024808405.1 | Salinibacter_ruber | Bacteria>Pseudomonadati>FCB_group>Bacteroidota/Chlorobiota_group>Rhodothermota>Rhodothermia>Rhodothermales>Salinibacteraceae>Salinibacter | 597 | GGP77_002962 | 24201..25994 | 146919 | putative-ATP-dependent-endonuclease-of-OLD-family-[Salinibacter-ruber] |
| WP_259165676.1 | OLD-ABC+TOPRIM | GCF_024806515.1 | Salinibacter_ruber | Bacteria>Pseudomonadati>FCB_group>Bacteroidota/Chlorobiota_group>Rhodothermota>Rhodothermia>Rhodothermales>Salinibacteraceae>Salinibacter | 578 | GGP90_RS03900 | 954066..955802 | 146919 | ATP-dependent-endonuclease-[Salinibacter-ruber] |
| WP_259048231.1 | OLD-ABC+TOPRIM | GCF_024808395.1 | Salinibacter_ruber | Bacteria>Pseudomonadati>FCB_group>Bacteroidota/Chlorobiota_group>Rhodothermota>Rhodothermia>Rhodothermales>Salinibacteraceae>Salinibacter | 577 | GGP52_RS13875 | 21156..22889 | 146919 | ATP-dependent-endonuclease-[Salinibacter-ruber] |
| WP_259086345.1 | OLD-ABC+TOPRIM | GCF_024808115.1 | Salinibacter_ruber | Bacteria>Pseudomonadati>FCB_group>Bacteroidota/Chlorobiota_group>Rhodothermota>Rhodothermia>Rhodothermales>Salinibacteraceae>Salinibacter | 577 | GGQ04_RS13425 | 52258..53991 | 146919 | ATP-dependent-endonuclease-[Salinibacter-ruber] |
| WP_259107790.1 | OLD-ABC+TOPRIM | GCF_024808295.1 | Salinibacter_ruber | Bacteria>Pseudomonadati>FCB_group>Bacteroidota/Chlorobiota_group>Rhodothermota>Rhodothermia>Rhodothermales>Salinibacteraceae>Salinibacter | 577 | GGP63_RS14305 | 40923..42656 | 146919 | ATP-dependent-endonuclease-[Salinibacter-ruber] |
| WP_259120990.1 | OLD-ABC+TOPRIM | GCF_024808535.1 | Salinibacter_ruber | Bacteria>Pseudomonadati>FCB_group>Bacteroidota/Chlorobiota_group>Rhodothermota>Rhodothermia>Rhodothermales>Salinibacteraceae>Salinibacter | 577 | GGP96_RS14220 | 74380..76113 | 146919 | ATP-dependent-endonuclease-[Salinibacter-ruber] |
| WP_259122110.1 | OLD-ABC+TOPRIM | GCF_024809285.1 | Salinibacter_ruber | Bacteria>Pseudomonadati>FCB_group>Bacteroidota/Chlorobiota_group>Rhodothermota>Rhodothermia>Rhodothermales>Salinibacteraceae>Salinibacter | 577 | GGQ10_RS14885 | 89623..91356 | 146919 | ATP-dependent-endonuclease-[Salinibacter-ruber] |
| WP_259127190.1 | OLD-ABC+TOPRIM | GCF_024807525.1 | Salinibacter_ruber | Bacteria>Pseudomonadati>FCB_group>Bacteroidota/Chlorobiota_group>Rhodothermota>Rhodothermia>Rhodothermales>Salinibacteraceae>Salinibacter | 577 | GGP44_RS04615 | 1105857..1107590 | 146919 | ATP-dependent-endonuclease-[Salinibacter-ruber] |
| WP_259220525.1 | OLD-ABC+TOPRIM | GCF_024807065.1 | Salinibacter_ruber | Bacteria>Pseudomonadati>FCB_group>Bacteroidota/Chlorobiota_group>Rhodothermota>Rhodothermia>Rhodothermales>Salinibacteraceae>Salinibacter | 545 | GGP94_RS12990 | 33318..34955 | 146919 | AAA-family-ATPase-[Salinibacter-ruber] |
| MCB5228478.1 | OLD-ABC+TOPRIM | GCA_020532325.1 | Candidatus_Cloacimonadota_bacterium | Bacteria>Pseudomonadati>FCB_group>Candidatus_Cloacimonadota>unclassified_Candidatus_Cloacimonadota | 634 | LHW44_00160 | 33990..35894 | 2030808 | AAA-family-ATPase-[Candidatus-Cloacimonadota-bacterium] |
| NUM53504.1 | OLD-ABC+TOPRIM | GCA_013359265.1 | Candidatus_Hydrogenedentota_bacterium | Bacteria>Pseudomonadati>FCB_group>Candidatus_Hydrogenedentota>unclassified_Candidatus_Hydrogenedentota | 578 | HUU46_07665 | 140981..142717 | 2030809 | AAA-family-ATPase-[Candidatus-Hydrogenedentes-bacterium] |
| MCF7837531.1 | OLD-ABC+TOPRIM | GCA_021734645.1 | Fidelibacterota_bacterium | Bacteria>Pseudomonadati>FCB_group>Fidelibacterota>unclassified_Fidelibacterota | 603 | K9N49_02785 | 16381..18192 | 2026760 | ATP-binding-protein-[Candidatus-Marinimicrobia-bacterium] |
| MDE3256979.1 | OLD-ABC+TOPRIM | GCA_028877215.1 | Gemmatimonadota_bacterium | Bacteria>Pseudomonadati>FCB_group>Gemmatimonadota>unclassified_Gemmatimonadota | 580 | OYM47_04005 | 45929..47671 | 2026742 | MAG:-AAA-family-ATPase-[Gemmatimonadota-bacterium] |
| MCD6198159.1 | OLD-ABC+TOPRIM | GCA_021159445.1 | Deltaproteobacteria_bacterium | Bacteria>Pseudomonadati>Myxococcota>Myxococcia>unclassified_Myxococcia | 672 | J7K15_06240 | 5418..7436 | 2026735 | AAA-family-ATPase-[Deltaproteobacteria-bacterium] |
| MBI4706261.1 | OLD-ABC+TOPRIM | GCA_016209145.1 | Deltaproteobacteria_bacterium | Bacteria>Pseudomonadati>Myxococcota>Myxococcia>unclassified_Myxococcia | 577 | HY744_34625 | 10745..12478 | 2026735 | AAA-family-ATPase-[Deltaproteobacteria-bacterium] |
| MCC7070412.1 | OLD-ABC+TOPRIM | GCA_020850995.1 | Deltaproteobacteria_bacterium | Bacteria>Pseudomonadati>Myxococcota>Myxococcia>unclassified_Myxococcia | 531 | IT383_03760 | 34275..35870 | 2026735 | DUF2813-domain-containing-protein-[Deltaproteobacteria-bacterium] |
| MDE0331735.1 | OLD-ABC+TOPRIM | GCA_028820795.1 | Nitrospinota_bacterium | Bacteria>Pseudomonadati>Nitrospinota/Tectimicrobiota_group>Nitrospinota>unclassified_Nitrospinota | 675 | OXL41_07690 | 650..2677 | 2026769 | MAG:-AAA-family-ATPase-[Nitrospinae-bacterium] |
| NOS82781.1 | OLD-ABC+TOPRIM | GCA_013140535.1 | Nitrospira_sp. | Bacteria>Pseudomonadati>Nitrospirota>Nitrospiria>Nitrospirales>Nitrospiraceae>Nitrospira>unclassified_Nitrospira | 642 | HOP32_14485 | 3800..5728 | 70125 | AAA-family-ATPase-[Nitrospira-sp.] |
| WP_215861192.1 | OLD-ABC+TOPRIM | GCF_018854875.1 | Acidithiobacillus_ferriphilus | Bacteria>Pseudomonadati>Pseudomonadota>Acidithiobacillia>Acidithiobacillales>Acidithiobacillaceae>Acidithiobacillus | 611 | HF928_RS01660 | 15052..16887 | 1689834 | ATP-binding-protein-[Acidithiobacillus-ferriphilus] |
| GIX11745.1 | OLD-ABC+TOPRIM | GCA_026004895.1 | Elioraea_sp. | Bacteria>Pseudomonadati>Pseudomonadota>Alphaproteobacteria>Acetobacterales>Elioraeaceae>Elioraea>unclassified_Elioraea | 569 | KatS3mg116_3455 | 529419..531128 | 2185103 | MAG:-hypothetical-protein-KatS3mg116_3455-[Elioraea-sp.] |
| WP_114817808.1 | OLD-ABC+TOPRIM | GCF_034424665.1 | Brevundimonas_bullata | Bacteria>Pseudomonadati>Pseudomonadota>Alphaproteobacteria>Caulobacterales>Caulobacteraceae>Brevundimonas | 674 | DXK21_RS10915 | 48770..50794 | 13160 | AAA-family-ATPase-[Brevundimonas-bullata] |
| WP_206455642.1 | OLD-ABC+TOPRIM | GCF_017183135.1 | Aurantimonas_marina | Bacteria>Pseudomonadati>Pseudomonadota>Alphaproteobacteria>Hyphomicrobiales>Aurantimonadaceae>Aurantimonas | 711 | ING15_RS14955 | 2116..4251 | 2780508 | AAA-family-ATPase-[Aurantimonas-marina] |
| WP_071923270.1 | OLD-ABC+TOPRIM | GCF_001887265.1 | Chelatococcus_daeguensis | Bacteria>Pseudomonadati>Pseudomonadota>Alphaproteobacteria>Hyphomicrobiales>Chelatococcaceae>Chelatococcus | 637 | BOQ54_RS03325 | 715169..717082 | 444444 | AAA-family-ATPase-[Chelatococcus-daeguensis] |
| MBX3547059.1 | OLD-ABC+TOPRIM | GCA_019635905.1 | Chelatococcus_sp. | Bacteria>Pseudomonadati>Pseudomonadota>Alphaproteobacteria>Hyphomicrobiales>Chelatococcaceae>Chelatococcus>unclassified_Chelatococcus | 650 | KF818_30260 | 34806..36758 | 1953771 | AAA-family-ATPase-[Chelatococcus-sp.] |
| WP_246513576.1 | OLD-ABC+TOPRIM | GCF_016801025.1 | Paradevosia_shaoguanensis | Bacteria>Pseudomonadati>Pseudomonadota>Alphaproteobacteria>Hyphomicrobiales>Devosiaceae>Paradevosia | 600 | JNE37_RS07700 | 1642673..1644475 | 1335043 | ATP-binding-protein-[Paradevosia-shaoguanensis] |
| WP_130224759.1 | OLD-ABC+TOPRIM | GCF_004212675.1 | Bradyrhizobium_sp._Leo121 | Bacteria>Pseudomonadati>Pseudomonadota>Alphaproteobacteria>Hyphomicrobiales>Nitrobacteraceae>Bradyrhizobium>unclassified_Bradyrhizobium | 652 | CWO90_RS07125 | 62079..64037 | 1571195 | AAA-family-ATPase-[Bradyrhizobium-sp.-Leo121] |
| MAM94523.1 | OLD-ABC+TOPRIM | GCA_002694985.1 | Parvibaculum_sp. | Bacteria>Pseudomonadati>Pseudomonadota>Alphaproteobacteria>Hyphomicrobiales>Parvibaculaceae>Parvibaculum>unclassified_Parvibaculum | 618 | CMI61_07825 | 156951..158807 | 2024848 | ATP-dependent-endonuclease-[Parvibaculum-sp.] |
| WP_077987194.1 | OLD-ABC+TOPRIM | GCF_030440405.1 | Agrobacterium_pusense | Bacteria>Pseudomonadati>Pseudomonadota>Alphaproteobacteria>Hyphomicrobiales>Rhizobiaceae>Rhizobium/Agrobacterium_group>Agrobacterium | 795 | BTE56_RS17170 | 6358..8745 | 648995 | AAA-family-ATPase-[Rhizobium-pusense] |
| WP_222362944.1 | OLD-ABC+TOPRIM | GCF_019798305.1 | Rhizobium_leguminosarum | Bacteria>Pseudomonadati>Pseudomonadota>Alphaproteobacteria>Hyphomicrobiales>Rhizobiaceae>Rhizobium/Agrobacterium_group>Rhizobium | 755 | HFO99_RS19340 | 180669..182936 | 384 | ATP-dependent-endonuclease-[Rhizobium-leguminosarum] |
| WP_012755923.1 | OLD-ABC+TOPRIM | - | Rhizobium_leguminosarum | Bacteria>Pseudomonadati>Pseudomonadota>Alphaproteobacteria>Hyphomicrobiales>Rhizobiaceae>Rhizobium/Agrobacterium_group>Rhizobium | 654 | RLEG_RS00995 | 197189..199153 | 384 | AAA-family-ATPase-[Rhizobium-leguminosarum] |
| MCZ7659370.1 | OLD-ABC+TOPRIM | GCA_027493405.1 | Xanthobacteraceae_bacterium | Bacteria>Pseudomonadati>Pseudomonadota>Alphaproteobacteria>Hyphomicrobiales>Xanthobacteraceae>unclassified_Xanthobacteraceae | 441 | M5U07_16610 | 12726..14051 | 2021379 | MAG:-YjbD-family-protein-[Xanthobacteraceae-bacterium] |
| WP_076528444.1 | OLD-ABC+TOPRIM | GCF_014636395.1 | Gemmobacter_megaterium | Bacteria>Pseudomonadati>Pseudomonadota>Alphaproteobacteria>Rhodobacterales>Paracoccaceae>Gemmobacter | 617 | IEV79_RS02785 | 594344..596197 | 1086013 | ATP-binding-protein-[Gemmobacter-megaterium] |
| WP_271692173.1 | OLD-ABC+TOPRIM | GCF_028201715.1 | Sulfitobacter_faviae | Bacteria>Pseudomonadati>Pseudomonadota>Alphaproteobacteria>Rhodobacterales>Roseobacteraceae>Sulfitobacter | 564 | PL335_RS12855 | 2591117..2592811 | 1775881 | ATP-binding-protein-[Sulfitobacter-faviae] |
| WP_147041136.1 | OLD-ABC+TOPRIM | GCF_007991775.1 | Skermanella_aerolata | Bacteria>Pseudomonadati>Pseudomonadota>Alphaproteobacteria>Rhodospirillales>Azospirillaceae>Skermanella | 655 | SAE02_RS33465 | 19572..21539 | 393310 | AAA-family-ATPase-[Skermanella-aerolata] |
| MDD9991560.1 | OLD-ABC+TOPRIM | GCA_028818565.1 | Rhodospirillales_bacterium | Bacteria>Pseudomonadati>Pseudomonadota>Alphaproteobacteria>Rhodospirillales>unclassified_Rhodospirillales | 585 | OXP75_07155 | 953..2710 | 2026786 | MAG:-AAA-family-ATPase-[Rhodospirillales-bacterium] |
| TVR79631.1 | OLD-ABC+TOPRIM | GCA_007695045.1 | Rhodospirillales_bacterium | Bacteria>Pseudomonadati>Pseudomonadota>Alphaproteobacteria>Rhodospirillales>unclassified_Rhodospirillales | 568 | EA405_12445 | 8494..10200 | 2026786 | AAA-family-ATPase-[Rhodospirillales-bacterium] |
| MDD3182284.1 | OLD-ABC+TOPRIM | GCA_028691835.1 | Alphaproteobacteria_bacterium | Bacteria>Pseudomonadati>Pseudomonadota>Alphaproteobacteria>unclassified_Alphaproteobacteria | 669 | PHD48_05735 | 4763..6772 | 1913988 | MAG:-AAA-family-ATPase-[Alphaproteobacteria-bacterium] |
| WP_125277464.1 | OLD-ABC+TOPRIM | GCF_003938225.2 | Burkholderiales | Bacteria>Pseudomonadati>Pseudomonadota>Betaproteobacteria | 651 | EGK70_RS02755 | 604972..606927 | 80840 | AAA-family-ATPase-[Alcaligenes-aquatilis] |
| HCN68768.1 | OLD-ABC+TOPRIM | GCA_003535635.1 | Accumulibacter_sp. | Bacteria>Pseudomonadati>Pseudomonadota>Betaproteobacteria>Betaproteobacteria_incertae_sedis>Candidatus_Accumulibacter>unclassified_Candidatus_Accumulibacter | 575 | DIT03_10985 | 39643..41370 | 2053492 | hypothetical-protein-[Candidatus-Accumulibacter-sp.] |
| HBT33988.1 | OLD-ABC+TOPRIM | GCA_003509985.1 | Pusillimonas_sp. | Bacteria>Pseudomonadati>Pseudomonadota>Betaproteobacteria>Burkholderiales>Alcaligenaceae>Pusillimonas>unclassified_Pusillimonas | 651 | DEB15_14770 | 26918..28873 | 3040095 | ATP-dependent-endonuclease-[Pusillimonas-sp.-(ex-Stolz-et-al.-2005)] |
| WP_059567312.1 | OLD-ABC+TOPRIM | GCF_001523965.1 | Burkholderia_cepacia_complex | Bacteria>Pseudomonadati>Pseudomonadota>Betaproteobacteria>Burkholderiales>Burkholderiaceae>Burkholderia | 576 | C6T69_RS26895 | 6838..8568 | 87882 | MULTISPECIES:-DUF2813-domain-containing-protein-[Burkholderia-cepacia-complex] |
| WP_186089003.1 | OLD-ABC+TOPRIM | GCF_902831595.1 | Burkholderia_gladioli | Bacteria>Pseudomonadati>Pseudomonadota>Betaproteobacteria>Burkholderiales>Burkholderiaceae>Burkholderia | 575 | H9D22_RS09660 | 581634..583361 | 28095 | DUF2813-domain-containing-protein-[Burkholderia-gladioli] |
| WP_186109345.1 | OLD-ABC+TOPRIM | GCF_902831275.1 | Burkholderia_gladioli | Bacteria>Pseudomonadati>Pseudomonadota>Betaproteobacteria>Burkholderiales>Burkholderiaceae>Burkholderia | 575 | H9D39_RS35045 | 63440..65167 | 28095 | DUF2813-domain-containing-protein-[Burkholderia-gladioli] |
| WP_186146943.1 | OLD-ABC+TOPRIM | GCF_902831935.1 | Burkholderia_gladioli | Bacteria>Pseudomonadati>Pseudomonadota>Betaproteobacteria>Burkholderiales>Burkholderiaceae>Burkholderia | 575 | H9E56_RS05555 | 441821..443548 | 28095 | DUF2813-domain-containing-protein-[Burkholderia-gladioli] |
| WP_081068806.1 | OLD-ABC+TOPRIM | GCF_001528605.1 | Burkholderia_multivorans | Bacteria>Pseudomonadati>Pseudomonadota>Betaproteobacteria>Burkholderiales>Burkholderiaceae>Burkholderia>Burkholderia_cepacia_complex | 616 | WK33_RS03210 | 6056..7906 | 87883 | AAA-family-ATPase-[Burkholderia-multivorans] |
| WP_269509112.1 | OLD-ABC+TOPRIM | GCF_027257875.1 | Burkholderia_sp._IMCC1007 | Bacteria>Pseudomonadati>Pseudomonadota>Betaproteobacteria>Burkholderiales>Burkholderiaceae>Burkholderia>unclassified_Burkholderia | 575 | O1Q03_RS33575 | 18927..20654 | 3004104 | DUF2813-domain-containing-protein-[Burkholderia-sp.-IMCC1007] |
| WP_062175041.1 | OLD-ABC+TOPRIM | GCF_001557535.2 | Burkholderia_sp._PAMC_26561 | Bacteria>Pseudomonadati>Pseudomonadota>Betaproteobacteria>Burkholderiales>Burkholderiaceae>Burkholderia>unclassified_Burkholderia | 522 | AXG89_RS34135 | 125152..126720 | 1795043 | AAA-family-ATPase-[Burkholderia-sp.-PAMC-26561] |
| SAK86095.1 | OLD-ABC+TOPRIM | GCA_001545075.2 | Caballeronia_ptereochthonis | Bacteria>Pseudomonadati>Pseudomonadota>Betaproteobacteria>Burkholderiales>Burkholderiaceae>Caballeronia | 575 | AWB83_04704 | 118299..120026 | 1777144 | ATP-dependent-OLD-family-endonuclease-[Caballeronia-ptereochthonis] |
| WP_250522807.1 | OLD-ABC+TOPRIM | GCF_023631205.1 | unclassified_Caballeronia | Bacteria>Pseudomonadati>Pseudomonadota>Betaproteobacteria>Burkholderiales>Burkholderiaceae>Caballeronia | 575 | NCS65_RS00825 | 178297..180024 | 2646786 | MULTISPECIES:-DUF2813-domain-containing-protein-[unclassified-Caballeronia] |
| WP_143968628.1 | OLD-ABC+TOPRIM | GCF_007556755.1 | Tepidimonas_fonticaldi | Bacteria>Pseudomonadati>Pseudomonadota>Betaproteobacteria>Burkholderiales>Burkholderiales_genera_incertae_sedis>Tepidimonas | 616 | Tfont_RS05080 | 100753..102603 | 1101373 | AAA-family-ATPase-[Tepidimonas-fonticaldi] |
| WP_086929122.1 | OLD-ABC+TOPRIM | GCF_002157185.1 | Acidovorax_carolinensis | Bacteria>Pseudomonadati>Pseudomonadota>Betaproteobacteria>Burkholderiales>Comamonadaceae>Acidovorax | 656 | CBP35_RS20920 | 154359..156329 | 553814 | AAA-family-ATPase-[Acidovorax-carolinensis] |
| WP_086927317.1 | OLD-ABC+TOPRIM | GCF_002157185.1 | Acidovorax_carolinensis | Bacteria>Pseudomonadati>Pseudomonadota>Betaproteobacteria>Burkholderiales>Comamonadaceae>Acidovorax | 575 | CBP35_RS08890 | 1922781..1924508 | 553814 | DUF2813-domain-containing-protein-[Acidovorax-carolinensis] |
| MBN9628763.1 | OLD-ABC+TOPRIM | GCA_017307415.1 | Acidovorax_sp. | Bacteria>Pseudomonadati>Pseudomonadota>Betaproteobacteria>Burkholderiales>Comamonadaceae>Acidovorax>unclassified_Acidovorax | 643 | JOG38_21235 | 4468..6399 | 1872122 | AAA-family-ATPase-[Acidovorax-sp.] |
| ODS67620.1 | OLD-ABC+TOPRIM | GCA_001724795.1 | Acidovorax_sp._SCN_65-28 | Bacteria>Pseudomonadati>Pseudomonadota>Betaproteobacteria>Burkholderiales>Comamonadaceae>Acidovorax>unclassified_Acidovorax | 643 | ABS39_19770 | 17431..19362 | 1660086 | MAG:-ATP-dependent-endonuclease-[Acidovorax-sp.-SCN-65-28] |
| WP_101103327.1 | OLD-ABC+TOPRIM | GCF_002837135.1 | Macromonas_bipunctata | Bacteria>Pseudomonadati>Pseudomonadota>Betaproteobacteria>Burkholderiales>Comamonadaceae>Macromonas | 575 | CON76_RS10600 | 11927..13654 | 183670 | DUF2813-domain-containing-protein-[Macromonas-bipunctata] |
| WP_011797931.1 | OLD-ABC+TOPRIM | GCF_000015505.1 | Polaromonas_naphthalenivorans | Bacteria>Pseudomonadati>Pseudomonadota>Betaproteobacteria>Burkholderiales>Comamonadaceae>Polaromonas | 645 | PNAP_RS21260 | 168340..170277 | 216465 | AAA-family-ATPase-[Polaromonas-naphthalenivorans] |
| MBA3056687.1 | OLD-ABC+TOPRIM | GCA_013791765.1 | Rhodoferax_sp. | Bacteria>Pseudomonadati>Pseudomonadota>Betaproteobacteria>Burkholderiales>Comamonadaceae>Rhodoferax>unclassified_Rhodoferax | 660 | FP823_02090 | 55274..57256 | 50421 | AAA-family-ATPase-[Rhodoferax-sp.] |
| WP_265282595.1 | OLD-ABC+TOPRIM | GCF_026127365.1 | Verminephrobacter_aporrectodeae | Bacteria>Pseudomonadati>Pseudomonadota>Betaproteobacteria>Burkholderiales>Comamonadaceae>Verminephrobacter | 654 | D5039_RS14595 | 1241273..1243237 | 1110389 | AAA-family-ATPase-[Verminephrobacter-aporrectodeae] |
| WP_265654213.1 | OLD-ABC+TOPRIM | GCF_026192715.1 | Verminephrobacter_aporrectodeae | Bacteria>Pseudomonadati>Pseudomonadota>Betaproteobacteria>Burkholderiales>Comamonadaceae>Verminephrobacter | 654 | D8B27_RS10355 | 19495..21459 | 1110389 | AAA-family-ATPase-[Verminephrobacter-aporrectodeae] |
| WP_182661246.1 | OLD-ABC+TOPRIM | GCF_014145335.1 | Aquariibacter_albus | Bacteria>Pseudomonadati>Pseudomonadota>Betaproteobacteria>Burkholderiales>Sphaerotilaceae>Aquariibacter | 605 | H4F90_RS02730 | 594945..596762 | 2759899 | AAA-family-ATPase-[Aquariibacter-albus] |
| MBL0086263.1 | OLD-ABC+TOPRIM | GCA_016721465.1 | Ideonella_sp. | Bacteria>Pseudomonadati>Pseudomonadota>Betaproteobacteria>Burkholderiales>Sphaerotilaceae>Ideonella>unclassified_Ideonella | 605 | IPP44_06035 | 513094..514911 | 1929293 | AAA-family-ATPase-[Ideonella-sp.] |
| MBK7531538.1 | OLD-ABC+TOPRIM | GCA_016706565.1 | Piscinibacter_sp. | Bacteria>Pseudomonadati>Pseudomonadota>Betaproteobacteria>Burkholderiales>Sphaerotilaceae>Piscinibacter>unclassified_Piscinibacter | 674 | IPI47_13310 | 101550..103574 | 1903157 | AAA-family-ATPase-[Piscinibacter-sp.] |
| WP_164962558.1 | OLD-ABC+TOPRIM | GCF_011065555.1 | Rubrivivax_sp._JA1026 | Bacteria>Pseudomonadati>Pseudomonadota>Betaproteobacteria>Burkholderiales>Sphaerotilaceae>Rubrivivax>unclassified_Rubrivivax | 576 | G5B56_RS05555 | 397378..399108 | 2710888 | DUF2813-domain-containing-protein-[Rubrivivax-sp.-JA1026] |
| WP_179636387.1 | OLD-ABC+TOPRIM | GCF_013410775.1 | Sphaerotilus_montanus | Bacteria>Pseudomonadati>Pseudomonadota>Betaproteobacteria>Burkholderiales>Sphaerotilaceae>Sphaerotilus | 712 | HZU83_RS20735 | 14132..16270 | 522889 | AAA-family-ATPase-[Sphaerotilus-montanus] |
| NNM65797.1 | OLD-ABC+TOPRIM | GCA_013044035.1 | Burkholderiales_bacterium | Bacteria>Pseudomonadati>Pseudomonadota>Betaproteobacteria>Burkholderiales>unclassified_Burkholderiales | 656 | HKL99_14540 | 60574..62544 | 1891238 | AAA-family-ATPase-[Burkholderiales-bacterium] |
| NNM65794.1 | OLD-ABC+TOPRIM | GCA_013044035.1 | Burkholderiales_bacterium | Bacteria>Pseudomonadati>Pseudomonadota>Betaproteobacteria>Burkholderiales>unclassified_Burkholderiales | 624 | HKL99_14525 | 56426..58300 | 1891238 | AAA-family-ATPase-[Burkholderiales-bacterium] |
| NDU85860.1 | OLD-ABC+TOPRIM | GCA_010436015.1 | Ferrovum_sp. | Bacteria>Pseudomonadati>Pseudomonadota>Betaproteobacteria>Ferrovales>Ferrovaceae>Ferrovum>unclassified_Ferrovum | 613 | G3H99_04515 | 19773..21614 | 2609467 | AAA-family-ATPase-[Ferrovum-sp.] |
| WP_239902640.1 | OLD-ABC+TOPRIM | GCF_022394475.1 | Laribacter_hongkongensis | Bacteria>Pseudomonadati>Pseudomonadota>Betaproteobacteria>Neisseriales>Aquaspirillaceae>Laribacter | 574 | LH449_RS00395 | 85427..87151 | 168471 | DUF2813-domain-containing-protein-[Laribacter-hongkongensis] |
| TXI44181.1 | OLD-ABC+TOPRIM | GCA_008015755.1 | Methylophilus_sp. | Bacteria>Pseudomonadati>Pseudomonadota>Betaproteobacteria>Nitrosomonadales>Methylophilaceae>Methylophilus>unclassified_Methylophilus | 627 | E6Q52_09980 | 21234..23117 | 29541 | MAG:-DUF2813-domain-containing-protein-[Methylophilus-sp.] |
| WP_072293653.1 | OLD-ABC+TOPRIM | GCF_900119085.1 | Nitrosovibrio_sp._Nv17 | Bacteria>Pseudomonadati>Pseudomonadota>Betaproteobacteria>Nitrosomonadales>Nitrosomonadaceae>Nitrosovibrio>unclassified_Nitrosovibrio | 651 | BT108_RS06125 | 112956..114911 | 1855339 | AAA-family-ATPase-[Nitrosovibrio-sp.-Nv17] |
| WP_153146316.1 | OLD-ABC+TOPRIM | GCF_009469615.1 | Dechloromonas_sp._H13 | Bacteria>Pseudomonadati>Pseudomonadota>Betaproteobacteria>Rhodocyclales>Azonexaceae>Dechloromonas>unclassified_Dechloromonas | 645 | GH660_RS08915 | 119268..121205 | 2570193 | AAA-family-ATPase-[Dechloromonas-sp.-H13] |
| MBT9520404.1 | OLD-ABC+TOPRIM | GCA_018780195.1 | Dechloromonas_sp. | Bacteria>Pseudomonadati>Pseudomonadota>Betaproteobacteria>Rhodocyclales>Azonexaceae>Dechloromonas>unclassified_Dechloromonas | 619 | IV101_05870 | 39596..41455 | 1917218 | ATP-dependent-endonuclease-[Dechloromonas-sp.] |
| WP_205230200.1 | OLD-ABC+TOPRIM | GCF_905120365.1 | Azospira_oryzae | Bacteria>Pseudomonadati>Pseudomonadota>Betaproteobacteria>Rhodocyclales>Rhodocyclaceae>Azospira | 576 | JYJ48_RS05800 | 10267..11997 | 146939 | DUF2813-domain-containing-protein-[Azospira-oryzae] |
| WP_198363324.1 | OLD-ABC+TOPRIM | GCF_002354895.1 | Thauera | Bacteria>Pseudomonadati>Pseudomonadota>Betaproteobacteria>Rhodocyclales>Zoogloeaceae | 571 | CCZ27_RS10615 | 2438091..2439806 | 33057 | ATP-binding-protein-[Thauera-sp.-K11] |
| MBI3370953.1 | OLD-ABC+TOPRIM | GCA_016195915.1 | Betaproteobacteria_bacterium | Bacteria>Pseudomonadati>Pseudomonadota>Betaproteobacteria>unclassified_Betaproteobacteria | 594 | HY017_04225 | 109631..111415 | 1891241 | AAA-family-ATPase-[Betaproteobacteria-bacterium] |
| PKO27555.1 | OLD-ABC+TOPRIM | GCA_002842215.1 | Betaproteobacteria_bacterium_HGW-Betaproteobacteria-9 | Bacteria>Pseudomonadati>Pseudomonadota>Betaproteobacteria>unclassified_Betaproteobacteria | 575 | CVU36_19690 | 60059..61786 | 2013721 | hypothetical-protein-CVU36_19690-[Betaproteobacteria-bacterium-HGW-Betaproteobacteria-9] |
| BBR09768.1 | OLD-ABC+TOPRIM | GCA_014158455.1 | Aeromonas_caviae | Bacteria>Pseudomonadati>Pseudomonadota>Gammaproteobacteria>Aeromonadales>Aeromonadaceae>Aeromonas | 651 | WP3S18E02_14290 | 1544041..1545996 | 648 | hypothetical-protein-WP3S18E02_14290-[Aeromonas-caviae] |
| WP_242470461.1 | OLD-ABC+TOPRIM | GCF_016584135.1 | Allochromatium_vinosum | Bacteria>Pseudomonadati>Pseudomonadota>Gammaproteobacteria>Chromatiales>Chromatiaceae>Allochromatium | 575 | CKO29_RS16460 | 11976..13703 | 1049 | DUF2813-domain-containing-protein-[Allochromatium-vinosum] |
| WP_200346463.1 | OLD-ABC+TOPRIM | GCF_016584085.1 | Halochromatium_glycolicum | Bacteria>Pseudomonadati>Pseudomonadota>Gammaproteobacteria>Chromatiales>Chromatiaceae>Halochromatium | 578 | CKO40_RS12045 | 35195..36931 | 85075 | DUF2813-domain-containing-protein-[Halochromatium-glycolicum] |
| WP_242518914.1 | OLD-ABC+TOPRIM | GCF_016653315.1 | Halochromatium_roseum | Bacteria>Pseudomonadati>Pseudomonadota>Gammaproteobacteria>Chromatiales>Chromatiaceae>Halochromatium | 575 | CCR96_RS10460 | 45727..47454 | 391920 | DUF2813-domain-containing-protein-[Halochromatium-roseum] |
| WP_062271489.1 | OLD-ABC+TOPRIM | GCF_001583505.1 | Marichromatium_gracile | Bacteria>Pseudomonadati>Pseudomonadota>Gammaproteobacteria>Chromatiales>Chromatiaceae>Marichromatium | 561 | AY586_RS00790 | 15432..17117 | 1048 | DUF2813-domain-containing-protein-[Marichromatium-gracile] |
| MCG7871838.1 | OLD-ABC+TOPRIM | GCA_016848925.1 | Candidatus_Thiodiazotropha_lotti | Bacteria>Pseudomonadati>Pseudomonadota>Gammaproteobacteria>Chromatiales>Sedimenticolaceae>Candidatus_Thiodiazotropha | 641 | JAZ11_07010 | 11599..13524 | 2792787 | AAA-family-ATPase-[Candidatus-Thiodiazotropha-lotti] |
| KAB0815081.1 | OLD-ABC+TOPRIM | GCA_008805175.1 | Cronobacter_sakazakii | Bacteria>Pseudomonadati>Pseudomonadota>Gammaproteobacteria>Enterobacterales>Enterobacteriaceae>Cronobacter | 550 | FZI24_17875 | 24337..25989 | 28141 | AAA-family-ATPase-[Cronobacter-sakazakii] |
| WP_104677906.1 | OLD-ABC+TOPRIM | GCF_002942265.1 | Cronobacter_sakazakii | Bacteria>Pseudomonadati>Pseudomonadota>Gammaproteobacteria>Enterobacterales>Enterobacteriaceae>Cronobacter | 550 | C3D67_RS10985 | 287137..288789 | 28141 | AAA-family-ATPase-[Cronobacter-sakazakii] |
| WP_182062515.1 | OLD-ABC+TOPRIM | GCF_013927005.1 | Enterobacter_sp._RHBSTW-00175 | Bacteria>Pseudomonadati>Pseudomonadota>Gammaproteobacteria>Enterobacterales>Enterobacteriaceae>Enterobacter>unclassified_Enterobacter | 552 | HV107_RS06390 | 1360783..1362441 | 2742639 | AAA-family-ATPase-[Enterobacter-sp.-RHBSTW-00175] |
| WP_195869509.1 | OLD-ABC+TOPRIM | GCF_015645305.1 | Serratia | Bacteria>Pseudomonadati>Pseudomonadota>Gammaproteobacteria>Enterobacterales>Yersiniaceae | 608 | I1A46_RS14775 | 223531..225357 | 613 | ATP-binding-protein-[Serratia-liquefaciens] |
| CNI93823.1 | OLD-ABC+TOPRIM | GCA_001319685.1 | Yersinia_pseudotuberculosis | Bacteria>Pseudomonadati>Pseudomonadota>Gammaproteobacteria>Enterobacterales>Yersiniaceae>Yersinia>Yersinia_pseudotuberculosis_complex | 422 | ERS008465_02110 | 82648..83916 | 633 | cobalt-transporter-ATP-binding-subunit-[Yersinia-pseudotuberculosis] |
| WP_057532440.1 | OLD-ABC+TOPRIM | GCF_001319705.1 | Yersinia_pseudotuberculosis | Bacteria>Pseudomonadati>Pseudomonadota>Gammaproteobacteria>Enterobacterales>Yersiniaceae>Yersinia>Yersinia_pseudotuberculosis_complex | 422 | EGX52_RS11295 | 2332580..2333848 | 633 | AAA-family-ATPase-[Yersinia-pseudotuberculosis] |
| MBH2008930.1 | OLD-ABC+TOPRIM | GCA_015999665.1 | Lysobacteraceae_bacterium | Bacteria>Pseudomonadati>Pseudomonadota>Gammaproteobacteria>Lysobacterales>Lysobacteraceae>unclassified_Lysobacteraceae | 624 | I8H71_04425 | 2295..4169 | 1926873 | AAA-family-ATPase-[Xanthomonadaceae-bacterium] |
| TAN08685.1 | OLD-ABC+TOPRIM | GCA_004299555.1 | Rhodanobacteraceae_bacterium | Bacteria>Pseudomonadati>Pseudomonadota>Gammaproteobacteria>Lysobacterales>Rhodanobacteraceae>unclassified_Rhodanobacteraceae | 463 | EPN38_01825 | 136280..137671 | 2052182 | hypothetical-protein-EPN38_01825-[Rhodanobacteraceae-bacterium] |
| WP_005369542.1 | OLD-ABC+TOPRIM | GCF_000214275.2 | Methylomicrobium_album | Bacteria>Pseudomonadati>Pseudomonadota>Gammaproteobacteria>Methylococcales>Methylococcaceae>Methylomicrobium | 606 | METAL_RS03110 | 745837..747657 | 39775 | ATP-binding-protein-[Methylomicrobium-album] |
| WP_000593543.1 | OLD-ABC+TOPRIM | GCF_000069245.1 | Acinetobacter | Bacteria>Pseudomonadati>Pseudomonadota>Gammaproteobacteria>Moraxellales>Moraxellaceae | 651 | J687_RS15345 | 32204..34159 | 469 | MULTISPECIES:-AAA-family-ATPase-[Acinetobacter] |
| WP_040118384.1 | OLD-ABC+TOPRIM | GCF_018139165.1 | Acinetobacter_nosocomialis | Bacteria>Pseudomonadati>Pseudomonadota>Gammaproteobacteria>Moraxellales>Moraxellaceae>Acinetobacter>Acinetobacter_calcoaceticus/baumannii_complex | 645 | KC103_RS15120 | 89055..90989 | 106654 | AAA-family-ATPase,-partial-[Acinetobacter-nosocomialis] |

| Protein-ID | Domain | Assembly-ID | Strain | Taxonomy | Length | Locus-Tag | Genomic-coordinates | TaxID | Fasta-defline |
|---|---|---|---|---|---|---|---|---|---|
| WP_226789418.1 | OLD-ABC+TOPRIM | GCF_020535225.1 | Acinetobacter_pittii | Bacteria>Pseudomonadati>Pseudomonadota>Gammaproteobacteria>Moraxellales>Moraxellaceae>Acinetobacter>Acinetobacter_calcoaceticus/baumannii_complex | 561 | LG653_RS19205 | 17305..18990 | 48296 | AAA-family-ATPase-[Acinetobacter-pittii] |
| CAM84754.1 | OLD-ABC+TOPRIM | GCA_000069245.1 | Acinetobacter_baumannii_AYE | Bacteria>Pseudomonadati>Pseudomonadota>Gammaproteobacteria>Moraxellales>Moraxellaceae>Acinetobacter>Acinetobacter_calcoaceticus/baumannii_complex>Acinetobacter_baumannii | 651 | p3ABAYE0061 | 45667..47622 | 509173 | conserved-hypothetical-protein-(plasmid)-[Acinetobacter-baumannii-AYE] |
| EXR99986.1 | OLD-ABC+TOPRIM | GCA_000588415.1 | Acinetobacter_sp._225588 | Bacteria>Pseudomonadati>Pseudomonadota>Gammaproteobacteria>Moraxellales>Moraxellaceae>Acinetobacter>Acinetobacter_calcoaceticus/baumannii_complex>unclassified_Acinetobacter_calcoaceticus/baumannii_complex | 651 | J687_2213 | 32204..34159 | 1310788 | AAA-ATPase-domain-protein-[Acinetobacter-sp.-225588] |
| WP_159177825.1 | OLD-ABC+TOPRIM | GCF_902506335.1 | Halomonadaceae | Bacteria>Pseudomonadati>Pseudomonadota>Gammaproteobacteria>Oceanospirillales | 686 | JMX33_RS24505 | 5294004..5296064 | 28256 | MULTISPECIES:-ATP-dependent-endonuclease-[unclassified-Halomonas] |
| WP_008928711.1 | OLD-ABC+TOPRIM | GCF_000300995.1 | Alcanivorax_hongdengensis | Bacteria>Pseudomonadati>Pseudomonadota>Gammaproteobacteria>Oceanospirillales>Alcanivoracaceae>Alcanivorax | 651 | A11A3_RS07675 | 79342..81297 | 519051 | AAA-family-ATPase-[Alcanivorax-hongdengensis] |
| WP_272975232.1 | OLD-ABC+TOPRIM | GCF_963971865.1 | Alcanivorax_jadensis | Bacteria>Pseudomonadati>Pseudomonadota>Gammaproteobacteria>Oceanospirillales>Alcanivoracaceae>Alcanivorax | 610 | B2Z32_RS11815 | 41642..43474 | 64988 | ATP-binding-protein-[Alcanivorax-jadensis] |
| WP_161382588.1 | OLD-ABC+TOPRIM | - | Alcanivorax_sp._ZXX171 | Bacteria>Pseudomonadati>Pseudomonadota>Gammaproteobacteria>Oceanospirillales>Alcanivoracaceae>Alcanivorax>unclassified_Alcanivorax | 686 | GN155_RS00685 | 149271..151331 | 2695858 | ATP-dependent-endonuclease-[Alcanivorax-sp.-ZXX171] |
| MCH2557386.1 | OLD-ABC+TOPRIM | GCA_022449455.1 | Alcanivorax_sp. | Bacteria>Pseudomonadati>Pseudomonadota>Gammaproteobacteria>Oceanospirillales>Alcanivoracaceae>Alcanivorax>unclassified_Alcanivorax | 651 | MK005_08805 | 142611..144566 | 1872427 | ATP-binding-protein-[Alcanivorax-sp.] |
| WP_213229323.1 | OLD-ABC+TOPRIM | - | Halomonas_litopenaei | Bacteria>Pseudomonadati>Pseudomonadota>Gammaproteobacteria>Oceanospirillales>Halomonadaceae>Halomonas | 669 | DYI26_RS12510 | 34541..36550 | 2109328 | AAA-family-ATPase-[Halomonas-litopenaei] |
| KXS54420.1 | OLD-ABC+TOPRIM | GCA_001577575.1 | Marinobacter_sp._T13-3 | Bacteria>Pseudomonadati>Pseudomonadota>Gammaproteobacteria>Pseudomonadales>Marinobacteraceae>Marinobacter>unclassified_Marinobacter | 669 | AWU57_1202 | 10858..12867 | 1794906 | ATP-dependent-OLD-family-endonuclease-[Marinobacter-sp.-T13-3] |
| WP_269819148.1 | OLD-ABC+TOPRIM | GCF_027498055.1 | Denitrificimonas_caeni | Bacteria>Pseudomonadati>Pseudomonadota>Gammaproteobacteria>Pseudomonadales>Pseudomonadaceae>Denitrificimonas | 460 | O6P33_RS05170 | 1080183..1081565 | 521720 | AAA-family-ATPase-[Denitrificimonas-caeni] |
| MBM2633273.1 | OLD-ABC+TOPRIM | GCA_016836445.1 | Pseudomonas_aeruginosa | Bacteria>Pseudomonadati>Pseudomonadota>Gammaproteobacteria>Pseudomonadales>Pseudomonadaceae>Pseudomonas>Pseudomonas_aeruginosa_group | 712 | JQ480_07730 | 1625189..1627327 | 287 | AAA-family-ATPase-[Pseudomonas-aeruginosa] |
| WP_023113868.1 | OLD-ABC+TOPRIM | GCF_036232585.1 | Pseudomonas_aeruginosa | Bacteria>Pseudomonadati>Pseudomonadota>Gammaproteobacteria>Pseudomonadales>Pseudomonadaceae>Pseudomonas>Pseudomonas_aeruginosa_group | 712 | JQ482_RS07420 | 1570253..1572391 | 287 | AAA-family-ATPase-[Pseudomonas-aeruginosa] |
| HCF1890562.1 | OLD-ABC+TOPRIM | GCA_023942415.1 | Pseudomonas_aeruginosa | Bacteria>Pseudomonadati>Pseudomonadota>Gammaproteobacteria>Pseudomonadales>Pseudomonadaceae>Pseudomonas>Pseudomonas_aeruginosa_group | 653 | NH920_001196 | 103026..104987 | 287 | AAA-family-ATPase-[Pseudomonas-aeruginosa] |
| HBO7582680.1 | OLD-ABC+TOPRIM | GCA_021673555.1 | Pseudomonas_aeruginosa | Bacteria>Pseudomonadati>Pseudomonadota>Gammaproteobacteria>Pseudomonadales>Pseudomonadaceae>Pseudomonas>Pseudomonas_aeruginosa_group | 651 | L4920_006244 | 2937..4892 | 287 | AAA-family-ATPase-[Pseudomonas-aeruginosa] |
| RTS56723.1 | OLD-ABC+TOPRIM | GCA_003968225.1 | Pseudomonas_aeruginosa | Bacteria>Pseudomonadati>Pseudomonadota>Gammaproteobacteria>Pseudomonadales>Pseudomonadaceae>Pseudomonas>Pseudomonas_aeruginosa_group | 651 | DY943_06680 | 159257..161212 | 287 | ATP-dependent-endonuclease-[Pseudomonas-aeruginosa] |
| WP_060853049.1 | OLD-ABC+TOPRIM | GCF_001548335.1 | Pseudomonas_aeruginosa | Bacteria>Pseudomonadati>Pseudomonadota>Gammaproteobacteria>Pseudomonadales>Pseudomonadaceae>Pseudomonas>Pseudomonas_aeruginosa_group | 651 | IOMTU133_RS11690 | 2535903..2537858 | 287 | AAA-family-ATPase-[Pseudomonas-aeruginosa] |
| WP_077565919.1 | OLD-ABC+TOPRIM | GCF_002002765.1 | Pseudomonas_aeruginosa | Bacteria>Pseudomonadati>Pseudomonadota>Gammaproteobacteria>Pseudomonadales>Pseudomonadaceae>Pseudomonas>Pseudomonas_aeruginosa_group | 651 | BM548_RS18920 | 102795..104750 | 287 | AAA-family-ATPase-[Pseudomonas-aeruginosa] |
| WP_126583670.1 | OLD-ABC+TOPRIM | GCF_003968225.1 | Pseudomonas_aeruginosa | Bacteria>Pseudomonadati>Pseudomonadota>Gammaproteobacteria>Pseudomonadales>Pseudomonadaceae>Pseudomonas>Pseudomonas_aeruginosa_group | 651 | DY943_RS06680 | 159257..161212 | 287 | AAA-family-ATPase-[Pseudomonas-aeruginosa] |
| ERV97551.1 | OLD-ABC+TOPRIM | GCA_000481225.1 | Pseudomonas_aeruginosa_BWHPSA024 | Bacteria>Pseudomonadati>Pseudomonadota>Gammaproteobacteria>Pseudomonadales>Pseudomonadaceae>Pseudomonas>Pseudomonas_aeruginosa_group>Pseudomonas_aeruginosa | 712 | Q037_05161 | 19676..21814 | 1402524 | hypothetical-protein-Q037_05161-[Pseudomonas-aeruginosa-BWHPSA024] |
| WP_007159181.1 | OLD-ABC+TOPRIM | GCF_000236825.1 | Pseudomonas_oryzihabitans | Bacteria>Pseudomonadati>Pseudomonadota>Gammaproteobacteria>Pseudomonadales>Pseudomonadaceae>Pseudomonas>Pseudomonas_putida_group | 720 | PPL19_RS03935 | 160374..162536 | 47885 | AAA-family-ATPase-[Pseudomonas-psychrotolerans] |
| MXY51428.1 | OLD-ABC+TOPRIM | GCA_009837725.1 | Gammaproteobacteria_bacterium | Bacteria>Pseudomonadati>Pseudomonadota>Gammaproteobacteria>unclassified_Gammaproteobacteria | 680 | F4Y86_02710 | 37640..39682 | 1913989 | AAA-family-ATPase-[Gammaproteobacteria-bacterium] |
| MDE0453910.1 | OLD-ABC+TOPRIM | GCA_028821145.1 | Gammaproteobacteria_bacterium | Bacteria>Pseudomonadati>Pseudomonadota>Gammaproteobacteria>unclassified_Gammaproteobacteria | 638 | OXJ63_01220 | 26008..27924 | 1913989 | MAG:-AAA-family-ATPase-[Gammaproteobacteria-bacterium] |
| HDH16016.1 | OLD-ABC+TOPRIM | GCA_011040115.1 | Gammaproteobacteria_bacterium | Bacteria>Pseudomonadati>Pseudomonadota>Gammaproteobacteria>unclassified_Gammaproteobacteria | 570 | ENG90_06020 | 2596..4308 | 1913989 | DUF2813-domain-containing-protein-[Gammaproteobacteria-bacterium] |
| MDD9824623.1 | OLD-ABC+TOPRIM | GCA_028817255.1 | Gammaproteobacteria_bacterium | Bacteria>Pseudomonadati>Pseudomonadota>Gammaproteobacteria>unclassified_Gammaproteobacteria | 552 | OXU43_05580 | 4988..6646 | 1913989 | MAG:-AAA-family-ATPase-[Gammaproteobacteria-bacterium] |
| MDD9822475.1 | OLD-ABC+TOPRIM | GCA_028816685.1 | Gammaproteobacteria_bacterium | Bacteria>Pseudomonadati>Pseudomonadota>Gammaproteobacteria>unclassified_Gammaproteobacteria | 550 | OXU98_06285 | 1035..2687 | 1913989 | MAG:-AAA-family-ATPase-[Gammaproteobacteria-bacterium] |
| MDE0129993.1 | OLD-ABC+TOPRIM | GCA_028819355.1 | Gammaproteobacteria_bacterium | Bacteria>Pseudomonadati>Pseudomonadota>Gammaproteobacteria>unclassified_Gammaproteobacteria | 530 | OXQ86_10565 | 54700..56292 | 1913989 | MAG:-AAA-family-ATPase-[Gammaproteobacteria-bacterium] |
| MDE0129999.1 | OLD-ABC+TOPRIM | GCA_028819355.1 | Gammaproteobacteria_bacterium | Bacteria>Pseudomonadati>Pseudomonadota>Gammaproteobacteria>unclassified_Gammaproteobacteria | 405 | OXQ86_10595 | 67697..68914 | 1913989 | MAG:-AAA-family-ATPase-[Gammaproteobacteria-bacterium] |
| MBK8482431.1 | OLD-ABC+TOPRIM | GCA_016711215.1 | Pseudomonadota_bacterium | Bacteria>Pseudomonadati>Pseudomonadota>unclassified_Pseudomonadota | 609 | IPL40_14910 | 139244..141073 | 1977087 | AAA-family-ATPase-[Pseudomonadota-bacterium] |
| MDD4099139.1 | OLD-ABC+TOPRIM | GCA_028701535.1 | Lentisphaeria_bacterium | Bacteria>Pseudomonadati>PVC_group>Lentisphaerota>Lentisphaeria>unclassified_Lentisphaeria | 404 | PHC30_10240 | 1805..3017 | 2053569 | MAG:-AAA-family-ATPase,-partial-[Lentisphaeria-bacterium] |
| MBT3379344.1 | OLD-ABC+TOPRIM | GCA_018648355.1 | Lentisphaerota_bacterium | Bacteria>Pseudomonadati>PVC_group>Lentisphaerota>unclassified_Lentisphaerota | 571 | HN406_27335 | 15771..17486 | 1932692 | DUF2813-domain-containing-protein-[Lentisphaerota-bacterium] |
| MBT4818759.1 | OLD-ABC+TOPRIM | GCA_018664235.1 | Lentisphaerota_bacterium | Bacteria>Pseudomonadati>PVC_group>Lentisphaerota>unclassified_Lentisphaerota | 571 | HON70_23825 | 2079..3794 | 1932692 | DUF2813-domain-containing-protein-[Lentisphaerota-bacterium] |
| MBT5612039.1 | OLD-ABC+TOPRIM | GCA_018666605.1 | Lentisphaerota_bacterium | Bacteria>Pseudomonadati>PVC_group>Lentisphaerota>unclassified_Lentisphaerota | 571 | HOJ57_39235 | 5716..7431 | 1932692 | DUF2813-domain-containing-protein-[Lentisphaerota-bacterium] |
| MBX3394307.1 | OLD-ABC+TOPRIM | GCA_019636475.1 | Phycisphaerae_bacterium | Bacteria>Pseudomonadati>PVC_group>Planctomycetota>Phycisphaerae>unclassified_Phycisphaerae | 601 | KF841_02965 | 760407..762212 | 2026778 | AAA-family-ATPase-[Phycisphaerae-bacterium] |
| WP_184301674.1 | OLD-ABC+TOPRIM | GCF_014192335.1 | Aporhodopirellula_rubra | Bacteria>Pseudomonadati>PVC_group>Planctomycetota>Planctomycetia>Pirellulales>Pirellulaceae>Aporhodopirellula | 620 | FHS27_RS03230 | 259106..260968 | 980271 | AAA-family-ATPase-[Rhodopirellula-rubra] |
| WP_235989928.1 | OLD-ABC+TOPRIM | GCF_013618625.1 | Bremerella_alba | Bacteria>Pseudomonadati>PVC_group>Planctomycetota>Planctomycetia>Pirellulales>Pirellulaceae>Bremerella | 600 | HOV93_RS05920 | 51357..53159 | 980252 | AAA-family-ATPase-[Bremerella-alba] |
| MCF7962595.1 | OLD-ABC+TOPRIM | GCA_021736575.1 | Pirellula_sp. | Bacteria>Pseudomonadati>PVC_group>Planctomycetota>Planctomycetia>Pirellulales>Pirellulaceae>Pirellula>unclassified_Pirellula | 658 | K9M08_17795 | 5229..7205 | 117 | AAA-family-ATPase-[Pirellula-sp.] |
| MBQ7111127.1 | OLD-ABC+TOPRIM | GCA_017436025.1 | Thermoguttaceae_bacterium | Bacteria>Pseudomonadati>PVC_group>Planctomycetota>Planctomycetia>Pirellulales>Thermoguttaceae>unclassified_Thermoguttaceae | 686 | IJO06_07895 | 24751..26811 | 2691551 | AAA-family-ATPase-[Thermoguttaceae-bacterium] |
| MBP3531677.1 | OLD-ABC+TOPRIM | GCA_017937485.1 | Thermoguttaceae_bacterium | Bacteria>Pseudomonadati>PVC_group>Planctomycetota>Planctomycetia>Pirellulales>Thermoguttaceae>unclassified_Thermoguttaceae | 615 | J6K25_10930 | 4777..6624 | 2691551 | AAA-family-ATPase-[Thermoguttaceae-bacterium] |
| OJW08013.1 | OLD-ABC+TOPRIM | GCA_001898385.1 | Planctomycetales_bacterium_71-10 | Bacteria>Pseudomonadati>PVC_group>Planctomycetota>Planctomycetia>Planctomycetales>unclassified_Planctomycetales | 622 | BGO49_20705 | 30639..32507 | 1895807 | MAG:-ATP-dependent-endonuclease-[Planctomycetales-bacterium-71-10] |
| MCX5669696.1 | OLD-ABC+TOPRIM | GCA_026387055.1 | Planctomycetota_bacterium | Bacteria>Pseudomonadati>PVC_group>Planctomycetota>unclassified_Planctomycetota | 682 | NTU94_00025 | 2150..4198 | 2026780 | MAG:-AAA-family-ATPase-[Planctomycetota-bacterium] |
| NMD35176.1 | OLD-ABC+TOPRIM | GCA_012797405.1 | Planctomycetota_bacterium | Bacteria>Pseudomonadati>PVC_group>Planctomycetota>unclassified_Planctomycetota | 657 | GYA73_04765 | 6599..8572 | 2026780 | AAA-family-ATPase-[Planctomycetota-bacterium] |
| MBL8860738.1 | OLD-ABC+TOPRIM | GCA_016795205.1 | Planctomycetota_bacterium | Bacteria>Pseudomonadati>PVC_group>Planctomycetota>unclassified_Planctomycetota | 618 | JNK02_01905 | 3757..5613 | 2026780 | AAA-family-ATPase-[Planctomycetota-bacterium] |
| MBK9387521.1 | OLD-ABC+TOPRIM | GCA_016719295.1 | Planctomycetota_bacterium | Bacteria>Pseudomonadati>PVC_group>Planctomycetota>unclassified_Planctomycetota | 588 | IPN34_22130 | 5827316..5829082 | 2026780 | ATP-dependent-endonuclease-[Planctomycetota-bacterium] |
| MBA3936143.1 | OLD-ABC+TOPRIM | GCA_013821615.1 | Planctomycetota_bacterium | Bacteria>Pseudomonadati>PVC_group>Planctomycetota>unclassified_Planctomycetota | 560 | HOX38_01685 | 532..2214 | 2026780 | AAA-family-ATPase-[Planctomycetota-bacterium] |
| MCH8539500.1 | OLD-ABC+TOPRIM | GCA_022565675.1 | Opitutales_bacterium | Bacteria>Pseudomonadati>PVC_group>Verrucomicrobiota>Opitutia>Opitutales>unclassified_Opitutales | 606 | LAT58_01880 | 15066..16886 | 2026772 | ATP-binding-protein-[Opitutales-bacterium] |
| MCI0539267.1 | OLD-ABC+TOPRIM | GCA_022615925.1 | Verrucomicrobiales_bacterium | Bacteria>Pseudomonadati>PVC_group>Verrucomicrobiota>Verrucomicrobiia>Verrucomicrobiales>unclassified_Verrucomicrobiales | 695 | LOZ50_29015 | 2064..4151 | 2026801 | AAA-family-ATPase-[Verrucomicrobiales-bacterium] |
| AEJ20207.1 | OLD-ABC+TOPRIM | GCA_000219725.1 | Gracilinema_caldarium_DSM_7334 | Bacteria>Pseudomonadati>Spirochaetota>Spirochaetia>Spirochaetales>Breznakiellaceae>Gracilinema>Gracilinema_caldarium | 557 | Spica_2080 | 2339623..2341296 | 744872 | SMC-domain-protein-[Gracilinema-caldarium-DSM-7334] |
| SLM14725.1 | OLD-ABC+TOPRIM | GCA_900163985.1 | uncultured_spirochete | Bacteria>Pseudomonadati>Spirochaetota>Spirochaetia>Spirochaetales>environmental_samples | 554 | SPIROBIBN47_350051 | 60171..61835 | 156406 | SMC-domain-protein-[uncultured-spirochete] |
| MCK9286307.1 | OLD-ABC+TOPRIM | GCA_023230445.1 | Sphaerochaetaceae_bacterium | Bacteria>Pseudomonadati>Spirochaetota>Spirochaetia>Spirochaetales>Sphaerochaetaceae>unclassified_Sphaerochaetaceae | 603 | MOP29_01915 | 120052..121863 | 2791017 | AAA-family-ATPase-[Sphaerochaetaceae-bacterium] |
| WP_010252930.1 | OLD-ABC+TOPRIM | GCF_000297095.1 | Treponema_primitia | Bacteria>Pseudomonadati>Spirochaetota>Spirochaetia>Spirochaetales>Treponemataceae>Treponema | 541 | TPRIMZ1_RS0100105 | 25205..26830 | 88058 | AAA-family-ATPase-[Treponema-primitia] |
| MCA1950499.1 | OLD-ABC+TOPRIM | GCA_020216655.1 | Treponema_sp. | Bacteria>Pseudomonadati>Spirochaetota>Spirochaetia>Spirochaetales>Treponemataceae>Treponema>unclassified_Treponema | 555 | LDL24_07990 | 13100..14767 | 166 | AAA-family-ATPase-[Treponema-sp.] |
| WP_132781618.1 | OLD-ABC+TOPRIM | GCF_004343725.1 | Treponema_sp._J25 | Bacteria>Pseudomonadati>Spirochaetota>Spirochaetia>Spirochaetales>Treponemataceae>Treponema>unclassified_Treponema | 554 | C5O22_RS10545 | 18861..20525 | 2094121 | ATP-binding-protein-[Treponema-sp.-J25] |
| KMY65635.1 | OLD-ABC+TOPRIM | GCA_001184205.1 | Desulfocarbo_indianensis | Bacteria>Pseudomonadati>Thermodesulfobacteriota>Desulfarculia>Desulfarculales>Desulfarculaceae>Desulfocarbo | 571 | AAU61_21225 | 20473..22188 | 1348163 | hypothetical-protein-AAU61_21225-[Desulfocarbo-indianensis] |
| KGO32869.1 | OLD-ABC+TOPRIM | GCA_000769715.1 | Desulfobulbus_sp._Tol-SR | Bacteria>Pseudomonadati>Thermodesulfobacteriota>Desulfobulbia>Desulfobulbales>Desulfobulbaceae>Desulfobulbus>unclassified_Desulfobulbus | 614 | JT06_17855 | 7836..9680 | 1536652 | hypothetical-protein-JT06_17855-[Desulfobulbus-sp.-Tol-SR] |
| MBA3013970.1 | OLD-ABC+TOPRIM | GCA_013792005.1 | Desulfobulbaceae_bacterium | Bacteria>Pseudomonadati>Thermodesulfobacteriota>Desulfobulbia>Desulfobulbales>Desulfobulbaceae>unclassified_Desulfobulbaceae | 645 | FP815_03335 | 5005..6942 | 2053307 | AAA-family-ATPase-[Desulfobulbaceae-bacterium] |
| WP_009368943.1 | OLD-ABC+TOPRIM | GCF_020592975.1 | Bilophila | Bacteria>Pseudomonadati>Thermodesulfobacteriota>Desulfovibrionia>Desulfovibrionales>Desulfovibrionaceae | 612 | LJE06_RS05500 | 43222..45060 | 35832 | MULTISPECIES:-ATP-binding-protein-[Bilophila] |
| WP_144304547.1 | OLD-ABC+TOPRIM | GCF_007625075.1 | Oceanidesulfovibrio_indonesiensis | Bacteria>Pseudomonadati>Thermodesulfobacteriota>Desulfovibrionia>Desulfovibrionales>Desulfovibrionaceae>Oceanidesulfovibrio | 595 | DPQ33_RS17555 | 7826..9613 | 54767 | AAA-family-ATPase-[Oceanidesulfovibrio-indonesiensis] |
| WP_011735761.1 | OLD-ABC+TOPRIM | GCF_000015045.1 | Pelobacter_propionicus | Bacteria>Pseudomonadati>Thermodesulfobacteriota>Desulfuromonadia>Desulfuromonadales>Desulfuromonadaceae>Pelobacter | 703 | PPRO_RS09325 | 2015460..2017571 | 29543 | AAA-family-ATPase-[Pelobacter-propionicus] |
| OIP92315.1 | OLD-ABC+TOPRIM | GCA_001873745.1 | Syntrophaceae_bacterium_CG2_30_58_14 | Bacteria>Pseudomonadati>Thermodesulfobacteriota>Syntrophia>Syntrophales>Syntrophaceae>unclassified_Syntrophaceae | 657 | AUK26_04960 | 2268..4241 | 1805378 | hypothetical-protein-AUK26_04960-[Syntrophaceae-bacterium-CG2_30_58_14] |
| GKT09453.1 | OLD-ABC+TOPRIM | GCA_022836495.1 | Desulforhabdus_sp._TSK | Bacteria>Pseudomonadati>Thermodesulfobacteriota>Syntrophobacteria>Syntrophobacterales>Syntrophobacteraceae>Desulforhabdus>unclassified_Desulforhabdus | 674 | DSTSK_27580 | 38598..40622 | 2925014 | hypothetical-protein-DSTSK_27580-[Desulforhabdus-sp.-TSK] |
| WP_244084751.1 | OLD-ABC+TOPRIM | GCF_022836495.1 | Desulforhabdus_sp._TSK | Bacteria>Pseudomonadati>Thermodesulfobacteriota>Syntrophobacteria>Syntrophobacterales>Syntrophobacteraceae>Desulforhabdus>unclassified_Desulforhabdus | 674 | MWC30_RS13655 | 38598..40622 | 2925014 | ATP-dependent-endonuclease-[Desulforhabdus-sp.-TSK] |
| MCU4137449.1 | OLD-ABC+TOPRIM | GCA_024699695.1 | Thermodesulfobacteriota_bacterium | Bacteria>Pseudomonadati>Thermodesulfobacteriota>unclassified_Thermodesulfobacteriota | 613 | MW689_001020 | 114950..116791 | 2202153 | putative-ATP-dependent-endonuclease-of-the-OLD-family-[Thermodesulfobacteriota-bacteirum] |
| WP_015716389.1 | OLD-ABC+TOPRIM | GCF_000187005.1 | Thermus | Bacteria>Thermotogati>Deinococcota>Deinococci>Thermales>Thermaceae | 528 | TSC_RS02325 | 417341..418927 | 270 | AAA-family-ATPase-[Thermus-scotoductus] |
| WP_267262113.1 | OLD-ABC+TOPRIM | GCF_026427635.1 | Thermus_sp._PS18 | Bacteria>Thermotogati>Deinococcota>Deinococci>Thermales>Thermaceae>Thermus>unclassified_Thermus | 621 | KQ693_RS05010 | 874654..876519 | 2849039 | ATP-dependent-endonuclease-[Thermus-sp.-PS18] |
| MBN2383964.1 | OLD-ABC+TOPRIM | GCA_016934675.1 | bacterium | Bacteria>unclassified_Bacteria | 656 | JXQ70_13895 | 22432..24402 | 1869227 | ATP-dependent-endonuclease-[bacterium] |
| MDE0416421.1 | OLD-ABC+TOPRIM | GCA_028821455.1 | bacterium | Bacteria>unclassified_Bacteria | 653 | OXI95_05720 | 20177..22138 | 1869227 | MAG:-AAA-family-ATPase-[bacterium] |
| NUO19885.1 | OLD-ABC+TOPRIM | GCA_013360195.1 | bacterium | Bacteria>unclassified_Bacteria | 630 | HUU59_10585 | 4894..6786 | 1869227 | AAA-family-ATPase-[bacterium] |
| MCY3637185.1 | OLD-ABC+TOPRIM | GCA_026705625.1 | bacterium | Bacteria>unclassified_Bacteria | 626 | OXH23_16410 | 19895..21775 | 1869227 | MAG:-ATP-dependent-endonuclease-[bacterium] |
| MDE2480823.1 | OLD-ABC+TOPRIM | GCA_028869385.1 | bacterium | Bacteria>unclassified_Bacteria | 559 | KGN02_01365 | 61264..62943 | 1869227 | MAG:-DUF2813-domain-containing-protein-[bacterium] |
| --TOPRIM | | | | | | | | | |
| RLI77770.1 | TOPRIM | GCA_003662925.1 | Archaeoglobales_archaeon | Archaea>Methanobacteriati>Methanobacteriota>Archaeoglobi>Archaeoglobales>unclassified_Archaeoglobales | 171 | DRP04_10875 | 387..902 | 2250258 | hypothetical-protein-DRP04_10875-[Archaeoglobales-archaeon] |
| WP_250543618.1 | TOPRIM | - | Methanocaldococcus_lauensis | Archaea>Methanobacteriati>Methanobacteriota>Methanomada_group>Methanococci>Methanococcales>Methanocaldococcaceae>Methanocaldococcus | 122 | KMP69_RS08090 | 457584..457952 | 2546128 | hypothetical-protein-[Methanocaldococcus-lauensis] |
| WP_250543619.1 | TOPRIM | GCF_902827225.1 | Methanocaldococcus_lauensis | Archaea>Methanobacteriati>Methanobacteriota>Methanomada_group>Methanococci>Methanococcales>Methanocaldococcaceae>Methanocaldococcus | 118 | KMP69_RS08095 | 458018..458374 | 2546128 | hypothetical-protein-[Methanocaldococcus-lauensis] |
| MCW3129355.1 | TOPRIM | GCA_025966515.1 | Candidatus_Methanophagales_archaeon | Archaea>Methanobacteriati>Methanobacteriota>Stenosarchaea_group>Methanomicrobia>Candidatus_Methanophagales>unclassified_Candidatus_Methanophagales | 156 | N2V75_04560 | 3185..3655 | 2056316 | MAG:-hypothetical-protein-N2V75_04560-[Methanophagales-archaeon] |
| MCW3134162.1 | TOPRIM | GCA_025966595.1 | Candidatus_Methanophagales_archaeon | Archaea>Methanobacteriati>Methanobacteriota>Stenosarchaea_group>Methanomicrobia>Candidatus_Methanophagales>unclassified_Candidatus_Methanophagales | 156 | N2V78_07545 | 1301..1771 | 2056316 | MAG:-hypothetical-protein-N2V78_07545-[Methanophagales-archaeon] |
| MBR1368480.1 | TOPRIM | GCA_018132105.1 | Methanocalculus_chunghsingensis | Archaea>Methanobacteriati>Methanobacteriota>Stenosarchaea_group>Methanomicrobia>Methanomicrobiales>Methanocalculaceae>Methanocalculus | 224 | RJ53_02755 | 15859..16533 | 156457 | hypothetical-protein-[Methanocalculus-chunghsingensis] |
| WP_211530108.1 | TOPRIM | GCF_018132105.1 | Methanocalculus_chunghsingensis | Archaea>Methanobacteriati>Methanobacteriota>Stenosarchaea_group>Methanomicrobia>Methanomicrobiales>Methanocalculaceae>Methanocalculus | 224 | RJ53_RS02820 | 15859..16533 | 156457 | DUF4276-family-protein-[Methanocalculus-chunghsingensis] |
| WP_253488282.1 | TOPRIM | GCF_024170505.1 | Methanocalculus_alkaliphilus | Archaea>Methanobacteriati>Methanobacteriota>Stenosarchaea_group>Methanomicrobia>Methanomicrobiales>Methanocalculaceae>Methanocalculus | 224 | J2T58_RS06395 | 4468..5142 | 768730 | DUF4276-family-protein-[Methanocalculus-alkaliphilus] |
| WP_167816001.1 | TOPRIM | - | Methanocorpusculum_sp._GPch4 | Archaea>Methanobacteriati>Methanobacteriota>Stenosarchaea_group>Methanomicrobia>Methanomicrobiales>Methanocorpusculaceae>Methanocorpusculum>unclassified_Methanocorpusculum | 188 | EYA77_RS04460 | 15333..15899 | 2527877 | DUF4276-family-protein-[Methanocorpusculum-sp.-GPch4] |
| WP_011448870.1 | TOPRIM | GCF_000013445.1 | Methanospirillum | Archaea>Methanobacteriati>Methanobacteriota>Stenosarchaea_group>Methanomicrobia>Methanomicrobiales>Methanospirillaceae | 197 | MHUN_RS09830 | 2133889..2134482 | 2202 | hypothetical-protein-[Methanospirillum-hungatei] |
| WP_011448106.1 | TOPRIM | GCF_000013445.1 | Methanospirillum_hungatei | Archaea>Methanobacteriati>Methanobacteriota>Stenosarchaea_group>Methanomicrobia>Methanomicrobiales>Methanospirillaceae>Methanospirillum | 226 | MHUN_RS05645 | 1218177..1218857 | 2203 | DUF4276-family-protein-[Methanospirillum-hungatei] |
| WP_011448740.1 | TOPRIM | GCF_000013445.1 | Methanospirillum_hungatei | Archaea>Methanobacteriati>Methanobacteriota>Stenosarchaea_group>Methanomicrobia>Methanomicrobiales>Methanospirillaceae>Methanospirillum | 205 | MHUN_RS09165 | 1984129..1984746 | 2203 | DUF4276-family-protein-[Methanospirillum-hungatei] |
| ABD40828.1 | TOPRIM | GCA_000013445.1 | Methanospirillum_hungatei_JF-1 | Archaea>Methanobacteriati>Methanobacteriota>Stenosarchaea_group>Methanomicrobia>Methanomicrobiales>Methanospirillaceae>Methanospirillum>Methanospirillum_hungatei | 226 | Mhun_1079 | 1218177..1218857 | 323259 | conserved-hypothetical-protein-[Methanospirillum-hungatei-JF-1] |
| ABD41476.1 | TOPRIM | GCA_000013445.1 | Methanospirillum_hungatei_JF-1 | Archaea>Methanobacteriati>Methanobacteriota>Stenosarchaea_group>Methanomicrobia>Methanomicrobiales>Methanospirillaceae>Methanospirillum>Methanospirillum_hungatei | 205 | Mhun_1755 | 1984129..1984746 | 323259 | conserved-hypothetical-protein-[Methanospirillum-hungatei-JF-1] |
| NLL10044.1 | TOPRIM | GCA_012520015.1 | Methanomicrobiales_archaeon | Archaea>Methanobacteriati>Methanobacteriota>Stenosarchaea_group>Methanomicrobia>Methanomicrobiales>unclassified_Methanomicrobiales | 197 | GX268_03950 | 1802..2395 | 2184052 | DUF4276-family-protein-[Methanomicrobiales-archaeon] |
| QNO47558.1 | TOPRIM | - | Candidatus_Methanogaster_sp._ANME-2c_ERB4 | Archaea>Methanobacteriati>Methanobacteriota>Stenosarchaea_group>Methanomicrobia>Methanosarcinales>ANME-2_cluster>Candidatus_Methanogasteraceae>Candidatus_Methanogaster>unclassified_Candidatus_Methanogaster | 154 | GGGHDLIA_00048 | 32676..33140 | 2759911 | hypothetical-protein-GGGHDLIA_00048-[Methanosarcinales-archaeon-ANME-2c-ERB4] |
| OFV65718.1 | TOPRIM | GCA_001766825.1 | Candidatus_Syntrophoarchaeum_butanivorans | Archaea>Methanobacteriati>Methanobacteriota>Stenosarchaea_group>Methanomicrobia>Methanosarcinales>ANME-2_cluster>Candidatus_Syntrophoarchaeum | 153 | SBU_001301 | 3381..3842 | 1839936 | hypothetical-protein-SBU_001301-[Candidatus-Syntrophoarchaeum-butanivorans] |
| WP_270096524.1 | TOPRIM | GCF_027594145.1 | Methanococcoides_alaskense | Archaea>Methanobacteriati>Methanobacteriota>Stenosarchaea_group>Methanomicrobia>Methanosarcinales>Methanosarcinaceae>Methanococcoides | 199 | PBV53_RS07465 | 729956..730555 | 325778 | RloB-family-protein-[Methanococcoides-alaskense] |
| NPE28177.1 | TOPRIM | GCA_013166765.1 | Methanococcoides_sp._SA1 | Archaea>Methanobacteriati>Methanobacteriota>Stenosarchaea_group>Methanomicrobia>Methanosarcinales>Methanosarcinaceae>Methanococcoides>unclassified_Methanococcoides | 198 | HNV12_09440 | 4852..5448 | 2735871 | RloB-domain-containing-protein-[Methanococcoides-sp.-SA1] |
| WP_091932396.1 | TOPRIM | GCF_900114835.1 | Methanolobus_profundi | Archaea>Methanobacteriati>Methanobacteriota>Stenosarchaea_group>Methanomicrobia>Methanosarcinales>Methanosarcinaceae>Methanolobus | 199 | BM008_RS01860 | 360893..361492 | 487685 | RloB-family-protein-[Methanolobus-profundi] |
| KXS44091.1 | TOPRIM | GCA_001577565.1 | Methanolobus_sp._T82-4 | Archaea>Methanobacteriati>Methanobacteriota>Stenosarchaea_group>Methanomicrobia>Methanosarcinales>Methanosarcinaceae>Methanolobus>unclassified_Methanolobus | 199 | AWU59_724 | 6709..7308 | 1794908 | MAG:-hypothetical-protein-AWU59_724-[Methanolobus-sp.-T82-4] |
| WP_197073949.1 | TOPRIM | GCF_000969965.1 | Methanosarcina_sp._WWM596 | Archaea>Methanobacteriati>Methanobacteriota>Stenosarchaea_group>Methanomicrobia>Methanosarcinales>Methanosarcinaceae>Methanosarcina>unclassified_Methanosarcina | 219 | MSWHS_RS14815 | 3359869..3360528 | 1434103 | hypothetical-protein-[Methanosarcina-sp.-WWM596] |
| WP_273278520.1 | TOPRIM | GCF_913757535.1 | Methanothrix_soehngenii | Archaea>Methanobacteriati>Methanobacteriota>Stenosarchaea_group>Methanomicrobia>Methanotrichales>Methanotrichaceae>Methanothrix | 209 | NW278_RS08465 | 1664494..1665123 | 2223 | hypothetical-protein-[Methanothrix-soehngenii] |
| MBK7385731.1 | TOPRIM | GCA_016706325.1 | Methanothrix_sp. | Archaea>Methanobacteriati>Methanobacteriota>Stenosarchaea_group>Methanomicrobia>Methanotrichales>Methanotrichaceae>Methanothrix>unclassified_Methanothrix | 212 | IPI63_02960 | 571700..572338 | 90426 | DUF4276-family-protein-[Methanothrix-sp.] |
| MBA3046425.1 | TOPRIM | GCA_013791785.1 | Methanobacteriota_archaeon | Archaea>Methanobacteriati>Methanobacteriota>unclassified_Methanobacteriota | 216 | FP824_09470 | 33012..33662 | 2026739 | RloB-domain-containing-protein-[Euryarchaeota-archaeon] |
| MBU4220745.1 | TOPRIM | GCA_018898145.1 | Methanobacteriota_archaeon | Archaea>Methanobacteriati>Methanobacteriota>unclassified_Methanobacteriota | 155 | KKA10_03840 | 4243..4710 | 2026739 | hypothetical-protein-[Euryarchaeota-archaeon] |
| OQA55745.1 | TOPRIM | GCA_002069575.1 | Euryarchaeota_archaeon_ADurb.Bin294 | Archaea>Methanobacteriati>Methanobacteriota>unclassified_Methanobacteriota | 133 | BWY45_02072 | 3938..4339 | 1852878 | hypothetical-protein-BWY45_02072-[Euryarchaeota-archaeon-ADurb.Bin294] |
| RLF71962.1 | TOPRIM | GCA_003663625.1 | Thermoplasmata_archaeon | Archaea>Methanobacteriati>Thermoplasmatota>Thermoplasmata>unclassified_Thermoplasmata | 163 | DRN40_01335 | 2071..2562 | 1906666 | MAG:-hypothetical-protein-DRN40_01335-[Thermoplasmata-archaeon] |
| MCY3892499.1 | TOPRIM | GCA_026707105.1 | Acidimicrobiaceae_bacterium | Bacteria>Bacillati>Actinomycetota>Acidimicrobiia>Acidimicrobiales>Acidimicrobiaceae>unclassified_Acidimicrobiaceae | 224 | OXF65_04290 | 2486..3160 | 2024894 | MAG:-hypothetical-protein-OXF65_04290-[Acidimicrobiaceae-bacterium] |
| MXW95269.1 | TOPRIM | GCA_009836165.1 | Acidimicrobiaceae_bacterium | Bacteria>Bacillati>Actinomycetota>Acidimicrobiia>Acidimicrobiales>Acidimicrobiaceae>unclassified_Acidimicrobiaceae | 211 | F4Z64_07155 | 1577..2212 | 2024894 | RloB-domain-containing-protein-[Acidimicrobiaceae-bacterium] |
| MYH43041.1 | TOPRIM | GCA_009843995.1 | Acidimicrobiaceae_bacterium | Bacteria>Bacillati>Actinomycetota>Acidimicrobiia>Acidimicrobiales>Acidimicrobiaceae>unclassified_Acidimicrobiaceae | 211 | F4131_04990 | 20354..20989 | 2024894 | RloB-domain-containing-protein-[Acidimicrobiaceae-bacterium] |
| MXW97853.1 | TOPRIM | GCA_009835735.1 | Acidimicrobiaceae_bacterium | Bacteria>Bacillati>Actinomycetota>Acidimicrobiia>Acidimicrobiales>Acidimicrobiaceae>unclassified_Acidimicrobiaceae | 207 | F4Z89_04470 | 11300..11923 | 2024894 | hypothetical-protein-[Acidimicrobiaceae-bacterium] |
| MBF0967810.1 | TOPRIM | GCA_015259445.1 | Actinomyces_sp. | Bacteria>Bacillati>Actinomycetota>Actinomycetes>Actinomycetales>Actinomycetaceae>Actinomyces>unclassified_Actinomyces | 207 | HXK08_00805 | 9292..9915 | 29317 | hypothetical-protein-[Actinomyces-sp.] |
| MBS6968673.1 | TOPRIM | GCA_018381115.1 | Actinomyces_sp. | Bacteria>Bacillati>Actinomycetota>Actinomycetes>Actinomycetales>Actinomycetaceae>Actinomyces>unclassified_Actinomyces | 204 | KH140_06970 | 7994..8608 | 29317 | hypothetical-protein-[Actinomyces-sp.] |
| OFQ22230.1 | TOPRIM | GCA_001813985.1 | Actinomyces_sp._HMSC062G12 | Bacteria>Bacillati>Actinomycetota>Actinomycetes>Actinomycetales>Actinomycetaceae>Actinomyces>unclassified_Actinomyces | 204 | HMPREF2946_04750 | 254483..255097 | 1739450 | hypothetical-protein-HMPREF2946_04750-[Actinomyces-sp.-HMSC062G12] |
| AOS47542.1 | TOPRIM | GCA_001746855.1 | Pauljensenia_hongkongensis | Bacteria>Bacillati>Actinomycetota>Actinomycetes>Actinomycetales>Actinomycetaceae>Pauljensenia | 214 | BH719_06535 | 1592747..1593391 | 178339 | hypothetical-protein-BH719_06535-[Pauljensenia-hongkongensis] |
| WP_009744025.1 | TOPRIM | GCF_001746855.1 | Pauljensenia_hongkongensis | Bacteria>Bacillati>Actinomycetota>Actinomycetes>Actinomycetales>Actinomycetaceae>Pauljensenia | 214 | BH719_RS06535 | 1592747..1593391 | 178339 | hypothetical-protein-[Pauljensenia-hongkongensis] |
| WP_081966325.1 | TOPRIM | GCF_000770965.1 | Bifidobacterium_pullorum | Bacteria>Bacillati>Actinomycetota>Actinomycetes>Bifidobacteriales>Bifidobacteriaceae>Bifidobacterium | 216 | OU28_RS09680 | 27878..28528 | 78448 | RloB-family-protein-[Bifidobacterium-pullorum] |
| WP_204389274.1 | TOPRIM | GCF_016900615.1 | Bifidobacterium_pullorum | Bacteria>Bacillati>Actinomycetota>Actinomycetes>Bifidobacteriales>Bifidobacteriaceae>Bifidobacterium | 216 | H6A14_RS02715 | 191377..192027 | 78448 | RloB-family-protein-[Bifidobacterium-pullorum] |
| PJM72969.1 | TOPRIM | GCA_002802875.1 | Bifidobacterium_primatium | Bacteria>Bacillati>Actinomycetota>Actinomycetes>Bifidobacteriales>Bifidobacteriaceae>Bifidobacterium | 215 | CS006_06865 | 166309..166956 | 2045438 | hypothetical-protein-CS006_06865-[Bifidobacterium-primatium] |
| WP_100511072.1 | TOPRIM | GCF_002802875.1 | Bifidobacterium_primatium | Bacteria>Bacillati>Actinomycetota>Actinomycetes>Bifidobacteriales>Bifidobacteriaceae>Bifidobacterium | 215 | CS006_RS06845 | 166309..166956 | 2045438 | RloB-family-protein-[Bifidobacterium-primatium] |
| WP_248824791.1 | TOPRIM | GCF_023242945.1 | Frankia_umida | Bacteria>Bacillati>Actinomycetota>Actinomycetes>Frankiales>Frankiaceae>Frankia | 205 | MXD59_RS12305 | 102827..103444 | 573489 | DUF4276-family-protein-[Frankia-sp.-Ag45/Mut15] |

| Protein-ID | Domain | Assembly-ID | Strain | Taxonomy | Length | Locus-Tag | Genomic-coordinates | TaxID | Fasta-defline |
|---|---|---|---|---|---|---|---|---|---|
| WP_242606587.1 | TOPRIM | GCF_900634745.1 | Protofrankia_symbiont_of_Coriaria_ruscifolia | Bacteria>Bacillati>Actinomycetota>Actinomycetes>Frankiales>Frankiaceae>Protofrankia>unclassified_Protofrankia | 175 | CV1CTNOD_RS11000 | 7231..7758 | 1306542 | RloB-family-protein-[Frankia-symbiont-of-Coriaria-ruscifolia] |
| WP_123459804.1 | TOPRIM | GCF_044743135.1 | Streptomyces | Bacteria>Bacillati>Actinomycetota>Actinomycetes>Kitasatosporales>Streptomycetaceae | 222 | EDD95_RS00805 | 188003..188671 | 1883 | DUF4276-family-protein-[Streptomyces-sp.-CEV-2-1] |
| WP_233477892.1 | TOPRIM | - | Streptomyces | Bacteria>Bacillati>Actinomycetota>Actinomycetes>Kitasatosporales>Streptomycetaceae | 185 | BJ962_RS25970 | 2063446..2064003 | 1883 | MULTISPECIES:-hypothetical-protein-[Streptomyces] |
| WP_229869911.1 | TOPRIM | GCF_014648815.1 | Streptomyces_flaveolus | Bacteria>Bacillati>Actinomycetota>Actinomycetes>Kitasatosporales>Streptomycetaceae>Streptomyces | 219 | IEZ05_RS29285 | 129907..130566 | 67297 | DUF4276-family-protein-[Streptomyces-flaveolus] |
| WP_043500389.1 | TOPRIM | GCF_000761215.1 | Streptomyces_glaucescens | Bacteria>Bacillati>Actinomycetota>Actinomycetes>Kitasatosporales>Streptomycetaceae>Streptomyces | 212 | SGLAU_RS10270 | 2378473..2379111 | 1907 | hypothetical-protein-[Streptomyces-glaucescens] |
| WP_077964583.1 | TOPRIM | GCF_009296025.1 | Streptomyces_tsukubensis | Bacteria>Bacillati>Actinomycetota>Actinomycetes>Kitasatosporales>Streptomycetaceae>Streptomyces | 212 | GBW32_RS05170 | 1160649..1161287 | 83656 | hypothetical-protein-[Streptomyces-tsukubensis] |
| WP_105973299.1 | TOPRIM | GCF_002954775.1 | Streptomyces_geranii | Bacteria>Bacillati>Actinomycetota>Actinomycetes>Kitasatosporales>Streptomycetaceae>Streptomyces | 211 | CW359_RS29875 | 51091..51726 | 2058923 | hypothetical-protein-[Streptomyces-geranii] |
| WP_245226720.1 | TOPRIM | - | Streptomyces_smyrnaeus | Bacteria>Bacillati>Actinomycetota>Actinomycetes>Kitasatosporales>Streptomycetaceae>Streptomyces | 195 | JW613_RS32965 | 12659..13246 | 1387713 | DUF4276-family-protein-[Streptomyces-smyrnaeus] |
| ALO95601.1 | TOPRIM | GCA_001447075.1 | Streptomyces_hygroscopicus_subsp._limoneus | Bacteria>Bacillati>Actinomycetota>Actinomycetes>Kitasatosporales>Streptomycetaceae>Streptomyces>Streptomyces_violaceusniger_group>Streptomyces_hygroscopicus | 210 | SHL15_4511 | 5050786..5051418 | 264445 | RloB-[Streptomyces-hygroscopicus-subsp.-limoneus] |
| WP_214971905.1 | TOPRIM | GCF_018614615.1 | Streptomyces_sp._ISL-100 | Bacteria>Bacillati>Actinomycetota>Actinomycetes>Kitasatosporales>Streptomycetaceae>Streptomyces>unclassified_Streptomyces | 222 | J7E89_RS10775 | 54169..54837 | 2819173 | DUF4276-family-protein-[Streptomyces-sp.-ISL-100] |
| WP_225252716.1 | TOPRIM | GCF_002879675.3 | Streptomyces_sp._SMS_SU21 | Bacteria>Bacillati>Actinomycetota>Actinomycetes>Kitasatosporales>Streptomycetaceae>Streptomyces>unclassified_Streptomyces | 219 | C1C94_RS24655 | 1680341..1681000 | 2069440 | DUF4276-family-protein-[Streptomyces-sp.-SMS_SU21] |
| WP_037951085.1 | TOPRIM | GCF_000590515.1 | Streptomyces_sp._PRh5 | Bacteria>Bacillati>Actinomycetota>Actinomycetes>Kitasatosporales>Streptomycetaceae>Streptomyces>unclassified_Streptomyces | 215 | Z951_RS11090 | 75911..76558 | 1158056 | hypothetical-protein-[Streptomyces-sp.-PRh5] |
| WP_215167461.1 | TOPRIM | GCF_018614415.1 | Streptomyces_sp._ISL-12 | Bacteria>Bacillati>Actinomycetota>Actinomycetes>Kitasatosporales>Streptomycetaceae>Streptomyces>unclassified_Streptomyces | 214 | J7I94_RS14405 | 80422..81066 | 2819177 | DUF4276-family-protein-[Streptomyces-sp.-ISL-12] |
| WP_086823276.1 | TOPRIM | GCF_002156145.1 | Streptomyces_sp._NRRL_B-24572 | Bacteria>Bacillati>Actinomycetota>Actinomycetes>Kitasatosporales>Streptomycetaceae>Streptomyces>unclassified_Streptomyces | 212 | CCN56_RS03175 | 27794..28432 | 1962156 | hypothetical-protein-[Streptomyces-sp.-NRRL-B-24572] |
| WP_235479566.1 | TOPRIM | GCF_021545335.1 | Streptomyces_sp._GQFP | Bacteria>Bacillati>Actinomycetota>Actinomycetes>Kitasatosporales>Streptomycetaceae>Streptomyces>unclassified_Streptomyces | 211 | LUX31_RS36420 | 8081996..8082631 | 2907545 | hypothetical-protein-[Streptomyces-sp.-GQFP] |
| WP_266686662.1 | TOPRIM | GCF_026342335.1 | Streptomyces_sp._NBC_00268 | Bacteria>Bacillati>Actinomycetota>Actinomycetes>Kitasatosporales>Streptomycetaceae>Streptomyces>unclassified_Streptomyces | 211 | OOK47_RS16775 | 3612072..3612707 | 2975695 | hypothetical-protein-[Streptomyces-sp.-NBC_00268] |
| WP_251075135.1 | TOPRIM | - | Streptomyces_sp._ISL-12 | Bacteria>Bacillati>Actinomycetota>Actinomycetes>Kitasatosporales>Streptomycetaceae>Streptomyces>unclassified_Streptomyces | 162 | J7I94_RS14455 | 92244..92732 | 2819177 | RloB-family-protein-[Streptomyces-sp.-ISL-12] |
| MBX3116433.1 | TOPRIM | GCA_019638875.1 | Cryobacterium_sp. | Bacteria>Bacillati>Actinomycetota>Actinomycetes>Micrococcales>Microbacteriaceae>Cryobacterium>unclassified_Cryobacterium | 195 | KF808_03695 | 33096..33683 | 1926290 | hypothetical-protein-[Cryobacterium-sp.] |
| WP_090804559.1 | TOPRIM | GCF_016862535.1 | Asanoa_ishikariensis | Bacteria>Bacillati>Actinomycetota>Actinomycetes>Micromonosporales>Micromonosporaceae>Asanoa | 279 | BLV39_RS40680 | 267514..268353 | 137265 | hypothetical-protein-[Asanoa-ishikariensis] |
| WP_088962574.1 | TOPRIM | GCF_900091515.1 | Micromonospora_purpureochromogenes | Bacteria>Bacillati>Actinomycetota>Actinomycetes>Micromonosporales>Micromonosporaceae>Micromonospora | 317 | GA0074696_RS20360 | 4407493..4408446 | 47872 | hypothetical-protein-[Micromonospora-purpureochromogenes] |
| WP_076473574.1 | TOPRIM | GCF_900156065.1 | Micromonospora_avicenniae | Bacteria>Bacillati>Actinomycetota>Actinomycetes>Micromonosporales>Micromonosporaceae>Micromonospora | 283 | BXA10_RS26495 | 28410..29261 | 1198245 | hypothetical-protein-[Micromonospora-avicenniae] |
| SCF44511.1 | TOPRIM | GCA_900091555.1 | Micromonospora_mirobrigensis | Bacteria>Bacillati>Actinomycetota>Actinomycetes>Micromonosporales>Micromonosporaceae>Micromonospora | 279 | GA0070564_110102 | 118752..119591 | 262898 | hypothetical-protein-GA0070564_110102-[Micromonospora-mirobrigensis] |
| WP_244254910.1 | TOPRIM | GCF_017599015.1 | Micromonospora_tulbaghiae | Bacteria>Bacillati>Actinomycetota>Actinomycetes>Micromonosporales>Micromonosporaceae>Micromonospora | 268 | J5U46_RS17355 | 28223..29029 | 479978 | hypothetical-protein-[Micromonospora-tulbaghiae] |
| WP_179779351.1 | TOPRIM | GCF_013410645.1 | Micromonospora_jinlongensis | Bacteria>Bacillati>Actinomycetota>Actinomycetes>Micromonosporales>Micromonosporaceae>Micromonospora | 214 | HNR22_RS05335 | 1261236..1261880 | 1287877 | hypothetical-protein-[Micromonospora-jinlongensis] |
| WP_123809687.1 | TOPRIM | GCF_003176745.1 | Micromonospora_globispora | Bacteria>Bacillati>Actinomycetota>Actinomycetes>Micromonosporales>Micromonosporaceae>Micromonospora | 204 | DLE60_RS16070 | 6780..7394 | 1450148 | hypothetical-protein-[Micromonospora-globispora] |
| WP_203807951.1 | TOPRIM | GCF_016862435.1 | Paractinoplanes_tereljensis | Bacteria>Bacillati>Actinomycetota>Actinomycetes>Micromonosporales>Micromonosporaceae>Paractinoplanes | 281 | Ate02nite_RS20660 | 72712..73557 | 571912 | hypothetical-protein-[Actinoplanes-tereljensis] |
| WP_270679268.1 | TOPRIM | GCF_027706745.1 | Nocardiopsis_suaedae | Bacteria>Bacillati>Actinomycetota>Actinomycetes>Streptosporangiales>Nocardiopsidaceae>Nocardiopsis | 227 | O4U47_RS19130 | 27620..28303 | 3018444 | hypothetical-protein-[Nocardiopsis-sp.-LSu2-4] |
| WP_067482822.1 | TOPRIM | GCF_001552635.1 | Actinomadura_hibisca | Bacteria>Bacillati>Actinomycetota>Actinomycetes>Streptosporangiales>Thermomonosporaceae>Actinomadura | 324 | AH6_RS22060 | 114615..115589 | 68565 | hypothetical-protein-[Actinomadura-hibisca] |
| WP_182841315.1 | TOPRIM | GCF_039528335.1 | Actinomadura_namibiensis | Bacteria>Bacillati>Actinomycetota>Actinomycetes>Streptosporangiales>Thermomonosporaceae>Actinomadura | 324 | HNR61_RS01550 | 335809..336783 | 182080 | hypothetical-protein-[Actinomadura-namibiensis] |
| WP_067795479.1 | TOPRIM | GCF_001552155.1 | Actinomadura_formosensis | Bacteria>Bacillati>Actinomycetota>Actinomycetes>Streptosporangiales>Thermomonosporaceae>Actinomadura | 323 | AF1_RS09870 | 43669..44640 | 60706 | hypothetical-protein-[Actinomadura-formosensis] |
| TMR42116.1 | TOPRIM | GCA_005889745.1 | Actinomadura_geliboluensis | Bacteria>Bacillati>Actinomycetota>Actinomycetes>Streptosporangiales>Thermomonosporaceae>Actinomadura | 220 | ETD96_02085 | 28615..29277 | 882440 | RloB-domain-containing-protein-[Actinomadura-geliboluensis] |
| WP_225960999.1 | TOPRIM | - | Actinomadura_algeriensis | Bacteria>Bacillati>Actinomycetota>Actinomycetes>Streptosporangiales>Thermomonosporaceae>Actinomadura | 207 | H4W34_RS03565 | 876461..877084 | 1679523 | RloB-family-protein-[Actinomadura-algeriensis] |
| WP_242884438.1 | TOPRIM | GCF_022751195.1 | Actinomadura_litoris | Bacteria>Bacillati>Actinomycetota>Actinomycetes>Streptosporangiales>Thermomonosporaceae>Actinomadura | 205 | K2F48_RS09340 | 531784..532401 | 2678616 | RloB-family-protein-[Actinomadura-litoris] |
| GFP42416.1 | TOPRIM | GCA_013282075.1 | Candidatus_Hakubella_thermalkaliphila | Bacteria>Bacillati>Actinomycetota>Actinomycetota_incertae_sedis>Candidatus_Hakubellales>Candidatus_Hakubellaceae>Candidatus_Hakubella | 152 | HKBW3C_01542 | 7603..8061 | 2754717 | hypothetical-protein-HKBW3C_01542-[Candidatus-Hakubanella-thermoalkaliphilus] |
| WP_176233420.1 | TOPRIM | GCF_013281955.1 | Candidatus_Hakubella_thermalkaliphila | Bacteria>Bacillati>Actinomycetota>Actinomycetota_incertae_sedis>Candidatus_Hakubellales>Candidatus_Hakubellaceae>Candidatus_Hakubella | 152 | HW440_RS05445 | 7887..8345 | 2754717 | hypothetical-protein-[Candidatus-Hakubanella-thermoalkaliphilus] |
| GIU94955.1 | TOPRIM | GCA_026002995.1 | Gaiellaceae_bacterium | Bacteria>Bacillati>Actinomycetota>Thermoleophilia>Gaiellales>Gaiellaceae>unclassified_Gaiellaceae | 199 | KatS3mg012_1412 | 1400388..1400987 | 2838783 | MAG:-hypothetical-protein-KatS3mg012_1412-[Gaiellaceae-bacterium] |
| MDA8117680.1 | TOPRIM | GCA_028064745.1 | Actinomycetota_bacterium | Bacteria>Bacillati>Actinomycetota>unclassified_Actinomycetota | 193 | M0000_09965 | 5428..6009 | 2900548 | MAG:-DUF4276-family-protein-[Actinomycetota-bacterium] |
| NLH98009.1 | TOPRIM | GCA_012517255.1 | Chthonomonadales_bacterium | Bacteria>Bacillati>Armatimonadota>Chthonomonadia>Chthonomonadales>unclassified_Chthonomonadales | 202 | GX446_00795 | 19999..20607 | 2282151 | DUF4276-family-protein-[Chthonomonadales-bacterium] |
| NCO91845.1 | TOPRIM | GCA_009993145.1 | Armatimonadota_bacterium | Bacteria>Bacillati>Armatimonadota>unclassified_Armatimonadota | 208 | GW880_11040 | 78..704 | 2033014 | MAG:-hypothetical-protein-GW880_11040-[Armatimonadetes-bacterium] |
| OIO95802.1 | TOPRIM | GCA_001871675.1 | Armatimonadetes_bacterium_CG2_30_66_41 | Bacteria>Bacillati>Armatimonadota>unclassified_Armatimonadota | 208 | AUJ96_25910 | 10470..11096 | 1805028 | hypothetical-protein-AUJ96_25910-[Armatimonadetes-bacterium-CG2_30_66_41] |
| PIU89757.1 | TOPRIM | GCA_002779825.1 | Armatimonadetes_bacterium_CG06_land_8_20_14_3_00_66_21 | Bacteria>Bacillati>Armatimonadota>unclassified_Armatimonadota | 208 | COS65_27515 | 732..1358 | 1973915 | hypothetical-protein-COS65_27515-[Armatimonadetes-bacterium-CG06_land_8_20_14_3_00_66_21] |
| PIZ50108.1 | TOPRIM | GCA_002791775.1 | Armatimonadetes_bacterium_CG_4_10_14_0_8_um_filter_66_14 | Bacteria>Bacillati>Armatimonadota>unclassified_Armatimonadota | 208 | COY42_02300 | 752..1378 | 1973908 | hypothetical-protein-COY42_02300-[Armatimonadetes-bacterium-CG_4_10_14_0_8_um_filter_66_14] |
| NDK16275.1 | TOPRIM | GCA_010119115.1 | Armatimonadota_bacterium | Bacteria>Bacillati>Armatimonadota>unclassified_Armatimonadota | 205 | GW911_29965 | 1..614 | 2033014 | hypothetical-protein-[Armatimonadota-bacterium] |
| PJB73374.1 | TOPRIM | GCA_002789735.1 | Armatimonadetes_bacterium_CG_4_9_14_3_um_filter_66_14 | Bacteria>Bacillati>Armatimonadota>unclassified_Armatimonadota | 200 | CO096_06120 | 37183..37782 | 1973914 | hypothetical-protein-CO096_06120,-partial-[Armatimonadetes-bacterium-CG_4_9_14_3_um_filter_66_14] |
| WP_163530733.1 | TOPRIM | GCF_902726645.1 | Halobacillus_ihumii | Bacteria>Bacillati>Bacillota>Bacilli>Bacillales>Bacillaceae>Halobacillus | 307 | G6R08_RS20415 | 4078892..4079815 | 2686092 | toprim-domain-containing-protein-[Halobacillus-ihumii] |
| WP_028784623.1 | TOPRIM | GCF_000496835.1 | Thalassobacillus_devorans | Bacteria>Bacillati>Bacillota>Bacilli>Bacillales>Bacillaceae>Thalassobacillus | 306 | P855_RS0116620 | 83879..84799 | 279813 | toprim-domain-containing-protein-[Thalassobacillus-devorans] |
| AQQ52247.1 | TOPRIM | GCA_001999905.1 | Planococcus_lenghuensis | Bacteria>Bacillati>Bacillota>Bacilli>Bacillales>Caryophanaceae>Planococcus | 253 | BOX71_03385 | 630725..631486 | 2213202 | hypothetical-protein-BOX71_03385-[Planococcus-lenghuensis] |
| WP_241559443.1 | TOPRIM | - | Lacticaseibacillus_hulanensis | Bacteria>Bacillati>Bacillota>Bacilli>Lactobacillales>Lactobacillaceae>Lacticaseibacillus | 271 | EKN00_RS02435 | 68327..69142 | 2493111 | DUF2399-domain-containing-protein-[Lacticaseibacillus-hulanensis] |
| WP_179394504.1 | TOPRIM | GCF_013407595.1 | Lacticaseibacillus_absianus | Bacteria>Bacillati>Bacillota>Bacilli>Lactobacillales>Lactobacillaceae>Lacticaseibacillus | 268 | HII33_RS04060 | 944956..945762 | 2729623 | DUF2399-domain-containing-protein-[Lacticaseibacillus-absianus] |
| WP_087716720.1 | TOPRIM | GCF_002179515.1 | Levilactobacillus_brevis | Bacteria>Bacillati>Bacillota>Bacilli>Lactobacillales>Lactobacillaceae>Levilactobacillus | 267 | CA846_RS04715 | 153810..154613 | 1580 | DUF2399-domain-containing-protein-[Levilactobacillus-brevis] |
| WP_251420884.1 | TOPRIM | GCF_042849115.1 | Lactococcus | Bacteria>Bacillati>Bacillota>Bacilli>Lactobacillales>Streptococcaceae | 186 | NFR64_RS02410 | 37110..37670 | 1357 | MULTISPECIES:-RloB-family-protein-[Lactococcus] |
| WP_058211092.1 | TOPRIM | GCF_001456725.1 | Lactococcus_lactis | Bacteria>Bacillati>Bacillota>Bacilli>Lactobacillales>Streptococcaceae>Lactococcus | 179 | KF134_RS08790 | 319144..319683 | 1358 | RloB-domain-containing-protein-[Lactococcus-lactis] |
| WP_191672701.1 | TOPRIM | GCF_014830225.1 | Lactococcus_petauri | Bacteria>Bacillati>Bacillota>Bacilli>Lactobacillales>Streptococcaceae>Lactococcus | 179 | DXX96_RS03205 | 89039..89578 | 1940789 | RloB-family-protein-[Lactococcus-petauri] |
| WP_164950927.1 | TOPRIM | GCF_011065955.2 | Clostridium_sp._FP2 | Bacteria>Bacillati>Bacillota>Clostridia>Eubacteriales>Clostridiaceae>Clostridium>unclassified_Clostridium | 270 | G9F71_RS16300 | 3392170..3392982 | 2724481 | hypothetical-protein-[Clostridium-sp.-FP2] |
| WP_035381187.1 | TOPRIM | GCF_000601455.1 | Fervidicella_metallireducens | Bacteria>Bacillati>Bacillota>Clostridia>Eubacteriales>Clostridiaceae>Fervidicella | 243 | Q428_RS11725 | 11729..12460 | 655338 | hypothetical-protein-[Fervidicella-metallireducens] |
| KGP76500.1 | TOPRIM | GCA_000770645.1 | Desulfosporosinus_sp._Tol-M | Bacteria>Bacillati>Bacillota>Clostridia>Eubacteriales>Desulfitobacteriaceae>Desulfosporosinus>unclassified_Desulfosporosinus | 221 | JT05_04465 | 16100..16765 | 1536651 | hypothetical-protein-JT05_04465-[Desulfosporosinus-sp.-Tol-M] |
| WP_091728836.1 | TOPRIM | GCF_900107485.1 | Proteiniborus | Bacteria>Bacillati>Bacillota>Clostridia>Eubacteriales>Eubacteriales_incertae_sedis | 243 | BLV37_RS06300 | 19469..20200 | 415014 | DUF3226-domain-containing-protein-[Proteiniborus-ethanoligenes] |
| WP_073163774.1 | TOPRIM | GCF_900129285.1 | Desulfofundulus_australicus | Bacteria>Bacillati>Bacillota>Clostridia>Eubacteriales>Peptococcaceae>Desulfofundulus | 206 | BUB44_RS05255 | 44672..45292 | 1566 | hypothetical-protein-[Desulfofundulus-australicus] |
| SHE91832.1 | TOPRIM | GCA_900129285.1 | Desulfofundulus_australicus_DSM_11792 | Bacteria>Bacillati>Bacillota>Clostridia>Eubacteriales>Peptococcaceae>Desulfofundulus>Desulfofundulus_australicus | 206 | SAMN02745218_01065 | 44672..45292 | 1121425 | hypothetical-protein-SAMN02745218_01065-[Desulfofundulus-australicus-DSM-11792] |
| WP_138262158.1 | TOPRIM | GCF_015550395.1 | [Clostridium]_hylemonae | Bacteria>Bacillati>Bacillota>Clostridia>Lachnospirales>Lachnospiraceae>Lachnoclostridium | 232 | I2C33_RS01890 | 48587..49285 | 89153 | RloB-family-protein-[[Clostridium] |
| NBJ03624.1 | TOPRIM | GCA_009911175.1 | Lachnospiraceae_bacterium | Bacteria>Bacillati>Bacillota>Clostridia>Lachnospirales>Lachnospiraceae>unclassified_Lachnospiraceae | 249 | D3Z62_26945 | 5317..6066 | 1898203 | hypothetical-protein-[Lachnospiraceae-bacterium] |
| MBR4608211.1 | TOPRIM | GCA_017536085.1 | Lachnospiraceae_bacterium | Bacteria>Bacillati>Bacillota>Clostridia>Lachnospirales>Lachnospiraceae>unclassified_Lachnospiraceae | 224 | IKO41_18540 | 26191..26865 | 1898203 | DUF4276-family-protein-[Lachnospiraceae-bacterium] |
| WP_255228678.1 | TOPRIM | GCF_024363565.1 | Lutispora_saccharofermentans | Bacteria>Bacillati>Bacillota>Clostridia>Lutisporales>Lutisporaceae>Lutispora | 225 | LJD61_RS16665 | 80141..80818 | 3024236 | DUF4276-family-protein-[Lutispora-saccharofermentans] |
| WP_071520602.1 | TOPRIM | GCF_001874065.1 | Moorella_thermoacetica | Bacteria>Bacillati>Bacillota>Clostridia>Moorellales>Moorellaceae>Moorella | 205 | MOOR_RS04465 | 137905..138522 | 1525 | hypothetical-protein-[Moorella-thermoacetica] |
| WP_099191958.1 | TOPRIM | GCF_002724055.1 | Tepidibacter_mesophilus | Bacteria>Bacillati>Bacillota>Clostridia>Peptostreptococcales>Peptostreptococcaceae>Tepidibacter | 246 | Tmeso_RS13180 | 42381..43121 | 655607 | hypothetical-protein-[Tepidibacter-mesophilus] |
| MBP3656731.1 | TOPRIM | GCA_017938795.1 | Clostridia_bacterium | Bacteria>Bacillati>Bacillota>Clostridia>unclassified_Clostridia | 214 | J6K32_08555 | 21687..22331 | 2044939 | DUF4276-family-protein-[Clostridia-bacterium] |
| MCR4419671.1 | TOPRIM | GCA_024653205.1 | Clostridia_bacterium | Bacteria>Bacillati>Bacillota>Clostridia>unclassified_Clostridia | 210 | NUV99_05985 | 81131..81763 | 2044939 | hypothetical-protein-[Clostridia-bacterium] |
| WP_021772224.1 | TOPRIM | GCF_000469545.1 | Mitsuokella_sp._oral_taxon_131 | Bacteria>Bacillati>Bacillota>Negativicutes>Selenomonadales>Selenomonadaceae>Mitsuokella>unclassified_Mitsuokella | 388 | HMPREF1985_RS08705 | 184100..185266 | 1321780 | hypothetical-protein-[Mitsuokella-sp.-oral-taxon-131] |
| NCB76906.1 | TOPRIM | GCA_009929485.1 | Negativicutes_bacterium | Bacteria>Bacillati>Bacillota>Negativicutes>unclassified_Negativicutes | 227 | EOM42_07420 | 8063..8746 | 2499140 | DUF4276-family-protein-[Negativicutes-bacterium] |
| WP_072743896.1 | TOPRIM | GCF_900130025.1 | Sporanaerobacter_acetigenes | Bacteria>Bacillati>Bacillota>Tissierellia>Tissierellales>Sporanaerobacteraceae>Sporanaerobacter | 243 | BUA21_RS05955 | 81541..82272 | 165813 | hypothetical-protein-[Sporanaerobacter-acetigenes] |
| WP_054870027.1 | TOPRIM | GCF_001316435.1 | Caloranaerobacter_sp._TR13 | Bacteria>Bacillati>Bacillota>Tissierellia>Tissierellales>Thermohalobacteraceae>Caloranaerobacter>unclassified_Caloranaerobacter | 243 | TR13x_RS01060 | 231272..232003 | 1302151 | hypothetical-protein-[Caloranaerobacter-sp.-TR13] |
| MBU4541891.1 | TOPRIM | GCA_018903255.1 | Bacillota_bacterium | Bacteria>Bacillati>Bacillota>unclassified_Bacillota | 252 | KJ774_11800 | 609..1367 | 1879010 | DUF3226-domain-containing-protein-[Bacillota-bacterium] |
| MBT9164709.1 | TOPRIM | GCA_018725485.1 | Candidatus_Lithacetigena_glycinireducens | Bacteria>Bacillati>Candidatus_Lithacetigenota>Candidatus_Lithacetigena | 152 | DDT23_00713 | 3881..4339 | 2986055 | hypothetical-protein-[candidate-division-WS2-bacterium] |
| MBQ6502302.1 | TOPRIM | GCA_017432685.1 | Flexilinea_sp. | Bacteria>Bacillati>Chloroflexota>Anaerolineae>Anaerolineales>Anaerolineaceae>Flexilinea>unclassified_Flexilinea | 208 | IJI57_00135 | 13873..14499 | 2017390 | RloB-domain-containing-protein-[Flexilinea-sp.] |
| WP_062416897.1 | TOPRIM | GCF_001050255.2 | Levilinea_saccharolytica | Bacteria>Bacillati>Chloroflexota>Anaerolineae>Anaerolineales>Anaerolineaceae>Levilinea | 220 | LSAC_RS05150 | 237043..237705 | 229921 | DUF4276-family-protein-[Levilinea-saccharolytica] |
| WP_062419289.1 | TOPRIM | GCF_001050255.2 | Levilinea_saccharolytica | Bacteria>Bacillati>Chloroflexota>Anaerolineae>Anaerolineales>Anaerolineaceae>Levilinea | 208 | LSAC_RS18185 | 3296895..3297521 | 229921 | RloB-family-protein-[Levilinea-saccharolytica] |
| MBN1317035.1 | TOPRIM | GCA_016927355.1 | Anaerolineales_bacterium | Bacteria>Bacillati>Chloroflexota>Anaerolineae>Anaerolineales>unclassified_Anaerolineales | 226 | JXA42_16255 | 34080..34760 | 2073117 | DUF4276-family-protein-[Anaerolineales-bacterium] |
| MCL4868347.1 | TOPRIM | GCA_023384415.1 | Anaerolineae_bacterium | Bacteria>Bacillati>Chloroflexota>Anaerolineae>unclassified_Anaerolineae | 205 | KJ063_05215 | 195188..195805 | 2052143 | RloB-domain-containing-protein-[Anaerolineae-bacterium] |
| MBN2007578.1 | TOPRIM | GCA_016931835.1 | Anaerolineae_bacterium | Bacteria>Bacillati>Chloroflexota>Anaerolineae>unclassified_Anaerolineae | 193 | JXA21_29805 | 7225..7806 | 2052143 | hypothetical-protein-[Anaerolineae-bacterium] |
| MCR4407532.1 | TOPRIM | GCA_024653325.1 | Anaerolineae_bacterium | Bacteria>Bacillati>Chloroflexota>Anaerolineae>unclassified_Anaerolineae | 186 | NUW24_11535 | 6662..7222 | 2052143 | DUF4276-family-protein-[Anaerolineae-bacterium] |
| MCX7854726.1 | TOPRIM | GCA_026417235.1 | Anaerolineae_bacterium | Bacteria>Bacillati>Chloroflexota>Anaerolineae>unclassified_Anaerolineae | 143 | N2556_01895 | 7683..8114 | 2052143 | MAG:-DUF4276-family-protein-[Anaerolineae-bacterium] |
| MCZ7568050.1 | TOPRIM | GCA_027493995.1 | Ardenticatenaceae_bacterium | Bacteria>Bacillati>Chloroflexota>Ardenticatenia>Ardenticatenales>Ardenticatenaceae>unclassified_Ardenticatenaceae | 109 | M5U01_05525 | 1317129..1317458 | 2614247 | MAG:-hypothetical-protein-M5U01_05525-[Ardenticatenaceae-bacterium] |
| MCB0025413.1 | TOPRIM | GCA_020433165.1 | Caldilinea_sp. | Bacteria>Bacillati>Chloroflexota>Caldilineae>Caldilineales>Caldilineaceae>Caldilinea>unclassified_Caldilinea | 196 | KDD91_20415 | 454..1044 | 2293560 | DUF4276-family-protein-[Caldilinea-sp.] |
| MXX27765.1 | TOPRIM | GCA_009835815.1 | Caldilineaceae_bacterium_SB0668_bin_21 | Bacteria>Bacillati>Chloroflexota>Caldilineae>Caldilineales>Caldilineaceae>unclassified_Caldilineaceae | 216 | F4Z82_20250 | 24643..25293 | 2605264 | hypothetical-protein-[Caldilineaceae-bacterium-SB0668_bin_21] |
| MCY4079347.1 | TOPRIM | GCA_026707715.1 | Caldilineaceae_bacterium | Bacteria>Bacillati>Chloroflexota>Caldilineae>Caldilineales>Caldilineaceae>unclassified_Caldilineaceae | 214 | OXF54_03835 | 15794..16438 | 2614250 | MAG:-hypothetical-protein-OXF54_03835-[Caldilineaceae-bacterium] |
| MCY4117636.1 | TOPRIM | GCA_026708105.1 | Caldilineaceae_bacterium | Bacteria>Bacillati>Chloroflexota>Caldilineae>Caldilineales>Caldilineaceae>unclassified_Caldilineaceae | 214 | OXF55_12185 | 33834..34478 | 2614250 | MAG:-hypothetical-protein-OXF55_12185-[Caldilineaceae-bacterium] |
| MDE0200337.1 | TOPRIM | GCA_028819675.1 | Caldilineaceae_bacterium | Bacteria>Bacillati>Chloroflexota>Caldilineae>Caldilineales>Caldilineaceae>unclassified_Caldilineaceae | 214 | OXK78_19180 | 4408..5052 | 2614250 | MAG:-hypothetical-protein-OXK78_19180-[Caldilineaceae-bacterium] |
| MDE0431260.1 | TOPRIM | GCA_028821555.1 | Caldilineaceae_bacterium | Bacteria>Bacillati>Chloroflexota>Caldilineae>Caldilineales>Caldilineaceae>unclassified_Caldilineaceae | 214 | OXH98_15970 | 80319..80963 | 2614250 | MAG:-hypothetical-protein-OXH98_15970-[Caldilineaceae-bacterium] |
| MDE0464747.1 | TOPRIM | GCA_028821465.1 | Caldilineaceae_bacterium | Bacteria>Bacillati>Chloroflexota>Caldilineae>Caldilineales>Caldilineaceae>unclassified_Caldilineaceae | 214 | OXH93_20180 | 5534..6178 | 2614250 | MAG:-hypothetical-protein-OXH93_20180-[Caldilineaceae-bacterium] |
| MXX28116.1 | TOPRIM | GCA_009835815.1 | Caldilineaceae_bacterium_SB0668_bin_21 | Bacteria>Bacillati>Chloroflexota>Caldilineae>Caldilineales>Caldilineaceae>unclassified_Caldilineaceae | 214 | F4Z82_22080 | 3446..4090 | 2605264 | hypothetical-protein-[Caldilineaceae-bacterium-SB0668_bin_21] |
| MDE0631426.1 | TOPRIM | GCA_028824195.1 | Caldilineaceae_bacterium | Bacteria>Bacillati>Chloroflexota>Caldilineae>Caldilineales>Caldilineaceae>unclassified_Caldilineaceae | 212 | OXH73_07975 | 22584..23222 | 2614250 | MAG:-DUF4276-family-protein-[Caldilineaceae-bacterium] |
| MCB9118709.1 | TOPRIM | GCA_020634765.1 | Caldilineaceae_bacterium | Bacteria>Bacillati>Chloroflexota>Caldilineae>Caldilineales>Caldilineaceae>unclassified_Caldilineaceae | 196 | H6640_03150 | 17621..18211 | 2614250 | DUF4276-family-protein-[Caldilineaceae-bacterium] |
| MCB9124680.1 | TOPRIM | GCA_020634715.1 | Caldilineaceae_bacterium | Bacteria>Bacillati>Chloroflexota>Caldilineae>Caldilineales>Caldilineaceae>unclassified_Caldilineaceae | 196 | H6643_09240 | 35860..36450 | 2614250 | MAG:-DUF4276-family-protein-[Caldilineaceae-bacterium] |
| MBK9096488.1 | TOPRIM | GCA_016719785.1 | Candidatus_Amarolinea_dominans | Bacteria>Bacillati>Chloroflexota>Caldilineae>Caldilineales>Candidatus_Amarolineaceae>Candidatus_Amarolinea | 250 | IPM84_27770 | 243010..243762 | 3140696 | hypothetical-protein-[Anaerolineae-bacterium] |
| MBK6429946.1 | TOPRIM | GCA_016704685.1 | Candidatus_Amarolinea_dominans | Bacteria>Bacillati>Chloroflexota>Caldilineae>Caldilineales>Candidatus_Amarolineaceae>Candidatus_Amarolinea | 210 | IPF85_04085 | 213897..214529 | 3140696 | hypothetical-protein-[Anaerolineae-bacterium] |
| MBK9093187.1 | TOPRIM | GCA_016719785.1 | Candidatus_Amarolinea_dominans | Bacteria>Bacillati>Chloroflexota>Caldilineae>Caldilineales>Candidatus_Amarolineaceae>Candidatus_Amarolinea | 210 | IPM84_10460 | 141622..142254 | 3140696 | hypothetical-protein-[Anaerolineae-bacterium] |
| WP_129673428.1 | TOPRIM | GCF_004138165.1 | Candidatus_Chloroploca_sp._Khr17 | Bacteria>Bacillati>Chloroflexota>Chloroflexia>Chloroflexales>Chloroflexineae>Oscillochloridaceae>Candidatus_Chloroploca>unclassified_Candidatus_Chloroploca | 186 | EKD05_RS13140 | 8408..8968 | 2496869 | DUF4276-family-protein-[Candidatus-Chloroploca-sp.-Khr17] |
| WP_088570294.1 | TOPRIM | GCF_900187885.1 | Thermoflexus_hugenholtzii | Bacteria>Bacillati>Chloroflexota>Thermoflexia>Thermoflexales>Thermoflexaceae>Thermoflexus | 219 | CFB18_RS02795 | 21111..21770 | 1495650 | DUF4276-family-protein-[Thermoflexus-hugenholtzii] |
| MCE7937789.1 | TOPRIM | GCA_021404695.1 | Chloroflexi_bacterium_CFX6 | Bacteria>Bacillati>Chloroflexota>unclassified_Chloroflexota | 273 | DYG90_04150 | 3251..4072 | 2293710 | DUF4276-family-protein-[Chloroflexi-bacterium-CFX6] |
| OGN72035.1 | TOPRIM | GCA_001796335.1 | Chloroflexi_bacterium_GWB2_49_20 | Bacteria>Bacillati>Chloroflexota>unclassified_Chloroflexota | 231 | A2X25_12870 | 145214..145909 | 1797612 | hypothetical-protein-A2X25_12870-[Chloroflexi-bacterium-GWB2_49_20] |
| MBI4327840.1 | TOPRIM | GCA_016210465.1 | Chloroflexota_bacterium | Bacteria>Bacillati>Chloroflexota>unclassified_Chloroflexota | 210 | HY674_21630 | 2100..2732 | 2026724 | hypothetical-protein-[Chloroflexota-bacterium] |
| MBM3127991.1 | TOPRIM | GCA_016865935.1 | Chloroflexota_bacterium | Bacteria>Bacillati>Chloroflexota>unclassified_Chloroflexota | 209 | FJ009_05065 | 19361..19990 | 2026724 | hypothetical-protein-[Chloroflexota-bacterium] |
| MYC29509.1 | TOPRIM | GCA_009840135.1 | Chloroflexota_bacterium | Bacteria>Bacillati>Chloroflexota>unclassified_Chloroflexota | 202 | F4X65_05400 | 5756..6364 | 2026724 | hypothetical-protein-[Chloroflexota-bacterium] |
| MYD49642.1 | TOPRIM | GCA_009841575.1 | Chloroflexota_bacterium | Bacteria>Bacillati>Chloroflexota>unclassified_Chloroflexota | 202 | F4W95_14350 | 8800..9408 | 2026724 | hypothetical-protein-[Chloroflexota-bacterium] |
| MBI2863488.1 | TOPRIM | GCA_016190625.1 | Chloroflexota_bacterium | Bacteria>Bacillati>Chloroflexota>unclassified_Chloroflexota | 195 | HYX94_02880 | 6274..6861 | 2026724 | hypothetical-protein-[Chloroflexota-bacterium] |
| MXZ89878.1 | TOPRIM | GCA_009837355.1 | Chloroflexota_bacterium | Bacteria>Bacillati>Chloroflexota>unclassified_Chloroflexota | 138 | F4Z05_00145 | 21675..22091 | 2026724 | hypothetical-protein-[Chloroflexota-bacterium] |
| PZV08801.1 | TOPRIM | GCA_003249105.1 | Leptolyngbya_sp. | Bacteria>Bacillati>Cyanobacteriota/Melainabacteria_group>Cyanobacteriota>Cyanophyceae>Leptolyngbyales>Leptolyngbyaceae>Leptolyngbya_group>Leptolyngbya>unclassified_Leptolyngbya | 207 | DCF32_03935 | 19363..19986 | 47254 | hypothetical-protein-DCF32_03935-[Leptolyngbya-sp.] |
| WP_127084954.1 | TOPRIM | GCF_003991905.1 | Dulcicalothrix_desertica | Bacteria>Bacillati>Cyanobacteriota/Melainabacteria_group>Cyanobacteriota>Cyanophyceae>Nostocales>Calotrichaceae>Dulcicalothrix | 215 | DSM106972_RS34640 | 64960..65607 | 32056 | DUF4276-family-protein-[Dulcicalothrix-desertica] |
| MBR8836044.1 | TOPRIM | GCA_016903675.1 | Stigonema_ocellatum_SAG_48.90_=_DSM_106950 | Bacteria>Bacillati>Cyanobacteriota/Melainabacteria_group>Cyanobacteriota>Cyanophyceae>Nostocales>Stigonemataceae>Stigonema>Stigonema_ocellatum | 203 | DSM106950_18975 | 43967..44578 | 221294 | DUF4276-family-protein-[Stigonema-ocellatum-SAG-48.90-=-DSM-106950] |
| MCA6584620.1 | TOPRIM | GCA_020404925.1 | Pseudanabaena_sp._M34BS1SP1A06MG | Bacteria>Bacillati>Cyanobacteriota/Melainabacteria_group>Cyanobacteriota>Cyanophyceae>Pseudanabaenales>Pseudanabaenaceae>Pseudanabaena>unclassified_Pseudanabaena | 211 | IM536_21890 | 14274..14909 | 2771240 | DUF4276-family-protein-[Pseudanabaena-sp.-M34BS1SP1A06MG] |
| MCA6593930.1 | TOPRIM | GCA_020404945.1 | Pseudanabaena_sp._M38BS1SP1A06MG | Bacteria>Bacillati>Cyanobacteriota/Melainabacteria_group>Cyanobacteriota>Cyanophyceae>Pseudanabaenales>Pseudanabaenaceae>Pseudanabaena>unclassified_Pseudanabaena | 211 | IM535_17910 | 12113..12748 | 2771238 | DUF4276-family-protein-[Pseudanabaena-sp.-M38BS1SP1A06MG] |
| MCA6597706.1 | TOPRIM | GCA_020405005.1 | Pseudanabaena_sp._M046S1SP1A06QC | Bacteria>Bacillati>Cyanobacteriota/Melainabacteria_group>Cyanobacteriota>Cyanophyceae>Pseudanabaenales>Pseudanabaenaceae>Pseudanabaena>unclassified_Pseudanabaena | 211 | IM539_15750 | 14266..14901 | 2771236 | hypothetical-protein-[Pseudanabaena-sp.-M046S1SP1A06QC] |
| MCF8140151.1 | TOPRIM | GCA_021737825.1 | Cyanobium_usitatum_Tobar12.5m-G36 | Bacteria>Bacillati>Cyanobacteriota/Melainabacteria_group>Cyanobacteriota>Cyanophyceae>Synechococcales>Prochlorococcaceae>Cyanobium>Cyanobium_usitatum | 188 | K9J75_03465 | 35381..35947 | 2874848 | DUF4276-family-protein-[Cyanobium-usitatum-Tobar12.5m-G36] |

| Protein-ID | Domain | Assembly-ID | Strain | Taxonomy | Length | Locus-Tag | Genomic-coordinates | TaxID | Fasta-defline |
|---|---|---|---|---|---|---|---|---|---|
| MCX6842951.1 | TOPRIM | GCA_026397595.1 | candidate_division_WOR-3_bacterium | Bacteria>Bacteria_incertae_sedis>Bacteria_candidate_phyla>candidate_division_WOR-3 | 184 | NTX53_11790 | 8004..8558 | 2052148 | MAG:-hypothetical-protein-NTX53_11790-[candidate-division-WOR-3-bacterium] |
| MCG2812993.1 | TOPRIM | GCA_022013915.1 | Candidatus_Aminicenantes_bacterium | Bacteria>Bacteria_incertae_sedis>Bacteria_candidate_phyla>Candidatus_Aminicenantota>unclassified_Candidatus_Aminicenantes | 208 | L6428_16305 | 2918..3544 | 2052149 | DUF4276-family-protein-[Candidatus-Aminicenantes-bacterium] |
| QAA75941.1 | TOPRIM | GCA_004102645.1 | Candidatus_Bipolaricaulis_sibiricus | Bacteria>Bacteria_incertae_sedis>Bacteria_candidate_phyla>Candidatus_Bipolaricaulota>Candidatus_Bipolaricaulia>Candidatus_Bipolaricaulales>Candidatus_Bipolaricaulaceae>Candidatus_Bipolaricaulis | 212 | BIP78_0173 | 165516..166154 | 2501609 | hypothetical-protein-BIP78_0173-[Candidatus-Bipolaricaulis-sibiricus] |
| MDD5559007.1 | TOPRIM | GCA_028716865.1 | Candidatus_Methylomirabilis_sp. | Bacteria>Bacteria_incertae_sedis>Bacteria_candidate_phyla>Candidatus_Methylomirabilota>Candidatus_Methylomirabilia>Candidatus_Methylomirabilales>Candidatus_Methylomirabilaceae>Candidatus_Methylomirabilis>unclassified_Candidatus_Methylomirabilis | 206 | PHV01_05035 | 52251..52871 | 2032687 | MAG:-hypothetical-protein-PHV01_05035-[Candidatus-Methylomirabilis-sp.] |
| MDE2058886.1 | TOPRIM | GCA_028864815.1 | Candidatus_Methylomirabilota_bacterium | Bacteria>Bacteria_incertae_sedis>Bacteria_candidate_phyla>Candidatus_Methylomirabilota>unclassified_Candidatus_Methylomirabilota | 213 | KGJ27_04055 | 12454..13095 | 2072417 | MAG:-RloB-domain-containing-protein-[candidate-division-NC10-bacterium] |
| MDE2058202.1 | TOPRIM | GCA_028864815.1 | Candidatus_Methylomirabilota_bacterium | Bacteria>Bacteria_incertae_sedis>Bacteria_candidate_phyla>Candidatus_Methylomirabilota>unclassified_Candidatus_Methylomirabilota | 201 | KGJ27_00520 | 28897..29502 | 2072417 | MAG:-hypothetical-protein-KGJ27_00520-[candidate-division-NC10-bacterium] |
| MBI4536777.1 | TOPRIM | GCA_016209735.1 | Candidatus_Methylomirabilota_bacterium | Bacteria>Bacteria_incertae_sedis>Bacteria_candidate_phyla>Candidatus_Methylomirabilota>unclassified_Candidatus_Methylomirabilota | 199 | HY712_02335 | 64973..65572 | 2072417 | hypothetical-protein-[candidate-division-NC10-bacterium] |
| MDE0183897.1 | TOPRIM | GCA_028819845.1 | Candidatus_Poribacteria_bacterium | Bacteria>Bacteria_incertae_sedis>Bacteria_candidate_phyla>Candidatus_Poribacteria>unclassified_Candidatus_Poribacteria | 216 | OXP71_00405 | 95292..95942 | 2026781 | MAG:-DUF4276-family-protein-[Candidatus-Poribacteria-bacterium] |
| MBF0500209.1 | TOPRIM | GCA_015233155.1 | Candidatus_Rifleibacteriota_bacterium | Bacteria>Bacteria_incertae_sedis>Bacteria_candidate_phyla>Candidatus_Rifleibacteriota>unclassified_Candidatus_Rifleibacteriota | 223 | HQM09_08750 | 72239..72910 | 2053586 | DUF4276-family-protein-[Candidatus-Riflebacteria-bacterium] |
| MBI2567006.1 | TOPRIM | GCA_016188575.1 | Candidatus_Schekmanbacteria_bacterium | Bacteria>Bacteria_incertae_sedis>Bacteria_candidate_phyla>Candidatus_Schekmaniibacteriota>unclassified_Candidatus_Schekmanbacteria | 233 | HYV63_08260 | 61280..61981 | 2419984 | hypothetical-protein-[Candidatus-Schekmanbacteria-bacterium] |
| MBN1883903.1 | TOPRIM | GCA_016931295.1 | Candidatus_Krumholzibacteriota_bacterium | Bacteria>Candidatus_Krumholzibacteriota>unclassified_Candidatus_Krumholzibacteriota | 190 | JW876_00095 | 28124..28696 | 2725313 | DUF4276-family-protein-[Candidatus-Krumholzibacteriota-bacterium] |
| EFK06075.1 | TOPRIM | - | delta_proteobacterium_NaphS2 | Bacteria>delta/epsilon_subdivisions>Deltaproteobacteria>unclassified_Deltaproteobacteria | 231 | NPH_2624 | 40617..41312 | 88274 | conserved-hypothetical-protein-[delta-proteobacterium-NaphS2] |
| OGQ97433.1 | TOPRIM | GCA_001799275.1 | Deltaproteobacteria_bacterium_RIFOXYD12_FULL_57_12 | Bacteria>delta/epsilon_subdivisions>Deltaproteobacteria>unclassified_Deltaproteobacteria | 227 | A2521_08590 | 17632..18315 | 1797906 | hypothetical-protein-A2521_08590-[Deltaproteobacteria-bacterium-RIFOXYD12_FULL_57_12] |
| PJB45676.1 | TOPRIM | GCA_002789955.1 | Deltaproteobacteria_bacterium_CG_4_9_14_3_um_filter_63_12 | Bacteria>delta/epsilon_subdivisions>Deltaproteobacteria>unclassified_Deltaproteobacteria | 197 | CO108_06830 | 10847..11440 | 1973967 | hypothetical-protein-CO108_06830-[Deltaproteobacteria-bacterium-CG_4_9_14_3_um_filter_63_12] |
| WP_023036015.1 | TOPRIM | GCF_024140205.1 | Fusobacterium | Bacteria>Fusobacteriati>Fusobacteriota>Fusobacteriia>Fusobacteriales>Fusobacteriaceae | 259 | BKN39_RS05485 | 8906..9685 | 848 | MULTISPECIES:-hypothetical-protein-[Fusobacterium] |
| WP_008700608.1 | TOPRIM | GCF_000273625.1 | Fusobacterium | Bacteria>Fusobacteriati>Fusobacteriota>Fusobacteriia>Fusobacteriales>Fusobacteriaceae | 256 | YWH7199_RS06525 | 20774..21544 | 848 | MULTISPECIES:-hypothetical-protein-[Fusobacterium] |
| WP_238972695.1 | TOPRIM | GCF_037911825.1 | Fusobacterium | Bacteria>Fusobacteriati>Fusobacteriota>Fusobacteriia>Fusobacteriales>Fusobacteriaceae | 256 | K5I19_RS02515 | 132934..133704 | 848 | hypothetical-protein-[Fusobacterium-nucleatum] |
| WP_032836036.1 | TOPRIM | GCF_045159995.1 | Fusobacterium | Bacteria>Fusobacteriati>Fusobacteriota>Fusobacteriia>Fusobacteriales>Fusobacteriaceae | 255 | HMPREF1501_RS01110 | 61461..62228 | 848 | hypothetical-protein-[Fusobacterium-sp.-OBRC1] |
| WP_249525222.1 | TOPRIM | GCF_023383525.1 | Fusobacterium_nucleatum | Bacteria>Fusobacteriati>Fusobacteriota>Fusobacteriia>Fusobacteriales>Fusobacteriaceae>Fusobacterium | 280 | YWH7054_RS02300 | 19627..20469 | 851 | hypothetical-protein-[Fusobacterium-nucleatum] |
| WP_008797393.1 | TOPRIM | GCF_000163915.2 | Fusobacterium_vincentii | Bacteria>Fusobacteriati>Fusobacteriota>Fusobacteriia>Fusobacteriales>Fusobacteriaceae>Fusobacterium | 259 | HMPREF0405_RS05630 | 1190901..1191680 | 155615 | hypothetical-protein-[Fusobacterium-vincentii] |
| WP_060676305.1 | TOPRIM | GCF_001296085.1 | Fusobacterium_animalis | Bacteria>Fusobacteriati>Fusobacteriota>Fusobacteriia>Fusobacteriales>Fusobacteriaceae>Fusobacterium | 258 | RN98_RS07260 | 1459417..1460193 | 76859 | hypothetical-protein-[Fusobacterium-animalis] |
| WP_099972119.1 | TOPRIM | GCF_963546905.1 | Fusobacterium_pseudoperiodonticum | Bacteria>Fusobacteriati>Fusobacteriota>Fusobacteriia>Fusobacteriales>Fusobacteriaceae>Fusobacterium | 258 | CTM72_RS01095 | 204877..205653 | 2663009 | hypothetical-protein-[Fusobacterium-pseudoperiodonticum] |
| WP_124794844.1 | TOPRIM | GCF_003859915.1 | Fusobacterium_canifelinum | Bacteria>Fusobacteriati>Fusobacteriota>Fusobacteriia>Fusobacteriales>Fusobacteriaceae>Fusobacterium | 258 | EII27_RS00790 | 178024..178800 | 285729 | hypothetical-protein-[Fusobacterium-canifelinum] |
| WP_158410487.1 | TOPRIM | GCF_050219895.1 | Fusobacterium_animalis | Bacteria>Fusobacteriati>Fusobacteriota>Fusobacteriia>Fusobacteriales>Fusobacteriaceae>Fusobacterium | 258 | CI111_RS06860 | 4524..5300 | 76859 | hypothetical-protein-[Fusobacterium-animalis] |
| WP_220308655.1 | TOPRIM | GCF_019552125.1 | Fusobacterium_animalis | Bacteria>Fusobacteriati>Fusobacteriota>Fusobacteriia>Fusobacteriales>Fusobacteriaceae>Fusobacterium | 258 | JY401_RS04540 | 952784..953560 | 76859 | hypothetical-protein-[Fusobacterium-animalis] |
| WP_008692037.1 | TOPRIM | GCF_000218655.1 | Fusobacterium_animalis | Bacteria>Fusobacteriati>Fusobacteriota>Fusobacteriia>Fusobacteriales>Fusobacteriaceae>Fusobacterium | 256 | HMPREF0401_RS10110 | 562326..563096 | 76859 | hypothetical-protein-[Fusobacterium-animalis] |
| WP_099987860.1 | TOPRIM | GCF_002763695.1 | Fusobacterium_pseudoperiodonticum | Bacteria>Fusobacteriati>Fusobacteriota>Fusobacteriia>Fusobacteriales>Fusobacteriaceae>Fusobacterium | 255 | CTM74_RS08995 | 1719039..1719806 | 2663009 | hypothetical-protein-[Fusobacterium-pseudoperiodonticum] |
| WP_088387745.1 | TOPRIM | GCF_002204435.1 | Fusobacterium_polymorphum | Bacteria>Fusobacteriati>Fusobacteriota>Fusobacteriia>Fusobacteriales>Fusobacteriaceae>Fusobacterium | 254 | CA839_RS02160 | 472102..472866 | 76857 | hypothetical-protein-[Fusobacterium-polymorphum] |
| WP_098702330.1 | TOPRIM | GCF_002573625.1 | Fusobacterium_polymorphum | Bacteria>Fusobacteriati>Fusobacteriota>Fusobacteriia>Fusobacteriales>Fusobacteriaceae>Fusobacterium | 254 | RN96_RS03480 | 674524..675288 | 76857 | hypothetical-protein-[Fusobacterium-polymorphum] |
| WP_098973856.1 | TOPRIM | GCF_037897565.1 | Fusobacterium_polymorphum | Bacteria>Fusobacteriati>Fusobacteriota>Fusobacteriia>Fusobacteriales>Fusobacteriaceae>Fusobacterium | 254 | CBG58_RS00350 | 82643..83407 | 76857 | hypothetical-protein-[Fusobacterium-polymorphum] |
| WP_220302803.1 | TOPRIM | GCF_019552045.1 | Fusobacterium_polymorphum | Bacteria>Fusobacteriati>Fusobacteriota>Fusobacteriia>Fusobacteriales>Fusobacteriaceae>Fusobacterium | 254 | JY397_RS08310 | 1736487..1737251 | 76857 | hypothetical-protein-[Fusobacterium-polymorphum] |
| WP_233486204.1 | TOPRIM | GCF_002761955.1 | Fusobacterium_pseudoperiodonticum | Bacteria>Fusobacteriati>Fusobacteriota>Fusobacteriia>Fusobacteriales>Fusobacteriaceae>Fusobacterium | 204 | CTM71_RS09945 | 1965330..1965944 | 2663009 | hypothetical-protein-[Fusobacterium-pseudoperiodonticum] |
| WP_023036663.1 | TOPRIM | GCF_048313155.1 | Fusobacterium_polymorphum | Bacteria>Fusobacteriati>Fusobacteriota>Fusobacteriia>Fusobacteriales>Fusobacteriaceae>Fusobacterium | 144 | HMPREF1767_RS12910 | 31388..31822 | 76857 | hypothetical-protein-[Fusobacterium-nucleatum] |
| WP_265152270.1 | TOPRIM | GCF_026089295.1 | Fusobacterium_simiae | Bacteria>Fusobacteriati>Fusobacteriota>Fusobacteriia>Fusobacteriales>Fusobacteriaceae>Fusobacterium | 125 | OCK72_RS06720 | 35400..35777 | 855 | hypothetical-protein-[Fusobacterium-simiae] |
| ALF25294.1 | TOPRIM | GCA_001296185.1 | Fusobacterium_nucleatum_subsp._nucleatum | Bacteria>Fusobacteriati>Fusobacteriota>Fusobacteriia>Fusobacteriales>Fusobacteriaceae>Fusobacterium>Fusobacterium_nucleatum | 255 | RN95_02160 | 460194..460961 | 76856 | hypothetical-protein-RN95_02160-[Fusobacterium-nucleatum-subsp.-nucleatum] |
| WP_032846248.1 | TOPRIM | GCF_000524395.1 | Fusobacterium_sp._CM22 | Bacteria>Fusobacteriati>Fusobacteriota>Fusobacteriia>Fusobacteriales>Fusobacteriaceae>Fusobacterium>unclassified_Fusobacterium | 255 | HMPREF1500_RS09200 | 122421..123188 | 936563 | hypothetical-protein-[Fusobacterium-sp.-CM22] |
| WP_070766194.1 | TOPRIM | GCF_001815715.1 | Fusobacterium_sp._HMSC064B11 | Bacteria>Fusobacteriati>Fusobacteriota>Fusobacteriia>Fusobacteriales>Fusobacteriaceae>Fusobacterium>unclassified_Fusobacterium | 255 | HMPREF3051_RS04310 | 256840..257607 | 1739543 | hypothetical-protein-[Fusobacterium-sp.-HMSC064B11] |
| CDA09058.1 | TOPRIM | GCA_000433695.1 | Fusobacterium_sp._CAG:649 | Bacteria>Fusobacteriati>Fusobacteriota>Fusobacteriia>Fusobacteriales>Fusobacteriaceae>Fusobacterium>unclassified_Fusobacterium | 186 | BN748_00882 | 21261..21821 | 1262900 | putative-uncharacterized-protein-[Fusobacterium-sp.-CAG:649] |
| WP_270932598.1 | TOPRIM | GCF_045055315.1 | Pseudomonadota | Bacteria>Pseudomonadati | 217 | JR315_RS22295 | 18312..18965 | 1224 | DUF4276-family-protein-[Pseudomonas-aeruginosa] |
| WP_079541820.1 | TOPRIM | GCF_963530725.1 | Pseudomonadota | Bacteria>Pseudomonadati | 180 | CHUV0807_RS10770 | 36968..37510 | 1224 | DUF4276-family-protein-[Cardiobacterium-hominis] |
| QQR73411.1 | TOPRIM | GCA_016699405.1 | Holophagales_bacterium | Bacteria>Pseudomonadati>Acidobacteriota>Holophagae>Holophagales>unclassified_Holophagales | 174 | IPJ17_18305 | 4434757..4435281 | 2478486 | DUF4276-family-protein-[Holophagales-bacterium] |
| MCY4597197.1 | TOPRIM | GCA_026713455.1 | Bryobacterales_bacterium | Bacteria>Pseudomonadati>Acidobacteriota>Terriglobia>Bryobacterales>unclassified_Bryobacterales | 218 | OXC19_20645 | 11729..12385 | 2026791 | MAG:-hypothetical-protein-OXC19_20645-[Bryobacterales-bacterium] |
| MXY68845.1 | TOPRIM | GCA_009838515.1 | Terriglobia_bacterium | Bacteria>Pseudomonadati>Acidobacteriota>Terriglobia>unclassified_Terriglobia | 186 | F4Y47_09835 | 225922..226482 | 2081523 | hypothetical-protein-[Acidobacteriia-bacterium] |
| MYK11749.1 | TOPRIM | GCA_009845805.1 | Terriglobia_bacterium | Bacteria>Pseudomonadati>Acidobacteriota>Terriglobia>unclassified_Terriglobia | 186 | F4041_19180 | 4181..4741 | 2081523 | hypothetical-protein-[Acidobacteriia-bacterium] |
| MCU0292532.1 | TOPRIM | GCA_025456225.1 | Thermoanaerobaculaceae_bacterium | Bacteria>Pseudomonadati>Acidobacteriota>Thermoanaerobaculia>Thermoanaerobaculales>Thermoanaerobaculaceae>unclassified_Thermoanaerobaculaceae | 223 | MUF10_11195 | 8519..9190 | 2699758 | hypothetical-protein-[Thermoanaerobaculaceae-bacterium] |
| MBP7799238.1 | TOPRIM | GCA_018000955.1 | Thermoanaerobaculaceae_bacterium | Bacteria>Pseudomonadati>Acidobacteriota>Thermoanaerobaculia>Thermoanaerobaculales>Thermoanaerobaculaceae>unclassified_Thermoanaerobaculaceae | 203 | KA072_11470 | 13719..14330 | 2699758 | hypothetical-protein-[Thermoanaerobaculaceae-bacterium] |
| MBP7799280.1 | TOPRIM | GCA_018000955.1 | Thermoanaerobaculaceae_bacterium | Bacteria>Pseudomonadati>Acidobacteriota>Thermoanaerobaculia>Thermoanaerobaculales>Thermoanaerobaculaceae>unclassified_Thermoanaerobaculaceae | 201 | KA072_11715 | 31180..31785 | 2699758 | hypothetical-protein-[Thermoanaerobaculaceae-bacterium] |
| MCG3192273.1 | TOPRIM | GCA_022071945.1 | Thermoanaerobaculia_bacterium | Bacteria>Pseudomonadati>Acidobacteriota>Thermoanaerobaculia>unclassified_Thermoanaerobaculia | 208 | DIJKHBIC_01511 | 133408..134034 | 2651171 | hypothetical-protein-[Thermoanaerobaculia-bacterium] |
| MCG3190933.1 | TOPRIM | GCA_022071945.1 | Thermoanaerobaculia_bacterium | Bacteria>Pseudomonadati>Acidobacteriota>Thermoanaerobaculia>unclassified_Thermoanaerobaculia | 204 | DIJKHBIC_00156 | 192051..192665 | 2651171 | hypothetical-protein-[Thermoanaerobaculia-bacterium] |
| MCG3192295.1 | TOPRIM | GCA_022071945.1 | Thermoanaerobaculia_bacterium | Bacteria>Pseudomonadati>Acidobacteriota>Thermoanaerobaculia>unclassified_Thermoanaerobaculia | 187 | DIJKHBIC_01533 | 16201..16764 | 2651171 | hypothetical-protein-[Thermoanaerobaculia-bacterium] |
| MYN66271.1 | TOPRIM | GCA_009861545.1 | Acidobacteriota_bacterium | Bacteria>Pseudomonadati>Acidobacteriota>unclassified_Acidobacteriota | 239 | F4X11_14770 | 7932..8651 | 1978231 | hypothetical-protein-[Acidobacteriota-bacterium] |
| MDE2979141.1 | TOPRIM | GCA_028873735.1 | Acidobacteriota_bacterium | Bacteria>Pseudomonadati>Acidobacteriota>unclassified_Acidobacteriota | 215 | OXU63_16715 | 12557..13204 | 1978231 | MAG:-DUF4276-family-protein-[Acidobacteriota-bacterium] |
| MYN68212.1 | TOPRIM | GCA_009861545.1 | Acidobacteriota_bacterium | Bacteria>Pseudomonadati>Acidobacteriota>unclassified_Acidobacteriota | 210 | F4X11_24875 | 55924..56556 | 1978231 | hypothetical-protein-[Acidobacteriota-bacterium] |
| MCC6131986.1 | TOPRIM | GCA_020845555.1 | Acidobacteriota_bacterium | Bacteria>Pseudomonadati>Acidobacteriota>unclassified_Acidobacteriota | 208 | IT186_18850 | 12786..13412 | 1978231 | hypothetical-protein-[Acidobacteriota-bacterium] |
| MCY3965601.1 | TOPRIM | GCA_026707545.1 | Acidobacteriota_bacterium | Bacteria>Pseudomonadati>Acidobacteriota>unclassified_Acidobacteriota | 206 | OXG83_11230 | 2821940..2822560 | 1978231 | MAG:-hypothetical-protein-OXG83_11230-[Acidobacteria-bacterium] |
| MBI4902186.1 | TOPRIM | GCA_016213775.1 | Acidobacteriota_bacterium | Bacteria>Pseudomonadati>Acidobacteriota>unclassified_Acidobacteriota | 205 | HY820_01035 | 16922..17539 | 1978231 | DUF4276-family-protein-[Acidobacteriota-bacterium] |
| MYN67291.1 | TOPRIM | GCA_009861545.1 | Acidobacteriota_bacterium | Bacteria>Pseudomonadati>Acidobacteriota>unclassified_Acidobacteriota | 194 | F4X11_20050 | 69393..69977 | 1978231 | hypothetical-protein-[Acidobacteriota-bacterium] |
| MCY3965604.1 | TOPRIM | GCA_026707545.1 | Acidobacteriota_bacterium | Bacteria>Pseudomonadati>Acidobacteriota>unclassified_Acidobacteriota | 190 | OXG83_11245 | 2827057..2827629 | 1978231 | MAG:-hypothetical-protein-OXG83_11245-[Acidobacteria-bacterium] |
| MCC6130028.1 | TOPRIM | GCA_020845555.1 | Acidobacteriota_bacterium | Bacteria>Pseudomonadati>Acidobacteriota>unclassified_Acidobacteriota | 187 | IT186_08865 | 24849..25412 | 1978231 | DUF4276-family-protein-[Acidobacteriota-bacterium] |
| MCY4659245.1 | TOPRIM | GCA_026713985.1 | Acidobacteriota_bacterium | Bacteria>Pseudomonadati>Acidobacteriota>unclassified_Acidobacteriota | 167 | OXF93_05450 | 11081..11584 | 1978231 | MAG:-hypothetical-protein-OXF93_05450-[Acidobacteria-bacterium] |
| MBI5244007.1 | TOPRIM | GCA_016212885.1 | Elusimicrobiota_bacterium | Bacteria>Pseudomonadati>Elusimicrobiota>unclassified_Elusimicrobiota | 211 | HY922_10080 | 1021..1656 | 2030800 | hypothetical-protein-[Elusimicrobiota-bacterium] |
| MBI4676021.1 | TOPRIM | GCA_016209035.1 | Elusimicrobiota_bacterium | Bacteria>Pseudomonadati>Elusimicrobiota>unclassified_Elusimicrobiota | 202 | HY748_00375 | 76926..77534 | 2030800 | DUF4276-family-protein-[Elusimicrobiota-bacterium] |
| MBI4676019.1 | TOPRIM | GCA_016209035.1 | Elusimicrobiota_bacterium | Bacteria>Pseudomonadati>Elusimicrobiota>unclassified_Elusimicrobiota | 197 | HY748_00365 | 75102..75695 | 2030800 | DUF4276-family-protein-[Elusimicrobiota-bacterium] |
| WP_241209994.1 | TOPRIM | GCF_963928575.1 | Bacteroidales | Bacteria>Pseudomonadati>FCB_group>Bacteroidota/Chlorobiota_group>Bacteroidota>Bacteroidia | 208 | M3087_RS02160 | 168668..169294 | 171549 | MULTISPECIES:-RloB-family-protein-[Bacteroidales] |
| WP_217745555.1 | TOPRIM | GCF_019127365.1 | Prevotellaceae | Bacteria>Pseudomonadati>FCB_group>Bacteroidota/Chlorobiota_group>Bacteroidota>Bacteroidia>Bacteroidales | 279 | KSW85_RS06185 | 45470..46309 | 171552 | DUF4435-domain-containing-protein-[Prevotella-copri] |
| WP_072542574.1 | TOPRIM | GCF_900128455.1 | Bacteroides_mediterraneensis | Bacteria>Pseudomonadati>FCB_group>Bacteroidota/Chlorobiota_group>Bacteroidota>Bacteroidia>Bacteroidales>Bacteroidaceae>Bacteroides | 282 | BQ7480_RS08750 | 2127450..2128298 | 1841856 | DUF4435-domain-containing-protein-[Bacteroides-mediterraneensis] |
| WP_204433171.1 | TOPRIM | - | Bacteroides_mediterraneensis | Bacteria>Pseudomonadati>FCB_group>Bacteroidota/Chlorobiota_group>Bacteroidota>Bacteroidia>Bacteroidales>Bacteroidaceae>Bacteroides | 282 | H6B28_RS04785 | 131090..131938 | 1841856 | DUF4435-domain-containing-protein-[Bacteroides-mediterraneensis] |
| CDD33288.1 | TOPRIM | GCA_000436055.1 | Bacteroides_sp._CAG:714 | Bacteria>Pseudomonadati>FCB_group>Bacteroidota/Chlorobiota_group>Bacteroidota>Bacteroidia>Bacteroidales>Bacteroidaceae>Bacteroides>unclassified_Bacteroides | 282 | BN762_00979 | 2038..2886 | 1262749 | putative-uncharacterized-protein-[Bacteroides-sp.-CAG:714] |
| CDD50106.1 | TOPRIM | GCA_000435415.1 | Bacteroides_sp._CAG:875 | Bacteria>Pseudomonadati>FCB_group>Bacteroidota/Chlorobiota_group>Bacteroidota>Bacteroidia>Bacteroidales>Bacteroidaceae>Bacteroides>unclassified_Bacteroides | 282 | BN800_01991 | 22486..23334 | 1262752 | putative-uncharacterized-protein-[Bacteroides-sp.-CAG:875] |
| HJF82221.1 | TOPRIM | GCA_020742805.1 | Phocaeicola_plebeius | Bacteria>Pseudomonadati>FCB_group>Bacteroidota/Chlorobiota_group>Bacteroidota>Bacteroidia>Bacteroidales>Bacteroidaceae>Phocaeicola | 282 | K8V40_11355 | 8447..9295 | 310297 | DUF4435-domain-containing-protein-[Phocaeicola-plebeius] |
| WP_118441920.1 | TOPRIM | GCF_003474535.1 | Phocaeicola_plebeius | Bacteria>Pseudomonadati>FCB_group>Bacteroidota/Chlorobiota_group>Bacteroidota>Bacteroidia>Bacteroidales>Bacteroidaceae>Phocaeicola | 282 | DW041_RS13385 | 8904..9752 | 310297 | DUF4435-domain-containing-protein-[Phocaeicola-plebeius] |
| WP_148330683.1 | TOPRIM | GCF_025567125.1 | Phocaeicola_fibrisolvens | Bacteria>Pseudomonadati>FCB_group>Bacteroidota/Chlorobiota_group>Bacteroidota>Bacteroidia>Bacteroidales>Bacteroidaceae>Phocaeicola | 282 | OCV81_RS14575 | 2381..3229 | 2981793 | DUF4435-domain-containing-protein-[Phocaeicola-fibrisolvens] |
| MCR4768678.1 | TOPRIM | GCA_024682395.1 | Bacteroidaceae_bacterium | Bacteria>Pseudomonadati>FCB_group>Bacteroidota/Chlorobiota_group>Bacteroidota>Bacteroidia>Bacteroidales>Bacteroidaceae>unclassified_Bacteroidaceae | 191 | K5874_00450 | 902..1477 | 2212467 | RloB-family-protein-[Bacteroidaceae-bacterium] |
| HAH91108.1 | TOPRIM | GCA_003450425.1 | Prevotella_sp. | Bacteria>Pseudomonadati>FCB_group>Bacteroidota/Chlorobiota_group>Bacteroidota>Bacteroidia>Bacteroidales>Prevotellaceae>Prevotella>unclassified_Prevotella | 296 | DCL96_05240 | 12086..12976 | 59823 | hypothetical-protein-[Prevotella-sp.] |
| MCI6336760.1 | TOPRIM | GCA_022777825.1 | Prevotella_sp. | Bacteria>Pseudomonadati>FCB_group>Bacteroidota/Chlorobiota_group>Bacteroidota>Bacteroidia>Bacteroidales>Prevotellaceae>Prevotella>unclassified_Prevotella | 288 | MR802_02270 | 3371..4237 | 59823 | DUF4435-domain-containing-protein-[Prevotella-sp.] |
| MBN2915286.1 | TOPRIM | GCA_016939035.1 | Prevotella_sp. | Bacteria>Pseudomonadati>FCB_group>Bacteroidota/Chlorobiota_group>Bacteroidota>Bacteroidia>Bacteroidales>Prevotellaceae>Prevotella>unclassified_Prevotella | 279 | JTJ26_04205 | 23431..24270 | 59823 | DUF4435-domain-containing-protein-[Prevotella-sp.] |
| WP_176452272.1 | TOPRIM | GCF_002265625.1 | Prevotella_sp._885 | Bacteria>Pseudomonadati>FCB_group>Bacteroidota/Chlorobiota_group>Bacteroidota>Bacteroidia>Bacteroidales>Prevotellaceae>Prevotella>unclassified_Prevotella | 279 | CHL74_RS14180 | 2519..3358 | 2022527 | DUF4435-domain-containing-protein-[Prevotella-sp.-885] |
| MCI6195662.1 | TOPRIM | GCA_022773545.1 | Prevotella_sp. | Bacteria>Pseudomonadati>FCB_group>Bacteroidota/Chlorobiota_group>Bacteroidota>Bacteroidia>Bacteroidales>Prevotellaceae>Prevotella>unclassified_Prevotella | 277 | MR670_00295 | 48005..48838 | 59823 | DUF4435-domain-containing-protein-[Prevotella-sp.] |
| WP_181992711.1 | TOPRIM | GCF_003465445.1 | Segatella_copri | Bacteria>Pseudomonadati>FCB_group>Bacteroidota/Chlorobiota_group>Bacteroidota>Bacteroidia>Bacteroidales>Prevotellaceae>Segatella | 279 | DWV60_RS07155 | 36228..37067 | 165179 | DUF4435-domain-containing-protein-[Prevotella-copri] |
| MBQ6275507.1 | TOPRIM | GCA_017428605.1 | Bacteroidales_bacterium | Bacteria>Pseudomonadati>FCB_group>Bacteroidota/Chlorobiota_group>Bacteroidota>Bacteroidia>Bacteroidales>unclassified_Bacteroidales | 228 | IJK62_02220 | 86427..87113 | 2030927 | hypothetical-protein-[Bacteroidales-bacterium] |
| MBR6930694.1 | TOPRIM | GCA_017519875.1 | Bacteroidales_bacterium | Bacteria>Pseudomonadati>FCB_group>Bacteroidota/Chlorobiota_group>Bacteroidota>Bacteroidia>Bacteroidales>unclassified_Bacteroidales | 218 | IKH61_10800 | 58136..58792 | 2030927 | DUF4276-family-protein-[Bacteroidales-bacterium] |
| MBD5181934.1 | TOPRIM | GCA_014800235.1 | Bacteroidales_bacterium | Bacteria>Pseudomonadati>FCB_group>Bacteroidota/Chlorobiota_group>Bacteroidota>Bacteroidia>Bacteroidales>unclassified_Bacteroidales | 216 | HDS98_05225 | 29001..29651 | 2030927 | DUF4276-family-protein-[Bacteroidales-bacterium] |
| MCQ2228646.1 | TOPRIM | GCA_024401335.1 | Bacteroidales_bacterium | Bacteria>Pseudomonadati>FCB_group>Bacteroidota/Chlorobiota_group>Bacteroidota>Bacteroidia>Bacteroidales>unclassified_Bacteroidales | 198 | MJZ13_02740 | 170259..170855 | 2030927 | RloB-family-protein-[Bacteroidales-bacterium] |
| MCL2726874.1 | TOPRIM | GCA_009781275.1 | Bacteroidales_bacterium | Bacteria>Pseudomonadati>FCB_group>Bacteroidota/Chlorobiota_group>Bacteroidota>Bacteroidia>Bacteroidales>unclassified_Bacteroidales | 196 | FWD56_00655 | 17122..17712 | 2030927 | DUF4276-family-protein-[Bacteroidales-bacterium] |
| WP_201982523.1 | TOPRIM | GCF_016734815.1 | Hymenobacter_rubidus | Bacteria>Pseudomonadati>FCB_group>Bacteroidota/Chlorobiota_group>Bacteroidota>Cytophagia>Cytophagales>Hymenobacteraceae>Hymenobacter | 774 | JKL48_RS11455 | 8298..10622 | 1441626 | type-I-DNA-topoisomerase-[Hymenobacter-rubidus] |
| MBX7052697.1 | TOPRIM | GCA_019694805.1 | Flavobacteriales_bacterium | Bacteria>Pseudomonadati>FCB_group>Bacteroidota/Chlorobiota_group>Bacteroidota>Flavobacteriia>Flavobacteriales>unclassified_Flavobacteriales | 213 | K1X54_11725 | 18442..19083 | 2021391 | hypothetical-protein-[Flavobacteriales-bacterium] |
| WP_020534910.1 | TOPRIM | GCF_000379805.1 | Lewinella_cohaerens | Bacteria>Pseudomonadati>FCB_group>Bacteroidota/Chlorobiota_group>Bacteroidota>Saprospiria>Saprospirales>Lewinellaceae>Lewinella | 296 | A3EU_RS0104380 | 126194..127084 | 70995 | hypothetical-protein-[Lewinella-cohaerens] |
| MCB9053305.1 | TOPRIM | GCA_020636405.1 | Lewinellaceae_bacterium | Bacteria>Pseudomonadati>FCB_group>Bacteroidota/Chlorobiota_group>Bacteroidota>Saprospiria>Saprospirales>Lewinellaceae>unclassified_Lewinellaceae | 199 | H6556_28040 | 308377..308976 | 2760807 | RloB-domain-containing-protein-[Lewinellaceae-bacterium] |
| MCB9053321.1 | TOPRIM | GCA_020636405.1 | Lewinellaceae_bacterium | Bacteria>Pseudomonadati>FCB_group>Bacteroidota/Chlorobiota_group>Bacteroidota>Saprospiria>Saprospirales>Lewinellaceae>unclassified_Lewinellaceae | 187 | H6556_28120 | 340638..341201 | 2760807 | RloB-domain-containing-protein-[Lewinellaceae-bacterium] |
| MBK8920120.1 | TOPRIM | GCA_016719895.1 | Saprospirales_bacterium | Bacteria>Pseudomonadati>FCB_group>Bacteroidota/Chlorobiota_group>Bacteroidota>Saprospiria>Saprospirales>unclassified_Saprospirales | 213 | IPM81_01190 | 287326..287967 | 2026790 | hypothetical-protein-[Saprospirales-bacterium] |
| MCC5934982.1 | TOPRIM | GCA_020832405.1 | Balneolales_bacterium | Bacteria>Pseudomonadati>FCB_group>Bacteroidota/Chlorobiota_group>Balneolota>Balneolia>Balneolales>unclassified_Balneolales | 219 | JJU35_12120 | 7666..8325 | 2497613 | hypothetical-protein-[Balneolales-bacterium] |
| MBV5328346.1 | TOPRIM | GCA_019162945.1 | Chlorobium_sp. | Bacteria>Pseudomonadati>FCB_group>Bacteroidota/Chlorobiota_group>Chlorobiota>Chlorobiia>Chlorobiales>Chlorobiaceae>Chlorobium/Pelodictyon_group>Chlorobium>unclassified_Chlorobium | 227 | JZU65_12025 | 4706..5389 | 1095 | DUF4276-family-protein-[Chlorobium-sp.] |
| TNJ34081.1 | TOPRIM | GCA_006265245.1 | Prosthecochloris_vibrioformis | Bacteria>Pseudomonadati>FCB_group>Bacteroidota/Chlorobiota_group>Chlorobiota>Chlorobiia>Chlorobiales>Chlorobiaceae>Prosthecochloris | 295 | FGF68_10420 | 6349..7236 | 1098 | hypothetical-protein-FGF68_10420-[Prosthecochloris-vibrioformis] |
| WP_139626999.1 | TOPRIM | GCF_006265245.1 | Prosthecochloris_vibrioformis | Bacteria>Pseudomonadati>FCB_group>Bacteroidota/Chlorobiota_group>Chlorobiota>Chlorobiia>Chlorobiales>Chlorobiaceae>Prosthecochloris | 295 | FGF68_RS10415 | 6349..7236 | 1098 | hypothetical-protein-[Prosthecochloris-vibrioformis] |
| MBK7265054.1 | TOPRIM | GCA_016707285.1 | Ignavibacteriales_bacterium | Bacteria>Pseudomonadati>FCB_group>Bacteroidota/Chlorobiota_group>Ignavibacteriota>Ignavibacteria>Ignavibacteriales>unclassified_Ignavibacteriales | 233 | IPI12_01705 | 360727..361428 | 2049428 | RloB-domain-containing-protein-[Ignavibacteriales-bacterium] |
| MCB5285391.1 | TOPRIM | GCA_020532085.1 | Candidatus_Cloacimonadota_bacterium | Bacteria>Pseudomonadati>FCB_group>Candidatus_Cloacimonadota>unclassified_Candidatus_Cloacimonadota | 213 | LHW45_07370 | 26486..27127 | 2030808 | DUF4276-family-protein-[Candidatus-Cloacimonadota-bacterium] |
| MCB5285395.1 | TOPRIM | GCA_020532085.1 | Candidatus_Cloacimonadota_bacterium | Bacteria>Pseudomonadati>FCB_group>Candidatus_Cloacimonadota>unclassified_Candidatus_Cloacimonadota | 204 | LHW45_07390 | 32087..32701 | 2030808 | DUF4276-family-protein-[Candidatus-Cloacimonadota-bacterium] |
| MBP7719544.1 | TOPRIM | GCA_017999735.1 | Candidatus_Fermentibacter_sp. | Bacteria>Pseudomonadati>FCB_group>Candidatus_Fermentibacterota>Candidatus_Fermentibacteria_(class)>Candidatus_Fermentibacterales>Candidatus_Fermentibacteraceae>Candidatus_Fermentibacter>unclassified_Candidatus_Fermentibacter | 194 | KA108_02785 | 13857..14441 | 2779163 | DUF4276-family-protein-[Candidatus-Fermentibacter-sp.] |
| NOQ23465.1 | TOPRIM | GCA_013138965.1 | Candidatus_Aegiribacteria_sp. | Bacteria>Pseudomonadati>FCB_group>FCB_group_incertae_sedis>Candidatus_Aegiribacteria>unclassified_Candidatus_Aegiribacteria | 297 | GQ565_12570 | 6676..7569 | 2528271 | hypothetical-protein-[Candidatus-Aegiribacteria-sp.] |
| NLD91455.1 | TOPRIM | GCA_012513915.1 | Fibrobacter_sp. | Bacteria>Pseudomonadati>FCB_group>Fibrobacterota>Fibrobacteria>Fibrobacterales>Fibrobacteraceae>Fibrobacter>unclassified_Fibrobacter | 208 | GX639_02180 | 60734..61360 | 35828 | DUF4276-family-protein-[Fibrobacter-sp.] |
| MXX77943.1 | TOPRIM | GCA_009836785.1 | Gemmatimonadales_bacterium | Bacteria>Pseudomonadati>FCB_group>Gemmatimonadota>Gemmatimonadia>Gemmatimonadales>unclassified_Gemmatimonadales | 190 | F4Z33_02940 | 635..1207 | 2448054 | DUF4276-family-protein-[Gemmatimonadales-bacterium] |
| MYA34702.1 | TOPRIM | GCA_009838765.1 | Gemmatimonadales_bacterium | Bacteria>Pseudomonadati>FCB_group>Gemmatimonadota>Gemmatimonadia>Gemmatimonadales>unclassified_Gemmatimonadales | 188 | F4Y33_13790 | 67763..68329 | 2448054 | hypothetical-protein-[Gemmatimonadales-bacterium] |
| MYB67637.1 | TOPRIM | GCA_009840005.1 | Gemmatimonadota_bacterium | Bacteria>Pseudomonadati>FCB_group>Gemmatimonadota>unclassified_Gemmatimonadota | 218 | F4X75_03960 | 9978..10634 | 2026742 | DUF4276-family-protein-[Gemmatimonadota-bacterium] |
| MCY4603566.1 | TOPRIM | GCA_026713655.1 | Gemmatimonadota_bacterium | Bacteria>Pseudomonadati>FCB_group>Gemmatimonadota>unclassified_Gemmatimonadota | 207 | OXE49_04980 | 6010..6633 | 2026742 | MAG:-hypothetical-protein-OXE49_04980-[Gemmatimonadetes-bacterium] |
| MBX6362425.1 | TOPRIM | GCA_019689775.1 | Gemmatimonadota_bacterium | Bacteria>Pseudomonadati>FCB_group>Gemmatimonadota>unclassified_Gemmatimonadota | 203 | IRZ00_01045 | 22629..23240 | 2026742 | hypothetical-protein-[Gemmatimonadota-bacterium] |
| MDE2723063.1 | TOPRIM | GCA_028875455.1 | Gemmatimonadota_bacterium | Bacteria>Pseudomonadati>FCB_group>Gemmatimonadota>unclassified_Gemmatimonadota | 193 | OXI59_06775 | 7221..7802 | 2026742 | MAG:-DUF4276-family-protein-[Gemmatimonadota-bacterium] |
| MXX38677.1 | TOPRIM | GCA_009835785.1 | Gemmatimonadota_bacterium | Bacteria>Pseudomonadati>FCB_group>Gemmatimonadota>unclassified_Gemmatimonadota | 193 | F4Z85_11555 | 5877..6458 | 2026742 | DUF4276-family-protein-[Gemmatimonadota-bacterium] |
| MXX38674.1 | TOPRIM | GCA_009835785.1 | Gemmatimonadota_bacterium | Bacteria>Pseudomonadati>FCB_group>Gemmatimonadota>unclassified_Gemmatimonadota | 190 | F4Z85_11540 | 617..1189 | 2026742 | hypothetical-protein-[Gemmatimonadota-bacterium] |
| MDE2654307.1 | TOPRIM | GCA_028875795.1 | Gemmatimonadota_bacterium | Bacteria>Pseudomonadati>FCB_group>Gemmatimonadota>unclassified_Gemmatimonadota | 187 | OXI71_10905 | 86590..87153 | 2026742 | MAG:-DUF4276-family-protein-[Gemmatimonadota-bacterium] |
| MDE2723059.1 | TOPRIM | GCA_028875455.1 | Gemmatimonadota_bacterium | Bacteria>Pseudomonadati>FCB_group>Gemmatimonadota>unclassified_Gemmatimonadota | 187 | OXI59_06755 | 1988..2551 | 2026742 | MAG:-hypothetical-protein-OXI59_06755-[Gemmatimonadota-bacterium] |
| MYD59775.1 | TOPRIM | GCA_009841665.1 | Gemmatimonadota_bacterium | Bacteria>Pseudomonadati>FCB_group>Gemmatimonadota>unclassified_Gemmatimonadota | 187 | F4W91_01920 | 34436..34999 | 2026742 | hypothetical-protein-[Gemmatimonadota-bacterium] |
| MYA24523.1 | TOPRIM | GCA_009836855.1 | Gemmatimonadota_bacterium | Bacteria>Pseudomonadati>FCB_group>Gemmatimonadota>unclassified_Gemmatimonadota | 104 | F4Z30_17855 | 7863..8179 | 2026742 | DUF4276-family-protein-[Gemmatimonadota-bacterium] |
| HCF56389.1 | TOPRIM | GCA_003534605.1 | Myxococcales_bacterium | Bacteria>Pseudomonadati>Myxococcota>Myxococcia>Myxococcales>unclassified_Myxococcales | 234 | DFS52_00130 | 4349..5053 | 2026763 | hypothetical-protein-[Myxococcales-bacterium] |

| Protein-ID | Domain | Assembly-ID | Strain | Taxonomy | Length | Locus-Tag | Genomic-coordinates | TaxID | Fasta-defline |
|---|---|---|---|---|---|---|---|---|---|
| MCB9544270.1 | TOPRIM | GCA_020633435.1 | Myxococcales_bacterium | Bacteria>Pseudomonadati>Myxococcota>Myxococcia>Myxococcales>unclassified_Myxococcales | 208 | H6706_00075 | 19380..20006 | 2026763 | hypothetical-protein-[Myxococcales-bacterium] |
| MBL9045675.1 | TOPRIM | GCA_016794465.1 | Myxococcales_bacterium | Bacteria>Pseudomonadati>Myxococcota>Myxococcia>Myxococcales>unclassified_Myxococcales | 193 | JNM83_28960 | 8994..9575 | 2026763 | DUF4276-family-protein-[Myxococcales-bacterium] |
| MCG5052487.1 | TOPRIM | GCA_022184915.1 | Myxococcales_bacterium | Bacteria>Pseudomonadati>Myxococcota>Myxococcia>Myxococcales>unclassified_Myxococcales | 191 | KA712_05970 | 1446128..1446703 | 2026763 | DUF4276-family-protein-[Myxococcales-bacterium] |
| MBI4706188.1 | TOPRIM | GCA_016209145.1 | Deltaproteobacteria_bacterium | Bacteria>Pseudomonadati>Myxococcota>Myxococcia>unclassified_Myxococcia | 369 | HY744_34260 | 8566..9675 | 2026735 | DUF4276-family-protein-[Deltaproteobacteria-bacterium] |
| MCC6811539.1 | TOPRIM | GCA_020848545.1 | Deltaproteobacteria_bacterium | Bacteria>Pseudomonadati>Myxococcota>Myxococcia>unclassified_Myxococcia | 258 | IT381_29180 | 87778..88554 | 2026735 | RloB-domain-containing-protein-[Deltaproteobacteria-bacterium] |
| MBI4818891.1 | TOPRIM | GCA_016208175.1 | Deltaproteobacteria_bacterium | Bacteria>Pseudomonadati>Myxococcota>Myxococcia>unclassified_Myxococcia | 221 | HY791_21650 | 833..1498 | 2026735 | DUF4276-family-protein-[Deltaproteobacteria-bacterium] |
| PXF54131.1 | TOPRIM | GCA_003194485.1 | Deltaproteobacteria_bacterium | Bacteria>Pseudomonadati>Myxococcota>Myxococcia>unclassified_Myxococcia | 214 | C4B57_08255 | 22962..23606 | 2026735 | hypothetical-protein-C4B57_08255-[Deltaproteobacteria-bacterium] |
| PXF57192.1 | TOPRIM | GCA_003194495.1 | Deltaproteobacteria_bacterium | Bacteria>Pseudomonadati>Myxococcota>Myxococcia>unclassified_Myxococcia | 213 | C4B58_10700 | 13345..13986 | 2026735 | hypothetical-protein-C4B58_10700-[Deltaproteobacteria-bacterium] |
| MBO6936776.1 | TOPRIM | GCA_017644045.1 | Deltaproteobacteria_bacterium | Bacteria>Pseudomonadati>Myxococcota>Myxococcia>unclassified_Myxococcia | 212 | JJ863_17535 | 236806..237444 | 2026735 | DUF4276-family-protein-[Deltaproteobacteria-bacterium] |
| MCX5910051.1 | TOPRIM | GCA_026388955.1 | Deltaproteobacteria_bacterium | Bacteria>Pseudomonadati>Myxococcota>Myxococcia>unclassified_Myxococcia | 209 | NTY64_23460 | 379..1008 | 2026735 | MAG:-hypothetical-protein-NTY64_23460-[Deltaproteobacteria-bacterium] |
| MBT3255202.1 | TOPRIM | GCA_018644785.1 | Deltaproteobacteria_bacterium | Bacteria>Pseudomonadati>Myxococcota>Myxococcia>unclassified_Myxococcia | 205 | HN366_01915 | 15223..15840 | 2026735 | DUF4276-family-protein-[Deltaproteobacteria-bacterium] |
| MCC6811536.1 | TOPRIM | GCA_020848545.1 | Deltaproteobacteria_bacterium | Bacteria>Pseudomonadati>Myxococcota>Myxococcia>unclassified_Myxococcia | 205 | IT381_29165 | 81581..82198 | 2026735 | hypothetical-protein-[Deltaproteobacteria-bacterium] |
| MBW1911017.1 | TOPRIM | GCA_019307765.1 | Deltaproteobacteria_bacterium | Bacteria>Pseudomonadati>Myxococcota>Myxococcia>unclassified_Myxococcia | 204 | JRJ11_16020 | 7763..8377 | 2026735 | hypothetical-protein-[Deltaproteobacteria-bacterium] |
| MBW1922688.1 | TOPRIM | GCA_019307945.1 | Deltaproteobacteria_bacterium | Bacteria>Pseudomonadati>Myxococcota>Myxococcia>unclassified_Myxococcia | 202 | JRJ35_04380 | 4994..5602 | 2026735 | hypothetical-protein-[Deltaproteobacteria-bacterium] |
| NLN59462.1 | TOPRIM | GCA_012522415.1 | Deltaproteobacteria_bacterium | Bacteria>Pseudomonadati>Myxococcota>Myxococcia>unclassified_Myxococcia | 202 | GX147_01900 | 35641..36249 | 2026735 | hypothetical-protein-[Deltaproteobacteria-bacterium] |
| RLC08399.1 | TOPRIM | GCA_003647435.1 | Deltaproteobacteria_bacterium | Bacteria>Pseudomonadati>Myxococcota>Myxococcia>unclassified_Myxococcia | 202 | DRH43_10120 | 1070..1678 | 2026735 | hypothetical-protein-DRH43_10120-[Deltaproteobacteria-bacterium] |
| TRZ75306.1 | TOPRIM | GCA_007279985.1 | Deltaproteobacteria_bacterium | Bacteria>Pseudomonadati>Myxococcota>Myxococcia>unclassified_Myxococcia | 200 | D4R93_05185 | 3466..4068 | 2026735 | DUF4276-family-protein-[Deltaproteobacteria-bacterium] |
| MBI4706263.1 | TOPRIM | GCA_016209145.1 | Deltaproteobacteria_bacterium | Bacteria>Pseudomonadati>Myxococcota>Myxococcia>unclassified_Myxococcia | 198 | HY744_34635 | 15417..16013 | 2026735 | hypothetical-protein-[Deltaproteobacteria-bacterium] |
| MDD3553582.1 | TOPRIM | GCA_028695735.1 | Deltaproteobacteria_bacterium | Bacteria>Pseudomonadati>Myxococcota>Myxococcia>unclassified_Myxococcia | 197 | PHC35_03570 | 35351..35944 | 2026735 | MAG:-DUF4276-family-protein-[Deltaproteobacteria-bacterium] |
| MBN1905586.1 | TOPRIM | GCA_016931115.1 | Deltaproteobacteria_bacterium | Bacteria>Pseudomonadati>Myxococcota>Myxococcia>unclassified_Myxococcia | 196 | JW927_10855 | 36348..36938 | 2026735 | RloB-domain-containing-protein-[Deltaproteobacteria-bacterium] |
| MBI3795503.1 | TOPRIM | GCA_016200595.1 | Deltaproteobacteria_bacterium | Bacteria>Pseudomonadati>Myxococcota>Myxococcia>unclassified_Myxococcia | 195 | HY268_00835 | 5464..6051 | 2026735 | hypothetical-protein-[Deltaproteobacteria-bacterium] |
| MBM4396668.1 | TOPRIM | GCA_016875475.1 | Deltaproteobacteria_bacterium | Bacteria>Pseudomonadati>Myxococcota>Myxococcia>unclassified_Myxococcia | 192 | FJ087_13390 | 4990..5568 | 2026735 | hypothetical-protein-[Deltaproteobacteria-bacterium] |
| MDD9873632.1 | TOPRIM | GCA_028817295.1 | Deltaproteobacteria_bacterium | Bacteria>Pseudomonadati>Myxococcota>Myxococcia>unclassified_Myxococcia | 184 | OXU65_09085 | 8871..9425 | 2026735 | MAG:-hypothetical-protein-OXU65_09085-[Deltaproteobacteria-bacterium] |
| MAG32382.1 | TOPRIM | GCA_002688255.1 | Deltaproteobacteria_bacterium | Bacteria>Pseudomonadati>Myxococcota>Myxococcia>unclassified_Myxococcia | 180 | CL908_15985 | 9086..9628 | 2026735 | hypothetical-protein-[Deltaproteobacteria-bacterium] |
| MBK8262772.1 | TOPRIM | GCA_016712315.1 | Nannocystis_sp. | Bacteria>Pseudomonadati>Myxococcota>Polyangia>Nannocystales>Nannocystaceae>Nannocystis>unclassified_Nannocystis | 198 | IPK80_15735 | 181771..182367 | 1962667 | hypothetical-protein-[Nannocystis-sp.] |
| WP_129573700.1 | TOPRIM | GCF_004135755.1 | Sorangium | Bacteria>Pseudomonadati>Myxococcota>Polyangia>Polyangiales>Polyangiaceae | 208 | SOCE836_RS08205 | 2234660..2235286 | 39643 | MULTISPECIES:-DUF4276-family-protein-[Sorangium] |
| WP_129345941.1 | TOPRIM | GCF_004135735.1 | Sorangium_cellulosum | Bacteria>Pseudomonadati>Myxococcota>Polyangia>Polyangiales>Polyangiaceae>Sorangium | 208 | SOCEGT47_RS05510 | 1455367..1455993 | 56 | DUF4276-family-protein-[Sorangium-cellulosum] |
| MBN2191665.1 | TOPRIM | GCA_016933275.1 | Polyangiaceae_bacterium | Bacteria>Pseudomonadati>Myxococcota>Polyangia>Polyangiales>Polyangiaceae>unclassified_Polyangiaceae | 227 | JW751_02520 | 18132..18815 | 2268199 | DUF4276-family-protein-[Polyangiaceae-bacterium] |
| MBK6517526.1 | TOPRIM | GCA_016704335.1 | Polyangiaceae_bacterium | Bacteria>Pseudomonadati>Myxococcota>Polyangia>Polyangiales>Polyangiaceae>unclassified_Polyangiaceae | 223 | IPG04_26275 | 278392..279063 | 2268199 | DUF4276-family-protein-[Polyangiaceae-bacterium] |
| MBK8937973.1 | TOPRIM | GCA_016719945.1 | Polyangiaceae_bacterium | Bacteria>Pseudomonadati>Myxococcota>Polyangia>Polyangiales>Polyangiaceae>unclassified_Polyangiaceae | 223 | IPM79_10095 | 432545..433216 | 2268199 | DUF4276-family-protein-[Polyangiaceae-bacterium] |
| MBP7125458.1 | TOPRIM | GCA_017994695.1 | Myxococcota_bacterium | Bacteria>Pseudomonadati>Myxococcota>unclassified_Myxococcota | 217 | KBD49_03720 | 63489..64142 | 2818507 | hypothetical-protein-[Myxococcota-bacterium] |
| MCS6797215.1 | TOPRIM | GCA_025054695.1 | Myxococcota_bacterium | Bacteria>Pseudomonadati>Myxococcota>unclassified_Myxococcota | 199 | NZ898_01575 | 56611..57210 | 2818507 | hypothetical-protein-[Myxococcota-bacterium] |
| MCK6575221.1 | TOPRIM | GCA_023150895.1 | Myxococcota_bacterium | Bacteria>Pseudomonadati>Myxococcota>unclassified_Myxococcota | 192 | L6V77_29495 | 386..964 | 2818507 | RloB-family-protein-[Myxococcota-bacterium] |
| MCU0671308.1 | TOPRIM | GCA_025453555.1 | Myxococcota_bacterium | Bacteria>Pseudomonadati>Myxococcota>unclassified_Myxococcota | 84 | MUE69_00800 | 1..252 | 2818507 | hypothetical-protein-[Myxococcota-bacterium] |
| MCC6543977.1 | TOPRIM | GCA_020846775.1 | Nitrospirota_bacterium | Bacteria>Pseudomonadati>Nitrospirota>unclassified_Nitrospirota | 214 | IT392_05665 | 170536..171180 | 2026887 | DUF4276-family-protein-[Nitrospirota-bacterium] |
| PIX82646.1 | TOPRIM | GCA_002783945.1 | Nitrospirae_bacterium_CG_4_10_14_3_um_filter_70_108 | Bacteria>Pseudomonadati>Nitrospirota>unclassified_Nitrospirota | 190 | COZ33_09560 | 3583..4155 | 1974054 | cytoplasmic-protein-[Nitrospirae-bacterium-CG_4_10_14_3_um_filter_70_108] |
| WP_000605305.1 | TOPRIM | GCF_000299535.2 | Gammaproteobacteria | Bacteria>Pseudomonadati>Pseudomonadota | 252 | B934_RS0100880 | 10388..11146 | 1236 | MULTISPECIES:-hypothetical-protein-[Gammaproteobacteria] |
| WP_014998458.1 | TOPRIM | - | Gammaproteobacteria | Bacteria>Pseudomonadati>Pseudomonadota | 225 | AMBLS11_RS09805 | 2292064..2292741 | 1236 | DUF4276-family-protein-[Alteromonas-macleodii] |
| WP_022988842.1 | TOPRIM | GCF_000475255.1 | Gammaproteobacteria | Bacteria>Pseudomonadati>Pseudomonadota | 225 | DFO68_RS01955 | 8525..9202 | 1236 | MULTISPECIES:-DUF4276-family-protein-[Gammaproteobacteria] |
| WP_065846577.1 | TOPRIM | GCF_011762225.1 | Alphaproteobacteria | Bacteria>Pseudomonadati>Pseudomonadota | 209 | FHS82_RS15740 | 28592..29221 | 28211 | MULTISPECIES:-DUF4276-family-protein-[Alphaproteobacteria] |
| WP_215861191.1 | TOPRIM | GCF_018854875.1 | Acidithiobacillus_ferriphilus | Bacteria>Pseudomonadati>Pseudomonadota>Acidithiobacillia>Acidithiobacillales>Acidithiobacillaceae>Acidithiobacillus | 297 | HF928_RS01655 | 14061..14954 | 1689834 | DUF4435-domain-containing-protein-[Acidithiobacillus-ferriphilus] |
| WP_014029722.1 | TOPRIM | GCF_000214095.2 | Acidithiobacillus_ferrivorans | Bacteria>Pseudomonadati>Pseudomonadota>Acidithiobacillia>Acidithiobacillales>Acidithiobacillaceae>Acidithiobacillus | 218 | ACIFE_RS11340 | 2372457..2373113 | 160808 | DUF4276-family-protein-[Acidithiobacillus-ferrivorans] |
| WP_123104884.1 | TOPRIM | GCF_003721225.2 | Acidithiobacillus_sulfuriphilus | Bacteria>Pseudomonadati>Pseudomonadota>Acidithiobacillia>Acidithiobacillales>Acidithiobacillaceae>Acidithiobacillus | 205 | EC580_RS10650 | 6720..7337 | 1867749 | DUF4276-family-protein-[Acidithiobacillus-sulfuriphilus] |
| WP_218285165.1 | TOPRIM | GCF_019203965.1 | Elioraea_tepida | Bacteria>Pseudomonadati>Pseudomonadota>Alphaproteobacteria>Acetobacterales>Elioraeaceae>Elioraea | 172 | KO353_RS13035 | 2714065..2714583 | 2843330 | DUF4276-family-protein-[Elioraea-tepida] |
| MCB1832523.1 | TOPRIM | GCA_020445435.1 | Geminicoccaceae_bacterium | Bacteria>Pseudomonadati>Pseudomonadota>Alphaproteobacteria>Geminicoccales>Geminicoccaceae>unclassified_Geminicoccaceae | 209 | KDH19_03615 | 17611..18240 | 2448052 | DUF4276-family-protein-[Geminicoccaceae-bacterium] |
| VXB71474.1 | TOPRIM | GCA_902506225.1 | Enhydrobacter_sp._8BJ | Bacteria>Pseudomonadati>Pseudomonadota>Alphaproteobacteria>Hyphomicrobiales>Hyphomicrobiales_incertae_sedis>Enhydrobacter>unclassified_Enhydrobacter | 244 | ENHYD8BJ_80436 | 466404..467138 | 2653137 | conserved-hypothetical-protein-[Enhydrobacter-sp.-8BJ] |
| WP_115156542.1 | TOPRIM | GCF_004329915.1 | Rhizobium_leguminosarum | Bacteria>Pseudomonadati>Pseudomonadota>Alphaproteobacteria>Hyphomicrobiales>Rhizobiaceae>Rhizobium/Agrobacterium_group>Rhizobium | 209 | EOH64_RS12780 | 332589..333218 | 384 | DUF4276-family-protein-[Rhizobium-leguminosarum] |
| MBF0262094.1 | TOPRIM | GCA_015231775.1 | Magnetococcales_bacterium | Bacteria>Pseudomonadati>Pseudomonadota>Alphaproteobacteria>Magnetococcales>unclassified_Magnetococcales | 208 | HQL97_09710 | 7864..8490 | 2026759 | hypothetical-protein-[Magnetococcales-bacterium] |
| MBF0181808.1 | TOPRIM | GCA_015231235.1 | Magnetococcales_bacterium | Bacteria>Pseudomonadati>Pseudomonadota>Alphaproteobacteria>Magnetococcales>unclassified_Magnetococcales | 205 | HQM03_17460 | 10479..11096 | 2026759 | hypothetical-protein-[Magnetococcales-bacterium] |
| OOY26624.1 | TOPRIM | GCA_002020275.1 | Thioclava_sp._L04-15 | Bacteria>Pseudomonadati>Pseudomonadota>Alphaproteobacteria>Rhodobacterales>Paracoccaceae>Thioclava>unclassified_Thioclava | 343 | BMI90_16655 | 33137..34168 | 1915318 | DNA-primase-[Thioclava-sp.-L04-15] |
| WP_078530616.1 | TOPRIM | GCF_002020275.1 | Thioclava_sp._L04-15 | Bacteria>Pseudomonadati>Pseudomonadota>Alphaproteobacteria>Rhodobacterales>Paracoccaceae>Thioclava>unclassified_Thioclava | 343 | BMI90_RS16585 | 33137..34168 | 1915318 | toprim-domain-containing-protein-[Thioclava-sp.-L04-15] |
| OOY26630.1 | TOPRIM | GCA_002020275.1 | Thioclava_sp._L04-15 | Bacteria>Pseudomonadati>Pseudomonadota>Alphaproteobacteria>Rhodobacterales>Paracoccaceae>Thioclava>unclassified_Thioclava | 211 | BMI90_16690 | 42246..42881 | 1915318 | hypothetical-protein-BMI90_16690-[Thioclava-sp.-L04-15] |
| WP_078530622.1 | TOPRIM | GCF_002020275.1 | Thioclava_sp._L04-15 | Bacteria>Pseudomonadati>Pseudomonadota>Alphaproteobacteria>Rhodobacterales>Paracoccaceae>Thioclava>unclassified_Thioclava | 211 | BMI90_RS16620 | 42246..42881 | 1915318 | DUF4276-family-protein-[Thioclava-sp.-L04-15] |
| QCO03443.1 | TOPRIM | GCA_005222185.1 | Azospirillum_argentinense | Bacteria>Pseudomonadati>Pseudomonadota>Alphaproteobacteria>Rhodospirillales>Azospirillaceae>Azospirillum | 231 | D3867_15335 | 174553..175248 | 2970906 | DUF4276-family-protein-[Azospirillum-argentinense] |
| MDE0384392.1 | TOPRIM | GCA_028821395.1 | Defluviicoccus_sp. | Bacteria>Pseudomonadati>Pseudomonadota>Alphaproteobacteria>Rhodospirillales>Rhodospirillaceae>Defluviicoccus>unclassified_Defluviicoccus | 187 | OXI22_10950 | 42322..42885 | 2682137 | MAG:-hypothetical-protein-OXI22_10950-[Defluviicoccus-sp.] |
| MDA8051500.1 | TOPRIM | GCA_028063655.1 | Rhodospirillales_bacterium | Bacteria>Pseudomonadati>Pseudomonadota>Alphaproteobacteria>Rhodospirillales>unclassified_Rhodospirillales | 201 | M0002_16115 | 4752..5357 | 2026786 | MAG:-hypothetical-protein-M0002_16115-[Rhodospirillales-bacterium] |
| MCB9746494.1 | TOPRIM | GCA_020632815.1 | Alphaproteobacteria_bacterium | Bacteria>Pseudomonadati>Pseudomonadota>Alphaproteobacteria>unclassified_Alphaproteobacteria | 227 | H6740_28220 | 8726..9409 | 1913988 | hypothetical-protein-[Alphaproteobacteria-bacterium] |
| MCB9797659.1 | TOPRIM | GCA_020632765.1 | Alphaproteobacteria_bacterium | Bacteria>Pseudomonadati>Pseudomonadota>Alphaproteobacteria>unclassified_Alphaproteobacteria | 227 | H6741_33635 | 15741..16424 | 1913988 | hypothetical-protein-[Alphaproteobacteria-bacterium] |
| MCY3753478.1 | TOPRIM | GCA_026706245.1 | Alphaproteobacteria_bacterium | Bacteria>Pseudomonadati>Pseudomonadota>Alphaproteobacteria>unclassified_Alphaproteobacteria | 187 | OXG99_05235 | 12449..13012 | 1913988 | MAG:-hypothetical-protein-OXG99_05235-[Alphaproteobacteria-bacterium] |
| WP_071038553.1 | TOPRIM | GCF_001854305.1 | Burkholderiales | Bacteria>Pseudomonadati>Pseudomonadota>Betaproteobacteria | 209 | BKK79_RS15565 | 3509322..3509951 | 80840 | MULTISPECIES:-DUF4276-family-protein-[Burkholderiales] |
| WP_138677189.1 | TOPRIM | GCF_016790805.1 | Candidatus_Accumulibacter | Bacteria>Pseudomonadati>Pseudomonadota>Betaproteobacteria>Betaproteobacteria_incertae_sedis | 209 | ACCUM_RS00295 | 70141..70770 | 327159 | hypothetical-protein-[Candidatus-Accumulibacter-phosphatis] |
| RDE49749.1 | TOPRIM | GCA_003332265.1 | Candidatus_Accumulibacter_meliphilus | Bacteria>Pseudomonadati>Pseudomonadota>Betaproteobacteria>Betaproteobacteria_incertae_sedis>Candidatus_Accumulibacter | 248 | DVS81_14910 | 15657..16403 | 2211374 | MAG:-hypothetical-protein-DVS81_14910-[Candidatus-Accumulibacter-meliphilus] |
| WP_169070207.1 | TOPRIM | GCF_036625285.1 | Candidatus_Accumulibacter_contiguus | Bacteria>Pseudomonadati>Pseudomonadota>Betaproteobacteria>Betaproteobacteria_incertae_sedis>Candidatus_Accumulibacter | 248 | E4Q08_RS09475 | 42655..43401 | 2954381 | hypothetical-protein-[Candidatus-Accumulibacter-contiguus] |
| KFB72163.1 | TOPRIM | GCA_000585035.2 | Candidatus_Accumulibacter_phosphatis | Bacteria>Pseudomonadati>Pseudomonadota>Betaproteobacteria>Betaproteobacteria_incertae_sedis>Candidatus_Accumulibacter | 221 | AW09_002655 | 8716..9381 | 327160 | MAG:-hypothetical-protein-AW09_002655-[Candidatus-Accumulibacter-phosphatis] |
| QLH49279.1 | TOPRIM | GCA_013414765.1 | Candidatus_Accumulibacter_cognatus | Bacteria>Pseudomonadati>Pseudomonadota>Betaproteobacteria>Betaproteobacteria_incertae_sedis>Candidatus_Accumulibacter | 213 | HWD57_05385 | 1166481..1167122 | 2954383 | MAG:-RloB-domain-containing-protein-[Candidatus-Accumulibacter-cognatus] |
| QLH49281.1 | TOPRIM | GCA_013414765.1 | Candidatus_Accumulibacter_cognatus | Bacteria>Pseudomonadati>Pseudomonadota>Betaproteobacteria>Betaproteobacteria_incertae_sedis>Candidatus_Accumulibacter | 209 | HWD57_05400 | 1168514..1169143 | 2954383 | MAG:-hypothetical-protein-HWD57_05400-[Candidatus-Accumulibacter-cognatus] |
| KFB72165.1 | TOPRIM | GCA_000585035.2 | Candidatus_Accumulibacter_phosphatis | Bacteria>Pseudomonadati>Pseudomonadota>Betaproteobacteria>Betaproteobacteria_incertae_sedis>Candidatus_Accumulibacter | 207 | AW09_002657 | 10739..11362 | 327160 | MAG:-hypothetical-protein-AW09_002657-[Candidatus-Accumulibacter-phosphatis] |
| TLD46496.1 | TOPRIM | GCA_005524045.1 | Accumulibacter_sp. | Bacteria>Pseudomonadati>Pseudomonadota>Betaproteobacteria>Betaproteobacteria_incertae_sedis>Candidatus_Accumulibacter>unclassified_Candidatus_Accumulibacter | 224 | FAZ92_01233 | 11569..12243 | 2053492 | MAG:-hypothetical-protein-FAZ92_01233-[Candidatus-Accumulibacter-sp.] |
| MBK8115028.1 | TOPRIM | GCA_016712935.1 | Accumulibacter_sp. | Bacteria>Pseudomonadati>Pseudomonadota>Betaproteobacteria>Betaproteobacteria_incertae_sedis>Candidatus_Accumulibacter>unclassified_Candidatus_Accumulibacter | 213 | IPK44_11020 | 655031..655672 | 2053492 | RloB-domain-containing-protein-[Candidatus-Accumulibacter-sp.] |
| MBN8517034.1 | TOPRIM | GCA_017302385.1 | Accumulibacter_sp. | Bacteria>Pseudomonadati>Pseudomonadota>Betaproteobacteria>Betaproteobacteria_incertae_sedis>Candidatus_Accumulibacter>unclassified_Candidatus_Accumulibacter | 213 | JOL71_04000 | 49485..50126 | 2053492 | RloB-domain-containing-protein-[Candidatus-Accumulibacter-sp.] |
| MBN8517036.1 | TOPRIM | GCA_017302385.1 | Accumulibacter_sp. | Bacteria>Pseudomonadati>Pseudomonadota>Betaproteobacteria>Betaproteobacteria_incertae_sedis>Candidatus_Accumulibacter>unclassified_Candidatus_Accumulibacter | 209 | JOL71_04015 | 51518..52147 | 2053492 | hypothetical-protein-[Candidatus-Accumulibacter-sp.] |
| MBN8451981.1 | TOPRIM | GCA_017302415.1 | Accumulibacter_sp. | Bacteria>Pseudomonadati>Pseudomonadota>Betaproteobacteria>Betaproteobacteria_incertae_sedis>Candidatus_Accumulibacter>unclassified_Candidatus_Accumulibacter | 204 | JOM31_02655 | 90681..91295 | 2053492 | DUF4276-family-protein-[Candidatus-Accumulibacter-sp.] |
| MBK8387655.1 | TOPRIM | GCA_016711725.1 | Accumulibacter_sp. | Bacteria>Pseudomonadati>Pseudomonadota>Betaproteobacteria>Betaproteobacteria_incertae_sedis>Candidatus_Accumulibacter>unclassified_Candidatus_Accumulibacter | 163 | IPL11_19655 | 166..657 | 2053492 | DUF4276-family-protein-[Candidatus-Accumulibacter-sp.] |
| MCX8099547.1 | TOPRIM | GCA_026414825.1 | Casimicrobiaceae_bacterium | Bacteria>Pseudomonadati>Pseudomonadota>Betaproteobacteria>Betaproteobacteria_incertae_sedis>Casimicrobiaceae>unclassified_Casimicrobiaceae | 199 | N3F11_11210 | 12010..12609 | 2906524 | MAG:-hypothetical-protein-N3F11_11210-[Casimicrobiaceae-bacterium] |
| MCH1991585.1 | TOPRIM | GCA_022444305.1 | Achromobacter_xylosoxidans | Bacteria>Pseudomonadati>Pseudomonadota>Betaproteobacteria>Burkholderiales>Alcaligenaceae>Achromobacter | 217 | L7Q78_02315 | 21786..22439 | 85698 | DUF4276-family-protein-[Achromobacter-xylosoxidans] |
| SUA58549.1 | TOPRIM | GCA_900454315.1 | Oligella_ureolytica | Bacteria>Pseudomonadati>Pseudomonadota>Betaproteobacteria>Burkholderiales>Alcaligenaceae>Oligella | 141 | NCTC11998_02424 | 2465198..2465623 | 90244 | Uncharacterised-protein-[Oligella-ureolytica] |
| NYT60765.1 | TOPRIM | GCA_013416445.1 | Alcaligenaceae_bacterium | Bacteria>Pseudomonadati>Pseudomonadota>Betaproteobacteria>Burkholderiales>Alcaligenaceae>unclassified_Alcaligenaceae | 227 | HOA66_00270 | 64829..65512 | 2021368 | DUF4276-family-protein-[Alcaligenaceae-bacterium] |
| WP_254215275.1 | TOPRIM | - | Burkholderia_multivorans | Bacteria>Pseudomonadati>Pseudomonadota>Betaproteobacteria>Burkholderiales>Burkholderiaceae>Burkholderia>Burkholderia_cepacia_complex | 196 | KTD09_RS06775 | 73482..74072 | 87883 | DUF4276-family-protein-[Burkholderia-multivorans] |
| KIX33649.1 | TOPRIM | GCA_000932075.1 | Burkholderia_pseudomallei | Bacteria>Pseudomonadati>Pseudomonadota>Betaproteobacteria>Burkholderiales>Burkholderiaceae>Burkholderia>pseudomallei_group | 209 | SY87_33350 | 9932..10561 | 28450 | hypothetical-protein-SY87_33350-[Burkholderia-pseudomallei] |
| WP_044359464.1 | TOPRIM | GCF_000932075.1 | Burkholderia_pseudomallei | Bacteria>Pseudomonadati>Pseudomonadota>Betaproteobacteria>Burkholderiales>Burkholderiaceae>Burkholderia>pseudomallei_group | 209 | SY87_RS33400 | 9932..10561 | 28450 | DUF4276-family-protein-[Burkholderia-pseudomallei] |
| AOY94228.1 | TOPRIM | GCA_001854305.1 | Cupriavidus_sp._USMAA2-4 | Bacteria>Pseudomonadati>Pseudomonadota>Betaproteobacteria>Burkholderiales>Burkholderiaceae>Cupriavidus>unclassified_Cupriavidus | 209 | BKK79_15570 | 3509322..3509951 | 876364 | hypothetical-protein-BKK79_15570-[Cupriavidus-sp.-USMAA2-4] |
| WP_017772489.1 | TOPRIM | GCF_000341045.1 | Paraburkholderia_kururiensis | Bacteria>Pseudomonadati>Pseudomonadota>Betaproteobacteria>Burkholderiales>Burkholderiaceae>Paraburkholderia | 209 | G118_RS0102335 | 508655..509284 | 984307 | DUF4276-family-protein-[Paraburkholderia-kururiensis] |
| WP_208110744.1 | TOPRIM | GCF_004362855.1 | Aquabacterium_commune | Bacteria>Pseudomonadati>Pseudomonadota>Betaproteobacteria>Burkholderiales>Burkholderiales_genera_incertae_sedis>Aquabacterium | 212 | EV672_RS09265 | 192755..193393 | 70586 | hypothetical-protein-[Aquabacterium-commune] |
| WP_143969785.1 | TOPRIM | GCF_007556755.1 | Tepidimonas_fonticaldi | Bacteria>Pseudomonadati>Pseudomonadota>Betaproteobacteria>Burkholderiales>Burkholderiales_genera_incertae_sedis>Tepidimonas | 195 | Tfont_RS13270 | 8019..8606 | 1101373 | DUF4276-family-protein-[Tepidimonas-fonticaldi] |
| WP_231572172.1 | TOPRIM | GCF_000807215.1 | Tepidimonas_taiwanensis | Bacteria>Pseudomonadati>Pseudomonadota>Betaproteobacteria>Burkholderiales>Burkholderiales_genera_incertae_sedis>Tepidimonas | 191 | PL21_RS04255 | 84409..84984 | 307486 | DUF4276-family-protein-[Tepidimonas-taiwanensis] |
| WP_260682384.1 | TOPRIM | GCF_007556755.1 | Tepidimonas_fonticaldi | Bacteria>Pseudomonadati>Pseudomonadota>Betaproteobacteria>Burkholderiales>Burkholderiales_genera_incertae_sedis>Tepidimonas | 189 | Tfont_RS12355 | 12335..12904 | 1101373 | DUF4276-family-protein-[Tepidimonas-fonticaldi] |
| WP_272568619.1 | TOPRIM | GCF_030159035.1 | unclassified_Acidovorax | Bacteria>Pseudomonadati>Pseudomonadota>Betaproteobacteria>Burkholderiales>Comamonadaceae>Acidovorax | 247 | M5C96_RS08965 | 1970169..1970912 | 2684926 | hypothetical-protein-[Acidovorax-sp.-GBBC-1281] |
| WP_157896445.1 | TOPRIM | GCF_002157145.1 | Acidovorax_carolinensis | Bacteria>Pseudomonadati>Pseudomonadota>Betaproteobacteria>Burkholderiales>Comamonadaceae>Acidovorax | 223 | CBP34_RS19385 | 1390398..1391069 | 553814 | hypothetical-protein-[Acidovorax-carolinensis] |
| WP_008906817.1 | TOPRIM | GCF_000238595.1 | Acidovorax_sp._NO-1 | Bacteria>Pseudomonadati>Pseudomonadota>Betaproteobacteria>Burkholderiales>Comamonadaceae>Acidovorax>unclassified_Acidovorax | 253 | KYG_RS20350 | 15302..16063 | 512030 | hypothetical-protein-[Acidovorax-sp.-NO-1] |
| GAO25243.1 | TOPRIM | GCA_000974645.1 | Alicycliphilus_sp._B1 | Bacteria>Pseudomonadati>Pseudomonadota>Betaproteobacteria>Burkholderiales>Comamonadaceae>Alicycliphilus>unclassified_Alicycliphilus | 222 | ALISP_5063 | 1531335..1532003 | 1603291 | hypothetical-protein-ALISP_5063-[Alicycliphilus-sp.-B1] |
| WP_091817124.1 | TOPRIM | GCF_900110225.1 | Brachymonas_denitrificans | Bacteria>Pseudomonadati>Pseudomonadota>Betaproteobacteria>Burkholderiales>Comamonadaceae>Brachymonas | 217 | BMW28_RS09515 | 74246..74899 | 28220 | hypothetical-protein-[Brachymonas-denitrificans] |
| SEN73507.1 | TOPRIM | GCA_900110225.1 | Brachymonas_denitrificans_DSM_15123 | Bacteria>Pseudomonadati>Pseudomonadota>Betaproteobacteria>Burkholderiales>Comamonadaceae>Brachymonas>Brachymonas_denitrificans | 217 | SAMN02745977_01907 | 74246..74899 | 1121117 | hypothetical-protein-SAMN02745977_01907-[Brachymonas-denitrificans-DSM-15123] |
| WP_054075979.1 | TOPRIM | GCF_001294075.1 | Comamonas_testosteroni | Bacteria>Pseudomonadati>Pseudomonadota>Betaproteobacteria>Burkholderiales>Comamonadaceae>Comamonas | 188 | KF712C_RS25805 | 24094..24658 | 285 | hypothetical-protein,-partial-[Comamonas-testosteroni] |
| MBF5004208.1 | TOPRIM | GCA_015354245.1 | Diaphorobacter_caeni | Bacteria>Pseudomonadati>Pseudomonadota>Betaproteobacteria>Burkholderiales>Comamonadaceae>Diaphorobacter | 209 | ISG32_08045 | 70216..70845 | 2784387 | DUF4276-family-protein-[Diaphorobacter-caeni] |
| QIL80933.1 | TOPRIM | GCA_011305995.1 | Diaphorobacter_sp._HDW4A | Bacteria>Pseudomonadati>Pseudomonadota>Betaproteobacteria>Burkholderiales>Comamonadaceae>Diaphorobacter>unclassified_Diaphorobacter | 223 | G7047_14260 | 3129635..3130306 | 2714924 | DUF4276-family-protein-[Diaphorobacter-sp.-HDW4A] |
| WP_166306536.1 | TOPRIM | GCF_011305995.1 | Diaphorobacter_sp._HDW4A | Bacteria>Pseudomonadati>Pseudomonadota>Betaproteobacteria>Burkholderiales>Comamonadaceae>Diaphorobacter>unclassified_Diaphorobacter | 223 | G7047_RS14255 | 3129635..3130306 | 2714924 | DUF4276-family-protein-[Diaphorobacter-sp.-HDW4A] |
| WP_104799763.1 | TOPRIM | GCF_002943475.1 | Limnohabitans_sp._TS-CS-82 | Bacteria>Pseudomonadati>Pseudomonadota>Betaproteobacteria>Burkholderiales>Comamonadaceae>Limnohabitans>unclassified_Limnohabitans | 250 | C5F52_RS18140 | 193040..193792 | 2094193 | hypothetical-protein-[Limnohabitans-sp.-TS-CS-82] |
| WP_146187722.1 | TOPRIM | GCF_003063315.1 | Limnohabitans_sp._T6-5 | Bacteria>Pseudomonadati>Pseudomonadota>Betaproteobacteria>Burkholderiales>Comamonadaceae>Limnohabitans>unclassified_Limnohabitans | 250 | B9Z51_RS07540 | 403771..404523 | 1100724 | hypothetical-protein-[Limnohabitans-sp.-T6-5] |
| WP_108373345.1 | TOPRIM | GCF_003063315.1 | Limnohabitans_sp._T6-5 | Bacteria>Pseudomonadati>Pseudomonadota>Betaproteobacteria>Burkholderiales>Comamonadaceae>Limnohabitans>unclassified_Limnohabitans | 222 | B9Z51_RS07550 | 405725..406393 | 1100724 | hypothetical-protein-[Limnohabitans-sp.-T6-5] |
| WP_108373334.1 | TOPRIM | GCF_003063315.1 | Limnohabitans_sp._T6-5 | Bacteria>Pseudomonadati>Pseudomonadota>Betaproteobacteria>Burkholderiales>Comamonadaceae>Limnohabitans>unclassified_Limnohabitans | 194 | B9Z51_RS07520 | 400517..401101 | 1100724 | DUF4276-family-protein-[Limnohabitans-sp.-T6-5] |
| WP_209423132.1 | TOPRIM | GCF_021249385.1 | Melaminivora_jejuensis | Bacteria>Pseudomonadati>Pseudomonadota>Betaproteobacteria>Burkholderiales>Comamonadaceae>Melaminivora | 210 | LVC68_RS01460 | 288691..289323 | 1267217 | hypothetical-protein-[Melaminivora-jejuensis] |
| WP_265288961.1 | TOPRIM | GCF_026127525.1 | Verminephrobacter_eiseniae | Bacteria>Pseudomonadati>Pseudomonadota>Betaproteobacteria>Burkholderiales>Comamonadaceae>Verminephrobacter | 245 | D5047_RS18230 | 3906525..3907262 | 364317 | hypothetical-protein-[Verminephrobacter-eiseniae] |
| WP_208624500.1 | TOPRIM | GCF_002327145.1 | Janthinobacterium_svalbardensis | Bacteria>Pseudomonadati>Pseudomonadota>Betaproteobacteria>Burkholderiales>Oxalobacteraceae>Janthinobacterium | 202 | CNX70_RS06895 | 1573014..1573622 | 368607 | hypothetical-protein-[Janthinobacterium-svalbardensis] |
| WP_251969636.1 | TOPRIM | GCF_023734135.1 | Sphaerotilus_microaerophilus | Bacteria>Pseudomonadati>Pseudomonadota>Betaproteobacteria>Burkholderiales>Sphaerotilaceae>Sphaerotilus | 254 | NGK70_RS16745 | 3932408..3933172 | 2914710 | hypothetical-protein-[Sphaerotilus-sp.-FB-5] |
| MCK6404268.1 | TOPRIM | GCA_023150255.1 | Sphaerotilus_sulfidivorans | Bacteria>Pseudomonadati>Pseudomonadota>Betaproteobacteria>Burkholderiales>Sphaerotilaceae>Sphaerotilus | 185 | L6Q74_20555 | 8773..9330 | 639200 | DUF4276-family-protein-[Sphaerotilus-sulfidivorans] |
| NRT58385.1 | TOPRIM | GCA_013294065.1 | Sphaerotilus_uruguayifluvii | Bacteria>Pseudomonadati>Pseudomonadota>Betaproteobacteria>Burkholderiales>Sphaerotilaceae>Sphaerotilus | 185 | HNQ01_004153 | 21410..21967 | 2735897 | hypothetical-protein-[Leptothrix-sp.-C29] |
| WP_173807404.1 | TOPRIM | GCF_013294065.1 | Sphaerotilus_uruguayifluvii | Bacteria>Pseudomonadati>Pseudomonadota>Betaproteobacteria>Burkholderiales>Sphaerotilaceae>Sphaerotilus | 185 | HNQ01_RS20660 | 21410..21967 | 2735897 | DUF4276-family-protein-[Leptothrix-sp.-C29] |
| WP_273473500.1 | TOPRIM | GCF_023150255.1 | Sphaerotilus_sulfidivorans | Bacteria>Pseudomonadati>Pseudomonadota>Betaproteobacteria>Burkholderiales>Sphaerotilaceae>Sphaerotilus | 185 | L6Q74_RS20515 | 8773..9330 | 639200 | DUF4276-family-protein-[Sphaerotilus-sulfidivorans] |
| MBP7846547.1 | TOPRIM | GCA_018001735.1 | Burkholderiales_bacterium | Bacteria>Pseudomonadati>Pseudomonadota>Betaproteobacteria>Burkholderiales>unclassified_Burkholderiales | 262 | KAZ94_00265 | 62247..63035 | 1891238 | RloB-domain-containing-protein-[Burkholderiales-bacterium] |
| MBX9598970.1 | TOPRIM | GCA_019748155.1 | Burkholderiales_bacterium | Bacteria>Pseudomonadati>Pseudomonadota>Betaproteobacteria>Burkholderiales>unclassified_Burkholderiales | 262 | K2X04_10430 | 11887..12675 | 1891238 | RloB-family-protein-[Burkholderiales-bacterium] |
| MBK6715973.1 | TOPRIM | GCA_016703325.1 | Burkholderiales_bacterium | Bacteria>Pseudomonadati>Pseudomonadota>Betaproteobacteria>Burkholderiales>unclassified_Burkholderiales | 258 | IPG57_12975 | 78148..78924 | 1891238 | hypothetical-protein-[Burkholderiales-bacterium] |
| MCZ2419566.1 | TOPRIM | GCA_026931465.1 | Burkholderiales_bacterium | Bacteria>Pseudomonadati>Pseudomonadota>Betaproteobacteria>Burkholderiales>unclassified_Burkholderiales | 217 | LC123_06985 | 88907..89560 | 1891238 | MAG:-RloB-family-protein-[Burkholderiales-bacterium] |
| MBE0627112.1 | TOPRIM | GCA_014860355.1 | Burkholderiales_bacterium | Bacteria>Pseudomonadati>Pseudomonadota>Betaproteobacteria>Burkholderiales>unclassified_Burkholderiales | 216 | IH606_20120 | 4574..5224 | 1891238 | RloB-domain-containing-protein-[Burkholderiales-bacterium] |

| Protein-ID | Domain | Assembly-ID | Strain | Taxonomy | Length | Locus-Tag | Genomic-coordinates | TaxID | Fasta-defline |
|---|---|---|---|---|---|---|---|---|---|
| MBX9598968.1 | TOPRIM | GCA_019748155.1 | Burkholderiales_bacterium | Bacteria>Pseudomonadati>Pseudomonadota>Betaproteobacteria>Burkholderiales>unclassified_Burkholderiales | 202 | K2X04_10420 | 9904..10512 | 1891238 | DUF4276-family-protein-[Burkholderiales-bacterium] |
| NMA26875.1 | TOPRIM | GCA_012799975.1 | Burkholderiales_bacterium | Bacteria>Pseudomonadati>Pseudomonadota>Betaproteobacteria>Burkholderiales>unclassified_Burkholderiales | 188 | GX934_03855 | 2000..2566 | 1891238 | DUF4276-family-protein-[Burkholderiales-bacterium] |
| QKE41379.1 | TOPRIM | GCA_013181415.1 | Ferrovum_myxofaciens | Bacteria>Pseudomonadati>Pseudomonadota>Betaproteobacteria>Ferrovales>Ferrovaceae>Ferrovum | 99 | HO274_08655 | 1791264..1791563 | 416213 | hypothetical-protein-HO274_08655-[Ferrovum-myxofaciens] |
| WP_031597355.1 | TOPRIM | GCF_000735045.1 | Ferrovum_myxofaciens | Bacteria>Pseudomonadati>Pseudomonadota>Betaproteobacteria>Ferrovales>Ferrovaceae>Ferrovum | 99 | IV91_RS0107685 | 4583..4882 | 416213 | RloB-domain-containing-protein-[Ferrovum-myxofaciens] |
| WP_273118815.1 | TOPRIM | GCF_013181415.1 | Ferrovum_myxofaciens | Bacteria>Pseudomonadati>Pseudomonadota>Betaproteobacteria>Ferrovales>Ferrovaceae>Ferrovum | 99 | HO274_RS08685 | 1791264..1791563 | 416213 | RloB-domain-containing-protein-[Ferrovum-myxofaciens] |
| NDU85856.1 | TOPRIM | GCA_010436015.1 | Ferrovum_sp. | Bacteria>Pseudomonadati>Pseudomonadota>Betaproteobacteria>Ferrovales>Ferrovaceae>Ferrovum>unclassified_Ferrovum | 210 | G3H99_04495 | 12264..12896 | 2609467 | RloB-domain-containing-protein-[Ferrovum-sp.] |
| WP_022652886.1 | TOPRIM | GCF_000420525.1 | Aquaspirillum_serpens | Bacteria>Pseudomonadati>Pseudomonadota>Betaproteobacteria>Neisseriales>Aquaspirillaceae>Aquaspirillum | 217 | G465_RS0102385 | 19242..19895 | 190 | DUF4276-family-protein-[Aquaspirillum-serpens] |
| MCG9000248.1 | TOPRIM | GCA_022393975.1 | Laribacter_hongkongensis | Bacteria>Pseudomonadati>Pseudomonadota>Betaproteobacteria>Neisseriales>Aquaspirillaceae>Laribacter | 247 | LH454_02010 | 64245..64988 | 168471 | hypothetical-protein-[Laribacter-hongkongensis] |
| MCG9059781.1 | TOPRIM | GCA_022394335.1 | Laribacter_hongkongensis | Bacteria>Pseudomonadati>Pseudomonadota>Betaproteobacteria>Neisseriales>Aquaspirillaceae>Laribacter | 247 | LH452_12735 | 23349..24092 | 168471 | hypothetical-protein-[Laribacter-hongkongensis] |
| WP_239891832.1 | TOPRIM | GCF_022394335.1 | Laribacter_hongkongensis | Bacteria>Pseudomonadati>Pseudomonadota>Betaproteobacteria>Neisseriales>Aquaspirillaceae>Laribacter | 247 | LH452_RS12725 | 23349..24092 | 168471 | hypothetical-protein-[Laribacter-hongkongensis] |
| MCG9052523.1 | TOPRIM | GCA_022394235.1 | Laribacter_hongkongensis | Bacteria>Pseudomonadati>Pseudomonadota>Betaproteobacteria>Neisseriales>Aquaspirillaceae>Laribacter | 223 | LH447_05305 | 10305..10976 | 168471 | hypothetical-protein-[Laribacter-hongkongensis] |
| WP_239900338.1 | TOPRIM | GCF_022394275.1 | Laribacter_hongkongensis | Bacteria>Pseudomonadati>Pseudomonadota>Betaproteobacteria>Neisseriales>Aquaspirillaceae>Laribacter | 215 | LH442_RS13355 | 13608..14255 | 168471 | hypothetical-protein-[Laribacter-hongkongensis] |
| WP_047248548.1 | TOPRIM | GCF_001020585.1 | Chromobacterium_subtsugae | Bacteria>Pseudomonadati>Pseudomonadota>Betaproteobacteria>Neisseriales>Chromobacteriaceae>Chromobacterium | 272 | AA094_RS04295 | 49663..50481 | 251747 | DUF4435-domain-containing-protein-[Chromobacterium-subtsugae] |
| WP_013292865.1 | TOPRIM | GCF_000145255.1 | Gallionella_capsiferriformans | Bacteria>Pseudomonadati>Pseudomonadota>Betaproteobacteria>Nitrosomonadales>Gallionellaceae>Gallionella | 246 | GALF_RS04455 | 942015..942755 | 370405 | hypothetical-protein-[Gallionella-capsiferriformans] |
| MBP9798942.1 | TOPRIM | GCA_018062105.1 | Sterolibacterium_sp. | Bacteria>Pseudomonadati>Pseudomonadota>Betaproteobacteria>Nitrosomonadales>Sterolibacteriaceae>Sterolibacterium>unclassified_Sterolibacterium | 209 | KBD39_02335 | 26937..27566 | 2819261 | DUF4276-family-protein-[Sterolibacterium-sp.] |
| NOT17075.1 | TOPRIM | GCA_013140765.1 | Sulfuriferula_sp. | Bacteria>Pseudomonadati>Pseudomonadota>Betaproteobacteria>Nitrosomonadales>Sulfuricellaceae>Sulfuriferula>unclassified_Sulfuriferula | 245 | HOP20_03260 | 14032..14769 | 2025307 | hypothetical-protein-[Sulfuriferula-sp.] |
| MCX7674152.1 | TOPRIM | GCA_026419045.1 | Thiobacillaceae_bacterium | Bacteria>Pseudomonadati>Pseudomonadota>Betaproteobacteria>Nitrosomonadales>Thiobacillaceae>unclassified_Thiobacillaceae | 193 | N2Z63_11265 | 18153..18734 | 2528409 | MAG:-DUF4276-family-protein-[Thiobacillaceae-bacterium] |
| MBI5658239.1 | TOPRIM | GCA_016223025.1 | Nitrosomonadales_bacterium | Bacteria>Pseudomonadati>Pseudomonadota>Betaproteobacteria>Nitrosomonadales>unclassified_Nitrosomonadales | 232 | HZC43_01525 | 59526..60224 | 2026761 | RloB-domain-containing-protein-[Nitrosomonadales-bacterium] |
| MBI3902218.1 | TOPRIM | GCA_016199835.1 | Nitrosomonadales_bacterium | Bacteria>Pseudomonadati>Pseudomonadota>Betaproteobacteria>Nitrosomonadales>unclassified_Nitrosomonadales | 222 | HY306_04650 | 126134..126802 | 2026761 | RloB-domain-containing-protein-[Nitrosomonadales-bacterium] |
| MBI1887195.1 | TOPRIM | GCA_016177795.1 | Nitrosomonadales_bacterium | Bacteria>Pseudomonadati>Pseudomonadota>Betaproteobacteria>Nitrosomonadales>unclassified_Nitrosomonadales | 220 | HYS19_02315 | 93421..94083 | 2026761 | RloB-domain-containing-protein-[Nitrosomonadales-bacterium] |
| WP_153146307.1 | TOPRIM | GCF_009469615.1 | Dechloromonas_sp._H13 | Bacteria>Pseudomonadati>Pseudomonadota>Betaproteobacteria>Rhodocyclales>Azonexaceae>Dechloromonas>unclassified_Dechloromonas | 210 | GH660_RS08855 | 101887..102519 | 2570193 | DUF4276-family-protein-[Dechloromonas-sp.-H13] |
| MCK0506218.1 | TOPRIM | GCA_012910705.2 | Aromatoleum_anaerobium | Bacteria>Pseudomonadati>Pseudomonadota>Betaproteobacteria>Rhodocyclales>Rhodocyclaceae>Aromatoleum | 271 | GO606_004920 | 98270..99085 | 182180 | DUF4435-domain-containing-protein-[Aromatoleum-anaerobium] |
| WP_011238775.1 | TOPRIM | GCF_000025965.1 | Aromatoleum_aromaticum | Bacteria>Pseudomonadati>Pseudomonadota>Betaproteobacteria>Rhodocyclales>Rhodocyclaceae>Aromatoleum | 271 | EBN1_RS14790 | 3103986..3104801 | 551760 | DUF4435-domain-containing-protein-[Aromatoleum-aromaticum] |
| WP_169117246.1 | TOPRIM | GCF_012910705.2 | Aromatoleum_anaerobium | Bacteria>Pseudomonadati>Pseudomonadota>Betaproteobacteria>Rhodocyclales>Rhodocyclaceae>Aromatoleum | 271 | GO606_RS04920 | 98270..99085 | 182180 | DUF4435-domain-containing-protein-[Aromatoleum-anaerobium] |
| WP_153111720.1 | TOPRIM | GCF_009469695.1 | Propionivibrio_limicola | Bacteria>Pseudomonadati>Pseudomonadota>Betaproteobacteria>Rhodocyclales>Rhodocyclaceae>Propionivibrio | 217 | GH659_RS12235 | 3771..4424 | 167645 | DUF4276-family-protein-[Propionivibrio-limicola] |
| AUM01341.1 | TOPRIM | GCA_002863805.1 | Rhodocyclaceae_bacterium | Bacteria>Pseudomonadati>Pseudomonadota>Betaproteobacteria>Rhodocyclales>Rhodocyclaceae>unclassified_Rhodocyclaceae | 208 | B4966_15130 | 3174259..3174885 | 1898103 | hypothetical-protein-B4966_15130-[Rhodocyclaceae-bacterium] |
| TRZ91909.1 | TOPRIM | GCA_007280105.1 | Rhodocyclaceae_bacterium | Bacteria>Pseudomonadati>Pseudomonadota>Betaproteobacteria>Rhodocyclales>Rhodocyclaceae>unclassified_Rhodocyclaceae | 204 | D4R84_13300 | 1066..1680 | 1898103 | DUF4276-family-protein-[Rhodocyclaceae-bacterium] |
| TRZ91924.1 | TOPRIM | GCA_007280105.1 | Rhodocyclaceae_bacterium | Bacteria>Pseudomonadati>Pseudomonadota>Betaproteobacteria>Rhodocyclales>Rhodocyclaceae>unclassified_Rhodocyclaceae | 193 | D4R84_13315 | 4890..5471 | 1898103 | DUF4276-family-protein-[Rhodocyclaceae-bacterium] |
| MCX8086914.1 | TOPRIM | GCA_026415155.1 | Rhodocyclaceae_bacterium | Bacteria>Pseudomonadati>Pseudomonadota>Betaproteobacteria>Rhodocyclales>Rhodocyclaceae>unclassified_Rhodocyclaceae | 189 | N3C63_08445 | 31421..31990 | 1898103 | MAG:-DUF4276-family-protein-[Rhodocyclaceae-bacterium] |
| CAH1748217.1 | TOPRIM | GCA_932126575.1 | Thauera_humireducens | Bacteria>Pseudomonadati>Pseudomonadota>Betaproteobacteria>Rhodocyclales>Zoogloeaceae>Thauera | 249 | C4PIVTH_3077 | 3150983..3151732 | 1134435 | conserved-protein-of-unknown-function-[Thauera-sp.-PIV-1] |
| WP_247725475.1 | TOPRIM | GCF_023093595.1 | Thauera_aromatica | Bacteria>Pseudomonadati>Pseudomonadota>Betaproteobacteria>Rhodocyclales>Zoogloeaceae>Thauera | 249 | MXC99_RS10885 | 47042..47791 | 59405 | hypothetical-protein-[Thauera-aromatica] |
| AVZ79924.1 | TOPRIM | GCA_003054515.1 | Zoogloeaceae_bacteirum_Par-f-2 | Bacteria>Pseudomonadati>Pseudomonadota>Betaproteobacteria>Rhodocyclales>Zoogloeaceae>unclassified_Zoogloeaceae | 210 | C3497_11120 | 2341766..2342398 | 2080469 | hypothetical-protein-C3497_11120-[Zoogloeaceae-bacteirum-Par-f-2] |
| MBI3372696.1 | TOPRIM | GCA_016195915.1 | Betaproteobacteria_bacterium | Bacteria>Pseudomonadati>Pseudomonadota>Betaproteobacteria>unclassified_Betaproteobacteria | 214 | HY017_13205 | 57501..58145 | 1891241 | DUF4276-family-protein-[Betaproteobacteria-bacterium] |
| GHT91732.1 | TOPRIM | GCA_026000515.1 | Betaproteobacteria_bacterium | Bacteria>Pseudomonadati>Pseudomonadota>Betaproteobacteria>unclassified_Betaproteobacteria | 203 | FACS1894116_00770 | 35198..35809 | 1891241 | hypothetical-protein-FACS1894116_00770-[Betaproteobacteria-bacterium] |
| MBI3917900.1 | TOPRIM | GCA_016199545.1 | Betaproteobacteria_bacterium | Bacteria>Pseudomonadati>Pseudomonadota>Betaproteobacteria>unclassified_Betaproteobacteria | 197 | HY322_12935 | 27528..28121 | 1891241 | hypothetical-protein-[Betaproteobacteria-bacterium] |
| WP_099755073.1 | TOPRIM | GCF_902168565.1 | Enterobacterales | Bacteria>Pseudomonadati>Pseudomonadota>Gammaproteobacteria | 277 | FS908_RS01790 | 62034..62867 | 91347 | MULTISPECIES:-DUF4435-domain-containing-protein-[Enterobacterales] |
| WP_112020506.1 | TOPRIM | GCF_013624605.1 | Enterobacterales | Bacteria>Pseudomonadati>Pseudomonadota>Gammaproteobacteria | 277 | HV094_RS19980 | 4136016..4136849 | 91347 | MULTISPECIES:-DUF4435-domain-containing-protein-[Enterobacterales] |
| WP_112781821.1 | TOPRIM | GCF_902387755.1 | Enterobacterales | Bacteria>Pseudomonadati>Pseudomonadota>Gammaproteobacteria | 277 | FYC20_RS23050 | 3905..4738 | 91347 | MULTISPECIES:-DUF4435-domain-containing-protein-[Enterobacteriaceae] |
| WP_129328055.1 | TOPRIM | GCF_036708225.1 | Enterobacterales | Bacteria>Pseudomonadati>Pseudomonadota>Gammaproteobacteria | 277 | DD604_RS20950 | 24203..25036 | 91347 | DUF4435-domain-containing-protein-[Enterobacter-cloacae] |
| WP_237558162.1 | TOPRIM | GCF_963882275.1 | Enterobacterales | Bacteria>Pseudomonadati>Pseudomonadota>Gammaproteobacteria | 200 | HV083_RS19910 | 4150139..4150741 | 91347 | MULTISPECIES:-DUF4276-family-protein-[Enterobacterales] |
| WP_139392559.1 | TOPRIM | GCF_038524185.1 | Aeromonas | Bacteria>Pseudomonadati>Pseudomonadota>Gammaproteobacteria>Aeromonadales>Aeromonadaceae | 273 | OKC32_RS09730 | 19414..20235 | 642 | MULTISPECIES:-DUF4435-domain-containing-protein-[Aeromonas] |
| WP_159144194.1 | TOPRIM | GCF_009834065.1 | Aeromonas | Bacteria>Pseudomonadati>Pseudomonadota>Gammaproteobacteria>Aeromonadales>Aeromonadaceae | 272 | GRF56_RS18070 | 3923964..3924782 | 642 | MULTISPECIES:-DUF4435-domain-containing-protein-[Aeromonas] |
| WP_183149506.1 | TOPRIM | GCF_014158455.1 | Aeromonas | Bacteria>Pseudomonadati>Pseudomonadota>Gammaproteobacteria>Aeromonadales>Aeromonadaceae | 254 | J4G65_RS17595 | 85426..86190 | 642 | MULTISPECIES:-hypothetical-protein-[Aeromonas] |
| WP_059169683.1 | TOPRIM | GCF_963869605.1 | Aeromonas | Bacteria>Pseudomonadati>Pseudomonadota>Gammaproteobacteria>Aeromonadales>Aeromonadaceae | 253 | OKB81_RS14895 | 5979..6740 | 642 | MULTISPECIES:-hypothetical-protein-[Aeromonas] |
| WP_128297390.1 | TOPRIM | GCF_040908055.1 | Aeromonas | Bacteria>Pseudomonadati>Pseudomonadota>Gammaproteobacteria>Aeromonadales>Aeromonadaceae | 253 | DN612_RS19430 | 7451..8212 | 642 | hypothetical-protein-[Aeromonas-caviae] |
| WP_160837713.1 | TOPRIM | GCF_013085725.1 | Aeromonas | Bacteria>Pseudomonadati>Pseudomonadota>Gammaproteobacteria>Aeromonadales>Aeromonadaceae | 253 | NG896_RS17515 | 26170..26931 | 642 | MULTISPECIES:-hypothetical-protein-[Aeromonas] |
| WP_219254716.1 | TOPRIM | GCF_038524185.1 | Aeromonas | Bacteria>Pseudomonadati>Pseudomonadota>Gammaproteobacteria>Aeromonadales>Aeromonadaceae | 253 | GPV00_RS13990 | 284379..285140 | 642 | hypothetical-protein-[Aeromonas-hydrophila] |
| WP_234605107.1 | TOPRIM | GCF_026173065.1 | Aeromonas | Bacteria>Pseudomonadati>Pseudomonadota>Gammaproteobacteria>Aeromonadales>Aeromonadaceae | 253 | ORM39_RS00825 | 19448..20209 | 642 | hypothetical-protein-[Aeromonas-salmonicida] |
| WP_236766293.1 | TOPRIM | GCF_021654355.1 | Aeromonas | Bacteria>Pseudomonadati>Pseudomonadota>Gammaproteobacteria>Aeromonadales>Aeromonadaceae | 253 | L6424_RS03525 | 748148..748909 | 642 | hypothetical-protein-[Aeromonas-hydrophila] |
| WP_064341062.1 | TOPRIM | GCF_014169835.1 | Aeromonas | Bacteria>Pseudomonadati>Pseudomonadota>Gammaproteobacteria>Aeromonadales>Aeromonadaceae | 185 | EHZ86_RS11605 | 15248..15805 | 642 | MULTISPECIES:-DUF4276-family-protein-[Aeromonas] |
| WP_103858987.1 | TOPRIM | GCF_019048725.1 | Aeromonas | Bacteria>Pseudomonadati>Pseudomonadota>Gammaproteobacteria>Aeromonadales>Aeromonadaceae | 185 | I6L37_RS11485 | 2395267..2395824 | 642 | MULTISPECIES:-DUF4276-family-protein-[Aeromonas] |
| WP_219296381.1 | TOPRIM | GCF_019971655.1 | Aeromonas | Bacteria>Pseudomonadati>Pseudomonadota>Gammaproteobacteria>Aeromonadales>Aeromonadaceae | 185 | LDN21_RS02450 | 70972..71529 | 642 | MULTISPECIES:-DUF4276-family-protein-[Aeromonas] |
| WP_244812703.1 | TOPRIM | GCF_022869435.1 | Aeromonas | Bacteria>Pseudomonadati>Pseudomonadota>Gammaproteobacteria>Aeromonadales>Aeromonadaceae | 185 | MUW49_RS10490 | 2526..3083 | 642 | MULTISPECIES:-DUF4276-family-protein-[Aeromonas] |
| BBR09399.1 | TOPRIM | GCA_014158455.1 | Aeromonas_caviae | Bacteria>Pseudomonadati>Pseudomonadota>Gammaproteobacteria>Aeromonadales>Aeromonadaceae>Aeromonas | 273 | WP3S18E02_10600 | 1130802..1131623 | 648 | hypothetical-protein-WP3S18E02_10600-[Aeromonas-caviae] |
| MBS4697261.1 | TOPRIM | GCA_018359965.1 | Aeromonas_allosaccharophila | Bacteria>Pseudomonadati>Pseudomonadota>Gammaproteobacteria>Aeromonadales>Aeromonadaceae>Aeromonas | 273 | J4G65_17580 | 77926..78747 | 656 | DUF4435-domain-containing-protein-[Aeromonas-allosaccharophila] |
| MBW3845319.1 | TOPRIM | GCA_019348915.1 | Aeromonas_hydrophila | Bacteria>Pseudomonadati>Pseudomonadota>Gammaproteobacteria>Aeromonadales>Aeromonadaceae>Aeromonas | 273 | GPV00_14025 | 275132..275953 | 644 | DUF4435-domain-containing-protein-[Aeromonas-hydrophila] |
| WP_042011549.1 | TOPRIM | GCF_000819885.1 | Aeromonas_fluvialis | Bacteria>Pseudomonadati>Pseudomonadota>Gammaproteobacteria>Aeromonadales>Aeromonadaceae>Aeromonas | 273 | BN1125_RS07610 | 69553..70374 | 591962 | DUF4435-domain-containing-protein-[Aeromonas-fluvialis] |
| WP_183149510.1 | TOPRIM | GCF_014158455.1 | Aeromonas_caviae | Bacteria>Pseudomonadati>Pseudomonadota>Gammaproteobacteria>Aeromonadales>Aeromonadaceae>Aeromonas | 273 | H7X73_RS05565 | 1130802..1131623 | 648 | DUF4435-domain-containing-protein-[Aeromonas-caviae] |
| WP_202047003.1 | TOPRIM | GCF_016730065.1 | Aeromonas_media | Bacteria>Pseudomonadati>Pseudomonadota>Gammaproteobacteria>Aeromonadales>Aeromonadaceae>Aeromonas | 273 | JD497_RS12350 | 523449..524270 | 651 | DUF4435-domain-containing-protein-[Aeromonas-media] |
| WP_265783516.1 | TOPRIM | GCF_026230585.1 | Aeromonas_veronii | Bacteria>Pseudomonadati>Pseudomonadota>Gammaproteobacteria>Aeromonadales>Aeromonadaceae>Aeromonas | 272 | OD810_RS03275 | 98261..99079 | 654 | hypothetical-protein-[Aeromonas-veronii] |
| WP_182788687.1 | TOPRIM | GCF_014161425.1 | Aeromonas_media | Bacteria>Pseudomonadati>Pseudomonadota>Gammaproteobacteria>Aeromonadales>Aeromonadaceae>Aeromonas | 257 | H7K58_RS09760 | 2088911..2089684 | 651 | hypothetical-protein-[Aeromonas-media] |
| BBR09395.1 | TOPRIM | GCA_014158455.1 | Aeromonas_caviae | Bacteria>Pseudomonadati>Pseudomonadota>Gammaproteobacteria>Aeromonadales>Aeromonadaceae>Aeromonas | 254 | WP3S18E02_10560 | 1123347..1124111 | 648 | hypothetical-protein-WP3S18E02_10560-[Aeromonas-caviae] |
| MBS4697265.1 | TOPRIM | GCA_018359965.1 | Aeromonas_allosaccharophila | Bacteria>Pseudomonadati>Pseudomonadota>Gammaproteobacteria>Aeromonadales>Aeromonadaceae>Aeromonas | 254 | J4G65_17600 | 85426..86190 | 656 | hypothetical-protein-[Aeromonas-allosaccharophila] |
| WP_087756128.1 | TOPRIM | GCF_963873515.1 | Aeromonas_salmonicida | Bacteria>Pseudomonadati>Pseudomonadota>Gammaproteobacteria>Aeromonadales>Aeromonadaceae>Aeromonas | 254 | O23A_RS10580 | 2335104..2335868 | 645 | hypothetical-protein-[Aeromonas-salmonicida] |
| MBW3845325.1 | TOPRIM | GCA_019348915.1 | Aeromonas_hydrophila | Bacteria>Pseudomonadati>Pseudomonadota>Gammaproteobacteria>Aeromonadales>Aeromonadaceae>Aeromonas | 253 | GPV00_14055 | 284379..285140 | 644 | hypothetical-protein-[Aeromonas-hydrophila] |
| WP_042011556.1 | TOPRIM | GCF_000819885.1 | Aeromonas_fluvialis | Bacteria>Pseudomonadati>Pseudomonadota>Gammaproteobacteria>Aeromonadales>Aeromonadaceae>Aeromonas | 253 | BN1125_RS07635 | 77273..78034 | 591962 | hypothetical-protein-[Aeromonas-fluvialis] |
| WP_050664998.1 | TOPRIM | GCF_000820105.1 | Aeromonas_schubertii | Bacteria>Pseudomonadati>Pseudomonadota>Gammaproteobacteria>Aeromonadales>Aeromonadaceae>Aeromonas | 253 | BN1133_RS02625 | 43548..44309 | 652 | hypothetical-protein-[Aeromonas-schubertii] |
| WP_076362314.1 | TOPRIM | GCF_900155975.1 | Aeromonas_hydrophila | Bacteria>Pseudomonadati>Pseudomonadota>Gammaproteobacteria>Aeromonadales>Aeromonadaceae>Aeromonas | 253 | BW963_RS17630 | 140157..140918 | 644 | hypothetical-protein-[Aeromonas-hydrophila] |
| WP_101617902.1 | TOPRIM | GCF_002850695.3 | Aeromonas_hydrophila | Bacteria>Pseudomonadati>Pseudomonadota>Gammaproteobacteria>Aeromonadales>Aeromonadaceae>Aeromonas | 253 | C1A23_RS07655 | 1220103..1220864 | 644 | hypothetical-protein-[Aeromonas-hydrophila] |
| WP_123173887.1 | TOPRIM | GCF_003722175.1 | Aeromonas_veronii | Bacteria>Pseudomonadati>Pseudomonadota>Gammaproteobacteria>Aeromonadales>Aeromonadaceae>Aeromonas | 253 | EFI48_RS19260 | 4193012..4193773 | 654 | hypothetical-protein-[Aeromonas-veronii] |
| WP_139392565.1 | TOPRIM | GCF_006243275.1 | Aeromonas_hydrophila | Bacteria>Pseudomonadati>Pseudomonadota>Gammaproteobacteria>Aeromonadales>Aeromonadaceae>Aeromonas | 253 | CF138_RS14415 | 157721..158482 | 644 | hypothetical-protein-[Aeromonas-hydrophila] |
| WP_190283728.1 | TOPRIM | GCF_014679585.1 | Aeromonas_hydrophila | Bacteria>Pseudomonadati>Pseudomonadota>Gammaproteobacteria>Aeromonadales>Aeromonadaceae>Aeromonas | 253 | IFA22_RS17950 | 3925988..3926749 | 644 | hypothetical-protein-[Aeromonas-hydrophila] |
| WP_193000844.1 | TOPRIM | GCF_014892695.1 | Aeromonas_simiae | Bacteria>Pseudomonadati>Pseudomonadota>Gammaproteobacteria>Aeromonadales>Aeromonadaceae>Aeromonas | 253 | FE240_RS10735 | 2270886..2271647 | 218936 | hypothetical-protein-[Aeromonas-simiae] |
| WP_201932777.1 | TOPRIM | GCF_016729935.1 | Aeromonas_caviae | Bacteria>Pseudomonadati>Pseudomonadota>Gammaproteobacteria>Aeromonadales>Aeromonadaceae>Aeromonas | 253 | JD546_RS01985 | 62680..63441 | 648 | hypothetical-protein-[Aeromonas-caviae] |
| WP_202047006.1 | TOPRIM | GCF_016730065.1 | Aeromonas_media | Bacteria>Pseudomonadati>Pseudomonadota>Gammaproteobacteria>Aeromonadales>Aeromonadaceae>Aeromonas | 253 | JD497_RS12365 | 529197..529958 | 651 | hypothetical-protein-[Aeromonas-media] |
| WP_234578016.1 | TOPRIM | GCF_963898355.1 | Aeromonas_salmonicida | Bacteria>Pseudomonadati>Pseudomonadota>Gammaproteobacteria>Aeromonadales>Aeromonadaceae>Aeromonas | 253 | LZ632_RS15260 | 21301..22062 | 645 | hypothetical-protein-[Aeromonas-salmonicida] |
| WP_236274795.1 | TOPRIM | GCF_021609985.1 | Aeromonas_caviae | Bacteria>Pseudomonadati>Pseudomonadota>Gammaproteobacteria>Aeromonadales>Aeromonadaceae>Aeromonas | 253 | L1871_RS17015 | 3636041..3636802 | 648 | hypothetical-protein-[Aeromonas-caviae] |
| WP_257688090.1 | TOPRIM | GCF_049175235.1 | Aeromonas_veronii | Bacteria>Pseudomonadati>Pseudomonadota>Gammaproteobacteria>Aeromonadales>Aeromonadaceae>Aeromonas | 253 | NUK45_RS19500 | 1588..2349 | 654 | hypothetical-protein-[Aeromonas-veronii] |
| WP_257712002.1 | TOPRIM | GCF_024649525.1 | Aeromonas_caviae | Bacteria>Pseudomonadati>Pseudomonadota>Gammaproteobacteria>Aeromonadales>Aeromonadaceae>Aeromonas | 253 | NUK31_RS14220 | 15482..16243 | 648 | hypothetical-protein-[Aeromonas-caviae] |
| WP_265455063.1 | TOPRIM | GCF_026174125.1 | Aeromonas_salmonicida | Bacteria>Pseudomonadati>Pseudomonadota>Gammaproteobacteria>Aeromonadales>Aeromonadaceae>Aeromonas | 253 | ORM31_RS04460 | 130275..131036 | 645 | hypothetical-protein-[Aeromonas-salmonicida] |
| WP_265776108.1 | TOPRIM | GCF_049168655.1 | Aeromonas_veronii | Bacteria>Pseudomonadati>Pseudomonadota>Gammaproteobacteria>Aeromonadales>Aeromonadaceae>Aeromonas | 253 | OD804_RS12715 | 1824..2585 | 654 | hypothetical-protein-[Aeromonas-veronii] |
| WP_265783521.1 | TOPRIM | GCF_026230585.1 | Aeromonas_veronii | Bacteria>Pseudomonadati>Pseudomonadota>Gammaproteobacteria>Aeromonadales>Aeromonadaceae>Aeromonas | 253 | OD810_RS03310 | 105590..106351 | 654 | hypothetical-protein-[Aeromonas-veronii] |
| WP_270793603.1 | TOPRIM | GCF_027705385.1 | unclassified_Aeromonas | Bacteria>Pseudomonadati>Pseudomonadota>Gammaproteobacteria>Aeromonadales>Aeromonadaceae>Aeromonas | 253 | OKB76_RS13255 | 6915..7676 | 257493 | MULTISPECIES:-hypothetical-protein-[unclassified-Aeromonas] |
| WP_275056845.1 | TOPRIM | GCF_029024985.1 | Aeromonas_allosaccharophila | Bacteria>Pseudomonadati>Pseudomonadota>Gammaproteobacteria>Aeromonadales>Aeromonadaceae>Aeromonas | 253 | PYU98_RS18270 | 3927213..3927974 | 656 | hypothetical-protein-[Aeromonas-allosaccharophila] |
| WP_201944274.1 | TOPRIM | GCF_016729595.1 | Aeromonas_veronii | Bacteria>Pseudomonadati>Pseudomonadota>Gammaproteobacteria>Aeromonadales>Aeromonadaceae>Aeromonas | 247 | JD494_RS16535 | 20611..21354 | 654 | hypothetical-protein-[Aeromonas-veronii] |
| WP_270793607.1 | TOPRIM | GCF_027705385.1 | unclassified_Aeromonas | Bacteria>Pseudomonadati>Pseudomonadota>Gammaproteobacteria>Aeromonadales>Aeromonadaceae>Aeromonas | 191 | OKB76_RS13275 | 12962..13537 | 257493 | MULTISPECIES:-hypothetical-protein-[unclassified-Aeromonas] |
| MCF3097927.1 | TOPRIM | GCA_021538905.1 | Aeromonas_australiensis | Bacteria>Pseudomonadati>Pseudomonadota>Gammaproteobacteria>Aeromonadales>Aeromonadaceae>Aeromonas | 185 | EHZ86_11635 | 15248..15805 | 1114880 | DUF4276-family-protein-[Aeromonas-australiensis] |
| WP_064338618.1 | TOPRIM | GCF_001634325.1 | Aeromonas_veronii | Bacteria>Pseudomonadati>Pseudomonadota>Gammaproteobacteria>Aeromonadales>Aeromonadaceae>Aeromonas | 185 | WM43_RS07830 | 1776803..1777360 | 654 | DUF4276-family-protein-[Aeromonas-veronii] |
| WP_257709588.1 | TOPRIM | GCF_024649925.1 | Aeromonas_caviae | Bacteria>Pseudomonadati>Pseudomonadota>Gammaproteobacteria>Aeromonadales>Aeromonadaceae>Aeromonas | 185 | NUK49_RS00055 | 11714..12271 | 648 | DUF4276-family-protein-[Aeromonas-caviae] |
| WP_264787583.1 | TOPRIM | GCF_026000055.1 | Aeromonas_caviae | Bacteria>Pseudomonadati>Pseudomonadota>Gammaproteobacteria>Aeromonadales>Aeromonadaceae>Aeromonas | 185 | OPI30_RS08275 | 1646267..1646824 | 648 | DUF4276-family-protein-[Aeromonas-caviae] |
| WP_270698351.1 | TOPRIM | GCF_027705185.1 | unclassified_Aeromonas | Bacteria>Pseudomonadati>Pseudomonadota>Gammaproteobacteria>Aeromonadales>Aeromonadaceae>Aeromonas | 185 | OKC33_RS16315 | 8484..9041 | 257493 | MULTISPECIES:-DUF4276-family-protein-[unclassified-Aeromonas] |
| WP_270698109.1 | TOPRIM | GCF_027705185.1 | Aeromonas_sp._QDB05 | Bacteria>Pseudomonadati>Pseudomonadota>Gammaproteobacteria>Aeromonadales>Aeromonadaceae>Aeromonas>unclassified_Aeromonas | 253 | OKC32_RS09700 | 9929..10690 | 2990478 | hypothetical-protein-[Aeromonas-sp.-QDB05] |
| WP_163133167.1 | TOPRIM | GCF_900537085.1 | Agarivorans_sp._Alg241-V36 | Bacteria>Pseudomonadati>Pseudomonadota>Gammaproteobacteria>Alteromonadales>Alteromonadaceae>Agarivorans>unclassified_Agarivorans | 225 | G6R11_RS11255 | 296740..297417 | 2305992 | DUF4276-family-protein-[Agarivorans-sp.-Alg241-V36] |
| WP_100972009.1 | TOPRIM | GCF_002831605.1 | Alteromonas_sp._MB-3u-76 | Bacteria>Pseudomonadati>Pseudomonadota>Gammaproteobacteria>Alteromonadales>Alteromonadaceae>Alteromonas/Salinimonas_group>Alteromonas>unclassified_Alteromonas | 225 | CW735_RS10465 | 2333388..2334065 | 2058133 | DUF4276-family-protein-[Alteromonas-sp.-MB-3u-76] |
| WP_146799744.1 | TOPRIM | GCF_007954305.1 | Colwellia_hornerae | Bacteria>Pseudomonadati>Pseudomonadota>Gammaproteobacteria>Alteromonadales>Colwelliaceae>Colwellia | 225 | ESZ26_RS11970 | 110532..111209 | 89402 | DUF4276-family-protein-[Colwellia-hornerae] |
| WP_165723734.1 | TOPRIM | GCF_943789085.1 | Pseudoalteromonas | Bacteria>Pseudomonadati>Pseudomonadota>Gammaproteobacteria>Alteromonadales>Pseudoalteromonadaceae | 245 | QDP19_RS17630 | 3063445..3064182 | 53246 | MULTISPECIES:-hypothetical-protein-[Pseudoalteromonas] |
| WP_130146689.1 | TOPRIM | GCF_004208955.1 | Pseudoalteromonas_sp._CO325X | Bacteria>Pseudomonadati>Pseudomonadota>Gammaproteobacteria>Alteromonadales>Pseudoalteromonadaceae>Pseudoalteromonas>unclassified_Pseudoalteromonas | 245 | EXT46_RS08640 | 68834..69571 | 1777262 | hypothetical-protein-[Pseudoalteromonas-sp.-CO325X] |
| WP_208162986.1 | TOPRIM | GCF_017580535.1 | Shewanella_algae | Bacteria>Pseudomonadati>Pseudomonadota>Gammaproteobacteria>Alteromonadales>Shewanellaceae>Shewanella | 252 | I6M49_RS22470 | 8364..9122 | 38313 | hypothetical-protein-[Shewanella-algae] |
| WP_208251009.1 | TOPRIM | GCF_017580085.1 | Shewanella_algae | Bacteria>Pseudomonadati>Pseudomonadota>Gammaproteobacteria>Alteromonadales>Shewanellaceae>Shewanella | 248 | I6M29_RS02115 | 14278..15024 | 38313 | hypothetical-protein-[Shewanella-algae] |
| WP_006086469.1 | TOPRIM | GCF_000231345.1 | Shewanella_baltica | Bacteria>Pseudomonadati>Pseudomonadota>Gammaproteobacteria>Alteromonadales>Shewanellaceae>Shewanella | 245 | SBAL625DRAFT_RS15540 | 219626..220363 | 62322 | hypothetical-protein-[Shewanella-baltica] |
| WP_076500073.1 | TOPRIM | GCF_900156405.1 | Shewanella_morhuae | Bacteria>Pseudomonadati>Pseudomonadota>Gammaproteobacteria>Alteromonadales>Shewanellaceae>Shewanella | 223 | BW947_RS12475 | 15057..15728 | 365591 | DUF4276-family-protein-[Shewanella-morhuae] |
| WP_261732938.1 | TOPRIM | GCF_025398315.1 | Shewanella_phaeophyticola | Bacteria>Pseudomonadati>Pseudomonadota>Gammaproteobacteria>Alteromonadales>Shewanellaceae>Shewanella | 223 | N4T56_RS08720 | 1862822..1863493 | 2978345 | DUF4276-family-protein-[Shewanella-sp.-KJ10-1] |
| WP_143247599.1 | TOPRIM | GCF_002312815.1 | Agaribacterium_haliotis | Bacteria>Pseudomonadati>Pseudomonadota>Gammaproteobacteria>Cellvibrionales>Cellvibrionaceae>Agaribacterium | 248 | CFK45_RS18355 | 10057..10803 | 2013869 | hypothetical-protein-[Agaribacterium-haliotis] |
| WP_050977049.1 | TOPRIM | GCF_000263355.1 | Cellvibrio_sp._BR | Bacteria>Pseudomonadati>Pseudomonadota>Gammaproteobacteria>Cellvibrionales>Cellvibrionaceae>Cellvibrio>unclassified_Cellvibrio | 251 | O59_RS11275 | 228289..229044 | 1134474 | hypothetical-protein-[Cellvibrio-sp.-BR] |
| WP_054114049.1 | TOPRIM | GCF_001296225.1 | Marinagarivorans_algicola | Bacteria>Pseudomonadati>Pseudomonadota>Gammaproteobacteria>Cellvibrionales>Cellvibrionaceae>Marinagarivorans | 248 | AC526_RS14080 | 28952..29698 | 1513270 | hypothetical-protein-[Marinagarivorans-algicola] |
| WP_226986432.1 | TOPRIM | GCF_000013665.1 | Saccharophagus_degradans | Bacteria>Pseudomonadati>Pseudomonadota>Gammaproteobacteria>Cellvibrionales>Cellvibrionaceae>Saccharophagus | 236 | SDE_RS14410 | 3486955..3487665 | 86304 | DUF4276-family-protein-[Saccharophagus-degradans] |
| WP_135445664.1 | TOPRIM | GCF_004745945.1 | Mangrovimicrobium_sediminis | Bacteria>Pseudomonadati>Pseudomonadota>Gammaproteobacteria>Cellvibrionales>Halieaceae>Mangrovimicrobium | 185 | E4634_RS16030 | 158833..159390 | 2562682 | DUF4276-family-protein-[Haliea-sp.-SAOS-164] |
| WP_176976533.1 | TOPRIM | GCF_013385175.1 | Allochromatium_humboldtianum | Bacteria>Pseudomonadati>Pseudomonadota>Gammaproteobacteria>Chromatiales>Chromatiaceae>Allochromatium | 255 | HW932_RS10980 | 215865..216632 | 504901 | hypothetical-protein-[Allochromatium-humboldtianum] |
| BCU06240.1 | TOPRIM | GCA_018409545.1 | Allochromatium_tepidum | Bacteria>Pseudomonadati>Pseudomonadota>Gammaproteobacteria>Chromatiales>Chromatiaceae>Allochromatium | 214 | Atep_09170 | 997164..997808 | 553982 | hypothetical-protein-Atep_09170-[Allochromatium-tepidum] |
| WP_213380487.1 | TOPRIM | GCF_018409545.1 | Allochromatium_tepidum | Bacteria>Pseudomonadati>Pseudomonadota>Gammaproteobacteria>Chromatiales>Chromatiaceae>Allochromatium | 214 | Atep_RS04615 | 997164..997808 | 553982 | hypothetical-protein-[Allochromatium-tepidum] |
| MTW22276.1 | TOPRIM | GCA_009720725.1 | Allochromatium_palmeri | Bacteria>Pseudomonadati>Pseudomonadota>Gammaproteobacteria>Chromatiales>Chromatiaceae>Allochromatium | 196 | GJ668_14450 | 38292..38882 | 231048 | DUF4276-family-protein-[Allochromatium-palmeri] |
| WP_155450844.1 | TOPRIM | GCF_009720725.1 | Allochromatium_palmeri | Bacteria>Pseudomonadati>Pseudomonadota>Gammaproteobacteria>Chromatiales>Chromatiaceae>Allochromatium | 196 | GJ668_RS14455 | 38292..38882 | 231048 | DUF4276-family-protein-[Allochromatium-palmeri] |
| WP_200346139.1 | TOPRIM | GCF_016584085.1 | Halochromatium_glycolicum | Bacteria>Pseudomonadati>Pseudomonadota>Gammaproteobacteria>Chromatiales>Chromatiaceae>Halochromatium | 232 | CKO40_RS10405 | 39242..39940 | 85075 | hypothetical-protein-[Halochromatium-glycolicum] |

| Protein-ID | Domain | Assembly-ID | Strain | Taxonomy | Length | Locus-Tag | Genomic-coordinates | TaxID | Fasta-defline |
|---|---|---|---|---|---|---|---|---|---|
| WP_201223301.1 | TOPRIM | GCF_016653315.1 | Halochromatium_roseum | Bacteria>Pseudomonadati>Pseudomonadota>Gammaproteobacteria>Chromatiales>Chromatiaceae>Halochromatium | 207 | CCR96_RS21085 | 10494..11117 | 391920 | hypothetical-protein-[Halochromatium-roseum] |
| WP_020507738.1 | TOPRIM | GCF_000379525.1 | Lamprocystis_purpurea | Bacteria>Pseudomonadati>Pseudomonadota>Gammaproteobacteria>Chromatiales>Chromatiaceae>Lamprocystis | 229 | A390_RS0126775 | 74891..75580 | 61598 | RloB-family-protein-[Lamprocystis-purpurea] |
| WP_211211616.1 | TOPRIM | GCF_000379525.1 | Lamprocystis_purpurea | Bacteria>Pseudomonadati>Pseudomonadota>Gammaproteobacteria>Chromatiales>Chromatiaceae>Lamprocystis | 226 | A390_RS0116900 | 21364..22044 | 61598 | hypothetical-protein-[Lamprocystis-purpurea] |
| WP_020507363.1 | TOPRIM | GCF_000379525.1 | Lamprocystis_purpurea | Bacteria>Pseudomonadati>Pseudomonadota>Gammaproteobacteria>Chromatiales>Chromatiaceae>Lamprocystis | 64 | A390_RS0124820 | 127457..127651 | 61598 | DUF4276-family-protein-[Lamprocystis-purpurea] |
| WP_200387140.1 | TOPRIM | GCF_016583985.1 | Thiocapsa_imhoffii | Bacteria>Pseudomonadati>Pseudomonadota>Gammaproteobacteria>Chromatiales>Chromatiaceae>Thiocapsa | 281 | CK025_RS06620 | 40907..41752 | 382777 | DUF4435-domain-containing-protein-[Thiocapsa-imhoffii] |
| QVL49526.1 | TOPRIM | GCA_018399035.1 | Thiocapsa_sp. | Bacteria>Pseudomonadati>Pseudomonadota>Gammaproteobacteria>Chromatiales>Chromatiaceae>Thiocapsa>unclassified_Thiocapsa | 348 | KFB96_03160 | 686804..687850 | 2024551 | MAG:-toprim-domain-containing-protein-[Thiocapsa-sp.] |
| CRI67898.1 | TOPRIM | GCA_001499735.1 | Thiocapsa_sp._KS1 | Bacteria>Pseudomonadati>Pseudomonadota>Gammaproteobacteria>Chromatiales>Chromatiaceae>Thiocapsa>unclassified_Thiocapsa | 305 | THIOKS1890031 | 39934..40851 | 610332 | hypothetical-protein-THIOKS1890031-[Thiocapsa-sp.-KS1] |
| WP_093186248.1 | TOPRIM | - | Thiocapsa_sp._KS1 | Bacteria>Pseudomonadati>Pseudomonadota>Gammaproteobacteria>Chromatiales>Chromatiaceae>Thiocapsa>unclassified_Thiocapsa | 305 | THIOKS1_RS05385 | 1205531..1206448 | 610332 | hypothetical-protein-[Thiocapsa-sp.-KS1] |
| QVL48949.1 | TOPRIM | GCA_018399035.1 | Thiocapsa_sp. | Bacteria>Pseudomonadati>Pseudomonadota>Gammaproteobacteria>Chromatiales>Chromatiaceae>Thiocapsa>unclassified_Thiocapsa | 281 | KFB96_26070 | 5692906..5693751 | 2024551 | MAG:-DUF4435-domain-containing-protein-[Thiocapsa-sp.] |
| QVL49516.1 | TOPRIM | GCA_018399035.1 | Thiocapsa_sp. | Bacteria>Pseudomonadati>Pseudomonadota>Gammaproteobacteria>Chromatiales>Chromatiaceae>Thiocapsa>unclassified_Thiocapsa | 217 | KFB96_03100 | 672243..672896 | 2024551 | MAG:-hypothetical-protein-KFB96_03100-[Thiocapsa-sp.] |
| QVL49212.1 | TOPRIM | GCA_018399035.1 | Thiocapsa_sp. | Bacteria>Pseudomonadati>Pseudomonadota>Gammaproteobacteria>Chromatiales>Chromatiaceae>Thiocapsa>unclassified_Thiocapsa | 208 | KFB96_01390 | 297361..297987 | 2024551 | MAG:-hypothetical-protein-KFB96_01390-[Thiocapsa-sp.] |
| QVL47945.1 | TOPRIM | GCA_018399035.1 | Thiocapsa_sp. | Bacteria>Pseudomonadati>Pseudomonadota>Gammaproteobacteria>Chromatiales>Chromatiaceae>Thiocapsa>unclassified_Thiocapsa | 207 | KFB96_20145 | 4422897..4423520 | 2024551 | MAG:-hypothetical-protein-KFB96_20145-[Thiocapsa-sp.] |
| QVL47238.1 | TOPRIM | GCA_018399035.1 | Thiocapsa_sp. | Bacteria>Pseudomonadati>Pseudomonadota>Gammaproteobacteria>Chromatiales>Chromatiaceae>Thiocapsa>unclassified_Thiocapsa | 198 | KFB96_16115 | 3546241..3546837 | 2024551 | MAG:-DUF4276-family-protein-[Thiocapsa-sp.] |
| WP_200375565.1 | TOPRIM | GCF_016583575.1 | Thiocystis_violacea | Bacteria>Pseudomonadati>Pseudomonadota>Gammaproteobacteria>Chromatiales>Chromatiaceae>Thiocystis | 202 | CK027_RS14230 | 27360..27968 | 13725 | hypothetical-protein-[Thiocystis-violacea] |
| WP_100919681.1 | TOPRIM | GCF_002813775.1 | Candidatus_Thiodictyon_syntrophicum | Bacteria>Pseudomonadati>Pseudomonadota>Gammaproteobacteria>Chromatiales>Chromatiaceae>Thiodictyon | 281 | THSYN_RS13920 | 3312490..3313335 | 1166950 | DUF4435-domain-containing-protein-[Candidatus-Thiodictyon-syntrophicum] |
| WP_100917908.1 | TOPRIM | GCF_002813775.1 | Candidatus_Thiodictyon_syntrophicum | Bacteria>Pseudomonadati>Pseudomonadota>Gammaproteobacteria>Chromatiales>Chromatiaceae>Thiodictyon | 195 | THSYN_RS03435 | 787921..788508 | 1166950 | hypothetical-protein-[Candidatus-Thiodictyon-syntrophicum] |
| AUB82574.1 | TOPRIM | GCA_002813775.1 | Candidatus_Thiodictyon_syntrophicum | Bacteria>Pseudomonadati>Pseudomonadota>Gammaproteobacteria>Chromatiales>Chromatiaceae>Thiodictyon | 90 | THSYN_17570 | 4229872..4230144 | 1166950 | hypothetical-protein-THSYN_17570-[Candidatus-Thiodictyon-syntrophicum] |
| WP_100920296.1 | TOPRIM | GCF_002813775.1 | Candidatus_Thiodictyon_syntrophicum | Bacteria>Pseudomonadati>Pseudomonadota>Gammaproteobacteria>Chromatiales>Chromatiaceae>Thiodictyon | 90 | THSYN_RS17705 | 4229872..4230144 | 1166950 | hypothetical-protein-[Candidatus-Thiodictyon-syntrophicum] |
| WP_150092067.1 | TOPRIM | GCF_008632335.1 | Thiohalocapsa_marina | Bacteria>Pseudomonadati>Pseudomonadota>Gammaproteobacteria>Chromatiales>Chromatiaceae>Thiohalocapsa | 207 | F2Q65_RS07750 | 162869..163492 | 424902 | hypothetical-protein-[Thiohalocapsa-marina] |
| WP_058557218.1 | TOPRIM | GCF_001469165.1 | Thiohalocapsa_sp._ML1 | Bacteria>Pseudomonadati>Pseudomonadota>Gammaproteobacteria>Chromatiales>Chromatiaceae>Thiohalocapsa>unclassified_Thiohalocapsa | 258 | ML01_RS20305 | 8080..8856 | 1431688 | hypothetical-protein-[Thiohalocapsa-sp.-ML1] |
| WP_164452828.1 | TOPRIM | GCF_010820565.1 | Thiorhodococcus_minor | Bacteria>Pseudomonadati>Pseudomonadota>Gammaproteobacteria>Chromatiales>Chromatiaceae>Thiorhodococcus | 206 | G3446_RS10735 | 66322..66942 | 57489 | DUF4276-family-protein-[Thiorhodococcus-minor] |
| MBP8282507.1 | TOPRIM | GCA_018007135.1 | Chromatiaceae_bacterium | Bacteria>Pseudomonadati>Pseudomonadota>Gammaproteobacteria>Chromatiales>Chromatiaceae>unclassified_Chromatiaceae | 217 | KAX46_01100 | 746..1399 | 2049432 | hypothetical-protein-[Chromatiaceae-bacterium] |
| MBP6806941.1 | TOPRIM | GCA_017991215.1 | Chromatiaceae_bacterium | Bacteria>Pseudomonadati>Pseudomonadota>Gammaproteobacteria>Chromatiales>Chromatiaceae>unclassified_Chromatiaceae | 214 | KA125_01450 | 6534..7178 | 2049432 | DUF4276-family-protein-[Chromatiaceae-bacterium] |
| MBK8637651.1 | TOPRIM | GCA_016714645.1 | Chromatiaceae_bacterium | Bacteria>Pseudomonadati>Pseudomonadota>Gammaproteobacteria>Chromatiales>Chromatiaceae>unclassified_Chromatiaceae | 209 | IPN92_04960 | 1069213..1069842 | 2049432 | hypothetical-protein-[Chromatiaceae-bacterium] |
| MBK8637138.1 | TOPRIM | GCA_016714645.1 | Chromatiaceae_bacterium | Bacteria>Pseudomonadati>Pseudomonadota>Gammaproteobacteria>Chromatiales>Chromatiaceae>unclassified_Chromatiaceae | 207 | IPN92_02245 | 475824..476447 | 2049432 | hypothetical-protein-[Chromatiaceae-bacterium] |
| MBP6733369.1 | TOPRIM | GCA_017990755.1 | Chromatiaceae_bacterium | Bacteria>Pseudomonadati>Pseudomonadota>Gammaproteobacteria>Chromatiales>Chromatiaceae>unclassified_Chromatiaceae | 98 | KA142_01515 | 6047..6341 | 2049432 | hypothetical-protein-[Chromatiaceae-bacterium] |
| MBP8282512.1 | TOPRIM | GCA_018007135.1 | Chromatiaceae_bacterium | Bacteria>Pseudomonadati>Pseudomonadota>Gammaproteobacteria>Chromatiales>Chromatiaceae>unclassified_Chromatiaceae | 98 | KAX46_01125 | 7788..8082 | 2049432 | hypothetical-protein-[Chromatiaceae-bacterium] |
| MBP6582543.1 | TOPRIM | GCA_017989555.1 | Chromatiaceae_bacterium | Bacteria>Pseudomonadati>Pseudomonadota>Gammaproteobacteria>Chromatiales>Chromatiaceae>unclassified_Chromatiaceae | 87 | KA204_03590 | 3395..3660 | 2049432 | hypothetical-protein-[Chromatiaceae-bacterium] |
| MBP6733368.1 | TOPRIM | GCA_017990755.1 | Chromatiaceae_bacterium | Bacteria>Pseudomonadati>Pseudomonadota>Gammaproteobacteria>Chromatiales>Chromatiaceae>unclassified_Chromatiaceae | 87 | KA142_01510 | 5681..5946 | 2049432 | hypothetical-protein-[Chromatiaceae-bacterium] |
| MBP8282511.1 | TOPRIM | GCA_018007135.1 | Chromatiaceae_bacterium | Bacteria>Pseudomonadati>Pseudomonadota>Gammaproteobacteria>Chromatiales>Chromatiaceae>unclassified_Chromatiaceae | 87 | KAX46_01120 | 7422..7687 | 2049432 | hypothetical-protein-[Chromatiaceae-bacterium] |
| MBP6582544.1 | TOPRIM | GCA_017989555.1 | Chromatiaceae_bacterium | Bacteria>Pseudomonadati>Pseudomonadota>Gammaproteobacteria>Chromatiales>Chromatiaceae>unclassified_Chromatiaceae | 61 | KA204_03595 | 3761..3944 | 2049432 | hypothetical-protein-[Chromatiaceae-bacterium] |
| WP_255705564.1 | TOPRIM | - | unclassified_Ectothiorhodospira | Bacteria>Pseudomonadati>Pseudomonadota>Gammaproteobacteria>Chromatiales>Ectothiorhodospiraceae>Ectothiorhodospira | 222 | LPW28_RS03030 | 33451..34119 | 2684909 | MULTISPECIES:-DUF4276-family-protein-[unclassified-Ectothiorhodospira] |
| WP_238621057.1 | TOPRIM | GCF_022227785.1 | Ectothiorhodospira_lacustris | Bacteria>Pseudomonadati>Pseudomonadota>Gammaproteobacteria>Chromatiales>Ectothiorhodospiraceae>Ectothiorhodospira | 217 | LQ952_RS02365 | 43650..44303 | 2899127 | DUF4276-family-protein-[Ectothiorhodospira-sp.-B14B] |
| WP_202906241.1 | TOPRIM | GCF_900110965.1 | Ectothiorhodospira_magna | Bacteria>Pseudomonadati>Pseudomonadota>Gammaproteobacteria>Chromatiales>Ectothiorhodospiraceae>Ectothiorhodospira | 189 | BMX62_RS09255 | 44069..44638 | 867345 | hypothetical-protein-[Ectothiorhodospira-magna] |
| WP_238617952.1 | TOPRIM | GCF_022227765.1 | Ectothiorhodospira_shaposhnikovii | Bacteria>Pseudomonadati>Pseudomonadota>Gammaproteobacteria>Chromatiales>Ectothiorhodospiraceae>Ectothiorhodospira | 130 | LQD11_RS13155 | 105942..106334 | 1054 | hypothetical-protein-[Ectothiorhodospira-shaposhnikovii] |
| WP_156776974.1 | TOPRIM | GCF_000153205.1 | Nitrococcus_mobilis | Bacteria>Pseudomonadati>Pseudomonadota>Gammaproteobacteria>Chromatiales>Ectothiorhodospiraceae>Nitrococcus | 218 | NB231_RS09970 | 2187352..2188008 | 35797 | hypothetical-protein-[Nitrococcus-mobilis] |
| QEP43014.1 | TOPRIM | GCA_008375315.1 | Ectothiorhodospiraceae_bacterium_BW-2 | Bacteria>Pseudomonadati>Pseudomonadota>Gammaproteobacteria>Chromatiales>Ectothiorhodospiraceae>unclassified_Ectothiorhodospiraceae | 226 | D5085_07730 | 1604990..1605670 | 947515 | DUF4276-family-protein-[Ectothiorhodospiraceae-bacterium-BW-2] |
| MBF2761743.1 | TOPRIM | GCA_015278165.1 | Ectothiorhodospiraceae_bacterium_AqS1 | Bacteria>Pseudomonadati>Pseudomonadota>Gammaproteobacteria>Chromatiales>Ectothiorhodospiraceae>unclassified_Ectothiorhodospiraceae | 204 | ISN28_16030 | 9451..10065 | 1725326 | hypothetical-protein-[Ectothiorhodospiraceae-bacterium-AqS1] |
| MBF2761737.1 | TOPRIM | GCA_015278165.1 | Ectothiorhodospiraceae_bacterium_AqS1 | Bacteria>Pseudomonadati>Pseudomonadota>Gammaproteobacteria>Chromatiales>Ectothiorhodospiraceae>unclassified_Ectothiorhodospiraceae | 202 | ISN28_16000 | 216..824 | 1725326 | hypothetical-protein-[Ectothiorhodospiraceae-bacterium-AqS1] |
| PLY12611.1 | TOPRIM | GCA_002869585.1 | Sedimenticola_sp. | Bacteria>Pseudomonadati>Pseudomonadota>Gammaproteobacteria>Chromatiales>Sedimenticolaceae>Sedimenticola>unclassified_Sedimenticola | 220 | C0631_16550 | 13994..14656 | 1940285 | MAG:-hypothetical-protein-C0631_16550-[Sedimenticola-sp.] |
| WP_177417922.1 | TOPRIM | GCF_011947365.2 | endosymbiont_of_Lamellibrachia_barhami | Bacteria>Pseudomonadati>Pseudomonadota>Gammaproteobacteria>Chromatiales>unclassified_Chromatiales | 241 | G9Q73_RS00035 | 9566..10291 | 205975 | hypothetical-protein-[endosymbiont-of-Lamellibrachia-barhami] |
| WP_023157614.1 | TOPRIM | GCF_033235525.1 | Enterobacteriaceae | Bacteria>Pseudomonadati>Pseudomonadota>Gammaproteobacteria>Enterobacterales | 277 | HMPREF1620_RS03660 | 13710..14543 | 543 | DUF4435-domain-containing-protein-[Escherichia-coli] |
| WP_057055969.1 | TOPRIM | GCF_014169295.1 | Enterobacteriaceae | Bacteria>Pseudomonadati>Pseudomonadota>Gammaproteobacteria>Enterobacterales | 277 | H7R61_RS03355 | 679172..680005 | 543 | MULTISPECIES:-DUF4435-domain-containing-protein-[Enterobacter-cloacae-complex] |
| WP_100126888.1 | TOPRIM | GCF_014103055.1 | Enterobacteriaceae | Bacteria>Pseudomonadati>Pseudomonadota>Gammaproteobacteria>Enterobacterales | 277 | M4W13_RS12115 | 115925..116758 | 543 | MULTISPECIES:-DUF4435-domain-containing-protein-[Enterobacteriaceae] |
| WP_163363283.1 | TOPRIM | GCF_046630575.1 | Enterobacteriaceae | Bacteria>Pseudomonadati>Pseudomonadota>Gammaproteobacteria>Enterobacterales | 277 | G4556_RS12895 | 11307..12140 | 543 | DUF4435-domain-containing-protein-[Enterobacter-cloacae] |
| WP_241501909.1 | TOPRIM | GCF_963899015.1 | Morganellaceae | Bacteria>Pseudomonadati>Pseudomonadota>Gammaproteobacteria>Enterobacterales | 245 | ORM28_RS01815 | 14227..14964 | 1903414 | MULTISPECIES:-hypothetical-protein-[Morganellaceae] |
| WP_210849181.1 | TOPRIM | GCF_022349905.1 | Morganellaceae | Bacteria>Pseudomonadati>Pseudomonadota>Gammaproteobacteria>Enterobacterales | 199 | PSE52_RS01010 | 230917..231516 | 1903414 | MULTISPECIES:-hypothetical-protein-[Morganellaceae] |
| WP_006812022.1 | TOPRIM | GCF_013635755.1 | Enterobacteriaceae | Bacteria>Pseudomonadati>Pseudomonadota>Gammaproteobacteria>Enterobacterales | 189 | K4X45_RS16765 | 78906..79475 | 543 | MULTISPECIES:-hypothetical-protein-[Enterobacteriaceae] |
| WP_117108867.1 | TOPRIM | GCF_903684385.1 | Enterobacteriaceae | Bacteria>Pseudomonadati>Pseudomonadota>Gammaproteobacteria>Enterobacterales | 189 | GQQ45_RS24730 | 5063..5632 | 543 | MULTISPECIES:-hypothetical-protein-[Enterobacteriaceae] |
| WP_148977797.1 | TOPRIM | GCF_902807175.1 | Enterobacteriaceae | Bacteria>Pseudomonadati>Pseudomonadota>Gammaproteobacteria>Enterobacterales | 189 | PBS85_RS02765 | 577522..578091 | 543 | MULTISPECIES:-hypothetical-protein-[Enterobacteriaceae] |
| WP_222691534.1 | TOPRIM | GCF_019841255.1 | Enterobacteriaceae | Bacteria>Pseudomonadati>Pseudomonadota>Gammaproteobacteria>Enterobacterales | 189 | K7A35_RS15420 | 9916..10485 | 543 | MULTISPECIES:-DUF4276-family-protein-[Enterobacteriaceae] |
| WP_100165588.1 | TOPRIM | GCF_003186415.1 | Enterobacter | Bacteria>Pseudomonadati>Pseudomonadota>Gammaproteobacteria>Enterobacterales>Enterobacteriaceae | 277 | NJA11_RS17915 | 126405..127238 | 547 | MULTISPECIES:-DUF4435-domain-containing-protein-[Enterobacter] |
| WP_010989100.1 | TOPRIM | GCF_000006945.2 | Salmonella | Bacteria>Pseudomonadati>Pseudomonadota>Gammaproteobacteria>Enterobacterales>Enterobacteriaceae | 273 | NMU37_RS20795 | 4235976..4236797 | 590 | MULTISPECIES:-DUF4435-domain-containing-protein-[Salmonella] |
| WP_000238714.1 | TOPRIM | GCF_000264195.1 | Escherichia | Bacteria>Pseudomonadati>Pseudomonadota>Gammaproteobacteria>Enterobacterales>Enterobacteriaceae | 189 | PP93_RS24135 | 21345..21914 | 561 | MULTISPECIES:-hypothetical-protein-[Escherichia] |
| WP_023224628.1 | TOPRIM | GCF_001569545.1 | Salmonella | Bacteria>Pseudomonadati>Pseudomonadota>Gammaproteobacteria>Enterobacterales>Enterobacteriaceae | 189 | NOQ95_RS21575 | 4456909..4457478 | 590 | MULTISPECIES:-hypothetical-protein-[Salmonella] |
| WP_149691149.1 | TOPRIM | GCF_008373795.1 | Citrobacter_pasteurii | Bacteria>Pseudomonadati>Pseudomonadota>Gammaproteobacteria>Enterobacterales>Enterobacteriaceae>Citrobacter | 277 | DXF85_RS00945 | 189718..190551 | 1563222 | DUF4435-domain-containing-protein-[Citrobacter-pasteurii] |
| HAU4366730.1 | TOPRIM | GCA_016081455.1 | Citrobacter_amalonaticus | Bacteria>Pseudomonadati>Pseudomonadota>Gammaproteobacteria>Enterobacterales>Enterobacteriaceae>Citrobacter | 273 | F6W26_02160 | 127309..128130 | 35703 | DUF4435-domain-containing-protein-[Citrobacter-amalonaticus] |
| WP_080721694.1 | TOPRIM | GCF_000731055.1 | Citrobacter_amalonaticus | Bacteria>Pseudomonadati>Pseudomonadota>Gammaproteobacteria>Enterobacterales>Enterobacteriaceae>Citrobacter | 273 | DQ02_RS05330 | 406987..407808 | 35703 | DUF4435-domain-containing-protein-[Citrobacter-amalonaticus] |
| WP_203457738.1 | TOPRIM | GCF_016859695.1 | Citrobacter_braakii | Bacteria>Pseudomonadati>Pseudomonadota>Gammaproteobacteria>Enterobacterales>Enterobacteriaceae>Citrobacter>Citrobacter_freundii_complex | 275 | GF710_RS09110 | 93435..94262 | 57706 | DUF4435-domain-containing-protein-[Citrobacter-braakii] |
| WP_241511102.1 | TOPRIM | GCF_022585175.1 | Citrobacter_freundii | Bacteria>Pseudomonadati>Pseudomonadota>Gammaproteobacteria>Enterobacterales>Enterobacteriaceae>Citrobacter>Citrobacter_freundii_complex | 271 | MMQ01_RS10390 | 138499..139314 | 546 | DUF4435-domain-containing-protein-[Citrobacter-freundii] |
| HBC2890666.1 | TOPRIM | GCA_018089985.1 | Citrobacter_freundii | Bacteria>Pseudomonadati>Pseudomonadota>Gammaproteobacteria>Enterobacterales>Enterobacteriaceae>Citrobacter>Citrobacter_freundii_complex | 189 | JD321_004758 | 9883..10452 | 546 | hypothetical-protein-[Citrobacter-freundii] |
| WP_269506673.1 | TOPRIM | GCF_027277125.1 | Cronobacter_sakazakii | Bacteria>Pseudomonadati>Pseudomonadota>Gammaproteobacteria>Enterobacterales>Enterobacteriaceae>Cronobacter | 277 | O6D31_RS12690 | 372554..373387 | 28141 | DUF4435-domain-containing-protein-[Cronobacter-sakazakii] |
| WP_257246763.1 | TOPRIM | GCF_024607805.1 | unclassified_Enterobacter | Bacteria>Pseudomonadati>Pseudomonadota>Gammaproteobacteria>Enterobacterales>Enterobacteriaceae>Enterobacter | 271 | NR293_RS15380 | 78188..79003 | 2608935 | MULTISPECIES:-DUF4435-domain-containing-protein-[unclassified-Enterobacter] |
| WP_032634487.1 | TOPRIM | GCF_019974415.1 | Enterobacter_cloacae_complex | Bacteria>Pseudomonadati>Pseudomonadota>Gammaproteobacteria>Enterobacterales>Enterobacteriaceae>Enterobacter | 189 | CI630_RS14020 | 10373..10942 | 354276 | MULTISPECIES:-hypothetical-protein-[Enterobacter-cloacae-complex] |
| MCE1563405.1 | TOPRIM | GCA_021285765.1 | Enterobacter_hormaechei | Bacteria>Pseudomonadati>Pseudomonadota>Gammaproteobacteria>Enterobacterales>Enterobacteriaceae>Enterobacter>Enterobacter_cloacae_complex | 277 | LWT81_04615 | 332261..333094 | 158836 | DUF4435-domain-containing-protein-[Enterobacter-hormaechei] |
| WP_032621384.1 | TOPRIM | GCF_023734955.1 | Enterobacter_hormaechei | Bacteria>Pseudomonadati>Pseudomonadota>Gammaproteobacteria>Enterobacterales>Enterobacteriaceae>Enterobacter>Enterobacter_cloacae_complex | 277 | NVR78_RS04770 | 320614..321447 | 158836 | DUF4435-domain-containing-protein-[Enterobacter-hormaechei] |
| WP_069597815.1 | TOPRIM | GCF_001729805.1 | Enterobacter_roggenkampii | Bacteria>Pseudomonadati>Pseudomonadota>Gammaproteobacteria>Enterobacterales>Enterobacteriaceae>Enterobacter>Enterobacter_cloacae_complex | 277 | BFV67_RS02555 | 551759..552592 | 1812935 | DUF4435-domain-containing-protein-[Enterobacter-roggenkampii] |
| WP_219280526.1 | TOPRIM | GCF_019351595.1 | Enterobacter_cloacae | Bacteria>Pseudomonadati>Pseudomonadota>Gammaproteobacteria>Enterobacterales>Enterobacteriaceae>Enterobacter>Enterobacter_cloacae_complex | 277 | JW319_RS20575 | 182268..183101 | 550 | DUF4435-domain-containing-protein-[Enterobacter-cloacae] |
| WP_248241153.1 | TOPRIM | GCF_023153155.1 | Enterobacter_roggenkampii | Bacteria>Pseudomonadati>Pseudomonadota>Gammaproteobacteria>Enterobacterales>Enterobacteriaceae>Enterobacter>Enterobacter_cloacae_complex | 277 | L8Q49_RS17925 | 120750..121583 | 1812935 | DUF4435-domain-containing-protein-[Enterobacter-roggenkampii] |
| WP_058675373.1 | TOPRIM | GCF_030324195.1 | Enterobacter_hormaechei | Bacteria>Pseudomonadati>Pseudomonadota>Gammaproteobacteria>Enterobacterales>Enterobacteriaceae>Enterobacter>Enterobacter_cloacae_complex | 275 | NUJ48_RS00165 | 37581..38408 | 158836 | DUF4435-domain-containing-protein-[Enterobacter-hormaechei] |
| WP_274416989.1 | TOPRIM | GCF_028765355.1 | Enterobacter_kobei | Bacteria>Pseudomonadati>Pseudomonadota>Gammaproteobacteria>Enterobacterales>Enterobacteriaceae>Enterobacter>Enterobacter_cloacae_complex | 275 | PVM09_RS00205 | 40829..41656 | 208224 | hypothetical-protein-[Enterobacter-kobei] |
| WP_095453566.1 | TOPRIM | GCF_002278355.1 | Enterobacter_cloacae | Bacteria>Pseudomonadati>Pseudomonadota>Gammaproteobacteria>Enterobacterales>Enterobacteriaceae>Enterobacter>Enterobacter_cloacae_complex | 273 | CIW57_RS16425 | 144340..145161 | 550 | DUF4435-domain-containing-protein-[Enterobacter-cloacae] |
| WP_063851025.1 | TOPRIM | GCF_902160625.1 | Enterobacter_hormaechei | Bacteria>Pseudomonadati>Pseudomonadota>Gammaproteobacteria>Enterobacterales>Enterobacteriaceae>Enterobacter>Enterobacter_cloacae_complex | 271 | H0848_RS18570 | 36155..36970 | 158836 | DUF4435-domain-containing-protein-[Enterobacter-hormaechei] |
| WP_248053566.1 | TOPRIM | GCF_023154935.1 | Enterobacter_chengduensis | Bacteria>Pseudomonadati>Pseudomonadota>Gammaproteobacteria>Enterobacterales>Enterobacteriaceae>Enterobacter>Enterobacter_cloacae_complex | 189 | L8P32_RS20975 | 14889..15458 | 2494701 | DUF4276-family-protein-[Enterobacter-chengduensis] |
| MBT2005127.1 | TOPRIM | GCA_018597745.1 | Enterobacter_hormaechei_subsp._xiangfangensis | Bacteria>Pseudomonadati>Pseudomonadota>Gammaproteobacteria>Enterobacterales>Enterobacteriaceae>Enterobacter>Enterobacter_cloacae_complex>Enterobacter_hormaechei | 277 | KKZ39_07400 | 335060..335893 | 1296536 | DUF4435-domain-containing-protein-[Enterobacter-hormaechei-subsp.-xiangfangensis] |
| MCI9498940.1 | TOPRIM | GCA_022800955.1 | Enterobacter_hormaechei_subsp._steigerwaltii | Bacteria>Pseudomonadati>Pseudomonadota>Gammaproteobacteria>Enterobacterales>Enterobacteriaceae>Enterobacter>Enterobacter_cloacae_complex>Enterobacter_hormaechei | 189 | HFE69_15005 | 337478..338047 | 299766 | DUF4276-family-protein-[Enterobacter-hormaechei-subsp.-steigerwaltii] |
| WP_182063471.1 | TOPRIM | GCF_013927005.1 | Enterobacter_sp._RHBSTW-00175 | Bacteria>Pseudomonadati>Pseudomonadota>Gammaproteobacteria>Enterobacterales>Enterobacteriaceae>Enterobacter>unclassified_Enterobacter | 189 | HV107_RS06450 | 1377100..1377669 | 2742639 | hypothetical-protein-[Enterobacter-sp.-RHBSTW-00175] |
| WP_060616119.1 | TOPRIM | GCF_034719175.1 | Escherichia_coli | Bacteria>Pseudomonadati>Pseudomonadota>Gammaproteobacteria>Enterobacterales>Enterobacteriaceae>Escherichia | 277 | DT570_RS09695 | 5120..5953 | 562 | DUF4435-domain-containing-protein-[Escherichia-coli] |
| MCN5345819.1 | TOPRIM | GCA_023869305.1 | Escherichia_coli | Bacteria>Pseudomonadati>Pseudomonadota>Gammaproteobacteria>Enterobacterales>Enterobacteriaceae>Escherichia | 233 | MLU97_13825 | 78202..78903 | 562 | DUF4435-domain-containing-protein-[Escherichia-coli] |
| EFA6389815.1 | TOPRIM | - | Escherichia_coli | Bacteria>Pseudomonadati>Pseudomonadota>Gammaproteobacteria>Enterobacterales>Enterobacteriaceae>Escherichia | 189 | CIW09_001010 | 346561..347130 | 562 | DUF4276-family-protein-[Escherichia-coli] |
| EFC0631246.1 | TOPRIM | - | Escherichia_coli | Bacteria>Pseudomonadati>Pseudomonadota>Gammaproteobacteria>Enterobacterales>Enterobacteriaceae>Escherichia | 189 | FA817_13725 | 294988..295557 | 562 | hypothetical-protein-FA817_13725-[Escherichia-coli] |
| HAG7757103.1 | TOPRIM | GCA_012297605.1 | Escherichia_coli | Bacteria>Pseudomonadati>Pseudomonadota>Gammaproteobacteria>Enterobacterales>Enterobacteriaceae>Escherichia | 189 | HGR18_09660 | 346516..347085 | 562 | TPA:-hypothetical-protein-HGR18_09660-[Escherichia-coli] |
| HAX4880264.1 | TOPRIM | GCA_017016975.1 | Escherichia_coli | Bacteria>Pseudomonadati>Pseudomonadota>Gammaproteobacteria>Enterobacterales>Enterobacteriaceae>Escherichia | 189 | JZF83_001469 | 103863..104432 | 562 | TPA:-DUF4276-family-protein-[Escherichia-coli] |
| HCJ9453759.1 | TOPRIM | GCA_024682335.1 | Escherichia_coli | Bacteria>Pseudomonadati>Pseudomonadota>Gammaproteobacteria>Enterobacterales>Enterobacteriaceae>Escherichia | 189 | NV934_004007 | 17681..18250 | 562 | TPA:-hypothetical-protein-NV934_004007-[Escherichia-coli] |
| WP_181716350.1 | TOPRIM | - | Escherichia_coli | Bacteria>Pseudomonadati>Pseudomonadota>Gammaproteobacteria>Enterobacterales>Enterobacteriaceae>Escherichia | 189 | H5E40_RS01440 | 287176..287745 | 562 | hypothetical-protein-[Escherichia-coli] |
| WP_154062669.1 | TOPRIM | GCF_902159535.1 | Klebsiella | Bacteria>Pseudomonadati>Pseudomonadota>Gammaproteobacteria>Enterobacterales>Enterobacteriaceae>Klebsiella/Raoultella_group | 277 | FN635_RS25515 | 5350637..5351470 | 570 | DUF4435-domain-containing-protein-[Klebsiella-michiganensis] |
| WP_196588768.1 | TOPRIM | GCF_024917295.1 | Klebsiella | Bacteria>Pseudomonadati>Pseudomonadota>Gammaproteobacteria>Enterobacterales>Enterobacteriaceae>Klebsiella/Raoultella_group | 271 | M5R35_RS24855 | 5273695..5274510 | 570 | MULTISPECIES:-DUF4435-domain-containing-protein-[Klebsiella] |
| WP_182409421.1 | TOPRIM | GCF_014129405.1 | Klebsiella_michiganensis | Bacteria>Pseudomonadati>Pseudomonadota>Gammaproteobacteria>Enterobacterales>Enterobacteriaceae>Klebsiella/Raoultella_group>Klebsiella | 277 | H0753_RS11135 | 92566..93399 | 1134687 | DUF4435-domain-containing-protein-[Klebsiella-michiganensis] |
| WP_217074480.1 | TOPRIM | GCF_019048885.1 | Klebsiella_aerogenes | Bacteria>Pseudomonadati>Pseudomonadota>Gammaproteobacteria>Enterobacterales>Enterobacteriaceae>Klebsiella/Raoultella_group>Klebsiella | 277 | KSL85_RS22170 | 4647362..4648195 | 548 | DUF4435-domain-containing-protein-[Klebsiella-aerogenes] |
| WP_224260390.1 | TOPRIM | GCF_026224455.1 | Klebsiella_michiganensis | Bacteria>Pseudomonadati>Pseudomonadota>Gammaproteobacteria>Enterobacterales>Enterobacteriaceae>Klebsiella/Raoultella_group>Klebsiella | 277 | ORL69_RS11455 | 326440..327273 | 1134687 | DUF4435-domain-containing-protein-[Klebsiella-michiganensis] |
| WP_264975994.1 | TOPRIM | GCF_026008315.1 | Klebsiella_pneumoniae_complex | Bacteria>Pseudomonadati>Pseudomonadota>Gammaproteobacteria>Enterobacterales>Enterobacteriaceae>Klebsiella/Raoultella_group>Klebsiella | 277 | OPC12_RS09050 | 182069..182902 | 3390273 | DUF4435-domain-containing-protein-[Klebsiella-quasipneumoniae] |
| GKP62592.1 | TOPRIM | GCA_026008315.1 | Klebsiella_quasipneumoniae | Bacteria>Pseudomonadati>Pseudomonadota>Gammaproteobacteria>Enterobacterales>Enterobacteriaceae>Klebsiella/Raoultella_group>Klebsiella>Klebsiella_pneumoniae_complex | 277 | NUKP47_17660 | 182069..182902 | 1463165 | hypothetical-protein-NUKP47_17660-[Klebsiella-quasipneumoniae] |
| HBR1207679.1 | TOPRIM | GCA_021909055.1 | Klebsiella_pneumoniae | Bacteria>Pseudomonadati>Pseudomonadota>Gammaproteobacteria>Enterobacterales>Enterobacteriaceae>Klebsiella/Raoultella_group>Klebsiella>Klebsiella_pneumoniae_complex | 277 | L9K92_004774 | 35072..35905 | 573 | DUF4435-domain-containing-protein-[Klebsiella-pneumoniae] |
| HBS6654369.1 | TOPRIM | GCA_021975495.1 | Klebsiella_pneumoniae | Bacteria>Pseudomonadati>Pseudomonadota>Gammaproteobacteria>Enterobacterales>Enterobacteriaceae>Klebsiella/Raoultella_group>Klebsiella>Klebsiella_pneumoniae_complex | 277 | MAN10_004111 | 47965..48798 | 573 | TPA:-DUF4435-domain-containing-protein-[Klebsiella-pneumoniae] |
| HBS7652077.1 | TOPRIM | GCA_021979555.1 | Klebsiella_pneumoniae | Bacteria>Pseudomonadati>Pseudomonadota>Gammaproteobacteria>Enterobacterales>Enterobacteriaceae>Klebsiella/Raoultella_group>Klebsiella>Klebsiella_pneumoniae_complex | 277 | MAR21_002851 | 325922..326755 | 573 | TPA:-DUF4435-domain-containing-protein-[Klebsiella-pneumoniae] |
| HBS9599069.1 | TOPRIM | GCA_021986995.1 | Klebsiella_pneumoniae | Bacteria>Pseudomonadati>Pseudomonadota>Gammaproteobacteria>Enterobacterales>Enterobacteriaceae>Klebsiella/Raoultella_group>Klebsiella>Klebsiella_pneumoniae_complex | 277 | MAW98_001663 | 47964..48797 | 573 | TPA:-DUF4435-domain-containing-protein-[Klebsiella-pneumoniae] |
| HBX7751794.1 | TOPRIM | GCA_022258155.1 | Klebsiella_pneumoniae | Bacteria>Pseudomonadati>Pseudomonadota>Gammaproteobacteria>Enterobacterales>Enterobacteriaceae>Klebsiella/Raoultella_group>Klebsiella>Klebsiella_pneumoniae_complex | 277 | MIG23_15235 | 9214..10047 | 573 | TPA:-DUF4435-domain-containing-protein-[Klebsiella-pneumoniae] |
| HCF8566124.1 | TOPRIM | GCA_023980665.1 | Klebsiella_pneumoniae | Bacteria>Pseudomonadati>Pseudomonadota>Gammaproteobacteria>Enterobacterales>Enterobacteriaceae>Klebsiella/Raoultella_group>Klebsiella>Klebsiella_pneumoniae_complex | 277 | NIV99_001795 | 26697..27530 | 573 | TPA:-DUF4435-domain-containing-protein-[Klebsiella-pneumoniae] |
| TWY76108.1 | TOPRIM | GCA_007955295.1 | Klebsiella_pneumoniae | Bacteria>Pseudomonadati>Pseudomonadota>Gammaproteobacteria>Enterobacterales>Enterobacteriaceae>Klebsiella/Raoultella_group>Klebsiella>Klebsiella_pneumoniae_complex | 277 | FTO41_25005 | 26682..27515 | 573 | DUF4435-domain-containing-protein-[Klebsiella-pneumoniae] |
| WP_040153454.1 | TOPRIM | GCF_900775295.1 | Klebsiella_pneumoniae | Bacteria>Pseudomonadati>Pseudomonadota>Gammaproteobacteria>Enterobacterales>Enterobacteriaceae>Klebsiella/Raoultella_group>Klebsiella>Klebsiella_pneumoniae_complex | 277 | JEK00_RS18425 | 14177..15010 | 573 | DUF4435-domain-containing-protein-[Klebsiella-pneumoniae] |
| WP_048289720.1 | TOPRIM | GCF_934247105.1 | Klebsiella_pneumoniae | Bacteria>Pseudomonadati>Pseudomonadota>Gammaproteobacteria>Enterobacterales>Enterobacteriaceae>Klebsiella/Raoultella_group>Klebsiella>Klebsiella_pneumoniae_complex | 277 | A8T29_RS25595 | 19955..20788 | 573 | DUF4435-domain-containing-protein-[Klebsiella-pneumoniae] |
| WP_064143071.1 | TOPRIM | GCF_900085185.1 | Klebsiella_variicola | Bacteria>Pseudomonadati>Pseudomonadota>Gammaproteobacteria>Enterobacterales>Enterobacteriaceae>Klebsiella/Raoultella_group>Klebsiella>Klebsiella_pneumoniae_complex | 277 | A8S52_RS23970 | 28172..29005 | 244366 | DUF4435-domain-containing-protein-[Klebsiella-variicola] |
| WP_104443230.1 | TOPRIM | GCF_018156605.1 | Klebsiella_pneumoniae | Bacteria>Pseudomonadati>Pseudomonadota>Gammaproteobacteria>Enterobacterales>Enterobacteriaceae>Klebsiella/Raoultella_group>Klebsiella>Klebsiella_pneumoniae_complex | 277 | J5D66_RS24900 | 10702..11535 | 573 | DUF4435-domain-containing-protein-[Klebsiella-pneumoniae] |
| WP_147075819.1 | TOPRIM | GCF_019927485.1 | Klebsiella_pneumoniae | Bacteria>Pseudomonadati>Pseudomonadota>Gammaproteobacteria>Enterobacterales>Enterobacteriaceae>Klebsiella/Raoultella_group>Klebsiella>Klebsiella_pneumoniae_complex | 277 | PXW47_RS20405 | 333083..333916 | 573 | DUF4435-domain-containing-protein-[Klebsiella-pneumoniae] |
| WP_117268633.1 | TOPRIM | GCF_900509065.1 | Klebsiella_pneumoniae | Bacteria>Pseudomonadati>Pseudomonadota>Gammaproteobacteria>Enterobacterales>Enterobacteriaceae>Klebsiella/Raoultella_group>Klebsiella>Klebsiella_pneumoniae_complex | 271 | D2728_RS15280 | 262624..263439 | 573 | DUF4435-domain-containing-protein-[Klebsiella-pneumoniae] |
| EIV3902453.1 | TOPRIM | GCA_021871195.1 | Klebsiella_pneumoniae | Bacteria>Pseudomonadati>Pseudomonadota>Gammaproteobacteria>Enterobacterales>Enterobacteriaceae>Klebsiella/Raoultella_group>Klebsiella>Klebsiella_pneumoniae_complex | 189 | L8E75_001736 | 320630..321199 | 573 | hypothetical-protein-[Klebsiella-pneumoniae] |
| EIY5117085.1 | TOPRIM | GCA_022503905.1 | Klebsiella_quasipneumoniae | Bacteria>Pseudomonadati>Pseudomonadota>Gammaproteobacteria>Enterobacterales>Enterobacteriaceae>Klebsiella/Raoultella_group>Klebsiella>Klebsiella_pneumoniae_complex | 189 | ML076_004999 | 28394..28963 | 1463165 | hypothetical-protein-[Klebsiella-quasipneumoniae] |
| HBR1207026.1 | TOPRIM | GCA_021909055.1 | Klebsiella_pneumoniae | Bacteria>Pseudomonadati>Pseudomonadota>Gammaproteobacteria>Enterobacterales>Enterobacteriaceae>Klebsiella/Raoultella_group>Klebsiella>Klebsiella_pneumoniae_complex | 189 | L9K92_004100 | 88909..89478 | 573 | hypothetical-protein-[Klebsiella-pneumoniae] |
| HBX5876370.1 | TOPRIM | GCA_022249175.1 | Klebsiella_pneumoniae | Bacteria>Pseudomonadati>Pseudomonadota>Gammaproteobacteria>Enterobacterales>Enterobacteriaceae>Klebsiella/Raoultella_group>Klebsiella>Klebsiella_pneumoniae_complex | 189 | MH570_26230 | 10413..10982 | 573 | TPA:-hypothetical-protein-MH570_26230-[Klebsiella-pneumoniae] |
| WP_048999819.1 | TOPRIM | GCF_040365915.1 | Klebsiella_pneumoniae | Bacteria>Pseudomonadati>Pseudomonadota>Gammaproteobacteria>Enterobacterales>Enterobacteriaceae>Klebsiella/Raoultella_group>Klebsiella>Klebsiella_pneumoniae_complex | 189 | LF666_RS16210 | 19147..19716 | 573 | hypothetical-protein-[Klebsiella-pneumoniae] |
| WP_135732748.1 | TOPRIM | GCF_900775595.1 | Klebsiella_quasipneumoniae | Bacteria>Pseudomonadati>Pseudomonadota>Gammaproteobacteria>Enterobacterales>Enterobacteriaceae>Klebsiella/Raoultella_group>Klebsiella>Klebsiella_pneumoniae_complex | 187 | ICC30_RS25235 | 20120..20683 | 1463165 | hypothetical-protein-[Klebsiella-quasipneumoniae] |
| HCI4589557.1 | TOPRIM | GCA_024379455.1 | Klebsiella_quasipneumoniae_subsp._quasipneumoniae | Bacteria>Pseudomonadati>Pseudomonadota>Gammaproteobacteria>Enterobacterales>Enterobacteriaceae>Klebsiella/Raoultella_group>Klebsiella>Klebsiella_pneumoniae_complex>Klebsiella_quasipneumoniae | 277 | N0899_004585 | 24616..25449 | 1667327 | DUF4435-domain-containing-protein-[Klebsiella-quasipneumoniae-subsp.-quasipneumoniae] |
| HCB0569862.1 | TOPRIM | GCA_023030625.1 | Klebsiella_quasipneumoniae_subsp._similipneumoniae | Bacteria>Pseudomonadati>Pseudomonadota>Gammaproteobacteria>Enterobacterales>Enterobacteriaceae>Klebsiella/Raoultella_group>Klebsiella>Klebsiella_pneumoniae_complex>Klebsiella_quasipneumoniae | 257 | MYL58_003259 | 42187..42960 | 1463164 | hypothetical-protein-[Klebsiella-quasipneumoniae-subsp.-similipneumoniae] |

| Protein-ID | Domain | Assembly-ID | Strain | Taxonomy | Length | Locus-Tag | Genomic-coordinates | TaxID | Fasta-defline |
|---|---|---|---|---|---|---|---|---|---|
| HCI5938464.1 | TOPRIM | GCA_024431715.1 | Klebsiella_quasipneumoniae_subsp._quasipneumoniae | Bacteria>Pseudomonadati>Pseudomonadota>Gammaproteobacteria>Enterobacterales>Enterobacteriaceae>Klebsiella/Raoultella_group>Klebsiella>Klebsiella_pneumoniae_complex>Klebsiella_quasipneumoniae | 189 | NPN45_004675 | 10606..11175 | 1667327 | hypothetical-protein-[Klebsiella-quasipneumoniae-subsp.-quasipneumoniae] |
| WP_102104495.1 | TOPRIM | GCF_002870005.1 | unclassified_Lelliottia | Bacteria>Pseudomonadati>Pseudomonadota>Gammaproteobacteria>Enterobacterales>Enterobacteriaceae>Lelliottia | 189 | F159LOC_RS08765 | 162805..163374 | 2642424 | MULTISPECIES:-hypothetical-protein-[unclassified-Lelliottia] |
| WP_233672186.1 | TOPRIM | GCF_021368255.1 | Plesiomonas_sp._PI-19 | Bacteria>Pseudomonadati>Pseudomonadota>Gammaproteobacteria>Enterobacterales>Enterobacteriaceae>Plesiomonas>unclassified_Plesiomonas | 273 | LQ901_RS09565 | 2137177..2137998 | 2898798 | DUF4435-domain-containing-protein-[Plesiomonas-sp.-PI-19] |
| WP_233672191.1 | TOPRIM | GCF_021368255.1 | Plesiomonas_sp._PI-19 | Bacteria>Pseudomonadati>Pseudomonadota>Gammaproteobacteria>Enterobacterales>Enterobacteriaceae>Plesiomonas>unclassified_Plesiomonas | 253 | LQ901_RS09590 | 2143460..2144221 | 2898798 | hypothetical-protein-[Plesiomonas-sp.-PI-19] |
| EAA3385006.1 | TOPRIM | GCA_004174495.1 | Salmonella_enterica | Bacteria>Pseudomonadati>Pseudomonadota>Gammaproteobacteria>Enterobacterales>Enterobacteriaceae>Salmonella | 273 | DMV54_10725 | 140340..141161 | 28901 | hypothetical-protein-[Salmonella-enterica] |
| EAA7604357.1 | TOPRIM | GCA_004218255.1 | Salmonella_enterica | Bacteria>Pseudomonadati>Pseudomonadota>Gammaproteobacteria>Enterobacterales>Enterobacteriaceae>Salmonella | 273 | ADQ66_03570 | 138886..139707 | 28901 | DUF4435-domain-containing-protein-[Salmonella-enterica] |
| EAA8317757.1 | TOPRIM | GCA_004221325.1 | Salmonella_enterica | Bacteria>Pseudomonadati>Pseudomonadota>Gammaproteobacteria>Enterobacterales>Enterobacteriaceae>Salmonella | 273 | AOS09_04980 | 171040..171861 | 28901 | DUF4435-domain-containing-protein-[Salmonella-enterica] |
| EAA8875708.1 | TOPRIM | GCA_004223005.1 | Salmonella_enterica | Bacteria>Pseudomonadati>Pseudomonadota>Gammaproteobacteria>Enterobacterales>Enterobacteriaceae>Salmonella | 273 | CIU45_09640 | 390257..391078 | 28901 | DUF4435-domain-containing-protein-[Salmonella-enterica] |
| EAA9319739.1 | TOPRIM | GCA_004226405.1 | Salmonella_enterica | Bacteria>Pseudomonadati>Pseudomonadota>Gammaproteobacteria>Enterobacterales>Enterobacteriaceae>Salmonella | 273 | CRQ39_09915 | 175803..176624 | 28901 | DUF4435-domain-containing-protein-[Salmonella-enterica] |
| EAB1768966.1 | TOPRIM | GCA_004236005.1 | Salmonella_enterica | Bacteria>Pseudomonadati>Pseudomonadota>Gammaproteobacteria>Enterobacterales>Enterobacteriaceae>Salmonella | 273 | EIS13_11235 | 26702..27523 | 28901 | DUF4435-domain-containing-protein-[Salmonella-enterica] |
| EAB2831581.1 | TOPRIM | GCA_004240605.1 | Salmonella_enterica | Bacteria>Pseudomonadati>Pseudomonadota>Gammaproteobacteria>Enterobacterales>Enterobacteriaceae>Salmonella | 273 | EAV48_05285 | 166334..167155 | 28901 | DUF4435-domain-containing-protein-[Salmonella-enterica] |
| EAB2910639.1 | TOPRIM | GCA_004240965.1 | Salmonella_enterica | Bacteria>Pseudomonadati>Pseudomonadota>Gammaproteobacteria>Enterobacterales>Enterobacteriaceae>Salmonella | 273 | EHO38_04040 | 126995..127816 | 28901 | DUF4435-domain-containing-protein-[Salmonella-enterica] |
| EAM3013485.1 | TOPRIM | GCA_005412095.1 | Salmonella_enterica | Bacteria>Pseudomonadati>Pseudomonadota>Gammaproteobacteria>Enterobacterales>Enterobacteriaceae>Salmonella | 273 | CBJ93_01880 | 403047..403868 | 28901 | DUF4435-domain-containing-protein-[Salmonella-enterica] |
| EAM5567467.1 | TOPRIM | GCA_005423155.1 | Salmonella_enterica | Bacteria>Pseudomonadati>Pseudomonadota>Gammaproteobacteria>Enterobacterales>Enterobacteriaceae>Salmonella | 273 | EK556_21765 | 5492..6313 | 28901 | DUF4435-domain-containing-protein-[Salmonella-enterica] |
| EAM5877055.1 | TOPRIM | GCA_005424775.1 | Salmonella_enterica | Bacteria>Pseudomonadati>Pseudomonadota>Gammaproteobacteria>Enterobacterales>Enterobacteriaceae>Salmonella | 273 | EPF42_07730 | 340744..341565 | 28901 | DUF4435-domain-containing-protein-[Salmonella-enterica] |
| EAM6081346.1 | TOPRIM | GCA_005424535.1 | Salmonella_enterica | Bacteria>Pseudomonadati>Pseudomonadota>Gammaproteobacteria>Enterobacterales>Enterobacteriaceae>Salmonella | 273 | ERR94_15335 | 11821..12642 | 28901 | DUF4435-domain-containing-protein-[Salmonella-enterica] |
| EAO2684617.1 | TOPRIM | GCA_005496785.1 | Salmonella_enterica | Bacteria>Pseudomonadati>Pseudomonadota>Gammaproteobacteria>Enterobacterales>Enterobacteriaceae>Salmonella | 273 | E2X65_05665 | 75960..76781 | 28901 | DUF4435-domain-containing-protein-[Salmonella-enterica] |
| EAO8797694.1 | TOPRIM | GCA_005530015.1 | Salmonella_enterica | Bacteria>Pseudomonadati>Pseudomonadota>Gammaproteobacteria>Enterobacterales>Enterobacteriaceae>Salmonella | 273 | E0391_14550 | 34305..35126 | 28901 | DUF4435-domain-containing-protein-[Salmonella-enterica] |
| EAP0128504.1 | TOPRIM | GCA_005535515.1 | Salmonella_enterica | Bacteria>Pseudomonadati>Pseudomonadota>Gammaproteobacteria>Enterobacterales>Enterobacteriaceae>Salmonella | 273 | DKP73_00830 | 169790..170611 | 28901 | DUF4435-domain-containing-protein-[Salmonella-enterica] |
| EAP0644640.1 | TOPRIM | GCA_005538295.1 | Salmonella_enterica | Bacteria>Pseudomonadati>Pseudomonadota>Gammaproteobacteria>Enterobacterales>Enterobacteriaceae>Salmonella | 273 | C6277_11590 | 102450..103271 | 28901 | DUF4435-domain-containing-protein-[Salmonella-enterica] |
| EAQ0497030.1 | TOPRIM | GCA_005582475.1 | Salmonella_enterica | Bacteria>Pseudomonadati>Pseudomonadota>Gammaproteobacteria>Enterobacterales>Enterobacteriaceae>Salmonella | 273 | EW683_07770 | 45316..46137 | 28901 | DUF4435-domain-containing-protein-[Salmonella-enterica] |
| EAQ6130546.1 | TOPRIM | GCA_005607075.1 | Salmonella_enterica | Bacteria>Pseudomonadati>Pseudomonadota>Gammaproteobacteria>Enterobacterales>Enterobacteriaceae>Salmonella | 273 | AXA91_18745 | 70785..71606 | 28901 | DUF4435-domain-containing-protein-[Salmonella-enterica] |
| EAQ7293578.1 | TOPRIM | GCA_005612115.1 | Salmonella_enterica | Bacteria>Pseudomonadati>Pseudomonadota>Gammaproteobacteria>Enterobacterales>Enterobacteriaceae>Salmonella | 273 | CRQ46_08460 | 415145..415966 | 28901 | DUF4435-domain-containing-protein-[Salmonella-enterica] |
| EAS9365413.1 | TOPRIM | GCA_005712675.1 | Salmonella_enterica | Bacteria>Pseudomonadati>Pseudomonadota>Gammaproteobacteria>Enterobacterales>Enterobacteriaceae>Salmonella | 273 | EPA30_12285 | 179275..180096 | 28901 | DUF4435-domain-containing-protein-[Salmonella-enterica] |
| EAT1639066.1 | TOPRIM | GCA_005720135.1 | Salmonella_enterica | Bacteria>Pseudomonadati>Pseudomonadota>Gammaproteobacteria>Enterobacterales>Enterobacteriaceae>Salmonella | 273 | EKF56_11755 | 33451..34272 | 28901 | DUF4435-domain-containing-protein-[Salmonella-enterica] |
| EAT1838144.1 | TOPRIM | GCA_005717435.1 | Salmonella_enterica | Bacteria>Pseudomonadati>Pseudomonadota>Gammaproteobacteria>Enterobacterales>Enterobacteriaceae>Salmonella | 273 | EJR21_08330 | 111418..112239 | 28901 | DUF4435-domain-containing-protein-[Salmonella-enterica] |
| EAT2430710.1 | TOPRIM | GCA_005726115.1 | Salmonella_enterica | Bacteria>Pseudomonadati>Pseudomonadota>Gammaproteobacteria>Enterobacterales>Enterobacteriaceae>Salmonella | 273 | EI207_04340 | 185575..186396 | 28901 | DUF4435-domain-containing-protein-[Salmonella-enterica] |
| EAU0116270.1 | TOPRIM | GCA_005756415.1 | Salmonella_enterica | Bacteria>Pseudomonadati>Pseudomonadota>Gammaproteobacteria>Enterobacterales>Enterobacteriaceae>Salmonella | 273 | ASR58_09445 | 34317..35138 | 28901 | DUF4435-domain-containing-protein-[Salmonella-enterica] |
| EAU0152694.1 | TOPRIM | GCA_005756695.1 | Salmonella_enterica | Bacteria>Pseudomonadati>Pseudomonadota>Gammaproteobacteria>Enterobacterales>Enterobacteriaceae>Salmonella | 273 | AZG01_01880 | 399164..399985 | 28901 | DUF4435-domain-containing-protein-[Salmonella-enterica] |
| EAV3792367.1 | TOPRIM | GCA_005840905.1 | Salmonella_enterica | Bacteria>Pseudomonadati>Pseudomonadota>Gammaproteobacteria>Enterobacterales>Enterobacteriaceae>Salmonella | 273 | E2X64_08250 | 34040..34861 | 28901 | DUF4435-domain-containing-protein-[Salmonella-enterica] |
| EAV4850771.1 | TOPRIM | GCA_005847515.1 | Salmonella_enterica | Bacteria>Pseudomonadati>Pseudomonadota>Gammaproteobacteria>Enterobacterales>Enterobacteriaceae>Salmonella | 273 | EGE73_10375 | 31417..32238 | 28901 | DUF4435-domain-containing-protein-[Salmonella-enterica] |
| EAW1006427.1 | TOPRIM | GCA_005898145.1 | Salmonella_enterica | Bacteria>Pseudomonadati>Pseudomonadota>Gammaproteobacteria>Enterobacterales>Enterobacteriaceae>Salmonella | 273 | EA086_11675 | 165491..166312 | 28901 | DUF4435-domain-containing-protein-[Salmonella-enterica] |
| EAW5830200.1 | TOPRIM | GCA_005917905.1 | Salmonella_enterica | Bacteria>Pseudomonadati>Pseudomonadota>Gammaproteobacteria>Enterobacterales>Enterobacteriaceae>Salmonella | 273 | FF074_06120 | 179145..179966 | 28901 | DUF4435-domain-containing-protein-[Salmonella-enterica] |
| EAW8145858.1 | TOPRIM | GCA_005934015.1 | Salmonella_enterica | Bacteria>Pseudomonadati>Pseudomonadota>Gammaproteobacteria>Enterobacterales>Enterobacteriaceae>Salmonella | 273 | HM10_03840 | 171926..172747 | 28901 | DUF4435-domain-containing-protein-[Salmonella-enterica] |
| EAW9404812.1 | TOPRIM | GCA_005940875.1 | Salmonella_enterica | Bacteria>Pseudomonadati>Pseudomonadota>Gammaproteobacteria>Enterobacterales>Enterobacteriaceae>Salmonella | 273 | QD58_16775 | 31935..32756 | 28901 | DUF4435-domain-containing-protein-[Salmonella-enterica] |
| EAX4456713.1 | TOPRIM | GCA_005963445.1 | Salmonella_enterica | Bacteria>Pseudomonadati>Pseudomonadota>Gammaproteobacteria>Enterobacterales>Enterobacteriaceae>Salmonella | 273 | AC467_14835 | 26471..27292 | 28901 | DUF4435-domain-containing-protein-[Salmonella-enterica] |
| EAX8471830.1 | TOPRIM | GCA_005980785.1 | Salmonella_enterica | Bacteria>Pseudomonadati>Pseudomonadota>Gammaproteobacteria>Enterobacterales>Enterobacteriaceae>Salmonella | 273 | AH421_04155 | 254948..255769 | 28901 | DUF4435-domain-containing-protein-[Salmonella-enterica] |
| EAZ5773631.1 | TOPRIM | GCA_006057755.1 | Salmonella_enterica | Bacteria>Pseudomonadati>Pseudomonadota>Gammaproteobacteria>Enterobacterales>Enterobacteriaceae>Salmonella | 273 | CCZ51_07190 | 102451..103272 | 28901 | DUF4435-domain-containing-protein-[Salmonella-enterica] |
| EAZ9311964.1 | TOPRIM | GCA_006073635.1 | Salmonella_enterica | Bacteria>Pseudomonadati>Pseudomonadota>Gammaproteobacteria>Enterobacterales>Enterobacteriaceae>Salmonella | 273 | CBJ28_03040 | 167120..167941 | 28901 | DUF4435-domain-containing-protein-[Salmonella-enterica] |
| EBA4497720.1 | TOPRIM | GCA_006097275.1 | Salmonella_enterica | Bacteria>Pseudomonadati>Pseudomonadota>Gammaproteobacteria>Enterobacterales>Enterobacteriaceae>Salmonella | 273 | A7G53_12035 | 217312..218133 | 28901 | DUF4435-domain-containing-protein-[Salmonella-enterica] |
| EBB0684612.1 | TOPRIM | GCA_006125875.1 | Salmonella_enterica | Bacteria>Pseudomonadati>Pseudomonadota>Gammaproteobacteria>Enterobacterales>Enterobacteriaceae>Salmonella | 273 | CG859_01595 | 340077..340898 | 28901 | DUF4435-domain-containing-protein-[Salmonella-enterica] |
| EBB1941187.1 | TOPRIM | GCA_006131775.1 | Salmonella_enterica | Bacteria>Pseudomonadati>Pseudomonadota>Gammaproteobacteria>Enterobacterales>Enterobacteriaceae>Salmonella | 273 | CJP21_07745 | 47112..47933 | 28901 | DUF4435-domain-containing-protein-[Salmonella-enterica] |
| EBB3832832.1 | TOPRIM | GCA_006140415.1 | Salmonella_enterica | Bacteria>Pseudomonadati>Pseudomonadota>Gammaproteobacteria>Enterobacterales>Enterobacteriaceae>Salmonella | 273 | CJ140_05535 | 158169..158990 | 28901 | DUF4435-domain-containing-protein-[Salmonella-enterica] |
| EBC0774148.1 | TOPRIM | GCA_006172705.1 | Salmonella_enterica | Bacteria>Pseudomonadati>Pseudomonadota>Gammaproteobacteria>Enterobacterales>Enterobacteriaceae>Salmonella | 273 | CS696_13030 | 17930..18751 | 28901 | DUF4435-domain-containing-protein-[Salmonella-enterica] |
| EBC4250054.1 | TOPRIM | GCA_006188785.1 | Salmonella_enterica | Bacteria>Pseudomonadati>Pseudomonadota>Gammaproteobacteria>Enterobacterales>Enterobacteriaceae>Salmonella | 273 | C2B00_07705 | 222282..223103 | 28901 | DUF4435-domain-containing-protein-[Salmonella-enterica] |
| EBD9210541.1 | TOPRIM | GCA_006272795.1 | Salmonella_enterica | Bacteria>Pseudomonadati>Pseudomonadota>Gammaproteobacteria>Enterobacterales>Enterobacteriaceae>Salmonella | 273 | DA198_05740 | 181389..182210 | 28901 | DUF4435-domain-containing-protein-[Salmonella-enterica] |
| EBE1252192.1 | TOPRIM | GCA_006282055.1 | Salmonella_enterica | Bacteria>Pseudomonadati>Pseudomonadota>Gammaproteobacteria>Enterobacterales>Enterobacteriaceae>Salmonella | 273 | DDQ39_02655 | 132850..133671 | 28901 | DUF4435-domain-containing-protein-[Salmonella-enterica] |
| EBE5553329.1 | TOPRIM | GCA_006301715.1 | Salmonella_enterica | Bacteria>Pseudomonadati>Pseudomonadota>Gammaproteobacteria>Enterobacterales>Enterobacteriaceae>Salmonella | 273 | CP597_01385 | 304318..305139 | 28901 | DUF4435-domain-containing-protein-[Salmonella-enterica] |
| EBE9623451.1 | TOPRIM | GCA_006320265.1 | Salmonella_enterica | Bacteria>Pseudomonadati>Pseudomonadota>Gammaproteobacteria>Enterobacterales>Enterobacteriaceae>Salmonella | 273 | DA967_03440 | 179396..180217 | 28901 | DUF4435-domain-containing-protein-[Salmonella-enterica] |
| EBF0840959.1 | TOPRIM | GCA_006325425.1 | Salmonella_enterica | Bacteria>Pseudomonadati>Pseudomonadota>Gammaproteobacteria>Enterobacterales>Enterobacteriaceae>Salmonella | 273 | DDQ86_07965 | 158701..159522 | 28901 | DUF4435-domain-containing-protein-[Salmonella-enterica] |
| EBG3411776.1 | TOPRIM | GCA_006423935.1 | Salmonella_enterica | Bacteria>Pseudomonadati>Pseudomonadota>Gammaproteobacteria>Enterobacterales>Enterobacteriaceae>Salmonella | 273 | FIW39_06220 | 179461..180282 | 28901 | DUF4435-domain-containing-protein-[Salmonella-enterica] |
| EBH0811992.1 | TOPRIM | GCA_006475165.1 | Salmonella_enterica | Bacteria>Pseudomonadati>Pseudomonadota>Gammaproteobacteria>Enterobacterales>Enterobacteriaceae>Salmonella | 273 | FJ896_02685 | 179365..180186 | 28901 | DUF4435-domain-containing-protein-[Salmonella-enterica] |
| EBH4970959.1 | TOPRIM | GCA_006499395.1 | Salmonella_enterica | Bacteria>Pseudomonadati>Pseudomonadota>Gammaproteobacteria>Enterobacterales>Enterobacteriaceae>Salmonella | 273 | CKR35_08940 | 178780..179601 | 28901 | DUF4435-domain-containing-protein-[Salmonella-enterica] |
| EBI8027933.1 | TOPRIM | GCA_006571625.1 | Salmonella_enterica | Bacteria>Pseudomonadati>Pseudomonadota>Gammaproteobacteria>Enterobacterales>Enterobacteriaceae>Salmonella | 273 | CHM23_07405 | 179352..180173 | 28901 | DUF4435-domain-containing-protein-[Salmonella-enterica] |
| EBK5803038.1 | TOPRIM | GCA_006648525.1 | Salmonella_enterica | Bacteria>Pseudomonadati>Pseudomonadota>Gammaproteobacteria>Enterobacterales>Enterobacteriaceae>Salmonella | 273 | DPM88_04080 | 166848..167669 | 28901 | DUF4435-domain-containing-protein-[Salmonella-enterica] |
| EBK7108453.1 | TOPRIM | GCA_006653645.1 | Salmonella_enterica | Bacteria>Pseudomonadati>Pseudomonadota>Gammaproteobacteria>Enterobacterales>Enterobacteriaceae>Salmonella | 273 | DSC93_06700 | 179362..180183 | 28901 | DUF4435-domain-containing-protein-[Salmonella-enterica] |
| EBK9871842.1 | TOPRIM | GCA_006668805.1 | Salmonella_enterica | Bacteria>Pseudomonadati>Pseudomonadota>Gammaproteobacteria>Enterobacterales>Enterobacteriaceae>Salmonella | 273 | DYD89_19165 | 34317..35138 | 28901 | DUF4435-domain-containing-protein-[Salmonella-enterica] |
| EBL1260732.1 | TOPRIM | GCA_006672445.1 | Salmonella_enterica | Bacteria>Pseudomonadati>Pseudomonadota>Gammaproteobacteria>Enterobacterales>Enterobacteriaceae>Salmonella | 273 | D0790_07485 | 179563..180384 | 28901 | DUF4435-domain-containing-protein-[Salmonella-enterica] |
| EBL2857389.1 | TOPRIM | GCA_006679365.1 | Salmonella_enterica | Bacteria>Pseudomonadati>Pseudomonadota>Gammaproteobacteria>Enterobacterales>Enterobacteriaceae>Salmonella | 273 | D3Q95_12485 | 126778..127599 | 28901 | DUF4435-domain-containing-protein-[Salmonella-enterica] |
| EBL7150921.1 | TOPRIM | GCA_006698245.1 | Salmonella_enterica | Bacteria>Pseudomonadati>Pseudomonadota>Gammaproteobacteria>Enterobacterales>Enterobacteriaceae>Salmonella | 273 | CFL44_11215 | 74563..75384 | 28901 | DUF4435-domain-containing-protein-[Salmonella-enterica] |
| EBM6187523.1 | TOPRIM | GCA_006738745.1 | Salmonella_enterica | Bacteria>Pseudomonadati>Pseudomonadota>Gammaproteobacteria>Enterobacterales>Enterobacteriaceae>Salmonella | 273 | D1943_04245 | 109009..109830 | 28901 | DUF4435-domain-containing-protein-[Salmonella-enterica] |
| EBR4438816.1 | TOPRIM | GCA_006955825.1 | Salmonella_enterica | Bacteria>Pseudomonadati>Pseudomonadota>Gammaproteobacteria>Enterobacterales>Enterobacteriaceae>Salmonella | 273 | BSE18_11490 | 102451..103272 | 28901 | DUF4435-domain-containing-protein-[Salmonella-enterica] |
| EBS7702175.1 | TOPRIM | GCA_007022165.1 | Salmonella_enterica | Bacteria>Pseudomonadati>Pseudomonadota>Gammaproteobacteria>Enterobacterales>Enterobacteriaceae>Salmonella | 273 | CD603_08295 | 58771..59592 | 28901 | DUF4435-domain-containing-protein-[Salmonella-enterica] |
| EBT4520739.1 | TOPRIM | GCA_007056145.1 | Salmonella_enterica | Bacteria>Pseudomonadati>Pseudomonadota>Gammaproteobacteria>Enterobacterales>Enterobacteriaceae>Salmonella | 273 | CKQ08_17665 | 102451..103272 | 28901 | DUF4435-domain-containing-protein-[Salmonella-enterica] |
| EBU3750857.1 | TOPRIM | GCA_007101645.1 | Salmonella_enterica | Bacteria>Pseudomonadati>Pseudomonadota>Gammaproteobacteria>Enterobacterales>Enterobacteriaceae>Salmonella | 273 | CVX24_10940 | 34314..35135 | 28901 | DUF4435-domain-containing-protein-[Salmonella-enterica] |
| ECG9327784.1 | TOPRIM | GCA_007700105.1 | Salmonella_enterica | Bacteria>Pseudomonadati>Pseudomonadota>Gammaproteobacteria>Enterobacterales>Enterobacteriaceae>Salmonella | 273 | FO199_03400 | 126236..127057 | 28901 | DUF4435-domain-containing-protein-[Salmonella-enterica] |
| ECH3013707.1 | TOPRIM | GCA_007716945.1 | Salmonella_enterica | Bacteria>Pseudomonadati>Pseudomonadota>Gammaproteobacteria>Enterobacterales>Enterobacteriaceae>Salmonella | 273 | FOV18_07405 | 337485..338306 | 28901 | DUF4435-domain-containing-protein-[Salmonella-enterica] |
| ECJ6092449.1 | TOPRIM | GCA_007834095.1 | Salmonella_enterica | Bacteria>Pseudomonadati>Pseudomonadota>Gammaproteobacteria>Enterobacterales>Enterobacteriaceae>Salmonella | 273 | GR78_06775 | 380777..381598 | 28901 | DUF4435-domain-containing-protein-[Salmonella-enterica] |
| ECO5284317.1 | TOPRIM | GCA_008153185.1 | Salmonella_enterica | Bacteria>Pseudomonadati>Pseudomonadota>Gammaproteobacteria>Enterobacterales>Enterobacteriaceae>Salmonella | 273 | FYE58_11860 | 406384..407205 | 28901 | DUF4435-domain-containing-protein-[Salmonella-enterica] |
| ECQ9432771.1 | TOPRIM | GCA_008312495.1 | Salmonella_enterica | Bacteria>Pseudomonadati>Pseudomonadota>Gammaproteobacteria>Enterobacterales>Enterobacteriaceae>Salmonella | 273 | F0371_05180 | 11240..12061 | 28901 | DUF4435-domain-containing-protein-[Salmonella-enterica] |
| EDC2377954.1 | TOPRIM | - | Salmonella_enterica | Bacteria>Pseudomonadati>Pseudomonadota>Gammaproteobacteria>Enterobacterales>Enterobacteriaceae>Salmonella | 273 | GAA61_09320 | 34317..35138 | 28901 | DUF4435-domain-containing-protein-[Salmonella-enterica] |
| EDH2237973.1 | TOPRIM | - | Salmonella_enterica | Bacteria>Pseudomonadati>Pseudomonadota>Gammaproteobacteria>Enterobacterales>Enterobacteriaceae>Salmonella | 273 | GC580_06325 | 178946..179767 | 28901 | DUF4435-domain-containing-protein-[Salmonella-enterica] |
| EDI9669680.1 | TOPRIM | - | Salmonella_enterica | Bacteria>Pseudomonadati>Pseudomonadota>Gammaproteobacteria>Enterobacterales>Enterobacteriaceae>Salmonella | 273 | GEV75_12820 | 47101..47922 | 28901 | hypothetical-protein-GEV75_12820-[Salmonella-enterica] |
| EDL6685017.1 | TOPRIM | - | Salmonella_enterica | Bacteria>Pseudomonadati>Pseudomonadota>Gammaproteobacteria>Enterobacterales>Enterobacteriaceae>Salmonella | 273 | CXC07_02685 | 179346..180167 | 28901 | hypothetical-protein-CXC07_02685-[Salmonella-enterica] |
| EDL7421469.1 | TOPRIM | - | Salmonella_enterica | Bacteria>Pseudomonadati>Pseudomonadota>Gammaproteobacteria>Enterobacterales>Enterobacteriaceae>Salmonella | 273 | CU840_19770 | 17582..18403 | 28901 | hypothetical-protein-CU840_19770-[Salmonella-enterica] |
| EDV2155935.1 | TOPRIM | - | Salmonella_enterica | Bacteria>Pseudomonadati>Pseudomonadota>Gammaproteobacteria>Enterobacterales>Enterobacteriaceae>Salmonella | 273 | JR65_004158 | 33526..34347 | 28901 | DUF4435-domain-containing-protein-[Salmonella-enterica] |
| EDW7722491.1 | TOPRIM | - | Salmonella_enterica | Bacteria>Pseudomonadati>Pseudomonadota>Gammaproteobacteria>Enterobacterales>Enterobacteriaceae>Salmonella | 273 | GTR06_001874 | 406342..407163 | 28901 | DUF4435-domain-containing-protein-[Salmonella-enterica] |
| EDX0369806.1 | TOPRIM | - | Salmonella_enterica | Bacteria>Pseudomonadati>Pseudomonadota>Gammaproteobacteria>Enterobacterales>Enterobacteriaceae>Salmonella | 273 | GTL21_000364 | 394973..395794 | 28901 | DUF4435-domain-containing-protein-[Salmonella-enterica] |
| EEB8473039.1 | TOPRIM | - | Salmonella_enterica | Bacteria>Pseudomonadati>Pseudomonadota>Gammaproteobacteria>Enterobacterales>Enterobacteriaceae>Salmonella | 273 | GN471_06060 | 179151..179972 | 28901 | DUF4435-domain-containing-protein-[Salmonella-enterica] |
| EEH0007754.1 | TOPRIM | - | Salmonella_enterica | Bacteria>Pseudomonadati>Pseudomonadota>Gammaproteobacteria>Enterobacterales>Enterobacteriaceae>Salmonella | 273 | G3P26_001651 | 363358..364179 | 28901 | DUF4435-domain-containing-protein-[Salmonella-enterica] |
| EEH3158633.1 | TOPRIM | - | Salmonella_enterica | Bacteria>Pseudomonadati>Pseudomonadota>Gammaproteobacteria>Enterobacterales>Enterobacteriaceae>Salmonella | 273 | GZB32_002493 | 102451..103272 | 28901 | DUF4435-domain-containing-protein-[Salmonella-enterica] |
| EEI0202358.1 | TOPRIM | - | Salmonella_enterica | Bacteria>Pseudomonadati>Pseudomonadota>Gammaproteobacteria>Enterobacterales>Enterobacteriaceae>Salmonella | 273 | GNM15_17105 | 102451..103272 | 28901 | DUF4435-domain-containing-protein-[Salmonella-enterica] |
| EEL0135866.1 | TOPRIM | - | Salmonella_enterica | Bacteria>Pseudomonadati>Pseudomonadota>Gammaproteobacteria>Enterobacterales>Enterobacteriaceae>Salmonella | 273 | G4L63_001977 | 120507..121328 | 28901 | DUF4435-domain-containing-protein-[Salmonella-enterica] |
| EEL7097506.1 | TOPRIM | - | Salmonella_enterica | Bacteria>Pseudomonadati>Pseudomonadota>Gammaproteobacteria>Enterobacterales>Enterobacteriaceae>Salmonella | 273 | GKH84_08590 | 167316..168137 | 28901 | DUF4435-domain-containing-protein-[Salmonella-enterica] |
| EEP2127995.1 | TOPRIM | - | Salmonella_enterica | Bacteria>Pseudomonadati>Pseudomonadota>Gammaproteobacteria>Enterobacterales>Enterobacteriaceae>Salmonella | 273 | HA780_002413 | 119698..120519 | 28901 | DUF4435-domain-containing-protein-[Salmonella-enterica] |
| EEP5746978.1 | TOPRIM | - | Salmonella_enterica | Bacteria>Pseudomonadati>Pseudomonadota>Gammaproteobacteria>Enterobacterales>Enterobacteriaceae>Salmonella | 273 | HBB29_002272 | 102451..103272 | 28901 | DUF4435-domain-containing-protein-[Salmonella-enterica] |
| EFP5236620.1 | TOPRIM | - | Salmonella_enterica | Bacteria>Pseudomonadati>Pseudomonadota>Gammaproteobacteria>Enterobacterales>Enterobacteriaceae>Salmonella | 273 | H1F05_002030 | 178781..179602 | 28901 | DUF4435-domain-containing-protein-[Salmonella-enterica] |
| EFQ1578020.1 | TOPRIM | - | Salmonella_enterica | Bacteria>Pseudomonadati>Pseudomonadota>Gammaproteobacteria>Enterobacterales>Enterobacteriaceae>Salmonella | 273 | H1L81_001835 | 47105..47926 | 28901 | DUF4435-domain-containing-protein-[Salmonella-enterica] |
| EFQ9845219.1 | TOPRIM | - | Salmonella_enterica | Bacteria>Pseudomonadati>Pseudomonadota>Gammaproteobacteria>Enterobacterales>Enterobacteriaceae>Salmonella | 273 | HZL63_001576 | 144404..145225 | 28901 | DUF4435-domain-containing-protein-[Salmonella-enterica] |
| EFR0716068.1 | TOPRIM | - | Salmonella_enterica | Bacteria>Pseudomonadati>Pseudomonadota>Gammaproteobacteria>Enterobacterales>Enterobacteriaceae>Salmonella | 273 | HYQ91_004639 | 6956..7777 | 28901 | DUF4435-domain-containing-protein-[Salmonella-enterica] |
| EFS3421935.1 | TOPRIM | - | Salmonella_enterica | Bacteria>Pseudomonadati>Pseudomonadota>Gammaproteobacteria>Enterobacterales>Enterobacteriaceae>Salmonella | 273 | HX201_001608 | 158570..159391 | 28901 | DUF4435-domain-containing-protein-[Salmonella-enterica] |
| EFS5965941.1 | TOPRIM | - | Salmonella_enterica | Bacteria>Pseudomonadati>Pseudomonadota>Gammaproteobacteria>Enterobacterales>Enterobacteriaceae>Salmonella | 273 | HW654_000975 | 203749..204570 | 28901 | DUF4435-domain-containing-protein-[Salmonella-enterica] |
| EFS7669871.1 | TOPRIM | - | Salmonella_enterica | Bacteria>Pseudomonadati>Pseudomonadota>Gammaproteobacteria>Enterobacterales>Enterobacteriaceae>Salmonella | 273 | HWZ11_001256 | 8031..8852 | 28901 | DUF4435-domain-containing-protein-[Salmonella-enterica] |
| EFU4335449.1 | TOPRIM | - | Salmonella_enterica | Bacteria>Pseudomonadati>Pseudomonadota>Gammaproteobacteria>Enterobacterales>Enterobacteriaceae>Salmonella | 273 | HUC58_001816 | 178782..179603 | 28901 | DUF4435-domain-containing-protein-[Salmonella-enterica] |
| EGB5523465.1 | TOPRIM | - | Salmonella_enterica | Bacteria>Pseudomonadati>Pseudomonadota>Gammaproteobacteria>Enterobacterales>Enterobacteriaceae>Salmonella | 273 | H7074_002107 | 167484..168305 | 28901 | DUF4435-domain-containing-protein-[Salmonella-enterica] |
| EGC3767045.1 | TOPRIM | - | Salmonella_enterica | Bacteria>Pseudomonadati>Pseudomonadota>Gammaproteobacteria>Enterobacterales>Enterobacteriaceae>Salmonella | 273 | H9A68_002333 | 47105..47926 | 28901 | DUF4435-domain-containing-protein-[Salmonella-enterica] |
| EGD1472164.1 | TOPRIM | - | Salmonella_enterica | Bacteria>Pseudomonadati>Pseudomonadota>Gammaproteobacteria>Enterobacterales>Enterobacteriaceae>Salmonella | 273 | IAV68_002847 | 102451..103272 | 28901 | DUF4435-domain-containing-protein-[Salmonella-enterica] |
| EGF1249456.1 | TOPRIM | - | Salmonella_enterica | Bacteria>Pseudomonadati>Pseudomonadota>Gammaproteobacteria>Enterobacterales>Enterobacteriaceae>Salmonella | 273 | IB037_001324 | 179352..180173 | 28901 | DUF4435-domain-containing-protein-[Salmonella-enterica] |
| EGG4120339.1 | TOPRIM | - | Salmonella_enterica | Bacteria>Pseudomonadati>Pseudomonadota>Gammaproteobacteria>Enterobacterales>Enterobacteriaceae>Salmonella | 273 | SI35_20755 | 165758..166579 | 28901 | DUF4435-domain-containing-protein-[Salmonella-enterica] |
| EGH1765317.1 | TOPRIM | - | Salmonella_enterica | Bacteria>Pseudomonadati>Pseudomonadota>Gammaproteobacteria>Enterobacterales>Enterobacteriaceae>Salmonella | 273 | ID459_000692 | 179120..179941 | 28901 | DUF4435-domain-containing-protein-[Salmonella-enterica] |
| EGH2238324.1 | TOPRIM | - | Salmonella_enterica | Bacteria>Pseudomonadati>Pseudomonadota>Gammaproteobacteria>Enterobacterales>Enterobacteriaceae>Salmonella | 273 | ID764_000357 | 399247..400068 | 28901 | DUF4435-domain-containing-protein-[Salmonella-enterica] |
| EGH8508056.1 | TOPRIM | - | Salmonella_enterica | Bacteria>Pseudomonadati>Pseudomonadota>Gammaproteobacteria>Enterobacterales>Enterobacteriaceae>Salmonella | 273 | IFE76_001739 | 34526..35347 | 28901 | DUF4435-domain-containing-protein-[Salmonella-enterica] |
| EGI8941865.1 | TOPRIM | - | Salmonella_enterica | Bacteria>Pseudomonadati>Pseudomonadota>Gammaproteobacteria>Enterobacterales>Enterobacteriaceae>Salmonella | 273 | IG687_002356 | 281431..282252 | 28901 | DUF4435-domain-containing-protein-[Salmonella-enterica] |
| EGK3406183.1 | TOPRIM | - | Salmonella_enterica | Bacteria>Pseudomonadati>Pseudomonadota>Gammaproteobacteria>Enterobacterales>Enterobacteriaceae>Salmonella | 273 | IOS83_004390 | 19590..20411 | 28901 | DUF4435-domain-containing-protein-[Salmonella-enterica] |
| EGL7108187.1 | TOPRIM | - | Salmonella_enterica | Bacteria>Pseudomonadati>Pseudomonadota>Gammaproteobacteria>Enterobacterales>Enterobacteriaceae>Salmonella | 273 | IMT80_001264 | 179362..180183 | 28901 | DUF4435-domain-containing-protein-[Salmonella-enterica] |
| EGP1242558.1 | TOPRIM | - | Salmonella_enterica | Bacteria>Pseudomonadati>Pseudomonadota>Gammaproteobacteria>Enterobacterales>Enterobacteriaceae>Salmonella | 273 | IV995_001918 | 68573..69394 | 28901 | DUF4435-domain-containing-protein-[Salmonella-enterica] |
| EGT1190055.1 | TOPRIM | - | Salmonella_enterica | Bacteria>Pseudomonadati>Pseudomonadota>Gammaproteobacteria>Enterobacterales>Enterobacteriaceae>Salmonella | 273 | I3G58_002372 | 102451..103272 | 28901 | DUF4435-domain-containing-protein-[Salmonella-enterica] |
| EGU3680636.1 | TOPRIM | - | Salmonella_enterica | Bacteria>Pseudomonadati>Pseudomonadota>Gammaproteobacteria>Enterobacterales>Enterobacteriaceae>Salmonella | 273 | HOR40_002398 | 35829..36650 | 28901 | DUF4435-domain-containing-protein-[Salmonella-enterica] |
| EGU5502540.1 | TOPRIM | - | Salmonella_enterica | Bacteria>Pseudomonadati>Pseudomonadota>Gammaproteobacteria>Enterobacterales>Enterobacteriaceae>Salmonella | 273 | HGK88_001888 | 337683..338504 | 28901 | DUF4435-domain-containing-protein-[Salmonella-enterica] |
| EGV8384251.1 | TOPRIM | - | Salmonella_enterica | Bacteria>Pseudomonadati>Pseudomonadota>Gammaproteobacteria>Enterobacterales>Enterobacteriaceae>Salmonella | 273 | EHN43_00845 | 179354..180175 | 28901 | DUF4435-domain-containing-protein-[Salmonella-enterica] |
| EHA2523076.1 | TOPRIM | - | Salmonella_enterica | Bacteria>Pseudomonadati>Pseudomonadota>Gammaproteobacteria>Enterobacterales>Enterobacteriaceae>Salmonella | 273 | JKI82_003472 | 174321..175142 | 28901 | DUF4435-domain-containing-protein-[Salmonella-enterica] |
| EHD7670521.1 | TOPRIM | - | Salmonella_enterica | Bacteria>Pseudomonadati>Pseudomonadota>Gammaproteobacteria>Enterobacterales>Enterobacteriaceae>Salmonella | 273 | JQD35_001757 | 179015..179836 | 28901 | DUF4435-domain-containing-protein-[Salmonella-enterica] |
| EHH9023837.1 | TOPRIM | GCA_017748765.1 | Salmonella_enterica | Bacteria>Pseudomonadati>Pseudomonadota>Gammaproteobacteria>Enterobacterales>Enterobacteriaceae>Salmonella | 273 | J8439_003632 | 26750..27571 | 28901 | DUF4435-domain-containing-protein-[Salmonella-enterica] |

| Protein-ID | Domain | Assembly-ID | Strain | Taxonomy | Length | Locus-Tag | Genomic-coordinates | TaxID | Fasta-defline |
|---|---|---|---|---|---|---|---|---|---|
| EHI1197049.1 | TOPRIM | GCA_017785955.1 | Salmonella_enterica | Bacteria>Pseudomonadati>Pseudomonadota>Gammaproteobacteria>Enterobacterales>Enterobacteriaceae>Salmonella | 273 | J8551_000467 | 102451..103272 | 28901 | DUF4435-domain-containing-protein-[Salmonella-enterica] |
| EHJ0877872.1 | TOPRIM | GCA_017846115.1 | Salmonella_enterica | Bacteria>Pseudomonadati>Pseudomonadota>Gammaproteobacteria>Enterobacterales>Enterobacteriaceae>Salmonella | 273 | KAB51_002700 | 26558..27379 | 28901 | DUF4435-domain-containing-protein-[Salmonella-enterica] |
| EHL7445942.1 | TOPRIM | GCA_018171815.1 | Salmonella_enterica | Bacteria>Pseudomonadati>Pseudomonadota>Gammaproteobacteria>Enterobacterales>Enterobacteriaceae>Salmonella | 273 | KEH66_000995 | 392538..393359 | 28901 | DUF4435-domain-containing-protein-[Salmonella-enterica] |
| EHN4465871.1 | TOPRIM | GCA_018387865.1 | Salmonella_enterica | Bacteria>Pseudomonadati>Pseudomonadota>Gammaproteobacteria>Enterobacterales>Enterobacteriaceae>Salmonella | 273 | KIZ21_001041 | 342787..343608 | 28901 | DUF4435-domain-containing-protein-[Salmonella-enterica] |
| EHO5101497.1 | TOPRIM | GCA_018532585.1 | Salmonella_enterica | Bacteria>Pseudomonadati>Pseudomonadota>Gammaproteobacteria>Enterobacterales>Enterobacteriaceae>Salmonella | 273 | KLP74_002742 | 6808..7629 | 28901 | DUF4435-domain-containing-protein-[Salmonella-enterica] |
| EHO5627404.1 | TOPRIM | GCA_018535225.1 | Salmonella_enterica | Bacteria>Pseudomonadati>Pseudomonadota>Gammaproteobacteria>Enterobacterales>Enterobacteriaceae>Salmonella | 273 | KLQ90_002310 | 39200..40021 | 28901 | DUF4435-domain-containing-protein-[Salmonella-enterica] |
| EHQ7935668.1 | TOPRIM | GCA_018976465.1 | Salmonella_enterica | Bacteria>Pseudomonadati>Pseudomonadota>Gammaproteobacteria>Enterobacterales>Enterobacteriaceae>Salmonella | 273 | KSM73_002314 | 102451..103272 | 28901 | DUF4435-domain-containing-protein-[Salmonella-enterica] |
| EHS4662098.1 | TOPRIM | GCA_019144395.1 | Salmonella_enterica | Bacteria>Pseudomonadati>Pseudomonadota>Gammaproteobacteria>Enterobacterales>Enterobacteriaceae>Salmonella | 273 | KV604_003671 | 67928..68749 | 28901 | DUF4435-domain-containing-protein-[Salmonella-enterica] |
| EHT1034284.1 | TOPRIM | GCA_019186465.1 | Salmonella_enterica | Bacteria>Pseudomonadati>Pseudomonadota>Gammaproteobacteria>Enterobacterales>Enterobacteriaceae>Salmonella | 273 | KW750_001287 | 157547..158368 | 28901 | DUF4435-domain-containing-protein-[Salmonella-enterica] |
| EHT6941869.1 | TOPRIM | GCA_019250575.1 | Salmonella_enterica | Bacteria>Pseudomonadati>Pseudomonadota>Gammaproteobacteria>Enterobacterales>Enterobacteriaceae>Salmonella | 273 | KYH33_001190 | 179351..180172 | 28901 | DUF4435-domain-containing-protein-[Salmonella-enterica] |
| EHU5044811.1 | TOPRIM | GCA_019291225.1 | Salmonella_enterica | Bacteria>Pseudomonadati>Pseudomonadota>Gammaproteobacteria>Enterobacterales>Enterobacteriaceae>Salmonella | 273 | KY920_001888 | 94120..94941 | 28901 | DUF4435-domain-containing-protein-[Salmonella-enterica] |
| EHU5815177.1 | TOPRIM | GCA_019298295.1 | Salmonella_enterica | Bacteria>Pseudomonadati>Pseudomonadota>Gammaproteobacteria>Enterobacterales>Enterobacteriaceae>Salmonella | 273 | KZE80_000352 | 392778..393599 | 28901 | DUF4435-domain-containing-protein-[Salmonella-enterica] |
| EHU7518555.1 | TOPRIM | GCA_019318345.1 | Salmonella_enterica | Bacteria>Pseudomonadati>Pseudomonadota>Gammaproteobacteria>Enterobacterales>Enterobacteriaceae>Salmonella | 273 | KZL07_001986 | 281411..282232 | 28901 | DUF4435-domain-containing-protein-[Salmonella-enterica] |
| EHV2030462.1 | TOPRIM | GCA_019356775.1 | Salmonella_enterica | Bacteria>Pseudomonadati>Pseudomonadota>Gammaproteobacteria>Enterobacterales>Enterobacteriaceae>Salmonella | 273 | KZ138_001534 | 398126..398947 | 28901 | DUF4435-domain-containing-protein-[Salmonella-enterica] |
| EHW6237844.1 | TOPRIM | GCA_019485205.1 | Salmonella_enterica | Bacteria>Pseudomonadati>Pseudomonadota>Gammaproteobacteria>Enterobacterales>Enterobacteriaceae>Salmonella | 273 | K2495_002340 | 39184..40005 | 28901 | DUF4435-domain-containing-protein-[Salmonella-enterica] |
| EHZ3783991.1 | TOPRIM | GCA_019697575.1 | Salmonella_enterica | Bacteria>Pseudomonadati>Pseudomonadota>Gammaproteobacteria>Enterobacterales>Enterobacteriaceae>Salmonella | 273 | K5U30_001275 | 179352..180173 | 28901 | DUF4435-domain-containing-protein-[Salmonella-enterica] |
| EHZ9862033.1 | TOPRIM | GCA_019744275.1 | Salmonella_enterica | Bacteria>Pseudomonadati>Pseudomonadota>Gammaproteobacteria>Enterobacterales>Enterobacteriaceae>Salmonella | 273 | K6Z50_000359 | 404324..405145 | 28901 | DUF4435-domain-containing-protein-[Salmonella-enterica] |
| EIA3194370.1 | TOPRIM | GCA_019788365.1 | Salmonella_enterica | Bacteria>Pseudomonadati>Pseudomonadota>Gammaproteobacteria>Enterobacterales>Enterobacteriaceae>Salmonella | 273 | K7D38_001958 | 181388..182209 | 28901 | DUF4435-domain-containing-protein-[Salmonella-enterica] |
| EIA3335814.1 | TOPRIM | GCA_019789135.1 | Salmonella_enterica | Bacteria>Pseudomonadati>Pseudomonadota>Gammaproteobacteria>Enterobacterales>Enterobacteriaceae>Salmonella | 273 | K7D78_001301 | 408562..409383 | 28901 | DUF4435-domain-containing-protein-[Salmonella-enterica] |
| EIB1064375.1 | TOPRIM | GCA_019855255.1 | Salmonella_enterica | Bacteria>Pseudomonadati>Pseudomonadota>Gammaproteobacteria>Enterobacterales>Enterobacteriaceae>Salmonella | 273 | K7907_001962 | 102451..103272 | 28901 | DUF4435-domain-containing-protein-[Salmonella-enterica] |
| EIC4097435.1 | TOPRIM | GCA_019951695.1 | Salmonella_enterica | Bacteria>Pseudomonadati>Pseudomonadota>Gammaproteobacteria>Enterobacterales>Enterobacteriaceae>Salmonella | 273 | K9796_003006 | 30591..31412 | 28901 | DUF4435-domain-containing-protein-[Salmonella-enterica] |
| EIF9427098.1 | TOPRIM | GCA_020283115.1 | Salmonella_enterica | Bacteria>Pseudomonadati>Pseudomonadota>Gammaproteobacteria>Enterobacterales>Enterobacteriaceae>Salmonella | 273 | HW154_002106 | 34317..35138 | 28901 | DUF4435-domain-containing-protein-[Salmonella-enterica] |
| EIF9623263.1 | TOPRIM | GCA_020284415.1 | Salmonella_enterica | Bacteria>Pseudomonadati>Pseudomonadota>Gammaproteobacteria>Enterobacterales>Enterobacteriaceae>Salmonella | 273 | HQ981_002278 | 34317..35138 | 28901 | DUF4435-domain-containing-protein-[Salmonella-enterica] |
| EIJ0643395.1 | TOPRIM | GCA_020492735.1 | Salmonella_enterica | Bacteria>Pseudomonadati>Pseudomonadota>Gammaproteobacteria>Enterobacterales>Enterobacteriaceae>Salmonella | 273 | LH177_001936 | 66685..67506 | 28901 | DUF4435-domain-containing-protein-[Salmonella-enterica] |
| EIJ2988435.1 | TOPRIM | GCA_020513955.1 | Salmonella_enterica | Bacteria>Pseudomonadati>Pseudomonadota>Gammaproteobacteria>Enterobacterales>Enterobacteriaceae>Salmonella | 273 | LIS40_000164 | 179352..180173 | 28901 | DUF4435-domain-containing-protein-[Salmonella-enterica] |
| EIM3792266.1 | TOPRIM | GCA_020780095.1 | Salmonella_enterica | Bacteria>Pseudomonadati>Pseudomonadota>Gammaproteobacteria>Enterobacterales>Enterobacteriaceae>Salmonella | 273 | LM721_003557 | 35781..36602 | 28901 | DUF4435-domain-containing-protein-[Salmonella-enterica] |
| EIO7765746.1 | TOPRIM | GCA_020997775.1 | Salmonella_enterica | Bacteria>Pseudomonadati>Pseudomonadota>Gammaproteobacteria>Enterobacterales>Enterobacteriaceae>Salmonella | 273 | LQR80_000702 | 179352..180173 | 28901 | DUF4435-domain-containing-protein-[Salmonella-enterica] |
| EIQ4281224.1 | TOPRIM | GCA_021189835.1 | Salmonella_enterica | Bacteria>Pseudomonadati>Pseudomonadota>Gammaproteobacteria>Enterobacterales>Enterobacteriaceae>Salmonella | 273 | LU454_001108 | 111263..112084 | 28901 | DUF4435-domain-containing-protein-[Salmonella-enterica] |
| EIR8875062.1 | TOPRIM | GCA_021305935.1 | Salmonella_enterica | Bacteria>Pseudomonadati>Pseudomonadota>Gammaproteobacteria>Enterobacterales>Enterobacteriaceae>Salmonella | 273 | LXE31_003495 | 11212..12033 | 28901 | DUF4435-domain-containing-protein-[Salmonella-enterica] |
| EIU6597717.1 | TOPRIM | GCA_021692515.1 | Salmonella_enterica | Bacteria>Pseudomonadati>Pseudomonadota>Gammaproteobacteria>Enterobacterales>Enterobacteriaceae>Salmonella | 273 | L5W02_001571 | 385687..386508 | 28901 | DUF4435-domain-containing-protein-[Salmonella-enterica] |
| EJB4514211.1 | TOPRIM | GCA_022945175.1 | Salmonella_enterica | Bacteria>Pseudomonadati>Pseudomonadota>Gammaproteobacteria>Enterobacterales>Enterobacteriaceae>Salmonella | 273 | MT088_002309 | 388573..389394 | 28901 | DUF4435-domain-containing-protein-[Salmonella-enterica] |
| EJC3638309.1 | TOPRIM | GCA_023047235.1 | Salmonella_enterica | Bacteria>Pseudomonadati>Pseudomonadota>Gammaproteobacteria>Enterobacterales>Enterobacteriaceae>Salmonella | 273 | MYX88_003396 | 159533..160354 | 28901 | DUF4435-domain-containing-protein-[Salmonella-enterica] |
| EJF4683724.1 | TOPRIM | GCA_023476405.1 | Salmonella_enterica | Bacteria>Pseudomonadati>Pseudomonadota>Gammaproteobacteria>Enterobacterales>Enterobacteriaceae>Salmonella | 273 | M8N25_002790 | 102451..103272 | 28901 | DUF4435-domain-containing-protein-[Salmonella-enterica] |
| EJF5860349.1 | TOPRIM | GCA_023495345.1 | Salmonella_enterica | Bacteria>Pseudomonadati>Pseudomonadota>Gammaproteobacteria>Enterobacterales>Enterobacteriaceae>Salmonella | 273 | M8X04_001884 | 178947..179768 | 28901 | DUF4435-domain-containing-protein-[Salmonella-enterica] |
| EJK0228889.1 | TOPRIM | GCA_024068175.1 | Salmonella_enterica | Bacteria>Pseudomonadati>Pseudomonadota>Gammaproteobacteria>Enterobacterales>Enterobacteriaceae>Salmonella | 273 | NKK90_001816 | 317562..318383 | 28901 | DUF4435-domain-containing-protein-[Salmonella-enterica] |
| EJM7985886.1 | TOPRIM | GCA_024373005.1 | Salmonella_enterica | Bacteria>Pseudomonadati>Pseudomonadota>Gammaproteobacteria>Enterobacterales>Enterobacteriaceae>Salmonella | 273 | NO486_001115 | 377516..378337 | 28901 | DUF4435-domain-containing-protein-[Salmonella-enterica] |
| EJN9121679.1 | TOPRIM | GCA_024535515.1 | Salmonella_enterica | Bacteria>Pseudomonadati>Pseudomonadota>Gammaproteobacteria>Enterobacterales>Enterobacteriaceae>Salmonella | 273 | NR521_001806 | 179362..180183 | 28901 | DUF4435-domain-containing-protein-[Salmonella-enterica] |
| EJO5979500.1 | TOPRIM | GCA_024573615.1 | Salmonella_enterica | Bacteria>Pseudomonadati>Pseudomonadota>Gammaproteobacteria>Enterobacterales>Enterobacteriaceae>Salmonella | 273 | NRR85_002628 | 167484..168305 | 28901 | DUF4435-domain-containing-protein-[Salmonella-enterica] |
| EJS8182323.1 | TOPRIM | GCA_024988615.1 | Salmonella_enterica | Bacteria>Pseudomonadati>Pseudomonadota>Gammaproteobacteria>Enterobacterales>Enterobacteriaceae>Salmonella | 273 | NWJ04_001333 | 34317..35138 | 28901 | DUF4435-domain-containing-protein-[Salmonella-enterica] |
| HAC8921682.1 | TOPRIM | GCA_010188005.1 | Salmonella_enterica | Bacteria>Pseudomonadati>Pseudomonadota>Gammaproteobacteria>Enterobacterales>Enterobacteriaceae>Salmonella | 273 | G0J09_01275 | 53018..53839 | 28901 | TPA_asm:-DUF4435-domain-containing-protein-[Salmonella-enterica] |
| HAF0812800.1 | TOPRIM | GCA_011125695.1 | Salmonella_enterica | Bacteria>Pseudomonadati>Pseudomonadota>Gammaproteobacteria>Enterobacterales>Enterobacteriaceae>Salmonella | 273 | G9G17_001780 | 102451..103272 | 28901 | TPA:-DUF4435-domain-containing-protein-[Salmonella-enterica] |
| HAF9991659.1 | TOPRIM | GCA_011779145.1 | Salmonella_enterica | Bacteria>Pseudomonadati>Pseudomonadota>Gammaproteobacteria>Enterobacterales>Enterobacteriaceae>Salmonella | 273 | G8Q07_002651 | 179177..179998 | 28901 | TPA:-DUF4435-domain-containing-protein-[Salmonella-enterica] |
| HAG2842213.1 | TOPRIM | GCA_011791385.1 | Salmonella_enterica | Bacteria>Pseudomonadati>Pseudomonadota>Gammaproteobacteria>Enterobacterales>Enterobacteriaceae>Salmonella | 273 | G8Z41_002589 | 109360..110181 | 28901 | TPA:-DUF4435-domain-containing-protein-[Salmonella-enterica] |
| HAG5552353.1 | TOPRIM | GCA_011798725.1 | Salmonella_enterica | Bacteria>Pseudomonadati>Pseudomonadota>Gammaproteobacteria>Enterobacterales>Enterobacteriaceae>Salmonella | 273 | G8Q25_001509 | 181157..181978 | 28901 | TPA:-DUF4435-domain-containing-protein-[Salmonella-enterica] |
| HAK8092291.1 | TOPRIM | GCA_013643495.1 | Salmonella_enterica | Bacteria>Pseudomonadati>Pseudomonadota>Gammaproteobacteria>Enterobacterales>Enterobacteriaceae>Salmonella | 273 | H2V91_002195 | 161525..162346 | 28901 | DUF4435-domain-containing-protein-[Salmonella-enterica] |
| HAK8810850.1 | TOPRIM | GCA_013646795.1 | Salmonella_enterica | Bacteria>Pseudomonadati>Pseudomonadota>Gammaproteobacteria>Enterobacterales>Enterobacteriaceae>Salmonella | 273 | H2X42_001965 | 166798..167619 | 28901 | DUF4435-domain-containing-protein-[Salmonella-enterica] |
| MBH5277081.1 | TOPRIM | GCA_016019715.1 | Salmonella_enterica | Bacteria>Pseudomonadati>Pseudomonadota>Gammaproteobacteria>Enterobacterales>Enterobacteriaceae>Salmonella | 273 | I3A81_01635 | 353020..353841 | 28901 | DUF4435-domain-containing-protein-[Salmonella-enterica] |
| MCM9949028.1 | TOPRIM | GCA_023843835.1 | Salmonella_enterica | Bacteria>Pseudomonadati>Pseudomonadota>Gammaproteobacteria>Enterobacterales>Enterobacteriaceae>Salmonella | 273 | M5G05_07265 | 342710..343531 | 28901 | DUF4435-domain-containing-protein-[Salmonella-enterica] |
| MDG46366.1 | TOPRIM | GCA_003733245.1 | Salmonella_enterica | Bacteria>Pseudomonadati>Pseudomonadota>Gammaproteobacteria>Enterobacterales>Enterobacteriaceae>Salmonella | 273 | EB289_07935 | 47105..47926 | 28901 | DUF4435-domain-containing-protein-[Salmonella-enterica] |
| WP_077248484.1 | TOPRIM | GCF_027943645.1 | Salmonella_enterica | Bacteria>Pseudomonadati>Pseudomonadota>Gammaproteobacteria>Enterobacterales>Enterobacteriaceae>Salmonella | 273 | F1E41_RS17175 | 159107..159928 | 28901 | DUF4435-domain-containing-protein-[Salmonella-enterica] |
| WP_077906189.1 | TOPRIM | GCF_967854935.1 | Salmonella_enterica | Bacteria>Pseudomonadati>Pseudomonadota>Gammaproteobacteria>Enterobacterales>Enterobacteriaceae>Salmonella | 273 | J3T40_RS14920 | 388823..389644 | 28901 | DUF4435-domain-containing-protein-[Salmonella-enterica] |
| WP_077908842.1 | TOPRIM | GCF_000412855.1 | Salmonella_enterica | Bacteria>Pseudomonadati>Pseudomonadota>Gammaproteobacteria>Enterobacterales>Enterobacteriaceae>Salmonella | 273 | GX95_RS20920 | 4063515..4064336 | 28901 | DUF4435-domain-containing-protein-[Salmonella-enterica] |
| WP_077917342.1 | TOPRIM | GCF_943114325.1 | Salmonella_enterica | Bacteria>Pseudomonadati>Pseudomonadota>Gammaproteobacteria>Enterobacterales>Enterobacteriaceae>Salmonella | 273 | OCU44_RS01000 | 220444..221265 | 28901 | DUF4435-domain-containing-protein-[Salmonella-enterica] |
| WP_077945597.1 | TOPRIM | GCF_003324795.1 | Salmonella_enterica | Bacteria>Pseudomonadati>Pseudomonadota>Gammaproteobacteria>Enterobacterales>Enterobacteriaceae>Salmonella | 273 | DOE60_RS06785 | 1372763..1373584 | 28901 | DUF4435-domain-containing-protein-[Salmonella-enterica] |
| WP_077949404.1 | TOPRIM | GCF_001570285.1 | Salmonella_enterica | Bacteria>Pseudomonadati>Pseudomonadota>Gammaproteobacteria>Enterobacterales>Enterobacteriaceae>Salmonella | 273 | AW153_RS18930 | 166786..167607 | 28901 | DUF4435-domain-containing-protein-[Salmonella-enterica] |
| WP_077958470.1 | TOPRIM | GCF_040956775.1 | Salmonella_enterica | Bacteria>Pseudomonadati>Pseudomonadota>Gammaproteobacteria>Enterobacterales>Enterobacteriaceae>Salmonella | 273 | SES26_RS19330 | 4017896..4018717 | 28901 | DUF4435-domain-containing-protein-[Salmonella-enterica] |
| WP_078056646.1 | TOPRIM | GCF_001975325.1 | Salmonella_enterica | Bacteria>Pseudomonadati>Pseudomonadota>Gammaproteobacteria>Enterobacterales>Enterobacteriaceae>Salmonella | 273 | CHD02_RS05520 | 1133822..1134643 | 28901 | DUF4435-domain-containing-protein-[Salmonella-enterica] |
| WP_078061208.1 | TOPRIM | GCF_001975445.1 | Salmonella_enterica | Bacteria>Pseudomonadati>Pseudomonadota>Gammaproteobacteria>Enterobacterales>Enterobacteriaceae>Salmonella | 273 | LFZ24_RS24365 | 4837473..4838294 | 28901 | DUF4435-domain-containing-protein-[Salmonella-enterica] |
| WP_079808685.1 | TOPRIM | GCF_002035875.1 | Salmonella_enterica | Bacteria>Pseudomonadati>Pseudomonadota>Gammaproteobacteria>Enterobacterales>Enterobacteriaceae>Salmonella | 273 | Y985_RS08715 | 114436..115257 | 28901 | DUF4435-domain-containing-protein-[Salmonella-enterica] |
| WP_079902379.1 | TOPRIM | GCF_003325275.1 | Salmonella_enterica | Bacteria>Pseudomonadati>Pseudomonadota>Gammaproteobacteria>Enterobacterales>Enterobacteriaceae>Salmonella | 273 | CHE40_RS23165 | 4537910..4538731 | 28901 | DUF4435-domain-containing-protein-[Salmonella-enterica] |
| WP_079944399.1 | TOPRIM | GCF_037083205.1 | Salmonella_enterica | Bacteria>Pseudomonadati>Pseudomonadota>Gammaproteobacteria>Enterobacterales>Enterobacteriaceae>Salmonella | 273 | CCO43_RS23240 | 7265..8086 | 28901 | DUF4435-domain-containing-protein-[Salmonella-enterica] |
| WP_080162868.1 | TOPRIM | GCF_002062115.1 | Salmonella_enterica | Bacteria>Pseudomonadati>Pseudomonadota>Gammaproteobacteria>Enterobacterales>Enterobacteriaceae>Salmonella | 273 | SO66_RS04070 | 167204..168025 | 28901 | DUF4435-domain-containing-protein-[Salmonella-enterica] |
| WP_080166835.1 | TOPRIM | GCF_002208255.1 | Salmonella_enterica | Bacteria>Pseudomonadati>Pseudomonadota>Gammaproteobacteria>Enterobacterales>Enterobacteriaceae>Salmonella | 273 | LFZ35_RS21105 | 4070415..4071236 | 28901 | DUF4435-domain-containing-protein-[Salmonella-enterica] |
| WP_080213761.1 | TOPRIM | GCF_025176825.1 | Salmonella_enterica | Bacteria>Pseudomonadati>Pseudomonadota>Gammaproteobacteria>Enterobacterales>Enterobacteriaceae>Salmonella | 273 | AS04_RS02265 | 52828..53649 | 28901 | DUF4435-domain-containing-protein-[Salmonella-enterica] |
| WP_094946979.1 | TOPRIM | GCF_002266315.1 | Salmonella_enterica | Bacteria>Pseudomonadati>Pseudomonadota>Gammaproteobacteria>Enterobacterales>Enterobacteriaceae>Salmonella | 273 | CCO36_RS01895 | 395409..396230 | 28901 | DUF4435-domain-containing-protein-[Salmonella-enterica] |
| WP_094964982.1 | TOPRIM | GCF_002266055.1 | Salmonella_enterica | Bacteria>Pseudomonadati>Pseudomonadota>Gammaproteobacteria>Enterobacterales>Enterobacteriaceae>Salmonella | 273 | CCO49_RS14350 | 407509..408330 | 28901 | DUF4435-domain-containing-protein-[Salmonella-enterica] |
| WP_097368037.1 | TOPRIM | GCF_002507625.1 | Salmonella_enterica | Bacteria>Pseudomonadati>Pseudomonadota>Gammaproteobacteria>Enterobacterales>Enterobacteriaceae>Salmonella | 273 | CBI66_RS11195 | 179492..180313 | 28901 | DUF4435-domain-containing-protein-[Salmonella-enterica] |
| WP_097369132.1 | TOPRIM | GCF_002507575.1 | Salmonella_enterica | Bacteria>Pseudomonadati>Pseudomonadota>Gammaproteobacteria>Enterobacterales>Enterobacteriaceae>Salmonella | 273 | CBI65_RS08840 | 179325..180146 | 28901 | DUF4435-domain-containing-protein-[Salmonella-enterica] |
| WP_111761034.1 | TOPRIM | GCF_019173605.1 | Salmonella_enterica | Bacteria>Pseudomonadati>Pseudomonadota>Gammaproteobacteria>Enterobacterales>Enterobacteriaceae>Salmonella | 273 | KXD81_RS02505 | 16375..17196 | 28901 | DUF4435-domain-containing-protein-[Salmonella-enterica] |
| WP_111772031.1 | TOPRIM | GCF_025292575.1 | Salmonella_enterica | Bacteria>Pseudomonadati>Pseudomonadota>Gammaproteobacteria>Enterobacterales>Enterobacteriaceae>Salmonella | 273 | DQN83_RS20195 | 3955107..3955928 | 28901 | DUF4435-domain-containing-protein-[Salmonella-enterica] |
| WP_140237024.1 | TOPRIM | GCF_006379925.2 | Salmonella_enterica | Bacteria>Pseudomonadati>Pseudomonadota>Gammaproteobacteria>Enterobacterales>Enterobacteriaceae>Salmonella | 273 | FG677_RS21925 | 11286..12107 | 28901 | DUF4435-domain-containing-protein-[Salmonella-enterica] |
| WP_183072300.1 | TOPRIM | GCF_903931105.1 | Salmonella_enterica | Bacteria>Pseudomonadati>Pseudomonadota>Gammaproteobacteria>Enterobacterales>Enterobacteriaceae>Salmonella | 273 | H7T17_RS07625 | 62601..63422 | 28901 | DUF4435-domain-containing-protein-[Salmonella-enterica] |
| WP_223365096.1 | TOPRIM | GCF_002106425.1 | Salmonella_enterica | Bacteria>Pseudomonadati>Pseudomonadota>Gammaproteobacteria>Enterobacterales>Enterobacteriaceae>Salmonella | 273 | R537_RS06015 | 148837..149658 | 28901 | DUF4435-domain-containing-protein-[Salmonella-enterica] |
| WP_267272779.1 | TOPRIM | GCF_026460795.1 | Salmonella_enterica | Bacteria>Pseudomonadati>Pseudomonadota>Gammaproteobacteria>Enterobacterales>Enterobacteriaceae>Salmonella | 273 | OWO74_RS14180 | 5746..6567 | 28901 | hypothetical-protein-[Salmonella-enterica] |
| EAB2013165.1 | TOPRIM | GCA_004236885.1 | Salmonella_enterica | Bacteria>Pseudomonadati>Pseudomonadota>Gammaproteobacteria>Enterobacterales>Enterobacteriaceae>Salmonella | 189 | D7J64_23105 | 13563..14132 | 28901 | hypothetical-protein-[Salmonella-enterica] |
| EAP9950627.1 | TOPRIM | GCA_005576875.1 | Salmonella_enterica | Bacteria>Pseudomonadati>Pseudomonadota>Gammaproteobacteria>Enterobacterales>Enterobacteriaceae>Salmonella | 189 | D3G27_06350 | 184544..185113 | 28901 | hypothetical-protein-[Salmonella-enterica] |
| EAW3206016.1 | TOPRIM | GCA_005905825.1 | Salmonella_enterica | Bacteria>Pseudomonadati>Pseudomonadota>Gammaproteobacteria>Enterobacterales>Enterobacteriaceae>Salmonella | 189 | FEL99_04735 | 227440..228009 | 28901 | hypothetical-protein-[Salmonella-enterica] |
| EAW9396390.1 | TOPRIM | GCA_005940865.1 | Salmonella_enterica | Bacteria>Pseudomonadati>Pseudomonadota>Gammaproteobacteria>Enterobacterales>Enterobacteriaceae>Salmonella | 189 | QE38_19750 | 56840..57409 | 28901 | hypothetical-protein-QE38_19750-[Salmonella-enterica] |
| EAY5508621.1 | TOPRIM | GCA_006012725.1 | Salmonella_enterica | Bacteria>Pseudomonadati>Pseudomonadota>Gammaproteobacteria>Enterobacterales>Enterobacteriaceae>Salmonella | 189 | NE47_11030 | 57826..58395 | 28901 | hypothetical-protein-NE47_11030-[Salmonella-enterica] |
| EBF4455782.1 | TOPRIM | GCA_006343165.1 | Salmonella_enterica | Bacteria>Pseudomonadati>Pseudomonadota>Gammaproteobacteria>Enterobacterales>Enterobacteriaceae>Salmonella | 189 | FH443_09920 | 420549..421118 | 28901 | hypothetical-protein-FH443_09920-[Salmonella-enterica] |
| EBH5838352.1 | TOPRIM | GCA_006503215.1 | Salmonella_enterica | Bacteria>Pseudomonadati>Pseudomonadota>Gammaproteobacteria>Enterobacterales>Enterobacteriaceae>Salmonella | 189 | CQL58_06360 | 299812..300381 | 28901 | hypothetical-protein-[Salmonella-enterica] |
| EBR5149896.1 | TOPRIM | GCA_006958505.1 | Salmonella_enterica | Bacteria>Pseudomonadati>Pseudomonadota>Gammaproteobacteria>Enterobacterales>Enterobacteriaceae>Salmonella | 189 | B5W64_09860 | 359243..359812 | 28901 | hypothetical-protein-[Salmonella-enterica] |
| EDZ0760565.1 | TOPRIM | - | Salmonella_enterica | Bacteria>Pseudomonadati>Pseudomonadota>Gammaproteobacteria>Enterobacterales>Enterobacteriaceae>Salmonella | 189 | GPN34_002363 | 419081..419650 | 28901 | DUF4276-family-protein-[Salmonella-enterica] |
| WP_064013332.1 | TOPRIM | GCF_002211845.1 | Salmonella_enterica | Bacteria>Pseudomonadati>Pseudomonadota>Gammaproteobacteria>Enterobacterales>Enterobacteriaceae>Salmonella | 189 | CE137_RS21065 | 4104206..4104775 | 28901 | hypothetical-protein-[Salmonella-enterica] |
| EIC8525258.1 | TOPRIM | GCA_019993965.1 | Salmonella_enterica | Bacteria>Pseudomonadati>Pseudomonadota>Gammaproteobacteria>Enterobacterales>Enterobacteriaceae>Salmonella | 187 | LAR10_002218 | 38840..39403 | 28901 | hypothetical-protein-[Salmonella-enterica] |
| EAA5451234.1 | TOPRIM | GCA_004184965.1 | Salmonella_enterica_subsp._enterica | Bacteria>Pseudomonadati>Pseudomonadota>Gammaproteobacteria>Enterobacterales>Enterobacteriaceae>Salmonella>Salmonella_enterica | 273 | DQC86_00815 | 168958..169779 | 59201 | DUF4435-domain-containing-protein-[Salmonella-enterica-subsp.-enterica] |
| EAA9627985.1 | TOPRIM | GCA_004227565.1 | Salmonella_enterica_subsp._enterica | Bacteria>Pseudomonadati>Pseudomonadota>Gammaproteobacteria>Enterobacterales>Enterobacteriaceae>Salmonella>Salmonella_enterica | 273 | DV01_07270 | 179352..180173 | 59201 | DUF4435-domain-containing-protein-[Salmonella-enterica-subsp.-enterica] |
| EAB6416576.1 | TOPRIM | GCA_004256685.1 | Salmonella_enterica_subsp._enterica | Bacteria>Pseudomonadati>Pseudomonadota>Gammaproteobacteria>Enterobacterales>Enterobacteriaceae>Salmonella>Salmonella_enterica | 273 | D6S13_10925 | 60727..61548 | 59201 | DUF4435-domain-containing-protein-[Salmonella-enterica-subsp.-enterica] |
| EAB6843528.1 | TOPRIM | GCA_004258725.1 | Salmonella_enterica_subsp._salamae | Bacteria>Pseudomonadati>Pseudomonadota>Gammaproteobacteria>Enterobacterales>Enterobacteriaceae>Salmonella>Salmonella_enterica | 273 | XT66_14200 | 59365..60186 | 59202 | DUF4435-domain-containing-protein-[Salmonella-enterica-subsp.-salamae] |
| EAB9078799.1 | TOPRIM | GCA_004265565.1 | Salmonella_enterica_subsp._enterica | Bacteria>Pseudomonadati>Pseudomonadota>Gammaproteobacteria>Enterobacterales>Enterobacteriaceae>Salmonella>Salmonella_enterica | 273 | AAL03_01525 | 60799..61620 | 59201 | DUF4435-domain-containing-protein-[Salmonella-enterica-subsp.-enterica] |
| EAC1236048.1 | TOPRIM | GCA_004276135.1 | Salmonella_enterica_subsp._enterica | Bacteria>Pseudomonadati>Pseudomonadota>Gammaproteobacteria>Enterobacterales>Enterobacteriaceae>Salmonella>Salmonella_enterica | 273 | DTG84_00825 | 166364..167185 | 59201 | DUF4435-domain-containing-protein-[Salmonella-enterica-subsp.-enterica] |
| EBG5655088.1 | TOPRIM | GCA_006430455.1 | Salmonella_enterica_subsp._enterica | Bacteria>Pseudomonadati>Pseudomonadota>Gammaproteobacteria>Enterobacterales>Enterobacteriaceae>Salmonella>Salmonella_enterica | 273 | FI236_11960 | 102451..103272 | 59201 | DUF4435-domain-containing-protein-[Salmonella-enterica-subsp.-enterica] |
| ECC0743073.1 | TOPRIM | GCA_007457215.1 | Salmonella_enterica_subsp._enterica | Bacteria>Pseudomonadati>Pseudomonadota>Gammaproteobacteria>Enterobacterales>Enterobacteriaceae>Salmonella>Salmonella_enterica | 273 | FMV06_07380 | 102451..103272 | 59201 | DUF4435-domain-containing-protein-[Salmonella-enterica-subsp.-enterica] |
| ECC1667302.1 | TOPRIM | GCA_007463295.1 | Salmonella_enterica_subsp._salamae | Bacteria>Pseudomonadati>Pseudomonadota>Gammaproteobacteria>Enterobacterales>Enterobacteriaceae>Salmonella>Salmonella_enterica | 273 | FNN95_01270 | 276275..277096 | 59202 | DUF4435-domain-containing-protein-[Salmonella-enterica-subsp.-salamae] |
| ECC3171800.1 | TOPRIM | GCA_007472855.1 | Salmonella_enterica_subsp._enterica | Bacteria>Pseudomonadati>Pseudomonadota>Gammaproteobacteria>Enterobacterales>Enterobacteriaceae>Salmonella>Salmonella_enterica | 273 | R914_02610 | 86040..86861 | 59201 | DUF4435-domain-containing-protein-[Salmonella-enterica-subsp.-enterica] |
| ECD1705584.1 | TOPRIM | GCA_007509785.1 | Salmonella_enterica_subsp._enterica | Bacteria>Pseudomonadati>Pseudomonadota>Gammaproteobacteria>Enterobacterales>Enterobacteriaceae>Salmonella>Salmonella_enterica | 273 | EWD17_08935 | 102451..103272 | 59201 | DUF4435-domain-containing-protein-[Salmonella-enterica-subsp.-enterica] |
| ECD9608665.1 | TOPRIM | GCA_007542335.1 | Salmonella_enterica_subsp._salamae | Bacteria>Pseudomonadati>Pseudomonadota>Gammaproteobacteria>Enterobacterales>Enterobacteriaceae>Salmonella>Salmonella_enterica | 273 | FNZ18_04575 | 264949..265770 | 59202 | DUF4435-domain-containing-protein-[Salmonella-enterica-subsp.-salamae] |
| ECG0705018.1 | TOPRIM | GCA_007641515.1 | Salmonella_enterica_subsp._salamae | Bacteria>Pseudomonadati>Pseudomonadota>Gammaproteobacteria>Enterobacterales>Enterobacteriaceae>Salmonella>Salmonella_enterica | 273 | DL122_04715 | 43975..44796 | 59202 | hypothetical-protein-DL122_04715-[Salmonella-enterica-subsp.-salamae] |
| ECG1483125.1 | TOPRIM | GCA_007649655.1 | Salmonella_enterica_subsp._enterica | Bacteria>Pseudomonadati>Pseudomonadota>Gammaproteobacteria>Enterobacterales>Enterobacteriaceae>Salmonella>Salmonella_enterica | 273 | DT144_20835 | 60846..61667 | 59201 | DUF4435-domain-containing-protein-[Salmonella-enterica-subsp.-enterica] |
| ECG1893120.1 | TOPRIM | GCA_007650535.1 | Salmonella_enterica_subsp._enterica | Bacteria>Pseudomonadati>Pseudomonadota>Gammaproteobacteria>Enterobacterales>Enterobacteriaceae>Salmonella>Salmonella_enterica | 273 | EI460_05120 | 164069..164890 | 59201 | DUF4435-domain-containing-protein-[Salmonella-enterica-subsp.-enterica] |
| ECG5099192.1 | TOPRIM | GCA_007668905.1 | Salmonella_enterica_subsp._enterica | Bacteria>Pseudomonadati>Pseudomonadota>Gammaproteobacteria>Enterobacterales>Enterobacteriaceae>Salmonella>Salmonella_enterica | 273 | E0547_20060 | 68345..69166 | 59201 | DUF4435-domain-containing-protein-[Salmonella-enterica-subsp.-enterica] |
| ECI3716683.1 | TOPRIM | GCA_007768905.1 | Salmonella_enterica_subsp._enterica | Bacteria>Pseudomonadati>Pseudomonadota>Gammaproteobacteria>Enterobacterales>Enterobacteriaceae>Salmonella>Salmonella_enterica | 273 | DK077_11335 | 52833..53654 | 59201 | hypothetical-protein-[Salmonella-enterica-subsp.-enterica] |
| ECI6492270.1 | TOPRIM | GCA_007781245.1 | Salmonella_enterica_subsp._enterica | Bacteria>Pseudomonadati>Pseudomonadota>Gammaproteobacteria>Enterobacterales>Enterobacteriaceae>Salmonella>Salmonella_enterica | 273 | EIL18_20710 | 64226..65047 | 59201 | DUF4435-domain-containing-protein-[Salmonella-enterica-subsp.-enterica] |
| ECI6612374.1 | TOPRIM | GCA_007781585.1 | Salmonella_enterica_subsp._enterica | Bacteria>Pseudomonadati>Pseudomonadota>Gammaproteobacteria>Enterobacterales>Enterobacteriaceae>Salmonella>Salmonella_enterica | 273 | EU408_19970 | 71179..72000 | 59201 | DUF4435-domain-containing-protein-[Salmonella-enterica-subsp.-enterica] |
| ECI6683604.1 | TOPRIM | GCA_007781825.1 | Salmonella_enterica_subsp._enterica | Bacteria>Pseudomonadati>Pseudomonadota>Gammaproteobacteria>Enterobacterales>Enterobacteriaceae>Salmonella>Salmonella_enterica | 273 | EHE65_20740 | 60537..61358 | 59201 | DUF4435-domain-containing-protein-[Salmonella-enterica-subsp.-enterica] |
| ECI7333733.1 | TOPRIM | GCA_007783945.1 | Salmonella_enterica_subsp._enterica | Bacteria>Pseudomonadati>Pseudomonadota>Gammaproteobacteria>Enterobacterales>Enterobacteriaceae>Salmonella>Salmonella_enterica | 273 | EZ611_10900 | 102451..103272 | 59201 | DUF4435-domain-containing-protein-[Salmonella-enterica-subsp.-enterica] |
| ECR2528671.1 | TOPRIM | GCA_008341925.1 | Salmonella_enterica_subsp._enterica | Bacteria>Pseudomonadati>Pseudomonadota>Gammaproteobacteria>Enterobacterales>Enterobacteriaceae>Salmonella>Salmonella_enterica | 273 | F1J42_15850 | 102451..103272 | 59201 | DUF4435-domain-containing-protein-[Salmonella-enterica-subsp.-enterica] |
| EDS5421361.1 | TOPRIM | - | Salmonella_enterica_subsp._enterica | Bacteria>Pseudomonadati>Pseudomonadota>Gammaproteobacteria>Enterobacterales>Enterobacteriaceae>Salmonella>Salmonella_enterica | 273 | ZH17_000159 | 166677..167498 | 59201 | DUF4435-domain-containing-protein-[Salmonella-enterica-subsp.-enterica] |
| EDT2838943.1 | TOPRIM | - | Salmonella_enterica_subsp._enterica | Bacteria>Pseudomonadati>Pseudomonadota>Gammaproteobacteria>Enterobacterales>Enterobacteriaceae>Salmonella>Salmonella_enterica | 273 | S958_003410 | 9204..10025 | 59201 | DUF4435-domain-containing-protein-[Salmonella-enterica-subsp.-enterica] |
| EDU7470134.1 | TOPRIM | - | Salmonella_enterica_subsp._enterica | Bacteria>Pseudomonadati>Pseudomonadota>Gammaproteobacteria>Enterobacterales>Enterobacteriaceae>Salmonella>Salmonella_enterica | 273 | CQO18_002337 | 102451..103272 | 59201 | DUF4435-domain-containing-protein-[Salmonella-enterica-subsp.-enterica] |
| EDU9161209.1 | TOPRIM | - | Salmonella_enterica_subsp._enterica | Bacteria>Pseudomonadati>Pseudomonadota>Gammaproteobacteria>Enterobacterales>Enterobacteriaceae>Salmonella>Salmonella_enterica | 273 | IW54_004326 | 5967..6788 | 59201 | DUF4435-domain-containing-protein-[Salmonella-enterica-subsp.-enterica] |
| EDU9624698.1 | TOPRIM | - | Salmonella_enterica_subsp._enterica | Bacteria>Pseudomonadati>Pseudomonadota>Gammaproteobacteria>Enterobacterales>Enterobacteriaceae>Salmonella>Salmonella_enterica | 273 | XW65_002880 | 68238..69059 | 59201 | DUF4435-domain-containing-protein-[Salmonella-enterica-subsp.-enterica] |
| EDV4604928.1 | TOPRIM | - | Salmonella_enterica_subsp._enterica | Bacteria>Pseudomonadati>Pseudomonadota>Gammaproteobacteria>Enterobacterales>Enterobacteriaceae>Salmonella>Salmonella_enterica | 273 | ZP62_000970 | 66809..67630 | 59201 | DUF4435-domain-containing-protein-[Salmonella-enterica-subsp.-enterica] |

| Protein-ID | Domain | Assembly-ID | Strain | Taxonomy | Length | Locus-Tag | Genomic-coordinates | TaxID | Fasta-defline |
|---|---|---|---|---|---|---|---|---|---|
| EDV5116993.1 | TOPRIM | - | Salmonella_enterica_subsp._enterica | Bacteria>Pseudomonadati>Pseudomonadota>Gammaproteobacteria>Enterobacterales>Enterobacteriaceae>Salmonella>Salmonella_enterica | 273 | ZJ36_000828 | 37348..38169 | 59201 | DUF4435-domain-containing-protein-[Salmonella-enterica-subsp.-enterica] |
| EDV9109401.1 | TOPRIM | - | Salmonella_enterica_subsp._enterica | Bacteria>Pseudomonadati>Pseudomonadota>Gammaproteobacteria>Enterobacterales>Enterobacteriaceae>Salmonella>Salmonella_enterica | 273 | ZK68_001603 | 60425..61246 | 59201 | DUF4435-domain-containing-protein-[Salmonella-enterica-subsp.-enterica] |
| EDW0696248.1 | TOPRIM | - | Salmonella_enterica_subsp._enterica | Bacteria>Pseudomonadati>Pseudomonadota>Gammaproteobacteria>Enterobacterales>Enterobacteriaceae>Salmonella>Salmonella_enterica | 273 | KV01_000538 | 158795..159616 | 59201 | DUF4435-domain-containing-protein-[Salmonella-enterica-subsp.-enterica] |
| EDW4488618.1 | TOPRIM | - | Salmonella_enterica_subsp._enterica | Bacteria>Pseudomonadati>Pseudomonadota>Gammaproteobacteria>Enterobacterales>Enterobacteriaceae>Salmonella>Salmonella_enterica | 273 | YV39_000490 | 174499..175320 | 59201 | DUF4435-domain-containing-protein-[Salmonella-enterica-subsp.-enterica] |
| EDZ7084223.1 | TOPRIM | - | Salmonella_enterica_subsp._enterica | Bacteria>Pseudomonadati>Pseudomonadota>Gammaproteobacteria>Enterobacterales>Enterobacteriaceae>Salmonella>Salmonella_enterica | 273 | GS086_003559 | 11248..12069 | 59201 | DUF4435-domain-containing-protein-[Salmonella-enterica-subsp.-enterica] |
| EEC0292874.1 | TOPRIM | - | Salmonella_enterica_subsp._enterica | Bacteria>Pseudomonadati>Pseudomonadota>Gammaproteobacteria>Enterobacterales>Enterobacteriaceae>Salmonella>Salmonella_enterica | 273 | YV76_000182 | 194816..195637 | 59201 | DUF4435-domain-containing-protein-[Salmonella-enterica-subsp.-enterica] |
| EEC1070529.1 | TOPRIM | - | Salmonella_enterica_subsp._enterica | Bacteria>Pseudomonadati>Pseudomonadota>Gammaproteobacteria>Enterobacterales>Enterobacteriaceae>Salmonella>Salmonella_enterica | 273 | AH545_000141 | 148422..149243 | 59201 | DUF4435-domain-containing-protein-[Salmonella-enterica-subsp.-enterica] |
| EEJ3565318.1 | TOPRIM | - | Salmonella_enterica_subsp._enterica | Bacteria>Pseudomonadati>Pseudomonadota>Gammaproteobacteria>Enterobacterales>Enterobacteriaceae>Salmonella>Salmonella_enterica | 273 | AA437_001847 | 134139..134960 | 59201 | DUF4435-domain-containing-protein-[Salmonella-enterica-subsp.-enterica] |
| EEJ7235531.1 | TOPRIM | - | Salmonella_enterica_subsp._salamae | Bacteria>Pseudomonadati>Pseudomonadota>Gammaproteobacteria>Enterobacterales>Enterobacteriaceae>Salmonella>Salmonella_enterica | 273 | S483_003522 | 160826..161647 | 59202 | DUF4435-domain-containing-protein-[Salmonella-enterica-subsp.-salamae] |
| EHA8894996.1 | TOPRIM | - | Salmonella_enterica_subsp._enterica | Bacteria>Pseudomonadati>Pseudomonadota>Gammaproteobacteria>Enterobacterales>Enterobacteriaceae>Salmonella>Salmonella_enterica | 273 | JHW15_001257 | 52807..53628 | 59201 | DUF4435-domain-containing-protein-[Salmonella-enterica-subsp.-enterica] |
| HAT0208693.1 | TOPRIM | GCA_015882745.1 | Salmonella_enterica_subsp._enterica | Bacteria>Pseudomonadati>Pseudomonadota>Gammaproteobacteria>Enterobacterales>Enterobacteriaceae>Salmonella>Salmonella_enterica | 273 | I8Q75_000457 | 69452..70273 | 59201 | TPA:-DUF4435-domain-containing-protein-[Salmonella-enterica-subsp.-enterica] |
| HAV0565024.1 | TOPRIM | GCA_016249375.1 | Salmonella_enterica_subsp._enterica | Bacteria>Pseudomonadati>Pseudomonadota>Gammaproteobacteria>Enterobacterales>Enterobacteriaceae>Salmonella>Salmonella_enterica | 273 | JF300_000745 | 83760..84581 | 59201 | TPA:-DUF4435-domain-containing-protein-[Salmonella-enterica-subsp.-enterica] |
| MLQ35365.1 | TOPRIM | GCA_003910355.1 | Salmonella_enterica_subsp._enterica | Bacteria>Pseudomonadati>Pseudomonadota>Gammaproteobacteria>Enterobacterales>Enterobacteriaceae>Salmonella>Salmonella_enterica | 273 | DRM12_04725 | 256320..257141 | 59201 | hypothetical-protein-[Salmonella-enterica-subsp.-enterica] |
| SUG73763.1 | TOPRIM | GCA_900456435.1 | Salmonella_enterica_subsp._enterica | Bacteria>Pseudomonadati>Pseudomonadota>Gammaproteobacteria>Enterobacterales>Enterobacteriaceae>Salmonella>Salmonella_enterica | 197 | NCTC5798_05052 | 4081035..4081628 | 59201 | putative-cytoplasmic-protein-[Salmonella-enterica-subsp.-enterica] |
| ECI8263214.1 | TOPRIM | GCA_007791395.1 | Salmonella_enterica_subsp._salamae | Bacteria>Pseudomonadati>Pseudomonadota>Gammaproteobacteria>Enterobacterales>Enterobacteriaceae>Salmonella>Salmonella_enterica | 189 | S481_13090 | 130438..131007 | 59202 | hypothetical-protein-S481_13090-[Salmonella-enterica-subsp.-salamae] |
| EDT7223676.1 | TOPRIM | - | Salmonella_enterica_subsp._enterica | Bacteria>Pseudomonadati>Pseudomonadota>Gammaproteobacteria>Enterobacterales>Enterobacteriaceae>Salmonella>Salmonella_enterica | 189 | AH268_003337 | 177720..178289 | 59201 | DUF4276-family-protein-[Salmonella-enterica-subsp.-enterica] |
| EGB5697878.1 | TOPRIM | - | Salmonella_enterica_subsp._enterica | Bacteria>Pseudomonadati>Pseudomonadota>Gammaproteobacteria>Enterobacterales>Enterobacteriaceae>Salmonella>Salmonella_enterica | 189 | H7531_003068 | 11924..12493 | 59201 | DUF4276-family-protein-[Salmonella-enterica-subsp.-enterica] |
| EAA1506271.1 | TOPRIM | GCA_004160295.1 | Salmonella_enterica_subsp._enterica_serovar_Agama | Bacteria>Pseudomonadati>Pseudomonadota>Gammaproteobacteria>Enterobacterales>Enterobacteriaceae>Salmonella>Salmonella_enterica>Salmonella_enterica_subsp._enterica | 273 | DL104_05745 | 37654..38475 | 399581 | DUF4435-domain-containing-protein-[Salmonella-enterica-subsp.-enterica] |
| EAA3988069.1 | TOPRIM | GCA_004177075.1 | Salmonella_enterica_subsp._enterica_serovar_Typhimurium | Bacteria>Pseudomonadati>Pseudomonadota>Gammaproteobacteria>Enterobacterales>Enterobacteriaceae>Salmonella>Salmonella_enterica>Salmonella_enterica_subsp._enterica | 273 | DO557_10280 | 127065..127886 | 90371 | DUF4435-domain-containing-protein-[Salmonella-enterica-subsp.-enterica] |
| EAA4842521.1 | TOPRIM | GCA_004183005.1 | Salmonella_enterica_subsp._enterica_serovar_Typhimurium | Bacteria>Pseudomonadati>Pseudomonadota>Gammaproteobacteria>Enterobacterales>Enterobacteriaceae>Salmonella>Salmonella_enterica>Salmonella_enterica_subsp._enterica | 273 | DPP49_10720 | 68549..69370 | 90371 | DUF4435-domain-containing-protein-[Salmonella-enterica-subsp.-enterica] |
| EAA5580613.1 | TOPRIM | GCA_004185805.1 | Salmonella_enterica_subsp._enterica_serovar_Glostrup | Bacteria>Pseudomonadati>Pseudomonadota>Gammaproteobacteria>Enterobacterales>Enterobacteriaceae>Salmonella>Salmonella_enterica>Salmonella_enterica_subsp._enterica | 273 | DQD90_06855 | 67364..68185 | 1151180 | hypothetical-protein-[Salmonella-enterica-subsp.-enterica] |
| EAA5642509.1 | TOPRIM | GCA_004185395.1 | Salmonella_enterica_subsp._enterica_serovar_Javiana | Bacteria>Pseudomonadati>Pseudomonadota>Gammaproteobacteria>Enterobacterales>Enterobacteriaceae>Salmonella>Salmonella_enterica>Salmonella_enterica_subsp._enterica | 273 | DQJ71_10405 | 157500..158321 | 363569 | DUF4435-domain-containing-protein-[Salmonella-enterica-subsp.-enterica] |
| EAA6071873.1 | TOPRIM | GCA_004187475.1 | Salmonella_enterica_subsp._enterica_serovar_Corvallis | Bacteria>Pseudomonadati>Pseudomonadota>Gammaproteobacteria>Enterobacterales>Enterobacteriaceae>Salmonella>Salmonella_enterica>Salmonella_enterica_subsp._enterica | 273 | DQZ25_06125 | 166691..167512 | 593905 | hypothetical-protein-[Salmonella-enterica-subsp.-enterica] |
| EAA6168127.1 | TOPRIM | GCA_004188635.1 | Salmonella_enterica_subsp._enterica_serovar_Ohio | Bacteria>Pseudomonadati>Pseudomonadota>Gammaproteobacteria>Enterobacterales>Enterobacteriaceae>Salmonella>Salmonella_enterica>Salmonella_enterica_subsp._enterica | 273 | DQZ68_14450 | 19043..19864 | 117541 | DUF4435-domain-containing-protein-[Salmonella-enterica-subsp.-enterica] |
| EAA7109618.1 | TOPRIM | GCA_004192185.1 | Salmonella_enterica_subsp._enterica_serovar_Ouagadougou | Bacteria>Pseudomonadati>Pseudomonadota>Gammaproteobacteria>Enterobacterales>Enterobacteriaceae>Salmonella>Salmonella_enterica>Salmonella_enterica_subsp._enterica | 273 | DSE94_16645 | 60572..61393 | 2564899 | hypothetical-protein-[Salmonella-enterica-subsp.-enterica] |
| EAA7310020.1 | TOPRIM | GCA_004192645.1 | Salmonella_enterica_subsp._enterica_serovar_Duesseldorf | Bacteria>Pseudomonadati>Pseudomonadota>Gammaproteobacteria>Enterobacterales>Enterobacteriaceae>Salmonella>Salmonella_enterica>Salmonella_enterica_subsp._enterica | 273 | TL61_10440 | 164426..165247 | 322832 | DUF4435-domain-containing-protein-[Salmonella-enterica-subsp.-enterica] |
| EAA8523364.1 | TOPRIM | GCA_004222265.1 | Salmonella_enterica_subsp._enterica_serovar_Cerro | Bacteria>Pseudomonadati>Pseudomonadota>Gammaproteobacteria>Enterobacterales>Enterobacteriaceae>Salmonella>Salmonella_enterica>Salmonella_enterica_subsp._enterica | 273 | ADQ53_00830 | 169424..170245 | 340188 | DUF4435-domain-containing-protein-[Salmonella-enterica-subsp.-enterica] |
| EAB6380444.1 | TOPRIM | GCA_004255205.1 | Salmonella_enterica_subsp._enterica_serovar_Emek | Bacteria>Pseudomonadati>Pseudomonadota>Gammaproteobacteria>Enterobacterales>Enterobacteriaceae>Salmonella>Salmonella_enterica>Salmonella_enterica_subsp._enterica | 273 | CWG97_20705 | 24534..25355 | 286784 | DUF4435-domain-containing-protein-[Salmonella-enterica-subsp.-enterica] |
| EAB7776929.1 | TOPRIM | GCA_004262645.1 | Salmonella_enterica_subsp._enterica_serovar_Cannstatt | Bacteria>Pseudomonadati>Pseudomonadota>Gammaproteobacteria>Enterobacterales>Enterobacteriaceae>Salmonella>Salmonella_enterica>Salmonella_enterica_subsp._enterica | 273 | D6413_08730 | 60763..61584 | 2564307 | DUF4435-domain-containing-protein-[Salmonella-enterica-subsp.-enterica] |
| EAB9374478.1 | TOPRIM | GCA_004270415.1 | Salmonella_enterica_subsp._enterica_serovar_Llandoff | Bacteria>Pseudomonadati>Pseudomonadota>Gammaproteobacteria>Enterobacterales>Enterobacteriaceae>Salmonella>Salmonella_enterica>Salmonella_enterica_subsp._enterica | 273 | D7N03_15215 | 69116..69937 | 2564701 | DUF4435-domain-containing-protein-[Salmonella-enterica-subsp.-enterica] |
| EAC0120240.1 | TOPRIM | GCA_004271765.1 | Salmonella_enterica_subsp._enterica_serovar_Typhimurium | Bacteria>Pseudomonadati>Pseudomonadota>Gammaproteobacteria>Enterobacterales>Enterobacteriaceae>Salmonella>Salmonella_enterica>Salmonella_enterica_subsp._enterica | 273 | EBO89_06940 | 66624..67445 | 90371 | DUF4435-domain-containing-protein-[Salmonella-enterica-subsp.-enterica] |
| EAC1009598.1 | TOPRIM | GCA_004275715.1 | Salmonella_enterica_subsp._enterica_serovar_Jangwani | Bacteria>Pseudomonadati>Pseudomonadota>Gammaproteobacteria>Enterobacterales>Enterobacteriaceae>Salmonella>Salmonella_enterica>Salmonella_enterica_subsp._enterica | 273 | YR14_13815 | 158840..159661 | 2564577 | DUF4435-domain-containing-protein-[Salmonella-enterica-subsp.-enterica] |
| EAO4363252.1 | TOPRIM | GCA_005505735.1 | Salmonella_enterica_subsp._enterica_serovar_Bere | Bacteria>Pseudomonadati>Pseudomonadota>Gammaproteobacteria>Enterobacterales>Enterobacteriaceae>Salmonella>Salmonella_enterica>Salmonella_enterica_subsp._enterica | 273 | E5197_09145 | 183791..184612 | 2024271 | DUF4435-domain-containing-protein-[Salmonella-enterica] |
| EBB4495258.1 | TOPRIM | GCA_006143655.1 | Salmonella_enterica_subsp._enterica_serovar_Typhimurium | Bacteria>Pseudomonadati>Pseudomonadota>Gammaproteobacteria>Enterobacterales>Enterobacteriaceae>Salmonella>Salmonella_enterica>Salmonella_enterica_subsp._enterica | 273 | CET45_10245 | 179351..180172 | 90371 | DUF4435-domain-containing-protein-[Salmonella-enterica-subsp.-enterica-serovar-Typhimurium] |
| EBF8311520.1 | TOPRIM | GCA_006394955.1 | Salmonella_enterica_subsp._enterica_serovar_Tamberma | Bacteria>Pseudomonadati>Pseudomonadota>Gammaproteobacteria>Enterobacterales>Enterobacteriaceae>Salmonella>Salmonella_enterica>Salmonella_enterica_subsp._enterica | 273 | FGE90_18595 | 61649..62470 | 2565079 | DUF4435-domain-containing-protein-[Salmonella-enterica-subsp.-enterica] |
| EBG0770115.1 | TOPRIM | GCA_006406395.1 | Salmonella_enterica_subsp._enterica_serovar_Derby | Bacteria>Pseudomonadati>Pseudomonadota>Gammaproteobacteria>Enterobacterales>Enterobacteriaceae>Salmonella>Salmonella_enterica>Salmonella_enterica_subsp._enterica | 273 | FH890_08340 | 11491..12312 | 28144 | DUF4435-domain-containing-protein-[Salmonella-enterica-subsp.-enterica-serovar-Derby] |
| EBL4178056.1 | TOPRIM | GCA_006685525.1 | Salmonella_enterica_subsp._enterica_serovar_Typhimurium | Bacteria>Pseudomonadati>Pseudomonadota>Gammaproteobacteria>Enterobacterales>Enterobacteriaceae>Salmonella>Salmonella_enterica>Salmonella_enterica_subsp._enterica | 273 | E5L78_02840 | 26597..27418 | 90371 | DUF4435-domain-containing-protein-[Salmonella-enterica-subsp.-enterica-serovar-Typhimurium] |
| EBL6294788.1 | TOPRIM | GCA_006694585.1 | Salmonella_enterica_subsp._enterica_serovar_Typhimurium | Bacteria>Pseudomonadati>Pseudomonadota>Gammaproteobacteria>Enterobacterales>Enterobacteriaceae>Salmonella>Salmonella_enterica>Salmonella_enterica_subsp._enterica | 273 | AC864_13980 | 176200..177021 | 90371 | DUF4435-domain-containing-protein-[Salmonella-enterica-subsp.-enterica-serovar-Typhimurium] |
| EBR9945164.1 | TOPRIM | GCA_006983125.1 | Salmonella_enterica_subsp._enterica_serovar_Krefeld | Bacteria>Pseudomonadati>Pseudomonadota>Gammaproteobacteria>Enterobacterales>Enterobacteriaceae>Salmonella>Salmonella_enterica>Salmonella_enterica_subsp._enterica | 273 | DTF03_11160 | 161839..162660 | 1242084 | DUF4435-domain-containing-protein-[Salmonella-enterica-subsp.-enterica-serovar-Krefeld] |
| EBS4429085.1 | TOPRIM | GCA_007008305.1 | Salmonella_enterica_subsp._enterica_serovar_Poona | Bacteria>Pseudomonadati>Pseudomonadota>Gammaproteobacteria>Enterobacterales>Enterobacteriaceae>Salmonella>Salmonella_enterica>Salmonella_enterica_subsp._enterica | 273 | DQJ97_07380 | 34327..35148 | 436295 | DUF4435-domain-containing-protein-[Salmonella-enterica-subsp.-enterica-serovar-Poona] |
| EBU6515332.1 | TOPRIM | GCA_007137465.1 | Salmonella_enterica_subsp._enterica_serovar_Typhimurium | Bacteria>Pseudomonadati>Pseudomonadota>Gammaproteobacteria>Enterobacterales>Enterobacteriaceae>Salmonella>Salmonella_enterica>Salmonella_enterica_subsp._enterica | 273 | DKJ21_01485 | 13247..14068 | 90371 | DUF4435-domain-containing-protein-[Salmonella-enterica-subsp.-enterica-serovar-Typhimurium] |
| EBU6749122.1 | TOPRIM | GCA_007137985.1 | Salmonella_enterica_subsp._enterica_serovar_Typhimurium | Bacteria>Pseudomonadati>Pseudomonadota>Gammaproteobacteria>Enterobacterales>Enterobacteriaceae>Salmonella>Salmonella_enterica>Salmonella_enterica_subsp._enterica | 273 | DKS55_01050 | 7419..8240 | 90371 | DUF4435-domain-containing-protein-[Salmonella-enterica-subsp.-enterica-serovar-Typhimurium] |
| EBU7641091.1 | TOPRIM | GCA_007142065.1 | Salmonella_enterica_subsp._enterica_serovar_Brevik | Bacteria>Pseudomonadati>Pseudomonadota>Gammaproteobacteria>Enterobacterales>Enterobacteriaceae>Salmonella>Salmonella_enterica>Salmonella_enterica_subsp._enterica | 273 | DK709_13730 | 67825..68646 | 2564275 | hypothetical-protein-[Salmonella-enterica-subsp.-enterica-serovar-Brevik] |
| EBU8677657.1 | TOPRIM | GCA_007146445.1 | Salmonella_enterica_subsp._enterica_serovar_Parkroyal | Bacteria>Pseudomonadati>Pseudomonadota>Gammaproteobacteria>Enterobacterales>Enterobacteriaceae>Salmonella>Salmonella_enterica>Salmonella_enterica_subsp._enterica | 273 | DLR30_20700 | 34413..35234 | 2564909 | hypothetical-protein-[Salmonella-enterica-subsp.-enterica-serovar-Parkroyal] |
| EBU9775081.1 | TOPRIM | GCA_007151525.1 | Salmonella_enterica_subsp._enterica_serovar_Kumasi | Bacteria>Pseudomonadati>Pseudomonadota>Gammaproteobacteria>Enterobacterales>Enterobacteriaceae>Salmonella>Salmonella_enterica>Salmonella_enterica_subsp._enterica | 273 | DM304_21660 | 11501..12322 | 2493647 | hypothetical-protein-[Salmonella-enterica-subsp.-enterica-serovar-Kumasi] |
| EBU9878492.1 | TOPRIM | GCA_007152025.1 | Salmonella_enterica_subsp._enterica_serovar_Typhimurium | Bacteria>Pseudomonadati>Pseudomonadota>Gammaproteobacteria>Enterobacterales>Enterobacteriaceae>Salmonella>Salmonella_enterica>Salmonella_enterica_subsp._enterica | 273 | DM361_04545 | 68648..69469 | 90371 | DUF4435-domain-containing-protein-[Salmonella-enterica-subsp.-enterica-serovar-Typhimurium] |
| EBV0268833.1 | TOPRIM | GCA_007152865.1 | Salmonella_enterica_subsp._enterica_serovar_Typhimurium | Bacteria>Pseudomonadati>Pseudomonadota>Gammaproteobacteria>Enterobacterales>Enterobacteriaceae>Salmonella>Salmonella_enterica>Salmonella_enterica_subsp._enterica | 273 | DNH74_05360 | 68434..69255 | 90371 | DUF4435-domain-containing-protein-[Salmonella-enterica-subsp.-enterica-serovar-Typhimurium] |
| EBV2735695.1 | TOPRIM | GCA_007164145.1 | Salmonella_enterica_subsp._enterica_serovar_Stourbridge | Bacteria>Pseudomonadati>Pseudomonadota>Gammaproteobacteria>Enterobacterales>Enterobacteriaceae>Salmonella>Salmonella_enterica>Salmonella_enterica_subsp._enterica | 273 | DOA65_09460 | 67371..68192 | 1160766 | hypothetical-protein-[Salmonella-enterica-subsp.-enterica-serovar-Stourbridge] |
| EBV4609759.1 | TOPRIM | GCA_007170635.1 | Salmonella_enterica_subsp._enterica_serovar_Solt | Bacteria>Pseudomonadati>Pseudomonadota>Gammaproteobacteria>Enterobacterales>Enterobacteriaceae>Salmonella>Salmonella_enterica>Salmonella_enterica_subsp._enterica | 273 | DOX24_02190 | 11497..12318 | 2565040 | hypothetical-protein-[Salmonella-enterica-subsp.-enterica-serovar-Solt] |
| EBV4672046.1 | TOPRIM | GCA_007170935.1 | Salmonella_enterica_subsp._enterica_serovar_Tamberma | Bacteria>Pseudomonadati>Pseudomonadota>Gammaproteobacteria>Enterobacterales>Enterobacteriaceae>Salmonella>Salmonella_enterica>Salmonella_enterica_subsp._enterica | 273 | DOW22_12535 | 157605..158426 | 2565079 | hypothetical-protein-[Salmonella-enterica-subsp.-enterica-serovar-Tamberma] |
| EBV5450234.1 | TOPRIM | GCA_007175875.1 | Salmonella_enterica_subsp._enterica_serovar_Typhimurium | Bacteria>Pseudomonadati>Pseudomonadota>Gammaproteobacteria>Enterobacterales>Enterobacteriaceae>Salmonella>Salmonella_enterica>Salmonella_enterica_subsp._enterica | 273 | DO810_06285 | 68652..69473 | 90371 | DUF4435-domain-containing-protein-[Salmonella-enterica-subsp.-enterica-serovar-Typhimurium] |
| EBV7110757.1 | TOPRIM | GCA_007180535.1 | Salmonella_enterica_subsp._enterica_serovar_Ohio | Bacteria>Pseudomonadati>Pseudomonadota>Gammaproteobacteria>Enterobacterales>Enterobacteriaceae>Salmonella>Salmonella_enterica>Salmonella_enterica_subsp._enterica | 273 | YY60_02805 | 163136..163957 | 117541 | DUF4435-domain-containing-protein-[Salmonella-enterica-subsp.-enterica-serovar-Ohio] |
| EBW1953459.1 | TOPRIM | GCA_007201375.1 | Salmonella_enterica_subsp._enterica_serovar_Cerro | Bacteria>Pseudomonadati>Pseudomonadota>Gammaproteobacteria>Enterobacterales>Enterobacteriaceae>Salmonella>Salmonella_enterica>Salmonella_enterica_subsp._enterica | 273 | C3N96_11425 | 52492..53313 | 340188 | DUF4435-domain-containing-protein-[Salmonella-enterica-subsp.-enterica-serovar-Cerro] |
| EBW2731419.1 | TOPRIM | GCA_007205375.1 | Salmonella_enterica_subsp._enterica_serovar_Typhimurium | Bacteria>Pseudomonadati>Pseudomonadota>Gammaproteobacteria>Enterobacterales>Enterobacteriaceae>Salmonella>Salmonella_enterica>Salmonella_enterica_subsp._enterica | 273 | DPA65_05350 | 179368..180189 | 90371 | DUF4435-domain-containing-protein-[Salmonella-enterica-subsp.-enterica-serovar-Typhimurium] |
| EBW3295111.1 | TOPRIM | GCA_007207035.1 | Salmonella_enterica_subsp._enterica_serovar_Cerro | Bacteria>Pseudomonadati>Pseudomonadota>Gammaproteobacteria>Enterobacterales>Enterobacteriaceae>Salmonella>Salmonella_enterica>Salmonella_enterica_subsp._enterica | 273 | DPE09_02375 | 52491..53312 | 340188 | DUF4435-domain-containing-protein-[Salmonella-enterica-subsp.-enterica-serovar-Cerro] |
| EBW5570925.1 | TOPRIM | GCA_007216555.1 | Salmonella_enterica_subsp._enterica_serovar_Emek | Bacteria>Pseudomonadati>Pseudomonadota>Gammaproteobacteria>Enterobacterales>Enterobacteriaceae>Salmonella>Salmonella_enterica>Salmonella_enterica_subsp._enterica | 273 | DPY54_19330 | 24526..25347 | 286784 | hypothetical-protein-DPY54_19330-[Salmonella-enterica-subsp.-enterica-serovar-Emek] |
| EBW8394991.1 | TOPRIM | GCA_007228915.1 | Salmonella_enterica_subsp._enterica_serovar_Florida | Bacteria>Pseudomonadati>Pseudomonadota>Gammaproteobacteria>Enterobacterales>Enterobacteriaceae>Salmonella>Salmonella_enterica>Salmonella_enterica_subsp._enterica | 273 | A3X38_19930 | 11498..12319 | 2564436 | DUF4435-domain-containing-protein-[Salmonella-enterica-subsp.-enterica-serovar-Florida] |
| EBX4839080.1 | TOPRIM | GCA_007256215.1 | Salmonella_enterica_subsp._enterica_serovar_Tamberma | Bacteria>Pseudomonadati>Pseudomonadota>Gammaproteobacteria>Enterobacterales>Enterobacteriaceae>Salmonella>Salmonella_enterica>Salmonella_enterica_subsp._enterica | 273 | DSB42_14915 | 171998..172819 | 2565079 | hypothetical-protein-[Salmonella-enterica-subsp.-enterica-serovar-Tamberma] |
| EBX6018290.1 | TOPRIM | GCA_007258805.1 | Salmonella_enterica_subsp._enterica_serovar_Dortmund | Bacteria>Pseudomonadati>Pseudomonadota>Gammaproteobacteria>Enterobacterales>Enterobacteriaceae>Salmonella>Salmonella_enterica>Salmonella_enterica_subsp._enterica | 273 | DSR74_22230 | 6430..7251 | 2564368 | hypothetical-protein-[Salmonella-enterica-subsp.-enterica-serovar-Dortmund] |
| EBX6302554.1 | TOPRIM | GCA_007260765.1 | Salmonella_enterica_subsp._enterica_serovar_Enugu | Bacteria>Pseudomonadati>Pseudomonadota>Gammaproteobacteria>Enterobacterales>Enterobacteriaceae>Salmonella>Salmonella_enterica>Salmonella_enterica_subsp._enterica | 273 | DS191_04285 | 34232..35053 | 2564400 | hypothetical-protein-[Salmonella-enterica-subsp.-enterica-serovar-Enugu] |
| EBX8836175.1 | TOPRIM | GCA_007270555.1 | Salmonella_enterica_subsp._enterica_serovar_Urbana | Bacteria>Pseudomonadati>Pseudomonadota>Gammaproteobacteria>Enterobacterales>Enterobacteriaceae>Salmonella>Salmonella_enterica>Salmonella_enterica_subsp._enterica | 273 | DTU76_05385 | 60656..61477 | 260677 | DUF4435-domain-containing-protein-[Salmonella-enterica-subsp.-enterica-serovar-Urbana] |
| EBY1390421.1 | TOPRIM | GCA_007281935.1 | Salmonella_enterica_subsp._enterica_serovar_Wagenia | Bacteria>Pseudomonadati>Pseudomonadota>Gammaproteobacteria>Enterobacterales>Enterobacteriaceae>Salmonella>Salmonella_enterica>Salmonella_enterica_subsp._enterica | 273 | DUW06_08890 | 147275..148096 | 322833 | DUF4435-domain-containing-protein-[Salmonella-enterica-subsp.-enterica-serovar-Wagenia] |
| EBY1767818.1 | TOPRIM | GCA_007283275.1 | Salmonella_enterica_subsp._enterica_serovar_Georgia | Bacteria>Pseudomonadati>Pseudomonadota>Gammaproteobacteria>Enterobacterales>Enterobacteriaceae>Salmonella>Salmonella_enterica>Salmonella_enterica_subsp._enterica | 273 | DU102_16120 | 59051..59872 | 2564467 | DUF4435-domain-containing-protein-[Salmonella-enterica-subsp.-enterica-serovar-Georgia] |
| EBY2665623.1 | TOPRIM | GCA_007287115.1 | Salmonella_enterica_subsp._enterica_serovar_Teko | Bacteria>Pseudomonadati>Pseudomonadota>Gammaproteobacteria>Enterobacterales>Enterobacteriaceae>Salmonella>Salmonella_enterica>Salmonella_enterica_subsp._enterica | 273 | DVF23_02330 | 25546..26367 | 2079198 | DUF4435-domain-containing-protein-[Salmonella-enterica-subsp.-enterica-serovar-Teko] |
| EBZ6440067.1 | TOPRIM | GCA_007344875.1 | Salmonella_enterica_subsp._enterica_serovar_Typhimurium | Bacteria>Pseudomonadati>Pseudomonadota>Gammaproteobacteria>Enterobacterales>Enterobacteriaceae>Salmonella>Salmonella_enterica>Salmonella_enterica_subsp._enterica | 273 | ED482_12680 | 7488..8309 | 90371 | DUF4435-domain-containing-protein-[Salmonella-enterica-subsp.-enterica-serovar-Typhimurium] |
| EBZ6491795.1 | TOPRIM | GCA_007345015.1 | Salmonella_enterica_subsp._enterica_serovar_Dugbe | Bacteria>Pseudomonadati>Pseudomonadota>Gammaproteobacteria>Enterobacterales>Enterobacteriaceae>Salmonella>Salmonella_enterica>Salmonella_enterica_subsp._enterica | 273 | ED480_20075 | 25274..26095 | 2564376 | DUF4435-domain-containing-protein-[Salmonella-enterica-subsp.-enterica-serovar-Dugbe] |
| ECA8969865.1 | TOPRIM | GCA_007399965.1 | Salmonella_enterica_subsp._enterica_serovar_Omuna | Bacteria>Pseudomonadati>Pseudomonadota>Gammaproteobacteria>Enterobacterales>Enterobacteriaceae>Salmonella>Salmonella_enterica>Salmonella_enterica_subsp._enterica | 273 | ETB55_05740 | 66177..66998 | 2564887 | DUF4435-domain-containing-protein-[Salmonella-enterica-subsp.-enterica-serovar-Omuna] |
| ECB3805267.1 | TOPRIM | GCA_007422765.1 | Salmonella_enterica_subsp._enterica_serovar_Fufu | Bacteria>Pseudomonadati>Pseudomonadota>Gammaproteobacteria>Enterobacterales>Enterobacteriaceae>Salmonella>Salmonella_enterica>Salmonella_enterica_subsp._enterica | 273 | EW367_11375 | 60509..61330 | 2564449 | DUF4435-domain-containing-protein-[Salmonella-enterica-subsp.-enterica-serovar-Fufu] |
| ECD2821738.1 | TOPRIM | GCA_007513645.1 | Salmonella_enterica_subsp._enterica_serovar_Fanti | Bacteria>Pseudomonadati>Pseudomonadota>Gammaproteobacteria>Enterobacterales>Enterobacteriaceae>Salmonella>Salmonella_enterica>Salmonella_enterica_subsp._enterica | 273 | EXA45_11165 | 60745..61566 | 2564418 | DUF4435-domain-containing-protein-[Salmonella-enterica-subsp.-enterica-serovar-Fanti] |
| ECD3895177.1 | TOPRIM | GCA_007520185.1 | Salmonella_enterica_subsp._enterica_serovar_Gloucester | Bacteria>Pseudomonadati>Pseudomonadota>Gammaproteobacteria>Enterobacterales>Enterobacteriaceae>Salmonella>Salmonella_enterica>Salmonella_enterica_subsp._enterica | 273 | EOU70_05965 | 173184..174005 | 2564474 | DUF4435-domain-containing-protein-[Salmonella-enterica-subsp.-enterica-serovar-Gloucester] |
| ECE0383740.1 | TOPRIM | GCA_007548695.1 | Salmonella_enterica_subsp._enterica_serovar_Aba | Bacteria>Pseudomonadati>Pseudomonadota>Gammaproteobacteria>Enterobacterales>Enterobacteriaceae>Salmonella>Salmonella_enterica>Salmonella_enterica_subsp._enterica | 273 | XZ45_12025 | 34122..34943 | 2564103 | DUF4435-domain-containing-protein-[Salmonella-enterica-subsp.-enterica] |
| ECF4176038.1 | TOPRIM | GCA_007613175.1 | Salmonella_enterica_subsp._enterica_serovar_Typhimurium | Bacteria>Pseudomonadati>Pseudomonadota>Gammaproteobacteria>Enterobacterales>Enterobacteriaceae>Salmonella>Salmonella_enterica>Salmonella_enterica_subsp._enterica | 273 | E5U03_12940 | 102451..103272 | 90371 | DUF4435-domain-containing-protein-[Salmonella-enterica-subsp.-enterica-serovar-Typhimurium] |
| ECG3781076.1 | TOPRIM | GCA_007660485.1 | Salmonella_enterica_subsp._enterica_serovar_Breukelen | Bacteria>Pseudomonadati>Pseudomonadota>Gammaproteobacteria>Enterobacterales>Enterobacteriaceae>Salmonella>Salmonella_enterica>Salmonella_enterica_subsp._enterica | 273 | EXA83_17835 | 60538..61359 | 2564274 | DUF4435-domain-containing-protein-[Salmonella-enterica-subsp.-enterica-serovar-Breukelen] |
| ECG8301817.1 | TOPRIM | GCA_007694625.1 | Salmonella_enterica_subsp._enterica_serovar_Glostrup | Bacteria>Pseudomonadati>Pseudomonadota>Gammaproteobacteria>Enterobacterales>Enterobacteriaceae>Salmonella>Salmonella_enterica>Salmonella_enterica_subsp._enterica | 273 | E1853_02150 | 67362..68183 | 1151180 | DUF4435-domain-containing-protein-[Salmonella-enterica-subsp.-enterica-serovar-Glostrup] |
| ECI2073655.1 | TOPRIM | GCA_007762095.1 | Salmonella_enterica_subsp._enterica_serovar_Kentucky | Bacteria>Pseudomonadati>Pseudomonadota>Gammaproteobacteria>Enterobacterales>Enterobacteriaceae>Salmonella>Salmonella_enterica>Salmonella_enterica_subsp._enterica | 273 | AUD11_09530 | 102451..103272 | 192955 | DUF4435-domain-containing-protein-[Salmonella-enterica-subsp.-enterica-serovar-Kentucky] |
| ECK5773203.1 | TOPRIM | GCA_007886645.1 | Salmonella_enterica_subsp._enterica_serovar_Typhimurium | Bacteria>Pseudomonadati>Pseudomonadota>Gammaproteobacteria>Enterobacterales>Enterobacteriaceae>Salmonella>Salmonella_enterica>Salmonella_enterica_subsp._enterica | 273 | AXS32_00830 | 181010..181831 | 90371 | DUF4435-domain-containing-protein-[Salmonella-enterica-subsp.-enterica-serovar-Typhimurium] |
| ECM3029731.1 | TOPRIM | GCA_008020915.1 | Salmonella_enterica_subsp._enterica_serovar_Typhimurium | Bacteria>Pseudomonadati>Pseudomonadota>Gammaproteobacteria>Enterobacterales>Enterobacteriaceae>Salmonella>Salmonella_enterica>Salmonella_enterica_subsp._enterica | 273 | QB31_08420 | 68660..69481 | 90371 | DUF4435-domain-containing-protein-[Salmonella-enterica-subsp.-enterica-serovar-Typhimurium] |
| ECM3363966.1 | TOPRIM | GCA_008021815.1 | Salmonella_enterica_subsp._enterica_serovar_Typhimurium | Bacteria>Pseudomonadati>Pseudomonadota>Gammaproteobacteria>Enterobacterales>Enterobacteriaceae>Salmonella>Salmonella_enterica>Salmonella_enterica_subsp._enterica | 273 | PW85_07005 | 102451..103272 | 90371 | DUF4435-domain-containing-protein-[Salmonella-enterica-subsp.-enterica-serovar-Typhimurium] |
| ECN0902567.1 | TOPRIM | GCA_008061715.1 | Salmonella_enterica_subsp._enterica_serovar_Typhimurium | Bacteria>Pseudomonadati>Pseudomonadota>Gammaproteobacteria>Enterobacterales>Enterobacteriaceae>Salmonella>Salmonella_enterica>Salmonella_enterica_subsp._enterica | 273 | YR82_02420 | 68435..69256 | 90371 | DUF4435-domain-containing-protein-[Salmonella-enterica-subsp.-enterica-serovar-Typhimurium] |
| ECO0332139.1 | TOPRIM | GCA_008112305.1 | Salmonella_enterica_subsp._enterica_serovar_Typhimurium | Bacteria>Pseudomonadati>Pseudomonadota>Gammaproteobacteria>Enterobacterales>Enterobacteriaceae>Salmonella>Salmonella_enterica>Salmonella_enterica_subsp._enterica | 273 | ZS66_00460 | 102451..103272 | 90371 | DUF4435-domain-containing-protein-[Salmonella-enterica-subsp.-enterica-serovar-Typhimurium] |
| ECT8081317.1 | TOPRIM | GCA_008481645.1 | Salmonella_enterica_subsp._enterica_serovar_Carrau | Bacteria>Pseudomonadati>Pseudomonadota>Gammaproteobacteria>Enterobacterales>Enterobacteriaceae>Salmonella>Salmonella_enterica>Salmonella_enterica_subsp._enterica | 273 | BLY96_09875 | 158521..159342 | 1160739 | DUF4435-domain-containing-protein-[Salmonella-enterica-subsp.-enterica-serovar-Carrau] |
| ECU0288390.1 | TOPRIM | GCA_008492025.1 | Salmonella_enterica_subsp._enterica_serovar_Typhimurium | Bacteria>Pseudomonadati>Pseudomonadota>Gammaproteobacteria>Enterobacterales>Enterobacteriaceae>Salmonella>Salmonella_enterica>Salmonella_enterica_subsp._enterica | 273 | C7B88_00725 | 148458..149279 | 90371 | DUF4435-domain-containing-protein-[Salmonella-enterica-subsp.-enterica-serovar-Typhimurium] |
| ECU9351203.1 | TOPRIM | GCA_008530195.1 | Salmonella_enterica_subsp._enterica_serovar_Typhimurium | Bacteria>Pseudomonadati>Pseudomonadota>Gammaproteobacteria>Enterobacterales>Enterobacteriaceae>Salmonella>Salmonella_enterica>Salmonella_enterica_subsp._enterica | 273 | CIH13_12055 | 179462..180283 | 90371 | DUF4435-domain-containing-protein-[Salmonella-enterica-subsp.-enterica-serovar-Typhimurium] |
| ECV7778492.1 | TOPRIM | GCA_008576505.1 | Salmonella_enterica_subsp._enterica_serovar_Typhimurium | Bacteria>Pseudomonadati>Pseudomonadota>Gammaproteobacteria>Enterobacterales>Enterobacteriaceae>Salmonella>Salmonella_enterica>Salmonella_enterica_subsp._enterica | 273 | ZX52_02605 | 68435..69256 | 90371 | DUF4435-domain-containing-protein-[Salmonella-enterica-subsp.-enterica-serovar-Typhimurium] |
| ECY8229164.1 | TOPRIM | GCA_008752395.1 | Salmonella_enterica_subsp._enterica_serovar_Stanley | Bacteria>Pseudomonadati>Pseudomonadota>Gammaproteobacteria>Enterobacterales>Enterobacteriaceae>Salmonella>Salmonella_enterica>Salmonella_enterica_subsp._enterica | 273 | AU695_08295 | 169100..169921 | 192953 | DUF4435-domain-containing-protein-[Salmonella-enterica-subsp.-enterica-serovar-Stanley] |
| ECZ9372114.1 | TOPRIM | GCA_008834045.1 | Salmonella_enterica_subsp._enterica_serovar_Typhimurium | Bacteria>Pseudomonadati>Pseudomonadota>Gammaproteobacteria>Enterobacterales>Enterobacteriaceae>Salmonella>Salmonella_enterica>Salmonella_enterica_subsp._enterica | 273 | A3U95_10230 | 102451..103272 | 90371 | DUF4435-domain-containing-protein-[Salmonella-enterica-subsp.-enterica-serovar-Typhimurium] |
| EDA5898039.1 | TOPRIM | GCA_008864045.1 | Salmonella_enterica_subsp._enterica_serovar_Poona | Bacteria>Pseudomonadati>Pseudomonadota>Gammaproteobacteria>Enterobacterales>Enterobacteriaceae>Salmonella>Salmonella_enterica>Salmonella_enterica_subsp._enterica | 273 | F9X73_05950 | 34317..35138 | 436295 | DUF4435-domain-containing-protein-[Salmonella-enterica-subsp.-enterica] |
| EDA7420103.1 | TOPRIM | - | Salmonella_enterica_subsp._enterica_serovar_Typhimurium | Bacteria>Pseudomonadati>Pseudomonadota>Gammaproteobacteria>Enterobacterales>Enterobacteriaceae>Salmonella>Salmonella_enterica>Salmonella_enterica_subsp._enterica | 273 | A3U84_08775 | 68652..69473 | 90371 | DUF4435-domain-containing-protein-[Salmonella-enterica-subsp.-enterica-serovar-Typhimurium] |
| EDB9656971.1 | TOPRIM | - | Salmonella_enterica_subsp._enterica_serovar_Typhimurium | Bacteria>Pseudomonadati>Pseudomonadota>Gammaproteobacteria>Enterobacterales>Enterobacteriaceae>Salmonella>Salmonella_enterica>Salmonella_enterica_subsp._enterica | 273 | BEO39_09560 | 35888..36709 | 90371 | DUF4435-domain-containing-protein-[Salmonella-enterica-subsp.-enterica-serovar-Typhimurium] |
| EDE2293301.1 | TOPRIM | - | Salmonella_enterica_subsp._enterica_serovar_Typhimurium | Bacteria>Pseudomonadati>Pseudomonadota>Gammaproteobacteria>Enterobacterales>Enterobacteriaceae>Salmonella>Salmonella_enterica>Salmonella_enterica_subsp._enterica | 273 | GBE85_07240 | 102451..103272 | 90371 | DUF4435-domain-containing-protein-[Salmonella-enterica] |
| EDH5631748.1 | TOPRIM | - | Salmonella_enterica_subsp._enterica_serovar_Claibornei | Bacteria>Pseudomonadati>Pseudomonadota>Gammaproteobacteria>Enterobacterales>Enterobacteriaceae>Salmonella>Salmonella_enterica>Salmonella_enterica_subsp._enterica | 273 | CB172_15095 | 166845..167666 | 2572709 | hypothetical-protein-[Salmonella-enterica-subsp.-enterica-serovar-Claibornei] |
| EDI5281733.1 | TOPRIM | - | Salmonella_enterica_subsp._enterica_serovar_Typhimurium | Bacteria>Pseudomonadati>Pseudomonadota>Gammaproteobacteria>Enterobacterales>Enterobacteriaceae>Salmonella>Salmonella_enterica>Salmonella_enterica_subsp._enterica | 273 | CEM90_10190 | 179251..180072 | 90371 | hypothetical-protein-CEM90_10190-[Salmonella-enterica-subsp.-enterica-serovar-Typhimurium] |
| EDL1929628.1 | TOPRIM | - | Salmonella_enterica_subsp._enterica_serovar_Typhimurium | Bacteria>Pseudomonadati>Pseudomonadota>Gammaproteobacteria>Enterobacterales>Enterobacteriaceae>Salmonella>Salmonella_enterica>Salmonella_enterica_subsp._enterica | 273 | CQD92_11435 | 102451..103272 | 90371 | hypothetical-protein-CQD92_11435-[Salmonella-enterica-subsp.-enterica-serovar-Typhimurium] |
| EDL2092520.1 | TOPRIM | - | Salmonella_enterica_subsp._enterica_serovar_Typhimurium | Bacteria>Pseudomonadati>Pseudomonadota>Gammaproteobacteria>Enterobacterales>Enterobacteriaceae>Salmonella>Salmonella_enterica>Salmonella_enterica_subsp._enterica | 273 | CQF25_02840 | 66800..67621 | 90371 | hypothetical-protein-CQF25_02840-[Salmonella-enterica-subsp.-enterica-serovar-Typhimurium] |
| EDM4126308.1 | TOPRIM | - | Salmonella_enterica_subsp._enterica_serovar_Typhimurium | Bacteria>Pseudomonadati>Pseudomonadota>Gammaproteobacteria>Enterobacterales>Enterobacteriaceae>Salmonella>Salmonella_enterica>Salmonella_enterica_subsp._enterica | 273 | CR810_06750 | 181011..181832 | 90371 | hypothetical-protein-CR810_06750-[Salmonella-enterica-subsp.-enterica-serovar-Typhimurium] |
| EDM6640283.1 | TOPRIM | - | Salmonella_enterica_subsp._enterica_serovar_Typhimurium | Bacteria>Pseudomonadati>Pseudomonadota>Gammaproteobacteria>Enterobacterales>Enterobacteriaceae>Salmonella>Salmonella_enterica>Salmonella_enterica_subsp._enterica | 273 | CUM00_00825 | 179350..180171 | 90371 | hypothetical-protein-CUM00_00825-[Salmonella-enterica-subsp.-enterica-serovar-Typhimurium] |
| EDR5087375.1 | TOPRIM | - | Salmonella_enterica_subsp._enterica_serovar_Kingston | Bacteria>Pseudomonadati>Pseudomonadota>Gammaproteobacteria>Enterobacterales>Enterobacteriaceae>Salmonella>Salmonella_enterica>Salmonella_enterica_subsp._enterica | 273 | GRZ33_003682 | 32712..33533 | 2161663 | DUF4435-domain-containing-protein-[Salmonella-enterica-subsp.-enterica] |
| EDS5116135.1 | TOPRIM | - | Salmonella_enterica_subsp._enterica_serovar_Agoueve | Bacteria>Pseudomonadati>Pseudomonadota>Gammaproteobacteria>Enterobacterales>Enterobacteriaceae>Salmonella>Salmonella_enterica>Salmonella_enterica_subsp._enterica | 273 | CBJ63_000888 | 173463..174284 | 1505961 | DUF4435-domain-containing-protein-[Salmonella-enterica-subsp.-enterica-serovar-Agoueve] |
| EDT1456938.1 | TOPRIM | - | Salmonella_enterica_subsp._enterica_serovar_Ramatgan | Bacteria>Pseudomonadati>Pseudomonadota>Gammaproteobacteria>Enterobacterales>Enterobacteriaceae>Salmonella>Salmonella_enterica>Salmonella_enterica_subsp._enterica | 273 | CE894_001673 | 161590..162411 | 2564942 | DUF4435-domain-containing-protein-[Salmonella-enterica-subsp.-enterica-serovar-Ramatgan] |
| EDT5820085.1 | TOPRIM | - | Salmonella_enterica_subsp._enterica_serovar_4,[5],12:i:- | Bacteria>Pseudomonadati>Pseudomonadota>Gammaproteobacteria>Enterobacterales>Enterobacteriaceae>Salmonella>Salmonella_enterica>Salmonella_enterica_subsp._enterica | 273 | B7921_002247 | 102451..103272 | 440524 | DUF4435-domain-containing-protein-[Salmonella-enterica-subsp.-enterica-serovar-4,[5] |
| EDT7239827.1 | TOPRIM | - | Salmonella_enterica_subsp._enterica_serovar_Warragul | Bacteria>Pseudomonadati>Pseudomonadota>Gammaproteobacteria>Enterobacterales>Enterobacteriaceae>Salmonella>Salmonella_enterica>Salmonella_enterica_subsp._enterica | 273 | AHZ96_001970 | 101809..102630 | 2565180 | DUF4435-domain-containing-protein-[Salmonella-enterica-subsp.-enterica] |
| EDU3766995.1 | TOPRIM | - | Salmonella_enterica_subsp._enterica_serovar_Minnesota | Bacteria>Pseudomonadati>Pseudomonadota>Gammaproteobacteria>Enterobacterales>Enterobacteriaceae>Salmonella>Salmonella_enterica>Salmonella_enterica_subsp._enterica | 273 | C4O09_000360 | 399192..400013 | 70803 | DUF4435-domain-containing-protein-[Salmonella-enterica-subsp.-enterica-serovar-Minnesota] |
| EDV2418009.1 | TOPRIM | - | Salmonella_enterica_subsp._enterica_serovar_Ohio | Bacteria>Pseudomonadati>Pseudomonadota>Gammaproteobacteria>Enterobacterales>Enterobacteriaceae>Salmonella>Salmonella_enterica>Salmonella_enterica_subsp._enterica | 273 | CQ222_000230 | 220139..220960 | 117541 | DUF4435-domain-containing-protein-[Salmonella-enterica-subsp.-enterica-serovar-Ohio] |
| EDV9644207.1 | TOPRIM | - | Salmonella_enterica_subsp._enterica_serovar_Carno | Bacteria>Pseudomonadati>Pseudomonadota>Gammaproteobacteria>Enterobacterales>Enterobacteriaceae>Salmonella>Salmonella_enterica>Salmonella_enterica_subsp._enterica | 273 | HM25_004000 | 172943..173764 | 2024268 | DUF4435-domain-containing-protein-[Salmonella-enterica-subsp.-enterica-serovar-Carno] |
| EDW2626495.1 | TOPRIM | - | Salmonella_enterica_subsp._enterica_serovar_Sangalkam | Bacteria>Pseudomonadati>Pseudomonadota>Gammaproteobacteria>Enterobacterales>Enterobacteriaceae>Salmonella>Salmonella_enterica>Salmonella_enterica_subsp._enterica | 273 | XX60_002385 | 165130..165951 | 2564992 | DUF4435-domain-containing-protein-[Salmonella-enterica-subsp.-enterica] |
| EDX7895852.1 | TOPRIM | - | Salmonella_enterica_subsp._enterica_serovar_4,[5],12:i:- | Bacteria>Pseudomonadati>Pseudomonadota>Gammaproteobacteria>Enterobacterales>Enterobacteriaceae>Salmonella>Salmonella_enterica>Salmonella_enterica_subsp._enterica | 273 | C2Y98_001771 | 47095..47916 | 440524 | DUF4435-domain-containing-protein-[Salmonella-enterica-subsp.-enterica-serovar-4,[5] |
| EED3636778.1 | TOPRIM | - | Salmonella_enterica_subsp._enterica_serovar_Sundsvall | Bacteria>Pseudomonadati>Pseudomonadota>Gammaproteobacteria>Enterobacterales>Enterobacteriaceae>Salmonella>Salmonella_enterica>Salmonella_enterica_subsp._enterica | 273 | S719_003284 | 166006..166827 | 2024273 | DUF4435-domain-containing-protein-[Salmonella-enterica-subsp.-enterica] |
| EEE1152711.1 | TOPRIM | - | Salmonella_enterica_subsp._enterica_serovar_Teko | Bacteria>Pseudomonadati>Pseudomonadota>Gammaproteobacteria>Enterobacterales>Enterobacteriaceae>Salmonella>Salmonella_enterica>Salmonella_enterica_subsp._enterica | 273 | CAE49_001972 | 325206..326027 | 2079198 | DUF4435-domain-containing-protein-[Salmonella-enterica-subsp.-enterica-serovar-Teko] |
| EEE6742965.1 | TOPRIM | - | Salmonella_enterica_subsp._enterica_serovar_Westhampton | Bacteria>Pseudomonadati>Pseudomonadota>Gammaproteobacteria>Enterobacterales>Enterobacteriaceae>Salmonella>Salmonella_enterica>Salmonella_enterica_subsp._enterica | 273 | CBJ05_002023 | 417228..418049 | 356621 | DUF4435-domain-containing-protein-[Salmonella-enterica-subsp.-enterica-serovar-Westhampton] |
| EEH2707249.1 | TOPRIM | - | Salmonella_enterica_subsp._enterica_serovar_Lille | Bacteria>Pseudomonadati>Pseudomonadota>Gammaproteobacteria>Enterobacterales>Enterobacteriaceae>Salmonella>Salmonella_enterica>Salmonella_enterica_subsp._enterica | 273 | GYE71_000440 | 62436..63257 | 486997 | DUF4435-domain-containing-protein-[Salmonella-enterica-subsp.-enterica] |
| EEH9995097.1 | TOPRIM | - | Salmonella_enterica_subsp._enterica_serovar_4,[5],12:i:- | Bacteria>Pseudomonadati>Pseudomonadota>Gammaproteobacteria>Enterobacterales>Enterobacteriaceae>Salmonella>Salmonella_enterica>Salmonella_enterica_subsp._enterica | 273 | G7H85_000162 | 178852..179673 | 440524 | DUF4435-domain-containing-protein-[Salmonella-enterica-subsp.-enterica-serovar-4,[5] |
| EEJ7298181.1 | TOPRIM | - | Salmonella_enterica_subsp._enterica_serovar_Livingstone | Bacteria>Pseudomonadati>Pseudomonadota>Gammaproteobacteria>Enterobacterales>Enterobacteriaceae>Salmonella>Salmonella_enterica>Salmonella_enterica_subsp._enterica | 273 | DV24_002302 | 306506..307327 | 260680 | DUF4435-domain-containing-protein-[Salmonella-enterica-subsp.-enterica-serovar-Livingstone] |
| EEJ9771969.1 | TOPRIM | - | Salmonella_enterica_subsp._enterica_serovar_Typhimurium | Bacteria>Pseudomonadati>Pseudomonadota>Gammaproteobacteria>Enterobacterales>Enterobacteriaceae>Salmonella>Salmonella_enterica>Salmonella_enterica_subsp._enterica | 273 | HL79_09645 | 179364..180185 | 90371 | DUF4435-domain-containing-protein-[Salmonella-enterica-subsp.-enterica-serovar-Typhimurium] |

| Protein-ID | Domain | Assembly-ID | Strain | Taxonomy | Length | Locus-Tag | Genomic-coordinates | TaxID | Fasta-defline |
|---|---|---|---|---|---|---|---|---|---|
| EEK0538628.1 | TOPRIM | - | Salmonella_enterica_subsp._enterica_serovar_Typhimurium | Bacteria>Pseudomonadati>Pseudomonadota>Gammaproteobacteria>Enterobacterales>Enterobacteriaceae>Salmonella>Salmonella_enterica>Salmonella_enterica_subsp._enterica | 273 | IH96_05495 | 144410..145231 | 90371 | DUF4435-domain-containing-protein-[Salmonella-enterica-subsp.-enterica-serovar-Typhimurium] |
| EEO3564963.1 | TOPRIM | - | Salmonella_enterica_subsp._enterica_serovar_Poona | Bacteria>Pseudomonadati>Pseudomonadota>Gammaproteobacteria>Enterobacterales>Enterobacteriaceae>Salmonella>Salmonella_enterica>Salmonella_enterica_subsp._enterica | 273 | DEI68_09010 | 60813..61634 | 436295 | DUF4435-domain-containing-protein-[Salmonella-enterica-subsp.-enterica-serovar-Poona] |
| EEP8318666.1 | TOPRIM | - | Salmonella_enterica_subsp._enterica_serovar_Livingstone | Bacteria>Pseudomonadati>Pseudomonadota>Gammaproteobacteria>Enterobacterales>Enterobacteriaceae>Salmonella>Salmonella_enterica>Salmonella_enterica_subsp._enterica | 273 | HCM12_000957 | 60840..61661 | 260680 | DUF4435-domain-containing-protein-[Salmonella-enterica-subsp.-enterica] |
| EEP9788444.1 | TOPRIM | - | Salmonella_enterica_subsp._enterica_serovar_Saintpaul | Bacteria>Pseudomonadati>Pseudomonadota>Gammaproteobacteria>Enterobacterales>Enterobacteriaceae>Salmonella>Salmonella_enterica>Salmonella_enterica_subsp._enterica | 273 | HCP67_002108 | 73349..74170 | 90105 | DUF4435-domain-containing-protein-[Salmonella-enterica-subsp.-enterica-serovar-Saintpaul] |
| EGA9157243.1 | TOPRIM | - | Salmonella_enterica_subsp._enterica_serovar_Kirkee | Bacteria>Pseudomonadati>Pseudomonadota>Gammaproteobacteria>Enterobacterales>Enterobacteriaceae>Salmonella>Salmonella_enterica>Salmonella_enterica_subsp._enterica | 273 | H6M98_002634 | 60734..61555 | 2564624 | DUF4435-domain-containing-protein-[Salmonella-enterica-subsp.-enterica-serovar-Kirkee] |
| EGH9456391.1 | TOPRIM | - | Salmonella_enterica_subsp._enterica_serovar_Leeuwarden | Bacteria>Pseudomonadati>Pseudomonadota>Gammaproteobacteria>Enterobacterales>Enterobacteriaceae>Salmonella>Salmonella_enterica>Salmonella_enterica_subsp._enterica | 273 | HKU92_000350 | 397764..398585 | 2564678 | DUF4435-domain-containing-protein-[Salmonella-enterica-subsp.-enterica-serovar-Leeuwarden] |
| EGT0819114.1 | TOPRIM | - | Salmonella_enterica_subsp._enterica_serovar_Ohio | Bacteria>Pseudomonadati>Pseudomonadota>Gammaproteobacteria>Enterobacterales>Enterobacteriaceae>Salmonella>Salmonella_enterica>Salmonella_enterica_subsp._enterica | 273 | I3I53_003215 | 19508..20329 | 117541 | DUF4435-domain-containing-protein-[Salmonella-enterica-subsp.-enterica-serovar-Ohio] |
| EGZ8568478.1 | TOPRIM | - | Salmonella_enterica_subsp._enterica_serovar_Typhimurium | Bacteria>Pseudomonadati>Pseudomonadota>Gammaproteobacteria>Enterobacterales>Enterobacteriaceae>Salmonella>Salmonella_enterica>Salmonella_enterica_subsp._enterica | 273 | JJM06_003395 | 21018..21839 | 90371 | DUF4435-domain-containing-protein-[Salmonella-enterica-subsp.-enterica-serovar-Typhimurium] |
| EHC3781324.1 | TOPRIM | - | Salmonella_enterica_subsp._enterica_serovar_Typhimurium | Bacteria>Pseudomonadati>Pseudomonadota>Gammaproteobacteria>Enterobacterales>Enterobacteriaceae>Salmonella>Salmonella_enterica>Salmonella_enterica_subsp._enterica | 273 | JS923_000753 | 181477..182298 | 90371 | DUF4435-domain-containing-protein-[Salmonella-enterica-subsp.-enterica-serovar-Typhimurium] |
| EHD9459648.1 | TOPRIM | - | Salmonella_enterica_subsp._enterica_serovar_Typhimurium | Bacteria>Pseudomonadati>Pseudomonadota>Gammaproteobacteria>Enterobacterales>Enterobacteriaceae>Salmonella>Salmonella_enterica>Salmonella_enterica_subsp._enterica | 273 | JYL48_000534 | 179365..180186 | 90371 | DUF4435-domain-containing-protein-[Salmonella-enterica-subsp.-enterica-serovar-Typhimurium] |
| EHJ1242125.1 | TOPRIM | GCA_017851875.1 | Salmonella_enterica_subsp._enterica_serovar_Typhimurium | Bacteria>Pseudomonadati>Pseudomonadota>Gammaproteobacteria>Enterobacterales>Enterobacteriaceae>Salmonella>Salmonella_enterica>Salmonella_enterica_subsp._enterica | 273 | KAC06_001463 | 75745..76566 | 90371 | DUF4435-domain-containing-protein-[Salmonella-enterica-subsp.-enterica-serovar-Typhimurium] |
| EHN4525916.1 | TOPRIM | GCA_018388245.1 | Salmonella_enterica_subsp._enterica_serovar_Ohio | Bacteria>Pseudomonadati>Pseudomonadota>Gammaproteobacteria>Enterobacterales>Enterobacteriaceae>Salmonella>Salmonella_enterica>Salmonella_enterica_subsp._enterica | 273 | KI018_001084 | 100452..101273 | 117541 | DUF4435-domain-containing-protein-[Salmonella-enterica-subsp.-enterica-serovar-Ohio] |
| EHO4506763.1 | TOPRIM | GCA_018524125.1 | Salmonella_enterica_subsp._enterica_serovar_Carrau | Bacteria>Pseudomonadati>Pseudomonadota>Gammaproteobacteria>Enterobacterales>Enterobacteriaceae>Salmonella>Salmonella_enterica>Salmonella_enterica_subsp._enterica | 273 | KD666_001069 | 157707..158528 | 1160739 | DUF4435-domain-containing-protein-[Salmonella-enterica-subsp.-enterica-serovar-Carrau] |
| EHP4799943.1 | TOPRIM | GCA_018776965.1 | Salmonella_enterica_subsp._enterica_serovar_Sangera | Bacteria>Pseudomonadati>Pseudomonadota>Gammaproteobacteria>Enterobacterales>Enterobacteriaceae>Salmonella>Salmonella_enterica>Salmonella_enterica_subsp._enterica | 273 | KM018_002018 | 53101..53922 | 1173574 | DUF4435-domain-containing-protein-[Salmonella-enterica-subsp.-enterica-serovar-Sangera] |
| EHS3976103.1 | TOPRIM | GCA_019140515.1 | Salmonella_enterica_subsp._enterica_serovar_Poona | Bacteria>Pseudomonadati>Pseudomonadota>Gammaproteobacteria>Enterobacterales>Enterobacteriaceae>Salmonella>Salmonella_enterica>Salmonella_enterica_subsp._enterica | 273 | KV430_003535 | 10409..11230 | 436295 | DUF4435-domain-containing-protein-[Salmonella-enterica-subsp.-enterica-serovar-Poona] |
| EHS5456438.1 | TOPRIM | GCA_019151305.1 | Salmonella_enterica_subsp._enterica_serovar_Corvallis | Bacteria>Pseudomonadati>Pseudomonadota>Gammaproteobacteria>Enterobacterales>Enterobacteriaceae>Salmonella>Salmonella_enterica>Salmonella_enterica_subsp._enterica | 273 | KV802_001389 | 173625..174446 | 593905 | DUF4435-domain-containing-protein-[Salmonella-enterica-subsp.-enterica-serovar-Corvallis] |
| EIE3342239.1 | TOPRIM | GCA_020155375.1 | Salmonella_enterica_subsp._enterica_serovar_Typhimurium | Bacteria>Pseudomonadati>Pseudomonadota>Gammaproteobacteria>Enterobacterales>Enterobacteriaceae>Salmonella>Salmonella_enterica>Salmonella_enterica_subsp._enterica | 273 | LDC84_002301 | 68642..69463 | 90371 | DUF4435-domain-containing-protein-[Salmonella-enterica-subsp.-enterica-serovar-Typhimurium] |
| EII7462487.1 | TOPRIM | GCA_020462505.1 | Salmonella_enterica_subsp._enterica_serovar_Typhimurium | Bacteria>Pseudomonadati>Pseudomonadota>Gammaproteobacteria>Enterobacterales>Enterobacteriaceae>Salmonella>Salmonella_enterica>Salmonella_enterica_subsp._enterica | 273 | LHR57_000985 | 7391..8212 | 90371 | DUF4435-domain-containing-protein-[Salmonella-enterica-subsp.-enterica-serovar-Typhimurium] |
| EIJ0372102.1 | TOPRIM | GCA_020490715.1 | Salmonella_enterica_subsp._enterica_serovar_Typhimurium | Bacteria>Pseudomonadati>Pseudomonadota>Gammaproteobacteria>Enterobacterales>Enterobacteriaceae>Salmonella>Salmonella_enterica>Salmonella_enterica_subsp._enterica | 273 | LH081_001562 | 179352..180173 | 90371 | DUF4435-domain-containing-protein-[Salmonella-enterica-subsp.-enterica-serovar-Typhimurium] |
| EIJ6143995.1 | TOPRIM | GCA_020548845.1 | Salmonella_enterica_subsp._enterica_serovar_4:a:- | Bacteria>Pseudomonadati>Pseudomonadota>Gammaproteobacteria>Enterobacterales>Enterobacteriaceae>Salmonella>Salmonella_enterica>Salmonella_enterica_subsp._enterica | 273 | LI678_001017 | 403155..403976 | 2885150 | DUF4435-domain-containing-protein-[Salmonella-enterica-subsp.-enterica-serovar-4:a:-] |
| EIR7569903.1 | TOPRIM | GCA_021281445.1 | Salmonella_enterica_subsp._enterica_serovar_Agbeni | Bacteria>Pseudomonadati>Pseudomonadota>Gammaproteobacteria>Enterobacterales>Enterobacteriaceae>Salmonella>Salmonella_enterica>Salmonella_enterica_subsp._enterica | 273 | LXI61_003345 | 24741..25562 | 1967642 | DUF4435-domain-containing-protein-[Salmonella-enterica-subsp.-enterica-serovar-Agbeni] |
| EJN2869331.1 | TOPRIM | GCA_024442015.1 | Salmonella_enterica_subsp._enterica_serovar_Derby | Bacteria>Pseudomonadati>Pseudomonadota>Gammaproteobacteria>Enterobacterales>Enterobacteriaceae>Salmonella>Salmonella_enterica>Salmonella_enterica_subsp._enterica | 273 | NPT71_003581 | 173451..174272 | 28144 | DUF4435-domain-containing-protein-[Salmonella-enterica-subsp.-enterica-serovar-Derby] |
| EJW7283565.1 | TOPRIM | GCA_025474975.1 | Salmonella_enterica_subsp._enterica_serovar_Typhimurium | Bacteria>Pseudomonadati>Pseudomonadota>Gammaproteobacteria>Enterobacterales>Enterobacteriaceae>Salmonella>Salmonella_enterica>Salmonella_enterica_subsp._enterica | 273 | OCR56_001318 | 75654..76475 | 90371 | DUF4435-domain-containing-protein-[Salmonella-enterica-subsp.-enterica-serovar-Typhimurium] |
| HAD0448191.1 | TOPRIM | GCA_010193865.1 | Salmonella_enterica_subsp._enterica_serovar_Typhimurium | Bacteria>Pseudomonadati>Pseudomonadota>Gammaproteobacteria>Enterobacterales>Enterobacteriaceae>Salmonella>Salmonella_enterica>Salmonella_enterica_subsp._enterica | 273 | G0N57_00695 | 144413..145234 | 90371 | TPA_asm:-DUF4435-domain-containing-protein-[Salmonella-enterica-subsp.-enterica-serovar-Typhimurium] |
| HAD1283635.1 | TOPRIM | GCA_010197825.1 | Salmonella_enterica_subsp._enterica_serovar_Typhimurium | Bacteria>Pseudomonadati>Pseudomonadota>Gammaproteobacteria>Enterobacterales>Enterobacteriaceae>Salmonella>Salmonella_enterica>Salmonella_enterica_subsp._enterica | 273 | GTH76_16760 | 102451..103272 | 90371 | TPA_asm:-DUF4435-domain-containing-protein-[Salmonella-enterica-subsp.-enterica-serovar-Typhimurium] |
| HAE3790826.1 | TOPRIM | GCA_010485735.1 | Salmonella_enterica_subsp._enterica_serovar_Typhimurium | Bacteria>Pseudomonadati>Pseudomonadota>Gammaproteobacteria>Enterobacterales>Enterobacteriaceae>Salmonella>Salmonella_enterica>Salmonella_enterica_subsp._enterica | 273 | G4A23_000146 | 155135..155956 | 90371 | TPA_asm:-DUF4435-domain-containing-protein-[Salmonella-enterica-subsp.-enterica-serovar-Typhimurium] |
| HAS9574204.1 | TOPRIM | GCA_015871885.1 | Salmonella_enterica_subsp._enterica_serovar_Typhimurium | Bacteria>Pseudomonadati>Pseudomonadota>Gammaproteobacteria>Enterobacterales>Enterobacteriaceae>Salmonella>Salmonella_enterica>Salmonella_enterica_subsp._enterica | 273 | I8M64_002286 | 102452..103273 | 90371 | TPA:-DUF4435-domain-containing-protein-[Salmonella-enterica-subsp.-enterica-serovar-Typhimurium] |
| HBJ3502654.1 | TOPRIM | GCA_020048005.1 | Salmonella_enterica_subsp._enterica_serovar_Typhimurium | Bacteria>Pseudomonadati>Pseudomonadota>Gammaproteobacteria>Enterobacterales>Enterobacteriaceae>Salmonella>Salmonella_enterica>Salmonella_enterica_subsp._enterica | 273 | LB282_001660 | 179136..179957 | 90371 | TPA:-DUF4435-domain-containing-protein-[Salmonella-enterica-subsp.-enterica-serovar-Typhimurium] |
| HBL9882172.1 | TOPRIM | GCA_021103155.1 | Salmonella_enterica_subsp._enterica_serovar_Jangwani | Bacteria>Pseudomonadati>Pseudomonadota>Gammaproteobacteria>Enterobacterales>Enterobacteriaceae>Salmonella>Salmonella_enterica>Salmonella_enterica_subsp._enterica | 273 | LT029_003493 | 163644..164465 | 2564577 | TPA:-DUF4435-domain-containing-protein-[Salmonella-enterica-subsp.-enterica-serovar-Jangwani] |
| HCB4841067.1 | TOPRIM | GCA_023082205.1 | Salmonella_enterica_subsp._enterica_serovar_Bredeney | Bacteria>Pseudomonadati>Pseudomonadota>Gammaproteobacteria>Enterobacterales>Enterobacteriaceae>Salmonella>Salmonella_enterica>Salmonella_enterica_subsp._enterica | 273 | M0F44_000354 | 398830..399651 | 134047 | DUF4435-domain-containing-protein-[Salmonella-enterica-subsp.-enterica-serovar-Bredeney] |
| MBT0981088.1 | TOPRIM | GCA_018531305.1 | Salmonella_enterica_subsp._enterica_serovar_Typhimurium | Bacteria>Pseudomonadati>Pseudomonadota>Gammaproteobacteria>Enterobacterales>Enterobacteriaceae>Salmonella>Salmonella_enterica>Salmonella_enterica_subsp._enterica | 273 | KJ032_08035 | 179382..180203 | 90371 | DUF4435-domain-containing-protein-[Salmonella-enterica-subsp.-enterica-serovar-Typhimurium] |
| MBZ4898654.1 | TOPRIM | GCA_020070625.1 | Salmonella_enterica_subsp._enterica_serovar_Typhimurium | Bacteria>Pseudomonadati>Pseudomonadota>Gammaproteobacteria>Enterobacterales>Enterobacteriaceae>Salmonella>Salmonella_enterica>Salmonella_enterica_subsp._enterica | 273 | HU117_01200 | 53335..54156 | 90371 | DUF4435-domain-containing-protein-[Salmonella-enterica-subsp.-enterica-serovar-Typhimurium] |
| MCL8965755.1 | TOPRIM | GCA_023608175.1 | Salmonella_enterica_subsp._enterica_serovar_Enteritidis | Bacteria>Pseudomonadati>Pseudomonadota>Gammaproteobacteria>Enterobacterales>Enterobacteriaceae>Salmonella>Salmonella_enterica>Salmonella_enterica_subsp._enterica | 273 | MU637_15190 | 178982..179803 | 149539 | DUF4435-domain-containing-protein-[Salmonella-enterica-subsp.-enterica-serovar-Enteritidis] |
| MIX28207.1 | TOPRIM | GCA_003876625.1 | Salmonella_enterica_subsp._enterica_serovar_Livingstone | Bacteria>Pseudomonadati>Pseudomonadota>Gammaproteobacteria>Enterobacterales>Enterobacteriaceae>Salmonella>Salmonella_enterica>Salmonella_enterica_subsp._enterica | 273 | CRQ34_16025 | 167609..168430 | 260680 | DUF4435-domain-containing-protein-[Salmonella-enterica-subsp.-enterica-serovar-Livingstone] |
| QSG19453.1 | TOPRIM | GCA_017094545.1 | Salmonella_enterica_subsp._enterica_serovar_Typhimurium | Bacteria>Pseudomonadati>Pseudomonadota>Gammaproteobacteria>Enterobacterales>Enterobacteriaceae>Salmonella>Salmonella_enterica>Salmonella_enterica_subsp._enterica | 273 | IRY60_23140 | 4738531..4739352 | 90371 | DUF4435-domain-containing-protein-[Salmonella-enterica-subsp.-enterica-serovar-Typhimurium] |
| QSG25300.1 | TOPRIM | GCA_017094585.1 | Salmonella_enterica_subsp._enterica_serovar_Typhimurium | Bacteria>Pseudomonadati>Pseudomonadota>Gammaproteobacteria>Enterobacterales>Enterobacteriaceae>Salmonella>Salmonella_enterica>Salmonella_enterica_subsp._enterica | 273 | IRY58_23100 | 4744085..4744906 | 90371 | DUF4435-domain-containing-protein-[Salmonella-enterica-subsp.-enterica-serovar-Typhimurium] |
| OFB20553.1 | TOPRIM | GCA_001756205.1 | Salmonella_enterica_subsp._enterica_serovar_Typhimurium | Bacteria>Pseudomonadati>Pseudomonadota>Gammaproteobacteria>Enterobacterales>Enterobacteriaceae>Salmonella>Salmonella_enterica>Salmonella_enterica_subsp._enterica | 270 | BG487_15370 | 8407..9219 | 90371 | hypothetical-protein-BG487_15370-[Salmonella-enterica-subsp.-enterica-serovar-Typhimurium] |
| SUH94773.1 | TOPRIM | GCA_900456925.1 | Salmonella_enterica_subsp._enterica_serovar_Typhimurium | Bacteria>Pseudomonadati>Pseudomonadota>Gammaproteobacteria>Enterobacterales>Enterobacteriaceae>Salmonella>Salmonella_enterica>Salmonella_enterica_subsp._enterica | 270 | NCTC13347_04216 | 4181440..4182252 | 90371 | putative-cytoplasmic-protein-[Salmonella-enterica-subsp.-enterica-serovar-Typhimurium] |
| SUI30169.1 | TOPRIM | GCA_900456825.1 | Salmonella_enterica_subsp._enterica_serovar_Derby | Bacteria>Pseudomonadati>Pseudomonadota>Gammaproteobacteria>Enterobacterales>Enterobacteriaceae>Salmonella>Salmonella_enterica>Salmonella_enterica_subsp._enterica | 270 | NCTC5722_04073 | 4070906..4071718 | 28144 | putative-cytoplasmic-protein-[Salmonella-enterica-subsp.-enterica-serovar-Derby] |
| AZT73894.1 | TOPRIM | GCA_003994735.1 | Salmonella_enterica_subsp._enterica_serovar_Waycross | Bacteria>Pseudomonadati>Pseudomonadota>Gammaproteobacteria>Enterobacterales>Enterobacteriaceae>Salmonella>Salmonella_enterica>Salmonella_enterica_subsp._enterica | 189 | ELZ91_19535 | 3959832..3960401 | 58101 | hypothetical-protein-ELZ91_19535-[Salmonella-enterica-subsp.-enterica-serovar-Waycross] |
| EBU9347430.1 | TOPRIM | GCA_007149485.1 | Salmonella_enterica_subsp._enterica_serovar_Angoda | Bacteria>Pseudomonadati>Pseudomonadota>Gammaproteobacteria>Enterobacterales>Enterobacteriaceae>Salmonella>Salmonella_enterica>Salmonella_enterica_subsp._enterica | 189 | DMU95_04675 | 157854..158423 | 2564159 | hypothetical-protein-[Salmonella-enterica-subsp.-enterica-serovar-Angoda] |
| ECY3255618.1 | TOPRIM | GCA_008731355.1 | Salmonella_enterica_subsp._enterica_serovar_Alachua | Bacteria>Pseudomonadati>Pseudomonadota>Gammaproteobacteria>Enterobacterales>Enterobacteriaceae>Salmonella>Salmonella_enterica>Salmonella_enterica_subsp._enterica | 189 | AU577_03155 | 25493..26062 | 913240 | hypothetical-protein-[Salmonella-enterica-subsp.-enterica-serovar-Alachua] |
| EDN6133536.1 | TOPRIM | - | Salmonella_enterica_subsp._enterica_serovar_Kentucky | Bacteria>Pseudomonadati>Pseudomonadota>Gammaproteobacteria>Enterobacterales>Enterobacteriaceae>Salmonella>Salmonella_enterica>Salmonella_enterica_subsp._enterica | 189 | GJQ78_12625 | 70812..71381 | 192955 | hypothetical-protein-GJQ78_12625-[Salmonella-enterica-subsp.-enterica-serovar-Kentucky] |
| EDS3575402.1 | TOPRIM | - | Salmonella_enterica_subsp._enterica_serovar_Sangera | Bacteria>Pseudomonadati>Pseudomonadota>Gammaproteobacteria>Enterobacterales>Enterobacteriaceae>Salmonella>Salmonella_enterica>Salmonella_enterica_subsp._enterica | 189 | GR179_002521 | 11833..12402 | 1173574 | DUF4276-family-protein-[Salmonella-enterica-subsp.-enterica] |
| EEI5362888.1 | TOPRIM | - | Salmonella_enterica_subsp._enterica_serovar_Kentucky | Bacteria>Pseudomonadati>Pseudomonadota>Gammaproteobacteria>Enterobacterales>Enterobacteriaceae>Salmonella>Salmonella_enterica>Salmonella_enterica_subsp._enterica | 189 | GXM95_001566 | 181807..182376 | 192955 | DUF4276-family-protein-[Salmonella-enterica-subsp.-enterica-serovar-Kentucky] |
| EEM2455578.1 | TOPRIM | - | Salmonella_enterica_subsp._enterica_serovar_Kentucky | Bacteria>Pseudomonadati>Pseudomonadota>Gammaproteobacteria>Enterobacterales>Enterobacteriaceae>Salmonella>Salmonella_enterica>Salmonella_enterica_subsp._enterica | 189 | CWG70_02790 | 182140..182709 | 192955 | hypothetical-protein-CWG70_02790-[Salmonella-enterica-subsp.-enterica-serovar-Kentucky] |
| EHJ7117736.1 | TOPRIM | GCA_017926475.1 | Salmonella_enterica_subsp._enterica_serovar_Kentucky | Bacteria>Pseudomonadati>Pseudomonadota>Gammaproteobacteria>Enterobacterales>Enterobacteriaceae>Salmonella>Salmonella_enterica>Salmonella_enterica_subsp._enterica | 189 | KAP12_002968 | 420763..421332 | 192955 | hypothetical-protein-KAP12_002968-[Salmonella-enterica-subsp.-enterica-serovar-Kentucky] |
| UTZ95785.1 | TOPRIM | GCA_024442275.1 | Salmonella_enterica_subsp._enterica_serovar_Kentucky | Bacteria>Pseudomonadati>Pseudomonadota>Gammaproteobacteria>Enterobacterales>Enterobacteriaceae>Salmonella>Salmonella_enterica>Salmonella_enterica_subsp._enterica | 189 | NOQ95_21575 | 4456909..4457478 | 192955 | DUF4276-family-protein-[Salmonella-enterica-subsp.-enterica-serovar-Kentucky] |
| ECF0142192.1 | TOPRIM | GCA_007595955.1 | Salmonella_enterica_subsp._enterica_serovar_Monschaui | Bacteria>Pseudomonadati>Pseudomonadota>Gammaproteobacteria>Enterobacterales>Enterobacteriaceae>Salmonella>Salmonella_enterica>Salmonella_enterica_subsp._enterica | 187 | EX931_14170 | 78110..78673 | 29480 | hypothetical-protein-[Salmonella-enterica-subsp.-enterica-serovar-Monschaui] |
| ESG75018.1 | TOPRIM | GCA_000487055.1 | Salmonella_enterica_subsp._enterica_serovar_Kentucky_str._ATCC_9263 | Bacteria>Pseudomonadati>Pseudomonadota>Gammaproteobacteria>Enterobacterales>Enterobacteriaceae>Salmonella>Salmonella_enterica>Salmonella_enterica_subsp._enterica>Salmonella_enterica_subsp._enterica_serovar_Kentucky | 189 | SEEK9263_16877 | 27652..28221 | 984236 | hypothetical-protein-SEEK9263_16877-[Salmonella-enterica-subsp.-enterica-serovar-Kentucky-str.-ATCC-9263] |
| EDH9995456.1 | TOPRIM | - | Salmonella_enterica_subsp._enterica_serovar_Java | Bacteria>Pseudomonadati>Pseudomonadota>Gammaproteobacteria>Enterobacterales>Enterobacteriaceae>Salmonella>Salmonella_enterica>Salmonella_enterica_subsp._enterica>Salmonella_enterica_subsp._enterica_serovar_Paratyphi_B | 273 | CC627_22220 | 7285..8106 | 224729 | hypothetical-protein-CC627_22220-[Salmonella-enterica-subsp.-enterica-serovar-Java] |
| ASE02937.1 | TOPRIM | GCA_002208315.1 | Salmonella_enterica_subsp._enterica_serovar_Quebec_str._S-1267 | Bacteria>Pseudomonadati>Pseudomonadota>Gammaproteobacteria>Enterobacterales>Enterobacteriaceae>Salmonella>Salmonella_enterica>Salmonella_enterica_subsp._enterica>Salmonella_enterica_subsp._enterica_serovar_Quebec | 189 | LFZ40_19390 | 3804579..3805148 | 1243591 | hypothetical-protein-LFZ40_19390-[Salmonella-enterica-subsp.-enterica-serovar-Quebec-str.-S-1267] |
| ECS4740322.1 | TOPRIM | GCA_008423865.1 | Salmonella_enterica_subsp._enterica_serovar_Typhimurium_var._5- | Bacteria>Pseudomonadati>Pseudomonadota>Gammaproteobacteria>Enterobacterales>Enterobacteriaceae>Salmonella>Salmonella_enterica>Salmonella_enterica_subsp._enterica>Salmonella_enterica_subsp._enterica_serovar_Typhimurium | 273 | A9R80_15945 | 80385..81206 | 1620419 | DUF4435-domain-containing-protein-[Salmonella-enterica-subsp.-enterica-serovar-Typhimurium-var.-5-] |
| EDV6600963.1 | TOPRIM | - | Salmonella_enterica_subsp._enterica_serovar_Typhimurium_var._5- | Bacteria>Pseudomonadati>Pseudomonadota>Gammaproteobacteria>Enterobacterales>Enterobacteriaceae>Salmonella>Salmonella_enterica>Salmonella_enterica_subsp._enterica>Salmonella_enterica_subsp._enterica_serovar_Typhimurium | 273 | BH535_001149 | 75730..76551 | 1620419 | DUF4435-domain-containing-protein-[Salmonella-enterica-subsp.-enterica-serovar-Typhimurium-var.-5-] |
| HCM2417895.1 | TOPRIM | GCA_025074775.1 | Salmonella_enterica_subsp._enterica_serovar_Typhimurium_var._monophasic_4,[5],12:i:- | Bacteria>Pseudomonadati>Pseudomonadota>Gammaproteobacteria>Enterobacterales>Enterobacteriaceae>Salmonella>Salmonella_enterica>Salmonella_enterica_subsp._enterica>Salmonella_enterica_subsp._enterica_serovar_Typhimurium | 273 | N3D22_002713 | 102451..103272 | 1461749 | TPA:-DUF4435-domain-containing-protein-[Salmonella-enterica-subsp.-enterica-serovar-Typhimurium-var.-monophasic-4,[5] |
| EJU7772066.1 | TOPRIM | GCA_025208445.1 | Salmonella_enterica_subsp._salamae_serovar_4,12:e,n,x:1,6 | Bacteria>Pseudomonadati>Pseudomonadota>Gammaproteobacteria>Enterobacterales>Enterobacteriaceae>Salmonella>Salmonella_enterica>Salmonella_enterica_subsp._salamae | 273 | KAB52_003011 | 287851..288672 | 2978379 | DUF4435-domain-containing-protein-[Salmonella-enterica-subsp.-salamae-serovar-4,12:e,n,x:1,6] |
| VUC90729.1 | TOPRIM | GCA_902109495.1 | Salmonella_sp._NCTC_11881 | Bacteria>Pseudomonadati>Pseudomonadota>Gammaproteobacteria>Enterobacterales>Enterobacteriaceae>Salmonella>unclassified_Salmonella | 103 | NCTC11881_05568 | 4066879..4067190 | 2583579 | putative-cytoplasmic-protein-[Salmonella-sp.-NCTC-11881] |
| WP_154209855.1 | TOPRIM | GCF_009668105.1 | Pantoea_agglomerans | Bacteria>Pseudomonadati>Pseudomonadota>Gammaproteobacteria>Enterobacterales>Erwiniaceae>Pantoea>Pantoea_agglomerans_group | 277 | GJV05_RS04765 | 294048..294881 | 549 | DUF4435-domain-containing-protein-[Pantoea-agglomerans] |
| WP_245115613.1 | TOPRIM | GCF_022966105.1 | Pantoea_agglomerans | Bacteria>Pseudomonadati>Pseudomonadota>Gammaproteobacteria>Enterobacterales>Erwiniaceae>Pantoea>Pantoea_agglomerans_group | 277 | LZ609_RS16070 | 2640923..2641756 | 549 | DUF4435-domain-containing-protein-[Pantoea-agglomerans] |
| WP_137241126.1 | TOPRIM | GCF_005233495.2 | Pantoea_agglomerans | Bacteria>Pseudomonadati>Pseudomonadota>Gammaproteobacteria>Enterobacterales>Erwiniaceae>Pantoea>Pantoea_agglomerans_group | 271 | PagCFBP13505_RS08755 | 49274..50089 | 549 | DUF4435-domain-containing-protein-[Pantoea-agglomerans] |
| WP_007890766.1 | TOPRIM | GCF_000282675.1 | Pantoea_sp._GM01 | Bacteria>Pseudomonadati>Pseudomonadota>Gammaproteobacteria>Enterobacterales>Erwiniaceae>Pantoea>unclassified_Pantoea | 271 | PMI17_RS16920 | 190779..191594 | 1144320 | DUF4435-domain-containing-protein-[Pantoea-sp.-GM01] |
| WP_222233107.1 | TOPRIM | GCF_019795975.1 | Pantoea_sp._DY-17 | Bacteria>Pseudomonadati>Pseudomonadota>Gammaproteobacteria>Enterobacterales>Erwiniaceae>Pantoea>unclassified_Pantoea | 186 | K5Y33_RS01210 | 246804..247364 | 2871490 | DUF4435-domain-containing-protein-[Pantoea-sp.-DY-17] |
| WP_226228489.1 | TOPRIM | - | Hafnia_paralvei | Bacteria>Pseudomonadati>Pseudomonadota>Gammaproteobacteria>Enterobacterales>Hafniaceae>Hafnia | 271 | LA328_RS17875 | 3841712..3842527 | 546367 | DUF4435-domain-containing-protein-[Hafnia-paralvei] |
| WP_241543891.1 | TOPRIM | GCF_963899445.1 | Moellerella_wisconsensis | Bacteria>Pseudomonadati>Pseudomonadota>Gammaproteobacteria>Enterobacterales>Morganellaceae>Moellerella | 249 | MNY64_RS16765 | 125677..126426 | 158849 | hypothetical-protein-[Moellerella-wisconsensis] |
| WP_209036128.1 | TOPRIM | GCF_017693395.1 | Proteus_mirabilis | Bacteria>Pseudomonadati>Pseudomonadota>Gammaproteobacteria>Enterobacterales>Morganellaceae>Proteus | 245 | IGB04_RS13140 | 87474..88211 | 584 | hypothetical-protein-[Proteus-mirabilis] |
| HAT2164889.1 | TOPRIM | GCA_015896365.1 | Proteus_mirabilis | Bacteria>Pseudomonadati>Pseudomonadota>Gammaproteobacteria>Enterobacterales>Morganellaceae>Proteus | 199 | I8052_002914 | 20311..20910 | 584 | DUF4276-family-protein-[Proteus-mirabilis] |
| MBI6417156.1 | TOPRIM | GCA_016238675.1 | Proteus_mirabilis | Bacteria>Pseudomonadati>Pseudomonadota>Gammaproteobacteria>Enterobacterales>Morganellaceae>Proteus | 199 | JEP37_15635 | 62009..62608 | 584 | hypothetical-protein-[Proteus-mirabilis] |
| WP_175403158.1 | TOPRIM | GCF_005157325.1 | Providencia_heimbachae | Bacteria>Pseudomonadati>Pseudomonadota>Gammaproteobacteria>Enterobacterales>Morganellaceae>Providencia | 275 | C9446_RS17905 | 3862088..3862915 | 333962 | DUF4435-domain-containing-protein-[Providencia-heimbachae] |
| WP_045967683.1 | TOPRIM | GCF_963866745.1 | Xenorhabdus_doucetiae | Bacteria>Pseudomonadati>Pseudomonadota>Gammaproteobacteria>Enterobacterales>Morganellaceae>Xenorhabdus | 191 | LY16_RS12970 | 18801..19376 | 351671 | hypothetical-protein-[Xenorhabdus-doucetiae] |
| WP_103160992.1 | TOPRIM | GCF_002904045.1 | Pectobacterium_odoriferum | Bacteria>Pseudomonadati>Pseudomonadota>Gammaproteobacteria>Enterobacterales>Pectobacteriaceae>Pectobacterium | 277 | BVY05_RS19625 | 82346..83179 | 78398 | DUF4435-domain-containing-protein-[Pectobacterium-odoriferum] |
| WP_116155990.1 | TOPRIM | GCF_003382585.2 | Pectobacterium_aquaticum | Bacteria>Pseudomonadati>Pseudomonadota>Gammaproteobacteria>Enterobacterales>Pectobacteriaceae>Pectobacterium | 277 | DMB81_RS10750 | 13325..14158 | 2204145 | DUF4435-domain-containing-protein-[Pectobacterium-aquaticum] |
| WP_181847882.1 | TOPRIM | GCF_013913645.1 | Pectobacterium_aroidearum | Bacteria>Pseudomonadati>Pseudomonadota>Gammaproteobacteria>Enterobacterales>Pectobacteriaceae>Pectobacterium | 275 | H2Y54_RS11270 | 231022..231849 | 1201031 | DUF4435-domain-containing-protein-[Pectobacterium-aroidearum] |
| WP_005974425.1 | TOPRIM | GCF_000291725.1 | Pectobacterium_wasabiae | Bacteria>Pseudomonadati>Pseudomonadota>Gammaproteobacteria>Enterobacterales>Pectobacteriaceae>Pectobacterium | 271 | Y17_RS17675 | 16711..17526 | 55208 | DUF4435-domain-containing-protein-[Pectobacterium-wasabiae] |
| WP_103199121.1 | TOPRIM | GCF_002904085.1 | Pectobacterium_odoriferum | Bacteria>Pseudomonadati>Pseudomonadota>Gammaproteobacteria>Enterobacterales>Pectobacteriaceae>Pectobacterium | 271 | BV921_RS15375 | 108366..109181 | 78398 | DUF4435-domain-containing-protein-[Pectobacterium-odoriferum] |
| WP_268198599.1 | TOPRIM | GCF_026723725.1 | Pectobacterium_brasiliense | Bacteria>Pseudomonadati>Pseudomonadota>Gammaproteobacteria>Enterobacterales>Pectobacteriaceae>Pectobacterium | 271 | OWC53_RS08215 | 1776792..1777607 | 180957 | hypothetical-protein-[Pectobacterium-brasiliense] |
| GKX39592.1 | TOPRIM | GCA_024343445.1 | Pectobacterium_carotovorum_subsp._carotovorum | Bacteria>Pseudomonadati>Pseudomonadota>Gammaproteobacteria>Enterobacterales>Pectobacteriaceae>Pectobacterium>Pectobacterium_carotovorum | 271 | SOASR014_33310 | 119310..120125 | 555 | hypothetical-protein-SOASR014_33310-[Pectobacterium-carotovorum-subsp.-carotovorum] |
| WP_112197430.1 | TOPRIM | GCF_003263525.1 | Rahnella_sp._NRRL_B-41462 | Bacteria>Pseudomonadati>Pseudomonadota>Gammaproteobacteria>Enterobacterales>Yersiniaceae>Rahnella>unclassified_Rahnella | 277 | UC14_RS06685 | 39908..40741 | 1610579 | DUF4435-domain-containing-protein-[Rahnella-sp.-NRRL-B-41462] |
| HBE9089663.1 | TOPRIM | GCA_018627075.1 | Serratia_fonticola | Bacteria>Pseudomonadati>Pseudomonadota>Gammaproteobacteria>Enterobacterales>Yersiniaceae>Serratia | 277 | KNG84_001687 | 42995..43828 | 47917 | DUF4435-domain-containing-protein-[Serratia-fonticola] |
| WP_195869523.1 | TOPRIM | - | Serratia_liquefaciens | Bacteria>Pseudomonadati>Pseudomonadota>Gammaproteobacteria>Enterobacterales>Yersiniaceae>Serratia | 189 | I1A46_RS14710 | 208373..208942 | 614 | hypothetical-protein-[Serratia-liquefaciens] |
| WP_053011790.1 | TOPRIM | GCF_001244675.1 | Yersinia_intermedia | Bacteria>Pseudomonadati>Pseudomonadota>Gammaproteobacteria>Enterobacterales>Yersiniaceae>Yersinia | 275 | ALC37_RS11590 | 468329..469156 | 631 | DUF4435-domain-containing-protein-[Yersinia-intermedia] |
| WP_050124853.1 | TOPRIM | GCF_001115185.1 | Yersinia_aleksiciae | Bacteria>Pseudomonadati>Pseudomonadota>Gammaproteobacteria>Enterobacterales>Yersiniaceae>Yersinia | 271 | AEB18_RS00880 | 225044..225859 | 263819 | DUF4435-domain-containing-protein-[Yersinia-aleksiciae] |
| WP_145532659.1 | TOPRIM | GCF_902170345.1 | Yersinia_alsatica | Bacteria>Pseudomonadati>Pseudomonadota>Gammaproteobacteria>Enterobacterales>Yersiniaceae>Yersinia | 271 | FTB07_RS01255 | 31043..31858 | 2890317 | DUF4435-domain-containing-protein-[Yersinia-alsatica] |
| WP_145580445.1 | TOPRIM | GCF_902170795.1 | Yersinia_mollaretii | Bacteria>Pseudomonadati>Pseudomonadota>Gammaproteobacteria>Enterobacterales>Yersiniaceae>Yersinia | 271 | FTB33_RS16520 | 15131..15946 | 33060 | DUF4435-domain-containing-protein-[Yersinia-mollaretii] |
| WP_038251186.1 | TOPRIM | GCF_000173755.1 | Yersinia_ruckeri | Bacteria>Pseudomonadati>Pseudomonadota>Gammaproteobacteria>Enterobacterales>Yersiniaceae>Yersinia | 189 | YRUCK0001_RS14095 | 23754..24323 | 29486 | hypothetical-protein-[Yersinia-ruckeri] |
| QDW34852.1 | TOPRIM | GCA_005937895.2 | Yersinia_sp._KBS0713 | Bacteria>Pseudomonadati>Pseudomonadota>Gammaproteobacteria>Enterobacterales>Yersiniaceae>Yersinia>unclassified_Yersinia_(in:_enterobacteria) | 277 | FFE93_018525 | 3902026..3902859 | 1179669 | DUF4435-domain-containing-protein-[Yersinia-sp.-KBS0713] |
| BAL53120.1 | TOPRIM | - | uncultured_Gammaproteobacteria_bacterium | Bacteria>Pseudomonadati>Pseudomonadota>Gammaproteobacteria>environmental_samples | 175 | HGMM_F04D09C17 | 15299..15826 | 86473 | hypothetical-conserved-protein-[uncultured-Gammaproteobacteria-bacterium] |
| MBK8754676.1 | TOPRIM | GCA_016710535.1 | Candidatus_Competibacteraceae_bacterium | Bacteria>Pseudomonadati>Pseudomonadota>Gammaproteobacteria>Gammaproteobacteria_incertae_sedis>Candidatus_Competibacteraceae>unclassified_Candidatus_Competibacteraceae | 206 | IPL99_24905 | 44238..44858 | 2053538 | DUF4276-family-protein-[Candidatus-Competibacteraceae-bacterium] |
| MBK8754770.1 | TOPRIM | GCA_016710535.1 | Candidatus_Competibacteraceae_bacterium | Bacteria>Pseudomonadati>Pseudomonadota>Gammaproteobacteria>Gammaproteobacteria_incertae_sedis>Candidatus_Competibacteraceae>unclassified_Candidatus_Competibacteraceae | 200 | IPL99_25410 | 32203..32805 | 2053538 | DUF4276-family-protein-[Candidatus-Competibacteraceae-bacterium] |
| WP_248566601.1 | TOPRIM | GCF_023195885.1 | Methylonatrum_kenyense | Bacteria>Pseudomonadati>Pseudomonadota>Gammaproteobacteria>Gammaproteobacteria_incertae_sedis>Methylonatrum | 217 | M0534_RS01325 | 280474..281127 | 455253 | DUF4276-family-protein-[Methylonatrum-kenyense] |
| MAZ66824.1 | TOPRIM | GCA_002707725.1 | Kangiellaceae_bacterium | Bacteria>Pseudomonadati>Pseudomonadota>Gammaproteobacteria>Kangiellales>Kangiellaceae>unclassified_Kangiellaceae | 200 | CMF25_06915 | 5851..6453 | 2026751 | hypothetical-protein-[Kangiellaceae-bacterium] |
| WP_054661692.1 | TOPRIM | GCF_002739955.1 | Stenotrophomonas | Bacteria>Pseudomonadati>Pseudomonadota>Gammaproteobacteria>Lysobacterales>Lysobacteraceae | 217 | BRB81_RS09935 | 111434..112087 | 40323 | MULTISPECIES:-DUF4276-family-protein-[Stenotrophomonas] |
| PZO64464.1 | TOPRIM | GCA_003240695.1 | Pseudoxanthomonas_suwonensis | Bacteria>Pseudomonadati>Pseudomonadota>Gammaproteobacteria>Lysobacterales>Lysobacteraceae>Pseudoxanthomonas | 211 | DI635_05300 | 3522..4157 | 314722 | hypothetical-protein-DI635_05300-[Pseudoxanthomonas-suwonensis] |
| ALJ26656.1 | TOPRIM | GCA_001314305.1 | Stenotrophomonas_acidaminiphila | Bacteria>Pseudomonadati>Pseudomonadota>Gammaproteobacteria>Lysobacterales>Lysobacteraceae>Stenotrophomonas | 217 | AOT14_01950 | 239076..239729 | 128780 | enolase-[Stenotrophomonas-acidaminiphila] |
| MTI74443.1 | TOPRIM | GCA_009711865.1 | Stenotrophomonas_sp. | Bacteria>Pseudomonadati>Pseudomonadota>Gammaproteobacteria>Lysobacterales>Lysobacteraceae>Stenotrophomonas>unclassified_Stenotrophomonas | 220 | FH747_12430 | 111600..112262 | 69392 | RloB-domain-containing-protein-[Stenotrophomonas-sp.] |
| WP_230434264.1 | TOPRIM | GCF_020990895.1 | Xanthomonas_melonis | Bacteria>Pseudomonadati>Pseudomonadota>Gammaproteobacteria>Lysobacterales>Lysobacteraceae>Xanthomonas | 209 | JWH05_RS03155 | 13099..13728 | 56456 | DUF4276-family-protein-[Xanthomonas-melonis] |
| WP_259393302.1 | TOPRIM | - | Dokdonella_fugitiva | Bacteria>Pseudomonadati>Pseudomonadota>Gammaproteobacteria>Lysobacterales>Rhodanobacteraceae>Dokdonella | 187 | FHW13_RS05510 | 303474..304037 | 328517 | DUF4276-family-protein-[Dokdonella-fugitiva] |
| WP_262965893.1 | TOPRIM | GCF_025583945.1 | Methylobacter_psychrophilus | Bacteria>Pseudomonadati>Pseudomonadota>Gammaproteobacteria>Methylococcales>Methylococcaceae>Methylobacter | 225 | N7974_RS12255 | 23519..24196 | 96941 | hypothetical-protein-[Methylobacter-psychrophilus] |
| WP_031438514.1 | TOPRIM | GCF_000733835.1 | Methylobacter_tundripaludum | Bacteria>Pseudomonadati>Pseudomonadota>Gammaproteobacteria>Methylococcales>Methylococcaceae>Methylobacter | 224 | T449_RS0121500 | 4648531..4649205 | 173365 | hypothetical-protein-[Methylobacter-tundripaludum] |
| WP_218800687.1 | TOPRIM | GCF_019285575.1 | Methylococcus_capsulatus | Bacteria>Pseudomonadati>Pseudomonadota>Gammaproteobacteria>Methylococcales>Methylococcaceae>Methylococcus | 209 | KW113_RS00565 | 121055..121684 | 414 | DUF4276-family-protein-[Methylococcus-capsulatus] |
| WP_218796634.1 | TOPRIM | GCF_019285555.1 | Methylococcus_capsulatus | Bacteria>Pseudomonadati>Pseudomonadota>Gammaproteobacteria>Methylococcales>Methylococcaceae>Methylococcus | 195 | KW114_RS08325 | 1837595..1838182 | 414 | DUF4276-family-protein-[Methylococcus-capsulatus] |
| WP_255189602.1 | TOPRIM | GCF_024360925.2 | Methylomonas_rapida | Bacteria>Pseudomonadati>Pseudomonadota>Gammaproteobacteria>Methylococcales>Methylococcaceae>Methylomonas | 274 | NM686_RS20140 | 4255687..4256511 | 2963939 | DUF4435-domain-containing-protein-[Methylomonas-rapida] |
| WP_255188310.1 | TOPRIM | GCF_024360925.2 | Methylomonas_rapida | Bacteria>Pseudomonadati>Pseudomonadota>Gammaproteobacteria>Methylococcales>Methylococcaceae>Methylomonas | 244 | NM686_RS13210 | 2819088..2819822 | 2963939 | hypothetical-protein-[Methylomonas-rapida] |
| WP_255187196.1 | TOPRIM | GCF_024360925.2 | Methylomonas_rapida | Bacteria>Pseudomonadati>Pseudomonadota>Gammaproteobacteria>Methylococcales>Methylococcaceae>Methylomonas | 223 | NM686_RS07170 | 1513925..1514596 | 2963939 | DUF4276-family-protein-[Methylomonas-rapida] |
| MBS3964852.1 | TOPRIM | GCA_018333915.1 | Methylomonas_sp. | Bacteria>Pseudomonadati>Pseudomonadota>Gammaproteobacteria>Methylococcales>Methylococcaceae>Methylomonas>unclassified_Methylomonas | 272 | KGZ80_10235 | 15622..16440 | 418 | DUF4435-domain-containing-protein-[Methylomonas-sp.] |
| MBE0471465.1 | TOPRIM | GCA_014859465.1 | Methyloprofundus_sp. | Bacteria>Pseudomonadati>Pseudomonadota>Gammaproteobacteria>Methylococcales>Methylococcaceae>Methyloprofundus>unclassified_Methyloprofundus | 168 | IBX55_18390 | 2008..2514 | 2020875 | hypothetical-protein-[Methyloprofundus-sp.] |
| NOT11400.1 | TOPRIM | GCA_013140705.1 | Methylococcaceae_bacterium | Bacteria>Pseudomonadati>Pseudomonadota>Gammaproteobacteria>Methylococcales>Methylococcaceae>unclassified_Methylococcaceae | 225 | HOP23_06150 | 42128..42805 | 1933926 | RloB-domain-containing-protein-[Methylococcaceae-bacterium] |
| NOT11403.1 | TOPRIM | GCA_013140705.1 | Methylococcaceae_bacterium | Bacteria>Pseudomonadati>Pseudomonadota>Gammaproteobacteria>Methylococcales>Methylococcaceae>unclassified_Methylococcaceae | 188 | HOP23_06165 | 46937..47503 | 1933926 | DUF4276-family-protein-[Methylococcaceae-bacterium] |
| WP_087528531.1 | TOPRIM | GCF_002165305.2 | Acinetobacter | Bacteria>Pseudomonadati>Pseudomonadota>Gammaproteobacteria>Moraxellales>Moraxellaceae | 245 | CDG62_RS00800 | 29932..30669 | 469 | hypothetical-protein-[Acinetobacter-sp.-WCHA55] |
| WP_108675354.1 | TOPRIM | GCF_003072585.2 | Acinetobacter | Bacteria>Pseudomonadati>Pseudomonadota>Gammaproteobacteria>Moraxellales>Moraxellaceae | 245 | C0119_RS05590 | 1125568..1126305 | 469 | MULTISPECIES:-hypothetical-protein-[Acinetobacter] |
| WP_109590313.1 | TOPRIM | GCF_904845945.1 | Psychrobacter | Bacteria>Pseudomonadati>Pseudomonadota>Gammaproteobacteria>Moraxellales>Moraxellaceae | 169 | KF936_RS05010 | 117539..118048 | 497 | MULTISPECIES:-hypothetical-protein-[Psychrobacter] |
| WP_204509372.1 | TOPRIM | GCF_003051055.1 | Agitococcus_lubricus | Bacteria>Pseudomonadati>Pseudomonadota>Gammaproteobacteria>Moraxellales>Moraxellaceae>Agitococcus | 136 | C8N29_RS17075 | 14639..15049 | 1077255 | hypothetical-protein-[Agitococcus-lubricus] |
| WP_201558736.1 | TOPRIM | - | Psychrobacter_immobilis | Bacteria>Pseudomonadati>Pseudomonadota>Gammaproteobacteria>Moraxellales>Moraxellaceae>Psychrobacter | 172 | JMW74_RS07745 | 30152..30670 | 498 | hypothetical-protein-[Psychrobacter-immobilis] |
| OLF41366.1 | TOPRIM | GCA_001921755.1 | Psychrobacter_sp._Rd_27.2 | Bacteria>Pseudomonadati>Pseudomonadota>Gammaproteobacteria>Moraxellales>Moraxellaceae>Psychrobacter>unclassified_Psychrobacter | 228 | BTV99_04805 | 272907..273593 | 1926479 | hypothetical-protein-BTV99_04805-[Psychrobacter-sp.-Rd-27.2] |

| Protein-ID | Domain | Assembly-ID | Strain | Taxonomy | Length | Locus-Tag | Genomic-coordinates | TaxID | Fasta-defline |
|---|---|---|---|---|---|---|---|---|---|
| WP_075101227.1 | TOPRIM | GCF_052263635.1 | Psychrobacter_sp._Rd_27.2 | Bacteria>Pseudomonadati>Pseudomonadota>Gammaproteobacteria>Moraxellales>Moraxellaceae>Psychrobacter>unclassified_Psychrobacter | 228 | BTV99_RS04800 | 272907..273593 | 1926479 | DUF4276-family-protein-[Psychrobacter-sp.-Rd-27.2] |
| MCP5177612.1 | TOPRIM | GCA_024234655.1 | Moraxellaceae_bacterium | Bacteria>Pseudomonadati>Pseudomonadota>Gammaproteobacteria>Moraxellales>Moraxellaceae>unclassified_Moraxellaceae | 229 | H6997_08685 | 42024..42713 | 1889775 | DUF4276-family-protein-[Moraxellaceae-bacterium] |
| TQC95075.1 | TOPRIM | GCA_006546595.1 | Moraxellaceae_bacterium_AER2_44_116 | Bacteria>Pseudomonadati>Pseudomonadota>Gammaproteobacteria>Moraxellales>Moraxellaceae>unclassified_Moraxellaceae | 229 | FK216_14935 | 6690..7379 | 2591038 | DUF4276-family-protein-[Moraxellaceae-bacterium-AER2_44_116] |
| WP_093286150.1 | TOPRIM | GCF_900111015.1 | Solimonas_aquatica | Bacteria>Pseudomonadati>Pseudomonadota>Gammaproteobacteria>Nevskiales>Nevskiaceae>Solimonas | 224 | BMY12_RS11545 | 202751..203425 | 489703 | DUF4276-family-protein-[Solimonas-aquatica] |
| WP_062362840.1 | TOPRIM | GCF_011399095.1 | Halomonadaceae | Bacteria>Pseudomonadati>Pseudomonadota>Gammaproteobacteria>Oceanospirillales | 274 | HHSLTHF2_RS10280 | 2145983..2146807 | 28256 | MULTISPECIES:-DUF4435-domain-containing-protein-[Halomonas] |
| WP_052475611.1 | TOPRIM | GCF_900099955.1 | Halomonadaceae | Bacteria>Pseudomonadati>Pseudomonadota>Gammaproteobacteria>Oceanospirillales | 240 | BLR10_RS07675 | 63181..63903 | 28256 | MULTISPECIES:-hypothetical-protein-[Halomonas] |
| HIK75725.1 | TOPRIM | GCA_015660495.1 | Alcanivorax_sp. | Bacteria>Pseudomonadati>Pseudomonadota>Gammaproteobacteria>Oceanospirillales>Alcanivoracaceae>Alcanivorax>unclassified_Alcanivorax | 225 | EYG00_13650 | 6364..7041 | 1872427 | DUF4276-family-protein-[Alcanivorax-sp.] |
| MAD69923.1 | TOPRIM | GCA_002685555.1 | Alcanivorax_sp. | Bacteria>Pseudomonadati>Pseudomonadota>Gammaproteobacteria>Oceanospirillales>Alcanivoracaceae>Alcanivorax>unclassified_Alcanivorax | 225 | CL538_05300 | 41162..41839 | 1872427 | hypothetical-protein-[Alcanivorax-sp.] |
| MBE8233361.1 | TOPRIM | GCA_015163765.1 | Endozoicomonadaceae_bacterium | Bacteria>Pseudomonadati>Pseudomonadota>Gammaproteobacteria>Oceanospirillales>Endozoicomonadaceae>unclassified_Endozoicomonadaceae | 239 | HAW67_06455 | 10418..11137 | 2590577 | hypothetical-protein-[Endozoicomonadaceae-bacterium] |
| WP_041598694.1 | TOPRIM | GCF_000012985.1 | Hahella_chejuensis | Bacteria>Pseudomonadati>Pseudomonadota>Gammaproteobacteria>Oceanospirillales>Hahellaceae>Hahella | 206 | HCH_RS15590 | 3600878..3601498 | 158327 | DUF4276-family-protein-[Hahella-chejuensis] |
| WP_101147213.1 | TOPRIM | GCF_037070215.1 | unclassified_Halomonas | Bacteria>Pseudomonadati>Pseudomonadota>Gammaproteobacteria>Oceanospirillales>Halomonadaceae>Halomonas | 272 | CXF87_RS15380 | 112921..113739 | 2609666 | DUF4435-domain-containing-protein-[Halomonas-sp.-MES3-P3E] |
| WP_275437441.1 | TOPRIM | - | unclassified_Halomonas | Bacteria>Pseudomonadati>Pseudomonadota>Gammaproteobacteria>Oceanospirillales>Halomonadaceae>Halomonas | 222 | MHM35_RS10440 | 131038..131706 | 2609666 | MULTISPECIES:-DUF4276-family-protein-[unclassified-Halomonas] |
| WP_169956454.1 | TOPRIM | GCF_013000985.1 | Halomonas_sp._PGE1 | Bacteria>Pseudomonadati>Pseudomonadota>Gammaproteobacteria>Oceanospirillales>Halomonadaceae>Halomonas>unclassified_Halomonas | 225 | HIR79_RS02130 | 431842..432519 | 2730360 | DUF4276-family-protein-[Halomonas-sp.-PGE1] |
| WP_218314910.1 | TOPRIM | GCF_019211725.1 | Halomonas_sp._18071143 | Bacteria>Pseudomonadati>Pseudomonadota>Gammaproteobacteria>Oceanospirillales>Halomonadaceae>Halomonas>unclassified_Halomonas | 224 | KVY02_RS07745 | 1672034..1672708 | 2855441 | hypothetical-protein-[Halomonas-sp.-18071143] |
| WP_231664590.1 | TOPRIM | GCF_021151495.1 | Halomonas_sp._IOP_31 | Bacteria>Pseudomonadati>Pseudomonadota>Gammaproteobacteria>Oceanospirillales>Halomonadaceae>Halomonas>unclassified_Halomonas | 217 | K7I12_RS15000 | 16130..16783 | 2876584 | DUF4276-family-protein-[Halomonas-sp.-IOP_31] |
| WP_171051221.1 | TOPRIM | GCF_005937175.1 | Halomonas_sp._ATBC28 | Bacteria>Pseudomonadati>Pseudomonadota>Gammaproteobacteria>Oceanospirillales>Halomonadaceae>Halomonas>unclassified_Halomonas | 197 | E0L35_RS22990 | 7954..8547 | 2545264 | DUF4276-family-protein-[Halomonas-sp.-ATBC28] |
| WP_139788404.1 | TOPRIM | - | Halomonas_sp._BC1 | Bacteria>Pseudomonadati>Pseudomonadota>Gammaproteobacteria>Oceanospirillales>Halomonadaceae>Halomonas>unclassified_Halomonas | 123 | B7L72_RS18045 | 30065..30436 | 1670448 | hypothetical-protein-[Halomonas-sp.-BC1] |
| WP_079550438.1 | TOPRIM | GCF_900142895.1 | Vreelandella_subglaciescola | Bacteria>Pseudomonadati>Pseudomonadota>Gammaproteobacteria>Oceanospirillales>Halomonadaceae>Vreelandella | 250 | B5495_RS00625 | 141713..142465 | 29571 | hypothetical-protein-[Halomonas-subglaciescola] |
| WP_193059744.1 | TOPRIM | GCF_052263255.1 | Vreelandella_alkaliphila | Bacteria>Pseudomonadati>Pseudomonadota>Gammaproteobacteria>Oceanospirillales>Halomonadaceae>Vreelandella | 248 | INT28_RS17810 | 14169..14915 | 272774 | MULTISPECIES:-hypothetical-protein-[unclassified-Halomonas] |
| WP_027853085.1 | TOPRIM | GCF_000428985.1 | Marinobacterium_litorale | Bacteria>Pseudomonadati>Pseudomonadota>Gammaproteobacteria>Oceanospirillales>Oceanospirillaceae>Marinobacterium | 225 | G403_RS0105365 | 184712..185389 | 404770 | DUF4276-family-protein-[Marinobacterium-litorale] |
| WP_206838937.1 | TOPRIM | GCF_017310015.1 | Marinobacterium_iners | Bacteria>Pseudomonadati>Pseudomonadota>Gammaproteobacteria>Oceanospirillales>Oceanospirillaceae>Marinobacterium | 225 | CFI10_RS02425 | 490002..490679 | 48076 | DUF4276-family-protein-[Marinobacterium-iners] |
| WP_254579409.1 | TOPRIM | GCF_024267675.1 | Marinobacterium_sedimentorum | Bacteria>Pseudomonadati>Pseudomonadota>Gammaproteobacteria>Oceanospirillales>Oceanospirillaceae>Marinobacterium | 204 | MRS74_RS06195 | 159291..159905 | 2927804 | DUF4276-family-protein-[Marinobacterium-sedimentorum] |
| WP_226752734.1 | TOPRIM | GCF_020532605.1 | Marinomonas_algarum | Bacteria>Pseudomonadati>Pseudomonadota>Gammaproteobacteria>Oceanospirillales>Oceanospirillaceae>Marinomonas | 252 | LG368_RS00215 | 44490..45248 | 2883105 | hypothetical-protein-[Marinomonas-algarum] |
| WP_067206353.1 | TOPRIM | GCF_900089755.1 | Marinomonas_aquimarina | Bacteria>Pseudomonadati>Pseudomonadota>Gammaproteobacteria>Oceanospirillales>Oceanospirillaceae>Marinomonas | 245 | MAQ5080_RS17365 | 82005..82742 | 295068 | hypothetical-protein-[Marinomonas-aquimarina] |
| WP_036514527.1 | TOPRIM | GCF_000585235.1 | Nitrincola_nitratireducens | Bacteria>Pseudomonadati>Pseudomonadota>Gammaproteobacteria>Oceanospirillales>Oceanospirillaceae>Nitrincola | 215 | D791_RS17630 | 5270..5917 | 1229521 | DUF4276-family-protein-[Nitrincola-nitratireducens] |
| QEW05878.1 | TOPRIM | GCA_008727795.1 | Nitrincola_iocasae | Bacteria>Pseudomonadati>Pseudomonadota>Gammaproteobacteria>Oceanospirillales>Oceanospirillaceae>Nitrincola | 204 | F5I99_04900 | 1041348..1041962 | 2614693 | DUF4276-family-protein-[Nitrincola-iocasae] |
| WP_151053917.1 | TOPRIM | GCF_008727795.1 | Nitrincola_iocasae | Bacteria>Pseudomonadati>Pseudomonadota>Gammaproteobacteria>Oceanospirillales>Oceanospirillaceae>Nitrincola | 204 | F5I99_RS04900 | 1041348..1041962 | 2614693 | DUF4276-family-protein-[Nitrincola-iocasae] |
| WP_260997647.1 | TOPRIM | GCF_025345565.1 | Thalassolituus_hydrocarboniclasticus | Bacteria>Pseudomonadati>Pseudomonadota>Gammaproteobacteria>Oceanospirillales>Oceanospirillaceae>Thalassolituus | 212 | HUF19_RS16695 | 3716061..3716699 | 2742796 | DUF4276-family-protein-[Thalassolituus-hydrocarbonoclasticus] |
| MBM94537.1 | TOPRIM | GCA_002723385.1 | Oceanospirillaceae_bacterium | Bacteria>Pseudomonadati>Pseudomonadota>Gammaproteobacteria>Oceanospirillales>Oceanospirillaceae>unclassified_Oceanospirillaceae | 252 | CMI09_01625 | 45908..46666 | 1899355 | hypothetical-protein-[Oceanospirillaceae-bacterium] |
| MBM94534.1 | TOPRIM | GCA_002723385.1 | Oceanospirillaceae_bacterium | Bacteria>Pseudomonadati>Pseudomonadota>Gammaproteobacteria>Oceanospirillales>Oceanospirillaceae>unclassified_Oceanospirillaceae | 199 | CMI09_01610 | 40465..41064 | 1899355 | hypothetical-protein-[Oceanospirillaceae-bacterium] |
| WP_116688146.1 | TOPRIM | GCF_963878705.1 | Pelagibaculum_spongiae | Bacteria>Pseudomonadati>Pseudomonadota>Gammaproteobacteria>Oceanospirillales>Oceanospirillales_incertae_sedis>Pelagibaculum | 242 | DC094_RS16070 | 170555..171283 | 2080658 | hypothetical-protein-[Pelagibaculum-spongiae] |
| WP_116688144.1 | TOPRIM | GCF_963878705.1 | Pelagibaculum_spongiae | Bacteria>Pseudomonadati>Pseudomonadota>Gammaproteobacteria>Oceanospirillales>Oceanospirillales_incertae_sedis>Pelagibaculum | 198 | DC094_RS16060 | 168744..169340 | 2080658 | hypothetical-protein-[Pelagibaculum-spongiae] |
| MBR9867108.1 | TOPRIM | GCA_018223265.1 | Oceanospirillales_bacterium | Bacteria>Pseudomonadati>Pseudomonadota>Gammaproteobacteria>Oceanospirillales>unclassified_Oceanospirillales | 199 | GYB18_08395 | 14000..14599 | 1914300 | DUF4276-family-protein-[Oceanospirillales-bacterium] |
| WP_023127683.1 | TOPRIM | GCF_905479555.1 | Pseudomonadaceae | Bacteria>Pseudomonadati>Pseudomonadota>Gammaproteobacteria>Pseudomonadales | 227 | ACP89_RS12350 | 15816..16499 | 135621 | MULTISPECIES:-DUF4276-family-protein-[Pseudomonas-aeruginosa-group] |
| WP_092021236.1 | TOPRIM | GCF_900114775.1 | Marinobacter_zhejiangensis | Bacteria>Pseudomonadati>Pseudomonadota>Gammaproteobacteria>Pseudomonadales>Marinobacteraceae>Marinobacter | 283 | BMZ67_RS07000 | 353221..354072 | 488535 | hypothetical-protein-[Marinobacter-zhejiangensis] |
| WP_139214341.1 | TOPRIM | GCF_900114775.1 | Marinobacter_zhejiangensis | Bacteria>Pseudomonadati>Pseudomonadota>Gammaproteobacteria>Pseudomonadales>Marinobacteraceae>Marinobacter | 247 | BMZ67_RS06975 | 344742..345485 | 488535 | hypothetical-protein-[Marinobacter-zhejiangensis] |
| BBJ05058.1 | TOPRIM | - | Marinobacter_nauticus | Bacteria>Pseudomonadati>Pseudomonadota>Gammaproteobacteria>Pseudomonadales>Marinobacteraceae>Marinobacter | 225 | YBY_29070 | 3198969..3199646 | 2743 | hypothetical-protein-YBY_29070-[Marinobacter-nauticus] |
| WP_011784107.1 | TOPRIM | GCF_000015365.1 | Marinobacter_nauticus | Bacteria>Pseudomonadati>Pseudomonadota>Gammaproteobacteria>Pseudomonadales>Marinobacteraceae>Marinobacter | 225 | MAQU_RS02835 | 642481..643158 | 2743 | DUF4276-family-protein-[Marinobacter-nauticus] |
| WP_104416049.1 | TOPRIM | GCF_002934485.1 | Marinobacter_persicus | Bacteria>Pseudomonadati>Pseudomonadota>Gammaproteobacteria>Pseudomonadales>Marinobacteraceae>Marinobacter | 225 | BY455_RS09010 | 77340..78017 | 930118 | DUF4276-family-protein-[Marinobacter-persicus] |
| EHJ03981.1 | TOPRIM | GCA_000235625.2 | Marinobacter_manganoxydans_MnI7-9 | Bacteria>Pseudomonadati>Pseudomonadota>Gammaproteobacteria>Pseudomonadales>Marinobacteraceae>Marinobacter>Marinobacter_manganoxydans | 176 | KYE_13710 | 52243..52769 | 1094979 | hypothetical-protein-KYE_13710,-partial-[Marinobacter-manganoxydans-MnI7-9] |
| MBC7192645.1 | TOPRIM | GCA_014359475.1 | Marinobacter_sp. | Bacteria>Pseudomonadati>Pseudomonadota>Gammaproteobacteria>Pseudomonadales>Marinobacteraceae>Marinobacter>unclassified_Marinobacter | 217 | H5U31_09865 | 12327..12980 | 50741 | DUF4276-family-protein-[Marinobacter-sp.] |
| WP_094987830.1 | TOPRIM | GCF_918697895.1 | Pseudomonas | Bacteria>Pseudomonadati>Pseudomonadota>Gammaproteobacteria>Pseudomonadales>Pseudomonadaceae | 227 | LQ224_RS08505 | 120251..120934 | 286 | MULTISPECIES:-DUF4276-family-protein-[Pseudomonas] |
| WP_152390154.1 | TOPRIM | GCF_009363155.1 | Azotobacter_salinestris | Bacteria>Pseudomonadati>Pseudomonadota>Gammaproteobacteria>Pseudomonadales>Pseudomonadaceae>Azotobacter_group>Azotobacter | 244 | GCU53_RS24405 | 143454..144188 | 69964 | hypothetical-protein-[Azotobacter-salinestris] |
| WP_161892248.1 | TOPRIM | GCF_009932375.1 | Pseudomonas_juntendi | Bacteria>Pseudomonadati>Pseudomonadota>Gammaproteobacteria>Pseudomonadales>Pseudomonadaceae>Pseudomonas | 227 | GYA94_RS10100 | 84702..85385 | 2666183 | DUF4276-family-protein-[Pseudomonas-juntendi] |
| WP_150549233.1 | TOPRIM | GCF_900682675.2 | Pseudomonas_marincola | Bacteria>Pseudomonadati>Pseudomonadota>Gammaproteobacteria>Pseudomonadales>Pseudomonadaceae>Pseudomonas | 226 | PMYSY11_RS11195 | 2409085..2409765 | 437900 | DUF4276-family-protein-[Pseudomonas-marincola] |
| MBA1269667.1 | TOPRIM | GCA_013522615.1 | Pseudomonas_carnis | Bacteria>Pseudomonadati>Pseudomonadota>Gammaproteobacteria>Pseudomonadales>Pseudomonadaceae>Pseudomonas | 209 | G7029_19565 | 128805..129434 | 2487355 | DUF4276-family-protein-[Pseudomonas-carnis] |
| WP_181127555.1 | TOPRIM | GCF_013522615.1 | Pseudomonas_carnis | Bacteria>Pseudomonadati>Pseudomonadota>Gammaproteobacteria>Pseudomonadales>Pseudomonadaceae>Pseudomonas | 209 | G7029_RS19475 | 128805..129434 | 2487355 | DUF4276-family-protein-[Pseudomonas-carnis] |
| HBO1619175.1 | TOPRIM | GCA_021598085.1 | Pseudomonas_aeruginosa | Bacteria>Pseudomonadati>Pseudomonadota>Gammaproteobacteria>Pseudomonadales>Pseudomonadaceae>Pseudomonas>Pseudomonas_aeruginosa_group | 227 | L4F62_005708 | 9922..10605 | 287 | TPA:-DUF4276-family-protein-[Pseudomonas-aeruginosa] |
| WP_059319047.1 | TOPRIM | GCF_001516265.1 | Pseudomonas_aeruginosa | Bacteria>Pseudomonadati>Pseudomonadota>Gammaproteobacteria>Pseudomonadales>Pseudomonadaceae>Pseudomonas>Pseudomonas_aeruginosa_group | 227 | HV84_RS13895 | 320931..321614 | 287 | DUF4276-family-protein-[Pseudomonas-aeruginosa] |
| RRI70512.1 | TOPRIM | GCA_003863585.1 | Pseudomonas_aeruginosa | Bacteria>Pseudomonadati>Pseudomonadota>Gammaproteobacteria>Pseudomonadales>Pseudomonadaceae>Pseudomonas>Pseudomonas_aeruginosa_group | 219 | EIM10_18030 | 72381..73040 | 287 | DUF4276-family-protein-[Pseudomonas-aeruginosa] |
| WP_124988195.1 | TOPRIM | GCF_032334355.1 | Pseudomonas_aeruginosa | Bacteria>Pseudomonadati>Pseudomonadota>Gammaproteobacteria>Pseudomonadales>Pseudomonadaceae>Pseudomonas>Pseudomonas_aeruginosa_group | 219 | MX104_RS19085 | 62914..63573 | 287 | DUF4276-family-protein-[Pseudomonas-aeruginosa] |
| HBO7921140.1 | TOPRIM | GCA_021678155.1 | Pseudomonas_aeruginosa | Bacteria>Pseudomonadati>Pseudomonadota>Gammaproteobacteria>Pseudomonadales>Pseudomonadaceae>Pseudomonas>Pseudomonas_aeruginosa_group | 217 | L5C37_005434 | 27206..27859 | 287 | DUF4276-family-protein-[Pseudomonas-aeruginosa] |
| MBX6030767.1 | TOPRIM | GCA_019688625.1 | Pseudomonas_aeruginosa | Bacteria>Pseudomonadati>Pseudomonadota>Gammaproteobacteria>Pseudomonadales>Pseudomonadaceae>Pseudomonas>Pseudomonas_aeruginosa_group | 217 | ISE67_08710 | 280817..281470 | 287 | DUF4276-family-protein-[Pseudomonas-aeruginosa] |
| MBG4863331.1 | TOPRIM | GCA_015735985.1 | Pseudomonas_aeruginosa | Bacteria>Pseudomonadati>Pseudomonadota>Gammaproteobacteria>Pseudomonadales>Pseudomonadaceae>Pseudomonas>Pseudomonas_aeruginosa_group | 216 | GHU23_29005 | 12179..12829 | 287 | DUF4276-family-protein-[Pseudomonas-aeruginosa] |
| MBG6730511.1 | TOPRIM | GCA_015764285.1 | Pseudomonas_aeruginosa | Bacteria>Pseudomonadati>Pseudomonadota>Gammaproteobacteria>Pseudomonadales>Pseudomonadaceae>Pseudomonas>Pseudomonas_aeruginosa_group | 216 | I5I92_11350 | 14314..14964 | 287 | DUF4276-family-protein-[Pseudomonas-aeruginosa] |
| HBO1294564.1 | TOPRIM | GCA_021597005.1 | Pseudomonas_aeruginosa | Bacteria>Pseudomonadati>Pseudomonadota>Gammaproteobacteria>Pseudomonadales>Pseudomonadaceae>Pseudomonas>Pseudomonas_aeruginosa_group | 209 | L4E86_006700 | 3495..4124 | 287 | DUF4276-family-protein-[Pseudomonas-aeruginosa] |
| WP_094776640.1 | TOPRIM | GCF_002239565.1 | Pseudomonas_aeruginosa | Bacteria>Pseudomonadati>Pseudomonadota>Gammaproteobacteria>Pseudomonadales>Pseudomonadaceae>Pseudomonas>Pseudomonas_aeruginosa_group | 207 | CF337_RS09110 | 69983..70606 | 287 | DUF4276-family-protein-[Pseudomonas-aeruginosa] |
| HCK4430923.1 | TOPRIM | GCA_024830635.1 | Pseudomonas_aeruginosa | Bacteria>Pseudomonadati>Pseudomonadota>Gammaproteobacteria>Pseudomonadales>Pseudomonadaceae>Pseudomonas>Pseudomonas_aeruginosa_group | 191 | N0G72_005468 | 45581..46156 | 287 | TPA:-DUF4276-family-protein-[Pseudomonas-aeruginosa] |
| HCL3757804.1 | TOPRIM | GCA_025003865.1 | Pseudomonas_aeruginosa | Bacteria>Pseudomonadati>Pseudomonadota>Gammaproteobacteria>Pseudomonadales>Pseudomonadaceae>Pseudomonas>Pseudomonas_aeruginosa_group | 191 | N2B17_005732 | 10294..10869 | 287 | TPA:-DUF4276-family-protein-[Pseudomonas-aeruginosa] |
| WP_049269902.1 | TOPRIM | GCF_038431925.1 | Pseudomonas_aeruginosa | Bacteria>Pseudomonadati>Pseudomonadota>Gammaproteobacteria>Pseudomonadales>Pseudomonadaceae>Pseudomonas>Pseudomonas_aeruginosa_group | 191 | EQH74_RS32315 | 3311..3886 | 287 | DUF4276-family-protein-[Pseudomonas-aeruginosa] |
| WP_124127282.1 | TOPRIM | GCF_022551795.1 | Pseudomonas_aeruginosa | Bacteria>Pseudomonadati>Pseudomonadota>Gammaproteobacteria>Pseudomonadales>Pseudomonadaceae>Pseudomonas>Pseudomonas_aeruginosa_group | 188 | IPC776_RS25340 | 81767..82333 | 287 | DUF4276-family-protein-[Pseudomonas-aeruginosa] |
| WP_047283131.1 | TOPRIM | GCF_018449005.1 | Pseudomonas_lundensis | Bacteria>Pseudomonadati>Pseudomonadota>Gammaproteobacteria>Pseudomonadales>Pseudomonadaceae>Pseudomonas>Pseudomonas_chlororaphis_group | 227 | KIY13_RS15510 | 3393412..3394095 | 86185 | DUF4276-family-protein-[Pseudomonas-lundensis] |
| ARQ76271.1 | TOPRIM | GCA_002128325.1 | Pseudomonas_fragi | Bacteria>Pseudomonadati>Pseudomonadota>Gammaproteobacteria>Pseudomonadales>Pseudomonadaceae>Pseudomonas>Pseudomonas_chlororaphis_group | 224 | B6D87_19570 | 4220090..4220764 | 296 | hypothetical-protein-B6D87_19570-[Pseudomonas-fragi] |
| WP_086798820.1 | TOPRIM | GCF_002128325.1 | Pseudomonas_fragi | Bacteria>Pseudomonadati>Pseudomonadota>Gammaproteobacteria>Pseudomonadales>Pseudomonadaceae>Pseudomonas>Pseudomonas_chlororaphis_group | 224 | B6D87_RS19590 | 4220090..4220764 | 296 | DUF4276-family-protein-[Pseudomonas-fragi] |
| WP_106118418.1 | TOPRIM | GCF_003001555.1 | Pseudomonas_fluorescens | Bacteria>Pseudomonadati>Pseudomonadota>Gammaproteobacteria>Pseudomonadales>Pseudomonadaceae>Pseudomonas>Pseudomonas_fluorescens_group | 250 | C7A10_RS23720 | 135601..136353 | 294 | hypothetical-protein-[Pseudomonas-fluorescens] |
| WP_236310038.1 | TOPRIM | GCF_021605645.1 | Pseudomonas_gessardii | Bacteria>Pseudomonadati>Pseudomonadota>Gammaproteobacteria>Pseudomonadales>Pseudomonadaceae>Pseudomonas>Pseudomonas_fluorescens_group | 229 | GIW56_RS19750 | 21253..21942 | 78544 | DUF4276-family-protein-[Pseudomonas-gessardii] |
| WP_261753790.1 | TOPRIM | GCF_025397875.1 | Pseudomonas_veronii | Bacteria>Pseudomonadati>Pseudomonadota>Gammaproteobacteria>Pseudomonadales>Pseudomonadaceae>Pseudomonas>Pseudomonas_fluorescens_group | 189 | MN113_RS13510 | 503091..503660 | 76761 | DUF4276-family-protein-[Pseudomonas-veronii] |
| CAB5576354.1 | TOPRIM | GCA_903684465.1 | Pseudomonas_putida | Bacteria>Pseudomonadati>Pseudomonadota>Gammaproteobacteria>Pseudomonadales>Pseudomonadaceae>Pseudomonas>Pseudomonas_putida_group | 189 | GHA_00150 | 171767..172336 | 303 | Uncharacterised-protein-[Pseudomonas-putida] |
| WP_122701524.1 | TOPRIM | - | Pseudomonas_viridiflava | Bacteria>Pseudomonadati>Pseudomonadota>Gammaproteobacteria>Pseudomonadales>Pseudomonadaceae>Pseudomonas>Pseudomonas_syringae_group | 189 | EFA20_RS09790 | 78842..79411 | 33069 | DUF4276-family-protein-[Pseudomonas-viridiflava] |
| WP_222018158.1 | TOPRIM | GCF_019775695.1 | Pseudomonas_sp._AO-1 | Bacteria>Pseudomonadati>Pseudomonadota>Gammaproteobacteria>Pseudomonadales>Pseudomonadaceae>Pseudomonas>unclassified_Pseudomonas | 250 | KVQ82_RS04455 | 957125..957877 | 2855434 | hypothetical-protein-[Pseudomonas-sp.-AO-1] |
| WP_273917576.1 | TOPRIM | GCF_028656895.1 | Pseudomonas_sp._TNT2022_ID1044 | Bacteria>Pseudomonadati>Pseudomonadota>Gammaproteobacteria>Pseudomonadales>Pseudomonadaceae>Pseudomonas>unclassified_Pseudomonas | 250 | M5G20_RS01560 | 345543..346295 | 2942636 | hypothetical-protein-[Pseudomonas-sp.-TNT2022-ID1044] |
| MBB59700.1 | TOPRIM | GCA_002709345.1 | Pseudomonas_sp. | Bacteria>Pseudomonadati>Pseudomonadota>Gammaproteobacteria>Pseudomonadales>Pseudomonadaceae>Pseudomonas>unclassified_Pseudomonas | 227 | CML00_00825 | 31709..32392 | 306 | DUF4276-domain-containing-protein-[Pseudomonas-sp.] |
| MCK9534934.1 | TOPRIM | GCA_023228505.1 | Pseudomonas_sp. | Bacteria>Pseudomonadati>Pseudomonadota>Gammaproteobacteria>Pseudomonadales>Pseudomonadaceae>Pseudomonas>unclassified_Pseudomonas | 225 | M0Q98_09615 | 15771..16448 | 306 | DUF4276-family-protein-[Pseudomonas-sp.] |
| TXH86523.1 | TOPRIM | GCA_008015405.1 | Pseudomonas_sp. | Bacteria>Pseudomonadati>Pseudomonadota>Gammaproteobacteria>Pseudomonadales>Pseudomonadaceae>Pseudomonas>unclassified_Pseudomonas | 191 | E6Q71_06450 | 3588..4163 | 306 | DUF4276-family-protein-[Pseudomonas-sp.] |
| MCQ4319998.1 | TOPRIM | GCA_024448935.1 | Stutzerimonas_stutzeri | Bacteria>Pseudomonadati>Pseudomonadota>Gammaproteobacteria>Pseudomonadales>Pseudomonadaceae>Stutzerimonas>Stutzerimonas_stutzeri_group>Stutzerimonas_stutzeri_subgroup | 250 | NA637_07225 | 22542..23294 | 316 | hypothetical-protein-[Stutzerimonas-stutzeri] |
| PAO90332.1 | TOPRIM | GCA_002282995.1 | Stutzerimonas_stutzeri | Bacteria>Pseudomonadati>Pseudomonadota>Gammaproteobacteria>Pseudomonadales>Pseudomonadaceae>Stutzerimonas>Stutzerimonas_stutzeri_group>Stutzerimonas_stutzeri_subgroup | 227 | BV581_21210 | 15238..15921 | 316 | hypothetical-protein-BV581_21210-[Stutzerimonas-stutzeri] |
| WP_231692716.1 | TOPRIM | GCF_001267275.1 | Thiopseudomonas_alkaliphila | Bacteria>Pseudomonadati>Pseudomonadota>Gammaproteobacteria>Pseudomonadales>Pseudomonadaceae>Thiopseudomonas | 236 | AKN90_RS00635 | 133039..133749 | 1697053 | DUF4276-family-protein-[Thiopseudomonas-alkaliphila] |
| MAB42107.1 | TOPRIM | GCA_002684155.1 | Pseudomonadales_bacterium | Bacteria>Pseudomonadati>Pseudomonadota>Gammaproteobacteria>Pseudomonadales>unclassified_Pseudomonadales | 226 | CMK77_07470 | 6559..7239 | 1891229 | hypothetical-protein-[Pseudomonadales-bacterium] |
| MCP5320938.1 | TOPRIM | GCA_024234165.1 | Pseudomonadales_bacterium | Bacteria>Pseudomonadati>Pseudomonadota>Gammaproteobacteria>Pseudomonadales>unclassified_Pseudomonadales | 206 | H7A12_08970 | 736203..736823 | 1891229 | DUF4276-family-protein-[Pseudomonadales-bacterium] |
| MCG8614351.1 | TOPRIM | GCA_022361325.1 | Pseudomonadales_bacterium | Bacteria>Pseudomonadati>Pseudomonadota>Gammaproteobacteria>Pseudomonadales>unclassified_Pseudomonadales | 186 | MI864_27890 | 12036..12596 | 1891229 | DUF4276-family-protein-[Pseudomonadales-bacterium] |
| MAS10375.1 | TOPRIM | GCA_002702145.1 | Salinisphaera_sp. | Bacteria>Pseudomonadati>Pseudomonadota>Gammaproteobacteria>Salinisphaerales>Salinisphaeraceae>Salinisphaera>unclassified_Salinisphaera | 184 | CMN26_09990 | 8097..8651 | 1914330 | hypothetical-protein-[Salinisphaera-sp.] |
| WP_022954805.1 | TOPRIM | GCF_000419525.1 | Leucothrix_mucor | Bacteria>Pseudomonadati>Pseudomonadota>Gammaproteobacteria>Thiotrichales>Thiotrichaceae>Leucothrix | 247 | LEUMU_RS0123795 | 73738..74481 | 45248 | hypothetical-protein-[Leucothrix-mucor] |
| OQX14374.1 | TOPRIM | GCA_002083875.1 | Thiothrix_lacustris | Bacteria>Pseudomonadati>Pseudomonadota>Gammaproteobacteria>Thiotrichales>Thiotrichaceae>Thiothrix | 307 | BWK73_09675 | 14122..15045 | 525917 | hypothetical-protein-BWK73_09675-[Thiothrix-lacustris] |
| WP_002710594.1 | TOPRIM | GCF_000260135.1 | Thiothrix_nivea | Bacteria>Pseudomonadati>Pseudomonadota>Gammaproteobacteria>Thiotrichales>Thiotrichaceae>Thiothrix | 240 | THINI_RS21435 | 4306911..4307633 | 1031 | hypothetical-protein-[Thiothrix-nivea] |
| QLQ31995.1 | TOPRIM | GCA_013693955.1 | Candidatus_Thiothrix_singaporensis | Bacteria>Pseudomonadati>Pseudomonadota>Gammaproteobacteria>Thiotrichales>Thiotrichaceae>Thiothrix | 215 | HZT40_10795 | 2140338..2140985 | 2799669 | hypothetical-protein-HZT40_10795-[Candidatus-Thiothrix-singaporensis] |
| WP_210226005.1 | TOPRIM | GCF_017901155.1 | Candidatus_Thiothrix_anitrata | Bacteria>Pseudomonadati>Pseudomonadota>Gammaproteobacteria>Thiotrichales>Thiotrichaceae>Thiothrix | 206 | J8380_RS12845 | 2552451..2553071 | 2823902 | DUF4276-family-protein-[Candidatus-Thiothrix-anitrata] |
| MCY4286578.1 | TOPRIM | GCA_026709935.1 | Thiotrichales_bacterium | Bacteria>Pseudomonadati>Pseudomonadota>Gammaproteobacteria>Thiotrichales>unclassified_Thiotrichales | 199 | OXC65_14720 | 7127..7726 | 2026796 | MAG:-hypothetical-protein-OXC65_14720-[Thiotrichales-bacterium] |
| MYA37127.1 | TOPRIM | GCA_009838725.1 | Gammaproteobacteria_bacterium | Bacteria>Pseudomonadati>Pseudomonadota>Gammaproteobacteria>unclassified_Gammaproteobacteria | 297 | F4Y34_10765 | 14037..14930 | 1913989 | hypothetical-protein-[Gammaproteobacteria-bacterium] |
| MCP5451260.1 | TOPRIM | GCA_024235115.1 | Gammaproteobacteria_bacterium | Bacteria>Pseudomonadati>Pseudomonadota>Gammaproteobacteria>unclassified_Gammaproteobacteria | 224 | H6972_12095 | 50379..51053 | 1913989 | DUF4276-family-protein-[Gammaproteobacteria-bacterium] |
| MCP4410536.1 | TOPRIM | GCA_024222475.1 | Gammaproteobacteria_bacterium | Bacteria>Pseudomonadati>Pseudomonadota>Gammaproteobacteria>unclassified_Gammaproteobacteria | 221 | GY807_22890 | 8848..9513 | 1913989 | DUF4276-family-protein-[Gammaproteobacteria-bacterium] |
| NLY14761.1 | TOPRIM | GCA_012728415.1 | Gammaproteobacteria_bacterium | Bacteria>Pseudomonadati>Pseudomonadota>Gammaproteobacteria>unclassified_Gammaproteobacteria | 220 | GXZ05_00055 | 5726..6388 | 1913989 | DUF4276-family-protein-[Gammaproteobacteria-bacterium] |
| MXY90448.1 | TOPRIM | GCA_009837675.1 | Gammaproteobacteria_bacterium | Bacteria>Pseudomonadati>Pseudomonadota>Gammaproteobacteria>unclassified_Gammaproteobacteria | 209 | F4Y89_07910 | 37501..38130 | 1913989 | hypothetical-protein-[Gammaproteobacteria-bacterium] |
| MDD9864538.1 | TOPRIM | GCA_028817305.1 | Gammaproteobacteria_bacterium | Bacteria>Pseudomonadati>Pseudomonadota>Gammaproteobacteria>unclassified_Gammaproteobacteria | 208 | OXU54_08200 | 3690..4316 | 1913989 | MAG:-hypothetical-protein-OXU54_08200-[Gammaproteobacteria-bacterium] |
| MDE0157709.1 | TOPRIM | GCA_028819495.1 | Gammaproteobacteria_bacterium | Bacteria>Pseudomonadati>Pseudomonadota>Gammaproteobacteria>unclassified_Gammaproteobacteria | 204 | OXS28_19165 | 2856..3470 | 1913989 | MAG:-hypothetical-protein-OXS28_19165-[Gammaproteobacteria-bacterium] |
| MDE0512493.1 | TOPRIM | GCA_028823865.1 | Gammaproteobacteria_bacterium | Bacteria>Pseudomonadati>Pseudomonadota>Gammaproteobacteria>unclassified_Gammaproteobacteria | 204 | OXI88_11975 | 8468..9082 | 1913989 | MAG:-hypothetical-protein-OXI88_11975-[Gammaproteobacteria-bacterium] |
| NPU94899.1 | TOPRIM | GCA_013178375.1 | Gammaproteobacteria_bacterium | Bacteria>Pseudomonadati>Pseudomonadota>Gammaproteobacteria>unclassified_Gammaproteobacteria | 204 | HPY82_23590 | 10374..10988 | 1913989 | DUF4276-family-protein-[Gammaproteobacteria-bacterium] |
| MCZ0943262.1 | TOPRIM | GCA_026914145.1 | Gammaproteobacteria_bacterium | Bacteria>Pseudomonadati>Pseudomonadota>Gammaproteobacteria>unclassified_Gammaproteobacteria | 194 | OXJ53_09415 | 7263..7847 | 1913989 | MAG:-DUF4276-family-protein-[Gammaproteobacteria-bacterium] |
| MDE0073870.1 | TOPRIM | GCA_028819075.1 | Gammaproteobacteria_bacterium | Bacteria>Pseudomonadati>Pseudomonadota>Gammaproteobacteria>unclassified_Gammaproteobacteria | 188 | OXU32_07800 | 2726..3292 | 1913989 | MAG:-DUF4276-family-protein-[Gammaproteobacteria-bacterium] |
| MDE0038694.1 | TOPRIM | GCA_028818795.1 | Gammaproteobacteria_bacterium | Bacteria>Pseudomonadati>Pseudomonadota>Gammaproteobacteria>unclassified_Gammaproteobacteria | 186 | OXU77_14245 | 15231..15791 | 1913989 | MAG:-hypothetical-protein-OXU77_14245-[Gammaproteobacteria-bacterium] |
| TAK54307.1 | TOPRIM | GCA_004297335.1 | Gammaproteobacteria_bacterium | Bacteria>Pseudomonadati>Pseudomonadota>Gammaproteobacteria>unclassified_Gammaproteobacteria | 181 | EPO25_07525 | 12564..13109 | 1913989 | DUF4276-family-protein-[Gammaproteobacteria-bacterium] |
| MDD9817276.1 | TOPRIM | GCA_028817495.1 | Gammaproteobacteria_bacterium | Bacteria>Pseudomonadati>Pseudomonadota>Gammaproteobacteria>unclassified_Gammaproteobacteria | 167 | OXU61_03935 | 3753..4256 | 1913989 | MAG:-hypothetical-protein-OXU61_03935-[Gammaproteobacteria-bacterium] |
| WP_170894127.1 | TOPRIM | GCF_033441435.1 | Vibrio | Bacteria>Pseudomonadati>Pseudomonadota>Gammaproteobacteria>Vibrionales>Vibrionaceae | 252 | G3220_RS19465 | 64487..65245 | 662 | MULTISPECIES:-hypothetical-protein-[Vibrio] |
| WP_116869875.1 | TOPRIM | GCF_050888945.2 | Vibrio | Bacteria>Pseudomonadati>Pseudomonadota>Gammaproteobacteria>Vibrionales>Vibrionaceae | 245 | B5C31_RS21485 | 48945..49682 | 662 | MULTISPECIES:-hypothetical-protein-[Vibrio] |
| WP_113594182.1 | TOPRIM | GCF_043352035.1 | Vibrio | Bacteria>Pseudomonadati>Pseudomonadota>Gammaproteobacteria>Vibrionales>Vibrionaceae | 221 | DLR66_RS03575 | 65223..65888 | 662 | DUF4276-family-protein-[Vibrio-paracholerae] |
| WP_054775223.1 | TOPRIM | GCF_014647395.1 | Aliivibrio_fischeri | Bacteria>Pseudomonadati>Pseudomonadota>Gammaproteobacteria>Vibrionales>Vibrionaceae>Aliivibrio | 227 | AVFI_RS07665 | 1640666..1641349 | 668 | DUF4276-family-protein-[Aliivibrio-fischeri] |
| WP_190395656.1 | TOPRIM | GCF_014694495.1 | Aliivibrio_sp._S10_S31 | Bacteria>Pseudomonadati>Pseudomonadota>Gammaproteobacteria>Vibrionales>Vibrionaceae>Aliivibrio>unclassified_Aliivibrio | 227 | HC728_RS13695 | 66053..66736 | 2720224 | DUF4276-family-protein-[Aliivibrio-sp.-S10_S31] |
| PLC56102.1 | TOPRIM | GCA_002849605.1 | Photobacterium_carnosum | Bacteria>Pseudomonadati>Pseudomonadota>Gammaproteobacteria>Vibrionales>Vibrionaceae>Photobacterium | 252 | CIK00_20175 | 15344..16102 | 2023717 | hypothetical-protein-CIK00_20175-[Photobacterium-carnosum] |
| WP_176285153.1 | TOPRIM | GCF_013371045.1 | Photobacterium_damselae | Bacteria>Pseudomonadati>Pseudomonadota>Gammaproteobacteria>Vibrionales>Vibrionaceae>Photobacterium | 221 | HU831_RS17620 | 65199..65864 | 38293 | DUF4276-family-protein-[Photobacterium-damselae] |
| WP_197573430.1 | TOPRIM | GCF_014930975.1 | Photobacterium_damselae | Bacteria>Pseudomonadati>Pseudomonadota>Gammaproteobacteria>Vibrionales>Vibrionaceae>Photobacterium | 221 | IL982_RS10180 | 2136822..2137487 | 38293 | DUF4276-family-protein-[Photobacterium-damselae] |
| CAQ34955.1 | TOPRIM | - | Photobacterium_damselae_subsp._piscicida | Bacteria>Pseudomonadati>Pseudomonadota>Gammaproteobacteria>Vibrionales>Vibrionaceae>Photobacterium>Photobacterium_damselae | 249 | . | 26030..26779 | 38294 | hypothetical-protein-[Photobacterium-damselae-subsp.-piscicida] |
| QOQ68158.1 | TOPRIM | GCA_014930975.1 | Photobacterium_damselae_subsp._damselae | Bacteria>Pseudomonadati>Pseudomonadota>Gammaproteobacteria>Vibrionales>Vibrionaceae>Photobacterium>Photobacterium_damselae | 221 | IL982_10180 | 2136822..2137487 | 85581 | DUF4276-family-protein-[Photobacterium-damselae-subsp.-damselae] |
| WP_077485091.1 | TOPRIM | GCF_001995745.1 | Salinivibrio_sp._ML323 | Bacteria>Pseudomonadati>Pseudomonadota>Gammaproteobacteria>Vibrionales>Vibrionaceae>Salinivibrio>unclassified_Salinivibrio | 221 | BZG13_RS03570 | 133457..134122 | 1909474 | DUF4276-family-protein-[Salinivibrio-sp.-ML323] |
| TNF18466.1 | TOPRIM | GCA_006226665.1 | Vibrionaceae_bacterium | Bacteria>Pseudomonadati>Pseudomonadota>Gammaproteobacteria>Vibrionales>Vibrionaceae>unclassified_Vibrionaceae | 224 | EP325_05040 | 59860..60534 | 1889773 | DUF4276-family-protein-[Vibrionaceae-bacterium] |

| Protein-ID | Domain | Assembly-ID | Strain | Taxonomy | Length | Locus-Tag | Genomic-coordinates | TaxID | Fasta-defline |
|---|---|---|---|---|---|---|---|---|---|
| WP_054541290.1 | TOPRIM | GCF_963869775.1 | Vibrio_splendidus | Bacteria>Pseudomonadati>Pseudomonadota>Gammaproteobacteria>Vibrionales>Vibrionaceae>Vibrio | 254 | AN167_RS02945 | 112340..113104 | 29497 | hypothetical-protein-[Vibrio-splendidus] |
| AAL59729.1 | TOPRIM | - | Vibrio_cholerae | Bacteria>Pseudomonadati>Pseudomonadota>Gammaproteobacteria>Vibrionales>Vibrionaceae>Vibrio | 252 | . | 33237..33995 | 666 | unknown-[Vibrio-cholerae] |
| EJB8347938.1 | TOPRIM | GCA_022966375.1 | Vibrio_cholerae | Bacteria>Pseudomonadati>Pseudomonadota>Gammaproteobacteria>Vibrionales>Vibrionaceae>Vibrio | 252 | MW328_000280 | 38660..39418 | 666 | hypothetical-protein-[Vibrio-cholerae] |
| EJB8581802.1 | TOPRIM | GCA_022967455.1 | Vibrio_cholerae | Bacteria>Pseudomonadati>Pseudomonadota>Gammaproteobacteria>Vibrionales>Vibrionaceae>Vibrio | 252 | MW361_003121 | 33035..33793 | 666 | hypothetical-protein-[Vibrio-cholerae] |
| WP_274695777.1 | TOPRIM | GCF_028858205.1 | Vibrio_aestuarianus | Bacteria>Pseudomonadati>Pseudomonadota>Gammaproteobacteria>Vibrionales>Vibrionaceae>Vibrio | 252 | MEG05_RS14955 | 3092878..3093636 | 28171 | hypothetical-protein-[Vibrio-aestuarianus] |
| WP_261872481.1 | TOPRIM | GCF_024347075.1 | Vibrio_rarus | Bacteria>Pseudomonadati>Pseudomonadota>Gammaproteobacteria>Vibrionales>Vibrionaceae>Vibrio | 248 | OCU56_RS06800 | 1450526..1451272 | 413403 | hypothetical-protein-[Vibrio-rarus] |
| EGQ8649253.1 | TOPRIM | - | Vibrio_cholerae | Bacteria>Pseudomonadati>Pseudomonadota>Gammaproteobacteria>Vibrionales>Vibrionaceae>Vibrio | 245 | GNS53_08420 | 55264..56001 | 666 | hypothetical-protein-[Vibrio-cholerae] |
| EJK2098982.1 | TOPRIM | GCA_024103575.1 | Vibrio_cholerae | Bacteria>Pseudomonadati>Pseudomonadota>Gammaproteobacteria>Vibrionales>Vibrionaceae>Vibrio | 245 | NK421_001791 | 212144..212881 | 666 | hypothetical-protein-[Vibrio-cholerae] |
| WP_210447445.1 | TOPRIM | GCF_017917465.1 | Vibrio_crassostreae | Bacteria>Pseudomonadati>Pseudomonadota>Gammaproteobacteria>Vibrionales>Vibrionaceae>Vibrio | 227 | KB624_RS13675 | 50492..51175 | 246167 | DUF4276-family-protein-[Vibrio-crassostreae] |
| EGR0286975.1 | TOPRIM | - | Vibrio_cholerae | Bacteria>Pseudomonadati>Pseudomonadota>Gammaproteobacteria>Vibrionales>Vibrionaceae>Vibrio | 221 | FK206_11685 | 80034..80699 | 666 | DUF4276-family-protein-[Vibrio-cholerae] |
| EGR2430330.1 | TOPRIM | - | Vibrio_cholerae | Bacteria>Pseudomonadati>Pseudomonadota>Gammaproteobacteria>Vibrionales>Vibrionaceae>Vibrio | 221 | DYB83_10155 | 19173..19838 | 666 | DUF4276-family-protein-[Vibrio-cholerae] |
| EJL6578341.1 | TOPRIM | GCA_024217525.1 | Vibrio_cholerae | Bacteria>Pseudomonadati>Pseudomonadota>Gammaproteobacteria>Vibrionales>Vibrionaceae>Vibrio | 221 | NMS50_002627 | 66115..66780 | 666 | DUF4276-family-protein-[Vibrio-cholerae] |
| WP_045588643.1 | TOPRIM | GCF_000959745.1 | Vibrio_vulnificus | Bacteria>Pseudomonadati>Pseudomonadota>Gammaproteobacteria>Vibrionales>Vibrionaceae>Vibrio | 221 | WG98_RS00465 | 99776..100441 | 672 | DUF4276-family-protein-[Vibrio-vulnificus] |
| EII3725631.1 | TOPRIM | GCA_020414365.1 | Vibrio_cholerae | Bacteria>Pseudomonadati>Pseudomonadota>Gammaproteobacteria>Vibrionales>Vibrionaceae>Vibrio | 199 | LG468_000344 | 122032..122631 | 666 | DUF4276-family-protein-[Vibrio-cholerae] |
| HAS6259591.1 | TOPRIM | GCA_015777345.1 | Vibrio_vulnificus | Bacteria>Pseudomonadati>Pseudomonadota>Gammaproteobacteria>Vibrionales>Vibrionaceae>Vibrio | 199 | I7226_03350 | 256748..257347 | 672 | DUF4276-family-protein-[Vibrio-vulnificus] |
| WP_032482148.1 | TOPRIM | GCF_000736875.1 | Vibrio_cholerae | Bacteria>Pseudomonadati>Pseudomonadota>Gammaproteobacteria>Vibrionales>Vibrionaceae>Vibrio | 199 | DN38_RS05525 | 19040..19639 | 666 | DUF4276-family-protein-[Vibrio-cholerae] |
| WP_142738827.1 | TOPRIM | GCF_006803025.1 | Vibrio_cholerae | Bacteria>Pseudomonadati>Pseudomonadota>Gammaproteobacteria>Vibrionales>Vibrionaceae>Vibrio | 199 | FLL79_RS01430 | 46255..46854 | 666 | DUF4276-family-protein-[Vibrio-cholerae] |
| WP_102376353.1 | TOPRIM | GCF_002876115.1 | Vibrio_breoganii | Bacteria>Pseudomonadati>Pseudomonadota>Gammaproteobacteria>Vibrionales>Vibrionaceae>Vibrio | 189 | BCT61_RS02985 | 187181..187750 | 553239 | hypothetical-protein-[Vibrio-breoganii] |
| WP_108187611.1 | TOPRIM | GCF_003049965.1 | Vibrio_splendidus | Bacteria>Pseudomonadati>Pseudomonadota>Gammaproteobacteria>Vibrionales>Vibrionaceae>Vibrio | 189 | CWO07_RS09200 | 38792..39361 | 29497 | hypothetical-protein-[Vibrio-splendidus] |
| WP_134990668.1 | TOPRIM | - | Vibrio_parahaemolyticus | Bacteria>Pseudomonadati>Pseudomonadota>Gammaproteobacteria>Vibrionales>Vibrionaceae>Vibrio>Vibrio_harveyi_group | 252 | DA442_RS13690 | 2689187..2689945 | 670 | hypothetical-protein-[Vibrio-parahaemolyticus] |
| MBE3881972.1 | TOPRIM | GCA_014919165.1 | Vibrio_parahaemolyticus | Bacteria>Pseudomonadati>Pseudomonadota>Gammaproteobacteria>Vibrionales>Vibrionaceae>Vibrio>Vibrio_harveyi_group | 248 | HJ168_09875 | 231118..231864 | 670 | hypothetical-protein-[Vibrio-parahaemolyticus] |
| EJE4207360.1 | TOPRIM | GCA_023313315.1 | Vibrio_parahaemolyticus | Bacteria>Pseudomonadati>Pseudomonadota>Gammaproteobacteria>Vibrionales>Vibrionaceae>Vibrio>Vibrio_harveyi_group | 245 | M3899_001771 | 265184..265921 | 670 | hypothetical-protein-[Vibrio-parahaemolyticus] |
| HCE3362110.1 | TOPRIM | GCA_023821665.1 | Vibrio_parahaemolyticus | Bacteria>Pseudomonadati>Pseudomonadota>Gammaproteobacteria>Vibrionales>Vibrionaceae>Vibrio>Vibrio_harveyi_group | 245 | NG589_000242 | 270534..271271 | 670 | hypothetical-protein-[Vibrio-parahaemolyticus] |
| WP_274056405.1 | TOPRIM | GCF_028616275.1 | Vibrio_parahaemolyticus | Bacteria>Pseudomonadati>Pseudomonadota>Gammaproteobacteria>Vibrionales>Vibrionaceae>Vibrio>Vibrio_harveyi_group | 245 | O8051_RS20170 | 62518..63255 | 670 | hypothetical-protein-[Vibrio-parahaemolyticus] |
| EGR1144245.1 | TOPRIM | - | Vibrio_parahaemolyticus | Bacteria>Pseudomonadati>Pseudomonadota>Gammaproteobacteria>Vibrionales>Vibrionaceae>Vibrio>Vibrio_harveyi_group | 221 | EC562_15490 | 35823..36488 | 670 | DUF4276-family-protein-[Vibrio-parahaemolyticus] |
| EGR1548463.1 | TOPRIM | - | Vibrio_parahaemolyticus | Bacteria>Pseudomonadati>Pseudomonadota>Gammaproteobacteria>Vibrionales>Vibrionaceae>Vibrio>Vibrio_harveyi_group | 221 | D6Y11_16935 | 156024..156689 | 670 | DUF4276-family-protein-[Vibrio-parahaemolyticus] |
| EGR3302283.1 | TOPRIM | - | Vibrio_parahaemolyticus | Bacteria>Pseudomonadati>Pseudomonadota>Gammaproteobacteria>Vibrionales>Vibrionaceae>Vibrio>Vibrio_harveyi_group | 221 | DMJ21_13145 | 16119..16784 | 670 | DUF4276-domain-containing-protein-[Vibrio-parahaemolyticus] |
| EHH2507628.1 | TOPRIM | GCA_017637525.1 | Vibrio_parahaemolyticus | Bacteria>Pseudomonadati>Pseudomonadota>Gammaproteobacteria>Vibrionales>Vibrionaceae>Vibrio>Vibrio_harveyi_group | 221 | J7X10_003791 | 104315..104980 | 670 | DUF4276-family-protein-[Vibrio-parahaemolyticus] |
| EII2403784.1 | TOPRIM | GCA_020407245.1 | Vibrio_parahaemolyticus | Bacteria>Pseudomonadati>Pseudomonadota>Gammaproteobacteria>Vibrionales>Vibrionaceae>Vibrio>Vibrio_harveyi_group | 221 | LG159_002757 | 111606..112271 | 670 | DUF4276-family-protein-[Vibrio-parahaemolyticus] |
| EIS4855659.1 | TOPRIM | GCA_021362915.1 | Vibrio_parahaemolyticus | Bacteria>Pseudomonadati>Pseudomonadota>Gammaproteobacteria>Vibrionales>Vibrionaceae>Vibrio>Vibrio_harveyi_group | 221 | LZX80_001785 | 101452..102117 | 670 | DUF4276-family-protein-[Vibrio-parahaemolyticus] |
| EJC1210120.1 | TOPRIM | GCA_022997325.1 | Vibrio_parahaemolyticus | Bacteria>Pseudomonadati>Pseudomonadota>Gammaproteobacteria>Vibrionales>Vibrionaceae>Vibrio>Vibrio_harveyi_group | 221 | MX350_002597 | 13737..14402 | 670 | DUF4276-family-protein-[Vibrio-parahaemolyticus] |
| EJG0936608.1 | TOPRIM | GCA_015814185.2 | Vibrio_parahaemolyticus | Bacteria>Pseudomonadati>Pseudomonadota>Gammaproteobacteria>Vibrionales>Vibrionaceae>Vibrio>Vibrio_harveyi_group | 221 | C4G97_RS14660 | 113058..113723 | 670 | DUF4276-family-protein-[Vibrio-parahaemolyticus] |
| EJG0981290.1 | TOPRIM | GCA_015813375.2 | Vibrio_parahaemolyticus | Bacteria>Pseudomonadati>Pseudomonadota>Gammaproteobacteria>Vibrionales>Vibrionaceae>Vibrio>Vibrio_harveyi_group | 221 | C4G85_RS03485 | 771446..772111 | 670 | DUF4276-family-protein-[Vibrio-parahaemolyticus] |
| HCM0920434.1 | TOPRIM | GCA_025068375.1 | Vibrio_parahaemolyticus | Bacteria>Pseudomonadati>Pseudomonadota>Gammaproteobacteria>Vibrionales>Vibrionaceae>Vibrio>Vibrio_harveyi_group | 221 | N2828_003372 | 101453..102118 | 670 | TPA:-DUF4276-family-protein-[Vibrio-parahaemolyticus] |
| KOY19346.1 | TOPRIM | GCA_001280635.1 | Vibrio_parahaemolyticus | Bacteria>Pseudomonadati>Pseudomonadota>Gammaproteobacteria>Vibrionales>Vibrionaceae>Vibrio>Vibrio_harveyi_group | 221 | ACX06_21945 | 162226..162891 | 670 | hypothetical-protein-ACX06_21945-[Vibrio-parahaemolyticus] |
| ODY54468.1 | TOPRIM | GCA_001727265.1 | Vibrio_parahaemolyticus | Bacteria>Pseudomonadati>Pseudomonadota>Gammaproteobacteria>Vibrionales>Vibrionaceae>Vibrio>Vibrio_harveyi_group | 221 | BBM25_06255 | 11475..12140 | 670 | hypothetical-protein-BBM25_06255-[Vibrio-parahaemolyticus] |
| WP_029846593.1 | TOPRIM | GCF_000491375.1 | Vibrio_parahaemolyticus | Bacteria>Pseudomonadati>Pseudomonadota>Gammaproteobacteria>Vibrionales>Vibrionaceae>Vibrio>Vibrio_harveyi_group | 221 | PO77_RS19435 | 162628..163293 | 670 | DUF4276-family-protein-[Vibrio-parahaemolyticus] |
| WP_053816407.1 | TOPRIM | GCF_001280635.1 | Vibrio_parahaemolyticus | Bacteria>Pseudomonadati>Pseudomonadota>Gammaproteobacteria>Vibrionales>Vibrionaceae>Vibrio>Vibrio_harveyi_group | 221 | ACX06_RS21945 | 162226..162891 | 670 | DUF4276-family-protein-[Vibrio-parahaemolyticus] |
| WP_062854376.1 | TOPRIM | GCF_001608855.1 | Vibrio_parahaemolyticus | Bacteria>Pseudomonadati>Pseudomonadota>Gammaproteobacteria>Vibrionales>Vibrionaceae>Vibrio>Vibrio_harveyi_group | 221 | AWQ17_RS02390 | 101860..102525 | 670 | DUF4276-family-protein-[Vibrio-parahaemolyticus] |
| WP_065970498.1 | TOPRIM | GCF_001704915.1 | Vibrio_parahaemolyticus | Bacteria>Pseudomonadati>Pseudomonadota>Gammaproteobacteria>Vibrionales>Vibrionaceae>Vibrio>Vibrio_harveyi_group | 221 | BBM25_RS06255 | 11475..12140 | 670 | DUF4276-family-protein-[Vibrio-parahaemolyticus] |
| WP_102780434.1 | TOPRIM | GCF_030238805.1 | Vibrio_parahaemolyticus | Bacteria>Pseudomonadati>Pseudomonadota>Gammaproteobacteria>Vibrionales>Vibrionaceae>Vibrio>Vibrio_harveyi_group | 221 | CSR03_RS22365 | 129654..130319 | 670 | DUF4276-family-protein-[Vibrio-parahaemolyticus] |
| WP_238812499.1 | TOPRIM | GCF_022290755.1 | Vibrio_parahaemolyticus | Bacteria>Pseudomonadati>Pseudomonadota>Gammaproteobacteria>Vibrionales>Vibrionaceae>Vibrio>Vibrio_harveyi_group | 221 | K6U44_RS14245 | 15859..16524 | 670 | DUF4276-family-protein-[Vibrio-parahaemolyticus] |
| MBE3824869.1 | TOPRIM | GCA_014918955.1 | Vibrio_parahaemolyticus | Bacteria>Pseudomonadati>Pseudomonadota>Gammaproteobacteria>Vibrionales>Vibrionaceae>Vibrio>Vibrio_harveyi_group | 199 | HJ179_16450 | 248782..249381 | 670 | DUF4276-family-protein-[Vibrio-parahaemolyticus] |
| WP_104969989.1 | TOPRIM | GCF_002953335.1 | Vibrio_diabolicus | Bacteria>Pseudomonadati>Pseudomonadota>Gammaproteobacteria>Vibrionales>Vibrionaceae>Vibrio>Vibrio_harveyi_group>Vibrio_diabolicus_subgroup | 191 | AL552_RS16295 | 3107096..3107671 | 50719 | hypothetical-protein-[Vibrio-diabolicus] |
| OEE71515.1 | TOPRIM | GCA_000287075.2 | Vibrio_ordalii_FF-167 | Bacteria>Pseudomonadati>Pseudomonadota>Gammaproteobacteria>Vibrionales>Vibrionaceae>Vibrio>Vibrio_ordalii | 252 | A1QQ_08595 | 35598..36356 | 617131 | hypothetical-protein-A1QQ_08595-[Vibrio-ordalii-FF-167] |
| BBD77928.1 | TOPRIM | GCA_003574215.1 | Hydrogenophilus_thermoluteolus | Bacteria>Pseudomonadati>Pseudomonadota>Hydrogenophilia>Hydrogenophilales>Hydrogenophilaceae>Hydrogenophilus | 207 | HPTL_1670 | 1768660..1769283 | 297 | hypothetical-protein-HPTL_1670-[Hydrogenophilus-thermoluteolus] |
| NPU98923.1 | TOPRIM | GCA_013177805.1 | Candidatus_Omnitrophota_bacterium | Bacteria>Pseudomonadati>PVC_group>Candidatus_Omnitrophota>unclassified_Candidatus_Omnitrophota | 208 | HPY51_17140 | 17547..18173 | 2035772 | DUF4276-family-protein-[Candidatus-Omnitrophota-bacterium] |
| MCF7790536.1 | TOPRIM | GCA_021733985.1 | Victivallales_bacterium | Bacteria>Pseudomonadati>PVC_group>Lentisphaerota>Lentisphaeria>Victivallales>unclassified_Victivallales | 205 | K9M56_00935 | 7122..7739 | 2212475 | RloB-family-protein-[Victivallales-bacterium] |
| NLF62182.1 | TOPRIM | GCA_012515215.1 | Lentisphaerota_bacterium | Bacteria>Pseudomonadati>PVC_group>Lentisphaerota>unclassified_Lentisphaerota | 206 | GX574_13645 | 36122..36742 | 1932692 | DUF4276-family-protein-[Lentisphaerota-bacterium] |
| OQC12436.1 | TOPRIM | GCA_002070945.1 | Lentisphaerae_bacterium_ADurb.Bin082 | Bacteria>Pseudomonadati>PVC_group>Lentisphaerota>unclassified_Lentisphaerota | 189 | BWX73_02929 | 53102..53671 | 1852899 | MAG:-hypothetical-protein-BWX73_02929-[Lentisphaerae-bacterium-ADurb.Bin082] |
| MBP7937488.1 | TOPRIM | GCA_018003015.1 | Phycisphaerae_bacterium | Bacteria>Pseudomonadati>PVC_group>Planctomycetota>Phycisphaerae>unclassified_Phycisphaerae | 211 | KA354_22840 | 7708..8343 | 2026778 | DUF4276-family-protein-[Phycisphaerae-bacterium] |
| MCK6483012.1 | TOPRIM | GCA_023150595.1 | Phycisphaerae_bacterium | Bacteria>Pseudomonadati>PVC_group>Planctomycetota>Phycisphaerae>unclassified_Phycisphaerae | 189 | L6R00_02620 | 134417..134986 | 2026778 | hypothetical-protein-[Phycisphaerae-bacterium] |
| MBY0527366.1 | TOPRIM | GCA_019752375.1 | Gemmataceae_bacterium | Bacteria>Pseudomonadati>PVC_group>Planctomycetota>Planctomycetia>Gemmatales>Gemmataceae>unclassified_Gemmataceae | 223 | K2R98_28480 | 32835..33506 | 2052164 | hypothetical-protein-[Gemmataceae-bacterium] |
| GIW78772.1 | TOPRIM | GCA_026004675.1 | Gemmatales_bacterium | Bacteria>Pseudomonadati>PVC_group>Planctomycetota>Planctomycetia>Gemmatales>unclassified_Gemmatales | 210 | KatS3mg105_0579 | 674069..674701 | 2838789 | MAG:-hypothetical-protein-KatS3mg105_0579-[Gemmatales-bacterium] |
| WP_011117923.1 | TOPRIM | GCF_000196115.1 | Rhodopirellula_baltica | Bacteria>Pseudomonadati>PVC_group>Planctomycetota>Planctomycetia>Pirellulales>Pirellulaceae>Rhodopirellula | 224 | DDX19_RS10400 | 130951..131625 | 265606 | DUF4276-family-protein-[Rhodopirellula-baltica] |
| WP_218933730.1 | TOPRIM | GCF_007741535.1 | Rubripirellula_lacrimiformis | Bacteria>Pseudomonadati>PVC_group>Planctomycetota>Planctomycetia>Pirellulales>Pirellulaceae>Rubripirellula | 217 | K227x_RS02715 | 778163..778816 | 1930273 | DUF4276-family-protein-[Rubripirellula-lacrimiformis] |
| NLY99305.1 | TOPRIM | GCA_012744395.1 | Pirellulaceae_bacterium | Bacteria>Pseudomonadati>PVC_group>Planctomycetota>Planctomycetia>Pirellulales>Pirellulaceae>unclassified_Pirellulaceae | 221 | GXY25_02075 | 5864..6529 | 2699754 | hypothetical-protein-[Pirellulaceae-bacterium] |
| GIW94556.1 | TOPRIM | GCA_026004775.1 | Pirellulaceae_bacterium | Bacteria>Pseudomonadati>PVC_group>Planctomycetota>Planctomycetia>Pirellulales>Pirellulaceae>unclassified_Pirellulaceae | 207 | KatS3mg110_2597 | 16999..17622 | 2699754 | MAG:-hypothetical-protein-KatS3mg110_2597-[Pirellulaceae-bacterium] |
| MDD4270715.1 | TOPRIM | GCA_028703175.1 | Thermoguttaceae_bacterium | Bacteria>Pseudomonadati>PVC_group>Planctomycetota>Planctomycetia>Pirellulales>Thermoguttaceae>unclassified_Thermoguttaceae | 221 | PHN77_21020 | 3146..3811 | 2691551 | MAG:-hypothetical-protein-PHN77_21020-[Thermoguttaceae-bacterium] |
| WP_075094280.1 | TOPRIM | GCF_001610835.1 | Planctomyces_sp._SH-PL14 | Bacteria>Pseudomonadati>PVC_group>Planctomycetota>Planctomycetia>Planctomycetales>Planctomycetaceae>Planctomyces>unclassified_Planctomyces | 256 | VT03_RS18110 | 4715854..4716624 | 1632864 | hypothetical-protein-[Planctomyces-sp.-SH-PL14] |
| MBA4188927.1 | TOPRIM | GCA_013823135.1 | Planctomycetaceae_bacterium | Bacteria>Pseudomonadati>PVC_group>Planctomycetota>Planctomycetia>Planctomycetales>Planctomycetaceae>unclassified_Planctomycetaceae | 248 | C0467_13100 | 104983..105729 | 2026779 | hypothetical-protein-[Planctomycetaceae-bacterium] |
| HCS52254.1 | TOPRIM | GCA_003531135.1 | Planctomycetaceae_bacterium | Bacteria>Pseudomonadati>PVC_group>Planctomycetota>Planctomycetia>Planctomycetales>Planctomycetaceae>unclassified_Planctomycetaceae | 246 | DIW81_11790 | 6655..7395 | 2026779 | hypothetical-protein-[Planctomycetaceae-bacterium] |
| HBI47066.1 | TOPRIM | GCA_003508275.1 | Planctomycetales_bacterium | Bacteria>Pseudomonadati>PVC_group>Planctomycetota>Planctomycetia>Planctomycetales>unclassified_Planctomycetales | 291 | DDY78_30065 | 16167..17042 | 2053591 | hypothetical-protein-[Planctomycetales-bacterium] |
| HBI47075.1 | TOPRIM | GCA_003508275.1 | Planctomycetales_bacterium | Bacteria>Pseudomonadati>PVC_group>Planctomycetota>Planctomycetia>Planctomycetales>unclassified_Planctomycetales | 247 | DDY78_30110 | 29660..30403 | 2053591 | hypothetical-protein-[Planctomycetales-bacterium] |
| OJW08010.1 | TOPRIM | GCA_001898385.1 | Planctomycetales_bacterium_71-10 | Bacteria>Pseudomonadati>PVC_group>Planctomycetota>Planctomycetia>Planctomycetales>unclassified_Planctomycetales | 228 | BGO49_20690 | 23159..23845 | 1895807 | MAG:-hypothetical-protein-BGO49_20690-[Planctomycetales-bacterium-71-10] |
| MCA9126644.1 | TOPRIM | GCA_020426625.1 | Planctomycetales_bacterium | Bacteria>Pseudomonadati>PVC_group>Planctomycetota>Planctomycetia>Planctomycetales>unclassified_Planctomycetales | 202 | KDB22_06155 | 18648..19256 | 2053591 | DUF4276-family-protein-[Planctomycetales-bacterium] |
| MCA9232429.1 | TOPRIM | GCA_020427155.1 | Planctomycetales_bacterium | Bacteria>Pseudomonadati>PVC_group>Planctomycetota>Planctomycetia>Planctomycetales>unclassified_Planctomycetales | 195 | KDA57_17400 | 7874..8461 | 2053591 | RloB-domain-containing-protein-[Planctomycetales-bacterium] |
| QDU83133.1 | TOPRIM | GCA_007750655.1 | Rohdeia_mirabilis | Bacteria>Pseudomonadati>PVC_group>Planctomycetota>Planctomycetia>Planctomycetia_incertae_sedis>Rohdeia | 190 | Pla163_02300 | 293287..293859 | 2528008 | hypothetical-protein-Pla163_02300-[Planctomycetes-bacterium-Pla163] |
| NQW48682.1 | TOPRIM | GCA_013205375.1 | Planctomycetota_bacterium | Bacteria>Pseudomonadati>PVC_group>Planctomycetota>unclassified_Planctomycetota | 219 | HQ464_13055 | 10863..11522 | 2026780 | hypothetical-protein-[Planctomycetota-bacterium] |
| OQA03649.1 | TOPRIM | GCA_002068635.1 | Planctomycetes_bacterium_ADurb.Bin401 | Bacteria>Pseudomonadati>PVC_group>Planctomycetota>unclassified_Planctomycetota | 218 | BWY69_00334 | 11943..12599 | 1852904 | hypothetical-protein-BWY69_00334-[Planctomycetes-bacterium-ADurb.Bin401] |
| MCG3135900.1 | TOPRIM | GCA_022072415.1 | Planctomycetota_bacterium | Bacteria>Pseudomonadati>PVC_group>Planctomycetota>unclassified_Planctomycetota | 217 | HMLKMBBP_03676 | 11785..12438 | 2026780 | hypothetical-protein-[Planctomycetota-bacterium] |
| RIK67121.1 | TOPRIM | GCA_003576905.1 | Planctomycetota_bacterium | Bacteria>Pseudomonadati>PVC_group>Planctomycetota>unclassified_Planctomycetota | 211 | DCC65_07835 | 149670..150305 | 2026780 | MAG:-hypothetical-protein-DCC65_07835-[Planctomycetota-bacterium] |
| MBI1826654.1 | TOPRIM | GCA_016178855.1 | Planctomycetota_bacterium | Bacteria>Pseudomonadati>PVC_group>Planctomycetota>unclassified_Planctomycetota | 209 | HYR83_09755 | 17344..17973 | 2026780 | DUF4276-family-protein-[Planctomycetota-bacterium] |
| MBW3538715.1 | TOPRIM | GCA_019347905.1 | Planctomycetota_bacterium | Bacteria>Pseudomonadati>PVC_group>Planctomycetota>unclassified_Planctomycetota | 208 | KY476_00445 | 26705..27331 | 2026780 | DUF4276-family-protein-[Planctomycetota-bacterium] |
| GIW70820.1 | TOPRIM | GCA_026004615.1 | Planctomycetota_bacterium | Bacteria>Pseudomonadati>PVC_group>Planctomycetota>unclassified_Planctomycetota | 207 | KatS3mg102_0362 | 454538..455161 | 2026780 | MAG:-hypothetical-protein-KatS3mg102_0362-[Planctomycetota-bacterium] |
| MBM4083198.1 | TOPRIM | GCA_016873055.1 | Planctomycetota_bacterium | Bacteria>Pseudomonadati>PVC_group>Planctomycetota>unclassified_Planctomycetota | 207 | FJ272_00235 | 2648..3271 | 2026780 | hypothetical-protein-[Planctomycetota-bacterium] |
| MBI4584118.1 | TOPRIM | GCA_016209605.1 | Planctomycetota_bacterium | Bacteria>Pseudomonadati>PVC_group>Planctomycetota>unclassified_Planctomycetota | 197 | HY717_08855 | 13074..13667 | 2026780 | hypothetical-protein-[Planctomycetota-bacterium] |
| MBK6942984.1 | TOPRIM | GCA_016709145.1 | Planctomycetota_bacterium | Bacteria>Pseudomonadati>PVC_group>Planctomycetota>unclassified_Planctomycetota | 191 | IPH13_22665 | 260313..260888 | 2026780 | DUF4276-family-protein-[Planctomycetota-bacterium] |
| MBI4586097.1 | TOPRIM | GCA_016209605.1 | Planctomycetota_bacterium | Bacteria>Pseudomonadati>PVC_group>Planctomycetota>unclassified_Planctomycetota | 141 | HY717_18970 | 25354..25779 | 2026780 | DUF4276-family-protein-[Planctomycetota-bacterium] |
| AHF94667.1 | TOPRIM | GCA_000242935.3 | Opitutaceae_bacterium_TAV5 | Bacteria>Pseudomonadati>PVC_group>Verrucomicrobiota>Opitutia>Opitutales>Opitutaceae>unclassified_Opitutaceae | 278 | OPIT5_23235 | 5474827..5475663 | 794903 | hypothetical-protein-OPIT5_23235-[Opitutaceae-bacterium-TAV5] |
| MCH8539503.1 | TOPRIM | GCA_022565675.1 | Opitutales_bacterium | Bacteria>Pseudomonadati>PVC_group>Verrucomicrobiota>Opitutia>Opitutales>unclassified_Opitutales | 253 | LAT58_01895 | 21262..22023 | 2026772 | RloB-family-protein-[Opitutales-bacterium] |
| MBE2203057.1 | TOPRIM | GCA_014879575.1 | Chthoniobacterales_bacterium | Bacteria>Pseudomonadati>PVC_group>Verrucomicrobiota>Spartobacteria>Chthoniobacterales>unclassified_Chthoniobacterales | 231 | IAE94_01780 | 25135..25830 | 2201154 | hypothetical-protein-[Chthoniobacterales-bacterium] |
| MBM3880205.1 | TOPRIM | GCA_016871715.1 | Verrucomicrobiota_bacterium | Bacteria>Pseudomonadati>PVC_group>Verrucomicrobiota>unclassified_Verrucomicrobiota | 277 | FJ387_10880 | 27227..28060 | 2026799 | hypothetical-protein-[Verrucomicrobiota-bacterium] |
| MBP7947741.1 | TOPRIM | GCA_018003215.1 | Verrucomicrobiota_bacterium | Bacteria>Pseudomonadati>PVC_group>Verrucomicrobiota>unclassified_Verrucomicrobiota | 221 | KA191_17000 | 9242..9907 | 2026799 | hypothetical-protein-[Verrucomicrobiota-bacterium] |
| NMD22050.1 | TOPRIM | GCA_012797305.1 | Verrucomicrobiota_bacterium | Bacteria>Pseudomonadati>PVC_group>Verrucomicrobiota>unclassified_Verrucomicrobiota | 221 | GYA76_17655 | 3103..3768 | 2026799 | hypothetical-protein-[Verrucomicrobiota-bacterium] |
| MCF7707714.1 | TOPRIM | GCA_021733835.1 | Verrucomicrobiota_bacterium | Bacteria>Pseudomonadati>PVC_group>Verrucomicrobiota>unclassified_Verrucomicrobiota | 220 | K9N52_02295 | 31387..32049 | 2026799 | DUF4276-family-protein-[Verrucomicrobiota-bacterium] |
| MCX6966134.1 | TOPRIM | GCA_026398255.1 | Verrucomicrobiota_bacterium | Bacteria>Pseudomonadati>PVC_group>Verrucomicrobiota>unclassified_Verrucomicrobiota | 218 | NTW41_12415 | 33959..34615 | 2026799 | MAG:-hypothetical-protein-NTW41_12415-[Verrucomicrobia-bacterium] |
| MBU4201309.1 | TOPRIM | GCA_018898455.1 | Verrucomicrobiota_bacterium | Bacteria>Pseudomonadati>PVC_group>Verrucomicrobiota>unclassified_Verrucomicrobiota | 217 | KJ919_15485 | 16857..17510 | 2026799 | hypothetical-protein-[Verrucomicrobiota-bacterium] |
| TAE90782.1 | TOPRIM | GCA_004293285.1 | Verrucomicrobiota_bacterium | Bacteria>Pseudomonadati>PVC_group>Verrucomicrobiota>unclassified_Verrucomicrobiota | 215 | EAZ81_08660 | 9420..10067 | 2026799 | hypothetical-protein-EAZ81_08660-[Verrucomicrobiota-bacterium] |
| NLH73612.1 | TOPRIM | GCA_012517065.1 | Verrucomicrobiota_bacterium | Bacteria>Pseudomonadati>PVC_group>Verrucomicrobiota>unclassified_Verrucomicrobiota | 209 | GX456_11210 | 19236..19865 | 2026799 | hypothetical-protein-[Verrucomicrobiota-bacterium] |
| PWU13197.1 | TOPRIM | GCA_003176045.1 | Verrucomicrobiota_bacterium | Bacteria>Pseudomonadati>PVC_group>Verrucomicrobiota>unclassified_Verrucomicrobiota | 209 | C5B50_19755 | 18402..19031 | 2026799 | hypothetical-protein-C5B50_19755-[Verrucomicrobiota-bacterium] |
| RME93004.1 | TOPRIM | GCA_003695015.1 | Verrucomicrobiota_bacterium | Bacteria>Pseudomonadati>PVC_group>Verrucomicrobiota>unclassified_Verrucomicrobiota | 206 | D6766_09045 | 8644..9264 | 2026799 | hypothetical-protein-D6766_09045-[Verrucomicrobiota-bacterium] |
| MBP7947739.1 | TOPRIM | GCA_018003215.1 | Verrucomicrobiota_bacterium | Bacteria>Pseudomonadati>PVC_group>Verrucomicrobiota>unclassified_Verrucomicrobiota | 205 | KA191_16990 | 7321..7938 | 2026799 | hypothetical-protein-[Verrucomicrobiota-bacterium] |
| NMD22052.1 | TOPRIM | GCA_012797305.1 | Verrucomicrobiota_bacterium | Bacteria>Pseudomonadati>PVC_group>Verrucomicrobiota>unclassified_Verrucomicrobiota | 205 | GYA76_17665 | 5072..5689 | 2026799 | hypothetical-protein-[Verrucomicrobiota-bacterium] |
| MBI4624650.1 | TOPRIM | GCA_016209255.1 | Verrucomicrobiota_bacterium | Bacteria>Pseudomonadati>PVC_group>Verrucomicrobiota>unclassified_Verrucomicrobiota | 203 | HY736_15705 | 14532..15143 | 2026799 | DUF4276-family-protein-[Verrucomicrobiota-bacterium] |
| MBU6399159.1 | TOPRIM | GCA_018969165.1 | Verrucomicrobiota_bacterium | Bacteria>Pseudomonadati>PVC_group>Verrucomicrobiota>unclassified_Verrucomicrobiota | 203 | KGS61_02475 | 2095..2706 | 2026799 | DUF4276-family-protein-[Verrucomicrobiota-bacterium] |
| MBM3880421.1 | TOPRIM | GCA_016871715.1 | Verrucomicrobiota_bacterium | Bacteria>Pseudomonadati>PVC_group>Verrucomicrobiota>unclassified_Verrucomicrobiota | 197 | FJ387_12005 | 8561..9154 | 2026799 | DUF4276-family-protein-[Verrucomicrobiota-bacterium] |
| MBP7947572.1 | TOPRIM | GCA_018003215.1 | Verrucomicrobiota_bacterium | Bacteria>Pseudomonadati>PVC_group>Verrucomicrobiota>unclassified_Verrucomicrobiota | 192 | KA191_16100 | 4544..5122 | 2026799 | hypothetical-protein-[Verrucomicrobiota-bacterium] |
| MCF7668202.1 | TOPRIM | GCA_021733785.1 | Verrucomicrobiota_bacterium | Bacteria>Pseudomonadati>PVC_group>Verrucomicrobiota>unclassified_Verrucomicrobiota | 190 | K9N48_00290 | 28966..29538 | 2026799 | DUF4276-family-protein-[Verrucomicrobiota-bacterium] |
| MCC5848506.1 | TOPRIM | GCA_020831635.1 | Verrucomicrobiota_bacterium | Bacteria>Pseudomonadati>PVC_group>Verrucomicrobiota>unclassified_Verrucomicrobiota | 183 | JJU29_10450 | 22691..23242 | 2026799 | DUF4276-family-protein-[Verrucomicrobiota-bacterium] |
| WP_165106900.1 | TOPRIM | GCF_011044775.1 | Limisphaera_ngatamarikiensis | Bacteria>Pseudomonadati>PVC_group>Verrucomicrobiota>Verrucomicrobiia>Limisphaerales>Limisphaeraceae>Limisphaera | 213 | G4L39_RS06740 | 85332..85973 | 1324935 | hypothetical-protein-[Limisphaera-ngatamarikiensis] |
| MBE7503148.1 | TOPRIM | GCA_015075445.1 | Verrucomicrobiales_bacterium | Bacteria>Pseudomonadati>PVC_group>Verrucomicrobiota>Verrucomicrobiia>Verrucomicrobiales>unclassified_Verrucomicrobiales | 209 | HS113_23260 | 55115..55744 | 2026801 | hypothetical-protein-[Verrucomicrobiales-bacterium] |
| MCI0536694.1 | TOPRIM | GCA_022615925.1 | Verrucomicrobiales_bacterium | Bacteria>Pseudomonadati>PVC_group>Verrucomicrobiota>Verrucomicrobiia>Verrucomicrobiales>unclassified_Verrucomicrobiales | 201 | LOZ50_15865 | 8109..8714 | 2026801 | DUF4276-family-protein-[Verrucomicrobiales-bacterium] |
| MBE7502033.1 | TOPRIM | GCA_015075445.1 | Verrucomicrobiales_bacterium | Bacteria>Pseudomonadati>PVC_group>Verrucomicrobiota>Verrucomicrobiia>Verrucomicrobiales>unclassified_Verrucomicrobiales | 173 | HS113_17415 | 120778..121299 | 2026801 | RloB-domain-containing-protein-[Verrucomicrobiales-bacterium] |
| MBO7485541.1 | TOPRIM | GCA_017649985.1 | Spirochaetaceae_bacterium | Bacteria>Pseudomonadati>Spirochaetota>Spirochaetia>Spirochaetales>Spirochaetaceae>unclassified_Spirochaetaceae | 216 | J6T84_05720 | 12567..13217 | 1898206 | DUF4276-family-protein-[Spirochaetaceae-bacterium] |
| MDD9989002.1 | TOPRIM | GCA_028818385.1 | Spirochaetaceae_bacterium | Bacteria>Pseudomonadati>Spirochaetota>Spirochaetia>Spirochaetales>Spirochaetaceae>unclassified_Spirochaetaceae | 191 | OXQ31_22215 | 6417..6992 | 1898206 | MAG:-hypothetical-protein-OXQ31_22215-[Spirochaetaceae-bacterium] |
| MDE0447652.1 | TOPRIM | GCA_028821475.1 | Spirochaetaceae_bacterium | Bacteria>Pseudomonadati>Spirochaetota>Spirochaetia>Spirochaetales>Spirochaetaceae>unclassified_Spirochaetaceae | 191 | OXH96_13345 | 28157..28732 | 1898206 | MAG:-hypothetical-protein-OXH96_13345-[Spirochaetaceae-bacterium] |
| OQY41118.1 | TOPRIM | GCA_002084785.1 | Spirochaetaceae_bacterium_4572_7 | Bacteria>Pseudomonadati>Spirochaetota>Spirochaetia>Spirochaetales>Spirochaetaceae>unclassified_Spirochaetaceae | 191 | B6229_00030 | 6838..7413 | 1972456 | hypothetical-protein-B6229_00030-[Spirochaetaceae-bacterium-4572_7] |
| GHU24468.1 | TOPRIM | GCA_026002265.1 | Spirochaetia_bacterium | Bacteria>Pseudomonadati>Spirochaetota>Spirochaetia>unclassified_Spirochaetia | 214 | FACS1894164_11180 | 3980..4624 | 2053615 | hypothetical-protein-FACS1894164_11180-[Spirochaetia-bacterium] |
| GHV83241.1 | TOPRIM | GCA_025997755.1 | Spirochaetia_bacterium | Bacteria>Pseudomonadati>Spirochaetota>Spirochaetia>unclassified_Spirochaetia | 213 | AGMMS50212_05810 | 5581..6222 | 2053615 | hypothetical-protein-AGMMS50212_05810-[Spirochaetia-bacterium] |
| TXT44737.1 | TOPRIM | GCA_008080685.1 | Spirochaetota_bacterium | Bacteria>Pseudomonadati>Spirochaetota>unclassified_Spirochaetota | 210 | FD137_1512 | 8247..8879 | 2202144 | hypothetical-protein-FD137_1512-[Spirochaetota-bacterium] |
| OQB96945.1 | TOPRIM | GCA_002070755.1 | Spirochaetes_bacterium_ADurb.Bin110 | Bacteria>Pseudomonadati>Spirochaetota>unclassified_Spirochaetota | 209 | BWX81_01214 | 13653..14282 | 1852908 | hypothetical-protein-BWX81_01214-[Spirochaetes-bacterium-ADurb.Bin110] |
| HEC67880.1 | TOPRIM | GCA_011052795.1 | Candidatus_Desulfofervidus_auxilii | Bacteria>Pseudomonadati>Thermodesulfobacteriota>Candidatus_Desulfofervidia>Candidatus_Desulfofervidales>Candidatus_Desulfofervidaceae>Candidatus_Desulfofervidus | 157 | ENI35_03590 | 7079..7552 | 1621989 | hypothetical-protein-[Candidatus-Desulfofervidus-auxilii] |

| Protein-ID | Domain | Assembly-ID | Strain | Taxonomy | Length | Locus-Tag | Genomic-coordinates | TaxID | Fasta-defline |
|---|---|---|---|---|---|---|---|---|---|
| WP_073476322.1 | TOPRIM | GCF_900142135.1 | Desulfatibacillum_alkenivorans | Bacteria>Pseudomonadati>Thermodesulfobacteriota>Desulfobacteria>Desulfobacterales>Desulfatibacillaceae>Desulfatibacillum | 230 | BUE18_RS12860 | 65677..66369 | 259354 | DUF4276-family-protein-[Desulfatibacillum-alkenivorans] |
| WP_028324420.1 | TOPRIM | GCF_000429925.1 | Desulfatirhabdium_butyrativorans | Bacteria>Pseudomonadati>Thermodesulfobacteriota>Desulfobacteria>Desulfobacterales>Desulfatirhabdiaceae>Desulfatirhabdium | 226 | G492_RS0109405 | 93715..94395 | 340467 | DUF4276-family-protein-[Desulfatirhabdium-butyrativorans] |
| WP_028324161.1 | TOPRIM | GCF_000429925.1 | Desulfatirhabdium_butyrativorans | Bacteria>Pseudomonadati>Thermodesulfobacteriota>Desulfobacteria>Desulfobacterales>Desulfatirhabdiaceae>Desulfatirhabdium | 205 | G492_RS0107595 | 72019..72636 | 340467 | DUF4276-family-protein-[Desulfatirhabdium-butyrativorans] |
| WP_080806590.1 | TOPRIM | - | Desulfamplus_magnetovallimortis | Bacteria>Pseudomonadati>Thermodesulfobacteriota>Desulfobacteria>Desulfobacterales>Desulfobacteraceae>Desulfamplus | 220 | MTBBW1_RS20360 | 77271..77933 | 1246637 | DUF4276-family-protein-[Desulfamplus-magnetovallimortis] |
| MBF0468018.1 | TOPRIM | GCA_015232935.1 | Desulfamplus_sp. | Bacteria>Pseudomonadati>Thermodesulfobacteriota>Desulfobacteria>Desulfobacterales>Desulfobacteraceae>Desulfamplus>unclassified_Desulfamplus | 238 | HQK61_03900 | 2867..3583 | 2006184 | hypothetical-protein-[Desulfamplus-sp.] |
| HBT88453.1 | TOPRIM | GCA_003510365.1 | Desulfobacter_sp. | Bacteria>Pseudomonadati>Thermodesulfobacteriota>Desulfobacteria>Desulfobacterales>Desulfobacteraceae>Desulfobacter>unclassified_Desulfobacter | 219 | DEB50_08310 | 26676..27335 | 2294 | hypothetical-protein-[Desulfobacter-sp.] |
| MCF8089784.1 | TOPRIM | GCA_021737465.1 | Desulfotignum_sp. | Bacteria>Pseudomonadati>Thermodesulfobacteriota>Desulfobacteria>Desulfobacterales>Desulfobacteraceae>Desulfotignum>unclassified_Desulfotignum | 237 | K9L23_17205 | 7806..8519 | 2049434 | DUF4276-family-protein-[Desulfotignum-sp.] |
| MBC8432679.1 | TOPRIM | GCA_014383215.1 | Candidatus_Desulfatibia_vada | Bacteria>Pseudomonadati>Thermodesulfobacteriota>Desulfobacteria>Desulfobacterales>Desulfobacterales_incertae_sedis>Candidatus_Desulfatibia | 218 | H8D96_12275 | 9976..10632 | 2841696 | DUF4276-family-protein-[Desulfobacterales-bacterium] |
| WP_207691045.1 | TOPRIM | GCF_017377355.1 | Desulfonema_limicola | Bacteria>Pseudomonadati>Thermodesulfobacteriota>Desulfobacteria>Desulfobacterales>Desulfococcaceae>Desulfonema | 298 | dnl_RS07070 | 1636651..1637547 | 45656 | hypothetical-protein-[Desulfonema-limicola] |
| WP_207678194.1 | TOPRIM | GCF_017377395.1 | Desulfonema_magnum | Bacteria>Pseudomonadati>Thermodesulfobacteriota>Desulfobacteria>Desulfobacterales>Desulfococcaceae>Desulfonema | 134 | dnm_RS17445 | 4556429..4556833 | 45655 | RloB-family-protein-[Desulfonema-magnum] |
| WP_155316142.1 | TOPRIM | GCF_009688945.1 | Desulfosarcina_alkanivorans | Bacteria>Pseudomonadati>Thermodesulfobacteriota>Desulfobacteria>Desulfobacterales>Desulfosarcinaceae>Desulfosarcina | 185 | DSCA_RS09155 | 2076587..2077144 | 571177 | DUF4276-family-protein-[Desulfosarcina-alkanivorans] |
| MBC2710213.1 | TOPRIM | GCA_014237405.1 | Desulfosarcina_sp. | Bacteria>Pseudomonadati>Thermodesulfobacteriota>Desulfobacteria>Desulfobacterales>Desulfosarcinaceae>Desulfosarcina>unclassified_Desulfosarcina | 208 | HGJ94_04180 | 15554..16180 | 2027861 | hypothetical-protein-[Desulfosarcina-sp.] |
| MBF0101460.1 | TOPRIM | GCA_015228675.1 | Desulfobacterales_bacterium | Bacteria>Pseudomonadati>Thermodesulfobacteriota>Desulfobacteria>Desulfobacterales>unclassified_Desulfobacterales | 225 | HQK77_11180 | 5617..6294 | 2044940 | DUF4276-family-protein-[Desulfobacterales-bacterium] |
| MBL0712136.1 | TOPRIM | GCA_016735415.1 | Desulfobacterales_bacterium | Bacteria>Pseudomonadati>Thermodesulfobacteriota>Desulfobacteria>Desulfobacterales>unclassified_Desulfobacterales | 220 | JJV98_00415 | 8780..9442 | 2044940 | hypothetical-protein-[Desulfosarcina-sp.] |
| MCG2778952.1 | TOPRIM | GCA_022013615.1 | Desulfobacterales_bacterium | Bacteria>Pseudomonadati>Thermodesulfobacteriota>Desulfobacteria>Desulfobacterales>unclassified_Desulfobacterales | 204 | L6406_25025 | 8280..8894 | 2044940 | DUF4276-family-protein-[Desulfobacterales-bacterium] |
| MDD2316835.1 | TOPRIM | GCA_028683495.1 | Desulfobacterales_bacterium | Bacteria>Pseudomonadati>Thermodesulfobacteriota>Desulfobacteria>Desulfobacterales>unclassified_Desulfobacterales | 188 | PHD57_05755 | 12416..12982 | 2044940 | MAG:-DUF4276-family-protein-[Desulfobacterales-bacterium] |
| MCU0558985.1 | TOPRIM | GCA_025454255.1 | Desulfobacterales_bacterium | Bacteria>Pseudomonadati>Thermodesulfobacteriota>Desulfobacteria>Desulfobacterales>unclassified_Desulfobacterales | 80 | MUD16_02170 | 69903..70145 | 2044940 | hypothetical-protein-[Desulfobacterales-bacterium] |
| WP_051554418.1 | TOPRIM | GCF_000621145.1 | Desulfobulbus_elongatus | Bacteria>Pseudomonadati>Thermodesulfobacteriota>Desulfobulbia>Desulfobulbales>Desulfobulbaceae>Desulfobulbus | 211 | Q362_RS21225 | 6684..7319 | 53332 | hypothetical-protein-[Desulfobulbus-elongatus] |
| WP_205241969.1 | TOPRIM | GCF_016918545.1 | Desulfobulbus_alkaliphilus | Bacteria>Pseudomonadati>Thermodesulfobacteriota>Desulfobulbia>Desulfobulbales>Desulfobulbaceae>Desulfobulbus | 183 | JWG43_RS06295 | 125254..125805 | 869814 | hypothetical-protein-[Desulfobulbus-alkaliphilus] |
| WP_013164347.1 | TOPRIM | GCF_000092205.1 | Desulfurivibrio_alkaliphilus | Bacteria>Pseudomonadati>Thermodesulfobacteriota>Desulfobulbia>Desulfobulbales>Desulfobulbaceae>Desulfurivibrio | 213 | DAAHT2_RS10965 | 2517357..2517998 | 427923 | DUF4276-family-protein-[Desulfurivibrio-alkaliphilus] |
| MBE0584989.1 | TOPRIM | GCA_014860245.1 | Desulfofustis_sp. | Bacteria>Pseudomonadati>Thermodesulfobacteriota>Desulfobulbia>Desulfobulbales>Desulfocapsaceae>Desulfofustis>unclassified_Desulfofustis | 185 | IH612_14680 | 884..1441 | 2053546 | DUF4276-family-protein-[Desulfofustis-sp.] |
| BDV01250.1 | TOPRIM | GCA_027925145.1 | Thermodesulfomicrobium_sp._WS | Bacteria>Pseudomonadati>Thermodesulfobacteriota>Desulfovibrionia>Desulfovibrionales>Desulfomicrobiaceae>Thermodesulfomicrobium>unclassified_Thermodesulfomicrobium | 204 | TDMWS_13350 | 1436069..1436683 | 3004129 | hypothetical-protein-TDMWS_13350-[Thermodesulfomicrobium-sp.-WS] |
| HCF05839.1 | TOPRIM | GCA_003530935.1 | Desulfomicrobiaceae_bacterium | Bacteria>Pseudomonadati>Thermodesulfobacteriota>Desulfovibrionia>Desulfovibrionales>Desulfomicrobiaceae>unclassified_Desulfomicrobiaceae | 194 | DEU72_06350 | 12171..12755 | 2053547 | cytoplasmic-protein-[Desulfomicrobiaceae-bacterium] |
| MBZ4685272.1 | TOPRIM | GCA_020063575.1 | Desulfomicrobiaceae_bacterium | Bacteria>Pseudomonadati>Thermodesulfobacteriota>Desulfovibrionia>Desulfovibrionales>Desulfomicrobiaceae>unclassified_Desulfomicrobiaceae | 194 | JG774_1017 | 62696..63280 | 2053547 | Uncharacterized-protein-[Desulfomicrobiaceae-bacterium] |
| MBC7355964.1 | TOPRIM | GCA_014359945.1 | Desulfomicrobiaceae_bacterium | Bacteria>Pseudomonadati>Thermodesulfobacteriota>Desulfovibrionia>Desulfovibrionales>Desulfomicrobiaceae>unclassified_Desulfomicrobiaceae | 128 | H5U09_07175 | 27894..28280 | 2053547 | hypothetical-protein-[Desulfomicrobiaceae-bacterium] |
| WP_008869232.1 | TOPRIM | GCF_000174435.1 | Desulfonatronospira | Bacteria>Pseudomonadati>Thermodesulfobacteriota>Desulfovibrionia>Desulfovibrionales>Desulfonatronovibrionaceae | 223 | DTHIO_RS04840 | 1064428..1065099 | 488937 | DUF4276-family-protein-[Desulfonatronospira-thiodismutans] |
| WP_045217258.1 | TOPRIM | GCF_000934755.1 | Desulfonatronovibrio_magnus | Bacteria>Pseudomonadati>Thermodesulfobacteriota>Desulfovibrionia>Desulfovibrionales>Desulfonatronovibrionaceae>Desulfonatronovibrio | 220 | LZ23_RS20530 | 18744..19406 | 698827 | hypothetical-protein-[Desulfonatronovibrio-magnus] |
| WP_270570095.1 | TOPRIM | - | Bilophila_wadsworthia | Bacteria>Pseudomonadati>Thermodesulfobacteriota>Desulfovibrionia>Desulfovibrionales>Desulfovibrionaceae>Bilophila | 220 | PG042_RS02155 | 64144..64806 | 35833 | hypothetical-protein-[Bilophila-wadsworthia] |
| MCI7405011.1 | TOPRIM | GCA_022783735.1 | Desulfovibrio_piger | Bacteria>Pseudomonadati>Thermodesulfobacteriota>Desulfovibrionia>Desulfovibrionales>Desulfovibrionaceae>Desulfovibrio | 221 | MSS64_01025 | 29647..30312 | 901 | DUF4276-family-protein-[Desulfovibrio-piger] |
| HJB66435.1 | TOPRIM | GCA_019119045.1 | Candidatus_Mailhella_merdavium | Bacteria>Pseudomonadati>Thermodesulfobacteriota>Desulfovibrionia>Desulfovibrionales>Desulfovibrionaceae>Mailhella | 214 | H9768_09165 | 45490..46134 | 2838657 | hypothetical-protein-[Candidatus-Mailhella-merdavium] |
| PLX83383.1 | TOPRIM | GCA_002868845.1 | Desulfuromonas_sp. | Bacteria>Pseudomonadati>Thermodesulfobacteriota>Desulfuromonadia>Desulfuromonadales>Desulfuromonadaceae>Desulfuromonas>unclassified_Desulfuromonas | 177 | C0617_12100 | 151924..152457 | 892 | MAG:-abortive-phage-resistance-protein-[Desulfuromonas-sp.] |
| WP_207162786.1 | TOPRIM | GCF_017338855.1 | Geobacter_benzoatilyticus | Bacteria>Pseudomonadati>Thermodesulfobacteriota>Desulfuromonadia>Geobacterales>Geobacteraceae>Geobacter | 193 | JZM60_RS12585 | 2734246..2734827 | 2815309 | RloB-family-protein-[Geobacter-benzoatilyticus] |
| WP_148202434.1 | TOPRIM | GCF_000013405.1 | Syntrophus_aciditrophicus | Bacteria>Pseudomonadati>Thermodesulfobacteriota>Syntrophia>Syntrophales>Syntrophaceae>Syntrophus | 200 | SYN_RS00155 | 34295..34897 | 316277 | DUF4276-family-protein-[Syntrophus-aciditrophicus] |
| ABC75910.1 | TOPRIM | GCA_000013405.1 | Syntrophus_aciditrophicus_SB | Bacteria>Pseudomonadati>Thermodesulfobacteriota>Syntrophia>Syntrophales>Syntrophaceae>Syntrophus>Syntrophus_aciditrophicus | 203 | SYN_00009 | 34295..34906 | 56780 | hypothetical-cytosolic-protein-[Syntrophus-aciditrophicus-SB] |
| OPY81710.1 | TOPRIM | GCA_002067745.1 | Syntrophus_sp._PtaU1.Bin005 | Bacteria>Pseudomonadati>Thermodesulfobacteriota>Syntrophia>Syntrophales>Syntrophaceae>Syntrophus>unclassified_Syntrophus_(in:_bacteria) | 198 | A4E70_01076 | 23145..23741 | 1811712 | MAG:-hypothetical-protein-A4E70_01076-[Syntrophus-sp.-PtaU1.Bin005] |
| OIP91483.1 | TOPRIM | GCA_001873745.1 | Syntrophaceae_bacterium_CG2_30_58_14 | Bacteria>Pseudomonadati>Thermodesulfobacteriota>Syntrophia>Syntrophales>Syntrophaceae>unclassified_Syntrophaceae | 214 | AUK26_06770 | 2541..3185 | 1805378 | hypothetical-protein-AUK26_06770-[Syntrophaceae-bacterium-CG2_30_58_14] |
| MCK9230423.1 | TOPRIM | GCA_023230845.1 | Syntrophales_bacterium | Bacteria>Pseudomonadati>Thermodesulfobacteriota>Syntrophia>Syntrophales>unclassified_Syntrophales | 117 | MOP18_07075 | 1960..2313 | 2928017 | DUF4276-family-protein-[Syntrophales-bacterium] |
| MCK9528170.1 | TOPRIM | GCA_023228425.1 | Syntrophales_bacterium | Bacteria>Pseudomonadati>Thermodesulfobacteriota>Syntrophia>Syntrophales>unclassified_Syntrophales | 117 | MOQ23_05910 | 66500..66853 | 2928017 | DUF4276-family-protein-[Syntrophales-bacterium] |
| WP_137424066.1 | TOPRIM | GCF_005377625.1 | Desulfoglaeba_alkanexedens | Bacteria>Pseudomonadati>Thermodesulfobacteriota>Syntrophobacteria>Syntrophobacterales>Syntrophobacteraceae>Desulfoglaeba | 202 | FDQ92_RS07905 | 1745466..1746074 | 361111 | DUF4276-family-protein-[Desulfoglaeba-alkanexedens] |
| WP_137423744.1 | TOPRIM | GCF_005377625.1 | Desulfoglaeba_alkanexedens | Bacteria>Pseudomonadati>Thermodesulfobacteriota>Syntrophobacteria>Syntrophobacterales>Syntrophobacteraceae>Desulfoglaeba | 199 | FDQ92_RS06005 | 1284292..1284891 | 361111 | hypothetical-protein-[Desulfoglaeba-alkanexedens] |
| WP_170161766.1 | TOPRIM | GCF_003751385.1 | Desulfosoma_caldarium | Bacteria>Pseudomonadati>Thermodesulfobacteriota>Syntrophobacteria>Syntrophobacterales>Syntrophobacteraceae>Desulfosoma | 202 | EDC27_RS10785 | 104772..105380 | 610254 | DUF4276-family-protein-[Desulfosoma-caldarium] |
| WP_123290642.1 | TOPRIM | GCF_003751385.1 | Desulfosoma_caldarium | Bacteria>Pseudomonadati>Thermodesulfobacteriota>Syntrophobacteria>Syntrophobacterales>Syntrophobacteraceae>Desulfosoma | 201 | EDC27_RS10770 | 99515..100120 | 610254 | hypothetical-protein-[Desulfosoma-caldarium] |
| PIV05609.1 | TOPRIM | GCA_002779905.1 | Syntrophobacterales_bacterium_CG03_land_8_20_14_0_80_58_14 | Bacteria>Pseudomonadati>Thermodesulfobacteriota>Syntrophobacteria>Syntrophobacterales>unclassified_Syntrophobacterales | 232 | COS57_06995 | 11279..11977 | 1974100 | hypothetical-protein-COS57_06995-[Syntrophobacterales-bacterium-CG03_land_8_20_14_0_80_58_14] |
| MBW6486652.1 | TOPRIM | GCA_019429105.1 | Syntrophobacterales_bacterium | Bacteria>Pseudomonadati>Thermodesulfobacteriota>Syntrophobacteria>Syntrophobacterales>unclassified_Syntrophobacterales | 185 | K0B01_10945 | 10808..11365 | 2282155 | DUF4276-family-protein-[Syntrophobacterales-bacterium] |
| WP_051408773.1 | TOPRIM | GCF_000512235.1 | Syntrophorhabdus_aromaticivorans | Bacteria>Pseudomonadati>Thermodesulfobacteriota>Syntrophorhabdia>Syntrophorhabdales>Syntrophorhabdaceae>Syntrophorhabdus | 208 | SYNAR_RS19505 | 128178..128804 | 328301 | FGGY-family-carbohydrate-kinase-[Syntrophorhabdus-aromaticivorans] |
| OPY73255.1 | TOPRIM | GCA_002067415.1 | Syntrophorhabdus_sp._PtaU1.Bin002 | Bacteria>Pseudomonadati>Thermodesulfobacteriota>Syntrophorhabdia>Syntrophorhabdales>Syntrophorhabdaceae>Syntrophorhabdus>unclassified_Syntrophorhabdus | 216 | A4E62_00571 | 22979..23629 | 1811702 | MAG:-hypothetical-protein-A4E62_00571-[Syntrophorhabdus-sp.-PtaU1.Bin002] |
| MBX3143489.1 | TOPRIM | GCA_019637455.1 | Trueperaceae_bacterium | Bacteria>Thermotogati>Deinococcota>Deinococci>Trueperales>Trueperaceae>unclassified_Trueperaceae | 233 | KF813_07025 | 11318..12019 | 2026798 | hypothetical-protein-[Trueperaceae-bacterium] |
| MCR9098962.1 | TOPRIM | GCA_024742755.1 | bacterium | Bacteria>unclassified_Bacteria | 223 | NXI25_03360 | 18909..19580 | 1869227 | hypothetical-protein-[bacterium] |
| MCY3637187.1 | TOPRIM | GCA_026705625.1 | bacterium | Bacteria>unclassified_Bacteria | 219 | OXH23_16420 | 22533..23192 | 1869227 | MAG:-RloB-family-protein-[bacterium] |
| MBD5384948.1 | TOPRIM | GCA_014802625.1 | bacterium | Bacteria>unclassified_Bacteria | 216 | HDR69_00780 | 102154..102804 | 1869227 | DUF4276-family-protein-[bacterium] |
| MCP4662278.1 | TOPRIM | GCA_024231315.1 | bacterium | Bacteria>unclassified_Bacteria | 208 | GY856_43305 | 7076..7702 | 1869227 | DUF4276-family-protein-[bacterium] |
| KAF0250486.1 | TOPRIM | GCA_009773835.1 | bacterium | Bacteria>unclassified_Bacteria | 198 | FD167_112 | 10413..11009 | 1869227 | hypothetical-protein-FD167_112-[bacterium] |
| MBU1261807.1 | TOPRIM | GCA_018819855.1 | bacterium | Bacteria>unclassified_Bacteria | 157 | KJ640_02500 | 1481..1954 | 1869227 | hypothetical-protein-[bacterium] |
| MBU1598555.1 | TOPRIM | GCA_018822205.1 | bacterium | Bacteria>unclassified_Bacteria | 157 | KKG61_00345 | 2877..3350 | 1869227 | hypothetical-protein-[bacterium] |
| MBU2462379.1 | TOPRIM | GCA_018828965.1 | bacterium | Bacteria>unclassified_Bacteria | 157 | KKH65_05850 | 2287..2760 | 1869227 | hypothetical-protein-[bacterium] |

```
# ; HEPN-EndoRNase

                   1---------11--------21--------31--------41--------51--------61--------71--------81--------91--------101-------111--
OrigSeq            MDELTGAILEKIVYRADRIIRYVTEHTQETFLADEMVQDAVCMNLLNIGELVKQLPIDIRTQYPLVPWRKLAGLRDITAHRYDSLRMVDIWTAANKEVSALKTQVEEILEQREMK

Jnet               -----HHHHHHHHHHHHHHHHHHHHH--HHHHH--HHHHHHHHHHHHHHHHHHHH---HHHHHH-----HHHHHH----EE-------HHHHHHHHHH--HHHHHHHHHHHHHH----
jhmm               -----HHHHHHHHHHHHHHHHHHHHH--HH-----HHHHHHHHHHHHHHHHHHHH---HHHHHH-----HHHHHH----HEE------HHHHHHHHHH-H-HHHHHHHHHHH-----
jpssm              -----HHHHHHHHHHHHHHHHHHHHH--HHHHHH-HHHHHHHHHHHHHHHHHHH----HHHHHH-----HHHHHH---EEE-------HHHHHHHHHHH-HHHHHHHHHHHHHHH---
GHU66456.1         MDELTGAILEKIVYRADRIIRYVTEHTQETFLADEMVQDAVCMNLLNIGELVKQLPIDIRTQYPLVPWRKLAGLRDITAHRYDSLRMVDIWTAANKEVSALKTQVEEILEQREMK
GIW33265.1         MRPDDRVRLQHMVEAAESIATFVAGRTRGDLDTDRMLLFAVVRAIEIIGEAASQISDETRSRAEDVPWRAMVGMRNRIVHAYFDIDADIVWKTAVEEVPGLLPRLRELLEQAKKG
GIW47697.1         MRRGDREFLMDILEACRRIKNYTKGLSYEVFLRSEEKQDAVIRNMEVIGEAVKHISYNLRNKYPEVDWKKVAGIRDKLIHFYFGISLEIVWTVIEKEIPKLEESVKRIMEKEGWS
HAA34482.1         MLKHDLVRVKHMRDGAEEIQKFIKGKTRNDLEHDRMLSLAIVQLLEIIGEAANKVTPEFCAKYTQIPWKSVVGMRNRLIHGYFDIDLNIVWKTAVDEIPSLNKALTEIIKTEEFA
HAF70682.1         FKKDWRLFAQDILESIARIESYVADLAYEDFLADNKTQDAVVRNLEIIGEAARHIPQDVQSQHPGIPWAQLIGLRNRLIHGYFVVDYEIVWDIVKNELPGLKRKLAAILEELRGK
MBA1445771.1       MSKRDQLYLEHIEQAMDRIQRYLVEFDHAAFLENEEKQDAVIRNLEIIGEASGKIQPDFASEHPEFPLKAAYGTRNVLAHGYFKVDLDVVWKTVEKDLPELRARLAAVLSSLKQM
MBC8359967.1       MQKSDIIRLRHMFDAAKEAASFIQEEERASLDVDRKLVLALMKSIEIIGEAATKITKECQEDLSQIPWPNIIGMRNRLIHAYFDVNLEILWKTVTEDLPGLIDELEKILPSAKV-
MBC8458332.1       RHKSPLPFIKDIVDSIEKIEKYCQNITYEQFLQDDKTKDAVVRNLEIIGEAAKNIPDHIKKISSKTAWKSMTGMRDKLIHEYFGVSYSTVWETVKGDLPLLKESIRELYDTIKAK
MBE7444887.1       MPRDFRLYLDDILEAIHQIRTYMVDMDEEAFCADRKTQDAVIRNLEVIGEAAGNLPEQIQKAAPEVDWRKIKGLRNILIHEYFGINLPIVWDVIQNKLDPLETVCRKLVEIAIET
MBE7469122.1       -MRDDRQRLLDVQEAIERIEKYTT-QGRAAFEQDELIQTWVIHHLQIIGEACRAISQPLKDQYPTLPWTKIIGMRHILVHTYFGIDEEIVWSVIEQDLSGLKRDIEAILQQLGHD
MBF0120189.1       MKIQDKVRFEHISEAILEIESYVKGISEGDFVRNSLIRSATVRQFEIIGEAVKAISEELQGQYPKIPWRLWANFRNVLIHQYFGVDYEEVYKTVQNDLPTLKRQIDEILSKA---
MBI2569496.1       KQRDARVYLYDALQACVLVCQFTAGKGHTDYSGDPFLRSAVERQLEIVGEAMNRALKCDPSLANELPSSKIIAFRNRLVHAYFAISDAVVWRIVESDVPILRDKLRRLLPPDEVA
MBK8639648.1       SDREWRFYIDDMVDFAEHVLAYTQGLDQAGLVADRLRYDATLRNLELIGEAATHLPREVRMANPDIPWRLVIAMRNRLVHGYLGIDNDTLWSVVQQDVPALLPKLRALRDAVQTT
MBQ6961346.1       MKHSDQVILQKIIYYCNEIDMLMNGKSQEAYEQDRAYQYAVSTCIVQIGELVARLTEETLAANDSIPWRAIRGMRNIYVHDYEKTKYKTMWQTMTEDIPTLRSQLLDLLDHDEDS
MBU2055999.1       MKKDPRVYLAQILERIDRILEYTAE-GKDAFFTDARTQDAVIRNFEVIGEAAKRIPGDYRENHPLIPWRALSGFRDVLIHQYEGVSVAEIWQIIERDLETLREAIIKILPPLKEL
MBW7864499.1       -MRTDPILLQDMLDAIGEVIENTPA-DRDTYNADKFLRSHLVRNIQIIGEAAVRLSKELRDANPQVPWRQIAGMRHAIVHDYFQIDWNEVYDTAVRDIPPLNSLIADIIAALPET
MBX6329368.1       MKPDDRIRLRHMAEALTSLGRFVSGRRRQDLDNDQMLAYALVHALQIVGEAGSKISAETRAMNPAIPWAEIIGMRHRLVHAYVDVDLDLVWTTATEAAPALLAQIAPLLD----G
MBZ4682112.1       PNRDWKLLILDILESINKIERYTEGMDYELFEKAEQTKDAVVRNLEIIGEASSKIPKDIQQKYNHIPWGKIIAMRNRMIHGYFTIDYRIVWKIIKEDLPELKEQLEQTLKKEKG-
MCI1262622.1       FMRRELLLLREMRDAALAARELVAGLSVEQVEADRMRRSALLWHFTVLGEAASQVPPETKSSEPAIAWRASTRMRNRIVHGYWDIDVATLVATAADDLPRMIAQLEALIMVLDVD
MCL4543925.1       QPREWTLRIEDMLEAAAHCRQYVAGMDFAAFTSDQKTIDAVVRNLEIIGEAARYVPPEIELRYADIPWVDIRGMRHRIAHGYFAVDLDVVWGTATTSLNTLVPLLEQLLQQESQP
MCQ3930199.1       MSRDYKLYLRDILAAIQNIERFTAGVDEETFSADELRLHSVLYNLMVIGEAVKSLPDEMRARVPEVPWRDIKRFRDKLVHHYFSIKTSIVWQIIQEDLATLKTAAETLLTNLEEE
MCR4416750.1       MKRKSSLYLKDILTAIQKIDEFIEGMNFEQFLEDDKTSSAVIRKLEIIGEAVKQLPPEIRERFSEIPWAAMAKSRDKIIHFYHGIDYEIIWQIIKEDLPRVKEKLQQIYFLIVKE
MCS4056783.1       STSDDLLYLDHIEERIQRIEDCAEE-GQSAFEKSHVLQDAVIRNFEVIGEAVKQLSPKLRAQYPEVPWRRIAGFRDVLIHDYMGVDLQEVWNVIENDLPALKEIVGAMREDLGEE
MCS7301718.1       MSRELRLFLKDILDAMDAIEEFTHGMDYDAFLADRRTRNAVERNLEIIGEAVRHVPEEMRAQFPEVEWRKIAGMRDRIIHAYFGVDYTLVWETVRTHIPATRPQIERVLQTIEQQ
MCU0836360.1       MTKQWQPYASHILDTIGKIRRIQ---ARGDLMQDEVLYDAPLRNLQTLSEATQHLPEELKQQHPGIPWKQISGFRNILVHQYLEIDPQTVRQVVEQHLTPLEYAIQSMLTGLGE-
MCX7997939.1       SSRNYKYFIADIIEAINRIETYTNECSFENFQKDIKTQDAVIKNLEVIGEASNKIPTVIKEKYLEIPWKEISGLRNHLIHAYFTIDFNIIWNIIQQELPKLKIEFEKIRNDIQE-
MCZ7663781.1       --MRDLLLLLNIRESIESIEEYVRD-GRSAFLGDKKTRDAVLRNLHTLSESALRVSPELKARYPEVAWRDIAAFRNVVVHDYLALDLAVIWDVVVADLPVLRSQVVGVLEDLSTR
MDE2692296.1       DRGTDLVLLEHIRDRISRIGEYTGG-DSSKFFGSTLVQDAVVRNLQTIAESTQRLSETLRATEPEIPWRAIAGFRNVLVHDYFGIDVEAVWTVVEQDLPSLAAAVERMAALAAGT
MXY29379.1         PMRNYRLYLKDIFEAMGTAQIFVEGMDFEAFVADDKTASAVVRKLEIIGEATKNVPKTVRQKYSQVPWRQMAGMRDRIIHAYFDVNYTIVWEVVTEMIPPLQPIIAQILKDLEEE
NBK98919.1         SQRDWTLYVKDIFRECERIETFVYGITYEQFTDNIEKVYAVAKAFENIGEAVKQLPQELTSKYPNIPWSEIAKMRDVLTHHYFGLDDKVLWDTLDEDFKLFKLTIEQMLKEIG--
NUQ24581.1         SKRSNHELLVDILECIDKIKRYTQDMSFEQFRGDEKSIDAVVRNFEIIGEAANRVDETIKRKYASVEWRRIIGLRNRIVHEYFGIDLQIIWTIIEENMDAFKFEIEEIIDEVS--
OYT64650.1         SKRDIKLYVEGILELIKAIEKFVEGADFESFKEDEKASSAVIRKFEIIGEATKNIPQSIKEKYLHISRREILGLCDRLIRSHFKIQYDVVWDTIKLKLPELRNNIERVPKDLEGE
PFG73562.1         -MRDPRERLRDMLEAIAAIERYL-GRGQAAFEQDELLQGWFVRNLQVIGEAARALPQETRDLAPDVPWPAIIGMRNVLVHGYFDIDTAIVWEAASRDLPALRPRLEALLQRLEEQ
PMB78269.1         MKRDYRLFLKDIIDAMESIERFVEGMELTDFMKDDKTSSAVIRKFEIIGEAVKYLPDHIKEKHARIPWKKMAGMRDKLIHAYFGVDYLLIWETIKSEIPKLKPELRKVLQELEKK
RMH48111.1         -MKSDAVYLHHVLDALRQIEDYMDGISLDVFLSDRMRQDAVVRQLEIIGEACRQLSDALRARHPDVPWRAIIGMRNRIAHDYLNIDLHVIWDVVQVDLPSLKTSIERILG-----
WP_011958257.1     MSRNVLVYIKDILQNMRDAEQFIQGMTYEQFVADKKTVNAILRSIEVIGEAAKRVPDDVRAQYPQIPWKEMAGMRDKVIHLYFSVDNETVWLVVKERIPSLQPLIEQIVRDLEDQ
WP_015227929.1     MSRSPEDYLQHILDEAEYLFQSSQNLSKEDFLKDETLKRAYIRSLEVIGEATKNLPNSFKAEHPEVAWRRMAGMRDKLIHNYFGVDYDIVWNAIIDEIPNLKVKIENILKEIKGE
WP_119292280.1     MSPDDLIRLRHMADAAALAMEFARGRERAELDRDAMLRLALTRAVEIVGEAASKVSAEARQGLEKVAWAAIVGMRNRLVHAYFDVDVDQLWMVVIRDLPELLKQLQPFIEKPEEG
WP_156999266.1     PSRDDPARVDAILADIRRFGVTAATRGRSRFVDDDDDQRRIARSITDLSTAADRLPAAFRASHPDIDWAGIRATRNFIAHDYDGTDDEVLWQAIAVQLPAILARLTDA-------
WP_201244805.1     NSKTLRVYLGHILQAIDRIQAYTDQASKQAFVDNAQLQDAVIRNLEIIGEASRNVQPDLIAQHQDIPWSALYAMRNRISHGYWSVDLEIVWTVVQRDLPDLRVQLQSLVSQLKA-
WP_207689225.1     MSNLHLDYLRHILDETVFLLKESKGISEDEFLEDEKLKRAFVRSFEIIGEASKNIPDDEKKKYRSIDWKAMARMRDRLIHHYFGVDYHIVWDTVQENIPSLQKEIEKIIASNASE
consensus/100%     .......bh..h......h..........h..s..........h..lup....h..................pp...+.a...p...h...h...h......h..........
consensus/95%      .......bh.ph...h..h..........h..s..........h..luEus..l..........lsh..h...Rp.hhH.Y..lp...la..h..phs.h...h..h.......
consensus/90%      ...p...bl.ch.p.h..h..h....sb..h..s.....uhhb.hp.lGEAs..ls.p.b.....lsW..h..hRs.lhH.Yb.ls...lWp.h..pls.lb..l..h.......
consensus/85%      ..pp...bl.ch.p.h..h.ph....sb..h..spb...Alh+php.lGEAsppls.p.b...splsW..h.uhRs.lhH.Yhsls...lWp.h.pcls.Lb..l..hh......
consensus/80%      ..cp.bhbl.chhp.h.plbpa....sb..a..Dpb..pAll+phplIGEAsppls.phb...splsW..h.uhRs.llH.Yhslc.p.lWpshppcls.Lb..l..lh......
consensus/75%      .p+p.bhhlpchh-sh.plbpas.s.sbp.a..Dpb.bpAllRslplIGEAsppls.phb.p.splsWp.hhuhRsbllH.Yaslchp.lWpslppclP.Lb..lppllp.....
consensus/70%      .p+s.blhlpclh-shpplbpas.sbsbpsF..Dcb.bsAllRslElIGEAsppls.ch+.p.splPWp.lhGhRsbllH.YaslDhphlWpslpp-lPsLb.plppllpph...

# ; HEPN-EndoRNase
                   1---------11--------21--------31--------41--------51--------61--------71--------81--------91--------101-------111-------121-------131
OrigSeq            MKKKEFVRQWLAKAEGNLRASRHLLDGGPGFAAAAAFHAQQAAEKSLKSVLVWHQVEFTKTRDIGAILELLARVEGELVGRLADANVLTPYAARYRYPGEVPEPTISEAGHAADIAETVFREVVLRLASELHP

Jnet               ---HHHHHHHHHHHHHHHHHHHHHHH------HHHHHHHHHHHHHHHHHHHHHHEE-------EE-HHHHHHHHH----------HHHHHH----EEEE----------HHHHHHHHHHHHHHHHHHHHHHHHHH----
jhmm               ---HHHHHHHHHHHHHHHHHHHHHHH------HHHHHHHHHHHHHHHHHHHHHEEE-------EE-HHHHHHHHH------------HHHH----EEEE----------HHHHHHHHHHHHHHHHHHHHHHHHHH----
jpssm              ---HHHHHHHHHHHHHHHHHHHHHHH------HHHHHHHHHHHHHHHHHHHHHHHE---------HHHHHHHHHH----------HHHHHH-----EE----------HHHHHHHHHHHHHHHHHHHHHHHHHH----
HAZ27920.1         MKKKEFVRQWLAKAEGNLRASRHLLDGGPGFAAAAAFHAQQAAEKSLKSVLVWHQVEFTKTRDIGAILELLARVEGELVGRLADANVLTPYAARYRYPGEVPEPTISEAGHAADIAETVFREVVLRLASELHP
GBD39154.1         M-KRKVVVQWLIKANDDLKTSEILLKSEDIPTGAVCFHAQQAVEKFLKAYLTNIDVRTGKTHDIAKLLNMCIQHDKDFKLNREKLEKLTFYAVEIRYPEEFYTPTTEEAKEAFELALTVKEFISKKLQIDIKE
GFP32492.1         MRNKELVNDWIKRAKSNLERAKAGRISQDVLYEDLCFDAQQCVEKSLKSLLVSLDVEFPWKHDIDVLFGLISKSGIKIPDDLKSAVILTRYAVHARYPGLAEPVSEEDYQEALALAGKVFNWALKILREDKE-
MBA4158335.1       MTALQAAREWFDIAEGDLRSAKLLLEADPPEIRSATFHCQQAAEKYLKGFLVVHGEEPPKIHILEVLLERAEQHDPSLPNLLGDASVLTTFAVEPRYPLGGTSPRLSDARDALARAGRIRDAIQAAIPELSQP
MBF0261193.1       MIVQTQVDYWRTGSDEDWEAALLLVEKG--KIHHGLFFLHLALEKLLKAHVCAIHDHPPRIHDLVRLTRRAELELT--EEWLLLLGKVNAFCQIGRYPGNPMS-SLEVAHRYIDNVQEVRAWLQQKLTES---
MBF0565079.1       MDTVKYMNNWLESAERDLKTADSLFLNA--KYDWCLFIGHLVIEKALKALYMKKNIAPPKTHNLIGLTENLEIHLT--EEQIVLLSRISDFNIDARYPNEKDS-TKEFTREYLGKIKEIYRWLLLQIKQ----
MBI2357705.1       --MAQPPEEWLRQADYDMDTADYMFSGG--RYFYAVFMCHLSIEKALKGLYRKLRQEPPRVHNLIYLLNEIGIKPP--EAIGRFIVKLNEASVATRYPEEIRK-TQGIVKGILSEGRKALEWIKKQL------
MBI2357706.1       MSGTPVADEWMQIARKDWRRAERSLRDR--DSEAAGFFLQQSLEKYLKAFLLKHDWNLRKIHELDALLDEASKCQKDLETYRDLCERVSGYYIAERYPPLSAL-SSADIRKDLKEAR---EFVKLMFPREKLP
MBI4730025.1       MRLIAVCAEWVAKAENDLTNAAHALTLGEMPTDTVCFHAQQCVEKYLKAILVLEGIDFPKTHDLETLVVRVPARLRP-GLTPEEQARLTEYATGARYPGWEEI-PLAAARRAVAVARRVRRDARRTLPRNALR
MBI4774969.1       --MNAPGDRWLEFARQDLQMAELALKEG--IYNQVCFHSQQCVEKCIKGLIANIGKTPPRTHSIVDLLGLLPTGYL--SHLQDDLAQLDIFYIPTRYPDALPGPGEEDARDAIEAAQACWKKIGSKELGGR--
MBI5740567.1       MQITEVIREWINKAENDLINAANTLKMGDDPTDTVCFHAQQCAEKYVKALLVWKGIPFPKTHNLSLLMALLPLKTPAL-LSVAEQKLLTDYAVVTRYPGDEEEPTLGEARKAVSLARRVRKNIRGMLPKEALT
MBK7674931.1       MTPLDHARALLARARDDFYVVRHLRGEADAPLWVLGFHAQQAVEKALKSVLSSRGIEYPRTHNLVMLVEMLRRAQIGLPPDADALGLLVPFGVLLRYEDVEGAPQHIDPERFDGFVVRTIDWAVEHLKKAD--
MBL8540232.1       ----------MRKAADDLYVARRLAQDVDAPAWILGFHAQQAVEKSLKAVLSLHRIEFPRTHNLSMLLELLRRAALALPPDGDELARLIPFGVIARYETALDAEVEIDRVWAVDVATRTFECGEGAVGLDRDP
MBN1472031.1       --MNELVREWIDKAEGDYHSALREYRARKFNYDAAGFHAQQCAEKYLKAILQLKNIRFEKVHDLLALLTLCLPSAPELELHKELLAYLNPFAVAFRYPGES--ATRDQARQAIQIIKKLRLILRETLHIEG--
MBO9362158.1       MNRAGEALRWLRFAGEDLREAEQLAAQTQSIPRHICWFAQQAAEKALKAALILEDMEVPFWHNLDALRNLLPPGWSV-KQEHPDLAELTAWAVEARYPGDWPEATWEDAHRAVALARAVYDCIVRDFEQRGVK
MBU1261809.1       MTNQTLVEKWLKRAKSNLEIARAKIVSEEILYEDLCFDCQQVVEKSLKALLVHINVVFPWTHSITKLIELIEEKGIDVPEEVKESISLTTYAVTTRYPGDYESVDEEEYNEAVEIAEKVLKWVERWIQEVI--
MBV9397052.1       MPIPGSPQEWLVRAKGDLAMARIALPEGA-HYEDLCFHAQQAAEKAVKAVYRARKLKFAYIHDIAELLAGLEREGIQVPQSVLESAVLSDFAWQARYPGTAEPVDEAEYRDALMVAEATVKWAEDLVNRGT--
MBW2329464.1       MKNLDKANEWLKRAKSNMARAKSGRVSPDILYEDLCYDAQQAVEKAFKSACIIHEIVFPKTHDIAYLIELLEKGGVEVPINLQNAKILTGYAVETRYPGDYEPVDEEDIRKAIEIAKEVLKWVKKEIDKSQRP
MBW8002949.1       MTREQYAREWLQYAENDLAAARFLFGMKPLPPEIICFHCQQAAEKALKAYLAYNGTNVPKTHDLTNLNELCAVHEKDIESLVEQCIALNDYAVEVRYPDESPI-EERDAHQALKDADIVFNYIRERLKHE---
MCA1947579.1       --MFEQARRLLRKAQQDGEAAQRLAEAGTTALWIVGFHAQQAIEKAFKAVLTARGVPHARTHNLVMLFTLLKQQGATPPGSSDEHALLTPFGVAFRYDDTWDPDVRLDMAAALSLVAKVLAWAQHEVTGGEES
MCA6523508.1       MDKDELVQNWLIKAQHDLLAAKKLGIEPETYADIAIYHCQQSAEKAVKGFLILHNQEFPRTHDIRLLIQLAIRINPNFEHYQETSEILTPYATEFRYPSDVMEPSLEELEDALIKAEELFDFVVSLLPDEIKT
MCC6048505.1       --MSHKVKLWLIKAFEDYMGLKFLSEVENILRSIICFHAQQMVEKLLKAFLSFHNIPFPRTHILEIILKQKCIELDEDF--QNLDFKKLSKYAVEIRYPEEFKIPSIEETKECIRIALQVKDFILEKLNITENE
MCC6629588.1       MPLVADARAWLVRAAADLRAADVDLAALPPLLEDALFHSQQAVEKAFKAFLTFHLQPFRRTHSLEEIGAACLALDASLRPTVGETAPLSEYAWAYRHPGPAMAPDRVEAEQTRAIAQRSYQAILARLPEVVHP
```

```
MCD6257886.1    M-SIEAAKKWILKAENDLKIAKDEITMENPATDAICFHAQQCAEKYLKAYLIFNGKEVRKTHDIAELISLCSEVNIEFKLNQVDIVSLTSYAVEVRYAEDFYFPSVEEAKYAIELAEKVKNFVLKSLRKAGFR
MCP9469843.1    MSLEDLVRSWVDKAEHDLLNIDNNLAAKEIPWDTVGFHAQQCAEKYLKALLVFHQIDFPKIHDLTELYAFLPKDFRP-PIDPRLLDRLNPYAIEGRYPGVWEPVERDEALRAVEAARAVRDAVRRALPSSCLE
MCS6883593.1    M---NEGETWFQFAREDLAMAELARQAG--ILNQVCFHAQLCVEKVLKGLLSTQGVTPPRAHRLGDLLPLLPSNPF--DDIALDLQLLDRFYIPARYPDALPGPNESDADEALLLARQVMQRIASSSPAATNG
MCX5669699.1    MRLHTILAEWVEKAEADLASAVRLLAEKEAAAETICFHAQQCVEKYLKTMLVLARIDFPKTHDVRRLMGMLPSASRP-DLSQAERDTLTPYATVTRYPTGLGPLHICQARQAVAMARRVRKHARSVLPRPALR
MCX7789964.1    M-SPPSVQAWVRKAESDLKVARDELATSEPATDAVCFHSQQCAEKYLKAYLICHQQETPRTHNIADLLRRCMDIDIRFQLLESDVPLLTSYAVAMRYPGDELFPPVEEARDAVRLAEQVKEFVWARLTEAG--
MCY3559221.1    MNEPQDPERWLGYAEEDLRAARMFHERGDFSPRHACWYAQQSAEKALKAVVLTVQPLARKTHNMADL---LPSPAGSI-VSTADLNHLTDLVETARYPDATPTPTEADARRAVDVAGTIYDFVAAEFERRGAI
MCY4393098.1    MSPAEDALRWLRFARADLGAARQFLGDAAIPPHHACLHSQQAAEKALKALLRAHGIAFPPIHDLEQLHDLLPATSAL-PASRPHLDRLTDWHIEARYPGPWAEPTMADAEQAASIAAAVNDYVVGEFGRRGIA
MDA8063429.1    MRNYDEARRWLANTDDDLRALHALARESEAPGRIICFMAHLAVEKALKATLIDAGVPFKKTHDLVALHAMCVGSGRLTDVDTGQLVLLSPWGVDGRYADDLVDAERALASRLADFAERLVEAVHQILEETDGG
MDA8316553.1    MDAHGQARRWLRWADEDLVAARHSAADTEVAPRVACSLAQQAAEKAIKALVVSSDLDPPKTHNLLRLARMLAEETAD-RLLDVDLEDLTRWAIEGRYPGDLDEATSRDAARAVELADQVLAIAQAALEAADGD
MDD9986486.1    M--------LLEAAERDVGALRVMRRSHEVSDEVFGFHVQQAAEKSLKAWLALLGKVYPLTHNLEALLELLSERNDV-SCRFRKLIDYTPYAVEFRYEGVESGTEPIDRKRALTLVEALLAQVRRQLPDTDSS
MDE0200335.1    MNVAQEARRWLRFSVEDLEVAQRLLADGLSSSRFVCWHSQQAAEKALKAALILEEIEFPFTHDLNALHNLLPDSWPL-PVDHSELADLTEWAVEARYPGEWPQATESDAIRAESQARATRDSVVAEFQRRGIN
MDE0688676.1    MQRLTDTAGWLRHAEEDLTTAEIFLGHTHVPPRQSCWCAQQAAEKALKAALIFLQIDFQRTHDLNVLQDMLPERWKPLKTALPNLRDLNRWAIEARYPEEGQEPTQTDASTAVEHARSVWTSLSTELIQHGFL
MXW25934.1      MSAAAEARRWLRYALDDLEIAGQMAREAG-RPRYVCWFSQQAAEKALKAALVLEDIEFPYTHDLRRLRNLVPGAWPA-GAGPDILARLTEWGAESRYPGDWPEPTTQEAAQAQADAWAVYDSIAAEFGRRGLS
MXX86605.1      MKPPDDPREWLNRARSNLVRAKNRIPGA--YLEDLCFDAQQAVEKAVKAVMLDRGIEFPYVHDLARLLSILEADGEPIPPEVWSASKLTRYAVHSRYPDLEEAVLENEYEETIATAEEVVGWAEKRL------
OGV65470.1      --MNAVVEEWLAKAEGDFATATREFAAAETNYDAVCFHAQQCIEKHIKAILIARGVIPPKTHDLLELDKLLVAACPTWHWAVEELRLLSRAAVVFRYPGES--AGHEEAAALDVCTRIREKLRGLIGA----
PMB78263.1      M-SKETARKWYRQALHDLEMAEKTIEIG--GYDVSAFLSHQCVEKLLKAGFILDGKEVPKTHYIDELAKKLNLP----NELIDEIYELTLDYTLSRYPDVSETPSEEIAKEKLEIARKVLNYFKKRWGEL---
QBI18518.1      MSDDAEFHRWQRHAHRALDTARLAANGE--RYEWACFLAEQAAQLSVKGLLHGIGEE-AWGHDLTVLASLLSPHWPDAPDAADAAARLTRHYIPTRYPDAHPGARRSDADAAIADAERLIDNAARAWAALG--
SPE35561.1      MTLLDETRRWVQRAQGDLRAAEILATE---LPATALFHCQQAAEKYLKAYLTWNQTAFRKTHELQELAAACAVIDPALQSALDPAMALSKYAWQFRYPGAPYEPDAQEAALGRALAEQVRADISSRLPQAALS
TEU12998.1      M-TTEEFEDWLRRAESNLHIARRGRGEG-VFLEDLCFEAQQAAEKALKALLIHLSGDYPRVHAFTLLLERLQQH-VAVPEPIREVVELSDYAVQIRYPGDYYPVSEEEYERALELAERVLRWVRSQTGQKR--
WP_045165445.1  --MNDIVNNWIKKALEDLKVAEHELSLEENVTSAICFHSQQAVEKLLKAYLTSKNVKVGKSHDIAYLLQLCIHQDSDFKLKELNVEQLTFYAVEIRYPDDFYIPSIEEAKASYELAVLVKEFILKKIEV----
WP_070065820.1  MSLPASPEEWVTHARSDLTLAQIGRDSSKVLHEQICFHAQQAAEKAIKAVLLFYKVDFPLTHDLEELVDILDKNGISIPKEILDAGILTPYAVEARYPGYWDEITEHDVDDALQLAEKIVIWAGKCIKKN---
WP_085440418.1  MNVQRQIDHWRTGSDEDLEAAEIMITAG--KRRHGLFFLHLSVEKILKAHVWVTQEHPPKIHNLLRLVQIAALSLG--QEQSALLVKLNRHCLEGRYAEEVAAPDAAETEILLQQAKEVQTWLRNQL------
WP_132542049.1  MTQREEAARLLRLARRDQAALQALLGAPGVGIAVAFFHAQQAAEKALKAAMCLHALEYRRTHDLEELAGQLADAGHPPSVAEADLRLLTPYAVEYRYDDEA--PELLTAAQALAIVAAVLAWSDEHMQVQP--
WP_134213464.1  --MRAQASVWIDIAYDDLDAAEHCFQGD--RFLWAMFMCQQAVEKAIKAVYFKTGLTPPKKHDLVSLAGSAGLLEKCTKELRDLLRRLSLYYIETRYPDKRAV-TKEHTKEILERSKGAVEWLRSMLK-----
consensus/100%  ...........s..s......................pb.hbb.hKs.h..........+.h..l...h................h.......Ra.................................
consensus/95%   .........h...u..sh..h..............a..pbshEK.hKu.h........b.H.l..L...h...............ls......RY.................h......h.........
consensus/90%   .........hh..A..sh..h............h.a.spbshEK.lKuhh........b.Hpl..L...h...............Ls.a....RYs................h......h...h......
consensus/85%   .........Wh..A..Dh..up...........h.a.sQQshEKhlKuhh........b.Hsl..L..bh...............Ls.ash..RYss.....p..ph..hh..h..hb..h...h......
consensus/80%   M.....sppWl.bAp.Dh..Ap.........p.hhF.sQQshEKhlKuhl.......sb.Hsl..L..bh.............h..Ls.ash..RYPs.....p..ch.phh..u..lbp.h...h......
consensus/75%   M.....sppWl.bAppDL..Ac..h.......p.hhF.uQQshEKhlKAhl.......s+sHsl..L..bh.............h..Lo.aul..RYPs....ssb.-hpphl..Ap.lbp.h...h......
consensus/70%   M.....sppWlpbAcpDL.sAc..h.......c.hhFauQQshEKhLKAhLh.p..phs+sHsl..Lhphh...........ph.bLo.aul..RYPs....ssb.-hcpAl.hApplbphl.pbl......

# ; HEPN-EndoRNase
                1---------11--------21--------31--------41--------51--------61--------71--------81--------91--------101-------111-------121-------131-------141----
OrigSeq         MKSQEHHYAHAALRHFKDAVFLLDNSRLPNADYHFGFAVECALKSLLLRFTAAPKPNGKPSKSPWVVAHSYSHLPWVATDVALLTHGRSATRLSAALGGLTAFDTWSVDHRYLDGSSVVEADVRQRRTVAENILRLHEQALITGRLP

Jnet            -------HHHHHHHHHHHHHHHHHHH--------HHHHHHHHHHHHHHHHH------------------------HHHHHHHHHHHHH---------------------------------------HHHHHHHHHHHHHHHHHHHHHHHH----
jhmm            -------HHHHHHHHHHHHHHHHHHH--------HHHHHHHHHHHHHHHHH-------------------------HHHHHHHHHHHH---------------------------------------HHHHHHHHHHHHHHHHHHHHHHHHHH----
jpssm           -------HHHHHHHHHHHHHHHHHHH-------HHHHHHHHHHHHHHHHHHHH-----------------------HHHHHHHHHHHH----------E-----------------------------HHHHHHHHHHHHHHHHHHHHHHH-----
AL095594.1      MKSQEHHYAHAALRHFKDAVFLLDNSRLPNADYHFGFAVECALKSLLLRFTAAPKPNGKPSKSPWVVAHSYSHLPWVATDVALLTHGRSATRLSAALGGLTAFDTWSVDHRYLDGSSVVEADVRQRRTVAENILRLHEQALITGRLP
EXI73735.1      M----INFRGAARRHFADAKLLEQQVRTANAGQLYGFCVECGLKALLVAHGLPTESNGDIQRKP--KTGYREHMPSLGSGLTAFPDGRQATRYVAMLPGIAAFSDWSVDHRYYDESRLPLASLPGWRGAATEVMSMLDQAIVDGVMQ
MBI2566522.1    MRSNGDDHPDASGKHLADAQALLAAGRFDGAAYLAGYVVECALKSLIQVE---TKSQPRFT-----------HKKALHAVAYLVAGAATAKYVTPAVLGSAQISGWTEQLRYRGPSFSSTD-AQAWVAEASQIADTIAQMIHDGVI-
MBK6972677.1    M---STDFRDASDRHFEDGGLLEAQGRFANADHLYGFSAECSLKAAMLGVGHPKAAKGDWPD------GHCDHVNALWVGFQSFAHGLLDAKYAAHVPPTNPFSTWHANQRYWGRHHFTAGSVAPHKAAADGCRRLLRELILDGVF-
MBK7955384.1    MVTPRVDYLLCARRHIRDAHFLLDRGRKANAGQLFGFSVECGLKALLVRSGAPVDGDGNILKAS----GLREHLPKLGNAMTTLPDGQAANIIQSQLPHLIDTHDWHTDHRYWRASAVPLASLDNWEMAALEMDVLLDDVISKGLL-
MBL8465278.1    M----LDYADATRRHWDDGLFLLEDRRLANADHLFGLAAECALKAVMLPLGMKLRPSGAPED------RRHGHIDRLWDEFGAFVSSRGGARYGALLSAQNPFASWKVDQRYDSRSGFTLAAVASHRDGAQTAKRCLDAAVLDGVIA
MBM3879663.1    MIATGEDYPAAASKHCTDARALLKANRADGAAYLSGYGIECVLKSLVQVE---TRATGRVAGR----------HNQDLGLDLATLPTGRTWRYLKSDPVSGVPPAEWSESLRYHPEFTVPPGIAAQWLQEADRLSEVIGGMAKDGVIT
MBM3880841.1    M---ATASISAAQKHLTDSSLLLEQQRFDNAAYLSGYVVECCLKALLDIDGMP--------------SAKELGHEPLMALMLGYAAPSRSRYNLVPTPDLDRLINDWKPDSRYEKENTTSQALAESRLKGAQAAEQIGIAMILDGTIQ
MBT2380837.1    MAVMEQQYALAGARHYHDAVYLCGDQRWPTADHLFGFAAECALKSLLLRFAVHTYRNGNAMNRARWKEDWLGHINQVLGAAPLLAYGRGGPAFVEAVDRLNVFASWDLADRYRDGRQVSDSVVASRQEAARLALSLYELYELTGRTS
MCF7977305.1    M---STAFEDAAQRHWDDAELLSDHQRWPNADHLYGFAAECALKAVMVSLGMQLGPTGKPPK------PYAVHIHQLWDQFLTFASGQGQAGYAAMMQQSNPFSAWHESQRYWDSTTIGPADARDHRAAALQALGVLSQARQDGRVK
MCL4821148.1    MPGSSDDHPDAARKHLLDAAALLHESRPDGAAYLSGYVVECALKSLWLLG-LPSLPPKA----PW-PSGSKGHESYLGSGLAAFAGPQAGRYLGAASRNVSAIAAWNPAMRYESESRTQAD-AVAWWALADAVMETIAQMQLDGVL-
MDD5374990.1    M---QADFMDAHERHWDDAERLLQAQRWANADHLYGIAAECGLKRMMLAFGMPYDATRDRPASG----QDRVHADGVWARFESYRSGHIQGT-GYVLPANNPYYDWNVSDRYAHQSNFDQARAQGHQAGAQTVRNLIAKAQREGLLI
QFY91198.1      M---QEDFLDAHRRHLSDAGYLFQAERWANADHLYGISAECGLKVLIEKLKGS------SLDR-----GDRLHVNNAWLKYQTYLSGHAKAA-KLALPTLNPFADWLVSQRYANQSNFDHVRVQSHQEGAKAVAALLKKDAVEGFK-
WP_096450981.1  M---STRFLDASDRHFHDAELLLAQARGANADHLFGLSAECSLKAVMLALGMAVSPDGAPQDK-----GHRVHMPELWAAFQSFAEGGLASRYLEPLDKSNPFADWSIDQRYWARDAITASAIPSHRAAANQCRVSLEKLIFDGVTT
WP_149501938.1  MPARRVDYVASARRHHRDAQLLWAEGRAANASQLWGLAMECGLKVVLLAAGVVPEADGSLPDRR----RFREHLPTLADLLGHLPDGRLAHRYLALLPGRAAFANWAVDQRYWHEDALPAATLPAWQAAADEVARMLDEALLDGVLR
WP_152627977.1  MKPVEDHYASAAQRHFEDGVFLHDDGRLPNADHHYGFAAECALKSLLLRLVPEKKPGKPPATKPWLRALDLGHIQGMLDVLPWLLRGRAGAAVLAGLASPAEFRAWSESDRYTDGSEVLEGTVVRRRLAAEEIVKLHNQQ-LGGVRP
WP_164382632.1  MAAPPPAFRAASGHHYHSARLAAADRLTIAADHSAGLAAECAIKAILLDYLGSYNSALELSSQP--KEYLHEHLPDLWGQLTAILDRRRARETGVLFLRHNPFHDWDVSGRYCDGTAISEADLDRHLRAAYDLLAAHEQARILGTGA
WP_193814302.1  M---STDFCDAAKRHMADAELLMEQSHWANADHLFGIASECAIKQVMISLGMATHPDGNPVDR-----AHRVHMPDIWAAFHSFAQGRLASRYLEPLSPGCPYGNWSVDQRYAHRGDFSEAQANDHQLAADECMASLMMTLLDGGMN
WP_196191500.1  MATPPPAFATASRNHYRAATLAAERDLRISADHLAGLAAECAVKAILIDFLGSEHPEVKYLDRS--WTNKHGHLPELWEHLTAVAHRRRGGGSGALFTQNNPFAGWDIGDRYCDGTAITDTVIGRHLKAAYDLIAAHEQASILGTRT
WP_223780483.1  MKTANTDYACAGERHYEDASYLHGDGRLLNADHLFGLAVECLMKALLLRFGGSRNSGGSESDRLWWRRRALGHINEMIQALPLLLDGRPGPALTEALTRSADFEEWRVDERYADGTHLDPALLSRRQEAATLAQELHLHVQITGKLP
WP_334662745.1  MPTQQQHFASAASRHFLDGDYLRKGSRLPSADHLYGFAAECAVKSLLLYFTEVLEGEEKPSAKPRARIIEFGHVNELVREIQLFASGRSGAPLHALLDSLRAFRRWSVAHRYRDGGWLQADVVDKRREAAHRILALHETAMIRKRLP
WSN45801.1      MPSVEHHYGSAAERHFHDAVYLHDDQRLPTADHLFGFAAECAVKSLLLRFTAETDSKGRTSRKPWWKLEPLGHIANVLPAVSLLTHGRTGADLLQAMGSLSDFASWTEADRYTDGTSVEASVVACRRKAATLVLALHEQAVITGKLS
WTK31128.1      MMTTEDHFASAARRHYDDARYLCKRKRWPNADHHLGFAAECALKSLLLRFTEAKPPKGKQGNKPWI-VQEYGHLPWSETELAVLATGPLGQHLLTALGDLSAFARWTEQDRYHHCDHITQETVTALHDATQFILALHESARITGTLE
consensus/100%  M.........s..pH..su..h.......sAs...Ghs.EC.hK.hh.........................H.......h..h......................W....RY..........h......s..h...............
consensus/95%   M.......a..u..+Hh.su.hh....b..sAs...GhshECslK.hh.........................H...h....h..h......................W....RY........s.h.......A..h...........G...
consensus/90%   M.......a..As.+Hh.Du.hL....+..sAs...GhshECslKslh.........................H...h....h..h..s...............h..Wp.s.RY........s.h....b..A..h...........G...
consensus/85%   M......a.sAu.+Hh.Du.hL....R.ssAsa.hGhshECulKslh........s................H...h....h..h..u.............h...ssh..Wp.s.RY...s.h..s.h....b.sA..h.......h..pG...
consensus/80%   M.....pa.sAup+Hh.Du.hL....R.ssAsaLhGhuhECulKulhl.......s................Hhs.h...h..h.pup....h...hs..ssh.sWp.spRY.p.s.hs.s.h....b.uAp.h..h....h.bpG.b.
consensus/75%   M.....pa.sAup+HhpDA.hL..p.R.ssAsaLhGhuhECulKulhl.......s................Hhs.l...h..hspup.s..h...hs..ssa.sWp.spRY.p.s.hs.s.h....b.uAp.h..h...ph.hpG.b.
consensus/70%   M.....ca.sAupRHhpDA.hL..p.RhssAsaLhGhuhECuLKulhl.hs....sp.p..p..........Hhs.lhs.h..hspGp.u..h...hs..ssassWp.spRY.s.s.hs.u.hs..b.uAp.hb.h..phblsG.h.

# ; HEPN-EndoRNase
                1---------11--------21--------31--------41--------51--------61--------71--------81--------91--------101-------111-------121-------131
OrigSeq         MADAEREIKELMRKAQESLEAAKLLLQRGYYDFAASRGYYAMFYAAEAALLSKGLSFSKHSAVVAAFGQHLAAPGELPTHLHRYLLDAFDLRMVGDYDAPGAVGESRASQVLGWAEEFLQEIERFLAASGYR

Jnet            ---HHHHHHHHHHHHHHHHHHHHHHHHHHH---HHHHHHHHHHHHHHHHHHHHHHH-------HHHHHHHHH----------HHHHHHHHHHHHHH--------EEEE--HHHHHHHHHHHHHHHHHHHHHHHH----
jhmm            ---HHHHHHHHHHHHHHHHHHHHHHHHHHH---HHHHHHHHHHHHHHHHHHHHHHH-------HHHHHHHH------------HHHHHHHHHHHHHH--------EEEE--HHHHHHHHHHHHHHHHHHHHHHHH----
jpssm           ---HHHHHHHHHHHHHHHHHHHHHHHHHH----HHHHHHHHHHHHHHHHHHHHHHH-------HHHHHHHHHHH--------HHHHHHHHHHHHHHHH-------EEE----HHHHHHHHHHHHHHHHHHHHHH-----
KUK26301.1      MADAEREIKELMRKAQESLEAAKLLLQRGYYDFAASRGYYAMFYAAEAALLSKGLSFSKHSAVVAAFGQHLAAPGELPTHLHRYLLDAFDLRMVGDYDAPGAVGESRASQVLGWAEEFLQEIERFLAASGYR
MBC7332926.1    KQEFIDLAKYRLEKARNTLSDAKKYINNATLESTVNRIYYAMFYAVNALLITKGLASSKHSGVLALFNGEIVKKGLIERRFGKFYSDMLNIRQEGDYKDFVKFEREDVEEWLKKAEEFIAKIEEITLKMIE-
MBW1922079.1    PAQKEALVRYWFEKARESLESARDEYRSGRLSFAMNRIYYACFYALTGVFQDRNKTFKKHKGLRSALHRDLVKGGIIDAQWGKFFDEVFESRQRGDYMPIAVFESAQVEESLEEARQFLNEMEKLIVK----
MBW2077702.1    KAHKDALIRYWMEKARESIEAAQSEYDSGRYATAVRDLYYACFYALTAVLLKEGHSFKKHTSVKAALHKDLIRTGIVQPEWGKFYNKIFDSRHEGDYQPLRVFEAEEVKMFLDQGAGFIANMEEILAK----
MCI9358066.1    DEKRKDLCRYRLEKAETCLSSARILVQLKDYCGAANRSYYSIFHCIRSLLALEGVDFSKHSGVMAYFQKNYIKPGVFGKEYSKILTEAFQIRNDSDYDDYYILSKEDVDVQIQNAQIFFNGIVSYVNRTL--
MCL1489689.1    ------EIKQLLSLAIEDLETAKILLELQRYRPCLSRAYYAMYYAAQSLLLSAGLETSTHKGVIKLINFHFENTGKISKDIAKLLKSTYDLRQSGDYNTAFIADQATAETAIANAQKFIDEVQRIMNLDE--
MCL2124496.1    MITKTDLMRYRLEQARECLHTAEATVDVSM-KNAANRSYYCIFHAMRAVLAXDEFDSKKHSGIISYFRQNYIKTGKFXADFSKTIAKAFEVRGGSDYQDFYVVSKPEVLEQIENAKLFLEAVEAYINTLPE-
PIV20447.1      -MSQQAIIDYWFEKANQDIASAQENFLAGRYVNAVRDAYFACFHAFSSVLLKSGKAFRKHKEVRSALHRDFIRMQKIDRKWGKHYDWLYDNRQKADYRPLVEFESEQVKEIIEQSQAFVEEIKRLLNRGV--
QVL47098.1      -----------MDKAKRASASAELLLNAGDPEGACNRAYYAMFDAARAALIASRTKVRTHSGLITAFGLRLVKTGRLPVELGRALNRASELRLAADYTG-GPVTKGQAEWMVNQSKAFIAEVHRAFTSHEG-
RJP35072.1      NEEKLDLIKYRLEQADDSLMDAKVLLDSQRTQSIVNRAYYAMFYAALALLQKIRKVPRKHRGVLSLFDTEFVLKGLFPKYLSKDFHEIFELRQVSDYQITEKITMEKAEKTLISARNFVQEVKTYLLSDQT-
RLE33860.1      ------DIASLLEKAERSLSAARSLLDEGHHDFAVSRAYYAMFYVAEALLASLGEAYSSHGAVIGAFGRIFAKSGKLDPKFHRWLIDAQDLRNVGDYGVGRTITAEQVKEVITQAQEFIAAGRAYLISNRR-
```

```
WP_081364174.1    KEEIKQLILYRMERAKEAIAEAELLFSEGHIRTFVNRLYYACFYAVSAILLAKDYSSAKHSGIRSLFHQKIVKAGLVNPSAGTLYNRLFDARQKADYADLVKFEADIVAPWFDEVKSLVDQIETLVVKEIR-
WP_199302091.1    MNIPNEAVQYRLERAEEAYHDALTLAESGGWNSCVNRLYYSCFYAVSALLLKDNLSSPKHTGVRGIFNKDYVRTGIISKDQARLYNDLFESRQEADYGDFVYFEESEISPLLPQVRDLLDSITKVLAQD---
WP_238455596.1    DVNFRDLSRYRIEMAERKISAAEVLIKEGMYADAVSRIYYGIFHAARAMLALKGLDSKKHSGIISLFNQHFVKTGLLPKEFGRIIKNAKDLREESDYNEFYLVSIKDAEELLKDGTRFIIGIKKRIQEILNK
consensus/100%    ...........hpbA.p....Ab..h........hpc.Yasha.hh.uhh........pH..l..hh...h.....h.....p.h....p.R..uDY.....h....h...h..s..hh.............
consensus/95%     ...........hpbA.p....Ab..h........hpc.Yasha.hh.uhh........pH..l..hh...h.....h.....p.h....p.R..uDY.....h....h...h..s..hh.............
consensus/90%     ..........bhcbAppsh.sAb..hp......hhpc.YYuhFahhpuhL........pHpulbshh..phh..Gbh..ph.+.h.phbp.R..uDY.....hp..ph...l.psp.hl..hp.hh......
consensus/85%     .......h.bbh-bAppsl.sAb.bhp.....hhsR.YYuhFaAhpuhLh....s.ppHpulbuhh.bphhp.Gbh..phu+.h.phb-.R..uDY.s...hp..php..l.pup.Fl..hpphh.p....
consensus/80%     .......h.bbh-bAppsl.sAb.bhp.....hhsR.YYuhFaAhpuhLh....s.ppHpulbuhh.bphhp.Gbh..phu+.h.phb-.R..uDY.s...hp..php..l.pup.Fl..hpphh.p....
consensus/75%     ......bhpbbhEbAccsl.sAc.bhp.u..pshhsRhYYuhFaAhpulLh.psbs.pKHpulbuhhpbchl+sGbls.chu+hhpchh-.Rp.uDY.s...hp..php..l.pup.Fl..lcphl.p....
consensus/70%     ..p...hhcYbhE+AccslpsAc.hhp.uphpshhsRhYYAhFaAhpAlLh.psbs.pKHsGlbuhFpbchl+sGbls.chu+hhpcha-hRp.uDYpsh..hp.pphcbhlppup.Fl.plcphlsp....

# ; CBS+HEPN
                  1--------11--------21--------31--------41--------51--------61--------71--------81--------91--------101-------111-------121-------131-------141-------151-------161-------171-------181-------191-------201-------211-------221-------231-------241-------251-------261-------271-------281-------291-------301-------311-------321-------331-------341-
OrigSeq           MMSHTPSEEQLRELRGKLIPVSELLALFGTRVRNDQSVLFITQTLKDVGLSTIPYFAHCSLRADVQIVPEESVAPPEPGGEDEEAEQDALSRGTLPQHSFRIGDLPAARNGVDSVPSTAQVKTATHLMLVKNYSQVPVIDGLSTLRGVVTWSSVARMYERNEDTALANAMVEDPPTADVHGDFFSLLPMISEYGYLLVRDMSGRITGIVTAADVTERFEATAWPFFVIGEIEFRLRKCLGAALEPDAIRAVQRRNKQTGQITDLMFNGYVKLLDGHQENPTLCQKADQNWQALGWSLADRVQFVRQLDRVREIRNMIAHFDPEPLSQKLCQELRQFLGLLRLYT

Jnet              -------HHHHHHHH-----HHHHHHH-----------HHHHHHHHH----E-----------EEEEE------------------------------------------EEEE------HHHHHHHHH----EEEEE-----EEEEEE-HHHHHHHH-----------------HHHH---HHHHHHHHH---EEEEE----EEEEEEE---HHHHH------EEEEEHHHHHHHHH------HHHHHHH-------------H-HHHHH------HHHHHHHHHHHHHH-------HHHHHHHHHHHHHHHHHH---------HHHHHHHHHHHHHHHHH--
jhmm              -------HHHHHHHH-----HHHHHH------------HHHHHHHHH---EE------------EEEEE------------------------------------------EEEE------HHEEEEEHH----EEEEE----EEEEEEE-HHHHHHHH-----------------HHHH---HHHH--HHH---EEEEE----EEEEEE--HHHHHH-------EEEEEHHHHHHH-------HHHHHH---------EE-------HHHH------HHHHHHHHHHHHHH-------HHHHHHHHHHHHHHH-------------HHHHHHHHHHHHHHHHH--
jpssm             -------HHHHHHHH-----HHHHHHH-----------HHHHHHHHH------E---------EEEEEE-----------------------HHHHH----------EEEEE-----HHHHHHHHH------EEE-----EEEEEEEHHHHHHHH------H---------HHHH---HHHHHHHHH---EEEEE---EEEEEEEEE--HHHHH------EEEEHHHHHHHHHH--------HHHHHE----------HHHHHHHHHHH-----HHHHHHH----HHH-------HHHHHHHHHHHHHHHHHH---------HHHHHHHHHHHHHHHHH--
GAA0539971.1      MMSHTPSEEQLRELRGKLIPVSELLALFGTRVRNDQSVLFITQTLKDVGLSTIPYFAHCSLRADVQIVPEESVAPPEPGGEDEEAEQDALSRGTLPQHSFRIGDLPAARNGVDSVPSTAQVKTATHLMLVKNYSQVPVIDGLSTLRGVVTWSSVARMYERNEDTALANAMVEDPPTADVHGDFFSLLPMISEYGYLLVRDMSGRITGIVTAADVTERFEATAWPFFVIGEIEFRLRKCLGAALEPDAIRAVQRRNKQTGQITDLMFNGYVKLLDGHQENPTLCQKADQNWQALGWSLADRVQFVRQLDRVREIRNMIAHFDPEPLSQKLCQELRQFLGLLRLYT
MBB5890598.1      -MTDNSLRQQLLERRGASMALSELLALFDVRVRTYQTVSLIRDTLQEIGLATVPSFATCGLKANLLIVGEEEGVAEAV---DEAGDGEL-APGALPQQSFKIGDIPAASAGLISVTSSDPLTRATYLMRANNYSQVPVIDGQSDLRGVVTWSSVAARYETGLAPTLANAMVTDLPVAEVHQELFAQLPLVAEHGYVLVRSNSGVVCGIVTAADIADRFHETALPFFLVGEIEFRLRKCLG-ELPPERIKAVQPNRTQTGNIADLMFGDYVKLLDAHQQDRRLRENADGNWQALGWSGVDRTQFVHHLKEVKSTRNMIAHFDSKRLTPERITELRQFSGLLRQLM
WP_026343754.1    -M---PTIEHIYALKGRDLTVSKLLAFFGFRYRNFESVPTIDAALENAGLTTEPSFATCKLNMTVKVVDLNQATEDPV---DEAEDPQL---DSLPTRPFLVSDLPSATAGLTSITFADGLPYALHLMQTEGYSQIPVIDGTSSLKGIVTWRSVAAVFLRGVAHTLSNA-MEKAETVELHLDLFRELSRLCEHGYLLVRNNDGELTGIITPTDITNQFHDTALPFFLVGEIEFRLRACLRDRIGPDSIREVQRY-KKTGDIDDLVFGDYLSLLRTDQQKVSLNASAEANWQNLGWHGVDRNMFIDHLIQVKNIRNSIAHFDAKPTSLEDITVLRKFSALLKALG
WP_039986433.1    -MTTRPQERDVASLKGRSVSVQEILDLFQVRVRDHRTVHRISQALTDAGLTTLPDFAVCGLRSTVDVVPLAAVPQRTPPESEAEEADEALPSHALPQR-LLLGDIPSARRGLVSVGPGTPLSQTTFLMRTKGLSQVPVTTGMAQVHGVVTWGSVAKMYEAGKEATLDNAMEKDLPVADARQEFFTTLPVIREHGYLLVRGDDGCLSGIVTAADVTERFEGAARPFFIVGEIESLLRRCLGAALDAEAIKAVQTNKQRTGQVSDLMFGDYLRLLDGDQTKQSFAERADRNWEALKWPNMPREQFVGLLKRVKDIRNRIAHFDEKPLPQEMINELTTFAKVLRAFA
WP_081522441.1    -MTQTPSTTDIAALKGKSIYVRELLTLFGLSNRDDQAVMVVKQALKDAGLTTVPSFATCGHGIPLHIQVQEAVAPADAAESDEGEEEDL-PAGALPQQAMRVGDMLVGSSGFVSLEPTAQLVQATYLMREHGFSQLPVVES-TTLHGVVTWKSVVKLYETNKPLTLANALETDVPVAEEQHDFFSILPRISEKGYVLVRGSDGAFKGIVTASDVTERFTVTARPFFMVGEIELRLRKCLGAVIAPEAIKAVQTR-KKTGLVADLTLGAYVVLLDCQQSKDQLRLRAEQNWADLKWDGVDRVMFVRQLAQVRDIRNRIAHFDPQPLPETKAEDLRRFLQLLRDYV
WP_164373103.1    -MTTPPSDDSLLALKGTRISVPDLLALFGTRRRDDQSVLAIAQALRDVGLSTVPSFATCGRGIEVHVVALETVAAPD-SASDEESEEEP-TPGLLPQHPLRIGDIPAARAGLESLPSSAQLSEATYLMRTRDYSQLPIIDGTTTLCGAVTWNSVARMYEKNVDRVLANAMVDDPPVVEEHQNFFAALPLVCEKGYVLVRRYDGRFSGIVTSADITERFDATARPFFIVGEIEFLLRKCLGAAIDPEAIKAVQPKKRQTGKISDLMFGDYVSLLDGQQSKEALRPKADQNWQDLGWPMVNRVQFVYQLDQVRRIRNKIAHFAPEPLDQRSLDELGQFLRVLRQLV
WP_169086812.1    -MAKTPSAQQLLDLKGQDVPVLELLDLFDERVRDNPTCLRIQQTLNAAGLSTVPYFGTCARRASVHMVALDSATEPDPEDEEDAEEPED-PAGSLPERPLSIGDVLNG-DGVDSLPPNSPLNTVTHVMRHNNYSQLPIIEGKSTLRGVVTWRSVARMYEQGSVPTLANAAPSDWAVAQESQDFFALLPMISEYGYALIRRNDGTIGGIVTAADITERFAGTARPFFLVGEIELRLRKCLGTAIGPDAIRAVQRKYQQTGQISDLTFGEYVMLIDGEQRKESLRPLADQNWLALKWSGADRVQFVRRLKRVKDIRNKIAHFDPQPLPDHMHEELHRFCLLLRQYV
WP_195024435.1    -MTVAPPQQDVRALLNTSIPVADLLEYFGFRTRNYDTVPVITKALQAAGLTTTPSFATCRRDARVMIVAADDAPNQPI---EDPEGDQF-LPGTLPQRPFLVGDLPAARGRLISVTSDVSLTKAVHLMRSHDISQIPVIDGSSDLKGVVTWQSVAALREKHVTPTLAAATVGTVDSAETHHELFPRLPIIQKQGFLLVRERTGEFSGIITTADVTAHFHQSALPFFLIGEIEGKLRKLLG-SLSPQAIALVQTGRNKTGDIHDLMFGQYELLLRTNHTKEELRASADANWQRLGWTTVDRTLFVAQLGRVRAIRNAIAHFSPKQLPESDLDELRSFAQLLNHLL
WP_250297837.1    -MSQNPTEEQLLALKGRQIPVLELLALFGTRLRNELTVPRVEQALKDVGLATLPYFGNCNNGADVHIVPYQQAASEEPSGDETAEDAEGLPSGGLPARPLRIGDIPSAVGGLFSVTPDTPLTEVTWLMRTKKYSQIPVIEGGSTLHGVVTWSSVAKVYETTSAAVLAEAMVDDVPVVDVRHDFFSLLPRVSDDDFLLVRANNGTLCGIVTAADITDRFGTTAWPFFVVGEIESRLRKCLGAKLDEDAIRAVQGPSMKTGLITDLMFNGYVKLLDGDQQNPALRALSDDNWTALGWPGVDRGQFVRQLDRVREIRNSIAHFEAEPLGAKRSGELREFVGLLRQIT
WP_254895758.1    MVLRTPPEEELLALKGRRIPLVDLLACFNTRFRNDQAILHIERILKDTGLSTVPYFATCGSRTEVLVVARESTVSESAADGDQEDEAGL-LPGALPQHPLRIGDLPCAHAGLASVTPAEDLTDATYIMHTHNYSQVPVLEGQYTVVGVVTWRSLAKVHATGGEAALANAIVEDPPVVHAHDDFFSVLPAICEHGYVLVRANSGQISGIVTAADITQRFDATAWPFFVVGEIEFRLRKCLGAKLGGDAIRAVQRNDQKTGNIADLMFGQYVMLLDGDQRKEALCTAADQNWTTLGWTGVNRVQFVRQLDRVRDIRNHIAHFDPEPLSPQRSEELRQFVGLLRQLT
WP_267842812.1    -MNRKPSAEELLALKGTEVPLGELLDLFSVRVRRPWNVHDIAQALTDAGLATLPDFANCAPGERVQIVAFVTAVQARPPDDADAAERDPLPSASLPQR-LRIGDLPSARGGVRCVSLTTELSQATYLMRRRGYAQIPVTTGMTTLHGVITWRSLALMYERGKESSLENAMQQELPVADARQEFFACLPMLSEHGYMLVRGEDGSLSGIVTQAHVVHRFESAARPFFLLGEIESLLRRWLGARLTQDAVKAVQTNKHRSGRVEDLTFGDYVYLLNGEQNKVAMAQQADLNWAALQCESLDRGQFVRHLRRVKDIRNRIAHFDAEPLPSEAMRELTAFTTLLRDYV
WP_322753656.1    -M---MDEAALTALRGTELTIRELLTRWGYRIRNADSVSRIRFDLERVGLVTSPTFDIGSMNITIRVVPITAPEEPEPPTDETPADH---AAEELPAAAIRVDDLPSARTPVVCVTPGDTVAYATSLMMKLDYSQIPVVEGDSSLHGVVTWQSVARMYANQSERTLANTPASDVTIVHTGDNLLAVLPTIAACDYVLVRNQHGHICGIVTTADLAVQFGTVARPFFVIGEIERRLRGFLTPVFDHDAVKKATR--KRTDRVGEMMFGDYVKLL-----------SSPENWQRLGWPFVDRDLFIELLDEVRKVRNTVAHFRPTTLTTVENQQVDQLLLMLKNVG
consensus/100%    .h........l..bbs..h.l.clL..a..p.Rp..sh..l...Lp..GLsT.P.Fs.s.....l.h.s...s......................LP...h.lsDh.su...h.sl..s..l..s..lM....huQlPl.pu..pl.GhlTW.Slh.........sL..s...p...hp...phh..Ls.l...sahLlR..pG.h.GIlT.scls.pF..sAbPFFhlGEIE.bLR.hL...h..p.l..hp....pos.l.-h.hs.Y..Ll.............u..NW..L.h..hsR.bFl..L.pV+..RN.lAHF..p..s.......l..h..hLp.h.
consensus/95%     .h........l..bbs..h.l.clL..a..p.Rp..sh..l...Lp..GLsT.P.Fs.s.....l.h.s...s......................LP...h.lsDh.su...h.sl..s..l..s..lM....huQlPl.pu..pl.GhlTW.Slh.........sL..s...p...hp...phh..Ls.l...sahLlR..pG.h.GIlT.scls.pF..sAbPFFhlGEIE.bLR.hL...h..p.l..hp....pos.l.-h.hs.Y..Ll.............u..NW..L.h..hsR.bFl..L.pV+..RN.lAHF..p..s.......l..h..hLp.h.
consensus/90%     .M...s..ppl..L+Gpplsl.-LLshFs.R.Rs..sl..l..sLp.hGLsT.P.Fu.C....pl.lVsb..ss.......-p...........LP.p.hbluDl.su.ssl.sls.ss.ls.ssalMb.pshSQlPl.pG.sslpGlVTWpSlA.ha.p....sLssA..pc.sshc.p.phFs.LP.l.c.sahLVR..sG.hsGIlTsuDls.pF..sAbPFFllGEIE.bLR.hLs..ls.-ul+.VQ...ppTG.l.DL.Fs.Yl.LLpspppp..h...u-.NW..L.W..hsRsbFl..L.pV+plRN.IAHFpsp.Ls.p..ppLppF..lL+.h.
consensus/85%     .M...s..ppl..L+Gpplsl.-LLshFs.R.Rs..sl..l..sLp.hGLsT.P.Fu.C....pl.lVsb..ss.......-p...........LP.p.hbluDl.su.ssl.sls.ss.ls.ssalMb.pshSQlPl.pG.sslpGlVTWpSlA.ha.p....sLssA..pc.sshc.p.phFs.LP.l.c.sahLVR..sG.hsGIlTsuDls.pF..sAbPFFllGEIE.bLR.hLs..ls.-ul+.VQ...ppTG.l.DL.Fs.Yl.LLpspppp..h...u-.NW..L.W..hsRsbFl..L.pV+plRN.IAHFpsp.Ls.p..ppLppF..lL+.h.
consensus/80%     .M...Ps.ppl.sL+Gpplsl.-LLshFs.R.Rs..sV..IppsLpshGLsT.P.FusCs...pl.lVsbp.ss...s...-p..-.p...sssLPppshblGDlPsApsGl.Slssss.LspsTaLMbppsaSQlPVl-G.osL+GVVTWpSVA.habps...sLuNA..p-.sshc.pp-hFs.LP.ls-pGYlLVR.psGphsGIVTsADlsp+F.ssAbPFFllGEIE.bLR+hLGs.ls.-AI+hVQ...ppTGplsDL.FG.Yl.LLcspQpp..h...A-.NW.sL.Ws.hsRsbFV.pL.pV+pIRN.IAHFcspsLs.p..p-LppF..lL+.hs
consensus/75%     .Msp.PsppplhuL+Gpplsl.-LLshFs.RhRs.poV..IppsLpchGLsTlP.FAsCu..hpVplVsbpsss..ps...-p..-.p..ssusLPp+shblGDlPuApsGl.Slssss.LspATaLM+pcsYSQlPVl-G.ooL+GVVTWpSVAbhYEps...sLANAh.pDhslh-.+p-hFu.LPblsEpGYlLVRspsGplsGIVTsADlTpRFpsoAbPFFllGEIE.+LR+CLGs.lss-AI+AVQp..ppTGplsDLhFGsYVbLLcspQpc..Lp.pAD.NWpsL.WsslDRsbFVcpLccV+cIRNpIAHF-scPLs.p..pELcpFh.LLRphs
consensus/70%     .Msp.PsppplhuL+Gpplsl.-LLshFs.RhRs.poV..IppsLpchGLsTlP.FAsCu..hpVplVsbpsss..ps...-p..-.p..ssusLPp+shblGDlPuApsGl.Slssss.LspATaLM+pcsYSQlPVl-G.ooL+GVVTWpSVAbhYEps...sLANAh.pDhslh-.+p-hFu.LPblsEpGYlLVRspsGplsGIVTsADlTpRFpsoAbPFFllGEIE.+LR+CLGs.lss-AI+AVQp..ppTGplsDLhFGsYVbLLcspQpc..Lp.pAD.NWpsL.WsslDRsbFVcpLccV+cIRNpIAHF-scPLs.p..pELcpFh.LLRphs
```

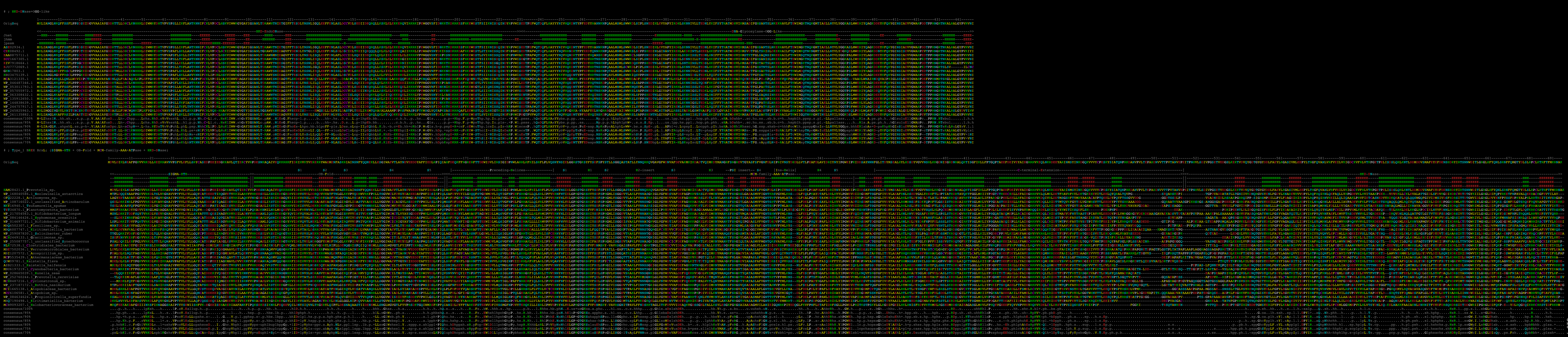

### ; HKD-DNase+OGG-like

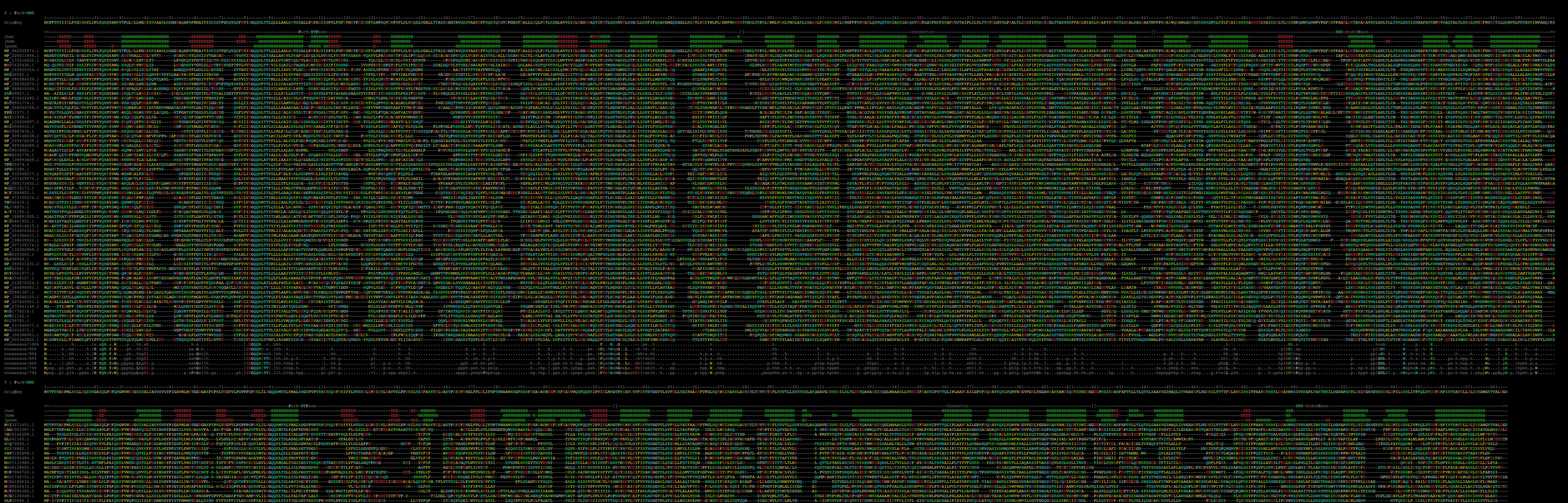

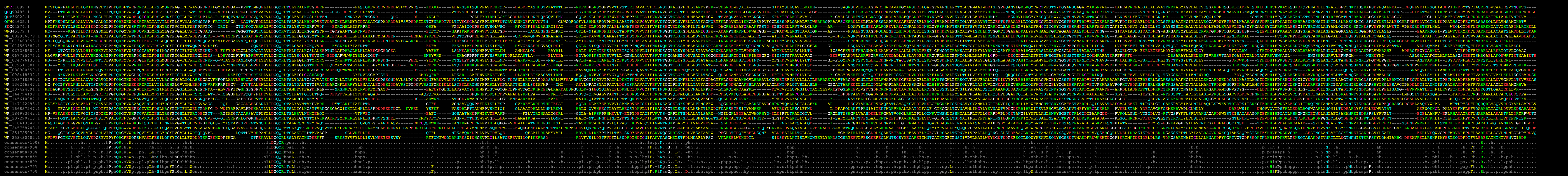

### ; wHTH(SBD)+HNH

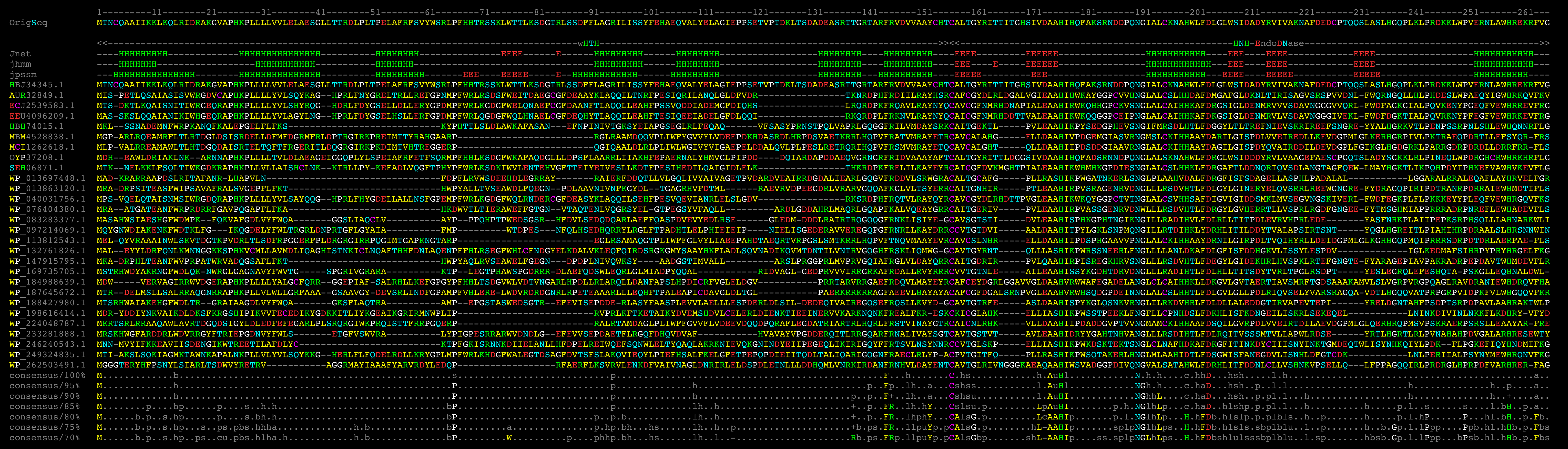

### ; SAD(SRA)+wHTH+HNH

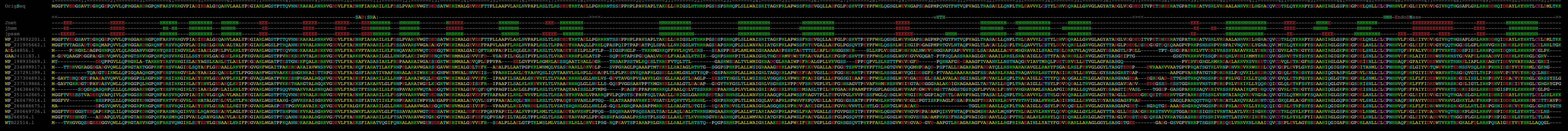

```
consensus/100%  M...................hlLQ.bG.....u..pa.+oh..Gl.lsp....l......L..la..G.ubhWGu..s....p.K..Al..pbsGDpVhFYA...FhupApl...hps..hA..lW.......saEahhALs.h..h.....A............lb.h.b..s.p......bh...................h....hh..l..l..............slshLW.lub.................l..hl.c.G.............h..o.hWp.p.......................sh...s..bh.....b.phl..........-...lh.ph.Lssa.pAsG.........................R..s...b..Rs...s..lK.ha...CQlC...l........bAAHlbGlu.PH.GsD...NhLCLCPNpH.bFD.h.halD.p..Vbps.s...h..h.R...H.ls...l..HR.bh........
consensus/95%   M...................hlLQ.bG.....u..pa.+oh..Gl.lsp....l......L..la..G.ubhWGu..s....p.K..Al..pbsGDpVhFYA...FhupApl...hps..hA..lW.......saEahhALs.h..h.....A............lb.h.b..s.p......bh...................h....hh..l..l..............slshLW.lub.................l..hl.c.G.............h..o.hWp.p.......................sh...s..bh.....b.phl..........-...lh.ph.Lssa.pAsG.........................R..s...b..Rs...s..lK.ha...CQlC...l........bAAHlbGlu.PH.GsD...NhLCLCPNpH.bFD.h.halD.p..Vbps.s...h..h.R...H.ls...l..HR.bh........
consensus/90%   M.................sbhlLQ.RG....pG.bpa.+ShbpGl.lsph...lG.ph..L..laPcG.A+hWGu.ss.p..p.Ks.Alp.p+sGDpVLFYA...FhApAplh.hhps..hAc.lW....p..sWEHhhALsph..h....h.A..hL..L.....lb.h.L..u.p..p...bls...........s......h...phh..l.sL..........ss.p.sLslLW.lu+hsss..Rbs....hp..ls.lL.caG.....s...sh..L..o.lW-hpu.......s..........uGh...sh.bhpps..R.phls.L.......hDb..LL.plGLsGY.pAsG.....................ss..s.Rb.spspc..Rs..hs..lK.ha...CQlC...Lb.....aSEAAHIbGlG.PHpGsD.l.NhLCLCPNHH.bFD.h.lalD.c..V+psps.p.l..L.Rp..H.Is...lpaHR.hC..s.....
consensus/85%   M.................sQhlLQPRGs...pGPbpa.+SlbpGl.lu-h...LG.ph..L..laP-G.ARLWGussspp.pp.Ks.Alpsp+VGDcVLFYA..pFhApApll.lhps..lA+.lW...p-..sWEHhhALu-l.bh...hsA..hL..Ls....lb.LpL.pA.pb.p...hls.........ss.s......hs..pLh..l.sL.........ssbp.sLslLWsluRhsss..Rhs..p.hp.plssLL.caG..s...s.c.sh..Lp.oslWElcu........s.........s.s.uGh...sh.lLpps.sR.cAls.Lp.phh.chDb..LL.plGLuGY.sAsG.....................ss.ss.Rb.spspR..Rs..hsp.lK.ha.p.CQlCs.pLb.b...aSEAAHIbGlG.PH-GPD.l.NhLCLCPNHHsbFD.h.lalD.-..V+popssp.l.pLbRcs.H.Isp..lpaHR.hC..s.....
consensus/80%   M.................sQhlLQPRGssp.+GPbpF.+Sl+pGl.lu-h...LG.ph..L..laP-GlARLWGuoPspp.sp.KspAlcsp+VGDcVLFYAcppFhA+ApIl.LFcs..lA+.lWs..p-s.TWEHhhALu-l.bFp..hsA..lL..Ls....LRsLpL.pA.pb.p...Lls.........ss.s....s.hs..-Llp.l.sL.s.....s..ssbc.sLsLLWuIuRlssub.RLs.bp.Fp.plusLL.-FG..s.phsPc.sh.pLpsoGlWElcGh.....s.s....h..s.s.AGhs..sAbLLpps.sR.cAlsbLp.pahs-hDb..LL.+lGLuGYsoAsG..s..........ussss.R+.ssssR.sRss.lspplK.la.p.CQlCs.pLb.b...YSEAAHIbGlGpPH-GPDcl.NLLCLCPNHHsbFD.h.lalD.-..VRpopssc.lupLbRcspH.Isp..lcYHR.hC.bs.....
consensus/75%   M................ssQhlLQPRGssp.+GPbpFp+SlRpGl.Lu-h.p.LGppu..L..LaPDGlARLWGuTPspp.ss.KspALRsp+VGDcVLFYA-ppFlA+ApIL.LFcs..lA+.lWGspc-u.TWEHlMALG-l.EFs..lsA..lL..Lsh.s.LRuLTL.sAppb.+hh.LLs.....s.s..ss.s...ssphspp-Llp.lssLpsp....s.sssRabsLsLLWuIuRlssGb.RLsPWp.FcsplGsLL.-FG.ss.phsPE.PFbpLpsSGlWEVcGh...s.s.s.ssshsps.s.AGhs..uAcLLpps.sRhcAlsbLpspahs-lDb..LL.RVGLAGYsoAsG..s.s.......s....ussss.R+.ssuoR.sRssplsppVK.la.p.CQVCs.pLb.+...YSEAAHIbGlGpPHDGPDcLpNLLCLCPNHHsbFD.h.lYlD.-..VRcocssc.lupL+RcspH.ID-palcYHR.LC.bs.....
consensus/70%   M................ssQhlLQPRGssp.+GPbpFp+SlRpGl.Lu-h.p.LGppu..L..LaPDGlARLWGuTPspp.ss.KspALRsp+VGDcVLFYA-ppFlA+ApIL.LFcs..lA+.lWGspc-u.TWEHlMALG-l.EFs..lsA..lL..Lsh.s.LRuLTL.sAppb.+hh.LLs.....s.s..ss.s...ssphspp-Llp.lssLpsp....s.sssRabsLsLLWuIuRlssGb.RLsPWp.FcsplGsLL.-FG.ss.phsPE.PFbpLpsSGlWEVcGh...s.s.s.ssshsps.s.AGhs..uAcLLpps.sRhcAlsbLpspahs-lDb..LL.RVGLAGYsoAsG..s.s.......s....ussss.R+.ssuoR.sRssplsppVK.la.p.CQVCs.pLb.+...YSEAAHIbGlGpPHDGPDcLpNLLCLCPNHHsbFD.h.lYlD.-..VRcocssc.lupL+RcspH.ID-palcYHR.LC.bs.....

# ; Standalone HNH

                1---------11--------21--------31--------41--------51--------61--------71--------81--------91--------101-------111-------121-------131-------141-------151-------161-------171-------181-------191-------201-------211-------221-----
OrigSeq         MRHVLPTGAGPVALSQAHQTPARRWHGFARHKQTVINSLLEDQYMLCCYSELRADWYGLGYHIEHIENKSQYPQLTFDWPNLAASALSSEDGLPEVNFGGHAKQNQVDLAHFVSIRDPQCARYFAYLSNGEMGPNLNRSPADQARAEYTIRVLNLNSPYLVTLRRNLWDELDALLAEHLDKGWSIEHLCQGEITPHGRPALNSFFSLARQFFGPVAEQTLRQYAPGLV

Jnet            --EE-------HHHHHHH------------HHHHHHHHHHHHH---EEEE----------EEEEEEE--------------EEEEE------------------------------------EEEE----EEEE--------HHHHHHHHHHH----HHHHHHHHHHHHHHHHHHHHH----HHHHHHHHHHH---------HHHHHHHHHHHHHHHHHHHHH-----
jhmm            --EEE------HHHHHHH------------HHHHHHHHHHHHH---EEEE----------EEEEEEE--------------EEEEE------------------------------------EEEE----EEEE--------HHHHHHHHHHH----HHHHHHHHHHHHHHHHHHHHH---HHHHHHHHHHHHHH----------HHHHHHHHHHHHHHHHHH-----
jpssm           -----------HHHHHHH-----------HHHHHHHHHHHHHH----EEEEEE-------EEEEEEEE-------------EEEEE------------------------------------EEEE----EEEE-------HHHHHHHHHHH----HHHHHHHHHHHHHHHHHHHHHHHH----HHHHHHHHH---------HHHHHHHHHHHHHHHHHHHHHH----
MBN8437007.1    MRHVLPTGAGPVALSQAHQTPARRWHGFARHKQTVINSLLEDQYMLCCYSELRADWYGLGYHIEHIENKSQYPQLTFDWPNLAASALSSEDGLPEVNFGGHAKQNQVDLAHFVSIRDPQCARYFAYLSNGEMGPNLNRSPADQARAEYTIRVLNLNSPYLVTLRRNLWDELDALLAEHLDKGWSIEHLCQGEITPHGRPALNSFFSLARQFFGPVAEQTLRQYAPGLV
WP_081344814.1  MKNVTKG-SPPQSFEAWKML-NPDWGNLQPQKRALHESLLEEQGWVCCYCGRDVTIRD--SHIEHFRPQTQYEDLALSYENLFASCIRETAPG-MPLHCGHAKDDDFDEERYISPLDTTCEGRFLYTLLGEVSPTD----LTDERAAYIVDLVQLDIQFLRNRREEALQRTFDADFLAS----ELRKLQDAFQRHDNAGHSENFGHVLARYAGQRLADRRGGEDVAAQ
TXH92073.1      MKHISKQ-PEPQALLDWKAQ-NADWELLSAPKQALKQALMVEQGFICCYCEQRLRDDD--SHIEHFRPQSDPACDALDYTNLLCSCQNRIERG-APRHCGNLKEDWFDEQLLVSPLSVGCEKHFAFAGDGSIRPAN----FDDQAAANTIEKLGLNIRKLQDLRKAAIEPFLDPELSD-----DLQLFTQAYLTRDADNRLGEFWSTVAHLFGRDEQGVYGA------
WP_057786318.1  MLQIHKQ-AEPQEVINFKAL-QDNWKGLTEAKTTLRKALVTEQGGVCCYCNRHIGGPANEFHIEHFRPQHAFTHLELDYQNLHASCFKNDSVK-GRRHCGPAKDKWFDTALTLSPL--NNSGIIRYGQNGTIFSNS------SPAAAETIKQLALDHPMLIAERAEAIAGKLDA---------TNDDLVKLYRAHQQMGRFQPFTLAVMQCIAALLPDPLKAT-----
MBQ6473653.1    MHKLIRT-EEPVELTEAREAQSLDWEGFSQNKHIVQDALLLMQNFRCAYCECALNEHE--GHIEHFRRKNIHPELTFVWSNLFYSCQIPGNKT-SHIRCGISKDEGFNYGLLIDPCVDNPEDFLVFTKKGYVYPKTNLSDTDRLRAETTIKMFNLNEEHLVVSRKNAIKKYKWLKDIIDKNKIEIDQYLDSI------KNSEPFVTAIFHYFGRRVVE---------
MCY4606376.1    MKRILAR-NKPTLLAEYLAA-DANWNGFRASYNELIEKLIDVQHGLCGYCEINLTELD--RQIEHIIPRRDGASKEVDITNMIACCKGRTSPVFDNMSCGQAKED---DADFLEPRLPALPSLTKVLVDGRIEVDEESEDIPTTRVTRTIEILNLNAERLKLAREKQWNNLEEEADQIGDPDNIMEAWIRSVLMPDDAGQLVSFFTTTRSYFGPLAERVLEQQPQAWI
TGX97922.1      MIVIKRG-KEPNSLLKFRKDNDADYEDMPDVAKDIREQMWEEQRCLCAYCMKKIDAPAV-VRIEHCRPRHEHKKATLDFKWMLGVCYGNSLVRGEDKTCDAHRGN---TELTVNPFDELSVRKIKYKADGSIYSDD------ADINKDVTETLNLNCQALPQTRKNVLMAEKNRIMRKCKGKSFIHELERTYASLVQERNLTPYCGIIISWLESKLK-----IS----
WP_020614468.1  MIKINKG-SSPSELDQYKRECTTKLDDFS-RKDDLRISLAEEQGYLCAYCMQRINPENGDTKIEHFHPQTEYPTEQLDYSNLLLCCNGNQGD--ANQHCDTKKKN---AEIKYNPASPSIEQTIRYERDGKIVSDD------EDWNNHIDNVLNLNYSRLRENRESVLRAVDQALAKNSGMRSQIDTIIQNWKAKNSSGELKEYCGAAIYRLQKHPAYKRG-------
RGH23085.1      MILIKKF-SAPPELEKLKKDASEGYDLLRPLKDQVRESLMKEQGHLCAYCMRRIPDSRIKVYIEHWKARESTENKGLDYNNLFAVCSGNEKAPGRHFTCDKRRDS---ESLKVNPLDESTIETIYYSMDGTMMSTD------SQIEDDINRRLNLNCNKLPQYRKEVLDTVMEYLAEQDGDYYNCEEQLQMWENEEDP--KTPYIGIAIWWLKKELAQRNEDGNR---
OQY33157.1      MVNINKK-TPPQSLIDYKRQQKDIFDDFK-EKSDLRTSLIEEQGYICAYCMQKIENDPLKCKIEHCISQDESSKKKLDYDNLLLCCKGNEGH--VDQHCDTKKGE---KSFHFNPSEQDMERLIQYNSQGKILSTD------AELNDQLVQTLNLNFKMLKMHRLGVLQEIERVLSKKKGTRTEIQKLINRWEALGSKGQHKPFYGVAVYKLKKHQSYCRA-------
MBR2506168.1    MVQIIKK-TAPKSLVEYKIA-NTSYNDFS-NKDDIRISLVQEQNELCAYCMGKIAADSTKMKIEHFKCQTTYPSLQLEYSNMLGCCKGNEGF--KMQTCDTYKGD---NILSLNPSNPFSKMQIIYSDDGTIKSLN------SQFDKELNDVLNLNTNVLKANRKTMIESAKMALSSKKGTRSEIQKLIQKFE-----AAHKPYYGAAIYYLEKKLKGAK--------
WP_168935609.1  MKLINKK-IEPRRLRKYRRKH-GTWQKVPRLRRHCRYKCMKQQGNICCYCECSLHHQTIFCRVEHFHQKSDGKNWHLDWQNMLAACNGKTDKNKNNLSCDSYKNYIQCEGLIMNPLLAAFPNLFKVGGSGELKPDDNKLATTKDLVQNTIDSLNLNCDRLRANRRTVLLFLNQRIAQEKKAGKALPAIVHQYFS----KKWPAYFTTIRCRLGKYAEDYLHSIAFQG-
WP_211114072.1  VKHCRKS-PEPPKLTEFRAAN-AKWDELRACYGDIRDATRRDQGGLCAYCELSLTDDN--EQIAHFHPKSDTVNWALKWENLWLACKGGTQTHHDNRSCDENKTK---DGHVLSPEVPLFPRIFRFEQNPRIVPDEEEAGIPIEKVQRTIDVFNLNCQRLAEARLSVYRPIESTLKKLRESSFKQRQFAARHLSIDAGHRWPRFFTLLRWQFGMVAETHLNSLPYQG-
WP_232474707.1  MKPVRGL-SGPPPLKRFCDD-PQNWETFGAAKQEVANALARSQRGLCCYCEATLVSGD--RQVEHVVPRSDGSARAVDATNMLASCKGGTERNFDNMSCGQRKGE----TIVMDPRLPPARSIFVVAADGKILTDPIEHGIDANEADKAIIDLGLNVERLRIQRAAILASLSSV--------PVWDALARGTLLPDEGGQLRPWFTTTRSFFRELAESILAQPPQDWI
MBX3668834.1    MKRVLRQ-PEPIQLTNYRNAN-STWEQMRAAYEESRKKLVADQGGICAFCEIDIRDNDPKCRVEHFHPKSDAHNWALDWNNLLAVCAGGSYRFAENLSCDAHKDREQCEGWILNPATPASPSLFRLMSTGRLLPDSNQHADVQTLVQHTIDMLNLNCDRLCNARLAIIRDIERNKKKQRDAGFGLGNLAVHYLR----QRWPGFFSTIRLCLGTAAETHLANLGYQG-
MBK9093497.1    MKKCRKL-SEPPTLEAYRLAQ-GSWDQMRQAYTDVKRTLVRGQRCLCAYCEIEIAGGMDEQRVEHFHPKEDTLNWALHWSNLWAVCPGGSQRPPANLSCDAFKDHASPEGWLLASDVPAFPLLFQFAPNGTPEPHPSHYPNTTTLVSKTIEHLNLGCPRLNRNRCIAKAQLEKQIRKAQQSAPVMLDLARRLLSQDADSPWPQYFSLVRWRLGDPAEAHLQSLNYAG-
MXY53198.1      M-------ETPTPLAAYKEQ-SATWEGFGDAKRELMAALEELQHGLCGYCESAFPSQG--RQLEHVVARSDGPERALDHTNIIACCGGGQRSSSGHLSCGQRKGE---DADFLDPRLPSSPSLMRVRSDGEIVPDEAEAAIETARVAKTIRILGLNVARLREARRVQRGLALDDHDGCS----RIRATAQSWLLLDRGGRLPAFFTTVRGVFGRTGEQILAAPGQPWT
WP_168963707.1  MKKIPSK-TEPLSFRNYRLANERNWDSFRQAYNDLKRQLFQEQGEICAYCEEHVPGNR--RRIEHFHPKSDSHNWTLDWCNMLGVCMGGEYEAYANLSCDSYKNHEQCDHDLLNPYIVSSENLFCFPVTGEITPNEGQTKTTLDLVRDTICFLNLNCARLKDKRKKIHQELQRWLRTQRQKHRAKRELAKRWFS----QRWPSFFTTRRCLLANYAEEFLRNSNYNG-
PZN82191.1      MKKISTLGNSPQGLQHYLQN-DRNWESFRYAYKELMLALTRKQHNLCAYCEIELKDTD--RQIEHFHPKSDGVEWAFLPSNLLAACKGGDNPNEENRSCGASHGN---LQNDQHPALPAMPSIFLVYRHGEITVNHSKNAEFQKIAQSTIVSLNLNCRRLKNARSGLFERLLKEFNQRTQEQTAMSEMALASLLPDDSGKLPPFFTTRRSFLVEFAEELLQHHPEVWI
MCB6991566.1    MIHIKKR--EPIEVQKELRE-SLEWDAMRTCKQEIRQCLLEEQHYLCAYCMSAIENKELHMSIEHFVPLSKCKEKVLDYRNYLGVCKGGADLPRKCICCDASKGN---ELQCINPFNEAQMHQIKYLSDGVIYYERRESDSEEKAIQKDIETLKLNGEKLVKNRKDAYEECESYIAELYANEMQLESDLQNIMERDHY-----AYDGVRLFLLKMAYENLQQIPNM--
NLX95049.1      MRTITRN-PNPPALTQ-----GQDWHAFMPCHQVVGESLLTEQHHVCCYCEVEIAADD--SHIEHMVPRSVTQGHDYDYTNLAASCDGG-----AVEHCGRFKDDRYDPNRFCPPQDPDTTRLLQYLPSGSVTPALGIGPQEAQKAQYMIGYLGLQCARLTGRRRTHARQLIDTLGPT-----LIAWAIGYYLNPDPQGKLRQFTSLSRKILNQ------------
WP_262567767.1  MRHIKKS-AAPAAFNAYKGC-GATYKNFVPLYNEVKQQLIGDQLGQCCYCGIRVELDS--THIEHVLDQNKHKKLQLEFSNLLASCNGG-----RQEHCGHAKGS---RPLPVTPLQDDCEERFVFRSNGYVASVE-----DDQDADTSIDVLNLNIVKLRNMRKAALSGSGLFEDD------DIEEYIELYNSADDNGQLEPFSQVVVNRLRQEQALLLQAERVSG-
MCI8605655.1    MRQIKKS-PEPGHFIKWKKDFDAEYEDLNNEYLLLKKSLVEEQGYICCYCEKRIGKQQYDCDIEHFMPRHDSPKAQMDYANLFASCKGEIGE--SADHCNHKKDNWFSFRSCISPSDPRIGGVFGFGLDGRLFVKD-------PAGQEMARHLNLNSYVLQEQRKAAFDAMMEVEFEDEDLLGYVEAVIGDYESMEG-GEYTEFCSMIVYCLRENWGGGVSWCNLAKE
WP_159471210.1  MITIRKG-REPREWTAKKKT-----EGFKEPIPKLRKALLKEQGNICAYCMREIPLSKRTSKIEHIKSREFHSDEQLDYDNMVICCPGCID---GNDHCDKSKKS---QEISFSPFDPNVQNSITYSKDGSIRSNR-------EDWDRDINQILCLNNKRLKANRTSVLKGIIELLDKKH----VIQQIIEEWKTPDKDGKMKPYCGIIIAFLEKRNRAI---------
WP_119395056.1  MHKIVKR-CNRPELRTAQQSPASRWRRM--RKDLLSKFLWQEQRGLCAYSEIQLKIEGGGYHIDHVQPKSAYPSRTFDEYNLVLSAFAQTDLARDAIFGGDPKDNHYDEHRFISPLQDNCRGYFLFLPNGEMTPHPSKSITDQQNAQYTIDLLNLNAEPLKLEREKWYDEIEQAFDVHIENGDCLTSLMGCDLVPRG-GQLSPFFSLTSQIYGDLAHTFVTEVETTSA
WP_078924152.1  MIFIRKRG-----LEQAHVNPTSRWSCFG-HKPAVTCFLEDEQYGLCAYTELRPDQAGLGTHIEHVQPKSRYPQRTFDFSNLVLCALADADLQGADLFGGHAKLSEYNAALFVSCLQSDCLRFFAYLSDGRVVAAANLSVNETQQAQYTIDLLNLNAPYLLVQRKNWLDELDKLIDEHLDNDNSLEDLAAIDLLPTS-GKLSRFFSATRQRFGRIAEPLLQEHAPELL
MBK9497785.1    MIKVEKK-PNPDNLIDQKAKANKSPEGISYGQADVKKKLYELYHSKCGYCEIKEHNDV--LQIDHYRPKNRKSNLALEWSNLVLVCG----------YCNRYKSEKQENPLIVNPEVDNPDNSLYFKNDGTIHAKD--------NKGIYTIKVFHLYDDRLVKKRKEIIDHYIEDFLNKYKAFKEFKNFLMIQVKATINKRQKPFIDYVIPYFETPLQDVIKNVYNQLK
HBL44679.1      MIRIDID------LEQQIET-KPGWDELESIWSPIKQVFTDLQFSKCVFCEKEIEDQPIEQDVEHFRPKNNKPNLAYKFLNYAAACK----------TCNSRKRDKSEKAYLIYPISDDPEDLIEF----GLSPQAKKSGFGRKRALVTIEMFQLDDERLFKRRAMLIYLLFTALEKVNNGNTEGQAGVDFLTSSKCE-----HTNCLRCFKKLYDTDFPKAEIVSQE
MBE6357837.1    MHKFIRS-KAPVNFTRVEQK-YQRWDELIEEHTIIGNALYEQQDHYCAYCEIKIEEMQGGGHIEHLERRSDNPQRTFDWNNMFFSCN-------NLDSCGKFKDNKYDVKDIVDPSREDPADFFTFDALGNVSPLN---HSVEHRALETIRIFNLQDSVLKRKRAEIAETVKYFLESS-----DITEFLTSLTGQD--------YISVYYNMLNRRMA-----------
MBI5767833.1    MHKLDRS-SAPVCLAGYDYR-TQTWADLDGCKRQVRLALQRIQGQRCAYCEGAIRHG---GHLEHFRRKNRYPELTFAWENLFLACA-------SQAHCGHYKDRAYDPDDLVKPDIHDPDAFFYFHSTGEVMVRNGMTDDERRRASETIRVFNLDCSTLRGARSRALKTYSDRNRGILEFLMEVADREAFIRGEIEATCWEPYAITIKHFFEKAA-----------
MBL8657368.1    MRQLDRA-QAPACLANYNHG-AQTWDDVSADKSGIRASLEVMQGRRCSYCEGGIDELG--QHIEHFRRKRDYPALTFDWSNLFWSCD-------QTDSCGHFKDRAYNPDDLVDPCGDDPDEYFLFRADGTISVREGLPANREHRAVETLRVFSLNPQWLRNMRQAAVAGYIRDADEAASAGLTPDEIRAYFADCLEYVKEFPFYTAIRHVLTER------------
MBX3408156.1    MRKLDRA-TCPTCLGEFVPG--SNYGQLRHEKQEIRDALLNMQGHRCAYCERRTGEERDEGHIEHFRKQADQPALQLAWPNMFWSCL-------DERSCGKHKDKSFNVDELLNPCDDDPDGFLEFLADGTVRARDGLSAHDLRRAQESLRVFQLDANALRQSRKDAVSPYMSAVDALLKHGVAVKGYVASVQSQINA---APFSAAIKHYLRGLQ------------
MCG9130915.1    MIRLKRIGKEAFELNQRRIKR----ENFKDRWKPAKEQLLAETGGKCAYCEAPTSPVAF-GDVEHYRPKSSY--LAYCYDNYLASCQ----------LCNQRKRDQQERPLLLNPYFDDPAKYFAWRADDEVTPNSRVDNTEEPIATASIEHYGLNRMELKNRRYNIYGHYRTYKQMLEYEQILIRHSIEKMKAPRAE-----FAGMIRYF---------------
MCP9859838.1    MKLLYRT-LNPNFLTQYNPN-TDKW-SVKDHRNQIWIALLSIQGGVCAYCEAGLQNG---RHIEHFANRGKRPDLTFEWTNLFGSCN-------RNDCCGHYKDSGYNLADLIKPDREDPWHFLVFGSDGRVSVRDGLSVNDKIRGQTTIDVFNLNAPH-VPERLSKVQGHLGLIEEDAE-----EAIVDLILDEIRSPHLDTYSSAALQPLELILGPENYSTP----
OYP37213.1      MKKLDRS-QAPECLGNYCHK-THTWDDFTEDKKSVWIQLDLMQKGKCAYCESFLGRT---RHIEHFRRKSQHQSLTFAWSNLLGCCG-------ANTSCGHHKDLPYQANDLIDPSTEDPDHFFHFQTDGSIKLREGLNPLEEHRAKETLRVLNLDEDRLRRRRAKVVQLYFQLEPGIVQALMEPADAKAYAQEELARIEDDELSAPLRHLFIDFVAA------N---
WP_010033157.1  MIRIVVT------LRARIEE-KAGWTALRNEWSDIKEAYIEFQHSKCAFCEKPLEGKI-EQDVEHFRPKGAPPGLAYHPLNYLTACK----------TCNSRKKNAGEQAYLIYPLSDDPEHLIEF----GITPRPGLTVFERLRARVTIEFFGLDNERLLIDRFETLEKVYLALKGRTEFATRADQVVTAYQTSRRR-----HTNCIKHFVALFTRHPEQAEQLYQD
WP_020558460.1  MHKLTRP-EAPDCLSRYRHG-RNQWDEVSAEKSEIWEKLDKMQQGRCAYCENALSRSGEDAHIEHFRQRGRYPQGIFEWTNLFGSCN-------RKDSCGKHKDEVYPHEDLIKPDVEDPEHFFLFVVDGSIAIRQGLSAVEQHRASETLWILNLDAEHLRQMRSSVVKGYIETAEYLAAMYADEDEWLELLQQELAAVEHLPFVTAIKHILKSES-----------
WP_061531640.1  MYRLTDA------IKEYEYE----WDKLKYLKYKIKSSLIQNPLNICYYCKSPLDCGTTPGDIEHIVHKSKYEIFTYEPINLTLACD----------RCNTAKGS---EDILITDLDSYTEEDYPLHSDAYIQIQDFVGIDQNNKGENTIKCCNLNRSKIKQVKSENSSPVKKMINGAVDSEKTLKEIEKIFEKPSHEQQLKEFQKRSKTVLLTLLEELLDSYDLSNV
WP_061808763.1  MIHIKRI-SNPDSLQEAIKH-DRNF-EFKYKKGNVKEILETMFNYKCAYCESRISPVTY-GDIEHFRPKKAYHSLAMNWENLLLACD----------TCNRTKKNIIEDPKLIDPCDEDPQDYMYFTKEGIVKYIG----EEGSKGDISIHVYGLYNPKLTENRKELILGYLKRISKLTKASNELKEVIESDVKDLEINNLTAYGPIIEKLLSRFNQLTDNN------
WP_077851366.1  MHKVKRN-KAPLKLIEKDKEWEPDWAKFSALKKKTIAALEEMYKGCCCYCESKIKSTSY-PEIEHFKPKSKHKELCFDYNNLHYSCK----------RCNIAKGS---NDNLINPSEDNPEEYIRYIGELAISIDD------NKRGVETIDVIKLNERALKEERIRYLNEFRKNYELLISAMEIVKPFIANFIENVKTNHGESYCTMIKHNFSDKLDMLMEVLEEVK-
WP_101103330.1  MKTVVKG-VEPKELRDWKRSNNRHYDNLSAVKKAMLEKLAAEQGWLCAYTMKQITDKDGQAHIEHILPRSAHPDQSLEWQNLIACVP-------PGVSCEYKKDADPATQPFVNPTMGGVSAQFRFRENGEVDGLT-----EAARVTVSQEVLHLNHPNLTNDRKGKIQGAIAHRPTA-----QARQRAQLLRKTDQNGMLEPYCEAVAQVLEAYATRLEKRASRTSK
WP_164949143.1  MHKVNRG-IKPKSLNDNDLEWNNTFTDIPYKQEDVKQVLIEMYEGLCCYCESPIGVQTY-EQIEHLKPKSIFPQFCFEWTNLHLSCQ----------VCNIKKTNWDKDNPILDPTVDKIEDFLEILESGEIVPIN-----NNARANTTIEHTGLNRDNLVKRRKSIIITMKKLLIIAVNANEDFKEILEIMSEDMG------YKLLINTFIKKL------------
WP_200158834.1  MRSIVKG-AEPRALIQWKAENNLVYGGF--PADAVRAALLAEQFHLCAYTMKRLKTRAESCHIEHLLPQARKPGEDIDYRNMVACYP-------SRIACEFAKGDDPMVGAFVSPLLPKVERHFEFDARGGITGR-------TPEGSATIQVLKLNHPALVNDRAAVIRGALHPKGKPLSA--AARRLATVMTQPNRDGCLPAYCVAVATAALAWAERAERRAARRRK
WP_312777483.1  MRKLSRP-EGPECLEAVPAG--TPWSSFGPCKPKVRQALEAMNMDLCSYCESLLVENSG-SHIEHLFPKSEYPDRTYRWDNLYLSCQ-------NTDHCGHYKDSDYDPADLIRPDIDSPDGYLHFHSSGEVTVRGGLSDAERHRAQTTIDVLNLNAPSLIASRHAVASQVQAMILPDLELLDDEDELSELFSAEIESFRAGAHPTTARHFFA--------------
consensus/100%  h..........................h..................C.as............l.Hh..b............h..sh............s...+......................h............................L........+................................................................
consensus/95%   M..........................h......h...h.......Csas............l-Hh..b.......h...Nh..sh............ss..+.........h.s........h.h.....h.............s......h.L........R........................................a.......................
consensus/90%   M..h.........h.............h......h...h...b...CsYsb..h.......plEHh..b.......h...Nhh.sh............Cs..K.p.......h.P........h.h...s.l.............s......h.Ls...L...R........................................a......h................
consensus/85%   M..l......s..h..........a..h...b..lb..L...b..bCsYCb..l.......plEHh..bp....b.hpapNhh.sC...........pCs..K.p.......hpP........h.h..pG.l.s...........s...hp.h.Ls...L...R...h..h..............p.h................a.s.....h...............
consensus/80%   M..l.p....P..l..........ap.h...b..lb..L..bQ..bCsYCb..l.......plEHh.sbp....b.hpapNhhhsC...........pCs..K.p......hlpP...s....h.a..pG.l.s..........bs...lp.hsLs...L...R...h..hbp............p.hh...............a.s..b..h...............
consensus/75%   Mbpl.+....P..L.php.....pap.h...bp.lb..LbpbQ..hCsYCc..l.......cIEHh.s+sp..pb.hcapNhhhsC...........pCs..Ksp......hlsP..sss.pbh.a..sGpl.sp........pbsp.slc.hsLs...L...Rp.hhp.hbp.b..........hpphh..h............a.s.hb.hh...............
consensus/70%   Mbpl.+....P..L.php.....papphp..bpplbp.LhpbQ..hCuYCE..l..p....cIEHhps+sp..pbshcapNhhhsC...........pCsp.Ksp....s.hlsP..sss.phh.a..sGpl.spp......ppbsppoIchhsLs..pL...Rp.hhp.hbp.b.........hpphhp.hb.......p...ahshhbphh...............
```

```
# ; PIN-EndoRNase

                 1---------11--------21--------31--------41--------51--------61--------71--------81--------91--------101-------111-------121----
OrigSeq          MNGFLLDTTILIDFLRGRKEAIDMLNRLVEAGPLACCPVTVAEVFSGARPEELPKIEEFFEALLFYPVGYKTARRAGLYRREYGKKGITLSISDTIIAAVAVENSLTLVTKNVRHFPMPELAVIEHG

Jnet             ---EEE--HHHHHH----HHHHHHHHH------EEEEHHHHHHHHHHHHHHHHHHHHHHHH---------HHHHHHHHHHHHHHHH------HHHHHHHHHHHH---EEEE-----------------
jhmm             --EEEEHHHHHHHH----HHHHHHHHH------EEEEHHHHHHHHHHHHHHHHHHHHHHHH---------HHHHHHHHHHHHHHHH------HHHHHHHHHHHH--EEEEE-----------------
jpssm            ---EEE--HHHHHHH---HHHHHHHHH------EEEEEHHHHHHHHHHHHHHHHHHHHHHH---------HHHHHHHHHHHHHHH-------HHHHHHHHHHHH---EEEE------------E----
AEG13740.1       MNGFLLDTTILIDFLRGRKEAIDMLNRLVEAGPLACCPVTVAEVFSGARPEELPKIEEFFEALLFYPVGYKTARRAGLYRREYGKKGITLSISDTIIAAVAVENSLTLVTKNVRHFPMPELAVIEHG
AUB85623.1       -MGFLIDTNVLSELQKRRRGVVAWYANTRSRD-IYLSVLVLGEVRQGIERL-ERRMALIQVGDRILPVTRAIAERWGNNNVP-----DALPVIDGLLAATALEHDLILVTRNVRDVERCGVRLLNP-
BAQ63188.1       -MTYLLDTCVISEYIKKKPLVIEWLDNQAKDN-LWLSILTIAELKKGIFKI-SQWLEFVKTKGHILPLNEEVLNTWAEITGESEACGKKLSVIDSLIGATAQQYNLIIVTRNINDFNFSSLPIFNP-
EXI76475.1       --MILLDTNICIYLINTRPEVLARFHQYRLGD-IGLSSVVAAELAFGVAKS--QALEMFLAPLEILPFDTAAMWAYGDLRADLERRGEPIGSLDTLIAAHALSVHAVLVTNNTREFAKVRGLQLEN-
HAL93439.1       --MYYLDTNTCIYFLNGKYSIKEKVLSIPPNE-IGIPSIVKAELFFGAYKS--EKVEMFLEPFEIIAFDDLMTYVYADIREKMEKTGALIGPNDLLIASIVKFYDGILVTNNEKEFGKVEVLRIEN-
HAL93443.1       ---MILDTDVLIWYLRGNEKAKNAVETAL---PFSISVVTYIELVQGMKNKEYRLFQKQIQDTDIIQIDREISSRAMFYIQEYVFSHS-MYLADALIAATVVQRGEMLFTANDKHYKFIPTIQCKR-
KAF0185694.1     MARFLLDTNIVSDLVRNPQRTSAKIAELGEDA-VATSIIVAAELRYGAAKKLAEQLETILAAMEVIPLEAPADATYGATRVALEAAGTPIGANDLLIAAQTLALDMVLVTNNEREFERVSGLKVEN-
MBI2378667.1     -MGALIDTSVLVHVERGNLRVAAALAA-LEDS-PVVSAITASELLHGVHRA-AAYVERVLAGVPVIPFDLAAAREHARVWAEVKKAGISVGERDLMIAACALAHGHRVVTCDARSFGLVPGLTVET-
MBI2378673.1     -----MSSSIATPTRHNAEGVLERLLAQSPRD-VTVSVVSVGELRTGAAKS--ARVESFLQPLTVLDFVRDDASVYAEVRARLERSGTPIGPLDTLIAAHALARKLTLVSNNEKELRRVKGLRVEN-
MBI5243144.1     MPELLLDSDVIIEALRGSPDTLAELAEVEAAGLISYTPIAKAEIYHGLRRGEEKATEGFFAACRCLTITGDVGEKAGRYLAAYHAGHG-TEIADALVAASAFVHKAALFTLNRRHYPMTDIAFHKR-
MBM5817551.1     --MFLIDTNVISEARKGSRGVRAFFVAASEQDPLFVTSITIGELRRGVDLI-EHWLAVLLEADRVLALDAEAAQLWGQLRVP-----HPEHALDKQIAAIALLHDLTVVTGNTGDFLATGVALLNP-
MBM5817553.1     MALLVVDTDVLIDYLRDQSDAVAFLEGCTQ--PLALSVVSVAELYVGVRDGERHRLDAFVAAFDALPLERKAAVQAGLWRRQYGPSHG-TGLADALIAASVATAGATLVSLNRRHFPMLADVLVPY-
MBR1880070.1     MKQFLLDTNILIELMCDNPNVAQRLLDVGFDN-CRMSAISLHELNFGAFNAEMVRIKKLLEKIKVLSLPAEAAEYYGKIKYTLERQGKRVDEFDMLIGGHALATGMTVVTDNIKHFENMPDVDIEN-
MBV5272981.1     -MGYLLDTNICIHLIDRHPQVLARLAPLPLGD-AVMSVISYAELWHGVERY-ARALKRLIARVPVLDYPALAAEHYGELAAAVKDRRRDV--MDRLIAAHARSFGLTLVTNNEADFVGYPGLTVEN-
MBV8611621.1     -MSFLLDTDICSAFLKGERNVGNRFLQ-YGGR-LHVSTVTVGELYTGVLRS--PLMLGMLNNLTIFDVTYDVAYKFGTIRAPLLDRGLPAPGMDLLIAATAIVHGLTLVTHNTPHYAHVPSLTLAD-
MCA9013094.1     --MFILDSDHLSLLQQTSTRLAIRISAYSADL-FFVTIVSFHEQVNGWTAYGYTELDGILSDAQVLPFSAAAADIYDDL----KKRKIRVGAMDLRIASIAIANQMTVLTRNTIDFERIPNLSFED-
MCD6306395.1     -MAFLFDTDAISELLRPHPKYIQWLLSVPRED-QFTSAVVIGELYKGAYRSHLKNIEQVLPAVIVLPYDVATAKVFGQIRARLEESGNILPDADIQIAATAVYHDIELVTGNLHHFSRIDGLKVNTG
MCE2718754.1     MLIYLLDTNVCIMYLKGKSNINHHIDNLKPEK-IAVCSVVKSELFYGSMGS--NIQKSFLRQFVSLPFDDNCAEVYGKIRADLAKAGTPIGLNDIQIATIALANNLILVTHNVKEFSRIEGLKFED-
MCY4474613.1     -MNYLVDSDWAIHYLRGAQPVIDRLQPLTHEG-VGISIISLAELYHGVFRS--ENLNGFLSQLTVVGLNHEVCKLFGQERAKLQLRRIRVPDFDLMIGCTAVYHDLTLLTNNRRDFENIEAINIET-
MCY4597194.1     --MIPVGTNVLSELIRRRPRELAWLEARDP--LLAAPATALLELRYGISRP-LRFLQMTRDHARILLFDESATEAYGEIAAEAERSRGRLNVAAGQIPAISLVHGTTIQTRKLIAFKASGASLVNP-
MDE2076669.1     MKGYLLDTSVVSALAPGREAALAEWLQAHHQE-LFLPAIAVAEMAQGICKL-DRWLDGLLAADRILPLDAHAARLAGQLSDAAMAQGCHPGFADVAIAALAQHAGLLLLTCNLRHFQPLGVACADPG
MPZ42200.1       -MSWLLDTCVISELVRPEPQVVAWVDSCEEET-LFLSAITLGELEKGIAKL-ESWIRRLPERGRILPVDETVATRWGAVSGAAEIRGTPLPVIDALIAATGMAHDLTVVTRNTADFARCGARCFDP-
MXZ78954.1       --MYVLDTSIVSEIRKSKTAVVEWSSRVPTGL-TYISAITIQELEHGVLLA-RHWLDGVASAQRILPVDTTVARLAASLHVP-----DPAPIIDALIAATALANGMTVVTLNTRDFERFDQLQV---
PXF57195.1       MQLVIFDTDVLIDAGRRIDDAISCLQRAEQKATLGVSIITQMELMVGCRNKEFRALEKFLCRFQILGLNENIGNTSVDLIRHYRLSHG-LLIADALIAATALTTRVPFVTKNRRDYRFIEGLELLSS
QVL49521.1       -MTWLIDTNALSELKKRVPGVADWFAERPAGS-LHLSVLTLGELRKGIHAM-SAWLESVPVYGRILDIDGTIAERWGRLCADAR---RPLPAIDSLLAATALVHGLTLVTRNLADFRVPSLRVVNP-
WP_008932881.1   --MIVLDTDVISELWKSQPNVLAWVDAQAIET-LYLSIVTVAELRYGIAAM-QTRLEHVLPAGRVLAFDLEASQAYAELMARARAGGKAIGMADGYIAATAAVRGLVVATRDISPYQAAGLVTINP-
WP_015245483.1   -MNFLLDTDTCSAHMRRPAQLAHRFIQ-HIGQ-LAISSVTLAELYAGAYKH--ALVADLLQEVHVVDFDPACAHTFGQVRGTLWQQGISVPTADLMIASAALVHNLTLVTHNTADYQYIPGLRLDD-
WP_017842530.1   MLIYLLDTNTISEPLKATPQVIENIKR-YHAQ-IAIAAMVFYELVRGAELL-TDYIESVLIKFPILPYTEKAAKWHGQTAAHLQQQGKTPPFVDAQIAAVAKTNNLILVTRNVDDFANFSDLLIEN-
WP_019463149.1   MLRYLLDTNLCIRVLRDRPGLRGRFNA-EAAA-LCTSDVVLYELLYGAERS--REVERFVARLSVLPFDGDAAAHTAEIRASLERRGGVIGPYDLMIVGQARSRGLVVVTGNLSEFGRVEGLRCED-
WP_020565138.1   MISYLLDSNILSEPARLKPKVLQQLAD-HDGE-YATAAIVWHELAYGCELL-QSYLAMLLANLIVLPYDQAAADWYAKERARLQRHGLTCAYADGEIAAIAATQQLILVTRNTLDFGNFRNLALQN-
WP_152828753.1   --MILLDTNVISELMRVEPIVLDWFGQHEAAD-LFISAITEAELRTGVAIL-QLAIDAIDQDTRVLPFDSSAARSYAEIATARRAKGKPIMDADCQIAAIARACGAAIATRNVKDFEGPGISVINP-
WP_177417919.1   --MYMLDTDTCIYVLKNRSRLRHKFKA--IKN-ICISSITYAELCFGIENG--EQLELFTRRLLIDPLDEDAARHYGYIRALLKKQGTPIGNNDLLISAHARSMNAVLVTNNLREFQRVPDLSVEN-
WP_179052053.1   MEQVLLDTDTLSAIMRQHPVVIPKAQAYLAQHSFTFSIITRYEILRGLKAKQLQAFEHFCANNIITPLTDEIIVQASDIYAELRKKGITIGDADILIAASAMVNRIAVITNNQSHFCHISGLTIYN-
WP_201066874.1   -MAYLLDTNAAIALMKHHPTVQSHVRRVGRQQ-LRICATVESELWFGVAKS--ERLLRLLDWLPSLPFSGEATRRFGEIRVYLNRQGTPIGPYDLQIAAIALAAELTLVTNNTSEFERVPGLRCED-
WP_235751991.1   --MYLLDTNICIALLKNNPLAVSEFTV-KTEE-SYLSTIVVSELYKGVYCS--KTLNEFIALMSIMEFDLQAALEFGKIQAELRKMGKPTGELDAMIAAVARSRQLILVTDNIRHFENIANLQLEN-
WP_241191480.1   MVRVLLDTNVISELVRPRRAVVAYLSGLEPDQ-SYLSLISVGEIERGIAQV-RNWLDGVVPQGRLLPLDEPVIRRWGELMALPEVRARSGIAIEALIAATASVHRLTLVTRNSRDFAVFPVASYDP-
WP_242470923.1   -----MDTDVCIEILRGNPRVLEQRRA-TLDH-VVTTWITACELAYGAANS--ILVTEFLSTLPILDFNLPSALLFGRHKARLRGLGKTVADADLMIAAMALAQGATLVTGNTRHYERFDDLRLEDG
WP_247725416.1   -MNYLLDTNVLSDLLRGQEPVKSRFTRVRGHVRFILSPVVDFEIRRYLLLKNQAQYDALVGAWFEPAFDAAHWQQAAELWAARHRIGKPIADADLLIAVTALRQNATLVTNNMDHFLDL-GLSLED-
WP_254428054.1   MPVIVLDTNVISELMRPQPRVLAWANSLDPES-AAITAMNEAEILHGLARL-RESWDAAADLGRVWAFTSEAAHWYGELVSHRERLARPIATADAVIAAIGLAHGAPLATRNTSDFADLGLQLINP-
consensus/100%   .....hsos.h.......................s..s..E......................h...........................lss.s......h.o.p...h..........
consensus/95%    .....hsos.h....p..................s..s..Eh..Gh.................h.......s................-..lss.s......lho.s...h.......h...
consensus/90%    ...hhhDos.h....p....h...h..........s.hs..El..Gh.......h..h.....hhshs.......u................D.bIuu.s......lhT.N..ca.......h...
consensus/85%    ...hllDTs.h..h.c....h...h........h.hs.ls..El..Gh.......h..h.....hhshs..hs..hu................D.bIuu.u....h.lhT.N..ca.......hps.
consensus/80%    ...hllDTshh..hb+.p..h..bh........h.hs.ls..El..Gh.......hp.hh....llshs..hs..huph.s........hs.hD.bIAu.A....h.llT.N..ca..h..h.hps.
consensus/75%    ...hllDTshh..hh+.p..h..bh........h.hoslsh.ELb.Gh.......lp.hh....llshs..hsp.huph.s.h......hs.hD.hIAA.Ab.p.h.llTpN.pcF..h.sh.hps.
consensus/70%    ...hlLDTslh..hh+sp..lh.bh........h.loslshsELb.Gh.....pblc.hh....llshs..hup.aupl.s.hp.....ls.hDhhIAA.AhspshslVTpN.pcF..hssh.lps.
```

```
# ; PIN-EndoRNase

                  1---------11--------21--------31--------41--------51--------61--------71--------81--------91--------101-------111-------121-------131-------141-------151-------161-------171-------181-------191-------201-
OrigSeq           METTKRRTTVYIDGFNAYNGMKAAGLRNCRWLDWVAYVSGLVGDDVLTVRYFTAVVNSPLDARERQSHYLRALRANGGLDIIEGSFMARSMKCPDCGHSWKRPKEQFTDVNIAIAMTHDAREGSLDRAMLISGDSDLVPAVRYVQDLGIEVVNVAPPRRKGDELASACDGLLHMDDVAMRKAQLPNPVMESFRGRPLHAPEGWT

Jnet              ------EEEEEE----HHHHHHH----------HHHHHHHHHHHHHHEEEEEEEEE-------HHHHHHHHHHHH----EEEEEE-----EE------EEEEEEE---HHHHHHHHHHHHHH---EEEEEE------HHHHHHHHH---EEEEEE------HHHHHH----EHHHHHHHHH---------------------
jhmm              ------EEEEEE----HHHHHH-----------HHHHHHHHHHHHHHHEEEEEEEE-------HHHHHHHHHHHH----EEEEEE------E------EEEEEEE---HHHHHHHHHHHHH----EEEEEE------HHHHHHHHH---EEEEE-------HHHHHH----HHHHHHHHHH---------------------
jpssm             ------EEEEEE----HHHHHHH----------HHHHHHHHHHHHHEEEEEEEEE-------HHHHHHHHHHHHH----EEEEEE---EEEE------EEE----EE-HHHHHHHHHHHHHH---EEEEEE------HHHHHHHHHH--EEEEEE-----HHHHHH----EEEE-HHHHHH-------------E---------
BAN04396.1        METTKRRTTVYIDGFNAYNGMKAAGLRNCRWLDWVAYVSGLVGDDVLTVRYFTAVVNSPLDARERQSHYLRALRANGGLDIIEGSFMARSMKCPDCGHSWKRPKEQFTDVNIAIAMTHDAREGSLDRAMLISGDSDLVPAVRYVQDLGIEVVNVAPPRRKGDELASACDGLLHMDDVAMRKAQLPNPVMESFRGRPLHAPEGWT
EPL60618.1        M----QRVIAYVDGFNLYFGLKDSQFKKYYWLDLPAMAAALLDQQLVATHYFTARIRRNADDAKRQTTYIDALAAQPNLTVHEGHYLEKTQRCKSCGASWKVYEEKMTDVNIAAEMLADAYEDRFDTAFVISGDSDLTTPIQQVRK-NKRLIVVFPPNRQSAQLKKAANGYLSIGEDKLRQNQLPNTV-MTSSGFALHRPTHWK
EXI65854.1        M----TRVISYIDGFNLYFGLRSKGWRKYYWLDLVALSTSLLGQELQAVHYFTTRIRHNSQDIQRQNMYLEALGMRPALSLHFGHYLEKPKQCRKCGATWMDFEEKMTDVNIAVQLLADAFDDRFDTALLVSADSVLTSPVRQVRN-LKRIVVAQPPGRHSSELTKAATGFVTVGEDKLRQSQLPPLV-ARSDGFVLQRPTYWT
GHU44760.1        M----NRVKVYVDGFNFYFGLRSKGWRKYYWLDLNALSTALMDQTLAGVHYFTSRIRDNVPDMQRQSLYLEALGTLSDVQLHFGHYLQKNRRCFECGAKWKNYEEKMTDVNIAVQLLADAYDNSFDTALLISADSDLTTPVRKLLA-DKRVIVVFPPDRRSADLAKAATATFLINETALRQSQLPAEV-QRADGFILQRPAYWQ
GII23649.1        M----ASVIAYVDGFNLYHGLRERYQHRYLWLDLAKLVGRLRNDTIIAVRYFTAAVRNDPAAEARQHTYLEALKAGNSIEVILGRYQAKSRSCRSCGAVWTSYEEKETDVNIAVSLVADAASAAADIALIVSADSDLCPAVRKARTVKMGMIAAFPPRRNSNELRSLIPASFQLAHADIRNSLLPDTVIDAATGRAHQRPAKWH
GIW54144.1        MEPFPKRAVAFVDGQNLYHAVREAFGYTYPNYDVQSLAQRICGWDLVQVRFYTGVPTDNPFWHGFWANKLRALSWQ-HVHVYARRYRNKRVVLPDGTHTFLAGEEKGIDVRIAIDVIRMAHHKEYDVALLFSQDQDLSEVAEEIRVIWIKVACAFPNITNRRGIEK--TDWVKIDRATYEACLDRRDYRPKSTGLGGT-P----
HCF05835.1        M----SRVIAYIDGFNLYFGLKSKGWKRHYWLDVVGLSRSLLGQTLQAVHYFTARIRNNYADMQRQADYLDALATSPLLTIQFGHYLPKGRQCCRCGNTWTDYEEKMTDVNIAVQLLADAFDDRFDTALLISADSDLTTPVQQVRA-AKRVIVAQPPGRNSVQLAAAATAAFTISETKVRQNQLPAAV-TTASGYVINRPEHWR
MAS10378.1        M----TRVASYIDGFNLYFGLKSKHWKRYYWLDLWAMPQALLGQQLVASTYFTSRIRINQADMDRQSDYLDALSTNPNLSIREGHFLEKPRRCRACGATWTTYEEKMTDVNIATALLSDAHRDEYDTALIISGDSDLTPPIREVRR-HKRLIVAFPPNRHSQDLRRAAHGFIAIGADTLRQNQLPAYV-TMPNGHVIHRPATWK
MBC7252725.1      IRPSPLRVVVFIDGQNFYNDCKRNFGK--GETHPHLLGQELCDRRLVQVRFYTGIHTRNPRMHAIMTKRIEEMKRN-DVYVFTRKYSLEWVKLPDNSQKW-KGREKGVDVKLALDLVMMAVEDKYDVAVVVSTDTDLDEAVEEVLILWIAVENAIPPTGKRPPFKRNAGRLLYIDESIFRRIVDNTDYSG---------P----
MBF0143009.1      M----ARVIAYIDGFNLYFGLRSKGWRRFYWLDLVGLSRSLLGQELQTVHYFTARIRRNDEDLRRQNTCLEALSTQTDLYLHFGHYLEKKRERRRCGTQWLDYEKKMTDVNIAVRLLDNACEDRFDTALIVSGDSDLTAPLNLVRS-TKRVVVALPPGRHSVQLTRSAGGYFTIGEDKLRQNQLPETI-IRADGFILTRPAHWR
MBI1799724.1      MEPLHPRAMVFVDGQNLFHSARESFGYTYPNYDVRALAEAVCAWALSETRFYTGVPDDNAAWNHFWTHKLAMLGRQ-GVHTYSRRYRTRVIRLPDGNHSYLAGEEKGIDVRIAVDVIRLAHRQAYDVALVFSQDQDLSEVAEEIRVIWIKIACAYPLTRNRRGIQK--TDWIPIDKSTYDACLDRRDYRQKKAGEAEKGP----
MBK8644368.1      MEQKKERVIAYIDGFNLYFGMREAGFDSCRWLNIKLLVEKLLHQELIGVKYFTSRVSNNPDKQKRQSTYIDALES-TGAKAIYGNYQDVSEECLRCGHIWRTAKEKMTDVNIATAIMVDAFQDNYDMAMLISGDGDLTPPIREVHR-NKRVFIAFPPKRHNSSMALVAKGSEIIGRKKLVDAQFGEEV-ESRTGYMLKRPKEWV
MBL9184320.1      M----ERVITYIDGFNLYFGLRQKRWQRYYWLDVLALSRALLTQQLEMVRYFTSRVTSDPGQPRRQHIYLEALETLPQTTLHFGHYLSKPATCKSCGAQYTKYEEKMTDVNIAMELLTDAHQDAFDTAIVISGDSDLTGPVLKIRD-TKKVIVAFPPERSSDQLKRAAHGHFIIGRANLAASQLPDPV-SSKTGFPLKRPSSWA
MBN2361665.1      MEPQIKRAIAFFDGQNLFYAAREAFGYRYPNYDPIALAQVACGWQLAQVRFYTGVPDDDQFWNHFWVAKLGQMGRE-GVHTFSRRYRHQTVRLPDGSQVALIGQEKGIDIRIALDVVRLAHRREYDVGVIFSQDQDLSEVAEEIRVIYIKLCSAFPSSRNTRGINK--TDWLRIDRTTYTTCVDKRDYRPKQDQKPDR-P----
MBP7275756.1      M--TGTPTMIYVDGFNLYYGALKG--TPYKWLNFEALFARVFNNRIVGIRYFTAPINSDPDQPLRQQRLFRALRTLPLLTIIEGHYLCHPKSMPLVNAQVLFTEEKGSDVNLASYLLMDGFKNAYDCAIVVSGDSDLATPITMARQ-KTVGVLL-PQRRRSARLQQVASFFRDIRTGVLAASQFPSTL-TDAHG-TFTKPASW-
MBS0225818.1      M----NRVVGYIDGFNLFFGLRDSGLRRYYWLSPELLIQNLMWQTLAGVRYFSARISGDPDKHLRQQTYLEAVETLAGVEAIYGHYLSKPKQCRACGAQWQQAEEKMTDVNIAVRLLSDAMDDAFDTAMIVSADSDLVPPVEAVRA-AKRIIIASPPARHSAKLAAAANACFTIGRKKLQDSQLPATI-TKPDGFVLTRPASWN
MBU0673019.1      M----QRVIAYIDGFNLYFGLRENRWRKYYWLDLVKLVQSLLDQQFVAAHYFTARIRNNSSDSKRQSIYLDALATLPAITIHEGHYLQKKVQCRKCGVTWSTFEEKMTDVNIAAQLLVDAFDNRYDTALIISGDSDLTTPVQRVRE-DKRVVIALPPKRHSQQLTLAANGAFTIGEAKLRQSQLPDKV-VTTGGYILQRPEHWR
MBX3391866.1      -----MNVICYIDGFNLYHGLRSKGWRKYYWLDLWALAERFLGQTLVEVVYCTTLIKDDPPGQQRQLTFIDALKARPGIKVVYGHYLAKEMKCFKCANTYTRHEEKMTDVNIACHMLTDAMDCRFDVALVLSGDSDLVPPVKTVRD-QKRVIAVFPPERYSDALKRACHGHTWIGEEKLRQSQLPVVV-RTAAGLDLTRPTEWD
MCC6958029.1      -----MNTYVYVDGFNLYYGAVKG--TPYKWLNILDLSQRLLKNNIVKIKYFTALVTGDPDQPNRQQIYLRALRTLPNLEIIYGHFLQHPVKMPLVSANVMKTEEKGSDVNLAAHMVNDGYKSLYQVAVLVSNDSDLVEPIKIVRS-LPVIVLN-PRSTPSHELRKYASFVKPIRQSVLANSQFPDEL-TDAIG-TFHKPPGW-
MCY2953680.1      MEPPVKRAVAFFDGQSLFHCAKGAFGYTWANYDPVKLATAICAWKLAQTRFYTGIHTGNAYLHGLWKNKLSVLGRQPNIHIFSRRYRQKTVELPDGSHTFDTAEEKGVDVRLALDLIRLAMRAQYDVALVFSQDQDLSEAADEVRAIWIKIASAFPVGGNAHGINK--TDWIPIDKALYETCIDHRDYRPKPTGGAH--P----
MCY3902334.1      M----QRVAVYIDGFNLHYGLLDRGWRRYVWLNVRRLAENLLHQRFVLARYFTAKMRHNSGNSVRQETFLEATELLPDVHIHLGYYSRKRKICAGCGADFWVFEEKMTDVNIASSLLCDAQDDIFDTGIIVSADSDLVSPIREVLF-GKRMVVAFPPKRVSKHLRQVATGHFVIGRKKFEDSQLPSQV-IKPDGYVLTRPPAWN
MCY4278448.1      M----NRVLSYIDGLNLYHGLRSREWKKYYWLDLVGLTQRLLQQILEAVHYFTSTILGHDSAMRRQICYLDALRTLPLLNIHLGRFSEKSSQCRHCGAEWRKYEEKMTDVNIATQLLTDAFDDRFETALLISADSDLTTPVRVIRC-QKRVIVAWPPERRSEDLRRAATASTSISETALRLSQLPETV-TSESGHVLKRPKEWH
MCY4533096.1      M----NRVISYIDGFNFCFGLRAKGWRQYYWLDLAAMTRSFLGQQFEHCHYFTARIR-NGQSTKRQSLWLDALATRGEITCHFGHYLPKDQQCRTCGAKWVSHEEKMTDVNIATQLLVDAYDDHYDTAIIVSGDSDLTTPVRQIRN-NKRLIIAFPPTRRSAKLKKAAHGHFVIGADKLRQNLLPNEI-TTPRGYVLKRPARWY
MDD3182287.1      M----TRLITYIDGFNLYFGLREKAWKKYYWLDLQGLSERLLGQVLAATHYFTARIRHNSSDQQRQNTYLEALQTKPLITIHYGHYLEKTRTCRTCGAQWKDYEEKMTDVNIATQLLSDAFDDNFDVALIISGDSDLTTTIQSVLR-RKKIIVVYPPARHSSSLANAATASFVLGEANLRKSQFPQSV-TRQDGFQLLRPTRWR
MDE0079140.1      M----NKVIAYVDGFNLYFGLRDRGWRKYYWLDLEELLRRLLHQELEGIHYFTSRIQNNKASMDRQAIYLHALNTLDSVSLHFGHYLQKPRHCLNCGATWMSHEEKMTDVNIATQLLADAYEDRFDTALLMSADSDLTTPVRLVLD-NKRIVVVQPPRRRSVELGSAATACFTVGEANLRRSQFPDPL-TTKDGLILRRPEHWK
MPZ20363.1        M----ANVNVYVDGFNLYYGGLKR--TPYKWLDLAKLCQIMLQDVVQHIYYYTARVSHNPSAPMDQQVYLRALRTIPNLSVTYGHFMTHSVRMALSRVWVDKTEEKGSDVNLATHLVRDAFQKRFELAVLVTNDSDLAEPVRIVTQ-LPVGILN-PHQYHSQVLKRLATFVKRIRQSALHASQFPVTL-ADEKG-IFHKPSSW-
NLO26758.1        -----MRVGVYIDGYNLYYGGRKRGVPGWRWLDLRALATDLVSSTISRITYCTARIKSNPSGFRGQDIYLKALAVSGSVYIEYGRYVERVKRSPLATVSHSDREEKGSDVNVASHLLVDVLGKSVEAAIVISNDSDLRFPIQEARK----ICTVNPSAQTAGDLRGGGHWWLRLTADDYKKHQLPDPC------RQYRKPHGW-
WP_015065388.1    M----ERVIAYVDGFNLYFGLREAKLKRYYWLDVSLLAKNLLQQHLVATHYFTARIQNNGPDRKRQNDYLEAL-TLVGVSCHFGHYLQKTQRCHKCGASWPYYEEKMTDVNIATQMLLDACDDLFDVALIVSGDSDLSTPIQRVRS-NKRVIVAFPPRRSSEQLKKHASGYLHIGKDKLRKSQLPDRL-TKPDGHVLVRPSTWR
WP_045536711.1    M----SRVVAYVDGFNLYFGLRSKGWKKYYWLDLCALATLLLGQTLEGVHYFTSRIRRNGADMQRQTSYLEALETRSKLRSHFGHYLEKPKQCHQCGAQWMDYEEKMTDVNIAIQLLTDAYDDRFDTALLISADSDLTTPVSVVRA-GKNIIVALPPNRRSHQLTQAASGSFIINETAYRRSLLPVQV-QREDGFVLQRPAHWR
WP_166837249.1    M----KRVMVYVDGFNLYFGLRSKQWRRYYWLDLVALGNALLDQQLTQVHYFTSRIRHNTSDVQRQNTYLEALATLPDVQIHYGHYLEKPKQCRSCGAQWMDYEEKMTDVNIAVQMLSDAFADRFDTALLISADSDLTTPVKQLRN-NKRVIAVFPPHRRSFDLGNAANGSFTLGEANLRQSQLPNNV-TRADGFVLQRPTHWN
WP_187288494.1    M----SRLIVYVDGFNFYYGAVKQ--TPYKWLNFNKLFKILFKDDLVLVRYFTARVKLNPGQRERQGTYLRALRTLPDFEIHEGFFLSHPVNMPVAPARVLKFEEKGSDVNLATYLMLDAIEDRFDTAVVVSHDSDLALPIQIVNE-KPVGVVGYDPPRTSKSLRKVAAFYKTLRESAFKRAQFPDVV-QSQDG-PLHKPSGW-
WP_211163379.1    M----QRVITYIDGFNLYFGLKQAGFKRYYWLDVAALGQQLLGQTLVATHYFTARIRRNVADQKRQNDYLEALAV-QGVRCQFGHYLEKVRECRACHATWPDYEEKMTDVNIAIQLMTDAFDDAFDTALVISGDSDLVTPIRRVRE-NKRVIVAFPPRRHSKALKQTANGHLAIGEDKLRASQLPEHI-AKPDGFVLHRPASWS
WP_219975496.1    M----KSTNVYVDGFNLYYGCLKG--TPFKWLDLSKLCGILLGNAVNRIRCFTARVRGDPNQQQRQQVYIRALETIPNLSVHYGHFLAHPVRMPLANVEAIKTEEKGSDVNLATYLLYDGFRGDYEVAVVVSNDSDLLEPIRIVRR-LTVGILN-PQKRPSRVLAQEADFLKQIRAGALKASQFPTTL-KDAHG-IITKPKEW-
WP_273009214.1    M----SRVISYIDGFNLYFGLKDKGWKRYYWLDLEALSAALMDQQLAATHYFTARLRNNQADKQRQDDYLDALLTRHSLHIHYGHYLRKNRTCRSCGASWPDYEEKMTDVNIAVQLIADAFDDSFDTAIIISADSDLTTPVQMVRQ-HKKIVIAQPPGRNSNQLRQAADGAFQISRTKLQQCQLPETI-TTAAGVTLQRPATWT
consensus/100%    .........ahDGbsh..s...........p...h...h....h....hhos..........b...h..............a........s.......bcb..DlplA..h...s....hp.uhlho.D.sL..sh..h...........s.......h.........h...h.......h...........P....
consensus/95%     ......p..sahDGbNha.s...........hs...h...hh...h....ahTs..p.s......b..bl..h.....h......a..p.......s...h...cEK..DVplA..hh..u.....h-.uhlhS.DpDL..shp.h.......h.s.P.......h.........l....h.......h...........P....
consensus/90%     h.....ps.salDGbNhaauhbp.....a..hs...h...hh...l...paaTu..p.s.....bb..blchl.....lph...pa..+..ph..ss..h...cEK.sDVplA..hh..u.p..a-.AhlhS.DpDLs.shp.lb.......h.s.P..p.s..l..........l....h..sb....h.....G.....P....
consensus/85%     M.....pshsYlDGhNLaaGhbp...p.a.WLsh..Lsp.lh.ppl...+aaTu.hp.s.....bQ..hlcAL.....lph.bG+ab.+..ph..ss.ph...EEK.oDVNlA..hl.sA.ps.aDsAlllSsDoDLs.slpblb.....hhhs.Ps.ppu..l.p.sp..b.lsps.hp.sbhs..h..p..G..bp.P..W.
consensus/80%     M.....+shsYlDGaNLYaGh+p...p.abWLch..Lsp.lh.ppl..s+YaTu.lp.ss....cQ..hLcAL...s.lph.bG+ab.+s.ph..ss.pa...EEK.oDVNlA.pll.DA.cs.aDsAlllSsDSDLs.slpblb...b.lhhs.Ps.Rpu..L.p.ss..b.lsps.hp.sbhP..l..p..G..hp+P..W.
consensus/75%     M....pRlhsYlDGFNLYaGh+p...ppYbWLDl..Lsp.Lh.ppl..s+YFTublppsss..pRQp.aLcAL.sbsslpl.bG+Ybp+s.ph..suspa.p.EEK.oDVNIAspll.DAhcs.aDsAlllSsDSDLs.Plpbl+...bpllhsbPs.RpS..L.p.Ass.h.Iscs.hcpsQhPp.l..p.sG..hp+P..W.
consensus/70%     M....pRVhsYlDGFNLYaGh+pp..ppYbWLDl..Lsp.LhsppL..s+YFTublppNss..pRQp.YLcAL.sbsslpl.hG+Yhp+s.ph.psuspa.p.EEK.oDVNIAspLl.DAhccpaDsAlllSuDSDLssPlpblRp..b+lllshPPpRpS.pLppsAss.h.Iscs.lcpsQhPspl..p.sG.slp+P..W.
```

```
# ; PIN-EndoRNase

                  1---------11--------21--------31--------41--------51--------61--------71--------81--------91--------101-------111-------121--
OrigSeq           MNLLLDTHILIWWLENNPRLPSEAQKAITEPSNLVYVSAITVWEIGIKKAIGKLTIPDNLEEALVVNRFQLLPITISHALVAGKLPLLHSDPFDRMLVAQAQVEILTIVTVDDRIKQYNVNYLKI

Jnet              --HHHHHHHEEEEE-------HHHHHHHH------EEEHHHHHHHHHHHH---------HHHHHH---EEEEE--HHHHHHHH----------HHHHHHHHHH---EEEE---------------
```

```
jhmm              -HHHHHHHHHHHHH-------HHHHHHHH------EEEHHHHHHHHHHHH---------HHHHH----EEEEE--HHHHHHHH-----------HHHHHHHHHH--EEE----------------
jpssm             -------EEEEEEE-------HHHHHHHH------EEEEHHHHHHHHHHH---EE----HHHHHH----EEEE--HHHHHHHH---------HHHHHHHHHHH---EEEEE---------EEEE-
KAF0250482.1      MNLLLDTHILIWWLENNPRLPSEAQKAITEPSNLVYVSAITVWEIGIKKAIGKLTIPDNLEEALVVNRFQLLPITISHALVAGKLPLLHSDPFDRMLVAQAQVEILTIVTVDDRIKQYNVNYLKI
KPA14791.1        MKILIDTHTLLWFLSNDVKLSEKAKKIIENSKNDVFVSIASFWEISIKLSLSKLVLDIPFVESENL-GIQILQIKKEHLIYLTKLPFFHKDPFDRIIICQGIIEDTAIISIDEKFDEYKVERIW-
MBF0143012.1      IMLLLDTHVWIWWAEQDRRLPRALKEKMEEHKGILAVSSATIYETLTLARRNRITLNRAIERAIYGADIEIIPLDGEIARQAGNLPFHHGDPIDRLIIATAICQKALLASMDSQFPHYEALAGRL
MBF0214026.1      DWLLLDTHIWIWWAKRDRQLAPSIAKRLDETRAGLAISSVSVYEALLQIRRGRVAIDLPLHAATVEAGIEVIAVNAEIAAKAALLPLHHGDPLDRIIIATALHHDVGVVSVDGQFPHYEALAGRL
MBK8255141.1      MRLLLDTHAFLWFVAGSSRLSTTARAAIEARENERFLSAASIFEMAVKMSLGKLTVGLPLGRELASGGFALLPIEIRHVGQLASLPFYHRDPFDRLLAAQALVDALMFVSADEAFDSYGVQRHW-
MBK8638070.1      -MIVLDTHIWHWWTNQDGKLTSGQIALIEE-ADDIAVSAISCFEMAWLHTHGRIDLGMPLSEVEASATVVFLPVTPWIAARAVALPEHHKDPQDRLIIATALVHQAHLMSFDGHFPSYQELDGRL
MBK8754772.1      -MLILDTHVWLWWIEQNARLPCWLAAQVADPATLIALSAVSIYELSLGVGRGRIVLNRPIQQATVEAEIQVLPVTAKIAGLAGQLPRHHLDPLDRLIIATALTHQAQLASLDDKFPLYAELNKTL
MBS3952429.1      MRVLLDTHAFLWFLLADPKLSTPAQEIISDAQTRVSVSPATYWEIAIKISLGKYALPEPMEQQLAINNFRILPILPGHTAILTSLPFHHRDPFDRLLIAQSIAEQVPLVSCDSAFDAYGISRIW-
MCI0624091.1      MRLLLDTHTFLWFLLNDAQLTETARSLIADSGNDIEISPASIWEIAIKISLGKYALTEPMEREMTANGFRLLALEIKHISVVTTLPFHHRDPFDRLLVAQAIAEQIPIISGDQMLDAYPVQRLW-
MCY3931993.1      KRLVLDTHVWVWSALDSSRLTAVARAAILD-ADAVHVSPISLYEVTQKARLGKWPEIAPIDALVAESETLPAPLTRAVAARAGMLDWVHRDPFDRFIAASALELRCRLVSKDDEFDALEGMPGWQ
MCZ2341132.1      MDVLLDTHTLLWAVDDPTKLGTTAAATLRDGNNTLQLSMASVWEIGIKYGKQKLPLSLPIDQAIAALDIRLLNITVDHIERHVQLAGYHGDPFDRMIAAQSLVRGIPLISVDPKLDAYGVTRIWN
MDB5306880.1      MRHLLDAHTLLWSQDDTGKLSAAATAALTEPAHDRLLSIVTVWEIGIKVALGKLPLSKPIQTAISDLVLSDLPITLDHVERQMSLPFHHRDPFDRLPAAQALVEGIPLISCDTIFDAYGVTRIWD
MDE0038235.1      MRYLLDTHVLLWSRASPERLSAEVVGLLQSTENTLYVSMATLWECAIKVCIGKLAVPEG----IVANDYEILDIELAHVEATLQLPLLHRDPFDRLLVAQARLGGLTLVTRDRNVIRYQVPVVEA
MXW42975.1        TILLLDTHVLQWWSAEPHRLSSNAVAVLED-ASELAVAAITWYELAWQAHQGSITVEQPLNTL--SSMVRTVALTPAIAAIAAFLPRFPGDPADRLIYATAIELGWQLVTKDDRMCKHRHSRSIV
QQS46396.1        MNLLLDTHAFLWFVLGDSRLSSEARQHIESLENKVYVSPAIYWELAIKISIGKYVLQDSIDRGMGQNGFITLPVEPRHTAILTTLPFHHRDPFDRLIIAQAIVEGLSVVSLDTSFDSYPVNRIW-
REJ55683.1        -MIILDTHIWLWWVDSSSRLTARLEQLILEHVSQLGVSIFSCWEVAKLVEKNRVELSYPLNYAISYPGVQLLPLTLPIIVQSTQLAGFHNDPADQIIVATAKVHNCPLLTVDEKILAYPDVNTLK
TSA38616.1        QKILLDTCTFLWLVRDDAALSSKARELIVDIDNTIFFSSVSAWEITVKHDLGRLPLADLIQNERQKHGIESLPLVEEAVAQLGKLPTYHNDPFDRMLICQAIYHGCVLLTPDYAIIRYPVNSQW-
WAS06094.1        MRLLLDTCTLLWFLTNDPKLSQNAKNAIEDPLSERWLSPISLLEIAVKYKLGKLPLPKPFPAALTAEDIHLLPLESAHIEPLSTMPLYHRDPFDRLIAATALVEGLCLVSADGTFDCYGLIRLW-
WP_005004505.1    MDILLDTHVLLWWLSDDPRLSAKARNLIADPANAITVSAATAWEIAIKQGLGKLMFDGDLEQAVREQDFQMLPITFAHTAQIGQLPEIHRDPFDRMLVAQARSENLALLSADPHVARYPVNVIEP
WP_009149688.1    -MIVLDTHIWLWWITRDRQLKQRWREKITA-ADQLGVAAISCFEVAWLVQHGRVELSEPFDKALNGSGIRLLAITPEIARIAAQLPEHHRDPQDRLIIATTLAHEATLISADGKFTLYHELQGRL
WP_073727328.1    MRLLLDTRVVMWWIADSAHLSASLKELLET-EPAVYVSAVSPWEITIKQFLGRLEGPGDLAEQVRDLQFRALPITADHGVRAGRLPMVHADPFDRILIAQAQAEGMVLVTRNRWIPQYDVQVMSA
WP_075888514.1    SSVIADTHTIIWYLSNNEKLSATAGTALSASGDNIFVSAISLIEITYLIEKGRLANEAILKAAIQAIGIELIPIDIEIATAIAQIDRTVSDMPDRIIAATALTLDLPLVTRDGKIQKALNIKTIW
WP_094343934.1    MSFLLDTHILLWFLENDSKLSDRVREVITNPENLIFVSAISAWEISIKQSLGKLIAPSNLEEALRFSRFEVLSMTLAHGIKVADLPLHHKDPFDRMLIAQALVEGLIIITVDEKFKFYDVPLFSD
WP_094560568.1    EAYLLDSHALLWWWFDPDRLSGAVRELLSDPATPVLVSAASVWELSLKHHQGKLPELNGLPGLLQADGFEALPISLAHGLRAGGYSQPHRDPFDRMLAAQAELDRLVLLSADPQLSTFPCQTLW-
WP_121969235.1    SAVVADTHAVIWFLRNAKQLSDNAATAMDGANQPIYIASISLVEMVYLTERDRIPAAAFLMATLDISTFEVVPLDLTITLELASIDRIVPEMPDRIIAATAQSLGLPLVTRDRSIQNLSTLKTIW
WP_194535995.1    PSLLLDTHAVVWYAQPSPRLSAKARHAIQQAGDPVQISAVSLVEITYLAEKRRLSADTFVVAIVQYSGLEVVAFDLAMADCLRQISVRVPDMPDRMIAATALYWNIPLVTADRQLHSLPNLSCIW
WP_213458102.1    MRLLLDTCILYDWMMD-ALADPETVALIQ--QTGAFVSAVAVWEMSIKHALGKMTLPSRIVEDVEAQGFGWLNITPSHAQAVLDLPAHHKDPFDRLLIAQAQCEGMQILTYDRLFRDYLPETRLV
WP_227382628.1    SAIVADTHAIVWYFRSPEKLAYDASIAFDHAGYPVHISAISIVEITYLVEKNRLPQVALLIENLADVGLVVVPMDILIARTLAKISRTVPEMGDRIIAATALYLGLPLVTKDHRIRELSVIQTIW
WP_242411966.1    DVIILDTHIWIWWITKHEKLTNKNLNLIEKYYQGLGISMISCWEIAKLVEKNRLQFQCSLNSAMKYPDIKILDLTLPIILQSTKLDGFHQDPADQIIVATAMVYDCPLITQDSKILKYPSVKILS
consensus/100%    ...lhDsp.h.........h.......h........hu.h..hE........ph.................h.h..........hs....-..Dph.hspsb.....hho.s..h..........
consensus/95%     ..hlhDop.h.W.......Ls......h........lu.hshhEh..b....+h.....h........h..lsh..........hs....-..Dphlhspub...h.lho.D..h..h.......
consensus/90%     ..hlhDTp.hhW....s.pLs......h........lu.hohhEhs.b..bs+h.....h........h..lshp.........ls....D..DRhlhApub...h.lho.D..h..h.......
consensus/85%     ..llLDTH.hlWhh.ps.pLs..h...l....s.l.lS.hohhEhshb..bs+h.....h........h..lslp.......s.ls....DP.DRhlhApAb...h.llo.D..h..h.....b.
consensus/80%     ..llLDTH.hlWah.ps.pLs..h...l.p..s.l.lS.hohaEhshb..bs+l.h...h........h.hlslp.....hspls..H.DP.DRhlhApAb...h.llo.D..h..a.....b.
consensus/75%     ..llLDTHhhlWah.ss.+Lssph..hlpp..s.l.lSshohaElshb.pbu+lshs.sl...h...shphlsls...s..hspLsb.H.DPhDRhlhApAbh..h.llo.D.ph..Y.s...b.
consensus/70%     ..llLDTHhhlWah.ss.+Losph..hlpp..s.l.lSshohaElshb.pbG+lshs.sl...h...shphLsls...s.bhspLsb.H.DPhDRllhApAhhp.hsLlo.D.ph..Y.s.p.b.
```

### ; Type-5 BREX and Type-3 BREX BrxHII-Helicase with C-terminal Active PDDEXK REase; (TUDOR+SF1-SF2-Helicase+iREase+CTD+REase)

```
               1---------11--------21--------31--------41--------51--------61--------71--------81--------91--------101-------111-------121-------131-------141-------151-------161-------171--
OrigSeq        MFHAGEVVYGPYFPENVEVKRCELLEDFYIIEAIGRETHQFYELMLEKEKIAELKRLKENASQQPVQGEDIQRYLQYLMFRHEMMYSRTRALGNKKLLPLPHQIEAVYGRMLQVPQIRFLLADDPGAGKTIMAGMLIKELKARYSIERILILTPPLVVKQWQEEMMEKFHEHFSI[illegible]

               <<-----------------------TUDOR------------------>><<----------------------------------------------------------------------------------Active-P-loop(Swi2/Snf2-Helicase)-------[illegible]
Jnet           ------EE-------EEEEEEE-----EEEEEEE-----HHHHHHH-HHHHHHHHH-----------HHHHHHHHHHHHHHHHHHHHHHHHHHH-----------HHHHH---------EEEE-------HHHHHHHHHHHHHHHHHHHH-HEEE-----HHHHHHHHHHH----EEE[illegible]
jhmm           ------EE-------EEEEEE------HEEHHHHH--HHHHHHHHH-HHHHHHHHH-----------HHHHHHHHHHHHHHHHHHHHHHHHHHH-----------HHHHH----------EE--------HHHHHHHHHHHHHHHHHHHH-EEEE------HHHHHHHHH----EEEE[illegible]
jpssm          ------E---------EEEEEEE----EEEEEEE-----HHHHHH--HHHHHHHHH-----------HHHHHHHHHHHHHHHHHHHHHHHHHHH---------HHHHHHHH--------EEEE-------HHHHHHHHHHHHHHHHHHHHHEE----HHHHHHHHHHHHH----EE[illegible]
AKM18688.1     MFHAGEVVYGPYFPENVEVKRCELLEDFYIIEAIGRETHQFYELMLEKEKIAELKRLKENASQQPVQGEDIQRYLQYLMFRHEMMYSRTRALGNKKLLPLPHQIEAVYGRMLQVPQIRFLLADDPGAGKTIMAGMLIKELKARYSIERILILTPPLVVKQWQEEMMEKFHEHFSI[illegible]
ASS85707.1     MIQVGDIVKGPLWPEVVEIKNCTMIQNFYQVEAYGKNTSKYYQVIILPEQLTNIEIINTSN-KKTYSSEELQDYLRYFEFVVDKRFSKTRALGNQHLIPLPHQIEAVYGRMLRVPSVRFLLADDPGAGKTIMAGMLIQELKARQSANRILILVPPLVLKQWQQELKEKFNEDFLI[illegible]
KJS50637.1     LIKVGDIIQGPYWAETVEIKKCEPIDKLFLVEAIGRQTREYYESYLELEQLESLELLQENL-NPVATLDQISHYLRYHVLKLDERYSKSRMRGNQQIIPLPHQIEAVYSRMLQTPQVRFLLADDPGAGKTIMAGMLIRELLARQSVERILILVPPLVLPQWQEELSEKFGEEFKI[illegible]
KKE80054.1     MLKKGEIVGASFWPEPVQIKHIETIDELYLVEAVGRESNQFFENYLESYQLHEVERYSEED-VSDNLKERFQHYLQYHVLKTEREYSQSRARGNKNMIPLPHQIEAVYSKMLQAPQVRYLLADDPGAGKTIMSGMLIRELKARNMVRKTLILVPPLVLKQWQAELKEKFDEDFL[illegible]
MBO8173522.1   MLRAGEIVKGSLFPESVEIKKCECISEMYLVEALGRQSNQFFELMLDENEINTLERLCVHNDKKELSAREELQHFLQYHTLAIEQKYSSSRALGNKNLIPLPHQIEAVYSRMLQVPKVRFLLADDPGAGKTIMSGMLIRELKARRSAERILILVPPLVVKQWQEELKEKFSEDFTI[illegible]
[illegible]
consensus/95%  .h..Gp.l.us.as[illegible]
consensus/90%  .hp.Gcllpusbas[illegible]
consensus/85%  .hp.G-llpusbasE.VclKph-.h..hh.lpuhGpposp aap.hlp..ph.php.hpp.......hp.pplQphlba..h..-.baSps+hbGspplhPLPHQIEAVYu+MLQssplRaLLADDPGAGKTIMuGMLl+ELbAR.phpRlLILVPPLVL+QWQpEhppKF.EpF.I[illegible]
consensus/80%  Mhp.G-IlpusbaPEsVEIK+h-.hspha.lEulGcpospaYc.hlpp.pl.plp.lppp.....hpspclQphlbY..lb.-.caSpsRAbGspplhPLPHQIEAVYu+MLQsPplRaLLADDPGAGKTIMuGMLI+ELbAR.shpRlLILVPPLVL+QWQEELppKF.EpF.I[illegible]
consensus/75%  MhphG-IlpGsbaPEsVEIK+hE.hschallEAlG+poNpaYE.hlppppl.plpblpppp.p..hsspclQphLbYb.lb.-.+aSpsRAbGNppllPLPHQIEAVYuRMLQsPplRaLLADDPGAGKTIMuGMLI+ELbARpshcRILILVPPLVL+QWQEELppKF.EpFpI[illegible]
consensus/70%  MlphG-IlpGshaPEsVEIK+hE.ls-hallEAlG+poNpaYE.hlpppplpplpblpppp.p..hsspclQpaLbYbhLb.-p+aSpsRAlGNppllPLPHQIEAVYuRMLQsPpVRaLLADDPGAGKTIMuGMLI+ELbARpshcRILILVPPLVL+QWQEELppKFsEpFpI[illegible]
```

### ; Type-2 BREX-Specific; REase+SF2-Helicase

```
               1---------11--------21--------31--------41--------51--------61--------71--------81--------91--------101-------111-------121-------131-------141-------151-------161-------171--
OrigSeq        MLAFQTSAKQILELTREAQLVEQLTAHLTRVGFRISKSERSSWQQSLPALAMELVGAGLGELDVLVECQLPRSSKRIDVILAGMEPLTRRPAYLVVELKQWSVATTYDEAEKLVQVPGMAEPQVHPAVQVENYCSYLTDFLPIFAESPDRVHGLAYLHNAEQGHIPAMTTKEDDG[illegible]

Jnet           --HHHHHHHHHHHHHH---HHHHHHHHHHH------HHHHHHHHHHHHHHHHHHHHH------EEEEEE-------EEEEEEEE--------EEEEEE-----EEEE------EEEE----------HHHHHHHHHHHHHH-----------EEEEEEE------------------[illegible]
jhmm           -HHHHHHHHHHHHHH----HHHHHHHHHHH------HHHHHHHHHHHHHHHHHHHHH------EEEEEE-------EEEEEEEE--------EEEEEE-----EEEE------EEEE----------HHHHHHHHHHHHHH------------EEEEEE------------------[illegible]
jpssm          --HHHHHHHHHHHHHH---HHHHHHHHHHH-------HHHHHHHHHHHHHHHHHHHH-----EEEEEEE--------EEEEEEE--------EEEEEEE----EEEE-----EEEEE---------HHHHHHHHHHHHHH----------EEEEEEEEE-----------------[illegible]
ADL44852.1     MLAFQTSAKQILELTREAQLVEQLTAHLTRVGFRISKSERSSWQQSLPALAMELVGAGLGELDVLVECQLPRSSKRIDVILAGMEPLTRRPAYLVVELKQWSVATTYDEAEKLVQVPGMAEPQVHPAVQVENYCSYLTDFLPIFAESPDRVHGLAYLHNAEQGHIPAMTTKEDDG[illegible]
ARF59783.1     MLLLKLSAQDLLTLDSRDRMVPHLAARWRYFGADASESERSAWAESLVSLANDLVAAGRGNVEMIVECAATGDPRLIDVVLVGRHPERHVPSVQLVELKRWSTVSRVERATALLHVPGTKKEKKHPALQLGAYYEAFTSARGPFQGADFECGGFAYLHNATDDSVRTLIDV-DAP[illegible]
EAZ58597.1     MIVYEATKKQFLQDSDNDDIEEVILQHFKAVGKSVGNSEINSWKGSLGYMAKVLRDEEVDDTGLAIELHLPQTSKRIDITLTGRDE-SGGKNAVLVELKQWSKASATGKDAIRTALGNGLREVVHPSYQAWSYATLLEGFNEAVYEASISVRPCAYLHNYPR--DGVIDSPHYRD[illegible]
[illegible]
```

```
consensus/75%   b.hh..os.phh......ph..pl.pph......supsE..SWppShs.hhp.L.puulsphphhlEhphP.sspRhDhlLhG.cs.s.p.shllVELKQWsphp...p...bs.hs...p...HPs.Ql.sYhphl.sa...h.p...pl.uhAaLHNh.p..s.slb...... [illegible]
consensus/70%   b.lh..os.phh.....splh.pl.pph...s..supsE..SWppSls.lhp.L.puuluphplhlEaplP.ospRhDhlLhG.cs.ssp.shllVELKQWsphp...p...bs.hss...p...HPs.Ql.sYhphlpsa...h.p...pl.uhAaLHNh.p.ps.slb..... [illegible]

# ; SF1-SF2-Helicase+REase

                1---------11--------21--------31--------41--------51--------61--------71--------81--------91--------101-------111-------121-------131-------141-------151-------161-------171-- [illegible]
OrigSeq         MMPTPTVQQQTAITTVDRALVVEAGAGTGKTWVLVERYLHLLALHPDWPIESIVAITFTEKAAREMRNRIRGGVEARARRHPHDVHWQERRRRLDQIQVSTIHSLCARILRENAIAATLDPKFDVLDEAAATLLAEEAIRQTLADLSQDADHPALALFASLSVFDVKNTMAELLS [illegible]

                <<----------------------------------------------------------------------------Active-P-loop-Domain(SF2-Helicase)--------------------------------------------------------- [illegible]
Jnet            ------HHHHHHHH-----EEEEE------HHHHHHHHHHHHH---------EEEEEHHHHHHHHHHHHHHHHHHHHHH-----HHHHHHHHHH-------HHHHHHHHHHHH-----------------HHHHHHHHHHHHHHHHHHH-----HHHHHHHH---HHHHHHHHHHHH [illegible]
jhmm            ------HHHHHHHH-----EEEEE------HHHHHHHHHHHHH---------EEEEEHHHHHHHHHHHHHHHHHHHHHH-----HHHHHHHHHH-------HHHHHHHHHHHH-----------------HHHHHHHHHHHHHHHHHHH-----HHHHHHHH---HHHHHHHHHHHH [illegible]
jpssm           ------HHHHHHHHH----EEEEE------HHHHHHHHHHHHH---------EEEEEEHHHHHHHHHHHHHHHHHHHHH------HHHHHHHHH----EEEEHHHHHHHHHHHH-----------EE--HHHHHHHHHHHHHHHHHHH-----HHHHHHHH---HHHHHHHHHHHH [illegible]
MBI1293580.1    MMPTPTVQQQTAITTVDRALVVEAGAGTGKTWVLVERYLHLLALHPDWPIESIVAITFTEKAAREMRNRIRGGVEARARRHPHDVHWQERRRRLDQIQVSTIHSLCARILRENAIAATLDPKFDVLDEAAATLLAEEAIRQTLADLSQDADHPALALFASLSVFDVKNTMAELLS [illegible]
MAD36289.1      MN--------------GPISVVSAGAGTGKTWRLSTEYVEAA--ESAMSTARIIATTFTVKAASELLERVRARLIQEG--------HADAAQSALAGLVGTVNSVCGRLVGDFAIDGGLSPVAEVIAPEMADSLFRLATEEAIQKFSDRI---------RLRQDDWPTTVLEIVR [illegible]
ELZ84494.1      MSEDPKPQQRTIIESERYPMRVLAGAGTGKTFTMVRKIENLID-EHGVSPDRILALTFTNNAADSMREKLNAKLGPAG----------------YDVDAYTYHSICNEILTDYAYDAGLDPDFEIATDAEKYVIVLDVLDEIDYRAVKPNVATKLLSFIGISPTEIEEYLGSPEQ [illegible]
MBP7846544.1    MTNKLHPLQQISFTQANQAYVVEASAGTGKTWTIERLFIKALLLHIPLAIENILVVTFTNDATDELKQRISEQIQTTINQIIYLNLLTRALQNFDQAAIYTIHGFCNRILQDYQFECQTPAEFELLAS--------QLIDELVLDFMRRE---------SIFSSQLELVINNLEG [illegible]
MBT4336381.1    MT-------NEIKTQVDKACVILAGAGTGKSYTIKQKAKYLVE-ELKYNPEEILCLTFSNEATNSLKKGMQEEINST------------------EDVDVKTFHAFCADILREEGHLVGVQEGFEILLPDDAKIFMHKYLE-----------SNRYINTISGISIDDIKGHLNKLKE [illegible]
MBZ5659869.1    MMIIDQSERLRVLTDLDSTLLVEAGAGTGKTALMAGRVAMLL--AAGVPTREIAAITFTEAAASELFARIFDYSSELREGRIPPENLLRASDSLDELICSTIHSFCQQLTKPYPVEANLDPGAVIMDEVQADLAYDVLFEEWLRSELSQSDDRPVAVLLVKGRRLIEQIVAAQQN [illegible]
MCC7380509.1    MS------KPRILSEIDRHAVIEASAGTGKTFTLESLLVELV--LAGAPIQSILVVTFTEKATAELRRRLRAKLQAMVRGEGAVERLDSALLDLDAASVSTIHAFCQRALTEQAFLN------RLFGEVDGGLTFTRMFTHLLRTELARDELQSYLEAWGQGLDELRALLWRVQA [illegible]
MCE9567299.1    MDKPLKPDQQEAATHEGTPYLLEAGPGTGKTQTLVGRVEYLL--TKGIEPSKILILTFSNKAAGELSERIGSKHRAAA----------------ASIWVGTFHGFGLDIIRRFHDHLKLPHNPKLLERTDAIGFLEDEYPKLGLKHFKNL-----------PSASIKSILDAISR [illegible]
MCG7876678.1    ML--------------PALTIIPAGAGSGKTYTIQQQLGEWVV-EGLVAPERIVAVTFTEAAAAELRERIRAKLLSLG--------RLDDALKLDQAYISTIHGFGLRLLTEFAFDAGLSPRPRLLNEDEENTLIRLALARTEKADVITV---------KSAEELFRDDLLRIVM [illegible]
MCR5727522.1    MEKKWTDAQKSVIDTKDKNILVSAAAGSGKTAVLVERIIKMIT-EKPVDIDRLVVITFTRAAAAQMKQKISEKIEELIEKEPENA-YRRQYHLLGNATISTIDSFCLNIVRSYFQKQELDPAFRVADETELGLIKSDCMDELMEKYHEEGDFYYLIEAYYKSDEKLTQYIFNLAR [illegible]
MCR9155220.1    M---------------SNLSIITAGAGSGKTYRITEEVVKAFG-DPSLEPRHLIVTTFTKVAAQELRERVAERLLQES--------MYLPLASLHEARIGTVHSVCQ [illegible]
MDD3564057.1    MI--------------DVDRFISASAGTGKTHTISKNYVDLFE-DEPLSIANVVAITFTRKAAAELKSRILKMISSRDDGNPA---WQQLRSSMAFAWISTIDSFAC [illegible]
MDE2877865.1    MIKNLTDAQSEAVNFDPSNLLILACAGSGKTETLACRAAQMV--QDGVPRGSIVAFTFTEHAAAELKHRIRRRVEEANPDEPA----------LGDMYVGTIHAYCI [illegible]
MYC78299.1      MR--LTPNQQQALDI-DRHICVTAGAGSGKTTVLVERYLKILR-EGNVNPQEIVAITFTDKAAAEMKERIIEELSVQEEREGSE-SLQGFREKMSTAHISTIHAFCS [illegible]
OBB45423.1      MSITDRSARDAVINDLDSTIFLEAGAGSGKTSCLVNRFVALV--ESGVPADHVAAITFTEKAAAELVDRIRAELRRRAPKNDA---CRDALLVLDSAAIGTLHSFAC [illegible]
OUL09461.1      MTEKFTETQWQSIYQTGDNILVSASAGSGKTTVLIQRIIEKV--KGGTNVDELLVVTFTESAAKEMKERMTGAIQTAINEAQSQEHLIKQLSLMPQANISTIHAFCL [illegible]
WDH83309.1      MTPADQADRDRILTDLDTTLLVEAGAGSGKTTSLVGRLLALI--ESGVNIGHIAAITFTNKAADEMKERFRLALEKAYRRSSEGSRLSEALGNLDLIFIGTIHSFCG [illegible]
WP_018248935.1  MGMVFTDQQAQAVLTLDRNLAINAGAGSGKTRVLTERYLEILLMYQKDALNKVIAITFTRKAAGEMKDRVRERFEKYLSENRDN-WIFYILDNLNQGRISTIHSFCS [illegible]
WP_019531411.1  M---------------DNITFISAGAGSGKTYNLMERLAALLL-NSGIKPEGVMATTFTVRASNELKERVQEKLLHEQ---------RPELAAQMESAFIGTVNGVCG [illegible]
WP_023554879.1  MNPSDQQARKRILEELNTTFLVEAGAGSGKTTSLVGRLLALIE-TGTAKMEHIAAITFTNKAADEMKERFRIALERKTADVEGE-RMEEALANLDQIFIGTIHAFCS [illegible]
WP_031304998.1  MMLKDDGARRDAISHHHRSILVEAGAGSGKTAVMAGRIALML--AQGVAPRAIAAVTFTELAASELLARVRDFVADLAAGHIAPENLIAATAAIDEITCSTIHGFCQ [illegible]
WP_055898782.1  MSFVDSAARERIDADIGTTLFVEAGAGVGKTHHLVTRILTLI--AADVPVSRIAAITFTEKAAGELRDRMRTALQVSTALTAAQ--RDAALDDLDAAPIGTIHSFAA [illegible]
WP_066478873.1  MT----------VNADQGLVIVPAGAGAGKTHRIKTQLSDWVK-RGIVRPERILAVTFTEAAAGELRERIRAGLLADG--------LVAEAMAVERAYVSTIHGLGL [illegible]
WP_070087174.1  MT----------LNPLHGSRLIEASAGTGKTFTIALLYVRLVLFKGPLTPPQILVVTFTDAATQELRDRIRARLTEAAQCFLEPDRLQLASEWMDEAAVSTIHGWCY [illegible]
WP_076118195.1  MV-DDHTARERIVEGLDRTLLVEAGAGSGKTTSLVGRMIALID-SGSAVAEQIAAITFTRKAADELKSRFSLELERQIRNAVEP-LLQRALKEIDRCYIGTIHSFCG [illegible]
WP_090826231.1  MNPNDWEARNRIEDDLETNLLVEAGAGSGKTTSLVGRMISLIR-TGNAQVNRITAITFTNKAADELRERFRLALEKSLSNAEDAERYAAAMEGLDQIFIGTIHAFCG [illegible]
WP_096390026.1  MTIQLTEEQQDALVQ-GRNVAITAGAGTGKTTTLTERYVTILA-NPSLTPENIVTITFTRKAAAELTERVREEVYDRLEAADSPARWRNVLDDLEDGYVHTIHAFCT [illegible]
WP_099255987.1  MSRTLRNAQATIRDRFSGLFAMNCVAGAGKSMTVQRIAAETII-GDRTPAQSVAVMSFTTNEAAAIGPAVCEHIREIVDHELIPEVAYLVTRVRHAPFIGTVDSVLR [illegible]
WP_129664631.1  MTLTDEHVRNRISTDLSSGLFVEAGAGSGKTTVLVARIVALV--ASGVPLRQIAAVTFTEKAATELRDRVRVELERAG--------ERAAVDELDGAAIGTLHSFAR [illegible]
WP_143068399.1  MSHTFTEAQARAIFS-EGSVAISAGAGSGKTRVLAERILNFL--ARGVRPAQIVAVTFTEAAAAELRERVTAVVGRRAEAEGG--HWPALVTDLALMTVGTIHSLCA [illegible]
WP_158203515.1  MVDDLTEEQQQAVDGLDRNVTLTAGAGTGKTTTLTARYLAMIEGDERLLPEHILTTTFTERAANELEESVREEITDRVGTLDAE-AWRTVADELEQGYIHTVHGFCA [illegible]
WP_167854939.1  MS--------------TAIHFISAGAGSGKTYRLTQKLNDLLS-QQAVSPAGVIATTFTKLAAGELQERVRGALIDAG--------QLVVANEMEQALIGTVNSVCG [illegible]
WP_190937389.1  MS--LTEQQQRAAYA-NGNVAVIAGAGTGKTYMLAERYLYHLR-VREFSPLEIVAVTYTEKAAAELRSRIRLLVSQELPERSD------LLAELEAAQISTIHALAM [illegible]
WP_213510442.1  ML-NDQAVRERILTELDTSFLVEAGAGSGKTTSLVGRMLAYIE-SGKAKVGQLAAITFTRKAADELRGRFRFELEKRIRTAKEP-RLEAAIQEIDQCYIGTIHAFCV [illegible]
WP_221091404.1  MT----TQEQTLITEYRGSVSIDAGAGSGKTFVLTRRLLGLL--THDLAAEELVAVTFTEAAAAELRGRLQALLDEEAQRRAQP-RVVAAAQALPLAQISTIHALCT [illegible]
WP_244705661.1  MTITPNDQQQELIDATEGTYLVDAGAGTGKTFTVTRRYANILD-QDDVEPEDLLLLTFTNNAAEEMKDRIVAHCDYSM--------------AALRDAPISTFHSLAH [illegible]
WP_248566597.1  MS--------RTLDPLADRHLIEASAGTGKTYTLAALYLRLVLERAPMLPPDILVVTFTEAATKELRDRIRGRLAEAARCFAGHAHLRAAADWMDEAAIYTIHGFCH [illegible]
WP_254767386.1  MT-EPNDQQRELIDSTEGIYLVDAGAGTGKTFAITRRYATIVE-QPDVEPDDVLLVTFTNNAATEMKERIVGRSDYGM--------------RELADAPIQTFHSLCH [illegible]
WP_267638592.1  MTITPTDEQKDALVL-DRNVTITAGAGTGKTTTLTHRYLEFLR-NPSATPKNIVTITFTRKAAAELETRVREEIYEELQSVEDPARWRAVLDELDDGYTHTIHAFCA [illegible]
WP_270096520.1  MTKGLNKGQSKAVDYTSGPLLILAGPGSGKTLTIAEKVVKLI--KDGTAPERTLALTFSEKAAGEMQERIEKQIGTDS------------------GITVSTFHSFCN [illegible]
consensus/100%  M...................h.sssGsGKo..h...h...h.............oao......h...h...........................Thpuh.. [illegible]
consensus/95%   M...................l.ussGsGKo..h..bh...h...........lh..TFo..As.ph...h..........................h.Thpuhs. [illegible]
consensus/90%   M...................l.AuAGoGKT..l..bh..hl.........plhh.TFop.As.-hb.ch...h.......................l.Thcuhs.phh.p [illegible]
consensus/85%   M.................hl.AuAGoGKT..l..bhh.hl.....h...plhhhTFTp.Au.Ehb.+h...l...................h....l.Thcuhs.cllpc.s [illegible]
consensus/80%   M.................hl.AuAGoGKT.sls.bhh.hl....sh...plhhlTFTp.AAsEhbpRl...l...................h..h.l.TlHuhs.cllpc.s....hs..hp.hsp [illegible]
consensus/75%   M.......b.......p..hhlpAuAGoGKT.sls.chh.hl..p.sh..pplhhlTFTp.AAsEhbpRlp..l...................h..h.luTlHuhs.cllpc.sh.s.hs..hplhsc [illegible]
consensus/70%   Ms......b.phh...s.shhlpAGAGoGKT.sls.+hh.hl..p.sh.sppllhlTFTppAAsEhcpRlp.bl.p..................hp.h.luTlHuhs.clLpc.shpssls..hplhsc [illegible]

# ; SF1-SF2-Helicase+REase

                1---------11--------21--------31--------41--------51--------61--------71--------81--------91--------101-------111-------121-------13 [illegible]
OrigSeq         MNSSNRQRNYVLCSIAENTFEHQQNAAKKAFAKLISGYKAVVIAAEMQSGKSGIALALACFQRLSLSDDLLSDRKYLKDTLYLVTMADLALQEQAKRDLDSCINVVVSNFSNFEAALKYNFKSHPPKLIII [illegible]

                <<-------------------------------------------------------------------------------------Active-P-loop-Domain(SF2-Helicase)------ [illegible]
Jnet            ---HHHHHHHHHHHH---HHHHHHHHHHHHHHH----EEEEEEEE------HHHHHHHHHH-----HHHH-HHHHH--EEEEEE----HHHHHHHHHHH----EEEE--HHHHHHHHHHH------EEEEE [illegible]
jhmm            ---HHHHHHHHHHH----HHHHHHHHHHHHHHH----EEEEEEEE------HHHHHHHHHH------------HH----EEEEEE-----HHHHHHHHHHH---EEEE--HHHHHHHHHHH------EEEEE [illegible]
jpssm           ---HHHHHHHHHHHH---HHHHHHHHHHHHHHHH---EEEEEEEEE-----HHHHHHHHHHH----HHHHHHHHHH---EEEE----HHHHHHHHHH-----EEEE--HHHHHHHHHHHH----EEEEEE [illegible]
ECJ2539584.1    MNSSNRQRNYVLCSIAENTFEHQQNAAKKAFAKLISGYKAVVIAAEMQSGKSGIALALACFQRLSLSDDLLSDRKYLKDTLYLVTMADLALQEQAKRDLDSCINVVVSNFSNFEAALKYNFKSHPPKLIII [illegible]
EGR0286973.1    MNNSNNHREHIAQALAKHTFEHQRSAAEKAFSKFNTGSKAVVVAAEMQSGKSGIALALACKQRLSLADEQLTSRKHLKDTLYLVTMADMALQEQAKRDLTPCKNVVVSNFTNFELALKADFKYQSPKLIII [illegible]
HAT7499720.1    MNIENKHRLYVKEILEAQTFEHQRIAANKTFSKLLSGSRAVVLAAEMQSGKSGIALALAGLQRLSLSDELIGNRKLLKDTLYLVTMGDISLQEQAKRDLSYCKNVVVSNFINFDITIKTEFRHQNPKLIII [illegible]
MCP5206313.1    ANTANIHRDAIEKVLSQNTFAHQRSAAQTAFEHFDNHNRAVVITAEMQSGKSGIALALSCIQRRKLNDADICDRKQLKDILYLVTMADLSLLEQAKQDLAPCNNVVVSNFTNYESALSSQFKNQQPKLIII [illegible]
NYA44687.1      KMMAKRHRERIGLVLNNYVFAHQKDAAYRAFNYFNGGARAVIVAAEMQSGKSGIALAMSCLQRDSLSDCDLCDRKKLKDTLYLVTMADISLLEQAKNDFSTSSNMVISNFANFRQSLETEFKHQPPKLIII [illegible]
NYA44688.1      TAKTDSLRQKVQSILNGQNLEHQRTAAQKTFDALNGNVRAVITAAEMQAGKSGVALALCCLQRLSMNDDDICDRLKLKDTLYLVTMPDVALLEQAEKDLADAKNVVVSNFIRFDQDLERSFKGSPPKLILI [illegible]
WP_010634557.1  TNQANAHRTRIQRVLQDTCFQHQLDAANKVNLALQSGARAVVIAAEMQSGKSGIALALACLQRLGLDDTTICDRKQLKDTLYLVTMADITLLEQAKEDLSLSPNLVVSNFSNFQMALASQFKTCPPKLIII [illegible]
WP_014998456.1  SLNAEQHRLIIDEILDANTFEHQRSAASKAHAKFAGGSRAVVIAAEMQSGKSGIALALACYQRNSLSDTDIVNRKLLKDTLYLVTMADVALQSQAKKDLEAAKNVVISNFTNYETALKAYFKHRPPKLIII [illegible]
WP_042889125.1  MSHADIMRQKVQEVLNSDKLEHQRSAAKKTFDALEGKVRAVVTAAEMQAGKSGVALALCCLQRLSMTEADICNRAKLKDTLYLVTMPDTALLEQAQDDLKDARNVVVSNFVRFEKDLEASFKGHPPKLIII [illegible]
WP_076500053.1  DSKAQNMRQKVQHILDSNALSHQINAAQKTFNALSSEVRAVVTAAEMQAGKSGIALALCCLQRLSLTVQQICNRAQLKDTLYLVTMPDTALKEQAEEDLKLAKNVVVSNFVNFEKALENEFKGNPPKLIII [illegible]
WP_076500065.1  MTALYKYRQQVENAISANTFEHQRNAAAKTFAKFSAGSRAVVLAAEMQSGKSGIALALSGLQRLSLTDDQISDRKRLKDTLYLITMADIALQEQAKKDLAPCKNVVVSNFTNFETALKVDFKHHQPQLIII [illegible]
WP_077485095.1  MKNANKHRKLTDVIITSNTFPHQRKAAQQAFAQFETGSRAVVIAAEMQSGKSGIALALSCLQRLSLSDEAISDRKYLKDTLYLVTMADLALQEQAKQDLAASTNVVVSNFTNFESTLKTHFRRQAPKLIII [illegible]
WP_173294712.1  TNKAIIHREKIAKVIEQETFEHQRQAANKAHIALNSGSRAVVIAAEMQSGKSGISLALACLQRQSLDDTVICDRKQLKDTLYLVTMADIALFDQAKHDLSACPNVVVSNFNNFRQALSLQFKNQQPKLIII [illegible]
WP_173294713.1  AKSAQELRNGVLKVLEGKSLDHQREAAKKTFEAFDKDTRAVVTAAEMQAGKSGVALALCCQQRLSLSDLDITSRKKLRDTLYLITMPDVALLEQAKDDLNGAKNVIVSNFVRFENDLENEFKENPPKLIII [illegible]
WP_242698654.1  LNNPETHRKLITNILSDSTLKHQIDAAQKTFSAFDGDIRAVVTTAEMQAGKSGIALALCCFQRLSLADTDICDRKQLKDTLYLVTMPDTALRDQAEDDLKLAKNVVVSNFVNFEKALENDFKGNPPKLIII [illegible]
consensus/100%  .......R......l....h.HQb.AA.ps...h....+AVl.sAEMQuGKSGluLAhssbQR..hs...lssR..L+D.LYLlTMsD.sLbpQAbpDh..s.NhllSNF.pap.sl...F+.p.PbLIlI [illegible]
consensus/95%   .......R......l....h.HQb.AA.ps...h....+AVl.sAEMQuGKSGluLAhssbQR..hs...lssR..L+D.LYLlTMsD.sLbpQAbpDh..s.NhllSNF.pap.sl...F+.p.PbLIlI [illegible]
consensus/90%   .......R..l..hl..psh.HQbpAA.psa..h.s..+AVl.sAEMQuGKSGlALALssbQR.uhs-..lssR.bLKDTLYLlTMsD.uLb-QAcpDL..s.NlVlSNF.pac.sLp.pF+.p.PKLIII [illegible]
consensus/85%   ...sp..Rp.l.phlp.psh.HQbpAAp+sa..h.s.sRAVV.AAEMQuGKSGlALALsChQRbSLsD..lssRbbLKDTLYLVTMsD.uLb-QAcpDL..s.NVVVSNFspFc.sLp.pFK.p.PKLIII [illegible]
consensus/80%   .ppsp..Rpbl.pllp.pshpHQbpAApKsF..hpsssRAVV.AAEMQuGKSGIALALsChQRbSLsDp.IssRKbLKDTLYLVTMsDhuLbEQAccDL..spNVVVSNFsNF-.sLcspFK.psPKLIII [illegible]
consensus/75%   .ppsp..Rpbl.pllp.pshpHQbpAApKsF..hpsssRAVV.AAEMQuGKSGIALALsChQRbSLsDp.IssRKbLKDTLYLVTMsDhuLbEQAccDL..spNVVVSNFsNF-.sLcspFK.psPKLIII [illegible]
consensus/70%   .spupp.Rpbl.plLsspshpHQ+sAApKsFs.hpussRAVV.AAEMQuGKSGIALALsChQRLSLsDppIssRKbLKDTLYLVTMsDlALbEQAKcDLsssNVVVSNFsNF-psLcspFK.psPKLIII [illegible]

# ; CR-ATPase+REase

                1---------11--------21--------31--------41--------51--------61--------71--------81--------91--------101-------111-------121-------13 [illegible]
OrigSeq         MIDRPMYTDKIMAYVDAPFIKILTGIRRCGKSTILKIIMEKLIISCRFDSMEYDDMTSKEIYDTLKSRLLSDRKTYVFLDEVQEIDGWEKLVNSLASDFDVDLYITGSNSRMTSSEISTYLTGRYISFRIFT [illegible]

                <<------------------------------------------------------------------------------------------------------CR-ATPase------------- [illegible]
Jnet            -----HHHHHHHHHH----EEEEEE----HHHHHHHHHHHHH-EEEE-----HHHHHHHHHHHHHHH-------EEEEEEEEE----HHHHHHEEEE----EEEEEE---HHHH---------EEEEEEE-- [illegible]
jhmm            -----HHHHHHHHHH----EEEEEE----HHHHHHHHHHHHH-EEEE-----HHHHHHHHHHHHHHH-------EEEEEEEEE----HHHHHHEEEE----EEEEEE----HH----------EEEEEEE-- [illegible]
jpssm           ----HHHHHHHHHHH----EEEEE---E--HHHHHHHHHHHH-EEEE-----HHHHHHHHHHHHHHH-------EEEEEEEEE-----HHHHEEEEE----EEEEEE--HHHHHH---------EEEEEEE-- [illegible]
GJM58068.1      MIDRPMYTDKIMAYVDAPFIKILTGIRRCGKSTILKIIMEKLIISCRFDSMEYDDMTSKEIYDTLKSRLLSDRKTYVFLDEVQEIDGWEKLVNSLASDFDVDLYITGSNSRMTSSEISTYLTGRYISFRIFT [illegible]
HBI14539.1      FINRAVS--LASFLVPGKAL-IIYGPRRAGKTTLLKSYLASCLHYRTGDDLRIRHLLGKQIVA------FAEGYDLVAIDEAQQIPDIGMGLKILVDHPDLRIIATGSSSFDLAGVIGEPLTGRKRTLTLFP [illegible]
KAA0252889.1    YLARQAEQEVLGKLRRNPVV-AIVGPRQCGKTTLARRVLERVGLYVDLERPSDAARLSEAFFRLHADE-------TVCLDEVQRIPGLFPVLRSVVDERNGQVLVLGSASPDLLRQSSETLAGRISFLELTP [illegible]
MBF0105430.1    PIKRNLNEK------KHKLL-FISGPRQSGKTYLVLHALKPA-WTLNMDIASERLLFKQAALDWLAVHNQGMEKPLIFVDEIHKVKGWRNIIKGTFDKNMVDCVASGSSAFEIRQDKGDSLAGRAVWLTLFP [illegible]
MBL8842554.1    PYRRPQLATLLARLREPRRFQVVAGARQVGKSTLVDQALAELLPALSADEPTLHDAAWWQRARALADSTAAEGGALLVVDEVQKVPAWSETVKRLWDERPLRVVLLGSAPLRLQQGLTESLAGRFELLHLPH [illegible]
```

```
MBP3741085.1      YKQRIADKLLEKKLEGTGAV-LIEGPKWCGKTTTAEQQAKSV-VYMPEEGTNWHQI[illegible]
MBQ6380319.1      YIERAITPILKKRVSESKCL-LLTGARQVGKSTLIKHVFPEY-TRANFDDRMTRLQ[illegible]
MBQ7062981.1      MKNRPIYTAKIEPFIDKPIVKVITGIRRSGKSGLLQLLQQHLIVYLNYESLRWSSY[illegible]
MBQ7238831.1      MYQRRIAQELKKRIAEQRDRQVISGPRQVGKSTMVKQVLEECVPYLSADNVPPTDV[illegible]
MBQ7946200.1      MTRRLYFNVLVTDLSIRRTI-ILMGPRRVGKSVMLYHTIEQLIIYASIDTPLYSGI[illegible]
MBQ8703690.1      MYRRIFD--IENKLDEGM---FLFGARQTGKSTLLRERFKEA-IYFDLLNPNVRKA[illegible]
MBR2835997.1      YKPRCIDQMISSSLEVFGAV-LIRGPKYCGKTWSGRNHANSE-ATLTLENPSPLEM[illegible]
MCB0520338.1      MIPRQLAPILRETCSLYPVL-SVTGPRQSGKTTLLQNLFPDY-GYVNFEEPNTRRE[illegible]
MCB0524511.1      MIKRILEQSIRDKLFKGKAI-LLFGPRQTGKTTLLNALFKNEVTWLNGDDPETRII[illegible]
MCK9435713.1      HILRKPELSKLSVLKDKNLIKVVTGVRRAGKSTLLIQFQELLIVSVNMDLPEFRFI[illegible]
NCP08804.1        MIIRDITEELVRSAEEYPVV-TILGPRQSGKTTLARMTFPDK-PYFSLEDPDVRMA[illegible]
NMA29240.1        LTRRDATLS--------NKVHAVTGMRRAGKTTFLHQLQAERAVFLGFDDDRLAGI[illegible]
NQX01744.1        PIPRHLKAVALRHLEGFPAI-AITGPRQSGKTTLARELGGDR-PYASLENPDVRRY[illegible]
UYJ11708.1        MVLRKSYLDKIIPFIDQDLIKVLVGIRRCGKTVLLGQIKDVLIIQVNFESMRFRNT[illegible]
WP_001255534.1    NIRRDGYLKFLNTWQDRDVIKVLSGVRRSGKSTLLAMFQQDLIIAINFEFMEFEEI[illegible]
WP_004078480.1    YLNRPKYQDKIELFRDSGEIIVLTGIRRSGKSTLLKLEMKALLLYVNFEDLRLYNC[illegible]
WP_010252916.1    LIERRPYMQQLRELRDEHIIKVITGVRRCGKSTLLEMFADELIQFYNFEDPDIFAI[illegible]
WP_013460553.1    FSPREKLTHALSFLKAKEVI-AILGARRVGKSSLARLMIRSLIFFINLEKTAFIPY[illegible]
WP_044088296.1    MIDRPQYLRKLDAWKDRDVVKVITGVRRCGKSTLMTLWKQRLVIHINLELLENEQE[illegible]
WP_193852558.1    LYARPRYQMLLERYRDIDQIKVLRGVRRCGKSAILTMFMRRLIFYKRFDEFGMPAM[illegible]
WP_200291402.1    HHPRYLRQQVLDALSDTPVV-CVLGPRQVGKTTLARLLQPER-TYVSFDDATLLAA[illegible]
WP_207635334.1    MIERKEYLDWLIRWKEKQIIKVVSGARRCGKSTMFDIFCNYLIIMLNFEDIDLEHI[illegible]
WP_224302002.1    LIRRRAEDLVAEALDDTRVV-LVNGARQAGKSTLTRLTAARRAPVLLLDDAATLRA[illegible]
WP_235402039.1    YLNRVFDLRLKAHLKAMGAV-LIEGPKWCGKTTTAKQLANSV-ISLTDHREEYLAT[illegible]
WP_250702184.1    MYKRAEYQVIKCRLEEQRRFQVVMGARQIGKSTVVKQVLKDLAPYQSADNVPATNS[illegible]
WP_251231673.1    YRSRYIESTIERALASSGAV-LVSGPKFCGKTTTCMRFQKSF----RLNNSQVISI[illegible]
WP_251851847.1    MKQRPYYLNKLIELKDFEQIKVITGVRRSGKSYLLQLFKDHLIVYMNFENLIYSDI[illegible]
consensus/100%    ...R....................l.G.+b.GKo.........................[illegible]
consensus/95%     ...R....................l.G.+bsGKo.................p.......[illegible]
consensus/90%     ...R.........h.....h.hl.G.+bsGKos.h...........p.-.........h...........hlDEhQ.h..h...lp.......phhh.GS..........p..sGR...h.h.shob.E..........h..h..G.hP[illegible]
consensus/85%     ...R.........h.....l.hl.G.RbsGKoshh..........hsh-.........h...........hlDEhQph..h...lp...sp...phhl.GSs........p..sGR...h.h.shoh.Eh.........h..hh..G.aP[illegible]
consensus/80%     hb.R.........h.....l.hl.GsRbsGKoshhp....p....hsh-p.....h....hhp..............hhlDEhQpls.hbphlp...sp...phhloGSss..h...hsp.LsGRh..hph.shSh.Eh.p........h..hh..GuaP[illegible]
consensus/75%     hb.R.....bb..h.....l.hl.GsRbsGKoTlhpbhb.ph...hsh-p.p...h....hhp..............hhlDElQpls.hbphlc.bhsp...chhloGSss.blpppisp.LsGRh..hph.shSh.Eh.p...........b.hsphh.bGuaP..h......[illegible]
consensus/70%     hh.R.b...bbp.hpp..hl.ll.GsRbsGKoTlhpbhbpph..hhsh-p.ph..h.pp.hhp..............lhlDElQpls.hbphlcsbhsp...chhloGSss.bLppplsp.LsGRh..hpl.PhSh.Eh.p...........b.hsphl.bGuaP..h......p[illegible]
```

### ; CR-ATPase+REase

```
                  1---------11--------21--------31--------41--------51--------61--------71--------81--------91--------101-------111-------121-------131-------141-------151-------161-------171--[illegible]
OrigSeq           MKKIPIGINDFKTLIENNYYYIDKTKHIEDILNDGSEVILFTRPRRFGKTLNMSMLKYFFDIENKEENKKLFNDLYIKNSEYISEQGKYPVIYISFKDLKAKNWESSIFKLKNQLKDLFNKIIYMKDNANYEYALKYLTEYLYKYYKQKVIVIIDEYDSPVVDSYENNYYDEAIM[illegible]

                  <<-------------------------------------------------------------------------------------------------------------------------------------------------------------------------------C[illegible]
Jnet              ---------HHHHHHH--EEEEEHHHHHHHHHH----EEEEEE------HHHHHHHHHHHHH------HHHHH-----------HHHH----EEEEE--------HHHHHHHHHHHHHHHHHHHHH--------HHHHHHHHHHHHHHH---EEEEEE----HHHH------HHHHHH[illegible]
jhmm              ---------HHHHHHH--EEEEEHHHHHHHHHH----EEEEEE------HHHHHHHHHHHHH------HHHH------------HHHH----EEEEE--------HHHHHHHHHHHHHHHHHHHHH--------HHHHHHHHHHHHHHH---EEEEEE----HHHH------HHHHHH[illegible]
jpssm             ----------HHHHHH--EEEEE---HHHHHH----EEEEEE------HHHHHHHHHHHHHH-----HHHHH-----------HHHH----EEEEE--------HHHHHHHHHHHHHHH-------------HHHHHHHHHHHHHHH---EEEEEE--HHHHHE------HHHHHH[illegible]
CDA09053.1        MKKIPIGINDFKTLIENNYYYIDKTKHIEDILNDGSEVILFTRPRRFGKTLNMSMLKYFFDIENKEENKKLFNDLYIKNSEYISEQGKYPVIYISFKDLKAKNWESSIFKLKNQLKDLFNKIIYMKDNANYEYALKYLTEYLYKYYKQKVIVIIDEYDSPVVDSYENNYYDEAIM[illegible]
MBD5489939.1      LQKCSVGITSYKE-ASTECYYVDKTRLIKELIDEHSKVFLFIRPRRFGKTLTMDMVKTFFEISES-DTSVYFKGREIWGCGDERYQGAYPVIYISFKDAHQKTWEDMLRSLKFTIKNEFNRILDMEADIEYQAALGELSYMLSAHYAAKTIIIIDEYDTPIQQGYSNGYYEQTTQ[illegible]
MBE6101271.1      IRRLPVGVQSFEDLREGGFVYVDKTKYIYELARRSKQY-FLSRPRRFGKSLFLSTLKAYWEGKRE-----LFEGLEIAELEKDKAFEAYPVFYFDFNGQNYQAA-ALEDVLDKMLSR-WEEQYDCKG-GTLSGRFQDLLIAAREKTGKKCVVLVDEYDKPLLEVLENNLEEHNKA[illegible]
MBF0120840.1      FKAIPYGIADFQKIQQNNLYYVDKTRFIPEIEKS-GDFLFLIRPRRFGKSLFLSILEGYYDISCEKEFDTLFQDTWIHEHPTE-NKNKYLILSFNFSVVNPDIK-ELAESFEDHVKNQYKNLLDESYFIEKSHSKLNFLLKYATIKGLKVYILIDEYDNFTNTILSASSYHDITH[illegible]
MBO4331561.1      LRKYPVGIQTFDKVIKGGYVYIDKTELIFNMVHTGEYY-FLSRPRRFGKSLLISTLQAYFEGRKD-----LFRGLAIERLETD--WEKYPVFRLDFSGNKYTEEGDLLHVANLFLHN-YEEIYGAKAEQTFGSRFQGLLRRAMEQTGRQCVVLIDEYDKPLTDTIGNTMQEKNRE[illegible]
MBP1566398.1      RKPIPVGVEDFKRLVDKEYYFIDKTLMIKDIIDCRDTISLITRPRRFGKTLNMSMIQRFFEKTDE-SNAYLFENLRISEEGEEKYQGRYPVITISLKSMKQPDYKQAFTEFKNIITNEFEAIISDEADSVYNTSLQLLSRCLKEFYKENVIVLIDEYDVPLENSYFCGYYDKMVS[illegible]
MBP1566411.1      ---FNPSNTGFKKSAEDE-LYVDKSGLLNLLNKKIKNCISVSHARRFGKSQAADMIDAYYSKGCD--SENLFTKLEISKSPDYKHLNQYNVIHIDISSFADYYKDKLVSKIVELIYNEYPDIL------GTDNPFSAILHQVYKKSEIPFVIIIDEWDCVVRNFSDKPLVHEYLQ[illegible]
MBQ7457437.1      LERLSLSSSNFHALRALNQIYIDKTDNIGDLAQVRGRF-FLSRPRRFGKTLLLSTFASLFANGIAD-----FAGLHLENRWKE---EAFPVISLNFASFRRESAEAFRDGLFCALQR-FTNVFPHLAGTNKDTPDSLLARYCVSIQTLNLVMLIDEYDAPIIHSLPDEKAQSILD[illegible]
MBR2836002.1      ---VNPGNIGFKTIRAGK--YVDKSDIIAVINNSLDKLTCVSRPRRFGKSFAAQMLSAYYDCSCD--SSELFDDLRISRSADYKYLNQFNVVYLDIASLKTATQDNLVSFISGQVANE-----------NHETPLISALTKAAEILQKQCIMIIDEWDAPIRETPE--LAEGYLE[illegible]
MCP5494231.1      MLQIAYGEANFENLRKDKSFFVDKTHFIPLVERI-KKF-FFIRPRRFGKSLWISVLISYYDINGEKKFDELFDGLYIKENPTE-YKNSYLVLKFDFSGLNTVNLESLRRDFNNRIRSSNDSIFGNEFYDMSASVMIDFLKKEVLNTRNKVYTFIDEYDHFANKLASEGFVRDIIS[illegible]
MCY3660413.1      LPPLHTNRADFSNIRSRREFYVDKTRFFRDLLAVTGHL-FLARPRRFGKTLLINTLETWFQGLPPESPAWLWDGLA----GIDGTHGWHPVIRLDMGQAVNPNLGSLQDKLQACLWGA--TLWRQRGVDAPDTILSSLIRALRQHYRRSPAVLVDEYDAPLVEFIGTDDPSRALA[illegible]
MCY3837524.1      YPRIPYGWADYAAIRQEGRMYVDKTRHLRALEE--ERYAVLIRPRRFGKSCWVSLMECYYDRNRADAFDALFGDTDIGRKPTR-RRNRYVILRFDFSAVSDVLE-TLEREFETYCESENPDVFSENAAPPSIEAKLNELFRHTRRNSIPLYVLIDEYDNFANTILAHRAYQSLTH[illegible]
UYJ11706.1        MKDFPVGISNFEKIREGGYYYIDKTNLISELLSGGAEVTLITRPRRFGKSLGMSTLANFLDITKD--SKQMFEGLAISKNTELKWMNQCPVVFFSFKDTDGLTFESAYGMLCMKLAFAFKRILGRTASDETKSCFLLLTRMLEIHFKKSAVVILDEYDVPIAKASSNGYYSQMLD[illegible]
WP_002710596.1    MKKLPIGISTLEKIIQDDFVYVDKTVHIQELTKKPGYY-FLSRPRRFGKSLLLDTLHQLFAGNEP-----LFRGLHIHPHWDW--SLAYPVIRLSFGSGIPKNKADLDRSIQRQLRR-NQQLLGVTCTNDSVGCFAELIQKAADHYGQPVVVLVDEYDKPILDNVTDDTATVMRN[illegible]
WP_005980881.1    MKELPIGIDDFKNIIDNDYFYADKSLFIEEISKNIGKTLLFTRPRRFGKTLNMSMLKYFYDIKGNKKNRELFKSLNIEKTSTMVEQGQYPVIFISFKEVKFNTYEKLFEQVKILISEVFINYLKKEVNVEAINSLKFLSQLLYDQYGKKVVVLIDEYDTAIVSAYENGYYKEAME[illegible]
WP_048067427.1    IPKFRVGTDDFKKLRDEGGYFVDKTLFIKEIIDG-NEVTLIPRPRRFGKTLNMTMLRYFFERSEE-SRAYLFDSCAIADYPMYEHQGQYPVIFLSLKDLKRDSYDHFIKATKAEIGRLYLRICREESPHELYNSLKELISHCYSFYKKPVIVLIDEYDTPMIEAFSKGYYNEMSD[illegible]
WP_118127446.1    LKGIPIGREDFKDIIENNYYYIDKTKFIEDLLLDGTHVKLFCRPRRFGKTLNMNTLKYFFDIKNGEENRKLFNGLYIENSPMIKEQGKYPVIFLTLKELKGSNLETIYLQIRTLISDLFDNIWKRKDE-DYSNSLKFLIRCLSEYYNQKVILLIDEYDAPFITAYEFGYYEEILQ[illegible]
WP_164949150.1    MKRIPYGISNFEVLREKNYLYVDKTSYIDILDTY-SPYQFFIRPRRFGKSLFISMLENYYDINKSDKFEKLFGDLYIGKNPTE-ERNNFLVWKISFAGVDAGHGEELRRSFNEKIILSYSNLLEEKVVIKSAEIAVQYISLLASKINKHVFVLIDEYDNFANELITGGTYKSIVR[illegible]
WP_235392640.1    --KFPIGIQDFKTIRKDGFVYVDKTALLYKLADEGQTY-FLSRPRRFGKSLLVSTI[illegible]
WP_235392649.1    IRKYPVGMQTFDKIREGNYVYIDKTKYIVKMMKNGSQYLFLSRPRRFGKSLFVSTI[illegible]
consensus/100%    ...h..s..sh.........ahDKo..h..h.........h.+sRRFGKo..hp.h[illegible]
consensus/95%     ...h..u..sap.......halDKo..l..h......h.hh.RPRRFGKoh.hs.h[illegible]
consensus/90%     ...hs.u..sap.l.p...halDKo..l..l.p....h.hh.RPRRFGKoh.hs.l[illegible]
consensus/85%     b..hshG..sFcplbp...hYlDKT..l..l.p...ph.hh.RPRRFGKoL.hs.l[illegible]
consensus/80%     h..hshG..sFcplbp...hYlDKT.hI..l.p...ph.hh.RPRRFGKoL.hs.l[illegible]
consensus/75%     h..hshG.psFcplbp..bhYlDKT.hI.pl.p...ph.hhsRPRRFGKoL.hs.I[illegible]
consensus/70%     hbphshGbpsFcplbpssbhYlDKTphI.cl.p..sph.hlsRPRRFGKoL.ho.I[illegible]
```

### ; (w)HTH+REase

```
                  1---------11--------21--------31--------41--------51----[illegible]
OrigSeq           MEQKSHETGDLFADTCEIVESAQSIAYAAVDTILVQRNWLIGKRIAMECLDEKGKA[illegible]

                  <<------------------------------------------------------[illegible]
Jnet              --------HHHHHHHHHHHHHHHHHHHHHHHHHHHHHHHHHHHHHHHHH--HHHH[illegible]
jhmm              -------HHHHHHHHHHHHHHHHHHHHHHHHHHHHHHHHHHHHHHHHHH---HHH[illegible]
jpssm             ----H-----HHHHHHHHHHHHHHHHHHHHHHHHHHHHHHHHHHHHHHH-HHHHH[illegible]
EOS42582.1        MEQKSHETGDLFADTCEIVESAQSIAYAAVDTILVQRNWLIGKRIAMECLDEKGKA[illegible]
EXI84707.1        MTDPAADYQDWLKTIKQRIHATRMKVALAANGELISLYYEIGAQIADRE----AHA[illegible]
GDY21842.1        MAAKRSALGQVLPEIQTLIESSRQHVVTTANLTLVWLYWNVGRVITQDIQKNAKRA[illegible]
KUO61375.1        MKK--VTLVSLYSQIRQILDTARSSSYKAVNFLMVQSYWQIGKLIVEEEQNGSNRA[illegible]
MAD46038.1        MNKPSRLSDRLLTDARSIITTAQSNAMRSVEFHRVEMYWQLGERIFVEQQQQDRA[illegible]
MBF0105030.1      MEG--SAKGNLFERIVFILEKARANVVRAVNSNMVIAYWLIGREIVLEVQQGKSRA[illegible]
MBI2505142.1      MKKRTADYGRLVAEVSGLLEQARRTAARAVNSVLTAAYWEIGRRIVEHEQGGKGRA[illegible]
MBK7550236.1      MKNESADYVELIRTLKDRVREARSQAVLSVNRQMVLLYWTIGQHILASQ----KHH[illegible]
MBL7172015.1      MNKHEEIPKSLFQDVRSILADARQRAYAAVNFIMVEAYWKIGRRIVKEEQGGKGRA[illegible]
MBN1601228.1      MQK--MQSTILLNQIRELIAEARVSVVRNIDFIQVWTNFHIGRMIVEHEQQGSHRA[illegible]
MBQ9588228.1      MVQDNNDYADWIADIKHRYRTAQVKAAVRVNAEKLLFNWMLGRDLVQRK----AEE[illegible]
MBS0213166.1      MKEKRPAQSDLFATVSGLIEEARRTLARQANSATVFLFWRIGQQVNSEI-LNNQRA[illegible]
MBV6627481.1      M-----NYEELIHDISNINNTSLTTVTRAVNKILTIRNWFIGAYIVEYEQNGVDRA[illegible]
MCC6178179.1      MHKKTESDDRGFGEVLEMIEVARGRALAEVNSALVDLYWRVGEYISKKI----ASA[illegible]
MCC8119637.1      MAE--QPLETQFSELLTIIQGHRSAASRAVNTEQLQMAWEVGAYVSSKL----KTE[illegible]
MCC8150844.1      MSELLEEYRCWIQELGNRYKVSQIKASIRVNNEMLMYYWSVGRDITERH----ANS[illegible]
MCE9554598.1      MKSKGADYRALLDNLKVRIRSAQVRAALSVNRELIQLYWDIGRAITVKQ----EAE[illegible]
MCL2053371.1      MLINDNSYFQALEDIKTRIHDAQHKAVLGMNREQIILFWNIGNVIIENT------[illegible]
MCP4577329.1      MDDERDGYANLLSSLKKRVQQERIKTVLSANCAMVMLYWDIGTSILKKQ----QTE[illegible]
MCP5421180.1      MSQNIDDYRNLIADLKTRVQAAQIKAAVTVNTQLMALYWDIGQQIAERQ----QAS[illegible]
MCP5560210.1      MPAKKGTSGVLFDRVASILEQARANVVRAVNSNMVLAYWLIGREIVQEIQGGKERA[illegible]
MCY3788285.1      MTEPVTQYRAIFNDVVGIVDAARSSAARSVNAVMTAAYWLIGRYIVEFEQGGEKRA[illegible]
NOQ23472.1        MDMIKWNFENLVNVIRQTHEYMAIQAGRAVNISLTMRNWVIGCYIREYEQNGEDRA[illegible]
TVL98973.1        MSKEVLTYKQLLVTFKQQVQSAQAKAALAVNSSLIQLYWNMGKMIADNQ----ALE[illegible]
TVL98974.1        MPD--EILINNFQQLYDLIRQARSRSIVFANREHLNLFWQVGAYIDAKL----AGG[illegible]
```

```
TXT43481.1          [illegible]
WP_004077397.1      [illegible]
WP_012508726.1      [illegible]
WP_019991367.1      [illegible]
WP_021798348.1      [illegible]
WP_070561015.1      [illegible]
WP_087287646.1      [illegible]
WP_172658767.1      [illegible]
WP_205241477.1      [illegible]
WP_217494871.1      [illegible]
CRL42136.1          [illegible]
consensus/100%      [illegible]
consensus/95%       [illegible]
consensus/90%       [illegible]
consensus/85%       [illegible]
consensus/80%       [illegible]
consensus/75%       [illegible]
consensus/70%       [illegible]
```

### ; HTH+REase

```
OrigSeq             [illegible]
Jnet                [illegible]
jhmm                [illegible]
jpssm               [illegible]
AOO75266.1          [illegible]
ARA42374.1          [illegible]
MAR01292.1          [illegible]
MBL3581664.1        [illegible]
MCH9867674.1        [illegible]
MCP6718635.1        [illegible]
MCQ4803589.1        [illegible]
NLA16131.1          [illegible]
NPV30264.1          [illegible]
QNI65419.1          [illegible]
SFW99816.1          [illegible]
THE19346.1          [illegible]
WP_004808453.1      [illegible]
WP_009555624.1      [illegible]
WP_016373141.1      [illegible]
WP_051006310.1      [illegible]
WP_068488201.1      [illegible]
WP_122014533.1      [illegible]
WP_126762074.1      [illegible]
WP_141142700.1      [illegible]
WP_152261654.1      [illegible]
WP_153832545.1      [illegible]
WP_179394512.1      [illegible]
WP_188891969.1      [illegible]
WP_233943242.1      [illegible]
WP_265072434.1      [illegible]
WP_270447888.1      [illegible]
WP_271847091.1      [illegible]
consensus/100%      [illegible]
consensus/95%       [illegible]
consensus/90%       [illegible]
consensus/85%       [illegible]
consensus/80%       [illegible]
consensus/75%       [illegible]
consensus/70%       [illegible]
```

### ; McrC-NTD+REase

```
OrigSeq             [illegible]
Jnet                [illegible]
jhmm                [illegible]
jpssm               [illegible]
WP_221091412.1      [illegible]
CCU85343.1          [illegible]
WP_006438850.1      [illegible]
WP_244713432.1      [illegible]
WP_247995975.1      [illegible]
WP_164594475.1      [illegible]
WP_254334240.1      [illegible]
WP_036689505.1      [illegible]
WP_075107045.1      [illegible]
WP_124569533.1      [illegible]
WP_170155634.1      [illegible]
WP_066770451.1      [illegible]
WP_058302017.1      [illegible]
MBN1921824.1        [illegible]
WP_036463105.1      [illegible]
GFM17531.1          [illegible]
WP_226023956.1      [illegible]
MBL7764606.1        [illegible]
WP_213510440.1      [illegible]
WP_260358536.1      [illegible]
MBZ4644072.1        [illegible]
WP_158531734.1      [illegible]
WP_189000187.1      [illegible]
WP_129664649.1      [illegible]
WP_195040820.1      [illegible]
WP_146078436.1      [illegible]
WP_036163551.1      [illegible]
HCG7393715.1        [illegible]
consensus/100%      [illegible]
consensus/95%       [illegible]
consensus/90%       [illegible]
consensus/85%       [illegible]
consensus/80%       [illegible]
```

```
consensus/75%   M....hplbE......c...l..h........lps.pbVGhl.h.s.plbl.PKls.pphl..hhb.......p...h..p...l.phhh.hahpbhpphhppGlbp.Y...ppsb..l+G+l.h.pphb+p....h.s.ap-hs.sp.bNphlb.slpbl.p.s.ss..ppph[illegible]
consensus/70%   M....hplbE.....sch..L.ph........lps.paVGhl.h.s.plblbPKls.pphLh.hhb.......p.sph..p...lb-hhh.hahpbhpphhcpGlbp.Y...ppsb..l+G+l.hspphb+p....ph.spap-hs.spsbNpllb.slpblbpbs.ss..ppph[illegible]
```

##### Standalone PDDEXK REase

### ; PDDEXK-REase

```
                1---------11--------21--------31--------41--------51--------61--------71--------81--------91--------101-------111-------121-------131-------141-------151-------161-------171-[illegible]
OrigSeq         MSARINTKLRERDIEIDFTDAIDTLVFDQMRSDQPNYHGIGEMHRVDFIVELDRAILFVEVKDPGNPKAQEKGLQKFMEELNNGTLSSTFASKFIDSFFYRWAEGRLSKDVYYLSLVTLEGEVLPNLSDEIAKKLSPVGKGSPRWQRHPVQNCQVFNIQTWNENFPKWPVTRLAA[illegible]

Jnet            ----EEEEEEE--EEEEE------EE-------------------EEEEEE---EEEEEEE-----------HHHHHHHH----HHHHHHHHHHHHHHHHHHHH-----EEEEEEEE---HHHHHHHHHHHHH-----------HHHHHHH---HHHHHHH-----EEEEE-[illegible]
jhmm            ----EEEEEEE--EEEE-------EE--------------------EEEEE---EEEEEEE-----------HHHHHHH-----HHHHHHHHHHHHHHHHHHHH-----EEEEEEEE---HHHHHHHHHHHHH-----------HHHHHHH---HHHHHHH-----EEEE--[illegible]
jpssm           ---EEEEEEEE--EEEEE-----EEEE-----------------EEEEE----EEEEEEE----------HHHHHHHHH---HHHHHHHHHHHHHHHHHHHHHH-----EEEEEEEE---HHHHHHHHHHHHH-----------HHHHHHHH--HHHHHH-----EEEEEEEE[illegible]
AUZ76296.1      MSARINTKLRERDIEIDFTDAIDTLVFDQMRSDQPNYHGIGEMHRVDFIVELDRAILFVEVKDPGNPKAQEKGLQKFMEELNNGTLSSTFASKFIDSFFYRWAEGRLSKDVYYLSLVTLEGEVLPNLSDEIAKKLSPVGKGSPRWQRHPVQNCQVFNIQTWNENFPKWPVTRLAA[illegible]
HCF96072.1      ------MILREGTLQFDFAGAIEAFKFDEQARELKTYHGLSHMKAVDFVVEYSDHYLFVEIKQPRTPERYGSDRDKF-------DLIQDLTKKFRDTFLYRWAEDKLDKPVYYHCLVELDNAQTSYLMKQLKNQLPTEK-QPQRWRRPLARSCSVANQATWNATFPNIQASRVAG[illegible]
HCG7393725.1    -----MSKFAEEDLEFDFSGARQAFRFDSDD-----LHGASSAQRVDFIAEYDDHYLFIEVKDPDNPAAS--NPEAFMAKLRNQKLEKSLAGKFRDTLFFRHLTSNQNKSIKYVVLLAIEPAMLMAKRDLLYREIPL---ENPAWWNNRVADCILVNLQQYKDLFGENSVRRISE[illegible]
HIJ32931.1      ------MEISEGNLSFTFNGALSGYKFDEQDQTSTHYHGLSHMKGVDLVIEQEDVWLFIEVKDPSHPDSTPEQIAAFAKKARSTELVSDLVHKYRDTFLYHWAEQKVEKPIHYLCLITLESALVLQLMNQLKRQLPEGRANNQRWQRAIADRCIVANLETWNSSFPHIPVQRVRG[illegible]
MBD5149895.1    MP--DVLIEEKRNYGIDCRNAV--WASDEIHAV---YHSCGLLCDADFVVETEDAIFLIEYKNANTPEARATAATEY--DPFQSEKFNKIVSKYYDSLHYLRLMGK-EKPIHYIFVLKGDSVSRKMLRNRLKKRLPFRLQEQFDTGIRLIDSVSVLNISEWNADYGQYPILPVSQ[illegible]
[illegible]
```

```
consensus/95%   [illegible]
consensus/90%   .........p..h.hphs.u...b..Dp........p...hp.lDhlhE..p...hlEhKs.s...........b..p.....b..[illegible]
consensus/85%   .........p..h.hshssA...b..Dp.p......p...hp.VDhlhE..p.hhhlEhKs.s...s.......b..p.....b.p.[illegible]
consensus/80%   ........b.p..hbhshssA...h..Dp.p....h+s...hp.VDhlhEb.p.hhhlEhKs.s.s.u.....p.h..p....ph.p.[illegible]
consensus/75%   .......bpp..hbhcFssA...h.hDp.p....hHs.s.hp.VDhlhEb.s.hhalEhKsss.s.A.....p.h..p....ph.psl[illegible]
consensus/70%   ......b.bpc.shbh-FssAh.sh.FDp.p....hHshsphp.VDFllEbpc.hlFlEhKsss.s.A...s.p.a.pp....pl.psl[illegible]
```

### ; PDDEXK-REase

```
                1---------11--------21--------31--------41--------51--------61--------71--------81--------[illegible]
OrigSeq         METPGERWKAAFMDIVHHHENALPLRDASLNGNLRKWTTELTSIVSSSCRALSWEVAALGHKLEKLPVSREEYLSLDVTAFEKKWENESG[illegible]

Jnet            ---HHHHHHHHHHHHH------HHHHHHHHHH-HHHHHHHHHHHHHHHHHH---EEEE-------------EEEEEEEEE----------[illegible]
jhmm            --HHHHHHHHHHHHHH------HHHHHHHHHH-HHHHHHHHHHHHHHHHHH---EEEE-------------EEE--EEEE----------[illegible]
jpssm           ---HHHHHHHHHHHHH------HHHHHHHHHH-HHHHHHHHHHHHHHHHHHHH-HHHHH--------------EEEEEEEE----------[illegible]
AKB47808.1      METPGERWKAAFMDIVHHHENALPLRDASLNGNLRKWTTELTSIVSSSCRALSWEVAALGHKLEKLPVSREEYLSLDVTAFEKKWENESG[illegible]
EGK85304.1      MQNIAYQWQTKFWQTLQQPENADPLKQAALNGQLTKWTTALTTTVVATSTAMGWQSSARGNSLGLFPVSTSEYLALDVMSFAP------G[illegible]
KUK43675.1      AVDIARRWQEAFLGILRPLEKSRPLKEAAASGNLGDWTCALTGLVVLSVESLGWQAAALGHPCRALPVSRKEYLSLDVLAFAPAPATDHN[illegible]
MAO24326.1      QAGFGLRWRNAFYRVIEETSVSDALREYADGAHLTEWTGHLTTSVIRSCELLGLKAAGKGHPLDLLPQSGQEYLGLDATAFLPESA---E[illegible]
MBE7497141.1    MNPTSQAFQTAFHSVVNGTEAGAMLREASLEGKLGPWTKHLTASCVQACEELSLRASAKGHKLDLLPVARSEYLSLDVVAFPE------T[illegible]
[illegible]
```

```
consensus/95%   [illegible]
consensus/90%   .......a...h...hp....s..Lb.su...pL..WT..hT..ls.shp.hsh.ssubs.....hP...pEYLuhDh.uF........[illegible]
consensus/85%   .......Wb..h...lp....u..L+.uu...pL..WT..LT.hVs.shp.huh.suA+G...p.lP...pEYLulDlhAF.......s[illegible]
consensus/80%   ..s..p.Wb..F...lp....u..L+pAu...pL..WT..LT.hVs.shp.hua.uuA+G...p.LP..ppEYLulDlhAF.......s[illegible]
consensus/75%   ..s.sp.WbpsF.p.lpp.p.u..L+pAuh..pL.pWTp.LT.hVl.oCc.hGWpAuA+Ga.hphLP.sppEYLulDVhAF.......s[illegible]
consensus/70%   ..s.sppWbpsFbp.lpp.p.u.sL+pAuh.upL.pWTp.LTshVV.oCc.hGWpAuA+Ga.hphLP.sppEYLulDVhAF.s......s[illegible]
```

### ; PDDEXK-REase

```
                1---------11--------21--------31--------41--------51--------61--------71--------81--------[illegible]
OrigSeq         MPVELQEGQLTFTFPDDWVASKYDAWAFYRNQFVKQLAGIQAVDFIAIDPRGTAYLIEVKDYRHPNTDKPSQLAQTIANKVLMTLAALLP[illegible]

Jnet            ---EEEE--EEEE-----EEEE----HHHHHHH-------EEEEEEEE-----EEEEEE-------------HHHHHHHHHHHHHHHHH[illegible]
jhmm            ---EEEE--EEEE-----EEEE----HHHHHHH--------EEEEEEE-----EEEEEE-------------HHHHHHHHHHHHHHHHH[illegible]
jpssm           --EEEEE--EEEE-----EEEE----HHHHHHHH------EEEEEEEE-----EEEEEEE------------HHHHHHHHHHHHHHHHH[illegible]
GIX36092.1      MPVELQEGQLTFTFPDDWVASKYDAWAFYRNQFVKQLAGIQAVDFIAIDPRGTAYLIEVKDYRHPNTDKPSQLAQTIANKVLMTLAALLP[illegible]
HBO5114777.1    -MTQLVEGGLRLTFPAGWQALKFDDTSWHRQRF----GHHQAMDVLATH-SGQHWWIEIKDCEGFEQDNRPRMSDELIEKLRSTLVSVLA[illegible]
```

```
KYF87241.1       -MPTIAADGLNFDFPSAWETSKYDEWAFHRKRFSKVRDGIKALDLLAVDPDGTAWLIEVKDYRVHARTKPSALDDEIAGKVLDTLAAMLPAKINANDAHEAEMAAAVLGAKKLRVVLHLEQPTKHSTAINPADVQQALRRRLKPIDAHPLVAETSRMG--SLAWKVT
MBE6357834.1     -MQNITVDSRTFCFDDSWLIMKYDETPFYKNHLSNKQKSMKGVDLIAFDPAKVLFLIESKDYRVHQRKKKLSPEKEFFEKVIDTFTGILPTVLNSRTNGEDVLNESVAKAKKIRLIYQFEQPIKHSKVYDTADMQSKLRQELRIFDPHLLVIDSTTQS--KVAWSVI
MBF0177024.1     -MTAITEGRLTFTFPGQWDAVKYDDWSFYRNQFNGCCGGTKAVDILAIGHGRTLWVIEVKDYRIRPRTKPSDLGDEVAAKVRDSLAGLVAAARNAHDSSEKSFARSALASASLRIVLHLEQPVRRSKLVDPSNVQMKLKRLLKSVDPRVRVLEVATHAR-SVSWTVA
MBP6559244.1     MSRTADVDGMRLTMPAGWWVWKYDDSDFHQTQFQNFAGGVKAMDAIALDPAQTLWLIELKDYRQNKRRKPSSVFAEVAGKARDTLAGLAVARVSAQAQQERALAQQALGSRQIRVALQLAQSARPHRVVEPADAGLQLRRAVKQVDAAPVCAVGSLGQHNMLPWNTV
MBP7545006.1     PTHSIYEGQLRFDFPQSWQALKFDDAPWYRNTIK---TRVKAMDLVACL-DATHWWIEVKDCKGFEADNLPRLSDEVAEKLEGTLLSLAAAVRQAEATAVAPFARVMDTTVQWSVVLLLTWARDFGR------LRDKLRQRLAAYNVQCFVLNESESAP-HQPWTLT
MBU0551511.1     -MAQVQVEHLVFDFSAGVAPFQYEQSGAVVAGWP---NGSKVVDVIATDAAVTAWFIEVKDFRIPKPANLQKLAQTVERKVRDSLSGVKAVAASSANCAAQKHAAVASRAAHRRVVLHLEPGGPHSFGF-AANVLMKLQSQVRDIDPKPLVLRVSTTPKAGVPWSVR
MBX3448930.1     -MPAYVEGNLTFSFPRYWKVLKYDDSVFHVSEFQSTCGGAKAIDFVALEPKACCWMIEVKDYRVHCQTNAVDLAAIVAHKVRDTLAGLAVARLYASQQHEKVHAQRALKCSGLRVVLHMENPVSLSAPINPANVLQELKQRLRVIDASLKVVSLTQSG--SVAWTVN
MCB5246761.1     -MIQIQEGQLTFVFSDSVMACKYDDWSFYRNQFQNGCRDNKAMDLLCHS-GSVAWMIEIKDYRCHHRTKAVDLADEIAIKVRDTLAGLAAAKTQANDRDEKLFAKKMLGATRLRVVCHIEQPAKSSRVIEPDKLKDRLRRELKAIDPHPIVMDKKNTAV-AVPWQVR
MCC6374736.1     -MPDIKEGLFTYQFPANWVISKFDELPFYLRHFQNFAGGCAAVDVWAFAPQDEVWLIEAKDYRAYPRLKSSDIFDEMALKALGTLACMVAARANASDGLSEVMATTVLRKPKLKIVLHLEQPQKPSKVVDPKSAKDKMRQKLRAVDPHAIVGDKALLNK-LMAWQVI
MCK9487879.1     MPSAIDVDGLRLTPPAAWQVTKYDEWAFYRRHFQSFAGGKKAVDAMGVSDGGTLWLIEVKDYRRNRRSKPSRIEHEIAGKVADSLSGLAVARVRATG-SEQQMAESGLRCSAIRVVLQLAQPVNPSRVVDPADVSIKLKRAVRAVDPHPMAVVGDISSA-RLPWKTS
MCL5961064.1     -MSTLNEGNLACTFPDSWKVTKYDDWEFYRRNRFAGSCGGNKAVDFLAHDPGRTLWLIELKDYRRFRRTKDDQLSDEVALKVRDTLAGLFAAKVDPAH-PEQSYAALALRVTSLRVVLHLEQPATGSKAYHPADVQQKLKQMLKPIDAHPRVIELNNMR--AVPWSAT
MCR7692358.1     IEQVINEDRLTFTFPAGAMATKYDDWAHYRRQFNSAFGGTKAIDLLYAE-DRVGWLVEIKDYRLHVRTKPTDIADEVAHKVRDTLAGLVSAQHHATDTDERRFARKMLRCQRLRVVLHLEQPRKTSRAMDPAAVQIRLRGLLKAIDPHVCVVDQHALKA-EMNWTVI
MCX6879290.1     -MATLQVDTLKFDFESTVSAQIYDQWQHYSSVWNAPPGGQKAVDVIAVNVESTAWLIEAKDFRVPKASNISGLAQFVTEKVQHTLAGLADASAKAALPDEKQLATDALACPTKRVVLHLEPTGVHTAGF-AASVLQRLRQLVTAIDSNPLVLNIANTPASGVPWTAS
SBT04364.1       APLAVEVDGQRFTFADGWRVLKYDDSRFHRNQFQSFAGGSKAVDAVALSPAGECWLIEVKDYRRNRRSKAGSVFSEVAGKVRATLAGLATARLQANDPDERDFCQQAMRANVLRIALHLDQPDHPSHIIDPRTAVTQLRHELHAVDPHALCYGTGVAIA-RLPWRVE
TQE99817.1       MDVDVKESRLRFSFPDSWTAEPYDEWAFYRNHFNGVC-KSKADDIVACDGTQALYLIEVKDYRRQRRTKPIDIADEIAHKCRDTLAGLAAARVRASNHNEQQAADQSMRCRTIRVVLHLEQPRRPSKSVDPKSALEKLRKRVRAIDPHPRIVDRAT----AMPWTVA
WP_022949218.1   -MNDFTEGGLTFSFANGCDATAFDKWSFIRNQFQSICGGSKAVDFLCID-NDVLWIIEVKDYRRPGSKKPSNLSLDVAKKVRDSIAGLVAAQCNATDQDEKKFARKALRATQLRVVLHVEQPCKASKAVDPSDLKSKLKKLLKAIDPHPEIVDQNNLKQ-NMNWTVT
WP_025282663.1   MTTTLMEQQLTFTFAEGVEATQYDDWAFYRNQFNAVCGGTKAVDFVCLD-GDQLWLIEVKDYRHHRRTKLVDLGDEVAAKVRDTLAGLVAASCNANEAEERRVARQAVGRSRLRVVLHLEQPRHPSRAVDPDDVMLKLKQRLKAVDAHPRVVDQSTLDA-RMPWAVR
WP_076345236.1   -MATLTEGKLRFTFAEDWTVSQYDDWGHYRNQFIKVCGSIKAVDVVAIEPNGCCWLLEIKDYREHVRTKTIHLADEMAEKVRDTLAGLVAAQFRADDESEKIAAKQALRATSLRVVLHLEQPTKPSKAINPATVLQRLRQLIKAIDPHPQVVETATSR--NLPWTIH
WP_077999169.1   -MTAITEGALTFTFPDEALVCKYDETAFYRNQFIKMAGGIKAMDLVFVD-NQTFWLIEVKDYRVHRRTKAIELADEIAGKLCNTLAGLWAAKCNANDQNERQFAAAAINTSKMCVVLHLEEPPQQGRKTLKTGVLQKMKQKLKSVDAHPCLVDQHSLKP-AMNWTVT
WP_132962624.1   -MSDVQVERLTFSFPSDWRVGKYDDWSFYRNQFSRQFDGIGAVDVIACSAHGETFLIEVKDYRHPDTTKPSDLPTIIANKVIMTLAALLPAALHANDAAEQALAKKALRCRSLRVVAHIEQPRAHRPAVDPADIKQKLCQLLRAVDPHPSVVSANAMQ--NLPWQVQ
WP_157118906.1   -MTVITEGTLKLTFAAGWCAIKIDDTDWYRDG-DAIANQVKAVDITAYGPPQEHWWIEIKDCFGHEQANRPRLVDEVVQKVAGTITTLAVASRESRAVAVQPYAKAFAPGVALKVVLFLTWGPDFGV------LKTAMEQRLHAFNVTCLVVNETVTAP-AQPWTVG
WP_165226835.1   -MISITEGRLTFDFPDGWNAMKYDAWTYYIRSFQSVGGGCKGVDLVAFPSRDCLWLIEIKDYEVAPRQKNLELVEEIVIKVRDTLAGLAAARWRATKEDEAGFASLALQCRDVKVVLHLEQPRNPTR-EDVSKLVLKLKERLRAIDNHPRIASLNDGR--SFGWTVR
WP_168935606.1   MPVKEDVDGLSFVFPTGWKTTKFDDLTFYRNHFLKMHDGIKAVDIVAISPGATLWLIEVKDFRRKRREKSLPLYEEIWQKVFATLAALLPTSVNASTDREQQFAKFALKVSKVRVVFQGEQPRKPSKSFTPADLQKKFKMIFKAIDPHALVVSSQDMPS-QIPWTVQ
WP_178366470.1   -MREIDEGQLRFFFPDNTLASKYDDWSFYRNQFSKAFDGIKAMDFIHVD-NGHTWLIEVKDYRVNRRTKPIDIGDEIALKARDTLAGLVAAKLNATDAQEQTIAENALRKNRLRIILHLEQPKKHSKAIDPSKVKQKLKQRLKSIDPHPRVVDKSDLEKDDMNWSVK
WP_189059187.1   -MTMLDVDGLSFTFPDDWDVSKFDDWAFCK-RFQNIVPKTGAVDLMAISPPITLWLIEVKDYRQHLRENPSTLDVVIAEKVRGTLAALLPAKLDAHDAGHRRVAGKALNSKRIRILLHLEQPAVRSAAINRANVLADLKTRVRSIDPHPRIVDRTT---PDLPWTVT
WP_230262564.1   -MTTFDEQEQAFDFPDEWDVSKFDEWSFYRGQFSSLAAGTKGIDFLAIESDHDCWQIEVKDYRKTLESTFLFLADEVALKVRDTLACLVVARTGSNNADEQRWAERAIGCSRQHVVLHLEQPTSNSRRKRAADVQQRLRQLVKCIDRRARVVDQTMNG--SVGWTVT
WP_231663148.1   -MPEVREGQLTFHFPEGWQLLKYDDSTWHRQRMK---SRSKGVDILAHMGGPTHWWVEVKDCEGYEPENRPRLSDEIEEKLRDTLAAIAVAEREAAP-ELAGFSPLHAPFPRLRIVLLLTWQRDFKR------LQTRLERALAPYGVQGLVVNEQTTGSAGLHCRVT
consensus/100%   ........b.h.........h-................s.Dh.........a.lE.KDhb.....p.......h..Kh..oh.sh.ss...s.......s.........plhh................h..hp..h..hs....h............h...
consensus/95%    ....h....b.h.hs..h....aD...a.....h........uhDhh.......ahlEhKDhc.....p....h...h..Kh..olsshhss...u.........s.........plhh.h..............h...hp..l..hs.p..hh.........shp..
consensus/90%    ....h....b.h.Fs..h...baDp..a.....h........uhDhhh.......ahlEhKDhc.....p....l...h..Kh..olsslhss..pu.........s.........bpllh.hp.s............h..phcp.l.shssps.hh.........hsWph.
consensus/85%    ....h..s.hphsFs.sh.h.baDc.saabppa....s..puhDhlh.......WhIElKDa+...ppp..pl.pplh.KlbsTLuulhsA..pup....p.hA..hh.s.h+lllplpbs...s.......hb.cL+p.l+shDspsblhp.........hsWpl.
consensus/80%    ....lp.s.LpasFsssh.h.baD-.saabppa....s..KAhDhlhh.....p.WhIElKDaR...ppp..pl.pplh.KlbsTLuulhsAp.pups...pp.hA..hh.s.ph+llLplpbs.p.s.......lb.cL+p.l+slDspsblhs.........hsWpl.
consensus/75%    ....lp.s.LpFsFsssh.h.KaD-.saa+ppFp...su.KAlDhlAhs.s.shWhIElKDYR...ppp..plspElh.KlbsTLAuLhsAp.pAss..Ep.hAp.hh.s.plRVVLaLEbP.p.up.hpsssLb.+L+pbl+slDs+sblls.ss.....hsWpVp
consensus/70%    .bsslp.s.LpFsFsssW.h.KYD-.saaRspFp...su.KAlDhlAhs.s.shWlIElKDYR...psp..plssElA.KVbsTLAuLhsAphpAss.pEp.hAp.hl.spplRVVLHLEQP.p.o+.hpssslb.+L+pbl+slDs+sbVls.ss.....hsWsVp

# ; PDDEXK-REase

                 1---------11--------21--------31--------41--------51--------61--------71--------81--------91--------101-------111-------121-------131-------141-------151--
OrigSeq          MNHEDLTAKSAKGAKEELLFKDECYAIQGAIFEVYREMGCGFLEAVYQECLEKEFRSRELPFEAQKELLLSYKSQPLSQTYKPDFICYGNIIVELKSVKEIASEHKAQLLNYLKATGLELGLLVNFGAYPKVEIVRIANSNFRDFSAFRGSRNLS

Jnet             ---------------HHHHHHHHHHHHHHHHHHHHHHHH----HHHHHHHHHHHHHHHHH-----EE--EEEEEE--EEE----EEEEEE---EEEEEEHH----HHHHHHHHHHHH----EEEEEE------EEEEEEEE--------------
jhmm             ---------------HHHHHHHHHHHHHHHHHHHHHHHH----HHHHHHHHHHHHHHHHH----------EEEEE--EE-----EEEEEE---EEEEEEEE-----HHHHHHHHHHHH---EEEEEE------EEEEEEEEE-------------
jpssm            ---------------HHHHHHHHHHHHHHHHHHHHHHHH-----HHHHHHHHHHHHHHHH----EEEEEEEEEEE-EEEE----EEEEEEEEEE---EEEHHHHH----HHHHHHHHHHHH-----EEEEEE------EEEEEEEEE-------------
AAS96496.1       MNHEDLTAKSAKGAKEELLFKDECYAIQGAIFEVYREMGCGFLEAVYQECLEKEFRSRELPFEAQKELLLSYKSQPLSQTYKPDFICYGNIIVELKSVKEIASEHKAQLLNYLKATGLELGLLVNFGAYPKVEIVRIANSNFRDFSAFRGSRNLS
ESQ16531.1       MTTENTEG-------TELEKDPLTGAVIGAAIEVHRHLGPGLLESAYEACLAHELGQQGVGYRLQVPLPVKYKGASLDCGYRIDMVVENRLLLELKAIEQVLPIHTAQILTYMRLASIDTGLLINFNV-TRLVVQRFKL--------------
GIV69802.1       -----------------MLHADITAEIIRAFYKVYNTLGYGFLEKVYENALAHELGRQGLSVRQQTPIQVYYEGRLVG-DYFADLMVAEKVIVELKAAEALRPEHEAQLLNYLKATEIEVGLLLNFGP-EPSYVRKIFTNDRKRKKQIRPCRSVS
GIX04292.1       M---RLPWNFPLGAMNEAELNRLTELIIGCAFTVSNTLRCGFLEKVYESALSHELRKNGLLVKQQEPIKVIYDGVIVG-EYFADILVKDSVIIELKAAKDISEAFAAQCLHYLKATGLPLCLLLNFGK-PKVEIKRFRL--------------
MBF0239770.1     -----------------MNENDLSRGIIGAAIEVHQELGPGLLESVYEEALCLELALRAFNVQRQVPVPISYKGQSLKTPLYLDVLVNGKIIVEVKATEKHNTIFEAQLLTYLRLTGLKLGLVINFGE-KRVVIHRVVNKL-------------
MBI2568436.1     MNHQGTKT--PRGTEVPESTDRVAHQVVDAAYRVHTTLGPGLLEAVYEVCLAHELTKRGLKVLRQVSLPVVYDGQQLDAGFRIDLLVEHCVIVELKAIESLLPVHKAQVLAYLKLSGHRLGLLINFNV-PLIKIVRLAL--------------
MBI4514359.1     MNSSVSSA-------QSVDRDPRTYAIIGAAMEVHRQLGCGFLEAVYQEALAIELESRAIPFRREVELPISYKGAQLSCSYRADFVCYDGVIVELKALGVLGGPEDAQVINYLKATGFSTGLLLNFGT-PRLEHKRFIYTKSTDDAD--GHR---
MBI4525151.1     ------------------MEDALTEKIIGAAIEVHRILGPGLLESIYEEALCVELDLRGVRCQRQIDVDVNYKGRTIK-GQRIDLLVAGEVVVELKAVATLPEVATAQVLSYLKATGLKRGLLVNFNA-PRLVIKRISL--------------
MBK6429950.1     -----------------MEDEALTEKIIGCAMKVHRTLGPGFLESVYENAFAHEMRKAGLRVECQVPIQVYYDGIVVG-DFSADMRVEGRILIENKAVLALNTTHEVQLVNYLTATGIEIGLLINFGA-ARLDVKRKYRTYRKAG----FGQDVQ
MBM3880409.1     M---DAN--------EKLLLKNEVYQIVGCAIEVLNELGHGLIEKPYENALVVEFGLKQIPFIQQPRFEVRYKGVAVG-EFVPDLIAFDALVVDTKVIEAITDHERGQMLNYLKITGHKVGVILNFKR-AKLEWERIVLERNR-----------
MBM3880844.1     M--------------NDLVYRDECYRIMGACFEVYKEKGCGFLEAVYQECLEIEFEMQGILFVPKQPLALSYKGRPLKQRYEPDFICMDKIIVELKAVSALADEHRAQVLNYLNATGYKLGLLVNFGHYPRVEWERIANTARETLPTARETRETH
MCI0624093.1     M-SFSTSGRGRVNSAKELIHKELSESIIGAAMTVLNTLKPGLDEKLYEKGLVIELRKRNHQVEQQRQFPVRYEGQLIG-TLTPDLIVDGLVIVDTKVVTAFNETHMAQMLGYLNITGLRLALLLNFKN-AKLEWKRVVR--------------
MCK6629897.1     M--------------AEILHKELAYAIVGAAMEVHRLLGPGFLEAVYQAALAHEFTLRNLPFEQFKKLPVTYKGVLVG-DYEVDFVVANQIILEIKAASSLHPKHEAQALNYLAVTGFRLAILLNFGA-DSLQHKRLVR--------------
MCL4302339.1     -----------------MDIEEVARNIVDSAIKVHRALGPGLLESAYQHCLIYELRKRSFEVKHELVLPVMYDGQQIDAGYRIDMLVEDCIIVENKAVDQLLPIHEAQLLTYLKLRNCQLGFLLNWNV-VLMKIKRMVNNLNR-----------
MCL5961068.1     MNHKDTKD-------TKARSDFLSSQVIGAAIEVHRLLGPGLLESAYEECLTRELTLRDVPFQRQVPLPVEYKGTLVDCAYRLDLLVADLVIVELKTVEKLEAIHDAQLLTYLRLKGLWLGLLINFSV-PVLRIKRLVNG-------------
OGH97201.1       -----------------MEFDELSNKVISLAIEVHKNLGPGLLESTYKQCLAYELAKADIKFIMEAELPVQYKDIKLSCGYKIDLFIEDTLIVELKSVEKMNPIYDAQILTYMKLSKVKVGLLLNFNN-KLLKIKRFVL--------------
OIP61732.1       ------------------MKEITEKIIGAAIEVHKTIGPGLLESAYEECLAHEMQLRGLKFDRQVPLPVTYKGVKLECGYRLDFLVEEAVVLELKVVDALQPIHEAQLLTYLKLGGWTVGLKNSPRS-LRLRGERSFYGTLVQGGN--TKRSSR
WP_120797040.1   -----------------MDENEIGTIVVDCAVKVHMRLGPGLLESVYETVLTYELVKRGLLVERQVVVPIVYDNLRFEEGFRADLIVGGKVILELKSVEHVTNAHKKQLLTYLRLTGLTVGYLLNFGE-ALMKITRTINGKLGGE----------
WP_258932063.1   M---NTNE-------REYIESDLTYKVVGCAMAVVNVLGHGLREKTYERALCVEFENQGLSFQQQHIYPIYYRNQHID-DYIPDLEVESRLIVEVKTVDRIVDEHIGQVLNYLRISDLEAGVILNFKH-PKLEWKKIVLQQGRKAK---------
consensus/100%   ....................s..l...hh.V...b..Gh.E..Yb.sh..Eh......h..b..h.l.Ycs..h...b..Dh.h....lll-.Ks..........Qhl.Yh.h.....sh..s..........+................
consensus/95%    ....................s..l...hh.V...b..Gh.E..Yb.sh..Eh......h..b..h.l.Ycs..h...b..Dh.h....lll-.Ks..........Qhl.Yh.h.....sh..s..........+................
consensus/90%    ...................p..s..lhsshhpV.p.h..GhbE..Yb.sL..Eh....h.h..b..h.l.Ycs..l...h..Dh.h....lll-.Ksh..h.....sQhlsYhph....uhllNa.....h...+...............
consensus/85%    ...................p.bs..llsshhpV.p.lG.GhlE.sYb.sL..Eh..psh.hb.h..h.l.Ycs..l...h..Dhhh....lllE.Ksh..h.....uQhLsYLphsshp.ulllNF.....h..pR...............
consensus/80%    ...............h..p.bs..llssAhcV.p.LGsGhLE.sYbpsLs.Eh.bpsl.hb.b..l.l.Ycu..l..sab.Dhhh.s.lllE.Kshp.l.s.apuQhLsYLphsshphsGlllNF....bl.hpRhh...............
consensus/75%    ...............h..p.ls..llusAhcVap.LGsGhLE.sYbpsLs.Eh.bpsl.hbbb..l.l.YcG..l...sab.Dhll.s.lllElKslp.l.s.apAQlLsYL+houhplGlLlNFs..sblbhcRhh...............
consensus/70%    ...............h..cpls..llGsAhcVappLGsGhLEusYcpsLs.Eh.bpsl.hbbQ..lsl.YcG..ls.sabhDhlV.splIlElKulp.l.s.acAQlLsYL+hoGhplGLLlNFs..sblchcRhh...............

# ; PDDEXK-REase

                 1---------11--------21--------31--------41--------51--------61--------71--------81--------91--------101-------111-------121-------131-------141-------151-------161-
OrigSeq          MNRTELQETQTPPAMVASSAQVSERMSRLPRKDTTPELALRRLLFASGLRYRVHERVTGMPRRTVDISFTRVRVAVFVDGCFWHGCPDHGMTPKANRDWWSRKLDGNRSRDAETSAHLSALGWVVLRFWSHEEPDAMAESVKACVLSRRLALGLKPVRRARGLT

Jnet             ------------------HHHHHHHHHHH------HHHHHHHHHHH----EEE---------EEEEEEE---EEEEEE---------------HHHHHHHHHHHHHHHHHHHHHHHHHHHHH--EEEEEEE----HHHHHHHHHHHHHHHH-----HHHH-----
jhmm             ----E-------------HHHHHHHHHH-------HHHHHHHHHHH----EEE---------EEEEEEE---EEEEEE---------------HHHHHHHHHHHHH-HHHHHHHHHHHHHH--EEEEEEE-----HHHHHHHHHHHH---------------
jpssm            -------------------HHHHHHHHHH-----HHHHHHHHHHHH----EEE---------EEEEEEE---EEEEEE-----------------HHHHHHHHHHHHHHHHHHHHHHHHH----EEEEE-----HHHHHHHHHHHHHHHHHH--HHHHHHH---
WP_239130011.1   MNRTELQETQTPPAMVASSAQVSERMSRLPRKDTTPELALRRLLFASGLRYRVHERVTGMPRRTVDISFTRVRVAVFVDGCFWHGCPDHGMTPKANRDWWSRKLDGNRSRDAETSAHLSALGWVVLRFWSHEEPDAMAESVKACVLSRRLALGLKPVRRARGLT
WP_234415357.1   SDAEIGAMTRADASSWASTPGVARSMRANTRRDTKPELAIRRLLHARGLRYRVDMAPWSNKRRRADIVFTRRRLAVFIDGCFWHGCPEHATVPVTNADYWRPKLARNIERDRETTAMAEAEGWRVLRIWEHVPPEDAVRLIVRALETGR-----------DG--
WP_240770227.1   TNGTLGGMSQQEPDRVSPGGALSRKMSTLARRDTAPELALRRALHARGLRYRVQMKVPGNRRRSIDVALTRVRLAVYVDGCFWHGCPEHGSTPRANSDWWRWKIERNRARDADTTASLEAAGWTVLRFWEHEDPSLAAVIVEAERRRL-----LEALTRPHGRG
WP_281063407.1   -----------------MNEGRARNMRANRRVDTKPEVQLRKRLHSMGFRFRKDYRIDSGTCVRPDIVFTARKVAVFVDGCFWHVCPVHGRQPTRNEWYWTPKLQRNVERDRRNDLALASGGWTVVRIWEHEALDYAAEVVAETLRSQSA----GQPHAQSP--
WP_101639483.1   --------------------------MSRQKAKNIAPKVELRRELRRLGLGYRIQLPLPGMPRRRCDVGFIGAQVAVFVNGCFWHTCRVHATVPSQNRDWLIDKLAANVTRDHESEERLSAASWLSIRVWEHEDMATAVSRIDEVVRRRRR-------------
WP_182766933.1   --------------------------MARQHRRDTKPELFLRRELHQLGLRYRVDHPLPGMPRRRADVLFTRAKVAVFVDGCFWHACPEHKTSPANNALWWTAKLARNVERDRETDAHLVALGWTVLRVWEHEDMPQAAVRIREAVRATGP-------RSVAGSS
WP_231559476.1   -------MSPEHAAGPALDAGVRKRMSRQRRDTVPELELRRALRALGLGYRVNAVLPGLPRRRCDVMFKGVKVVVFVDGCFWHGCPEHSTAPRNNADWWARKLSGNRARDRDTDRHMMALGWTVILVREHQDMTAAAGEISDLVRGLRG----EGPHKQVP--
MBI2825940.1     -------M-----DNLSQAQRSLTMARIRSRDTEPEMIVRRLIHHLGFRYRLHLSLPGRP----DIVLHRLKKVVLVNGCFWHRCKFGRPLPATNRTYWRAKLQRNADRDRLNRRRLRRQGWEVLVLWECWTRR-PD-VVAKRLHQF-----LQT-------
MBI3184671.1     -------MTAQRERTTFGALSRGELMARIRSSNLTTEQRLASLLRDAGLAGWRRFRVAGRP----DFAWRSVKVAVFVDGCFWHGCGRNL-SPRTNVEAWAEKIRRTKERDRKATQRLRQAGWTVVRVWECQLAKDPK-RCLDRIRGA-----LPD-------
MBY8951554.1     -------M-----DVVSKEVRSRMMAGIRGSNTMPEMKVRKLLHSHGFRYRLHRQLPGRP----DVVLPRYRLCLFIHGCFWHRCPYAT-NPKSREDFWRAKFEQNVKRDIRNRNELLQQGWRVFELWECGIRR-PE-AEIQWLLKA-----VPD-------
MCE9614799.1     -------MA-----DTISKAHRSWVMSRIRGSGTKPELMVRSLLHRAGYRFSLERALPGRP----DILLPRYRTAVLVHGCFFHGCRIAK-LPKSNTAFWRRKIAGNIERDQRVCRQLRHLGWQVAVVWECQVMADPK-KVLMHLAKH-----LER---AGGRP
MCK6484532.1     ------------------------MGRIKGRDTSPERIVRSILRGLGFKHRLHAALPGTP----DIVIDAAKTVILVHGCFWHRCPRGRSSPATRAAFWQAKFARNIRRDRQAARALRRSGWSVVIVWECQISRASRGRLTSRLRRF-----LDSRFRAAGAE
MCV7527921.1     -------MT----STWASTPGVRASMQSNKGRDTKPELLVRRLVHAAGLRYRVNARPEKDLRRTADLLFRPVKIAVLIDGCFWHGCPAHYTRPKANNAYWAEKVQRNRDRDAETTRVLTERGWLVLRFWEHEDPADVAARVISSVEARRL-------RRIAGAT
MDE2668028.1     -------MT-----DTLSPAQRSALMSRIRGRDTGPERALRRLLTDMGYRYRLQQRVPGRP----DIALPGRRKVIWVHGCFWHRCPKAT-LPKSRRDFWIPKLAANRHRDLVAQAEAEELGWDTLVVWECELRD-PN-RVVQELRNF-----LDE-------
MYE95019.1       -------MTSVSSLDTLTASERSARMALIRSTDTRPEMRVRRLLTHGMGYRYRLHKDLPGRP----DLVFRPRRKVVFVHGCFWHRCPRTR-MPKSRREYWRKKFEGNVMRDQQHRDALERMGWTTLVIWECETTD-AE-RLAALLRRF-----LDG-------
NLX98933.1       -------M-----DTFTKDKRRAIMQAVRRQRTRPEEALALLLRKAGVKFRRNRSLPGSP----DFYVPTQRLAVFIHGCFWHGCAKGRSRPKSNCSYWRRKITINQQRDQRVARQLRRLGYSVYTVWECQLTSNPP-RLLARLTVPLL-------------
OGW72083.1       -------MT-----DVLTKEQRSYNMSQIRSKWTKQEWVIHNYLKSQKIRHTMHPNIEGKP----DIILKDSNTAIFLDGCFWHKCPKCFKMPRTRTDFWKNKITKNITNDKKTTNRLRKNGYRVVRVWEHEIKKIEKSDLLRVFRIV--------------
OYP42210.1       -------MS-----DHLTKEQRHKNMAAIRSKDTKPEMIVRHGLWKRGFRYRLNKRLPGHP----DLVLRKYRTCIFVNGCFWHGCCK---IPKTNRDFWVEKIHRNKVRDKEEQQTLAKMGWHCITVWECELKPQKREVTLTDSIAFTLNHIWLQD----HGAK
RRJ96055.1       -------MS-----DIFTPSKRSEIMSKVRSKNRTTELRFIELMRRHGITGWRRSTLPGSP----DIIFPKQRVAVFIDGCFWHGCPRHFTLPENNMDFWLAKIGKNRTRDRRANAQLWKSGWRVIRIWEHALQKTTQDKLVARIRRA-----LEL-------
consensus/100%   ......................M..........c..h........h....p...........Dh.h...p.hlhlpGCFaH.C......P..p..hh..Kh..s..pD......h...ua.sh.hbpp......................................
consensus/95%    ......................M..........c..h........h....p...........Dh.h...p.hlhlpGCFaH.C......P..p..hh..Kh..s..pD......h...ua.sh.hbpp......................................
consensus/90%    ......................M.....p...sE..l.p.h...Gh.....p.....u.....Dh.h...p.hlhlpGCFWH.C......P..p..aW..Kh..N..RD......h...Ga.sh.hWEp.........h.......................
consensus/85%    ......................M...p.psp.sEb.l.p.lp..Gh.ab.p..l.u......Dlhh...+.hlhlpGCFWH.C......P.sp..aW..Kh..N..RD.psp..h...GW.sh.hWEp.........h...l...................
consensus/80%    ......................M...+.psT.PEb.l+pbl+..Ghpab.p..lsG.s....Dlhh...+.hlhlcGCFWH.Cs.....P.sp..aW..Kl..N..RD.psp..L...GWpsl.lWEpb....s...l...l...................
consensus/75%    ......................Mu..+.psT.PEh.lRphL+..Gh+aRhp..lsG.P....Dlhh...+.hlalcGCFWH.Csb....Ppsp.saW..Kl..N.pRDbcsp..L...GWpVl.lWEpb....s...l...lb...................
consensus/70%    ................hs.......Mu.b+.psTpPEh.lRchL+..Gh+aRlp..lsGpP....Dlhh...+.hValcGCFWH.Csb....PpsN.saW..Kl..N.pRDbcsp.pL...GWpVl.lWEpb..p.s...l...lc...................

# ; PDDEXK-REase
```

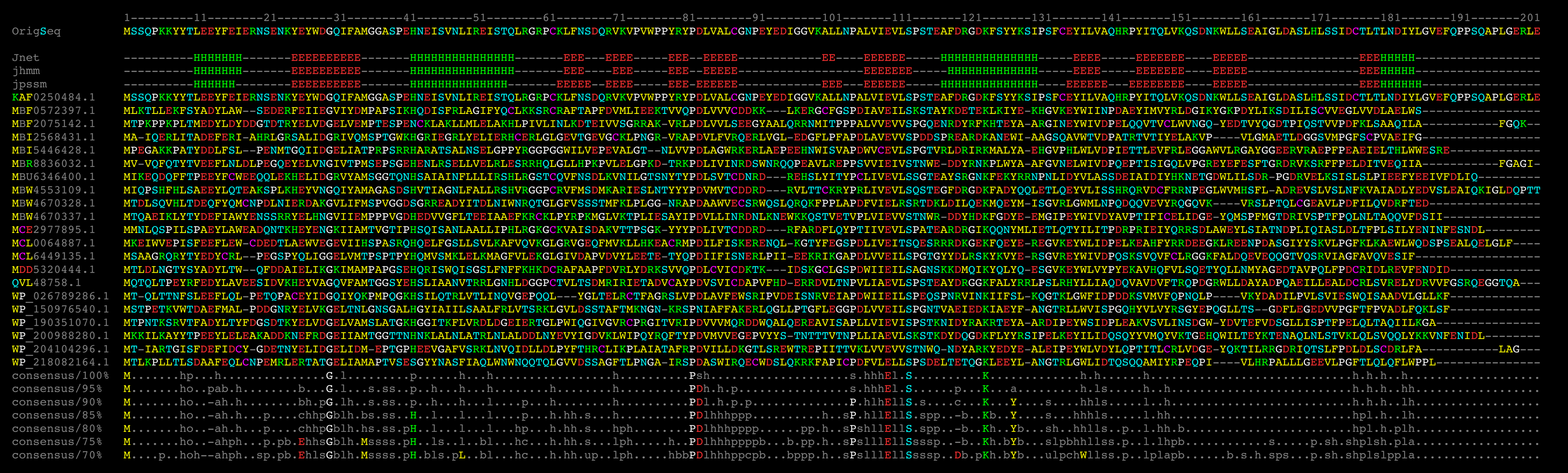

### ; Schlafen+HATPase+(w)HTH

[illegible]

### ; OLD+TOPRIM (Previously Classified OLD-ABCs, associated with ABC-ATPase centric Conflict Systems)

```
FINAL            --EEEEEEEE--EEEEEE----EEEEEEE-----HHHHHHHH---------------H-HH-------------------------EE-EEEE-EE------E---E-EE----------E----EEEE [illegible]
ALIGN            ----HHHHHHHH----------EEEEE--------EHEEHH------------------------------------------EE-EEE----------------H---------H-H----HHHH [illegible]
HMM              --EEEEEEEEEEEEEEEE----EEEEEEE----HHHHHHHHH---------------H-------------------------EE-EEEE-E--------E---E-EEEE------E-E----EEEE [illegible]
FREQ             --HHHHHH------EEEE----EEEEE-------HHHHHHHH---------------H-HHH-------------------------EE-EEEE-E--------E---E-EE [illegible]
PSSM             ----EEEE------EEEE-----EEEEE-------HHHHHHH---------------H-------------HH------EE-EEEE-EE---------------------------EEEE [illegible]
WP_052314664.1   LKHVTIHNFRGIIDASMN-FHDYALLVGANNGGKSSVIDCIRAFYE-----KDGFK----------FKK-DSD-F-PLKGSTD-EESWV-ELTF-GLI-DGEQS-S---LKEEY-QTP-E------MTLK [illegible]
WP_049228189.1   IKKITIDRLKQINNLTVE-FKPISIIAGVNRSGKTTILTTIA---------------C-SHENF--KKR-NIT-N-GILERHT-WQDFM-KFTV-NDC-QNDDY---K-YSIEY-KIG-S-N----IQTK [illegible]
OHA79738.1       LLSFRVENRRSVRLAEVSSVPPVMIIAGPNGVGKSTILYAIK---------------D-GAGLF------NPN-T-KILYQGP-NRVMR-KTNV-HRSWLTGAY---K-SFQDL-LSG-T-D----VSGY [illegible]
AFI05518.1       KRILRLHHFRNLTELNAD-FDGLVILVGENNVGKSNVLEGLKYF----NN-DNNFL-C-PIEDYEK-EV-YNN-D-NNQQQIE-KLSWV-ELKE-NQA-NKQAF---E-SFYPF-ENH-I-ESDKDIYLT [illegible]
AOS84045.1       LHAISVSQYRSISSKRIR-LGRSTVLIGPNNEGKSNILRGLMTILT-RDR-YRTIN---SARGYNWEKD-YPI-N-LQKKYPN-GETEI-ILEF-TLE-DKEYD-E---FYQEV-NSR-I-TGNLPIKIG [illegible]
CRH62638.1       VKSITINNHSRISNCTID-VRDNLILVGANGSGKSSIIRCVDLVLG-KTT-QQLYY---SIDNSDFENE-GQP-F-AVEVKLE-GLSED-ELSF-FPD-DYDVI-D-N-LTVRM-EAA-L--DKDDLTIR [illegible]
WP_026389839.1   ERVLKFKNYRNIQKLNHT-LDDVVILIGPNNSGKSNVLSGLKAFAN-KQI-NSEDIND-GFYELSDSFK-NNK-Y-KVNFKYE-KNEYS-LNKI-SDN-AEERI---K-SSKEY-KIA-W-E------EK [illegible]
KPL80624.1       LKSITIQKFGPFFEATLI-LESITVLTGRNDTGKSALLRAIQLLYS-----RKAIV-E-DDANIDW----LSE-N---TVSWK-DDPDI-RCNA-IFT-TKNMQ---Q-YVTGS-NIT-N--ETGDIELQ [illegible]
OGP68124.1       IKDITIENYRSITPQ-LT-LGKYNVIVGPNNAGKSNILRAFQLFFN-GHV---------DSLPYDASVD-FPK-A-TNLHNAQ-TRITV-TMQF-DPS-KDNKI-DRKVTESSQ-KRL-S-NNLLRLRLE [illegible]
[illegible]
consensus/90%    ...h.lp.hp.h.p...........hhG.N..GKoslh.sl..h. [illegible]
consensus/85%    h..h.lppac.h.p..........hhhG.N.sGKoslh.sl..hh.............................ph..........h. [illegible]
consensus/80%    lpplplppa+ph.p.......p.shlhG.NssGKosllpul..hh..................sh..................h...ph......p......h. [illegible]
consensus/75%    lpplplpsa+ph.p......sphshllG.NssGKosllpulp.hh..................sh..................h..hph......p......h.................h. [illegible]
consensus/70%    lpplplpsa+shpp.....hsphshllG.NssGKoslLpAlp.hh...............p..sh...pp..............p.....h..hph......p......hpp...p.........h.h. [illegible]
```

### ; BREX-systems OLD-ABC (TOPRIM-linked); Representative_1;

```
                 1---------11--------21--------31--------41--------51--------61--------71--------81--------91--------101-------111-------121 [illegible]
OrigSeq          MALIEGLKIQNYRSLQDISMPLTPLTVVIGKNGVGKSSIFDAFGFIADCMETDLETACNARGGYDKLISSGTDQPIMITIYYRESRDRPITYEVSIKEETDLPIVLSERLRQRASGRPRSFLHLENGKGT [illegible]

Jnet             ----EEEEE---------E-----EEEEE------HHHHHHHHHHHHHHH---HHHHHH-----------------EEEEEEEE-------EEEEEEEE----EEEEEEEEE-------EEEEEE----EE [illegible]
jhmm             ----EEEEE---------------EEEEE------HHHHHHHHHHHHHHH---HHHHHH-----H-----------EEEEEEEE-------EEEEEEEE----EEEEEEEEE-------EEEEEE----EE [illegible]
jpssm            ----EEEEE---------EEEE-----EEEEE-------HHHHHHHHHHHHH----HHHHH---------EE-----EEEEEEEEE-------EEEEEEEE----EEEEEEEEE-------EEEEEE----EE [illegible]
HBT88454.1       MALIEGLKIQNYRSLQDISMPLTPLTVVIGKNGVGKSSIFDAFGFIADCMETDLETACNARGGYDKLISSGTDQPIMITIYYRESRDRPITYEVSIKEETDLPIVLSERLRQRASGRPRSFLHLENGKGT [illegible]
ABD40829.1       -MKLKNICIKGFKSLEYVDLPLNHLTVCIGANGSGKSNLLSAFYFLHRIAEGRFSYAIAKAGGAHSLLYNGPDKKIEFTLTF-----EEFVYHAACEPSDDNLVFIQEDIKRPQ-----GVREVSHAKN- [illegible]
AVG41561.1       --MLTAITLEDFKSYRRATLPLSALTVLIGANASGKSNVIEGMRLLSWLAQGKLSAIRFAVQSGDQVVRGKLEESFGLGVHTNATE--WNHLSMRLSRRADGLHITAESLTHD-GAKVPLYTLDQPSSGQ [illegible]
CDD75189.1       --MIKCFTFENFKSFEKAELNLEALTTLIGTNSSGKSNAIEGIAILAKAATGELNTILDGQKSTGAVVRGGSRSAFKLGCLISLDD-YDLLYEMKVG-VGERVGIEEEGLYLVKGPKGNKIFKTKKVEDG [illegible]
EFK06062.1       M-RIKHIRIEGFKSIKNEPLMLGAINLLIGPNGSGKSNLISFFQLLNYVMSEGMQTYVATHGYANSFLYLGSKPRICGEVEFAT--GGTNRYCFTLVPANDILIFSEESIFFQA---HPWPIENTLDP-- [illegible]
GHT84069.1       M-LIKSIKLNNFLSFGDETVQLRPLNVIIGPNGSGKSNLLEAIDLLRNAPNE-LTKPIREGGGVLEWLWKGAKPNASLDVNIECLN-YSLDYGLVFCAADQRFFISAEKIKTDNKVSKFFYWSDRGKGAI [illegible]
[illegible]
```

[illegible]

### ; BREX-systems OLD-ABC (TOPRIM-linked); Representative_2;

[illegible]

### ; BREX-systems OLD-ABC (TOPRIM-linked); Representative_3;

[illegible]

[illegible]

consensus/90% M.lppl.lpsa+.h....h.h....slhhG.NusGKoslLcuhs..hs.h[illegible]

consensus/85% M.lpplplpNa+.h.p.ph.hp..hslhlG.NusGKoslL-uls..hu.h...h[illegible]

consensus/80% M.lpplplpNaRsh.phphphp.phslllG.NGsGKoslL-Alsh.hu.h...h[illegible]

consensus/75% MblpplplpNaRshcphplphp.phslllG.NGsGKoolL-Aluhhhushh..h.s[illegible]

consensus/70% MblpplplpNFRshcphplshp.phslllG.NGsGKTolL-Alulhlushh..h.s[illegible]

### ; BREX-related (DUF499 centered) systems OLD-ABC (Associated with TOPRIM/REase/HNH)

```
                 1--------11--------21--------31--------41--------51--------61--------71--------81--------91--------101-------111-------121-------131-------141
OrigSeq          MIEQITLKNFRCFELLDVTFHPQMTVLIAPNGAGKTTILDAVRIALWPFVKGFDLGSQIEDVRLERREQEMEPILPSQIAAQASGPFQWQQSRDQVKQSTQLLTKSAKSYQSVIYEAIDLPLVVYLTGRLWYQGRYASELDT[illegible]

Jnet             ---EEEEE------EEEEE----EEEEEE-----HHHHHHHHHHHHHHHH------------HHEE---------EEEEEEEE----EEEEEEE-----HHHHHHHHHHHHH-------HHHHHHHH[illegible]
jhmm             ---EEEEE-------EEEE----EEEEEE-----HHHHHHHHHHHHHHH------------HHHHH----------EEEEEEE----EEEEEEE----EEEHHHHHHHHH-----------HHHHH---EE[illegible]
jpssm            ---EEEEEE----EEEEEE----EEEEEE------HHHHHHHHHHHHHHH------------EEEE---------EEEEEEEEE----EEEEEE------HHHHHHHHHHHHHH------HHHHHHHH[illegible]
WP_201871425.1   MIEQITLKNFRCFELLDVTFHPQMTVLIAPNGAGKTTILDAVRIALWPFVKGFDLGSQIEDVRLERREQEMEPILPSQIAAQASGPFQWQQSRDQVKQSTQLLTKSAKSYQSVIYEAIDLPLVVYLTGRLWYQGRYASELDT[illegible]
MBI2833274.1     MIKQLVLKRFRSIPSERMTFDN-PTIFVGRNGSGKSNLVSVFSFLAEAMASPLQAVFDKAGISAVRNRTSYPPNLGLRVDFGRMNSGFYAFEVRALYVVREQCSVG---------VNGMKPA--LDPASLSLPL[illegible]
MDA8417185.1     LLKELKLYDYKKFEELRIHFEKNLTVLVGVNAAGKTSILDAVIKALTWIKAGLEAEGRDSDIRIADYSK-------TCTTFEFDGDGVYEYCLAKSGVRQLMYRRV---------YPIELPIFVYFVGRTWISSKV-DKTTL[illegible]
WP_129673426.1   MITRLTGRGFKGLREFDIQ-SK-INVLIGANGTGKTNFADLIDFFSLIARRGLVEAFGEYGLEQVRTRTPRAFTMRFRLELGEKDRVSYGFELSRAIIKHEVLYER---------IVRPSPNLEEITERLFLDE-------S[illegible]
MCW5668672.1     MLERLHLRNFRCFEELTIDFGKRLTVIIAENGAGKTAVLDAIAIGFGRYLTKLPGIAGDTDLRVADGER-----REMMLAWEATRPVVWASGRKRDVRGLKEL DDF---TLGLVQQAYFLPVIAYYTNRAIREEVQ-RRGFK[illegible]
```

[illegible]

consensus/90% hl.ph.l.sa+sh.p..l.h.....hhhu.Nu.GKoshhchh.h[illegible]

consensus/85% hl.ph.l.sa+sh.p.plph.....hhhu.Nu.GKoslhchh.h[illegible]

consensus/80% Ml.plplpsa+sh.phplph...hshhhu.NusGKosllchl.hh[illegible]

consensus/75% Mlpplplpsa+sh.phplsh...hshllu.NusGKosllcul.hh[illegible]

consensus/70% MlpplplpNa+sh.phplsh.s.hslllGsNGuGKoslL-Al.hh[illegible]

### ; OLD-ABC+TOPRIM

```
                 1--------11--------21--------31--------41--------51--------61--------71--------81--------91--------101-------111-------121-------131-------141
OrigSeq          MIIDQIQISNFRSFDNCTVDLNNIQAIVGRNNVGKSNLLKAIRLFMEASKRLLDEDCFHHHDTEEPIRIRVRFVDLNEWERQKFEEKTGEDDYSIDRFAIVREPEESEIALTVNGLDFGDELGTNPRGYKGVLKGGLPEFILV[illegible]

Jnet             -EEEEEEEE------EEEEE---EEEEEE-----HHHHHHHHHHHH-------------------EEEEEEEEE----HHHHHH-E----EEEEEEEEE----------HEEEEEE---------------HHH----EEEE[illegible]
jhmm             -EEEEEEEE-------EEEE---EEEEEE-----HHHHHHHHHHHH-------------------EEEEEEEEE----HHHHHH--------EEEEEEE-----------EEEEEE-----------------H------EEEE[illegible]
jpssm            -EEEEEEEE------EEEEE----EEEEE-----HHHHHHHHHHH--------------------EEEEEEEEE---HHHHHHH-EE--EEEEEEE---------H---HHH-----------------HHHHH----EEEE[illegible]
WP_259195585.1   MIIDQIQISNFRSFDNCTVDLNNIQAIVGRNNVGKSNLLKAIRLFMEASKRLLDEDCFHHHDTEEPIRIRVRFVDLNEWERQKFEEKTGEDDYSIDRFAIVREPEESEIALTVNGLDFGDELGTNPRGYKGVLKGGLPEFILV[illegible]
MBH2008930.1     IAITDVRVSNFRSLANVEVELNDLTVLIGANNAGKTSFLDAMFAAIGSGRKLLGQD-VRLAPKERKITIDLRIRPVPDGKAGSFWWGSGIALTPIRTSLLWSGS-KSEYVVERNFLKE------QTTPVRAAQ-LEPIALHL[illegible]
WP_007159181.1   VYLSALKIENFRQFGQLNIQFNGVTALVGENDAGKTTVIDAIRYVLQTRDARLQID-FHIATQAEAITLRCTLEGLSLAELGAFA-------LYVHWS-------AGRIKWVDISVRSGEYG--EGPSLDVGV-RQLLATAYI[illegible]
BBM69557.1       MRLKTLVLRNFRGYKEVRIPIHDLTAFVGKNDVGKSTILDALAIFFD----KFDTD-----AEDMEVRIGCVFDELATSTTKLQE--------PRVVARAMHPAAENSDLKRRLRQKQVDATG--IDLRSNPSI-RKALRL--[illegible]
```

[illegible]

[illegible]

```
# ; TOPRIM Nuclease (Associated with OLD-ABC)

                   1---------11--------21--------31--------41--------51--------61--------71--------81--------91--------101-------111-------121-------131-------141-------151-----
OrigSeq            MPKKCILIVCEGAETEPNYFESLRRYLKLSTVQIRKDRAGAPISVVDEAQNQIKKRERSFESVWCVFDVENPHHNQTFDRAVFLADQNNYQLAVSNPAFEFWYILHFERTTRPFANDELKEHLRHHIPRYVASTQTAVQHANSIHLLVQEMIEMSPSG

Jnet               ----EEEEEEE----HHHHHHHHHHH-----EEEEE------HHHHHHHHHHHHH------EEEEEEE----HHHHHHHHHHHH-----EEEEE---HHHHHHHHH---------HHHHHHHHHHHHHHHHHHHHHHHHHHHHHHHHHHHHHHHHH---
jhmm               ----EEEEEEE----HHHHHHHHHHH------EEEE------HHHHHHHHHHHHH------EEEEEEE---HHHHHHHHHHHHH-----EEEEEE---HHHHHH-----------HHHHHHHHHHHHHHHHHHHHHHHHH--HHHHHHHHHHHHHHHHH---
jpssm              ----EEEEEEE----HHHHHHHHHHHH----EEEE------HHHHHHHHHHHHH-----EEEEEEEE------HHHHHHHHHHH----EEEEE--HHHHHHHHHHH---------HHHHHHHHHHH-------HHHHHHHHHHHHHHHHHHHHHHHHH---
OGN72035.1         MPKKCILIVCEGAETEPNYFESLRRYLKLSTVQIRKDRAGAPISVVDEAQNQIKKRERSFESVWCVFDVENPHHNQTFDRAVFLADQNNYQLAVSNPAFEFWYILHFERTTRPFANDELKEHLRHHIPRYVASTQTAVQHANSIHLLVQEMIEMSPSG
MBR6930694.1       M-MSKIGIVSEGVSDY--ILKHILERYLKKDVYAIKQVGFGWQGVFDYISLIVEAVKEGCEFVVVQIDTDKEAFWNEVKAKLVESVE-KLVFAICIQEIECWLIPFVSSSESEQDTNIVNKHIA--QKLYSRKIGECSQYNYGFARFVEQMDAIAKAL
WP_270679268.1     MSKPIVVLAGEDSNDR-TSMKYLLQKICPRVVDILKAATPE-GALRKRVQTLVRMARVELACVFVHEDFDYEKTHTRVERELHKALG--GHYVLAVEEMEAWMLLFPDALTAV---KRLKEALMNERRKYFKAPEKAQGSNRSWERFSDHLG------
KFB72163.1         M---HIEILVEDSSGR-KLLQVLLPQLLGHTWRLTKADPAK-RILLDQLPRLLRGYGKTPGVVVVVLDADV--FLAELKAAADACHP--TLFRLAIEEIEAWYLGDQDALLAAYPRDS-AELLADAKAGWGPL---ERNLSPSFGKLRDGLRRLLSDA
MCH8539503.1       MPRSSILVVTEGENTEPVYFDFIRQRFAAPTVELVGAGRGDPRALVDAALKVQKDRKRDFDAIWIVFDTDL--DPQKLHDGIAYAQSKGIQIAHSEPCFEYWLLLHGTFTTAMENVPFLEETF---WRNYEPIIALAVKHAKRVDRLIHAINEAVSLA
CAQ34955.1         M-ITSIVVTGEGSSDMGGLLVRLLRNILPPTDWITKKLKKGFVEHANRATAGYAKNNG--HQLFYFHDTDFTNLHQSITFGFDGVDGVRGIPMIPKPTSEAWLICGRKSDPYG---ETQKKVLAQLEQQY---IDVTRINMPSYNQFKTDLTSAIEVI
NOT11400.1         MGSDKVLIVCEGEKTEPLYFKELIDHYEIHTANVRGQCGSDPVSVVDHGIQLYQDEKSPFDRVYCVFDRDN--YLQALDKIKHTKPDDTFFAIPSVPCFEYWLLLHFNYTTAPYTSNAVVKELQRYWPEYLQKLASAKQNAVRSHLVVDYLQKIK---
MCG3135900.1       MVRSISLIVCEGETEQ-EYFEAARIKYGLTTAEIINTIGPAPISVVECAEGKCAEGG--YDKVFCVFDRDR--ARKKIKTLADRKKKLAIREAISIPCFEFWVLLHHERTDAPFTKADVIDRVRNHMPGYMARVEDALGNADWIHHVLRHFAQVSNQK
MBQ6275507.1       M--ASIALICEGVSET-NILTHIINRYLGDSIVVNKQTTEGWLQVLNHCDDIIKDIMAANDYLVIQIDTDDDILYERVCDRIRRNISNKILFAICINETECWLLPLYNDVKKRSTTYILNQKLA--VKAYLKKIYRIAQYNYGFKMFIQQLDCISDDS
MBI5658239.1       MGSSRFLIVCEGMKTEPYYFDEFCQVHQLPRVRIAGEEGSSPDCVVAYAENLFEEDAPRYDQVFCVIDRDYQAAIQRIADL--NAEGKPFVVIPSVPCFEYWLLLHFTYTRQSFHAGTVIQELRKQFAGYKELTGTAIKHAQRAHHLVEKLQELAATH
WP_028324161.1     M--TDLFIASEDALSE-SVLERLVE-----DANEVRMGRKGNGYLKQRILEFIKTAHA--IPVLLMTDLDC--APVLIDDWCGRHPLKNMLFRVAVREVEAWLLADREGFSKYSGVES-KQMLMKLGFGYSRFPVKAAENADSLARTRNRLREIGKQG
OQB96945.1         M-KTIIGIACEGDVTEYRYFEEFKERYTDSRIHIEEASESAPEHVISSIDAFKKSAPD--DEFWVVIDYDR-WGEKKLSEIAQQASQKSYHVAVSRPCFEAWLLLHWTGIDAMLSEKS-IDEIRRIHGHYGEYAERAIKNARSLYQLCERIK------
CAH1748217.1       M---LLLLSGEGPSDLGRLVDQIIQDVLKVPTRLRKKQGQETGYFMKMAWGKLALQIADEDAVVFFRDTDWAEKWHSIERGFGRAGF--GVPMLPKPTSEAWLLCAAKGSPYQ---EDLKKEL---ARAFREWFEPSAMEMGSYRAFRERMVEVLTTV
WP_054541290.1     M-ITSIIVSGEGRTDIGGLIERMLARFVPNPLEVSKTLHQEHNGKYKHTRAGLATER---GQLVYFIDTDQQKIVQALNEGYIKGGF--GVAMVPKPTSEAWLICHCQTSPYA---ETRKKELERFGVGYFDKLDIDQLDMPSFNRFKDDLKSAIAAT
WP_008906817.1     M---RVIVSGEGPSDMGAWLQRLWAALLNVPGAVIRSKAAETGYYFNNALGLKAKELGPDEAVVLFHDTDWNTKWDSMMDGFGAAGF--GVPMVPKPVSEAWLLACTQQGHHS---EEIKALL---AQAFAEWENWPLQTMPSFKAFFDRFHEVAELI
OJW08010.1         M-STKVGIIAEGPIDY-HLIPALLERIARRPDDLVRKRGHG--GVLDMVRKMVEALSEVYDAFVILLDRKTRAVQEEVRQLISGHDR--FVMGVAIEEIEAWWLGDRRNTLAWTNLHCLKVTLDELDRFYAEELDDIRSQCSAYVPFTQHFRNAANLA
MBF2761743.1       M---TKLISAFGEDQYHQVIDGLIRRISMKNVEVRRNMYGGYGRVVSEAKKYLRNVDQAVDLVVVVTDANYNRRSKDIS--IDKNTI--VVCAIPDPHVERWLLLDGAAFKKVYGKEKKKKIL---SDAYVSELNRAARSDVSLKRFLDDLR------
MBX9598968.1       M--IRLVFWLEEYSAQ-VLLEGLIPRLFVANLNLEK------SDLERNIRKIHKYYSPDNLQVLILRDQDQ--VKQKLLNMVSDVDD--VLVRIACHELESFYLADLAAVETALQIDT-KDLLDKLAGKYGKVIKN--QRSPSFSHLVEAIQKLLVAC
WP_138262158.1     MPKERYYIFCEGEQTEPKYFEGFKNFIEDNLVLIDEPCAAETLRVIGQAEKYVKQNGIHKGQVWCVYDKDF--PAKDFNGVVERSEKENYYAAWSNECIEYWFILHFAYYESNNHRKEYVKFLNDKIGKYLNHPKQACKYAKKIYMLVEELAKYLPED
MCQ2228646.1       MAREIPTIIGAGITEQ--YFEHLKNLF-GYRLKIRRFFGNE---NIYYIEQRIRQVLDEGGRAICIFDADA--EKKKLEELKRKYGNPDVLICDSLPSIEYWFLLHFRKTAKFFTERELSSYIR----NYIQMQNDAITRAKEIYKAIEELKSK----
MBV5328346.1       M-SDEVIAIVEGKTEQ-VFIEKILAPYLGHKNIGVR-----FSRIKRDLEGHLKQRT---DTVTTFIDKEPAKIAKTINDAFSEQHACRFIPFVAVHEFEALLFSDAGILAEQLGIEAISKRLWSANGKFAKQIEKMRSACPLFNEWLNILESIPRGV
WP_099191958.1     M---RSILLCEGKTDA--ILSYYLGKVKGKKLNIIWAVGGK-DNFDYAISKIIDINKEVFNKIIILRDRDD--ISNDLDKSFSDKECDELTLIIPFDKLETFLLDAISELGDE------EKVLKSFLTKYLRNWEK-------YKSVFKKLEEI----
MCF8089784.1       M--PRLHITTEGQTEE-RFVKEVLAAYLGGKGVWRRESA--YAAVKKDICAWMKEDRN--PDVSTMFDKDPYRRVSLLEKALSADINTRFVSYIQLHEFEALLLADPEQLKWEFLEELISKRIIAEIHEYADKMETLCRRCRHFKEWIETLVALGKG-
PIV05609.1         M-GEKILILSGEFTEE-SFIRDVLHPHLGNKGVITS-----YGKFRKEIKMLLADTS---ACVTTMIDGDIFERANRLEQAFARDVDRRFVPYLSLHEFEALLFSSPEEIAGIAPDEEIAARITGLILAFVGRLTKIRNECRHFDAWLTKLEEFGE--
MBA3046425.1       MRDHKYVIVCEGEKDEIDYFDGFSTR---SGIKIMHGKCTDPKHIIEFAIEQKKKDNG--DGIWCVFDVDE-NQNRELQEAKANAKRKDIKTALSNPCFELWLLLHDEDINASFSRTEIKRKLKRYIPDYKIKLPVAIERAKRKYEVVEEIVKVSKKN
EFK06075.1         M-SHQIIFYVEGYAEL-EFVNEIIGPHLNAIGLVRK-----YGPIKKDLQRLLAQYGG--TDFTTLLDADPLAKVKAIEDAWRNDINYRFLPNLLLHEYETLILSNPASLLAGYPGEDISKRIIRANIRYVLELNTIRKECPKFDSWVKKLESLADN-
AOS47542.1         M-TVHPQVFVEGNTDE-PIVRALMT---ATGWRI--FCAKGSGNIIRSITKHAEAA----RQIILFLDSDPVDMRKDLEKELTHIPA--FVLRIVCTCIESWVLADCEGLASFCGV---KNELLKVSDDYADLAERAAQNNDSLRRAIARLKDLRARL
MCF7790536.1       MKNTTYYICVEGRATEKEYFNIFKS---DSTIDVQNKSKSHPKHVLDKLKKHLSENGSKYLKAWIVVDRDR--SEEQLNQVFEECGIKKYNFALSNPCFELWLLFHFESPSSK---RKCRNRLKKFLPNYKPEINNAIDNAEAKYRLVKEIYQE----
MXW95269.1         MPQPRFVIFCEGELTEPLYLKAFAALDEVGIATLDRGMGYEPRRLVEEARAVKRSERTGATEYWCVFDVEAPTQHARLKEAVQMARDNGINVAISNPCFELWLVLHHAEHERWIDNETCRGLLHEHGASYMPRRQQAIGRARRLYRLLEAVD------
TLD46496.1         M--TRLLMLVEGQSEE-VFVKRTLTPHLAQHGVFSN-----WSQIRRSLLPLTRDG----DAVSTLLDEDPREKVVALQERFAVELNQRFIPFLALHEFEAWLFSAPDAVEAHFGKELIKARLQSLVGGYLEKIAAVRAACPHFDCWLNRLEALCGGN
WP_062419289.1     MKSQRFLIVCEGTRTEPQYFERFRP-----GLVVREGVGMNTLRLVDEAINCCDED---YDQVWCVFDKDD-FPIENFENAIQRARERNMHVAYSNQAFELWYVLHFEYLNSAIDRKAYMEKLTQYGFKYQCKMDTAIRNAERLYQLVIALREQAKPL
MBK8920120.1       M--KDVIIVGEDPVTR-HVIKKLLQVTCPEGFNIIREDPVRGSEVKKLTPNYNALAAS--FPIILLADLDC--PPTEQAAWLNNQPKPHFIFRFAVDEAESWLLADRRGFAHFLGIYNLKTSLRELSSEYLPFVQTAIENSYSLQKMVRRLQEWCTQP
MBK6715973.1       M---KLMLSGEGPTDLGALVDKCCHMALDGGDVVVVKRPAGLGGITTQAWGILAVEERKSGRVILFHDSDWSEQVAAIKQGFARAEC--GVAMVPRPKSEGWLICALRQPHYQ---EESKAMLN--GGEYADWIDPRRIDMPSFAAFRRELDRACGVA
MBX7052697.1       M---DIYIVGEDPVTR-EVLRRIVQFCVPHEFNL-KDEPARGTEIIKKLDNYIRLAQV--HPVLMLTDLDC--AKIFVEELLHEKKV--FSFRVAVTEAESWLMSDRVAMARFLGVQDNSEMMMNIGPEYLPFIEAAMKNSYSLRRTVNRIRIILENL
MBN2191665.1       M---IIHALVEGASEK-AFLEPWAKRLLKHDLRAGPARRDQ-RGLLDQLPAKLAAFGSRQERVLVLVDADN--LERELEVLKARLDP--VIFRFAVEELEAFYLADQKALAAAFPDDS-IELFGRV----ASKIKAAESRSPSFKKLCNGLQELVASP
WP_226752734.1     M-ILHLIATGEGPTDIGILIIKLLENCLPQDFTHTKKAVKGHLEHAKRAAAEFTKENPETWSLVYFHDCDQANRVKAVLSGFKAVGYKHGVAMIPKPTSESWIICSVKKNPYC---EMDKVVLGNANQDYIAEVNR-LEDMPSFSQLKSNIIAVVKNA
WP_058557218.1     M---RLVLIVEDRADA-EAASMLADRVFIIDAETLHGRGLGPGAAARKALLLVIQTHDNIQALVLVRDMDE--RRASMESALADKLA--VAIAAPDPKREAWILHGFEPQNDNLGFERLKLVIERL--EYTTLLRERGQALTAYLAEVERLLPLLA--
MBK7385731.1       M--EPINLAYEDELSL-EVLLRLLNSPNVRCFS-VCFHGRGYGYLKKNITRFNEASKG--MPYLILTDLDR--APLLINEWLPQTRNPNLIFRIAVREVESWLLADDAGFARFLGIDD-KQTLIALGPAYIPFLEKARWHSHSLEKALRALETFEPQW
HCF56389.1         M-VQRLHFLVEGPADV-SLLENLLPRLIPHAWEIAKVDSSR-RGLLDNLPMKLRAWGVLDDRVVVLLDVDG--LLNRLKRMQQALAS--TIFRLAIEESESWYLGDRVAIREAFPKDS-RELFMQV----GKTVNEKANVSPSFVKFCRRVRELAGEH
MCE7937789.1       MDHRIVQPIVEGQGEA-SAFPLLLRRLAAGAWDIQQLVQKD------SLQTAVRVAGC--ASILVLFDADD--LAPRLEGWAREAAG--CHVVMANREYEAWLLASIESLRGTCGIET-KGELDARGNTYTAKLASAYRACRSFRKLVRAFGGVLQAG
ALO95601.1         MARPPILVVCGAKNTEPAYLRGLLASVDNRAVAVIKVCDQDPVSVVRHTVT--ERDRGHYHQAWCVLDVDE---FAHLDEALRLAADEDVEVALSNPCFELWLLLHHRDHRAEVAGQQAKRHLVAHHPGYRGGWPHAVERARELWRLVREIVPAHLQS
WP_208624500.1     M--VKIGFIVEGDTEK--LIESTAFRSWTKANNIINAKGGG-NLLPHHIEPMLAQFHAAPEHIVILTDLENEADIDSVKARITTTHT--EYIFVAVKAIEAWFLADTLALQRWLKIEN-TERLKALKPAFCRRIAQAANHPACAKLFHDTLLSLGSNQ
HCB0569862.1       M-LQTLLVFGEGPTDLGCLVHRLLLRHLPHAIPSVKKTKPGYLIHSKRAAGTLAQQAQRTHEIIYFHDTDPLDLEHAVNEGFRASGFEKGVAMIPQPTSEAWLICHCKPNSYQ---ETQKQMLGNYKPAYMTVIELDRLDMPTFNRFRHALCNAIDQI
MCY3637187.1       MARRTVLVFCEGRRTEADYLKALKQQPEVVAIEIDHAAGFDPLALVRRAAAAHKDAGSEVDEVWCIFDVEQPKKHSRLNEAMGLANDLGIQVAVSNPCFELWLLLHFEDKTAALTTSAAKRELARHGSMYMPSRHEAADRARSLYRFLSAVEGIREDL
OQA03649.1         M--IRVHIFCEGQTEE-TFVNNLLFLHFQNAGIIST-----YGKIKRQVDIKCKEDP---SSVTTLIDSDSMQKVIFLESKFDSDISKNFIPNIVLHEFEGLLFSDPTAFNRWFGEEDISKRILSICKNYAEKLERIRSECAHFNAWLNKMENIGGHT
WP_011117923.1     M--KKIVSFVEGEGDA-EAVPFLLRKILSQAFDAVNKLRKNFRDWHNKIRAAMKQPNV--GGILLLLDGDAMTVARELAAEALKVGA--VACVFACLEFESWLIAAAPGMGTLPDGDSLKAWLNTVPGGYTKALEVRAANLRSFRRLENALAELVAAV
WP_176976533.1     M---LILVSGEGPTDMGRLVDQIAETSEGDAELMFRRRGKETAYYFRNARAQIAAELKRWEPVILFRDGDWRDKWDSMCHGFAYEGC--GVPMIPNPKSEAWLLCALKSFQPY---EQAKAQL---EAALAELVNAARIEMSSFVKFRRRLQAVLTEP
MBD5181934.1       M--KRLIIVCEGPTEH-EFCVNVLAPELLKADIYVP-----WSSIKRQIEMHLHEK----DAVSLLVDKDLSTRLQFLCDGMKADIESRFIPYMQIHEFESLLFSDINVFKDNFDDEEISKRLIEAIPGYADELQKILDKCPLFSQWFLTLQSI----
WP_241209994.1     MGKQSFLIVCEGVRTEPDYFKAFRT-----AATVKVGQAMNTTSLVSKAISIRDADQRTYDQCWVVFDKDD-FPAKDFNEAITLAEKNGFKVAYSNQAFEYWFLLHFNLYKGALHRNSYADMLSRLGIPYLHLQSQAIKNASAVHLLVEELNKFI---
consensus/100%     M................h...............................h....D.p........................E.hhh.................h.......................h.......
consensus/95%      M......h.sbs..p...hh..h..........................h....D.-.......h.................E.hhh.................h......h...............h.......
consensus/90%      M......h.sEs..p...hh..h...............h..........h..h.D.-.......h............hs....E.hhh.................l......a.............h..h.p.h......
consensus/85%      M....h.h.sEs..pb..hhp.h.........h.........h.p.h..........hhhh.D.D......hp..h............hs..p.E.ahl...........p.....l......a.............h..h.p.h..h....
consensus/80%      M....h.hhsEG..pb..hhp.h.........h.p.........h.pph...........lhhh.D.D......lp..h............hs..p.E.Whl...........p.....l......Y.............h..h.p.l..h....
consensus/75%      M...pl.lhsEG.spb..hhp.hh......s..h.p........h.pph...h........lhhh.D.D.....p.lpp.h.......h..hhs..phE.Wll.p.p.........pp.ppbl......Y..bhp....ps.ph..h.p.l..h....
consensus/70%      M...plhlhsEG.ssb..hhp.hh......sh.l.p........l.cph..bhp........lhhhbD.D.....pplpp.h.......h.hhls..phEhWllhp.p........cp.ppbL......Y..bhpph.pps.ph.phhpplpph....
```
